# The physical form of microbial ligands bypasses the need for dendritic cell migration to stimulate adaptive immunity

**DOI:** 10.1101/2020.03.03.973727

**Authors:** Francesco Borriello, Roberto Spreafico, Valentina Poli, Janet Chou, Nora A. Barrett, Lucrezia Lacanfora, Marcella E Franco, Laura Marongiu, Yoichiro Iwakura, Ferdinando Pucci, Michael D Kruppa, Zuchao Ma, David L Wiliams, Ivan Zanoni

## Abstract

A central paradigm of immunology is that the innate immune system first detects infectious agents in peripheral tissues, shortly after a pathogen has breached an epithelial barrier. This detection event is mediated by pattern recognition receptors in phagocytes, which then migrate to draining lymph nodes (dLNs), where information of a microbial encounter is conveyed to T and B lymphocytes to generate adaptive immunity. Through the study of fungal moieties, we present data that challenge this model. We found that soluble fungal polysaccharides are immunosilent in the periphery, but become potent immunogens in the dLN. These ligands completely bypass the need of phagocyte migration and, instead, directly activate an immune response that is most similar to those that typify viral infections. These data establish a class of microbial products that violate a central tenet of the immunological lexicon and illustrate that the physical form (not just the chemical structure) impacts innate and adaptive immunity.

## Introduction

The dialogue between innate and adaptive branches of the immune system is a central paradigm of modern immunology and is vital for protection against infections as well as pathogenesis of autoimmune, allergic and inflammatory diseases (*1–4*). According to the current model, peripheral tissue infection and/or damage leads to activation and migration of innate immune cells to the draining lymph node (dLN) where they initiate an antigen-dependent adaptive immune response. The dLN has been thoroughly scrutinized for its capacity to host adaptive immune responses, but recent reports highlight that activation of immune cells in the peripheral tissue and migration of these cells to the dLN also elicit an antigen-independent lymph node innate response (LIR) (*5–13*). Albeit poorly characterized, LIR consists of at least two components: antigen-independent LN expansion and establishment of a pro-inflammatory milieu, which are both critical for the establishment of an effective adaptive immune response.

Innate immune cells recognize pathogen-associated molecular patterns (PAMPs) as well as damage-associated molecular patterns (DAMPs) through pattern recognition receptors (PRRs) (*2, 3*). Although PRRs were described for their ability to detect unique patterns associated with the chemical properties of PAMPs (*14*), some reports show that the physical form (e.g. dimension and solubility) of innate stimuli also impact recognition by, and activation of, innate immune cells (*15*). If and how these physical properties of PRR ligands influence development of the immune response in general, and of the LIR in particular, has been largely overlooked. The C-type lectin receptors (CLRs) Dectin-1 (*Clec7a*) and Dectin-2 (*Clec4n*) bind fungal polysaccharides in soluble as well as insoluble forms; but only the latter induce efficient receptor clustering and activation (*16, 17*). This model has also been extended to other CLRs, namely the mannose-binding receptor DC-SIGN (*18*). The differential responsiveness of Dectin receptors to fungal polysaccharides provides a model to investigate the impact of physical properties of innate stimuli on the LIR.

In this study, we focused our attention on soluble fungal polysaccharides, which are immunologically silent in the periphery. These microbial products do not cause any inflammation in the peripheral tissues and would therefore be expected to be ineffective at stimulating adaptive immunity. In contrast to this expectation, we found that these soluble fungal products can access the dLN and stimulate potent antigen-specific T and B cell responses. Remarkably, the responses we observed bypass the need of dendritic cell migration from the periphery to the dLN. The fungal molecules we identified therefore represent a novel class of PAMPs that uniquely stimulate innate and adaptive responses in the dLN - not in the periphery. Overall, we demonstrated that the physical form of fungal ligands dictates the location where the initial immune response takes place, and thereby determines the final outcome of the response. Our data yield novel insights into how physical properties of PAMPs affect the immune response and should help in the design of new vaccine strategies.

## Results

Dectin-1 and −2 are CLRs that respectively bind the fungal cell wall polysaccharides β-glucans and mannans and both activate the kinase Syk either directly (Dectin-1) or through the FcRγ chain (Dectin-2) (*19–21*). We employed preparations of β-glucans and mannans isolated from *Candida albicans* that exhibit distinct physical forms being respectively insoluble with a diameter of ~500 nm and soluble with a diameter of ~20 nm (**Fig. S1**). In keeping with the current model of dectin activation (*16, 17*), mannans were unable to elicit *in vitro* activation of GM-CSF-differentiated bone marrow-derived phagocytes while β-glucans induced cytokine production and co-stimulatory molecule expression in a dectin-1-depenent manner. On the other hand, immobilization of mannans onto microbeads resulted in Dectin-2 and FcRγ-dependent phagocyte activation (**Fig. S2**, lipopolysaccharide (LPS) and curdlan were used as controls). Accordingly, β-glucans elicited formation of skin abscesses and lesions upon *in vivo* intradermal injection whereas mannans did not induce signs of skin inflammation (Fig. 1A). These results were confirmed by transcriptomic analysis of skin samples injected with saline, β-glucans and mannans (Fig. 1B and **Table S1**). Pathway analysis of the cluster of differentially expressed genes (DEGs) upregulated by β-glucans revealed enrichment for pro-inflammatory and type II interferon (IFN) pathways, which is consistent with the innate immune response elicited by *C. albicans* skin infection (**Fig. S3**) (*22*). In contrast to the model of peripheral inflammation-elicited LIR, mannans induced dLN expansion and lymphocyte accrual as early as 6 hours post-injection (h.p.i.) which was sustained at 24 h.p.i. (Fig. 1C) and dependent on circulating leukocyte recruitment (Fig. 1D). β-glucans also elicited LN expansion but only at 24 h.p.i. (Fig. 1C). Considering the kinetics of response to mannans and their diameter (compatible with lymphatic drainage), we reasoned that mannans might activate LN-intrinsic circuits that eventually lead to dLN expansion. In keeping with this hypothesis, mannans rapidly accumulated to the dLN (within 6 h.p.i.) (Fig. 1E) and were able to induce a dLN expansion even in *Ccr7*^*−/−*^ mice (Fig. 1F) in which migration of immune cells from the periphery (i.e. skin) to the dLNs is abolished (*23*). Collectively, these data demonstrate that mannans activate a unique program that induces a potent LIR in the absence of peripheral (skin) inflammation and/or phagocyte migration to the dLN. To further characterize the molecular events associated with LN expansion we performed transcriptomic analysis of dLNs isolated from saline-, β-glucan- and mannan-injected mice and found a completely opposite profile compared to the skin, with mannans eliciting an earlier and more pronounced transcriptional response compared to β-glucans (Fig. 1G and **I**, **Table S2)**. Pathway analysis revealed that the cluster of DEGs upregulated by mannans is enriched for type I and II IFN pathways (Fig. 1H). The soluble nature of mannans was required for the induction of IFN-stimulated genes (ISGs) since their immobilization onto microbeads (which are retained in the skin based on their diameter) led to proinflammatory gene expression in the skin while no ISG expression in the dLN could be detected (**Fig. S4**). Underscoring the functional relevance of the IFN pathways, we found that simultaneous blockade of type I and type II IFNs (using IFNAR KO mice and an anti-IFNγ blocking antibody) prevented mannan-elicited LN expansion and induction of ISGs (Fig. 1J). Therefore, our results demonstrate that soluble mannans isolated from *C. albicans* elicit a LIR that differs from the one induced by β-glucans, namely in the induction of ISGs. To further establish whether these differences are due to distinct physical forms of fungal polysaccharides, we evaluated the response to whole glucan particles (WGP) in dispersible (D) or soluble (S) forms which have been respectively characterized as Dectin-1 agonists and antagonists (*16*). Intriguingly, WGP-S did not induce skin inflammation but elicited LN expansion and ISG expression (Fig. 1K and **L**). Altogether, these results support a model in which the physical form of fungal polysaccharides drive a LIR characterized by dLN expansion and ISG induction.

**Fig. 1.**
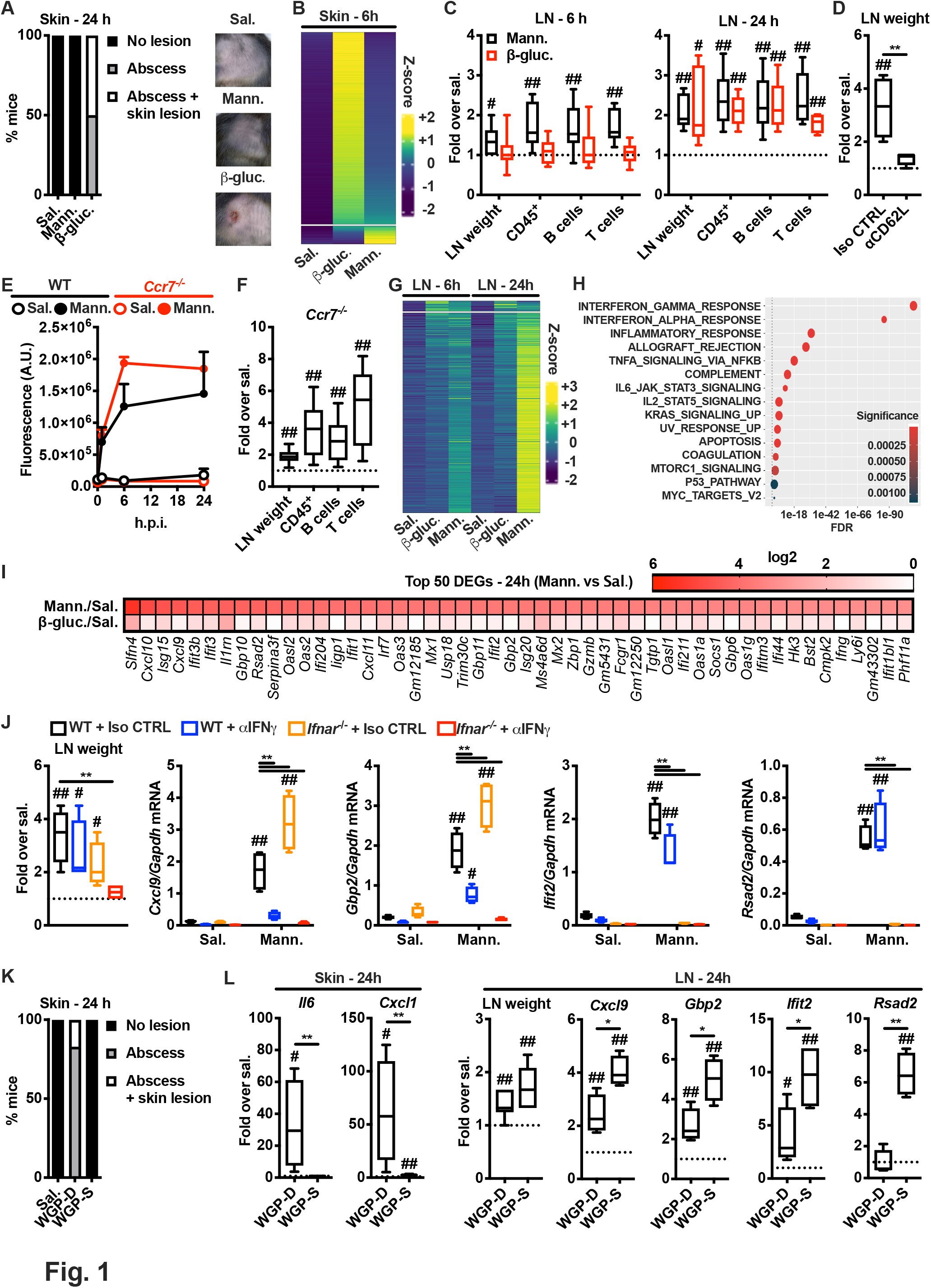
Soluble fungal polysaccharides induce a LN-restricted innate response. (**A**) Mice were injected intradermally with saline, mannans (Mann.) or β-glucans (β-gluc.). 24 hours later the injection site was assessed for the presence of an abscess with or without skin lesion. The graph shows percentages of mice in each of the indicated categories. Representative pictures of skin appearance at injection sites of saline, mannans and β-glucans are also shown. N = 5 mice per group. (**B**) Transcriptomic analysis of skin samples collected 6 hours after injection of saline, mannans or β-glucans. Heatmap of abundance (z-scored log2 normalized counts) of genes induced by β-glucans and/or mannans compared to saline control, ranked by abundance difference between β-glucans and mannans. Gap splits genes in two clusters, one that is highly upregulated by β-glucans and one that is highly upregulated by mannans. N = 3 mice per group. (**C**) Mice were injected intradermally with saline, mannans or β-glucans. 6 or 24 hours later dLNs were collected and analyzed for weight as well as absolute numbers of CD45^+^, B and T cells. Results are expressed as fold over contralateral, saline-injected LN. N = 5 - 9 mice per group. (**D**) Mice were injected intravenously with a blocking anti-CD62L antibody (αCD62L) or the same dose of an isotype control (Iso CTRL) one day before intradermal injections of saline or mannans. 24 hours later dLNs were collected and their weights were measured. Results are expressed as fold over contralateral, saline-injected LN. N = 4 mice per group. (**E**) WT and *Ccr7*^*−/−*^ mice were intradermally injected with saline or fluorescently labelled mannans. 1, 6 and 24 hours later dLNs were collected and homogenized to measure total fluorescence. Results are expressed as arbitrary units (A.U.) of fluorescence and shown as mean + SEM. N = 3 mice per timepoint. (**F**) *Ccr7*^*−/−*^ mice were injected intradermally with saline or mannans. 24 hours later dLNs were collected and analyzed as indicated in **B**. N = 6 mice. (**G**) Transcriptomic analysis of dLNs collected 6 and 24 hours after intradermal injection of saline, mannans or β-glucans. Heatmap of abundance (z-scored log2 normalized counts) of genes induced by β-glucans and/or mannans compared to saline control, ranked by abundance difference between β-glucans and mannans. Gap splits genes in two clusters, one that is highly upregulated by β-glucans and one that is highly upregulated by mannans. N = 4-5 mice per group. (**H**) Pathway enrichment analysis of genes belonging to the cluster upregulated by mannans as depicted in **G**. (**I**) Heatmap representation of the average expression levels of the top 50 genes upregulated in mannan-treated dLNs 24 hours after the injection compared to the saline control. N = 4-5 mice per group. (**J**) WT and *Ifnar*^*−/−*^ were intravenously injected with an anti-IFNγ blocking antibody or the same dose of an isotype control (Iso CTRL) on day −1 and 0. On day 0 mice were also intradermally injected with saline or mannans. 24 hours later dLNs were collected, their weights were measured and RNA was extracted for gene expression analysis. Results are expressed as fold over contralateral, saline-injected LN (weight) or as relative expression compared to *Gapdh*. N = 4 mice per group. (**K** and **L**) Mice were injected intradermally with saline, WGP-D or WGP-S. 24 hours later injection sites were assessed as indicated in **A** (**K**), skin samples and dLNs were collected, LN weights were measured and RNA extracted for gene expression analysis (**L**). Results are expressed as fold over the median value of saline injected skin samples or LNs. N = 4 mice per group. # and ## respectively indicate *p* ≤ 0.05 and 0.01 when comparing each group against the value 1 (which represent the contralateral control sample expressed as fold). * and ** respectively indicate *p* ≤ 0.05 and 0.01 when comparing among different experimental groups.

Dectin-2 is the major receptor for mannans (*24*) and, in agreement with *in vitro* data, we found that Dectin-2 and FcRγ were also required for mannan-elicited LIR (Fig. 2A). Dectin-2 is expressed mainly by myeloid cells (*25*). Therefore, we employed fluorescently labelled mannans to perform an immunophenotypic analysis of the non-lymphoid (CD3/CD19/NK1.1^−^) compartment of dLNs to identify cellular targets of mannans. The vast majority of mannan-laden cells were CD45^+^ cells (Fig. 2B). Imaging cytometry analysis confirmed that these cells internalized mannans (Fig. 2C). In addition, confocal microscopy analysis of dLNs 1 h.p.i. of mannans showed colocalization of phospho-Syk and mannans (Fig. 2D), which is indicative of Dectin-2/FcRγ-mediated activation. Accordingly, mannan-laden cells exhibited the highest levels of CD86 expression (marker of innate immune cell activation) in an FcRγ-dependent manner (Fig. 2E). We further characterized the phenotype of mannan-laden cells and found that more than 50% were Ly6G^−^ (CD11b^+^ Ly6C^+^)^−^ CD11c^+^ cells, while less abundant cell subsets were neutrophils (CD11b^+^ Ly6G^+^) and monocytes/monocyte-derived cells (MoCs, Ly6G^−^ CD11b^+^ Ly6C^+^) (Fig. 2F). Interestingly, using diphtheria toxin (DT)-mediated depletion of CD11c^+^ cells in CD11c-DT receptor (DTR) mice we were able to completely abolish mannan-elicited LIR (Fig. 2G). On the other hand, mannan-elicited LIR was not affected in *Ccr2*^*−/−*^ mice (in which monocyte egress from the bone marrow is impaired) (Fig. 2H) or by antibody-mediated depletion of neutrophils (Fig. 2I). CD11c^+^ cells can be further distinguished based on the expression of CD11b (Fig. 2F). Since Dectin-2 is critical for mannan-elicited LIR, we assessed its expression on CD11b^−^ CD11c^+^ and CD11b^+^ CD11c^+^ cells at steady state. Dectin-2 was mainly expressed by CD11b^+^ CD11c^+^ cells (Fig. 2J and **Fig. S5**), and the majority of CD11b^+^ CD11c^+^ Dectin-2^+^ cells expressed the macrophage marker CD64 (**Fig. S6**).

**Fig. 2.**
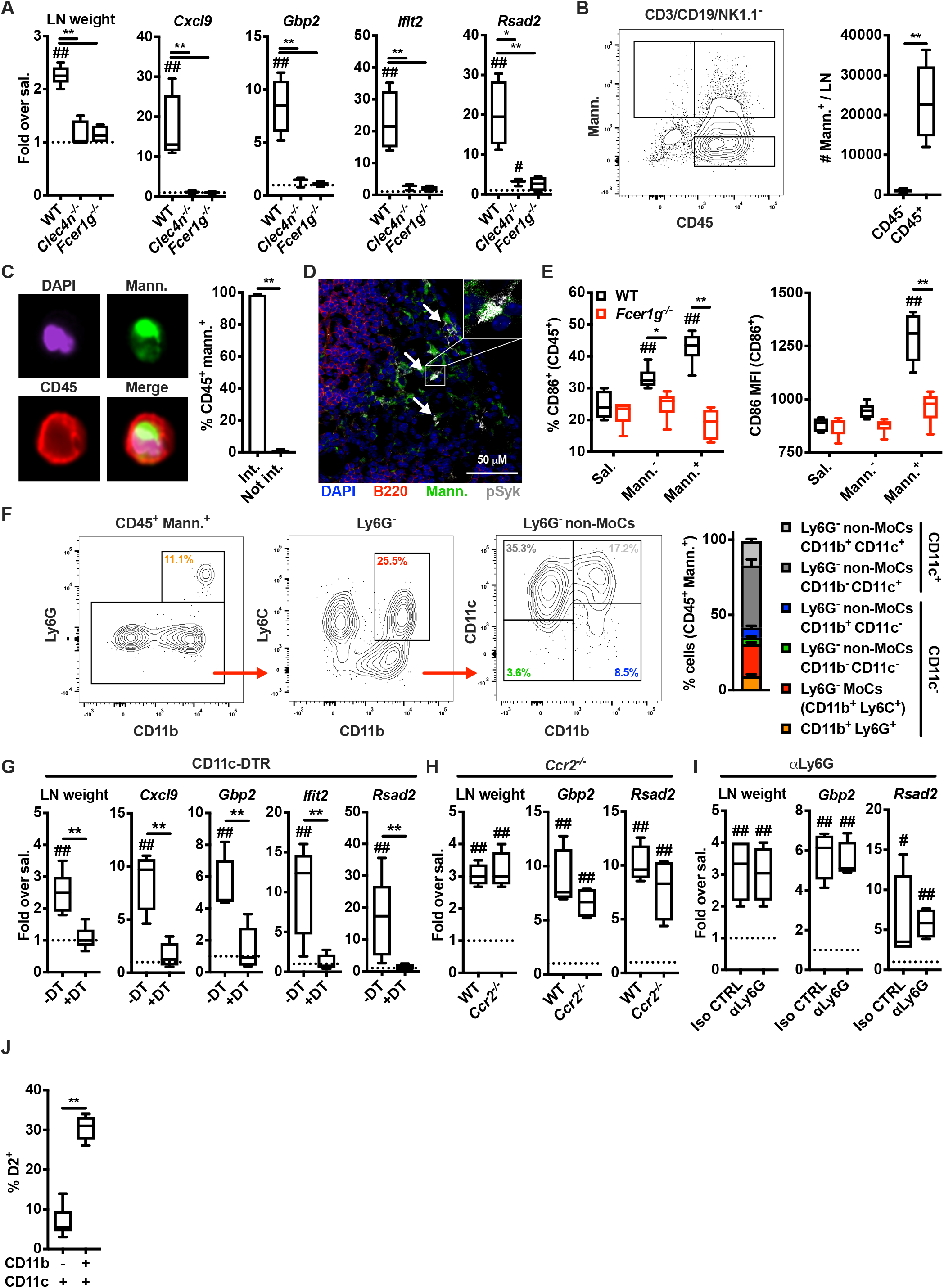
Dectin-2^+^ macrophages are required for mannan-elicited LIR. (**A**) WT, *Clec4n*^*−/−*^ and *Fcer1g*^*−/−*^ mice were intradermally injected with saline or mannans. 24 hours later dLNs were collected, their weights were measured and RNA was extracted for gene expression analysis. Results are expressed as fold over contralateral, saline-injected LN. N = 3-5 mice per genotype. (**B**) WT were intradermally injected with fluorescently labelled mannans. 6 hours later dLNs were collected and the absolute numbers of mannan-laden (Mann.^+^) CD3/CD19/NK1.1^−^ cells were quantified by flow cytometry. N = 6 mice. (**C**) Mice were treated as in **B**. Imaging cytometry analysis and quantification of mannan internalization was performed on CD3/CD19/NK1.1-depeted, CD45^+^ mannan-laden (Mann.^+^) cells. N = 4 mice. (**D**) WT mice were intradermally injected with fluorescently labelled mannans. 1 hour later dLNs were collected for confocal microscopy analysis using antibodies against B220 and phospho-Syk (pSyk). DAPI was used for nuclear counterstaining. One representative image is shown. (**E**) WT and *Fcer1g*^*−/−*^ mice were injected with saline or fluorescently labelled mannans. 6 hours later dLNs were collected and CD86 expression levels were assessed by flow cytometry on CD3/CD19/NK1.1^−^ CD45^+^ mannan-laden (Mann.^+^) cells, CD45^+^ cells that did not capture mannans (Mann.^−^) and CD45^+^ cells from saline-injected dLNs. N = 6 mice per genotype. (**F**) WT mice were intradermally injected with fluorescently labelled mannans. 6 hours later dLNs were collected and the phenotype of CD3/CD19/NK1.1^−^ CD45^+^ mannan-laden (Mann.^+^) cells was assessed by flow cytometry. N = 6 mice. (**G** - **I**) DT-treated CD11c-DTR, *Ccr2*^*−/−*^ and isotype control (Iso CTRL)- or anti-Ly6G (αLy6G)-treated mice were treated and analyzed as in **A**. N = 4 mice per group. (**J**) LNs were isolated from untreated WT mice and the expression of Dectin-2 (D2) was evaluated by flow cytometry as percentage of expression in the indicated CD3/CD19/NK1.1^−^ CD45^+^ cell subsets. N = 6. # and ## respectively indicate *p ≤* 0.05 and 0.01 when comparing each group against the value 1 (which represent the contralateral control sample expressed as fold) or saline control. * and ** respectively indicate *p ≤* 0.05 and 0.01 when comparing among different experimental groups.

The evidence of Syk phosphorylation in LN-resident, mannan-laden cells as early as 1 h.p.i. points to activation of signaling pathways downstream of Dectin-2. A key step in the Dectin receptor-Syk pathway is the activation of CARD9 which in turn regulates the activity of several signaling molecules and transcription factors, including MAPKs and canonical NF-κB (*19*). Therefore, we employed *Card9*^*−/−*^ mice to characterize the molecular events associated with mannan-elicited LIR. Surprisingly, mannan-elicited LN expansion was comparable between wild type (WT) and *Card9*^*−/−*^ mice (Fig. 3A). In keeping with the relevance of the IFN signature in driving the LIR, ISG induction was completely abrogated in *Fcer1g*^*−/−*^ mice (Fig. 2A), but it was largely maintained in *Card9*^−/−^ mice (Fig. 3A). In particular, type I IFN-dependent genes were unchanged in *Card9*^−/−^ compared to WT mice, while type II IFN-dependent ISGs, although significantly decreased compared to WT mice, were still partially induced (Fig. 3A). In addition, upregulation of CD86 on mannan-laden CD45^+^ cells was largely maintained in *Card9*^−/−^ mice compared to WT mice (Fig. 3B). Overall, these data demonstrate that LN expansion, immune cell accrual, as well as the type I IFN (and to a lesser extent type II IFN) signature and the upregulation of co-stimulatory molecules are regulated in the LN independently of CARD9.

**Fig. 3.**
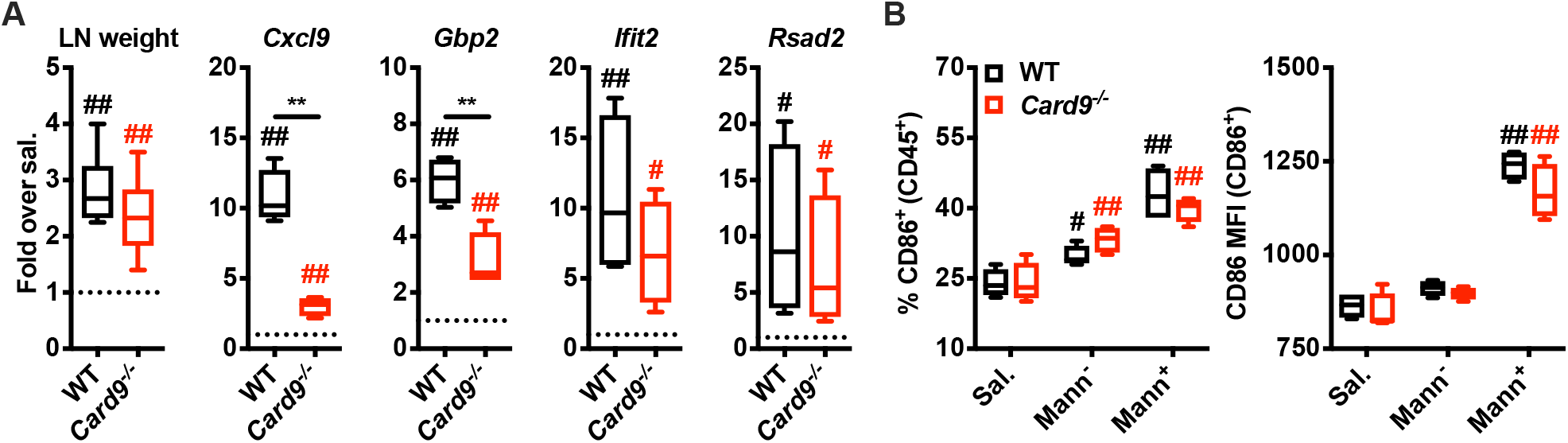
Mannans drive a LN-intrinsic Dectin-2/FcRγ-dependent, CARD9-independent transcriptional program. (**A**) WT and *Card9*^*−/−*^ mice were treated and analyzed as in **Fig. 2A**. N = 9 (for LN weight) or 4 (for gene expression analysis) mice per genotype. (**B**) WT and *Card9*^*−/−*^ mice were treated and analyzed as in **Fig. 2E**. N = 4 mice per genotype. (# and ## respectively indicate *p* ≤ 0.05 and 0.01 when comparing each group against the value 1 (which represent the contralateral control sample expressed as fold) or saline control. * and ** respectively indicate *p* ≤ 0.05 and 0.01 when comparing among different experimental groups.

There is evidence suggesting that the LIR is essential to control the development of adaptive immune responses (*7, 10, 26*). We reasoned that lymphocyte accrual and IFN signatures induced by mannan may favor the encounter of T and B cells with their cognate antigen and its efficient presentation by the innate immune compartment, thereby improving the adaptive immune response. Therefore, we investigated the impact of mannan-elicited LIR on adaptive immunity. First, we employed a model of adoptive transfer of CFSE-labelled, OVA-specific OT-II CD4^+^ T cells to assess modulation of T cell proliferation. Combining mannans with OVA resulted in a strong increase in the numbers of OT-II cells in the dLN compared to mice injected with saline as well as OVA or OVA combined with alum (a commonly used adjuvant) (Fig. 4A). This increase was likely due to higher OT-II CD4^+^ T cell recruitment to the dLN and more efficient antigen presentation/co-stimulation by LN-resident innate immune cells. Indeed, we detected a lower percentage of non-proliferating T cells as well as a higher percentage of T cells undergoing 6 or 7 divisions in mice treated with OVA and mannans compared to OVA or OVA and alum (Fig. 4B and **Fig. S7**). Of note, the effect of mannans on T cell proliferation was abrogated in *Fcer1g*^*−/−*^ mice while it was only attenuated in *Card9*^*−/−*^ mice (Fig. 4C and **D**). Then, to explore the impact of mannans on the antigen-specific antibody response we used the recombinant hemagglutinin (rHA) influenza vaccine as clinically relevant model antigen since induction of antibodies against HA plays a critical role in establishing protection from influenza infection (*27*). Anti-rHA IgG serum titers (Fig. 4E) and absolute numbers of germinal center (GC) B cells (Fig. 4F) in the dLNs of mice immunized with mannans and rHA were higher than mice immunized with rHA alone and comparable to mice immunized with rHA and alum. Interestingly, mannans were more efficient than alum at inducing type 1 polarization of the rHA-specific antibody response, as assessed by anti-rHA IgG2c serum titers (an antibody isotype induced by IFNγ) (*28*) (Fig. 4G) and anti-rHA IgG2c/IgG1 ratio (Fig. 4H). As observed for the T cell response, induction of anti-rHA IgG2c was reverted in *Fcer1g*^*−/−*^ mice and only reduced in *Card9*^*−/−*^ mice (Fig. 4I), therefore correlating with mannan-induced innate type II IFN signature. Overall, our results show that mannans act as adjuvant by enhancing antigen-specific T and B cell responses whose magnitudes correlate with mannan-elicited LIR.

**Fig. 4.**
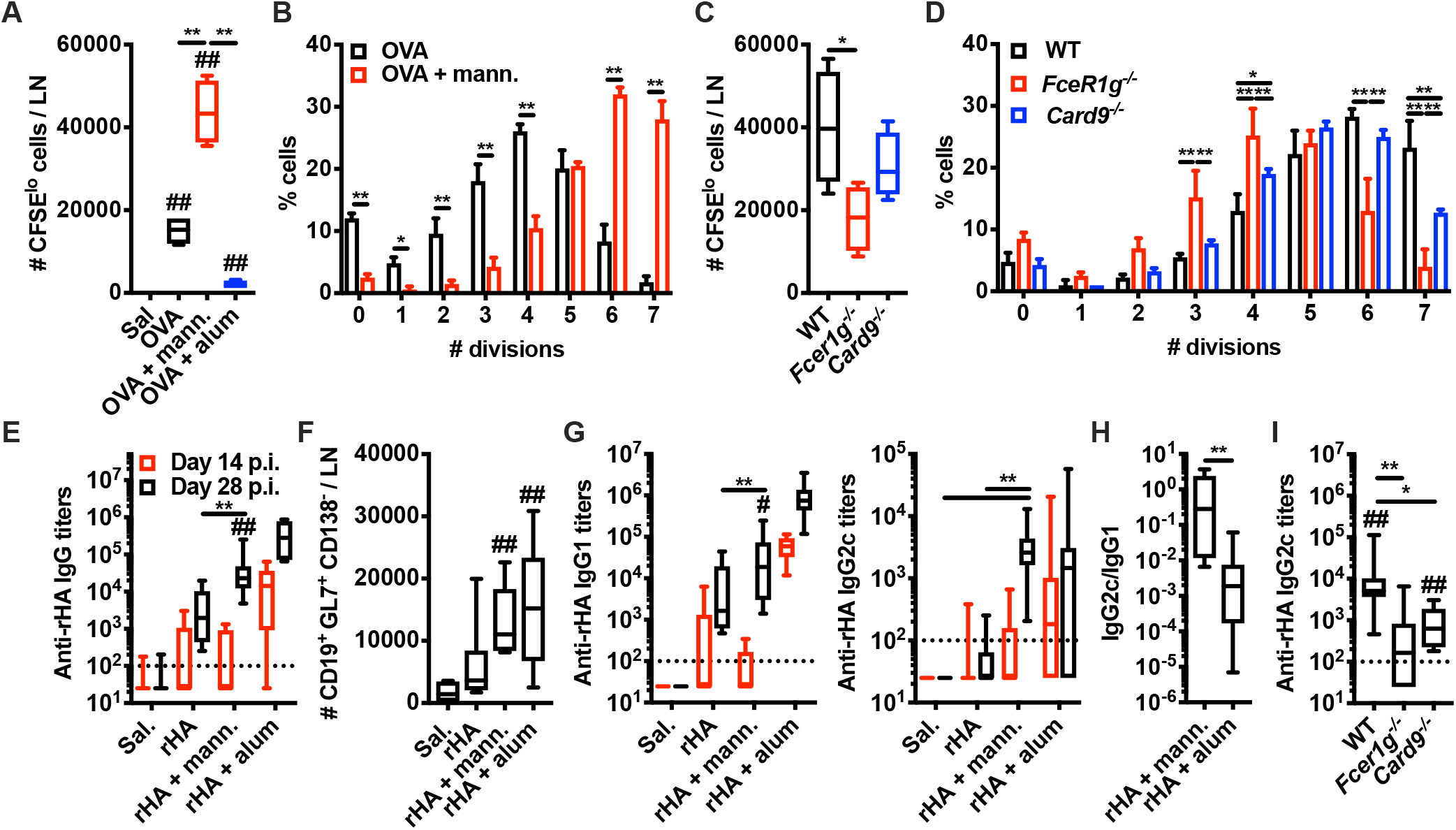
Mannan-elicited LIR directs the adaptive immune response. (**A**, **B**) CFSE-labelled OT-II CD4^+^ T cells were intravenously injected in WT mice on day −1. On day 0 the mice were intradermally injected with saline (Sal.), ovalbumin (OVA), OVA combined with mannan (OVA + mann.) or alum alhydrogel (OVA + alum). 3 days later dLNs were isolated and the absolute numbers of CFSE^lo^ cells (i.e. cells that underwent at least one cycle of cell division) (**A**) or the percentages of cells in each division peak (**B**) were quantified by flow cytometry. N = 4 mice per group. (**C**, **D**) WT, *Fcer1g*^*−/−*^ and *Card9*^*−/−*^ mice were treated and analyzed as in **A** and **B** (with the exception that all mice received OVA+mann.) N = 4 mice per genotype. (**E**, **G**, **H**) WT mice were intradermally injected with saline (Sal.), recombinant hemagglutinin (rHA), rHA combined with mannan (rHA + mann.) or alum alhydrogel (rHA + alum) on day 0 (prime) and day +14 (boost). Anti-rHA IgG (**E**), IgG1 and IgG2c (**G**) antibody titers were measured in plasma samples collected on day +14 (pre-boost) and day +28. The dotted line in the graphs indicates the limit of detection of the ELISA. Ratios of anti-rHA IgG2c and IgG1 titers were calculated for the experimental conditions rHA + mann. and rHA + alum (**H**). N = 10-11 mice per group. (**F**) WT mice were treated as in **E**, **G**. On day +28 dLNs were collected and absolute numbers of germinal center B cells (CD19^+^ GL7^+^ CD138^−^) were assessed by flow cytometry. N = 6 mice per group. (**I**) WT, *Fcer1g*^*−/−*^ and *Card9*^*−/−*^ mice were intradermally injected with rHA + mann. on day 0 and day +14. Anti-rHA IgG2c antibody titers were measured in plasma samples collected on day +28. The dotted line in the graph indicates the limit of detection of the ELISA. N = 13 (WT), 7 (*Fcer1g*^*−/−*^) or 6 (*Card9*^*−/−*^) mice. # and ## respectively indicate *p* ≤ 0.05 and 0.01 when comparing each group against the value 100 (which represent the limit of detection of the ELISA) or saline control. * and ** respectively indicate *p* ≤ 0.05 and 0.01 when comparing among different experimental groups.

## Discussion

Activation of innate immune cells by PAMPs or DAMPs is regarded as a critical step for the initiation of the adaptive immune response (*1–4*). Dendritic cells dispersed throughout peripheral tissues sense the presence of microbial clues released during an infection, are activated and migrate to the draining lymph node (dLN) - enabling a transfer of “information” from peripheral tissue to the dLN - where the antigen-dependent adaptive immune response against the pathogen is initiated. As such, it is assumed that migratory dendritic cells are an essential component of any strategy to stimulate adaptive immune responses in the dLN. Interestingly, another proposed model posits that localization, kinetics and dose of the antigen as well as its structure determine the activation of the adaptive immune response (*29*). These parameters are all influenced by the antigen physical form, and recent evidence has further proved that their modulation determines the outcome of vaccine immunotherapy (*30–32*). Even though the impact of the localization of inflammatory ligands at cellular and organismal levels in activating innate immune cells is being increasingly appreciated (*33, 34*), if the physical form of inflammatory moieties modulates their localization and in turn affects the activation of the innate immune system has been largely overlooked. In this study we report that soluble fungal polysaccharides, while largely devoid of pro-inflammatory activity *in vitro* and *in vivo* at the injection site, elicited a unique LN phenotype which was unpredictable based on the current model of Dectin-mediated pro-inflammatory activation but rather explained by their localization to the dLN. This anatomically restricted response also unraveled novel and unexpected properties of Dectin signaling. Indeed, Dectin-induced responses rely on the signaling molecule CARD9. In our model we found that soluble mannan-elicited LIR is Dectin-2/FcRγ-dependent but largely CARD9-independent. The transcriptional programs elicited upon mannan capture in LN-resident cells are also Dectin-2/FcRγ-dependent but largely CARD9-independent. This might be due to activation of a specific cell subset in which Dectin signaling does not rely on CARD9 or local accumulation of mannans at a dose that activates the Dectin-2/FcRγ/Syk axis but bypasses CARD9 requirement. Regardless of the specific mechanism, in both cases it is the localization of the fungal polysaccharide to the dLN that dictates the outcome of the innate immune response. For this reason we dubbed this class of inflammatory stimuli “locaPAMPs”.

Another novel finding of our work is the requirement for both type I and II IFNs to sustain mannan-induced lymphocyte accrual and LN expansion. This signature strikingly resembles antiviral responses and is not the type of response typically associated with anti-fungal immunity (*11, 13*). Thus, we have not only identified a novel means to stimulate dLN immune responses, but the nature of the immune response induced by locaPAMPs is also unique. It is still unclear whether in this model type I and II IFNs act on the same cell subset or on different ones, a likely occurrence due to the ubiquitous distribution of their receptors. IFNs can induce chemokine expression by myeloid cells and modulate vascular permeability (*35, 36*). Hence, it is conceivable that mannan-elicited IFN signatures affect LN-resident myeloid and stromal compartments eventually leading to lymphocyte recruitment. Of note, a combined type I and II IFN signature has recently been described in a range of infectious and inflammatory disease models (*37*), suggesting that its mechanistic dissection can provide insights into the pathophysiology of several immune-related disorders.

It is worth noting that increased plasma levels of soluble fungal polysaccharides, namely β-glucans and galcatomannans, have been associated with invasive fungal infections in humans (*38*). Our results suggest that along with being used as biomarkers, soluble polysaccharides might participate in the pathophysiology of fungal infections by activating innate immune cells in a Dectin-dependent manner.

Our results show that locaPAMP-elicited LIR can be exploited in the context of vaccine immunotherapy to enhance and polarize the adaptive immune response. This is in line with previous reports showing that LIR promotes adaptive immunity (*7, 10, 26*). In addition, mannans have been previously investigated as adjuvant candidates (*39*), but their mechanism of action has been poorly characterized so far. Our work provides the mechanistic foundation for targeting Dectin-2 (and possibly other CLRs) on LN-resident immune cells to enhance vaccine immunotherapy.

Overall, we demonstrate that the *in vivo* activity of PAMPs is defined not only by their receptor specificity but also by their physical form which in turn determines the categories of location, kinetics and dose of activity. As shown by our data, accounting for the physical form of PAMPs can uncover *in vivo* signaling pathways and immune signatures that might be unpredictable based on the current model of innate immune cell activation. Ultimately, we propose that extending the model of location, time, dose and structure (*29*) to the analysis of the innate immune system is poised to provide quantitative insights into tissue-specific mechanisms of innate immune cell activation as well as define novel properties of innate stimuli that can be harnessed in the context of vaccine immunotherapy.

## Acknowledgments

We thank Drs. JC Kagan, RS Geha, and F Granucci for discussion, help and support.

## Authors contributions

FB designed, performed, analyzed the experiments and wrote the paper; RS performed the analysis of the sequencing data; JC contributed to the design of the sequencing analysis; NAB provided *Clec4n* mice and contributed to the design of the experiments; VP, LL, MEF, LM participated to *in vitro* and *in vivo* experiments; YI provided *Clec4n* mice; FP provided mice; MDK, ZM and DLW provided fungal ligands and contributed to the design of the experiments; IZ conceived the project, designed the experiments, supervised the study and wrote the paper.

## Data and materials availability

all data is available in the manuscript or the supplementary materials.

## Materials and methods

### Mice

C57BL/6J (Jax 00664) (wild-type), B6.129P2(C)-*Ccr7*^*tm1Rfor*^/J (*Ccr7*^*−/−*^, Jax 006621), B6.129S2-*Ifnar1*^*tm1Agt*^/Mmjax (*Ifnar*^*−/−*^, Jax 32045-JAX), B6.FVB-*1700016L21Rik*^*Tg(Itgax-DTR/EGFP)57Lan*^/J (CD11c-DTR, Jax 004509), B6.129S4-*Ccr2*^*tm1Ifc*^/J (*Ccr2*^*−/−*^, Jax 004999), B6.129-*Card9*^*tm1Xlin*^/J (*Card9*^*−/−*^, Jax 028652) and B6.129S6-*Clec7a*^*tm1Gdb*^/J (*Clec7a*^*−/−*^, Jax 012337) were purchased from Jackson Labs. B6.129P2-*Fcer1g*^*tm1Rav*^ N12 (*Fcer1g*^*−/−*^, Model 583) were purchased from Taconic. *Clec4n*^*−/−*^ mice were kindly provided by Drs. Nora A. Barrett and Yoichiro Iwakura. B6.Cg-Tg(TcraTcrb)425Cbn/J (OT-II, Jax 004194) were kindly provided by Juan Manuel Leyva-Castillo. Mice were housed under specific pathogen-free conditions at Boston Children’s Hospital, and all the procedures were approved under the Institutional Animal Care and Use Committee (IACUC) and operated under the supervision of the department of Animal Resources at Children’s Hospital (ARCH).

### Reagents and antibodies

for flow cytometry, imaging cytometry, fluorescence-activated cell sorting (FACS) and confocal microscopy experiments the following reagents and antibodies were used: anti-CD45 BV510 (30-F11), anti-CD45 Alexa Fluor 700 (30-F11), anti-CD45 APC (30-F11), anti-CD3 PE/Dazzle 594 (17A2), anti-CD3 BV510 (17A2), anti-CD19 PE/Dazzle 594 (6D5), anti-CD19 BV650 (6D5), anti-NK1.1 PE/Dazzle 594 (PK136), anti-Ter119 PE/Dazzle 594 (TER-119), anti-I-A/I-E PE/Cy7 (M5/114.15.2), anti-Ly6G PerCP/Cy5.5 (1A8), anti-CD11b Pacific Blue (M1/70), anti-Ly6C BV711 (HK1.4), anti-CD11c BV785 (N418), anti-CD11c APC (N418), anti-CD86 APC/Cy7 (GL-1), anti-CD86 APC (GL-1), anti-OX40L PE (RM134L), anti-CD4 APC/Fire 750 (GK1.5), anti-GL7 PE (GL7), anti-CD138 BV421 (281-2), anti-CD45R/B220 Alexa Fluor 594 (RA3-6B2), TrueStain FcX (93), True-Stain Monocyte Blocker and Zombie Red Fixable Viability Kit were purchased from Biolegend; anti-Dectin-2 PE (REA1001) and anti-CD64 APC (REA286) were purchased from Miltenyi Biotec; rat anti-Dectin-2 (D2.11E4) was purchased from GeneTex; anti-phopsho-Syk (Tyr525/526) (C87C1) was purchased from Cell Signaling Technology; CellTrace CFSE Cell Proliferation Kit, Alexa Fluor 488 NHS Ester (Succinimidyl Ester) and DAPI were purchased from ThermoFisher Scientific.

For *in vitro* and *in vivo* experiments the following reagents were used: Iscove’s Modified Dubecco’s Medium (IMDM), Phosphate Buffer Saline (PBS), penicillin/streptomycin (pen/strep) and L-Glutamine (L-Gln) were purchased from Lonza; Fetal Bovine Serum (FBS) was purchased from ThermoFisher Scientific; collagenase from *Clostridium histolyticum*, deoxyribonuclease (DNase) I from bovine pancreas and dispase II were purchased from MilliporeSigma; TLRGrade *Escherichia coli* LPS (Serotype O555:B5, 1 μg/ml) was purchased from Enzo Life Sciences; curdlan (10 μg/ml) was purchased from Wako Chemicals; mannans, Alexa Fluor 488-conjugated mannans (AF488-mannans) and β-glucans (10 μg/ml for *in vitro* experiments, 500 μg/mouse for *in vivo* experiments) were provided by Michael D Kruppa, Zuchao Ma and David L Wiliams (East Tennessee State University); carboxyl latex beads 3 μm were purchased from ThermoFisher Scientific and used directly (cell:bead ratio 1:10 for *in vitro* experiments, 5 × 10^6^ beads/mouse for *in vivo* experiments) or after coating with diaminopropane derivatized mannans provided by Michael D Kruppa, Zuchao Ma and David L Wiliams (East Tennessee State University); WGP-S and WGP-D (500 μg/mouse for *in vivo* experiments) were purchased from Invivogen; diphtheria toxin (unnicked) from *Corynebacterium diphtheriae* (200 ng/mouse for CD11-DTR mice, 500 ng/mouse for CD169-DTR mice) was purchased form Cayman Chamical; ovalbumin (OVA) EndoFit (5 μg/mouse) and Alhydrogel adjuvant 2% (alum, 100 μg/mouse) were purchased from Invivogen; recombinant hemagglutinin (rHA) influenza vaccine FluBlok (1 μg/mouse) was purchased from the Boston Children’s Hospital pharmacy; anti-CD62L (Mel-14, 100 μg/mouse), anti-IFNγ (XMG1.2, 100 μg/mouse), anti-Ly6G (1A8, 50 μg/mouse) and their respective isotype controls rat IgG2a (2A3) and rat IgG1 (HRPN) were purchased from Bio X Cell.

### Isolation of mannan from *C. albicans*

*Candida albicans* strain SC5314 was maintained on blood agar (Remel) plates grown at 37°C. For mannan isolation, *C. albicans* was inoculated into 15 l of YPD (1% yeast extract, 2% peptone, 2% dextrose) and grown for 20 hours at 37°C. Cells were harvested by centrifugation at 5,000 x *g* for 5 min. This resulted in a 100 g pellet from 15 l of media. We used a standard protocol for isolation and NMR characterization of the mannan (*42, 43*). Essentially, the cell pellets were suspended in 200 ml of acetone to delipidate the cells for 20min. The cells were then centrifuged at 5,000x *g* for 5 min, the acetone was discarded and the pellet was allowed to dry for an hour. The dried pellet was, broken up and transferred to a beadbeater. An equivalent volume of acid-washed glass beads were added and 200mL of dH_2_O was added to the mixture. The cells were subjected to bead beating for three 30 second pulses before the entire mixture was transferred to a 1 l flask. The material was autoclaved for 2 hours, allowed to cool and then centrifuged for 5 min at 5000 x *g*. The supernatant was retained and the cell pellet discarded. Pronase (500 mg in 20 ml dH_2_O), which had been filter sterilized and heat treated for 20 min at 65°C (to remove any glycosidic acitivity) was added to the supernatant along with sodium azide to a concentration of 1mM. The mixture was then incubated overnight (20 hours) at 37°C to allow for degradation of any proteins in the solution. Mannans were extracted by addition of an equal volume of Fehling’s solution to the protease treated mannan solution and allowed to mix for one hour at RT. After mixing the solution was allowed to stand for 20 min to allow for the mannan to precipitate. The supernatant was decanted and the precipitate was dissolved in 10ml of 3M HCl, this allows for release of copper from the reducing ends of the mannans. To the solution of dissolved mannans we added 500mL of an 8:1 mixture of methanol:acetic acid, stirred the mixture and allowed the mannan to precipitate overnight. After the material had settled, the supernatant was decanted, washed again with 500mL of methanol, allowing six hours for the mannans to settle. The supernatant was decanted and the remaining precipitate was dissolved in 200 ml dH_2_O. The mannans were dialyzed against a 200-fold change of dH_2_O over 48 hours using a 2000MW cutoff membrane to remove residual acid, methanol and other contaminants from the extraction process. The dialysate was then subjected to lyophilization for seven days at stored at −20°C until needed. A small sample (10 mg) of the material was subjected to NMR and confirmed for its purity of N-linked mannans (*43*) and assessment of molecular weight determination (*42*). Prior to *in vitro* or *in vivo* use the mannan is depyrogenated and filter sterilized.

### Preparation of the diaminopropane derivatized mannan for fluorescent labeling

Mannan (100 mg) was dissolved in 1 ml of dimethyl sulfoxide (DMSO) in 4 ml vial after one hour of stirring. 1,3-Diaminopropane (100 μL) was added and stirred at ambient temperature for 3 hours. Sodium cyanoborohydride (100 mg) was added and the reaction mixture was stirred for 48 hours, followed by addition of sodium borohydride (50 mg) and stirring for 24 hours. Acetic acid (200.0 μl) was added dropwise at 0 °C to quench the reaction and the reaction mixture was stirred at ambient temperature for 3 hours, then dialyzed with a 1000 MWCO RC membrane against ultrapure water (1000 ml x 4). The retentate was harvested and lyophilized to yield the DAP attached mannan. The recovery was 88.5 mg, ~88%. The mannan-DAP was characterized by 1H-NMR to confirm the identity of the compound.

For conjugation with Alexa Fluor 488 NHS Ester (Succinimidyl Ester), ~15 mg of mannan-DAP were resuspended in 1 ml of sodium borate conjugation buffer (100 mM, pH 8.5) and allowed to solvate for at least 24 hours. Then, 1 mg of Alexa Fluor 488 NHS Ester resuspended in 35 μl of DMSO was added to the solution and incubated overnight in the dark at room temperature with gentle agitation. The reaction mixture was dialyzed with a 6000-8000 MWCO RC membrane against saline (1000 mL x 4) and the retentate was filter sterilized.

For conjugation with carboxyl latex beads 3 μm, mannan-DAP was resuspended at a concentration of 10 mg/1 ml of BupH MES conjugation buffer pH 4.5 (ThermoFisher Scientific) and allowed to solvate for at least 24 hours. 1 ml of mannan-DAP was added to 50 × 10^6^ beads and then mixed with 4 mg/1 ml of EDC (ThermoFisher Scientific) resuspended in pure water. The reaction mixture was incubated for 4 hours in the dark at room temperature with gentle agitation. Then, the beads were washed twice (4000 *g* for 10 minutes) with saline and resuspended in saline at a concentration of 10^8^ beads/ml.

### Preparation of *Candida albicans* glucan particles

Glucan particles were isolated from *Candida albicans* SC5314 as previously described by our laboratory (*44*). Briefly, glucan was isolated from *C.albicans* using a base/acid extraction approach with provides water insoluble glucan particles that are ≥ 95% pure. The structure and purity of the glucan was determined by 1H-NMR in DMSO-d6. Prior to *in vitro* or *in vivo* use the glucan particles are depyrogenated and filter sterilized.

### Analysis of skin and LN responses

To assess skin and LN innate responses, mice were intradermally injected on day 0 with the indicated compounds in a volume of 50 μl on each side of the back (one side for the compound and the contralateral side for saline of vehicle control). 6 or 24 hours post-injection (h.p.i.) skin samples at the injection sites and draining (brachial) LNs were collected for subsequent analysis.

Skin samples were transferred to a beadbeater and homogenized in 1 ml of TRI Reagent (Zymo Research). Then, samples were centrifuged 12000 *g* for 10 minutes and 800 μl of cleared supernatant were transferred to a new tube for subsequent RNA isolation.

LNs were weighted on an analytic balance before being processed to generate a LN cell suspension by modification of a previously published protocol. Briefly, individual LNs were incubated at 37°C for 20 minutes in 400 μl of digestion mix (IMDM + pen/strep + FBS 2% + collagenase 100 mg/ml + dispase II 100 mg/ml + DNase 10 mg/ml). Then, LNs were grinded by pipetting with a 1000 μl tip, supernatants were transferred to new tubes and kept at 4°C while 200 μl of digestion mix were added to the pellets and incubated at 37°C for 10 minutes. This cycle was repeated one more time, then pooled supernatants of individual LNs were divided into two aliquots: one for flow cytometry analysis, another one was centrifuged at 300 *g* for 5 minutes and the cell pellet was resuspended in 800 μl of TRI Reagent for subsequent RNA isolation.

For specific experiments mice were treated with: anti-CD62L blocking antibody or isotype control, intravenous injections on day −1; anti-IFNγ blocking antibody or isotype control, intravenous injections on day −1 and 0; anti-Ly6G depleting antibody or isotype control, intraperitoneal injections on day −1 and 0; diphtheria toxin, intravenous injections on day −1 and intradermal injections (co-injected with mannans) on day 0 for CD11c-DTR mice, intraperitoneal injection on day −2 for CD169-DTR mice.

### *In vitro* stimulation of GM-CSF-differentiated, bone marrow-derived phagocytes

Bone marrow-derived phagocytes were differentiated from bone marrow in IMDM + 10% B16-GM-CSF derived supernatant + 10% FBS + pen/strep + L-Gln and used after 7 days of culture. Then, cells were harvested, plated in flat bottom 96 well plates at a density of 10^5^ cells/200 μl/well in IMDM + 10% FBS + pen/strep + L-Gln and stimulated with the indicated compounds for 18-21 hours. At the end of stimulation, supernatants were harvested and TNF and IL-2 levels were measured by ELISA (Biolegend) according to the manufacturer’s protocol. Cells were detached with PBS + EDTA 2 mM and transferred to a round bottom 96 well plate for subsequent flow cytometry staining and analysis.

### *In vivo* quantification of fluorescently labelled mannans

WT and *Ccr7*^*−/−*^ mice were intradermally injected with AF488-mannans and saline. After 1, 6 and 24 hours dLNs were collected, transferred to a beadbeater and homogenized in 400 μl of deionized water. Then, samples were centrifuged (12000 *g* for 10 minutes) and cleared supernatants were transferred to a 96 well clear bottom black plate. Fluorescence values were measured with SpectraMax i3x microplate reader (Molecular Devices) and expressed as arbitrary units after background (deionized water) subtraction.

### Flow cytometry, fluorescence-activated cell sorting (FACS), imaging cytometry and confocal microscopy

For flow cytometry analysis, cells were first stained with Zombie Red Fixable Viability in PBS for 5 minutes at 4°C, washed once with PBS + BSA 0.2% + NaN_3_ 0.05% (300 *g* for 5 minutes) and then stained with antibodies against surface antigens diluted in PBS + BSA 0.2% + NaN_3_ 0.05% for 20 minutes at 4°C. Cells were then washed, fixed with 2% paraformaldehyde for 10 minutes at room temperature, washed again and resuspended in PBS + BSA 0.2% + NaN_3_ 0.05%. Samples were acquired on a BD LSRFortessa flow cytometer and data were analyzed using FlowJo v.10 software (BD Biosciences). CountBright Absolute Counting Beads were used to quantify absolute cell numbers.

For FACS and imaging cytometry, mice were intradermally injected with AF488-mannans and 6 hours later dLNs were harvested to obtain LN cell suspensions. For FACS, cells were stained with antibodies against surface antigens diluted in PBS + BSA 0.2% for 20 minutes at 4°C. Cells were then washed once, resuspended in 1 ml of PBS + BSA 0.2%, filtered through 70 μm cell strainers (Fisher Scientific) and sorted with a Sony MA900 cell sorter directly into 1 ml of TRI Reagent. The following cell subset was sorted: CD3^−^ CD19^−^ NK1.1^−^ Ter119^−^ CD45^+^ AF488-mannan^+^ Ly6G^−^ (CD11b^+^ Ly6C^+^)^−^ CD11b^+^ CD11c^+^. For imaging cytometry, cells were depleted of lymphoid and erythroid cells by sequential staining with biotinylated antibodies against anti-CD3, anti-CD19, anti-NK1.1, anti-Ter119 and Streptavidin Microbeads (Miltenyi Biotec) according to the manufacturer’s protocol. The remaining cells were stained with anti-CD45 APC, fixed with BD FACSLyse, washed once and resuspended in 60 μl of PBS + DAPI (0.2 μg/ml). Samples were then acquired on an Amnis ImageStream X Mark II (Luminex Corporation). Mannan internalization was analyzed with Amnis Ideas Software and calculated with Internalization Feature as AF488 signal within the APC mask.

For confocal microscopy, dLNs were isolated at steady state or 1 h.p.i. of AF488-mannans and fixed with 4% paraformaldehyde overnight. Slides were imaged on a Zeiss 880 laser scanning confocal microscope.

### RNA isolation, qPCR, transcriptomic and pathway analyses

RNA was isolated from TRI Reagent samples using phenol-chloroform extraction or column-based extraction systems (Direct-zol RNA Microprep and Miniprep, Zymo Research) according to the manufacturer’s protocol. RNA concentration and purity (260/280 and 260/230 ratios) were measured by NanoDrop (ThermoFisher Scientific).

Purified RNA was analyzed for gene expression by qPCR on a CFX384 real time cycler (Bio-rad) using pre-designed KiCqStart SYBR Green Primers (MilliporeSigma) specific for *Cxcl9* (RM1_Cxcl9 and FM1_Cxcl9), *Gbp2* (RM1_Gpb2 and FM1_Gbp2), *Ifit2* (RM1_Ifit2 and FM1_Ifit2), *Rsad2* (RM1_Rsad2 and FM1_Rsad2), *Il6* (RM1_Il6 and FM1_Il6), *Cxcl1* (RM1_Cxcl1 and FM1_Cxcl1) or pre-designed PrimeTime qPCR Primers (Integrated DNA Technologies) specific for *Gapdh* (Mm.PT.39a.1).

For bulk RNAseq analysis, RNA isolated from LN samples was submitted to Genewiz. RNA samples were quantified using Qubit 2.0 Fluorometer (ThermoFisher Scientific) and RNA integrity was checked with RNA Screen Tape on Agilent 2200 TapeStation (Agilent Technologies). RNA sequencing library preparation was prepared using TruSeq Stranded mRNA library Prep kit following manufacturer’s protocol (Illumina, Cat# RS-122-2101). Briefly, mRNAs were first enriched with Oligod(T) beads. Enriched mRNAs were fragmented for 8 minutes at 94°C. First strand and second strand cDNA were subsequently synthesized. The second strand of cDNA was marked by incorporating dUTP during the synthesis. cDNA fragments were adenylated at 3’ends, and indexed adapter was ligated to cDNA fragments. Limited cycle PCR was used for library enrichment. The incorporated dUTP in second strand cDNA quenched the amplification of second strand, which helped to preserve the strand specificity. Sequencing libraries were validated using DNA Analysis Screen Tape on the Agilent 2200 TapeStation (Agilent Technologies), and quantified by using Qubit 2.0 Fluorometer (ThermoFisher Scientific) as well as by quantitative PCR (Applied Biosystems). The sequencing libraries were multiplexed and clustered on 1 lanes of flowcell. After clustering, the flowcell was loaded on the Illumina HiSeq instrument according to manufacturer’s instructions. The samples were sequenced using a 2×150 Pair-End (PE) High Output configuration. Image analysis and base calling were conducted by the HiSeq Control Software (HCS) on the HiSeq instrument. Raw sequence data (.bcl files) generated from Illumina HiSeq was converted into fastq files and de-multiplexed using Illumina bcl2fastq program version 2.17. One mismatch was allowed for index sequence identification. Reads were quality-controlled using FastQC. Illumina adapters were removed using cutadapt. Trimmed reads were mapped to the mouse transcriptome (GRCm38) based on Ensembl annotations using Kallisto (*45*). Transcript counts were imported and aggregated to gene counts using tximport (*46*). Gene counts were analyzed using the R package DESeq2 (*47*). When applicable, batch was used as a blocking factor in the statistical model. Differentially expressed genes (DEGs) were identified as those passing a threshold of FDR significance threshold (0.05 for skin; 0.01 for lymph nodes, because of the higher power due higher number of replicates) where the alternate hypothesis was that the absolute log2 FC was greater than 0. Genes induced by mannan or glucan treatment over saline were plotted in heatmaps using the R package ComplexHeatmap, using Z-scored log2 normalized abundance. Genes were arranged by abundance delta between glucan and mannan (aggregated from multiple time points when appropriate), with a gap delimiting two clusters: genes more highly expressed upon mannan stimulation vs genes more highly expressed upon glucan stimulation. Pathway analysis was performed with the R package hypeR (*48*), using hypergeometric enrichment tests of genes belonging to a cluster of interest and the Hallmark gene set collection from the Broad Institute’s MSigDB collection.

### *In vivo* T cell proliferation assay

Spleens were isolated from OT-II mice and meshed with the plunger end of a syringe. Then, splenocyte cell suspensions were treated with ACK lysis buffer (2 minutes at room temperature), washed with PBS (300 *g* for 5 minutes) and filtered through 70 μm cell strainers. CD4^+^ T cells were purified using CD4 (L3T4) MicroBeads (Miltenyi Biotech) according to the manufacturer’s protocol and stained with CellTrace CFSE (5 μM in PBS + FBS 2.5% for 20 minutes in the dark). At the end of incubation, cells were washed twice with PBS, resuspended at a concentration of 5 × 10^5^ cells/100 μl saline and 100 μl of cell suspension was intravenously injected into each mouse. 24 hours later (day 0) mice were intradermally injected with OVA combined with mannans or alum. Saline-injected mice were used as control. On day +3 dLNs were harvested and LN cell suspension were stained with anti-CD19, anti-Ter119, anti-CD3 and anti-CD4 antibodies. Adoptively transferred, CFSE-labelled cells were detected in the CD19^−^ Ter119^−^ CD3^+^ CD4^+^ gate. Results are expressed as absolute number of CD19^−^ Ter119^−^ CD3^+^ CD4^+^ CFSE^lo^ cells (i.e. cells undergoing at least one division cycle) of percentage of each division peak within the CD19^−^ Ter119^−^ CD3^+^ CD4^+^ gate.

### Immunization and antibody quantification

Mice were immunized by intradermal injection of rHA alone or combined with mannans or alum on day 0 and day +14. Saline-injected mice were used as control. Blood samples were collected by retroorbital bleeding on day +14 (pre-boost) and day +28, and plasma was isolated after centrifugation of blood samples at 500 *g* for 20 minutes. rHA-specific IgG, IgG1, IgG2c antibody titers were quantified in plasma samples by ELISA as previously described (*49*). Briefly, high binding flat bottom 96-well plates were coated with 1 µg/ml rHA in carbonate buffer pH 9.6, incubated overnight at 4°C and blocked with PBS + BSA 1% for 1 h at room temperature. Then, plasma samples were added with an initial dilution of 1:100 and 1:4 serial dilutions in PBS + BSA 1% and incubated for 2 h at room temperature. Plates were then washed and incubated for 1 h at room temperature with HRP-conjugated anti-mouse IgG, IgG1 or IgG2c (Southern Biotech). At the end of the incubation, plates were washed again and developed with tetramethylbenzidine (BD Biosciences) for 5 min, then stopped with 2 N H_2_SO_4_. The optical density was read at 450 nm with SpectraMax i3x microplate reader (Molecular Devices) and endpoint titers were calculated using as cutoff three times the optical density of the background.

### Statistics

Data were log-transformed to approximate normal distributions. One-sample t test was used to compare each group against the value 1 (0 after log-transformation, which represent the contralateral control sample expressed as fold). Statistical differences between groups in datasets with one categorical variable were evaluated by two sample t test (2 groups) of one-way ANOVA (more than 2 groups). Statistical differences between groups in datasets with two categorical variables were evaluated by two-way ANOVA. # or * and ** or ## respectively indicate *p* ≤ 0.05 and 0.01.

