## Supplementary Figures and Tables for "The physical form of microbial ligands bypasses the need for dendritic cell migration to stimulate adaptive immunity"

**Supplementary Material**

**Fig. S1. Mannans and β-glucans exhibit different diameters.** Hydrodynamic diameters of mannan (Mann.) and β-glucan (β-gluc.) preparations were measured by dynamic light scattering. Results from 1 representative experiment are shown.

**Fig. S2. Soluble mannans are inactive *in vitro*.** GM-CSF differentiated, bone marrow-derived phagocytes were generated from WT, *Clec7a*$^{-/-}$, *Clec4n*$^{-/-}$ and *Fcer1g*$^{-/-}$ mice and stimulated with LPS, curdlan, β-glucans (β-gluc.), soluble mannans (Mann.), uncoated microbeads (B only) and microbeads covalently linked to mannans (B:Mann.). After 18-21 hours supernatants were collected, and TNF and IL-2 protein levels were measured by ELISA while cells were harvested and expression levels of CD86 and OX40L were measured by flow cytometry. ## indicate $p \leq 0.01$ when comparing each group against its untreated control (CTRL).

**Fig. S3.** Pathway enrichment analysis of genes belonging to the cluster upregulated by β-glucans as depicted in **Fig. 1B**.

**Fig. S4. Mannans immobilized onto microbeads do not elicit a LN-specific IFN signature.** WT mice were intradermally injected with uncoated microbeads (B only) or microbeads covalently linked to mannans (5 x $10^6$ per mouse). 24 hours later skin samples and dLNs were collected and RNA was extracted for gene expression analysis. Results are expressed as fold over contralateral, B only-injected skin or LN. N = 4 mice. # indicate $p \leq 0.05$ when comparing each group against the value 1 (which represent the contralateral control sample expressed as fold).

**Fig. S5. LN-resident CD11b$^+$ CD11c$^+$ cells express higher levels of Dectin-2.** LNs were isolated from untreated WT mice and the expression of Dectin-2 (D2) was evaluated by flow cytometry as MFI in the indicated CD3/CD19/NK1.1$^-$ CD45$^+$ cell subsets. N = 6 mice. ** indicate $p \leq 0.01$ when comparing among different experimental groups.

**Fig. S6. LN-resident CD11b$^+$ CD11c$^+$ Dectin-2$^+$ cells exhibit a macrophage phenotype.** LNs were isolated from untreated WT mice and the expression of the macrophage marker CD64 was evaluated by flow cytometry as MFI in the indicated CD3/CD19/NK1.1$^-$ CD45$^+$ CD11b$^+$ CD11c$^+$ Dectin-2$^-$ and Dectin-2$^+$ cells. N = 6 mice.

**Fig. S7. OVA combined with mannan boosts proliferation of CFSE-labelled OT-II CD4$^+$ T cells.** Representative histograms of the results presented in **Fig. 4A** and **B**.

**Table S1.** List of genes detected by bulk RNAseq analysis that are expressed in the skin 6 hours after the injection of saline, β-glucans or mannans.

**Table S2.** List of genes detected by bulk RNAseq analysis that are expressed in the dLN 6 and 24 hours after the injection of saline, β-glucans or mannans.

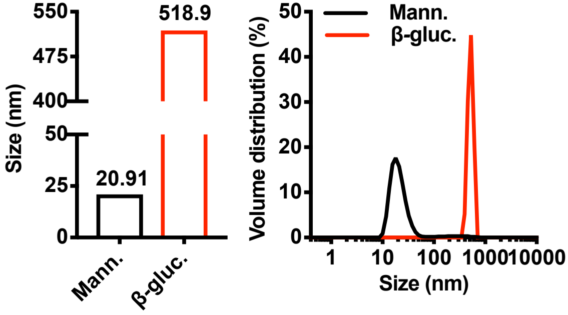
550
475
400
50
25
0
Size (nm)
518.9
20.91
Mann.
β-gluc.
Volume distribution (%)
50
40
30
20
10
0
1
10
100
1000
10000
Size (nm)
Mann.
β-gluc.

### Fig. S1

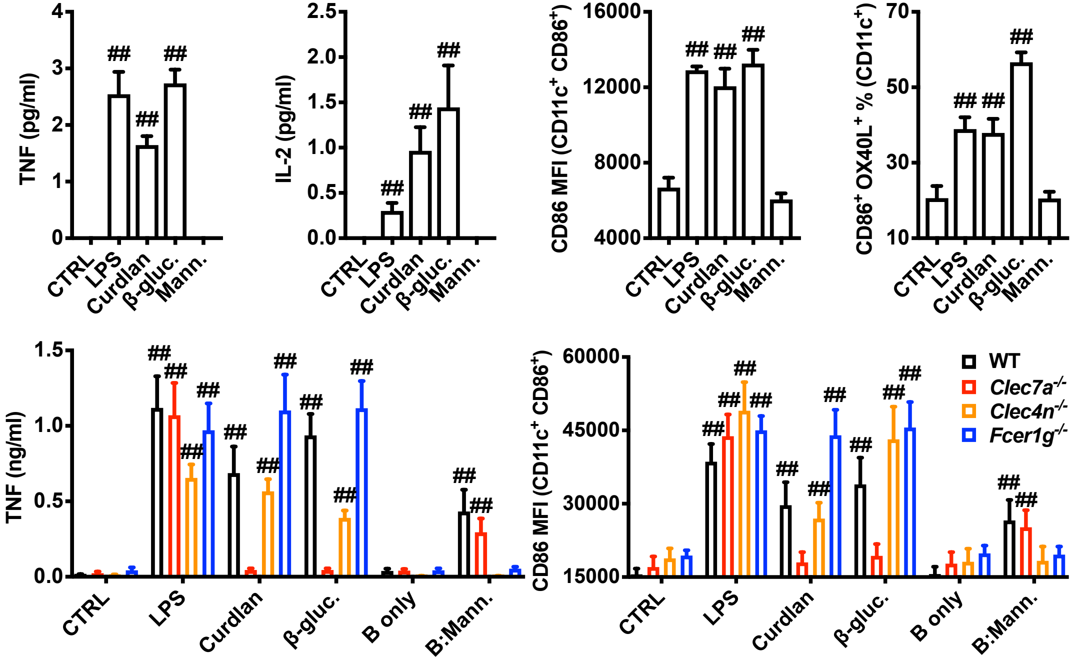
TNF (pg/ml)
IL-2 (pg/ml)
CD86 MFI (CD11c+ CD86+)
CD86+ OX40L+ % (CD11c+)
CTRL
LPS
Curdlan
β-gluc.
Mann.
##
TNF (ng/ml)
B only
B:Mann.
WT
Clec7a-/-
Clec4n-/-
Fcer1g-/-

Fig. S2

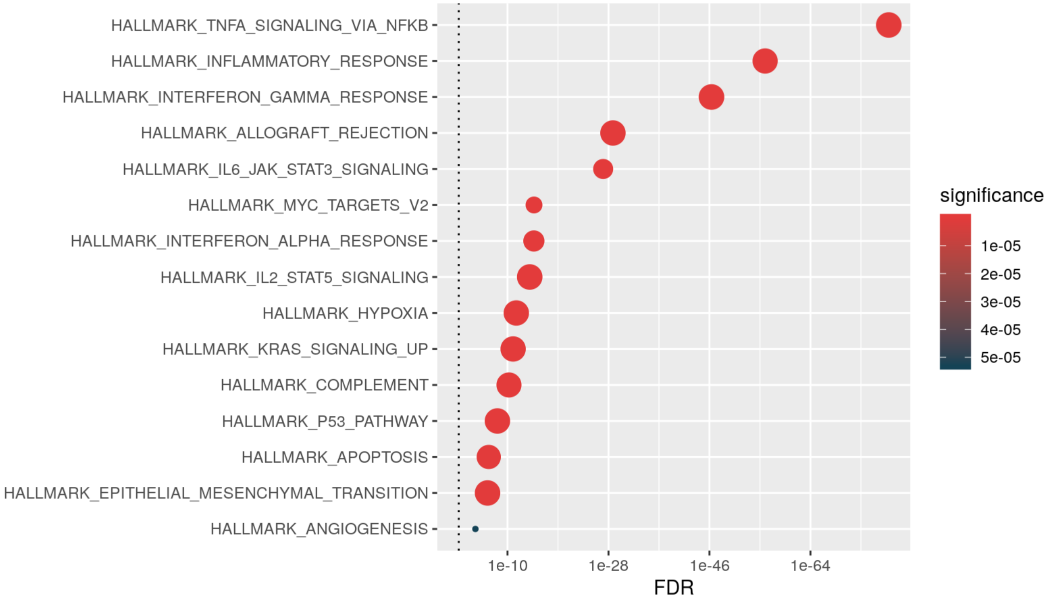
HALLMARK_TNFA_SIGNALING_VIA_NFKB
HALLMARK_INFLAMMATORY_RESPONSE
HALLMARK_INTERFERON_GAMMA_RESPONSE
HALLMARK_ALLOGRAFT_REJECTION
HALLMARK_IL6_JAK_STAT3_SIGNALING
HALLMARK_MYC_TARGETS_V2
HALLMARK_INTERFERON_ALPHA_RESPONSE
HALLMARK_IL2_STAT5_SIGNALING
HALLMARK_HYPOXIA
HALLMARK_KRAS_SIGNALING_UP
HALLMARK_COMPLEMENT
HALLMARK_P53_PATHWAY
HALLMARK_APOPTOSIS
HALLMARK_EPITHELIAL_MESENCHYMAL_TRANSITION
HALLMARK_ANGIOGENESIS
1e-10
1e-28
1e-46
1e-64
FDR
significance
1e-05
2e-05
3e-05
4e-05
5e-05

**Fig. S3**

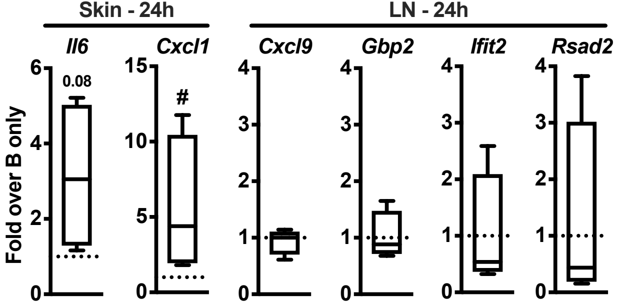

Skin - 24h
LN - 24h
Il6
Cxcl1
Cxcl9
Gbp2
Ifit2
Rsad2
Fold over B only
0.08
#
0
2
4
6
0
5
10
15
0
1
2
3
4

### Fig. S4

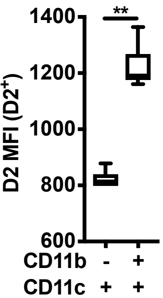

### Fig. S5

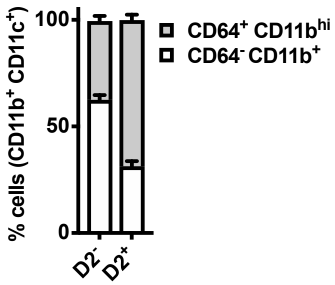

### Fig. S6

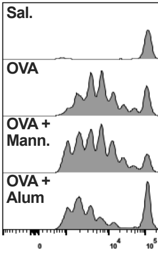

### Fig. S7

**Table S1**

| GeneSymbol | b-gluc._6h | Mann._6h | Sal._6h | Mann./Sal._6h | b-gluc./Sal._6h |
|---|---|---|---|---|---|
| Cxcl2 | 13.27124631 | 9.218731659 | 7.194096679 | 2.02463498 | 6.077149628 |
| Cxcl3 | 11.29377768 | 7.579321042 | 6.159446058 | 1.419874985 | 5.134331626 |
| Ccl3 | 11.97144507 | 8.56675652 | 6.864523117 | 1.702233403 | 5.10692195 |
| Ccl4 | 12.18385131 | 8.580089382 | 7.184234491 | 1.39585489 | 4.999616822 |
| Acod1 | 12.64037033 | 9.159329796 | 7.774152019 | 1.385177776 | 4.866218311 |
| Osm | 10.78928784 | 7.719293684 | 6.173781125 | 1.545512559 | 4.615506715 |
| Il1b | 13.58816159 | 11.18198472 | 9.017277374 | 2.164707342 | 4.570884212 |
| Cd14 | 12.77476697 | 9.92464453 | 8.410451807 | 1.514192723 | 4.364315166 |
| S100a9 | 13.46465867 | 10.88637598 | 9.26178557 | 1.624590407 | 4.202873103 |
| Gm5483 | 9.576950841 | 7.232036735 | 5.597070231 | 1.634966505 | 3.979880611 |
| Hdc | 12.45665206 | 9.625777607 | 8.489212337 | 1.13656527 | 3.967439723 |
| Retnlg | 10.49716285 | 7.732081988 | 6.54079755 | 1.191284437 | 3.956365296 |
| Ankrd1 | 10.77855394 | 11.98857459 | 6.845698617 | 5.142875977 | 3.932855327 |
| S100a8 | 12.56230535 | 10.23026422 | 8.63692267 | 1.59334155 | 3.925382684 |
| Spp1 | 12.32845372 | 9.605482128 | 8.40758376 | 1.197898369 | 3.92086996 |
| Trem1 | 9.980438353 | 7.447134243 | 6.100908468 | 1.346225774 | 3.879529885 |
| Rrad | 11.00721108 | 9.808440117 | 7.219437549 | 2.589002568 | 3.787773531 |
| Nlrp3 | 10.90140527 | 8.636405541 | 7.173532082 | 1.462873458 | 3.727873188 |
| Clec4d | 11.72079802 | 9.789933704 | 8.048307084 | 1.741626619 | 3.672490932 |
| Ptx3 | 13.23530858 | 10.37349581 | 9.602985436 | 0.770510377 | 3.632323147 |
| Cxcl5 | 11.84956725 | 9.793316065 | 8.279655695 | 1.51366037 | 3.569911559 |
| Slc7a11 | 11.38905721 | 9.566645991 | 7.904643528 | 1.662002463 | 3.484413682 |
| Plaur | 11.64315721 | 9.687026662 | 8.196604295 | 1.490422367 | 3.446552919 |
| Clec4e | 11.47398666 | 9.81970874 | 8.04843184 | 1.7712769 | 3.42555482 |
| Adamts4 | 10.59955132 | 8.912832248 | 7.224288029 | 1.688544219 | 3.375263291 |
| Thbs1 | 15.25749342 | 13.30289424 | 11.88882447 | 1.414069777 | 3.36866895 |
| Ptgs2 | 11.49748986 | 8.751841136 | 8.156009868 | 0.595831267 | 3.341479988 |
| Slpi | 11.24349618 | 9.2920217 | 7.913146096 | 1.378875604 | 3.33035008 |
| Cxcl1 | 10.71553478 | 8.756278003 | 7.437970214 | 1.318307789 | 3.27756457 |
| Cyp4f18 | 9.913065334 | 7.448158048 | 6.696107699 | 0.752050349 | 3.216957635 |

| | | | | | |
|---|---|---|---|---|---|
| Mrgpra2b | 8.570957025 | 6.324534089 | 5.463178258 | 0.861355831 | 3.107778767 |
| Slfn4 | 13.42571073 | 11.40840635 | 10.34068551 | 1.067720833 | 3.08502522 |
| Srgn | 12.9601352 | 11.29603368 | 9.91111394 | 1.384919736 | 3.04902126 |
| Fgf23 | 8.725683275 | 7.183757592 | 5.697639114 | 1.486118478 | 3.028044162 |
| Ccr1 | 11.41923113 | 10.13684077 | 8.436257264 | 1.700583507 | 2.982973863 |
| Inhba | 10.72140807 | 8.220575759 | 7.769114987 | 0.451460772 | 2.95229308 |
| Nfkbid | 9.585559504 | 7.759291597 | 6.651253053 | 1.108038544 | 2.934306452 |
| Ccrl2 | 9.657334725 | 7.3745059 | 6.753637704 | 0.620868196 | 2.903697021 |
| Csf3 | 7.88968386 | 5.89918312 | 5.005272124 | 0.893910996 | 2.884411736 |
| Il6 | 9.743802745 | 7.554500596 | 6.873930171 | 0.680570425 | 2.869872574 |
| Plek | 13.05753062 | 11.49854853 | 10.2336691 | 1.264879433 | 2.823861523 |
| Lilr4b | 13.03777989 | 11.64476398 | 10.23604069 | 1.408723296 | 2.801739199 |
| Cd300lf | 10.35227179 | 9.52875008 | 7.588037002 | 1.940713078 | 2.764234788 |
| Lilrb4a | 12.86973803 | 11.78731654 | 10.14413878 | 1.643177757 | 2.72559925 |
| Fpr2 | 8.897125929 | 7.502166786 | 6.259601711 | 1.242565076 | 2.637524218 |
| Myod1 | 8.519377625 | 8.204503219 | 5.889554391 | 2.314948829 | 2.629823234 |
| Slc15a3 | 11.0804778 | 9.45672375 | 8.485618702 | 0.971105048 | 2.594859098 |
| Clec7a | 11.04363364 | 9.919855257 | 8.488234387 | 1.43162087 | 2.555399256 |
| Serpine1 | 11.92217966 | 10.97805598 | 9.368667413 | 1.609388564 | 2.553512251 |
| Samsn1 | 10.0875543 | 8.558061168 | 7.548153186 | 1.009907982 | 2.539401113 |
| Socs3 | 12.66902843 | 11.39176402 | 10.13103444 | 1.260729583 | 2.537993993 |
| Stfa2l1 | 7.862648202 | 6.163543484 | 5.341872393 | 0.821671091 | 2.520775809 |
| Ccl2 | 12.72393993 | 11.2770677 | 10.21384005 | 1.063227642 | 2.510099879 |
| Ch25h | 8.897375897 | 7.091713917 | 6.399876039 | 0.691837878 | 2.497499858 |
| Nr4a3 | 10.55290975 | 9.615831858 | 8.062720572 | 1.553111286 | 2.490189181 |
| Mt2 | 14.36771381 | 13.40971534 | 11.88039188 | 1.52932346 | 2.487321923 |
| Mmp8 | 8.77133484 | 7.134660711 | 6.298626357 | 0.836034355 | 2.472708484 |
| Mcoln2 | 10.31818184 | 9.116406082 | 7.847924406 | 1.268481676 | 2.470257437 |
| Mcemp1 | 9.252006677 | 7.582117465 | 6.792075861 | 0.790041605 | 2.459930816 |
| Olr1 | 7.950523437 | 5.905238944 | 5.491344606 | 0.413894339 | 2.459178831 |
| Cxcr2 | 11.01995712 | 10.1224583 | 8.562722392 | 1.559735905 | 2.457234728 |
| Tnfaip6 | 10.74924861 | 9.381446632 | 8.294959399 | 1.086487232 | 2.454289214 |

| | | | | | |
|---|---|---|---|---|---|
| Tnf | 8.887652136 | 7.480002489 | 6.454466133 | 1.025536356 | 2.433186004 |
| Has3 | 10.76604938 | 9.461692677 | 8.343769366 | 1.11792331 | 2.42228001 |
| Rnd1 | 9.312634064 | 7.341641188 | 6.897338916 | 0.444302272 | 2.415295148 |
| Csf3r | 11.28836038 | 10.79884903 | 8.877181037 | 1.92166799 | 2.411179346 |
| Xirp1 | 11.65006293 | 13.43129167 | 9.245516386 | 4.185775288 | 2.404546541 |
| Ccl7 | 12.54577914 | 11.25427238 | 10.16189971 | 1.092372674 | 2.383879428 |
| Fpr1 | 7.953899027 | 6.392290038 | 5.573325969 | 0.818964069 | 2.380573058 |
| Cd80 | 9.119354844 | 7.661681043 | 6.779291024 | 0.882390019 | 2.34006382 |
| Tnc | 13.8369703 | 13.17673535 | 11.51671402 | 1.660021333 | 2.320256287 |
| Timp1 | 11.77530112 | 10.72330501 | 9.463182449 | 1.260122565 | 2.312118668 |
| Rsad2 | 12.78316374 | 10.13015552 | 10.4713589 | -0.341203376 | 2.311804836 |
| Slc11a1 | 10.32434956 | 9.172750813 | 8.025278653 | 1.14747216 | 2.299070904 |
| Csrnp1 | 11.28389703 | 10.11383586 | 9.011588387 | 1.102247472 | 2.27230864 |
| Lcn2 | 10.3420601 | 9.137927729 | 8.073423975 | 1.064503754 | 2.26863612 |
| Ccn1 | 12.07444034 | 11.45929416 | 9.834595663 | 1.6246985 | 2.239844674 |
| Trim30b | 9.210892529 | 8.3256128 | 6.987785438 | 1.337827362 | 2.223107091 |
| Adam8 | 12.03423704 | 10.95548581 | 9.815941477 | 1.139544336 | 2.218295563 |
| Sh2d5 | 8.841973394 | 7.910116499 | 6.63138897 | 1.278727529 | 2.210584425 |
| Rab20 | 9.112493417 | 7.830646164 | 6.918727543 | 0.911918621 | 2.193765874 |
| Tarm1 | 8.521753294 | 7.510524243 | 6.347518029 | 1.163006215 | 2.174235265 |
| Slc2a3 | 9.613451211 | 7.59357165 | 7.441077266 | 0.152494384 | 2.172373945 |
| Ankrd33b | 9.979642925 | 9.233386424 | 7.812184664 | 1.421201761 | 2.167458261 |
| Trpm2 | 8.207763606 | 7.279560655 | 6.048135455 | 1.2314252 | 2.159628151 |
| Rab44 | 10.28652385 | 8.924436451 | 8.128238335 | 0.796198116 | 2.158285515 |
| Tlr2 | 10.51561355 | 9.500917332 | 8.358721406 | 1.142195927 | 2.156892148 |
| Lif | 7.885982708 | 6.304005666 | 5.737029976 | 0.56697569 | 2.148952732 |
| Gm13889 | 8.638212277 | 7.859712063 | 6.508716265 | 1.350995798 | 2.129496012 |
| Mt1 | 13.49145294 | 12.80134889 | 11.36436465 | 1.436984243 | 2.127088297 |
| Slc39a14 | 11.7198673 | 10.95562534 | 9.596374833 | 1.359250507 | 2.123492467 |
| Sell | 10.70582446 | 10.06193648 | 8.590568757 | 1.471367727 | 2.115255706 |
| Chil1 | 10.36593075 | 9.633960735 | 8.254043243 | 1.379917491 | 2.111887504 |
| Cxcl10 | 11.56362321 | 8.623377777 | 9.487005316 | -0.863627539 | 2.076617897 |

| | | | | | |
|---|---|---|---|---|---|
| Il1rn | 13.34718802 | 11.73039544 | 11.2730472 | 0.45734824 | 2.074140823 |
| Gm45551 | 10.13001494 | 9.143363762 | 8.063677005 | 1.079686756 | 2.066337931 |
| Klra2 | 8.841889857 | 7.690545922 | 6.776828915 | 0.913717008 | 2.065060942 |
| Dusp5 | 10.42559986 | 9.099591329 | 8.467361633 | 0.632229696 | 1.958238231 |
| Ier3 | 11.01287293 | 9.825281285 | 9.059899951 | 0.765381334 | 1.952972976 |
| Ifitm1 | 9.787157016 | 8.619413838 | 7.835490345 | 0.783923493 | 1.951666671 |
| Bcl2a1b | 8.831961095 | 7.946318199 | 6.883369806 | 1.062948393 | 1.948591289 |
| Arg2 | 8.363286555 | 7.179794872 | 6.426722671 | 0.753072201 | 1.936563884 |
| Adm | 9.221384495 | 7.739497474 | 7.286683565 | 0.452813909 | 1.93470093 |
| Cxcl9 | 9.103927595 | 6.783551423 | 7.177958369 | -0.394406946 | 1.925969226 |
| Sp140 | 10.25471325 | 8.21435057 | 8.350896556 | -0.136545986 | 1.903816691 |
| Rnf122 | 9.179787199 | 8.091662445 | 7.276301296 | 0.815361149 | 1.903485904 |
| Cd33 | 9.789736084 | 9.061023665 | 7.892001597 | 1.169022068 | 1.897734487 |
| Il4ra | 12.62255872 | 12.05572841 | 10.73027036 | 1.325458057 | 1.892288367 |
| Il18rap | 9.277195728 | 8.422186763 | 7.395010113 | 1.02717665 | 1.882185615 |
| Csf2rb | 12.13760204 | 11.24419505 | 10.25648589 | 0.987709157 | 1.881116147 |
| Bcl3 | 10.57450724 | 9.531068668 | 8.693486613 | 0.837582055 | 1.881020627 |
| Sgms2 | 9.327015054 | 8.30461535 | 7.448742868 | 0.855872482 | 1.878272186 |
| Olfr1033 | 12.34869406 | 11.38710079 | 10.47283611 | 0.914264677 | 1.875857947 |
| C5ar1 | 10.54731803 | 9.525997285 | 8.676305998 | 0.849691287 | 1.871012032 |
| Isg15 | 11.26453873 | 9.471694836 | 9.396726972 | 0.074967864 | 1.867811758 |
| Smim3 | 9.908859372 | 8.478864745 | 8.041157133 | 0.437707613 | 1.867702239 |
| Gm49339 | 10.5928598 | 10.25191866 | 8.728956772 | 1.522961891 | 1.863903028 |
| Selp | 9.701898413 | 8.661852658 | 7.838976501 | 0.822876158 | 1.862921913 |
| Slc16a3 | 10.74768979 | 9.49735564 | 8.885089345 | 0.612266295 | 1.862600442 |
| A530064D06Rik | 8.445710193 | 6.921129696 | 6.588114246 | 0.333015449 | 1.857595946 |
| Csf1 | 12.87726795 | 11.75506522 | 11.02722253 | 0.727842687 | 1.850045423 |
| Clec4n | 10.07999621 | 9.538842737 | 8.23197549 | 1.306867247 | 1.848020717 |
| Mefv | 9.511206532 | 8.763598878 | 7.663330988 | 1.10026789 | 1.847875544 |
| Iigp1 | 12.07738251 | 10.02481574 | 10.2605561 | -0.235740359 | 1.816826409 |
| Loxl4 | 9.745309886 | 8.974770144 | 7.930095008 | 1.044675136 | 1.815214878 |
| Mrgpra2a | 6.198420315 | 4.553180536 | 4.384321352 | 0.168859184 | 1.814098963 |

| | | | | | |
|---|---|---|---|---|---|
| Thbd | 12.22155138 | 11.04769483 | 10.41265038 | 0.635044453 | 1.808900997 |
| Itgal | 9.968003088 | 9.310949246 | 8.167552232 | 1.143397014 | 1.800450856 |
| Adamts9 | 10.63643955 | 9.574948646 | 8.836741896 | 0.738206751 | 1.799697654 |
| Gpr35 | 10.19708264 | 9.207330736 | 8.397408717 | 0.809922019 | 1.799673926 |
| Gadd45b | 10.29435018 | 9.3422076 | 8.49718053 | 0.84502707 | 1.797169647 |
| Hbegf | 10.09932995 | 9.330819007 | 8.310350277 | 1.02046873 | 1.788979676 |
| Junb | 11.57281619 | 11.0890226 | 9.784052023 | 1.304970574 | 1.78876417 |
| Rgs16 | 8.426990786 | 7.464828819 | 6.639141204 | 0.825687615 | 1.787849582 |
| Isg20 | 9.029382135 | 7.568627989 | 7.257773037 | 0.310854952 | 1.771609098 |
| Stx11 | 9.781205263 | 8.566946217 | 8.020020816 | 0.546925401 | 1.761184447 |
| Cxcr4 | 9.135988886 | 8.086719011 | 7.385413324 | 0.701305687 | 1.750575562 |
| Chil3 | 10.9283306 | 10.08666163 | 9.189596846 | 0.89706478 | 1.73873375 |
| Btg2 | 12.15306724 | 11.24708947 | 10.42178929 | 0.825300183 | 1.731277947 |
| Snx20 | 8.999911483 | 8.134331373 | 7.271789134 | 0.862542239 | 1.728122348 |
| Fgr | 10.26891062 | 9.68032981 | 8.561307067 | 1.119022743 | 1.70760355 |
| Tagap | 7.930086763 | 6.350913812 | 6.22759844 | 0.123315372 | 1.702488323 |
| Bcl2l11 | 9.70129041 | 8.700353204 | 8.004579447 | 0.695773757 | 1.696710963 |
| Marcksl1 | 11.04489837 | 9.882546927 | 9.351550891 | 0.530996037 | 1.693347476 |
| Cd53 | 12.01938527 | 11.03275755 | 10.32893625 | 0.7038213 | 1.69044902 |
| Cass4 | 8.82476804 | 7.632007977 | 7.13792107 | 0.494086907 | 1.686846971 |
| Pdpn | 11.06072314 | 10.57501908 | 9.378838573 | 1.19618051 | 1.68188457 |
| Tnfrsf1b | 11.4097754 | 10.65136836 | 9.731647259 | 0.919721101 | 1.678128144 |
| Dusp1 | 12.56498699 | 11.27313274 | 10.88949301 | 0.38363973 | 1.67549398 |
| F5 | 8.280116818 | 7.28643601 | 6.608154025 | 0.678281985 | 1.671962793 |
| Oas3 | 10.04645933 | 9.067181127 | 8.376302043 | 0.690879084 | 1.670157283 |
| Phlda1 | 9.571631468 | 8.73473152 | 7.912178311 | 0.822553209 | 1.659453156 |
| Pik3ap1 | 10.54152093 | 9.297245736 | 8.882304489 | 0.414941248 | 1.659216438 |
| Tnfaip2 | 11.83946167 | 10.75360013 | 10.1885096 | 0.565090526 | 1.65095207 |
| Traf1 | 9.207496011 | 8.261492789 | 7.561633566 | 0.699859223 | 1.645862445 |
| Slfn2 | 11.91225348 | 11.02621804 | 10.26958573 | 0.756632317 | 1.64266775 |
| Zfp36 | 12.35769548 | 11.61255401 | 10.72041696 | 0.892137047 | 1.637278513 |
| Car4 | 8.140636877 | 6.853552595 | 6.505063961 | 0.348488633 | 1.635572916 |

| | | | | | |
|---|---|---|---|---|---|
| Cd300ld | 9.604590221 | 8.311835268 | 7.98121217 | 0.330623098 | 1.623378051 |
| F10 | 8.964950759 | 8.987599529 | 7.345179411 | 1.642420118 | 1.619771347 |
| Saa3 | 9.716218187 | 8.671517661 | 8.112098937 | 0.559418725 | 1.604119251 |
| Cmpk2 | 10.60317322 | 8.540859835 | 9.002036171 | -0.461176336 | 1.601137049 |
| Arid5a | 10.34690638 | 9.796641262 | 8.749371338 | 1.047269924 | 1.597535038 |
| Hcar2 | 11.75995801 | 10.35282848 | 10.1647158 | 0.18811268 | 1.59524221 |
| Gsr | 11.19691007 | 10.3770548 | 9.602021241 | 0.775033559 | 1.594888833 |
| Ppp1r3b | 9.091241968 | 7.75680346 | 7.503021132 | 0.253782327 | 1.588220836 |
| Smox | 11.49266586 | 10.48294693 | 9.921338106 | 0.561608824 | 1.571327751 |
| Gadd45a | 9.720840249 | 8.583649923 | 8.16379609 | 0.419853833 | 1.557044159 |
| Angptl4 | 11.19214211 | 10.27359518 | 9.638410265 | 0.635184915 | 1.553731848 |
| Lmnb1 | 11.01319218 | 9.765165877 | 9.462956766 | 0.302209111 | 1.550235411 |
| Cd274 | 10.49782277 | 9.247566639 | 8.948498117 | 0.299068522 | 1.549324656 |
| Slc29a2 | 8.654348128 | 7.882163649 | 7.105537558 | 0.776626091 | 1.54881057 |
| Cd177 | 7.820656116 | 7.21547381 | 6.273156074 | 0.942317736 | 1.547500042 |
| Ptges | 12.49057195 | 11.410509 | 10.9438087 | 0.4667003 | 1.546763243 |
| Gbp5 | 10.01006839 | 8.452255936 | 8.46994937 | -0.017693434 | 1.540119017 |
| Ifit1 | 11.85787028 | 9.9633609 | 10.31796068 | -0.35459978 | 1.539909596 |
| Tubb6 | 11.05303849 | 10.65511975 | 9.51472595 | 1.1403938 | 1.538312537 |
| Sh2d3c | 8.574625887 | 7.907847204 | 7.04843522 | 0.859411985 | 1.526190667 |
| Ripor2 | 9.089138903 | 8.544021614 | 7.563107494 | 0.980914119 | 1.526031409 |
| Sbno2 | 12.4276998 | 12.05698541 | 10.90213725 | 1.154848163 | 1.525562553 |
| Apold1 | 8.452092295 | 7.59476447 | 6.929633483 | 0.665130987 | 1.522458812 |
| Entpd1 | 11.61441701 | 10.308689 | 10.09290003 | 0.215788971 | 1.521516981 |
| Gbp2 | 11.97569062 | 10.3596465 | 10.45564612 | -0.09599962 | 1.5200445 |
| Pde4b | 11.0713496 | 9.840146267 | 9.556531453 | 0.283614814 | 1.51481815 |
| Mmp9 | 9.789238102 | 8.842342105 | 8.2755522 | 0.566789905 | 1.513685902 |
| Upp1 | 7.939919483 | 7.03696643 | 6.427337165 | 0.609629264 | 1.512582318 |
| Il10ra | 10.02644028 | 8.884262349 | 8.514786157 | 0.369476191 | 1.511654119 |
| Alpk2 | 9.621737131 | 9.670573548 | 8.110970011 | 1.559603537 | 1.510767119 |
| Mxd1 | 12.14109552 | 11.15576257 | 10.63582542 | 0.519937147 | 1.505270103 |
| Gpr132 | 8.770399384 | 8.109821558 | 7.267899395 | 0.841922163 | 1.502499989 |

| | | | | | |
|---|---|---|---|---|---|
| Nfkbia | 12.59179858 | 11.58690161 | 11.09918306 | 0.487718553 | 1.492615523 |
| Fmnl1 | 10.80740318 | 10.00933737 | 9.319013959 | 0.690323409 | 1.488389221 |
| Ptafr | 10.37914826 | 9.92337366 | 8.891737815 | 1.031635845 | 1.487410449 |
| Rcan1 | 12.34889488 | 11.66156218 | 10.86330007 | 0.798262113 | 1.48559481 |
| Fosl1 | 8.916992532 | 8.464152901 | 7.434299797 | 1.029853104 | 1.482692735 |
| Pglyrp1 | 7.793080429 | 6.820298505 | 6.313729495 | 0.50656901 | 1.479350934 |
| Tnfaip3 | 11.05377458 | 9.882024662 | 9.583883527 | 0.298141136 | 1.469891053 |
| Nr4a1 | 10.99804089 | 10.51141824 | 9.528583419 | 0.982834818 | 1.469457468 |
| Gm6361 | 5.626279976 | 4.345572059 | 4.163543351 | 0.182028708 | 1.462736625 |
| Tm4sf1 | 10.54875827 | 9.779142374 | 9.087208636 | 0.691933739 | 1.461549634 |
| Srxn1 | 11.55851015 | 11.36954636 | 10.09718202 | 1.272364337 | 1.461328127 |
| Tlr6 | 7.711917439 | 6.971678334 | 6.253272987 | 0.718405347 | 1.458644452 |
| Krt6b | 14.30150983 | 14.01290221 | 12.84308225 | 1.169819963 | 1.458427587 |
| AB124611 | 8.374541843 | 7.525184911 | 6.921658289 | 0.603526622 | 1.452883554 |
| Klhl40 | 9.898659394 | 10.8564126 | 8.446844514 | 2.409568083 | 1.45181488 |
| Akap12 | 10.33405795 | 9.168340031 | 8.889386922 | 0.27895311 | 1.444671025 |
| Alpl | 9.402580562 | 8.671959898 | 7.965832815 | 0.706127082 | 1.436747747 |
| Lcp2 | 9.839051727 | 9.253482801 | 8.404317057 | 0.849165744 | 1.43473467 |
| Il1a | 10.08397128 | 8.719859207 | 8.649265662 | 0.070593545 | 1.434705615 |
| Helz2 | 11.23954604 | 10.29236518 | 9.809455167 | 0.482910013 | 1.430090876 |
| Csf2ra | 9.616375474 | 9.141562073 | 8.189342594 | 0.952219478 | 1.42703288 |
| P2ry2 | 8.994457366 | 8.194068217 | 7.568251113 | 0.625817104 | 1.426206253 |
| B4galt5 | 10.47669584 | 10.14320536 | 9.051736075 | 1.091469282 | 1.424959769 |
| Trib1 | 11.55012116 | 10.70159709 | 10.12856852 | 0.573028576 | 1.421552643 |
| Ly6c2 | 9.600171845 | 9.043367145 | 8.180790807 | 0.862576338 | 1.419381038 |
| Ncf4 | 8.966324809 | 8.49144532 | 7.551855948 | 0.939589372 | 1.414468861 |
| Sh2b2 | 8.571486193 | 7.972995963 | 7.158619127 | 0.814376836 | 1.412867066 |
| Epgn | 10.27827002 | 9.424127159 | 8.867458257 | 0.556668902 | 1.41081176 |
| Basp1 | 9.061402859 | 8.413496109 | 7.651273142 | 0.762222967 | 1.410129717 |
| Nfil3 | 9.619780538 | 9.261038766 | 8.222114806 | 1.03892396 | 1.397665733 |
| Tnfsf14 | 7.141761935 | 6.578719426 | 5.746964687 | 0.831754738 | 1.394797248 |
| Procr | 9.123355707 | 8.575645223 | 7.730619115 | 0.845026108 | 1.392736592 |

| | | | | | |
|---|---|---|---|---|---|
| Oas2 | 10.05927388 | 9.357361283 | 8.670136249 | 0.687225034 | 1.389137634 |
| Rhoh | 7.709439655 | 6.900623082 | 6.320464375 | 0.580158707 | 1.38897528 |
| Ppp1r15a | 11.02474809 | 10.50697971 | 9.64130038 | 0.865679333 | 1.383447713 |
| Siglece | 7.887293053 | 7.502812271 | 6.504216331 | 0.998595941 | 1.383076722 |
| Myd88 | 11.66742858 | 10.80501439 | 10.28703491 | 0.517979483 | 1.38039367 |
| Krt16 | 15.05801476 | 14.47924163 | 13.67803355 | 0.80120808 | 1.379981213 |
| Trex1 | 10.49583061 | 9.716303568 | 9.122934127 | 0.593369441 | 1.372896486 |
| Tlr13 | 9.221519437 | 9.084677875 | 7.85762122 | 1.227056656 | 1.363898217 |
| Ncf2 | 10.51755766 | 9.713665754 | 9.154147726 | 0.559518028 | 1.36340993 |
| Ifit3 | 12.5728024 | 10.25395579 | 11.21140316 | -0.957447371 | 1.361399243 |
| Myog | 6.863487465 | 6.62152782 | 5.508098921 | 1.113428899 | 1.355388544 |
| Glipr2 | 9.741489134 | 9.037679154 | 8.38770203 | 0.649977124 | 1.353787104 |
| Slfn1 | 8.310822901 | 7.89804172 | 6.957772786 | 0.940268934 | 1.353050114 |
| Ifi211 | 11.55629523 | 10.5685558 | 10.20472386 | 0.363831939 | 1.351571365 |
| G0s2 | 10.05841469 | 8.352381618 | 8.706912479 | -0.354530861 | 1.351502207 |
| Hk3 | 8.93701962 | 8.830415816 | 7.586297821 | 1.244117995 | 1.350721799 |
| Pim1 | 11.86222446 | 10.97630336 | 10.51298695 | 0.463316403 | 1.349237503 |
| Cebpd | 9.459336217 | 9.157101543 | 8.110417175 | 1.046684367 | 1.348919041 |
| Trim30a | 11.20852668 | 10.24852784 | 9.86026457 | 0.38826327 | 1.348262113 |
| Crispld2 | 12.15054351 | 11.3636052 | 10.80272122 | 0.560883987 | 1.347822293 |
| Syk | 10.73720211 | 10.14031277 | 9.394816056 | 0.745496714 | 1.34238605 |
| Lilra6 | 8.713765323 | 8.206486726 | 7.37416114 | 0.832325586 | 1.339604183 |
| Lyn | 10.91904466 | 10.26617599 | 9.581325121 | 0.684850869 | 1.337719542 |
| Hcls1 | 10.63819633 | 9.835036634 | 9.300889472 | 0.534147162 | 1.337306861 |
| Crem | 9.016136819 | 7.935737912 | 7.682021186 | 0.253716726 | 1.334115633 |
| Clic4 | 13.67189803 | 12.83602433 | 12.34068779 | 0.495336533 | 1.33121024 |
| Pram1 | 7.161847607 | 6.915884541 | 5.836023908 | 1.079860633 | 1.325823699 |
| Itgam | 11.57608104 | 10.99224774 | 10.25150322 | 0.740744514 | 1.324577817 |
| Ccdc88b | 8.38245866 | 8.238440587 | 7.059877787 | 1.1785628 | 1.322580873 |
| Ccl12 | 8.291255968 | 7.403806609 | 6.96886261 | 0.434943999 | 1.322393358 |
| Slc2a6 | 8.178983248 | 7.650724432 | 6.856651674 | 0.794072759 | 1.322331574 |
| Oasl2 | 11.52561335 | 10.56901527 | 10.2040671 | 0.36494817 | 1.321546253 |

| | | | | | |
|---|---|---|---|---|---|
| Il17ra | 11.16605093 | 10.70916529 | 9.846843912 | 0.862321378 | 1.319207018 |
| Ccl5 | 7.023972483 | 5.767719938 | 5.705362002 | 0.062357936 | 1.318610481 |
| Ccr5 | 10.20511643 | 9.957114715 | 8.894436848 | 1.062677867 | 1.310679579 |
| Hck | 9.393143351 | 8.836861652 | 8.083722842 | 0.75313881 | 1.309420509 |
| Nrg1 | 7.224623747 | 6.55657397 | 5.915850024 | 0.640723945 | 1.308773722 |
| Gbp6 | 10.4777061 | 8.91116628 | 9.170161562 | -0.258995282 | 1.307544538 |
| Egr1 | 12.33988097 | 12.34674513 | 11.0365995 | 1.310145623 | 1.303281463 |
| Ier5 | 11.18281722 | 10.56947919 | 9.880502837 | 0.688976353 | 1.302314379 |
| Vcam1 | 9.553227421 | 8.175699387 | 8.252912952 | -0.077213565 | 1.300314469 |
| Ifi209 | 10.49187836 | 9.769404202 | 9.196726816 | 0.572677386 | 1.295151541 |
| Zbp1 | 10.49460022 | 9.496554688 | 9.200616392 | 0.295938297 | 1.293983832 |
| Prss27 | 8.792268434 | 7.982516764 | 7.499412523 | 0.48310424 | 1.292855911 |
| Stom | 11.98949907 | 11.22930786 | 10.69776218 | 0.531545683 | 1.29173689 |
| Sirpb1b | 7.875379458 | 7.392353162 | 6.586472648 | 0.805880514 | 1.28890681 |
| Dusp2 | 7.322692242 | 7.061226437 | 6.038005288 | 1.023221148 | 1.284686954 |
| Ifi204 | 12.07374798 | 11.12144861 | 10.79291429 | 0.328534323 | 1.280833697 |
| Tmprss11g | 7.755695292 | 6.688026553 | 6.475750227 | 0.212276326 | 1.279945066 |
| Itgb2 | 11.25072431 | 10.78868673 | 9.971283793 | 0.817402941 | 1.279440517 |
| Cd300lb | 8.743677707 | 7.876109637 | 7.464611108 | 0.411498528 | 1.279066599 |
| Ccl17 | 6.852474796 | 6.486581631 | 5.575356423 | 0.911225208 | 1.277118373 |
| Tyrobp | 11.30375695 | 10.62430601 | 10.02745996 | 0.596846052 | 1.276296995 |
| Chst11 | 9.143667538 | 8.278865957 | 7.868862367 | 0.410003589 | 1.274805171 |
| Rac2 | 9.884831793 | 9.324382711 | 8.614688618 | 0.709694093 | 1.270143175 |
| Igtp | 10.90325869 | 9.662891909 | 9.635647371 | 0.027244537 | 1.267611319 |
| H2-Q10 | 9.502449202 | 8.720007868 | 8.236196337 | 0.483811531 | 1.266252865 |
| Icam1 | 11.00801738 | 10.44602582 | 9.750502601 | 0.695523215 | 1.257514779 |
| Sod2 | 12.42859707 | 11.59633294 | 11.17383524 | 0.4224977 | 1.254761827 |
| Ccr3 | 6.839678966 | 6.060705718 | 5.587800973 | 0.472904745 | 1.251877993 |
| Runx1 | 9.973792738 | 9.624361486 | 8.722417394 | 0.901944092 | 1.251375344 |
| Ddah1 | 9.637588231 | 9.25083992 | 8.387205317 | 0.863634603 | 1.250382914 |
| Irf7 | 11.09870174 | 9.986541267 | 9.851862138 | 0.134679129 | 1.246839605 |
| Tmem173 | 9.849090072 | 9.310134377 | 8.6049746 | 0.705159777 | 1.244115472 |

| | | | | | |
|---|---|---|---|---|---|
| Akap2 | 11.56900978 | 10.92695381 | 10.32547625 | 0.60147756 | 1.24353353 |
| Padi4 | 7.609276768 | 6.94573973 | 6.368734074 | 0.577005657 | 1.240542694 |
| Tnfrsf12a | 10.30930177 | 10.34466116 | 9.071819366 | 1.272841791 | 1.237482408 |
| Mustn1 | 10.20639429 | 11.38921387 | 8.978759443 | 2.41045443 | 1.227634846 |
| Ifi205 | 12.40255405 | 11.19314685 | 11.17761724 | 0.015529603 | 1.224936807 |
| Oasl1 | 9.616606052 | 8.345078808 | 8.393326405 | -0.048247598 | 1.223279647 |
| Atp1a3 | 7.948442154 | 7.168773276 | 6.725679203 | 0.443094073 | 1.222762951 |
| Ifit3b | 10.4342284 | 8.481405692 | 9.21276482 | -0.731359128 | 1.221463584 |
| Gk | 10.23459193 | 9.554271257 | 9.013385178 | 0.540886078 | 1.221206748 |
| Nfkbie | 9.06732826 | 8.520179514 | 7.846577846 | 0.673601668 | 1.220750414 |
| S1pr3 | 8.719383138 | 7.880821829 | 7.500093056 | 0.380728774 | 1.219290082 |
| 5430427O19Rik | 7.047854972 | 6.107612974 | 5.829516612 | 0.278096362 | 1.21833836 |
| Dennd1c | 8.608410549 | 8.08994729 | 7.392614145 | 0.697333145 | 1.215796404 |
| Trim30c | 7.35687943 | 6.896490873 | 6.142619251 | 0.753871622 | 1.214260179 |
| Cd44 | 13.73049116 | 12.95881482 | 12.51844719 | 0.440367627 | 1.21204397 |
| Cebpb | 10.45659546 | 9.895419426 | 9.244960082 | 0.650459344 | 1.211635375 |
| Gch1 | 9.444275711 | 8.519050926 | 8.234177364 | 0.284873561 | 1.210098347 |
| Gm10108 | 6.426797771 | 5.455519205 | 5.217583757 | 0.237935448 | 1.209214014 |
| Kdm6b | 11.21606295 | 10.7286942 | 10.00945602 | 0.719238178 | 1.206606932 |
| Samhd1 | 12.73087926 | 11.80209119 | 11.52433999 | 0.277751197 | 1.20653927 |
| Calhm6 | 6.695933241 | 5.412312212 | 5.492384344 | -0.080072132 | 1.203548897 |
| Pilra | 8.167504964 | 7.577224844 | 6.96871055 | 0.608514294 | 1.198794414 |
| Ipcef1 | 7.082301531 | 6.03828589 | 5.885255612 | 0.153030278 | 1.19704592 |
| Mndal | 10.9891573 | 9.685585499 | 9.80082445 | -0.115238951 | 1.188332847 |
| Nfkbiz | 11.07449575 | 10.25130191 | 9.890382425 | 0.360919482 | 1.184113322 |
| Phf11d | 9.954611439 | 8.654710934 | 8.772741682 | -0.118030748 | 1.181869757 |
| Myc | 10.78539639 | 10.44103223 | 9.60567907 | 0.83535316 | 1.17971732 |
| Alox15 | 7.720023689 | 7.042805688 | 6.541458733 | 0.501346956 | 1.178564956 |
| Dennd4a | 10.66399467 | 9.747729985 | 9.485881921 | 0.261848064 | 1.178112749 |
| Fos | 10.89519861 | 10.75379503 | 9.71716517 | 1.036629857 | 1.17803344 |
| Hmox1 | 12.52036667 | 12.88497792 | 11.34389151 | 1.541086413 | 1.176475157 |
| Rasip1 | 9.222211286 | 8.461895039 | 8.046154778 | 0.415740261 | 1.176056508 |

| | | | | | |
|---|---|---|---|---|---|
| Plk3 | 10.78320446 | 9.911579181 | 9.611490825 | 0.300088356 | 1.171713635 |
| Fcgr4 | 7.75277995 | 7.055752069 | 6.581198865 | 0.474553204 | 1.171581086 |
| Rnf213 | 12.57286639 | 11.67280165 | 11.40353346 | 0.269268193 | 1.16933293 |
| Hbb-bs | 12.27092146 | 10.81774635 | 11.10202052 | -0.284274166 | 1.168900944 |
| Mx1 | 9.995724941 | 8.28101483 | 8.830312241 | -0.549297411 | 1.165412699 |
| AA467197 | 5.957306422 | 5.044874383 | 4.794935159 | 0.249939224 | 1.162371263 |
| Mmp25 | 7.760113132 | 7.122641785 | 6.599882552 | 0.522759233 | 1.16023058 |
| Lmcd1 | 10.16643954 | 10.83319756 | 9.0062325 | 1.826965057 | 1.160207044 |
| Trim30d | 9.973574817 | 9.060721002 | 8.813688748 | 0.247032255 | 1.159886069 |
| Tgm2 | 12.68473739 | 11.9575832 | 11.52518727 | 0.43239593 | 1.15955012 |
| Cd244a | 8.789585342 | 8.43987501 | 7.631501561 | 0.808373449 | 1.158083781 |
| Ptprc | 11.72076214 | 11.03389324 | 10.56460044 | 0.469292803 | 1.1561617 |
| Vcan | 11.25901968 | 10.89716301 | 10.10332356 | 0.793839446 | 1.155696116 |
| Itgax | 9.344893724 | 8.868570001 | 8.191436616 | 0.677133385 | 1.153457108 |
| Maff | 9.579529028 | 9.194276533 | 8.426619128 | 0.767657404 | 1.1529099 |
| Bhlhe40 | 11.85998456 | 11.03558069 | 10.70853221 | 0.32704848 | 1.151452353 |
| Cyth4 | 10.4444589 | 9.826942437 | 9.293779833 | 0.533162604 | 1.150679071 |
| Aqp3 | 13.67469808 | 13.01437034 | 12.52409121 | 0.490279127 | 1.15060687 |
| Lcp1 | 12.7743119 | 12.18967467 | 11.62399103 | 0.565683647 | 1.150320873 |
| Slc7a5 | 10.96673446 | 10.3300233 | 9.818155736 | 0.511867568 | 1.148578724 |
| Themis2 | 9.830992438 | 8.894780541 | 8.683602269 | 0.211178272 | 1.147390169 |
| Apobr | 9.30101844 | 8.788534316 | 8.15435688 | 0.634177436 | 1.14666156 |
| Gm12250 | 8.802701405 | 7.506002238 | 7.657205264 | -0.151203026 | 1.145496141 |
| Gsta4 | 9.954388702 | 9.432638267 | 8.809977164 | 0.622661103 | 1.144411537 |
| Adamts1 | 11.89121826 | 11.3832832 | 10.74735745 | 0.635925747 | 1.143860803 |
| Krt6a | 14.79940961 | 14.49037659 | 13.65667047 | 0.83370612 | 1.142739133 |
| Fgl2 | 12.86265209 | 12.26495825 | 11.72382492 | 0.541133327 | 1.13882717 |
| Zyx | 11.58145178 | 11.19982409 | 10.44351271 | 0.756311377 | 1.137939067 |
| Gm14548 | 8.287961496 | 7.692168488 | 7.150649906 | 0.541518582 | 1.13731159 |
| Csf2rb2 | 10.57957452 | 10.20461146 | 9.442427363 | 0.762184101 | 1.137147154 |
| Zfp729a | 11.70702539 | 10.8871997 | 10.57137071 | 0.315828993 | 1.135654683 |
| Flnc | 12.6326158 | 13.70392523 | 11.49716945 | 2.20675578 | 1.135446353 |

| | | | | | |
|---|---|---|---|---|---|
| Bcl2a1d | 7.814985035 | 7.369462998 | 6.68033475 | 0.689128248 | 1.134650285 |
| Galnt6 | 10.29361769 | 9.522099791 | 9.159759687 | 0.362340104 | 1.133858006 |
| Hif1a | 12.72544648 | 11.89434382 | 11.59311749 | 0.301226333 | 1.132328993 |
| Il33 | 11.47511588 | 10.76440754 | 10.34362426 | 0.42078328 | 1.13149162 |
| Slfn8 | 9.303812565 | 8.51615614 | 8.174364639 | 0.341791502 | 1.129447926 |
| Odc1 | 12.59319709 | 11.92516158 | 11.46541837 | 0.45974321 | 1.127778727 |
| Agap2 | 7.21023056 | 6.499497096 | 6.084771389 | 0.414725707 | 1.12545917 |
| Atf3 | 9.880861204 | 10.99898779 | 8.756497478 | 2.242490308 | 1.124363726 |
| Ifi203 | 11.48021183 | 10.28872322 | 10.35630597 | -0.067582743 | 1.123905863 |
| Olfm4 | 5.593208183 | 4.782086225 | 4.46946048 | 0.312625745 | 1.123747704 |
| Herc6 | 9.568889579 | 8.586825891 | 8.451670968 | 0.135154923 | 1.117218611 |
| Gadd45g | 8.82175442 | 9.091539156 | 7.706125738 | 1.385413417 | 1.115628682 |
| Fgd3 | 8.904041991 | 8.1547766 | 7.789552897 | 0.365223703 | 1.114489094 |
| Pde1b | 8.994189098 | 8.303318778 | 7.880658926 | 0.422659853 | 1.113530172 |
| Ms4a4c | 9.550711565 | 8.799774875 | 8.438859431 | 0.360915444 | 1.111852134 |
| Ctps | 10.49567768 | 9.999872439 | 9.38545927 | 0.614413169 | 1.11021841 |
| Rtp4 | 10.1523623 | 9.246910147 | 9.043117092 | 0.203793055 | 1.109245208 |
| Serpina3f | 6.111525013 | 5.227978746 | 5.002386714 | 0.225592032 | 1.109138299 |
| Parp14 | 11.73181061 | 10.85881296 | 10.62343016 | 0.2353828 | 1.108380453 |
| B3gnt5 | 6.656758741 | 5.865461072 | 5.549056471 | 0.3164046 | 1.10770227 |
| Gvin1 | 11.62889347 | 10.34735632 | 10.52237279 | -0.17501647 | 1.106520677 |
| Prg4 | 10.13969998 | 9.444297826 | 9.033790418 | 0.410507409 | 1.105909559 |
| Nek6 | 9.605513283 | 9.225583673 | 8.500011275 | 0.725572398 | 1.105502008 |
| Litaf | 12.5706724 | 11.95640339 | 11.46838654 | 0.488016857 | 1.10228586 |
| Tslp | 7.20314548 | 5.987661359 | 6.101508785 | -0.113847426 | 1.101636695 |
| Itga5 | 11.05941327 | 10.58603151 | 9.959408891 | 0.626622619 | 1.100004379 |
| Irak2 | 10.17069188 | 9.451074806 | 9.071277074 | 0.379797732 | 1.099414808 |
| Ngf | 7.766761768 | 7.212300153 | 6.669426468 | 0.542873685 | 1.0973353 |
| Cfb | 11.86286625 | 10.49838004 | 10.77010755 | -0.27172751 | 1.092758703 |
| I830077J02Rik | 7.995465078 | 7.466669914 | 6.906466921 | 0.560202993 | 1.088998158 |
| Adgrg3 | 7.277239483 | 6.815675581 | 6.193297751 | 0.62237783 | 1.083941732 |
| Psg16 | 10.40093019 | 9.661736104 | 9.317611489 | 0.344124616 | 1.083318698 |

| | | | | | |
|---|---|---|---|---|---|
| Plekho2 | 11.13802517 | 10.77049261 | 10.06073329 | 0.709759315 | 1.077291879 |
| Selenon | 10.49254582 | 9.588019295 | 9.415599855 | 0.17241944 | 1.076945968 |
| Dusp6 | 10.80195833 | 9.937710905 | 9.726110311 | 0.211600595 | 1.075848023 |
| Casp4 | 9.917756997 | 9.127403984 | 8.842576589 | 0.284827395 | 1.075180408 |
| Gm4841 | 7.566478616 | 6.181434195 | 6.493054804 | -0.311620609 | 1.073423813 |
| Mpeg1 | 12.01330798 | 11.67861221 | 10.93995332 | 0.73865889 | 1.07335466 |
| Oas1a | 9.802219603 | 9.051833682 | 8.72931683 | 0.322516852 | 1.072902773 |
| Relt | 7.8554328 | 7.751163725 | 6.783233344 | 0.967930381 | 1.072199456 |
| Urah | 12.31606838 | 11.7895894 | 11.24419179 | 0.545397613 | 1.07187659 |
| Plac8 | 10.24989499 | 9.519888677 | 9.178405057 | 0.34148362 | 1.071489937 |
| Cd52 | 10.60609213 | 9.966458649 | 9.535637744 | 0.430820905 | 1.07045439 |
| Gsdmc | 10.57715691 | 10.4583261 | 9.506793927 | 0.951532176 | 1.070362983 |
| Cytip | 10.03804428 | 9.45087158 | 8.968146552 | 0.482725028 | 1.069897723 |
| Trmt61a | 8.910336028 | 8.521856641 | 7.840998683 | 0.680857958 | 1.069337345 |
| Pvr | 9.537577023 | 9.340147575 | 8.470084409 | 0.870063166 | 1.067492614 |
| Xylt1 | 9.467609473 | 8.877293613 | 8.401273564 | 0.476020049 | 1.066335909 |
| Gm4070 | 11.52187427 | 10.51287591 | 10.45735389 | 0.05552202 | 1.064520373 |
| Srm | 10.61412117 | 10.14726014 | 9.549637247 | 0.597622897 | 1.064483927 |
| Rrp1b | 9.671543757 | 9.303223101 | 8.608233339 | 0.694989762 | 1.063310417 |
| Clec5a | 8.15628095 | 7.576758572 | 7.098216607 | 0.478541966 | 1.058064343 |
| Bcat1 | 7.7348823 | 7.20217687 | 6.677744889 | 0.524431981 | 1.05713741 |
| Sp110 | 9.671085416 | 8.635321803 | 8.614976231 | 0.020345571 | 1.056109184 |
| Slc10a6 | 9.454812981 | 8.732950854 | 8.398841511 | 0.334109343 | 1.055971469 |
| Tes | 9.657492104 | 8.909540364 | 8.602572421 | 0.306967943 | 1.054919683 |
| Nop2 | 10.64607901 | 10.14483441 | 9.598281686 | 0.546552729 | 1.047797321 |
| Dhrs9 | 9.511691439 | 8.708595497 | 8.465508914 | 0.243086583 | 1.046182525 |
| Rtn4rl2 | 6.628336095 | 6.139958188 | 5.582397907 | 0.557560281 | 1.045938188 |
| Ackr1 | 9.736676602 | 9.081810834 | 8.691286649 | 0.390524186 | 1.045389954 |
| Rrp9 | 9.499019675 | 8.855151906 | 8.453648599 | 0.401503307 | 1.045371076 |
| Ankrd2 | 8.835881209 | 8.85062214 | 7.79056678 | 1.060055361 | 1.045314429 |
| Arhgap4 | 8.278712057 | 7.740540595 | 7.233804366 | 0.506736229 | 1.04490769 |
| Lrrc32 | 9.673695837 | 9.356314746 | 8.628967411 | 0.727347335 | 1.044728426 |

| | | | | | |
|---|---|---|---|---|---|
| Il1r2 | 11.62228475 | 10.7665516 | 10.57794089 | 0.188610713 | 1.044343867 |
| Nop58 | 11.55729224 | 10.99886381 | 10.51305143 | 0.485812383 | 1.04424081 |
| Gm8995 | 10.46445804 | 9.84364875 | 9.423164546 | 0.420484205 | 1.041293491 |
| Rnf149 | 11.14976915 | 10.38824852 | 10.11088804 | 0.277360483 | 1.038881113 |
| Vav1 | 9.463312985 | 9.3088122 | 8.42807467 | 0.88073753 | 1.035238315 |
| Jak2 | 11.49587763 | 10.8494587 | 10.46119778 | 0.38826092 | 1.034679847 |
| Ltb4r1 | 9.460160196 | 9.007353183 | 8.426896998 | 0.580456185 | 1.033263198 |
| Osmr | 11.63411788 | 11.07908546 | 10.60496119 | 0.47412427 | 1.029156687 |
| Shb | 9.587131776 | 8.953076758 | 8.559904729 | 0.393172029 | 1.027227047 |
| Rab31 | 11.20639789 | 10.53234252 | 10.17958286 | 0.352759653 | 1.026815027 |
| Usp18 | 9.941367348 | 8.660244617 | 8.914677411 | -0.254432794 | 1.026689937 |
| Fosl2 | 12.62873123 | 11.99383364 | 11.60264434 | 0.391189303 | 1.026086897 |
| Gsap | 8.555362566 | 8.006434183 | 7.529598693 | 0.47683549 | 1.025763873 |
| Gm18853 | 7.487489681 | 6.648052876 | 6.462778462 | 0.185274415 | 1.02471122 |
| Grwd1 | 9.654741852 | 9.126899082 | 8.632176119 | 0.494722963 | 1.022565733 |
| Ms4a6d | 10.64225192 | 10.25057307 | 9.620227792 | 0.630345275 | 1.022024125 |
| Krt17 | 14.81427642 | 14.55477367 | 13.79232738 | 0.76244629 | 1.021949037 |
| B430306N03Rik | 7.04693867 | 6.734553705 | 6.028367398 | 0.706186307 | 1.018571272 |
| Sdc4 | 13.52307157 | 12.72107357 | 12.5061833 | 0.21489027 | 1.016888273 |
| Vmn1r46 | 8.425168178 | 7.690282074 | 7.408890215 | 0.281391859 | 1.016277963 |
| Tmem252 | 7.561405832 | 6.918558577 | 6.54611933 | 0.372439247 | 1.015286502 |
| Osgin1 | 8.432914906 | 9.319890734 | 7.418452182 | 1.901438552 | 1.014462724 |
| Ifitm6 | 9.822492643 | 8.896810736 | 8.809954677 | 0.086856059 | 1.012537967 |
| Siglecg | 6.959261174 | 6.231157831 | 5.94749044 | 0.283667392 | 1.011770734 |
| Pirb | 11.02467861 | 10.71033771 | 10.01295009 | 0.697387615 | 1.011728519 |
| Pla2g7 | 10.08537961 | 9.569307603 | 9.07478495 | 0.494522654 | 1.010594658 |
| Hdac7 | 11.00769599 | 10.46757215 | 9.999177541 | 0.468394606 | 1.008518446 |
| Ifi207 | 11.51749323 | 10.95223748 | 10.50917941 | 0.443058077 | 1.008313823 |
| Dgat1 | 11.06335412 | 10.03926735 | 10.05676062 | -0.017493269 | 1.006593498 |
| Ifih1 | 10.93952731 | 9.679123265 | 9.93699664 | -0.257873375 | 1.002530667 |
| Nod2 | 9.2088935 | 8.782901205 | 8.208622913 | 0.574278292 | 1.000270587 |
| Evi2b | 9.165220329 | 8.702961512 | 8.165651107 | 0.537310405 | 0.999569222 |

| | | | | | |
|---|---|---|---|---|---|
| Fcgr3 | 10.64187607 | 10.07345452 | 9.642488766 | 0.430965751 | 0.999387301 |
| Enc1 | 10.65416986 | 10.09851178 | 9.655178391 | 0.443333389 | 0.998991466 |
| Gm6377 | 7.281379216 | 6.808636745 | 6.286732809 | 0.521903937 | 0.994646408 |
| Tiparp | 10.683201 | 9.769267209 | 9.68999848 | 0.079268729 | 0.993202516 |
| Ifit2 | 10.19146193 | 8.843474399 | 9.198552743 | -0.355078344 | 0.992909185 |
| Atp8b4 | 8.164774893 | 7.712771994 | 7.172549465 | 0.540222529 | 0.992225428 |
| Slc25a48 | 10.65731252 | 9.989352123 | 9.665289191 | 0.324062931 | 0.992023329 |
| Stat3 | 13.22723506 | 12.77762665 | 12.23609846 | 0.541528183 | 0.9911366 |
| Sorl1 | 10.84831217 | 10.31218483 | 9.85750831 | 0.45467652 | 0.990803864 |
| Hivep3 | 10.38897633 | 9.743822744 | 9.398570726 | 0.345252019 | 0.990405604 |
| Gla | 8.836238892 | 8.679902439 | 7.846429773 | 0.833472665 | 0.989809119 |
| Ddx21 | 12.27364428 | 11.78557333 | 11.28406999 | 0.501503337 | 0.98957429 |
| Fas | 9.1418296 | 8.387968152 | 8.154421341 | 0.233546811 | 0.987408259 |
| Has1 | 8.277511824 | 7.866182099 | 7.29058864 | 0.575593459 | 0.986923184 |
| Ier2 | 10.07218454 | 9.97308814 | 9.086080495 | 0.887007645 | 0.98610404 |
| Efhd2 | 11.50682271 | 11.21829212 | 10.52198245 | 0.696309677 | 0.984840263 |
| Sla | 9.320819305 | 8.838069999 | 8.336003729 | 0.50206627 | 0.984815576 |
| Ifi213 | 8.465065257 | 7.690892818 | 7.480315925 | 0.210576893 | 0.984749332 |
| Prss12 | 8.833596694 | 8.256815502 | 7.849910609 | 0.406904893 | 0.983686084 |
| Stk17b | 10.97711823 | 10.26224392 | 9.994202975 | 0.268040945 | 0.982915258 |
| Rin3 | 9.499161765 | 9.284213454 | 8.519962577 | 0.764250877 | 0.979199188 |
| Mlkl | 8.919386796 | 8.11156846 | 7.940353351 | 0.171215109 | 0.979033445 |
| Fcgr1 | 9.920955975 | 9.54322139 | 8.942347829 | 0.600873561 | 0.978608146 |
| Elfn1 | 5.84622511 | 5.422867678 | 4.868901252 | 0.553966427 | 0.977323859 |
| Gcnt2 | 11.59877313 | 10.922374 | 10.62264297 | 0.29973103 | 0.97613016 |
| Heatr1 | 10.88909411 | 10.46643919 | 9.914698052 | 0.551741141 | 0.974396061 |
| Ampd3 | 10.2361907 | 9.875086874 | 9.262345276 | 0.612741598 | 0.973845421 |
| Gnl3 | 11.04111663 | 10.48969778 | 10.0676904 | 0.422007377 | 0.973426227 |
| St3gal1 | 11.39302981 | 10.8450311 | 10.4202672 | 0.4247639 | 0.97276261 |
| Noc4l | 9.664217165 | 9.191254082 | 8.692051678 | 0.499202404 | 0.972165487 |
| Emilin2 | 11.28159049 | 10.93030327 | 10.31101924 | 0.619284027 | 0.97057125 |
| Pwp2 | 9.946144804 | 9.533217434 | 8.975999182 | 0.557218252 | 0.970145622 |

| | | | | | |
|---|---|---|---|---|---|
| Treml2 | 6.250113787 | 5.980664164 | 5.2803835 | 0.700280664 | 0.969730288 |
| Fxyd5 | 11.53213343 | 10.86553348 | 10.56327388 | 0.302259607 | 0.968859553 |
| Rnasel | 10.33978482 | 9.601211495 | 9.372653647 | 0.228557848 | 0.96713117 |
| Ifi206 | 8.37800997 | 7.423540368 | 7.411083534 | 0.012456834 | 0.966926435 |
| Spi1 | 9.757427172 | 9.424911045 | 8.790957826 | 0.633953219 | 0.966469346 |
| Gbp3 | 10.25497575 | 8.813097725 | 9.28862342 | -0.475525695 | 0.966352334 |
| Zc3h12a | 10.19177544 | 9.632433528 | 9.22610272 | 0.406330808 | 0.965672717 |
| Arl4d | 9.258992927 | 8.482497405 | 8.29345507 | 0.189042334 | 0.965537856 |
| Sp100 | 10.94911443 | 9.975610989 | 9.984460105 | -0.008849116 | 0.964654325 |
| Ifi47 | 9.995103606 | 8.942839511 | 9.03105734 | -0.088217829 | 0.964046266 |
| Gpsm3 | 8.605175153 | 7.949973229 | 7.642448104 | 0.307525125 | 0.962727049 |
| Rrp12 | 10.32330384 | 9.85752797 | 9.36058044 | 0.49694753 | 0.9627234 |
| Tet1 | 6.701302714 | 5.898574104 | 5.739258935 | 0.159315169 | 0.962043779 |
| Coro1a | 11.14792778 | 10.61979072 | 10.18966966 | 0.430121063 | 0.95825812 |
| Stat5a | 10.63781005 | 10.15416866 | 9.682752997 | 0.471415666 | 0.955057053 |
| Nop56 | 11.37999331 | 10.86652121 | 10.42570774 | 0.44081347 | 0.95428557 |
| Nolc1 | 11.3052677 | 10.78900518 | 10.35100342 | 0.43800176 | 0.95426428 |
| Gda | 12.29454197 | 11.61684125 | 11.34039189 | 0.276449367 | 0.95415008 |
| Glipr1 | 7.808663371 | 7.34160785 | 6.854767994 | 0.486839856 | 0.953895377 |
| Pstpip1 | 8.847912801 | 8.446924331 | 7.897645319 | 0.549279012 | 0.950267483 |
| Vps37b | 9.521965276 | 9.150694701 | 8.573964374 | 0.576730328 | 0.948000902 |
| Slc2a1 | 11.35272183 | 10.9316498 | 10.40710925 | 0.524540553 | 0.94561258 |
| Neurl3 | 9.093249277 | 8.263974342 | 8.147999587 | 0.115974756 | 0.945249691 |
| Isy1 | 10.15000426 | 9.500629895 | 9.206877382 | 0.293752513 | 0.943126876 |
| Ppan | 9.737562074 | 9.457194304 | 8.79530026 | 0.661894043 | 0.942261813 |
| Acap1 | 7.919769771 | 7.276000589 | 6.978053875 | 0.297946714 | 0.941715895 |
| Ptgir | 8.495360372 | 7.802643261 | 7.55444155 | 0.24820171 | 0.940918821 |
| Samd9l | 10.76839842 | 9.785955576 | 9.827601916 | -0.04164634 | 0.940796501 |
| Trpv2 | 9.15151073 | 8.662339321 | 8.211982403 | 0.450356918 | 0.939528327 |
| Tnip1 | 10.61485729 | 9.984729226 | 9.675386345 | 0.309342881 | 0.939470945 |
| Lgals9 | 11.23046267 | 10.48593347 | 10.29106259 | 0.194870873 | 0.939400077 |
| Ltb4r2 | 10.021497 | 9.53174289 | 9.083227107 | 0.448515783 | 0.938269896 |

| | | | | | |
|---|---|---|---|---|---|
| Fam107b | 11.45245244 | 10.53193048 | 10.51563449 | 0.016295993 | 0.936817947 |
| Jdp2 | 8.420008783 | 7.684965138 | 7.484115221 | 0.200849917 | 0.935893562 |
| Kri1 | 10.07947673 | 9.408878889 | 9.143584518 | 0.265294371 | 0.935892212 |
| Gm2564 | 9.977329042 | 9.824947042 | 9.0416488 | 0.783298243 | 0.935680242 |
| Tnfrsf26 | 7.139145644 | 6.489665791 | 6.204187855 | 0.285477935 | 0.934957788 |
| Slc7a1 | 10.48099046 | 10.01053453 | 9.546576007 | 0.463958524 | 0.93441445 |
| Hilpda | 8.682207904 | 7.891246401 | 7.747949759 | 0.143296642 | 0.934258145 |
| Pprc1 | 11.02734062 | 10.56214234 | 10.09337102 | 0.468771315 | 0.933969595 |
| Slco2a1 | 12.00393439 | 11.2999185 | 11.07045892 | 0.22945958 | 0.933475463 |
| Ptprj | 10.70643444 | 9.964321584 | 9.775437504 | 0.188884079 | 0.930996939 |
| Ccl9 | 12.35300706 | 12.03354323 | 11.42230697 | 0.611236263 | 0.93070009 |
| I830127L07Rik | 6.449158138 | 5.982709796 | 5.518778503 | 0.463931293 | 0.930379636 |
| Gm4951 | 8.584214763 | 7.377287129 | 7.653971974 | -0.276684845 | 0.93024279 |
| Alox8 | 6.226726986 | 5.351721858 | 5.297478082 | 0.054243777 | 0.929248904 |
| Xaf1 | 9.593220029 | 8.67109296 | 8.664211925 | 0.006881036 | 0.929008104 |
| Sema4d | 10.49197224 | 9.944273101 | 9.56425197 | 0.380021131 | 0.92772027 |
| Cybb | 10.59726122 | 10.03468424 | 9.669580381 | 0.365103856 | 0.927680843 |
| Cgas | 9.06598583 | 8.399819947 | 8.139304172 | 0.260515775 | 0.926681658 |
| Plscr1 | 7.368509523 | 6.666259059 | 6.441992274 | 0.224266785 | 0.926517249 |
| Bcl2a1a | 6.679432248 | 6.41288744 | 5.754668759 | 0.658218681 | 0.924763489 |
| Gpatch4 | 9.817350685 | 9.22083043 | 8.893149347 | 0.327681083 | 0.924201338 |
| Gm1966 | 8.511064324 | 8.141080229 | 7.587283431 | 0.553796798 | 0.923780893 |
| Malt1 | 10.37121299 | 9.615011608 | 9.449607247 | 0.165404361 | 0.92160574 |
| Ccl6 | 12.1977957 | 11.55423019 | 11.27917322 | 0.27505697 | 0.91862248 |
| Bnc1 | 9.775299027 | 9.210348105 | 8.860551153 | 0.349796952 | 0.914747874 |
| Dhx58 | 9.198766886 | 8.501340071 | 8.284939807 | 0.216400264 | 0.913827079 |
| Slc19a1 | 8.602607605 | 8.166950277 | 7.688817236 | 0.478133041 | 0.913790369 |
| Pira2 | 8.367425864 | 8.219670894 | 7.453936318 | 0.765734576 | 0.913489546 |
| Gfpt2 | 9.860006775 | 9.433059168 | 8.947989543 | 0.485069625 | 0.912017233 |
| Nrros | 10.20322969 | 10.04149743 | 9.292483771 | 0.749013656 | 0.910745916 |
| Atp10a | 9.982950626 | 9.205302917 | 9.072730809 | 0.132572108 | 0.910219817 |
| Nfe2 | 6.790415271 | 6.331621311 | 5.881311446 | 0.450309864 | 0.909103825 |

| | | | | | |
|---|---|---|---|---|---|
| Soat2 | 6.105935105 | 5.728881924 | 5.197098174 | 0.53178375 | 0.908836931 |
| Nfkb2 | 10.66875738 | 10.31081942 | 9.761427752 | 0.549391665 | 0.907329628 |
| Gmfg | 8.59072827 | 7.814626012 | 7.684933713 | 0.129692299 | 0.905794557 |
| Rin1 | 10.21596498 | 9.740990351 | 9.310702191 | 0.430288161 | 0.905262786 |
| Inhbb | 9.791222052 | 9.306540928 | 8.88615261 | 0.420388318 | 0.905069442 |
| Wfdc17 | 10.03736943 | 9.489755509 | 9.134962312 | 0.354793197 | 0.902407113 |
| Stat4 | 5.982809507 | 4.997190588 | 5.080464126 | -0.083273538 | 0.902345381 |
| Cd209a | 10.34068236 | 9.885362618 | 9.438980277 | 0.446382341 | 0.901702086 |
| Wdr43 | 11.10175077 | 10.54216733 | 10.20069506 | 0.34147227 | 0.901055717 |
| Mafb | 11.35033428 | 11.12204863 | 10.45238228 | 0.669666353 | 0.897951997 |
| Abcc8 | 6.106005162 | 5.597107699 | 5.208768294 | 0.388339405 | 0.897236868 |
| Vmp1 | 11.7169307 | 11.30343073 | 10.82228392 | 0.48114681 | 0.894646773 |
| Il1r1 | 11.38890791 | 10.97147939 | 10.49486847 | 0.476610913 | 0.894039433 |
| Irgm1 | 10.70808887 | 9.829881649 | 9.814841372 | 0.015040278 | 0.893247498 |
| Actb | 15.76945248 | 15.2306164 | 14.87636079 | 0.35425561 | 0.89309169 |
| Phf11b | 7.78511923 | 6.839348241 | 6.892045317 | -0.052697076 | 0.893073913 |
| Uck2 | 11.47259168 | 11.15328567 | 10.58050134 | 0.572784333 | 0.892090337 |
| Myf6 | 9.102743201 | 8.398773133 | 8.210775163 | 0.18799797 | 0.891968038 |
| Slc38a1 | 9.141163874 | 8.630289222 | 8.249361222 | 0.380928 | 0.891802651 |
| Prr13 | 11.670452 | 11.11530297 | 10.77866463 | 0.336638333 | 0.891787363 |
| Cd84 | 9.659418467 | 9.171232398 | 8.767684438 | 0.40354796 | 0.891734028 |
| Pqlc1 | 10.91817067 | 10.49134597 | 10.02659618 | 0.46474979 | 0.891574486 |
| Adh6a | 8.853152948 | 8.206191212 | 7.964655129 | 0.241536082 | 0.888497818 |
| Rrp15 | 9.298450221 | 8.826038151 | 8.410384855 | 0.415653296 | 0.888065366 |
| Myh9 | 13.47794839 | 13.22571385 | 12.59007069 | 0.635643167 | 0.8878777 |
| Txnrd1 | 12.35276841 | 12.24040261 | 11.46564118 | 0.77476143 | 0.88712723 |
| Aldh18a1 | 10.05364701 | 9.601865629 | 9.167033986 | 0.434831643 | 0.886613028 |
| Mrto4 | 10.04033745 | 9.570373308 | 9.153857324 | 0.416515984 | 0.886480128 |
| Fst | 10.86477934 | 10.06860953 | 9.978810166 | 0.089799364 | 0.885969174 |
| Bst1 | 8.72049367 | 8.207507475 | 7.834567698 | 0.372939777 | 0.885925972 |
| Fes | 9.646887849 | 9.387755664 | 8.761080804 | 0.62667486 | 0.885807045 |
| Cd101 | 6.365449875 | 5.727539552 | 5.481203905 | 0.246335647 | 0.88424597 |

| | | | | | |
|---|---|---|---|---|---|
| Msr1 | 10.08954079 | 10.0030793 | 9.205602159 | 0.797477138 | 0.883938634 |
| Grina | 12.25003174 | 11.63969797 | 11.36628224 | 0.273415737 | 0.883749503 |
| Mvp | 11.58297837 | 11.40382314 | 10.69943447 | 0.704388673 | 0.883543903 |
| Adora2b | 9.524881102 | 9.075279666 | 8.642132802 | 0.433146864 | 0.8827483 |
| Rnf19b | 11.04582481 | 10.5185631 | 10.16365246 | 0.354910643 | 0.882172357 |
| Uap1 | 11.17878349 | 10.46094407 | 10.2975338 | 0.16341027 | 0.881249687 |
| Rrs1 | 9.962474708 | 9.488396165 | 9.083273194 | 0.405122971 | 0.879201514 |
| Ddx58 | 10.57525945 | 9.745882092 | 9.696153591 | 0.049728501 | 0.879105859 |
| Scimp | 6.909747593 | 6.19482031 | 6.031521668 | 0.163298643 | 0.878225925 |
| Ppp1r18 | 10.74714925 | 10.35100916 | 9.869201453 | 0.481807704 | 0.877947797 |
| Tgfa | 10.44226571 | 9.835896298 | 9.564329375 | 0.271566922 | 0.877936331 |
| Tinf2 | 9.719681485 | 8.822600179 | 8.842272336 | -0.019672157 | 0.87740915 |
| Olfr56 | 6.669413271 | 5.774594788 | 5.793802875 | -0.019208088 | 0.875610395 |
| Nip7 | 9.882928159 | 9.330660311 | 9.009130827 | 0.321529484 | 0.873797333 |
| Ifitm3 | 13.28019211 | 12.69957253 | 12.40951248 | 0.290060047 | 0.87067963 |
| Mybbp1a | 12.16549025 | 11.63218768 | 11.29538236 | 0.33680532 | 0.870107893 |
| Errfi1 | 11.80888132 | 11.41914926 | 10.93900743 | 0.480141823 | 0.869873883 |
| Stk40 | 11.84450501 | 11.44265168 | 10.97534863 | 0.467303047 | 0.869156377 |
| Tcof1 | 10.37256075 | 10.09877916 | 9.504520992 | 0.594258168 | 0.868039755 |
| Birc3 | 10.70348688 | 10.16045681 | 9.836351449 | 0.324105361 | 0.867135431 |
| Wdr46 | 9.631411134 | 9.335632416 | 8.76617651 | 0.569455905 | 0.865234624 |
| Slfn5 | 12.3106297 | 11.60909342 | 11.44693819 | 0.162155227 | 0.863691507 |
| Eif1a | 11.33992823 | 10.83778002 | 10.47655961 | 0.361220413 | 0.863368627 |
| Cxcl14 | 11.16634843 | 10.50561091 | 10.30336098 | 0.202249937 | 0.862987457 |
| Arhgap30 | 10.29728159 | 10.04571064 | 9.434577925 | 0.611132714 | 0.862703662 |
| Zfp281 | 9.614940592 | 8.981938961 | 8.758176307 | 0.223762654 | 0.856764285 |
| A530032D15Rik | 4.888004722 | 4.086987594 | 4.03169446 | 0.055293134 | 0.856310262 |
| Gm43302 | 5.370547862 | 4.521676784 | 4.514766655 | 0.006910129 | 0.855781207 |
| Arhgap45 | 9.98467722 | 9.761596646 | 9.12971836 | 0.631878286 | 0.85495886 |
| Msn | 13.27797787 | 12.91941762 | 12.42369015 | 0.495727463 | 0.85428772 |
| Sectm1a | 11.06477424 | 10.81644183 | 10.21147801 | 0.604963823 | 0.85329623 |
| Ifi35 | 10.34638951 | 9.741304674 | 9.495140677 | 0.246163997 | 0.851248836 |

| | | | | | |
|---|---|---|---|---|---|
| Cav3 | 8.678433071 | 8.018113055 | 7.828510628 | 0.189602427 | 0.849922443 |
| Actg1 | 15.63954501 | 15.03194864 | 14.7896261 | 0.24232254 | 0.84991891 |
| Il1f9 | 11.47606777 | 10.79246964 | 10.62758577 | 0.16488387 | 0.848481997 |
| Socs1 | 7.158394343 | 6.863412761 | 6.310376656 | 0.553036105 | 0.848017687 |
| Arc | 8.19208634 | 9.206000129 | 7.344162728 | 1.8618374 | 0.847923611 |
| C920009B18Rik | 6.901792874 | 6.330967931 | 6.054886148 | 0.276081783 | 0.846906727 |
| Dkc1 | 10.77931154 | 10.18113709 | 9.933132511 | 0.248004579 | 0.846179029 |
| Rhoc | 10.26584415 | 9.601429094 | 9.42047096 | 0.180958134 | 0.845373194 |
| Pilrb2 | 6.350916765 | 5.914239297 | 5.50590454 | 0.408334757 | 0.845012225 |
| Selplg | 9.921225086 | 9.613305809 | 9.076739753 | 0.536566056 | 0.844485333 |
| Gm9733 | 6.551952683 | 6.219764691 | 5.71017777 | 0.509586921 | 0.841774912 |
| Csrp3 | 9.580296813 | 10.81796042 | 8.738555182 | 2.079405235 | 0.841741631 |
| Ehd1 | 10.42452989 | 10.02452635 | 9.584193196 | 0.440333157 | 0.840336697 |
| Nol10 | 9.679886297 | 9.058126289 | 8.841923103 | 0.216203186 | 0.837963195 |
| Arhgap26 | 8.877956736 | 8.474277398 | 8.040243215 | 0.434034183 | 0.837713521 |
| Cpne2 | 9.724190859 | 9.350396364 | 8.887008175 | 0.463388189 | 0.837182684 |
| Rasgrp4 | 9.216110576 | 9.02290694 | 8.379793357 | 0.643113584 | 0.83631722 |
| Myo1f | 9.908309563 | 9.832960561 | 9.07326556 | 0.759695 | 0.835044003 |
| Tgfb1 | 10.07508785 | 9.502353425 | 9.240045404 | 0.262308021 | 0.835042446 |
| Gm27029 | 7.058232834 | 7.020524956 | 6.223667763 | 0.796857192 | 0.83456507 |
| Rassf1 | 10.22003209 | 9.788286611 | 9.385935716 | 0.402350894 | 0.834096374 |
| Utp20 | 10.76835996 | 10.45686305 | 9.934964827 | 0.521898223 | 0.833395136 |
| Steap4 | 12.36384289 | 11.65065095 | 11.53133832 | 0.119312623 | 0.832504567 |
| Fcer1g | 11.17761883 | 10.54178247 | 10.34570136 | 0.19608111 | 0.83191747 |
| Slc20a1 | 10.13164653 | 9.917687691 | 9.301371761 | 0.616315929 | 0.830274765 |
| Bst2 | 10.02333562 | 9.316986355 | 9.19307458 | 0.123911775 | 0.830261039 |
| Cryab | 12.34180648 | 13.61939533 | 11.51163254 | 2.10776279 | 0.830173943 |
| Sirpb1c | 7.245841583 | 7.157241863 | 6.415694054 | 0.741547809 | 0.830147529 |
| Trem3 | 5.276066275 | 4.816318843 | 4.447317257 | 0.369001586 | 0.828749018 |
| Ccdc86 | 9.287302606 | 8.917156959 | 8.45948392 | 0.457673039 | 0.827818687 |
| Tlr4 | 9.305584522 | 8.897790033 | 8.478244623 | 0.41954541 | 0.827339899 |
| Cyp7b1 | 8.490675087 | 8.073087422 | 7.663550098 | 0.409537325 | 0.827124989 |

| | | | | | |
|---|---|---|---|---|---|
| Dennd3 | 7.503662584 | 6.972715821 | 6.681476736 | 0.291239085 | 0.822185848 |
| Slamf8 | 7.993661914 | 7.85869013 | 7.171941281 | 0.686748849 | 0.821720633 |
| Lyz2 | 14.10173404 | 13.5913412 | 13.28067776 | 0.31066344 | 0.821056277 |
| Ms4a4a | 9.614023943 | 8.876757606 | 8.794757786 | 0.08199982 | 0.819266157 |
| Dsg3 | 13.04382566 | 12.49959448 | 12.22599164 | 0.27360284 | 0.81783402 |
| 2610528A11Rik | 8.029444694 | 7.306142253 | 7.211763808 | 0.094378445 | 0.817680886 |
| Gm45799 | 6.303799581 | 5.97810296 | 5.486237649 | 0.491865311 | 0.817561932 |
| Slc16a1 | 11.43993285 | 10.91070562 | 10.62278855 | 0.28791707 | 0.817144303 |
| Cdkn2d | 9.018211681 | 8.310431977 | 8.201227421 | 0.109204556 | 0.81698426 |
| Fcgr2b | 11.53679801 | 11.1319052 | 10.7219239 | 0.409981297 | 0.81487411 |
| Tpd52 | 12.05278874 | 11.53659388 | 11.23839775 | 0.298196133 | 0.81439099 |
| Stap1 | 6.887750967 | 6.323387625 | 6.073680527 | 0.249707098 | 0.81407044 |
| Parp9 | 10.69540613 | 9.915448257 | 9.881701006 | 0.03374725 | 0.81370512 |
| Siglecf | 6.011557351 | 5.363262197 | 5.198642678 | 0.164619519 | 0.812914673 |
| Tgif1 | 10.02757948 | 9.534136489 | 9.215069305 | 0.319067184 | 0.81251018 |
| Serpina3n | 13.64116537 | 13.33615066 | 12.828907 | 0.50724366 | 0.81225837 |
| Bach1 | 11.62826066 | 11.27299465 | 10.81676539 | 0.456229257 | 0.811495273 |
| Jaml | 8.429547714 | 8.073670203 | 7.618219474 | 0.455450729 | 0.81132824 |
| Sema4c | 9.783679202 | 9.266101342 | 8.97591171 | 0.290189632 | 0.807767492 |
| Dhrs13 | 6.948463901 | 6.445670102 | 6.140718755 | 0.304951347 | 0.807745146 |
| Midn | 11.70316147 | 11.64335189 | 10.89757975 | 0.745772137 | 0.805581723 |
| 1700017B05Rik | 9.413404756 | 9.095450839 | 8.608534345 | 0.486916494 | 0.804870411 |
| Lst1 | 7.869914925 | 7.404480932 | 7.065371068 | 0.339109864 | 0.804543857 |
| Pgs1 | 11.22423977 | 10.89349044 | 10.42008244 | 0.473407997 | 0.804157333 |
| Zfp593 | 8.519571652 | 8.248770536 | 7.715916265 | 0.532854271 | 0.803655387 |
| Pik3r5 | 9.231875921 | 9.444420085 | 8.428589296 | 1.015830789 | 0.803286625 |
| Adgre5 | 10.59065224 | 10.21913751 | 9.787365779 | 0.431771732 | 0.803286465 |
| Lyar | 9.525223374 | 9.01690141 | 8.722435342 | 0.294466068 | 0.802788032 |
| Sycp2 | 5.674103368 | 5.141315946 | 4.871651539 | 0.269664407 | 0.802451829 |
| Vasp | 11.43661319 | 11.00905782 | 10.63571657 | 0.373341243 | 0.800896617 |
| Cd40 | 7.653023912 | 7.403387423 | 6.853092284 | 0.550295139 | 0.799931628 |
| Bdkrb1 | 5.710204894 | 5.283732469 | 4.910432371 | 0.373300098 | 0.799772523 |

| | | | | | |
|---|---|---|---|---|---|
| Utp4 | 10.11643312 | 9.628697757 | 9.317106633 | 0.311591124 | 0.799326488 |
| Pus7l | 7.863949385 | 7.371412885 | 7.065015703 | 0.306397182 | 0.798933683 |
| Abcc3 | 8.164110341 | 7.775284938 | 7.366460377 | 0.408824561 | 0.797649964 |
| Smyd5 | 9.356513302 | 8.955446987 | 8.558958268 | 0.396488719 | 0.797555034 |
| Ppp1r3d | 7.519056291 | 6.954903773 | 6.721591786 | 0.233311988 | 0.797464505 |
| Sema7a | 9.363787749 | 8.911379552 | 8.566819201 | 0.344560351 | 0.796968547 |
| Xdh | 13.08539767 | 12.49059175 | 12.2887088 | 0.201882947 | 0.79668887 |
| Dok2 | 9.424851392 | 9.163234985 | 8.628744669 | 0.534490316 | 0.796106722 |
| Slc3a2 | 11.89995704 | 11.49720158 | 11.10529592 | 0.39190566 | 0.794661127 |
| Cdk17 | 9.471452597 | 8.927196065 | 8.677654401 | 0.249541664 | 0.793798196 |
| Gpr84 | 5.092381755 | 4.74646323 | 4.300808668 | 0.445654562 | 0.791573087 |
| Bop1 | 10.33154456 | 10.15291882 | 9.540437888 | 0.612480932 | 0.791106672 |
| Glt1d1 | 7.521867717 | 7.079902153 | 6.730981639 | 0.348920514 | 0.790886078 |
| Ripk3 | 9.258449597 | 8.914050568 | 8.468343101 | 0.445707467 | 0.790106496 |
| Cd24a | 12.01426547 | 11.41218031 | 11.22556863 | 0.186611683 | 0.788696837 |
| Fyb | 9.239408911 | 9.056274609 | 8.451250243 | 0.605024366 | 0.788158668 |
| Daxx | 10.48090702 | 9.63142508 | 9.693374934 | -0.061949855 | 0.787532089 |
| Napsa | 9.174143192 | 8.784002926 | 8.387316641 | 0.396686285 | 0.786826551 |
| Rdh10 | 9.52714848 | 8.920112279 | 8.740651214 | 0.179461064 | 0.786497266 |
| Ifrd1 | 11.75333498 | 11.37762765 | 10.96704034 | 0.410587307 | 0.786294637 |
| Fth1 | 14.80967718 | 14.29052545 | 14.02404684 | 0.26647861 | 0.78563034 |
| Gbp10 | 6.334893463 | 5.509576025 | 5.550299516 | -0.040723491 | 0.784593947 |
| Hbb-bt | 9.436246056 | 7.981733443 | 8.652257332 | -0.670523889 | 0.783988724 |
| Hspa5 | 13.7606879 | 13.91370784 | 12.97716341 | 0.93654443 | 0.783524483 |
| Stat1 | 11.82901617 | 11.2413633 | 11.04568961 | 0.195673687 | 0.78332656 |
| Nos3 | 8.598956158 | 8.461998142 | 7.817227434 | 0.644770708 | 0.781728724 |
| Ccr2 | 11.28060322 | 10.97282141 | 10.49899627 | 0.473825143 | 0.78160695 |
| Mmp3 | 11.15045967 | 10.80111095 | 10.36990909 | 0.431201853 | 0.780550577 |
| Resf1 | 10.90377758 | 10.43734531 | 10.12340902 | 0.313936297 | 0.78036856 |
| Cad | 10.45357672 | 10.33962737 | 9.675397226 | 0.664230141 | 0.778179494 |
| Rhog | 10.77768825 | 10.36767683 | 9.999725831 | 0.367950995 | 0.777962422 |
| Chd7 | 11.19575795 | 10.48255238 | 10.41968375 | 0.062868633 | 0.776074207 |

| | | | | | |
|---|---|---|---|---|---|
| Cxcr1 | 5.020371413 | 4.468100441 | 4.244610517 | 0.223489923 | 0.775760896 |
| Asb5 | 9.930571501 | 10.35503648 | 9.159120352 | 1.195916125 | 0.771451149 |
| Ets2 | 12.55969197 | 12.02529537 | 11.78851342 | 0.236781953 | 0.771178547 |
| Igsf6 | 8.391027919 | 8.420530332 | 7.620717051 | 0.799813281 | 0.770310868 |
| Nfe2l2 | 11.73496345 | 11.26058281 | 10.96494276 | 0.295640047 | 0.77002069 |
| Adamts8 | 6.187315895 | 5.590184032 | 5.417395721 | 0.172788311 | 0.769920174 |
| Map2k1 | 10.90361878 | 10.29806714 | 10.13378651 | 0.16428063 | 0.769832277 |
| Fam49b | 10.64425603 | 10.15386115 | 9.874424601 | 0.279436546 | 0.769831432 |
| Polr1b | 9.827993074 | 9.373089546 | 9.058423226 | 0.31466632 | 0.769569848 |
| Nlrp12 | 5.574395809 | 5.281175048 | 4.804848663 | 0.476326385 | 0.769547145 |
| Ifi44 | 7.820691809 | 6.535593681 | 7.051414424 | -0.515820743 | 0.769277385 |
| Fbl | 11.50295523 | 11.24123464 | 10.73375707 | 0.50747757 | 0.76919816 |
| Noc2l | 11.31130341 | 10.93193061 | 10.54327742 | 0.388653183 | 0.76802599 |
| Impdh1 | 9.394180006 | 9.067469549 | 8.626432199 | 0.44103735 | 0.767747807 |
| Prelp | 11.94598815 | 11.49178841 | 11.17849329 | 0.313295117 | 0.767494857 |
| Spsb1 | 8.734133533 | 8.278303576 | 7.966833593 | 0.311469982 | 0.76729994 |
| Nifk | 10.55159813 | 10.03332496 | 9.784807319 | 0.248517636 | 0.766790811 |
| Tcirg1 | 9.964181094 | 9.82873174 | 9.197621616 | 0.631110124 | 0.766559478 |
| Cox17 | 9.381782997 | 9.007739692 | 8.616255961 | 0.391483731 | 0.765527036 |
| Scn8a | 6.380698779 | 5.952716357 | 5.615183738 | 0.337532619 | 0.765515041 |
| Adora3 | 7.695892475 | 7.082667234 | 6.930549744 | 0.15211749 | 0.765342732 |
| Irgm2 | 10.76705558 | 9.846335354 | 10.00197286 | -0.155637507 | 0.765082718 |
| Ackr3 | 10.43074961 | 9.604182656 | 9.665781983 | -0.061599327 | 0.764967627 |
| Irx1 | 8.598569551 | 8.482542185 | 7.833765462 | 0.648776724 | 0.764804089 |
| Gapt | 6.912867833 | 6.274708664 | 6.148467724 | 0.12624094 | 0.764400109 |
| Eef1e1 | 9.163020464 | 8.540655588 | 8.39877539 | 0.141880198 | 0.764245074 |
| Hk2 | 11.86437977 | 12.14418361 | 11.10014502 | 1.04403859 | 0.764234747 |
| Mdn1 | 10.60204518 | 10.43321918 | 9.838022718 | 0.595196465 | 0.764022465 |
| Bysl | 9.750631488 | 9.3584562 | 8.986837858 | 0.371618342 | 0.76379363 |
| Ftl1 | 14.17123507 | 14.27786618 | 13.40801235 | 0.869853833 | 0.76322272 |
| Cdh3 | 10.48086271 | 10.02489339 | 9.718513544 | 0.306379846 | 0.762349169 |
| Mapkapk2 | 12.19200989 | 11.57955516 | 11.42980208 | 0.149753083 | 0.762207817 |

| | | | | | |
|---|---|---|---|---|---|
| Lipg | 9.207101477 | 8.612283099 | 8.445540443 | 0.166742656 | 0.761561034 |
| Kcnh1 | 7.450525012 | 7.048119819 | 6.689592406 | 0.358527413 | 0.760932607 |
| Ruvbl2 | 9.459199055 | 9.190958599 | 8.698588994 | 0.492369605 | 0.760610061 |
| Ms4a4b | 6.896753819 | 5.999139555 | 6.136954498 | -0.137814942 | 0.759799322 |
| Id1 | 9.344761449 | 9.194568193 | 8.585335838 | 0.609232355 | 0.759425611 |
| Irf9 | 10.83083606 | 10.47072556 | 10.07405448 | 0.396671083 | 0.756781583 |
| Eif2ak2 | 10.62663654 | 9.881936813 | 9.870659774 | 0.011277039 | 0.755976766 |
| Ptpn12 | 11.15081519 | 10.64349375 | 10.39518325 | 0.248310503 | 0.755631937 |
| Dok3 | 7.613659901 | 7.314192558 | 6.858756777 | 0.455435781 | 0.754903124 |
| Tuba1c | 12.80361623 | 12.3748092 | 12.04875091 | 0.326058283 | 0.754865313 |
| Cyba | 9.89884095 | 9.787034641 | 9.144246997 | 0.642787644 | 0.754593953 |
| Tnfrsf1a | 12.70966483 | 12.2613412 | 11.95660938 | 0.304731827 | 0.75305545 |
| Parp10 | 10.08501018 | 9.65517138 | 9.33203681 | 0.32313457 | 0.752973372 |
| Cyp26b1 | 12.84033257 | 12.5967646 | 12.08778929 | 0.508975313 | 0.75254328 |
| Nfkb1 | 11.94460288 | 11.43801788 | 11.19255344 | 0.245464443 | 0.752049437 |
| Osbpl8 | 10.9951 | 10.44609593 | 10.24358828 | 0.20250765 | 0.75151172 |
| Stat2 | 10.96420109 | 10.54032999 | 10.21323148 | 0.327098506 | 0.75096961 |
| Mdfi | 7.299952302 | 6.934729586 | 6.551450357 | 0.383279229 | 0.748501945 |
| Pop1 | 8.429262327 | 8.218155599 | 7.681059272 | 0.537096327 | 0.748203054 |
| Hrh2 | 6.763109231 | 6.420467182 | 6.015272517 | 0.405194666 | 0.747836714 |
| Nmi | 9.144036832 | 8.438647915 | 8.396296835 | 0.04235108 | 0.747739997 |
| Synpo | 10.57120874 | 10.09927418 | 9.823692614 | 0.275581566 | 0.747516126 |
| Avpi1 | 10.9299161 | 10.42929192 | 10.18324635 | 0.246045567 | 0.74666975 |
| Fpgs | 9.549020865 | 9.112799483 | 8.803549846 | 0.309249637 | 0.745471019 |
| Ebna1bp2 | 10.67976566 | 10.24789401 | 9.934542247 | 0.313351763 | 0.745223409 |
| Hlx | 7.138843641 | 6.800088733 | 6.394769598 | 0.405319136 | 0.744074043 |
| Pitpnm1 | 7.952254345 | 7.77254874 | 7.208655191 | 0.563893549 | 0.743599153 |
| Il2rb | 8.013986449 | 7.505540413 | 7.271399083 | 0.23414133 | 0.742587366 |
| Ninj1 | 9.911461192 | 9.541715425 | 9.169686634 | 0.372028791 | 0.741774559 |
| Stfa1 | 9.844240243 | 9.537067396 | 9.103579504 | 0.433487892 | 0.740660739 |
| Fosb | 8.566356837 | 8.338911585 | 7.826211887 | 0.512699698 | 0.74014495 |
| 1810055G02Rik | 9.108174484 | 8.653047477 | 8.368154167 | 0.28489331 | 0.740020317 |

| | | | | | |
|---|---|---|---|---|---|
| Ms4a6c | 10.55381598 | 10.02727327 | 9.813957316 | 0.213315952 | 0.739858667 |
| Gar1 | 9.478245268 | 9.051391719 | 8.738540503 | 0.312851216 | 0.739704765 |
| Cd83 | 8.314530089 | 8.256334698 | 7.575227589 | 0.68110711 | 0.7393025 |
| Nol8 | 9.636828675 | 9.144978845 | 8.89765243 | 0.247326415 | 0.739176245 |
| Il15ra | 10.05343524 | 9.492129918 | 9.314681108 | 0.17744881 | 0.738754134 |
| Lrrc59 | 11.79351173 | 11.49271996 | 11.05539171 | 0.43732825 | 0.738120023 |
| Aqp9 | 9.778124517 | 9.21890384 | 9.040332609 | 0.178571231 | 0.737791908 |
| Parp12 | 10.57911113 | 9.857445708 | 9.842621331 | 0.014824378 | 0.736489799 |
| Nop16 | 9.927426241 | 9.524801401 | 9.190949707 | 0.333851694 | 0.736476534 |
| Sh3bp2 | 10.1688834 | 9.999529009 | 9.434231367 | 0.565297642 | 0.73465203 |
| Sphk1 | 10.90668533 | 10.58093968 | 10.17217172 | 0.40876796 | 0.73451361 |
| Hsp90aa1 | 13.24536839 | 13.94392076 | 12.51086408 | 1.433056683 | 0.73450431 |
| F830016B08Rik | 8.021780533 | 6.78666889 | 7.28734967 | -0.50068078 | 0.734430863 |
| Pilrb1 | 6.355022501 | 6.054444091 | 5.62080357 | 0.433640522 | 0.734218931 |
| Ptpn1 | 11.44605502 | 11.27663006 | 10.71215459 | 0.564475473 | 0.73390043 |
| Nedd9 | 9.576191565 | 8.864149997 | 8.842685419 | 0.021464578 | 0.733506146 |
| Hyou1 | 11.7362982 | 11.70821822 | 11.0029819 | 0.70523632 | 0.733316307 |
| Adcy4 | 8.883251872 | 8.5887385 | 8.149947121 | 0.438791379 | 0.733304751 |
| Map3k14 | 9.026095857 | 9.113727283 | 8.293557734 | 0.82016955 | 0.732538123 |
| Mcl1 | 13.35122654 | 12.87948667 | 12.61885792 | 0.260628747 | 0.732368617 |
| Itga3 | 11.54885143 | 11.2053423 | 10.8164997 | 0.3888426 | 0.732351727 |
| Tsku | 8.732452187 | 8.275391856 | 8.000414336 | 0.27497752 | 0.732037851 |
| Cntfr | 7.412270672 | 7.203255915 | 6.680728587 | 0.522527329 | 0.731542085 |
| Nfam1 | 10.3663636 | 10.15190659 | 9.635410541 | 0.516496052 | 0.730953059 |
| Ncf1 | 9.513538557 | 9.002193769 | 8.78299754 | 0.219196229 | 0.730541016 |
| Rab8b | 11.74152816 | 11.16699471 | 11.01118093 | 0.155813787 | 0.730347233 |
| Nckap1l | 10.59263993 | 10.22711508 | 9.862313392 | 0.364801688 | 0.730326541 |
| Ptk2b | 10.64909281 | 10.01804302 | 9.920204214 | 0.097838801 | 0.728888599 |
| Brca2 | 8.471895927 | 7.811780699 | 7.743544192 | 0.068236507 | 0.728351735 |
| Ogfr | 10.48541078 | 10.03171796 | 9.757209113 | 0.274508847 | 0.72820167 |
| Mak16 | 10.09592563 | 9.61177162 | 9.368017568 | 0.243754052 | 0.727908063 |
| Pum3 | 10.81857348 | 10.37760745 | 10.0911083 | 0.286499143 | 0.72746518 |

| | | | | | |
|---|---|---|---|---|---|
| Plbd1 | 12.51442457 | 12.13847568 | 11.78738509 | 0.35109059 | 0.72703948 |
| Tsr1 | 10.68057711 | 10.32356203 | 9.954092229 | 0.369469804 | 0.726484881 |
| Fam217b | 8.66601116 | 7.988496498 | 7.940645139 | 0.047851359 | 0.725366021 |
| Lrp8 | 7.22338493 | 7.044916251 | 6.500822734 | 0.544093517 | 0.722562196 |
| Tnfrsf23 | 8.436561489 | 8.053899215 | 7.714535667 | 0.339363549 | 0.722025823 |
| Id3 | 9.927338447 | 9.807653877 | 9.205527079 | 0.602126798 | 0.721811368 |
| Ftsj3 | 11.27249416 | 10.84169212 | 10.55155448 | 0.290137647 | 0.72093968 |
| Gm18787 | 5.033537674 | 4.453244062 | 4.312628256 | 0.140615806 | 0.720909418 |
| Snai1 | 7.11638382 | 6.808673156 | 6.395629599 | 0.413043557 | 0.720754221 |
| Shisa5 | 12.09344132 | 11.70048747 | 11.3727223 | 0.327765167 | 0.720719017 |
| Camkk2 | 10.48098914 | 10.16531556 | 9.760460337 | 0.404855227 | 0.7205288 |
| Pdcd1lg2 | 6.013245167 | 5.692737957 | 5.293158302 | 0.399579654 | 0.720086865 |
| C130050O18Rik | 5.743518888 | 5.396920479 | 5.02359883 | 0.373321649 | 0.719920058 |
| Prex1 | 9.49537902 | 9.29094879 | 8.775762183 | 0.515186607 | 0.719616837 |
| Ncl | 14.09412515 | 13.7039456 | 13.37504807 | 0.328897527 | 0.719077077 |
| Ptger2 | 5.882108918 | 5.521152772 | 5.165953109 | 0.355199663 | 0.716155809 |
| Ier5l | 6.529477135 | 6.313722392 | 5.81547883 | 0.498243562 | 0.713998304 |
| Il2rg | 9.912219021 | 9.503124167 | 9.198262646 | 0.304861521 | 0.713956375 |
| Ceacam1 | 8.350839356 | 7.882790057 | 7.637415895 | 0.245374162 | 0.713423461 |
| Plod3 | 10.80660506 | 10.42060871 | 10.09318591 | 0.327422797 | 0.713419147 |
| Polr3d | 9.003643383 | 8.654596507 | 8.29122056 | 0.363375947 | 0.712422824 |
| Ptpn23 | 10.24943698 | 10.26688726 | 9.537181722 | 0.729705541 | 0.712255255 |
| Plk2 | 10.12229731 | 9.800838603 | 9.410485685 | 0.390352917 | 0.711811628 |
| Pmaip1 | 8.918132628 | 8.755476823 | 8.208251298 | 0.547225525 | 0.70988133 |
| 9830107B12Rik | 4.332508222 | 3.72473312 | 3.623626735 | 0.101106385 | 0.708881487 |
| Ccl19 | 9.053226201 | 9.00288926 | 8.34439598 | 0.65849328 | 0.708830221 |
| Nat10 | 10.17786705 | 9.628302853 | 9.469040822 | 0.159262031 | 0.708826225 |
| Ifi208 | 7.38175229 | 6.383834602 | 6.673535254 | -0.289700652 | 0.708217037 |
| Klf2 | 8.707596335 | 8.164475559 | 7.999551746 | 0.164923813 | 0.708044589 |
| Card19 | 9.068965092 | 8.802391688 | 8.361490704 | 0.440900983 | 0.707474388 |
| Milr1 | 7.294345741 | 6.648900001 | 6.586917182 | 0.061982818 | 0.707428559 |
| Mmp19 | 11.02141515 | 10.933045 | 10.31405598 | 0.618989013 | 0.707359167 |

| | | | | | |
|---|---|---|---|---|---|
| Havcr2 | 7.803653935 | 7.50845956 | 7.09665456 | 0.411805 | 0.706999375 |
| Rara | 9.580795487 | 9.584098346 | 8.874503389 | 0.709594957 | 0.706292098 |
| Gzma | 5.423724298 | 4.754192797 | 4.717656246 | 0.036536551 | 0.706068052 |
| Dyrk3 | 5.96181913 | 5.660971692 | 5.256497528 | 0.404474165 | 0.705321602 |
| Ecscr | 8.722363621 | 8.438023173 | 8.017652879 | 0.420370294 | 0.704710742 |
| Rap1gap2 | 8.146937448 | 7.608576218 | 7.442408359 | 0.166167859 | 0.704529089 |
| Snx10 | 10.09675703 | 9.498878223 | 9.39271484 | 0.106163383 | 0.704042186 |
| Spry4 | 8.13644648 | 7.611874664 | 7.432429484 | 0.179445181 | 0.704016996 |
| Epop | 7.343684591 | 7.004023018 | 6.639758634 | 0.364264384 | 0.703925957 |
| Uba7 | 9.572075316 | 8.853611935 | 8.870277629 | -0.016665694 | 0.701797687 |
| Gpx2 | 7.245151359 | 6.952275235 | 6.544886727 | 0.407388508 | 0.700264632 |
| Smad3 | 10.06377343 | 9.346285524 | 9.364230034 | -0.01794451 | 0.6995434 |
| Psmb10 | 9.591182467 | 9.212525603 | 8.891912129 | 0.320613474 | 0.699270338 |
| Akr1b8 | 9.892719768 | 9.399459434 | 9.193880608 | 0.205578826 | 0.69883916 |
| Mafk | 10.04912364 | 10.25246892 | 9.350442739 | 0.902026185 | 0.698680899 |
| Cadm3 | 11.18257682 | 10.82133108 | 10.48392002 | 0.337411067 | 0.698656807 |
| Ets1 | 10.0477225 | 9.464274861 | 9.349256907 | 0.115017955 | 0.698465592 |
| Inpp5j | 5.571894376 | 5.138920282 | 4.873570552 | 0.26534973 | 0.698323824 |
| Tpm3 | 13.03932499 | 12.68760597 | 12.34126443 | 0.34634154 | 0.698060563 |
| Slc9a1 | 11.49126959 | 11.43284064 | 10.79371006 | 0.639130583 | 0.697559533 |
| Inpp5d | 9.977808785 | 9.619187832 | 9.280385004 | 0.338802828 | 0.697423781 |
| Mx2 | 7.998975219 | 7.07816196 | 7.301563359 | -0.223401399 | 0.69741186 |
| 3110082I17Rik | 8.158192356 | 7.741347169 | 7.461921647 | 0.279425522 | 0.696270709 |
| Nat8l | 10.27360548 | 9.92935023 | 9.577499528 | 0.351850701 | 0.696105952 |
| Prss53 | 8.551248655 | 8.726421309 | 7.855820238 | 0.870601071 | 0.695428417 |
| Zbtb21 | 9.083545568 | 8.753164764 | 8.388635056 | 0.364529708 | 0.694910512 |
| Pnp | 11.62636602 | 10.99293877 | 10.93173329 | 0.061205483 | 0.694632727 |
| Thap6 | 7.956694949 | 7.351043442 | 7.262406955 | 0.088636487 | 0.694287994 |
| Wdr77 | 10.80690895 | 10.41855621 | 10.11334719 | 0.305209022 | 0.693561762 |
| Slc25a22 | 9.769248243 | 9.354987147 | 9.076601812 | 0.278385335 | 0.692646431 |
| Cd69 | 6.390951512 | 5.767371063 | 5.698420731 | 0.068950332 | 0.692530781 |
| Bin2 | 8.093691702 | 7.724059234 | 7.401850654 | 0.32220858 | 0.691841048 |

| | | | | | |
|---|---|---|---|---|---|
| Nfkbib | 9.234938503 | 8.97212277 | 8.543165875 | 0.428956895 | 0.691772627 |
| Scn5a | 6.00163292 | 6.092809207 | 5.310024507 | 0.7827847 | 0.691608413 |
| Apobec3 | 9.360177628 | 8.934299879 | 8.668854319 | 0.26544556 | 0.691323309 |
| Tomm40 | 10.21161037 | 9.967838149 | 9.522035986 | 0.445802163 | 0.689574381 |
| Irf1 | 10.49162538 | 9.867135878 | 9.803238292 | 0.063897586 | 0.688387091 |
| Ptpre | 9.502951492 | 9.143873401 | 8.814592451 | 0.32928095 | 0.688359041 |
| Dph2 | 8.431910829 | 8.158688481 | 7.744123411 | 0.414565071 | 0.687787419 |
| Sema6b | 7.652050912 | 7.493719376 | 6.964507068 | 0.529212308 | 0.687543844 |
| Apbb3 | 8.691963563 | 8.071797375 | 8.004639128 | 0.067158248 | 0.687324436 |
| Gm18852 | 5.804198905 | 5.381017128 | 5.116954265 | 0.264062863 | 0.68724464 |
| Ptpn2 | 9.784751274 | 9.288031039 | 9.099245995 | 0.188785044 | 0.685505279 |
| Nle1 | 8.487299493 | 8.447648574 | 7.801985289 | 0.645663285 | 0.685314204 |
| Rilpl2 | 7.939008779 | 7.523570552 | 7.253702133 | 0.269868419 | 0.685306646 |
| Ctsz | 11.31228639 | 11.20979437 | 10.62744642 | 0.582347953 | 0.68483997 |
| Fbxo10 | 8.414379868 | 7.817817659 | 7.729615557 | 0.088202102 | 0.684764311 |
| Rras2 | 9.860790492 | 9.337251912 | 9.176181004 | 0.161070908 | 0.684609488 |
| Akna | 9.059525097 | 8.89839496 | 8.375879174 | 0.522515786 | 0.683645922 |
| Farsa | 10.07690683 | 9.767888234 | 9.393307204 | 0.374581031 | 0.683599626 |
| Oas1g | 7.796262494 | 7.202433367 | 7.113442939 | 0.088990427 | 0.682819555 |
| Arap3 | 9.171085665 | 8.700116146 | 8.48922096 | 0.210895186 | 0.681864705 |
| Laptm5 | 11.93051792 | 11.62876976 | 11.24899414 | 0.379775617 | 0.681523777 |
| Sdcbp | 12.34163648 | 11.93496986 | 11.66028927 | 0.274680597 | 0.681347217 |
| Dmxl2 | 9.077329653 | 8.837193928 | 8.396661732 | 0.440532196 | 0.680667921 |
| Ctu2 | 8.6458894 | 8.295346166 | 7.965871458 | 0.329474707 | 0.680017942 |
| Polr3h | 8.830906747 | 8.314336489 | 8.150945783 | 0.163390706 | 0.679960964 |
| Csf2 | 4.46064406 | 3.836718068 | 3.781197416 | 0.055520651 | 0.679446643 |
| Slc41a1 | 10.82654926 | 10.2993847 | 10.14766007 | 0.151724627 | 0.67888919 |
| Rps6ka2 | 9.370922028 | 8.518029349 | 8.692290777 | -0.174261429 | 0.678631251 |
| Il1rl1 | 9.429044406 | 9.076728097 | 8.751197276 | 0.325530821 | 0.67784713 |
| Wdr74 | 9.632941487 | 9.203145453 | 8.955632312 | 0.247513141 | 0.677309175 |
| Syde1 | 9.04310764 | 8.671259206 | 8.365814999 | 0.305444207 | 0.677292641 |
| Nfyb | 9.910147357 | 9.34738692 | 9.232911036 | 0.114475884 | 0.677236321 |

| | | | | | |
|---|---|---|---|---|---|
| Urb1 | 9.35593591 | 9.040049472 | 8.679452955 | 0.360596517 | 0.676482955 |
| Nol6 | 10.67369939 | 10.37656887 | 9.997660025 | 0.378908842 | 0.676039365 |
| Npm3 | 9.844798958 | 9.592347021 | 9.168848421 | 0.4234986 | 0.675950536 |
| Tnfrsf9 | 6.877347798 | 6.768593492 | 6.202045186 | 0.566548306 | 0.675302611 |
| Gimap3 | 5.882243286 | 5.187999384 | 5.208973808 | -0.020974424 | 0.673269477 |
| Taf4b | 8.206540213 | 7.859547193 | 7.53432157 | 0.325225623 | 0.672218643 |
| Ass1 | 9.799597843 | 9.346045655 | 9.127520042 | 0.218525613 | 0.672077801 |
| Crlf2 | 6.851256677 | 6.77172655 | 6.17945881 | 0.592267741 | 0.671797867 |
| Ppp4r1 | 11.03466758 | 10.73385963 | 10.36365923 | 0.370200397 | 0.671008347 |
| Myrfl | 4.777367965 | 4.2217279 | 4.106459082 | 0.115268818 | 0.670908883 |
| Abi3 | 7.216757573 | 6.858572353 | 6.547036497 | 0.311535856 | 0.669721076 |
| Cd209e | 9.088978597 | 8.543746371 | 8.419337506 | 0.124408865 | 0.669641091 |
| Gzmb | 5.55118455 | 4.878559897 | 4.881917779 | -0.003357883 | 0.669266771 |
| Chka | 8.647925249 | 8.880458497 | 7.979557137 | 0.90090136 | 0.668368112 |
| Noc3l | 9.403630688 | 9.038382579 | 8.735858385 | 0.302524194 | 0.667772304 |
| Imp4 | 10.11079847 | 9.824004435 | 9.443123627 | 0.380880808 | 0.667674847 |
| Gripap1 | 10.11045609 | 9.799905938 | 9.443277701 | 0.356628238 | 0.667178386 |
| Hap1 | 7.014899905 | 6.371645338 | 6.349533148 | 0.02211219 | 0.665366756 |
| D11Wsu47e | 7.674850388 | 7.245380512 | 7.010135225 | 0.235245287 | 0.664715163 |
| N4bp1 | 11.07831855 | 10.70725569 | 10.41423018 | 0.293025503 | 0.664088367 |
| Mrpl19 | 9.339843573 | 8.837858196 | 8.676901731 | 0.160956465 | 0.662941842 |
| Gart | 10.80265581 | 10.4609298 | 10.1398163 | 0.321113501 | 0.662839504 |
| Itga2b | 7.513390805 | 7.174269433 | 6.851063711 | 0.323205722 | 0.662327094 |
| Il13ra1 | 12.6384126 | 12.13207954 | 11.9766579 | 0.155421643 | 0.6617547 |
| Gbp7 | 10.48970948 | 9.574579834 | 9.82840078 | -0.253820946 | 0.661308703 |
| Asns | 10.36911807 | 10.13507662 | 9.708443521 | 0.426633095 | 0.660674552 |
| Homer1 | 9.698951511 | 10.30809594 | 9.039129443 | 1.268966501 | 0.659822068 |
| Gramd1a | 10.28763511 | 10.11928156 | 9.628456907 | 0.490824653 | 0.659178206 |
| Alpk1 | 8.298207012 | 7.897214905 | 7.639457735 | 0.25775717 | 0.658749277 |
| Cxcl11 | 5.072813715 | 4.279983711 | 4.414390013 | -0.134406301 | 0.658423702 |
| Ell2 | 10.82002995 | 10.40715034 | 10.16279305 | 0.244357293 | 0.6572369 |
| Serpinb1a | 10.37804161 | 9.728078195 | 9.721786377 | 0.006291818 | 0.65625523 |

| | | | | | |
|---|---|---|---|---|---|
| D1Ertd622e | 9.621609115 | 9.25581679 | 8.965455295 | 0.290361495 | 0.65615382 |
| Gclm | 9.963468654 | 10.02802746 | 9.30836972 | 0.719657735 | 0.655098934 |
| Has2 | 7.274275145 | 6.707935106 | 6.619823279 | 0.088111827 | 0.654451866 |
| Ltbr | 10.92799555 | 10.6966159 | 10.27363388 | 0.422982017 | 0.654361667 |
| Ddx18 | 10.36756735 | 9.938947983 | 9.715285014 | 0.223662969 | 0.652282333 |
| Relb | 9.410364008 | 9.168705774 | 8.758291407 | 0.410414367 | 0.652072601 |
| Wdr18 | 9.785192547 | 9.702246765 | 9.133533365 | 0.568713399 | 0.651659182 |
| Wdr12 | 9.673334375 | 9.250672655 | 9.022852484 | 0.227820171 | 0.650481891 |
| Was | 8.020797188 | 7.77900583 | 7.370409204 | 0.408596626 | 0.650387983 |
| Sh3pxd2b | 11.39804793 | 11.13006383 | 10.74851619 | 0.38154764 | 0.64953174 |
| Cotl1 | 10.57110981 | 10.30061752 | 9.922689773 | 0.377927751 | 0.648420041 |
| Wnt16 | 8.334446841 | 8.019737767 | 7.686959649 | 0.332778119 | 0.647487193 |
| Tma16 | 8.631296898 | 8.299995698 | 7.984286142 | 0.315709557 | 0.647010756 |
| Pml | 10.08975762 | 9.644029369 | 9.442993977 | 0.201035392 | 0.646763641 |
| Tnip2 | 9.565507174 | 9.218784928 | 8.918902708 | 0.29988222 | 0.646604466 |
| Map3k6 | 10.20768093 | 9.894727645 | 9.561139128 | 0.333588517 | 0.646541805 |
| Fermt3 | 9.279815204 | 9.107233743 | 8.633586148 | 0.473647595 | 0.646229056 |
| Cd86 | 8.903844909 | 8.477639277 | 8.257893828 | 0.219745449 | 0.64595108 |
| Kti12 | 8.986295881 | 8.707950019 | 8.340739285 | 0.367210734 | 0.645556596 |
| St3gal4 | 10.00911974 | 9.842705525 | 9.364689289 | 0.478016237 | 0.644430455 |
| Sele | 7.989218312 | 7.704987219 | 7.344847063 | 0.360140157 | 0.644371249 |
| Orai2 | 8.403690735 | 8.04297764 | 7.759565386 | 0.283412254 | 0.64412535 |
| Cct3 | 12.2938657 | 11.97478475 | 11.64987931 | 0.324905437 | 0.643986383 |
| Ppp1r27 | 7.50574701 | 7.941898168 | 6.862469596 | 1.079428572 | 0.643277414 |
| Pdcd11 | 10.28116058 | 9.979337145 | 9.63878887 | 0.340548274 | 0.642371706 |
| Wdr55 | 9.106590058 | 8.776830461 | 8.464919235 | 0.311911226 | 0.641670823 |
| Adgra2 | 10.16664853 | 10.21515489 | 9.525504871 | 0.689650016 | 0.641143656 |
| Slc41a2 | 8.026699553 | 7.688304722 | 7.385754446 | 0.302550276 | 0.640945107 |
| Aatf | 9.439109001 | 8.986158798 | 8.798329049 | 0.187829749 | 0.640779952 |
| Cdk6 | 11.17397851 | 10.77161659 | 10.53367479 | 0.2379418 | 0.640303717 |
| Gm11427 | 8.034153108 | 7.707769798 | 7.394224749 | 0.313545049 | 0.639928359 |
| Cyp1b1 | 10.85116999 | 10.43376488 | 10.2112815 | 0.22248338 | 0.639888497 |

| | | | | | |
|---|---|---|---|---|---|
| Slc12a7 | 10.04172755 | 9.5864967 | 9.401947009 | 0.184549691 | 0.639780543 |
| Serpina3i | 8.725266119 | 8.456844742 | 8.085515745 | 0.371328997 | 0.639750374 |
| Irak4 | 8.974996382 | 8.585803835 | 8.335951687 | 0.249852148 | 0.639044695 |
| Ifit1bl2 | 6.406862156 | 5.978811335 | 5.768409791 | 0.210401544 | 0.638452365 |
| Celsr3 | 5.030600609 | 4.673907766 | 4.39262325 | 0.281284516 | 0.637977359 |
| Efna5 | 10.34016291 | 9.667163855 | 9.702729114 | -0.035565258 | 0.6374338 |
| Gbp9 | 9.277582819 | 8.486474306 | 8.640743438 | -0.154269132 | 0.63683938 |
| Epsti1 | 8.521152177 | 7.706479303 | 7.884345152 | -0.177865849 | 0.636807024 |
| Nsmce1 | 9.176499147 | 8.708051922 | 8.539736595 | 0.168315327 | 0.636762552 |
| Arhgap8 | 6.907784742 | 6.590571853 | 6.271163159 | 0.319408695 | 0.636621583 |
| Qtrt2 | 9.078109197 | 8.795983542 | 8.442154229 | 0.353829313 | 0.635954968 |
| Zfp608 | 8.284409847 | 7.784561937 | 7.648548697 | 0.136013239 | 0.63586115 |
| Lrg1 | 10.59495386 | 10.63473856 | 9.95930944 | 0.67542912 | 0.635644416 |
| Steap1 | 5.487663692 | 5.076027191 | 4.852623399 | 0.223403792 | 0.635040293 |
| Arpc1b | 12.24359273 | 12.1789617 | 11.60870312 | 0.570258587 | 0.634889613 |
| Cln3 | 9.227970423 | 9.171163027 | 8.593115625 | 0.578047402 | 0.634854798 |
| Eps8 | 9.959624708 | 9.891401089 | 9.325012781 | 0.566388308 | 0.634611928 |
| Nabp1 | 10.76126914 | 10.42421141 | 10.12697154 | 0.297239862 | 0.634297595 |
| Etv6 | 10.56002498 | 10.13083317 | 9.926027811 | 0.204805362 | 0.633997172 |
| Ifrd2 | 9.984388814 | 9.563156477 | 9.35046392 | 0.212692557 | 0.633924894 |
| Mthfd1l | 8.739550927 | 8.520936161 | 8.105632928 | 0.415303234 | 0.633917999 |
| F3 | 10.38627005 | 9.659551856 | 9.752879964 | -0.093328107 | 0.633390083 |
| Pfkfb4 | 9.795765569 | 9.610932182 | 9.16279638 | 0.448135802 | 0.632969189 |
| Dab2 | 11.98739448 | 11.71724402 | 11.35446766 | 0.362776363 | 0.63292682 |
| Pus1 | 9.195005016 | 8.866628391 | 8.562203632 | 0.304424759 | 0.632801384 |
| Sde2 | 10.05195543 | 9.760425323 | 9.419350577 | 0.341074746 | 0.632604856 |
| Furin | 11.68323008 | 11.60771042 | 11.05105145 | 0.55665897 | 0.63217863 |
| Tbl3 | 9.883279635 | 9.503300247 | 9.252038659 | 0.251261588 | 0.631240976 |
| Esd | 11.23235491 | 11.07251887 | 10.60129119 | 0.47122768 | 0.631063713 |
| Prmt1 | 11.04703742 | 10.65685095 | 10.41631589 | 0.240535057 | 0.630721527 |
| Atf4 | 12.40399639 | 12.20586487 | 11.77343726 | 0.432427613 | 0.630559133 |
| Sec24b | 10.94048931 | 10.66951776 | 10.31059637 | 0.358921387 | 0.629892937 |

| | | | | | |
|---|---|---|---|---|---|
| Emp1 | 14.01628927 | 13.59377909 | 13.38654958 | 0.207229507 | 0.629739687 |
| Ip6k3 | 8.464378782 | 7.537153892 | 7.834887593 | -0.297733702 | 0.629491188 |
| Alox5ap | 10.46364516 | 9.652396204 | 9.834479856 | -0.182083652 | 0.629165304 |
| Slc5a8 | 9.514891979 | 9.081124656 | 8.886142848 | 0.194981807 | 0.628749131 |
| Runx2 | 7.94586919 | 7.327978806 | 7.317173934 | 0.010804872 | 0.628695256 |
| Smad1 | 10.34515913 | 10.07177042 | 9.71678764 | 0.35498278 | 0.62837149 |
| Etv5 | 9.457167801 | 8.739674837 | 8.828831022 | -0.089156185 | 0.628336779 |
| Naip5 | 7.319177583 | 7.087099021 | 6.691204646 | 0.395894376 | 0.627972937 |
| Cish | 9.857681732 | 9.940146979 | 9.22987472 | 0.710272259 | 0.627807012 |
| Fam83a | 6.836364477 | 6.495919132 | 6.208848033 | 0.287071099 | 0.627516444 |
| Pgm2 | 8.913911331 | 8.43683783 | 8.287371877 | 0.149465952 | 0.626539453 |
| Snx18 | 10.6866888 | 10.21378787 | 10.06039636 | 0.153391515 | 0.626292442 |
| Nhp2 | 10.72894823 | 10.31864517 | 10.10287316 | 0.21577201 | 0.62607507 |
| Antxrl | 4.46317775 | 3.957448258 | 3.837638727 | 0.119809531 | 0.625539023 |
| Atp6v0c | 12.30517141 | 12.4171467 | 11.6796766 | 0.7374701 | 0.625494817 |
| Batf2 | 6.338046037 | 5.754991157 | 5.714131993 | 0.040859163 | 0.623914044 |
| Tle3 | 11.46522575 | 11.20044866 | 10.84196022 | 0.358488437 | 0.623265527 |
| Cpne8 | 10.66173175 | 10.09980548 | 10.03993975 | 0.059865728 | 0.621791997 |
| Aen | 10.14160139 | 9.928312525 | 9.520123257 | 0.408189268 | 0.621478136 |
| Chchd4 | 8.803939509 | 8.407294981 | 8.182925579 | 0.224369402 | 0.62101393 |
| Fubp1 | 11.64982009 | 11.52640025 | 11.0292207 | 0.49717955 | 0.620599393 |
| Creld2 | 9.52558079 | 9.532123536 | 8.905305062 | 0.626818474 | 0.620275728 |
| Mphosph10 | 9.949753469 | 9.505716914 | 9.330020439 | 0.175696475 | 0.61973303 |
| Rpf2 | 8.866054591 | 8.554574054 | 8.247277269 | 0.307296785 | 0.618777322 |
| Ubash3b | 9.344263416 | 8.962999278 | 8.726660858 | 0.23633842 | 0.617602558 |
| Rcc1 | 9.750407286 | 9.573659968 | 9.133532646 | 0.440127322 | 0.61687464 |
| Mtap | 9.577051364 | 9.08957995 | 8.960628023 | 0.128951927 | 0.616423341 |
| Myo1g | 9.653078764 | 9.849515879 | 9.037155622 | 0.812360256 | 0.615923141 |
| Atp1a1 | 13.65614667 | 13.4546778 | 13.04049638 | 0.41418142 | 0.615650287 |
| Vars | 10.66521224 | 10.68328182 | 10.04956862 | 0.633713203 | 0.61564362 |
| Fam78b | 7.212961775 | 6.974790511 | 6.597365177 | 0.377425333 | 0.615596597 |
| Trim25 | 11.33426189 | 10.84473337 | 10.71878868 | 0.125944693 | 0.61547321 |

| | | | | | |
|---|---|---|---|---|---|
| Timm10 | 8.409417767 | 7.886391675 | 7.795241229 | 0.091150446 | 0.614176539 |
| St6galnac4 | 8.14937656 | 8.305764085 | 7.535339305 | 0.77042478 | 0.614037255 |
| Rhov | 9.595930506 | 9.087893172 | 8.982086423 | 0.105806749 | 0.613844083 |
| Tor3a | 10.41574977 | 9.642359265 | 9.802030613 | -0.159671348 | 0.613719154 |
| Naa25 | 11.28655224 | 10.92873745 | 10.67343107 | 0.255306377 | 0.613121167 |
| Arrdc4 | 10.28115314 | 9.813065595 | 9.668709226 | 0.144356369 | 0.612443917 |
| Arl5c | 6.961248975 | 6.853318272 | 6.348849975 | 0.504468297 | 0.612399 |
| Entpd3 | 6.501115961 | 6.318113214 | 5.889847699 | 0.428265514 | 0.611268261 |
| Timm9 | 8.852714534 | 8.397061739 | 8.241507032 | 0.155554706 | 0.611207502 |
| Cd37 | 8.618191684 | 8.213039067 | 8.007127345 | 0.205911721 | 0.611064339 |
| Tox2 | 5.874007601 | 5.664672943 | 5.263125356 | 0.401547586 | 0.610882245 |
| Gnb4 | 8.700661021 | 8.079542685 | 8.089999748 | -0.010457064 | 0.610661272 |
| Wdr75 | 10.11775904 | 9.656760874 | 9.507739635 | 0.14902124 | 0.610019403 |
| Rnf40 | 10.96024704 | 10.79288021 | 10.35061898 | 0.44226123 | 0.609628063 |
| Gm9797 | 6.772570892 | 6.869795207 | 6.163083458 | 0.706711749 | 0.609487434 |
| Ddx39 | 10.81284076 | 10.61258876 | 10.20394604 | 0.408642722 | 0.608894728 |
| Igf2bp2 | 7.528478126 | 7.873735675 | 6.919741502 | 0.953994173 | 0.608736624 |
| Tmem185b | 8.938656838 | 8.614490917 | 8.330489418 | 0.284001499 | 0.608167421 |
| Ly6g | 4.098365847 | 3.583540875 | 3.491492978 | 0.092047897 | 0.606872869 |
| Ngp | 3.75930904 | 3.190712637 | 3.153159797 | 0.03755284 | 0.606149243 |
| Rnf31 | 9.807284116 | 9.343962231 | 9.201615057 | 0.142347173 | 0.605669058 |
| Htr7 | 5.862482981 | 5.696678553 | 5.25711085 | 0.439567703 | 0.605372131 |
| Qtrt1 | 7.778276426 | 7.542190236 | 7.173416116 | 0.36877412 | 0.604860309 |
| Hspa8 | 15.38561215 | 15.64467309 | 14.78078618 | 0.863886903 | 0.60482597 |
| Yars | 10.70049024 | 10.31755717 | 10.09571273 | 0.221844435 | 0.604777505 |
| Rela | 10.78796977 | 10.60569852 | 10.18331276 | 0.422385753 | 0.604657007 |
| Gm49368 | 4.957703887 | 4.788439125 | 4.353084948 | 0.435354176 | 0.604618939 |
| Ube2l6 | 9.36088911 | 8.592603296 | 8.756680487 | -0.164077192 | 0.604208622 |
| Ehbp1l1 | 11.184055 | 10.95110793 | 10.58034529 | 0.370762633 | 0.603709703 |
| Cd3eap | 9.895499935 | 9.522261928 | 9.291910939 | 0.23035099 | 0.603588996 |
| 2410002F23Rik | 10.17435073 | 9.738443913 | 9.571983942 | 0.166459971 | 0.602366791 |
| Ddx31 | 8.398633414 | 7.991542773 | 7.796665767 | 0.194877006 | 0.601967647 |

| | | | | | |
|---|---|---|---|---|---|
| Hs3st3b1 | 8.961662858 | 8.729921248 | 8.359985155 | 0.369936093 | 0.601677704 |
| Naip2 | 7.909798385 | 7.52729862 | 7.308175156 | 0.219123464 | 0.601623229 |
| Ehf | 9.888213091 | 9.441954472 | 9.287190531 | 0.154763941 | 0.60102256 |
| Togaram2 | 7.727794029 | 7.270404819 | 7.127509722 | 0.142895097 | 0.600284307 |
| Flot1 | 10.53269226 | 10.08782903 | 9.932514475 | 0.155314555 | 0.600177788 |
| Pxdc1 | 9.175868745 | 8.82448233 | 8.578474595 | 0.246007735 | 0.59739415 |
| Gm28048 | 9.751454215 | 8.985084351 | 9.154747041 | -0.16966269 | 0.596707174 |
| Ttc27 | 9.238401253 | 8.873592672 | 8.641833709 | 0.231758963 | 0.596567544 |
| Bid | 8.160403683 | 7.971169345 | 7.564133844 | 0.4070355 | 0.596269838 |
| Pycr2 | 9.119685714 | 8.802321503 | 8.523443749 | 0.278877754 | 0.596241965 |
| Cd300a | 8.801328071 | 8.731938587 | 8.205760712 | 0.526177875 | 0.595567359 |
| Rgs14 | 7.487732572 | 7.178105557 | 6.892252111 | 0.285853446 | 0.595480462 |
| Sertad1 | 9.924711004 | 9.774501836 | 9.329889904 | 0.444611932 | 0.594821099 |
| Gemin5 | 9.618478593 | 9.333126342 | 9.023756019 | 0.309370323 | 0.594722574 |
| Ruvbl1 | 9.463803114 | 9.026113659 | 8.869303469 | 0.15681019 | 0.594499645 |
| Il4i1 | 7.425413181 | 8.046867414 | 6.831364943 | 1.215502471 | 0.594048238 |
| Dctd | 8.197136946 | 8.030113621 | 7.603101926 | 0.427011695 | 0.59403502 |
| Lrrc25 | 8.276867195 | 8.257550015 | 7.683631793 | 0.573918221 | 0.593235401 |
| Ppm1h | 8.524537277 | 8.117840997 | 7.932103663 | 0.185737334 | 0.592433614 |
| Tpm4 | 13.33762683 | 12.9499771 | 12.74534469 | 0.204632403 | 0.592282133 |
| Lpar1 | 10.08631509 | 9.71313922 | 9.494243223 | 0.218895997 | 0.592071862 |
| Mthfd2 | 8.759087432 | 8.453077998 | 8.167248712 | 0.285829285 | 0.591838719 |
| Nup98 | 11.03671494 | 10.7824517 | 10.44513418 | 0.33731752 | 0.591580763 |
| Pnpt1 | 9.820870582 | 9.352136728 | 9.229538324 | 0.122598404 | 0.591332258 |
| Ikzf3 | 5.787394641 | 5.323973409 | 5.196106386 | 0.127867022 | 0.591288255 |
| Exoc3l4 | 5.984831593 | 5.54771953 | 5.393883695 | 0.153835835 | 0.590947899 |
| Slc30a7 | 10.28010117 | 10.06653719 | 9.689673012 | 0.376864175 | 0.590428155 |
| Timm8a1 | 9.356662818 | 8.779224035 | 8.766662227 | 0.012561808 | 0.590000591 |
| Lpin2 | 9.063323655 | 8.892658049 | 8.47344674 | 0.419211309 | 0.589876915 |
| Ttc37 | 10.69445462 | 10.17576371 | 10.10477619 | 0.070987525 | 0.589678431 |
| Pik3cd | 9.512835658 | 9.447067083 | 8.925551185 | 0.521515899 | 0.587284473 |
| Rbm12 | 9.298353108 | 9.120585402 | 8.711139847 | 0.409445555 | 0.587213261 |

| | | | | | |
|---|---|---|---|---|---|
| Itga7 | 10.54785688 | 10.26605036 | 9.960707811 | 0.305342553 | 0.587149066 |
| Pak6 | 10.61752022 | 10.42208519 | 10.03085416 | 0.391231028 | 0.586666058 |
| Armc6 | 7.797060044 | 7.702312392 | 7.210905223 | 0.491407169 | 0.586154821 |
| Layn | 7.660229698 | 7.478812276 | 7.074153868 | 0.404658408 | 0.58607583 |
| Socs2 | 8.857899471 | 8.919084048 | 8.272966343 | 0.646117705 | 0.584933128 |
| Gjb1 | 5.705657376 | 5.514193076 | 5.12097671 | 0.393216366 | 0.584680666 |
| Rdh16 | 6.123658289 | 5.740313232 | 5.53950646 | 0.200806772 | 0.584151829 |
| Gpr171 | 6.964799959 | 6.625236742 | 6.381266725 | 0.243970017 | 0.583533234 |
| Bag3 | 10.88648157 | 11.76524436 | 10.3031077 | 1.462136667 | 0.58337387 |
| Arhgdib | 11.77502834 | 11.39928603 | 11.1920744 | 0.207211633 | 0.582953947 |
| Hspb7 | 10.86073318 | 11.13471295 | 10.2780208 | 0.856692153 | 0.58271238 |
| Gm9531 | 10.04991151 | 9.765441145 | 9.467260144 | 0.298181001 | 0.582651362 |
| Ampd2 | 9.757568916 | 9.435755197 | 9.175582128 | 0.260173069 | 0.581986788 |
| Clip2 | 9.063553798 | 9.013685084 | 8.48281915 | 0.530865934 | 0.580734648 |
| Engase | 7.880684984 | 7.355073638 | 7.299986829 | 0.055086809 | 0.580698155 |
| Rnf114 | 10.6221382 | 10.17486846 | 10.04155813 | 0.133310325 | 0.580580069 |
| Trim12c | 9.644530144 | 9.179695345 | 9.064032955 | 0.115662389 | 0.580497189 |
| Ccl24 | 8.545261916 | 8.283837838 | 7.965102125 | 0.318735712 | 0.580159791 |
| Slc39a1 | 12.06530314 | 11.67919551 | 11.48528754 | 0.193907967 | 0.5800156 |
| Tmem71 | 6.841970809 | 6.379105933 | 6.262109199 | 0.116996734 | 0.57986161 |
| Plscr2 | 6.743270871 | 5.845589705 | 6.163450966 | -0.317861261 | 0.579819905 |
| Gtpbp4 | 11.0031666 | 10.63027086 | 10.42339504 | 0.206875817 | 0.57977156 |
| Naf1 | 8.941411118 | 8.53043527 | 8.362411163 | 0.168024107 | 0.578999955 |
| Tars | 11.05655547 | 10.60652601 | 10.47784604 | 0.128679973 | 0.578709437 |
| Atad3a | 10.12848065 | 9.7698922 | 9.550095214 | 0.219796985 | 0.57838544 |
| Bend3 | 8.468408326 | 8.079694059 | 7.890028721 | 0.189665338 | 0.578379605 |
| Rsl1d1 | 11.04474492 | 10.74682875 | 10.46689948 | 0.279929267 | 0.57784544 |
| Isyna1 | 8.963544454 | 8.896715726 | 8.386714898 | 0.510000829 | 0.576829556 |
| Samd4b | 10.7848234 | 10.67893054 | 10.20803864 | 0.470891903 | 0.57678476 |
| Bdkrb2 | 7.292585817 | 6.871002672 | 6.715829055 | 0.155173617 | 0.576756762 |
| Jak3 | 8.375721743 | 8.267499257 | 7.799151209 | 0.468348048 | 0.576570534 |
| H2-T24 | 8.37023715 | 7.810835 | 7.793769697 | 0.017065303 | 0.576467453 |

| | | | | | |
|---|---|---|---|---|---|
| Rbpj | 11.61220442 | 11.264794 | 11.03592843 | 0.22886557 | 0.576275993 |
| Slc7a8 | 9.110466848 | 8.768259542 | 8.534254237 | 0.234005305 | 0.576212611 |
| Manf | 10.82822594 | 10.83715441 | 10.25221097 | 0.584943443 | 0.576014977 |
| Srfbp1 | 8.512886384 | 8.166815271 | 7.937348876 | 0.229466395 | 0.575537508 |
| B4galt1 | 11.86381511 | 11.4530593 | 11.28833297 | 0.16472633 | 0.57548214 |
| Flot2 | 10.46967111 | 10.00695265 | 9.894219329 | 0.11273332 | 0.575451784 |
| Exosc2 | 9.392955259 | 9.02817107 | 8.819012891 | 0.209158179 | 0.573942368 |
| Gm5970 | 5.083478743 | 4.48281433 | 4.510598615 | -0.027784285 | 0.572880128 |
| Cmip | 10.19587682 | 10.06582395 | 9.623186428 | 0.442637519 | 0.572690392 |
| Farsb | 10.57440845 | 10.19817637 | 10.00177433 | 0.196402039 | 0.572634126 |
| Ccdc137 | 9.063566921 | 8.877451025 | 8.491303644 | 0.386147381 | 0.572263276 |
| Ccl20 | 6.976309139 | 6.595424032 | 6.404274783 | 0.191149249 | 0.572034356 |
| Epb41l3 | 9.707945521 | 9.57196281 | 9.136127889 | 0.435834921 | 0.571817633 |
| Map1s | 7.4327525 | 7.465081057 | 6.861571068 | 0.603509989 | 0.571181432 |
| Klf7 | 10.01272217 | 9.605484722 | 9.441884891 | 0.16359983 | 0.570837277 |
| Irf5 | 10.08234934 | 9.908267423 | 9.512049879 | 0.396217543 | 0.570299464 |
| Ran | 12.45703532 | 12.15453787 | 11.88688159 | 0.267656277 | 0.57015373 |
| Picalm | 12.59707216 | 12.15071684 | 12.0275026 | 0.123214247 | 0.56956956 |
| Gad1-ps | 5.289889867 | 5.150782365 | 4.720475667 | 0.430306698 | 0.5694142 |
| Cd209c | 8.301583759 | 7.982413943 | 7.732626456 | 0.249787487 | 0.568957304 |
| Ccl11 | 10.40318504 | 9.808684752 | 9.835134644 | -0.026449892 | 0.568050399 |
| Cmtr1 | 10.6149099 | 10.22988936 | 10.04720256 | 0.182686796 | 0.567707336 |
| Pak1ip1 | 10.42689199 | 10.05747673 | 9.859703063 | 0.197773667 | 0.567188924 |
| Panx1 | 9.418843616 | 9.165404848 | 8.851710283 | 0.313694564 | 0.567133333 |
| Limd2 | 8.745885856 | 8.278554492 | 8.178820975 | 0.099733516 | 0.56706488 |
| Mettl1 | 9.320056237 | 9.066247328 | 8.753112229 | 0.313135099 | 0.566944008 |
| Lsp1 | 11.48294267 | 11.14178526 | 10.91628346 | 0.225501807 | 0.566659213 |
| Mrpl12 | 10.49091816 | 10.21175207 | 9.924463579 | 0.287288491 | 0.566454585 |
| AW011738 | 6.95164588 | 6.430451338 | 6.386036924 | 0.044414413 | 0.565608955 |
| Coq10b | 9.849827802 | 9.43517291 | 9.284752843 | 0.150420067 | 0.565074959 |
| Skil | 10.06362888 | 9.766083486 | 9.498675499 | 0.267407987 | 0.564953384 |
| Rcc1l | 8.862915832 | 8.616368427 | 8.299212721 | 0.317155706 | 0.563703111 |

| | | | | | |
|---|---|---|---|---|---|
| Nts | 4.979533343 | 4.660121702 | 4.416095224 | 0.244026478 | 0.563438119 |
| Sorcs2 | 7.908219245 | 7.847736446 | 7.344897577 | 0.50283887 | 0.563321668 |
| Tfec | 6.348716692 | 6.389761518 | 5.78552838 | 0.604233138 | 0.563188312 |
| Il11ra2 | 6.579152809 | 6.37967288 | 6.016297901 | 0.363374979 | 0.562854908 |
| Brix1 | 10.20934871 | 9.885318745 | 9.646688933 | 0.238629812 | 0.562659774 |
| Map2k3 | 11.88822224 | 11.48360139 | 11.3259578 | 0.15764359 | 0.562264443 |
| Ikzf1 | 9.089788702 | 8.943931184 | 8.527952274 | 0.41597891 | 0.561836428 |
| Mettl13 | 9.192784022 | 8.899397596 | 8.631311097 | 0.268086499 | 0.561472925 |
| Ldlr | 11.71575133 | 11.54564858 | 11.15433327 | 0.391315303 | 0.561418057 |
| Ralgds | 11.39265016 | 11.03301741 | 10.83158342 | 0.20143399 | 0.56106674 |
| Snrpa | 9.902155221 | 9.734463377 | 9.341501565 | 0.392961811 | 0.560653656 |
| Dock2 | 9.628529335 | 9.276795789 | 9.068254911 | 0.208540878 | 0.560274425 |
| Tut1 | 9.247079003 | 8.874362513 | 8.686979442 | 0.187383071 | 0.560099561 |
| Antxr2 | 10.46011248 | 9.635662553 | 9.900044084 | -0.264381531 | 0.560068399 |
| Slurp1 | 9.047182608 | 8.708331296 | 8.487269534 | 0.221061762 | 0.559913074 |
| Mrps18b | 9.772799413 | 9.458301476 | 9.212963559 | 0.245337917 | 0.559835854 |
| Hba-a2 | 10.96721048 | 9.974997543 | 10.40765662 | -0.432659073 | 0.559553864 |
| Tnfrsf18 | 9.850207504 | 9.474839882 | 9.290793294 | 0.184046588 | 0.55941421 |
| Zmynd19 | 9.353732419 | 9.077742468 | 8.794725938 | 0.28301653 | 0.559006481 |
| E2f4 | 10.48070507 | 10.2019172 | 9.922087354 | 0.27982985 | 0.558617716 |
| Il10rb | 10.09514103 | 9.778195109 | 9.53668648 | 0.241508628 | 0.558454553 |
| Grk6 | 10.39116444 | 10.07031022 | 9.8329217 | 0.23738852 | 0.558242737 |
| Adar | 10.5937634 | 10.25814753 | 10.03557599 | 0.222571543 | 0.558187416 |
| Cyth3 | 11.00173228 | 10.50850961 | 10.44372756 | 0.064782053 | 0.558004723 |
| Znfx1 | 11.16670232 | 10.56454856 | 10.60892653 | -0.044377967 | 0.557775797 |
| Rfng | 10.03849416 | 9.828896833 | 9.481001971 | 0.347894862 | 0.55749219 |
| 1810037I17Rik | 10.83221219 | 10.49182733 | 10.27475622 | 0.217071107 | 0.557455967 |
| Prag1 | 7.581481275 | 7.471387952 | 7.024891158 | 0.446496794 | 0.556590117 |
| Nus1 | 10.83784602 | 10.54426312 | 10.28135135 | 0.262911767 | 0.556494673 |
| Gm5150 | 5.531887282 | 5.391469759 | 4.977011145 | 0.414458614 | 0.554876137 |
| Trim8 | 10.17092747 | 9.990925889 | 9.616203784 | 0.374722105 | 0.554723682 |
| Car9 | 6.411046679 | 6.089375433 | 5.857005696 | 0.232369737 | 0.554040983 |

| | | | | | |
|---|---|---|---|---|---|
| C1qbp | 10.86006505 | 10.48830155 | 10.30685177 | 0.181449777 | 0.55321328 |
| Gpr65 | 7.012984001 | 6.77400362 | 6.459923572 | 0.314080048 | 0.553060429 |
| Spry2 | 9.267409295 | 8.954007139 | 8.714898255 | 0.239108884 | 0.55251104 |
| Trim12a | 9.303784522 | 8.579502382 | 8.751614643 | -0.172112261 | 0.552169879 |
| Rsl24d1 | 10.17475122 | 9.793588667 | 9.622732851 | 0.170855816 | 0.552018373 |
| Apln | 6.724990046 | 6.853178464 | 6.173429355 | 0.679749109 | 0.551560691 |
| Ndufaf4 | 9.12130567 | 8.728709744 | 8.570262245 | 0.158447499 | 0.551043424 |
| Gm20390 | 10.98664112 | 10.99437822 | 10.43578247 | 0.558595743 | 0.55085865 |
| Por | 11.6552473 | 11.47966789 | 11.10450503 | 0.37516286 | 0.550742267 |
| Prpf31 | 10.05029489 | 9.747566255 | 9.500161109 | 0.247405146 | 0.550133783 |
| Ddx54 | 10.3315737 | 10.29982481 | 9.781447555 | 0.518377258 | 0.550126148 |
| Cebpz | 10.6270122 | 10.17287603 | 10.07702211 | 0.095853917 | 0.549990083 |
| Cyth1 | 10.56542141 | 9.98289968 | 10.01630353 | -0.033403848 | 0.549117878 |
| Dhx37 | 8.895453665 | 8.568118492 | 8.347380209 | 0.220738283 | 0.548073457 |
| 40057 | 11.52266729 | 11.22540916 | 10.97489606 | 0.250513097 | 0.547771227 |
| Capn2 | 12.09891704 | 11.8223022 | 11.55136099 | 0.270941213 | 0.547556057 |
| Cfl1 | 13.59668302 | 13.34168965 | 13.04965328 | 0.292036373 | 0.54702974 |
| Tsr2 | 9.060380749 | 8.697290327 | 8.51425781 | 0.183032517 | 0.546122939 |
| Surf2 | 9.110231744 | 8.768400148 | 8.564143188 | 0.204256961 | 0.546088556 |
| Tgfbi | 13.88965531 | 13.74538934 | 13.34381154 | 0.401577793 | 0.545843763 |
| Flt1 | 10.03147786 | 9.486409697 | 9.485959669 | 0.000450028 | 0.545518188 |
| Zdhhc18 | 9.940824013 | 9.927605719 | 9.395812756 | 0.531792963 | 0.545011257 |
| Lsm7 | 8.752383955 | 8.66388897 | 8.207501057 | 0.456387913 | 0.544882898 |
| Pde12 | 9.373178994 | 9.048443286 | 8.828315329 | 0.220127958 | 0.544863665 |
| Il18r1 | 8.133161444 | 7.62151733 | 7.589139606 | 0.032377724 | 0.544021838 |
| Dcstamp | 5.591310493 | 5.251489546 | 5.047591847 | 0.203897699 | 0.543718646 |
| Lpxn | 8.023927778 | 7.994425848 | 7.480451699 | 0.513974149 | 0.543476079 |
| Tnfrsf11b | 8.650343645 | 7.814077981 | 8.10741855 | -0.293340568 | 0.542925096 |
| Sco2 | 7.714842581 | 7.666581859 | 7.17202645 | 0.49455541 | 0.542816131 |
| Map3k15 | 5.605160481 | 5.196304792 | 5.062359496 | 0.133945295 | 0.542800984 |
| Car12 | 11.40950289 | 11.03651354 | 10.8667049 | 0.169808643 | 0.542797997 |
| Ifitm2 | 12.52346928 | 12.07560692 | 11.98072728 | 0.094879643 | 0.542741997 |

| | | | | | |
|---|---|---|---|---|---|
| Itprip | 10.51613702 | 10.07236629 | 9.974063987 | 0.098302303 | 0.542073033 |
| Gmppb | 9.259690111 | 9.123041433 | 8.717780864 | 0.405260569 | 0.541909247 |
| Gsdmd | 9.263782281 | 8.959424583 | 8.722843969 | 0.236580614 | 0.540938312 |
| Rffl | 9.353803352 | 8.942142456 | 8.813276684 | 0.128865772 | 0.540526668 |
| Cndp2 | 11.54609435 | 11.32442726 | 11.00594471 | 0.31848255 | 0.540149637 |
| Timm23 | 10.88395172 | 10.66980636 | 10.34394324 | 0.325863117 | 0.540008477 |
| Pwp1 | 9.713274172 | 9.354655865 | 9.173543553 | 0.181112312 | 0.539730619 |
| Prmt3 | 9.539598673 | 9.127899962 | 9.000172949 | 0.127727013 | 0.539425724 |
| Zfp263 | 9.864033973 | 9.489623423 | 9.325225124 | 0.164398298 | 0.538808849 |
| Trim16 | 10.14996376 | 9.921259523 | 9.611580834 | 0.309678688 | 0.538382929 |
| Pxn | 10.70295596 | 10.63859996 | 10.16479606 | 0.473803897 | 0.5381599 |
| Hpx | 5.266964136 | 4.814841618 | 4.729093784 | 0.085747835 | 0.537870353 |
| Pfkl | 10.53477738 | 10.3278897 | 9.99710619 | 0.330783513 | 0.53767119 |
| Card9 | 7.506978077 | 7.248287056 | 6.969465752 | 0.278821304 | 0.537512325 |
| Dnajb11 | 11.12943066 | 10.87858287 | 10.5919267 | 0.286656167 | 0.537503957 |
| Psmb8 | 10.59379804 | 10.21100311 | 10.05667078 | 0.154332335 | 0.537127258 |
| Casp6 | 8.889983715 | 8.628456806 | 8.352924968 | 0.275531838 | 0.537058747 |
| Msrb1 | 11.25241653 | 10.83563378 | 10.71546189 | 0.12017189 | 0.536954637 |
| Rhoj | 10.15510318 | 9.559381843 | 9.618608249 | -0.059226406 | 0.536494931 |
| Rrp8 | 9.674219111 | 9.318652159 | 9.137810221 | 0.180841938 | 0.53640889 |
| Itgb7 | 8.872323481 | 8.484170661 | 8.336067883 | 0.148102778 | 0.536255599 |
| Mfsd2a | 8.846981027 | 8.387866068 | 8.312044678 | 0.07582139 | 0.534936349 |
| Rabgef1 | 10.80428078 | 10.54659206 | 10.26961869 | 0.276973363 | 0.534662087 |
| Trim11 | 9.712786374 | 9.467965844 | 9.178264932 | 0.289700913 | 0.534521442 |
| Rcl1 | 9.406518978 | 8.9415289 | 8.872410564 | 0.069118336 | 0.534108414 |
| Tyro3 | 10.93184947 | 10.58169313 | 10.39777636 | 0.183916763 | 0.534073107 |
| Hba-a1 | 10.54676492 | 9.815132576 | 10.01282986 | -0.197697283 | 0.53393506 |
| Tomm5 | 9.735763225 | 9.388611425 | 9.201897178 | 0.186714248 | 0.533866048 |
| Mfsd4b1 | 5.138893522 | 4.524942571 | 4.605159631 | -0.080217059 | 0.533733892 |
| Clec12a | 7.987011493 | 7.59197081 | 7.453801088 | 0.138169722 | 0.533210406 |
| Hspbp1 | 8.975747114 | 8.913011602 | 8.443045481 | 0.469966121 | 0.532701634 |
| Fem1c | 10.00228656 | 9.497488594 | 9.469903901 | 0.027584693 | 0.532382656 |

| | | | | | |
|---|---|---|---|---|---|
| Sf3b4 | 9.905392335 | 9.930960982 | 9.373067006 | 0.557893976 | 0.532325329 |
| Sdad1 | 10.24982303 | 9.923430967 | 9.717525934 | 0.205905033 | 0.532297096 |
| Fam43a | 9.581112819 | 9.373602851 | 9.049319361 | 0.32428349 | 0.531793459 |
| Wdr1 | 12.33330429 | 12.02994568 | 11.80155882 | 0.22838686 | 0.53174547 |
| Nob1 | 9.456518693 | 9.213384669 | 8.92493473 | 0.288449939 | 0.531583963 |
| Eef1akmt4 | 7.390768334 | 7.058245759 | 6.859448323 | 0.198797436 | 0.531320011 |
| Gnl2 | 10.47961189 | 10.15387588 | 9.94883847 | 0.20503741 | 0.53077342 |
| Pfas | 9.379113373 | 9.139151503 | 8.848728544 | 0.290422959 | 0.530384829 |
| Map3k5 | 9.584600769 | 8.903026352 | 9.054366858 | -0.151340506 | 0.530233912 |
| Etv4 | 9.161209494 | 8.811341548 | 8.631008458 | 0.18033309 | 0.530201036 |
| Aprt | 9.938712611 | 9.803087433 | 9.408535405 | 0.394552028 | 0.530177206 |
| Ltv1 | 9.970994968 | 9.55534331 | 9.440984888 | 0.114358422 | 0.53001008 |
| Atxn7l1 | 9.243648435 | 8.840842302 | 8.714394893 | 0.126447409 | 0.529253542 |
| Abce1 | 11.70718247 | 11.4034499 | 11.17844697 | 0.225002927 | 0.528735497 |
| Slc27a3 | 10.8438793 | 10.85083832 | 10.31524422 | 0.5355941 | 0.528635077 |
| Nudcd1 | 9.571118289 | 9.27635069 | 9.042702023 | 0.233648667 | 0.528416266 |
| Mall | 11.00569331 | 10.62943071 | 10.47729352 | 0.152137183 | 0.52839979 |
| Ppard | 10.27842136 | 10.35555392 | 9.75025324 | 0.605300684 | 0.528168124 |
| Rab5c | 12.20607837 | 11.94347079 | 11.6783737 | 0.265097097 | 0.52770467 |
| Ptpn7 | 7.928595369 | 7.524227647 | 7.401009486 | 0.12321816 | 0.527585883 |
| Ddx27 | 10.18095364 | 9.93636399 | 9.653528764 | 0.282835226 | 0.52742488 |
| Plekha4 | 8.813748156 | 8.384969182 | 8.286713653 | 0.098255529 | 0.527034503 |
| Zdhhc13 | 9.148426143 | 8.855862081 | 8.622064067 | 0.233798014 | 0.526362076 |
| Snu13 | 11.18547121 | 10.81485342 | 10.65930533 | 0.155548083 | 0.526165877 |
| Acot11 | 8.245068037 | 7.917519332 | 7.719347295 | 0.198172037 | 0.525720742 |
| Rhbdf2 | 10.49223352 | 10.38951036 | 9.966987381 | 0.422522976 | 0.525246136 |
| Slc5a1 | 9.626840115 | 9.149875326 | 9.10193501 | 0.047940316 | 0.524905106 |
| Sipa1l3 | 11.06122962 | 11.02751682 | 10.53634321 | 0.49117361 | 0.524886407 |
| Cdhr1 | 5.288091615 | 4.780702424 | 4.763414012 | 0.017288411 | 0.524677603 |
| Ciita | 9.255910825 | 9.150303387 | 8.731525599 | 0.418777788 | 0.524385226 |
| Cmtm6 | 12.03278969 | 11.64047648 | 11.50949896 | 0.13097752 | 0.523290727 |
| B4galnt4 | 5.170325746 | 5.023302646 | 4.647324338 | 0.375978307 | 0.523001407 |

| | | | | | |
|---|---|---|---|---|---|
| Sssca1 | 8.069960831 | 8.010331031 | 7.547095801 | 0.46323523 | 0.52286503 |
| Mgam | 4.614694041 | 4.098433692 | 4.091905187 | 0.006528505 | 0.522788854 |
| Abcg2 | 9.68900007 | 9.303233078 | 9.166505706 | 0.136727372 | 0.522494364 |
| Champ1 | 9.499542165 | 9.256501583 | 8.977600942 | 0.278900642 | 0.521941223 |
| Impdh2 | 11.88168472 | 11.63821937 | 11.36016898 | 0.278050397 | 0.52151574 |
| Elac2 | 9.827107375 | 9.433783722 | 9.305675048 | 0.128108674 | 0.521432328 |
| Eif4a1 | 14.03763918 | 13.74182589 | 13.51647249 | 0.2253534 | 0.521166683 |
| Itgb4 | 12.32819066 | 12.31615429 | 11.80766137 | 0.508492913 | 0.520529283 |
| Mrpl52 | 9.37635083 | 9.201781364 | 8.855955134 | 0.34582623 | 0.520395696 |
| Slc25a37 | 9.471531783 | 9.156143848 | 8.951666864 | 0.204476984 | 0.519864919 |
| Ddx60 | 8.679297838 | 7.514320046 | 8.159543271 | -0.645223225 | 0.519754566 |
| Large2 | 9.629407212 | 9.6167995 | 9.110000741 | 0.506798759 | 0.519406471 |
| Cnksr1 | 9.493222393 | 9.812947334 | 8.973823892 | 0.839123441 | 0.519398501 |
| Polr1a | 10.24798274 | 9.947935106 | 9.728811938 | 0.219123167 | 0.519170805 |
| Ciao2b | 8.485485643 | 8.226889824 | 7.96657835 | 0.260311474 | 0.518907293 |
| Gm47283 | 8.912982953 | 9.249834575 | 8.395038939 | 0.854795636 | 0.517944014 |
| Nkrf | 8.235374685 | 7.824358068 | 7.717530462 | 0.106827606 | 0.517844223 |
| Chmp4b | 12.23687521 | 11.87826297 | 11.71907585 | 0.159187117 | 0.517799357 |
| Uhrf1bp1l | 10.33384104 | 10.15364977 | 9.816117187 | 0.337532583 | 0.517723853 |
| Sf3a2 | 9.655942142 | 9.660747168 | 9.139524455 | 0.521222713 | 0.516417687 |
| Cd200r1 | 8.159567869 | 8.117869262 | 7.643153209 | 0.474716054 | 0.51641466 |
| Il24 | 4.57824903 | 4.447294038 | 4.062040634 | 0.385253404 | 0.516208396 |
| Apex1 | 10.34966135 | 10.06788049 | 9.833646651 | 0.234233835 | 0.516014702 |
| Mrps25 | 9.272784275 | 9.050724641 | 8.757149506 | 0.293575134 | 0.515634769 |
| Tet2 | 10.15176458 | 9.798842002 | 9.636682844 | 0.162159158 | 0.515081732 |
| Lsg1 | 9.62047458 | 9.377760214 | 9.105456452 | 0.272303762 | 0.515018128 |
| Ext1 | 11.07490935 | 10.56670851 | 10.5603215 | 0.006387013 | 0.514587847 |
| Itpk1 | 9.062193935 | 8.628482974 | 8.547770064 | 0.08071291 | 0.514423871 |
| Prrc2a | 12.59300112 | 12.65488958 | 12.07896667 | 0.575922907 | 0.51403445 |
| Ciapin1 | 9.228136696 | 8.987236726 | 8.714143062 | 0.273093664 | 0.513993634 |
| Casp3 | 9.374962476 | 8.94782096 | 8.860984634 | 0.086836327 | 0.513977842 |
| Il21r | 5.673725474 | 5.310409673 | 5.159777805 | 0.150631867 | 0.513947669 |

| | | | | | |
|---|---|---|---|---|---|
| Flna | 13.81579878 | 13.65348144 | 13.30189152 | 0.35158992 | 0.513907267 |
| Rbm28 | 10.36620945 | 10.1231262 | 9.853115487 | 0.270010713 | 0.513093959 |
| Myo1e | 11.15433512 | 10.82756034 | 10.64134398 | 0.186216357 | 0.512991137 |
| Slc28a2 | 8.534910431 | 8.196854302 | 8.022052419 | 0.174801883 | 0.512858012 |
| Xpo5 | 10.65601591 | 10.39391409 | 10.14333098 | 0.250583105 | 0.512684929 |
| Pgam5 | 9.6180114 | 9.223909707 | 9.10664986 | 0.117259847 | 0.51136154 |
| Mat2a | 11.44721766 | 11.09504327 | 10.93662552 | 0.158417747 | 0.510592133 |
| Wdr36 | 10.59819273 | 10.09078207 | 10.08789218 | 0.002889889 | 0.51030055 |
| Bhlhe22 | 6.186008271 | 6.122372676 | 5.676164312 | 0.446208365 | 0.50984396 |
| Zc3h12c | 7.938307647 | 7.738614856 | 7.428816306 | 0.30979855 | 0.509491341 |
| Zswim4 | 10.50804048 | 10.32644066 | 9.998840957 | 0.327599703 | 0.509199526 |
| Map7d1 | 11.48290178 | 11.35056952 | 10.97381904 | 0.37675048 | 0.509082737 |
| Ms4a8a | 6.491149632 | 5.943938675 | 5.98273552 | -0.038796846 | 0.508414112 |
| Psme3 | 11.48498949 | 11.26160364 | 10.97667234 | 0.284931303 | 0.508317157 |
| Utp18 | 9.709859206 | 9.330808591 | 9.201606136 | 0.129202456 | 0.50825307 |
| Gemin6 | 7.656974675 | 7.293444625 | 7.148742045 | 0.14470258 | 0.50823263 |
| Apol9a | 6.936159928 | 6.26305033 | 6.428747181 | -0.165696851 | 0.507412747 |
| Ddx10 | 9.802870869 | 9.388744155 | 9.296205727 | 0.092538427 | 0.506665141 |
| Tgfbr2 | 12.41670194 | 12.13667294 | 11.9102478 | 0.22642514 | 0.50645414 |
| Acvrl1 | 9.658339249 | 9.570857355 | 9.152073225 | 0.41878413 | 0.506266024 |
| Eif2b3 | 8.9586728 | 8.513484417 | 8.452509469 | 0.060974948 | 0.506163331 |
| Pinx1 | 7.937306086 | 7.648245457 | 7.431330621 | 0.216914836 | 0.505975465 |
| Ptpn6 | 10.35413715 | 10.26367753 | 9.848398902 | 0.415278631 | 0.505738244 |
| Erh | 10.69127114 | 10.38575445 | 10.18681103 | 0.198943417 | 0.50446011 |
| Ifit1bl1 | 6.862420328 | 6.130624447 | 6.358072043 | -0.227447596 | 0.504348285 |
| Arid3a | 7.713726591 | 7.588668644 | 7.209923677 | 0.378744966 | 0.503802913 |
| Cdh5 | 10.34683595 | 9.988878039 | 9.843251851 | 0.145626188 | 0.503584103 |
| Nr2c2ap | 8.798616131 | 8.574416991 | 8.295598315 | 0.278818676 | 0.503017816 |
| Fmnl2 | 9.584852159 | 9.234832863 | 9.082535417 | 0.152297446 | 0.502316741 |
| Alg3 | 8.601999108 | 8.407980579 | 8.099772562 | 0.308208017 | 0.502226546 |
| Klra17 | 4.61068518 | 4.24488377 | 4.108677801 | 0.136205969 | 0.502007379 |
| Tifa | 8.118564932 | 7.339299049 | 7.616927422 | -0.277628373 | 0.501637509 |

| | | | | | |
|---|---|---|---|---|---|
| Nop14 | 9.930730112 | 9.678431739 | 9.429118903 | 0.249312836 | 0.501611209 |
| Skap2 | 10.95458516 | 10.63733338 | 10.45306675 | 0.18426663 | 0.50151841 |
| Bax | 9.404707837 | 9.335129496 | 8.903420488 | 0.431709007 | 0.501287348 |
| Rpl7l1 | 11.05657093 | 10.77316393 | 10.55589978 | 0.217264147 | 0.500671147 |
| Diexf | 8.842696448 | 8.481554181 | 8.34235142 | 0.13920276 | 0.500345028 |
| G3bp1 | 12.26809384 | 11.99820702 | 11.76789529 | 0.23031173 | 0.500198547 |
| Nol9 | 9.86705534 | 9.598527481 | 9.367304521 | 0.23122296 | 0.499750818 |
| Igf1r | 10.893946 | 10.65429267 | 10.39488628 | 0.259406387 | 0.49905972 |
| Stk19 | 9.138867647 | 9.004140049 | 8.640177592 | 0.363962457 | 0.498690055 |
| Gcc1 | 9.517736338 | 9.348352203 | 9.019111758 | 0.329240444 | 0.498624579 |
| Gprc5a | 5.445221704 | 5.351775811 | 4.94708546 | 0.404690351 | 0.498136244 |
| Atic | 10.71221155 | 10.46860271 | 10.21424498 | 0.25435773 | 0.497966567 |
| Npl | 10.71459438 | 10.41882047 | 10.21667062 | 0.20214985 | 0.49792376 |
| Pmepa1 | 9.470522438 | 9.829997272 | 8.972614406 | 0.857382866 | 0.497908032 |
| Tmem156 | 5.456660287 | 5.037508613 | 4.958931011 | 0.078577602 | 0.497729276 |
| Slc25a28 | 9.619221752 | 9.350599755 | 9.121707316 | 0.228892439 | 0.497514436 |
| Rps19bp1 | 8.317577012 | 8.122814561 | 7.821582186 | 0.301232375 | 0.495994826 |
| Cenpw | 7.752959408 | 7.283654956 | 7.257565648 | 0.026089308 | 0.49539376 |
| Mid1ip1 | 11.44419121 | 11.04349153 | 10.94883844 | 0.094653097 | 0.495352777 |
| Agfg1 | 11.06384637 | 10.70185398 | 10.56896794 | 0.132886043 | 0.494878437 |
| Smn1 | 8.87893433 | 8.63011766 | 8.384440071 | 0.245677589 | 0.494494258 |
| Auts2 | 9.516087269 | 9.568791823 | 9.021624091 | 0.547167731 | 0.494463177 |
| Rap1b | 12.26403676 | 11.85023118 | 11.76960867 | 0.080622507 | 0.494428087 |
| Chsy1 | 8.714668888 | 8.451772995 | 8.220806672 | 0.230966323 | 0.493862216 |
| Dtx3l | 10.10881585 | 9.637469733 | 9.61502871 | 0.022441024 | 0.493787137 |
| Tnfsf8 | 5.548077419 | 5.183747235 | 5.054582744 | 0.129164491 | 0.493494675 |
| Spout1 | 8.969834471 | 8.769180992 | 8.476386577 | 0.292794414 | 0.493447894 |
| Spata5 | 8.63610814 | 8.279185413 | 8.142781875 | 0.136403537 | 0.493326264 |
| Sfn | 14.0314794 | 13.85184735 | 13.53820357 | 0.313643783 | 0.493275833 |
| Elavl1 | 10.73313625 | 10.50485214 | 10.23991646 | 0.264935683 | 0.49321979 |
| Pisd | 10.4054047 | 10.16062554 | 9.912350521 | 0.248275016 | 0.493054176 |
| Xbp1 | 12.42049898 | 12.27332372 | 11.92750774 | 0.345815977 | 0.49299124 |

| | | | | | |
|---|---|---|---|---|---|
| Adpgk | 9.789484851 | 9.603447903 | 9.296845439 | 0.306602463 | 0.492639412 |
| Mogs | 10.15790057 | 10.09165999 | 9.66562191 | 0.426038081 | 0.492278657 |
| Gcn1l1 | 10.38504357 | 10.45016499 | 9.892778409 | 0.557386578 | 0.492265158 |
| Hgh1 | 8.12998117 | 7.750633722 | 7.637850744 | 0.112782978 | 0.492130426 |
| Ston2 | 10.37893414 | 10.07486326 | 9.887075002 | 0.187788261 | 0.491859141 |
| Dnajc2 | 10.25894021 | 9.817913273 | 9.76728739 | 0.050625883 | 0.491652823 |
| U2af2 | 11.85603572 | 11.81398443 | 11.36475583 | 0.4492286 | 0.491279893 |
| Jpt2 | 10.50479446 | 10.33577511 | 10.01354972 | 0.322225393 | 0.49124474 |
| Nars | 11.80773173 | 11.62099021 | 11.31655824 | 0.304431967 | 0.491173487 |
| Amer2 | 4.616108767 | 4.413902204 | 4.125125791 | 0.288776413 | 0.490982976 |
| Irf8 | 8.681028888 | 8.491744669 | 8.190518687 | 0.301225982 | 0.490510201 |
| Tpst2 | 8.56675041 | 8.274554963 | 8.076806595 | 0.197748369 | 0.489943816 |
| Wipf1 | 9.903681009 | 9.625719768 | 9.413753963 | 0.211965805 | 0.489927046 |
| Exosc1 | 8.89454608 | 8.643577268 | 8.404750238 | 0.238827031 | 0.489795842 |
| Cd163l1 | 6.491707441 | 6.433611349 | 6.002268415 | 0.431342933 | 0.489439026 |
| Nsun2 | 11.93310082 | 11.65788192 | 11.44390905 | 0.21397287 | 0.489191767 |
| Kcnn4 | 6.688454577 | 6.622519414 | 6.199305601 | 0.423213813 | 0.489148977 |
| Drd1 | 4.334353407 | 3.895164484 | 3.845380263 | 0.049784221 | 0.488973145 |
| Znhit6 | 8.660144931 | 8.394913177 | 8.171187304 | 0.223725873 | 0.488957628 |
| Sirpa | 11.66839235 | 11.61769023 | 11.18035359 | 0.437336643 | 0.488038757 |
| Pno1 | 9.695484847 | 9.392188746 | 9.207725153 | 0.184463594 | 0.487759695 |
| Pom121 | 10.07100604 | 10.0547481 | 9.583305156 | 0.471442944 | 0.487700884 |
| Polr1e | 8.336675036 | 7.964560589 | 7.848989343 | 0.115571246 | 0.487685693 |
| Osgin2 | 9.06874862 | 8.853640023 | 8.581572886 | 0.272067137 | 0.487175734 |
| Ipo4 | 10.42605613 | 10.44749122 | 9.938899389 | 0.508591828 | 0.487156745 |
| Set | 12.48458502 | 12.15285079 | 11.99755059 | 0.1553002 | 0.487034423 |
| Gm21188 | 9.12080705 | 8.894887589 | 8.634240147 | 0.260647442 | 0.486566903 |
| Slc4a8 | 8.898495781 | 8.650732059 | 8.411998773 | 0.238733286 | 0.486497008 |
| Pfdn2 | 9.764945384 | 9.391726483 | 9.278555599 | 0.113170885 | 0.486389786 |
| Ell | 9.683153306 | 9.456045101 | 9.196826271 | 0.25921883 | 0.486327036 |
| Apobec1 | 8.899317295 | 8.859284126 | 8.413157063 | 0.446127063 | 0.486160232 |
| Il4 | 3.984789581 | 3.523113311 | 3.498862625 | 0.024250686 | 0.485926956 |

| | | | | | |
|---|---|---|---|---|---|
| Ms4a6b | 9.376355159 | 8.96329914 | 8.891031934 | 0.072267206 | 0.485323225 |
| Ehd3 | 8.288164362 | 8.031674555 | 7.803027231 | 0.228647324 | 0.485137131 |
| Lbp | 10.35864507 | 9.846243705 | 9.87359677 | -0.027353065 | 0.485048303 |
| Pfn1 | 12.84211625 | 12.59956527 | 12.3574558 | 0.24210947 | 0.48466045 |
| Tagln2 | 12.94074368 | 12.67167455 | 12.45611007 | 0.215564483 | 0.48463361 |
| Ece1 | 12.41530217 | 12.18728286 | 11.93085904 | 0.256423817 | 0.484443123 |
| Gm45140 | 9.668877526 | 9.117573782 | 9.18450573 | -0.066931948 | 0.484371796 |
| Ccdc71l | 9.179304318 | 8.891666113 | 8.69497084 | 0.196695273 | 0.484333478 |
| Hnrnph2 | 11.63681131 | 11.23269359 | 11.15274913 | 0.079944453 | 0.48406218 |
| Alkbh1 | 8.925316056 | 8.681478059 | 8.441966712 | 0.239511346 | 0.483349344 |
| Adora2a | 5.89359613 | 5.631763722 | 5.410270775 | 0.221492947 | 0.483325355 |
| Areg | 6.480278261 | 6.163638434 | 5.997189909 | 0.166448525 | 0.483088351 |
| Klhdc4 | 9.443655349 | 9.274205101 | 8.960685882 | 0.313519219 | 0.482969467 |
| Olfr1175-ps | 5.513689614 | 5.112164515 | 5.031140858 | 0.081023657 | 0.482548756 |
| Pdgfa | 9.085455738 | 9.015390624 | 8.603030968 | 0.412359656 | 0.48242477 |
| Gpr146 | 9.947799165 | 9.552323992 | 9.465383894 | 0.086940098 | 0.482415271 |
| Fastkd2 | 8.879075585 | 8.419423827 | 8.396882009 | 0.022541818 | 0.482193576 |
| Nmrk2 | 6.558875882 | 7.402561158 | 6.077031151 | 1.325530007 | 0.481844731 |
| Etf1 | 12.29674417 | 11.99012236 | 11.81539663 | 0.174725737 | 0.481347547 |
| Rpl27 | 10.62199789 | 10.70604284 | 10.14072926 | 0.565313579 | 0.481268625 |
| Ino80e | 9.366337192 | 9.312318769 | 8.885143137 | 0.427175632 | 0.481194054 |
| Riox1 | 8.247123729 | 7.970408289 | 7.766603579 | 0.20380471 | 0.48052015 |
| Esf1 | 9.983085311 | 9.593852856 | 9.502587568 | 0.091265288 | 0.480497743 |
| Hcst | 5.354376028 | 5.020559259 | 4.874057261 | 0.146501998 | 0.480318767 |
| Rbmxl1 | 10.06039805 | 9.809162457 | 9.580314181 | 0.228848275 | 0.480083869 |
| Sik3 | 10.46700145 | 10.14950525 | 9.987105191 | 0.162400056 | 0.479896262 |
| Tor1aip1 | 11.47979636 | 11.06532367 | 11.00007352 | 0.065250143 | 0.479722833 |
| Ccnd3 | 10.81989935 | 10.64051441 | 10.34036677 | 0.30014764 | 0.479532583 |
| Tent5b | 9.418290131 | 9.285869557 | 8.938938787 | 0.34693077 | 0.479351344 |
| Myo9b | 11.08655088 | 10.9050199 | 10.60732543 | 0.297694463 | 0.479225447 |
| Utp14a | 10.23565149 | 9.965163026 | 9.757117362 | 0.208045664 | 0.478534128 |
| Larp1 | 12.57580648 | 12.32743862 | 12.09771057 | 0.229728047 | 0.478095907 |

| | | | | | |
|---|---|---|---|---|---|
| Rel | 10.2701048 | 9.901028957 | 9.792017508 | 0.10901145 | 0.478087292 |
| Znrf1 | 10.01770333 | 9.752874236 | 9.539634973 | 0.213239264 | 0.478068355 |
| Rbm19 | 9.057626108 | 8.900439859 | 8.579689643 | 0.320750216 | 0.477936465 |
| Utp15 | 9.562076815 | 9.31054296 | 9.084210194 | 0.226332766 | 0.477866621 |
| Susd6 | 11.87680153 | 11.65206892 | 11.39901054 | 0.253058387 | 0.477790997 |
| Dgkg | 5.367404529 | 4.862034467 | 4.889858427 | -0.02782396 | 0.477546102 |
| Gm6548 | 8.263237096 | 7.699371419 | 7.785813973 | -0.086442554 | 0.477423123 |
| Jakmip2 | 6.124363688 | 5.661097924 | 5.646961202 | 0.014136723 | 0.477402486 |
| Slc4a7 | 9.110147751 | 8.655449768 | 8.632813391 | 0.022636377 | 0.47733436 |
| Rsc1a1 | 7.760491543 | 7.40901192 | 7.283219359 | 0.125792561 | 0.477272184 |
| Srsf7 | 11.27593335 | 11.11017306 | 10.79869325 | 0.311479807 | 0.477240093 |
| Lrrfip1 | 11.28942026 | 10.89804642 | 10.81219714 | 0.08584928 | 0.47722312 |
| Haus8 | 8.620861299 | 8.333859282 | 8.143711339 | 0.190147943 | 0.47714996 |
| Nmd3 | 9.727563696 | 9.47420257 | 9.250422237 | 0.223780333 | 0.477141458 |
| Dhx30 | 10.31063396 | 9.961009048 | 9.833541241 | 0.127467807 | 0.477092716 |
| Neu1 | 9.072575328 | 8.972002985 | 8.595663429 | 0.376339556 | 0.4769119 |
| Itgb6 | 8.329007548 | 8.048357403 | 7.852290075 | 0.196067328 | 0.476717473 |
| Dhx33 | 9.67039681 | 9.398918826 | 9.193698353 | 0.205220473 | 0.476698457 |
| Fjx1 | 7.059337219 | 6.767825624 | 6.582676285 | 0.185149339 | 0.476660934 |
| Runx3 | 7.977885362 | 7.654924413 | 7.501496392 | 0.153428021 | 0.47638897 |
| Lypla2 | 9.957173344 | 9.917441835 | 9.480823471 | 0.436618364 | 0.476349873 |
| Adamts5 | 11.33610488 | 10.55001974 | 10.85977226 | -0.30975252 | 0.47633262 |
| Rnh1 | 12.81381359 | 12.7310503 | 12.33784049 | 0.393209813 | 0.475973103 |
| Dhx38 | 9.951038395 | 9.912036781 | 9.475584585 | 0.436452197 | 0.47545381 |
| Marcks | 11.96051027 | 11.56561458 | 11.48514909 | 0.080465497 | 0.47536118 |
| Nufip1 | 8.68930344 | 8.27817 | 8.214188959 | 0.063981041 | 0.475114481 |
| Slc23a2 | 10.16896807 | 9.773316372 | 9.694141195 | 0.079175177 | 0.474826872 |
| Tmem51 | 7.509064682 | 7.337016202 | 7.034619423 | 0.302396779 | 0.474445259 |
| Tmem251 | 8.361694967 | 8.156546581 | 7.887435132 | 0.269111449 | 0.474259835 |
| Npm1 | 13.31797834 | 13.0606581 | 12.84375803 | 0.21690007 | 0.474220313 |
| C130026I21Rik | 4.598411718 | 4.006086698 | 4.124429983 | -0.118343285 | 0.473981736 |
| Riok3 | 11.95042806 | 11.58361084 | 11.47646108 | 0.107149767 | 0.473966987 |

| | | | | | |
|---|---|---|---|---|---|
| Glrx | 11.4594978 | 10.9801702 | 10.9856917 | -0.005521503 | 0.4738061 |
| Jmjd6 | 9.669169328 | 9.576328164 | 9.195544531 | 0.380783633 | 0.473624797 |
| H13 | 11.11061829 | 10.92647405 | 10.63702124 | 0.289452807 | 0.473597053 |
| Klk6 | 10.81251652 | 10.69330139 | 10.33893592 | 0.354365467 | 0.473580597 |
| Ydjc | 6.268605723 | 6.102942837 | 5.795228456 | 0.307714381 | 0.473377267 |
| Arhgap9 | 8.489603655 | 8.073191284 | 8.016272141 | 0.056919143 | 0.473331514 |
| Fam206a | 8.685550952 | 8.494095354 | 8.212318556 | 0.281776797 | 0.473232396 |
| Gtse1 | 7.413019561 | 7.422238434 | 6.940253943 | 0.481984491 | 0.472765618 |
| Btla | 6.334746584 | 6.03864375 | 5.86204831 | 0.17659544 | 0.472698274 |
| Cct8 | 12.20972675 | 11.97262227 | 11.73727155 | 0.23535072 | 0.4724552 |
| Spred1 | 11.19140568 | 10.807116 | 10.71930464 | 0.087811357 | 0.47210104 |
| Raver1 | 10.13355547 | 10.13178007 | 9.661699674 | 0.470080399 | 0.471855793 |
| Zfp385a | 11.24386604 | 11.13575077 | 10.77223274 | 0.363518027 | 0.471633297 |
| Snrpa1 | 9.502530061 | 9.278430937 | 9.031902881 | 0.246528056 | 0.47062718 |
| Kif1a | 7.705175824 | 7.643696913 | 7.234683454 | 0.409013459 | 0.47049237 |
| Map4k1 | 7.165883575 | 7.180676216 | 6.696193513 | 0.484482703 | 0.469690062 |
| Klrb1b | 8.289086452 | 7.75018071 | 7.8198193 | -0.069638589 | 0.469267152 |
| Il6st | 13.1666561 | 12.90283956 | 12.69756743 | 0.205272127 | 0.46908867 |
| Ranbp1 | 9.518279986 | 9.319025817 | 9.049520391 | 0.269505426 | 0.468759595 |
| Snrpd1 | 10.12707132 | 9.640697307 | 9.658320365 | -0.017623058 | 0.468750958 |
| Cnr2 | 6.925056675 | 6.867043212 | 6.456383246 | 0.410659966 | 0.46867343 |
| Simc1 | 7.717173039 | 7.592542752 | 7.248858377 | 0.343684375 | 0.468314662 |
| Ptges3 | 11.82703271 | 11.55749985 | 11.35891547 | 0.19858438 | 0.468117243 |
| Hspd1 | 12.67534448 | 12.54429007 | 12.20730768 | 0.33698239 | 0.468036807 |
| Stambpl1 | 8.187847357 | 7.719311292 | 7.719868328 | -0.000557036 | 0.467979029 |
| Arid3b | 6.562125169 | 6.250277437 | 6.094302528 | 0.155974909 | 0.467822641 |
| Exo1 | 7.420521953 | 7.122858129 | 6.952792452 | 0.170065677 | 0.467729501 |
| Ddx24 | 11.01008817 | 10.67982727 | 10.54250086 | 0.13732641 | 0.467587313 |
| Eif3g | 11.02526021 | 10.80746325 | 10.55824318 | 0.249220077 | 0.467017033 |
| Polr2l | 9.470446513 | 9.507966708 | 9.003738418 | 0.504228291 | 0.466708095 |
| Pa2g4 | 12.47704385 | 12.17402302 | 12.01037966 | 0.163643363 | 0.466664193 |
| Elf5 | 7.233665308 | 6.904659357 | 6.767492446 | 0.137166912 | 0.466172862 |

| | | | | | |
|---|---|---|---|---|---|
| Ddx51 | 8.92900519 | 8.46643298 | 8.462981092 | 0.003451889 | 0.466024098 |
| Col12a1 | 11.16688878 | 11.08537153 | 10.70123231 | 0.38413922 | 0.465656467 |
| Cdk2ap2 | 8.932137095 | 8.829850774 | 8.46690601 | 0.362944764 | 0.465231085 |
| Heatr3 | 10.11840807 | 9.808609368 | 9.653195774 | 0.155413594 | 0.465212296 |
| Sf3b3 | 11.87449502 | 11.61986633 | 11.41003133 | 0.209835003 | 0.46446369 |
| Nop9 | 9.385345344 | 9.135622678 | 8.921030847 | 0.214591831 | 0.464314497 |
| Pelp1 | 9.769244964 | 9.583953134 | 9.304980554 | 0.278972579 | 0.46426441 |
| Pgam1 | 12.23578217 | 11.9502509 | 11.77179285 | 0.178458047 | 0.46398932 |
| Nol11 | 9.71661273 | 9.378997266 | 9.252782912 | 0.126214354 | 0.463829818 |
| Eif6 | 11.75215236 | 11.36267626 | 11.28913544 | 0.07354082 | 0.46301692 |
| Hnrnpd | 11.62296439 | 11.3010463 | 11.16001479 | 0.141031507 | 0.462949597 |
| Il7r | 8.375858571 | 8.410615951 | 7.913114673 | 0.497501279 | 0.462743898 |
| Syngr2 | 11.57657807 | 11.36771389 | 11.11401124 | 0.253702647 | 0.46256683 |
| Llph | 9.983739202 | 9.576947097 | 9.521426222 | 0.055520875 | 0.462312979 |
| Btg1 | 12.693858 | 12.33575987 | 12.23163671 | 0.104123153 | 0.462221287 |
| Nepro | 8.58756401 | 8.451843122 | 8.125458727 | 0.326384396 | 0.462105284 |
| H2-K2 | 6.905537321 | 6.616132084 | 6.444179244 | 0.17195284 | 0.461358077 |
| Clcf1 | 6.934087098 | 6.773079912 | 6.472752354 | 0.300327559 | 0.461334744 |
| Usp36 | 10.34913801 | 10.16774061 | 9.887945177 | 0.279795429 | 0.46119283 |
| Cflar | 11.30927228 | 11.09822722 | 10.84816339 | 0.250063837 | 0.46110889 |
| Eif5a | 14.04537702 | 13.81850337 | 13.58446616 | 0.234037207 | 0.460910857 |
| Gm49342 | 9.655932659 | 9.168273447 | 9.195070225 | -0.026796778 | 0.460862434 |
| Rab7 | 12.95098482 | 12.63318156 | 12.49022579 | 0.14295577 | 0.460759027 |
| Spata13 | 9.108174384 | 8.636927102 | 8.64746952 | -0.010542418 | 0.460704864 |
| Gpr176 | 6.079675798 | 5.806841423 | 5.618975292 | 0.187866131 | 0.460700506 |
| Bud23 | 8.928073148 | 8.815752684 | 8.467575286 | 0.348177398 | 0.460497862 |
| Arhgap15 | 7.479617151 | 6.940438394 | 7.019665907 | -0.079227513 | 0.459951244 |
| Baz1a | 10.97913296 | 10.63318402 | 10.51929203 | 0.113891997 | 0.459840933 |
| Wars | 10.0357019 | 9.803338416 | 9.57623311 | 0.227105306 | 0.459468793 |
| Grrp1 | 6.26572929 | 6.064671559 | 5.806383053 | 0.258288507 | 0.459346237 |
| Vav3 | 10.72930062 | 10.42962069 | 10.26996443 | 0.159656263 | 0.459336193 |
| Serpinb3b | 9.213352802 | 8.779731877 | 8.754040333 | 0.025691544 | 0.459312468 |

| | | | | | |
|---|---|---|---|---|---|
| Evl | 8.779836418 | 8.783860788 | 8.320583342 | 0.463277447 | 0.459253076 |
| Ckap4 | 10.80916349 | 10.64650067 | 10.35006629 | 0.296434377 | 0.459097193 |
| Slc43a1 | 9.01010127 | 8.41429272 | 8.551182248 | -0.136889528 | 0.458919023 |
| Arhgdia | 12.85029388 | 12.75536412 | 12.39144542 | 0.363918703 | 0.458848463 |
| Sqstm1 | 12.24865966 | 12.43001527 | 11.78983018 | 0.640185087 | 0.458829473 |
| Dph3 | 9.316733379 | 9.03559231 | 8.858427859 | 0.177164451 | 0.45830552 |
| Atp8b1 | 9.871730494 | 9.569288168 | 9.413436905 | 0.155851263 | 0.458293589 |
| Serinc2 | 10.39065304 | 10.20434587 | 9.932562075 | 0.271783792 | 0.458090965 |
| Rab23 | 8.313732838 | 8.076421614 | 7.855658218 | 0.220763396 | 0.45807462 |
| Gm21992 | 6.698905045 | 6.522677011 | 6.240921636 | 0.281755375 | 0.457983409 |
| Rnps1 | 11.03480317 | 10.8259123 | 10.57683603 | 0.249076263 | 0.45796714 |
| Syt12 | 8.513765446 | 8.184997982 | 8.05585494 | 0.129143042 | 0.457910506 |
| Anxa11 | 11.14652454 | 10.84849505 | 10.68886919 | 0.159625853 | 0.45765535 |
| Pdlim7 | 9.586602708 | 9.725609155 | 9.129051508 | 0.596557647 | 0.4575512 |
| Ddx3x | 13.67532868 | 13.33264648 | 13.21820623 | 0.114440247 | 0.457122447 |
| Ppa1 | 10.90655703 | 10.63212929 | 10.44947817 | 0.182651117 | 0.457078857 |
| Gtf2f1 | 10.4627497 | 10.24810616 | 10.00573611 | 0.242370053 | 0.45701359 |
| Ppid | 10.42569953 | 10.28597897 | 9.968852171 | 0.317126796 | 0.456847356 |
| Trmt10c | 8.962909267 | 8.66652892 | 8.50641393 | 0.160114989 | 0.456495337 |
| Mitd1 | 8.930157563 | 8.616929537 | 8.473706749 | 0.143222787 | 0.456450813 |
| Gja1 | 12.9947894 | 12.60840322 | 12.53863043 | 0.06977279 | 0.45615897 |
| Abtb2 | 9.703311384 | 9.351846916 | 9.247868158 | 0.103978759 | 0.455443226 |
| Eef2kmt | 8.823050093 | 8.577993548 | 8.367716844 | 0.210276704 | 0.455333248 |
| Spn | 7.767405335 | 7.41750434 | 7.312557499 | 0.104946842 | 0.454847837 |
| Il19 | 5.34052868 | 4.856580503 | 4.885899148 | -0.029318645 | 0.454629532 |
| C5ar2 | 6.203592764 | 6.023319513 | 5.749072366 | 0.274247147 | 0.454520399 |
| Zfp36l1 | 11.90126714 | 11.69428256 | 11.44688739 | 0.247395173 | 0.454379753 |
| Fam208b | 9.977983038 | 9.515860361 | 9.523634439 | -0.007774078 | 0.454348599 |
| Saa1 | 5.509678313 | 5.097788469 | 5.055432751 | 0.042355717 | 0.454245561 |
| Elp1 | 9.624150002 | 9.379208963 | 9.169973441 | 0.209235522 | 0.454176561 |
| Pou2f2 | 7.106693449 | 7.054390627 | 6.652529278 | 0.401861348 | 0.454164171 |
| Hsh2d | 5.224866615 | 5.070530636 | 4.770783978 | 0.299746658 | 0.454082637 |

| | | | | | |
|---|---|---|---|---|---|
| Hnrnpab | 12.90120735 | 12.62299119 | 12.44718364 | 0.17580755 | 0.45402371 |
| Ltb | 6.359583128 | 6.639064226 | 5.905567215 | 0.733497011 | 0.454015913 |
| Eng | 10.29079755 | 10.19635905 | 9.837390028 | 0.358969025 | 0.453407518 |
| Gm6545 | 5.655142387 | 5.267447542 | 5.201855923 | 0.065591619 | 0.453286464 |
| Shmt2 | 10.38169687 | 10.44175137 | 9.928609068 | 0.513142306 | 0.453087799 |
| Gbp4 | 8.477456689 | 7.853124269 | 8.024745468 | -0.171621199 | 0.452711221 |
| Dop1b | 10.01970794 | 10.02374508 | 9.567052594 | 0.456692484 | 0.452655347 |
| Id2 | 9.986696461 | 9.695356858 | 9.534431986 | 0.160924872 | 0.452264475 |
| Cemip2 | 10.005896 | 9.724606625 | 9.554002385 | 0.17060424 | 0.451893611 |
| Snrpb | 11.0844692 | 11.08307825 | 10.63290151 | 0.45017674 | 0.451567693 |
| Btk | 8.136511933 | 7.829699781 | 7.685607313 | 0.144092468 | 0.45090462 |
| Tbc1d9 | 8.394565608 | 8.280010625 | 7.943888938 | 0.336121688 | 0.45067667 |
| Gm48786 | 5.988821608 | 5.606778108 | 5.538360661 | 0.068417447 | 0.450460947 |
| Hnrnpf | 13.39873677 | 13.1070887 | 12.94852096 | 0.158567743 | 0.450215813 |
| Mid1 | 9.836785357 | 10.09295924 | 9.38669762 | 0.706261619 | 0.450087737 |
| Timm13 | 9.671246294 | 9.614776425 | 9.221609957 | 0.393166468 | 0.449636337 |
| Prkg1 | 8.203655355 | 7.568732515 | 7.754105289 | -0.185372774 | 0.449550066 |
| Sh2d2a | 5.652554588 | 5.253354453 | 5.203132361 | 0.050222092 | 0.449422227 |
| Magohb | 7.11418883 | 6.840044336 | 6.66476936 | 0.175274976 | 0.44941947 |
| Treml4 | 4.610097421 | 4.467523193 | 4.161321788 | 0.306201405 | 0.448775633 |
| Gm12407 | 5.67037215 | 5.542514542 | 5.22204754 | 0.320467002 | 0.44832461 |
| Atxn2l | 11.29329296 | 11.27068339 | 10.84503761 | 0.42564578 | 0.44825535 |
| Gfra2 | 9.121669621 | 8.609316154 | 8.673414311 | -0.064098157 | 0.44825531 |
| Diaph1 | 11.93654129 | 11.7421052 | 11.48837788 | 0.253727327 | 0.44816341 |
| Kif21b | 8.100531335 | 7.711329859 | 7.652410586 | 0.058919273 | 0.448120749 |
| Nol12 | 9.300279057 | 9.031623838 | 8.852304081 | 0.179319757 | 0.447974976 |
| Dph5 | 8.226773916 | 7.920599905 | 7.778944132 | 0.141655772 | 0.447829784 |
| Ddx56 | 9.962933883 | 9.74471436 | 9.515381911 | 0.229332448 | 0.447551972 |
| Srsf2 | 12.44247184 | 12.16557695 | 11.99497356 | 0.170603393 | 0.447498283 |
| Trmt6 | 9.475411281 | 9.133208212 | 9.027933806 | 0.105274406 | 0.447477474 |
| Sema6d | 9.254635351 | 8.969129903 | 8.807221664 | 0.161908239 | 0.447413687 |
| Nubp2 | 9.305669645 | 9.08021418 | 8.858266922 | 0.221947258 | 0.447402723 |

| | | | | | |
|---|---|---|---|---|---|
| Cx3cr1 | 7.136240511 | 6.704891122 | 6.689077926 | 0.015813195 | 0.447162585 |
| Nlgn2 | 9.100742826 | 8.972143985 | 8.653932137 | 0.318211849 | 0.44681069 |
| Il18bp | 7.351048719 | 7.005570797 | 6.904451113 | 0.101119684 | 0.446597607 |
| Xrcc6 | 9.47317019 | 9.055031398 | 9.026740635 | 0.028290763 | 0.446429555 |
| Ramp3 | 6.696051867 | 6.153826187 | 6.249805678 | -0.095979491 | 0.446246189 |
| M6pr | 11.74481566 | 11.41687122 | 11.29860039 | 0.118270827 | 0.446215267 |
| Prickle2 | 9.791905908 | 9.477526319 | 9.345725141 | 0.131801178 | 0.446180766 |
| Mrpl33 | 10.00995499 | 9.641789495 | 9.564206067 | 0.077583428 | 0.445748924 |
| Batf | 6.178417893 | 6.080036176 | 5.732705292 | 0.347330884 | 0.445712601 |
| Rpp38 | 7.793553373 | 7.511530024 | 7.348009362 | 0.163520662 | 0.445544011 |
| Gmip | 9.482915424 | 9.554790704 | 9.037608382 | 0.517182322 | 0.445307042 |
| Ifi202b | 8.926870889 | 8.703356446 | 8.481590029 | 0.221766417 | 0.44528086 |
| Vegfa | 10.15502405 | 10.05297338 | 9.709778322 | 0.343195055 | 0.445245731 |
| Zc3hav1 | 11.2733967 | 10.91328159 | 10.82824592 | 0.08503567 | 0.445150777 |
| Eif3d | 11.69486052 | 11.40722868 | 11.24975798 | 0.1574707 | 0.44510254 |
| Cers2 | 10.69353865 | 10.52934014 | 10.24846974 | 0.280870407 | 0.44506891 |
| Ncln | 10.42498931 | 10.3145478 | 9.980060067 | 0.33448773 | 0.44492924 |
| Sh2b3 | 9.992997217 | 9.791380388 | 9.548298019 | 0.243082368 | 0.444699197 |
| Rrbp1 | 12.4506367 | 12.11201272 | 12.0062705 | 0.105742217 | 0.4443662 |
| Slfn10-ps | 6.276857739 | 5.99673827 | 5.832561827 | 0.164176443 | 0.444295912 |
| Gm48499 | 4.277062517 | 4.461682911 | 3.832792319 | 0.628890591 | 0.444270197 |
| Pag1 | 8.654925741 | 8.347264788 | 8.210801356 | 0.136463432 | 0.444124385 |
| Wsb1 | 10.6491724 | 10.43016254 | 10.20568049 | 0.224482047 | 0.443491913 |
| Pold2 | 8.874889279 | 8.830081463 | 8.431655019 | 0.398426444 | 0.44323426 |
| Tfpi2 | 6.198530183 | 5.965129184 | 5.755348922 | 0.209780262 | 0.443181261 |
| Nup62 | 10.39425138 | 10.27609159 | 9.951099906 | 0.324991684 | 0.443151471 |
| Lmnb2 | 8.683564884 | 8.684608995 | 8.240460994 | 0.444148001 | 0.44310389 |
| Rpl39l | 7.191130177 | 6.953492258 | 6.748300237 | 0.205192021 | 0.44282994 |
| CR974586.5 | 6.2782833 | 6.256158065 | 5.835635754 | 0.42052231 | 0.442647546 |
| Cyp27b1 | 4.792785241 | 4.652958667 | 4.350156687 | 0.30280198 | 0.442628554 |
| Hspb1 | 12.87014899 | 13.33827713 | 12.42768684 | 0.910590297 | 0.44246215 |
| Ly6a | 12.95583388 | 12.53518917 | 12.51342851 | 0.02176066 | 0.442405367 |

| | | | | | |
|---|---|---|---|---|---|
| Eda | 7.571002324 | 7.280079751 | 7.128778555 | 0.151301197 | 0.44222377 |
| Hmga1 | 11.50506699 | 11.3938088 | 11.06313825 | 0.33067055 | 0.44192874 |
| Acsl4 | 10.89294314 | 10.36520978 | 10.45143033 | -0.086220547 | 0.441512813 |
| Sec61a1 | 11.89841934 | 11.93533595 | 11.45705167 | 0.478284277 | 0.441367663 |
| Trafd1 | 11.32255143 | 10.99505268 | 10.88119312 | 0.113859563 | 0.441358307 |
| Map4k4 | 12.0359397 | 11.79590939 | 11.59488661 | 0.201022787 | 0.441053093 |
| Eif2s1 | 10.95709734 | 10.67445397 | 10.51607376 | 0.15838021 | 0.441023573 |
| Ubac2 | 10.09758594 | 9.869929513 | 9.657078329 | 0.212851185 | 0.440507611 |
| Tnfrsf21 | 9.931127992 | 9.616143359 | 9.49071906 | 0.125424299 | 0.440408932 |
| H2-Q4 | 11.65159937 | 11.27268387 | 11.21192287 | 0.060760997 | 0.439676503 |
| Snap23 | 10.27894822 | 10.00372314 | 9.83957068 | 0.164152458 | 0.43937754 |
| BC037034 | 9.093018028 | 9.25541896 | 8.653837635 | 0.601581325 | 0.439180394 |
| Tbk1 | 9.976168429 | 9.679894034 | 9.536989236 | 0.142904798 | 0.439179193 |
| Tspan4 | 10.44528952 | 10.23947658 | 10.00659644 | 0.232880141 | 0.438693078 |
| Ippk | 9.17686159 | 8.990481197 | 8.738337654 | 0.252143543 | 0.438523936 |
| Ncbp1 | 10.81040993 | 10.51991918 | 10.37206443 | 0.14785475 | 0.438345503 |
| Ifitm5 | 4.481782814 | 4.273352337 | 4.043534031 | 0.229818306 | 0.438248783 |
| Gpr55 | 5.615438068 | 5.381979577 | 5.177478748 | 0.204500829 | 0.43795932 |
| Rgl1 | 10.45520201 | 10.28169296 | 10.0173953 | 0.264297652 | 0.437806702 |
| Depdc7 | 7.135987125 | 6.764954195 | 6.698312676 | 0.066641519 | 0.437674449 |
| Eif4a-ps4 | 11.40496314 | 11.43445199 | 10.96755339 | 0.466898607 | 0.437409757 |
| Mettl16 | 9.80728651 | 9.504679175 | 9.370054959 | 0.134624216 | 0.437231551 |
| Ugcg | 11.33660339 | 11.07875785 | 10.89961029 | 0.179147553 | 0.436993093 |
| Cyb561 | 10.64415077 | 10.26872391 | 10.20744869 | 0.061275213 | 0.436702077 |
| Arhgef2 | 9.617324021 | 9.594794388 | 9.180749929 | 0.41404446 | 0.436574092 |
| Gypc | 9.030006104 | 8.174415889 | 8.593780718 | -0.419364829 | 0.436225387 |
| Smndc1 | 9.969189647 | 9.695750444 | 9.533752176 | 0.161998269 | 0.435437471 |
| Mon1a | 8.911363537 | 8.835528896 | 8.475981587 | 0.359547309 | 0.43538195 |
| Tpk1 | 7.746367779 | 7.399063142 | 7.311106226 | 0.087956916 | 0.435261553 |
| Apol9b | 6.44231201 | 5.754810349 | 6.007236368 | -0.252426019 | 0.435075642 |
| Urb2 | 9.309222729 | 8.990888012 | 8.87453568 | 0.116352332 | 0.434687049 |
| Rbms1 | 12.05311749 | 11.67991451 | 11.61879993 | 0.06111458 | 0.434317563 |

| | | | | | |
|---|---|---|---|---|---|
| Mzt2 | 7.893282352 | 7.513695852 | 7.459526226 | 0.054169626 | 0.433756126 |
| Rpl12 | 9.282792125 | 9.194975707 | 8.8493888 | 0.345586907 | 0.433403325 |
| Gm42715 | 10.43099623 | 9.956911107 | 9.997877406 | -0.040966298 | 0.433118821 |
| Aars | 11.29265625 | 11.25572214 | 10.85998061 | 0.395741527 | 0.432675637 |
| Pdk4 | 10.3649679 | 10.61851378 | 9.932367339 | 0.686146441 | 0.432600565 |
| Syvn1 | 10.09710824 | 10.41717053 | 9.665077277 | 0.75209325 | 0.432030963 |
| Rps6ka4 | 10.93376642 | 10.89078013 | 10.5018382 | 0.388941927 | 0.43192822 |
| Tapbp | 11.72640354 | 11.39497919 | 11.29449219 | 0.100486993 | 0.431911343 |
| Smpd5 | 4.633715096 | 4.298165216 | 4.201806227 | 0.09635899 | 0.431908869 |
| Klf10 | 10.78237393 | 10.50829622 | 10.35060961 | 0.157686603 | 0.431764317 |
| Emg1 | 10.42434661 | 10.19951579 | 9.992838636 | 0.20667715 | 0.431507977 |
| Amotl2 | 11.07070446 | 10.86344234 | 10.63933623 | 0.224106113 | 0.431368233 |
| Arrb2 | 9.391840653 | 9.294310056 | 8.960768224 | 0.333541832 | 0.431072429 |
| Tbrg4 | 10.03796655 | 9.918048687 | 9.60718465 | 0.310864037 | 0.430781904 |
| Rufy4 | 6.345186634 | 5.912118152 | 5.914744495 | -0.002626342 | 0.43044214 |
| Kars | 11.19934572 | 10.99676642 | 10.76949358 | 0.227272847 | 0.429852143 |
| Eif1 | 13.33663722 | 13.06522461 | 12.90708464 | 0.15813997 | 0.429552577 |
| Tap1 | 11.49131827 | 11.18988917 | 11.06217717 | 0.127712003 | 0.429141107 |
| Apbb1ip | 9.659460509 | 9.535797528 | 9.230323425 | 0.305474103 | 0.429137084 |
| Nampt | 11.38628705 | 10.93674269 | 10.9572412 | -0.020498503 | 0.429045857 |
| Pik3r6 | 7.497305694 | 7.295106634 | 7.068520732 | 0.226585902 | 0.428784962 |
| Bmpr1b | 6.420406485 | 6.422769597 | 5.991826577 | 0.430943019 | 0.428579908 |
| Dok1 | 7.998110987 | 7.855225218 | 7.56966047 | 0.285564748 | 0.428450517 |
| Pdcd2 | 8.228479955 | 8.021315435 | 7.800034752 | 0.221280682 | 0.428445202 |
| Isg20l2 | 10.03802175 | 9.771660591 | 9.60964824 | 0.162012352 | 0.428373507 |
| Sprr1a | 12.54237726 | 12.90174936 | 12.11407055 | 0.787678807 | 0.42830671 |
| Pus7 | 10.18233286 | 9.965807096 | 9.754134063 | 0.211673033 | 0.428198797 |
| Dis3 | 9.557026563 | 9.422858445 | 9.129238567 | 0.293619878 | 0.427787996 |
| A230050P20Rik | 7.281209566 | 7.182754513 | 6.853653729 | 0.329100784 | 0.427555837 |
| Dgkh | 8.521286739 | 8.042680267 | 8.09403068 | -0.051350413 | 0.42725606 |
| Preb | 10.69082992 | 10.48751689 | 10.263609 | 0.223907883 | 0.42722092 |
| Cpxm1 | 10.64201291 | 10.64900803 | 10.21493757 | 0.434070453 | 0.42707534 |

| | | | | | |
|---|---|---|---|---|---|
| Ccdc134 | 8.47109283 | 8.538788893 | 8.044118199 | 0.494670695 | 0.426974632 |
| Nxf1 | 11.35525277 | 11.17202391 | 10.92842774 | 0.24359617 | 0.426825027 |
| Kras | 10.73021865 | 10.30907278 | 10.30369379 | 0.005378993 | 0.42652486 |
| Cdh11 | 8.941017924 | 8.63523508 | 8.514576987 | 0.120658093 | 0.426440937 |
| Pygl | 10.11757577 | 9.26284674 | 9.691450766 | -0.428604026 | 0.426125003 |
| Peg10 | 7.095453744 | 7.587367767 | 6.669381329 | 0.917986438 | 0.426072415 |
| Zpr1 | 9.997185268 | 9.813092654 | 9.571131761 | 0.241960893 | 0.426053507 |
| Phb2 | 11.45451818 | 11.24846666 | 11.02847283 | 0.21999383 | 0.42604535 |
| Wfs1 | 8.401770587 | 8.467485883 | 7.975919991 | 0.491565892 | 0.425850596 |
| Ube2f | 10.25484556 | 10.09290536 | 9.82912803 | 0.26377733 | 0.425717527 |
| Slc35c2 | 9.111410278 | 9.093083519 | 8.685950441 | 0.407133078 | 0.425459837 |
| Ube2q2 | 10.66451698 | 10.26011541 | 10.2391037 | 0.021011707 | 0.425413277 |
| Trim34a | 8.641772638 | 8.234340264 | 8.216523902 | 0.017816362 | 0.425248735 |
| Socs5 | 10.3116676 | 10.08006871 | 9.886531996 | 0.193536717 | 0.4251356 |
| Usb1 | 8.84503095 | 8.553374408 | 8.420023659 | 0.13335075 | 0.425007291 |
| Thap7 | 7.871135844 | 7.812041259 | 7.446342657 | 0.365698602 | 0.424793187 |
| Trp53i11 | 9.485379095 | 9.328313783 | 9.060693798 | 0.267619985 | 0.424685297 |
| Strap | 11.4458336 | 11.12360511 | 11.02161709 | 0.101988027 | 0.424216517 |
| Arpc3 | 12.06898428 | 11.70933486 | 11.64491891 | 0.064415953 | 0.42406537 |
| Tfap4 | 8.143427841 | 7.820561256 | 7.719569859 | 0.100991397 | 0.423857982 |
| Dennd4b | 9.463031099 | 9.258129029 | 9.03934233 | 0.2187867 | 0.42368877 |
| Med12 | 10.10434271 | 10.15566948 | 9.680830865 | 0.474838615 | 0.42351184 |
| Plcb2 | 7.610559808 | 7.398700984 | 7.187327113 | 0.211373871 | 0.423232695 |
| Slc16a6 | 9.325424491 | 9.134828328 | 8.902334849 | 0.232493479 | 0.423089643 |
| Trim27 | 9.901185361 | 9.556219373 | 9.478131044 | 0.078088329 | 0.423054317 |
| Cdk5r1 | 6.818806651 | 6.577293095 | 6.395955278 | 0.181337816 | 0.422851373 |
| Scube2 | 7.890356232 | 7.861741051 | 7.467852092 | 0.39388896 | 0.42250414 |
| Arih2 | 10.56726821 | 10.24552059 | 10.14515521 | 0.100365383 | 0.422113003 |
| Ksr1 | 10.2879486 | 9.990669188 | 9.866140387 | 0.124528801 | 0.421808213 |
| Traf3ip3 | 5.991640441 | 5.702719232 | 5.570665192 | 0.13205404 | 0.420975249 |
| Aimp2 | 9.565105447 | 9.342398254 | 9.144295515 | 0.19810274 | 0.420809933 |
| Jun | 10.33276858 | 11.10564475 | 9.912340107 | 1.193304639 | 0.420428476 |

| | | | | | |
|---|---|---|---|---|---|
| Eva1c | 6.961720803 | 6.765636597 | 6.541772006 | 0.223864592 | 0.419948797 |
| Mfsd10 | 9.091082843 | 9.068374272 | 8.671569958 | 0.396804314 | 0.419512885 |
| Cd68 | 10.73966819 | 10.85745283 | 10.32018138 | 0.537271457 | 0.419486817 |
| Pnrc1 | 10.29792542 | 9.975657542 | 9.878727593 | 0.09692995 | 0.419197827 |
| Ptp4a1 | 11.948705 | 11.57955476 | 11.52956342 | 0.04999134 | 0.419141573 |
| Taf5l | 9.734507373 | 9.539810243 | 9.315522438 | 0.224287805 | 0.418984936 |
| Slfn9 | 10.052738 | 9.721032525 | 9.633978245 | 0.08705428 | 0.418759751 |
| Eif2s2 | 12.09438623 | 11.80241324 | 11.67573866 | 0.126674577 | 0.418647567 |
| Ahsa1 | 10.99976385 | 11.0249411 | 10.58132846 | 0.44361264 | 0.418435387 |
| Naa15 | 11.66759405 | 11.46612108 | 11.24953689 | 0.21658419 | 0.418057163 |
| Morf4l1-ps1 | 8.567892261 | 8.43031489 | 8.150227268 | 0.280087622 | 0.417664993 |
| Prr5l | 6.958163113 | 6.705971737 | 6.540536555 | 0.165435182 | 0.417626558 |
| Gtpbp3 | 8.010859864 | 7.999103864 | 7.59328294 | 0.405820924 | 0.417576924 |
| Shq1 | 7.64581086 | 7.30081251 | 7.228650087 | 0.072162424 | 0.417160773 |
| Traf4 | 7.856579042 | 7.67996926 | 7.439483143 | 0.240486116 | 0.417095899 |
| Pkp4 | 10.96578372 | 10.60008596 | 10.54881796 | 0.051267993 | 0.416965753 |
| Ero1l | 10.39720871 | 10.17304685 | 9.98042201 | 0.192624843 | 0.416786697 |
| Il23a | 3.951246694 | 3.596256085 | 3.534461109 | 0.061794976 | 0.416785585 |
| Suco | 10.40672797 | 10.06245879 | 9.990266693 | 0.0721921 | 0.416461274 |
| Cap1 | 12.79967051 | 12.57814142 | 12.3835708 | 0.194570623 | 0.416099713 |
| Rpain | 8.062460337 | 7.729635585 | 7.646420362 | 0.083215223 | 0.416039975 |
| Atg7 | 9.37418154 | 9.101008129 | 8.95819105 | 0.142817079 | 0.41599049 |
| Acy1 | 7.86546978 | 7.705299518 | 7.449489393 | 0.255810125 | 0.415980387 |
| Gm15922 | 6.598599136 | 6.638652997 | 6.183457882 | 0.455195115 | 0.415141254 |
| Srf | 9.619617537 | 9.605563939 | 9.204647597 | 0.400916342 | 0.41496994 |
| Kcnip3 | 6.78262664 | 6.545559033 | 6.367813908 | 0.177745125 | 0.414812732 |
| Clpp | 8.826288075 | 8.691389718 | 8.411525893 | 0.279863825 | 0.414762182 |
| Fbxl5 | 11.02893606 | 10.70864858 | 10.61434643 | 0.094302147 | 0.414589623 |
| Tgm1 | 11.40686694 | 11.01265865 | 10.99230363 | 0.020355013 | 0.414563307 |
| Gm49388 | 6.607172291 | 6.082582252 | 6.192756192 | -0.11017394 | 0.414416099 |
| Hspbap1 | 8.14579142 | 7.844614682 | 7.731398907 | 0.113215775 | 0.414392513 |
| Tor2a | 9.105314454 | 8.909774356 | 8.691291679 | 0.218482677 | 0.414022775 |

| | | | | | |
|---|---|---|---|---|---|
| Nlrc5 | 11.47288941 | 11.26892092 | 11.05889712 | 0.2100238 | 0.413992283 |
| Atp2a2 | 12.83849012 | 12.6985855 | 12.42474172 | 0.273843777 | 0.413748403 |
| Gm47036 | 4.955273121 | 4.843219653 | 4.541597967 | 0.301621686 | 0.413675154 |
| Ms4a4d | 9.082844951 | 8.452451519 | 8.669211449 | -0.216759931 | 0.413633502 |
| Ppp1r15b | 11.34674948 | 11.03760927 | 10.93311826 | 0.10449101 | 0.413631217 |
| Cyp4f16 | 8.099039444 | 7.891200043 | 7.68563667 | 0.205563373 | 0.413402774 |
| Kat2a | 9.928468782 | 9.884047007 | 9.515307667 | 0.36873934 | 0.413161115 |
| Rpl28 | 11.88238487 | 12.05231613 | 11.46927125 | 0.583044883 | 0.413113623 |
| Dot1l | 10.28623451 | 10.50061873 | 9.873451368 | 0.627167362 | 0.412783146 |
| Prg2 | 3.321247131 | 2.983332012 | 2.9091086 | 0.074223412 | 0.412138531 |
| Cdca7 | 10.05431962 | 9.852069257 | 9.64256413 | 0.209505127 | 0.411755487 |
| Ccdc115 | 9.304517631 | 8.986916925 | 8.892874239 | 0.094042686 | 0.411643392 |
| Pusl1 | 8.385143641 | 8.340784929 | 7.973546864 | 0.367238065 | 0.411596777 |
| Lama1 | 7.025539295 | 7.39959269 | 6.613986776 | 0.785605915 | 0.411552519 |
| Plcg2 | 10.07499983 | 9.949638476 | 9.663502571 | 0.286135904 | 0.411497256 |
| Enoph1 | 8.36792465 | 8.171416485 | 7.956878 | 0.214538485 | 0.41104665 |
| Exosc3 | 8.606607896 | 8.322503117 | 8.195926647 | 0.12657647 | 0.410681249 |
| Iars | 10.69032981 | 10.45791413 | 10.27996296 | 0.177951173 | 0.41036685 |
| Atrip | 8.529989955 | 8.350826967 | 8.11993382 | 0.230893147 | 0.410056135 |
| Mtrr | 8.427011123 | 8.269278952 | 8.017681404 | 0.251597548 | 0.409329719 |
| Gpaa1 | 9.693802536 | 9.819000006 | 9.284480005 | 0.534520001 | 0.409322531 |
| Sap18b | 9.934951594 | 9.956042503 | 9.525720893 | 0.43032161 | 0.409230701 |
| Ccdc116 | 5.170374628 | 4.923078037 | 4.761419377 | 0.16165866 | 0.408955251 |
| Tmem125 | 8.897968395 | 8.668091093 | 8.489896051 | 0.178195041 | 0.408072344 |
| A630001G21Rik | 6.294656244 | 5.951513337 | 5.886641623 | 0.064871714 | 0.408014622 |
| Gpr4 | 6.311091589 | 6.326298935 | 5.903143468 | 0.423155467 | 0.407948121 |
| Rps2-ps10 | 6.130213727 | 5.973296015 | 5.722389858 | 0.250906157 | 0.407823869 |
| Dctpp1 | 8.348711479 | 8.178379979 | 7.941289562 | 0.237090417 | 0.407421917 |
| Eif3j2 | 10.46529456 | 10.24612678 | 10.05801694 | 0.188109845 | 0.407277622 |
| Lrrc47 | 10.48733711 | 10.36191295 | 10.08054006 | 0.281372891 | 0.406797051 |
| Actr3 | 13.28912013 | 12.999495 | 12.88239808 | 0.117096917 | 0.406722043 |
| Trmt1 | 9.75412452 | 9.732175282 | 9.34744909 | 0.384726191 | 0.40667543 |

| | | | | | |
|---|---|---|---|---|---|
| Slc25a25 | 9.248659929 | 8.944114541 | 8.842154571 | 0.10195997 | 0.406505358 |
| Slco4a1 | 7.207251903 | 7.021936779 | 6.800853092 | 0.221083687 | 0.406398811 |
| Sox11 | 4.544464861 | 4.361387924 | 4.138239368 | 0.223148556 | 0.406225494 |
| Cox10 | 8.732507757 | 8.50249857 | 8.326309113 | 0.176189457 | 0.406198644 |
| Elob | 10.40836009 | 10.48801482 | 10.00243304 | 0.485581781 | 0.405927051 |
| Mus81 | 8.023392793 | 7.932023135 | 7.617574011 | 0.314449124 | 0.405818782 |
| Dock5 | 8.942819561 | 8.781674478 | 8.537128875 | 0.244545603 | 0.405690686 |
| Mrm3 | 7.67599897 | 7.382345303 | 7.27049138 | 0.111853922 | 0.405507589 |
| Ptk6 | 9.262218733 | 8.890513142 | 8.856775075 | 0.033738067 | 0.405443658 |
| Tirap | 8.673474896 | 8.383614262 | 8.268196481 | 0.115417781 | 0.405278415 |
| Rab4b | 9.012483567 | 8.887224204 | 8.607475303 | 0.279748901 | 0.405008264 |
| Ints5 | 9.650302021 | 9.577127655 | 9.245559805 | 0.33156785 | 0.404742216 |
| Fam111a | 10.16571937 | 9.921604739 | 9.761005597 | 0.160599142 | 0.404713776 |
| Hsph1 | 10.81473651 | 12.2256097 | 10.41045257 | 1.815157133 | 0.40428394 |
| Spred3 | 6.713353832 | 6.675967162 | 6.309087013 | 0.366880149 | 0.404266819 |
| Cdc42se1 | 11.95100328 | 11.78471654 | 11.54686876 | 0.23784778 | 0.40413452 |
| Mrpl17 | 10.01840889 | 9.719714726 | 9.61445635 | 0.105258376 | 0.403952535 |
| Crtc2 | 9.968932498 | 9.954052485 | 9.565070878 | 0.388981607 | 0.40386162 |
| Ahi1 | 8.170681729 | 7.695998538 | 7.766839483 | -0.070840945 | 0.403842246 |
| Enah | 10.65312047 | 10.99827353 | 10.2493592 | 0.74891433 | 0.403761267 |
| Abhd11 | 9.613356721 | 9.396689693 | 9.209717906 | 0.186971787 | 0.403638814 |
| Amigo2 | 7.084406282 | 6.674682725 | 6.681137121 | -0.006454396 | 0.403269162 |
| Wwc1 | 9.742216931 | 9.515288539 | 9.338967646 | 0.176320893 | 0.403249285 |
| Mical1 | 8.911459511 | 8.595198233 | 8.508475558 | 0.086722675 | 0.402983953 |
| Twist2 | 7.52426625 | 7.151308606 | 7.121732507 | 0.029576099 | 0.402533744 |
| Serpina3m | 6.263234287 | 6.008994122 | 5.860764367 | 0.148229755 | 0.402469919 |
| Tsc22d1 | 11.05004222 | 10.62062478 | 10.64819678 | -0.027572003 | 0.40184544 |
| Aldh3b3 | 7.053137987 | 6.79635216 | 6.651508785 | 0.144843375 | 0.401629202 |
| Sntb2 | 10.21605031 | 9.871963705 | 9.814493759 | 0.057469946 | 0.401556548 |
| Nsun5 | 8.150510185 | 7.998468748 | 7.749238291 | 0.249230456 | 0.401271893 |
| Ubap1 | 10.4737025 | 10.19004243 | 10.07247101 | 0.11757142 | 0.401231493 |
| Tmed5 | 11.78141918 | 11.30766962 | 11.38019406 | -0.07252444 | 0.40122512 |

| | | | | | |
|---|---|---|---|---|---|
| Piwil2 | 7.378959907 | 7.246775155 | 6.977935561 | 0.268839593 | 0.401024346 |
| Arid5b | 11.11924774 | 11.04929996 | 10.71852487 | 0.330775087 | 0.40072287 |
| Wdr5 | 10.2917545 | 10.09320068 | 9.891221396 | 0.201979287 | 0.400533101 |
| Oas1b | 7.344493113 | 7.052534662 | 6.9439742 | 0.108560462 | 0.400518913 |
| Yrdc | 9.623594029 | 9.346517292 | 9.22350963 | 0.123007662 | 0.400084399 |
| Ankrd28 | 10.46089339 | 10.09703188 | 10.06109408 | 0.035937799 | 0.399799314 |
| Fxyd4 | 5.57951425 | 5.266533815 | 5.179996082 | 0.086537732 | 0.399518168 |
| Rpl36 | 11.05951191 | 11.32505555 | 10.66011987 | 0.664935683 | 0.399392043 |
| Lrrc42 | 9.4519875 | 9.103241154 | 9.052811564 | 0.05042959 | 0.399175935 |
| Naip6 | 6.575868009 | 6.434910137 | 6.176777262 | 0.258132875 | 0.399090747 |
| Cacybp | 10.54174641 | 10.40247554 | 10.14270674 | 0.259768807 | 0.399039677 |
| Apaf1 | 9.05744148 | 9.009994543 | 8.658489651 | 0.351504891 | 0.398951829 |
| Ngdn | 9.301515148 | 9.197432126 | 8.902792588 | 0.294639538 | 0.398722561 |
| H3f3b | 13.53374351 | 13.15886394 | 13.13515339 | 0.02371055 | 0.398590113 |
| Gm12918 | 5.896214794 | 9.439376401 | 5.497638675 | 3.941737726 | 0.398576119 |
| Arhgap25 | 8.375403079 | 8.175686995 | 7.976925991 | 0.198761004 | 0.398477088 |
| Itga4 | 9.277811208 | 9.05310289 | 8.879350203 | 0.173752687 | 0.398461005 |
| Snapc4 | 8.94162792 | 8.777917846 | 8.543293908 | 0.234623938 | 0.398334012 |
| Cct7 | 12.19650312 | 11.97518517 | 11.7982681 | 0.17691707 | 0.39823502 |
| Top1 | 11.63572356 | 11.38154185 | 11.23773272 | 0.143809133 | 0.397990843 |
| Cnn2 | 11.32051844 | 11.19431541 | 10.92276434 | 0.27155107 | 0.397754103 |
| Epb41l4a | 8.519275788 | 8.166693118 | 8.121683521 | 0.045009598 | 0.397592267 |
| Ppp4c | 10.52955661 | 10.31220836 | 10.13240425 | 0.17980411 | 0.397152357 |
| Slc25a30 | 9.379107936 | 9.09870763 | 8.98245638 | 0.11625125 | 0.396651556 |
| Hirip3 | 8.404005553 | 8.322786022 | 8.007522734 | 0.315263288 | 0.396482819 |
| Dnah10 | 4.938399047 | 4.68335283 | 4.541926133 | 0.141426697 | 0.396472915 |
| Taf7 | 8.565176635 | 8.614389786 | 8.168861941 | 0.445527845 | 0.396314693 |
| D16Ertd472e | 10.08770507 | 9.774423125 | 9.691503636 | 0.082919489 | 0.396201437 |
| Tatdn2 | 10.64575447 | 10.29185521 | 10.24961238 | 0.04224283 | 0.396142087 |
| Hnrnpa3 | 12.89793004 | 12.77885608 | 12.50212133 | 0.276734743 | 0.39580871 |
| Ccdc25 | 10.05237161 | 9.709670175 | 9.656619853 | 0.053050322 | 0.395751755 |
| Btbd19 | 8.504233065 | 8.34276785 | 8.108681588 | 0.234086261 | 0.395551477 |

| | | | | | |
|---|---|---|---|---|---|
| Gfm1 | 10.21223964 | 9.896958461 | 9.817170492 | 0.079787969 | 0.395069144 |
| Stn1 | 8.404107629 | 8.291267099 | 8.009054631 | 0.282212468 | 0.395052998 |
| Sec23b | 10.45796119 | 10.28251834 | 10.0629868 | 0.21953154 | 0.394974387 |
| Med29 | 8.132055945 | 7.943027424 | 7.737965785 | 0.205061639 | 0.39409016 |
| Psmd3 | 11.47627157 | 11.4933673 | 11.08274025 | 0.410627047 | 0.393531317 |
| Timeless | 9.203611165 | 8.818324924 | 8.810088618 | 0.008236306 | 0.393522547 |
| Fblim1 | 7.82703044 | 7.71619089 | 7.433930169 | 0.282260721 | 0.393100271 |
| Hspa4 | 12.37554587 | 12.25699352 | 11.98260942 | 0.2743841 | 0.392936453 |
| Polr2h | 8.83611244 | 8.564380726 | 8.443351319 | 0.121029407 | 0.392761121 |
| Gm2260 | 8.062276282 | 7.442847285 | 7.669569008 | -0.226721723 | 0.392707274 |
| Etnk1 | 10.85623549 | 10.47838406 | 10.46364626 | 0.0147378 | 0.392589233 |
| Parvg | 7.101183232 | 6.931197148 | 6.708859306 | 0.222337841 | 0.392323926 |
| Gpr183 | 7.118891435 | 7.08582207 | 6.726658118 | 0.359163952 | 0.392233317 |
| Abcf2 | 11.07436635 | 10.88642224 | 10.68213573 | 0.204286513 | 0.392230627 |
| Rae1 | 9.711490461 | 9.449565851 | 9.319399836 | 0.130166014 | 0.392090624 |
| Snapc1 | 8.827651257 | 8.488496702 | 8.435790513 | 0.052706188 | 0.391860744 |
| Cltb | 12.13138492 | 12.01046843 | 11.73957015 | 0.270898287 | 0.391814777 |
| Gm14303 | 6.436566126 | 6.463018359 | 6.044762563 | 0.418255796 | 0.391803563 |
| Azin1 | 11.21078864 | 11.17589314 | 10.81939093 | 0.356502203 | 0.39139771 |
| Pdlim1 | 10.43722937 | 10.29222718 | 10.04600139 | 0.246225788 | 0.391227985 |
| Pitpna | 12.16267643 | 11.89040112 | 11.77154145 | 0.118859663 | 0.391134977 |
| Trim56 | 10.53782743 | 10.08582881 | 10.14673764 | -0.06090883 | 0.39108979 |
| Medag | 10.66735911 | 10.43500531 | 10.27648559 | 0.15851972 | 0.390873517 |
| Cfp | 10.10625191 | 9.714492309 | 9.715381306 | -0.000888997 | 0.3908706 |
| Zeb2 | 11.10199426 | 10.79571439 | 10.71145052 | 0.084263863 | 0.39054374 |
| Itga9 | 8.935098263 | 8.625099175 | 8.545031188 | 0.080067987 | 0.390067075 |
| Ddx39b | 11.98208741 | 11.84360122 | 11.592224 | 0.25137722 | 0.389863403 |
| Rgs20 | 8.045603362 | 7.610463343 | 7.656115465 | -0.045652122 | 0.389487896 |
| Mtcl1 | 5.083983261 | 4.923245652 | 4.694547897 | 0.228697755 | 0.389435364 |
| Rab32 | 9.280878987 | 9.033182192 | 8.891629794 | 0.141552398 | 0.389249193 |
| Fus | 12.76941258 | 12.44173933 | 12.38031351 | 0.061425817 | 0.389099067 |
| Dcaf13 | 8.889445876 | 8.552445871 | 8.500613236 | 0.051832635 | 0.38883264 |

| | | | | | |
|---|---|---|---|---|---|
| Rrp7a | 10.44523414 | 10.3664044 | 10.05692185 | 0.309482555 | 0.388312291 |
| Smg9 | 8.714463225 | 8.694040845 | 8.326174545 | 0.367866301 | 0.38828868 |
| Gpc1 | 11.00934907 | 10.91365651 | 10.6210877 | 0.292568807 | 0.388261367 |
| Lamc2 | 11.35309315 | 11.34596692 | 10.96493597 | 0.381030947 | 0.388157177 |
| Dnaaf5 | 8.601123339 | 8.408846052 | 8.213161772 | 0.19568428 | 0.387961567 |
| Zkscan6 | 8.881957197 | 8.588793237 | 8.494235814 | 0.094557423 | 0.387721383 |
| Mif | 10.58681434 | 10.74103943 | 10.19926476 | 0.541774677 | 0.38754958 |
| Phf11a | 5.714961317 | 5.195105045 | 5.327710854 | -0.132605809 | 0.387250462 |
| Nqo1 | 8.433412863 | 8.083370774 | 8.046164426 | 0.037206348 | 0.387248437 |
| Pla2g4e | 9.374418763 | 9.040129711 | 8.98722544 | 0.052904271 | 0.387193323 |
| Rangap1 | 11.72378707 | 11.5749081 | 11.33693223 | 0.23797587 | 0.386854843 |
| Gal | 3.7682696 | 3.6162906 | 3.381602004 | 0.234688596 | 0.386667595 |
| Irf2 | 10.08608509 | 9.727012155 | 9.699431411 | 0.027580744 | 0.386653675 |
| Ppm1j | 6.960150072 | 6.303864892 | 6.573700867 | -0.269835975 | 0.386449205 |
| Fcrlb | 3.999665799 | 3.838620577 | 3.613388709 | 0.225231868 | 0.38627709 |
| Rgs19 | 8.889179166 | 8.70022604 | 8.5029378 | 0.19728824 | 0.386241366 |
| Kbtbd8 | 7.166767443 | 6.733474303 | 6.780547524 | -0.047073221 | 0.386219919 |
| Sap30 | 7.740899859 | 7.839913345 | 7.355012153 | 0.484901192 | 0.385887706 |
| Vill | 7.336912845 | 7.174572112 | 6.951242156 | 0.223329955 | 0.385670689 |
| Txn1 | 11.58740598 | 11.3614767 | 11.20192067 | 0.159556033 | 0.385485307 |
| Ralb | 9.917175673 | 9.644005406 | 9.532545251 | 0.111460155 | 0.384630422 |
| Snrpf | 8.843435038 | 8.698555792 | 8.458896836 | 0.239658956 | 0.384538202 |
| Ccnl1 | 11.25481438 | 10.92802674 | 10.87036004 | 0.0576667 | 0.384454343 |
| Bak1 | 9.295301339 | 9.268558183 | 8.911026611 | 0.357531572 | 0.384274728 |
| Gm28285 | 6.839994024 | 6.622312036 | 6.455849899 | 0.166462137 | 0.384144125 |
| Nrcam | 7.763772668 | 7.771804537 | 7.380063789 | 0.391740749 | 0.38370888 |
| Gm9320 | 4.511052083 | 4.184786919 | 4.127839646 | 0.056947273 | 0.383212436 |
| Pfdn6 | 9.360947752 | 9.131159394 | 8.977842978 | 0.153316416 | 0.383104775 |
| Anp32b | 12.0244677 | 11.65519259 | 11.64137747 | 0.013815113 | 0.383090227 |
| Gabpb1 | 9.09127008 | 8.775509324 | 8.708245121 | 0.067264204 | 0.38302496 |
| Il17rc | 10.8301697 | 10.84962903 | 10.44722161 | 0.402407417 | 0.38294809 |
| Fam71f2 | 3.821838265 | 3.420682089 | 3.438991134 | -0.018309045 | 0.382847131 |

| | | | | | |
|---|---|---|---|---|---|
| Hmgb1-ps8 | 4.814758671 | 4.574757131 | 4.432204985 | 0.142552146 | 0.382553685 |
| Slc25a33 | 7.327575516 | 7.14316398 | 6.945025452 | 0.198138528 | 0.382550064 |
| Sart1 | 10.26574889 | 10.15130854 | 9.883202409 | 0.268106131 | 0.382546484 |
| Moxd1 | 11.42054121 | 11.15644079 | 11.03808349 | 0.118357303 | 0.382457723 |
| Alyref | 10.02363115 | 9.811071567 | 9.641274712 | 0.169796855 | 0.382356434 |
| Pcgf5 | 9.696639853 | 9.243337045 | 9.314474162 | -0.071137117 | 0.382165691 |
| Dll1 | 8.974412827 | 9.034882341 | 8.592345704 | 0.442536637 | 0.382067123 |
| Esyt1 | 11.21909234 | 11.05935527 | 10.83782507 | 0.2215302 | 0.38126727 |
| Tln1 | 12.15715704 | 12.16775222 | 11.77630295 | 0.39144927 | 0.380854097 |
| Taf1d | 10.48634314 | 10.07071918 | 10.10566663 | -0.034947453 | 0.380676503 |
| Dusp16 | 10.62912085 | 10.33065734 | 10.24850547 | 0.082151873 | 0.380615387 |
| Ssbp4 | 8.649163128 | 8.69385349 | 8.268608609 | 0.425244881 | 0.38055452 |
| Ctdp1 | 9.40111479 | 9.51769256 | 9.020626407 | 0.497066153 | 0.380488383 |
| Tomm20 | 11.2478416 | 11.01791654 | 10.86738892 | 0.150527613 | 0.38045268 |
| Dimt1 | 8.183785039 | 7.891507753 | 7.803343621 | 0.088164132 | 0.380441418 |
| Dlk2 | 7.699285751 | 8.01288098 | 7.318975365 | 0.693905616 | 0.380310387 |
| Rbm42 | 9.685558701 | 9.756553522 | 9.305717642 | 0.45083588 | 0.379841059 |
| Cdk11b | 10.99681101 | 10.79966347 | 10.61698606 | 0.182677407 | 0.379824947 |
| Ntmt1 | 8.035825812 | 7.89119476 | 7.656203053 | 0.234991708 | 0.37962276 |
| Gmps | 10.63657152 | 10.35128589 | 10.25719076 | 0.09409513 | 0.379380767 |
| Dnaja1 | 11.84808898 | 12.56678576 | 11.4688932 | 1.097892557 | 0.379195777 |
| Gm10600 | 8.816692262 | 8.808619001 | 8.437944594 | 0.370674407 | 0.378747668 |
| Il1rap | 9.960787035 | 9.726630286 | 9.582194339 | 0.144435947 | 0.378592695 |
| Ifi203-ps | 6.573162725 | 5.994584379 | 6.194689411 | -0.200105032 | 0.378473314 |
| S1pr2 | 9.33717435 | 9.111228214 | 8.958773719 | 0.152454495 | 0.378400631 |
| Phc2 | 10.72060688 | 10.64703085 | 10.34232536 | 0.30470549 | 0.378281523 |
| Hspa9 | 13.06238613 | 12.91485996 | 12.68425926 | 0.2306007 | 0.37812687 |
| Tap2 | 10.35373902 | 10.28244183 | 9.975990182 | 0.306451651 | 0.377748841 |
| Ccdc59 | 9.466098837 | 9.116572521 | 9.088353203 | 0.028219318 | 0.377745634 |
| Cd300c2 | 7.723125775 | 7.470784107 | 7.345747751 | 0.125036356 | 0.377378024 |
| Pkm | 14.4948649 | 14.44995061 | 14.117863 | 0.332087617 | 0.377001907 |
| Ung | 8.490184362 | 8.37185126 | 8.113368389 | 0.258482871 | 0.376815973 |

| | | | | | |
|---|---|---|---|---|---|
| Prep | 10.72870226 | 10.55612438 | 10.35193064 | 0.204193737 | 0.37677162 |
| Snrnp40 | 8.619199728 | 8.364434232 | 8.242458491 | 0.121975741 | 0.376741237 |
| Atp13a1 | 10.37290557 | 10.42065765 | 9.996187554 | 0.424470099 | 0.376718012 |
| Plec | 13.24967935 | 13.37809046 | 12.87331783 | 0.504772627 | 0.376361517 |
| Slc12a9 | 8.197906788 | 8.245512874 | 7.821875187 | 0.423637687 | 0.376031601 |
| Tcerg1 | 10.69919916 | 10.54266804 | 10.32352057 | 0.219147467 | 0.375678593 |
| Xpo4 | 9.701540967 | 9.555099448 | 9.325932061 | 0.229167387 | 0.375608906 |
| Sash3 | 8.040873479 | 7.926411692 | 7.665354353 | 0.261057338 | 0.375519125 |
| Mars2 | 8.698084975 | 8.400195213 | 8.322616776 | 0.077578437 | 0.375468199 |
| Ddr2 | 12.12075735 | 11.8932931 | 11.74529707 | 0.14799603 | 0.375460273 |
| Phf5a | 9.75173045 | 9.542139064 | 9.376494201 | 0.165644863 | 0.375236248 |
| Ptbp1 | 12.22295425 | 12.25411273 | 11.84771944 | 0.406393287 | 0.375234807 |
| Foxp4 | 8.502945358 | 8.462490599 | 8.127786014 | 0.334704585 | 0.375159344 |
| Fam98a | 9.822174407 | 9.652384685 | 9.447081928 | 0.205302757 | 0.375092479 |
| Ppat | 8.972980353 | 8.646827969 | 8.598154306 | 0.048673663 | 0.374826047 |
| U2af1 | 10.87180997 | 10.66290339 | 10.49722598 | 0.165677413 | 0.374583993 |
| Capn10 | 9.312030478 | 9.114888465 | 8.937663017 | 0.177225448 | 0.374367461 |
| Cactin | 8.909002854 | 9.053990566 | 8.534648008 | 0.519342558 | 0.374354846 |
| Sat1 | 11.66595213 | 11.33679326 | 11.2918797 | 0.044913563 | 0.374072437 |
| Adcy3 | 9.499394062 | 9.346874302 | 9.12598764 | 0.220886662 | 0.373406421 |
| Gata2 | 7.398463347 | 7.093139348 | 7.025057776 | 0.068081571 | 0.373405571 |
| Snx8 | 9.231846384 | 9.345890282 | 8.858458336 | 0.487431946 | 0.373388048 |
| Dnttip2 | 10.20052238 | 9.995018467 | 9.827153089 | 0.167865378 | 0.373369291 |
| Ppm1g | 10.88675915 | 10.65272877 | 10.51347689 | 0.139251883 | 0.373282263 |
| Siah2 | 8.770856409 | 8.754565842 | 8.397822834 | 0.356743008 | 0.373033575 |
| Nsd3 | 11.08437858 | 10.78323488 | 10.71151425 | 0.07172063 | 0.372864333 |
| Zc3h7a | 10.96994364 | 10.90357767 | 10.59755782 | 0.306019847 | 0.37238582 |
| Cybc1 | 10.06948322 | 9.907472134 | 9.697230978 | 0.210241157 | 0.372252242 |
| Thy1 | 12.24839863 | 12.19285369 | 11.87616086 | 0.316692827 | 0.372237767 |
| Rinl | 7.699313126 | 7.416045744 | 7.327118245 | 0.088927499 | 0.372194881 |
| Shmt1 | 9.789393999 | 9.557739869 | 9.417285395 | 0.140454475 | 0.372108604 |
| Gm5859 | 7.958884109 | 8.062977465 | 7.586824691 | 0.476152774 | 0.372059418 |

| | | | | | |
|---|---|---|---|---|---|
| Alg8 | 8.21529902 | 8.206808158 | 7.843465484 | 0.363342674 | 0.371833536 |
| Mrpl21 | 8.959324014 | 8.872628352 | 8.587587673 | 0.285040679 | 0.371736341 |
| Nckap5l | 6.613172105 | 6.451241884 | 6.241508479 | 0.209733405 | 0.371663626 |
| Rybp | 10.20869842 | 10.08807603 | 9.837188204 | 0.250887823 | 0.371510213 |
| Ccnyl1 | 9.007568067 | 8.864668244 | 8.636103787 | 0.228564457 | 0.37146428 |
| Stt3a | 11.59380155 | 11.58291219 | 11.22262289 | 0.360289303 | 0.371178667 |
| Cbr3 | 9.207121721 | 9.136423626 | 8.836149654 | 0.300273971 | 0.370972066 |
| Ctse | 4.357935801 | 4.132966184 | 3.987114593 | 0.145851591 | 0.370821208 |
| Dll4 | 6.615799125 | 6.542084282 | 6.245000317 | 0.297083965 | 0.370798809 |
| Fnip2 | 10.03950754 | 10.18397368 | 9.668709767 | 0.515263917 | 0.370797776 |
| Sra1 | 9.951938674 | 9.591593198 | 9.581870858 | 0.00972234 | 0.370067816 |
| Ubxn8 | 8.162961835 | 8.061438834 | 7.79293518 | 0.268503655 | 0.370026656 |
| Ubqln4 | 10.65070061 | 10.54406962 | 10.28069245 | 0.263377167 | 0.370008153 |
| Pelo | 8.287020617 | 8.26273273 | 7.917387453 | 0.345345277 | 0.369633164 |
| Grb2 | 10.91219505 | 10.81912706 | 10.54264677 | 0.276480293 | 0.369548277 |
| Ifi214 | 5.176775579 | 4.833354345 | 4.807234149 | 0.026120196 | 0.36954143 |
| Nadk | 11.99647388 | 11.84382092 | 11.62711771 | 0.21670321 | 0.369356167 |
| P4ha1 | 10.13067867 | 10.09953652 | 9.761428476 | 0.338108048 | 0.369250194 |
| Tnfsf9 | 5.768944075 | 5.640567137 | 5.399717678 | 0.240849459 | 0.369226397 |
| Kpnb1 | 12.62837428 | 12.50732318 | 12.25944844 | 0.247874733 | 0.368925833 |
| Gata1 | 4.716554693 | 4.42296446 | 4.347769039 | 0.075195421 | 0.368785654 |
| Adrm1 | 10.69653176 | 10.81332176 | 10.32788735 | 0.485434413 | 0.36864441 |
| Zfp142 | 8.82753615 | 9.00166881 | 8.459355417 | 0.542313393 | 0.368180733 |
| Eif3a | 13.07342917 | 12.79747137 | 12.70530188 | 0.092169487 | 0.36812729 |
| Dnajc11 | 10.45062373 | 10.32002288 | 10.08263546 | 0.237387415 | 0.367988265 |
| Sfxn5 | 8.340710141 | 8.253170883 | 7.97295504 | 0.280215843 | 0.367755101 |
| Cd3d | 6.611120847 | 6.284981018 | 6.243546134 | 0.041434884 | 0.367574713 |
| B3galt6 | 7.688012764 | 7.509987136 | 7.320498381 | 0.189488755 | 0.367514383 |
| Gnl1 | 10.48421439 | 10.31892978 | 10.11684706 | 0.202082721 | 0.367367337 |
| Gpr137b-ps | 8.710469405 | 8.64825795 | 8.34336955 | 0.3048884 | 0.367099855 |
| Polr3e | 9.651666308 | 9.553655057 | 9.284966839 | 0.268688218 | 0.366699468 |
| Eif3b | 12.41489896 | 12.1835629 | 12.04829991 | 0.135262997 | 0.366599057 |

| | | | | | |
|---|---|---|---|---|---|
| B230303A05Rik | 4.146584604 | 3.864258684 | 3.780099022 | 0.084159662 | 0.366485582 |
| 36951 | 8.03531946 | 7.695164968 | 7.668959814 | 0.026205154 | 0.366359646 |
| Grap2 | 7.469440167 | 7.182363048 | 7.103286479 | 0.079076569 | 0.366153688 |
| Git1 | 10.44615662 | 10.48186814 | 10.08034093 | 0.401527212 | 0.365815688 |
| Polr2e | 9.892489388 | 9.854514818 | 9.526771596 | 0.327743223 | 0.365717792 |
| Bzw1 | 12.76438038 | 12.54275034 | 12.39872252 | 0.14402782 | 0.365657863 |
| Psmg3 | 8.260316806 | 8.118194289 | 7.894698788 | 0.223495501 | 0.365618018 |
| Nme1 | 10.88581984 | 10.48663212 | 10.52022804 | -0.03359592 | 0.365591797 |
| Egfr | 11.56545796 | 11.25653616 | 11.19994473 | 0.056591427 | 0.36551323 |
| Surf6 | 9.399518111 | 9.286968623 | 9.03401266 | 0.252955963 | 0.365505451 |
| Zfp330 | 9.68054457 | 9.491086406 | 9.315296589 | 0.175789817 | 0.36524798 |
| Brpf1 | 10.02936588 | 9.986086517 | 9.664237608 | 0.321848909 | 0.365128273 |
| Ints2 | 8.963437401 | 8.779658783 | 8.598452183 | 0.181206599 | 0.364985218 |
| Mob3a | 9.681014249 | 9.515900318 | 9.316066869 | 0.199833449 | 0.36494738 |
| Pgf | 5.964182177 | 5.817036929 | 5.599465745 | 0.217571184 | 0.364716432 |
| Aff1 | 11.72997476 | 11.60157641 | 11.3653633 | 0.23621311 | 0.364611463 |
| Irak3 | 9.564504874 | 9.001544328 | 9.199907847 | -0.198363519 | 0.364597027 |
| Fndc3a | 10.38117249 | 10.0001351 | 10.01683537 | -0.016700265 | 0.364337122 |
| Gtpbp6 | 7.898065287 | 8.028369359 | 7.534022993 | 0.494346366 | 0.364042293 |
| Steap2 | 6.986796492 | 6.608181276 | 6.622804615 | -0.014623339 | 0.363991877 |
| Sys1 | 8.905221411 | 8.796251281 | 8.541240573 | 0.255010708 | 0.363980837 |
| Tlr3 | 8.980319236 | 8.155771862 | 8.616386225 | -0.460614363 | 0.363933011 |
| Mex3a | 6.880354564 | 6.641232895 | 6.516521848 | 0.124711047 | 0.363832716 |
| Cpsf3 | 10.0251465 | 9.771579257 | 9.661365548 | 0.11021371 | 0.363780949 |
| Def6 | 9.091453883 | 8.983781784 | 8.727680093 | 0.256101691 | 0.363773789 |
| Ist1 | 11.70409484 | 11.52679101 | 11.34044065 | 0.186350363 | 0.363654187 |
| E2f7 | 7.514604101 | 7.198743315 | 7.150974477 | 0.047768838 | 0.363629623 |
| H6pd | 10.9496424 | 10.55791893 | 10.58640839 | -0.028489457 | 0.36323401 |
| Mapk1ip1l | 10.93071759 | 10.71696609 | 10.56752012 | 0.14944597 | 0.36319747 |
| Lrrfip2 | 10.97195337 | 10.59850951 | 10.6089272 | -0.01041769 | 0.363026177 |
| Lima1 | 11.96600358 | 11.77013716 | 11.60301814 | 0.16711902 | 0.36298544 |
| Hspe1 | 10.63695349 | 10.90826071 | 10.27400359 | 0.634257123 | 0.362949903 |

| | | | | | |
|---|---|---|---|---|---|
| Smchd1 | 10.3638336 | 9.962643513 | 10.00108971 | -0.038446201 | 0.362743883 |
| Vsir | 10.56215764 | 10.36933348 | 10.19949155 | 0.169841934 | 0.362666091 |
| Serpinb2 | 12.25819439 | 11.71736251 | 11.89588927 | -0.17852676 | 0.362305113 |
| Nomo1 | 11.28442533 | 11.14189263 | 10.92223379 | 0.21965884 | 0.362191543 |
| Fam169b | 6.369978582 | 6.024578075 | 6.00787708 | 0.016700996 | 0.362101503 |
| Arpc2 | 12.768574 | 12.5344346 | 12.40666377 | 0.127770823 | 0.361910223 |
| Rbm34 | 8.94608417 | 8.521707852 | 8.584486802 | -0.06277895 | 0.361597368 |
| Dram1 | 10.54435513 | 10.17366424 | 10.1827754 | -0.009111163 | 0.36157973 |
| Prok2 | 3.902039711 | 3.823687061 | 3.540472713 | 0.283214348 | 0.361566998 |
| Kcna3 | 4.670732312 | 4.521223392 | 4.309284519 | 0.211938873 | 0.361447793 |
| Mcub | 8.229783069 | 8.013823592 | 7.868361 | 0.145462592 | 0.361422069 |
| Tmem268 | 8.286104358 | 8.090953326 | 7.924765672 | 0.166187654 | 0.361338686 |
| Ptgdr | 6.478856514 | 6.420401893 | 6.117721535 | 0.302680359 | 0.361134979 |
| Rlim | 11.15617531 | 10.86588289 | 10.79546175 | 0.07042114 | 0.360713567 |
| Iqsec1 | 10.26744642 | 9.986763314 | 9.906768909 | 0.079994405 | 0.360677511 |
| Cyp4b1 | 11.09427983 | 10.85886985 | 10.73367192 | 0.125197937 | 0.36060791 |
| Ascc3 | 10.48994922 | 10.25428499 | 10.12953482 | 0.12475017 | 0.360414407 |
| Nckipsd | 7.994755941 | 7.940042822 | 7.634345537 | 0.305697285 | 0.360410404 |
| Xrn2 | 10.97472972 | 10.64743703 | 10.61456737 | 0.032869657 | 0.360162353 |
| Arhgef16 | 8.308655084 | 8.275047191 | 7.94863512 | 0.326412071 | 0.360019964 |
| Igsf9b | 7.167364476 | 7.287029436 | 6.80774127 | 0.479288167 | 0.359623206 |
| Msl2 | 10.63682766 | 10.44036873 | 10.27724499 | 0.163123733 | 0.359582667 |
| Cyp2b10 | 10.3261729 | 10.07927157 | 9.966714028 | 0.112557539 | 0.359458872 |
| Stip1 | 11.56450012 | 11.66166549 | 11.2052073 | 0.456458183 | 0.35929282 |
| Nuak2 | 8.721607549 | 8.616263239 | 8.362372012 | 0.253891227 | 0.359235537 |
| Supv3l1 | 8.725242723 | 8.4909648 | 8.366221147 | 0.124743653 | 0.359021576 |
| Tmsb10 | 11.35211964 | 11.07193858 | 10.99323949 | 0.078699093 | 0.358880157 |
| Zrsr2 | 8.985869118 | 8.90137271 | 8.627218348 | 0.274154362 | 0.35865077 |
| St3gal6 | 9.359988959 | 8.898946101 | 9.001545763 | -0.102599662 | 0.358443196 |
| B3gnt3 | 7.631018797 | 7.419716509 | 7.273174841 | 0.146541667 | 0.357843955 |
| Trip6 | 9.410181711 | 9.332269196 | 9.052365732 | 0.279903464 | 0.35781598 |
| Tle1 | 9.034912478 | 8.794448558 | 8.677325503 | 0.117123055 | 0.357586975 |

| | | | | | |
|---|---|---|---|---|---|
| Nudc | 10.6418982 | 10.52009165 | 10.28436271 | 0.235728947 | 0.357535497 |
| Lsm2 | 8.693760503 | 8.472956071 | 8.336302957 | 0.136653114 | 0.357457546 |
| Cycs | 10.63560501 | 10.47947668 | 10.27821222 | 0.201264457 | 0.35739279 |
| Cdc42ep4 | 10.71733923 | 10.56885027 | 10.35995196 | 0.20889831 | 0.357387273 |
| Rbm8a | 10.34538305 | 10.09229214 | 9.988011222 | 0.104280915 | 0.357371828 |
| Bccip | 10.40974452 | 10.14430518 | 10.0525278 | 0.091777383 | 0.357216717 |
| Adsl | 10.01543623 | 9.790521237 | 9.65829832 | 0.132222918 | 0.357137913 |
| Ddx49 | 9.477283586 | 9.321441302 | 9.120185357 | 0.201255945 | 0.357098229 |
| Glo1 | 11.53820689 | 11.27863137 | 11.18159749 | 0.097033883 | 0.356609397 |
| Zfp628 | 7.666533482 | 7.751735603 | 7.309989762 | 0.441745841 | 0.35654372 |
| Exosc4 | 8.350923555 | 8.238362376 | 7.994410597 | 0.243951778 | 0.356512958 |
| Anxa2 | 15.11208425 | 14.94249091 | 14.75612862 | 0.186362287 | 0.35595563 |
| Dhps | 8.630801493 | 8.39666894 | 8.274850077 | 0.121818864 | 0.355951416 |
| Tomm70a | 11.19420824 | 10.91996882 | 10.8383315 | 0.081637323 | 0.35587674 |
| Col4a2 | 12.18717857 | 11.91695313 | 11.83132339 | 0.085629743 | 0.355855177 |
| Taldo1 | 11.84676473 | 11.62684247 | 11.49112753 | 0.135714937 | 0.355637193 |
| Phf23 | 10.32002172 | 10.18754832 | 9.964691906 | 0.222856411 | 0.355329814 |
| Znhit2 | 7.256240396 | 7.384239907 | 6.900953103 | 0.483286804 | 0.355287292 |
| Gm5641 | 13.12392294 | 13.00118103 | 12.7690731 | 0.232107933 | 0.354849847 |
| Aste1 | 7.729213872 | 7.478262671 | 7.374741169 | 0.103521502 | 0.354472703 |
| Abcg4 | 7.547872592 | 7.394645045 | 7.193670321 | 0.200974724 | 0.354202271 |
| Cherp | 10.14287815 | 10.26347413 | 9.788741551 | 0.474732579 | 0.354136599 |
| Arpc5 | 12.01727622 | 11.77897743 | 11.66327196 | 0.115705473 | 0.354004263 |
| Gtf2e2 | 9.224934656 | 8.903872658 | 8.870961311 | 0.032911347 | 0.353973345 |
| Gnai2 | 12.37861844 | 12.15737574 | 12.02468064 | 0.132695093 | 0.353937793 |
| Ilf3 | 10.31806126 | 10.17372151 | 9.964267392 | 0.209454118 | 0.353793864 |
| Bms1 | 10.30944267 | 10.16758689 | 9.955865615 | 0.211721272 | 0.353577052 |
| Tgif2 | 6.507728311 | 6.175290594 | 6.154172674 | 0.021117919 | 0.353555637 |
| Atp2b1 | 11.74285148 | 11.52719664 | 11.38931949 | 0.137877147 | 0.353531987 |
| Prdx6 | 12.07126482 | 11.64205243 | 11.71804271 | -0.075990273 | 0.35322211 |
| Ppig | 11.25294251 | 10.87280685 | 10.89988643 | -0.027079583 | 0.35305608 |
| Srprb | 10.12622024 | 10.05823101 | 9.773178536 | 0.28505247 | 0.353041707 |

| | | | | | |
|---|---|---|---|---|---|
| Gm29288 | 5.294978708 | 4.890387262 | 4.942251192 | -0.051863931 | 0.352727516 |
| Vrk3 | 8.827025219 | 8.656805043 | 8.474313241 | 0.182491801 | 0.352711978 |
| Rbm47 | 9.058994519 | 8.95970794 | 8.706385102 | 0.253322838 | 0.352609417 |
| Psd4 | 10.4950249 | 10.11133944 | 10.1425371 | -0.031197659 | 0.352487805 |
| Foxp3 | 5.282626353 | 4.897144679 | 4.930207475 | -0.033062796 | 0.352418877 |
| Egr2 | 9.208223713 | 9.135738507 | 8.856203074 | 0.279535433 | 0.352020639 |
| Foxk1 | 10.63353967 | 10.51416083 | 10.28157856 | 0.232582267 | 0.351961113 |
| Wdr3 | 9.919002159 | 9.80045519 | 9.567194735 | 0.233260455 | 0.351807424 |
| Akt1 | 11.58138611 | 11.45158789 | 11.22967574 | 0.22191215 | 0.351710363 |
| Cmtr2 | 7.616312623 | 7.364837381 | 7.264680777 | 0.100156604 | 0.351631846 |
| Ifngr1 | 11.53629182 | 11.10496393 | 11.18509616 | -0.080132237 | 0.35119566 |
| Gmeb1 | 9.083636714 | 8.941938587 | 8.732598343 | 0.209340245 | 0.351038371 |
| Fdx1l | 8.54159929 | 8.301924998 | 8.190608072 | 0.111316926 | 0.350991218 |
| Prmt7 | 9.655552158 | 9.4322801 | 9.30457256 | 0.127707539 | 0.350979597 |
| Gnpnat1 | 8.60498311 | 8.282468409 | 8.254272361 | 0.028196048 | 0.350710749 |
| Dnph1 | 8.238038723 | 8.072492606 | 7.887574886 | 0.184917719 | 0.350463836 |
| Klre1 | 4.081035198 | 3.868325466 | 3.730626041 | 0.137699426 | 0.350409158 |
| Prss22 | 6.807753491 | 6.670020404 | 6.457528883 | 0.21249152 | 0.350224607 |
| Pop7 | 7.519594824 | 7.492130852 | 7.169419505 | 0.322711347 | 0.350175319 |
| Gm21541 | 8.764427434 | 8.588627572 | 8.414322127 | 0.174305445 | 0.350105307 |
| Phrf1 | 10.34019804 | 10.30257757 | 9.990248162 | 0.312329411 | 0.349949881 |
| Nup93 | 9.94729521 | 9.70815164 | 9.597404581 | 0.110747058 | 0.349890629 |
| Gnal | 6.470488694 | 6.177006138 | 6.12080102 | 0.056205119 | 0.349687675 |
| Mapk6 | 11.41211684 | 11.18061262 | 11.06266959 | 0.117943033 | 0.349447253 |
| Lrpprc | 10.39456047 | 10.22789179 | 10.04514643 | 0.182745363 | 0.349414043 |
| Eif4ebp1 | 9.89452269 | 9.647608802 | 9.545248056 | 0.102360746 | 0.349274633 |
| Pxk | 9.928397966 | 9.62906852 | 9.579511536 | 0.049556985 | 0.348886431 |
| Brd4 | 11.19973446 | 11.31171044 | 10.85097398 | 0.460736457 | 0.348760473 |
| Sf3b5 | 8.626347305 | 8.69583905 | 8.277755834 | 0.418083215 | 0.348591471 |
| Efemp2 | 9.400728921 | 9.271884808 | 9.052170316 | 0.219714492 | 0.348558605 |
| Rps12-ps3 | 13.50169182 | 13.74432931 | 13.15333228 | 0.590997033 | 0.34835954 |
| Gm10309 | 3.758955281 | 3.411176804 | 3.410733879 | 0.000442925 | 0.348221402 |

| | | | | | |
|---|---|---|---|---|---|
| Ugdh | 10.89272241 | 10.65846626 | 10.54454054 | 0.11392572 | 0.348181867 |
| Ssr1 | 12.2077211 | 12.04743691 | 11.85961462 | 0.187822283 | 0.348106477 |
| Glis1 | 4.314956856 | 4.204452785 | 3.966878391 | 0.237574394 | 0.348078465 |
| Eprs | 11.99809466 | 11.81062166 | 11.65001937 | 0.16060229 | 0.348075287 |
| Alcam | 10.26676478 | 9.985246644 | 9.919455837 | 0.065790807 | 0.347308947 |
| Gm10693 | 4.566718652 | 4.38592134 | 4.219464836 | 0.166456504 | 0.347253816 |
| Pin1 | 9.294228832 | 9.272728036 | 8.947108227 | 0.32561981 | 0.347120606 |
| Arf4 | 10.70278571 | 10.43641042 | 10.35611248 | 0.080297943 | 0.346673227 |
| P2ry10 | 7.315290166 | 6.9710026 | 6.968717459 | 0.002285141 | 0.346572707 |
| Arpc4 | 11.50526192 | 11.29095433 | 11.15872593 | 0.1322284 | 0.346535983 |
| Pdia6 | 11.9661473 | 12.06402881 | 11.61978812 | 0.444240687 | 0.34635918 |
| Jarid2 | 10.41775504 | 10.26432502 | 10.07141658 | 0.192908443 | 0.34633846 |
| CT010583.1 | 4.319316165 | 4.155134029 | 3.972981135 | 0.182152894 | 0.34633503 |
| Slc39a7 | 11.28808318 | 11.21419975 | 10.94181995 | 0.2723798 | 0.346263223 |
| Gm20425 | 6.982876142 | 6.516579958 | 6.636691885 | -0.120111928 | 0.346184257 |
| Eif3j1 | 10.48232248 | 10.27539839 | 10.13646929 | 0.138929097 | 0.345853187 |
| Riox2 | 9.625432918 | 9.401598292 | 9.279852636 | 0.121745656 | 0.345580282 |
| Bcar1 | 9.7380514 | 9.684998116 | 9.392471491 | 0.292526625 | 0.345579909 |
| Mars | 10.46288689 | 10.37763788 | 10.11757557 | 0.260062309 | 0.345311322 |
| Mb21d2 | 9.776121391 | 9.492627713 | 9.430955261 | 0.061672452 | 0.34516613 |
| Prmt5 | 9.995091954 | 9.810348307 | 9.650332688 | 0.160015619 | 0.344759266 |
| Gm28043 | 7.764990759 | 7.017510471 | 7.420241257 | -0.402730786 | 0.344749502 |
| Llph-ps2 | 8.624493013 | 8.287597314 | 8.279776142 | 0.007821171 | 0.34471687 |
| Ago2 | 11.53508829 | 11.44039285 | 11.19049794 | 0.24989491 | 0.344590343 |
| Lsm12 | 10.22047073 | 9.908934073 | 9.876083088 | 0.032850985 | 0.344387642 |
| Gm11346 | 5.694372702 | 5.476534871 | 5.349994709 | 0.126540162 | 0.344377993 |
| Gca | 6.689125914 | 6.322823929 | 6.344897106 | -0.022073177 | 0.344228808 |
| Smpdl3b | 7.605389581 | 7.435507894 | 7.261216915 | 0.174290978 | 0.344172666 |
| 2010005H15Rik | 4.028295994 | 3.660920524 | 3.684138827 | -0.023218302 | 0.344157167 |
| Kctd1 | 9.926447755 | 9.666021681 | 9.582415841 | 0.08360584 | 0.344031914 |
| Ssr2 | 11.21383755 | 11.05646915 | 10.87006912 | 0.18640003 | 0.343768433 |
| Eif4a3 | 10.73567766 | 10.45263141 | 10.39200037 | 0.06063104 | 0.34367729 |

| | | | | | |
|---|---|---|---|---|---|
| Max | 10.80438378 | 10.60811125 | 10.46091217 | 0.14719908 | 0.34347161 |
| Ifnar2 | 9.554585278 | 9.172877806 | 9.211185608 | -0.038307803 | 0.34339967 |
| Abcb1b | 8.862109767 | 8.748899165 | 8.518797644 | 0.230101521 | 0.343312124 |
| Pcgf6 | 8.316447215 | 8.154307142 | 7.973207396 | 0.181099746 | 0.34323982 |
| Tnfrsf10b | 7.679947791 | 7.620328205 | 7.336743221 | 0.283584984 | 0.343204569 |
| Ly6i | 4.486587901 | 4.301847207 | 4.143788227 | 0.158058981 | 0.342799674 |
| Rars2 | 9.304174463 | 9.014837909 | 8.961556108 | 0.053281801 | 0.342618355 |
| Nipa2 | 9.977385182 | 9.693192822 | 9.634886505 | 0.058306316 | 0.342498677 |
| Gsdmc2 | 6.320452229 | 6.669327401 | 5.978067199 | 0.691260202 | 0.34238503 |
| Pik3r1 | 11.96421555 | 11.59257122 | 11.62197482 | -0.029403603 | 0.342240727 |
| Dph1 | 8.058241674 | 7.892614969 | 7.716181659 | 0.17643331 | 0.342060014 |
| Igsf3 | 9.610041203 | 9.45154293 | 9.268032592 | 0.183510338 | 0.342008611 |
| 2900026A02Rik | 11.08049355 | 10.94696669 | 10.73860422 | 0.208362467 | 0.341889333 |
| Hpcal1 | 9.337879469 | 9.030690214 | 8.996002959 | 0.034687254 | 0.34187651 |
| Nup214 | 10.06942758 | 9.802163878 | 9.727721618 | 0.07444226 | 0.341705958 |
| Cald1 | 11.47222779 | 11.05328607 | 11.13075561 | -0.07746954 | 0.34147218 |
| Tdrd7 | 9.691519257 | 9.276454619 | 9.350066259 | -0.07361164 | 0.341452998 |
| Nlrx1 | 9.157213052 | 9.035999705 | 8.815760413 | 0.220239291 | 0.341452638 |
| Nom1 | 9.828306799 | 9.691375943 | 9.486885987 | 0.204489956 | 0.341420812 |
| Ankrd50 | 10.5579042 | 10.42469203 | 10.21650626 | 0.208185769 | 0.341397932 |
| Met | 10.35759742 | 10.04601068 | 10.01643674 | 0.029573944 | 0.341160676 |
| Dlgap4 | 11.03860954 | 10.93371339 | 10.69751772 | 0.236195667 | 0.34109182 |
| Fam185a | 7.494586387 | 7.216313033 | 7.153540515 | 0.062772518 | 0.341045872 |
| Med13l | 11.26894636 | 11.08847192 | 10.92791517 | 0.160556747 | 0.341031183 |
| Knop1 | 10.44371393 | 10.2474538 | 10.10271876 | 0.14473504 | 0.34099517 |
| Map3k8 | 8.708977703 | 8.582512992 | 8.368026316 | 0.214486676 | 0.340951387 |
| Cpsf2 | 10.42829656 | 10.16288392 | 10.08738785 | 0.07549607 | 0.340908707 |
| Pcbp1 | 11.8964192 | 11.77700283 | 11.55551242 | 0.221490403 | 0.340906777 |
| Tuba8 | 10.97534738 | 10.79948042 | 10.63452607 | 0.164954343 | 0.34082131 |
| Zcchc2 | 10.14373838 | 9.997459241 | 9.803536384 | 0.193922857 | 0.340201993 |
| Fndc4 | 7.582869743 | 7.47344529 | 7.242802065 | 0.230643225 | 0.340067678 |
| Noa1 | 8.83799336 | 8.594998394 | 8.49816906 | 0.096829334 | 0.339824299 |

| | | | | | |
|---|---|---|---|---|---|
| Slc35c1 | 9.303695612 | 9.198659556 | 8.964111981 | 0.234547574 | 0.339583631 |
| Pex14 | 9.681670069 | 9.60157302 | 9.342157079 | 0.259415941 | 0.33951299 |
| Fbxl6 | 8.925862724 | 8.930619481 | 8.586395898 | 0.344223582 | 0.339466826 |
| Tent5a | 11.31271193 | 10.91768435 | 10.97331486 | -0.05563051 | 0.33939707 |
| 2810004N23Rik | 9.422214888 | 9.110415624 | 9.083159782 | 0.027255842 | 0.339055107 |
| Lgals3bp | 11.16208041 | 10.43329907 | 10.82309206 | -0.38979299 | 0.338988357 |
| Gm16867 | 7.539331626 | 7.342682328 | 7.200374662 | 0.142307666 | 0.338956964 |
| Tgfbr1 | 10.11469287 | 9.90526885 | 9.776013417 | 0.129255433 | 0.33867945 |
| Larp4 | 10.84935465 | 10.67781004 | 10.51071054 | 0.167099503 | 0.33864411 |
| Mrps2 | 9.177922516 | 8.952540488 | 8.839431685 | 0.113108804 | 0.338490831 |
| Eno1 | 13.22163545 | 13.11838315 | 12.88321591 | 0.235167247 | 0.338419547 |
| P2rx1 | 6.014175883 | 5.562874855 | 5.67584583 | -0.112970975 | 0.338330053 |
| Exosc10 | 10.17978118 | 10.05286476 | 9.841765731 | 0.211099029 | 0.338015445 |
| Hars | 11.24115993 | 11.01797538 | 10.90326556 | 0.114709823 | 0.337894373 |
| Txk | 4.434307533 | 4.192106371 | 4.096547888 | 0.095558482 | 0.337759644 |
| Umps | 8.779112911 | 8.68155986 | 8.441395933 | 0.240163927 | 0.337716978 |
| Tmem248 | 10.32892453 | 10.27222584 | 9.991316855 | 0.280908982 | 0.337607678 |
| Klra8 | 3.705754956 | 3.441170408 | 3.368274246 | 0.072896163 | 0.33748071 |
| Gfra1 | 9.303225233 | 9.18654425 | 8.965751135 | 0.220793116 | 0.337474098 |
| Tbc1d10c | 5.582034116 | 5.287656755 | 5.245056026 | 0.042600729 | 0.33697809 |
| Tlr8 | 7.290858045 | 6.984326758 | 6.953988153 | 0.030338605 | 0.336869892 |
| Slc52a2 | 7.547228432 | 7.54061935 | 7.210392063 | 0.330227287 | 0.336836369 |
| Sdf2l1 | 8.570850588 | 8.877247383 | 8.234022822 | 0.643224561 | 0.336827766 |
| Scamp3 | 9.553750735 | 9.780971432 | 9.217033649 | 0.563937783 | 0.336717086 |
| Gm20604 | 7.425674279 | 7.297969057 | 7.08897892 | 0.208990136 | 0.336695359 |
| Gm49336 | 10.70780235 | 10.42199263 | 10.37115451 | 0.050838123 | 0.336647837 |
| Aldh1a3 | 7.263296014 | 6.684213472 | 6.927892722 | -0.24367925 | 0.335403292 |
| Gars | 11.61766245 | 11.36135145 | 11.2823285 | 0.079022953 | 0.335333957 |
| Il3ra | 6.811491938 | 6.793925162 | 6.476217431 | 0.317707731 | 0.335274507 |
| Casp8 | 9.903982667 | 9.634561481 | 9.568828345 | 0.065733137 | 0.335154322 |
| Tmem11 | 9.459102888 | 9.276316939 | 9.124025719 | 0.15229122 | 0.33507717 |
| Rrn3 | 10.48806458 | 10.22267395 | 10.15308004 | 0.069593907 | 0.334984537 |

| | | | | | |
|---|---|---|---|---|---|
| Ercc8 | 7.823702944 | 7.567923075 | 7.489023999 | 0.078899076 | 0.334678945 |
| Calr | 13.5157753 | 13.54878826 | 13.18120258 | 0.367585683 | 0.334572723 |
| Rgcc | 9.917313817 | 9.557484118 | 9.58320681 | -0.025722692 | 0.334107007 |
| Dna2 | 8.004428488 | 8.038179901 | 7.670368626 | 0.367811275 | 0.334059862 |
| Pnpo | 9.21576596 | 8.947807072 | 8.881731465 | 0.066075607 | 0.334034495 |
| Serp1 | 12.07839639 | 11.82877803 | 11.74476892 | 0.08400911 | 0.33362747 |
| Rars | 11.11463275 | 10.87889544 | 10.78123204 | 0.097663403 | 0.33340071 |
| Cdh23 | 3.973163025 | 3.795575253 | 3.640077682 | 0.155497571 | 0.333085343 |
| Timm50 | 9.386386288 | 9.287663677 | 9.053453639 | 0.234210038 | 0.332932649 |
| Arf2 | 9.785899451 | 9.587714232 | 9.45296976 | 0.134744472 | 0.332929691 |
| Gfi1 | 3.999775195 | 3.687356743 | 3.667141038 | 0.020215705 | 0.332634158 |
| Sf3a1 | 10.71349707 | 10.66157465 | 10.3810442 | 0.28053045 | 0.33245287 |
| Gm21748 | 5.103558077 | 5.329421534 | 4.771230445 | 0.558191088 | 0.332327631 |
| Gm21860 | 5.103558077 | 5.329421534 | 4.771230445 | 0.558191088 | 0.332327631 |
| Ebi3 | 5.873097462 | 5.731757233 | 5.540805648 | 0.190951585 | 0.332291814 |
| Cand1 | 11.18738958 | 11.13236765 | 10.8551675 | 0.27720015 | 0.332222077 |
| Ssb | 11.51591974 | 11.27319518 | 11.18369821 | 0.08949697 | 0.332221537 |
| Exosc9 | 9.41204546 | 9.092423893 | 9.08013391 | 0.012289983 | 0.33191155 |
| Akap11 | 10.24753545 | 9.91838327 | 9.915739515 | 0.002643754 | 0.331795931 |
| Chtf8 | 11.01460617 | 10.99808605 | 10.68309255 | 0.314993497 | 0.33151362 |
| Masp1 | 10.50517306 | 10.61146715 | 10.1737489 | 0.43771825 | 0.33142416 |
| Ptprn | 3.28973863 | 2.975581195 | 2.958415729 | 0.017165466 | 0.331322901 |
| Pigw | 7.709655738 | 7.50376318 | 7.378455999 | 0.125307181 | 0.33119974 |
| Zfp276 | 8.717330083 | 8.873732674 | 8.386359053 | 0.48737362 | 0.330971029 |
| Twistnb | 9.410086873 | 8.896718176 | 9.079812953 | -0.183094777 | 0.330273921 |
| Tinagl1 | 9.252807173 | 9.286993523 | 8.922645904 | 0.36434762 | 0.33016127 |
| Hsf1 | 9.440243268 | 9.502676778 | 9.110133309 | 0.392543469 | 0.330109959 |
| Ddit3 | 8.168399895 | 8.896422911 | 7.83843258 | 1.057990331 | 0.329967315 |
| Nup160 | 9.430370566 | 9.278559789 | 9.101091566 | 0.177468223 | 0.329279001 |
| Rabggta | 9.250786207 | 9.07737724 | 8.92186801 | 0.15550923 | 0.328918197 |
| Eaf1 | 10.52522807 | 10.37754795 | 10.19675791 | 0.180790033 | 0.328470157 |
| Sae1 | 10.83837775 | 10.65372128 | 10.50997843 | 0.143742843 | 0.328399317 |

| | | | | | |
|---|---|---|---|---|---|
| Riok2 | 9.490668759 | 9.17029577 | 9.162285059 | 0.00801071 | 0.328383699 |
| Jagn1 | 9.307167001 | 9.008097755 | 8.978832728 | 0.029265027 | 0.328334272 |
| Asap3 | 8.80483753 | 8.743981309 | 8.47662192 | 0.267359389 | 0.32821561 |
| Ugt1a7c | 8.03696372 | 7.905973731 | 7.708843823 | 0.197129908 | 0.328119897 |
| P2ry6 | 7.613081018 | 7.505169161 | 7.285146166 | 0.220022995 | 0.327934852 |
| Alox5 | 8.775914479 | 8.133654296 | 8.44800036 | -0.314346063 | 0.327914119 |
| Mafg | 10.77129107 | 10.91667506 | 10.44348795 | 0.473187107 | 0.327803113 |
| Myo1c | 12.52563956 | 12.37161043 | 12.19793152 | 0.17367891 | 0.32770804 |
| Arsi | 8.49931066 | 8.597256418 | 8.171725651 | 0.425530767 | 0.327585009 |
| Npc2 | 12.50801525 | 12.08058722 | 12.18045497 | -0.099867753 | 0.32756028 |
| Prkab1 | 9.050384191 | 8.83245634 | 8.72289268 | 0.10956366 | 0.327491511 |
| Apoc2 | 5.001031746 | 4.842356232 | 4.673609482 | 0.16874675 | 0.327422265 |
| Fam241a | 8.954697078 | 8.530692402 | 8.627363837 | -0.096671435 | 0.327333241 |
| Orc2 | 9.351610488 | 9.128328001 | 9.024325346 | 0.104002655 | 0.327285142 |
| Rasal3 | 6.294060733 | 6.230456288 | 5.966798503 | 0.263657785 | 0.32726223 |
| Arhgef1 | 11.2716832 | 11.36580448 | 10.94447234 | 0.42133214 | 0.327210863 |
| Zc3h18 | 10.07931177 | 10.13958137 | 9.752107758 | 0.387473615 | 0.327204015 |
| Olfm1 | 10.16501271 | 9.869259376 | 9.837888243 | 0.031371133 | 0.327124467 |
| Arf5 | 10.98947909 | 10.89899082 | 10.6624658 | 0.236525023 | 0.327013293 |
| Lgals8 | 10.40410858 | 10.15266165 | 10.07747323 | 0.075188417 | 0.326635343 |
| Orai1 | 9.227309951 | 9.333126772 | 8.900777584 | 0.432349189 | 0.326532367 |
| Rubcnl | 5.999263774 | 5.591995097 | 5.672803943 | -0.080808847 | 0.32645983 |
| Trim63 | 9.798167762 | 11.27189062 | 9.471880259 | 1.800010357 | 0.326287503 |
| Psma5 | 11.14173595 | 10.92191502 | 10.81555322 | 0.106361803 | 0.32618273 |
| Top1mt | 8.065360662 | 7.934757846 | 7.73940344 | 0.195354406 | 0.325957222 |
| Prpf38a | 9.522136913 | 9.18818816 | 9.196474076 | -0.008285916 | 0.325662837 |
| Rplp2 | 12.43988745 | 12.57468391 | 12.11427415 | 0.460409757 | 0.325613293 |
| Rpl15-ps3 | 11.77609543 | 11.45233644 | 11.4505368 | 0.001799643 | 0.32555863 |
| Ear6 | 4.369525831 | 4.062871584 | 4.043987042 | 0.018884542 | 0.325538789 |
| Klrb1c | 4.439864499 | 4.069083072 | 4.114514446 | -0.045431374 | 0.325350052 |
| Bmp7 | 10.01736863 | 9.892666227 | 9.692018636 | 0.200647591 | 0.325349997 |
| Lrfn4 | 7.744986938 | 7.963699775 | 7.419825102 | 0.543874673 | 0.325161836 |

| | | | | | |
|---|---|---|---|---|---|
| Casp1 | 9.184666935 | 8.857674544 | 8.859598741 | -0.001924196 | 0.325068194 |
| Sptlc2 | 10.77221706 | 10.49943688 | 10.44720924 | 0.052227643 | 0.325007817 |
| Ipo5 | 11.92289812 | 11.82605699 | 11.59795369 | 0.228103297 | 0.324944427 |
| Pak2 | 11.51470721 | 11.32634645 | 11.18982149 | 0.13652496 | 0.32488572 |
| Mrpl47 | 8.394683528 | 8.227464245 | 8.070072381 | 0.157391864 | 0.324611147 |
| Metap1 | 10.43281001 | 10.14281791 | 10.1089706 | 0.033847303 | 0.32383941 |
| Alg1 | 8.648995632 | 8.574938071 | 8.325400849 | 0.249537223 | 0.323594784 |
| Tram2 | 8.770990975 | 8.627689757 | 8.447840486 | 0.179849271 | 0.323150489 |
| Mob3c | 8.418852839 | 8.331541711 | 8.095736707 | 0.235805004 | 0.323116131 |
| Txn2 | 11.40611216 | 11.24474877 | 11.08300456 | 0.161744207 | 0.3231076 |
| Ppp1r9b | 11.61227899 | 11.51374496 | 11.28921461 | 0.22453035 | 0.323064377 |
| Col4a1 | 13.01564994 | 12.75077915 | 12.69273657 | 0.05804258 | 0.322913377 |
| Fkbp11 | 6.183071156 | 6.140910403 | 5.860284226 | 0.280626177 | 0.32278693 |
| Myl12a | 12.14530919 | 11.95750532 | 11.82253366 | 0.134971653 | 0.32277553 |
| Gadd45gip1 | 8.192378659 | 8.23669783 | 7.869625876 | 0.367071954 | 0.322752783 |
| Ap1ar | 10.08001781 | 9.931262078 | 9.757350449 | 0.173911629 | 0.322667358 |
| Ndufb7 | 9.809039033 | 9.857945594 | 9.486518723 | 0.371426871 | 0.32252031 |
| Ptgis | 7.340848756 | 6.977668229 | 7.018487704 | -0.040819475 | 0.322361052 |
| Med7 | 8.573288331 | 8.375333232 | 8.251328479 | 0.124004753 | 0.321959852 |
| Krr1 | 9.853750237 | 9.469315431 | 9.531892708 | -0.062577277 | 0.321857528 |
| Ntng2 | 7.35953421 | 7.223857154 | 7.037885089 | 0.185972065 | 0.321649121 |
| Sub1 | 11.0702292 | 10.79230149 | 10.74874503 | 0.043556463 | 0.321484173 |
| Tgoln1 | 12.16272121 | 11.90516625 | 11.8412584 | 0.063907847 | 0.32146281 |
| Scd3 | 11.0503231 | 10.89922557 | 10.72887663 | 0.170348937 | 0.32144647 |
| Nab2 | 9.706112748 | 9.701994649 | 9.384686744 | 0.317307905 | 0.321426004 |
| Hnrnpc | 12.07079501 | 11.91089272 | 11.74953628 | 0.161356443 | 0.321258733 |
| Timm10b | 8.522937472 | 8.480721919 | 8.2017732 | 0.278948719 | 0.321164271 |
| Acvr1b | 11.10454355 | 10.98757449 | 10.7834869 | 0.20408759 | 0.32105665 |
| Crtc3 | 9.71734328 | 9.648715858 | 9.396552816 | 0.252163042 | 0.320790464 |
| Smad7 | 7.981175502 | 7.818383782 | 7.66059846 | 0.157785321 | 0.320577042 |
| Ccdc97 | 9.799236942 | 9.751450458 | 9.478733843 | 0.272716615 | 0.320503098 |
| Rap1gds1 | 10.64542613 | 10.38850095 | 10.32503282 | 0.063468123 | 0.320393303 |

| | | | | | |
|---|---|---|---|---|---|
| Nedd4l | 9.579230936 | 9.228106009 | 9.25887217 | -0.030766161 | 0.320358765 |
| Mbd3 | 10.65344552 | 10.65828649 | 10.33311106 | 0.325175427 | 0.32033446 |
| Ergic1 | 10.66647774 | 10.54620408 | 10.34617375 | 0.200030327 | 0.320303987 |
| Slc52a3 | 8.332441766 | 7.928359559 | 8.012172357 | -0.083812798 | 0.320269409 |
| Pim3 | 10.49019417 | 10.05833654 | 10.16993449 | -0.111597952 | 0.320259673 |
| Slc25a32 | 8.351865455 | 7.983091842 | 8.031841263 | -0.048749421 | 0.320024192 |
| Gm16286 | 10.74891949 | 10.56334039 | 10.42889701 | 0.13444338 | 0.320022477 |
| Klf16 | 9.095168497 | 8.971497196 | 8.775281258 | 0.196215937 | 0.319887239 |
| Rbm18 | 9.711539141 | 9.50054624 | 9.391712048 | 0.108834192 | 0.319827093 |
| Wdr4 | 9.016979039 | 8.910099125 | 8.69717384 | 0.212925285 | 0.319805199 |
| Xpo6 | 10.65924365 | 10.48315017 | 10.33954766 | 0.14360251 | 0.319695997 |
| Tsen54 | 8.09677156 | 8.032459297 | 7.77713824 | 0.255321057 | 0.31963332 |
| Tnrc18 | 10.55769953 | 10.86662974 | 10.23829535 | 0.628334393 | 0.31940418 |
| Slc35f6 | 9.996800185 | 10.06456867 | 9.677401391 | 0.387167282 | 0.319398794 |
| Arhgef10l | 10.25295325 | 10.33527764 | 9.933613547 | 0.401664089 | 0.319339699 |
| Plekhg2 | 9.256760184 | 9.35696844 | 8.93768922 | 0.41927922 | 0.319070964 |
| Rcc2 | 11.54133266 | 11.51581659 | 11.22240862 | 0.29340797 | 0.318924037 |
| Mrpl20 | 9.951126377 | 9.666952569 | 9.632500947 | 0.034451621 | 0.318625429 |
| Mapk11 | 8.366765584 | 8.254315772 | 8.048156502 | 0.20615927 | 0.318609082 |
| Phox2a | 3.109697023 | 2.868595875 | 2.791105545 | 0.07749033 | 0.318591478 |
| Nup153 | 10.81948535 | 10.75831976 | 10.50089649 | 0.257423273 | 0.318588857 |
| Focad | 9.276326039 | 9.206125798 | 8.957768801 | 0.248356997 | 0.318557238 |
| Sec24d | 10.07489984 | 9.830236761 | 9.756364806 | 0.073871954 | 0.318535034 |
| Zcchc10 | 7.570924814 | 7.377596494 | 7.252461141 | 0.125135352 | 0.318463672 |
| Nup155 | 9.667014086 | 9.579315521 | 9.348610455 | 0.230705067 | 0.318403632 |
| Eif3c | 12.37067328 | 12.32529589 | 12.05245982 | 0.27283607 | 0.31821346 |
| Trim21 | 8.738874334 | 8.354184996 | 8.421185266 | -0.06700027 | 0.317689067 |
| Adgrd1 | 10.41276405 | 10.11729786 | 10.09519915 | 0.022098711 | 0.317564901 |
| Tnfrsf14 | 6.826977356 | 6.627869475 | 6.509696498 | 0.118172977 | 0.317280858 |
| Tha1 | 6.005024192 | 5.900047785 | 5.687811108 | 0.212236677 | 0.317213084 |
| Rbm17 | 10.48005198 | 10.18110423 | 10.16304004 | 0.01806419 | 0.317011937 |
| Ftl1-ps1 | 5.858518324 | 5.706824122 | 5.541965747 | 0.164858375 | 0.316552577 |

| | | | | | |
|---|---|---|---|---|---|
| Cmtm7 | 8.248352547 | 8.024263749 | 7.931883756 | 0.092379994 | 0.316468792 |
| Bdh1 | 9.157436188 | 8.877779172 | 8.841350227 | 0.036428945 | 0.316085961 |
| Pdlim4 | 8.31787689 | 8.195612163 | 8.001853283 | 0.19375888 | 0.316023607 |
| Vmn2r97 | 7.965218956 | 7.957919636 | 7.649254222 | 0.308665413 | 0.315964734 |
| Map1lc3b | 12.25944355 | 12.04064785 | 11.94364307 | 0.097004787 | 0.31580048 |
| Denr | 10.33425011 | 10.09320269 | 10.01848842 | 0.07471427 | 0.315761687 |
| Slc22a3 | 8.325033278 | 7.998591619 | 8.009313816 | -0.010722197 | 0.315719462 |
| 5830411N06Rik | 4.994587376 | 4.934274012 | 4.679056515 | 0.255217497 | 0.315530861 |
| Mgat2 | 10.38768792 | 10.19082636 | 10.07216041 | 0.11866595 | 0.315527503 |
| Lbr | 10.34837606 | 10.08363801 | 10.03298798 | 0.050650027 | 0.315388073 |
| Adam9 | 11.06422061 | 10.82887466 | 10.74883304 | 0.08004162 | 0.31538757 |
| Tubb5 | 12.82688202 | 12.72728717 | 12.51155246 | 0.215734703 | 0.315329557 |
| Ammecr1 | 10.19626129 | 9.927197146 | 9.881153578 | 0.046043568 | 0.315107708 |
| Cct2 | 12.43138036 | 12.27383134 | 12.116536 | 0.157295343 | 0.31484436 |
| Tgs1 | 9.388795547 | 9.269768634 | 9.074289888 | 0.195478746 | 0.314505659 |
| Ephb1 | 5.637153835 | 5.427079636 | 5.323019011 | 0.104060625 | 0.314134824 |
| Slamf6 | 5.496508167 | 5.395737547 | 5.182571382 | 0.213166165 | 0.313936785 |
| Gm15542 | 7.839781037 | 8.052495882 | 7.525870402 | 0.52662548 | 0.313910635 |
| Prpf40a | 11.19285958 | 10.97752363 | 10.87897414 | 0.098549493 | 0.313885443 |
| Ppargc1b | 6.794749259 | 6.6772003 | 6.480958354 | 0.196241946 | 0.313790905 |
| Arfrp1 | 9.678512151 | 9.51115037 | 9.364736003 | 0.146414366 | 0.313776148 |
| Rangrf | 8.056294007 | 7.872622291 | 7.742901768 | 0.129720522 | 0.313392239 |
| Cdk9 | 10.41115391 | 10.31871593 | 10.09800368 | 0.220712254 | 0.313150234 |
| Gjb2 | 9.724421013 | 9.464718034 | 9.411292034 | 0.053426 | 0.313128979 |
| Cacna2d2 | 7.724635664 | 7.570413303 | 7.411539888 | 0.158873415 | 0.313095777 |
| B9d2 | 6.812756816 | 6.580290498 | 6.499834828 | 0.08045567 | 0.312921988 |
| Batf3 | 4.915710481 | 4.755913502 | 4.602857229 | 0.153056274 | 0.312853252 |
| Ccnt1 | 10.56553052 | 10.27511279 | 10.25280317 | 0.02230962 | 0.312727353 |
| Azi2 | 10.51188399 | 10.35771366 | 10.19943138 | 0.158282277 | 0.312452607 |
| Slamf1 | 4.469508427 | 4.359299063 | 4.157058735 | 0.202240328 | 0.312449692 |
| Cdh4 | 8.995259622 | 8.844017328 | 8.683234772 | 0.160782556 | 0.312024849 |
| Atf6b | 10.12160124 | 10.24919329 | 9.809622067 | 0.439571227 | 0.311979177 |

| | | | | | |
|---|---|---|---|---|---|
| Cyp4f37 | 6.75474356 | 6.564072443 | 6.442806573 | 0.12126587 | 0.311936987 |
| Slc35g2 | 4.914528781 | 4.657115271 | 4.602799531 | 0.05431574 | 0.31172925 |
| Xkr8 | 6.948077378 | 6.597590392 | 6.636410544 | -0.038820152 | 0.311666834 |
| Xpot | 10.73151726 | 10.69117232 | 10.41992473 | 0.271247587 | 0.311592533 |
| Adprh | 9.760999381 | 9.622966444 | 9.449433131 | 0.173533313 | 0.31156625 |
| 9130008F23Rik | 7.210911449 | 7.076890102 | 6.899386289 | 0.177503813 | 0.31152516 |
| Bcl10 | 10.11664369 | 9.821477438 | 9.805286673 | 0.016190765 | 0.311357017 |
| Gm7666 | 3.761228895 | 3.672442304 | 3.44996118 | 0.222481124 | 0.311267715 |
| Rexo4 | 9.70986562 | 9.630626894 | 9.398623108 | 0.232003786 | 0.311242512 |
| Frk | 7.45607717 | 7.093405237 | 7.145260466 | -0.051855229 | 0.310816704 |
| Fam167a | 9.888504291 | 9.50713758 | 9.577886835 | -0.070749255 | 0.310617456 |
| Ccno | 4.123029343 | 3.895017963 | 3.812498628 | 0.082519335 | 0.310530715 |
| Jrk | 6.763914139 | 6.628725649 | 6.45351234 | 0.17521331 | 0.310401799 |
| Bri3bp | 9.043302522 | 8.849977649 | 8.732917442 | 0.117060207 | 0.31038508 |
| Rnf4 | 11.37671913 | 11.15129313 | 11.06653052 | 0.084762613 | 0.31018861 |
| Klhl6 | 6.366968144 | 6.028072234 | 6.056820782 | -0.028748549 | 0.310147362 |
| Prpf4 | 9.335346434 | 9.111404655 | 9.025202302 | 0.086202354 | 0.310144132 |
| Bzw2 | 10.57349686 | 10.289221 | 10.26336746 | 0.02585354 | 0.3101294 |
| Aldh1b1 | 5.792094107 | 5.642864435 | 5.482069183 | 0.160795252 | 0.310024924 |
| Asb6 | 8.78578017 | 8.92282738 | 8.475957103 | 0.446870277 | 0.309823066 |
| Amd1 | 11.96254211 | 11.83170717 | 11.65295212 | 0.17875505 | 0.309589993 |
| Angpt2 | 7.242612617 | 7.070652017 | 6.933029391 | 0.137622626 | 0.309583226 |
| Mul1 | 9.396805497 | 9.377023559 | 9.087366632 | 0.289656927 | 0.309438865 |
| Zfp566 | 6.049670204 | 5.975361047 | 5.740342587 | 0.23501846 | 0.309327617 |
| Muc1 | 5.766013371 | 5.594592945 | 5.456828454 | 0.137764491 | 0.309184918 |
| Ube2j2 | 9.702825296 | 9.613642511 | 9.393758972 | 0.219883539 | 0.309066324 |
| E130309D02Rik | 8.558838908 | 8.599230416 | 8.249920824 | 0.349309592 | 0.308918084 |
| Cct5 | 12.46035222 | 12.30048749 | 12.1515085 | 0.148978993 | 0.308843723 |
| Atg4d | 9.421769586 | 9.373768671 | 9.113101985 | 0.260666686 | 0.308667601 |
| Zfp655 | 9.995421416 | 9.95871713 | 9.687025624 | 0.271691507 | 0.308395792 |
| Msantd3 | 7.487699462 | 7.148290064 | 7.179536399 | -0.031246334 | 0.308163064 |
| Pofut2 | 9.276075146 | 9.3337522 | 8.967988408 | 0.365763791 | 0.308086738 |

| | | | | | |
|---|---|---|---|---|---|
| Nbeal2 | 10.16413887 | 10.21350609 | 9.856164219 | 0.357341867 | 0.307974655 |
| Shank3 | 8.202104092 | 8.19786012 | 7.894174682 | 0.303685438 | 0.30792941 |
| Twnk | 9.065812053 | 8.813870879 | 8.757918753 | 0.055952126 | 0.3078933 |
| Eml4 | 9.631653896 | 9.480924917 | 9.323784687 | 0.15714023 | 0.307869209 |
| Drg2 | 9.740910136 | 9.548076097 | 9.433052163 | 0.115023934 | 0.307857973 |
| Eef1d | 11.96236317 | 11.78962744 | 11.65458257 | 0.135044877 | 0.307780603 |
| 37135 | 7.086563387 | 6.824230373 | 6.778819101 | 0.045411272 | 0.307744286 |
| Aqr | 10.35620837 | 10.14981636 | 10.04861689 | 0.101199471 | 0.307591487 |
| Pes1 | 11.00391422 | 10.81168474 | 10.69648048 | 0.115204257 | 0.30743374 |
| Tyw3 | 6.578290696 | 6.665409452 | 6.271017136 | 0.394392316 | 0.30727356 |
| Sart3 | 9.58192872 | 9.583732032 | 9.274664111 | 0.309067921 | 0.307264609 |
| P2ry13 | 6.561772293 | 6.291779024 | 6.254562029 | 0.037216995 | 0.307210265 |
| Dhx29 | 9.336370736 | 9.372507126 | 9.029216608 | 0.343290518 | 0.307154127 |
| Utp11 | 9.517989749 | 9.2996898 | 9.210907996 | 0.088781803 | 0.307081752 |
| Amdhd2 | 7.938262731 | 8.012727043 | 7.631333565 | 0.381393478 | 0.306929165 |
| Oaf | 8.808463156 | 8.379733692 | 8.501537111 | -0.121803419 | 0.306926045 |
| Dusp9 | 4.91773273 | 4.744297236 | 4.610936223 | 0.133361013 | 0.306796507 |
| Crygn | 4.794604 | 4.565243788 | 4.487939511 | 0.077304277 | 0.306664489 |
| Rac1 | 12.86427023 | 12.63346979 | 12.55767736 | 0.075792433 | 0.306592873 |
| Stk39 | 8.479242213 | 8.348390571 | 8.172778164 | 0.175612407 | 0.306464049 |
| Cd2ap | 11.77082483 | 11.46926618 | 11.46436254 | 0.004903647 | 0.30646229 |
| Trp53 | 10.5047886 | 10.43563044 | 10.19834064 | 0.237289793 | 0.30644796 |
| Polr2d | 8.972952205 | 8.703993521 | 8.666647148 | 0.037346373 | 0.306305057 |
| Hspa14 | 9.864194206 | 9.545149149 | 9.558167343 | -0.013018194 | 0.306026863 |
| Htra2 | 8.992093337 | 8.783777393 | 8.686184435 | 0.097592957 | 0.305908902 |
| Snrpd2 | 10.00487767 | 9.879785355 | 9.699038637 | 0.180746718 | 0.30583903 |
| Coro2a | 8.333175603 | 8.110835168 | 8.027474961 | 0.083360207 | 0.305700642 |
| Ripk1 | 10.28527329 | 10.17573036 | 9.979675462 | 0.196054898 | 0.305597831 |
| Tox4 | 10.73686968 | 10.52054363 | 10.43146913 | 0.0890745 | 0.305400543 |
| Ddx41 | 9.303242678 | 9.160255688 | 8.997895522 | 0.162360166 | 0.305347156 |
| Gm18445 | 6.539134339 | 6.509138069 | 6.233793579 | 0.27534449 | 0.30534076 |
| Mrpl57 | 8.105841726 | 8.015035701 | 7.800554735 | 0.214480966 | 0.305286992 |

| | | | | | |
|---|---|---|---|---|---|
| Ric8a | 10.34778848 | 10.20719755 | 10.04252445 | 0.164673098 | 0.305264031 |
| Ctla2a | 8.923090679 | 8.564648237 | 8.617915518 | -0.053267281 | 0.305175161 |
| Lcmt2 | 7.76409058 | 7.699837453 | 7.458980809 | 0.240856644 | 0.305109771 |
| Pfkp | 11.7083944 | 11.56550395 | 11.40336736 | 0.162136593 | 0.305027043 |
| Tmc8 | 4.97346009 | 4.919484893 | 4.668500746 | 0.250984146 | 0.304959344 |
| Scamp4 | 10.16977379 | 10.11897053 | 9.864908626 | 0.254061904 | 0.304865164 |
| Tardbp | 12.39723034 | 12.48202557 | 12.09281023 | 0.38921534 | 0.30442011 |
| Galnt2 | 8.295526748 | 8.623929343 | 7.991204241 | 0.632725103 | 0.304322507 |
| Zfand5 | 11.48798952 | 11.45971479 | 11.18371263 | 0.276002163 | 0.30427689 |
| Gfod1 | 9.318526137 | 9.058283485 | 9.014387533 | 0.043895952 | 0.304138604 |
| Tm6sf1 | 8.939739454 | 8.555804227 | 8.635688083 | -0.079883856 | 0.304051372 |
| Ptcd3 | 10.17762852 | 9.8675196 | 9.873586993 | -0.006067393 | 0.304041523 |
| Rab11fip5 | 10.38232118 | 10.19135004 | 10.07834409 | 0.113005949 | 0.303977093 |
| Dbr1 | 8.486316 | 8.253882052 | 8.182511303 | 0.071370749 | 0.303804697 |
| Cerk | 10.73583131 | 10.60573832 | 10.43204685 | 0.17369147 | 0.303784457 |
| Atad1 | 10.86635097 | 10.5874798 | 10.56259783 | 0.024881973 | 0.30375314 |
| Abi3bp | 11.09837709 | 10.84310509 | 10.79463675 | 0.048468337 | 0.303740337 |
| Bola2 | 8.748598801 | 8.499090704 | 8.444931091 | 0.054159613 | 0.30366771 |
| Slc35a4 | 10.97898217 | 10.79236071 | 10.67531515 | 0.117045557 | 0.303667017 |
| Actn1 | 11.77196842 | 11.70164497 | 11.46841659 | 0.233228377 | 0.303551827 |
| Ikbkb | 10.72821759 | 10.59792153 | 10.42467362 | 0.17324791 | 0.303543973 |
| Zfp783 | 7.06765135 | 7.04031838 | 6.764281594 | 0.276036786 | 0.303369756 |
| Capza1 | 12.01328371 | 11.82514719 | 11.71012376 | 0.115023433 | 0.303159953 |
| Synj1 | 9.711752675 | 9.673304987 | 9.408634416 | 0.264670571 | 0.303118259 |
| Brd2 | 12.20973101 | 12.08664209 | 11.90662087 | 0.18002122 | 0.303110143 |
| Cdk20 | 7.192962866 | 7.252321637 | 6.890321944 | 0.361999693 | 0.302640923 |
| Ptges2 | 8.656522971 | 8.436457678 | 8.354228664 | 0.082229014 | 0.302294307 |
| Gm10521 | 5.072993537 | 4.983378398 | 4.770714531 | 0.212663868 | 0.302279006 |
| Aup1 | 10.36477321 | 10.35608893 | 10.0625068 | 0.293582133 | 0.302266406 |
| Nutf2-ps1 | 9.885479777 | 9.617024743 | 9.583240393 | 0.03378435 | 0.302239385 |
| Anxa7 | 12.636922 | 12.37468453 | 12.33491323 | 0.039771297 | 0.302008767 |
| B4galt3 | 9.054494087 | 8.877592407 | 8.752587093 | 0.125005314 | 0.301906994 |

| | | | | | |
|---|---|---|---|---|---|
| Mllt6 | 10.60883847 | 10.58124782 | 10.307089 | 0.27415882 | 0.301749473 |
| Leo1 | 8.910174907 | 8.707099349 | 8.608444827 | 0.098654521 | 0.30173008 |
| Ak4 | 9.7014227 | 9.300743457 | 9.399724354 | -0.098980897 | 0.301698346 |
| Als2 | 9.53472666 | 9.244621447 | 9.233152881 | 0.011468566 | 0.301573779 |
| Gm13301 | 8.305293271 | 8.318147382 | 8.003807303 | 0.314340079 | 0.301485968 |
| Chl1 | 12.05857995 | 11.91117046 | 11.75718864 | 0.153981827 | 0.30139131 |
| Gm7332 | 9.0049033 | 9.021218466 | 8.703550295 | 0.317668171 | 0.301353005 |
| Spidr | 8.633288256 | 8.360085358 | 8.332080999 | 0.028004358 | 0.301207257 |
| Snrpc | 9.218565153 | 9.198712172 | 8.917437485 | 0.281274687 | 0.301127669 |
| Klf6 | 12.27741119 | 12.05516789 | 11.97629917 | 0.07886872 | 0.30111202 |
| Neto2 | 6.140455493 | 6.05118034 | 5.839347218 | 0.211833122 | 0.301108275 |
| Vps18 | 8.867848434 | 9.126879465 | 8.566748778 | 0.560130687 | 0.301099656 |
| Vcl | 11.3357019 | 11.16303492 | 11.03463721 | 0.12839771 | 0.301064693 |
| Rpusd2 | 6.723553486 | 6.547679385 | 6.422516786 | 0.125162599 | 0.3010367 |
| Scrg1 | 3.720637778 | 3.373796544 | 3.419660954 | -0.04586441 | 0.300976823 |
| Bcar3 | 8.478528183 | 8.400974557 | 8.177713709 | 0.223260848 | 0.300814474 |
| Ctss | 11.76770666 | 11.66626011 | 11.4669097 | 0.199350407 | 0.30079696 |
| Hgf | 6.31971491 | 5.983784821 | 6.01892999 | -0.035145169 | 0.30078492 |
| 38412 | 9.637871595 | 9.191399531 | 9.337089327 | -0.145689796 | 0.300782269 |
| Zmiz2 | 10.92710855 | 11.08236036 | 10.62647566 | 0.4558847 | 0.300632893 |
| Thbs2 | 12.50881946 | 12.25532653 | 12.20853388 | 0.046792647 | 0.300285577 |
| Grk5 | 9.322067039 | 9.22614309 | 9.021906262 | 0.204236828 | 0.300160777 |
| 2310057M21Rik | 8.645926081 | 8.536818925 | 8.346093872 | 0.190725053 | 0.299832209 |
| Donson | 7.78696451 | 7.607012762 | 7.487161227 | 0.119851535 | 0.299803283 |
| Arap1 | 10.85834843 | 10.71429773 | 10.55869229 | 0.155605433 | 0.299656137 |
| Tra2b | 11.68943154 | 11.64319451 | 11.38986649 | 0.25332802 | 0.299565043 |
| Tfe3 | 10.46561846 | 10.41495706 | 10.16622068 | 0.248736387 | 0.299397783 |
| Pkp2 | 7.434395322 | 6.945615699 | 7.135086607 | -0.189470908 | 0.299308715 |
| Cd63 | 12.26571368 | 12.13308251 | 11.96665264 | 0.166429873 | 0.299061043 |
| Unc93b1 | 9.897149991 | 9.714449511 | 9.598232773 | 0.116216738 | 0.298917218 |
| Amigo3 | 6.76197076 | 6.6390872 | 6.463139397 | 0.175947803 | 0.298831363 |
| Rps6ka3 | 11.67123527 | 11.39038687 | 11.37240395 | 0.017982923 | 0.29883132 |

| | | | | | |
|---|---|---|---|---|---|
| Cdipt | 10.18449997 | 10.13029781 | 9.885775923 | 0.244521884 | 0.298724043 |
| Gbp11 | 5.156189425 | 4.84680701 | 4.857730272 | -0.010923262 | 0.298459153 |
| 40787 | 10.85997932 | 10.74209908 | 10.56167521 | 0.180423867 | 0.298304107 |
| Cdk4 | 11.2697504 | 11.15272188 | 10.97151973 | 0.181202147 | 0.298230673 |
| Golt1b | 10.07339795 | 9.839089172 | 9.775399441 | 0.06368973 | 0.297998509 |
| Afg3l2 | 10.59817494 | 10.44227892 | 10.30025735 | 0.14202157 | 0.29791759 |
| Ppp1r11 | 9.620559202 | 9.401628006 | 9.322755704 | 0.078872303 | 0.297803498 |
| Atp6v1h | 10.88796489 | 10.91696478 | 10.59049723 | 0.326467543 | 0.29746766 |
| Ern1 | 10.01712194 | 9.873993942 | 9.719706721 | 0.15428722 | 0.297415223 |
| 9130401M01Rik | 7.892308281 | 7.772393693 | 7.594915887 | 0.177477806 | 0.297392394 |
| Uqcr11 | 10.22148422 | 10.28571452 | 9.924409395 | 0.361305125 | 0.297074826 |
| Evi2 | 6.174896929 | 5.894064197 | 5.878213616 | 0.015850581 | 0.296683313 |
| Bcor | 9.657643977 | 9.526801411 | 9.361205379 | 0.165596032 | 0.296438599 |
| Pcdhgb6 | 8.071125931 | 8.164611206 | 7.77486297 | 0.389748236 | 0.296262961 |
| Hnrnpa0 | 11.05407519 | 10.85991166 | 10.75811932 | 0.101792343 | 0.295955873 |
| Sfpq | 12.1081506 | 12.05506646 | 11.81226687 | 0.24279959 | 0.295883733 |
| Grk3 | 8.902237476 | 9.128670787 | 8.606474672 | 0.522196115 | 0.295762804 |
| Sf3a3 | 9.942945332 | 9.7606418 | 9.647356317 | 0.113285483 | 0.295589015 |
| Gpm6b | 10.24659489 | 10.07034433 | 9.951108936 | 0.119235397 | 0.295485957 |
| Ccn3 | 7.458774545 | 7.203687638 | 7.163496253 | 0.040191386 | 0.295278292 |
| Pold1 | 8.431769513 | 8.737416269 | 8.136673083 | 0.600743186 | 0.29509643 |
| Ndufaf8 | 6.883502169 | 6.903513846 | 6.588433684 | 0.315080162 | 0.295068485 |
| Cdc42ep2 | 8.332025314 | 8.115599182 | 8.037021454 | 0.078577727 | 0.29500386 |
| Patl1 | 9.723588244 | 9.658586074 | 9.428720864 | 0.22986521 | 0.294867379 |
| Abt1 | 8.361212899 | 8.245172681 | 8.066474164 | 0.178698517 | 0.294738735 |
| Ccdc12 | 9.463920615 | 9.357079936 | 9.169360567 | 0.187719369 | 0.294560048 |
| Gucy2c | 3.666288944 | 3.379982358 | 3.371870286 | 0.008112072 | 0.294418658 |
| Mrps5 | 9.441606102 | 9.33877685 | 9.147378028 | 0.191398822 | 0.294228074 |
| S1pr1 | 9.320447965 | 9.035864678 | 9.026855323 | 0.009009355 | 0.293592642 |
| Ttll12 | 9.51836041 | 9.468813663 | 9.224854005 | 0.243959658 | 0.293506405 |
| Trmt2a | 8.38883033 | 8.193430763 | 8.095403234 | 0.098027529 | 0.293427096 |
| Syncrip | 12.0533011 | 11.93232819 | 11.7598907 | 0.172437487 | 0.293410397 |

| | | | | | |
|---|---|---|---|---|---|
| Rabggtb | 10.25853383 | 10.00890409 | 9.965130049 | 0.043774044 | 0.293403781 |
| Sprr2a1 | 7.232093786 | 7.183124566 | 6.938709941 | 0.244414625 | 0.293383845 |
| D10Wsu102e | 11.1916045 | 11.00534617 | 10.89837685 | 0.106969317 | 0.29322765 |
| Adgrg2 | 8.069606929 | 7.528583081 | 7.776409734 | -0.247826653 | 0.293197195 |
| Tfrc | 11.05713752 | 10.72858383 | 10.76403533 | -0.035451507 | 0.29310219 |
| Cd200r4 | 5.812335212 | 5.750878121 | 5.519419846 | 0.231458275 | 0.292915366 |
| Gls | 11.25548685 | 10.91000919 | 10.96261096 | -0.052601773 | 0.29287589 |
| Bloc1s4 | 7.159391174 | 6.908125564 | 6.866526993 | 0.041598571 | 0.29286418 |
| Arf6 | 11.1051398 | 11.01282154 | 10.81247441 | 0.200347127 | 0.29266539 |
| Itga6 | 13.04764401 | 12.84668004 | 12.75502343 | 0.091656613 | 0.292620577 |
| Mrps22 | 8.506644862 | 8.286179913 | 8.214059012 | 0.072120901 | 0.292585851 |
| Rgs1 | 5.9853233 | 5.830426685 | 5.69315033 | 0.137276355 | 0.29217297 |
| Rpia | 9.36011916 | 9.257094703 | 9.068053645 | 0.189041059 | 0.292065516 |
| Msra | 8.619234031 | 8.204053737 | 8.327208052 | -0.123154315 | 0.292025979 |
| Atp6v0d2 | 5.359272636 | 5.688086959 | 5.067379747 | 0.620707212 | 0.291892889 |
| Kcmf1 | 11.01033886 | 10.90205403 | 10.71845439 | 0.183599633 | 0.291884463 |
| Nup85 | 10.02171398 | 9.889765189 | 9.730117395 | 0.159647794 | 0.291596589 |
| Magoh | 9.636883711 | 9.36227922 | 9.345432913 | 0.016846307 | 0.291450799 |
| Utp6 | 10.46020781 | 10.17669221 | 10.16887087 | 0.007821333 | 0.291336933 |
| Thada | 8.793669958 | 8.652725921 | 8.50248001 | 0.15024591 | 0.291189947 |
| Psmb5 | 10.95524114 | 10.90318004 | 10.6641216 | 0.239058443 | 0.291119537 |
| Rbm14 | 9.380242156 | 9.604662391 | 9.089193875 | 0.515468516 | 0.291048281 |
| Trim28 | 11.60805072 | 11.45739618 | 11.31704347 | 0.140352707 | 0.291007247 |
| Zfp131 | 9.692790394 | 9.64121487 | 9.401816527 | 0.239398343 | 0.290973866 |
| Dad1 | 10.33769519 | 10.13755986 | 10.04680135 | 0.090758512 | 0.290893842 |
| Gspt1 | 11.61109656 | 11.39480762 | 11.32034791 | 0.07445971 | 0.290748657 |
| Mydgf | 9.86140482 | 9.738561889 | 9.570786578 | 0.167775311 | 0.290618242 |
| Prrx2 | 6.510615 | 6.56958602 | 6.220021017 | 0.349565003 | 0.290593983 |
| A430005L14Rik | 8.607606797 | 8.64501754 | 8.317027831 | 0.327989709 | 0.290578966 |
| Ak2 | 11.4618669 | 11.27154346 | 11.1714546 | 0.100088857 | 0.2904123 |
| Lrrk2 | 8.304008634 | 7.87762349 | 8.013596587 | -0.135973096 | 0.290412047 |
| Carhsp1 | 11.24022381 | 10.86197876 | 10.94988395 | -0.087905193 | 0.290339857 |

| | | | | | |
|---|---|---|---|---|---|
| Gm15448 | 5.162311591 | 5.13259495 | 4.872034449 | 0.260560502 | 0.290277142 |
| Gcsh | 9.443760183 | 9.228736066 | 9.153683927 | 0.075052139 | 0.290076256 |
| Gm48493 | 7.264193588 | 7.047119374 | 6.974177693 | 0.07294168 | 0.290015895 |
| Psmg1 | 8.476335712 | 8.346434806 | 8.186341834 | 0.160092972 | 0.289993879 |
| Dck | 9.202639293 | 8.961808946 | 8.913030243 | 0.048778703 | 0.28960905 |
| Nudt4 | 11.79808885 | 11.28109981 | 11.50864773 | -0.22754792 | 0.289441123 |
| Gm12551 | 8.403732261 | 8.256777558 | 8.114320753 | 0.142456805 | 0.289411508 |
| Khsrp | 10.70428868 | 10.83265147 | 10.41509476 | 0.41755671 | 0.289193917 |
| Prrc2c | 12.11984415 | 11.95387808 | 11.83068564 | 0.123192447 | 0.28915851 |
| E2f3 | 8.499321503 | 8.34777364 | 8.210280762 | 0.137492877 | 0.289040741 |
| Rps16-ps2 | 12.08085941 | 11.99976405 | 11.79190163 | 0.207862417 | 0.288957773 |
| Irf2bp2 | 10.78691528 | 10.62055157 | 10.49796628 | 0.12258529 | 0.288949003 |
| Cinp | 9.253996639 | 9.15593836 | 8.965076385 | 0.190861975 | 0.288920254 |
| Col23a1 | 10.74166782 | 10.78705368 | 10.45286618 | 0.334187497 | 0.288801643 |
| Usp39 | 8.390065632 | 8.172568558 | 8.101264049 | 0.071304509 | 0.288801583 |
| Bcr | 10.05010259 | 10.04383504 | 9.761434018 | 0.282401019 | 0.288668576 |
| Il1f10 | 6.856912228 | 6.475287629 | 6.568340626 | -0.093052996 | 0.288571602 |
| Moap1 | 6.473373759 | 6.401444272 | 6.184830789 | 0.216613483 | 0.28854297 |
| Txndc11 | 9.516891147 | 9.533329735 | 9.228505831 | 0.304823905 | 0.288385317 |
| Srsf3 | 12.59306371 | 12.44432058 | 12.30471202 | 0.139608567 | 0.288351697 |
| Dhx8 | 9.819832647 | 9.839896515 | 9.531532228 | 0.308364287 | 0.288300419 |
| Ltbp1 | 10.59358049 | 10.41220708 | 10.30528294 | 0.10692414 | 0.28829755 |
| Irf2bpl | 9.96126883 | 10.07918285 | 9.673010497 | 0.406172353 | 0.288258333 |
| Ccdc124 | 9.202077447 | 9.259661942 | 8.913842845 | 0.345819097 | 0.288234601 |
| Rps6ka1 | 9.832106425 | 9.790329006 | 9.543907473 | 0.246421533 | 0.288198952 |
| Cwc25 | 9.076202226 | 9.037098926 | 8.788009138 | 0.249089788 | 0.288193088 |
| Utp23 | 8.261697009 | 8.163236091 | 7.97369545 | 0.189540641 | 0.28800156 |
| Hoxa1 | 6.099085442 | 6.010914305 | 5.811256655 | 0.19965765 | 0.287828787 |
| Mrpl16 | 8.78981099 | 8.646900214 | 8.50247436 | 0.144425855 | 0.28733663 |
| Susd1 | 6.467474666 | 6.03011929 | 6.180302433 | -0.150183143 | 0.287172233 |
| Tfip11 | 9.373463384 | 9.366130985 | 9.086306692 | 0.279824293 | 0.287156692 |
| Dda1 | 9.613214215 | 9.62227213 | 9.326058181 | 0.296213949 | 0.287156034 |

| | | | | | |
|---|---|---|---|---|---|
| Mthfd1 | 10.19733579 | 10.23081104 | 9.910203471 | 0.320607569 | 0.287132316 |
| Ogfod1 | 9.468187448 | 9.306842895 | 9.181097446 | 0.125745449 | 0.287090002 |
| Eloc | 10.45405218 | 10.19543233 | 10.16697879 | 0.028453543 | 0.287073393 |
| Emp3 | 9.987896783 | 10.03083582 | 9.700890081 | 0.329945739 | 0.287006702 |
| Metap2 | 11.86934328 | 11.6563814 | 11.5824465 | 0.073934907 | 0.286896787 |
| Prl8a1 | 7.645865385 | 7.545098839 | 7.359159034 | 0.185939805 | 0.286706351 |
| Gpatch3 | 6.916253594 | 6.951993467 | 6.629813786 | 0.322179681 | 0.286439809 |
| Nr4a2 | 7.453824143 | 7.009869331 | 7.167566132 | -0.157696801 | 0.286258011 |
| Mocs3 | 6.79911379 | 6.733678048 | 6.512917342 | 0.220760706 | 0.286196448 |
| Gm13092 | 8.091112208 | 7.921769639 | 7.804996319 | 0.11677332 | 0.286115889 |
| Slc25a13 | 8.517108425 | 8.418828394 | 8.231086793 | 0.187741601 | 0.286021632 |
| Alas1 | 10.67322864 | 10.73103146 | 10.38723858 | 0.343792877 | 0.28599006 |
| F2rl2 | 6.10088003 | 5.786695268 | 5.814935579 | -0.028240311 | 0.28594445 |
| Dhodh | 7.728027183 | 7.690720225 | 7.442558219 | 0.248162006 | 0.285468964 |
| Lrwd1 | 8.353875763 | 8.385826642 | 8.068466563 | 0.317360079 | 0.2854092 |
| Bpnt1 | 9.026081898 | 8.877074226 | 8.74076407 | 0.136310157 | 0.285317829 |
| Itgb1 | 12.92318677 | 12.83817758 | 12.63788166 | 0.200295927 | 0.285305113 |
| Las1l | 9.885370699 | 9.676909252 | 9.600096593 | 0.076812659 | 0.285274106 |
| Mov10 | 7.948418287 | 7.621861544 | 7.663224108 | -0.041362564 | 0.285194179 |
| Plagl2 | 9.732159938 | 9.663493937 | 9.447051618 | 0.21644232 | 0.28510832 |
| Slc9a8 | 9.069813527 | 8.8685439 | 8.784713736 | 0.083830164 | 0.285099791 |
| Pgd | 11.56520771 | 11.45202509 | 11.28015027 | 0.17187482 | 0.285057433 |
| Rpp25l | 8.242957329 | 8.199018926 | 7.95793078 | 0.241088146 | 0.285026548 |
| Usp16 | 10.40050004 | 10.22475399 | 10.11595863 | 0.108795367 | 0.284541413 |
| Setdb2 | 8.055759649 | 7.79397444 | 7.77137878 | 0.02259566 | 0.28438087 |
| Mrps35 | 8.771102852 | 8.515420119 | 8.486760552 | 0.028659567 | 0.284342299 |
| Med17 | 9.464345373 | 9.37973482 | 9.180108736 | 0.199626084 | 0.284236636 |
| Bud31 | 9.995105546 | 9.847587063 | 9.710915329 | 0.136671734 | 0.284190216 |
| Mctp2 | 8.010325739 | 7.607872612 | 7.726169231 | -0.118296618 | 0.284156508 |
| Clint1 | 11.09316173 | 10.94170967 | 10.80907482 | 0.132634857 | 0.284086917 |
| Cluh | 11.56563098 | 11.50433212 | 11.28154594 | 0.22278618 | 0.284085043 |
| Dohh | 8.907158361 | 8.862650751 | 8.623118958 | 0.239531793 | 0.284039403 |

| | | | | | |
|---|---|---|---|---|---|
| Rbm45 | 9.051292808 | 8.79964975 | 8.76732959 | 0.03232016 | 0.283963218 |
| Klra4 | 3.423313885 | 3.173415983 | 3.139498477 | 0.033917505 | 0.283815408 |
| Ccr7 | 5.982848683 | 6.242410458 | 5.699198495 | 0.543211963 | 0.283650189 |
| Nxt1 | 8.650166624 | 8.452069945 | 8.366697769 | 0.085372176 | 0.283468855 |
| Supt16 | 11.19462386 | 10.99493769 | 10.91143754 | 0.08350015 | 0.283186317 |
| Ubr4 | 11.85947182 | 11.85286601 | 11.57628593 | 0.276580077 | 0.283185883 |
| Ipo7 | 11.96236332 | 11.88001677 | 11.67932754 | 0.20068923 | 0.28303578 |
| Atp5g1 | 11.35777329 | 11.35409484 | 11.07492377 | 0.279171063 | 0.282849513 |
| Gm15459 | 4.099820908 | 3.94726255 | 3.816972002 | 0.130290548 | 0.282848906 |
| Sigmar1 | 10.39301828 | 10.29712547 | 10.1102076 | 0.186917876 | 0.282810682 |
| Zfp513 | 8.497277228 | 8.506150606 | 8.214546671 | 0.291603935 | 0.282730557 |
| Pip5k1b | 4.581727634 | 4.31624144 | 4.299258462 | 0.016982978 | 0.282469173 |
| Cacna1b | 3.512920894 | 3.472686734 | 3.230486116 | 0.242200618 | 0.282434778 |
| H2-Eb1 | 12.7807684 | 12.68748258 | 12.49838135 | 0.189101237 | 0.282387053 |
| Taf5 | 7.598664202 | 7.370958478 | 7.316314282 | 0.054644196 | 0.282349919 |
| Ap1m1 | 9.650672916 | 9.674686446 | 9.36834015 | 0.306346296 | 0.282332766 |
| Atp6v1e1 | 11.58746194 | 11.30714498 | 11.30528346 | 0.001861523 | 0.282178483 |
| Klrk1 | 7.290184993 | 6.777559321 | 7.008016017 | -0.230456696 | 0.282168976 |
| Lrp10 | 11.45247585 | 11.38374247 | 11.17045064 | 0.213291833 | 0.28202521 |
| Usp12 | 10.13939826 | 10.0046884 | 9.857415695 | 0.147272707 | 0.281982568 |
| Carm1 | 10.3554706 | 10.23843514 | 10.0737209 | 0.164714241 | 0.281749701 |
| Usp42 | 8.679306787 | 8.513150936 | 8.397559145 | 0.115591792 | 0.281747642 |
| Sntg2 | 5.106824335 | 4.955456761 | 4.825079123 | 0.130377638 | 0.281745211 |
| Timm17a | 10.12594252 | 9.909199855 | 9.844205999 | 0.064993856 | 0.281736524 |
| Vnn3 | 6.621444718 | 6.195963564 | 6.339823765 | -0.143860201 | 0.281620953 |
| Ccdc58 | 8.195806624 | 8.040201297 | 7.914196033 | 0.126005264 | 0.281610591 |
| Riok1 | 9.180503587 | 8.897377508 | 8.898907984 | -0.001530476 | 0.281595602 |
| Tubb4b | 13.18282173 | 13.14671122 | 12.90129029 | 0.245420927 | 0.28153144 |
| S100a11 | 13.72949487 | 13.47407617 | 13.44829881 | 0.025777363 | 0.281196063 |
| Gorasp2 | 11.17468318 | 11.09357644 | 10.89359535 | 0.199981093 | 0.281087833 |
| Gramd1b | 8.24771027 | 8.161668828 | 7.966623971 | 0.195044857 | 0.281086299 |
| Tti2 | 8.427415078 | 8.209235373 | 8.146478574 | 0.062756798 | 0.280936504 |

| | | | | | |
|---|---|---|---|---|---|
| Thumpd1 | 10.35660301 | 10.12631482 | 10.07567747 | 0.050637343 | 0.28092554 |
| Rwdd4a | 9.123638639 | 8.737369973 | 8.842854333 | -0.10548436 | 0.280784306 |
| Fastkd5 | 8.302068426 | 8.191952741 | 8.02130093 | 0.170651811 | 0.280767496 |
| Bend4 | 6.597285887 | 6.441782161 | 6.316602932 | 0.125179229 | 0.280682955 |
| Mrpl3 | 9.511152313 | 9.270281031 | 9.23061175 | 0.039669281 | 0.280540563 |
| Dus1l | 9.581290688 | 9.514917572 | 9.300784015 | 0.214133556 | 0.280506672 |
| Smg8 | 8.484719798 | 8.332870363 | 8.204303317 | 0.128567046 | 0.280416481 |
| Safb2 | 10.33152688 | 10.33510646 | 10.05112723 | 0.283979235 | 0.280399655 |
| Ndufs7 | 9.568710665 | 9.642447226 | 9.288317172 | 0.354130054 | 0.280393493 |
| Tcp1 | 12.4356035 | 12.32510234 | 12.15527649 | 0.169825853 | 0.28032701 |
| Psrc1 | 8.045495092 | 7.903822057 | 7.765248182 | 0.138573875 | 0.280246911 |
| Rbm43 | 8.169448731 | 7.920768077 | 7.88924269 | 0.031525386 | 0.280206041 |
| Tm2d3 | 8.202145058 | 8.106480885 | 7.921957639 | 0.184523245 | 0.280187418 |
| Chd1 | 10.48451429 | 10.28342748 | 10.20432745 | 0.07910003 | 0.280186847 |
| Mt3 | 5.336089876 | 5.710490561 | 5.055923241 | 0.654567319 | 0.280166635 |
| Slc35b1 | 10.16439162 | 10.15145404 | 9.884892375 | 0.266561665 | 0.279499242 |
| Mmp12 | 7.812795676 | 7.542097014 | 7.533438719 | 0.008658296 | 0.279356958 |
| Srsf9 | 9.682394083 | 9.536518165 | 9.403149627 | 0.133368538 | 0.279244456 |
| Fastkd3 | 8.024676763 | 7.75153635 | 7.745532352 | 0.006003998 | 0.279144411 |
| Artn | 3.723729432 | 3.466020708 | 3.44477871 | 0.021241998 | 0.278950722 |
| Ncdn | 9.078029839 | 9.132207084 | 8.799080501 | 0.333126583 | 0.278949338 |
| Nt5dc3 | 9.237241258 | 9.450693943 | 8.958545619 | 0.492148323 | 0.278695639 |
| Igsf8 | 9.927202269 | 9.791874266 | 9.64852081 | 0.143353456 | 0.278681459 |
| Hsd17b2 | 7.253534463 | 7.176673382 | 6.974945935 | 0.201727447 | 0.278588528 |
| Gpr182 | 6.474011238 | 6.30844157 | 6.195548315 | 0.112893255 | 0.278462923 |
| Cd93 | 9.305724144 | 9.285322922 | 9.02729611 | 0.258026812 | 0.278428034 |
| Cd38 | 7.590832828 | 6.954849943 | 7.312426817 | -0.357576874 | 0.278406011 |
| Supt6 | 11.45772338 | 11.41030479 | 11.17935617 | 0.23094862 | 0.278367217 |
| Cpsf1 | 10.23515668 | 10.36875638 | 9.956855704 | 0.411900673 | 0.27830098 |
| 2310022A10Rik | 9.82109245 | 9.921832668 | 9.542890229 | 0.378942439 | 0.278202222 |
| Slit1 | 5.352454245 | 5.113674545 | 5.074380509 | 0.039294037 | 0.278073737 |
| Zcchc17 | 8.212305463 | 7.984483966 | 7.934461674 | 0.050022292 | 0.277843788 |

| | | | | | |
|---|---|---|---|---|---|
| Eme2 | 8.357151557 | 8.440642687 | 8.079346537 | 0.36129615 | 0.27780502 |
| Puf60 | 10.94372701 | 10.85349605 | 10.66614038 | 0.18735567 | 0.277586627 |
| Eya3 | 10.13408293 | 9.914428357 | 9.856539636 | 0.057888721 | 0.277543298 |
| Pcdhgc5 | 5.681030326 | 5.738111152 | 5.403553827 | 0.334557326 | 0.277476499 |
| Thoc1 | 9.750056439 | 9.580447284 | 9.472761709 | 0.107685575 | 0.27729473 |
| Larp7 | 9.264982623 | 9.105580937 | 8.987778753 | 0.117802184 | 0.27720387 |
| Ubd | 4.451211292 | 4.362075111 | 4.174050805 | 0.188024305 | 0.277160487 |
| Uspl1 | 9.691264699 | 9.722021518 | 9.414252039 | 0.307769479 | 0.27701266 |
| Ces2c | 3.415565559 | 3.178747731 | 3.1385547 | 0.040193031 | 0.277010859 |
| Gm49325 | 5.029892402 | 5.4471178 | 4.75302355 | 0.694094251 | 0.276868852 |
| Ppp2r3d | 6.814631971 | 6.963239553 | 6.537912253 | 0.425327299 | 0.276719718 |
| C3ar1 | 9.347370846 | 9.47321216 | 9.070767274 | 0.402444885 | 0.276603571 |
| Dhx9 | 11.87816529 | 11.80301063 | 11.60162409 | 0.201386543 | 0.2765412 |
| Sln | 6.441393411 | 6.151453549 | 6.165173318 | -0.013719769 | 0.276220093 |
| Pum1 | 9.753436389 | 9.631542914 | 9.477280055 | 0.15426286 | 0.276156335 |
| Tbkbp1 | 7.814481952 | 7.869791019 | 7.538328466 | 0.331462553 | 0.276153486 |
| Rps4l | 7.802682285 | 7.786885118 | 7.52657579 | 0.260309328 | 0.276106494 |
| Cdca4 | 9.722818581 | 9.579147802 | 9.446726896 | 0.132420907 | 0.276091685 |
| Gm48774 | 5.48818417 | 5.277249273 | 5.212443394 | 0.064805879 | 0.275740776 |
| Rrp36 | 8.67918097 | 8.616184951 | 8.403526605 | 0.212658346 | 0.275654364 |
| Srsf4 | 9.989371683 | 9.932729853 | 9.713788431 | 0.218941422 | 0.275583252 |
| Rab11a | 12.18442595 | 11.89554274 | 11.90886901 | -0.013326273 | 0.27555694 |
| Gm48348 | 8.631345567 | 8.591445902 | 8.355797056 | 0.235648847 | 0.275548511 |
| Bcl6b | 6.437880773 | 6.381175813 | 6.162344241 | 0.218831572 | 0.275536532 |
| Smyd2 | 9.706953572 | 9.590163986 | 9.43176732 | 0.158396665 | 0.275186252 |
| Vdr | 11.67762456 | 11.55522247 | 11.40271819 | 0.15250428 | 0.27490637 |
| Fance | 8.459825233 | 8.424267633 | 8.185110642 | 0.239156991 | 0.274714591 |
| Gstcd | 7.922720894 | 7.678813114 | 7.648066516 | 0.030746598 | 0.274654378 |
| Glis3 | 7.811716284 | 7.830180918 | 7.537198821 | 0.292982097 | 0.274517463 |
| Gm18294 | 4.17286394 | 3.969223297 | 3.898394206 | 0.070829091 | 0.274469734 |
| Zfand2a | 9.248917438 | 9.811936352 | 8.974469191 | 0.83746716 | 0.274448247 |
| Nasp | 10.63619636 | 10.44886802 | 10.36175822 | 0.087109797 | 0.27443814 |

| | | | | | |
|---|---|---|---|---|---|
| Rbm22 | 9.798431243 | 9.59859383 | 9.524059696 | 0.074534134 | 0.274371548 |
| Mettl2 | 8.455897246 | 8.377320077 | 8.181764389 | 0.195555687 | 0.274132857 |
| Trmt5 | 7.744562572 | 7.346170211 | 7.470451178 | -0.124280966 | 0.274111394 |
| Gtf2f2 | 9.218027997 | 8.945099592 | 8.943921728 | 0.001177864 | 0.274106269 |
| Zfp597 | 8.302060806 | 8.263941818 | 8.028090167 | 0.235851651 | 0.273970639 |
| Uchl5 | 9.682992921 | 9.422330334 | 9.409162353 | 0.013167981 | 0.273830568 |
| Abhd17b | 9.334318884 | 9.038269979 | 9.060555561 | -0.022285582 | 0.273763323 |
| Dzank1 | 4.880194498 | 4.380499775 | 4.606530123 | -0.226030348 | 0.273664375 |
| Slc7a7 | 7.204690519 | 7.101870289 | 6.931077373 | 0.170792916 | 0.273613146 |
| Dnajc25 | 8.319082679 | 8.241335665 | 8.045496631 | 0.195839034 | 0.273586048 |
| Lrrc8d | 9.672639874 | 9.378597197 | 9.399500505 | -0.020903308 | 0.273139369 |
| Fau | 12.51746473 | 12.29548054 | 12.24433995 | 0.051140593 | 0.273124777 |
| Cers6 | 10.31901847 | 10.24289151 | 10.0459419 | 0.19694961 | 0.273076567 |
| Alkbh4 | 7.247462697 | 7.138256901 | 6.974435377 | 0.163821524 | 0.273027321 |
| AC124502.3 | 4.838363258 | 4.748419963 | 4.565418229 | 0.183001734 | 0.272945029 |
| Olfr1372-ps1 | 8.660137168 | 8.646930761 | 8.387412805 | 0.259517956 | 0.272724364 |
| Dysf | 9.111646666 | 9.154364611 | 8.838957821 | 0.31540679 | 0.272688845 |
| Lat2 | 8.293200019 | 8.540145742 | 8.020516727 | 0.519629015 | 0.272683292 |
| Ube2s | 10.41744072 | 10.21924985 | 10.14493077 | 0.074319073 | 0.272509947 |
| Xab2 | 9.652606214 | 9.687607869 | 9.380108995 | 0.307498874 | 0.272497219 |
| Gm3608 | 5.582326352 | 5.288292423 | 5.309914869 | -0.021622446 | 0.272411483 |
| Ptges3-ps | 8.271902009 | 8.428204434 | 7.999517521 | 0.428686912 | 0.272384488 |
| Cnot3 | 10.08416669 | 10.10411135 | 9.811941068 | 0.292170285 | 0.272225619 |
| Parvb | 8.907857549 | 8.74153333 | 8.635722438 | 0.105810891 | 0.27213511 |
| Ube2n | 11.16896026 | 10.97416286 | 10.8970011 | 0.077161753 | 0.27195916 |
| Nup54 | 9.362816512 | 9.136119733 | 9.090883192 | 0.045236541 | 0.27193332 |
| Csprs | 7.206663649 | 6.797045762 | 6.934881513 | -0.137835752 | 0.271782135 |
| Nelfa | 9.115396646 | 8.989454698 | 8.843627816 | 0.145826882 | 0.271768831 |
| Snrpd3 | 10.40508395 | 10.16722949 | 10.13339263 | 0.033836863 | 0.27169132 |
| Ecd | 9.856750355 | 9.672212815 | 9.585059468 | 0.087153347 | 0.271690886 |
| Faap100 | 8.590452957 | 8.568303337 | 8.318868993 | 0.249434344 | 0.271583964 |
| Csrp2 | 8.280729182 | 7.904963889 | 8.009265217 | -0.104301327 | 0.271463965 |

| | | | | | |
|---|---|---|---|---|---|
| Necap1 | 9.05989247 | 8.93127691 | 8.788525097 | 0.142751813 | 0.271367373 |
| Ythdf2 | 10.27774338 | 10.11856526 | 10.00651001 | 0.112055253 | 0.271233373 |
| Prkcsh | 10.25578296 | 10.28984591 | 9.98456418 | 0.30528173 | 0.27121878 |
| Arl8a | 9.795775238 | 9.755486128 | 9.524564802 | 0.230921327 | 0.271210436 |
| Apex2 | 7.811968251 | 7.740932904 | 7.540885691 | 0.200047212 | 0.27108256 |
| Gm11331 | 3.970628159 | 3.781987062 | 3.699856168 | 0.082130895 | 0.270771991 |
| Rwdd1 | 9.822310107 | 9.628976021 | 9.551590545 | 0.077385476 | 0.270719562 |
| Atf1 | 10.06798827 | 9.997466118 | 9.797396099 | 0.200070018 | 0.270592173 |
| Fut4 | 5.411815479 | 5.359310578 | 5.14129549 | 0.218015088 | 0.270519989 |
| Rplp1 | 13.49139875 | 13.80168685 | 13.22134169 | 0.580345153 | 0.270057057 |
| Zfp574 | 8.08065304 | 8.221590557 | 7.810711368 | 0.410879189 | 0.269941672 |
| Kdf1 | 7.975970949 | 8.091013486 | 7.706548962 | 0.384464523 | 0.269421987 |
| Rlf | 9.94210588 | 9.670688938 | 9.672840574 | -0.002151636 | 0.269265307 |
| Ftsj1 | 8.551152378 | 8.398021272 | 8.282023378 | 0.115997895 | 0.269129 |
| Dus3l | 8.658633244 | 8.916227854 | 8.389619845 | 0.52660801 | 0.269013399 |
| D17Wsu92e | 12.27907265 | 12.15383725 | 12.01038688 | 0.143450367 | 0.268685767 |
| Prkca | 9.086030326 | 8.921469932 | 8.817550776 | 0.103919156 | 0.26847955 |
| Srsf6 | 12.19199947 | 12.09468332 | 11.92355604 | 0.17112728 | 0.26844343 |
| Rnf126 | 9.603325047 | 9.786388505 | 9.335276054 | 0.451112451 | 0.268048993 |
| Tsfm | 8.277874666 | 8.165409214 | 8.01005588 | 0.155353334 | 0.267818785 |
| Dclre1c | 8.564511359 | 8.374966107 | 8.296829057 | 0.07813705 | 0.267682301 |
| Nptx1 | 7.175628316 | 7.017271142 | 6.907990199 | 0.109280943 | 0.267638117 |
| Rbm25 | 11.90006808 | 11.72450252 | 11.63252774 | 0.091974777 | 0.267540333 |
| Ints12 | 8.697706976 | 8.424316011 | 8.430765134 | -0.006449123 | 0.266941842 |
| Bfsp1 | 6.026401326 | 5.980526099 | 5.759665296 | 0.220860803 | 0.26673603 |
| Med27 | 8.131261251 | 7.951424705 | 7.864528523 | 0.086896181 | 0.266732728 |
| Adgre4 | 7.053932266 | 6.697901835 | 6.787226359 | -0.089324524 | 0.266705907 |
| Pym1 | 8.581636494 | 8.41071639 | 8.314947532 | 0.095768858 | 0.266688963 |
| Edem1 | 11.06208114 | 11.0211666 | 10.79549837 | 0.225668223 | 0.26658277 |
| Gm9973 | 5.142343343 | 4.991219982 | 4.875761621 | 0.115458361 | 0.266581722 |
| Prmt6 | 7.81370691 | 7.401423171 | 7.547272479 | -0.145849308 | 0.266434431 |
| Bag1 | 11.9636814 | 11.76573776 | 11.69736964 | 0.068368117 | 0.266311757 |

| | | | | | |
|---|---|---|---|---|---|
| Szrd1 | 11.02408107 | 10.9982386 | 10.75816455 | 0.240074053 | 0.26591652 |
| Prkcd | 11.28423539 | 11.40311583 | 11.01834146 | 0.38477437 | 0.265893933 |
| Dtwd1 | 7.132010513 | 6.789896528 | 6.866324988 | -0.07642846 | 0.265685525 |
| Ccdc51 | 7.120725082 | 6.943620494 | 6.855090669 | 0.088529826 | 0.265634414 |
| Lpcat4 | 6.615814917 | 6.684037671 | 6.350251054 | 0.333786616 | 0.265563863 |
| Pcdhgc4 | 10.29185305 | 9.76308425 | 10.02636745 | -0.263283204 | 0.265485596 |
| Tent4a | 9.681779982 | 9.54715923 | 9.416430857 | 0.130728373 | 0.265349125 |
| Mbd2 | 11.06440457 | 10.80033711 | 10.79909394 | 0.00124317 | 0.265310637 |
| Nppb | 5.332497517 | 6.602781103 | 5.067234714 | 1.535546389 | 0.265262803 |
| Stoml2 | 9.999010755 | 9.826292162 | 9.733877145 | 0.092415017 | 0.26513361 |
| Tmco6 | 7.774922478 | 7.534585837 | 7.5098009 | 0.024784938 | 0.265121579 |
| Kdelr2 | 10.36741373 | 10.31979358 | 10.10241248 | 0.2173811 | 0.265001243 |
| Mtf1 | 9.560243765 | 9.398495822 | 9.295432444 | 0.103063378 | 0.264811321 |
| Mcrip1 | 9.940125511 | 9.866019379 | 9.675336883 | 0.190682496 | 0.264788627 |
| Ubxn4 | 11.3196961 | 11.23208738 | 11.05498373 | 0.177103643 | 0.264712363 |
| Med15 | 9.997346671 | 10.12337469 | 9.732679737 | 0.390694956 | 0.264666934 |
| Taf3 | 8.820110877 | 8.746043235 | 8.55560969 | 0.190433545 | 0.264501187 |
| Tdh | 8.147627809 | 8.013010206 | 7.883302925 | 0.129707281 | 0.264324884 |
| Smarca5 | 11.33031291 | 11.12227555 | 11.06607897 | 0.05619658 | 0.264233943 |
| Mllt1 | 9.955199712 | 10.04448504 | 9.691060917 | 0.353424123 | 0.264138795 |
| Arhgap39 | 7.996715383 | 7.942373584 | 7.732608251 | 0.209765333 | 0.264107132 |
| Gab2 | 9.323347598 | 9.158671447 | 9.059297212 | 0.099374235 | 0.264050386 |
| Ears2 | 7.867659131 | 7.695476885 | 7.603688574 | 0.091788311 | 0.263970557 |
| Rab22a | 11.35722387 | 11.18823326 | 11.09333276 | 0.0949005 | 0.26389111 |
| Thg1l | 8.020624814 | 7.863336131 | 7.756913181 | 0.10642295 | 0.263711633 |
| Dpp9 | 9.726333329 | 9.807238469 | 9.462662725 | 0.344575744 | 0.263670604 |
| Atp11b | 10.84030671 | 10.53832787 | 10.57667362 | -0.038345747 | 0.26363309 |
| Lgmn | 11.70970582 | 11.65318369 | 11.44608762 | 0.20709607 | 0.263618197 |
| Psma3 | 11.01473775 | 10.8556623 | 10.75124686 | 0.104415437 | 0.26349089 |
| Rpl37rt | 8.819506949 | 8.823483775 | 8.556064704 | 0.267419071 | 0.263442245 |
| Actr2 | 12.64214397 | 12.42835785 | 12.37879395 | 0.049563903 | 0.263350023 |
| Egln3 | 10.11556321 | 9.782985811 | 9.852222375 | -0.069236564 | 0.263340835 |

| | | | | | |
|---|---|---|---|---|---|
| Slc1a1 | 8.151500819 | 8.072671161 | 7.88821854 | 0.184452621 | 0.263282279 |
| Ankrd49 | 9.143364787 | 8.966312409 | 8.880093189 | 0.08621922 | 0.263271598 |
| Ubr5 | 12.18782172 | 11.96582825 | 11.92460293 | 0.04122532 | 0.263218797 |
| Tfdp1 | 10.91857591 | 10.82954423 | 10.65547562 | 0.174068603 | 0.26310029 |
| Emd | 10.32730129 | 10.08735265 | 10.06438494 | 0.022967717 | 0.262916357 |
| Chpf | 7.648084619 | 7.751601385 | 7.385184227 | 0.366417158 | 0.262900393 |
| Impa1 | 9.387179003 | 9.249908383 | 9.124314601 | 0.125593782 | 0.262864402 |
| Surf4 | 12.17044335 | 12.12477178 | 11.90759148 | 0.217180297 | 0.262851873 |
| Eif4e | 11.46577225 | 11.26531009 | 11.20310718 | 0.062202917 | 0.262665073 |
| Npepps | 12.13355713 | 11.9622426 | 11.8710713 | 0.0911713 | 0.262485827 |
| Evi2a | 9.192742462 | 8.795202356 | 8.930508354 | -0.135305999 | 0.262234108 |
| Arl4c | 10.23886964 | 10.04430927 | 9.976713367 | 0.067595901 | 0.262156269 |
| 2310011J03Rik | 9.330807548 | 9.30093822 | 9.068737976 | 0.232200244 | 0.262069573 |
| Pcna | 11.51385892 | 11.38997107 | 11.25180316 | 0.138167913 | 0.262055757 |
| Col27a1 | 7.665951169 | 7.745662502 | 7.4039419 | 0.341720602 | 0.262009269 |
| Hes1 | 9.589464184 | 9.339559936 | 9.32754459 | 0.012015346 | 0.261919594 |
| Eif2b5 | 10.80017915 | 10.55435952 | 10.53833687 | 0.016022653 | 0.261842283 |
| Sgpp2 | 7.690300698 | 7.367776231 | 7.428560608 | -0.060784377 | 0.261740091 |
| Imp3 | 9.532812314 | 9.325651973 | 9.271241818 | 0.054410155 | 0.261570496 |
| Lrrc75a | 7.386138016 | 7.367100908 | 7.124823445 | 0.242277463 | 0.261314571 |
| Kazn | 8.657233123 | 8.591013768 | 8.396017009 | 0.194996758 | 0.261216113 |
| Dus4l | 7.271368633 | 7.068917115 | 7.010202532 | 0.058714583 | 0.261166101 |
| Pias4 | 8.629891096 | 8.578133393 | 8.369235316 | 0.208898077 | 0.260655781 |
| Rec8 | 8.063761946 | 7.747496527 | 7.803116102 | -0.055619575 | 0.260645844 |
| Pitrm1 | 10.30175181 | 10.20803556 | 10.04119201 | 0.166843545 | 0.260559802 |
| Shkbp1 | 8.835877581 | 8.779594787 | 8.575391247 | 0.20420354 | 0.260486334 |
| Fam133b | 9.125410861 | 8.930329103 | 8.865072873 | 0.065256231 | 0.260337988 |
| Mrpl15 | 9.552589783 | 9.291609054 | 9.292750766 | -0.001141713 | 0.259839017 |
| Entpd7 | 8.16576374 | 8.228810957 | 7.905960034 | 0.322850924 | 0.259803706 |
| Lrrc8a | 9.690468541 | 9.688592432 | 9.430680842 | 0.25791159 | 0.259787699 |
| Qpctl | 8.420780061 | 8.369638511 | 8.161273917 | 0.208364594 | 0.259506144 |
| 4930453N24Rik | 9.779963797 | 9.661149973 | 9.520695155 | 0.140454818 | 0.259268643 |

| | | | | | |
|---|---|---|---|---|---|
| Yae1d1 | 9.385904187 | 9.179497545 | 9.126671704 | 0.052825841 | 0.259232483 |
| Mrpl36 | 8.978696498 | 8.876711208 | 8.719530925 | 0.157180283 | 0.259165573 |
| Kcnc3 | 6.776911563 | 6.986599427 | 6.517763455 | 0.468835972 | 0.259148108 |
| Pcdhga10 | 7.258021101 | 7.209677996 | 6.998951109 | 0.210726887 | 0.259069992 |
| Rpp14 | 9.173346867 | 8.835492775 | 8.91442743 | -0.078934656 | 0.258919436 |
| U2surp | 10.76942869 | 10.533584 | 10.51060388 | 0.022980127 | 0.258824813 |
| Gm14494 | 7.231167109 | 7.13988248 | 6.972410721 | 0.167471759 | 0.258756388 |
| Rps29 | 12.54067141 | 12.63826011 | 12.28194204 | 0.356318073 | 0.258729367 |
| Zscan22 | 7.922077092 | 7.963227201 | 7.663392922 | 0.299834279 | 0.25868417 |
| Elp5 | 9.699318929 | 9.432139079 | 9.440951066 | -0.008811986 | 0.258367863 |
| Lypd5 | 12.31454661 | 12.25312626 | 12.0562483 | 0.19687796 | 0.25829831 |
| Mrpl50 | 9.477776621 | 9.189397583 | 9.219486946 | -0.030089362 | 0.258289675 |
| Lars | 10.74387699 | 10.5485453 | 10.48566965 | 0.06287565 | 0.258207347 |
| Fam160a2 | 9.208485986 | 9.153037893 | 8.950357637 | 0.202680256 | 0.258128349 |
| Pold4 | 7.846631814 | 7.748852796 | 7.588579281 | 0.160273515 | 0.258052533 |
| Fam162a | 10.56132747 | 10.38201006 | 10.30339191 | 0.07861815 | 0.25793556 |
| Atp6v1f | 9.992732995 | 9.890930637 | 9.734970164 | 0.155960473 | 0.257762831 |
| Kpna2 | 11.12362056 | 10.91980096 | 10.8658895 | 0.053911463 | 0.257731067 |
| Rock2 | 11.36576692 | 11.18138238 | 11.10810537 | 0.07327701 | 0.257661553 |
| Elovl7 | 11.77896862 | 11.53786857 | 11.52133588 | 0.01653269 | 0.25763274 |
| Zfp36l2 | 12.54129039 | 12.44605871 | 12.28374351 | 0.162315203 | 0.257546877 |
| Col18a1 | 11.5709528 | 11.78386305 | 11.31354259 | 0.470320463 | 0.257410207 |
| Srrt | 11.16427064 | 11.09507257 | 10.90702305 | 0.188049513 | 0.257247583 |
| Slc10a3 | 8.753536988 | 8.671255331 | 8.496394918 | 0.174860412 | 0.257142069 |
| Rps6kc1 | 8.764802012 | 8.937315874 | 8.507807034 | 0.42950884 | 0.256994978 |
| Trak2 | 10.71781334 | 10.58881109 | 10.46082849 | 0.127982597 | 0.256984853 |
| Senp2 | 10.44868863 | 10.28623755 | 10.19180743 | 0.094430113 | 0.256881197 |
| Ccm2 | 8.966494372 | 8.829114167 | 8.709713001 | 0.119401165 | 0.256781371 |
| Rbm48 | 7.331412134 | 7.217573457 | 7.074670176 | 0.142903281 | 0.256741958 |
| Hyal2 | 9.36407287 | 9.34835373 | 9.107383577 | 0.240970153 | 0.256689293 |
| Ubc | 13.86528574 | 14.30816934 | 13.60870032 | 0.699469023 | 0.256585423 |
| Ing3 | 8.275798787 | 8.084457478 | 8.019255017 | 0.065202461 | 0.25654377 |

| | | | | | |
|---|---|---|---|---|---|
| Snrnp48 | 9.506794363 | 9.366403883 | 9.250282203 | 0.11612168 | 0.256512159 |
| Fbxl14 | 9.146318961 | 9.128962601 | 8.889870189 | 0.239092412 | 0.256448772 |
| Gtf2h1 | 10.22900574 | 10.25099169 | 9.972701038 | 0.278290655 | 0.256304705 |
| Tac1 | 4.380858551 | 4.354856639 | 4.12459221 | 0.230264428 | 0.25626634 |
| Il10 | 4.793641564 | 4.625723549 | 4.53743884 | 0.088284709 | 0.256202724 |
| Dock10 | 9.445120702 | 9.202059241 | 9.189077921 | 0.012981319 | 0.25604278 |
| Trub1 | 8.42099614 | 8.346471315 | 8.164985752 | 0.181485563 | 0.256010388 |
| Gfap | 4.610072625 | 4.675243491 | 4.354101166 | 0.321142325 | 0.255971459 |
| 37865 | 8.210016053 | 7.98513015 | 7.954133319 | 0.030996831 | 0.255882734 |
| Fastkd1 | 7.951337801 | 7.840978406 | 7.69561421 | 0.145364196 | 0.255723591 |
| Tifab | 8.144944838 | 8.078385441 | 7.889252378 | 0.189133063 | 0.25569246 |
| Ggta1 | 10.90970127 | 10.8030375 | 10.65409551 | 0.148941993 | 0.255605767 |
| Txnrd3 | 9.186345785 | 8.949427012 | 8.930779721 | 0.018647292 | 0.255566065 |
| Psmd7 | 11.48742919 | 11.34459845 | 11.23201335 | 0.112585097 | 0.255415837 |
| Gpn1 | 9.111467416 | 8.966984906 | 8.856124301 | 0.110860605 | 0.255343115 |
| H2-D1 | 14.22997071 | 14.06875531 | 13.97472576 | 0.094029553 | 0.25524495 |
| Sec13 | 10.70018105 | 10.56563555 | 10.44496863 | 0.120666917 | 0.25521242 |
| Mtx1 | 8.878804552 | 8.756960486 | 8.623722381 | 0.133238104 | 0.25508217 |
| Btf3 | 12.71761305 | 12.52790189 | 12.46259854 | 0.065303343 | 0.25501451 |
| Mapre1 | 11.92219354 | 11.83178695 | 11.66721331 | 0.164573637 | 0.254980227 |
| Tor1aip2 | 11.78822023 | 11.54799487 | 11.53328496 | 0.014709917 | 0.254935273 |
| Olfr539 | 6.073939913 | 5.922557333 | 5.81915307 | 0.103404263 | 0.254786843 |
| Xrcc5 | 9.319201787 | 9.111006044 | 9.064868187 | 0.046137857 | 0.2543336 |
| Gpx1 | 12.09953431 | 12.00811102 | 11.84537964 | 0.16273138 | 0.254154673 |
| Kmo | 7.137447999 | 6.698534955 | 6.883342368 | -0.184807412 | 0.254105631 |
| Serpinb3a | 5.250660036 | 5.215399803 | 4.996597759 | 0.218802044 | 0.254062276 |
| Uqcrq | 10.1439588 | 10.19481579 | 9.889972633 | 0.304843157 | 0.253986167 |
| Mybpc3 | 4.180033278 | 4.05672893 | 3.926323832 | 0.130405097 | 0.253709446 |
| Stard3 | 8.883400763 | 8.773424554 | 8.629735635 | 0.143688919 | 0.253665128 |
| Rnd3 | 10.73606614 | 10.61133923 | 10.482423 | 0.128916237 | 0.25364314 |
| Ezr | 12.20204339 | 12.0882705 | 11.94855244 | 0.139718053 | 0.25349095 |
| Ptbp3 | 12.62186854 | 12.40077458 | 12.36840749 | 0.03236709 | 0.25346105 |

| | | | | | |
|---|---|---|---|---|---|
| Sash1 | 10.77219502 | 10.52911406 | 10.51882451 | 0.010289547 | 0.253370507 |
| Ccdc85c | 8.913062235 | 8.842225275 | 8.659892866 | 0.182332408 | 0.253169369 |
| Myadm | 11.33346583 | 10.93335133 | 11.08037267 | -0.147021343 | 0.25309316 |
| Clp1 | 8.21864321 | 7.933925384 | 7.965738592 | -0.031813208 | 0.252904618 |
| Zfp456 | 5.590438137 | 5.752546554 | 5.337663413 | 0.41488314 | 0.252774723 |
| Fam136a | 9.025243852 | 8.722711496 | 8.772548501 | -0.049837005 | 0.252695351 |
| Mettl14 | 9.155252556 | 8.906287396 | 8.902665368 | 0.003622027 | 0.252587188 |
| Eif4g1 | 13.59694306 | 13.37506478 | 13.34444796 | 0.030616823 | 0.252495097 |
| Nkain1 | 5.20758977 | 5.125380145 | 4.955095803 | 0.170284342 | 0.252493967 |
| Ddx47 | 10.03913431 | 9.80726582 | 9.786705821 | 0.020559999 | 0.252428488 |
| Gm20594 | 11.86695615 | 12.2013052 | 11.61466524 | 0.586639957 | 0.252290913 |
| Dpagt1 | 9.401042498 | 9.266038946 | 9.148768673 | 0.117270274 | 0.252273826 |
| Hspa2 | 8.228382976 | 8.080757249 | 7.976166603 | 0.104590645 | 0.252216373 |
| Tmem214 | 10.14079342 | 10.3054804 | 9.888577904 | 0.4169025 | 0.252215516 |
| Dnmt3b | 6.892936042 | 6.596984021 | 6.640730457 | -0.043746436 | 0.252205585 |
| Gpat3 | 8.515014372 | 8.196786795 | 8.263003229 | -0.066216435 | 0.252011142 |
| Mcm3 | 10.1205585 | 10.04165318 | 9.868730995 | 0.172922185 | 0.251827509 |
| Ly6e | 13.30173521 | 12.90161997 | 13.04995948 | -0.148339513 | 0.25177573 |
| Mcmbp | 10.59291267 | 10.52271947 | 10.34117658 | 0.181542887 | 0.25173609 |
| Ndufa2 | 9.908660068 | 9.863865005 | 9.656954259 | 0.206910746 | 0.251705809 |
| Dmxl1 | 10.11318612 | 9.67124142 | 9.861529679 | -0.190288259 | 0.251656437 |
| Creb3l1 | 8.923981727 | 8.91037208 | 8.672424268 | 0.237947812 | 0.251557459 |
| Mrps18a | 9.195868067 | 9.031384427 | 8.944545854 | 0.086838573 | 0.251322212 |
| Gadl1 | 10.04158918 | 9.982608321 | 9.790368 | 0.192240321 | 0.251221179 |
| Rpp25 | 5.69928121 | 5.511005681 | 5.448176741 | 0.06282894 | 0.25110447 |
| 1110002E22Rik | 9.580488261 | 11.11664645 | 9.32941678 | 1.787229673 | 0.25107148 |
| Tuba4a | 12.77994617 | 12.71642679 | 12.52891116 | 0.187515627 | 0.25103501 |
| Zfp202 | 7.264070514 | 7.006586002 | 7.01320745 | -0.006621448 | 0.250863064 |
| Klrc2 | 3.670487625 | 3.513696466 | 3.419839309 | 0.093857157 | 0.250648316 |
| Ankrd17 | 11.56948347 | 11.44008736 | 11.31895789 | 0.12112947 | 0.25052558 |
| Zfp703 | 10.39171353 | 10.29769568 | 10.14126022 | 0.156435453 | 0.250453307 |
| L3mbtl2 | 8.592335378 | 8.502380715 | 8.342051413 | 0.160329302 | 0.250283965 |

| | | | | | |
|---|---|---|---|---|---|
| Fbrsl1 | 9.157635127 | 9.169457435 | 8.907372185 | 0.26208525 | 0.250262942 |
| Bdp1 | 10.13754315 | 10.03607018 | 9.887444921 | 0.14862526 | 0.250098229 |
| Ddx28 | 8.36463529 | 8.130121687 | 8.114780625 | 0.015341062 | 0.249854665 |
| Nras | 11.58922406 | 11.42789341 | 11.33940488 | 0.088488533 | 0.249819177 |
| Rest | 9.74922866 | 9.582039956 | 9.499695685 | 0.082344271 | 0.249532975 |
| Grpel1 | 10.01614615 | 9.841853282 | 9.766621388 | 0.075231894 | 0.249524758 |
| Pik3cg | 8.300354475 | 8.138199463 | 8.051160845 | 0.087038618 | 0.24919363 |
| Bok | 9.698725065 | 9.600353803 | 9.449647725 | 0.150706078 | 0.24907734 |
| Mgat1 | 10.51736395 | 10.37340634 | 10.26838514 | 0.1050212 | 0.248978813 |
| Hmga1b | 2.946603336 | 2.609394896 | 2.697651026 | -0.08825613 | 0.248952309 |
| BC117090 | 4.46885501 | 4.238251805 | 4.220073962 | 0.018177843 | 0.248781048 |
| Spink4 | 3.744873118 | 3.59795753 | 3.496263844 | 0.101693686 | 0.248609274 |
| B4galnt1 | 9.725954242 | 9.86838617 | 9.477557126 | 0.390829045 | 0.248397116 |
| Clcn5 | 9.417816508 | 9.369686916 | 9.169426322 | 0.200260594 | 0.248390186 |
| Ovca2 | 8.743230365 | 8.522529584 | 8.49496843 | 0.027561154 | 0.248261936 |
| Znhit1 | 8.549225621 | 8.566105733 | 8.300965874 | 0.265139859 | 0.248259747 |
| Mdm2 | 10.23360584 | 10.24611763 | 9.985397389 | 0.260720241 | 0.248208448 |
| Srrd | 7.077389343 | 6.822623458 | 6.829233356 | -0.006609898 | 0.248155987 |
| Sf1 | 10.6706816 | 10.63921884 | 10.42299266 | 0.216226183 | 0.247688937 |
| Gad1 | 4.202332184 | 4.256632145 | 3.954647927 | 0.301984217 | 0.247684257 |
| Aox4 | 10.03916365 | 9.752688923 | 9.791576315 | -0.038887392 | 0.247587336 |
| Ahrr | 7.261235505 | 7.039402263 | 7.013651601 | 0.025750662 | 0.247583904 |
| Ifnar1 | 10.16068011 | 10.1389589 | 9.913138244 | 0.225820659 | 0.247541863 |
| Pcdhga12 | 6.459521739 | 6.349738246 | 6.212018178 | 0.137720069 | 0.247503562 |
| Tlnrd1 | 8.594886663 | 8.600214875 | 8.34744307 | 0.252771806 | 0.247443593 |
| Psmb9 | 9.318212775 | 8.920918582 | 9.070801374 | -0.149882792 | 0.247411401 |
| Irf2bp1 | 8.759159613 | 8.834903818 | 8.511988717 | 0.322915101 | 0.247170896 |
| Fcf1 | 9.179039466 | 8.974514456 | 8.9319972 | 0.042517256 | 0.247042266 |
| A430078G23Rik | 4.993788263 | 4.890460009 | 4.746824196 | 0.143635814 | 0.246964067 |
| Tgtp2 | 7.614074393 | 6.969136704 | 7.367235815 | -0.398099111 | 0.246838578 |
| Kctd10 | 11.29747081 | 11.19110986 | 11.05073848 | 0.140371373 | 0.246732327 |
| Chst13 | 4.201713611 | 4.013855827 | 3.95507176 | 0.058784066 | 0.246641851 |

| | | | | | |
|---|---|---|---|---|---|
| Pnkp | 8.944281885 | 8.902755487 | 8.697827358 | 0.204928129 | 0.246454527 |
| Ppie | 9.054388193 | 8.922759092 | 8.807943744 | 0.114815348 | 0.246444449 |
| Mrps27 | 9.181235115 | 9.128720768 | 8.934907779 | 0.193812989 | 0.246327336 |
| Gbf1 | 10.66490167 | 10.69266966 | 10.41860067 | 0.274068983 | 0.246300997 |
| Polr2f | 9.355065578 | 9.248994047 | 9.108829006 | 0.140165041 | 0.246236572 |
| H2-Ab1 | 13.02595791 | 12.93589052 | 12.77973129 | 0.15615923 | 0.246226623 |
| Crk | 11.5729714 | 11.45138061 | 11.32676165 | 0.124618967 | 0.246209753 |
| Dgkd | 10.30961149 | 10.05003143 | 10.06349778 | -0.013466352 | 0.24611371 |
| Spen | 10.4289386 | 10.50385599 | 10.18291882 | 0.32093717 | 0.246019774 |
| Dnajb9 | 9.790045084 | 10.41465016 | 9.544030078 | 0.870620079 | 0.246015006 |
| Sec31a | 11.55157624 | 11.56116276 | 11.30566669 | 0.255496073 | 0.245909557 |
| Tmem147 | 10.2351703 | 10.05779693 | 9.989292608 | 0.068504318 | 0.245877692 |
| Dhx16 | 9.955984282 | 9.811010969 | 9.710111516 | 0.100899453 | 0.245872766 |
| Fam220a | 9.790195949 | 9.619592604 | 9.544346675 | 0.07524593 | 0.245849274 |
| Capg | 11.79359879 | 11.83348204 | 11.54779768 | 0.285684363 | 0.245801107 |
| Dio2 | 8.831585579 | 8.502784911 | 8.58578803 | -0.083003119 | 0.24579755 |
| Pdcd1 | 3.306862453 | 3.109017487 | 3.061088191 | 0.047929295 | 0.245774262 |
| Ifnlr1 | 8.70925479 | 8.506963222 | 8.463499853 | 0.043463369 | 0.245754937 |
| Gm10053 | 11.04526929 | 10.8757964 | 10.79964736 | 0.076149033 | 0.245621923 |
| Klhl30 | 7.772378443 | 8.397132838 | 7.526902803 | 0.870230035 | 0.24547564 |
| Dnajb6 | 11.30905587 | 11.03610662 | 11.06373652 | -0.0276299 | 0.245319347 |
| Kpna4 | 11.45624338 | 11.29623979 | 11.21096783 | 0.085271957 | 0.245275547 |
| Tmed9 | 11.75145217 | 11.69936367 | 11.506179 | 0.193184673 | 0.245273173 |
| Musk | 5.451173797 | 5.369568681 | 5.206015861 | 0.163552819 | 0.245157936 |
| Slc25a39 | 10.51441348 | 10.5716654 | 10.26942382 | 0.30224158 | 0.244989663 |
| Ddx23 | 10.62785362 | 10.5212552 | 10.38288552 | 0.138369687 | 0.244968107 |
| Gm826 | 4.328255591 | 4.152319889 | 4.083384428 | 0.068935461 | 0.244871163 |
| Tamm41 | 8.029204822 | 7.842551379 | 7.78443136 | 0.058120019 | 0.244773462 |
| Tra2a | 10.99353268 | 10.90797462 | 10.74909543 | 0.158879187 | 0.244437247 |
| Gcnt1 | 8.234863025 | 8.114384038 | 7.990486194 | 0.123897844 | 0.244376832 |
| Vps37c | 9.776355438 | 9.813899969 | 9.531989106 | 0.281910863 | 0.244366333 |
| P2ry14 | 6.653257355 | 6.229681133 | 6.409005163 | -0.17932403 | 0.244252192 |

| | | | | | |
|---|---|---|---|---|---|
| Mief1 | 9.399402338 | 9.433390667 | 9.155177873 | 0.278212794 | 0.244224464 |
| Il5ra | 4.435651561 | 4.320927156 | 4.191453159 | 0.129473997 | 0.244198402 |
| Rpl29 | 12.55409777 | 12.61819449 | 12.3101712 | 0.308023293 | 0.243926567 |
| Galnt4 | 9.027518348 | 8.732771834 | 8.783634573 | -0.050862739 | 0.243883774 |
| Gm46223 | 10.96679357 | 10.78575793 | 10.72329022 | 0.062467703 | 0.243503347 |
| Tango6 | 7.663799423 | 7.491598244 | 7.420483455 | 0.071114789 | 0.243315968 |
| Cxcl12 | 11.94624932 | 11.82608589 | 11.70301814 | 0.12306775 | 0.24323118 |
| Pkd1l2 | 4.031827137 | 3.897163211 | 3.788700271 | 0.10846294 | 0.243126867 |
| Sgf29 | 8.330739445 | 8.121591189 | 8.087747526 | 0.033843663 | 0.242991918 |
| Ppp1r14b | 10.68309675 | 10.68904228 | 10.44050511 | 0.24853717 | 0.242591643 |
| Tspan31 | 10.53794312 | 10.3032993 | 10.29535698 | 0.00794232 | 0.24258614 |
| Elf4 | 10.15079962 | 10.21030436 | 9.908238338 | 0.302066019 | 0.242561285 |
| Nup205 | 10.36331622 | 10.27161658 | 10.12090416 | 0.150712427 | 0.242412063 |
| Tmsb4x | 14.17872272 | 13.83427709 | 13.93634133 | -0.102064233 | 0.242381397 |
| Abhd17a | 9.047744462 | 9.18115275 | 8.805369007 | 0.375783743 | 0.242375455 |
| Nkg7 | 6.210856347 | 6.026139145 | 5.968489464 | 0.057649681 | 0.242366883 |
| Psmb4 | 11.55769184 | 11.40130904 | 11.31539741 | 0.085911633 | 0.24229443 |
| Hipk1 | 11.60247312 | 11.48105362 | 11.36031369 | 0.120739927 | 0.242159427 |
| Bcl2l1 | 10.43978091 | 10.42347693 | 10.19772684 | 0.225750093 | 0.24205407 |
| Zfp1 | 7.545234449 | 7.351317421 | 7.303293536 | 0.048023885 | 0.241940913 |
| Pus3 | 7.877536172 | 7.768887508 | 7.635763657 | 0.133123851 | 0.241772516 |
| Tmem8 | 8.050821941 | 7.958227712 | 7.809109004 | 0.149118709 | 0.241712938 |
| Gm10131 | 8.615510314 | 8.410064125 | 8.373831278 | 0.036232847 | 0.241679036 |
| Oaz1-ps | 8.604234769 | 8.471637993 | 8.362739405 | 0.108898588 | 0.241495363 |
| Prdx1 | 13.3750715 | 13.24728784 | 13.13359046 | 0.113697383 | 0.241481037 |
| Lin28a | 3.045696678 | 2.927160872 | 2.804267363 | 0.122893509 | 0.241429314 |
| Tesk1 | 9.056776687 | 9.264454854 | 8.815367163 | 0.449087691 | 0.241409523 |
| Cntn1 | 7.097724937 | 7.022815858 | 6.856369344 | 0.166446514 | 0.241355593 |
| Srp68 | 10.73283785 | 10.64877146 | 10.4915474 | 0.15722406 | 0.24129045 |
| Cd74 | 14.38153753 | 14.27015662 | 14.14025741 | 0.12989921 | 0.241280127 |
| Eif2b2 | 9.218746513 | 9.19590608 | 8.97754525 | 0.21836083 | 0.241201263 |
| Armc7 | 7.705212861 | 7.675885669 | 7.464059359 | 0.21182631 | 0.241153502 |

| | | | | | |
|---|---|---|---|---|---|
| Ctnnbl1 | 8.825012247 | 8.713978087 | 8.583897191 | 0.130080896 | 0.241115056 |
| Tfap2a | 10.35431543 | 10.38308591 | 10.11326996 | 0.269815943 | 0.241045463 |
| Cuzd1 | 4.800819654 | 4.729801834 | 4.559791873 | 0.170009962 | 0.241027782 |
| Gm13722 | 6.277559811 | 6.242990333 | 6.036653486 | 0.206336848 | 0.240906326 |
| Unc93a | 3.284523558 | 3.107697563 | 3.043619147 | 0.064078416 | 0.240904412 |
| Eed | 9.569449087 | 9.547120964 | 9.32855438 | 0.218566584 | 0.240894707 |
| Yap1 | 12.15474804 | 11.99825697 | 11.91389771 | 0.084359267 | 0.24085033 |
| Ltf | 3.278729999 | 3.028086232 | 3.037977105 | -0.009890872 | 0.240752895 |
| Rpl18 | 12.99538606 | 12.88908551 | 12.75465637 | 0.134429143 | 0.24072969 |
| Rfc2 | 9.308444116 | 9.305582527 | 9.067826323 | 0.237756204 | 0.240617794 |
| Gpr27 | 6.396351504 | 6.130605126 | 6.155757174 | -0.025152047 | 0.24059433 |
| Zcchc9 | 8.803545214 | 8.755769621 | 8.563236225 | 0.192533396 | 0.240308989 |
| Atp6v0a1 | 11.05601355 | 11.03317085 | 10.81590493 | 0.217265917 | 0.240108623 |
| Nnmt | 9.325845677 | 8.925222213 | 9.08581103 | -0.160588817 | 0.240034647 |
| Aadacl4 | 10.00826383 | 9.788906154 | 9.768589377 | 0.020316777 | 0.23967445 |
| Tdg | 10.57465718 | 10.46027773 | 10.33500777 | 0.125269967 | 0.239649413 |
| Smim15 | 9.965175469 | 9.738291758 | 9.72556111 | 0.012730647 | 0.239614359 |
| Elk3 | 9.25118433 | 8.958571524 | 9.011627418 | -0.053055894 | 0.239556911 |
| Emc10 | 10.30576364 | 10.42034732 | 10.06620944 | 0.354137873 | 0.239554197 |
| Vdac2 | 11.40764844 | 11.25399278 | 11.16835316 | 0.08563962 | 0.239295273 |
| Cdyl | 8.978018149 | 8.729051359 | 8.738736678 | -0.00968532 | 0.239281471 |
| Zfp384 | 10.11376845 | 10.11001618 | 9.874581707 | 0.235434473 | 0.23918674 |
| Acot7 | 8.404372672 | 8.417374941 | 8.165332029 | 0.252042912 | 0.239040643 |
| Rps9 | 13.94648532 | 13.85116701 | 13.70747436 | 0.14369265 | 0.23901096 |
| Rbsn | 10.1845936 | 10.06794318 | 9.945616196 | 0.122326981 | 0.238977404 |
| Rnf135 | 7.302165711 | 6.952181576 | 7.063390245 | -0.111208668 | 0.238775467 |
| Zswim8 | 10.69829701 | 10.68608372 | 10.45960609 | 0.22647763 | 0.23869092 |
| Gm10925 | 14.32487054 | 14.29976959 | 14.08627179 | 0.213497807 | 0.238598757 |
| Efl1 | 9.210770093 | 9.079070613 | 8.97229699 | 0.106773622 | 0.238473102 |
| Vps13d | 11.05204141 | 10.86968081 | 10.81360932 | 0.05607149 | 0.238432087 |
| Nudt5 | 10.11909528 | 10.03772512 | 9.880783529 | 0.156941594 | 0.238311755 |
| Exosc8 | 9.338237831 | 9.218904733 | 9.099972703 | 0.11893203 | 0.238265128 |

| | | | | | |
|---|---|---|---|---|---|
| Foxn2 | 10.3086057 | 10.17128955 | 10.07046304 | 0.100826513 | 0.238142667 |
| Hps4 | 7.958530771 | 8.095285528 | 7.720411025 | 0.374874503 | 0.238119746 |
| Caml | 8.642007038 | 8.497699133 | 8.403896556 | 0.093802578 | 0.238110482 |
| Slc39a6 | 10.67525846 | 10.60564333 | 10.43722798 | 0.168415347 | 0.238030473 |
| Polr2c | 9.570256258 | 9.397523851 | 9.332719314 | 0.064804537 | 0.237536944 |
| Nap1l1 | 12.06350474 | 11.91274755 | 11.82606128 | 0.086686267 | 0.23744346 |
| Vmn1r65 | 5.241484328 | 4.896971203 | 5.004134634 | -0.107163431 | 0.237349694 |
| Flvcr1 | 9.211651797 | 8.939821384 | 8.974370063 | -0.034548679 | 0.237281734 |
| Cstb | 11.31538319 | 11.28885984 | 11.07814082 | 0.210719017 | 0.237242373 |
| Kansl2 | 10.24167601 | 10.07878375 | 10.00448001 | 0.074303735 | 0.237195992 |
| Mrps17 | 9.819758659 | 9.684919195 | 9.582584028 | 0.102335167 | 0.237174631 |
| Trp53rka | 8.376173532 | 8.124277304 | 8.139105099 | -0.014827794 | 0.237068434 |
| Abcf1 | 11.59688014 | 11.35884045 | 11.35981645 | -0.000976007 | 0.23706369 |
| Polr2a | 11.62669516 | 11.6167267 | 11.38983769 | 0.226889013 | 0.23685747 |
| Dbnl | 11.58395728 | 11.51920197 | 11.34715777 | 0.172044193 | 0.236799503 |
| Eif2s3x | 12.59414516 | 12.36892609 | 12.35745811 | 0.011467983 | 0.23668705 |
| Skp1a | 12.34448664 | 12.11698364 | 12.107849 | 0.009134637 | 0.236637637 |
| Ufm1 | 10.27234077 | 10.21878005 | 10.03576425 | 0.183015793 | 0.236576513 |
| Esam | 8.900651779 | 8.620829405 | 8.664164362 | -0.043334957 | 0.236487417 |
| 1600014C10Rik | 9.605136913 | 9.373750234 | 9.368700069 | 0.005050165 | 0.236436844 |
| Hgsnat | 9.812756791 | 9.525060973 | 9.576346439 | -0.051285466 | 0.236410351 |
| Sbno1 | 11.30326442 | 11.23922604 | 11.0669744 | 0.172251643 | 0.236290027 |
| Lefty1 | 6.862761732 | 6.744671261 | 6.626484765 | 0.118186496 | 0.236276966 |
| Hopx | 10.67601555 | 10.44563843 | 10.43979504 | 0.005843393 | 0.236220517 |
| Lck | 6.373034538 | 6.174806299 | 6.136886699 | 0.0379196 | 0.236147839 |
| Trim47 | 8.914268558 | 8.982981995 | 8.678278506 | 0.304703489 | 0.235990053 |
| Ncr1 | 3.273179373 | 3.078366113 | 3.037245572 | 0.041120541 | 0.235933801 |
| Mybl2 | 7.470784438 | 7.525429966 | 7.234861579 | 0.290568387 | 0.235922859 |
| Pim2 | 7.447270113 | 7.462144546 | 7.211436632 | 0.250707913 | 0.235833481 |
| Cdc37 | 11.39048076 | 11.38216598 | 11.15471805 | 0.227447923 | 0.235762703 |
| Zc3h12d | 6.129009564 | 6.043435052 | 5.893485018 | 0.149950034 | 0.235524546 |
| Atg16l1 | 9.811469187 | 9.680028818 | 9.575970047 | 0.104058771 | 0.23549914 |

| | | | | | |
|---|---|---|---|---|---|
| Nrtn | 5.629788071 | 5.86751609 | 5.39442563 | 0.473090461 | 0.235362442 |
| Gm7694 | 7.804690055 | 7.664381217 | 7.569336108 | 0.095045109 | 0.235353947 |
| Map4k2 | 8.279493476 | 8.205785726 | 8.044459987 | 0.161325739 | 0.235033489 |
| Il17re | 8.924625416 | 9.04329063 | 8.689611145 | 0.353679485 | 0.235014271 |
| Prpf3 | 9.325142347 | 9.206420703 | 9.090130754 | 0.116289949 | 0.235011593 |
| Esrra | 9.421362751 | 9.69055792 | 9.186534153 | 0.504023767 | 0.234828597 |
| Tmem167 | 10.45548419 | 10.35893171 | 10.22074442 | 0.13818729 | 0.234739773 |
| Eid3 | 5.408989333 | 6.095079403 | 5.17437235 | 0.920707054 | 0.234616983 |
| Cnbd2 | 5.645686273 | 5.631311893 | 5.41121696 | 0.220094933 | 0.234469313 |
| Serbp1 | 12.96034075 | 12.83496154 | 12.72606279 | 0.108898743 | 0.234277957 |
| Napa | 11.28512366 | 11.40164186 | 11.05102677 | 0.35061509 | 0.234096893 |
| Cdk7 | 9.188327442 | 9.116982527 | 8.954238807 | 0.162743719 | 0.234088635 |
| CAAA01118383.1 | 7.585633145 | 7.648516967 | 7.351557618 | 0.296959349 | 0.234075527 |
| Gm8325 | 5.994296805 | 5.431095997 | 5.760362982 | -0.329266985 | 0.233933823 |
| Adrb2 | 8.701227845 | 8.624835552 | 8.46734963 | 0.157485922 | 0.233878215 |
| Med28 | 9.656679997 | 9.544752952 | 9.422811195 | 0.121941757 | 0.233868802 |
| Trmt11 | 7.843985776 | 7.648928008 | 7.610286876 | 0.038641132 | 0.2336989 |
| Prrc1 | 10.45493455 | 10.37312094 | 10.22126897 | 0.151851975 | 0.233665588 |
| Ndel1 | 11.80491528 | 11.69860634 | 11.57133616 | 0.12727018 | 0.23357912 |
| Dcp2 | 10.08768664 | 9.833468412 | 9.85419254 | -0.020724128 | 0.233494096 |
| Taf10 | 10.24356073 | 10.01818841 | 10.01007585 | 0.008112564 | 0.233484881 |
| Nubp1 | 9.2259726 | 9.195286365 | 8.992518326 | 0.202768039 | 0.233454274 |
| Taf12 | 9.104376153 | 9.009222888 | 8.871055812 | 0.138167075 | 0.233320341 |
| Begain | 5.471275904 | 5.528901967 | 5.237965521 | 0.290936446 | 0.233310383 |
| Herc4 | 10.64666188 | 10.40276764 | 10.4134092 | -0.010641563 | 0.233252683 |
| Zbtb48 | 7.509035926 | 7.534621733 | 7.276014305 | 0.258607428 | 0.233021621 |
| Khdrbs3 | 7.888774851 | 7.663429678 | 7.655840249 | 0.007589429 | 0.232934602 |
| Il27ra | 5.654057609 | 5.608546905 | 5.421138488 | 0.187408416 | 0.232919121 |
| Ccdc85b | 7.304447178 | 7.272522314 | 7.071532393 | 0.200989921 | 0.232914785 |
| Gatad2a | 11.17531959 | 11.25324365 | 10.94258218 | 0.31066147 | 0.232737413 |
| Nfya | 8.899403275 | 8.952927144 | 8.666701274 | 0.28622587 | 0.232702002 |
| Spcs3 | 11.06591113 | 10.90788037 | 10.83321754 | 0.07466283 | 0.23269359 |

| | | | | | |
|---|---|---|---|---|---|
| Btg3 | 9.385506366 | 9.251592412 | 9.152874504 | 0.098717908 | 0.232631862 |
| Rps6 | 13.4206841 | 13.37179342 | 13.18810293 | 0.18369049 | 0.23258117 |
| Nufip2 | 10.15268932 | 10.12753781 | 9.92026514 | 0.20727267 | 0.23242418 |
| Mrps26 | 9.110949099 | 8.984422579 | 8.878579533 | 0.105843046 | 0.232369566 |
| Gosr2 | 11.11634995 | 10.92932932 | 10.88409142 | 0.045237903 | 0.232258533 |
| Cct4 | 11.95108873 | 11.84171466 | 11.7188378 | 0.122876857 | 0.232250933 |
| Leprotl1 | 10.0542952 | 9.868942827 | 9.822087828 | 0.046854999 | 0.232207371 |
| E030030I06Rik | 5.436866533 | 5.38495304 | 5.204790338 | 0.180162702 | 0.232076195 |
| Pcdhga11 | 7.446427863 | 7.232006033 | 7.214434731 | 0.017571302 | 0.231993132 |
| Nsfl1c | 10.73928857 | 10.55669248 | 10.50734543 | 0.049347053 | 0.23194314 |
| Rrp1 | 11.3107728 | 11.1370584 | 11.07885881 | 0.058199583 | 0.231913983 |
| Ankrd13b | 8.099440775 | 8.218477878 | 7.867631904 | 0.350845973 | 0.231808871 |
| Tomm6 | 10.23059312 | 10.14238883 | 9.998795252 | 0.143593574 | 0.231797864 |
| Rexo1 | 9.267862897 | 9.450070967 | 9.036149274 | 0.413921694 | 0.231713624 |
| Snx15 | 8.542851608 | 8.562491342 | 8.311392724 | 0.251098618 | 0.231458884 |
| Tuba1b | 13.57387089 | 13.46471118 | 13.34241246 | 0.122298723 | 0.23145843 |
| Uxt | 8.156640273 | 8.028272697 | 7.925369843 | 0.102902854 | 0.23127043 |
| Cnpy3 | 9.393004457 | 9.414034431 | 9.161964403 | 0.252070028 | 0.231040054 |
| Cst7 | 5.849551399 | 5.689767814 | 5.618523981 | 0.071243833 | 0.231027418 |
| En1 | 5.840409089 | 5.796995316 | 5.609399703 | 0.187595613 | 0.231009387 |
| Mtpn | 11.71160364 | 11.5659408 | 11.48062192 | 0.085318877 | 0.230981713 |
| Lbhd1 | 7.827746544 | 7.744437885 | 7.596832614 | 0.147605271 | 0.23091393 |
| Wdr83os | 9.736228746 | 9.661776308 | 9.505322411 | 0.156453897 | 0.230906336 |
| Spire1 | 9.674368976 | 9.433719125 | 9.44355204 | -0.009832915 | 0.230816936 |
| Vps28 | 10.26072678 | 10.04066941 | 10.03011357 | 0.010555834 | 0.230613209 |
| Ypel2 | 9.775614284 | 9.320641029 | 9.545077788 | -0.224436759 | 0.230536496 |
| Apol6 | 10.17092096 | 9.508661251 | 9.94038814 | -0.431726888 | 0.23053282 |
| Gna13 | 11.24186387 | 11.18726458 | 11.01139948 | 0.1758651 | 0.230464393 |
| Brms1 | 9.052098623 | 8.973822464 | 8.821719808 | 0.152102656 | 0.230378815 |
| Slc39a10 | 10.2117076 | 10.2775198 | 9.981370117 | 0.296149683 | 0.230337483 |
| Frmd4a | 9.544601116 | 9.511485107 | 9.31429618 | 0.197188927 | 0.230304936 |
| Ndufb6 | 9.508639924 | 9.496776701 | 9.278345886 | 0.218430815 | 0.230294038 |

| | | | | | |
|---|---|---|---|---|---|
| Spty2d1 | 9.996401006 | 9.956301046 | 9.766145315 | 0.190155731 | 0.230255691 |
| Tubb2a | 11.16428776 | 11.29943397 | 10.9342442 | 0.365189777 | 0.230043563 |
| Tmem208 | 9.115426237 | 9.058003557 | 8.885473542 | 0.172530015 | 0.229952695 |
| Ranbp3 | 10.12452651 | 10.16148235 | 9.894594006 | 0.266888347 | 0.2299325 |
| Awat1 | 10.71624964 | 10.55448604 | 10.48631802 | 0.068168017 | 0.229931617 |
| Sco1 | 7.864987723 | 7.638098312 | 7.63509095 | 0.003007362 | 0.229896773 |
| Ccnl2 | 11.38774189 | 11.45242133 | 11.15786451 | 0.294556827 | 0.229877387 |
| Gm10698 | 8.506483446 | 8.745407522 | 8.276659896 | 0.468747626 | 0.22982355 |
| Med24 | 9.967135243 | 9.987696936 | 9.737328102 | 0.250368835 | 0.229807141 |
| AI467606 | 7.058159322 | 6.853222578 | 6.828461682 | 0.024760896 | 0.22969764 |
| Mrap | 8.311521011 | 7.522720821 | 8.081881602 | -0.559160781 | 0.229639409 |
| Pnn | 10.73882083 | 10.72299154 | 10.50926148 | 0.213730057 | 0.22955935 |
| Dusp7 | 11.29893237 | 11.28968555 | 11.06938122 | 0.22030433 | 0.229551153 |
| Naaa | 10.61180728 | 10.32244874 | 10.38226422 | -0.05981548 | 0.229543057 |
| Sp2 | 8.467867084 | 8.472508079 | 8.23833478 | 0.234173299 | 0.229532303 |
| Soga1 | 8.289493935 | 8.394876295 | 8.059968427 | 0.334907868 | 0.229525508 |
| Krtcap2 | 9.211219993 | 9.207506816 | 8.981853991 | 0.225652825 | 0.229366002 |
| Mcm4 | 10.41187835 | 10.31320263 | 10.1825285 | 0.130674127 | 0.229349847 |
| Pkn1 | 9.644781224 | 9.710093893 | 9.415441359 | 0.294652534 | 0.229339865 |
| Nup50 | 10.16674544 | 10.14855308 | 9.937666383 | 0.210886701 | 0.229079061 |
| Aoah | 6.694313453 | 6.53126774 | 6.465272447 | 0.065995293 | 0.229041006 |
| Rin2 | 11.11613155 | 10.8770649 | 10.88716391 | -0.010099017 | 0.22896764 |
| Lsm5 | 7.886780964 | 7.729253735 | 7.657946922 | 0.071306813 | 0.228834042 |
| Arv1 | 6.783432657 | 6.78084176 | 6.554631672 | 0.226210088 | 0.228800986 |
| Tmem132e | 4.058315541 | 3.778551693 | 3.8296438 | -0.051092107 | 0.228671742 |
| Morc3 | 9.934357995 | 9.818739177 | 9.705762635 | 0.112976543 | 0.228595361 |
| Fam104a | 9.320896936 | 9.215549546 | 9.092349027 | 0.123200519 | 0.22854791 |
| Tmem128 | 8.851851474 | 8.702851845 | 8.623395026 | 0.079456818 | 0.228456448 |
| Slc35e4 | 9.450779588 | 9.546414451 | 9.222518317 | 0.323896135 | 0.228261271 |
| Cklf | 7.299969171 | 7.169329928 | 7.071732051 | 0.097597878 | 0.22823712 |
| Gm13124 | 6.259218419 | 6.059193164 | 6.031272525 | 0.027920639 | 0.227945894 |
| Icos | 5.756054546 | 5.547715984 | 5.528345099 | 0.019370886 | 0.227709447 |

| | | | | | |
|---|---|---|---|---|---|
| Adgrb2 | 6.933475984 | 7.230965798 | 6.705882268 | 0.52508353 | 0.227593716 |
| Wdr89 | 6.772979619 | 6.621126374 | 6.545530594 | 0.07559578 | 0.227449025 |
| Fnbp4 | 10.56505314 | 10.47359826 | 10.33765724 | 0.135941023 | 0.227395903 |
| Nfx1 | 10.68771692 | 10.49871709 | 10.46062002 | 0.038097063 | 0.2270969 |
| Rbmx2 | 7.292840085 | 7.109787509 | 7.065814708 | 0.043972802 | 0.227025378 |
| Arf1 | 12.81173303 | 12.74042982 | 12.58489614 | 0.15553368 | 0.226836893 |
| Pkig | 8.786154653 | 8.744360782 | 8.559379987 | 0.184980795 | 0.226774666 |
| Ctdnep1 | 10.84729004 | 10.74598869 | 10.62067695 | 0.125311743 | 0.226613097 |
| Mgp | 8.947883154 | 8.469775203 | 8.721275311 | -0.251500109 | 0.226607843 |
| Tsr3 | 8.218196384 | 8.054340355 | 7.991662316 | 0.062678039 | 0.226534068 |
| Malsu1 | 7.921460159 | 7.704885121 | 7.694964605 | 0.009920516 | 0.226495554 |
| Rmc1 | 9.525009913 | 9.322024658 | 9.298560803 | 0.023463854 | 0.22644911 |
| Ggnbp2 | 11.03463776 | 10.87449757 | 10.8083282 | 0.06616937 | 0.22630956 |
| Slirp | 8.879448344 | 8.750205142 | 8.653175724 | 0.097029417 | 0.22627262 |
| Ythdf1 | 10.47441218 | 10.32127216 | 10.24821448 | 0.07305768 | 0.226197703 |
| 1810009A15Rik | 8.554257051 | 8.455676614 | 8.328096538 | 0.127580076 | 0.226160513 |
| Tmem201 | 8.533273539 | 8.572888323 | 8.307144678 | 0.265743645 | 0.226128861 |
| Dgat2l6 | 8.844581032 | 8.735121254 | 8.618469957 | 0.116651296 | 0.226111074 |
| Plaa | 10.79771943 | 10.7041478 | 10.57182593 | 0.13232187 | 0.225893507 |
| Cebpe | 3.368266866 | 3.228172543 | 3.142425653 | 0.08574689 | 0.225841212 |
| Mrpl35 | 9.080095435 | 8.770937139 | 8.854421044 | -0.083483905 | 0.225674392 |
| Smc1a | 11.50927342 | 11.34653499 | 11.28360032 | 0.062934663 | 0.225673093 |
| Rnmt | 9.479405392 | 9.198541225 | 9.253815287 | -0.055274061 | 0.225590105 |
| Zfp598 | 10.43677122 | 10.5848203 | 10.21122594 | 0.373594363 | 0.225545283 |
| Atp2a3 | 8.836969202 | 8.60924684 | 8.611458877 | -0.002212037 | 0.225510325 |
| Med31 | 7.401765931 | 7.217313452 | 7.176335197 | 0.040978255 | 0.225430734 |
| Trmt13 | 7.777163918 | 7.562221781 | 7.551760719 | 0.010461062 | 0.225403199 |
| Sinhcaf | 9.244201866 | 9.032213016 | 9.018816761 | 0.013396255 | 0.225385104 |
| Snd1 | 11.33614209 | 11.3273453 | 11.11078318 | 0.216562127 | 0.22535891 |
| Gm14654 | 4.328787924 | 4.255270705 | 4.103437069 | 0.151833636 | 0.225350855 |
| H2-DMb1 | 10.01975713 | 9.835226285 | 9.794519138 | 0.040707147 | 0.22523799 |
| Strn4 | 10.61509448 | 10.68243647 | 10.38991295 | 0.292523527 | 0.22518153 |

| | | | | | |
|---|---|---|---|---|---|
| Plekhj1 | 8.847435009 | 8.68507126 | 8.622279702 | 0.062791558 | 0.225155307 |
| Gm48269 | 6.175405497 | 5.974873131 | 5.950269985 | 0.024603146 | 0.225135512 |
| Gm21994 | 5.708149045 | 5.517294146 | 5.483014603 | 0.034279543 | 0.225134442 |
| Nlrp1a | 5.044291229 | 4.906666329 | 4.819170679 | 0.08749565 | 0.22512055 |
| Mrps7 | 9.75319453 | 9.682172407 | 9.528074635 | 0.154097772 | 0.225119895 |
| Col8a1 | 7.398675561 | 7.378884838 | 7.173599439 | 0.205285399 | 0.225076122 |
| Gm28036 | 8.999056706 | 8.787595467 | 8.774233007 | 0.01336246 | 0.224823699 |
| Thoc5 | 9.536356353 | 9.427394228 | 9.311535691 | 0.115858537 | 0.224820663 |
| Gm16464 | 3.340566054 | 3.270552554 | 3.11574901 | 0.154803544 | 0.224817044 |
| Ssrp1 | 11.7984327 | 11.61645864 | 11.57382006 | 0.04263858 | 0.22461264 |
| 1110038F14Rik | 8.290244531 | 8.197740132 | 8.065691676 | 0.132048455 | 0.224552854 |
| Polr1d | 11.00269524 | 10.84623621 | 10.77818124 | 0.068054977 | 0.224514 |
| Rnf125 | 6.878487315 | 6.572107955 | 6.654014733 | -0.081906778 | 0.224472582 |
| Phf11c | 6.038096573 | 5.915444359 | 5.813641802 | 0.101802557 | 0.224454771 |
| Ocstamp | 4.69005579 | 4.600723007 | 4.465623518 | 0.13509949 | 0.224432272 |
| Prkrip1 | 8.191867211 | 8.070478562 | 7.967460624 | 0.103017938 | 0.224406587 |
| Dpm3 | 6.877220856 | 7.13101878 | 6.652880769 | 0.478138011 | 0.224340087 |
| Romo1 | 8.168242285 | 8.262253379 | 7.943983148 | 0.318270231 | 0.224259137 |
| Dlst | 11.6654188 | 11.4566917 | 11.44117668 | 0.015515027 | 0.22424212 |
| E2f5 | 7.569645232 | 7.261184907 | 7.345455373 | -0.084270466 | 0.22418986 |
| Yju2 | 7.500448534 | 7.414724982 | 7.276306011 | 0.138418971 | 0.224142523 |
| Slc7a6os | 9.184680725 | 9.161582732 | 8.960632616 | 0.200950116 | 0.224048109 |
| Msto1 | 8.583659569 | 8.495273218 | 8.359681646 | 0.135591572 | 0.223977924 |
| Hmces | 8.06632049 | 7.875959434 | 7.842545433 | 0.033414001 | 0.223775057 |
| B3gnt2 | 8.210309755 | 8.062158899 | 7.98655541 | 0.07560349 | 0.223754345 |
| Sdc1 | 13.1810893 | 13.10485578 | 12.95745306 | 0.147402713 | 0.223636237 |
| Vamp4 | 9.804495868 | 9.495473051 | 9.580862779 | -0.085389727 | 0.22363309 |
| Sh2d6 | 3.598265065 | 3.504118136 | 3.374658925 | 0.129459211 | 0.22360614 |
| Rpp30 | 7.735711042 | 7.510469902 | 7.512218585 | -0.001748683 | 0.223492457 |
| 1700037C18Rik | 5.674165186 | 5.552115927 | 5.450675189 | 0.101440738 | 0.223489997 |
| Foxred1 | 9.144972833 | 8.996860459 | 8.921519301 | 0.075341158 | 0.223453532 |
| Pcdhgb7 | 7.092389026 | 6.967094677 | 6.86895432 | 0.098140357 | 0.223434706 |

| | | | | | |
|---|---|---|---|---|---|
| Ppp4r2 | 11.26292455 | 11.12653064 | 11.03949379 | 0.087036857 | 0.223430763 |
| Iws1 | 10.09149498 | 9.971418578 | 9.868270701 | 0.103147877 | 0.223224282 |
| 9430038I01Rik | 6.819028841 | 6.64251454 | 6.595928442 | 0.046586098 | 0.223100398 |
| Il11 | 4.549686196 | 4.496404558 | 4.326625817 | 0.169778741 | 0.223060378 |
| Snip1 | 8.877236606 | 8.833459549 | 8.654330614 | 0.179128935 | 0.222905993 |
| Hnrnpu | 12.10933239 | 12.02846988 | 11.88653371 | 0.141936177 | 0.22279868 |
| Adap2 | 7.269995602 | 7.000692689 | 7.0472803 | -0.046587611 | 0.222715302 |
| Matk | 6.054021025 | 5.975327634 | 5.831389655 | 0.143937979 | 0.22263137 |
| Fam98b | 9.291469427 | 9.133127103 | 9.068939514 | 0.064187589 | 0.222529912 |
| Olfr1388 | 7.389424995 | 7.367966528 | 7.166947847 | 0.201018681 | 0.222477148 |
| Rpl8 | 13.64401636 | 13.63410226 | 13.42173723 | 0.212365027 | 0.222279127 |
| Adnp2 | 9.104886487 | 9.188694857 | 8.882775844 | 0.305919013 | 0.222110643 |
| Osbpl3 | 9.780539124 | 9.634168812 | 9.558440056 | 0.075728756 | 0.222099067 |
| Mtmr10 | 9.770891045 | 9.769710568 | 9.54887738 | 0.220833188 | 0.222013665 |
| Pfdn4 | 7.989857229 | 7.815501643 | 7.767873578 | 0.047628065 | 0.221983651 |
| Dpysl3 | 9.891488079 | 9.644391259 | 9.669520643 | -0.025129384 | 0.221967436 |
| Chd9 | 12.11514991 | 11.94196001 | 11.89322846 | 0.048731557 | 0.221921457 |
| Atp5d | 11.26318153 | 11.3170345 | 11.0412772 | 0.275757307 | 0.221904337 |
| Bub3 | 10.33959093 | 10.27763456 | 10.11771031 | 0.159924253 | 0.221880626 |
| Gm15501 | 4.102112992 | 3.895280285 | 3.880253411 | 0.015026874 | 0.221859581 |
| 2510039O18Rik | 9.8007446 | 9.89042069 | 9.57892526 | 0.311495429 | 0.221819339 |
| Fkbp1a | 11.59849065 | 11.41876359 | 11.37678655 | 0.041977037 | 0.221704093 |
| Bend6 | 5.067909478 | 4.780029124 | 4.846303039 | -0.066273916 | 0.221606439 |
| Sapcd2 | 5.739720897 | 5.638561864 | 5.51812869 | 0.120433173 | 0.221592207 |
| Cenpm | 5.861767394 | 5.848331969 | 5.640183734 | 0.208148236 | 0.22158366 |
| Hsp90b1 | 13.31696944 | 13.48430788 | 13.09547226 | 0.388835617 | 0.221497177 |
| Dctn3 | 9.405579989 | 9.33792732 | 9.184128057 | 0.153799263 | 0.221451932 |
| Suv39h2 | 6.951477055 | 6.77481341 | 6.730186018 | 0.044627392 | 0.221291037 |
| Zfp366 | 7.406282177 | 7.626647158 | 7.185017667 | 0.441629491 | 0.22126451 |
| Rnase2b | 9.713748294 | 9.559194054 | 9.492593212 | 0.066600843 | 0.221155082 |
| Pdia4 | 11.25468315 | 11.37346224 | 11.03363267 | 0.339829567 | 0.22105048 |
| Eln | 10.37671676 | 10.68477456 | 10.15567822 | 0.529096339 | 0.221038532 |

| | | | | | |
|---|---|---|---|---|---|
| 3110001I22Rik | 5.823089358 | 5.745572095 | 5.602331638 | 0.143240457 | 0.22075772 |
| Olfr1216 | 3.621941229 | 3.458716395 | 3.401192802 | 0.057523593 | 0.220748428 |
| Olfr1122 | 4.986990686 | 5.002329675 | 4.766327882 | 0.236001793 | 0.220662803 |
| Prkx | 10.58625979 | 10.44113956 | 10.36570608 | 0.075433483 | 0.22055371 |
| Psmg4 | 6.81296652 | 6.716449649 | 6.592616217 | 0.123833432 | 0.220350303 |
| Pigt | 10.59640564 | 10.57060262 | 10.37610196 | 0.194500657 | 0.22030368 |
| Trmo | 6.728556183 | 6.694081011 | 6.50830065 | 0.185780361 | 0.220255533 |
| Nfatc4 | 9.082957588 | 9.101036914 | 8.862845242 | 0.238191672 | 0.220112346 |
| Eif2b1 | 9.693960428 | 9.519698376 | 9.473900468 | 0.045797908 | 0.22005996 |
| Sf3b6 | 10.36248251 | 10.19535409 | 10.14243685 | 0.052917243 | 0.22004566 |
| Dock8 | 9.818480515 | 9.739409147 | 9.598438377 | 0.14097077 | 0.220042138 |
| Mrpl30 | 9.814881895 | 9.614242739 | 9.594841655 | 0.019401084 | 0.22004024 |
| Cd47 | 11.98651115 | 11.63105456 | 11.7665432 | -0.135488643 | 0.21996795 |
| Sec61b | 9.346577895 | 9.380728591 | 9.126991858 | 0.253736733 | 0.219586037 |
| Ppp1r8 | 9.539013603 | 9.384646812 | 9.319648099 | 0.064998713 | 0.219365504 |
| Fads3 | 10.26125191 | 9.920040185 | 10.04193946 | -0.121899279 | 0.219312443 |
| Abcf3 | 10.20440006 | 10.20600286 | 9.98511712 | 0.22088574 | 0.219282937 |
| Podxl | 9.908708622 | 9.797387843 | 9.689508691 | 0.107879152 | 0.219199931 |
| Ehd4 | 10.7896761 | 10.75965701 | 10.57050105 | 0.189155967 | 0.21917505 |
| Srsf10 | 11.06846219 | 10.93320627 | 10.84931483 | 0.08389144 | 0.219147357 |
| Cxcl16 | 10.3614163 | 10.31182943 | 10.14227336 | 0.169556077 | 0.219142947 |
| Myo19 | 9.734938298 | 9.806578301 | 9.515870091 | 0.290708211 | 0.219068207 |
| Ciao2a | 9.300054688 | 9.156972056 | 9.08111777 | 0.075854286 | 0.218936917 |
| Atp6v0b | 10.97475838 | 10.97660055 | 10.75593569 | 0.220664863 | 0.218822687 |
| Mlx | 9.627985062 | 9.523543148 | 9.409246337 | 0.114296811 | 0.218738725 |
| Ripk2 | 8.932003857 | 8.667601028 | 8.713284174 | -0.045683146 | 0.218719682 |
| Uqcr10 | 10.10835181 | 10.20015257 | 9.889910077 | 0.310242496 | 0.218441735 |
| Vapa | 11.29388039 | 11.14981668 | 11.07544445 | 0.074372227 | 0.218435933 |
| Trnau1ap | 8.897681791 | 8.620474153 | 8.679267697 | -0.058793544 | 0.218414094 |
| BC023105 | 7.091774578 | 6.088467232 | 6.873481406 | -0.785014174 | 0.218293172 |
| Gps1 | 10.62572977 | 10.52801099 | 10.4075422 | 0.120468787 | 0.21818757 |
| Lgals7 | 14.47000628 | 14.45727444 | 14.25185119 | 0.20542325 | 0.218155093 |

| | | | | | |
|---|---|---|---|---|---|
| Tespa1 | 5.09088287 | 4.848255949 | 4.872745571 | -0.024489622 | 0.218137299 |
| Snx5 | 11.63517894 | 11.47350265 | 11.41717349 | 0.056329157 | 0.218005443 |
| Irs2 | 9.054396159 | 9.541302326 | 8.836641118 | 0.704661208 | 0.217755041 |
| Ppm1f | 9.505604826 | 9.428904401 | 9.287892695 | 0.141011706 | 0.21771213 |
| Slc12a4 | 10.2612544 | 10.43592104 | 10.04356765 | 0.392353387 | 0.217686747 |
| Arl6ip5 | 11.01620167 | 10.93761823 | 10.79864094 | 0.138977287 | 0.217560727 |
| Gtf2e1 | 8.510258904 | 8.33089603 | 8.29294778 | 0.03794825 | 0.217311124 |
| Gdf15 | 4.015585115 | 4.363984139 | 3.798428952 | 0.565555187 | 0.217156162 |
| Dnajc1 | 9.761301669 | 9.711843992 | 9.544283256 | 0.167560736 | 0.217018413 |
| Ankle2 | 10.04893681 | 9.672809635 | 9.831998381 | -0.159188746 | 0.216938432 |
| Atp6v1c1 | 9.178814638 | 9.125762527 | 8.961920769 | 0.163841758 | 0.216893868 |
| Tmem222 | 9.452723322 | 9.395879845 | 9.235975129 | 0.159904716 | 0.216748193 |
| Tlr9 | 6.900550376 | 6.875836955 | 6.683882132 | 0.191954823 | 0.216668244 |
| Selenok | 10.5215186 | 10.26562665 | 10.3049436 | -0.039316953 | 0.216574993 |
| Gnb1l | 6.937451838 | 6.759173593 | 6.720972554 | 0.03820104 | 0.216479285 |
| Ppp1r16b | 7.085554939 | 6.713927845 | 6.869145139 | -0.155217294 | 0.2164098 |
| H2afy | 11.17247843 | 11.07291297 | 10.95618641 | 0.116726557 | 0.21629202 |
| Eftud2 | 10.79865091 | 10.77031906 | 10.58238559 | 0.187933473 | 0.216265327 |
| Ldha | 14.30988688 | 14.08928468 | 14.09366932 | -0.004384637 | 0.216217563 |
| Mboat7 | 10.59163754 | 10.49951739 | 10.37557002 | 0.123947373 | 0.21606752 |
| Fndc3b | 10.49719335 | 10.30957772 | 10.28119144 | 0.02838628 | 0.21600191 |
| Trim15 | 3.010712765 | 2.787210798 | 2.794723687 | -0.007512889 | 0.215989078 |
| Mrps12 | 8.209185071 | 8.182017306 | 7.993198251 | 0.188819054 | 0.215986819 |
| Wbp11 | 10.42046837 | 10.22391467 | 10.20449426 | 0.019420413 | 0.215974113 |
| Dgcr8 | 8.563375116 | 8.530308648 | 8.347449989 | 0.18285866 | 0.215925127 |
| A630089N07Rik | 7.113241319 | 6.902662385 | 6.897641507 | 0.005020878 | 0.215599812 |
| Atf6 | 10.62764637 | 10.52498593 | 10.412065 | 0.112920927 | 0.21558137 |
| Kmt2d | 11.39098625 | 11.62483839 | 11.17554266 | 0.449295727 | 0.215443587 |
| Mcfd2 | 10.69873294 | 10.53835318 | 10.48344232 | 0.054910863 | 0.21529062 |
| Gm1821 | 6.538493451 | 6.724733125 | 6.323224002 | 0.401509123 | 0.215269449 |
| Sgms1 | 10.65723541 | 10.4528741 | 10.44209877 | 0.01077533 | 0.215136643 |
| Ppbp | 4.19422685 | 4.203164321 | 3.979222626 | 0.223941695 | 0.215004224 |

| | | | | | |
|---|---|---|---|---|---|
| Lyrm9 | 6.996133842 | 6.790962084 | 6.7811992 | 0.009762884 | 0.214934642 |
| Smpd4 | 9.20759983 | 9.324921963 | 8.992770126 | 0.332151837 | 0.214829704 |
| Aim2 | 6.920241616 | 6.641056091 | 6.705478693 | -0.064422602 | 0.214762923 |
| Tbrg1 | 11.07490646 | 10.85809041 | 10.86025304 | -0.002162633 | 0.214653417 |
| Naa50 | 12.00304918 | 11.91290077 | 11.78854906 | 0.124351713 | 0.214500123 |
| Stc1 | 8.738203639 | 8.433587401 | 8.523811068 | -0.090223667 | 0.214392571 |
| Unc13d | 8.657544846 | 8.810021466 | 8.443165813 | 0.366855653 | 0.214379033 |
| Spcs2 | 11.04319638 | 10.9328449 | 10.8288831 | 0.103961793 | 0.21431328 |
| Morf4l2 | 12.02700977 | 12.00009991 | 11.81288838 | 0.18721153 | 0.214121393 |
| Card11 | 5.161286736 | 5.080404872 | 4.947281096 | 0.133123776 | 0.21400564 |
| Mgat4a | 9.042597632 | 8.832241582 | 8.828612159 | 0.003629422 | 0.213985473 |
| Clptm1l | 11.37314475 | 11.20668407 | 11.15922389 | 0.047460173 | 0.213920853 |
| R3hcc1l | 9.338696432 | 9.165093747 | 9.12487966 | 0.040214088 | 0.213816772 |
| Zfp335 | 9.007811196 | 9.273154783 | 8.794006737 | 0.479148046 | 0.213804459 |
| Ppp2ca | 12.58964428 | 12.41128683 | 12.37589211 | 0.03539472 | 0.213752173 |
| Znrd1 | 8.702475376 | 8.537536119 | 8.488784516 | 0.048751604 | 0.21369086 |
| Ccnk | 9.467803851 | 9.445915466 | 9.254119496 | 0.19179597 | 0.213684354 |
| Snrnp35 | 7.494895888 | 7.257025065 | 7.281279809 | -0.024254744 | 0.213616079 |
| Cfap298 | 8.056401705 | 7.843956981 | 7.842884839 | 0.001072142 | 0.213516866 |
| Mrpl38 | 9.176699698 | 9.154550817 | 8.963267076 | 0.191283741 | 0.213432622 |
| Olfr366 | 4.925579707 | 4.834662069 | 4.712169046 | 0.122493023 | 0.213410661 |
| Mex3c | 10.5382817 | 10.44308718 | 10.32499267 | 0.11809451 | 0.213289027 |
| Fam69a | 8.940583772 | 8.667949216 | 8.727373437 | -0.059424221 | 0.213210335 |
| Ahctf1 | 10.8023879 | 10.7006612 | 10.58920043 | 0.11146077 | 0.21318747 |
| Zbtb9 | 8.222490972 | 8.195447305 | 8.00937407 | 0.186073235 | 0.213116902 |
| Tmem104 | 8.481761706 | 8.675045874 | 8.268795858 | 0.406250016 | 0.212965848 |
| Cyp11a1 | 7.481130062 | 7.043729677 | 7.268164851 | -0.224435174 | 0.212965211 |
| Bet1l | 9.408096254 | 9.428538691 | 9.195280418 | 0.233258273 | 0.212815837 |
| CAAA01147332.1 | 6.388431559 | 6.897164956 | 6.175728592 | 0.721436364 | 0.212702967 |
| Tnfrsf22 | 7.742674184 | 7.53848264 | 7.530116547 | 0.008366093 | 0.212557637 |
| Tsen2 | 7.732424908 | 7.378926015 | 7.519958282 | -0.141032267 | 0.212466626 |
| Ttc22 | 7.597990002 | 7.538267416 | 7.385603979 | 0.152663438 | 0.212386023 |

| | | | | | |
|---|---|---|---|---|---|
| Lime1 | 7.606497968 | 7.714111675 | 7.394371343 | 0.319740333 | 0.212126625 |
| Birc6 | 11.44533404 | 11.30730394 | 11.23321466 | 0.074089283 | 0.212119377 |
| Ddx5 | 14.31305521 | 14.20344365 | 14.10112502 | 0.102318627 | 0.21193019 |
| Mettl6 | 9.132769637 | 8.968203502 | 8.920856982 | 0.047346519 | 0.211912655 |
| Gclc | 9.488507039 | 9.597541351 | 9.276634201 | 0.32090715 | 0.211872839 |
| Prpf19 | 11.35382502 | 11.22245209 | 11.14230069 | 0.080151403 | 0.21152433 |
| Saal1 | 8.068767377 | 7.875504762 | 7.85739068 | 0.018114082 | 0.211376697 |
| Ptk2 | 9.907675311 | 9.852104407 | 9.696334402 | 0.155770005 | 0.211340909 |
| Slc35e1 | 10.72163927 | 10.58632438 | 10.51041036 | 0.07591402 | 0.21122891 |
| Ddx20 | 8.708050848 | 8.607556517 | 8.496837488 | 0.110719029 | 0.21121336 |
| Wfdc5 | 10.0103035 | 9.742754807 | 9.799303789 | -0.056548982 | 0.210999709 |
| Rab11fip1 | 10.89579138 | 10.71075275 | 10.68480226 | 0.02595049 | 0.210989113 |
| Eif2b4 | 9.775206865 | 9.659999882 | 9.564237077 | 0.095762805 | 0.210969788 |
| Snf8 | 9.954527733 | 9.765666987 | 9.743575003 | 0.022091984 | 0.210952731 |
| Fhod1 | 8.448083878 | 8.557419787 | 8.237501073 | 0.319918714 | 0.210582805 |
| Caprin1 | 12.28565578 | 12.17795861 | 12.07514053 | 0.102818083 | 0.21051525 |
| Sec11a | 10.19621209 | 10.019848 | 9.985757466 | 0.034090537 | 0.210454621 |
| Ptrh2 | 8.874626178 | 8.601704888 | 8.664268522 | -0.062563634 | 0.210357656 |
| Rps27 | 6.574006498 | 7.406485148 | 6.363795175 | 1.042689974 | 0.210211323 |
| Bag5 | 9.1754394 | 8.827116445 | 8.965229312 | -0.138112867 | 0.210210088 |
| Vrk1 | 9.175324316 | 8.891851933 | 8.965222252 | -0.073370319 | 0.210102064 |
| Mamld1 | 7.216478724 | 7.025660815 | 7.006395384 | 0.019265431 | 0.21008334 |
| Hdgfl2 | 9.218552958 | 9.315648591 | 9.008669906 | 0.306978685 | 0.209883052 |
| Cstf1 | 9.139285402 | 9.072329683 | 8.929418587 | 0.142911096 | 0.209866814 |
| Rps15 | 13.13303893 | 13.06366177 | 12.92319408 | 0.140467693 | 0.209844847 |
| Ctbp2 | 8.710553056 | 8.652449818 | 8.500753836 | 0.151695982 | 0.209799219 |
| Trip10 | 9.960081104 | 9.856925267 | 9.7503034 | 0.106621867 | 0.209777704 |
| Pigf | 7.951682522 | 7.786769301 | 7.741957401 | 0.0448119 | 0.209725121 |
| Lmx1a | 4.069113932 | 3.88255133 | 3.859498229 | 0.023053101 | 0.209615703 |
| Tmem161a | 8.969901432 | 9.102565693 | 8.760295856 | 0.342269837 | 0.209605577 |
| Psmd12 | 10.88197448 | 10.71793124 | 10.67255042 | 0.045380817 | 0.20942406 |
| Ssh3 | 10.56266395 | 10.65325945 | 10.35327915 | 0.299980297 | 0.209384803 |

| | | | | | |
|---|---|---|---|---|---|
| Stk10 | 9.582758092 | 9.492986642 | 9.3733936 | 0.119593042 | 0.209364492 |
| Taok3 | 9.309751878 | 9.174356332 | 9.100504853 | 0.073851479 | 0.209247025 |
| Capzb | 12.39591728 | 12.24968931 | 12.18677625 | 0.062913063 | 0.209141033 |
| Stx12 | 11.54458334 | 11.34727172 | 11.33552149 | 0.011750237 | 0.20906185 |
| Mios | 8.751231087 | 8.508613333 | 8.542201545 | -0.033588212 | 0.209029542 |
| Dnajc5 | 11.44046224 | 11.27241843 | 11.23144346 | 0.040974973 | 0.20901878 |
| Snrpe | 9.983679818 | 9.875826328 | 9.774673442 | 0.101152886 | 0.209006376 |
| Flii | 11.32615723 | 11.26207528 | 11.11718651 | 0.144888767 | 0.208970713 |
| Bcam | 11.63254319 | 11.67731891 | 11.42377328 | 0.253545627 | 0.208769913 |
| A730049H05Rik | 2.786193481 | 2.659948888 | 2.577437516 | 0.082511372 | 0.208755964 |
| Lasp1 | 11.16884513 | 11.10897048 | 10.96012195 | 0.148848533 | 0.208723183 |
| Mup2 | 6.082180522 | 6.248397928 | 5.873524447 | 0.374873481 | 0.208656075 |
| Aar2 | 9.356818036 | 9.312314938 | 9.148234391 | 0.164080547 | 0.208583645 |
| Cacng2 | 3.320057041 | 3.14642935 | 3.111603633 | 0.034825716 | 0.208453408 |
| Zc3h7b | 10.60445882 | 10.45938607 | 10.39619344 | 0.063192627 | 0.20826538 |
| Ilf2 | 10.37130202 | 10.19112155 | 10.16311497 | 0.028006583 | 0.208187053 |
| Rell2 | 4.416006905 | 4.296897293 | 4.207830344 | 0.08906695 | 0.208176561 |
| Syt8 | 6.159264262 | 6.040743822 | 5.951323644 | 0.089420179 | 0.207940619 |
| Rnf24 | 8.230877064 | 8.040774125 | 8.02294712 | 0.017827005 | 0.207929944 |
| Maip1 | 8.258738969 | 8.075021356 | 8.050877756 | 0.0241436 | 0.207861213 |
| CR974586.4 | 2.680471013 | 2.554140034 | 2.472942263 | 0.081197771 | 0.20752875 |
| Gm3893 | 2.680471013 | 2.554140034 | 2.472942263 | 0.081197771 | 0.20752875 |
| Tex10 | 9.146810086 | 9.03681625 | 8.939306043 | 0.097510207 | 0.207504043 |
| Plpbp | 10.24410873 | 10.45391612 | 10.03665927 | 0.417256855 | 0.207449465 |
| Cnbp | 13.02340391 | 12.76928283 | 12.81598206 | -0.046699223 | 0.207421857 |
| Tbc1d2 | 8.148143712 | 8.399797454 | 7.941026667 | 0.458770787 | 0.207117045 |
| Adprhl2 | 9.546853803 | 9.527705547 | 9.339824844 | 0.187880703 | 0.207028958 |
| Rad9a | 7.29068739 | 7.185185842 | 7.083757145 | 0.101428697 | 0.206930245 |
| Man2b1 | 10.7387504 | 10.5433576 | 10.53184232 | 0.011515283 | 0.206908087 |
| Dnajb5 | 9.561006877 | 9.382214533 | 9.354188596 | 0.028025937 | 0.206818281 |
| Otud7a | 3.503470093 | 3.369609334 | 3.29677272 | 0.072836614 | 0.206697374 |
| Gpr68 | 7.140417281 | 7.199190818 | 6.93381273 | 0.265378087 | 0.206604551 |

| | | | | | |
|---|---|---|---|---|---|
| 9930111J21Rik1 | 6.583124511 | 6.230357269 | 6.376545234 | -0.146187966 | 0.206579276 |
| Luc7l | 10.32245647 | 10.42138436 | 10.11596418 | 0.305420183 | 0.20649229 |
| Mrtfa | 9.308370038 | 9.503545497 | 9.101887144 | 0.401658353 | 0.206482894 |
| Eif4g2 | 14.62215444 | 14.42765019 | 14.41581121 | 0.011838977 | 0.20634323 |
| Sirpb1a | 5.747815534 | 5.769747246 | 5.541546148 | 0.228201099 | 0.206269387 |
| Akap1 | 9.764507642 | 9.689019357 | 9.558546772 | 0.130472585 | 0.205960869 |
| Cdv3 | 11.90303052 | 11.78364423 | 11.69721116 | 0.086433073 | 0.20581936 |
| Hp | 14.25398666 | 13.80407962 | 14.04823422 | -0.244154593 | 0.205752443 |
| Muc13 | 3.53627862 | 3.421156792 | 3.330648225 | 0.090508567 | 0.205630395 |
| Mfsd14a | 10.35476465 | 10.22251035 | 10.14915103 | 0.073359317 | 0.20561362 |
| Ndufaf6 | 6.674185221 | 6.372662941 | 6.468627492 | -0.095964551 | 0.205557729 |
| Itpa | 9.526392713 | 9.367302919 | 9.320871921 | 0.046430998 | 0.205520791 |
| Syn1 | 4.747431105 | 4.716181263 | 4.541925526 | 0.174255737 | 0.205505579 |
| H2-Ob | 6.205203956 | 5.963883068 | 5.99972156 | -0.035838492 | 0.205482395 |
| Tarbp1 | 8.02151308 | 7.927640655 | 7.816087197 | 0.111553458 | 0.205425883 |
| Ablim2 | 7.733120828 | 7.650560467 | 7.527807171 | 0.122753297 | 0.205313658 |
| Gm13502 | 6.488538531 | 6.448294921 | 6.283266563 | 0.165028357 | 0.205271968 |
| Rps28 | 11.68905646 | 11.73051287 | 11.48378856 | 0.246724313 | 0.205267903 |
| Xxylt1 | 9.12878349 | 8.97711341 | 8.923520332 | 0.053593078 | 0.205263158 |
| Psmd9 | 8.777511103 | 8.771920082 | 8.572386145 | 0.199533936 | 0.205124958 |
| Tspo | 10.59023977 | 10.52535995 | 10.38519586 | 0.140164093 | 0.205043913 |
| Parp1 | 10.11241501 | 9.997016928 | 9.907413226 | 0.089603702 | 0.20500178 |
| Celf4 | 7.025040365 | 6.929349833 | 6.820081663 | 0.10926817 | 0.204958702 |
| Etnppl | 4.569795718 | 4.391360533 | 4.364920947 | 0.026439586 | 0.204874771 |
| Aldh16a1 | 8.730587321 | 8.889499917 | 8.525789149 | 0.363710769 | 0.204798173 |
| Akap13 | 11.26738919 | 10.95550106 | 11.06265271 | -0.10715165 | 0.204736477 |
| Map1b | 9.25793878 | 9.292554533 | 9.053342588 | 0.239211945 | 0.204596192 |
| Ube2d3 | 12.93102707 | 12.82340445 | 12.72643735 | 0.0969671 | 0.204589723 |
| Hdac6 | 9.001049056 | 8.949433724 | 8.796518362 | 0.152915362 | 0.204530695 |
| Srp72 | 11.33578751 | 11.20378346 | 11.13131034 | 0.072473117 | 0.204477163 |
| Hpf1 | 8.957369428 | 8.833868303 | 8.753000882 | 0.080867421 | 0.204368546 |
| Mfsd5 | 10.5081116 | 10.50116135 | 10.30379888 | 0.197362467 | 0.204312717 |

| | | | | | |
|---|---|---|---|---|---|
| Mphosph6 | 8.207214441 | 7.940880346 | 8.002938564 | -0.062058219 | 0.204275876 |
| Ecsit | 9.273716531 | 9.227905728 | 9.069446538 | 0.15845919 | 0.204269992 |
| Tubb2b | 8.711107907 | 8.751891667 | 8.506891097 | 0.24500057 | 0.20421681 |
| Exosc5 | 8.437757642 | 8.52678261 | 8.233701568 | 0.293081042 | 0.204056074 |
| Wdfy4 | 9.008862296 | 8.949447455 | 8.804885187 | 0.144562268 | 0.203977108 |
| Hax1 | 9.886931651 | 9.888284041 | 9.683007256 | 0.205276786 | 0.203924395 |
| Mest | 7.779927353 | 7.619712045 | 7.576011914 | 0.043700131 | 0.203915439 |
| Rnaseh2a | 9.317996469 | 9.304264291 | 9.11413226 | 0.190132031 | 0.20386421 |
| Plekha3 | 8.873406043 | 8.747511894 | 8.669615781 | 0.077896113 | 0.203790262 |
| Kbtbd2 | 9.808907706 | 9.751694991 | 9.60518147 | 0.146513521 | 0.203726236 |
| 1810022K09Rik | 9.387880074 | 9.204337197 | 9.184275224 | 0.020061973 | 0.20360485 |
| Derl3 | 3.682722068 | 3.699927086 | 3.479187931 | 0.220739155 | 0.203534137 |
| Taf15 | 10.56704286 | 10.43964232 | 10.36351246 | 0.076129867 | 0.203530403 |
| Slc30a5 | 10.11624851 | 9.922967817 | 9.912727662 | 0.010240155 | 0.203520848 |
| Usp27x | 6.480683 | 6.285838049 | 6.277175597 | 0.008662452 | 0.203507403 |
| Hnrnpm | 11.91542438 | 11.75775286 | 11.71204692 | 0.045705943 | 0.203377467 |
| Tbccd1 | 8.508387793 | 8.36398481 | 8.305049374 | 0.058935436 | 0.203338418 |
| Tpd52l2 | 10.91608811 | 10.75813571 | 10.71277521 | 0.0453605 | 0.2033129 |
| Phf1 | 8.708219822 | 8.640418487 | 8.505043038 | 0.135375448 | 0.203176783 |
| Mcts2 | 6.956321639 | 6.820216643 | 6.753195005 | 0.067021638 | 0.203126634 |
| Axdnd1 | 4.25568091 | 4.051802819 | 4.05257818 | -0.000775361 | 0.20310273 |
| Pet100 | 8.831433733 | 8.657533588 | 8.628353667 | 0.029179922 | 0.203080066 |
| Polr2j | 9.058200131 | 8.965952568 | 8.855161538 | 0.11079103 | 0.203038592 |
| Gimap9 | 6.598659845 | 6.476204807 | 6.395639022 | 0.080565785 | 0.203020823 |
| Eif3i | 11.55605887 | 11.459502 | 11.35334302 | 0.106158977 | 0.202715853 |
| Ahcy | 10.26344264 | 10.2750847 | 10.06073962 | 0.214345073 | 0.202703016 |
| Lap3 | 10.80737022 | 10.70218705 | 10.60468858 | 0.09749847 | 0.202681643 |
| Rundc3b | 4.446685149 | 4.309699415 | 4.24400595 | 0.065693465 | 0.202679199 |
| Cdr2l | 7.522499791 | 7.548394996 | 7.319860016 | 0.22853498 | 0.202639775 |
| Gnb2 | 11.89334162 | 11.92644653 | 11.69083459 | 0.23561194 | 0.202507023 |
| Nup43 | 6.97446075 | 6.877962058 | 6.771987731 | 0.105974327 | 0.202473018 |
| Nrip3 | 8.935273768 | 8.62748355 | 8.732810782 | -0.105327232 | 0.202462986 |

| | | | | | |
|---|---|---|---|---|---|
| Pcdhgb8 | 5.484786493 | 5.433305593 | 5.282466772 | 0.15083882 | 0.202319721 |
| Arl11 | 5.987499801 | 5.797265746 | 5.785297683 | 0.011968063 | 0.202202118 |
| Mrps10 | 7.878949451 | 7.730129516 | 7.676769637 | 0.053359879 | 0.202179814 |
| Eif5b | 11.79432576 | 11.58244245 | 11.5921938 | -0.009751347 | 0.202131963 |
| Yif1a | 9.70121653 | 9.621411894 | 9.499084699 | 0.122327195 | 0.202131831 |
| Aldh3b1 | 7.214162362 | 7.058612945 | 7.01208493 | 0.046528015 | 0.202077432 |
| Gm17455 | 3.222921901 | 3.090869955 | 3.020850727 | 0.070019228 | 0.202071174 |
| Stx6 | 10.10365374 | 10.139588 | 9.901613744 | 0.237974253 | 0.20204 |
| Gm19040 | 3.346756813 | 3.212984839 | 3.144729505 | 0.068255334 | 0.202027308 |
| Clns1a | 10.12251188 | 9.963553788 | 9.920517928 | 0.043035861 | 0.201993952 |
| Orc1 | 6.573980858 | 6.581902134 | 6.372007271 | 0.209894862 | 0.201973587 |
| Gm340 | 6.916453084 | 6.753362467 | 6.714669676 | 0.038692791 | 0.201783407 |
| Jpt1 | 10.52606425 | 10.47480208 | 10.3244292 | 0.15037288 | 0.201635053 |
| Psmc4 | 11.03503023 | 10.97375482 | 10.83353244 | 0.140222383 | 0.201497793 |
| Sfxn1 | 10.2615456 | 10.01591229 | 10.06022051 | -0.044308216 | 0.201325095 |
| Cd2bp2 | 10.3410438 | 10.22231129 | 10.13992076 | 0.082390527 | 0.20112304 |
| Ino80 | 9.940898897 | 9.786824601 | 9.739899701 | 0.046924899 | 0.200999195 |
| Atp13a3 | 11.06930469 | 10.96136293 | 10.86832697 | 0.093035963 | 0.200977723 |
| Tal1 | 6.755000647 | 6.380088944 | 6.554070499 | -0.173981555 | 0.200930147 |
| Leng1 | 7.8353417 | 7.803929005 | 7.634501606 | 0.169427399 | 0.200840094 |
| Unc5c | 4.570944925 | 4.527810582 | 4.370129083 | 0.157681499 | 0.200815842 |
| Psmd4 | 11.33294419 | 11.29987715 | 11.1321377 | 0.167739443 | 0.200806487 |
| Gpn2 | 7.852872408 | 7.798987717 | 7.652160109 | 0.146827608 | 0.200712298 |
| Sdc3 | 10.75387903 | 10.54466364 | 10.55323402 | -0.008570387 | 0.20064501 |
| Rnf34 | 9.052246056 | 8.886984138 | 8.851643423 | 0.035340715 | 0.200602633 |
| Mcm5 | 10.18791445 | 10.28846785 | 9.987322059 | 0.301145791 | 0.200592394 |
| BC003965 | 8.913857933 | 8.826300489 | 8.713348326 | 0.112952163 | 0.200509607 |
| Nudt22 | 7.156718441 | 7.094020144 | 6.956226792 | 0.137793352 | 0.20049165 |
| Sf3b1 | 12.68843204 | 12.54453355 | 12.48796632 | 0.056567233 | 0.200465727 |
| Armc3 | 3.063174225 | 2.937238455 | 2.862744128 | 0.074494327 | 0.200430097 |
| Trnt1 | 9.539578081 | 9.340188572 | 9.339153875 | 0.001034697 | 0.200424206 |
| Usp25 | 11.60083368 | 11.25583002 | 11.40043971 | -0.144609693 | 0.200393963 |

| | | | | | |
|---|---|---|---|---|---|
| Wsb2 | 10.91955164 | 10.8435094 | 10.71919032 | 0.124319077 | 0.200361317 |
| Zmat2 | 10.58795763 | 10.32521648 | 10.38767945 | -0.062462967 | 0.20027818 |
| Taf1a | 8.148292388 | 8.102826501 | 7.948042632 | 0.154783869 | 0.200249756 |
| Mrpl37 | 9.855839614 | 9.819345047 | 9.655622024 | 0.163723023 | 0.20021759 |
| Nars2 | 7.55356432 | 7.280608004 | 7.353390588 | -0.072782584 | 0.200173732 |
| Jak1 | 12.48284589 | 12.38518014 | 12.28274237 | 0.102437767 | 0.200103517 |
| Cox7c | 10.61034153 | 10.58620972 | 10.41024736 | 0.175962363 | 0.200094173 |
| Plat | 8.926717241 | 8.762566473 | 8.726679409 | 0.035887063 | 0.200037831 |
| Pphln1 | 9.516651025 | 9.37477578 | 9.316712012 | 0.058063768 | 0.199939013 |
| Pdgfra | 11.39408667 | 11.34380653 | 11.19441047 | 0.149396053 | 0.1996762 |
| Gm3636 | 5.037741062 | 4.925014905 | 4.838171736 | 0.086843169 | 0.199569326 |
| Mdc1 | 9.731997157 | 9.66132434 | 9.532427836 | 0.128896504 | 0.199569321 |
| Atp5e | 10.75238662 | 10.68853837 | 10.55300811 | 0.135530253 | 0.199378503 |
| Cpeb4 | 10.34161128 | 10.45620769 | 10.14233671 | 0.313870977 | 0.199274573 |
| Itga1 | 9.58486174 | 9.379761267 | 9.385684812 | -0.005923545 | 0.199176927 |
| Nup188 | 10.32593427 | 10.31028317 | 10.1268122 | 0.183470967 | 0.199122073 |
| Eomes | 2.952196634 | 2.785615636 | 2.753091238 | 0.032524398 | 0.199105396 |
| Psmd11 | 11.41967811 | 11.4536449 | 11.22061479 | 0.23303011 | 0.199063313 |
| Notch2 | 11.67293577 | 11.60668186 | 11.47391316 | 0.132768707 | 0.199022613 |
| Aldh8a1 | 4.124269055 | 3.979494497 | 3.925293291 | 0.054201206 | 0.198975764 |
| Pbdc1 | 9.209083055 | 9.246360367 | 9.010159588 | 0.236200779 | 0.198923467 |
| Ube2o | 9.499658188 | 9.598927425 | 9.300898453 | 0.298028972 | 0.198759735 |
| Qrsl1 | 7.77131053 | 7.558238464 | 7.572896687 | -0.014658223 | 0.198413843 |
| Plekhm2 | 9.787105561 | 10.03562414 | 9.588705882 | 0.446918263 | 0.198399679 |
| Kdm7a | 11.08049007 | 10.81870667 | 10.88222343 | -0.06351676 | 0.19826664 |
| Tcp10b | 3.296514601 | 3.156363533 | 3.098327846 | 0.058035687 | 0.198186755 |
| Dtymk | 9.027848455 | 8.989589762 | 8.829840453 | 0.15974931 | 0.198008002 |
| Tlr1 | 7.419210601 | 7.736225451 | 7.221209216 | 0.515016234 | 0.198001384 |
| Mpzl2 | 10.68955785 | 10.69565248 | 10.49161456 | 0.20403792 | 0.197943293 |
| Tcp11l1 | 7.224454624 | 7.13202635 | 7.02658303 | 0.105443321 | 0.197871595 |
| Ppia | 13.74518622 | 13.80940226 | 13.54740039 | 0.26200187 | 0.197785833 |
| Smarcd1 | 8.828321855 | 8.654110654 | 8.630557447 | 0.023553207 | 0.197764407 |

| | | | | | |
|---|---|---|---|---|---|
| Tipin | 8.857104746 | 8.597873012 | 8.659358235 | -0.061485223 | 0.197746511 |
| Polg | 10.38171751 | 10.38080279 | 10.18399632 | 0.196806467 | 0.19772119 |
| Uchl3 | 9.926701439 | 9.766756083 | 9.729112753 | 0.03764333 | 0.197588686 |
| Lmna | 13.05107007 | 13.27171131 | 12.85357783 | 0.41813348 | 0.197492243 |
| Dnmt3l | 5.218767246 | 5.198334203 | 5.021275475 | 0.177058728 | 0.197491771 |
| Btbd10 | 9.134252845 | 8.990238307 | 8.936812212 | 0.053426095 | 0.197440633 |
| Tmem14c | 10.29168486 | 10.23511108 | 10.09425671 | 0.14085437 | 0.197428147 |
| Lrch4 | 9.07277403 | 9.090127592 | 8.875441649 | 0.214685943 | 0.197332381 |
| Palld | 11.11040877 | 10.97335678 | 10.91326356 | 0.060093217 | 0.197145207 |
| Arel1 | 9.89940794 | 9.755746695 | 9.702309094 | 0.053437601 | 0.197098846 |
| Sec61g | 7.274339894 | 7.003974137 | 7.077373414 | -0.073399277 | 0.196966479 |
| Atf5 | 10.61389057 | 10.79321541 | 10.41712283 | 0.376092573 | 0.196767733 |
| Srsf5 | 12.70760451 | 12.51493285 | 12.51085751 | 0.004075343 | 0.196747003 |
| Sirt1 | 9.357747964 | 9.371838582 | 9.161170059 | 0.210668523 | 0.196577905 |
| Rpl5 | 9.859566309 | 9.796932447 | 9.663005199 | 0.133927248 | 0.19656111 |
| Rnf44 | 10.57902384 | 10.7093608 | 10.38257132 | 0.326789477 | 0.19645252 |
| Gm8080 | 4.541208949 | 4.341416873 | 4.344882828 | -0.003465955 | 0.196326121 |
| Comt | 9.997513386 | 9.798930646 | 9.801260789 | -0.002330143 | 0.196252597 |
| Trac | 4.952516525 | 4.735768096 | 4.756279339 | -0.020511243 | 0.196237186 |
| Ssr3 | 12.38652437 | 12.26617383 | 12.19047907 | 0.07569476 | 0.196045303 |
| Asb4 | 4.960299114 | 4.758576725 | 4.764384769 | -0.005808044 | 0.195914345 |
| Fem1b | 10.81606748 | 10.83860721 | 10.62021566 | 0.218391543 | 0.19585182 |
| Rbck1 | 10.3115728 | 10.37585831 | 10.11578199 | 0.260076318 | 0.195790804 |
| Hmga2 | 5.144427666 | 4.995779018 | 4.948718145 | 0.047060872 | 0.195709521 |
| Pabpc1 | 14.01146616 | 13.99242907 | 13.81582606 | 0.176603007 | 0.195640103 |
| Serpina10 | 3.620226974 | 3.60085189 | 3.424673235 | 0.176178655 | 0.195553739 |
| Khdrbs1 | 11.42135302 | 11.20314403 | 11.22583778 | -0.022693743 | 0.195515247 |
| Lrp3 | 7.192214204 | 7.238207709 | 6.996726521 | 0.241481188 | 0.195487683 |
| Traf2 | 9.352817647 | 9.431941199 | 9.157377471 | 0.274563728 | 0.195440176 |
| Gins1 | 7.296185099 | 7.220690031 | 7.10082558 | 0.119864452 | 0.195359519 |
| Ripor1 | 8.914194034 | 8.882553824 | 8.719057009 | 0.163496814 | 0.195137024 |
| Entr1 | 10.30318383 | 10.30813914 | 10.10810857 | 0.200030575 | 0.195075258 |

| | | | | | |
|---|---|---|---|---|---|
| Stx7 | 10.21419577 | 10.13551414 | 10.01915643 | 0.116357707 | 0.19503934 |
| Abracl | 9.933181193 | 9.752848743 | 9.738156602 | 0.01469214 | 0.19502459 |
| Upf1 | 10.68164877 | 10.85150635 | 10.48672037 | 0.364785987 | 0.1949284 |
| Hamp2 | 5.338785071 | 5.13887692 | 5.144042117 | -0.005165197 | 0.194742954 |
| Lfng | 9.251919844 | 9.614407557 | 9.057191975 | 0.557215583 | 0.19472787 |
| Iqgap1 | 13.35493626 | 13.2454215 | 13.16027987 | 0.085141627 | 0.194656383 |
| Uba5 | 9.903724082 | 9.826425509 | 9.709121926 | 0.117303583 | 0.194602156 |
| Ticam1 | 8.538434002 | 8.732411331 | 8.343877148 | 0.388534183 | 0.194556854 |
| Nt5dc2 | 8.923472767 | 8.932800417 | 8.728974547 | 0.20382587 | 0.19449822 |
| Omp | 3.608517553 | 3.406809565 | 3.41403392 | -0.007224356 | 0.194483633 |
| Ranbp2 | 11.76200348 | 11.74081816 | 11.56760506 | 0.1732131 | 0.194398423 |
| Uba52 | 10.53618904 | 10.6466441 | 10.34179781 | 0.304846283 | 0.194391223 |
| Saa2 | 4.266671713 | 4.099714447 | 4.07240757 | 0.027306877 | 0.194264143 |
| Nacc1 | 10.98616223 | 11.06440846 | 10.79193697 | 0.272471493 | 0.19422526 |
| Prf1 | 6.336263755 | 6.218942353 | 6.142056262 | 0.076886091 | 0.194207493 |
| Ccdc126 | 6.715311135 | 6.544315584 | 6.521341627 | 0.022973957 | 0.193969508 |
| Ly75 | 7.525824311 | 7.856647205 | 7.331904033 | 0.524743172 | 0.193920278 |
| Tmc6 | 9.391555225 | 9.313924749 | 9.197701087 | 0.116223662 | 0.193854139 |
| Celf1 | 10.49063571 | 10.41596696 | 10.29681519 | 0.119151773 | 0.19382052 |
| Mmp1b | 3.396645529 | 3.217455159 | 3.202868002 | 0.014587157 | 0.193777527 |
| Rock1 | 11.35602844 | 11.16523105 | 11.16230939 | 0.002921663 | 0.193719053 |
| Ccdc93 | 9.336132356 | 9.318001022 | 9.142424507 | 0.175576516 | 0.193707849 |
| Sec23ip | 10.12278731 | 10.01966295 | 9.929122332 | 0.090540613 | 0.193664974 |
| Rap2c | 10.58416459 | 10.43896556 | 10.3905402 | 0.04842536 | 0.19362439 |
| Wdr70 | 8.828271339 | 8.76406129 | 8.634673593 | 0.129387697 | 0.193597746 |
| Otub1 | 10.0845993 | 10.08974211 | 9.891303179 | 0.198438934 | 0.193296124 |
| Twf2 | 9.256420255 | 9.918930372 | 9.063210725 | 0.855719648 | 0.19320953 |
| 0610037L13Rik | 9.309293123 | 9.191181521 | 9.116111568 | 0.075069953 | 0.193181555 |
| Olfr510 | 3.233368494 | 3.112651077 | 3.040281444 | 0.072369633 | 0.19308705 |
| Mto1 | 8.779162285 | 8.518650525 | 8.586306807 | -0.067656283 | 0.192855477 |
| A4galt | 8.164724729 | 7.726095527 | 7.971888859 | -0.245793332 | 0.19283587 |
| Tmem87b | 10.19593394 | 10.00709889 | 10.00316827 | 0.003930617 | 0.192765665 |

| | | | | | |
|---|---|---|---|---|---|
| Naa20 | 10.01564232 | 9.858507349 | 9.822889949 | 0.0356174 | 0.19275237 |
| Rpl36al | 12.0257185 | 11.85453136 | 11.83297792 | 0.02155344 | 0.192740577 |
| Psme2 | 11.48455388 | 11.32757907 | 11.29183939 | 0.03573968 | 0.192714493 |
| Slk | 11.80639257 | 11.69158425 | 11.61383107 | 0.07775318 | 0.192561503 |
| 9130230L23Rik | 5.608748799 | 5.520136882 | 5.4162751 | 0.103861782 | 0.192473699 |
| Plpp3 | 11.78739562 | 11.38166872 | 11.59492846 | -0.213259743 | 0.192467163 |
| Gsk3a | 10.94225206 | 10.93644753 | 10.74979196 | 0.18665557 | 0.1924601 |
| Ube2e1 | 10.45528154 | 10.20774429 | 10.26283047 | -0.055086183 | 0.192451067 |
| Banf1 | 10.64585869 | 10.51949359 | 10.45350866 | 0.065984933 | 0.192350037 |
| Ada | 6.553097414 | 6.43106995 | 6.360788275 | 0.070281675 | 0.192309139 |
| Trmt112 | 9.318137953 | 9.247628384 | 9.125895875 | 0.121732508 | 0.192242078 |
| Ces2f | 8.020509502 | 8.047143645 | 7.828300049 | 0.218843596 | 0.192209453 |
| Cdc42ep3 | 8.956273706 | 8.721706217 | 8.764065263 | -0.042359047 | 0.192208443 |
| Med26 | 7.750718561 | 7.763639571 | 7.558513356 | 0.205126216 | 0.192205205 |
| Pdzk1ip1 | 11.28289909 | 10.99990377 | 11.09072302 | -0.090819243 | 0.19217607 |
| Mex3d | 7.64434206 | 7.640843238 | 7.452241564 | 0.188601674 | 0.192100496 |
| Lonrf1 | 8.304940402 | 8.047539203 | 8.112843935 | -0.065304732 | 0.192096467 |
| Sarnp | 9.967383673 | 9.831352683 | 9.775323362 | 0.056029321 | 0.19206031 |
| Lamtor4 | 9.578544658 | 9.489593993 | 9.386694722 | 0.102899271 | 0.191849936 |
| Tufm | 10.56488454 | 10.48054071 | 10.37304367 | 0.107497037 | 0.19184087 |
| Dnajc8 | 10.54453137 | 10.325356 | 10.35270554 | -0.027349543 | 0.191825823 |
| Arhgef7 | 10.46837942 | 10.46191139 | 10.27657925 | 0.185332133 | 0.191800167 |
| Gm7609 | 4.05620253 | 3.910594368 | 3.864525998 | 0.04606837 | 0.191676532 |
| Tfg | 10.56422008 | 10.55297909 | 10.37258335 | 0.18039574 | 0.191636733 |
| Ube2m | 10.74864267 | 10.63768628 | 10.5570242 | 0.080662077 | 0.191618467 |
| Klra13-ps | 4.357751408 | 4.040238553 | 4.166179983 | -0.125941429 | 0.191571425 |
| Pdzd8 | 10.06783008 | 9.934132409 | 9.876288439 | 0.05784397 | 0.19154164 |
| Ubtf | 11.4151562 | 11.35314125 | 11.22376499 | 0.129376267 | 0.19139121 |
| A830018L16Rik | 4.307697193 | 4.15353915 | 4.116307527 | 0.037231623 | 0.191389665 |
| Sec24c | 11.7086494 | 11.63919981 | 11.51728312 | 0.121916687 | 0.191366283 |
| Kmt2b | 10.48408547 | 10.63942364 | 10.292763 | 0.34666064 | 0.191322463 |
| Coq8b | 9.656433171 | 9.689043602 | 9.465171016 | 0.223872586 | 0.191262155 |

| | | | | | |
|---|---|---|---|---|---|
| Chsy3 | 4.513887411 | 4.303321993 | 4.322657726 | -0.019335732 | 0.191229685 |
| Fam110c | 7.115454719 | 6.815264147 | 6.924383196 | -0.109119049 | 0.191071523 |
| Cnot9 | 10.08283739 | 9.917136833 | 9.891853767 | 0.025283067 | 0.190983627 |
| Sv2c | 5.382284757 | 5.36388566 | 5.191425 | 0.17246066 | 0.190859757 |
| Gdnf | 7.234480083 | 7.25360232 | 7.043624512 | 0.209977808 | 0.190855571 |
| Tmem115 | 9.313875098 | 9.381647918 | 9.123021709 | 0.258626209 | 0.190853389 |
| Parp4 | 9.399716152 | 9.17119474 | 9.208916589 | -0.037721849 | 0.190799563 |
| Tph1 | 5.620428497 | 5.289015146 | 5.429712398 | -0.140697251 | 0.190716099 |
| Txnl1 | 10.85387655 | 10.77467058 | 10.66321275 | 0.111457823 | 0.190663797 |
| Itk | 6.332192488 | 5.993473798 | 6.141552064 | -0.148078266 | 0.190640424 |
| Prps1 | 9.456925588 | 9.22531617 | 9.266598332 | -0.041282163 | 0.190327256 |
| Pi4k2a | 10.76637993 | 10.72533847 | 10.57615857 | 0.149179897 | 0.19022136 |
| Bambi | 6.05925105 | 5.965464301 | 5.869035883 | 0.096428418 | 0.190215167 |
| Acox2 | 8.628172822 | 8.421002869 | 8.438033283 | -0.017030414 | 0.190139539 |
| Mei4 | 4.821802464 | 4.658652555 | 4.631768336 | 0.026884219 | 0.190034129 |
| Hnrnpa2b1 | 13.81377308 | 13.78302337 | 13.62380434 | 0.159219027 | 0.189968733 |
| Lox | 9.618929241 | 9.049051093 | 9.429194721 | -0.380143628 | 0.189734521 |
| Kif5c | 5.956429153 | 5.760978063 | 5.766768307 | -0.005790243 | 0.189660846 |
| Adam19 | 8.857685873 | 8.77218569 | 8.668098401 | 0.104087289 | 0.189587472 |
| Psenen | 10.19032037 | 9.986740137 | 10.00091782 | -0.014177686 | 0.189402547 |
| Rpl14 | 13.06612588 | 13.01654789 | 12.87672432 | 0.139823563 | 0.189401557 |
| Canx | 13.01756328 | 12.96296115 | 12.82830968 | 0.134651467 | 0.1892536 |
| Prkcb | 8.488373634 | 8.061095665 | 8.299216801 | -0.238121136 | 0.189156834 |
| Wbp1l | 11.51484615 | 11.33360608 | 11.32584113 | 0.007764943 | 0.189005013 |
| Lims1 | 10.74608243 | 10.62159504 | 10.55711973 | 0.064475317 | 0.188962707 |
| Trir | 9.805170463 | 9.706745279 | 9.616307622 | 0.090437657 | 0.188862841 |
| Fam171a2 | 7.489899623 | 7.178815991 | 7.301094271 | -0.12227828 | 0.188805352 |
| Dusp27 | 7.756606091 | 8.609885577 | 7.567926802 | 1.041958775 | 0.188679289 |
| Psmb2 | 10.9757172 | 10.90847281 | 10.7870463 | 0.121426507 | 0.188670903 |
| Atp11a | 10.97930176 | 10.95567346 | 10.7906443 | 0.16502916 | 0.188657463 |
| Npdc1 | 9.63600345 | 9.665479768 | 9.447408815 | 0.218070953 | 0.188594635 |
| Fam222b | 7.984879976 | 8.051807646 | 7.796293345 | 0.255514301 | 0.188586631 |

| | | | | | |
|---|---|---|---|---|---|
| AC125149.3 | 2.991578518 | 2.794084191 | 2.802995969 | -0.008911779 | 0.188582549 |
| Hdac4 | 9.761600843 | 9.725127145 | 9.573108716 | 0.152018429 | 0.188492127 |
| Gm10408 | 2.495610693 | 2.423265088 | 2.307229449 | 0.116035639 | 0.188381244 |
| Uxs1 | 8.989905112 | 8.664718013 | 8.801556135 | -0.136838122 | 0.188348977 |
| Rabepk | 8.144754971 | 8.024493862 | 7.956463325 | 0.068030537 | 0.188291646 |
| Col9a3 | 4.978082631 | 5.084515897 | 4.789864218 | 0.294651679 | 0.188218413 |
| Josd2 | 8.51582796 | 8.764546137 | 8.327710739 | 0.436835399 | 0.188117222 |
| Nup37 | 7.864359988 | 7.653268979 | 7.676641055 | -0.023372077 | 0.187718933 |
| Gm8459 | 2.997775536 | 2.904449574 | 2.810199677 | 0.094249896 | 0.187575859 |
| AI837181 | 8.772760417 | 8.792516158 | 8.585193696 | 0.207322462 | 0.187566721 |
| 2210016F16Rik | 9.413669818 | 9.259805057 | 9.226287522 | 0.033517535 | 0.187382296 |
| Tbcb | 9.587250404 | 9.561666541 | 9.399930848 | 0.161735693 | 0.187319555 |
| Spryd4 | 8.524065837 | 8.295496626 | 8.336891199 | -0.041394572 | 0.187174638 |
| Sulf1 | 8.586164269 | 8.17223217 | 8.399034559 | -0.22680239 | 0.18712971 |
| Zfp831 | 4.945740385 | 4.797620326 | 4.758747137 | 0.038873189 | 0.186993248 |
| Ret | 7.039133156 | 6.987118585 | 6.852147028 | 0.134971557 | 0.186986128 |
| Gsdma3 | 6.613794079 | 6.589883799 | 6.426827616 | 0.163056184 | 0.186966463 |
| Sh3bgrl3 | 11.83870874 | 11.75915081 | 11.65176861 | 0.107382193 | 0.186940123 |
| Ibtk | 10.58632658 | 10.3605965 | 10.39940757 | -0.038811067 | 0.186919013 |
| Psmc2 | 11.45341609 | 11.336242 | 11.26677986 | 0.069462137 | 0.186636227 |
| Phb | 10.81934461 | 10.60139472 | 10.63280376 | -0.031409047 | 0.18654085 |
| Gm13305 | 7.712389657 | 7.687017062 | 7.525890894 | 0.161126168 | 0.186498763 |
| Polr1c | 9.277212642 | 9.149216651 | 9.090738765 | 0.058477886 | 0.186473877 |
| Ankrd66 | 3.643980127 | 3.52811106 | 3.457602231 | 0.070508829 | 0.186377896 |
| Rtcb | 11.22504992 | 11.00193803 | 11.03874375 | -0.036805713 | 0.186306177 |
| Tnfsf11 | 3.149349206 | 2.970414558 | 2.963287725 | 0.007126833 | 0.186061481 |
| Pdap1 | 10.88438527 | 10.76189909 | 10.69835205 | 0.063547037 | 0.186033213 |
| Lamtor3 | 10.05876931 | 10.07881739 | 9.87301437 | 0.205803017 | 0.185754937 |
| Hist2h2aa2 | 6.320506046 | 6.474668066 | 6.134752579 | 0.339915488 | 0.185753467 |
| Myl12b | 11.80318602 | 11.66422169 | 11.61768515 | 0.04653654 | 0.185500867 |
| Tma7 | 10.27126484 | 10.03115959 | 10.08619784 | -0.055038253 | 0.185067003 |
| Psma7 | 11.25796043 | 11.08894565 | 11.07292293 | 0.016022717 | 0.185037497 |

| | | | | | |
|---|---|---|---|---|---|
| Zcchc7 | 9.782429672 | 9.65271461 | 9.597609741 | 0.055104869 | 0.184819931 |
| H2-T23 | 11.4495647 | 11.24772061 | 11.26475157 | -0.017030963 | 0.184813127 |
| Lins1 | 7.781842056 | 7.639421472 | 7.597056264 | 0.042365208 | 0.184785791 |
| Smarce1 | 10.84202182 | 10.56453777 | 10.65727651 | -0.092738743 | 0.18474531 |
| Tmtc2 | 7.245786877 | 6.884279058 | 7.061051463 | -0.176772404 | 0.184735414 |
| Tcfl5 | 3.434027835 | 3.272915056 | 3.249305072 | 0.023609984 | 0.184722764 |
| Rasl10b | 3.708667949 | 3.585739512 | 3.524126583 | 0.061612929 | 0.184541366 |
| Atp6ap2 | 10.78839208 | 10.72311933 | 10.6039383 | 0.11918103 | 0.18445378 |
| Mcm2 | 9.870791909 | 10.01546265 | 9.686343837 | 0.329118811 | 0.184448072 |
| Lamp2 | 13.17014875 | 12.99448147 | 12.98570612 | 0.008775357 | 0.18444263 |
| Ttll3 | 5.961289219 | 5.887860214 | 5.776934813 | 0.110925401 | 0.184354406 |
| Slc25a5 | 13.03405539 | 12.96304156 | 12.84971528 | 0.113326283 | 0.184340117 |
| Sema3a | 8.152474142 | 7.965616723 | 7.968287482 | -0.002670758 | 0.184186661 |
| Rad18 | 7.885240777 | 7.79898584 | 7.701096881 | 0.097888959 | 0.184143896 |
| Ostc | 10.06191619 | 9.984840367 | 9.877830752 | 0.107009615 | 0.184085435 |
| Samd11 | 4.306439602 | 4.57881767 | 4.12241138 | 0.456406291 | 0.184028223 |
| Sec22b | 10.35879471 | 10.27411507 | 10.17479472 | 0.09932035 | 0.18399999 |
| Ncoa5 | 9.468932784 | 9.391893238 | 9.284952586 | 0.106940652 | 0.183980198 |
| Fbxl22 | 6.05064644 | 6.003548617 | 5.866714063 | 0.136834554 | 0.183932377 |
| Ttc39c | 9.028346154 | 8.659383568 | 8.844454188 | -0.185070619 | 0.183891967 |
| Serpinb6e | 8.591537796 | 8.412233628 | 8.407693608 | 0.00454002 | 0.183844188 |
| Pld4 | 8.088465383 | 7.917282032 | 7.904759687 | 0.012522345 | 0.183705696 |
| Senp3 | 10.30899666 | 10.34626222 | 10.12532247 | 0.220939753 | 0.18367419 |
| Fam84b | 9.050364371 | 8.844542116 | 8.866758387 | -0.022216271 | 0.183605984 |
| Api5 | 11.40669157 | 11.264819 | 11.22309653 | 0.04172247 | 0.18359504 |
| Pafah1b2 | 10.90718555 | 10.8256634 | 10.72374905 | 0.10191435 | 0.183436503 |
| Tgm3 | 8.127429937 | 8.021731304 | 7.944070368 | 0.077660936 | 0.183359569 |
| Mmp13 | 6.85285696 | 6.911461189 | 6.669511998 | 0.241949192 | 0.183344963 |
| Gramd3 | 10.46878994 | 10.22849537 | 10.2855457 | -0.057050327 | 0.183244243 |
| Tmem86b | 5.921466889 | 5.967909054 | 5.738228815 | 0.229680239 | 0.183238074 |
| Itgav | 10.88227963 | 10.6302832 | 10.69912992 | -0.06884672 | 0.183149707 |
| Rab3il1 | 9.543360399 | 9.738165409 | 9.360228994 | 0.377936415 | 0.183131405 |

| | | | | | |
|---|---|---|---|---|---|
| Pld1 | 9.507047793 | 9.340401189 | 9.323968429 | 0.01643276 | 0.183079364 |
| Cenpl | 7.212042066 | 7.006653809 | 7.029151978 | -0.022498168 | 0.182890088 |
| Svbp | 7.976826709 | 7.808810095 | 7.793969158 | 0.014840937 | 0.182857551 |
| Zgpat | 9.147513224 | 9.376186944 | 8.964688407 | 0.411498537 | 0.182824817 |
| Mrps30 | 9.065955658 | 8.78154772 | 8.883136788 | -0.101589067 | 0.182818871 |
| Rab5if | 10.34411096 | 10.16914415 | 10.16130982 | 0.007834337 | 0.182801143 |
| Tigd5 | 5.908896428 | 5.820681163 | 5.72609577 | 0.094585394 | 0.182800659 |
| Gm14440 | 6.138194168 | 6.36371978 | 5.955450449 | 0.408269331 | 0.182743719 |
| Pak1 | 9.886148419 | 9.702924346 | 9.703478981 | -0.000554635 | 0.182669438 |
| Ndst1 | 11.66865565 | 11.6144395 | 11.4860635 | 0.128375993 | 0.182592147 |
| Rraga | 10.40394711 | 10.20301475 | 10.22148673 | -0.01847198 | 0.18246038 |
| Sla2 | 4.769221016 | 4.618826157 | 4.586767714 | 0.032058443 | 0.182453302 |
| Hccs | 8.770392699 | 8.52235638 | 8.588118105 | -0.065761725 | 0.182274594 |
| Ces4a | 10.95983066 | 10.89169105 | 10.77760699 | 0.114084063 | 0.182223673 |
| Scly | 8.052945654 | 8.029542004 | 7.870727324 | 0.158814681 | 0.182218331 |
| Exoc3l2 | 8.244508861 | 8.259709683 | 8.062345043 | 0.19736464 | 0.182163817 |
| Sars | 11.2148923 | 11.17573516 | 11.03284836 | 0.142886803 | 0.182043943 |
| Snrpg | 9.54943417 | 9.340286056 | 9.367408277 | -0.027122222 | 0.182025892 |
| P4hb | 13.38795114 | 13.41090218 | 13.20594498 | 0.2049572 | 0.182006157 |
| Pcdhga9 | 5.433813913 | 5.365582879 | 5.252225553 | 0.113357325 | 0.18158836 |
| Irx3 | 8.543386435 | 8.792480744 | 8.361825403 | 0.430655342 | 0.181561033 |
| Rnf20 | 10.5599771 | 10.44558821 | 10.3784569 | 0.06713131 | 0.181520203 |
| Acat3 | 4.352634081 | 4.362554825 | 4.171124234 | 0.19143059 | 0.181509846 |
| Traf7 | 10.77014944 | 10.85581908 | 10.58874995 | 0.26706913 | 0.181399483 |
| Med18 | 6.708349715 | 6.658275779 | 6.527019253 | 0.131256526 | 0.181330461 |
| Ndufa8 | 10.21685995 | 10.11782897 | 10.03555343 | 0.08227554 | 0.181306527 |
| Uso1 | 11.30395675 | 11.16687084 | 11.12273789 | 0.044132957 | 0.181218867 |
| Khdc4 | 10.47962641 | 10.53862997 | 10.29844375 | 0.240186217 | 0.18118266 |
| Dbndd2 | 8.826973477 | 8.757798384 | 8.645794127 | 0.112004257 | 0.18117935 |
| Tut7 | 11.21716413 | 11.02530194 | 11.03600867 | -0.01070673 | 0.181155463 |
| Ube2d2a | 11.35871217 | 11.09909104 | 11.17762356 | -0.078532523 | 0.181088607 |
| Ywhag | 12.15555371 | 12.05640197 | 11.97461326 | 0.08178871 | 0.18094045 |

| | | | | | |
|---|---|---|---|---|---|
| Cd276 | 8.135294631 | 8.235671642 | 7.95437288 | 0.281298762 | 0.180921751 |
| Tpp2 | 10.433153 | 10.29974525 | 10.25248061 | 0.047264633 | 0.180672387 |
| Med19 | 8.824770535 | 8.748918167 | 8.644118486 | 0.104799681 | 0.180652049 |
| Ppp6c | 10.74274155 | 10.59773453 | 10.56209148 | 0.035643057 | 0.18065007 |
| Dnaja3 | 10.33689863 | 10.11900813 | 10.15637536 | -0.037367223 | 0.180523277 |
| Ppp2r1a | 12.21650628 | 12.05388792 | 12.03607988 | 0.017808037 | 0.180426403 |
| Braf | 9.813163576 | 9.617508446 | 9.632737731 | -0.015229285 | 0.180425845 |
| Mvb12a | 8.921793625 | 8.977953117 | 8.741629138 | 0.236323979 | 0.180164486 |
| Bola1 | 6.081833917 | 6.299089601 | 5.901693034 | 0.397396567 | 0.180140882 |
| Atg2b | 10.2091164 | 10.04094175 | 10.02901706 | 0.011924693 | 0.180099343 |
| Gm49377 | 3.770439104 | 3.758340323 | 3.590374293 | 0.16796603 | 0.180064811 |
| Scnn1a | 9.843877871 | 9.854270122 | 9.663940525 | 0.190329597 | 0.179937347 |
| Cfdp1 | 10.4418886 | 10.19448339 | 10.26201498 | -0.067531597 | 0.179873613 |
| Etv3 | 11.01210474 | 11.13869791 | 10.83225116 | 0.30644675 | 0.17985358 |
| Gpbp1 | 10.8986115 | 10.80164719 | 10.71878281 | 0.082864377 | 0.179828693 |
| Tmem259 | 10.58052271 | 10.75927944 | 10.4007876 | 0.35849184 | 0.179735107 |
| Cdc42ep1 | 9.769159868 | 9.775741827 | 9.589443396 | 0.186298431 | 0.179716472 |
| Rpp21 | 7.263383019 | 7.304157587 | 7.083736525 | 0.220421062 | 0.179646495 |
| Get4 | 9.863292242 | 9.746053234 | 9.683739748 | 0.062313486 | 0.179552493 |
| Stk16 | 9.494112433 | 9.349551018 | 9.314607529 | 0.034943488 | 0.179504903 |
| Dxo | 8.335532264 | 8.228877185 | 8.156030307 | 0.072846878 | 0.179501957 |
| Sec63 | 11.11588452 | 11.05189456 | 10.93645676 | 0.1154378 | 0.17942776 |
| Tomm34 | 9.187851211 | 9.040152199 | 9.008460202 | 0.031691996 | 0.179391008 |
| Maml1 | 9.570724012 | 9.587285663 | 9.391453183 | 0.19583248 | 0.179270829 |
| Slc35g1 | 7.875921027 | 7.735229133 | 7.69669536 | 0.038533773 | 0.179225667 |
| Dcun1d5 | 9.905599194 | 9.716221975 | 9.726400294 | -0.01017832 | 0.1791989 |
| Tmem106a | 9.812030487 | 9.802200588 | 9.632847756 | 0.169352832 | 0.17918273 |
| Cox6a1 | 10.88506945 | 10.90860674 | 10.70597614 | 0.202630607 | 0.179093313 |
| Psmb3 | 10.80605875 | 10.84482847 | 10.62699257 | 0.217835897 | 0.179066177 |
| Col26a1 | 3.664413655 | 3.688925038 | 3.485426907 | 0.203498132 | 0.178986748 |
| Utp3 | 9.696344649 | 9.574200822 | 9.517499492 | 0.05670133 | 0.178845157 |
| Klri2 | 5.351078309 | 5.216407964 | 5.172275803 | 0.044132161 | 0.178802506 |

| | | | | | |
|---|---|---|---|---|---|
| Cstf3 | 8.937324218 | 8.919741166 | 8.758612318 | 0.161128848 | 0.1787119 |
| Ttc1 | 9.250405661 | 9.035073843 | 9.071699328 | -0.036625484 | 0.178706333 |
| Dse | 10.04101577 | 9.734320708 | 9.862325388 | -0.12800468 | 0.178690383 |
| Ppme1 | 9.770820998 | 9.730251896 | 9.592176026 | 0.13807587 | 0.178644971 |
| Snrpb2 | 9.564364665 | 9.407170833 | 9.385745163 | 0.021425671 | 0.178619503 |
| Limk1 | 8.281164068 | 8.444139953 | 8.10264735 | 0.341492603 | 0.178516718 |
| Psmg2 | 8.772299411 | 8.604699255 | 8.59379773 | 0.010901525 | 0.178501681 |
| Rbfa | 9.038812192 | 8.790771678 | 8.860334528 | -0.069562849 | 0.178477665 |
| Piezo1 | 10.80787057 | 10.85708983 | 10.62939387 | 0.22769596 | 0.1784767 |
| Gfi1b | 3.034753019 | 2.871837066 | 2.856313874 | 0.015523193 | 0.178439145 |
| Ywhaz | 13.947747 | 13.75845737 | 13.76934776 | -0.010890387 | 0.178399237 |
| Lpar5 | 7.239999083 | 7.324391195 | 7.061601155 | 0.26279004 | 0.178397928 |
| Bcl2l12 | 7.455366485 | 7.595006055 | 7.277002367 | 0.318003688 | 0.178364118 |
| Edf1 | 10.25572521 | 10.36941107 | 10.07740366 | 0.292007407 | 0.178321553 |
| Peak1 | 9.974553825 | 9.857401094 | 9.796417338 | 0.060983756 | 0.178136487 |
| Cmas | 9.218643927 | 8.880415033 | 9.040520443 | -0.16010541 | 0.178123484 |
| Depdc1b | 5.706167791 | 5.483551493 | 5.528054307 | -0.044502815 | 0.178113483 |
| Ak6 | 10.3817962 | 10.13604174 | 10.20387651 | -0.067834767 | 0.177919687 |
| Zfyve27 | 9.439897452 | 9.511821296 | 9.26211784 | 0.249703456 | 0.177779612 |
| Dok7 | 5.452231294 | 6.21782554 | 5.274636062 | 0.943189478 | 0.177595232 |
| Tmem189 | 10.60886109 | 10.55542445 | 10.43126947 | 0.124154983 | 0.177591623 |
| Safb | 10.664261 | 10.86248766 | 10.48674787 | 0.37573979 | 0.17751313 |
| Pgk1 | 12.99335888 | 12.85265731 | 12.81602096 | 0.036636347 | 0.17733792 |
| Inip | 8.951680211 | 8.854136452 | 8.774384955 | 0.079751498 | 0.177295256 |
| Btbd9 | 9.659111864 | 9.505163143 | 9.481891386 | 0.023271757 | 0.177220478 |
| Tspan8 | 10.13746978 | 9.849833804 | 9.960334158 | -0.110500354 | 0.177135625 |
| Ier3ip1 | 9.783391841 | 9.916752464 | 9.606271476 | 0.310480988 | 0.177120366 |
| Prelid3b | 10.58538791 | 10.35295928 | 10.40841392 | -0.05545464 | 0.176973997 |
| Ceacam10 | 2.21829326 | 2.108063109 | 2.041543409 | 0.066519701 | 0.176749851 |
| Nt5dc1 | 8.936416485 | 8.646353966 | 8.7596691 | -0.113315134 | 0.176747385 |
| Mmadhc | 10.88213648 | 10.61828051 | 10.70538921 | -0.087108703 | 0.176747263 |
| Cd200r3 | 5.662223254 | 5.461216174 | 5.485537295 | -0.024321121 | 0.176685959 |

| | | | | | |
|---|---|---|---|---|---|
| AW822252 | 4.634095587 | 4.455769509 | 4.457516495 | -0.001746986 | 0.176579092 |
| Epha2 | 9.626169911 | 9.803958825 | 9.449699274 | 0.354259551 | 0.176470637 |
| Foxk2 | 10.29202717 | 10.34922759 | 10.11573602 | 0.233491567 | 0.176291147 |
| Ostf1 | 10.85696082 | 10.6929342 | 10.68079916 | 0.01213504 | 0.176161657 |
| Mcpt-ps1 | 3.460289338 | 3.266835735 | 3.284396652 | -0.017560917 | 0.175892687 |
| Seh1l | 10.76045298 | 10.58381406 | 10.58460044 | -0.000786373 | 0.175852543 |
| Cks2 | 8.244408988 | 8.114424564 | 8.06855855 | 0.045866014 | 0.175850438 |
| Aamp | 11.36739001 | 11.35448496 | 11.19157585 | 0.162909113 | 0.175814163 |
| Pam16 | 5.679180217 | 5.708777398 | 5.503466718 | 0.20531068 | 0.175713499 |
| Ppp1r10 | 10.21401742 | 10.40879772 | 10.03865749 | 0.370140234 | 0.175359934 |
| Hnrnpa1 | 13.15934269 | 13.04333067 | 12.98405007 | 0.0592806 | 0.175292617 |
| Erich2 | 2.871931408 | 2.804722952 | 2.696719497 | 0.108003454 | 0.17521191 |
| Tnfrsf4 | 7.528672444 | 7.573955222 | 7.353568778 | 0.220386443 | 0.175103665 |
| Morc2a | 10.18048521 | 10.2365657 | 10.00539338 | 0.231172314 | 0.175091827 |
| Mecr | 8.742890175 | 8.733047681 | 8.567815719 | 0.165231962 | 0.175074456 |
| Aasdhppt | 9.121746161 | 8.953857366 | 8.946957387 | 0.006899979 | 0.174788774 |
| Dbn1 | 7.323678087 | 7.142742647 | 7.149076319 | -0.006333672 | 0.174601767 |
| Uhrf1 | 10.02562194 | 10.18868584 | 9.851052063 | 0.33763378 | 0.174569878 |
| Tmem258 | 9.193096218 | 9.044433501 | 9.018620852 | 0.025812649 | 0.174475366 |
| Cnnm1 | 4.634789156 | 4.62406947 | 4.460397798 | 0.163671672 | 0.174391358 |
| Sh3rf1 | 9.037114718 | 8.909306733 | 8.862804302 | 0.046502432 | 0.174310417 |
| Zfp365 | 8.014923316 | 7.793836618 | 7.840643627 | -0.046807009 | 0.174279688 |
| Ripk4 | 10.16252226 | 10.14910288 | 9.988249882 | 0.160852994 | 0.174272381 |
| Psmd6 | 10.66680028 | 10.56113503 | 10.49265088 | 0.06848415 | 0.174149393 |
| Pcdhga7 | 5.83510325 | 5.902251716 | 5.661031082 | 0.241220635 | 0.174072168 |
| Ddx46 | 11.13651607 | 11.11630468 | 10.96249149 | 0.153813187 | 0.174024573 |
| Tnfsf18 | 4.947157883 | 4.840610329 | 4.773144968 | 0.067465362 | 0.174012915 |
| Skap1 | 5.385677821 | 5.213123112 | 5.211678169 | 0.001444943 | 0.173999652 |
| Cars | 10.19973911 | 10.06045787 | 10.02586569 | 0.034592181 | 0.173873421 |
| Snx17 | 10.45540681 | 10.34872928 | 10.28153818 | 0.067191103 | 0.173868627 |
| D8Ertd738e | 10.47482711 | 10.46682272 | 10.30096171 | 0.165861003 | 0.173865393 |
| Selenos | 10.47464315 | 10.34771163 | 10.30095165 | 0.046759973 | 0.173691497 |

| | | | | | |
|---|---|---|---|---|---|
| Kctd13 | 6.904056874 | 6.900584656 | 6.730585271 | 0.169999385 | 0.173471603 |
| Enkd1 | 5.902348597 | 5.916694623 | 5.728877646 | 0.187816977 | 0.173470951 |
| Psmb6 | 11.1215359 | 11.08031798 | 10.94806963 | 0.132248347 | 0.173466263 |
| Wfdc3 | 8.633286795 | 8.580886335 | 8.459828954 | 0.121057381 | 0.173457841 |
| Per1 | 10.81055756 | 11.03758635 | 10.63712198 | 0.400464373 | 0.17343558 |
| Gm14735 | 3.941599686 | 3.928191107 | 3.768192851 | 0.159998256 | 0.173406835 |
| Cox18 | 7.275091066 | 7.178819865 | 7.101735868 | 0.077083997 | 0.173355198 |
| Tedc2 | 7.656334643 | 7.700669576 | 7.483038204 | 0.217631372 | 0.173296439 |
| Ccnd1 | 10.99246342 | 10.93104693 | 10.81917379 | 0.11187314 | 0.173289627 |
| Foxj3 | 9.94911995 | 9.948528257 | 9.775836051 | 0.172692206 | 0.173283899 |
| Lyst | 10.35561514 | 10.26677052 | 10.18237893 | 0.084391587 | 0.173236207 |
| Gm42417 | 3.507457151 | 3.395155639 | 3.334290021 | 0.060865618 | 0.17316713 |
| Mpp1 | 10.71605773 | 10.48634185 | 10.54289386 | -0.056552003 | 0.17316387 |
| Camta2 | 9.56039324 | 9.838807981 | 9.387250037 | 0.451557944 | 0.173143203 |
| Rps6kb1 | 10.23857759 | 10.05786338 | 10.06551435 | -0.007650972 | 0.173063232 |
| Cdyl2 | 5.161050644 | 4.967378673 | 4.988001765 | -0.020623093 | 0.173048879 |
| Osbpl7 | 7.459991563 | 7.231355176 | 7.286975761 | -0.055620585 | 0.173015802 |
| Tmem267 | 7.936626533 | 8.50122416 | 7.763722442 | 0.737501717 | 0.17290409 |
| Pgm3 | 7.855705718 | 7.733505547 | 7.68286334 | 0.050642208 | 0.172842379 |
| Nupr1 | 10.61306499 | 10.47325448 | 10.44026059 | 0.032993887 | 0.1728044 |
| Asxl1 | 10.19191605 | 10.07941439 | 10.01913185 | 0.060282534 | 0.172784194 |
| Mrpl13 | 9.418862555 | 9.220006769 | 9.246155494 | -0.026148724 | 0.172707061 |
| Sprtn | 9.524889083 | 9.383468026 | 9.352256717 | 0.031211309 | 0.172632366 |
| Cul2 | 9.859449925 | 9.756120494 | 9.686840116 | 0.069280378 | 0.172609809 |
| 2410004B18Rik | 8.336258344 | 8.296710664 | 8.163666848 | 0.133043816 | 0.172591496 |
| Zc3h10 | 8.114594076 | 8.116353324 | 7.942003425 | 0.174349899 | 0.172590651 |
| Mrpl55 | 8.251376776 | 8.378249104 | 8.078828559 | 0.299420545 | 0.172548217 |
| Insrr | 3.723665732 | 3.578811658 | 3.551180369 | 0.027631289 | 0.172485363 |
| Rif1 | 10.00232488 | 9.831812433 | 9.829941367 | 0.001871066 | 0.172383511 |
| B3gnt7 | 4.571902383 | 4.601055069 | 4.399519446 | 0.201535623 | 0.172382937 |
| Stk32c | 4.316418193 | 4.238830336 | 4.144087877 | 0.094742459 | 0.172330316 |
| Hbs1l | 10.50746616 | 10.42471308 | 10.33538566 | 0.089327423 | 0.172080503 |

| | | | | | |
|---|---|---|---|---|---|
| Havcr1 | 2.884847756 | 2.871067772 | 2.71279523 | 0.158272541 | 0.172052526 |
| 9930111J21Rik2 | 9.287530747 | 8.87071685 | 9.11557551 | -0.24485866 | 0.171955236 |
| Smg5 | 10.81081156 | 10.81679464 | 10.6388778 | 0.17791684 | 0.17193376 |
| Rbm3 | 13.10603142 | 12.88560521 | 12.93419965 | -0.048594443 | 0.17183177 |
| Aebp2 | 10.86886266 | 10.83037655 | 10.69713007 | 0.133246477 | 0.171732593 |
| Ndufs8 | 9.878184016 | 9.885433176 | 9.706494495 | 0.178938681 | 0.171689521 |
| Limk2 | 12.07708882 | 12.10032683 | 11.90543957 | 0.194887263 | 0.171649257 |
| Lama5 | 10.23256974 | 10.28933122 | 10.06092062 | 0.2284106 | 0.171649124 |
| Galnt9 | 4.206250916 | 4.076877422 | 4.034657171 | 0.042220251 | 0.171593744 |
| Tmem50b | 10.24403711 | 10.08735241 | 10.07244927 | 0.014903135 | 0.171587839 |
| Ncor2 | 11.02944403 | 11.28450346 | 10.8578694 | 0.42663406 | 0.171574623 |
| Fzd5 | 8.276525704 | 8.323260748 | 8.105004315 | 0.218256433 | 0.171521389 |
| Polr2i | 8.067268431 | 8.048623529 | 7.895776928 | 0.152846601 | 0.171491502 |
| Nhej1 | 6.540802409 | 6.666770301 | 6.369354769 | 0.297415533 | 0.17144764 |
| Prkar1a | 12.16939474 | 12.00136197 | 11.99806514 | 0.00329683 | 0.171329603 |
| Zfp511 | 7.878893544 | 7.904303843 | 7.707575127 | 0.196728716 | 0.171318417 |
| Rdh12 | 10.67682482 | 10.44630886 | 10.50557188 | -0.059263023 | 0.171252943 |
| Smu1 | 10.73710903 | 10.53676064 | 10.5658692 | -0.02910856 | 0.171239827 |
| Hcfc1 | 11.07989067 | 11.10673388 | 10.90869925 | 0.19803463 | 0.171191423 |
| Clec2d | 11.25649591 | 11.05311142 | 11.08533912 | -0.032227703 | 0.171156783 |
| Agk | 8.281250678 | 8.178780598 | 8.110096077 | 0.068684521 | 0.171154601 |
| Fip1l1 | 10.16174563 | 10.1550881 | 9.990700508 | 0.164387595 | 0.171045122 |
| Rps26 | 12.51662303 | 12.60408782 | 12.34560574 | 0.258482083 | 0.171017287 |
| Rnaseh1 | 7.816889627 | 7.778655474 | 7.645925161 | 0.132730312 | 0.170964466 |
| C2cd2l | 8.621173914 | 8.545906374 | 8.45041703 | 0.095489344 | 0.170756885 |
| Ednrb | 9.389471026 | 9.491207517 | 9.218762832 | 0.272444685 | 0.170708194 |
| Tmem97 | 9.425205645 | 9.253752277 | 9.254504014 | -0.000751738 | 0.170701631 |
| Tmem39a | 9.036870657 | 9.226747465 | 8.86619332 | 0.360554144 | 0.170677337 |
| Ankrd10 | 9.585061627 | 9.913121411 | 9.414522692 | 0.498598719 | 0.170538935 |
| Gm29397 | 5.553438096 | 5.080065494 | 5.383023041 | -0.302957547 | 0.170415056 |
| Ptpn22 | 7.378451957 | 7.291849454 | 7.208057508 | 0.083791946 | 0.170394449 |
| Rpl19 | 13.56853664 | 13.50064966 | 13.39818602 | 0.10246364 | 0.170350623 |

| | | | | | |
|---|---|---|---|---|---|
| Emc6 | 9.495745482 | 9.332827136 | 9.325439169 | 0.007387967 | 0.170306313 |
| Naa10 | 9.622305923 | 9.627778353 | 9.45207934 | 0.175699013 | 0.170226583 |
| Sin3b | 10.33999404 | 10.24386097 | 10.16993206 | 0.073928917 | 0.17006198 |
| Prickle1 | 7.845014488 | 7.825100486 | 7.674987054 | 0.150113432 | 0.170027433 |
| Tmem266 | 3.955737085 | 3.825819753 | 3.785780079 | 0.040039674 | 0.169957006 |
| 9930104L06Rik | 8.330835097 | 8.254952901 | 8.160948554 | 0.094004346 | 0.169886543 |
| Serf2 | 11.98438138 | 11.87692294 | 11.81461896 | 0.06230398 | 0.16976242 |
| Dclre1b | 8.295065646 | 8.078122061 | 8.125429221 | -0.047307161 | 0.169636425 |
| Mrps28 | 8.345848457 | 8.382550793 | 8.176271985 | 0.206278808 | 0.169576472 |
| Pcnx | 10.72387218 | 10.58781346 | 10.55447927 | 0.033334187 | 0.169392907 |
| Grip2 | 6.74988379 | 6.730165079 | 6.580537411 | 0.149627668 | 0.169346379 |
| Vps4b | 11.34905151 | 11.17920826 | 11.17972808 | -0.00051982 | 0.169323437 |
| Bicd2 | 11.64798816 | 11.62570109 | 11.47866617 | 0.147034923 | 0.169321993 |
| Prr14 | 9.943473765 | 9.896145144 | 9.774198665 | 0.121946479 | 0.1692751 |
| Prr5 | 6.768486722 | 6.781789348 | 6.599236297 | 0.18255305 | 0.169250424 |
| Pomk | 8.3634006 | 8.176778347 | 8.194238963 | -0.017460615 | 0.169161637 |
| Psma6 | 11.51232616 | 11.28197645 | 11.34322199 | -0.061245533 | 0.169104177 |
| Rnf138 | 8.760910621 | 8.460351543 | 8.592004913 | -0.13165337 | 0.168905708 |
| Ankrd44 | 9.643936656 | 9.541145353 | 9.475097921 | 0.066047432 | 0.168838735 |
| Yars2 | 7.981342095 | 7.854123134 | 7.812544895 | 0.041578239 | 0.1687972 |
| Cd7 | 7.498319046 | 7.419179894 | 7.329630451 | 0.089549442 | 0.168688595 |
| Gdap2 | 8.687982016 | 8.596147378 | 8.519342581 | 0.076804797 | 0.168639435 |
| Serpinb6b | 5.553923693 | 5.508585154 | 5.385317567 | 0.123267587 | 0.168606126 |
| Zdhhc17 | 8.878489022 | 8.53739024 | 8.70990852 | -0.17251828 | 0.168580502 |
| Lpcat3 | 11.33898483 | 11.23936816 | 11.17044857 | 0.068919583 | 0.16853626 |
| Dars | 10.69227681 | 10.48048201 | 10.52376598 | -0.043283963 | 0.16851083 |
| Cdkn1a | 12.17849055 | 12.15675809 | 12.01002651 | 0.146731587 | 0.168464047 |
| Nes | 8.633657282 | 8.921706593 | 8.465207355 | 0.456499238 | 0.168449927 |
| Sft2d1 | 8.733874401 | 8.611420034 | 8.565425881 | 0.045994153 | 0.16844852 |
| Lrfn1 | 5.300373303 | 5.271419059 | 5.131976689 | 0.13944237 | 0.168396614 |
| Mrpl4 | 9.503072995 | 9.517721396 | 9.334676659 | 0.183044737 | 0.168396336 |
| Luzp1 | 11.14839844 | 11.15629486 | 10.98010445 | 0.176190417 | 0.168293993 |

| | | | | | |
|---|---|---|---|---|---|
| Rbks | 5.837136665 | 5.695090968 | 5.668850352 | 0.026240616 | 0.168286313 |
| Fmnl3 | 8.259118306 | 8.018390273 | 8.09101399 | -0.072623717 | 0.168104316 |
| Gm614 | 2.711703117 | 2.594240489 | 2.543881329 | 0.05035916 | 0.167821788 |
| Ppfia1 | 10.64195056 | 10.42155782 | 10.47436756 | -0.05280974 | 0.167583 |
| Traf3 | 9.54009496 | 9.764445872 | 9.372528785 | 0.391917087 | 0.167566175 |
| Cldn12 | 9.294512466 | 9.173392862 | 9.127026312 | 0.04636655 | 0.167486154 |
| Smarcc1 | 10.7958294 | 10.73924359 | 10.62844615 | 0.11079744 | 0.16738325 |
| Trp53rkb | 6.401644714 | 6.380459487 | 6.234484681 | 0.145974805 | 0.167160032 |
| Nf1 | 9.438064825 | 9.335020377 | 9.270908789 | 0.064111589 | 0.167156037 |
| Emc8 | 9.392487003 | 9.450595767 | 9.225378514 | 0.225217254 | 0.16710849 |
| Cbl | 11.24653458 | 11.38163418 | 11.07946702 | 0.302167163 | 0.16706756 |
| Baiap3 | 3.757100035 | 3.567837544 | 3.590063857 | -0.022226313 | 0.167036178 |
| P2rx4 | 9.57547256 | 9.507665698 | 9.408440507 | 0.099225191 | 0.167032053 |
| Snrnp27 | 9.122621732 | 8.962721619 | 8.95563545 | 0.007086169 | 0.166986282 |
| Srarp | 4.449785591 | 4.498647837 | 4.282968582 | 0.215679255 | 0.166817009 |
| Adat1 | 6.993190964 | 6.950328199 | 6.826464003 | 0.123864197 | 0.166726961 |
| Rps13 | 12.03923552 | 12.06465845 | 11.87263261 | 0.192025843 | 0.166602917 |
| Stau1 | 10.59960217 | 10.4476685 | 10.43301136 | 0.014657143 | 0.166590813 |
| Gna14 | 7.959113221 | 7.910970779 | 7.792612423 | 0.118358356 | 0.166500798 |
| Klrd1 | 6.253801371 | 6.100854778 | 6.08744979 | 0.013404988 | 0.166351581 |
| Slco4c1 | 3.13889117 | 3.0787286 | 2.972619698 | 0.106108902 | 0.166271472 |
| Itgb1bp1 | 9.198728975 | 9.04973793 | 9.032521309 | 0.017216621 | 0.166207666 |
| Dynll1 | 12.36726921 | 12.2992344 | 12.20117108 | 0.09806332 | 0.166098137 |
| Atp6v1g1 | 11.10864483 | 10.97502801 | 10.94255028 | 0.03247773 | 0.16609455 |
| Ssh1 | 8.47223107 | 8.416643639 | 8.30622077 | 0.110422869 | 0.1660103 |
| Zdhhc23 | 8.030939491 | 7.889797689 | 7.864952101 | 0.024845588 | 0.165987391 |
| Trps1 | 9.244189948 | 8.981378585 | 9.078404926 | -0.097026342 | 0.165785021 |
| Cpne3 | 10.92965553 | 10.82261754 | 10.76388498 | 0.058732563 | 0.16577055 |
| Txndc5 | 11.19890302 | 11.08948174 | 11.03321041 | 0.05627133 | 0.165692607 |
| Mesd | 9.774628928 | 9.590307618 | 9.608990874 | -0.018683256 | 0.165638054 |
| Lzic | 8.807652258 | 8.352723954 | 8.642023179 | -0.289299226 | 0.165629078 |
| Aftph | 11.16984176 | 10.8797843 | 11.00433506 | -0.124550763 | 0.165506697 |

| | | | | | |
|---|---|---|---|---|---|
| Tank | 9.853981457 | 9.731070791 | 9.688560496 | 0.042510295 | 0.165420961 |
| Drg1 | 10.27196757 | 10.11339295 | 10.10661283 | 0.006780127 | 0.165354743 |
| Btaf1 | 10.37650784 | 10.35863079 | 10.21119197 | 0.147438823 | 0.165315873 |
| Rps5 | 13.27809613 | 13.30127014 | 13.11280089 | 0.188469257 | 0.16529524 |
| Spns3 | 4.18578068 | 4.077905635 | 4.020712164 | 0.057193471 | 0.165068516 |
| Dagla | 7.399865515 | 7.377649354 | 7.234866038 | 0.142783316 | 0.164999478 |
| Trabd | 10.13684246 | 10.32295123 | 9.971891485 | 0.351059742 | 0.164950978 |
| Ctsb | 14.39455995 | 14.3748331 | 14.22963973 | 0.145193373 | 0.164920223 |
| Reps1 | 9.876686121 | 9.796823568 | 9.711781788 | 0.085041779 | 0.164904332 |
| Nsrp1 | 9.413606989 | 9.174623261 | 9.248732453 | -0.074109192 | 0.164874536 |
| Gm5416 | 4.940139455 | 4.819966113 | 4.775312787 | 0.044653326 | 0.164826668 |
| Hat1 | 9.588926103 | 9.436114798 | 9.424190683 | 0.011924115 | 0.16473542 |
| Samd4 | 8.392110513 | 8.68518168 | 8.227402912 | 0.457778768 | 0.164707601 |
| Msx3 | 2.821215377 | 2.692024639 | 2.656522112 | 0.035502527 | 0.164693265 |
| Mrps34 | 8.600230501 | 8.506046264 | 8.435572906 | 0.070473358 | 0.164657595 |
| Otud6b | 10.09023561 | 9.920056055 | 9.925809263 | -0.005753208 | 0.164426344 |
| Tigit | 3.871707898 | 3.730021713 | 3.707367215 | 0.022654499 | 0.164340684 |
| Tbc1d1 | 9.522851273 | 9.337053311 | 9.358560165 | -0.021506854 | 0.164291108 |
| Apba3 | 9.057060019 | 9.185530291 | 8.892832573 | 0.292697718 | 0.164227447 |
| Rhbdd2 | 8.386406659 | 8.476397778 | 8.222218078 | 0.2541797 | 0.164188581 |
| Ercc3 | 9.556553971 | 9.466166546 | 9.392395204 | 0.073771342 | 0.164158767 |
| Ywhae | 13.06962324 | 12.93360226 | 12.90548102 | 0.028121237 | 0.164142217 |
| Fam122a | 7.22027194 | 7.108047906 | 7.056187291 | 0.051860615 | 0.164084649 |
| Cracr2a | 5.901623794 | 5.634678336 | 5.737597634 | -0.102919299 | 0.164026159 |
| Caap1 | 7.287837415 | 7.151007462 | 7.123852358 | 0.027155104 | 0.163985057 |
| Pebp1 | 11.51742645 | 11.45670844 | 11.35366415 | 0.10304429 | 0.163762307 |
| Babam2 | 9.079518862 | 8.864899693 | 8.915925398 | -0.051025705 | 0.163593463 |
| Rpusd4 | 7.851527763 | 7.836947102 | 7.687978305 | 0.148968797 | 0.163549458 |
| Necap2 | 10.06203897 | 10.11684753 | 9.898506271 | 0.218341256 | 0.163532699 |
| Gm45717 | 3.774335688 | 3.776920098 | 3.610885855 | 0.166034243 | 0.163449833 |
| Ero1lb | 7.724825211 | 7.814292199 | 7.561419048 | 0.252873151 | 0.163406163 |
| Rnf219 | 8.175638008 | 7.995913938 | 8.012271139 | -0.0163572 | 0.163366869 |

| | | | | | |
|---|---|---|---|---|---|
| Olfr839-ps1 | 5.617343461 | 5.492945568 | 5.453982926 | 0.038962642 | 0.163360535 |
| Nptx2 | 7.179821053 | 7.089263106 | 7.016466904 | 0.072796202 | 0.163354148 |
| Arhgdig | 3.665770036 | 3.598024494 | 3.502474009 | 0.095550485 | 0.163296027 |
| Gpatch2l | 9.122039343 | 9.007603675 | 8.958869349 | 0.048734326 | 0.163169994 |
| Cables1 | 8.585556839 | 8.433342666 | 8.422444769 | 0.010897897 | 0.16311207 |
| Lrrc14 | 8.509146761 | 8.424436918 | 8.346060299 | 0.078376619 | 0.163086462 |
| Psmd1 | 12.04839524 | 12.00691978 | 11.88531832 | 0.12160146 | 0.16307692 |
| Tnpo3 | 11.19808366 | 11.0770888 | 11.03512691 | 0.041961893 | 0.162956747 |
| Dstn | 13.77389352 | 13.63042401 | 13.61108399 | 0.019340013 | 0.162809523 |
| Rnase6 | 6.653072891 | 6.301585604 | 6.490377151 | -0.188791547 | 0.16269574 |
| Htr1b | 4.464030543 | 4.288911766 | 4.301347023 | -0.012435257 | 0.162683519 |
| Nrarp | 10.69710413 | 10.82904301 | 10.53442678 | 0.294616227 | 0.162677347 |
| 2610001J05Rik | 10.17214179 | 9.936087252 | 10.00946623 | -0.07337898 | 0.162675562 |
| Rps8 | 13.79535298 | 13.6600306 | 13.63273079 | 0.027299803 | 0.162622183 |
| Drosha | 9.899737329 | 9.677483626 | 9.73724315 | -0.059759524 | 0.162494179 |
| Srebf2 | 11.80534543 | 11.92573839 | 11.64293601 | 0.282802383 | 0.16240942 |
| Zfp516 | 9.31684913 | 9.641120754 | 9.154481317 | 0.486639437 | 0.162367813 |
| Rassf8 | 9.030427972 | 8.85434721 | 8.868122328 | -0.013775118 | 0.162305643 |
| Ago3 | 9.399588845 | 9.514765497 | 9.237345844 | 0.277419653 | 0.162243001 |
| Prrx1 | 11.49224135 | 11.13716005 | 11.33001514 | -0.192855093 | 0.162226207 |
| Arfip2 | 9.504959951 | 9.586231828 | 9.342754317 | 0.243477511 | 0.162205634 |
| Ankfy1 | 11.21859976 | 11.12709216 | 11.05651831 | 0.07057385 | 0.16208145 |
| Prnp | 12.62008372 | 12.42490637 | 12.4580243 | -0.033117933 | 0.162059417 |
| Smarca4 | 11.54949177 | 11.57540863 | 11.38754758 | 0.187861047 | 0.161944193 |
| Emb | 11.45601603 | 11.2695744 | 11.29410102 | -0.024526617 | 0.161915007 |
| Thop1 | 10.14089328 | 10.19965829 | 9.979109264 | 0.220549029 | 0.161784016 |
| Wrnip1 | 8.954888564 | 9.054616051 | 8.793127349 | 0.261488702 | 0.161761215 |
| Krt75 | 10.8896141 | 10.74038987 | 10.7278857 | 0.012504173 | 0.161728407 |
| Kdm5c | 11.56492261 | 11.61479427 | 11.40322154 | 0.21157273 | 0.161701067 |
| Prkcz | 7.428372146 | 7.221240235 | 7.26668716 | -0.045446925 | 0.161684986 |
| Mthfsl | 7.619164566 | 7.550101744 | 7.457518874 | 0.09258287 | 0.161645692 |
| Ghdc | 7.841187244 | 7.887022995 | 7.679541846 | 0.207481149 | 0.161645398 |

| | | | | | |
|---|---|---|---|---|---|
| Arl3 | 8.787906233 | 8.640349893 | 8.6263763 | 0.013973593 | 0.161529933 |
| Tssc4 | 8.558892283 | 8.578372739 | 8.397385335 | 0.180987404 | 0.161506948 |
| Plekhm1 | 9.998980413 | 10.13802643 | 9.837611712 | 0.300414718 | 0.161368701 |
| Dhx15 | 11.8655784 | 11.8381436 | 11.70422955 | 0.133914047 | 0.161348847 |
| Nmt1 | 11.17043191 | 11.04528158 | 11.00910378 | 0.036177803 | 0.161328137 |
| Wdr73 | 8.28626549 | 8.180372656 | 8.124955128 | 0.055417528 | 0.161310362 |
| Ubiad1 | 8.174685797 | 8.052594394 | 8.013419592 | 0.039174802 | 0.161266205 |
| Tmem199 | 8.348013748 | 8.241055388 | 8.186999879 | 0.054055509 | 0.161013869 |
| Ube2w | 9.453988586 | 9.213322952 | 9.29303756 | -0.079714608 | 0.160951026 |
| Oaz1 | 12.22836531 | 12.19286137 | 12.06741598 | 0.12544539 | 0.160949327 |
| Tmem39b | 7.496355918 | 7.42426811 | 7.335474595 | 0.088793515 | 0.160881323 |
| Prdm5 | 5.730818087 | 5.651102506 | 5.569943665 | 0.08115884 | 0.160874422 |
| Bahd1 | 9.305017831 | 9.383221758 | 9.144171753 | 0.239050005 | 0.160846078 |
| Srsf11 | 11.30971552 | 11.2878988 | 11.14891915 | 0.13897965 | 0.160796373 |
| Sidt1 | 4.507696001 | 4.393286712 | 4.347077785 | 0.046208927 | 0.160618216 |
| Snupn | 7.699078855 | 7.460079315 | 7.538501726 | -0.078422411 | 0.160577129 |
| Srrm1 | 11.27092604 | 11.24996227 | 11.11037936 | 0.13958291 | 0.16054668 |
| Prr7 | 2.833718357 | 2.774988147 | 2.673326048 | 0.101662099 | 0.160392309 |
| Ncbp2 | 9.827067752 | 9.789868874 | 9.666887804 | 0.12298107 | 0.160179948 |
| Ndufa11 | 9.820575657 | 9.844737727 | 9.660493045 | 0.184244682 | 0.160082611 |
| Ryk | 11.47553218 | 11.39126479 | 11.31545656 | 0.07580823 | 0.16007562 |
| Ddx52 | 9.25208634 | 9.097155915 | 9.092073867 | 0.005082048 | 0.160012473 |
| Col11a2 | 4.845760675 | 4.9356534 | 4.685802193 | 0.249851207 | 0.159958481 |
| Spopl | 8.867746982 | 8.649815012 | 8.707814515 | -0.057999503 | 0.159932467 |
| Slc9a3r1 | 10.12102114 | 10.16704108 | 9.961144963 | 0.205896113 | 0.159876173 |
| Ssbp1 | 8.823165459 | 8.655792504 | 8.663292294 | -0.00749979 | 0.159873165 |
| Praf2 | 7.983584531 | 7.902238651 | 7.823876484 | 0.078362167 | 0.159708047 |
| Rpp40 | 7.077104999 | 7.10563594 | 6.917511362 | 0.188124578 | 0.159593637 |
| Psmd2 | 12.65928979 | 12.61474707 | 12.4998043 | 0.114942767 | 0.159485493 |
| Stx18 | 8.509174662 | 8.287882334 | 8.349759201 | -0.061876867 | 0.15941546 |
| Tmbim6 | 13.15759005 | 13.13219329 | 12.99840226 | 0.133791023 | 0.159187787 |
| Bnip2 | 10.82122051 | 10.76780579 | 10.66203999 | 0.1057658 | 0.159180523 |

| | | | | | |
|---|---|---|---|---|---|
| Rpn1 | 11.86771498 | 11.80832646 | 11.70856545 | 0.09976101 | 0.15914953 |
| Clk2 | 9.46159136 | 9.570307626 | 9.302575117 | 0.267732509 | 0.159016243 |
| Exoc7 | 10.10033452 | 9.837311879 | 9.941364435 | -0.104052557 | 0.158970088 |
| Ptpn9 | 9.524887553 | 9.405850203 | 9.366090482 | 0.039759721 | 0.158797071 |
| Kif17 | 3.738909183 | 3.740921323 | 3.580178863 | 0.16074246 | 0.15873032 |
| Pomgnt1 | 10.07838058 | 9.934036256 | 9.919677036 | 0.01435922 | 0.158703547 |
| Lsm4 | 8.245877967 | 8.110521715 | 8.087352985 | 0.02316873 | 0.158524982 |
| Smg7 | 10.89582488 | 10.77524542 | 10.7373749 | 0.037870527 | 0.15844998 |
| Snrnp70 | 11.84413286 | 11.8879203 | 11.6856925 | 0.2022278 | 0.158440363 |
| Plxnb1 | 11.12660359 | 11.32304073 | 10.9682589 | 0.354781823 | 0.158344687 |
| Nop10 | 9.418193304 | 9.218472245 | 9.259854107 | -0.041381862 | 0.158339197 |
| Cabin1 | 9.94804739 | 10.12213475 | 9.789741224 | 0.332393526 | 0.158306166 |
| Pcdhga3 | 5.739632209 | 5.726701153 | 5.58147628 | 0.145224873 | 0.158155929 |
| Eif4e2 | 10.42770472 | 10.31400407 | 10.26957267 | 0.044431403 | 0.158132053 |
| Edrf1 | 9.184996628 | 9.145472585 | 9.026948822 | 0.118523762 | 0.158047805 |
| Lrrc8c | 8.541660674 | 8.243780602 | 8.383620702 | -0.1398401 | 0.158039972 |
| Ntf5 | 6.187421333 | 6.103440259 | 6.029468557 | 0.073971701 | 0.157952776 |
| Gm8741 | 3.902581933 | 3.722577606 | 3.744799517 | -0.022221911 | 0.157782416 |
| Mrpl54 | 8.719134052 | 8.722130351 | 8.561389099 | 0.160741252 | 0.157744952 |
| Plekhf1 | 7.338792976 | 7.368023872 | 7.181064787 | 0.186959085 | 0.157728189 |
| Tgtp1 | 6.935691718 | 6.507384736 | 6.77801279 | -0.270628054 | 0.157678928 |
| Stk11 | 10.48667502 | 10.52975612 | 10.32906492 | 0.2006912 | 0.157610103 |
| Mvk | 9.852797797 | 9.936611192 | 9.695319579 | 0.241291613 | 0.157478217 |
| Banp | 7.873844329 | 8.318469952 | 7.716459675 | 0.602010278 | 0.157384654 |
| Nipsnap3b | 8.291956911 | 8.096479925 | 8.134606989 | -0.038127064 | 0.157349923 |
| Pdcd6 | 10.36096419 | 10.15741207 | 10.20369991 | -0.04628784 | 0.157264277 |
| Dcp1a | 9.578249524 | 9.512320994 | 9.42103003 | 0.091290964 | 0.157219494 |
| Gtpbp10 | 8.416301764 | 8.183969867 | 8.259087351 | -0.075117484 | 0.157214412 |
| Pqbp1 | 9.112708936 | 8.996348572 | 8.955666299 | 0.040682274 | 0.157042637 |
| Dph6 | 8.313574466 | 8.098748165 | 8.156552263 | -0.057804098 | 0.157022203 |
| Lman2 | 11.74071004 | 11.79944949 | 11.58377795 | 0.215671543 | 0.156932087 |
| Syt13 | 6.543731294 | 6.385549617 | 6.386809418 | -0.001259801 | 0.156921876 |

| | | | | | |
|---|---|---|---|---|---|
| Ddx55 | 8.490438974 | 8.459065008 | 8.333576568 | 0.12548844 | 0.156862406 |
| Bmp2k | 10.10045744 | 9.968698273 | 9.943661832 | 0.025036441 | 0.156795605 |
| Dmap1 | 8.234943798 | 8.164512098 | 8.078149561 | 0.086362537 | 0.156794237 |
| Drap1 | 10.75211234 | 10.62538111 | 10.59542424 | 0.029956873 | 0.156688103 |
| Jund | 9.028075779 | 9.180749141 | 8.871671759 | 0.309077382 | 0.15640402 |
| Mrps11 | 7.383901338 | 7.438489517 | 7.227510838 | 0.210978679 | 0.1563905 |
| Gm48850 | 2.860084974 | 2.729093473 | 2.70372604 | 0.025367434 | 0.156358934 |
| Dedd2 | 9.792573941 | 10.07489105 | 9.636270628 | 0.438620425 | 0.156303313 |
| Sox9 | 10.04726831 | 10.13098131 | 9.891006827 | 0.239974479 | 0.156261485 |
| Gmppa | 9.462460324 | 9.434759678 | 9.306202258 | 0.12855742 | 0.156258066 |
| Chst2 | 5.970377898 | 5.905308167 | 5.81417744 | 0.091130727 | 0.156200458 |
| Mocs1 | 9.522267003 | 9.361732596 | 9.366139408 | -0.004406813 | 0.156127594 |
| Adm2 | 2.652924279 | 2.499842442 | 2.496854641 | 0.002987801 | 0.156069638 |
| Psmc1 | 11.33237164 | 11.27229752 | 11.17634277 | 0.095954747 | 0.15602887 |
| Fam207a | 9.125428216 | 9.169413285 | 8.969427787 | 0.199985498 | 0.156000429 |
| Cse1l | 10.69185339 | 10.57767593 | 10.5358691 | 0.041806823 | 0.155984283 |
| Gpkow | 9.778382928 | 9.5889891 | 9.622590854 | -0.033601753 | 0.155792074 |
| Ccdc174 | 8.791733619 | 8.631149905 | 8.63629645 | -0.005146546 | 0.155437169 |
| Dnttip1 | 9.120235756 | 8.908383324 | 8.964804577 | -0.056421254 | 0.155431179 |
| Olfr12 | 3.611485844 | 3.578558073 | 3.456059044 | 0.12249903 | 0.1554268 |
| Eif1b | 9.119447821 | 9.046855417 | 8.964035984 | 0.082819433 | 0.155411837 |
| Shf | 7.055098096 | 6.929037728 | 6.899724483 | 0.029313244 | 0.155373613 |
| Foxf1 | 2.732345831 | 2.588911288 | 2.577057272 | 0.011854016 | 0.155288559 |
| Zfat | 6.782924487 | 6.835162401 | 6.627646073 | 0.207516328 | 0.155278414 |
| Rbm7 | 10.08563427 | 9.963918536 | 9.930357968 | 0.033560568 | 0.155276299 |
| Lsm1 | 8.102192857 | 7.950974451 | 7.946996177 | 0.003978274 | 0.15519668 |
| BC147527 | 5.948463701 | 5.641658974 | 5.793317338 | -0.151658364 | 0.155146363 |
| Acin1 | 11.38856422 | 11.45487907 | 11.23350951 | 0.221369557 | 0.15505471 |
| Prdx3 | 10.18188032 | 9.950113936 | 10.02683021 | -0.076716278 | 0.155050109 |
| 4931406B18Rik | 2.997129661 | 2.833761096 | 2.842108758 | -0.008347663 | 0.155020903 |
| Gnb1 | 12.62241838 | 12.50409918 | 12.46743689 | 0.03666229 | 0.154981483 |
| Eny2 | 9.606993341 | 9.444831241 | 9.452055232 | -0.007223992 | 0.154938109 |

| | | | | | |
|---|---|---|---|---|---|
| Trip13 | 7.385916137 | 7.284044028 | 7.230986594 | 0.053057434 | 0.154929543 |
| Polr2b | 10.61381448 | 10.43209858 | 10.45901537 | -0.026916797 | 0.154799107 |
| Gtf3c4 | 9.717655933 | 9.653786906 | 9.562923961 | 0.090862945 | 0.154731973 |
| Vmn2r115 | 5.338086216 | 5.215860006 | 5.183403522 | 0.032456484 | 0.154682694 |
| Cd27 | 4.179556068 | 4.063082197 | 4.024893162 | 0.038189035 | 0.154662906 |
| Ebp | 10.25510899 | 10.15082937 | 10.10044754 | 0.050381833 | 0.154661453 |
| Kcne4 | 6.127932273 | 5.900829075 | 5.973279439 | -0.072450363 | 0.154652834 |
| Pycrl | 7.794463962 | 7.889341073 | 7.639824159 | 0.249516914 | 0.154639803 |
| Ttc7 | 11.07419719 | 11.01881978 | 10.91956056 | 0.09925922 | 0.154636633 |
| Zbtb11 | 9.357369555 | 9.437937523 | 9.202749256 | 0.235188267 | 0.154620299 |
| Gm10282 | 6.074546255 | 5.898318837 | 5.919928994 | -0.021610156 | 0.154617262 |
| Tyw5 | 7.109776139 | 6.961392255 | 6.955208279 | 0.006183976 | 0.15456786 |
| Ppp1r35 | 6.098966747 | 6.171127365 | 5.944432684 | 0.226694681 | 0.154534063 |
| Mrc1 | 11.94619252 | 11.87524587 | 11.79174585 | 0.083500027 | 0.15444667 |
| Adam17 | 11.04743575 | 11.01215037 | 10.89307767 | 0.119072697 | 0.15435808 |
| Mrpl41 | 9.049379939 | 8.937152014 | 8.895094023 | 0.042057991 | 0.154285916 |
| Gm1604a | 1.969159072 | 1.801617641 | 1.815048533 | -0.013430892 | 0.154110539 |
| Slc39a13 | 9.02438305 | 9.183261586 | 8.870288547 | 0.312973039 | 0.154094504 |
| Tgfb1i1 | 8.313312824 | 8.289756507 | 8.159330639 | 0.130425869 | 0.153982185 |
| Eefsec | 7.941976047 | 7.850870953 | 7.788049735 | 0.062821218 | 0.153926312 |
| Nos1ap | 5.653428864 | 5.590406789 | 5.499518214 | 0.090888576 | 0.15391065 |
| Tfpt | 7.951280464 | 7.85486431 | 7.7973812 | 0.05748311 | 0.153899264 |
| Mcrip2 | 6.78665641 | 6.646102716 | 6.632758971 | 0.013343746 | 0.153897439 |
| Nudt21 | 9.95365864 | 9.880925826 | 9.799806911 | 0.081118915 | 0.153851728 |
| Zfp326 | 9.118900624 | 9.034161817 | 8.965080353 | 0.069081464 | 0.153820272 |
| Zc3h15 | 10.96052018 | 10.80711557 | 10.80675536 | 0.00036021 | 0.153764823 |
| Cmah | 7.602099316 | 7.373101414 | 7.448403206 | -0.075301792 | 0.15369611 |
| Cldn3 | 5.1296235 | 5.159250709 | 4.975998148 | 0.18325256 | 0.153625351 |
| Rundc1 | 8.489718657 | 8.454209226 | 8.336181562 | 0.118027663 | 0.153537095 |
| Chmp2a | 11.41016364 | 11.23517682 | 11.25680895 | -0.021632123 | 0.15335469 |
| Cwh43 | 10.08567146 | 9.973590176 | 9.932517673 | 0.041072503 | 0.153153791 |
| Golph3 | 10.96984052 | 10.80669887 | 10.81672922 | -0.010030357 | 0.153111293 |

| | | | | | |
|---|---|---|---|---|---|
| Bloc1s2 | 8.683911445 | 8.564815984 | 8.530867516 | 0.033948468 | 0.15304393 |
| Lipt2 | 6.0511367 | 5.971350348 | 5.898197577 | 0.073152771 | 0.152939123 |
| Tmem200a | 4.550505935 | 4.875603728 | 4.397587291 | 0.478016437 | 0.152918645 |
| Ppp1r12c | 9.910820458 | 9.922339806 | 9.758026725 | 0.164313081 | 0.152793733 |
| Pigs | 9.539849364 | 9.464401275 | 9.387061669 | 0.077339606 | 0.152787694 |
| Dmtf1 | 9.610159224 | 9.48940824 | 9.457461375 | 0.031946865 | 0.152697849 |
| Timm8b | 8.910381194 | 8.799562804 | 8.757741013 | 0.041821791 | 0.152640182 |
| Srsf1 | 11.27404419 | 11.20815453 | 11.12153518 | 0.086619357 | 0.152509013 |
| Hnrnph1 | 12.29971914 | 12.31417405 | 12.1473961 | 0.166777957 | 0.15232304 |
| Sephs2 | 9.675129414 | 9.623974644 | 9.522822748 | 0.101151896 | 0.152306665 |
| Bmx | 5.332122894 | 4.979704183 | 5.179889074 | -0.200184891 | 0.15223382 |
| Aph1a | 11.12078727 | 11.05905556 | 10.96855558 | 0.09049998 | 0.152231693 |
| Car2 | 5.007054688 | 4.944645411 | 4.854872305 | 0.089773105 | 0.152182383 |
| Uba1 | 12.69759384 | 12.57819789 | 12.54547452 | 0.032723367 | 0.15211932 |
| Lnx2 | 8.743827666 | 8.844064564 | 8.591779682 | 0.252284882 | 0.152047984 |
| Thap1 | 7.478164756 | 7.429709644 | 7.326144683 | 0.103564962 | 0.152020073 |
| Anp32e | 11.01791127 | 10.87176381 | 10.86589137 | 0.005872443 | 0.152019903 |
| Rnf121 | 9.632478764 | 9.456821903 | 9.480608734 | -0.023786831 | 0.151870029 |
| Rab35 | 10.41968193 | 10.4698274 | 10.26781507 | 0.202012333 | 0.151866863 |
| Fkbp8 | 11.08994936 | 11.27555324 | 10.93815639 | 0.337396843 | 0.151792963 |
| Cnn3 | 11.07215516 | 10.81590469 | 10.92042845 | -0.104523757 | 0.151726717 |
| Rchy1 | 9.698478372 | 9.587478908 | 9.546804293 | 0.040674615 | 0.151674078 |
| Mob1a | 12.11675601 | 12.04982366 | 11.96511275 | 0.084710917 | 0.151643263 |
| Rab6a | 11.35692073 | 11.12060985 | 11.20529876 | -0.08468891 | 0.151621977 |
| Diaph2 | 9.854909846 | 9.697440128 | 9.703433074 | -0.005992946 | 0.151476772 |
| Erf | 9.846591009 | 10.11488136 | 9.695148834 | 0.419732526 | 0.151442175 |
| Nog | 6.041284221 | 5.88865927 | 5.889873974 | -0.001214704 | 0.151410248 |
| Lmbrd1 | 10.06845546 | 10.05491573 | 9.917070894 | 0.137844835 | 0.151384562 |
| Actl6a | 9.723082069 | 9.684314968 | 9.571760088 | 0.11255488 | 0.151321981 |
| Cdc42 | 13.05447855 | 12.84502949 | 12.90328427 | -0.058254777 | 0.151194277 |
| Pspc1 | 8.792726225 | 8.585889944 | 8.641575852 | -0.055685908 | 0.151150373 |
| Dync1li1 | 10.2326722 | 10.14807143 | 10.08162075 | 0.066450683 | 0.151051457 |

| | | | | | |
|---|---|---|---|---|---|
| Cry1 | 8.658117649 | 8.65874893 | 8.507277535 | 0.151471395 | 0.150840114 |
| Chtop | 11.36861728 | 11.27850158 | 11.21781952 | 0.06068206 | 0.150797753 |
| Eif4h | 12.62846962 | 12.65208358 | 12.47786038 | 0.1742232 | 0.150609243 |
| Cops8 | 10.23976537 | 10.03809229 | 10.08940976 | -0.051317469 | 0.150355608 |
| Cep85 | 8.950464814 | 8.844276667 | 8.80016942 | 0.044107246 | 0.150295393 |
| Cpsf4 | 8.454700101 | 8.376829403 | 8.30458781 | 0.072241593 | 0.150112291 |
| St8sia4 | 8.144440448 | 7.770493383 | 7.994336365 | -0.223842982 | 0.150104083 |
| Hspa4l | 9.410144503 | 9.402182162 | 9.260217621 | 0.141964541 | 0.149926882 |
| Rhob | 10.33863579 | 10.34688473 | 10.1887423 | 0.158142423 | 0.149893487 |
| Dok5 | 4.900712178 | 4.865628406 | 4.750907119 | 0.114721287 | 0.149805059 |
| Scaf4 | 9.367910321 | 9.407446496 | 9.218108271 | 0.189338225 | 0.14980205 |
| St14 | 10.68127603 | 10.70944547 | 10.53154358 | 0.177901887 | 0.149732447 |
| Chuk | 9.927162857 | 9.962545123 | 9.777438614 | 0.185106509 | 0.149724242 |
| Ndc1 | 8.879206882 | 8.833975 | 8.729492757 | 0.104482243 | 0.149714125 |
| Mta1 | 9.53119811 | 9.615502281 | 9.381504744 | 0.233997537 | 0.149693366 |
| Nfkbil1 | 6.816498496 | 6.873348619 | 6.666809692 | 0.206538927 | 0.149688803 |
| Hcfc2 | 9.092356618 | 8.973951134 | 8.942709461 | 0.031241673 | 0.149647157 |
| Tial1 | 10.42099071 | 10.36578752 | 10.27136213 | 0.094425393 | 0.149628577 |
| Smagp | 9.888351222 | 9.701547944 | 9.738744395 | -0.03719645 | 0.149606827 |
| Snrnp200 | 11.29830484 | 11.22976139 | 11.14871157 | 0.081049813 | 0.149593267 |
| Slc22a15 | 8.732498435 | 8.564090204 | 8.582920345 | -0.018830141 | 0.14957809 |
| Rgs6 | 5.68147329 | 5.79927197 | 5.531988594 | 0.267283375 | 0.149484695 |
| Ptpa | 11.65881772 | 11.65549048 | 11.50940849 | 0.146081997 | 0.14940923 |
| Arih1 | 11.49724157 | 11.42293292 | 11.34785335 | 0.07507957 | 0.149388227 |
| Ptchd4 | 2.792365763 | 2.695394647 | 2.643074001 | 0.052320646 | 0.149291762 |
| Srp19 | 9.432094476 | 9.324831395 | 9.282809938 | 0.042021456 | 0.149284537 |
| Sem1 | 11.15751593 | 11.07507026 | 11.00826626 | 0.066803997 | 0.149249667 |
| Nrf1 | 9.144636604 | 9.102974194 | 8.995402306 | 0.107571888 | 0.149234299 |
| B4galt7 | 7.860253829 | 7.869116816 | 7.711095741 | 0.158021075 | 0.149158088 |
| Raly | 11.09974426 | 11.08265608 | 10.95074057 | 0.131915513 | 0.149003697 |
| Dpm2 | 9.322721079 | 9.155973732 | 9.173790575 | -0.017816843 | 0.148930504 |
| Prss16 | 5.319583732 | 5.462482463 | 5.170681243 | 0.29180122 | 0.14890249 |

| | | | | | |
|---|---|---|---|---|---|
| Zfp592 | 9.7028266 | 9.820177004 | 9.554062142 | 0.266114862 | 0.148764457 |
| Arpin | 8.959973254 | 8.794859165 | 8.811230527 | -0.016371362 | 0.148742727 |
| Grin1 | 3.317616019 | 3.261787363 | 3.168896301 | 0.092891062 | 0.148719718 |
| Degs1 | 11.548066 | 11.27129367 | 11.39936452 | -0.12807085 | 0.148701483 |
| Clcn6 | 8.427764546 | 8.569407134 | 8.279466764 | 0.28994037 | 0.148297782 |
| Slc39a12 | 3.201017783 | 3.0934066 | 3.052821353 | 0.040585247 | 0.14819643 |
| Pabpc4 | 10.76637478 | 10.87035646 | 10.6182659 | 0.252090563 | 0.148108883 |
| Cox5a | 11.31103903 | 11.18285757 | 11.16293273 | 0.019924837 | 0.1481063 |
| Rala | 10.30790414 | 10.13140527 | 10.15980349 | -0.02839822 | 0.14810065 |
| Dnajc21 | 9.684556797 | 9.475383683 | 9.536558373 | -0.06117469 | 0.147998424 |
| Commd7 | 9.261075698 | 9.185107487 | 9.113080393 | 0.072027093 | 0.147995305 |
| Edn1 | 6.996653016 | 7.075281396 | 6.848868969 | 0.226412427 | 0.147784047 |
| Crabp2 | 10.3137691 | 10.14845677 | 10.16599419 | -0.017537417 | 0.14777491 |
| Sh3bp4 | 8.613451208 | 8.48338973 | 8.465692995 | 0.017696735 | 0.147758213 |
| Slc8a1 | 7.924969457 | 7.592810654 | 7.77725682 | -0.184446166 | 0.147712637 |
| Gm42791 | 3.796465472 | 3.588884492 | 3.648752904 | -0.059868412 | 0.147712568 |
| Smug1 | 8.788782036 | 8.733034933 | 8.64121001 | 0.091824923 | 0.147572026 |
| Abl2 | 10.24477853 | 10.21663042 | 10.09726742 | 0.119363 | 0.147511103 |
| Plscr3 | 10.20525666 | 10.24691646 | 10.05780383 | 0.189112629 | 0.147452829 |
| Map2k2 | 10.53077202 | 10.78072604 | 10.38332999 | 0.39739605 | 0.14744203 |
| Myo1b | 9.70820808 | 9.568832658 | 9.5608102 | 0.008022458 | 0.14739788 |
| Lrrc4 | 6.421198105 | 6.107059156 | 6.273862141 | -0.166802985 | 0.147335964 |
| Zup1 | 8.538407814 | 8.256148467 | 8.391092358 | -0.134943891 | 0.147315456 |
| Gm49321 | 2.473001122 | 2.414835046 | 2.325698737 | 0.089136309 | 0.147302385 |
| Sik1 | 10.39734174 | 10.39942483 | 10.25006857 | 0.149356257 | 0.147273163 |
| Zfp408 | 8.059677771 | 8.23512561 | 7.912450468 | 0.322675142 | 0.147227303 |
| AW551984 | 7.761968595 | 7.439326715 | 7.61474808 | -0.175421365 | 0.147220515 |
| Fastk | 9.852536168 | 9.956140097 | 9.705337346 | 0.250802751 | 0.147198822 |
| Rbx1 | 10.46979387 | 10.36839793 | 10.32269886 | 0.045699063 | 0.147095003 |
| Cope | 10.71951139 | 10.75013211 | 10.57243259 | 0.17769952 | 0.147078807 |
| Gm21750 | 5.664506621 | 5.455371341 | 5.517483658 | -0.062112317 | 0.147022963 |
| Msh6 | 9.918768258 | 9.843197592 | 9.771948542 | 0.07124905 | 0.146819716 |

| | | | | | |
|---|---|---|---|---|---|
| Ccnq | 7.183773588 | 7.070487395 | 7.036968441 | 0.033518953 | 0.146805147 |
| Bcap29 | 9.518846994 | 9.191089443 | 9.372059804 | -0.18097036 | 0.146787191 |
| Dtx4 | 9.396531469 | 9.307823461 | 9.249765212 | 0.05805825 | 0.146766257 |
| Naa40 | 9.353038891 | 9.276431816 | 9.206287216 | 0.0701446 | 0.146751675 |
| Slc35b2 | 9.584495408 | 9.445490506 | 9.437839122 | 0.007651384 | 0.146656285 |
| Gng2 | 8.611582175 | 8.594734484 | 8.464973009 | 0.129761475 | 0.146609166 |
| Eef1g | 13.76418684 | 13.68455842 | 13.6175797 | 0.066978713 | 0.146607137 |
| Galnt18 | 8.358225518 | 8.587984785 | 8.211683757 | 0.376301028 | 0.146541761 |
| Cxxc1 | 9.584242297 | 9.768992433 | 9.437769138 | 0.331223296 | 0.146473159 |
| Rpl11 | 12.9244259 | 12.94366217 | 12.77798926 | 0.165672907 | 0.146436633 |
| Uqcc1 | 9.562890423 | 9.267045762 | 9.416486849 | -0.149441087 | 0.146403573 |
| Rasl11a | 5.224624708 | 5.167658831 | 5.078284075 | 0.089374755 | 0.146340632 |
| Rpl38 | 11.92437208 | 11.77900913 | 11.77803164 | 0.000977493 | 0.14634044 |
| Krt14 | 16.58854399 | 16.77355606 | 16.44222722 | 0.331328843 | 0.146316773 |
| Inpp5a | 9.954253418 | 9.813460692 | 9.808107562 | 0.00535313 | 0.146145855 |
| Ring1 | 8.535699496 | 8.66704547 | 8.389790682 | 0.277254788 | 0.145908813 |
| Prr3 | 7.854739364 | 7.715808767 | 7.708831743 | 0.006977024 | 0.145907622 |
| Rnf185 | 9.609226526 | 9.546458041 | 9.463377827 | 0.083080214 | 0.145848699 |
| Gap43 | 4.85845201 | 4.782105366 | 4.712696043 | 0.069409323 | 0.145755967 |
| Ahsa2 | 8.772364297 | 9.101307928 | 8.626726498 | 0.47458143 | 0.145637799 |
| Trim46 | 4.696726002 | 4.691947383 | 4.551130878 | 0.140816505 | 0.145595124 |
| Pdxp | 6.319598967 | 6.31319672 | 6.17406256 | 0.13913416 | 0.145536407 |
| Far1 | 10.90893897 | 10.83729431 | 10.76341315 | 0.073881167 | 0.145525823 |
| Ttc39b | 11.15241664 | 10.91348915 | 11.0069662 | -0.093477057 | 0.145450437 |
| Qrich1 | 10.5505345 | 10.52522917 | 10.40512497 | 0.120104197 | 0.145409527 |
| Sap30bp | 9.438712662 | 9.392717524 | 9.293344586 | 0.099372937 | 0.145368076 |
| Socs6 | 10.11837836 | 10.01384174 | 9.973033252 | 0.040808484 | 0.145345108 |
| Tmem184b | 9.738088974 | 9.850376157 | 9.592810867 | 0.25756529 | 0.145278106 |
| Mns1 | 5.081997744 | 5.064422564 | 4.936828848 | 0.127593716 | 0.145168896 |
| Il27 | 2.536213838 | 2.472146017 | 2.391115185 | 0.081030832 | 0.145098654 |
| Hdx | 4.672206575 | 4.520034089 | 4.527115504 | -0.007081415 | 0.145091071 |
| Cp | 10.39033117 | 9.922908978 | 10.24539326 | -0.322484282 | 0.144937907 |

| | | | | | |
|---|---|---|---|---|---|
| Csnk1d | 11.64622882 | 11.58220603 | 11.50133553 | 0.080870497 | 0.14489329 |
| Psmc3ip | 6.37432055 | 6.380541496 | 6.2295031 | 0.151038396 | 0.14481745 |
| Ap2m1 | 12.53294529 | 12.4991003 | 12.38819408 | 0.110906223 | 0.14475121 |
| Rpl13 | 13.48101334 | 13.36056561 | 13.33627085 | 0.024294763 | 0.144742497 |
| Pitpnb | 10.91100309 | 10.78268974 | 10.76633057 | 0.016359173 | 0.144672527 |
| Mfsd6l | 2.825657739 | 2.759224507 | 2.681183373 | 0.078041134 | 0.144474366 |
| Plekho1 | 8.829685104 | 8.926026383 | 8.685233335 | 0.240793048 | 0.144451769 |
| Coa4 | 6.765120566 | 6.65916023 | 6.620688931 | 0.038471299 | 0.144431634 |
| Serpina11 | 2.280653055 | 2.206736868 | 2.136258903 | 0.070477965 | 0.144394152 |
| Elmo3 | 9.267451825 | 9.178764666 | 9.123106273 | 0.055658393 | 0.144345552 |
| Rpf1 | 8.870951996 | 8.805502841 | 8.726613686 | 0.078889155 | 0.14433831 |
| G3bp2 | 11.45783664 | 11.10098069 | 11.31354274 | -0.212562047 | 0.144293903 |
| Slit2 | 9.062794132 | 9.077114157 | 8.918558273 | 0.158555884 | 0.144235859 |
| Cox6a2 | 9.909332338 | 10.16631536 | 9.765138329 | 0.401177033 | 0.144194009 |
| Arap2 | 10.74074899 | 10.50197094 | 10.59664493 | -0.09467399 | 0.144104057 |
| Aip | 8.291808031 | 8.296738311 | 8.147785128 | 0.148953184 | 0.144022903 |
| C1galt1c1 | 8.774652659 | 8.522479823 | 8.630693089 | -0.108213266 | 0.143959569 |
| Kif5b | 12.40284779 | 12.24197444 | 12.25895753 | -0.016983093 | 0.143890253 |
| Svep1 | 10.91118171 | 10.74856623 | 10.76733821 | -0.018771983 | 0.1438435 |
| Hs6st1 | 9.811910957 | 9.809736495 | 9.668114678 | 0.141621817 | 0.143796279 |
| Fscn1 | 9.193882386 | 9.531776613 | 9.050086796 | 0.481689817 | 0.143795591 |
| Psme2b | 6.209773462 | 6.062706265 | 6.065980697 | -0.003274432 | 0.143792765 |
| Lrp1 | 13.05060898 | 13.34939453 | 12.90684682 | 0.442547713 | 0.14376216 |
| Ror1 | 7.720151978 | 7.734919658 | 7.576409095 | 0.158510563 | 0.143742882 |
| Pcif1 | 9.926007319 | 9.9105477 | 9.782278842 | 0.128268858 | 0.143728477 |
| Fgf2 | 9.345919524 | 8.840557598 | 9.202321191 | -0.361763593 | 0.143598333 |
| Rbm10 | 9.654620939 | 9.771094357 | 9.511024561 | 0.260069796 | 0.143596378 |
| Sema4a | 10.60288755 | 10.60074609 | 10.45930453 | 0.141441553 | 0.14358302 |
| Nap1l4 | 11.08046569 | 10.95182624 | 10.93699015 | 0.014836097 | 0.14347554 |
| Lyrm4 | 7.01581242 | 6.90276001 | 6.872357026 | 0.030402984 | 0.143455394 |
| 3110009E18Rik | 5.247124356 | 5.162490301 | 5.103743543 | 0.058746758 | 0.143380813 |
| Gm13033 | 4.096334118 | 3.995070506 | 3.953130914 | 0.041939592 | 0.143203205 |

| | | | | | |
|---|---|---|---|---|---|
| Myef2 | 7.642879161 | 7.59646515 | 7.499753445 | 0.096711704 | 0.143125716 |
| Fbxo28 | 10.04276659 | 9.925727652 | 9.899663319 | 0.026064333 | 0.143103271 |
| Dtnbp1 | 9.532434904 | 9.4307265 | 9.389450795 | 0.041275705 | 0.142984109 |
| Fam102a | 9.901349181 | 10.08167446 | 9.758378548 | 0.323295908 | 0.142970632 |
| Zfp710 | 9.466561972 | 9.515616466 | 9.323645768 | 0.191970698 | 0.142916204 |
| 2410131K14Rik | 8.669427604 | 8.645273719 | 8.526655252 | 0.118618467 | 0.142772352 |
| Glrx5 | 9.487702491 | 9.45133765 | 9.344999105 | 0.106338544 | 0.142703385 |
| Alkbh6 | 7.817549359 | 7.840840252 | 7.674881476 | 0.165958776 | 0.142667883 |
| Scand1 | 5.039917627 | 5.208480951 | 4.897475485 | 0.311005466 | 0.142442141 |
| Ppp1cc | 11.66693038 | 11.49360705 | 11.52452033 | -0.03091328 | 0.142410043 |
| Stac2 | 6.584105942 | 6.625825778 | 6.441772795 | 0.184052983 | 0.142333146 |
| Cd59b | 6.993901983 | 6.982142633 | 6.851659229 | 0.130483404 | 0.142242754 |
| Eloa | 10.76365142 | 10.64571832 | 10.62150882 | 0.0242095 | 0.142142603 |
| Tagap1 | 8.414511069 | 8.219341004 | 8.272378758 | -0.053037755 | 0.14213231 |
| Scyl2 | 10.13830616 | 10.07841597 | 9.996182721 | 0.082233245 | 0.142123442 |
| Ankrd13a | 10.79128843 | 10.81880958 | 10.64924383 | 0.169565747 | 0.142044603 |
| Gm16976 | 2.77894317 | 2.638295024 | 2.636934821 | 0.001360203 | 0.142008349 |
| Cdt1 | 8.11370161 | 8.25122217 | 7.971693326 | 0.279528844 | 0.142008284 |
| Slc39a8 | 10.14338311 | 9.873728804 | 10.00149003 | -0.127761223 | 0.141893086 |
| Taf9 | 5.575772845 | 5.744410771 | 5.433886026 | 0.310524745 | 0.141886819 |
| Rapgef1 | 11.1191966 | 11.22462733 | 10.9773284 | 0.24729893 | 0.1418682 |
| Fam168b | 11.75183052 | 11.67843004 | 11.61007633 | 0.068353707 | 0.141754187 |
| Rad1 | 7.660273394 | 7.55709867 | 7.518544829 | 0.038553841 | 0.141728565 |
| Fbxo42 | 9.490136734 | 9.568972525 | 9.348430674 | 0.220541851 | 0.141706059 |
| Raet1d | 3.031751205 | 2.972194261 | 2.890100186 | 0.082094075 | 0.141651019 |
| Pgp | 8.992375411 | 8.832046437 | 8.850838322 | -0.018791886 | 0.141537088 |
| Cic | 11.05799 | 11.12249938 | 10.91646265 | 0.206036733 | 0.141527347 |
| Nsun4 | 8.196860022 | 8.027009558 | 8.055340347 | -0.028330789 | 0.141519675 |
| Bfar | 9.694297071 | 9.532020666 | 9.552811851 | -0.020791185 | 0.141485221 |
| Znhit3 | 7.494870635 | 7.217061742 | 7.353397861 | -0.136336119 | 0.141472773 |
| Chrna7 | 2.495383378 | 2.415874078 | 2.353934648 | 0.061939429 | 0.141448729 |
| Sgip1 | 5.984451149 | 5.84403713 | 5.843044919 | 0.000992211 | 0.14140623 |

| | | | | | |
|---|---|---|---|---|---|
| Adgrg1 | 10.96311234 | 10.91240858 | 10.82171954 | 0.090689043 | 0.141392803 |
| Nradd | 8.185377036 | 8.210007702 | 8.043993866 | 0.166013836 | 0.14138317 |
| Galnt3 | 7.897535698 | 7.595342354 | 7.756251487 | -0.160909133 | 0.141284211 |
| Rad23b | 12.07655441 | 12.11193601 | 11.93530818 | 0.17662783 | 0.141246233 |
| Supt5 | 11.36391455 | 11.41217528 | 11.22276927 | 0.189406013 | 0.141145277 |
| Csnk1a1 | 13.37931666 | 13.19604061 | 13.23823998 | -0.04219937 | 0.141076677 |
| Lrpap1 | 10.14745609 | 9.896842703 | 10.00643666 | -0.109593954 | 0.141019433 |
| Myl6 | 13.24963679 | 13.1060045 | 13.10861856 | -0.002614057 | 0.141018233 |
| Atg3 | 10.26804335 | 10.11479787 | 10.12705336 | -0.012255497 | 0.140989987 |
| Htr2b | 3.848727944 | 3.743985765 | 3.707851782 | 0.036133982 | 0.140876162 |
| Naga | 9.855098631 | 9.674588172 | 9.714321847 | -0.039733675 | 0.140776784 |
| Prelid1 | 11.63473252 | 11.63803856 | 11.49406439 | 0.143974173 | 0.14066813 |
| Hnrnpk | 13.36128945 | 13.19397299 | 13.22062273 | -0.026649743 | 0.140666717 |
| Gemin2 | 7.835092716 | 7.727543268 | 7.694481485 | 0.033061783 | 0.140611231 |
| Tmc4 | 6.809042471 | 6.863251561 | 6.668456086 | 0.194795475 | 0.140586385 |
| Comtd1 | 6.265185184 | 6.390690066 | 6.124656494 | 0.266033572 | 0.14052869 |
| Cyb561d2 | 7.435883538 | 7.405600902 | 7.295367484 | 0.110233417 | 0.140516054 |
| BC048403 | 7.821831323 | 7.627773918 | 7.681402473 | -0.053628555 | 0.140428851 |
| Fbxo30 | 9.082812216 | 9.090819885 | 8.94240813 | 0.148411755 | 0.140404086 |
| Inka1 | 6.984243218 | 6.846313099 | 6.843972404 | 0.002340695 | 0.140270815 |
| Dctn4 | 10.73187966 | 10.60385959 | 10.59164485 | 0.012214747 | 0.14023481 |
| Gfer | 8.315457452 | 8.125647858 | 8.175249411 | -0.049601553 | 0.140208041 |
| Gpr18 | 4.230745438 | 4.053232053 | 4.090622128 | -0.037390075 | 0.14012331 |
| Phf8 | 9.682018014 | 9.555588274 | 9.542000856 | 0.013587418 | 0.140017158 |
| Tmem54 | 10.17927428 | 10.20438763 | 10.03933736 | 0.165050271 | 0.139936924 |
| Ttc41 | 4.731318566 | 4.711048422 | 4.591391865 | 0.119656557 | 0.139926702 |
| Tnn | 5.979776179 | 5.823739248 | 5.839885248 | -0.016146 | 0.13989093 |
| Txnl4a | 9.06390565 | 8.962867033 | 8.924052672 | 0.038814361 | 0.139852979 |
| Pcdhgb4 | 5.468258895 | 5.509903785 | 5.328419337 | 0.181484448 | 0.139839558 |
| Stxbp2 | 10.05517508 | 10.30576278 | 9.91536536 | 0.39039742 | 0.139809723 |
| Polrmt | 8.550450387 | 8.585639497 | 8.410665985 | 0.174973512 | 0.139784403 |
| Slx4 | 8.322539621 | 8.410269417 | 8.182798798 | 0.227470619 | 0.139740823 |

| | | | | | |
|---|---|---|---|---|---|
| Rex1bd | 8.021535844 | 8.180729835 | 7.881827404 | 0.298902431 | 0.13970844 |
| Nudt3 | 10.71266152 | 10.57511951 | 10.57296109 | 0.002158423 | 0.139700433 |
| Copz1 | 11.2763816 | 11.12213145 | 11.13677561 | -0.014644157 | 0.13960599 |
| Hnrnpdl | 11.84767326 | 11.79989635 | 11.70812049 | 0.091775857 | 0.139552773 |
| Hells | 9.074305924 | 9.03918739 | 8.934839271 | 0.104348118 | 0.139466653 |
| Mrps18c | 8.886207696 | 8.596424659 | 8.746805039 | -0.15038038 | 0.139402658 |
| Rab29 | 8.167710808 | 8.041418902 | 8.028333078 | 0.013085824 | 0.13937773 |
| Abca7 | 9.520164295 | 9.742039402 | 9.380863139 | 0.361176263 | 0.139301157 |
| Arcn1 | 11.9615332 | 11.87240199 | 11.82225166 | 0.050150327 | 0.13928154 |
| Mapk8ip3 | 9.956941953 | 10.17670083 | 9.817683823 | 0.35901701 | 0.13925813 |
| Nedd8 | 10.56639581 | 10.43521355 | 10.42716497 | 0.008048583 | 0.139230843 |
| Chchd2 | 11.90607881 | 11.9545424 | 11.76689404 | 0.18764836 | 0.139184777 |
| Nup107 | 9.034741407 | 8.727954587 | 8.895617965 | -0.167663378 | 0.139123442 |
| Mast4 | 10.53549609 | 10.53125989 | 10.39642832 | 0.13483157 | 0.13906777 |
| Ppp2cb | 10.19440507 | 10.05458348 | 10.05543724 | -0.000853757 | 0.13896783 |
| Zglp1 | 2.846522236 | 2.67053643 | 2.707585991 | -0.037049561 | 0.138936245 |
| Bicra | 8.734661706 | 8.871078432 | 8.595850062 | 0.27522837 | 0.138811644 |
| Cadm4 | 7.721404745 | 7.788025173 | 7.582595178 | 0.205429996 | 0.138809567 |
| Rogdi | 8.144609195 | 8.34991963 | 8.005824782 | 0.344094848 | 0.138784414 |
| Atp9b | 10.17207294 | 10.08939906 | 10.03329707 | 0.056101992 | 0.138775865 |
| Skint11 | 9.050613013 | 8.770703957 | 8.911904538 | -0.141200581 | 0.138708474 |
| Sh3gl1 | 10.26040263 | 10.3980043 | 10.12170985 | 0.276294449 | 0.138692786 |
| Sirt7 | 9.43878762 | 9.478465573 | 9.300252954 | 0.178212619 | 0.138534666 |
| Zc3hc1 | 8.325917076 | 8.284753056 | 8.187405104 | 0.097347952 | 0.138511973 |
| 2310039H08Rik | 6.585529356 | 6.629382619 | 6.447058251 | 0.182324368 | 0.138471105 |
| Toe1 | 8.930701305 | 8.845850311 | 8.792338594 | 0.053511716 | 0.138362711 |
| Ppp4r3a | 10.65444302 | 10.60623903 | 10.51608056 | 0.09015847 | 0.13836246 |
| Ccdc63 | 5.236982119 | 5.321250549 | 5.098628858 | 0.222621691 | 0.138353262 |
| Mtln | 7.829500754 | 7.893350794 | 7.691270815 | 0.202079979 | 0.138229939 |
| Med10 | 8.917612781 | 8.801910592 | 8.779501707 | 0.022408884 | 0.138111073 |
| Mdfic | 9.795366026 | 9.873464102 | 9.657468329 | 0.215995773 | 0.137897697 |
| Trap1 | 10.82258143 | 10.78547426 | 10.68469361 | 0.100780653 | 0.137887827 |

| | | | | | |
|---|---|---|---|---|---|
| Faap24 | 7.254027106 | 7.238889071 | 7.116158825 | 0.122730246 | 0.137868281 |
| Tbcc | 7.200799573 | 7.25097772 | 7.063151441 | 0.187826279 | 0.137648132 |
| Zfp622 | 9.081795893 | 8.986197665 | 8.944153533 | 0.042044132 | 0.137642359 |
| Ubtd1 | 9.060180462 | 9.187129726 | 8.922721407 | 0.264408319 | 0.137459055 |
| Usp31 | 10.03918766 | 10.03056388 | 9.901753241 | 0.128810636 | 0.137434418 |
| Cnih4 | 9.742362893 | 9.551342072 | 9.604935471 | -0.053593399 | 0.137427423 |
| Trappc4 | 9.774098813 | 9.584918977 | 9.636690464 | -0.051771487 | 0.137408349 |
| Tspan3 | 10.69769859 | 10.64503109 | 10.56031303 | 0.08471806 | 0.137385557 |
| Adss | 10.48079276 | 10.34688412 | 10.34343789 | 0.003446233 | 0.137354873 |
| Ppef2 | 4.203973868 | 4.072265416 | 4.066783548 | 0.005481868 | 0.137190321 |
| Gm10443 | 10.34313734 | 10.26309214 | 10.20599837 | 0.057093763 | 0.13713897 |
| Ik | 11.07508635 | 10.90961922 | 10.93797256 | -0.02835334 | 0.137113783 |
| Med8 | 9.92251292 | 9.812124904 | 9.785485468 | 0.026639436 | 0.137027452 |
| Fbrs | 10.55626143 | 10.86096328 | 10.41931944 | 0.441643833 | 0.136941983 |
| Tbc1d10b | 9.943345671 | 10.07999775 | 9.806440109 | 0.273557644 | 0.136905562 |
| Hprt | 10.66054174 | 10.49521097 | 10.52377219 | -0.028561223 | 0.13676955 |
| Zfp747 | 7.378219961 | 7.412391823 | 7.241465379 | 0.170926444 | 0.136754582 |
| Ttc9 | 9.335079311 | 9.212965277 | 9.198388094 | 0.014577182 | 0.136691217 |
| Rpl7a | 13.65314909 | 13.57971453 | 13.51646372 | 0.06325081 | 0.136685373 |
| Vmn1r58 | 4.604061602 | 4.51805133 | 4.467656717 | 0.050394613 | 0.136404885 |
| Sharpin | 9.009127764 | 8.945151303 | 8.872747368 | 0.072403935 | 0.136380396 |
| Parp3 | 9.956087823 | 9.913802095 | 9.819772658 | 0.094029438 | 0.136315165 |
| Mbd6 | 9.719955274 | 9.95176954 | 9.583676561 | 0.368092979 | 0.136278713 |
| Sytl3 | 5.495771712 | 5.330312007 | 5.359494468 | -0.029182461 | 0.136277244 |
| Ppp1r2 | 11.29906924 | 11.17017417 | 11.16292298 | 0.007251193 | 0.136146263 |
| Smcr8 | 10.04306986 | 10.12950781 | 9.907122897 | 0.222384909 | 0.135946958 |
| Cxadr | 11.86883821 | 11.97502719 | 11.73298985 | 0.24203734 | 0.135848363 |
| Ndor1 | 9.489424967 | 9.56707779 | 9.353611613 | 0.213466176 | 0.135813353 |
| Sphk2 | 10.21130047 | 10.47341375 | 10.07549892 | 0.397914833 | 0.13580155 |
| Ccl19-ps1 | 4.59609227 | 4.642501133 | 4.460308756 | 0.182192377 | 0.135783514 |
| Trmt112-ps2 | 5.97286353 | 6.201067872 | 5.837137105 | 0.363930767 | 0.135726425 |
| Nif3l1 | 8.813748811 | 8.698335256 | 8.678117361 | 0.020217895 | 0.13563145 |

| | | | | | |
|---|---|---|---|---|---|
| Scap | 11.20779891 | 11.33807817 | 11.07220696 | 0.265871207 | 0.13559195 |
| 6030458C11Rik | 9.384284055 | 9.146999962 | 9.24890505 | -0.101905088 | 0.135379006 |
| Rbm15 | 8.634295778 | 8.600868518 | 8.498918064 | 0.101950453 | 0.135377714 |
| Zbed5 | 4.881230084 | 4.831104509 | 4.7458979 | 0.085206609 | 0.135332183 |
| Tor1b | 10.11581006 | 10.05472759 | 9.980492971 | 0.074234616 | 0.135317086 |
| Cd320 | 7.846562238 | 7.84386046 | 7.711488045 | 0.132372415 | 0.135074193 |
| Hmgn1 | 12.3970972 | 12.2988401 | 12.26203104 | 0.03680906 | 0.135066153 |
| Ptpn11 | 11.60616682 | 11.54384619 | 11.47120987 | 0.072636323 | 0.134956957 |
| 2310009B15Rik | 6.275101448 | 6.282386596 | 6.140201893 | 0.142184703 | 0.134899555 |
| Gjc1 | 7.079863373 | 6.945213839 | 6.945006147 | 0.000207692 | 0.134857226 |
| Gpr137b | 8.520549647 | 8.759704624 | 8.385881057 | 0.373823568 | 0.13466859 |
| Rspry1 | 8.963555982 | 8.915606729 | 8.829001008 | 0.086605721 | 0.134554974 |
| Cdc26 | 9.35437669 | 9.215032996 | 9.219890655 | -0.004857659 | 0.134486035 |
| Ndst2 | 9.721203929 | 9.49666124 | 9.586787716 | -0.090126476 | 0.134416213 |
| Lonp1 | 10.53185004 | 10.53180983 | 10.39745243 | 0.134357403 | 0.134397617 |
| Spats2 | 8.009740165 | 7.891000247 | 7.875357388 | 0.015642859 | 0.134382777 |
| Zfp429 | 6.745074747 | 6.537978493 | 6.610843567 | -0.072865074 | 0.13423118 |
| Enthd1 | 2.165164887 | 2.034288463 | 2.030959038 | 0.003329424 | 0.134205849 |
| Mrpl51 | 10.1512541 | 10.0574994 | 10.01704887 | 0.040450534 | 0.134205228 |
| Klhdc8a | 4.084069646 | 4.012145012 | 3.949882302 | 0.06226271 | 0.134187344 |
| Bad | 8.347382034 | 8.466284754 | 8.21321564 | 0.253069114 | 0.134166393 |
| Spry1 | 9.816963539 | 9.484310192 | 9.682805657 | -0.198495465 | 0.134157882 |
| Rab21 | 10.83787316 | 10.7156105 | 10.70377827 | 0.01183223 | 0.13409489 |
| H2-Q7 | 11.36127952 | 11.40670188 | 11.2272005 | 0.179501387 | 0.134079023 |
| Pdf | 7.694059861 | 7.54788216 | 7.559987145 | -0.012104984 | 0.134072717 |
| Atxn7l3 | 10.45308666 | 10.52446108 | 10.31903999 | 0.205421097 | 0.13404667 |
| Kcnc4 | 7.615423598 | 7.345375317 | 7.481502857 | -0.136127541 | 0.133920741 |
| Nme6 | 7.619080996 | 7.603126144 | 7.485200516 | 0.117925628 | 0.13388048 |
| Tiam2 | 7.472480348 | 7.533728331 | 7.338635471 | 0.19509286 | 0.133844877 |
| Cblb | 10.19426911 | 10.10775901 | 10.06047798 | 0.047281033 | 0.13379113 |
| Pcdhgc3 | 10.749393 | 10.83557948 | 10.61560433 | 0.21997515 | 0.13378867 |
| Rbbp8 | 9.850692188 | 9.580438845 | 9.716914887 | -0.136476042 | 0.133777301 |

| | | | | | |
|---|---|---|---|---|---|
| Parp11 | 8.516084463 | 8.324148858 | 8.382319364 | -0.058170506 | 0.133765098 |
| Med30 | 8.036506947 | 7.885012282 | 7.903024106 | -0.018011824 | 0.133482841 |
| Senp5 | 9.820461424 | 9.816159028 | 9.686990135 | 0.129168893 | 0.133471288 |
| Micall2 | 7.56005136 | 7.242015523 | 7.426582088 | -0.184566565 | 0.133469272 |
| Adprm | 7.399270275 | 7.187144394 | 7.26586294 | -0.078718546 | 0.133407335 |
| Tchh | 6.789481351 | 6.629220867 | 6.656077545 | -0.026856678 | 0.133403806 |
| Gm2033 | 2.266335296 | 2.1692092 | 2.132948117 | 0.036261082 | 0.133387178 |
| Teddm3 | 10.16138364 | 9.968643359 | 10.02816924 | -0.059525881 | 0.133214396 |
| Tmem67 | 6.998641346 | 6.74396113 | 6.865507515 | -0.121546385 | 0.133133831 |
| Slc25a29 | 6.192557726 | 6.07870618 | 6.059428287 | 0.019277893 | 0.133129439 |
| 2700062C07Rik | 7.511125494 | 7.376790328 | 7.378076618 | -0.00128629 | 0.133048876 |
| Wfdc12 | 6.112244348 | 5.997066673 | 5.979315367 | 0.017751307 | 0.132928981 |
| Pmpcb | 9.990999118 | 9.638148329 | 9.858072321 | -0.219923992 | 0.132926797 |
| Il15 | 7.77785166 | 7.186956237 | 7.644945655 | -0.457989418 | 0.132906005 |
| Crb3 | 7.856440367 | 7.947845927 | 7.723606227 | 0.224239699 | 0.13283414 |
| B3gnt8 | 7.423040504 | 7.651085841 | 7.290317958 | 0.360767883 | 0.132722546 |
| Alg13 | 7.640876351 | 7.617187819 | 7.508163892 | 0.109023927 | 0.132712459 |
| Zbed4 | 8.859810494 | 8.887696999 | 8.727138713 | 0.160558286 | 0.132671781 |
| Ccnj | 7.541268565 | 7.338631367 | 7.408788101 | -0.070156734 | 0.132480464 |
| Aida | 10.45752853 | 10.16105954 | 10.32524005 | -0.164180517 | 0.13228848 |
| Lrrc66 | 3.113259252 | 3.007424202 | 2.981173738 | 0.026250464 | 0.132085513 |
| Rhof | 7.395253601 | 7.468862762 | 7.263176009 | 0.205686753 | 0.132077592 |
| Opa3 | 10.12165137 | 10.08058096 | 9.989639618 | 0.090941339 | 0.132011749 |
| Klra1 | 2.749651494 | 2.65403044 | 2.617647911 | 0.036382529 | 0.132003583 |
| Rab37 | 5.207558938 | 5.141266494 | 5.075563955 | 0.065702539 | 0.131994983 |
| Ankrd54 | 8.481805812 | 8.590899571 | 8.349814254 | 0.241085317 | 0.131991558 |
| Sesn2 | 9.316760949 | 9.499140602 | 9.184923122 | 0.314217479 | 0.131837827 |
| 6430548M08Rik | 11.04005463 | 10.93111221 | 10.90827652 | 0.022835687 | 0.131778107 |
| Dnlz | 8.866965477 | 8.884911826 | 8.73519313 | 0.149718695 | 0.131772347 |
| Sbds | 10.1756346 | 10.14118116 | 10.04389908 | 0.097282079 | 0.131735519 |
| Thumpd3 | 9.771877385 | 9.75097288 | 9.640168386 | 0.110804494 | 0.131708999 |
| Utp14b | 7.351589711 | 7.666569861 | 7.219930245 | 0.446639616 | 0.131659465 |

| | | | | | |
|---|---|---|---|---|---|
| Mctp1 | 6.809933553 | 6.467409011 | 6.678281014 | -0.210872003 | 0.131652539 |
| Pacs1 | 9.170400883 | 9.068956348 | 9.038852542 | 0.030103806 | 0.131548341 |
| Tnfrsf8 | 3.802582332 | 3.845480565 | 3.671114038 | 0.174366527 | 0.131468294 |
| Eif1ad | 10.25807751 | 10.08809715 | 10.12668138 | -0.038584227 | 0.13139613 |
| Mphosph8 | 9.263576044 | 9.099116776 | 9.132276458 | -0.033159683 | 0.131299586 |
| Catip | 3.483950603 | 3.361284788 | 3.352660643 | 0.008624145 | 0.13128996 |
| Rxylt1 | 8.631900307 | 8.433243101 | 8.500753601 | -0.0675105 | 0.131146706 |
| Gja5 | 6.357029065 | 6.457926784 | 6.225948547 | 0.231978236 | 0.131080518 |
| Gm48417 | 5.108443917 | 5.254363332 | 4.977489916 | 0.276873416 | 0.130954001 |
| Med1 | 10.22432711 | 10.1397781 | 10.09338504 | 0.046393057 | 0.130942067 |
| Bcl6 | 10.81639517 | 10.74030994 | 10.68545981 | 0.054850127 | 0.13093536 |
| Diablo | 9.834794548 | 9.711752364 | 9.703910449 | 0.007841915 | 0.130884099 |
| Vac14 | 9.39329755 | 9.474984012 | 9.262421993 | 0.212562019 | 0.130875557 |
| Dus2 | 8.169024526 | 8.042827194 | 8.038151852 | 0.004675342 | 0.130872674 |
| Ercc6 | 9.227665546 | 9.237851398 | 9.09680098 | 0.141050418 | 0.130864566 |
| Tmed7 | 11.20087928 | 11.10726118 | 11.07002925 | 0.037231927 | 0.130850033 |
| Ube2i | 11.29130211 | 11.27853333 | 11.16058911 | 0.117944223 | 0.130713 |
| Ube2l3 | 11.51098312 | 11.35655466 | 11.38029494 | -0.023740277 | 0.13068818 |
| Taf1 | 10.26271512 | 10.22869139 | 10.13204822 | 0.09664317 | 0.130666903 |
| Gls2 | 6.068638464 | 5.793381612 | 5.93805673 | -0.144675118 | 0.130581734 |
| Xcr1 | 5.531662099 | 5.391957015 | 5.401085705 | -0.00912869 | 0.130576394 |
| Alg5 | 9.274022333 | 9.127113969 | 9.143473334 | -0.016359365 | 0.130548999 |
| Cuta | 9.405322592 | 9.465382189 | 9.274827575 | 0.190554614 | 0.130495017 |
| Ctsw | 6.143571401 | 5.952782365 | 6.013114279 | -0.060331915 | 0.130457121 |
| Tprn | 8.092107195 | 8.205289304 | 7.961769474 | 0.24351983 | 0.130337721 |
| Cryaa | 4.485557453 | 4.557057029 | 4.355257086 | 0.201799943 | 0.130300367 |
| Rce1 | 9.047758248 | 9.146749416 | 8.917524249 | 0.229225167 | 0.130233999 |
| Mtpap | 9.037066627 | 8.889630393 | 8.90689499 | -0.017264597 | 0.130171637 |
| Lats2 | 10.06136692 | 9.973500081 | 9.931296228 | 0.042203852 | 0.130070691 |
| Pde7b | 7.859574601 | 7.41372187 | 7.72965431 | -0.31593244 | 0.129920291 |
| Ap2s1 | 10.26602684 | 10.07565147 | 10.1361681 | -0.060516631 | 0.12985874 |
| Coq5 | 8.769536027 | 8.658223056 | 8.639744233 | 0.018478823 | 0.129791794 |

| | | | | | |
|---|---|---|---|---|---|
| Cpsf7 | 10.59040617 | 10.56505377 | 10.46062006 | 0.104433713 | 0.129786113 |
| Ankib1 | 9.880449007 | 9.744153589 | 9.75078874 | -0.006635151 | 0.129660267 |
| Pttg1 | 9.061112896 | 8.719629911 | 8.931490757 | -0.211860846 | 0.129622139 |
| Dhtkd1 | 7.161018732 | 7.021273674 | 7.031421725 | -0.01014805 | 0.129597007 |
| Cd6 | 4.311772146 | 4.110976978 | 4.182195436 | -0.071218458 | 0.12957671 |
| Fam13b | 9.059187517 | 9.151557717 | 8.929612691 | 0.221945026 | 0.129574826 |
| Pdia3 | 13.13318974 | 13.22098223 | 13.00365153 | 0.2173307 | 0.129538217 |
| Zfp951 | 7.107909252 | 7.075052224 | 6.978442998 | 0.096609225 | 0.129466254 |
| Pomgnt2 | 6.642592871 | 6.651624136 | 6.513151745 | 0.138472391 | 0.129441126 |
| Clpb | 9.314144936 | 9.251151614 | 9.18473842 | 0.066413194 | 0.129406517 |
| Gm14434 | 5.685903959 | 5.821103973 | 5.556499474 | 0.264604499 | 0.129404485 |
| Btnl2 | 4.452440176 | 4.322317477 | 4.323065236 | -0.000747759 | 0.12937494 |
| BC031181 | 9.948525038 | 9.735184199 | 9.819158717 | -0.083974517 | 0.129366321 |
| Mcrs1 | 9.630184676 | 9.532326764 | 9.500842623 | 0.031484141 | 0.129342053 |
| Nutf2 | 6.24303068 | 6.55203537 | 6.113788852 | 0.438246518 | 0.129241827 |
| Nfu1 | 9.039683896 | 8.760802589 | 8.910443891 | -0.149641302 | 0.129240005 |
| Dusp12 | 8.016352375 | 7.884799666 | 7.887115001 | -0.002315335 | 0.129237374 |
| Ndnf | 7.300896677 | 7.288631399 | 7.171717998 | 0.116913401 | 0.129178678 |
| Mrpl43 | 9.181986212 | 9.123163727 | 9.052922561 | 0.070241167 | 0.129063652 |
| Becn1 | 11.0525009 | 10.84113693 | 10.92344364 | -0.082306703 | 0.12905726 |
| Capns1 | 12.07221684 | 11.98459128 | 11.94318937 | 0.041401907 | 0.12902747 |
| Eif3m | 11.20616387 | 11.04416431 | 11.07713696 | -0.032972653 | 0.129026913 |
| Ndufa13 | 10.2291497 | 10.27803155 | 10.10014764 | 0.177883908 | 0.129002057 |
| Zfp143 | 8.805680765 | 8.71531937 | 8.676785359 | 0.038534011 | 0.128895406 |
| Gpr85 | 4.397322428 | 4.298119893 | 4.268481184 | 0.029638709 | 0.128841243 |
| Casp12 | 8.666639529 | 8.360790912 | 8.537876137 | -0.177085226 | 0.128763392 |
| Psme1 | 11.05796816 | 10.86564124 | 10.92927517 | -0.063633923 | 0.12869299 |
| Grid2ip | 4.866906203 | 5.016706047 | 4.738255361 | 0.278450686 | 0.128650842 |
| Dnm2 | 11.10046936 | 11.1927977 | 10.97185077 | 0.220946933 | 0.128618597 |
| Trmt44 | 7.83010014 | 7.911486721 | 7.701596465 | 0.209890256 | 0.128503675 |
| Kcne3 | 5.294393696 | 4.962835876 | 5.165915024 | -0.203079147 | 0.128478673 |
| Gramd4 | 10.17183678 | 10.33937443 | 10.04365117 | 0.295723259 | 0.128185609 |

| | | | | | |
|---|---|---|---|---|---|
| Gtf3c2 | 10.72374646 | 10.65172229 | 10.59560267 | 0.056119623 | 0.128143793 |
| Mrps9 | 8.974112263 | 8.906653937 | 8.846019687 | 0.06063425 | 0.128092576 |
| Acbd6 | 9.564315418 | 9.509063633 | 9.436243537 | 0.072820096 | 0.128071881 |
| BC005537 | 12.56220478 | 12.60177325 | 12.43415916 | 0.167614097 | 0.128045627 |
| Afap1 | 9.341358433 | 9.124970351 | 9.213425329 | -0.088454978 | 0.127933104 |
| Klhl2 | 9.762748457 | 9.563776229 | 9.634840138 | -0.07106391 | 0.127908319 |
| Rps2 | 14.31799447 | 14.32786944 | 14.19016279 | 0.137706653 | 0.12783168 |
| Hypk | 8.207720343 | 7.966201662 | 8.079925907 | -0.113724245 | 0.127794435 |
| Camp | 2.563923579 | 2.441982464 | 2.436130286 | 0.005852178 | 0.127793294 |
| Abcc1 | 11.60507075 | 11.57877477 | 11.47729646 | 0.101478313 | 0.127774293 |
| F11r | 10.87213744 | 10.79460111 | 10.74436852 | 0.050232587 | 0.127768917 |
| Sfmbt2 | 3.653769783 | 3.502985652 | 3.526011063 | -0.023025411 | 0.12775872 |
| Gins2 | 7.598385234 | 7.794849173 | 7.470629758 | 0.324219415 | 0.127755476 |
| Dnd1 | 4.624013415 | 4.696236275 | 4.496412734 | 0.199823541 | 0.127600682 |
| Ppif | 9.829771326 | 9.746436038 | 9.702203562 | 0.044232476 | 0.127567764 |
| Zmpste24 | 10.01012679 | 9.960349986 | 9.882643483 | 0.077706503 | 0.127483304 |
| 4930486L24Rik | 2.437663784 | 2.327668414 | 2.310185898 | 0.017482516 | 0.127477887 |
| Vma21 | 9.944580611 | 9.850291148 | 9.817102881 | 0.033188267 | 0.12747773 |
| Mgat5b | 6.163559176 | 6.270522813 | 6.03610968 | 0.234413133 | 0.127449496 |
| Tada2b | 8.937379772 | 8.786955935 | 8.81011469 | -0.023158754 | 0.127265082 |
| Itga2 | 8.650691973 | 8.527154462 | 8.523453533 | 0.003700929 | 0.127238439 |
| Mfhas1 | 9.867784752 | 9.922060279 | 9.740564194 | 0.181496085 | 0.127220558 |
| Letm1 | 10.47265354 | 10.64722105 | 10.34549586 | 0.301725187 | 0.127157683 |
| Gm18517 | 5.411114492 | 5.268835605 | 5.283965771 | -0.015130165 | 0.127148721 |
| Cops7a | 10.76785224 | 10.80951402 | 10.6407398 | 0.168774217 | 0.127112443 |
| Vcpip1 | 9.999366409 | 9.870869325 | 9.872298936 | -0.001429611 | 0.127067473 |
| Tead4 | 8.769682097 | 8.78790123 | 8.64262642 | 0.14527481 | 0.127055677 |
| Snx1 | 10.44566969 | 10.49619846 | 10.31861668 | 0.17758178 | 0.12705301 |
| Tiprl | 9.358467907 | 9.252921275 | 9.231458086 | 0.021463189 | 0.127009821 |
| Eif1ax | 11.19875949 | 11.02353836 | 11.07188565 | -0.04834729 | 0.126873843 |
| Uqcc2 | 9.881687912 | 9.902279734 | 9.75493433 | 0.147345404 | 0.126753583 |
| Nup88 | 10.00099028 | 9.850885244 | 9.874237221 | -0.023351977 | 0.126753057 |

| | | | | | |
|---|---|---|---|---|---|
| Gm5148 | 4.966796996 | 4.93622061 | 4.840084127 | 0.096136484 | 0.12671287 |
| Enpp4 | 8.72949436 | 8.110574824 | 8.602799051 | -0.492224227 | 0.126695309 |
| Mtdh | 11.96256546 | 11.77729828 | 11.8359162 | -0.05861792 | 0.12664926 |
| Foxs1 | 4.897883641 | 4.856982419 | 4.771549344 | 0.085433075 | 0.126334297 |
| Rasa3 | 11.0004091 | 11.05124116 | 10.8741398 | 0.177101363 | 0.126269303 |
| Gm43064 | 2.229197758 | 2.242059415 | 2.103072311 | 0.138987104 | 0.126125446 |
| H2-Aa | 13.1221631 | 13.06141434 | 12.99610535 | 0.065308993 | 0.126057753 |
| Zzef1 | 10.28362758 | 10.27282815 | 10.15765699 | 0.115171153 | 0.125970583 |
| Cstf2t | 9.33882986 | 9.163678441 | 9.212876445 | -0.049198004 | 0.125953415 |
| Trappc13 | 9.802773994 | 9.725914121 | 9.677056645 | 0.048857475 | 0.125717349 |
| Cluap1 | 8.52374437 | 8.301583043 | 8.398166985 | -0.096583943 | 0.125577385 |
| Tnpo1 | 11.46023587 | 11.47603637 | 11.33466199 | 0.141374383 | 0.12557388 |
| Apod | 11.79467177 | 11.29458597 | 11.66914229 | -0.374556323 | 0.12552948 |
| Rpl41 | 13.63626686 | 13.56053089 | 13.51082502 | 0.049705877 | 0.125441843 |
| Pip5k1a | 10.01015082 | 10.09698937 | 9.884760618 | 0.212228752 | 0.125390207 |
| Cystm1 | 8.232422147 | 8.428297576 | 8.107074378 | 0.321223199 | 0.125347769 |
| Tlk2 | 9.971925666 | 9.783474891 | 9.846623806 | -0.063148915 | 0.12530186 |
| Otulin | 9.201208185 | 9.030521363 | 9.076096486 | -0.045575122 | 0.125111699 |
| Zfp146 | 9.43755974 | 9.367803308 | 9.312482188 | 0.05532112 | 0.125077552 |
| Txndc9 | 10.3807127 | 10.31438391 | 10.25568639 | 0.05869752 | 0.12502631 |
| Usp1 | 10.3041861 | 10.15347652 | 10.17917221 | -0.02569569 | 0.125013897 |
| Nemf | 10.46089742 | 10.35445321 | 10.33593902 | 0.018514197 | 0.1249584 |
| Cadm1 | 9.545556358 | 9.452025062 | 9.420726789 | 0.031298273 | 0.12482957 |
| Ptgdr2 | 2.531586371 | 2.506826551 | 2.406782421 | 0.10004413 | 0.12480395 |
| Hnrnpr | 10.91116843 | 10.77516544 | 10.78643031 | -0.011264867 | 0.124738117 |
| Kifc3 | 9.443183202 | 9.556448718 | 9.318513474 | 0.237935245 | 0.124669728 |
| Csnk1g2 | 10.56097947 | 10.6531923 | 10.43635302 | 0.216839287 | 0.12462645 |
| Gtf2h4 | 8.130956479 | 8.022915328 | 8.00635718 | 0.016558149 | 0.124599299 |
| Hspg2 | 11.68910752 | 11.99513528 | 11.56454684 | 0.43058844 | 0.12456068 |
| Fibp | 9.372752659 | 9.303196369 | 9.248222585 | 0.054973785 | 0.124530074 |
| Ccdc136 | 6.967331815 | 7.049693446 | 6.842815339 | 0.206878107 | 0.124516475 |
| Uchl1 | 4.338889123 | 4.514560627 | 4.214377815 | 0.300182812 | 0.124511308 |

| | | | | | |
|---|---|---|---|---|---|
| Pnpla7 | 8.99463387 | 8.785036654 | 8.870157129 | -0.085120476 | 0.12447674 |
| Ddhd1 | 8.898555973 | 9.004595194 | 8.774197165 | 0.230398029 | 0.124358808 |
| Slamf7 | 8.072537687 | 8.390843713 | 7.948191802 | 0.442651912 | 0.124345886 |
| Wfdc18 | 3.485880131 | 3.436009718 | 3.361591332 | 0.074418386 | 0.124288799 |
| Amfr | 11.41222369 | 11.55303549 | 11.28795304 | 0.26508245 | 0.124270657 |
| Gm10038 | 2.315748389 | 2.248081906 | 2.191498757 | 0.056583149 | 0.124249631 |
| Gm4956 | 2.440866734 | 2.301927624 | 2.316637498 | -0.014709874 | 0.124229236 |
| Nudt19 | 8.98598263 | 8.822688204 | 8.861812792 | -0.039124587 | 0.124169838 |
| Stk4 | 10.55062554 | 10.49730599 | 10.42647656 | 0.070829427 | 0.12414898 |
| H2afx | 8.14281529 | 8.152036458 | 8.01875172 | 0.133284738 | 0.12406357 |
| Slc26a6 | 6.27307315 | 6.309078185 | 6.149013284 | 0.160064901 | 0.124059866 |
| Helq | 7.606274142 | 7.550131523 | 7.48222897 | 0.067902553 | 0.124045173 |
| Nup133 | 9.187451927 | 9.112360506 | 9.063460562 | 0.048899944 | 0.123991365 |
| Man2a1 | 11.3803044 | 11.26167497 | 11.25631428 | 0.005360693 | 0.12399012 |
| Tmem68 | 8.977854544 | 8.921258905 | 8.853908687 | 0.067350218 | 0.123945857 |
| Rarres2 | 11.47784307 | 11.28689957 | 11.35390699 | -0.067007423 | 0.12393608 |
| Dcaf1 | 9.715349625 | 9.834836362 | 9.591421814 | 0.243414547 | 0.123927811 |
| Fhad1 | 4.270675306 | 4.476932165 | 4.146781879 | 0.330150286 | 0.123893427 |
| Naa16 | 8.778615198 | 8.695729235 | 8.65472402 | 0.041005215 | 0.123891178 |
| Gm8818 | 4.509973846 | 6.249496951 | 4.386122702 | 1.863374249 | 0.123851144 |
| Copg1 | 11.060166 | 11.10757497 | 10.93637661 | 0.171198363 | 0.123789393 |
| Ska2 | 7.791493139 | 7.642584812 | 7.667722961 | -0.025138149 | 0.123770178 |
| Clu | 11.02699138 | 11.30085718 | 10.90325821 | 0.39759897 | 0.123733163 |
| Me3 | 8.097655883 | 7.98351794 | 7.973953992 | 0.009563949 | 0.123701891 |
| Tmem203 | 6.781666555 | 6.673496203 | 6.658003215 | 0.015492988 | 0.12366334 |
| Fis1 | 10.43662089 | 10.29418096 | 10.31304129 | -0.018860327 | 0.123579603 |
| BC024978 | 7.745559732 | 7.85859115 | 7.622025514 | 0.236565636 | 0.123534217 |
| Plcb3 | 11.19445928 | 11.20242853 | 11.07100805 | 0.131420483 | 0.123451237 |
| Ndufa12 | 9.951603189 | 9.891752722 | 9.828153206 | 0.063599516 | 0.123449983 |
| Gm44126 | 5.814867943 | 5.679869719 | 5.691443498 | -0.011573779 | 0.123424445 |
| Cmpk1 | 10.86343877 | 10.66265542 | 10.74021601 | -0.07756059 | 0.123222753 |
| Larp1b | 9.888173496 | 9.677724024 | 9.765010977 | -0.087286953 | 0.123162519 |

| | | | | | |
|---|---|---|---|---|---|
| Plin2 | 12.37642636 | 12.35592819 | 12.25330426 | 0.102623927 | 0.123122097 |
| Nbas | 9.455118413 | 9.434475294 | 9.332035372 | 0.102439921 | 0.123083041 |
| Rps21 | 11.5605636 | 11.59595633 | 11.43761477 | 0.158341557 | 0.122948827 |
| Nsf | 10.41697015 | 10.44077991 | 10.29403519 | 0.146744713 | 0.122934953 |
| Kcnh4 | 2.808541062 | 2.691628115 | 2.685622352 | 0.006005763 | 0.12291871 |
| Arntl2 | 8.788377418 | 8.823021965 | 8.665535796 | 0.157486169 | 0.122841623 |
| Rbm27 | 10.21780496 | 10.16433528 | 10.09504893 | 0.069286343 | 0.122756027 |
| Adck1 | 8.60787453 | 8.603509182 | 8.485220003 | 0.11828918 | 0.122654527 |
| Usp14 | 10.45310485 | 10.38523845 | 10.33048062 | 0.05475783 | 0.12262423 |
| 4930578G10Rik | 4.724371704 | 4.669545532 | 4.601785928 | 0.067759604 | 0.122585776 |
| Nup210l | 4.736354373 | 4.811111849 | 4.613854616 | 0.197257234 | 0.122499757 |
| Irx2 | 10.26260363 | 10.47403311 | 10.14011576 | 0.333917357 | 0.122487873 |
| Rab11b | 11.13352958 | 11.12436407 | 11.01118709 | 0.113176977 | 0.122342483 |
| Nudcd2 | 8.939178725 | 8.804332887 | 8.816837202 | -0.012504315 | 0.122341522 |
| Nvl | 9.920070207 | 9.831324285 | 9.797780458 | 0.033543827 | 0.122289749 |
| Bank1 | 4.773226535 | 4.420589628 | 4.650963286 | -0.230373659 | 0.122263249 |
| Ldlrap1 | 10.40987732 | 10.62168089 | 10.28762837 | 0.334052513 | 0.122248943 |
| Dedd | 9.19137755 | 9.281774625 | 9.06918184 | 0.212592785 | 0.12219571 |
| Rpl35 | 11.97650734 | 11.81512027 | 11.85437414 | -0.039253877 | 0.1221332 |
| Atp5g3 | 12.05017988 | 11.97333784 | 11.9280545 | 0.04528334 | 0.12212538 |
| Topbp1 | 10.4621695 | 10.5890087 | 10.34027872 | 0.248729977 | 0.121890783 |
| Pole4 | 9.371624137 | 9.24558794 | 9.249754524 | -0.004166585 | 0.121869613 |
| Fermt2 | 10.48229679 | 10.1950879 | 10.36043228 | -0.165344383 | 0.121864507 |
| Mmp1a | 3.04578614 | 2.974231779 | 2.923936609 | 0.05029517 | 0.121849531 |
| Thoc3 | 9.348996077 | 9.086994059 | 9.227191951 | -0.140197892 | 0.121804126 |
| Sac3d1 | 7.22247585 | 7.398665147 | 7.100676645 | 0.297988502 | 0.121799204 |
| Mrpl11 | 9.006846141 | 8.814360248 | 8.885067165 | -0.070706917 | 0.121778976 |
| Zfp451 | 9.648441227 | 9.544753211 | 9.526669256 | 0.018083955 | 0.121771971 |
| 2210016L21Rik | 8.161137792 | 8.163217274 | 8.039383921 | 0.123833354 | 0.121753871 |
| Gm12240 | 3.899971402 | 3.924451388 | 3.778250316 | 0.146201072 | 0.121721086 |
| Col13a1 | 4.768584607 | 4.802272438 | 4.646888384 | 0.155384055 | 0.121696224 |
| Mre11a | 8.992613817 | 8.935691051 | 8.870985619 | 0.064705431 | 0.121628197 |

| | | | | | |
|---|---|---|---|---|---|
| Psmc5 | 11.65283648 | 11.53427722 | 11.53135439 | 0.002922833 | 0.12148209 |
| Orm2 | 4.344540861 | 4.328782064 | 4.223093049 | 0.105689015 | 0.121447813 |
| Tradd | 7.746744556 | 7.851755562 | 7.625314382 | 0.22644118 | 0.121430175 |
| Aldh1a2 | 5.057979016 | 4.942176205 | 4.936553953 | 0.005622252 | 0.121425063 |
| Atp23 | 6.52040565 | 6.378726564 | 6.399002849 | -0.020276285 | 0.121402801 |
| Slc15a4 | 9.048730052 | 8.990765147 | 8.927331615 | 0.063433531 | 0.121398437 |
| Krt18 | 3.05119803 | 3.003426408 | 2.92985467 | 0.073571738 | 0.12134336 |
| Sec14l4 | 10.4139886 | 10.31966975 | 10.29292504 | 0.026744713 | 0.121063563 |
| Tbc1d2b | 10.90448347 | 10.93237234 | 10.78343398 | 0.148938353 | 0.121049483 |
| Ubap2l | 11.48083871 | 11.5140432 | 11.35987602 | 0.15416718 | 0.12096269 |
| Tnr | 2.650525536 | 2.613212711 | 2.529610744 | 0.083601967 | 0.120914792 |
| B3gat3 | 9.120494923 | 9.265881647 | 8.999640034 | 0.266241613 | 0.120854889 |
| Prdm4 | 9.465625143 | 9.375943527 | 9.344787518 | 0.031156009 | 0.120837625 |
| Prkce | 8.842413721 | 8.490684016 | 8.72159849 | -0.230914475 | 0.120815231 |
| Yy2 | 6.092105575 | 5.925495482 | 5.971328707 | -0.045833225 | 0.120776868 |
| Olfr1039 | 3.137210488 | 3.062462306 | 3.016511556 | 0.04595075 | 0.120698932 |
| Dbx2 | 2.744185626 | 2.672082473 | 2.623499376 | 0.048583097 | 0.12068625 |
| Faim | 7.865709095 | 7.692924063 | 7.745026522 | -0.052102459 | 0.120682573 |
| Slc25a3 | 12.85699961 | 12.73797083 | 12.73639988 | 0.00157095 | 0.120599733 |
| Gga1 | 10.31786058 | 10.51574613 | 10.19727167 | 0.318474453 | 0.120588907 |
| Irgc1 | 2.52020782 | 2.465797304 | 2.399718223 | 0.066079081 | 0.120489597 |
| Zc3h13 | 10.32477277 | 10.22665533 | 10.20435086 | 0.022304467 | 0.12042191 |
| Zfp64 | 7.979083425 | 8.044072064 | 7.858697197 | 0.185374867 | 0.120386228 |
| Rreb1 | 10.56033782 | 10.64649746 | 10.43999318 | 0.206504283 | 0.120344643 |
| Pip5k1c | 10.11787386 | 10.37956273 | 9.997534475 | 0.382028255 | 0.120339385 |
| Smarcad1 | 9.126930333 | 9.026050596 | 9.006598994 | 0.019451602 | 0.120331338 |
| Meaf6 | 8.991993003 | 9.064928072 | 8.871677797 | 0.193250276 | 0.120315206 |
| Tacc1 | 10.50463375 | 10.3068974 | 10.38432284 | -0.077425437 | 0.12031091 |
| Ctla4 | 5.476541404 | 5.54269463 | 5.356231516 | 0.186463114 | 0.120309888 |
| Eif4g3 | 11.06453284 | 10.92900473 | 10.94423901 | -0.015234287 | 0.12029383 |
| Gm4813 | 4.713875946 | 4.629812377 | 4.593589218 | 0.036223159 | 0.120286728 |
| Gm11518 | 11.033654 | 10.87417814 | 10.9134736 | -0.039295457 | 0.120180397 |

| | | | | | |
|---|---|---|---|---|---|
| Rpl27-ps3 | 11.033654 | 10.87417814 | 10.9134736 | -0.039295457 | 0.120180397 |
| Pts | 8.722901443 | 8.440435389 | 8.602743222 | -0.162307833 | 0.120158221 |
| Ubtd2 | 7.158134414 | 7.120809248 | 7.037983724 | 0.082825524 | 0.12015069 |
| Pcsk5 | 8.152857929 | 8.1284156 | 8.03272568 | 0.09568992 | 0.120132249 |
| Col17a1 | 14.75116955 | 15.00477732 | 14.63104684 | 0.373730483 | 0.120122717 |
| C1rl | 6.60760242 | 6.292087036 | 6.487527728 | -0.195440691 | 0.120074692 |
| Dtwd2 | 6.263416856 | 6.057314722 | 6.143368408 | -0.086053686 | 0.120048449 |
| Pecam1 | 10.60155861 | 10.37682623 | 10.48151106 | -0.10468483 | 0.120047553 |
| Yif1b | 8.538086487 | 8.598417005 | 8.418198302 | 0.180218703 | 0.119888185 |
| B3galt5 | 3.714664602 | 3.579757136 | 3.594800423 | -0.015043287 | 0.119864179 |
| Zfp428 | 4.060992714 | 3.978420254 | 3.941176711 | 0.037243543 | 0.119816003 |
| Gpr141 | 8.172548987 | 8.002309773 | 8.052774083 | -0.05046431 | 0.119774904 |
| Tars2 | 9.393723228 | 9.394073958 | 9.274092249 | 0.119981709 | 0.119630979 |
| Clec2l | 4.521020273 | 4.540467961 | 4.401493733 | 0.138974228 | 0.119526539 |
| Rngtt | 8.609996565 | 8.526134352 | 8.490562216 | 0.035572135 | 0.119434349 |
| Mcpt8 | 3.746819833 | 3.711167693 | 3.627418877 | 0.083748816 | 0.119400956 |
| Prdm2 | 10.43334163 | 10.50946873 | 10.31409821 | 0.195370517 | 0.119243417 |
| Anapc10 | 7.837290064 | 7.547976316 | 7.718058727 | -0.170082411 | 0.119231337 |
| Fam53c | 9.63704465 | 9.746336863 | 9.51785274 | 0.228484123 | 0.11919191 |
| Serpinh1 | 11.78570295 | 12.15128553 | 11.66653341 | 0.484752123 | 0.119169537 |
| Scyl1 | 9.95560744 | 10.07546659 | 9.83644048 | 0.23902611 | 0.11916696 |
| Hint1 | 10.74651464 | 10.56683736 | 10.62736896 | -0.0605316 | 0.119145673 |
| Fcrl1 | 3.734181957 | 3.725719243 | 3.61503801 | 0.110681233 | 0.119143947 |
| Cwc22 | 9.564236963 | 9.435074773 | 9.445148404 | -0.010073632 | 0.119088559 |
| Asna1 | 9.766378072 | 9.787886863 | 9.647298155 | 0.140588709 | 0.119079918 |
| Tysnd1 | 8.196882192 | 8.398235331 | 8.078252368 | 0.319982963 | 0.118629824 |
| Rsl1 | 6.735464783 | 6.782755439 | 6.616854808 | 0.165900631 | 0.118609976 |
| Wtip | 8.418606799 | 8.401512701 | 8.299997001 | 0.1015157 | 0.118609798 |
| Ube2q1 | 10.41767881 | 10.31212641 | 10.29915984 | 0.012966573 | 0.11851897 |
| F7 | 4.989781073 | 5.309564044 | 4.871324835 | 0.438239209 | 0.118456238 |
| Pcdhga4 | 5.574394222 | 5.536639534 | 5.456030679 | 0.080608856 | 0.118363543 |
| Itfg2 | 8.386393145 | 8.334606889 | 8.268128275 | 0.066478614 | 0.11826487 |

| | | | | | |
|---|---|---|---|---|---|
| Hsp90ab1 | 15.13586417 | 15.22018418 | 15.01764869 | 0.202535497 | 0.11821548 |
| Gabarap | 12.22464182 | 12.08555692 | 12.10642878 | -0.020871857 | 0.11821304 |
| Lig4 | 7.706564407 | 7.604342416 | 7.58835933 | 0.015983086 | 0.118205077 |
| Psma2 | 11.36919355 | 11.21742015 | 11.25100616 | -0.033586017 | 0.118187383 |
| Scaf11 | 11.22251741 | 11.09678193 | 11.10439548 | -0.007613553 | 0.11812193 |
| Son | 13.18205749 | 13.17789436 | 13.06397081 | 0.11392355 | 0.118086677 |
| Alkbh2 | 5.853136732 | 5.888078895 | 5.73511681 | 0.152962085 | 0.118019922 |
| Phf12 | 9.498112313 | 9.457274637 | 9.38011103 | 0.077163607 | 0.118001283 |
| Tomm22 | 10.84842731 | 10.68898848 | 10.73045193 | -0.04146345 | 0.117975377 |
| Rnf187 | 10.84049514 | 10.81740283 | 10.7226968 | 0.094706027 | 0.117798343 |
| Pmm2 | 9.936584795 | 9.869425377 | 9.818978823 | 0.050446554 | 0.117605972 |
| Uba6 | 9.203251967 | 9.123240541 | 9.085764772 | 0.037475769 | 0.117487195 |
| Trip11 | 10.92499277 | 10.87820728 | 10.80752619 | 0.070681083 | 0.117466573 |
| Srpr | 11.71909692 | 11.5317861 | 11.60163624 | -0.069850133 | 0.11746068 |
| Tbc1d25 | 7.861697346 | 7.797891969 | 7.74428006 | 0.053611908 | 0.117417286 |
| Ddx4 | 2.607030632 | 2.485180086 | 2.489741925 | -0.004561839 | 0.117288706 |
| Chic2 | 9.691190696 | 9.49491227 | 9.573954438 | -0.079042168 | 0.117236258 |
| Ctnnd2 | 7.239252854 | 7.134973069 | 7.12208557 | 0.012887499 | 0.117167283 |
| Mcpt1 | 1.426877458 | 1.339356913 | 1.309816696 | 0.029540217 | 0.117060762 |
| Lhcgr | 4.66752917 | 4.562578848 | 4.550512632 | 0.012066216 | 0.117016538 |
| Adgrg5 | 4.882592834 | 4.765015596 | 4.765618713 | -0.000603118 | 0.11697412 |
| Meis2 | 7.458860984 | 7.378213043 | 7.341887872 | 0.03632517 | 0.116973112 |
| Whamm | 8.308041098 | 8.315353642 | 8.191090643 | 0.124262999 | 0.116950455 |
| Pdrg1 | 9.962740789 | 9.962746159 | 9.845904279 | 0.116841879 | 0.11683651 |
| Gpank1 | 8.229333641 | 8.252102365 | 8.112548387 | 0.139553978 | 0.116785255 |
| Rem1 | 5.403200918 | 5.322696932 | 5.286426147 | 0.036270784 | 0.11677477 |
| Ptpn18 | 6.797256281 | 6.641178718 | 6.680495091 | -0.039316373 | 0.11676119 |
| Ubb | 14.14381432 | 14.3367375 | 14.02707863 | 0.30965887 | 0.116735697 |
| Zap70 | 4.699161813 | 4.671112457 | 4.582438034 | 0.088674422 | 0.116723778 |
| Sugp1 | 9.227388704 | 9.19240872 | 9.110693232 | 0.081715488 | 0.116695472 |
| Fbxl15 | 5.364515879 | 5.332994674 | 5.247908435 | 0.085086239 | 0.116607444 |
| Cep83 | 8.880477159 | 8.678033022 | 8.763933351 | -0.085900329 | 0.116543808 |

| | | | | | |
|---|---|---|---|---|---|
| Vwa1 | 8.700684736 | 8.672217731 | 8.584203059 | 0.088014672 | 0.116481677 |
| Wrb | 7.806445834 | 7.542164729 | 7.690020213 | -0.147855484 | 0.116425621 |
| Srp54b | 8.819628173 | 9.062712612 | 8.703202752 | 0.35950986 | 0.11642542 |
| Larp4b | 11.28799247 | 11.13397594 | 11.17158132 | -0.037605383 | 0.116411153 |
| Med22 | 10.65893816 | 10.60259322 | 10.5425295 | 0.06006372 | 0.11640866 |
| Tspoap1 | 5.851939005 | 5.908424482 | 5.735551782 | 0.1728727 | 0.116387223 |
| 6430531B16Rik | 2.526041133 | 2.485255026 | 2.409769187 | 0.075485838 | 0.116271946 |
| Gm13835 | 10.17754219 | 10.0429678 | 10.06131281 | -0.018345007 | 0.11622938 |
| Coa7 | 7.52255804 | 7.375916624 | 7.406388964 | -0.03047234 | 0.116169076 |
| Olfr2 | 3.400884004 | 3.299536061 | 3.284790407 | 0.014745653 | 0.116093597 |
| Tpgs1 | 6.362114648 | 6.530918094 | 6.246033488 | 0.284884605 | 0.116081159 |
| Sumf2 | 8.784865289 | 8.844323981 | 8.668860856 | 0.175463125 | 0.116004433 |
| Rabif | 9.480257288 | 9.364835814 | 9.36428159 | 0.000554224 | 0.115975698 |
| Nlrp1b | 6.550436978 | 6.457945263 | 6.434468281 | 0.023476982 | 0.115968696 |
| Plekhg4 | 2.624499182 | 2.631611144 | 2.508610386 | 0.123000758 | 0.115888797 |
| Psmf1 | 9.072489899 | 8.910578361 | 8.956608567 | -0.046030206 | 0.115881332 |
| Dcps | 8.22723397 | 8.114988463 | 8.11140384 | 0.003584623 | 0.115830129 |
| Sec1 | 3.768502283 | 3.772466588 | 3.652702125 | 0.119764463 | 0.115800157 |
| Vmn2r-ps54 | 2.586249306 | 2.508372262 | 2.470545877 | 0.037826384 | 0.115703428 |
| Lamtor1 | 10.60452488 | 10.52747784 | 10.48882654 | 0.038651297 | 0.115698337 |
| Samd15 | 2.567492341 | 2.449554091 | 2.451868964 | -0.002314873 | 0.115623376 |
| Gpr151 | 1.239321652 | 1.120224673 | 1.123711022 | -0.00348635 | 0.115610629 |
| Xlr4b | 5.170175075 | 4.649595533 | 5.054574636 | -0.404979103 | 0.115600439 |
| Crls1 | 8.765659085 | 8.662534897 | 8.650097877 | 0.01243702 | 0.115561208 |
| Ezh2 | 9.508916811 | 9.542027787 | 9.393386221 | 0.148641566 | 0.115530591 |
| Bud13 | 8.080436574 | 7.635404614 | 7.964923902 | -0.329519288 | 0.115512672 |
| Prpf4b | 11.07734675 | 11.01324967 | 10.96184609 | 0.051403577 | 0.11550066 |
| Cib1 | 8.89581061 | 8.753989721 | 8.780482568 | -0.026492847 | 0.115328042 |
| Ndufab1 | 10.07812007 | 10.0331398 | 9.962822922 | 0.070316881 | 0.115297146 |
| Gm48592 | 3.708028613 | 3.650192796 | 3.592772113 | 0.057420682 | 0.115256499 |
| Dgkz | 10.23508879 | 10.42549915 | 10.11985466 | 0.305644496 | 0.11523413 |
| Hrh1 | 5.305786868 | 5.14986169 | 5.190650193 | -0.040788503 | 0.115136674 |

| | | | | | |
|---|---|---|---|---|---|
| Nelfe | 8.983168691 | 8.956193708 | 8.868039502 | 0.088154206 | 0.115129189 |
| Itpr1 | 9.062136938 | 8.865187003 | 8.947047902 | -0.081860899 | 0.115089037 |
| Card14 | 10.13762054 | 10.03004045 | 10.02265121 | 0.007389236 | 0.114969325 |
| Slc19a3 | 3.801983533 | 3.779966523 | 3.687332011 | 0.092634512 | 0.114651521 |
| Mpp7 | 9.849910306 | 9.775844908 | 9.735265355 | 0.040579553 | 0.114644951 |
| Il13 | 2.133082938 | 2.005007926 | 2.018504066 | -0.013496139 | 0.114578873 |
| Pomp | 11.22730267 | 11.09426379 | 11.11274925 | -0.018485467 | 0.114553413 |
| Zbtb7b | 11.39448583 | 11.58648321 | 11.27996875 | 0.306514453 | 0.114517077 |
| Ano6 | 11.27643593 | 11.26285839 | 11.16194075 | 0.100917643 | 0.11449518 |
| Carmil2 | 4.580825363 | 4.586987559 | 4.466337262 | 0.120650297 | 0.114488101 |
| Eif5 | 12.78173884 | 12.87849415 | 12.66725557 | 0.211238577 | 0.114483273 |
| Sprr2i | 5.754752814 | 5.795704502 | 5.640317565 | 0.155386937 | 0.114435248 |
| Fbxw7 | 9.81895475 | 9.731552646 | 9.704599883 | 0.026952763 | 0.114354867 |
| Gtpbp1 | 9.99335399 | 9.88886605 | 9.879002715 | 0.009863335 | 0.114351275 |
| Gm11549 | 2.799927734 | 2.669338229 | 2.68565891 | -0.016320681 | 0.114268824 |
| Pecr | 7.533118016 | 7.408885859 | 7.418870227 | -0.009984368 | 0.114247789 |
| Fam3b | 7.454304016 | 7.153772927 | 7.34016581 | -0.186392883 | 0.114138206 |
| Tmem171 | 5.161930958 | 5.120198879 | 5.047819257 | 0.072379622 | 0.114111701 |
| Gstp1 | 9.868104041 | 9.891638626 | 9.753997999 | 0.137640627 | 0.114106042 |
| Nxpe5 | 8.000464404 | 7.987574866 | 7.886589603 | 0.100985263 | 0.113874801 |
| Kdm5a | 10.61197422 | 10.4447546 | 10.49815675 | -0.053402157 | 0.11381747 |
| Jakmip1 | 4.123404723 | 3.993684625 | 4.00960615 | -0.015921525 | 0.113798573 |
| Ppp6r1 | 11.18323243 | 11.22813807 | 11.06951729 | 0.15862078 | 0.113715137 |
| Lrch1 | 9.761244021 | 9.681364307 | 9.647560954 | 0.033803353 | 0.113683066 |
| Fkbp4 | 11.59169439 | 11.58469795 | 11.47810521 | 0.10659274 | 0.113589177 |
| Rtn4ip1 | 8.177317496 | 7.949814635 | 8.063755012 | -0.113940377 | 0.113562484 |
| Klra9 | 2.940093824 | 2.794295434 | 2.826579002 | -0.032283568 | 0.113514822 |
| Sdhaf2 | 9.578850651 | 9.509742225 | 9.46534197 | 0.044400255 | 0.113508681 |
| Pmpca | 10.82841956 | 10.75304202 | 10.71513882 | 0.037903207 | 0.11328074 |
| Akirin1 | 10.17147878 | 9.995743767 | 10.0582765 | -0.062532732 | 0.113202284 |
| Gm13340 | 4.28368527 | 4.267201121 | 4.170485467 | 0.096715653 | 0.113199802 |
| Tmem243 | 7.923620693 | 7.557513225 | 7.810499064 | -0.252985839 | 0.113121629 |

| | | | | | |
|---|---|---|---|---|---|
| Fasl | 2.189398558 | 2.047860904 | 2.076306978 | -0.028446073 | 0.11309158 |
| Serpinb13 | 4.644810218 | 4.365943386 | 4.531857431 | -0.165914044 | 0.112952787 |
| Dapp1 | 8.69195326 | 8.669928282 | 8.579015335 | 0.090912947 | 0.112937924 |
| Skiv2l | 10.41141623 | 10.407178 | 10.29857469 | 0.108603303 | 0.11284154 |
| Rnf113a2 | 7.712144016 | 7.628052019 | 7.599357525 | 0.028694494 | 0.11278649 |
| Ywhah | 11.45150424 | 11.31210389 | 11.3387419 | -0.02663801 | 0.112762337 |
| Nbn | 9.079088628 | 9.037693717 | 8.966349055 | 0.071344662 | 0.112739573 |
| Ces2g | 9.289548674 | 9.331594408 | 9.176821348 | 0.15477306 | 0.112727326 |
| Erlin1 | 10.66480111 | 10.52350253 | 10.55227028 | -0.028767747 | 0.112530833 |
| Pex26 | 8.901868616 | 8.903654803 | 8.789360666 | 0.114294137 | 0.11250795 |
| Fgfbp1 | 11.3392039 | 11.24689156 | 11.22686332 | 0.02002824 | 0.112340583 |
| Lad1 | 11.98447894 | 11.95873641 | 11.87217923 | 0.086557173 | 0.112299707 |
| Olfr1487 | 2.809301222 | 2.702945144 | 2.697016029 | 0.005929115 | 0.112285193 |
| Uimc1 | 9.634625671 | 9.503320492 | 9.52245975 | -0.019139258 | 0.112165921 |
| Gm11847 | 5.232149911 | 4.989243359 | 5.119991448 | -0.130748089 | 0.112158463 |
| Wdr33 | 9.994755774 | 9.998519787 | 9.882802947 | 0.11571684 | 0.111952827 |
| Shc1 | 10.85327917 | 10.76931643 | 10.74133281 | 0.027983617 | 0.11194636 |
| Jade2 | 9.16632384 | 9.138382836 | 9.054405646 | 0.08397719 | 0.111918195 |
| Gm7897 | 3.501680785 | 3.379013581 | 3.389784284 | -0.010770703 | 0.111896501 |
| Epcam | 11.36238449 | 11.26536714 | 11.25049331 | 0.014873833 | 0.11189118 |
| Ythdc1 | 10.82947406 | 10.65689566 | 10.7175854 | -0.060689747 | 0.111888657 |
| Zfp35 | 8.797751193 | 8.657296743 | 8.686042561 | -0.028745818 | 0.111708632 |
| Setd1a | 10.07979696 | 10.38590453 | 9.96811034 | 0.417794186 | 0.11168662 |
| Jmjd1c | 10.07576232 | 9.982598987 | 9.964159084 | 0.018439903 | 0.111603239 |
| Rpn2 | 12.20130034 | 12.15067676 | 12.08979167 | 0.060885087 | 0.11150867 |
| Tm9sf3 | 12.48565943 | 12.32546369 | 12.37415837 | -0.048694677 | 0.111501063 |
| Tm9sf4 | 10.69161322 | 10.6343798 | 10.58011312 | 0.054266683 | 0.111500103 |
| Cers5 | 11.15729885 | 11.09073723 | 11.04593494 | 0.04480229 | 0.11136391 |
| Psma4 | 11.45808146 | 11.35841475 | 11.34689946 | 0.011515293 | 0.111182 |
| Tarbp2 | 9.033260794 | 9.134039303 | 8.922173394 | 0.211865909 | 0.1110874 |
| Gm5526 | 3.25964637 | 3.305885692 | 3.148681336 | 0.157204356 | 0.110965034 |
| Cggbp1 | 11.07149816 | 10.93791857 | 10.96065291 | -0.02273434 | 0.11084525 |

| | | | | | |
|---|---|---|---|---|---|
| Tcam1 | 2.488928587 | 2.440780687 | 2.378149432 | 0.062631255 | 0.110779155 |
| Nppc | 3.03144039 | 2.984499309 | 2.920718571 | 0.063780738 | 0.110721819 |
| Gga3 | 9.106794033 | 9.161631127 | 8.996137632 | 0.165493495 | 0.110656401 |
| Tmem98 | 6.57900285 | 6.465134866 | 6.468380386 | -0.00324552 | 0.110622464 |
| Polr3b | 9.155009411 | 9.033511396 | 9.044428278 | -0.010916881 | 0.110581133 |
| Sugp2 | 9.162839452 | 9.21694214 | 9.052368928 | 0.164573212 | 0.110470524 |
| Crtam | 2.892185172 | 2.885763424 | 2.781731704 | 0.10403172 | 0.110453469 |
| Rer1 | 11.37749582 | 11.20103101 | 11.26705231 | -0.0660213 | 0.110443513 |
| Mrgprb1 | 7.564223414 | 7.065608317 | 7.453820929 | -0.388212613 | 0.110402484 |
| Sdhd | 10.9307294 | 10.82764847 | 10.82038419 | 0.00726428 | 0.11034521 |
| Rexo2 | 10.28859195 | 10.24658871 | 10.17825086 | 0.068337853 | 0.11034109 |
| S100a10 | 12.52663159 | 12.29108637 | 12.41630481 | -0.12521844 | 0.11032678 |
| C030006K11Rik | 7.553387356 | 7.629454979 | 7.44310865 | 0.186346329 | 0.110278706 |
| Lsm6 | 9.276950559 | 9.214664915 | 9.166689394 | 0.04797552 | 0.110261165 |
| Ube2v2 | 10.05668417 | 9.871776609 | 9.946496544 | -0.074719935 | 0.110187622 |
| Ddx1 | 11.33248192 | 11.18254603 | 11.22230582 | -0.039759787 | 0.1101761 |
| Eea1 | 11.386398 | 11.25566002 | 11.27629258 | -0.020632563 | 0.110105413 |
| Tns3 | 10.52033391 | 10.45604308 | 10.4103609 | 0.04568218 | 0.109973017 |
| Gss | 8.352632123 | 8.394204659 | 8.242661614 | 0.151543044 | 0.109970509 |
| Bcl7b | 8.939821172 | 8.948626296 | 8.829948359 | 0.118677937 | 0.109872813 |
| Zcrb1 | 9.61791354 | 9.455977784 | 9.508052099 | -0.052074315 | 0.109861441 |
| Ldlrad4 | 6.766025247 | 6.63345734 | 6.656188568 | -0.022731228 | 0.109836679 |
| Map1lc3a | 11.42455222 | 11.40064222 | 11.31476158 | 0.08588064 | 0.109790637 |
| Srp14 | 9.91633233 | 9.73966281 | 9.806553112 | -0.066890303 | 0.109779218 |
| Mettl4 | 8.641003015 | 8.657938491 | 8.531233371 | 0.12670512 | 0.109769644 |
| P3h1 | 8.188320429 | 8.186255378 | 8.078575918 | 0.10767946 | 0.109744511 |
| Dap3 | 9.897742631 | 9.852408025 | 9.788024306 | 0.064383719 | 0.109718325 |
| Gm15925 | 2.360284513 | 2.312324936 | 2.250566942 | 0.061757994 | 0.109717571 |
| Cpsf6 | 10.71675731 | 10.65170747 | 10.60709378 | 0.04461369 | 0.109663523 |
| Mcmdc2 | 3.436948779 | 3.408036466 | 3.327313876 | 0.08072259 | 0.109634903 |
| Triml1 | 2.566397292 | 2.481584991 | 2.456851726 | 0.024733264 | 0.109545565 |
| Phactr4 | 9.894174928 | 9.863930364 | 9.784645108 | 0.079285256 | 0.10952982 |

| | | | | | |
|---|---|---|---|---|---|
| Zfr | 10.66076621 | 10.58537241 | 10.55125741 | 0.034115 | 0.109508797 |
| Gng10 | 8.839811819 | 8.653949037 | 8.730384757 | -0.076435719 | 0.109427062 |
| Mtfr2 | 5.754615343 | 5.694677407 | 5.645237581 | 0.049439826 | 0.109377762 |
| Wdr91 | 8.926161553 | 9.117443462 | 8.816791112 | 0.300652351 | 0.109370441 |
| Wnt10a | 9.024737363 | 9.270042843 | 8.915383574 | 0.354659268 | 0.109353789 |
| Cpz | 7.188980458 | 7.320322558 | 7.079717246 | 0.240605312 | 0.109263212 |
| Cops9 | 9.317752472 | 9.115104228 | 9.208553683 | -0.093449456 | 0.109198789 |
| Pcdhb21 | 4.232407293 | 4.115137518 | 4.123279665 | -0.008142147 | 0.109127628 |
| Cd9 | 13.20611002 | 12.99009316 | 13.09703904 | -0.106945877 | 0.109070983 |
| 1110004F10Rik | 10.50665214 | 10.3690368 | 10.39764712 | -0.02861032 | 0.10900502 |
| Vmn2r26 | 3.100830659 | 2.993815181 | 2.991834915 | 0.001980266 | 0.108995744 |
| Wdcp | 6.865909075 | 6.929377529 | 6.756985002 | 0.172392527 | 0.108924073 |
| Khdrbs2 | 2.756059684 | 2.627008266 | 2.647165045 | -0.020156779 | 0.108894639 |
| Ambp | 2.083783863 | 2.042818309 | 1.974962528 | 0.06785578 | 0.108821335 |
| Ndufa5 | 9.665970138 | 9.498592572 | 9.557164211 | -0.058571639 | 0.108805927 |
| Nudt1 | 8.010959125 | 7.935299662 | 7.90221987 | 0.033079792 | 0.108739255 |
| Kcnd3 | 5.162740521 | 4.934426597 | 5.054018395 | -0.119591798 | 0.108722126 |
| a | 3.174507948 | 3.171850822 | 3.065924058 | 0.105926765 | 0.10858389 |
| Cox14 | 8.970154582 | 8.986355445 | 8.861590411 | 0.124765033 | 0.108564171 |
| Smarcb1 | 10.11845554 | 10.17167791 | 10.00993854 | 0.161739368 | 0.108516998 |
| Zfp623 | 7.50671344 | 7.526485404 | 7.398258082 | 0.128227322 | 0.108455358 |
| Ddost | 11.35611429 | 11.29565518 | 11.24769225 | 0.04796293 | 0.108422037 |
| Dazap1 | 9.951495333 | 10.13149437 | 9.843103808 | 0.288390565 | 0.108391525 |
| Ccl22 | 8.903445004 | 10.54464162 | 8.795194701 | 1.749446923 | 0.108250304 |
| Spic | 4.702004186 | 4.913613658 | 4.593758894 | 0.319854763 | 0.108245291 |
| Fzd9 | 4.378109674 | 4.339733649 | 4.269898729 | 0.06983492 | 0.108210945 |
| Optn | 8.789856228 | 8.619022291 | 8.681680828 | -0.062658537 | 0.1081754 |
| Rpl10a-ps1 | 12.46216564 | 12.41315614 | 12.35401642 | 0.059139723 | 0.108149223 |
| Pdcd7 | 8.752662553 | 8.523953249 | 8.644549407 | -0.120596158 | 0.108113146 |
| Rab8a | 10.52181145 | 10.46452307 | 10.41373961 | 0.05078346 | 0.108071833 |
| Klk10 | 10.1318678 | 10.01579189 | 10.02379728 | -0.008005387 | 0.108070515 |
| Gm11808 | 12.8529771 | 12.85658444 | 12.74498895 | 0.111595487 | 0.107988153 |

| | | | | | |
|---|---|---|---|---|---|
| Rgs17 | 2.982465277 | 2.901900835 | 2.874518451 | 0.027382384 | 0.107946826 |
| Fn1 | 12.6276907 | 12.41900619 | 12.51980218 | -0.10079599 | 0.107888513 |
| Nabp2 | 9.343052981 | 9.299959648 | 9.235182995 | 0.064776653 | 0.107869987 |
| Cnot11 | 9.294240541 | 9.031959051 | 9.186374689 | -0.154415638 | 0.107865852 |
| Gm8203 | 8.351526229 | 8.573664082 | 8.243842965 | 0.329821118 | 0.107683265 |
| Slc39a11 | 9.715978819 | 9.785867167 | 9.60835361 | 0.177513557 | 0.107625209 |
| Fancf | 5.756417668 | 5.684610672 | 5.648844042 | 0.03576663 | 0.107573627 |
| Rpa2 | 8.759794782 | 8.700697332 | 8.652323033 | 0.048374299 | 0.107471748 |
| Rnpep | 10.22273246 | 10.27826114 | 10.11533561 | 0.162925525 | 0.107396849 |
| Arl2bp | 9.511748196 | 9.346439477 | 9.40449372 | -0.058054243 | 0.107254476 |
| Zhx2 | 8.786600824 | 8.707539046 | 8.679353699 | 0.028185347 | 0.107247126 |
| Ap2b1 | 11.31070851 | 11.2721762 | 11.20354054 | 0.068635653 | 0.10716797 |
| Nme2 | 11.7633695 | 11.62376771 | 11.65623399 | -0.03246628 | 0.107135507 |
| Sox15 | 6.574414728 | 6.677208653 | 6.467307244 | 0.209901409 | 0.107107484 |
| Olfr1309 | 4.338490975 | 4.233573191 | 4.23143113 | 0.002142061 | 0.107059846 |
| Vps16 | 10.04268773 | 9.79129164 | 9.935694413 | -0.144402773 | 0.106993319 |
| Prss23 | 11.34562751 | 11.06964597 | 11.23863824 | -0.16899227 | 0.10698927 |
| Kirrel3 | 2.813051732 | 2.745050471 | 2.706075388 | 0.038975083 | 0.106976344 |
| Rps7 | 12.79826575 | 12.85446173 | 12.6912953 | 0.163166433 | 0.106970453 |
| Thoc6 | 8.246109428 | 8.555371659 | 8.139230215 | 0.416141444 | 0.106879213 |
| Ercc1 | 8.50718156 | 8.561459579 | 8.400370627 | 0.161088952 | 0.106810933 |
| Gm3755 | 2.78995686 | 2.683694164 | 2.683218804 | 0.00047536 | 0.106738056 |
| Ybx1 | 12.5633298 | 12.51511318 | 12.45663055 | 0.058482637 | 0.10669925 |
| Rnf166 | 8.33877542 | 8.392494941 | 8.232098384 | 0.160396557 | 0.106677036 |
| Ercc2 | 8.764802271 | 8.866287306 | 8.658219742 | 0.208067564 | 0.106582529 |
| Snx11 | 9.365601653 | 9.460911644 | 9.259028991 | 0.201882653 | 0.106572662 |
| Cmss1 | 7.345238895 | 7.340739186 | 7.23869909 | 0.102040096 | 0.106539805 |
| E230025N22Rik | 1.162698699 | 1.07187823 | 1.05617765 | 0.01570058 | 0.106521049 |
| Msantd1 | 3.859417661 | 3.959368382 | 3.752906076 | 0.206462306 | 0.106511585 |
| Fancl | 6.990172428 | 6.923354035 | 6.883734165 | 0.03961987 | 0.106438263 |
| Selenot | 11.87855748 | 11.6990551 | 11.77215504 | -0.073099943 | 0.106402437 |
| Psmd13 | 11.19227719 | 11.10132799 | 11.08590634 | 0.015421643 | 0.106370843 |

| | | | | | |
|---|---|---|---|---|---|
| Wdr24 | 8.600623828 | 8.617051875 | 8.494332754 | 0.122719121 | 0.106291075 |
| Fen1 | 8.856295223 | 8.825107149 | 8.750011073 | 0.075096077 | 0.106284151 |
| Clec1b | 1.945378313 | 1.889295808 | 1.839132264 | 0.050163544 | 0.106246049 |
| Mcm10 | 7.54262884 | 7.584083306 | 7.436411861 | 0.147671445 | 0.10621698 |
| Eif3l | 11.56418043 | 11.54678208 | 11.45797207 | 0.088810007 | 0.10620836 |
| Tmem186 | 8.450711377 | 8.31405141 | 8.344570573 | -0.030519163 | 0.106140804 |
| Tmco4 | 8.845897425 | 8.768054287 | 8.739798717 | 0.02825557 | 0.106098707 |
| Spryd3 | 9.818840573 | 9.782055121 | 9.712800992 | 0.069254129 | 0.106039581 |
| Piga | 9.043341281 | 9.145523427 | 8.937378423 | 0.208145004 | 0.105962858 |
| Pole3 | 9.958704126 | 9.828672364 | 9.852897995 | -0.024225631 | 0.105806131 |
| Clec9a | 4.727842853 | 4.691106712 | 4.622046579 | 0.069060133 | 0.105796274 |
| Gm19001 | 3.400738185 | 3.399956491 | 3.294956436 | 0.105000055 | 0.105781749 |
| Nox3 | 1.991644782 | 1.887500834 | 1.885900938 | 0.001599896 | 0.105743844 |
| Trim14 | 6.01852681 | 5.711317945 | 5.912800107 | -0.201482162 | 0.105726703 |
| Gm49356 | 3.333767741 | 3.235378139 | 3.228057041 | 0.007321098 | 0.1057107 |
| Txlna | 10.6568311 | 10.75082173 | 10.55113929 | 0.199682433 | 0.105691807 |
| Taf1c | 8.173709977 | 8.539579916 | 8.068030475 | 0.471549441 | 0.105679502 |
| Pgap3 | 7.999068815 | 8.010089633 | 7.893389459 | 0.116700174 | 0.105679356 |
| Cdk8 | 9.294605664 | 9.251827776 | 9.188957478 | 0.062870299 | 0.105648186 |
| Spata2 | 9.249475869 | 9.22918704 | 9.143828906 | 0.085358134 | 0.105646963 |
| Elmo1 | 8.79956844 | 8.807695276 | 8.694010791 | 0.113684485 | 0.10555765 |
| Ndufs6 | 9.874481457 | 9.814520646 | 9.768926649 | 0.045593997 | 0.105554808 |
| Gm6505 | 3.49571237 | 3.437194624 | 3.390316349 | 0.046878275 | 0.105396021 |
| Usmg5 | 10.41079387 | 10.28816695 | 10.3054868 | -0.017319847 | 0.10530707 |
| Tsnax | 10.80985165 | 10.67281354 | 10.70454798 | -0.031734433 | 0.10530367 |
| Phykpl | 7.860614945 | 7.77602145 | 7.755333888 | 0.020687561 | 0.105281057 |
| Heatr6 | 8.818674299 | 8.846074368 | 8.713413667 | 0.132660701 | 0.105260632 |
| Pced1a | 7.52282697 | 7.618253629 | 7.417645055 | 0.200608574 | 0.105181915 |
| Rab3b | 3.866602238 | 3.792684189 | 3.761444178 | 0.031240011 | 0.10515806 |
| Cttn | 11.65991684 | 11.59406918 | 11.55476982 | 0.03929936 | 0.10514702 |
| Klhl15 | 8.137249848 | 7.827274151 | 8.032109507 | -0.204835356 | 0.105140341 |
| Erp44 | 10.40923572 | 10.4397031 | 10.3041063 | 0.135596807 | 0.105129423 |

| | | | | | |
|---|---|---|---|---|---|
| Appbp2 | 10.29167257 | 10.23301526 | 10.18655661 | 0.04645865 | 0.105115957 |
| Wtap | 10.6253495 | 10.51345254 | 10.52032195 | -0.006869413 | 0.105027547 |
| Tmem38b | 9.402434749 | 8.918323696 | 9.297451326 | -0.37912763 | 0.104983423 |
| Cyp1a1 | 9.465529731 | 9.519572816 | 9.360589891 | 0.158982925 | 0.10493984 |
| Crnkl1 | 9.38254419 | 9.299159137 | 9.277630311 | 0.021528826 | 0.104913879 |
| Rpl37 | 11.94949986 | 11.86112339 | 11.84462448 | 0.016498917 | 0.104875387 |
| Zfp821 | 7.990831715 | 8.023482287 | 7.886089874 | 0.137392413 | 0.104741841 |
| Meox1 | 5.974317933 | 5.972794086 | 5.869617875 | 0.103176211 | 0.104700057 |
| Mtg2 | 7.938676799 | 7.943648224 | 7.834124159 | 0.109524065 | 0.10455264 |
| Rps27a-ps2 | 1.969490093 | 1.993514918 | 1.86497467 | 0.128540247 | 0.104515423 |
| Serpinb9 | 7.819425205 | 7.656929027 | 7.714947033 | -0.058018006 | 0.104478172 |
| Tnip3 | 6.781446731 | 6.724583946 | 6.676998963 | 0.047584983 | 0.104447767 |
| H2-K1 | 14.21983254 | 14.04741061 | 14.11540294 | -0.06799233 | 0.104429593 |
| Chaf1b | 8.0860723 | 8.080853406 | 7.981658713 | 0.099194693 | 0.104413586 |
| Tmed2 | 12.53340363 | 12.31696413 | 12.42899908 | -0.112034947 | 0.10440455 |
| Nkd2 | 8.388904711 | 8.394128904 | 8.284568966 | 0.109559937 | 0.104335744 |
| Fbxo4 | 7.739129029 | 7.571000687 | 7.634884079 | -0.063883392 | 0.10424495 |
| Pum2 | 12.16084306 | 11.99141863 | 12.056599 | -0.065180367 | 0.104244067 |
| Ccdc146 | 4.413389888 | 4.314044879 | 4.309197447 | 0.004847432 | 0.104192441 |
| N4bp2l2 | 10.2269241 | 10.10525559 | 10.12273596 | -0.017480363 | 0.104188147 |
| Bcas1 | 5.885959042 | 5.654435703 | 5.781811879 | -0.127376176 | 0.104147163 |
| Abhd17c | 10.09570974 | 9.993839164 | 9.991566506 | 0.002272658 | 0.104143238 |
| Elmod2 | 9.026261779 | 8.641982416 | 8.922335887 | -0.280353471 | 0.103925893 |
| Lpgat1 | 11.39517592 | 11.14036271 | 11.29129033 | -0.150927627 | 0.103885587 |
| Cox5b | 10.905664 | 10.80298724 | 10.80178175 | 0.001205493 | 0.10388225 |
| Colgalt1 | 10.89208243 | 10.89475303 | 10.78828438 | 0.106468647 | 0.10379805 |
| Atp5l | 11.52122938 | 11.36970663 | 11.4174702 | -0.047763577 | 0.10375918 |
| Rab39b | 2.442278741 | 2.314266973 | 2.338540467 | -0.024273494 | 0.103738274 |
| Krt86 | 2.238944827 | 2.125465977 | 2.135245411 | -0.009779433 | 0.103699416 |
| Nucb2 | 10.0846889 | 9.951429623 | 9.981007964 | -0.029578341 | 0.103680937 |
| Vps54 | 10.71877631 | 10.62400872 | 10.61518663 | 0.008822083 | 0.103589673 |
| Gm4824 | 2.071179706 | 1.982527954 | 1.967621228 | 0.014906726 | 0.103558478 |

| | | | | | |
|---|---|---|---|---|---|
| Btf3l4 | 10.1997496 | 9.997317217 | 10.09622367 | -0.09890645 | 0.103525933 |
| Pop5 | 8.776975665 | 8.656750135 | 8.673487768 | -0.016737632 | 0.103487898 |
| Dnmt1 | 10.98186579 | 11.09160519 | 10.87840466 | 0.213200527 | 0.103461127 |
| Hira | 9.84058108 | 9.80756519 | 9.737161324 | 0.070403867 | 0.103419756 |
| 9330161L09Rik | 3.206726747 | 3.130840456 | 3.10331722 | 0.027523237 | 0.103409527 |
| Srp9 | 10.1741595 | 10.08415198 | 10.07090454 | 0.013247438 | 0.10325496 |
| Gm48216 | 6.027823055 | 5.906879983 | 5.92466153 | -0.017781546 | 0.103161526 |
| Zkscan2 | 3.721153204 | 3.646348438 | 3.618029435 | 0.028319003 | 0.103123769 |
| Hk1 | 10.91967436 | 10.90380053 | 10.81655324 | 0.087247293 | 0.103121123 |
| Ints1 | 9.667034971 | 9.836012706 | 9.564057225 | 0.271955481 | 0.102977746 |
| Zdhhc7 | 9.554533744 | 9.568251222 | 9.451575594 | 0.116675628 | 0.10295815 |
| Ccdc180 | 3.202411376 | 3.194611484 | 3.099559861 | 0.095051623 | 0.102851515 |
| Larp6 | 6.586198201 | 6.375834312 | 6.483392075 | -0.107557763 | 0.102806126 |
| Zfp869 | 8.28447586 | 8.294561905 | 8.18173288 | 0.112829026 | 0.10274298 |
| Sympk | 10.67407796 | 10.86332235 | 10.57135663 | 0.291965713 | 0.10272133 |
| Aven | 8.587441416 | 8.51787074 | 8.484736503 | 0.033134236 | 0.102704912 |
| Cnot4 | 9.351065368 | 9.365793571 | 9.248391457 | 0.117402114 | 0.10267391 |
| Uox | 3.29043103 | 3.267054337 | 3.187760489 | 0.079293848 | 0.102670541 |
| Atg4b | 9.978742625 | 9.794933844 | 9.876089703 | -0.081155859 | 0.102652922 |
| Aurkaip1 | 9.669152251 | 9.573320735 | 9.566608515 | 0.00671222 | 0.102543736 |
| Ing5 | 8.921377471 | 8.724164486 | 8.818860483 | -0.094695997 | 0.102516988 |
| Ctu1 | 7.415775745 | 7.509737426 | 7.313267967 | 0.196469459 | 0.102507778 |
| Yipf5 | 10.01447648 | 9.946537561 | 9.911989209 | 0.034548352 | 0.102487271 |
| Zscan25 | 7.559841648 | 7.686218799 | 7.457459958 | 0.22875884 | 0.10238169 |
| Nsmce3 | 8.57525838 | 8.447856467 | 8.47289104 | -0.025034573 | 0.10236734 |
| Pds5a | 11.32729229 | 11.20330455 | 11.22493357 | -0.021629027 | 0.10235872 |
| Tmem241 | 6.591941744 | 6.453922471 | 6.489664705 | -0.035742234 | 0.102277039 |
| Tubb4a | 6.708130134 | 6.688864957 | 6.605858322 | 0.083006636 | 0.102271813 |
| Ascc2 | 9.888075432 | 9.961185869 | 9.785803625 | 0.175382244 | 0.102271807 |
| Gm20431 | 3.121374206 | 3.102204944 | 3.019105593 | 0.083099352 | 0.102268613 |
| BC100530 | 12.09729288 | 12.14336104 | 11.99502676 | 0.148334287 | 0.10226612 |
| Tbx21 | 3.126375287 | 2.99562898 | 3.024194379 | -0.028565399 | 0.102180908 |

| | | | | | |
|---|---|---|---|---|---|
| Ggt5 | 8.146205276 | 8.044881591 | 8.044063612 | 0.000817979 | 0.102141664 |
| Ints7 | 9.416544036 | 9.414555761 | 9.314419476 | 0.100136285 | 0.10212456 |
| Inafm2 | 9.14390714 | 9.033027782 | 9.041882906 | -0.008855124 | 0.102024233 |
| Cdcp1 | 10.33982257 | 10.32328073 | 10.23807612 | 0.085204613 | 0.101746453 |
| Gnai3 | 12.32429223 | 12.14433079 | 12.22262446 | -0.078293677 | 0.101667763 |
| Mrpl10 | 9.652309259 | 9.539020227 | 9.550679842 | -0.011659615 | 0.101629416 |
| Cd226 | 6.136014779 | 6.069555897 | 6.034403416 | 0.035152481 | 0.101611364 |
| Tmprss5 | 3.647844024 | 3.615099026 | 3.546245855 | 0.068853171 | 0.101598169 |
| Otulinl | 8.715009325 | 8.901949806 | 8.613432645 | 0.288517161 | 0.10157668 |
| Ube2k | 10.75042202 | 10.66555623 | 10.64884631 | 0.01670992 | 0.101575703 |
| Rab19 | 4.919870082 | 4.863657938 | 4.818334117 | 0.045323821 | 0.101535965 |
| Gm17230 | 1.313746941 | 1.229004245 | 1.212271012 | 0.016733233 | 0.10147593 |
| Pcdhgb1 | 4.682943814 | 4.643983362 | 4.581628995 | 0.062354368 | 0.10131482 |
| Tlr7 | 7.506502426 | 7.538492602 | 7.405267717 | 0.133224885 | 0.101234709 |
| Nln | 9.037349916 | 8.826007905 | 8.936238131 | -0.110230226 | 0.101111785 |
| Mrps6 | 8.685078321 | 8.963850714 | 8.584041068 | 0.379809646 | 0.101037253 |
| Mpp6 | 9.420463912 | 9.305950988 | 9.319512261 | -0.013561274 | 0.100951651 |
| Erg | 7.285264658 | 7.108716343 | 7.184503213 | -0.075786869 | 0.100761446 |
| Hmgxb3 | 9.442925228 | 9.535100176 | 9.342164162 | 0.192936014 | 0.100761066 |
| Frrs1 | 10.17811301 | 10.22165986 | 10.07744858 | 0.144211279 | 0.100664425 |
| Cavin3 | 10.24603828 | 10.26648919 | 10.14537765 | 0.121111547 | 0.10066063 |
| Hc | 2.111144349 | 2.072529648 | 2.010514984 | 0.062014664 | 0.100629365 |
| Psmd14 | 10.87142697 | 10.75149542 | 10.77086141 | -0.019365993 | 0.100565553 |
| Ogfrl1 | 9.332699575 | 9.103809291 | 9.232184855 | -0.128375565 | 0.10051472 |
| Phf10 | 10.26621708 | 10.18127089 | 10.16574215 | 0.01552874 | 0.100474933 |
| 4930503L19Rik | 7.195010132 | 7.18438534 | 7.094598729 | 0.089786611 | 0.100411404 |
| Pole2 | 7.609084907 | 7.698398614 | 7.508698309 | 0.189700306 | 0.100386599 |
| Rpl24 | 12.78356901 | 12.62475216 | 12.68318789 | -0.058435733 | 0.100381117 |
| 1810013L24Rik | 10.60637206 | 10.54086762 | 10.50602211 | 0.03484551 | 0.10034995 |
| Gm48119 | 6.892258834 | 6.870741272 | 6.791923533 | 0.078817738 | 0.1003353 |
| Mdh2 | 12.05331837 | 11.98915183 | 11.95302395 | 0.036127873 | 0.100294417 |
| Ankrd37 | 5.882993308 | 5.684117672 | 5.782790462 | -0.098672791 | 0.100202846 |

| | | | | | |
|---|---|---|---|---|---|
| Ttc30a1 | 6.513828666 | 6.441490102 | 6.413656544 | 0.027833558 | 0.100172122 |
| Rbm15b | 10.26839976 | 10.35200985 | 10.16824484 | 0.183765017 | 0.100154923 |
| Camsap2 | 10.75997929 | 10.77279936 | 10.65987134 | 0.11292802 | 0.10010795 |
| Tmem160 | 6.842489678 | 6.975959731 | 6.742450191 | 0.23350954 | 0.100039488 |
| Rmi2 | 6.321903536 | 6.494270863 | 6.221875167 | 0.272395696 | 0.100028369 |
| Rbm6 | 10.43980684 | 10.40601697 | 10.33983373 | 0.066183247 | 0.099973117 |
| AC126438.1 | 3.378056001 | 3.370362711 | 3.278085074 | 0.092277638 | 0.099970927 |
| Noxred1 | 3.740845303 | 3.791735531 | 3.640879364 | 0.150856167 | 0.09996594 |
| Sfrp1 | 10.47655886 | 10.3246012 | 10.37660922 | -0.052008023 | 0.099949643 |
| Plrg1 | 9.660622827 | 9.548449913 | 9.560701729 | -0.012251816 | 0.099921099 |
| Gimap7 | 1.83910108 | 1.76600348 | 1.73918424 | 0.02681924 | 0.09991684 |
| AC166110.3 | 3.766057305 | 3.978851487 | 3.66615391 | 0.312697577 | 0.099903396 |
| Ctrc | 2.689343752 | 2.588642649 | 2.589492501 | -0.000849852 | 0.099851251 |
| Gpcpd1 | 10.91917691 | 10.47985812 | 10.8193377 | -0.339479573 | 0.099839213 |
| Ccnh | 8.555184051 | 8.382008807 | 8.455354015 | -0.073345208 | 0.099830036 |
| Lmo2 | 7.907485679 | 7.795410237 | 7.80767183 | -0.012261593 | 0.09981385 |
| Coa6 | 6.603491656 | 6.583750192 | 6.503696016 | 0.080054176 | 0.09979564 |
| Fam129a | 9.620382608 | 9.476262852 | 9.520627209 | -0.044364357 | 0.099755399 |
| Bahcc1 | 8.679061149 | 8.82149471 | 8.579310156 | 0.242184554 | 0.099750993 |
| Olfr509 | 4.234480402 | 4.242893133 | 4.13473278 | 0.108160353 | 0.099747622 |
| Dmac1 | 8.36702292 | 8.371588781 | 8.267299534 | 0.104289247 | 0.099723386 |
| Zmat5 | 7.427514192 | 7.450350111 | 7.327807487 | 0.122542624 | 0.099706705 |
| Gpr174 | 3.997288141 | 3.877612465 | 3.897634648 | -0.020022183 | 0.099653493 |
| Phlda3 | 10.73562435 | 10.672434 | 10.63597478 | 0.036459223 | 0.099649573 |
| Vhl | 8.61712893 | 8.44548181 | 8.517547024 | -0.072065214 | 0.099581906 |
| Ighg1 | 4.389879848 | 4.415129724 | 4.290307202 | 0.124822522 | 0.099572646 |
| Zc3h8 | 6.936089829 | 6.954515049 | 6.83667158 | 0.117843469 | 0.099418249 |
| Vta1 | 9.443284862 | 9.271897341 | 9.34391208 | -0.072014739 | 0.099372782 |
| Zeb1 | 9.457945728 | 9.374751937 | 9.358637324 | 0.016114613 | 0.099308404 |
| Rapgef5 | 9.422358903 | 9.261722312 | 9.323052835 | -0.061330523 | 0.099306068 |
| H2-T22 | 11.55668272 | 11.29226808 | 11.45740095 | -0.165132867 | 0.09928177 |
| Tbc1d10a | 9.856111457 | 10.09426232 | 9.756831245 | 0.337431071 | 0.099280211 |

| | | | | | |
|---|---|---|---|---|---|
| Serpina3g | 8.655138302 | 8.817042219 | 8.555924986 | 0.261117233 | 0.099213316 |
| Dppa5a | 3.187711202 | 3.185401682 | 3.088509139 | 0.096892543 | 0.099202064 |
| Gm43498 | 4.183017304 | 4.051372695 | 4.083819705 | -0.03244701 | 0.099197599 |
| Lag3 | 5.492347173 | 5.503059843 | 5.393168759 | 0.109891084 | 0.099178414 |
| Capza2 | 12.08777225 | 11.85306188 | 11.98859502 | -0.13553314 | 0.099177227 |
| Grem2 | 8.972433366 | 8.729477675 | 8.873269517 | -0.143791842 | 0.099163849 |
| Tm9sf1 | 10.50637827 | 10.45380123 | 10.4072196 | 0.046581627 | 0.099158667 |
| Gm2000 | 11.32332798 | 11.2490913 | 11.22420958 | 0.024881717 | 0.0991184 |
| Gfpt1 | 9.900723088 | 9.880881496 | 9.801664429 | 0.079217066 | 0.099058659 |
| Fer1l4 | 2.879119402 | 2.737547462 | 2.780092548 | -0.042545087 | 0.099026853 |
| Afmid | 6.276905991 | 6.073673958 | 6.178019188 | -0.10434523 | 0.098886803 |
| Vcp | 13.37126702 | 13.42556022 | 13.27243439 | 0.153125827 | 0.098832633 |
| Exoc5 | 10.50107336 | 10.43648505 | 10.40226826 | 0.034216793 | 0.098805097 |
| Erap1 | 10.48542227 | 10.44232485 | 10.3867379 | 0.055586943 | 0.098684363 |
| Gm49383 | 5.916980983 | 5.784712342 | 5.818316532 | -0.03360419 | 0.098664451 |
| Lat | 6.309097112 | 6.154570443 | 6.210451198 | -0.055880755 | 0.098645914 |
| Nudt2 | 7.931990441 | 7.79228056 | 7.833470266 | -0.041189707 | 0.098520174 |
| Hpdl | 6.111146868 | 6.13812925 | 6.012702368 | 0.125426882 | 0.0984445 |
| Il2ra | 5.92565265 | 5.854182121 | 5.827253174 | 0.026928947 | 0.098399477 |
| Golga7 | 10.93887597 | 10.81768272 | 10.84049313 | -0.022810417 | 0.09838284 |
| Secisbp2 | 8.793437893 | 8.882511906 | 8.695218292 | 0.187293614 | 0.098219602 |
| Strip1 | 10.52318555 | 10.45795604 | 10.42498174 | 0.032974303 | 0.09820381 |
| Zfp697 | 5.445264647 | 5.470490845 | 5.347080789 | 0.123410055 | 0.098183858 |
| Cxcr6 | 5.938324719 | 5.672624029 | 5.840160917 | -0.167536888 | 0.098163801 |
| Zfp786 | 4.271074532 | 4.087085041 | 4.172996904 | -0.085911863 | 0.098077629 |
| Lypd6b | 9.248439938 | 8.883153605 | 9.150411676 | -0.267258071 | 0.098028262 |
| Olfr1385 | 6.687426096 | 6.657148253 | 6.589429283 | 0.06771897 | 0.097996813 |
| Anapc7 | 9.297719751 | 9.281372524 | 9.199740872 | 0.081631652 | 0.097978879 |
| Slc2a5 | 7.698642581 | 7.796002062 | 7.600665542 | 0.19533652 | 0.09797704 |
| Trdc | 7.782544888 | 7.500725868 | 7.684591295 | -0.183865427 | 0.097953593 |
| Ssu72 | 9.994522991 | 9.814990922 | 9.896728719 | -0.081737797 | 0.097794272 |
| Ptgfrn | 11.67874364 | 11.66695382 | 11.58096208 | 0.085991737 | 0.09778156 |

| | | | | | |
|---|---|---|---|---|---|
| Pabpn1 | 10.58473734 | 10.59531435 | 10.48704438 | 0.10826997 | 0.09769296 |
| Rpl15 | 12.41172921 | 12.37953251 | 12.31409231 | 0.065440197 | 0.097636897 |
| Prpf8 | 12.13727921 | 12.13966206 | 12.03968154 | 0.099980523 | 0.09759767 |
| Krt84 | 5.428588899 | 5.166105369 | 5.331094125 | -0.164988756 | 0.097494774 |
| Blcap | 9.735373709 | 9.629928159 | 9.638088017 | -0.008159858 | 0.097285691 |
| Cfap69 | 2.765478349 | 2.67526408 | 2.668219236 | 0.007044844 | 0.097259113 |
| Cdc25a | 8.560888026 | 8.46973425 | 8.463676846 | 0.006057404 | 0.09721118 |
| Gm9855 | 1.574602835 | 1.468673697 | 1.477441896 | -0.008768199 | 0.097160939 |
| Phactr2 | 9.927824513 | 9.724361635 | 9.830692158 | -0.106330522 | 0.097132355 |
| Gm48270 | 4.955117327 | 4.817004993 | 4.858014083 | -0.04100909 | 0.097103244 |
| Coil | 7.174618551 | 7.073081333 | 7.077547547 | -0.004466214 | 0.097071003 |
| Cops6 | 10.53898964 | 10.42189674 | 10.44197455 | -0.020077807 | 0.09701509 |
| Terb1 | 3.863959175 | 3.850538339 | 3.767049015 | 0.083489325 | 0.096910161 |
| Plekhb2 | 10.20405459 | 10.20866834 | 10.10716475 | 0.101503593 | 0.096889847 |
| Lcor | 7.347335961 | 7.208071964 | 7.250512185 | -0.042440221 | 0.096823776 |
| Manbal | 8.797706914 | 8.724024524 | 8.700902435 | 0.023122088 | 0.096804479 |
| Egr3 | 10.89494187 | 11.01146396 | 10.79814791 | 0.213316047 | 0.096793957 |
| Samd3 | 2.28724776 | 2.214651773 | 2.190513131 | 0.024138642 | 0.096734629 |
| Rpap3 | 8.407542678 | 8.2715215 | 8.310808551 | -0.039287051 | 0.096734127 |
| Cfap54 | 2.800348247 | 2.821623705 | 2.703813255 | 0.117810449 | 0.096534992 |
| Stat6 | 11.41341445 | 11.3989982 | 11.31706967 | 0.081928533 | 0.096344783 |
| Cpne1 | 9.930438458 | 9.888320378 | 9.834105364 | 0.054215014 | 0.096333094 |
| Mrpl32 | 8.80774193 | 8.577273708 | 8.711457953 | -0.134184244 | 0.096283977 |
| Gtf3c6 | 9.269044279 | 9.134397126 | 9.172804621 | -0.038407496 | 0.096239658 |
| Iqgap2 | 8.358868996 | 8.091775593 | 8.262635257 | -0.170859663 | 0.09623374 |
| Gm10020 | 2.780775897 | 2.76879021 | 2.684555342 | 0.084234868 | 0.096220555 |
| Vmn1r216 | 4.167158731 | 4.149597111 | 4.070967324 | 0.078629787 | 0.096191407 |
| Gnaz | 4.859019643 | 4.859944687 | 4.762843488 | 0.097101198 | 0.096176155 |
| Isca2 | 8.566332053 | 8.341769802 | 8.470283258 | -0.128513456 | 0.096048795 |
| Tnfsf13b | 5.070855591 | 4.983618461 | 4.974840724 | 0.008777737 | 0.096014867 |
| Pdcd6ip | 11.99906447 | 12.02168343 | 11.90305526 | 0.118628173 | 0.096009213 |
| Tnks1bp1 | 12.0026238 | 12.0908092 | 11.90672615 | 0.184083047 | 0.095897643 |

| | | | | | |
|---|---|---|---|---|---|
| Fam50a | 9.507672395 | 9.391947146 | 9.411777245 | -0.019830099 | 0.09589515 |
| Dbt | 9.619904878 | 9.359030893 | 9.524052869 | -0.165021975 | 0.095852009 |
| Mamdc4 | 4.93157406 | 5.077287588 | 4.835737866 | 0.241549722 | 0.095836194 |
| Ttll4 | 9.690892433 | 9.812029947 | 9.595117803 | 0.216912144 | 0.09577463 |
| Plcl2 | 8.337967315 | 8.144976644 | 8.242230754 | -0.09725411 | 0.095736561 |
| Mtrex | 10.26789217 | 10.16865668 | 10.172165 | -0.003508313 | 0.095727173 |
| H2-Q6 | 10.84972316 | 10.72484951 | 10.75407177 | -0.029222263 | 0.095651383 |
| Actr1b | 10.76091933 | 10.79732797 | 10.66527489 | 0.13205308 | 0.095644447 |
| Slc18a2 | 5.562719931 | 5.334263843 | 5.467107993 | -0.132844149 | 0.095611939 |
| Ltbp2 | 9.034441083 | 8.989265115 | 8.938837296 | 0.050427819 | 0.095603787 |
| Pola2 | 8.626082224 | 8.619919493 | 8.530489601 | 0.089429892 | 0.095592623 |
| Fam126a | 10.05383996 | 10.00816725 | 9.958254749 | 0.049912498 | 0.095585207 |
| Gm3052 | 4.593168227 | 4.586305527 | 4.49764133 | 0.088664197 | 0.095526897 |
| Mettl3 | 9.005833654 | 8.997634912 | 8.910333629 | 0.087301283 | 0.095500025 |
| Hook2 | 9.492168083 | 9.663340083 | 9.396682559 | 0.266657524 | 0.095485524 |
| Pak4 | 9.321169076 | 9.667910331 | 9.225729391 | 0.44218094 | 0.095439685 |
| Olfr393 | 2.578506224 | 2.589951448 | 2.4831624 | 0.106789048 | 0.095343824 |
| Elp6 | 7.146884219 | 7.106954005 | 7.051807738 | 0.055146268 | 0.095076481 |
| Afg3l1 | 10.10706508 | 9.996821343 | 10.01205298 | -0.01523164 | 0.095012093 |
| Sars2 | 7.445204281 | 7.729282934 | 7.350221984 | 0.37906095 | 0.094982297 |
| Sfr1 | 11.21550887 | 11.10607855 | 11.12061182 | -0.014533267 | 0.094897057 |
| Rdh13 | 9.414956544 | 9.334738998 | 9.32009594 | 0.014643058 | 0.094860604 |
| Fam174b | 7.507886452 | 7.415088556 | 7.413037771 | 0.002050785 | 0.094848681 |
| Zfp830 | 7.9929305 | 7.84198854 | 7.898122977 | -0.056134437 | 0.094807523 |
| Kin | 8.064526492 | 8.030403005 | 7.969809355 | 0.06059365 | 0.094717137 |
| Dr1 | 10.15713192 | 10.09307141 | 10.06246259 | 0.03060882 | 0.094669333 |
| Eral1 | 8.536535314 | 8.413972696 | 8.441866036 | -0.027893339 | 0.094669279 |
| Nktr | 11.36256362 | 11.50797624 | 11.26793557 | 0.24004067 | 0.094628047 |
| Ppp1r3e | 7.112419244 | 7.082295 | 7.017854808 | 0.064440192 | 0.094564436 |
| Nsg2 | 3.120178193 | 3.022167428 | 3.025641277 | -0.003473849 | 0.094536916 |
| Ep400 | 10.76515166 | 11.05943623 | 10.67064843 | 0.3887878 | 0.094503227 |
| Gm47331 | 4.255143299 | 4.200112586 | 4.160668787 | 0.039443799 | 0.094474512 |

| | | | | | |
|---|---|---|---|---|---|
| Tub | 4.184128901 | 4.174030508 | 4.089692443 | 0.084338065 | 0.094436458 |
| Ceacam16 | 2.250629216 | 2.234986592 | 2.156225954 | 0.078760639 | 0.094403262 |
| Cst6 | 12.97578473 | 13.03989506 | 12.88139673 | 0.158498333 | 0.094388 |
| Invs | 7.159720252 | 7.000390277 | 7.065346204 | -0.064955926 | 0.094374048 |
| Cchcr1 | 8.287181374 | 8.391346424 | 8.192815992 | 0.198530432 | 0.094365382 |
| Capn15 | 9.87564994 | 10.30146277 | 9.781291385 | 0.520171381 | 0.094358554 |
| Misp | 2.448753797 | 2.372509636 | 2.354530028 | 0.017979608 | 0.094223769 |
| Hist1h4i | 5.16248511 | 5.116429817 | 5.06828061 | 0.048149207 | 0.0942045 |
| Gsto1 | 10.43690865 | 10.39271167 | 10.34276676 | 0.049944903 | 0.094141883 |
| Ttf1 | 8.581392315 | 8.60714295 | 8.487252715 | 0.119890234 | 0.0941396 |
| Cc2d1a | 9.30160229 | 9.372295933 | 9.20746294 | 0.164832993 | 0.09413935 |
| Gm5831 | 1.439249926 | 1.332130572 | 1.345142231 | -0.013011659 | 0.094107695 |
| Lipc | 2.758384295 | 2.662942209 | 2.664314324 | -0.001372115 | 0.094069971 |
| Phgdh | 10.42865988 | 10.39538478 | 10.33460929 | 0.06077549 | 0.094050593 |
| Ergic2 | 10.02330897 | 10.0164757 | 9.929342268 | 0.087133429 | 0.093966703 |
| 1700001O22Rik | 4.946528686 | 4.995950088 | 4.852563487 | 0.143386601 | 0.093965199 |
| Hps1 | 8.110868926 | 8.192254503 | 8.016923942 | 0.175330561 | 0.093944983 |
| Scoc | 9.098063207 | 8.808119255 | 9.004153098 | -0.196033843 | 0.093910109 |
| LTO1 | 8.528956957 | 8.377230148 | 8.435052578 | -0.05782243 | 0.093904379 |
| Vmn1r85 | 4.783445312 | 4.723528961 | 4.689609504 | 0.033919457 | 0.093835808 |
| Tpsab1 | 4.631149089 | 4.28220917 | 4.537350873 | -0.255141702 | 0.093798216 |
| Rnasek | 9.781500704 | 9.915687003 | 9.687725152 | 0.227961851 | 0.093775551 |
| Mboat1 | 8.846194344 | 8.681445732 | 8.752460003 | -0.071014272 | 0.093734341 |
| Stra6 | 4.083291656 | 3.99808979 | 3.989623682 | 0.008466108 | 0.093667974 |
| Micall1 | 10.98102829 | 11.21671837 | 10.88742177 | 0.3292966 | 0.09360652 |
| Cux1 | 11.01197942 | 11.02402964 | 10.91864499 | 0.105384653 | 0.093334437 |
| Vmn2r108 | 3.7381517 | 3.640095763 | 3.644875496 | -0.004779733 | 0.093276203 |
| Gm7582 | 1.373294215 | 1.289164513 | 1.280078002 | 0.00908651 | 0.093216213 |
| Ndrg1 | 11.39347242 | 11.04048781 | 11.30030855 | -0.259820743 | 0.093163863 |
| H2-T10 | 7.826910114 | 7.69346326 | 7.733806899 | -0.040343639 | 0.093103215 |
| Gm47951 | 5.125585698 | 5.19318613 | 5.03255971 | 0.16062642 | 0.093025989 |
| Slitrk5 | 2.802830802 | 2.780742312 | 2.709844429 | 0.070897883 | 0.092986373 |

| | | | | | |
|---|---|---|---|---|---|
| Gm10175 | 11.38325047 | 11.37218162 | 11.29027711 | 0.081904513 | 0.092973363 |
| Ccdc57 | 5.828580884 | 5.975024434 | 5.735677374 | 0.23934706 | 0.092903511 |
| Babam1 | 9.830830293 | 9.791797557 | 9.738005954 | 0.053791603 | 0.092824339 |
| Gm21244 | 5.090120627 | 5.044760001 | 4.997338525 | 0.047421475 | 0.092782102 |
| Nlk | 8.237673436 | 8.104011423 | 8.144894066 | -0.040882643 | 0.092779371 |
| Cxcr5 | 1.742594895 | 1.752930686 | 1.649914707 | 0.103015978 | 0.092680187 |
| Bex6 | 3.645275938 | 3.600676556 | 3.552601235 | 0.048075321 | 0.092674702 |
| Rnf7 | 10.23337678 | 10.09110543 | 10.1407764 | -0.049670971 | 0.092600383 |
| Gpd2 | 10.13694422 | 9.931361252 | 10.0443612 | -0.112999944 | 0.092583025 |
| Plp2 | 11.80473049 | 11.70801932 | 11.71222291 | -0.004203583 | 0.092507583 |
| Matn4 | 3.692181774 | 3.685206982 | 3.59967835 | 0.085528632 | 0.092503424 |
| Lats1 | 9.670813259 | 9.587600641 | 9.578338286 | 0.009262355 | 0.092474973 |
| Alyref2 | 5.496471418 | 5.40206035 | 5.404002295 | -0.001941945 | 0.092469123 |
| Got2 | 11.69149658 | 11.7138323 | 11.59904433 | 0.11478797 | 0.092452247 |
| Olfr907 | 2.563316033 | 2.487492542 | 2.47088115 | 0.016611392 | 0.092434883 |
| 4930415O20Rik | 2.648373473 | 2.588770026 | 2.555949572 | 0.032820454 | 0.092423901 |
| Metrnl | 10.36484189 | 10.37623987 | 10.27241806 | 0.103821813 | 0.09242383 |
| 0610010K14Rik | 8.637702506 | 8.671411654 | 8.545291559 | 0.126120096 | 0.092410948 |
| Gm4366 | 1.792643838 | 1.808764756 | 1.700278139 | 0.108486617 | 0.092365699 |
| Epn1 | 11.10722337 | 11.28386926 | 11.01493754 | 0.268931717 | 0.092285827 |
| Mterf3 | 9.072397784 | 8.941850762 | 8.980134584 | -0.038283822 | 0.0922632 |
| Mri1 | 8.463066762 | 8.631713979 | 8.370864531 | 0.260849447 | 0.09220223 |
| Itgb3 | 8.074725125 | 7.955163307 | 7.982552153 | -0.027388847 | 0.092172972 |
| Slc35b3 | 8.281106023 | 8.266933402 | 8.189033209 | 0.077900193 | 0.092072814 |
| Fam32a | 10.42095703 | 10.28434841 | 10.3289128 | -0.04456439 | 0.092044227 |
| Adh5 | 10.05687517 | 9.935817072 | 9.964870717 | -0.029053645 | 0.092004452 |
| Ly86 | 8.413770668 | 8.104359897 | 8.321854962 | -0.217495065 | 0.091915707 |
| Ccdc22 | 8.737126033 | 8.857896445 | 8.645219755 | 0.21267669 | 0.091906278 |
| Kcnh7 | 2.359000791 | 2.272787967 | 2.267105912 | 0.005682055 | 0.09189488 |
| Gm6863 | 2.279858954 | 2.254986575 | 2.18799219 | 0.066994386 | 0.091866764 |
| Mea1 | 9.512476827 | 9.45091474 | 9.42068102 | 0.03023372 | 0.091795807 |
| Pnpla5 | 10.61910612 | 10.60115557 | 10.52736876 | 0.073786817 | 0.09173736 |

| | | | | | |
|---|---|---|---|---|---|
| Rfxap | 8.068273901 | 8.016527668 | 7.97654686 | 0.039980808 | 0.091727041 |
| Gm8399 | 2.717368437 | 2.600442229 | 2.625658271 | -0.025216042 | 0.091710166 |
| Ndufs4 | 10.09904779 | 9.992482607 | 10.00736153 | -0.014878921 | 0.091686257 |
| Tmem238 | 7.060989423 | 6.962507469 | 6.969326487 | -0.006819018 | 0.091662936 |
| B4gat1 | 8.868786668 | 8.61866556 | 8.777135577 | -0.158470016 | 0.091651091 |
| Cox8a | 10.83194481 | 10.72670735 | 10.74029873 | -0.01359138 | 0.091646077 |
| Tmx1 | 10.14010035 | 10.07848449 | 10.0484655 | 0.030018986 | 0.091634845 |
| Oit3 | 1.971061266 | 1.946137412 | 1.879465277 | 0.066672134 | 0.091595989 |
| Osr1 | 6.390503029 | 6.183475792 | 6.298925731 | -0.115449939 | 0.091577298 |
| Chid1 | 8.987463872 | 9.043068907 | 8.896017121 | 0.147051786 | 0.091446751 |
| Akap8 | 10.8788941 | 10.92611586 | 10.78745184 | 0.13866402 | 0.09144226 |
| Dock3 | 5.065736758 | 4.885429402 | 4.974319884 | -0.088890483 | 0.091416874 |
| Tmem181b-ps | 7.383192386 | 7.317410312 | 7.291789887 | 0.025620425 | 0.091402499 |
| Gm29094 | 2.143559519 | 2.094197852 | 2.05217412 | 0.042023732 | 0.091385398 |
| Gm6741 | 3.405887827 | 3.313275313 | 3.314523214 | -0.001247901 | 0.091364613 |
| Cgref1 | 7.239192518 | 7.495771202 | 7.147906367 | 0.347864835 | 0.091286151 |
| Glp2r | 2.085793264 | 1.980277423 | 1.994517622 | -0.014240199 | 0.091275642 |
| Gm18537 | 6.993468958 | 7.109556621 | 6.902246011 | 0.207310609 | 0.091222947 |
| Pstpip2 | 7.717116471 | 7.589954542 | 7.625909854 | -0.035955312 | 0.091206617 |
| Elf1 | 10.47683377 | 10.45612953 | 10.38578909 | 0.07034044 | 0.09104468 |
| Ppp2r2d | 9.207309762 | 9.026079618 | 9.116286489 | -0.090206871 | 0.091023273 |
| Gm44245 | 2.805219886 | 2.747210311 | 2.714201791 | 0.03300852 | 0.091018095 |
| Hmbs | 9.388426062 | 9.404776259 | 9.297436467 | 0.107339792 | 0.090989595 |
| Poc5 | 8.267517338 | 7.975742606 | 8.176531636 | -0.20078903 | 0.090985702 |
| Mfsd13a | 6.371438726 | 6.248421161 | 6.280517921 | -0.03209676 | 0.090920805 |
| Kif5a | 5.917054747 | 6.038105011 | 5.826141001 | 0.21196401 | 0.090913746 |
| Trit1 | 8.15057313 | 8.104772459 | 8.059672129 | 0.045100331 | 0.090901002 |
| Gm8091 | 2.02513127 | 2.009524694 | 1.934263022 | 0.075261672 | 0.090868248 |
| Rpl31 | 12.23693998 | 12.10720762 | 12.14610112 | -0.038893507 | 0.090838853 |
| Il20ra | 8.839877198 | 8.661543694 | 8.74904778 | -0.087504086 | 0.090829418 |
| Zfp207 | 11.74988575 | 11.83617109 | 11.65909238 | 0.17707871 | 0.090793377 |
| Eif3k | 11.26160492 | 11.20370227 | 11.17082677 | 0.0328755 | 0.090778147 |

| | | | | | |
|---|---|---|---|---|---|
| 5430419D17Rik | 5.528736645 | 5.395811192 | 5.438005642 | -0.04219445 | 0.090731003 |
| Rasgef1c | 4.993259966 | 4.981742679 | 4.902543457 | 0.079199222 | 0.090716508 |
| Atp6v1b2 | 11.67164342 | 11.75487721 | 11.58098916 | 0.173888043 | 0.090654257 |
| Grm1 | 3.01395527 | 2.954938389 | 2.923366804 | 0.031571586 | 0.090588466 |
| Mapk4 | 3.364423061 | 3.419090415 | 3.273840704 | 0.145249711 | 0.090582357 |
| Sergef | 7.455688641 | 7.335329988 | 7.365194736 | -0.029864748 | 0.090493905 |
| Schip1 | 8.993184609 | 8.46888842 | 8.90270414 | -0.433815719 | 0.090480469 |
| Gak | 11.08843937 | 11.00922559 | 10.99805353 | 0.01117206 | 0.09038584 |
| Sdhaf1 | 7.83080403 | 7.65733165 | 7.74044533 | -0.08311368 | 0.0903587 |
| Timm17b | 8.458255442 | 8.383794362 | 8.3679425 | 0.015851861 | 0.090312942 |
| Cox16 | 8.337811474 | 8.183718393 | 8.247519334 | -0.063800941 | 0.09029214 |
| Mrpl39 | 8.782434279 | 8.655553849 | 8.692328243 | -0.036774394 | 0.090106037 |
| Rasa2 | 9.204221999 | 8.95472084 | 9.114183017 | -0.159462176 | 0.090038983 |
| Prodh2 | 2.327215617 | 2.340378602 | 2.237237853 | 0.103140749 | 0.089977765 |
| Timmdc1 | 8.562605089 | 8.456972361 | 8.472633691 | -0.01566133 | 0.089971398 |
| Trim39 | 9.098012519 | 9.173718206 | 9.00805126 | 0.165666946 | 0.089961259 |
| Gm43960 | 4.576656551 | 4.530669959 | 4.486847481 | 0.043822478 | 0.08980907 |
| Ewsr1 | 12.055158 | 12.05740061 | 11.96548004 | 0.091920577 | 0.08967796 |
| Sod3 | 10.96357887 | 10.93947523 | 10.87396628 | 0.065508943 | 0.08961259 |
| Wdfy1 | 8.652652624 | 8.610208702 | 8.563094405 | 0.047114297 | 0.089558219 |
| Pcyox1 | 12.18661835 | 12.08174479 | 12.09706353 | -0.01531874 | 0.089554817 |
| Zmym1 | 7.742841274 | 7.868373646 | 7.653349778 | 0.215023867 | 0.089491496 |
| Fam204a | 8.550807716 | 8.150455583 | 8.461324249 | -0.310868666 | 0.089483467 |
| Cyc1 | 11.22866329 | 11.21795233 | 11.13918269 | 0.078769633 | 0.0894806 |
| Mrpl53 | 8.252950327 | 8.358020827 | 8.163564323 | 0.194456504 | 0.089386004 |
| Rps16 | 12.38089352 | 12.25416277 | 12.29152984 | -0.037367073 | 0.089363677 |
| Nptxr | 7.555292067 | 7.442441654 | 7.4659792 | -0.023537545 | 0.089312868 |
| Mbd1 | 9.801568954 | 9.969288341 | 9.71233651 | 0.256951831 | 0.089232444 |
| Zfp407 | 8.439605628 | 8.49148515 | 8.350471466 | 0.141013684 | 0.089134162 |
| Copb2 | 11.69393998 | 11.61926069 | 11.60484612 | 0.014414563 | 0.089093853 |
| Il12rb1 | 3.171022951 | 3.198581287 | 3.081967386 | 0.116613901 | 0.089055566 |
| Olfr318 | 3.511934142 | 3.384652077 | 3.422882215 | -0.038230138 | 0.089051927 |

| | | | | | |
|---|---|---|---|---|---|
| Gm10447 | 3.009613891 | 2.989136891 | 2.920606468 | 0.068530423 | 0.089007423 |
| Fam20c | 7.102518935 | 7.273207419 | 7.013542707 | 0.259664711 | 0.088976228 |
| Hsf4 | 5.035834049 | 4.980762834 | 4.94686918 | 0.033893654 | 0.088964869 |
| Gm13423 | 2.863051595 | 2.837147004 | 2.774214096 | 0.062932907 | 0.088837499 |
| Med16 | 8.48674505 | 8.678090763 | 8.397911389 | 0.280179374 | 0.088833661 |
| Slc30a6 | 8.0234208 | 7.965479127 | 7.934644404 | 0.030834723 | 0.088776397 |
| Exoc8 | 8.745645477 | 8.729236314 | 8.65689951 | 0.072336804 | 0.088745967 |
| Rell1 | 10.31888556 | 10.17390005 | 10.23019818 | -0.056298133 | 0.08868738 |
| 2610021A01Rik | 7.840000473 | 7.836934715 | 7.751319889 | 0.085614826 | 0.088680584 |
| Rbbp7 | 11.28882835 | 11.20067144 | 11.20017069 | 0.000500743 | 0.088657653 |
| Cdk18 | 5.575065039 | 5.595457162 | 5.486622913 | 0.108834249 | 0.088442126 |
| Gstp3 | 5.59646164 | 5.579561248 | 5.508039879 | 0.071521369 | 0.088421761 |
| Hoxb4 | 7.053219502 | 7.067890733 | 6.964836418 | 0.103054315 | 0.088383084 |
| Srp54a | 10.90874439 | 10.86755399 | 10.82040348 | 0.047150503 | 0.088340907 |
| Sirt6 | 7.274496186 | 7.523412263 | 7.186156611 | 0.337255652 | 0.088339575 |
| Epha5 | 4.677949871 | 4.529259243 | 4.589640759 | -0.060381516 | 0.088309112 |
| Mapk13 | 10.85008301 | 10.93544874 | 10.76179296 | 0.17365578 | 0.088290047 |
| Ankrd52 | 10.44787409 | 10.61874806 | 10.35966034 | 0.259087723 | 0.08821375 |
| Hras | 11.22670627 | 11.26833747 | 11.13850021 | 0.12983726 | 0.088206057 |
| Ppp1r37 | 10.13976594 | 10.36285059 | 10.05157489 | 0.311275703 | 0.08819105 |
| Casc3 | 10.35061619 | 10.38467354 | 10.26244696 | 0.122226573 | 0.088169223 |
| Wdr95 | 1.657714836 | 1.577203641 | 1.569569087 | 0.007634554 | 0.088145749 |
| Wnt5a | 8.926265648 | 8.838403502 | 8.83829043 | 0.000113072 | 0.087975218 |
| Tac4 | 6.018559394 | 5.951933166 | 5.930587312 | 0.021345854 | 0.087972082 |
| Zfyve26 | 9.361880801 | 9.290727212 | 9.273919735 | 0.016807477 | 0.087961067 |
| Rpl28-ps1 | 12.32598133 | 12.20695376 | 12.23803897 | -0.031085207 | 0.087942367 |
| Fam169a | 2.08369285 | 2.06130312 | 1.995847147 | 0.065455973 | 0.087845703 |
| Zbtb25 | 6.958911843 | 6.763979013 | 6.871104321 | -0.107125307 | 0.087807523 |
| N6amt1 | 8.276370727 | 8.121079928 | 8.188620483 | -0.067540555 | 0.087750245 |
| Rpe | 9.800645861 | 9.617052296 | 9.712905559 | -0.095853262 | 0.087740302 |
| Ddx42 | 10.59067962 | 10.42335249 | 10.5029876 | -0.079635117 | 0.087692013 |
| Sec16a | 11.27635572 | 11.19591328 | 11.18869 | 0.007223277 | 0.087665717 |

| | | | | | |
|---|---|---|---|---|---|
| Znrf2 | 11.33296844 | 11.18489998 | 11.24532039 | -0.06042041 | 0.087648053 |
| Swt1 | 8.468663984 | 8.811608641 | 8.381024915 | 0.430583726 | 0.087639069 |
| Parp8 | 8.566858581 | 8.411705223 | 8.479303616 | -0.067598392 | 0.087554966 |
| E130114P18Rik | 2.156085563 | 2.102298972 | 2.068556247 | 0.033742725 | 0.087529316 |
| Acaa1a | 10.68160635 | 10.74629823 | 10.59408681 | 0.152211427 | 0.087519543 |
| Grm6 | 2.359556693 | 2.342508174 | 2.27204172 | 0.070466454 | 0.087514973 |
| Mindy1 | 10.72606242 | 10.60285802 | 10.63860445 | -0.03574643 | 0.087457967 |
| Ctsc | 11.56162802 | 11.21055639 | 11.47417977 | -0.263623383 | 0.087448247 |
| Nod1 | 10.27565835 | 10.29259311 | 10.18822256 | 0.104370543 | 0.087435783 |
| R3hdm4 | 10.70467589 | 10.59946737 | 10.61726788 | -0.01780051 | 0.08740801 |
| Zbtb2 | 8.063530405 | 8.21660477 | 7.976126203 | 0.240478567 | 0.087404202 |
| Tmem87a | 10.15521536 | 9.974077696 | 10.06785266 | -0.093774964 | 0.087362703 |
| Casp7 | 8.192951802 | 8.030354535 | 8.105654182 | -0.075299647 | 0.087297619 |
| Kdm8 | 6.796329076 | 6.755330978 | 6.709038292 | 0.046292686 | 0.087290785 |
| Ppp1r14c | 10.64058506 | 10.49448236 | 10.55331165 | -0.058829287 | 0.087273417 |
| Polr3f | 8.778427638 | 8.616264357 | 8.691163093 | -0.074898735 | 0.087264546 |
| Scfd2 | 8.41498573 | 8.371301715 | 8.327747011 | 0.043554704 | 0.087238719 |
| Insl3 | 2.694746697 | 2.687229961 | 2.607559477 | 0.079670484 | 0.087187219 |
| Tmigd3 | 2.092059822 | 2.023030301 | 2.004939115 | 0.018091186 | 0.087120707 |
| Arpp19 | 10.74369077 | 10.58456397 | 10.65661801 | -0.072054037 | 0.087072757 |
| Zfp930 | 7.775916087 | 7.729940327 | 7.688951999 | 0.040988328 | 0.086964088 |
| Gosr1 | 9.903832202 | 9.820548434 | 9.816869522 | 0.003678912 | 0.08696268 |
| Dctn6 | 9.458391717 | 9.384485603 | 9.371444099 | 0.013041504 | 0.086947619 |
| Ddx6 | 12.26753687 | 12.20501742 | 12.18061472 | 0.0244027 | 0.086922153 |
| Vmn2r112 | 5.303182566 | 5.312591962 | 5.216305107 | 0.096286855 | 0.086877459 |
| Tcf25 | 12.1849043 | 12.12422319 | 12.09803297 | 0.026190213 | 0.086871323 |
| Dcbld1 | 7.85908838 | 7.899078146 | 7.772239274 | 0.126838872 | 0.086849107 |
| Hgs | 9.953907263 | 10.20147301 | 9.867113048 | 0.334359965 | 0.086794215 |
| Rai1 | 8.562676063 | 8.786164424 | 8.475980625 | 0.310183799 | 0.086695438 |
| Rnf41 | 9.165797849 | 9.333050455 | 9.079115393 | 0.253935062 | 0.086682456 |
| Ppp5c | 9.65232994 | 9.857618753 | 9.565681685 | 0.291937068 | 0.086648254 |
| Mcm3ap | 9.420251707 | 9.339649054 | 9.333648056 | 0.006000998 | 0.086603652 |

| | | | | | |
|---|---|---|---|---|---|
| Timm29 | 9.970126318 | 9.893714956 | 9.883641505 | 0.010073451 | 0.086484813 |
| Gm47968 | 4.432103114 | 4.390910513 | 4.345648263 | 0.04526225 | 0.086454852 |
| Naca | 13.23884068 | 13.12876662 | 13.15246155 | -0.02369493 | 0.08637913 |
| Ola1 | 10.46130817 | 10.40302606 | 10.3749324 | 0.028093663 | 0.086375767 |
| Usp10 | 10.37632591 | 10.16601868 | 10.28995125 | -0.12393257 | 0.086374663 |
| Smap2 | 10.35157319 | 10.28532504 | 10.26524042 | 0.020084617 | 0.08633277 |
| S100a16 | 12.39162734 | 12.40271132 | 12.30546128 | 0.097250043 | 0.086166063 |
| Senp6 | 10.57594907 | 10.55790167 | 10.48980448 | 0.068097197 | 0.086144597 |
| Dtd2 | 7.201729954 | 7.009835347 | 7.115675545 | -0.105840199 | 0.086054408 |
| Prpf6 | 10.3073939 | 10.26556704 | 10.22135312 | 0.044213923 | 0.086040777 |
| Ppil4 | 9.619618486 | 9.498698428 | 9.533582333 | -0.034883905 | 0.086036153 |
| Esco1 | 9.472404328 | 9.399912875 | 9.386433591 | 0.013479284 | 0.085970737 |
| Bcs1l | 8.177902329 | 8.02500822 | 8.091953562 | -0.066945342 | 0.085948767 |
| Smim12 | 8.098178663 | 8.024640353 | 8.01223533 | 0.012405023 | 0.085943332 |
| Adat3 | 3.682687914 | 3.893240943 | 3.596789657 | 0.296451286 | 0.085898257 |
| Med20 | 9.447338969 | 9.548671527 | 9.36145238 | 0.187219147 | 0.085886589 |
| Otud4 | 10.89707014 | 10.69748871 | 10.81124019 | -0.113751483 | 0.085829943 |
| Myo1d | 10.07013576 | 9.953394303 | 9.984320598 | -0.030926295 | 0.085815167 |
| 9530077C05Rik | 4.111305793 | 4.109682049 | 4.025628453 | 0.084053597 | 0.085677341 |
| Podnl1 | 2.225765307 | 2.176184398 | 2.140092982 | 0.036091417 | 0.085672325 |
| Cemip | 5.601634914 | 5.537135436 | 5.515969992 | 0.021165444 | 0.085664922 |
| Zfp397 | 9.441480123 | 9.475651287 | 9.355826821 | 0.119824467 | 0.085653303 |
| Rnf26 | 8.97474684 | 8.938879319 | 8.889095447 | 0.049783872 | 0.085651393 |
| Patj | 8.703675719 | 8.706224256 | 8.618067047 | 0.088157209 | 0.085608672 |
| Trim9 | 2.869181146 | 2.709052999 | 2.783591831 | -0.074538833 | 0.085589315 |
| Gm5421 | 2.550801481 | 2.467896045 | 2.465257007 | 0.002639038 | 0.085544474 |
| Rasgrp2 | 6.718886287 | 6.639023905 | 6.633398662 | 0.005625243 | 0.085487625 |
| Nans | 9.310706218 | 9.259538615 | 9.225347771 | 0.034190844 | 0.085358448 |
| Mark3 | 10.48678677 | 10.55606351 | 10.40144094 | 0.15462257 | 0.085345827 |
| Pcdhb7 | 4.009693297 | 4.009557524 | 3.924389631 | 0.085167893 | 0.085303666 |
| Hhat | 6.183530686 | 6.160189235 | 6.098228754 | 0.061960481 | 0.085301932 |
| Sec14l2 | 9.594620629 | 9.631611396 | 9.509334919 | 0.122276477 | 0.08528571 |

| | | | | | |
|---|---|---|---|---|---|
| Paf1 | 10.2535085 | 10.0473413 | 10.16833227 | -0.120990965 | 0.08517623 |
| Timm44 | 9.852355016 | 9.742740265 | 9.767272276 | -0.024532011 | 0.085082741 |
| Nudt18 | 8.689939256 | 8.491492888 | 8.604975203 | -0.113482314 | 0.084964053 |
| Gm16385 | 2.620088795 | 2.602676501 | 2.535184056 | 0.067492445 | 0.084904739 |
| Nos2 | 3.68384046 | 3.710892531 | 3.598946744 | 0.111945788 | 0.084893717 |
| Mnt | 9.214818684 | 9.240973229 | 9.129944967 | 0.111028262 | 0.084873717 |
| Asb7 | 8.268341951 | 7.99624924 | 8.183512947 | -0.187263707 | 0.084829004 |
| Zfp236 | 9.301897428 | 9.492455133 | 9.217257306 | 0.275197827 | 0.084640122 |
| Tpm3-rs7 | 2.787652816 | 2.797765688 | 2.703021543 | 0.094744145 | 0.084631274 |
| Dmrta2 | 1.729207019 | 1.706956587 | 1.644611305 | 0.062345282 | 0.084595714 |
| Tpcn2 | 8.152201022 | 8.303660457 | 8.067609456 | 0.236051001 | 0.084591566 |
| Nkiras2 | 9.649890884 | 9.724316371 | 9.565329576 | 0.158986795 | 0.084561308 |
| Sult4a1 | 2.864896033 | 2.808179965 | 2.780368788 | 0.027811177 | 0.084527245 |
| Wdr45b | 9.926617026 | 9.909696422 | 9.842154281 | 0.067542141 | 0.084462745 |
| Ikzf5 | 9.492524678 | 9.395363847 | 9.408066952 | -0.012703105 | 0.084457725 |
| Rpl18a | 13.12920378 | 13.21402298 | 13.0447533 | 0.169269677 | 0.084450477 |
| Pnldc1 | 4.614984216 | 4.404082494 | 4.530641244 | -0.126558751 | 0.084342972 |
| Mroh1 | 9.198258713 | 9.274739056 | 9.114017417 | 0.160721639 | 0.084241296 |
| Rps10 | 12.65091334 | 12.5867879 | 12.56670181 | 0.02008609 | 0.08421153 |
| Lhfp | 9.570998156 | 9.328829549 | 9.486809942 | -0.157980393 | 0.084188215 |
| Lman1 | 10.80187163 | 10.82944034 | 10.71779765 | 0.111642687 | 0.084073973 |
| Tmem101 | 7.979054776 | 7.863763317 | 7.895010173 | -0.031246856 | 0.084044602 |
| Atp13a4 | 7.124723696 | 7.100304791 | 7.040686016 | 0.059618774 | 0.08403768 |
| Grik3 | 5.339356342 | 5.445585176 | 5.255334391 | 0.190250785 | 0.084021952 |
| Rps11 | 13.28415075 | 13.15169355 | 13.20013803 | -0.048444483 | 0.08401272 |
| Emc4 | 9.813636375 | 9.751228222 | 9.729648258 | 0.021579964 | 0.083988116 |
| Pqlc2 | 8.075250417 | 8.278129701 | 7.991309296 | 0.286820406 | 0.083941121 |
| Cd151 | 11.2368481 | 11.15074605 | 11.15290985 | -0.0021638 | 0.08393825 |
| Gm47103 | 4.037779472 | 3.835735968 | 3.953891594 | -0.118155627 | 0.083887878 |
| Mad2l2 | 7.795157196 | 7.682306932 | 7.711286528 | -0.028979595 | 0.083870669 |
| Olfr77 | 4.308571323 | 4.221793301 | 4.224736935 | -0.002943634 | 0.083834388 |
| Irf3 | 9.668592418 | 9.76348224 | 9.584842224 | 0.178640016 | 0.083750194 |

| | | | | | |
|---|---|---|---|---|---|
| Aarsd1 | 8.76836016 | 8.746976388 | 8.684642354 | 0.062334033 | 0.083717806 |
| Ints14 | 9.165780932 | 9.051070613 | 9.082125889 | -0.031055276 | 0.083655043 |
| Ppp2r2a | 10.80396358 | 10.76537907 | 10.72032217 | 0.045056893 | 0.083641407 |
| Psmc6 | 11.16194884 | 11.16728503 | 11.07836032 | 0.088924707 | 0.083588513 |
| Sgta | 10.64820769 | 10.79668898 | 10.56463609 | 0.232052883 | 0.0835716 |
| Chmp1a | 11.33813806 | 11.30212735 | 11.25457611 | 0.04755124 | 0.083561947 |
| Exosc7 | 9.443187278 | 9.36452861 | 9.359643962 | 0.004884648 | 0.083543316 |
| Ext2 | 10.02299729 | 9.919234503 | 9.939621047 | -0.020386543 | 0.083376242 |
| Cyfip2 | 8.963450977 | 8.814587135 | 8.88008785 | -0.065500715 | 0.083363127 |
| Loxl1 | 9.687986193 | 9.955956596 | 9.604625565 | 0.351331031 | 0.083360628 |
| Gm16238 | 2.570056507 | 2.537839327 | 2.486741433 | 0.051097893 | 0.083315073 |
| 1190007I07Rik | 6.250714735 | 6.189318411 | 6.167458095 | 0.021860316 | 0.08325664 |
| Vps11 | 9.883195717 | 9.88474399 | 9.800000248 | 0.084743742 | 0.083195468 |
| Gm8991 | 1.522562075 | 1.427470177 | 1.439462789 | -0.011992612 | 0.083099286 |
| Tyw1 | 8.132005646 | 8.082578872 | 8.048935352 | 0.03364352 | 0.083070294 |
| Jtb | 9.481258815 | 9.479848196 | 9.398223096 | 0.0816251 | 0.083035719 |
| Gm8738 | 3.584291024 | 3.576371879 | 3.501377081 | 0.074994798 | 0.082913942 |
| Rps14 | 13.12265851 | 13.01414824 | 13.03975329 | -0.025605047 | 0.08290522 |
| Ipo11 | 9.209652743 | 9.169432505 | 9.126770436 | 0.042662069 | 0.082882307 |
| Magt1 | 10.60235297 | 10.59310191 | 10.51953271 | 0.073569203 | 0.082820263 |
| Gimap4 | 6.702355777 | 6.407330045 | 6.619540347 | -0.212210302 | 0.08281543 |
| Igflr1 | 4.291359444 | 4.532446125 | 4.208550502 | 0.323895624 | 0.082808942 |
| Soga3 | 1.681009507 | 1.615312659 | 1.598291857 | 0.017020802 | 0.08271765 |
| Mthfr | 9.446891856 | 9.612395353 | 9.364208285 | 0.248187069 | 0.082683572 |
| Polr2g | 9.476615173 | 9.308066934 | 9.393969877 | -0.085902943 | 0.082645296 |
| Nkap | 8.942446975 | 8.789145719 | 8.859804628 | -0.07065891 | 0.082642347 |
| Bbs12 | 6.248497138 | 6.183528443 | 6.165943182 | 0.017585261 | 0.082553956 |
| Usp8 | 10.43260668 | 10.33777232 | 10.35007694 | -0.012304617 | 0.08252974 |
| Btnl10 | 3.225379344 | 3.203226702 | 3.142883023 | 0.060343679 | 0.082496321 |
| Snx2 | 11.46923847 | 11.25780504 | 11.38677612 | -0.128971087 | 0.08246235 |
| Gngt2 | 7.56539151 | 7.734119665 | 7.482945009 | 0.251174657 | 0.082446501 |
| Gpr89 | 9.069296989 | 9.089271002 | 8.986859997 | 0.102411005 | 0.082436992 |

| | | | | | |
|---|---|---|---|---|---|
| Aurka | 7.568419377 | 7.417211401 | 7.485983813 | -0.068772412 | 0.082435564 |
| Rtf2 | 10.88415343 | 10.72476024 | 10.80175472 | -0.07699448 | 0.082398703 |
| Rbm26 | 10.08135195 | 9.955530579 | 9.99901617 | -0.043485591 | 0.082335777 |
| Tmem120a | 9.529659569 | 9.34169647 | 9.447324941 | -0.105628471 | 0.082334628 |
| Vmn1r181 | 2.020016974 | 1.975430908 | 1.937718581 | 0.037712327 | 0.082298393 |
| Spatc1 | 1.934272261 | 1.835865011 | 1.852018733 | -0.016153722 | 0.082253528 |
| Hhex | 5.607197522 | 5.357552047 | 5.524966883 | -0.167414836 | 0.082230639 |
| Ddx19a | 9.857760778 | 9.871712037 | 9.775608405 | 0.096103631 | 0.082152372 |
| Fez1 | 7.13212134 | 6.962810251 | 7.050010766 | -0.087200515 | 0.082110574 |
| Pglyrp3 | 7.266229824 | 7.359827628 | 7.184144893 | 0.175682735 | 0.082084931 |
| Proser2 | 6.460082924 | 6.441040026 | 6.378029222 | 0.063010804 | 0.082053702 |
| Mboat4 | 2.378079091 | 2.314151078 | 2.296087147 | 0.018063931 | 0.081991944 |
| Flrt3 | 9.699993327 | 9.790539318 | 9.618006106 | 0.172533212 | 0.081987221 |
| Rps19 | 12.99691159 | 12.90210832 | 12.91501756 | -0.01290924 | 0.08189403 |
| Twf1 | 11.06395854 | 10.96708103 | 10.98208422 | -0.015003193 | 0.08187432 |
| Zfyve16 | 9.029726581 | 8.930889126 | 8.947928159 | -0.017039033 | 0.081798422 |
| Vmn1r193 | 3.500474191 | 3.474088512 | 3.418677432 | 0.05541108 | 0.081796759 |
| Clba1 | 5.363051726 | 5.360077023 | 5.281297251 | 0.078779772 | 0.081754475 |
| Ppil1 | 9.154829024 | 9.170265576 | 9.073121799 | 0.097143777 | 0.081707226 |
| Erbb2 | 11.3941567 | 11.47485204 | 11.31244993 | 0.162402117 | 0.081706773 |
| Hikeshi | 8.45245875 | 8.445124069 | 8.370777681 | 0.074346388 | 0.081681069 |
| Galk1 | 7.022866346 | 7.108573102 | 6.94119575 | 0.167377352 | 0.081670596 |
| Slc6a20a | 8.667604789 | 8.68963754 | 8.585969423 | 0.103668117 | 0.081635365 |
| Tshz3 | 7.167209505 | 7.231831115 | 7.085637307 | 0.146193808 | 0.081572198 |
| Gpr165 | 2.281145949 | 2.21220455 | 2.199610279 | 0.01259427 | 0.08153567 |
| Tlcd2 | 9.597192151 | 9.509418219 | 9.515684656 | -0.006266436 | 0.081507496 |
| Dis3l | 9.269629208 | 9.182676074 | 9.188218575 | -0.005542501 | 0.081410633 |
| Tmem219 | 8.406948316 | 8.17700215 | 8.32559896 | -0.14859681 | 0.081349356 |
| Hddc2 | 7.826571578 | 7.68695101 | 7.745251605 | -0.058300595 | 0.081319974 |
| Cnot2 | 10.480109 | 10.35710414 | 10.39878988 | -0.041685737 | 0.081319127 |
| Rhbdl3 | 8.744301421 | 8.751293644 | 8.662988045 | 0.0883056 | 0.081313376 |
| Actn4 | 12.79832112 | 13.01950664 | 12.71701846 | 0.302488183 | 0.08130266 |

| | | | | | |
|---|---|---|---|---|---|
| Krt90 | 4.54188094 | 4.784507146 | 4.460633897 | 0.323873249 | 0.081247043 |
| Vps53 | 9.873818755 | 9.86519131 | 9.792621149 | 0.072570161 | 0.081197606 |
| Chchd10 | 9.077474454 | 9.133934133 | 8.996323491 | 0.137610642 | 0.081150964 |
| Alas2 | 6.625329602 | 6.006645635 | 6.544204075 | -0.53755844 | 0.081125527 |
| Gm47356 | 3.522132514 | 3.576568804 | 3.44101765 | 0.135551154 | 0.081114864 |
| Car13 | 11.11103826 | 10.95062722 | 11.02995079 | -0.07932357 | 0.08108747 |
| Vmn1r227 | 4.530392295 | 4.480723991 | 4.449321653 | 0.031402338 | 0.081070642 |
| Gm43587 | 6.145048992 | 6.070996441 | 6.06397881 | 0.007017631 | 0.081070182 |
| Mrps24 | 8.393418074 | 8.426707866 | 8.312368035 | 0.114339831 | 0.081050039 |
| Zfp462 | 7.939167458 | 7.944737694 | 7.858121207 | 0.086616487 | 0.081046251 |
| Rnf141 | 10.39673252 | 10.29773884 | 10.31582154 | -0.018082693 | 0.080910983 |
| Nxpe3 | 5.562877605 | 5.495256571 | 5.482058984 | 0.013197587 | 0.08081862 |
| Gm8773 | 2.923278151 | 2.851646522 | 2.842550042 | 0.00909648 | 0.080728109 |
| Ccn4 | 6.296417448 | 6.472177747 | 6.215745997 | 0.25643175 | 0.080671452 |
| Rassf2 | 9.929503466 | 9.968742892 | 9.848857134 | 0.119885759 | 0.080646332 |
| Saraf | 10.89272341 | 10.88430823 | 10.81212411 | 0.072184113 | 0.0805993 |
| Kctd3 | 9.434963795 | 9.491494607 | 9.354367714 | 0.137126893 | 0.080596081 |
| Bclaf1 | 11.2201672 | 11.14016922 | 11.13958862 | 0.000580603 | 0.08057858 |
| Gm12191 | 2.489893779 | 2.300531133 | 2.409321874 | -0.108790741 | 0.080571905 |
| Sike1 | 9.698098384 | 9.758136582 | 9.6175768 | 0.140559782 | 0.080521585 |
| Tfeb | 8.400743899 | 8.653336066 | 8.320225881 | 0.333110185 | 0.080518018 |
| Olfr1120 | 3.836824022 | 3.795114504 | 3.756310502 | 0.038804002 | 0.08051352 |
| C87414 | 3.461462844 | 3.460626358 | 3.38097226 | 0.079654098 | 0.080490584 |
| Tpr | 11.75187448 | 11.6371298 | 11.67139718 | -0.034267377 | 0.080477307 |
| Polh | 9.106889717 | 8.992283371 | 9.026517679 | -0.034234308 | 0.080372038 |
| Zfp423 | 6.902365374 | 6.715852319 | 6.822042836 | -0.106190517 | 0.080322538 |
| Svil | 11.10646122 | 11.00273368 | 11.02614757 | -0.02341389 | 0.080313647 |
| Zranb2 | 10.6272897 | 10.69282645 | 10.5470427 | 0.14578375 | 0.080247 |
| Fhl2 | 5.797850599 | 5.662168219 | 5.717736157 | -0.055567938 | 0.080114441 |
| Adamts3 | 2.990384767 | 2.913507733 | 2.910310946 | 0.003196787 | 0.080073821 |
| Wdr61 | 10.19441106 | 10.10533347 | 10.11440744 | -0.009073963 | 0.08000362 |
| Rnpepl1 | 11.00907706 | 11.20516639 | 10.92914075 | 0.27602564 | 0.07993631 |

| | | | | | |
|---|---|---|---|---|---|
| Ganab | 11.53825992 | 11.57827983 | 11.4583522 | 0.11992763 | 0.079907727 |
| Tmprss2 | 5.085478093 | 5.101073542 | 5.005582167 | 0.095491375 | 0.079895926 |
| Hsd17b7 | 10.3492474 | 10.3174965 | 10.26937857 | 0.048117923 | 0.079868826 |
| Clcc1 | 9.792365188 | 9.843795147 | 9.71251729 | 0.131277857 | 0.079847898 |
| Rnf111 | 9.871894036 | 9.91131451 | 9.792106707 | 0.119207804 | 0.079787329 |
| Pip4k2a | 9.831974414 | 9.711171753 | 9.75222637 | -0.041054617 | 0.079748044 |
| Srms | 6.126718678 | 6.203933402 | 6.046972919 | 0.156960483 | 0.079745759 |
| Ndufaf2 | 7.244076146 | 7.218983874 | 7.164332356 | 0.054651518 | 0.07974379 |
| Ccsap | 5.324943542 | 5.138021176 | 5.245218439 | -0.107197263 | 0.079725103 |
| Gm12033 | 2.232597458 | 2.246356786 | 2.152874512 | 0.093482273 | 0.079722946 |
| Nelfb | 10.47175161 | 10.38140803 | 10.39207794 | -0.010669913 | 0.07967367 |
| Vcpkmt | 7.455824281 | 7.37211776 | 7.376208902 | -0.004091142 | 0.079615379 |
| Atn1 | 10.24439357 | 10.47964651 | 10.16479286 | 0.314853653 | 0.079600707 |
| Gcsam | 5.589269195 | 5.505799279 | 5.509668829 | -0.00386955 | 0.079600365 |
| Nlrp6 | 1.512306297 | 1.447206467 | 1.432930017 | 0.014276451 | 0.07937628 |
| Mtmr2 | 10.12253049 | 9.967234615 | 10.04316453 | -0.075929913 | 0.079365965 |
| Tm9sf2 | 11.49402075 | 11.39609928 | 11.41467137 | -0.018572087 | 0.079349377 |
| Teddm1b | 1.426366143 | 1.347549985 | 1.347029827 | 0.000520158 | 0.079336316 |
| Gm20695 | 3.197867874 | 3.161093754 | 3.118667396 | 0.042426358 | 0.079200478 |
| Gm20482 | 4.08948492 | 3.960303681 | 4.010285389 | -0.049981708 | 0.079199531 |
| Mrpl44 | 8.850310987 | 8.739382995 | 8.771187039 | -0.031804044 | 0.079123949 |
| Fbxw8 | 10.08543521 | 10.09683514 | 10.00645891 | 0.090376232 | 0.078976308 |
| Fkbp15 | 10.22945617 | 10.15008213 | 10.15052879 | -0.000446663 | 0.07892738 |
| Pi16 | 11.59773727 | 11.52373003 | 11.51883 | 0.004900033 | 0.07890727 |
| Cisd2 | 10.01335334 | 9.999063165 | 9.934459957 | 0.064603208 | 0.078893386 |
| Psmd5 | 9.69040011 | 9.653222411 | 9.611566648 | 0.041655763 | 0.078833462 |
| Zbtb32 | 4.732798983 | 4.756019372 | 4.654009975 | 0.102009398 | 0.078789008 |
| Srp54c | 10.38320481 | 10.24910761 | 10.30441669 | -0.05530908 | 0.078788117 |
| Ubox5 | 7.585619418 | 7.429521465 | 7.506846973 | -0.077325507 | 0.078772446 |
| Pcdh15 | 1.689942669 | 1.660646522 | 1.611212982 | 0.04943354 | 0.078729688 |
| Rad17 | 8.658627486 | 8.670704591 | 8.580124147 | 0.090580444 | 0.078503339 |
| Psap | 14.37413468 | 14.53023297 | 14.29569525 | 0.23453772 | 0.078439423 |

| | | | | | |
|---|---|---|---|---|---|
| Gm18186 | 3.338306309 | 3.275729323 | 3.259868431 | 0.015860892 | 0.078437878 |
| Slc7a4 | 5.923594656 | 5.943310891 | 5.845183246 | 0.098127645 | 0.07841141 |
| Nploc4 | 11.06237057 | 11.15417927 | 10.98401872 | 0.17016055 | 0.078351857 |
| Pla2g12a | 8.851672808 | 8.702543018 | 8.773345217 | -0.070802199 | 0.078327591 |
| Nsmf | 8.686384115 | 8.82272834 | 8.608062653 | 0.214665687 | 0.078321461 |
| Apon | 2.732933221 | 2.605547067 | 2.654623705 | -0.049076637 | 0.078309516 |
| Gm19345 | 1.558283201 | 1.512200253 | 1.48002424 | 0.032176013 | 0.078258961 |
| Ap1g1 | 11.19906643 | 11.17943298 | 11.12085548 | 0.058577497 | 0.07821095 |
| Gm47678 | 5.124336049 | 5.12642346 | 5.046145985 | 0.080277475 | 0.078190063 |
| St7 | 7.755821968 | 7.682386402 | 7.677632042 | 0.00475436 | 0.078189926 |
| Cnot1 | 11.65263433 | 11.58206447 | 11.57446042 | 0.00760405 | 0.078173907 |
| Cdk2ap1 | 10.83813072 | 10.78061164 | 10.75996162 | 0.020650023 | 0.078169103 |
| Cacna2d3 | 5.041560931 | 5.042238596 | 4.963397769 | 0.078840828 | 0.078163163 |
| L2hgdh | 8.651338497 | 8.443351042 | 8.573204334 | -0.129853292 | 0.078134163 |
| Slc44a5 | 1.647386643 | 1.567551705 | 1.569270068 | -0.001718363 | 0.078116575 |
| Gm49395 | 2.763467495 | 2.655139091 | 2.685386159 | -0.030247067 | 0.078081336 |
| Olfr705 | 1.542932552 | 1.498877722 | 1.464868567 | 0.034009155 | 0.078063985 |
| Rpl37a | 12.7036295 | 12.59707588 | 12.62559734 | -0.02852146 | 0.078032157 |
| Acot5 | 6.568527079 | 6.344988962 | 6.490538099 | -0.145549138 | 0.07798898 |
| Uggt1 | 11.03304224 | 11.01247144 | 10.95507821 | 0.057393227 | 0.077964033 |
| Scrib | 10.69719241 | 11.08897642 | 10.6192419 | 0.469734517 | 0.077950507 |
| Emc1 | 9.934773233 | 9.855461603 | 9.856907681 | -0.001446079 | 0.077865551 |
| Tjp2 | 10.59637081 | 10.58908206 | 10.51853475 | 0.07054731 | 0.07783606 |
| Camk2d | 10.42000319 | 10.30530808 | 10.34216792 | -0.03685984 | 0.07783527 |
| Ankrd39 | 6.403567794 | 6.40503779 | 6.325739618 | 0.079298172 | 0.077828176 |
| Ahcyl1 | 11.5382373 | 11.4816134 | 11.46041864 | 0.021194753 | 0.07781866 |
| 2200002D01Rik | 6.703682539 | 6.711209557 | 6.625883904 | 0.085325653 | 0.077798635 |
| 2610318N02Rik | 2.120997724 | 2.133419549 | 2.043210257 | 0.090209292 | 0.077787466 |
| Dmwd | 8.021884685 | 8.122637291 | 7.944136108 | 0.178501183 | 0.077748578 |
| Pkp3 | 12.54115493 | 12.65080383 | 12.46341133 | 0.187392497 | 0.077743597 |
| Rmdn3 | 9.941263848 | 9.913370959 | 9.863561075 | 0.049809884 | 0.077702772 |
| 0610010F05Rik | 8.50060666 | 8.34943282 | 8.422932293 | -0.073499473 | 0.077674367 |

| | | | | | |
|---|---|---|---|---|---|
| Tsc22d4 | 10.08091981 | 10.32432806 | 10.00338876 | 0.320939301 | 0.077531053 |
| Fam118b | 8.212206093 | 8.180997829 | 8.134681153 | 0.046316676 | 0.07752494 |
| Itpkc | 10.51050404 | 10.57826721 | 10.4329918 | 0.145275403 | 0.077512233 |
| Cars2 | 8.404166464 | 8.579608731 | 8.326781443 | 0.252827288 | 0.077385021 |
| Sestd1 | 9.013033968 | 8.845275789 | 8.935675427 | -0.090399638 | 0.077358541 |
| Grpr | 2.402514883 | 2.419890784 | 2.325287855 | 0.094602929 | 0.077227028 |
| Trmt12 | 7.223178264 | 7.113041819 | 7.145952906 | -0.032911087 | 0.077225358 |
| Tcte2 | 3.63078649 | 3.626212831 | 3.553589621 | 0.072623209 | 0.077196869 |
| Tmem256 | 9.131468892 | 8.944072043 | 9.054357409 | -0.110285367 | 0.077111483 |
| Rps15a-ps7 | 1.963292842 | 1.882064461 | 1.886198809 | -0.004134348 | 0.077094034 |
| Lgals3 | 13.60279901 | 13.7175152 | 13.52573264 | 0.191782557 | 0.077066363 |
| Brk1 | 11.06304167 | 10.95084899 | 10.98597955 | -0.03513056 | 0.077062117 |
| Clic1 | 12.10221086 | 11.96560185 | 12.02515071 | -0.059548867 | 0.077060143 |
| Gm45871 | 6.895038687 | 6.79649583 | 6.818163522 | -0.021667692 | 0.076875165 |
| Elmo2 | 9.912947289 | 9.975386448 | 9.836082874 | 0.139303574 | 0.076864414 |
| Olfr825 | 3.189125905 | 3.116622737 | 3.112275762 | 0.004346975 | 0.076850143 |
| Ap3s2 | 9.829542167 | 9.818096399 | 9.752694465 | 0.065401934 | 0.076847702 |
| Akt2 | 11.95558739 | 11.88748371 | 11.87888712 | 0.00859659 | 0.07670027 |
| Tmem88b | 5.46449833 | 5.358451382 | 5.387811802 | -0.029360421 | 0.076686528 |
| Dusp8 | 8.398243455 | 8.420849403 | 8.321627806 | 0.099221597 | 0.076615649 |
| Olfr642 | 2.187746807 | 2.112629095 | 2.111217392 | 0.001411703 | 0.076529415 |
| Yipf3 | 9.912886395 | 10.07501325 | 9.836405049 | 0.238608205 | 0.076481345 |
| Tcf7 | 8.270645603 | 8.145695788 | 8.194171745 | -0.048475957 | 0.076473858 |
| Ttc32 | 6.166514669 | 6.104954539 | 6.090309798 | 0.014644741 | 0.07620487 |
| Ssr4 | 10.55037057 | 10.47687057 | 10.47419253 | 0.00267804 | 0.076178037 |
| Gm48541 | 6.224356173 | 6.263373218 | 6.148196145 | 0.115177073 | 0.076160028 |
| Dicer1 | 10.5012879 | 10.43702792 | 10.42512898 | 0.011898947 | 0.07615892 |
| Apeh | 9.375411115 | 9.255749793 | 9.299263091 | -0.043513298 | 0.076148024 |
| Olfr667 | 3.64756776 | 3.653477397 | 3.571437395 | 0.082040002 | 0.076130365 |
| Gm3362 | 3.068342521 | 3.02355387 | 2.992277596 | 0.031276274 | 0.076064925 |
| Cul5 | 10.27794237 | 10.07232369 | 10.20188506 | -0.129561367 | 0.076057307 |
| Gm47495 | 2.356076361 | 2.358570994 | 2.280041732 | 0.078529261 | 0.076034629 |

| | | | | | |
|---|---|---|---|---|---|
| Dnah14 | 2.713417235 | 2.543864231 | 2.637471098 | -0.093606866 | 0.075946138 |
| Chst7 | 5.298791389 | 5.205024203 | 5.222971741 | -0.017947538 | 0.075819648 |
| Clvs1 | 3.254205421 | 3.235021737 | 3.178392265 | 0.056629473 | 0.075813157 |
| Mfap3 | 9.708114537 | 9.692852956 | 9.632314353 | 0.060538603 | 0.075800183 |
| Sppl2a | 11.52581632 | 11.3118543 | 11.45007813 | -0.138223837 | 0.07573819 |
| Tmem70 | 9.3430078 | 9.22992922 | 9.267292793 | -0.037363572 | 0.075715007 |
| Slc26a11 | 7.459425048 | 7.436244798 | 7.383742771 | 0.052502026 | 0.075682277 |
| Mia2 | 11.18016555 | 11.05777736 | 11.10453633 | -0.04675897 | 0.075629217 |
| Tc2n | 5.058758456 | 4.985226674 | 4.983159233 | 0.002067442 | 0.075599224 |
| Fuca2 | 9.966456951 | 9.906902001 | 9.890866701 | 0.016035301 | 0.075590251 |
| Stat5b | 10.25226332 | 10.24105071 | 10.1766804 | 0.06437031 | 0.075582913 |
| Pcdhga8 | 5.482526633 | 5.420919759 | 5.406963569 | 0.013956189 | 0.075563063 |
| Dpep2 | 8.092459464 | 7.845052839 | 8.017005378 | -0.171952539 | 0.075454086 |
| BC035044 | 3.734340749 | 3.75708766 | 3.659077338 | 0.098010322 | 0.075263411 |
| Bcas2 | 10.01278154 | 9.87336934 | 9.937555808 | -0.064186468 | 0.075225732 |
| Gtf2b | 9.219592163 | 9.18830717 | 9.144369239 | 0.043937931 | 0.075222924 |
| Nrp2 | 10.38444687 | 10.46576838 | 10.30923647 | 0.156531907 | 0.075210403 |
| Zfp62 | 9.054592786 | 8.978901838 | 8.979382949 | -0.000481111 | 0.075209838 |
| Rhoa | 12.86892703 | 12.71453237 | 12.79372972 | -0.079197353 | 0.07519731 |
| Adam15 | 10.93097305 | 10.73044904 | 10.85577856 | -0.125329523 | 0.075194487 |
| Crcp | 8.892790463 | 8.807279146 | 8.817617904 | -0.010338759 | 0.075172558 |
| Katnb1 | 7.866135864 | 7.986654176 | 7.790968001 | 0.195686175 | 0.075167864 |
| Higd1a | 10.55644797 | 10.37197463 | 10.48133441 | -0.109359783 | 0.075113557 |
| Ndufa1 | 9.505944067 | 9.475236972 | 9.430864918 | 0.044372054 | 0.075079149 |
| Gm5559 | 1.483320032 | 1.423103219 | 1.408246093 | 0.014857126 | 0.075073938 |
| Slc26a3 | 3.257599919 | 3.203349935 | 3.182535124 | 0.020814811 | 0.075064795 |
| Klf13 | 11.80113163 | 11.67350926 | 11.72613095 | -0.05262169 | 0.075000673 |
| Dtd1 | 7.660418155 | 7.454160183 | 7.585457016 | -0.131296833 | 0.074961139 |
| Nectin1 | 11.51072486 | 11.59239015 | 11.43585542 | 0.15653473 | 0.07486944 |
| Grm2 | 1.637428976 | 1.565213669 | 1.56257198 | 0.002641689 | 0.074856995 |
| Ogfod2 | 7.655934601 | 7.688030666 | 7.581121955 | 0.106908711 | 0.074812646 |
| Fxr2 | 10.35624493 | 10.48880305 | 10.28150411 | 0.207298947 | 0.074740823 |

| | | | | | |
|---|---|---|---|---|---|
| Vmn1r179 | 2.947622548 | 2.962294457 | 2.872932552 | 0.089361905 | 0.074689995 |
| Spink10 | 2.982852702 | 3.000602915 | 2.908186687 | 0.092416228 | 0.074666015 |
| Cnot7 | 9.87507441 | 9.612122272 | 9.800437394 | -0.188315122 | 0.074637017 |
| Tspan14 | 10.31207503 | 10.30272646 | 10.2374663 | 0.065260157 | 0.07460873 |
| Cox4i1 | 12.18128561 | 12.0927006 | 12.1066847 | -0.0139841 | 0.074600913 |
| AC126807.1 | 2.295145091 | 2.191423116 | 2.220561567 | -0.029138451 | 0.074583524 |
| Ube2cbp | 6.178542634 | 6.127370881 | 6.104002364 | 0.023368517 | 0.07454027 |
| Kcnk13 | 5.487180468 | 5.357482989 | 5.412730544 | -0.055247555 | 0.074449923 |
| Ttc17 | 9.326336793 | 9.174095433 | 9.251932347 | -0.077836914 | 0.074404446 |
| Gm6055 | 5.833286691 | 5.805116199 | 5.758888092 | 0.046228107 | 0.074398599 |
| Sptbn4 | 3.657584906 | 3.797680786 | 3.583234642 | 0.214446144 | 0.074350264 |
| Psen2 | 8.270888536 | 8.071672535 | 8.19654485 | -0.124872315 | 0.074343687 |
| Cd8b1 | 2.5625123 | 2.491804153 | 2.488327998 | 0.003476155 | 0.074184302 |
| Ficd | 7.374398013 | 7.390812678 | 7.300213735 | 0.090598943 | 0.074184278 |
| Fan1 | 7.397521946 | 7.263924524 | 7.323382869 | -0.059458345 | 0.074139077 |
| Trappc9 | 9.295314253 | 9.334323122 | 9.221226674 | 0.113096447 | 0.074087578 |
| Asap2 | 8.87366357 | 8.924213808 | 8.79958769 | 0.124626118 | 0.07407588 |
| Gm10705 | 6.607159613 | 6.805634342 | 6.533104517 | 0.272529825 | 0.074055096 |
| Dgkk | 2.03453226 | 1.972247859 | 1.960501451 | 0.011746408 | 0.074030809 |
| Dnaja2 | 11.86313424 | 11.75474965 | 11.78912718 | -0.03437753 | 0.074007063 |
| Adamdec1 | 1.469404164 | 1.420752561 | 1.395415071 | 0.025337491 | 0.073989094 |
| Prpf38b | 10.96524263 | 11.03813441 | 10.89129962 | 0.146834793 | 0.073943007 |
| Kif3b | 9.610950595 | 9.510078477 | 9.53705698 | -0.026978503 | 0.073893615 |
| Pcnx3 | 10.04694175 | 10.3557281 | 9.9730979 | 0.382630197 | 0.073843854 |
| Gm17971 | 3.719507659 | 3.709594535 | 3.645699137 | 0.063895398 | 0.073808522 |
| Josd1 | 9.788727114 | 9.697136488 | 9.714918692 | -0.017782204 | 0.073808422 |
| Cenpq | 7.093422344 | 7.073966907 | 7.019678167 | 0.05428874 | 0.073744177 |
| Spag9 | 11.89763504 | 11.6985091 | 11.82391193 | -0.125402837 | 0.07372311 |
| Usp38 | 10.02550774 | 10.01010811 | 9.951811085 | 0.058297027 | 0.073696657 |
| Ube2g2 | 10.56322311 | 10.56986454 | 10.48954524 | 0.080319307 | 0.073677873 |
| Gm5617 | 4.782563365 | 4.88375144 | 4.708926754 | 0.174824686 | 0.073636611 |
| Chrnd | 2.734203092 | 2.734247038 | 2.660606681 | 0.073640357 | 0.073596411 |

| | | | | | |
|---|---|---|---|---|---|
| Itpripl1 | 9.085188429 | 8.958086096 | 9.011691435 | -0.053605338 | 0.073496994 |
| Sh3tc1 | 7.904665245 | 8.060976198 | 7.831187174 | 0.229789024 | 0.073478071 |
| Sass6 | 7.280621732 | 7.164866593 | 7.207186364 | -0.042319771 | 0.073435369 |
| Gm12366 | 2.786131133 | 2.721322479 | 2.712702969 | 0.00861951 | 0.073428163 |
| Pcdhb22 | 6.290289245 | 6.392793437 | 6.216883966 | 0.175909471 | 0.073405279 |
| Tfap2c | 9.935305388 | 9.934798009 | 9.861969894 | 0.072828115 | 0.073335493 |
| Gm45492 | 2.045211937 | 2.001311877 | 1.97196558 | 0.029346298 | 0.073246357 |
| Lmf2 | 9.365332756 | 9.594216801 | 9.292091311 | 0.302125491 | 0.073241446 |
| Naprt | 8.597442906 | 8.634986882 | 8.524221676 | 0.110765206 | 0.07322123 |
| Coch | 11.03380598 | 10.97329142 | 10.96059582 | 0.012695593 | 0.07321016 |
| Psph | 9.951637415 | 9.899757908 | 9.878472934 | 0.021284974 | 0.07316448 |
| Zzz3 | 10.5320238 | 10.41569719 | 10.45885995 | -0.04316276 | 0.07316385 |
| Slc48a1 | 10.95631985 | 10.96509807 | 10.88320065 | 0.081897417 | 0.073119197 |
| Prima1 | 3.711735507 | 3.740734733 | 3.63862892 | 0.102105812 | 0.073106587 |
| Ppih | 8.354601478 | 8.318639495 | 8.28153705 | 0.037102445 | 0.073064427 |
| Tmed10 | 11.93758045 | 11.77501586 | 11.86453937 | -0.08952351 | 0.07304108 |
| Gas8 | 7.738375735 | 7.710148729 | 7.665335509 | 0.04481322 | 0.073040226 |
| Gm9762 | 7.238036118 | 7.251272887 | 7.165034291 | 0.086238596 | 0.073001827 |
| Zfp523 | 7.742305422 | 7.841921945 | 7.669427195 | 0.17249475 | 0.072878228 |
| Olfr716 | 2.293679159 | 2.25943836 | 2.220843863 | 0.038594497 | 0.072835296 |
| Satb1 | 9.470660409 | 9.340049161 | 9.397921363 | -0.057872202 | 0.072739046 |
| Foxq1 | 7.364828927 | 7.392926378 | 7.292107691 | 0.100818687 | 0.072721236 |
| Ap2a1 | 10.12676554 | 10.29326703 | 10.05414613 | 0.239120908 | 0.072619415 |
| Fzd8 | 7.125124689 | 7.177961217 | 7.052724597 | 0.125236619 | 0.072400092 |
| Actr10 | 10.47418687 | 10.36173785 | 10.40190289 | -0.040165033 | 0.07228398 |
| Gnmt | 3.871324367 | 3.914806765 | 3.799072809 | 0.115733956 | 0.072251557 |
| Arid4a | 9.551494857 | 9.546045224 | 9.479266726 | 0.066778498 | 0.072228131 |
| Ccar2 | 9.820501329 | 9.810980208 | 9.748286875 | 0.062693333 | 0.072214454 |
| Ankrd26 | 7.52068887 | 7.700563502 | 7.448582804 | 0.251980698 | 0.072106066 |
| Olfr1278 | 3.395746513 | 3.341858464 | 3.32368305 | 0.018175414 | 0.072063464 |
| Ccnjl | 1.029393084 | 0.998969114 | 0.957337761 | 0.041631353 | 0.072055323 |
| Ovgp1 | 5.495888501 | 5.374561856 | 5.423844241 | -0.049282385 | 0.07204426 |

| | | | | | |
|---|---|---|---|---|---|
| Cstf2 | 9.65257762 | 9.594282416 | 9.580559314 | 0.013723102 | 0.072018306 |
| Cops5 | 10.74558672 | 10.67945349 | 10.67359704 | 0.00585645 | 0.071989677 |
| Gm8502 | 2.388335817 | 2.370567287 | 2.316376602 | 0.054190685 | 0.071959215 |
| Efcab14 | 10.4948434 | 10.29732248 | 10.42291449 | -0.125592013 | 0.071928913 |
| Rnf25 | 8.842710005 | 8.807875188 | 8.770795543 | 0.037079645 | 0.071914462 |
| Papss1 | 10.9200034 | 10.87935151 | 10.84812256 | 0.031228947 | 0.071880843 |
| AC134576.3 | 1.679812006 | 1.640224008 | 1.607944538 | 0.03227947 | 0.071867468 |
| Gm29427 | 2.240168003 | 2.374192165 | 2.16833305 | 0.205859115 | 0.071834953 |
| Tm4sf19 | 2.872801275 | 2.913700582 | 2.800979413 | 0.11272117 | 0.071821862 |
| Inpp1 | 7.97240304 | 7.540570171 | 7.900611443 | -0.360041272 | 0.071791597 |
| Itgad | 1.973845188 | 1.895837066 | 1.902086065 | -0.006248999 | 0.071759123 |
| Faim2 | 2.794013831 | 2.724932474 | 2.722294181 | 0.002638293 | 0.07171965 |
| Gpr155 | 8.82317668 | 8.563898354 | 8.751466579 | -0.187568225 | 0.071710101 |
| Ptdss1 | 10.86300186 | 10.72969204 | 10.79134965 | -0.061657603 | 0.071652213 |
| Ptrh1 | 8.213808256 | 8.265789529 | 8.142209026 | 0.123580503 | 0.07159923 |
| Atg101 | 8.604664163 | 8.704731869 | 8.533073618 | 0.171658251 | 0.071590545 |
| 1700010I14Rik | 2.174718063 | 2.09262594 | 2.103263798 | -0.010637859 | 0.071454264 |
| Rpl23a-ps3 | 2.550807339 | 2.522033627 | 2.479367621 | 0.042666006 | 0.071439718 |
| Gmeb2 | 9.035531916 | 8.970590708 | 8.96413322 | 0.006457488 | 0.071398696 |
| Csrp1 | 11.75383374 | 11.51906479 | 11.68243761 | -0.163372817 | 0.071396137 |
| Samd13 | 3.188290194 | 3.199692342 | 3.116901054 | 0.082791288 | 0.071389141 |
| Dennd5a | 10.47169006 | 10.4138879 | 10.40032097 | 0.01356693 | 0.07136909 |
| Xlr4a | 4.148927353 | 3.964766098 | 4.077577655 | -0.112811556 | 0.071349698 |
| AW209491 | 7.855520794 | 7.721589302 | 7.784203878 | -0.062614576 | 0.071316916 |
| Serpind1 | 3.31595323 | 3.201664566 | 3.244717596 | -0.04305303 | 0.071235634 |
| Jag2 | 10.68954234 | 11.18097872 | 10.61833904 | 0.562639687 | 0.0712033 |
| Anxa5 | 12.24455421 | 12.27308798 | 12.17344603 | 0.099641953 | 0.07110818 |
| Slc35d1 | 8.46783932 | 8.450956585 | 8.396733087 | 0.054223498 | 0.071106232 |
| Trappc5 | 8.756439741 | 8.784420595 | 8.685343878 | 0.099076717 | 0.071095863 |
| Gpr45 | 2.642678559 | 2.593448979 | 2.571608185 | 0.021840794 | 0.071070374 |
| Sumo2 | 11.77217855 | 11.69548146 | 11.70111543 | -0.005633967 | 0.071063123 |
| Rnf113a1 | 6.202798751 | 6.171771779 | 6.131758596 | 0.040013183 | 0.071040155 |

| | | | | | |
|---|---|---|---|---|---|
| Gm6524 | 2.746377158 | 2.991897343 | 2.675368999 | 0.316528343 | 0.071008159 |
| Med6 | 8.198518357 | 8.117325371 | 8.127514618 | -0.010189247 | 0.071003739 |
| Dcaf7 | 10.94273838 | 11.03182111 | 10.87174811 | 0.160073003 | 0.070990273 |
| Gm7511 | 2.13855369 | 2.098242709 | 2.06757822 | 0.03066449 | 0.07097547 |
| Zfp800 | 8.87992313 | 8.848582011 | 8.809019361 | 0.039562651 | 0.070903769 |
| Setd6 | 8.434540213 | 8.348991478 | 8.363651089 | -0.014659611 | 0.070889124 |
| Serpina3k | 2.595458412 | 2.579554375 | 2.524604553 | 0.054949822 | 0.070853859 |
| Upk1a | 2.649568841 | 2.636082523 | 2.578716578 | 0.057365945 | 0.070852263 |
| Gemin8 | 6.597837704 | 6.5892498 | 6.526999554 | 0.062250246 | 0.07083815 |
| Narfl | 8.76540369 | 8.684271933 | 8.694581022 | -0.010309089 | 0.070822668 |
| Crhr2 | 7.596827309 | 6.854549932 | 7.526118611 | -0.671568679 | 0.070708698 |
| Cd300c | 1.303701095 | 1.243630279 | 1.233058391 | 0.010571888 | 0.070642705 |
| Plekhg1 | 8.913034842 | 8.737474218 | 8.842440234 | -0.104966016 | 0.070594608 |
| Armc5 | 8.375943065 | 8.709591138 | 8.305387548 | 0.40420359 | 0.070555517 |
| Ctdsp1 | 11.10488735 | 11.1489865 | 11.03434154 | 0.114644957 | 0.070545803 |
| Ptger4 | 8.830371368 | 8.989797791 | 8.759880228 | 0.229917563 | 0.070491139 |
| Pdgfb | 8.874268741 | 9.053664752 | 8.803828082 | 0.24983667 | 0.070440659 |
| Gm5905 | 2.319161122 | 2.335351256 | 2.248721916 | 0.08662934 | 0.070439206 |
| Hmgcl | 9.43315871 | 9.44043425 | 9.362761811 | 0.077672439 | 0.0703969 |
| Gcnt4 | 6.755923825 | 6.592351333 | 6.685594084 | -0.09324275 | 0.070329741 |
| Eif4e3 | 9.541231331 | 9.239673819 | 9.470903063 | -0.231229244 | 0.070328268 |
| Gm4891 | 1.219025765 | 1.153082294 | 1.148811152 | 0.004271142 | 0.070214613 |
| Nceh1 | 8.971007397 | 9.187634231 | 8.900798978 | 0.286835253 | 0.070208419 |
| Rpsa | 14.46922095 | 14.44320183 | 14.39903865 | 0.044163187 | 0.0701823 |
| Slc35e3 | 8.826815128 | 8.679106377 | 8.756642126 | -0.077535749 | 0.070173002 |
| Gm49333 | 3.253844533 | 3.292123205 | 3.183729924 | 0.108393281 | 0.070114608 |
| Gapdhs | 3.126859834 | 3.104187895 | 3.056749309 | 0.047438586 | 0.070110525 |
| Cks1b | 8.250142466 | 8.135001284 | 8.180038714 | -0.045037429 | 0.070103752 |
| Gm10177 | 9.999973006 | 9.840692935 | 9.929872592 | -0.089179657 | 0.070100413 |
| Bnip1 | 8.274835675 | 8.29622686 | 8.204798038 | 0.091428821 | 0.070037636 |
| Nepn | 1.811672039 | 1.716433467 | 1.741655119 | -0.025221652 | 0.07001692 |
| Eef1akmt3 | 5.659453015 | 5.66790205 | 5.589465369 | 0.078436681 | 0.069987647 |

| | | | | | |
|---|---|---|---|---|---|
| Tmbim1 | 11.61623944 | 11.55740567 | 11.54627204 | 0.011133637 | 0.0699674 |
| Med21 | 8.7787285 | 8.685978417 | 8.708776857 | -0.02279844 | 0.069951643 |
| Gm37348 | 1.659033765 | 1.639697286 | 1.589099681 | 0.050597605 | 0.069934084 |
| Med9 | 8.193567953 | 8.154417654 | 8.123655545 | 0.030762109 | 0.069912409 |
| Lair1 | 6.109359611 | 6.34496784 | 6.039454091 | 0.305513748 | 0.06990552 |
| Dnal4 | 8.28512583 | 8.369540282 | 8.215235477 | 0.154304805 | 0.069890353 |
| Cdca7l | 9.119699857 | 9.131485088 | 9.049847289 | 0.0816378 | 0.069852568 |
| Jmjd4 | 7.97089393 | 7.99074537 | 7.901063327 | 0.089682043 | 0.069830603 |
| Papola | 11.56277076 | 11.59709286 | 11.4930934 | 0.10399946 | 0.069677363 |
| Coq7 | 8.1513623 | 8.126555678 | 8.081722328 | 0.044833351 | 0.069639972 |
| Agtpbp1 | 9.41611857 | 9.308811342 | 9.346518049 | -0.037706707 | 0.069600521 |
| Homer2 | 11.028974 | 10.94328103 | 10.95937714 | -0.01609611 | 0.069596863 |
| H2-DMb2 | 7.929420212 | 7.774022461 | 7.859921212 | -0.085898751 | 0.069499001 |
| Poln | 3.799745417 | 3.641700069 | 3.730253959 | -0.088553889 | 0.069491459 |
| Syt11 | 7.44888486 | 7.43083873 | 7.379401281 | 0.051437449 | 0.069483579 |
| Commd5 | 7.883780344 | 7.777859113 | 7.814336954 | -0.036477841 | 0.06944339 |
| Rbl1 | 8.98652464 | 8.935661023 | 8.917132816 | 0.018528207 | 0.069391824 |
| Exoc3 | 10.43171009 | 10.35073053 | 10.36238175 | -0.01165122 | 0.069328333 |
| Mipol1 | 5.75289653 | 5.75205901 | 5.683579355 | 0.068479654 | 0.069317175 |
| Pcdhb14 | 5.647928811 | 5.591441772 | 5.578650555 | 0.012791217 | 0.069278256 |
| Ppp2r3c | 8.573391538 | 8.432723335 | 8.504159109 | -0.071435773 | 0.069232429 |
| Vmn1r18 | 4.682086909 | 4.747953678 | 4.612883591 | 0.135070087 | 0.069203318 |
| Tmem269 | 5.240921078 | 5.232252662 | 5.171732426 | 0.060520236 | 0.069188652 |
| Swap70 | 9.251546721 | 9.26684765 | 9.182362495 | 0.084485156 | 0.069184227 |
| Dnajb1 | 10.90662232 | 12.01848759 | 10.83747744 | 1.181010153 | 0.069144883 |
| Taok2 | 10.32560256 | 10.52882115 | 10.25646194 | 0.272359213 | 0.06914062 |
| Chchd1 | 8.904022394 | 8.808080512 | 8.835058138 | -0.026977626 | 0.068964256 |
| Ank2 | 8.305866009 | 8.240208701 | 8.236931803 | 0.003276898 | 0.068934206 |
| Prorsd1 | 7.80497551 | 7.641743214 | 7.73608137 | -0.094338156 | 0.06889414 |
| Eno1b | 2.531409949 | 2.663016943 | 2.462516615 | 0.200500327 | 0.068893334 |
| Gpr15 | 2.744752585 | 2.696327718 | 2.675885356 | 0.020442362 | 0.068867229 |
| Cog3 | 9.729480646 | 9.733631785 | 9.660714547 | 0.072917238 | 0.068766099 |

| | | | | | |
|---|---|---|---|---|---|
| Gm10171 | 7.422146604 | 6.911969836 | 7.353408507 | -0.441438671 | 0.068738097 |
| Gm10233 | 7.422146604 | 6.911969836 | 7.353408507 | -0.441438671 | 0.068738097 |
| Rps6kl1 | 2.675290883 | 2.602643177 | 2.606584881 | -0.003941704 | 0.068706002 |
| Tmem168 | 9.952332098 | 9.904749138 | 9.883643841 | 0.021105297 | 0.068688256 |
| Ighd | 3.680572537 | 3.656145629 | 3.611933424 | 0.044212205 | 0.068639113 |
| Prr16 | 7.503114663 | 7.327754953 | 7.434476537 | -0.106721584 | 0.068638126 |
| Mrpl2 | 8.856997433 | 8.926634779 | 8.788378666 | 0.138256114 | 0.068618768 |
| Spsb4 | 3.689150477 | 3.679234214 | 3.620571055 | 0.058663159 | 0.068579421 |
| Itpripl2 | 10.4173636 | 10.37099951 | 10.34879898 | 0.022200523 | 0.068564613 |
| Tmem121 | 2.991947014 | 2.958771767 | 2.92339623 | 0.035375537 | 0.068550784 |
| Gng5 | 10.58317175 | 10.53436565 | 10.51474154 | 0.019624107 | 0.06843021 |
| Rps12 | 9.529743356 | 9.363558585 | 9.461351413 | -0.097792828 | 0.068391942 |
| Pou5f2 | 2.685636906 | 2.70654975 | 2.61724689 | 0.08930286 | 0.068390016 |
| Lyrm1 | 7.158791521 | 7.062989331 | 7.090454273 | -0.027464942 | 0.068337248 |
| B2m | 14.39984274 | 14.24289096 | 14.33151898 | -0.088628017 | 0.06832376 |
| Naa35 | 9.669188794 | 9.50514453 | 9.600915757 | -0.095771227 | 0.068273037 |
| Alkbh7 | 6.338337879 | 6.373191606 | 6.270289617 | 0.10290199 | 0.068048262 |
| Aimp1 | 10.0796893 | 9.931792353 | 10.01164243 | -0.079850074 | 0.068046873 |
| Ppp1ccb | 9.472607252 | 9.29105344 | 9.404652735 | -0.113599295 | 0.067954517 |
| Proser1 | 8.547279326 | 8.718373482 | 8.479330314 | 0.239043168 | 0.067949012 |
| Zfp213 | 7.003667647 | 7.089919752 | 6.93572644 | 0.154193312 | 0.067941206 |
| Krt83 | 2.488557699 | 2.428050772 | 2.42076215 | 0.007288621 | 0.067795549 |
| Orc4 | 9.402064243 | 9.35727267 | 9.334322607 | 0.022950063 | 0.067741636 |
| Pgghg | 9.019656246 | 8.938333025 | 8.951988655 | -0.01365563 | 0.067667591 |
| Galns | 7.239199162 | 7.365688194 | 7.171647388 | 0.194040806 | 0.067551773 |
| D10Jhu81e | 9.853166496 | 9.854985189 | 9.785630475 | 0.069354714 | 0.067536021 |
| Agpat1 | 9.399766433 | 9.495314843 | 9.332270384 | 0.16304446 | 0.067496049 |
| Sqor | 9.19794468 | 8.879734168 | 9.130462067 | -0.250727899 | 0.067482613 |
| Hnrnpll | 9.809146464 | 9.810746841 | 9.741837869 | 0.068908972 | 0.067308595 |
| Ctrl | 2.888114365 | 2.864884534 | 2.820827063 | 0.044057471 | 0.067287302 |
| Lef1 | 5.595238119 | 5.430832512 | 5.527955122 | -0.09712261 | 0.067282998 |
| Agps | 9.650960658 | 9.56814888 | 9.583730136 | -0.015581256 | 0.067230522 |

| | | | | | |
|---|---|---|---|---|---|
| Hs6st2 | 6.009530497 | 5.77366717 | 5.942318135 | -0.168650965 | 0.067212362 |
| Pi4kb | 10.01671206 | 9.838664592 | 9.949517274 | -0.110852682 | 0.06719479 |
| Ubr7 | 9.454815364 | 9.387554766 | 9.387651648 | -9.68813E-05 | 0.067163717 |
| Syt5 | 1.837185032 | 1.770011242 | 1.770040345 | -2.91037E-05 | 0.067144687 |
| Cdc123 | 10.33761826 | 10.27611787 | 10.2705174 | 0.005600477 | 0.06710086 |
| Rheb | 10.23807373 | 10.23213517 | 10.17100869 | 0.061126473 | 0.067065033 |
| 9530050K03Rik | 2.998369182 | 2.96666475 | 2.931437944 | 0.035226806 | 0.066931238 |
| Vmn1r76 | 4.615392237 | 4.598789011 | 4.548505238 | 0.050283773 | 0.066886999 |
| Tmem151b | 1.254687915 | 1.218183438 | 1.1878363 | 0.030347138 | 0.066851616 |
| Tubgcp6 | 9.132731526 | 9.268281078 | 9.065902657 | 0.202378421 | 0.066828869 |
| Gm43802 | 3.064397001 | 3.225208261 | 2.997612252 | 0.227596009 | 0.066784749 |
| Fam129b | 13.17130298 | 13.33064933 | 13.10454304 | 0.226106283 | 0.066759933 |
| Hesx1 | 2.741373413 | 2.63615605 | 2.674719983 | -0.038563933 | 0.06665343 |
| D130007C19Rik | 3.285617665 | 3.255202139 | 3.218998607 | 0.036203532 | 0.066619058 |
| Recql4 | 6.819420672 | 7.027388657 | 6.752813883 | 0.274574775 | 0.066606789 |
| Kat5 | 9.195687452 | 9.132254445 | 9.129172224 | 0.003082221 | 0.066515228 |
| Siah1b | 6.573048218 | 6.555044434 | 6.506581164 | 0.04846327 | 0.066467054 |
| Gm17084 | 5.666778318 | 5.641381656 | 5.600514573 | 0.040867083 | 0.066263745 |
| Il11ra1 | 11.00942106 | 10.9773913 | 10.94317531 | 0.03421599 | 0.066245753 |
| Gsk3b | 10.97726313 | 10.96472956 | 10.91107391 | 0.05365565 | 0.066189217 |
| Wdr92 | 8.038355124 | 7.973108123 | 7.972220346 | 0.000887777 | 0.066134778 |
| Ado | 8.699190029 | 8.684784564 | 8.633100836 | 0.051683728 | 0.066089193 |
| Dnaaf2 | 6.456278309 | 6.516920811 | 6.390240683 | 0.126680128 | 0.066037626 |
| Olfr57 | 3.631691899 | 3.561382183 | 3.5656766 | -0.004294417 | 0.0660153 |
| Tkt | 12.59269143 | 12.31953765 | 12.52669199 | -0.207154333 | 0.065999447 |
| Csnk1g3 | 9.152684755 | 9.047143154 | 9.086688037 | -0.039544883 | 0.065996718 |
| Gpr37l1 | 5.656176591 | 5.778660633 | 5.590254242 | 0.188406391 | 0.065922349 |
| Miip | 8.361493285 | 8.302312944 | 8.295666421 | 0.006646523 | 0.065826864 |
| Awat2 | 9.416075114 | 9.589116776 | 9.350260845 | 0.238855931 | 0.065814269 |
| Eif4ebp3 | 6.295321931 | 6.365798875 | 6.229656239 | 0.136142636 | 0.065665692 |
| Gm15772 | 1.739115991 | 1.699264364 | 1.673468527 | 0.025795836 | 0.065647464 |
| Cask | 10.20369742 | 10.18035143 | 10.13811693 | 0.042234497 | 0.065580487 |

| | | | | | |
|---|---|---|---|---|---|
| Olfr243 | 2.105705393 | 2.109259233 | 2.040139525 | 0.069119708 | 0.065565868 |
| Olfr829 | 2.616613164 | 2.554803322 | 2.551060589 | 0.003742733 | 0.065552574 |
| Bcl11b | 11.00822211 | 11.0610339 | 10.94267516 | 0.118358737 | 0.06554695 |
| Arglu1 | 10.78533672 | 10.84084173 | 10.71980786 | 0.12103387 | 0.065528857 |
| Gm10638 | 3.034368777 | 2.995509886 | 2.968841189 | 0.026668697 | 0.065527588 |
| Rps27l | 10.22549153 | 10.1822137 | 10.15999144 | 0.02222226 | 0.06550009 |
| Pms2 | 7.636948064 | 7.654413567 | 7.571454308 | 0.082959259 | 0.065493756 |
| Gm13002 | 2.495859924 | 2.501956763 | 2.430373374 | 0.071583389 | 0.06548655 |
| Trp73 | 8.499966486 | 8.976547781 | 8.434507728 | 0.542040053 | 0.065458758 |
| Sh2d1a | 1.278676552 | 1.240846837 | 1.213265411 | 0.027581427 | 0.065411141 |
| Purb | 12.42423935 | 12.29852501 | 12.35887988 | -0.06035487 | 0.06535947 |
| Gm44821 | 3.686426654 | 3.625717804 | 3.621067967 | 0.004649837 | 0.065358687 |
| Pdcl3 | 9.854646823 | 9.746699491 | 9.789312697 | -0.042613206 | 0.065334125 |
| Cdh17 | 2.692482432 | 2.705313285 | 2.627172598 | 0.078140687 | 0.065309834 |
| Gsg1 | 1.758125214 | 1.730624058 | 1.692830714 | 0.037793344 | 0.0652945 |
| Derl2 | 9.792177597 | 9.790536684 | 9.726887249 | 0.063649435 | 0.065290348 |
| Coa3 | 9.286809736 | 9.243311045 | 9.221564642 | 0.021746402 | 0.065245093 |
| Vmn2r22 | 1.930075998 | 1.879080021 | 1.864831795 | 0.014248227 | 0.065244203 |
| Kcnd1 | 5.19350447 | 5.223690296 | 5.128305514 | 0.095384782 | 0.065198956 |
| Bend5 | 4.367961218 | 4.288923569 | 4.302774975 | -0.013851406 | 0.065186243 |
| Tnk1 | 9.128639962 | 9.122501533 | 9.063495724 | 0.05900581 | 0.065144238 |
| Gm10132 | 2.315612349 | 2.301340941 | 2.250544203 | 0.050796738 | 0.065068146 |
| Crlf3 | 8.966528378 | 8.925668698 | 8.901486083 | 0.024182615 | 0.065042295 |
| Rfc4 | 7.804586672 | 7.636000377 | 7.739669396 | -0.103669019 | 0.064917276 |
| Kdm2a | 11.55982475 | 11.52099487 | 11.49493156 | 0.026063313 | 0.064893197 |
| Dock4 | 8.499262839 | 8.379814537 | 8.434383539 | -0.054569001 | 0.0648793 |
| Tigd3 | 4.173198505 | 4.166302045 | 4.108458857 | 0.057843188 | 0.064739648 |
| Adat2 | 6.449659496 | 6.301451594 | 6.384944702 | -0.083493108 | 0.064714794 |
| Gpat4 | 10.42393001 | 10.30911573 | 10.35931809 | -0.050202353 | 0.064611923 |
| Arf3 | 11.96176325 | 11.87121839 | 11.89718069 | -0.025962307 | 0.064582553 |
| Itm2c | 10.51533041 | 10.3376259 | 10.45075891 | -0.113133003 | 0.0645715 |
| Msh5 | 4.575141288 | 4.633738049 | 4.510660548 | 0.123077501 | 0.06448074 |

| | | | | | |
|---|---|---|---|---|---|
| Plppr4 | 3.566843748 | 3.524004302 | 3.502449171 | 0.021555132 | 0.064394577 |
| Bhlha15 | 3.001669919 | 2.997331135 | 2.937291354 | 0.060039782 | 0.064378566 |
| Espn | 4.533093345 | 4.542037453 | 4.468724405 | 0.073313048 | 0.06436894 |
| C1galt1 | 10.53412097 | 10.55593766 | 10.46975793 | 0.086179727 | 0.064363033 |
| Hdac3 | 10.70358702 | 10.5905978 | 10.63923998 | -0.048642177 | 0.064347037 |
| Prodh | 7.810661662 | 7.688153872 | 7.746321789 | -0.058167917 | 0.064339873 |
| Ubfd1 | 9.664884542 | 9.875589159 | 9.600587318 | 0.27500184 | 0.064297224 |
| Nucks1 | 11.44952856 | 11.23820944 | 11.38527078 | -0.147061337 | 0.06425778 |
| Dazap2 | 11.98750508 | 11.85768961 | 11.92325438 | -0.06556477 | 0.064250697 |
| Ube3c | 10.59290034 | 10.46547817 | 10.52865299 | -0.06317482 | 0.064247353 |
| Olfr729 | 1.679548313 | 1.644305039 | 1.615327689 | 0.02897735 | 0.064220624 |
| Gm2619 | 1.630594911 | 1.587790707 | 1.566421481 | 0.021369226 | 0.064173431 |
| Cma2 | 2.018592181 | 1.968719146 | 1.954438102 | 0.014281044 | 0.064154079 |
| Arhgap23 | 10.52453448 | 10.48366151 | 10.46039186 | 0.023269653 | 0.064142623 |
| Ccer2 | 5.293471422 | 5.285331141 | 5.229342725 | 0.055988416 | 0.064128696 |
| Gm49391 | 6.170388794 | 6.125856422 | 6.106272363 | 0.019584059 | 0.064116431 |
| Pigm | 9.755282123 | 9.706031888 | 9.691225509 | 0.014806379 | 0.064056614 |
| Klra7 | 3.097488856 | 3.062888183 | 3.033527113 | 0.02936107 | 0.063961743 |
| Gm11027 | 5.512470965 | 5.333635993 | 5.448648245 | -0.115012252 | 0.06382272 |
| Ly96 | 6.826696977 | 6.792360236 | 6.762919708 | 0.029440528 | 0.063777269 |
| Emc7 | 10.18444423 | 10.09843778 | 10.1206974 | -0.022259623 | 0.063746823 |
| Gm42420 | 2.242002324 | 2.270585064 | 2.178343299 | 0.092241765 | 0.063659025 |
| Cdc5l | 10.3913881 | 10.29811393 | 10.32775838 | -0.02964445 | 0.063629717 |
| AI661453 | 10.93288096 | 11.05116189 | 10.86926603 | 0.181895853 | 0.063614923 |
| Pcsk4 | 5.379481366 | 5.484708109 | 5.31595366 | 0.168754449 | 0.063527706 |
| Dlx2 | 2.376008003 | 2.3799582 | 2.31248261 | 0.06747559 | 0.063525393 |
| Pcid2 | 9.080732742 | 9.008809759 | 9.017251395 | -0.008441636 | 0.063481347 |
| Gsdme | 6.248846539 | 6.099583991 | 6.185374697 | -0.085790705 | 0.063471842 |
| Gm4759 | 2.438540187 | 2.385951606 | 2.375178102 | 0.010773504 | 0.063362085 |
| Pex5 | 9.957281445 | 10.02177973 | 9.893967242 | 0.127812491 | 0.063314203 |
| Klri1 | 3.65697307 | 3.542584107 | 3.593702228 | -0.051118121 | 0.063270842 |
| Gm6749 | 1.472388499 | 1.447668194 | 1.409229412 | 0.038438781 | 0.063159086 |

| | | | | | |
|---|---|---|---|---|---|
| Olfr506 | 1.56822732 | 1.539134078 | 1.505171263 | 0.033962815 | 0.063056057 |
| Stx3 | 8.753495379 | 8.752404016 | 8.690453048 | 0.061950968 | 0.063042331 |
| 8030462N17Rik | 8.372191515 | 8.317247282 | 8.309241123 | 0.00800616 | 0.062950393 |
| Mcm6 | 10.79059285 | 10.82523887 | 10.72768942 | 0.09754945 | 0.06290343 |
| Mis12 | 9.312666177 | 9.424184386 | 9.249783194 | 0.174401192 | 0.062882983 |
| Gm6158 | 7.224400875 | 7.268903938 | 7.161533702 | 0.107370236 | 0.062867172 |
| Pign | 9.590753716 | 9.426581723 | 9.527899375 | -0.101317652 | 0.062854341 |
| Selenow | 12.33143449 | 12.24540625 | 12.26861643 | -0.023210173 | 0.062818067 |
| Tram1 | 11.09016235 | 11.14171825 | 11.02734672 | 0.114371527 | 0.062815633 |
| Fcrla | 3.169415821 | 3.215574809 | 3.106622117 | 0.108952691 | 0.062793703 |
| 6330403K07Rik | 1.837469 | 1.782546209 | 1.774710305 | 0.007835904 | 0.062758695 |
| D030056L22Rik | 7.480186505 | 7.297107703 | 7.417529002 | -0.120421299 | 0.062657503 |
| Calca | 3.3619658 | 3.294579858 | 3.299325957 | -0.004746099 | 0.062639843 |
| Cth | 5.91350484 | 5.914692629 | 5.850903844 | 0.063788785 | 0.062600996 |
| Olfr545 | 1.445105211 | 1.383283208 | 1.382519686 | 0.000763522 | 0.062585525 |
| Gm3468 | 3.471030915 | 3.398156169 | 3.408529748 | -0.010373579 | 0.062501167 |
| Alg11 | 8.568929665 | 8.450953504 | 8.506446008 | -0.055492504 | 0.062483657 |
| Nova2 | 5.468374359 | 5.525039943 | 5.40589691 | 0.119143033 | 0.062477449 |
| Slc1a3 | 9.850781472 | 9.758841924 | 9.788329612 | -0.029487688 | 0.06245186 |
| Nab1 | 10.88225805 | 10.83343813 | 10.81981567 | 0.013622457 | 0.062442383 |
| Itgb2l | 1.598241041 | 1.501717042 | 1.535824638 | -0.034107596 | 0.062416403 |
| Keap1 | 10.02294064 | 10.03409178 | 9.960529809 | 0.073561971 | 0.062410832 |
| Zfp668 | 7.600051228 | 7.755658539 | 7.537680016 | 0.217978523 | 0.062371211 |
| Prdx5 | 12.32407429 | 12.16970946 | 12.26172383 | -0.092014363 | 0.062350463 |
| Flt3 | 6.858130222 | 6.832416584 | 6.795816889 | 0.036599695 | 0.062313333 |
| Kif2a | 9.455466416 | 9.454790637 | 9.393154605 | 0.061636032 | 0.062311811 |
| Vmn1r-ps144 | 4.953853045 | 4.876886282 | 4.891600316 | -0.014714034 | 0.062252729 |
| Dlx6 | 3.154700629 | 3.13035857 | 3.09245224 | 0.03790633 | 0.062248389 |
| Ubl4a | 9.997556096 | 10.01811164 | 9.935308347 | 0.082803292 | 0.062247749 |
| Unc80 | 3.219895047 | 3.267524952 | 3.157649632 | 0.10987532 | 0.062245415 |
| Katna1 | 8.964864023 | 8.854130016 | 8.902638654 | -0.048508638 | 0.062225369 |
| Ndufa7 | 9.57564828 | 9.555288794 | 9.513542975 | 0.041745819 | 0.062105305 |

| | | | | | |
|---|---|---|---|---|---|
| Trim34b | 2.332192016 | 2.287856131 | 2.270120119 | 0.017736012 | 0.062071897 |
| Tonsl | 6.932080718 | 7.312921413 | 6.870070178 | 0.442851235 | 0.06201054 |
| Lsm14a | 11.08614325 | 11.04859863 | 11.02414183 | 0.0244568 | 0.06200142 |
| Iscu | 9.629643221 | 9.449313238 | 9.567648112 | -0.118334874 | 0.061995109 |
| Tnfaip8l2 | 7.31903446 | 7.183317367 | 7.257070563 | -0.073753197 | 0.061963896 |
| Rgs22 | 2.88053782 | 2.819575857 | 2.818609713 | 0.000966145 | 0.061928108 |
| Brat1 | 8.164447873 | 8.37574294 | 8.102549598 | 0.273193343 | 0.061898275 |
| Gm37108 | 3.403375267 | 3.375306457 | 3.341481181 | 0.033825276 | 0.061894086 |
| Krt4 | 4.186864373 | 4.208577547 | 4.125005354 | 0.083572193 | 0.061859019 |
| Slc50a1 | 8.90171488 | 8.795752368 | 8.839904018 | -0.04415165 | 0.061810862 |
| Nek3 | 5.273204839 | 5.296874863 | 5.211419535 | 0.085455328 | 0.061785304 |
| Tmem126a | 8.511329199 | 8.370954993 | 8.449562931 | -0.078607938 | 0.061766268 |
| Myl2 | 1.556501856 | 1.49689853 | 1.494744544 | 0.002153987 | 0.061757313 |
| Gm43110 | 6.09653883 | 5.976824959 | 6.034792488 | -0.057967529 | 0.061746341 |
| Arid1a | 11.16665245 | 11.34281783 | 11.10491859 | 0.237899243 | 0.061733867 |
| Wfdc21 | 9.891711075 | 9.498949676 | 9.82997879 | -0.331029113 | 0.061732286 |
| Bik | 3.887094307 | 3.881675247 | 3.825409351 | 0.056265896 | 0.061684956 |
| AC148019.3 | 2.005940194 | 1.97852596 | 1.944289871 | 0.034236088 | 0.061650323 |
| Abcc10 | 7.485551016 | 7.700748667 | 7.423930108 | 0.27681856 | 0.061620908 |
| Zfp280c | 8.416559056 | 8.243073924 | 8.354956495 | -0.111882571 | 0.061602561 |
| Gm10130 | 2.314133196 | 2.300908526 | 2.252605557 | 0.048302969 | 0.061527639 |
| Zdhhc16 | 8.81389739 | 8.916449761 | 8.752381286 | 0.164068475 | 0.061516104 |
| Rhox8 | 2.055320563 | 1.987520538 | 1.993838782 | -0.006318244 | 0.061481781 |
| Mag | 5.502119336 | 5.641314732 | 5.440692271 | 0.200622461 | 0.061427065 |
| 4933434E20Rik | 10.09175267 | 9.984317034 | 10.03032562 | -0.046008587 | 0.061427045 |
| Mrpl18 | 9.451324145 | 9.398913846 | 9.390012658 | 0.008901187 | 0.061311487 |
| Ube2ql1 | 3.353551676 | 3.386086077 | 3.292243232 | 0.093842845 | 0.061308445 |
| Mtfr1l | 10.08586076 | 9.907295083 | 10.02456734 | -0.117272252 | 0.061293427 |
| Mindy3 | 10.1440632 | 10.04769323 | 10.0827762 | -0.035082967 | 0.061287007 |
| Duox1 | 7.888827739 | 7.873553668 | 7.827561477 | 0.045992191 | 0.061266261 |
| Tent2 | 9.806414328 | 9.615242618 | 9.745151795 | -0.129909176 | 0.061262534 |
| Creg1 | 10.39631425 | 10.22434969 | 10.3350682 | -0.11071851 | 0.06124605 |

| | | | | | |
|---|---|---|---|---|---|
| Commd4 | 8.99865282 | 8.970724466 | 8.937430767 | 0.033293699 | 0.061222053 |
| Zfp583 | 4.616373696 | 4.603341835 | 4.555168727 | 0.048173108 | 0.061204968 |
| Olfr887 | 1.84465393 | 1.79893296 | 1.783458679 | 0.015474281 | 0.061195251 |
| Ankrd63 | 1.478902419 | 1.425204586 | 1.417730305 | 0.007474281 | 0.061172114 |
| S1pr4 | 5.497220697 | 5.421744339 | 5.436158601 | -0.014414262 | 0.061062096 |
| Gm7336 | 7.930193887 | 7.856403841 | 7.869133106 | -0.012729265 | 0.061060781 |
| Sdhb | 10.7602386 | 10.73837805 | 10.69921504 | 0.03916301 | 0.061023563 |
| Gm11425 | 3.488218701 | 3.459246498 | 3.427198727 | 0.032047771 | 0.061019974 |
| 4930562C15Rik | 2.854003379 | 2.903660191 | 2.792984724 | 0.110675467 | 0.061018655 |
| Cwf19l1 | 7.96227756 | 7.966005127 | 7.901283245 | 0.064721882 | 0.060994315 |
| Ctnnal1 | 10.04048321 | 10.11687603 | 9.979595649 | 0.137280377 | 0.060887562 |
| Rprd2 | 9.924631625 | 9.88422157 | 9.863751425 | 0.020470145 | 0.060880199 |
| Vmn1r77 | 2.125604256 | 2.063822841 | 2.064724866 | -0.000902025 | 0.06087939 |
| Rnf157 | 6.22185079 | 6.191428512 | 6.160992791 | 0.030435721 | 0.060857999 |
| Slc33a1 | 9.344046425 | 9.269526953 | 9.283226967 | -0.013700014 | 0.060819457 |
| Gm4045 | 3.798994749 | 3.698752626 | 3.738177621 | -0.039424995 | 0.060817128 |
| Wnk1 | 12.72474338 | 12.6301726 | 12.66396035 | -0.033787753 | 0.060783027 |
| Pgap2 | 10.09356711 | 10.14864207 | 10.03287922 | 0.115762848 | 0.06068789 |
| Slc16a2 | 9.059698036 | 9.091678619 | 8.999057754 | 0.092620866 | 0.060640282 |
| Crybb3 | 3.851786873 | 3.996791544 | 3.791197615 | 0.205593929 | 0.060589258 |
| Gm10226 | 2.756107407 | 2.7153245 | 2.695568325 | 0.019756175 | 0.060539082 |
| 4930447C04Rik | 2.128235267 | 2.075423766 | 2.06774749 | 0.007676276 | 0.060487777 |
| Mospd4 | 1.101037758 | 1.064530748 | 1.040583876 | 0.023946872 | 0.060453882 |
| Slc39a4 | 4.471848998 | 4.385576578 | 4.411450521 | -0.025873943 | 0.060398477 |
| Nt5c1a | 3.266503978 | 3.232886343 | 3.206118836 | 0.026767507 | 0.060385142 |
| Mcm7 | 10.11037308 | 10.16964126 | 10.04999644 | 0.119644823 | 0.060376643 |
| Hdac2 | 10.80947068 | 10.64080973 | 10.74912947 | -0.108319737 | 0.060341217 |
| Gm29666 | 6.148746702 | 6.038413302 | 6.088473954 | -0.050060652 | 0.060272748 |
| Maea | 10.3399992 | 10.48219981 | 10.2797818 | 0.202418017 | 0.0602174 |
| Trmt10a | 8.213537401 | 7.987676364 | 8.153354635 | -0.165678271 | 0.060182766 |
| Pnp2 | 5.929911101 | 5.842068656 | 5.869743912 | -0.027675256 | 0.060167189 |
| Nr1h2 | 9.604677515 | 9.799279425 | 9.544586377 | 0.254693048 | 0.060091138 |

| | | | | | |
|---|---|---|---|---|---|
| Tmem33 | 10.48694227 | 10.63022499 | 10.42685399 | 0.203371 | 0.060088283 |
| 1110008P14Rik | 8.450865337 | 8.548765911 | 8.390790541 | 0.15797537 | 0.060074796 |
| Ylpm1 | 10.18792226 | 10.12204448 | 10.12789077 | -0.005846287 | 0.06003149 |
| Il13ra2 | 6.878453188 | 6.760836495 | 6.818444524 | -0.057608029 | 0.060008664 |
| Mapk8 | 9.140221159 | 9.157081673 | 9.080220725 | 0.076860949 | 0.060000434 |
| Sdf4 | 11.53039753 | 11.37904984 | 11.47041226 | -0.091362417 | 0.059985267 |
| Zbtb17 | 8.869377656 | 9.089721346 | 8.809433846 | 0.280287501 | 0.05994381 |
| Ssh2 | 10.24353597 | 10.1183079 | 10.18364073 | -0.06533283 | 0.059895237 |
| Sar1a | 11.89206524 | 11.80742551 | 11.83217412 | -0.024748607 | 0.059891117 |
| Mrgbp | 8.609653748 | 8.56311973 | 8.549794279 | 0.013325452 | 0.059859469 |
| Boc | 9.588652986 | 9.589033661 | 9.528807136 | 0.060226525 | 0.05984585 |
| Tubg1 | 8.763646563 | 8.879327863 | 8.70381284 | 0.175515023 | 0.059833723 |
| 0610040J01Rik | 4.888396962 | 4.640409664 | 4.8285874 | -0.188177736 | 0.059809562 |
| Gm8369 | 3.080477711 | 3.05041999 | 3.020710501 | 0.029709489 | 0.05976721 |
| Il17b | 2.228872899 | 2.228234878 | 2.169143505 | 0.059091373 | 0.059729393 |
| Setd1b | 9.701489372 | 10.07576851 | 9.64176695 | 0.434001563 | 0.059722422 |
| Dapk3 | 8.429187398 | 8.594641413 | 8.369478805 | 0.225162608 | 0.059708593 |
| Dnah17 | 4.461884154 | 4.55970476 | 4.402201284 | 0.157503476 | 0.05968287 |
| Klra14-ps | 1.539520924 | 1.459459281 | 1.47989312 | -0.02043384 | 0.059627804 |
| Cbx4 | 8.909988214 | 9.129355763 | 8.850437597 | 0.278918166 | 0.059550617 |
| Usp17la | 3.580845133 | 3.569719891 | 3.521316084 | 0.048403808 | 0.059529049 |
| Rap2b | 9.779358808 | 9.908744212 | 9.719844811 | 0.188899401 | 0.059513997 |
| Immt | 11.56157938 | 11.45023365 | 11.50207658 | -0.05184293 | 0.059502797 |
| Edc4 | 9.831788974 | 9.901298224 | 9.772327038 | 0.128971186 | 0.059461936 |
| Mad1l1 | 8.696186834 | 8.985302193 | 8.636747249 | 0.348554944 | 0.059439585 |
| Psmb7 | 11.53639596 | 11.39871046 | 11.47710243 | -0.078391977 | 0.059293523 |
| Agpat4 | 10.57440933 | 10.36160654 | 10.51512575 | -0.153519207 | 0.05928358 |
| Dhx36 | 9.766497463 | 9.531111782 | 9.70723161 | -0.176119828 | 0.059265854 |
| Phlda2 | 3.594052231 | 3.648825377 | 3.534795769 | 0.114029608 | 0.059256462 |
| Bod1 | 10.40150292 | 10.23110037 | 10.34226533 | -0.111164953 | 0.059237597 |
| Gm7232 | 1.029163974 | 0.986153885 | 0.970005149 | 0.016148736 | 0.059158825 |
| Tsen15 | 7.692631513 | 7.545067467 | 7.633522978 | -0.088455511 | 0.059108535 |

| | | | | | |
|---|---|---|---|---|---|
| S100a6 | 13.21120168 | 12.99912288 | 13.15209581 | -0.15297293 | 0.05910587 |
| Dnah6 | 2.138535075 | 2.122999736 | 2.079588373 | 0.043411363 | 0.058946701 |
| Cdc20 | 8.678185709 | 8.54683022 | 8.619250609 | -0.072420389 | 0.0589351 |
| Zdhhc6 | 9.64640288 | 9.61120971 | 9.58753508 | 0.02367463 | 0.0588678 |
| Rpsa-ps2 | 1.45112445 | 1.398882843 | 1.392321147 | 0.006561696 | 0.058803303 |
| Gm5176 | 2.192923615 | 2.179861092 | 2.134129259 | 0.045731833 | 0.058794356 |
| Vasn | 9.565106618 | 9.931424408 | 9.506348696 | 0.425075712 | 0.058757922 |
| Vps9d1 | 8.900993549 | 9.110495914 | 8.842255908 | 0.268240006 | 0.058737641 |
| Ubqln1 | 11.9031518 | 11.97407049 | 11.84443587 | 0.12963462 | 0.05871593 |
| Flnb | 11.54932752 | 11.67485434 | 11.49066867 | 0.184185667 | 0.05865885 |
| Ash2l | 9.929029953 | 9.78704695 | 9.870388993 | -0.083342043 | 0.058640959 |
| Rps18 | 13.49804842 | 13.4653874 | 13.43958322 | 0.025804173 | 0.0584652 |
| Jag1 | 10.43287295 | 10.33763637 | 10.37445602 | -0.03681965 | 0.058416937 |
| Hsd17b14 | 7.107298628 | 7.080913305 | 7.048925478 | 0.031987827 | 0.05837315 |
| Ubash3a | 4.639747917 | 4.645278726 | 4.581408942 | 0.063869784 | 0.058338975 |
| Trp63 | 12.60023006 | 12.43291023 | 12.54189536 | -0.108985133 | 0.058334697 |
| Nefh | 2.164481349 | 2.1734292 | 2.106146996 | 0.067282203 | 0.058334353 |
| Plxnb2 | 12.16821646 | 12.2036748 | 12.10988671 | 0.093788087 | 0.05832975 |
| Tctn3 | 7.226658935 | 7.164694843 | 7.168351658 | -0.003656815 | 0.058307278 |
| Hspa1a | 9.702928867 | 12.90919604 | 9.644666408 | 3.264529629 | 0.058262459 |
| Cdc23 | 9.795564836 | 9.710370414 | 9.737315599 | -0.026945185 | 0.058249237 |
| Vars2 | 8.251451513 | 8.270048116 | 8.193212984 | 0.076835132 | 0.058238528 |
| Zfp595 | 7.202506044 | 7.158523573 | 7.144290695 | 0.014232878 | 0.058215349 |
| Ncbp3 | 9.734542957 | 9.634497328 | 9.676332933 | -0.041835605 | 0.058210024 |
| Cplx1 | 2.038558404 | 2.101015835 | 1.98038617 | 0.120629665 | 0.058172234 |
| Cep152 | 7.530068022 | 7.601945568 | 7.471898205 | 0.130047362 | 0.058169817 |
| Uvrag | 9.438676809 | 9.558391623 | 9.380545959 | 0.177845664 | 0.05813085 |
| Gtf2h3 | 9.117887823 | 9.096579653 | 9.059765067 | 0.036814586 | 0.058122756 |
| Chchd5 | 7.441222808 | 7.365090208 | 7.383100482 | -0.018010274 | 0.058122326 |
| Ctnna1 | 12.64956369 | 12.59537937 | 12.59146119 | 0.00391818 | 0.058102497 |
| Plekhs1 | 2.440382982 | 2.431929947 | 2.382293975 | 0.049635973 | 0.058089007 |
| Ccdc162 | 4.372713451 | 4.416672464 | 4.314652397 | 0.102020067 | 0.058061054 |

| | | | | | |
|---|---|---|---|---|---|
| Card6 | 6.997481034 | 6.759719703 | 6.939437538 | -0.179717834 | 0.058043497 |
| Gnl3l | 9.878088497 | 9.757642112 | 9.820070952 | -0.06242884 | 0.058017545 |
| Clpx | 10.39648741 | 10.23246318 | 10.3384788 | -0.10601562 | 0.058008603 |
| Sugt1 | 10.11016755 | 9.970676147 | 10.05216098 | -0.081484828 | 0.058006572 |
| Rpap1 | 9.105186168 | 9.223558036 | 9.047194655 | 0.17636338 | 0.057991513 |
| Nrde2 | 8.138055677 | 8.138918474 | 8.080068188 | 0.058850287 | 0.05798749 |
| Plekha8 | 8.562217438 | 8.410770609 | 8.504230541 | -0.093459932 | 0.057986897 |
| Ythdf3 | 10.82961254 | 10.78777221 | 10.77171541 | 0.016056793 | 0.05789713 |
| Igsf21 | 3.007855199 | 2.977448467 | 2.9499803 | 0.027468167 | 0.057874898 |
| Arsj | 4.99726197 | 4.933208757 | 4.939420905 | -0.006212148 | 0.057841065 |
| Rap1a | 10.47537415 | 10.35256841 | 10.41755103 | -0.064982617 | 0.05782312 |
| Zbtb18 | 9.597385473 | 9.80400219 | 9.539577095 | 0.264425095 | 0.057808379 |
| Wac | 10.85522239 | 10.89061566 | 10.79746596 | 0.093149703 | 0.05775643 |
| Gad2 | 1.501552791 | 1.467514823 | 1.443808974 | 0.023705849 | 0.057743818 |
| Gpha2 | 3.161759861 | 3.114131177 | 3.10410468 | 0.010026497 | 0.057655181 |
| Guca2b | 2.251356039 | 2.249503082 | 2.193702512 | 0.05580057 | 0.057653527 |
| Btbd11 | 9.078351718 | 9.122017624 | 9.020728354 | 0.10128927 | 0.057623364 |
| Ssna1 | 9.31975006 | 9.34441893 | 9.262135368 | 0.082283562 | 0.057614692 |
| Gspt2 | 5.584348682 | 5.469598185 | 5.52682203 | -0.057223845 | 0.057526652 |
| Olfr767 | 4.719116401 | 4.75213728 | 4.661647108 | 0.090490172 | 0.057469294 |
| Gm17195 | 3.165784964 | 3.152640301 | 3.108382532 | 0.044257769 | 0.057402432 |
| Vps26a | 10.98939543 | 10.99260655 | 10.93216406 | 0.06044249 | 0.057231367 |
| Zscan12 | 7.123511564 | 7.064115611 | 7.06636485 | -0.002249239 | 0.057146714 |
| Gpatch2 | 8.458951056 | 8.375620336 | 8.401850809 | -0.026230473 | 0.057100247 |
| Armc1 | 9.503957161 | 9.371464925 | 9.446886777 | -0.075421852 | 0.057070384 |
| B4galnt2 | 3.220112018 | 3.160196457 | 3.163045146 | -0.002848689 | 0.057066872 |
| Tmem192 | 8.519819932 | 8.37386416 | 8.462763969 | -0.088899809 | 0.057055963 |
| Amn1 | 8.06889004 | 7.620840965 | 8.011858498 | -0.391017533 | 0.057031542 |
| Sox12 | 6.809930217 | 6.80095845 | 6.752935564 | 0.048022886 | 0.056994653 |
| Slit3 | 10.57818459 | 10.49226418 | 10.52125062 | -0.028986443 | 0.056933973 |
| Glrx3 | 11.18939127 | 11.05471802 | 11.13254571 | -0.077827693 | 0.056845557 |
| Gm18113 | 2.671782177 | 2.681880208 | 2.614938154 | 0.066942053 | 0.056844023 |

| | | | | | |
|---|---|---|---|---|---|
| Map3k20 | 11.47633842 | 11.38144423 | 11.41951787 | -0.038073637 | 0.056820553 |
| Olfr1349 | 1.744270225 | 1.71083342 | 1.687451792 | 0.023381628 | 0.056818433 |
| Olfr118 | 2.289715886 | 2.270112572 | 2.232928261 | 0.037184311 | 0.056787625 |
| Rab1a | 12.4289324 | 12.3590046 | 12.37220551 | -0.01320091 | 0.056726893 |
| Ptprcap | 4.012158103 | 3.962983608 | 3.955468279 | 0.007515328 | 0.056689824 |
| Ube2n-ps1 | 8.522904554 | 8.478752142 | 8.466216298 | 0.012535844 | 0.056688256 |
| Ptk7 | 10.33376933 | 10.46761002 | 10.27708576 | 0.190524257 | 0.056683567 |
| Grasp | 6.509992462 | 6.461193827 | 6.453309701 | 0.007884126 | 0.056682761 |
| Ddx43 | 2.549908241 | 2.554591515 | 2.493267126 | 0.061324389 | 0.056641115 |
| Acan | 1.59018868 | 1.554590181 | 1.53355578 | 0.021034401 | 0.0566329 |
| Cd8a | 3.649217952 | 3.578177856 | 3.592589004 | -0.014411149 | 0.056628948 |
| Prps1l3 | 8.178658581 | 8.157372842 | 8.122034221 | 0.035338622 | 0.05662436 |
| Dlgap1 | 4.583886203 | 4.579036822 | 4.527276792 | 0.05176003 | 0.056609411 |
| Chordc1 | 10.34074926 | 10.68021958 | 10.2841865 | 0.396033077 | 0.05656276 |
| Pip4k2b | 9.980090513 | 9.939389417 | 9.923568783 | 0.015820634 | 0.056521729 |
| BC005624 | 9.429016236 | 9.336514202 | 9.372551401 | -0.036037199 | 0.056464835 |
| Zdhhc5 | 11.57399959 | 11.57504685 | 11.51760438 | 0.057442463 | 0.056395207 |
| Gpalpp1 | 9.872533062 | 9.712364153 | 9.816150878 | -0.103786726 | 0.056382184 |
| Nagpa | 8.599581845 | 8.634580185 | 8.54324649 | 0.091333696 | 0.056335355 |
| Gm20498 | 5.819228743 | 5.648849076 | 5.762905324 | -0.114056248 | 0.056323419 |
| Rad23a | 10.58163831 | 10.54150189 | 10.52533791 | 0.016163977 | 0.056300403 |
| Rnase12 | 2.034493805 | 1.993638127 | 1.978253511 | 0.015384615 | 0.056240293 |
| H2afj | 9.261426145 | 9.327367996 | 9.205223157 | 0.122144839 | 0.056202988 |
| Gm14324 | 3.613625786 | 3.562250041 | 3.557442241 | 0.0048078 | 0.056183545 |
| Gm2606 | 2.246300708 | 2.220227214 | 2.190203369 | 0.030023845 | 0.056097339 |
| Fkbp2 | 9.348331202 | 9.363004142 | 9.292237223 | 0.070766919 | 0.056093979 |
| Mrpl23-ps1 | 9.187983318 | 9.070410221 | 9.131895325 | -0.061485104 | 0.056087994 |
| Trpm6 | 5.581806851 | 5.434534672 | 5.525726489 | -0.091191817 | 0.056080362 |
| Klrb1a | 2.551211266 | 2.48348517 | 2.495220314 | -0.011735144 | 0.055990953 |
| Myg1 | 8.561604108 | 8.506624964 | 8.50561953 | 0.001005434 | 0.055984579 |
| Lrrc41 | 9.908196975 | 10.04885478 | 9.852221604 | 0.196633172 | 0.05597537 |
| Mtx2 | 9.586239979 | 9.483477364 | 9.530274186 | -0.046796822 | 0.055965794 |

| | | | | | |
|---|---|---|---|---|---|
| Padi2 | 9.21181989 | 8.975479298 | 9.155876772 | -0.180397475 | 0.055943117 |
| Qsox1 | 11.22558663 | 11.1968417 | 11.16965731 | 0.027184387 | 0.055929317 |
| Ubxn1 | 10.79356697 | 10.78865226 | 10.7376423 | 0.051009963 | 0.05592467 |
| Vps35l | 9.809488994 | 9.706440837 | 9.753592453 | -0.047151617 | 0.05589654 |
| Gm44041 | 1.510599762 | 1.476706731 | 1.454712853 | 0.021993878 | 0.055886909 |
| Gsdmcl2 | 1.669332465 | 1.617366143 | 1.613450628 | 0.003915515 | 0.055881837 |
| Rab3gap2 | 10.13847731 | 10.22527095 | 10.08263552 | 0.142635423 | 0.055841787 |
| Zfp553 | 8.017653142 | 7.962456936 | 7.961816938 | 0.000639998 | 0.055836203 |
| Smdt1 | 9.968320074 | 9.840941432 | 9.912496921 | -0.071555489 | 0.055823153 |
| Rabl3 | 7.473277334 | 7.396273587 | 7.417458535 | -0.021184948 | 0.055818799 |
| Olfr152 | 3.424445218 | 3.448389587 | 3.368646535 | 0.079743052 | 0.055798683 |
| A630023A22Rik | 3.999169186 | 4.008559715 | 3.943399801 | 0.065159914 | 0.055769385 |
| Gm48054 | 4.26778433 | 4.211063608 | 4.21202252 | -0.000958912 | 0.05576181 |
| Tomm7 | 10.11587008 | 9.973943726 | 10.06011116 | -0.08616743 | 0.055758922 |
| E430018J23Rik | 6.499881584 | 6.346213606 | 6.444123512 | -0.097909906 | 0.055758072 |
| Gm48048 | 1.488214385 | 1.424365441 | 1.432469404 | -0.008103963 | 0.055744981 |
| Scarf1 | 6.32979394 | 6.238552096 | 6.274057741 | -0.035505644 | 0.055736199 |
| Sag | 5.320790592 | 5.215727303 | 5.265055306 | -0.049328004 | 0.055735286 |
| Fetub | 4.768938422 | 4.638341234 | 4.713203262 | -0.074862028 | 0.055735161 |
| Usp19 | 11.16931554 | 11.31361717 | 11.11359354 | 0.20002363 | 0.055722007 |
| Olfr351 | 1.690516611 | 1.589876903 | 1.634807724 | -0.044930821 | 0.055708888 |
| Flt4 | 7.98095145 | 7.991339091 | 7.925306646 | 0.066032446 | 0.055644804 |
| Slbp | 9.613813058 | 9.605672684 | 9.558188513 | 0.047484171 | 0.055624544 |
| Cnfn | 10.91346669 | 11.15384488 | 10.85785444 | 0.295990447 | 0.055612253 |
| Gm18190 | 4.45995216 | 4.605951508 | 4.404404647 | 0.201546861 | 0.055547513 |
| C2cd5 | 8.812445277 | 8.76024575 | 8.757079534 | 0.003166216 | 0.055365743 |
| Phf6 | 8.731101466 | 8.701607433 | 8.675777693 | 0.02582974 | 0.055323773 |
| Jrkl | 8.144022914 | 7.926101915 | 8.088706865 | -0.16260495 | 0.05531605 |
| Tmem69 | 7.993955763 | 7.857528528 | 7.938688365 | -0.081159837 | 0.055267398 |
| Ino80c | 9.648740976 | 9.57192694 | 9.593488589 | -0.021561649 | 0.055252387 |
| Dppa3 | 2.333077667 | 2.274252017 | 2.277840389 | -0.003588372 | 0.055237278 |
| Gm47773 | 1.440193101 | 1.429226496 | 1.384971921 | 0.044254576 | 0.05522118 |

| | | | | | |
|---|---|---|---|---|---|
| Ap4s1 | 8.684206401 | 8.548861711 | 8.628989105 | -0.080127394 | 0.055217296 |
| Gorab | 7.807510552 | 7.600898261 | 7.7522993 | -0.151401039 | 0.055211251 |
| Ndufb9 | 11.17797596 | 11.0386012 | 11.12277629 | -0.084175083 | 0.055199673 |
| Spr-ps1 | 2.522604944 | 2.506169789 | 2.467522884 | 0.038646905 | 0.05508206 |
| Dhx35 | 7.835884374 | 7.942082433 | 7.780905878 | 0.161176555 | 0.054978496 |
| Spib | 4.387237199 | 4.500043873 | 4.332333653 | 0.16771022 | 0.054903546 |
| AA414768 | 4.865529927 | 4.762619823 | 4.810644277 | -0.048024454 | 0.05488565 |
| Hist1h2ap | 5.383117389 | 5.812919111 | 5.328257528 | 0.484661583 | 0.054859861 |
| Lepr | 9.144266492 | 8.819165839 | 9.089424471 | -0.270258631 | 0.054842021 |
| Ifngr2 | 10.29908368 | 10.17197816 | 10.24426113 | -0.072282967 | 0.054822557 |
| Tnxb | 12.64443233 | 12.71622855 | 12.58963845 | 0.1265901 | 0.054793883 |
| Ociad1 | 11.3041304 | 11.12854486 | 11.24936228 | -0.120817423 | 0.05476812 |
| Cd300e | 2.442750647 | 2.391516552 | 2.388097701 | 0.00341885 | 0.054652946 |
| Pgls | 9.099848374 | 9.265605272 | 9.045196061 | 0.220409211 | 0.054652313 |
| Nup35 | 8.327933766 | 8.372645328 | 8.273289483 | 0.099355845 | 0.054644283 |
| Vps33b | 8.575157927 | 8.457918103 | 8.520525602 | -0.062607499 | 0.054632325 |
| Npy | 1.811039687 | 1.800722401 | 1.75644654 | 0.044275861 | 0.054593148 |
| Gm12789 | 2.010705408 | 1.967704838 | 1.956121377 | 0.011583461 | 0.054584032 |
| Scamp2 | 10.75399287 | 10.76671434 | 10.69942089 | 0.067293457 | 0.054571983 |
| Olfr107 | 2.339687337 | 2.31074869 | 2.285269577 | 0.025479113 | 0.05441776 |
| Gm19684 | 3.710771448 | 3.661569415 | 3.656424963 | 0.005144451 | 0.054346485 |
| Atr | 8.946449526 | 8.78723197 | 8.892148662 | -0.104916692 | 0.054300864 |
| Ilkap | 9.585506493 | 9.639080653 | 9.531294803 | 0.10778585 | 0.05421169 |
| D17H6S53E | 8.421642786 | 8.377027873 | 8.367470449 | 0.009557424 | 0.054172337 |
| Snx3 | 11.02423778 | 10.99144133 | 10.97008098 | 0.021360353 | 0.054156807 |
| Ddx19b | 9.697944053 | 9.490233324 | 9.643793076 | -0.153559752 | 0.054150976 |
| Syndig1 | 2.354686203 | 2.308109438 | 2.300536888 | 0.00757255 | 0.054149315 |
| Fiz1 | 8.640724492 | 8.788522779 | 8.586633656 | 0.201889123 | 0.054090837 |
| Lrtm2 | 2.71001502 | 2.658305796 | 2.655950146 | 0.00235565 | 0.054064874 |
| Asl | 8.044080873 | 8.120426655 | 7.990095116 | 0.130331539 | 0.053985757 |
| Zfp809 | 8.006435513 | 8.059727357 | 7.952465119 | 0.107262238 | 0.053970395 |
| Mrpl9 | 9.725459557 | 9.648163476 | 9.671502756 | -0.02333928 | 0.053956801 |

| | | | | | |
|---|---|---|---|---|---|
| Dars2 | 8.363755228 | 8.206620037 | 8.309873883 | -0.103253846 | 0.053881345 |
| Lct | 1.965185936 | 1.858580105 | 1.911334574 | -0.052754469 | 0.053851362 |
| mt-Co2 | 16.16276705 | 16.03626489 | 16.1089348 | -0.07266991 | 0.053832253 |
| Kpna1 | 10.95706551 | 10.98175174 | 10.90329763 | 0.078454107 | 0.053767877 |
| Maz | 10.55281999 | 10.72979393 | 10.49907455 | 0.230719377 | 0.053745443 |
| Nox1 | 2.73877289 | 2.731956142 | 2.685122628 | 0.046833514 | 0.053650263 |
| Gm20388 | 11.0397524 | 11.01655773 | 10.98611775 | 0.03043998 | 0.053634643 |
| 4932438H23Rik | 3.01030573 | 2.850195016 | 2.95668738 | -0.106492364 | 0.05361835 |
| Poldip3 | 10.88653829 | 10.92471682 | 10.83293765 | 0.091779177 | 0.05360064 |
| Mid1-ps1 | 3.052355495 | 3.307403663 | 2.998890159 | 0.308513504 | 0.053465337 |
| Gm18194 | 4.858698578 | 4.882676105 | 4.805258241 | 0.077417864 | 0.053440337 |
| AC124512.1 | 2.135758181 | 2.184947465 | 2.082322215 | 0.10262525 | 0.053435966 |
| Commd2 | 8.801981912 | 8.658912866 | 8.748553605 | -0.089640739 | 0.053428307 |
| Bco2 | 5.903438247 | 5.546474495 | 5.850013431 | -0.303538936 | 0.053424816 |
| Casc1 | 2.940794175 | 2.889078449 | 2.887385797 | 0.001692652 | 0.053408377 |
| Iffo2 | 12.6115301 | 12.55031726 | 12.55815108 | -0.00783382 | 0.053379023 |
| Msl1 | 11.12519658 | 11.02179496 | 11.07186209 | -0.050067133 | 0.053334487 |
| Ncoa6 | 10.22187135 | 10.20736274 | 10.16854058 | 0.038822163 | 0.053330767 |
| Pigx | 8.186675878 | 8.112205635 | 8.133356182 | -0.021150547 | 0.053319696 |
| Mrps14 | 9.53224343 | 9.398070343 | 9.478925471 | -0.080855129 | 0.053317958 |
| Ube2v1 | 11.40763407 | 11.44107458 | 11.35437611 | 0.086698463 | 0.053257957 |
| Cdc73 | 9.900889511 | 9.825335989 | 9.84767126 | -0.022335271 | 0.053218251 |
| Impg2 | 3.995631389 | 3.987249755 | 3.942430592 | 0.044819163 | 0.053200797 |
| Rpgrip1 | 4.303821252 | 4.169169243 | 4.250638689 | -0.081469446 | 0.053182564 |
| Tmem209 | 9.181247452 | 9.053646731 | 9.128073706 | -0.074426975 | 0.053173746 |
| Gm7645 | 2.089095237 | 2.000039824 | 2.035982506 | -0.035942681 | 0.053112731 |
| Emilin1 | 7.976818392 | 8.256538504 | 7.923737294 | 0.33280121 | 0.053081098 |
| Pced1b | 5.899867052 | 5.689487045 | 5.846854739 | -0.157367694 | 0.053012313 |
| Eif3e | 11.70502377 | 11.60415326 | 11.65201626 | -0.047862993 | 0.05300751 |
| Snurf | 2.479384344 | 2.396922484 | 2.426399211 | -0.029476727 | 0.052985133 |
| Wrap73 | 7.594178218 | 7.728597967 | 7.541200661 | 0.187397305 | 0.052977557 |
| Rpl35a | 12.69467863 | 12.52021692 | 12.6417532 | -0.12153628 | 0.052925433 |

| | | | | | |
|---|---|---|---|---|---|
| Olfr720 | 1.384935156 | 1.331102099 | 1.332061254 | -0.000959155 | 0.052873902 |
| Atxn1l | 10.08526963 | 9.983184592 | 10.03251771 | -0.04933312 | 0.052751916 |
| Cpox | 9.923994363 | 10.03817184 | 9.871303889 | 0.166867951 | 0.052690474 |
| Lrrc40 | 8.525351367 | 8.306167363 | 8.472692042 | -0.166524679 | 0.052659325 |
| Gm14049 | 1.512590362 | 1.501155751 | 1.459943722 | 0.041212029 | 0.05264664 |
| Zfp345 | 2.06429572 | 2.019723949 | 2.011663434 | 0.008060515 | 0.052632286 |
| Setdb1 | 10.09009244 | 10.04641399 | 10.03748479 | 0.008929203 | 0.052607651 |
| Slc30a2 | 3.589070967 | 3.467106684 | 3.536503611 | -0.069396928 | 0.052567356 |
| Polr2k | 8.114424531 | 7.955697694 | 8.061976618 | -0.106278924 | 0.052447913 |
| Gm48132 | 6.703126536 | 6.456285369 | 6.650687211 | -0.194401842 | 0.052439325 |
| Mrfap1 | 12.15671072 | 12.1256062 | 12.10433126 | 0.02127494 | 0.05237946 |
| Fgfr3 | 11.00300503 | 11.16927703 | 10.95063233 | 0.218644707 | 0.052372707 |
| Hs2st1 | 9.555660269 | 9.507028796 | 9.503442538 | 0.003586258 | 0.052217731 |
| Garem2 | 2.368041517 | 2.336103144 | 2.315828676 | 0.020274468 | 0.052212841 |
| Hsd17b10 | 9.823882139 | 9.6663473 | 9.771740758 | -0.105393458 | 0.052141381 |
| Cep104 | 9.343749629 | 9.406490794 | 9.291629031 | 0.114861763 | 0.052120598 |
| Necab1 | 3.045372069 | 3.019855747 | 2.993274766 | 0.026580981 | 0.052097304 |
| Ndufs3 | 9.555667221 | 9.440988431 | 9.503647117 | -0.062658686 | 0.052020104 |
| Urgcp | 9.122104309 | 9.281844199 | 9.070104029 | 0.21174017 | 0.05200028 |
| Cops7b | 8.387715278 | 8.395857971 | 8.335720363 | 0.060137608 | 0.051994914 |
| Gm21897 | 7.205669653 | 7.062039279 | 7.153714661 | -0.091675382 | 0.051954992 |
| Glyr1 | 11.2916106 | 11.29141515 | 11.23966963 | 0.051745513 | 0.05194097 |
| Rbmx | 9.735831934 | 9.65934874 | 9.683903151 | -0.024554411 | 0.051928783 |
| Ube2a | 9.285336226 | 9.294644305 | 9.233411048 | 0.061233256 | 0.051925178 |
| Gps2 | 9.182828722 | 9.245156489 | 9.130956921 | 0.114199568 | 0.0518718 |
| Gbp8 | 5.229317063 | 5.201996832 | 5.17745247 | 0.024544361 | 0.051864593 |
| Mrpl28 | 9.160211811 | 9.144112274 | 9.108379611 | 0.035732663 | 0.0518322 |
| Olfr1260 | 2.927090855 | 2.914115383 | 2.875286708 | 0.038828676 | 0.051804148 |
| Cdkl1 | 2.48862768 | 2.425797797 | 2.436851783 | -0.011053986 | 0.051775897 |
| Pacsin2 | 10.59969078 | 10.73051211 | 10.54795359 | 0.182558523 | 0.05173719 |
| Scnn1b | 7.129839194 | 7.170983365 | 7.078152579 | 0.092830786 | 0.051686615 |
| Tmem202 | 3.685170671 | 3.649834662 | 3.633489696 | 0.016344966 | 0.051680975 |

| | | | | | |
|---|---|---|---|---|---|
| Celsr1 | 10.6381622 | 10.89840289 | 10.58650749 | 0.311895397 | 0.05165471 |
| Vmn1r-ps14 | 1.9722189 | 1.982070018 | 1.920585156 | 0.061484863 | 0.051633745 |
| Cast | 12.45004571 | 12.37269706 | 12.39842162 | -0.025724563 | 0.051624083 |
| Gmpr2 | 7.901190508 | 7.813294177 | 7.849602309 | -0.036308131 | 0.051588199 |
| Trpm1 | 5.568133265 | 5.48121835 | 5.516579979 | -0.035361629 | 0.051553286 |
| Gm12070 | 4.237420805 | 4.292125895 | 4.185946397 | 0.106179497 | 0.051474408 |
| Zfp382 | 4.933087016 | 4.921484692 | 4.881627614 | 0.039857078 | 0.051459402 |
| Garem1 | 8.732198904 | 8.66856873 | 8.680770694 | -0.012201964 | 0.051428211 |
| Aunip | 5.589594839 | 5.572769547 | 5.538247116 | 0.03452243 | 0.051347723 |
| Rfc3 | 8.28530718 | 8.312538083 | 8.234113482 | 0.078424601 | 0.051193697 |
| Amn | 2.521812488 | 2.561225149 | 2.470687218 | 0.090537931 | 0.05112527 |
| Sgsm3 | 8.18740461 | 8.332898467 | 8.136311471 | 0.196586996 | 0.051093139 |
| Dph7 | 7.495027245 | 7.316267854 | 7.44395308 | -0.127685226 | 0.051074165 |
| Gm20900 | 1.085409895 | 1.054486533 | 1.034341313 | 0.02014522 | 0.051068581 |
| Mgl2 | 10.49331953 | 10.01344086 | 10.4422811 | -0.428840236 | 0.051038437 |
| Erp27 | 2.243597307 | 2.216644836 | 2.192570313 | 0.024074523 | 0.051026994 |
| Arhgap31 | 9.671967231 | 9.523252892 | 9.620975697 | -0.097722805 | 0.050991535 |
| Zfpl1 | 8.418510732 | 8.507673348 | 8.367525173 | 0.140148175 | 0.050985559 |
| Gabra2 | 2.305047588 | 2.269585485 | 2.254070949 | 0.015514536 | 0.050976639 |
| Mycbp2 | 11.95170505 | 11.96323606 | 11.90079525 | 0.062440803 | 0.050909793 |
| Clec4a2 | 8.555236424 | 8.488741697 | 8.504374455 | -0.015632758 | 0.050861969 |
| Dpf2 | 10.06882522 | 10.09229348 | 10.0180048 | 0.07428868 | 0.050820417 |
| Vmn1r201 | 2.978948537 | 2.97244597 | 2.928131099 | 0.044314871 | 0.050817437 |
| Oacyl | 2.576201498 | 2.453359053 | 2.525442579 | -0.072083526 | 0.050758919 |
| Rab3c | 3.350664385 | 3.277690425 | 3.299935354 | -0.02224493 | 0.050729031 |
| Prpsap1 | 9.627614016 | 9.727449513 | 9.576938069 | 0.150511444 | 0.050675947 |
| Unc119b | 9.467572918 | 9.515406019 | 9.416960478 | 0.098445541 | 0.050612439 |
| Rtca | 8.325060155 | 8.184875448 | 8.274476647 | -0.089601199 | 0.050583508 |
| Rpl21 | 13.28703669 | 13.11406951 | 13.23646416 | -0.122394653 | 0.05057253 |
| Hmgb1-ps5 | 2.669941649 | 2.691829729 | 2.619370025 | 0.072459704 | 0.050571624 |
| Unc45a | 10.54247582 | 10.7501491 | 10.49193451 | 0.258214583 | 0.05054131 |
| Armc8 | 9.545022769 | 9.344413375 | 9.494546549 | -0.150133174 | 0.05047622 |

| | | | | | |
|---|---|---|---|---|---|
| Gtf2a2 | 9.444430953 | 9.404288155 | 9.394057053 | 0.010231102 | 0.0503739 |
| 9130019O22Rik | 6.351363329 | 6.299372245 | 6.301056637 | -0.001684392 | 0.050306692 |
| Gm48800 | 2.860876963 | 2.854065106 | 2.81057851 | 0.043486596 | 0.050298453 |
| Olfr1220 | 2.554031028 | 2.504724614 | 2.503745494 | 0.000979121 | 0.050285534 |
| Bag2 | 7.492169978 | 7.488357787 | 7.441957479 | 0.046400308 | 0.050212499 |
| Syt4 | 2.021042768 | 2.062648082 | 1.970885793 | 0.09176229 | 0.050156975 |
| Zfp97 | 7.921963506 | 7.786904019 | 7.871855212 | -0.084951193 | 0.050108294 |
| Zdhhc21 | 10.4533031 | 10.38150567 | 10.40321684 | -0.021711177 | 0.050086253 |
| Pde1a | 7.785436611 | 7.38624079 | 7.73535934 | -0.349118549 | 0.050077272 |
| Slc7a6 | 10.80468867 | 10.77118295 | 10.75463716 | 0.016545787 | 0.050051503 |
| Cpa6 | 4.665651126 | 4.692351911 | 4.615605281 | 0.07674663 | 0.050045845 |
| Coq2 | 9.841146781 | 9.730335065 | 9.791113779 | -0.060778715 | 0.050033002 |
| Sav1 | 10.29076298 | 10.20941726 | 10.24073567 | -0.03131841 | 0.050027307 |
| Prom2 | 11.80076223 | 11.89649808 | 11.75075941 | 0.14573867 | 0.05000282 |
| Atxn2 | 9.609288969 | 9.756783298 | 9.559311441 | 0.197471857 | 0.049977527 |
| Tctex1d2 | 6.733169731 | 6.609948765 | 6.683231434 | -0.073282669 | 0.049938297 |
| Pygo2 | 9.943055525 | 10.10063124 | 9.893119521 | 0.207511716 | 0.049936004 |
| Ints8 | 9.354148778 | 9.294101534 | 9.304216827 | -0.010115293 | 0.049931951 |
| Rundc3a | 4.09673612 | 4.108235874 | 4.046830016 | 0.061405858 | 0.049906104 |
| Gm14685 | 1.460513403 | 1.515911432 | 1.410629092 | 0.105282339 | 0.049884311 |
| Gm7143 | 4.254633049 | 4.31382037 | 4.204792036 | 0.109028334 | 0.049841013 |
| Gm12693 | 3.578374376 | 3.354160894 | 3.528628126 | -0.174467232 | 0.04974625 |
| Naa80 | 8.771212051 | 9.055940634 | 8.721471649 | 0.334468985 | 0.049740402 |
| Gm49273 | 6.793055713 | 7.574011707 | 6.743324245 | 0.830687462 | 0.049731469 |
| Tmem138 | 6.744822876 | 6.824349732 | 6.695117439 | 0.129232293 | 0.049705437 |
| Trbj2-3 | 1.746597691 | 1.742953073 | 1.696907454 | 0.046045618 | 0.049690236 |
| Nudcd3 | 10.81076668 | 10.61718086 | 10.76107946 | -0.1438986 | 0.049687223 |
| Pdhx | 9.096802289 | 8.968878053 | 9.047162034 | -0.078283981 | 0.049640255 |
| Rpl6 | 13.7411979 | 13.67110685 | 13.69158101 | -0.02047416 | 0.049616883 |
| Dnajc3 | 11.24757905 | 11.34689662 | 11.19796732 | 0.1489293 | 0.04961173 |
| Amz1 | 7.194320467 | 7.442760981 | 7.144709359 | 0.298051622 | 0.049611107 |
| Chst5 | 1.426527991 | 1.407891459 | 1.377005008 | 0.03088645 | 0.049522983 |

| | | | | | |
|---|---|---|---|---|---|
| 1700123O20Rik | 8.863092289 | 8.772803112 | 8.813575236 | -0.040772124 | 0.049517053 |
| A3galt2 | 3.42617596 | 3.422664703 | 3.37668899 | 0.045975713 | 0.04948697 |
| Zbtb24 | 7.981810184 | 8.015917815 | 7.932376604 | 0.083541211 | 0.04943358 |
| Dhrs2 | 1.537539293 | 1.526559702 | 1.488108998 | 0.038450704 | 0.049430295 |
| D6Ertd527e | 5.583284447 | 5.628224338 | 5.534026909 | 0.09419743 | 0.049257539 |
| Smim26 | 6.790287939 | 6.718070336 | 6.741036273 | -0.022965937 | 0.049251666 |
| Suv39h1 | 8.479497198 | 8.482916497 | 8.430263835 | 0.052652663 | 0.049233363 |
| Zbtb45 | 7.598084258 | 7.677203738 | 7.548883034 | 0.128320704 | 0.049201224 |
| Spag7 | 9.560274048 | 9.459716938 | 9.511077886 | -0.051360948 | 0.049196162 |
| Vps8 | 8.791616759 | 8.833727726 | 8.742428049 | 0.091299677 | 0.04918871 |
| Sptlc1 | 10.50155391 | 10.36201763 | 10.45239079 | -0.090373163 | 0.04916312 |
| Zfp560 | 6.720020694 | 6.719666838 | 6.670903874 | 0.048762964 | 0.04911682 |
| Gm42517 | 3.354798143 | 3.367555234 | 3.30571583 | 0.061839404 | 0.049082314 |
| Trim26 | 9.797505694 | 9.694292227 | 9.748432829 | -0.054140602 | 0.049072866 |
| Bcdin3d | 6.975392267 | 6.924810599 | 6.926335244 | -0.001524645 | 0.049057023 |
| Cd200 | 9.947779653 | 9.688788736 | 9.898736877 | -0.209948142 | 0.049042776 |
| Brf2 | 6.997696634 | 6.890358712 | 6.948699913 | -0.0583412 | 0.048996722 |
| Cep41 | 6.686953686 | 6.687956219 | 6.637987361 | 0.049968857 | 0.048966324 |
| Rbbp5 | 9.522904612 | 9.506911745 | 9.474029668 | 0.032882076 | 0.048874943 |
| Csk | 10.21576243 | 10.18864419 | 10.16689063 | 0.021753553 | 0.048871797 |
| AC122861.1 | 1.930167461 | 1.934470781 | 1.881350433 | 0.053120347 | 0.048817027 |
| Gm9974 | 1.985754562 | 1.966973942 | 1.936947699 | 0.030026243 | 0.048806862 |
| Rpl34 | 12.5332189 | 12.42463139 | 12.48443154 | -0.059800153 | 0.048787357 |
| Cog8 | 9.430318087 | 9.40705288 | 9.381574966 | 0.025477913 | 0.048743121 |
| Gm6682 | 2.78258579 | 2.827490754 | 2.733885302 | 0.093605452 | 0.048700488 |
| Pacsin1 | 2.478022012 | 2.418005097 | 2.429365575 | -0.011360478 | 0.048656437 |
| Gm18997 | 1.840864419 | 1.783964259 | 1.792213817 | -0.008249558 | 0.048650603 |
| Zswim5 | 2.641829575 | 2.63127089 | 2.593229157 | 0.038041733 | 0.048600418 |
| Psmc3 | 11.54930074 | 11.58905839 | 11.50070778 | 0.088350613 | 0.04859296 |
| Limd1 | 11.3425236 | 11.39066665 | 11.29394674 | 0.09671991 | 0.048576863 |
| Samd8 | 9.895359215 | 9.825697531 | 9.84681052 | -0.021112989 | 0.048548695 |
| Ormdl2 | 9.675961502 | 9.57472707 | 9.627518659 | -0.052791589 | 0.048442844 |

| | | | | | |
|---|---|---|---|---|---|
| Mkln1 | 10.63505323 | 10.55780734 | 10.58662329 | -0.02881595 | 0.048429943 |
| Cyp2b23 | 4.867119649 | 4.817873871 | 4.818721869 | -0.000847998 | 0.048397781 |
| Pcyt2 | 9.753772305 | 9.916015276 | 9.705435168 | 0.210580108 | 0.048337137 |
| Gm7514 | 1.057131761 | 1.076943885 | 1.008809042 | 0.068134843 | 0.048322718 |
| Ceacam3 | 2.718516422 | 2.6634769 | 2.670295276 | -0.006818375 | 0.048221146 |
| Gm6904 | 2.865549241 | 2.746863298 | 2.817402426 | -0.070539127 | 0.048146815 |
| Ttc9c | 9.445427823 | 9.531134663 | 9.39728968 | 0.133844984 | 0.048138144 |
| Psg25 | 2.525419436 | 2.490114763 | 2.477324524 | 0.012790239 | 0.048094911 |
| Timd4 | 5.950979136 | 5.629129527 | 5.902920712 | -0.273791185 | 0.048058424 |
| Fam193a | 9.368019437 | 9.419849613 | 9.3199652 | 0.099884413 | 0.048054237 |
| Sos2 | 9.978483422 | 9.756629473 | 9.930434303 | -0.17380483 | 0.04804912 |
| Gm19426 | 2.740159735 | 2.698422597 | 2.692126542 | 0.006296055 | 0.048033193 |
| Cd247 | 6.425173627 | 6.251362512 | 6.377143395 | -0.125780884 | 0.048030231 |
| Prcc | 9.63026134 | 9.632008336 | 9.582265594 | 0.049742742 | 0.047995747 |
| Pigl | 6.762981346 | 6.821414629 | 6.715003404 | 0.106411225 | 0.047977941 |
| Trappc2l | 8.552177775 | 8.378122742 | 8.504382185 | -0.126259443 | 0.04779559 |
| Dclk2 | 5.277294106 | 5.409048671 | 5.229498807 | 0.179549865 | 0.0477953 |
| Apoc4 | 3.92748505 | 3.871123065 | 3.879737058 | -0.008613993 | 0.047747992 |
| Stt3b | 11.8408558 | 11.7894426 | 11.79312152 | -0.00367892 | 0.04773428 |
| Topors | 9.455891091 | 9.366847155 | 9.408231963 | -0.041384808 | 0.047659128 |
| Kif18b | 7.209914072 | 7.306482873 | 7.162320285 | 0.144162588 | 0.047593787 |
| Rap1gap | 7.005247668 | 7.148781923 | 6.957666235 | 0.191115689 | 0.047581433 |
| Psmb1 | 11.55464204 | 11.44888631 | 11.50707525 | -0.058188937 | 0.04756679 |
| Dctn1 | 10.85034086 | 11.10913906 | 10.80281728 | 0.30632178 | 0.04752358 |
| Acaa1b | 7.492215422 | 7.632391417 | 7.44470591 | 0.187685506 | 0.047509512 |
| Ost4 | 9.920621003 | 9.898306546 | 9.873125163 | 0.025181383 | 0.04749584 |
| Lipt1 | 5.974703924 | 5.966118562 | 5.927216907 | 0.038901654 | 0.047487017 |
| Cyren | 6.963971087 | 7.024640719 | 6.916540814 | 0.108099905 | 0.047430273 |
| Vmn2r94 | 2.46929114 | 2.407190324 | 2.42186655 | -0.014676226 | 0.047424591 |
| Olfr145 | 1.24646024 | 1.210461548 | 1.199035717 | 0.011425831 | 0.047424522 |
| Olfr1124 | 1.791359055 | 1.748382433 | 1.7439662 | 0.004416234 | 0.047392856 |
| Rpl10-ps3 | 13.57821818 | 13.52426662 | 13.53083125 | -0.006564623 | 0.04738693 |

| | | | | | |
|---|---|---|---|---|---|
| Gm10420 | 2.258407547 | 2.199877914 | 2.211030607 | -0.011152693 | 0.047376939 |
| Psma8 | 2.131310709 | 2.10573618 | 2.083942402 | 0.021793778 | 0.047368307 |
| Gm6451 | 3.186738751 | 3.102174492 | 3.139423703 | -0.037249211 | 0.047315047 |
| Gatc | 8.794830426 | 8.727264336 | 8.747572108 | -0.020307772 | 0.047258318 |
| Rabl6 | 10.72362241 | 10.78622054 | 10.67640643 | 0.109814103 | 0.047215977 |
| Gm11991 | 1.507197855 | 1.486194063 | 1.45999026 | 0.026203802 | 0.047207595 |
| Nol4l | 6.520303258 | 6.660271072 | 6.473118746 | 0.187152325 | 0.047184512 |
| Lemd2 | 9.898749886 | 10.04529564 | 9.851579288 | 0.193716352 | 0.047170598 |
| Tcea1 | 11.24775846 | 11.16450777 | 11.2006284 | -0.03612063 | 0.047130063 |
| Rusc1 | 8.430739133 | 8.543072809 | 8.383627353 | 0.159445456 | 0.04711178 |
| Gm14730 | 2.363255357 | 2.367358301 | 2.316158351 | 0.051199951 | 0.047097006 |
| Atl1 | 6.250916994 | 5.969473115 | 6.203881399 | -0.234408284 | 0.047035595 |
| Loxl3 | 6.791107023 | 6.736339322 | 6.744082987 | -0.007743665 | 0.047024037 |
| Gm17018 | 7.521077641 | 7.333764665 | 7.474101501 | -0.140336836 | 0.04697614 |
| Agtrap | 9.081221017 | 9.203727892 | 9.034295353 | 0.169432539 | 0.046925664 |
| Urm1 | 9.177273762 | 9.21837033 | 9.130359862 | 0.088010468 | 0.0469139 |
| Wdr31 | 6.552186358 | 6.482424954 | 6.505283062 | -0.022858109 | 0.046903295 |
| Zc3h11a | 12.07961375 | 12.08776118 | 12.03280789 | 0.054953287 | 0.04680586 |
| Acd | 8.571608734 | 8.646405715 | 8.524826326 | 0.121579389 | 0.046782408 |
| Catsper2 | 5.445878664 | 5.484792441 | 5.399104311 | 0.08568813 | 0.046774353 |
| Tmbim4 | 9.354003418 | 9.219658017 | 9.307234789 | -0.087576772 | 0.046768629 |
| Ywhaq | 11.52272423 | 11.44519257 | 11.47596764 | -0.030775077 | 0.046756583 |
| Ndufa4 | 12.01540865 | 11.87801507 | 11.96868905 | -0.09067398 | 0.0467196 |
| Gm6061 | 2.564102219 | 2.516186547 | 2.517383298 | -0.001196751 | 0.046718921 |
| Lsm8 | 9.171614519 | 9.022604371 | 9.124916633 | -0.102312262 | 0.046697886 |
| Gm2897 | 2.033080793 | 2.065729019 | 1.986386707 | 0.079342312 | 0.046694086 |
| Ak5 | 3.713016892 | 3.631624282 | 3.66641411 | -0.034789828 | 0.046602782 |
| Vmn1r218 | 5.375618938 | 5.474626488 | 5.329046712 | 0.145579776 | 0.046572226 |
| Crmp1 | 3.250719895 | 3.119205725 | 3.204185936 | -0.084980211 | 0.046533958 |
| Lpcat2 | 9.277800137 | 9.137871019 | 9.23132164 | -0.093450621 | 0.046478497 |
| Ap5b1 | 6.246402377 | 6.496586748 | 6.199936399 | 0.296650349 | 0.046465978 |
| Cox6c | 11.43834184 | 11.34885461 | 11.39194628 | -0.043091673 | 0.046395563 |

| | | | | | |
|---|---|---|---|---|---|
| Cngb3 | 2.931690947 | 2.768600724 | 2.885345407 | -0.116744683 | 0.04634554 |
| Gm11960 | 4.022218292 | 3.948222981 | 3.975875154 | -0.027652173 | 0.046343138 |
| Gm11225 | 1.813213619 | 1.778260232 | 1.766901093 | 0.011359139 | 0.046312526 |
| Hsbp1 | 10.54141976 | 10.45391295 | 10.49516436 | -0.041251413 | 0.046255397 |
| Mst1r | 9.756496241 | 10.02431391 | 9.710245606 | 0.314068307 | 0.046250635 |
| Dynlrb1 | 10.62051678 | 10.49781812 | 10.57439305 | -0.076574923 | 0.04612373 |
| Lrrc7 | 3.257526993 | 3.149462907 | 3.211495239 | -0.062032332 | 0.046031754 |
| Dvl3 | 9.800827846 | 9.948561114 | 9.754799991 | 0.193761123 | 0.046027854 |
| Dcakd | 8.833898637 | 8.928093323 | 8.787924094 | 0.140169229 | 0.045974542 |
| 2310034G01Rik | 2.448304626 | 2.412391011 | 2.402342121 | 0.010048891 | 0.045962505 |
| Zfp385c | 2.216999095 | 2.182298129 | 2.171062164 | 0.011235966 | 0.045936932 |
| 2610528J11Rik | 7.46155096 | 7.544993427 | 7.415632377 | 0.129361049 | 0.045918583 |
| Farp1 | 8.744479502 | 8.768815754 | 8.698577255 | 0.070238499 | 0.045902247 |
| Zbtb41 | 9.414683682 | 9.194420591 | 9.368803555 | -0.174382964 | 0.045880127 |
| Gm2004 | 3.871768206 | 3.940218291 | 3.825938206 | 0.114280085 | 0.04583 |
| Mrps33 | 8.99806556 | 8.82097621 | 8.952265606 | -0.131289396 | 0.045799954 |
| Cd207 | 8.769511937 | 9.119771455 | 8.723735724 | 0.396035731 | 0.045776213 |
| Tnfaip1 | 11.15399536 | 11.09738306 | 11.10824014 | -0.010857083 | 0.04575522 |
| Serpinb5 | 13.35683965 | 13.21767094 | 13.31117848 | -0.093507543 | 0.045661167 |
| Atp5b | 14.18748096 | 14.09072179 | 14.14183948 | -0.051117683 | 0.045641487 |
| Pemt | 5.22195162 | 5.279358097 | 5.176358803 | 0.102999294 | 0.045592817 |
| Mycbpap | 2.366652982 | 2.432480218 | 2.321063727 | 0.11141649 | 0.045589254 |
| Hspb6 | 11.18860864 | 11.35108962 | 11.14304617 | 0.208043447 | 0.045562473 |
| Dpp10 | 4.525100631 | 4.539793153 | 4.479539125 | 0.060254028 | 0.045561507 |
| Gm15079 | 1.681515017 | 1.717051396 | 1.635957872 | 0.081093524 | 0.045557145 |
| Rack1 | 14.08553088 | 13.98767226 | 14.04006139 | -0.05238913 | 0.04546949 |
| Gm45902 | 6.478678493 | 6.398424179 | 6.433299086 | -0.034874907 | 0.045379407 |
| Lrmda | 5.436474438 | 5.296425578 | 5.391095258 | -0.09466968 | 0.04537918 |
| Vwa3a | 2.893058203 | 2.864411637 | 2.847685106 | 0.016726531 | 0.045373097 |
| Rptor | 9.853891375 | 10.03935526 | 9.808540038 | 0.230815226 | 0.045351337 |
| Fam216a | 7.156256177 | 7.115486642 | 7.110982802 | 0.00450384 | 0.045273376 |
| Pcdhb10 | 2.511194934 | 2.486761123 | 2.465965942 | 0.020795181 | 0.045228992 |

| | | | | | |
|---|---|---|---|---|---|
| Gm47355 | 5.209650392 | 5.149179322 | 5.164443491 | -0.015264169 | 0.045206901 |
| Sufu | 9.025255272 | 9.049527993 | 8.980098668 | 0.069429325 | 0.045156604 |
| Gm3953 | 1.795413159 | 1.760971342 | 1.750260777 | 0.010710565 | 0.045152383 |
| Vmn2r92 | 3.574491768 | 3.582624867 | 3.529360552 | 0.053264315 | 0.045131216 |
| Rftn1 | 8.190211495 | 8.407423295 | 8.145083557 | 0.262339738 | 0.045127938 |
| Accsl | 1.793776552 | 1.787660338 | 1.748660282 | 0.039000056 | 0.04511627 |
| Slc25a34 | 5.457817157 | 5.649350079 | 5.412702763 | 0.236647316 | 0.045114394 |
| Rhou | 10.56389844 | 10.41321614 | 10.51881407 | -0.10559793 | 0.04508437 |
| Olfr1310 | 3.204366655 | 3.170205497 | 3.159298336 | 0.010907161 | 0.045068319 |
| Pcdhb4 | 3.907673495 | 3.862637475 | 3.86263182 | 5.65533E-06 | 0.045041675 |
| Olfr1271 | 2.631671726 | 2.669923705 | 2.586646248 | 0.083277457 | 0.045025478 |
| Gprin2 | 10.47603001 | 10.38095072 | 10.43101835 | -0.050067627 | 0.045011657 |
| Gm47630 | 2.498256698 | 2.516074434 | 2.453255909 | 0.062818525 | 0.04500079 |
| Rap2a | 9.409574968 | 9.506999946 | 9.364664137 | 0.142335808 | 0.044910831 |
| Ralbp1 | 11.95275809 | 11.98630987 | 11.90793775 | 0.078372117 | 0.044820343 |
| Glyctk | 5.795735135 | 5.962750197 | 5.750939255 | 0.211810942 | 0.044795881 |
| Prr22 | 2.407123508 | 2.42383157 | 2.362373425 | 0.061458145 | 0.044750083 |
| Gpi1 | 12.6055983 | 12.54017307 | 12.56086913 | -0.020696063 | 0.044729167 |
| Recql | 8.926460813 | 9.082997946 | 8.881736571 | 0.201261375 | 0.044724243 |
| Olfr873 | 1.624504003 | 1.62708926 | 1.579808314 | 0.047280946 | 0.044695689 |
| Calm1 | 13.15459299 | 13.13378491 | 13.10993399 | 0.023850917 | 0.044659003 |
| Kndc1 | 4.018920776 | 4.059904469 | 3.974271365 | 0.085633104 | 0.044649411 |
| Crkl | 10.23936763 | 10.24031794 | 10.19473526 | 0.04558268 | 0.044632363 |
| Fam192a | 9.624640987 | 9.478268094 | 9.580013417 | -0.101745323 | 0.04462757 |
| Pigu | 9.268627458 | 9.272246827 | 9.224090577 | 0.04815625 | 0.044536881 |
| Dnajc30 | 7.998150731 | 7.998708809 | 7.953638839 | 0.04506997 | 0.044511892 |
| H2afz | 11.28458717 | 11.36123735 | 11.24009463 | 0.121142723 | 0.04449254 |
| Pcbp2 | 12.50689128 | 12.46156894 | 12.46246002 | -0.000891077 | 0.044431257 |
| Stk24 | 10.59429878 | 10.45947629 | 10.54991893 | -0.090442633 | 0.044379857 |
| Gm5478 | 1.061534066 | 1.032898389 | 1.017154653 | 0.015743736 | 0.044379414 |
| Tmem81 | 6.697123186 | 6.621396858 | 6.652759541 | -0.031362683 | 0.044363645 |
| Rccd1 | 7.021123077 | 6.995044409 | 6.976796369 | 0.01824804 | 0.044326708 |

| | | | | | |
|---|---|---|---|---|---|
| Aacs | 11.23660479 | 11.30683001 | 11.19232268 | 0.11450733 | 0.04428211 |
| Ppp1r7 | 9.720873232 | 9.669998191 | 9.67659728 | -0.00659909 | 0.044275952 |
| Olfr398 | 3.127660695 | 3.10177845 | 3.083401572 | 0.018376878 | 0.044259123 |
| Fam83e | 1.70907637 | 1.660571298 | 1.664902366 | -0.004331067 | 0.044174004 |
| Ffar1 | 2.30864372 | 2.255289069 | 2.264490026 | -0.009200957 | 0.044153694 |
| Olfr333-ps1 | 1.962750351 | 1.928652846 | 1.918615461 | 0.010037385 | 0.044134889 |
| Adamts15 | 7.898076255 | 7.938770069 | 7.853997518 | 0.084772551 | 0.044078737 |
| Acta2 | 11.51766549 | 11.35735476 | 11.47360612 | -0.11625136 | 0.044059373 |
| Nfyc | 9.616449608 | 9.638366886 | 9.572410174 | 0.065956711 | 0.044039434 |
| Xpnpep1 | 10.65066891 | 10.64225891 | 10.60663144 | 0.035627473 | 0.044037473 |
| Trim69 | 2.510676407 | 2.542278081 | 2.466650919 | 0.075627162 | 0.044025488 |
| Rps11-ps1 | 2.244068345 | 2.216675839 | 2.200054635 | 0.016621204 | 0.04401371 |
| Rps27a | 13.34145906 | 13.15565278 | 13.29755636 | -0.141903573 | 0.043902707 |
| Cpne7 | 5.26738137 | 5.268344257 | 5.223483825 | 0.044860432 | 0.043897546 |
| Gm4691 | 1.478482702 | 1.493243115 | 1.434666356 | 0.05857676 | 0.043816346 |
| Tmem154 | 10.69624119 | 10.67075986 | 10.65249507 | 0.018264797 | 0.04374612 |
| Gm3379 | 2.722488746 | 2.659385131 | 2.678767339 | -0.019382208 | 0.043721407 |
| Ppp3cc | 7.410860222 | 7.261822743 | 7.367139468 | -0.105316725 | 0.043720754 |
| Zfp991 | 8.516786632 | 8.437471847 | 8.473076947 | -0.0356051 | 0.043709684 |
| Gm16147 | 2.360845635 | 2.335267595 | 2.317231998 | 0.018035598 | 0.043613637 |
| Jchain | 1.963915437 | 1.922683741 | 1.920343875 | 0.002339865 | 0.043571561 |
| Ttc39a | 9.104179027 | 8.962961439 | 9.060617247 | -0.097655808 | 0.04356178 |
| Ajm1 | 3.07701999 | 3.220886762 | 3.033479594 | 0.187407169 | 0.043540397 |
| Il23r | 2.541096707 | 2.461330566 | 2.497666523 | -0.036335957 | 0.043430184 |
| Lvrn | 7.444049326 | 7.201061204 | 7.40064764 | -0.199586437 | 0.043401686 |
| Xpr1 | 9.96751129 | 9.939851901 | 9.924151636 | 0.015700265 | 0.043359653 |
| Dpy30 | 8.71445889 | 8.585127209 | 8.671115128 | -0.085987919 | 0.043343763 |
| Kcnq3 | 3.505817233 | 3.397854687 | 3.46247821 | -0.064623523 | 0.043339023 |
| Tbxas1 | 6.484055913 | 6.5652034 | 6.440752082 | 0.124451318 | 0.043303831 |
| Tpbg | 9.332181575 | 9.339222174 | 9.288889097 | 0.050333077 | 0.043292478 |
| Gm2574 | 2.890517408 | 2.890173457 | 2.847307021 | 0.042866437 | 0.043210387 |
| Gm5138 | 2.890517408 | 2.890173457 | 2.847307021 | 0.042866437 | 0.043210387 |

| | | | | | |
|---|---|---|---|---|---|
| Gm12917 | 1.864330895 | 1.808169598 | 1.821166914 | -0.012997316 | 0.043163981 |
| Ybx3 | 13.3403717 | 13.38660618 | 13.29722984 | 0.089376333 | 0.043141857 |
| Cacna2d4 | 3.053498182 | 2.946490119 | 3.010390982 | -0.063900863 | 0.0431072 |
| Camk1d | 9.713810147 | 9.561575403 | 9.670792807 | -0.109217405 | 0.043017339 |
| Me2 | 8.500164422 | 8.325645178 | 8.457165216 | -0.131520039 | 0.042999206 |
| Ctbp1 | 10.98449425 | 11.14734436 | 10.94156384 | 0.205780517 | 0.042930413 |
| Eif4enif1 | 10.1925129 | 10.21015547 | 10.14960328 | 0.060552197 | 0.042909627 |
| A130010J15Rik | 7.91732583 | 7.877661289 | 7.87458564 | 0.00307565 | 0.04274019 |
| Gm12338 | 8.634333807 | 8.445583427 | 8.591597971 | -0.146014544 | 0.042735836 |
| Naif1 | 6.935079351 | 7.027618486 | 6.892354793 | 0.135263693 | 0.042724557 |
| Gm3883 | 4.013471267 | 4.078378226 | 3.970753262 | 0.107624964 | 0.042718005 |
| Ttyh3 | 9.212350305 | 9.525064345 | 9.169767546 | 0.355296799 | 0.042582759 |
| Bex2 | 1.569190298 | 1.568725241 | 1.526702877 | 0.042022364 | 0.04248742 |
| Sf3b2 | 11.76271878 | 11.76258104 | 11.72027747 | 0.042303567 | 0.04244131 |
| Selenoh | 7.699447463 | 7.827657445 | 7.657024127 | 0.170633318 | 0.042423335 |
| Frrs1l | 2.245771174 | 2.191586404 | 2.203377837 | -0.011791433 | 0.042393337 |
| Paxip1 | 9.1529336 | 9.359107071 | 9.110604006 | 0.248503065 | 0.042329594 |
| Mthfs | 6.921425143 | 6.838844659 | 6.879109702 | -0.040265043 | 0.042315441 |
| Fam205a4 | 3.233718105 | 2.975542638 | 3.191456967 | -0.215914329 | 0.042261138 |
| Vps50 | 9.686014071 | 9.713991167 | 9.643799081 | 0.070192086 | 0.042214989 |
| Pcsk7 | 9.541187533 | 9.64040258 | 9.49898531 | 0.14141727 | 0.042202223 |
| Cox8b | 8.734688279 | 8.818671879 | 8.692493862 | 0.126178017 | 0.042194417 |
| Tcrg-C1 | 6.888598433 | 6.719415151 | 6.846418373 | -0.127003222 | 0.042180059 |
| Olfr1322 | 1.948116348 | 1.990853686 | 1.905973671 | 0.084880014 | 0.042142677 |
| Tmem116 | 4.412136946 | 4.354505165 | 4.370001369 | -0.015496204 | 0.042135576 |
| Hyal3 | 3.517869739 | 3.426666087 | 3.475775564 | -0.049109476 | 0.042094175 |
| Pear1 | 7.489300917 | 7.493877542 | 7.44721362 | 0.046663922 | 0.042087297 |
| Olfr1438-ps1 | 2.181074008 | 2.150784124 | 2.138990052 | 0.011794071 | 0.042083956 |
| Tchp | 7.016410858 | 7.24456849 | 6.974332049 | 0.270236441 | 0.042078809 |
| Tstd1 | 3.931957546 | 3.911314236 | 3.88998025 | 0.021333987 | 0.041977297 |
| Csmd2 | 2.036413213 | 2.065189786 | 1.99445296 | 0.070736826 | 0.041960253 |
| Rpl17-ps10 | 1.400796498 | 1.417828508 | 1.358857063 | 0.058971446 | 0.041939435 |

| | | | | | |
|---|---|---|---|---|---|
| Gm6477 | 1.85751013 | 1.831923138 | 1.81565914 | 0.016263998 | 0.04185099 |
| Setd4 | 6.976494239 | 6.910224379 | 6.934652582 | -0.024428203 | 0.041841657 |
| Akr1b3 | 10.54210655 | 10.50252411 | 10.50030183 | 0.00222228 | 0.041804723 |
| Igsf11 | 4.699696951 | 4.737942935 | 4.657904969 | 0.080037965 | 0.041791982 |
| Zcchc4 | 7.869968682 | 7.738567305 | 7.828254039 | -0.089686735 | 0.041714643 |
| Rnf8 | 8.664941474 | 8.620724053 | 8.623248331 | -0.002524277 | 0.041693143 |
| Evpl | 12.10438998 | 12.43788912 | 12.06276293 | 0.375126197 | 0.041627053 |
| Gm15185 | 2.297459736 | 2.230743262 | 2.255892103 | -0.025148841 | 0.041567632 |
| Asgr2 | 4.044060625 | 3.921889432 | 4.002502671 | -0.080613239 | 0.041557954 |
| Uba2 | 11.06708133 | 11.00381757 | 11.02558153 | -0.021763967 | 0.041499793 |
| Gm7488 | 7.913469209 | 7.768060894 | 7.872027153 | -0.103966259 | 0.041442056 |
| Trim44 | 10.54598348 | 10.47524062 | 10.50459226 | -0.02935164 | 0.041391223 |
| Rps12-ps4 | 2.13352651 | 2.112842433 | 2.092171797 | 0.020670636 | 0.041354713 |
| Ap1s3 | 8.651682691 | 8.706220076 | 8.610416102 | 0.095803974 | 0.041266588 |
| Thap4 | 9.237046277 | 9.212342065 | 9.195799694 | 0.016542372 | 0.041246583 |
| Gm15696 | 2.699344991 | 2.764027167 | 2.658125895 | 0.105901272 | 0.041219096 |
| Zfp719 | 7.398730426 | 7.518821187 | 7.357552601 | 0.161268586 | 0.041177825 |
| E330009J07Rik | 9.719875998 | 9.580100119 | 9.678782245 | -0.098682126 | 0.041093753 |
| Gm49196 | 1.53587431 | 1.528051327 | 1.494807595 | 0.033243731 | 0.041066714 |
| Tmem41b | 10.38367312 | 10.62688195 | 10.34260687 | 0.284275083 | 0.041066247 |
| BC030867 | 6.279306878 | 6.299504594 | 6.238264691 | 0.061239903 | 0.041042188 |
| Gm13302 | 5.276382821 | 5.373063074 | 5.235402434 | 0.13766064 | 0.040980387 |
| Olfr975 | 3.164501649 | 3.191058012 | 3.123574925 | 0.067483087 | 0.040926724 |
| Ift57 | 7.42202923 | 7.190537488 | 7.381109106 | -0.190571618 | 0.040920124 |
| Clec4b2 | 1.719496762 | 1.77756122 | 1.678609381 | 0.098951839 | 0.040887381 |
| Egln2 | 9.62917594 | 9.834337615 | 9.588392513 | 0.245945103 | 0.040783428 |
| Blm | 7.122693974 | 7.140938919 | 7.081920119 | 0.0590188 | 0.040773855 |
| Raet1e | 2.657414971 | 2.691352736 | 2.616678059 | 0.074674677 | 0.040736912 |
| Zdhhc12 | 6.134288967 | 6.263737506 | 6.09356491 | 0.170172596 | 0.040724057 |
| Dusp11 | 11.40651797 | 11.48039597 | 11.36580879 | 0.11458718 | 0.040709177 |
| Ccdc153 | 3.066589093 | 3.086312071 | 3.025901475 | 0.060410596 | 0.040687618 |
| Slc12a8 | 4.054650949 | 4.003478749 | 4.013983513 | -0.010504764 | 0.040667436 |

| | | | | | |
|---|---|---|---|---|---|
| Gm3045 | 3.383814613 | 3.374762089 | 3.34322217 | 0.031539919 | 0.040592444 |
| Nono | 11.94319796 | 11.9331125 | 11.90265596 | 0.030456537 | 0.040541993 |
| Gm47102 | 5.709208283 | 5.721152978 | 5.668669401 | 0.052483576 | 0.040538882 |
| Txnl4b | 7.287699144 | 7.210376844 | 7.247185856 | -0.036809012 | 0.040513288 |
| Gabrb1 | 1.511681344 | 1.487072136 | 1.471197059 | 0.015875077 | 0.040484285 |
| Zfp105 | 5.166963387 | 5.05700461 | 5.12648468 | -0.06948007 | 0.040478708 |
| Tmem132d | 2.267103191 | 2.175804233 | 2.226663896 | -0.050859664 | 0.040439295 |
| Gjb3 | 11.08458815 | 11.20815837 | 11.04415539 | 0.16400298 | 0.040432767 |
| Cnot10 | 9.841735068 | 9.788175895 | 9.801338424 | -0.013162529 | 0.040396644 |
| Guf1 | 8.766756303 | 8.702277002 | 8.72636626 | -0.024089258 | 0.040390043 |
| 1700102P08Rik | 4.247576788 | 4.428203983 | 4.207195507 | 0.221008476 | 0.040381281 |
| Gm49274 | 3.556901043 | 3.542448575 | 3.516522918 | 0.025925657 | 0.040378125 |
| Gas2l1 | 10.21498126 | 10.4016574 | 10.17462109 | 0.227036303 | 0.04036017 |
| Gm49384 | 6.25866705 | 6.0947138 | 6.218320555 | -0.123606754 | 0.040346495 |
| Tmem179 | 1.789288067 | 1.763774297 | 1.748949656 | 0.014824641 | 0.040338411 |
| Olfr102 | 3.671644613 | 3.556250378 | 3.631407708 | -0.07515733 | 0.040236906 |
| Adam4 | 3.625962362 | 3.704075651 | 3.585734992 | 0.118340659 | 0.04022737 |
| Atoh8 | 5.712076731 | 5.752537257 | 5.671896164 | 0.080641094 | 0.040180568 |
| Fth-ps2 | 2.672815136 | 2.598799149 | 2.632656467 | -0.033857319 | 0.040158669 |
| Dmp1 | 2.725109296 | 2.788650791 | 2.684972814 | 0.103677977 | 0.040136482 |
| Sirt2 | 10.2999232 | 10.36416195 | 10.25983772 | 0.10432423 | 0.040085483 |
| Olfr212 | 1.628444428 | 1.577714627 | 1.58838066 | -0.010666033 | 0.040063767 |
| Calu | 12.46624923 | 12.43810982 | 12.42618874 | 0.011921087 | 0.040060493 |
| Rprd1b | 9.793236414 | 9.775849035 | 9.75321283 | 0.022636205 | 0.040023585 |
| Gabrb2 | 1.597385835 | 1.524837417 | 1.557398476 | -0.032561059 | 0.039987359 |
| Rpl13a | 13.82929578 | 13.78139186 | 13.78931261 | -0.00792075 | 0.039983167 |
| Dennd6b | 7.088663358 | 7.147422327 | 7.04877693 | 0.098645398 | 0.039886428 |
| Tcte3 | 2.180917743 | 2.117027815 | 2.141069895 | -0.02404208 | 0.039847848 |
| Gm14150 | 2.840584106 | 2.753089166 | 2.800761516 | -0.04767235 | 0.03982259 |
| Olfr607 | 5.115051239 | 5.136735403 | 5.07526447 | 0.061470933 | 0.039786769 |
| Gm6967 | 4.124790295 | 4.148796087 | 4.085071787 | 0.0637243 | 0.039718508 |
| Olfr740 | 2.690030581 | 2.668767154 | 2.650334916 | 0.018432238 | 0.039695664 |

| | | | | | |
|---|---|---|---|---|---|
| Bbip1 | 8.135983202 | 8.124513767 | 8.096298614 | 0.028215154 | 0.039684588 |
| Rab39 | 3.805671046 | 3.646756443 | 3.76603529 | -0.119278847 | 0.039635756 |
| Mef2d | 10.79593023 | 10.91431208 | 10.75631053 | 0.158001553 | 0.0396197 |
| Vmn1r68 | 4.071349898 | 4.101151002 | 4.031745963 | 0.069405039 | 0.039603935 |
| Fra10ac1 | 8.044692936 | 7.97942886 | 8.005103465 | -0.025674605 | 0.039589471 |
| Rprm | 1.61269061 | 1.542391835 | 1.573127552 | -0.030735717 | 0.039563058 |
| Lsamp | 6.849519642 | 6.724636687 | 6.809962295 | -0.085325609 | 0.039557347 |
| Fndc11 | 1.604032904 | 1.623890446 | 1.564497998 | 0.059392447 | 0.039534906 |
| Zbtb6 | 8.959034148 | 8.870214319 | 8.919510025 | -0.049295706 | 0.039524122 |
| Padi3 | 1.298278218 | 1.255800011 | 1.258811255 | -0.003011244 | 0.039466964 |
| Apom | 1.979580746 | 1.964996755 | 1.940158759 | 0.024837995 | 0.039421987 |
| Rbpms | 8.540313966 | 8.485047754 | 8.500978937 | -0.015931184 | 0.039335029 |
| Hipk2 | 9.798708559 | 9.704573367 | 9.75940774 | -0.054834373 | 0.039300819 |
| Senp1 | 10.09558817 | 10.01080824 | 10.05630994 | -0.045501696 | 0.039278233 |
| Gm45665 | 3.999385672 | 4.123884857 | 3.960128197 | 0.163756661 | 0.039257475 |
| Ina | 2.432449791 | 2.438086151 | 2.393195225 | 0.044890926 | 0.039254566 |
| P2rx5 | 6.753420081 | 6.656536671 | 6.71418349 | -0.057646819 | 0.039236591 |
| Ubald2 | 9.403988623 | 9.661577608 | 9.36476447 | 0.296813138 | 0.039224154 |
| Klk1 | 1.342514178 | 1.313290334 | 1.303300832 | 0.009989502 | 0.039213346 |
| Gm20721 | 2.234507883 | 2.261431219 | 2.195391073 | 0.066040146 | 0.03911681 |
| Crp | 2.556851291 | 2.514106593 | 2.517773326 | -0.003666733 | 0.039077965 |
| Mrgprx1 | 1.007500895 | 1.004474625 | 0.968435196 | 0.036039429 | 0.039065699 |
| Rgs3 | 9.007816487 | 8.748563699 | 8.968752807 | -0.220189108 | 0.03906368 |
| Acsf3 | 7.928955202 | 7.871599831 | 7.88989707 | -0.018297239 | 0.039058132 |
| Trbv13-3 | 1.034830675 | 0.981685775 | 0.995819407 | -0.014133632 | 0.039011268 |
| Olfr397 | 2.020364286 | 1.994748491 | 1.981452847 | 0.013295644 | 0.038911439 |
| Endov | 8.228108524 | 8.42040617 | 8.189206084 | 0.231200085 | 0.03890244 |
| Khnyn | 9.475624082 | 9.395969435 | 9.436791609 | -0.040822174 | 0.038832473 |
| Ltk | 2.182966586 | 2.124693268 | 2.144207694 | -0.019514426 | 0.038758891 |
| Phyhipl | 3.164532551 | 3.133690792 | 3.125820191 | 0.007870601 | 0.03871236 |
| Rab14 | 11.9557667 | 11.86494734 | 11.91706058 | -0.052113247 | 0.03870612 |
| Gskip | 8.977502858 | 8.9687402 | 8.93883615 | 0.02990405 | 0.038666708 |

| | | | | | |
|---|---|---|---|---|---|
| Fdx1 | 7.939690157 | 7.806778153 | 7.901031057 | -0.094252904 | 0.0386591 |
| Hist1h2br | 1.574645154 | 1.570654976 | 1.536067939 | 0.034587038 | 0.038577216 |
| 5031439G07Rik | 11.34655727 | 11.7780101 | 11.30799915 | 0.470010947 | 0.038558123 |
| Suz12 | 10.07717545 | 9.983464765 | 10.03865247 | -0.055187706 | 0.038522979 |
| Pigq | 10.48310494 | 10.60933887 | 10.44460276 | 0.164736113 | 0.038502187 |
| Ppp1r13l | 10.4526658 | 10.90835128 | 10.41417229 | 0.494178987 | 0.03849351 |
| Washc4 | 9.889954246 | 9.728253454 | 9.851518289 | -0.123264835 | 0.038435957 |
| Crx | 2.089885864 | 2.086533194 | 2.051536547 | 0.034996647 | 0.038349317 |
| Gm49380 | 2.085912117 | 2.045634547 | 2.047582569 | -0.001948022 | 0.038329548 |
| Disc1 | 3.383670351 | 3.377710228 | 3.345345976 | 0.032364252 | 0.038324375 |
| Epha10 | 3.972244523 | 3.915960075 | 3.933920938 | -0.017960863 | 0.038323585 |
| Gm42372 | 2.920210119 | 2.966848922 | 2.88190503 | 0.084943892 | 0.038305089 |
| Nlrc4 | 5.235541933 | 5.153308468 | 5.197313124 | -0.044004656 | 0.038228809 |
| Ccdc13 | 2.496849165 | 2.47173005 | 2.458658821 | 0.013071228 | 0.038190344 |
| Dennd1b | 9.455904512 | 9.282922498 | 9.417726755 | -0.134804257 | 0.038177757 |
| Gm8822 | 2.51023819 | 2.46402634 | 2.472088439 | -0.008062099 | 0.038149751 |
| Gm48976 | 2.339144865 | 2.364237257 | 2.301049509 | 0.063187748 | 0.038095355 |
| Guk1 | 9.141760745 | 9.195779044 | 9.103728265 | 0.092050779 | 0.03803248 |
| Cd164 | 12.69406695 | 12.5682095 | 12.65605782 | -0.08784832 | 0.03800913 |
| Glod4 | 10.30179283 | 10.16552203 | 10.26384487 | -0.098322843 | 0.037947957 |
| Prelid2 | 4.365570898 | 4.236336575 | 4.327629492 | -0.091292916 | 0.037941406 |
| Gm12286 | 4.108166728 | 4.027162307 | 4.070253776 | -0.043091469 | 0.037912952 |
| Eif2ak1 | 10.13320918 | 10.04249798 | 10.09529895 | -0.052800976 | 0.037910227 |
| Gm15737 | 2.058566095 | 2.039535653 | 2.02072565 | 0.018810003 | 0.037840445 |
| Hist1h3d | 2.49412159 | 2.521743082 | 2.456301115 | 0.065441967 | 0.037820476 |
| Ankrd55 | 4.770778769 | 4.865866973 | 4.733001556 | 0.132865417 | 0.037777213 |
| Zw10 | 9.34468041 | 9.343219267 | 9.30690782 | 0.036311447 | 0.03777259 |
| Vmn2r89 | 3.566814087 | 3.505101557 | 3.529064016 | -0.023962459 | 0.037750071 |
| Mrps16 | 9.006785695 | 8.89849024 | 8.969051747 | -0.070561507 | 0.037733948 |
| Cenps | 6.550169917 | 6.598209185 | 6.512436596 | 0.085772589 | 0.03773332 |
| Gsdma2 | 4.354634444 | 4.332326184 | 4.31697109 | 0.015355094 | 0.037663354 |
| Tbc1d5 | 9.257247225 | 9.345048149 | 9.219593475 | 0.125454674 | 0.03765375 |

| | | | | | |
|---|---|---|---|---|---|
| Rab3ip | 7.846029158 | 7.89390186 | 7.80837975 | 0.08552211 | 0.037649407 |
| Atxn7l3b | 11.31722238 | 11.21303153 | 11.27964641 | -0.066614883 | 0.037575967 |
| Scrn2 | 6.715585646 | 6.925794669 | 6.678077172 | 0.247717498 | 0.037508474 |
| Flad1 | 9.288280411 | 9.283071787 | 9.250784511 | 0.032287276 | 0.037495899 |
| Otop2 | 2.746318945 | 2.783529127 | 2.708879701 | 0.074649426 | 0.037439244 |
| Vwa5b2 | 2.584629392 | 2.665386509 | 2.547267355 | 0.118119154 | 0.037362037 |
| Vps33a | 9.928207282 | 9.937183023 | 9.890887593 | 0.04629543 | 0.037319689 |
| Hfe | 9.173569281 | 8.878395442 | 9.136259797 | -0.257864355 | 0.037309484 |
| Gm11517 | 2.682326295 | 2.699619892 | 2.645069269 | 0.054550623 | 0.037257026 |
| Lonrf3 | 6.09004727 | 7.418411264 | 6.052790715 | 1.36562055 | 0.037256555 |
| Tbpl1 | 8.628104917 | 8.60489057 | 8.590901486 | 0.013989084 | 0.037203431 |
| Kit | 8.053970986 | 7.6069997 | 8.016795307 | -0.409795608 | 0.037175679 |
| Tex38 | 1.848037846 | 1.834703016 | 1.810864293 | 0.023838723 | 0.037173553 |
| Gm38317 | 1.718588302 | 1.632915295 | 1.68142824 | -0.048512945 | 0.037160062 |
| Dhx34 | 7.55868038 | 7.773297077 | 7.521561627 | 0.251735451 | 0.037118753 |
| Mrpl40 | 8.778434009 | 8.648692679 | 8.741368377 | -0.092675697 | 0.037065632 |
| Mrpl58 | 8.390974297 | 8.268056261 | 8.353909059 | -0.085852798 | 0.037065239 |
| Src | 9.150411089 | 9.357375473 | 9.1133478 | 0.244027673 | 0.037063289 |
| Exo5 | 7.366924726 | 7.364121436 | 7.329952189 | 0.034169247 | 0.036972537 |
| Gimap1 | 5.321511349 | 5.093022113 | 5.28454853 | -0.191526417 | 0.036962819 |
| Gm5466 | 1.795316984 | 1.774116491 | 1.758354533 | 0.015761958 | 0.036962451 |
| Usp37 | 9.321134773 | 9.359954921 | 9.284280413 | 0.075674507 | 0.03685436 |
| Gkn1 | 4.496628085 | 4.478968346 | 4.459822089 | 0.019146257 | 0.036805996 |
| Olfr535 | 1.622161971 | 1.572197405 | 1.585366669 | -0.013169264 | 0.036795302 |
| Plet1 | 9.756469247 | 9.743665853 | 9.719712359 | 0.023953495 | 0.036756888 |
| Gm29036 | 2.237172622 | 2.165557698 | 2.20044938 | -0.034891682 | 0.036723242 |
| Vps51 | 9.017485789 | 8.980488571 | 8.980775684 | -0.000287114 | 0.036710105 |
| Grpel2 | 9.555178138 | 9.398490581 | 9.518499984 | -0.120009403 | 0.036678155 |
| AC154369.1 | 2.657505017 | 2.625687274 | 2.620855362 | 0.004831912 | 0.036649655 |
| R3hdm1 | 9.842854058 | 9.8207172 | 9.80632485 | 0.01439235 | 0.036529208 |
| Olfr1152 | 4.018514503 | 4.005197819 | 3.98200175 | 0.02319607 | 0.036512753 |
| Dhh | 5.995068077 | 6.062884623 | 5.95856495 | 0.104319673 | 0.036503127 |

| | | | | | |
|---|---|---|---|---|---|
| Gm17374 | 2.976497045 | 2.961128391 | 2.940018503 | 0.021109887 | 0.036478542 |
| Mlf2 | 11.50984889 | 11.55391664 | 11.47338294 | 0.0805337 | 0.036465943 |
| Nrgn | 3.67263432 | 3.704412833 | 3.63617746 | 0.068235373 | 0.03645686 |
| Rps23 | 11.71951399 | 11.58157922 | 11.68306942 | -0.101490193 | 0.036444577 |
| Spsb2 | 7.415771596 | 7.587163319 | 7.37935193 | 0.207811389 | 0.036419666 |
| B630019K06Rik | 5.371738451 | 5.341683628 | 5.335327358 | 0.00635627 | 0.036411093 |
| Gpatch11 | 8.505596204 | 8.378582028 | 8.469186356 | -0.090604328 | 0.036409848 |
| Gm12481 | 8.666489939 | 8.566312933 | 8.630097002 | -0.063784069 | 0.036392937 |
| Vmn2r3 | 3.353970228 | 3.408249843 | 3.317668048 | 0.090581795 | 0.03630218 |
| Desi1 | 10.95816073 | 10.86258085 | 10.92186004 | -0.05927919 | 0.03630069 |
| Foxf2 | 1.458456496 | 1.504016988 | 1.422162954 | 0.081854034 | 0.036293542 |
| Klrg1 | 2.714813796 | 2.601157447 | 2.678600496 | -0.077443049 | 0.0362133 |
| Ubtfl1 | 2.103094328 | 2.098048368 | 2.06688978 | 0.031158588 | 0.036204548 |
| Ipo13 | 10.24887909 | 10.25444222 | 10.21274676 | 0.041695454 | 0.036132327 |
| Wdfy2 | 10.5759752 | 10.51593136 | 10.53985141 | -0.023920047 | 0.036123793 |
| Prrg2 | 8.915202582 | 9.06772101 | 8.879085167 | 0.188635843 | 0.036117415 |
| Ammecr1l | 9.425500699 | 9.511881941 | 9.389483765 | 0.122398176 | 0.036016933 |
| Cltc | 13.1312914 | 13.19134744 | 13.0952968 | 0.096050643 | 0.0359946 |
| Lamtor5 | 9.248854786 | 9.212369898 | 9.212878261 | -0.000508363 | 0.035976525 |
| Trmt10b | 7.107402502 | 6.91505801 | 7.071460731 | -0.156402721 | 0.035941771 |
| Gm12439 | 4.392692649 | 4.35488987 | 4.356771859 | -0.001881989 | 0.03592079 |
| Xrcc1 | 8.794506201 | 8.887801324 | 8.758608858 | 0.129192466 | 0.035897343 |
| Rps6kb2 | 8.707242879 | 8.721782101 | 8.671405031 | 0.050377069 | 0.035837848 |
| Kbtbd7 | 7.907686608 | 7.836989991 | 7.871867349 | -0.034877358 | 0.035819259 |
| Ice1 | 9.916004743 | 9.857216169 | 9.880186454 | -0.022970285 | 0.035818288 |
| Ppp1ca | 11.24541149 | 11.18168323 | 11.20960412 | -0.027920887 | 0.035807373 |
| Gm9442 | 2.007574656 | 2.019588001 | 1.97180917 | 0.047778832 | 0.035765486 |
| Gm12184 | 4.598768808 | 4.692885132 | 4.563005258 | 0.129879874 | 0.03576355 |
| Gm38356 | 5.709086175 | 5.619707852 | 5.673394827 | -0.053686975 | 0.035691347 |
| Nccrp1 | 11.50604724 | 11.67596003 | 11.47038056 | 0.20557947 | 0.03566668 |
| Adgb | 3.67540301 | 3.710672611 | 3.63980878 | 0.070863831 | 0.035594231 |
| Fabp6 | 4.86414131 | 4.855920957 | 4.828627835 | 0.027293122 | 0.035513475 |

| | | | | | |
|---|---|---|---|---|---|
| Gmfb | 11.36428676 | 11.32180419 | 11.32880165 | -0.006997467 | 0.03548511 |
| Alkbh5 | 11.0626679 | 10.93327742 | 11.02719408 | -0.093916663 | 0.035473817 |
| Gm16580 | 3.625988362 | 3.627372963 | 3.590524816 | 0.036848147 | 0.035463546 |
| Ndufb4c | 9.36284023 | 9.334428422 | 9.327381954 | 0.007046468 | 0.035458276 |
| Med11 | 8.242673453 | 8.19622605 | 8.207235903 | -0.011009853 | 0.03543755 |
| Atp5f1 | 12.06534354 | 11.94683586 | 12.02992219 | -0.083086333 | 0.035421347 |
| Zfp488 | 6.351846914 | 6.431920786 | 6.316444789 | 0.115475996 | 0.035402125 |
| Olfr354 | 3.222851661 | 3.226724732 | 3.187457289 | 0.039267443 | 0.035394372 |
| Scfd1 | 9.766356698 | 9.623485718 | 9.730975574 | -0.107489855 | 0.035381124 |
| Frmpd3 | 2.810461436 | 2.839645571 | 2.77511452 | 0.064531051 | 0.035346916 |
| Bap1 | 9.940027942 | 9.858322743 | 9.90469733 | -0.046374587 | 0.035330612 |
| Mterf1a | 6.436594025 | 6.406353825 | 6.40134648 | 0.005007345 | 0.035247545 |
| Ly6d | 12.70729155 | 12.74767692 | 12.67206892 | 0.075607997 | 0.035222627 |
| Zcchc3 | 6.513486694 | 6.4949256 | 6.478266391 | 0.016659208 | 0.035220303 |
| Odf3b | 2.408226031 | 2.341330019 | 2.373057938 | -0.03172792 | 0.035168093 |
| Olfr76 | 2.72443881 | 2.729822785 | 2.68927189 | 0.040550896 | 0.03516692 |
| Azin2 | 6.34646929 | 6.410236667 | 6.311354356 | 0.098882311 | 0.035114934 |
| Pou2af1 | 1.417937191 | 1.415626177 | 1.382852887 | 0.03277329 | 0.035084304 |
| Uhmk1 | 11.03781615 | 11.02134782 | 11.00282706 | 0.018520763 | 0.034989097 |
| Scd4 | 4.416258254 | 4.332335399 | 4.381284826 | -0.048949427 | 0.034973428 |
| Cyb5r4 | 10.59677094 | 10.46627447 | 10.56180528 | -0.09553081 | 0.034965663 |
| Gm44845 | 1.656695126 | 1.606012791 | 1.621749413 | -0.015736622 | 0.034945713 |
| Vmn1r175 | 2.680240918 | 2.636011257 | 2.645325288 | -0.009314031 | 0.03491563 |
| Cdc16 | 9.962179794 | 9.758080695 | 9.927268994 | -0.1691883 | 0.0349108 |
| Dytn | 1.855071047 | 1.850482658 | 1.820199746 | 0.030282912 | 0.034871301 |
| Gm20708 | 1.735598811 | 1.725133872 | 1.700735917 | 0.024397955 | 0.034862895 |
| Cib3 | 1.865750287 | 1.817199557 | 1.830911333 | -0.013711776 | 0.034838954 |
| Cntn2 | 3.650352784 | 3.771944947 | 3.61558973 | 0.156355217 | 0.034763054 |
| Cept1 | 10.37134625 | 10.34980807 | 10.33667657 | 0.013131503 | 0.03466968 |
| Atp6v0d1 | 11.15784067 | 11.15667587 | 11.12317493 | 0.033500943 | 0.03466574 |
| Vmn1r220 | 2.078419225 | 2.07622365 | 2.043804033 | 0.032419617 | 0.034615193 |
| Magef1 | 5.895344401 | 5.857277345 | 5.860750787 | -0.003473441 | 0.034593615 |

| | | | | | |
|---|---|---|---|---|---|
| Tulp1 | 3.825414071 | 3.901487519 | 3.79082763 | 0.11065989 | 0.034586441 |
| Ap3d1 | 10.93394674 | 11.1395501 | 10.89936687 | 0.24018323 | 0.03457987 |
| Gm2808 | 1.421421214 | 1.467304408 | 1.386891056 | 0.080413352 | 0.034530158 |
| Mecp2 | 9.930564123 | 9.876247783 | 9.896067796 | -0.019820014 | 0.034496327 |
| Abhd14a | 7.721892325 | 7.898682322 | 7.6874089 | 0.211273422 | 0.034483424 |
| Gm40020 | 2.024041683 | 2.024524349 | 1.989565431 | 0.034958918 | 0.034476252 |
| Gm40881 | 1.307611986 | 1.343321809 | 1.273166651 | 0.070155157 | 0.034445334 |
| Gm18829 | 1.759684605 | 1.734333711 | 1.725250887 | 0.009082824 | 0.034433718 |
| Smco4 | 7.732636944 | 7.711284178 | 7.698216147 | 0.013068031 | 0.034420797 |
| Stub1 | 10.54689789 | 10.59051625 | 10.51260318 | 0.077913077 | 0.034294717 |
| Wipf2 | 9.99119673 | 10.09149737 | 9.956926558 | 0.134570815 | 0.034270172 |
| E2f6 | 9.004481129 | 9.0418071 | 8.970223352 | 0.071583748 | 0.034257777 |
| Hdlbp | 12.89548733 | 12.97547066 | 12.86127685 | 0.11419381 | 0.034210483 |
| Peg12 | 3.567616503 | 3.513485989 | 3.5334565 | -0.01997051 | 0.034160003 |
| Vmn1r60 | 3.027681859 | 2.969947999 | 2.993624067 | -0.023676068 | 0.034057793 |
| Gm15433 | 3.491923667 | 3.475819303 | 3.45788205 | 0.017937253 | 0.034041616 |
| Vmn1r70 | 1.671198438 | 1.679940466 | 1.637162626 | 0.04277784 | 0.034035813 |
| Mfap1b | 9.884843911 | 9.848465141 | 9.850813389 | -0.002348248 | 0.034030522 |
| Fam53a | 9.162379159 | 9.177230489 | 9.128370667 | 0.048859822 | 0.034008492 |
| Ptma | 14.42727078 | 14.27536251 | 14.39331875 | -0.117956243 | 0.033952027 |
| Trim43b | 1.624050179 | 1.625194036 | 1.590100841 | 0.035093196 | 0.033949339 |
| Ppip5k2 | 9.437832404 | 9.394147499 | 9.403962638 | -0.009815138 | 0.033869766 |
| B230307C23Rik | 6.888224581 | 6.715097797 | 6.854365266 | -0.139267469 | 0.033859315 |
| Nipal4 | 10.84439677 | 10.73996414 | 10.81056681 | -0.07060267 | 0.03382996 |
| Gm10293 | 2.361433062 | 2.347104322 | 2.327610845 | 0.019493477 | 0.033822218 |
| Cab39 | 11.80947839 | 11.69905925 | 11.7756957 | -0.076636447 | 0.033782693 |
| Ooep | 1.793613664 | 1.727156904 | 1.759865641 | -0.032708737 | 0.033748023 |
| Ankdd1b | 1.843741158 | 1.852054408 | 1.810041267 | 0.042013141 | 0.033699891 |
| Ifng | 1.247083702 | 1.178321415 | 1.213509164 | -0.035187749 | 0.033574538 |
| Gm15427 | 3.495951232 | 3.529382833 | 3.462391094 | 0.066991739 | 0.033560137 |
| Nfe2l1 | 12.89019199 | 12.86572434 | 12.85665385 | 0.009070487 | 0.033538133 |
| Ccdc61 | 6.931557147 | 6.864806945 | 6.898020548 | -0.033213603 | 0.033536599 |

| | | | | | |
|---|---|---|---|---|---|
| Gm9964 | 2.033337591 | 1.989984924 | 1.999835277 | -0.009850354 | 0.033502314 |
| Unc119 | 7.274305519 | 7.112042627 | 7.240855003 | -0.128812376 | 0.033450516 |
| Bex4 | 1.816691667 | 1.78554346 | 1.783268502 | 0.002274958 | 0.033423165 |
| Mapkapk5 | 9.634249355 | 9.633097808 | 9.600929108 | 0.0321687 | 0.033320246 |
| Prss55 | 3.190818926 | 3.203684596 | 3.157551862 | 0.046132734 | 0.033267064 |
| Gm17081 | 4.682939513 | 4.741432561 | 4.649704832 | 0.091727729 | 0.033234681 |
| CR974586.1 | 3.596433843 | 3.621876506 | 3.56320229 | 0.058674216 | 0.033231554 |
| Gm48663 | 2.860156062 | 2.76902546 | 2.827004007 | -0.057978546 | 0.033152055 |
| Zscan18 | 6.025692349 | 6.029535705 | 5.992591269 | 0.036944435 | 0.03310108 |
| Gon4l | 10.20840763 | 10.33898514 | 10.17530879 | 0.163676347 | 0.03309884 |
| 2210408I21Rik | 5.779586763 | 5.810117075 | 5.746535328 | 0.063581747 | 0.033051436 |
| Tomm40l | 9.355259037 | 9.422794723 | 9.322220082 | 0.100574642 | 0.033038955 |
| Tmem79 | 10.69020549 | 10.73890895 | 10.65717806 | 0.081730893 | 0.033027427 |
| AC116841.1 | 5.538384782 | 5.587904612 | 5.505402581 | 0.082502031 | 0.032982201 |
| Gm5616 | 2.577431891 | 2.483935861 | 2.544496099 | -0.060560238 | 0.032935792 |
| Mdk | 4.098646193 | 4.202752766 | 4.065713796 | 0.13703897 | 0.032932397 |
| Btbd1 | 10.625011 | 10.45268978 | 10.59208468 | -0.139394897 | 0.03292632 |
| Dnaic2 | 2.806057313 | 2.782755166 | 2.773239824 | 0.009515342 | 0.032817488 |
| Cwc27 | 8.262858307 | 8.115602259 | 8.230063876 | -0.114461617 | 0.032794431 |
| Fbxw11 | 10.70379358 | 10.76502528 | 10.67100018 | 0.094025107 | 0.0327934 |
| Hsd17b1 | 1.903859957 | 1.943246579 | 1.871093555 | 0.072153023 | 0.032766402 |
| Pcbp3 | 7.047528158 | 7.006509084 | 7.01482224 | -0.008313156 | 0.032705918 |
| Ap1s1 | 10.21966411 | 10.05564965 | 10.18697243 | -0.131322784 | 0.032691683 |
| Nlrp5 | 2.079884865 | 2.131161979 | 2.047207129 | 0.08395485 | 0.032677736 |
| Gm5697 | 3.464133906 | 3.45652176 | 3.431489412 | 0.025032348 | 0.032644493 |
| Atp6ap1 | 11.95209952 | 11.96437804 | 11.91950261 | 0.04487543 | 0.032596903 |
| Fgf9 | 4.188463077 | 4.027375514 | 4.155871554 | -0.12849604 | 0.032591523 |
| Lhb | 1.502020154 | 1.507957896 | 1.469468329 | 0.038489567 | 0.032551826 |
| Rnf220 | 10.06078251 | 10.05122211 | 10.02828045 | 0.022941663 | 0.03250206 |
| Atp5k | 10.65828734 | 10.56146238 | 10.62578588 | -0.0643235 | 0.03250146 |
| Kcnj13 | 1.605684251 | 1.556119534 | 1.57320619 | -0.017086656 | 0.032478062 |
| Dupd1 | 4.404680913 | 4.426580563 | 4.372215273 | 0.054365289 | 0.032465639 |

| | | | | | |
|---|---|---|---|---|---|
| Olfr656 | 1.919299155 | 1.884000786 | 1.886839867 | -0.002839081 | 0.032459288 |
| 6030408B16Rik | 2.179456425 | 2.082942249 | 2.147107998 | -0.064165749 | 0.032348427 |
| AC154766.1 | 1.253098671 | 1.272241843 | 1.220819031 | 0.051422812 | 0.032279639 |
| Pate2 | 1.82081194 | 1.732578154 | 1.78857691 | -0.055998756 | 0.03223503 |
| Pars2 | 6.680891027 | 6.653457171 | 6.648707778 | 0.004749393 | 0.032183249 |
| Gm12689 | 2.433389906 | 2.423671862 | 2.401230821 | 0.022441041 | 0.032159085 |
| Olfr1031 | 1.420517282 | 1.42980657 | 1.388365218 | 0.041441352 | 0.032152063 |
| Pdzd3 | 2.89411831 | 2.861696387 | 2.861966704 | -0.000270317 | 0.032151606 |
| Tmem82 | 4.444663246 | 4.618690251 | 4.412540854 | 0.206149398 | 0.032122393 |
| Ndufaf5 | 7.600724121 | 7.519861299 | 7.568752834 | -0.048891535 | 0.031971287 |
| Rims2 | 2.608731435 | 2.655144775 | 2.576915185 | 0.078229591 | 0.03181625 |
| Fabp7 | 1.545889441 | 1.534662379 | 1.514108063 | 0.020554316 | 0.031781377 |
| Rnf19a | 9.812027029 | 9.720951097 | 9.780278498 | -0.059327401 | 0.031748531 |
| Herpud2 | 9.852484436 | 9.776252221 | 9.820790704 | -0.044538482 | 0.031693732 |
| Washc5 | 9.822772421 | 9.934465162 | 9.791090125 | 0.143375037 | 0.031682296 |
| Gm47853 | 3.279074282 | 3.233998844 | 3.247422766 | -0.013423922 | 0.031651516 |
| Olfr137 | 3.395975452 | 3.294717595 | 3.364344952 | -0.069627357 | 0.0316305 |
| Ube4b | 10.54241964 | 10.79296213 | 10.51079013 | 0.282172003 | 0.031629517 |
| Tbx6 | 4.034499893 | 4.23207864 | 4.00287563 | 0.229203011 | 0.031624263 |
| Snapc2 | 8.126216161 | 8.218344515 | 8.094614452 | 0.123730063 | 0.031601709 |
| Vmn1r47 | 1.706773225 | 1.665177815 | 1.67521805 | -0.010040235 | 0.031555175 |
| Usp20 | 9.619974717 | 9.661322303 | 9.588497396 | 0.072824907 | 0.031477321 |
| Olfr1250 | 3.210750964 | 3.156807628 | 3.179315958 | -0.02250833 | 0.031435006 |
| Gm3512 | 1.338325272 | 1.345608888 | 1.306905143 | 0.038703745 | 0.031420129 |
| Cdk10 | 8.959264213 | 9.076925113 | 8.927853924 | 0.149071188 | 0.031410288 |
| Ndufc2 | 9.387935199 | 9.40979668 | 9.356566414 | 0.053230265 | 0.031368784 |
| Frmd8 | 9.128607619 | 9.532502814 | 9.097318932 | 0.435183882 | 0.031288687 |
| Adgra1 | 1.792203248 | 1.731325066 | 1.760919907 | -0.029594841 | 0.031283341 |
| Usp30 | 8.353205622 | 8.384789201 | 8.321936599 | 0.062852602 | 0.031269023 |
| Ugt1a1 | 10.62180867 | 10.45022912 | 10.59054616 | -0.140317047 | 0.031262507 |
| Pex12 | 7.854125636 | 7.859747092 | 7.822867811 | 0.036879281 | 0.031257825 |
| Ptger1 | 4.494431696 | 4.566257161 | 4.463182763 | 0.103074399 | 0.031248933 |

| | | | | | |
|---|---|---|---|---|---|
| Gm8355 | 2.572413412 | 2.503421893 | 2.541165674 | -0.037743781 | 0.031247737 |
| Fcer1a | 4.249625704 | 3.944459687 | 4.218508593 | -0.274048906 | 0.031117111 |
| Calr4 | 2.138736467 | 2.131645762 | 2.107624186 | 0.024021576 | 0.031112281 |
| Brd1 | 10.58718718 | 10.72210954 | 10.55607773 | 0.16603181 | 0.031109443 |
| Gm8899 | 2.416005895 | 2.465950132 | 2.384931103 | 0.081019029 | 0.031074792 |
| Phf20l1 | 10.68321363 | 10.39229033 | 10.65216525 | -0.25987492 | 0.031048383 |
| 1500009C09Rik | 2.327493354 | 2.29966892 | 2.296456843 | 0.003212077 | 0.031036511 |
| Kdm1a | 10.23501965 | 10.26080056 | 10.20401052 | 0.056790033 | 0.03100913 |
| Gm20878 | 3.692971788 | 3.807413618 | 3.662064832 | 0.145348786 | 0.030906956 |
| Gm13071 | 3.057296136 | 3.030672042 | 3.026395766 | 0.004276276 | 0.03090037 |
| Igkc | 2.976729301 | 2.944235636 | 2.945835573 | -0.001599938 | 0.030893727 |
| Shd | 2.388919263 | 2.372729661 | 2.358048228 | 0.014681433 | 0.030871036 |
| Wdr34 | 7.240583401 | 7.367213346 | 7.209763237 | 0.157450109 | 0.030820164 |
| Gm12960 | 2.278719525 | 2.256090749 | 2.247947838 | 0.008142911 | 0.030771687 |
| Creb3l2 | 9.772245757 | 9.809998747 | 9.741493846 | 0.068504901 | 0.030751911 |
| Gm12276 | 3.692337717 | 3.760156499 | 3.661592066 | 0.098564433 | 0.030745651 |
| Nrbf2 | 7.935811724 | 7.769501104 | 7.905114302 | -0.135613198 | 0.030697422 |
| Gm9544 | 3.574502386 | 3.586681769 | 3.543820649 | 0.04286112 | 0.030681737 |
| Krt13 | 1.733843667 | 1.713237453 | 1.703191551 | 0.010045902 | 0.030652116 |
| Gm10509 | 4.197319902 | 4.261227252 | 4.166722369 | 0.094504883 | 0.030597533 |
| Olfr691 | 1.911569978 | 1.838544797 | 1.880983371 | -0.042438574 | 0.030586607 |
| Msi1 | 3.283959267 | 3.219879741 | 3.253405812 | -0.033526071 | 0.030553455 |
| Tsn | 10.41063244 | 10.49881298 | 10.38012753 | 0.118685457 | 0.03050491 |
| Klrc1 | 4.845712144 | 4.725855336 | 4.815213362 | -0.089358026 | 0.030498782 |
| Vax2 | 4.119121701 | 4.094708929 | 4.088633531 | 0.006075398 | 0.03048817 |
| Gm14327 | 5.365448949 | 5.233044927 | 5.334983037 | -0.10193811 | 0.030465912 |
| Dolpp1 | 9.472855481 | 9.675598996 | 9.442414676 | 0.23318432 | 0.030440805 |
| Kctd20 | 10.88647713 | 10.85043236 | 10.85607123 | -0.00563887 | 0.030405903 |
| Myrf | 3.569214365 | 3.782057674 | 3.538853158 | 0.243204516 | 0.030361208 |
| Brsk2 | 1.867076552 | 1.873594304 | 1.836745597 | 0.036848708 | 0.030330955 |
| Fam173b | 7.931396135 | 7.976406076 | 7.901067874 | 0.075338202 | 0.030328261 |
| Ikzf4 | 6.163733765 | 6.213128734 | 6.133442277 | 0.079686457 | 0.030291488 |

| | | | | | |
|---|---|---|---|---|---|
| Vmn1r214 | 1.915602048 | 1.8721325 | 1.885325299 | -0.013192799 | 0.030276749 |
| Cops4 | 10.17550038 | 10.10725248 | 10.14525094 | -0.037998463 | 0.03024944 |
| BC030499 | 3.921237292 | 3.877261701 | 3.891013284 | -0.013751583 | 0.030224008 |
| Zfp709 | 7.355265077 | 7.265964734 | 7.325053053 | -0.059088319 | 0.030212024 |
| Nhsl2 | 8.302295873 | 8.203209682 | 8.27209171 | -0.068882027 | 0.030204163 |
| Fbxo11 | 10.68478684 | 10.71988994 | 10.65459209 | 0.06529785 | 0.030194753 |
| Gon7 | 6.377220708 | 6.388158324 | 6.347057183 | 0.041101141 | 0.030163525 |
| Olfr1028 | 4.235760758 | 4.223624841 | 4.205660588 | 0.017964253 | 0.03010017 |
| Colq | 3.516877167 | 3.413873676 | 3.486823462 | -0.072949786 | 0.030053705 |
| Olfr313 | 1.293408412 | 1.243807271 | 1.2633591 | -0.019551829 | 0.030049312 |
| Vmn1r203 | 5.960729056 | 5.93696 | 5.930683294 | 0.006276707 | 0.030045763 |
| Oxa1l | 9.770175462 | 9.635185517 | 9.740144214 | -0.104958698 | 0.030031248 |
| Tagln3 | 2.028817366 | 2.070643985 | 1.998799887 | 0.071844098 | 0.030017479 |
| Mgarp | 2.641873322 | 2.585617965 | 2.611906866 | -0.026288901 | 0.029966456 |
| Rpl17-ps9 | 4.914047063 | 4.737487185 | 4.88408541 | -0.146598225 | 0.029961653 |
| Madd | 10.07600324 | 10.0814774 | 10.04605708 | 0.035420322 | 0.029946165 |
| Ddx50 | 10.49764686 | 10.4416789 | 10.46771023 | -0.026031333 | 0.029936623 |
| Mark4 | 10.72659726 | 10.94338193 | 10.69671099 | 0.246670937 | 0.029886263 |
| Pcdhb3 | 2.855777607 | 2.840392075 | 2.825904565 | 0.01448751 | 0.029873042 |
| Cbln3 | 2.36398397 | 2.312833806 | 2.334119877 | -0.021286071 | 0.029864093 |
| Snx4 | 9.719770989 | 9.642801208 | 9.689939362 | -0.047138154 | 0.029831627 |
| BC049762 | 1.835918389 | 1.813740679 | 1.806088477 | 0.007652202 | 0.029829912 |
| Zfp217 | 10.06446814 | 10.07281065 | 10.03465158 | 0.038159073 | 0.029816565 |
| Krt5 | 16.74043301 | 16.9585547 | 16.71062004 | 0.247934667 | 0.02981297 |
| Ddx11 | 7.140433372 | 7.486070885 | 7.110659759 | 0.375411127 | 0.029773613 |
| Polr2m | 11.39548157 | 11.23145002 | 11.36571569 | -0.134265667 | 0.029765877 |
| Slc47a2 | 2.680972431 | 2.618169088 | 2.651208554 | -0.033039466 | 0.029763877 |
| Tead1 | 11.2094527 | 11.06110239 | 11.17969588 | -0.118593493 | 0.029756813 |
| Fv1 | 6.810160082 | 6.779049198 | 6.780426127 | -0.00137693 | 0.029733955 |
| Ireb2 | 10.42841322 | 10.34213511 | 10.39872291 | -0.056587803 | 0.029690313 |
| Gm9803 | 7.517611168 | 7.327113826 | 7.48800487 | -0.160891044 | 0.029606299 |
| Olfr536 | 2.951201911 | 2.922916099 | 2.921596995 | 0.001319104 | 0.029604915 |

| | | | | | |
|---|---|---|---|---|---|
| Upk3b | 4.570758379 | 4.541679964 | 4.541214166 | 0.000465798 | 0.029544213 |
| Xkr9 | 4.232987317 | 4.155113199 | 4.203449849 | -0.04833665 | 0.029537468 |
| Tcrg-C2 | 4.600053239 | 4.647167912 | 4.570540275 | 0.076627637 | 0.029512964 |
| H2-M9 | 4.554229896 | 4.497957738 | 4.524832855 | -0.026875117 | 0.029397041 |
| Gm5620 | 3.128558634 | 3.171257418 | 3.099172907 | 0.072084511 | 0.029385727 |
| Gm49051 | 1.967417119 | 1.920353819 | 1.938060717 | -0.017706898 | 0.029356402 |
| Gm14117 | 5.903284221 | 5.778933348 | 5.873978338 | -0.09504499 | 0.029305884 |
| Cdh10 | 2.037298476 | 2.032274332 | 2.008084267 | 0.024190065 | 0.029214209 |
| Olfr733 | 1.421407085 | 1.398582344 | 1.392268673 | 0.006313671 | 0.029138412 |
| Tmem181c-ps | 2.739596472 | 2.820477589 | 2.71047936 | 0.109998228 | 0.029117112 |
| Polr3a | 8.8575266 | 8.809240089 | 8.828428675 | -0.019188585 | 0.029097926 |
| Lrrc51 | 4.825145423 | 4.79583969 | 4.796058488 | -0.000218799 | 0.029086935 |
| Elp3 | 9.818408915 | 9.718427277 | 9.789330179 | -0.070902901 | 0.029078736 |
| Vps29 | 9.783006341 | 9.724945812 | 9.753936908 | -0.028991096 | 0.029069432 |
| Gm36128 | 2.345229619 | 2.322061475 | 2.316233695 | 0.00582778 | 0.028995924 |
| Gm6397 | 1.152047241 | 1.119263694 | 1.123081564 | -0.00381787 | 0.028965677 |
| Ucp2 | 12.52654677 | 12.5626956 | 12.49759905 | 0.065096557 | 0.028947723 |
| Rtl10 | 5.452766812 | 5.372397112 | 5.423830707 | -0.051433596 | 0.028936104 |
| Olfr109 | 2.51600793 | 2.527076488 | 2.4871378 | 0.039938688 | 0.028870129 |
| Gm11231 | 2.242281584 | 2.204600391 | 2.213422798 | -0.008822407 | 0.028858786 |
| Plekhm3 | 8.857097958 | 8.909195414 | 8.828242767 | 0.080952646 | 0.028855191 |
| Cdc34 | 9.96392767 | 9.870177293 | 9.93510906 | -0.064931767 | 0.02881861 |
| Pros1 | 11.20284244 | 11.07084709 | 11.17405076 | -0.10320367 | 0.02879168 |
| Olfr1254 | 1.765916466 | 1.722357613 | 1.737126691 | -0.014769077 | 0.028789775 |
| Zbtb46 | 7.02211618 | 6.903059992 | 6.993459584 | -0.090399592 | 0.028656596 |
| Rps13-ps2 | 11.08244602 | 10.41744177 | 11.05383285 | -0.636391077 | 0.028613173 |
| Fgfr4 | 6.759942848 | 7.183841619 | 6.731334785 | 0.452506835 | 0.028608064 |
| Trbj1-6 | 1.905478481 | 1.924963413 | 1.876881497 | 0.048081916 | 0.028596984 |
| Gm21762 | 0.757454967 | 0.706456973 | 0.728860202 | -0.02240323 | 0.028594765 |
| Dnah2 | 5.200790508 | 5.12742275 | 5.172237412 | -0.044814662 | 0.028553096 |
| Gm8648 | 1.270918325 | 1.240230038 | 1.242397653 | -0.002167615 | 0.028520672 |
| Agt | 8.766053882 | 8.766950336 | 8.737535924 | 0.029414412 | 0.028517958 |

| | | | | | |
|---|---|---|---|---|---|
| Zdhhc9 | 10.97502574 | 10.93153311 | 10.94651247 | -0.01497936 | 0.02851327 |
| Rasgef1a | 4.545859852 | 4.63796011 | 4.517373 | 0.12058711 | 0.028486852 |
| Olfr914 | 3.672528728 | 3.697083587 | 3.644046938 | 0.053036649 | 0.028481789 |
| Ltn1 | 10.12543389 | 10.04314673 | 10.0969835 | -0.053836767 | 0.02845039 |
| Zfp324 | 7.623934844 | 7.684817681 | 7.595508165 | 0.089309515 | 0.028426678 |
| Ttc19 | 8.175158661 | 7.990180829 | 8.146737372 | -0.156556542 | 0.02842129 |
| Gm21811 | 6.507505908 | 6.386312185 | 6.479094245 | -0.09278206 | 0.028411664 |
| Dusp10 | 7.226328828 | 7.39025749 | 7.197939213 | 0.192318277 | 0.028389615 |
| Dnajc7 | 10.95514426 | 10.93011102 | 10.92676854 | 0.003342477 | 0.028375713 |
| Hdgf | 12.81205499 | 12.82630835 | 12.78371167 | 0.042596677 | 0.028343317 |
| Rassf5 | 10.20118742 | 10.23370796 | 10.17285274 | 0.060855223 | 0.028334683 |
| Rps3a1 | 14.22104171 | 14.12553983 | 14.19271604 | -0.067176217 | 0.028325663 |
| Gm10931 | 1.925307977 | 1.991878329 | 1.896998775 | 0.094879554 | 0.028309202 |
| Nfrkb | 8.997119051 | 9.040742276 | 8.968813933 | 0.071928343 | 0.028305117 |
| Acnat1 | 2.014671081 | 1.975847642 | 1.986371576 | -0.010523934 | 0.028299505 |
| Slc35f2 | 7.010519398 | 6.807930674 | 6.982237531 | -0.174306858 | 0.028281867 |
| Mmgt1 | 10.46852692 | 10.38629543 | 10.44026221 | -0.053966783 | 0.028264707 |
| Hnrnpul1 | 11.60315768 | 11.61897012 | 11.57495124 | 0.044018877 | 0.028206433 |
| Lilra5 | 1.308928493 | 1.230256778 | 1.280747987 | -0.050491209 | 0.028180506 |
| Qars | 11.78625893 | 11.75543373 | 11.75807876 | -0.002645027 | 0.028180173 |
| CT868723.1 | 3.344449304 | 3.32199528 | 3.316287375 | 0.005707905 | 0.028161929 |
| BC107364 | 3.197622075 | 3.208286037 | 3.169482295 | 0.038803743 | 0.02813978 |
| Gpr3 | 2.117263343 | 2.23608553 | 2.089171522 | 0.146914008 | 0.028091821 |
| Btbd17 | 1.392117144 | 1.350016692 | 1.364048863 | -0.014032171 | 0.028068281 |
| Alg14 | 8.979082001 | 8.97295436 | 8.951060363 | 0.021893997 | 0.028021638 |
| Hsf2bp | 3.675790504 | 3.999972868 | 3.647785788 | 0.35218708 | 0.028004716 |
| Trim40 | 3.008168416 | 3.037423774 | 2.980173065 | 0.057250708 | 0.027995351 |
| Olfr1392 | 2.097651333 | 2.108404493 | 2.069656957 | 0.038747536 | 0.027994377 |
| N4bp2 | 8.326077118 | 8.111935544 | 8.298146314 | -0.18621077 | 0.027930805 |
| Ykt6 | 11.40783361 | 11.35023417 | 11.38000684 | -0.029772663 | 0.027826777 |
| Tprg | 9.886509215 | 9.840804594 | 9.858688093 | -0.017883499 | 0.027821122 |
| Zmat4 | 2.475166322 | 2.551703924 | 2.447377018 | 0.104326906 | 0.027789304 |

| | | | | | |
|---|---|---|---|---|---|
| Nenf | 9.019255779 | 8.869442906 | 8.991482773 | -0.122039868 | 0.027773006 |
| Unc13a | 3.304354428 | 3.219393252 | 3.276582105 | -0.057188852 | 0.027772323 |
| Erv3 | 1.568520136 | 1.527389899 | 1.540754393 | -0.013364494 | 0.027765743 |
| Tectb | 1.549243469 | 1.550015192 | 1.521489148 | 0.028526044 | 0.027754321 |
| Gdf5 | 3.391506395 | 3.460958557 | 3.363780605 | 0.097177952 | 0.02772579 |
| Spag4 | 3.211126819 | 3.221924566 | 3.183448379 | 0.038476188 | 0.02767844 |
| Ttll11 | 3.557630342 | 3.535631971 | 3.529980977 | 0.005650994 | 0.027649366 |
| Rpl27-ps1 | 5.36822004 | 5.188384165 | 5.340576118 | -0.152191953 | 0.027643922 |
| Stk32a | 2.335405646 | 2.347453827 | 2.30776888 | 0.039684947 | 0.027636766 |
| Gng12 | 11.85932078 | 11.85013567 | 11.83169188 | 0.01844379 | 0.0276289 |
| Olfr259 | 1.745782711 | 1.704744983 | 1.718162889 | -0.013417906 | 0.027619823 |
| Slc25a15 | 8.965677429 | 8.864841882 | 8.938098283 | -0.073256401 | 0.027579146 |
| Zfp965 | 5.609575126 | 5.594293392 | 5.582010266 | 0.012283126 | 0.02756486 |
| Tmem185a | 8.617140882 | 8.508061278 | 8.589599587 | -0.081538308 | 0.027541295 |
| Cdk12 | 9.887645545 | 9.909879487 | 9.860184953 | 0.049694534 | 0.027460592 |
| Rbm20 | 7.086224787 | 6.842464157 | 7.058779993 | -0.216315836 | 0.027444794 |
| Tle4 | 9.840680278 | 9.822134036 | 9.813261999 | 0.008872037 | 0.02741828 |
| Gm42742 | 8.638680017 | 8.680455877 | 8.611309949 | 0.069145928 | 0.027370068 |
| Atp5h | 11.45454672 | 11.3167744 | 11.42722186 | -0.110447457 | 0.027324863 |
| Pcdhb18 | 4.765798729 | 4.748835785 | 4.738482212 | 0.010353573 | 0.027316517 |
| Siva1 | 7.671254405 | 7.920885764 | 7.643951344 | 0.27693442 | 0.027303061 |
| Tbl2 | 9.398932812 | 9.255055821 | 9.371648687 | -0.116592866 | 0.027284125 |
| Olfr715b | 2.713494611 | 2.690173096 | 2.686226309 | 0.003946787 | 0.027268301 |
| Gm12922 | 1.655089989 | 1.655930541 | 1.627833976 | 0.028096566 | 0.027256013 |
| Sectm1b | 6.709456267 | 6.656556656 | 6.682229855 | -0.025673199 | 0.027226412 |
| Polr3k | 9.452108891 | 9.414584898 | 9.424921398 | -0.010336499 | 0.027187493 |
| 1700030C10Rik | 2.396300022 | 2.401218402 | 2.369126681 | 0.032091721 | 0.027173341 |
| Snca | 5.68390219 | 5.50632406 | 5.656730692 | -0.150406632 | 0.027171499 |
| Rsph1 | 3.877955929 | 3.896182576 | 3.850818743 | 0.045363833 | 0.027137186 |
| Coro7 | 9.005935647 | 9.161312983 | 8.97880951 | 0.182503473 | 0.027126137 |
| C1d | 9.763182188 | 9.469547912 | 9.736094176 | -0.266546264 | 0.027088012 |
| Olfr629 | 1.339048142 | 1.337781285 | 1.312049738 | 0.025731547 | 0.026998404 |

| | | | | | |
|---|---|---|---|---|---|
| Gm28041 | 1.91907141 | 1.857027715 | 1.892185795 | -0.035158079 | 0.026885616 |
| Dgcr2 | 10.89912391 | 11.02071842 | 10.87228142 | 0.148437003 | 0.026842493 |
| Taf2 | 9.405144328 | 9.349637409 | 9.378349993 | -0.028712584 | 0.026794335 |
| Pagr1a | 4.884142594 | 5.030651861 | 4.857384849 | 0.173267012 | 0.026757745 |
| Gm6641 | 1.893099705 | 1.889616583 | 1.866348936 | 0.023267647 | 0.026750769 |
| Pcnp | 10.9824128 | 10.88635051 | 10.95567684 | -0.069326327 | 0.026735957 |
| Olfr160 | 1.631455631 | 1.625069296 | 1.60478378 | 0.020285516 | 0.026671851 |
| Gm13577 | 2.414332431 | 2.315243164 | 2.38768082 | -0.072437655 | 0.026651612 |
| Erfe | 4.538537576 | 4.676048473 | 4.511891728 | 0.164156745 | 0.026645848 |
| 4933427I04Rik | 1.683713484 | 1.691884797 | 1.657112931 | 0.034771866 | 0.026600553 |
| Ccdc96 | 2.514425879 | 2.49045688 | 2.487828617 | 0.002628263 | 0.026597261 |
| Ywhab | 12.6491251 | 12.48040634 | 12.62259729 | -0.142190953 | 0.02652781 |
| Slain1 | 4.389558331 | 4.321087126 | 4.363062879 | -0.041975753 | 0.026495453 |
| Icam4 | 2.070350185 | 2.01892499 | 2.043897088 | -0.024972098 | 0.026453097 |
| Olfr1316 | 1.904314371 | 1.939046866 | 1.877921855 | 0.061125012 | 0.026392516 |
| Ufc1 | 9.978201958 | 9.856598476 | 9.951825841 | -0.095227366 | 0.026376117 |
| Cog1 | 9.132791965 | 9.177456933 | 9.106454593 | 0.07100234 | 0.026337372 |
| Ttpa | 6.462170961 | 6.386202414 | 6.435905821 | -0.049703408 | 0.02626514 |
| Mtmr6 | 10.71141772 | 10.51647275 | 10.68517679 | -0.168704037 | 0.026240927 |
| Ythdc2 | 8.267034706 | 8.110908391 | 8.240797177 | -0.129888786 | 0.026237529 |
| Gm9817 | 3.179406158 | 3.226554207 | 3.153206522 | 0.073347685 | 0.026199636 |
| Gm44715 | 4.855443595 | 4.893879244 | 4.829248548 | 0.064630696 | 0.026195047 |
| Ttpal | 10.78606321 | 10.86092151 | 10.75988483 | 0.101036677 | 0.02617838 |
| Bhmt2 | 1.462621062 | 1.446503008 | 1.436489743 | 0.010013265 | 0.026131319 |
| Gm47364 | 1.51817454 | 1.518531015 | 1.492054861 | 0.026476154 | 0.026119679 |
| 4932438A13Rik | 10.53057174 | 10.45367788 | 10.50448599 | -0.05080811 | 0.026085747 |
| 1700030J22Rik | 4.084041342 | 4.040634076 | 4.058014537 | -0.017380462 | 0.026026805 |
| Best1 | 3.698891995 | 3.746919952 | 3.67287614 | 0.074043812 | 0.026015855 |
| Gm47611 | 5.222991023 | 5.192002754 | 5.196989273 | -0.004986519 | 0.02600175 |
| Slc4a1ap | 8.921361231 | 8.6789131 | 8.895378673 | -0.216465573 | 0.025982559 |
| Dcp1b | 7.318228001 | 7.317925406 | 7.29228193 | 0.025643475 | 0.025946071 |
| Vmn2r93 | 1.599986029 | 1.580415612 | 1.574080278 | 0.006335334 | 0.025905751 |

| | | | | | |
|---|---|---|---|---|---|
| 38047 | 1.655207583 | 1.600159108 | 1.6293179 | -0.029158792 | 0.025889683 |
| Plk5 | 3.358922975 | 3.381667416 | 3.333046361 | 0.048621055 | 0.025876614 |
| Pin4 | 8.373234903 | 8.232099618 | 8.347381853 | -0.115282235 | 0.02585305 |
| Pcdh12 | 5.518028957 | 5.340960668 | 5.492187834 | -0.151227166 | 0.025841123 |
| Ikbkg | 10.02271168 | 10.23912248 | 9.996872623 | 0.242249857 | 0.025839053 |
| Erich3 | 1.792693089 | 1.82799905 | 1.766900625 | 0.061098425 | 0.025792464 |
| Gsto2 | 3.270715349 | 3.238254303 | 3.244937422 | -0.00668312 | 0.025777927 |
| Rhod | 9.144806889 | 9.221221032 | 9.119055118 | 0.102165914 | 0.025751771 |
| Fam131c | 3.289953504 | 3.384080435 | 3.264271841 | 0.119808594 | 0.025681663 |
| Grk2 | 11.52879232 | 11.61528741 | 11.50313373 | 0.112153683 | 0.02565859 |
| Zfp653 | 6.384715509 | 6.649684914 | 6.359074875 | 0.290610039 | 0.025640634 |
| Myo1a | 2.002700069 | 1.983837346 | 1.977084833 | 0.006752513 | 0.025615236 |
| Zkscan17 | 8.804396911 | 8.813394595 | 8.778784209 | 0.034610386 | 0.025612703 |
| Olfr1196 | 2.995217857 | 2.955219219 | 2.969616806 | -0.014397587 | 0.025601051 |
| Olfr843 | 1.886901411 | 1.885731086 | 1.861328306 | 0.02440278 | 0.025573105 |
| Rps15a-ps4 | 3.025132881 | 3.060577911 | 2.999569243 | 0.061008668 | 0.025563638 |
| Got1 | 10.45287364 | 10.28769026 | 10.42740973 | -0.13971947 | 0.025463907 |
| Gm10157 | 2.042053754 | 2.012292924 | 2.016628038 | -0.004335114 | 0.025425716 |
| RPS10-NUDT3 | 1.102857809 | 1.033256741 | 1.077478338 | -0.044221597 | 0.025379471 |
| Haus2 | 8.666352903 | 8.474466264 | 8.64100734 | -0.166541076 | 0.025345563 |
| Ggt7 | 4.760888563 | 4.862657715 | 4.735543311 | 0.127114404 | 0.025345252 |
| Il16 | 8.525282862 | 8.448865131 | 8.499941161 | -0.051076031 | 0.025341701 |
| Tm2d2 | 9.437753051 | 9.382267171 | 9.412445921 | -0.03017875 | 0.02530713 |
| Sobp | 6.607155569 | 6.405253899 | 6.581963593 | -0.176709693 | 0.025191976 |
| Gm3531 | 1.914192563 | 1.863122414 | 1.889029115 | -0.025906701 | 0.025163448 |
| Arfgap1 | 9.69005254 | 9.760778496 | 9.664918624 | 0.095859872 | 0.025133917 |
| Enkur | 2.914615484 | 2.90049325 | 2.889491517 | 0.011001732 | 0.025123967 |
| Olfr127 | 2.429816323 | 2.377774164 | 2.404705398 | -0.026931234 | 0.025110925 |
| Olfr1167 | 3.696684401 | 3.753988197 | 3.671608423 | 0.082379774 | 0.025075978 |
| Dip2c | 9.009090714 | 8.853085094 | 8.984042749 | -0.130957655 | 0.025047966 |
| Arpc1a | 11.10764788 | 11.06370476 | 11.08260531 | -0.018900547 | 0.025042577 |
| 9430025C20Rik | 2.165223075 | 2.172734378 | 2.140212132 | 0.032522245 | 0.025010943 |

| | | | | | |
|---|---|---|---|---|---|
| Olfr1307 | 6.404542045 | 6.365460685 | 6.379648506 | -0.014187822 | 0.024893538 |
| Tmem161b | 8.361083631 | 8.212468292 | 8.336319614 | -0.123851322 | 0.024764017 |
| Lmtk3 | 3.255849034 | 3.332504329 | 3.23110508 | 0.10139925 | 0.024743954 |
| Slc1a7 | 4.397913645 | 4.533810375 | 4.373255023 | 0.160555351 | 0.024658621 |
| Six6 | 1.776193813 | 1.79942369 | 1.751536544 | 0.047887147 | 0.02465727 |
| Aars2 | 7.58061412 | 7.841483451 | 7.555999582 | 0.285483869 | 0.024614538 |
| Pnma2 | 1.786777517 | 1.773082022 | 1.762182409 | 0.010899613 | 0.024595108 |
| Mal2 | 10.43908929 | 10.19278716 | 10.41453301 | -0.221745847 | 0.02455628 |
| Zyg11a | 4.491632522 | 4.677896312 | 4.467088911 | 0.210807401 | 0.024543612 |
| Tbc1d23 | 9.60171113 | 9.646503839 | 9.577171116 | 0.069332722 | 0.024540014 |
| Btbd16 | 3.336077469 | 3.450797457 | 3.311539347 | 0.13925811 | 0.024538122 |
| Slc15a5 | 1.588753107 | 1.527593029 | 1.564219586 | -0.036626557 | 0.024533521 |
| Tent5c | 7.285376578 | 8.534748111 | 7.260891546 | 1.273856565 | 0.024485031 |
| Gm9512 | 1.61587203 | 1.553339679 | 1.591399705 | -0.038060026 | 0.024472325 |
| Pkd1l3 | 2.66373366 | 2.67308612 | 2.63935233 | 0.03373379 | 0.02438133 |
| Tcf3 | 9.355684066 | 9.618346399 | 9.331322704 | 0.287023696 | 0.024361362 |
| Elk4 | 10.47988851 | 10.38683781 | 10.45558451 | -0.068746693 | 0.024304 |
| Sugct | 4.155525093 | 4.113327681 | 4.131227571 | -0.01789989 | 0.024297522 |
| Tbp | 8.710472651 | 8.652278675 | 8.686186073 | -0.033907398 | 0.024286578 |
| Pebp4 | 1.736124691 | 1.715588262 | 1.711868686 | 0.003719576 | 0.024256005 |
| Gm5590 | 3.486211127 | 3.643589966 | 3.461973951 | 0.181616015 | 0.024237176 |
| Gm10639 | 1.931161604 | 1.989944085 | 1.906933423 | 0.083010662 | 0.024228181 |
| Pou4f1 | 4.31706676 | 4.325709206 | 4.292840557 | 0.032868649 | 0.024226203 |
| Psma1 | 11.20676817 | 11.1605451 | 11.18263138 | -0.02208628 | 0.02413679 |
| Rbm4 | 8.603724214 | 8.531930514 | 8.579590015 | -0.047659501 | 0.024134199 |
| Ildr1 | 5.45935717 | 5.410745634 | 5.43524775 | -0.024502116 | 0.02410942 |
| Tlr12 | 2.48635078 | 2.55483366 | 2.462253598 | 0.092580062 | 0.024097181 |
| Grcc10 | 9.004466872 | 9.014612704 | 8.980407867 | 0.034204837 | 0.024059005 |
| Zfp738 | 8.52195596 | 8.493762529 | 8.497954207 | -0.004191679 | 0.024001753 |
| Gm20449 | 2.888184001 | 2.825047574 | 2.864193892 | -0.039146319 | 0.023990108 |
| Zcchc14 | 9.984759084 | 10.02576601 | 9.960770775 | 0.064995238 | 0.02398831 |
| Ndufa9 | 10.84629415 | 10.73513681 | 10.82232176 | -0.087184957 | 0.023972387 |

| | | | | | |
|---|---|---|---|---|---|
| Spin1 | 10.55695325 | 10.48376579 | 10.53299637 | -0.049230583 | 0.02395688 |
| Klhl32 | 2.793917099 | 2.785562646 | 2.769970884 | 0.015591762 | 0.023946215 |
| Pde6h | 3.986579737 | 3.92197923 | 3.962635585 | -0.040656355 | 0.023944152 |
| Creb1 | 10.31748405 | 10.35248534 | 10.29354486 | 0.058940487 | 0.023939197 |
| Ly6k | 2.206969763 | 2.145868503 | 2.183045456 | -0.037176954 | 0.023924307 |
| Hspd1-ps4 | 6.709753023 | 6.455676057 | 6.685850071 | -0.230174013 | 0.023902953 |
| Ptrhd1 | 6.917725619 | 6.83360514 | 6.893870204 | -0.060265064 | 0.023855415 |
| Pcdhga5 | 6.360188372 | 6.426893376 | 6.336346415 | 0.090546961 | 0.023841957 |
| Gm8979 | 2.587751731 | 2.574798688 | 2.563910778 | 0.01088791 | 0.023840952 |
| Ndufs5 | 9.901853685 | 9.760465672 | 9.878094436 | -0.117628765 | 0.023759249 |
| Rps19-ps4 | 1.171246528 | 1.159684432 | 1.147557505 | 0.012126926 | 0.023689023 |
| Synb | 1.829528311 | 1.832728443 | 1.805842316 | 0.026886128 | 0.023685995 |
| Rps25 | 13.11610049 | 12.99335252 | 13.09242161 | -0.099069097 | 0.02367888 |
| Olfr906 | 2.142166421 | 2.153214232 | 2.118499374 | 0.034714859 | 0.023667047 |
| Gm7696 | 3.033158834 | 3.040788248 | 3.009520819 | 0.031267429 | 0.023638015 |
| Gm47924 | 7.014584261 | 6.874047814 | 6.991003882 | -0.116956068 | 0.023580379 |
| Pdss1 | 9.032226492 | 8.88658456 | 9.008670835 | -0.122086275 | 0.023555657 |
| Haao | 5.021346674 | 4.947085261 | 4.997810731 | -0.05072547 | 0.023535943 |
| Ercc6l2 | 9.108394534 | 8.961403738 | 9.084869561 | -0.123465823 | 0.023524973 |
| Cox11 | 7.054483055 | 6.932855828 | 7.030966469 | -0.098110642 | 0.023516586 |
| Gm7701 | 1.428307825 | 1.395378321 | 1.404834977 | -0.009456656 | 0.023472848 |
| Anks1 | 9.623204461 | 9.811874521 | 9.59973336 | 0.212141161 | 0.023471101 |
| Gm26920 | 2.754003361 | 2.657502944 | 2.730562917 | -0.073059973 | 0.023440444 |
| Scgb1a1 | 3.530107974 | 3.45411873 | 3.506668664 | -0.052549934 | 0.02343931 |
| Phc1 | 7.6881452 | 7.716109754 | 7.664711497 | 0.051398257 | 0.023433703 |
| Cd160 | 2.371245717 | 2.382792518 | 2.347822328 | 0.034970191 | 0.023423389 |
| H2-T-ps | 4.748107235 | 4.817814272 | 4.724730643 | 0.093083628 | 0.023376592 |
| Pde6a | 2.279854054 | 2.249673181 | 2.256481036 | -0.006807855 | 0.023373017 |
| Med25 | 9.71570287 | 9.870285357 | 9.692334953 | 0.177950404 | 0.023367916 |
| Tdp1 | 7.365875733 | 7.383446266 | 7.342545503 | 0.040900764 | 0.023330231 |
| Mettl17 | 7.807586958 | 8.001813598 | 7.784319452 | 0.217494146 | 0.023267506 |
| Mrgpra4 | 2.904499646 | 2.86842307 | 2.881247546 | -0.012824476 | 0.0232521 |

| | | | | | |
|---|---|---|---|---|---|
| Stoml3 | 1.801281478 | 1.887986139 | 1.778118456 | 0.109867683 | 0.023163022 |
| Gm6882 | 3.10931398 | 3.14648765 | 3.086156688 | 0.060330962 | 0.023157292 |
| Vmn1r184 | 2.820364636 | 2.835351887 | 2.797219028 | 0.038132859 | 0.023145607 |
| Synpr | 3.746869127 | 3.754388458 | 3.723739872 | 0.030648587 | 0.023129255 |
| Gfra3 | 5.61770626 | 5.610113179 | 5.594585661 | 0.015527518 | 0.023120599 |
| Vmn2r73 | 3.228067466 | 3.185039267 | 3.204964563 | -0.019925296 | 0.023102902 |
| Gm11688 | 1.209876361 | 1.244234321 | 1.186783877 | 0.057450444 | 0.023092484 |
| Hmgn2-ps1 | 1.816873651 | 1.790720428 | 1.793788062 | -0.003067634 | 0.023085589 |
| Stam2 | 10.09437338 | 10.07681548 | 10.07129203 | 0.005523443 | 0.023081343 |
| Olfr1341 | 2.016155493 | 1.999006765 | 1.993088707 | 0.005918058 | 0.023066786 |
| Olfr250 | 2.631517602 | 2.61734967 | 2.608461076 | 0.008888594 | 0.023056527 |
| Gm9903 | 1.908349471 | 1.907036068 | 1.885303405 | 0.021732663 | 0.023046065 |
| Birc5 | 8.155645187 | 8.115587767 | 8.132606692 | -0.017018925 | 0.023038495 |
| Npy4r | 4.621603828 | 4.578806338 | 4.598575921 | -0.019769582 | 0.023027908 |
| Muc2 | 2.251441144 | 2.272292292 | 2.228428559 | 0.043863733 | 0.023012585 |
| Irf6 | 12.16942272 | 12.21441646 | 12.14643146 | 0.067984993 | 0.022991253 |
| Zwint | 10.04044417 | 10.22679076 | 10.01745649 | 0.209334264 | 0.022987677 |
| Acbd3 | 10.24252424 | 10.21782714 | 10.21953774 | -0.001710597 | 0.0229865 |
| Mff | 10.35874093 | 10.25314719 | 10.33576035 | -0.08261316 | 0.02298058 |
| Slc12a3 | 3.053355271 | 3.029796712 | 3.030411553 | -0.000614841 | 0.022943719 |
| Ms4a2 | 6.420195305 | 6.226210376 | 6.397264696 | -0.17105432 | 0.02293061 |
| Olfr380 | 2.843126991 | 2.777732611 | 2.820202932 | -0.04247032 | 0.022924059 |
| Tpd52l1 | 6.854209848 | 6.821546825 | 6.831285793 | -0.009738968 | 0.022924055 |
| Ccdc43 | 8.908406153 | 8.791126846 | 8.885493026 | -0.09436618 | 0.022913127 |
| Rtn1 | 5.720385103 | 5.667671041 | 5.697489266 | -0.029818226 | 0.022895837 |
| Olfr904 | 2.256703649 | 2.221004272 | 2.233810116 | -0.012805844 | 0.022893533 |
| Plekhg3 | 10.10366757 | 10.26546846 | 10.08079557 | 0.184672883 | 0.022872 |
| Sdsl | 2.890786212 | 2.840822564 | 2.8679517 | -0.027129137 | 0.022834512 |
| Lrrc73 | 2.967205902 | 3.021225967 | 2.944394646 | 0.07683132 | 0.022811256 |
| Cd22 | 4.129038812 | 4.183958305 | 4.106229733 | 0.077728571 | 0.022809078 |
| Gm17082 | 1.430048641 | 1.396837631 | 1.40726633 | -0.010428699 | 0.022782311 |
| Gm13498 | 1.777921921 | 1.725905655 | 1.755148297 | -0.029242642 | 0.022773625 |

| | | | | | |
|---|---|---|---|---|---|
| Baz2b | 10.30296385 | 10.18374672 | 10.28023 | -0.09648328 | 0.022733843 |
| Gm6793 | 2.117962154 | 2.120036467 | 2.09526231 | 0.024774157 | 0.022699844 |
| Hapln3 | 3.051588906 | 3.168596986 | 3.02890658 | 0.139690406 | 0.022682326 |
| Uqcrfs1 | 10.95958102 | 10.82452744 | 10.93695819 | -0.112430753 | 0.022622823 |
| Erich6 | 1.943014201 | 1.958532701 | 1.9204452 | 0.038087501 | 0.022569001 |
| Snx12 | 9.654901313 | 9.753370219 | 9.632369686 | 0.121000533 | 0.022531627 |
| Aak1 | 10.15939061 | 10.19241905 | 10.13686868 | 0.05555037 | 0.022521927 |
| Tnfrsf13c | 2.742035662 | 2.768551015 | 2.719535876 | 0.049015139 | 0.022499786 |
| Gm49392 | 4.237593597 | 4.108402476 | 4.215141932 | -0.106739456 | 0.022451665 |
| Tg | 1.47739851 | 1.499926433 | 1.454954509 | 0.044971924 | 0.022444001 |
| Usp43 | 6.217884845 | 6.218234524 | 6.195485547 | 0.022748977 | 0.022399298 |
| Gm8213 | 2.109659654 | 2.103743193 | 2.087293541 | 0.016449652 | 0.022366113 |
| Tsg101 | 10.54941074 | 10.48897454 | 10.52706511 | -0.038090567 | 0.022345637 |
| 1700066B19Rik | 1.711515826 | 1.689483348 | 1.689217923 | 0.000265424 | 0.022297902 |
| Gm12816 | 2.090605936 | 2.094250606 | 2.068440074 | 0.025810533 | 0.022165863 |
| Sh2b1 | 9.872607596 | 10.03450236 | 9.850448326 | 0.184054037 | 0.02215927 |
| Fbxo46 | 7.624925026 | 7.913764758 | 7.602793014 | 0.310971744 | 0.022132012 |
| Ubl5 | 10.65976561 | 10.47464505 | 10.6376494 | -0.16300435 | 0.02211621 |
| Olfr1284 | 2.847884774 | 2.845033953 | 2.825782572 | 0.019251381 | 0.022102202 |
| Tmem102 | 7.527335252 | 7.692591923 | 7.505250844 | 0.187341079 | 0.022084408 |
| Sh3bp1 | 10.08938937 | 10.49093535 | 10.06734009 | 0.423595256 | 0.022049279 |
| Vmn2r65 | 1.612300975 | 1.602119884 | 1.590265219 | 0.011854665 | 0.022035756 |
| Rgs4 | 6.140604864 | 6.015085809 | 6.118572874 | -0.103487065 | 0.02203199 |
| Kat14 | 9.29988953 | 9.359980201 | 9.277864805 | 0.082115396 | 0.022024725 |
| Acr | 1.353049638 | 1.324448585 | 1.331059689 | -0.006611104 | 0.021989949 |
| Gm17132 | 7.447047194 | 7.644002367 | 7.425059309 | 0.218943058 | 0.021987885 |
| Prss30 | 1.829981447 | 1.834704881 | 1.808018591 | 0.026686289 | 0.021962856 |
| Usf1 | 9.326336618 | 9.233343589 | 9.304460524 | -0.071116935 | 0.021876094 |
| Mslnl | 1.349492408 | 1.453117117 | 1.327693025 | 0.125424093 | 0.021799383 |
| Zbtb8os | 8.599438776 | 8.598779661 | 8.577695146 | 0.021084515 | 0.021743631 |
| Smad2 | 10.20820637 | 10.18822514 | 10.18646778 | 0.001757357 | 0.02173859 |
| Gsdmc3 | 1.610561274 | 1.629741603 | 1.588829507 | 0.040912096 | 0.021731767 |

| | | | | | |
|---|---|---|---|---|---|
| Sgk2 | 1.776258026 | 1.757136724 | 1.75453066 | 0.002606063 | 0.021727365 |
| Alkal2 | 3.62404388 | 3.518012414 | 3.602358203 | -0.084345789 | 0.021685677 |
| Pias2 | 9.132721192 | 9.024310028 | 9.111037542 | -0.086727515 | 0.021683649 |
| Mroh2a | 3.06314763 | 3.092979038 | 3.041498072 | 0.051480966 | 0.021649558 |
| Sctr | 3.955734946 | 3.927198826 | 3.934127249 | -0.006928422 | 0.021607697 |
| Cwf19l2 | 9.429226298 | 9.216427284 | 9.407624771 | -0.191197487 | 0.021601528 |
| Gm3194 | 1.559979676 | 1.494528469 | 1.538391781 | -0.043863312 | 0.021587895 |
| Dop1a | 9.512295741 | 9.42351995 | 9.490729818 | -0.067209868 | 0.021565923 |
| Olfr769 | 5.202327937 | 5.194362392 | 5.180780983 | 0.013581408 | 0.021546954 |
| Megf11 | 3.454159855 | 3.487405226 | 3.432619009 | 0.054786218 | 0.021540846 |
| Elf3 | 4.939353472 | 5.009999992 | 4.917912034 | 0.092087958 | 0.021441438 |
| Gm49530 | 3.256676222 | 3.252677503 | 3.235239992 | 0.017437511 | 0.021436229 |
| Tmem196 | 1.169372163 | 1.106085019 | 1.147991381 | -0.041906362 | 0.021380783 |
| Commd9 | 8.50424528 | 8.602146473 | 8.482888222 | 0.119258251 | 0.021357058 |
| Vkorc1l1 | 10.68362063 | 10.55473786 | 10.66227086 | -0.107533 | 0.02134977 |
| 4930431P19Rik | 4.977753343 | 4.938862326 | 4.956405128 | -0.017542802 | 0.021348215 |
| Epas1 | 11.14369033 | 10.85416192 | 11.12235392 | -0.268192003 | 0.021336413 |
| Zbtb34 | 8.228084103 | 8.081725521 | 8.206755728 | -0.125030206 | 0.021328375 |
| Nrk | 2.494776851 | 2.468794673 | 2.473490968 | -0.004696295 | 0.021285883 |
| Dscam | 2.362143841 | 2.305999915 | 2.340885774 | -0.034885859 | 0.021258067 |
| Smim4 | 6.303733747 | 6.194263472 | 6.28249835 | -0.088234878 | 0.021235397 |
| Gsdma | 11.25953343 | 11.27737577 | 11.23830954 | 0.039066233 | 0.021223893 |
| Hmox2 | 9.892279442 | 9.700029074 | 9.87106526 | -0.171036186 | 0.021214183 |
| Gm3222 | 1.913991761 | 1.865467518 | 1.892843623 | -0.027376105 | 0.021148138 |
| Slc9a4 | 3.769679619 | 3.852751895 | 3.748558798 | 0.104193097 | 0.021120821 |
| Mia | 2.340790368 | 2.451157322 | 2.319697099 | 0.131460223 | 0.02109327 |
| Tm2d1 | 8.129021316 | 7.998127146 | 8.107932217 | -0.109805071 | 0.021089099 |
| Nek11 | 2.503222485 | 2.505662767 | 2.482147778 | 0.023514989 | 0.021074707 |
| Rps19-ps3 | 3.995336913 | 3.964407871 | 3.974295678 | -0.009887807 | 0.021041235 |
| 2010003K11Rik | 5.686905008 | 5.594391272 | 5.665890967 | -0.071499695 | 0.02101404 |
| Os9 | 11.08271186 | 11.03294922 | 11.06175019 | -0.02880097 | 0.020961677 |
| Mfsd4b5 | 3.251051369 | 3.238329759 | 3.230136968 | 0.008192791 | 0.020914401 |

| | | | | | |
|---|---|---|---|---|---|
| Tmem45a2 | 2.244360592 | 2.238690448 | 2.223495276 | 0.015195171 | 0.020865316 |
| Tedc1 | 6.82343013 | 6.932169509 | 6.802579866 | 0.129589643 | 0.020850264 |
| Mdm4-ps | 1.862609301 | 1.798345721 | 1.841779927 | -0.043434206 | 0.020829374 |
| Gm12889 | 2.882287289 | 2.873158842 | 2.861461269 | 0.011697573 | 0.02082602 |
| Ncan | 2.941178908 | 2.953240671 | 2.920357942 | 0.032882728 | 0.020820966 |
| Foxd2 | 2.718019268 | 2.7195138 | 2.697199971 | 0.022313829 | 0.020819297 |
| Rprml | 2.375926063 | 2.329024164 | 2.355114393 | -0.026090229 | 0.02081167 |
| Csn1s2b | 1.630011537 | 1.618189264 | 1.609264059 | 0.008925205 | 0.020747478 |
| Gm8850 | 1.966238933 | 1.951227463 | 1.945535956 | 0.005691507 | 0.020702977 |
| Gdf3 | 4.167507835 | 4.24450399 | 4.146819748 | 0.097684241 | 0.020688087 |
| Gm5917 | 2.401904231 | 2.408560552 | 2.381255155 | 0.027305396 | 0.020649076 |
| Xpo1 | 11.22636735 | 11.26347931 | 11.20580348 | 0.057675837 | 0.02056387 |
| Mrrf | 8.356688832 | 8.372330659 | 8.336146574 | 0.036184085 | 0.020542258 |
| Snx16 | 7.745059869 | 7.800154037 | 7.724574648 | 0.075579388 | 0.020485221 |
| Sdr39u1 | 8.946174946 | 8.884396389 | 8.925742033 | -0.041345643 | 0.020432914 |
| Prkag1 | 10.20199647 | 10.0821502 | 10.18157061 | -0.099420413 | 0.020425863 |
| Gdpgp1 | 6.570406058 | 6.651620898 | 6.550030369 | 0.101590529 | 0.020375689 |
| Pax5 | 3.141745872 | 3.045541521 | 3.121376558 | -0.075835038 | 0.020369313 |
| Sap18 | 9.676319119 | 8.895781035 | 9.655950841 | -0.760169806 | 0.020368278 |
| Olfr943 | 1.778209077 | 1.808379995 | 1.757952834 | 0.050427161 | 0.020256243 |
| Fbxo16 | 2.962005065 | 3.062314086 | 2.941749543 | 0.120564543 | 0.020255522 |
| Bag6 | 11.31133675 | 11.40968284 | 11.29108718 | 0.118595663 | 0.020249573 |
| Tstd2 | 9.018994428 | 9.132463876 | 8.998751793 | 0.133712084 | 0.020242635 |
| AC122306.1 | 2.556767775 | 2.513098277 | 2.536554234 | -0.023455958 | 0.020213541 |
| Chchd6 | 7.346164889 | 7.258705945 | 7.326021236 | -0.067315291 | 0.020143653 |
| Eif2ak4 | 8.053021522 | 7.903961946 | 8.032891026 | -0.128929081 | 0.020130496 |
| Olfr714 | 1.499682362 | 1.53201681 | 1.479554728 | 0.052462081 | 0.020127634 |
| Usp4 | 11.55892116 | 11.51452747 | 11.53880255 | -0.02427508 | 0.020118613 |
| Fam69c | 3.832865172 | 3.908678853 | 3.812747691 | 0.095931162 | 0.020117481 |
| Rps6-ps3 | 1.548705506 | 1.603310007 | 1.52865978 | 0.074650226 | 0.020045726 |
| Dpf1 | 5.404878629 | 6.693040953 | 5.384842122 | 1.308198831 | 0.020036507 |
| Ppp1r1b | 1.558651922 | 1.550853954 | 1.538645101 | 0.012208853 | 0.020006821 |

| | | | | | |
|---|---|---|---|---|---|
| Adarb2 | 4.553308682 | 4.545374513 | 4.533307263 | 0.01206725 | 0.020001418 |
| Saysd1 | 7.068075069 | 6.957993087 | 7.048091072 | -0.090097985 | 0.019983997 |
| Trbv13-2 | 1.505369533 | 1.45981947 | 1.48543993 | -0.025620461 | 0.019929602 |
| Arfgef1 | 11.1010351 | 11.05305193 | 11.08114043 | -0.028088493 | 0.01989467 |
| Gm13456 | 3.239109674 | 3.120798719 | 3.219238306 | -0.098439587 | 0.019871368 |
| Gm2163 | 5.500833466 | 5.540283522 | 5.481047172 | 0.05923635 | 0.019786294 |
| Klhl36 | 5.860972356 | 5.998427716 | 5.841190069 | 0.157237647 | 0.019782287 |
| Mettl22 | 7.602759261 | 7.728463605 | 7.582980746 | 0.145482858 | 0.019778514 |
| Rpl22l1 | 10.72021912 | 10.58188013 | 10.70049545 | -0.118615313 | 0.019723677 |
| 0610012G03Rik | 8.377011625 | 8.381081828 | 8.357334274 | 0.023747553 | 0.019677351 |
| Traf3ip2 | 9.294330971 | 9.26543934 | 9.274667462 | -0.009228122 | 0.019663509 |
| Gm3272 | 3.760046875 | 3.826824747 | 3.740392323 | 0.086432424 | 0.019654551 |
| Rpl27a | 11.80662163 | 11.5800675 | 11.78697906 | -0.206911563 | 0.019642563 |
| Lypd2 | 2.553997438 | 2.486742377 | 2.534368689 | -0.047626312 | 0.019628749 |
| Gm14740 | 1.813222074 | 1.810361185 | 1.79360319 | 0.016757996 | 0.019618884 |
| Ttll9 | 1.773780085 | 1.793455692 | 1.754162863 | 0.039292828 | 0.019617222 |
| Shoc2 | 10.48866965 | 10.37446302 | 10.46907356 | -0.09461054 | 0.019596087 |
| Pcf11 | 10.7832716 | 10.85473415 | 10.76369226 | 0.091041897 | 0.019579347 |
| Ccdc107 | 7.473096889 | 7.38827699 | 7.453531899 | -0.065254909 | 0.01956499 |
| AC154652.2 | 3.781024866 | 3.662094125 | 3.761531496 | -0.099437371 | 0.01949337 |
| Gm38392 | 6.384645277 | 6.345831277 | 6.365153441 | -0.019322164 | 0.019491835 |
| Atp5a1 | 12.8408479 | 12.77143209 | 12.82144797 | -0.050015877 | 0.01939993 |
| Gm8093 | 1.777771571 | 1.778148588 | 1.758378335 | 0.019770252 | 0.019393236 |
| Gm7600 | 1.710915133 | 1.7792766 | 1.691557415 | 0.087719185 | 0.019357718 |
| Alg9 | 8.748959866 | 8.654192358 | 8.729620045 | -0.075427687 | 0.019339821 |
| Zfp961 | 7.737206516 | 7.838233515 | 7.717952936 | 0.120280578 | 0.019253579 |
| Gm19046 | 2.110736543 | 2.158431691 | 2.091511953 | 0.066919738 | 0.01922459 |
| Homer3 | 8.11155668 | 8.300689689 | 8.092345147 | 0.208344542 | 0.019211533 |
| Gm14822 | 2.275401606 | 2.22729212 | 2.256199363 | -0.028907243 | 0.019202243 |
| Syce2 | 6.321901618 | 6.279348325 | 6.302745218 | -0.023396892 | 0.0191564 |
| Mpi | 8.418551761 | 8.332912833 | 8.39941914 | -0.066506307 | 0.019132621 |
| Tspyl3 | 6.640247331 | 6.673438628 | 6.621134008 | 0.05230462 | 0.019113323 |

| | | | | | |
|---|---|---|---|---|---|
| Gm10182 | 5.426413044 | 5.46539046 | 5.407352386 | 0.058038074 | 0.019060658 |
| 39326 | 11.68132655 | 11.58293949 | 11.66235131 | -0.079411817 | 0.018975247 |
| Mylk3 | 3.603016667 | 3.564299642 | 3.584076591 | -0.019776949 | 0.018940076 |
| D6Wsu163e | 8.514401722 | 8.519634365 | 8.495475754 | 0.024158611 | 0.018925968 |
| Draxin | 3.028302175 | 3.062880198 | 3.009429781 | 0.053450418 | 0.018872394 |
| Scaf8 | 10.14062685 | 10.1382559 | 10.12176646 | 0.01648944 | 0.018860393 |
| Sdk1 | 5.964500462 | 6.179409185 | 5.945666693 | 0.233742492 | 0.018833769 |
| Vmn1r12 | 2.922094178 | 2.849922851 | 2.903292377 | -0.053369526 | 0.018801801 |
| Olfr889 | 2.319889425 | 2.377245526 | 2.301206571 | 0.076038955 | 0.018682854 |
| Gm35315 | 5.873402539 | 5.727946949 | 5.854728736 | -0.126781787 | 0.018673802 |
| Efnb2 | 11.2277672 | 11.18061363 | 11.2091373 | -0.02852367 | 0.018629897 |
| Vps52 | 10.0454064 | 10.11971099 | 10.02677728 | 0.09293371 | 0.018629112 |
| Cntn3 | 1.861545005 | 1.86648473 | 1.842921067 | 0.023563662 | 0.018623937 |
| Mroh4 | 2.799126725 | 2.884223678 | 2.780511985 | 0.103711693 | 0.01861474 |
| Wdr44 | 8.499725583 | 8.3947896 | 8.481116265 | -0.086326665 | 0.018609318 |
| Gm48793 | 4.611357051 | 4.591536956 | 4.592836057 | -0.001299101 | 0.018520993 |
| Rpl7a-ps3 | 0.641987443 | 0.631072806 | 0.623479882 | 0.007592924 | 0.018507561 |
| Anapc15 | 8.352491895 | 8.404221873 | 8.333986396 | 0.070235477 | 0.018505499 |
| Stxbp1 | 7.449935657 | 7.503496134 | 7.431474171 | 0.072021963 | 0.018461487 |
| Cdh13 | 10.66760671 | 10.58633082 | 10.64917838 | -0.06284756 | 0.018428327 |
| Hist1h3e | 1.68778792 | 1.750300299 | 1.669360731 | 0.080939568 | 0.018427189 |
| B3gnt6 | 1.762325563 | 1.761263455 | 1.74394622 | 0.017317235 | 0.018379343 |
| Vim | 13.71548644 | 13.68119158 | 13.69716676 | -0.01597518 | 0.01831968 |
| Olfr1116 | 3.924011024 | 3.867110435 | 3.905697381 | -0.038586945 | 0.018313643 |
| Yeats4 | 9.037460504 | 8.99252841 | 9.01916485 | -0.02663644 | 0.018295654 |
| Pknox1 | 8.795924282 | 8.949912112 | 8.777637628 | 0.172274484 | 0.018286655 |
| Atp5j2 | 10.98393458 | 10.90117702 | 10.96564846 | -0.064471447 | 0.018286117 |
| Olfr802 | 2.976944036 | 2.964045706 | 2.958662245 | 0.005383462 | 0.018281792 |
| Bdnf | 5.802610218 | 5.78709229 | 5.78434454 | 0.00274775 | 0.018265677 |
| Olfr628 | 1.859418412 | 1.815782556 | 1.841201187 | -0.025418631 | 0.018217225 |
| Rhox4b | 1.811936494 | 1.796126126 | 1.793719607 | 0.002406519 | 0.018216887 |
| Osbpl11 | 9.611407852 | 9.59288064 | 9.593215723 | -0.000335083 | 0.018192129 |

| | | | | | |
|---|---|---|---|---|---|
| Naaladl1 | 1.988688449 | 1.96481912 | 1.970620007 | -0.005800887 | 0.018068442 |
| Galnt16 | 8.84896527 | 8.745708916 | 8.830927722 | -0.085218806 | 0.018037548 |
| Rhbdd1 | 9.259245217 | 9.236242988 | 9.241225973 | -0.004982985 | 0.018019244 |
| Fbxo17 | 5.551490907 | 5.54839654 | 5.533477233 | 0.014919307 | 0.018013674 |
| Alg12 | 7.845853027 | 7.963416944 | 7.827863488 | 0.135553456 | 0.017989538 |
| D330045A20Rik | 2.291879701 | 2.256942219 | 2.273893177 | -0.016950958 | 0.017986524 |
| Mon1b | 9.108414247 | 9.267615414 | 9.090466065 | 0.177149349 | 0.017948182 |
| Tec | 9.154909731 | 9.102302349 | 9.136971752 | -0.034669403 | 0.017937978 |
| Slc24a5 | 1.740747488 | 1.759719432 | 1.722844229 | 0.036875203 | 0.017903259 |
| Vmn2r59 | 5.204458773 | 5.207838508 | 5.186562436 | 0.021276072 | 0.017896336 |
| Ctla2b | 5.855525073 | 5.753836423 | 5.837664384 | -0.083827961 | 0.01786069 |
| Cacna1i | 2.430508648 | 2.430871411 | 2.412684958 | 0.018186453 | 0.01782369 |
| Gm8189 | 1.819517388 | 1.869410549 | 1.801785462 | 0.067625087 | 0.017731926 |
| Ccdc24 | 2.543858826 | 2.655066244 | 2.526195992 | 0.128870252 | 0.017662834 |
| Mcpt4 | 11.06417904 | 10.49126037 | 11.0465167 | -0.555256327 | 0.01766234 |
| Prkcq | 9.03759926 | 8.865974715 | 9.019951567 | -0.153976852 | 0.017647693 |
| Sspo | 3.492374227 | 3.513550204 | 3.474754113 | 0.038796092 | 0.017620115 |
| 9530068E07Rik | 11.1191714 | 10.96617014 | 11.10157736 | -0.135407217 | 0.01759404 |
| Gm47860 | 2.830698321 | 2.822890356 | 2.813106012 | 0.009784344 | 0.017592309 |
| Nefm | 2.578517716 | 2.844500426 | 2.560931378 | 0.283569048 | 0.017586338 |
| Slc7a3 | 3.370334875 | 3.437454 | 3.35276935 | 0.084684649 | 0.017565525 |
| Kat7 | 10.33372478 | 10.26475184 | 10.31619628 | -0.051444437 | 0.017528507 |
| Anxa8 | 14.59603659 | 14.46937716 | 14.57851129 | -0.10913413 | 0.0175253 |
| Rnf180 | 4.32507755 | 4.304123083 | 4.3075656 | -0.003442516 | 0.017511951 |
| Elf2 | 10.30871895 | 10.3732779 | 10.29122803 | 0.08204987 | 0.01749092 |
| Mocos | 8.870293525 | 8.90520187 | 8.852804147 | 0.052397723 | 0.017489378 |
| Hoxc9 | 2.430268736 | 2.464340592 | 2.412781429 | 0.051559163 | 0.017487306 |
| Olfr352 | 1.439725322 | 1.429414301 | 1.422245521 | 0.00716878 | 0.017479801 |
| Wrap53 | 7.560458592 | 7.586879231 | 7.543009065 | 0.043870166 | 0.017449528 |
| St8sia5 | 5.856384146 | 5.946343053 | 5.838951465 | 0.107391588 | 0.017432681 |
| Nphs1 | 1.759542137 | 1.726230809 | 1.742128498 | -0.015897688 | 0.017413639 |
| Bard1 | 7.038328086 | 6.924577998 | 7.020927866 | -0.096349868 | 0.01740022 |

| | | | | | |
|---|---|---|---|---|---|
| Otop1 | 2.057452758 | 2.073507989 | 2.040091304 | 0.033416686 | 0.017361454 |
| Kcnk4 | 2.596079083 | 2.62102329 | 2.578811812 | 0.042211478 | 0.017267271 |
| Pcdhga1 | 4.657263291 | 4.679401885 | 4.640021464 | 0.039380421 | 0.017241827 |
| Thyn1 | 7.651837332 | 7.644225335 | 7.634619429 | 0.009605906 | 0.017217904 |
| Vmn2r69 | 3.053309922 | 3.054682796 | 3.036105391 | 0.018577405 | 0.017204531 |
| Prn | 2.79094726 | 2.750304049 | 2.77376938 | -0.023465331 | 0.017177879 |
| Olfr465-ps1 | 1.897637065 | 1.91875521 | 1.880462626 | 0.038292584 | 0.017174439 |
| Vmn2r56 | 4.291598295 | 4.169800335 | 4.274433544 | -0.104633209 | 0.017164751 |
| AL731706.2 | 2.137664405 | 2.082745031 | 2.120519616 | -0.037774585 | 0.017144789 |
| Olfr323 | 2.574889904 | 2.576939881 | 2.557747049 | 0.019192833 | 0.017142856 |
| Park7 | 10.68134034 | 10.66798447 | 10.66421818 | 0.00376629 | 0.01712216 |
| Gm4969 | 1.843495262 | 1.854146286 | 1.826392433 | 0.027753853 | 0.017102829 |
| Sfswap | 9.839929207 | 9.971078488 | 9.822859343 | 0.148219145 | 0.017069864 |
| Cacfd1 | 9.305456449 | 9.359817349 | 9.288393461 | 0.071423888 | 0.017062988 |
| Vmn2r79 | 5.789007478 | 5.932027637 | 5.771998438 | 0.160029199 | 0.01700904 |
| Hnrnpl | 12.12434176 | 12.09859818 | 12.107334 | -0.008735817 | 0.017007767 |
| Sft2d3 | 6.388984625 | 6.241057345 | 6.372056539 | -0.130999194 | 0.016928086 |
| Arl14epl | 1.438616387 | 1.389808194 | 1.421712392 | -0.031904198 | 0.016903994 |
| Serping1 | 12.49941563 | 12.25467783 | 12.48252069 | -0.227842857 | 0.016894937 |
| Rps19-ps6 | 1.536929187 | 1.52918154 | 1.520107609 | 0.009073932 | 0.016821578 |
| Pdcd5-ps | 7.736456589 | 7.525999521 | 7.719662329 | -0.193662808 | 0.01679426 |
| Ggn | 3.216109315 | 3.223549892 | 3.199320831 | 0.02422906 | 0.016788483 |
| Pramel4 | 1.09835246 | 1.080783883 | 1.081564915 | -0.000781032 | 0.016787545 |
| 1700028K03Rik | 2.023933595 | 2.006069449 | 2.007159361 | -0.001089912 | 0.016774234 |
| Mrpl23 | 5.631007416 | 6.015444773 | 5.614245755 | 0.401199018 | 0.016761661 |
| 39142 | 11.13547926 | 10.82233541 | 11.11872391 | -0.2963885 | 0.016755347 |
| Otud5 | 10.26397871 | 10.36241249 | 10.24734275 | 0.115069733 | 0.016635957 |
| Fubp3 | 9.756443504 | 9.767674279 | 9.739854968 | 0.02781931 | 0.016588536 |
| Morf4l1 | 12.31877192 | 12.22777479 | 12.3022086 | -0.074433813 | 0.01656332 |
| Cox7a2 | 10.63745502 | 10.57008441 | 10.62098956 | -0.05090515 | 0.016465453 |
| Vmn2r20 | 3.221528937 | 3.156101233 | 3.205064365 | -0.048963132 | 0.016464572 |
| Gm12057 | 2.130985286 | 2.117237156 | 2.114552342 | 0.002684814 | 0.016432944 |

| | | | | | |
|---|---|---|---|---|---|
| Gm10481 | 1.816297315 | 1.832723205 | 1.799867629 | 0.032855576 | 0.016429687 |
| Usp50 | 3.843267938 | 3.823823898 | 3.826850958 | -0.00302706 | 0.01641698 |
| Rpl5-ps2 | 1.659743342 | 1.683969761 | 1.643379765 | 0.040589996 | 0.016363578 |
| Gm17906 | 1.885344776 | 1.840229419 | 1.869025459 | -0.02879604 | 0.016319317 |
| Map2k4 | 11.66848004 | 11.55280499 | 11.65217038 | -0.099365383 | 0.01630966 |
| Frmd3 | 4.673919919 | 4.822126462 | 4.657629494 | 0.164496968 | 0.016290425 |
| Eef1akmt1 | 8.019480588 | 7.958487798 | 8.003241328 | -0.04475353 | 0.01623926 |
| Vmn1r188 | 2.915014255 | 2.880487157 | 2.898779211 | -0.018292054 | 0.016235045 |
| Adamtsl2 | 5.341514233 | 5.667880956 | 5.325285545 | 0.342595411 | 0.016228689 |
| Gm6092 | 1.899207259 | 1.882206699 | 1.882983398 | -0.000776698 | 0.016223861 |
| Clta | 11.06753705 | 11.0068849 | 11.05132563 | -0.04444073 | 0.016211417 |
| Hs3st3a1 | 4.367977137 | 4.222661259 | 4.351787972 | -0.129126714 | 0.016189165 |
| Wdr27 | 3.923990003 | 3.828860482 | 3.907814784 | -0.078954302 | 0.016175219 |
| Ubap1l | 2.074780774 | 2.118269295 | 2.058607038 | 0.059662256 | 0.016173736 |
| Chrdl2 | 2.075322264 | 2.043936306 | 2.059174903 | -0.015238598 | 0.016147361 |
| Vmn2r63 | 4.015139162 | 4.027797743 | 3.998992748 | 0.028804994 | 0.016146414 |
| Dlg1 | 10.55013206 | 10.47893605 | 10.53402075 | -0.055084697 | 0.01611131 |
| Amhr2 | 4.093273504 | 4.146639527 | 4.077167119 | 0.069472408 | 0.016106385 |
| 9530053A07Rik | 3.375364958 | 3.364325139 | 3.359279244 | 0.005045895 | 0.016085714 |
| Rpl9-ps4 | 1.893456876 | 1.865824238 | 1.877374508 | -0.01155027 | 0.016082368 |
| Gm48862 | 1.874188425 | 1.877009976 | 1.858143005 | 0.018866971 | 0.016045419 |
| Pik3c2a | 9.854852214 | 9.776697075 | 9.838808495 | -0.06211142 | 0.016043719 |
| Cox7b | 11.22360781 | 11.12963804 | 11.20756552 | -0.077927477 | 0.016042293 |
| Gm13178 | 9.144021694 | 8.99130234 | 9.128028409 | -0.136726068 | 0.015993285 |
| Bmp8b | 2.012968395 | 1.994278008 | 1.997005053 | -0.002727045 | 0.015963342 |
| Olfr287 | 1.263424347 | 1.240431737 | 1.247465461 | -0.007033725 | 0.015958886 |
| Gm49331 | 3.921410578 | 3.9955611 | 3.905517159 | 0.090043941 | 0.015893418 |
| Hcn4 | 5.020371071 | 5.018478846 | 5.004485665 | 0.013993181 | 0.015885406 |
| Mmp24 | 1.802012145 | 1.789703588 | 1.786151576 | 0.003552012 | 0.015860569 |
| Zfp568 | 8.01204786 | 8.316155005 | 7.996206672 | 0.319948333 | 0.015841188 |
| Ido1 | 1.290797561 | 1.288329161 | 1.275000665 | 0.013328496 | 0.015796896 |
| Aoc2 | 6.203511736 | 6.388927472 | 6.187722162 | 0.20120531 | 0.015789574 |

| | | | | | |
|---|---|---|---|---|---|
| Hgfac | 2.823397797 | 2.887099811 | 2.807681727 | 0.079418084 | 0.015716071 |
| Bora | 7.409335909 | 7.381330223 | 7.393658622 | -0.012328399 | 0.015677288 |
| Spata19 | 0.90328427 | 0.846443159 | 0.887609086 | -0.041165926 | 0.015675185 |
| Rs1 | 2.09508867 | 2.13533498 | 2.079460391 | 0.055874589 | 0.01562828 |
| Bod1l | 10.55295906 | 10.50414633 | 10.53735435 | -0.033208023 | 0.015604707 |
| C1qtnf4 | 2.413552931 | 2.448964023 | 2.39805083 | 0.050913193 | 0.015502101 |
| Mtrf1l | 6.899836674 | 6.838407463 | 6.884356467 | -0.045949004 | 0.015480207 |
| Iqch | 2.033752206 | 1.962707571 | 2.018281814 | -0.055574243 | 0.015470392 |
| Ajap1 | 2.828758389 | 2.844821945 | 2.813317218 | 0.031504727 | 0.015441172 |
| Slc25a14 | 6.417248388 | 6.447563529 | 6.401813814 | 0.045749715 | 0.015434575 |
| Bloc1s1 | 9.367443155 | 9.30466289 | 9.352029834 | -0.047366944 | 0.015413321 |
| 1700021F05Rik | 8.601178557 | 8.558583514 | 8.585780709 | -0.027197196 | 0.015397848 |
| Neil1 | 7.22759434 | 7.292365112 | 7.212230406 | 0.080134706 | 0.015363934 |
| Cd79a | 1.500418683 | 1.506050873 | 1.485076843 | 0.020974031 | 0.015341841 |
| Rps8-ps4 | 1.117773733 | 1.101177841 | 1.102449717 | -0.001271876 | 0.015324016 |
| Eif2d | 9.515798754 | 9.47657287 | 9.500505364 | -0.023932494 | 0.01529339 |
| Olfr1221 | 5.230129768 | 5.287721103 | 5.214902635 | 0.072818468 | 0.015227133 |
| Ormdl3 | 10.83954088 | 10.89555359 | 10.82431461 | 0.07123898 | 0.01522627 |
| Gipc1 | 9.812890114 | 9.938346903 | 9.797683152 | 0.140663751 | 0.015206962 |
| Hoxb2 | 7.284721942 | 7.266515772 | 7.269540303 | -0.003024532 | 0.015181639 |
| Wnt7a | 1.982544015 | 1.986787392 | 1.967433314 | 0.019354077 | 0.015110701 |
| Ap1b1 | 11.59705243 | 11.69536568 | 11.58194695 | 0.113418727 | 0.015105477 |
| Adgrg6 | 9.879506296 | 9.841437101 | 9.864439179 | -0.023002077 | 0.015067118 |
| Hacd3 | 11.09379061 | 11.04178504 | 11.07874771 | -0.03696267 | 0.015042907 |
| Gm49355 | 2.767052608 | 2.812085772 | 2.752020036 | 0.060065736 | 0.015032572 |
| Shc4 | 6.15055625 | 6.110858726 | 6.135540342 | -0.024681617 | 0.015015908 |
| Gpr63 | 3.793176838 | 3.80618344 | 3.778181257 | 0.028002182 | 0.01499558 |
| AC169506.2 | 1.819235408 | 1.852042968 | 1.804259522 | 0.047783446 | 0.014975886 |
| Zic4 | 4.565652175 | 4.549789045 | 4.550678484 | -0.000889438 | 0.014973691 |
| Cd302 | 9.23172509 | 8.918028626 | 9.216759918 | -0.298731292 | 0.014965172 |
| Cfap61 | 2.334526891 | 2.379585168 | 2.319622872 | 0.059962296 | 0.014904019 |
| P2rx3 | 3.507346196 | 3.568190494 | 3.492447886 | 0.075742608 | 0.01489831 |

| | | | | | |
|---|---|---|---|---|---|
| Ube2j1 | 10.01152755 | 10.02083883 | 9.996692886 | 0.024145944 | 0.014834666 |
| Mrgprc2-ps | 3.350998071 | 3.350913114 | 3.33620224 | 0.014710874 | 0.01479583 |
| Exog | 7.434357977 | 7.545418102 | 7.419563762 | 0.12585434 | 0.014794215 |
| Olfr905 | 2.080206562 | 2.052433311 | 2.065418781 | -0.012985471 | 0.01478778 |
| C530008M17Rik | 4.333739497 | 4.386227862 | 4.318975453 | 0.067252409 | 0.014764044 |
| Rmdn2 | 6.039484726 | 5.843982556 | 6.024744758 | -0.180762202 | 0.014739968 |
| Slc38a10 | 10.82145517 | 10.87833087 | 10.80674466 | 0.071586213 | 0.014710517 |
| Il9r | 2.991134987 | 3.014780618 | 2.976467146 | 0.038313472 | 0.014667841 |
| Rpsa-ps9 | 2.280727626 | 2.245615 | 2.266081846 | -0.020466845 | 0.01464578 |
| Gm4247 | 2.057097792 | 2.095171221 | 2.042483082 | 0.052688139 | 0.01461471 |
| Vps37a | 10.05128201 | 10.00029434 | 10.03666953 | -0.036375186 | 0.014612486 |
| Zfp524 | 7.381716347 | 7.532583595 | 7.367151098 | 0.165432497 | 0.014565249 |
| Adam18 | 1.978817281 | 1.967407035 | 1.964265783 | 0.003141252 | 0.014551498 |
| Entpd4b | 9.695593667 | 9.675007961 | 9.681076242 | -0.006068281 | 0.014517425 |
| Zc3h3 | 8.246035473 | 8.355407749 | 8.231575973 | 0.123831776 | 0.0144595 |
| Kank4 | 5.948620696 | 5.89160476 | 5.934163559 | -0.042558799 | 0.014457137 |
| Mrm2 | 6.965933211 | 7.038873212 | 6.951548108 | 0.087325103 | 0.014385102 |
| Vmn1r20 | 2.598504713 | 2.609622531 | 2.58418079 | 0.025441741 | 0.014323923 |
| Zfp787 | 7.297882547 | 7.369534901 | 7.283573767 | 0.085961134 | 0.01430878 |
| Hsd3b7 | 9.466486804 | 9.549099077 | 9.452180911 | 0.096918166 | 0.014305894 |
| Mettl25 | 6.581213593 | 6.567416486 | 6.566907904 | 0.000508582 | 0.014305689 |
| Abcc2 | 3.72859522 | 3.581458372 | 3.714293949 | -0.132835577 | 0.01430127 |
| Rpl13-ps6 | 1.722927994 | 1.760997796 | 1.708635428 | 0.052362367 | 0.014292566 |
| Luc7l3 | 11.2793819 | 11.33552305 | 11.26510696 | 0.070416093 | 0.01427494 |
| Gm6034 | 1.605403767 | 1.596952053 | 1.59114627 | 0.005805783 | 0.014257497 |
| Cpn1 | 2.006551582 | 2.046467253 | 1.992319034 | 0.054148219 | 0.014232548 |
| Olfr1233 | 2.268064192 | 2.336134585 | 2.253852401 | 0.082282184 | 0.014211791 |
| Cyp19a1 | 1.818057501 | 1.833753273 | 1.80384977 | 0.029903502 | 0.01420773 |
| Apc-ps1 | 1.953303683 | 1.990371687 | 1.939109682 | 0.051262005 | 0.014194001 |
| Derl1 | 11.16594888 | 11.15179445 | 11.15176427 | 3.01833E-05 | 0.014184617 |
| Atxn3 | 10.09483228 | 9.937718343 | 10.08068978 | -0.142971434 | 0.014142507 |
| Ints13 | 9.548913667 | 9.622324269 | 9.534777416 | 0.087546853 | 0.014136251 |

| | | | | | |
|---|---|---|---|---|---|
| Nsl1 | 6.60290934 | 6.545656893 | 6.588838123 | -0.04318123 | 0.014071217 |
| Speer6-ps1 | 2.248540148 | 2.299754385 | 2.234479585 | 0.065274799 | 0.014060563 |
| Gm14219 | 2.925686691 | 2.875388064 | 2.911627127 | -0.036239063 | 0.014059564 |
| Gm867 | 1.356187399 | 1.409592197 | 1.342133324 | 0.067458872 | 0.014054075 |
| Trbj2-4 | 1.602589504 | 1.615305672 | 1.58855891 | 0.026746762 | 0.014030594 |
| Wwp2 | 10.30148168 | 10.27807765 | 10.28749238 | -0.00941473 | 0.013989303 |
| Man1b1 | 10.49475448 | 10.46803351 | 10.48076837 | -0.01273486 | 0.013986113 |
| Smg1 | 11.85393988 | 12.00676004 | 11.83996691 | 0.16679313 | 0.013972973 |
| Zfp948 | 8.483149418 | 8.530476114 | 8.469248535 | 0.061227579 | 0.013900882 |
| Mrps15 | 8.921612896 | 8.901283777 | 8.907736282 | -0.006452505 | 0.013876614 |
| Vmn2r6 | 3.584404037 | 3.493879628 | 3.570535494 | -0.076655867 | 0.013868543 |
| Htt | 10.60454442 | 10.70164175 | 10.59071371 | 0.110928033 | 0.013830707 |
| Mrpl22 | 8.311354057 | 8.153739042 | 8.297530297 | -0.143791256 | 0.01382376 |
| Olfr1555-ps1 | 1.792151575 | 1.783975336 | 1.778344822 | 0.005630514 | 0.013806753 |
| Kcnk10 | 3.420733049 | 3.37260797 | 3.406963864 | -0.034355894 | 0.013769185 |
| Gm10012 | 2.160970789 | 2.157732437 | 2.147208631 | 0.010523806 | 0.013762158 |
| Mrpl14 | 8.375757492 | 8.47468065 | 8.362004489 | 0.112676161 | 0.013753003 |
| H2-Oa | 4.035286184 | 3.922036843 | 4.02157172 | -0.099534878 | 0.013714463 |
| Them6 | 5.561546899 | 5.553516192 | 5.547863248 | 0.005652944 | 0.013683651 |
| Eef1akmt2 | 7.462148819 | 7.452227694 | 7.448486671 | 0.003741023 | 0.013662148 |
| Gm438 | 7.061958815 | 6.995659361 | 7.048298832 | -0.052639471 | 0.013659984 |
| Ucp1 | 0.988097575 | 1.036675985 | 0.974486165 | 0.06218982 | 0.01361141 |
| Edc3 | 9.330609405 | 9.401035254 | 9.317008917 | 0.084026337 | 0.013600488 |
| AC121135.2 | 1.311978388 | 1.290171399 | 1.298444781 | -0.008273383 | 0.013533606 |
| Ap2a2 | 11.22538442 | 11.23728116 | 11.21187045 | 0.025410713 | 0.013513973 |
| Mmgt2 | 7.25316244 | 7.309502019 | 7.239653816 | 0.069848204 | 0.013508624 |
| Fyb2 | 4.575142926 | 4.517984943 | 4.561645252 | -0.043660309 | 0.013497674 |
| Zcchc12 | 2.541441501 | 2.50321937 | 2.527949484 | -0.024730114 | 0.013492017 |
| Prr18 | 1.315463676 | 1.320680859 | 1.30198625 | 0.018694609 | 0.013477426 |
| Gm5182 | 2.505819074 | 2.413561137 | 2.492349731 | -0.078788595 | 0.013469343 |
| Cwc15 | 10.35156496 | 10.24889237 | 10.33810151 | -0.08920914 | 0.013463447 |
| Pcdhb8 | 2.530145358 | 2.52875612 | 2.516708637 | 0.012047483 | 0.013436722 |

| | | | | | |
|---|---|---|---|---|---|
| Arhgap17 | 9.535656374 | 9.596135599 | 9.52225992 | 0.073875679 | 0.013396454 |
| Clnk | 2.790613975 | 2.771836112 | 2.777239766 | -0.005403654 | 0.013374209 |
| Lrmp | 8.250445817 | 8.180490854 | 8.237073373 | -0.056582519 | 0.013372444 |
| Tom1 | 10.2598849 | 10.42079415 | 10.24653279 | 0.174261363 | 0.01335211 |
| Ppib | 11.74183875 | 11.72500536 | 11.72848673 | -0.003481367 | 0.013352023 |
| Cryba4 | 2.432458303 | 2.484549531 | 2.419147578 | 0.065401953 | 0.013310725 |
| Hic2 | 5.82114974 | 5.752193381 | 5.80788838 | -0.055694999 | 0.01326136 |
| Fbp1 | 3.089931858 | 3.069921307 | 3.07668529 | -0.006763982 | 0.013246568 |
| Tcrg-C4 | 4.554075834 | 4.603689143 | 4.540838619 | 0.062850523 | 0.013237215 |
| Camk2n2 | 4.29680989 | 4.309870831 | 4.283581136 | 0.026289695 | 0.013228754 |
| Traf3ip1 | 6.945113951 | 7.043467907 | 6.931888119 | 0.111579788 | 0.013225832 |
| Ints11 | 9.784937447 | 9.853063228 | 9.771733484 | 0.081329744 | 0.013203962 |
| Gpr39 | 2.241284667 | 2.244272947 | 2.22810482 | 0.016168127 | 0.013179847 |
| Gm1848 | 2.249398182 | 2.26382764 | 2.236244775 | 0.027582865 | 0.013153406 |
| 2410004P03Rik | 3.851192732 | 3.793171147 | 3.838044944 | -0.044873797 | 0.013147788 |
| Lpcat2b | 1.600164201 | 1.639995038 | 1.587021167 | 0.052973871 | 0.013143034 |
| Slc35a2 | 8.666383723 | 8.710702339 | 8.65326357 | 0.057438769 | 0.013120153 |
| Olfr814 | 2.171689471 | 2.172181761 | 2.158592056 | 0.013589705 | 0.013097414 |
| Fadd | 8.092413521 | 8.090106087 | 8.079322596 | 0.010783491 | 0.013090924 |
| Chn1 | 5.32889372 | 5.193861716 | 5.315854578 | -0.121992862 | 0.013039142 |
| Ncaph2 | 11.1607096 | 11.12089102 | 11.14768169 | -0.026790673 | 0.01302791 |
| Axin1 | 10.12682677 | 10.15737443 | 10.11380841 | 0.043566027 | 0.013018363 |
| Olfr675 | 1.655590995 | 1.636362877 | 1.642582619 | -0.006219742 | 0.013008376 |
| Atp1b3 | 12.92258634 | 12.91644475 | 12.90960295 | 0.006841807 | 0.012983397 |
| Zfp7 | 7.618369947 | 7.610958763 | 7.605432035 | 0.005526728 | 0.012937912 |
| Gm13237 | 1.10712059 | 1.12814104 | 1.094182998 | 0.033958042 | 0.012937592 |
| Clcn7 | 10.13495302 | 10.32099059 | 10.12202859 | 0.198962007 | 0.01292443 |
| Gm9770 | 1.115182275 | 1.136418885 | 1.102268505 | 0.03415038 | 0.01291377 |
| Ncstn | 11.03222989 | 11.01804156 | 11.01933625 | -0.00129469 | 0.012893643 |
| Tcrg-V6 | 1.254547548 | 1.25417238 | 1.241662146 | 0.012510234 | 0.012885402 |
| Olfr646 | 1.719169759 | 1.67698323 | 1.706368789 | -0.029385559 | 0.01280097 |
| Gm6012 | 1.836194737 | 1.854430846 | 1.823415691 | 0.031015155 | 0.012779046 |

| | | | | | |
|---|---|---|---|---|---|
| Hmgb2 | 10.12904297 | 10.06095466 | 10.11628065 | -0.055325989 | 0.01276232 |
| Thpo | 4.622277955 | 4.443413703 | 4.609568653 | -0.16615495 | 0.012709302 |
| Arnt | 10.4711647 | 10.42159342 | 10.45846475 | -0.03687133 | 0.01269995 |
| Cabyr | 2.250045129 | 2.24998554 | 2.23736538 | 0.01262016 | 0.012679749 |
| Hsd17b12 | 12.27887393 | 12.1620904 | 12.2662282 | -0.104137803 | 0.01264573 |
| Pcdhgb2 | 6.294081866 | 6.353563831 | 6.281445698 | 0.072118134 | 0.012636168 |
| Zfp746 | 8.98466836 | 9.029964555 | 8.97203336 | 0.057931196 | 0.012635001 |
| Il12a | 3.215872272 | 3.140263881 | 3.2032519 | -0.062988019 | 0.012620372 |
| Gm7729 | 1.379577867 | 1.392013817 | 1.367006206 | 0.025007612 | 0.012571661 |
| Zfp457 | 3.004721825 | 2.971251469 | 2.992154688 | -0.020903219 | 0.012567137 |
| Hist1h4h | 3.319003867 | 3.397145215 | 3.306444179 | 0.090701036 | 0.012559688 |
| Dtl | 8.19362925 | 8.221677323 | 8.181159126 | 0.040518197 | 0.012470123 |
| Stam | 9.142639405 | 9.197838118 | 9.130203354 | 0.067634764 | 0.012436051 |
| Nrm | 8.19946093 | 8.154948932 | 8.187041072 | -0.03209214 | 0.012419858 |
| Rps27a-ps1 | 1.4144251 | 1.407529335 | 1.40200931 | 0.005520025 | 0.012415789 |
| Bloc1s6 | 8.808583561 | 8.755097552 | 8.796173481 | -0.041075929 | 0.01241008 |
| Cd72 | 7.286701293 | 7.11770596 | 7.274327612 | -0.156621652 | 0.012373682 |
| Nupr1l | 4.034095264 | 4.023478441 | 4.021723045 | 0.001755395 | 0.012372219 |
| Lrrc24 | 1.834232336 | 1.877406913 | 1.821893552 | 0.055513361 | 0.012338785 |
| Tfcp2l1 | 4.531034148 | 4.605778889 | 4.518718074 | 0.087060815 | 0.012316074 |
| Fam234b | 9.500513506 | 9.524345762 | 9.488237203 | 0.036108559 | 0.012276303 |
| Gm14057 | 2.196011355 | 2.21465879 | 2.183754923 | 0.030903867 | 0.012256433 |
| Khdc3 | 2.456866118 | 2.497937283 | 2.444657764 | 0.053279519 | 0.012208355 |
| Lbx1 | 5.274792521 | 5.101344101 | 5.262680099 | -0.161335998 | 0.012112422 |
| Tax1bp1 | 12.13016561 | 12.07911132 | 12.11810784 | -0.038996513 | 0.012057773 |
| Pyy | 1.56570986 | 1.590996901 | 1.553659465 | 0.037337435 | 0.012050395 |
| Adgrb3 | 2.11658702 | 2.06932378 | 2.104567855 | -0.035244075 | 0.012019165 |
| Fbxl13 | 1.937336037 | 2.021653123 | 1.925330022 | 0.096323101 | 0.012006015 |
| Osgep | 9.408189017 | 9.343483937 | 9.396185664 | -0.052701727 | 0.012003353 |
| Olfr1245 | 2.231026855 | 2.230211348 | 2.219102663 | 0.011108686 | 0.011924193 |
| D630039A03Rik | 2.426384567 | 2.389771335 | 2.414466306 | -0.024694971 | 0.011918261 |
| Gm7240 | 4.039984383 | 3.919992708 | 4.028079558 | -0.10808685 | 0.011904825 |

| | | | | | |
|---|---|---|---|---|---|
| Olfr149 | 3.310856552 | 3.339580891 | 3.298956899 | 0.040623992 | 0.011899653 |
| 2810039B14Rik | 4.527227124 | 4.506621362 | 4.515335994 | -0.008714632 | 0.01189113 |
| Zfand3 | 10.49579393 | 10.43532695 | 10.48392643 | -0.04859948 | 0.0118675 |
| Enho | 4.647044485 | 4.688694348 | 4.635189291 | 0.053505057 | 0.011855194 |
| Bace2 | 8.798261646 | 8.748630295 | 8.786420061 | -0.037789766 | 0.011841585 |
| Polb | 10.04627211 | 10.04012578 | 10.03447802 | 0.005647763 | 0.011794094 |
| Cln5 | 9.629009894 | 9.630937415 | 9.617262629 | 0.013674786 | 0.011747265 |
| Olfr780 | 1.506254411 | 1.534000336 | 1.494545404 | 0.039454932 | 0.011709007 |
| Ralgps1 | 7.304088472 | 7.386448879 | 7.292467423 | 0.093981456 | 0.011621049 |
| Fam57b | 7.771988338 | 7.664966853 | 7.760395841 | -0.095428989 | 0.011592497 |
| Kdr | 8.854136384 | 8.762426789 | 8.842598314 | -0.080171526 | 0.01153807 |
| Jup | 14.05730062 | 14.23593854 | 14.04577065 | 0.19016789 | 0.01152997 |
| Pop4 | 8.640021612 | 8.585189263 | 8.628514024 | -0.043324761 | 0.011507588 |
| Rps12-ps26 | 1.423307396 | 1.425957009 | 1.41180288 | 0.014154129 | 0.011504516 |
| Gm29216 | 15.48182712 | 15.33829453 | 15.47035373 | -0.132059197 | 0.011473393 |
| Gm15807 | 2.490961995 | 2.495213773 | 2.479565449 | 0.015648324 | 0.011396546 |
| Olfr1269 | 2.502899225 | 2.524266569 | 2.491508791 | 0.032757777 | 0.011390433 |
| Ten1 | 8.068223771 | 8.083038238 | 8.056836901 | 0.026201337 | 0.01138687 |
| Grin2d | 5.050939511 | 5.197880876 | 5.039580414 | 0.158300462 | 0.011359098 |
| AL731706.1 | 7.097445146 | 7.257737326 | 7.0861002 | 0.171637127 | 0.011344946 |
| Gm15538 | 2.040876588 | 2.044954884 | 2.029629522 | 0.015325362 | 0.011247066 |
| Kansl3 | 10.1596101 | 10.30559025 | 10.1484011 | 0.157189143 | 0.011209 |
| Mdga2 | 4.552318626 | 4.581586785 | 4.541110708 | 0.040476077 | 0.011207918 |
| Cldnd1 | 10.58129676 | 10.65916897 | 10.57014328 | 0.089025697 | 0.011153483 |
| Krt8 | 2.554984261 | 2.566601436 | 2.543875379 | 0.022726057 | 0.011108882 |
| Gtf3c5 | 8.963387829 | 8.958358749 | 8.952331957 | 0.006026792 | 0.011055872 |
| Pcdhgb5 | 5.999469659 | 6.105331703 | 5.988432871 | 0.116898832 | 0.011036788 |
| Sned1 | 10.42398149 | 10.71959912 | 10.41298889 | 0.306610233 | 0.010992603 |
| Tex15 | 3.860420795 | 3.823708529 | 3.849438375 | -0.025729846 | 0.01098242 |
| Ndufv2 | 11.050117 | 10.92937813 | 11.03914269 | -0.109764567 | 0.010974303 |
| Olfr381 | 1.588136444 | 1.592382971 | 1.577162235 | 0.015220736 | 0.01097421 |
| Gm13573 | 1.423010499 | 1.406199556 | 1.41206532 | -0.005865765 | 0.010945179 |

| | | | | | |
|---|---|---|---|---|---|
| Blk | 4.20399543 | 4.248469837 | 4.193092055 | 0.055377782 | 0.010903375 |
| Olfr859 | 3.269721493 | 3.162782736 | 3.258862513 | -0.096079777 | 0.01085898 |
| 37500 | 11.91612285 | 11.85538193 | 11.90528519 | -0.049903263 | 0.010837653 |
| Ap3m1 | 10.44534556 | 10.47762686 | 10.43459375 | 0.04303311 | 0.010751807 |
| Olfr1109 | 2.099590465 | 2.052027268 | 2.088858965 | -0.036831697 | 0.010731501 |
| Apoc3 | 1.85558042 | 1.826435948 | 1.844895787 | -0.018459838 | 0.010684633 |
| Chrna4 | 2.122707695 | 2.155437195 | 2.112069996 | 0.043367199 | 0.010637699 |
| Msc | 2.869514956 | 2.924517372 | 2.858934415 | 0.065582957 | 0.010580541 |
| Gm5619 | 2.899734435 | 2.858967453 | 2.889189013 | -0.030221561 | 0.010545422 |
| Gtf2h5 | 9.589532143 | 9.51221457 | 9.579024406 | -0.066809836 | 0.010507736 |
| Oxsr1 | 10.41832376 | 10.33347259 | 10.40783784 | -0.074365247 | 0.01048592 |
| Gm45268 | 2.274685612 | 2.279957395 | 2.264210903 | 0.015746492 | 0.010474709 |
| Gm5345 | 2.890097029 | 2.850899737 | 2.879637141 | -0.028737404 | 0.010459888 |
| Cenpb | 11.94440974 | 11.98362935 | 11.9339579 | 0.049671457 | 0.010451847 |
| Ptcd1 | 8.32047995 | 8.36030474 | 8.31007604 | 0.0502287 | 0.01040391 |
| Krt42 | 2.753409582 | 2.7877435 | 2.743007813 | 0.044735686 | 0.010401769 |
| Gm32687 | 6.419914164 | 6.330529934 | 6.409521475 | -0.078991541 | 0.010392689 |
| C8a | 2.158896764 | 2.136014412 | 2.148515028 | -0.012500616 | 0.010381736 |
| Ribc1 | 3.999959462 | 4.134432032 | 3.989589242 | 0.144842791 | 0.010370221 |
| Dner | 2.092262523 | 2.085593432 | 2.081903142 | 0.00369029 | 0.010359381 |
| Vmn1r180 | 1.647180435 | 1.648839801 | 1.636832726 | 0.012007074 | 0.010347709 |
| Gm21319 | 2.368023454 | 2.410531333 | 2.357716035 | 0.052815298 | 0.010307419 |
| B4galt2 | 6.816791593 | 6.767231601 | 6.806507363 | -0.039275762 | 0.01028423 |
| Gm6480 | 1.297467197 | 1.257747734 | 1.287196584 | -0.02944885 | 0.010270613 |
| Lrrc27 | 4.498657503 | 4.442079882 | 4.488432839 | -0.046352957 | 0.010224663 |
| Vmn2r117 | 3.787397998 | 3.778728606 | 3.777207266 | 0.00152134 | 0.010190733 |
| Klra3 | 2.836790158 | 2.827521649 | 2.826638053 | 0.000883597 | 0.010152105 |
| Kif26b | 7.455785667 | 7.666481402 | 7.445683299 | 0.220798104 | 0.010102369 |
| Pcdha4 | 1.109307694 | 1.115521458 | 1.099207763 | 0.016313695 | 0.010099931 |
| Catsper4 | 3.887526162 | 4.161378101 | 3.877577344 | 0.283800757 | 0.009948819 |
| Cyp2ab1 | 3.71657241 | 3.629286228 | 3.706660584 | -0.077374356 | 0.009911825 |
| Ttc30a2 | 2.573411356 | 2.613769696 | 2.563531337 | 0.050238359 | 0.009880019 |

| | | | | | |
|---|---|---|---|---|---|
| Tcaf2 | 8.630201629 | 8.556722547 | 8.620334188 | -0.063611642 | 0.009867441 |
| Gm6637 | 1.315467449 | 1.275205727 | 1.30560754 | -0.030401813 | 0.00985991 |
| Slc25a26 | 7.745569755 | 7.710718605 | 7.735717481 | -0.024998877 | 0.009852274 |
| Olfr695 | 2.503528916 | 2.498846061 | 2.493680802 | 0.005165259 | 0.009848115 |
| Rnf217 | 9.659807718 | 9.443284904 | 9.64996162 | -0.206676716 | 0.009846098 |
| Pcdhb20 | 5.466725338 | 5.452157948 | 5.456882366 | -0.004724418 | 0.009842973 |
| Cep128 | 7.212958898 | 7.059168516 | 7.203129233 | -0.143960717 | 0.009829665 |
| Tbca | 9.84353857 | 9.734451281 | 9.833734762 | -0.099283482 | 0.009803807 |
| Hmgxb4 | 8.921487693 | 8.835154851 | 8.911684977 | -0.076530125 | 0.009802717 |
| R3hcc1 | 8.123381571 | 8.081252782 | 8.113600289 | -0.032347507 | 0.009781282 |
| Lrfn2 | 2.751027323 | 2.762635415 | 2.741261235 | 0.02137418 | 0.009766088 |
| Ccr6 | 3.25050694 | 3.222483201 | 3.24075415 | -0.01827095 | 0.009752789 |
| Tent4b | 9.58419138 | 9.677534873 | 9.574446566 | 0.103088307 | 0.009744815 |
| Slc10a2 | 1.281172432 | 1.310591775 | 1.271441209 | 0.039150566 | 0.009731223 |
| Itih4 | 4.134746408 | 4.675772153 | 4.125023018 | 0.550749135 | 0.00972339 |
| Cuedc2 | 8.937856684 | 8.885822057 | 8.928145687 | -0.04232363 | 0.009710997 |
| Cdh22 | 1.809239919 | 1.789915397 | 1.79963646 | -0.009721064 | 0.009603459 |
| Ldhd | 7.719882879 | 7.769316703 | 7.710284035 | 0.059032668 | 0.009598845 |
| Fanci | 6.867063098 | 6.958646076 | 6.857467459 | 0.101178617 | 0.009595639 |
| Ly6g5b | 4.641955851 | 4.585339275 | 4.632391799 | -0.047052523 | 0.009564052 |
| Cd3e | 7.192057874 | 6.986481756 | 7.182531768 | -0.196050012 | 0.009526106 |
| Vmn2r95 | 4.15402592 | 4.184259665 | 4.144513207 | 0.039746457 | 0.009512713 |
| Gm13547 | 1.490520403 | 1.516749651 | 1.481012733 | 0.035736918 | 0.00950767 |
| Synj2bp | 9.798932285 | 9.739376717 | 9.789473398 | -0.050096681 | 0.009458887 |
| Prnd | 4.406348689 | 4.644011428 | 4.39691288 | 0.247098548 | 0.009435809 |
| Rabep1 | 9.833630893 | 9.830591186 | 9.824332782 | 0.006258405 | 0.009298111 |
| Nlrp5-ps | 1.710432087 | 1.708300571 | 1.701137975 | 0.007162596 | 0.009294112 |
| Meis3 | 6.966545459 | 7.156554626 | 6.957274376 | 0.199280251 | 0.009271084 |
| Hpse2 | 2.620956713 | 2.641813663 | 2.611743934 | 0.03006973 | 0.009212779 |
| Aggf1 | 10.46507961 | 10.57493573 | 10.45592909 | 0.119006643 | 0.009150517 |
| Mms22l | 7.872649448 | 7.851008127 | 7.863504892 | -0.012496765 | 0.009144556 |
| Olfr446 | 2.796806266 | 2.751503232 | 2.787710542 | -0.03620731 | 0.009095724 |

| | | | | | |
|---|---|---|---|---|---|
| Crabp1 | 8.742482222 | 8.58706598 | 8.733389533 | -0.146323554 | 0.009092689 |
| Olfr774 | 2.536064723 | 2.490672654 | 2.526978532 | -0.036305878 | 0.009086191 |
| Sec11c | 10.08067981 | 9.995243029 | 10.07170488 | -0.076461851 | 0.008974927 |
| Ndufaf7 | 8.426786847 | 8.310623735 | 8.417823418 | -0.107199683 | 0.008963428 |
| Crym | 3.959418563 | 3.973250456 | 3.950469616 | 0.022780841 | 0.008948947 |
| Tmem91 | 1.941989463 | 1.914569758 | 1.933080992 | -0.018511235 | 0.008908471 |
| Gm17823 | 1.781283068 | 1.746060137 | 1.772403707 | -0.02634357 | 0.008879361 |
| Armt1 | 8.640915698 | 8.674691227 | 8.632060699 | 0.042630528 | 0.008854999 |
| Plxnc1 | 7.836028055 | 8.028723826 | 7.827173235 | 0.201550591 | 0.00885482 |
| Dscaml1 | 1.612022972 | 1.641485146 | 1.60320477 | 0.038280376 | 0.008818202 |
| Fam19a3 | 4.167855178 | 4.059815612 | 4.159061863 | -0.099246251 | 0.008793315 |
| Atp5mpl | 10.0027942 | 9.920533107 | 9.99405302 | -0.073519913 | 0.008741184 |
| Zfp563 | 6.965692326 | 7.131801495 | 6.956986791 | 0.174814704 | 0.008705535 |
| Myf5 | 1.997400618 | 2.042843842 | 1.98870861 | 0.054135232 | 0.008692008 |
| Vgll2 | 4.856020519 | 4.851871632 | 4.847336514 | 0.004535118 | 0.008684005 |
| Cdc6 | 7.254837425 | 7.340795675 | 7.246193344 | 0.094602331 | 0.008644081 |
| Cyp20a1 | 8.35069468 | 8.311216071 | 8.342125396 | -0.030909325 | 0.008569283 |
| Atf7ip2 | 1.33615845 | 1.349169646 | 1.327638249 | 0.021531397 | 0.008520201 |
| Caps2 | 2.253099799 | 2.213512236 | 2.244668361 | -0.031156125 | 0.008431438 |
| 5730480H06Rik | 7.280934292 | 7.317169691 | 7.272543079 | 0.044626612 | 0.008391213 |
| Mia3 | 10.72977849 | 10.7205192 | 10.72139392 | -0.000874727 | 0.00838457 |
| Uqcrb | 11.08667352 | 11.00353882 | 11.07830061 | -0.07476179 | 0.008372907 |
| Pcdhb15 | 3.304845933 | 3.335506847 | 3.296486984 | 0.039019863 | 0.008358949 |
| Vmn1r49 | 2.080123086 | 2.062806096 | 2.071778323 | -0.008972227 | 0.008344763 |
| Rgs12 | 10.84882847 | 11.0246926 | 10.84048908 | 0.184203517 | 0.00833939 |
| Gm3383 | 1.547285648 | 1.48711641 | 1.538956415 | -0.051840005 | 0.008329233 |
| Clec12b | 4.01299102 | 4.015770092 | 4.004662103 | 0.011107988 | 0.008328917 |
| Atg2a | 8.662285473 | 9.017674957 | 8.65407525 | 0.363599706 | 0.008210223 |
| Kiss1r | 3.675997738 | 3.718633661 | 3.667807877 | 0.050825784 | 0.00818986 |
| Gm15931 | 3.537378436 | 3.628651691 | 3.529215614 | 0.099436077 | 0.008162822 |
| Gpr82 | 2.290376294 | 2.210920753 | 2.282215272 | -0.071294519 | 0.008161022 |
| Slc5a7 | 3.076384232 | 3.019981795 | 3.068413743 | -0.048431948 | 0.007970489 |

| | | | | | |
|---|---|---|---|---|---|
| BC053393 | 2.407585493 | 2.428640225 | 2.399617641 | 0.029022584 | 0.007967852 |
| Ly9 | 6.527654692 | 6.632670971 | 6.519687857 | 0.112983114 | 0.007966835 |
| Gm48365 | 5.347761826 | 5.408087842 | 5.339807352 | 0.06828049 | 0.007954474 |
| Cfap74 | 6.094638374 | 6.391864744 | 6.086698871 | 0.305165873 | 0.007939503 |
| Wars2 | 7.028102953 | 7.073696144 | 7.020167731 | 0.053528414 | 0.007935223 |
| 1110008L16Rik | 7.973755369 | 7.878996755 | 7.965842091 | -0.086845335 | 0.007913278 |
| Gm43235 | 2.810734614 | 2.860662472 | 2.802822733 | 0.057839739 | 0.007911881 |
| Dhrs1 | 10.26237625 | 10.11310727 | 10.25447239 | -0.141365123 | 0.007903857 |
| Fgd2 | 7.441040904 | 7.510916811 | 7.43316401 | 0.077752801 | 0.007876894 |
| Dync1h1 | 12.72341336 | 13.00589004 | 12.71554663 | 0.290343417 | 0.007866733 |
| Gm10357 | 1.409714045 | 1.383667281 | 1.40189093 | -0.018223649 | 0.007823115 |
| Gast | 1.611459067 | 1.623825746 | 1.603641705 | 0.020184041 | 0.007817362 |
| Atp6v1a | 11.5417582 | 11.6722507 | 11.53397661 | 0.13827409 | 0.007781597 |
| Mapk7 | 8.295553827 | 8.48567142 | 8.287844442 | 0.197826978 | 0.007709385 |
| Pikfyve | 10.05543883 | 9.995061716 | 10.04777216 | -0.052710444 | 0.007666673 |
| Ints6 | 9.376046692 | 9.433798913 | 9.368382785 | 0.065416128 | 0.007663907 |
| Zfp536 | 5.942008833 | 6.27701463 | 5.934433501 | 0.342581129 | 0.007575332 |
| Ppp1r36 | 3.801174956 | 3.67857657 | 3.79363217 | -0.1150556 | 0.007542786 |
| Gm12098 | 1.528520417 | 1.536674666 | 1.521028656 | 0.015646009 | 0.007491761 |
| Tcp11 | 2.474851217 | 2.533189533 | 2.467408033 | 0.0657815 | 0.007443184 |
| Olfr652 | 1.861172313 | 1.880198238 | 1.853733017 | 0.026465221 | 0.007439296 |
| Serpinb9b | 4.007498906 | 3.946761213 | 4.000063243 | -0.05330203 | 0.007435663 |
| Olfr1198 | 1.668740562 | 1.660836275 | 1.661381033 | -0.000544757 | 0.007359529 |
| Ptgs1 | 12.14433173 | 12.13569392 | 12.13702263 | -0.00132871 | 0.007309103 |
| Ect2l | 2.597301858 | 2.597152339 | 2.590183177 | 0.006969163 | 0.007118682 |
| Yes1 | 9.40894399 | 9.302485863 | 9.401940855 | -0.099454993 | 0.007003134 |
| Gm10490 | 1.922186429 | 1.859977771 | 1.915187023 | -0.055209252 | 0.006999406 |
| Zfand2b | 8.858879692 | 8.880509152 | 8.851887599 | 0.028621554 | 0.006992093 |
| Gm15280 | 3.521713264 | 3.499027908 | 3.514756487 | -0.015728579 | 0.006956777 |
| Fxyd6 | 9.72128042 | 9.556977458 | 9.714333224 | -0.157355766 | 0.006947196 |
| D630023F18Rik | 2.300919088 | 2.358496663 | 2.293979468 | 0.064517196 | 0.006939621 |
| Impdh2-ps | 0.856564443 | 0.854066272 | 0.849632964 | 0.004433309 | 0.006931479 |

| | | | | | |
|---|---|---|---|---|---|
| Efcab12 | 2.068259953 | 2.080732705 | 2.061374591 | 0.019358114 | 0.006885362 |
| Olfr678 | 2.003998863 | 2.022661932 | 1.997123282 | 0.02553865 | 0.00687558 |
| Anxa4 | 11.15001857 | 11.19879641 | 11.143151 | 0.05564541 | 0.00686757 |
| Gm47842 | 4.586905416 | 4.619845161 | 4.580051585 | 0.039793575 | 0.006853831 |
| Appl1 | 10.70896203 | 10.63671665 | 10.70218907 | -0.065472417 | 0.006772967 |
| Gm20517 | 3.601192741 | 3.548248024 | 3.594448394 | -0.046200369 | 0.006744347 |
| 2310030G06Rik | 7.08340431 | 7.268882273 | 7.076667895 | 0.192214378 | 0.006736415 |
| Slc25a44 | 9.371869167 | 9.360711517 | 9.365169116 | -0.004457599 | 0.006700051 |
| Gm2274 | 3.404854931 | 3.567905421 | 3.398161775 | 0.169743646 | 0.006693156 |
| Bbs5 | 6.26621973 | 6.289887034 | 6.259528734 | 0.0303583 | 0.006690996 |
| Ikbke | 9.057343707 | 9.370603086 | 9.050666493 | 0.319936593 | 0.006677214 |
| Pcsk9 | 3.536182611 | 3.553182138 | 3.529527082 | 0.023655055 | 0.006655528 |
| Lancl3 | 2.082117357 | 2.03830467 | 2.075594047 | -0.037289378 | 0.006523309 |
| AC129217.1 | 1.705606616 | 1.761354834 | 1.699085528 | 0.062269306 | 0.006521088 |
| Thap12 | 11.05672291 | 10.91801874 | 11.05020686 | -0.132188113 | 0.00651605 |
| Olfr177 | 1.411542982 | 1.39814931 | 1.405122296 | -0.006972986 | 0.006420686 |
| Iqck | 3.878663135 | 3.814142773 | 3.872242628 | -0.058099855 | 0.006420507 |
| Abcc6 | 2.323516496 | 2.279335598 | 2.317119921 | -0.037784323 | 0.006396575 |
| Fgf4 | 1.399611088 | 1.385342632 | 1.393236017 | -0.007893385 | 0.006375071 |
| Prr15l | 4.727060796 | 4.76209061 | 4.720701636 | 0.041388974 | 0.00635916 |
| Nhlrc4 | 1.689616299 | 1.72463522 | 1.683265005 | 0.041370216 | 0.006351294 |
| Gm7862 | 3.111913247 | 3.190146562 | 3.105564311 | 0.084582251 | 0.006348935 |
| Vwce | 2.131531587 | 2.138582763 | 2.12524071 | 0.013342053 | 0.006290877 |
| Rps17 | 12.88149107 | 12.83534246 | 12.87520508 | -0.039862617 | 0.006285987 |
| Nhlrc3 | 8.820115692 | 8.786040118 | 8.813905944 | -0.027865825 | 0.006209749 |
| Fermt1 | 9.462988134 | 9.543068462 | 9.45679647 | 0.086271992 | 0.006191664 |
| Naa30 | 9.937887147 | 9.781434285 | 9.931728103 | -0.150293818 | 0.006159044 |
| 4932411N23Rik | 1.797275092 | 1.821110104 | 1.791129203 | 0.029980901 | 0.006145889 |
| Bri3 | 9.386946933 | 9.397063401 | 9.380830905 | 0.016232496 | 0.006116028 |
| Arl1 | 11.23297503 | 11.08591739 | 11.22689935 | -0.140981963 | 0.006075673 |
| Pramel7 | 1.996116426 | 2.003788076 | 1.9900816 | 0.013706476 | 0.006034826 |
| Gm12573 | 4.873222356 | 4.715284964 | 4.867196605 | -0.151911641 | 0.006025751 |

| | | | | | |
|---|---|---|---|---|---|
| Gm21887 | 3.844435673 | 3.92320923 | 3.838443987 | 0.084765243 | 0.005991687 |
| Prpf18 | 9.861971351 | 9.824420915 | 9.85601732 | -0.031596405 | 0.005954031 |
| Myo5b | 8.71615691 | 8.518169474 | 8.710222804 | -0.192053329 | 0.005934106 |
| Gen1 | 7.049620678 | 7.064744254 | 7.043686821 | 0.021057433 | 0.005933857 |
| Tmpo | 11.09254918 | 11.14482833 | 11.08664523 | 0.0581831 | 0.00590395 |
| Olfr1454 | 1.301518662 | 1.29345113 | 1.295624088 | -0.002172958 | 0.005894574 |
| Gm5075 | 2.15128002 | 2.16629914 | 2.145389379 | 0.020909761 | 0.005890641 |
| Prdm15 | 8.232881378 | 8.502811859 | 8.227021296 | 0.275790563 | 0.005860082 |
| Vmn2r86 | 1.85812094 | 1.886963051 | 1.852263702 | 0.034699349 | 0.005857239 |
| Matr3 | 11.46260896 | 11.38416365 | 11.45677511 | -0.072611463 | 0.00583385 |
| Olfr1191-ps1 | 3.763781332 | 3.761220529 | 3.75798372 | 0.003236809 | 0.005797612 |
| Olfr1156 | 3.412794988 | 3.365003609 | 3.40702116 | -0.042017551 | 0.005773828 |
| Gm10260 | 2.138682456 | 2.070959302 | 2.13291302 | -0.061953719 | 0.005769436 |
| Tsta3 | 8.444702864 | 8.465192131 | 8.438949729 | 0.026242402 | 0.005753135 |
| Npffr1 | 1.97536574 | 1.993880598 | 1.969636272 | 0.024244326 | 0.005729468 |
| Cyb5d1 | 7.720807056 | 7.624559741 | 7.715096112 | -0.090536371 | 0.005710944 |
| Sfmbt1 | 8.554077383 | 8.708624967 | 8.54843379 | 0.160191177 | 0.005643593 |
| Bmp15 | 1.568430139 | 1.58642414 | 1.562791531 | 0.023632609 | 0.005638608 |
| 6030468B19Rik | 1.265907375 | 1.229626613 | 1.260292038 | -0.030665425 | 0.005615337 |
| 5830417I10Rik | 9.559142071 | 9.617334955 | 9.553536562 | 0.063798393 | 0.00560551 |
| Bclaf3 | 8.207339999 | 8.138578939 | 8.201750497 | -0.063171558 | 0.005589502 |
| Slc45a3 | 8.310528328 | 8.330774084 | 8.304966323 | 0.025807761 | 0.005562005 |
| Bcl2l10 | 2.2268862 | 2.301735321 | 2.221327187 | 0.080408134 | 0.005559013 |
| Mtmr7 | 3.97412906 | 4.001556799 | 3.96858709 | 0.032969709 | 0.00554197 |
| Serinc4 | 1.656450935 | 1.687003469 | 1.650965354 | 0.036038115 | 0.005485581 |
| Gm17981 | 1.87349076 | 1.891663486 | 1.868009803 | 0.023653683 | 0.005480957 |
| Haus3 | 7.914888232 | 7.986779279 | 7.909460914 | 0.077318365 | 0.005427319 |
| B430305J03Rik | 1.998983145 | 1.999203743 | 1.993597495 | 0.005606247 | 0.00538565 |
| Pcdhb13 | 3.110025925 | 3.054030087 | 3.104676042 | -0.050645956 | 0.005349883 |
| Gm19000 | 3.444229635 | 3.460443358 | 3.438898336 | 0.021545021 | 0.005331299 |
| Atcay | 1.459824524 | 1.491050395 | 1.454513068 | 0.036537327 | 0.005311456 |
| Rnf11 | 11.75512706 | 11.56196763 | 11.74983132 | -0.18786369 | 0.005295743 |

| | | | | | |
|---|---|---|---|---|---|
| Gm4468 | 1.756510286 | 1.760258094 | 1.751221721 | 0.009036372 | 0.005288565 |
| Aph1c | 6.839957623 | 6.852363232 | 6.834695221 | 0.017668011 | 0.005262402 |
| Ager | 3.955527045 | 3.998692624 | 3.950267418 | 0.048425206 | 0.005259627 |
| Olfr1537 | 2.32361184 | 2.316576438 | 2.318417736 | -0.001841298 | 0.005194104 |
| Copa | 12.25377417 | 12.32245605 | 12.24860421 | 0.073851843 | 0.00516996 |
| Rwdd2b | 7.105175754 | 7.004671261 | 7.100025892 | -0.095354631 | 0.005149862 |
| Olfr713 | 3.80631131 | 3.787705579 | 3.801185976 | -0.013480397 | 0.005125334 |
| Gm8172 | 1.34860027 | 1.289429873 | 1.343485992 | -0.054056119 | 0.005114278 |
| Gm10563 | 2.654181829 | 2.680132423 | 2.64908564 | 0.031046782 | 0.005096189 |
| Gm10031 | 0.804440241 | 0.832898914 | 0.799369779 | 0.033529135 | 0.005070462 |
| Gm49358 | 1.453199916 | 1.443251714 | 1.448178363 | -0.004926649 | 0.005021553 |
| BC049715 | 2.862345179 | 2.967559812 | 2.857326923 | 0.110232889 | 0.005018256 |
| Skp2 | 7.296784633 | 7.395094561 | 7.291797532 | 0.10329703 | 0.004987102 |
| Olfr851 | 2.743757741 | 2.698214065 | 2.738783318 | -0.040569253 | 0.004974423 |
| Abhd13 | 9.91973265 | 9.784880641 | 9.914786112 | -0.129905471 | 0.004946538 |
| Cdc42se2 | 10.01417235 | 9.990340401 | 10.00927244 | -0.018932035 | 0.004899918 |
| Ccdc60 | 3.372789117 | 3.357846968 | 3.367930417 | -0.010083449 | 0.0048587 |
| Tert | 4.053862958 | 4.007918791 | 4.049027547 | -0.041108756 | 0.004835411 |
| Fbxo27 | 4.494481084 | 4.565555626 | 4.489648429 | 0.075907197 | 0.004832655 |
| Chpf2 | 9.101002438 | 9.497725043 | 9.09617431 | 0.401550733 | 0.004828128 |
| Olfr279 | 3.876624606 | 3.840875346 | 3.871853343 | -0.030977997 | 0.004771263 |
| Cyp2j12 | 7.444424175 | 7.298646611 | 7.439702706 | -0.141056095 | 0.00472147 |
| 4930548H24Rik | 2.171596103 | 2.224041593 | 2.166889894 | 0.057151699 | 0.004706209 |
| Amph | 3.654915584 | 3.597450461 | 3.650213424 | -0.052762963 | 0.00470216 |
| Arl5b | 10.0725218 | 10.19395194 | 10.0678233 | 0.126128636 | 0.004698495 |
| Dnajb3 | 5.502586775 | 5.344741764 | 5.497902374 | -0.15316061 | 0.004684401 |
| Gm13769 | 2.339128436 | 2.302169598 | 2.334473657 | -0.032304059 | 0.00465478 |
| A130006I12Rik | 1.657328461 | 1.688669154 | 1.65267926 | 0.035989894 | 0.004649201 |
| Gm12161 | 1.921100468 | 2.005992086 | 1.916468756 | 0.08952333 | 0.004631712 |
| Frmd5 | 3.013697096 | 3.034484089 | 3.009067725 | 0.025416364 | 0.004629371 |
| Actr6 | 7.079481493 | 6.972838759 | 7.074903714 | -0.102064955 | 0.004577779 |
| Gm12166 | 0.846961023 | 0.796820998 | 0.842429588 | -0.04560859 | 0.004531435 |

| | | | | | |
|---|---|---|---|---|---|
| Mro | 2.643939003 | 2.631150809 | 2.639417056 | -0.008266247 | 0.004521947 |
| Cisd3 | 8.463043205 | 8.362477843 | 8.458549827 | -0.096071984 | 0.004493378 |
| Spns1 | 9.16441974 | 9.396962023 | 9.159960387 | 0.237001637 | 0.004459353 |
| Zfp748 | 7.899854427 | 7.984520306 | 7.895425324 | 0.089094982 | 0.004429103 |
| Gm6710 | 6.262009778 | 6.192412835 | 6.257617787 | -0.065204952 | 0.004391991 |
| Ppp2r1b | 10.1233615 | 10.06482848 | 10.11897799 | -0.054149513 | 0.004383507 |
| Ehmt2 | 10.59487943 | 10.64362506 | 10.59051636 | 0.0531087 | 0.00436307 |
| Tdrd9 | 1.430609188 | 1.444286742 | 1.426246242 | 0.0180405 | 0.004362946 |
| Gm48177 | 3.263570949 | 3.288980446 | 3.259301167 | 0.02967928 | 0.004269783 |
| Hspa13 | 9.380474655 | 9.409842325 | 9.376264141 | 0.033578184 | 0.004210514 |
| Ywhaq-ps3 | 0.928214841 | 0.922503399 | 0.924019084 | -0.001515685 | 0.004195757 |
| Olfr307 | 1.256347054 | 1.221375332 | 1.252155798 | -0.030780466 | 0.004191256 |
| 2410015M20Rik | 10.00408248 | 9.894339215 | 9.999916171 | -0.105576955 | 0.004166308 |
| Vwa3b | 3.0169101 | 3.021032178 | 3.012765254 | 0.008266924 | 0.004144845 |
| Prdm11 | 7.033702836 | 7.070128708 | 7.029653758 | 0.04047495 | 0.004049078 |
| Epha7 | 4.630753506 | 4.64025987 | 4.626731302 | 0.013528568 | 0.004022205 |
| Gm14569 | 2.077387794 | 2.054314069 | 2.073371812 | -0.019057743 | 0.004015982 |
| Vti1b | 10.13632118 | 9.999183778 | 10.13234402 | -0.133160246 | 0.003977157 |
| Pid1 | 9.208884869 | 9.078872593 | 9.204920514 | -0.126047921 | 0.003964355 |
| Tmx4 | 8.741674299 | 8.557427665 | 8.737719427 | -0.180291762 | 0.003954872 |
| Acvr1 | 9.1959824 | 9.154505808 | 9.192052471 | -0.037546663 | 0.003929929 |
| Zfp91 | 11.85860897 | 11.71631176 | 11.85471277 | -0.138401007 | 0.003896207 |
| Gm3200 | 1.937107268 | 2.004376654 | 1.933252006 | 0.071124648 | 0.003855262 |
| Cd300ld2 | 2.894781841 | 2.88500198 | 2.890935814 | -0.005933834 | 0.003846026 |
| Gm42421 | 0.583224324 | 0.599524022 | 0.579379507 | 0.020144514 | 0.003844817 |
| Pogk | 10.01195962 | 9.906138814 | 10.00812103 | -0.101982219 | 0.003838585 |
| Tyms | 8.811061737 | 8.951277854 | 8.807252335 | 0.144025519 | 0.003809401 |
| Dnajc13 | 10.6790097 | 11.10985181 | 10.67521956 | 0.434632243 | 0.003790133 |
| Olfr689 | 1.853262986 | 1.826463388 | 1.849473534 | -0.023010146 | 0.003789452 |
| Gm13717 | 3.510208725 | 3.547097118 | 3.506420448 | 0.040676669 | 0.003788277 |
| E330034G19Rik | 2.335250088 | 2.310505998 | 2.331504478 | -0.02099848 | 0.00374561 |
| Platr17 | 2.438085023 | 2.394605495 | 2.434370262 | -0.039764767 | 0.003714761 |

| | | | | | |
|---|---|---|---|---|---|
| Ankhd1 | 10.74371364 | 10.72653223 | 10.74000987 | -0.01347764 | 0.00370377 |
| Olfr1268-ps1 | 2.948132701 | 2.95028359 | 2.944441607 | 0.005841983 | 0.003691094 |
| Glul | 13.23360281 | 12.9266425 | 13.22992169 | -0.303279187 | 0.003681127 |
| Slc23a4 | 4.619465648 | 4.656640535 | 4.615786929 | 0.040853607 | 0.003678719 |
| Ap5z1 | 8.008314473 | 8.305652632 | 8.004704821 | 0.300947812 | 0.003609652 |
| Olfr1440 | 2.096131642 | 2.083451659 | 2.092527058 | -0.009075399 | 0.003604584 |
| Hist1h2ac | 2.948291032 | 2.982030621 | 2.944755704 | 0.037274917 | 0.003535328 |
| Olfr1178 | 3.169147656 | 3.217299598 | 3.165720191 | 0.051579407 | 0.003427465 |
| Scn3b | 4.583315993 | 4.384285397 | 4.579907811 | -0.195622414 | 0.003408182 |
| BC052040 | 6.750206057 | 6.75623484 | 6.74679989 | 0.00943495 | 0.003406167 |
| Pcdhb17 | 6.903489675 | 6.888269181 | 6.900084911 | -0.01181573 | 0.003404765 |
| Mss51 | 5.161176636 | 5.22538526 | 5.157806007 | 0.067579253 | 0.003370629 |
| Vmn1r6 | 4.183624142 | 4.219733207 | 4.180258048 | 0.03947516 | 0.003366095 |
| P2ry1 | 9.534449258 | 9.682925824 | 9.531090261 | 0.151835563 | 0.003358997 |
| Nol7 | 9.39802457 | 9.336858965 | 9.394756212 | -0.057897247 | 0.003268358 |
| Stag3 | 4.977014914 | 5.19055773 | 4.973747368 | 0.216810362 | 0.003267546 |
| Pax6 | 2.655785453 | 2.564250851 | 2.652534367 | -0.088283516 | 0.003251086 |
| Slc26a4 | 2.507224782 | 2.553540442 | 2.504013494 | 0.049526948 | 0.003211288 |
| Nup210 | 7.984106638 | 8.071737822 | 7.980908968 | 0.090828854 | 0.00319767 |
| Smc3 | 10.78182677 | 10.79285364 | 10.77867451 | 0.01417913 | 0.003152267 |
| Catsper3 | 1.618256446 | 1.641731945 | 1.615106401 | 0.026625545 | 0.003150045 |
| Zdhhc22 | 1.491410838 | 1.512481642 | 1.488276585 | 0.024205057 | 0.003134253 |
| 2210010C04Rik | 1.528600963 | 1.481196486 | 1.525482388 | -0.044285902 | 0.003118575 |
| Hvcn1 | 7.456499751 | 7.877394849 | 7.453417563 | 0.423977286 | 0.003082188 |
| 1110065P20Rik | 7.220413391 | 7.175491398 | 7.217341598 | -0.0418502 | 0.003071793 |
| Gm47632 | 3.489081566 | 3.560500557 | 3.48601707 | 0.074483488 | 0.003064496 |
| Rplp0 | 14.71111606 | 14.80201666 | 14.70806288 | 0.09395378 | 0.00305318 |
| Wdr82 | 10.76132507 | 10.79516964 | 10.75832397 | 0.036845667 | 0.003001097 |
| Hist1h4d | 1.959376419 | 2.019068932 | 1.956453892 | 0.06261504 | 0.002922527 |
| Borcs5 | 7.954845343 | 7.858911214 | 7.951925682 | -0.093014468 | 0.00291966 |
| Aptx | 8.790393084 | 8.686703218 | 8.787480678 | -0.100777459 | 0.002912406 |
| Olfr1264 | 1.324325029 | 1.355132907 | 1.321427415 | 0.033705491 | 0.002897613 |

| | | | | | |
|---|---|---|---|---|---|
| Efcab6 | 4.412648654 | 5.118964908 | 4.409752666 | 0.709212241 | 0.002895988 |
| Tor4a | 8.82951206 | 8.905099392 | 8.826620119 | 0.078479273 | 0.002891941 |
| Tmem9b | 10.32283647 | 10.17116768 | 10.31995585 | -0.148788173 | 0.00288062 |
| Sprr1b | 10.90107086 | 11.21120404 | 10.8982102 | 0.312993833 | 0.00286066 |
| Tmem230 | 8.538162842 | 8.455209834 | 8.535342709 | -0.080132875 | 0.002820133 |
| Asxl3 | 3.051982915 | 3.044915286 | 3.049172349 | -0.004257063 | 0.002810566 |
| Espnl | 1.289520426 | 1.358698507 | 1.286713976 | 0.071984531 | 0.00280645 |
| Loxhd1 | 1.787114161 | 1.827066584 | 1.784344353 | 0.042722231 | 0.002769808 |
| Crtac1 | 3.213548692 | 3.17013524 | 3.210795124 | -0.040659883 | 0.002753568 |
| Tusc1 | 5.477493156 | 5.403063427 | 5.474779037 | -0.07171561 | 0.002714118 |
| Vmn2r30 | 2.32125712 | 2.264083164 | 2.318583503 | -0.05450034 | 0.002673617 |
| Nrl | 1.246059982 | 1.221064328 | 1.243419734 | -0.022355406 | 0.002640248 |
| Ren1 | 1.100901669 | 1.11108749 | 1.098337022 | 0.012750468 | 0.002564647 |
| Vmn1r42 | 1.972516659 | 1.957454897 | 1.969984634 | -0.012529736 | 0.002532025 |
| Bptf | 10.94863979 | 11.025525 | 10.94610786 | 0.07941714 | 0.002531937 |
| L3mbtl1 | 2.317284472 | 2.386963288 | 2.314800283 | 0.072163005 | 0.002484189 |
| Ube3a | 11.09116059 | 10.88017233 | 11.08868571 | -0.208513387 | 0.002474873 |
| G6pc3 | 8.434322898 | 8.507735456 | 8.431864202 | 0.075871255 | 0.002458696 |
| Hoxa6 | 3.343567887 | 3.421428968 | 3.341144513 | 0.080284456 | 0.002423375 |
| Pip4k2c | 10.34038997 | 10.3838444 | 10.33797348 | 0.045870923 | 0.00241649 |
| Rpl23a | 13.35771088 | 13.32559098 | 13.35531384 | -0.029722863 | 0.00239704 |
| Gm5864 | 0.844996553 | 0.873853716 | 0.842607636 | 0.03124608 | 0.002388917 |
| Rrh | 1.428809223 | 1.440839634 | 1.426427918 | 0.014411715 | 0.002381305 |
| Gm6169 | 2.904427936 | 2.880201165 | 2.90205014 | -0.021848975 | 0.002377795 |
| Gipc3 | 2.70944647 | 2.720011468 | 2.707079562 | 0.012931905 | 0.002366907 |
| Gm20763 | 1.962470515 | 1.992523319 | 1.960104674 | 0.032418645 | 0.002365841 |
| Slc5a5 | 1.709806668 | 1.701540725 | 1.707471597 | -0.005930871 | 0.002335071 |
| Pank4 | 8.485275068 | 8.608855068 | 8.482944736 | 0.125910332 | 0.002330332 |
| Satb2 | 5.029822319 | 5.047265629 | 5.027493724 | 0.019771905 | 0.002328595 |
| Ap4b1 | 8.520818459 | 8.458619182 | 8.518491334 | -0.059872151 | 0.002327126 |
| Hcn3 | 2.133820634 | 2.188637425 | 2.131504008 | 0.057133417 | 0.002316626 |
| Gm7536 | 12.60885776 | 12.6134111 | 12.60657539 | 0.006835713 | 0.002282367 |

| | | | | | |
|---|---|---|---|---|---|
| Gm12577 | 1.585492528 | 1.603749095 | 1.583230798 | 0.020518296 | 0.00226173 |
| Vmn2r4 | 3.951742854 | 3.904859076 | 3.949481415 | -0.044622339 | 0.002261439 |
| Fam98c | 7.792277416 | 8.112944646 | 7.790048995 | 0.322895651 | 0.002228421 |
| Gcm1 | 2.339051677 | 2.323567287 | 2.336839618 | -0.01327233 | 0.00221206 |
| Stk35 | 9.34680329 | 9.488720377 | 9.344672521 | 0.144047856 | 0.002130769 |
| Armcx6 | 4.893788909 | 4.903801203 | 4.891696377 | 0.012104827 | 0.002092532 |
| Gapdh | 14.6846706 | 14.74994824 | 14.68258133 | 0.067366903 | 0.00208927 |
| 4921524J17Rik | 7.495382199 | 7.387174527 | 7.493373437 | -0.10619891 | 0.002008762 |
| Cr2 | 1.109640298 | 1.282697984 | 1.107706294 | 0.174991691 | 0.001934004 |
| Yipf1 | 9.112536127 | 8.98831134 | 9.110603262 | -0.122291922 | 0.001932865 |
| Lipo2 | 1.781508234 | 1.812321849 | 1.779674677 | 0.032647171 | 0.001833556 |
| Ccl28 | 3.403027201 | 3.412994754 | 3.40119969 | 0.011795064 | 0.00182751 |
| Gm20784 | 1.508337178 | 1.557942058 | 1.506641127 | 0.051300931 | 0.00169605 |
| Gm3488 | 1.380394021 | 1.399093848 | 1.378703025 | 0.020390823 | 0.001690996 |
| Acsm2 | 1.69506826 | 1.687860078 | 1.693391197 | -0.005531119 | 0.001677063 |
| 1810024B03Rik | 4.201349854 | 4.186248605 | 4.199692089 | -0.013443484 | 0.001657765 |
| Cyp2a5 | 4.298574388 | 4.265730054 | 4.296927172 | -0.031197118 | 0.001647216 |
| Gm7909 | 2.680941766 | 2.704985664 | 2.6793147 | 0.025670964 | 0.001627065 |
| Ccdc36 | 3.826207671 | 3.791885683 | 3.824620886 | -0.032735203 | 0.001586785 |
| Fbxo39 | 1.429930236 | 1.435282855 | 1.428353866 | 0.00692899 | 0.00157637 |
| Ccl1 | 2.817580758 | 2.859395555 | 2.816039079 | 0.043356476 | 0.001541679 |
| 2610028H24Rik | 1.683242146 | 1.660379029 | 1.681716584 | -0.021337555 | 0.001525562 |
| 1110059G10Rik | 7.898959666 | 7.710884353 | 7.897435079 | -0.186550726 | 0.001524587 |
| Gm47967 | 3.844655737 | 3.894805288 | 3.843143283 | 0.051662005 | 0.001512454 |
| Cenpt | 8.041434118 | 8.213841212 | 8.03998728 | 0.173853932 | 0.001446838 |
| Pitx3 | 1.717676378 | 1.715324042 | 1.716242142 | -0.0009181 | 0.001434236 |
| Naa38 | 8.342395059 | 8.301512566 | 8.340964499 | -0.039451933 | 0.00143056 |
| Trbj2-7 | 3.677161075 | 3.666166395 | 3.675772941 | -0.009606546 | 0.001388133 |
| Olfr6 | 2.217667761 | 2.227332055 | 2.216307923 | 0.011024132 | 0.001359838 |
| Eri1 | 9.414561471 | 9.458047524 | 9.413225838 | 0.044821686 | 0.001335633 |
| Rasef | 3.406816947 | 3.332136172 | 3.405481824 | -0.073345652 | 0.001335123 |
| Dera | 8.547590183 | 8.506790769 | 8.546274206 | -0.039483437 | 0.001315977 |

| | | | | | |
|---|---|---|---|---|---|
| Chil5 | 1.65509368 | 1.592174968 | 1.653809817 | -0.061634849 | 0.001283863 |
| Borcs6 | 8.232220867 | 8.209460476 | 8.230941227 | -0.021480751 | 0.00127964 |
| Atp7b | 2.002661652 | 2.001386586 | 2.001419258 | -3.26717E-05 | 0.001242394 |
| AC115631.1 | 9.18723888 | 9.239915082 | 9.1860029 | 0.053912182 | 0.001235979 |
| Sel1l2 | 2.189412134 | 2.17000465 | 2.188180932 | -0.018176283 | 0.001231202 |
| H1fx | 4.004857345 | 4.053337295 | 4.003649725 | 0.04968757 | 0.00120762 |
| Fzd10 | 9.559740785 | 9.72868632 | 9.558596669 | 0.170089651 | 0.001144116 |
| Ccdc117 | 9.191623431 | 9.514015795 | 9.19048785 | 0.323527945 | 0.001135581 |
| Rtl4 | 1.012170264 | 1.016871123 | 1.011042357 | 0.005828766 | 0.001127907 |
| Chd1l | 8.786977146 | 8.737103375 | 8.785861648 | -0.048758274 | 0.001115498 |
| Virma | 10.0563076 | 10.05910751 | 10.05519843 | 0.003909086 | 0.001109173 |
| Ptcd2 | 9.24245473 | 9.233262558 | 9.241428503 | -0.008165945 | 0.001026227 |
| mt-Nd6 | 10.37701554 | 10.11426664 | 10.37602713 | -0.26176049 | 0.000988413 |
| Aig1 | 8.370840265 | 8.361932978 | 8.369890827 | -0.007957849 | 0.000949438 |
| Gm48789 | 1.477594223 | 1.478442808 | 1.476670274 | 0.001772535 | 0.000923949 |
| C330021F23Rik | 2.518509947 | 2.49281967 | 2.517599738 | -0.024780068 | 0.000910209 |
| Macf1 | 12.49566648 | 12.53101059 | 12.49476097 | 0.036249617 | 0.00090551 |
| Rhbdf1 | 9.923370385 | 10.34349158 | 9.922511822 | 0.420979758 | 0.000858563 |
| Gm12097 | 2.954045055 | 3.015903414 | 2.953190979 | 0.062712435 | 0.000854077 |
| Hipk3 | 11.83519921 | 11.68741902 | 11.83435039 | -0.146931377 | 0.00084882 |
| Olfr147 | 4.737980941 | 4.839394202 | 4.737150542 | 0.102243659 | 0.000830398 |
| Rpl9 | 13.4252911 | 13.28629807 | 13.42449541 | -0.138197337 | 0.000795687 |
| Ofcc1 | 4.481967801 | 4.43891352 | 4.481176582 | -0.042263062 | 0.000791218 |
| Apba2 | 3.168833182 | 3.22187943 | 3.168088035 | 0.053791396 | 0.000745147 |
| Apol7e | 4.244937384 | 4.167184773 | 4.244209047 | -0.077024274 | 0.000728337 |
| Star | 4.768614789 | 4.929897468 | 4.767923167 | 0.161974301 | 0.000691623 |
| Vmn1r81 | 6.445173621 | 6.521622095 | 6.44448518 | 0.077136915 | 0.000688441 |
| Lonp2 | 10.77457061 | 10.607741 | 10.77388599 | -0.166144993 | 0.000684617 |
| Gm5575 | 1.885898595 | 1.940523691 | 1.885241139 | 0.055282552 | 0.000657457 |
| Rabep2 | 8.48775992 | 8.820873312 | 8.487103603 | 0.333769709 | 0.000656317 |
| Gm7972 | 2.0632246 | 2.097738423 | 2.062571235 | 0.035167189 | 0.000653366 |
| Ntn3 | 3.698445934 | 3.800103417 | 3.697821877 | 0.102281539 | 0.000624057 |

| | | | | | |
|---|---|---|---|---|---|
| Inhbe | 2.776430981 | 2.817799652 | 2.775828316 | 0.041971336 | 0.000602664 |
| Gm10499 | 3.342126586 | 3.364501286 | 3.341538992 | 0.022962294 | 0.000587594 |
| Tenm1 | 3.748718687 | 3.719702701 | 3.748155896 | -0.028453195 | 0.000562791 |
| Fam71a | 1.217580203 | 1.360825829 | 1.217062837 | 0.143762992 | 0.000517366 |
| Tph2 | 2.430394458 | 2.454439602 | 2.429883531 | 0.02455607 | 0.000510926 |
| Gm42814 | 1.726953972 | 1.865697176 | 1.726480853 | 0.139216323 | 0.000473119 |
| Rapgef3 | 7.783024667 | 7.75692852 | 7.782555692 | -0.025627173 | 0.000468975 |
| Ccr9 | 6.689656027 | 6.790268192 | 6.689198059 | 0.101070133 | 0.000457969 |
| Prpf40b | 8.505229753 | 8.561506842 | 8.504800926 | 0.056705916 | 0.000428827 |
| Ajuba | 10.709331 | 10.95850032 | 10.70890649 | 0.249593827 | 0.000424507 |
| Cd40lg | 1.747654338 | 1.748247061 | 1.747232652 | 0.001014409 | 0.000421686 |
| Trgv2 | 2.133404933 | 2.164302482 | 2.133003311 | 0.03129917 | 0.000401622 |
| Wnt2b | 4.461495685 | 4.399761412 | 4.461103242 | -0.06134183 | 0.000392443 |
| Gm5431 | 6.635459647 | 6.376314874 | 6.635126842 | -0.258811967 | 0.000332805 |
| Aaas | 8.747437858 | 8.816434405 | 8.747139822 | 0.069294583 | 0.000298037 |
| Six2 | 1.525126876 | 1.523826871 | 1.524833723 | -0.001006852 | 0.000293153 |
| Fasn | 13.87258084 | 13.53051458 | 13.87230041 | -0.341785833 | 0.00028043 |
| Nrg2 | 5.67628703 | 5.995685417 | 5.676088789 | 0.319596629 | 0.000198241 |
| Rpl19-ps11 | 1.787560192 | 1.847601472 | 1.787363661 | 0.060237811 | 0.000196532 |
| Pcsk1 | 2.265250237 | 2.299937476 | 2.26506775 | 0.034869726 | 0.000182487 |
| Aldoc | 7.341788291 | 7.179674391 | 7.341633374 | -0.161958983 | 0.000154917 |
| Rdh1 | 10.41792264 | 10.36106758 | 10.4177842 | -0.05671662 | 0.000138443 |
| Taf6l | 7.020454919 | 7.150257205 | 7.020336877 | 0.129920328 | 0.000118042 |
| Sbf2 | 9.859117839 | 9.817253836 | 9.859005514 | -0.041751678 | 0.000112324 |
| Olfr432 | 3.167223144 | 3.137965002 | 3.167169524 | -0.029204522 | 5.36197E-05 |
| Gm7450 | 4.153403609 | 4.132110984 | 4.153352001 | -0.021241017 | 5.1608E-05 |
| Zfp607a | 6.941277126 | 6.982236181 | 6.94122612 | 0.041010061 | 5.1006E-05 |
| Piwil4 | 2.748686344 | 2.763864328 | 2.748646165 | 0.015218162 | 4.01783E-05 |
| H3f3c | 0.536262242 | 0.54578705 | 0.536329161 | 0.009457889 | -6.6919E-05 |
| Nf2 | 11.00343064 | 11.07086388 | 11.00352616 | 0.067337723 | -9.55233E-05 |
| Mrpl46 | 8.224300544 | 8.116057357 | 8.224412923 | -0.108355566 | -0.000112379 |
| Gm13081 | 2.302667251 | 2.283085203 | 2.302823111 | -0.019737908 | -0.00015586 |

| | | | | | |
|---|---|---|---|---|---|
| Sbf1 | 10.83301787 | 11.01643613 | 10.83322165 | 0.183214483 | -0.000203783 |
| Abcc4 | 9.572271541 | 9.579450441 | 9.572496946 | 0.006953495 | -0.000225405 |
| Gm10364 | 1.083546078 | 1.069292754 | 1.083773876 | -0.014481122 | -0.000227798 |
| Map4k5 | 10.1826276 | 10.14829878 | 10.18287632 | -0.03457754 | -0.000248727 |
| Vpreb1 | 1.837383677 | 1.868892028 | 1.837675007 | 0.031217022 | -0.000291329 |
| Thrap3 | 11.75191233 | 11.72500702 | 11.75221934 | -0.027212313 | -0.00030701 |
| Stk3 | 8.548748896 | 8.479757684 | 8.549057932 | -0.069300248 | -0.000309036 |
| Actr5 | 7.226338009 | 7.46101497 | 7.22666148 | 0.23435349 | -0.000323471 |
| Otx1 | 3.80772885 | 3.821418735 | 3.808058488 | 0.013360247 | -0.000329638 |
| Nudt9 | 9.30064268 | 9.22740024 | 9.30099457 | -0.07359433 | -0.00035189 |
| Coq6 | 8.950003373 | 8.927738406 | 8.950391578 | -0.022653172 | -0.000388205 |
| Cdkl4 | 4.478666599 | 4.396977504 | 4.479078171 | -0.082100666 | -0.000411572 |
| Serp2 | 2.238402878 | 2.226570726 | 2.238849179 | -0.012278454 | -0.000446301 |
| Gm5637 | 1.772179327 | 1.825447277 | 1.772716773 | 0.052730504 | -0.000537446 |
| Gm11879 | 3.014444709 | 3.0545645 | 3.014996998 | 0.039567502 | -0.000552289 |
| Gm9892 | 1.31054115 | 1.350265228 | 1.31118387 | 0.039081357 | -0.00064272 |
| Irak1 | 10.68198389 | 10.74665622 | 10.68265843 | 0.06399779 | -0.000674547 |
| Rtf1 | 10.4408602 | 10.38573936 | 10.44154921 | -0.055809847 | -0.000689013 |
| Ptprh | 1.545758474 | 1.572984942 | 1.546499847 | 0.026485095 | -0.000741372 |
| Erich1 | 7.28076234 | 7.271832303 | 7.281536701 | -0.009704398 | -0.000774362 |
| 1110032F04Rik | 2.697688688 | 2.692966907 | 2.698477284 | -0.005510377 | -0.000788596 |
| Gm49373 | 2.616897483 | 2.856794022 | 2.617733576 | 0.239060446 | -0.000836093 |
| Gm45105 | 2.441140142 | 2.514932856 | 2.442024462 | 0.072908395 | -0.000884319 |
| Gm6254 | 3.382840717 | 3.481786605 | 3.383752444 | 0.098034161 | -0.000911727 |
| Rhox4d | 1.39047455 | 1.408076239 | 1.391403785 | 0.016672454 | -0.000929235 |
| Tspan13 | 8.619030178 | 8.248614895 | 8.619964807 | -0.371349912 | -0.000934629 |
| Hrh4 | 1.714924916 | 1.7599928 | 1.715886167 | 0.044106633 | -0.000961251 |
| AC148174.1 | 1.614358441 | 1.613134781 | 1.615346702 | -0.002211921 | -0.000988261 |
| Apol10b | 4.800184877 | 4.769621851 | 4.80122012 | -0.031598269 | -0.001035243 |
| Pcp4 | 1.96687485 | 1.953978647 | 1.967934198 | -0.013955552 | -0.001059348 |
| Gm11127 | 3.154340779 | 3.104132865 | 3.155402736 | -0.051269871 | -0.001061957 |
| Lta4h | 10.09554947 | 10.15702111 | 10.09661496 | 0.060406147 | -0.00106549 |

| | | | | | |
|---|---|---|---|---|---|
| Wdr83 | 8.2757704 | 8.228739789 | 8.276840407 | -0.048100619 | -0.001070007 |
| Olfr341 | 2.920660934 | 2.928796522 | 2.921735207 | 0.007061315 | -0.001074273 |
| Gm7285 | 3.066870482 | 3.15745942 | 3.067968706 | 0.089490714 | -0.001098224 |
| Gm37568 | 2.455963855 | 2.451187713 | 2.457107876 | -0.005920163 | -0.001144021 |
| Crybb1 | 1.907559151 | 1.917984971 | 1.908716362 | 0.009268609 | -0.001157211 |
| Gm13785 | 1.480408716 | 1.466420406 | 1.481602807 | -0.015182401 | -0.001194092 |
| Agbl2 | 2.939777465 | 2.890582029 | 2.941046662 | -0.050464633 | -0.001269198 |
| Gm49181 | 2.220714671 | 2.243435103 | 2.222011988 | 0.021423115 | -0.001297317 |
| Gm12879 | 0.987501686 | 0.985151366 | 0.988810471 | -0.003659105 | -0.001308785 |
| A030005L19Rik | 1.777450567 | 1.812081885 | 1.778792877 | 0.033289008 | -0.00134231 |
| Rab33a | 1.401330088 | 1.362443989 | 1.402698385 | -0.040254397 | -0.001368297 |
| D630033A02Rik | 5.058734961 | 4.958056935 | 5.060124325 | -0.10206739 | -0.001389364 |
| Pcdhb5 | 4.060178673 | 4.079395651 | 4.061572479 | 0.017823172 | -0.001393806 |
| Foxi3 | 2.066221895 | 2.083407952 | 2.067636522 | 0.01577143 | -0.001414627 |
| Olfr155 | 1.705582396 | 1.708522695 | 1.70700718 | 0.001515514 | -0.001424785 |
| Snap29 | 9.859050491 | 9.91551753 | 9.860476897 | 0.055040633 | -0.001426406 |
| Acot8 | 7.661707356 | 7.887627401 | 7.663137987 | 0.224489414 | -0.001430631 |
| Cyp8b1 | 1.481123195 | 1.477056909 | 1.482560445 | -0.005503536 | -0.00143725 |
| Trub2 | 8.978392037 | 8.897299052 | 8.979835001 | -0.082535949 | -0.001442964 |
| Tekt5 | 2.752931416 | 2.78157205 | 2.754412762 | 0.027159288 | -0.001481346 |
| Ctnna2 | 2.48568599 | 2.473466644 | 2.487192549 | -0.013725905 | -0.001506559 |
| Vav2 | 8.212544112 | 8.065802348 | 8.214054736 | -0.148252388 | -0.001510624 |
| Hps5 | 9.34144292 | 9.450833193 | 9.342963687 | 0.107869505 | -0.001520767 |
| Ino80b | 7.185386832 | 7.390054247 | 7.186938454 | 0.203115792 | -0.001551623 |
| Ndufv1 | 11.07685024 | 11.03526077 | 11.07841052 | -0.04314975 | -0.001560287 |
| Trdv4 | 5.409704836 | 5.396966871 | 5.411278733 | -0.014311862 | -0.001573897 |
| Xylb | 5.484329245 | 5.564498919 | 5.485921138 | 0.078577781 | -0.001591893 |
| Cul4b | 9.342729417 | 9.266902766 | 9.344408187 | -0.077505421 | -0.001678771 |
| Slc15a2 | 6.977056793 | 6.817368824 | 6.978757762 | -0.161388938 | -0.001700969 |
| Gm17509 | 1.786070336 | 1.765271063 | 1.787781984 | -0.022510921 | -0.001711648 |
| Slc7a14 | 3.081907536 | 3.076366589 | 3.083628171 | -0.007261581 | -0.001720634 |
| Gm13736 | 6.290147627 | 6.154674034 | 6.291937133 | -0.137263099 | -0.001789507 |

| | | | | | |
|---|---|---|---|---|---|
| Uty | 2.98973526 | 3.108408314 | 2.991565427 | 0.116842887 | -0.001830167 |
| Gm20458 | 0.948244084 | 0.897258888 | 0.950081551 | -0.052822664 | -0.001837467 |
| Plod2 | 7.410259642 | 7.589792349 | 7.412136087 | 0.177656261 | -0.001876446 |
| Gm3550 | 1.978831522 | 1.917062724 | 1.980708501 | -0.063645777 | -0.001876979 |
| Thoc7 | 9.719825874 | 9.62480809 | 9.721766238 | -0.096958148 | -0.001940365 |
| Slc35f5 | 10.4544095 | 10.47286975 | 10.45635742 | 0.016512327 | -0.00194792 |
| Eid2 | 5.538951823 | 5.422064167 | 5.540922779 | -0.118858612 | -0.001970956 |
| Ndufa3 | 9.927431595 | 9.851109962 | 9.929429299 | -0.078319337 | -0.001997703 |
| Pmp2 | 3.774283469 | 3.880171227 | 3.776361101 | 0.103810126 | -0.002077632 |
| Sap30l | 8.333060862 | 8.272831224 | 8.335178332 | -0.062347108 | -0.00211747 |
| Olfr373 | 2.714614762 | 2.737870794 | 2.716744082 | 0.021126712 | -0.00212932 |
| Gpx4 | 11.50595817 | 11.37750118 | 11.50811854 | -0.13061736 | -0.00216037 |
| Faxc | 5.126050564 | 5.154060875 | 5.128221973 | 0.025838902 | -0.00217141 |
| Memo1 | 9.894103092 | 9.885980494 | 9.896347216 | -0.010366722 | -0.002244125 |
| Sec22a | 8.850620536 | 8.714426812 | 8.852883899 | -0.138457087 | -0.002263363 |
| Gm15304 | 1.10141472 | 1.104705867 | 1.103680507 | 0.00102536 | -0.002265787 |
| Tex30 | 7.31660855 | 7.232165463 | 7.318888849 | -0.086723386 | -0.002280298 |
| Pura | 9.820521573 | 9.770714722 | 9.822807771 | -0.052093049 | -0.002286198 |
| Npm2 | 1.992536808 | 1.989234195 | 1.994823902 | -0.005589707 | -0.002287095 |
| Trim5 | 6.666935459 | 6.50123847 | 6.669255889 | -0.168017419 | -0.00232043 |
| Morn1 | 4.674274449 | 4.627272481 | 4.676646853 | -0.049374371 | -0.002372404 |
| Gm17655 | 5.71797209 | 5.685554333 | 5.720386353 | -0.03483202 | -0.002414262 |
| Olfr1048 | 2.08161979 | 2.080245027 | 2.084058498 | -0.003813472 | -0.002438709 |
| Whrn | 5.140319963 | 5.243113413 | 5.1427677 | 0.100345713 | -0.002447737 |
| Dio3 | 1.856218799 | 1.97131494 | 1.858686007 | 0.112628933 | -0.002467208 |
| Gm13357 | 2.08425539 | 2.097879267 | 2.086724036 | 0.011155231 | -0.002468646 |
| Vat1 | 12.34863332 | 12.40822142 | 12.35112827 | 0.057093153 | -0.00249495 |
| Nprl2 | 8.303531242 | 8.314378442 | 8.3060282 | 0.008350242 | -0.002496959 |
| Pabpc1l | 3.825669227 | 3.765494998 | 3.828203897 | -0.062708899 | -0.002534669 |
| Siglech | 4.883527728 | 4.73070941 | 4.886095268 | -0.155385857 | -0.00256754 |
| Gm15694 | 1.973833704 | 1.976395067 | 1.976403756 | -8.68933E-06 | -0.002570053 |
| Metrn | 5.9882149 | 6.157755289 | 5.990799503 | 0.166955785 | -0.002584603 |

| | | | | | |
|---|---|---|---|---|---|
| Ptprn2 | 2.085855484 | 2.122235594 | 2.088454679 | 0.033780915 | -0.002599195 |
| Cend1 | 1.411240663 | 1.411124186 | 1.413882827 | -0.002758641 | -0.002642164 |
| Dis3l2 | 8.512672525 | 8.647989849 | 8.515340181 | 0.132649668 | -0.002667657 |
| Zfp26 | 9.285324035 | 9.094327663 | 9.288040818 | -0.193713155 | -0.002716783 |
| Atp6v0e | 10.62981549 | 10.54850135 | 10.63257796 | -0.084076617 | -0.00276247 |
| Dcaf12l1 | 2.829124485 | 2.829136615 | 2.831907444 | -0.002770829 | -0.002782959 |
| Baz1b | 11.29241587 | 11.37454431 | 11.29526487 | 0.079279443 | -0.002848993 |
| Hist2h2aa1 | 2.071943577 | 2.078600696 | 2.074793183 | 0.003807513 | -0.002849606 |
| Kcnn1 | 4.929641991 | 4.922258955 | 4.932499438 | -0.010240482 | -0.002857447 |
| Rad54l2 | 9.523635472 | 9.631101573 | 9.526525771 | 0.104575802 | -0.002890299 |
| 1700057G04Rik | 1.551495011 | 1.591181481 | 1.554401935 | 0.036779547 | -0.002906924 |
| Slc6a8 | 11.82763917 | 11.92775121 | 11.83055238 | 0.09719883 | -0.00291321 |
| Gm5949 | 3.010520553 | 2.998665573 | 3.01344171 | -0.014776136 | -0.002921157 |
| Psd | 3.27506847 | 3.282906323 | 3.278044652 | 0.004861672 | -0.002976182 |
| Apol8 | 6.302854505 | 6.350540486 | 6.305835692 | 0.044704794 | -0.002981187 |
| 5930422O12Rik | 1.201389588 | 1.250259006 | 1.204375534 | 0.045883472 | -0.002985946 |
| Gm45147 | 1.863526105 | 1.826823599 | 1.866517928 | -0.039694329 | -0.002991823 |
| Cep170 | 9.106216494 | 9.087772758 | 9.109211002 | -0.021438244 | -0.002994508 |
| Cox19 | 8.626625866 | 8.698379569 | 8.629671174 | 0.068708395 | -0.003045308 |
| Prkar1b | 5.2980373 | 5.494900536 | 5.301100708 | 0.193799828 | -0.003063408 |
| Fgf12 | 3.124303561 | 3.125998398 | 3.127368187 | -0.001369788 | -0.003064626 |
| Vmn1r90 | 3.941781987 | 4.049594476 | 3.944853087 | 0.104741388 | -0.0030711 |
| Fbn2 | 5.025850733 | 5.114770556 | 5.028925757 | 0.085844799 | -0.003075025 |
| Smarca5-ps | 1.512795779 | 1.493647191 | 1.515877178 | -0.022229987 | -0.003081399 |
| Gm10478 | 2.660430846 | 2.634512007 | 2.663549481 | -0.029037474 | -0.003118635 |
| St3gal5 | 7.092565697 | 7.023946102 | 7.095696929 | -0.071750826 | -0.003131231 |
| S100a14 | 13.15882534 | 13.09580725 | 13.16204999 | -0.06624274 | -0.003224653 |
| Spg20 | 9.944057219 | 9.821184692 | 9.947311264 | -0.126126572 | -0.003254045 |
| Rnf6 | 10.10915096 | 10.1022604 | 10.11241589 | -0.010155487 | -0.003264925 |
| Haus6 | 8.47134897 | 8.403697153 | 8.47462191 | -0.070924757 | -0.00327294 |
| Ccdc40 | 1.736520048 | 1.725024126 | 1.739808603 | -0.014784477 | -0.003288555 |
| Snai3 | 4.565438623 | 4.903235653 | 4.568737897 | 0.334497755 | -0.003299274 |

| | | | | | |
|---|---|---|---|---|---|
| Tmprss11a | 4.222346432 | 4.220965439 | 4.225649188 | -0.004683749 | -0.003302756 |
| Ilk | 10.59976924 | 10.4405549 | 10.60309505 | -0.16254015 | -0.003325803 |
| Gm6483 | 4.186114791 | 4.297867252 | 4.189451571 | 0.10841568 | -0.00333678 |
| Gm6850 | 2.669866632 | 2.634355181 | 2.6732057 | -0.03885052 | -0.003339068 |
| Vmn2r13 | 2.258014375 | 2.270629672 | 2.261364282 | 0.009265391 | -0.003349906 |
| 1600002K03Rik | 6.954307574 | 7.091961763 | 6.957667979 | 0.134293785 | -0.003360405 |
| Nhlrc1 | 6.223556115 | 6.2720262 | 6.226918252 | 0.045107948 | -0.003362137 |
| Tmc3 | 5.398198289 | 5.299358294 | 5.401579248 | -0.102220953 | -0.003380958 |
| Gimap5 | 5.417815244 | 5.333737056 | 5.421254727 | -0.087517672 | -0.003439483 |
| Coprs | 5.944089219 | 6.05785087 | 5.947547484 | 0.110303386 | -0.003458265 |
| Npat | 8.586428444 | 8.586933142 | 8.589895298 | -0.002962156 | -0.003466854 |
| Gm8909 | 3.078022526 | 3.066772706 | 3.081511812 | -0.014739106 | -0.003489286 |
| Gm8624 | 2.93968816 | 2.982321103 | 2.943180811 | 0.039140292 | -0.003492651 |
| Map3k7cl | 5.556042571 | 5.568924188 | 5.559540478 | 0.00938371 | -0.003497907 |
| Gm14667 | 1.739509624 | 1.752229841 | 1.743009641 | 0.0092202 | -0.003500016 |
| A930017M01Rik | 2.23008452 | 2.276742374 | 2.233601567 | 0.043140806 | -0.003517047 |
| Vmn2r5 | 2.50307225 | 2.521083882 | 2.506594868 | 0.014489014 | -0.003522618 |
| Gm20537 | 1.304392745 | 1.289736438 | 1.307978693 | -0.018242255 | -0.003585948 |
| Mrgprx2 | 5.213071175 | 4.866468463 | 5.216725175 | -0.350256712 | -0.003654001 |
| Grik4 | 2.95505201 | 2.932341496 | 2.958742112 | -0.026400616 | -0.003690103 |
| Gm1141 | 2.00562785 | 2.022802612 | 2.009328424 | 0.013474187 | -0.003700574 |
| Rab10 | 12.25349979 | 12.18492106 | 12.25721677 | -0.072295707 | -0.00371698 |
| AC133509.1 | 2.966206282 | 2.959244203 | 2.969935052 | -0.01069085 | -0.00372877 |
| Ptchd1 | 2.013288782 | 2.087266183 | 2.017110706 | 0.070155477 | -0.003821924 |
| Pam | 11.39434885 | 11.26374921 | 11.39820046 | -0.13445125 | -0.003851607 |
| Parp2 | 8.945146668 | 8.871044059 | 8.949017126 | -0.077973068 | -0.003870458 |
| Rdh11 | 10.75994625 | 10.6445908 | 10.7638173 | -0.119226497 | -0.003871047 |
| Prcd | 1.573207427 | 1.557819174 | 1.577120681 | -0.019301507 | -0.003913255 |
| Sprr2j-ps | 3.320275875 | 3.302720676 | 3.324202274 | -0.021481598 | -0.003926399 |
| Zfp985 | 2.114218769 | 2.15604341 | 2.118157926 | 0.037885484 | -0.003939157 |
| Mtor | 9.989068291 | 10.14720224 | 9.993049011 | 0.154153229 | -0.00398072 |
| 9230102O04Rik | 2.096478108 | 2.113498003 | 2.10047884 | 0.013019163 | -0.004000732 |

| | | | | | |
|---|---|---|---|---|---|
| Kif22 | 7.775508597 | 7.845269646 | 7.779602122 | 0.065667523 | -0.004093525 |
| Vmn1r9 | 2.867038353 | 2.819052187 | 2.871140943 | -0.052088756 | -0.004102591 |
| Hist1h4c | 1.438495213 | 1.53281241 | 1.442667889 | 0.090144522 | -0.004172676 |
| Vmn2r100 | 2.66500882 | 2.632124846 | 2.669226442 | -0.037101596 | -0.004217621 |
| Gm2398 | 2.292148646 | 2.208107535 | 2.296411874 | -0.088304339 | -0.004263227 |
| Vmn2r124 | 4.56797097 | 4.512505225 | 4.572244355 | -0.05973913 | -0.004273385 |
| Kcnk7 | 7.061501665 | 7.168667448 | 7.065790604 | 0.102876844 | -0.004288939 |
| Gm20481 | 1.847082902 | 2.04351257 | 1.851373891 | 0.192138679 | -0.004290989 |
| Cbll1 | 8.702529421 | 8.779542553 | 8.706841404 | 0.07270115 | -0.004311982 |
| Obox3 | 1.371803035 | 1.355471757 | 1.376158431 | -0.020686674 | -0.004355397 |
| Morc2b | 2.848093445 | 2.812789276 | 2.852470105 | -0.039680829 | -0.00437666 |
| Xpo7 | 10.57269888 | 10.61553933 | 10.57709779 | 0.038441543 | -0.004398913 |
| Gm10709 | 1.428350463 | 1.398123476 | 1.432770989 | -0.034647513 | -0.004420526 |
| Man2c1 | 9.751947928 | 9.955031398 | 9.75640883 | 0.198622568 | -0.004460902 |
| Des | 13.13909463 | 13.37795697 | 13.14359218 | 0.23436479 | -0.004497557 |
| Gm14849 | 2.555714008 | 2.580125248 | 2.560216056 | 0.019909192 | -0.004502048 |
| Cttnbp2nl | 9.68144671 | 9.788690789 | 9.685965416 | 0.102725373 | -0.004518706 |
| Htr2a | 5.204273905 | 5.226959549 | 5.20879804 | 0.018161509 | -0.004524135 |
| Eapp | 9.493969998 | 9.337313596 | 9.498503177 | -0.161189581 | -0.004533179 |
| Dpysl4 | 2.059555858 | 2.066122274 | 2.064127683 | 0.001994591 | -0.004571825 |
| Ndufb8 | 10.72770414 | 10.65573201 | 10.73230177 | -0.076569763 | -0.004597637 |
| Ccdc158 | 1.973308345 | 1.981638791 | 1.977912297 | 0.003726494 | -0.004603951 |
| Adgrg4 | 1.249185254 | 1.250385871 | 1.253792432 | -0.003406561 | -0.004607178 |
| Vmn1r202 | 2.635404138 | 2.650427889 | 2.640025136 | 0.010402753 | -0.004620998 |
| Sri | 10.04066941 | 9.875721618 | 10.04529353 | -0.169571912 | -0.004624118 |
| Ranbp17 | 4.73688488 | 4.849256935 | 4.741521215 | 0.10773572 | -0.004636335 |
| Angpt1 | 7.61200141 | 6.970253822 | 7.616654627 | -0.646400805 | -0.004653218 |
| Otud3 | 7.022258483 | 7.166158877 | 7.026913885 | 0.139244992 | -0.004655402 |
| Fgf7 | 8.023189275 | 7.685956225 | 8.027908816 | -0.341952591 | -0.004719541 |
| Gm48391 | 4.043255696 | 4.017874047 | 4.047988459 | -0.030114412 | -0.004732763 |
| Zbtb44 | 9.816328066 | 9.773825188 | 9.821175048 | -0.04734986 | -0.004846982 |
| Hipk4 | 3.694417672 | 3.773479354 | 3.699267475 | 0.074211879 | -0.004849803 |

| | | | | | |
|---|---|---|---|---|---|
| Tcea2 | 5.607717077 | 5.644951961 | 5.612649512 | 0.032302449 | -0.004932435 |
| Olfr418 | 1.860951991 | 1.856095657 | 1.865902353 | -0.009806696 | -0.004950361 |
| Gm49760 | 3.534249395 | 3.555466694 | 3.53926765 | 0.016199044 | -0.005018255 |
| 2300009A05Rik | 5.834817168 | 5.770263629 | 5.839836564 | -0.069572935 | -0.005019395 |
| Olfr167 | 1.776554722 | 1.796316076 | 1.781578412 | 0.014737663 | -0.00502369 |
| Vmn1r26 | 4.077393627 | 4.061439034 | 4.08245911 | -0.021020076 | -0.005065484 |
| Tcrg-V1 | 1.23272728 | 1.222949441 | 1.237815975 | -0.014866534 | -0.005088694 |
| Blnk | 9.385050635 | 9.365709169 | 9.39017892 | -0.024469751 | -0.005128285 |
| Gm9057 | 2.161475587 | 2.180844801 | 2.166617369 | 0.014227432 | -0.005141782 |
| Shank2 | 2.586230335 | 2.683132657 | 2.591393497 | 0.09173916 | -0.005163162 |
| Dnm1l | 10.62931527 | 10.66082355 | 10.63448802 | 0.026335533 | -0.00517275 |
| Rasgef1b | 8.436770731 | 8.504280086 | 8.441953408 | 0.062326677 | -0.005182678 |
| Aasdh | 7.490336786 | 7.659699341 | 7.495609817 | 0.164089525 | -0.005273031 |
| Kynu | 4.922461874 | 4.895756102 | 4.927741129 | -0.031985027 | -0.005279255 |
| Olfr479 | 2.425255109 | 2.415548576 | 2.430536105 | -0.014987529 | -0.005280996 |
| 2310061I04Rik | 8.369028663 | 8.338013527 | 8.374320845 | -0.036307318 | -0.005292182 |
| Gm49328 | 2.784641728 | 2.728581147 | 2.789938529 | -0.061357382 | -0.0052968 |
| Sh3glb1 | 12.01792673 | 11.86861176 | 12.02327046 | -0.1546587 | -0.005343737 |
| Prim2 | 7.874914835 | 7.848083593 | 7.880290871 | -0.032207278 | -0.005376036 |
| Gm7592 | 3.573878236 | 3.584864518 | 3.57930078 | 0.005563737 | -0.005422544 |
| Fndc3c1 | 1.496899561 | 1.441979757 | 1.502341674 | -0.060361917 | -0.005442113 |
| Pon2 | 10.96688142 | 10.88227897 | 10.97232747 | -0.090048493 | -0.005446043 |
| Ctr9 | 9.942045955 | 10.02058647 | 9.947513726 | 0.073072747 | -0.00546777 |
| Fcer2a | 4.695639284 | 4.617867277 | 4.701175053 | -0.083307777 | -0.00553577 |
| Vmn2r27 | 2.139309388 | 2.123933393 | 2.144874031 | -0.020940638 | -0.005564643 |
| Liph | 8.996516065 | 9.02446969 | 9.002096064 | 0.022373626 | -0.005579998 |
| Mta2 | 11.77703534 | 11.85396091 | 11.78263177 | 0.071329137 | -0.005596433 |
| Slc6a20b | 4.791047013 | 4.855881895 | 4.796668819 | 0.059213076 | -0.005621806 |
| Gemin7 | 8.651910088 | 8.665327961 | 8.657557407 | 0.007770554 | -0.005647319 |
| Epb42 | 2.963209369 | 2.938695345 | 2.968901619 | -0.030206275 | -0.005692251 |
| Snw1 | 10.56048647 | 10.43566149 | 10.56618633 | -0.13052484 | -0.005699857 |
| Palb2 | 6.296614499 | 6.520496352 | 6.302339137 | 0.218157216 | -0.005724638 |

| | | | | | |
|---|---|---|---|---|---|
| Fam120a | 12.1604454 | 12.29621633 | 12.16618318 | 0.130033147 | -0.005737787 |
| Atp7a | 8.912803605 | 8.987695164 | 8.918551347 | 0.069143817 | -0.005747742 |
| Tmem108 | 3.630656502 | 3.715802092 | 3.636409456 | 0.079392636 | -0.005752954 |
| Vmn1r168 | 2.834874969 | 2.819367964 | 2.840635508 | -0.021267543 | -0.005760539 |
| Itsn2 | 11.26683958 | 11.30670775 | 11.27260508 | 0.034102663 | -0.0057655 |
| Tbc1d22b | 9.302383535 | 9.310717416 | 9.308160569 | 0.002556846 | -0.005777034 |
| Olfr1157 | 2.820058499 | 2.796974444 | 2.825845813 | -0.028871369 | -0.005787314 |
| Gna11 | 9.813644593 | 10.01117735 | 9.819446955 | 0.191730394 | -0.005802363 |
| Tor1a | 8.71736497 | 8.578991852 | 8.723177074 | -0.144185222 | -0.005812104 |
| Klk14 | 2.118526319 | 2.104990365 | 2.124386225 | -0.01939586 | -0.005859906 |
| Olfr808 | 1.331745173 | 1.328323267 | 1.337606601 | -0.009283333 | -0.005861428 |
| Tmem179b | 8.345456946 | 8.323052021 | 8.351334864 | -0.028282843 | -0.005877918 |
| Cldn7 | 1.979274363 | 1.980769308 | 1.98517363 | -0.004404322 | -0.005899268 |
| Mettl8 | 7.572840455 | 7.589459979 | 7.578748296 | 0.010711683 | -0.005907841 |
| Fcrl5 | 0.807228755 | 0.801569092 | 0.813164324 | -0.011595233 | -0.005935569 |
| Gm28362 | 1.081566569 | 1.097115445 | 1.087513828 | 0.009601617 | -0.005947259 |
| Slc35a1 | 8.274410986 | 8.23101893 | 8.280392522 | -0.049373593 | -0.005981536 |
| Orc3 | 8.914439917 | 8.932360149 | 8.920437938 | 0.011922211 | -0.005998021 |
| Lrfn5 | 1.476555272 | 1.485372665 | 1.482582555 | 0.00279011 | -0.006027283 |
| Gm15666 | 2.53127408 | 2.550412458 | 2.537340332 | 0.013072126 | -0.006066252 |
| Cfap53 | 2.038101361 | 2.045891681 | 2.044176151 | 0.00171553 | -0.00607479 |
| Dctn5 | 9.8650738 | 9.836747201 | 9.871157136 | -0.034409935 | -0.006083337 |
| Hsd11b1 | 8.461241786 | 7.895597607 | 8.467326456 | -0.57172885 | -0.00608467 |
| Ccdc189 | 4.811205941 | 4.857569615 | 4.817315452 | 0.040254163 | -0.006109512 |
| Scx | 5.223534489 | 5.335061673 | 5.229649195 | 0.105412479 | -0.006114706 |
| Dpep3 | 2.786137794 | 2.767282016 | 2.792303229 | -0.025021213 | -0.006165435 |
| Pank2 | 9.767746828 | 9.618868419 | 9.773943893 | -0.155075474 | -0.006197065 |
| Trbc1 | 6.82880467 | 6.943817153 | 6.835030085 | 0.108787068 | -0.006225415 |
| Tmem167b | 10.15852863 | 10.11534624 | 10.16478325 | -0.049437013 | -0.006254617 |
| Gm47932 | 4.39765964 | 4.355229697 | 4.40392893 | -0.048699232 | -0.00626929 |
| Olfr204 | 3.313309725 | 3.358656599 | 3.319581984 | 0.039074615 | -0.006272258 |
| Glyat | 1.734313922 | 1.789304618 | 1.740644266 | 0.048660352 | -0.006330344 |

| | | | | | |
|---|---|---|---|---|---|
| Aldh1l2 | 6.508126768 | 6.854745649 | 6.51447111 | 0.340274539 | -0.006344342 |
| Fat2 | 11.35794908 | 11.61629018 | 11.36437011 | 0.25192007 | -0.00642103 |
| Arl8b | 11.71234268 | 11.74870806 | 11.7188168 | 0.02989126 | -0.006474123 |
| Txndc12 | 8.95035388 | 8.808771412 | 8.956904623 | -0.148133211 | -0.006550743 |
| Iba57 | 7.518176862 | 7.709962225 | 7.524739033 | 0.185223192 | -0.006562171 |
| Pdss2 | 7.256748382 | 7.172083108 | 7.263318548 | -0.09123544 | -0.006570166 |
| Coq4 | 7.621604962 | 7.693640962 | 7.628189449 | 0.065451513 | -0.006584487 |
| Zfp92 | 3.315752702 | 3.328319548 | 3.322377905 | 0.005941642 | -0.006625203 |
| Zfp941 | 4.413914112 | 4.362197987 | 4.420594596 | -0.058396609 | -0.006680484 |
| Mgat4c | 2.560087866 | 2.540467378 | 2.566795586 | -0.026328209 | -0.006707721 |
| Cd82 | 10.83831172 | 10.88040317 | 10.84507376 | 0.035329403 | -0.006762043 |
| P4ha2 | 8.599214684 | 8.671532735 | 8.60600119 | 0.065531545 | -0.006786506 |
| Lta | 2.976341874 | 3.019879894 | 2.983159868 | 0.036720026 | -0.006817994 |
| Serpinb3c | 1.828953354 | 1.808814532 | 1.835777464 | -0.026962932 | -0.00682411 |
| Duoxa2 | 2.691779652 | 2.683133165 | 2.698626074 | -0.015492909 | -0.006846421 |
| Llgl1 | 9.958679139 | 10.19330443 | 9.965530368 | 0.227774062 | -0.006851229 |
| Dpm1 | 8.614575574 | 8.334053063 | 8.621436686 | -0.287383623 | -0.006861112 |
| Olfr846 | 2.229091256 | 2.219059153 | 2.23599952 | -0.016940367 | -0.006908264 |
| Sox30 | 1.007332001 | 0.995332038 | 1.014264006 | -0.018931967 | -0.006932004 |
| Vash1 | 5.188961433 | 5.426856144 | 5.195910807 | 0.230945337 | -0.006949374 |
| Rcn1 | 11.34925813 | 11.30009444 | 11.35624742 | -0.056152983 | -0.006989293 |
| Gm14411 | 3.330286099 | 3.326606658 | 3.337281198 | -0.010674541 | -0.006995099 |
| Cdkn2aipnl | 9.178237412 | 9.160525676 | 9.185246347 | -0.024720671 | -0.007008936 |
| Commd1 | 9.357425455 | 9.301294863 | 9.364453587 | -0.063158724 | -0.007028132 |
| Themis | 4.213816221 | 4.263544247 | 4.220871844 | 0.042672403 | -0.007055623 |
| Pnmt | 2.051568894 | 2.034288703 | 2.058633045 | -0.024344342 | -0.007064152 |
| Vopp1 | 9.65401076 | 9.610544887 | 9.661097934 | -0.050553047 | -0.007087174 |
| Gm48401 | 3.380051894 | 3.419658784 | 3.387139599 | 0.032519185 | -0.007087705 |
| Tacr3 | 3.665362072 | 3.602655562 | 3.672476821 | -0.069821259 | -0.007114749 |
| Pafah1b3 | 7.825011709 | 7.811018494 | 7.832132846 | -0.021114352 | -0.007121136 |
| Clec2f | 6.101319474 | 5.969867526 | 6.108471391 | -0.138603864 | -0.007151917 |
| Cacng4 | 3.056508186 | 3.058155017 | 3.063662065 | -0.005507048 | -0.007153879 |

| | | | | | |
|---|---|---|---|---|---|
| Pbx2 | 9.417183676 | 9.544625735 | 9.424384918 | 0.120240818 | -0.007201242 |
| Wapl | 11.36755133 | 11.43365154 | 11.37479996 | 0.058851577 | -0.007248633 |
| Lemd3 | 8.976822114 | 8.953472141 | 8.984071806 | -0.030599666 | -0.007249692 |
| 7530416G11Rik | 2.012779439 | 2.017294057 | 2.020042037 | -0.002747981 | -0.007262598 |
| Vps4a | 9.95713744 | 9.883749398 | 9.964413638 | -0.08066424 | -0.007276198 |
| Kansl1l | 8.941881222 | 8.885915666 | 8.949192948 | -0.063277282 | -0.007311726 |
| 1700020L24Rik | 1.62981654 | 1.598282857 | 1.637138385 | -0.038855528 | -0.007321845 |
| Stk11ip | 8.741345277 | 8.853735518 | 8.748686946 | 0.105048572 | -0.007341669 |
| Ccz1 | 9.774392128 | 9.676032135 | 9.78176803 | -0.105735895 | -0.007375902 |
| Gm13864 | 1.951299217 | 1.950871857 | 1.958678515 | -0.007806657 | -0.007379297 |
| Olfr1202 | 2.086331101 | 2.095702347 | 2.093726439 | 0.001975908 | -0.007395338 |
| Zbtb8b | 2.664123637 | 2.659500618 | 2.671582159 | -0.012081542 | -0.007458523 |
| Tmem245 | 10.69254948 | 10.63129848 | 10.70001172 | -0.06871324 | -0.00746224 |
| Orc6 | 8.771697274 | 8.730005548 | 8.77919132 | -0.049185772 | -0.007494046 |
| Gm32742 | 2.754512652 | 2.800253224 | 2.762040844 | 0.03821238 | -0.007528192 |
| Gm43028 | 1.725657432 | 1.777612825 | 1.733241286 | 0.044371539 | -0.007583854 |
| Kcng2 | 2.179009916 | 2.321754562 | 2.186594368 | 0.135160193 | -0.007584453 |
| Spo11 | 1.843023089 | 1.853691735 | 1.850645872 | 0.003045863 | -0.007622782 |
| Ccnb1ip1 | 2.079654857 | 2.011511663 | 2.087286548 | -0.075774885 | -0.007631691 |
| Slc2a4rg-ps | 7.310121058 | 7.620205471 | 7.317761305 | 0.302444166 | -0.007640247 |
| Arpc5l | 10.26767276 | 10.10701088 | 10.27536756 | -0.16835668 | -0.007694797 |
| Wiz | 10.02069633 | 10.2502562 | 10.02839741 | 0.221858795 | -0.007701079 |
| Golga3 | 10.03313241 | 10.04468707 | 10.04089433 | 0.003792743 | -0.007761919 |
| Gm47974 | 3.711269246 | 3.812199394 | 3.719096694 | 0.0931027 | -0.007827448 |
| Cenpn | 6.456135675 | 6.51063463 | 6.463980037 | 0.046654593 | -0.007844362 |
| Snx19 | 9.491282807 | 9.344040188 | 9.499159395 | -0.155119207 | -0.007876588 |
| Krt28 | 4.693670537 | 4.774966203 | 4.701550124 | 0.073416079 | -0.007879587 |
| Mast1 | 2.070251039 | 2.055567049 | 2.078180031 | -0.022612982 | -0.007928992 |
| Eipr1 | 8.6191711 | 8.571397008 | 8.627104769 | -0.055707761 | -0.007933668 |
| Faap20 | 8.21136893 | 8.31271196 | 8.219311753 | 0.093400207 | -0.007942823 |
| Mon2 | 11.02443286 | 10.97736129 | 11.03238011 | -0.055018823 | -0.007947253 |
| Gm8730 | 1.70768965 | 1.71828984 | 1.715694885 | 0.002594955 | -0.008005235 |

| | | | | | |
|---|---|---|---|---|---|
| Jsrp1 | 8.274741712 | 8.847623514 | 8.282763176 | 0.564860338 | -0.008021464 |
| Dnajc17 | 7.154611124 | 7.109584291 | 7.162668427 | -0.053084135 | -0.008057303 |
| 6430550D23Rik | 3.146893587 | 3.159217552 | 3.154975928 | 0.004241625 | -0.008082341 |
| Zfp341 | 7.13980218 | 7.13535092 | 7.147939346 | -0.012588426 | -0.008137166 |
| Kcnmb1 | 5.030612431 | 5.079811902 | 5.038800501 | 0.0410114 | -0.00818807 |
| Psd2 | 2.510224006 | 2.474477954 | 2.518422467 | -0.043944513 | -0.008198461 |
| Cnih1 | 10.52399752 | 10.43018702 | 10.53222337 | -0.102036353 | -0.008225857 |
| Foxd3 | 2.462817546 | 2.47720056 | 2.471046711 | 0.006153849 | -0.008229165 |
| 3110053B16Rik | 4.122789573 | 4.183129079 | 4.131034968 | 0.052094111 | -0.008245395 |
| Creg2 | 2.546912671 | 2.544253265 | 2.555178737 | -0.010925472 | -0.008266066 |
| Msantd4 | 9.326795082 | 9.260983391 | 9.335134971 | -0.07415158 | -0.008339889 |
| Rgma | 9.943437486 | 10.00609148 | 9.951793521 | 0.054297962 | -0.008356035 |
| AU041133 | 5.482633251 | 5.42168524 | 5.49102358 | -0.06933834 | -0.00839033 |
| Vmn1r206 | 1.926301516 | 1.927978339 | 1.934727589 | -0.00674925 | -0.008426072 |
| Mei1 | 1.179991452 | 1.197440076 | 1.188424066 | 0.00901601 | -0.008432614 |
| Baalc | 3.941582713 | 4.01877891 | 3.950023609 | 0.068755301 | -0.008440897 |
| Cdca5 | 6.859915061 | 6.792444827 | 6.86840392 | -0.075959093 | -0.008488859 |
| Gucy2g | 3.166216929 | 3.169497252 | 3.174721049 | -0.005223797 | -0.00850412 |
| Olfr600 | 3.332937559 | 3.443800687 | 3.341463111 | 0.102337577 | -0.008525552 |
| A330044P14Rik | 3.350868958 | 3.368221485 | 3.359420191 | 0.008801294 | -0.008551234 |
| Pef1 | 9.853083345 | 9.923293214 | 9.861647593 | 0.061645621 | -0.008564249 |
| Guca1b | 2.472281624 | 2.503807588 | 2.480856806 | 0.022950782 | -0.008575182 |
| Spcs1 | 10.56604625 | 10.50333266 | 10.57462628 | -0.071293617 | -0.00858003 |
| Gigyf2 | 10.39275883 | 10.4589768 | 10.40136369 | 0.057613107 | -0.008604863 |
| Nek10 | 1.684403438 | 1.673700964 | 1.693015173 | -0.019314209 | -0.008611735 |
| Ces2b | 2.601647167 | 2.567467176 | 2.610295113 | -0.042827937 | -0.008647946 |
| Icmt | 10.44896188 | 10.58896228 | 10.45761253 | 0.131349757 | -0.008650647 |
| Rorb | 2.991540305 | 2.990190517 | 3.000223117 | -0.0100326 | -0.008682812 |
| Gm7902 | 1.30454836 | 1.319143854 | 1.313273526 | 0.005870328 | -0.008725166 |
| Adap1 | 7.680943381 | 7.632776795 | 7.689681841 | -0.056905046 | -0.00873846 |
| C2cd4c | 2.524028124 | 2.520212672 | 2.532788471 | -0.012575799 | -0.008760346 |
| Arl6ip4 | 8.781541351 | 8.765964944 | 8.790303134 | -0.02433819 | -0.008761783 |

| | | | | | |
|---|---|---|---|---|---|
| Gm11478 | 1.307173664 | 1.295901596 | 1.31594407 | -0.020042474 | -0.008770406 |
| Ttn | 14.28002176 | 14.2912931 | 14.2888045 | 0.002488593 | -0.008782747 |
| Slc25a45 | 7.647147317 | 7.566157847 | 7.655943623 | -0.089785776 | -0.008796305 |
| 4931440P22Rik | 3.378985429 | 3.353930972 | 3.38778936 | -0.033858388 | -0.008803931 |
| Olfr586 | 2.654066341 | 2.68997144 | 2.662892455 | 0.027078985 | -0.008826114 |
| Mkrn2os | 4.126327245 | 4.073135946 | 4.135195004 | -0.062059058 | -0.008867759 |
| Tvp23a | 4.463708592 | 4.419243177 | 4.472606924 | -0.053363746 | -0.008898331 |
| Serpinb1b | 2.15897011 | 2.172967583 | 2.167905964 | 0.005061619 | -0.008935854 |
| Golga5 | 9.739636339 | 9.702119692 | 9.748678846 | -0.046559155 | -0.009042508 |
| Gm13341 | 3.039249512 | 3.052060259 | 3.048292784 | 0.003767475 | -0.009043272 |
| Gm21960 | 1.863749423 | 1.93419449 | 1.872853653 | 0.061340836 | -0.00910423 |
| Dpp7 | 8.691646654 | 8.960336733 | 8.700859968 | 0.259476765 | -0.009213314 |
| Zfp518a | 8.45667895 | 8.517787539 | 8.465893057 | 0.051894482 | -0.009214107 |
| Hapln4 | 2.529446231 | 2.519536734 | 2.538713135 | -0.019176401 | -0.009266904 |
| Asic2 | 1.954538689 | 1.918070096 | 1.9638296 | -0.045759505 | -0.009290911 |
| Vmn2r66 | 10.23075925 | 10.19600076 | 10.24005923 | -0.044058473 | -0.009299977 |
| Tesmin | 2.126576784 | 2.108880944 | 2.135883488 | -0.027002545 | -0.009306705 |
| Vmn2r31 | 3.022667287 | 3.037914192 | 3.031991587 | 0.005922606 | -0.0093243 |
| Sema5a | 10.27932587 | 10.57602163 | 10.28865243 | 0.287369207 | -0.009326557 |
| Gnaq | 10.24493402 | 10.16479687 | 10.25430551 | -0.08950864 | -0.009371487 |
| 1110017D15Rik | 1.033656405 | 1.053087928 | 1.043036861 | 0.010051067 | -0.009380456 |
| Pms1 | 6.488564175 | 6.58401293 | 6.497955187 | 0.086057743 | -0.009391012 |
| Hcn2 | 3.847662508 | 3.892275508 | 3.857062386 | 0.035213122 | -0.009399878 |
| T2 | 1.807584969 | 1.832687899 | 1.816984928 | 0.015702971 | -0.009399959 |
| Got2-ps1 | 1.185682538 | 1.208402236 | 1.195145211 | 0.013257025 | -0.009462673 |
| Olfr63 | 4.508420999 | 4.499448898 | 4.517889553 | -0.018440655 | -0.009468554 |
| Hspb3 | 4.954090657 | 5.188524738 | 4.963566962 | 0.224957775 | -0.009476305 |
| Top2a | 10.86865691 | 10.74662272 | 10.87814682 | -0.131524093 | -0.009489907 |
| Fbxo6 | 9.154907773 | 9.020984237 | 9.164405467 | -0.14342123 | -0.009497695 |
| Vkorc1 | 9.240306642 | 9.164947786 | 9.249830254 | -0.084882467 | -0.009523612 |
| Rnaseh2c | 6.768175025 | 6.774853043 | 6.777704054 | -0.002851011 | -0.009529029 |
| Yy1 | 10.67052937 | 10.74432808 | 10.6800901 | 0.064237983 | -0.009560723 |

| | | | | | |
|---|---|---|---|---|---|
| Fkrp | 8.542100155 | 8.572186963 | 8.551670494 | 0.02051647 | -0.009570339 |
| Vmn2r24 | 4.620314405 | 4.556012389 | 4.629926191 | -0.073913802 | -0.009611786 |
| Med23 | 8.984772151 | 9.100844824 | 8.994451436 | 0.106393388 | -0.009679285 |
| Gm8956 | 2.442882714 | 2.386122943 | 2.4525789 | -0.066455957 | -0.009696187 |
| Gm15645 | 1.713193383 | 1.729580992 | 1.722918508 | 0.006662484 | -0.009725125 |
| Elac1 | 8.546351106 | 8.357066831 | 8.556189899 | -0.199123068 | -0.009838794 |
| Psen1 | 11.18865141 | 11.23692264 | 11.19850004 | 0.0384226 | -0.009848627 |
| Tmem132b | 2.88779819 | 2.925501514 | 2.897650054 | 0.02785146 | -0.009851864 |
| Gm7899 | 1.549012029 | 1.553849515 | 1.558949834 | -0.00510032 | -0.009937806 |
| Cfd | 13.29938444 | 13.30419632 | 13.30932227 | -0.005125957 | -0.00993783 |
| Gm3336 | 4.591752247 | 4.60427356 | 4.601709151 | 0.002564409 | -0.009956904 |
| Tm6sf2 | 3.115425254 | 3.168753897 | 3.125400604 | 0.043353294 | -0.009975349 |
| Olfr270 | 2.451648363 | 2.54018392 | 2.461686489 | 0.078497432 | -0.010038126 |
| Hmgb1-ps2 | 2.156619185 | 2.188310952 | 2.166663111 | 0.02164784 | -0.010043926 |
| Chga | 6.062898711 | 6.275326638 | 6.072945007 | 0.202381632 | -0.010046295 |
| BC029722 | 8.65089351 | 8.670034345 | 8.661002318 | 0.009032027 | -0.010108808 |
| Troap | 5.693985124 | 5.841875159 | 5.704117819 | 0.13775734 | -0.010132695 |
| 9030612E09Rik | 2.291405265 | 2.257241005 | 2.301569829 | -0.044328823 | -0.010164563 |
| Gm18492 | 0.979201225 | 1.009710341 | 0.989378151 | 0.020332191 | -0.010176925 |
| Rfc5 | 8.388733662 | 8.521359734 | 8.39892548 | 0.122434254 | -0.010191818 |
| Cdc42bpb | 10.13821051 | 10.56808514 | 10.14841884 | 0.419666292 | -0.010208338 |
| Fbxo47 | 2.88427694 | 2.812947353 | 2.894502902 | -0.081555549 | -0.010225962 |
| Tbx18 | 6.027693778 | 6.044403124 | 6.037920924 | 0.0064822 | -0.010227146 |
| Commd10 | 8.625417995 | 8.561142534 | 8.63564932 | -0.074506786 | -0.010231325 |
| Emc3 | 10.7739065 | 10.74163076 | 10.78413895 | -0.042508187 | -0.01023245 |
| Gabrg3 | 1.798984553 | 1.846178997 | 1.809227173 | 0.036951824 | -0.01024262 |
| Hpn | 4.803135375 | 4.891622308 | 4.813379093 | 0.078243215 | -0.010243719 |
| Pla2g6 | 7.980779704 | 8.119514494 | 7.991053797 | 0.128460697 | -0.010274094 |
| Gm15382 | 1.273786776 | 1.254929702 | 1.284098324 | -0.029168623 | -0.010311549 |
| Gne | 7.72082881 | 8.068480199 | 7.731190636 | 0.337289563 | -0.010361827 |
| Impa2 | 8.239675159 | 8.272353898 | 8.250070026 | 0.022283872 | -0.010394867 |
| Gm45647 | 2.382817935 | 2.403882172 | 2.393234536 | 0.010647636 | -0.010416602 |

| | | | | | |
|---|---|---|---|---|---|
| Ranbp3l | 2.962208821 | 2.923539951 | 2.972641006 | -0.049101054 | -0.010432185 |
| Hmgb1-rs16 | 2.303410452 | 2.324482652 | 2.313855509 | 0.010627143 | -0.010445057 |
| Abtb1 | 8.370423315 | 8.536396895 | 8.380874482 | 0.155522412 | -0.010451168 |
| Dab2ip | 10.49248178 | 10.58892389 | 10.50293699 | 0.085986903 | -0.010455207 |
| Gp6 | 2.490745249 | 2.551396468 | 2.50121945 | 0.050177018 | -0.010474201 |
| Stc2 | 7.27283744 | 7.17651948 | 7.283320661 | -0.10680118 | -0.01048322 |
| Fam209 | 1.799964415 | 1.810777408 | 1.810453081 | 0.000324327 | -0.010488666 |
| Ccnb2-ps | 1.788191525 | 1.811881392 | 1.798687772 | 0.013193621 | -0.010496246 |
| Taf11 | 8.710093182 | 8.744037396 | 8.720761078 | 0.023276318 | -0.010667896 |
| Iqsec3 | 3.649713809 | 3.8141523 | 3.660434221 | 0.153718079 | -0.010720412 |
| Kcnj10 | 2.71873107 | 2.663628145 | 2.729475717 | -0.065847573 | -0.010744648 |
| Olfr1023 | 1.70167966 | 1.699732726 | 1.712487665 | -0.012754939 | -0.010808005 |
| Trmt1l | 8.754763058 | 9.001316256 | 8.765573969 | 0.235742287 | -0.010810911 |
| Rhot1 | 9.97769888 | 9.910363984 | 9.988536951 | -0.078172968 | -0.010838072 |
| Tomt | 1.119107471 | 1.141853002 | 1.129946771 | 0.011906231 | -0.0108393 |
| Ifitm10 | 6.088987039 | 6.285093667 | 6.09985801 | 0.185235657 | -0.010870971 |
| Gm8226 | 2.473361581 | 2.465776365 | 2.484234793 | -0.018458427 | -0.010873212 |
| Gm11870 | 3.145912436 | 3.187733358 | 3.156815132 | 0.030918226 | -0.010902696 |
| H2-M3 | 7.985479955 | 7.785990263 | 7.996383896 | -0.210393632 | -0.010903941 |
| Anapc11 | 8.793668707 | 8.922733091 | 8.804589008 | 0.118144083 | -0.010920302 |
| Asap1 | 10.16015294 | 10.3179127 | 10.17112065 | 0.146792053 | -0.010967708 |
| Tmem191c | 3.998130802 | 4.055982049 | 4.009156033 | 0.046826016 | -0.011025231 |
| Pdcd10 | 9.69468913 | 9.612852062 | 9.705734008 | -0.092881947 | -0.011044878 |
| Slc13a1 | 1.872657665 | 1.860515278 | 1.883742079 | -0.023226801 | -0.011084414 |
| Cnnm2 | 7.600757522 | 7.316002367 | 7.611862357 | -0.29585999 | -0.011104836 |
| Cab39l | 10.64156398 | 10.61663067 | 10.65269958 | -0.036068913 | -0.0111356 |
| Gm13238 | 1.251455356 | 1.26276158 | 1.262595492 | 0.000166088 | -0.011140136 |
| Olfr237-ps1 | 2.375185229 | 2.418216553 | 2.386337024 | 0.031879529 | -0.011151794 |
| Rem2 | 2.658004674 | 2.675576286 | 2.669160696 | 0.00641559 | -0.011156023 |
| Slc30a4 | 9.617760103 | 9.460776683 | 9.628949439 | -0.168172756 | -0.011189336 |
| Gm8173 | 2.771390494 | 2.765423 | 2.782623386 | -0.017200385 | -0.011232892 |
| Tnxa | 2.513586409 | 2.541802263 | 2.524829981 | 0.016972282 | -0.011243572 |

| | | | | | |
|---|---|---|---|---|---|
| Mprip | 12.39567548 | 12.44166981 | 12.40694203 | 0.034727777 | -0.01126655 |
| Slc31a1 | 10.40548214 | 10.4269347 | 10.41678545 | 0.010149253 | -0.01130331 |
| Gm12403 | 1.950096745 | 1.970629771 | 1.961408705 | 0.009221066 | -0.011311961 |
| Gm45844 | 5.827016185 | 5.820496509 | 5.838341385 | -0.017844876 | -0.0113252 |
| Gpr179 | 1.300291185 | 1.296154731 | 1.311616477 | -0.015461746 | -0.011325292 |
| Arhgef15 | 7.635222736 | 7.61180805 | 7.646561584 | -0.034753535 | -0.011338849 |
| Srbd1 | 8.403943405 | 8.461003855 | 8.415332846 | 0.045671009 | -0.011389441 |
| Hspa1b | 11.02252268 | 13.48404572 | 11.03395161 | 2.45009411 | -0.01142893 |
| Gm10073 | 2.336993676 | 2.365132406 | 2.348428824 | 0.016703582 | -0.011435149 |
| Lrrd1 | 1.56743356 | 1.564325451 | 1.578870944 | -0.014545493 | -0.011437384 |
| Gm10548 | 3.010157518 | 2.998312273 | 3.021654161 | -0.023341888 | -0.011496643 |
| Rasa4 | 8.487170058 | 8.412334766 | 8.498689197 | -0.08635443 | -0.011519139 |
| St3gal3 | 8.143081324 | 8.06266969 | 8.154604543 | -0.091934854 | -0.011523219 |
| Opalin | 2.296407418 | 2.310405343 | 2.307957959 | 0.002447384 | -0.011550541 |
| BC048671 | 1.873592569 | 1.897526688 | 1.885163574 | 0.012363114 | -0.011571006 |
| Prr15 | 4.199248935 | 4.167565036 | 4.21082588 | -0.043260844 | -0.011576945 |
| Lysmd3 | 8.8719882 | 8.922837555 | 8.883665371 | 0.039172184 | -0.011677171 |
| Csde1 | 13.13200801 | 13.01309225 | 13.14375826 | -0.130666007 | -0.011750247 |
| 2010109A12Rik | 2.83225506 | 3.068971641 | 2.844046796 | 0.224924845 | -0.011791736 |
| Ube2t | 6.052995672 | 6.060801391 | 6.064803103 | -0.004001712 | -0.011807431 |
| Col5a3 | 10.28742574 | 10.10079929 | 10.29923678 | -0.198437494 | -0.011811043 |
| Gm10045 | 0.655554897 | 0.658648988 | 0.66739896 | -0.008749972 | -0.011844063 |
| Vmn2r7 | 1.014217915 | 1.015393845 | 1.02614397 | -0.010750125 | -0.011926055 |
| Atp5o | 11.47860191 | 11.42838099 | 11.49053491 | -0.062153917 | -0.011933003 |
| 2310033P09Rik | 7.736276843 | 7.791749228 | 7.7482572 | 0.043492028 | -0.011980357 |
| Palm2 | 3.673333786 | 3.471138764 | 3.685357289 | -0.214218526 | -0.012023503 |
| Timm22 | 8.659644691 | 8.73915396 | 8.67174855 | 0.067405409 | -0.012103859 |
| Nudt15 | 6.613876814 | 6.741232921 | 6.626002942 | 0.115229979 | -0.012126128 |
| Slc5a11 | 2.951301864 | 2.945060427 | 2.963429274 | -0.018368847 | -0.01212741 |
| Dlx4 | 2.498848252 | 2.575751064 | 2.511039193 | 0.064711871 | -0.012190941 |
| Rpl9-ps7 | 1.938649345 | 1.998854671 | 1.950851143 | 0.048003527 | -0.012201798 |
| Gm20589 | 3.074314019 | 3.197950311 | 3.086521031 | 0.11142928 | -0.012207012 |

| | | | | | |
|---|---|---|---|---|---|
| Crebzf | 9.553060309 | 9.528973625 | 9.565276879 | -0.036303254 | -0.01221657 |
| Vmn1r224 | 1.661048524 | 1.673150796 | 1.673367935 | -0.000217139 | -0.012319411 |
| 40238 | 1.608175782 | 1.58587205 | 1.620499465 | -0.034627415 | -0.012323683 |
| Vmn2r-ps117 | 2.122061801 | 2.12262714 | 2.134402432 | -0.011775292 | -0.012340631 |
| 9030025P20Rik | 6.903194983 | 6.849459366 | 6.915536834 | -0.066077468 | -0.012341851 |
| Hmgn2 | 11.51291415 | 11.41678994 | 11.52530313 | -0.10851319 | -0.01238898 |
| BC005561 | 8.053355215 | 8.000857131 | 8.065755167 | -0.064898036 | -0.012399952 |
| Gm49405 | 5.87682891 | 6.115625567 | 5.889248058 | 0.226377509 | -0.012419149 |
| Capn11 | 2.540111719 | 2.805875787 | 2.55257296 | 0.253302827 | -0.012461241 |
| Gm2099 | 2.805693958 | 2.83235654 | 2.818216343 | 0.014140197 | -0.012522385 |
| Csrnp3 | 2.314048329 | 2.286615587 | 2.32657086 | -0.039955274 | -0.012522531 |
| Gm8186 | 6.851780785 | 6.831867141 | 6.864317241 | -0.0324501 | -0.012536456 |
| Henmt1 | 1.03442272 | 1.016221503 | 1.046967301 | -0.030745798 | -0.012544581 |
| Gm4791 | 1.88129852 | 1.877762167 | 1.893847029 | -0.016084862 | -0.012548509 |
| Adipor1 | 12.02588317 | 11.94775013 | 12.03846807 | -0.09071794 | -0.012584903 |
| Slc27a2 | 3.188885685 | 3.192431125 | 3.201497977 | -0.009066852 | -0.012612292 |
| Sez6l | 3.423232309 | 3.40906501 | 3.435846571 | -0.026781561 | -0.012614262 |
| Olfr1262 | 1.438877711 | 1.46143472 | 1.451514855 | 0.009919865 | -0.012637144 |
| Uts2r | 2.305403324 | 2.315076542 | 2.318075785 | -0.002999242 | -0.01267246 |
| Smad4 | 10.19627585 | 10.19394485 | 10.20896304 | -0.01501819 | -0.012687193 |
| Fcgbp | 13.7212702 | 14.0483443 | 13.73396138 | 0.314382923 | -0.012691183 |
| Brox | 10.19480041 | 10.30980771 | 10.20749966 | 0.10230805 | -0.01269925 |
| Shcbp1l | 2.787428423 | 2.872639491 | 2.800140998 | 0.072498493 | -0.012712575 |
| 4932414N04Rik | 2.56599561 | 2.548716897 | 2.578722815 | -0.030005919 | -0.012727206 |
| Gm14296 | 8.369449761 | 8.489038319 | 8.382225869 | 0.10681245 | -0.012776108 |
| Nyx | 3.92586607 | 3.947689467 | 3.938646378 | 0.009043089 | -0.012780308 |
| Gm8113 | 8.334906018 | 8.302824847 | 8.347699546 | -0.044874699 | -0.012793528 |
| Olfr1564 | 1.611937974 | 1.626310946 | 1.624737438 | 0.001573507 | -0.012799465 |
| Ccdc78 | 2.665992147 | 2.682409091 | 2.678822881 | 0.003586209 | -0.012830734 |
| Gm42785 | 6.847880905 | 6.837931457 | 6.860748556 | -0.022817098 | -0.012867651 |
| Zfp639 | 8.584605146 | 8.48118148 | 8.597488741 | -0.116307261 | -0.012883594 |
| Mrpl49 | 8.964176416 | 8.885536847 | 8.977061026 | -0.091524179 | -0.01288461 |

| | | | | | |
|---|---|---|---|---|---|
| Syngr3 | 1.393152087 | 1.427498087 | 1.406082591 | 0.021415496 | -0.012930504 |
| Tmem86a | 9.245800623 | 9.262237588 | 9.258780269 | 0.003457319 | -0.012979646 |
| Ppp1r32 | 2.681693733 | 2.649880018 | 2.694749852 | -0.044869834 | -0.013056119 |
| Tasp1 | 7.335469205 | 7.303213839 | 7.348585241 | -0.045371403 | -0.013116037 |
| Creb3 | 9.428929805 | 9.487199122 | 9.442104618 | 0.045094504 | -0.013174813 |
| Vmn1r230 | 2.432601018 | 2.448333815 | 2.445788325 | 0.00254549 | -0.013187307 |
| Gm49776 | 1.827121398 | 1.73924898 | 1.840327995 | -0.101079015 | -0.013206597 |
| Irak1bp1 | 6.078425362 | 5.940623547 | 6.091636789 | -0.151013242 | -0.013211428 |
| Gm14414 | 1.124164741 | 1.147876407 | 1.137493208 | 0.010383199 | -0.013328467 |
| Uqcc3 | 7.899614727 | 7.83619651 | 7.912950698 | -0.076754188 | -0.013335971 |
| Olfr1205 | 2.613036538 | 2.607764593 | 2.626377283 | -0.01861269 | -0.013340744 |
| Ccdc103 | 2.16441005 | 2.184295392 | 2.177751563 | 0.00654383 | -0.013341512 |
| Slc13a3 | 6.860406869 | 7.328366557 | 6.873754474 | 0.454612083 | -0.013347605 |
| Gm28455 | 3.130994997 | 3.072493405 | 3.144363152 | -0.071869748 | -0.013368156 |
| Gm12291 | 1.428861197 | 1.372957265 | 1.442252848 | -0.069295583 | -0.01339165 |
| Vmn1r197 | 4.086725871 | 4.129530767 | 4.100119233 | 0.029411533 | -0.013393363 |
| Npas3 | 1.034865393 | 1.045944665 | 1.048259864 | -0.002315199 | -0.013394471 |
| Psmd10 | 7.319804725 | 7.235435883 | 7.333221883 | -0.097786001 | -0.013417158 |
| Ttll13 | 2.669659478 | 2.695171186 | 2.683078975 | 0.012092211 | -0.013419497 |
| Kir3dl1 | 3.646352211 | 3.714444984 | 3.659787083 | 0.054657901 | -0.013434872 |
| Mchr1 | 2.71795685 | 2.69045587 | 2.73142723 | -0.04097136 | -0.01347038 |
| Cpne9 | 2.015604666 | 2.04282327 | 2.029088829 | 0.013734441 | -0.013484163 |
| Icam2 | 6.372160674 | 6.306115661 | 6.385658121 | -0.07954246 | -0.013497447 |
| Xrra1 | 1.973881195 | 2.092626366 | 1.98738434 | 0.105242026 | -0.013503145 |
| Mup13 | 1.54134808 | 1.543510429 | 1.554860249 | -0.011349821 | -0.01351217 |
| Vps25 | 10.19283406 | 10.15973428 | 10.20634705 | -0.046612763 | -0.01351299 |
| Gm11627 | 2.304470533 | 2.348917298 | 2.318026669 | 0.030890629 | -0.013556136 |
| Gm4858 | 1.616457295 | 1.60160715 | 1.630027439 | -0.028420289 | -0.013570144 |
| Vmn1r207-ps | 1.10817749 | 1.077489116 | 1.121750184 | -0.044261069 | -0.013572695 |
| Reg2 | 3.1416674 | 3.191863909 | 3.155262262 | 0.036601647 | -0.013594862 |
| Sfxn2 | 8.961448622 | 8.887098434 | 8.975082291 | -0.087983857 | -0.013633669 |
| Gm13233 | 1.806280435 | 1.900065683 | 1.819933348 | 0.080132335 | -0.013652913 |

| | | | | | |
|---|---|---|---|---|---|
| Zfp711 | 5.447237701 | 5.515088361 | 5.460896668 | 0.054191693 | -0.013658967 |
| Gm45880 | 1.673865818 | 1.65560979 | 1.687540255 | -0.031930465 | -0.013674437 |
| Gm47626 | 6.871172548 | 6.961700361 | 6.884865516 | 0.076834844 | -0.013692968 |
| Il17d | 3.982937408 | 3.968588134 | 3.996636168 | -0.028048034 | -0.01369876 |
| Ttyh1 | 5.539951879 | 5.755315667 | 5.553664748 | 0.201650919 | -0.013712869 |
| Tcf24 | 4.028413689 | 4.042461028 | 4.042158046 | 0.000302982 | -0.013744357 |
| Gm47476 | 5.221987795 | 5.197554467 | 5.235749603 | -0.038195137 | -0.013761808 |
| Ces1e | 4.15623597 | 4.042739855 | 4.170022576 | -0.127282722 | -0.013786607 |
| Gcm2 | 1.544701121 | 1.513970626 | 1.558531287 | -0.044560662 | -0.013830166 |
| Lsm3 | 8.809799788 | 8.748910132 | 8.82364013 | -0.074729998 | -0.013840341 |
| Trpv1 | 4.064391808 | 4.108563549 | 4.078232781 | 0.030330768 | -0.013840974 |
| AC132444.6 | 4.768283693 | 4.730952105 | 4.782153894 | -0.051201789 | -0.013870201 |
| Zic5 | 1.401466681 | 1.422666691 | 1.415379521 | 0.00728717 | -0.01391284 |
| Pex2 | 9.248108918 | 9.046122563 | 9.262027396 | -0.215904833 | -0.013918478 |
| Gm47759 | 2.242034849 | 2.314185553 | 2.255966487 | 0.058219066 | -0.013931638 |
| Cilp2 | 2.92712013 | 2.978716639 | 2.94105888 | 0.037657758 | -0.01393875 |
| Mpg | 8.154764661 | 8.108158558 | 8.168703488 | -0.06054493 | -0.013938828 |
| Rnf115 | 9.919491367 | 10.19840129 | 9.933430708 | 0.264970585 | -0.013939341 |
| Cyb561d1 | 8.937826285 | 8.820537205 | 8.951794526 | -0.131257321 | -0.013968242 |
| Ndufb2 | 9.291259727 | 9.45082544 | 9.305238682 | 0.145586758 | -0.013978955 |
| Ahsg | 1.828846251 | 1.805434107 | 1.84284492 | -0.037410813 | -0.01399867 |
| Mpp4 | 2.593118627 | 2.646426449 | 2.607119035 | 0.039307415 | -0.014000408 |
| Chrna5 | 4.005544264 | 3.962468498 | 4.019556581 | -0.057088084 | -0.014012317 |
| Zfp638 | 10.43835017 | 10.40161607 | 10.45236588 | -0.05074981 | -0.014015703 |
| Gabrd | 1.852838491 | 1.857895618 | 1.866911993 | -0.009016376 | -0.014073502 |
| Ppp2r2c | 9.895222194 | 10.02017165 | 9.909300322 | 0.110871324 | -0.014078128 |
| Pcgf1 | 7.160082968 | 7.18324554 | 7.174196204 | 0.009049336 | -0.014113236 |
| 4930524J08Rik | 1.983895857 | 1.991329958 | 1.998078265 | -0.006748307 | -0.014182408 |
| Nek5 | 1.159748772 | 1.145006213 | 1.173943584 | -0.028937371 | -0.014194812 |
| Prr36 | 2.866210549 | 2.869965066 | 2.880418599 | -0.010453533 | -0.01420805 |
| Ccdc183 | 1.890020983 | 1.972716948 | 1.904229578 | 0.06848737 | -0.014208595 |
| Msh4 | 3.880327168 | 3.868847814 | 3.894552854 | -0.02570504 | -0.014225686 |

| | | | | | |
|---|---|---|---|---|---|
| Ndufb10 | 10.46327528 | 10.40102403 | 10.47752432 | -0.076500287 | -0.014249037 |
| Tspan2 | 8.008180954 | 7.851463941 | 8.022432257 | -0.170968316 | -0.014251303 |
| Coro1c | 11.51567922 | 11.55699624 | 11.52996277 | 0.027033473 | -0.01428355 |
| Olfr824 | 0.51812873 | 0.526759966 | 0.53244988 | -0.005689913 | -0.014321149 |
| Tmx2 | 9.750163029 | 9.693709874 | 9.764485517 | -0.070775643 | -0.014322487 |
| Defb42 | 2.35133474 | 2.346088946 | 2.365755187 | -0.019666241 | -0.014420447 |
| Sapcd1 | 1.430674917 | 1.441795432 | 1.445112562 | -0.00331713 | -0.014437644 |
| Gm48208 | 1.393702063 | 1.380896003 | 1.408171563 | -0.027275561 | -0.0144695 |
| 3830403N18Rik | 2.465976283 | 2.531178299 | 2.480460896 | 0.050717403 | -0.014484613 |
| Gm4149 | 2.610882506 | 2.583952045 | 2.62547297 | -0.041520926 | -0.014590465 |
| Mpdu1 | 9.926752137 | 9.83770437 | 9.94136946 | -0.10366509 | -0.014617323 |
| Mcam | 10.30426734 | 10.22331018 | 10.31890194 | -0.095591767 | -0.014634607 |
| Ppp1r12a | 11.18806287 | 11.22605164 | 11.20272031 | 0.023331333 | -0.014657437 |
| Mtx3 | 8.429206347 | 8.501889863 | 8.443887382 | 0.05800248 | -0.014681035 |
| Parpbp | 6.011997399 | 5.889038504 | 6.026698811 | -0.137660307 | -0.014701412 |
| Gm12736 | 3.514837062 | 3.508082442 | 3.529540208 | -0.021457766 | -0.014703145 |
| 1700061G19Rik | 2.344812678 | 2.363849617 | 2.35952239 | 0.004327227 | -0.014709713 |
| Rpl3 | 14.10978691 | 14.08862306 | 14.12452221 | -0.03589915 | -0.0147353 |
| Car10 | 2.118723565 | 2.14814054 | 2.133465136 | 0.014675404 | -0.014741571 |
| Lpcat1 | 7.901128111 | 7.877094688 | 7.915893601 | -0.038798913 | -0.01476549 |
| Gm48552 | 2.929428918 | 3.069121199 | 2.944206804 | 0.124914395 | -0.014777887 |
| Nanos1 | 3.911747229 | 4.020434068 | 3.926584225 | 0.093849843 | -0.014836996 |
| Uqcrh | 11.60609328 | 11.45207942 | 11.62098973 | -0.168910303 | -0.014896447 |
| Hmgb1-ps3 | 2.161042502 | 2.231867494 | 2.176014711 | 0.055852783 | -0.014972209 |
| Gm17205 | 2.792552221 | 2.81138073 | 2.80752629 | 0.00385444 | -0.014974069 |
| Cog2 | 9.27440587 | 9.364104489 | 9.289397969 | 0.074706521 | -0.014992099 |
| Rnaset2a | 7.011285542 | 6.954645137 | 7.026277657 | -0.07163252 | -0.014992115 |
| Iars2 | 10.71403472 | 10.60789451 | 10.72907007 | -0.121175553 | -0.015035347 |
| Tns2 | 10.34859047 | 10.33146576 | 10.36364149 | -0.032175723 | -0.01505102 |
| Maoa | 9.748819512 | 9.669806703 | 9.763888538 | -0.094081835 | -0.015069026 |
| Zfp414 | 7.505225158 | 7.768190447 | 7.520294506 | 0.247895942 | -0.015069347 |
| Vmn1r198 | 3.509229798 | 3.569508333 | 3.524330996 | 0.045177337 | -0.015101198 |

| | | | | | |
|---|---|---|---|---|---|
| Olfr671 | 1.82016896 | 1.79454065 | 1.835281963 | -0.040741314 | -0.015113004 |
| AI593442 | 1.521143595 | 1.502925814 | 1.536270792 | -0.033344978 | -0.015127197 |
| Fam19a1 | 1.661344016 | 1.73519889 | 1.676476298 | 0.058722591 | -0.015132283 |
| Slc11a2 | 10.11033474 | 10.32283166 | 10.12551423 | 0.197317427 | -0.015179497 |
| Gm5380 | 2.484960822 | 2.451405552 | 2.500168982 | -0.048763429 | -0.01520816 |
| Zxdc | 8.980401321 | 8.866514755 | 8.995613165 | -0.129098409 | -0.015211843 |
| Rnf10 | 12.07668702 | 12.13120338 | 12.09189892 | 0.03930446 | -0.0152119 |
| G6pdx | 11.2256369 | 11.13719817 | 11.24088998 | -0.103691817 | -0.015253087 |
| Gm11675 | 1.878583551 | 1.885613551 | 1.893842549 | -0.008228998 | -0.015258998 |
| Gm48990 | 1.976249045 | 2.028104411 | 1.991508515 | 0.036595896 | -0.01525947 |
| Fam205a2 | 3.825584323 | 3.786873851 | 3.840861571 | -0.05398772 | -0.015277248 |
| Gm21981 | 3.027754054 | 3.04221507 | 3.043034532 | -0.000819462 | -0.015280478 |
| Wasf3 | 3.548040571 | 3.599597461 | 3.563333459 | 0.036264003 | -0.015292887 |
| Cdkn2b | 6.957812137 | 7.018416375 | 6.973130549 | 0.045285826 | -0.015318411 |
| Myh14 | 10.42923989 | 10.51098955 | 10.44458953 | 0.066400027 | -0.01534964 |
| Triap1 | 9.112147816 | 9.092612819 | 9.127505457 | -0.034892638 | -0.015357641 |
| 6030445D17Rik | 1.798341952 | 1.793191505 | 1.813700357 | -0.020508852 | -0.015358405 |
| Rnpc3 | 8.512769672 | 8.50971721 | 8.528138043 | -0.018420833 | -0.015368371 |
| 9130213A22Rik | 1.599891202 | 1.615442543 | 1.615296585 | 0.000145958 | -0.015405383 |
| Cntnap2 | 2.962533431 | 2.933858322 | 2.977950637 | -0.044092315 | -0.015417207 |
| Gmds | 7.225643457 | 7.295187356 | 7.241065462 | 0.054121894 | -0.015422005 |
| Mief2 | 8.278797808 | 8.154644184 | 8.294254639 | -0.139610454 | -0.015456831 |
| Cela2a | 1.760444294 | 1.766859365 | 1.775902746 | -0.009043381 | -0.015458452 |
| Ttbk1 | 2.982031832 | 3.017775155 | 2.997575167 | 0.020199988 | -0.015543334 |
| Klhl12 | 9.118368535 | 9.040514048 | 9.133936208 | -0.09342216 | -0.015567673 |
| Hist1h2bh | 2.594694207 | 2.600048968 | 2.61030961 | -0.010260642 | -0.015615402 |
| Gm18180 | 1.165008484 | 1.239350326 | 1.180695667 | 0.058654659 | -0.015687183 |
| Oscar | 1.300666471 | 1.387376895 | 1.316361371 | 0.071015524 | -0.015694899 |
| Gm13292 | 1.878825673 | 1.960982787 | 1.8945238 | 0.066458987 | -0.015698127 |
| Pcdhb9 | 3.06153265 | 3.065827993 | 3.077231759 | -0.011403766 | -0.015699109 |
| Chtf18 | 6.646734137 | 6.602922504 | 6.66247286 | -0.059550356 | -0.015738723 |
| Gprin1 | 2.298130842 | 2.264323902 | 2.313884786 | -0.049560883 | -0.015753943 |

| | | | | | |
|---|---|---|---|---|---|
| Hist1h2bq | 3.875407984 | 3.924254618 | 3.891164193 | 0.033090425 | -0.01575621 |
| Ndufs2 | 11.73335569 | 11.60511041 | 11.74911576 | -0.144005347 | -0.015760063 |
| 1700067K01Rik | 1.972491201 | 2.016545096 | 1.988251521 | 0.028293575 | -0.01576032 |
| Pold3 | 9.066784366 | 9.11343213 | 9.082556574 | 0.030875556 | -0.015772208 |
| Sowahc | 10.34749545 | 10.3192059 | 10.36328347 | -0.044077573 | -0.015788027 |
| Trdmt1 | 7.03199442 | 6.977492201 | 7.047792619 | -0.070300418 | -0.015798199 |
| Olfr822 | 1.420126749 | 1.407607228 | 1.435931052 | -0.028323824 | -0.015804303 |
| Pigb | 8.119954676 | 8.088292174 | 8.135838674 | -0.0475465 | -0.015883998 |
| Thumpd2 | 6.007784478 | 5.869839818 | 6.023695047 | -0.153855228 | -0.015910569 |
| Phex | 3.389070222 | 3.3499425 | 3.404996811 | -0.055054311 | -0.015926588 |
| Slf1 | 7.670218654 | 7.689050772 | 7.686175673 | 0.002875099 | -0.015957019 |
| Slc6a15 | 2.93701838 | 2.983163127 | 2.953043497 | 0.03011963 | -0.016025117 |
| Dnali1 | 1.337988577 | 1.337175093 | 1.354071226 | -0.016896133 | -0.01608265 |
| Cdc27 | 10.1766779 | 10.21979908 | 10.19277307 | 0.027026013 | -0.016095173 |
| Trat1 | 6.882190615 | 7.1149261 | 6.89837921 | 0.21654689 | -0.016188595 |
| Elmsan1 | 10.43343689 | 10.47703042 | 10.44965405 | 0.02737637 | -0.016217163 |
| Heatr5a | 9.830669686 | 9.87641627 | 9.846915699 | 0.029500571 | -0.016246013 |
| Gm12394 | 1.13751617 | 1.145416193 | 1.153769314 | -0.008353121 | -0.016253144 |
| Cisd1 | 9.99772677 | 9.92330853 | 10.01400243 | -0.090693895 | -0.016275655 |
| 1500011B03Rik | 7.284068256 | 7.173526478 | 7.300395213 | -0.126868735 | -0.016326957 |
| Slc29a4 | 1.836891832 | 1.839968696 | 1.853235196 | -0.0132665 | -0.016343364 |
| Asb17 | 1.565546786 | 1.589097427 | 1.581908789 | 0.007188639 | -0.016362003 |
| 1700028J19Rik | 3.106272101 | 3.080336531 | 3.122639246 | -0.042302715 | -0.016367145 |
| Rps3 | 13.80740039 | 13.68931256 | 13.82377269 | -0.134460127 | -0.016372297 |
| Jakmip3 | 1.767809725 | 1.771005951 | 1.784195566 | -0.013189615 | -0.016385841 |
| Plk1 | 7.323351411 | 7.34344731 | 7.339817739 | 0.003629572 | -0.016466328 |
| Krt24 | 8.346916995 | 8.219192093 | 8.36347485 | -0.144282757 | -0.016557856 |
| Atox1 | 10.55577605 | 10.61064563 | 10.57233417 | 0.038311457 | -0.016558117 |
| Aff2 | 4.693775403 | 4.713027945 | 4.710355881 | 0.002672064 | -0.016580477 |
| Pdzd11 | 9.493386792 | 9.377615586 | 9.509989151 | -0.132373565 | -0.016602359 |
| Prickle3 | 8.947972451 | 9.18070148 | 8.964579605 | 0.216121875 | -0.016607154 |
| Arl9 | 1.620146905 | 1.647197567 | 1.636758573 | 0.010438994 | -0.016611668 |

| | | | | | |
|---|---|---|---|---|---|
| Lrrc57 | 8.127488018 | 8.048095453 | 8.144158685 | -0.096063232 | -0.016670667 |
| Card10 | 9.970176079 | 10.48209211 | 9.986850942 | 0.495241171 | -0.016674864 |
| Ncoa7 | 8.209619879 | 7.796661178 | 8.226360868 | -0.42969969 | -0.016740989 |
| Rragc | 10.86086818 | 10.8952484 | 10.87764907 | 0.017599323 | -0.01678089 |
| Akap4 | 1.661627137 | 1.684045397 | 1.678444799 | 0.005600598 | -0.016817662 |
| Eaf2 | 3.673569243 | 3.710827052 | 3.690487977 | 0.020339076 | -0.016918733 |
| Cfap97d2 | 3.300462684 | 3.298134572 | 3.317428667 | -0.019294095 | -0.016965983 |
| Dhx40 | 9.259246893 | 9.346818158 | 9.276215341 | 0.070602816 | -0.016968448 |
| Rpl32 | 13.24690264 | 13.12323035 | 13.2639375 | -0.14070715 | -0.017034857 |
| Ubr2 | 10.64110536 | 10.76474279 | 10.65814618 | 0.10659661 | -0.01704082 |
| Tpt1 | 14.95340692 | 14.81801226 | 14.97050461 | -0.15249235 | -0.017097683 |
| Spata24 | 4.639798201 | 4.536070924 | 4.656905539 | -0.120834615 | -0.017107337 |
| Selenoi | 10.47749379 | 10.48940751 | 10.49460921 | -0.0052017 | -0.01711542 |
| 5430437J10Rik | 2.755134032 | 2.847385426 | 2.772252953 | 0.075132473 | -0.017118921 |
| Dnhd1 | 5.711099011 | 5.496353498 | 5.728223039 | -0.231869541 | -0.017124028 |
| Gpr33 | 1.544047533 | 1.573160849 | 1.561214718 | 0.011946131 | -0.017167185 |
| Gm8798 | 1.993018704 | 1.998191981 | 2.010238057 | -0.012046076 | -0.017219352 |
| Gm6988 | 1.588582382 | 1.557105031 | 1.605811386 | -0.048706355 | -0.017229004 |
| Rpl26 | 13.18526105 | 13.07873781 | 13.2025313 | -0.12379349 | -0.017270253 |
| Pdpk1 | 10.70610404 | 10.65185822 | 10.72338524 | -0.071527017 | -0.017281203 |
| Ccdc130 | 7.232236642 | 7.265426842 | 7.249533379 | 0.015893462 | -0.017296738 |
| Trhr | 1.660730931 | 1.683106248 | 1.678029723 | 0.005076525 | -0.017298792 |
| Angptl8 | 3.616825547 | 3.457222352 | 3.634135568 | -0.176913217 | -0.017310021 |
| Gm8221 | 2.38945747 | 2.504173033 | 2.406797134 | 0.097375899 | -0.017339664 |
| Shisal1 | 2.033398815 | 2.070721237 | 2.05081093 | 0.019910307 | -0.017412115 |
| Gm8584 | 2.844325921 | 2.866673943 | 2.861749184 | 0.004924759 | -0.017423263 |
| Itgae | 7.440630956 | 7.481254836 | 7.458065608 | 0.023189228 | -0.017434652 |
| H2-DMa | 9.444405374 | 9.319212121 | 9.461841603 | -0.142629483 | -0.017436229 |
| Gm49619 | 1.676426561 | 1.688926107 | 1.693879075 | -0.004952968 | -0.017452514 |
| Ppp1r1c | 2.583371765 | 2.51122557 | 2.600828438 | -0.089602868 | -0.017456673 |
| Ubn1 | 10.86724689 | 10.98472657 | 10.88471612 | 0.100010447 | -0.01746923 |
| Gm12106 | 2.736753397 | 2.801064688 | 2.754230746 | 0.046833942 | -0.01747735 |

| | | | | | |
|---|---|---|---|---|---|
| Pmf1 | 8.161847235 | 8.364194633 | 8.179332337 | 0.184862296 | -0.017485102 |
| Pola1 | 8.821844516 | 8.897658337 | 8.839352213 | 0.058306124 | -0.017507697 |
| Kl | 3.57562307 | 3.618405154 | 3.593132676 | 0.025272479 | -0.017509606 |
| Cdh15 | 5.496226251 | 5.526697683 | 5.513758984 | 0.012938699 | -0.017532734 |
| Dync1i2 | 11.22502586 | 11.12193333 | 11.24256313 | -0.120629797 | -0.017537273 |
| Lrif1 | 8.361951938 | 8.248769065 | 8.379517763 | -0.130748698 | -0.017565826 |
| Gm8251 | 1.954480533 | 1.951012014 | 1.972055262 | -0.021043248 | -0.017574728 |
| Fam229b | 4.185927561 | 4.240089937 | 4.20353231 | 0.036557627 | -0.017604749 |
| Klrb1f | 4.308881065 | 4.319689021 | 4.326534894 | -0.006845873 | -0.01765383 |
| Gm47933 | 5.430652009 | 5.446424528 | 5.448327026 | -0.001902498 | -0.017675017 |
| Ica1l | 2.777069442 | 2.773634663 | 2.794762832 | -0.021128169 | -0.017693389 |
| Sh2d1b1 | 4.063600926 | 3.997987556 | 4.081297004 | -0.083309448 | -0.017696078 |
| Fam45a | 8.594297483 | 8.5085043 | 8.612006152 | -0.103501852 | -0.017708669 |
| Tube1 | 5.708401555 | 5.697771743 | 5.726136096 | -0.028364353 | -0.017734542 |
| Mc1r | 1.700788194 | 1.692619987 | 1.718530256 | -0.025910269 | -0.017742062 |
| Qrich2 | 2.809836675 | 2.903057041 | 2.827579059 | 0.075477982 | -0.017742384 |
| Lrp2 | 2.507937788 | 2.54055617 | 2.525685919 | 0.014870251 | -0.017748131 |
| Nxf7 | 1.579395436 | 1.610433759 | 1.597186297 | 0.013247462 | -0.017790861 |
| Krt9 | 2.436909317 | 2.476016104 | 2.454704454 | 0.021311651 | -0.017795136 |
| Gm16574 | 2.933613206 | 2.955267816 | 2.951422258 | 0.003845557 | -0.017809053 |
| Prkcg | 3.259875773 | 3.411140459 | 3.277701251 | 0.133439208 | -0.017825478 |
| Rasl2-9 | 1.959397741 | 1.944106329 | 1.977238097 | -0.033131768 | -0.017840356 |
| Gm16487 | 2.559724437 | 2.569322254 | 2.577627794 | -0.00830554 | -0.017903358 |
| Fbxo24 | 2.28479816 | 2.349053717 | 2.302712292 | 0.046341425 | -0.017914132 |
| Rtbdn | 0.872557413 | 0.900161518 | 0.890481053 | 0.009680465 | -0.01792364 |
| Rps13-ps4 | 2.747632648 | 2.797383332 | 2.765582388 | 0.031800944 | -0.01794974 |
| Celsr2 | 11.02480044 | 11.40420324 | 11.04281973 | 0.361383503 | -0.018019297 |
| Dalrd3 | 8.770834161 | 8.799080456 | 8.788883143 | 0.010197313 | -0.018048982 |
| Ahr | 10.56348128 | 10.67186205 | 10.58153088 | 0.09033117 | -0.018049603 |
| Gm49369 | 4.750702343 | 4.736527076 | 4.768774169 | -0.032247093 | -0.018071826 |
| Sdhc | 11.07404519 | 11.07982136 | 11.09219237 | -0.012371017 | -0.018147187 |
| Olfr31 | 3.612071619 | 3.56388306 | 3.630231368 | -0.066348308 | -0.018159749 |

| | | | | | |
|---|---|---|---|---|---|
| Tecta | 1.603261608 | 1.573271457 | 1.621432859 | -0.048161402 | -0.018171251 |
| Brinp1 | 1.100852008 | 1.061510105 | 1.119051722 | -0.057541616 | -0.018199714 |
| Nop53 | 11.24599077 | 11.32336013 | 11.26420897 | 0.059151157 | -0.018218197 |
| Abr | 10.74569858 | 10.9406038 | 10.7639634 | 0.176640393 | -0.018264827 |
| Trappc3 | 10.07166562 | 10.00203613 | 10.0899349 | -0.08789877 | -0.018269273 |
| Fuom | 7.012215971 | 7.000999562 | 7.030492696 | -0.029493133 | -0.018276725 |
| Slc18a1 | 2.861120253 | 2.878507471 | 2.879483276 | -0.000975805 | -0.018363023 |
| Mtf2 | 9.337572037 | 9.241655304 | 9.355940269 | -0.114284964 | -0.018368232 |
| Kpna3 | 10.55636857 | 10.48807684 | 10.57475141 | -0.086674573 | -0.01838284 |
| 5730507C01Rik | 4.446039241 | 4.347588771 | 4.464423552 | -0.116834782 | -0.018384311 |
| Cul4a | 10.73569692 | 10.6332245 | 10.75412435 | -0.120899853 | -0.018427433 |
| Ubald1 | 9.214293783 | 9.553150795 | 9.232779932 | 0.320370863 | -0.018486149 |
| Zbtb12 | 6.439349456 | 6.477076135 | 6.45787344 | 0.019202695 | -0.018523984 |
| Vmn2r67 | 3.423107227 | 3.447927794 | 3.441638618 | 0.006289176 | -0.01853139 |
| Vmn1r208 | 1.092553078 | 1.113538588 | 1.111149198 | 0.00238939 | -0.01859612 |
| Tmem30b | 9.17838078 | 9.042756273 | 9.197014693 | -0.15425842 | -0.018633912 |
| Bhlhe41 | 8.474132943 | 8.417451454 | 8.492779948 | -0.075328494 | -0.018647005 |
| Dhrs11 | 6.836260607 | 6.907616522 | 6.854922232 | 0.05269429 | -0.018661626 |
| Axl | 11.95104869 | 11.79130579 | 11.96972806 | -0.178422267 | -0.018679363 |
| Gm12818 | 1.739636022 | 1.785399974 | 1.758329936 | 0.027070038 | -0.018693914 |
| Gm8797 | 3.018355688 | 3.044935546 | 3.037069396 | 0.00786615 | -0.018713708 |
| Gm28693 | 2.198018753 | 2.230256718 | 2.216742541 | 0.013514177 | -0.018723788 |
| 2700097O09Rik | 7.586700705 | 7.630526286 | 7.605432825 | 0.025093461 | -0.01873212 |
| Vmn1r59 | 1.090548928 | 1.107570657 | 1.109334646 | -0.001763989 | -0.018785719 |
| Gm43951 | 2.521201265 | 2.541035409 | 2.539990425 | 0.001044984 | -0.01878916 |
| Stk25 | 10.57720942 | 10.65031037 | 10.59603141 | 0.054278957 | -0.018821987 |
| Spata33 | 4.43501802 | 4.40838885 | 4.453842823 | -0.045453973 | -0.018824803 |
| Cacnb2 | 4.287829059 | 4.354244487 | 4.306678767 | 0.047565719 | -0.018849708 |
| Gucy2e | 2.118398125 | 2.201469734 | 2.137285808 | 0.064183926 | -0.018887683 |
| Rpsa-ps4 | 1.44713953 | 1.452987226 | 1.466038229 | -0.013051003 | -0.0188987 |
| Gm17420 | 3.44719342 | 3.4911891 | 3.466112168 | 0.025076932 | -0.018918748 |
| Spink6 | 1.740440644 | 1.785413482 | 1.759361712 | 0.02605177 | -0.018921068 |

| | | | | | |
|---|---|---|---|---|---|
| 3110070M22Rik | 3.347547007 | 3.411164204 | 3.366478875 | 0.044685329 | -0.018931868 |
| Trpc4ap | 10.93811782 | 11.04339114 | 10.95705656 | 0.08633458 | -0.01893874 |
| Chia1 | 1.603618832 | 1.630677287 | 1.6226081 | 0.008069187 | -0.018989268 |
| Cfap43 | 2.634251604 | 2.609825087 | 2.653246489 | -0.043421402 | -0.018994884 |
| Zfp473 | 5.126883234 | 5.311064475 | 5.145942019 | 0.165122456 | -0.019058785 |
| Epha4 | 9.736320818 | 9.706145703 | 9.755385994 | -0.049240291 | -0.019065175 |
| Borcs7 | 7.640068814 | 7.596491483 | 7.659180972 | -0.06268949 | -0.019112158 |
| Cacna1d | 5.943554558 | 5.99767832 | 5.962722686 | 0.034955634 | -0.019168128 |
| Hspd1-ps3 | 2.362934292 | 2.452878788 | 2.38219558 | 0.070683208 | -0.019261289 |
| Gm5812 | 2.964862376 | 2.919216275 | 2.984137201 | -0.064920926 | -0.019274825 |
| Efr3a | 11.12949496 | 11.02856485 | 11.14880033 | -0.120235483 | -0.019305377 |
| Pttg1ip | 11.13165203 | 11.08793343 | 11.15097054 | -0.06303711 | -0.019318517 |
| Olfr73 | 1.619145919 | 1.606651991 | 1.63851416 | -0.03186217 | -0.019368241 |
| Susd3 | 5.12913898 | 5.136707078 | 5.148508075 | -0.011800997 | -0.019369095 |
| Gm14122 | 1.622063649 | 1.580631539 | 1.641434812 | -0.060803273 | -0.019371163 |
| Pcdhb16 | 5.983629987 | 6.067916667 | 6.00300543 | 0.064911237 | -0.019375442 |
| Rps23-ps1 | 12.8130611 | 12.691232 | 12.83244396 | -0.14121196 | -0.019382863 |
| Gpr161 | 8.542029413 | 8.611570554 | 8.561414424 | 0.05015613 | -0.019385011 |
| Tubb4b-ps1 | 1.372746375 | 1.379202949 | 1.392131946 | -0.012928997 | -0.019385571 |
| Olfr1423 | 2.259125105 | 2.252227287 | 2.278525833 | -0.026298546 | -0.019400728 |
| 3110067C02Rik | 2.345883393 | 2.396089506 | 2.36529668 | 0.030792826 | -0.019413287 |
| Trappc6a | 7.819184388 | 7.993471172 | 7.838680148 | 0.154791023 | -0.01949576 |
| Psors1c2 | 8.740888174 | 8.848313526 | 8.76041486 | 0.087898666 | -0.019526686 |
| H2-Q1 | 1.502894625 | 1.532650687 | 1.52244604 | 0.010204647 | -0.019551415 |
| Wdr72 | 2.014603278 | 2.015857689 | 2.034164286 | -0.018306598 | -0.019561008 |
| Gopc | 9.747387206 | 9.671135756 | 9.766969568 | -0.095833812 | -0.019582362 |
| Nmu | 3.285834571 | 3.763682408 | 3.305437923 | 0.458244486 | -0.019603351 |
| Marveld1 | 10.69119194 | 10.51581691 | 10.71081557 | -0.194998663 | -0.01962363 |
| Gm12941 | 1.4850653 | 1.545250341 | 1.504698933 | 0.040551408 | -0.019633632 |
| Dnajc15 | 8.683838106 | 8.447864007 | 8.703473411 | -0.255609404 | -0.019635305 |
| Zdhhc19 | 1.38537829 | 1.378550288 | 1.405022257 | -0.026471969 | -0.019643967 |
| Fam210a | 9.647382735 | 9.670792656 | 9.667160865 | 0.003631791 | -0.01977813 |

| | | | | | |
|---|---|---|---|---|---|
| Ptpro | 7.976594388 | 7.860919527 | 7.996422692 | -0.135503165 | -0.019828304 |
| Abcb11 | 1.68616193 | 1.80320865 | 1.705992398 | 0.097216252 | -0.019830468 |
| Gm3650 | 6.08875387 | 6.151271391 | 6.108607773 | 0.042663618 | -0.019853903 |
| Map6 | 8.140899802 | 8.149453889 | 8.160771594 | -0.011317705 | -0.019871792 |
| Tmem63a | 10.73551131 | 10.81213644 | 10.75539033 | 0.05674611 | -0.019879023 |
| Fgfr1op2 | 11.00131349 | 10.97415456 | 11.02120952 | -0.047054957 | -0.019896027 |
| Angptl6 | 5.12011371 | 5.234937693 | 5.140035715 | 0.094901978 | -0.019922005 |
| Arfgap3 | 8.821471847 | 8.965179742 | 8.841424338 | 0.123755403 | -0.019952491 |
| Olfr46 | 5.997069156 | 6.036110428 | 6.017037894 | 0.019072534 | -0.019968738 |
| Rabac1 | 9.826412877 | 9.880513123 | 9.846407107 | 0.034106016 | -0.01999423 |
| Minpp1 | 9.566779728 | 9.581578011 | 9.586856636 | -0.005278625 | -0.020076907 |
| Gzmm | 4.426077687 | 4.427624952 | 4.446191181 | -0.018566229 | -0.020113494 |
| Krit1 | 9.550071695 | 9.634145431 | 9.570200296 | 0.063945135 | -0.020128601 |
| Olfr1312 | 2.548803889 | 2.556552882 | 2.568953644 | -0.012400762 | -0.020149755 |
| E4f1 | 8.469573718 | 8.506276637 | 8.489750038 | 0.0165266 | -0.02017632 |
| Dand5 | 5.392694537 | 5.39625472 | 5.412915221 | -0.016660501 | -0.020220684 |
| Cd2 | 5.934656018 | 5.812581904 | 5.954915469 | -0.142333565 | -0.02025945 |
| Idnk | 8.09285194 | 8.125815253 | 8.113112396 | 0.012702857 | -0.020260456 |
| Poc1a | 6.177252387 | 6.154236134 | 6.197530224 | -0.04329409 | -0.020277837 |
| Reep3 | 11.64806658 | 11.65256134 | 11.66837526 | -0.015813923 | -0.020308687 |
| Mapkap1 | 9.705814074 | 9.706889071 | 9.72614742 | -0.019258349 | -0.020333345 |
| Gm18905 | 2.962318236 | 2.941095822 | 2.982665 | -0.041569178 | -0.020346764 |
| Pglyrp2 | 5.331650926 | 5.463718057 | 5.352058647 | 0.111659411 | -0.020407721 |
| Gm12346 | 4.249349185 | 4.288386219 | 4.269785301 | 0.018600917 | -0.020436117 |
| Olfr1297 | 5.692609758 | 5.569218985 | 5.713061562 | -0.143842578 | -0.020451804 |
| C1s2 | 1.641353898 | 1.685961722 | 1.661859071 | 0.024102651 | -0.020505174 |
| Klkb1 | 2.350473265 | 2.333158194 | 2.370990114 | -0.037831919 | -0.020516848 |
| Egfl7 | 8.485289691 | 8.563576409 | 8.505869243 | 0.057707166 | -0.020579552 |
| Macc1 | 9.336021376 | 9.311097794 | 9.356607998 | -0.045510204 | -0.020586622 |
| Rab2a | 11.57570911 | 11.47731402 | 11.59630095 | -0.11898693 | -0.02059184 |
| Olfr1484 | 2.503876576 | 2.529088097 | 2.52447273 | 0.004615368 | -0.020596154 |
| Syf2 | 10.11607459 | 10.03580665 | 10.1366951 | -0.100888449 | -0.020620503 |

| | | | | | |
|---|---|---|---|---|---|
| Cda | 3.232455242 | 3.265140092 | 3.253098321 | 0.012041771 | -0.020643079 |
| 1700056E22Rik | 1.498722579 | 1.517103625 | 1.519379544 | -0.002275919 | -0.020656965 |
| Mknk1 | 9.135898762 | 9.170620389 | 9.156566936 | 0.014053453 | -0.020668175 |
| Brsk1 | 5.180794974 | 5.256690469 | 5.201487913 | 0.055202556 | -0.020692939 |
| Rbm44 | 1.826756493 | 1.830253809 | 1.847450993 | -0.017197184 | -0.020694499 |
| Ddx3y | 1.352580819 | 1.38077026 | 1.373327647 | 0.007442614 | -0.020746828 |
| Mrps21 | 9.459946159 | 9.411024954 | 9.480721218 | -0.069696265 | -0.02077506 |
| Tmem74 | 1.159072854 | 1.208100605 | 1.179887712 | 0.028212893 | -0.020814858 |
| Olfr173 | 2.554125934 | 2.530326624 | 2.574946083 | -0.044619459 | -0.020820149 |
| Heatr5b | 8.9585635 | 9.106428814 | 8.979386669 | 0.127042145 | -0.020823169 |
| Uap1l1 | 8.870791636 | 9.164777136 | 8.891616484 | 0.273160653 | -0.020824848 |
| Copb1 | 11.26275537 | 11.29358709 | 11.28359659 | 0.009990493 | -0.020841227 |
| Gm45208 | 1.322747656 | 1.331177162 | 1.343600998 | -0.012423837 | -0.020853343 |
| Mlh1 | 8.174626199 | 8.276543389 | 8.19559324 | 0.080950149 | -0.020967041 |
| Vmn2r82 | 2.037898719 | 2.037365825 | 2.058867945 | -0.021502121 | -0.020969227 |
| Spaca9 | 2.177809479 | 2.118130089 | 2.198784907 | -0.080654819 | -0.020975428 |
| C1ql3 | 2.673930903 | 2.717564836 | 2.69494531 | 0.022619526 | -0.021014406 |
| Phyh | 10.44792089 | 10.32942235 | 10.46900456 | -0.139582213 | -0.021083673 |
| Lamb3 | 10.81198044 | 10.99788745 | 10.83312986 | 0.164757597 | -0.021149417 |
| Gm37988 | 1.096685815 | 1.158344236 | 1.117869344 | 0.040474892 | -0.021183529 |
| Gata6 | 7.170989887 | 7.147348046 | 7.192177607 | -0.044829561 | -0.02118772 |
| Vamp3 | 11.38149309 | 11.28321944 | 11.4026996 | -0.119480163 | -0.021206517 |
| Catsperd | 2.789186175 | 2.808713031 | 2.81047666 | -0.001763628 | -0.021290485 |
| Ddb1 | 12.77233413 | 12.78587686 | 12.79363805 | -0.00776119 | -0.021303917 |
| Pnma1 | 2.311414522 | 2.334893521 | 2.332728877 | 0.002164644 | -0.021314354 |
| Nae1 | 9.365502244 | 9.320875005 | 9.386824903 | -0.065949898 | -0.021322659 |
| C2cd4d | 1.873207525 | 1.885992878 | 1.89453629 | -0.008543412 | -0.021328765 |
| CT025731.2 | 1.88414844 | 1.933275897 | 1.905487605 | 0.027788292 | -0.021339165 |
| Gnptg | 8.795873698 | 8.813637992 | 8.817220632 | -0.00358264 | -0.021346934 |
| Phactr3 | 1.176220409 | 1.164749021 | 1.197633507 | -0.032884486 | -0.021413098 |
| Mlana | 2.843424476 | 2.898753509 | 2.864874606 | 0.033878903 | -0.02145013 |
| Cpsf4l | 4.431541988 | 4.453488221 | 4.453000031 | 0.00048819 | -0.021458043 |

| | | | | | |
|---|---|---|---|---|---|
| Ccdc184 | 1.716015176 | 1.726751906 | 1.737504756 | -0.010752851 | -0.02148958 |
| Vmn2r99 | 3.700606735 | 3.775400227 | 3.722150464 | 0.053249763 | -0.021543728 |
| Gm6210 | 1.782828862 | 1.846814856 | 1.804385978 | 0.042428878 | -0.021557116 |
| Olfr853 | 2.074510685 | 2.071787074 | 2.09609494 | -0.024307866 | -0.021584255 |
| Zc3h4 | 9.991461752 | 10.10646549 | 10.01306272 | 0.093402776 | -0.021600966 |
| Nrxn2 | 3.555772184 | 3.747390641 | 3.577425918 | 0.169964723 | -0.021653734 |
| Vmn1r39 | 1.740379028 | 1.768521328 | 1.762071063 | 0.006450264 | -0.021692035 |
| Rhot2 | 9.866304615 | 10.01391722 | 9.888029102 | 0.125888117 | -0.021724486 |
| Cd79b | 3.102888914 | 3.129838704 | 3.124642415 | 0.00519629 | -0.021753501 |
| Zfp94 | 5.933566045 | 5.896294152 | 5.955325801 | -0.059031649 | -0.021759756 |
| Hdac1 | 10.63144754 | 10.57774828 | 10.65321656 | -0.07546828 | -0.021769017 |
| Adcy2 | 7.905457658 | 7.7405778 | 7.927252238 | -0.186674438 | -0.02179458 |
| Gm21988 | 6.349642692 | 6.629267298 | 6.371550698 | 0.2577166 | -0.021908006 |
| Eif5a2 | 7.307273115 | 7.247173157 | 7.329182015 | -0.082008858 | -0.021908901 |
| Gm49594 | 2.008077062 | 2.043863906 | 2.030011456 | 0.01385245 | -0.021934394 |
| Mkrn1 | 10.17438593 | 10.09784707 | 10.19635126 | -0.098504183 | -0.021965327 |
| Olfr622 | 2.035751851 | 2.065462375 | 2.057735849 | 0.007726526 | -0.021983998 |
| Gm14276 | 4.438971751 | 4.485165328 | 4.460958285 | 0.024207043 | -0.021986534 |
| Ms4a12 | 3.374080548 | 3.335166486 | 3.396095808 | -0.060929323 | -0.02201526 |
| Gm5781 | 1.714938213 | 1.751069319 | 1.737055761 | 0.014013558 | -0.022117548 |
| Gm3854 | 4.172607937 | 4.283025346 | 4.194737601 | 0.088287745 | -0.022129663 |
| Nav3 | 8.162735232 | 7.787340318 | 8.184866932 | -0.397526614 | -0.022131699 |
| Gabrb3 | 1.84690596 | 1.870584886 | 1.869065783 | 0.001519103 | -0.022159823 |
| Tagln | 10.58675256 | 10.47587577 | 10.60893091 | -0.13305514 | -0.022178347 |
| 1700003E16Rik | 2.6012773 | 2.66969293 | 2.623493223 | 0.046199707 | -0.022215923 |
| Krtap3-3 | 1.879135671 | 1.869576333 | 1.901394719 | -0.031818386 | -0.022259048 |
| Phf2 | 9.907453682 | 10.12839283 | 9.929712892 | 0.198679935 | -0.02225921 |
| Cdk5rap1 | 7.588622046 | 7.355098105 | 7.61091949 | -0.255821384 | -0.022297443 |
| Cox6b1 | 11.59486071 | 11.47525729 | 11.61716678 | -0.141909497 | -0.022306073 |
| Cpa2 | 2.146515334 | 2.206560311 | 2.168853103 | 0.037707208 | -0.022337769 |
| Smad5 | 11.01456344 | 11.00081903 | 11.03694466 | -0.036125627 | -0.022381217 |
| Srsf12 | 1.246386343 | 1.287444064 | 1.268806697 | 0.018637367 | -0.022420354 |

| | | | | | |
|---|---|---|---|---|---|
| Adra2b | 2.834712713 | 2.893950607 | 2.857151184 | 0.036799423 | -0.022438472 |
| Kpna6 | 10.93133804 | 10.99040888 | 10.95380003 | 0.03660885 | -0.02246199 |
| Tmem139 | 6.148126699 | 6.097214112 | 6.170654034 | -0.073439922 | -0.022527335 |
| Osbp | 10.79061644 | 10.84570298 | 10.81314401 | 0.032558967 | -0.022527577 |
| Olfr578 | 1.543998186 | 1.536499745 | 1.566528913 | -0.030029169 | -0.022530728 |
| 2810459M11Rik | 2.031266921 | 2.045690818 | 2.053811955 | -0.008121137 | -0.022545034 |
| AC116766.1 | 2.714338542 | 2.746523653 | 2.736906226 | 0.009617427 | -0.022567684 |
| Efna1 | 9.724629306 | 9.773828217 | 9.747201658 | 0.026626559 | -0.022572352 |
| Fem1a | 10.86650698 | 10.72500411 | 10.88910456 | -0.16410045 | -0.02259758 |
| Zfyve28 | 4.187384378 | 4.218090386 | 4.210076519 | 0.008013868 | -0.022692141 |
| Kat6a | 11.06603716 | 11.11493546 | 11.08874291 | 0.026192553 | -0.022705743 |
| Cblc | 8.27388866 | 8.22843853 | 8.296615631 | -0.068177101 | -0.022726971 |
| Optc | 3.418151135 | 3.376674638 | 3.440968727 | -0.064294089 | -0.022817592 |
| Mmp17 | 6.41690036 | 6.555457363 | 6.439720564 | 0.115736799 | -0.022820203 |
| Zfp959 | 6.93196693 | 6.997178451 | 6.95479032 | 0.042388131 | -0.02282339 |
| Gm5740 | 2.707175019 | 2.718633352 | 2.730041533 | -0.011408181 | -0.022866514 |
| Zcchc18 | 2.488666858 | 2.471272035 | 2.511533466 | -0.040261432 | -0.022866608 |
| Rpap3-ps2 | 1.765687609 | 1.822773862 | 1.788561663 | 0.034212199 | -0.022874054 |
| Pot1a | 8.366121393 | 8.109823521 | 8.3890472 | -0.279223679 | -0.022925807 |
| Cenpx | 8.601119598 | 8.685129542 | 8.624055697 | 0.061073845 | -0.022936099 |
| Spr | 9.038381994 | 9.121963302 | 9.061322661 | 0.060640641 | -0.022940667 |
| Upk2 | 1.740256937 | 1.750818032 | 1.763237694 | -0.012419662 | -0.022980758 |
| Prss35 | 3.67869691 | 3.664800345 | 3.70171215 | -0.036911805 | -0.02301524 |
| Zfp418 | 5.794753092 | 5.816972503 | 5.817772206 | -0.000799703 | -0.023019114 |
| Olfr734 | 2.697437224 | 2.724806764 | 2.72045936 | 0.004347404 | -0.023022136 |
| Olfr75-ps1 | 2.569771394 | 2.571660953 | 2.592803056 | -0.021142103 | -0.023031662 |
| Hmgcr | 11.79306414 | 11.72131567 | 11.8161343 | -0.094818627 | -0.023070163 |
| Rpl36-ps12 | 0.641638582 | 0.688496228 | 0.664732348 | 0.02376388 | -0.023093766 |
| Fntb | 8.459808036 | 8.539794874 | 8.482940379 | 0.056854496 | -0.023132343 |
| Fam196b | 1.618842093 | 1.617463999 | 1.641993881 | -0.024529882 | -0.023151788 |
| Hip1r | 10.61378851 | 10.97584834 | 10.63694807 | 0.338900273 | -0.02315956 |
| Gm4034 | 2.807123015 | 2.800434894 | 2.830282611 | -0.029847718 | -0.023159596 |

| | | | | | |
|---|---|---|---|---|---|
| Csgalnact1 | 7.793598287 | 7.946900342 | 7.816773444 | 0.130126898 | -0.023175158 |
| Stmn4 | 2.530345287 | 2.681047683 | 2.553525373 | 0.12752231 | -0.023180086 |
| Grsf1 | 10.37101242 | 10.2598492 | 10.39420088 | -0.13435168 | -0.023188467 |
| Gm16588 | 2.407050553 | 2.408866558 | 2.430245639 | -0.021379081 | -0.023195085 |
| Otoa | 3.942240181 | 4.157008337 | 3.965437242 | 0.191571095 | -0.023197061 |
| Catsperz | 3.382379338 | 3.372249643 | 3.405596147 | -0.033346504 | -0.023216809 |
| Ctsg | 2.659716309 | 2.6593438 | 2.682952333 | -0.023608533 | -0.023236024 |
| Nap1l2 | 2.660869901 | 2.625742908 | 2.684106146 | -0.058363238 | -0.023236245 |
| Prrt4 | 3.933330031 | 3.918933619 | 3.956611879 | -0.03767826 | -0.023281848 |
| Dnajc24 | 6.746095613 | 6.753628828 | 6.769382674 | -0.015753846 | -0.02328706 |
| Nelfcd | 8.900759827 | 8.922823568 | 8.924077118 | -0.001253549 | -0.02331729 |
| Olfr731 | 5.07409007 | 5.009298112 | 5.097575276 | -0.088277164 | -0.023485206 |
| Pip4p1 | 9.654090834 | 9.694725731 | 9.677596852 | 0.017128879 | -0.023506018 |
| Trappc12 | 9.204814078 | 9.304270383 | 9.22835794 | 0.075912443 | -0.023543862 |
| Atxn7 | 9.878003699 | 9.871698361 | 9.901569742 | -0.029871381 | -0.023566043 |
| Olfr918 | 1.410290742 | 1.391250472 | 1.433867594 | -0.042617122 | -0.023576852 |
| Ppp1cb | 12.67398269 | 12.62257026 | 12.69756479 | -0.07499453 | -0.023582097 |
| Clec4a3 | 8.240865849 | 8.353672038 | 8.264499487 | 0.089172551 | -0.023633639 |
| Mup-ps19 | 1.021840072 | 1.019253159 | 1.045541955 | -0.026288796 | -0.023701883 |
| Olfr1183 | 3.110802399 | 3.14072059 | 3.134540683 | 0.006179907 | -0.023738284 |
| Ube3b | 11.01531756 | 11.19614906 | 11.03915442 | 0.15699464 | -0.02383686 |
| Gpr162 | 4.463421069 | 4.677902383 | 4.48726505 | 0.190637333 | -0.023843982 |
| Rpl22 | 12.32170859 | 12.23893485 | 12.34556645 | -0.106631603 | -0.023857867 |
| Arhgap11a | 8.773679404 | 8.832310465 | 8.797553559 | 0.034756906 | -0.023874154 |
| Gm13762 | 2.098543071 | 2.132168248 | 2.122424644 | 0.009743604 | -0.023881573 |
| Bsdc1 | 9.987933114 | 9.943803492 | 10.01182719 | -0.068023695 | -0.023894072 |
| Klhdc3 | 9.972974259 | 10.07994432 | 9.996884286 | 0.083060031 | -0.023910027 |
| Grm5 | 3.632722922 | 3.642212207 | 3.656683476 | -0.014471269 | -0.023960553 |
| Syt6 | 2.819515175 | 2.844836826 | 2.843479951 | 0.001356876 | -0.023964776 |
| Bin3 | 9.857173664 | 9.822226257 | 9.881168197 | -0.058941941 | -0.023994534 |
| Rnls | 4.758930754 | 4.688731435 | 4.78293637 | -0.094204935 | -0.024005616 |
| Map3k7 | 10.63383924 | 10.67422619 | 10.65785826 | 0.016367927 | -0.02401902 |

| | | | | | |
|---|---|---|---|---|---|
| 4933408B17Rik | 3.556270158 | 3.605695758 | 3.58031319 | 0.025382569 | -0.024043031 |
| Tbc1d7 | 7.41543608 | 7.310912762 | 7.439480016 | -0.128567255 | -0.024043936 |
| Sil1 | 8.430407039 | 8.459919877 | 8.454463529 | 0.005456347 | -0.024056491 |
| Masp2 | 1.668366327 | 1.707560935 | 1.692446042 | 0.015114893 | -0.024079715 |
| Cnn1 | 8.110161954 | 8.171510761 | 8.134290681 | 0.03722008 | -0.024128727 |
| Cbfa2t3 | 8.003627378 | 8.292455735 | 8.027774797 | 0.264680938 | -0.024147418 |
| Chic1 | 5.88145547 | 5.881558327 | 5.905621878 | -0.024063551 | -0.024166408 |
| Trbc2 | 8.776758867 | 8.664974615 | 8.800927685 | -0.135953071 | -0.024168818 |
| Tmem56 | 10.64396756 | 10.53216931 | 10.6681661 | -0.135996783 | -0.02419854 |
| Cenpu | 6.307299187 | 6.38523561 | 6.331552873 | 0.053682737 | -0.024253687 |
| Trpm4 | 8.536834123 | 8.974901386 | 8.561088172 | 0.413813215 | -0.024254048 |
| Otos | 1.789029924 | 1.72886757 | 1.813284695 | -0.084417125 | -0.024254771 |
| Atp6v0a2 | 10.72944632 | 10.82885746 | 10.75371596 | 0.075141507 | -0.024269637 |
| Ddx59 | 7.443825846 | 7.284739842 | 7.468100351 | -0.183360509 | -0.024274505 |
| Myo16 | 3.462552664 | 3.505616546 | 3.486829869 | 0.018786676 | -0.024277205 |
| Kif24 | 6.840858774 | 7.002670299 | 6.865153934 | 0.137516365 | -0.02429516 |
| Ipmk | 10.78997645 | 10.71465912 | 10.8142959 | -0.099636777 | -0.024319447 |
| Gm12080 | 1.29816933 | 1.314041294 | 1.322489341 | -0.008448046 | -0.024320011 |
| Spint2 | 11.22323518 | 11.26062351 | 11.2475593 | 0.01306421 | -0.024324113 |
| Gm14176 | 1.145791962 | 1.146914061 | 1.170126314 | -0.023212253 | -0.024334352 |
| Tmed1 | 8.072708324 | 8.103066379 | 8.097101493 | 0.005964886 | -0.02439317 |
| Chrm3 | 4.179074176 | 4.187269536 | 4.203494874 | -0.016225338 | -0.024420698 |
| Olfr1321 | 2.357685456 | 2.390952891 | 2.382112382 | 0.008840509 | -0.024426926 |
| Mgea5 | 11.55496029 | 11.51106923 | 11.57942891 | -0.068359683 | -0.024468623 |
| Vmn1r86 | 2.879666831 | 2.987203562 | 2.904188805 | 0.083014757 | -0.024521974 |
| Gldn | 8.069769415 | 8.187951214 | 8.094330801 | 0.093620413 | -0.024561386 |
| Tmem158 | 6.222389835 | 6.176240064 | 6.246984077 | -0.070744013 | -0.024594242 |
| Vps72 | 9.161187352 | 9.304728072 | 9.185784352 | 0.11894372 | -0.024597001 |
| Gp1ba | 1.152873005 | 1.145620811 | 1.177488025 | -0.031867214 | -0.024615021 |
| Esr2 | 3.113134683 | 3.119247077 | 3.137763517 | -0.01851644 | -0.024628834 |
| Gm16409 | 3.665989066 | 3.723075431 | 3.690655903 | 0.032419529 | -0.024666837 |
| Nostrin | 5.242737144 | 5.23480542 | 5.267404118 | -0.032598698 | -0.024666974 |

| | | | | | |
|---|---|---|---|---|---|
| Nsmce2 | 8.348351453 | 8.424568047 | 8.373030717 | 0.051537329 | -0.024679264 |
| Stox1 | 2.253833241 | 2.278579254 | 2.27855536 | 2.38937E-05 | -0.024722119 |
| Ccdc148 | 2.759212507 | 2.777657844 | 2.783943906 | -0.006286062 | -0.024731399 |
| Dpp6 | 3.716609106 | 3.7589708 | 3.741347257 | 0.017623542 | -0.024738152 |
| Col6a4 | 2.59754224 | 2.611440916 | 2.622325373 | -0.010884457 | -0.024783133 |
| Gm5105 | 2.063987736 | 2.034743105 | 2.088778945 | -0.05403584 | -0.02479121 |
| Fkbpl | 7.012808264 | 7.048574656 | 7.037602641 | 0.010972015 | -0.024794377 |
| Rph3al | 6.103075911 | 6.18644115 | 6.127888693 | 0.058552457 | -0.024812782 |
| Miga2 | 9.896436669 | 10.07828237 | 9.921254876 | 0.157027498 | -0.024818207 |
| Tmem30c | 2.180400287 | 2.207756136 | 2.205222979 | 0.002533157 | -0.024822692 |
| Slc30a9 | 10.0459559 | 9.972527843 | 10.07078257 | -0.098254731 | -0.024826675 |
| Zpbp2 | 0.854351381 | 0.832877887 | 0.879187171 | -0.046309284 | -0.02483579 |
| Iqcd | 1.377517333 | 1.408290082 | 1.402372751 | 0.005917332 | -0.024855418 |
| Brd3 | 10.18646883 | 10.4158491 | 10.2113346 | 0.204514497 | -0.024865767 |
| Fam19a5 | 5.050972909 | 5.185982824 | 5.07586381 | 0.110119014 | -0.024890901 |
| Filip1l | 10.36043252 | 10.41660505 | 10.38533557 | 0.031269483 | -0.024903043 |
| Gm6030 | 2.505584839 | 2.544032959 | 2.530508772 | 0.013524187 | -0.024923933 |
| Trp53bp2 | 9.965806978 | 10.00482603 | 9.990734967 | 0.014091063 | -0.024927989 |
| Tmem107 | 7.002896694 | 7.094420715 | 7.027832422 | 0.066588294 | -0.024935727 |
| Dcaf15 | 8.486668454 | 8.633580773 | 8.511656219 | 0.121924554 | -0.024987765 |
| Gm10110 | 3.70126356 | 3.780069426 | 3.726260108 | 0.053809318 | -0.024996548 |
| Traf5 | 7.630261925 | 7.580229431 | 7.65526321 | -0.075033779 | -0.025001285 |
| Mfge8 | 11.24562865 | 11.47407038 | 11.27071547 | 0.203354913 | -0.02508682 |
| Gm45038 | 3.218205539 | 3.175365381 | 3.243304531 | -0.06793915 | -0.025098991 |
| Rsph3b | 9.043505791 | 9.012231411 | 9.068613287 | -0.056381876 | -0.025107496 |
| Defb3 | 2.465669478 | 2.486682396 | 2.490794885 | -0.004112489 | -0.025125407 |
| Duxbl3 | 1.793500001 | 1.851268537 | 1.818667834 | 0.032600702 | -0.025167833 |
| Gm26712 | 4.531331288 | 4.533328793 | 4.55651489 | -0.023186097 | -0.025183602 |
| AC117239.2 | 1.055809292 | 1.080455365 | 1.081003708 | -0.000548343 | -0.025194416 |
| Rpl10-ps5 | 3.329995865 | 3.343949383 | 3.355209533 | -0.011260149 | -0.025213668 |
| Atf2 | 9.689700563 | 9.672470466 | 9.714927846 | -0.04245738 | -0.025227283 |
| Vmn1r213 | 3.754373194 | 3.765121146 | 3.779603477 | -0.014482331 | -0.025230283 |

| | | | | | |
|---|---|---|---|---|---|
| Kdm3b | 10.22130275 | 10.14792758 | 10.24654385 | -0.09861627 | -0.025241102 |
| Dffa | 10.53885987 | 10.56354995 | 10.56410718 | -0.000557233 | -0.025247313 |
| Hdac10 | 8.336283988 | 8.4998404 | 8.361535918 | 0.138304482 | -0.025251931 |
| Unc50 | 9.383927202 | 9.343736055 | 9.409200332 | -0.065464277 | -0.02527313 |
| Gm20605 | 8.34280802 | 8.960795327 | 8.368105545 | 0.592689782 | -0.025297525 |
| Gm49376 | 3.407393742 | 3.406862397 | 3.43274858 | -0.025886182 | -0.025354838 |
| Capsl | 4.36985999 | 4.331474106 | 4.395234716 | -0.06376061 | -0.025374726 |
| Nat2 | 7.024348182 | 6.907221685 | 7.049730518 | -0.142508833 | -0.025382336 |
| Nme3 | 6.623728408 | 6.596656245 | 6.649164158 | -0.052507912 | -0.02543575 |
| Gm4767 | 3.146974011 | 3.229447666 | 3.172465674 | 0.056981992 | -0.025491663 |
| Lrrc75b | 3.586572342 | 3.62773386 | 3.612072738 | 0.015661121 | -0.025500396 |
| Tmod3 | 11.13270689 | 11.08036459 | 11.15822859 | -0.077864007 | -0.0255217 |
| Gm6333 | 1.467903922 | 1.484205013 | 1.493477273 | -0.00927226 | -0.025573351 |
| Cyp2r1 | 4.966110283 | 5.068918368 | 4.991698445 | 0.077219923 | -0.025588162 |
| Pigyl | 6.637439281 | 6.905663302 | 6.66305441 | 0.242608892 | -0.02561513 |
| Ldhal6b | 1.670041758 | 1.666275913 | 1.695715644 | -0.029439731 | -0.025673886 |
| Mettl21c | 3.423349931 | 3.348707671 | 3.449056258 | -0.100348587 | -0.025706327 |
| Cers3 | 10.46831011 | 10.37407136 | 10.49406994 | -0.119998583 | -0.025759837 |
| Dpy19l1 | 9.297745387 | 9.2387496 | 9.323597892 | -0.084848292 | -0.025852504 |
| Gm12416 | 3.389967585 | 3.429675797 | 3.415823056 | 0.013852741 | -0.025855471 |
| Olfr1105 | 1.281472146 | 1.317913005 | 1.30735628 | 0.010556725 | -0.025884134 |
| Gm12482 | 2.484880298 | 2.537477871 | 2.51078173 | 0.026696141 | -0.025901432 |
| Gm15443 | 3.465772181 | 3.414054898 | 3.491681394 | -0.077626496 | -0.025909213 |
| Arl6ip1 | 11.48506555 | 11.39236327 | 11.5109777 | -0.118614423 | -0.02591215 |
| Rbbp4 | 11.49904907 | 11.40503468 | 11.52504565 | -0.12001097 | -0.02599658 |
| Slc25a18 | 1.396527553 | 1.399009329 | 1.422536767 | -0.023527439 | -0.026009214 |
| Gm16072 | 1.430159905 | 1.431153617 | 1.45617219 | -0.025018573 | -0.026012285 |
| Cops3 | 10.10116544 | 10.05865855 | 10.12718301 | -0.068524466 | -0.026017573 |
| Tex19.1 | 3.042031933 | 3.288449173 | 3.068059257 | 0.220389917 | -0.026027323 |
| Fam199x | 8.509629843 | 8.325904727 | 8.535666923 | -0.209762196 | -0.02603708 |
| Gm4735 | 1.565725716 | 1.652017068 | 1.591805853 | 0.060211215 | -0.026080137 |
| Sdk2 | 6.991384428 | 7.276739576 | 7.017473916 | 0.25926566 | -0.026089488 |

| | | | | | |
|---|---|---|---|---|---|
| Ccdc127 | 9.839082273 | 9.679137402 | 9.865174258 | -0.186036856 | -0.026091985 |
| Gm12185 | 5.440573448 | 4.876537502 | 5.466674514 | -0.590137012 | -0.026101066 |
| Olfr984 | 1.628655743 | 1.629186603 | 1.654768275 | -0.025581672 | -0.026112532 |
| 3300002I08Rik | 3.43417069 | 3.453522796 | 3.460351332 | -0.006828536 | -0.026180643 |
| Mettl18 | 6.004582209 | 5.977955155 | 6.030785507 | -0.052830352 | -0.026203297 |
| Rpl36a | 12.43238264 | 12.34707139 | 12.4586106 | -0.11153921 | -0.02622796 |
| Gjc2 | 3.101626441 | 3.228691707 | 3.12790377 | 0.100787936 | -0.026277329 |
| Pih1d2 | 4.461161831 | 4.447064853 | 4.487464761 | -0.040399909 | -0.02630293 |
| Rpl30 | 12.80147549 | 12.76186223 | 12.82785547 | -0.065993237 | -0.026379983 |
| Coa5 | 10.95781578 | 10.9175441 | 10.98440479 | -0.066860683 | -0.026589003 |
| Ddc | 3.189606424 | 3.14828896 | 3.216302992 | -0.068014031 | -0.026696567 |
| Galc | 7.434934451 | 7.426498885 | 7.461670368 | -0.035171483 | -0.026735916 |
| Fkbp5 | 11.1974489 | 11.14867893 | 11.22420752 | -0.075528587 | -0.02675862 |
| Pcdhb12 | 3.547506572 | 3.563481205 | 3.574267966 | -0.010786761 | -0.026761394 |
| Nudt16l1 | 9.202042726 | 9.22057125 | 9.228813436 | -0.008242186 | -0.02677071 |
| Nrbp1 | 10.96565212 | 10.94424671 | 10.99247335 | -0.048226643 | -0.026821237 |
| Gm9985 | 2.250589497 | 2.236596183 | 2.277463678 | -0.040867495 | -0.026874181 |
| Asic4 | 1.803163303 | 1.964947986 | 1.830040496 | 0.134907489 | -0.026877193 |
| Chrm1 | 2.728688193 | 2.761722666 | 2.755584932 | 0.006137734 | -0.026896738 |
| Ntrk1 | 1.756548323 | 1.897350755 | 1.783459938 | 0.113890817 | -0.026911614 |
| Olfr66 | 3.093631125 | 3.0538463 | 3.120567585 | -0.066721285 | -0.026936461 |
| Nit2 | 8.768605146 | 8.81215098 | 8.795585312 | 0.016565668 | -0.026980166 |
| Chrac1 | 8.481150879 | 8.334476921 | 8.508132177 | -0.173655256 | -0.026981299 |
| Gm15712 | 2.13584929 | 2.112215906 | 2.16291606 | -0.050700154 | -0.027066771 |
| Gm9480 | 2.273001333 | 2.285085288 | 2.300080807 | -0.014995518 | -0.027079474 |
| Trim17 | 1.554525214 | 1.577211602 | 1.581616225 | -0.004404623 | -0.027091011 |
| Gm45713 | 3.434116274 | 3.565380044 | 3.461284211 | 0.104095833 | -0.027167937 |
| Urod | 9.62015644 | 9.663289522 | 9.647359554 | 0.015929968 | -0.027203114 |
| Olfr1336 | 5.263582873 | 5.313708616 | 5.290820678 | 0.022887938 | -0.027237805 |
| Olfr753-ps1 | 2.214484953 | 2.237743939 | 2.241724052 | -0.003980113 | -0.027239099 |
| Nfxl1 | 8.543258187 | 8.496695082 | 8.570534663 | -0.073839581 | -0.027276476 |
| Minos1 | 10.22716223 | 10.15873273 | 10.25462633 | -0.095893597 | -0.027464093 |

| | | | | | |
|---|---|---|---|---|---|
| Zfp661 | 6.97529556 | 7.088560731 | 7.002796089 | 0.085764642 | -0.027500529 |
| Xcl1 | 4.509400615 | 4.482258747 | 4.536911042 | -0.054652295 | -0.027510427 |
| Fam222a | 3.074591066 | 3.204987985 | 3.102101498 | 0.102886487 | -0.027510433 |
| Adgrg7 | 3.404231401 | 3.48732313 | 3.431756395 | 0.055566735 | -0.027524993 |
| Fbxw4 | 8.211928502 | 8.327664034 | 8.239460987 | 0.088203047 | -0.027532485 |
| Grik2 | 4.246108159 | 4.359775332 | 4.273710905 | 0.086064427 | -0.027602746 |
| Rab36 | 4.469909944 | 4.525217917 | 4.497518659 | 0.027699258 | -0.027608715 |
| Dnajb13 | 3.332869866 | 3.227268228 | 3.360497115 | -0.133228886 | -0.027627248 |
| Gucy2f | 1.422056404 | 1.424628239 | 1.449803027 | -0.025174788 | -0.027746623 |
| Gm13409 | 2.070616499 | 2.110070111 | 2.098437567 | 0.011632544 | -0.027821068 |
| Gm38247 | 1.532802579 | 1.509831839 | 1.560676012 | -0.050844173 | -0.027873433 |
| Ngb | 2.347837978 | 2.353867979 | 2.375750188 | -0.021882209 | -0.02791221 |
| 5530401A14Rik | 1.175486645 | 1.226679197 | 1.203496924 | 0.023182273 | -0.028010278 |
| Akr1a1 | 12.55274301 | 12.51125195 | 12.58085902 | -0.06960707 | -0.02811601 |
| Efcab9 | 1.748931479 | 1.75124146 | 1.777053698 | -0.025812238 | -0.028122219 |
| Prmt9 | 8.368493738 | 8.349938316 | 8.396620937 | -0.046682621 | -0.028127199 |
| Rad50 | 9.461904891 | 9.448597492 | 9.4900541 | -0.041456608 | -0.028149209 |
| Elp4 | 7.312105188 | 7.634503126 | 7.340262037 | 0.294241089 | -0.028156849 |
| Ccdc71 | 9.297842369 | 9.387506841 | 9.326083399 | 0.061423443 | -0.028241029 |
| Gm48501 | 4.528380571 | 4.564423423 | 4.556679871 | 0.007743552 | -0.0282993 |
| Bcan | 1.752815524 | 1.678051204 | 1.781143603 | -0.103092399 | -0.028328079 |
| Rhpn1 | 2.123442833 | 2.131173216 | 2.151772156 | -0.02059894 | -0.028329323 |
| Slc16a4 | 1.873787229 | 1.910871467 | 1.902124132 | 0.008747335 | -0.028336903 |
| Rfk | 9.580080019 | 9.4099518 | 9.608474594 | -0.198522794 | -0.028394575 |
| Gm15710 | 1.665948917 | 1.661754426 | 1.694349774 | -0.032595348 | -0.028400856 |
| Ap1m2 | 9.955575904 | 10.04563585 | 9.983997195 | 0.061638653 | -0.028421291 |
| Bsg | 12.32648687 | 12.29300924 | 12.3549336 | -0.061924357 | -0.028446727 |
| 9130208D14Rik | 3.153216831 | 3.102239883 | 3.181671756 | -0.079431873 | -0.028454925 |
| Lrrk1 | 9.803801871 | 9.851384546 | 9.832264976 | 0.01911957 | -0.028463104 |
| Dnajc19 | 8.727901889 | 8.755865899 | 8.756375392 | -0.000509493 | -0.028473503 |
| Pla2g15 | 8.968093143 | 9.094855888 | 8.996579896 | 0.098275992 | -0.028486753 |
| Gm19220 | 6.335213981 | 6.469031959 | 6.363728244 | 0.105303715 | -0.028514263 |

| | | | | | |
|---|---|---|---|---|---|
| Ctnnb1 | 13.24247156 | 13.24569701 | 13.27098974 | -0.02529273 | -0.028518173 |
| Rrm2 | 9.467804047 | 9.54447413 | 9.496323291 | 0.04815084 | -0.028519244 |
| Crip2 | 11.42333642 | 11.57756296 | 11.45188571 | 0.125677253 | -0.02854929 |
| Gm14914 | 1.747313168 | 1.750615922 | 1.775884476 | -0.025268554 | -0.028571307 |
| Agbl4 | 1.980459511 | 1.9626907 | 2.009035381 | -0.046344681 | -0.02857587 |
| Tsen34 | 9.758495703 | 9.909494554 | 9.78709088 | 0.122403674 | -0.028595177 |
| Fstl4 | 3.57242199 | 3.671966361 | 3.601060255 | 0.070906105 | -0.028638265 |
| B930094E09Rik | 1.904673249 | 1.889518374 | 1.933314909 | -0.043796535 | -0.02864166 |
| Hectd2 | 6.333364618 | 6.34065647 | 6.362039986 | -0.021383516 | -0.028675368 |
| Olfr896-ps1 | 1.735604973 | 1.777444202 | 1.764284063 | 0.013160139 | -0.02867909 |
| Gm8162 | 1.978141058 | 2.02392534 | 2.006827184 | 0.017098156 | -0.028686125 |
| Gm9794 | 10.17873043 | 10.21637208 | 10.20741962 | 0.00895246 | -0.028689193 |
| Foxp1 | 10.20219493 | 10.12288256 | 10.2308859 | -0.108003333 | -0.028690963 |
| Abcb8 | 8.910245176 | 8.986168172 | 8.938969656 | 0.047198515 | -0.02872448 |
| Edem2 | 9.103451481 | 9.277454007 | 9.13225113 | 0.145202877 | -0.02879965 |
| Tfpi | 8.241247241 | 7.978201724 | 8.270102225 | -0.2919005 | -0.028854984 |
| Anapc13 | 8.848443353 | 8.74050579 | 8.877326448 | -0.136820658 | -0.028883095 |
| Haspin | 5.990021427 | 5.971535154 | 6.018910268 | -0.047375114 | -0.028888841 |
| Gm48040 | 1.676047408 | 1.673381884 | 1.704948357 | -0.031566473 | -0.028900948 |
| Ppp2r5c | 11.22342352 | 11.26188611 | 11.25234334 | 0.00954277 | -0.02891982 |
| Ccdc154 | 0.956352615 | 0.985056523 | 0.985409335 | -0.000352812 | -0.02905672 |
| Hoxb1 | 2.274960128 | 2.279778812 | 2.304023947 | -0.024245134 | -0.029063819 |
| Xylt2 | 8.820773449 | 8.978495879 | 8.849844015 | 0.128651864 | -0.029070566 |
| Lingo3 | 4.160412594 | 4.189815411 | 4.189543555 | 0.000271855 | -0.029130961 |
| Setmar | 6.188858629 | 6.087257555 | 6.218021734 | -0.13076418 | -0.029163105 |
| Ttyh2 | 8.80282398 | 8.811635728 | 8.832010646 | -0.020374919 | -0.029186667 |
| Gm20696 | 2.124315175 | 2.212985796 | 2.153508578 | 0.059477217 | -0.029193404 |
| Gm43720 | 4.396441089 | 4.299035205 | 4.425634748 | -0.126599543 | -0.029193659 |
| Sit1 | 2.760329169 | 2.723826536 | 2.789529487 | -0.065702952 | -0.029200318 |
| 4930426L09Rik | 2.610981505 | 2.612032134 | 2.640215231 | -0.028183097 | -0.029233726 |
| Zfp882 | 4.710545154 | 4.726683187 | 4.739787668 | -0.013104481 | -0.029242514 |
| Yaf2 | 7.883307679 | 7.88768733 | 7.912550423 | -0.024863093 | -0.029242744 |

| | | | | | |
|---|---|---|---|---|---|
| Cdk3-ps | 2.955111511 | 3.072232105 | 2.9843659 | 0.087866205 | -0.029254389 |
| Eif4a2 | 12.36599972 | 12.37447665 | 12.39525576 | -0.020779113 | -0.029256043 |
| Pafah1b1 | 12.08499627 | 12.07222394 | 12.11425293 | -0.04202899 | -0.029256657 |
| Ccdc47 | 11.14844034 | 11.1125706 | 11.17771344 | -0.065142837 | -0.029273097 |
| Fbxo34 | 8.738283308 | 8.696598339 | 8.767604235 | -0.071005895 | -0.029320927 |
| Zfp846 | 7.744798191 | 7.813719489 | 7.774131652 | 0.039587837 | -0.029333462 |
| Tmprss11e | 5.93328815 | 5.71857871 | 5.962633606 | -0.244054895 | -0.029345456 |
| Gfod2 | 7.908568029 | 8.224578678 | 7.93792417 | 0.286654508 | -0.029356141 |
| Gm18806 | 2.009481869 | 2.04410811 | 2.038878954 | 0.005229156 | -0.029397085 |
| Ss18l2 | 8.115951932 | 8.00086086 | 8.14538873 | -0.144527869 | -0.029436798 |
| Chit1 | 11.62623192 | 11.67597899 | 11.6556885 | 0.020290487 | -0.029456583 |
| Nlrp9c | 1.786143724 | 1.812067278 | 1.815652128 | -0.00358485 | -0.029508404 |
| Dvl2 | 8.077959179 | 8.443883654 | 8.107468715 | 0.336414939 | -0.029509536 |
| Zdhhc3 | 11.5630807 | 11.54480928 | 11.59268904 | -0.047879757 | -0.029608333 |
| Gdap1l1 | 1.113277162 | 1.13265974 | 1.142921946 | -0.010262205 | -0.029644783 |
| Zfp472 | 7.053820136 | 7.0983142 | 7.083483862 | 0.014830338 | -0.029663726 |
| Vmn1r13 | 4.428518839 | 4.396633113 | 4.458212994 | -0.061579881 | -0.029694155 |
| Efnb3 | 5.018701145 | 5.024547668 | 5.048399676 | -0.023852007 | -0.029698531 |
| Spin2c | 4.335454524 | 4.367857444 | 4.365164471 | 0.002692973 | -0.029709948 |
| Xkr4 | 3.551828177 | 3.531201136 | 3.581564875 | -0.050363739 | -0.029736698 |
| 5330417C22Rik | 2.114241409 | 2.191546897 | 2.143996712 | 0.047550185 | -0.029755303 |
| Renbp | 7.706840324 | 8.133007904 | 7.736614446 | 0.396393458 | -0.029774122 |
| Ism2 | 2.470706819 | 2.502646432 | 2.500523479 | 0.002122953 | -0.02981666 |
| Gm14584 | 4.051581856 | 3.943070147 | 4.081399334 | -0.138329188 | -0.029817479 |
| Ebf4 | 3.925403669 | 3.898490376 | 3.955222491 | -0.056732115 | -0.029818822 |
| Ly6g6d | 1.911534023 | 1.90292381 | 1.941355426 | -0.038431616 | -0.029821404 |
| Lrrc49 | 7.592963931 | 7.464492705 | 7.622789619 | -0.158296914 | -0.029825688 |
| Slc22a19 | 1.753083541 | 1.786506993 | 1.782984586 | 0.003522407 | -0.029901045 |
| Gria2 | 2.492111056 | 2.529480928 | 2.522028228 | 0.0074527 | -0.029917171 |
| Mnat1 | 8.18995064 | 8.051984804 | 8.219911566 | -0.167926762 | -0.029960926 |
| Fam83g | 11.21000814 | 11.49717157 | 11.23997323 | 0.257198343 | -0.029965093 |
| Rab42 | 2.146884644 | 2.161058511 | 2.176863461 | -0.01580495 | -0.029978817 |

| | | | | | |
|---|---|---|---|---|---|
| Gm14267 | 2.706771573 | 2.674856556 | 2.73690166 | -0.062045104 | -0.030130087 |
| Aox2 | 3.266888681 | 3.30954699 | 3.297021238 | 0.012525752 | -0.030132557 |
| Gm4462 | 2.43886958 | 2.484328414 | 2.469017378 | 0.015311036 | -0.030147798 |
| Mcm8 | 6.67674951 | 6.664079978 | 6.706907568 | -0.04282759 | -0.030158059 |
| Rrnad1 | 8.306768337 | 8.456919943 | 8.336936394 | 0.119983549 | -0.030168057 |
| Slc12a5 | 3.708625487 | 3.842163105 | 3.738816931 | 0.103346173 | -0.030191444 |
| Gm48035 | 4.460331592 | 4.4217675 | 4.490548702 | -0.068781203 | -0.03021711 |
| Rps6-ps1 | 2.023049878 | 1.9712087 | 2.053421944 | -0.082213244 | -0.030372066 |
| Gm3839 | 3.382735954 | 3.453117696 | 3.413114379 | 0.040003317 | -0.030378425 |
| Olfr1402 | 2.521045372 | 2.523394333 | 2.551488508 | -0.028094175 | -0.030443136 |
| Gpr108 | 9.832942455 | 9.885619503 | 9.863420888 | 0.022198615 | -0.030478432 |
| Fh1 | 10.58053376 | 10.37582212 | 10.61102916 | -0.235207043 | -0.030495403 |
| Srrm2 | 12.93413749 | 13.17746424 | 12.96464754 | 0.212816703 | -0.030510053 |
| Trim43a | 1.487729036 | 1.536647604 | 1.518242039 | 0.018405566 | -0.030513003 |
| Gm49286 | 1.03017995 | 1.041243784 | 1.06069578 | -0.019451996 | -0.03051583 |
| Lyl1 | 6.421724399 | 6.274984549 | 6.452252209 | -0.17726766 | -0.03052781 |
| Olfr39 | 1.412824455 | 1.416612247 | 1.443355207 | -0.026742961 | -0.030530752 |
| Zscan20 | 6.505844411 | 6.77894645 | 6.536377038 | 0.242569412 | -0.030532627 |
| Galnt10 | 9.306796883 | 9.381510706 | 9.337421266 | 0.04408944 | -0.030624383 |
| Mup16 | 7.194692924 | 7.07092348 | 7.225334147 | -0.154410667 | -0.030641223 |
| Fgd4 | 8.46120931 | 8.335503346 | 8.491885626 | -0.156382279 | -0.030676316 |
| Glmp | 10.40988049 | 10.32974718 | 10.44058401 | -0.11083683 | -0.030703517 |
| Tmem200b | 5.201138075 | 5.241217714 | 5.231847197 | 0.009370517 | -0.030709122 |
| Pacs2 | 10.07929423 | 10.24302515 | 10.11001682 | 0.133008337 | -0.030722587 |
| Atg10 | 6.916789937 | 6.756278885 | 6.947517913 | -0.191239027 | -0.030727976 |
| Usp15 | 10.73460413 | 10.6771164 | 10.76533496 | -0.08821856 | -0.030730823 |
| Rhno1 | 8.562251377 | 8.512477555 | 8.593016307 | -0.080538752 | -0.03076493 |
| Adra1b | 2.930905134 | 2.954285858 | 2.961762651 | -0.007476793 | -0.030857517 |
| Olfr344 | 4.796178933 | 4.794204184 | 4.827111408 | -0.032907224 | -0.030932475 |
| Rnf216 | 9.649207178 | 9.700641157 | 9.680164412 | 0.020476745 | -0.030957234 |
| Lin7c | 10.89187911 | 10.81132563 | 10.92288641 | -0.111560787 | -0.031007303 |
| Cage1 | 4.38175895 | 4.413517471 | 4.412769181 | 0.00074829 | -0.031010231 |

| | | | | | |
|---|---|---|---|---|---|
| Znf41-ps | 7.571012116 | 7.627757526 | 7.602032819 | 0.025724706 | -0.031020703 |
| Wnt3 | 9.248232109 | 9.52810208 | 9.279273799 | 0.248828281 | -0.031041689 |
| Stx4a | 10.42218714 | 10.3962362 | 10.45322986 | -0.05699366 | -0.031042723 |
| Fignl1 | 7.589636004 | 7.562899362 | 7.620702204 | -0.057802842 | -0.0310662 |
| Cript | 9.441307472 | 9.423522152 | 9.472375964 | -0.048853812 | -0.031068493 |
| Olfr1305 | 2.816667918 | 2.883767103 | 2.84774757 | 0.036019533 | -0.031079652 |
| Tmem30a | 11.64469875 | 11.50475292 | 11.67580573 | -0.17105281 | -0.031106983 |
| Fopnl | 9.437514484 | 9.297262568 | 9.468641036 | -0.171378468 | -0.031126553 |
| Ccdc169 | 1.727044106 | 1.766107833 | 1.758176785 | 0.007931048 | -0.031132679 |
| Chmp7 | 9.705492188 | 9.742342565 | 9.736682485 | 0.00566008 | -0.031190298 |
| H2-Eb2 | 4.639530563 | 5.07784518 | 4.670722746 | 0.407122435 | -0.031192183 |
| 4930523C07Rik | 9.581940841 | 9.397443278 | 9.613141148 | -0.215697869 | -0.031200306 |
| Mmd2 | 2.245079096 | 2.253054847 | 2.276282764 | -0.023227917 | -0.031203668 |
| Gm29228 | 1.720642383 | 1.70866531 | 1.751989496 | -0.043324186 | -0.031347113 |
| Ciao1 | 9.362990236 | 9.38162322 | 9.3943771 | -0.012753879 | -0.031386864 |
| Gm10046 | 1.07238468 | 1.282366853 | 1.103793685 | 0.178573168 | -0.031409005 |
| Ndufv3 | 9.89731359 | 9.869470212 | 9.92873163 | -0.059261418 | -0.03141804 |
| Ttc13 | 9.456431017 | 9.593007973 | 9.487857119 | 0.105150854 | -0.031426102 |
| Yme1l1 | 11.12112466 | 11.08291832 | 11.15256285 | -0.069644527 | -0.03143819 |
| Ttc4 | 9.225554771 | 9.149663949 | 9.257001765 | -0.107337816 | -0.031446994 |
| Zfp322a | 8.788124545 | 8.744227876 | 8.819645629 | -0.075417753 | -0.031521085 |
| Zfp672 | 9.130353449 | 9.043546297 | 9.161904395 | -0.118358098 | -0.031550946 |
| Kptn | 7.774038786 | 7.864396031 | 7.805616232 | 0.058779799 | -0.031577446 |
| Gm2962 | 1.27528173 | 1.284709682 | 1.306921201 | -0.022211519 | -0.031639472 |
| Rpl4 | 14.58994673 | 14.53853095 | 14.62160265 | -0.0830717 | -0.031655927 |
| Pde8b | 4.539287999 | 4.510435867 | 4.570959397 | -0.06052353 | -0.031671398 |
| Fbxo15 | 1.981666475 | 1.985384817 | 2.013363172 | -0.027978355 | -0.031696698 |
| Exoc2 | 9.39334636 | 9.446361963 | 9.425072059 | 0.021289904 | -0.031725699 |
| Cep63 | 8.981665691 | 9.021188375 | 9.013423404 | 0.007764971 | -0.031757713 |
| Rpl19-ps3 | 3.537382921 | 3.500448222 | 3.569141821 | -0.068693599 | -0.0317589 |
| Zfp735 | 1.12224007 | 1.148936685 | 1.154017343 | -0.005080657 | -0.031777273 |
| Slitrk2 | 3.241415349 | 3.274371374 | 3.273235817 | 0.001135556 | -0.031820468 |

| | | | | | |
|---|---|---|---|---|---|
| Fam25c | 12.13519126 | 12.25985568 | 12.16702896 | 0.092826717 | -0.0318377 |
| Mylf-ps | 0.822515134 | 0.810913782 | 0.854354308 | -0.043440526 | -0.031839174 |
| M1ap | 1.740594047 | 1.744519883 | 1.772493492 | -0.027973609 | -0.031899445 |
| Gm5941 | 3.761337128 | 3.73302627 | 3.793238434 | -0.060212164 | -0.031901307 |
| Kctd19 | 2.031310436 | 2.165147946 | 2.063216296 | 0.10193165 | -0.031905859 |
| Hist3h2ba | 2.216808822 | 2.245706928 | 2.248756913 | -0.003049985 | -0.031948092 |
| Krt71 | 2.449306461 | 2.456914562 | 2.481278228 | -0.024363666 | -0.031971767 |
| Sowaha | 2.589110865 | 2.637120418 | 2.621089303 | 0.016031115 | -0.031978438 |
| Map2k7 | 10.08164558 | 10.24852076 | 10.11366431 | 0.134856447 | -0.03201873 |
| Slc35d3 | 1.914212891 | 1.899988576 | 1.946308929 | -0.046320353 | -0.032096038 |
| Krt87 | 1.832332661 | 1.887383404 | 1.864431516 | 0.022951888 | -0.032098855 |
| Tcf20 | 10.21721073 | 10.25088505 | 10.24931634 | 0.00156871 | -0.032105613 |
| Gm10136 | 1.996610575 | 2.03201961 | 2.02872431 | 0.0032953 | -0.032113735 |
| Mup18 | 8.371426744 | 8.09343295 | 8.403588029 | -0.310155079 | -0.032161285 |
| Eps15l1 | 10.23614719 | 10.45839494 | 10.26833193 | 0.190063007 | -0.032184747 |
| Dcst2 | 4.467399577 | 4.50805657 | 4.499587724 | 0.008468847 | -0.032188147 |
| Rnf123 | 9.796313522 | 9.876677118 | 9.828592487 | 0.048084631 | -0.032278965 |
| Ces5a | 2.897208812 | 2.971701487 | 2.929502791 | 0.042198696 | -0.032293979 |
| Poldip2 | 10.26334112 | 10.18638401 | 10.29564634 | -0.10926233 | -0.03230522 |
| Gm16523 | 2.653498357 | 2.661067629 | 2.685807522 | -0.024739893 | -0.032309165 |
| Dbh | 1.80841739 | 1.810376417 | 1.840745086 | -0.03036867 | -0.032327696 |
| Pcmt1 | 9.117312852 | 9.07132516 | 9.149674961 | -0.078349801 | -0.032362109 |
| Atg5 | 9.323477376 | 9.398116323 | 9.355875875 | 0.042240448 | -0.032398499 |
| Sptan1 | 12.37116099 | 12.56835403 | 12.40355965 | 0.164794383 | -0.032398653 |
| Gm21092 | 5.468598028 | 5.479182368 | 5.501013407 | -0.021831039 | -0.032415379 |
| Gm44511 | 1.145826274 | 1.153447575 | 1.178248672 | -0.024801097 | -0.032422398 |
| Olfr1410 | 1.350263488 | 1.382047756 | 1.382700716 | -0.00065296 | -0.032437228 |
| Tk2 | 8.520365242 | 8.402913896 | 8.552817454 | -0.149903558 | -0.032452212 |
| Thap8 | 2.525945135 | 2.580651638 | 2.558405404 | 0.022246233 | -0.032460269 |
| Vmn1r1 | 1.870785953 | 1.898275832 | 1.903284049 | -0.005008218 | -0.032498097 |
| Lsm14b | 9.538357569 | 9.611611435 | 9.570860537 | 0.040750898 | -0.032502968 |
| Rbbp6 | 10.95999484 | 10.94568669 | 10.99250485 | -0.046818167 | -0.03251001 |

| | | | | | |
|---|---|---|---|---|---|
| Gm10275 | 13.26775245 | 13.12692398 | 13.30028255 | -0.173358577 | -0.032530103 |
| Spink5 | 13.57510493 | 13.46926001 | 13.60764457 | -0.138384557 | -0.03253964 |
| Uqcrc1 | 11.01581133 | 11.03996631 | 11.04835597 | -0.008389663 | -0.032544643 |
| Gpr156 | 2.002708153 | 2.054429782 | 2.035391056 | 0.019038726 | -0.032682903 |
| Gm47056 | 5.809825219 | 5.713080983 | 5.842551393 | -0.12947041 | -0.032726174 |
| Crebbp | 10.60104727 | 10.42516047 | 10.63380328 | -0.20864281 | -0.03275601 |
| Hcn1 | 4.113801449 | 4.063451287 | 4.146570349 | -0.083119062 | -0.0327689 |
| Sipa1 | 8.222085332 | 8.491502244 | 8.254921023 | 0.236581221 | -0.032835691 |
| Fam83h | 10.59839667 | 10.99949648 | 10.63125291 | 0.36824357 | -0.032856247 |
| Eef1b2 | 13.05527903 | 12.94863268 | 13.08813579 | -0.13950311 | -0.032856767 |
| Gm15421 | 2.579637495 | 2.546988208 | 2.612533145 | -0.065544937 | -0.03289565 |
| Cntnap4 | 2.116330144 | 2.121972514 | 2.149229385 | -0.027256872 | -0.032899242 |
| Stx16 | 9.752397106 | 9.959076639 | 9.785338821 | 0.173737818 | -0.032941715 |
| Rab26 | 1.382687271 | 1.39091625 | 1.415672388 | -0.024756139 | -0.032985117 |
| Dazl | 1.439350552 | 1.461254149 | 1.472432289 | -0.01117814 | -0.033081737 |
| Fam91a1 | 10.0756406 | 10.07122278 | 10.1087263 | -0.037503515 | -0.033085701 |
| Oprd1 | 1.381303255 | 1.425713168 | 1.414446858 | 0.01126631 | -0.033143603 |
| Gm9616 | 2.986584868 | 2.946530603 | 3.019764833 | -0.07323423 | -0.033179965 |
| Gchfr | 3.220571516 | 3.172128113 | 3.253784054 | -0.081655941 | -0.033212538 |
| Zmynd10 | 2.443907429 | 2.522953907 | 2.47718801 | 0.045765897 | -0.033280581 |
| Srcap | 11.04816669 | 11.28860786 | 11.08147397 | 0.207133893 | -0.033307273 |
| Vbp1 | 9.532771547 | 9.474968256 | 9.566104545 | -0.091136289 | -0.033332998 |
| Rnf208 | 8.252716857 | 8.518347311 | 8.286066359 | 0.232280952 | -0.033349502 |
| Spin4 | 5.338775921 | 5.219020085 | 5.372126338 | -0.153106253 | -0.033350417 |
| Olfr986 | 6.523356114 | 6.562666201 | 6.556712946 | 0.005953255 | -0.033356832 |
| Kpna2-ps | 2.389721319 | 2.468024681 | 2.423087024 | 0.044937658 | -0.033365705 |
| Inf2 | 10.81836718 | 11.112356 | 10.8518288 | 0.260527197 | -0.03346162 |
| 9030624G23Rik | 4.236548764 | 4.318139608 | 4.270011757 | 0.048127852 | -0.033462993 |
| Vmn2r113 | 3.205643614 | 3.197034032 | 3.239117323 | -0.042083291 | -0.03347371 |
| Olfr1226 | 2.370832437 | 2.343506851 | 2.404327625 | -0.060820774 | -0.033495188 |
| Gm15760 | 3.554400371 | 3.483581526 | 3.587905618 | -0.104324093 | -0.033505247 |
| Sytl5 | 1.479008715 | 1.526225497 | 1.51252966 | 0.013695838 | -0.033520945 |

| | | | | | |
|---|---|---|---|---|---|
| Dpp3 | 11.00498265 | 11.13063643 | 11.03851034 | 0.092126087 | -0.03352769 |
| 4930550C14Rik | 5.201942087 | 5.24132331 | 5.235484042 | 0.005839268 | -0.033541955 |
| Atxn7l2 | 6.43189984 | 6.709901036 | 6.46548913 | 0.244411906 | -0.03358929 |
| Ube2z | 11.17362356 | 11.17945217 | 11.20729107 | -0.0278389 | -0.033667517 |
| Usp5 | 11.05717647 | 11.20990533 | 11.09087257 | 0.119032763 | -0.0336961 |
| Grhl2 | 9.853959993 | 9.976129391 | 9.887662933 | 0.088466457 | -0.03370294 |
| Rnf17 | 2.386978477 | 2.467620156 | 2.420710879 | 0.046909277 | -0.033732402 |
| Klhl42 | 8.556750064 | 8.626321476 | 8.590496599 | 0.035824877 | -0.033746535 |
| Gm29394 | 2.781016745 | 2.7827337 | 2.814781287 | -0.032047587 | -0.033764541 |
| Olfr1356 | 2.320086544 | 2.322033294 | 2.35392481 | -0.031891516 | -0.033838266 |
| Pex10 | 7.258780589 | 7.439241962 | 7.292623112 | 0.14661885 | -0.033842523 |
| Mta3 | 9.966994938 | 10.06395881 | 10.00084894 | 0.06310987 | -0.033854 |
| Peli1 | 10.45004964 | 10.36596685 | 10.48392709 | -0.117960247 | -0.033877453 |
| Mysm1 | 9.128630836 | 9.044519934 | 9.162525868 | -0.118005934 | -0.033895032 |
| Gm15387 | 10.85647927 | 10.66276774 | 10.89038191 | -0.22761417 | -0.03390264 |
| Gm17893 | 2.809015012 | 2.808393125 | 2.842935163 | -0.034542038 | -0.033920151 |
| Tmem176b | 10.10946537 | 9.945947461 | 10.14340348 | -0.197456022 | -0.033938118 |
| Nxpe1-ps | 3.550740513 | 3.482334537 | 3.584686155 | -0.102351617 | -0.033945642 |
| Pbx1 | 10.76882388 | 10.76029329 | 10.80277768 | -0.042484387 | -0.033953797 |
| Cacul1 | 10.23676133 | 10.346435 | 10.27078697 | 0.075648027 | -0.03402564 |
| Gm40466 | 2.465197824 | 2.454175955 | 2.499227923 | -0.045051968 | -0.034030099 |
| Rapgef6 | 9.85753381 | 9.98994454 | 9.891618595 | 0.098325945 | -0.034084785 |
| Zfp977 | 2.773868765 | 2.7152633 | 2.808005553 | -0.092742253 | -0.034136788 |
| Gm14322 | 4.469761098 | 4.600951696 | 4.503988253 | 0.096963444 | -0.034227155 |
| Rmnd5b | 9.246984789 | 9.274553496 | 9.281248798 | -0.006695302 | -0.034264009 |
| Tspan33 | 4.996381002 | 5.220622717 | 5.030663239 | 0.189959478 | -0.034282237 |
| Uchl4 | 2.242633183 | 2.253312225 | 2.276919459 | -0.023607234 | -0.034286276 |
| Olfr613 | 3.528398056 | 3.59207413 | 3.562781632 | 0.029292498 | -0.034383576 |
| Gapvd1 | 10.94371207 | 10.98459604 | 10.97813181 | 0.006464233 | -0.034419737 |
| Creld1 | 8.702790406 | 8.981673477 | 8.737286549 | 0.244386928 | -0.034496143 |
| Fanca | 6.988966257 | 7.16382188 | 7.023487307 | 0.140334573 | -0.03452105 |
| Brd7 | 10.37446772 | 10.24477831 | 10.40901113 | -0.164232817 | -0.034543407 |

| | | | | | |
|---|---|---|---|---|---|
| Pgr | 1.880036837 | 1.932940463 | 1.914594494 | 0.018345969 | -0.034557656 |
| Dynlt1-ps1 | 4.930238045 | 4.798539392 | 4.964818876 | -0.166279484 | -0.034580831 |
| Tubgcp3 | 9.296058632 | 9.280583704 | 9.330643879 | -0.050060175 | -0.034585247 |
| Fbxo38 | 10.17524955 | 10.19594979 | 10.20983814 | -0.013888357 | -0.034588593 |
| Pou3f2 | 1.968986163 | 1.966872692 | 2.00359435 | -0.036721658 | -0.034608187 |
| 4931428L18Rik | 4.491307177 | 4.485462471 | 4.525964396 | -0.040501925 | -0.034657218 |
| Ufsp2 | 9.317607107 | 9.250960597 | 9.352314447 | -0.101353851 | -0.03470734 |
| Ascl4 | 1.335422476 | 1.364192287 | 1.370162102 | -0.005969815 | -0.034739626 |
| Tapt1 | 10.18015829 | 10.14928636 | 10.21491969 | -0.06563333 | -0.034761397 |
| Pde6g | 2.084223639 | 2.144089283 | 2.11905541 | 0.025033873 | -0.03483177 |
| Slc35a5 | 8.574823096 | 8.718228021 | 8.609662585 | 0.108565436 | -0.034839489 |
| Btn1a1 | 0.84284972 | 0.812163951 | 0.877692203 | -0.065528252 | -0.034842484 |
| Olfr414 | 2.005137761 | 2.014910144 | 2.0399875 | -0.025077356 | -0.034849739 |
| Dvl1 | 9.951700267 | 10.21187014 | 9.986578997 | 0.22529114 | -0.034878729 |
| Acpp | 12.20648017 | 12.21619044 | 12.24139815 | -0.025207707 | -0.03491798 |
| Emilin3 | 4.168731615 | 4.194114777 | 4.203680744 | -0.009565967 | -0.034949129 |
| Gabrr2 | 2.977222133 | 2.940305516 | 3.012247987 | -0.071942472 | -0.035025854 |
| Esrrb | 3.075533453 | 3.146716976 | 3.11058441 | 0.036132565 | -0.035050957 |
| Myo1h | 4.483079758 | 4.554642931 | 4.518204939 | 0.036437992 | -0.035125181 |
| Gm9944 | 5.465847792 | 5.444871314 | 5.500975665 | -0.056104351 | -0.035127873 |
| Traf6 | 10.22587367 | 10.26640561 | 10.26104713 | 0.00535848 | -0.035173463 |
| Edn2 | 2.133525063 | 2.166635913 | 2.168703624 | -0.002067711 | -0.035178561 |
| Aebp1 | 11.64813707 | 11.73314568 | 11.68337182 | 0.049773857 | -0.03523475 |
| Wnk4 | 7.449093822 | 7.376508263 | 7.484347779 | -0.107839516 | -0.035253957 |
| Fgl1 | 3.472663962 | 3.376221926 | 3.508007478 | -0.131785551 | -0.035343516 |
| Ankrd65 | 1.762611705 | 1.776004809 | 1.797993269 | -0.021988461 | -0.035381564 |
| Tmem150b | 3.782245797 | 3.674137563 | 3.81765999 | -0.143522427 | -0.035414193 |
| 1700012P22Rik | 2.263509977 | 2.28216096 | 2.298949884 | -0.016788924 | -0.035439907 |
| Lmbr1l | 9.129664133 | 9.251267638 | 9.165107056 | 0.086160582 | -0.035442923 |
| Tiam1 | 12.00020908 | 12.08766869 | 12.0356582 | 0.052010487 | -0.03544912 |
| Aire | 1.797581727 | 1.782574002 | 1.833030887 | -0.050456885 | -0.03544916 |
| Mrps31 | 8.107191445 | 8.106941642 | 8.142648776 | -0.035707134 | -0.035457331 |

| | | | | | |
|---|---|---|---|---|---|
| Gm45739 | 1.798763467 | 1.849374057 | 1.834233152 | 0.015140905 | -0.035469684 |
| Gm17334 | 1.525235435 | 1.515645214 | 1.560725888 | -0.045080674 | -0.035490453 |
| Chd8 | 10.91100844 | 10.89083049 | 10.94650046 | -0.05566997 | -0.035492023 |
| Nav2 | 8.800231611 | 8.919222577 | 8.835752037 | 0.08347054 | -0.035520426 |
| Olfr1173 | 5.285625573 | 5.386014628 | 5.321150325 | 0.064864304 | -0.035524752 |
| Zfp57 | 5.254928154 | 5.25190631 | 5.290520166 | -0.038613856 | -0.035592011 |
| Lrrc10b | 1.785323467 | 1.817889268 | 1.820922459 | -0.003033191 | -0.035598992 |
| Diras1 | 1.649992663 | 1.748888536 | 1.6856136 | 0.063274936 | -0.035620937 |
| Gprc5c | 8.385951239 | 7.875095348 | 8.42160891 | -0.546513562 | -0.035657672 |
| Nudt8 | 7.027021604 | 7.029191524 | 7.062694454 | -0.03350293 | -0.035672851 |
| Olfr1416 | 3.434391369 | 3.44911972 | 3.470066149 | -0.02094643 | -0.03567478 |
| 4930590J08Rik | 3.237897692 | 3.25370473 | 3.27362697 | -0.01992224 | -0.035729278 |
| Stmnd1 | 2.45078166 | 2.470608406 | 2.486516019 | -0.015907613 | -0.035734359 |
| Rps27rt | 12.98973671 | 12.81200517 | 13.02547352 | -0.213468343 | -0.035736803 |
| Txndc17 | 10.95481716 | 10.98215897 | 10.99058127 | -0.0084223 | -0.035764107 |
| Ctsa | 11.76612385 | 12.031623 | 11.80190212 | 0.229720883 | -0.03577827 |
| Gm20547 | 6.353395233 | 5.850212727 | 6.389252727 | -0.53904 | -0.035857494 |
| Ankfn1 | 1.721967438 | 1.746287427 | 1.757895154 | -0.011607727 | -0.035927716 |
| Cstad | 1.571125886 | 1.592737783 | 1.607057886 | -0.014320103 | -0.035932 |
| Stac | 3.963566759 | 3.977508238 | 3.999518807 | -0.022010569 | -0.035952048 |
| Sprn | 2.616108788 | 2.535636443 | 2.652080684 | -0.116444241 | -0.035971895 |
| Rtraf | 10.71093843 | 10.62442286 | 10.74693809 | -0.12251523 | -0.03599966 |
| Muc3a | 2.059800836 | 2.063857089 | 2.095807211 | -0.031950122 | -0.036006376 |
| Vmn2r111 | 1.413836455 | 1.456597952 | 1.449933279 | 0.006664673 | -0.036096824 |
| Gm4922 | 1.752539489 | 1.788747609 | 1.788683163 | 6.44463E-05 | -0.036143674 |
| Gja8 | 2.364603238 | 2.39219394 | 2.40075267 | -0.00855873 | -0.036149432 |
| Dcaf10 | 9.378901572 | 9.436317006 | 9.415159632 | 0.021157374 | -0.03625806 |
| Zfp629 | 8.796397037 | 8.886209316 | 8.832690958 | 0.053518359 | -0.036293921 |
| Necab3 | 3.560509076 | 3.709252209 | 3.596864427 | 0.112387781 | -0.036355351 |
| Gdi1 | 10.84755692 | 10.93190108 | 10.88392064 | 0.047980433 | -0.036363727 |
| 9330182L06Rik | 4.458289192 | 4.488588908 | 4.494710156 | -0.006121248 | -0.036420965 |
| Smim7 | 11.10282574 | 11.07591801 | 11.13932514 | -0.063407127 | -0.0364994 |

| | | | | | |
|---|---|---|---|---|---|
| Sgpl1 | 11.99106982 | 12.13157447 | 12.02758573 | 0.103988747 | -0.036515907 |
| Traip | 6.777890558 | 6.935751301 | 6.814445713 | 0.121305588 | -0.036555155 |
| Gm48416 | 7.411716882 | 7.45497847 | 7.448293091 | 0.006685379 | -0.036576209 |
| Nectin2 | 9.10087278 | 9.362143245 | 9.137480991 | 0.224662254 | -0.036608211 |
| Kdm3a | 10.03728636 | 10.03874719 | 10.07389624 | -0.035149041 | -0.03660988 |
| Dnah8 | 5.655875137 | 5.719531678 | 5.692503621 | 0.027028057 | -0.036628484 |
| Aifm3 | 2.043419436 | 2.054953672 | 2.080069187 | -0.025115515 | -0.036649751 |
| Gm4032 | 2.049110062 | 2.069677695 | 2.085764818 | -0.016087123 | -0.036654756 |
| Mtfp1 | 6.503157344 | 6.488506864 | 6.539819674 | -0.05131281 | -0.03666233 |
| Tmprss11b | 2.534152156 | 2.53367999 | 2.570826888 | -0.037146899 | -0.036674732 |
| Baiap2l2 | 3.905002166 | 4.010835353 | 3.941688514 | 0.069146838 | -0.036686349 |
| Mms19 | 9.829702833 | 9.789974813 | 9.866390793 | -0.07641598 | -0.03668796 |
| Tuba1a | 12.27312183 | 12.19841609 | 12.30984546 | -0.11142937 | -0.036723627 |
| Triobp | 9.33748015 | 9.625663617 | 9.374234374 | 0.251429243 | -0.036754224 |
| Mtss1 | 10.06673918 | 10.01378558 | 10.10350253 | -0.089716949 | -0.036763343 |
| Vsig2 | 4.698562254 | 4.644263258 | 4.735348345 | -0.091085087 | -0.036786091 |
| Paxx | 7.399203754 | 7.380508651 | 7.435994017 | -0.055485366 | -0.036790263 |
| Top3a | 7.650559502 | 7.763838051 | 7.687357284 | 0.076480767 | -0.036797782 |
| Actg2 | 8.687763724 | 8.673780918 | 8.724567525 | -0.050786607 | -0.036803801 |
| Plin5 | 4.851796775 | 4.913062968 | 4.888609756 | 0.024453212 | -0.03681298 |
| Cpeb2 | 10.77030223 | 10.64139752 | 10.80715005 | -0.165752527 | -0.036847817 |
| BC051019 | 3.446989583 | 3.437756083 | 3.483845181 | -0.046089098 | -0.036855598 |
| Fxn | 6.490427004 | 6.50533311 | 6.527296677 | -0.021963567 | -0.036869673 |
| Cyb5b | 11.57549882 | 11.50932987 | 11.61237774 | -0.103047877 | -0.03687892 |
| Tada2a | 8.146737833 | 8.173438953 | 8.183644301 | -0.010205348 | -0.036906468 |
| Olfr95 | 2.745918167 | 2.700059904 | 2.78284444 | -0.082784536 | -0.036926273 |
| Zfp84 | 8.378231945 | 8.280701371 | 8.415160892 | -0.13445952 | -0.036928947 |
| Gm7434 | 2.247542448 | 2.317060011 | 2.284489715 | 0.032570296 | -0.036947267 |
| Olfr433 | 2.430626033 | 2.41392325 | 2.467609084 | -0.053685834 | -0.036983051 |
| Gpat2 | 2.29395556 | 2.307903284 | 2.330975533 | -0.023072249 | -0.037019974 |
| Selenoo | 8.664641163 | 8.644077157 | 8.701700925 | -0.057623768 | -0.037059762 |
| Map3k11 | 9.779437347 | 10.13629912 | 9.816498392 | 0.319800728 | -0.037061045 |

| | | | | | |
|---|---|---|---|---|---|
| Gm47325 | 2.600472019 | 2.595213976 | 2.637544487 | -0.042330511 | -0.037072468 |
| Fgfr1op | 8.404933921 | 8.451495028 | 8.442027926 | 0.009467103 | -0.037094005 |
| Clcn4 | 10.97586649 | 10.99057039 | 11.01304191 | -0.022471517 | -0.037175417 |
| Ccdc106 | 3.360401814 | 3.426406031 | 3.397712861 | 0.028693171 | -0.037311047 |
| Abhd2 | 10.6701959 | 10.76065813 | 10.70751067 | 0.053147457 | -0.03731477 |
| Gm266 | 2.123398078 | 2.143801506 | 2.160722334 | -0.016920828 | -0.037324256 |
| Gm45855 | 3.238170346 | 3.242852748 | 3.275521142 | -0.032668394 | -0.037350797 |
| Gnat1 | 4.138711914 | 4.214180807 | 4.17607797 | 0.038102837 | -0.037366056 |
| Pwwp2b | 7.182350207 | 7.360004618 | 7.219744595 | 0.140260023 | -0.037394388 |
| Wdr20rt | 2.143102769 | 2.176859808 | 2.180521684 | -0.003661875 | -0.037418915 |
| Vmn2r16 | 2.696089675 | 2.745711628 | 2.733545481 | 0.012166147 | -0.037455806 |
| Nphp4 | 4.574299059 | 4.525412384 | 4.611761822 | -0.086349438 | -0.037462764 |
| Arhgef39 | 5.438788382 | 5.511670179 | 5.476309306 | 0.035360873 | -0.037520924 |
| Cspg4 | 9.677581198 | 9.714122363 | 9.715123674 | -0.001001311 | -0.037542476 |
| Stk33 | 1.319550415 | 1.343938934 | 1.357101061 | -0.013162127 | -0.037550646 |
| Nckap5 | 6.149992794 | 6.240787253 | 6.187562447 | 0.053224806 | -0.037569654 |
| Gm13689 | 3.378949759 | 3.404475173 | 3.416530084 | -0.012054911 | -0.037580325 |
| Nek7 | 11.22711041 | 11.07216332 | 11.26485247 | -0.19268915 | -0.037742063 |
| Gm47659 | 1.911159207 | 1.946140122 | 1.94895459 | -0.002814468 | -0.037795383 |
| 4632433K11Rik | 2.929054594 | 2.899096481 | 2.966866054 | -0.067769573 | -0.03781146 |
| Gm28053 | 1.937543664 | 1.897418344 | 1.975355965 | -0.077937622 | -0.037812301 |
| Gpr26 | 2.640309933 | 2.74674641 | 2.678156086 | 0.068590324 | -0.037846153 |
| Slc26a8 | 2.784408493 | 2.835863429 | 2.822310415 | 0.013553014 | -0.037901921 |
| Tbck | 8.685287358 | 8.711769083 | 8.723321644 | -0.011552561 | -0.038034286 |
| Dbi | 11.92473043 | 11.57188616 | 11.96282774 | -0.390941577 | -0.038097307 |
| Fxyd7 | 3.467969125 | 3.502507775 | 3.506080688 | -0.003572913 | -0.038111563 |
| Anapc16 | 9.083680763 | 9.229763773 | 9.121795932 | 0.107967841 | -0.038115169 |
| Bglap3 | 7.221863423 | 7.34716895 | 7.259994908 | 0.087174043 | -0.038131485 |
| Cdhr3 | 2.135258526 | 2.170411681 | 2.173403554 | -0.002991874 | -0.038145028 |
| Actr1a | 11.93431387 | 11.92015387 | 11.97246364 | -0.052309773 | -0.03814977 |
| Cdsn | 13.64874617 | 13.79310286 | 13.6869095 | 0.106193357 | -0.038163333 |
| Vmn1r-ps15 | 2.76695844 | 2.772945962 | 2.805125944 | -0.032179982 | -0.038167504 |

| | | | | | |
|---|---|---|---|---|---|
| Guca1a | 2.595266552 | 2.610233167 | 2.633438933 | -0.023205766 | -0.038172381 |
| Klk12 | 3.241251393 | 3.286282179 | 3.2794892 | 0.006792978 | -0.038237807 |
| Ckap2 | 8.039171215 | 7.956997385 | 8.077443271 | -0.120445886 | -0.038272056 |
| Pgbd5 | 3.315772677 | 3.450933157 | 3.354066354 | 0.096866803 | -0.038293677 |
| Ifi27l2a | 10.77165243 | 10.44016556 | 10.80996353 | -0.369797977 | -0.038311103 |
| Slc2a10 | 6.100182385 | 6.179685836 | 6.138495737 | 0.041190099 | -0.038313352 |
| Tmco3 | 8.342069143 | 8.405003461 | 8.380393245 | 0.024610216 | -0.038324102 |
| Cfap45 | 5.351986002 | 5.279424191 | 5.390330217 | -0.110906026 | -0.038344215 |
| Olfr1195 | 2.999818683 | 3.065789337 | 3.038179671 | 0.027609667 | -0.038360987 |
| Gbe1 | 9.261284245 | 8.867438067 | 9.299678515 | -0.432240448 | -0.03839427 |
| Gm12017 | 2.134699898 | 2.169772603 | 2.173114384 | -0.003341781 | -0.038414486 |
| Clrn1 | 2.97474986 | 2.980134165 | 3.013190732 | -0.033056568 | -0.038440873 |
| Spats1 | 2.20051399 | 2.243406928 | 2.239010022 | 0.004396906 | -0.038496032 |
| Aspscr1 | 9.369390494 | 9.572454081 | 9.407911991 | 0.16454209 | -0.038521497 |
| Eif2a | 10.38120818 | 10.30470289 | 10.41974545 | -0.115042567 | -0.03853727 |
| Cyp4f41-ps | 2.619457984 | 2.692577686 | 2.657999241 | 0.034578445 | -0.038541257 |
| Gm21962 | 1.922132783 | 2.002146219 | 1.960677518 | 0.041468701 | -0.038544735 |
| C1qtnf1 | 9.845457892 | 9.845216852 | 9.884019857 | -0.038803005 | -0.038561965 |
| Hdhd2 | 9.146235543 | 9.038421817 | 9.184828477 | -0.14640666 | -0.038592934 |
| Dbil5 | 2.316087857 | 2.384660505 | 2.354757538 | 0.029902967 | -0.038669681 |
| Lrguk | 3.524752908 | 3.395293092 | 3.563424543 | -0.168131451 | -0.038671635 |
| Vmn1r80 | 2.342488888 | 2.379279873 | 2.381201633 | -0.00192176 | -0.038712745 |
| Gm4734 | 2.362927983 | 2.548286342 | 2.401668787 | 0.146617555 | -0.038740804 |
| Gm48418 | 5.113003766 | 5.146073348 | 5.151747483 | -0.005674134 | -0.038743717 |
| Gm9493 | 1.957622961 | 1.959684507 | 1.996417185 | -0.036732678 | -0.038794224 |
| Uba1y | 2.169571656 | 2.251821809 | 2.208378413 | 0.043443396 | -0.038806757 |
| Nanos3 | 2.484369873 | 2.503210138 | 2.523199234 | -0.019989095 | -0.038829361 |
| Gm9812 | 1.859683444 | 1.879815209 | 1.898520756 | -0.018705546 | -0.038837312 |
| Unkl | 8.843929619 | 9.002497376 | 8.88280962 | 0.119687756 | -0.038880001 |
| Ighmbp2 | 7.308703832 | 7.356240339 | 7.347588297 | 0.008652041 | -0.038884466 |
| Gm47147 | 4.146899421 | 4.146369658 | 4.185816118 | -0.03944646 | -0.038916697 |
| Ppp1r3f | 4.906646584 | 4.918283514 | 4.945572523 | -0.027289009 | -0.03892594 |

| | | | | | |
|---|---|---|---|---|---|
| Olfr575 | 2.784145718 | 2.815186104 | 2.823094585 | -0.007908481 | -0.038948867 |
| Antxr1 | 11.16747985 | 11.20920632 | 11.20644591 | 0.002760413 | -0.03896606 |
| Ccdc69 | 6.307881006 | 6.389110295 | 6.34685083 | 0.042259465 | -0.038969824 |
| Crybg1 | 12.02540566 | 12.12880086 | 12.06439049 | 0.06441037 | -0.038984827 |
| Scml4 | 4.711065961 | 4.65488725 | 4.750094702 | -0.095207452 | -0.039028741 |
| Olfr466 | 2.720103418 | 2.733176553 | 2.759180061 | -0.026003508 | -0.039076642 |
| Sppl2b | 8.49624557 | 8.803151272 | 8.535328279 | 0.267822993 | -0.039082709 |
| Fitm1 | 7.90849677 | 7.999192413 | 7.947616268 | 0.051576145 | -0.039119497 |
| Tdrkh | 6.168463231 | 5.939119928 | 6.207645541 | -0.268525614 | -0.03918231 |
| Olfr775 | 2.78089787 | 2.804940943 | 2.82021984 | -0.015278897 | -0.03932197 |
| Cmtm8 | 6.425183309 | 6.443377293 | 6.464514501 | -0.021137208 | -0.039331192 |
| Slc29a3 | 9.310475305 | 9.496365578 | 9.349848744 | 0.146516834 | -0.039373439 |
| Yipf6 | 9.718319773 | 9.648945913 | 9.757705763 | -0.10875985 | -0.03938599 |
| Kdm4a | 10.08525985 | 10.25328327 | 10.12467827 | 0.128604997 | -0.039418427 |
| Krt36 | 1.318719688 | 1.301421458 | 1.358156565 | -0.056735107 | -0.039436877 |
| Cacna1f | 2.125757949 | 2.20432386 | 2.165210376 | 0.039113484 | -0.039452427 |
| Caln1 | 1.958097611 | 2.10827051 | 1.997584349 | 0.110686161 | -0.039486738 |
| Nkapd1 | 8.874802319 | 8.803673887 | 8.914295341 | -0.110621454 | -0.039493023 |
| Mup1 | 4.751669348 | 4.71689323 | 4.79116925 | -0.07427602 | -0.039499903 |
| Slc7a13 | 1.367448416 | 1.357946156 | 1.406976282 | -0.049030127 | -0.039527866 |
| Pla2g2d | 6.965776735 | 7.08479068 | 7.005355464 | 0.079435216 | -0.039578729 |
| Vmn2r88 | 2.909559082 | 2.786644483 | 2.949191816 | -0.162547333 | -0.039632735 |
| Vmn2r44 | 1.950756562 | 1.980450254 | 1.990396499 | -0.009946245 | -0.039639937 |
| Nsun7 | 3.378743488 | 3.364454739 | 3.418421113 | -0.053966374 | -0.039677625 |
| Gns | 11.22738004 | 11.16324435 | 11.26709525 | -0.103850903 | -0.039715213 |
| Wbp4 | 9.664973703 | 9.693087486 | 9.704728097 | -0.011640611 | -0.039754394 |
| Ubxn10 | 2.176204142 | 2.284920258 | 2.216012708 | 0.06890755 | -0.039808566 |
| Bmp1 | 10.36091143 | 10.43709614 | 10.40072685 | 0.036369287 | -0.03981542 |
| Yjefn3 | 2.188950735 | 2.286829032 | 2.22881397 | 0.058015062 | -0.039863235 |
| Gm7061 | 2.543696673 | 2.609363251 | 2.583575348 | 0.025787903 | -0.039878675 |
| AI413582 | 7.837954413 | 7.9428569 | 7.877839908 | 0.065016992 | -0.039885495 |
| Mrps23 | 8.18205231 | 8.244480382 | 8.221944432 | 0.02253595 | -0.039892122 |

| | | | | | |
|---|---|---|---|---|---|
| Msh2 | 9.303803989 | 9.287463378 | 9.343764592 | -0.056301213 | -0.039960602 |
| Fam214b | 9.806894529 | 10.20125344 | 9.846894258 | 0.354359182 | -0.039999729 |
| Ndufaf1 | 8.042143761 | 7.940989401 | 8.082180931 | -0.14119153 | -0.04003717 |
| Fgf16 | 2.899683545 | 2.985390182 | 2.939775672 | 0.04561451 | -0.040092127 |
| Ubxn11 | 4.563665575 | 4.663704966 | 4.603773263 | 0.059931703 | -0.040107688 |
| Atp6ap1l | 1.647580145 | 1.655086253 | 1.687697826 | -0.032611573 | -0.040117681 |
| Mzf1 | 4.032991589 | 4.079496386 | 4.073119188 | 0.006377198 | -0.040127599 |
| Prelid3a | 3.719519922 | 3.791630948 | 3.75965109 | 0.031979857 | -0.040131168 |
| Scnm1 | 8.375731801 | 8.291462202 | 8.415909758 | -0.124447556 | -0.040177957 |
| Gm31166 | 5.615275203 | 5.517187657 | 5.655462232 | -0.138274576 | -0.040187029 |
| Gm47265 | 1.35279472 | 1.401457961 | 1.393085738 | 0.008372223 | -0.040291019 |
| Lrrc55 | 3.351296891 | 3.388988067 | 3.391595061 | -0.002606995 | -0.04029817 |
| C9 | 3.972443276 | 4.053660719 | 4.01275812 | 0.040902599 | -0.040314844 |
| Tmem92 | 2.18720059 | 2.213794809 | 2.227546662 | -0.013751853 | -0.040346072 |
| Trim3 | 8.911057271 | 9.099829354 | 8.951442748 | 0.148386606 | -0.040385477 |
| Vamp8 | 9.923695309 | 9.854956855 | 9.964104987 | -0.109148132 | -0.040409679 |
| Trappc10 | 10.79492948 | 10.92770221 | 10.83537419 | 0.092328017 | -0.040444717 |
| Caskin1 | 2.171548492 | 2.226676046 | 2.21204703 | 0.014629017 | -0.040498537 |
| AC121967.1 | 2.567052446 | 2.55927622 | 2.607566662 | -0.048290442 | -0.040514216 |
| F8a | 6.868827131 | 6.891370669 | 6.909371979 | -0.01800131 | -0.040544848 |
| Klhdc10 | 10.08313838 | 10.18755598 | 10.1236938 | 0.063862173 | -0.04055542 |
| Tacc2 | 12.51264688 | 12.35833309 | 12.55324934 | -0.194916257 | -0.04060246 |
| Sap130 | 9.336303604 | 9.56915411 | 9.376906201 | 0.192247909 | -0.040602596 |
| Usp7 | 12.00764892 | 12.04443041 | 12.04829289 | -0.00386248 | -0.040643967 |
| Luzp2 | 2.41110712 | 2.401125722 | 2.451753749 | -0.050628026 | -0.040646629 |
| Pde11a | 3.511309345 | 3.530172241 | 3.551978091 | -0.02180585 | -0.040668746 |
| Ntn5 | 2.705064811 | 2.755237199 | 2.745784119 | 0.00945308 | -0.040719308 |
| Mapkapk3 | 11.18823657 | 11.25608927 | 11.22901038 | 0.027078883 | -0.040773813 |
| Exoc1 | 9.594655198 | 9.702993474 | 9.635450473 | 0.067543002 | -0.040795275 |
| Mrpl42 | 9.886165774 | 9.919340212 | 9.926978619 | -0.007638407 | -0.040812845 |
| Iffo1 | 7.315674018 | 7.293428242 | 7.356534686 | -0.063106444 | -0.040860667 |
| Gorasp1 | 8.784733528 | 8.878394988 | 8.825614192 | 0.052780796 | -0.040880664 |

| | | | | | |
|---|---|---|---|---|---|
| Zfp575 | 2.211626548 | 2.296570864 | 2.252591164 | 0.0439797 | -0.040964616 |
| Dut | 8.606559292 | 8.595303599 | 8.647539919 | -0.05223632 | -0.040980628 |
| Eml2 | 9.366154655 | 9.501410437 | 9.407146505 | 0.094263932 | -0.04099185 |
| Gja6 | 1.579458067 | 1.608980379 | 1.620490074 | -0.011509695 | -0.041032007 |
| Gm20716 | 11.21085021 | 11.22619711 | 11.25192273 | -0.025725623 | -0.041072523 |
| Mtmr12 | 9.728827049 | 9.769005579 | 9.77002684 | -0.001021261 | -0.041199791 |
| Nxph1 | 1.900705721 | 1.938160104 | 1.941910175 | -0.00375007 | -0.041204453 |
| Slc9b2 | 4.603632741 | 4.622077494 | 4.644851693 | -0.022774199 | -0.041218952 |
| Mpv17l2 | 6.870381836 | 6.991978487 | 6.911604963 | 0.080373523 | -0.041223127 |
| Dcc | 1.474334648 | 1.498534206 | 1.515619976 | -0.017085771 | -0.041285328 |
| BC025920 | 5.629058052 | 5.614525217 | 5.670360837 | -0.05583562 | -0.041302785 |
| Gal3st2b | 2.596233216 | 2.658091822 | 2.637553397 | 0.020538425 | -0.041320181 |
| Clec2e | 2.530327079 | 2.537374177 | 2.571684014 | -0.034309837 | -0.041356935 |
| Gm10557 | 4.687135572 | 4.779134105 | 4.728509485 | 0.05062462 | -0.041373913 |
| Trim54 | 8.487819039 | 8.983913327 | 8.529221716 | 0.454691611 | -0.041402677 |
| Rnft1 | 9.700672908 | 9.690665635 | 9.742130261 | -0.051464625 | -0.041457352 |
| Olfr920 | 5.354686397 | 5.213006482 | 5.396196156 | -0.183189674 | -0.041509759 |
| Magi2 | 5.579989469 | 5.477604657 | 5.621564792 | -0.143960135 | -0.041575323 |
| Adgrv1 | 5.614121436 | 5.405226203 | 5.655741181 | -0.250514978 | -0.041619745 |
| Kng2 | 2.864531523 | 2.912086925 | 2.906192301 | 0.005894624 | -0.041660778 |
| Klc1 | 10.63191678 | 10.7468091 | 10.67358902 | 0.07322008 | -0.041672243 |
| Zfp420 | 6.31836781 | 6.42574776 | 6.360040102 | 0.065707657 | -0.041672293 |
| Dnaaf1 | 2.141122469 | 2.189131722 | 2.182814134 | 0.006317588 | -0.041691665 |
| Gm48641 | 5.586434333 | 5.270850564 | 5.628136034 | -0.35728547 | -0.041701701 |
| St13 | 11.39320617 | 11.52734046 | 11.43492011 | 0.092420347 | -0.041713947 |
| Col22a1 | 4.55814349 | 4.584299951 | 4.599864739 | -0.015564788 | -0.041721249 |
| Rps15a | 13.06433463 | 13.0527865 | 13.10606849 | -0.053281997 | -0.04173386 |
| Serpinf2 | 4.12975957 | 4.124303024 | 4.171535672 | -0.047232648 | -0.041776102 |
| Pdia5 | 6.871546899 | 6.900135246 | 6.913343483 | -0.013208237 | -0.041796584 |
| Mier3 | 8.962563531 | 9.143559668 | 9.004382039 | 0.139177629 | -0.041818507 |
| Scn9a | 2.319905659 | 2.336257166 | 2.361724978 | -0.025467812 | -0.041819319 |
| Zdhhc20 | 10.14596437 | 10.28674522 | 10.1877883 | 0.09895692 | -0.041823933 |

| | | | | | |
|---|---|---|---|---|---|
| Gm15539 | 1.956868593 | 1.981786116 | 1.998717933 | -0.016931817 | -0.04184934 |
| Stfa2 | 6.541999087 | 6.660659118 | 6.583885488 | 0.076773631 | -0.041886401 |
| Prss57 | 1.532421584 | 1.594597747 | 1.574326261 | 0.020271486 | -0.041904677 |
| Nlrc3 | 5.985573648 | 6.028090367 | 6.027515502 | 0.000574865 | -0.041941854 |
| Glp1r | 1.399820263 | 1.373894207 | 1.441845292 | -0.067951085 | -0.042025029 |
| Pigr | 2.610114944 | 2.600392454 | 2.652148129 | -0.051755675 | -0.042033185 |
| Ints4 | 9.376550313 | 9.396564907 | 9.418595071 | -0.022030163 | -0.042044758 |
| Thap11 | 8.74570338 | 8.830685989 | 8.787822112 | 0.042863877 | -0.042118732 |
| Slc22a13 | 1.539310963 | 1.587281112 | 1.581442364 | 0.005838747 | -0.042131402 |
| Fam69b | 5.248498389 | 5.367562764 | 5.290637929 | 0.076924835 | -0.042139539 |
| Gm6853 | 2.53370363 | 2.53885755 | 2.575865674 | -0.037008124 | -0.042162044 |
| Gm2a | 11.06388093 | 10.94884359 | 11.10609072 | -0.157247123 | -0.042209787 |
| Chkb | 8.827976342 | 8.823754302 | 8.870219624 | -0.046465322 | -0.042243283 |
| Gm16503 | 1.736894489 | 1.758298696 | 1.779193392 | -0.020894696 | -0.042298904 |
| Gm14226 | 2.423075701 | 2.521088195 | 2.465388603 | 0.055699592 | -0.042312902 |
| Ccdc9 | 9.873435861 | 9.846493883 | 9.915757025 | -0.069263141 | -0.042321163 |
| Zfp459 | 5.467943227 | 5.547058888 | 5.5102899 | 0.036768987 | -0.042346674 |
| Gm14416 | 3.698727772 | 3.670200601 | 3.741101085 | -0.070900484 | -0.042373313 |
| Sipa1l2 | 9.491817531 | 9.669385508 | 9.534222157 | 0.135163351 | -0.042404626 |
| Gm6627 | 2.250117043 | 2.281087396 | 2.292526487 | -0.011439091 | -0.042409444 |
| Rpl9-ps3 | 3.251702547 | 3.247846221 | 3.294115453 | -0.046269232 | -0.042412906 |
| Lrrc23 | 1.131207879 | 1.15644864 | 1.173625794 | -0.017177154 | -0.042417915 |
| Efcab8 | 1.918580696 | 1.987821005 | 1.961002913 | 0.026818091 | -0.042422218 |
| Tmco1 | 9.751543797 | 9.658911467 | 9.793966955 | -0.135055488 | -0.042423157 |
| Pi4ka | 10.50623366 | 10.64026526 | 10.54866852 | 0.091596743 | -0.042434857 |
| Gm16223 | 1.556955399 | 1.5597136 | 1.599417574 | -0.039703974 | -0.042462175 |
| C87436 | 8.191435329 | 8.283641396 | 8.23391673 | 0.049724665 | -0.042481401 |
| Ofd1 | 8.145624439 | 8.030218151 | 8.188108272 | -0.15789012 | -0.042483833 |
| Bsn | 5.984968522 | 6.219365233 | 6.027469014 | 0.191896219 | -0.042500492 |
| Rps2-ps6 | 1.39487799 | 1.395070418 | 1.437392375 | -0.042321957 | -0.042514385 |
| Gm9670 | 2.460570764 | 2.454460122 | 2.503090104 | -0.048629982 | -0.04251934 |
| Klhl20 | 8.816373285 | 8.881770916 | 8.858892729 | 0.022878187 | -0.042519444 |

| | | | | | |
|---|---|---|---|---|---|
| Sh3rf3 | 5.329509665 | 5.598938178 | 5.372058385 | 0.226879793 | -0.04254872 |
| Gm17040 | 2.125225812 | 2.196273115 | 2.167827069 | 0.028446046 | -0.042601257 |
| Zfp972 | 3.241425627 | 3.279365296 | 3.284030779 | -0.004665484 | -0.042605153 |
| Ccdc42 | 2.933387032 | 2.929991753 | 2.976019817 | -0.046028064 | -0.042632785 |
| Olfr1459 | 4.602167356 | 4.628151673 | 4.644828725 | -0.016677052 | -0.042661369 |
| Usp24 | 11.2158387 | 11.25209646 | 11.2586144 | -0.006517937 | -0.0427757 |
| Gm45808 | 1.955091577 | 1.93021992 | 1.997887189 | -0.067667269 | -0.042795611 |
| Olfr1505 | 2.602525665 | 2.638836204 | 2.645391142 | -0.006554938 | -0.042865477 |
| Wnt4 | 10.62563834 | 10.86928984 | 10.66852455 | 0.200765297 | -0.042886207 |
| Slc14a1 | 4.431059083 | 4.360749894 | 4.473968726 | -0.113218832 | -0.042909643 |
| Nmnat1 | 6.922217669 | 6.884399918 | 6.965152468 | -0.080752551 | -0.042934799 |
| Tulp3 | 8.546677228 | 8.598310858 | 8.589629958 | 0.0086809 | -0.04295273 |
| Hunk | 5.081436124 | 5.046148686 | 5.124390504 | -0.078241818 | -0.04295438 |
| Rps4x | 14.22893033 | 14.15059361 | 14.27189271 | -0.121299103 | -0.042962377 |
| Ehmt1 | 9.635094606 | 9.732353066 | 9.678061443 | 0.054291622 | -0.042966837 |
| Tmem273 | 3.084556874 | 3.151855514 | 3.127525477 | 0.024330037 | -0.042968603 |
| Nisch | 11.37485805 | 11.63049128 | 11.41783055 | 0.212660723 | -0.042972503 |
| Ercc4 | 8.304783887 | 8.247966664 | 8.347793421 | -0.099826757 | -0.043009534 |
| Csmd1 | 5.56477729 | 5.681587685 | 5.607839558 | 0.073748127 | -0.043062268 |
| Gm12216 | 4.415298217 | 4.493838217 | 4.458420591 | 0.035417626 | -0.043122374 |
| Mynn | 8.891874935 | 8.800143088 | 8.935082294 | -0.134939206 | -0.043207359 |
| Ankle1 | 5.582124931 | 5.774036198 | 5.625342218 | 0.14869398 | -0.043217288 |
| Ogfod3 | 6.981524842 | 6.824455605 | 7.0247752 | -0.200319595 | -0.043250358 |
| Mfrp | 3.44043375 | 3.557966217 | 3.483711191 | 0.074255026 | -0.043277441 |
| Cux2 | 3.670035604 | 3.58119993 | 3.713323576 | -0.132123647 | -0.043287973 |
| Tmem164 | 11.93878097 | 11.94053652 | 11.98208105 | -0.041544527 | -0.043300077 |
| Zfp790 | 8.105784427 | 8.051137669 | 8.149135513 | -0.097997844 | -0.043351086 |
| Arfgap2 | 11.18497058 | 11.23366195 | 11.22835626 | 0.005305683 | -0.04338568 |
| Scgb1b27 | 3.168874075 | 3.161415216 | 3.21226198 | -0.050846764 | -0.043387904 |
| Slc5a6 | 8.309726739 | 8.353789867 | 8.353130501 | 0.000659366 | -0.043403762 |
| Nat8f4 | 4.489304643 | 4.529349134 | 4.532715409 | -0.003366275 | -0.043410766 |
| Olfr1150-ps1 | 2.421093588 | 2.401537005 | 2.464538685 | -0.06300168 | -0.043445097 |

| | | | | | |
|---|---|---|---|---|---|
| Lgi3 | 4.340135131 | 4.344838047 | 4.383586836 | -0.038748789 | -0.043451705 |
| Gm14921 | 2.56379603 | 2.547452264 | 2.607248649 | -0.059796385 | -0.04345262 |
| Kir3dl2 | 2.912493487 | 2.960088755 | 2.9559561 | 0.004132655 | -0.043462613 |
| Tmem183a | 10.22610067 | 10.13050397 | 10.26960366 | -0.139099693 | -0.043502987 |
| Angel1 | 8.553703516 | 8.568821934 | 8.597208431 | -0.028386497 | -0.043504916 |
| Pkn2 | 10.39569747 | 10.39783088 | 10.43925348 | -0.041422597 | -0.043556003 |
| 1700001L19Rik | 0.886461418 | 0.86697607 | 0.930028659 | -0.063052589 | -0.043567241 |
| Prdm6 | 2.016299155 | 2.027441217 | 2.059892682 | -0.032451465 | -0.043593527 |
| Smim17 | 1.972160158 | 2.029063727 | 2.015813518 | 0.013250209 | -0.04365336 |
| Tnni3k | 3.036112606 | 3.017515235 | 3.079791921 | -0.062276686 | -0.043679315 |
| Zfp455 | 5.184609843 | 5.109305451 | 5.228298732 | -0.118993281 | -0.043688889 |
| Tbcd | 9.700649543 | 9.743092808 | 9.744346661 | -0.001253853 | -0.043697119 |
| Gm29201 | 1.436469481 | 1.454802184 | 1.480182181 | -0.025379997 | -0.0437127 |
| Arg1 | 9.037556746 | 8.778679972 | 9.08129849 | -0.302618519 | -0.043741744 |
| Impact | 9.476973453 | 9.973350825 | 9.52072645 | 0.452624375 | -0.043752997 |
| Ess2 | 8.213256845 | 8.379426712 | 8.257037954 | 0.122388758 | -0.043781109 |
| Eif4e1b | 1.617327561 | 1.629106748 | 1.661116618 | -0.03200987 | -0.043789057 |
| Gm21833 | 3.89773708 | 3.85920706 | 3.941544584 | -0.082337523 | -0.043807503 |
| Suclg1 | 10.43219977 | 10.47984698 | 10.47602355 | 0.00382343 | -0.043823783 |
| Gm12248 | 3.982156556 | 4.13864506 | 4.025994575 | 0.112650485 | -0.043838019 |
| Dnmbp | 8.95595632 | 8.936846307 | 8.999903563 | -0.063057256 | -0.043947243 |
| Terf2ip | 8.692539449 | 8.768833525 | 8.736513799 | 0.032319727 | -0.04397435 |
| Sprr3 | 3.243345922 | 3.387560524 | 3.287327989 | 0.100232535 | -0.043982067 |
| Mesp2 | 6.382321235 | 6.425021187 | 6.426342448 | -0.001321261 | -0.044021213 |
| Ankrd6 | 6.689218044 | 6.719313325 | 6.733241779 | -0.013928454 | -0.044023735 |
| Hepacam | 2.885891085 | 2.871234373 | 2.929970263 | -0.05873589 | -0.044079178 |
| Nuggc | 4.241407925 | 4.292964446 | 4.285491826 | 0.00747262 | -0.0440839 |
| Stard6 | 3.023865599 | 3.012795493 | 3.067964524 | -0.055169031 | -0.044098925 |
| Olfr1286 | 1.49987183 | 1.497478385 | 1.543980114 | -0.046501729 | -0.044108284 |
| Slc17a7 | 1.735038187 | 1.835683124 | 1.779175924 | 0.0565072 | -0.044137737 |
| Kxd1 | 9.313844946 | 9.194326511 | 9.358007559 | -0.163681049 | -0.044162614 |
| Vmn1r-ps47 | 2.046875553 | 2.112561602 | 2.09103895 | 0.021522652 | -0.044163398 |

| | | | | | |
|---|---|---|---|---|---|
| Zfp493 | 5.074583976 | 5.098663055 | 5.118817116 | -0.020154062 | -0.04423314 |
| Ppp6r3 | 11.06116421 | 11.08487106 | 11.10540725 | -0.020536183 | -0.044243033 |
| Olfr1276 | 2.141409626 | 2.116885839 | 2.185668535 | -0.068782695 | -0.044258909 |
| Rflna | 3.705217293 | 3.700991389 | 3.749560748 | -0.048569359 | -0.044343455 |
| Extl2 | 7.810790612 | 7.576713671 | 7.855148318 | -0.278434647 | -0.044357706 |
| Sdhaf3 | 6.747163497 | 6.605581667 | 6.791525661 | -0.185943994 | -0.044362164 |
| Dmc1 | 1.892528065 | 1.910462799 | 1.936898033 | -0.026435234 | -0.044369967 |
| 4930516K23Rik | 2.64104318 | 2.708726128 | 2.685427389 | 0.023298739 | -0.044384209 |
| A630010A05Rik | 3.719290318 | 3.831347567 | 3.763687285 | 0.067660282 | -0.044396967 |
| Prokr2 | 3.250127824 | 3.279709643 | 3.294529189 | -0.014819546 | -0.044401365 |
| Igfl3 | 4.280069956 | 4.324120962 | 4.324516262 | -0.000395299 | -0.044446306 |
| Raf1 | 11.32219786 | 11.31003926 | 11.3666684 | -0.056629137 | -0.044470533 |
| Slc28a1 | 4.556230617 | 4.755264668 | 4.600722135 | 0.154542533 | -0.044491518 |
| Ltbp4 | 11.10507363 | 11.15872347 | 11.14962447 | 0.009098993 | -0.044550847 |
| Vmn1r72 | 2.960579469 | 2.965623187 | 3.005133949 | -0.039510762 | -0.04455448 |
| Mzt1 | 9.379797496 | 9.349365456 | 9.424362084 | -0.074996628 | -0.044564587 |
| Zscan29 | 8.757988091 | 8.892125578 | 8.802561667 | 0.089563911 | -0.044573576 |
| Cyth2 | 9.634176998 | 9.617625961 | 9.678758526 | -0.061132564 | -0.044581528 |
| Rab15 | 8.153899981 | 8.071160114 | 8.19849207 | -0.127331956 | -0.044592089 |
| Car1 | 1.051146453 | 1.095005591 | 1.095749343 | -0.000743752 | -0.04460289 |
| Atp4a | 3.93739596 | 3.946940404 | 3.982023667 | -0.035083264 | -0.044627708 |
| Supt7l | 8.27281057 | 8.367623771 | 8.317468561 | 0.05015521 | -0.044657991 |
| Nub1 | 10.13053408 | 10.04654325 | 10.17523876 | -0.128695507 | -0.04470468 |
| Tbx5 | 1.704830587 | 1.662283919 | 1.74963687 | -0.087352951 | -0.044806284 |
| Chrna2 | 1.820240148 | 1.820933407 | 1.865058611 | -0.044125204 | -0.044818463 |
| Qdpr | 9.667714385 | 9.575416509 | 9.712565036 | -0.137148528 | -0.044850652 |
| Adh6-ps1 | 1.732382551 | 1.696338213 | 1.77726311 | -0.080924897 | -0.044880559 |
| Rbm12b2 | 8.571955164 | 8.6317669 | 8.61685829 | 0.01490861 | -0.044903126 |
| Vmn1r191 | 3.285946136 | 3.240286563 | 3.330885916 | -0.090599352 | -0.04493978 |
| 4930432K21Rik | 2.507640743 | 2.504257213 | 2.552724695 | -0.048467482 | -0.045083952 |
| Tmem236 | 2.747979333 | 2.957840515 | 2.79307222 | 0.164768295 | -0.045092887 |
| Mak | 1.473701891 | 1.45637892 | 1.518837371 | -0.062458451 | -0.04513548 |

| | | | | | |
|---|---|---|---|---|---|
| Vil1 | 1.162605631 | 1.251352775 | 1.207754411 | 0.043598364 | -0.04514878 |
| Zfp952 | 7.667878951 | 7.62143729 | 7.713097918 | -0.091660628 | -0.045218967 |
| Repin1 | 7.705857421 | 7.760624173 | 7.751089974 | 0.0095342 | -0.045232552 |
| Pfdn5 | 10.94143497 | 10.85102435 | 10.98673408 | -0.135709737 | -0.045299117 |
| Parl | 10.03502173 | 9.989511039 | 10.08033244 | -0.090821401 | -0.04531071 |
| Rnf139 | 9.593285716 | 9.431173614 | 9.638604321 | -0.207430708 | -0.045318605 |
| Onecut2 | 2.36561862 | 2.414818087 | 2.410941974 | 0.003876114 | -0.045323354 |
| Ticrr | 7.147105349 | 7.185944386 | 7.192466955 | -0.006522569 | -0.045361606 |
| Osbp2 | 4.415543318 | 4.360508442 | 4.46104428 | -0.100535837 | -0.045500961 |
| Coasy | 9.288117233 | 9.292062728 | 9.333635946 | -0.041573218 | -0.045518713 |
| Tecpr1 | 9.138075054 | 9.468867276 | 9.183594473 | 0.285272803 | -0.045519419 |
| Folh1 | 3.387540386 | 3.231262255 | 3.433063582 | -0.201801326 | -0.045523196 |
| E130208F15Rik | 3.047034545 | 2.96642282 | 3.092562193 | -0.126139373 | -0.045527648 |
| Dld | 11.05054237 | 10.95294269 | 11.09609043 | -0.143147743 | -0.045548067 |
| Slc6a12 | 3.290465979 | 3.330121917 | 3.336114025 | -0.005992108 | -0.045648046 |
| Wasf2 | 11.46069186 | 11.53999195 | 11.50638301 | 0.033608937 | -0.045691147 |
| Mrpl48 | 9.541441006 | 9.521979297 | 9.587169109 | -0.065189812 | -0.045728103 |
| Tada1 | 8.88788092 | 8.947870174 | 8.933613892 | 0.014256281 | -0.045732973 |
| Gm47191 | 3.054202398 | 3.026388806 | 3.099942127 | -0.073553321 | -0.045739729 |
| Nprl3 | 8.401240738 | 8.247313962 | 8.446992889 | -0.199678927 | -0.04575215 |
| Gm47145 | 1.673599804 | 1.738235711 | 1.719381564 | 0.018854147 | -0.04578176 |
| Olfr190 | 0.912312935 | 0.914902366 | 0.958110038 | -0.043207672 | -0.045797103 |
| Rims3 | 2.841958682 | 2.867199536 | 2.88776002 | -0.020560484 | -0.045801338 |
| Stag1 | 10.23662714 | 10.26245087 | 10.28245922 | -0.02000835 | -0.045832087 |
| Slc25a20 | 9.873592915 | 9.784328228 | 9.919474439 | -0.135146211 | -0.045881523 |
| Ccp110 | 7.794795322 | 7.840442432 | 7.840682854 | -0.000240422 | -0.045887533 |
| Fbln1 | 10.36978019 | 10.45002237 | 10.41567156 | 0.034350807 | -0.045891367 |
| Acot9 | 10.27071012 | 10.19399727 | 10.31661109 | -0.122613823 | -0.045900967 |
| Spx | 5.036396565 | 5.121746633 | 5.082328385 | 0.039418248 | -0.04593182 |
| Mmel1 | 1.843472347 | 1.903564114 | 1.889410067 | 0.014154047 | -0.04593772 |
| Flcn | 9.367453146 | 9.547246581 | 9.413393288 | 0.133853293 | -0.045940142 |
| Zfp708 | 5.775998105 | 5.676086606 | 5.821947074 | -0.145860467 | -0.045948969 |

| | | | | | |
|---|---|---|---|---|---|
| Htra4 | 4.543062732 | 4.587013745 | 4.589016665 | -0.00200292 | -0.045953933 |
| Irx5 | 8.221475404 | 8.450927493 | 8.267502186 | 0.183425307 | -0.046026782 |
| Pdcd5 | 8.739182072 | 8.770542555 | 8.785223598 | -0.014681043 | -0.046041526 |
| Xrcc2 | 6.683812663 | 6.844519512 | 6.729880901 | 0.114638611 | -0.046068237 |
| Vtcn1 | 3.701583525 | 3.639270633 | 3.747651972 | -0.108381339 | -0.046068447 |
| Gm7331 | 4.317192946 | 4.243807015 | 4.363267183 | -0.119460168 | -0.046074237 |
| Olfr1012 | 1.505240681 | 1.501466848 | 1.551341025 | -0.049874176 | -0.046100344 |
| H2-Ke6 | 8.801973173 | 8.716075436 | 8.848077131 | -0.132001695 | -0.046103958 |
| Spink1 | 2.443789324 | 2.485111747 | 2.489905276 | -0.004793529 | -0.046115952 |
| Stx1a | 5.911003542 | 6.091343912 | 5.95717238 | 0.134171533 | -0.046168838 |
| Rsrc2 | 10.50641793 | 10.53911764 | 10.55260105 | -0.013483407 | -0.04618312 |
| AC102437.1 | 1.766828711 | 1.743545293 | 1.813031509 | -0.069486216 | -0.046202799 |
| Afap1l1 | 8.438873323 | 8.413956757 | 8.485078571 | -0.071121814 | -0.046205248 |
| Ppt1 | 10.27048371 | 10.21042766 | 10.31673604 | -0.10630838 | -0.046252327 |
| Tmem17 | 5.309859791 | 5.242766685 | 5.356166706 | -0.113400021 | -0.046306916 |
| Otof | 2.487652088 | 2.480295019 | 2.533962644 | -0.053667625 | -0.046310555 |
| Mrpl27 | 8.915295353 | 8.840892563 | 8.961607296 | -0.120714733 | -0.046311943 |
| Zfp292 | 9.416247672 | 9.421395688 | 9.462640384 | -0.041244696 | -0.046392712 |
| Adcy10 | 2.596520124 | 2.544847309 | 2.642938049 | -0.09809074 | -0.046417925 |
| Ksr2 | 3.216379296 | 3.191691945 | 3.262832108 | -0.071140162 | -0.046452811 |
| Gm2792 | 4.04679891 | 4.23822105 | 4.093284682 | 0.144936368 | -0.046485772 |
| Dlx1 | 2.351513065 | 2.422920179 | 2.398007679 | 0.0249125 | -0.046494614 |
| Gm12819 | 1.512852872 | 1.519392685 | 1.559369224 | -0.039976539 | -0.046516351 |
| Crtap | 9.766629369 | 9.622328719 | 9.813188396 | -0.190859678 | -0.046559027 |
| Cenpa | 8.399204114 | 8.438397307 | 8.445776122 | -0.007378815 | -0.046572008 |
| Vmn1r236 | 3.061826666 | 3.123422761 | 3.108410089 | 0.015012672 | -0.046583423 |
| Atp2b2 | 3.636619686 | 3.619581588 | 3.683213485 | -0.063631897 | -0.046593799 |
| Igf2bp3 | 2.995242709 | 3.000305238 | 3.041885455 | -0.041580217 | -0.046642746 |
| Cd3g | 7.749965799 | 7.655824332 | 7.79665473 | -0.140830398 | -0.046688931 |
| Cog7 | 8.739110127 | 8.691364424 | 8.78588979 | -0.094525367 | -0.046779663 |
| Amy2a5 | 1.210729677 | 1.260503471 | 1.257542652 | 0.002960818 | -0.046812976 |
| Gtf2ird2 | 6.960422112 | 6.981657335 | 7.007237632 | -0.025580298 | -0.04681552 |

| | | | | | |
|---|---|---|---|---|---|
| Sh2d4b | 3.981175301 | 4.137323005 | 4.028059392 | 0.109263613 | -0.046884091 |
| Trio | 9.830417955 | 10.07694372 | 9.87733102 | 0.199612695 | -0.046913065 |
| Apopt1 | 8.251535136 | 8.218559588 | 8.298526874 | -0.079967286 | -0.046991738 |
| Efhc2 | 1.756024843 | 1.729286195 | 1.803017869 | -0.073731675 | -0.046993027 |
| Deptor | 11.48612431 | 11.28001424 | 11.5331175 | -0.25310326 | -0.04699319 |
| Mterf4 | 7.751025389 | 7.745648082 | 7.79806723 | -0.052419148 | -0.047041841 |
| Kcns1 | 2.970784772 | 2.988327545 | 3.017845493 | -0.029517948 | -0.04706072 |
| Atp6v1d | 10.60782365 | 10.59051958 | 10.65488515 | -0.06436557 | -0.047061503 |
| Pax7 | 2.487607259 | 2.511426268 | 2.534688572 | -0.023262304 | -0.047081313 |
| Tapbpl | 9.334124537 | 9.381412389 | 9.38125242 | 0.000159969 | -0.047127883 |
| Vwc2 | 1.185734912 | 1.170825963 | 1.232904884 | -0.06207892 | -0.047169972 |
| Htr1d | 3.487733403 | 3.565982186 | 3.534945593 | 0.031036593 | -0.04721219 |
| Insig1 | 10.83256581 | 11.09155848 | 10.87978234 | 0.21177614 | -0.047216523 |
| Ddt | 8.134307218 | 8.209898017 | 8.181560534 | 0.028337484 | -0.047253315 |
| Gm3365 | 3.269455912 | 3.309512677 | 3.316745242 | -0.007232565 | -0.04728933 |
| Cpeb1 | 8.511314561 | 8.729246912 | 8.558608768 | 0.170638144 | -0.047294207 |
| Chmp3 | 11.30146986 | 11.22875662 | 11.34889182 | -0.120135207 | -0.04742196 |
| Det1 | 7.791969075 | 7.879855694 | 7.839398878 | 0.040456816 | -0.047429803 |
| Stk32b | 3.422433611 | 3.373699747 | 3.469864737 | -0.09616499 | -0.047431125 |
| Endou | 10.6976636 | 10.54017514 | 10.74513379 | -0.20495865 | -0.047470183 |
| Car5a | 1.884531822 | 1.925380507 | 1.932014982 | -0.006634475 | -0.047483159 |
| Gxylt1 | 9.528233974 | 9.577681741 | 9.575746175 | 0.001935566 | -0.047512201 |
| 4931414P19Rik | 6.256474978 | 6.208676043 | 6.304017208 | -0.095341164 | -0.04754223 |
| Elk1 | 8.269371757 | 8.36756337 | 8.316937102 | 0.050626268 | -0.047565345 |
| Top3b | 9.487395491 | 9.478815056 | 9.53498013 | -0.056165074 | -0.047584639 |
| Dnajc4 | 7.855424078 | 7.787468137 | 7.903032137 | -0.115564 | -0.047608059 |
| Gm11032 | 2.018474328 | 2.061176798 | 2.066122087 | -0.004945289 | -0.047647759 |
| Szt2 | 9.164013907 | 9.915736571 | 9.211662516 | 0.704074055 | -0.047648609 |
| Olfr1204 | 1.57154594 | 1.566316914 | 1.619197985 | -0.052881071 | -0.047652045 |
| Tbc1d20 | 11.51851099 | 11.56600761 | 11.56616312 | -0.000155507 | -0.047652123 |
| Snx27 | 10.61092201 | 10.71276296 | 10.65857798 | 0.054184973 | -0.04765597 |
| Sox10 | 6.964635395 | 7.349632088 | 7.012300695 | 0.337331393 | -0.047665301 |

| | | | | | |
|---|---|---|---|---|---|
| Brcc3 | 8.905228733 | 8.731662672 | 8.952942813 | -0.221280141 | -0.04771408 |
| Pcdhb19 | 5.629506551 | 5.702342539 | 5.677232578 | 0.025109962 | -0.047726027 |
| Vmn1r17 | 1.41049709 | 1.460620202 | 1.458225616 | 0.002394586 | -0.047728526 |
| Ccr8 | 3.150714079 | 3.153255802 | 3.198467706 | -0.045211904 | -0.047753627 |
| Nfe2l3 | 8.666397436 | 8.621551693 | 8.714157051 | -0.092605358 | -0.047759616 |
| Sez6 | 2.829380856 | 2.841163302 | 2.877182543 | -0.036019242 | -0.047801688 |
| Rbpj-ps3 | 0.497290574 | 0.530032048 | 0.54509674 | -0.015064691 | -0.047806166 |
| Dhdds | 9.621244776 | 9.727480223 | 9.669116698 | 0.058363525 | -0.047871922 |
| Ciz1 | 9.616821742 | 9.855608579 | 9.664726739 | 0.190881841 | -0.047904996 |
| Commd6 | 8.869549775 | 8.766952351 | 8.91747463 | -0.15052228 | -0.047924855 |
| Fndc8 | 1.83685915 | 1.840539953 | 1.884812069 | -0.044272116 | -0.047952919 |
| Gpr173 | 4.160838409 | 4.249351879 | 4.208812127 | 0.040539752 | -0.047973718 |
| Clec11a | 6.464703632 | 6.582974927 | 6.512685808 | 0.070289119 | -0.047982176 |
| Nr1h3 | 8.106966864 | 8.252729592 | 8.154993325 | 0.097736267 | -0.048026461 |
| Vmn1r226 | 1.40503875 | 1.43740169 | 1.453071997 | -0.015670307 | -0.048033248 |
| Rnft2 | 5.005581547 | 5.152183724 | 5.053637162 | 0.098546563 | -0.048055615 |
| Atp5g2 | 6.741221274 | 6.83102158 | 6.789297271 | 0.041724309 | -0.048075997 |
| Lhpp | 9.806029437 | 9.773151206 | 9.854184666 | -0.08103346 | -0.048155229 |
| Tfb2m | 9.26730638 | 9.12449136 | 9.315476121 | -0.190984761 | -0.048169741 |
| Lrrn4 | 1.58791303 | 1.610375703 | 1.636102428 | -0.025726725 | -0.048189398 |
| Pcdh11x | 1.338081616 | 1.358191809 | 1.386296987 | -0.028105178 | -0.048215371 |
| Hps6 | 7.016429817 | 7.250964188 | 7.064691142 | 0.186273046 | -0.048261325 |
| Erlin2 | 10.38471559 | 10.34549337 | 10.43300005 | -0.087506687 | -0.048284467 |
| Mrpl1 | 8.437877108 | 8.370638697 | 8.486183567 | -0.11554487 | -0.048306459 |
| Clasrp | 8.940236008 | 9.379631665 | 8.988553674 | 0.39107799 | -0.048317667 |
| Rpl23 | 13.4696251 | 13.37914124 | 13.51796162 | -0.138820377 | -0.048336517 |
| Fcamr | 5.417181174 | 5.618778834 | 5.465562423 | 0.15321641 | -0.04838125 |
| Dnaaf3 | 5.848847384 | 5.708690022 | 5.897234945 | -0.188544923 | -0.048387561 |
| Vmn1r233 | 3.420113337 | 3.495415691 | 3.468520559 | 0.026895132 | -0.048407222 |
| Dscc1 | 5.728161237 | 5.773826663 | 5.776569179 | -0.002742516 | -0.048407942 |
| Bcl2 | 8.718495968 | 8.626822405 | 8.766980781 | -0.140158375 | -0.048484812 |
| Zfp868 | 7.989635019 | 7.94867711 | 8.038152692 | -0.089475582 | -0.048517673 |

| | | | | | |
|---|---|---|---|---|---|
| Asf1a | 8.076090501 | 7.978087192 | 8.124649676 | -0.146562484 | -0.048559176 |
| Tmem109 | 12.00174939 | 12.02603424 | 12.05032663 | -0.024292387 | -0.048577243 |
| Sowahb | 6.321401129 | 6.128758068 | 6.370027508 | -0.241269439 | -0.048626379 |
| Orc5 | 8.394694211 | 8.349211513 | 8.443352187 | -0.094140674 | -0.048657976 |
| Pih1d1 | 8.755457176 | 8.838045527 | 8.804232025 | 0.033813502 | -0.048774849 |
| 9030619P08Rik | 4.04966445 | 4.249491825 | 4.098461269 | 0.151030556 | -0.048796819 |
| Olfr912 | 2.932289745 | 2.975316743 | 2.981111719 | -0.005794976 | -0.048821974 |
| Sbk3 | 3.893679233 | 3.989125985 | 3.942536968 | 0.046589017 | -0.048857736 |
| Upf3a | 9.193391863 | 9.155704539 | 9.242260071 | -0.086555533 | -0.048868209 |
| Ttc21a | 3.691968973 | 3.802927356 | 3.740918732 | 0.062008624 | -0.048949759 |
| Spc24 | 6.961546881 | 7.292666497 | 7.010525994 | 0.282140503 | -0.048979113 |
| Vmn1r61 | 1.761577344 | 1.803061766 | 1.81060256 | -0.007540795 | -0.049025217 |
| Krtcap3 | 7.235115331 | 7.326966865 | 7.284196647 | 0.042770219 | -0.049081316 |
| Ndufb11 | 10.54442855 | 10.58995259 | 10.59353007 | -0.003577483 | -0.049101517 |
| Rgs2 | 9.350304846 | 9.319757236 | 9.3994522 | -0.079694964 | -0.049147354 |
| Cct6b | 2.115668878 | 2.095103062 | 2.164879833 | -0.069776771 | -0.049210955 |
| Dlk1 | 2.980506346 | 3.090719761 | 3.029724056 | 0.060995705 | -0.04921771 |
| Inava | 9.068113687 | 9.447748999 | 9.117350745 | 0.330398253 | -0.049237059 |
| Ptprk | 9.21317356 | 9.050453721 | 9.262467823 | -0.212014101 | -0.049294262 |
| Gm5786 | 2.402409591 | 2.357052453 | 2.45170411 | -0.094651657 | -0.049294519 |
| Olfr338 | 4.436923732 | 4.33290913 | 4.486239407 | -0.153330277 | -0.049315675 |
| Tex14 | 3.294541463 | 3.331925323 | 3.34389032 | -0.011964997 | -0.049348857 |
| 4930444P10Rik | 2.032831783 | 2.021316042 | 2.082206528 | -0.060890486 | -0.049374745 |
| Elovl5 | 12.03412282 | 11.95230028 | 12.08352235 | -0.131222073 | -0.049399533 |
| Gucy1a2 | 2.786656168 | 2.812243037 | 2.83606445 | -0.023821413 | -0.049408282 |
| Mettl23 | 8.735971171 | 8.731391791 | 8.785409337 | -0.054017545 | -0.049438166 |
| Tmf1 | 9.966207452 | 9.957262254 | 10.01566291 | -0.05840066 | -0.049455462 |
| Gm48349 | 2.462285213 | 2.495293502 | 2.51177974 | -0.016486238 | -0.049494527 |
| Vmn2r25 | 2.256063536 | 2.308453794 | 2.305590612 | 0.002863181 | -0.049527076 |
| Vps37d | 3.447017509 | 3.539693575 | 3.496565917 | 0.043127658 | -0.049548408 |
| Shld1 | 6.898537676 | 6.753918057 | 6.948091203 | -0.194173146 | -0.049553527 |
| Gm6296 | 2.281752589 | 2.314192189 | 2.331341069 | -0.01714888 | -0.04958848 |

| | | | | | |
|---|---|---|---|---|---|
| BC028528 | 6.267729168 | 6.192172117 | 6.317389012 | -0.125216895 | -0.049659843 |
| Crybg2 | 10.44078793 | 10.57764665 | 10.49045668 | 0.087189977 | -0.04966875 |
| Abca17 | 1.642640747 | 1.668221086 | 1.692325739 | -0.024104652 | -0.049684992 |
| Mocs2 | 10.31903354 | 10.26195215 | 10.36872531 | -0.10677316 | -0.04969177 |
| Tpsg1 | 5.05787613 | 4.966299081 | 5.107586127 | -0.141287046 | -0.049709997 |
| Ccdc142 | 2.159068511 | 2.245700158 | 2.20878922 | 0.036910938 | -0.04972071 |
| Ly6c1 | 11.207731 | 10.93174923 | 11.25749421 | -0.32574498 | -0.049763217 |
| Olfr635 | 5.807957372 | 5.754203814 | 5.857810207 | -0.103606393 | -0.049852835 |
| Sin3a | 10.19665975 | 10.14360532 | 10.24652503 | -0.102919707 | -0.049865273 |
| Vti1a | 9.532005704 | 9.58376166 | 9.581967476 | 0.001794185 | -0.049961772 |
| Mien1 | 9.244199575 | 9.253520342 | 9.294161498 | -0.040641156 | -0.049961923 |
| Cox15 | 8.891125054 | 8.929374471 | 8.941196181 | -0.01182171 | -0.050071127 |
| Ing1 | 9.085181373 | 9.196335973 | 9.135279827 | 0.061056146 | -0.050098454 |
| Ryr2 | 3.514788105 | 3.527194833 | 3.564976621 | -0.037781788 | -0.050188516 |
| Gckr | 1.713873175 | 1.824355206 | 1.764078329 | 0.060276876 | -0.050205155 |
| Rnf112 | 1.640929733 | 1.694571449 | 1.69118247 | 0.003388978 | -0.050252737 |
| Tmem238l | 3.225647427 | 3.252133977 | 3.275906289 | -0.023772312 | -0.050258862 |
| Enpp6 | 3.692228411 | 3.665919687 | 3.742499825 | -0.076580138 | -0.050271414 |
| Gm45233 | 2.521965471 | 2.649077354 | 2.572293487 | 0.076783867 | -0.050328016 |
| Vwde | 2.687833161 | 2.691765451 | 2.738217729 | -0.046452278 | -0.050384568 |
| Gm12013 | 0.792339207 | 0.780812489 | 0.842762452 | -0.061949963 | -0.050423244 |
| Pomt1 | 8.267504874 | 8.396565577 | 8.317934689 | 0.078630888 | -0.050429815 |
| Cdhr4 | 2.338563375 | 2.309560744 | 2.389035687 | -0.079474943 | -0.050472312 |
| Zdhhc4 | 8.722290409 | 8.799735495 | 8.772792365 | 0.02694313 | -0.050501956 |
| Ift172 | 8.825170587 | 8.974839531 | 8.87571803 | 0.0991215 | -0.050547443 |
| Rp2 | 8.65614885 | 8.612859514 | 8.706719001 | -0.093859487 | -0.050570151 |
| Minar1 | 1.312681122 | 1.38774425 | 1.363251891 | 0.024492359 | -0.050570769 |
| Fads2 | 7.072189121 | 6.985073292 | 7.122767949 | -0.137694657 | -0.050578828 |
| Olfr718-ps1 | 4.644354074 | 4.726388593 | 4.69493292 | 0.031455674 | -0.050578845 |
| Men1 | 7.937557132 | 8.156530055 | 7.988169386 | 0.168360669 | -0.050612254 |
| Odf3l2 | 4.300474189 | 4.033290517 | 4.351096855 | -0.317806338 | -0.050622667 |
| Iqcg | 3.816264886 | 3.731394831 | 3.866929335 | -0.135534504 | -0.050664449 |

| | | | | | |
|---|---|---|---|---|---|
| Gm5316 | 2.951476452 | 2.931955742 | 3.002145231 | -0.070189489 | -0.050668779 |
| Dtnb | 7.053098346 | 7.400290087 | 7.103868529 | 0.296421558 | -0.050770183 |
| Bmp8a | 1.873910454 | 1.89345008 | 1.924691602 | -0.031241522 | -0.050781148 |
| Ppil3 | 8.258930043 | 8.233481633 | 8.309739858 | -0.076258225 | -0.050809814 |
| Folr1 | 3.267502287 | 3.374646941 | 3.318370468 | 0.056276472 | -0.050868182 |
| Prr29 | 2.883597952 | 2.910725552 | 2.934468684 | -0.023743132 | -0.050870732 |
| Lingo4 | 2.879627417 | 3.007550188 | 2.930520515 | 0.077029673 | -0.050893098 |
| Slc22a21 | 6.390225222 | 6.504091953 | 6.441212013 | 0.06287994 | -0.050986792 |
| Klf14 | 3.493915387 | 3.48859025 | 3.544908778 | -0.056318527 | -0.05099339 |
| Hist1h2bg | 2.564414652 | 2.541764076 | 2.615414416 | -0.073650339 | -0.050999764 |
| Spock3 | 2.312734921 | 2.3335925 | 2.363753232 | -0.030160732 | -0.051018311 |
| Zfp770 | 7.912231694 | 7.838206758 | 7.963305533 | -0.125098775 | -0.051073839 |
| Aifm1 | 9.149471663 | 9.061539578 | 9.200560786 | -0.139021208 | -0.051089123 |
| A4gnt | 6.703831511 | 6.776402467 | 6.754950462 | 0.021452005 | -0.051118952 |
| Arid4b | 10.13853605 | 10.10791545 | 10.18967449 | -0.081759037 | -0.051138443 |
| Myl4 | 4.882990835 | 4.851673735 | 4.934151541 | -0.082477805 | -0.051160706 |
| G430095P16Rik | 2.146463767 | 2.171345925 | 2.197632818 | -0.026286893 | -0.051169051 |
| Gm8430 | 10.40281012 | 10.43085507 | 10.45399993 | -0.023144863 | -0.05118981 |
| Unc5d | 2.786674944 | 2.870426929 | 2.837902979 | 0.03252395 | -0.051228035 |
| Hdhd3 | 6.240460026 | 6.446197378 | 6.29173411 | 0.154463268 | -0.051274084 |
| Fam166b | 3.712861131 | 3.739268777 | 3.764147098 | -0.024878321 | -0.051285967 |
| Adam28 | 2.634180697 | 2.613844544 | 2.685481814 | -0.071637271 | -0.051301117 |
| Akr7a5 | 8.056402606 | 8.288991685 | 8.107708881 | 0.181282804 | -0.051306275 |
| Gm6981 | 2.060548404 | 2.035641926 | 2.111872681 | -0.076230755 | -0.051324277 |
| Gm20541 | 2.763928803 | 2.75929061 | 2.815293212 | -0.056002602 | -0.051364409 |
| Rsph3a | 7.566036607 | 7.565519333 | 7.617431601 | -0.051912268 | -0.051394994 |
| Lsr | 10.74503717 | 10.85993692 | 10.79649806 | 0.06343886 | -0.051460883 |
| Zfp644 | 9.312534763 | 9.194271313 | 9.364024159 | -0.169752846 | -0.051489396 |
| Pradc1 | 7.488908522 | 7.514759051 | 7.540401215 | -0.025642164 | -0.051492693 |
| Hemk1 | 6.533341299 | 6.659418401 | 6.584840814 | 0.074577587 | -0.051499515 |
| Gbx1 | 2.932373743 | 2.973841028 | 2.983903093 | -0.010062066 | -0.05152935 |
| Gm15009 | 2.808258185 | 2.840738845 | 2.859792862 | -0.019054017 | -0.051534677 |

| | | | | | |
|---|---|---|---|---|---|
| Def8 | 10.40019987 | 10.44141368 | 10.45183265 | -0.010418977 | -0.051632787 |
| Olfr1489 | 3.402263634 | 3.37239753 | 3.453947898 | -0.081550368 | -0.051684264 |
| Mmp14 | 11.50327134 | 11.60176067 | 11.5549685 | 0.046792177 | -0.05169716 |
| Plb1 | 7.410424939 | 7.618120455 | 7.4621554 | 0.155965055 | -0.05173046 |
| Lamtor2 | 9.468393578 | 9.420435418 | 9.520150028 | -0.09971461 | -0.05175645 |
| Snx25 | 8.216087191 | 8.22771323 | 8.267891773 | -0.040178542 | -0.051804581 |
| Irgq | 9.423880665 | 9.550991479 | 9.475687718 | 0.075303761 | -0.051807053 |
| Galnt14 | 4.232390576 | 4.172683931 | 4.28420307 | -0.111519139 | -0.051812493 |
| Vmn1r234 | 4.046281215 | 4.104192628 | 4.098121063 | 0.006071565 | -0.051839848 |
| Slc3a1 | 1.936280361 | 1.958446788 | 1.988189223 | -0.029742435 | -0.051908862 |
| Eva1a | 6.826804982 | 7.002087409 | 6.878719858 | 0.123367551 | -0.051914876 |
| Lce1b | 11.66551591 | 11.91682538 | 11.71744769 | 0.19937769 | -0.051931777 |
| Nmnat2 | 4.160744837 | 4.254806041 | 4.212679791 | 0.042126251 | -0.051934954 |
| Faf2 | 10.46272954 | 10.50537144 | 10.5146795 | -0.009308063 | -0.051949967 |
| Zfp689 | 5.672598975 | 5.76250324 | 5.724569311 | 0.037933929 | -0.051970336 |
| Nlgn1 | 1.501098614 | 1.533650993 | 1.553096357 | -0.019445364 | -0.051997742 |
| 4930404N11Rik | 1.376340073 | 1.416624498 | 1.428403477 | -0.011778979 | -0.052063403 |
| Cipc | 10.2589095 | 10.41762006 | 10.31101023 | 0.10660983 | -0.05210073 |
| Myo15 | 2.538149115 | 2.575921362 | 2.590253702 | -0.01433234 | -0.052104587 |
| Mettl5 | 7.601666707 | 7.499188427 | 7.653778849 | -0.154590422 | -0.052112143 |
| Plekha2 | 10.01811362 | 9.815246316 | 10.07028892 | -0.2550426 | -0.0521753 |
| Olfr735 | 4.49756598 | 4.507640428 | 4.549770206 | -0.042129778 | -0.052204226 |
| Cnnm4 | 8.161917534 | 8.293495662 | 8.214124349 | 0.079371313 | -0.052206815 |
| Igsf1 | 1.533925813 | 1.55152567 | 1.586142888 | -0.034617218 | -0.052217075 |
| Setd2 | 10.65026544 | 10.69766052 | 10.70250465 | -0.004844133 | -0.052239213 |
| Xlr3a | 3.375709211 | 3.250464974 | 3.428000646 | -0.177535672 | -0.052291435 |
| Akp-ps1 | 3.20411923 | 3.330021503 | 3.256412208 | 0.073609295 | -0.052292978 |
| Gm49378 | 3.699554182 | 3.382510596 | 3.751882808 | -0.369372212 | -0.052328626 |
| Ccne1 | 6.949974575 | 7.195967843 | 7.002303268 | 0.193664575 | -0.052328693 |
| 4921507P07Rik | 2.811518139 | 2.85499481 | 2.86386668 | -0.00887187 | -0.052348541 |
| Gm45380 | 2.763812977 | 2.777839013 | 2.816177176 | -0.038338163 | -0.052364199 |
| Slco3a1 | 11.89065777 | 11.94363718 | 11.94302398 | 0.0006132 | -0.052366203 |

| | | | | | |
|---|---|---|---|---|---|
| Gba | 10.7483495 | 10.7725334 | 10.80073329 | -0.028199893 | -0.05238379 |
| Vmn2r102 | 4.345485264 | 4.476995369 | 4.397913777 | 0.079081592 | -0.052428512 |
| Syne2 | 11.73738093 | 11.82444415 | 11.78984333 | 0.03460082 | -0.052462403 |
| Dcaf12 | 10.53642471 | 10.52862343 | 10.58890287 | -0.060279437 | -0.052478163 |
| Ppp2r2b | 3.029557591 | 3.012304901 | 3.082048327 | -0.069743426 | -0.052490736 |
| Idua | 7.951395669 | 8.049930825 | 8.003915396 | 0.046015429 | -0.052519727 |
| Ak8 | 3.487827377 | 3.529632582 | 3.540366717 | -0.010734136 | -0.05253934 |
| Gm13394 | 2.270738729 | 2.357638523 | 2.323305153 | 0.034333371 | -0.052566423 |
| Rgs9 | 5.548628684 | 5.628839044 | 5.601204874 | 0.02763417 | -0.05257619 |
| Slc27a4 | 12.24671328 | 12.19249807 | 12.29930284 | -0.106804767 | -0.052589557 |
| Hoxb7 | 6.615780339 | 6.686362483 | 6.668519548 | 0.017842936 | -0.052739208 |
| Gm14760 | 3.120302054 | 3.228521464 | 3.173050583 | 0.055470881 | -0.052748529 |
| Chmp2b | 10.6835093 | 10.63150588 | 10.73628979 | -0.104783913 | -0.052780487 |
| Zfp777 | 8.602507063 | 8.482041154 | 8.655317939 | -0.173276785 | -0.052810876 |
| Asphd2 | 3.757188118 | 3.784320007 | 3.810001409 | -0.025681401 | -0.052813291 |
| Susd5 | 4.04602474 | 4.081240579 | 4.098845433 | -0.017604854 | -0.052820693 |
| Tnni1 | 2.75065232 | 2.762476862 | 2.803483353 | -0.041006491 | -0.052831033 |
| Coq3 | 8.56745355 | 8.474149958 | 8.620287396 | -0.146137438 | -0.052833846 |
| Gm3417 | 1.729565329 | 1.856858412 | 1.782403085 | 0.074455327 | -0.052837756 |
| Gm9294 | 2.416845598 | 2.38181662 | 2.469736136 | -0.087919517 | -0.052890538 |
| Tmem132a | 8.297301892 | 8.599137546 | 8.350231054 | 0.248906492 | -0.052929162 |
| Atad2b | 9.790159952 | 9.76770793 | 9.843135886 | -0.075427956 | -0.052975934 |
| Cbwd1 | 7.900740434 | 7.744891181 | 7.953779589 | -0.208888408 | -0.053039155 |
| Gcat | 9.973286036 | 10.07510408 | 10.026327 | 0.048777077 | -0.053040967 |
| Unk | 8.088216381 | 8.104436129 | 8.141292574 | -0.036856446 | -0.053076194 |
| Vmn2r114 | 6.388982946 | 6.458516792 | 6.442071597 | 0.016445195 | -0.053088651 |
| L3mbtl4 | 2.475084254 | 2.491715887 | 2.528231734 | -0.036515847 | -0.05314748 |
| Aqp1 | 11.49017082 | 11.22424068 | 11.5433307 | -0.31909002 | -0.053159877 |
| Gm47605 | 0.814734607 | 0.862494362 | 0.867948967 | -0.005454605 | -0.05321436 |
| Kdsr | 10.18098922 | 10.06945009 | 10.23422214 | -0.16477205 | -0.05323292 |
| Morc1 | 2.154162945 | 2.15152897 | 2.207422991 | -0.05589402 | -0.053260045 |
| Esyt2 | 11.16463895 | 11.07637753 | 11.21796937 | -0.141591837 | -0.053330413 |

| | | | | | |
|---|---|---|---|---|---|
| Paip1 | 10.14962925 | 10.14649848 | 10.20301648 | -0.056518003 | -0.053387237 |
| Rft1 | 8.141477826 | 7.971864739 | 8.194871023 | -0.223006284 | -0.053393197 |
| Pfdn1 | 9.755953124 | 9.765316646 | 9.80937494 | -0.044058294 | -0.053421816 |
| Cited4 | 7.842032307 | 7.937902789 | 7.895551096 | 0.042351693 | -0.053518789 |
| Nfasc | 6.460365864 | 6.800928008 | 6.513907786 | 0.287020222 | -0.053541922 |
| Idh3a | 11.16358421 | 11.04579733 | 11.21718027 | -0.17138294 | -0.05359606 |
| Slc25a17 | 10.50875218 | 10.44575282 | 10.56245851 | -0.116705693 | -0.05370633 |
| Gale | 8.515343349 | 8.729681001 | 8.569099127 | 0.160581874 | -0.053755778 |
| Sepsecs | 8.898696534 | 8.845250859 | 8.952490876 | -0.107240017 | -0.053794342 |
| Gm6542 | 1.673445219 | 1.703770751 | 1.727256721 | -0.023485969 | -0.053811502 |
| 4930519F16Rik | 1.809140424 | 1.802273703 | 1.863007107 | -0.060733404 | -0.053866682 |
| Stard3nl | 8.24619287 | 8.20591309 | 8.300083676 | -0.094170586 | -0.053890806 |
| Erbb4 | 2.007045585 | 2.015664949 | 2.060942555 | -0.045277606 | -0.05389697 |
| Gm14435 | 4.33129835 | 4.413438691 | 4.385227425 | 0.028211266 | -0.053929075 |
| Arpp21 | 5.533317973 | 5.402694903 | 5.587259734 | -0.184564831 | -0.053941761 |
| Pex6 | 9.243249082 | 9.57742122 | 9.297241807 | 0.280179413 | -0.053992725 |
| Cxcr3 | 4.328560298 | 4.287026287 | 4.382558082 | -0.095531795 | -0.053997784 |
| Snph | 2.531788279 | 2.681904202 | 2.585873484 | 0.096030718 | -0.054085205 |
| Tmcc1 | 9.890019774 | 9.987252793 | 9.944139852 | 0.043112941 | -0.054120078 |
| Gm48260 | 4.114256752 | 4.14303867 | 4.168418222 | -0.025379552 | -0.05416147 |
| Mkrn3 | 3.663998647 | 3.628869117 | 3.718182239 | -0.089313122 | -0.054183592 |
| B330016D10Rik | 4.561488809 | 4.647948552 | 4.615765715 | 0.032182838 | -0.054276905 |
| Dnajb4 | 10.1791902 | 10.60618105 | 10.23360859 | 0.37257246 | -0.05441839 |
| Gm28802 | 3.469853869 | 3.436052089 | 3.524272271 | -0.088220182 | -0.054418401 |
| Acad10 | 7.017663452 | 7.190302666 | 7.072115948 | 0.118186718 | -0.054452496 |
| Olfr117 | 2.399300191 | 2.357507753 | 2.453789676 | -0.096281923 | -0.054489485 |
| Ngly1 | 9.712988645 | 9.599517796 | 9.767479964 | -0.167962168 | -0.054491318 |
| Hspb11 | 6.477903456 | 6.459310942 | 6.532505832 | -0.07319489 | -0.054602375 |
| Churc1 | 8.496017576 | 8.362341413 | 8.550651175 | -0.188309762 | -0.054633598 |
| Zswim1 | 8.484171282 | 8.634700811 | 8.538843376 | 0.095857435 | -0.054672094 |
| Atp8b5 | 5.134255151 | 5.022677229 | 5.188932372 | -0.166255143 | -0.054677221 |
| Cxxc5 | 7.695771886 | 7.685341636 | 7.75054685 | -0.065205215 | -0.054774965 |

| | | | | | |
|---|---|---|---|---|---|
| 1500009L16Rik | 3.672873297 | 3.692770049 | 3.727651207 | -0.034881157 | -0.05477791 |
| Tmub1 | 6.89705997 | 7.084995823 | 6.951860987 | 0.133134837 | -0.054801016 |
| Chrne | 4.001970401 | 3.988356478 | 4.056782979 | -0.068426501 | -0.054812578 |
| Hacd4 | 8.008722738 | 7.636692588 | 8.063536318 | -0.426843729 | -0.054813579 |
| Abcb6 | 8.152833268 | 8.215193588 | 8.207673523 | 0.007520065 | -0.054840255 |
| Gm42718 | 3.689635224 | 3.732083896 | 3.744483541 | -0.012399644 | -0.054848316 |
| Mrpl34 | 7.215226935 | 7.403158338 | 7.270101144 | 0.133057194 | -0.054874209 |
| Gm11400 | 3.770252067 | 3.857034784 | 3.825126604 | 0.031908179 | -0.054874537 |
| Jmjd7 | 5.887774814 | 6.041737806 | 5.942688381 | 0.099049425 | -0.054913568 |
| Clk3 | 10.35302355 | 10.38742106 | 10.40802837 | -0.02060731 | -0.05500482 |
| Ndufb5 | 10.19271826 | 10.18491672 | 10.24773718 | -0.062820453 | -0.055018917 |
| Plekhd1 | 2.332860964 | 2.352831898 | 2.387897719 | -0.035065821 | -0.055036756 |
| Leng8 | 11.11443855 | 11.55669172 | 11.16955655 | 0.387135167 | -0.055118007 |
| Lig3 | 10.26239653 | 10.31691006 | 10.31756336 | -0.0006533 | -0.05516683 |
| Fam71e1 | 2.303245798 | 2.303530589 | 2.358414245 | -0.054883656 | -0.055168446 |
| Trabd2b | 9.264550378 | 9.102464533 | 9.319719888 | -0.217255354 | -0.05516951 |
| Zfp280b | 7.970926396 | 8.034719573 | 8.026124068 | 0.008595505 | -0.055197672 |
| Zscan30 | 2.499212666 | 2.548062301 | 2.554457977 | -0.006395676 | -0.055245311 |
| Psg23 | 3.693060802 | 3.710910522 | 3.748389345 | -0.037478823 | -0.055328543 |
| Pianp | 6.402689354 | 7.097407572 | 6.458068472 | 0.6393391 | -0.055379118 |
| Ccdc39 | 2.295980193 | 2.333177512 | 2.351391769 | -0.018214257 | -0.055411576 |
| Vmn2r116 | 2.447940308 | 2.431623095 | 2.503417722 | -0.071794627 | -0.055477414 |
| Rpl7 | 13.96329193 | 13.93087003 | 14.0187975 | -0.087927477 | -0.055505573 |
| Olfr111 | 3.374584601 | 3.500680802 | 3.430096989 | 0.070583813 | -0.055512388 |
| Tmem47 | 8.628890013 | 8.44017451 | 8.68443195 | -0.24425744 | -0.055541937 |
| Fam219b | 9.174747744 | 9.096796456 | 9.230345688 | -0.133549232 | -0.055597945 |
| Vmn1r87 | 5.358409538 | 5.40222057 | 5.414054922 | -0.011834352 | -0.055645384 |
| Gm20040 | 3.781806845 | 3.86806647 | 3.83746453 | 0.03060194 | -0.055657685 |
| Slc39a9 | 10.40770465 | 10.42791031 | 10.46340008 | -0.035489763 | -0.05569543 |
| Atpaf2 | 8.97323613 | 8.935469537 | 9.028964082 | -0.093494545 | -0.055727953 |
| Fam205a3 | 2.090413921 | 2.303068873 | 2.146226005 | 0.156842868 | -0.055812084 |
| Slc25a43 | 4.851019294 | 4.803908725 | 4.906837687 | -0.102928961 | -0.055818393 |

| | | | | | |
|---|---|---|---|---|---|
| Robo4 | 7.519214125 | 7.534294225 | 7.575044565 | -0.04075034 | -0.05583044 |
| Gm18336 | 3.059160677 | 3.093713284 | 3.114996179 | -0.021282895 | -0.055835502 |
| Emsy | 9.382598203 | 9.702007721 | 9.43843691 | 0.263570811 | -0.055838707 |
| Gm17909 | 0.939717744 | 0.973769119 | 0.995562326 | -0.021793207 | -0.055844581 |
| Rab24 | 9.917675955 | 9.830357289 | 9.973540901 | -0.143183612 | -0.055864946 |
| Mansc1 | 7.74807097 | 7.612169839 | 7.803967116 | -0.191797277 | -0.055896146 |
| Zfp558 | 3.92573993 | 3.974939417 | 3.981642978 | -0.006703561 | -0.055903048 |
| Myo15b | 2.363704363 | 2.424496865 | 2.419627704 | 0.00486916 | -0.055923341 |
| Obox8 | 2.106076525 | 2.117270336 | 2.162059168 | -0.044788833 | -0.055982643 |
| Cntrl | 9.421455692 | 9.502356968 | 9.477451349 | 0.024905619 | -0.055995657 |
| Dach2 | 2.6826691 | 2.647562781 | 2.73870052 | -0.091137739 | -0.05603142 |
| Olfr552 | 5.441148476 | 5.440452709 | 5.49719403 | -0.056741321 | -0.056045554 |
| Cebpzos | 7.917617898 | 7.785665134 | 7.973674735 | -0.188009601 | -0.056056837 |
| Gdf9 | 3.736062388 | 4.032112138 | 3.792130225 | 0.239981913 | -0.056067837 |
| Plbd2 | 10.16967377 | 10.30318382 | 10.22574797 | 0.077435853 | -0.056074197 |
| Cd19 | 2.825056795 | 2.90316269 | 2.881132812 | 0.022029877 | -0.056076017 |
| Vps26b | 10.27677037 | 10.28507626 | 10.33286732 | -0.047791057 | -0.05609695 |
| Cyp4f39 | 10.73264949 | 10.79027819 | 10.78874765 | 0.001530543 | -0.056098163 |
| Isca1 | 10.28790261 | 10.28500007 | 10.34401407 | -0.059013993 | -0.056111457 |
| Styxl1 | 1.813808966 | 1.815903136 | 1.869942801 | -0.054039665 | -0.056133835 |
| Papolb | 1.531160975 | 1.586223236 | 1.587302054 | -0.001078818 | -0.056141079 |
| Ces2e | 3.872891757 | 3.937996796 | 3.929034405 | 0.008962391 | -0.056142648 |
| Chd2 | 10.50300872 | 10.49067182 | 10.55915652 | -0.0684847 | -0.0561478 |
| Cabp1 | 3.522083095 | 3.624216729 | 3.578254405 | 0.045962323 | -0.056171311 |
| Fgf1 | 8.642867974 | 8.508062238 | 8.699051695 | -0.190989457 | -0.056183721 |
| Arl2 | 8.057378696 | 8.263446733 | 8.113730023 | 0.14971671 | -0.056351327 |
| Gm37601 | 2.704560292 | 2.743995106 | 2.760923269 | -0.016928163 | -0.056362978 |
| Gpatch1 | 8.205651949 | 8.231717535 | 8.262018215 | -0.03030068 | -0.056366266 |
| Tdrd1 | 3.082792225 | 3.101162485 | 3.13920139 | -0.038038905 | -0.056409165 |
| Veph1 | 2.970873531 | 2.972618655 | 3.02731611 | -0.054697455 | -0.056442579 |
| Zfp426 | 8.547127493 | 8.558244845 | 8.603598307 | -0.045353462 | -0.056470814 |
| Unc13c | 1.898789188 | 1.966376583 | 1.95534019 | 0.011036393 | -0.056551003 |

| | | | | | |
|---|---|---|---|---|---|
| Gm9194 | 1.841524302 | 1.833507835 | 1.898102159 | -0.064594324 | -0.056577857 |
| Dnpep | 9.850315387 | 10.0302305 | 9.906897042 | 0.123333454 | -0.056581656 |
| Cds2 | 10.66387986 | 10.60380014 | 10.72046399 | -0.116663857 | -0.056584133 |
| Prkar2a | 11.49270501 | 11.37260916 | 11.5493022 | -0.176693037 | -0.056597193 |
| Telo2 | 8.358487201 | 8.67196829 | 8.415090197 | 0.256878093 | -0.056602997 |
| Sp5 | 2.386002308 | 2.429548675 | 2.442610182 | -0.013061507 | -0.056607874 |
| Spg11 | 9.371256899 | 9.484545433 | 9.427896625 | 0.056648808 | -0.056639726 |
| Chm | 9.2239833 | 9.230790408 | 9.280732021 | -0.049941613 | -0.056748721 |
| Fat3 | 2.886978874 | 2.929855667 | 2.943729183 | -0.013873516 | -0.056750309 |
| Defb14 | 6.499937025 | 6.486978715 | 6.556724294 | -0.069745579 | -0.056787269 |
| Pou3f3 | 1.252195958 | 1.275376412 | 1.309007263 | -0.033630851 | -0.056811305 |
| Dsg4 | 4.863775244 | 4.931865204 | 4.920636417 | 0.011228787 | -0.056861173 |
| Tll1 | 6.71842079 | 6.662205298 | 6.775313876 | -0.113108579 | -0.056893086 |
| AI182371 | 3.499156017 | 3.52696781 | 3.556051912 | -0.029084102 | -0.056895895 |
| Ogdh | 12.60694423 | 12.56562277 | 12.66384789 | -0.098225123 | -0.056903663 |
| Olfr898 | 2.965330153 | 2.975482068 | 3.022237758 | -0.04675569 | -0.056907605 |
| Vipas39 | 9.308852877 | 9.371489635 | 9.365773375 | 0.00571626 | -0.056920498 |
| Tada3 | 8.196300718 | 8.523596656 | 8.253284058 | 0.270312598 | -0.05698334 |
| Nupl2 | 7.064424303 | 7.094527461 | 7.121419755 | -0.026892294 | -0.056995452 |
| Zfp346 | 8.453687011 | 8.439390355 | 8.510714892 | -0.071324537 | -0.057027881 |
| Plpp1 | 8.966836274 | 8.814697534 | 9.023888885 | -0.209191351 | -0.057052611 |
| Gm14443 | 5.961849247 | 6.028429313 | 6.018918348 | 0.009510965 | -0.057069102 |
| Usp44 | 1.921281545 | 1.95689471 | 1.978402958 | -0.021508248 | -0.057121413 |
| Mzb1 | 2.861450079 | 2.923752688 | 2.918587317 | 0.005165371 | -0.057137238 |
| Zfp69 | 3.613425152 | 3.66348719 | 3.670584596 | -0.007097406 | -0.057159444 |
| Gm33707 | 2.127274878 | 2.177992015 | 2.184481084 | -0.006489069 | -0.057206206 |
| Cetn4 | 4.526583219 | 4.608259687 | 4.583803818 | 0.024455869 | -0.057220599 |
| Nr5a2 | 3.477913768 | 3.509681721 | 3.535167563 | -0.025485842 | -0.057253794 |
| Mmp10 | 4.210644316 | 4.443412212 | 4.267957999 | 0.175454213 | -0.057313683 |
| Ttc25 | 4.429904982 | 4.466673827 | 4.4872191 | -0.020545273 | -0.057314118 |
| Zfp992 | 7.017753217 | 7.065694407 | 7.075094473 | -0.009400067 | -0.057341257 |
| Helz | 9.451765513 | 9.461455151 | 9.509119932 | -0.047664782 | -0.057354419 |

| | | | | | |
|---|---|---|---|---|---|
| Zmynd15 | 5.684199832 | 5.883934126 | 5.741559063 | 0.142375063 | -0.057359231 |
| Epg5 | 8.620198897 | 8.660035462 | 8.677559415 | -0.017523953 | -0.057360518 |
| Igfbp6 | 9.306196754 | 9.357585812 | 9.363560513 | -0.005974701 | -0.057363759 |
| Gm49327 | 4.462492142 | 4.523350667 | 4.519885017 | 0.00346565 | -0.057392875 |
| Rsg1 | 4.072585144 | 4.035364967 | 4.130018248 | -0.094653281 | -0.057433105 |
| Dusp26 | 6.960554408 | 6.819843982 | 7.01802773 | -0.198183748 | -0.057473322 |
| Capn12 | 4.760480258 | 4.999548972 | 4.817970331 | 0.181578641 | -0.057490073 |
| Magi3 | 7.639746751 | 7.573974343 | 7.697277487 | -0.123303144 | -0.057530735 |
| Armcx5 | 7.379666515 | 7.222310687 | 7.437200242 | -0.214889554 | -0.057533727 |
| Dcun1d4 | 8.118363494 | 8.16491896 | 8.175924825 | -0.011005864 | -0.057561331 |
| Tab1 | 8.376029251 | 8.457639163 | 8.433595504 | 0.024043659 | -0.057566252 |
| Slc1a6 | 1.4913014 | 1.522631077 | 1.548874178 | -0.026243101 | -0.057572778 |
| Mreg | 10.55121675 | 10.66522395 | 10.60880233 | 0.05642162 | -0.057585587 |
| Gm48836 | 4.568939175 | 4.568684534 | 4.626528244 | -0.05784371 | -0.057589069 |
| Pard6a | 5.988541025 | 6.129951451 | 6.046142063 | 0.083809389 | -0.057601038 |
| Chmp6 | 9.006371014 | 8.873926885 | 9.064118027 | -0.190191142 | -0.057747013 |
| Fmc1 | 7.135873448 | 6.980835876 | 7.193632133 | -0.212796257 | -0.057758685 |
| A930004D18Rik | 5.944890574 | 6.225130952 | 6.002668955 | 0.222461997 | -0.057778381 |
| Ergic3 | 10.39303752 | 10.40925417 | 10.45083272 | -0.041578547 | -0.057795203 |
| Wscd2 | 6.221475966 | 6.27381556 | 6.279282975 | -0.005467415 | -0.057807009 |
| Snai2 | 8.873629037 | 8.983558759 | 8.93145231 | 0.052106449 | -0.057823273 |
| Ppp1r16a | 8.489827438 | 8.751383311 | 8.547664312 | 0.203718998 | -0.057836874 |
| Rnf38 | 10.29367257 | 10.26276581 | 10.35152495 | -0.088759137 | -0.057852383 |
| Zfp853 | 2.25361948 | 2.247841908 | 2.311520816 | -0.063678908 | -0.057901335 |
| Gm13241 | 1.674265838 | 1.704762202 | 1.732195314 | -0.027433112 | -0.057929475 |
| Ipo8 | 10.12182882 | 10.19135742 | 10.17982072 | 0.011536693 | -0.0579919 |
| Lrrc36 | 2.124815987 | 2.175536988 | 2.182875416 | -0.007338428 | -0.05805943 |
| 2610507B11Rik | 12.32866585 | 12.17878571 | 12.38679444 | -0.208008737 | -0.05812859 |
| Olfr922 | 2.097641159 | 2.128684698 | 2.15577397 | -0.027089271 | -0.058132811 |
| Hist1h1d | 3.341154701 | 3.349593971 | 3.399317376 | -0.049723405 | -0.058162675 |
| 6820431F20Rik | 11.53101153 | 11.54105272 | 11.58922949 | -0.048176767 | -0.058217963 |
| Edn3 | 4.253785617 | 4.260948726 | 4.312054329 | -0.051105603 | -0.058268712 |

| | | | | | |
|---|---|---|---|---|---|
| Cacnb4 | 2.883329108 | 2.978527286 | 2.941629908 | 0.036897378 | -0.0583008 |
| Rgsl1 | 3.800476725 | 3.755709813 | 3.858784374 | -0.103074561 | -0.058307649 |
| Mrln | 5.787341185 | 5.704795281 | 5.845649249 | -0.140853968 | -0.058308064 |
| Uhrf1bp1 | 9.183784864 | 9.211232204 | 9.242157664 | -0.03092546 | -0.0583728 |
| Nudt17 | 9.141359983 | 9.18165015 | 9.199857179 | -0.01820703 | -0.058497196 |
| Ctcfl | 2.437925956 | 2.448678893 | 2.496440518 | -0.047761625 | -0.058514562 |
| Olfr1378 | 2.72764194 | 2.744265988 | 2.786182725 | -0.041916737 | -0.058540784 |
| Cetn3 | 9.788812835 | 9.808246996 | 9.847378452 | -0.039131457 | -0.058565618 |
| Ptpra | 11.01983372 | 10.98720559 | 11.07844423 | -0.091238637 | -0.058610507 |
| Nanog | 2.54450605 | 2.556990231 | 2.603157071 | -0.04616684 | -0.058651021 |
| Olfr11 | 1.964487129 | 1.996024304 | 2.023204146 | -0.027179842 | -0.058717017 |
| Zfp239 | 4.002957808 | 4.073560505 | 4.061675947 | 0.011884557 | -0.05871814 |
| C2cd3 | 8.773133228 | 8.557934915 | 8.831868956 | -0.273934042 | -0.058735729 |
| Wdr41 | 9.198488538 | 9.229838355 | 9.257235815 | -0.02739746 | -0.058747277 |
| Mogat1 | 3.976233621 | 4.139880644 | 4.03505682 | 0.104823824 | -0.058823199 |
| Ethe1 | 7.8036793 | 7.962272912 | 7.862612179 | 0.099660732 | -0.058932879 |
| Usp29 | 3.462346675 | 3.509851468 | 3.521288656 | -0.011437188 | -0.058941981 |
| Arntl | 8.515973884 | 8.852118292 | 8.574928854 | 0.277189438 | -0.058954971 |
| C730034F03Rik | 4.307010581 | 4.424913032 | 4.365972047 | 0.058940985 | -0.058961466 |
| Slc6a18 | 3.071493133 | 3.116216969 | 3.130485887 | -0.014268919 | -0.058992754 |
| Csnk2a1 | 11.71087868 | 11.73584957 | 11.76988268 | -0.034033107 | -0.059003997 |
| Ap1g2 | 9.507727865 | 9.554837399 | 9.566757411 | -0.011920012 | -0.059029546 |
| Stk38l | 9.362503628 | 9.377241798 | 9.42153583 | -0.044294032 | -0.059032201 |
| B130006D01Rik | 3.086904729 | 3.124075035 | 3.146010268 | -0.021935233 | -0.059105539 |
| Lztr1 | 10.05467604 | 9.977525419 | 10.11378809 | -0.136262668 | -0.059112044 |
| Gm13244 | 2.137014721 | 2.046928029 | 2.196154116 | -0.149226088 | -0.059139395 |
| Msln | 3.329680972 | 3.452244122 | 3.388842395 | 0.063401727 | -0.059161424 |
| Alx4 | 4.993412749 | 4.78241585 | 5.052585117 | -0.270169267 | -0.059172367 |
| Pxmp4 | 10.8084838 | 10.81207965 | 10.86769465 | -0.055615003 | -0.05921085 |
| Chmp1b | 10.19484172 | 10.18933207 | 10.25406101 | -0.064728937 | -0.059219287 |
| Atg9a | 10.25727967 | 10.52024465 | 10.31652217 | 0.203722477 | -0.059242497 |
| Catsperg2 | 2.225035432 | 2.266587643 | 2.284299819 | -0.017712176 | -0.059264387 |

| | | | | | |
|---|---|---|---|---|---|
| Bbx | 9.249389489 | 9.187771683 | 9.308672193 | -0.12090051 | -0.059282704 |
| Defb12 | 2.028171148 | 2.036976524 | 2.087473079 | -0.050496555 | -0.059301931 |
| Prpsap2 | 8.416375167 | 8.455037632 | 8.475755821 | -0.020718189 | -0.059380653 |
| Kdm4c | 9.083568659 | 9.074923462 | 9.142957733 | -0.068034271 | -0.059389074 |
| Insc | 4.315939006 | 4.377831619 | 4.375335324 | 0.002496296 | -0.059396318 |
| Sel1l | 11.35432395 | 11.42429194 | 11.41373669 | 0.010555253 | -0.059412743 |
| Zbtb7a | 10.85864266 | 11.17977884 | 10.91807713 | 0.26170171 | -0.059434473 |
| Hist2h4 | 2.128049602 | 2.213610307 | 2.187499283 | 0.026111024 | -0.059449681 |
| Zfp133-ps | 1.967186128 | 1.989185641 | 2.026643813 | -0.037458172 | -0.059457685 |
| Olfr982 | 1.087003445 | 1.089390958 | 1.146535133 | -0.057144176 | -0.059531688 |
| Pla2g4a | 9.538908579 | 9.510983747 | 9.598440782 | -0.087457035 | -0.059532202 |
| Kcnk5 | 5.29302394 | 5.325431726 | 5.352574126 | -0.0271424 | -0.059550187 |
| Dnajc14 | 10.77256936 | 10.6955255 | 10.83222199 | -0.136696487 | -0.059652633 |
| Slc22a13b-ps | 3.077181963 | 3.174348545 | 3.136841856 | 0.037506689 | -0.059659893 |
| Ufd1 | 10.14201917 | 10.19740841 | 10.20170093 | -0.004292513 | -0.059681753 |
| Mtr | 8.756578074 | 8.874682696 | 8.81627078 | 0.058411916 | -0.059692706 |
| Cyp2j13 | 1.795449514 | 1.839482008 | 1.855155623 | -0.015673615 | -0.059706109 |
| Akirin2 | 9.499966802 | 9.477265208 | 9.559675756 | -0.082410548 | -0.059708954 |
| Pirt | 1.25561382 | 1.287345571 | 1.315332334 | -0.027986763 | -0.059718514 |
| Gm2237 | 4.546171782 | 4.506262265 | 4.605904837 | -0.099642572 | -0.059733055 |
| Tsc22d2 | 9.995841722 | 10.25577706 | 10.05558429 | 0.200192769 | -0.059742569 |
| Astn1 | 2.870508369 | 2.755287164 | 2.930255213 | -0.174968049 | -0.059746845 |
| Adam1a | 6.420363685 | 6.493210217 | 6.48014098 | 0.013069237 | -0.059777295 |
| Esm1 | 1.714377576 | 1.735451391 | 1.774171371 | -0.038719981 | -0.059793795 |
| Vsx2 | 2.097557956 | 2.210143424 | 2.157355764 | 0.05278766 | -0.059797808 |
| Htatsf1 | 10.30224851 | 10.23875364 | 10.36206294 | -0.123309307 | -0.059814433 |
| Leng9 | 6.736750719 | 6.691046338 | 6.796589625 | -0.105543287 | -0.059838906 |
| Defb6 | 11.69867435 | 11.68578475 | 11.7585643 | -0.072779547 | -0.059889947 |
| Tmem42 | 6.765556062 | 6.75685788 | 6.825448318 | -0.068590438 | -0.059892256 |
| Slc29a1 | 10.21729339 | 10.21411711 | 10.27722464 | -0.063107527 | -0.059931253 |
| Cfap46 | 1.233512968 | 1.298187879 | 1.293463509 | 0.00472437 | -0.059950541 |
| Sun3 | 3.260670215 | 3.281678023 | 3.320630838 | -0.038952815 | -0.059960623 |

| | | | | | |
|---|---|---|---|---|---|
| Xrcc4 | 7.257545981 | 7.179142196 | 7.317515395 | -0.138373199 | -0.059969414 |
| Ankub1 | 2.356444695 | 2.309458308 | 2.416448075 | -0.106989767 | -0.06000338 |
| Cd96 | 5.456746544 | 5.384751943 | 5.51676615 | -0.132014207 | -0.060019606 |
| Gm47628 | 4.710857285 | 4.787816534 | 4.770889543 | 0.016926992 | -0.060032258 |
| AI481877 | 4.183292066 | 4.33494912 | 4.243325855 | 0.091623265 | -0.060033788 |
| Ap3b1 | 10.31688915 | 10.29825917 | 10.37693958 | -0.078680407 | -0.06005043 |
| Ralgapa1 | 10.61105243 | 10.50959504 | 10.6711363 | -0.161541253 | -0.06008387 |
| Rtn4r | 5.763017608 | 6.093473855 | 5.823120762 | 0.270353093 | -0.060103154 |
| Itsn1 | 10.07445918 | 10.17130601 | 10.13456671 | 0.0367393 | -0.060107532 |
| Poli | 6.588851036 | 6.624670153 | 6.648986791 | -0.024316638 | -0.060135755 |
| Vamp5 | 6.483451515 | 6.425379114 | 6.543595546 | -0.118216432 | -0.060144031 |
| Etfb | 10.53376115 | 10.48794271 | 10.59391288 | -0.105970177 | -0.060151733 |
| Kcnf1 | 3.948214939 | 4.0714628 | 4.008454101 | 0.063008699 | -0.060239163 |
| Trim36 | 6.126255687 | 6.395506031 | 6.186508953 | 0.208997078 | -0.060253266 |
| Gm4739 | 9.640015009 | 9.4766993 | 9.700309607 | -0.223610307 | -0.060294598 |
| Wfikkn1 | 4.115331053 | 4.335898052 | 4.175632386 | 0.160265666 | -0.060301333 |
| Tnfaip8l1 | 6.242474103 | 6.236892772 | 6.302791448 | -0.065898676 | -0.060317345 |
| Blvra | 8.516096506 | 8.471671121 | 8.57641586 | -0.104744739 | -0.060319354 |
| Csf1r | 11.23931553 | 11.06990194 | 11.29964542 | -0.229743483 | -0.060329897 |
| Etv3l | 2.499378402 | 2.553480038 | 2.559711736 | -0.006231699 | -0.060333334 |
| Rpl39 | 12.55261172 | 12.44707802 | 12.61300667 | -0.165928657 | -0.060394957 |
| Lsm10 | 7.505485206 | 7.536117157 | 7.565894036 | -0.029776879 | -0.060408831 |
| Nkain3 | 1.896760911 | 1.930572532 | 1.957174455 | -0.026601923 | -0.060413545 |
| Rpusd3 | 6.019487704 | 6.176992196 | 6.079903398 | 0.097088798 | -0.060415694 |
| Amdhd1 | 2.29405987 | 2.301052964 | 2.354536067 | -0.053483103 | -0.060476197 |
| Rhobtb3 | 8.017684485 | 7.890251656 | 8.078184198 | -0.187932542 | -0.060499713 |
| Olig1 | 2.541070061 | 2.641731811 | 2.601603802 | 0.04012801 | -0.06053374 |
| Gpr75 | 2.930233491 | 2.970664241 | 2.990781436 | -0.020117194 | -0.060547944 |
| Tceal9 | 10.02819746 | 9.869669924 | 10.08887038 | -0.21920046 | -0.060672927 |
| Gm10662 | 2.057480459 | 2.070085396 | 2.118175348 | -0.048089951 | -0.060694888 |
| Gm5891 | 2.057480459 | 2.070085396 | 2.118175348 | -0.048089951 | -0.060694888 |
| Prrt1 | 4.238387162 | 4.273033295 | 4.299092423 | -0.026059128 | -0.06070526 |

| | | | | | |
|---|---|---|---|---|---|
| Fam49a | 8.72348987 | 9.105913532 | 8.784245154 | 0.321668378 | -0.060755284 |
| Bbc3 | 5.446993737 | 5.566602752 | 5.507765987 | 0.058836765 | -0.06077225 |
| Ambra1 | 9.909683414 | 9.977641658 | 9.970466799 | 0.007174859 | -0.060783385 |
| Kbtbd11 | 7.503046967 | 7.296575476 | 7.563868554 | -0.267293078 | -0.060821587 |
| Olfr1330 | 2.833302328 | 2.822533944 | 2.894128935 | -0.071594991 | -0.060826607 |
| Nipbl | 11.28769274 | 11.29610945 | 11.3486117 | -0.052502243 | -0.060918953 |
| Hemgn | 1.249018519 | 1.27355008 | 1.310095313 | -0.036545233 | -0.061076794 |
| Pdhb | 11.04227721 | 10.89746158 | 11.10340821 | -0.20594663 | -0.061130997 |
| Gm16297 | 1.78532969 | 1.812084202 | 1.846464242 | -0.034380041 | -0.061134552 |
| Tfb1m | 6.747522759 | 6.718058575 | 6.808693765 | -0.09063519 | -0.061171006 |
| Gm35339 | 5.518054259 | 5.676953113 | 5.579231264 | 0.097721849 | -0.061177005 |
| 4921536K21Rik | 5.121274116 | 5.191720198 | 5.182488307 | 0.009231891 | -0.061214191 |
| Smim10l2a | 3.480889474 | 3.49851564 | 3.542224913 | -0.043709273 | -0.061335439 |
| Pkdrej | 2.881762571 | 2.817527373 | 2.943126599 | -0.125599225 | -0.061364028 |
| Fam160b2 | 9.585727628 | 9.783206686 | 9.64716791 | 0.136038777 | -0.061440281 |
| Ccpg1 | 10.30868693 | 10.10827136 | 10.37016942 | -0.261898063 | -0.06148249 |
| Olfr1328 | 2.059721286 | 2.103877541 | 2.121207901 | -0.01733036 | -0.061486615 |
| Plekha7 | 8.90696408 | 9.150714564 | 8.968480357 | 0.182234207 | -0.061516277 |
| Maats1 | 2.74607107 | 2.797336454 | 2.807606177 | -0.010269723 | -0.061535108 |
| Cysltr2 | 3.211128958 | 3.184038462 | 3.272664794 | -0.088626332 | -0.061535836 |
| Ppp2r5e | 9.966656568 | 10.0125987 | 10.02819357 | -0.015594875 | -0.061537004 |
| Itpka | 4.643080401 | 4.647031971 | 4.704636319 | -0.057604348 | -0.061555918 |
| Ccdc88c | 9.365959205 | 9.398634768 | 9.427534507 | -0.028899739 | -0.061575302 |
| Gm44578 | 3.303816077 | 3.379478082 | 3.365402433 | 0.014075649 | -0.061586356 |
| Elovl3 | 11.73856834 | 11.56490949 | 11.80015634 | -0.235246843 | -0.061587997 |
| Csgalnact2 | 9.459373247 | 9.335821989 | 9.520978779 | -0.18515679 | -0.061605532 |
| Amd2 | 3.791996609 | 3.782417195 | 3.853606339 | -0.071189145 | -0.06160973 |
| Cep192 | 8.715049237 | 8.716196575 | 8.776680939 | -0.060484365 | -0.061631702 |
| Ppp2r5b | 8.074889207 | 8.275238678 | 8.136527686 | 0.138710992 | -0.061638479 |
| Vwa5a | 10.91231375 | 10.83277647 | 10.97396561 | -0.14118914 | -0.061651853 |
| Gpc4 | 9.603055915 | 9.533978255 | 9.664708248 | -0.130729993 | -0.061652333 |
| Camk4 | 4.236326573 | 4.240346716 | 4.297987473 | -0.057640757 | -0.0616609 |

| | | | | | |
|---|---|---|---|---|---|
| Rxrb | 9.513215839 | 9.783768246 | 9.574889987 | 0.20887826 | -0.061674148 |
| Tnnc1 | 3.196345737 | 3.177137532 | 3.258020001 | -0.080882469 | -0.061674264 |
| Mroh8 | 1.959483853 | 2.007202621 | 2.021191185 | -0.013988564 | -0.061707331 |
| Foxc2 | 4.551196532 | 4.704299112 | 4.612931936 | 0.091367177 | -0.061735403 |
| Phax | 9.094988293 | 9.005331465 | 9.156741906 | -0.151410441 | -0.061753612 |
| Cnot8 | 9.662009986 | 9.557429804 | 9.723765835 | -0.16633603 | -0.061755848 |
| Cdk13 | 10.14783237 | 10.26175895 | 10.20961848 | 0.052140473 | -0.061786107 |
| Hoxa2 | 5.716413654 | 5.80414749 | 5.778257439 | 0.025890051 | -0.061843785 |
| Gm19241 | 1.893723017 | 1.922607996 | 1.955648273 | -0.033040277 | -0.061925256 |
| Tssk4 | 4.409534324 | 4.488062871 | 4.471469929 | 0.016592942 | -0.061935604 |
| Asic1 | 1.743429002 | 1.771611621 | 1.805428939 | -0.033817318 | -0.061999937 |
| Gm49396 | 7.290396487 | 7.503443947 | 7.35239804 | 0.151045907 | -0.062001552 |
| Mrpl24 | 10.13546909 | 10.13877211 | 10.19749701 | -0.058724907 | -0.062027927 |
| Car7 | 3.141197707 | 3.2178298 | 3.203247667 | 0.014582133 | -0.062049961 |
| Ctsd | 14.21458328 | 14.34771927 | 14.2766601 | 0.071059177 | -0.062076817 |
| Vmn1r209 | 2.56785963 | 2.589229792 | 2.629959733 | -0.040729941 | -0.062100104 |
| Serpini1 | 3.554424462 | 3.546363401 | 3.616531485 | -0.070168084 | -0.062107022 |
| Pole | 8.407061464 | 8.51667803 | 8.469174368 | 0.047503662 | -0.062112904 |
| Plau | 9.822692992 | 9.890878437 | 9.884852172 | 0.006026265 | -0.06215918 |
| Olfr49 | 3.307370699 | 3.338889532 | 3.369530397 | -0.030640865 | -0.062159698 |
| Ak9 | 2.188849999 | 2.215142788 | 2.251060444 | -0.035917656 | -0.062210445 |
| Ccdc144b | 4.626318196 | 4.537963582 | 4.688557751 | -0.150594168 | -0.062239555 |
| Slc16a13 | 8.871362488 | 9.151415762 | 8.933623783 | 0.21779198 | -0.062261295 |
| Casp8ap2 | 8.958184409 | 9.008814166 | 9.020614681 | -0.011800515 | -0.062430271 |
| Idi1 | 11.8615983 | 11.74552929 | 11.92405913 | -0.17852984 | -0.06246083 |
| Pbld2 | 3.058846668 | 3.053781164 | 3.12131721 | -0.067536046 | -0.062470542 |
| Gabbr2 | 1.48313145 | 1.53249778 | 1.545612523 | -0.013114742 | -0.062481073 |
| C4bp-ps1 | 0.728613405 | 0.744892122 | 0.79111703 | -0.046224908 | -0.062503625 |
| Ropn1l | 2.983733011 | 3.023330434 | 3.046242386 | -0.022911953 | -0.062509376 |
| Rbm46 | 1.380543481 | 1.423630244 | 1.443053594 | -0.01942335 | -0.062510113 |
| Tab2 | 11.36085172 | 11.38216981 | 11.42336473 | -0.041194927 | -0.062513013 |
| Ap4e1 | 8.32360958 | 8.340850966 | 8.386181381 | -0.045330415 | -0.062571801 |

| | | | | | |
|---|---|---|---|---|---|
| Clec4g | 4.059717229 | 3.945736065 | 4.122340292 | -0.176604227 | -0.062623063 |
| Sycp3 | 3.464460575 | 3.416364974 | 3.527120392 | -0.110755418 | -0.062659817 |
| Mfsd13b | 2.91565018 | 2.917592765 | 2.978339914 | -0.060747149 | -0.062689734 |
| Gm18716 | 3.654071497 | 3.732378514 | 3.716792392 | 0.015586121 | -0.062720896 |
| Fbxo7 | 8.766621826 | 8.743311408 | 8.829384358 | -0.08607295 | -0.062762533 |
| Vmn1r43 | 3.103795079 | 3.060149957 | 3.166575336 | -0.106425379 | -0.062780258 |
| Rnf182 | 2.061675745 | 2.07378053 | 2.124457243 | -0.050676714 | -0.062781498 |
| Eya2 | 5.323823182 | 5.229598724 | 5.386638626 | -0.157039902 | -0.062815444 |
| 1700029I15Rik | 5.629549915 | 5.743779498 | 5.692370638 | 0.05140886 | -0.062820723 |
| Olfr1238 | 5.114032606 | 5.146050583 | 5.176866714 | -0.030816131 | -0.062834108 |
| Med13 | 10.87301514 | 11.03621411 | 10.93584996 | 0.10036415 | -0.062834823 |
| Bex1 | 4.545905034 | 4.902025492 | 4.608762635 | 0.293262857 | -0.062857601 |
| Wnk3 | 3.603822007 | 3.645930651 | 3.66673141 | -0.020800759 | -0.062909403 |
| Myo18a | 12.05763193 | 12.23819559 | 12.12059452 | 0.117601073 | -0.062962583 |
| Wdr11 | 9.830566174 | 9.778299505 | 9.893554144 | -0.115254639 | -0.06298797 |
| Samm50 | 10.87634303 | 10.87800081 | 10.93937961 | -0.0613788 | -0.06303658 |
| Slc39a2 | 7.08300488 | 7.13660329 | 7.146162268 | -0.009558978 | -0.063157389 |
| Lmo4 | 10.40136339 | 10.25908898 | 10.46453463 | -0.20544565 | -0.06317124 |
| 37681 | 8.647510326 | 8.69910508 | 8.710688905 | -0.011583825 | -0.063178579 |
| Fhl4 | 1.587853332 | 1.585418738 | 1.651037656 | -0.065618918 | -0.063184324 |
| Smim6 | 4.244697825 | 4.144533406 | 4.307891029 | -0.163357623 | -0.063193204 |
| Cd28 | 4.141714364 | 4.023446723 | 4.204925177 | -0.181478454 | -0.063210813 |
| Vmn2r105 | 1.954463949 | 1.957493641 | 2.017700942 | -0.060207301 | -0.063236993 |
| Syna | 3.338598284 | 3.369760078 | 3.401839067 | -0.032078989 | -0.063240783 |
| Igfn1 | 1.765224764 | 1.797748877 | 1.828482358 | -0.030733481 | -0.063257594 |
| Arsg | 6.110732105 | 6.188631651 | 6.17402191 | 0.014609741 | -0.063289805 |
| Cnmd | 1.761649678 | 1.758138393 | 1.824959682 | -0.066821289 | -0.063310004 |
| Vmn1r55 | 6.532949825 | 6.561935405 | 6.59626979 | -0.034334385 | -0.063319965 |
| Ppm1e | 4.827622386 | 4.656385776 | 4.890947045 | -0.234561269 | -0.063324659 |
| Hps3 | 8.744354357 | 8.707526726 | 8.807681769 | -0.100155043 | -0.063327412 |
| Gm38699 | 4.338189146 | 4.336915009 | 4.401534963 | -0.064619954 | -0.063345817 |
| Reln | 8.371379386 | 8.500286999 | 8.434740376 | 0.065546622 | -0.06336099 |

| | | | | | |
|---|---|---|---|---|---|
| Zbtb49 | 6.791784278 | 6.883272408 | 6.855150739 | 0.028121669 | -0.063366461 |
| Slc39a3 | 8.777847104 | 9.063119922 | 8.841289483 | 0.221830439 | -0.063442379 |
| Shtn1 | 10.19793263 | 10.30099547 | 10.26138764 | 0.03960783 | -0.063455007 |
| Olfr639 | 5.5511735 | 5.635970096 | 5.614668644 | 0.021301452 | -0.063495144 |
| Nwd1 | 2.512323461 | 2.634093926 | 2.575832234 | 0.058261692 | -0.063508773 |
| Syne4 | 2.975028088 | 3.027900888 | 3.038589 | -0.010688112 | -0.063560913 |
| Cfi | 2.208824003 | 2.230884934 | 2.272387445 | -0.04150251 | -0.063563442 |
| Trappc2 | 7.950990589 | 7.918776702 | 8.014565324 | -0.095788622 | -0.063574735 |
| Nectin4 | 11.3250065 | 11.36134842 | 11.38859608 | -0.02724766 | -0.063589577 |
| Cacna1s | 10.79181668 | 11.09105879 | 10.85543536 | 0.235623427 | -0.063618679 |
| Ncor1 | 11.55528744 | 11.62378553 | 11.61896223 | 0.004823307 | -0.063674783 |
| Creb3l4 | 4.910442775 | 5.039349757 | 4.97414309 | 0.065206667 | -0.063700315 |
| Ccdc84 | 7.464277021 | 7.604924923 | 7.528012502 | 0.076912421 | -0.063735481 |
| Stard13 | 8.264800065 | 8.296527558 | 8.328554335 | -0.032026777 | -0.06375427 |
| Rbm12b1 | 7.430753947 | 7.588159778 | 7.494513567 | 0.093646211 | -0.06375962 |
| Mbip | 7.625516857 | 7.662151771 | 7.689317865 | -0.027166095 | -0.063801008 |
| Olfr1044 | 2.088920679 | 2.13366641 | 2.15287205 | -0.01920564 | -0.063951371 |
| Ano9 | 11.91186687 | 11.9851947 | 11.9758537 | 0.009341003 | -0.063986833 |
| Mup12 | 1.544546192 | 1.586922229 | 1.608553527 | -0.021631298 | -0.064007335 |
| Acox3 | 10.44185371 | 10.59417865 | 10.50594069 | 0.088237967 | -0.06408698 |
| Mrs2 | 10.01370572 | 10.18150626 | 10.0777986 | 0.103707657 | -0.064092886 |
| Fabp3 | 7.9743561 | 8.52652882 | 8.03848729 | 0.48804153 | -0.06413119 |
| Slc23a1 | 3.744852468 | 3.885062195 | 3.808987068 | 0.076075127 | -0.064134599 |
| Gtf2h2 | 8.104352729 | 8.127275993 | 8.168491182 | -0.041215188 | -0.064138453 |
| Gm13868 | 2.241330444 | 2.36443573 | 2.305543489 | 0.05889224 | -0.064213045 |
| 5730596B20Rik | 2.921680775 | 3.07958756 | 2.985932199 | 0.093655361 | -0.064251425 |
| Rab1b | 11.13168788 | 11.23068134 | 11.19594953 | 0.034731813 | -0.064261643 |
| Adgre1 | 9.658095921 | 9.526745667 | 9.722389067 | -0.195643399 | -0.064293145 |
| Arl14ep | 8.719234779 | 8.764688202 | 8.78352796 | -0.018839759 | -0.064293181 |
| Serpina1a | 1.407861955 | 1.406648662 | 1.472162323 | -0.065513661 | -0.064300368 |
| Pcdhga2 | 4.156183765 | 4.181096258 | 4.220497887 | -0.039401629 | -0.064314122 |
| Usp48 | 10.21525322 | 10.34377695 | 10.27963308 | 0.064143877 | -0.064379853 |

| | | | | | |
|---|---|---|---|---|---|
| Hoxa4 | 5.594942597 | 5.502594218 | 5.65935416 | -0.156759942 | -0.064411563 |
| Tceal8 | 9.338652646 | 9.34534994 | 9.403225295 | -0.057875355 | -0.06457265 |
| Zbtb42 | 8.173476062 | 8.300069409 | 8.238059661 | 0.062009748 | -0.064583599 |
| Ccdc74a | 4.090252612 | 4.174653487 | 4.154851208 | 0.019802279 | -0.064598597 |
| 38777 | 10.94676522 | 10.84852179 | 11.01138047 | -0.162858683 | -0.064615253 |
| Rps24 | 13.82117199 | 13.67308158 | 13.88581274 | -0.21273116 | -0.064640747 |
| Ttr | 4.390889481 | 4.467569384 | 4.455561307 | 0.012008077 | -0.064671826 |
| Nrbp2 | 10.67603393 | 10.77964335 | 10.74071659 | 0.03892676 | -0.06468266 |
| Foxr1 | 1.903687013 | 1.893689645 | 1.968397758 | -0.074708113 | -0.064710745 |
| Vmn2r10 | 1.749743766 | 1.771787495 | 1.814471444 | -0.042683948 | -0.064727678 |
| Snx9 | 10.37479796 | 10.33672387 | 10.43957085 | -0.10284698 | -0.06477289 |
| Hykk | 5.767270969 | 6.033477678 | 5.832110382 | 0.201367296 | -0.064839413 |
| Tmem178 | 1.866513259 | 1.880174325 | 1.931355406 | -0.051181081 | -0.064842147 |
| Cdc25c | 6.220016825 | 6.181146852 | 6.285058598 | -0.103911745 | -0.065041773 |
| Gm11841 | 1.503080216 | 1.508234819 | 1.568136017 | -0.059901198 | -0.065055801 |
| Lce1m | 12.8898876 | 13.04397539 | 12.95503563 | 0.088939767 | -0.065148027 |
| Vstm2a | 3.61630618 | 3.534586482 | 3.681515256 | -0.146928774 | -0.065209076 |
| Gm15833 | 2.290578119 | 2.353223365 | 2.355799146 | -0.002575781 | -0.065221027 |
| Zfp11 | 6.203359065 | 6.130803421 | 6.268580598 | -0.137777176 | -0.065221532 |
| Armcx3 | 9.394385025 | 9.453741926 | 9.459618925 | -0.005877 | -0.065233901 |
| Tbc1d13 | 9.996866362 | 10.04606975 | 10.06216544 | -0.016095695 | -0.065299078 |
| Gm14496 | 4.577479849 | 4.576881874 | 4.642819043 | -0.065937169 | -0.065339194 |
| Rgs7 | 5.483570916 | 5.418315989 | 5.548913238 | -0.130597249 | -0.065342322 |
| Tefm | 7.183531729 | 7.042347922 | 7.248875836 | -0.206527914 | -0.065344107 |
| Lingo1 | 4.832571147 | 5.056203814 | 4.897918497 | 0.158285317 | -0.065347349 |
| Siah1a | 8.774312828 | 8.769049016 | 8.839691927 | -0.070642912 | -0.065379099 |
| Pmp22 | 11.54304506 | 11.48352864 | 11.60844124 | -0.124912597 | -0.06539618 |
| Cd48 | 8.13769062 | 8.273009455 | 8.203088445 | 0.06992101 | -0.065397825 |
| Zfp955a | 7.817746779 | 7.787117817 | 7.883156417 | -0.0960386 | -0.065409638 |
| Mapk15 | 3.096945813 | 3.352594815 | 3.162371476 | 0.190223339 | -0.065425664 |
| Pde9a | 4.508262254 | 4.479678987 | 4.573714998 | -0.094036011 | -0.065452744 |
| 2210011C24Rik | 1.992050986 | 2.03898559 | 2.057552404 | -0.018566814 | -0.065501418 |

| | | | | | |
|---|---|---|---|---|---|
| Mtch2 | 9.980969359 | 9.947863835 | 10.04650764 | -0.098643803 | -0.065538279 |
| Gm5177 | 1.354774447 | 1.414703919 | 1.420339852 | -0.005635934 | -0.065565406 |
| Gm14401 | 2.115316204 | 2.121922024 | 2.180884981 | -0.058962957 | -0.065568777 |
| Gm21056 | 3.175581371 | 3.328679352 | 3.241220543 | 0.087458809 | -0.065639172 |
| Foxd1 | 3.521924544 | 3.600633971 | 3.587609394 | 0.013024578 | -0.06568485 |
| Ryr3 | 3.787110441 | 3.813679944 | 3.852851754 | -0.03917181 | -0.065741313 |
| Msx2 | 5.479529381 | 5.609766991 | 5.545317345 | 0.064449646 | -0.065787963 |
| Umad1 | 6.896608735 | 6.942150413 | 6.96242467 | -0.020274257 | -0.065815935 |
| St6gal2 | 2.002493925 | 2.017527642 | 2.068332092 | -0.05080445 | -0.065838167 |
| Eci3 | 3.090790316 | 3.07382352 | 3.156694615 | -0.082871095 | -0.065904299 |
| Cnot6 | 11.11179287 | 11.11598055 | 11.17778031 | -0.061799763 | -0.065987443 |
| Msl3l2 | 5.581303622 | 5.583491968 | 5.64735678 | -0.063864811 | -0.066053158 |
| Gm5851 | 1.898150529 | 1.948353083 | 1.964207735 | -0.015854652 | -0.066057206 |
| Tacr1 | 6.060749476 | 6.132192811 | 6.126819998 | 0.005372813 | -0.066070522 |
| Olfr1192-ps1 | 4.388632939 | 4.342266811 | 4.45471825 | -0.112451439 | -0.066085311 |
| Slc36a1 | 9.261951852 | 9.40592598 | 9.328077712 | 0.077848269 | -0.06612586 |
| N4bp3 | 8.316103957 | 8.507380401 | 8.382233295 | 0.125147105 | -0.066129339 |
| Gm7435 | 2.616876584 | 2.532754687 | 2.68300855 | -0.150253863 | -0.066131966 |
| Slc20a2 | 9.681192577 | 9.706333467 | 9.747360081 | -0.041026614 | -0.066167504 |
| Galnt1 | 11.12502583 | 11.2608361 | 11.19119541 | 0.06964069 | -0.066169573 |
| Olfr1009 | 1.456379038 | 1.445331446 | 1.522599847 | -0.077268401 | -0.066220809 |
| Syt17 | 2.167483126 | 2.157670382 | 2.233746155 | -0.076075773 | -0.066263029 |
| Kcnt1 | 2.509946058 | 2.510038265 | 2.576246116 | -0.066207851 | -0.066300059 |
| Swi5 | 10.83475553 | 10.75966048 | 10.90113157 | -0.141471083 | -0.066376037 |
| Kitl | 9.9882812 | 9.80042013 | 10.05466294 | -0.254242814 | -0.066381745 |
| Ephb4 | 10.05676852 | 10.16104751 | 10.12315644 | 0.03789107 | -0.066387918 |
| Pggt1b | 10.04442951 | 9.988867263 | 10.11081885 | -0.121951591 | -0.066389347 |
| Gm48841 | 3.912603781 | 3.95498775 | 3.979040138 | -0.024052388 | -0.066436357 |
| Ccny | 10.80146165 | 10.80626599 | 10.86790111 | -0.06163512 | -0.066439457 |
| Itgb3bp | 5.332684827 | 5.30281198 | 5.39914493 | -0.09633295 | -0.066460103 |
| Oxnad1 | 7.24195052 | 7.254117276 | 7.30841422 | -0.054296944 | -0.066463701 |
| Pcyox1l | 5.863352589 | 6.188171316 | 5.929876105 | 0.258295211 | -0.066523516 |

| | | | | | |
|---|---|---|---|---|---|
| Slc37a1 | 6.978155975 | 7.074049548 | 7.044686845 | 0.029362703 | -0.06653087 |
| Ddrgk1 | 9.711452932 | 9.777650513 | 9.77801562 | -0.000365108 | -0.066562689 |
| Clec2i | 4.763164603 | 4.70097256 | 4.8297925 | -0.128819941 | -0.066627898 |
| Prps2 | 9.368847511 | 9.288222097 | 9.435509579 | -0.147287482 | -0.066662069 |
| Ensa | 11.06405841 | 11.13605612 | 11.13074621 | 0.005309903 | -0.0666878 |
| Rcn2 | 9.282590767 | 9.192962881 | 9.349293168 | -0.156330287 | -0.066702401 |
| Tmprss11d | 4.850372002 | 5.136285234 | 4.917076294 | 0.21920894 | -0.066704292 |
| Flywch2 | 4.977892412 | 5.096549756 | 5.044692512 | 0.051857244 | -0.066800099 |
| Vmn2r17 | 3.557596635 | 3.613700282 | 3.624459153 | -0.010758872 | -0.066862518 |
| Gng8 | 3.275680001 | 3.281768248 | 3.34256093 | -0.060792682 | -0.066880928 |
| Abhd11os | 2.314297863 | 2.277754927 | 2.381262871 | -0.103507944 | -0.066965009 |
| Kif15 | 7.798675092 | 7.843752882 | 7.865650658 | -0.021897776 | -0.066975567 |
| Plgrkt | 9.444863517 | 9.509264962 | 9.511840389 | -0.002575427 | -0.066976872 |
| Rxfp1 | 2.660398667 | 2.678034069 | 2.727413016 | -0.049378947 | -0.067014349 |
| 1500015O10Rik | 1.710703703 | 1.686120073 | 1.777768247 | -0.091648173 | -0.067064544 |
| Krcc1 | 10.54769851 | 10.4703067 | 10.61479128 | -0.14448458 | -0.06709277 |
| Nmral1 | 7.072300977 | 6.999861575 | 7.139418666 | -0.139557091 | -0.067117689 |
| Pctp | 10.67776789 | 10.72212658 | 10.74490218 | -0.022775607 | -0.06713429 |
| Vmn2r1 | 2.900532354 | 2.957072899 | 2.967736136 | -0.010663237 | -0.067203782 |
| Pidd1 | 6.437108001 | 6.724971591 | 6.50433771 | 0.220633881 | -0.067229709 |
| Cyp3a57 | 7.806104573 | 7.905523147 | 7.87341425 | 0.032108897 | -0.067309677 |
| Cmc1 | 7.739620636 | 7.547201557 | 7.806974262 | -0.259772705 | -0.067353625 |
| Fsip1 | 2.452355707 | 2.526628022 | 2.519929576 | 0.006698447 | -0.067573869 |
| Gdf6 | 1.625696078 | 1.626564933 | 1.693301043 | -0.06673611 | -0.067604965 |
| Ephb2 | 8.024288573 | 8.204736682 | 8.091968472 | 0.11276821 | -0.067679899 |
| Psmb11 | 3.178942126 | 3.181781924 | 3.246638557 | -0.064856633 | -0.067696431 |
| Trip4 | 9.501115808 | 9.474398511 | 9.568819179 | -0.094420668 | -0.067703371 |
| Olfr45 | 4.07731582 | 4.057306348 | 4.145047701 | -0.087741353 | -0.067731881 |
| Ttk | 8.371695151 | 8.419355859 | 8.439549655 | -0.020193796 | -0.067854504 |
| Spint1 | 9.998251463 | 9.998224947 | 10.06611305 | -0.067888098 | -0.067861582 |
| Samd1 | 8.835754427 | 9.046773124 | 8.903652679 | 0.143120445 | -0.067898251 |
| Tnnc2 | 13.6885653 | 13.70415833 | 13.756589 | -0.052430667 | -0.068023697 |

| | | | | | |
|---|---|---|---|---|---|
| Gm10037 | 2.829482412 | 2.843141223 | 2.897550969 | -0.054409746 | -0.068068557 |
| Tram1l1 | 2.357284134 | 2.382728088 | 2.425373351 | -0.042645263 | -0.068089217 |
| 37316 | 9.840649316 | 9.807444326 | 9.908755788 | -0.101311462 | -0.068106472 |
| Adgrb1 | 2.028903033 | 2.138068456 | 2.097030143 | 0.041038313 | -0.06812711 |
| Fignl2 | 6.700001048 | 6.719546554 | 6.768170187 | -0.048623634 | -0.06816914 |
| E330021D16Rik | 3.112450734 | 3.183508305 | 3.180630539 | 0.002877766 | -0.068179806 |
| Rp1 | 2.001300072 | 1.995052105 | 2.069512051 | -0.074459946 | -0.068211979 |
| Sec62 | 11.78260999 | 11.87800632 | 11.85082586 | 0.027180463 | -0.06821587 |
| Nat1 | 1.116469515 | 1.141616272 | 1.184692082 | -0.04307581 | -0.068222567 |
| Bean1 | 3.09331993 | 3.143658224 | 3.161569563 | -0.017911339 | -0.068249633 |
| AC154486.3 | 3.660505926 | 3.718130989 | 3.728802043 | -0.010671054 | -0.068296117 |
| Tmem123 | 12.00849904 | 11.9946104 | 12.07680174 | -0.08219134 | -0.068302693 |
| Hnrnpul2 | 12.0781772 | 12.14666247 | 12.14651687 | 0.0001456 | -0.06833967 |
| Zfp865 | 8.379268623 | 8.695587532 | 8.447638633 | 0.247948899 | -0.06837001 |
| Upf3b | 9.079114667 | 9.200759596 | 9.147500404 | 0.053259192 | -0.068385737 |
| Rbm11 | 2.631881332 | 2.674066275 | 2.70032131 | -0.026255035 | -0.068439978 |
| Clptm1 | 10.95186696 | 11.00244323 | 11.0203952 | -0.017951977 | -0.068528247 |
| Ccdc173 | 4.724569832 | 4.826968824 | 4.793213675 | 0.033755148 | -0.068643843 |
| Vmn1r38 | 3.387160283 | 3.450018895 | 3.455845727 | -0.005826832 | -0.068685444 |
| Fam149a | 7.312721237 | 7.171483103 | 7.381415609 | -0.209932505 | -0.068694372 |
| Tubgcp2 | 9.090447459 | 9.187944207 | 9.159158976 | 0.028785231 | -0.068711517 |
| Map3k13 | 5.358941191 | 5.419341605 | 5.427686285 | -0.00834468 | -0.068745094 |
| Gm14326 | 7.309912091 | 7.305508858 | 7.37866305 | -0.073154192 | -0.068750959 |
| Slc8a2 | 4.067005208 | 4.584099684 | 4.135808051 | 0.448291633 | -0.068802843 |
| Snapc3 | 8.291416722 | 8.375646342 | 8.360252586 | 0.015393756 | -0.068835864 |
| Immp2l | 5.306885098 | 5.318776651 | 5.375751708 | -0.056975056 | -0.06886661 |
| Susd4 | 3.440995958 | 3.492780776 | 3.509882743 | -0.017101968 | -0.068886785 |
| Tro | 1.655137738 | 1.723710384 | 1.72402797 | -0.000317587 | -0.068890233 |
| Gm9403 | 3.402847631 | 3.421940035 | 3.471756942 | -0.049816907 | -0.068909311 |
| Gm14681 | 1.911794925 | 2.032889811 | 1.980742206 | 0.052147605 | -0.068947281 |
| Cst3 | 12.43131738 | 12.55799717 | 12.50026703 | 0.05773014 | -0.068949647 |
| Ccndbp1 | 9.285725422 | 9.201646221 | 9.354708996 | -0.153062775 | -0.068983575 |

| | | | | | |
|---|---|---|---|---|---|
| Ankrd22 | 8.80218816 | 8.695580912 | 8.871221806 | -0.175640894 | -0.069033646 |
| Abraxas2 | 9.188576855 | 9.311994075 | 9.257635232 | 0.054358844 | -0.069058376 |
| Gm6430 | 2.140501271 | 2.211065561 | 2.209580263 | 0.001485298 | -0.069078991 |
| Pakap | 5.87780306 | 5.871449249 | 5.946897849 | -0.0754486 | -0.069094789 |
| Rgl2 | 9.987119248 | 10.36017601 | 10.05621575 | 0.303960263 | -0.069096502 |
| Plcb1 | 7.450377238 | 7.331509598 | 7.519476611 | -0.187967012 | -0.069099373 |
| Hacd1 | 7.405399427 | 7.323823554 | 7.474500821 | -0.150677267 | -0.069101394 |
| Kbtbd6 | 4.194216538 | 4.197783819 | 4.263353395 | -0.065569576 | -0.069136856 |
| Rbms3 | 9.469743527 | 9.409894916 | 9.538899315 | -0.129004399 | -0.069155789 |
| Tanc2 | 10.05893327 | 9.981360843 | 10.12809721 | -0.146736363 | -0.069163933 |
| Gm3791 | 2.950788796 | 2.989482224 | 3.01996751 | -0.030485286 | -0.069178714 |
| Dkk1 | 1.536471388 | 1.561699712 | 1.605736854 | -0.044037142 | -0.069265466 |
| Itih2 | 2.462123112 | 2.514852761 | 2.531392933 | -0.016540173 | -0.069269821 |
| Neurl2 | 5.207219097 | 5.338623322 | 5.276525163 | 0.062098159 | -0.069306066 |
| Tubgcp4 | 9.304730492 | 9.531090841 | 9.374054824 | 0.157036017 | -0.069324332 |
| Tmem37 | 5.954044002 | 5.853544137 | 6.023373412 | -0.169829275 | -0.06932941 |
| Upf2 | 10.05747361 | 9.9384471 | 10.12683374 | -0.18838664 | -0.06936013 |
| BC024139 | 3.791481756 | 3.804046829 | 3.86085579 | -0.056808961 | -0.069374033 |
| Usp3 | 9.720891366 | 9.661655857 | 9.790343435 | -0.128687578 | -0.069452069 |
| Xlr4c | 3.371841172 | 3.317926478 | 3.441300477 | -0.123374 | -0.069459305 |
| Gm44954 | 1.83379194 | 1.888071524 | 1.90332636 | -0.015254836 | -0.069534421 |
| Rsu1 | 9.611061721 | 9.541336264 | 9.680613573 | -0.139277309 | -0.069551852 |
| Ndufb3 | 9.57044223 | 9.436563152 | 9.640012124 | -0.203448972 | -0.069569894 |
| Cables2 | 8.624523146 | 8.726720652 | 8.694094501 | 0.032626151 | -0.069571355 |
| Rpl10-ps6 | 4.337777811 | 4.406593008 | 4.407364613 | -0.000771605 | -0.069586802 |
| Sox7 | 9.145238519 | 9.245218464 | 9.214835621 | 0.030382842 | -0.069597102 |
| Gm13323 | 3.914269205 | 3.922812071 | 3.983875863 | -0.061063791 | -0.069606658 |
| Vmn2r52 | 3.144126798 | 3.234345776 | 3.213782826 | 0.02056295 | -0.069656028 |
| Gtpbp8 | 7.537081227 | 7.658624144 | 7.60679197 | 0.051832174 | -0.069710742 |
| Pigk | 9.906506317 | 9.722651726 | 9.976220171 | -0.253568446 | -0.069713855 |
| Dleu7 | 1.78133366 | 1.860768307 | 1.851067328 | 0.009700979 | -0.069733668 |
| Cr1l | 10.3807212 | 10.24505988 | 10.45058963 | -0.205529753 | -0.069868433 |

| | | | | | |
|---|---|---|---|---|---|
| Sidt2 | 12.00073421 | 12.15150316 | 12.07061172 | 0.08089144 | -0.069877503 |
| Tgds | 7.644271858 | 7.562177437 | 7.714169845 | -0.151992409 | -0.069897987 |
| Trip12 | 12.35178795 | 12.40815301 | 12.42173299 | -0.013579987 | -0.06994504 |
| Git2 | 9.785949163 | 9.907214883 | 9.8559532 | 0.051261683 | -0.070004038 |
| Disp2 | 2.213894145 | 2.25466247 | 2.283965419 | -0.029302949 | -0.070071274 |
| Pmm1 | 8.838500168 | 8.982784246 | 8.908595206 | 0.074189041 | -0.070095037 |
| Mblac1 | 4.289434439 | 4.270987184 | 4.359551226 | -0.088564042 | -0.070116788 |
| Fhit | 5.513574217 | 5.392504611 | 5.583800791 | -0.19129618 | -0.070226574 |
| Grin2b | 2.407772394 | 2.438449189 | 2.478011923 | -0.039562734 | -0.070239529 |
| Mrgprh | 1.634225665 | 1.662961996 | 1.70450187 | -0.041539874 | -0.070276205 |
| Kcng1 | 3.280981013 | 3.297396771 | 3.351279607 | -0.053882837 | -0.070298594 |
| Acrbp | 6.261407759 | 6.529474277 | 6.331722857 | 0.19775142 | -0.070315098 |
| Frat1 | 6.025736459 | 5.979352054 | 6.096072451 | -0.116720397 | -0.070335992 |
| Vmn2r90 | 3.110769513 | 3.182651427 | 3.181114642 | 0.001536786 | -0.070345129 |
| Rpl10a | 11.90396669 | 12.03209595 | 11.97432295 | 0.057772997 | -0.070356263 |
| A830005F24Rik | 2.300188883 | 2.358376297 | 2.370598739 | -0.012222442 | -0.070409856 |
| Gm12739 | 1.146851361 | 1.170379049 | 1.217276079 | -0.04689703 | -0.070424718 |
| Arl6ip6 | 8.308460044 | 8.483791819 | 8.37892092 | 0.104870899 | -0.070460876 |
| Pmel | 4.585630609 | 4.783439164 | 4.656188977 | 0.127250186 | -0.070558368 |
| Cyb5r2 | 2.306837432 | 2.317958338 | 2.37739954 | -0.059441201 | -0.070562107 |
| Mfsd2b | 4.448534381 | 4.546238481 | 4.519097802 | 0.02714068 | -0.07056342 |
| BC024063 | 4.35932888 | 4.330907029 | 4.429917525 | -0.099010496 | -0.070588645 |
| Gm5270 | 2.32584286 | 2.447680992 | 2.396472723 | 0.051208269 | -0.070629863 |
| Miga1 | 6.762480229 | 6.84383534 | 6.833142078 | 0.010693261 | -0.070661849 |
| Zfp52 | 7.744935403 | 7.756736231 | 7.815662498 | -0.058926268 | -0.070727095 |
| Fam217a | 2.1221025 | 2.140205012 | 2.19285353 | -0.052648518 | -0.07075103 |
| Inpp4a | 9.084296926 | 9.206286166 | 9.155086266 | 0.0511999 | -0.07078934 |
| Lce1a2 | 11.81976781 | 12.08592175 | 11.89057637 | 0.195345373 | -0.070808563 |
| Cnksr3 | 8.080780648 | 8.135429926 | 8.151650471 | -0.016220545 | -0.070869823 |
| Abhd10 | 7.489095484 | 7.558007276 | 7.559971527 | -0.001964251 | -0.070876043 |
| Gm5320 | 1.043305752 | 1.090783304 | 1.114186572 | -0.023403268 | -0.07088082 |
| Gm15653 | 2.832311733 | 2.891429779 | 2.903214205 | -0.011784426 | -0.070902472 |

| | | | | | |
|---|---|---|---|---|---|
| Tmem88 | 5.547058408 | 5.548227112 | 5.618014795 | -0.069787683 | -0.070956386 |
| Minar2 | 4.04636773 | 4.197474029 | 4.117340393 | 0.080133636 | -0.070972663 |
| Clec4a1 | 8.870044729 | 8.957277996 | 8.941017838 | 0.016260158 | -0.070973108 |
| Lrrc74b | 3.183092604 | 3.199236988 | 3.254130505 | -0.054893517 | -0.071037901 |
| Ccdc89 | 2.273926338 | 2.34465485 | 2.344969122 | -0.000314272 | -0.071042784 |
| Klhl14 | 1.900479279 | 1.921748812 | 1.971535117 | -0.049786305 | -0.071055838 |
| Itga10 | 5.602678303 | 5.794774361 | 5.673760655 | 0.121013706 | -0.071082352 |
| Ccl25 | 6.466244409 | 6.512570796 | 6.537416711 | -0.024845914 | -0.071172302 |
| 2810021J22Rik | 5.969334696 | 5.877566688 | 6.040539765 | -0.162973077 | -0.071205069 |
| Alg2 | 8.570456273 | 8.624962268 | 8.641677948 | -0.01671568 | -0.071221675 |
| Gm6551 | 8.916348816 | 8.682563011 | 8.98757209 | -0.305009079 | -0.071223274 |
| Psmd8 | 11.6418822 | 11.70897534 | 11.71311415 | -0.004138813 | -0.07123195 |
| Wdr59 | 8.270593532 | 8.35939136 | 8.341929733 | 0.017461627 | -0.071336201 |
| Emc2 | 9.800692301 | 9.701451553 | 9.872084776 | -0.170633223 | -0.071392475 |
| Fbxo2 | 3.469803408 | 3.528047959 | 3.541242219 | -0.013194259 | -0.07143881 |
| Fgf5 | 3.26389613 | 3.36935752 | 3.335381912 | 0.033975608 | -0.071485782 |
| Erbin | 11.63174839 | 11.66969622 | 11.70328616 | -0.033589937 | -0.071537763 |
| Sod1 | 10.84212801 | 10.76436602 | 10.91366763 | -0.149301607 | -0.07153962 |
| Gm15500 | 14.15279439 | 14.05377315 | 14.224347 | -0.170573843 | -0.07155261 |
| Pglyrp4 | 9.090395988 | 9.176807681 | 9.161987126 | 0.014820555 | -0.071591138 |
| Cmklr1 | 9.62301088 | 9.700668092 | 9.694633492 | 0.0060346 | -0.071622613 |
| Paics | 11.09886302 | 11.09096521 | 11.17049104 | -0.079525827 | -0.07162802 |
| Klhl21 | 10.11906286 | 10.54915347 | 10.1907052 | 0.358448267 | -0.071642347 |
| Mmachc | 8.011192785 | 7.921707298 | 8.082845331 | -0.161138033 | -0.071652546 |
| Exoc6b | 10.38812495 | 10.34771276 | 10.45980118 | -0.112088427 | -0.071676237 |
| Rpl31-ps4 | 2.011784494 | 2.141924193 | 2.08346583 | 0.058458363 | -0.071681336 |
| Efr3b | 6.62415911 | 9.190933054 | 6.695888948 | 2.495044106 | -0.071729838 |
| Gm5269 | 1.707622949 | 1.771562005 | 1.779394196 | -0.00783219 | -0.071771247 |
| Gm14138 | 2.00752137 | 2.040337691 | 2.07935106 | -0.039013369 | -0.071829691 |
| Celf5 | 3.168483874 | 3.34451231 | 3.240345193 | 0.104167116 | -0.071861319 |
| Gm37102 | 4.916301969 | 4.995773917 | 4.988200854 | 0.007573063 | -0.071898885 |
| Rpl10l | 2.316068114 | 2.307204858 | 2.387982133 | -0.080777276 | -0.071914019 |

| | | | | | |
|---|---|---|---|---|---|
| Rergl | 1.074534628 | 1.103196995 | 1.146457587 | -0.043260592 | -0.071922959 |
| Rpl34-ps1 | 1.500967623 | 1.583720772 | 1.572913207 | 0.010807565 | -0.071945583 |
| Ccn2 | 10.05904666 | 10.53423586 | 10.13102734 | 0.403208513 | -0.071980679 |
| Rbm39 | 12.10067577 | 12.13458451 | 12.17267105 | -0.038086543 | -0.071995287 |
| Serpinb6d | 8.031060871 | 7.705217615 | 8.103073026 | -0.397855411 | -0.072012156 |
| Col25a1 | 2.896061124 | 2.908912063 | 2.968115063 | -0.059203 | -0.072053939 |
| Gys2 | 4.022052129 | 3.757882344 | 4.094159069 | -0.336276725 | -0.07210694 |
| Fbxl2 | 5.331653122 | 5.373363933 | 5.40386076 | -0.030496827 | -0.072207638 |
| Ppfia2 | 2.214470806 | 2.298927391 | 2.286682894 | 0.012244497 | -0.072212088 |
| Piezo2 | 5.70146861 | 5.662972562 | 5.773702001 | -0.110729439 | -0.072233391 |
| Gabpa | 10.12049706 | 10.22888219 | 10.19273284 | 0.036149343 | -0.07223578 |
| 2410137M14Rik | 1.439884085 | 1.420612418 | 1.512142018 | -0.091529599 | -0.072257933 |
| Fam213b | 9.923347862 | 9.879291699 | 9.99561981 | -0.11632811 | -0.072271947 |
| 1700037H04Rik | 8.526501377 | 8.520190238 | 8.598802198 | -0.07861196 | -0.072300821 |
| Epyc | 1.401291045 | 1.405505145 | 1.473620981 | -0.068115837 | -0.072329937 |
| Col20a1 | 6.203796136 | 6.498360632 | 6.276234396 | 0.222126236 | -0.07243826 |
| Kcnd2 | 2.508252682 | 2.564012983 | 2.580758872 | -0.016745889 | -0.07250619 |
| Carnmt1 | 10.18033394 | 10.13656098 | 10.2528506 | -0.116289617 | -0.072516661 |
| Cyp4a31 | 4.046358392 | 4.096340198 | 4.118894214 | -0.022554016 | -0.072535822 |
| Atp2c1 | 10.57688365 | 10.7776249 | 10.64942542 | 0.128199483 | -0.072541763 |
| Ccdc112 | 6.539336825 | 6.616501453 | 6.61189948 | 0.004601973 | -0.072562655 |
| Daam2 | 7.404810911 | 7.484182502 | 7.477375849 | 0.006806653 | -0.072564938 |
| Gm49450 | 1.092032561 | 1.145516577 | 1.164679278 | -0.019162701 | -0.072646716 |
| Pramef8 | 7.284504265 | 7.304818091 | 7.357155329 | -0.052337238 | -0.072651064 |
| Gm21182 | 1.252996816 | 1.322768857 | 1.325663912 | -0.002895055 | -0.072667096 |
| Morc4 | 7.628991799 | 7.695500817 | 7.701666591 | -0.006165774 | -0.072674792 |
| Eqtn | 2.425287079 | 2.448242073 | 2.497970973 | -0.0497289 | -0.072683893 |
| Adam32 | 0.621777472 | 0.640728765 | 0.694472257 | -0.053743493 | -0.072694785 |
| Megf8 | 9.892791295 | 10.14334257 | 9.96550766 | 0.177834907 | -0.072716365 |
| Igfbp7 | 11.25044212 | 11.10853951 | 11.32318287 | -0.214643353 | -0.072740747 |
| Dcun1d3 | 9.34323897 | 9.35809112 | 9.415985943 | -0.057894822 | -0.072746973 |
| Gatad1 | 9.80203575 | 9.715778912 | 9.874807688 | -0.159028776 | -0.072771938 |

| | | | | | |
|---|---|---|---|---|---|
| Nudt10 | 3.027039033 | 3.13517654 | 3.099821708 | 0.035354832 | -0.072782675 |
| Srd5a3 | 9.20430922 | 9.195554453 | 9.277095635 | -0.081541182 | -0.072786415 |
| Gm8292 | 0.786156681 | 0.813246721 | 0.858961835 | -0.045715113 | -0.072805154 |
| Cep89 | 7.831922877 | 7.939462968 | 7.904786123 | 0.034676846 | -0.072863246 |
| Hinfp | 8.865835911 | 8.986738423 | 8.938737821 | 0.048000602 | -0.07290191 |
| Rsbn1 | 9.282226076 | 9.248969147 | 9.355149461 | -0.106180313 | -0.072923385 |
| App | 12.94842881 | 12.9465231 | 13.02135808 | -0.074834987 | -0.072929273 |
| Rfxank | 7.913958653 | 7.947961215 | 7.986909302 | -0.038948088 | -0.072950649 |
| Zfp27 | 6.565441369 | 6.660250156 | 6.638446018 | 0.021804139 | -0.073004649 |
| Abi1 | 11.29904201 | 11.27731911 | 11.37207958 | -0.09476047 | -0.073037563 |
| 3100002H09Rik | 1.352676121 | 1.406410339 | 1.425719523 | -0.019309183 | -0.073043401 |
| Wwox | 6.872099942 | 6.912569462 | 6.945238918 | -0.032669456 | -0.073138976 |
| Sephs1 | 9.482927867 | 9.506158417 | 9.556082779 | -0.049924363 | -0.073154912 |
| Sertad3 | 7.710485934 | 7.739965169 | 7.783644202 | -0.043679033 | -0.073158268 |
| Oat | 11.80083387 | 11.81411817 | 11.87402332 | -0.059905153 | -0.07318945 |
| Phpt1 | 8.453570742 | 8.460244098 | 8.526800191 | -0.066556092 | -0.073229449 |
| Npas4 | 3.177024812 | 3.244784143 | 3.250283869 | -0.005499726 | -0.073259057 |
| Gzf1 | 9.324846563 | 9.330280723 | 9.39813101 | -0.067850287 | -0.073284446 |
| Brf1 | 9.340751688 | 9.437488023 | 9.414044639 | 0.023443384 | -0.073292951 |
| Vmn2r23 | 4.258186236 | 4.337168072 | 4.33154091 | 0.005627162 | -0.073354674 |
| Pcyt1a | 10.95753892 | 11.02659149 | 11.03091856 | -0.00432707 | -0.073379647 |
| Olfr596 | 2.437014245 | 2.494285285 | 2.510398894 | -0.016113609 | -0.073384649 |
| Psca | 4.623639589 | 4.68372618 | 4.697049846 | -0.013323667 | -0.073410257 |
| Glmn | 7.293961945 | 7.224476904 | 7.367417608 | -0.142940704 | -0.073455663 |
| Klhl11 | 6.986556875 | 7.126864061 | 7.06004754 | 0.066816521 | -0.073490664 |
| Sltm | 11.13058757 | 11.22955953 | 11.20408127 | 0.025478267 | -0.0734937 |
| Tjap1 | 8.912558331 | 9.08225681 | 8.986135633 | 0.096121177 | -0.073577302 |
| Gal3st2c | 3.850835949 | 3.911824528 | 3.924436001 | -0.012611473 | -0.073600052 |
| Mob4 | 10.46282586 | 10.47552368 | 10.53643434 | -0.06091066 | -0.07360848 |
| Lce1c | 11.41754688 | 11.73487101 | 11.49115575 | 0.243715267 | -0.073608863 |
| Gm28230 | 4.753027679 | 4.777461847 | 4.826728588 | -0.049266741 | -0.073700909 |
| Npc1l1 | 4.432820135 | 4.643053551 | 4.506522221 | 0.13653133 | -0.073702086 |

| | | | | | |
|---|---|---|---|---|---|
| Gm45860 | 1.922618133 | 2.03125553 | 1.996423953 | 0.034831577 | -0.07380582 |
| mt-Nd3 | 13.27490093 | 13.17238895 | 13.34871249 | -0.176323543 | -0.073811563 |
| 1700024J04Rik | 2.41411538 | 2.444769746 | 2.487968044 | -0.043198298 | -0.073852664 |
| Mast2 | 10.31603175 | 10.37260368 | 10.38996755 | -0.017363877 | -0.073935803 |
| Fbxw9 | 7.690743596 | 7.867055061 | 7.764706326 | 0.102348735 | -0.07396273 |
| Mogat2 | 2.166214443 | 2.273776983 | 2.240193016 | 0.033583967 | -0.073978573 |
| Nap1l5 | 2.235921739 | 2.27468744 | 2.309936931 | -0.035249491 | -0.074015192 |
| Cldn14 | 3.263362317 | 3.24626321 | 3.337384069 | -0.091120859 | -0.074021752 |
| Kcnq2 | 3.476593258 | 3.767747087 | 3.550623829 | 0.217123258 | -0.074030571 |
| Snx6 | 10.46076814 | 10.41393596 | 10.53491804 | -0.120982073 | -0.0741499 |
| Ccdc102a | 6.16883893 | 6.26630233 | 6.242991873 | 0.023310456 | -0.074152944 |
| Ints10 | 9.069193385 | 9.137365659 | 9.143360893 | -0.005995234 | -0.074167508 |
| Lig1 | 9.50185866 | 9.627135889 | 9.576061505 | 0.051074384 | -0.074202845 |
| Ptp4a2 | 12.53771943 | 12.51701832 | 12.6119731 | -0.094954773 | -0.074253663 |
| 4933430I17Rik | 1.978165638 | 2.054490793 | 2.052509904 | 0.001980889 | -0.074344265 |
| H2-M6-ps | 1.679911775 | 1.680466472 | 1.754275551 | -0.073809079 | -0.074363776 |
| Cmc2 | 7.278485346 | 7.385012203 | 7.352907997 | 0.032104206 | -0.074422651 |
| Thsd7b | 2.752630793 | 2.797675541 | 2.82709002 | -0.02941448 | -0.074459227 |
| Gm45700 | 1.43121066 | 1.512667526 | 1.505673822 | 0.006993704 | -0.074463162 |
| Atp6v0a4 | 9.737562168 | 9.780891886 | 9.81214988 | -0.031257994 | -0.074587712 |
| Nell1 | 2.369226474 | 2.344262436 | 2.44384904 | -0.099586604 | -0.074622566 |
| Guca2a | 6.632248133 | 6.788746373 | 6.706877377 | 0.081868996 | -0.074629244 |
| Cd200r2 | 1.718291908 | 1.706759391 | 1.792929739 | -0.086170348 | -0.074637831 |
| Fbxo22 | 9.076251298 | 9.150619463 | 9.150899315 | -0.000279852 | -0.074648017 |
| Insyn1 | 5.658369305 | 5.739580282 | 5.73302768 | 0.006552602 | -0.074658375 |
| Bcap31 | 11.41150454 | 11.33412576 | 11.4861816 | -0.15205584 | -0.074677063 |
| Dnajc9 | 9.170186927 | 9.249017653 | 9.244912106 | 0.004105547 | -0.074725179 |
| Mepce | 9.556007314 | 9.727060985 | 9.630757313 | 0.096303672 | -0.074749999 |
| 3000002C10Rik | 5.239009413 | 5.250526224 | 5.313845531 | -0.063319308 | -0.074836118 |
| Spdl1 | 6.812824479 | 7.004271394 | 6.887668021 | 0.116603374 | -0.074843542 |
| Alkbh3 | 8.32572271 | 8.361163025 | 8.400566382 | -0.039403357 | -0.074843671 |
| Rab25 | 11.15958281 | 11.27811595 | 11.23443923 | 0.043676727 | -0.074856413 |

| | | | | | |
|---|---|---|---|---|---|
| Akt1s1 | 9.329216234 | 9.495279392 | 9.40414447 | 0.091134922 | -0.074928236 |
| Krt20 | 2.11397103 | 2.06532491 | 2.188914465 | -0.123589555 | -0.074943435 |
| Spa17 | 4.784198025 | 4.766795439 | 4.859220499 | -0.09242506 | -0.075022474 |
| Zbtb38 | 9.56322374 | 9.505261051 | 9.638254833 | -0.132993782 | -0.075031093 |
| Gnas | 12.90147449 | 12.93492039 | 12.97651859 | -0.041598203 | -0.075044107 |
| Ppil6 | 2.887383127 | 3.00608354 | 2.962454399 | 0.043629141 | -0.075071271 |
| Nt5m | 7.538289462 | 7.626759183 | 7.613449144 | 0.013310039 | -0.075159682 |
| AC154707.1 | 3.197972498 | 3.155638273 | 3.273134164 | -0.117495891 | -0.075161666 |
| Arhgap33 | 6.648318303 | 6.969153768 | 6.723645194 | 0.245508574 | -0.075326891 |
| Fabp5 | 13.96030996 | 13.92286084 | 14.03564669 | -0.11278585 | -0.075336733 |
| Aadacl3 | 7.9192537 | 7.991066308 | 7.994657356 | -0.003591048 | -0.075403656 |
| 4930518I15Rik | 4.523788925 | 4.633960852 | 4.599193083 | 0.034767769 | -0.075404158 |
| Qprt | 3.116630084 | 3.085224255 | 3.19208709 | -0.106862835 | -0.075457006 |
| Cnksr2 | 3.786410872 | 3.849418017 | 3.861922708 | -0.01250469 | -0.075511835 |
| Calm2 | 12.95033912 | 12.82632345 | 13.02585745 | -0.199534 | -0.07551833 |
| Fgf18 | 7.580628092 | 7.678090816 | 7.656219574 | 0.021871242 | -0.075591483 |
| Nme5 | 2.760952011 | 2.767511173 | 2.836545317 | -0.069034143 | -0.075593306 |
| Gm10025 | 2.441887491 | 2.561132962 | 2.517497943 | 0.043635019 | -0.075610452 |
| Vmn2r84 | 2.944555045 | 2.9694092 | 3.020192401 | -0.050783201 | -0.075637357 |
| AC170998.1 | 3.631350867 | 3.685981957 | 3.707009331 | -0.021027374 | -0.075658464 |
| Prdm10 | 8.241912748 | 8.370403447 | 8.317622159 | 0.052781288 | -0.075709411 |
| Zfp369 | 8.435764687 | 8.440581367 | 8.511495227 | -0.07091386 | -0.07573054 |
| Clmp | 9.846438491 | 9.881597108 | 9.922172952 | -0.040575844 | -0.075734461 |
| Ramac | 9.542733607 | 9.417526961 | 9.618469506 | -0.200942545 | -0.075735899 |
| Elof1 | 9.233874147 | 9.251149091 | 9.309788417 | -0.058639326 | -0.07591427 |
| Rnf145 | 10.37358101 | 10.28578799 | 10.44949679 | -0.163708803 | -0.075915783 |
| Krt7 | 4.754166455 | 4.773769317 | 4.830107618 | -0.056338301 | -0.075941163 |
| Vps13a | 9.909609784 | 9.968353759 | 9.985558919 | -0.01720516 | -0.075949135 |
| Ndufs1 | 11.22028358 | 11.14100093 | 11.29624855 | -0.155247617 | -0.075964967 |
| Rbp2 | 8.259114999 | 8.410938344 | 8.335093859 | 0.075844485 | -0.07597886 |
| Slurp2 | 6.297657036 | 6.194549505 | 6.373637684 | -0.17908818 | -0.075980649 |
| Astn2 | 3.452852656 | 3.499309535 | 3.528872275 | -0.02956274 | -0.076019619 |

| | | | | | |
|---|---|---|---|---|---|
| Tbc1d15 | 10.15386433 | 10.27960137 | 10.22988914 | 0.049712233 | -0.076024812 |
| Gm5628 | 1.834200147 | 1.855185177 | 1.910272679 | -0.055087502 | -0.076072532 |
| Rfx1 | 7.50461657 | 7.901973355 | 7.580714612 | 0.321258743 | -0.076098042 |
| Olfr1160 | 2.376963671 | 2.384540409 | 2.453089558 | -0.06854915 | -0.076125888 |
| Wdr37 | 8.976485126 | 8.914967918 | 9.052653562 | -0.137685644 | -0.076168437 |
| Zfp960 | 7.658311169 | 7.56519007 | 7.734553683 | -0.169363614 | -0.076242514 |
| Ccdc166 | 5.792757604 | 5.798163358 | 5.86905903 | -0.070895672 | -0.076301426 |
| Ccar1 | 10.81543023 | 10.91629219 | 10.89174295 | 0.02454924 | -0.076312717 |
| Wdr53 | 7.559720189 | 7.587317281 | 7.636051999 | -0.048734719 | -0.076331811 |
| Wnt9b | 1.954325967 | 2.049427421 | 2.030676136 | 0.018751285 | -0.076350169 |
| Ybey | 6.859962055 | 6.70814395 | 6.936324214 | -0.228180264 | -0.076362159 |
| Haus7 | 8.370944189 | 8.262558536 | 8.447344513 | -0.184785978 | -0.076400325 |
| Olfr1396 | 1.500491889 | 1.492567854 | 1.57695065 | -0.084382796 | -0.076458761 |
| Gm10702 | 3.775595317 | 3.752930412 | 3.8520559 | -0.099125488 | -0.076460583 |
| Vmn2r11 | 3.155532483 | 3.206231745 | 3.232002902 | -0.025771157 | -0.076470419 |
| Arfip1 | 10.20783294 | 10.19156744 | 10.28433113 | -0.092763693 | -0.076498193 |
| Scn2b | 5.307364995 | 5.288595153 | 5.383899015 | -0.095303862 | -0.07653402 |
| Nkapl | 3.627273972 | 3.665267325 | 3.703829157 | -0.038561832 | -0.076555185 |
| Eda2r | 7.425442921 | 7.428223034 | 7.502001173 | -0.073778139 | -0.076558252 |
| Vmn2r28 | 1.495700819 | 1.547234024 | 1.572260913 | -0.025026889 | -0.076560094 |
| Ccdc18 | 5.754956014 | 5.667289977 | 5.831529917 | -0.16423994 | -0.076573903 |
| Tom1l1 | 8.474118189 | 8.50200124 | 8.550695867 | -0.048694627 | -0.076577677 |
| Brd3os | 5.030653945 | 5.088031575 | 5.107269669 | -0.019238095 | -0.076615725 |
| Gm5422 | 1.414110618 | 1.457656653 | 1.490757378 | -0.033100725 | -0.07664676 |
| A530084C06Rik | 3.862552789 | 3.945108284 | 3.939237071 | 0.005871213 | -0.076684282 |
| Krt19 | 4.173384002 | 4.150514072 | 4.250163041 | -0.099648969 | -0.076779039 |
| Car15 | 4.452246618 | 4.623011136 | 4.529072233 | 0.093938903 | -0.076825615 |
| Zfp287 | 5.553456045 | 5.556823421 | 5.630359756 | -0.073536335 | -0.076903711 |
| Hoxd10 | 4.235861979 | 4.153879801 | 4.312803147 | -0.158923346 | -0.076941168 |
| Olfr784 | 5.112207022 | 5.229276435 | 5.189199781 | 0.040076654 | -0.076992759 |
| Prkd3 | 9.58992327 | 9.533667295 | 9.666924198 | -0.133256903 | -0.077000928 |
| Mark2 | 11.24579763 | 11.51661133 | 11.32290153 | 0.193709793 | -0.077103903 |

| | | | | | |
|---|---|---|---|---|---|
| Lias | 9.309774359 | 9.144811376 | 9.386899279 | -0.242087903 | -0.077124921 |
| Clasp1 | 10.49711025 | 10.57316226 | 10.5743441 | -0.001181833 | -0.07723385 |
| Irx6 | 4.122614058 | 4.151620814 | 4.199906566 | -0.048285752 | -0.077292508 |
| Tmem198 | 2.87648545 | 2.872915848 | 2.953780421 | -0.080864573 | -0.07729497 |
| Snx24 | 6.508093678 | 6.71368467 | 6.585417488 | 0.128267182 | -0.07732381 |
| Cfap126 | 4.358182571 | 4.348323923 | 4.435554286 | -0.087230364 | -0.077371715 |
| Hdgfl3 | 8.878855643 | 8.671924483 | 8.956252048 | -0.284327565 | -0.077396406 |
| Nsd2 | 10.02377567 | 10.1603704 | 10.10117466 | 0.05919574 | -0.07739899 |
| AC110262.4 | 3.315795229 | 3.413187615 | 3.393242336 | 0.019945279 | -0.077447107 |
| Cyld | 9.955052887 | 9.960966856 | 10.03254228 | -0.071575421 | -0.07748939 |
| Tmem127 | 10.74354637 | 10.79755265 | 10.82105919 | -0.023506547 | -0.077512827 |
| Rpusd1 | 6.825149785 | 7.161393903 | 6.902673568 | 0.258720335 | -0.077523783 |
| Abcb9 | 6.752356086 | 7.05071856 | 6.829904103 | 0.220814457 | -0.077548017 |
| Plekhh1 | 6.695279607 | 6.719700351 | 6.772933983 | -0.053233632 | -0.077654376 |
| Incenp | 8.923052594 | 8.963144625 | 9.000736094 | -0.037591469 | -0.077683499 |
| Gm12537 | 1.550628215 | 1.642343802 | 1.62837161 | 0.013972191 | -0.077743395 |
| Pex16 | 9.579395719 | 9.609064119 | 9.657199051 | -0.048134932 | -0.077803332 |
| Ugt3a1 | 2.68298457 | 2.70667055 | 2.76079376 | -0.05412321 | -0.07780919 |
| Olfr1280 | 2.575236092 | 2.622604829 | 2.653101342 | -0.030496514 | -0.07786525 |
| Mbp | 10.8432009 | 10.80619733 | 10.92121277 | -0.115015433 | -0.078011867 |
| Crisp2 | 1.31951162 | 1.301957636 | 1.397607926 | -0.09565029 | -0.078096306 |
| Mosmo | 8.992981182 | 9.01808864 | 9.071177464 | -0.053088824 | -0.078196283 |
| Wdr38 | 3.225138983 | 3.308206314 | 3.303337187 | 0.004869127 | -0.078198205 |
| Mplkip | 7.691963156 | 7.696091563 | 7.770241162 | -0.0741496 | -0.078278006 |
| Hexim1 | 9.453686057 | 9.812876487 | 9.532000801 | 0.280875686 | -0.078314744 |
| Ube2d-ps | 7.858313584 | 7.889463275 | 7.936636845 | -0.04717357 | -0.07832326 |
| Slc9a5 | 4.637215817 | 4.813183268 | 4.715596477 | 0.097586791 | -0.07838066 |
| Rassf4 | 9.932664486 | 10.05526343 | 10.01111421 | 0.044149214 | -0.078449728 |
| Smc6 | 10.45467307 | 10.45252294 | 10.53312933 | -0.080606393 | -0.078456257 |
| Adam10 | 11.00450555 | 11.00659886 | 11.08300825 | -0.07640939 | -0.078502697 |
| Dhrs7b | 9.017132954 | 9.067071953 | 9.095655127 | -0.028583174 | -0.078522173 |
| Tmem176a | 9.110681639 | 9.041601973 | 9.189245054 | -0.147643081 | -0.078563415 |

| | | | | | |
|---|---|---|---|---|---|
| K230010J24Rik | 5.305814433 | 5.450744213 | 5.384435365 | 0.066308848 | -0.078620932 |
| Tmed3 | 10.29895069 | 10.3240298 | 10.37765108 | -0.053621277 | -0.07870039 |
| Bola3 | 8.52027877 | 8.390894594 | 8.598993567 | -0.208098973 | -0.078714798 |
| Nudc-ps1 | 1.674250504 | 1.695249257 | 1.753118322 | -0.057869065 | -0.078867817 |
| Sumo3 | 10.72867831 | 10.7479235 | 10.80757761 | -0.05965411 | -0.0788993 |
| Smarcal1 | 8.338750666 | 8.418535425 | 8.41769093 | 0.000844495 | -0.078940265 |
| Srek1 | 10.15893379 | 10.18594866 | 10.237904 | -0.051955347 | -0.078970213 |
| Clca1 | 4.814915676 | 4.897761184 | 4.893891749 | 0.003869435 | -0.078976073 |
| Wdfy3 | 10.65128488 | 10.82080136 | 10.73026428 | 0.090537087 | -0.078979397 |
| Gm6525 | 2.230236786 | 2.289237248 | 2.309252122 | -0.020014874 | -0.079015335 |
| Nebl | 10.11420644 | 10.15447454 | 10.19322946 | -0.038754923 | -0.07902302 |
| Cdc45 | 7.679531902 | 7.916412239 | 7.758575244 | 0.157836995 | -0.079043342 |
| Rc3h2 | 10.14316815 | 10.10632944 | 10.2222155 | -0.115886056 | -0.07904735 |
| Zfp184 | 2.726166223 | 2.778618169 | 2.805301716 | -0.026683546 | -0.079135492 |
| Myo10 | 10.21910099 | 10.22735463 | 10.2982789 | -0.070924267 | -0.079177903 |
| Atpif1 | 10.53328169 | 10.57465087 | 10.61257234 | -0.037921467 | -0.079290647 |
| Zfp933 | 7.775651428 | 7.954753647 | 7.854973097 | 0.09978055 | -0.079321669 |
| Rab4a | 5.862740732 | 5.806800706 | 5.942087912 | -0.135287206 | -0.07934718 |
| Nfatc2ip | 7.884948836 | 7.980054114 | 7.964327279 | 0.015726835 | -0.079378443 |
| Tnks2 | 11.48826585 | 11.54016983 | 11.56767129 | -0.027501457 | -0.079405437 |
| Dqx1 | 5.342151236 | 5.39965561 | 5.421602214 | -0.021946604 | -0.079450978 |
| Styx | 9.562808798 | 9.498577569 | 9.642277894 | -0.143700325 | -0.079469096 |
| Gcfc2 | 7.310129144 | 7.278299599 | 7.38969887 | -0.111399271 | -0.079569726 |
| Gm5578 | 2.111642633 | 2.207374734 | 2.191214645 | 0.016160089 | -0.079572011 |
| Tvp23b | 9.040983203 | 9.019105124 | 9.120561315 | -0.101456191 | -0.079578112 |
| Acap3 | 9.710064591 | 9.964484726 | 9.78965665 | 0.174828076 | -0.079592059 |
| Mcm9 | 7.922600056 | 8.006814574 | 8.002210985 | 0.004603588 | -0.07961093 |
| 1700073E17Rik | 2.988036413 | 3.055604287 | 3.067662089 | -0.012057802 | -0.079625676 |
| Ppwd1 | 8.068342389 | 8.062093515 | 8.147975363 | -0.085881847 | -0.079632974 |
| 6330549D23Rik | 4.075712841 | 4.160075796 | 4.155351802 | 0.004723993 | -0.079638962 |
| Ttll2 | 3.202111076 | 3.263945895 | 3.281797881 | -0.017851986 | -0.079686805 |
| Cyyr1 | 8.506960006 | 8.318958439 | 8.586659977 | -0.267701538 | -0.07969997 |

| | | | | | |
|---|---|---|---|---|---|
| Arfgef3 | 3.570543398 | 3.679928071 | 3.650378065 | 0.029550006 | -0.079834667 |
| Ndufb4 | 7.883854251 | 7.726261944 | 7.963706466 | -0.237444522 | -0.079852215 |
| Rps20 | 12.5100535 | 12.5023549 | 12.58990752 | -0.087552613 | -0.079854013 |
| Vps36 | 10.06356061 | 10.14300237 | 10.1434718 | -0.000469433 | -0.079911187 |
| Zfp691 | 6.828484471 | 6.974041857 | 6.908571485 | 0.065470372 | -0.080087014 |
| Hivep1 | 10.21909009 | 10.14449215 | 10.2992458 | -0.154753647 | -0.080155703 |
| Nt5c3 | 10.87335949 | 10.79296256 | 10.95357948 | -0.16061692 | -0.08021999 |
| Pon1 | 5.955668586 | 5.830493818 | 6.035897497 | -0.205403678 | -0.08022891 |
| Csnk2b | 11.48060146 | 11.48027267 | 11.56083081 | -0.080558147 | -0.080229353 |
| Gm15446 | 7.92180297 | 7.803120722 | 8.002045868 | -0.198925146 | -0.080242898 |
| Mettl26 | 7.311786109 | 7.376007921 | 7.392037069 | -0.016029148 | -0.08025096 |
| Zfp410 | 9.713431621 | 9.815440535 | 9.79374168 | 0.021698854 | -0.08031006 |
| Hoxb6 | 5.987913571 | 6.17210308 | 6.068274336 | 0.103828745 | -0.080360764 |
| Uhrf2 | 9.470445976 | 9.631161177 | 9.550826987 | 0.08033419 | -0.08038101 |
| Tmsb15b2 | 3.252926136 | 3.24766179 | 3.333314127 | -0.085652337 | -0.080387991 |
| Ndufc1 | 9.29623216 | 9.322624985 | 9.376694599 | -0.054069614 | -0.080462439 |
| AC098880.2 | 6.928864358 | 6.973506173 | 7.009406061 | -0.035899888 | -0.080541702 |
| Slc17a8 | 2.869995625 | 2.910911159 | 2.950590494 | -0.039679335 | -0.08059487 |
| Zfp970 | 7.937306284 | 8.037563879 | 8.017907672 | 0.019656206 | -0.080601388 |
| Cma1 | 9.872314174 | 9.242943873 | 9.952955125 | -0.710011252 | -0.080640951 |
| Hus1 | 7.957590157 | 7.948704692 | 8.038368897 | -0.089664205 | -0.08077874 |
| C4bp | 3.078405014 | 3.118207949 | 3.159245133 | -0.041037184 | -0.080840119 |
| Tgfbr3l | 5.142433018 | 5.186096724 | 5.223275977 | -0.037179253 | -0.080842958 |
| Gm4707 | 3.974849381 | 3.989936748 | 4.055706247 | -0.065769498 | -0.080856866 |
| Rpsa-ps10 | 2.760657656 | 2.844134439 | 2.841552658 | 0.002581782 | -0.080895002 |
| Chadl | 3.55950382 | 3.645268058 | 3.64051064 | 0.004757418 | -0.08100682 |
| Ecpas | 11.5882722 | 11.62891511 | 11.66933604 | -0.04042093 | -0.081063847 |
| Mvb12b | 8.302710739 | 8.250061096 | 8.383775624 | -0.133714527 | -0.081064885 |
| Exoc3l | 3.908317982 | 3.979963801 | 3.989388856 | -0.009425055 | -0.081070874 |
| Setd5 | 11.24923495 | 11.27185119 | 11.33032122 | -0.058470033 | -0.081086277 |
| Actr3b | 4.500072396 | 4.522638762 | 4.581160883 | -0.058522121 | -0.081088487 |
| Cbx1 | 9.895500141 | 9.948055605 | 9.97661569 | -0.028560085 | -0.081115549 |

| | | | | | |
|---|---|---|---|---|---|
| Rpl36a-ps3 | 1.737510024 | 1.769933432 | 1.818631421 | -0.048697989 | -0.081121397 |
| Ostm1 | 9.36120812 | 9.252992426 | 9.44232969 | -0.189337263 | -0.08112157 |
| Chchd7 | 7.576088049 | 7.624828966 | 7.657321829 | -0.032492863 | -0.081233781 |
| Slc38a7 | 8.582891464 | 8.848584079 | 8.664227294 | 0.184356785 | -0.08133583 |
| Spaca1 | 2.694994427 | 2.734576216 | 2.776365498 | -0.041789282 | -0.081371071 |
| Itih3 | 1.985066625 | 2.045047502 | 2.066446547 | -0.021399045 | -0.081379922 |
| Xkr5 | 3.200171661 | 3.151824635 | 3.281607392 | -0.129782757 | -0.081435732 |
| Dido1 | 10.25476318 | 10.41872123 | 10.33620422 | 0.082517007 | -0.081441043 |
| Gm10134 | 3.650232121 | 3.628448714 | 3.731780611 | -0.103331897 | -0.081548491 |
| Pomt2 | 7.921740753 | 7.970059259 | 8.003313926 | -0.033254668 | -0.081573173 |
| Adam23 | 8.769239619 | 8.878401922 | 8.850874853 | 0.027527069 | -0.081635234 |
| Osbpl10 | 4.876571613 | 4.987801128 | 4.958216206 | 0.029584922 | -0.081644593 |
| Cbfb | 10.64623719 | 10.58422361 | 10.72790233 | -0.14367872 | -0.081665143 |
| Prdm1 | 9.669064438 | 9.830613513 | 9.750736759 | 0.079876754 | -0.081672321 |
| Gm33312 | 1.995008641 | 2.140330471 | 2.076693259 | 0.063637212 | -0.081684618 |
| Rai2 | 6.969983388 | 7.100673241 | 7.051686277 | 0.048986965 | -0.081702888 |
| 2900011O08Rik | 1.69678946 | 1.729428031 | 1.778557345 | -0.049129314 | -0.081767885 |
| Orm1 | 11.10639837 | 10.9123965 | 11.18816752 | -0.27577102 | -0.081769143 |
| Socs4 | 9.724095545 | 9.751185323 | 9.805912759 | -0.054727436 | -0.081817214 |
| Gm10330 | 3.756405951 | 3.859691378 | 3.838226666 | 0.021464712 | -0.081820715 |
| Tll2 | 1.639319911 | 1.694697911 | 1.721205069 | -0.026507158 | -0.081885158 |
| Cenpv | 7.327641365 | 7.430567731 | 7.409543513 | 0.021024218 | -0.081902148 |
| Vmn2r53 | 7.78465105 | 7.814738195 | 7.866581162 | -0.051842967 | -0.081930112 |
| Cep76 | 7.349265495 | 7.398012557 | 7.431227011 | -0.033214454 | -0.081961516 |
| Syt14 | 2.630650297 | 2.652950153 | 2.712655068 | -0.059704915 | -0.08200477 |
| Pla2g4c | 3.77649539 | 3.914917945 | 3.858510819 | 0.056407126 | -0.082015429 |
| Agap3 | 10.19122735 | 10.5464754 | 10.27324651 | 0.273228887 | -0.082019167 |
| 1810030O07Rik | 9.418115696 | 9.490723065 | 9.500183387 | -0.009460321 | -0.082067691 |
| Sigirr | 6.45784941 | 6.555550553 | 6.539979189 | 0.015571364 | -0.08212978 |
| Neurl4 | 9.26500707 | 9.37598056 | 9.347161871 | 0.028818689 | -0.082154801 |
| Faf1 | 10.22196458 | 10.29115656 | 10.30415348 | -0.012996923 | -0.0821889 |
| Pla1a | 6.686489014 | 6.296022904 | 6.768708128 | -0.472685224 | -0.082219114 |

| | | | | | |
|---|---|---|---|---|---|
| Tgfbrap1 | 9.81276365 | 9.880666569 | 9.894986158 | -0.014319589 | -0.082222508 |
| Sh3bp5l | 9.75016986 | 10.00168913 | 9.832433861 | 0.169255265 | -0.082264001 |
| Tead3 | 9.888093793 | 10.27169558 | 9.970361267 | 0.30133431 | -0.082267474 |
| Zfp68 | 9.626636089 | 9.716960555 | 9.7089181 | 0.008042455 | -0.082282011 |
| Olfr981 | 2.680655375 | 2.731459392 | 2.762961552 | -0.03150216 | -0.082306177 |
| Gpr19 | 5.691484761 | 5.669394653 | 5.773848984 | -0.104454331 | -0.082364224 |
| Tulp4 | 11.57772932 | 11.70767593 | 11.66009605 | 0.04757988 | -0.08236673 |
| Mtfmt | 7.45883263 | 7.59598806 | 7.541290137 | 0.054697923 | -0.082457507 |
| Fam184a | 3.904036162 | 3.95634741 | 3.986526201 | -0.030178791 | -0.082490039 |
| 4930452B06Rik | 1.383997831 | 1.394810895 | 1.466502233 | -0.071691338 | -0.082504401 |
| Hars2 | 8.904669366 | 9.074963249 | 8.987225402 | 0.087737847 | -0.082556036 |
| Fez2 | 8.911729566 | 8.836186789 | 8.99432102 | -0.158134231 | -0.082591454 |
| Anapc4 | 9.671758007 | 9.602773644 | 9.754362265 | -0.151588621 | -0.082604257 |
| Zfp706 | 11.89346192 | 11.92894261 | 11.97614711 | -0.047204493 | -0.082685187 |
| Uros | 7.813669806 | 7.717531769 | 7.896374502 | -0.178842733 | -0.082704696 |
| AC122270.2 | 2.053574492 | 2.062250826 | 2.13628714 | -0.074036314 | -0.082712648 |
| Gm5805 | 3.357476911 | 3.425134054 | 3.440237494 | -0.015103441 | -0.082760583 |
| Laptm4b | 10.35057775 | 10.38052903 | 10.43342066 | -0.052891623 | -0.082842903 |
| Kat8 | 8.385033417 | 8.410037964 | 8.467969485 | -0.057931521 | -0.082936068 |
| Kctd6 | 7.831009893 | 7.858453004 | 7.913958388 | -0.055505385 | -0.082948495 |
| Lmo1 | 7.701392718 | 7.784551119 | 7.784367764 | 0.000183355 | -0.082975046 |
| Mgme1 | 7.406854369 | 7.468901092 | 7.489865312 | -0.02096422 | -0.083010943 |
| 3110021N24Rik | 3.156511937 | 3.244058975 | 3.239537987 | 0.004520988 | -0.08302605 |
| Eef1a1 | 16.84827827 | 16.80872467 | 16.93145348 | -0.122728817 | -0.08317521 |
| Smim20 | 8.412758931 | 8.418065511 | 8.495938501 | -0.077872989 | -0.08317957 |
| Hey2 | 3.356724607 | 3.32024159 | 3.4399275 | -0.11968591 | -0.083202894 |
| Pkdcc | 8.051319763 | 8.298189338 | 8.134563443 | 0.163625894 | -0.08324368 |
| Fstl3 | 8.429391203 | 8.664422537 | 8.512704232 | 0.151718305 | -0.083313029 |
| Col4a3bp | 10.55853181 | 10.67956171 | 10.64187196 | 0.03768975 | -0.083340143 |
| Socs7 | 9.767322218 | 9.90895495 | 9.850671225 | 0.058283725 | -0.083349006 |
| Uprt | 6.598341024 | 6.624673284 | 6.681780466 | -0.057107183 | -0.083439442 |
| Carmil3 | 4.074430306 | 4.210445239 | 4.157947959 | 0.05249728 | -0.083517653 |

| | | | | | |
|---|---|---|---|---|---|
| 4930539E08Rik | 2.725059077 | 2.76499058 | 2.808601735 | -0.043611154 | -0.083542657 |
| Gfm2 | 9.434098065 | 9.458491801 | 9.517642879 | -0.059151078 | -0.083544814 |
| Olfr654 | 2.961198317 | 2.973315966 | 3.044800221 | -0.071484255 | -0.083601904 |
| P3h4 | 8.805527965 | 8.771999867 | 8.889182363 | -0.117182495 | -0.083654398 |
| Lgi1 | 5.292469399 | 5.110183354 | 5.376173933 | -0.265990579 | -0.083704535 |
| Sumo1 | 10.58658751 | 10.56480088 | 10.67030405 | -0.105503163 | -0.083716537 |
| P2rx7 | 8.224230479 | 9.07842419 | 8.307951871 | 0.770472319 | -0.083721392 |
| Pls3 | 12.00330206 | 12.04233963 | 12.08704886 | -0.04470923 | -0.0837468 |
| Gm13199 | 2.308238562 | 2.401372364 | 2.391992818 | 0.009379545 | -0.083754256 |
| Kcng4 | 4.405803985 | 4.526443746 | 4.489561394 | 0.036882352 | -0.083757409 |
| Arhgap1 | 10.77525087 | 10.97460703 | 10.85904054 | 0.115566487 | -0.083789677 |
| Sprydt7 | 9.650558256 | 9.491232444 | 9.734370645 | -0.243138201 | -0.083812389 |
| Rac3 | 7.162677283 | 7.23752704 | 7.246519888 | -0.008992849 | -0.083842605 |
| Nid1 | 11.43236508 | 11.09004515 | 11.51625637 | -0.426211223 | -0.083891293 |
| Olfr1352 | 3.854192641 | 3.877152616 | 3.938098508 | -0.060945892 | -0.083905867 |
| Eogt | 9.016790946 | 9.236021479 | 9.100712163 | 0.135309315 | -0.083921217 |
| Gm8864 | 1.385741994 | 1.451808521 | 1.469668634 | -0.017860113 | -0.08392664 |
| Pthlh | 5.232526526 | 5.125270302 | 5.316458668 | -0.191188365 | -0.083932142 |
| Zfp454 | 3.62888895 | 3.609148285 | 3.712856671 | -0.103708385 | -0.083967721 |
| Cyfip1 | 11.44475725 | 11.47638144 | 11.52874014 | -0.052358693 | -0.083982887 |
| Nosip | 8.985360256 | 9.184029017 | 9.069403798 | 0.114625219 | -0.084043542 |
| Gfra4 | 4.311901295 | 4.2773337 | 4.395949538 | -0.118615839 | -0.084048244 |
| Acp1 | 10.2988354 | 10.21925404 | 10.38288806 | -0.163634017 | -0.084052657 |
| Fsbp | 4.386411037 | 4.395644356 | 4.470486794 | -0.074842439 | -0.084075758 |
| Pacrg | 3.285946212 | 3.430996074 | 3.370061861 | 0.060934212 | -0.084115649 |
| Slc35a3 | 9.710742311 | 9.659379818 | 9.794876679 | -0.135496861 | -0.084134368 |
| Mipep | 8.64619425 | 8.635473714 | 8.730430205 | -0.094956491 | -0.084235954 |
| Rab7-ps1 | 1.488006678 | 1.522694969 | 1.572343205 | -0.049648236 | -0.084336526 |
| Fbxw2 | 10.19363488 | 10.0976268 | 10.2780108 | -0.180384003 | -0.08437592 |
| Thoc2 | 11.08424743 | 11.03951448 | 11.16867262 | -0.12915814 | -0.084425193 |
| Gm11703 | 1.761057448 | 1.76056627 | 1.845534233 | -0.084967964 | -0.084476785 |
| Gm9726 | 2.142966821 | 2.338021012 | 2.227445763 | 0.110575249 | -0.084478942 |

| | | | | | |
|---|---|---|---|---|---|
| Dlec1 | 2.629414835 | 2.707112969 | 2.713901044 | -0.006788075 | -0.084486209 |
| Trank1 | 4.189168752 | 4.114423769 | 4.273705642 | -0.159281872 | -0.08453689 |
| Olfr1246 | 2.752843945 | 2.870135874 | 2.837446411 | 0.032689463 | -0.084602466 |
| Zfp692 | 9.033507094 | 9.392329266 | 9.118150892 | 0.274178374 | -0.084643798 |
| Prtg | 3.020679655 | 2.994628516 | 3.105329422 | -0.110700907 | -0.084649767 |
| Dyrk1a | 10.35539957 | 10.55185893 | 10.44015484 | 0.111704087 | -0.08475527 |
| Zfp157 | 7.267542169 | 7.196370617 | 7.35231898 | -0.155948363 | -0.084776811 |
| Adamts19 | 3.766384576 | 3.771606889 | 3.851168338 | -0.079561449 | -0.084783762 |
| Slitrk3 | 2.023360176 | 2.060285998 | 2.10817754 | -0.047891542 | -0.084817364 |
| Gstm6 | 6.403235412 | 6.307217823 | 6.488150094 | -0.180932271 | -0.084914682 |
| Isoc2a | 8.590480402 | 8.55537759 | 8.675395208 | -0.120017618 | -0.084914806 |
| Cenpc1 | 8.177236546 | 8.222607601 | 8.2621742 | -0.039566599 | -0.084937654 |
| Zfp954 | 8.419259144 | 8.498711989 | 8.504236703 | -0.005524714 | -0.084977559 |
| Ap3s1 | 9.621414553 | 9.482969307 | 9.70641445 | -0.223445143 | -0.084999897 |
| Armc10 | 8.985543627 | 8.942224364 | 9.070593182 | -0.128368818 | -0.085049555 |
| Vmn2r60 | 4.610233146 | 4.666972547 | 4.695330916 | -0.02835837 | -0.08509777 |
| Atp8a1 | 9.826756351 | 9.843061493 | 9.911979165 | -0.068917672 | -0.085222814 |
| Xrn1 | 10.16611103 | 10.17416811 | 10.2513624 | -0.07719429 | -0.08525137 |
| Stx8 | 8.643066061 | 8.552630494 | 8.728328438 | -0.175697943 | -0.085262376 |
| Gm13378 | 2.864134319 | 2.986850587 | 2.949404258 | 0.037446329 | -0.085269939 |
| Gm42915 | 1.990451079 | 2.023450209 | 2.075861135 | -0.052410926 | -0.085410056 |
| Kif19a | 4.552789112 | 4.569825336 | 4.638231245 | -0.068405909 | -0.085442133 |
| Tmprss4 | 11.7220468 | 11.89810947 | 11.80750086 | 0.09060861 | -0.08545406 |
| Zbtb33 | 9.473215795 | 9.523748059 | 9.558689407 | -0.034941348 | -0.085473612 |
| Ctbs | 7.413934063 | 7.355713017 | 7.499466575 | -0.143753558 | -0.085532512 |
| Oaz3 | 2.765941445 | 2.778406402 | 2.851475045 | -0.073068643 | -0.0855336 |
| Pcsk2 | 2.409087734 | 2.465532725 | 2.494632517 | -0.029099792 | -0.085544783 |
| Rnf5 | 9.02876369 | 9.10341493 | 9.11436048 | -0.01094555 | -0.08559679 |
| Brd8 | 9.769418154 | 9.769487297 | 9.855144164 | -0.085656867 | -0.08572601 |
| Ctdspl2 | 8.760220472 | 8.780021664 | 8.845963348 | -0.065941683 | -0.085742875 |
| Mapre2 | 10.35429409 | 10.27135027 | 10.44010691 | -0.16875664 | -0.085812827 |
| Lrrc58 | 11.2231026 | 11.19640167 | 11.30892997 | -0.1125283 | -0.085827367 |

| | | | | | |
|---|---|---|---|---|---|
| Ankrd13c | 10.33454521 | 10.31010566 | 10.42038782 | -0.11028216 | -0.08584261 |
| Eef2 | 15.0440619 | 15.29441373 | 15.129957 | 0.164456727 | -0.085895103 |
| Etaa1 | 7.963013468 | 8.01355075 | 8.048983698 | -0.035432948 | -0.08597023 |
| Vps35 | 11.47635965 | 11.52844233 | 11.56239493 | -0.033952607 | -0.08603528 |
| 9430007A20Rik | 5.248320963 | 5.328216168 | 5.334357171 | -0.006141003 | -0.086036208 |
| Mtif2 | 8.862788683 | 8.855510326 | 8.948846962 | -0.093336635 | -0.086058279 |
| St7l | 8.48390293 | 8.551911818 | 8.569961788 | -0.01804997 | -0.086058858 |
| Brinp2 | 1.790783083 | 1.87586556 | 1.876949094 | -0.001083534 | -0.086166011 |
| Pfn4 | 2.593241239 | 2.621576052 | 2.679424491 | -0.057848439 | -0.086183252 |
| Zfp319 | 8.398027201 | 8.630644018 | 8.484276554 | 0.146367464 | -0.086249353 |
| Cdiptos | 3.364489914 | 3.348543393 | 3.45074543 | -0.102202036 | -0.086255516 |
| Tmem60 | 7.765835656 | 7.731780141 | 7.852133644 | -0.120353503 | -0.086297988 |
| Synpo2 | 11.19466094 | 10.9065858 | 11.28096953 | -0.37438373 | -0.08630859 |
| Gm49486 | 3.332464776 | 3.428146323 | 3.418774361 | 0.009371963 | -0.086309584 |
| Gm10040 | 5.683043303 | 5.744384399 | 5.769359732 | -0.024975333 | -0.086316429 |
| Cd163 | 10.90059322 | 10.61702586 | 10.98691659 | -0.36989073 | -0.08632337 |
| Smap1 | 10.56454713 | 10.59461912 | 10.65088596 | -0.056266843 | -0.086338827 |
| S100b | 6.060099588 | 5.990036955 | 6.146454126 | -0.156417171 | -0.086354538 |
| Spop | 11.73645512 | 11.67260294 | 11.82284622 | -0.150243287 | -0.0863911 |
| Gm39288 | 2.076562356 | 2.113836189 | 2.162959593 | -0.049123404 | -0.086397238 |
| Tax1bp3 | 9.89004451 | 9.95815383 | 9.976464686 | -0.018310856 | -0.086420176 |
| Gm11821 | 2.023727038 | 2.112618237 | 2.110152082 | 0.002466154 | -0.086425044 |
| Ece2 | 4.631068895 | 4.739252291 | 4.717503085 | 0.021749206 | -0.08643419 |
| Kcnn2 | 4.078283254 | 4.17497926 | 4.16473114 | 0.010248121 | -0.086447886 |
| Gpn3 | 7.985205182 | 8.037487756 | 8.07185385 | -0.034366093 | -0.086648667 |
| D830030K20Rik | 3.858474373 | 3.844566538 | 3.945153731 | -0.100587192 | -0.086679357 |
| Chfr | 9.813505425 | 9.947810676 | 9.900203992 | 0.047606684 | -0.086698567 |
| Olfr624 | 7.721321821 | 7.778850244 | 7.808060531 | -0.029210287 | -0.08673871 |
| Mllt10 | 9.827615186 | 10.00104306 | 9.914367332 | 0.086675731 | -0.086752147 |
| Gm4787 | 3.354063461 | 3.422226701 | 3.440818683 | -0.018591982 | -0.086755222 |
| Zfp458 | 5.477291914 | 5.469495004 | 5.564047247 | -0.094552243 | -0.086755333 |
| Zc3h12b | 4.18086867 | 4.174397857 | 4.26763157 | -0.093233713 | -0.0867629 |

| | | | | | |
|---|---|---|---|---|---|
| Spata6 | 7.934189122 | 7.711752064 | 8.020957642 | -0.309205578 | -0.086768519 |
| Patz1 | 7.209771703 | 7.238945802 | 7.296552901 | -0.057607099 | -0.086781198 |
| Plk4 | 7.002502124 | 7.049020835 | 7.089320451 | -0.040299616 | -0.086818327 |
| Rnf212 | 3.270342353 | 3.309549892 | 3.357174188 | -0.047624296 | -0.086831835 |
| Tnfsf13 | 5.966830099 | 5.809894363 | 6.053743718 | -0.243849355 | -0.086913619 |
| Gm10647 | 1.839460803 | 1.865454725 | 1.926375282 | -0.060920557 | -0.086914478 |
| Tbce | 9.221343309 | 9.291565932 | 9.308264831 | -0.016698899 | -0.086921522 |
| Cited1 | 2.012665931 | 2.076825391 | 2.099594988 | -0.022769597 | -0.086929057 |
| Lrrtm1 | 2.695569445 | 2.772220995 | 2.782511825 | -0.010290831 | -0.08694238 |
| AC102119.1 | 2.275354911 | 2.306473777 | 2.362309332 | -0.055835555 | -0.086954421 |
| Edem3 | 10.17271938 | 10.29177371 | 10.2597475 | 0.03202621 | -0.087028127 |
| Rbm38 | 8.002356655 | 8.453901078 | 8.089391621 | 0.364509457 | -0.087034966 |
| Gm28035 | 3.924206827 | 4.154776171 | 4.01125896 | 0.143517211 | -0.087052133 |
| Pagr1b | 2.410355824 | 2.398794869 | 2.497420323 | -0.098625453 | -0.087064499 |
| Serpina9 | 4.778491015 | 4.907368975 | 4.865577005 | 0.04179197 | -0.08708599 |
| Sox18 | 6.390367069 | 6.481967028 | 6.477472037 | 0.00449499 | -0.087104968 |
| Taf6 | 9.009235081 | 9.197095325 | 9.096372357 | 0.100722968 | -0.087137275 |
| Ermap | 4.37970333 | 4.266421989 | 4.466919404 | -0.200497415 | -0.087216074 |
| Tfr2 | 4.093181929 | 4.088758255 | 4.180401812 | -0.091643558 | -0.087219883 |
| Cul1 | 10.7528409 | 10.77453188 | 10.84009831 | -0.06556643 | -0.08725741 |
| Agrp | 3.467384446 | 3.575897893 | 3.554676773 | 0.02122112 | -0.087292327 |
| Blvrb | 8.52855682 | 8.686627387 | 8.615870996 | 0.070756391 | -0.087314176 |
| Gulo | 2.684665482 | 2.691986157 | 2.771988128 | -0.080001971 | -0.087322646 |
| Ptdss2 | 8.770337513 | 8.977795682 | 8.857671668 | 0.120124014 | -0.087334155 |
| Gabra4 | 3.98385515 | 4.148555409 | 4.071327012 | 0.077228397 | -0.087471862 |
| Plpp5 | 7.984523957 | 8.105002323 | 8.072070437 | 0.032931886 | -0.08754648 |
| Rsph10b | 2.579022584 | 2.676923468 | 2.666576408 | 0.01034706 | -0.087553824 |
| Farp2 | 7.514676963 | 7.625480599 | 7.60226026 | 0.023220339 | -0.087583298 |
| Tpm2 | 14.05205725 | 14.21180999 | 14.13965676 | 0.072153237 | -0.08759951 |
| AU022252 | 8.946162989 | 9.044178516 | 9.033804115 | 0.010374401 | -0.087641126 |
| Acadl | 11.12312746 | 10.93915052 | 11.21080758 | -0.271657053 | -0.087680113 |
| Naglu | 7.632785658 | 7.8574352 | 7.720485218 | 0.136949982 | -0.08769956 |

| | | | | | |
|---|---|---|---|---|---|
| Kctd7 | 6.023095635 | 6.187240046 | 6.110898292 | 0.076341754 | -0.087802657 |
| Kdm2b | 9.939409441 | 10.23407705 | 10.0272234 | 0.206853653 | -0.087813959 |
| Arsa | 9.328835513 | 9.416009413 | 9.416651816 | -0.000642403 | -0.087816303 |
| Hist2h3c2 | 3.561628563 | 3.698708468 | 3.649477562 | 0.049230906 | -0.087848999 |
| Lin37 | 7.507450798 | 7.619732282 | 7.595301185 | 0.024431097 | -0.087850387 |
| Zfp286 | 3.845954813 | 3.933417706 | 3.93384051 | -0.000422803 | -0.087885697 |
| Fam167b | 3.259681198 | 3.287248238 | 3.347570599 | -0.060322361 | -0.087889401 |
| Gtf3c1 | 10.15526748 | 10.45289972 | 10.24324884 | 0.20965088 | -0.08798136 |
| Rtel1 | 8.49467577 | 8.712953799 | 8.582692553 | 0.130261246 | -0.088016783 |
| Mindy4 | 5.882139946 | 5.915196685 | 5.970185473 | -0.054988788 | -0.088045527 |
| Efcab11 | 4.354313169 | 4.408351149 | 4.44244286 | -0.034091711 | -0.088129691 |
| Tatdn1 | 7.760941875 | 7.746799449 | 7.849072685 | -0.102273235 | -0.08813081 |
| Epc2 | 9.343221855 | 9.251107043 | 9.431399515 | -0.180292472 | -0.08817766 |
| Gm7049 | 2.524703351 | 2.569794566 | 2.612907081 | -0.043112515 | -0.088203731 |
| Mcph1 | 7.680838541 | 7.707440733 | 7.769074917 | -0.061634184 | -0.088236376 |
| Gm42927 | 8.492786859 | 8.732757463 | 8.581030398 | 0.151727065 | -0.08824354 |
| Cspg5 | 5.372831177 | 5.306576933 | 5.461093266 | -0.154516333 | -0.088262089 |
| Rsph9 | 4.226005813 | 4.203012079 | 4.314281201 | -0.111269122 | -0.088275387 |
| Erp29 | 10.71328014 | 10.82979842 | 10.80156827 | 0.02823015 | -0.08828813 |
| Bend7 | 5.328853271 | 5.278263462 | 5.417170394 | -0.138906932 | -0.088317123 |
| Prpf39 | 10.11383979 | 10.23802972 | 10.20217039 | 0.035859327 | -0.088330603 |
| Gm28530 | 2.515765309 | 2.56242658 | 2.604140802 | -0.041714222 | -0.088375493 |
| Ms4a14 | 5.757501244 | 5.897173419 | 5.845971455 | 0.051201964 | -0.088470211 |
| Stab2 | 3.404103102 | 3.422981637 | 3.49261767 | -0.069636032 | -0.088514568 |
| Cant1 | 9.644790599 | 9.786523497 | 9.733417189 | 0.053106308 | -0.08862659 |
| Gtf2a1 | 9.69891408 | 9.60476854 | 9.787568419 | -0.18279988 | -0.088654339 |
| AC163399.1 | 2.118385795 | 2.152722794 | 2.207041176 | -0.054318382 | -0.08865538 |
| Hes7 | 2.847287102 | 2.924816749 | 2.936030353 | -0.011213604 | -0.08874325 |
| Bscl2 | 10.63748579 | 10.65474729 | 10.72629022 | -0.071542923 | -0.088804427 |
| Cyb5r3 | 11.72156914 | 11.80411939 | 11.8104585 | -0.006339107 | -0.08888936 |
| Gngt1 | 3.07520786 | 3.08364244 | 3.164111491 | -0.080469051 | -0.088903631 |
| Wdr86 | 4.099014307 | 4.164312184 | 4.187949088 | -0.023636904 | -0.088934781 |

| | | | | | |
|---|---|---|---|---|---|
| Rtl8b | 7.770693274 | 7.984497499 | 7.859655278 | 0.124842222 | -0.088962003 |
| Zdhhc8 | 7.813773865 | 8.15957836 | 7.902755561 | 0.256822799 | -0.088981697 |
| Wdr81 | 8.912976465 | 9.334681456 | 9.001960726 | 0.332720729 | -0.088984261 |
| Syt3 | 4.614332392 | 4.603195688 | 4.70335164 | -0.100155952 | -0.089019247 |
| Abhd16a | 9.864949284 | 10.03900422 | 9.953975931 | 0.085028286 | -0.089026647 |
| Shisa7 | 7.181897554 | 7.164037954 | 7.270932803 | -0.106894849 | -0.089035249 |
| Hectd1 | 12.21264047 | 12.22840794 | 12.30168697 | -0.07327903 | -0.089046503 |
| Csnk1g1 | 9.581838061 | 9.547294188 | 9.67093638 | -0.123642193 | -0.089098319 |
| Spire2 | 5.747822257 | 5.90889762 | 5.836969357 | 0.071928264 | -0.0891471 |
| Trrap | 10.21229229 | 10.44196013 | 10.301452 | 0.14050813 | -0.089159703 |
| Lax1 | 5.731395913 | 5.869855354 | 5.820570815 | 0.04928454 | -0.089174901 |
| Dzip1l | 8.414090822 | 8.607517523 | 8.503271135 | 0.104246388 | -0.089180314 |
| Zfp300 | 3.742075116 | 3.716342414 | 3.831291379 | -0.114948966 | -0.089216263 |
| Pds5b | 9.536107355 | 9.579433031 | 9.625328385 | -0.045895354 | -0.08922103 |
| Ccdc114 | 4.498931476 | 4.623787547 | 4.588164152 | 0.035623395 | -0.089232676 |
| Bub1 | 7.038215874 | 7.05176963 | 7.127451342 | -0.075681712 | -0.089235468 |
| Gm48443 | 6.220610405 | 6.242481954 | 6.309898691 | -0.067416737 | -0.089288286 |
| Nucb1 | 11.29816005 | 11.38055086 | 11.38746233 | -0.006911473 | -0.08930228 |
| Cyb5r1 | 9.547408797 | 9.593742392 | 9.636715388 | -0.042972996 | -0.089306591 |
| Tmem59l | 2.89903312 | 2.929515021 | 2.988387364 | -0.058872343 | -0.089354244 |
| Mcts1 | 9.098172719 | 9.199905427 | 9.187543597 | 0.01236183 | -0.089370878 |
| 4930503B20Rik | 1.521854897 | 1.563755779 | 1.611236576 | -0.047480797 | -0.089381679 |
| Vmn1r4 | 2.629411146 | 2.66260669 | 2.718800159 | -0.056193469 | -0.089389013 |
| Fam71b | 2.287893529 | 2.326405267 | 2.377349138 | -0.050943872 | -0.089455609 |
| Ppa2 | 8.761474955 | 8.76835539 | 8.850940437 | -0.082585046 | -0.089465482 |
| Trim71 | 2.959031106 | 2.937226537 | 3.048499488 | -0.111272951 | -0.089468382 |
| Dnah1 | 4.365877876 | 4.591414976 | 4.455407777 | 0.136007199 | -0.089529901 |
| Numa1 | 11.38489787 | 11.60876236 | 11.47448005 | 0.134282307 | -0.089582183 |
| Atmin | 9.606793863 | 9.71183181 | 9.696462559 | 0.015369251 | -0.089668696 |
| 1110059E24Rik | 8.251223338 | 8.15142086 | 8.34091266 | -0.1894918 | -0.089689322 |
| Ccdc30 | 3.515431899 | 3.620223533 | 3.605170274 | 0.015053258 | -0.089738375 |
| Plxnd1 | 9.606978138 | 9.806138332 | 9.696837044 | 0.109301288 | -0.089858906 |

| | | | | | |
|---|---|---|---|---|---|
| Hdhd5 | 8.006115459 | 8.063855427 | 8.095985977 | -0.03213055 | -0.089870517 |
| Dchs2 | 3.419575693 | 3.492083713 | 3.509476296 | -0.017392584 | -0.089900604 |
| Ggct | 11.60218814 | 11.66404763 | 11.69212746 | -0.028079833 | -0.089939323 |
| Gm3219 | 7.64995534 | 7.625014146 | 7.740018274 | -0.115004128 | -0.090062934 |
| Dyx1c1 | 3.086805178 | 3.100102238 | 3.176883461 | -0.076781223 | -0.090078283 |
| Tacstd2 | 12.44085254 | 12.57862477 | 12.53097844 | 0.047646327 | -0.0901259 |
| Zfp715 | 8.382070298 | 8.464222309 | 8.472202225 | -0.007979917 | -0.090131927 |
| Cyp2j8 | 6.513384876 | 6.530555948 | 6.603533179 | -0.072977231 | -0.090148304 |
| Rsrc1 | 9.191940905 | 9.172329414 | 9.282114693 | -0.109785279 | -0.090173788 |
| Slc44a2 | 12.07380079 | 12.08838382 | 12.16399576 | -0.075611947 | -0.090194977 |
| Rnf169 | 9.25850889 | 9.461261042 | 9.348760434 | 0.112500608 | -0.090251544 |
| Rragb | 3.544130194 | 3.58038423 | 3.634392521 | -0.054008291 | -0.090262327 |
| Camta1 | 10.32765313 | 10.43078619 | 10.41794744 | 0.012838743 | -0.090294317 |
| Gm4118 | 2.592246912 | 2.589584562 | 2.682565128 | -0.092980565 | -0.090318216 |
| Gm44984 | 2.801263864 | 2.841487595 | 2.891617519 | -0.050129925 | -0.090353656 |
| Corin | 1.81864702 | 1.834070602 | 1.909031628 | -0.074961026 | -0.090384608 |
| CAAA01141682.1 | 4.025249853 | 4.098732622 | 4.115637926 | -0.016905304 | -0.090388073 |
| Rwdd2a | 3.437833003 | 3.398263525 | 3.528263295 | -0.12999977 | -0.090430293 |
| Pogz | 9.117764579 | 9.379483539 | 9.208203797 | 0.171279742 | -0.090439218 |
| Grm4 | 4.799120738 | 4.970614219 | 4.889648372 | 0.080965847 | -0.090527635 |
| Npepl1 | 9.520971182 | 9.647488113 | 9.611552278 | 0.035935835 | -0.090581096 |
| Tesc | 7.691905179 | 7.56454187 | 7.782539407 | -0.217997537 | -0.090634228 |
| Rad51 | 7.697674105 | 7.928646727 | 7.788352609 | 0.140294118 | -0.090678504 |
| Zfp955b | 7.85227692 | 7.765066109 | 7.942960574 | -0.177894466 | -0.090683654 |
| S100a1 | 9.373184399 | 9.272335275 | 9.463884592 | -0.191549316 | -0.090700193 |
| Eif3h | 11.71286952 | 11.66891223 | 11.80357747 | -0.134665247 | -0.09070795 |
| Zfr2 | 6.051275147 | 6.217292303 | 6.142029522 | 0.075262781 | -0.090754375 |
| Syt1 | 2.433959703 | 2.461461128 | 2.524716834 | -0.063255706 | -0.090757131 |
| Gm9687 | 3.293727491 | 3.400475361 | 3.384502224 | 0.015973137 | -0.090774733 |
| Gpr180 | 8.298285769 | 8.3328997 | 8.389065043 | -0.056165342 | -0.090779274 |
| Palmd | 10.31479883 | 9.972572981 | 10.40559131 | -0.433018329 | -0.09079248 |
| Ube2c | 7.793358951 | 7.825635424 | 7.884197142 | -0.058561718 | -0.090838191 |

| | | | | | |
|---|---|---|---|---|---|
| Tmem255b | 4.502442941 | 4.531571452 | 4.593309445 | -0.061737992 | -0.090866504 |
| Gm48368 | 8.020975186 | 8.288063985 | 8.111863545 | 0.17620044 | -0.090888359 |
| F2rl3 | 2.548394955 | 2.625917352 | 2.639292209 | -0.013374857 | -0.090897254 |
| Ltc4s | 5.345112596 | 5.373516666 | 5.436028368 | -0.062511701 | -0.090915772 |
| Mpv17l | 6.221749081 | 6.266801813 | 6.312889537 | -0.046087724 | -0.091140456 |
| Fndc10 | 6.974832734 | 6.985773782 | 7.06602172 | -0.080247938 | -0.091188987 |
| Slc38a2 | 12.54073751 | 12.93695986 | 12.63195207 | 0.305007793 | -0.091214557 |
| 0610009B22Rik | 7.681951847 | 7.73525306 | 7.77320835 | -0.03795529 | -0.091256503 |
| Tdrd5 | 2.429297934 | 2.442879006 | 2.520590634 | -0.077711628 | -0.0912927 |
| Ppox | 8.647833842 | 8.819167881 | 8.739142743 | 0.080025138 | -0.091308901 |
| Ccdc90b | 8.332192398 | 8.367983636 | 8.423571839 | -0.055588203 | -0.091379441 |
| Cyb5rl | 7.336416748 | 7.58825158 | 7.427799175 | 0.160452405 | -0.091382427 |
| Abcb7 | 9.56396095 | 9.506361332 | 9.655357734 | -0.148996402 | -0.091396784 |
| Gm5414 | 4.762829748 | 4.741701079 | 4.854235649 | -0.11253457 | -0.091405901 |
| Brwd3 | 9.151074055 | 9.064795897 | 9.242510141 | -0.177714244 | -0.091436087 |
| CT010433.1 | 3.213772494 | 3.327503224 | 3.305256574 | 0.02224665 | -0.09148408 |
| Snx7 | 8.821033087 | 8.758620384 | 8.912544951 | -0.153924566 | -0.091511863 |
| Ttc16 | 3.771770905 | 3.856479976 | 3.863313034 | -0.006833058 | -0.09154213 |
| Zfp148 | 10.02057749 | 10.02053618 | 10.11224468 | -0.091708504 | -0.091667188 |
| Pibf1 | 7.215342654 | 7.237684731 | 7.307112345 | -0.069427614 | -0.091769691 |
| Pex11b | 8.657142929 | 8.629552379 | 8.748999181 | -0.119446802 | -0.091856252 |
| Nsa2 | 11.34798253 | 11.29102524 | 11.43984336 | -0.14881812 | -0.09186083 |
| Hspa12a | 9.310175185 | 9.227999769 | 9.402052571 | -0.174052802 | -0.091877387 |
| Arl6 | 7.193875026 | 7.261910248 | 7.285776647 | -0.023866399 | -0.091901621 |
| Uqcrc2 | 11.75707479 | 11.67965951 | 11.84901574 | -0.169356223 | -0.09194095 |
| Mttp | 5.840275495 | 5.940042592 | 5.932241454 | 0.007801138 | -0.09196596 |
| Gm17511 | 2.558085319 | 2.619327455 | 2.650085001 | -0.030757546 | -0.091999681 |
| Drc1 | 5.235937882 | 5.461207075 | 5.327957132 | 0.133249943 | -0.09201925 |
| Gab3 | 4.931474599 | 5.008135569 | 5.023495909 | -0.01536034 | -0.092021309 |
| Scarb2 | 11.64952226 | 11.55677573 | 11.74160339 | -0.184827657 | -0.092081133 |
| Gm11839 | 3.427273815 | 3.540596355 | 3.519375948 | 0.021220407 | -0.092102133 |
| Pkmyt1 | 8.083220852 | 8.527650723 | 8.175378229 | 0.352272493 | -0.092157377 |

| | | | | | |
|---|---|---|---|---|---|
| Lrp11 | 5.327596965 | 5.264618401 | 5.41976506 | -0.155146659 | -0.092168094 |
| Tceanc2 | 8.581158673 | 8.341204082 | 8.673328054 | -0.332123972 | -0.092169381 |
| Pdgfc | 7.586863118 | 7.467184386 | 7.679059582 | -0.211875196 | -0.092196464 |
| P4htm | 4.5497545 | 4.477883919 | 4.641956455 | -0.164072536 | -0.092201955 |
| Gm48551 | 4.577122515 | 4.713230965 | 4.669352949 | 0.043878016 | -0.092230434 |
| Dlc1 | 10.50684033 | 10.30364286 | 10.59908099 | -0.29543813 | -0.092240663 |
| Rab3gap1 | 10.20283631 | 10.28941681 | 10.29516635 | -0.005749547 | -0.092330043 |
| Prr14l | 9.524084152 | 9.665811023 | 9.616456305 | 0.049354718 | -0.092372153 |
| Krt2 | 3.447410581 | 3.547472694 | 3.539817099 | 0.007655595 | -0.092406518 |
| Ecm1 | 11.53099775 | 11.56662561 | 11.623409 | -0.05678339 | -0.092411253 |
| Gng13 | 1.648248029 | 1.698441521 | 1.740705102 | -0.042263582 | -0.092457073 |
| Sarm1 | 4.092884277 | 4.239286548 | 4.185387354 | 0.053899194 | -0.092503077 |
| Grid1 | 2.495066965 | 2.497436779 | 2.587585174 | -0.090148395 | -0.092518209 |
| Washc2 | 10.29479468 | 10.59051111 | 10.38733699 | 0.20317412 | -0.092542313 |
| 37316 | 10.38514975 | 10.2400062 | 10.47770318 | -0.237696987 | -0.092553437 |
| Dap | 10.13356204 | 10.15489182 | 10.22612089 | -0.071229067 | -0.092558847 |
| Galnt7 | 8.950637122 | 9.037920627 | 9.043210944 | -0.005290317 | -0.092573822 |
| 9130023H24Rik | 7.286051437 | 7.262766822 | 7.378661288 | -0.115894466 | -0.092609851 |
| Ggact | 5.707054653 | 5.704212258 | 5.79966821 | -0.095455952 | -0.092613557 |
| Gm20422 | 3.019588059 | 3.023760493 | 3.112229809 | -0.088469316 | -0.092641749 |
| Pcdh1 | 10.59422919 | 10.88280902 | 10.68690548 | 0.195903537 | -0.092676287 |
| Gpsm1 | 9.151386848 | 9.398753728 | 9.244121769 | 0.15463196 | -0.092734921 |
| Rab27a | 9.23242941 | 9.279422288 | 9.325177637 | -0.045755348 | -0.092748226 |
| Porcn | 7.602288664 | 7.685047057 | 7.695049919 | -0.010002862 | -0.092761255 |
| Olfr709-ps1 | 2.564435504 | 2.662213934 | 2.657197022 | 0.005016912 | -0.092761518 |
| Ppp3r1 | 11.32428368 | 11.29510487 | 11.41706462 | -0.121959753 | -0.092780937 |
| Swsap1 | 7.454731533 | 7.413424886 | 7.5475167 | -0.134091814 | -0.092785167 |
| Aurkb | 7.640305479 | 7.808788598 | 7.733154844 | 0.075633753 | -0.092849366 |
| Nav1 | 9.702684388 | 9.766146938 | 9.795536904 | -0.029389967 | -0.092852517 |
| Kmt5a | 11.24665462 | 11.32833195 | 11.3395221 | -0.01119015 | -0.09286748 |
| Pbx4 | 4.192170562 | 4.364799235 | 4.28507634 | 0.079722895 | -0.092905779 |
| Lin54 | 8.648181825 | 8.727836682 | 8.741089556 | -0.013252874 | -0.09290773 |

| | | | | | |
|---|---|---|---|---|---|
| Phkg2 | 8.774209202 | 9.005314132 | 8.867142596 | 0.138171536 | -0.092933395 |
| Pcdhb11 | 3.639543322 | 3.698640486 | 3.73259114 | -0.033950654 | -0.093047818 |
| Mok | 3.731934746 | 3.897170804 | 3.825010033 | 0.072160771 | -0.093075288 |
| Slc35d2 | 7.101434133 | 7.004949955 | 7.194541431 | -0.189591475 | -0.093107297 |
| Gm16199 | 6.905855664 | 6.901501878 | 6.998965722 | -0.097463844 | -0.093110058 |
| Gm5514 | 2.001627802 | 2.026585282 | 2.094738604 | -0.068153323 | -0.093110802 |
| Grn | 12.07016825 | 12.05361831 | 12.16329415 | -0.109675833 | -0.0931259 |
| Abcb10 | 9.023280853 | 9.076694729 | 9.116423202 | -0.039728473 | -0.093142349 |
| R3hdm2 | 10.73954728 | 10.78030954 | 10.83270272 | -0.052393183 | -0.093155443 |
| Parm1 | 8.382178501 | 8.101064168 | 8.475417916 | -0.374353748 | -0.093239414 |
| Gm11875 | 1.646719676 | 1.694805759 | 1.740031532 | -0.045225773 | -0.093311856 |
| Pisd-ps1 | 6.36320387 | 6.508699598 | 6.456528041 | 0.052171556 | -0.093324172 |
| Kcna5 | 5.213089326 | 5.197294625 | 5.306430323 | -0.109135698 | -0.093340997 |
| Ticam2 | 6.494868777 | 6.355332222 | 6.588214083 | -0.232881861 | -0.093345307 |
| Kctd5 | 10.01230952 | 10.06557372 | 10.10569175 | -0.040118027 | -0.093382225 |
| F13a1 | 11.4633847 | 10.86399293 | 11.55685498 | -0.692862053 | -0.093470277 |
| Klc2 | 8.071078237 | 8.350552166 | 8.164567308 | 0.185984858 | -0.093489071 |
| Sema4b | 9.836207684 | 10.00252932 | 9.929703807 | 0.072825517 | -0.093496123 |
| Ccdc68 | 2.484285354 | 2.495611186 | 2.577842397 | -0.082231211 | -0.093557043 |
| Lipo3 | 8.526032992 | 8.588787091 | 8.619591513 | -0.030804423 | -0.093558521 |
| Vdac3 | 10.96055594 | 10.94705165 | 11.054122 | -0.10707035 | -0.093566053 |
| Cops2 | 11.04610891 | 10.9728377 | 11.1396825 | -0.1668448 | -0.093573587 |
| Alms1-ps2 | 2.955660673 | 3.117139574 | 3.04927035 | 0.067869224 | -0.093609677 |
| Poglut1 | 9.219218586 | 9.303469782 | 9.312832786 | -0.009363004 | -0.0936142 |
| Trappc6b | 9.224538511 | 9.07985412 | 9.31825577 | -0.23840165 | -0.093717259 |
| Vmn1r229 | 6.186640623 | 6.341149252 | 6.280377822 | 0.06077143 | -0.093737199 |
| Zc3h14 | 10.17569498 | 10.27312632 | 10.26946412 | 0.0036622 | -0.09376914 |
| Zfp112 | 5.555553192 | 5.622599454 | 5.649419339 | -0.026819885 | -0.093866147 |
| Serpina3e-ps | 3.086127907 | 3.204356854 | 3.180021531 | 0.024335323 | -0.093893624 |
| Tceal3 | 3.986129738 | 4.016245483 | 4.080045065 | -0.063799581 | -0.093915326 |
| Mtbp | 7.405259596 | 7.441323443 | 7.499211907 | -0.057888464 | -0.093952311 |
| Cspp1 | 8.214376926 | 8.362661538 | 8.308335117 | 0.05432642 | -0.093958192 |

| | | | | | |
|---|---|---|---|---|---|
| Mgll | 13.67618762 | 13.64761181 | 13.77017577 | -0.122563963 | -0.093988153 |
| Tmed8 | 8.462942417 | 8.432447083 | 8.55696654 | -0.124519457 | -0.094024123 |
| Lpp | 10.87423053 | 11.02811759 | 10.96825849 | 0.059859107 | -0.094027957 |
| Tmem234 | 10.47517304 | 10.53697826 | 10.56920791 | -0.03222965 | -0.094034877 |
| Gm5069 | 4.512226653 | 4.593142663 | 4.606273142 | -0.013130478 | -0.094046489 |
| Immp1l | 8.728692797 | 8.673899856 | 8.822772186 | -0.14887233 | -0.094079389 |
| Aqp5 | 5.400060372 | 5.614070524 | 5.494201526 | 0.119868998 | -0.094141154 |
| Ddi2 | 11.77069991 | 11.91265845 | 11.86487429 | 0.04778416 | -0.094174383 |
| Mettl21a | 7.782052225 | 7.759916181 | 7.876377538 | -0.116461357 | -0.094325313 |
| BC004004 | 10.09136476 | 9.993560366 | 10.18571426 | -0.192153894 | -0.094349497 |
| Asf1b | 7.510516463 | 7.76253636 | 7.604898588 | 0.157637772 | -0.094382125 |
| Sdr42e1 | 10.9277104 | 10.87921842 | 11.02209624 | -0.14287782 | -0.094385837 |
| Zfp65 | 7.554073034 | 7.65342516 | 7.64856546 | 0.0048597 | -0.094492426 |
| Gdi2 | 12.79486386 | 12.78665992 | 12.88940624 | -0.102746317 | -0.09454238 |
| AC158774.1 | 3.480112273 | 3.528218108 | 3.574657536 | -0.046439428 | -0.094545263 |
| Gm47135 | 3.809221346 | 3.900959568 | 3.903787565 | -0.002827997 | -0.094566219 |
| C2 | 9.134019283 | 8.782278336 | 9.228676362 | -0.446398026 | -0.094657079 |
| Cmtm5 | 4.068329804 | 4.133134133 | 4.163029908 | -0.029895774 | -0.094700104 |
| Ddias | 6.322777419 | 6.363351163 | 6.417479997 | -0.054128834 | -0.094702578 |
| Tnni2 | 13.60921221 | 13.62998984 | 13.70393897 | -0.07394913 | -0.094726763 |
| Acot12 | 2.761572439 | 2.792518355 | 2.85633065 | -0.063812295 | -0.094758211 |
| Kyat3 | 8.067630358 | 8.253769011 | 8.162524908 | 0.091244103 | -0.09489455 |
| Bmyc | 5.34565793 | 5.41989541 | 5.44062766 | -0.02073225 | -0.09496973 |
| Dnah7b | 3.421424908 | 3.413775536 | 3.516403422 | -0.102627885 | -0.094978514 |
| D730001G18Rik | 5.326124466 | 5.262112028 | 5.421121132 | -0.159009104 | -0.094996666 |
| 1600012H06Rik | 9.378810445 | 9.401275045 | 9.473999926 | -0.072724881 | -0.095189481 |
| Gm12390 | 2.044768927 | 2.130828021 | 2.139982148 | -0.009154127 | -0.095213221 |
| Mfsd4b3 | 4.262831706 | 4.219732327 | 4.358054844 | -0.138322517 | -0.095223138 |
| Ano4 | 2.936542066 | 3.009861332 | 3.031771445 | -0.021910113 | -0.095229379 |
| Cd46 | 5.822251637 | 5.962077662 | 5.917523747 | 0.044553916 | -0.095272109 |
| Fat1 | 11.07634487 | 11.38230238 | 11.1716373 | 0.210665073 | -0.095292437 |
| Rnf168 | 9.403287856 | 9.396550419 | 9.498615847 | -0.102065429 | -0.095327991 |

| | | | | | |
|---|---|---|---|---|---|
| Adra1d | 2.510425869 | 2.550090875 | 2.605791242 | -0.055700366 | -0.095365373 |
| Naa60 | 10.25373272 | 10.35940022 | 10.34910465 | 0.010295567 | -0.095371927 |
| Tnfrsf13b | 6.488677949 | 6.448375765 | 6.584055211 | -0.135679445 | -0.095377262 |
| Snx22 | 5.753837771 | 5.911528404 | 5.849297279 | 0.062231125 | -0.095459508 |
| Tprkb | 8.585605913 | 8.674783149 | 8.681224257 | -0.006441108 | -0.095618344 |
| Rpl17 | 13.30879326 | 13.18244998 | 13.40442192 | -0.221971947 | -0.09562866 |
| Pipox | 2.108881031 | 2.099895035 | 2.204585876 | -0.104690841 | -0.095704845 |
| Cdk5 | 8.672457043 | 8.838920035 | 8.768239091 | 0.070680943 | -0.095782048 |
| Syn3 | 2.944510138 | 2.965974297 | 3.040296032 | -0.074321735 | -0.095785895 |
| Vangl1 | 9.378268397 | 9.276840756 | 9.474082297 | -0.197241541 | -0.0958139 |
| U2af1l4 | 7.887180207 | 7.885933359 | 7.98302595 | -0.097092591 | -0.095845743 |
| Aldh2 | 11.74406198 | 11.70273723 | 11.83993208 | -0.137194847 | -0.0958701 |
| B3galt4 | 6.143157541 | 6.306485353 | 6.23914142 | 0.067343933 | -0.095983879 |
| Gm5678 | 7.114102133 | 7.163035412 | 7.210132633 | -0.047097222 | -0.0960305 |
| Gm10767 | 3.140830024 | 3.272272619 | 3.236863449 | 0.03540917 | -0.096033425 |
| Rasd1 | 7.714653028 | 7.560389857 | 7.81070069 | -0.250310833 | -0.096047661 |
| 2810408A11Rik | 4.071179706 | 4.014648184 | 4.167280672 | -0.152632487 | -0.096100966 |
| Grip1 | 6.417459586 | 6.38731954 | 6.513570993 | -0.126251453 | -0.096111407 |
| Nubpl | 6.399647814 | 6.450357158 | 6.495765199 | -0.045408041 | -0.096117385 |
| Gm9755 | 2.112385386 | 2.107005141 | 2.208539666 | -0.101534525 | -0.09615428 |
| Lmx1b | 4.285294443 | 4.291342057 | 4.381456855 | -0.090114798 | -0.096162413 |
| Gm6211 | 4.813217189 | 4.913376193 | 4.909399748 | 0.003976444 | -0.096182559 |
| Usp21 | 8.606350264 | 8.762650281 | 8.702571459 | 0.060078822 | -0.096221194 |
| Rnaseh2b | 8.506761622 | 8.502327505 | 8.602990958 | -0.100663452 | -0.096229336 |
| Lman2l | 8.668028965 | 8.739614921 | 8.76433483 | -0.024719909 | -0.096305865 |
| Ypel1 | 5.176959437 | 5.275469925 | 5.273299145 | 0.00217078 | -0.096339709 |
| Pdgfrb | 10.22653309 | 10.42430826 | 10.3228822 | 0.101426057 | -0.096349107 |
| Sphkap | 3.319639513 | 3.27903453 | 3.416011395 | -0.136976865 | -0.096371882 |
| Rfc1 | 10.21576592 | 10.19701042 | 10.31216046 | -0.115150043 | -0.096394543 |
| Chrm4 | 2.173535396 | 2.243066163 | 2.269947474 | -0.026881311 | -0.096412078 |
| 2210418O10Rik | 7.759192011 | 7.871114501 | 7.855712474 | 0.015402027 | -0.096520463 |
| Tbx1 | 7.122369197 | 7.25775265 | 7.218949413 | 0.038803236 | -0.096580216 |

| | | | | | |
|---|---|---|---|---|---|
| Sh3gl3 | 4.378554689 | 4.51575118 | 4.475212462 | 0.040538718 | -0.096657773 |
| Tex261 | 10.89372639 | 10.88996154 | 10.99038931 | -0.100427767 | -0.096662917 |
| 1700007K13Rik | 2.781561046 | 2.855730678 | 2.878234593 | -0.022503915 | -0.096673547 |
| Gm42669 | 6.389565899 | 6.401021599 | 6.486269948 | -0.085248349 | -0.096704049 |
| Gatb | 7.881465494 | 7.918940612 | 7.978176264 | -0.059235652 | -0.09671077 |
| Arfgef2 | 10.19501769 | 10.37905823 | 10.2917973 | 0.087260927 | -0.096779613 |
| Gm10123 | 12.65741426 | 12.53935169 | 12.75420884 | -0.21485715 | -0.096794577 |
| Itih5l-ps | 3.450918274 | 3.515121815 | 3.547762793 | -0.032640979 | -0.096844519 |
| Spdya | 1.522852242 | 1.547146167 | 1.619784726 | -0.072638559 | -0.096932485 |
| Slco1a5 | 3.068513123 | 3.144107303 | 3.165452761 | -0.021345458 | -0.096939638 |
| Chmp5 | 10.80368617 | 10.70318402 | 10.90064332 | -0.197459303 | -0.09695715 |
| Ebag9 | 7.961868557 | 7.841392583 | 8.058830049 | -0.217437466 | -0.096961492 |
| Spink2 | 3.844313711 | 3.816285797 | 3.941328838 | -0.125043042 | -0.097015127 |
| Vmn2r120 | 6.358173507 | 6.346556066 | 6.455203923 | -0.108647857 | -0.097030416 |
| Gm13597 | 6.822088174 | 6.777801078 | 6.919123039 | -0.141321962 | -0.097034865 |
| Pld3 | 10.74157861 | 11.06792208 | 10.83862581 | 0.22929627 | -0.0970472 |
| Zfp612 | 6.55120339 | 6.665414208 | 6.648343226 | 0.017070982 | -0.097139836 |
| Olfr161 | 3.842314082 | 3.797170842 | 3.939457483 | -0.142286641 | -0.097143401 |
| Ubap2 | 10.45818198 | 10.51963775 | 10.55533295 | -0.035695197 | -0.097150967 |
| Olfr648 | 4.131741638 | 4.206444551 | 4.228900857 | -0.022456306 | -0.097159219 |
| Suclg2 | 9.87141765 | 9.870864833 | 9.968669384 | -0.097804551 | -0.097251734 |
| Igha | 4.86525394 | 4.956359577 | 4.962523137 | -0.00616356 | -0.097269197 |
| Megf6 | 8.698907927 | 9.148205548 | 8.79620764 | 0.351997909 | -0.097299712 |
| Gm14295 | 7.119661964 | 7.359708859 | 7.216964328 | 0.142744531 | -0.097302364 |
| Ublcp1 | 9.467450815 | 9.433066771 | 9.564756236 | -0.131689466 | -0.097305421 |
| Rab17 | 3.416955711 | 3.381533243 | 3.514269009 | -0.132735766 | -0.097313297 |
| Nck2 | 8.561374228 | 8.742089271 | 8.658695042 | 0.083394229 | -0.097320814 |
| Lekr1 | 4.810080423 | 4.797433179 | 4.907481203 | -0.110048024 | -0.09740078 |
| Mrpl45 | 9.138558428 | 9.064681245 | 9.235984989 | -0.171303744 | -0.097426561 |
| Rtn4 | 12.6916995 | 12.66015837 | 12.78914779 | -0.12898942 | -0.09744829 |
| Hscb | 6.364551894 | 6.24765784 | 6.462054847 | -0.214397007 | -0.097502953 |
| Klhl18 | 9.564679128 | 9.773632351 | 9.662191939 | 0.111440412 | -0.097512811 |

| | | | | | |
|---|---|---|---|---|---|
| Eme1 | 5.253289543 | 5.482491779 | 5.350851448 | 0.131640332 | -0.097561904 |
| Ckb | 8.721560204 | 8.910334621 | 8.819168603 | 0.091166017 | -0.0976084 |
| Mgat4b | 9.601298017 | 9.624034374 | 9.698924812 | -0.074890439 | -0.097626795 |
| Ncapd3 | 8.969392642 | 9.082724773 | 9.067024344 | 0.015700429 | -0.097631703 |
| Caskin2 | 9.402614063 | 9.839847416 | 9.500257818 | 0.339589597 | -0.097643755 |
| Zcwpw2 | 2.116447164 | 2.136171646 | 2.214125042 | -0.077953396 | -0.097677878 |
| Klhl35 | 1.912197109 | 1.971871601 | 2.00988894 | -0.038017339 | -0.097691831 |
| Pitpnm2 | 9.373735428 | 9.748185472 | 9.471429183 | 0.276756289 | -0.097693755 |
| Dapk2 | 8.243742009 | 8.212810138 | 8.34149546 | -0.128685321 | -0.097753451 |
| Cox20 | 6.687007855 | 6.73613084 | 6.784841166 | -0.048710326 | -0.097833311 |
| Selenof | 11.59063313 | 11.4859094 | 11.68849591 | -0.202586517 | -0.09786278 |
| Slc9a6 | 9.863949549 | 9.922480736 | 9.961912816 | -0.03943208 | -0.097963267 |
| Vmn1r217 | 4.522334717 | 4.717578736 | 4.620298848 | 0.097279888 | -0.097964131 |
| Tnpo2 | 11.2772475 | 11.46720365 | 11.37526966 | 0.09193399 | -0.098022163 |
| Erc1 | 9.700987958 | 9.682423036 | 9.799018217 | -0.116595181 | -0.098030259 |
| Cd209d | 9.863001471 | 9.327173947 | 9.961073879 | -0.633899931 | -0.098072407 |
| Dkk3 | 9.081196857 | 8.996255351 | 9.17929632 | -0.18304097 | -0.098099463 |
| Gm13169 | 1.984424232 | 2.034257346 | 2.082529508 | -0.048272162 | -0.098105276 |
| Atf7 | 10.71566711 | 10.74140485 | 10.81381362 | -0.072408773 | -0.09814651 |
| Chek1 | 7.215590997 | 7.366407222 | 7.313787075 | 0.052620147 | -0.098196078 |
| Macrod1 | 8.556744735 | 8.682543625 | 8.655095856 | 0.02744777 | -0.09835112 |
| Uckl1 | 8.321092739 | 8.264588767 | 8.419468146 | -0.154879379 | -0.098375407 |
| Slc6a17 | 6.644738234 | 6.974202945 | 6.743170548 | 0.231032397 | -0.098432314 |
| Sall4 | 1.505225764 | 1.545863681 | 1.603745333 | -0.057881652 | -0.098519569 |
| Pnck | 4.642657839 | 4.616736094 | 4.741250318 | -0.124514224 | -0.098592479 |
| Gna12 | 9.745632943 | 9.972996091 | 9.844241895 | 0.128754195 | -0.098608952 |
| Gm15937 | 1.754658963 | 1.788478552 | 1.853293729 | -0.064815176 | -0.098634766 |
| Ssx2ip | 9.591830289 | 9.705402506 | 9.69047965 | 0.014922856 | -0.098649362 |
| Rimbp2 | 4.071043637 | 4.10923293 | 4.169697926 | -0.060464996 | -0.098654289 |
| Nrd1 | 12.46107623 | 12.44278877 | 12.55973264 | -0.116943873 | -0.098656413 |
| Gm3654 | 5.601925947 | 5.530398208 | 5.700609642 | -0.170211434 | -0.098683695 |
| Gm20150 | 2.168866662 | 2.285446286 | 2.267607602 | 0.017838683 | -0.09874094 |

| | | | | | |
|---|---|---|---|---|---|
| Fdxr | 6.928041931 | 7.237350491 | 7.026786631 | 0.210563861 | -0.098744699 |
| Trmu | 7.593923335 | 7.752728379 | 7.692669319 | 0.06005906 | -0.098745984 |
| Olfr523 | 2.62403706 | 2.708648852 | 2.722821982 | -0.01417313 | -0.098784922 |
| Scarb1 | 10.13123205 | 10.08937014 | 10.23001907 | -0.140648929 | -0.09878702 |
| Chrnb2 | 4.8617291 | 4.965974583 | 4.9605214 | 0.005453184 | -0.098792299 |
| Ppp3cb | 10.57162672 | 10.49985872 | 10.67056272 | -0.170703997 | -0.098936003 |
| Ep300 | 10.70980154 | 10.86983171 | 10.80885054 | 0.06098117 | -0.099049 |
| Frg2f1 | 5.475896257 | 5.688714545 | 5.574953385 | 0.11376116 | -0.099057128 |
| Chrna1 | 5.509147419 | 5.35146874 | 5.608253391 | -0.256784651 | -0.099105972 |
| Aplf | 7.729540518 | 7.810960579 | 7.828838208 | -0.017877629 | -0.099297689 |
| Gm17131 | 7.446889618 | 7.534205884 | 7.546302053 | -0.012096169 | -0.099412434 |
| Borcs8 | 8.164857268 | 8.29112449 | 8.264356278 | 0.026768212 | -0.099499009 |
| Nek9 | 10.85966872 | 10.84056866 | 10.95917552 | -0.118606867 | -0.0995068 |
| Fam131b | 2.973946194 | 2.962914003 | 3.073519702 | -0.110605699 | -0.099573508 |
| Abhd12b | 9.010877466 | 9.013731732 | 9.110451185 | -0.096719453 | -0.099573718 |
| Mbnl3 | 7.718220427 | 7.790900847 | 7.817803882 | -0.026903035 | -0.099583455 |
| Pgpep1 | 8.193501416 | 8.57816369 | 8.293099972 | 0.285063718 | -0.099598556 |
| Bbof1 | 5.585343676 | 5.66011886 | 5.685016859 | -0.024897999 | -0.099673183 |
| Prx | 8.868625348 | 8.893806242 | 8.968346162 | -0.07453992 | -0.099720814 |
| Csdc2 | 3.328725369 | 3.463845856 | 3.428522311 | 0.035323546 | -0.099796942 |
| Man1a2 | 10.58187269 | 10.76498355 | 10.6816738 | 0.083309757 | -0.09980111 |
| Dnajb14 | 9.726213141 | 9.686566561 | 9.82614814 | -0.139581579 | -0.099934999 |
| Fbxw10 | 4.043397142 | 4.144006071 | 4.143370452 | 0.000635619 | -0.09997331 |
| Zfp53 | 7.004344899 | 7.113335804 | 7.104343331 | 0.008992473 | -0.099998432 |
| Spindoc | 7.735179636 | 7.966696336 | 7.83522812 | 0.131468215 | -0.100048484 |
| Kbtbd3 | 5.993630001 | 5.887130729 | 6.093678578 | -0.206547849 | -0.100048578 |
| Ufsp1 | 6.736219623 | 6.827887476 | 6.836270916 | -0.00838344 | -0.100051293 |
| Tpx2 | 8.736871419 | 8.689641352 | 8.836937972 | -0.14729662 | -0.100066553 |
| Cep135 | 6.893908987 | 6.916258674 | 6.993991215 | -0.077732541 | -0.100082228 |
| Tchhl1 | 2.801023411 | 2.837232247 | 2.90119961 | -0.063967363 | -0.100176199 |
| Cdip1 | 9.84371868 | 9.929836723 | 9.943920378 | -0.014083654 | -0.100201698 |
| 2510002D24Rik | 8.118514449 | 8.305272724 | 8.218761468 | 0.086511256 | -0.100247018 |

| | | | | | |
|---|---|---|---|---|---|
| Zfp180 | 9.276771025 | 9.377885619 | 9.377030213 | 0.000855407 | -0.100259187 |
| P3h2 | 9.332273721 | 9.491981986 | 9.4325343 | 0.059447686 | -0.10026058 |
| Cep57 | 8.431853119 | 8.29282173 | 8.53216386 | -0.23934213 | -0.100310741 |
| Lypd3 | 11.75965566 | 11.89862799 | 11.85996999 | 0.038658003 | -0.100314327 |
| Fcho1 | 8.597777671 | 8.895219811 | 8.698169075 | 0.197050736 | -0.100391404 |
| Reep2 | 3.99631635 | 4.153653076 | 4.096708382 | 0.056944694 | -0.100392031 |
| Tlr11 | 4.859458147 | 4.96767767 | 4.959871713 | 0.007805957 | -0.100413566 |
| Acsl5 | 12.64349406 | 12.62769585 | 12.74391323 | -0.116217383 | -0.10041917 |
| Celf2 | 10.54077019 | 10.3023836 | 10.64122366 | -0.338840053 | -0.10045347 |
| Ash1l | 10.82017007 | 10.88860595 | 10.92068623 | -0.032080283 | -0.100516163 |
| Dnajb12 | 9.554273752 | 9.720766744 | 9.654834195 | 0.065932549 | -0.100560444 |
| Dnajc16 | 9.051297892 | 9.151779315 | 9.151910826 | -0.000131511 | -0.100612934 |
| Zmiz1 | 11.07926583 | 11.42408191 | 11.17990413 | 0.244177783 | -0.100638293 |
| Haus1 | 6.733783818 | 6.700605537 | 6.834471745 | -0.133866208 | -0.100687927 |
| Pcdhga6 | 6.65822448 | 6.615566653 | 6.758974671 | -0.143408018 | -0.10075019 |
| Prr32 | 3.019698992 | 3.003921852 | 3.12055899 | -0.116637138 | -0.100859998 |
| Clasp2 | 10.41192643 | 10.35986824 | 10.51293177 | -0.153063537 | -0.10100534 |
| Smurf1 | 9.733765965 | 9.940826533 | 9.834970135 | 0.105856397 | -0.10120417 |
| Zfp78 | 5.170718468 | 5.300303454 | 5.271923916 | 0.028379538 | -0.101205448 |
| Kcnh3 | 3.804620478 | 3.825871506 | 3.905849583 | -0.079978078 | -0.101229105 |
| Slc25a46 | 9.869603644 | 9.843909383 | 9.970844096 | -0.126934713 | -0.101240452 |
| Ttc7b | 8.295330627 | 8.321800399 | 8.396585187 | -0.074784788 | -0.10125456 |
| Egln1 | 10.56519399 | 10.50781427 | 10.66644916 | -0.15863489 | -0.101255163 |
| Gm11581 | 3.587575931 | 3.566106612 | 3.688871982 | -0.12276537 | -0.101296051 |
| Clip1 | 12.00782266 | 11.88289216 | 12.10912483 | -0.22623267 | -0.101302163 |
| Rbms2 | 10.01452278 | 9.928533416 | 10.11582761 | -0.187294198 | -0.101304838 |
| Wdr26 | 12.19476935 | 12.36187762 | 12.29608194 | 0.06579568 | -0.10131259 |
| Fyttd1 | 10.61397665 | 10.67698682 | 10.71530421 | -0.038317393 | -0.10132756 |
| Tmem150c | 5.523828356 | 5.522305701 | 5.625211019 | -0.102905318 | -0.101382664 |
| Mapt | 7.370939478 | 7.356381744 | 7.472333678 | -0.115951934 | -0.1013942 |
| Soat1 | 12.37292316 | 12.31979963 | 12.4743425 | -0.15454287 | -0.101419337 |
| Kifc1 | 7.503806273 | 7.570877179 | 7.605229542 | -0.034352362 | -0.101423269 |

| | | | | | |
|---|---|---|---|---|---|
| Pycr1 | 5.593463922 | 5.893651371 | 5.694896758 | 0.198754613 | -0.101432836 |
| Tcf7l2 | 8.361831781 | 8.351626766 | 8.463291432 | -0.111664666 | -0.101459651 |
| Gm20257 | 5.766717192 | 5.897536584 | 5.86820721 | 0.029329374 | -0.101490018 |
| 1810062G17Rik | 3.073016056 | 3.0580571 | 3.174549081 | -0.116491981 | -0.101533026 |
| Olfr804 | 4.411807486 | 4.435389811 | 4.51335089 | -0.077961078 | -0.101543404 |
| Tmub2 | 9.217764688 | 9.364319191 | 9.319369311 | 0.04494988 | -0.101604623 |
| Cmtm3 | 9.350316586 | 9.564405339 | 9.451957269 | 0.11244807 | -0.101640683 |
| Rfx5 | 8.437231278 | 8.476462187 | 8.538890251 | -0.062428064 | -0.101658973 |
| Epc1 | 9.280965903 | 9.459567821 | 9.382628991 | 0.07693883 | -0.101663088 |
| Cxxc4 | 2.290798453 | 2.35490877 | 2.392488038 | -0.037579268 | -0.101689585 |
| Rnf32 | 3.026237985 | 3.097442153 | 3.127939605 | -0.030497452 | -0.10170162 |
| Gm13464 | 2.938800956 | 2.958792776 | 3.040547801 | -0.081755025 | -0.101746845 |
| Tk1 | 8.63697752 | 8.827961909 | 8.738754253 | 0.089207656 | -0.101776733 |
| Prox2 | 5.315013801 | 5.337932936 | 5.416809504 | -0.078876569 | -0.101795704 |
| Sec24a | 9.693849964 | 9.877785473 | 9.795684628 | 0.082100846 | -0.101834663 |
| Gm4130 | 1.924086861 | 2.005144395 | 2.025923723 | -0.020779328 | -0.101836861 |
| Marco | 4.50644914 | 4.598671511 | 4.608297411 | -0.0096259 | -0.101848271 |
| Ddr1 | 11.09850437 | 11.53179294 | 11.20039716 | 0.331395777 | -0.10189279 |
| Rnf223 | 6.469258636 | 6.851607985 | 6.571166521 | 0.280441464 | -0.101907884 |
| Mbtps2 | 8.762842218 | 8.707034583 | 8.864813211 | -0.157778629 | -0.101970993 |
| Olfr472 | 6.175736512 | 6.234821796 | 6.277709866 | -0.042888071 | -0.101973354 |
| Fads6 | 7.288785252 | 7.285851482 | 7.390793936 | -0.104942454 | -0.102008684 |
| Tmigd1 | 6.972468283 | 7.140889855 | 7.074513378 | 0.066376477 | -0.102045095 |
| Pten | 11.86340397 | 11.92192297 | 11.96546973 | -0.04354676 | -0.102065757 |
| Taz | 9.853772446 | 9.885413335 | 9.955840941 | -0.070427606 | -0.102068494 |
| Fam114a2 | 10.71445423 | 10.74234535 | 10.81654173 | -0.07419638 | -0.102087507 |
| Phospho2 | 8.57779746 | 8.625842978 | 8.679899051 | -0.054056073 | -0.102101591 |
| P4ha3 | 4.14764169 | 4.260562169 | 4.249899186 | 0.010662983 | -0.102257497 |
| Insl6 | 3.931024812 | 3.998631915 | 4.03328674 | -0.034654825 | -0.102261928 |
| Gm48366 | 3.812596533 | 3.801204781 | 3.914979134 | -0.113774353 | -0.102382601 |
| Ldha-ps | 1.796121415 | 1.875126938 | 1.898599631 | -0.023472694 | -0.102478216 |
| Mad2l1 | 8.271154885 | 8.405308745 | 8.373639068 | 0.031669676 | -0.102484183 |

| | | | | | |
|---|---|---|---|---|---|
| Coro1b | 10.20856467 | 10.25290641 | 10.31109301 | -0.058186603 | -0.102528343 |
| Nedd1 | 8.552418794 | 8.658499548 | 8.654973715 | 0.003525833 | -0.102554921 |
| Nxnl1 | 2.458683808 | 2.449634833 | 2.561245303 | -0.11161047 | -0.102561494 |
| Gmnn | 7.704829035 | 7.865499118 | 7.807410773 | 0.058088345 | -0.102581738 |
| mt-Co3 | 16.27712536 | 16.21455881 | 16.37975619 | -0.165197383 | -0.102630833 |
| Ccdc138 | 5.540535165 | 5.632877779 | 5.643173566 | -0.010295787 | -0.102638401 |
| Taf13 | 9.953460729 | 10.13343277 | 10.05612139 | 0.077311373 | -0.102660665 |
| Gm38119 | 2.615288273 | 2.706462205 | 2.717991913 | -0.011529708 | -0.10270364 |
| Ndp | 3.653475838 | 3.738605562 | 3.756208211 | -0.017602649 | -0.102732373 |
| Vamp2 | 10.70624126 | 10.80541561 | 10.80898627 | -0.003570667 | -0.102745013 |
| Foxj2 | 9.750015727 | 9.920836052 | 9.852762592 | 0.06807346 | -0.102746865 |
| Bambi-ps1 | 3.293798638 | 3.350462397 | 3.396581966 | -0.046119568 | -0.102783328 |
| Slain2 | 10.90092628 | 10.98688659 | 11.00372029 | -0.016833703 | -0.10279401 |
| Mga | 10.35475675 | 10.36832805 | 10.45755124 | -0.08922319 | -0.102794497 |
| Vmn2r29 | 4.165025062 | 4.00306771 | 4.267860758 | -0.264793048 | -0.102835696 |
| Esrp2 | 11.16446406 | 11.43995565 | 11.26734387 | 0.17261178 | -0.102879813 |
| P2ry10b | 5.543257893 | 5.512135278 | 5.646223597 | -0.134088318 | -0.102965704 |
| Rab2b | 7.057754208 | 7.004255515 | 7.160754194 | -0.156498678 | -0.102999986 |
| Pm20d1 | 5.588822047 | 5.741126719 | 5.691824446 | 0.049302272 | -0.1030024 |
| Ubr1 | 9.593959143 | 9.534246411 | 9.696966081 | -0.16271967 | -0.103006938 |
| Zfhx2 | 6.739827655 | 7.022215942 | 6.842842509 | 0.179373433 | -0.103014854 |
| Strada | 8.270977025 | 8.334183227 | 8.374089261 | -0.039906035 | -0.103112237 |
| Dact2 | 5.226428734 | 5.044532476 | 5.329555021 | -0.285022545 | -0.103126287 |
| Dennd2d | 8.958190786 | 8.890802792 | 9.061322874 | -0.170520082 | -0.103132088 |
| Gm9780 | 7.340694188 | 7.7271419 | 7.443842837 | 0.283299063 | -0.103148649 |
| Igfbp2 | 9.416016709 | 9.546112854 | 9.51921094 | 0.026901914 | -0.103194231 |
| Prkch | 10.16393206 | 10.15549868 | 10.26717697 | -0.111678283 | -0.10324491 |
| Zfp37 | 5.549390879 | 5.454547048 | 5.652635902 | -0.198088854 | -0.103245022 |
| Hmgb1-ps6 | 2.188200384 | 2.182521719 | 2.291472626 | -0.108950907 | -0.103272242 |
| Scube3 | 4.955681098 | 5.010445056 | 5.058978708 | -0.048533652 | -0.10329761 |
| Sim1 | 1.688953616 | 1.717737358 | 1.79225537 | -0.074518012 | -0.103301753 |
| Adgrl2 | 11.09886266 | 11.10070656 | 11.20219767 | -0.101491113 | -0.103335007 |

| | | | | | |
|---|---|---|---|---|---|
| Cobll1 | 9.220107562 | 9.148616837 | 9.323488733 | -0.174871896 | -0.103381171 |
| Pou3f1 | 9.504507922 | 9.944728446 | 9.607905103 | 0.336823343 | -0.10339718 |
| Lin7a | 3.226933571 | 3.236608591 | 3.330355009 | -0.093746417 | -0.103421437 |
| Tmem231 | 7.366481005 | 7.472142337 | 7.470073465 | 0.002068872 | -0.10359246 |
| Cd5 | 5.630165306 | 5.524012437 | 5.733771489 | -0.209759052 | -0.103606183 |
| Dolk | 8.748223491 | 8.702833872 | 8.851834431 | -0.149000559 | -0.103610941 |
| Pcdhac2 | 2.654724178 | 2.653085134 | 2.758361471 | -0.105276337 | -0.103637293 |
| Col11a1 | 5.385711377 | 5.405296138 | 5.489357764 | -0.084061626 | -0.103646386 |
| Syap1 | 10.153996 | 10.00133787 | 10.25765362 | -0.256315744 | -0.103657617 |
| Akap10 | 8.892654123 | 8.97544956 | 8.996326603 | -0.020877043 | -0.103672481 |
| Vwf | 8.604092769 | 8.672932912 | 8.707775643 | -0.034842731 | -0.103682875 |
| Gprasp1 | 8.631069452 | 8.778295015 | 8.734783185 | 0.043511829 | -0.103713733 |
| Rpl17-ps8 | 2.542912577 | 2.591196812 | 2.646644181 | -0.055447369 | -0.103731604 |
| Gm4332 | 11.57354346 | 11.53984603 | 11.677332 | -0.137485977 | -0.103788543 |
| Lix1l | 9.534100634 | 9.576989211 | 9.637944286 | -0.060955075 | -0.103843652 |
| Gm9774 | 5.540556861 | 5.859745381 | 5.644475019 | 0.215270362 | -0.103918158 |
| Gtdc1 | 7.946502567 | 7.872714607 | 8.050472621 | -0.177758014 | -0.103970054 |
| Slc10a7 | 8.712717804 | 8.614572826 | 8.816719311 | -0.202146485 | -0.104001507 |
| Slc22a5 | 8.106837903 | 8.104893295 | 8.210881062 | -0.105987767 | -0.104043159 |
| Syngap1 | 6.707396571 | 7.12965764 | 6.811481004 | 0.318176636 | -0.104084433 |
| Serpina3b | 10.30668404 | 9.854478466 | 10.41079069 | -0.556312224 | -0.104106646 |
| Frat2 | 6.74467122 | 6.721433039 | 6.848835559 | -0.12740252 | -0.104164339 |
| Rnase1 | 5.948354687 | 5.931685323 | 6.052543065 | -0.120857743 | -0.104188379 |
| Mtmr1 | 10.39555679 | 10.39291738 | 10.49976706 | -0.10684968 | -0.104210273 |
| Gm15793 | 3.6099475 | 3.595907583 | 3.714175169 | -0.118267586 | -0.104227669 |
| Glrb | 4.171653399 | 4.116224006 | 4.275884008 | -0.159660001 | -0.104230609 |
| Camsap1 | 9.842396435 | 10.0029774 | 9.9466346 | 0.056342797 | -0.104238165 |
| Mfap1a | 9.74639 | 9.732393249 | 9.850636088 | -0.11824284 | -0.104246088 |
| Tspan9 | 11.0027614 | 11.13551357 | 11.1070264 | 0.028487173 | -0.104265003 |
| Tmsb15b1 | 3.80285465 | 3.837688394 | 3.907123157 | -0.069434763 | -0.104268507 |
| Scube1 | 4.812481391 | 4.822338684 | 4.916799952 | -0.094461267 | -0.104318561 |
| Gm5108 | 2.72605538 | 2.863838376 | 2.830406469 | 0.033431907 | -0.104351089 |

| | | | | | |
|---|---|---|---|---|---|
| Ap3b2 | 3.816745481 | 3.814510346 | 3.921102849 | -0.106592503 | -0.104357368 |
| Zfp512b | 9.606512693 | 9.847984864 | 9.710873036 | 0.137111828 | -0.104360343 |
| Dlat | 10.34710206 | 10.12624089 | 10.45146961 | -0.325228717 | -0.10436755 |
| Tmem100 | 7.640363942 | 7.038212089 | 7.744739986 | -0.706527897 | -0.104376044 |
| Lrrtm4 | 3.128854341 | 3.219698985 | 3.233238061 | -0.013539076 | -0.10438372 |
| Ptgds | 5.247309942 | 5.234870307 | 5.351762884 | -0.116892577 | -0.104452941 |
| Aldh9a1 | 9.587919663 | 9.687898094 | 9.692390324 | -0.00449223 | -0.104470661 |
| Lage3 | 8.438590118 | 8.392021 | 8.543180244 | -0.151159244 | -0.104590126 |
| Cnep1r1 | 9.22891851 | 9.275600736 | 9.333526211 | -0.057925474 | -0.1046077 |
| Fxyd1 | 9.716535594 | 9.725383646 | 9.821176717 | -0.095793072 | -0.104641124 |
| Fam76b | 9.442794121 | 9.448721837 | 9.547437246 | -0.098715409 | -0.104643125 |
| Tal2 | 3.678204558 | 3.590988888 | 3.782866967 | -0.191878079 | -0.10466241 |
| Rnf152 | 7.824236886 | 7.867721431 | 7.928982503 | -0.061261072 | -0.104745617 |
| Fam228b | 4.124401371 | 4.138733107 | 4.229167233 | -0.090434126 | -0.104765862 |
| Rbm4b | 7.785105634 | 7.954382479 | 7.889899224 | 0.064483255 | -0.104793591 |
| Ngrn | 8.155120436 | 8.184576427 | 8.25992436 | -0.075347933 | -0.104803924 |
| Samd10 | 8.01922027 | 8.049054562 | 8.124027935 | -0.074973373 | -0.104807665 |
| Plcxd1 | 8.68274124 | 9.229902488 | 8.787629786 | 0.442272702 | -0.104888546 |
| Sprr2f | 5.074498529 | 5.233463343 | 5.179392718 | 0.054070625 | -0.104894189 |
| Anks3 | 8.704869304 | 8.903922778 | 8.809800861 | 0.094121917 | -0.104931557 |
| Olfr487 | 2.740804058 | 2.768118609 | 2.845765707 | -0.077647098 | -0.10496165 |
| Olfr90 | 5.871660793 | 5.98111186 | 5.976652919 | 0.004458941 | -0.104992126 |
| Chn2 | 5.856618777 | 5.895645024 | 5.961612819 | -0.065967796 | -0.104994042 |
| Efhb | 2.948976628 | 2.950015048 | 3.054268224 | -0.104253176 | -0.105291597 |
| Adck5 | 7.722155913 | 7.966423692 | 7.82757472 | 0.138848972 | -0.105418806 |
| Bicd1 | 6.280025479 | 6.269367306 | 6.385525122 | -0.116157816 | -0.105499643 |
| Mnd1 | 6.63433064 | 6.596110252 | 6.739840099 | -0.143729847 | -0.105509459 |
| Use1 | 10.00686263 | 10.03947557 | 10.11237268 | -0.072897113 | -0.10551005 |
| Efcab7 | 4.173813615 | 4.307198081 | 4.279352404 | 0.027845678 | -0.105538788 |
| Tmem218 | 6.904499261 | 6.858933906 | 7.010074409 | -0.151140503 | -0.105575148 |
| Izumo1 | 3.775754515 | 3.94310828 | 3.881333641 | 0.061774639 | -0.105579126 |
| Lrrc46 | 3.573200633 | 3.705621961 | 3.678834987 | 0.026786974 | -0.105634354 |

| | | | | | |
|---|---|---|---|---|---|
| Gab1 | 10.07649634 | 10.20585435 | 10.18217566 | 0.023678693 | -0.105679323 |
| Mfn1 | 10.49981906 | 10.48875157 | 10.60550682 | -0.116755253 | -0.105687767 |
| Sntg1 | 1.692006246 | 1.809229074 | 1.797734497 | 0.011494577 | -0.105728251 |
| S100a4 | 12.52863037 | 12.5542476 | 12.63440602 | -0.080158417 | -0.105775647 |
| Samd14 | 6.178736791 | 6.376214071 | 6.284531848 | 0.091682223 | -0.105795057 |
| Mettl7b | 6.384840204 | 6.252740568 | 6.490661417 | -0.237920849 | -0.105821214 |
| Zbtb40 | 8.192321534 | 8.559076713 | 8.298206164 | 0.260870549 | -0.10588463 |
| Gm7312 | 2.876714297 | 2.941673433 | 2.982649212 | -0.040975778 | -0.105934914 |
| Wrn | 8.465908233 | 8.441503214 | 8.571870097 | -0.130366883 | -0.105961864 |
| Tmem129 | 8.424601876 | 8.50451397 | 8.530564941 | -0.026050971 | -0.105963065 |
| Slc25a24 | 9.549628205 | 9.569240374 | 9.655713511 | -0.086473137 | -0.106085306 |
| Zfp422 | 8.371735171 | 8.379279787 | 8.477828346 | -0.098548559 | -0.106093175 |
| Plekhg6 | 8.362919648 | 8.618427936 | 8.469116356 | 0.149311581 | -0.106196708 |
| Kdelr1 | 10.86809689 | 10.9704745 | 10.97435835 | -0.00388385 | -0.106261457 |
| Aldoa | 15.22038405 | 15.36584979 | 15.32675333 | 0.039096457 | -0.10636928 |
| Gm42688 | 5.985462931 | 6.138114857 | 6.091862346 | 0.046252511 | -0.106399415 |
| Cntf | 5.928367472 | 6.021710449 | 6.034881521 | -0.013171072 | -0.106514049 |
| S100a3 | 4.020222412 | 4.08124931 | 4.126806258 | -0.045556948 | -0.106583846 |
| Mtg1 | 7.974338657 | 7.990800928 | 8.080999897 | -0.090198968 | -0.106661239 |
| Camk2g | 10.00479987 | 10.04808309 | 10.11150851 | -0.06342542 | -0.106708634 |
| Aldh3a1 | 11.16129222 | 11.13154317 | 11.26804087 | -0.136497697 | -0.10674865 |
| Acat1 | 10.93238864 | 10.82722395 | 11.03921746 | -0.211993517 | -0.10682882 |
| Zbtb22 | 8.895432176 | 9.003282804 | 9.002283771 | 0.000999034 | -0.106851594 |
| Rbm5 | 11.0156533 | 11.21000497 | 11.12256018 | 0.087444783 | -0.10690688 |
| Ghitm | 11.94577263 | 11.97551481 | 12.0528712 | -0.077356393 | -0.107098577 |
| Tollip | 11.3814971 | 11.62480346 | 11.48860793 | 0.136195523 | -0.107110837 |
| Cthrc1 | 3.475232462 | 3.483921828 | 3.582493456 | -0.098571628 | -0.107260994 |
| Sez6l2 | 3.763748608 | 3.764940361 | 3.871026487 | -0.106086126 | -0.107277879 |
| Cenpi | 6.794654297 | 6.782573949 | 6.902027531 | -0.119453583 | -0.107373234 |
| Fam72a | 4.891011021 | 4.998517292 | 4.99841291 | 0.000104382 | -0.107401889 |
| Camkmt | 5.615933138 | 5.641248716 | 5.723341536 | -0.08209282 | -0.107408398 |
| Gcdh | 8.755658467 | 8.774666601 | 8.863139005 | -0.088472404 | -0.107480538 |

| | | | | | |
|---|---|---|---|---|---|
| Mesp1 | 2.94300929 | 2.98386831 | 3.050570437 | -0.066702127 | -0.107561146 |
| Styk1 | 5.44479366 | 5.667858361 | 5.552364728 | 0.115493633 | -0.107571068 |
| Btbd2 | 8.656669229 | 9.033024962 | 8.764289928 | 0.268735034 | -0.107620699 |
| Ccser1 | 6.14637211 | 6.101777154 | 6.254095926 | -0.152318771 | -0.107723816 |
| Pnkd | 9.217693168 | 9.45607592 | 9.325479043 | 0.130596877 | -0.107785875 |
| Duoxa1 | 7.008835871 | 7.143553866 | 7.116623146 | 0.02693072 | -0.107787274 |
| Cntd1 | 4.399422105 | 4.459298864 | 4.507227104 | -0.04792824 | -0.107804999 |
| Grin2a | 2.29459661 | 2.333557077 | 2.402402373 | -0.068845296 | -0.107805763 |
| Gdap1 | 3.94572761 | 3.84323406 | 4.053615356 | -0.210381296 | -0.107887746 |
| Fut1 | 8.571049245 | 8.574934804 | 8.678939925 | -0.104005121 | -0.107890681 |
| Dmrta1 | 2.679919683 | 2.769790796 | 2.787830174 | -0.018039378 | -0.107910492 |
| Mlec | 12.02943319 | 12.05719431 | 12.1373439 | -0.08014959 | -0.10791071 |
| Mrnip | 6.204137778 | 6.102142257 | 6.312067099 | -0.209924842 | -0.107929321 |
| Rnf207 | 4.887377429 | 4.866916473 | 4.995314212 | -0.128397739 | -0.107936783 |
| Gm13437 | 1.922548053 | 2.029950401 | 2.030532436 | -0.000582035 | -0.107984383 |
| Cdkn3 | 5.66437586 | 5.649241653 | 5.772365052 | -0.123123399 | -0.107989192 |
| Tecr | 12.2931215 | 12.27006427 | 12.40117621 | -0.131111937 | -0.10805471 |
| Zcchc24 | 10.5190883 | 10.29644093 | 10.62714685 | -0.330705923 | -0.108058557 |
| Abhd8 | 6.999721811 | 7.475097196 | 7.107806605 | 0.367290591 | -0.108084794 |
| Tfcp2 | 7.848300967 | 7.899679345 | 7.956399148 | -0.056719804 | -0.108098181 |
| Frg1 | 9.405019435 | 9.421055235 | 9.513117967 | -0.092062732 | -0.108098532 |
| Ddn | 3.571063738 | 3.656511178 | 3.679175436 | -0.022664258 | -0.108111698 |
| Dennd5b | 6.961551157 | 7.139927276 | 7.069797574 | 0.070129703 | -0.108246416 |
| Zfp810 | 6.746311481 | 6.668643187 | 6.854568721 | -0.185925534 | -0.10825724 |
| Mlxip | 10.20682706 | 10.35647347 | 10.31515754 | 0.041315923 | -0.108330483 |
| Mtmr9 | 9.551770898 | 9.614279415 | 9.660111927 | -0.045832512 | -0.108341029 |
| Tep1 | 9.955310978 | 9.905207502 | 10.06366503 | -0.158457524 | -0.108354049 |
| Pcdh9 | 3.7195299 | 3.658986642 | 3.828034619 | -0.169047977 | -0.108504719 |
| Pcbp4 | 8.200680018 | 8.375619345 | 8.309185963 | 0.066433382 | -0.108505945 |
| Sytl4 | 5.271482995 | 5.26631114 | 5.379997382 | -0.113686242 | -0.108514387 |
| Chd5 | 4.861890258 | 5.199343917 | 4.970486367 | 0.22885755 | -0.108596108 |
| Slc38a11 | 1.67398512 | 1.704677805 | 1.782583326 | -0.077905521 | -0.108598205 |

| | | | | | |
|---|---|---|---|---|---|
| Mtus2 | 3.965358249 | 3.92219267 | 4.074063021 | -0.151870351 | -0.108704772 |
| 4933411K16Rik | 2.472140546 | 2.52962141 | 2.580990998 | -0.051369588 | -0.108850452 |
| Sh3bgrl | 10.9376566 | 10.77508344 | 11.04655996 | -0.271476523 | -0.108903357 |
| Gm6802 | 4.108858804 | 4.298454738 | 4.217829916 | 0.080624822 | -0.108971112 |
| Lrrc45 | 8.760969334 | 9.24010406 | 8.870041411 | 0.37006265 | -0.109072076 |
| Zfp971 | 6.110622854 | 6.133701565 | 6.21982064 | -0.086119075 | -0.109197786 |
| Trem2 | 4.719884095 | 4.999265468 | 4.829089579 | 0.170175889 | -0.109205485 |
| Defb1 | 9.357465808 | 9.28274089 | 9.466783314 | -0.184042423 | -0.109317506 |
| Zfp141 | 7.545600167 | 7.664101963 | 7.654960852 | 0.009141112 | -0.109360685 |
| Ifi27 | 10.61345494 | 10.62634078 | 10.72284024 | -0.096499467 | -0.1093853 |
| Gm28344 | 3.456827357 | 3.594431475 | 3.566213207 | 0.028218268 | -0.10938585 |
| Rep15 | 3.616926816 | 3.670013011 | 3.726395597 | -0.056382586 | -0.109468781 |
| Aco2 | 12.24744142 | 12.29275835 | 12.35691654 | -0.064158197 | -0.109475123 |
| BC022960 | 3.36050776 | 3.452303027 | 3.470005896 | -0.017702869 | -0.109498136 |
| Mapkbp1 | 10.42270736 | 10.73240367 | 10.53223564 | 0.200168033 | -0.10952828 |
| Fli1 | 7.614604421 | 7.620026852 | 7.724157363 | -0.104130512 | -0.109552942 |
| Tmem223 | 7.974632976 | 8.036186076 | 8.084193171 | -0.048007094 | -0.109560195 |
| Zfp385b | 6.498830727 | 6.470554268 | 6.608393343 | -0.137839075 | -0.109562616 |
| Calr3 | 4.757031041 | 4.833130878 | 4.866661008 | -0.033530129 | -0.109629966 |
| St6galnac3 | 3.635188387 | 3.570604019 | 3.744821361 | -0.174217342 | -0.109632973 |
| Tnni3 | 4.621873154 | 4.690050643 | 4.731571971 | -0.041521328 | -0.109698817 |
| Serpina3h | 8.872215994 | 9.258174528 | 8.981973329 | 0.276201199 | -0.109757336 |
| Rrm1 | 10.65386269 | 10.68795218 | 10.76364415 | -0.07569197 | -0.109781463 |
| Tmem229b | 9.543415843 | 9.46817027 | 9.653207315 | -0.185037045 | -0.109791472 |
| Tmem72 | 8.415535746 | 8.497100962 | 8.525367967 | -0.028267004 | -0.109832221 |
| Rslcan18 | 5.013159965 | 5.041197764 | 5.123018962 | -0.081821198 | -0.109858997 |
| Zfp119b | 6.495973521 | 6.58395677 | 6.605835549 | -0.021878779 | -0.109862028 |
| Papolg | 8.099406482 | 8.193313814 | 8.209352287 | -0.016038472 | -0.109945804 |
| Acoxl | 8.902881531 | 8.800199233 | 9.012880326 | -0.212681093 | -0.109998794 |
| Pgbd1 | 4.354614959 | 4.41156208 | 4.464633524 | -0.053071444 | -0.110018565 |
| Slc6a7 | 3.059466392 | 3.235741819 | 3.169533706 | 0.066208113 | -0.110067315 |
| Zdbf2 | 4.144108019 | 4.402645118 | 4.254186068 | 0.14845905 | -0.110078049 |

| | | | | | |
|---|---|---|---|---|---|
| Znrd1as | 6.486160606 | 6.365240506 | 6.596293399 | -0.231052893 | -0.110132793 |
| Odf3l1 | 3.580868355 | 3.618090268 | 3.691020657 | -0.072930389 | -0.110152302 |
| Ccdc6 | 10.84912256 | 10.93182824 | 10.95928407 | -0.027455827 | -0.110161513 |
| Napg | 9.501162925 | 9.588555757 | 9.611339789 | -0.022784032 | -0.110176864 |
| Atp13a5 | 5.914891539 | 5.971976095 | 6.025150152 | -0.053174057 | -0.110258613 |
| Ap5s1 | 7.37118799 | 7.423869139 | 7.481526521 | -0.057657382 | -0.110338531 |
| Cep162 | 8.054678256 | 8.266360018 | 8.165140431 | 0.101219588 | -0.110462174 |
| Sema3c | 10.99502626 | 10.95436937 | 11.10551371 | -0.15114434 | -0.110487443 |
| Gde1 | 9.361646462 | 9.227960395 | 9.472136443 | -0.244176048 | -0.110489981 |
| Cutal | 4.179414591 | 4.292701288 | 4.289936788 | 0.0027645 | -0.110522197 |
| Nectin3 | 9.64743182 | 9.655989655 | 9.757987018 | -0.101997363 | -0.110555197 |
| Cdk5rap2 | 8.339799514 | 8.456694167 | 8.450382816 | 0.006311351 | -0.110583302 |
| Gm6161 | 2.150226906 | 2.11455374 | 2.260929436 | -0.146375696 | -0.11070253 |
| 4930544G11Rik | 1.932689477 | 1.980560838 | 2.043436825 | -0.062875987 | -0.110747348 |
| Rnf130 | 9.598800923 | 9.50248568 | 9.709580054 | -0.207094373 | -0.110779131 |
| Mettl9 | 10.16947957 | 10.09229025 | 10.28029119 | -0.18800094 | -0.110811617 |
| Mras | 7.509771468 | 7.695329185 | 7.620617514 | 0.074711671 | -0.110846046 |
| BC049352 | 3.145798688 | 3.073062064 | 3.256754566 | -0.183692502 | -0.110955878 |
| Chodl | 4.067198892 | 4.025223816 | 4.178182868 | -0.152959052 | -0.110983976 |
| Pias1 | 9.643032923 | 9.727645959 | 9.754038689 | -0.026392729 | -0.111005765 |
| Gabarapl2 | 10.91988336 | 10.99204541 | 11.03091002 | -0.038864607 | -0.111026653 |
| Homez | 7.107189555 | 7.184958015 | 7.218269152 | -0.033311137 | -0.111079597 |
| Pofut1 | 9.475484233 | 9.500278063 | 9.586632499 | -0.086354436 | -0.111148266 |
| Rims4 | 3.766510494 | 3.798495634 | 3.877702389 | -0.079206755 | -0.111191895 |
| Cradd | 6.289099588 | 6.363258203 | 6.400317247 | -0.037059044 | -0.111217659 |
| Gm2451 | 1.919389761 | 1.953174697 | 2.030799815 | -0.077625119 | -0.111410054 |
| Hacd2 | 11.43828155 | 11.45060655 | 11.54971194 | -0.099105387 | -0.11143039 |
| 1700025G04Rik | 10.21695946 | 10.21567465 | 10.32841196 | -0.112737313 | -0.1114525 |
| Cep126 | 4.968024061 | 4.851274105 | 5.079485533 | -0.228211428 | -0.111461472 |
| Kcnt2 | 3.920022391 | 3.936560617 | 4.031497439 | -0.094936821 | -0.111475048 |
| Grhpr | 8.192468629 | 8.343387549 | 8.304015527 | 0.039372022 | -0.111546898 |
| Cdkl3 | 5.43255233 | 5.603471749 | 5.544123767 | 0.059347982 | -0.111571437 |

| | | | | | |
|---|---|---|---|---|---|
| Orm3 | 4.603943073 | 4.635801713 | 4.715515604 | -0.07971389 | -0.111572531 |
| Smc5 | 9.22577674 | 9.330309084 | 9.337355419 | -0.007046335 | -0.111578678 |
| Chrnb1 | 6.840277016 | 7.056396184 | 6.951905842 | 0.104490342 | -0.111628826 |
| Cdk16 | 11.11559317 | 11.17361493 | 11.22728959 | -0.053674653 | -0.111696417 |
| Rhox10 | 4.489281677 | 4.550112898 | 4.601034077 | -0.050921179 | -0.111752401 |
| Vdac1 | 12.89475176 | 12.8512712 | 13.00657142 | -0.155300217 | -0.111819657 |
| H3f3a | 13.36800284 | 13.36959187 | 13.47982705 | -0.110235177 | -0.111824203 |
| Phf3 | 10.27588362 | 10.2926927 | 10.38773237 | -0.09503967 | -0.111848753 |
| Sap25 | 5.721428412 | 5.883142572 | 5.833337565 | 0.049805007 | -0.111909153 |
| Rnd2 | 7.536659222 | 7.637626294 | 7.648658811 | -0.011032517 | -0.111999589 |
| Rnf2 | 9.491964771 | 9.421375521 | 9.603973529 | -0.182598009 | -0.112008759 |
| Mettl15 | 5.908401944 | 5.904626894 | 6.020422428 | -0.115795533 | -0.112020484 |
| Wfdc1 | 6.661651479 | 6.519752432 | 6.773696553 | -0.253944121 | -0.112045074 |
| Pmvk | 10.11501655 | 10.0260927 | 10.22707941 | -0.200986704 | -0.11206286 |
| Clstn2 | 5.111081868 | 5.148123277 | 5.223388002 | -0.075264724 | -0.112306134 |
| Vmn1r183 | 4.608037817 | 4.669239369 | 4.720367597 | -0.051128228 | -0.11232978 |
| Zfx | 9.732663668 | 9.79256388 | 9.845086222 | -0.052522342 | -0.112422554 |
| Zfp599 | 4.90772074 | 4.952498855 | 5.02028122 | -0.067782364 | -0.112560479 |
| Zfp607b | 5.109491939 | 5.002594251 | 5.222055286 | -0.219461035 | -0.112563347 |
| Kcnu1 | 3.935873467 | 4.015424437 | 4.048458936 | -0.033034499 | -0.112585469 |
| Sgsh | 8.294308017 | 8.528089552 | 8.406900734 | 0.121188818 | -0.112592717 |
| Ankrd11 | 12.04163284 | 12.15358983 | 12.15424068 | -0.000650857 | -0.112607847 |
| Gm19196 | 3.154616734 | 3.160699459 | 3.267327418 | -0.106627959 | -0.112710684 |
| Lhfpl4 | 5.296971129 | 5.471567875 | 5.409760368 | 0.061807508 | -0.112789239 |
| Tatdn3 | 6.546106154 | 6.521767246 | 6.658916493 | -0.137149247 | -0.112810339 |
| Lrr1 | 5.02859923 | 5.109479744 | 5.141543272 | -0.032063527 | -0.112944042 |
| Ugt3a2 | 3.038661052 | 3.078419639 | 3.151622634 | -0.073202995 | -0.112961582 |
| Gm10335 | 2.098567048 | 2.124931085 | 2.211595067 | -0.086663982 | -0.113028019 |
| Msl3 | 9.347150869 | 9.340929673 | 9.46021359 | -0.119283917 | -0.113062722 |
| Galnt11 | 8.47609174 | 8.267947513 | 8.589191262 | -0.321243749 | -0.113099521 |
| Gm10642 | 3.743045598 | 3.752083258 | 3.856206139 | -0.10412288 | -0.11316054 |
| Pigo | 8.437696514 | 8.597009531 | 8.550887597 | 0.046121934 | -0.113191083 |

| | | | | | |
|---|---|---|---|---|---|
| Rho | 2.776831661 | 2.831799287 | 2.890049769 | -0.058250483 | -0.113218108 |
| Lyz1 | 9.239155093 | 8.978421515 | 9.352375642 | -0.373954126 | -0.113220549 |
| Rtn2 | 10.22600444 | 10.39064805 | 10.33924963 | 0.051398417 | -0.113245186 |
| Ndufa6 | 9.632905029 | 9.633944211 | 9.746167865 | -0.112223654 | -0.113262836 |
| Cerkl | 4.489023076 | 4.522115085 | 4.602340827 | -0.080225742 | -0.113317751 |
| Golph3l | 8.580234579 | 8.698520993 | 8.693597635 | 0.004923358 | -0.113363057 |
| Zfp953 | 6.840722647 | 6.885188907 | 6.95411256 | -0.068923654 | -0.113389914 |
| Galt | 8.19815778 | 8.345604279 | 8.311588075 | 0.034016204 | -0.113430295 |
| Shroom1 | 2.747340772 | 2.777796803 | 2.860819322 | -0.083022519 | -0.11347855 |
| Gm18848 | 1.723325036 | 1.784968782 | 1.836821274 | -0.051852492 | -0.113496238 |
| Rnase10 | 3.287277152 | 3.341678007 | 3.400815897 | -0.059137889 | -0.113538745 |
| Ncoa4 | 11.10143108 | 11.07993174 | 11.21499325 | -0.135061517 | -0.11356217 |
| Ccdc113 | 5.29117143 | 5.260332259 | 5.404765585 | -0.144433326 | -0.113594154 |
| Gm15720 | 2.045923727 | 2.069298656 | 2.159533777 | -0.090235121 | -0.11361005 |
| Gm436 | 4.501249524 | 4.376428805 | 4.614903358 | -0.238474553 | -0.113653834 |
| Opn1sw | 2.422567876 | 2.43068669 | 2.536222392 | -0.105535702 | -0.113654517 |
| Mdp1 | 9.120541926 | 9.217693612 | 9.234245561 | -0.016551949 | -0.113703635 |
| Atg12 | 9.484397067 | 9.58300456 | 9.598109238 | -0.015104678 | -0.113712171 |
| Gtf3a | 8.608644199 | 8.651054364 | 8.722381425 | -0.071327061 | -0.113737226 |
| P3h3 | 7.953232435 | 8.103604549 | 8.067125154 | 0.036479396 | -0.113892719 |
| Calhm5 | 7.362998637 | 7.328209534 | 7.476896871 | -0.148687337 | -0.113898235 |
| Wdr78 | 4.783276572 | 5.016914591 | 4.897187285 | 0.119727306 | -0.113910714 |
| Sh3bp5 | 8.931238364 | 8.871058668 | 9.045301409 | -0.174242741 | -0.114063045 |
| Lrrc3b | 2.639926043 | 2.642737392 | 2.753994745 | -0.111257353 | -0.114068702 |
| Sox2 | 3.051693567 | 3.089091842 | 3.165810676 | -0.076718833 | -0.114117109 |
| Fkbp3 | 9.841112657 | 9.769385464 | 9.955257578 | -0.185872114 | -0.114144921 |
| Lamp1 | 13.27164155 | 13.34371664 | 13.38579606 | -0.042079423 | -0.114154517 |
| Surf1 | 8.736992366 | 8.731108491 | 8.851152253 | -0.120043761 | -0.114159887 |
| Avil | 6.365232307 | 6.591544799 | 6.479410896 | 0.112133903 | -0.11417859 |
| Rcor2 | 4.916288731 | 5.046788846 | 5.030476273 | 0.016312574 | -0.114187541 |
| F8 | 4.293065699 | 4.384216089 | 4.407258802 | -0.023042713 | -0.114193103 |
| Pigv | 7.996646665 | 8.170451814 | 8.110883152 | 0.059568662 | -0.114236487 |

| | | | | | |
|---|---|---|---|---|---|
| Ccnc | 8.595535971 | 8.490549482 | 8.709840058 | -0.219290576 | -0.114304087 |
| Tbxa2r | 4.615983747 | 4.744632921 | 4.730326384 | 0.014306537 | -0.114342638 |
| Ppfia3 | 8.981070205 | 9.290865944 | 9.095451068 | 0.195414876 | -0.114380864 |
| Gm128 | 4.751122384 | 5.013553559 | 4.865563451 | 0.147990107 | -0.114441067 |
| Mup17 | 5.87951551 | 6.179970207 | 5.993983475 | 0.185986732 | -0.114467965 |
| Rdh16f1 | 10.66536015 | 10.62643908 | 10.77985262 | -0.153413543 | -0.114492467 |
| Tbc1d12 | 7.454247839 | 7.441907754 | 7.56875382 | -0.126846066 | -0.11450598 |
| Cldn11 | 3.265104393 | 3.386446212 | 3.379621073 | 0.006825139 | -0.11451668 |
| Zfp764 | 6.895670102 | 6.958463803 | 7.010293608 | -0.051829805 | -0.114623506 |
| AC169509.1 | 4.584060721 | 4.613769494 | 4.698701257 | -0.084931762 | -0.114640536 |
| Zswim7 | 6.1285369 | 5.980349805 | 6.243273352 | -0.262923547 | -0.114736452 |
| Tmx3 | 10.62057269 | 10.48630421 | 10.73531377 | -0.24900956 | -0.114741083 |
| Rxrg | 6.791780099 | 6.873703832 | 6.906579968 | -0.032876136 | -0.11479987 |
| Armcx2 | 7.391239508 | 7.569759619 | 7.506051354 | 0.063708265 | -0.114811846 |
| Idh3g | 10.70352881 | 10.69385157 | 10.81834611 | -0.124494543 | -0.1148173 |
| Pex13 | 9.683396764 | 9.532288512 | 9.798215342 | -0.26592683 | -0.114818578 |
| Cdpf1 | 7.590316012 | 7.815544331 | 7.705141143 | 0.110403188 | -0.114825131 |
| Cenph | 6.687989043 | 6.877131062 | 6.802863255 | 0.074267807 | -0.114874212 |
| Ubqln2 | 8.97086242 | 9.089463351 | 9.085746238 | 0.003717113 | -0.114883818 |
| Gm12183 | 2.363248723 | 2.366402146 | 2.478149833 | -0.111747686 | -0.11490111 |
| Rnf146 | 9.7424106 | 9.727584037 | 9.857318608 | -0.129734571 | -0.114908008 |
| Tyrp1 | 5.568369866 | 5.654084691 | 5.683443928 | -0.029359238 | -0.115074062 |
| Trdn | 11.68206221 | 11.66481798 | 11.79715685 | -0.13233887 | -0.11509464 |
| Dock9 | 10.70889719 | 10.6710656 | 10.82407803 | -0.15301243 | -0.11518084 |
| Dsc3 | 14.00613293 | 14.10676798 | 14.12132227 | -0.014554293 | -0.115189343 |
| Endog | 6.587102966 | 6.667403728 | 6.702295056 | -0.034891328 | -0.11519209 |
| Zfp827 | 6.905202189 | 6.840947501 | 7.020395736 | -0.179448236 | -0.115193547 |
| Pou6f1 | 7.914747372 | 7.995477472 | 8.029960995 | -0.034483523 | -0.115213623 |
| Luc7l2 | 11.69957607 | 11.69315767 | 11.81482334 | -0.121665673 | -0.11524727 |
| Mbnl2 | 11.81900171 | 11.86680846 | 11.93425692 | -0.067448467 | -0.115255213 |
| Poll | 7.331012337 | 7.337191968 | 7.446304638 | -0.10911267 | -0.115292302 |
| Zan | 3.171011232 | 3.236225861 | 3.286362307 | -0.050136446 | -0.115351075 |

| | | | | | |
|---|---|---|---|---|---|
| Cul7 | 8.420948579 | 8.571080016 | 8.536378343 | 0.034701673 | -0.115429764 |
| Rtkn2 | 3.607341804 | 3.600863527 | 3.722964007 | -0.12210048 | -0.115622203 |
| Atp5j | 11.06807329 | 11.08577801 | 11.18375572 | -0.097977713 | -0.115682433 |
| Ctsl | 12.83572484 | 12.90829611 | 12.95142637 | -0.043130267 | -0.115701533 |
| Naxd | 9.245246015 | 9.328162133 | 9.360984137 | -0.032822004 | -0.115738122 |
| Usp40 | 9.370051144 | 9.414685332 | 9.485803562 | -0.07111823 | -0.115752418 |
| Eps8l1 | 11.07728636 | 11.19040953 | 11.19313041 | -0.002720887 | -0.11584405 |
| Fbxo31 | 9.483657807 | 9.432408659 | 9.599556761 | -0.167148102 | -0.115898955 |
| Zswim9 | 7.05118412 | 7.140251213 | 7.167101424 | -0.026850211 | -0.115917304 |
| Nphp1 | 7.862027211 | 7.597228571 | 7.978018313 | -0.380789742 | -0.115991102 |
| Dync2h1 | 7.710924701 | 7.759306988 | 7.826971512 | -0.067664524 | -0.116046811 |
| Mup9 | 5.045995946 | 5.222201221 | 5.162080076 | 0.060121145 | -0.11608413 |
| Efna2 | 5.615616276 | 5.959005601 | 5.731705241 | 0.22730036 | -0.116088965 |
| Fars2 | 8.019572108 | 8.052782942 | 8.135727149 | -0.082944206 | -0.116155041 |
| Kifc5b | 5.869786937 | 6.074345387 | 5.985993329 | 0.088352058 | -0.116206392 |
| Zfp968 | 5.604702288 | 5.788472494 | 5.720920543 | 0.06755195 | -0.116218255 |
| Nat9 | 7.726758373 | 7.778448913 | 7.842978162 | -0.064529249 | -0.116219789 |
| Edar | 3.044483093 | 3.075436925 | 3.160703794 | -0.085266868 | -0.1162207 |
| Olfr1247 | 4.91475627 | 4.924160493 | 5.031131931 | -0.106971438 | -0.116375661 |
| Mad2l1bp | 7.661052717 | 7.840922343 | 7.777430604 | 0.063491739 | -0.116377888 |
| Adamts18 | 2.792864256 | 2.846893932 | 2.909283817 | -0.062389886 | -0.116419561 |
| Higd2a | 9.338720216 | 9.383372472 | 9.455204562 | -0.07183209 | -0.116484346 |
| Kansl1 | 9.934851998 | 10.1428585 | 10.05139207 | 0.091466433 | -0.116540069 |
| Gm16459 | 2.67299528 | 2.739401209 | 2.789554724 | -0.050153515 | -0.116559444 |
| Gm2539 | 6.51510965 | 6.417498224 | 6.631705136 | -0.214206912 | -0.116595486 |
| 1190005I06Rik | 4.221145971 | 4.36779269 | 4.337771776 | 0.030020913 | -0.116625805 |
| Stk26 | 7.832159696 | 7.890637606 | 7.948796978 | -0.058159373 | -0.116637283 |
| Tubb3 | 5.838032305 | 6.070596147 | 5.954684596 | 0.115911551 | -0.116652291 |
| Tcp11l2 | 9.372801391 | 9.256552952 | 9.489464435 | -0.232911483 | -0.116663044 |
| Zfp811 | 3.36690044 | 3.415375916 | 3.483577114 | -0.068201198 | -0.116676674 |
| C1qtnf2 | 6.798436707 | 6.902380889 | 6.915121096 | -0.012740207 | -0.116684389 |
| Shc2 | 7.454593505 | 7.778739471 | 7.571409351 | 0.20733012 | -0.116815846 |

| | | | | | |
|---|---|---|---|---|---|
| Pax3 | 4.652652533 | 4.695773782 | 4.769500001 | -0.073726219 | -0.116847469 |
| Bag4 | 9.69598608 | 9.673824745 | 9.812854794 | -0.139030048 | -0.116868713 |
| Krt80 | 13.15910098 | 13.29487564 | 13.27611755 | 0.018758087 | -0.117016577 |
| Tjp1 | 10.68916799 | 10.68058392 | 10.8062436 | -0.125659683 | -0.11707561 |
| Endod1 | 10.82507197 | 10.85768708 | 10.94223621 | -0.084549127 | -0.11716424 |
| Vwa7 | 5.642257577 | 5.829701786 | 5.759498902 | 0.070202883 | -0.117241325 |
| Kif14 | 6.954745661 | 6.987766747 | 7.072047007 | -0.08428026 | -0.117301346 |
| Mllt11 | 7.521159851 | 7.706362921 | 7.638521837 | 0.067841084 | -0.117361986 |
| Hnrnph3 | 8.921365877 | 9.002174409 | 9.038739004 | -0.036564594 | -0.117373127 |
| Rit1 | 9.367785299 | 9.400786329 | 9.48517647 | -0.084390141 | -0.117391172 |
| 0610030E20Rik | 8.816642978 | 8.916189891 | 8.934146144 | -0.017956253 | -0.117503166 |
| Ssc4d | 4.088031115 | 4.09366082 | 4.205586489 | -0.111925669 | -0.117555374 |
| Bcorl1 | 8.093779867 | 8.185785899 | 8.211345819 | -0.02555992 | -0.117565953 |
| Parg | 9.560459078 | 9.494572239 | 9.678084777 | -0.183512538 | -0.117625699 |
| Spc25 | 7.049126706 | 6.993025848 | 7.166826857 | -0.173801009 | -0.117700152 |
| Sgsm1 | 5.139878984 | 5.404350941 | 5.25759437 | 0.146756571 | -0.117715387 |
| Map3k21 | 4.054012381 | 4.158788132 | 4.17173374 | -0.012945607 | -0.117721359 |
| Rab12 | 9.876388072 | 9.753677371 | 9.994134844 | -0.240457472 | -0.117746772 |
| Lce3c | 8.555673407 | 8.792573751 | 8.673423293 | 0.119150458 | -0.117749887 |
| Zkscan1 | 9.571202641 | 9.602310581 | 9.689117505 | -0.086806925 | -0.117914865 |
| Speer5-ps1 | 4.475410801 | 4.389039101 | 4.59334164 | -0.204302539 | -0.117930839 |
| Tbc1d14 | 10.6946337 | 10.7298697 | 10.81259829 | -0.08272859 | -0.117964593 |
| Ogg1 | 6.798479251 | 6.955872453 | 6.916508501 | 0.039363952 | -0.11802925 |
| Cd4 | 6.293025494 | 6.231737263 | 6.411065352 | -0.179328089 | -0.118039857 |
| Brap | 10.66408053 | 10.85692061 | 10.78214452 | 0.07477609 | -0.11806399 |
| Plxna2 | 9.996888794 | 10.03179115 | 10.11498499 | -0.083193838 | -0.118096196 |
| Lrp2bp | 5.256858594 | 5.185311778 | 5.375023173 | -0.189711396 | -0.11816458 |
| Plekha1 | 10.78684127 | 10.90301438 | 10.9050159 | -0.00200152 | -0.118174627 |
| Gjb4 | 10.88752988 | 11.03350354 | 11.00570573 | 0.027797807 | -0.11817585 |
| Rab9b | 2.640467978 | 2.715558897 | 2.758653083 | -0.043094185 | -0.118185105 |
| Zxdb | 7.943631394 | 8.05498963 | 8.061849128 | -0.006859498 | -0.118217734 |
| Rpl10a-ps2 | 2.902166606 | 2.932072938 | 3.020398608 | -0.088325669 | -0.118232002 |

| | | | | | |
|---|---|---|---|---|---|
| Gm3226 | 3.979293979 | 4.068603023 | 4.097536006 | -0.028932983 | -0.118242027 |
| Gm2446 | 5.936521733 | 6.06826233 | 6.054892312 | 0.013370018 | -0.118370579 |
| Slc9a2 | 5.480381794 | 5.644932947 | 5.598761912 | 0.046171036 | -0.118380118 |
| Sh3tc2 | 7.8308826 | 7.904387827 | 7.949267067 | -0.04487924 | -0.118384467 |
| Trappc11 | 9.642800633 | 9.690061507 | 9.761199333 | -0.071137826 | -0.1183987 |
| Zfp654 | 8.97085234 | 8.930855088 | 9.089252558 | -0.15839747 | -0.118400218 |
| Zfp110 | 9.449430846 | 9.545183812 | 9.56784178 | -0.022657968 | -0.118410935 |
| Aldh1l1 | 9.583182602 | 9.251037072 | 9.701600195 | -0.450563123 | -0.118417593 |
| Ric1 | 10.28724409 | 10.41966825 | 10.40576158 | 0.013906677 | -0.11851749 |
| Supt20 | 9.73973688 | 9.877159316 | 9.85828806 | 0.018871256 | -0.11855118 |
| Bcl7c | 9.12807404 | 9.17535473 | 9.246628651 | -0.071273921 | -0.118554611 |
| Sdf2 | 9.803459059 | 9.80290191 | 9.922060353 | -0.119158443 | -0.118601294 |
| Morn2 | 5.621872011 | 5.70952026 | 5.740565401 | -0.031045141 | -0.11869339 |
| Eif3f | 11.83494656 | 11.82743728 | 11.95365261 | -0.126215323 | -0.11870605 |
| Gm48364 | 2.716741626 | 2.809594576 | 2.835454135 | -0.025859559 | -0.118712509 |
| Lzts2 | 9.087276356 | 9.502317303 | 9.206026261 | 0.296291042 | -0.118749905 |
| Ccnd2 | 12.35769904 | 12.26866762 | 12.47654092 | -0.2078733 | -0.11884188 |
| Wdr25 | 6.131840483 | 6.168555308 | 6.250729625 | -0.082174317 | -0.118889143 |
| Oas1c | 6.642578663 | 6.675169125 | 6.761492117 | -0.086322992 | -0.118913453 |
| Lama4 | 10.54679703 | 10.44623813 | 10.66579851 | -0.21956038 | -0.119001483 |
| Cop1 | 10.88230183 | 10.90596705 | 11.00134626 | -0.09537921 | -0.119044423 |
| Nacc2 | 9.005855587 | 8.995132032 | 9.124953819 | -0.129821787 | -0.119098232 |
| Rp9 | 8.743330775 | 8.686871445 | 8.862495884 | -0.175624438 | -0.119165108 |
| Dsc2 | 7.962672095 | 7.930625746 | 8.081866403 | -0.151240657 | -0.119194308 |
| Zfp446 | 6.74759927 | 6.965972976 | 6.866827498 | 0.099145478 | -0.119228228 |
| Fut7 | 3.3161853 | 3.344287303 | 3.43542233 | -0.091135026 | -0.11923703 |
| Galk2 | 8.697554258 | 8.785849839 | 8.816803471 | -0.030953632 | -0.119249213 |
| Rfesd | 7.271015118 | 7.135806602 | 7.390314459 | -0.254507857 | -0.119299341 |
| Lpin3 | 8.870564478 | 9.133746117 | 8.989886947 | 0.143859171 | -0.119322469 |
| Serac1 | 6.761849239 | 6.977659032 | 6.881209241 | 0.09644979 | -0.119360003 |
| Rpl12-ps2 | 2.223260817 | 2.288502045 | 2.342621425 | -0.05411938 | -0.119360608 |
| Akr1c12 | 1.56638038 | 1.573962356 | 1.685764567 | -0.111802212 | -0.119384188 |

| | | | | | |
|---|---|---|---|---|---|
| Mlst8 | 8.969138592 | 8.975932238 | 9.088528986 | -0.112596748 | -0.119390394 |
| Tmem181a | 9.637752383 | 9.719280388 | 9.757178526 | -0.037898139 | -0.119426144 |
| Lpar3 | 6.379966177 | 6.440816657 | 6.499435312 | -0.058618655 | -0.119469135 |
| Ubp1 | 10.63876704 | 10.7767017 | 10.75824164 | 0.01846006 | -0.119474593 |
| Sdha | 12.15824163 | 12.12515349 | 12.27773012 | -0.15257663 | -0.119488483 |
| 3110040N11Rik | 7.706228538 | 7.839906639 | 7.825747545 | 0.014159093 | -0.119519007 |
| Pla2g5 | 6.299313483 | 6.653819043 | 6.418838837 | 0.234980206 | -0.119525354 |
| Tmsb15l | 3.670826393 | 3.714239052 | 3.790426578 | -0.076187525 | -0.119600185 |
| Atp2b3 | 3.44260109 | 3.478810294 | 3.562203953 | -0.083393659 | -0.119602863 |
| Kcnab3 | 3.811609006 | 3.958801338 | 3.931253609 | 0.02754773 | -0.119644603 |
| Fa2h | 12.80985427 | 12.86126343 | 12.92949944 | -0.06823601 | -0.11964517 |
| Gm4617 | 2.380500416 | 2.432126962 | 2.500285753 | -0.06815879 | -0.119785337 |
| Nemp1 | 8.096513889 | 8.209625755 | 8.216321125 | -0.00669537 | -0.119807237 |
| Chrm2 | 3.193297612 | 3.273509809 | 3.313109487 | -0.039599678 | -0.119811875 |
| Cep290 | 7.836993251 | 7.899158284 | 7.95681634 | -0.057658057 | -0.11982309 |
| Tmem204 | 7.21386104 | 7.259157126 | 7.333769365 | -0.074612239 | -0.119908325 |
| Frmd7 | 2.649638627 | 2.723870153 | 2.76958678 | -0.045716626 | -0.119948152 |
| Flywch1 | 8.676707286 | 8.994652816 | 8.796657675 | 0.197995141 | -0.11995039 |
| Abraxas1 | 7.226598001 | 7.23237536 | 7.34659113 | -0.11421577 | -0.11999313 |
| Asb1 | 7.67181812 | 7.46438243 | 7.791814022 | -0.327431593 | -0.119995902 |
| Zfp879 | 3.417030873 | 3.418382264 | 3.537038413 | -0.118656149 | -0.120007539 |
| Aplp1 | 5.584498454 | 5.683949849 | 5.704521756 | -0.020571907 | -0.120023303 |
| Ephx4 | 3.95234872 | 4.007184992 | 4.0725127 | -0.065327708 | -0.12016398 |
| Rwdd3 | 4.631288713 | 4.628484216 | 4.751548305 | -0.123064089 | -0.120259592 |
| Nxpe2 | 4.290565266 | 4.391585052 | 4.410891018 | -0.019305966 | -0.120325752 |
| Ttc14 | 10.21441345 | 10.27521764 | 10.33473977 | -0.059522133 | -0.120326323 |
| Gm7002 | 2.117103719 | 2.235534495 | 2.237431997 | -0.001897502 | -0.120328278 |
| Akap5 | 4.63113537 | 4.699423983 | 4.751505051 | -0.052081068 | -0.120369681 |
| Znrf3 | 8.826045196 | 8.924539144 | 8.946419828 | -0.021880684 | -0.120374631 |
| Mier2 | 8.775148643 | 9.280617055 | 8.89553768 | 0.385079374 | -0.120389037 |
| Pfkfb3 | 10.47148847 | 10.3520673 | 10.59192654 | -0.239859237 | -0.120438063 |
| Prkn | 6.043838375 | 6.148740901 | 6.16427953 | -0.015538629 | -0.120441155 |

| | | | | | |
|---|---|---|---|---|---|
| Nsun6 | 6.974347651 | 6.983434467 | 7.094925543 | -0.111491076 | -0.120577892 |
| Lor | 13.96895436 | 14.18969557 | 14.08957761 | 0.10011796 | -0.120623243 |
| Smc4 | 10.70528821 | 10.68857254 | 10.8259717 | -0.137399163 | -0.120683493 |
| Ggps1 | 9.105241396 | 9.076458561 | 9.226056741 | -0.14959818 | -0.120815345 |
| Tpt1-ps3 | 12.85452482 | 12.85985281 | 12.97544483 | -0.11559202 | -0.12092001 |
| Nalcn | 3.085615181 | 3.105627119 | 3.206578297 | -0.100951178 | -0.120963117 |
| Mrvi1 | 7.635284911 | 7.867558718 | 7.756272116 | 0.111286602 | -0.120987205 |
| Primpol | 7.605149534 | 7.534708281 | 7.72622227 | -0.191513989 | -0.121072736 |
| Spata2l | 6.209138133 | 6.515364228 | 6.330220858 | 0.185143369 | -0.121082725 |
| Cpa3 | 9.926108731 | 9.470820428 | 10.0472714 | -0.576450967 | -0.121162664 |
| Dok6 | 4.160044815 | 4.252694081 | 4.281266669 | -0.028572588 | -0.121221854 |
| Thnsl1 | 6.261322589 | 6.138923139 | 6.38265823 | -0.24373509 | -0.121335641 |
| Slc37a3 | 9.178724905 | 9.357616418 | 9.300145319 | 0.057471099 | -0.121420414 |
| Ankrd12 | 10.70445675 | 10.95098196 | 10.82591992 | 0.12506204 | -0.12146317 |
| Psip1 | 9.572215264 | 9.589605848 | 9.693704026 | -0.104098178 | -0.121488762 |
| A430033K04Rik | 7.003807357 | 7.134007239 | 7.12533126 | 0.008675979 | -0.121523904 |
| Kbtbd4 | 7.46138829 | 7.52181609 | 7.582945693 | -0.061129603 | -0.121557403 |
| Rassf3 | 10.87690586 | 10.85211177 | 10.99847971 | -0.146367937 | -0.121573853 |
| Hhatl | 7.820426825 | 8.133241952 | 7.942001726 | 0.191240226 | -0.121574901 |
| Atp8a2 | 5.976646021 | 6.038646439 | 6.098229467 | -0.059583028 | -0.121583447 |
| Kcns3 | 6.346263319 | 6.102037268 | 6.467859451 | -0.365822183 | -0.121596132 |
| Zfp609 | 9.556624143 | 9.686243866 | 9.678323228 | 0.007920638 | -0.121699085 |
| Rabl2 | 6.470469044 | 6.799581166 | 6.592207404 | 0.207373762 | -0.12173836 |
| Sox5 | 5.362104504 | 5.339504549 | 5.483864733 | -0.144360183 | -0.121760229 |
| Washc1 | 8.940963441 | 9.195553266 | 9.062744908 | 0.132808358 | -0.121781467 |
| Vnn1 | 4.898333597 | 5.018013636 | 5.020126947 | -0.002113311 | -0.12179335 |
| Agap1 | 9.170015712 | 9.188134434 | 9.291883842 | -0.103749408 | -0.121868129 |
| Ccdc32 | 7.848597353 | 7.82007333 | 7.970479858 | -0.150406528 | -0.121882505 |
| Ly6h | 4.002018322 | 4.086797979 | 4.123937656 | -0.037139678 | -0.121919334 |
| Gnpda1 | 8.487793884 | 8.841999635 | 8.609754955 | 0.23224468 | -0.121961071 |
| Rtl8a | 7.76657131 | 7.910704378 | 7.88868361 | 0.022020768 | -0.122112299 |
| Hspb2 | 6.671638135 | 7.10352476 | 6.793786213 | 0.309738548 | -0.122148078 |

| | | | | | |
|---|---|---|---|---|---|
| Zmynd8 | 9.914045148 | 10.24260476 | 10.03620469 | 0.206400064 | -0.122159545 |
| Gm5513 | 2.512925544 | 2.63132796 | 2.635100324 | -0.003772364 | -0.12217478 |
| Srrm3 | 3.392361641 | 3.590615716 | 3.514582638 | 0.076033078 | -0.122220997 |
| Slf2 | 9.73261965 | 9.90742707 | 9.854888341 | 0.052538729 | -0.122268691 |
| Vmn2r76 | 6.132171301 | 6.29246563 | 6.254513096 | 0.037952534 | -0.122341795 |
| Gimap6 | 7.70674727 | 7.516093789 | 7.829138011 | -0.313044223 | -0.122390742 |
| Arl16 | 6.768467253 | 6.796092924 | 6.890864974 | -0.09477205 | -0.12239772 |
| Noct | 12.96715098 | 12.81802585 | 13.08957107 | -0.27154522 | -0.122420093 |
| Klc3 | 10.0715036 | 10.62793378 | 10.19392398 | 0.4340098 | -0.122420377 |
| Dynlt1b | 8.14134455 | 7.973162245 | 8.263820044 | -0.290657799 | -0.122475494 |
| Dtx2 | 9.240240119 | 9.822995558 | 9.362823924 | 0.460171634 | -0.122583805 |
| Ercc6l | 7.449868692 | 7.590771416 | 7.57248821 | 0.018283206 | -0.122619518 |
| Sox4 | 8.254660061 | 8.485265489 | 8.377290261 | 0.107975229 | -0.1226302 |
| Zfp12 | 8.469189149 | 8.743907066 | 8.59184322 | 0.152063846 | -0.122654071 |
| Cc2d1b | 10.04689704 | 10.05722439 | 10.1696593 | -0.112434906 | -0.122762262 |
| Fam181b | 3.178294485 | 3.201660694 | 3.301096027 | -0.099435333 | -0.122801541 |
| Pdlim2 | 10.33286032 | 10.54074232 | 10.4556999 | 0.085042423 | -0.122839577 |
| Trappc1 | 9.271189272 | 9.218326019 | 9.394040294 | -0.175714275 | -0.122851022 |
| Cyp4x1 | 3.204655921 | 3.293732535 | 3.327507865 | -0.03377533 | -0.122851944 |
| St8sia2 | 3.511569925 | 3.622686047 | 3.634479191 | -0.011793144 | -0.122909266 |
| Gm48672 | 2.541624564 | 2.62334789 | 2.664539843 | -0.041191954 | -0.12291528 |
| Rnaset2b | 9.965753076 | 9.939293195 | 10.08871273 | -0.149419532 | -0.122959651 |
| Tmem144 | 5.970455687 | 6.269115361 | 6.093417921 | 0.175697439 | -0.122962234 |
| Postn | 13.43361315 | 13.74807035 | 13.55659844 | 0.191471903 | -0.122985297 |
| Sypl | 11.51588902 | 11.44250324 | 11.63888374 | -0.1963805 | -0.122994723 |
| Gm8942 | 2.560368381 | 2.708793921 | 2.683423129 | 0.025370792 | -0.123054748 |
| Ctcf | 10.45684419 | 10.56214863 | 10.57997105 | -0.01782242 | -0.123126863 |
| Gimap8 | 6.425182303 | 6.336766994 | 6.548311393 | -0.211544399 | -0.12312909 |
| Gm11263 | 1.694335838 | 1.685666925 | 1.817470626 | -0.131803701 | -0.123134788 |
| Fam92a | 8.061278609 | 8.085006232 | 8.184413876 | -0.099407644 | -0.123135267 |
| Ispd | 6.840333575 | 6.822134255 | 6.963487219 | -0.141352964 | -0.123153644 |
| Ahdc1 | 8.75470983 | 9.079077672 | 8.877913808 | 0.201163864 | -0.123203979 |

| | | | | | |
|---|---|---|---|---|---|
| Hs3st1 | 6.196843185 | 6.269801675 | 6.320065778 | -0.050264103 | -0.123222592 |
| Wasl | 10.24208319 | 10.33865472 | 10.36533378 | -0.02667906 | -0.123250587 |
| Zic3 | 3.343895483 | 3.480009189 | 3.467148182 | 0.012861007 | -0.123252699 |
| Ctxn1 | 5.560414178 | 5.61001082 | 5.683682999 | -0.073672179 | -0.123268821 |
| Apc | 10.53611936 | 10.67511444 | 10.65941376 | 0.015700677 | -0.123294403 |
| Ssu2 | 2.011252942 | 2.037434225 | 2.134549837 | -0.097115613 | -0.123296895 |
| Gm3776 | 5.453301372 | 5.60845663 | 5.576640837 | 0.031815793 | -0.123339466 |
| Pnrc2 | 10.72755476 | 10.77555498 | 10.85096707 | -0.075412097 | -0.123412313 |
| Zfp507 | 9.044623217 | 8.951130296 | 9.168089289 | -0.216958994 | -0.123466072 |
| Tyr | 2.374972268 | 2.458069223 | 2.498480984 | -0.040411761 | -0.123508716 |
| Espl1 | 8.070912561 | 8.226113465 | 8.194480605 | 0.03163286 | -0.123568044 |
| Pabpc4l | 2.492999814 | 2.506990079 | 2.616579517 | -0.109589439 | -0.123579703 |
| Ncam2 | 4.278268405 | 4.369304267 | 4.401899137 | -0.03259487 | -0.123630732 |
| Gm14419 | 4.937839295 | 4.915880881 | 5.061481734 | -0.145600853 | -0.123642439 |
| Sp1 | 11.20601595 | 11.25034167 | 11.32966126 | -0.079319583 | -0.123645303 |
| Tet3 | 11.19729798 | 11.53867186 | 11.32113189 | 0.217539973 | -0.12383391 |
| Anp32a | 10.75412603 | 10.65942087 | 10.87798673 | -0.218565867 | -0.123860707 |
| Nacad | 3.856149643 | 3.967830136 | 3.980076005 | -0.012245869 | -0.123926362 |
| Gm6170 | 3.278846803 | 3.381400091 | 3.402953949 | -0.021553859 | -0.124107146 |
| Slc4a11 | 7.544950589 | 7.620392908 | 7.669061667 | -0.048668759 | -0.124111078 |
| Ngfr | 8.19005029 | 8.289003114 | 8.314184207 | -0.025181094 | -0.124133917 |
| Sema5b | 2.912893016 | 3.153351946 | 3.037095649 | 0.116256297 | -0.124202633 |
| Eci2 | 10.47783344 | 10.53902441 | 10.60206329 | -0.06303888 | -0.124229857 |
| Crocc2 | 3.780494169 | 3.805998086 | 3.904730649 | -0.098732563 | -0.12423648 |
| B3galnt2 | 9.847219046 | 9.883451935 | 9.971459476 | -0.08800754 | -0.12424043 |
| Gm46136 | 4.424638977 | 4.459978102 | 4.548893939 | -0.088915837 | -0.124254963 |
| Lrrc18 | 2.710007664 | 2.779217114 | 2.834467583 | -0.055250469 | -0.124459919 |
| Cpne5 | 6.81155126 | 6.855796403 | 6.936031872 | -0.080235468 | -0.124480611 |
| Tst | 7.880001607 | 7.966069429 | 8.00459832 | -0.038528891 | -0.124596713 |
| Abcg1 | 9.360094713 | 9.547247866 | 9.484733273 | 0.062514592 | -0.12463856 |
| Eif4ebp2 | 12.30875813 | 12.37109788 | 12.43343259 | -0.06233471 | -0.124674463 |
| Zfp979 | 5.385930962 | 5.512423104 | 5.510628857 | 0.001794246 | -0.124697896 |

| | | | | | |
|---|---|---|---|---|---|
| Gm15320 | 3.35992153 | 3.411529982 | 3.484660678 | -0.073130696 | -0.124739148 |
| Ppcs | 7.489555152 | 7.389307871 | 7.61435305 | -0.225045179 | -0.124797899 |
| Kdelc1 | 7.990965214 | 8.060887996 | 8.115770286 | -0.054882289 | -0.124805072 |
| Usp9x | 11.65662498 | 11.66256004 | 11.78146735 | -0.118907313 | -0.12484237 |
| Tldc1 | 7.4654831 | 7.628554336 | 7.590344493 | 0.038209843 | -0.124861392 |
| Mthfd2l | 7.101772037 | 7.046235588 | 7.226652084 | -0.180416496 | -0.124880047 |
| Strbp | 8.735780819 | 8.886734286 | 8.860679475 | 0.026054811 | -0.124898656 |
| Tex2 | 10.52855786 | 10.57518508 | 10.65347717 | -0.078292097 | -0.124919317 |
| Mgst3 | 9.658992743 | 9.662797407 | 9.783974192 | -0.121176785 | -0.124981449 |
| Ints3 | 10.14020995 | 10.4684266 | 10.26529734 | 0.203129267 | -0.125087387 |
| Papss2 | 8.83251826 | 8.625831672 | 8.957691607 | -0.331859935 | -0.125173347 |
| Mib1 | 10.8414419 | 10.8854153 | 10.96661618 | -0.08120088 | -0.12517428 |
| Rnf214 | 8.68516371 | 8.662175444 | 8.810385783 | -0.148210338 | -0.125222073 |
| Rgs11 | 5.907697177 | 5.993091722 | 6.032999159 | -0.039907437 | -0.125301983 |
| Nhs | 5.875637022 | 5.900875763 | 6.001068539 | -0.100192776 | -0.125431517 |
| Rgs18 | 6.759209604 | 6.425265047 | 6.884670382 | -0.459405335 | -0.125460778 |
| Wipi2 | 10.66805783 | 10.79325529 | 10.79355437 | -0.00029908 | -0.125496533 |
| Terf1 | 7.992295372 | 7.990112704 | 8.117832088 | -0.127719384 | -0.125536716 |
| Gm5835 | 3.269122891 | 3.382845303 | 3.394735325 | -0.011890022 | -0.125612434 |
| Chst8 | 2.951614549 | 3.060313249 | 3.077272076 | -0.016958826 | -0.125657527 |
| Fdps | 11.8034094 | 11.73074276 | 11.92914716 | -0.198404397 | -0.125737757 |
| Hoxb5 | 5.039842508 | 5.148007261 | 5.165614807 | -0.017607546 | -0.125772299 |
| Rpl3-ps1 | 2.040196999 | 2.096878109 | 2.166129531 | -0.069251422 | -0.125932532 |
| Rab18 | 11.86630358 | 11.83334478 | 11.99225586 | -0.158911083 | -0.12595228 |
| Camkv | 4.98459495 | 5.085779954 | 5.110553027 | -0.024773073 | -0.125958077 |
| Stx19 | 7.483832264 | 7.46756474 | 7.609791346 | -0.142226606 | -0.125959083 |
| Gnpat | 10.64873492 | 10.63918392 | 10.77470159 | -0.13551767 | -0.12596667 |
| Hmgb1 | 12.35789895 | 12.2940357 | 12.48391104 | -0.189875337 | -0.126012087 |
| Gm14387 | 4.049240916 | 4.250160155 | 4.175369633 | 0.074790522 | -0.126128717 |
| Gltp | 13.2942677 | 13.42857466 | 13.4204345 | 0.008140167 | -0.1261668 |
| Lrch3 | 9.497137399 | 9.606842629 | 9.623354359 | -0.01651173 | -0.126216961 |
| Gins3 | 6.605900247 | 6.71034548 | 6.732120927 | -0.021775447 | -0.12622068 |

| | | | | | |
|---|---|---|---|---|---|
| Slc8b1 | 9.293504887 | 9.48864775 | 9.419781338 | 0.068866412 | -0.126276451 |
| Grik5 | 6.351235839 | 6.547427385 | 6.477524439 | 0.069902946 | -0.1262886 |
| Nat8f1 | 3.752451634 | 3.807538494 | 3.878755386 | -0.071216892 | -0.126303752 |
| Lrriq3 | 2.844170621 | 2.908014131 | 2.970474931 | -0.0624608 | -0.126304311 |
| Sgk3 | 9.70928084 | 9.757365307 | 9.835721043 | -0.078355736 | -0.126440203 |
| Hsd3b2 | 5.465807581 | 5.561742973 | 5.592256959 | -0.030513986 | -0.126449378 |
| Ugp2 | 11.37694386 | 11.39722737 | 11.50340359 | -0.106176213 | -0.126459723 |
| Zfp518b | 6.854790893 | 6.732399314 | 6.981294527 | -0.248895213 | -0.126503634 |
| Ptar1 | 9.432221599 | 9.470360337 | 9.558760892 | -0.088400555 | -0.126539293 |
| Lrp12 | 8.542590737 | 8.698695332 | 8.669138146 | 0.029557186 | -0.12654741 |
| Gm9436 | 2.550235579 | 2.589408454 | 2.676786751 | -0.087378298 | -0.126551172 |
| Dennd1a | 8.86430262 | 9.058180736 | 8.990933887 | 0.067246849 | -0.126631266 |
| Gm44503 | 1.617209603 | 1.594681511 | 1.743850473 | -0.149168961 | -0.12664087 |
| Fam173a | 8.4220943 | 8.535470272 | 8.548742202 | -0.01327193 | -0.126647902 |
| Tigd2 | 7.833102669 | 7.662170354 | 7.959779396 | -0.297609042 | -0.126676727 |
| Gm8318 | 2.746015974 | 2.866242675 | 2.872700771 | -0.006458096 | -0.126684797 |
| Tyk2 | 9.475637911 | 9.79194302 | 9.60232813 | 0.189614889 | -0.126690219 |
| Stim2 | 9.353893163 | 9.194453988 | 9.480591575 | -0.286137587 | -0.126698413 |
| Mxi1 | 10.42579177 | 10.48873664 | 10.5526109 | -0.063874263 | -0.126819127 |
| Fchsd1 | 8.668984219 | 8.704154849 | 8.795809209 | -0.09165436 | -0.12682499 |
| E2f8 | 8.072595752 | 8.358993982 | 8.1994306 | 0.159563382 | -0.126834848 |
| Ska1 | 5.014703279 | 5.238088415 | 5.1415475 | 0.096540914 | -0.126844222 |
| D430042O09Rik | 8.502990117 | 8.637713269 | 8.629848551 | 0.007864718 | -0.126858434 |
| Esrrg | 5.049986973 | 5.059766494 | 5.176993233 | -0.117226738 | -0.12700626 |
| Tdp2 | 8.464639501 | 8.498451248 | 8.59174028 | -0.093289032 | -0.127100779 |
| Tnfrsf17 | 1.627037192 | 1.644815986 | 1.754144115 | -0.109328129 | -0.127106923 |
| Oard1 | 7.471798498 | 7.497911668 | 7.598976156 | -0.101064488 | -0.127177659 |
| Mfsd1 | 10.89188079 | 11.15725234 | 11.01909962 | 0.138152717 | -0.127218827 |
| Cs | 13.04077816 | 13.05969408 | 13.16806742 | -0.108373337 | -0.127289263 |
| Rbm33 | 10.21697671 | 10.27678491 | 10.34427184 | -0.06748693 | -0.127295127 |
| Fkbp1b | 4.306046068 | 4.305639507 | 4.433431083 | -0.127791576 | -0.127385015 |
| Ramp1 | 7.891543581 | 7.937690537 | 8.018929193 | -0.081238655 | -0.127385611 |

| | | | | | |
|---|---|---|---|---|---|
| Cdca8 | 8.060688685 | 8.211496697 | 8.188113428 | 0.023383269 | -0.127424743 |
| Trim45 | 5.725281885 | 5.86052356 | 5.852748345 | 0.007775216 | -0.12746646 |
| Hoxa3 | 6.027358275 | 6.326614938 | 6.154865602 | 0.171749336 | -0.127507327 |
| Ska3 | 6.241228659 | 6.242168281 | 6.368737621 | -0.12656934 | -0.127508962 |
| C8g | 2.878745757 | 2.883861871 | 3.006272761 | -0.12241089 | -0.127527004 |
| Gm5460 | 4.011027092 | 3.942429224 | 4.138630174 | -0.19620095 | -0.127603082 |
| Fbxw5 | 9.587724146 | 9.772112188 | 9.715360501 | 0.056751687 | -0.127636355 |
| Nat14 | 4.789895634 | 4.952044322 | 4.917626912 | 0.03441741 | -0.127731278 |
| Cck | 4.992884112 | 5.152990278 | 5.120622261 | 0.032368016 | -0.12773815 |
| Ss18 | 10.86242004 | 10.91470206 | 10.99022747 | -0.07552541 | -0.127807433 |
| Mutyh | 5.586872719 | 5.733974042 | 5.714689262 | 0.019284781 | -0.127816543 |
| Gpx4-ps2 | 2.294746403 | 2.349763002 | 2.422620105 | -0.072857103 | -0.127873702 |
| Stard5 | 10.87282092 | 10.97439266 | 11.00071703 | -0.026324373 | -0.127896117 |
| Tsga10 | 6.653580914 | 6.637690973 | 6.781484184 | -0.143793212 | -0.127903271 |
| Acer3 | 10.14510723 | 10.21719512 | 10.27301977 | -0.05582465 | -0.127912537 |
| Pdcd2l | 7.941331708 | 7.998220542 | 8.069367378 | -0.071146836 | -0.12803567 |
| Selenom | 7.306133326 | 7.409508128 | 7.434306161 | -0.024798033 | -0.128172835 |
| Zbtb39 | 8.565938132 | 8.597120462 | 8.694151109 | -0.097030647 | -0.128212977 |
| Lrrc8b | 9.261728804 | 9.314500936 | 9.390063646 | -0.07556271 | -0.128334841 |
| Unc13b | 8.546112803 | 8.833961971 | 8.674667897 | 0.159294075 | -0.128555094 |
| Blzf1 | 9.739321245 | 9.797408101 | 9.867989616 | -0.070581514 | -0.128668371 |
| Prkd2 | 8.781716734 | 9.103603669 | 8.910430854 | 0.193172815 | -0.12871412 |
| Nrxn1 | 4.654350641 | 4.635022242 | 4.783075991 | -0.148053749 | -0.12872535 |
| Fam135b | 4.862808053 | 4.928572759 | 4.991536702 | -0.062963942 | -0.128728649 |
| Mtfr1 | 9.901145088 | 9.896803937 | 10.02993328 | -0.133129339 | -0.128788188 |
| Sucla2 | 11.1124262 | 11.06875373 | 11.24129653 | -0.1725428 | -0.12887033 |
| Fam174a | 8.479133307 | 8.366891209 | 8.608010366 | -0.241119156 | -0.128877058 |
| Sh3kbp1 | 10.11168666 | 10.15774684 | 10.24059609 | -0.082849247 | -0.128909423 |
| Gpr37 | 5.993629327 | 6.094180756 | 6.122656151 | -0.028475395 | -0.129026824 |
| Tbc1d9b | 10.11063796 | 10.25279674 | 10.23966494 | 0.013131793 | -0.12902698 |
| Acadvl | 11.13702403 | 11.15899623 | 11.26609786 | -0.107101623 | -0.129073827 |
| Vstm5 | 7.626998852 | 7.604199571 | 7.756149098 | -0.151949528 | -0.129150247 |

| | | | | | |
|---|---|---|---|---|---|
| Mapk14 | 11.15801869 | 11.19813861 | 11.28718517 | -0.089046557 | -0.129166477 |
| Desi2 | 8.707060014 | 8.689059507 | 8.836246917 | -0.14718741 | -0.129186903 |
| Ppcdc | 8.150926789 | 8.223423048 | 8.280178112 | -0.056755064 | -0.129251323 |
| Ndst4 | 3.619648502 | 3.675018792 | 3.748969308 | -0.073950516 | -0.129320806 |
| Gm2011 | 5.006962631 | 5.103689557 | 5.136283685 | -0.032594129 | -0.129321054 |
| Fzr1 | 9.072502905 | 9.379019352 | 9.201850625 | 0.177168727 | -0.12934772 |
| Rfwd3 | 10.04245848 | 10.09004894 | 10.17181941 | -0.081770477 | -0.129360929 |
| Ankrd13d | 6.812854938 | 6.817882294 | 6.942236958 | -0.124354664 | -0.12938202 |
| Ppfia4 | 7.89475031 | 7.819673209 | 8.024254573 | -0.204581364 | -0.129504263 |
| Setx | 10.13779795 | 10.12394934 | 10.26731663 | -0.143367293 | -0.129518683 |
| Clic3 | 10.21087397 | 10.32457864 | 10.34044034 | -0.015861707 | -0.12956637 |
| Capn5 | 8.792141171 | 8.839884014 | 8.921743535 | -0.081859521 | -0.129602364 |
| Fam221b | 3.613149497 | 3.687602442 | 3.742794905 | -0.055192463 | -0.129645408 |
| Stk36 | 4.012149456 | 4.154682061 | 4.141811483 | 0.012870578 | -0.129662027 |
| Shank1 | 5.847043698 | 6.196417838 | 5.976840259 | 0.219577579 | -0.129796561 |
| Tead2 | 6.489022313 | 6.702348167 | 6.618884122 | 0.083464045 | -0.12986181 |
| Lrba | 10.20994789 | 10.18953859 | 10.33981544 | -0.15027685 | -0.129867547 |
| Gm960 | 4.316053759 | 4.457119157 | 4.44593553 | 0.011183628 | -0.12988177 |
| Hoxa10 | 3.900734146 | 3.895343842 | 4.030640973 | -0.135297132 | -0.129906827 |
| Cidea | 10.2864302 | 10.28321029 | 10.41634242 | -0.13313213 | -0.129912223 |
| Zfp956 | 7.171494535 | 7.258691022 | 7.301411928 | -0.042720906 | -0.129917393 |
| Adi1 | 9.182304689 | 9.254044663 | 9.312249237 | -0.058204574 | -0.129944548 |
| Birc2 | 9.725962624 | 9.72941116 | 9.855961583 | -0.126550423 | -0.12999896 |
| Sprr4 | 8.007214417 | 8.138517815 | 8.137219533 | 0.001298282 | -0.130005116 |
| Spata1 | 5.391805604 | 5.838990642 | 5.521813515 | 0.317177126 | -0.130007911 |
| Wdr93 | 3.541562889 | 3.559795796 | 3.671606584 | -0.111810788 | -0.130043695 |
| Eva1b | 6.074741176 | 6.406876318 | 6.20491509 | 0.201961228 | -0.130173914 |
| Kif27 | 2.453243061 | 2.566507419 | 2.583447397 | -0.016939978 | -0.130204337 |
| S100a13 | 9.23797603 | 9.040260332 | 9.368231152 | -0.32797082 | -0.130255123 |
| Creb3l3 | 3.356201092 | 3.441830644 | 3.486459803 | -0.044629159 | -0.130258711 |
| Parva | 10.37705231 | 10.28392155 | 10.50734384 | -0.22342229 | -0.130291533 |
| Itch | 10.45162666 | 10.51574515 | 10.58202129 | -0.066276137 | -0.130394627 |

| | | | | | |
|---|---|---|---|---|---|
| Atp5c1 | 11.97078426 | 11.91848733 | 12.10119817 | -0.182710847 | -0.130413913 |
| Maml3 | 9.05665779 | 9.144049423 | 9.187148317 | -0.043098894 | -0.130490526 |
| Rnf224 | 3.69370558 | 3.772592082 | 3.824268605 | -0.051676524 | -0.130563025 |
| Zfp750 | 11.13656085 | 11.32285996 | 11.26714722 | 0.05571274 | -0.130586367 |
| Krt79 | 13.98941925 | 14.05879816 | 14.12001361 | -0.06121545 | -0.13059436 |
| Dnah3 | 2.193418427 | 2.24782662 | 2.324058111 | -0.076231491 | -0.130639683 |
| Zfp58 | 6.850208584 | 6.858869948 | 6.980860639 | -0.121990692 | -0.130652056 |
| Comp | 3.65383426 | 3.744154729 | 3.78450155 | -0.040346821 | -0.130667291 |
| Scp2-ps2 | 3.034289393 | 3.162514466 | 3.164970417 | -0.002455951 | -0.130681024 |
| Bckdha | 9.661085378 | 9.848383923 | 9.791768321 | 0.056615602 | -0.130682944 |
| Gm14046 | 2.334735124 | 2.481011606 | 2.465426066 | 0.01558554 | -0.130690943 |
| Rufy1 | 9.625445254 | 9.742532248 | 9.756171682 | -0.013639434 | -0.130726428 |
| Gabbr1 | 8.564579095 | 8.579389431 | 8.695338989 | -0.115949558 | -0.130759894 |
| Elp2 | 10.00647145 | 10.04838276 | 10.13724442 | -0.08886166 | -0.130772963 |
| Acp6 | 9.460621555 | 9.600495999 | 9.591412165 | 0.009083834 | -0.13079061 |
| Egfl8 | 5.74368221 | 5.842098448 | 5.874500185 | -0.032401737 | -0.130817975 |
| Dbf4 | 7.931907109 | 8.059055987 | 8.062741729 | -0.003685742 | -0.13083462 |
| Zfp114 | 3.206593941 | 3.30792914 | 3.337539845 | -0.029610705 | -0.130945905 |
| Trpc6 | 3.764953567 | 3.788307138 | 3.895970357 | -0.107663219 | -0.13101679 |
| Trpt1 | 6.596551439 | 6.711833098 | 6.727575189 | -0.015742091 | -0.13102375 |
| Cetn2 | 8.595716928 | 8.615972434 | 8.726748357 | -0.110775923 | -0.131031429 |
| Tsacc | 4.771182942 | 4.82345051 | 4.902228569 | -0.078778058 | -0.131045627 |
| Kcnmb4 | 4.716245058 | 4.750288997 | 4.847311035 | -0.097022038 | -0.131065977 |
| Cbx3 | 10.76723355 | 10.79305789 | 10.8983192 | -0.105261303 | -0.13108565 |
| Dnajc27 | 8.250068793 | 8.166593708 | 8.381217177 | -0.214623469 | -0.131148384 |
| Rpgr | 5.647710798 | 5.74377317 | 5.778874917 | -0.035101747 | -0.131164119 |
| Lmf1 | 7.622412193 | 7.741712044 | 7.75363175 | -0.011919706 | -0.131219557 |
| Adh6b | 6.764054995 | 6.761339801 | 6.895333706 | -0.133993905 | -0.131278711 |
| Brwd1 | 10.61121267 | 10.85025255 | 10.74255379 | 0.10769876 | -0.13134112 |
| Sbsn | 15.28682173 | 15.40790627 | 15.41820109 | -0.01029482 | -0.131379353 |
| Rad9b | 5.610282401 | 5.634499698 | 5.741684848 | -0.10718515 | -0.131402447 |
| Fah | 8.023444859 | 7.807329424 | 8.154861517 | -0.347532094 | -0.131416659 |

| | | | | | |
|---|---|---|---|---|---|
| Setd3 | 10.87382812 | 10.84725054 | 11.00528131 | -0.158030767 | -0.13145319 |
| Unc5a | 6.27648457 | 6.086995394 | 6.407949879 | -0.320954485 | -0.131465309 |
| Lnpk | 8.91843384 | 8.908942658 | 9.049941782 | -0.140999124 | -0.131507942 |
| Iqsec2 | 8.897017727 | 8.925103433 | 9.02858935 | -0.103485918 | -0.131571623 |
| Fam151b | 6.045576631 | 6.029315688 | 6.177200654 | -0.147884966 | -0.131624023 |
| Nexmif | 3.414855272 | 3.486572204 | 3.546493044 | -0.059920841 | -0.131637773 |
| Pithd1 | 9.107836621 | 9.212574333 | 9.239489507 | -0.026915174 | -0.131652885 |
| Stap2 | 9.765812987 | 9.90348703 | 9.897476079 | 0.006010951 | -0.131663092 |
| Gm27023 | 2.064381154 | 2.124985464 | 2.196118418 | -0.071132954 | -0.131737265 |
| Nsun3 | 7.968900403 | 8.018371782 | 8.100694885 | -0.082323103 | -0.131794483 |
| Dcdc2b | 3.841978682 | 4.073772063 | 3.973830241 | 0.099941822 | -0.131851559 |
| Ints9 | 8.936650654 | 9.084017558 | 9.068557132 | 0.015460426 | -0.131906477 |
| Shprh | 9.324652816 | 9.305909805 | 9.456629663 | -0.150719858 | -0.131976846 |
| Prr9 | 3.679106462 | 3.837303212 | 3.811112449 | 0.026190763 | -0.132005987 |
| Chp1 | 11.90399766 | 11.82599384 | 12.03601557 | -0.210021727 | -0.132017903 |
| Ccdc167 | 7.672019391 | 7.848373855 | 7.804071332 | 0.044302523 | -0.132051941 |
| Pheta2 | 5.903058913 | 5.969592512 | 6.035120923 | -0.065528411 | -0.13206201 |
| Carmil1 | 8.922435429 | 9.050651246 | 9.054594909 | -0.003943663 | -0.13215948 |
| Tbx4 | 3.180561855 | 3.216789825 | 3.312737161 | -0.095947337 | -0.132175306 |
| Trappc8 | 10.29367147 | 10.37495908 | 10.42599771 | -0.051038633 | -0.13232624 |
| Ccdc9b | 9.138340783 | 9.208387495 | 9.270674369 | -0.062286873 | -0.132333586 |
| Dusp4 | 6.709085541 | 6.838960815 | 6.841433243 | -0.002472429 | -0.132347703 |
| Tspyl1 | 9.878424843 | 9.86008526 | 10.01077525 | -0.150689986 | -0.132350403 |
| Zfp59 | 6.981717629 | 6.967737483 | 7.114092401 | -0.146354917 | -0.132374772 |
| Mum1 | 9.332850905 | 9.550687596 | 9.465244228 | 0.085443367 | -0.132393323 |
| Dcxr | 7.263071769 | 7.289692178 | 7.395515683 | -0.105823505 | -0.132443914 |
| Gm19065 | 2.778748831 | 2.805081358 | 2.911201057 | -0.106119699 | -0.132452226 |
| Chaf1a | 8.810524843 | 9.061588679 | 8.943015525 | 0.118573154 | -0.132490682 |
| Th | 4.786500389 | 4.91577056 | 4.91899277 | -0.00322221 | -0.132492381 |
| Cdh1 | 12.68037008 | 12.74840944 | 12.81290991 | -0.06450047 | -0.132539827 |
| Adamts13 | 8.401309424 | 8.186940246 | 8.533872461 | -0.346932215 | -0.132563037 |
| Ift43 | 7.832495974 | 7.786572347 | 7.965063504 | -0.178491157 | -0.13256753 |

| | | | | | |
|---|---|---|---|---|---|
| Uri1 | 9.590690922 | 9.435476689 | 9.723283548 | -0.287806859 | -0.132592626 |
| Scgb2b27 | 2.861996669 | 2.958602051 | 2.994590537 | -0.035988486 | -0.132593869 |
| Pan2 | 9.567249207 | 9.722217945 | 9.69985233 | 0.022365615 | -0.132603123 |
| Igf2 | 6.772299743 | 6.866652557 | 6.904933018 | -0.038280461 | -0.132633275 |
| Sgpp1 | 12.41702315 | 12.43863081 | 12.54984514 | -0.11121433 | -0.132821997 |
| Sipa1l1 | 10.04859395 | 10.08844233 | 10.18142081 | -0.092978483 | -0.132826868 |
| Eml5 | 7.047468395 | 6.701850585 | 7.180348719 | -0.478498133 | -0.132880324 |
| Gm37515 | 7.441226501 | 7.567208036 | 7.574149632 | -0.006941596 | -0.132923131 |
| Pclo | 4.188625969 | 4.248394784 | 4.321580775 | -0.073185991 | -0.132954806 |
| Cbr1 | 9.082440741 | 9.078660556 | 9.215395949 | -0.136735393 | -0.132955208 |
| Dnajb2 | 10.13038811 | 10.54202611 | 10.26335527 | 0.27867084 | -0.132967163 |
| Osbpl9 | 11.25360801 | 11.11731144 | 11.38665141 | -0.269339967 | -0.133043403 |
| Map10 | 5.278016762 | 5.236448573 | 5.411112825 | -0.174664252 | -0.133096063 |
| Rpl10 | 11.1066855 | 11.34337399 | 11.23979596 | 0.103578027 | -0.133110457 |
| Dpp8 | 10.92813777 | 11.02991765 | 11.06129731 | -0.03137966 | -0.13315954 |
| Gm38394 | 10.28009518 | 10.35339084 | 10.41325508 | -0.059864233 | -0.133159893 |
| Uba3 | 10.21817937 | 10.20891362 | 10.35135663 | -0.142443017 | -0.13317726 |
| Prdx2 | 11.46720594 | 11.42444004 | 11.60040034 | -0.175960293 | -0.1331944 |
| Gcc2 | 8.291398983 | 8.096609611 | 8.424649064 | -0.328039453 | -0.133250081 |
| Tox3 | 3.819697904 | 3.90833907 | 3.953014518 | -0.044675448 | -0.133316614 |
| Ppp3ca | 11.92674255 | 11.88999058 | 12.06016245 | -0.170171873 | -0.133419897 |
| Zfp72 | 5.778501541 | 5.85104619 | 5.911930441 | -0.060884251 | -0.1334289 |
| Sms | 9.007080166 | 9.095430866 | 9.140697933 | -0.045267068 | -0.133617767 |
| Pdk3 | 9.024524425 | 8.991454683 | 9.158163888 | -0.166709205 | -0.133639463 |
| Eid2b | 5.796270646 | 5.792226448 | 5.929925324 | -0.137698876 | -0.133654677 |
| Tmppe | 7.969612899 | 8.281206541 | 8.103277417 | 0.177929125 | -0.133664518 |
| Ulk4 | 4.195093792 | 4.47592517 | 4.328805118 | 0.147120052 | -0.133711326 |
| Gm6274 | 2.321709847 | 2.445071018 | 2.455439034 | -0.010368016 | -0.133729187 |
| Idh3b | 10.99948708 | 11.12512941 | 11.13325261 | -0.008123197 | -0.13376553 |
| Hoxd3 | 3.667975079 | 4.031619255 | 3.80174703 | 0.229872225 | -0.133771952 |
| Mtif3 | 8.204669269 | 8.104712769 | 8.338464401 | -0.233751632 | -0.133795132 |
| Gm49322 | 6.703165811 | 6.747133869 | 6.836970019 | -0.08983615 | -0.133804208 |

| | | | | | |
|---|---|---|---|---|---|
| Pcna-ps2 | 1.958413693 | 2.059453271 | 2.092261112 | -0.032807841 | -0.133847419 |
| Macrod2 | 6.371817107 | 6.495441094 | 6.505671485 | -0.010230391 | -0.133854378 |
| Myh11 | 10.85201185 | 10.97004923 | 10.98588566 | -0.015836433 | -0.133873813 |
| Arhgap35 | 10.56383273 | 10.72874956 | 10.69771978 | 0.03102978 | -0.133887057 |
| Ipo9 | 10.17763153 | 10.23018106 | 10.31152062 | -0.08133956 | -0.13388909 |
| Agpat3 | 11.3725008 | 11.75913366 | 11.50646964 | 0.252664017 | -0.133968847 |
| Rexo5 | 5.617737221 | 5.657663174 | 5.75184478 | -0.094181606 | -0.134107559 |
| Tmlhe | 7.678309636 | 7.60647655 | 7.812469775 | -0.205993225 | -0.134160139 |
| Popdc2 | 7.324808542 | 7.779158568 | 7.459145187 | 0.32001338 | -0.134336646 |
| Pvalb | 13.89437771 | 13.87708834 | 14.02881931 | -0.151730963 | -0.1344416 |
| Nr2f6 | 8.58600497 | 8.850192923 | 8.720565945 | 0.129626978 | -0.134560975 |
| Map3k2 | 9.804549193 | 9.960788425 | 9.939115897 | 0.021672528 | -0.134566704 |
| Gsn | 15.50019592 | 15.44302888 | 15.6347861 | -0.191757213 | -0.134590173 |
| Csnk1e | 10.2611193 | 10.52806229 | 10.39577324 | 0.132289047 | -0.13465394 |
| Paxbp1 | 9.86348516 | 9.929372541 | 9.998154311 | -0.06878177 | -0.134669152 |
| Pla2g4d | 3.119080645 | 3.117858305 | 3.253788814 | -0.13593051 | -0.134708169 |
| Fbxl12 | 7.877929461 | 7.917024941 | 8.012645958 | -0.095621017 | -0.134716498 |
| Synrg | 10.15426519 | 10.27938901 | 10.2890233 | -0.00963429 | -0.134758107 |
| Rbm41 | 6.57989368 | 6.645084833 | 6.714692396 | -0.069607563 | -0.134798716 |
| Cyp2u1 | 4.308744345 | 4.304907054 | 4.443569361 | -0.138662307 | -0.134825016 |
| Dhrs4 | 8.809136363 | 8.774179491 | 8.943983 | -0.169803509 | -0.134846637 |
| Fbxl21 | 2.743703859 | 2.784253055 | 2.878555646 | -0.094302591 | -0.134851787 |
| 3830406C13Rik | 9.073710306 | 9.174692486 | 9.208569506 | -0.03387702 | -0.1348592 |
| Gm15732 | 2.696614874 | 2.792137054 | 2.831487376 | -0.039350323 | -0.134872503 |
| Polr3c | 9.245811967 | 9.332820226 | 9.380689714 | -0.047869488 | -0.134877748 |
| Entpd6 | 8.372996757 | 8.2597211 | 8.507883028 | -0.248161928 | -0.134886271 |
| Ankrd31 | 3.098979767 | 3.227510884 | 3.233897448 | -0.006386564 | -0.134917681 |
| Prune1 | 9.04296361 | 9.018240637 | 9.177948321 | -0.159707684 | -0.134984711 |
| Hapln1 | 3.707149901 | 3.754723236 | 3.842190635 | -0.087467398 | -0.135040734 |
| Zfp275 | 8.246250209 | 8.154180123 | 8.381323051 | -0.227142928 | -0.135072841 |
| Trim41 | 9.513681099 | 9.823071865 | 9.648843228 | 0.174228637 | -0.135162129 |
| B3gntl1 | 7.143592377 | 7.098763261 | 7.278764009 | -0.180000748 | -0.135171633 |

| | | | | | |
|---|---|---|---|---|---|
| Cog5 | 9.427833081 | 9.458847498 | 9.563037483 | -0.104189985 | -0.135204402 |
| Gabrq | 2.108254294 | 2.123973131 | 2.243474203 | -0.119501072 | -0.135219909 |
| Slc9a7 | 5.18478982 | 5.259687062 | 5.320011474 | -0.060324412 | -0.135221654 |
| Unc5b | 9.390168241 | 9.621602519 | 9.525404912 | 0.096197607 | -0.135236671 |
| Cdk1 | 8.606314012 | 8.627919547 | 8.741584444 | -0.113664898 | -0.135270432 |
| Slc25a47 | 5.235842072 | 5.472288348 | 5.371144143 | 0.101144206 | -0.135302071 |
| Fam20b | 11.03712106 | 11.02127734 | 11.17243605 | -0.151158713 | -0.135314993 |
| AI987944 | 7.478937033 | 7.514200889 | 7.614378618 | -0.100177729 | -0.135441585 |
| Jade3 | 7.645124536 | 7.910710048 | 7.780598414 | 0.130111634 | -0.135473878 |
| Ceacam19 | 10.16152829 | 10.26638874 | 10.29703379 | -0.030645053 | -0.1355055 |
| Lmtk2 | 9.804463004 | 9.96356349 | 9.939976008 | 0.023587482 | -0.135513004 |
| Serinc5 | 10.07385132 | 10.19670477 | 10.20938085 | -0.012676077 | -0.135529524 |
| Mis18a | 7.453648473 | 7.541421632 | 7.589184537 | -0.047762905 | -0.135536064 |
| Dctn2 | 11.42090456 | 11.50685704 | 11.5565581 | -0.04970106 | -0.135653537 |
| Slfn3 | 5.86465504 | 5.875639875 | 6.000314309 | -0.124674434 | -0.135659269 |
| Oser1 | 10.06867885 | 10.23529038 | 10.20436411 | 0.030926263 | -0.135685266 |
| Abca4 | 5.221829203 | 5.425355878 | 5.357526434 | 0.067829444 | -0.135697231 |
| Dip2b | 10.52894251 | 10.81606519 | 10.66469336 | 0.151371833 | -0.135750853 |
| Mroh3 | 3.523665922 | 3.751229892 | 3.659596785 | 0.091633106 | -0.135930864 |
| Xndc1 | 8.316076822 | 8.495379081 | 8.452050729 | 0.043328352 | -0.135973907 |
| Gm4724 | 6.547538083 | 6.478159193 | 6.683585191 | -0.205425998 | -0.136047109 |
| Suds3 | 10.54639072 | 10.63706803 | 10.68248685 | -0.045418817 | -0.13609613 |
| Wdr76 | 7.761840676 | 7.81171375 | 7.897967939 | -0.086254189 | -0.136127263 |
| Nmb | 4.472120841 | 4.437958103 | 4.608310104 | -0.170352002 | -0.136189263 |
| AI597479 | 8.819986177 | 8.825348537 | 8.956217097 | -0.13086856 | -0.136230921 |
| Serpinb1c | 7.431125895 | 7.535191124 | 7.567357472 | -0.032166348 | -0.136231577 |
| Hs3st5 | 2.859027869 | 2.873433889 | 2.99529909 | -0.121865202 | -0.136271221 |
| Prim1 | 8.716987788 | 8.915480729 | 8.853314481 | 0.062166249 | -0.136326693 |
| Rpl9-ps6 | 2.562031207 | 2.652756125 | 2.698408418 | -0.045652293 | -0.136377212 |
| Gm572 | 3.752770805 | 3.739665757 | 3.889187871 | -0.149522114 | -0.136417066 |
| Trim37 | 9.081721613 | 9.118136692 | 9.218266527 | -0.100129836 | -0.136544914 |
| Ninl | 6.447995038 | 6.379558318 | 6.584541764 | -0.204983446 | -0.136546726 |

| | | | | | |
|---|---|---|---|---|---|
| Herc2 | 10.8140675 | 10.83186127 | 10.95067825 | -0.118816977 | -0.13661075 |
| Tm7sf3 | 9.442168012 | 9.473206585 | 9.578782711 | -0.105576126 | -0.136614699 |
| Fbxl8 | 5.625798187 | 5.864368646 | 5.76243243 | 0.101936216 | -0.136634243 |
| Ttll5 | 7.578987616 | 7.664545638 | 7.715635292 | -0.051089654 | -0.136647676 |
| Nkain2 | 3.498451933 | 3.462432155 | 3.635177482 | -0.172745327 | -0.136725549 |
| Ift20 | 9.851268912 | 9.821095816 | 9.98799924 | -0.166903424 | -0.136730328 |
| Gm13508 | 2.601147601 | 2.776565118 | 2.737913767 | 0.038651351 | -0.136766165 |
| 4930451G09Rik | 3.280545321 | 3.319613356 | 3.417340052 | -0.097726696 | -0.136794731 |
| Serpinb6c | 6.982233011 | 6.874405492 | 7.119199396 | -0.244793904 | -0.136966385 |
| Zfp90 | 7.202329814 | 7.153700894 | 7.339298055 | -0.185597162 | -0.136968242 |
| Bloc1s5 | 7.661227745 | 7.792612441 | 7.798196457 | -0.005584016 | -0.136968712 |
| Gm14892 | 3.190299552 | 3.307855289 | 3.327338902 | -0.019483613 | -0.13703935 |
| Fam89a | 9.256373811 | 9.340828835 | 9.393549398 | -0.052720563 | -0.137175587 |
| Gas2l3 | 7.959052638 | 8.056240725 | 8.096243688 | -0.040002963 | -0.13719105 |
| Map7d2 | 2.589125172 | 2.666461676 | 2.726327694 | -0.059866018 | -0.137202522 |
| Tfam | 9.225152168 | 9.153737859 | 9.362360673 | -0.208622814 | -0.137208505 |
| Pip4p2 | 8.944649868 | 8.9581375 | 9.081870834 | -0.123733334 | -0.137220966 |
| Usp47 | 11.37444837 | 11.42579188 | 11.51172703 | -0.085935153 | -0.137278657 |
| Trib3 | 6.478220881 | 6.677433691 | 6.615515562 | 0.061918129 | -0.137294681 |
| Micu2 | 10.82136811 | 10.75659839 | 10.95872836 | -0.202129967 | -0.13736025 |
| Gm28551 | 2.117713009 | 2.146053558 | 2.255163709 | -0.109110151 | -0.1374507 |
| Pappa | 7.068606141 | 7.126587622 | 7.206171349 | -0.079583726 | -0.137565208 |
| Fam129c | 4.947955845 | 5.147081737 | 5.085632703 | 0.061449034 | -0.137676858 |
| Sel1l3 | 7.885189918 | 7.781547763 | 8.022931424 | -0.24138366 | -0.137741506 |
| Fryl | 10.74857099 | 10.85732383 | 10.88631647 | -0.028992637 | -0.137745473 |
| Ddx17 | 12.55388011 | 12.76844071 | 12.69186351 | 0.076577207 | -0.137983397 |
| Dhx32 | 9.783514987 | 9.710501723 | 9.921565505 | -0.211063781 | -0.138050517 |
| Cenpo | 7.072447629 | 7.284394466 | 7.210508868 | 0.073885598 | -0.138061238 |
| Agrn | 11.36428128 | 11.53538207 | 11.50235293 | 0.033029143 | -0.13807165 |
| Ift122 | 7.179663274 | 7.234392316 | 7.317754416 | -0.0833621 | -0.138091141 |
| Gm2115 | 3.795746108 | 3.860879249 | 3.933845661 | -0.072966412 | -0.138099553 |
| Cep350 | 10.56727779 | 10.64921648 | 10.70543394 | -0.05621746 | -0.138156157 |

| | | | | | |
|---|---|---|---|---|---|
| Doc2g | 5.135859335 | 5.345143016 | 5.274084396 | 0.07105862 | -0.138225061 |
| Tmem18 | 8.893302163 | 8.906599082 | 9.031534443 | -0.124935361 | -0.13823228 |
| Zfp637 | 8.075543231 | 8.167241166 | 8.21380795 | -0.046566784 | -0.138264719 |
| Pick1 | 8.631331536 | 8.82951622 | 8.769637809 | 0.059878411 | -0.138306274 |
| Zfp768 | 7.555926647 | 7.756755486 | 7.694263659 | 0.062491827 | -0.138337012 |
| Anapc1 | 10.92433755 | 11.04441417 | 11.06268816 | -0.01827399 | -0.138350607 |
| Commd8 | 9.541229596 | 9.407318938 | 9.679677375 | -0.272358437 | -0.138447779 |
| Gm42857 | 5.835122754 | 6.061130524 | 5.973614133 | 0.087516391 | -0.138491379 |
| Klhl28 | 8.535473759 | 8.520820145 | 8.674003808 | -0.153183663 | -0.138530049 |
| D230025D16Rik | 8.96102308 | 9.02077765 | 9.099583801 | -0.078806151 | -0.138560722 |
| Pbld1 | 3.834083196 | 3.785315223 | 3.972783755 | -0.187468532 | -0.138700558 |
| Nt5c | 8.724009135 | 8.696633521 | 8.862805268 | -0.166171747 | -0.138796133 |
| Btd | 8.510893657 | 8.509637221 | 8.64980005 | -0.140162829 | -0.138906393 |
| Ccs | 8.634403806 | 8.670761152 | 8.773366673 | -0.102605522 | -0.138962868 |
| Ccdc92b | 3.0551646 | 3.109558608 | 3.194170403 | -0.084611795 | -0.139005803 |
| Elmod3 | 8.107883847 | 8.303821723 | 8.246905153 | 0.05691657 | -0.139021306 |
| Atp6v1b1 | 3.741408543 | 3.885653751 | 3.880480479 | 0.005173272 | -0.139071936 |
| Gpr137c | 4.299187916 | 4.438219573 | 4.438294514 | -7.4941E-05 | -0.139106598 |
| Zwilch | 7.345946975 | 7.280868705 | 7.485071597 | -0.204202892 | -0.139124622 |
| Mucl1 | 4.602835396 | 4.604840287 | 4.742031903 | -0.137191615 | -0.139196506 |
| Kmt5b | 10.14474843 | 10.25812997 | 10.28397284 | -0.025842867 | -0.139224406 |
| Phf13 | 8.657969873 | 8.742005603 | 8.79727353 | -0.055267927 | -0.139303656 |
| Ror2 | 5.898172498 | 6.044678545 | 6.037483145 | 0.0071954 | -0.139310647 |
| Gm527 | 5.687001732 | 5.820952977 | 5.826382877 | -0.0054299 | -0.139381145 |
| Cdan1 | 8.063983527 | 8.245679463 | 8.203411215 | 0.042268248 | -0.139427688 |
| Armh3 | 8.771535718 | 8.776481031 | 8.911039475 | -0.134558444 | -0.139503757 |
| Fmn1 | 8.970406926 | 9.008059795 | 9.109925057 | -0.101865262 | -0.139518131 |
| Cyp17a1 | 9.177841245 | 9.111283837 | 9.31746563 | -0.206181793 | -0.139624385 |
| Slc28a3 | 7.818599202 | 7.823267278 | 7.958226778 | -0.1349595 | -0.139627576 |
| Hyal1 | 9.106444749 | 9.444620842 | 9.246074129 | 0.198546713 | -0.13962938 |
| Letmd1 | 8.725548222 | 8.91954035 | 8.865216069 | 0.054324281 | -0.139667847 |
| Cfap36 | 7.643188682 | 7.595167236 | 7.782879252 | -0.187712017 | -0.139690571 |

| | | | | | |
|---|---|---|---|---|---|
| Mgrn1 | 9.755549954 | 9.927830223 | 9.895308976 | 0.032521247 | -0.139759022 |
| Gm38393 | 3.190504182 | 3.23067923 | 3.330299706 | -0.099620476 | -0.139795525 |
| Dym | 9.016506175 | 9.065946872 | 9.156371663 | -0.090424791 | -0.139865488 |
| Aadacl2 | 8.445724074 | 8.406904419 | 8.585612062 | -0.178707643 | -0.139887988 |
| Pigz | 3.205974608 | 3.188494137 | 3.345864674 | -0.157370537 | -0.139890066 |
| Pex5l | 4.784665727 | 4.858488636 | 4.924602986 | -0.06611435 | -0.139937259 |
| Arl5a | 11.2353917 | 11.1761106 | 11.3753607 | -0.199250093 | -0.139968997 |
| Erg28 | 9.135953467 | 9.183526573 | 9.275928829 | -0.092402256 | -0.139975362 |
| Psme4 | 11.68085647 | 11.73251349 | 11.82085646 | -0.088342977 | -0.13999999 |
| Lgals1 | 12.13460194 | 12.09718144 | 12.27464482 | -0.177463377 | -0.14004288 |
| Pja1 | 9.559759527 | 9.676317548 | 9.699802597 | -0.023485049 | -0.14004307 |
| Arhgef4 | 9.573631539 | 9.90967457 | 9.713751654 | 0.195922916 | -0.140120115 |
| Syngr1 | 7.842859268 | 7.901804546 | 7.982998719 | -0.081194173 | -0.140139451 |
| Chd4 | 12.27444879 | 12.38048844 | 12.41459784 | -0.034109407 | -0.140149057 |
| Dpy19l3 | 7.579256108 | 7.883385899 | 7.719443035 | 0.163942864 | -0.140186927 |
| Entpd4 | 9.972115349 | 9.949730784 | 10.11235065 | -0.162619869 | -0.140235305 |
| Rad51d | 8.118341219 | 8.28235416 | 8.258606076 | 0.023748083 | -0.140264858 |
| Gm14676 | 2.382665707 | 2.491984317 | 2.523125425 | -0.031141108 | -0.140459717 |
| Gm6189 | 4.584562095 | 4.677025662 | 4.725057742 | -0.04803208 | -0.140495646 |
| Far2 | 13.17457805 | 13.19364592 | 13.3151278 | -0.12148188 | -0.140549747 |
| Pitpnm3 | 6.488157636 | 6.668032822 | 6.62878407 | 0.039248753 | -0.140626433 |
| Fahd1 | 7.531881797 | 7.607218788 | 7.672642804 | -0.065424016 | -0.140761007 |
| Vmn2r57 | 2.70125816 | 2.72877204 | 2.842060068 | -0.113288028 | -0.140801908 |
| BC017158 | 8.276324063 | 8.429597537 | 8.417150164 | 0.012447373 | -0.140826101 |
| Smim11 | 8.276658147 | 8.32253959 | 8.417502689 | -0.094963099 | -0.140844542 |
| Ggnbp1 | 7.172293391 | 7.298922783 | 7.313145049 | -0.014222265 | -0.140851657 |
| Lss | 10.56772187 | 10.52446469 | 10.70873236 | -0.18426767 | -0.14101049 |
| Hoxd4 | 6.143006148 | 6.278574126 | 6.284033187 | -0.005459061 | -0.141027039 |
| Fbxo44 | 5.503691351 | 5.670929746 | 5.644733367 | 0.026196378 | -0.141042016 |
| Trpv4 | 7.864595838 | 7.857326791 | 8.005700994 | -0.148374203 | -0.141105156 |
| Cenpk | 6.254441866 | 6.534115755 | 6.395684529 | 0.138431226 | -0.141242662 |
| 2310046K23Rik | 12.21622934 | 12.53055729 | 12.35758124 | 0.17297605 | -0.141351903 |

| | | | | | |
|---|---|---|---|---|---|
| Adamts2 | 10.4407731 | 10.65104099 | 10.58233087 | 0.06871012 | -0.14155777 |
| Phf14 | 9.827769149 | 9.76590518 | 9.969341201 | -0.203436022 | -0.141572052 |
| Cnga3 | 4.533155124 | 5.590456291 | 4.674747941 | 0.91570835 | -0.141592817 |
| Nek4 | 7.922248094 | 8.002756611 | 8.063851868 | -0.061095257 | -0.141603775 |
| Degs2 | 7.299919189 | 7.360363151 | 7.441557294 | -0.081194143 | -0.141638105 |
| Icam5 | 4.053864114 | 4.325040819 | 4.195531071 | 0.129509749 | -0.141666957 |
| Zranb3 | 6.589789244 | 6.570129492 | 6.731540565 | -0.161411072 | -0.141751321 |
| Gm9644 | 2.790590915 | 2.883046994 | 2.932451409 | -0.049404414 | -0.141860494 |
| Stoml1 | 7.781164356 | 7.860311693 | 7.923036748 | -0.062725055 | -0.141872392 |
| Vipr2 | 2.877094503 | 2.829345105 | 3.019025614 | -0.189680509 | -0.14193111 |
| Rpl31-ps8 | 11.89582182 | 11.88397646 | 12.03780068 | -0.153824227 | -0.141978867 |
| Gle1 | 9.487327676 | 9.674843559 | 9.629337136 | 0.045506422 | -0.14200946 |
| Klk11 | 9.764258619 | 10.02408145 | 9.906285586 | 0.117795862 | -0.142026967 |
| Slc38a6 | 8.826865849 | 8.828691933 | 8.969016155 | -0.140324222 | -0.142150306 |
| Elfn2 | 3.711355312 | 3.824307262 | 3.853604102 | -0.02929684 | -0.14224879 |
| Tmem131 | 10.88893786 | 11.13690078 | 11.0312854 | 0.105615387 | -0.142347533 |
| Ccdc15 | 4.980855211 | 5.047745614 | 5.123208757 | -0.075463143 | -0.142353546 |
| Gkn3 | 2.421699857 | 2.453441471 | 2.564123151 | -0.11068168 | -0.142423294 |
| Gphn | 9.547963494 | 9.479106206 | 9.69040951 | -0.211303304 | -0.142446016 |
| Lrch2 | 4.674252251 | 4.752070983 | 4.816702945 | -0.064631962 | -0.142450694 |
| Ttc5 | 8.80909873 | 8.826601894 | 8.951612942 | -0.125011048 | -0.142514213 |
| Ptpn5 | 4.454262572 | 4.546378035 | 4.596824122 | -0.050446087 | -0.14256155 |
| Ndfip2 | 10.26329608 | 10.3935459 | 10.40586506 | -0.012319157 | -0.14256898 |
| Spaca6 | 8.75790729 | 9.096745867 | 8.900527659 | 0.196218207 | -0.142620369 |
| Gm47685 | 3.50276467 | 3.654210518 | 3.645481305 | 0.008729212 | -0.142716635 |
| Apoe | 14.43406025 | 14.64192413 | 14.57678989 | 0.065134237 | -0.142729647 |
| Dcbld2 | 8.41466774 | 8.40092336 | 8.557446605 | -0.156523245 | -0.142778865 |
| Myb | 6.030881018 | 6.097701676 | 6.173683955 | -0.07598228 | -0.142802937 |
| Ctnnd1 | 12.34267325 | 12.45280074 | 12.48552648 | -0.03272574 | -0.14285323 |
| Cirbp | 9.538891402 | 9.550882155 | 9.681802851 | -0.130920697 | -0.142911449 |
| Nyap1 | 5.63597954 | 5.957138601 | 5.778915838 | 0.178222763 | -0.142936298 |
| Acot6 | 7.069767473 | 7.151790607 | 7.212703941 | -0.060913334 | -0.142936468 |

| | | | | | |
|---|---|---|---|---|---|
| Pyurf | 7.995087867 | 8.078364545 | 8.138072907 | -0.059708362 | -0.142985039 |
| Ahcyl2 | 10.61807036 | 10.84642391 | 10.76109254 | 0.085331373 | -0.14302218 |
| Fto | 10.47478882 | 10.47144852 | 10.61782492 | -0.146376403 | -0.143036097 |
| Lacc1 | 8.63800871 | 8.65056226 | 8.781141559 | -0.130579299 | -0.143132849 |
| Rtkn | 9.765769452 | 10.04761888 | 9.908944603 | 0.138674272 | -0.143175151 |
| Arhgap32 | 10.21814042 | 10.46890042 | 10.36133152 | 0.107568903 | -0.143191097 |
| Lrfn3 | 5.246005945 | 5.327926955 | 5.389312696 | -0.061385741 | -0.143306752 |
| Atrn | 10.10193938 | 10.10713655 | 10.24526316 | -0.138126606 | -0.14332378 |
| Acp5 | 9.682955031 | 9.675460691 | 9.826315052 | -0.150854361 | -0.143360021 |
| Ntf3 | 6.484846497 | 6.522956077 | 6.628254687 | -0.10529861 | -0.14340819 |
| Asxl2 | 9.703413051 | 9.796638352 | 9.846843637 | -0.050205286 | -0.143430587 |
| Zfp580 | 4.409342126 | 4.50137625 | 4.552844558 | -0.051468308 | -0.143502432 |
| Fut8 | 9.280733784 | 9.255839013 | 9.424271336 | -0.168432322 | -0.143537552 |
| Zfp334 | 6.141990331 | 6.286113994 | 6.285550312 | 0.000563682 | -0.143559981 |
| Apip | 7.493216999 | 7.430480593 | 7.636835447 | -0.206354854 | -0.143618447 |
| E2f1 | 8.256634355 | 8.49490847 | 8.400280967 | 0.094627502 | -0.143646612 |
| Zfp512 | 8.669570624 | 8.61772729 | 8.813220429 | -0.195493139 | -0.143649805 |
| Gm10643 | 4.669552159 | 4.656196067 | 4.81322741 | -0.157031343 | -0.143675251 |
| Zfp873 | 5.903302364 | 6.086758683 | 6.047003477 | 0.039755205 | -0.143701113 |
| Gm45716 | 6.39855203 | 6.199598347 | 6.542340653 | -0.342742306 | -0.143788623 |
| Pla2g4f | 10.50017527 | 10.62343469 | 10.64400454 | -0.020569847 | -0.143829263 |
| Asph | 10.7625706 | 10.47888804 | 10.90642093 | -0.42753289 | -0.143850337 |
| Emid1 | 5.890421312 | 5.980673405 | 6.034318323 | -0.053644918 | -0.143897012 |
| Rab33b | 8.303169431 | 8.233399292 | 8.447107657 | -0.213708365 | -0.143938226 |
| Lca5l | 3.56013865 | 3.635088845 | 3.704095118 | -0.069006273 | -0.143956468 |
| Adamts6 | 5.403811821 | 5.387283646 | 5.547794023 | -0.160510377 | -0.143982201 |
| Vps45 | 8.224990026 | 8.08239655 | 8.369047297 | -0.286650746 | -0.144057271 |
| Lce3a | 5.825149575 | 6.042879524 | 5.969272424 | 0.0736071 | -0.144122849 |
| Eci1 | 8.72505201 | 8.679423383 | 8.869186314 | -0.189762931 | -0.144134304 |
| Ube2e3 | 9.56949438 | 9.549183014 | 9.713647089 | -0.164464075 | -0.144152709 |
| Spink14 | 2.389344891 | 2.431196712 | 2.533586063 | -0.102389351 | -0.144241173 |
| Wfikkn2 | 4.745775539 | 4.841840083 | 4.890043618 | -0.048203535 | -0.144268078 |

| | | | | | |
|---|---|---|---|---|---|
| Tnfsfm13 | 6.4546145 | 6.532779937 | 6.598894581 | -0.066114644 | -0.144280081 |
| Tmem45b | 9.593113354 | 9.544279192 | 9.737449276 | -0.193170084 | -0.144335922 |
| Atp1b4 | 3.081954448 | 3.312703723 | 3.22631772 | 0.086386003 | -0.144363272 |
| Myocd | 4.251008123 | 4.276879572 | 4.395436519 | -0.118556947 | -0.144428396 |
| Spata7 | 5.703292069 | 5.767471731 | 5.847782539 | -0.080310808 | -0.14449047 |
| Fam193b | 9.632117503 | 9.798091247 | 9.776702748 | 0.021388499 | -0.144585245 |
| Sox8 | 3.333241258 | 3.397426016 | 3.477854357 | -0.080428341 | -0.144613099 |
| Dynlt1c | 6.05343765 | 6.192596336 | 6.198084935 | -0.005488599 | -0.144647285 |
| Adamts7 | 5.774324535 | 6.130741804 | 5.919127121 | 0.211614682 | -0.144802586 |
| Araf | 10.46055253 | 10.63046051 | 10.60538644 | 0.025074067 | -0.144833913 |
| Spg7 | 9.55032494 | 9.566703282 | 9.695161104 | -0.128457822 | -0.144836164 |
| Etnk2 | 2.142056248 | 2.187638212 | 2.286897799 | -0.099259587 | -0.14484155 |
| Man2b2 | 9.08656209 | 9.416245324 | 9.231409892 | 0.184835433 | -0.144847802 |
| Mvd | 9.528606671 | 9.836270411 | 9.673520346 | 0.162750065 | -0.144913675 |
| Mst1 | 7.428563177 | 7.751174889 | 7.573501855 | 0.177673034 | -0.144938677 |
| Atxn10 | 10.54547666 | 10.49091913 | 10.69048098 | -0.19956185 | -0.145004317 |
| Gm9745 | 2.051597366 | 2.084050232 | 2.196713057 | -0.112662825 | -0.145115691 |
| Capn1 | 11.94631132 | 12.13766877 | 12.09151376 | 0.04615501 | -0.145202443 |
| Tcrg-V5 | 4.651606497 | 4.767100626 | 4.796984635 | -0.029884009 | -0.145378138 |
| Cep78 | 7.296604104 | 7.366940907 | 7.441991427 | -0.07505052 | -0.145387323 |
| Otogl | 4.20490931 | 4.20816767 | 4.350304681 | -0.142137011 | -0.145395371 |
| Sc5d | 10.35866664 | 10.23590015 | 10.50412912 | -0.268228973 | -0.14546248 |
| Lypd1 | 3.08809159 | 3.135798062 | 3.233622289 | -0.097824227 | -0.145530699 |
| Rhoq | 10.3725768 | 10.22489477 | 10.51811864 | -0.293223867 | -0.145541843 |
| Ptpn4 | 9.420076432 | 9.430313096 | 9.565619411 | -0.135306315 | -0.145542978 |
| Atad2 | 8.276785624 | 8.46411818 | 8.422344733 | 0.041773448 | -0.145559108 |
| Rras | 9.599511488 | 9.580025518 | 9.745177607 | -0.165152089 | -0.145666118 |
| Lcorl | 7.886277871 | 8.010971269 | 8.031994946 | -0.021023677 | -0.145717075 |
| Ggcx | 7.261524333 | 7.202912153 | 7.407281209 | -0.204369056 | -0.145756875 |
| Hist1h1e | 4.57427896 | 4.769126422 | 4.7200789 | 0.049047523 | -0.14579994 |
| Slc35f1 | 5.173167027 | 5.249134379 | 5.319348374 | -0.070213995 | -0.146181347 |
| Mup7 | 8.731117958 | 8.856779384 | 8.877422633 | -0.020643248 | -0.146304675 |

| | | | | | |
|---|---|---|---|---|---|
| Tnks | 9.870294378 | 9.929692919 | 10.01668947 | -0.086996554 | -0.146395095 |
| Ppp4r1l-ps | 8.525861477 | 8.483841548 | 8.672408788 | -0.18856724 | -0.146547311 |
| Zfp51 | 7.661566151 | 7.699959375 | 7.808151336 | -0.108191961 | -0.146585185 |
| Polq | 6.131361502 | 6.153778611 | 6.278023519 | -0.124244908 | -0.146662017 |
| Tmem26 | 6.260171342 | 6.388461 | 6.406891599 | -0.018430599 | -0.146720256 |
| C1qtnf5 | 6.183084175 | 6.275670575 | 6.329934604 | -0.054264029 | -0.146850429 |
| Smc2 | 9.319230294 | 9.405677656 | 9.46622774 | -0.060550084 | -0.146997446 |
| Mast3 | 8.139425437 | 8.375186494 | 8.286435791 | 0.088750703 | -0.147010354 |
| Lrrc38 | 4.16657797 | 4.143153113 | 4.313603366 | -0.170450253 | -0.147025396 |
| Hsd3b6 | 9.022194344 | 9.026988642 | 9.169240858 | -0.142252216 | -0.147046513 |
| Dmtn | 6.597912777 | 6.416383168 | 6.744968706 | -0.328585538 | -0.14705593 |
| Mgmt | 5.645434437 | 5.708605633 | 5.792745224 | -0.084139592 | -0.147310788 |
| Dhrs7 | 11.0019482 | 10.88031371 | 11.14936653 | -0.269052823 | -0.147418327 |
| Gm14686 | 3.454709998 | 3.483444642 | 3.602178519 | -0.118733877 | -0.147468521 |
| Cox7a1 | 8.42417224 | 8.326459428 | 8.57164168 | -0.245182252 | -0.14746944 |
| Bet1 | 9.084212793 | 9.239341544 | 9.231690567 | 0.007650976 | -0.147477775 |
| Mylk | 11.02022005 | 11.03395946 | 11.16771855 | -0.13375909 | -0.147498503 |
| Gm3055 | 5.584240973 | 5.743057541 | 5.731791429 | 0.011266112 | -0.147550456 |
| Ercc5 | 9.303674484 | 9.359037133 | 9.451232541 | -0.092195408 | -0.147558057 |
| Padi1 | 2.803941659 | 2.834979589 | 2.951522891 | -0.116543302 | -0.147581232 |
| Dusp15 | 4.486660573 | 4.663925094 | 4.634255299 | 0.029669795 | -0.147594726 |
| Vezf1 | 9.105489367 | 9.089531893 | 9.253120747 | -0.163588854 | -0.14763138 |
| Med4 | 9.392250441 | 9.486690146 | 9.539917075 | -0.053226929 | -0.147666635 |
| Cyp51 | 12.02508297 | 12.04985472 | 12.1727636 | -0.122908873 | -0.147680623 |
| Gins4 | 8.212290875 | 8.265984193 | 8.359987379 | -0.094003186 | -0.147696504 |
| Gzmc | 6.160695698 | 6.329737199 | 6.308430549 | 0.02130665 | -0.147734851 |
| Slc26a7 | 8.563950002 | 7.683313609 | 8.71179718 | -1.028483572 | -0.147847178 |
| Slc24a2 | 4.743291348 | 4.72794766 | 4.891145547 | -0.163197887 | -0.147854199 |
| Arid1b | 10.14651687 | 10.46372796 | 10.29454877 | 0.169179193 | -0.148031897 |
| Gm11814 | 2.436888866 | 2.521009367 | 2.584933139 | -0.063923772 | -0.148044273 |
| Hyi | 7.719865426 | 7.863736706 | 7.86792435 | -0.004187644 | -0.148058924 |
| Cramp1l | 9.43008295 | 9.648952373 | 9.578165396 | 0.070786977 | -0.148082446 |

| | | | | | |
|---|---|---|---|---|---|
| 39873 | 7.750813872 | 7.965105429 | 7.898979369 | 0.06612606 | -0.148165497 |
| Gm48498 | 6.767210235 | 6.919888371 | 6.915467091 | 0.004421281 | -0.148256855 |
| Eif4b | 12.77647425 | 12.8025737 | 12.92475606 | -0.12218236 | -0.148281813 |
| Mdga1 | 6.921460261 | 6.787805651 | 7.069776389 | -0.281970738 | -0.148316128 |
| Zfp579 | 6.98413853 | 7.100583597 | 7.132526672 | -0.031943076 | -0.148388142 |
| Atp6v1g2 | 4.680221593 | 4.737170549 | 4.828625704 | -0.091455154 | -0.148404111 |
| Cnih2 | 4.106223366 | 4.187607765 | 4.254644115 | -0.06703635 | -0.148420749 |
| Pitpnc1 | 9.110675586 | 8.833404487 | 9.259098372 | -0.425693884 | -0.148422786 |
| Zhx1 | 9.084632425 | 9.174130477 | 9.233084914 | -0.058954437 | -0.148452489 |
| Rpa1 | 10.17442025 | 10.33429647 | 10.3230685 | 0.011227973 | -0.148648247 |
| Prss8 | 8.980174756 | 9.17649364 | 9.128826987 | 0.047666652 | -0.148652231 |
| Tmem265 | 7.724309001 | 7.825407851 | 7.872972986 | -0.047565134 | -0.148663985 |
| C330018D20Rik | 7.490931024 | 7.471366713 | 7.639665073 | -0.168298359 | -0.148734049 |
| Pbrm1 | 11.15507775 | 11.19938148 | 11.30382126 | -0.10443978 | -0.14874351 |
| Mab21l3 | 6.947734042 | 7.038516091 | 7.096482053 | -0.057965962 | -0.148748011 |
| Lrrn2 | 4.109297664 | 4.284930095 | 4.258078931 | 0.026851164 | -0.148781267 |
| Helb | 8.398285741 | 8.504354909 | 8.547083109 | -0.0427282 | -0.148797368 |
| Tmem205 | 7.987311825 | 8.114838273 | 8.136118713 | -0.021280439 | -0.148806888 |
| Erbb3 | 11.28094616 | 11.4331914 | 11.42984415 | 0.00334725 | -0.148897987 |
| Manba | 8.577359846 | 8.658846524 | 8.726289323 | -0.067442799 | -0.148929477 |
| Oxsm | 8.141039157 | 8.128906886 | 8.289979713 | -0.161072827 | -0.148940556 |
| Pot1b | 6.902950943 | 7.024931938 | 7.051918894 | -0.026986955 | -0.14896795 |
| Grk1 | 5.66833201 | 5.667217026 | 5.817302097 | -0.150085072 | -0.148970088 |
| Dynlt1f | 8.219830061 | 8.100513494 | 8.368880959 | -0.268367465 | -0.149050898 |
| Tmem255a | 3.028511126 | 3.107316749 | 3.177723452 | -0.070406703 | -0.149212326 |
| Tcf12 | 10.90526592 | 10.94570895 | 11.0545359 | -0.108826953 | -0.149269987 |
| Gm10778 | 6.706190178 | 6.675759118 | 6.855516241 | -0.179757123 | -0.149326063 |
| Adcy1 | 9.185176636 | 9.290592423 | 9.33466362 | -0.044071197 | -0.149486984 |
| Xiap | 10.82586542 | 10.87099019 | 10.9753973 | -0.104407107 | -0.149531877 |
| Edil3 | 4.524706863 | 4.502944618 | 4.674295684 | -0.171351066 | -0.149588821 |
| Tas1r3 | 4.072563551 | 4.166429387 | 4.222298985 | -0.055869598 | -0.149735434 |
| Ncmap | 6.444133252 | 6.39610115 | 6.593891822 | -0.197790672 | -0.14975857 |

| | | | | | |
|---|---|---|---|---|---|
| Vmn1r88 | 5.721470045 | 5.771316863 | 5.871236297 | -0.099919434 | -0.149766252 |
| Arnt2 | 6.213755073 | 6.293546442 | 6.363564407 | -0.070017965 | -0.149809334 |
| Mmab | 8.339824701 | 8.461625217 | 8.489656736 | -0.028031519 | -0.149832035 |
| Ankrd42 | 5.117837134 | 5.174315031 | 5.267699301 | -0.09338427 | -0.149862168 |
| Echdc2 | 8.161316753 | 8.249089561 | 8.311305862 | -0.062216301 | -0.149989109 |
| Prickle4 | 2.99019622 | 3.052735343 | 3.140221078 | -0.087485735 | -0.150024858 |
| Faah | 10.63650451 | 10.83151451 | 10.78655671 | 0.044957793 | -0.150052203 |
| Dmkn | 15.69689903 | 15.74555024 | 15.84699034 | -0.1014401 | -0.150091307 |
| Hr | 12.65287817 | 12.8349989 | 12.80297613 | 0.032022773 | -0.150097953 |
| Zfp87 | 7.484448531 | 7.485216015 | 7.634689401 | -0.149473386 | -0.15024087 |
| Mfap4 | 8.718906455 | 8.940741432 | 8.869148104 | 0.071593328 | -0.150241649 |
| Ncam1 | 6.590481288 | 6.532194722 | 6.740773028 | -0.208578307 | -0.15029174 |
| Mau2 | 10.22361838 | 10.49081205 | 10.3739273 | 0.116884747 | -0.15030892 |
| Wdr19 | 6.464815176 | 6.524484057 | 6.61513047 | -0.090646412 | -0.150315294 |
| Ttl | 8.520392464 | 8.637588828 | 8.670718933 | -0.033130105 | -0.150326469 |
| Zfp647 | 3.849430333 | 3.875405172 | 3.999810313 | -0.124405141 | -0.15037998 |
| Penk | 9.524792063 | 9.460192438 | 9.67520493 | -0.215012492 | -0.150412867 |
| 2010111I01Rik | 10.57341829 | 10.90298952 | 10.72400269 | 0.178986827 | -0.150584403 |
| Cers4 | 12.51160174 | 12.55090801 | 12.66233824 | -0.111430233 | -0.150736503 |
| Map2k5 | 8.072951079 | 8.160236238 | 8.223714323 | -0.063478085 | -0.150763244 |
| Fmo4 | 1.992415045 | 2.081688896 | 2.143198866 | -0.06150997 | -0.150783821 |
| Grtp1 | 6.227352453 | 6.162756681 | 6.378136811 | -0.215380131 | -0.150784358 |
| Pdcd4 | 10.71040191 | 10.70669698 | 10.8613926 | -0.15469562 | -0.15099069 |
| Kcnab1 | 4.346529475 | 4.35425185 | 4.497530775 | -0.143278925 | -0.151001301 |
| Zfand4 | 5.044122364 | 5.092322749 | 5.195144586 | -0.102821838 | -0.151022222 |
| Zmynd11 | 10.64749761 | 10.76160309 | 10.79858166 | -0.036978567 | -0.151084053 |
| Ccdc125 | 5.238696921 | 5.200626264 | 5.389804586 | -0.189178322 | -0.151107665 |
| Mrm1 | 7.504172549 | 7.662659399 | 7.655295442 | 0.007363957 | -0.151122893 |
| Fam120b | 9.412077484 | 9.526184895 | 9.563240173 | -0.037055278 | -0.151162688 |
| Fam118a | 7.230714172 | 7.048444893 | 7.382019378 | -0.333574485 | -0.151305206 |
| Nat8f7 | 4.04915683 | 4.318355741 | 4.20052127 | 0.117834472 | -0.151364439 |
| Avpr1a | 7.23727968 | 7.096854617 | 7.388682148 | -0.291827531 | -0.151402468 |

| | | | | | |
|---|---|---|---|---|---|
| Cenpj | 6.894133563 | 7.287226897 | 7.045566459 | 0.241660438 | -0.151432896 |
| Kcnip2 | 4.460640884 | 4.481057487 | 4.612112341 | -0.131054854 | -0.151471457 |
| Tmem170b | 9.293226384 | 9.28548895 | 9.444709319 | -0.159220369 | -0.151482935 |
| Gm14305 | 7.214696798 | 7.369489831 | 7.366250671 | 0.00323916 | -0.151553873 |
| Tmed4 | 9.246455736 | 9.292007145 | 9.398031227 | -0.106024082 | -0.151575491 |
| Pld5 | 3.025970461 | 3.07701417 | 3.177633472 | -0.100619302 | -0.151663012 |
| Lcat | 5.108523554 | 5.344227755 | 5.260271696 | 0.083956059 | -0.151748142 |
| Tpi1 | 12.91589691 | 12.86201447 | 13.06765546 | -0.205640993 | -0.15175855 |
| Vrk2 | 7.964497504 | 7.99275041 | 8.116380677 | -0.123630267 | -0.151883173 |
| Dusp3 | 9.822608703 | 10.01850964 | 9.974529632 | 0.043980009 | -0.151920929 |
| 4931406P16Rik | 9.64160468 | 9.781239964 | 9.793603198 | -0.012363234 | -0.151998517 |
| Rpa3 | 7.448489846 | 7.477767649 | 7.600518288 | -0.12275064 | -0.152028442 |
| Morn4 | 4.609415017 | 4.616552737 | 4.761531489 | -0.144978752 | -0.152116472 |
| Zfp119a | 6.394158788 | 6.421244186 | 6.546281094 | -0.125036909 | -0.152122306 |
| Zfp664 | 10.25376831 | 10.40789126 | 10.40591545 | 0.001975807 | -0.152147143 |
| Taco1 | 7.598456497 | 7.517070667 | 7.750621645 | -0.233550978 | -0.152165149 |
| Hhip | 4.854505851 | 5.006710119 | 5.00669741 | 1.27087E-05 | -0.152191559 |
| Iah1 | 8.535742382 | 8.475871973 | 8.688002558 | -0.212130585 | -0.152260177 |
| Sfrp5 | 6.982585118 | 7.117649214 | 7.134850152 | -0.017200938 | -0.152265034 |
| Lce1d | 10.96407206 | 11.29652459 | 11.1163738 | 0.180150787 | -0.152301737 |
| Txlng | 8.379416923 | 8.375334616 | 8.531735193 | -0.156400578 | -0.152318271 |
| Kcnab2 | 7.770386625 | 7.936485369 | 7.922714688 | 0.013770681 | -0.152328063 |
| Chchd3 | 10.53992752 | 10.47673147 | 10.69230263 | -0.21557116 | -0.152375107 |
| Tbc1d8b | 8.361140344 | 8.41218805 | 8.513526901 | -0.101338851 | -0.152386557 |
| Rad52 | 7.526988758 | 7.733405891 | 7.679440314 | 0.053965578 | -0.152451556 |
| Gm9372 | 2.493609626 | 2.594249856 | 2.646129281 | -0.051879425 | -0.152519655 |
| Snapc5 | 8.353126922 | 8.383897437 | 8.505678326 | -0.12178089 | -0.152551404 |
| 2010109I03Rik | 9.499053527 | 9.551022719 | 9.651605365 | -0.100582646 | -0.152551838 |
| Jam3 | 7.158747499 | 7.155733863 | 7.311303709 | -0.155569846 | -0.15255621 |
| Rapsn | 4.532781113 | 4.67663036 | 4.685345131 | -0.008714771 | -0.152564018 |
| Fgfbp3 | 7.007146889 | 6.829827792 | 7.159751141 | -0.329923349 | -0.152604252 |
| Gm44911 | 3.25121299 | 3.387915401 | 3.403838233 | -0.015922832 | -0.152625243 |

| | | | | | |
|---|---|---|---|---|---|
| Rasl10a | 5.065202942 | 5.088374422 | 5.217828517 | -0.129454094 | -0.152625575 |
| Elovl4 | 14.79004157 | 14.86431689 | 14.94278511 | -0.07846822 | -0.152743543 |
| Ano3 | 2.969765847 | 3.080108019 | 3.122530208 | -0.042422189 | -0.152764361 |
| Ccdc14 | 6.636673215 | 6.640846826 | 6.789468684 | -0.148621859 | -0.152795469 |
| Pwwp2a | 8.272827183 | 8.367061622 | 8.425633079 | -0.058571457 | -0.152805896 |
| Vezt | 9.018328444 | 9.033098763 | 9.171161806 | -0.138063044 | -0.152833363 |
| Myl6b | 4.217706413 | 4.27350149 | 4.370777809 | -0.09727632 | -0.153071397 |
| Smad6 | 5.058348738 | 5.112969517 | 5.21142638 | -0.098456862 | -0.153077642 |
| Kcnrg | 3.472495267 | 3.595734617 | 3.625595603 | -0.029860986 | -0.153100335 |
| 4930579G24Rik | 6.066235721 | 6.28998746 | 6.219337564 | 0.070649896 | -0.153101843 |
| Gpt2 | 10.95243783 | 10.6996524 | 11.10564604 | -0.405993637 | -0.153208207 |
| Nkpd1 | 7.854646945 | 8.29529646 | 8.007888415 | 0.287408045 | -0.15324147 |
| Gaa | 10.08559822 | 10.46749602 | 10.23884241 | 0.22865361 | -0.153244184 |
| 1700047I17Rik2 | 9.707860849 | 9.758330094 | 9.861143444 | -0.10281335 | -0.153282595 |
| Wbp2 | 10.62789149 | 10.8064112 | 10.78119329 | 0.025217907 | -0.1533018 |
| Lgi2 | 7.503774004 | 7.422684578 | 7.657135634 | -0.234451056 | -0.15336163 |
| Hacl1 | 8.068945556 | 7.976610726 | 8.222316187 | -0.245705462 | -0.153370631 |
| Gm9522 | 3.762310742 | 3.762042447 | 3.915730159 | -0.153687712 | -0.153419417 |
| Qsox2 | 9.379583223 | 9.401340954 | 9.533018597 | -0.131677643 | -0.153435374 |
| Lrat | 6.198167141 | 6.150981801 | 6.351628218 | -0.200646417 | -0.153461077 |
| Mpc1 | 10.72908118 | 10.58054405 | 10.88254333 | -0.301999283 | -0.15346215 |
| Brpf3 | 9.080704071 | 9.325938179 | 9.234170013 | 0.091768166 | -0.153465942 |
| Rps6-ps4 | 12.51569338 | 12.2485945 | 12.66925689 | -0.420662393 | -0.15356351 |
| Ubl3 | 10.81040436 | 10.78535206 | 10.96398629 | -0.17863423 | -0.15358193 |
| Abcd1 | 8.326353869 | 8.404347308 | 8.480038514 | -0.075691206 | -0.153684645 |
| 4931428F04Rik | 6.430992862 | 6.622707344 | 6.584727925 | 0.037979419 | -0.153735063 |
| Apoc1 | 10.89229577 | 10.75954992 | 11.04604277 | -0.286492853 | -0.153746997 |
| Apoo | 8.68840762 | 8.675714928 | 8.842155206 | -0.166440278 | -0.153747586 |
| Vat1l | 5.579473375 | 5.624398728 | 5.733299499 | -0.108900771 | -0.153826123 |
| Zfp24 | 9.216380437 | 9.227218049 | 9.370239357 | -0.143021308 | -0.15385892 |
| Plac9b | 9.438474217 | 9.194186844 | 9.592412997 | -0.398226153 | -0.15393878 |
| Myo7a | 7.205119265 | 7.439509243 | 7.35922192 | 0.080287324 | -0.154102655 |

| | | | | | |
|---|---|---|---|---|---|
| Zfp934 | 6.99004329 | 7.084669816 | 7.144165525 | -0.059495709 | -0.154122235 |
| Disp1 | 7.406382555 | 7.44963364 | 7.560520045 | -0.110886405 | -0.15413749 |
| Katnal1 | 6.931001521 | 6.886002497 | 7.085289585 | -0.199287088 | -0.154288065 |
| Pyroxd2 | 4.658905965 | 4.669540806 | 4.81325504 | -0.143714233 | -0.154349075 |
| Gm5538 | 9.076377191 | 9.23010766 | 9.230779507 | -0.000671847 | -0.154402316 |
| Arhgef26 | 7.909697853 | 7.89087044 | 8.064120378 | -0.173249938 | -0.154422526 |
| Mfn2 | 11.24748019 | 11.28177103 | 11.40190435 | -0.120133317 | -0.15442416 |
| Intu | 5.482666871 | 5.639035393 | 5.637489693 | 0.0015457 | -0.154822822 |
| Ccdc73 | 3.141830881 | 3.192056463 | 3.296724691 | -0.104668228 | -0.15489381 |
| Gm49320 | 3.173359789 | 3.297741631 | 3.328306178 | -0.030564547 | -0.154946389 |
| Tspyl2 | 8.456125106 | 8.621067443 | 8.611087273 | 0.00998017 | -0.154962167 |
| Mks1 | 6.486691339 | 6.631203409 | 6.641678514 | -0.010475105 | -0.154987175 |
| Icosl | 8.956227897 | 9.006395153 | 9.111272311 | -0.104877158 | -0.155044414 |
| C4b | 12.22631713 | 12.07408308 | 12.38139118 | -0.3073081 | -0.155074053 |
| Mpc2 | 9.921439322 | 9.748246499 | 10.07663949 | -0.328392989 | -0.155200166 |
| Myh7b | 3.062644105 | 3.087829817 | 3.217864518 | -0.130034701 | -0.155220412 |
| Glrx2 | 9.175819284 | 9.280240206 | 9.331070719 | -0.050830513 | -0.155251435 |
| Garnl3 | 5.298774349 | 5.393577264 | 5.454195949 | -0.060618685 | -0.155421599 |
| Prr12 | 9.179920689 | 9.738935871 | 9.335496796 | 0.403439075 | -0.155576107 |
| D430041D05Rik | 4.775711863 | 4.775883323 | 4.931406058 | -0.155522736 | -0.155694195 |
| Reep6 | 8.721758341 | 8.886240639 | 8.877537739 | 0.0087029 | -0.155779398 |
| Pnmal2 | 2.759337387 | 2.90948665 | 2.915150798 | -0.005664148 | -0.155813411 |
| Zfp712 | 5.783227955 | 5.823375656 | 5.939072816 | -0.11569716 | -0.155844861 |
| Ldhb | 10.39853189 | 10.4789114 | 10.55439111 | -0.07547971 | -0.15585922 |
| Tti1 | 8.768100371 | 8.899598696 | 8.923980461 | -0.024381765 | -0.15588009 |
| Lrrc1 | 10.35762707 | 10.53343789 | 10.51350963 | 0.019928257 | -0.155882557 |
| Ppdpf | 8.249981454 | 8.290538268 | 8.405912985 | -0.115374717 | -0.155931531 |
| Cep120 | 8.747999288 | 8.900592272 | 8.903941694 | -0.003349422 | -0.155942406 |
| Clec10a | 9.984899867 | 9.970203305 | 10.14110492 | -0.170901619 | -0.156205057 |
| Sgca | 7.866575661 | 7.986647887 | 8.022922824 | -0.036274937 | -0.156347163 |
| Mterf1b | 4.928306448 | 5.109984686 | 5.084677327 | 0.025307359 | -0.15637088 |
| Smtn | 9.625645578 | 9.940703324 | 9.78215579 | 0.158547534 | -0.156510212 |

| | | | | | |
|---|---|---|---|---|---|
| Mfsd8 | 7.527208192 | 7.547690053 | 7.683811674 | -0.136121621 | -0.156603483 |
| Mfsd12 | 8.37829754 | 8.892713879 | 8.534909426 | 0.357804453 | -0.156611886 |
| Dync2li1 | 6.447458215 | 6.455474175 | 6.604117598 | -0.148643424 | -0.156659383 |
| Tnfsf12 | 7.906485501 | 7.949160322 | 8.063240891 | -0.114080568 | -0.156755389 |
| Pif1 | 5.946013883 | 6.027267752 | 6.102845789 | -0.075578037 | -0.156831906 |
| C1qtnf12 | 9.375217911 | 9.346533516 | 9.532067611 | -0.185534095 | -0.1568497 |
| Gpr137 | 8.114622298 | 8.379253128 | 8.271520457 | 0.107732671 | -0.156898159 |
| Ptges3l | 6.92387768 | 6.999928902 | 7.080800637 | -0.080871735 | -0.156922957 |
| Atg16l2 | 6.932724194 | 7.017506832 | 7.089662059 | -0.072155227 | -0.156937865 |
| Sacm1l | 10.4883315 | 10.58884959 | 10.6453308 | -0.05648121 | -0.156999307 |
| Isoc1 | 10.80807504 | 10.82107032 | 10.96510291 | -0.14403259 | -0.15702787 |
| Gm3435 | 4.256327231 | 4.33329952 | 4.413362729 | -0.08006321 | -0.157035498 |
| Gm43540 | 4.146330385 | 4.101974308 | 4.303398007 | -0.201423699 | -0.157067622 |
| Klhl34 | 4.235270492 | 4.191804295 | 4.392348116 | -0.200543821 | -0.157077624 |
| Tcf15 | 3.321148313 | 3.34209506 | 3.478400826 | -0.136305766 | -0.157252513 |
| Inpp5f | 8.423850661 | 8.321398009 | 8.581113535 | -0.259715525 | -0.157262874 |
| Clcn3 | 10.5059786 | 10.5116577 | 10.66325439 | -0.151596697 | -0.157275797 |
| Haus5 | 7.435068333 | 7.73193758 | 7.592360376 | 0.139577205 | -0.157292043 |
| Ubl7 | 9.204678056 | 9.223504184 | 9.36200744 | -0.138503256 | -0.157329384 |
| Ttc26 | 5.879615123 | 6.019344366 | 6.037033677 | -0.01768931 | -0.157418553 |
| Fam172a | 9.280963792 | 9.350630954 | 9.438558915 | -0.087927961 | -0.157595123 |
| Pcdh17 | 4.986925702 | 4.917840391 | 5.144527079 | -0.226686688 | -0.157601377 |
| Vgll4 | 9.416966545 | 9.654486419 | 9.57458968 | 0.079896739 | -0.157623135 |
| Usp34 | 11.4689486 | 11.59151364 | 11.62666021 | -0.03514657 | -0.157711613 |
| Paqr7 | 11.74217938 | 11.92874602 | 11.89998568 | 0.028760347 | -0.157806293 |
| Hexim2 | 4.733308364 | 4.815933521 | 4.891199228 | -0.075265707 | -0.157890864 |
| Rdx | 11.32840361 | 11.34236599 | 11.4863867 | -0.14402071 | -0.15798309 |
| Jmjd8 | 9.219928599 | 9.339012507 | 9.377927583 | -0.038915077 | -0.157998984 |
| Pi15 | 7.642048627 | 7.610235607 | 7.800075491 | -0.189839884 | -0.158026864 |
| Accs | 5.945108378 | 5.927457524 | 6.103178354 | -0.17572083 | -0.158069976 |
| Efcab2 | 6.201639417 | 6.076720859 | 6.359756669 | -0.28303581 | -0.158117252 |
| Rtp3 | 3.960427438 | 3.9740008 | 4.118569872 | -0.144569072 | -0.158142435 |

| | | | | | |
|---|---|---|---|---|---|
| Plekhb1 | 7.997208384 | 7.958745224 | 8.155412111 | -0.196666887 | -0.158203727 |
| Actr8 | 8.653931039 | 8.734171591 | 8.812226007 | -0.078054415 | -0.158294968 |
| Nfs1 | 9.498931412 | 9.501457593 | 9.657266733 | -0.15580914 | -0.158335321 |
| Usf2 | 10.18813299 | 10.34834766 | 10.34647027 | 0.001877393 | -0.15833728 |
| Lrp5 | 10.48004103 | 10.8414346 | 10.63842359 | 0.203011003 | -0.158382563 |
| Smim13 | 9.849233257 | 9.774414624 | 10.00768679 | -0.233272166 | -0.158453533 |
| Hsd17b13 | 4.629124905 | 4.6344592 | 4.787583942 | -0.153124742 | -0.158459036 |
| Parp6 | 8.738737534 | 8.895515419 | 8.897261046 | -0.001745627 | -0.158523512 |
| Tubd1 | 6.964060851 | 6.977354699 | 7.122617828 | -0.14526313 | -0.158556978 |
| Acyp1 | 6.829793555 | 6.79492798 | 6.988356342 | -0.193428362 | -0.158562787 |
| Nck1 | 9.397267953 | 9.350713176 | 9.55591277 | -0.205199595 | -0.158644818 |
| Anxa3 | 8.193781349 | 7.938890946 | 8.352491576 | -0.41360063 | -0.158710227 |
| Rpl7a-ps12 | 2.741938472 | 2.871042169 | 2.900678543 | -0.029636374 | -0.158740071 |
| Zswim3 | 6.591213035 | 6.682523991 | 6.750036589 | -0.067512598 | -0.158823554 |
| Mtrf1 | 6.521595273 | 6.422140982 | 6.680438308 | -0.258297326 | -0.158843035 |
| Cdk5rap3 | 9.285786282 | 9.327769985 | 9.444660603 | -0.116890618 | -0.158874321 |
| Dnase2a | 10.08970165 | 10.27553313 | 10.24857718 | 0.02695595 | -0.158875537 |
| Myof | 10.99861515 | 11.12188915 | 11.1575136 | -0.03562445 | -0.15889845 |
| Rhpn2 | 5.615940162 | 5.635453424 | 5.774892906 | -0.139439483 | -0.158952744 |
| Mettl24 | 4.722932888 | 4.804195002 | 4.881967619 | -0.077772617 | -0.159034732 |
| Gper1 | 2.562803109 | 2.604274798 | 2.72186942 | -0.117594622 | -0.159066311 |
| Arl15 | 8.821659818 | 8.860141637 | 8.980793453 | -0.120651816 | -0.159133635 |
| D7Ertd443e | 7.254955956 | 7.544519351 | 7.414118378 | 0.130400973 | -0.159162422 |
| C9orf72 | 8.881491708 | 8.838165786 | 9.040693255 | -0.20252747 | -0.159201548 |
| Thap2 | 8.815625423 | 9.004739389 | 8.974828943 | 0.029910447 | -0.15920352 |
| Ankrd40 | 10.76488462 | 10.7731071 | 10.92410863 | -0.151001533 | -0.15922401 |
| Alad | 8.893940548 | 9.277857793 | 9.053201704 | 0.224656088 | -0.159261156 |
| Cacng1 | 7.471624097 | 7.7512323 | 7.63097177 | 0.12026053 | -0.159347673 |
| Zfp939 | 6.165179051 | 6.235042085 | 6.324542289 | -0.089500204 | -0.159363238 |
| Mical2 | 9.732635851 | 10.1002307 | 9.892028214 | 0.20820249 | -0.159392363 |
| Galnt5 | 3.342139178 | 3.391997297 | 3.501546837 | -0.10954954 | -0.159407659 |
| Lrrc31 | 3.677519573 | 3.729313219 | 3.836976845 | -0.107663626 | -0.159457271 |

| | | | | | |
|---|---|---|---|---|---|
| Fbxo48 | 3.432429009 | 3.557860689 | 3.591886408 | -0.034025719 | -0.159457399 |
| Klrg2 | 7.987328143 | 8.344310465 | 8.146809214 | 0.19750125 | -0.159481072 |
| Rab28 | 9.440292795 | 9.414142819 | 9.599787399 | -0.185644581 | -0.159494604 |
| Ttc21b | 7.042086968 | 7.194976548 | 7.201663203 | -0.006686655 | -0.159576235 |
| Tmem221 | 3.750496733 | 3.827544261 | 3.910089956 | -0.082545695 | -0.159593223 |
| Sprr2g | 6.479808766 | 6.642442667 | 6.639521644 | 0.002921022 | -0.159712879 |
| Zfp273 | 5.776673607 | 5.791967786 | 5.936438807 | -0.144471021 | -0.1597652 |
| Zfp780b | 7.838047846 | 8.0149427 | 7.997872644 | 0.017070056 | -0.159824798 |
| Drc3 | 5.198193223 | 5.318366082 | 5.358024784 | -0.039658702 | -0.159831561 |
| Ndufa10 | 10.70321846 | 10.73995318 | 10.86305108 | -0.1230979 | -0.15983262 |
| Rsbn1l | 9.115406186 | 8.988842338 | 9.275275768 | -0.28643343 | -0.159869582 |
| Ggt6 | 9.274001055 | 9.501668083 | 9.433882908 | 0.067785175 | -0.159881853 |
| Mtmr3 | 10.80939536 | 10.87152315 | 10.96930844 | -0.097785297 | -0.159913087 |
| Clock | 10.31081846 | 10.53400044 | 10.47075763 | 0.063242817 | -0.15993917 |
| CR974586.6 | 4.399792778 | 4.623505664 | 4.559736524 | 0.06376914 | -0.159943746 |
| Zfp687 | 8.787755644 | 8.888836806 | 8.94770847 | -0.058871663 | -0.159952826 |
| Rom1 | 6.004401588 | 6.158545752 | 6.164357839 | -0.005812087 | -0.159956251 |
| Pcgf3 | 10.34357865 | 10.45932819 | 10.50355059 | -0.044222397 | -0.159971933 |
| Ldah | 10.64332723 | 10.73537217 | 10.80337263 | -0.068000463 | -0.160045397 |
| Clstn3 | 8.876288763 | 8.876096483 | 9.036348329 | -0.160251846 | -0.160059566 |
| Kmt2c | 10.73527528 | 10.87083052 | 10.8953545 | -0.024523977 | -0.160079223 |
| Tmem35b | 6.075229005 | 6.192110195 | 6.235311257 | -0.043201063 | -0.160082252 |
| Gid8 | 10.5051602 | 10.50677995 | 10.66530722 | -0.158527273 | -0.160147017 |
| Slc6a1 | 3.036833619 | 3.107898296 | 3.197000192 | -0.089101896 | -0.160166573 |
| Zdhhc14 | 6.398403418 | 6.342229555 | 6.558731751 | -0.216502196 | -0.160328333 |
| Arhgap40 | 8.435245268 | 8.76786211 | 8.59569994 | 0.17216217 | -0.160454672 |
| Cdc40 | 9.094250511 | 9.116488071 | 9.254715294 | -0.138227223 | -0.160464783 |
| Twsg1 | 11.08680503 | 11.23682554 | 11.24728206 | -0.010456523 | -0.160477037 |
| Dhfr | 8.108855905 | 8.319321531 | 8.269426864 | 0.049894667 | -0.160570959 |
| Angel2 | 9.735163029 | 9.778858896 | 9.895750079 | -0.116891183 | -0.160587049 |
| Map3k3 | 9.86678419 | 10.00859015 | 10.02738826 | -0.018798115 | -0.160604072 |
| Zscan2 | 4.022132739 | 4.072585165 | 4.182736918 | -0.110151753 | -0.160604179 |

| | | | | | |
|---|---|---|---|---|---|
| Nme4 | 5.968524332 | 6.080509146 | 6.129156305 | -0.04864716 | -0.160631973 |
| Cul3 | 11.74687946 | 11.78417014 | 11.90755272 | -0.123382577 | -0.160673253 |
| Klhl25 | 8.554082778 | 8.69254802 | 8.714828575 | -0.022280554 | -0.160745796 |
| Adtrp | 10.29648137 | 10.37229143 | 10.45728141 | -0.084989973 | -0.160800037 |
| Plppr2 | 5.76246046 | 5.87905648 | 5.923367257 | -0.044310777 | -0.160906797 |
| Mgat5 | 9.418952346 | 9.572343487 | 9.579863648 | -0.007520161 | -0.160911302 |
| Cep164 | 8.697690093 | 8.847987623 | 8.858650926 | -0.010663302 | -0.160960833 |
| BC035947 | 4.235998082 | 4.362205437 | 4.396958961 | -0.034753524 | -0.160960879 |
| Wnt3a | 6.967967349 | 7.45463355 | 7.128932245 | 0.325701304 | -0.160964896 |
| Kif1bp | 8.748265327 | 8.902566323 | 8.909249746 | -0.006683423 | -0.160984419 |
| Tbc1d32 | 7.324289109 | 7.411019763 | 7.485283244 | -0.074263481 | -0.160994135 |
| Tmc7 | 6.573807964 | 6.670365832 | 6.734816701 | -0.064450869 | -0.161008737 |
| Nek2 | 6.301052271 | 6.258607413 | 6.462222461 | -0.203615048 | -0.16117019 |
| Mxd3 | 5.339464803 | 5.381032888 | 5.500640164 | -0.119607276 | -0.161175361 |
| St6galnac6 | 8.127089407 | 8.323308876 | 8.288382776 | 0.0349261 | -0.161293369 |
| Sirt4 | 6.810403644 | 6.995232792 | 6.971719072 | 0.023513721 | -0.161315427 |
| Rab40c | 9.238003858 | 9.455027125 | 9.39936391 | 0.055663215 | -0.161360052 |
| Tpst1 | 8.676823247 | 8.742862285 | 8.838229038 | -0.095366752 | -0.16140579 |
| Cela1 | 5.884891054 | 5.756033684 | 6.046301577 | -0.290267893 | -0.161410522 |
| Dffb | 6.950912713 | 7.156723652 | 7.112380183 | 0.044343469 | -0.16146747 |
| Tab3 | 8.750717134 | 8.776688049 | 8.912251436 | -0.135563387 | -0.161534302 |
| Nt5e | 9.466624533 | 9.354124565 | 9.628166152 | -0.274041587 | -0.161541619 |
| Ginm1 | 9.07757044 | 8.930401565 | 9.239126656 | -0.308725091 | -0.161556216 |
| Dlg4 | 7.288741744 | 7.679457088 | 7.450349634 | 0.229107454 | -0.161607891 |
| Stra6l | 4.563208209 | 5.619687626 | 4.724862235 | 0.894825391 | -0.161654026 |
| Adamtsl5 | 8.816177215 | 9.160677809 | 8.977872913 | 0.182804897 | -0.161695698 |
| Notch3 | 10.37247116 | 10.85667301 | 10.53418173 | 0.32249128 | -0.161710573 |
| Dhcr7 | 11.09314603 | 11.26498218 | 11.25486731 | 0.010114873 | -0.161721273 |
| Olfm2 | 7.871229838 | 7.99583188 | 8.032983628 | -0.037151748 | -0.16175379 |
| Tnrc6b | 10.65709327 | 10.82175354 | 10.8189312 | 0.002822337 | -0.161837927 |
| Gm5373 | 5.394270555 | 5.428042868 | 5.55615914 | -0.128116272 | -0.161888585 |
| Laptm4a | 12.4828914 | 12.41156993 | 12.64480516 | -0.23323523 | -0.16191376 |

| | | | | | |
|---|---|---|---|---|---|
| 1810010H24Rik | 3.508576653 | 3.499996969 | 3.670509821 | -0.170512852 | -0.161933168 |
| Rpl30-ps9 | 3.554513122 | 3.825881964 | 3.716508948 | 0.109373015 | -0.161995826 |
| Tecpr2 | 8.431000139 | 8.574584267 | 8.592997696 | -0.018413428 | -0.161997556 |
| Pgam2 | 11.23923603 | 10.9560452 | 11.40133522 | -0.445290027 | -0.162099197 |
| Lhfpl2 | 8.423821389 | 8.673212164 | 8.586005262 | 0.087206902 | -0.162183873 |
| Plpp2 | 9.39057998 | 9.479365906 | 9.552828686 | -0.073462781 | -0.162248707 |
| Usp28 | 8.637510393 | 9.408510952 | 8.799843416 | 0.608667536 | -0.162333023 |
| Snx29 | 6.849303351 | 7.036595622 | 7.011655891 | 0.024939731 | -0.16235254 |
| Mpzl3 | 10.70829151 | 10.62633047 | 10.87072665 | -0.244396183 | -0.162435147 |
| Rai14 | 10.39123601 | 10.51115577 | 10.55372246 | -0.042566693 | -0.162486453 |
| Ppp4r3b | 10.75867519 | 10.80927247 | 10.92120225 | -0.111929787 | -0.162527067 |
| Amer1 | 8.218311045 | 8.232521658 | 8.380870092 | -0.148348435 | -0.162559048 |
| Fam178b | 4.967096712 | 5.154653312 | 5.129679881 | 0.024973431 | -0.162583169 |
| Bhlhb9 | 7.461084851 | 7.560503437 | 7.623693132 | -0.063189695 | -0.162608281 |
| Enox1 | 6.239024732 | 6.340757094 | 6.401669588 | -0.060912494 | -0.162644856 |
| Rnf215 | 8.180692932 | 8.43080681 | 8.343349446 | 0.087457364 | -0.162656514 |
| Pdha1 | 12.21260066 | 12.06883873 | 12.37533441 | -0.306495687 | -0.16273375 |
| Cdk15 | 4.177564548 | 4.226780277 | 4.340299887 | -0.113519609 | -0.162735339 |
| Fktn | 8.848084427 | 8.868589363 | 9.010909699 | -0.142320336 | -0.162825272 |
| Sec14l1 | 10.48075394 | 10.72365541 | 10.64359213 | 0.08006328 | -0.16283819 |
| Gm48113 | 4.64827346 | 4.996582853 | 4.811244378 | 0.185338474 | -0.162970919 |
| Islr2 | 4.308774725 | 4.596005279 | 4.471773843 | 0.124231435 | -0.162999118 |
| Plcd3 | 7.196294051 | 7.508726457 | 7.359308718 | 0.149417739 | -0.163014667 |
| Foxc1 | 8.87062367 | 9.000097252 | 9.033645373 | -0.033548121 | -0.163021703 |
| Frs3 | 4.444888093 | 4.629536309 | 4.607913679 | 0.02162263 | -0.163025586 |
| Olfr365 | 4.750486491 | 4.957344022 | 4.913551065 | 0.043792957 | -0.163064574 |
| Trp53i13 | 6.835610804 | 7.058058657 | 6.998681909 | 0.059376748 | -0.163071105 |
| Kntc1 | 7.760598586 | 7.915753424 | 7.92371611 | -0.007962686 | -0.163117524 |
| Ccdc171 | 5.531275709 | 5.577659189 | 5.694459425 | -0.116800236 | -0.163183716 |
| Hip1 | 9.97083287 | 10.09407481 | 10.13406696 | -0.039992152 | -0.163234089 |
| Isl1 | 4.76457156 | 4.914913491 | 4.927809784 | -0.012896294 | -0.163238225 |
| Gm3448 | 4.414208163 | 4.530364788 | 4.577527332 | -0.047162544 | -0.163319169 |

| | | | | | |
|---|---|---|---|---|---|
| Cavin4 | 8.409547841 | 9.522465036 | 8.572953158 | 0.949511879 | -0.163405317 |
| Rnf14 | 10.22295125 | 10.23612052 | 10.38640272 | -0.150282207 | -0.163451473 |
| Bmpr2 | 11.58725728 | 11.41810736 | 11.75071401 | -0.332606643 | -0.163456723 |
| Uaca | 9.705319104 | 9.635220734 | 9.868809904 | -0.23358917 | -0.1634908 |
| Rgp1 | 10.09550818 | 10.45952438 | 10.25903472 | 0.20048966 | -0.163526537 |
| Fmn2 | 3.886452672 | 4.008329515 | 4.050000163 | -0.041670649 | -0.163547491 |
| Mycn | 3.815579148 | 3.838446084 | 3.979131419 | -0.140685335 | -0.163552272 |
| Fam76a | 9.776388406 | 9.818578757 | 9.939956701 | -0.121377944 | -0.163568295 |
| Zfp791 | 5.707508465 | 5.790306174 | 5.871078477 | -0.080772302 | -0.163570011 |
| Specc1 | 7.948112459 | 8.167921913 | 8.111703437 | 0.056218476 | -0.163590978 |
| Acsl6 | 4.933783577 | 5.057625468 | 5.097398852 | -0.039773385 | -0.163615276 |
| Armc9 | 6.894459474 | 7.069597828 | 7.058076909 | 0.011520919 | -0.163617435 |
| Noxo1 | 4.35626055 | 4.352908084 | 4.519940839 | -0.167032755 | -0.163680289 |
| Pln | 2.442360327 | 2.467633594 | 2.606115507 | -0.138481913 | -0.16375518 |
| Nmnat3 | 7.167991795 | 7.047258489 | 7.33181985 | -0.284561361 | -0.163828055 |
| Zscan21 | 8.049605994 | 8.158555607 | 8.213491647 | -0.05493604 | -0.163885654 |
| Slamf9 | 6.214262794 | 6.040741883 | 6.378151802 | -0.337409919 | -0.163889008 |
| Acer2 | 9.501093275 | 9.484543927 | 9.664991769 | -0.180447842 | -0.163898495 |
| Togaram1 | 8.789115035 | 9.042834043 | 8.953051277 | 0.089782766 | -0.163936242 |
| Adk | 9.865144963 | 9.810239894 | 10.02919588 | -0.218955985 | -0.164050916 |
| Rmi1 | 8.61191688 | 8.692401551 | 8.775996115 | -0.083594565 | -0.164079235 |
| Lbh | 9.054338796 | 9.010495618 | 9.218510021 | -0.208014403 | -0.164171226 |
| Daam1 | 11.00276494 | 11.11438599 | 11.16698246 | -0.05259647 | -0.164217523 |
| Plekhf2 | 9.550194309 | 9.598780428 | 9.714465378 | -0.11568495 | -0.16427107 |
| Paqr9 | 6.276150989 | 6.021689993 | 6.440452392 | -0.418762399 | -0.164301402 |
| Zbtb5 | 8.82431977 | 8.867425876 | 8.988634525 | -0.121208649 | -0.164314755 |
| Shroom2 | 7.830033705 | 7.997413671 | 7.994515159 | 0.002898512 | -0.164481454 |
| Insig2 | 10.5941678 | 10.59089009 | 10.75865341 | -0.16776332 | -0.16448561 |
| Kif1c | 13.07248089 | 13.24295144 | 13.23700098 | 0.00595046 | -0.164520097 |
| Mob1b | 10.32399255 | 10.34570031 | 10.48851778 | -0.142817477 | -0.16452523 |
| BC055324 | 6.544751233 | 6.880384208 | 6.70931244 | 0.171071767 | -0.164561208 |
| Nudt11 | 4.890745422 | 5.094001582 | 5.055348986 | 0.038652596 | -0.164603564 |

| | | | | | |
|---|---|---|---|---|---|
| Ccne2 | 7.702718468 | 7.983026631 | 7.867422555 | 0.115604076 | -0.164704087 |
| Dcun1d2 | 9.028955656 | 9.078929622 | 9.193665637 | -0.114736014 | -0.16470998 |
| Ccdc159 | 4.185291009 | 4.299984115 | 4.350101205 | -0.050117089 | -0.164810196 |
| Calml4 | 4.012354898 | 4.014108998 | 4.177266018 | -0.163157019 | -0.164911119 |
| Bex3 | 7.940603919 | 8.014503471 | 8.10555258 | -0.091049109 | -0.164948661 |
| Zfp646 | 9.645854437 | 9.923935412 | 9.810830652 | 0.113104761 | -0.164976214 |
| Slc1a5 | 11.95909206 | 11.73353497 | 12.12407152 | -0.39053655 | -0.164979453 |
| Dgkb | 2.874150216 | 2.889512751 | 3.03917342 | -0.149660669 | -0.165023204 |
| Btbd7 | 10.63673318 | 10.79152447 | 10.80177738 | -0.010252917 | -0.165044207 |
| Pllp | 5.585898748 | 5.604531115 | 5.750953884 | -0.146422769 | -0.165055136 |
| Bcl2l15 | 6.5018544 | 6.824573508 | 6.667007197 | 0.157566311 | -0.165152797 |
| Dusp28 | 6.851484656 | 6.896155706 | 7.016647067 | -0.120491361 | -0.165162411 |
| Notch1 | 11.23052061 | 11.93379686 | 11.39570185 | 0.53809501 | -0.165181243 |
| Nr2c1 | 7.765314242 | 7.889829611 | 7.930505493 | -0.040675882 | -0.165191252 |
| Zfp128 | 5.058118663 | 5.086909715 | 5.223312115 | -0.1364024 | -0.165193452 |
| C330007P06Rik | 9.796405956 | 9.775274061 | 9.961847854 | -0.186573793 | -0.165441898 |
| Slc18b1 | 7.559917765 | 7.540466369 | 7.725388225 | -0.184921855 | -0.16547046 |
| Gtf2ird1 | 8.683851797 | 8.926605717 | 8.849356528 | 0.077249189 | -0.165504731 |
| Zfp444 | 6.97688192 | 7.115064068 | 7.142465738 | -0.02740167 | -0.165583818 |
| Zfp872 | 6.105143093 | 6.1210668 | 6.270765193 | -0.149698393 | -0.1656221 |
| Dtx3 | 8.863829137 | 9.068103894 | 9.029690922 | 0.038412971 | -0.165861786 |
| Mettl21e | 3.182347597 | 3.131495529 | 3.348427663 | -0.216932134 | -0.166080066 |
| Kctd17 | 9.019293114 | 9.228638008 | 9.185450268 | 0.043187741 | -0.166157153 |
| Taf4 | 9.023953933 | 9.269147684 | 9.190164124 | 0.078983561 | -0.16621019 |
| Gm15159 | 3.208441212 | 3.335620303 | 3.374798211 | -0.039177908 | -0.166356999 |
| Cplx2 | 6.382921284 | 6.337445054 | 6.549463988 | -0.212018934 | -0.166542704 |
| Gm47406 | 4.487222058 | 4.461199618 | 4.653864151 | -0.192664533 | -0.166642093 |
| Cep55 | 7.406164717 | 7.334796839 | 7.572833886 | -0.238037047 | -0.166669169 |
| C1ra | 10.57033425 | 10.41122867 | 10.73701373 | -0.325785067 | -0.166679487 |
| Pde4c | 3.652273164 | 3.777106169 | 3.81895594 | -0.041849771 | -0.166682775 |
| Zbtb26 | 7.283203605 | 7.383626278 | 7.449895905 | -0.066269626 | -0.166692299 |
| Atp2c2 | 7.911846261 | 8.201464168 | 8.078545975 | 0.122918192 | -0.166699714 |

| | | | | | |
|---|---|---|---|---|---|
| Shcbp1 | 7.114351126 | 7.325117056 | 7.281069487 | 0.04404757 | -0.166718361 |
| Trim35 | 9.799492218 | 9.799109865 | 9.966226734 | -0.167116868 | -0.166734516 |
| Gpm6a | 6.430116227 | 6.34131634 | 6.596853563 | -0.255537223 | -0.166737336 |
| Strn3 | 10.05691991 | 10.05679288 | 10.22365793 | -0.166865053 | -0.166738024 |
| Stk38 | 10.58264444 | 10.73718228 | 10.74939739 | -0.01221511 | -0.166752957 |
| Nfatc3 | 10.58644866 | 10.76212144 | 10.75320647 | 0.008914973 | -0.166757813 |
| Tigar | 8.882157916 | 8.752256557 | 9.048946658 | -0.296690101 | -0.166788742 |
| Tacc3 | 8.755061517 | 8.892274865 | 8.921852076 | -0.029577211 | -0.166790559 |
| Gm43738 | 3.665320542 | 3.77297426 | 3.832155881 | -0.059181621 | -0.16683534 |
| Serpinc1 | 4.290658585 | 4.609235024 | 4.457523125 | 0.1517119 | -0.16686454 |
| Tmem151a | 3.386352343 | 3.705873407 | 3.553238985 | 0.152634422 | -0.166886642 |
| Zbtb3 | 5.579894846 | 5.793132676 | 5.746825077 | 0.046307599 | -0.16693023 |
| Osbpl5 | 11.06891774 | 11.34129241 | 11.23587034 | 0.105422077 | -0.166952597 |
| Ypel5 | 10.06505145 | 9.99117116 | 10.2320227 | -0.240851537 | -0.166971247 |
| Cldn22 | 2.813607999 | 2.834739859 | 2.980613518 | -0.145873658 | -0.167005519 |
| Slc13a4 | 4.327272726 | 4.29921503 | 4.494310649 | -0.195095619 | -0.167037923 |
| Efcc1 | 4.209546643 | 4.226979942 | 4.376590139 | -0.149610198 | -0.167043496 |
| Palm3 | 2.949406478 | 3.015349609 | 3.116499367 | -0.101149758 | -0.167092889 |
| Lama3 | 10.46521208 | 10.90089531 | 10.63236241 | 0.268532903 | -0.167150327 |
| Rad21 | 11.31764297 | 11.31299622 | 11.48484197 | -0.171845753 | -0.167199003 |
| Phtf1 | 9.176417309 | 9.258958526 | 9.343616349 | -0.084657823 | -0.16719904 |
| Hoga1 | 5.737882077 | 5.902142046 | 5.905183894 | -0.003041848 | -0.167301817 |
| D630045J12Rik | 6.63038006 | 6.87088378 | 6.797694072 | 0.073189708 | -0.167314012 |
| Mapk1 | 11.82826217 | 11.89923946 | 11.99560595 | -0.09636649 | -0.167343783 |
| Psapl1 | 13.54445091 | 13.71507031 | 13.71181916 | 0.003251157 | -0.167368247 |
| Dtx1 | 4.624268639 | 4.625712839 | 4.791688409 | -0.16597557 | -0.16741977 |
| Acat2 | 10.64378605 | 10.62655493 | 10.8114217 | -0.184866763 | -0.167635647 |
| Vamp7 | 8.713722917 | 8.750052923 | 8.881446477 | -0.131393554 | -0.167723559 |
| Capn7 | 10.15492989 | 10.16667999 | 10.32267726 | -0.15599727 | -0.16774737 |
| Pcbd1 | 4.826014076 | 4.767846988 | 4.993902109 | -0.226055121 | -0.167888033 |
| Huwe1 | 12.67091541 | 12.908173 | 12.83884213 | 0.06933087 | -0.167926713 |
| Dgcr6 | 8.779639303 | 8.954288845 | 8.947596603 | 0.006692243 | -0.167957299 |

| | | | | | |
|---|---|---|---|---|---|
| Apmap | 10.57259107 | 10.42212023 | 10.74063115 | -0.31851092 | -0.168040077 |
| Rad54l | 6.978031686 | 7.184311634 | 7.146095849 | 0.038215785 | -0.168064163 |
| Gm10288 | 11.23785665 | 10.81196725 | 11.40595126 | -0.59398401 | -0.168094607 |
| Lpar2 | 8.876823855 | 9.164342475 | 9.044939809 | 0.119402666 | -0.168115954 |
| Fam8a1 | 10.01286909 | 10.11793023 | 10.18100994 | -0.063079707 | -0.168140845 |
| Alg10b | 9.910478321 | 9.959471491 | 10.07864087 | -0.119169375 | -0.168162545 |
| Scyl3 | 8.97915752 | 9.072219194 | 9.147327673 | -0.075108479 | -0.168170153 |
| Cnppd1 | 10.13462393 | 10.36855252 | 10.30293698 | 0.065615537 | -0.16831305 |
| Zfp866 | 8.868747929 | 9.015024681 | 9.037109668 | -0.022084987 | -0.168361739 |
| Nptn | 11.53941105 | 11.49205931 | 11.70780015 | -0.215740837 | -0.1683891 |
| Ptprg | 9.669028536 | 9.803570394 | 9.837535965 | -0.033965571 | -0.168507429 |
| Kcnj14 | 3.275018074 | 3.337508292 | 3.443560952 | -0.10605266 | -0.168542878 |
| Thtpa | 8.009831965 | 8.006589017 | 8.178378354 | -0.171789338 | -0.168546389 |
| Pramef12 | 5.49106986 | 5.463188271 | 5.659630994 | -0.196442723 | -0.168561134 |
| Serinc3 | 12.59487103 | 12.46891253 | 12.76353517 | -0.294622643 | -0.168664143 |
| Tnfaip8l3 | 7.267169137 | 7.235433934 | 7.435933741 | -0.200499807 | -0.168764604 |
| Pgpep1l | 3.863629653 | 3.846037231 | 4.032431127 | -0.186393896 | -0.168801474 |
| Alkbh8 | 8.44863638 | 8.333742862 | 8.617474405 | -0.283731543 | -0.168838025 |
| Wdtc1 | 10.02012213 | 10.15820658 | 10.18896069 | -0.030754113 | -0.168838562 |
| Lrrc29 | 4.437515791 | 4.418817614 | 4.606377876 | -0.187560262 | -0.168862086 |
| Arhgap5 | 10.32725113 | 10.29739233 | 10.49613994 | -0.198747607 | -0.16888881 |
| Hmgcll1 | 4.060321949 | 4.166475214 | 4.229227246 | -0.062752032 | -0.168905297 |
| Akr1c19 | 4.699794123 | 4.757687377 | 4.86872681 | -0.111039432 | -0.168932687 |
| Rspo1 | 7.370044744 | 7.554088517 | 7.538994905 | 0.015093612 | -0.168950161 |
| Anks6 | 5.319560888 | 5.531988847 | 5.488547473 | 0.043441375 | -0.168986584 |
| Pdxk | 9.945194568 | 10.23136711 | 10.11420359 | 0.11716352 | -0.169009019 |
| Terf2 | 8.570965934 | 8.738044815 | 8.740006703 | -0.001961888 | -0.169040769 |
| Stambp | 8.809275797 | 8.773403304 | 8.978455437 | -0.205052133 | -0.169179639 |
| Hint3 | 7.462488218 | 7.545682879 | 7.631770129 | -0.08608725 | -0.169281911 |
| Colca2 | 3.728149137 | 3.703226058 | 3.8974315 | -0.194205442 | -0.169282363 |
| Rpap2 | 8.164065501 | 8.152057207 | 8.333349646 | -0.181292439 | -0.169284145 |
| Myh8 | 4.61029145 | 4.685106269 | 4.779598584 | -0.094492315 | -0.169307134 |

| | | | | | |
|---|---|---|---|---|---|
| Fbxl16 | 5.02590366 | 5.341450632 | 5.195314625 | 0.146136007 | -0.169410965 |
| Emc9 | 8.269653555 | 8.405406454 | 8.439136187 | -0.033729733 | -0.169482632 |
| C3 | 12.91412759 | 12.7207938 | 13.0836463 | -0.362852497 | -0.169518707 |
| Fam160b1 | 9.068277657 | 9.128973574 | 9.237959941 | -0.108986367 | -0.169682284 |
| Secisbp2l | 12.20757325 | 12.22121514 | 12.37726863 | -0.156053483 | -0.169695373 |
| Lin9 | 7.150551354 | 7.202729084 | 7.32033228 | -0.117603195 | -0.169780926 |
| Gpr21 | 2.566605487 | 2.577609275 | 2.736415847 | -0.158806571 | -0.16981036 |
| AW554918 | 7.723893186 | 7.760194712 | 7.893706682 | -0.13351197 | -0.169813496 |
| Adra2c | 4.20266633 | 4.297587604 | 4.37250577 | -0.074918166 | -0.16983944 |
| Slc22a18 | 6.192897751 | 6.371246248 | 6.362767394 | 0.008478854 | -0.169869643 |
| Slc40a1 | 9.103983055 | 9.154160512 | 9.273941617 | -0.119781105 | -0.169958562 |
| Bcl2l2 | 10.04168132 | 10.12098928 | 10.21167387 | -0.090684587 | -0.169992543 |
| Bnipl | 10.73591663 | 10.84397361 | 10.90594298 | -0.061969367 | -0.170026343 |
| Ei24 | 11.10868761 | 11.10299689 | 11.27877429 | -0.175777403 | -0.17008668 |
| Dhcr24 | 14.06775834 | 14.15594895 | 14.23789042 | -0.081941477 | -0.17013208 |
| Pafah2 | 9.692460562 | 9.833964889 | 9.862697418 | -0.028732529 | -0.170236856 |
| Ccl21a | 10.15722775 | 10.32534823 | 10.32747364 | -0.002125403 | -0.170245883 |
| Baiap2 | 10.18796502 | 10.64542086 | 10.35826984 | 0.287151023 | -0.170304823 |
| Mcat | 7.568315013 | 7.683103191 | 7.738650087 | -0.055546896 | -0.170335074 |
| Cdadc1 | 8.970439582 | 8.975581493 | 9.140796036 | -0.165214543 | -0.170356454 |
| Smyd3 | 7.353755954 | 7.244488599 | 7.524229593 | -0.279740995 | -0.170473639 |
| Sema3f | 10.3421547 | 10.81695128 | 10.51274053 | 0.30421075 | -0.170585827 |
| Gm48415 | 5.906442264 | 5.855779611 | 6.077117348 | -0.221337736 | -0.170675083 |
| Rdh5 | 5.863753386 | 6.01717667 | 6.034485742 | -0.017309072 | -0.170732356 |
| Spsb3 | 9.658990559 | 9.862463265 | 9.829913759 | 0.032549505 | -0.170923201 |
| Tbx3 | 5.874368994 | 5.656235292 | 6.045392925 | -0.389157633 | -0.171023932 |
| Crip1 | 10.97677027 | 10.89959104 | 11.14782048 | -0.248229437 | -0.17105021 |
| Adrb1 | 4.316035687 | 4.383497264 | 4.487149243 | -0.103651979 | -0.171113557 |
| Acly | 13.05324795 | 12.87442565 | 13.22448797 | -0.35006232 | -0.17124002 |
| Gpr107 | 10.46413765 | 10.51534172 | 10.63540746 | -0.12006574 | -0.171269803 |
| Depdc5 | 8.638563275 | 8.802305638 | 8.809960972 | -0.007655333 | -0.171397696 |
| 9930021J03Rik | 9.853868962 | 9.90433105 | 10.0252732 | -0.120942146 | -0.171404233 |

| | | | | | |
|---|---|---|---|---|---|
| Gm12461 | 3.678577787 | 3.862716495 | 3.850001383 | 0.012715111 | -0.171423596 |
| Ppp1r14a | 5.518696715 | 5.712924593 | 5.690126172 | 0.022798421 | -0.171429457 |
| Lpar6 | 7.948285283 | 7.863209481 | 8.119777995 | -0.256568514 | -0.171492712 |
| Rmdn1 | 7.004654858 | 7.041067844 | 7.176170292 | -0.135102448 | -0.171515434 |
| Pan3 | 10.20396873 | 10.33174679 | 10.37548483 | -0.043738033 | -0.171516097 |
| Mtmr4 | 9.871185422 | 10.31178019 | 10.0428294 | 0.268950791 | -0.171643978 |
| Arxes1 | 4.529243671 | 4.482077058 | 4.701092517 | -0.219015459 | -0.171848846 |
| Taf8 | 8.823723246 | 9.021567971 | 8.995580904 | 0.025987067 | -0.171857658 |
| Sprr2a3 | 7.044044158 | 7.728843471 | 7.215970303 | 0.512873168 | -0.171926145 |
| Aga | 7.947324266 | 7.944920342 | 8.119381779 | -0.174461436 | -0.172057513 |
| Mpig6b | 5.75145907 | 5.97462274 | 5.923576118 | 0.051046622 | -0.172117048 |
| Ttll10 | 5.27353268 | 5.457422102 | 5.445692442 | 0.01172966 | -0.172159762 |
| Cldn4 | 10.04140511 | 10.23604531 | 10.21366775 | 0.022377557 | -0.172262647 |
| Asb3 | 7.531084883 | 7.465895992 | 7.703353677 | -0.237457685 | -0.172268794 |
| Ptms | 11.93175689 | 12.00638613 | 12.10408362 | -0.097697493 | -0.172326733 |
| Hivep2 | 9.95288697 | 9.959874581 | 10.12522282 | -0.165348239 | -0.17233585 |
| Kctd11 | 10.613312 | 10.59487953 | 10.78573585 | -0.190856313 | -0.172423847 |
| Kirrel | 9.611941869 | 9.583569231 | 9.784430587 | -0.200861356 | -0.172488718 |
| Tbc1d22a | 9.088055788 | 9.262440786 | 9.26057334 | 0.001867446 | -0.172517552 |
| Zfp229 | 6.554990469 | 6.513753219 | 6.727511059 | -0.21375784 | -0.17252059 |
| Zdhhc24 | 7.072912983 | 7.298624726 | 7.245569839 | 0.053054887 | -0.172656857 |
| Ctnnbip1 | 10.23932366 | 10.42470391 | 10.41198171 | 0.0127222 | -0.172658047 |
| Bckdk | 10.25333648 | 10.2880637 | 10.4260197 | -0.137956003 | -0.17268322 |
| Sema3b | 8.029261596 | 7.981455474 | 8.202014058 | -0.220558583 | -0.172752462 |
| Riiad1 | 4.181419631 | 4.342441865 | 4.354180156 | -0.011738291 | -0.172760526 |
| Afg1l | 7.375210885 | 7.41500783 | 7.548181216 | -0.133173386 | -0.172970331 |
| Pcyt1b | 4.104259143 | 4.383643391 | 4.277252539 | 0.106390852 | -0.172993396 |
| Sdr16c6 | 12.13564548 | 12.17669438 | 12.30870427 | -0.132009883 | -0.17305879 |
| 1810043G02Rik | 6.823029904 | 7.027368564 | 6.996149874 | 0.03121869 | -0.17311997 |
| Gm17970 | 4.902169957 | 5.177932987 | 5.075371219 | 0.102561767 | -0.173201263 |
| Prkra | 8.012129764 | 8.057187746 | 8.185362526 | -0.128174779 | -0.173232761 |
| Hif1an | 10.81862286 | 10.85022951 | 10.99189303 | -0.14166352 | -0.17327017 |

| | | | | | |
|---|---|---|---|---|---|
| Eif2ak3 | 8.880243464 | 9.103238632 | 9.053577784 | 0.049660848 | -0.173334319 |
| Fam219a | 8.667541627 | 9.011825057 | 8.840887643 | 0.170937414 | -0.173346016 |
| Chat | 6.238941829 | 6.421540517 | 6.412353953 | 0.009186564 | -0.173412124 |
| Slc4a2 | 9.73009603 | 10.11651166 | 9.903584815 | 0.212926848 | -0.173488785 |
| Jph4 | 3.822990116 | 3.949299392 | 3.99651607 | -0.047216678 | -0.173525955 |
| Golga4 | 11.94894428 | 12.09531166 | 12.12247859 | -0.027166927 | -0.173534313 |
| Acad9 | 11.92142135 | 11.93742181 | 12.09495588 | -0.15753407 | -0.173534533 |
| Cep44 | 7.616177051 | 7.523550009 | 7.7898265 | -0.26627649 | -0.173649448 |
| Rapgef2 | 8.985858397 | 9.070456012 | 9.159517529 | -0.089061516 | -0.173659132 |
| Tlk1 | 10.22759171 | 10.33256811 | 10.40129966 | -0.068731553 | -0.17370795 |
| Gbgt1 | 4.673923737 | 4.63433801 | 4.847638611 | -0.2133006 | -0.173714874 |
| Slc25a12 | 10.3493357 | 10.34179783 | 10.52305528 | -0.18125745 | -0.17371958 |
| Trim67 | 3.869304558 | 3.884890888 | 4.043067226 | -0.158176338 | -0.173762668 |
| Zfp551 | 4.859337509 | 4.915857783 | 5.033161976 | -0.117304193 | -0.173824467 |
| Saxo2 | 3.677391127 | 3.708998664 | 3.85125383 | -0.142255167 | -0.173862704 |
| Castor1 | 7.636918778 | 7.872165932 | 7.810819113 | 0.061346819 | -0.173900334 |
| Oplah | 10.20966799 | 10.36218172 | 10.3837212 | -0.02153948 | -0.17405321 |
| Gm10036 | 3.763821447 | 3.969719762 | 3.937896167 | 0.031823595 | -0.17407472 |
| Msx1 | 3.794262551 | 3.859543757 | 3.968411903 | -0.108868146 | -0.174149352 |
| Cx3cl1 | 8.184820044 | 8.305509432 | 8.359023947 | -0.053514515 | -0.174203904 |
| Frmd4b | 10.50531612 | 10.52061241 | 10.67961666 | -0.15900425 | -0.174300533 |
| Rhobtb2 | 8.263072399 | 8.451311155 | 8.437389155 | 0.013922 | -0.174316756 |
| Grk4 | 5.993769429 | 5.847611113 | 6.168129922 | -0.320518809 | -0.174360493 |
| Tnnt1 | 4.103120432 | 4.120142054 | 4.277587865 | -0.157445811 | -0.174467433 |
| Casc4 | 7.542757397 | 7.462145634 | 7.717312586 | -0.255166952 | -0.174555188 |
| Apc2 | 3.18657305 | 3.242718011 | 3.361164315 | -0.118446304 | -0.174591265 |
| Lce1a1 | 12.2379961 | 12.49286569 | 12.41263893 | 0.08022676 | -0.174642833 |
| Cldn19 | 6.607170318 | 6.858149087 | 6.78186476 | 0.076284327 | -0.174694442 |
| Zfp354b | 3.860104799 | 3.928041117 | 4.034824228 | -0.106783111 | -0.174719429 |
| Gm14562 | 2.411177209 | 2.540655713 | 2.585924236 | -0.045268524 | -0.174747027 |
| Zfp219 | 9.086069615 | 9.498648465 | 9.260855125 | 0.23779334 | -0.17478551 |
| Sptbn1 | 12.87376299 | 12.81260327 | 13.04861653 | -0.236013257 | -0.174853533 |

| | | | | | |
|---|---|---|---|---|---|
| Herpud1 | 10.64385925 | 11.11741247 | 10.81873195 | 0.29868052 | -0.1748727 |
| Syt9 | 4.255989373 | 4.407005833 | 4.430866508 | -0.023860675 | -0.174877135 |
| Bmper | 6.611081297 | 6.335866639 | 6.785968401 | -0.450101762 | -0.174887104 |
| Dirc2 | 9.142501147 | 9.388180257 | 9.317458323 | 0.070721934 | -0.174957176 |
| Cenpe | 8.389290587 | 8.44424148 | 8.564252987 | -0.120011507 | -0.1749624 |
| Opa1 | 10.24721341 | 10.31211445 | 10.42220989 | -0.11009544 | -0.174996483 |
| Krtap3-2 | 6.690908373 | 6.800589545 | 6.86593028 | -0.065340735 | -0.175021906 |
| Susd2 | 12.29922409 | 12.4812901 | 12.47426837 | 0.007021733 | -0.17504428 |
| Pbk | 7.72214859 | 7.720805506 | 7.897199074 | -0.176393567 | -0.175050484 |
| Fndc5 | 6.552741786 | 6.690103734 | 6.727799638 | -0.037695905 | -0.175057853 |
| Mrps36 | 8.351540162 | 8.433332975 | 8.527214335 | -0.093881359 | -0.175674172 |
| Atg14 | 8.272099915 | 8.35287208 | 8.447779651 | -0.094907571 | -0.175679736 |
| Idi2 | 4.831673406 | 4.788972237 | 5.00738626 | -0.218414023 | -0.175712854 |
| Agbl5 | 6.588034476 | 6.891604657 | 6.763783883 | 0.127820774 | -0.175749407 |
| Crim1 | 10.14887661 | 10.35219312 | 10.32472516 | 0.027467957 | -0.175848553 |
| Rassf6 | 5.664299763 | 5.864258242 | 5.840167458 | 0.024090784 | -0.175867696 |
| Nol3 | 7.984102377 | 8.05839942 | 8.159984262 | -0.101584843 | -0.175881885 |
| Klhl5 | 9.371990206 | 9.355152931 | 9.548013757 | -0.192860825 | -0.176023551 |
| Nsmce4a | 9.56827897 | 9.587389961 | 9.744311304 | -0.156921343 | -0.176032333 |
| Hal | 12.40156457 | 12.44753695 | 12.57760186 | -0.13006491 | -0.176037297 |
| Tubgcp5 | 8.358201851 | 8.388448312 | 8.534269721 | -0.145821409 | -0.17606787 |
| Tceal5 | 5.036390898 | 5.140692335 | 5.21249154 | -0.071799205 | -0.176100642 |
| Cbx5 | 10.73867437 | 10.91158487 | 10.91492998 | -0.003345107 | -0.176255607 |
| Ptprr | 3.907576641 | 3.940134043 | 4.083838517 | -0.143704474 | -0.176261876 |
| Cdr2 | 10.16333463 | 10.2448286 | 10.33967886 | -0.09485026 | -0.176344227 |
| Tmprss7 | 4.173695237 | 4.397191008 | 4.350152339 | 0.047038669 | -0.176457101 |
| Hook1 | 9.54807657 | 9.607276171 | 9.724612967 | -0.117336796 | -0.176536397 |
| Crlf1 | 8.640799662 | 8.680943102 | 8.817390641 | -0.136447539 | -0.176590979 |
| Ntm | 4.266865536 | 4.319035436 | 4.443462418 | -0.124426982 | -0.176596882 |
| Zfp82 | 5.995688934 | 6.159820122 | 6.172299058 | -0.012478935 | -0.176610123 |
| Abhd5 | 11.00962036 | 11.0380042 | 11.18634974 | -0.148345543 | -0.176729377 |
| Usp45 | 8.549011473 | 8.666586946 | 8.725808207 | -0.059221261 | -0.176796734 |

| | | | | | |
|---|---|---|---|---|---|
| Ccdc66 | 7.458833931 | 7.588819379 | 7.635775035 | -0.046955656 | -0.176941105 |
| Uck1 | 9.181323107 | 9.354783956 | 9.358274823 | -0.003490867 | -0.176951716 |
| Glg1 | 11.41324336 | 11.55286641 | 11.59020773 | -0.03734132 | -0.176964367 |
| Deaf1 | 8.673864594 | 8.828583101 | 8.851079366 | -0.022496264 | -0.177214771 |
| Rint1 | 8.838139401 | 9.128429223 | 9.015368357 | 0.113060866 | -0.177228956 |
| Msantd2 | 7.737210281 | 7.845061698 | 7.914492146 | -0.069430448 | -0.177281865 |
| Gm10146 | 2.766449259 | 2.659382043 | 2.943736662 | -0.284354619 | -0.177287403 |
| Dynlt1a | 6.22103421 | 6.202177652 | 6.398459308 | -0.196281656 | -0.177425097 |
| Ntsr2 | 3.19276713 | 3.170342153 | 3.370195696 | -0.199853543 | -0.177428566 |
| Taok1 | 11.60928609 | 11.64416218 | 11.78673019 | -0.142568007 | -0.177444103 |
| Rad51ap1 | 7.031147155 | 7.132330087 | 7.208652665 | -0.076322577 | -0.177505509 |
| Fbxl17 | 8.892758384 | 8.893052735 | 9.07033268 | -0.177279946 | -0.177574296 |
| Cep68 | 8.578583789 | 8.591240991 | 8.756210571 | -0.16496958 | -0.177626781 |
| Ltbp3 | 9.982716227 | 10.40641708 | 10.1603608 | 0.24605628 | -0.177644576 |
| Rhbdl1 | 5.594436627 | 5.868343697 | 5.772086324 | 0.096257374 | -0.177649697 |
| 5730409E04Rik | 8.967594063 | 9.030364985 | 9.145423814 | -0.115058829 | -0.177829751 |
| Zfp784 | 5.524336804 | 5.580473118 | 5.702170786 | -0.121697668 | -0.177833982 |
| Mtmr14 | 8.500053901 | 8.600663297 | 8.677910877 | -0.077247581 | -0.177856976 |
| Taf9b | 7.660832441 | 7.585325953 | 7.838693396 | -0.253367444 | -0.177860955 |
| Bbs1 | 6.663070368 | 6.687773259 | 6.840967838 | -0.15319458 | -0.17789747 |
| Gm6472 | 11.78472078 | 11.57118599 | 11.96270012 | -0.39151413 | -0.177979333 |
| Klhl26 | 7.936478747 | 8.057447815 | 8.114496755 | -0.05704894 | -0.178018008 |
| Sprr2d | 6.987822186 | 7.355831946 | 7.165853858 | 0.189978088 | -0.178031672 |
| Foxm1 | 8.179410443 | 8.340773048 | 8.357539873 | -0.016766825 | -0.178129431 |
| Cryzl1 | 8.454045258 | 8.522048055 | 8.632219845 | -0.11017179 | -0.178174587 |
| Klhl9 | 10.22813384 | 10.29089336 | 10.40633867 | -0.115445317 | -0.17820483 |
| Zdhhc15 | 6.273465224 | 6.255022333 | 6.45172069 | -0.196698357 | -0.178255466 |
| Fbn1 | 11.39954257 | 11.64060577 | 11.57784132 | 0.062764443 | -0.178298753 |
| Chmp4c | 9.991018111 | 10.0233123 | 10.16932991 | -0.146017604 | -0.178311796 |
| Zfc3h1 | 10.38697699 | 10.64882384 | 10.56529475 | 0.083529087 | -0.178317763 |
| Gm20518 | 3.528247151 | 3.743213951 | 3.706581722 | 0.036632229 | -0.178334571 |
| Tmem110 | 8.608832453 | 8.727265579 | 8.787173554 | -0.059907975 | -0.178341101 |

| | | | | | |
|---|---|---|---|---|---|
| Hibadh | 10.59449132 | 10.54163954 | 10.77285271 | -0.23121317 | -0.178361387 |
| Zbtb37 | 8.850304369 | 8.83776323 | 9.028666369 | -0.190903139 | -0.178362 |
| Cacna1h | 3.369095761 | 3.498961511 | 3.547458342 | -0.048496831 | -0.178362581 |
| Fbxo9 | 9.544599541 | 9.658418502 | 9.723164968 | -0.064746466 | -0.178565427 |
| Maf1 | 9.699035142 | 9.787242922 | 9.877627875 | -0.090384954 | -0.178592734 |
| Armc2 | 4.498537519 | 4.774143555 | 4.677209932 | 0.096933623 | -0.178672413 |
| Txnrd2 | 6.267298089 | 6.228501122 | 6.446015155 | -0.217514033 | -0.178717066 |
| Commd3 | 9.539626953 | 9.659677163 | 9.718347616 | -0.058670453 | -0.178720663 |
| Drp2 | 8.649663784 | 8.632219079 | 8.828413337 | -0.196194258 | -0.178749552 |
| Zfp13 | 5.239015509 | 5.378415003 | 5.417767082 | -0.03935208 | -0.178751574 |
| Slc25a4 | 12.64886176 | 12.69087487 | 12.8276193 | -0.13674443 | -0.178757547 |
| Hspb8 | 11.68087471 | 13.32554156 | 11.85975119 | 1.465790373 | -0.178876477 |
| Mcu | 8.969390774 | 8.955864675 | 9.148327429 | -0.192462754 | -0.178936656 |
| 2810006K23Rik | 7.096808228 | 7.299471638 | 7.275764924 | 0.023706714 | -0.178956696 |
| Bicdl1 | 9.473157502 | 9.855685846 | 9.652116298 | 0.203569548 | -0.178958796 |
| 2610044O15Rik8 | 8.875766009 | 8.953764788 | 9.054849361 | -0.101084573 | -0.179083352 |
| Setbp1 | 8.191404776 | 8.154636302 | 8.370489175 | -0.215852873 | -0.179084399 |
| Sall2 | 6.770797263 | 6.815913951 | 6.94992329 | -0.134009339 | -0.179126026 |
| Tmem136 | 4.715010182 | 4.764754982 | 4.894174765 | -0.129419783 | -0.179164583 |
| Dgki | 4.45859771 | 4.613389212 | 4.637894234 | -0.024505022 | -0.179296524 |
| Maf | 11.44310732 | 11.48420526 | 11.62270482 | -0.138499567 | -0.179597507 |
| Rtl5 | 6.962211737 | 6.674796612 | 7.141834042 | -0.46703743 | -0.179622305 |
| Gna15 | 8.619260927 | 8.682264574 | 8.798915069 | -0.116650495 | -0.179654142 |
| Tstd3 | 6.432979581 | 6.379230433 | 6.612762409 | -0.233531975 | -0.179782827 |
| Qpct | 7.914479872 | 7.707089122 | 8.094350534 | -0.387261412 | -0.179870662 |
| Abhd3 | 7.612986483 | 7.701183223 | 7.792866543 | -0.09168332 | -0.17988006 |
| Retreg2 | 10.58281793 | 10.7105299 | 10.76275177 | -0.05222187 | -0.179933837 |
| Ak3 | 11.385756 | 11.36654303 | 11.56576569 | -0.199222657 | -0.180009687 |
| Echdc1 | 11.66032332 | 11.69879138 | 11.84039728 | -0.141605907 | -0.180073963 |
| Rab27b | 9.531739558 | 9.552934817 | 9.711875607 | -0.15894079 | -0.180136049 |
| Fam3c | 9.099808075 | 9.046187586 | 9.27996173 | -0.233774145 | -0.180153655 |
| Gm5643 | 2.590489181 | 2.721550831 | 2.770665551 | -0.04911472 | -0.18017637 |

| | | | | | |
|---|---|---|---|---|---|
| Haghl | 6.933885196 | 7.085124453 | 7.114067462 | -0.02894301 | -0.180182267 |
| Cutc | 7.561382936 | 7.585045009 | 7.741656367 | -0.156611359 | -0.180273432 |
| Dab1 | 8.708614788 | 8.612330802 | 8.888904638 | -0.276573835 | -0.18028985 |
| Clip4 | 11.77082002 | 11.73724501 | 11.95120972 | -0.21396471 | -0.180389703 |
| Gm3604 | 4.934388237 | 5.049664363 | 5.114881607 | -0.065217243 | -0.180493369 |
| Skida1 | 5.342274134 | 5.751525277 | 5.522775831 | 0.228749445 | -0.180501697 |
| Pdp2 | 7.650673566 | 7.672253557 | 7.831193174 | -0.158939617 | -0.180519608 |
| Msmo1 | 12.42151056 | 12.41766743 | 12.60206584 | -0.184398407 | -0.180555277 |
| Auh | 8.753147053 | 8.686214135 | 8.933725804 | -0.247511669 | -0.180578752 |
| Vgll3 | 10.57044497 | 10.58470427 | 10.75107937 | -0.166375097 | -0.180634403 |
| Fzd6 | 8.991045923 | 9.155694927 | 9.171742831 | -0.016047904 | -0.180696908 |
| Timm21 | 8.347818062 | 8.420068199 | 8.528533939 | -0.10846574 | -0.180715878 |
| Osbpl2 | 9.87208707 | 9.942039463 | 10.05285735 | -0.110817884 | -0.180770277 |
| Capn9 | 5.210838378 | 5.232134796 | 5.391742499 | -0.159607703 | -0.180904121 |
| Thap3 | 7.342158713 | 7.615927381 | 7.52306833 | 0.09285905 | -0.180909617 |
| Fbxw17 | 7.523233865 | 7.35872072 | 7.704147922 | -0.345427202 | -0.180914057 |
| Mpped2 | 3.848481113 | 3.909368122 | 4.029398979 | -0.120030857 | -0.180917866 |
| Cdh24 | 5.845710339 | 6.244129349 | 6.026666996 | 0.217462353 | -0.180956657 |
| Sertad2 | 9.951015699 | 10.06156486 | 10.13197612 | -0.07041126 | -0.180960424 |
| Rimklb | 3.391453236 | 3.674679465 | 3.572429088 | 0.102250378 | -0.180975852 |
| Pcgf2 | 7.965174587 | 8.359688998 | 8.146258483 | 0.213430516 | -0.181083895 |
| Dkkl1 | 9.450148206 | 9.505440727 | 9.631263574 | -0.125822847 | -0.181115368 |
| Aloxe3 | 10.10754444 | 10.48924816 | 10.28866472 | 0.20058344 | -0.181120286 |
| Arid2 | 10.20332721 | 10.35700859 | 10.3844481 | -0.02743951 | -0.181120883 |
| Cox4i2 | 4.449098204 | 4.548422586 | 4.630274015 | -0.081851429 | -0.181175811 |
| Trim33 | 9.715137842 | 9.826755333 | 9.896315484 | -0.069560151 | -0.181177642 |
| Sprr2e | 5.553612599 | 5.713456819 | 5.734821076 | -0.021364257 | -0.181208477 |
| Dmpk | 10.07069959 | 10.28951646 | 10.25196892 | 0.03754754 | -0.18126933 |
| Zfp619 | 5.801293975 | 5.884250177 | 5.982600356 | -0.098350179 | -0.181306382 |
| Tsc2 | 10.39003277 | 10.47015259 | 10.57135022 | -0.101197633 | -0.18131745 |
| Efhc1 | 4.621850599 | 4.801091797 | 4.803256275 | -0.002164478 | -0.181405676 |
| Lrrc28 | 9.69602144 | 9.635226279 | 9.877519282 | -0.242293003 | -0.181497843 |

| | | | | | |
|---|---|---|---|---|---|
| Gm5915 | 2.511132222 | 2.606040599 | 2.692631193 | -0.086590595 | -0.181498971 |
| Gja4 | 6.019687912 | 6.01223895 | 6.201280158 | -0.189041208 | -0.181592246 |
| Mbtd1 | 9.804431966 | 9.806994885 | 9.986158058 | -0.179163172 | -0.181726091 |
| Gm4924 | 5.185038298 | 5.155416636 | 5.366804359 | -0.211387723 | -0.181766062 |
| Als2cr12 | 4.966299304 | 4.971534744 | 5.14810712 | -0.176572376 | -0.181807816 |
| Hoxb9 | 4.266420989 | 4.323165117 | 4.448286895 | -0.125121778 | -0.181865906 |
| Olfr78 | 3.326501087 | 3.371007624 | 3.508435754 | -0.137428131 | -0.181934668 |
| Akap7 | 7.778445616 | 7.634954676 | 7.960404007 | -0.325449332 | -0.181958391 |
| C77080 | 10.66735408 | 11.07373021 | 10.8493556 | 0.22437461 | -0.182001517 |
| Nkiras1 | 7.984615732 | 7.980070644 | 8.166648687 | -0.186578043 | -0.182032955 |
| Gm9864 | 7.159019279 | 7.362178208 | 7.341052465 | 0.021125743 | -0.182033186 |
| Fbxl18 | 8.49656369 | 8.840301405 | 8.678624271 | 0.161677135 | -0.182060581 |
| Slc19a2 | 9.301886736 | 9.460844486 | 9.483973033 | -0.023128547 | -0.182086298 |
| Lamb2 | 10.39209694 | 10.6186945 | 10.57418609 | 0.044508417 | -0.18208915 |
| Gjb5 | 10.30137451 | 10.27031421 | 10.48346869 | -0.21315448 | -0.18209418 |
| Large1 | 8.348771714 | 8.524941857 | 8.530878657 | -0.0059368 | -0.182106943 |
| Vit | 6.445967946 | 6.377084432 | 6.628167214 | -0.251082782 | -0.182199268 |
| Ppt2 | 8.804708895 | 8.998524 | 8.986919552 | 0.011604448 | -0.182210657 |
| Igfals | 6.398737654 | 6.249137229 | 6.580963645 | -0.331826417 | -0.182225991 |
| Ndfip1 | 10.884332 | 10.77946533 | 11.06658031 | -0.287114977 | -0.182248307 |
| Inca1 | 4.446988091 | 4.472675309 | 4.629255173 | -0.156579863 | -0.182267082 |
| Slc1a4 | 7.751374973 | 7.966880118 | 7.933705026 | 0.033175092 | -0.182330053 |
| Zfp993 | 6.064329488 | 6.376204242 | 6.246742268 | 0.129461975 | -0.18241278 |
| Fbxo36 | 5.79820738 | 5.852786199 | 5.980657734 | -0.127871535 | -0.182450354 |
| Nhsl1 | 10.4213134 | 10.61567887 | 10.60385085 | 0.011828017 | -0.18253745 |
| Ppm1a | 11.20076126 | 11.21365858 | 11.38337769 | -0.16971911 | -0.182616433 |
| Adgrf5 | 9.541767428 | 9.391461223 | 9.724465524 | -0.333004301 | -0.182698096 |
| Crocc | 6.214192887 | 6.486495164 | 6.396988167 | 0.089506997 | -0.18279528 |
| Dennd4c | 10.43859757 | 10.57680495 | 10.62143767 | -0.044632713 | -0.182840097 |
| Ptprt | 2.938881575 | 2.815194211 | 3.121763268 | -0.306569057 | -0.182881693 |
| Unc79 | 5.264798285 | 5.36423838 | 5.447739796 | -0.083501415 | -0.182941511 |
| Ffar4 | 4.782119388 | 4.81141308 | 4.965079213 | -0.153666133 | -0.182959825 |

| | | | | | |
|---|---|---|---|---|---|
| Gabpb2 | 10.77420003 | 11.00105483 | 10.95716772 | 0.043887107 | -0.182967687 |
| Lysmd2 | 7.765134695 | 7.745002453 | 7.948109625 | -0.203107172 | -0.18297493 |
| Nudt14 | 7.278236042 | 7.617133777 | 7.461319426 | 0.155814351 | -0.183083384 |
| Cyb5d2 | 7.183749134 | 7.185943061 | 7.366879977 | -0.180936916 | -0.183130843 |
| Nxn | 10.21455118 | 10.42171874 | 10.39769824 | 0.0240205 | -0.183147063 |
| Kcnn3 | 5.104083598 | 5.114476458 | 5.287343392 | -0.172866934 | -0.183259794 |
| Gm3912 | 5.471293983 | 5.504713145 | 5.654589143 | -0.149875998 | -0.18329516 |
| B4galt6 | 8.377817843 | 8.33416866 | 8.561124195 | -0.226955535 | -0.183306352 |
| BC034090 | 7.467625472 | 7.711852013 | 7.651015128 | 0.060836885 | -0.183389655 |
| Smg6 | 9.486087217 | 9.68925824 | 9.669521003 | 0.019737237 | -0.183433786 |
| Angpt4 | 3.507707863 | 3.64847429 | 3.691198774 | -0.042724483 | -0.183490911 |
| Slc25a51 | 10.47169776 | 10.5851728 | 10.6552086 | -0.070035803 | -0.183510837 |
| Golga2 | 10.97647992 | 11.17777272 | 11.16005984 | 0.01771288 | -0.183579923 |
| Cln6 | 9.331561561 | 9.299924327 | 9.51515463 | -0.215230302 | -0.183593068 |
| Mpst | 8.283055672 | 8.512033766 | 8.466700207 | 0.045333559 | -0.183644535 |
| Fam131a | 6.865019522 | 6.966901218 | 7.048674725 | -0.081773506 | -0.183655202 |
| Tkfc | 7.170292037 | 7.338609866 | 7.353965656 | -0.015355791 | -0.18367362 |
| Zcchc8 | 9.389579417 | 9.5548345 | 9.573326947 | -0.018492447 | -0.18374753 |
| Baz2a | 11.2399049 | 11.47035046 | 11.42368943 | 0.046661023 | -0.18378453 |
| Fdxacb1 | 7.119238624 | 7.001027254 | 7.303061479 | -0.302034225 | -0.183822855 |
| Cdk2 | 9.310861859 | 9.500341449 | 9.494823785 | 0.005517664 | -0.183961926 |
| Spag17 | 2.54166125 | 2.643866846 | 2.72563628 | -0.081769434 | -0.18397503 |
| Otud7b | 9.972173215 | 9.960367213 | 10.15616211 | -0.195794894 | -0.183988891 |
| Cysrt1 | 9.896852306 | 10.05942034 | 10.08089339 | -0.021473058 | -0.184041088 |
| Ate1 | 9.496657807 | 9.545565272 | 9.680710536 | -0.135145264 | -0.184052729 |
| 1700109H08Rik | 5.261794916 | 5.312467062 | 5.445897181 | -0.133430119 | -0.184102265 |
| Bmt2 | 8.731361205 | 8.814558312 | 8.915486686 | -0.100928374 | -0.18412548 |
| Cyb5a | 11.47708144 | 11.46222278 | 11.66124858 | -0.1990258 | -0.184167137 |
| Gm13194 | 3.136204898 | 3.241963143 | 3.320487329 | -0.078524186 | -0.184282431 |
| Kdm5d | 5.491476991 | 5.686683112 | 5.675797907 | 0.010885205 | -0.184320916 |
| Eef2k | 10.91104109 | 11.07321957 | 11.09539531 | -0.022175733 | -0.184354213 |
| Zfp438 | 5.528463561 | 5.589641192 | 5.712822191 | -0.123180999 | -0.18435863 |

| | | | | | |
|---|---|---|---|---|---|
| Iqcb1 | 6.878168034 | 6.864640587 | 7.062543701 | -0.197903114 | -0.184375667 |
| Dynlt3 | 11.47850934 | 11.47811098 | 11.66289464 | -0.18478366 | -0.184385297 |
| Slc35e2 | 8.499572395 | 8.6380258 | 8.683965906 | -0.045940106 | -0.184393511 |
| Vapb | 10.47902046 | 10.59236212 | 10.66341863 | -0.07105651 | -0.18439817 |
| Rfx7 | 9.892187103 | 10.04480056 | 10.07658859 | -0.03178803 | -0.184401483 |
| Gm17555 | 4.255549968 | 4.310882056 | 4.44002554 | -0.129143483 | -0.184475572 |
| Ccdc27 | 4.33674124 | 4.497619362 | 4.521316236 | -0.023696873 | -0.184574996 |
| Gm9958 | 5.087135375 | 5.16184477 | 5.271764432 | -0.109919662 | -0.184629057 |
| Hint2 | 7.68646749 | 7.681130643 | 7.871325097 | -0.190194455 | -0.184857607 |
| Trp53bp1 | 9.268564308 | 9.235747689 | 9.453605521 | -0.217857832 | -0.185041213 |
| Tspan7 | 9.835027993 | 9.844271619 | 10.02007289 | -0.175801273 | -0.185044899 |
| Zfp944 | 7.810136145 | 7.81831599 | 7.995264445 | -0.176948455 | -0.1851283 |
| Purg | 6.042514997 | 6.101379045 | 6.227672802 | -0.126293757 | -0.185157804 |
| Lmln | 6.270970233 | 6.337922661 | 6.456173282 | -0.118250621 | -0.18520305 |
| Tmem198b | 7.657687741 | 7.745247326 | 7.842936104 | -0.097688778 | -0.185248363 |
| Oxld1 | 5.482286019 | 5.604959683 | 5.667548195 | -0.062588513 | -0.185262176 |
| Tbc1d30 | 4.214193761 | 4.194545764 | 4.399470205 | -0.204924441 | -0.185276444 |
| Zfp975 | 6.422768278 | 6.409482106 | 6.60812993 | -0.198647823 | -0.185361651 |
| Dnajc18 | 8.32126365 | 8.303764367 | 8.506681181 | -0.202916815 | -0.185417531 |
| Egfl6 | 6.064145192 | 6.021694383 | 6.249753594 | -0.22805921 | -0.185608401 |
| Spred2 | 9.106140099 | 9.297967705 | 9.29193515 | 0.006032555 | -0.185795051 |
| Ankrd16 | 9.116027118 | 9.19691177 | 9.301827852 | -0.104916082 | -0.185800734 |
| Mpnd | 9.36771408 | 9.489542168 | 9.553589235 | -0.064047066 | -0.185875154 |
| Rev1 | 9.029433605 | 9.130011954 | 9.215348586 | -0.085336633 | -0.185914981 |
| Gtf3c3 | 8.773758017 | 8.750593867 | 8.959816664 | -0.209222797 | -0.186058647 |
| Mif4gd | 9.154367395 | 9.305748313 | 9.340447912 | -0.0346996 | -0.186080517 |
| Extl3 | 10.6613002 | 10.69428508 | 10.84738288 | -0.153097797 | -0.18608268 |
| Hebp1 | 8.071765195 | 8.047791593 | 8.257900353 | -0.21010876 | -0.186135158 |
| Fcor | 6.32050386 | 6.095361618 | 6.506651071 | -0.411289453 | -0.18614721 |
| Pcnx4 | 7.927624415 | 8.046743156 | 8.113830308 | -0.067087151 | -0.186205892 |
| Cep112 | 5.171814948 | 5.119789807 | 5.358202472 | -0.238412665 | -0.186387524 |
| Lin52 | 7.62375771 | 7.711912387 | 7.810198259 | -0.098285872 | -0.186440549 |

| | | | | | |
|---|---|---|---|---|---|
| Kcnc2 | 2.214598268 | 2.249002015 | 2.401119054 | -0.152117038 | -0.186520786 |
| Stard10 | 9.541604246 | 9.64610014 | 9.728191337 | -0.082091198 | -0.186587092 |
| Ephx3 | 10.10812015 | 10.38515861 | 10.29474364 | 0.090414973 | -0.18662349 |
| Otud1 | 9.472590656 | 10.31631772 | 9.659287603 | 0.65703012 | -0.186696947 |
| Pcx | 11.44922732 | 11.13408002 | 11.63596444 | -0.501884427 | -0.186737127 |
| Polr3g | 9.697778395 | 9.774330537 | 9.884521961 | -0.110191424 | -0.186743567 |
| Rmnd1 | 8.07919437 | 8.21314704 | 8.266006538 | -0.052859498 | -0.186812168 |
| Zfp317 | 8.645308805 | 8.650658531 | 8.83215407 | -0.181495539 | -0.186845265 |
| Ppp2r5d | 9.734428293 | 9.93583306 | 9.921294543 | 0.014538517 | -0.18686625 |
| Fam83b | 9.908034152 | 10.14118821 | 10.0949472 | 0.04624101 | -0.186913048 |
| Zfp825 | 7.77414303 | 7.905786701 | 7.961105874 | -0.055319172 | -0.186962843 |
| Ccpg1os | 6.011433734 | 6.088552878 | 6.198682843 | -0.110129965 | -0.187249109 |
| Olfml3 | 9.809680464 | 9.908410038 | 9.997080569 | -0.088670531 | -0.187400104 |
| Rictor | 9.696804024 | 9.790350997 | 9.884284975 | -0.093933978 | -0.187480951 |
| Phlpp1 | 8.756201433 | 8.961476925 | 8.943684558 | 0.017792367 | -0.187483126 |
| Ntrk2 | 10.11353995 | 9.934960668 | 10.3010417 | -0.366081029 | -0.187501743 |
| Cxcl13 | 6.078728213 | 5.504261605 | 6.266245714 | -0.761984108 | -0.187517501 |
| Mypop | 5.803487449 | 6.049488087 | 5.991131229 | 0.058356858 | -0.187643779 |
| Tcn2 | 9.819844164 | 9.640120152 | 10.00751129 | -0.367391139 | -0.187667127 |
| Zbtb10 | 7.094819779 | 7.239300713 | 7.282499525 | -0.043198813 | -0.187679746 |
| Ift52 | 8.790728349 | 8.963911804 | 8.978412027 | -0.014500223 | -0.187683678 |
| Cited2 | 9.525543073 | 9.566759246 | 9.713277245 | -0.146517999 | -0.187734172 |
| Fbxl19 | 9.274137755 | 9.711335815 | 9.461884142 | 0.249451673 | -0.187746386 |
| Megf10 | 3.788867314 | 3.777451054 | 3.97667979 | -0.199228735 | -0.187812475 |
| Smurf2 | 9.950069114 | 10.17227494 | 10.13788742 | 0.034387513 | -0.187818309 |
| Il1rl2 | 11.26745865 | 11.34352284 | 11.45544333 | -0.11192049 | -0.187984673 |
| Ctif | 8.312798489 | 8.540404429 | 8.500814762 | 0.039589667 | -0.188016273 |
| Wdyhv1 | 7.072733366 | 7.216773933 | 7.260754274 | -0.04398034 | -0.188020908 |
| Shisa3 | 3.769118545 | 3.733005127 | 3.957195925 | -0.224190798 | -0.18807738 |
| Fdft1 | 11.04874625 | 11.11191908 | 11.23682431 | -0.124905227 | -0.188078053 |
| Apool | 8.116007763 | 8.151930687 | 8.304126702 | -0.152196015 | -0.188118938 |
| Ncoa3 | 10.45293763 | 10.80568221 | 10.64112087 | 0.16456134 | -0.188183233 |

| | | | | | |
|---|---|---|---|---|---|
| Magee1 | 6.844994331 | 6.992737735 | 7.03327412 | -0.040536385 | -0.188279789 |
| Myo9a | 10.26619816 | 10.44449103 | 10.45459517 | -0.010104137 | -0.188397003 |
| Hoxb8 | 5.498894234 | 5.687486289 | 5.687299311 | 0.000186977 | -0.188405077 |
| Kctd2 | 8.869152568 | 8.872521341 | 9.057662534 | -0.185141193 | -0.188509966 |
| Zfp982 | 4.40376864 | 4.480174717 | 4.5922824 | -0.112107683 | -0.18851376 |
| Notch4 | 6.676211265 | 6.958092576 | 6.864750738 | 0.093341837 | -0.188539474 |
| Dnajc10 | 10.11800683 | 10.12276128 | 10.30657686 | -0.18381558 | -0.188570027 |
| Zfp938 | 6.198538079 | 6.24641386 | 6.387135374 | -0.140721514 | -0.188597294 |
| Rnf165 | 6.466497207 | 6.751965059 | 6.655259787 | 0.096705272 | -0.18876258 |
| Gpr135 | 4.209322707 | 4.286601259 | 4.398092066 | -0.111490807 | -0.188769358 |
| Pccb | 9.98985015 | 9.892424504 | 10.17870052 | -0.286276019 | -0.188850374 |
| Ccdc120 | 9.598250026 | 9.699410512 | 9.787293994 | -0.087883482 | -0.189043968 |
| Gm13306 | 9.240780808 | 9.740702282 | 9.42983439 | 0.310867892 | -0.189053582 |
| Gstp2 | 6.747500993 | 7.039557384 | 6.936576244 | 0.10298114 | -0.189075251 |
| Ear2 | 8.192439752 | 8.066614932 | 8.381636495 | -0.315021563 | -0.189196743 |
| St6galnac5 | 8.370615426 | 8.264011103 | 8.559818382 | -0.295807279 | -0.189202956 |
| Tmem35a | 6.220423428 | 6.18841968 | 6.409668359 | -0.221248679 | -0.189244931 |
| Fndc7 | 3.882712997 | 3.961724695 | 4.072065955 | -0.11034126 | -0.189352957 |
| Tex264 | 10.81341966 | 10.91212482 | 11.00278456 | -0.090659747 | -0.189364903 |
| Mmaa | 8.371391807 | 8.497049109 | 8.560779527 | -0.063730418 | -0.18938772 |
| Rita1 | 6.279754236 | 6.58132198 | 6.469196823 | 0.112125157 | -0.189442587 |
| Ripor3 | 7.403260201 | 7.914648303 | 7.592717569 | 0.321930734 | -0.189457369 |
| Serinc1 | 11.46660896 | 11.47722014 | 11.65612549 | -0.178905347 | -0.18951653 |
| Cep295 | 9.000746119 | 9.091857247 | 9.19028121 | -0.098423963 | -0.189535091 |
| Adarb1 | 8.922542565 | 9.053746141 | 9.112090457 | -0.058344316 | -0.189547892 |
| Fam122b | 7.009626367 | 6.988703301 | 7.199216271 | -0.21051297 | -0.189589904 |
| Usp33 | 9.806854363 | 10.04236111 | 9.996464586 | 0.04589652 | -0.189610223 |
| Usp46 | 9.505999269 | 9.706572086 | 9.695621851 | 0.010950235 | -0.189622582 |
| Ttc12 | 5.347218718 | 5.509486715 | 5.536862107 | -0.027375392 | -0.18964339 |
| Ccnf | 7.785641146 | 7.987827711 | 7.975326112 | 0.012501598 | -0.189684967 |
| E130311K13Rik | 6.488008732 | 6.488308768 | 6.677702201 | -0.189393433 | -0.189693469 |
| Bcat2 | 10.46237958 | 10.55473168 | 10.65211758 | -0.0973859 | -0.189738003 |

| | | | | | |
|---|---|---|---|---|---|
| Tmem19 | 9.920129727 | 10.05328247 | 10.10988092 | -0.056598452 | -0.189751197 |
| Rab11fip3 | 9.209798213 | 9.412663155 | 9.39955584 | 0.013107315 | -0.189757628 |
| Ntpcr | 7.489760195 | 7.641011414 | 7.679574441 | -0.038563027 | -0.189814246 |
| Mybph | 9.397528064 | 9.714393999 | 9.587475887 | 0.126918112 | -0.189947823 |
| Zmat1 | 6.048718186 | 6.129165216 | 6.238783808 | -0.109618592 | -0.190065622 |
| Sumf1 | 9.113306833 | 9.196464953 | 9.303532661 | -0.107067707 | -0.190225828 |
| Etfrf1 | 8.607931778 | 8.579149254 | 8.798253354 | -0.2191041 | -0.190321575 |
| Ly6g6c | 12.90980718 | 13.04669917 | 13.10013533 | -0.053436157 | -0.190328143 |
| Ppp1r21 | 9.200946736 | 9.293031409 | 9.391330361 | -0.098298952 | -0.190383626 |
| Slc15a1 | 8.76086626 | 8.738282649 | 8.951257688 | -0.212975039 | -0.190391427 |
| Nim1k | 4.879288591 | 4.881948061 | 5.069691973 | -0.187743912 | -0.190403382 |
| Rcbtb1 | 8.414475256 | 8.748725959 | 8.604950521 | 0.143775438 | -0.190475265 |
| Cdkal1 | 7.883875064 | 7.84154428 | 8.074359772 | -0.232815493 | -0.190484709 |
| Hsbp1l1 | 6.91662968 | 6.925742229 | 7.107154507 | -0.181412278 | -0.190524827 |
| Gm20075 | 3.850021349 | 4.259415783 | 4.040558771 | 0.218857012 | -0.190537422 |
| Sema4f | 5.305741208 | 5.470344315 | 5.496305179 | -0.025960864 | -0.190563971 |
| Cngb1 | 2.852712658 | 3.03064847 | 3.043317337 | -0.012668867 | -0.190604678 |
| Lmbrd2 | 10.12997317 | 10.13980394 | 10.32061072 | -0.180806783 | -0.190637557 |
| Chst15 | 9.374306328 | 9.50978001 | 9.565051273 | -0.055271263 | -0.190744945 |
| Slc25a23 | 8.28058348 | 8.251257372 | 8.471333707 | -0.220076334 | -0.190750226 |
| Best2 | 4.498246681 | 4.707247799 | 4.688999378 | 0.018248421 | -0.190752697 |
| Nanp | 7.485121907 | 7.41385628 | 7.67589752 | -0.26204124 | -0.190775613 |
| Mis18bp1 | 6.772488483 | 6.699517066 | 6.963271337 | -0.263754271 | -0.190782854 |
| Ppp4r4 | 4.554415694 | 4.583005382 | 4.745230929 | -0.162225548 | -0.190815235 |
| Dclk3 | 4.143897926 | 4.206547484 | 4.334718542 | -0.128171058 | -0.190820616 |
| Sft2d2 | 11.14979785 | 11.10872053 | 11.34065905 | -0.231938517 | -0.1908612 |
| Nlgn3 | 4.860515508 | 4.832045164 | 5.051377948 | -0.219332784 | -0.19086244 |
| Chst14 | 7.644770405 | 7.78328803 | 7.835645833 | -0.052357803 | -0.190875428 |
| Zfp398 | 8.597643367 | 8.815402262 | 8.78853428 | 0.026867982 | -0.190890913 |
| Mamstr | 6.103720411 | 6.164283258 | 6.294643479 | -0.130360222 | -0.190923068 |
| Tob2 | 10.1563451 | 10.33938852 | 10.34737376 | -0.007985237 | -0.19102866 |
| Tmem59 | 11.61054429 | 11.58306856 | 11.80170966 | -0.2186411 | -0.191165377 |

| | | | | | |
|---|---|---|---|---|---|
| Pla2g2e | 6.498238957 | 6.823945193 | 6.689470654 | 0.134474539 | -0.191231697 |
| Gm5210 | 3.032760362 | 3.224669434 | 3.224025405 | 0.000644029 | -0.191265043 |
| Nt5c2 | 9.542116891 | 9.616467614 | 9.733386661 | -0.116919046 | -0.19126977 |
| Hmg20b | 9.058411895 | 9.324631638 | 9.249714751 | 0.074916887 | -0.191302856 |
| Perp | 15.27863857 | 15.32113295 | 15.47013441 | -0.149001463 | -0.191495837 |
| Ciart | 7.599752299 | 7.617197155 | 7.791344206 | -0.174147051 | -0.191591907 |
| Zbtb43 | 9.128733022 | 9.315207098 | 9.320404634 | -0.005197536 | -0.191671612 |
| Ip6k2 | 8.448752696 | 8.532552524 | 8.640537044 | -0.107984519 | -0.191784348 |
| Bloc1s3 | 7.880941716 | 8.069109581 | 8.072831943 | -0.003722362 | -0.191890227 |
| Rnf181 | 8.963497558 | 9.177017895 | 9.155387952 | 0.021629944 | -0.191890394 |
| Il6ra | 11.28698521 | 11.55984597 | 11.47888622 | 0.080959747 | -0.191901017 |
| Zfp277 | 8.599625335 | 8.729070747 | 8.79153261 | -0.062461863 | -0.191907275 |
| Atrnl1 | 10.28295146 | 10.24856683 | 10.47489484 | -0.22632801 | -0.19194338 |
| Gm37116 | 2.434674864 | 2.584741066 | 2.62677773 | -0.042036664 | -0.192102866 |
| Tnfsf15 | 4.379929823 | 4.464207234 | 4.572081599 | -0.107874365 | -0.192151776 |
| Rabgap1l | 9.963679529 | 9.822875143 | 10.15585499 | -0.332979843 | -0.192175458 |
| Nt5c3b | 8.525455719 | 8.715246249 | 8.717759394 | -0.002513146 | -0.192303676 |
| Prokr1 | 4.471465531 | 4.581156986 | 4.663857509 | -0.082700523 | -0.192391978 |
| 1700088E04Rik | 3.663833632 | 3.755715725 | 3.856235141 | -0.100519417 | -0.19240151 |
| Pawr | 7.919710469 | 8.000585992 | 8.11218723 | -0.111601238 | -0.192476761 |
| Cldn15 | 6.11543297 | 6.152402794 | 6.307938111 | -0.155535317 | -0.192505141 |
| Gsg1l | 6.344657134 | 6.468127804 | 6.537206515 | -0.06907871 | -0.192549381 |
| Clec4b1 | 5.772736969 | 5.952043538 | 5.965314086 | -0.013270548 | -0.192577117 |
| Ldlrad3 | 8.120387623 | 8.254674542 | 8.313025837 | -0.058351295 | -0.192638214 |
| Nhlrc2 | 9.479835961 | 9.621374543 | 9.672515522 | -0.051140979 | -0.192679561 |
| Glt8d1 | 8.212621663 | 8.286983772 | 8.405485643 | -0.118501872 | -0.192863981 |
| Pcdh10 | 4.431857383 | 4.570288549 | 4.624739105 | -0.054450556 | -0.192881722 |
| Fam187b | 5.346391749 | 5.512455537 | 5.5392899 | -0.026834363 | -0.19289815 |
| Fancm | 7.360325547 | 7.500183519 | 7.553308238 | -0.053124718 | -0.19298269 |
| Khk | 7.267523073 | 7.417918673 | 7.460567237 | -0.042648565 | -0.193044165 |
| Spaar | 5.239017243 | 5.279716295 | 5.432098444 | -0.152382149 | -0.1930812 |
| Agtr2 | 2.355609444 | 2.349287341 | 2.548718907 | -0.199431566 | -0.193109463 |

| | | | | | |
|---|---|---|---|---|---|
| Pter | 7.542736699 | 7.483707483 | 7.735921878 | -0.252214394 | -0.193185179 |
| Fam227a | 5.127446163 | 5.279071576 | 5.320638096 | -0.041566519 | -0.193191932 |
| Glb1l | 7.011302793 | 7.092561043 | 7.204681567 | -0.112120524 | -0.193378774 |
| Pygm | 12.23833656 | 12.36342242 | 12.43180055 | -0.06837813 | -0.193463993 |
| Heg1 | 10.27694893 | 10.27503085 | 10.47047652 | -0.19544567 | -0.193527598 |
| Best3 | 3.193084511 | 3.181057079 | 3.386725514 | -0.205668435 | -0.193641002 |
| Kctd14 | 5.357525474 | 5.494365065 | 5.551202145 | -0.056837081 | -0.193676672 |
| Ankdd1a | 4.149139984 | 4.340610864 | 4.342878753 | -0.002267889 | -0.193738769 |
| Lurap1 | 6.75932448 | 6.718342946 | 6.953071349 | -0.234728403 | -0.193746869 |
| Gsta1 | 5.62751142 | 5.907739272 | 5.821325639 | 0.086413634 | -0.193814219 |
| Fzd1 | 10.24452793 | 10.32269021 | 10.43834564 | -0.115655427 | -0.193817707 |
| Xpnpep3 | 8.281877808 | 8.577306499 | 8.475746433 | 0.101560066 | -0.193868625 |
| Specc1l | 10.03441759 | 10.17605293 | 10.22840275 | -0.052349827 | -0.193985163 |
| Dync1li2 | 11.00247279 | 11.09063138 | 11.19645928 | -0.1058279 | -0.19398649 |
| Paip2b | 10.08180521 | 10.19612366 | 10.27583191 | -0.07970825 | -0.194026693 |
| Phc3 | 10.04893985 | 10.19712016 | 10.24299332 | -0.045873157 | -0.194053467 |
| Srek1ip1 | 8.317756254 | 8.212821341 | 8.511917539 | -0.299096199 | -0.194161286 |
| Slu7 | 9.731727756 | 9.761727703 | 9.925890801 | -0.164163098 | -0.194163045 |
| Ptpn3 | 10.12778149 | 10.17849421 | 10.3219888 | -0.14349459 | -0.194207313 |
| Tmem242 | 9.141946637 | 9.14040351 | 9.33620875 | -0.19580524 | -0.194262114 |
| Csta1 | 10.42868894 | 10.39501914 | 10.62295585 | -0.227936703 | -0.194266907 |
| Carns1 | 6.125850762 | 6.240713615 | 6.320317518 | -0.079603902 | -0.194466755 |
| Gm49510 | 6.305155843 | 6.627509184 | 6.499734928 | 0.127774255 | -0.194579085 |
| Cntnap1 | 6.606618302 | 6.861448521 | 6.801211541 | 0.06023698 | -0.19459324 |
| 4930402H24Rik | 9.168931829 | 9.096484402 | 9.363525515 | -0.267041112 | -0.194593686 |
| 1700066M21Rik | 7.20841893 | 7.376938428 | 7.403015148 | -0.02607672 | -0.194596218 |
| Gm10644 | 3.437089749 | 3.550320926 | 3.63174593 | -0.081425004 | -0.194656182 |
| Mafa | 6.127203806 | 6.339362783 | 6.321994918 | 0.017367865 | -0.194791112 |
| Bach2 | 7.109842198 | 7.353344792 | 7.30484314 | 0.048501652 | -0.195000943 |
| Gdpd5 | 6.234811207 | 6.514190501 | 6.429846832 | 0.084343669 | -0.195035625 |
| Catsperg1 | 5.49292848 | 5.801217509 | 5.688189875 | 0.113027633 | -0.195261395 |
| Map3k1 | 10.56141412 | 10.72094287 | 10.7567061 | -0.03576323 | -0.19529198 |

| | | | | | |
|---|---|---|---|---|---|
| Supt4a | 8.810653593 | 8.802011585 | 9.005982019 | -0.203970435 | -0.195328426 |
| Gatad2b | 10.56331895 | 10.66858465 | 10.75867622 | -0.090091573 | -0.195357273 |
| 4833439L19Rik | 11.26257296 | 11.30824708 | 11.45798138 | -0.149734303 | -0.195408417 |
| Col4a3 | 5.41503825 | 5.323029403 | 5.61053699 | -0.287507587 | -0.19549874 |
| Stard7 | 10.87282773 | 10.87620873 | 11.06837957 | -0.19217084 | -0.195551837 |
| Slc8a3 | 7.025400028 | 6.961472021 | 7.220955218 | -0.259483197 | -0.195555191 |
| Trpc1 | 5.509095028 | 5.721662505 | 5.704807565 | 0.01685494 | -0.195712537 |
| Ift22 | 7.287113449 | 7.460208974 | 7.483092915 | -0.022883941 | -0.195979466 |
| Dusp23 | 7.386077356 | 7.411978704 | 7.582128273 | -0.170149568 | -0.196050917 |
| Acvr2b | 5.697104748 | 5.728623802 | 5.893201869 | -0.164578067 | -0.196097121 |
| Ptpdc1 | 7.466398281 | 7.608680994 | 7.662681797 | -0.054000803 | -0.196283516 |
| Ranbp6 | 8.669130157 | 8.754759417 | 8.865427847 | -0.11066843 | -0.19629769 |
| Aadac | 10.14215569 | 10.18736887 | 10.33848963 | -0.151120757 | -0.19633394 |
| Cadps | 4.166041628 | 4.304862357 | 4.362447176 | -0.05758482 | -0.196405548 |
| Snap91 | 7.086484554 | 7.046322766 | 7.282900769 | -0.236578003 | -0.196416216 |
| Zfp251 | 8.122577564 | 8.340640514 | 8.319077023 | 0.021563491 | -0.196499459 |
| Cpd | 11.66109312 | 11.72169138 | 11.8575988 | -0.135907413 | -0.19650568 |
| Adipor2 | 11.96553175 | 12.04780003 | 12.1621447 | -0.11434467 | -0.196612947 |
| Grb14 | 8.046930225 | 7.968414582 | 8.243601571 | -0.275186989 | -0.196671346 |
| Asnsd1 | 9.803137222 | 9.84145409 | 9.999852339 | -0.158398249 | -0.196715117 |
| Ids | 10.21578549 | 10.17834403 | 10.4126524 | -0.234308363 | -0.19686691 |
| Ubn2 | 10.83610861 | 11.10026398 | 11.03298744 | 0.067276533 | -0.196878837 |
| Nsmaf | 9.547100454 | 9.480089078 | 9.744011007 | -0.26392193 | -0.196910553 |
| Entpd2 | 8.411266558 | 8.596024378 | 8.608303629 | -0.012279251 | -0.197037071 |
| Lce1i | 10.04675068 | 10.53816273 | 10.24380034 | 0.29436239 | -0.197049663 |
| Slc25a38 | 8.387441381 | 8.684141913 | 8.58458009 | 0.099561823 | -0.197138709 |
| Akr1c18 | 10.95802214 | 11.21913984 | 11.15527736 | 0.06386248 | -0.197255223 |
| Nsdhl | 9.694159327 | 9.736562289 | 9.891442985 | -0.154880696 | -0.197283658 |
| Cep295nl | 4.836813404 | 4.995129314 | 5.034329927 | -0.039200613 | -0.197516523 |
| Rdh9 | 4.490890296 | 4.601307469 | 4.688415847 | -0.087108378 | -0.197525551 |
| Lrrc2 | 7.440458739 | 8.087806477 | 7.638045869 | 0.449760608 | -0.197587129 |
| Exoc4 | 9.817889871 | 9.797049733 | 10.01551219 | -0.218462457 | -0.197622319 |

| | | | | | |
|---|---|---|---|---|---|
| Zbtb1 | 9.653556854 | 9.689131191 | 9.851183603 | -0.162052412 | -0.197626748 |
| Ankrd46 | 9.347892598 | 9.401027637 | 9.545648378 | -0.144620741 | -0.19775578 |
| Ap3m2 | 7.51888574 | 7.517025017 | 7.716698732 | -0.199673715 | -0.197812992 |
| Trak1 | 11.1096768 | 11.23642451 | 11.30749735 | -0.071072843 | -0.197820547 |
| Snrnp25 | 7.535890648 | 7.616515375 | 7.733716871 | -0.117201497 | -0.197826223 |
| Wdhd1 | 8.496908147 | 8.541219673 | 8.694812009 | -0.153592336 | -0.197903861 |
| Dek | 11.16549741 | 11.1992711 | 11.36340225 | -0.16413115 | -0.19790484 |
| Akap8l | 8.918756119 | 9.177217454 | 9.116815221 | 0.060402233 | -0.198059103 |
| Slx4ip | 7.394919224 | 7.607409927 | 7.592996338 | 0.014413589 | -0.198077114 |
| Sncg | 9.917629217 | 9.631326555 | 10.11572635 | -0.484399797 | -0.198097136 |
| Ptp4a3 | 10.00313588 | 10.04515544 | 10.20126896 | -0.156113518 | -0.19813308 |
| Rtn3 | 12.11753724 | 12.16538059 | 12.31569406 | -0.15031347 | -0.198156823 |
| Gria3 | 5.391424505 | 5.510529453 | 5.589635812 | -0.079106359 | -0.198211306 |
| Cdc14a | 7.502274959 | 8.529146576 | 7.70049248 | 0.828654096 | -0.198217521 |
| Gm48111 | 5.885384678 | 6.201702673 | 6.08363275 | 0.118069924 | -0.198248071 |
| Odr4 | 9.306356551 | 9.20727007 | 9.504637153 | -0.297367083 | -0.198280602 |
| Pdik1l | 8.205048814 | 8.271415486 | 8.403437612 | -0.132022126 | -0.198388797 |
| Zfp976 | 6.376656367 | 6.459091658 | 6.575139759 | -0.116048102 | -0.198483392 |
| Dsp | 15.80494535 | 15.93734964 | 16.00349943 | -0.066149797 | -0.198554087 |
| Ppm1d | 8.944247074 | 9.128047088 | 9.142813206 | -0.014766118 | -0.198566132 |
| Sprr2a2 | 11.80269892 | 12.44946364 | 12.00128844 | 0.448175193 | -0.198589523 |
| Sfxn4 | 4.815366847 | 4.872593615 | 5.01401262 | -0.141419005 | -0.198645773 |
| Zmat3 | 9.882314236 | 10.09993884 | 10.08103347 | 0.018905367 | -0.198719234 |
| B4galnt3 | 6.974083868 | 7.151111512 | 7.172803157 | -0.021691644 | -0.198719289 |
| Slc17a5 | 7.720367838 | 7.786846313 | 7.919092827 | -0.132246515 | -0.19872499 |
| C2cd2 | 11.41313071 | 11.51496716 | 11.61185934 | -0.096892177 | -0.198728627 |
| Rabgap1 | 10.58352439 | 10.63447412 | 10.78237164 | -0.14789752 | -0.198847247 |
| Foxred2 | 7.946438849 | 7.882912135 | 8.145315991 | -0.262403856 | -0.198877142 |
| Sqle | 11.4486532 | 11.56818248 | 11.64753531 | -0.079352827 | -0.198882107 |
| Avl9 | 8.833976953 | 8.906780504 | 9.033072205 | -0.126291701 | -0.199095252 |
| Gm15335 | 2.699292426 | 2.791462057 | 2.898397718 | -0.10693566 | -0.199105291 |
| Dcn | 14.58544118 | 14.61096177 | 14.78456476 | -0.173602993 | -0.19912358 |

| | | | | | |
|---|---|---|---|---|---|
| Brd9 | 9.406029314 | 9.561358873 | 9.605171822 | -0.043812949 | -0.199142508 |
| Ckmt2 | 10.35199138 | 10.36343703 | 10.55117564 | -0.187738608 | -0.199184263 |
| 1810014B01Rik | 6.114718697 | 6.109363715 | 6.313918489 | -0.204554773 | -0.199199791 |
| 2700049A03Rik | 7.39195851 | 7.553093548 | 7.591283756 | -0.038190208 | -0.199325247 |
| Pde10a | 5.368497187 | 5.489985438 | 5.567892473 | -0.077907035 | -0.199395286 |
| Cdkn1c | 6.278952338 | 6.404552833 | 6.478348793 | -0.07379596 | -0.199396455 |
| Aco1 | 10.39031215 | 10.40464749 | 10.5897642 | -0.185116703 | -0.19945205 |
| Fkbp10 | 8.50590783 | 8.709282853 | 8.705366936 | 0.003915916 | -0.199459107 |
| Ralgapb | 10.66234081 | 10.82082445 | 10.86180861 | -0.040984163 | -0.199467797 |
| Mospd3 | 8.108093467 | 8.35830478 | 8.307678998 | 0.050625783 | -0.199585531 |
| Oip5 | 5.405691813 | 5.532998273 | 5.605308591 | -0.072310318 | -0.199616778 |
| Slc31a2 | 10.35628806 | 10.55372288 | 10.55591576 | -0.00219288 | -0.199627697 |
| Zfp942 | 7.923043769 | 8.084439783 | 8.122674424 | -0.038234641 | -0.199630655 |
| Efna4 | 7.986532623 | 8.202367069 | 8.186264707 | 0.016102362 | -0.199732084 |
| Gm14391 | 8.209847538 | 8.363952113 | 8.409627662 | -0.045675549 | -0.199780124 |
| Wdr54 | 4.849832911 | 4.895206913 | 5.049709751 | -0.154502838 | -0.19987684 |
| Dnase1l3 | 10.14098724 | 10.28539503 | 10.34090069 | -0.05550566 | -0.199913448 |
| Als2cl | 9.818714875 | 10.465292 | 10.01879045 | 0.44650155 | -0.200075572 |
| Ctsh | 10.72833986 | 10.82173276 | 10.92844681 | -0.10671405 | -0.200106953 |
| Sh3bgr | 9.236650724 | 9.261988021 | 9.436779744 | -0.174791724 | -0.20012902 |
| Eri3 | 8.844459484 | 8.905824684 | 9.044628921 | -0.138804237 | -0.200169437 |
| Gm9212 | 2.861764152 | 3.050014944 | 3.061948761 | -0.011933817 | -0.200184609 |
| Hist4h4 | 4.188870384 | 4.290302562 | 4.389134212 | -0.09883165 | -0.200263829 |
| Nudt6 | 5.937245632 | 6.007403897 | 6.137520713 | -0.130116816 | -0.200275081 |
| AI854703 | 5.694268809 | 6.031061789 | 5.894615756 | 0.136446033 | -0.200346947 |
| Dapl1 | 10.64465786 | 10.57296341 | 10.84517499 | -0.272211577 | -0.200517127 |
| Hexdc | 8.801552744 | 8.885976746 | 9.00207146 | -0.116094713 | -0.200518715 |
| Fam110b | 5.493308805 | 5.48846791 | 5.693835909 | -0.205367999 | -0.200527105 |
| Zfp316 | 7.303626047 | 7.309001573 | 7.504275254 | -0.195273681 | -0.200649207 |
| Casz1 | 11.0221464 | 11.35961488 | 11.22281798 | 0.1367969 | -0.200671587 |
| Nbdy | 7.717751555 | 7.717182026 | 7.918607533 | -0.201425507 | -0.200855978 |
| Gtpbp2 | 9.766112383 | 10.11988373 | 9.966999227 | 0.1528845 | -0.200886844 |

| | | | | | |
|---|---|---|---|---|---|
| Svip | 6.99743694 | 6.803927048 | 7.198453469 | -0.394526421 | -0.201016529 |
| Ube2g1 | 10.03100754 | 9.996038124 | 10.23207614 | -0.236038016 | -0.2010686 |
| Ncaph | 7.812991003 | 7.953792155 | 8.014182811 | -0.060390655 | -0.201191808 |
| Echs1 | 10.69656695 | 10.66597641 | 10.89787394 | -0.231897523 | -0.20130699 |
| Htatip2 | 9.118519996 | 9.146463167 | 9.319842201 | -0.173379034 | -0.201322205 |
| Golga7b | 5.170679372 | 5.390645573 | 5.372112741 | 0.018532833 | -0.201433368 |
| Nudt16 | 5.907940245 | 5.947396962 | 6.109528475 | -0.162131513 | -0.20158823 |
| Gnai1 | 10.90619146 | 10.8380624 | 11.10779393 | -0.269731527 | -0.201602467 |
| Plxna4 | 6.306883715 | 6.315075494 | 6.50853184 | -0.193456346 | -0.201648125 |
| Fundc2 | 10.08980451 | 9.97694984 | 10.29147002 | -0.314520183 | -0.20166551 |
| Rilp | 6.389936947 | 6.32128525 | 6.591624525 | -0.270339275 | -0.201687578 |
| Vcp-rs | 2.946069021 | 2.978441397 | 3.147773598 | -0.169332201 | -0.201704577 |
| Ube2e2 | 8.730932222 | 8.733861766 | 8.932647994 | -0.198786228 | -0.201715772 |
| Tspan11 | 7.986533081 | 8.329285915 | 8.188261272 | 0.141024643 | -0.201728191 |
| Rplp0-ps1 | 3.778752159 | 4.039733193 | 3.980497334 | 0.059235859 | -0.201745175 |
| Fer | 9.310522749 | 9.337374171 | 9.512383817 | -0.175009646 | -0.201861068 |
| Ivns1abp | 12.28221945 | 12.32083014 | 12.4841237 | -0.163293567 | -0.20190425 |
| St8sia1 | 5.971068369 | 5.942674895 | 6.173148459 | -0.230473564 | -0.20208009 |
| Arhgap27 | 8.919369945 | 9.25873424 | 9.121499753 | 0.137234487 | -0.202129808 |
| Sec61a2 | 8.553673926 | 8.789087601 | 8.755908182 | 0.033179419 | -0.202234256 |
| Rasd2 | 5.72620523 | 6.461986184 | 5.928641628 | 0.533344556 | -0.202436399 |
| Actn3 | 13.37328922 | 13.59715201 | 13.57574071 | 0.021411307 | -0.20245149 |
| Fnta | 10.3534714 | 10.39836839 | 10.55593194 | -0.157563547 | -0.202460537 |
| Dcun1d1 | 10.042215 | 10.19188263 | 10.24488127 | -0.052998637 | -0.202666266 |
| Orai3 | 8.104146675 | 8.139326904 | 8.306860388 | -0.167533484 | -0.202713714 |
| Brms1l | 8.524816029 | 8.553861753 | 8.727571848 | -0.173710095 | -0.202755819 |
| Galnt12 | 5.188238795 | 5.369615287 | 5.391018977 | -0.02140369 | -0.202780182 |
| Mtch1 | 11.34319371 | 11.41530079 | 11.54602046 | -0.130719667 | -0.202826743 |
| Tcf4 | 11.0464408 | 10.98148599 | 11.24931256 | -0.267826577 | -0.202871763 |
| Rtl8c | 4.876488475 | 5.010220955 | 5.079441943 | -0.069220988 | -0.202953468 |
| 2310050C09Rik | 12.43074414 | 12.5298105 | 12.6337248 | -0.103914297 | -0.20298066 |
| Dusp19 | 6.72774396 | 6.777982685 | 6.930746833 | -0.152764148 | -0.203002873 |

| | | | | | |
|---|---|---|---|---|---|
| Tusc3 | 8.658320998 | 8.680922885 | 8.861333 | -0.180410116 | -0.203012003 |
| Ino80d | 9.857769457 | 10.17523351 | 10.0608084 | 0.114425103 | -0.203038946 |
| Rasl12 | 7.341728863 | 7.541331091 | 7.54478479 | -0.003453698 | -0.203055927 |
| Acp2 | 9.278873628 | 9.473190406 | 9.481962738 | -0.008772332 | -0.203089111 |
| Zfp932 | 8.233312079 | 8.328101539 | 8.436404206 | -0.108302667 | -0.203092127 |
| Acsl3 | 8.415875394 | 8.747762527 | 8.619081247 | 0.128681279 | -0.203205853 |
| Tmem240 | 3.794958464 | 3.893710889 | 3.998206986 | -0.104496097 | -0.203248522 |
| Sox13 | 7.323405958 | 7.498821685 | 7.526709335 | -0.02788765 | -0.203303377 |
| Zbtb7c | 8.543695199 | 8.645340199 | 8.747032926 | -0.101692728 | -0.203337727 |
| Aatk | 7.889703375 | 8.130900893 | 8.09336587 | 0.037535023 | -0.203662495 |
| Tnnt2 | 6.678756395 | 6.615165793 | 6.882440478 | -0.267274684 | -0.203684083 |
| Gm48082 | 6.031226502 | 6.146227708 | 6.234975988 | -0.08874828 | -0.203749486 |
| Pard3b | 7.804805144 | 7.695961429 | 8.008562747 | -0.312601318 | -0.203757604 |
| Relch | 9.794174221 | 9.69910077 | 9.997944067 | -0.298843297 | -0.203769846 |
| Slc47a1 | 5.338052616 | 5.392622741 | 5.541829858 | -0.149207117 | -0.203777242 |
| Akip1 | 7.674517723 | 7.743252939 | 7.87836929 | -0.135116351 | -0.203851567 |
| Scrn1 | 6.398854675 | 6.721935089 | 6.602719997 | 0.119215092 | -0.203865322 |
| Tmem159 | 9.964586528 | 10.0098544 | 10.16845191 | -0.158597511 | -0.203865385 |
| Tbl1xr1 | 10.54062805 | 10.69902913 | 10.74452227 | -0.04549314 | -0.203894223 |
| Cebpg | 10.68192519 | 10.81595762 | 10.88587459 | -0.069916973 | -0.203949407 |
| Bpifc | 10.97786014 | 10.99183631 | 11.18184397 | -0.19000766 | -0.20398383 |
| Agpat5 | 10.46487485 | 10.54201381 | 10.6689955 | -0.126981693 | -0.204120653 |
| Aass | 2.73437782 | 2.877556305 | 2.938535091 | -0.060978786 | -0.204157271 |
| Arhgap21 | 10.13100558 | 10.16632179 | 10.33529501 | -0.16897322 | -0.204289429 |
| mt-Nd1 | 15.77507111 | 15.74755496 | 15.97940181 | -0.23184685 | -0.2043307 |
| CT030661.1 | 8.531689836 | 8.5019718 | 8.736178512 | -0.234206712 | -0.204488676 |
| Klhdc2 | 9.03340755 | 9.081910708 | 9.237916537 | -0.156005829 | -0.204508987 |
| Herc1 | 10.77110297 | 10.88318972 | 10.97568131 | -0.092491593 | -0.204578343 |
| Lgr4 | 10.38434554 | 10.35633162 | 10.58897722 | -0.232645607 | -0.204631683 |
| Prrg4 | 8.197207668 | 8.450560352 | 8.401868963 | 0.048691389 | -0.204661295 |
| Chrd | 5.59807065 | 5.667933962 | 5.802804119 | -0.134870156 | -0.204733469 |
| Cmtm4 | 9.542477301 | 9.677071954 | 9.747238309 | -0.070166354 | -0.204761007 |

| | | | | | |
|---|---|---|---|---|---|
| Fxyd3 | 11.65393167 | 11.71904779 | 11.85871642 | -0.139668633 | -0.204784753 |
| Hp1bp3 | 11.39804966 | 11.51688072 | 11.60285166 | -0.085970943 | -0.204801997 |
| Gstm4 | 7.116024348 | 7.231272263 | 7.32093186 | -0.089659596 | -0.204907512 |
| Cenpp | 5.328886743 | 5.439018965 | 5.533934282 | -0.094915316 | -0.205047538 |
| Gm38431 | 5.095004289 | 5.324031217 | 5.300060444 | 0.023970773 | -0.205056155 |
| Zgrf1 | 6.709570507 | 7.036103701 | 6.914658105 | 0.121445595 | -0.205087599 |
| Pxylp1 | 6.999541544 | 7.13341917 | 7.204663598 | -0.071244428 | -0.205122054 |
| Crct1 | 11.74098371 | 11.82424476 | 11.94624762 | -0.122002857 | -0.205263907 |
| Etfbkmt | 6.591635878 | 6.433967085 | 6.796963042 | -0.362995957 | -0.205327164 |
| Tmem95 | 4.097526107 | 4.205492877 | 4.302874938 | -0.097382061 | -0.205348832 |
| Kat6b | 9.29019211 | 9.331020717 | 9.495541466 | -0.164520749 | -0.205349356 |
| Colec11 | 2.909366589 | 2.979674426 | 3.114756597 | -0.135082171 | -0.205390008 |
| Radil | 5.456891393 | 5.591097377 | 5.662342557 | -0.07124518 | -0.205451164 |
| Zfp773 | 7.553848972 | 7.575442995 | 7.759352708 | -0.183909714 | -0.205503736 |
| Scaf1 | 9.74269207 | 10.1879489 | 9.948212724 | 0.239736176 | -0.205520654 |
| Pard6g | 10.40185201 | 10.6790629 | 10.60738223 | 0.07168067 | -0.205530227 |
| Krt222 | 3.536914378 | 3.545893263 | 3.742450898 | -0.196557635 | -0.20553652 |
| Dlg5 | 9.326374771 | 9.515721711 | 9.531934598 | -0.016212887 | -0.205559827 |
| Smarcd2 | 11.10725354 | 11.29928655 | 11.31284209 | -0.013555543 | -0.205588547 |
| Hmgn5 | 9.405004871 | 9.227951726 | 9.610639307 | -0.382687581 | -0.205634436 |
| Parn | 9.054492812 | 9.08521472 | 9.260277762 | -0.175063042 | -0.205784951 |
| St3gal2 | 9.328561681 | 9.630004605 | 9.534414433 | 0.095590172 | -0.205852752 |
| Gm10221 | 8.818950918 | 8.63708299 | 9.024843148 | -0.387760159 | -0.20589223 |
| Pex19 | 10.896719 | 10.85068216 | 11.10265937 | -0.251977207 | -0.205940373 |
| Phf21b | 5.340921948 | 5.59027158 | 5.546870781 | 0.043400799 | -0.205948833 |
| Fam161b | 5.769477348 | 6.099212168 | 5.975539072 | 0.123673096 | -0.206061724 |
| Serpinb11 | 9.163254234 | 9.204087246 | 9.36938536 | -0.165298113 | -0.206131125 |
| Evc2 | 8.024022734 | 8.223357398 | 8.230186139 | -0.006828741 | -0.206163405 |
| Fer1l6 | 4.354176509 | 4.348642823 | 4.560406211 | -0.211763388 | -0.206229702 |
| Rdh14 | 8.175701416 | 8.128113642 | 8.382051908 | -0.253938266 | -0.206350492 |
| Zfp212 | 8.246212412 | 8.306816016 | 8.452585137 | -0.145769122 | -0.206372725 |
| Abi2 | 8.425885979 | 8.488113911 | 8.632293904 | -0.144179993 | -0.206407925 |

| | | | | | |
|---|---|---|---|---|---|
| Nin | 9.623163974 | 9.787657312 | 9.829668691 | -0.042011379 | -0.206504717 |
| Vash2 | 4.58111928 | 5.115760296 | 4.787636023 | 0.328124273 | -0.206516742 |
| 1700029J07Rik | 4.754981531 | 4.83268339 | 4.96168778 | -0.12900439 | -0.206706249 |
| Gm3325 | 4.948301551 | 4.940365158 | 5.155094703 | -0.214729545 | -0.206793152 |
| Atpaf1 | 8.261547242 | 8.200063819 | 8.468344191 | -0.268280372 | -0.206796949 |
| Dlgap5 | 7.411720337 | 7.427265464 | 7.618523094 | -0.191257629 | -0.206802757 |
| Akap9 | 10.55210203 | 10.65882927 | 10.75907571 | -0.100246443 | -0.206973683 |
| Ubac1 | 9.531586776 | 9.590866735 | 9.738630278 | -0.147763543 | -0.207043502 |
| Gid4 | 10.2389339 | 10.28825238 | 10.44598591 | -0.157733527 | -0.207052003 |
| Fads1 | 9.012162592 | 8.923825113 | 9.219232657 | -0.295407544 | -0.207070065 |
| Gdpd2 | 10.74922753 | 10.88358551 | 10.95631059 | -0.072725083 | -0.207083063 |
| Pbx3 | 7.180990424 | 7.407344327 | 7.388094283 | 0.019250043 | -0.207103859 |
| Rarb | 3.890972935 | 3.903076444 | 4.098123423 | -0.19504698 | -0.207150488 |
| Kctd18 | 8.041610767 | 8.260400204 | 8.248771673 | 0.011628531 | -0.207160907 |
| Retn | 10.39283529 | 9.858890026 | 10.6000042 | -0.741114171 | -0.207168906 |
| Ccnt2 | 10.19852708 | 10.41507047 | 10.40574179 | 0.009328683 | -0.207214707 |
| Nckap1 | 12.17870977 | 12.27719484 | 12.3860073 | -0.108812457 | -0.20729753 |
| Slc6a9 | 9.55463449 | 9.773199569 | 9.761936181 | 0.011263388 | -0.207301691 |
| Nsd1 | 11.35504767 | 11.51185941 | 11.56238128 | -0.050521867 | -0.207333603 |
| Fam83c | 9.484293011 | 9.816574535 | 9.691707489 | 0.124867046 | -0.207414478 |
| Fuk | 7.370082182 | 7.663516173 | 7.577560517 | 0.085955656 | -0.207478335 |
| Apol7a | 5.518939962 | 5.715170603 | 5.726600406 | -0.011429802 | -0.207660444 |
| Tprgl | 10.78949974 | 10.9159446 | 10.99723925 | -0.08129465 | -0.20773951 |
| Zmym3 | 8.567383571 | 8.641598146 | 8.77526284 | -0.133664695 | -0.207879269 |
| Tceal7 | 5.392066139 | 5.595440141 | 5.599992011 | -0.00455187 | -0.207925872 |
| Rnf167 | 10.58820801 | 10.71505273 | 10.79629478 | -0.08124205 | -0.20808677 |
| Rnf144a | 7.326318815 | 7.040660521 | 7.534459412 | -0.493798891 | -0.208140597 |
| Golga1 | 9.686795124 | 9.925319584 | 9.894992679 | 0.030326904 | -0.208197555 |
| Camk1 | 8.964191369 | 8.980686663 | 9.172410142 | -0.19172348 | -0.208218773 |
| Zmym4 | 9.007929775 | 9.160728271 | 9.216167502 | -0.055439231 | -0.208237727 |
| Phf20 | 9.348449934 | 9.509070751 | 9.556965526 | -0.047894774 | -0.208515591 |
| Tmem184c | 7.960474611 | 8.062488857 | 8.169161916 | -0.10667306 | -0.208687305 |

| | | | | | |
|---|---|---|---|---|---|
| Sgo1 | 6.864698197 | 6.968013559 | 7.073503983 | -0.105490424 | -0.208805786 |
| Tmem62 | 8.187578638 | 8.489804898 | 8.396439413 | 0.093365485 | -0.208860775 |
| Rab3a | 6.701053012 | 6.762958936 | 6.909928805 | -0.146969869 | -0.208875792 |
| Gm29668 | 4.309930599 | 4.496056922 | 4.518904379 | -0.022847457 | -0.20897378 |
| Msrb3 | 10.04980657 | 9.896172666 | 10.25898354 | -0.362810874 | -0.209176966 |
| Prtn3 | 4.878560944 | 5.074161057 | 5.087764138 | -0.013603081 | -0.209203194 |
| Gm47692 | 2.618100375 | 2.782495492 | 2.827399396 | -0.044903904 | -0.209299021 |
| Bnc2 | 6.460792157 | 6.739721979 | 6.67009788 | 0.069624099 | -0.209305723 |
| Rida | 7.006814369 | 6.928405897 | 7.216209455 | -0.287803558 | -0.209395085 |
| Mfsd7a | 8.086974163 | 8.276346468 | 8.296370629 | -0.02002416 | -0.209396466 |
| Fahd2a | 6.697675818 | 6.873219896 | 6.907111309 | -0.033891414 | -0.209435491 |
| Ktn1 | 11.4458119 | 11.5195648 | 11.6554298 | -0.135865007 | -0.209617907 |
| Tdrp | 7.018111664 | 7.07664711 | 7.227754644 | -0.151107535 | -0.20964298 |
| Hyls1 | 5.929259337 | 6.041144954 | 6.138913488 | -0.097768534 | -0.209654152 |
| Zfp937 | 6.183013637 | 6.297580732 | 6.392713673 | -0.095132941 | -0.209700036 |
| Zfp990 | 2.422606748 | 2.605774341 | 2.632385021 | -0.02661068 | -0.209778273 |
| 4430402I18Rik | 3.896309225 | 3.879773924 | 4.106127886 | -0.226353963 | -0.209818661 |
| Cdca3 | 8.185875223 | 8.345804663 | 8.395840701 | -0.050036037 | -0.209965477 |
| Fgf22 | 6.462961472 | 6.722083994 | 6.673047689 | 0.049036306 | -0.210086216 |
| Vstm4 | 6.907681407 | 7.170986251 | 7.11780031 | 0.053185941 | -0.210118903 |
| Zfp740 | 10.5695965 | 10.74225797 | 10.77973467 | -0.0374767 | -0.21013817 |
| Ust | 9.307201913 | 9.39362387 | 9.517574345 | -0.123950475 | -0.210372432 |
| Rbak | 8.108094701 | 8.183588306 | 8.318501462 | -0.134913156 | -0.21040676 |
| Krt81 | 2.737202134 | 3.024113535 | 2.947628794 | 0.076484741 | -0.21042666 |
| Lyrm2 | 7.126392957 | 7.160598289 | 7.336854085 | -0.176255795 | -0.210461128 |
| Scamp1 | 10.95192394 | 10.98291743 | 11.16243403 | -0.179516597 | -0.210510087 |
| Flvcr2 | 6.668615258 | 6.893104636 | 6.879150854 | 0.013953782 | -0.210535596 |
| Rarres1 | 11.06604453 | 11.06780942 | 11.27669891 | -0.20888949 | -0.210654383 |
| Aldh3a2 | 9.858354851 | 9.924354567 | 10.06912586 | -0.144771297 | -0.210771012 |
| Rasgrf1 | 5.102747745 | 5.129091897 | 5.313540651 | -0.184448754 | -0.210792906 |
| Acot4 | 7.03592748 | 7.087785318 | 7.246798647 | -0.159013329 | -0.210871167 |
| Fig4 | 8.67828084 | 8.832191831 | 8.889177692 | -0.056985861 | -0.210896853 |

| | | | | | |
|---|---|---|---|---|---|
| Trib2 | 7.545536599 | 7.495967946 | 7.756447372 | -0.260479426 | -0.210910774 |
| Dusp14 | 8.96132378 | 9.100302875 | 9.172290747 | -0.071987872 | -0.210966967 |
| Reep5 | 12.06871583 | 11.948051 | 12.27970695 | -0.33165595 | -0.210991117 |
| Gm4821 | 3.587025841 | 3.678157061 | 3.798109238 | -0.119952177 | -0.211083397 |
| Adam11 | 5.562088823 | 5.533563051 | 5.773282 | -0.239718948 | -0.211193177 |
| Arhgef37 | 10.01146776 | 10.14983156 | 10.22269845 | -0.072866897 | -0.211230692 |
| Mapk9 | 9.259574949 | 9.358322405 | 9.470906194 | -0.112583788 | -0.211331245 |
| Wdr20 | 8.795336344 | 9.005692483 | 9.006771032 | -0.001078549 | -0.211434688 |
| Tenm3 | 7.545758826 | 7.709653107 | 7.7572365 | -0.047583393 | -0.211477675 |
| Mup3 | 5.117005274 | 5.172852828 | 5.328493141 | -0.155640314 | -0.211487867 |
| Tmem135 | 9.98435023 | 9.893663033 | 10.1958475 | -0.302184471 | -0.211497273 |
| Ugt8a | 5.256725217 | 5.32334356 | 5.468246991 | -0.144903431 | -0.211521774 |
| Atp9a | 10.33001184 | 10.4985979 | 10.54155617 | -0.042958277 | -0.21154433 |
| Gm4925 | 3.572351469 | 3.752547592 | 3.783898658 | -0.031351067 | -0.211547189 |
| Prkci | 10.70120263 | 10.91503627 | 10.91287947 | 0.0021568 | -0.211676847 |
| Ano1 | 10.28987815 | 10.41890507 | 10.50174101 | -0.082835943 | -0.211862863 |
| Wdr5b | 6.864019409 | 7.012070685 | 7.075962404 | -0.063891719 | -0.211942996 |
| Bspry | 8.17628843 | 8.359639275 | 8.388248673 | -0.028609397 | -0.211960242 |
| Cip2a | 8.067254134 | 8.054601071 | 8.279339623 | -0.224738551 | -0.212085489 |
| Tspan15 | 7.283958549 | 7.563640899 | 7.496112382 | 0.067528517 | -0.212153834 |
| Prrg1 | 6.787019281 | 6.709787376 | 6.999346288 | -0.289558912 | -0.212327006 |
| Anapc5 | 11.8785323 | 11.88676235 | 12.09088992 | -0.204127577 | -0.212357627 |
| Atg4a | 6.878174852 | 6.877194517 | 7.090550008 | -0.213355491 | -0.212375156 |
| Pcm1 | 10.16020023 | 10.2325428 | 10.37262369 | -0.140080893 | -0.212423467 |
| Lhx8 | 4.033869161 | 4.050518024 | 4.246463162 | -0.195945138 | -0.212594001 |
| Ormdl1 | 8.488454654 | 8.538736093 | 8.70116941 | -0.162433318 | -0.212714756 |
| Aes | 11.37758369 | 11.54948604 | 11.59031458 | -0.040828543 | -0.212730897 |
| Tceanc | 6.032077509 | 5.97656291 | 6.244902109 | -0.268339199 | -0.2128246 |
| Chac2 | 6.952553916 | 6.911761259 | 7.165431055 | -0.253669796 | -0.212877139 |
| Vsig10l | 9.444443939 | 9.819644226 | 9.657374852 | 0.162269374 | -0.212930913 |
| Gm13408 | 3.531290929 | 3.761978285 | 3.74432515 | 0.017653134 | -0.213034221 |
| Gstm5 | 9.769071786 | 9.694275909 | 9.982160965 | -0.287885056 | -0.213089179 |

| | | | | | |
|---|---|---|---|---|---|
| Sorcs3 | 3.771182193 | 3.922926071 | 3.984320531 | -0.061394461 | -0.213138338 |
| Kmt2a | 10.52461564 | 10.81218406 | 10.73777458 | 0.074409483 | -0.213158937 |
| Pm20d2 | 5.400552682 | 5.490902695 | 5.613722514 | -0.122819819 | -0.213169832 |
| Ubxn2a | 9.345286515 | 9.353121579 | 9.558706336 | -0.205584757 | -0.213419821 |
| Stmn2 | 6.442508054 | 6.363718958 | 6.655932199 | -0.292213241 | -0.213424145 |
| Frs2 | 9.723780781 | 9.874913972 | 9.937275559 | -0.062361587 | -0.213494778 |
| Hist1h2be | 5.972307069 | 6.103567818 | 6.185886095 | -0.082318278 | -0.213579026 |
| Lix1 | 3.633584436 | 3.622074384 | 3.847214128 | -0.225139744 | -0.213629692 |
| Zfp958 | 6.941301439 | 6.962283161 | 7.15493639 | -0.192653229 | -0.213634951 |
| Npbwr1 | 7.023524918 | 7.246439149 | 7.237193332 | 0.009245817 | -0.213668414 |
| Tmem165 | 9.972955796 | 10.12622564 | 10.1866789 | -0.060453263 | -0.213723104 |
| Gusb | 9.532925832 | 9.683354935 | 9.746655109 | -0.063300174 | -0.213729277 |
| Npas2 | 5.518356487 | 5.783472005 | 5.732087379 | 0.051384626 | -0.213730891 |
| Plcd1 | 9.37031475 | 9.597553938 | 9.584064429 | 0.013489508 | -0.213749679 |
| Rbfox2 | 10.44423771 | 10.52810497 | 10.65801286 | -0.129907897 | -0.213775157 |
| Crtc1 | 8.171302741 | 8.572887889 | 8.385194607 | 0.187693282 | -0.213891865 |
| Cadps2 | 4.131933942 | 4.217171345 | 4.345834326 | -0.128662981 | -0.213900384 |
| Brip1 | 6.819658537 | 7.034784913 | 7.033573349 | 0.001211564 | -0.213914812 |
| Nkain4 | 3.158763415 | 3.201375089 | 3.372722871 | -0.171347781 | -0.213959456 |
| Ranbp9 | 10.38466107 | 10.45022779 | 10.59862663 | -0.148398833 | -0.213965557 |
| Alg6 | 6.108642322 | 6.242014276 | 6.322643021 | -0.080628745 | -0.214000699 |
| Pbxip1 | 10.75531621 | 10.79285668 | 10.96944645 | -0.17658977 | -0.214130243 |
| Klhl41 | 10.87988125 | 11.56894829 | 11.09401298 | 0.474935307 | -0.214131733 |
| Gse1 | 10.05396113 | 10.4345256 | 10.26811997 | 0.166405637 | -0.214158835 |
| Rgl3 | 4.393404456 | 4.636954196 | 4.607568177 | 0.029386019 | -0.214163721 |
| Rbp7 | 4.371320866 | 4.479193955 | 4.585639629 | -0.106445674 | -0.214318762 |
| Glud1 | 11.88020785 | 11.92953434 | 12.09454311 | -0.165008773 | -0.21433526 |
| Sbk1 | 8.48913993 | 8.581048089 | 8.703531885 | -0.122483796 | -0.214391955 |
| Serpina3j | 10.53403168 | 10.45252149 | 10.74847168 | -0.295950197 | -0.214440007 |
| Higd1b | 2.624432146 | 2.719287833 | 2.838935886 | -0.119648053 | -0.21450374 |
| Abhd12 | 10.29515998 | 10.50130943 | 10.5096882 | -0.008378763 | -0.214528217 |
| Cnr1 | 5.165729729 | 6.042037973 | 5.380331278 | 0.661706695 | -0.214601549 |

| | | | | | |
|---|---|---|---|---|---|
| Lactb2 | 8.809413696 | 9.013192104 | 9.024050244 | -0.01085814 | -0.214636548 |
| Ice2 | 8.061192393 | 8.104863221 | 8.275839319 | -0.170976098 | -0.214646926 |
| Gm12901 | 3.0618014 | 3.292657042 | 3.2765531 | 0.016103941 | -0.214751701 |
| Mcee | 8.112412519 | 8.040244667 | 8.327179615 | -0.286934948 | -0.214767096 |
| Spry3 | 4.736196525 | 4.791911547 | 4.950987391 | -0.159075843 | -0.214790866 |
| Adgrl3 | 6.837002919 | 6.907931191 | 7.051942323 | -0.144011132 | -0.214939403 |
| Gm10762 | 4.884901921 | 4.852684261 | 5.099929873 | -0.247245612 | -0.215027952 |
| Rusc2 | 8.449018817 | 8.835725557 | 8.664113183 | 0.171612375 | -0.215094366 |
| Smim24 | 5.623500512 | 5.649329483 | 5.838600878 | -0.189271395 | -0.215100366 |
| Med12l | 6.425056926 | 6.384122764 | 6.640161254 | -0.25603849 | -0.215104328 |
| Pla2r1 | 7.67408388 | 7.760645535 | 7.889233908 | -0.128588373 | -0.215150028 |
| Tmem229a | 6.884650936 | 6.874967288 | 7.099820511 | -0.224853223 | -0.215169574 |
| B230219D22Rik | 10.28287545 | 10.40860417 | 10.49819387 | -0.089589693 | -0.215318413 |
| Etfdh | 10.20212937 | 10.22197284 | 10.41748926 | -0.195516423 | -0.215359897 |
| Gm8438 | 3.932507963 | 4.092687971 | 4.147968377 | -0.055280406 | -0.215460414 |
| Numbl | 8.152130749 | 8.595543135 | 8.367686629 | 0.227856506 | -0.21555588 |
| Gm13212 | 5.321242335 | 5.435583937 | 5.536843203 | -0.101259266 | -0.215600869 |
| Zfp445 | 10.5657492 | 10.67577136 | 10.7814021 | -0.105630743 | -0.2156529 |
| Fam102b | 10.49674613 | 10.58512365 | 10.71244405 | -0.1273204 | -0.21569792 |
| Fnip1 | 9.767054004 | 9.779412036 | 9.982912391 | -0.203500355 | -0.215858388 |
| Spast | 9.906833564 | 10.02216405 | 10.12281254 | -0.100648488 | -0.215978972 |
| Zfp729b | 7.685966965 | 7.804992651 | 7.901951896 | -0.096959245 | -0.215984931 |
| Wscd1 | 4.122609361 | 4.204456015 | 4.338617984 | -0.134161969 | -0.216008623 |
| Cd164l2 | 4.567237507 | 4.720346808 | 4.783305446 | -0.062958637 | -0.216067939 |
| Amy1 | 9.345243293 | 9.061510406 | 9.561415339 | -0.499904933 | -0.216172046 |
| Col28a1 | 6.679351226 | 6.771129057 | 6.895563028 | -0.124433972 | -0.216211803 |
| Gm14399 | 5.618806585 | 5.695814786 | 5.835157456 | -0.139342671 | -0.216350872 |
| Gria4 | 5.10693406 | 5.195624924 | 5.323367462 | -0.127742538 | -0.216433402 |
| Ntn4 | 7.993603902 | 8.051079445 | 8.210107955 | -0.15902851 | -0.216504054 |
| Eppk1 | 11.69895735 | 12.30437136 | 11.91559478 | 0.388776573 | -0.216637433 |
| Olah | 4.118077209 | 4.263311328 | 4.334760964 | -0.071449637 | -0.216683756 |
| Ick | 8.148885587 | 8.37838968 | 8.36557328 | 0.0128164 | -0.216687693 |

| | | | | | |
|---|---|---|---|---|---|
| Zfp658 | 5.521723749 | 5.709722779 | 5.738440724 | -0.028717945 | -0.216716975 |
| Bub1b | 8.499777502 | 8.633938536 | 8.716520303 | -0.082581768 | -0.216742802 |
| Ide | 12.09302451 | 12.07613451 | 12.30978037 | -0.23364586 | -0.216755857 |
| Zfp617 | 8.723540755 | 8.727637181 | 8.940378971 | -0.21274179 | -0.216838216 |
| Smim19 | 7.988624805 | 7.884498744 | 8.205541585 | -0.321042842 | -0.216916781 |
| Gas2 | 5.118456281 | 5.118403433 | 5.335422481 | -0.217019048 | -0.2169662 |
| Epha3 | 4.557618974 | 4.550739516 | 4.774672803 | -0.223933287 | -0.217053829 |
| Kcnq1 | 4.704324603 | 4.767814673 | 4.92146903 | -0.153654357 | -0.217144427 |
| Ski | 10.81769203 | 11.12640311 | 11.03486008 | 0.091543027 | -0.21716805 |
| Cdc7 | 7.530371151 | 7.763124368 | 7.747650535 | 0.015473834 | -0.217279384 |
| Map4k3 | 10.13485187 | 10.27539633 | 10.35223769 | -0.076841363 | -0.21738582 |
| Stxbp5 | 9.864783023 | 10.03828377 | 10.08220833 | -0.043924565 | -0.217425309 |
| Atf7ip | 10.78789286 | 11.09801972 | 11.00542876 | 0.092590963 | -0.217535897 |
| Pskh1 | 8.730193081 | 8.710109044 | 8.947798773 | -0.237689728 | -0.217605692 |
| Washc3 | 8.381808433 | 8.453487441 | 8.599427702 | -0.145940261 | -0.217619269 |
| Tmem29 | 7.649598061 | 7.735443914 | 7.867248264 | -0.13180435 | -0.217650203 |
| Olfr374 | 5.406178099 | 5.673542885 | 5.623992001 | 0.049550885 | -0.217813902 |
| Hoxa5 | 6.631040268 | 6.915774008 | 6.848925761 | 0.066848247 | -0.217885493 |
| Plvap | 9.320729174 | 9.524287077 | 9.538624649 | -0.014337571 | -0.217895474 |
| Slc4a4 | 7.35864259 | 7.344794194 | 7.576579815 | -0.231785621 | -0.217937225 |
| Syt15 | 6.160512509 | 6.220341808 | 6.378482624 | -0.158140816 | -0.217970115 |
| Usp53 | 8.728433517 | 8.778960235 | 8.946445429 | -0.167485194 | -0.218011912 |
| Adssl1 | 10.7099966 | 10.73941948 | 10.92801133 | -0.188591843 | -0.21801473 |
| Ing2 | 7.635454586 | 7.676154143 | 7.853543823 | -0.17738968 | -0.218089236 |
| Rtl6 | 6.416394143 | 6.460839362 | 6.634598787 | -0.173759425 | -0.218204644 |
| Sec23a | 9.769795152 | 9.740488885 | 9.988066542 | -0.247577657 | -0.21827139 |
| Maob | 9.21288782 | 9.176828473 | 9.431219349 | -0.254390876 | -0.218331529 |
| Tmem50a | 10.68010562 | 10.69193566 | 10.89852128 | -0.20658562 | -0.21841566 |
| Pkp1 | 14.30599234 | 14.51429475 | 14.52446272 | -0.01016797 | -0.21847038 |
| Nme7 | 7.487952454 | 7.636498074 | 7.706484091 | -0.069986016 | -0.218531637 |
| Isoc2b | 6.708185331 | 6.786944483 | 6.926728476 | -0.139783992 | -0.218543145 |
| Hpgd | 8.366999591 | 8.151165766 | 8.585543259 | -0.434377493 | -0.218543668 |

| | | | | | |
|---|---|---|---|---|---|
| Dennd6a | 8.03982438 | 8.140233582 | 8.258422229 | -0.118188647 | -0.21859785 |
| Sbspon | 5.300762575 | 5.401872311 | 5.519397096 | -0.117524786 | -0.218634522 |
| Ndst3 | 3.884134824 | 4.034915792 | 4.102869424 | -0.067953632 | -0.2187346 |
| Gm21926 | 7.873928893 | 7.995007474 | 8.092764126 | -0.097756652 | -0.218835232 |
| Tmem117 | 5.790401144 | 5.817641949 | 6.009256189 | -0.19161424 | -0.218855045 |
| Zfp605 | 5.743411517 | 5.740340371 | 5.962277598 | -0.221937226 | -0.218866081 |
| Srebf1 | 10.90817761 | 11.22429275 | 11.12713005 | 0.0971627 | -0.21895244 |
| Slc43a2 | 9.170085449 | 9.477412339 | 9.389133354 | 0.088278985 | -0.219047905 |
| Trf | 12.41296283 | 12.16304296 | 12.63201974 | -0.46897678 | -0.219056907 |
| Zfp108 | 4.706088556 | 4.85575341 | 4.925236521 | -0.069483111 | -0.219147965 |
| Lmntd2 | 4.692169816 | 4.902227535 | 4.911325223 | -0.009097688 | -0.219155406 |
| Nipsnap1 | 7.487340865 | 7.56929664 | 7.706558426 | -0.137261786 | -0.219217561 |
| St5 | 9.404542 | 9.671976778 | 9.623813088 | 0.048163689 | -0.219271089 |
| Kcnk1 | 5.631233687 | 5.710358354 | 5.850536423 | -0.140178069 | -0.219302735 |
| Ngef | 7.029660492 | 7.315130827 | 7.248965689 | 0.066165137 | -0.219305197 |
| Tpm1 | 13.94419072 | 14.09659216 | 14.16350623 | -0.06691407 | -0.219315517 |
| Arl10 | 6.314941893 | 6.593156956 | 6.53437325 | 0.058783705 | -0.219431357 |
| Rasal1 | 6.343166845 | 6.542600927 | 6.562610887 | -0.02000996 | -0.219444042 |
| G2e3 | 8.344150743 | 8.373968095 | 8.563596504 | -0.189628409 | -0.219445761 |
| Hexa | 10.21834272 | 10.27155269 | 10.43784238 | -0.166289687 | -0.21949966 |
| Ctps2 | 8.301222727 | 8.24193248 | 8.520751197 | -0.278818717 | -0.21952847 |
| Zfp931 | 6.065846076 | 6.250840491 | 6.285436238 | -0.034595747 | -0.219590162 |
| Cpt2 | 9.343635155 | 9.387033466 | 9.563243102 | -0.176209636 | -0.219607946 |
| Aamdc | 8.202121284 | 8.121011521 | 8.421795514 | -0.300783993 | -0.21967423 |
| B9d1 | 5.853351405 | 5.990589989 | 6.07304065 | -0.082450661 | -0.219689245 |
| Prkacb | 9.892276825 | 9.939101991 | 10.11204189 | -0.172939902 | -0.219765068 |
| Lce1l | 10.75455697 | 11.07940343 | 10.97434635 | 0.10505708 | -0.21978938 |
| Tmem41a | 8.578951888 | 8.813214207 | 8.79884279 | 0.014371416 | -0.219890902 |
| Akap17b | 6.934360083 | 7.063195314 | 7.154277584 | -0.09108227 | -0.219917501 |
| Amigo1 | 8.636653472 | 8.671052118 | 8.856575724 | -0.185523607 | -0.219922253 |
| Sbk2 | 6.994335762 | 7.077552766 | 7.214506465 | -0.136953699 | -0.220170703 |
| Tox | 4.667190096 | 4.668331816 | 4.887400646 | -0.21906883 | -0.22021055 |

| | | | | | |
|---|---|---|---|---|---|
| Trim59 | 8.316256573 | 8.468043941 | 8.536494647 | -0.068450705 | -0.220238073 |
| Rcor1 | 10.47625126 | 10.73660863 | 10.69653606 | 0.040072573 | -0.220284803 |
| Myl9 | 9.88132459 | 9.959785734 | 10.10162762 | -0.141841883 | -0.220303026 |
| Fnbp1l | 9.961660382 | 10.20299298 | 10.18197493 | 0.02101805 | -0.220314544 |
| Bicc1 | 9.20988923 | 9.269449805 | 9.430218614 | -0.160768809 | -0.220329384 |
| Adgrl1 | 9.306182233 | 9.760141932 | 9.52652492 | 0.233617012 | -0.220342687 |
| Tusc2 | 8.737374907 | 8.804970782 | 8.957779414 | -0.152808632 | -0.220404507 |
| Oma1 | 7.907802712 | 7.939455022 | 8.128213583 | -0.188758562 | -0.220410871 |
| Chml | 8.244936476 | 8.347467621 | 8.465417355 | -0.117949734 | -0.220480879 |
| Gm4217 | 3.10615015 | 3.241569072 | 3.326637709 | -0.085068637 | -0.22048756 |
| Slc2a9 | 5.977795258 | 6.166016919 | 6.198444457 | -0.032427538 | -0.220649199 |
| Smo | 10.0887415 | 10.36895089 | 10.30943787 | 0.059513027 | -0.220696366 |
| Lce3e | 6.415568919 | 6.707303676 | 6.636272876 | 0.0710308 | -0.220703957 |
| Calhm2 | 7.310325851 | 7.478850701 | 7.531112401 | -0.0522617 | -0.22078655 |
| Zbtb8a | 6.72677681 | 6.839102153 | 6.947586139 | -0.108483986 | -0.220809329 |
| Mut | 9.693823926 | 9.653463556 | 9.914752639 | -0.261289083 | -0.220928713 |
| Slc25a1 | 11.02793479 | 10.6601687 | 11.24894219 | -0.588773497 | -0.2210074 |
| Gmcl1 | 8.559686745 | 8.716396426 | 8.780697853 | -0.064301427 | -0.221011108 |
| Rufy3 | 9.131805869 | 9.409394492 | 9.352890115 | 0.056504377 | -0.221084246 |
| Zfp449 | 5.959621338 | 6.040240214 | 6.180868286 | -0.140628072 | -0.221246948 |
| Sox17 | 5.935272649 | 6.003074994 | 6.156604554 | -0.15352956 | -0.221331906 |
| Trim6 | 3.785530184 | 3.819085381 | 4.006963517 | -0.187878136 | -0.221433333 |
| Jazf1 | 6.306015209 | 6.251834597 | 6.527463307 | -0.27562871 | -0.221448098 |
| Zfp626 | 8.394205006 | 8.394306141 | 8.615689223 | -0.221383083 | -0.221484217 |
| Zfp160 | 8.478489687 | 8.516234369 | 8.700046817 | -0.183812449 | -0.22155713 |
| Fxr1 | 10.8507546 | 10.85892509 | 11.07235483 | -0.213429743 | -0.22160023 |
| Tmem28 | 3.435562896 | 3.506358805 | 3.657193669 | -0.150834864 | -0.221630773 |
| Alms1-ps1 | 2.594125663 | 2.743538825 | 2.815793659 | -0.072254834 | -0.221667996 |
| Mgst1 | 12.0860603 | 12.08225529 | 12.3077827 | -0.225527413 | -0.221722403 |
| Slc36a4 | 9.306241815 | 9.356169899 | 9.52811781 | -0.171947911 | -0.221875995 |
| Hspa1l | 6.141882795 | 7.038721322 | 6.363813119 | 0.674908203 | -0.221930324 |
| Itgb8 | 8.831275238 | 8.819260766 | 9.053300785 | -0.234040019 | -0.222025547 |

| | | | | | |
|---|---|---|---|---|---|
| Cpe | 11.21403147 | 11.27371021 | 11.43606948 | -0.162359267 | -0.222038007 |
| Ube4a | 10.41219559 | 10.6178176 | 10.6342515 | -0.016433907 | -0.222055913 |
| Bco1 | 4.725965129 | 4.768756663 | 4.948062657 | -0.179305993 | -0.222097527 |
| Rflnb | 9.363523262 | 9.355684454 | 9.585648251 | -0.229963797 | -0.222124988 |
| Trpc4 | 4.279353171 | 4.423520966 | 4.501506897 | -0.077985931 | -0.222153726 |
| Zfhx3 | 7.811144767 | 7.703372262 | 8.033396573 | -0.330024311 | -0.222251806 |
| Inha | 4.694087079 | 4.983147383 | 4.916353188 | 0.066794195 | -0.222266108 |
| Cdc37l1 | 10.11103928 | 10.19556886 | 10.33330937 | -0.137740513 | -0.222270087 |
| Ybx2 | 4.648195406 | 4.807046679 | 4.87056309 | -0.063516411 | -0.222367684 |
| Slc25a11 | 10.699071 | 10.77023422 | 10.92154975 | -0.15131553 | -0.22247875 |
| Cbarp | 7.958271063 | 8.309140566 | 8.180850583 | 0.128289983 | -0.22257952 |
| Pdzd4 | 7.767595591 | 7.931541019 | 7.990196731 | -0.058655712 | -0.22260114 |
| Mrgpre | 4.944836859 | 4.9876822 | 5.167737035 | -0.180054835 | -0.222900176 |
| Akr1c13 | 6.761322561 | 6.697182886 | 6.984330732 | -0.287147846 | -0.223008171 |
| Ctnna3 | 4.208449148 | 4.274043171 | 4.431470383 | -0.157427212 | -0.223021235 |
| Mapk3 | 11.30164517 | 11.37615348 | 11.52471799 | -0.14856451 | -0.223072813 |
| Arrdc1 | 8.98139503 | 9.312123942 | 9.204572756 | 0.107551186 | -0.223177726 |
| Recql5 | 8.675442641 | 8.852653921 | 8.898627421 | -0.0459735 | -0.22318478 |
| Numb | 9.428854021 | 9.622494433 | 9.652073817 | -0.029579384 | -0.223219796 |
| Kcnk6 | 9.298537973 | 9.585426009 | 9.521824449 | 0.06360156 | -0.223286476 |
| Xpnpep2 | 6.417512706 | 6.374388094 | 6.640879945 | -0.266491851 | -0.223367239 |
| Serpinb6a | 12.33008218 | 12.32661261 | 12.55348208 | -0.22686947 | -0.223399907 |
| Rnf39 | 9.106351828 | 9.413661008 | 9.329796918 | 0.08386409 | -0.22344509 |
| Upp2 | 4.782839452 | 4.956348238 | 5.006415531 | -0.050067294 | -0.223576079 |
| Foxo6 | 5.039597849 | 5.293461911 | 5.263223812 | 0.030238099 | -0.223625964 |
| Adcy7 | 9.608353303 | 9.734831339 | 9.832098737 | -0.097267398 | -0.223745434 |
| Mpz | 10.88253242 | 10.97508271 | 11.10637588 | -0.131293167 | -0.223843457 |
| Sorbs2 | 7.20767003 | 7.285462722 | 7.431531433 | -0.146068711 | -0.223861402 |
| Fam189b | 6.895580641 | 7.347629435 | 7.119482658 | 0.228146777 | -0.223902018 |
| Zfp940 | 5.240807574 | 5.342968141 | 5.464714651 | -0.12174651 | -0.223907076 |
| Adnp | 7.073601451 | 7.089658776 | 7.297876241 | -0.208217465 | -0.22427479 |
| Strn | 10.42487718 | 10.72808558 | 10.64916637 | 0.078919217 | -0.224289183 |

| | | | | | |
|---|---|---|---|---|---|
| Ppil2 | 9.947825858 | 10.1184006 | 10.17225286 | -0.053852263 | -0.224427002 |
| Efhd1 | 5.525795945 | 5.733757821 | 5.750394235 | -0.016636414 | -0.22459829 |
| Bbs10 | 4.838067323 | 4.900191689 | 5.062877021 | -0.162685333 | -0.224809699 |
| Doc2b | 3.8077998 | 3.851822461 | 4.032643464 | -0.180821003 | -0.224843664 |
| Gm6557 | 9.549927197 | 9.566103143 | 9.774864747 | -0.208761604 | -0.224937551 |
| Dnaja4 | 9.408432338 | 11.48844773 | 9.633383307 | 1.855064426 | -0.224950969 |
| Map3k4 | 9.631637987 | 9.952832734 | 9.856801126 | 0.096031609 | -0.225163139 |
| Ccl8 | 10.94141315 | 11.08479342 | 11.16662564 | -0.081832227 | -0.22521249 |
| Mpp5 | 9.454393214 | 9.733943681 | 9.679647915 | 0.054295766 | -0.225254701 |
| Pkd2l2 | 3.719092509 | 3.907305288 | 3.944447477 | -0.037142189 | -0.225354967 |
| Slc17a9 | 5.127492633 | 5.356916166 | 5.352870301 | 0.004045865 | -0.225377669 |
| Dock1 | 10.11782184 | 10.19775748 | 10.3432353 | -0.145477817 | -0.225413455 |
| B230217C12Rik | 3.840552634 | 3.980403305 | 4.065997233 | -0.085593928 | -0.225444598 |
| Dcaf4 | 8.840852895 | 9.086897303 | 9.066409833 | 0.02048747 | -0.225556938 |
| Cacna1c | 7.062332105 | 7.412041671 | 7.287915109 | 0.124126562 | -0.225583004 |
| Pigc | 8.857234479 | 8.87101688 | 9.082936307 | -0.211919427 | -0.225701828 |
| Golm1 | 7.607582074 | 7.763818817 | 7.833524848 | -0.069706031 | -0.225942773 |
| Mfsd3 | 6.333850649 | 6.423212189 | 6.559832375 | -0.136620186 | -0.225981726 |
| Cdkn2aip | 8.348623383 | 8.421644382 | 8.574656723 | -0.153012341 | -0.22603334 |
| Gm28557 | 5.425506231 | 5.407818115 | 5.651610318 | -0.243792203 | -0.226104087 |
| Fam89b | 8.138557311 | 8.378755941 | 8.364741274 | 0.014014667 | -0.226183964 |
| Dlg3 | 9.27712829 | 9.442248494 | 9.503320469 | -0.061071975 | -0.226192179 |
| Gng4 | 7.51090454 | 7.624388071 | 7.737183103 | -0.112795032 | -0.226278563 |
| Zfp871 | 10.52485913 | 10.69877203 | 10.75129435 | -0.052522323 | -0.226435227 |
| 2610301B20Rik | 6.656943324 | 6.752327718 | 6.883431976 | -0.131104258 | -0.226488652 |
| Teddm2 | 4.093027691 | 4.203163988 | 4.319516601 | -0.116352613 | -0.22648891 |
| Mlycd | 7.933215174 | 8.078393396 | 8.159724381 | -0.081330985 | -0.226509207 |
| Cfap97 | 8.349001787 | 8.400998861 | 8.57551982 | -0.174520959 | -0.226518033 |
| Hid1 | 6.023058945 | 6.255288688 | 6.249603474 | 0.005685215 | -0.226544528 |
| Klhl22 | 7.934943738 | 8.201128002 | 8.161553444 | 0.039574558 | -0.226609705 |
| Wwtr1 | 10.2884988 | 10.42073566 | 10.51518401 | -0.094448353 | -0.22668521 |
| Lrtm1 | 5.020546433 | 4.797168745 | 5.247292356 | -0.450123611 | -0.226745922 |

| | | | | | |
|---|---|---|---|---|---|
| Wdsub1 | 8.193403594 | 8.439602659 | 8.420180787 | 0.019421872 | -0.226777193 |
| Zfp125 | 6.277586048 | 6.402404108 | 6.504387427 | -0.10198332 | -0.22680138 |
| Acap2 | 11.11473774 | 11.21249909 | 11.34156428 | -0.12906519 | -0.226826543 |
| Cldn1 | 12.07582258 | 12.14219539 | 12.30273359 | -0.1605382 | -0.226911003 |
| Tmem206 | 6.262177132 | 6.612259363 | 6.48928647 | 0.122972893 | -0.227109338 |
| Cdh6 | 6.043697392 | 6.129440988 | 6.270882746 | -0.141441758 | -0.227185354 |
| Pstk | 7.599220445 | 7.556290255 | 7.826586094 | -0.270295839 | -0.227365649 |
| Gprasp2 | 4.289801546 | 4.43275081 | 4.517240569 | -0.084489759 | -0.227439023 |
| Pde7a | 8.725257465 | 8.888919241 | 8.952759061 | -0.063839821 | -0.227501597 |
| D5Ertd579e | 10.68616231 | 10.64894717 | 10.91381477 | -0.2648676 | -0.22765246 |
| Acads | 8.652262302 | 8.924434342 | 8.880041929 | 0.044392413 | -0.227779626 |
| Kera | 5.724072982 | 5.863866476 | 5.951940634 | -0.088074158 | -0.227867652 |
| Hcfc1r1 | 9.647218773 | 9.776475123 | 9.875382553 | -0.09890743 | -0.22816378 |
| Nbr1 | 11.03918491 | 11.20887732 | 11.26737937 | -0.05850205 | -0.22819446 |
| Tmem184a | 8.957224486 | 9.26146793 | 9.185455359 | 0.076012571 | -0.228230873 |
| Plod1 | 9.61262494 | 9.784223792 | 9.841157452 | -0.056933661 | -0.228532512 |
| Hadha | 11.78733703 | 11.88923341 | 12.01590581 | -0.1266724 | -0.228568773 |
| Dusp13 | 7.341571569 | 8.025796223 | 7.570243476 | 0.455552747 | -0.228671907 |
| Acvr2a | 9.990597327 | 9.927929304 | 10.21935097 | -0.291421663 | -0.22875364 |
| Lipa | 9.402102023 | 9.66325495 | 9.630883554 | 0.032371396 | -0.228781531 |
| Tgfb2 | 8.11304916 | 8.659707139 | 8.341901742 | 0.317805397 | -0.228852582 |
| Reep4 | 9.981417842 | 10.32953352 | 10.21037861 | 0.119154913 | -0.228960764 |
| Cd5l | 3.331923963 | 3.549429464 | 3.560892242 | -0.011462778 | -0.228968279 |
| Mllt3 | 9.059278593 | 9.180847588 | 9.288367407 | -0.107519819 | -0.229088814 |
| Smpdl3a | 10.31244755 | 10.28000627 | 10.54161027 | -0.261604003 | -0.229162723 |
| Ntan1 | 8.92850263 | 8.988519886 | 9.157683792 | -0.169163906 | -0.229181162 |
| Rassf7 | 9.021296662 | 9.428102086 | 9.250500373 | 0.177601713 | -0.229203711 |
| Cfl2 | 10.92137346 | 10.93942341 | 11.15061658 | -0.211193167 | -0.229243117 |
| Cfap100 | 4.207120714 | 4.402307131 | 4.436435042 | -0.034127911 | -0.229314328 |
| Cptp | 9.650824148 | 9.771324435 | 9.880143223 | -0.108818788 | -0.229319075 |
| Fbln7 | 5.294257487 | 5.420027799 | 5.523652786 | -0.103624986 | -0.229395299 |
| Fgf11 | 6.980490761 | 7.282940207 | 7.20995687 | 0.072983337 | -0.229466109 |

| | | | | | |
|---|---|---|---|---|---|
| Ociad2 | 8.70179704 | 8.774503429 | 8.931287009 | -0.15678358 | -0.229489969 |
| Zfp966 | 7.89053552 | 8.113062603 | 8.120039347 | -0.006976744 | -0.229503827 |
| H2afy2 | 6.295683211 | 6.437975908 | 6.52524282 | -0.087266912 | -0.229559609 |
| Gm4631 | 10.51334363 | 10.79749782 | 10.74301849 | 0.054479333 | -0.22967486 |
| Cnp | 8.938453215 | 8.88578503 | 9.168172741 | -0.282387711 | -0.229719526 |
| Tmem220 | 4.996422037 | 5.00437635 | 5.226248397 | -0.221872047 | -0.22982636 |
| Peli2 | 9.480622675 | 9.491651602 | 9.710454084 | -0.218802482 | -0.22983141 |
| Psat1 | 11.38670652 | 11.58572013 | 11.61655951 | -0.030839373 | -0.229852983 |
| S100a7a | 3.370287061 | 3.767962414 | 3.600241853 | 0.167720561 | -0.229954792 |
| Cav2 | 11.04149415 | 10.93528111 | 11.27145331 | -0.336172203 | -0.22995916 |
| Zkscan14 | 6.947045537 | 7.00693089 | 7.177151149 | -0.170220259 | -0.230105612 |
| Pdzrn3 | 9.483639834 | 9.676066343 | 9.713811 | -0.037744657 | -0.230171166 |
| Gin1 | 6.955999075 | 7.086828124 | 7.186280341 | -0.099452217 | -0.230281266 |
| Igf1 | 10.01618008 | 10.13265095 | 10.24672307 | -0.114072123 | -0.230542995 |
| Taf1b | 8.059688626 | 8.113442128 | 8.290361503 | -0.176919375 | -0.230672877 |
| Xrcc3 | 7.177388165 | 7.459644675 | 7.408137027 | 0.051507648 | -0.230748862 |
| Katnbl1 | 9.390225984 | 9.481910522 | 9.621168634 | -0.139258112 | -0.23094265 |
| Sema3e | 8.825437054 | 8.958060898 | 9.056396745 | -0.098335847 | -0.230959691 |
| Serf1 | 6.236000229 | 6.297136984 | 6.46701215 | -0.169875166 | -0.231011922 |
| Ascc1 | 8.1628145 | 8.345241749 | 8.393858248 | -0.048616498 | -0.231043748 |
| Nfat5 | 11.41434588 | 11.70139648 | 11.64554519 | 0.055851293 | -0.231199313 |
| Lgalsl | 11.79830124 | 11.74495241 | 12.02951689 | -0.28456448 | -0.231215647 |
| Clec16a | 9.626714127 | 9.719619903 | 9.857933378 | -0.138313475 | -0.231219251 |
| Tmem40 | 9.757434676 | 10.07109486 | 9.988669077 | 0.08242578 | -0.231234401 |
| Pde4d | 7.877201201 | 8.02984896 | 8.108489001 | -0.078640041 | -0.2312878 |
| Ocrl | 9.333332346 | 9.582541814 | 9.56463598 | 0.017905834 | -0.231303633 |
| Mup15 | 6.509339836 | 6.857581734 | 6.74070211 | 0.116879624 | -0.231362274 |
| Impad1 | 11.33763767 | 11.35147189 | 11.56900122 | -0.217529333 | -0.231363547 |
| Sp3 | 10.75779393 | 10.80349514 | 10.98926684 | -0.185771697 | -0.231472907 |
| Clk1 | 11.24721566 | 11.44475965 | 11.47883571 | -0.03407606 | -0.231620053 |
| Mpzl1 | 7.939842914 | 8.046175948 | 8.171516748 | -0.125340799 | -0.231673834 |
| Ccnb2 | 7.677704172 | 7.690028882 | 7.909441507 | -0.219412625 | -0.231737334 |

| | | | | | |
|---|---|---|---|---|---|
| Lypla1 | 10.44778874 | 10.47784609 | 10.67954459 | -0.201698497 | -0.23175585 |
| Cyb561a3 | 8.652843724 | 8.908544486 | 8.884608137 | 0.023936349 | -0.231764413 |
| Gm14325 | 6.811723945 | 6.963795041 | 7.043726601 | -0.07993156 | -0.232002656 |
| Prom1 | 4.960071011 | 4.917511174 | 5.192096363 | -0.274585188 | -0.232025352 |
| Hs1bp3 | 8.115028636 | 8.397430338 | 8.347059104 | 0.050371234 | -0.232030467 |
| L3mbtl3 | 8.5172546 | 8.670083562 | 8.749381102 | -0.07929754 | -0.232126502 |
| Vwa2 | 8.481201148 | 8.77994354 | 8.713345458 | 0.066598082 | -0.23214431 |
| Efcab1 | 5.733297622 | 5.846063574 | 5.965566509 | -0.119502935 | -0.232268887 |
| E130308A19Rik | 7.251248757 | 7.427130984 | 7.483556171 | -0.056425187 | -0.232307414 |
| Kif2c | 7.334799777 | 7.445475546 | 7.567139484 | -0.121663938 | -0.232339708 |
| Stx5a | 10.0230391 | 10.24265296 | 10.25545991 | -0.012806953 | -0.232420815 |
| Pla2g16 | 10.30894469 | 10.16750643 | 10.54144275 | -0.373936323 | -0.232498057 |
| Proca1 | 6.316630899 | 6.273487354 | 6.549198991 | -0.275711637 | -0.232568093 |
| Zfp111 | 7.799405582 | 7.963227218 | 8.032198168 | -0.06897095 | -0.232792586 |
| Llgl2 | 8.858150849 | 9.234868803 | 9.091009663 | 0.14385914 | -0.232858814 |
| Bmp4 | 7.858145291 | 8.135524426 | 8.091185472 | 0.044338953 | -0.233040181 |
| Leprot | 10.53265944 | 10.58828986 | 10.76580869 | -0.17751883 | -0.233149257 |
| Lzts3 | 6.379445509 | 6.646279298 | 6.612628862 | 0.033650436 | -0.233183353 |
| Efnb1 | 9.716948777 | 9.871429076 | 9.950138979 | -0.078709903 | -0.233190203 |
| Shroom3 | 8.749832174 | 9.071637475 | 8.983092781 | 0.088544694 | -0.233260607 |
| Epor | 2.976268312 | 3.070587599 | 3.209701693 | -0.139114093 | -0.233433381 |
| Tenm4 | 10.78427934 | 10.958259 | 11.01781487 | -0.05955587 | -0.233535533 |
| Cnrip1 | 6.445988572 | 6.765497668 | 6.679543359 | 0.085954308 | -0.233554787 |
| Rspo4 | 5.486898166 | 5.722481744 | 5.720482394 | 0.001999351 | -0.233584228 |
| Ube2d1 | 9.383350926 | 9.362910572 | 9.616951985 | -0.254041413 | -0.233601059 |
| Kmt2e | 11.32444097 | 11.46476101 | 11.55807985 | -0.093318837 | -0.23363888 |
| Tnmd | 6.748753858 | 6.929165675 | 6.982422272 | -0.053256596 | -0.233668414 |
| Map3k10 | 7.550244251 | 7.791376823 | 7.783925398 | 0.007451425 | -0.233681147 |
| Cd81 | 10.96786928 | 11.05209667 | 11.20159289 | -0.149496223 | -0.233723617 |
| Nlrp1c-ps | 4.726150586 | 4.903073964 | 4.959959321 | -0.056885357 | -0.233808735 |
| Nkd1 | 6.899174774 | 7.190842816 | 7.133011647 | 0.057831169 | -0.233836874 |
| Abl1 | 10.04694739 | 10.15238351 | 10.2808469 | -0.128463397 | -0.233899513 |

| | | | | | |
|---|---|---|---|---|---|
| Nagk | 9.109044648 | 9.280218101 | 9.342947528 | -0.062729426 | -0.233902879 |
| Aspg | 7.719095146 | 8.082871601 | 7.953056749 | 0.129814852 | -0.233961603 |
| Mbtps1 | 10.85657158 | 10.99839271 | 11.09058871 | -0.092196 | -0.23401713 |
| Cry2 | 9.705413401 | 9.765809409 | 9.939502684 | -0.173693275 | -0.234089283 |
| Xlr | 5.440005577 | 5.479276691 | 5.67414404 | -0.194867349 | -0.234138463 |
| Zfp688 | 6.20390741 | 6.15711448 | 6.438181259 | -0.281066779 | -0.234273849 |
| Kyat1 | 9.176106743 | 9.515080388 | 9.410391416 | 0.104688971 | -0.234284673 |
| Ppm1m | 8.60352784 | 8.624795555 | 8.837816082 | -0.213020527 | -0.234288242 |
| Sulf2 | 10.21759722 | 10.27386549 | 10.45190317 | -0.178037687 | -0.234305953 |
| 6720489N17Rik | 5.397987039 | 5.406010931 | 5.632344783 | -0.226333852 | -0.234357744 |
| Stxbp3 | 10.27305937 | 10.41427758 | 10.50746362 | -0.093186047 | -0.234404257 |
| Mcoln3 | 4.740432603 | 4.943273997 | 4.975025083 | -0.031751086 | -0.23459248 |
| Diras2 | 5.141246274 | 5.20586593 | 5.375883277 | -0.170017347 | -0.234637003 |
| Cysltr1 | 7.616094852 | 7.692333214 | 7.85074395 | -0.158410736 | -0.234649098 |
| Spg21 | 10.26796473 | 10.32659421 | 10.50271905 | -0.176124843 | -0.23475432 |
| Ctf1 | 6.248606786 | 6.277304928 | 6.48341627 | -0.206111342 | -0.234809484 |
| Ulk3 | 8.536083211 | 8.800200699 | 8.771041237 | 0.029159462 | -0.234958026 |
| Ampd1 | 10.44099241 | 10.38079832 | 10.67605492 | -0.295256597 | -0.235062508 |
| Slc16a11 | 5.222476509 | 5.384243149 | 5.45756861 | -0.073325461 | -0.235092101 |
| Gabarapl1 | 10.15827862 | 10.5024838 | 10.39346891 | 0.10901489 | -0.23519029 |
| Bdh2 | 3.771121885 | 3.92445541 | 4.006334927 | -0.081879517 | -0.235213041 |
| Cog4 | 9.856913269 | 9.979787774 | 10.09215326 | -0.112365489 | -0.235239995 |
| Rnf222 | 8.136810735 | 8.33087966 | 8.372073351 | -0.041193691 | -0.235262616 |
| Supt3 | 6.640960682 | 6.621437302 | 6.876376597 | -0.254939295 | -0.235415915 |
| Hlcs | 8.86197349 | 9.021239742 | 9.097609664 | -0.076369922 | -0.235636174 |
| Gm12504 | 3.053467242 | 3.237889347 | 3.28928171 | -0.051392363 | -0.235814468 |
| Gm21596 | 4.44906473 | 4.571152527 | 4.684923855 | -0.113771328 | -0.235859125 |
| Plekha6 | 9.400429307 | 9.551532824 | 9.636312791 | -0.084779967 | -0.235883484 |
| Klf9 | 11.03501067 | 10.97206599 | 11.27108436 | -0.299018367 | -0.236073693 |
| Mb | 11.12143196 | 11.129501 | 11.3576924 | -0.228191397 | -0.236260443 |
| Cyhr1 | 10.38206498 | 10.66148844 | 10.61834876 | 0.04313968 | -0.23628378 |
| Shld2 | 7.422657852 | 7.541574024 | 7.658976777 | -0.117402753 | -0.236318925 |

| | | | | | |
|---|---|---|---|---|---|
| Rnf128 | 7.138320432 | 7.050828401 | 7.37466269 | -0.323834288 | -0.236342257 |
| Mfsd6 | 10.91864843 | 10.973979 | 11.15502184 | -0.181042843 | -0.23637341 |
| Bcl11a | 8.526941287 | 8.70893921 | 8.763356492 | -0.054417282 | -0.236415205 |
| Mlph | 5.548823194 | 5.82924892 | 5.785262477 | 0.043986444 | -0.236439283 |
| Trim72 | 9.010704318 | 9.326752818 | 9.247221048 | 0.07953177 | -0.23651673 |
| Adra2a | 5.387814302 | 5.324957413 | 5.624376845 | -0.299419431 | -0.236562542 |
| Ntng1 | 5.297285416 | 5.282493738 | 5.534155918 | -0.25166218 | -0.236870502 |
| Dusp22 | 10.58907289 | 10.66412346 | 10.82594918 | -0.161825713 | -0.236876283 |
| Echdc3 | 6.318648285 | 6.303562836 | 6.555613931 | -0.252051095 | -0.236965646 |
| Ip6k1 | 10.73147531 | 11.01166771 | 10.96845606 | 0.04321165 | -0.23698075 |
| Pik3c3 | 9.1495485 | 9.138659075 | 9.386653232 | -0.247994157 | -0.237104732 |
| Gstm7 | 5.045054023 | 4.994834292 | 5.282204724 | -0.287370431 | -0.237150701 |
| Pacsin3 | 9.76499841 | 9.814732652 | 10.00224842 | -0.187515771 | -0.237250013 |
| Kif18a | 6.718514047 | 6.83251298 | 6.955792048 | -0.123279068 | -0.237278001 |
| Slx1b | 8.044237006 | 8.162838644 | 8.281798886 | -0.118960242 | -0.23756188 |
| Fstl1 | 11.82687741 | 11.81170318 | 12.06474314 | -0.253039963 | -0.237865733 |
| Sclt1 | 7.053566035 | 7.183365905 | 7.291459432 | -0.108093528 | -0.237893398 |
| Txndc15 | 8.815011661 | 8.931991582 | 9.05290806 | -0.120916478 | -0.237896399 |
| Igf2r | 9.849190311 | 10.1445553 | 10.08708891 | 0.057466393 | -0.237898599 |
| Duxbl1 | 4.504792492 | 4.422265649 | 4.742772348 | -0.320506699 | -0.237979856 |
| Fam135a | 10.3762086 | 10.43634569 | 10.61426535 | -0.177919653 | -0.238056743 |
| Slc5a9 | 8.585638587 | 8.809328182 | 8.823771892 | -0.014443711 | -0.238133305 |
| Plxna1 | 10.99034062 | 11.40108241 | 11.22852804 | 0.17255437 | -0.238187417 |
| Vegfb | 8.078873472 | 8.196304701 | 8.317114091 | -0.120809389 | -0.238240618 |
| Hddc3 | 6.978308727 | 7.12481374 | 7.216591316 | -0.091777576 | -0.238282589 |
| Fnbp1 | 10.94515617 | 11.21232045 | 11.18357619 | 0.028744267 | -0.238420013 |
| Lmbr1 | 8.365118478 | 8.550954318 | 8.603605847 | -0.052651529 | -0.238487369 |
| Ccdc82 | 9.678894095 | 9.812379912 | 9.917465179 | -0.105085267 | -0.238571084 |
| Fbh1 | 9.789248676 | 9.879510721 | 10.02794244 | -0.148431722 | -0.238693767 |
| Plppr3 | 5.232025149 | 5.424914437 | 5.470775136 | -0.045860699 | -0.238749987 |
| Tmem216 | 6.672468515 | 6.718207893 | 6.911251009 | -0.193043116 | -0.238782493 |
| Slc6a6 | 12.20443513 | 12.22346061 | 12.44329411 | -0.219833493 | -0.238858977 |

| | | | | | |
|---|---|---|---|---|---|
| Ncoa1 | 9.988230761 | 10.09453372 | 10.22709129 | -0.132557563 | -0.238860526 |
| Ccl27a | 8.743007686 | 9.120955831 | 8.981876288 | 0.139079544 | -0.238868602 |
| Naxe | 8.862004324 | 8.915746896 | 9.100873926 | -0.185127029 | -0.238869602 |
| 40422 | 8.71216765 | 8.719284876 | 8.951044075 | -0.231759199 | -0.238876426 |
| Raph1 | 9.752059036 | 9.875754834 | 9.991028828 | -0.115273994 | -0.238969792 |
| Sema6a | 7.600250423 | 7.637408004 | 7.839287431 | -0.201879428 | -0.239037008 |
| Bin1 | 10.33103406 | 10.56826707 | 10.57009661 | -0.00182954 | -0.239062547 |
| Plac9a | 6.442036073 | 6.313002426 | 6.681145141 | -0.368142715 | -0.239109068 |
| Dcst1 | 3.596438744 | 3.788596202 | 3.835605572 | -0.047009371 | -0.239166829 |
| Sult5a1 | 9.640301442 | 9.891114823 | 9.879487296 | 0.011627527 | -0.239185854 |
| Ppp1r26 | 5.391413942 | 5.525332831 | 5.630646393 | -0.105313562 | -0.239232451 |
| Thsd4 | 6.923512458 | 6.803474128 | 7.162798068 | -0.35932394 | -0.23928561 |
| Unc93a2 | 6.543628331 | 6.712810764 | 6.782942663 | -0.070131899 | -0.239314332 |
| AW146154 | 6.9926891 | 7.084046807 | 7.232020394 | -0.147973587 | -0.239331294 |
| Wwc2 | 9.608167551 | 9.863626478 | 9.847518614 | 0.016107864 | -0.239351063 |
| Inppl1 | 9.346890837 | 9.539373461 | 9.586285419 | -0.046911958 | -0.239394583 |
| Crip3 | 4.557556984 | 4.717152955 | 4.796951808 | -0.079798853 | -0.239394824 |
| Lpo | 5.70039898 | 5.595967317 | 5.939810151 | -0.343842835 | -0.239411172 |
| Adamtsl4 | 8.745214391 | 8.896508384 | 8.984658153 | -0.088149769 | -0.239443762 |
| Aif1 | 7.36857766 | 7.421355744 | 7.608076251 | -0.186720507 | -0.239498591 |
| Atp10d | 8.156164546 | 8.097491735 | 8.395668803 | -0.298177069 | -0.239504258 |
| Fam81a | 9.708372892 | 9.813804668 | 9.94803346 | -0.134228792 | -0.239660568 |
| Oaz2 | 10.03537832 | 10.15504039 | 10.27504071 | -0.120000317 | -0.239662388 |
| Ptch2 | 6.535505246 | 6.663498267 | 6.775180603 | -0.111682336 | -0.239675357 |
| Scd1 | 16.35513229 | 15.91418855 | 16.59490856 | -0.68072001 | -0.239776273 |
| Gramd2 | 7.992331434 | 8.436696723 | 8.232174532 | 0.204522191 | -0.239843098 |
| Tnrc6a | 10.43132102 | 10.68826989 | 10.67119044 | 0.017079443 | -0.239869427 |
| Slc25a19 | 7.973862003 | 8.366519947 | 8.213745663 | 0.152774284 | -0.23988366 |
| Fbxo5 | 7.25047821 | 7.362280875 | 7.490362092 | -0.128081217 | -0.239883882 |
| Lce3f | 6.915317013 | 7.263501618 | 7.155222265 | 0.108279353 | -0.239905252 |
| Ttf2 | 8.056037643 | 8.242294617 | 8.295943074 | -0.053648457 | -0.239905431 |
| mt-Cytb | 16.20404013 | 16.2276102 | 16.44398831 | -0.216378107 | -0.239948183 |

| | | | | | |
|---|---|---|---|---|---|
| Rnf43 | 8.193526552 | 8.549746727 | 8.433490691 | 0.116256036 | -0.239964138 |
| Sebox | 6.224825886 | 6.350568134 | 6.464930493 | -0.114362358 | -0.240104606 |
| Ubxn6 | 9.382395645 | 9.532495513 | 9.622527531 | -0.090032018 | -0.240131886 |
| Wnt11 | 5.800003544 | 6.053932914 | 6.040164416 | 0.013768499 | -0.240160871 |
| Mccc2 | 8.754258321 | 8.6576757 | 8.994576516 | -0.336900815 | -0.240318195 |
| C130079G13Rik | 9.67752759 | 9.805183613 | 9.917996465 | -0.112812852 | -0.240468876 |
| Adig | 8.499880385 | 8.443234596 | 8.740379327 | -0.297144731 | -0.240498942 |
| Lnpep | 11.60486642 | 11.61688238 | 11.84548666 | -0.228604283 | -0.24062024 |
| Nxpe4 | 6.501030526 | 6.575549605 | 6.741653834 | -0.166104229 | -0.240623308 |
| Pex11g | 7.477764212 | 7.644440026 | 7.718432666 | -0.073992639 | -0.240668454 |
| Cnnm3 | 8.594714857 | 8.910873317 | 8.835422779 | 0.075450538 | -0.240707922 |
| Tspan17 | 8.239007134 | 8.127480212 | 8.479722286 | -0.352242074 | -0.240715152 |
| Cdca2 | 6.994196777 | 6.919297824 | 7.234940818 | -0.315642994 | -0.240744041 |
| mt-Nd4 | 15.84911946 | 15.88217095 | 16.0901184 | -0.207947443 | -0.240998933 |
| Retsat | 8.626570284 | 8.696459695 | 8.867658784 | -0.171199089 | -0.241088499 |
| Zfp526 | 7.116060413 | 7.435143303 | 7.357172675 | 0.077970627 | -0.241112262 |
| Alb | 5.940195118 | 5.689826817 | 6.181362196 | -0.49153538 | -0.241167079 |
| Btrc | 7.555227566 | 7.572688111 | 7.79647456 | -0.223786449 | -0.241246994 |
| Pdlim5 | 11.49761758 | 11.41458173 | 11.7389169 | -0.324335163 | -0.241299313 |
| Hhipl1 | 4.586006284 | 4.738248724 | 4.827532003 | -0.08928328 | -0.24152572 |
| Hook3 | 10.84773653 | 10.95082627 | 11.08932003 | -0.138493753 | -0.241583497 |
| Tbl1x | 10.5910731 | 10.77500073 | 10.83266708 | -0.057666347 | -0.24159398 |
| Racgap1 | 8.808955441 | 8.972503104 | 9.050550147 | -0.078047043 | -0.241594706 |
| Lxn | 8.215342528 | 8.265680158 | 8.456947641 | -0.191267483 | -0.241605113 |
| 2610002M06Rik | 8.094690704 | 8.164111933 | 8.336339548 | -0.172227615 | -0.241648844 |
| Ddhd2 | 8.938513284 | 9.024294387 | 9.180166938 | -0.155872551 | -0.241653654 |
| Plcb4 | 8.792878136 | 8.783457599 | 9.034540263 | -0.251082663 | -0.241662127 |
| Rxra | 12.08807379 | 12.46683251 | 12.32985507 | 0.136977433 | -0.241781287 |
| Dennd2a | 8.030841627 | 8.150366698 | 8.272642101 | -0.122275404 | -0.241800474 |
| Rab5a | 11.11688775 | 11.21413654 | 11.35891842 | -0.144781877 | -0.242030673 |
| Trpm3 | 5.269270106 | 5.291183489 | 5.511442752 | -0.220259264 | -0.242172646 |
| Gm14403 | 4.537287094 | 4.719070692 | 4.779498745 | -0.060428052 | -0.242211651 |

| | | | | | |
|---|---|---|---|---|---|
| Ptov1 | 9.457816363 | 9.802597627 | 9.700065874 | 0.102531753 | -0.242249511 |
| Mageh1 | 6.366718256 | 6.420535428 | 6.60896858 | -0.188433152 | -0.242250324 |
| Tmem63b | 9.989869348 | 10.37426915 | 10.23229534 | 0.141973807 | -0.242425995 |
| Zfp781 | 4.823643272 | 4.829257756 | 5.066130814 | -0.236873057 | -0.242487542 |
| Pnpla1 | 9.384011296 | 9.566125058 | 9.626510186 | -0.060385128 | -0.24249889 |
| Gnb5 | 7.033295196 | 7.112304334 | 7.275804328 | -0.163499994 | -0.242509132 |
| Hecw2 | 6.327350562 | 6.508562556 | 6.569927471 | -0.061364915 | -0.242576909 |
| Spock1 | 4.534438577 | 4.590279531 | 4.77708534 | -0.18680581 | -0.242646763 |
| Aplp2 | 12.73890189 | 12.67053416 | 12.98159685 | -0.311062693 | -0.24269496 |
| 2210017I01Rik | 6.798349681 | 6.937760045 | 7.041175085 | -0.10341504 | -0.242825404 |
| Amacr | 7.67849202 | 7.666149186 | 7.921319486 | -0.2551703 | -0.242827466 |
| Mib2 | 8.094107527 | 8.607476728 | 8.336999992 | 0.270476737 | -0.242892464 |
| Bbs4 | 7.033931184 | 7.049825532 | 7.276834309 | -0.227008777 | -0.242903125 |
| Poc1b | 7.627455137 | 7.645182658 | 7.870396665 | -0.225214006 | -0.242941528 |
| Tarsl2 | 7.749927837 | 7.817205813 | 7.992962013 | -0.1757562 | -0.243034176 |
| Kcnj15 | 8.680916064 | 9.00442975 | 8.923961375 | 0.080468375 | -0.243045311 |
| AL732309.1 | 9.525911263 | 9.672625881 | 9.768990541 | -0.096364659 | -0.243079278 |
| Caprin2 | 7.038576557 | 7.404960033 | 7.281777569 | 0.123182463 | -0.243201012 |
| Prkaa1 | 9.752235721 | 9.902333874 | 9.99543952 | -0.093105645 | -0.243203798 |
| Stx17 | 9.294553487 | 9.396109455 | 9.53778546 | -0.141676005 | -0.243231973 |
| Slc26a9 | 7.021080015 | 7.209683531 | 7.264371914 | -0.054688382 | -0.243291899 |
| Pde1c | 3.50181815 | 3.593127923 | 3.745388551 | -0.152260628 | -0.2435704 |
| Hpgds | 10.57456894 | 10.79526236 | 10.81814079 | -0.022878427 | -0.24357185 |
| Ric3 | 5.990112867 | 6.061407621 | 6.23377013 | -0.172362509 | -0.243657263 |
| Wasf1 | 5.198913647 | 5.368708362 | 5.44257618 | -0.073867818 | -0.243662534 |
| Uggt2 | 6.555660074 | 6.525691738 | 6.799453044 | -0.273761306 | -0.24379297 |
| Cryzl2 | 7.187553328 | 7.357582036 | 7.431359329 | -0.073777293 | -0.243806001 |
| Cyp2g1 | 11.9851863 | 11.98038767 | 12.22900937 | -0.2486217 | -0.243823067 |
| Fn3k | 4.607210876 | 4.578254491 | 4.851077681 | -0.27282319 | -0.243866805 |
| Usp22 | 10.70322664 | 10.81910899 | 10.94710611 | -0.12799712 | -0.243879473 |
| Gabre | 5.279676236 | 5.568115817 | 5.523723913 | 0.044391904 | -0.244047676 |
| Wdr7 | 8.840046669 | 8.954838302 | 9.084168366 | -0.129330064 | -0.244121697 |

| | | | | | |
|---|---|---|---|---|---|
| Ccdc122 | 6.89380315 | 6.876217327 | 7.138171538 | -0.261954211 | -0.244368388 |
| Nphp3 | 7.157731848 | 7.03666726 | 7.40226821 | -0.36560095 | -0.244536362 |
| Qk | 11.63399064 | 11.70800123 | 11.87853213 | -0.170530903 | -0.244541493 |
| Rhbdl2 | 7.583553987 | 7.829468664 | 7.828095603 | 0.001373061 | -0.244541616 |
| Smarcc2 | 10.77567647 | 11.0360765 | 11.02025671 | 0.015819783 | -0.244580247 |
| Pla2g4b | 6.0815292 | 6.513306333 | 6.326131308 | 0.187175024 | -0.244602108 |
| Rhbdd3 | 7.442391128 | 7.671549003 | 7.687055199 | -0.015506196 | -0.244664071 |
| Atg9b | 9.484675812 | 9.895258882 | 9.729465235 | 0.165793647 | -0.244789423 |
| Agbl1 | 4.803895876 | 4.864998816 | 5.048763452 | -0.183764636 | -0.244867576 |
| Gm19680 | 7.299098622 | 7.367580315 | 7.543970295 | -0.176389981 | -0.244871673 |
| AU040320 | 9.399123716 | 9.637773822 | 9.643995972 | -0.00622215 | -0.244872256 |
| Magi1 | 8.287834961 | 8.435206347 | 8.532853697 | -0.09764735 | -0.245018735 |
| Fpgt | 8.389268274 | 8.426514683 | 8.634349048 | -0.207834365 | -0.245080774 |
| Ndufaf3 | 7.239113077 | 7.410687438 | 7.484204196 | -0.073516758 | -0.245091119 |
| P2rx6 | 5.534348729 | 5.405915344 | 5.779490291 | -0.373574947 | -0.245141562 |
| Etfa | 11.05608357 | 11.08879389 | 11.3012646 | -0.21247071 | -0.245181027 |
| Grem1 | 9.881006094 | 9.6895474 | 10.12623049 | -0.436683087 | -0.245224392 |
| Smim14 | 9.920751247 | 10.06430223 | 10.16609312 | -0.101790885 | -0.245341873 |
| Mindy2 | 10.63582871 | 10.6559738 | 10.88118283 | -0.225209023 | -0.24535412 |
| Fhl3 | 9.290947615 | 9.49063294 | 9.536353537 | -0.045720597 | -0.245405921 |
| Arhgef40 | 8.500343426 | 8.726190166 | 8.745827286 | -0.01963712 | -0.24548386 |
| Mdm4 | 10.89910851 | 11.19879918 | 11.14461758 | 0.0541816 | -0.24550907 |
| Lancl2 | 8.237154756 | 8.376383372 | 8.482669383 | -0.106286011 | -0.245514627 |
| Bicral | 8.944179396 | 9.086622187 | 9.18980248 | -0.103180293 | -0.245623084 |
| Ugt1a6b | 6.314379395 | 6.122179209 | 6.560050113 | -0.437870903 | -0.245670717 |
| Krtdap | 14.49792972 | 14.5527354 | 14.74365845 | -0.190923053 | -0.245728733 |
| Maneal | 4.951142457 | 5.056155506 | 5.196943519 | -0.140788013 | -0.245801062 |
| Duox2 | 5.278194706 | 5.352376867 | 5.524060313 | -0.171683446 | -0.245865608 |
| Fam110a | 8.790734493 | 9.092382 | 9.036644917 | 0.055737082 | -0.245910424 |
| 2700099C18Rik | 4.99132747 | 5.156963919 | 5.237260726 | -0.080296807 | -0.245933256 |
| Zfp28 | 6.176568639 | 6.175652247 | 6.422536941 | -0.246884694 | -0.245968301 |
| Gm9625 | 4.643120981 | 4.801877223 | 4.889191429 | -0.087314206 | -0.246070448 |

| | | | | | |
|---|---|---|---|---|---|
| Sfi1 | 8.503855314 | 8.674162168 | 8.75007513 | -0.075912962 | -0.246219817 |
| Tbcel | 8.4886859 | 8.549224403 | 8.734950935 | -0.185726532 | -0.246265035 |
| Pgrmc1 | 11.1241342 | 11.1910232 | 11.37046331 | -0.179440107 | -0.246329113 |
| Atat1 | 6.860584596 | 7.203136616 | 7.107087087 | 0.096049529 | -0.246502491 |
| Phf21a | 9.555254682 | 9.843240934 | 9.801757937 | 0.041482998 | -0.246503255 |
| Aldh3b2 | 10.7229355 | 10.96881216 | 10.9694443 | -0.000632143 | -0.2465088 |
| Trim66 | 5.444114978 | 5.608139628 | 5.690633001 | -0.082493373 | -0.246518023 |
| Kcnk3 | 5.087872837 | 5.080104472 | 5.334405023 | -0.254300551 | -0.246532187 |
| Mup11 | 11.31612368 | 11.41758803 | 11.56265851 | -0.145070477 | -0.24653483 |
| Tepsin | 8.274461536 | 8.775247371 | 8.521012233 | 0.254235138 | -0.246550697 |
| Tmem80 | 7.61326465 | 7.817051996 | 7.859858922 | -0.042806926 | -0.246594272 |
| Mbd5 | 8.387540087 | 8.492527136 | 8.634171185 | -0.141644049 | -0.246631098 |
| Zfp113 | 7.328638325 | 7.483176917 | 7.575361263 | -0.092184346 | -0.246722938 |
| Lrp6 | 10.86351145 | 11.04225556 | 11.11026461 | -0.068009047 | -0.246753153 |
| Rps3a3 | 3.342525637 | 3.520607212 | 3.58939612 | -0.068788908 | -0.246870483 |
| Neurl1b | 9.033675619 | 9.228401072 | 9.28054975 | -0.052148678 | -0.246874131 |
| Slco5a1 | 7.255705777 | 7.172311601 | 7.502710632 | -0.330399031 | -0.247004855 |
| Twist1 | 8.413342662 | 8.38708844 | 8.660496071 | -0.273407631 | -0.247153409 |
| Cd180 | 7.30221836 | 7.533484005 | 7.549372173 | -0.015888168 | -0.247153813 |
| Snap47 | 8.260211942 | 8.403784443 | 8.507389884 | -0.103605442 | -0.247177942 |
| Spta1 | 4.709857795 | 4.730199389 | 4.957036705 | -0.226837315 | -0.247178909 |
| A930033H14Rik | 4.700624699 | 4.779623144 | 4.94783582 | -0.168212676 | -0.247211121 |
| Ano10 | 9.85622782 | 9.796124199 | 10.10346003 | -0.307335834 | -0.247232213 |
| Lhx6 | 4.996038913 | 5.288655848 | 5.243357762 | 0.045298087 | -0.247318848 |
| Dgke | 8.292318368 | 8.29433012 | 8.539732193 | -0.245402073 | -0.247413825 |
| Kcnh2 | 4.415901755 | 4.542579605 | 4.663374273 | -0.120794668 | -0.247472518 |
| Insr | 10.17017405 | 10.1676893 | 10.41772283 | -0.250033523 | -0.247548777 |
| Napb | 6.642764242 | 6.789630348 | 6.890352685 | -0.100722337 | -0.247588443 |
| Sptbn2 | 11.58899018 | 12.06338885 | 11.83669707 | 0.22669178 | -0.247706893 |
| Klhl7 | 8.982126062 | 9.018184442 | 9.229897885 | -0.211713443 | -0.247771823 |
| Fam107a | 5.943244697 | 5.988734946 | 6.191043036 | -0.20230809 | -0.247798339 |
| Tdrd3 | 9.246190402 | 9.278602454 | 9.494151804 | -0.21554935 | -0.247961401 |

| | | | | | |
|---|---|---|---|---|---|
| Acbd5 | 10.08378844 | 10.12695268 | 10.33178397 | -0.204831283 | -0.247995529 |
| Jkamp | 8.60077397 | 8.660124153 | 8.84877909 | -0.188654937 | -0.24800512 |
| Zfp3 | 6.050065229 | 6.082118714 | 6.298124115 | -0.216005401 | -0.248058886 |
| Ccdc50 | 11.08585685 | 11.13514511 | 11.33393732 | -0.19879221 | -0.248080477 |
| AC122211.1 | 3.37842407 | 3.565891917 | 3.626508438 | -0.060616521 | -0.248084369 |
| Exoc6 | 8.375318808 | 8.267471789 | 8.623415642 | -0.355943852 | -0.248096834 |
| Abra | 7.713484678 | 8.941775696 | 7.961589916 | 0.98018578 | -0.248105238 |
| Arhgef11 | 10.32386183 | 10.52965415 | 10.57197963 | -0.042325477 | -0.248117803 |
| Ccnb1 | 8.662863899 | 8.784822055 | 8.91109436 | -0.126272305 | -0.24823046 |
| Fgd1 | 6.866836619 | 7.064031589 | 7.115077354 | -0.051045765 | -0.248240735 |
| Ldb1 | 10.70027049 | 11.10376868 | 10.94852575 | 0.155242923 | -0.24825526 |
| Tmem237 | 6.886409519 | 6.94260645 | 7.134673922 | -0.192067471 | -0.248264403 |
| Eml1 | 9.25933591 | 9.340800567 | 9.507702759 | -0.166902193 | -0.248366849 |
| Vsig10 | 8.304066063 | 8.562470486 | 8.552439793 | 0.010030693 | -0.24837373 |
| AC124237.1 | 5.582059072 | 5.796044702 | 5.830490929 | -0.034446227 | -0.248431857 |
| Zcwpw1 | 6.1267746 | 6.308863714 | 6.375273213 | -0.066409499 | -0.248498613 |
| Scarf2 | 7.07740321 | 7.358595698 | 7.325904067 | 0.032691631 | -0.248500857 |
| Cldn5 | 7.392585819 | 7.555084279 | 7.641174954 | -0.086090675 | -0.248589134 |
| Pigg | 8.946165755 | 9.103942028 | 9.194774553 | -0.090832525 | -0.248608798 |
| Prmt2 | 7.356614704 | 7.631676616 | 7.605251658 | 0.026424958 | -0.248636954 |
| Asprv1 | 14.39240863 | 14.55568046 | 14.64108099 | -0.085400527 | -0.248672353 |
| Cd55b | 3.222772381 | 3.422806638 | 3.471469591 | -0.048662953 | -0.24869721 |
| Pgrmc2 | 11.25553312 | 11.34959676 | 11.50431161 | -0.154714847 | -0.248778487 |
| Phip | 10.116144 | 10.14636999 | 10.36494254 | -0.21857255 | -0.248798537 |
| Klb | 5.256213712 | 4.982575879 | 5.505243206 | -0.522667327 | -0.249029493 |
| Ube2r2 | 11.21011067 | 11.3165145 | 11.45921218 | -0.142697673 | -0.249101507 |
| Tmem141 | 7.136589428 | 7.179858898 | 7.385716997 | -0.205858098 | -0.249127569 |
| Kcnq4 | 6.105264056 | 6.300463078 | 6.354443887 | -0.053980809 | -0.249179831 |
| Ube2b | 11.10980175 | 11.18447072 | 11.35905513 | -0.17458441 | -0.24925338 |
| Tpp1 | 11.07789067 | 11.2903749 | 11.32717238 | -0.03679748 | -0.249281713 |
| Mavs | 9.416651697 | 9.55882594 | 9.666055318 | -0.107229379 | -0.249403621 |
| Lce3b | 5.626842638 | 5.732519126 | 5.876248744 | -0.143729618 | -0.249406106 |

| | | | | | |
|---|---|---|---|---|---|
| Nek1 | 8.429834545 | 8.673449283 | 8.679296692 | -0.00584741 | -0.249462148 |
| Harbi1 | 7.28375221 | 7.316060203 | 7.533304528 | -0.217244326 | -0.249552319 |
| Mat2b | 10.12648978 | 10.0844899 | 10.37614376 | -0.29165386 | -0.249653977 |
| Nap1l3 | 3.649940853 | 3.754839489 | 3.899681512 | -0.144842022 | -0.249740659 |
| Evi5 | 9.916374729 | 9.868534672 | 10.16611694 | -0.297582265 | -0.249742207 |
| Pycard | 10.88341974 | 11.01470776 | 11.13322503 | -0.118517263 | -0.249805287 |
| Med14 | 10.32101416 | 10.4231997 | 10.57091179 | -0.147712083 | -0.249897623 |
| Cers1 | 4.172541976 | 4.206308899 | 4.422562681 | -0.216253781 | -0.250020705 |
| Acot1 | 9.332822302 | 9.56302676 | 9.582878126 | -0.019851365 | -0.250055823 |
| Clstn1 | 11.14051129 | 11.35397053 | 11.39062096 | -0.036650433 | -0.250109673 |
| Cacng7 | 4.677883308 | 4.699200902 | 4.928002981 | -0.228802079 | -0.250119673 |
| Lce1k | 9.355134011 | 9.678765652 | 9.605318096 | 0.073447556 | -0.250184085 |
| Dock7 | 9.234915328 | 9.416038687 | 9.485126786 | -0.069088099 | -0.250211458 |
| Zc3hav1l | 6.988337223 | 7.276343169 | 7.238692639 | 0.037650529 | -0.250355416 |
| Slc16a12 | 6.914223539 | 7.0589805 | 7.164717543 | -0.105737043 | -0.250494004 |
| Mark1 | 7.37663439 | 7.268741018 | 7.627144034 | -0.358403016 | -0.250509644 |
| Nmt2 | 9.462408537 | 9.487446229 | 9.712952247 | -0.225506018 | -0.25054371 |
| B4galt4 | 7.255610038 | 7.281378827 | 7.506167971 | -0.224789144 | -0.250557933 |
| Mkks | 8.127463893 | 8.051518301 | 8.378192615 | -0.326674314 | -0.250728722 |
| Fbxl4 | 8.111156757 | 7.948340402 | 8.361901216 | -0.413560814 | -0.250744459 |
| Plcl1 | 6.123306128 | 6.572013151 | 6.374096804 | 0.197916348 | -0.250790676 |
| Tie1 | 8.133173496 | 8.301164929 | 8.384179652 | -0.083014723 | -0.251006156 |
| Cyp4f13 | 7.914085049 | 8.015320812 | 8.165181847 | -0.149861035 | -0.251096798 |
| Ocel1 | 6.696539972 | 7.108389872 | 6.947638529 | 0.160751343 | -0.251098557 |
| Stbd1 | 6.733262529 | 6.812747196 | 6.984366284 | -0.171619088 | -0.251103755 |
| Lztfl1 | 8.709836477 | 8.778442358 | 8.960994999 | -0.182552642 | -0.251158522 |
| Cdh9 | 3.737469508 | 3.761962607 | 3.988639103 | -0.226676496 | -0.251169596 |
| Ssc5d | 7.588916644 | 7.762497942 | 7.840143676 | -0.077645734 | -0.251227032 |
| Ppm1b | 10.45835912 | 10.53114432 | 10.70963304 | -0.178488723 | -0.251273923 |
| Slc7a2 | 9.083105229 | 8.830899645 | 9.334420458 | -0.503520813 | -0.251315229 |
| Gm14308 | 4.999252315 | 4.958032768 | 5.250634183 | -0.292601415 | -0.251381868 |
| B3glct | 7.797828972 | 7.771440595 | 8.049238652 | -0.277798057 | -0.25140968 |

| | | | | | |
|---|---|---|---|---|---|
| D130040H23Rik | 5.044152754 | 5.25090942 | 5.295599297 | -0.044689877 | -0.251446543 |
| Msi2 | 9.421314368 | 9.439644686 | 9.672769716 | -0.23312503 | -0.251455348 |
| Zfyve9 | 10.51059814 | 10.58446242 | 10.76209808 | -0.17763566 | -0.251499937 |
| Zfp189 | 5.67968388 | 5.945997556 | 5.9312296 | 0.014767956 | -0.25154572 |
| Pdcl | 9.352657947 | 9.429491952 | 9.604269849 | -0.174777898 | -0.251611902 |
| Ing4 | 9.558192167 | 9.700268122 | 9.809817589 | -0.109549467 | -0.251625423 |
| Gnrh1 | 4.800798792 | 5.005216089 | 5.052481857 | -0.047265768 | -0.251683066 |
| Rassf10 | 7.865575413 | 7.983010547 | 8.117314205 | -0.134303658 | -0.251738792 |
| Gm4202 | 3.827165015 | 4.227260884 | 4.079246074 | 0.148014809 | -0.25208106 |
| Proser3 | 6.123664511 | 6.377407024 | 6.375760192 | 0.001646831 | -0.252095681 |
| Epb41l2 | 10.66352059 | 10.78638872 | 10.91562648 | -0.129237753 | -0.252105887 |
| Rasa1 | 10.0441511 | 10.06714941 | 10.29636646 | -0.229217047 | -0.252215363 |
| Ighm | 9.806990189 | 9.240346506 | 10.05922478 | -0.818878274 | -0.252234591 |
| Prepl | 8.45292776 | 8.64094738 | 8.705228981 | -0.064281601 | -0.252301221 |
| Gm14410 | 6.607123162 | 6.90689829 | 6.859580795 | 0.047317495 | -0.252457633 |
| Myoz2 | 6.553910555 | 6.520086729 | 6.806518677 | -0.286431948 | -0.252608122 |
| Gpatch8 | 9.970268514 | 10.16181532 | 10.22292947 | -0.061114153 | -0.252660956 |
| mt-Nd2 | 15.1770218 | 15.20530077 | 15.42971633 | -0.224415567 | -0.252694533 |
| Lctl | 5.766305321 | 5.613508621 | 6.019015466 | -0.405506845 | -0.252710145 |
| Tmem53 | 5.519696894 | 5.602653578 | 5.772474984 | -0.169821406 | -0.25277809 |
| Neil3 | 6.373793953 | 6.59321018 | 6.626661615 | -0.033451435 | -0.252867662 |
| Zfyve19 | 7.979855169 | 8.203487488 | 8.233000026 | -0.029512538 | -0.253144856 |
| Rttn | 7.223937349 | 7.456175765 | 7.47714093 | -0.020965165 | -0.253203581 |
| Evi5l | 9.042248914 | 9.241803528 | 9.295458792 | -0.053655264 | -0.253209878 |
| Srpk2 | 10.59475005 | 10.81438528 | 10.84820717 | -0.033821893 | -0.25345712 |
| Sult2b1 | 9.599738689 | 9.911413197 | 9.853450172 | 0.057963025 | -0.253711483 |
| Abhd18 | 5.944347298 | 6.030362706 | 6.198066742 | -0.167704036 | -0.253719443 |
| Zfp235 | 6.719030312 | 6.920257281 | 6.972777861 | -0.052520581 | -0.25374755 |
| Fchsd2 | 9.361769844 | 9.573209435 | 9.615563723 | -0.042354288 | -0.253793879 |
| Zfp39 | 6.778872069 | 6.793395447 | 7.032675761 | -0.239280315 | -0.253803692 |
| Krt32 | 6.131909531 | 6.289823026 | 6.385736012 | -0.095912987 | -0.253826481 |
| Melk | 7.294477643 | 7.366723876 | 7.548415001 | -0.181691125 | -0.253937358 |

| | | | | | |
|---|---|---|---|---|---|
| Adprhl1 | 6.13750378 | 6.939398509 | 6.39145267 | 0.547945839 | -0.25394889 |
| Zc4h2 | 4.97512058 | 5.109527033 | 5.229086577 | -0.119559545 | -0.253965997 |
| Cog6 | 9.268792716 | 9.330260378 | 9.522779331 | -0.192518953 | -0.253986615 |
| Foxo1 | 10.18786243 | 10.31872659 | 10.4420433 | -0.12331671 | -0.254180877 |
| Meis1 | 7.868825556 | 7.954228273 | 8.123086728 | -0.168858455 | -0.254261172 |
| Phospho1 | 9.884462669 | 10.35268565 | 10.1389112 | 0.213774452 | -0.254448529 |
| Rcsd1 | 8.967941932 | 8.812353123 | 9.222393511 | -0.410040388 | -0.254451579 |
| Bcl2l13 | 10.82295566 | 11.09899145 | 11.07741246 | 0.021578993 | -0.2544568 |
| Bckdhb | 9.145153695 | 9.281282653 | 9.399646315 | -0.118363662 | -0.25449262 |
| AI429214 | 6.09397205 | 6.257906827 | 6.348510173 | -0.090603346 | -0.254538123 |
| Man1c1 | 9.226445703 | 9.423859597 | 9.481000252 | -0.057140655 | -0.254554549 |
| Dmac2 | 8.035008608 | 8.221858057 | 8.289723748 | -0.067865691 | -0.25471514 |
| Zfp54 | 6.666061504 | 6.886736259 | 6.920988062 | -0.034251803 | -0.254926558 |
| Tango2 | 7.22697757 | 7.311589283 | 7.48193006 | -0.170340777 | -0.25495249 |
| 2810025M15Rik | 7.63216709 | 7.730153924 | 7.887133692 | -0.156979769 | -0.254966602 |
| Trim32 | 7.883143862 | 8.028644878 | 8.138141741 | -0.109496863 | -0.254997879 |
| Fitm2 | 10.37629182 | 10.44196961 | 10.63130188 | -0.18933227 | -0.255010053 |
| Tmem52 | 6.395218974 | 6.945672115 | 6.65029224 | 0.295379875 | -0.255073266 |
| Dkk2 | 8.396060536 | 8.221167309 | 8.651164482 | -0.429997173 | -0.255103946 |
| Tut4 | 8.409656011 | 8.20123917 | 8.66479731 | -0.46355814 | -0.255141299 |
| Ebpl | 7.424466728 | 7.464845477 | 7.679638681 | -0.214793204 | -0.255171953 |
| Map3k9 | 8.201574188 | 8.52451764 | 8.456825844 | 0.067691796 | -0.255251656 |
| Ces1f | 6.357396004 | 6.33422215 | 6.612771142 | -0.278548992 | -0.255375138 |
| Tekt2 | 4.79611403 | 5.059311323 | 5.051575178 | 0.007736145 | -0.255461148 |
| Barx2 | 9.648567295 | 9.755565165 | 9.904053239 | -0.148488074 | -0.255485944 |
| Dcaf5 | 9.440037515 | 9.518468411 | 9.695627366 | -0.177158956 | -0.255589851 |
| Prdx4 | 8.178069388 | 8.219223646 | 8.433687373 | -0.214463727 | -0.255617985 |
| Tmtc3 | 9.447615253 | 9.796193005 | 9.703350099 | 0.092842906 | -0.255734846 |
| Efemp1 | 11.31367421 | 11.26831808 | 11.56970301 | -0.301384937 | -0.256028803 |
| Rab9 | 9.299894452 | 9.32795292 | 9.556005013 | -0.228052093 | -0.256110561 |
| Gm21013 | 5.031803496 | 5.231910465 | 5.288064706 | -0.056154241 | -0.25626121 |
| Acaca | 11.84235984 | 11.60304055 | 12.09863562 | -0.495595073 | -0.25627578 |

| | | | | | |
|---|---|---|---|---|---|
| Ubxn2b | 7.558326104 | 7.734084409 | 7.814761139 | -0.08067673 | -0.256435035 |
| Gtf2i | 10.70283415 | 10.83163946 | 10.95930465 | -0.127665187 | -0.256470497 |
| Pyroxd1 | 8.268954495 | 8.414743587 | 8.525549038 | -0.110805451 | -0.256594543 |
| Ccna2 | 9.224059021 | 9.312602529 | 9.48085493 | -0.168252401 | -0.256795908 |
| Slc4a9 | 3.807957543 | 4.131402411 | 4.064939932 | 0.066462479 | -0.25698239 |
| Cpped1 | 9.170076972 | 9.346883685 | 9.427094506 | -0.080210821 | -0.257017534 |
| Mob2 | 9.473585658 | 9.68040388 | 9.73061764 | -0.05021376 | -0.257031982 |
| Apba1 | 6.950897863 | 6.94758463 | 7.207949551 | -0.260364921 | -0.257051688 |
| Zfp14 | 5.748072162 | 5.91428832 | 6.005296031 | -0.091007711 | -0.257223868 |
| Trim24 | 8.815157086 | 8.764296481 | 9.072401067 | -0.308104586 | -0.257243981 |
| Stau2 | 8.176540585 | 8.296959123 | 8.433812111 | -0.136852988 | -0.257271526 |
| Gm21451 | 3.98407968 | 4.079482786 | 4.241363595 | -0.161880809 | -0.257283914 |
| Pcbd2 | 6.448466413 | 6.510790122 | 6.705773445 | -0.194983324 | -0.257307032 |
| Smim8 | 7.403781558 | 7.497949531 | 7.661101803 | -0.163152272 | -0.257320245 |
| Popdc3 | 7.307781848 | 7.694899292 | 7.565337124 | 0.129562168 | -0.257555276 |
| Tns1 | 12.38211391 | 12.50566481 | 12.63969647 | -0.134031663 | -0.257582563 |
| Clybl | 8.065159934 | 8.020263159 | 8.322921428 | -0.30265827 | -0.257761494 |
| Actn2 | 10.42513004 | 10.76931151 | 10.68293876 | 0.086372747 | -0.257808728 |
| Hmmr | 7.963272232 | 8.011172994 | 8.221174543 | -0.210001548 | -0.257902311 |
| Igfbp4 | 11.25248553 | 11.27858308 | 11.51048311 | -0.231900023 | -0.257997573 |
| Nr2c2 | 10.04422781 | 10.1447333 | 10.30240119 | -0.15766789 | -0.258173376 |
| Apobec2 | 9.824626927 | 9.786048308 | 10.08281352 | -0.296765207 | -0.258186588 |
| F830045P16Rik | 5.971713298 | 5.700855959 | 6.22992071 | -0.529064752 | -0.258207413 |
| Kif7 | 5.768003914 | 5.998928116 | 6.026265578 | -0.027337462 | -0.258261664 |
| Smim5 | 7.869128165 | 7.995361097 | 8.127421257 | -0.13206016 | -0.258293092 |
| 36951 | 5.064813785 | 4.829917066 | 5.323483253 | -0.493566188 | -0.258669469 |
| Polm | 8.314713986 | 8.638244714 | 8.573471139 | 0.064773575 | -0.258757154 |
| Dpf3 | 4.581161028 | 4.614128128 | 4.839921223 | -0.225793095 | -0.258760195 |
| Pdxdc1 | 11.05523474 | 11.13855736 | 11.31420087 | -0.175643513 | -0.258966133 |
| Cacna1a | 5.816201547 | 6.128116688 | 6.07521822 | 0.052898468 | -0.259016673 |
| Gnptab | 10.18873428 | 10.41975493 | 10.44776694 | -0.02801201 | -0.259032657 |
| Prox1 | 6.030627472 | 6.123642362 | 6.289678251 | -0.166035889 | -0.25905078 |

| | | | | | |
|---|---|---|---|---|---|
| Pdgfrl | 6.287500273 | 6.381005302 | 6.546689933 | -0.165684631 | -0.25918966 |
| Trp53inp2 | 11.87342675 | 12.04860195 | 12.13262756 | -0.084025607 | -0.259200807 |
| Tbx20 | 2.10006757 | 2.15342263 | 2.359330874 | -0.205908244 | -0.259263304 |
| Wdr66 | 6.611016108 | 6.795939821 | 6.870359724 | -0.074419903 | -0.259343616 |
| Anapc2 | 10.49047871 | 10.66627063 | 10.74984177 | -0.08357114 | -0.25936306 |
| Gp1bb | 5.514694814 | 5.728170312 | 5.774103202 | -0.04593289 | -0.259408388 |
| Cep250 | 9.002354625 | 9.291951531 | 9.261872026 | 0.030079505 | -0.259517401 |
| Nr2f1 | 4.452870307 | 4.55712963 | 4.712404847 | -0.155275217 | -0.25953454 |
| Zfp109 | 5.538453802 | 5.794899072 | 5.798004436 | -0.003105364 | -0.259550635 |
| Rpl3l | 9.484607256 | 9.656510939 | 9.744216577 | -0.087705638 | -0.25960932 |
| Entpd5 | 10.11899521 | 10.13785483 | 10.37860479 | -0.240749967 | -0.25960958 |
| Zfp772 | 8.35691124 | 8.437104173 | 8.61660631 | -0.179502137 | -0.25969507 |
| Exd1 | 4.958354264 | 5.126119977 | 5.218151813 | -0.092031836 | -0.25979755 |
| Pinlyp | 6.880238556 | 7.056848751 | 7.140078307 | -0.083229556 | -0.259839752 |
| Gnao1 | 6.171318573 | 6.554296179 | 6.431180535 | 0.123115644 | -0.259861962 |
| Hltf | 8.871158078 | 9.085106127 | 9.131118801 | -0.046012675 | -0.259960723 |
| Npff | 5.342223028 | 5.597311934 | 5.602201094 | -0.004889161 | -0.259978067 |
| Gm12397 | 5.425768265 | 5.609244508 | 5.685784979 | -0.076540471 | -0.260016714 |
| Polk | 7.91558671 | 8.107168325 | 8.175607041 | -0.068438716 | -0.260020331 |
| Pisd-ps2 | 5.034935571 | 5.195888554 | 5.294959656 | -0.099071103 | -0.260024085 |
| Bgn | 11.81164803 | 12.00089744 | 12.07169786 | -0.070800427 | -0.260049833 |
| Sp4 | 7.454229706 | 7.394712013 | 7.714409831 | -0.319697819 | -0.260180125 |
| Arhgap29 | 10.41276029 | 10.39668093 | 10.67295948 | -0.276278547 | -0.260199187 |
| Stum | 5.320825074 | 5.310526124 | 5.581080953 | -0.270554829 | -0.260255879 |
| Nuf2 | 7.417803698 | 7.436160496 | 7.678130952 | -0.241970456 | -0.260327253 |
| Ttc33 | 8.388856828 | 8.384179568 | 8.649192623 | -0.265013055 | -0.260335795 |
| Rnf13 | 10.60579189 | 10.60585684 | 10.86618918 | -0.26033234 | -0.260397287 |
| Actc1 | 5.914683946 | 6.372379379 | 6.175189432 | 0.197189947 | -0.260505485 |
| Bicdl2 | 8.892208195 | 9.107273639 | 9.152920562 | -0.045646923 | -0.260712367 |
| Wdpcp | 5.858033563 | 6.022635002 | 6.118803688 | -0.096168686 | -0.260770125 |
| Mmp16 | 4.833455478 | 4.752497837 | 5.094266503 | -0.341768666 | -0.260811025 |
| Mlh3 | 8.012510721 | 8.071935791 | 8.273466794 | -0.201531003 | -0.260956073 |

| | | | | | |
|---|---|---|---|---|---|
| Asb8 | 9.026267949 | 8.913516263 | 9.287242283 | -0.37372602 | -0.260974333 |
| Mfsd4a | 5.924132792 | 5.988262731 | 6.185167086 | -0.196904354 | -0.261034293 |
| Senp8 | 6.459541679 | 6.671233184 | 6.720624349 | -0.049391165 | -0.26108267 |
| D630003M21Rik | 4.751270789 | 5.056039921 | 5.012713868 | 0.043326052 | -0.261443079 |
| D3Ertd254e | 8.002424172 | 8.059402296 | 8.263978981 | -0.204576685 | -0.26155481 |
| Scp2 | 12.13929879 | 12.10132236 | 12.40098297 | -0.29966061 | -0.261684173 |
| Colgalt2 | 4.272254267 | 4.352301777 | 4.534031031 | -0.181729254 | -0.261776764 |
| Rnf225 | 10.16878305 | 10.37540133 | 10.43057696 | -0.05517563 | -0.261793907 |
| Tm7sf2 | 7.709226623 | 7.787456765 | 7.971088329 | -0.183631564 | -0.261861707 |
| Inpp5e | 8.020590391 | 8.328641133 | 8.28245365 | 0.046187483 | -0.261863259 |
| Retreg1 | 10.55398694 | 11.13936357 | 10.8158972 | 0.323466373 | -0.261910263 |
| Klf12 | 6.62191323 | 6.807854389 | 6.883831318 | -0.075976929 | -0.261918088 |
| Gm45234 | 3.518418129 | 4.275958287 | 3.780389465 | 0.495568822 | -0.261971335 |
| Usp2 | 9.948816437 | 9.709888437 | 10.21086556 | -0.50097712 | -0.262049119 |
| Nfatc2 | 8.93949798 | 9.182786156 | 9.201743311 | -0.018957155 | -0.262245331 |
| Flrt2 | 9.724009218 | 9.786269615 | 9.986355362 | -0.200085747 | -0.262346144 |
| Gstk1 | 8.267841213 | 8.162561796 | 8.530298685 | -0.367736889 | -0.262457471 |
| Ttll1 | 5.851239759 | 6.103081573 | 6.113707848 | -0.010626274 | -0.262468089 |
| Ufl1 | 9.070866429 | 9.196417792 | 9.333473684 | -0.137055892 | -0.262607255 |
| Rere | 10.94194213 | 11.30027355 | 11.20456484 | 0.095708717 | -0.26262271 |
| Mecom | 5.767494681 | 5.877827988 | 6.030300264 | -0.152472276 | -0.262805583 |
| Glt8d2 | 6.120667788 | 6.061227341 | 6.383516724 | -0.322289382 | -0.262848935 |
| Polr3gl | 8.030979853 | 8.033951009 | 8.293910574 | -0.259959565 | -0.262930721 |
| Dusp18 | 9.162120236 | 9.350545334 | 9.425056119 | -0.074510785 | -0.262935884 |
| Tmem177 | 8.684364782 | 8.800591158 | 8.947378457 | -0.146787299 | -0.263013675 |
| Prdm16 | 6.590114478 | 7.185560744 | 6.853143239 | 0.332417504 | -0.263028761 |
| Lamc1 | 11.43821635 | 11.41380774 | 11.70127786 | -0.28747012 | -0.263061513 |
| Ephb6 | 11.05669391 | 11.36466314 | 11.31984533 | 0.04481781 | -0.263151423 |
| Ccdc157 | 7.140888644 | 7.422941924 | 7.404062661 | 0.018879263 | -0.263174017 |
| 9430015G10Rik | 7.625349332 | 7.996261751 | 7.888621699 | 0.107640051 | -0.263272367 |
| Bmp2 | 8.491650712 | 8.649064714 | 8.754940733 | -0.105876019 | -0.263290021 |
| Prkd1 | 7.068027669 | 7.020994196 | 7.331387642 | -0.310393445 | -0.263359972 |

| | | | | | |
|---|---|---|---|---|---|
| Zscan26 | 9.685782147 | 9.863689059 | 9.949468836 | -0.085779777 | -0.263686689 |
| Arhgef12 | 12.00792835 | 12.18692207 | 12.27163554 | -0.08471347 | -0.26370719 |
| Phlpp2 | 7.825718051 | 7.953533067 | 8.089456161 | -0.135923094 | -0.26373811 |
| Tnfrsf25 | 5.697589022 | 6.159515929 | 5.961345888 | 0.198170041 | -0.263756866 |
| Xntrpc | 4.768995565 | 4.960043707 | 5.032783895 | -0.072740188 | -0.26378833 |
| Sdr16c5 | 10.268102 | 10.30522687 | 10.53193466 | -0.22670779 | -0.263832667 |
| Pigh | 7.888096086 | 8.069100085 | 8.151939702 | -0.082839617 | -0.263843617 |
| Micu3 | 7.095708438 | 7.216825093 | 7.359587079 | -0.142761987 | -0.263878641 |
| Fign | 5.436981418 | 5.343222393 | 5.701029443 | -0.35780705 | -0.264048025 |
| Mfsd11 | 9.303340426 | 9.562213471 | 9.567684964 | -0.005471492 | -0.264344538 |
| Acadsb | 10.48437719 | 10.39275859 | 10.74873428 | -0.355975687 | -0.264357093 |
| Slc23a3 | 5.128075354 | 5.188840405 | 5.392446283 | -0.203605878 | -0.26437093 |
| Atg4c | 7.402974391 | 7.532825489 | 7.667354481 | -0.134528991 | -0.26438009 |
| Sardh | 7.490997447 | 7.754948557 | 7.755378538 | -0.000429981 | -0.264381091 |
| Pptc7 | 9.779100065 | 9.911259796 | 10.04365787 | -0.132398069 | -0.2645578 |
| Sirt5 | 7.73868698 | 7.86446484 | 8.00328296 | -0.138818121 | -0.26459598 |
| Nxt2 | 7.936162087 | 8.078518885 | 8.200777005 | -0.12225812 | -0.264614918 |
| Tmem106c | 7.937836122 | 8.045886887 | 8.202479486 | -0.156592599 | -0.264643365 |
| Gm36298 | 4.545984436 | 4.570720181 | 4.810692875 | -0.239972694 | -0.264708439 |
| Csad | 9.135293636 | 9.336436858 | 9.400012036 | -0.063575178 | -0.2647184 |
| Aldh7a1 | 9.144751788 | 9.073115113 | 9.409584665 | -0.336469552 | -0.264832876 |
| Myom1 | 11.53744595 | 11.59728898 | 11.80239193 | -0.205102953 | -0.264945977 |
| Kbtbd12 | 5.883946567 | 6.205216787 | 6.148991071 | 0.056225716 | -0.265044504 |
| Tmem260 | 8.605754719 | 8.777539811 | 8.870958779 | -0.093418968 | -0.265204059 |
| Tmem170 | 8.124322593 | 8.316666666 | 8.389553196 | -0.07288653 | -0.265230602 |
| Lyplal1 | 5.684551222 | 5.572613823 | 5.949877325 | -0.377263502 | -0.265326102 |
| Gm5141 | 5.523199176 | 5.819081228 | 5.788641269 | 0.030439959 | -0.265442093 |
| Usp6nl | 9.822689588 | 9.903950069 | 10.08820364 | -0.184253573 | -0.265514054 |
| Fbxl3 | 10.62458756 | 10.59940515 | 10.89015282 | -0.29074767 | -0.26556526 |
| Qser1 | 10.07289977 | 10.18772102 | 10.33850751 | -0.150786493 | -0.265607737 |
| Cmc4 | 6.220649949 | 6.473350504 | 6.486344654 | -0.01299415 | -0.265694705 |
| B230118H07Rik | 7.794064354 | 7.661547227 | 8.059938714 | -0.398391487 | -0.26587436 |

| | | | | | |
|---|---|---|---|---|---|
| Ovol2 | 5.361635505 | 5.705187779 | 5.627558701 | 0.077629078 | -0.265923196 |
| Dach1 | 7.231556565 | 7.171181467 | 7.497817472 | -0.326636005 | -0.266260908 |
| Eri2 | 7.348547627 | 7.273932302 | 7.615018152 | -0.34108585 | -0.266470525 |
| Ankrd9 | 7.363573046 | 7.231035919 | 7.630044501 | -0.399008582 | -0.266471455 |
| Clk4 | 9.779219221 | 10.00821435 | 10.04586797 | -0.037653621 | -0.266648752 |
| 38961 | 7.812010302 | 7.604465642 | 8.078857519 | -0.474391877 | -0.266847217 |
| Prrc2b | 11.82942947 | 12.16130898 | 12.09630249 | 0.065006497 | -0.26687302 |
| Col16a1 | 10.87268261 | 11.19807308 | 11.13957999 | 0.058493087 | -0.26689738 |
| Fam83d | 6.869773445 | 7.080565495 | 7.137031639 | -0.056466144 | -0.267258195 |
| Atp11c | 7.811572134 | 7.816671071 | 8.078965836 | -0.262294765 | -0.267393702 |
| Agbl3 | 5.341112265 | 5.48384323 | 5.608507152 | -0.124663921 | -0.267394887 |
| C1qtnf3 | 6.398089338 | 6.308312533 | 6.665576819 | -0.357264286 | -0.267487481 |
| Tmem140 | 8.562999161 | 8.791608046 | 8.830771383 | -0.039163337 | -0.267772223 |
| Cgrrf1 | 8.184178354 | 8.202933703 | 8.451957178 | -0.249023475 | -0.267778824 |
| Grap | 6.780686439 | 6.834377035 | 7.048467477 | -0.214090441 | -0.267781038 |
| Hmg20a | 9.390989407 | 9.549874164 | 9.658801928 | -0.108927764 | -0.267812521 |
| Hoxa9 | 4.140664093 | 4.175761171 | 4.408510262 | -0.232749091 | -0.267846169 |
| Acox1 | 11.52825654 | 11.56795147 | 11.79616886 | -0.22821739 | -0.267912317 |
| Ech1 | 10.96063377 | 11.03017483 | 11.2286398 | -0.19846497 | -0.26800603 |
| Ift80 | 7.738157027 | 7.822304278 | 8.006319385 | -0.184015107 | -0.268162358 |
| Lce3d | 7.743203843 | 8.117586223 | 8.011431192 | 0.106155031 | -0.268227349 |
| Htra1 | 10.9880818 | 11.04908442 | 11.25635954 | -0.20727512 | -0.268277743 |
| Zfp354a | 5.479224396 | 5.583095721 | 5.74771685 | -0.164621129 | -0.268492455 |
| Itfg1 | 10.23160554 | 10.263204 | 10.50017279 | -0.23696879 | -0.268567253 |
| Fancc | 7.797602404 | 7.860098718 | 8.066233628 | -0.20613491 | -0.268631224 |
| Xkr6 | 5.436486774 | 5.537731174 | 5.705125005 | -0.167393831 | -0.268638231 |
| Synj2 | 9.057093051 | 9.457578747 | 9.325747523 | 0.131831224 | -0.268654472 |
| Inpp5b | 9.791886618 | 10.02691199 | 10.06057217 | -0.033660175 | -0.268685549 |
| Ncs1 | 9.18796468 | 9.464947347 | 9.456718097 | 0.00822925 | -0.268753417 |
| Svopl | 5.504801658 | 5.566410428 | 5.773566934 | -0.207156506 | -0.268765276 |
| Cnot6l | 10.75937895 | 10.82379637 | 11.02815562 | -0.204359257 | -0.26877667 |
| Pex7 | 9.110856547 | 9.10252682 | 9.379753209 | -0.277226388 | -0.268896662 |

| | | | | | |
|---|---|---|---|---|---|
| Asb13 | 8.536751275 | 8.769254629 | 8.805677702 | -0.036423073 | -0.268926427 |
| Slmap | 10.91773509 | 11.06120262 | 11.18669662 | -0.125493993 | -0.268961527 |
| Pdzk1 | 6.932072266 | 7.262928745 | 7.201123483 | 0.061805262 | -0.269051217 |
| Hoxc8 | 7.509939448 | 7.666066741 | 7.77908778 | -0.113021039 | -0.269148332 |
| Rcbtb2 | 8.459384862 | 8.388761408 | 8.728597585 | -0.339836177 | -0.269212723 |
| Map7d3 | 4.486348075 | 4.542610967 | 4.755620983 | -0.213010016 | -0.269272909 |
| Sun2 | 11.65340598 | 11.77782685 | 11.92270775 | -0.144880903 | -0.269301767 |
| Zkscan3 | 9.643484332 | 9.898033008 | 9.912803208 | -0.0147702 | -0.269318876 |
| Akr1e1 | 8.157698363 | 8.244231491 | 8.427018707 | -0.182787216 | -0.269320344 |
| Rad51c | 6.25050136 | 6.646833803 | 6.519992112 | 0.126841692 | -0.269490752 |
| Fhl5 | 2.895469871 | 2.958441668 | 3.165007556 | -0.206565887 | -0.269537684 |
| Tmtc4 | 8.003957134 | 8.202347255 | 8.27350761 | -0.071160355 | -0.269550476 |
| Ss18l1 | 6.85408674 | 6.990223852 | 7.123725355 | -0.133501503 | -0.269638615 |
| Gyg | 9.875511956 | 9.894965895 | 10.14516851 | -0.250202619 | -0.269656558 |
| Cfap20 | 8.544773676 | 8.695627925 | 8.814440359 | -0.118812434 | -0.269666684 |
| Pik3r4 | 8.967479364 | 8.982715471 | 9.237273579 | -0.254558108 | -0.269794215 |
| Acaa2 | 10.9121448 | 11.03207163 | 11.18221401 | -0.150142373 | -0.270069203 |
| Pde6d | 7.512068837 | 7.712246177 | 7.782170396 | -0.069924219 | -0.27010156 |
| Them4 | 6.94512298 | 7.095569799 | 7.215235523 | -0.119665724 | -0.270112543 |
| Nit1 | 8.968297434 | 9.211644402 | 9.238501617 | -0.026857215 | -0.270204182 |
| Gemin4 | 7.06822735 | 7.448371736 | 7.338569909 | 0.109801827 | -0.270342558 |
| Pask | 7.035640192 | 7.302984342 | 7.306058218 | -0.003073876 | -0.270418026 |
| Smpd1 | 10.75418221 | 10.97412198 | 11.02469091 | -0.050568933 | -0.2705087 |
| Zc2hc1c | 3.699530142 | 3.834461095 | 3.970141559 | -0.135680463 | -0.270611417 |
| Cldn10 | 4.687147222 | 4.71526472 | 4.957762378 | -0.242497659 | -0.270615157 |
| Zfp282 | 9.168637124 | 9.511109744 | 9.43929053 | 0.071819213 | -0.270653406 |
| Tigd4 | 4.383129105 | 4.356832667 | 4.653826211 | -0.296993544 | -0.270697106 |
| Il17rb | 5.540067669 | 5.924353037 | 5.810820208 | 0.113532829 | -0.270752538 |
| Epm2aip1 | 9.801806349 | 9.830539274 | 10.07263345 | -0.242094178 | -0.270827103 |
| Lsm11 | 7.880911399 | 8.052630994 | 8.15175922 | -0.099128225 | -0.27084782 |
| Rtl3 | 2.71161718 | 2.746679331 | 2.982476169 | -0.235796838 | -0.270858989 |
| Eya4 | 6.806121934 | 6.759325455 | 7.07706395 | -0.317738495 | -0.270942015 |

| | | | | | |
|---|---|---|---|---|---|
| Napepld | 7.504429683 | 7.680169989 | 7.775384381 | -0.095214391 | -0.270954698 |
| Greb1 | 3.950248276 | 4.221201525 | 4.221205389 | -3.86467E-06 | -0.270957113 |
| Cby1 | 7.108412762 | 7.280121837 | 7.379427675 | -0.099305838 | -0.271014913 |
| Xpa | 7.087289755 | 7.169588758 | 7.358501413 | -0.188912655 | -0.271211658 |
| Marveld3 | 6.688963838 | 7.173319917 | 6.960212124 | 0.213107793 | -0.271248286 |
| Mthfsd | 8.073601636 | 8.268337162 | 8.344861429 | -0.076524267 | -0.271259793 |
| Rsf1 | 9.269229784 | 9.483926721 | 9.540552561 | -0.05662584 | -0.271322776 |
| Chac1 | 7.133436258 | 7.506268841 | 7.404904113 | 0.101364728 | -0.271467856 |
| Smarca1 | 5.032015782 | 5.156716113 | 5.303486462 | -0.146770349 | -0.27147068 |
| Gm37240 | 6.030477829 | 6.061690514 | 6.302022827 | -0.240332313 | -0.271544997 |
| Ptprz1 | 7.362807628 | 7.689929595 | 7.634422061 | 0.055507534 | -0.271614433 |
| Eml6 | 6.155165151 | 6.28897916 | 6.42678493 | -0.13780577 | -0.271619778 |
| Tmem150a | 5.924470867 | 6.215889268 | 6.196339182 | 0.019550085 | -0.271868315 |
| Smoc2 | 11.02980723 | 11.02589977 | 11.30178458 | -0.275884817 | -0.27197735 |
| Cpq | 8.776715603 | 8.655039618 | 9.048816966 | -0.393777347 | -0.272101363 |
| Car5b | 9.352421376 | 9.023768573 | 9.624667731 | -0.600899158 | -0.272246355 |
| Vps26c | 9.180687557 | 9.410373963 | 9.453001284 | -0.042627321 | -0.272313727 |
| Krt77 | 16.88220019 | 17.0399125 | 17.15456171 | -0.114649207 | -0.272361517 |
| Kdm6a | 10.04097742 | 10.14739185 | 10.31350172 | -0.166109873 | -0.272524304 |
| Zfp984 | 7.993249885 | 8.093654029 | 8.265896127 | -0.172242099 | -0.272646243 |
| Lyve1 | 8.751969486 | 8.281157229 | 9.024636988 | -0.743479759 | -0.272667501 |
| H2-M5 | 6.1688461 | 6.225320769 | 6.441517944 | -0.216197175 | -0.272671844 |
| Adamts17 | 4.590999212 | 4.667302945 | 4.86373134 | -0.196428395 | -0.272732128 |
| Gatm | 7.503888508 | 7.590671885 | 7.776646089 | -0.185974204 | -0.272757581 |
| Ndc80 | 6.850014055 | 6.911187069 | 7.122916283 | -0.211729215 | -0.272902229 |
| Grik1 | 3.531440655 | 3.654128237 | 3.804581734 | -0.150453496 | -0.273141079 |
| Wdr48 | 9.824963498 | 10.01531796 | 10.09829858 | -0.082980622 | -0.273335086 |
| B3galt1 | 4.336229942 | 4.512104582 | 4.609783516 | -0.097678934 | -0.273553574 |
| Zfp296 | 6.349344758 | 6.685478903 | 6.62303355 | 0.062445353 | -0.273688792 |
| Csnk2a2 | 11.43691024 | 11.57078678 | 11.71063911 | -0.139852327 | -0.273728863 |
| Ccdc149 | 6.527892259 | 6.708487419 | 6.801628251 | -0.093140832 | -0.273735991 |
| Tpmt | 6.13198759 | 6.184304778 | 6.405785257 | -0.221480479 | -0.273797667 |

| | | | | | |
|---|---|---|---|---|---|
| Tspan5 | 10.05929132 | 10.22996222 | 10.33319158 | -0.103229367 | -0.273900268 |
| Alms1 | 7.261398961 | 7.379525253 | 7.535304129 | -0.155778876 | -0.273905168 |
| Pcolce | 10.61812938 | 10.78656841 | 10.8920477 | -0.105479283 | -0.273918313 |
| Nr6a1 | 7.292727354 | 7.557065064 | 7.566783696 | -0.009718632 | -0.274056343 |
| Pard6b | 7.234538923 | 7.479220971 | 7.508660559 | -0.029439588 | -0.274121636 |
| Lyrm7 | 5.814315536 | 6.022192193 | 6.088450935 | -0.066258742 | -0.274135399 |
| Tgfbr3 | 11.11872207 | 11.16443253 | 11.39286859 | -0.22843606 | -0.274146517 |
| Obsl1 | 7.354675919 | 7.614591381 | 7.628940385 | -0.014349004 | -0.274264466 |
| Klhl29 | 7.238269642 | 7.539241146 | 7.51261533 | 0.026625816 | -0.274345688 |
| Casr | 4.687673302 | 4.887164569 | 4.962065461 | -0.074900892 | -0.274392159 |
| Oxr1 | 10.18764368 | 10.21787002 | 10.46214789 | -0.24427787 | -0.274504213 |
| Mier1 | 10.18346805 | 10.27894439 | 10.45798291 | -0.17903852 | -0.274514857 |
| Vps39 | 9.673556304 | 9.734114609 | 9.948091431 | -0.213976822 | -0.274535128 |
| Ilvbl | 8.859259641 | 9.041476268 | 9.133799371 | -0.092323103 | -0.27453973 |
| Rbpms2 | 6.100692802 | 6.169296938 | 6.375271975 | -0.205975037 | -0.274579173 |
| Hace1 | 9.197822124 | 9.375561961 | 9.472438304 | -0.096876343 | -0.27461618 |
| Pnpla8 | 10.97044581 | 10.97256102 | 11.24514803 | -0.272587007 | -0.274702217 |
| Epn2 | 10.33394908 | 10.39997434 | 10.60883773 | -0.208863393 | -0.274888653 |
| Sorcs1 | 4.773468946 | 4.824811921 | 5.048360941 | -0.22354902 | -0.274891995 |
| Slc46a2 | 9.08259682 | 9.271438652 | 9.357667911 | -0.086229259 | -0.275071091 |
| Nadsyn1 | 6.373749311 | 6.582281586 | 6.648962676 | -0.06668109 | -0.275213366 |
| Rdm1 | 7.72339675 | 7.725572263 | 7.998753716 | -0.273181453 | -0.275356966 |
| Scpep1 | 10.65272446 | 10.85727429 | 10.9282191 | -0.07094481 | -0.27549464 |
| Scmh1 | 8.686072099 | 8.912345041 | 8.96160006 | -0.049255019 | -0.275527961 |
| Ttbk2 | 8.13129348 | 8.312095439 | 8.406868562 | -0.094773123 | -0.275575082 |
| Ift88 | 7.271006106 | 7.399871887 | 7.54660634 | -0.146734453 | -0.275600233 |
| Tgm7 | 5.018508198 | 5.261412938 | 5.294158482 | -0.032745544 | -0.275650284 |
| Rprd1a | 8.771763364 | 8.953155433 | 9.047468072 | -0.09431264 | -0.275704709 |
| Rpl19-ps1 | 2.256471198 | 2.297216422 | 2.532184093 | -0.234967671 | -0.275712895 |
| Shox2 | 7.076062531 | 7.149599914 | 7.351787362 | -0.202187448 | -0.275724831 |
| Acta1 | 15.41776975 | 15.80967057 | 15.69352867 | 0.116141897 | -0.275758923 |
| Gm28042 | 6.532723462 | 6.42305122 | 6.80852261 | -0.385471389 | -0.275799148 |

| | | | | | |
|---|---|---|---|---|---|
| Cep19 | 7.21559821 | 7.451962745 | 7.491722842 | -0.039760097 | -0.276124632 |
| Gpc2 | 6.456148049 | 6.971490356 | 6.732388063 | 0.239102292 | -0.276240014 |
| Csrnp2 | 7.544690283 | 7.961465688 | 7.820941655 | 0.140524033 | -0.276251372 |
| Ccdc77 | 7.347461069 | 7.663962175 | 7.623721145 | 0.04024103 | -0.276260076 |
| Ccbe1 | 8.054120083 | 8.464812159 | 8.33040586 | 0.1344063 | -0.276285777 |
| Wbp1 | 8.004720077 | 8.241914039 | 8.281015302 | -0.039101263 | -0.276295225 |
| Pepd | 10.10196356 | 10.14924011 | 10.3789907 | -0.229750593 | -0.27702714 |
| Wif1 | 6.9574405 | 7.070517653 | 7.234495619 | -0.163977965 | -0.277055119 |
| Tspear | 4.10482533 | 4.211041154 | 4.381992114 | -0.17095096 | -0.277166784 |
| Cdkl2 | 8.432410055 | 8.580544681 | 8.70965865 | -0.129113969 | -0.277248595 |
| Igbp1 | 9.950154348 | 9.964120821 | 10.22742766 | -0.263306842 | -0.277273316 |
| Adal | 7.604426965 | 7.826439212 | 7.881703846 | -0.055264635 | -0.277276882 |
| Ptn | 10.11806793 | 10.2300627 | 10.39536194 | -0.165299243 | -0.277294007 |
| Dcaf17 | 8.385898938 | 8.45828428 | 8.663286268 | -0.205001989 | -0.27738733 |
| Nthl1 | 5.376291219 | 5.625951127 | 5.653791592 | -0.027840465 | -0.277500372 |
| Arhgef18 | 9.620940707 | 9.940141069 | 9.898477994 | 0.041663075 | -0.277537287 |
| Fam117b | 9.357599938 | 9.434783776 | 9.635144035 | -0.200360259 | -0.277544097 |
| She | 6.769875222 | 7.012705477 | 7.047633557 | -0.034928081 | -0.277758335 |
| Wdr35 | 7.942845729 | 8.058500844 | 8.220727876 | -0.162227032 | -0.277882146 |
| Gm7334 | 6.937580519 | 7.12981248 | 7.215479514 | -0.085667034 | -0.277898994 |
| 2310022B05Rik | 10.13108865 | 10.34612458 | 10.40908599 | -0.06296141 | -0.277997333 |
| Kalrn | 7.466795199 | 7.344018433 | 7.744822763 | -0.400804329 | -0.278027563 |
| Hsd17b4 | 11.01338471 | 11.04211191 | 11.29144402 | -0.24933211 | -0.278059313 |
| Itga8 | 7.269974896 | 7.378580082 | 7.54811463 | -0.169534548 | -0.278139734 |
| Cntrob | 7.348370854 | 7.727659738 | 7.626614035 | 0.101045703 | -0.278243181 |
| Hist3h2a | 7.295891557 | 7.555946881 | 7.57430861 | -0.018361729 | -0.278417053 |
| Fam177a | 4.743129027 | 5.679766019 | 5.021739809 | 0.65802621 | -0.278610782 |
| Bnip3l | 11.58368975 | 11.57597679 | 11.86231184 | -0.286335053 | -0.278622093 |
| Cep70 | 6.873149061 | 6.992611184 | 7.151838205 | -0.159227022 | -0.278689145 |
| Tmem143 | 7.538848905 | 7.917351569 | 7.817565223 | 0.099786346 | -0.278716318 |
| Ano8 | 8.936196722 | 9.501553257 | 9.214990824 | 0.286562433 | -0.278794101 |
| Amz2 | 9.3368405 | 9.508873632 | 9.615706704 | -0.106833072 | -0.278866204 |

| | | | | | |
|---|---|---|---|---|---|
| Triqk | 4.969922687 | 4.923297936 | 5.248854795 | -0.325556858 | -0.278932107 |
| Abcg3 | 5.973535803 | 6.269505303 | 6.252574711 | 0.016930592 | -0.279038908 |
| Gpd1l | 8.973203902 | 9.089439247 | 9.252256152 | -0.162816905 | -0.27905225 |
| Zfp839 | 7.511887641 | 7.778678514 | 7.791104458 | -0.012425944 | -0.279216816 |
| Zfp106 | 12.45200623 | 12.42054224 | 12.73167382 | -0.31113158 | -0.27966759 |
| Wnt2 | 4.084569293 | 4.216485225 | 4.36427228 | -0.147787055 | -0.279702987 |
| Klk7 | 13.36782481 | 13.5707462 | 13.647602 | -0.076855803 | -0.279777197 |
| Zfp120 | 9.17831841 | 9.26154011 | 9.458113159 | -0.196573049 | -0.279794749 |
| Dele1 | 9.222076042 | 9.531491097 | 9.501906758 | 0.029584339 | -0.279830716 |
| Gm21985 | 4.314907521 | 4.334382472 | 4.594804941 | -0.260422469 | -0.27989742 |
| Lysmd4 | 7.57156324 | 7.791414516 | 7.851514972 | -0.060100456 | -0.279951732 |
| Plag1 | 5.285333706 | 5.352883448 | 5.565296245 | -0.212412798 | -0.279962539 |
| Cep95 | 7.350996752 | 7.673322428 | 7.630960081 | 0.042362347 | -0.279963329 |
| Platr25 | 6.602282669 | 6.886852787 | 6.882253804 | 0.004598984 | -0.279971135 |
| Zfp248 | 5.761043639 | 5.981114221 | 6.04107578 | -0.059961558 | -0.28003214 |
| Pdzd7 | 4.291151554 | 4.508567467 | 4.571261895 | -0.062694429 | -0.280110341 |
| Tlr5 | 6.896956574 | 6.908209215 | 7.177124776 | -0.268915561 | -0.280168202 |
| Mfsd14b | 11.42430501 | 11.5280829 | 11.70449436 | -0.17641146 | -0.28018935 |
| Zfp651 | 8.564059473 | 9.029316454 | 8.844270997 | 0.185045457 | -0.280211525 |
| Stab1 | 10.12563708 | 10.08583317 | 10.40588845 | -0.32005528 | -0.280251368 |
| Zfp758 | 7.650923013 | 7.73717407 | 7.931202048 | -0.194027978 | -0.280279035 |
| Atp6v0e2 | 7.812495578 | 7.876538689 | 8.092897277 | -0.216358588 | -0.280401699 |
| Efcab5 | 3.44100589 | 3.667071399 | 3.721466898 | -0.054395499 | -0.280461008 |
| Dyrk2 | 9.644223056 | 9.687961022 | 9.924839613 | -0.236878591 | -0.280616557 |
| Chst10 | 4.547358314 | 4.857418002 | 4.828090561 | 0.029327441 | -0.280732247 |
| Col8a2 | 7.717267923 | 7.890890355 | 7.998025359 | -0.107135004 | -0.280757436 |
| Srgap1 | 8.644363555 | 8.890884559 | 8.925128739 | -0.03424418 | -0.280765184 |
| Man2a2 | 10.51977851 | 10.62795577 | 10.80072242 | -0.172766653 | -0.280943913 |
| Dgka | 10.42801391 | 10.69233042 | 10.70898532 | -0.016654897 | -0.28097141 |
| Ccdc170 | 3.78450609 | 3.821449016 | 4.065516587 | -0.244067571 | -0.281010497 |
| Prkdc | 8.554502393 | 8.784067943 | 8.835590179 | -0.051522237 | -0.281087786 |
| Ift27 | 7.457961782 | 7.485253091 | 7.73905669 | -0.253803599 | -0.281094908 |

| | | | | | |
|---|---|---|---|---|---|
| Slc38a4 | 7.695256894 | 8.026698668 | 7.976384439 | 0.050314228 | -0.281127546 |
| Zfp862-ps | 7.80282963 | 7.999556166 | 8.084239512 | -0.084683346 | -0.281409881 |
| Smim10l1 | 10.34077165 | 10.26251672 | 10.62225714 | -0.359740417 | -0.281485487 |
| 5430403G16Rik | 5.767117485 | 5.912102641 | 6.048607986 | -0.136505345 | -0.281490501 |
| Plxdc2 | 11.10099202 | 11.13493412 | 11.38258031 | -0.247646183 | -0.281588287 |
| Kif4 | 8.032155854 | 7.86905648 | 8.313930014 | -0.444873534 | -0.28177416 |
| Paip2 | 10.98472381 | 11.11763636 | 11.26650968 | -0.148873327 | -0.281785873 |
| Panx3 | 8.674488633 | 8.75492028 | 8.956313681 | -0.201393401 | -0.281825048 |
| Sgcb | 9.29768599 | 9.196998128 | 9.579540121 | -0.382541993 | -0.281854131 |
| Cox6b2 | 4.862076916 | 4.993267492 | 5.14399782 | -0.150730328 | -0.281920903 |
| Gpc6 | 6.378251491 | 6.341808014 | 6.660314964 | -0.31850695 | -0.282063473 |
| Fam149b | 8.603748983 | 8.859267573 | 8.88588405 | -0.026616477 | -0.282135067 |
| Cep131 | 6.351651044 | 6.793412342 | 6.633796761 | 0.159615582 | -0.282145717 |
| Snta1 | 8.788625077 | 8.877015198 | 9.070977753 | -0.193962554 | -0.282352676 |
| Zfp318 | 8.846704606 | 9.072415463 | 9.12914623 | -0.056730767 | -0.282441623 |
| Fyn | 9.209302015 | 9.185653806 | 9.491837589 | -0.306183782 | -0.282535573 |
| Syde2 | 7.562194148 | 7.543157624 | 7.844785691 | -0.301628066 | -0.282591543 |
| Map7 | 10.0487561 | 10.17347127 | 10.33134921 | -0.157877943 | -0.28259311 |
| Lrrc9 | 5.437025969 | 5.406090646 | 5.719672387 | -0.31358174 | -0.282646418 |
| Gucd1 | 9.059252589 | 9.132454579 | 9.341937655 | -0.209483076 | -0.282685066 |
| Meiob | 3.846248016 | 4.002957686 | 4.128969318 | -0.126011632 | -0.282721302 |
| Sos1 | 9.568635409 | 9.787519603 | 9.851537583 | -0.06401798 | -0.282902173 |
| Cd300ld3 | 7.217026714 | 7.250927086 | 7.499991257 | -0.249064171 | -0.282964543 |
| Zik1 | 5.001414478 | 5.218728198 | 5.284426079 | -0.065697881 | -0.283011601 |
| Lrrc61 | 8.183295528 | 8.215350153 | 8.466468301 | -0.251118149 | -0.283172773 |
| Fam210b | 9.422583421 | 9.49147616 | 9.705899006 | -0.214422845 | -0.283315585 |
| Npc1 | 10.29412442 | 10.66504449 | 10.57748141 | 0.087563077 | -0.283356987 |
| Bbs7 | 6.506478764 | 6.562080823 | 6.789865838 | -0.227785015 | -0.283387074 |
| Crat | 11.41170329 | 11.64709553 | 11.69522172 | -0.048126187 | -0.283518427 |
| Tnfrsf11a | 8.132931457 | 8.470715563 | 8.416675604 | 0.054039959 | -0.283744147 |
| Me1 | 12.53645094 | 12.38832763 | 12.82028392 | -0.431956293 | -0.28383298 |
| Trnp1 | 6.939344591 | 6.960371133 | 7.223262106 | -0.262890972 | -0.283917515 |

| | | | | | |
|---|---|---|---|---|---|
| Frmd6 | 10.71644562 | 10.96757476 | 11.00041364 | -0.032838873 | -0.28396802 |
| Calm3 | 10.87156969 | 10.99830492 | 11.15558118 | -0.157276263 | -0.28401149 |
| Alox12b | 10.60111125 | 10.76186049 | 10.88533294 | -0.123472447 | -0.28422169 |
| Wipf3 | 7.884336937 | 8.024631326 | 8.168636131 | -0.144004805 | -0.284299194 |
| Kif20a | 8.068755887 | 8.15902253 | 8.35319479 | -0.19417226 | -0.284438904 |
| Lamb1 | 10.17405376 | 10.19436061 | 10.45849592 | -0.264135317 | -0.28444216 |
| Pi4k2b | 8.647148181 | 8.826425193 | 8.931675467 | -0.105250275 | -0.284527286 |
| Srrm4 | 5.102765282 | 5.418724417 | 5.38743205 | 0.031292367 | -0.284666768 |
| Ccdc85a | 5.387452253 | 5.371997187 | 5.672157941 | -0.300160754 | -0.284705688 |
| Capn6 | 4.738522307 | 4.983835031 | 5.02332619 | -0.039491159 | -0.284803883 |
| Lrrc56 | 5.788250506 | 5.968784998 | 6.073097859 | -0.104312861 | -0.284847353 |
| Thbs4 | 8.463129788 | 8.837030573 | 8.74809351 | 0.088937063 | -0.284963722 |
| Gdpd3 | 8.56674933 | 8.709092422 | 8.851800926 | -0.142708504 | -0.285051596 |
| Kdm4b | 8.970319683 | 9.368403009 | 9.255379238 | 0.11302377 | -0.285059555 |
| Tmem25 | 3.715220778 | 3.717086458 | 4.000305452 | -0.283218994 | -0.285084674 |
| Yipf2 | 8.125524621 | 8.305421419 | 8.41068728 | -0.105265861 | -0.285162659 |
| Nxph4 | 5.109626457 | 5.217936359 | 5.394839102 | -0.176902742 | -0.285212645 |
| Zfp30 | 5.398131518 | 5.4028221 | 5.683383221 | -0.280561121 | -0.285251703 |
| Mup20 | 3.669184673 | 3.798024095 | 3.954451924 | -0.156427828 | -0.285267251 |
| 2010300C02Rik | 6.997327427 | 7.114937931 | 7.282670021 | -0.16773209 | -0.285342594 |
| Mup22 | 13.70692835 | 13.75891561 | 13.99229138 | -0.233375773 | -0.285363033 |
| Tpsb2 | 8.940144998 | 8.515226129 | 9.22555953 | -0.7103334 | -0.285414532 |
| Odf2 | 9.721403748 | 9.911432974 | 10.00693258 | -0.095499602 | -0.285528828 |
| Top2b | 11.68353568 | 11.91147528 | 11.9691941 | -0.057718817 | -0.28565842 |
| Lonrf2 | 4.109590231 | 4.127195677 | 4.395418128 | -0.268222451 | -0.285827898 |
| Plpp6 | 9.282715243 | 9.323581934 | 9.56854736 | -0.244965426 | -0.285832117 |
| Atp2a1 | 15.81432163 | 15.94067311 | 16.10019338 | -0.159520273 | -0.285871757 |
| Dpysl2 | 8.787065877 | 8.656194205 | 9.073069398 | -0.416875193 | -0.286003521 |
| Ube2h | 10.7408526 | 10.86537984 | 11.02724581 | -0.161865973 | -0.286393213 |
| Slc46a1 | 7.202530895 | 7.239335151 | 7.489030881 | -0.249695731 | -0.286499986 |
| Mfsd9 | 7.252866454 | 7.403854866 | 7.539396486 | -0.13554162 | -0.286530032 |
| Ptgr1 | 10.47092005 | 10.57704853 | 10.7574653 | -0.180416767 | -0.28654525 |

| | | | | | |
|---|---|---|---|---|---|
| Arhgap44 | 8.394321429 | 8.703199851 | 8.680987133 | 0.022212718 | -0.286665704 |
| Phf24 | 8.261910621 | 8.407063048 | 8.548590715 | -0.141527666 | -0.286680093 |
| Trim13 | 8.164080703 | 8.232561054 | 8.450785103 | -0.218224049 | -0.2867044 |
| Snx33 | 8.714477093 | 8.932155966 | 9.001218414 | -0.069062447 | -0.286741321 |
| Gpx8 | 8.19011795 | 8.312138583 | 8.476895291 | -0.164756708 | -0.286777341 |
| Asah1 | 10.35075759 | 10.51226821 | 10.63756753 | -0.125299317 | -0.286809943 |
| Zfp280d | 9.137609736 | 9.173346644 | 9.424429683 | -0.251083039 | -0.286819947 |
| Bcl7a | 8.341706926 | 8.666299777 | 8.628602496 | 0.037697281 | -0.28689557 |
| Gsta2 | 6.644455495 | 6.692712892 | 6.931514704 | -0.238801812 | -0.287059209 |
| Lin7b | 4.476968909 | 4.689214954 | 4.764037871 | -0.074822917 | -0.287068962 |
| Smad9 | 4.993419308 | 5.245536095 | 5.280502227 | -0.034966131 | -0.287082919 |
| Art4 | 7.865431651 | 8.171261986 | 8.15254834 | 0.018713646 | -0.287116689 |
| Tmem263 | 8.262914939 | 8.461735968 | 8.550052792 | -0.088316824 | -0.287137853 |
| Prkaca | 10.28446375 | 10.31436547 | 10.5716944 | -0.25732893 | -0.28723065 |
| Duxbl2 | 4.035404498 | 4.272931127 | 4.322673933 | -0.049742806 | -0.287269434 |
| Tspan32 | 5.909470486 | 5.852023708 | 6.19681436 | -0.344790652 | -0.287343873 |
| Tbc1d8 | 8.793454131 | 8.949127123 | 9.080912504 | -0.131785382 | -0.287458373 |
| Gm47654 | 4.91257062 | 5.007294449 | 5.200049066 | -0.192754617 | -0.287478446 |
| Dclk1 | 9.143076908 | 9.364665638 | 9.430737101 | -0.066071464 | -0.287660193 |
| Slc2a8 | 7.36566432 | 7.650364089 | 7.653327964 | -0.002963875 | -0.287663644 |
| Sh3bgrl2 | 10.21252735 | 10.47444849 | 10.50022526 | -0.025776773 | -0.287697907 |
| Zfp973 | 6.41952397 | 6.499312589 | 6.707302044 | -0.207989455 | -0.287778074 |
| D2hgdh | 8.026808064 | 8.309276668 | 8.314611529 | -0.00533486 | -0.287803464 |
| Cbx8 | 7.171734316 | 7.401020361 | 7.459635729 | -0.058615368 | -0.287901414 |
| Rfx3 | 6.569988306 | 6.717583473 | 6.857963523 | -0.140380051 | -0.287975217 |
| Mdm1 | 6.470655154 | 6.739871101 | 6.758673085 | -0.018801984 | -0.288017931 |
| Car8 | 4.295004296 | 4.328823975 | 4.583031514 | -0.254207539 | -0.288027218 |
| Dtna | 8.571570856 | 8.444079608 | 8.859623456 | -0.415543848 | -0.288052599 |
| Enox2 | 7.758291426 | 7.83843244 | 8.046368794 | -0.207936354 | -0.288077368 |
| Tnfaip8 | 10.35101351 | 10.48452471 | 10.63910385 | -0.15457914 | -0.288090343 |
| Tubg2 | 4.708838864 | 4.88773897 | 4.996998356 | -0.109259386 | -0.288159492 |
| Adck2 | 8.164434206 | 8.318602806 | 8.452628335 | -0.134025529 | -0.288194129 |

| | | | | | |
|---|---|---|---|---|---|
| Sh3pxd2a | 11.96185369 | 12.37067796 | 12.25012321 | 0.120554757 | -0.288269517 |
| Gpr153 | 8.454600686 | 8.580383161 | 8.743013314 | -0.162630154 | -0.288412628 |
| Myo18b | 9.709934301 | 10.47603939 | 9.998533937 | 0.477505456 | -0.288599636 |
| Cdc42bpa | 9.573561089 | 9.701052174 | 9.862184968 | -0.161132794 | -0.288623879 |
| Xk | 4.780667421 | 4.859003263 | 5.069424007 | -0.210420744 | -0.288756585 |
| Ppp1r13b | 9.326825373 | 9.614414942 | 9.615595184 | -0.001180242 | -0.288769811 |
| Ffar2 | 6.564552807 | 6.801837218 | 6.853330082 | -0.051492864 | -0.288777275 |
| Trim65 | 7.478629857 | 7.951227356 | 7.767857069 | 0.183370288 | -0.289227211 |
| 1110012L19Rik | 7.884680729 | 7.95378064 | 8.17395018 | -0.22016954 | -0.289269451 |
| Ccdc88a | 8.833697501 | 8.841828469 | 9.122987097 | -0.281158628 | -0.289289596 |
| Anxa6 | 11.20726772 | 11.2157066 | 11.49657315 | -0.280866553 | -0.289305433 |
| Hif3a | 4.754417529 | 4.785974737 | 5.043855351 | -0.257880614 | -0.289437823 |
| Aff4 | 11.49752848 | 11.77888177 | 11.78698612 | -0.008104353 | -0.289457637 |
| Nrip1 | 9.788358209 | 9.85565207 | 10.07795702 | -0.222304948 | -0.289598809 |
| Wdr60 | 7.567577974 | 7.693092773 | 7.857243673 | -0.1641509 | -0.289665699 |
| Daglb | 8.824800835 | 9.162758611 | 9.114497968 | 0.048260643 | -0.289697133 |
| Nipsnap2 | 10.70374551 | 10.61511968 | 10.99353408 | -0.3784144 | -0.289788573 |
| Ccdc34 | 7.368510322 | 7.479590389 | 7.658379755 | -0.178789366 | -0.289869433 |
| Notum | 6.590380884 | 6.697022952 | 6.880374634 | -0.183351682 | -0.289993749 |
| Fut10 | 7.019750462 | 7.149164861 | 7.309884561 | -0.1607197 | -0.290134099 |
| Tmc5 | 2.362216212 | 2.407430901 | 2.652466156 | -0.245035255 | -0.290249944 |
| Zfp2 | 6.318667623 | 6.385997207 | 6.608947327 | -0.22295012 | -0.290279704 |
| Cgnl1 | 9.585589402 | 9.759098964 | 9.875995443 | -0.116896479 | -0.290406041 |
| Zfp874a | 7.426963549 | 7.641832413 | 7.717548569 | -0.075716156 | -0.290585019 |
| Pkd2 | 9.626605479 | 9.810185307 | 9.917544018 | -0.107358711 | -0.29093854 |
| Cnpy2 | 9.446129383 | 9.484159759 | 9.737304856 | -0.253145096 | -0.291175473 |
| Mrgprf | 7.614168328 | 8.049050646 | 7.905428012 | 0.143622634 | -0.291259685 |
| Vps13b | 9.614629599 | 9.785707399 | 9.906037039 | -0.12032964 | -0.29140744 |
| Mapk8ip1 | 7.429803391 | 7.610105035 | 7.721568799 | -0.111463763 | -0.291765407 |
| Lce1h | 9.253561173 | 9.817797606 | 9.545427381 | 0.272370225 | -0.291866208 |
| Fundc1 | 8.866015479 | 9.00055905 | 9.158020571 | -0.157461521 | -0.292005092 |
| Rab13 | 7.331872267 | 7.302287133 | 7.624007167 | -0.321720035 | -0.2921349 |

| | | | | | |
|---|---|---|---|---|---|
| Retreg3 | 10.72804401 | 10.90330074 | 11.02024547 | -0.116944737 | -0.292201463 |
| Dpyd | 6.421930359 | 6.35805172 | 6.714136349 | -0.356084629 | -0.29220599 |
| Zkscan5 | 7.305792228 | 7.518284139 | 7.598012187 | -0.079728048 | -0.292219959 |
| Acot13 | 8.145268552 | 8.266509904 | 8.437570296 | -0.171060392 | -0.292301744 |
| Tcf19 | 8.449245541 | 8.759961533 | 8.741719731 | 0.018241802 | -0.29247419 |
| Tbx2 | 4.846912699 | 5.054398344 | 5.139389739 | -0.084991394 | -0.29247704 |
| Phf7 | 6.736836232 | 6.738086762 | 7.029321787 | -0.291235025 | -0.292485556 |
| Adra1a | 6.001772006 | 6.276122952 | 6.294261775 | -0.018138823 | -0.29248977 |
| Dram2 | 9.341974003 | 9.382338027 | 9.634514643 | -0.252176616 | -0.29254064 |
| Pank1 | 10.53049306 | 10.64346717 | 10.82304361 | -0.17957644 | -0.292550553 |
| Elovl1 | 12.24985109 | 12.51854401 | 12.54248556 | -0.02394155 | -0.292634463 |
| Smpd2 | 8.410531287 | 8.575303416 | 8.703232218 | -0.127928803 | -0.292700931 |
| Casd1 | 8.761376177 | 8.840060894 | 9.054086266 | -0.214025372 | -0.292710089 |
| Knl1 | 7.197652881 | 7.312334798 | 7.490385107 | -0.17805031 | -0.292732226 |
| Tmeff1 | 6.153973218 | 6.306585902 | 6.447071315 | -0.140485413 | -0.293098097 |
| Cndp1 | 3.972906412 | 4.016538605 | 4.266042217 | -0.249503612 | -0.293135805 |
| Ntn1 | 9.55518485 | 9.623637121 | 9.848325589 | -0.224688469 | -0.293140739 |
| Mtm1 | 8.271615705 | 8.216248719 | 8.564786731 | -0.348538012 | -0.293171026 |
| Trpc3 | 3.842724657 | 3.826050912 | 4.135949125 | -0.309898213 | -0.293224468 |
| Epb41 | 10.36327064 | 10.45918926 | 10.65655114 | -0.197361873 | -0.2932805 |
| Rgn | 4.849364567 | 4.899102648 | 5.142664042 | -0.243561394 | -0.293299475 |
| Metap1d | 8.481370226 | 8.670950403 | 8.774797268 | -0.103846864 | -0.293427042 |
| Kcna7 | 6.471600602 | 6.42035027 | 6.765061973 | -0.344711704 | -0.293461371 |
| Apbb2 | 8.846522437 | 9.008913486 | 9.14001076 | -0.131097273 | -0.293488322 |
| Btbd6 | 7.971355081 | 8.278600647 | 8.264875332 | 0.013725315 | -0.293520251 |
| Rufy2 | 7.77882592 | 8.029808928 | 8.072387088 | -0.04257816 | -0.293561168 |
| Zfp759 | 5.627171043 | 5.70948597 | 5.920848695 | -0.211362724 | -0.293677652 |
| Mbnl1 | 12.29624826 | 12.23376323 | 12.59003438 | -0.356271143 | -0.293786117 |
| Paox | 8.473351971 | 8.628722827 | 8.767464568 | -0.138741741 | -0.294112597 |
| Tns4 | 11.92054848 | 12.22399462 | 12.21467062 | 0.009324 | -0.294122137 |
| Haus4 | 7.377943562 | 7.658791911 | 7.672126126 | -0.013334215 | -0.294182564 |
| Acss2 | 10.36753959 | 9.933835671 | 10.66175837 | -0.727922695 | -0.294218779 |

| | | | | | |
|---|---|---|---|---|---|
| Pink1 | 11.4157134 | 11.36101088 | 11.71032062 | -0.349309733 | -0.29460722 |
| Mum1l1 | 3.980488011 | 4.19738589 | 4.275325107 | -0.077939217 | -0.294837096 |
| Srgap3 | 6.888751684 | 7.202168774 | 7.18370722 | 0.018461554 | -0.294955536 |
| Cyp46a1 | 3.355581605 | 3.548417989 | 3.650547916 | -0.102129927 | -0.294966311 |
| Habp4 | 8.533964765 | 8.748947515 | 8.828994335 | -0.08004682 | -0.29502957 |
| 4930430F08Rik | 7.543091338 | 7.640909831 | 7.838146681 | -0.197236851 | -0.295055344 |
| N4bp2l1 | 7.449389708 | 7.568326882 | 7.74445471 | -0.176127828 | -0.295065002 |
| Chrnb4 | 4.859446931 | 4.985383423 | 5.15452959 | -0.169146167 | -0.295082659 |
| Pcsk6 | 10.4315313 | 10.60558074 | 10.72682557 | -0.121244837 | -0.295294277 |
| Hpse | 7.76318757 | 7.96124732 | 8.058652311 | -0.09740499 | -0.29546474 |
| Gm42878 | 4.224037699 | 4.553147842 | 4.519620987 | 0.033526856 | -0.295583288 |
| Smyd4 | 6.914650433 | 7.280262108 | 7.210403033 | 0.069859074 | -0.2957526 |
| Pnpla2 | 11.58869784 | 11.55174177 | 11.88451267 | -0.332770893 | -0.295814827 |
| Anpep | 10.84412207 | 11.21317898 | 11.14009568 | 0.073083303 | -0.295973603 |
| Trpv3 | 8.684368489 | 8.926374647 | 8.980383511 | -0.054008864 | -0.296015022 |
| Tmem120b | 8.230474019 | 8.439983459 | 8.526831694 | -0.086848235 | -0.296357675 |
| Grhl1 | 11.68586883 | 11.81684162 | 11.98229673 | -0.16545511 | -0.296427897 |
| Mycbp | 8.126849756 | 8.232667605 | 8.423309652 | -0.190642047 | -0.296459896 |
| Capns2 | 10.55929286 | 10.632015 | 10.85576893 | -0.223753927 | -0.296476073 |
| Rbp1 | 7.639601498 | 7.700771875 | 7.936144005 | -0.23537213 | -0.296542507 |
| Ppm1k | 7.880401477 | 7.573512629 | 8.176984887 | -0.603472258 | -0.29658341 |
| Fn3krp | 7.554481102 | 7.63209779 | 7.851083558 | -0.218985768 | -0.296602455 |
| Ap5m1 | 8.774940625 | 8.976749516 | 9.071576313 | -0.094826796 | -0.296635688 |
| Zfp81 | 7.549003869 | 7.5546167 | 7.84570623 | -0.29108953 | -0.296702361 |
| Gdpd1 | 7.776494706 | 7.982057522 | 8.073370075 | -0.091312553 | -0.296875369 |
| Gstm2 | 9.910867344 | 9.887439881 | 10.20777842 | -0.320338536 | -0.296911072 |
| Hectd3 | 9.477273596 | 9.840434623 | 9.774226961 | 0.066207662 | -0.296953365 |
| Mup14 | 5.83949972 | 6.002749698 | 6.136520073 | -0.133770376 | -0.297020354 |
| Creb5 | 8.098791517 | 8.266590269 | 8.395923089 | -0.12933282 | -0.297131572 |
| Bmi1 | 9.366398005 | 9.461594346 | 9.663619916 | -0.20202557 | -0.297221911 |
| Atraid | 9.341165509 | 9.423167319 | 9.638432162 | -0.215264843 | -0.297266653 |
| Tmcc3 | 10.66167145 | 10.8033339 | 10.95902555 | -0.155691653 | -0.2973541 |

| | | | | | |
|---|---|---|---|---|---|
| Gm45351 | 8.08954537 | 7.911389902 | 8.387035138 | -0.475645236 | -0.297489768 |
| 5730455P16Rik | 8.394526449 | 8.358006123 | 8.692162034 | -0.334155911 | -0.297635585 |
| Krt1 | 16.02463322 | 16.13544591 | 16.32229159 | -0.186845683 | -0.297658373 |
| Gm49359 | 7.437813908 | 7.460089398 | 7.735477627 | -0.275388229 | -0.29766372 |
| Inpp5k | 9.49986942 | 9.681867039 | 9.797650841 | -0.115783803 | -0.297781421 |
| Nr1d1 | 10.64842416 | 10.73539433 | 10.94632187 | -0.21092754 | -0.297897707 |
| Kif9 | 5.698655154 | 5.758587311 | 5.996696906 | -0.238109594 | -0.298041752 |
| Kif3a | 7.948873066 | 8.076597876 | 8.247082048 | -0.170484172 | -0.298208982 |
| S100pbp | 8.42491431 | 8.773076284 | 8.723288436 | 0.049787848 | -0.298374126 |
| Paqr5 | 9.210764564 | 9.407992239 | 9.509141755 | -0.101149516 | -0.29837719 |
| Zfp943 | 8.352342157 | 8.467204536 | 8.650761745 | -0.183557209 | -0.298419588 |
| Rilpl1 | 8.451600593 | 8.2752432 | 8.750039564 | -0.474796364 | -0.298438971 |
| Tmem178b | 4.012556422 | 3.996716755 | 4.311023799 | -0.314307043 | -0.298467376 |
| Rrm2b | 8.188878712 | 8.364064819 | 8.487371278 | -0.123306459 | -0.298492566 |
| Lrig2 | 8.993193641 | 9.226569068 | 9.291690094 | -0.065121026 | -0.298496454 |
| Gm10033 | 7.181472064 | 7.289041768 | 7.479990659 | -0.190948891 | -0.298518595 |
| Ccdc91 | 9.109946959 | 9.093790556 | 9.40849416 | -0.314703604 | -0.2985472 |
| Slc36a2 | 7.897651237 | 7.821106003 | 8.196269785 | -0.375163782 | -0.298618548 |
| Atad5 | 7.816249888 | 8.115451848 | 8.114923601 | 0.000528247 | -0.298673713 |
| Gdf11 | 5.253940793 | 5.466468532 | 5.552687447 | -0.086218915 | -0.298746654 |
| Tspan6 | 7.264830913 | 7.314768559 | 7.56363173 | -0.24886317 | -0.298800817 |
| Erlec1 | 8.809209503 | 8.771327927 | 9.108090668 | -0.336762741 | -0.298881165 |
| Pheta1 | 7.32142129 | 7.695427343 | 7.620405089 | 0.075022254 | -0.298983799 |
| Bpgm | 8.642100511 | 8.562511032 | 8.941114518 | -0.378603486 | -0.299014008 |
| Tbc1d31 | 6.877689432 | 7.150762373 | 7.176891297 | -0.026128924 | -0.299201866 |
| Atp2b4 | 11.79889368 | 11.93241622 | 12.09831035 | -0.165894137 | -0.299416673 |
| Zfp442 | 5.86149454 | 6.048184011 | 6.16126398 | -0.113079969 | -0.299769439 |
| Gm6206 | 4.490844925 | 4.48725397 | 4.79072532 | -0.30347135 | -0.299880395 |
| Lrrc15 | 5.726795864 | 6.032442542 | 6.026749859 | 0.005692682 | -0.299953996 |
| Tbc1d19 | 7.243254721 | 7.452758971 | 7.543288597 | -0.090529626 | -0.300033876 |
| Sppl3 | 10.85874768 | 11.14153433 | 11.15883365 | -0.017299313 | -0.300085963 |
| Dsel | 9.169472689 | 9.273544585 | 9.469603601 | -0.196059016 | -0.300130911 |

| | | | | | |
|---|---|---|---|---|---|
| Yipf4 | 9.566864349 | 9.497905811 | 9.867116914 | -0.369211103 | -0.300252565 |
| Gm29609 | 4.620634404 | 4.998645255 | 4.920966846 | 0.077678409 | -0.300332442 |
| Ptprm | 7.923469283 | 7.862345711 | 8.223805626 | -0.361459915 | -0.300336343 |
| Ank | 11.36668702 | 11.51974357 | 11.66711167 | -0.147368103 | -0.300424657 |
| Ankzf1 | 8.522425778 | 8.794344983 | 8.822850955 | -0.028505972 | -0.300425177 |
| Slitrk6 | 9.351602081 | 9.349139767 | 9.652027814 | -0.302888047 | -0.300425733 |
| Zfp704 | 8.756532575 | 8.904019405 | 9.057192754 | -0.153173349 | -0.300660179 |
| Hsdl1 | 9.161155051 | 9.469617309 | 9.461823598 | 0.007793711 | -0.300668547 |
| Snx14 | 9.177699468 | 9.324236283 | 9.47841066 | -0.154174378 | -0.300711192 |
| Prkag2 | 9.257107339 | 9.472323636 | 9.557834117 | -0.08551048 | -0.300726778 |
| Plekhh3 | 8.99088217 | 9.411390575 | 9.291678373 | 0.119712202 | -0.300796203 |
| Gas1 | 10.21244551 | 10.37213571 | 10.51326807 | -0.141132363 | -0.30082256 |
| Slc44a1 | 10.91580887 | 11.00896967 | 11.21671831 | -0.207748637 | -0.300909443 |
| Spns2 | 10.87227198 | 11.13080679 | 11.17318582 | -0.04237903 | -0.300913837 |
| Cep72 | 5.868501813 | 6.153183713 | 6.169432612 | -0.016248899 | -0.300930799 |
| Acsm5 | 3.889857804 | 3.900805479 | 4.190848115 | -0.290042636 | -0.300990311 |
| 1110032A03Rik | 7.759940853 | 7.803403312 | 8.061111419 | -0.257708107 | -0.301170567 |
| Dzip3 | 6.886196704 | 7.034694053 | 7.187400534 | -0.152706481 | -0.30120383 |
| Pf4 | 9.010591007 | 8.744188258 | 9.311807898 | -0.56761964 | -0.301216891 |
| Wdr13 | 9.945351341 | 10.39217437 | 10.24657889 | 0.14559548 | -0.301227552 |
| Pkn3 | 8.186841735 | 8.516578848 | 8.488165849 | 0.028412998 | -0.301324115 |
| Zfp964 | 5.759558215 | 5.978228383 | 6.061019833 | -0.08279145 | -0.301461618 |
| Ankrd35 | 11.25249443 | 11.48237386 | 11.55396442 | -0.071590563 | -0.301469997 |
| Tcaim | 7.66134844 | 7.672087625 | 7.96290002 | -0.290812395 | -0.30155158 |
| Ckap5 | 10.18366616 | 10.4071814 | 10.48542794 | -0.078246543 | -0.301761783 |
| Pcca | 8.497178686 | 8.468163486 | 8.79907584 | -0.330912355 | -0.301897154 |
| Neurl1a | 7.919536327 | 7.889576621 | 8.22148336 | -0.331906739 | -0.301947033 |
| Cpxm2 | 9.2499658 | 9.191489824 | 9.5520065 | -0.360516676 | -0.302040701 |
| Klk9 | 9.738463712 | 10.10151353 | 10.04056252 | 0.060951004 | -0.302098811 |
| Ppp1r1a | 7.61959413 | 7.456606579 | 7.922016625 | -0.465410046 | -0.302422495 |
| Six5 | 6.443758139 | 6.947426506 | 6.746217536 | 0.20120897 | -0.302459397 |
| Vangl2 | 9.198798637 | 9.547353239 | 9.501398729 | 0.04595451 | -0.302600092 |

| | | | | | |
|---|---|---|---|---|---|
| Oscp1 | 6.185380273 | 5.890604194 | 6.488010808 | -0.597406614 | -0.302630535 |
| Rad54b | 6.265995956 | 6.43516229 | 6.568698501 | -0.133536211 | -0.302702545 |
| Lactb | 8.126746627 | 8.364612407 | 8.429548748 | -0.064936341 | -0.302802121 |
| Efna3 | 10.53030627 | 10.93143724 | 10.83323344 | 0.0982038 | -0.302927173 |
| Lgi4 | 8.509024565 | 8.835067171 | 8.812055863 | 0.023011308 | -0.303031299 |
| Tmem126b | 8.473678526 | 8.604304328 | 8.776752229 | -0.172447902 | -0.303073704 |
| Emx2 | 5.696552326 | 5.906047027 | 5.999641953 | -0.093594925 | -0.303089627 |
| Nnt | 7.709273375 | 7.743154556 | 8.012363353 | -0.269208797 | -0.303089978 |
| Enpp3 | 5.973453732 | 5.960199258 | 6.276628621 | -0.316429363 | -0.303174889 |
| Rbp4 | 8.758927473 | 8.176933882 | 9.062104094 | -0.885170211 | -0.303176621 |
| Mamdc2 | 4.821723434 | 4.845047973 | 5.125060912 | -0.280012939 | -0.303337478 |
| Yeats2 | 9.025584814 | 9.26339062 | 9.328940627 | -0.065550006 | -0.303355812 |
| Eps15 | 11.08191608 | 11.22105344 | 11.38535578 | -0.164302343 | -0.3034397 |
| Snhg11 | 7.650307249 | 7.882433659 | 7.953770644 | -0.071336985 | -0.303463394 |
| Plek2 | 9.538558004 | 9.759346509 | 9.842022902 | -0.082676392 | -0.303464897 |
| Itpr3 | 12.0985195 | 12.47679531 | 12.40199298 | 0.07480233 | -0.303473477 |
| Cybrd1 | 5.278544559 | 5.50489724 | 5.582119985 | -0.077222745 | -0.303575426 |
| Hadhb | 11.43462281 | 11.47624962 | 11.73844345 | -0.262193833 | -0.303820643 |
| Rnf103 | 9.100878368 | 9.311313353 | 9.40538171 | -0.094068358 | -0.304503342 |
| Atp8b2 | 10.11779479 | 10.47523329 | 10.42244428 | 0.052789017 | -0.304649483 |
| Vwa8 | 9.28478299 | 9.359417148 | 9.589439947 | -0.230022799 | -0.304656957 |
| Nek8 | 6.494619869 | 6.846258365 | 6.799474969 | 0.046783395 | -0.304855101 |
| Fuca1 | 10.06280261 | 10.11575408 | 10.36786025 | -0.25210617 | -0.305057643 |
| Fut2 | 9.610772432 | 9.889933936 | 9.915853502 | -0.025919565 | -0.30508107 |
| Cavin1 | 12.43778224 | 12.54538294 | 12.74295466 | -0.197571727 | -0.305172427 |
| Rbbp8nl | 5.366976689 | 5.823059115 | 5.672300934 | 0.150758181 | -0.305324245 |
| Ces1d | 11.11579889 | 10.91140594 | 11.42112513 | -0.509719183 | -0.305326237 |
| Pitx2 | 5.220611876 | 5.449394079 | 5.525954397 | -0.076560317 | -0.305342521 |
| Hist1h2bc | 8.523031771 | 8.749741715 | 8.82838749 | -0.078645775 | -0.305355719 |
| Stxbp6 | 7.516362504 | 7.643491528 | 7.821729204 | -0.178237676 | -0.305366701 |
| Dguok | 8.224226338 | 8.256605039 | 8.529737353 | -0.273132314 | -0.305511015 |
| Ddo | 5.940868406 | 5.881678173 | 6.246498592 | -0.36482042 | -0.305630186 |

| | | | | | |
|---|---|---|---|---|---|
| Tnrc6c | 9.878409961 | 10.17430173 | 10.18407945 | -0.00977772 | -0.305669493 |
| St8sia6 | 6.52144777 | 6.712285257 | 6.82721421 | -0.114928953 | -0.30576644 |
| Gm10297 | 4.596293551 | 4.614177783 | 4.902110925 | -0.287933142 | -0.305817374 |
| Wdr62 | 7.331345066 | 7.526504171 | 7.637298899 | -0.110794728 | -0.305953833 |
| Stim1 | 11.5045001 | 11.6379159 | 11.81056845 | -0.172652553 | -0.306068353 |
| Mc2r | 5.621588417 | 5.604762596 | 5.927726909 | -0.322964313 | -0.306138493 |
| BC021767 | 4.341777457 | 4.665417791 | 4.648231926 | 0.017185865 | -0.306454468 |
| Igdcc4 | 7.612010944 | 7.513216546 | 7.918499697 | -0.405283151 | -0.306488752 |
| Ccdc62 | 6.205514441 | 6.479061204 | 6.512080398 | -0.033019193 | -0.306565957 |
| Cds1 | 10.42020283 | 10.56181341 | 10.72709596 | -0.16528255 | -0.306893123 |
| Adam12 | 7.284465518 | 7.539685523 | 7.591363963 | -0.05167844 | -0.306898445 |
| Hexb | 9.729776402 | 9.766234754 | 10.03680411 | -0.270569354 | -0.307027706 |
| Nfatc1 | 10.10040071 | 10.25117839 | 10.40786138 | -0.156682997 | -0.307460671 |
| Zfp367 | 8.825431502 | 9.027171801 | 9.132971835 | -0.105800034 | -0.307540333 |
| Tmem254b | 9.833141114 | 9.997323911 | 10.14068911 | -0.143365195 | -0.307547993 |
| Tctn1 | 7.540892723 | 7.838809929 | 7.848619141 | -0.009809212 | -0.307726419 |
| St6gal1 | 7.956269124 | 7.973759506 | 8.264032178 | -0.290272672 | -0.307763054 |
| Cnpy4 | 8.22702375 | 8.437458903 | 8.534821045 | -0.097362142 | -0.307797295 |
| Usp32 | 10.28907141 | 10.52569307 | 10.5970307 | -0.07133763 | -0.30795929 |
| Slc25a40 | 7.157689685 | 7.134979477 | 7.465986849 | -0.331007372 | -0.308297164 |
| Klf8 | 6.608236096 | 6.625184801 | 6.916589465 | -0.291404664 | -0.308353369 |
| Zfp169 | 7.141548318 | 7.393265942 | 7.449975323 | -0.05670938 | -0.308427005 |
| Btbd8 | 6.216058007 | 6.166974155 | 6.524545698 | -0.357571543 | -0.308487691 |
| Cul9 | 7.545755671 | 7.730327745 | 7.854379061 | -0.124051316 | -0.30862339 |
| Zfp85 | 5.617925491 | 5.874452117 | 5.926878894 | -0.052426777 | -0.308953403 |
| Plekhg5 | 8.639482424 | 9.072258407 | 8.948447466 | 0.123810941 | -0.308965042 |
| Rgs9bp | 4.662106601 | 4.846416928 | 4.971155023 | -0.124738095 | -0.309048422 |
| Fcho2 | 10.10478947 | 10.21331567 | 10.413964 | -0.200648327 | -0.309174528 |
| Zfp182 | 7.424792889 | 7.717149164 | 7.734108068 | -0.016958905 | -0.309315179 |
| Sh2d4a | 8.893972046 | 9.067400754 | 9.203305789 | -0.135905035 | -0.309333743 |
| mt-Co1 | 17.45030573 | 17.4990837 | 17.75977886 | -0.26069516 | -0.30947313 |
| Trex2 | 9.527379394 | 9.946835458 | 9.836914485 | 0.109920973 | -0.309535091 |

| | | | | | |
|---|---|---|---|---|---|
| Snrpn | 6.947994675 | 7.044603229 | 7.257569912 | -0.212966683 | -0.309575237 |
| Stil | 6.443355697 | 6.588614786 | 6.753080548 | -0.164465761 | -0.30972485 |
| Pdlim3 | 10.31398148 | 10.51127522 | 10.62374381 | -0.112468587 | -0.30976233 |
| Esyt3 | 9.748822068 | 9.988452004 | 10.05858773 | -0.070135722 | -0.309765659 |
| Zmym2 | 9.852959474 | 10.13282779 | 10.16280505 | -0.029977267 | -0.30984558 |
| Olfr558 | 4.075032187 | 4.118453744 | 4.385075435 | -0.266621692 | -0.310043248 |
| Il31ra | 8.783013977 | 9.024982314 | 9.093142366 | -0.068160053 | -0.310128389 |
| Coq8a | 10.64783071 | 10.26678908 | 10.95803297 | -0.691243883 | -0.31020226 |
| Asb12 | 5.935598517 | 6.186895063 | 6.245825863 | -0.0589308 | -0.310227346 |
| Rab40b | 7.407207708 | 7.546628259 | 7.717468181 | -0.170839922 | -0.310260473 |
| Zfp652 | 10.48483918 | 10.66385025 | 10.79530533 | -0.13145508 | -0.31046615 |
| Snapin | 9.318367565 | 9.322481505 | 9.628971612 | -0.306490107 | -0.310604047 |
| Dcaf11 | 10.40763692 | 10.43596818 | 10.71830322 | -0.28233504 | -0.3106663 |
| Siglec1 | 8.318129306 | 7.764277013 | 8.628897306 | -0.864620293 | -0.310768 |
| Rhbg | 8.073549433 | 8.517132822 | 8.384320698 | 0.132812124 | -0.310771265 |
| Pex1 | 7.816064745 | 8.175837341 | 8.126838882 | 0.048998459 | -0.310774137 |
| Galnt17 | 7.287064915 | 7.317572827 | 7.597848703 | -0.280275876 | -0.310783788 |
| Copz2 | 8.731252515 | 8.712756982 | 9.04213234 | -0.329375359 | -0.310879825 |
| Pik3cb | 9.839228406 | 10.11827473 | 10.15018717 | -0.031912441 | -0.310958768 |
| Fsd1l | 7.059765232 | 6.931964519 | 7.370755507 | -0.438790988 | -0.310990275 |
| Pus10 | 8.957654351 | 9.081887196 | 9.269217103 | -0.187329907 | -0.311562752 |
| Fhdc1 | 10.04683876 | 10.33412951 | 10.3584406 | -0.02431109 | -0.311601834 |
| Zfp362 | 8.437346267 | 8.835068592 | 8.749061033 | 0.086007558 | -0.311714767 |
| Ctc1 | 8.329719883 | 8.638321597 | 8.641486501 | -0.003164904 | -0.311766618 |
| Coq9 | 9.992099636 | 10.02394668 | 10.30388049 | -0.279933813 | -0.311780857 |
| Ugt1a6a | 8.011810242 | 8.10689579 | 8.323649659 | -0.216753869 | -0.311839417 |
| Glb1 | 8.816070439 | 8.991189505 | 9.12798525 | -0.136795744 | -0.311914811 |
| Sgo2a | 6.785357562 | 6.92118062 | 7.097275696 | -0.176095077 | -0.311918134 |
| Stag2 | 11.22918966 | 11.32461472 | 11.54127718 | -0.21666246 | -0.31208752 |
| Ubr3 | 11.01655823 | 11.16776537 | 11.32865489 | -0.16088952 | -0.312096653 |
| Cpeb3 | 6.917729821 | 6.857207926 | 7.229990686 | -0.37278276 | -0.312260864 |
| Ucn2 | 5.614688293 | 6.186561342 | 5.927000351 | 0.259560991 | -0.312312058 |

| | | | | | |
|---|---|---|---|---|---|
| Sort1 | 11.09211119 | 11.06885416 | 11.40448229 | -0.335628127 | -0.312371097 |
| Prune2 | 7.305112911 | 7.402521236 | 7.617557352 | -0.215036116 | -0.312444441 |
| Lipk | 8.362240905 | 8.620585088 | 8.674783695 | -0.054198607 | -0.31254279 |
| Senp7 | 8.579884516 | 9.000385023 | 8.892528345 | 0.107856677 | -0.312643829 |
| Tcim | 8.91414334 | 9.027551998 | 9.226798128 | -0.19924613 | -0.312654788 |
| Rab34 | 8.807048783 | 8.894934967 | 9.119733834 | -0.224798867 | -0.312685051 |
| Serpina3c | 8.851938191 | 8.772402787 | 9.164780068 | -0.392377281 | -0.312841877 |
| Dhrs3 | 9.60330691 | 9.856917256 | 9.916435058 | -0.059517802 | -0.313128148 |
| Zadh2 | 9.524568308 | 9.469743626 | 9.837860758 | -0.368117132 | -0.313292451 |
| Arhgap12 | 9.652462628 | 9.847427522 | 9.965807685 | -0.118380163 | -0.313345057 |
| Dnal1 | 7.303046051 | 7.595557371 | 7.616445485 | -0.020888114 | -0.313399435 |
| Ric8b | 8.235066531 | 8.425696603 | 8.548510721 | -0.122814118 | -0.31344419 |
| Sypl2 | 9.578966197 | 9.504658957 | 9.892564877 | -0.38790592 | -0.31359868 |
| Fam208a | 9.875823176 | 9.964403859 | 10.18944133 | -0.225037468 | -0.313618151 |
| Tcea3 | 8.584808253 | 8.643079448 | 8.898492485 | -0.255413037 | -0.313684232 |
| Zfp40 | 6.502632853 | 6.538675411 | 6.816388156 | -0.277712745 | -0.313755303 |
| Tmem131l | 9.805590867 | 10.18746075 | 10.11936452 | 0.068096226 | -0.313773657 |
| Fmod | 8.706848801 | 8.695101798 | 9.020632907 | -0.325531109 | -0.313784106 |
| Strip2 | 6.134666571 | 6.231121374 | 6.448473919 | -0.217352545 | -0.313807348 |
| Add1 | 11.40550222 | 11.56075068 | 11.71933744 | -0.158586753 | -0.31383522 |
| Baiap2l1 | 7.844093979 | 8.235946459 | 8.157957573 | 0.077988886 | -0.313863594 |
| Kctd9 | 9.478380867 | 9.6539558 | 9.792263456 | -0.138307656 | -0.313882589 |
| Sat2 | 4.820132872 | 4.856221076 | 5.134032577 | -0.277811502 | -0.313899705 |
| Pcnt | 9.454732159 | 9.653999482 | 9.768721729 | -0.114722248 | -0.31398957 |
| Tle6 | 6.389390426 | 6.65156359 | 6.703449469 | -0.051885878 | -0.314059043 |
| Maco1 | 9.793268147 | 9.936752455 | 10.10737672 | -0.170624268 | -0.314108576 |
| Gm14421 | 5.685331864 | 5.821471623 | 5.999560806 | -0.178089183 | -0.314228942 |
| D930048N14Rik | 7.361277377 | 7.68674038 | 7.675593688 | 0.011146692 | -0.314316311 |
| Fmr1 | 10.33669595 | 10.36294614 | 10.65107338 | -0.28812724 | -0.314377423 |
| Hagh | 9.400275329 | 9.626644647 | 9.714808491 | -0.088163844 | -0.314533162 |
| AA986860 | 9.764608045 | 10.05337512 | 10.07917015 | -0.025795034 | -0.314562106 |
| Ptgr2 | 9.17448589 | 9.297948083 | 9.489076367 | -0.191128284 | -0.314590477 |

| | | | | | |
|---|---|---|---|---|---|
| Ipp | 7.153117875 | 7.208564382 | 7.467909268 | -0.259344887 | -0.314791394 |
| Spef1 | 7.79745655 | 8.048916359 | 8.112252595 | -0.063336237 | -0.314796045 |
| Ggh | 10.31853534 | 10.42634415 | 10.63336233 | -0.20701818 | -0.314826983 |
| Herc3 | 7.543540573 | 7.859780446 | 7.859057244 | 0.000723202 | -0.315516671 |
| Casp9 | 8.6372374 | 8.851986818 | 8.952776092 | -0.100789274 | -0.315538692 |
| AU021092 | 4.683924013 | 4.783250636 | 4.999641682 | -0.216391046 | -0.315717669 |
| Cntln | 7.067376496 | 7.082807217 | 7.383206994 | -0.300399777 | -0.315830499 |
| Ckmt1 | 11.16666761 | 11.42535664 | 11.48253684 | -0.0571802 | -0.31586923 |
| Sv2a | 4.202068292 | 4.57801046 | 4.518071474 | 0.059938986 | -0.316003182 |
| Tmem94 | 9.305015354 | 9.751508653 | 9.621035183 | 0.13047347 | -0.316019829 |
| Anxa1 | 13.54388802 | 13.5515453 | 13.86000413 | -0.30845883 | -0.31611611 |
| Fbxo25 | 7.339809269 | 7.551342781 | 7.656062969 | -0.104720188 | -0.316253701 |
| Golgb1 | 10.8850364 | 11.14832943 | 11.20149481 | -0.05316538 | -0.31645841 |
| Tcf23 | 4.065713012 | 4.153481246 | 4.382317926 | -0.22883668 | -0.316604915 |
| Arhgef5 | 11.40434188 | 11.6184405 | 11.72096166 | -0.102521153 | -0.316619773 |
| Mpv17 | 8.570248048 | 8.818560611 | 8.887012818 | -0.068452207 | -0.31676477 |
| Amt | 5.327910757 | 5.706737392 | 5.644762969 | 0.061974424 | -0.316852211 |
| Cryz | 7.126400786 | 7.177300236 | 7.443301634 | -0.266001398 | -0.316900848 |
| Greb1l | 4.589548022 | 4.829468239 | 4.906868522 | -0.077400283 | -0.317320501 |
| Podxl2 | 6.50531453 | 6.568457445 | 6.822914913 | -0.254457468 | -0.317600383 |
| Sms-ps | 6.095116102 | 6.172692453 | 6.41287637 | -0.240183917 | -0.317760268 |
| Pou2f1 | 8.900606676 | 8.949137926 | 9.218533942 | -0.269396017 | -0.317927267 |
| Zfp532 | 8.725437317 | 8.814076448 | 9.043409215 | -0.229332767 | -0.317971898 |
| Cfap57 | 7.364547023 | 7.538965216 | 7.682571991 | -0.143606775 | -0.318024969 |
| Atp5s | 6.791352114 | 6.800029146 | 7.109547304 | -0.309518158 | -0.31819519 |
| Zfp994 | 7.484864407 | 7.445607891 | 7.803118533 | -0.357510642 | -0.318254126 |
| Hspa12b | 7.403373951 | 7.673320268 | 7.721636228 | -0.04831596 | -0.318262277 |
| Rc3h1 | 10.19189614 | 10.46632004 | 10.51019421 | -0.043874177 | -0.318298076 |
| Trim23 | 8.416314147 | 8.413515093 | 8.734662105 | -0.321147011 | -0.318347958 |
| Rps6ka5 | 7.200319689 | 7.218704371 | 7.518825276 | -0.300120905 | -0.318505587 |
| Bmpr1a | 10.98985834 | 11.22767349 | 11.30843131 | -0.080757827 | -0.318572973 |
| Abca2 | 9.417622856 | 9.688083106 | 9.73622064 | -0.048137534 | -0.318597784 |

| | | | | | |
|---|---|---|---|---|---|
| E2f2 | 8.549249189 | 9.070724126 | 8.86786007 | 0.202864056 | -0.318610881 |
| Kcnma1 | 6.30049599 | 6.639796507 | 6.619209499 | 0.020587008 | -0.318713509 |
| Lce1j | 9.539082282 | 9.953280891 | 9.857796538 | 0.095484353 | -0.318714257 |
| Wdr6 | 8.707495484 | 8.896339068 | 9.026590688 | -0.13025162 | -0.319095204 |
| Pkd1 | 10.5039557 | 10.91093621 | 10.82310116 | 0.087835053 | -0.31914546 |
| Plxnb3 | 5.464946722 | 5.59038366 | 5.784167693 | -0.193784034 | -0.319220971 |
| Amotl1 | 12.05264511 | 12.36191158 | 12.37194644 | -0.010034867 | -0.319301337 |
| Nexn | 9.829111439 | 9.5557983 | 10.14841623 | -0.59261793 | -0.319304791 |
| Zfp963 | 6.062125151 | 6.286373036 | 6.38144424 | -0.095071203 | -0.319319089 |
| S1pr5 | 8.584086013 | 8.901297317 | 8.903463213 | -0.002165897 | -0.3193772 |
| Tpgs2 | 7.390208668 | 7.617144579 | 7.70964269 | -0.092498112 | -0.319434022 |
| Acyp2 | 8.011462878 | 7.963235853 | 8.331017281 | -0.367781428 | -0.319554403 |
| Rab3d | 11.44588852 | 11.52278951 | 11.76547805 | -0.24268854 | -0.31958953 |
| Arhgap10 | 9.704146722 | 9.976681454 | 10.02375602 | -0.047074568 | -0.319609299 |
| Eno2 | 6.239843001 | 6.8172393 | 6.559462283 | 0.257777017 | -0.319619281 |
| Slc45a4 | 8.442557136 | 8.924046772 | 8.762251593 | 0.16179518 | -0.319694457 |
| Ap1s2 | 8.812661074 | 8.819127593 | 9.13256204 | -0.313434446 | -0.319900966 |
| Mppe1 | 8.588010264 | 8.661388602 | 8.908106603 | -0.246718001 | -0.320096339 |
| Serpinf1 | 10.86806192 | 10.77426074 | 11.18818806 | -0.413927313 | -0.320126133 |
| Fam241b | 7.742513541 | 7.765006079 | 8.062747551 | -0.297741472 | -0.32023401 |
| Zfp174 | 5.947994002 | 6.138228551 | 6.2683293 | -0.130100749 | -0.320335298 |
| Kctd21 | 6.978081914 | 7.235329509 | 7.298602202 | -0.063272693 | -0.320520289 |
| Slc25a16 | 8.833461509 | 8.760997002 | 9.154148319 | -0.393151317 | -0.320686811 |
| Hdac5 | 10.32248767 | 10.65832972 | 10.64323527 | 0.01509445 | -0.320747597 |
| Slc35b4 | 9.282932364 | 9.548036582 | 9.603683563 | -0.055646981 | -0.320751199 |
| Gkap1 | 6.970880521 | 7.120847063 | 7.291694737 | -0.170847674 | -0.320814216 |
| Lypd6 | 4.715447006 | 4.859689383 | 5.036378562 | -0.176689179 | -0.320931557 |
| Gmpr | 7.674803234 | 7.764853267 | 7.995838208 | -0.230984941 | -0.321034974 |
| Rhebl1 | 5.755458967 | 5.857253005 | 6.07649526 | -0.219242255 | -0.321036293 |
| Apbb1 | 6.721979343 | 6.958472741 | 7.043083079 | -0.084610338 | -0.321103736 |
| Slc25a10 | 9.317208645 | 9.341040229 | 9.638341599 | -0.29730137 | -0.321132954 |
| Negr1 | 6.110748594 | 6.048289327 | 6.431992092 | -0.383702765 | -0.321243498 |

| | | | | | |
|---|---|---|---|---|---|
| Akr1cl | 5.919376094 | 6.118521072 | 6.240626729 | -0.122105657 | -0.321250635 |
| Tmprss13 | 10.43822925 | 10.78077824 | 10.75953442 | 0.02124382 | -0.321305177 |
| C7 | 4.388044071 | 4.325201171 | 4.70936581 | -0.384164639 | -0.321321739 |
| Kcp | 4.097931213 | 4.305754472 | 4.419294719 | -0.113540247 | -0.321363506 |
| Fgfr1 | 10.92188437 | 11.02126404 | 11.24336841 | -0.222104363 | -0.321484037 |
| Ppfibp1 | 10.44253239 | 10.52896485 | 10.76410986 | -0.235145013 | -0.321577477 |
| Dhrs7c | 8.503404373 | 8.611319727 | 8.824999853 | -0.213680126 | -0.32159548 |
| Raver2 | 5.778568799 | 5.993648755 | 6.100320164 | -0.106671408 | -0.321751364 |
| Per2 | 8.6938826 | 8.791442111 | 9.015700898 | -0.224258787 | -0.321818298 |
| Fbxo21 | 8.256169728 | 8.337169073 | 8.578007925 | -0.240838852 | -0.321838196 |
| Thsd7a | 6.197610293 | 6.207850203 | 6.519515211 | -0.311665008 | -0.321904918 |
| Pqlc3 | 9.00058182 | 9.090485782 | 9.322538754 | -0.232052972 | -0.321956933 |
| Ppic | 9.093220247 | 9.092153475 | 9.415504387 | -0.323350912 | -0.32228414 |
| Fam124b | 3.964731609 | 4.168756173 | 4.287056539 | -0.118300365 | -0.322324929 |
| Zfp850 | 6.571693815 | 6.751860885 | 6.894092607 | -0.142231722 | -0.322398792 |
| C1qb | 10.69805684 | 10.53289719 | 11.02052808 | -0.487630893 | -0.32247124 |
| Prr11 | 7.063745303 | 7.2238335 | 7.386323943 | -0.162490444 | -0.32257864 |
| Kcnb1 | 6.811941713 | 7.02314148 | 7.134567071 | -0.111425591 | -0.322625358 |
| Hoxc13 | 6.254780703 | 6.435237219 | 6.577440382 | -0.142203163 | -0.322659679 |
| Pias3 | 8.98763748 | 9.16931194 | 9.310332867 | -0.141020928 | -0.322695387 |
| Krba1 | 8.683727777 | 9.056409436 | 9.006435801 | 0.049973635 | -0.322708024 |
| Pja2 | 10.39949791 | 10.48245585 | 10.72240984 | -0.239953983 | -0.32291193 |
| Thsd1 | 8.203383941 | 8.418839712 | 8.526335491 | -0.10749578 | -0.322951551 |
| Camk2n1 | 7.926304401 | 7.912282287 | 8.249312281 | -0.337029994 | -0.32300788 |
| Acsf2 | 8.876138518 | 9.087610843 | 9.199281222 | -0.111670379 | -0.323142704 |
| Eya1 | 5.660874323 | 5.969104818 | 5.984074413 | -0.014969595 | -0.32320009 |
| Ica1 | 5.360261047 | 5.446448139 | 5.683948684 | -0.237500545 | -0.323687637 |
| Ednra | 8.001170227 | 7.921831334 | 8.325017406 | -0.403186072 | -0.323847179 |
| Kif13b | 10.12313359 | 10.42064328 | 10.44704406 | -0.026400787 | -0.323910471 |
| Lce1f | 10.75749165 | 11.23539243 | 11.08141355 | 0.153978877 | -0.323921907 |
| Ckm | 14.16152788 | 14.6013184 | 14.48546822 | 0.115850177 | -0.323940343 |
| Syne3 | 8.311804674 | 8.690826084 | 8.635794549 | 0.055031534 | -0.323989875 |

| | | | | | |
|---|---|---|---|---|---|
| Dhx57 | 8.401152687 | 8.619475072 | 8.725145617 | -0.105670545 | -0.32399293 |
| Cox7a2l | 11.09790189 | 11.15013026 | 11.42191699 | -0.271786733 | -0.324015097 |
| Arsk | 7.275768555 | 7.57455296 | 7.599846969 | -0.025294009 | -0.324078414 |
| Trim62 | 8.459699591 | 8.757343519 | 8.783802112 | -0.026458593 | -0.32410252 |
| Phldb3 | 8.682329825 | 9.291858396 | 9.006495782 | 0.285362614 | -0.324165957 |
| Fzd7 | 10.08143306 | 10.19679492 | 10.40574564 | -0.208950713 | -0.324312573 |
| Rcn3 | 8.520671204 | 8.742414743 | 8.845141329 | -0.102726586 | -0.324470125 |
| Mccc1 | 9.635786321 | 9.738029413 | 9.960281627 | -0.222252214 | -0.324495306 |
| Rab5b | 11.19257714 | 11.3657278 | 11.51717678 | -0.15144898 | -0.324599643 |
| Trpm7 | 10.51026845 | 10.80879004 | 10.83496608 | -0.026176047 | -0.32469763 |
| Abhd4 | 10.1721468 | 10.32308661 | 10.49706724 | -0.173980623 | -0.324920432 |
| Arrb1 | 10.70982872 | 10.91305915 | 11.03478071 | -0.121721553 | -0.324951983 |
| Nfix | 11.90967237 | 12.38197556 | 12.23473137 | 0.14724419 | -0.325059003 |
| Cldn34c1 | 5.108118375 | 5.144503297 | 5.433358765 | -0.288855468 | -0.325240391 |
| Thnsl2 | 6.287361445 | 6.380224281 | 6.612842179 | -0.232617898 | -0.325480734 |
| Syn2 | 5.29972179 | 5.489404923 | 5.625203764 | -0.135798841 | -0.325481973 |
| Zfp41 | 7.187013283 | 7.429568122 | 7.512559056 | -0.082990935 | -0.325545773 |
| Smoc1 | 8.587469806 | 8.357964124 | 8.913033992 | -0.555069869 | -0.325564187 |
| Ift140 | 8.371019114 | 8.618630486 | 8.696604007 | -0.077973521 | -0.325584892 |
| Sirt3 | 6.86039989 | 6.899340762 | 7.186302021 | -0.286961259 | -0.325902131 |
| Kif20b | 7.464685159 | 7.688255367 | 7.790597199 | -0.102341832 | -0.32591204 |
| Tom1l2 | 10.81768747 | 11.0914155 | 11.14375047 | -0.052334963 | -0.326062997 |
| Exd2 | 8.289116777 | 8.502316367 | 8.615360976 | -0.113044609 | -0.326244199 |
| Ctsf | 8.741630013 | 8.932665829 | 9.067926479 | -0.13526065 | -0.326296466 |
| Kbtbd13 | 4.964710643 | 4.994176413 | 5.291113692 | -0.29693728 | -0.32640305 |
| Lsmem1 | 5.775523853 | 5.726145223 | 6.101963302 | -0.375818079 | -0.326439449 |
| Nudt13 | 8.083762753 | 8.217551254 | 8.410461483 | -0.192910229 | -0.32669873 |
| Etl4 | 9.958916635 | 10.27468739 | 10.28564535 | -0.01095796 | -0.326728712 |
| Gm5127 | 5.55703543 | 5.769214237 | 5.883787981 | -0.114573744 | -0.326752551 |
| Otub2 | 8.286133632 | 8.416898974 | 8.612989333 | -0.196090359 | -0.326855701 |
| Hsd17b11 | 8.575208392 | 8.510515711 | 8.902122667 | -0.391606956 | -0.326914275 |
| Itpkb | 11.28450628 | 11.58205155 | 11.61144265 | -0.0293911 | -0.326936377 |

| | | | | | |
|---|---|---|---|---|---|
| Inafm1 | 8.479905998 | 8.689230461 | 8.806946662 | -0.117716201 | -0.327040664 |
| Atxn1 | 9.862356419 | 9.840800808 | 10.18953653 | -0.348735725 | -0.327180114 |
| Cfap300 | 6.172286196 | 6.345657797 | 6.499582771 | -0.153924974 | -0.327296575 |
| Zfp61 | 6.625221383 | 6.925307604 | 6.952524158 | -0.027216554 | -0.327302775 |
| Fam126b | 8.95731554 | 8.996922918 | 9.284664112 | -0.287741194 | -0.327348572 |
| Ptbp2 | 8.590295914 | 8.667867909 | 8.917703158 | -0.249835249 | -0.327407243 |
| Acp7 | 8.308082136 | 8.762021122 | 8.635509587 | 0.126511535 | -0.327427451 |
| Mbd4 | 6.642300773 | 7.026171319 | 6.969745528 | 0.056425792 | -0.327444755 |
| Fbxo3 | 10.48351658 | 10.52028155 | 10.81160032 | -0.291318777 | -0.328083747 |
| Dsn1 | 6.793650592 | 7.109717923 | 7.12173781 | -0.012019887 | -0.328087218 |
| Rab43 | 6.847040467 | 7.421338072 | 7.17515071 | 0.246187362 | -0.328110243 |
| Zfp775 | 5.72199907 | 5.649484036 | 6.050179113 | -0.400695077 | -0.328180043 |
| Gpbp1l1 | 10.33735252 | 10.49439407 | 10.66558396 | -0.171189887 | -0.328231443 |
| Dexi | 8.346958026 | 8.686237087 | 8.675224546 | 0.011012542 | -0.32826652 |
| Vps41 | 10.29581119 | 10.35856842 | 10.62446134 | -0.265892917 | -0.32865015 |
| Kremen2 | 6.361841656 | 6.682921539 | 6.690508362 | -0.007586824 | -0.328666706 |
| Copg2 | 10.39228999 | 10.54224864 | 10.72106309 | -0.178814447 | -0.328773093 |
| 1110051M20Rik | 7.569886347 | 7.743148469 | 7.89878731 | -0.15563884 | -0.328900962 |
| Sparc | 13.71157434 | 13.73355768 | 14.04048813 | -0.30693045 | -0.328913787 |
| Cacng6 | 6.590691624 | 6.707181248 | 6.919633427 | -0.21245218 | -0.328941803 |
| Mospd1 | 9.556342024 | 9.665305159 | 9.885669121 | -0.220363962 | -0.329327097 |
| Ckap2l | 8.592709111 | 8.660745721 | 8.922088727 | -0.261343006 | -0.329379616 |
| Zfp983 | 6.863790953 | 7.337319474 | 7.193224442 | 0.144095031 | -0.32943349 |
| Pde4dip | 12.71487876 | 12.7245543 | 13.04434675 | -0.319792453 | -0.32946799 |
| Gm14418 | 4.611957906 | 4.801553473 | 4.94143638 | -0.139882907 | -0.329478474 |
| Cat | 11.37504403 | 11.42415712 | 11.70457929 | -0.280422167 | -0.32953526 |
| Ghr | 11.58384596 | 11.40369375 | 11.91393994 | -0.510246197 | -0.33009398 |
| Lhfpl1 | 5.967699416 | 6.261647221 | 6.297901664 | -0.036254443 | -0.330202248 |
| 2010315B03Rik | 6.605682649 | 6.820324434 | 6.936019226 | -0.115694793 | -0.330336577 |
| Calml3 | 11.48031122 | 11.53245844 | 11.81076249 | -0.27830405 | -0.330451273 |
| Nmrk1 | 6.529492466 | 6.672274874 | 6.859969821 | -0.187694947 | -0.330477356 |
| Pde8a | 8.569603206 | 8.669077776 | 8.900317985 | -0.231240208 | -0.330714778 |

| | | | | | |
|---|---|---|---|---|---|
| Ccdc163 | 6.813862182 | 7.012196845 | 7.144641995 | -0.132445151 | -0.330779814 |
| Syp | 5.576849854 | 5.598799458 | 5.907789635 | -0.308990177 | -0.330939781 |
| 9930012K11Rik | 7.622587933 | 7.96075788 | 7.953660632 | 0.007097248 | -0.331072699 |
| Ccdc181 | 7.25378663 | 7.345834311 | 7.584875518 | -0.239041207 | -0.331088888 |
| Lcmt1 | 8.292255377 | 8.320048816 | 8.623387108 | -0.303338291 | -0.331131731 |
| Tmem45a | 12.35326766 | 12.58324419 | 12.68441659 | -0.101172407 | -0.331148937 |
| Fancg | 7.809137166 | 8.072591437 | 8.140380993 | -0.067789556 | -0.331243827 |
| Smtnl1 | 7.850295832 | 7.872467394 | 8.181805555 | -0.309338161 | -0.331509724 |
| Ubxn7 | 9.692948594 | 9.84212313 | 10.02449305 | -0.182369921 | -0.331544457 |
| Smpd3 | 10.12091325 | 10.51136103 | 10.45245924 | 0.058901797 | -0.331545987 |
| Cobl | 10.81860241 | 11.05252711 | 11.15015848 | -0.09763137 | -0.331556073 |
| 4933427D14Rik | 7.450021875 | 7.77587017 | 7.781642933 | -0.005772762 | -0.331621057 |
| Gm4294 | 7.6769034 | 8.472588775 | 8.008545636 | 0.464043139 | -0.331642236 |
| Hadh | 11.15984005 | 11.31199383 | 11.49171357 | -0.179719743 | -0.33187352 |
| Hjurp | 10.91271493 | 11.25451849 | 11.24465162 | 0.00986687 | -0.331936687 |
| Scg3 | 6.198786245 | 6.117850726 | 6.530766456 | -0.41291573 | -0.331980211 |
| Col6a2 | 11.43721739 | 11.6691417 | 11.769238 | -0.100096303 | -0.332020607 |
| Zfp329 | 8.530406262 | 8.783946625 | 8.862429717 | -0.078483092 | -0.332023455 |
| Rsad1 | 7.552187748 | 7.668549603 | 7.884244144 | -0.215694541 | -0.332056396 |
| Msh3 | 8.296182114 | 8.378303294 | 8.628420258 | -0.250116963 | -0.332238144 |
| Spag5 | 7.356810981 | 7.601525955 | 7.689086737 | -0.087560782 | -0.332275756 |
| Sfxn3 | 10.23730507 | 10.30937451 | 10.56982089 | -0.260446387 | -0.33251582 |
| Mpp3 | 5.559286336 | 5.481228464 | 5.891865432 | -0.410636968 | -0.332579096 |
| Ikbip | 8.138520864 | 8.304592022 | 8.471170009 | -0.166577987 | -0.332649145 |
| Zfp74 | 6.795037807 | 6.992592196 | 7.127871793 | -0.135279597 | -0.332833986 |
| Wdr90 | 7.192412407 | 7.706121259 | 7.525374985 | 0.180746274 | -0.332962578 |
| Cpt1c | 6.354066465 | 6.712774209 | 6.687387678 | 0.025386531 | -0.333321213 |
| Nipa1 | 7.194599521 | 7.311061408 | 7.528097364 | -0.217035956 | -0.333497843 |
| Mob3b | 8.050060548 | 8.242659406 | 8.383568942 | -0.140909536 | -0.333508394 |
| Stfa3 | 10.4228756 | 10.66989989 | 10.7564947 | -0.086594817 | -0.3336191 |
| Pnpla6 | 8.644970931 | 9.104394588 | 8.978596858 | 0.12579773 | -0.333625927 |
| Zfp677 | 6.663057263 | 6.764753871 | 6.996698026 | -0.231944155 | -0.333640763 |

| | | | | | |
|---|---|---|---|---|---|
| Tmem175 | 7.892825552 | 8.094164636 | 8.22657892 | -0.132414283 | -0.333753368 |
| Stradb | 8.319540271 | 8.32491974 | 8.653510013 | -0.328590273 | -0.333969742 |
| 1700019D03Rik | 5.873234857 | 5.845462522 | 6.207212763 | -0.36175024 | -0.333977906 |
| Trim68 | 7.800759085 | 7.957865293 | 8.134812998 | -0.176947705 | -0.334053913 |
| Cd59a | 8.892061108 | 8.933347535 | 9.226262458 | -0.292914923 | -0.334201351 |
| Mospd2 | 9.20337633 | 9.446046713 | 9.537740524 | -0.091693811 | -0.334364194 |
| Hmgcs1 | 12.90774624 | 13.10317058 | 13.24211586 | -0.13894528 | -0.334369623 |
| Zfp9 | 6.867589742 | 7.340899487 | 7.202097938 | 0.13880155 | -0.334508196 |
| Gpx3 | 12.54722155 | 12.37463367 | 12.8817377 | -0.507104033 | -0.334516153 |
| Iglon5 | 4.795270512 | 4.945386898 | 5.130094437 | -0.18470754 | -0.334823926 |
| Enpp1 | 8.240601779 | 8.413180617 | 8.575487661 | -0.162307044 | -0.334885882 |
| Tmem44 | 5.277853132 | 5.324484727 | 5.612873184 | -0.288388457 | -0.335020052 |
| Cracr2b | 8.028516018 | 8.401300007 | 8.363606434 | 0.037693573 | -0.335090416 |
| Acss1 | 7.832157094 | 7.882291165 | 8.167371467 | -0.285080302 | -0.335214372 |
| Stx1b | 5.772500514 | 5.935217502 | 6.107760313 | -0.172542811 | -0.335259799 |
| Hebp2 | 10.31129116 | 10.50088019 | 10.64671161 | -0.145831413 | -0.335420447 |
| Tspyl5 | 3.784778821 | 3.867472824 | 4.120224454 | -0.25275163 | -0.335445632 |
| Ereg | 10.4466597 | 10.66974399 | 10.78211122 | -0.112367227 | -0.33545152 |
| Ctns | 8.646509974 | 8.893468428 | 8.981987011 | -0.088518582 | -0.335477036 |
| Gm6136 | 4.090485865 | 4.346649475 | 4.42640316 | -0.079753685 | -0.335917294 |
| Zbed3 | 6.215740234 | 6.399826864 | 6.551877326 | -0.152050462 | -0.336137092 |
| Anln | 9.332817769 | 9.561006827 | 9.669050369 | -0.108043541 | -0.336232599 |
| Dnajc12 | 5.38128495 | 5.340079061 | 5.717688065 | -0.377609005 | -0.336403115 |
| Cdc42bpg | 9.815791887 | 10.48579195 | 10.15322376 | 0.332568187 | -0.337431877 |
| Dlx5 | 5.819527594 | 6.11372208 | 6.157327583 | -0.043605504 | -0.33779999 |
| Kif11 | 7.703118057 | 7.853357681 | 8.040976508 | -0.187618827 | -0.337858452 |
| Ctdsp2 | 12.34082499 | 12.5964262 | 12.67882316 | -0.082396963 | -0.337998177 |
| Aagab | 9.594942625 | 9.699055237 | 9.93298312 | -0.233927883 | -0.338040496 |
| Trmt2b | 8.445761285 | 8.687790889 | 8.783896705 | -0.096105816 | -0.33813542 |
| Pimreg | 5.146202724 | 5.289600559 | 5.484360723 | -0.194760164 | -0.338157999 |
| Cacnb1 | 8.339495537 | 8.577907891 | 8.677688932 | -0.09978104 | -0.338193394 |
| Arhgef17 | 8.382798133 | 8.638866913 | 8.72100798 | -0.082141068 | -0.338209848 |

| | | | | | |
|---|---|---|---|---|---|
| Rab6b | 8.423623531 | 8.346141543 | 8.761879192 | -0.415737649 | -0.338255661 |
| Iqce | 7.654010941 | 7.946385249 | 7.992386471 | -0.046001221 | -0.33837553 |
| Pclaf | 8.773397749 | 8.975693403 | 9.111802114 | -0.136108711 | -0.338404365 |
| AC149090.1 | 12.14981729 | 12.29596719 | 12.48833484 | -0.192367653 | -0.338517553 |
| Igsf10 | 9.527037024 | 9.640350066 | 9.865792487 | -0.225442421 | -0.338755463 |
| Sv2b | 4.21352448 | 4.577733492 | 4.552378863 | 0.025354629 | -0.338854383 |
| Manea | 8.07515763 | 8.07317171 | 8.414359098 | -0.341187387 | -0.339201468 |
| 4930412O13Rik | 4.836841053 | 5.088322368 | 5.176492523 | -0.088170155 | -0.33965147 |
| Sar1b | 10.72873805 | 10.72176265 | 11.06849406 | -0.34673141 | -0.33975601 |
| Zfp950 | 9.151273727 | 9.305486357 | 9.491045299 | -0.185558942 | -0.339771572 |
| Cdk14 | 8.494009689 | 8.704033338 | 8.833989118 | -0.12995578 | -0.339979429 |
| Mal | 7.417767756 | 7.490403043 | 7.757768969 | -0.267365926 | -0.340001212 |
| Fry | 10.43416342 | 10.43188027 | 10.77420728 | -0.34232701 | -0.34004386 |
| Akt3 | 8.246326791 | 8.214922601 | 8.586400551 | -0.37147795 | -0.34007376 |
| Ncapg | 7.60017038 | 7.66984287 | 7.940821566 | -0.270978696 | -0.340651186 |
| Mtcp1 | 6.839557581 | 7.001393325 | 7.18040646 | -0.179013135 | -0.340848879 |
| Mfap5 | 8.358924775 | 8.413192575 | 8.699774622 | -0.286582047 | -0.340849847 |
| Xpc | 8.548410571 | 8.757267085 | 8.88957846 | -0.132311375 | -0.341167889 |
| Cyp2s1 | 7.558315692 | 7.835438516 | 7.899722737 | -0.064284222 | -0.341407045 |
| Acacb | 10.92221221 | 11.11457613 | 11.2638503 | -0.14927417 | -0.34163809 |
| Smpx | 9.192025493 | 9.399717499 | 9.533777161 | -0.134059662 | -0.341751668 |
| BC064078 | 9.197466368 | 9.217636772 | 9.539276377 | -0.321639606 | -0.341810009 |
| Stard4 | 10.07151375 | 10.29381471 | 10.41346583 | -0.11965112 | -0.341952079 |
| Lnx1 | 8.005443689 | 8.277993881 | 8.347444243 | -0.069450362 | -0.342000554 |
| Myo5a | 10.44068188 | 10.78875579 | 10.78275499 | 0.0060008 | -0.342073107 |
| Nipal2 | 9.689762953 | 10.08245008 | 10.03199176 | 0.05045832 | -0.34222881 |
| Atl3 | 12.19419025 | 12.35243052 | 12.53646268 | -0.184032157 | -0.34227243 |
| Hoxb3 | 6.896369211 | 7.160940483 | 7.238652508 | -0.077712025 | -0.342283297 |
| Cep57l1 | 6.229216815 | 6.427652885 | 6.571541228 | -0.143888343 | -0.342324413 |
| Ift81 | 6.851016224 | 7.109451124 | 7.193541303 | -0.084090179 | -0.34252508 |
| Smtnl2 | 8.459343865 | 8.223554382 | 8.801914 | -0.578359618 | -0.342570135 |
| F2r | 8.155878885 | 8.407787004 | 8.498641059 | -0.090854055 | -0.342762174 |

| | | | | | |
|---|---|---|---|---|---|
| Ackr4 | 9.756956879 | 10.03054929 | 10.09982034 | -0.069271046 | -0.342863461 |
| Ganc | 8.116685786 | 8.137293279 | 8.459694305 | -0.322401026 | -0.343008519 |
| Fxyd2 | 5.691796352 | 5.742695245 | 6.035122017 | -0.292426772 | -0.343325665 |
| Ankrd27 | 9.808129776 | 10.03679695 | 10.15156387 | -0.114766927 | -0.343434098 |
| Pir | 7.182892241 | 7.290203793 | 7.526348087 | -0.236144294 | -0.343455846 |
| Tmem64 | 9.527436805 | 9.692187005 | 9.871180011 | -0.178993006 | -0.343743205 |
| Sh3glb2 | 9.73071762 | 10.10142087 | 10.07457164 | 0.026849237 | -0.343854016 |
| Epb41l5 | 8.022256416 | 8.094598249 | 8.366148086 | -0.271549836 | -0.34389167 |
| 4930449A18Rik | 7.814300692 | 7.999520833 | 8.158232147 | -0.158711315 | -0.343931455 |
| Aktip | 9.644534083 | 9.858777468 | 9.988501457 | -0.129723989 | -0.343967374 |
| Armcx4 | 7.46574892 | 7.469124889 | 7.809892033 | -0.340767144 | -0.344143113 |
| Rassf9 | 8.012384236 | 8.287163199 | 8.35658925 | -0.069426051 | -0.344205014 |
| Mmp27 | 6.116102867 | 6.155210359 | 6.460365708 | -0.305155349 | -0.344262841 |
| Prcp | 9.383696607 | 9.408451856 | 9.728013034 | -0.319561178 | -0.344316427 |
| Mcoln1 | 8.978970661 | 9.412718402 | 9.32344716 | 0.089271242 | -0.344476499 |
| Pde5a | 6.851844333 | 7.034181086 | 7.196413155 | -0.162232069 | -0.344568822 |
| Zfp820 | 5.805999127 | 6.045141879 | 6.151138909 | -0.105997029 | -0.345139782 |
| Crbn | 8.748222607 | 8.773583272 | 9.093479182 | -0.31989591 | -0.345256575 |
| Cbfa2t2 | 9.013522346 | 9.283708092 | 9.358836429 | -0.075128337 | -0.345314083 |
| Hmbox1 | 9.10023137 | 9.46023665 | 9.445577431 | 0.014659219 | -0.345346061 |
| Cacna1e | 6.691017798 | 6.667536145 | 7.036530211 | -0.368994066 | -0.345512413 |
| Zfp788 | 7.620975881 | 7.658763531 | 7.966610553 | -0.307847022 | -0.345634672 |
| Paqr4 | 8.869258063 | 9.025907925 | 9.215014467 | -0.189106542 | -0.345756404 |
| Vipr1 | 6.635543447 | 6.832879553 | 6.98138561 | -0.148506057 | -0.345842162 |
| Scn3a | 6.196537446 | 6.515603694 | 6.542688511 | -0.027084817 | -0.346151066 |
| Limch1 | 9.533547959 | 9.696551486 | 9.879878133 | -0.183326647 | -0.346330173 |
| Glt28d2 | 7.678184332 | 7.877213598 | 8.024548656 | -0.147335058 | -0.346364324 |
| Tnnt3 | 14.74052751 | 14.80765238 | 15.08699296 | -0.279340573 | -0.346465443 |
| Samd5 | 7.135783577 | 7.430891273 | 7.482414193 | -0.05152292 | -0.346630616 |
| Arhgef3 | 9.489674263 | 9.491603409 | 9.83638778 | -0.34478437 | -0.346713516 |
| Cryl1 | 7.602418241 | 7.602575541 | 7.949390275 | -0.346814734 | -0.346972035 |
| Kif23 | 8.225608512 | 8.357110318 | 8.572788854 | -0.215678536 | -0.347180342 |

| | | | | | |
|---|---|---|---|---|---|
| Magix | 5.187594914 | 5.369800785 | 5.534775458 | -0.164974673 | -0.347180544 |
| Cytl1 | 4.62117429 | 4.546514156 | 4.968562884 | -0.422048727 | -0.347388593 |
| Slc25a53 | 6.03794524 | 5.989188076 | 6.38536927 | -0.396181194 | -0.347424029 |
| Bbs9 | 6.922997672 | 6.897842574 | 7.270552841 | -0.372710266 | -0.347555169 |
| Foxn1 | 7.325894036 | 7.879499802 | 7.673473563 | 0.206026239 | -0.347579527 |
| Smim27 | 5.628439424 | 5.839806648 | 5.976226667 | -0.136420019 | -0.347787243 |
| Snn | 8.533608295 | 8.7419887 | 8.88152874 | -0.13954004 | -0.347920445 |
| Gck | 4.989236763 | 5.119118647 | 5.337178819 | -0.218060172 | -0.347942056 |
| Paqr6 | 5.867120738 | 6.161067055 | 6.215127091 | -0.054060037 | -0.348006353 |
| Rsrp1 | 12.15760241 | 12.42582299 | 12.50574708 | -0.07992409 | -0.34814467 |
| Depdc1a | 6.073082067 | 6.128296664 | 6.421325195 | -0.293028531 | -0.348243128 |
| Sema4g | 7.783237286 | 8.027423752 | 8.1314938 | -0.104070048 | -0.348256514 |
| Arsb | 8.72626784 | 8.722342973 | 9.074640937 | -0.352297964 | -0.348373097 |
| Acsm1 | 5.355309142 | 5.566875978 | 5.704323259 | -0.137447281 | -0.349014117 |
| Igip | 8.642413248 | 8.86917357 | 8.991477152 | -0.122303582 | -0.349063904 |
| Dennd2c | 9.549656655 | 9.803354414 | 9.898911798 | -0.095557384 | -0.349255143 |
| Itprid2 | 12.54103472 | 12.77890466 | 12.89059554 | -0.11169088 | -0.349560827 |
| Col15a1 | 10.22482509 | 10.19281377 | 10.57465775 | -0.38184398 | -0.34983266 |
| Decr1 | 10.0544678 | 10.00350407 | 10.40433452 | -0.400830453 | -0.349866723 |
| Gal3st1 | 7.095604185 | 7.517718742 | 7.445472866 | 0.072245875 | -0.349868681 |
| Bcl2l14 | 5.705822844 | 5.97721823 | 6.055761819 | -0.078543589 | -0.349938974 |
| Mfsd4b4 | 6.246936594 | 6.381221945 | 6.59702597 | -0.215804025 | -0.350089376 |
| Shisa2 | 8.731707189 | 8.790375328 | 9.081800175 | -0.291424846 | -0.350092985 |
| Snx13 | 10.05725092 | 10.13625819 | 10.40745561 | -0.271197417 | -0.350204686 |
| mt-Nd4l | 12.8634861 | 12.92164631 | 13.2138294 | -0.292183093 | -0.350343303 |
| Zfp60 | 7.942555199 | 8.270844485 | 8.29297164 | -0.022127155 | -0.350416441 |
| Fancb | 6.831093427 | 6.989865355 | 7.181551982 | -0.191686626 | -0.350458555 |
| Nde1 | 9.752600847 | 10.01086979 | 10.1031164 | -0.092246607 | -0.350515553 |
| Gas6 | 12.2658346 | 12.60479833 | 12.6163801 | -0.01158177 | -0.350545497 |
| Pde4a | 7.186704224 | 7.153756635 | 7.537451653 | -0.383695018 | -0.350747428 |
| As3mt | 6.798653295 | 6.897044224 | 7.149504279 | -0.252460055 | -0.350850984 |
| F2rl1 | 9.660862942 | 10.13689548 | 10.01172352 | 0.125171961 | -0.350860577 |

| | | | | | |
|---|---|---|---|---|---|
| Kctd4 | 8.879882154 | 9.094675895 | 9.230809389 | -0.136133493 | -0.350927235 |
| Hyal4 | 5.834549849 | 5.9261558 | 6.185654096 | -0.259498296 | -0.351104247 |
| Atp13a2 | 9.301993395 | 9.824092833 | 9.653251316 | 0.170841517 | -0.35125792 |
| Brca1 | 7.40715567 | 7.730269428 | 7.758555729 | -0.028286301 | -0.351400059 |
| Fzd2 | 8.249230911 | 8.663852176 | 8.600668233 | 0.063183943 | -0.351437322 |
| Kdm1b | 9.303115291 | 9.510023467 | 9.654593065 | -0.144569598 | -0.351477774 |
| Gk5 | 9.412057914 | 9.636822516 | 9.763576577 | -0.126754061 | -0.351518663 |
| Pcp4l1 | 7.363915618 | 7.585979927 | 7.715682023 | -0.129702096 | -0.351766406 |
| Sdhaf4 | 6.980805532 | 6.999524089 | 7.332635912 | -0.333111822 | -0.35183038 |
| Rerg | 6.070017423 | 6.120882917 | 6.422312641 | -0.301429724 | -0.352295217 |
| Mcur1 | 10.33071372 | 10.42302191 | 10.68304978 | -0.260027877 | -0.352336063 |
| Nsg1 | 7.948457726 | 7.976704254 | 8.300912424 | -0.32420817 | -0.352454697 |
| Prrg3 | 8.360769417 | 7.974191477 | 8.713230432 | -0.739038955 | -0.352461015 |
| Nbea | 7.164031625 | 7.269958386 | 7.516689299 | -0.246730913 | -0.352657674 |
| Dgkq | 8.980217367 | 9.408867235 | 9.332959546 | 0.07590769 | -0.352742178 |
| Uvssa | 8.525979804 | 8.709570993 | 8.878860016 | -0.169289024 | -0.352880212 |
| Marf1 | 10.46856524 | 10.77769156 | 10.82152774 | -0.04383618 | -0.352962493 |
| C1s1 | 10.96517354 | 10.89443317 | 11.31849086 | -0.424057687 | -0.353317317 |
| Hepacam2 | 4.731071721 | 4.690099202 | 5.084477839 | -0.394378637 | -0.353406119 |
| Kcnk2 | 7.337551185 | 7.394233778 | 7.690982307 | -0.296748529 | -0.353431122 |
| Plp1 | 8.542067254 | 8.551574827 | 8.895592016 | -0.344017189 | -0.353524762 |
| Slc1a2 | 4.001778717 | 4.403977524 | 4.355666759 | 0.048310765 | -0.353888042 |
| Zfpm2 | 4.544329939 | 4.684877628 | 4.89823429 | -0.213356662 | -0.353904351 |
| Mdh1 | 11.83315972 | 11.87438204 | 12.18707562 | -0.312693573 | -0.353915893 |
| Cdkl5 | 7.481531324 | 7.631890411 | 7.835629799 | -0.203739389 | -0.354098475 |
| Fbln5 | 9.726033943 | 9.838109388 | 10.0801565 | -0.242047112 | -0.354122557 |
| Tnfsf10 | 7.227054364 | 6.851510575 | 7.581488638 | -0.729978063 | -0.354434273 |
| Ablim1 | 12.26176273 | 12.44862187 | 12.61640553 | -0.16778366 | -0.354642807 |
| Klk5 | 11.00313463 | 11.27421545 | 11.35830241 | -0.084086967 | -0.355167783 |
| Slc37a4 | 8.359345947 | 8.499176261 | 8.71462363 | -0.21544737 | -0.355277684 |
| Apol7c | 6.704215275 | 7.735348587 | 7.059548435 | 0.675800152 | -0.35533316 |
| Srpk3 | 5.996361503 | 6.34085458 | 6.351815661 | -0.010961081 | -0.355454158 |

| | | | | | |
|---|---|---|---|---|---|
| Fbp2 | 7.857584475 | 7.885226146 | 8.213063027 | -0.327836881 | -0.355478552 |
| Mink1 | 10.98914174 | 11.47185958 | 11.344645 | 0.127214583 | -0.35550326 |
| Sp6 | 9.470100719 | 9.885438599 | 9.825716293 | 0.059722306 | -0.355615574 |
| Adam3 | 4.594035948 | 4.852472325 | 4.949776596 | -0.097304271 | -0.355740648 |
| Sdccag8 | 7.336646643 | 7.517827953 | 7.692461887 | -0.174633934 | -0.355815244 |
| Adamtsl1 | 7.849186088 | 7.86594795 | 8.205214758 | -0.339266807 | -0.356028669 |
| Dcaf8 | 10.96610788 | 11.12112127 | 11.32217475 | -0.20105348 | -0.356066873 |
| Kprp | 12.60996408 | 13.01073654 | 12.96644632 | 0.04429022 | -0.35648224 |
| Mef2a | 11.18706841 | 11.41330015 | 11.54360678 | -0.130306637 | -0.35653837 |
| Mical3 | 9.395151183 | 9.892701556 | 9.751809612 | 0.140891944 | -0.356658429 |
| Jmy | 9.769988361 | 9.937383955 | 10.12685845 | -0.189474498 | -0.356870092 |
| Sptssa | 11.88587615 | 12.05703239 | 12.24276421 | -0.185731817 | -0.356888057 |
| Runx1t1 | 6.369580111 | 6.485665628 | 6.726860902 | -0.241195274 | -0.35728079 |
| Gsta3 | 6.839448665 | 6.623239581 | 7.1967347 | -0.573495119 | -0.357286036 |
| Lce1g | 9.554489781 | 10.00841582 | 9.911829734 | 0.096586082 | -0.357339953 |
| Elovl6 | 13.96797955 | 14.19052807 | 14.32534332 | -0.13481525 | -0.35736377 |
| Lysmd1 | 8.961786504 | 9.081750968 | 9.319220993 | -0.237470025 | -0.357434489 |
| Slc38a9 | 8.76103197 | 9.110265603 | 9.118549767 | -0.008284164 | -0.357517797 |
| Pik3ca | 10.16937764 | 10.60128767 | 10.52696934 | 0.074318327 | -0.357591698 |
| Il7 | 5.848161735 | 6.130097623 | 6.205889116 | -0.075791493 | -0.357727381 |
| Fam53b | 9.615683988 | 10.04545499 | 9.97360322 | 0.071851768 | -0.357919232 |
| Gli3 | 7.882692817 | 8.217595816 | 8.240881131 | -0.023285315 | -0.358188314 |
| Maged2 | 7.669287547 | 7.765829436 | 8.027487559 | -0.261658123 | -0.358200012 |
| Pld2 | 9.452251365 | 9.824167466 | 9.810859346 | 0.01330812 | -0.35860798 |
| Gpld1 | 9.114164753 | 9.393059214 | 9.472831879 | -0.079772665 | -0.358667126 |
| Rpl14-ps1 | 10.72748984 | 10.55028569 | 11.08641007 | -0.536124383 | -0.358920237 |
| Arvcf | 8.973267408 | 9.37202183 | 9.332525249 | 0.039496581 | -0.359257841 |
| Vps13c | 9.225736185 | 9.553227393 | 9.584995904 | -0.03176851 | -0.359259719 |
| Tmem9 | 7.77706147 | 7.85208989 | 8.136480754 | -0.284390863 | -0.359419284 |
| Ogdhl | 4.614202014 | 4.82000724 | 4.973652775 | -0.153645535 | -0.359450761 |
| Cit | 6.890525541 | 7.398651875 | 7.250000427 | 0.148651448 | -0.359474886 |
| Bbs2 | 6.924786461 | 7.319820613 | 7.284328463 | 0.035492149 | -0.359542002 |

| | | | | | |
|---|---|---|---|---|---|
| Pla2g3 | 7.187490885 | 7.50535299 | 7.547397503 | -0.042044513 | -0.359906618 |
| Maged1 | 10.27332901 | 10.36055682 | 10.63349512 | -0.272938303 | -0.36016611 |
| Trmt9b | 5.216046613 | 5.449063158 | 5.576426318 | -0.12736316 | -0.360379705 |
| Art1 | 9.081466959 | 8.819601677 | 9.442226332 | -0.622624655 | -0.360759373 |
| Pex3 | 9.741677081 | 9.796269234 | 10.1024565 | -0.30618727 | -0.360779422 |
| Sybu | 7.152217451 | 7.386081725 | 7.513034564 | -0.126952839 | -0.360817113 |
| Ak1 | 11.10126912 | 11.19695503 | 11.46244119 | -0.26548616 | -0.36117207 |
| Itm2b | 14.23798594 | 14.28597109 | 14.59923366 | -0.313262567 | -0.361247723 |
| Ttc38 | 7.951728186 | 8.081127512 | 8.312986725 | -0.231859213 | -0.361258539 |
| Bcas3 | 8.648212614 | 8.846292009 | 9.00953285 | -0.163240841 | -0.361320236 |
| Slc12a6 | 9.809549351 | 10.0670592 | 10.17093267 | -0.103873472 | -0.361383319 |
| Atl2 | 10.78505789 | 10.88855842 | 11.14647353 | -0.257915113 | -0.361415637 |
| Tmem254c | 10.556007 | 10.63466909 | 10.91747538 | -0.282806297 | -0.361468383 |
| Atrx | 11.14233223 | 11.33611507 | 11.50390079 | -0.167785713 | -0.361568553 |
| Mboat2 | 11.10622962 | 11.22700199 | 11.46789473 | -0.240892733 | -0.361665107 |
| Kcnj8 | 7.588506359 | 7.722660648 | 7.950659295 | -0.227998646 | -0.362152936 |
| Eid1 | 10.28336342 | 10.43018521 | 10.6455826 | -0.215397387 | -0.36221918 |
| Tshz1 | 9.021295059 | 9.299858105 | 9.383524524 | -0.083666419 | -0.362229465 |
| Tmem14a | 4.914715541 | 4.957237301 | 5.277054496 | -0.319817195 | -0.362338955 |
| Pcdh7 | 10.43031887 | 10.6397273 | 10.79271996 | -0.152992663 | -0.362401093 |
| Klf5 | 11.07345731 | 11.33694677 | 11.43589583 | -0.098949063 | -0.36243852 |
| Plekhn1 | 11.14783397 | 11.53730348 | 11.51042129 | 0.026882183 | -0.36258732 |
| Zfp93 | 5.96707378 | 6.009125943 | 6.32970131 | -0.320575367 | -0.36262753 |
| Msrb2 | 6.342894763 | 6.559605585 | 6.705531573 | -0.145925988 | -0.36263681 |
| Lamc3 | 6.974377228 | 7.12432209 | 7.337047951 | -0.212725861 | -0.362670723 |
| Asb18 | 5.530068748 | 5.324350193 | 5.893506789 | -0.569156596 | -0.363438041 |
| Tbc1d4 | 9.193554278 | 9.356850742 | 9.557095278 | -0.200244536 | -0.363541001 |
| Dnm3 | 6.666204605 | 6.711899884 | 7.029907849 | -0.318007965 | -0.363703244 |
| Tfap2b | 9.609332456 | 9.937400148 | 9.973090414 | -0.035690267 | -0.363757958 |
| Robo2 | 7.797585742 | 7.699362279 | 8.161374746 | -0.462012467 | -0.363789005 |
| Dynll2 | 10.73334011 | 10.85400921 | 11.09717463 | -0.243165423 | -0.363834527 |
| Prss36 | 6.161831416 | 6.496228717 | 6.525741816 | -0.0295131 | -0.3639104 |

| | | | | | |
|---|---|---|---|---|---|
| Scnn1g | 6.096665901 | 6.474430987 | 6.460620438 | 0.013810549 | -0.363954536 |
| Mex3b | 5.389555818 | 5.665837672 | 5.753705323 | -0.087867651 | -0.364149505 |
| Gm14137 | 8.613037095 | 8.935836112 | 8.977254288 | -0.041418176 | -0.364217193 |
| Fggy | 6.294530752 | 6.482570773 | 6.65886542 | -0.176294647 | -0.364334668 |
| Fam83f | 9.428689837 | 9.960246942 | 9.79305103 | 0.167195912 | -0.364361193 |
| Idh2 | 10.06821623 | 10.15179393 | 10.43263489 | -0.280840957 | -0.364418654 |
| Arhgap6 | 6.505314345 | 6.571504901 | 6.869781284 | -0.298276383 | -0.36446694 |
| Prkab2 | 10.04239426 | 10.14444076 | 10.40688765 | -0.262446883 | -0.36449339 |
| Yipf7 | 6.775544733 | 6.912935371 | 7.140295738 | -0.227360367 | -0.364751005 |
| Aldh5a1 | 7.549387563 | 7.738432035 | 7.91422646 | -0.175794425 | -0.364838897 |
| Mmrn1 | 8.612536395 | 8.771237569 | 8.97744292 | -0.206205352 | -0.364906525 |
| Axin2 | 9.014378252 | 9.454723318 | 9.379323801 | 0.075399517 | -0.364945548 |
| Them5 | 10.77851403 | 10.84429132 | 11.1436012 | -0.299309883 | -0.365087173 |
| Aldh4a1 | 9.6851257 | 9.887626557 | 10.0502317 | -0.162605143 | -0.365106 |
| Clcn2 | 6.800862727 | 7.087484141 | 7.166258127 | -0.078773986 | -0.3653954 |
| Ppl | 12.95304572 | 13.32893629 | 13.31870642 | 0.010229867 | -0.365660697 |
| Fbxo45 | 9.461825955 | 9.573738329 | 9.8275686 | -0.25383027 | -0.365742645 |
| Acot2 | 6.796208588 | 7.068306284 | 7.161964817 | -0.093658533 | -0.365756229 |
| Dock11 | 9.093403772 | 9.063050871 | 9.459420148 | -0.396369277 | -0.366016376 |
| Fam57a | 11.27856889 | 11.48539853 | 11.64490474 | -0.15950621 | -0.36633585 |
| Sec31b | 6.104584923 | 6.221527889 | 6.470950472 | -0.249422583 | -0.366365549 |
| Fam160a1 | 9.668572057 | 9.981273666 | 10.03498252 | -0.053708854 | -0.366410462 |
| Ogt | 11.61953855 | 11.93945705 | 11.98595492 | -0.04649787 | -0.36641637 |
| Ppp6r2 | 9.294289194 | 9.47834503 | 9.661269014 | -0.182923985 | -0.366979821 |
| Srpk1 | 11.5705579 | 11.95539478 | 11.93756353 | 0.017831257 | -0.367005623 |
| Rb1cc1 | 10.92277745 | 11.15783554 | 11.28981625 | -0.131980713 | -0.367038797 |
| Art5 | 6.058589439 | 6.005171764 | 6.425645353 | -0.420473589 | -0.367055914 |
| Gba2 | 9.199612502 | 9.550499466 | 9.566826667 | -0.016327201 | -0.367214165 |
| Tmem132c | 7.338488602 | 7.391599158 | 7.706189063 | -0.314589905 | -0.367700461 |
| Abcd4 | 8.895198605 | 9.192941699 | 9.262904726 | -0.069963027 | -0.36770612 |
| Epn3 | 9.88155431 | 10.34545556 | 10.24954034 | 0.09591522 | -0.367986034 |
| Ndrg3 | 9.691137937 | 9.828647695 | 10.05912637 | -0.230478671 | -0.36798843 |

| | | | | | |
|---|---|---|---|---|---|
| Diaph3 | 7.88617788 | 7.882320757 | 8.25456408 | -0.372243323 | -0.3683862 |
| Skint10 | 9.259814753 | 9.311244962 | 9.628207377 | -0.316962415 | -0.368392623 |
| Phf19 | 6.223949282 | 6.216047249 | 6.592355998 | -0.376308748 | -0.368406716 |
| Rab30 | 7.898125979 | 8.346927588 | 8.266641553 | 0.080286036 | -0.368515574 |
| Klf3 | 11.7615656 | 12.00325858 | 12.13009199 | -0.126833407 | -0.368526387 |
| Wnt9a | 5.471533818 | 5.701434923 | 5.840303704 | -0.138868781 | -0.368769886 |
| Pck2 | 7.44879258 | 7.72709833 | 7.817719206 | -0.090620877 | -0.368926627 |
| Iqcc | 6.982574303 | 7.067302851 | 7.351633369 | -0.284330518 | -0.369059066 |
| Zfp641 | 5.971842058 | 5.98365186 | 6.341407295 | -0.357755435 | -0.369565238 |
| Klk13 | 7.055238017 | 7.328267762 | 7.424853482 | -0.096585719 | -0.369615465 |
| Slc25a42 | 7.528730917 | 7.826299741 | 7.898439387 | -0.072139647 | -0.36970847 |
| Ralgps2 | 9.734055476 | 9.983920326 | 10.10401142 | -0.120091092 | -0.369955942 |
| Art3 | 9.925110535 | 9.669527862 | 10.29511195 | -0.625584091 | -0.370001418 |
| Eno3 | 12.90748954 | 13.07263501 | 13.27754853 | -0.20491352 | -0.370058987 |
| Lclat1 | 8.629434693 | 8.737709072 | 8.999655335 | -0.261946263 | -0.370220642 |
| Zfp808 | 4.670664887 | 4.798665719 | 5.041082404 | -0.242416685 | -0.370417517 |
| Plcxd2 | 8.506824954 | 9.03391146 | 8.877540401 | 0.15637106 | -0.370715447 |
| Gamt | 6.709851038 | 6.727221595 | 7.080699107 | -0.353477511 | -0.370848068 |
| Gata3 | 11.29951498 | 11.79139416 | 11.67042482 | 0.120969337 | -0.37090984 |
| Gng7 | 6.139257893 | 6.337993687 | 6.510294274 | -0.172300587 | -0.371036381 |
| Pon3 | 8.71545945 | 8.704283623 | 9.086559957 | -0.382276334 | -0.371100507 |
| Adamts10 | 7.864840277 | 8.141106558 | 8.236139522 | -0.095032965 | -0.371299245 |
| Zfp358 | 7.420837263 | 7.676468587 | 7.792365777 | -0.11589719 | -0.371528514 |
| Tpra1 | 9.129561905 | 9.597200862 | 9.50136889 | 0.095831972 | -0.371806985 |
| Fbxo8 | 8.782651575 | 8.884205192 | 9.15446907 | -0.270263877 | -0.371817495 |
| Pik3c2b | 7.924902945 | 8.179921453 | 8.296725618 | -0.116804165 | -0.371822672 |
| Mmp15 | 8.347650508 | 8.883620023 | 8.719500836 | 0.164119186 | -0.371850328 |
| Aspm | 7.739961532 | 8.051368438 | 8.111845673 | -0.060477235 | -0.371884141 |
| Ptprd | 6.872310395 | 6.749267598 | 7.24424072 | -0.494973122 | -0.371930325 |
| Ankrd23 | 11.372975 | 11.57326803 | 11.74498252 | -0.17171449 | -0.37200752 |
| Scamp5 | 7.322718349 | 7.48578165 | 7.694867419 | -0.20908577 | -0.37214907 |
| Ect2 | 7.924029365 | 8.161546667 | 8.296261941 | -0.134715274 | -0.372232577 |

| | | | | | |
|---|---|---|---|---|---|
| Ddit4 | 9.399427404 | 9.786411332 | 9.772045376 | 0.014365957 | -0.372617972 |
| Ptprf | 12.28069363 | 12.77280544 | 12.65348496 | 0.119320477 | -0.37279133 |
| Ivd | 9.909553721 | 10.08024154 | 10.28249288 | -0.202251344 | -0.372939159 |
| Hephl1 | 9.650038709 | 9.957541262 | 10.02302479 | -0.065483525 | -0.372986078 |
| Gal3st4 | 8.005963375 | 8.182616689 | 8.379025309 | -0.19640862 | -0.373061934 |
| 4930522L14Rik | 6.835367664 | 6.995766181 | 7.20855759 | -0.212791409 | -0.373189926 |
| Lrrc4c | 4.470065302 | 4.442516069 | 4.843443696 | -0.400927627 | -0.373378394 |
| Npr3 | 9.761932771 | 9.156296399 | 10.1353877 | -0.979091297 | -0.373454925 |
| Ctsk | 10.95067991 | 11.188641 | 11.32437433 | -0.135733327 | -0.37369442 |
| Zfyve1 | 8.662256365 | 8.873807211 | 9.035958126 | -0.162150915 | -0.373701761 |
| Jade1 | 9.531347745 | 9.603338486 | 9.905104275 | -0.301765789 | -0.373756531 |
| Ttc30b | 7.032515637 | 7.129994731 | 7.406324899 | -0.276330168 | -0.373809262 |
| Gpr157 | 8.2307934 | 8.384366516 | 8.604639039 | -0.220272523 | -0.373845638 |
| Il17rd | 7.834367136 | 8.10654296 | 8.208331312 | -0.101788352 | -0.373964176 |
| Ccser2 | 10.37832057 | 10.41403739 | 10.75257987 | -0.338542477 | -0.374259303 |
| Cd209b | 8.118975691 | 7.530834699 | 8.493435684 | -0.962600986 | -0.374459994 |
| Cachd1 | 9.316969894 | 9.744139218 | 9.691505414 | 0.052633804 | -0.374535519 |
| Ankra2 | 8.2145749 | 8.410629963 | 8.589145333 | -0.17851537 | -0.374570433 |
| Snrk | 10.37740121 | 10.58247379 | 10.75225805 | -0.169784257 | -0.37485684 |
| Trim29 | 13.45234562 | 13.78750758 | 13.8272355 | -0.039727917 | -0.37488988 |
| Acad11 | 9.916777772 | 10.25241979 | 10.2917001 | -0.039280313 | -0.374922328 |
| Ccdc8 | 6.463115016 | 6.552853297 | 6.838209711 | -0.285356415 | -0.375094695 |
| Ulk1 | 10.2387345 | 10.73090608 | 10.61398602 | 0.116920057 | -0.37525152 |
| Ankrd24 | 6.128865418 | 6.228565677 | 6.504386831 | -0.275821155 | -0.375521413 |
| Slc26a2 | 10.46712472 | 10.7514139 | 10.8428015 | -0.0913876 | -0.375676777 |
| Ncoa2 | 10.05453674 | 10.29568791 | 10.43038013 | -0.134692227 | -0.375843398 |
| Irx4 | 7.21206296 | 7.447281754 | 7.58808036 | -0.140798606 | -0.3760174 |
| Pou2f3 | 10.79951844 | 11.07693521 | 11.17554225 | -0.09860704 | -0.376023807 |
| Sgtb | 5.402442471 | 5.548935567 | 5.778549189 | -0.229613622 | -0.376106718 |
| Mgat3 | 6.231335901 | 6.389416914 | 6.607595169 | -0.218178255 | -0.376259269 |
| Gga2 | 9.795604689 | 10.14912452 | 10.17191731 | -0.022792787 | -0.376312618 |
| Rapgef4 | 7.291055038 | 7.546721023 | 7.667465413 | -0.120744391 | -0.376410375 |

| | | | | | |
|---|---|---|---|---|---|
| Tsc1 | 9.07843986 | 9.370008577 | 9.454958385 | -0.084949809 | -0.376518526 |
| Kidins220 | 10.44465454 | 10.63435351 | 10.82120077 | -0.186847263 | -0.376546237 |
| Ccni | 11.40400477 | 11.48531145 | 11.78058926 | -0.29527781 | -0.376584483 |
| Rab11b-ps2 | 3.850872954 | 3.777756016 | 4.227544925 | -0.449788908 | -0.376671971 |
| Pdk1 | 9.72887534 | 9.722873105 | 10.10561352 | -0.382740413 | -0.376738178 |
| Gm5849 | 5.3892859 | 5.665104767 | 5.766112753 | -0.101007986 | -0.376826853 |
| Smim1 | 7.020420111 | 7.300790785 | 7.397255492 | -0.096464707 | -0.376835381 |
| Ascl2 | 5.123312947 | 5.595545557 | 5.500176286 | 0.095369271 | -0.376863339 |
| Dsg1c | 7.122376583 | 7.403009055 | 7.499418699 | -0.096409644 | -0.377042116 |
| Trove2 | 9.31440565 | 9.352224106 | 9.691460165 | -0.339236058 | -0.377054515 |
| Anxa9 | 10.36963422 | 10.59960361 | 10.74678321 | -0.147179597 | -0.377148987 |
| Mlip | 8.15919988 | 8.359180424 | 8.536568668 | -0.177388244 | -0.377368788 |
| 2610005L07Rik | 8.538023276 | 8.793938113 | 8.915481783 | -0.12154367 | -0.377458507 |
| Ermard | 7.715750224 | 8.065509918 | 8.093585824 | -0.028075906 | -0.3778356 |
| Dock6 | 8.897167652 | 9.356305382 | 9.275102991 | 0.08120239 | -0.377935339 |
| Dact3 | 5.940586008 | 6.164649728 | 6.318808013 | -0.154158285 | -0.378222004 |
| Ccdc80 | 12.05934049 | 11.82033546 | 12.43765852 | -0.617323063 | -0.37831803 |
| Hsdl2 | 10.48305132 | 10.6308295 | 10.86137143 | -0.230541937 | -0.378320113 |
| Gm20683 | 6.817099019 | 7.421249101 | 7.195619425 | 0.225629676 | -0.378520406 |
| Slc6a2 | 7.227951364 | 7.526666472 | 7.606533053 | -0.079866581 | -0.378581689 |
| Ncapd2 | 9.339607218 | 9.607898139 | 9.718191629 | -0.110293491 | -0.378584411 |
| Frmpd1 | 8.464967206 | 8.925734804 | 8.843662617 | 0.082072187 | -0.378695411 |
| Cplane1 | 8.210392569 | 8.153323673 | 8.589259482 | -0.43593581 | -0.378866913 |
| Col6a1 | 11.67511369 | 11.84434071 | 12.05404676 | -0.209706047 | -0.37893307 |
| Zfp867 | 6.84665023 | 6.946885698 | 7.225588461 | -0.278702763 | -0.378938231 |
| Egflam | 7.393980959 | 7.39781777 | 7.773048348 | -0.375230578 | -0.379067389 |
| Klhdc1 | 6.988838144 | 7.40852896 | 7.367956766 | 0.040572194 | -0.379118622 |
| Decr2 | 8.659860972 | 8.905395148 | 9.039191741 | -0.133796592 | -0.379330768 |
| B3gnt9 | 7.058415937 | 7.270281725 | 7.437973818 | -0.167692093 | -0.379557881 |
| Zkscan8 | 8.826565705 | 8.988948359 | 9.206193681 | -0.217245323 | -0.379627976 |
| Ankmy2 | 8.303427351 | 8.531979224 | 8.683152984 | -0.151173761 | -0.379725633 |
| Prc1 | 8.901239721 | 9.188162318 | 9.280979869 | -0.092817551 | -0.379740148 |

| | | | | | |
|---|---|---|---|---|---|
| Dchs1 | 6.966179267 | 7.627181789 | 7.346459548 | 0.280722241 | -0.380280281 |
| Gm10163 | 9.977011989 | 10.25758256 | 10.35762719 | -0.100044627 | -0.380615198 |
| Zfp386 | 8.141607736 | 8.221644922 | 8.522415371 | -0.300770449 | -0.380807635 |
| Trhde | 4.919641868 | 4.91573359 | 5.300554869 | -0.384821279 | -0.380913001 |
| Adhfe1 | 8.309775469 | 8.152506558 | 8.690810062 | -0.538303505 | -0.381034593 |
| mt-Atp8 | 13.48438621 | 13.61838255 | 13.86558764 | -0.247205093 | -0.38120143 |
| Ppp2r5a | 11.37094099 | 11.36187636 | 11.75313812 | -0.391261763 | -0.382197137 |
| Tfap2e | 7.045270496 | 7.488826858 | 7.427525415 | 0.061301443 | -0.382254919 |
| Calcb | 6.857714969 | 7.087357226 | 7.240157007 | -0.152799781 | -0.382442038 |
| Plcg1 | 10.49277841 | 10.77337768 | 10.87524922 | -0.10187154 | -0.382470813 |
| Acy3 | 5.543224843 | 5.639918368 | 5.925724617 | -0.285806249 | -0.382499774 |
| Il1f8 | 8.869006847 | 9.089157529 | 9.251779561 | -0.162622032 | -0.382772714 |
| Sorbs3 | 9.046164442 | 9.289467411 | 9.429116256 | -0.139648845 | -0.382951814 |
| Car14 | 5.984853915 | 5.990912045 | 6.367811292 | -0.376899247 | -0.382957377 |
| mt-Atp6 | 15.89720128 | 15.95717718 | 16.28022791 | -0.32305073 | -0.383026637 |
| Scaper | 8.531402835 | 8.689786859 | 8.914582417 | -0.224795558 | -0.383179582 |
| Phactr1 | 5.998994141 | 6.518380798 | 6.382444295 | 0.135936503 | -0.383450154 |
| Plch2 | 10.29867621 | 10.92880365 | 10.68239954 | 0.246404107 | -0.383723337 |
| Micu1 | 10.2289804 | 10.35119283 | 10.61285087 | -0.261658043 | -0.383870477 |
| Tmie | 5.309993168 | 5.473855071 | 5.693867204 | -0.220012133 | -0.383874035 |
| Dct | 6.672727309 | 6.957388737 | 7.056625185 | -0.099236449 | -0.383897877 |
| Sox21 | 8.858068528 | 9.333515832 | 9.242018659 | 0.091497173 | -0.383950131 |
| Mapk1ip1 | 6.910671981 | 7.19031014 | 7.294678483 | -0.104368343 | -0.384006502 |
| Spag1 | 8.700122782 | 9.082323866 | 9.08425249 | -0.001928625 | -0.384129708 |
| Aqp11 | 4.227350841 | 4.393119506 | 4.611661813 | -0.218542307 | -0.384310972 |
| H2-M2 | 8.170087461 | 9.09867587 | 8.554541911 | 0.544133959 | -0.38445445 |
| Akr1b10 | 7.351276755 | 7.383831766 | 7.735809513 | -0.351977747 | -0.384532758 |
| Hist1h1c | 9.224228236 | 9.279836037 | 9.608828952 | -0.328992915 | -0.384600716 |
| Serpinb10 | 8.865798516 | 9.196961842 | 9.250556612 | -0.05359477 | -0.384758096 |
| Adgrf2 | 7.759685959 | 8.109911288 | 8.144488982 | -0.034577694 | -0.384803023 |
| Ap4m1 | 8.859387612 | 9.129596557 | 9.244311564 | -0.114715007 | -0.384923952 |
| Rubcn | 9.649676229 | 10.00713467 | 10.03466267 | -0.027528 | -0.384986444 |

| | | | | | |
|---|---|---|---|---|---|
| Tshz2 | 8.454597362 | 8.377618137 | 8.83959965 | -0.461981514 | -0.385002289 |
| Prdm9 | 7.23418144 | 7.515401937 | 7.619270144 | -0.103868207 | -0.385088704 |
| Enpp5 | 8.312172518 | 8.219392871 | 8.697282193 | -0.477889322 | -0.385109675 |
| Slc30a1 | 8.883927244 | 9.233284454 | 9.269238434 | -0.03595398 | -0.38531119 |
| Rbfox1 | 7.946738784 | 7.912480046 | 8.332053631 | -0.419573585 | -0.385314847 |
| Il18 | 9.31504687 | 9.421000443 | 9.700570076 | -0.279569633 | -0.385523206 |
| Atm | 8.731561051 | 8.967861098 | 9.117179935 | -0.149318838 | -0.385618884 |
| Fam234a | 10.66253179 | 10.9555377 | 11.04850218 | -0.09296448 | -0.38597039 |
| Chd3 | 9.833759845 | 10.29380071 | 10.21985356 | 0.073947153 | -0.386093712 |
| Mmp28 | 8.463847171 | 8.749804264 | 8.849972514 | -0.10016825 | -0.386125343 |
| Glis2 | 7.217657566 | 7.563241077 | 7.603897274 | -0.040656197 | -0.386239708 |
| Cpa4 | 10.83163797 | 11.14118431 | 11.21795292 | -0.076768617 | -0.386314957 |
| Scin | 10.11508742 | 10.23373949 | 10.50147689 | -0.267737403 | -0.386389476 |
| Rb1 | 9.544445956 | 9.764365717 | 9.931173039 | -0.166807322 | -0.386727083 |
| Lancl1 | 10.23426246 | 10.37441482 | 10.6210688 | -0.246653983 | -0.386806343 |
| Casq1 | 12.36099489 | 12.4325127 | 12.74780598 | -0.315293283 | -0.38681109 |
| Ahnak | 15.16853328 | 15.39349515 | 15.55545473 | -0.161959573 | -0.386921447 |
| Hes6 | 6.699137171 | 6.957665623 | 7.086324897 | -0.128659275 | -0.387187727 |
| Coq10a | 9.270569847 | 9.392577664 | 9.657764851 | -0.265187186 | -0.387195004 |
| Ptprs | 10.0214097 | 10.42867325 | 10.40868987 | 0.019983377 | -0.387280171 |
| Snx30 | 7.789716705 | 7.940013413 | 8.177145014 | -0.237131601 | -0.387428309 |
| D3Ertd751e | 6.412640961 | 6.682101003 | 6.800149054 | -0.11804805 | -0.387508092 |
| Marveld2 | 7.98871656 | 8.348200663 | 8.376323364 | -0.028122701 | -0.387606804 |
| Pfkm | 12.28241137 | 12.43348203 | 12.67021976 | -0.23673773 | -0.38780839 |
| Calm4 | 14.5303194 | 14.53910717 | 14.91828336 | -0.379176187 | -0.387963957 |
| Cdh2 | 6.229463774 | 6.383192784 | 6.617445662 | -0.234252878 | -0.387981887 |
| Cilp | 11.20911885 | 11.39037599 | 11.59738676 | -0.207010777 | -0.388267917 |
| Fkbp9 | 11.09645236 | 11.15128128 | 11.48525327 | -0.333971993 | -0.388800917 |
| Osgepl1 | 7.633190263 | 7.748053429 | 8.022929551 | -0.274876122 | -0.389739288 |
| Ttc8 | 6.998436195 | 7.162867386 | 7.388206187 | -0.225338801 | -0.389769992 |
| Col6a3 | 12.11346383 | 12.17181016 | 12.5033174 | -0.331507237 | -0.389853567 |
| Zdhhc1 | 7.825612263 | 8.28442017 | 8.2155594 | 0.068860769 | -0.389947137 |

| | | | | | |
|---|---|---|---|---|---|
| Nbl1 | 10.32815257 | 10.64757123 | 10.71811096 | -0.07053973 | -0.389958387 |
| Arhgap28 | 5.08518589 | 5.416953341 | 5.475645152 | -0.058691811 | -0.390459262 |
| Fbxl7 | 5.333503577 | 5.54843463 | 5.724101667 | -0.175667037 | -0.390598091 |
| Lgals4 | 7.076691538 | 7.357554914 | 7.467404797 | -0.109849883 | -0.390713259 |
| Gpc3 | 9.163716164 | 9.006202611 | 9.554688984 | -0.548486373 | -0.39097282 |
| Zranb1 | 10.50023635 | 10.58624008 | 10.89124202 | -0.305001933 | -0.391005663 |
| Pea15a | 10.71731542 | 10.95929251 | 11.10878989 | -0.149497387 | -0.391474473 |
| 2700081O15Rik | 7.781822268 | 8.168259257 | 8.173396996 | -0.005137739 | -0.391574728 |
| Cadm2 | 6.09678538 | 6.309637674 | 6.488415234 | -0.17877756 | -0.391629854 |
| Camk2b | 8.141590637 | 8.456034745 | 8.533564689 | -0.077529945 | -0.391974053 |
| Zfand1 | 8.222941178 | 8.473519278 | 8.615070766 | -0.141551488 | -0.392129588 |
| Grb10 | 9.20006596 | 9.522707894 | 9.592232445 | -0.069524551 | -0.392166485 |
| Tln2 | 9.614176737 | 9.825912823 | 10.00663564 | -0.180722821 | -0.392458907 |
| Ccdc28b | 7.588086399 | 7.72888274 | 7.980805245 | -0.251922505 | -0.392718845 |
| Cecr2 | 8.460332423 | 8.689204544 | 8.853247231 | -0.164042687 | -0.392914808 |
| Asb11 | 8.129737262 | 8.16172347 | 8.522709484 | -0.360986014 | -0.392972222 |
| Myom3 | 7.716769643 | 7.873015591 | 8.10992509 | -0.236909499 | -0.393155447 |
| Ift74 | 7.314960451 | 7.315782066 | 7.708394667 | -0.392612601 | -0.393434216 |
| Ypel3 | 9.376688234 | 9.626798548 | 9.770263853 | -0.143465305 | -0.393575619 |
| Tnk2 | 10.09730549 | 10.73157274 | 10.4908966 | 0.240676147 | -0.39359111 |
| Scai | 7.707424685 | 7.69949187 | 8.101162877 | -0.401671007 | -0.393738192 |
| Gys1 | 10.12133549 | 10.12291268 | 10.51523966 | -0.392326977 | -0.393904171 |
| Prrt2 | 5.687477082 | 6.15099785 | 6.08147742 | 0.06952043 | -0.394000338 |
| Hist2h3c1 | 5.810764849 | 6.396207136 | 6.204777869 | 0.191429267 | -0.39401302 |
| Elmod1 | 9.966338022 | 10.18667543 | 10.36037989 | -0.17370446 | -0.394041865 |
| Stox2 | 9.160783661 | 9.524614325 | 9.554936272 | -0.030321947 | -0.394152611 |
| Pgm5 | 7.459658756 | 7.627523553 | 7.854505267 | -0.226981714 | -0.394846511 |
| Mmrn2 | 7.933563324 | 8.16456895 | 8.328486526 | -0.163917575 | -0.394923202 |
| 38231 | 5.758417357 | 5.866110139 | 6.153372935 | -0.287262796 | -0.394955578 |
| Rab11fip2 | 8.474904012 | 8.490676801 | 8.87011466 | -0.379437859 | -0.395210648 |
| Kdm5b | 10.3650855 | 10.65299821 | 10.7604871 | -0.107488893 | -0.395401603 |
| Clspn | 7.856929576 | 8.119363648 | 8.252376246 | -0.133012597 | -0.395446669 |

| | | | | | |
|---|---|---|---|---|---|
| Cep97 | 6.857516793 | 7.126821813 | 7.253143241 | -0.126321428 | -0.395626448 |
| Foxo3 | 10.42050271 | 10.77045288 | 10.81649915 | -0.04604627 | -0.395996447 |
| Slc27a6 | 7.617136708 | 7.822001366 | 8.013270252 | -0.191268886 | -0.396133545 |
| Wnt6 | 5.257638561 | 5.696088565 | 5.653847867 | 0.042240698 | -0.396209306 |
| Dpysl5 | 6.254718328 | 6.642455896 | 6.650976192 | -0.008520296 | -0.396257864 |
| Usp11 | 6.196759903 | 6.418686177 | 6.593066478 | -0.174380301 | -0.396306575 |
| Zhx3 | 9.5226185 | 9.833023316 | 9.918952662 | -0.085929346 | -0.396334162 |
| Aldh1a7 | 8.232110924 | 8.25226336 | 8.628583439 | -0.376320079 | -0.396472515 |
| Lrrc20 | 8.957344235 | 9.424617657 | 9.353904967 | 0.07071269 | -0.396560732 |
| Ndufa4l2 | 9.488962439 | 9.50712103 | 9.885526088 | -0.378405058 | -0.396563649 |
| Gpt | 9.574265877 | 9.743277727 | 9.970961674 | -0.227683947 | -0.396695797 |
| Clec1a | 5.721256409 | 5.871852778 | 6.118079664 | -0.246226886 | -0.396823255 |
| Gm12258 | 6.064508732 | 6.462981137 | 6.461352334 | 0.001628802 | -0.396843602 |
| Zfp946 | 6.976344533 | 7.38309515 | 7.373207024 | 0.009888126 | -0.39686249 |
| Pla2g2f | 12.97362051 | 13.29585258 | 13.37090328 | -0.0750507 | -0.397282767 |
| Slc43a3 | 9.105297047 | 8.807282981 | 9.502590506 | -0.695307525 | -0.397293459 |
| Pfkfb2 | 8.749634939 | 9.184333957 | 9.146973077 | 0.03736088 | -0.397338138 |
| Chst12 | 8.014833445 | 8.387955941 | 8.412248289 | -0.024292348 | -0.397414844 |
| Ntrk3 | 6.882639434 | 7.137955036 | 7.280071895 | -0.142116859 | -0.397432461 |
| Klc4 | 7.250215114 | 7.667222226 | 7.648135149 | 0.019087077 | -0.397920035 |
| Adamts20 | 4.98951905 | 5.149964791 | 5.387514223 | -0.237549432 | -0.397995173 |
| Pex11a | 8.802159709 | 8.928514291 | 9.200651331 | -0.272137039 | -0.398491622 |
| Tlcd1 | 9.050176961 | 9.132556534 | 9.448755908 | -0.316199374 | -0.398578947 |
| Colec12 | 9.257109685 | 9.271923074 | 9.655751837 | -0.383828763 | -0.398642152 |
| Dstyk | 9.889483423 | 10.00219924 | 10.28830688 | -0.286107636 | -0.398823457 |
| Lhx2 | 8.279834319 | 8.660069055 | 8.678692721 | -0.018623667 | -0.398858402 |
| Gng11 | 7.106366284 | 7.150671513 | 7.505324671 | -0.354653158 | -0.398958387 |
| Naaladl2 | 7.034742407 | 6.95085179 | 7.433814013 | -0.482962223 | -0.399071606 |
| Syt7 | 5.184894339 | 5.326018833 | 5.58401432 | -0.257995488 | -0.399119982 |
| Matn2 | 9.380555774 | 9.428755776 | 9.779969614 | -0.351213837 | -0.399413839 |
| Paqr8 | 8.329529085 | 8.751199387 | 8.729201731 | 0.021997656 | -0.399672647 |
| Ly6g6e | 8.704000566 | 9.104586427 | 9.103924593 | 0.000661834 | -0.399924027 |

| | | | | | |
|---|---|---|---|---|---|
| Epb41l4b | 9.888326305 | 10.18143954 | 10.28882121 | -0.107381673 | -0.400494908 |
| Omd | 4.696896226 | 4.63784549 | 5.097412512 | -0.459567022 | -0.400516286 |
| Mitf | 8.503875207 | 8.663730611 | 8.904501297 | -0.240770686 | -0.40062609 |
| Xkrx | 9.913671853 | 10.18382803 | 10.31432893 | -0.1305009 | -0.400657077 |
| Slc22a17 | 7.503347276 | 7.901854867 | 7.904113506 | -0.002258639 | -0.40076623 |
| Tgm5 | 9.976641423 | 10.30240229 | 10.37793086 | -0.075528563 | -0.401289433 |
| Rftn2 | 6.33833843 | 6.555402051 | 6.739974912 | -0.184572861 | -0.401636482 |
| C1qc | 9.912320212 | 9.916329235 | 10.31400289 | -0.397673658 | -0.401682682 |
| Lce1e | 9.836724969 | 10.28622856 | 10.23843151 | 0.04779705 | -0.401706537 |
| Fam3a | 8.786046775 | 8.994015741 | 9.187867975 | -0.193852234 | -0.4018212 |
| Osbpl1a | 10.25721811 | 10.39419778 | 10.65914412 | -0.26494634 | -0.401926013 |
| Ebf3 | 5.770536329 | 5.697314727 | 6.172752341 | -0.475437613 | -0.402216012 |
| Usf3 | 9.974955983 | 10.20567791 | 10.37787933 | -0.172201417 | -0.402923347 |
| Krt10 | 16.93303901 | 17.17187361 | 17.33614087 | -0.164267257 | -0.40310186 |
| Lrrn1 | 4.706902391 | 4.691675858 | 5.110308753 | -0.418632896 | -0.403406362 |
| Dlx3 | 8.240419366 | 8.723774939 | 8.644073301 | 0.079701638 | -0.403653935 |
| Zfp467 | 7.457081791 | 7.52449529 | 7.860859467 | -0.336364177 | -0.403777676 |
| Srd5a1 | 8.652921224 | 9.016917668 | 9.056803431 | -0.039885764 | -0.403882207 |
| Tmcc2 | 7.513977389 | 7.589057419 | 7.917870098 | -0.328812679 | -0.40389271 |
| Bcl9l | 10.53617761 | 11.19544287 | 10.94011789 | 0.255324983 | -0.403940273 |
| Atg13 | 10.10228985 | 10.42223122 | 10.50641833 | -0.08418711 | -0.404128477 |
| Ndn | 7.401146744 | 7.47897702 | 7.805286911 | -0.326309891 | -0.404140167 |
| Zfp874b | 7.408118545 | 7.856442206 | 7.812583249 | 0.043858957 | -0.404464704 |
| Gm14698 | 7.015631905 | 7.028972743 | 7.420341406 | -0.391368663 | -0.404709501 |
| Eef1a2 | 10.84347456 | 11.14616262 | 11.24824122 | -0.102078603 | -0.404766663 |
| Sync | 5.800293295 | 5.811407876 | 6.205211387 | -0.393803511 | -0.404918092 |
| Pcdh20 | 5.623473169 | 5.850741496 | 6.028421327 | -0.177679831 | -0.404948158 |
| Klk8 | 10.05906519 | 10.30578069 | 10.46416969 | -0.158388997 | -0.4051045 |
| Scel | 11.63020127 | 11.93926415 | 12.03539734 | -0.09613319 | -0.40519607 |
| 1700030K09Rik | 6.415011148 | 6.802539196 | 6.820395479 | -0.017856283 | -0.405384331 |
| Nfic | 10.98537008 | 11.37811878 | 11.39090663 | -0.01278785 | -0.405536547 |
| Tmem65 | 11.33651229 | 11.51695834 | 11.7424786 | -0.225520257 | -0.40596631 |

| | | | | | |
|---|---|---|---|---|---|
| Inpp4b | 8.279519221 | 8.41321745 | 8.685547126 | -0.272329676 | -0.406027905 |
| Nr3c1 | 11.30232728 | 11.41133708 | 11.70856466 | -0.297227583 | -0.40623738 |
| Serpine2 | 9.089449177 | 9.081943069 | 9.495930368 | -0.413987299 | -0.40648119 |
| Ikzf2 | 9.746076951 | 9.891104656 | 10.15302182 | -0.261917167 | -0.406944872 |
| Pknox2 | 7.433630297 | 7.482510504 | 7.840755188 | -0.358244684 | -0.407124891 |
| Frem1 | 5.614563221 | 5.904516359 | 6.022101362 | -0.117585003 | -0.407538141 |
| Rcor3 | 7.572138262 | 7.798737449 | 7.979701094 | -0.180963645 | -0.407562832 |
| Fkbp7 | 7.277249339 | 7.268816363 | 7.684863733 | -0.41604737 | -0.407614395 |
| Kank1 | 11.70423514 | 12.02729112 | 12.1118608 | -0.08456968 | -0.407625663 |
| Katnal2 | 5.101064803 | 5.181392684 | 5.508727071 | -0.327334387 | -0.407662268 |
| Pard3 | 10.45924505 | 10.76517 | 10.86693128 | -0.10176128 | -0.407686227 |
| Gm4977 | 4.312921673 | 4.599612835 | 4.72070061 | -0.121087776 | -0.407778937 |
| Abat | 5.98698107 | 5.973568289 | 6.394852797 | -0.421284508 | -0.407871726 |
| Abcd3 | 11.41260284 | 11.67639669 | 11.82084192 | -0.144445237 | -0.408239087 |
| Zic1 | 6.502571638 | 7.003578717 | 6.910960934 | 0.092617783 | -0.408389297 |
| Serpinb8 | 10.88546647 | 11.13795225 | 11.29403497 | -0.156082717 | -0.4085685 |
| Ackr2 | 7.953520555 | 8.268558908 | 8.362355091 | -0.093796183 | -0.408834536 |
| Dglucy | 6.808221307 | 7.024299732 | 7.217106528 | -0.192806796 | -0.408885221 |
| Gigyf1 | 9.107065753 | 9.532088248 | 9.51620477 | 0.015883477 | -0.409139018 |
| Ndrg2 | 12.27540019 | 12.29450696 | 12.68468794 | -0.390180977 | -0.409287743 |
| Lrrc3 | 5.463192822 | 5.591747158 | 5.872530038 | -0.28078288 | -0.409337216 |
| Kif26a | 6.409929251 | 6.876298011 | 6.81927958 | 0.057018431 | -0.40935033 |
| Arhgap22 | 5.878013056 | 6.429712082 | 6.287468056 | 0.142244027 | -0.409455 |
| Naalad2 | 6.054773339 | 6.123799594 | 6.464260296 | -0.340460702 | -0.409486956 |
| Cldn23 | 7.984918521 | 8.341749605 | 8.394612164 | -0.052862559 | -0.409693643 |
| Myh10 | 8.373415518 | 8.515704055 | 8.783128812 | -0.267424757 | -0.409713294 |
| Rbbp9 | 7.987488147 | 8.082925966 | 8.397266873 | -0.314340907 | -0.409778726 |
| Oxct1 | 12.32952211 | 12.39482996 | 12.73956532 | -0.344735353 | -0.41004321 |
| Plin3 | 9.943223223 | 10.33595372 | 10.35330217 | -0.01734845 | -0.410078951 |
| Odf2l | 7.707162923 | 8.05635735 | 8.11726744 | -0.06091009 | -0.410104517 |
| Zfp949 | 8.008072033 | 8.299236369 | 8.418395009 | -0.119158639 | -0.410322976 |
| Tesk2 | 6.453265442 | 6.712464313 | 6.864484585 | -0.152020272 | -0.411219143 |

| | | | | | |
|---|---|---|---|---|---|
| Arhgap42 | 8.50640501 | 8.910878948 | 8.917630821 | -0.006751873 | -0.41122581 |
| Dag1 | 11.57127155 | 11.79925789 | 11.98254046 | -0.183282573 | -0.411268913 |
| Tspan18 | 7.469226814 | 7.610371831 | 7.880531437 | -0.270159606 | -0.411304623 |
| Cir1 | 9.392085601 | 9.579236892 | 9.803427948 | -0.224191056 | -0.411342347 |
| Grb7 | 8.315134188 | 8.813973843 | 8.72661054 | 0.087363304 | -0.411476352 |
| Gm48128 | 7.174468477 | 7.397292411 | 7.586013065 | -0.188720655 | -0.411544588 |
| Sncaip | 4.989770461 | 5.112633041 | 5.401615535 | -0.288982494 | -0.411845074 |
| Scn7a | 9.404343713 | 9.352979049 | 9.816275384 | -0.463296334 | -0.41193167 |
| Jcad | 8.964468333 | 9.173069066 | 9.37657189 | -0.203502824 | -0.412103557 |
| Ccdc28a | 6.150536586 | 6.460178486 | 6.562665837 | -0.102487351 | -0.412129251 |
| Obscn | 11.40110686 | 11.77679389 | 11.8134386 | -0.03664471 | -0.412331737 |
| Lmo7 | 10.20935998 | 10.40044844 | 10.62170968 | -0.22126124 | -0.412349707 |
| Wipi1 | 8.788059423 | 8.908485747 | 9.20044594 | -0.291960193 | -0.412386517 |
| Asb16 | 7.353131183 | 7.412948663 | 7.766017159 | -0.353068496 | -0.412885976 |
| Parp16 | 6.920807729 | 7.086376114 | 7.333720832 | -0.247344718 | -0.412913103 |
| Vsnl1 | 6.52684538 | 6.921651129 | 6.939866031 | -0.018214902 | -0.413020651 |
| Tcf7l1 | 7.789733674 | 8.114679171 | 8.202772636 | -0.088093466 | -0.413038962 |
| Abca1 | 11.1789252 | 10.99111277 | 11.59214474 | -0.601031967 | -0.413219533 |
| Mmp23 | 6.507263047 | 6.72320124 | 6.920579788 | -0.197378547 | -0.41331674 |
| Eml3 | 9.238969078 | 9.742691242 | 9.652385067 | 0.090306175 | -0.413415989 |
| Natd1 | 8.918851135 | 9.346178468 | 9.332274838 | 0.01390363 | -0.413423703 |
| Acer1 | 11.2440721 | 11.41536501 | 11.6575601 | -0.242195093 | -0.413488 |
| Pik3r3 | 7.481976015 | 7.854483032 | 7.895475598 | -0.040992567 | -0.413499583 |
| Cercam | 7.00202738 | 7.106684603 | 7.415539337 | -0.308854735 | -0.413511957 |
| Zkscan4 | 4.542460046 | 4.769322317 | 4.956055183 | -0.186732866 | -0.413595137 |
| Plxdc1 | 7.757004491 | 8.174431599 | 8.17081839 | 0.003613209 | -0.413813899 |
| Slc34a2 | 7.826517197 | 8.302880379 | 8.240415247 | 0.062465132 | -0.41389805 |
| Olfml2b | 8.783382356 | 8.840246502 | 9.197450823 | -0.357204321 | -0.414068467 |
| Ctso | 8.33213978 | 8.378272544 | 8.746411545 | -0.368139001 | -0.414271765 |
| Smarcd3 | 8.284133308 | 8.44927702 | 8.698513943 | -0.249236923 | -0.414380635 |
| Yod1 | 10.21884045 | 10.42653728 | 10.63343516 | -0.20689788 | -0.414594717 |
| Arl13b | 8.81805813 | 8.994250677 | 9.232797758 | -0.238547081 | -0.414739628 |

| | | | | | |
|---|---|---|---|---|---|
| Shisa4 | 8.036105307 | 8.207472217 | 8.450950317 | -0.2434781 | -0.41484501 |
| Lurap1l | 6.467664416 | 6.716771501 | 6.882809416 | -0.166037915 | -0.415145 |
| Tshr | 7.204123072 | 7.043231225 | 7.619356905 | -0.57612568 | -0.415233833 |
| Gatd1 | 8.093651261 | 8.390254259 | 8.509004785 | -0.118750526 | -0.415353525 |
| Wnt5b | 6.161587312 | 6.37532055 | 6.577171651 | -0.201851101 | -0.415584339 |
| Zbtb14 | 8.540189675 | 8.665081095 | 8.955819026 | -0.290737931 | -0.415629351 |
| Lynx1 | 9.245980419 | 9.440235148 | 9.661666925 | -0.221431777 | -0.415686505 |
| Zfand6 | 10.71672752 | 10.92618572 | 11.13271334 | -0.206527617 | -0.41598582 |
| Efs | 9.357884006 | 9.959593777 | 9.774148388 | 0.18544539 | -0.416264382 |
| Zfp606 | 6.782100624 | 6.882655911 | 7.198503238 | -0.315847327 | -0.416402615 |
| Sertad4 | 8.617245146 | 8.74299726 | 9.033650038 | -0.290652778 | -0.416404892 |
| Gprin3 | 5.782542762 | 5.940777008 | 6.198961106 | -0.258184099 | -0.416418344 |
| Bphl | 7.344222058 | 7.484870504 | 7.760737309 | -0.275866805 | -0.416515251 |
| L3hypdh | 6.614460134 | 6.836046929 | 7.031035294 | -0.194988365 | -0.41657516 |
| Itpr2 | 10.36168549 | 10.37214053 | 10.77852183 | -0.406381297 | -0.416836337 |
| Slc25a36 | 9.271861292 | 9.394223939 | 9.689022335 | -0.294798395 | -0.417161043 |
| 39692 | 11.10212401 | 11.50902107 | 11.519678 | -0.01065693 | -0.41755399 |
| Knstrn | 7.386602575 | 7.739410825 | 7.804192191 | -0.064781366 | -0.417589616 |
| Phyhd1 | 6.713284365 | 6.999438002 | 7.131005389 | -0.131567387 | -0.417721024 |
| Rad51b | 5.563590679 | 5.815743676 | 5.981509312 | -0.165765636 | -0.417918633 |
| Tceal1 | 5.575989019 | 5.737323852 | 5.993929654 | -0.256605802 | -0.417940635 |
| Zrsr1 | 7.652249009 | 7.771247393 | 8.07033052 | -0.299083127 | -0.418081511 |
| Cep170b | 10.61731327 | 10.96812212 | 11.03551783 | -0.06739571 | -0.418204557 |
| Zfp995 | 6.80250906 | 7.095059302 | 7.220976994 | -0.125917692 | -0.418467933 |
| Podn | 9.056029612 | 9.201706125 | 9.474541707 | -0.272835582 | -0.418512094 |
| Gstm1 | 10.82564701 | 10.79739789 | 11.24464176 | -0.44724387 | -0.418994747 |
| Skint6 | 9.737379133 | 9.856072759 | 10.15657957 | -0.300506807 | -0.419200433 |
| Col4a6 | 7.317767269 | 7.576532153 | 7.737120563 | -0.16058841 | -0.419353294 |
| 4833420G17Rik | 9.164441139 | 9.445109256 | 9.583906233 | -0.138796977 | -0.419465094 |
| Bcl9 | 8.287434057 | 8.586354521 | 8.706942652 | -0.12058813 | -0.419508595 |
| Rasl11b | 6.457788804 | 6.854890969 | 6.877305156 | -0.022414187 | -0.419516352 |
| Cbr4 | 7.802247983 | 7.981827827 | 8.221798373 | -0.239970546 | -0.41955039 |

| | | | | | |
|---|---|---|---|---|---|
| Scn2a | 4.496858624 | 4.57043525 | 4.917148346 | -0.346713096 | -0.420289722 |
| Dsg1b | 10.89785232 | 11.26192863 | 11.3185435 | -0.05661487 | -0.420691173 |
| Siae | 7.991768658 | 8.132983313 | 8.412695388 | -0.279712075 | -0.42092673 |
| Nfib | 12.16817295 | 12.35836398 | 12.58938714 | -0.231023153 | -0.421214183 |
| Mtus1 | 10.57621962 | 10.5802133 | 10.99764034 | -0.417427037 | -0.421420717 |
| Hmgb3 | 8.068368696 | 8.157036566 | 8.490188231 | -0.333151665 | -0.421819535 |
| Col5a2 | 11.06508989 | 11.33874335 | 11.48694735 | -0.148203997 | -0.421857463 |
| Edaradd | 8.691490095 | 9.140441832 | 9.113353843 | 0.027087989 | -0.421863748 |
| Kdelr3 | 6.916904179 | 7.159281591 | 7.338998674 | -0.179717083 | -0.422094495 |
| Trpv6 | 5.674033856 | 6.055413674 | 6.096445721 | -0.041032047 | -0.422411865 |
| Myoz3 | 8.480522576 | 8.541589802 | 8.902991484 | -0.361401683 | -0.422468909 |
| Phka2 | 8.653660467 | 8.701806029 | 9.076497278 | -0.374691249 | -0.422836811 |
| Dpt | 12.22384239 | 12.11162557 | 12.64676395 | -0.53513838 | -0.42292156 |
| Myoz1 | 11.3931564 | 11.48601561 | 11.81652023 | -0.330504613 | -0.423363827 |
| Polg2 | 6.494590896 | 6.609626453 | 6.918243983 | -0.308617531 | -0.423653088 |
| Mterf2 | 5.847022426 | 5.940461077 | 6.270880142 | -0.330419065 | -0.423857716 |
| Man1a | 10.36863368 | 10.38059224 | 10.7928103 | -0.412218063 | -0.42417662 |
| Frzb | 5.703388994 | 5.390215226 | 6.1276334 | -0.737418174 | -0.424244406 |
| 2510009E07Rik | 8.846284058 | 9.035954412 | 9.270538578 | -0.234584165 | -0.42425452 |
| Ints6l | 9.529212398 | 9.837803665 | 9.953619837 | -0.115816172 | -0.424407439 |
| Hrnr | 14.03950629 | 14.30422313 | 14.46421938 | -0.159996247 | -0.42471309 |
| Stmn1 | 10.45509753 | 10.57542154 | 10.88002786 | -0.304606323 | -0.424930333 |
| Mybl1 | 5.898653732 | 6.130880009 | 6.323827275 | -0.192947266 | -0.425173543 |
| Acad8 | 8.427759883 | 8.633088422 | 8.852978681 | -0.219890259 | -0.425218798 |
| Ptch1 | 9.051231058 | 9.304669483 | 9.476564617 | -0.171895135 | -0.42533356 |
| Rorc | 8.896720008 | 9.129553526 | 9.322248566 | -0.192695039 | -0.425528558 |
| Sprr2h | 8.659336793 | 9.012998858 | 9.084880408 | -0.07188155 | -0.425543615 |
| Hoxd8 | 8.176697146 | 8.241296116 | 8.602713004 | -0.361416888 | -0.426015858 |
| Gm9696 | 7.657284977 | 7.922953242 | 8.083311303 | -0.160358061 | -0.426026326 |
| Letm2 | 6.027135582 | 6.208563341 | 6.453260253 | -0.244696912 | -0.426124671 |
| Dact1 | 6.389572769 | 6.337035474 | 6.815828271 | -0.478792797 | -0.426255501 |
| H2afv | 8.782853872 | 8.826947719 | 9.209189715 | -0.382241996 | -0.426335844 |

| | | | | | |
|---|---|---|---|---|---|
| Mapre3 | 8.237272472 | 8.657860246 | 8.663866058 | -0.006005812 | -0.426593585 |
| Fut11 | 8.050709716 | 8.262365003 | 8.477308855 | -0.214943853 | -0.426599139 |
| Itgb1bp2 | 8.454713398 | 8.458906588 | 8.881439063 | -0.422532475 | -0.426725665 |
| Pth1r | 7.887189053 | 8.272001971 | 8.3139343 | -0.041932329 | -0.426745246 |
| Il34 | 9.401339753 | 9.798882296 | 9.828141498 | -0.029259201 | -0.426801744 |
| Cand2 | 7.577516075 | 7.778920029 | 8.004380803 | -0.225460774 | -0.426864729 |
| Mfng | 8.116645044 | 8.062032677 | 8.543715568 | -0.481682891 | -0.427070524 |
| Ermp1 | 11.0621467 | 11.22762922 | 11.48958661 | -0.26195739 | -0.42743991 |
| Krt23 | 11.21635506 | 11.4486312 | 11.64416674 | -0.19553554 | -0.427811673 |
| Sgce | 8.197676017 | 8.183550683 | 8.625547419 | -0.441996736 | -0.427871402 |
| Adipoq | 11.6892598 | 11.36282779 | 12.11714172 | -0.754313923 | -0.427881917 |
| Zfp799 | 7.035768604 | 7.295494052 | 7.463761699 | -0.168267647 | -0.427993095 |
| Ephx2 | 9.003376765 | 9.044477236 | 9.431431338 | -0.386954102 | -0.428054573 |
| Dip2a | 9.011035714 | 9.57809176 | 9.439127463 | 0.138964297 | -0.428091748 |
| Fbxo33 | 8.631600028 | 8.968399323 | 9.05982467 | -0.091425347 | -0.428224642 |
| Opn3 | 5.81216783 | 5.725621911 | 6.240538451 | -0.51491654 | -0.428370621 |
| Mkrn2 | 8.67922412 | 8.825392087 | 9.107614531 | -0.282222444 | -0.428390411 |
| Dnajc6 | 6.451286643 | 6.686660813 | 6.88008932 | -0.193428507 | -0.428802677 |
| Hic1 | 5.751185735 | 6.070943322 | 6.180097138 | -0.109153816 | -0.428911404 |
| Ocln | 8.449354356 | 8.686196881 | 8.878369436 | -0.192172556 | -0.42901508 |
| Fam120c | 6.957894746 | 7.186103041 | 7.387063901 | -0.20096086 | -0.429169155 |
| Hoxa7 | 7.492930577 | 8.055729777 | 7.922145573 | 0.133584203 | -0.429214997 |
| Gm45062 | 8.729862114 | 9.020900189 | 9.159212448 | -0.138312259 | -0.429350334 |
| Spon1 | 8.791600111 | 8.96268307 | 9.221544088 | -0.258861018 | -0.429943976 |
| Pcdh19 | 7.576536259 | 7.729138854 | 8.00684475 | -0.277705896 | -0.430308491 |
| Map4 | 11.79707751 | 12.10184428 | 12.22774673 | -0.125902453 | -0.430669223 |
| Pltp | 9.836029383 | 9.852433413 | 10.26672429 | -0.414290877 | -0.430694907 |
| Iqgap3 | 7.158467619 | 7.536129844 | 7.589266544 | -0.053136699 | -0.430798924 |
| Slc25a35 | 7.070489172 | 7.23706433 | 7.501394199 | -0.264329869 | -0.430905026 |
| Pcmtd1 | 10.8976447 | 11.07756495 | 11.32857606 | -0.251011113 | -0.430931367 |
| Kat2b | 9.903313944 | 10.03899412 | 10.33427247 | -0.29527835 | -0.430958522 |
| Gpam | 11.42843637 | 11.64549993 | 11.85941169 | -0.213911763 | -0.43097532 |

| | | | | | |
|---|---|---|---|---|---|
| Kazald1 | 6.091390809 | 6.419425461 | 6.522414482 | -0.102989021 | -0.431023673 |
| Arhgef19 | 7.514473017 | 8.14347683 | 7.94551517 | 0.19796166 | -0.431042152 |
| Rnf144b | 10.15232835 | 10.30909564 | 10.58367666 | -0.274581023 | -0.431348305 |
| Zfp46 | 8.478219229 | 8.707145512 | 8.909700346 | -0.202554834 | -0.431481117 |
| Dclre1a | 7.605394004 | 8.029417639 | 8.036938451 | -0.007520812 | -0.431544446 |
| Tcap | 11.35557772 | 11.54489156 | 11.78714484 | -0.242253277 | -0.431567117 |
| Dsc1 | 12.33962383 | 12.60383182 | 12.77128265 | -0.167450833 | -0.43165882 |
| Sccpdh | 8.271198244 | 8.316120228 | 8.702910837 | -0.386790609 | -0.431712592 |
| Sgk1 | 10.78775834 | 11.25222982 | 11.21949511 | 0.03273471 | -0.431736773 |
| Ovol1 | 9.902332919 | 10.44130891 | 10.33411487 | 0.107194047 | -0.431781948 |
| Zfp763 | 6.241363537 | 6.205135045 | 6.673315356 | -0.468180311 | -0.431951819 |
| Lrp4 | 10.65171563 | 11.16278937 | 11.08376347 | 0.079025903 | -0.432047837 |
| Atp10b | 9.169598277 | 9.58990638 | 9.601678077 | -0.011771697 | -0.4320798 |
| Zfp433 | 6.183822568 | 6.214806432 | 6.615904573 | -0.401098141 | -0.432082005 |
| Nid2 | 8.83245337 | 8.583267205 | 9.264583827 | -0.681316622 | -0.432130457 |
| Ctxn3 | 5.869015829 | 5.899365305 | 6.301572022 | -0.402206717 | -0.432556193 |
| Jph2 | 10.41561032 | 10.24428736 | 10.84838051 | -0.604093143 | -0.432770183 |
| Ccn5 | 7.247774213 | 7.965221969 | 7.680903371 | 0.284318597 | -0.433129158 |
| Abhd15 | 6.543580962 | 6.686107149 | 6.976935191 | -0.290828042 | -0.433354229 |
| Dmd | 9.188541825 | 9.408124458 | 9.621940736 | -0.213816279 | -0.433398911 |
| Fam168a | 9.545960509 | 9.83251305 | 9.979501958 | -0.146988908 | -0.433541449 |
| Pde2a | 7.011867582 | 7.635603606 | 7.445550423 | 0.190053183 | -0.433682841 |
| Rab7b | 10.32578182 | 10.74623104 | 10.75979841 | -0.013567363 | -0.43401659 |
| Enpep | 7.947423494 | 7.364243743 | 8.381457566 | -1.017213823 | -0.434034071 |
| Gnpda2 | 7.925780237 | 8.039794346 | 8.359833829 | -0.320039483 | -0.434053592 |
| Tex9 | 6.042992335 | 6.250719715 | 6.477157104 | -0.22643739 | -0.434164769 |
| Fzd3 | 8.248742579 | 8.580744873 | 8.682933664 | -0.10218879 | -0.434191085 |
| Hrct1 | 5.493799391 | 5.66685632 | 5.928039587 | -0.261183267 | -0.434240197 |
| Esr1 | 8.846711424 | 8.937649268 | 9.281183403 | -0.343534135 | -0.43447198 |
| Mill2 | 7.80708945 | 7.827885024 | 8.241609946 | -0.413724922 | -0.434520496 |
| Pappa2 | 8.204321588 | 7.962201324 | 8.63890378 | -0.676702457 | -0.434582193 |
| Bmp5 | 3.75030074 | 3.86782674 | 4.185086341 | -0.317259601 | -0.434785601 |

| | | | | | |
|---|---|---|---|---|---|
| Gm14288 | 6.08698176 | 5.633379418 | 6.522429008 | -0.88904959 | -0.435447248 |
| Gem | 8.242844865 | 8.43242146 | 8.67840368 | -0.245982219 | -0.435558814 |
| Pfn2 | 9.107216217 | 9.117583676 | 9.542802858 | -0.425219182 | -0.43558664 |
| Ccdc191 | 6.712025412 | 6.871901343 | 7.14794726 | -0.276045916 | -0.435921848 |
| Mroh6 | 6.655531989 | 7.353578459 | 7.091593745 | 0.261984714 | -0.436061755 |
| Kremen1 | 10.60262465 | 11.00689089 | 11.03874502 | -0.031854127 | -0.436120363 |
| Rfx2 | 7.83017685 | 8.556412011 | 8.266343886 | 0.290068124 | -0.436167036 |
| Hrasls | 6.408978966 | 6.369980828 | 6.845350224 | -0.475369396 | -0.436371258 |
| Abca3 | 9.892459084 | 10.14259757 | 10.32887906 | -0.186281493 | -0.43641998 |
| Pnisr | 10.59549377 | 10.96178136 | 11.03192545 | -0.070144083 | -0.436431673 |
| Zfp383 | 6.616726013 | 6.820875089 | 7.053398186 | -0.232523096 | -0.436672172 |
| Uevld | 8.841705878 | 9.084984078 | 9.278407837 | -0.193423759 | -0.436701959 |
| Cacna1g | 6.834827141 | 7.479774189 | 7.27154043 | 0.208233759 | -0.436713289 |
| Fam20a | 5.382972821 | 5.856281462 | 5.819909763 | 0.036371699 | -0.436936942 |
| Dhdh | 8.578947274 | 8.488145996 | 9.016057982 | -0.527911986 | -0.437110708 |
| Rasal2 | 9.065776418 | 9.259040059 | 9.503045333 | -0.244005274 | -0.437268915 |
| Dnmt3a | 9.3005538 | 10.16343432 | 9.73784276 | 0.425591557 | -0.437288959 |
| Angptl2 | 10.72817073 | 10.88000737 | 11.16550689 | -0.285499523 | -0.43733616 |
| Ssbp3 | 9.633027163 | 9.931698242 | 10.07045971 | -0.138761464 | -0.437432544 |
| Crot | 9.567795945 | 9.776625072 | 10.00532449 | -0.228699418 | -0.437528545 |
| Timp2 | 12.32030152 | 12.34128022 | 12.75842261 | -0.417142387 | -0.438121087 |
| P2ry12 | 5.474879013 | 5.339446928 | 5.913143001 | -0.573696072 | -0.438263988 |
| Loxl2 | 9.217642584 | 9.478839869 | 9.656045976 | -0.177206107 | -0.438403392 |
| Myadml2 | 6.105286734 | 6.183852195 | 6.543891611 | -0.360039416 | -0.438604877 |
| Ptpru | 6.476773952 | 6.504390155 | 6.91573751 | -0.411347355 | -0.438963558 |
| Col7a1 | 10.50082717 | 11.2174583 | 10.9402725 | 0.2771858 | -0.439445327 |
| Pxmp2 | 7.189305665 | 7.152854536 | 7.62884562 | -0.475991083 | -0.439539954 |
| Pdzrn4 | 6.364915801 | 6.816518377 | 6.805087768 | 0.011430609 | -0.440171968 |
| Extl1 | 9.14022087 | 9.600263245 | 9.580604225 | 0.01965902 | -0.440383355 |
| Nxph3 | 5.465382273 | 5.827007004 | 5.90585394 | -0.078846935 | -0.440471667 |
| Zmym5 | 9.500099458 | 9.813271257 | 9.940636041 | -0.127364784 | -0.440536583 |
| Fam124a | 6.551349467 | 6.525200553 | 6.99231523 | -0.467114677 | -0.440965764 |

| | | | | | |
|---|---|---|---|---|---|
| Slc41a3 | 7.294168913 | 7.473444841 | 7.735262149 | -0.261817308 | -0.441093236 |
| Papln | 6.429862194 | 7.073614455 | 6.871424611 | 0.202189844 | -0.441562417 |
| Acsm3 | 6.075721007 | 6.065290435 | 6.517631765 | -0.45234133 | -0.441910758 |
| 1190002N15Rik | 9.048198188 | 9.261747216 | 9.490683728 | -0.228936512 | -0.44248554 |
| Cib2 | 9.012576413 | 9.050310671 | 9.455100564 | -0.404789893 | -0.442524152 |
| Abhd6 | 8.229203016 | 8.484056708 | 8.671850337 | -0.187793629 | -0.442647321 |
| Inka2 | 8.158291395 | 8.475905311 | 8.601208246 | -0.125302935 | -0.442916851 |
| H2-Q5 | 7.954889206 | 7.97055987 | 8.398105487 | -0.427545617 | -0.443216281 |
| Spats2l | 6.567454559 | 6.660785492 | 7.010860746 | -0.350075254 | -0.443406188 |
| Ncald | 6.219877018 | 6.364935769 | 6.663299988 | -0.298364219 | -0.44342297 |
| Hibch | 9.150177246 | 9.132320447 | 9.593624508 | -0.461304061 | -0.443447262 |
| Tnfrsf19 | 10.75618247 | 11.19189756 | 11.19977097 | -0.007873413 | -0.443588507 |
| Nemp2 | 7.21199612 | 7.337638809 | 7.65569193 | -0.318053121 | -0.44369581 |
| Hrc | 10.55639501 | 10.82493608 | 11.00020016 | -0.175264083 | -0.443805153 |
| Mkx | 5.989714504 | 6.380751505 | 6.433539407 | -0.052787902 | -0.443824903 |
| Rhobtb1 | 7.889096423 | 7.925836888 | 8.333098185 | -0.407261298 | -0.444001763 |
| Mr1 | 8.159917729 | 8.225779393 | 8.603948483 | -0.37816909 | -0.444030754 |
| Hdac8 | 6.787386486 | 7.002619171 | 7.232037895 | -0.229418724 | -0.444651409 |
| Lrrc8e | 8.668298354 | 9.008845961 | 9.113038106 | -0.104192144 | -0.444739751 |
| Gipc2 | 6.152196793 | 6.529878943 | 6.597470839 | -0.067591896 | -0.445274046 |
| Rarg | 11.24746813 | 11.88167238 | 11.69297477 | 0.188697603 | -0.445506647 |
| Kcna6 | 6.609685893 | 6.694042657 | 7.055242272 | -0.361199614 | -0.445556379 |
| Ago1 | 10.27490635 | 10.66953003 | 10.72050598 | -0.050975947 | -0.445599623 |
| Tbc1d16 | 8.399514473 | 8.721923718 | 8.845301922 | -0.123378203 | -0.445787449 |
| Epb41l1 | 9.462734414 | 9.663442694 | 9.908636489 | -0.245193794 | -0.445902075 |
| Net1 | 12.23735198 | 12.43050371 | 12.68326371 | -0.252760003 | -0.445911737 |
| Plcd4 | 6.414686443 | 6.437328565 | 6.860617538 | -0.423288973 | -0.445931095 |
| Sdcbp2 | 9.4378393 | 9.712225764 | 9.883868083 | -0.171642319 | -0.446028783 |
| Fech | 9.804717302 | 9.930329531 | 10.25074968 | -0.320420149 | -0.446032378 |
| Megf9 | 10.93978926 | 11.19785459 | 11.38588789 | -0.188033303 | -0.446098627 |
| Sema3g | 7.787190378 | 8.045878455 | 8.234013126 | -0.18813467 | -0.446822747 |
| Zfp771 | 5.599879214 | 5.895733994 | 6.046778735 | -0.151044741 | -0.446899522 |

| | | | | | |
|---|---|---|---|---|---|
| Mastl | 6.682785124 | 6.904335278 | 7.12971886 | -0.225383582 | -0.446933737 |
| Ralgapa2 | 9.458716416 | 9.510346881 | 9.906100747 | -0.395753867 | -0.447384332 |
| Neil2 | 5.428458796 | 5.59182703 | 5.875850667 | -0.284023637 | -0.447391871 |
| Sacs | 7.286606795 | 7.54142283 | 7.73415149 | -0.19272866 | -0.447544695 |
| Ranbp10 | 9.934486436 | 10.34058119 | 10.38223694 | -0.04165575 | -0.447750507 |
| Ccdc186 | 9.620308453 | 9.78976215 | 10.06806341 | -0.278301255 | -0.447754952 |
| Skint2 | 6.457944248 | 6.854129739 | 6.905940823 | -0.051811084 | -0.447996574 |
| Oas1f | 10.71987991 | 10.97635536 | 11.16796485 | -0.191609487 | -0.448084937 |
| Ang | 7.741662496 | 7.932843299 | 8.189772463 | -0.256929164 | -0.448109967 |
| Tmem182 | 9.511943092 | 9.565622511 | 9.960057901 | -0.39443539 | -0.448114809 |
| Klhl3 | 5.353542265 | 5.589662716 | 5.801686516 | -0.2120238 | -0.44814425 |
| Akr1d1 | 3.953704816 | 4.08136561 | 4.402024913 | -0.320659303 | -0.448320097 |
| Mc5r | 8.080647724 | 8.369048732 | 8.529272667 | -0.160223935 | -0.448624942 |
| Ddb2 | 7.751914605 | 8.044047977 | 8.200866554 | -0.156818577 | -0.448951949 |
| Npnt | 8.729161948 | 9.017098972 | 9.178286477 | -0.161187504 | -0.449124529 |
| Tmem43 | 11.6601853 | 11.94808339 | 12.1095602 | -0.161476807 | -0.449374897 |
| Neu3 | 7.17392975 | 7.697770686 | 7.623573012 | 0.074197673 | -0.449643262 |
| Txnip | 12.74512756 | 13.13024696 | 13.19481154 | -0.064564577 | -0.44968398 |
| Nipal1 | 8.667581513 | 8.983747462 | 9.117391619 | -0.133644157 | -0.449810106 |
| Ccdc92 | 6.248596311 | 6.680518103 | 6.698605258 | -0.018087155 | -0.450008947 |
| Rgs10 | 7.990505908 | 8.136005005 | 8.440674305 | -0.3046693 | -0.450168397 |
| Gabrp | 7.077478243 | 7.214342383 | 7.527961601 | -0.313619218 | -0.450483358 |
| Fgfr2 | 10.89440695 | 11.21713918 | 11.34499322 | -0.127854037 | -0.450586263 |
| Skint5 | 9.412384749 | 9.54142089 | 9.862991023 | -0.321570133 | -0.450606274 |
| Cenpf | 8.665398451 | 8.70324901 | 9.1161881 | -0.41293909 | -0.450789649 |
| Ldb3 | 11.28422867 | 11.34593698 | 11.73502789 | -0.389090903 | -0.450799217 |
| Abca5 | 9.348249196 | 9.510672305 | 9.799230286 | -0.288557981 | -0.45098109 |
| Mcc | 9.649867118 | 10.09928427 | 10.10085641 | -0.001572139 | -0.450989293 |
| Tpcn1 | 9.317613645 | 9.686901743 | 9.769021939 | -0.082120197 | -0.451408294 |
| Spon2 | 6.744737033 | 7.219694182 | 7.196231833 | 0.02346235 | -0.4514948 |
| Fbln2 | 10.58429721 | 10.83516479 | 11.03590163 | -0.200736837 | -0.451604417 |
| Zfp503 | 8.368998121 | 8.621591275 | 8.821138311 | -0.199547036 | -0.45214019 |

| | | | | | |
|---|---|---|---|---|---|
| Ncapg2 | 9.076247985 | 9.33812649 | 9.528486195 | -0.190359705 | -0.45223821 |
| Tbc1d17 | 9.012720197 | 9.362204635 | 9.465050198 | -0.102845563 | -0.452330001 |
| Fam114a1 | 8.114237054 | 8.176154666 | 8.566721478 | -0.390566812 | -0.452484425 |
| Fuz | 7.289087403 | 7.651775528 | 7.741603019 | -0.089827491 | -0.452515616 |
| Cubn | 5.069549924 | 5.329119446 | 5.522111791 | -0.192992346 | -0.452561868 |
| Zyg11b | 9.97695542 | 10.19497425 | 10.42976341 | -0.234789153 | -0.452807986 |
| Slc6a19 | 7.58247212 | 7.968371642 | 8.03538187 | -0.067010228 | -0.452909749 |
| Gpr160 | 8.653752035 | 8.799890537 | 9.106670357 | -0.30677982 | -0.452918322 |
| Klf11 | 7.880201495 | 8.134519804 | 8.333159953 | -0.198640148 | -0.452958458 |
| Zmym6 | 7.836531952 | 8.353596933 | 8.289508926 | 0.064088008 | -0.452976974 |
| Fbf1 | 8.686253512 | 9.115357155 | 9.139555251 | -0.024198096 | -0.453301739 |
| Flrt1 | 5.656244207 | 6.255543908 | 6.109673329 | 0.145870579 | -0.453429122 |
| Fgd5 | 7.538516555 | 7.789142565 | 7.992106092 | -0.202963527 | -0.453589537 |
| Hist2h2be | 6.626834955 | 6.902470438 | 7.080450914 | -0.177980476 | -0.453615959 |
| Car6 | 5.761087865 | 5.957937528 | 6.214719673 | -0.256782145 | -0.453631808 |
| Hectd4 | 9.647348285 | 10.23250488 | 10.10112598 | 0.131378892 | -0.453777699 |
| Asrgl1 | 7.581218075 | 7.858278367 | 8.035046058 | -0.176767691 | -0.453827983 |
| Cyp4f17 | 6.997168586 | 7.299737211 | 7.451306687 | -0.151569477 | -0.454138101 |
| Zbed6 | 8.240372359 | 8.278031344 | 8.695300791 | -0.417269447 | -0.454928432 |
| Prdm8 | 3.745886233 | 3.907668606 | 4.201106399 | -0.293437793 | -0.455220166 |
| Dpy19l4 | 8.882993361 | 9.076191261 | 9.338559391 | -0.26236813 | -0.45556603 |
| Slc6a13 | 5.05993058 | 5.194728744 | 5.515638114 | -0.32090937 | -0.455707534 |
| Bves | 6.980306815 | 7.263781317 | 7.436475487 | -0.17269417 | -0.456168672 |
| Glce | 9.198227798 | 9.487566516 | 9.654449238 | -0.166882722 | -0.45622144 |
| Fam198a | 5.195309115 | 5.569556765 | 5.651591976 | -0.082035211 | -0.456282861 |
| 2610008E11Rik | 8.286928694 | 8.497808819 | 8.743632642 | -0.245823823 | -0.456703948 |
| Mgst2 | 7.48784126 | 7.652119399 | 7.944648336 | -0.292528937 | -0.456807076 |
| Vamp1 | 8.181419185 | 8.458136166 | 8.638320126 | -0.180183961 | -0.456900941 |
| Fam78a | 6.164012651 | 6.174526457 | 6.621770664 | -0.447244207 | -0.457758013 |
| Gpr87 | 9.193449142 | 9.522060516 | 9.651259907 | -0.129199391 | -0.457810765 |
| Hsf2 | 9.284717419 | 9.680447681 | 9.742787405 | -0.062339724 | -0.458069987 |
| Cgn | 10.0592388 | 10.47821672 | 10.51770203 | -0.039485307 | -0.458463233 |

| | | | | | |
|---|---|---|---|---|---|
| Brdt | 5.73991495 | 5.980428124 | 6.198521289 | -0.218093165 | -0.458606339 |
| Gramd1c | 9.403519938 | 9.723387513 | 9.862388454 | -0.139000941 | -0.458868516 |
| Kctd15 | 10.12889591 | 10.48327868 | 10.58799706 | -0.104718377 | -0.459101147 |
| Gm12751 | 4.924521393 | 4.923555767 | 5.383750045 | -0.460194278 | -0.459228652 |
| Stxbp4 | 6.458499511 | 6.623585169 | 6.917756317 | -0.294171148 | -0.459256806 |
| Slc12a2 | 10.29037877 | 10.50898709 | 10.75017446 | -0.241187377 | -0.45979569 |
| Ephb3 | 9.0820856 | 9.697680998 | 9.542008007 | 0.155672991 | -0.459922407 |
| Sgcd | 6.806869812 | 6.848045825 | 7.266794691 | -0.418748866 | -0.459924879 |
| Rmnd5a | 11.47124306 | 11.70600541 | 11.93120587 | -0.22520046 | -0.459962813 |
| Ccdc17 | 6.158179285 | 6.432173514 | 6.618162841 | -0.185989327 | -0.459983555 |
| Zkscan7 | 7.27241801 | 7.543976276 | 7.732961221 | -0.188984944 | -0.460543211 |
| Gas7 | 11.28844367 | 11.60449268 | 11.74908895 | -0.144596273 | -0.46064528 |
| Hoxc5 | 6.326035264 | 6.762334608 | 6.787027311 | -0.024692703 | -0.460992047 |
| Slc46a3 | 7.554618821 | 7.925979515 | 8.015643779 | -0.089664264 | -0.461024959 |
| Zfyve21 | 7.8022746 | 8.128243045 | 8.26395689 | -0.135713844 | -0.461682289 |
| Zfp947 | 5.374205643 | 5.497884322 | 5.83609979 | -0.338215468 | -0.461894146 |
| Mknk2 | 11.42295463 | 12.13374107 | 11.88486148 | 0.24887959 | -0.46190685 |
| Tmem106b | 10.0952491 | 10.31561909 | 10.55716413 | -0.241545037 | -0.461915026 |
| Fkbp14 | 7.095270712 | 7.211107276 | 7.557193635 | -0.346086358 | -0.461922923 |
| Zfpm1 | 7.203937797 | 8.041157473 | 7.665928982 | 0.375228492 | -0.461991185 |
| Asb2 | 8.168828446 | 8.519448954 | 8.630849625 | -0.111400671 | -0.462021179 |
| Lipn | 8.309327308 | 8.627439517 | 8.771363453 | -0.143923935 | -0.462036145 |
| Slc38a3 | 7.035095534 | 6.927799241 | 7.497726524 | -0.569927283 | -0.46263099 |
| C1qa | 10.12415107 | 9.948703822 | 10.58697953 | -0.638275705 | -0.462828454 |
| Zfp101 | 8.140981608 | 8.515809921 | 8.603978857 | -0.088168936 | -0.462997249 |
| Zc2hc1a | 8.227362364 | 8.25700023 | 8.690473001 | -0.433472771 | -0.463110637 |
| Lce6a | 9.328769612 | 9.604408887 | 9.792259268 | -0.187850381 | -0.463489656 |
| Wdr45 | 9.09455939 | 9.411146912 | 9.558276616 | -0.147129703 | -0.463717226 |
| Mapk12 | 8.384701824 | 8.443047915 | 8.848671906 | -0.405623991 | -0.463970082 |
| Mettl27 | 6.059929078 | 6.334818737 | 6.523955613 | -0.189136876 | -0.464026535 |
| Ccdc3 | 9.770034134 | 9.985275736 | 10.23435393 | -0.24907819 | -0.464319793 |
| Zdhhc2 | 7.605517597 | 7.773036423 | 8.070103364 | -0.297066941 | -0.464585768 |

| | | | | | |
|---|---|---|---|---|---|
| Add3 | 10.26382593 | 10.52391542 | 10.72841948 | -0.204504057 | -0.464593553 |
| Opcml | 6.104683136 | 6.221894594 | 6.569501775 | -0.34760718 | -0.464818638 |
| Cebpa | 11.8031637 | 12.07335037 | 12.26802036 | -0.19466999 | -0.464856663 |
| Gabra3 | 5.386617399 | 5.349780362 | 5.85157069 | -0.501790328 | -0.464953291 |
| Ssbp2 | 8.824312118 | 9.05540144 | 9.29000101 | -0.234599571 | -0.465688892 |
| Tbc1d24 | 7.349504477 | 7.728311278 | 7.815263511 | -0.086952233 | -0.465759034 |
| Fam161a | 5.142990208 | 5.283857292 | 5.608980703 | -0.325123411 | -0.465990495 |
| Casp2 | 9.290848663 | 9.606183872 | 9.757302983 | -0.151119111 | -0.46645432 |
| Ccdc187 | 4.973490329 | 5.210067424 | 5.440266689 | -0.230199265 | -0.46677636 |
| Kank2 | 9.34999783 | 9.707759874 | 9.816801669 | -0.109041794 | -0.466803838 |
| Rnf170 | 9.073576656 | 9.397001372 | 9.540388672 | -0.1433873 | -0.466812016 |
| Dnajc28 | 6.399299347 | 6.475574358 | 6.866300772 | -0.390726414 | -0.467001425 |
| Ebf1 | 9.142933305 | 9.007870367 | 9.610001543 | -0.602131176 | -0.467068238 |
| Chrdl1 | 9.230156754 | 9.07749397 | 9.697766681 | -0.620272711 | -0.467609927 |
| Rnf227 | 10.23898787 | 10.6225611 | 10.70665075 | -0.084089653 | -0.467662886 |
| Itgb5 | 10.384675 | 10.71515117 | 10.85256258 | -0.137411417 | -0.467887583 |
| Serpina12 | 10.34622265 | 10.4684634 | 10.81426294 | -0.345799537 | -0.468040287 |
| Tmem254a | 7.155854784 | 7.911833487 | 7.623936844 | 0.287896643 | -0.46808206 |
| Ppip5k1 | 9.339578491 | 9.6213084 | 9.807961804 | -0.186653404 | -0.468383313 |
| Cd109 | 12.02522193 | 12.31531062 | 12.49380754 | -0.178496917 | -0.468585603 |
| Pdk2 | 9.79668422 | 10.02931447 | 10.26563795 | -0.236323485 | -0.468953734 |
| Kmt5c | 8.373472381 | 8.810234106 | 8.84274028 | -0.032506174 | -0.469267899 |
| Arhgap20 | 6.311466299 | 6.720996106 | 6.780817668 | -0.059821562 | -0.469351369 |
| AW549877 | 10.22128042 | 10.53628407 | 10.69066266 | -0.154378593 | -0.469382243 |
| Fras1 | 4.530547213 | 4.718969249 | 5.000027975 | -0.281058725 | -0.469480761 |
| Gpnmb | 8.744759785 | 9.1533446 | 9.214425389 | -0.06108079 | -0.469665604 |
| Cacnb3 | 8.494912867 | 9.349931479 | 8.964609301 | 0.385322178 | -0.469696435 |
| Arhgap18 | 8.501932782 | 8.695312817 | 8.971750249 | -0.276437432 | -0.469817467 |
| Golim4 | 10.80622274 | 11.09070009 | 11.27621769 | -0.1855176 | -0.469994943 |
| Sun1 | 10.77497669 | 11.13977359 | 11.24509448 | -0.10532089 | -0.470117793 |
| Krt15 | 14.24864851 | 14.54461485 | 14.71884411 | -0.17422926 | -0.4701956 |
| Utrn | 10.87831073 | 10.95636647 | 11.34870515 | -0.392338683 | -0.470394423 |

| | | | | | |
|---|---|---|---|---|---|
| Prob1 | 8.440940384 | 9.085787448 | 8.911491854 | 0.174295594 | -0.47055147 |
| Klhdc7a | 8.191001942 | 8.145598325 | 8.661686223 | -0.516087897 | -0.47068428 |
| Usp49 | 8.044849103 | 8.36868087 | 8.515580591 | -0.146899721 | -0.470731489 |
| Slc37a2 | 8.665569573 | 9.212108812 | 9.136420182 | 0.07568863 | -0.470850609 |
| Gpx7 | 6.597958781 | 6.776420002 | 7.068925976 | -0.292505974 | -0.470967195 |
| Heca | 10.55472419 | 10.86122441 | 11.02570309 | -0.164478673 | -0.470978897 |
| Prex2 | 8.037562825 | 8.036844507 | 8.508583719 | -0.471739212 | -0.471020894 |
| Skint3 | 10.62854037 | 10.87310044 | 11.099632 | -0.226531567 | -0.471091637 |
| Zfp760 | 7.345235807 | 7.568701838 | 7.81636719 | -0.247665352 | -0.471131383 |
| Serhl | 8.375559635 | 8.526695258 | 8.846752118 | -0.320056861 | -0.471192483 |
| Blmh | 12.20150889 | 12.43569947 | 12.672925 | -0.237225523 | -0.471416103 |
| Klhl8 | 8.518417886 | 8.636037789 | 8.990090723 | -0.354052934 | -0.471672837 |
| Gm45837 | 7.371686192 | 6.576383369 | 7.843381757 | -1.266998389 | -0.471695566 |
| Nox4 | 5.666495142 | 5.738635788 | 6.138193906 | -0.399558119 | -0.471698765 |
| Ccm2l | 5.387406507 | 5.731160596 | 5.859118731 | -0.127958134 | -0.471712224 |
| mt-Nd5 | 14.69900927 | 14.80933844 | 15.17116478 | -0.361826343 | -0.47215551 |
| Tmprss11f | 9.267964695 | 9.447640884 | 9.74012227 | -0.292481386 | -0.472157575 |
| Tmem8b | 6.972481737 | 7.190169505 | 7.444966373 | -0.254796868 | -0.472484636 |
| Cnst | 9.749680075 | 10.06992022 | 10.22216518 | -0.15224496 | -0.472485101 |
| Pls1 | 4.252239446 | 4.501520199 | 4.724795622 | -0.223275423 | -0.472556176 |
| Dapk1 | 9.325387495 | 9.790946117 | 9.798265881 | -0.007319765 | -0.472878387 |
| Fgd6 | 8.563428791 | 8.716836373 | 9.036432283 | -0.31959591 | -0.473003492 |
| Usp54 | 9.503763367 | 9.785174295 | 9.97680447 | -0.191630175 | -0.473041103 |
| Mmp2 | 11.74840107 | 11.93590518 | 12.22156926 | -0.285664077 | -0.47316819 |
| Tia1 | 9.281956082 | 9.434324279 | 9.755397182 | -0.321072903 | -0.4734411 |
| D430019H16Rik | 4.949457088 | 5.195433776 | 5.423167441 | -0.227733665 | -0.473710353 |
| Zbtb4 | 10.0074517 | 10.30874787 | 10.48121275 | -0.172464883 | -0.473761051 |
| Hdac9 | 6.377826119 | 6.41970913 | 6.851910676 | -0.432201546 | -0.474084557 |
| C6 | 4.33523709 | 4.42183818 | 4.809374464 | -0.387536284 | -0.474137374 |
| Nusap1 | 7.977036539 | 8.199599802 | 8.451332641 | -0.251732839 | -0.474296102 |
| Fibin | 7.701031928 | 7.831629132 | 8.175471824 | -0.343842693 | -0.474439897 |
| Ulbp1 | 5.062006393 | 5.499077115 | 5.537453796 | -0.038376681 | -0.475447403 |

| | | | | | |
|---|---|---|---|---|---|
| Myh7 | 3.492219298 | 3.511057788 | 3.967770549 | -0.456712761 | -0.475551251 |
| Cyp4v3 | 8.141702597 | 8.260319545 | 8.617776149 | -0.357456604 | -0.476073552 |
| Cln8 | 9.825333538 | 10.36703345 | 10.30179151 | 0.06524194 | -0.476457976 |
| Sec22c | 8.46442875 | 8.812025121 | 8.941108634 | -0.129083513 | -0.476679884 |
| Map9 | 7.302139954 | 7.731740506 | 7.77885134 | -0.047110834 | -0.476711387 |
| 39508 | 10.28517977 | 10.51336401 | 10.76248076 | -0.249116753 | -0.477300993 |
| Car11 | 5.517105116 | 5.928150115 | 5.994933158 | -0.066783043 | -0.477828042 |
| Asb10 | 6.488838095 | 6.612451158 | 6.966703216 | -0.354252058 | -0.477865121 |
| Phkb | 10.3767304 | 10.34795328 | 10.85478085 | -0.506827573 | -0.478050453 |
| BC067074 | 6.185843858 | 6.197133001 | 6.6638957 | -0.466762699 | -0.478051842 |
| Rcan3 | 7.817473377 | 8.043725204 | 8.295784404 | -0.252059199 | -0.478311027 |
| Eps8l2 | 10.35283439 | 10.91349087 | 10.83114676 | 0.08234411 | -0.478312367 |
| Nbeal1 | 9.104944183 | 9.149976393 | 9.583447662 | -0.433471269 | -0.478503478 |
| Phkg1 | 9.574923998 | 9.650916901 | 10.05497012 | -0.404053215 | -0.480046118 |
| Myo5c | 4.241581164 | 4.443430344 | 4.721746614 | -0.27831627 | -0.48016545 |
| Kcnj12 | 6.045190634 | 6.109944489 | 6.525601956 | -0.415657467 | -0.480411322 |
| P2rx2 | 5.383625236 | 5.343363379 | 5.864286683 | -0.520923305 | -0.480661447 |
| Lrrn3 | 6.467898125 | 6.761786277 | 6.949019352 | -0.187233075 | -0.481121227 |
| Bnip3 | 10.34145021 | 10.18020913 | 10.82292126 | -0.642712127 | -0.481471043 |
| 38596 | 10.80455952 | 11.29472676 | 11.28608828 | 0.00863848 | -0.481528753 |
| Slc25a27 | 6.891076776 | 7.289692433 | 7.372669135 | -0.082976702 | -0.481592359 |
| C1qtnf6 | 7.307056376 | 7.52860757 | 7.788838018 | -0.260230448 | -0.481781642 |
| Cyp2b19 | 9.478402466 | 10.02723265 | 9.960403183 | 0.066829468 | -0.482000716 |
| Kif16b | 9.339709721 | 9.668944804 | 9.821911659 | -0.152966855 | -0.482201937 |
| Gstz1 | 10.52152651 | 10.31954237 | 11.00381568 | -0.684273307 | -0.482289163 |
| Dmrt2 | 4.514931816 | 4.546345671 | 4.997506097 | -0.451160426 | -0.482574281 |
| BC016579 | 6.647673573 | 6.947792742 | 7.131071711 | -0.183278969 | -0.483398139 |
| Lrsam1 | 7.938413755 | 8.289832505 | 8.422342487 | -0.132509983 | -0.483928732 |
| Pdgfd | 5.409406468 | 5.581482648 | 5.893760088 | -0.31227744 | -0.48435362 |
| Mtmr11 | 9.26111633 | 9.521988339 | 9.745593211 | -0.223604872 | -0.484476881 |
| Lipm | 10.84652315 | 11.13026923 | 11.33101882 | -0.20074959 | -0.484495673 |
| Ddah2 | 7.478760979 | 7.793827457 | 7.96334609 | -0.169518633 | -0.484585111 |

| | | | | | |
|---|---|---|---|---|---|
| Phyhip | 10.26090584 | 10.90838174 | 10.74606976 | 0.16231198 | -0.48516392 |
| Timp3 | 12.15845373 | 12.18075718 | 12.64369988 | -0.462942697 | -0.485246143 |
| 6430571L13Rik | 5.691665477 | 5.779793397 | 6.177018182 | -0.397224785 | -0.485352705 |
| Gpd1 | 11.58270186 | 11.50444435 | 12.06860583 | -0.564161483 | -0.48590397 |
| Tcta | 8.519517102 | 8.611891908 | 9.005557906 | -0.393665998 | -0.486040804 |
| Sytl2 | 6.821571931 | 7.09630554 | 7.307690715 | -0.211385175 | -0.486118784 |
| Epdr1 | 8.201251152 | 8.067575267 | 8.687526909 | -0.619951642 | -0.486275757 |
| Il1f6 | 8.265389156 | 8.341453223 | 8.752192471 | -0.410739248 | -0.486803315 |
| Fhod3 | 6.965571142 | 7.261960769 | 7.452426947 | -0.190466178 | -0.486855805 |
| Pald1 | 7.621708516 | 7.813972585 | 8.108801767 | -0.294829182 | -0.487093251 |
| Gm8424 | 4.158383286 | 4.183284497 | 4.645875418 | -0.462590921 | -0.487492132 |
| Zer1 | 8.821349275 | 9.169073445 | 9.309241043 | -0.140167598 | -0.487891768 |
| Cdc42ep5 | 7.065839524 | 7.603121004 | 7.553999665 | 0.049121339 | -0.488160141 |
| Cdc14b | 8.362752415 | 8.947055746 | 8.85101339 | 0.096042356 | -0.488260974 |
| Hnmt | 6.680161904 | 6.691542818 | 7.168674064 | -0.477131246 | -0.48851216 |
| Clic5 | 9.230622814 | 9.301031357 | 9.719150604 | -0.418119247 | -0.48852779 |
| Arrdc3 | 11.31041284 | 11.60032356 | 11.7991233 | -0.198799743 | -0.48871046 |
| St6galnac2 | 6.760092214 | 7.027559413 | 7.249235661 | -0.221676248 | -0.489143447 |
| Col24a1 | 4.240694336 | 4.593055931 | 4.730093741 | -0.13703781 | -0.489399405 |
| Plce1 | 7.469654484 | 7.509690831 | 7.959385675 | -0.449694844 | -0.489731191 |
| Ezh1 | 9.655227865 | 9.982798946 | 10.14515927 | -0.162360328 | -0.489931408 |
| Spock2 | 8.153650263 | 8.078093549 | 8.643874134 | -0.565780585 | -0.490223871 |
| Lrig1 | 10.77288677 | 11.07721643 | 11.26323615 | -0.186019713 | -0.490349373 |
| Hey1 | 5.701235435 | 6.011497661 | 6.19197128 | -0.180473619 | -0.490735845 |
| Heyl | 5.625243606 | 5.730854479 | 6.116124395 | -0.385269916 | -0.490880789 |
| Mxra7 | 7.94476166 | 8.219686115 | 8.435656437 | -0.215970322 | -0.490894777 |
| Gli1 | 6.994390304 | 7.469132033 | 7.485789087 | -0.016657055 | -0.491398783 |
| Wls | 10.75931827 | 11.09663174 | 11.25080848 | -0.154176747 | -0.491490213 |
| Cpt1b | 8.680193301 | 8.845057004 | 9.171804172 | -0.326747169 | -0.491610871 |
| Mylpf | 13.30676812 | 13.49992791 | 13.7987914 | -0.298863493 | -0.49202328 |
| Bmp3 | 8.007845337 | 8.104237498 | 8.499874571 | -0.395637074 | -0.492029234 |
| Tspyl4 | 6.472014436 | 6.387851754 | 6.964216367 | -0.576364613 | -0.492201931 |

| | | | | | |
|---|---|---|---|---|---|
| Adh7 | 10.42040836 | 10.79074239 | 10.91275243 | -0.122010037 | -0.492344067 |
| Mrgprb3 | 7.072260892 | 7.37020155 | 7.564966807 | -0.194765256 | -0.492705914 |
| Rnase4 | 11.83202595 | 11.92389213 | 12.32478763 | -0.400895507 | -0.492761683 |
| Plagl1 | 7.764712119 | 8.146488517 | 8.257519715 | -0.111031198 | -0.492807596 |
| Fndc1 | 9.44107539 | 9.234014817 | 9.933887403 | -0.699872586 | -0.492812013 |
| Fam213a | 10.31164633 | 10.47305071 | 10.80474437 | -0.33169366 | -0.49309804 |
| Sh3yl1 | 7.857306075 | 8.068075541 | 8.350490855 | -0.282415314 | -0.493184779 |
| Carf | 7.233479634 | 7.582161107 | 7.726960217 | -0.14479911 | -0.493480583 |
| 4833423E24Rik | 9.254114403 | 9.4326654 | 9.747628827 | -0.314963427 | -0.493514424 |
| Zfp945 | 7.688719355 | 7.690460357 | 8.18266048 | -0.492200123 | -0.493941125 |
| Lrig3 | 9.079575439 | 9.562153584 | 9.573747355 | -0.011593771 | -0.494171916 |
| Kctd12 | 11.66285188 | 11.84291834 | 12.15748724 | -0.314568897 | -0.49463536 |
| Nrp1 | 9.024896124 | 9.105499438 | 9.519716517 | -0.41421708 | -0.494820393 |
| Klhdc8b | 7.602743636 | 8.189100298 | 8.097662426 | 0.091437872 | -0.49491879 |
| Izumo4 | 6.389683397 | 6.855076909 | 6.884695626 | -0.029618717 | -0.495012229 |
| Crybg3 | 10.04264936 | 10.63825874 | 10.53782254 | 0.100436203 | -0.495173185 |
| Mylk2 | 10.67448433 | 10.94589265 | 11.16981569 | -0.223923043 | -0.495331367 |
| Ccng1 | 11.63322846 | 11.78750441 | 12.1285708 | -0.341066393 | -0.495342343 |
| Muc15 | 8.754124077 | 9.031097028 | 9.249501995 | -0.218404967 | -0.495377918 |
| Trim2 | 10.55673414 | 10.94007222 | 11.05220655 | -0.112134323 | -0.495472407 |
| Armcx1 | 7.472840511 | 7.736103099 | 7.968490481 | -0.232387382 | -0.49564997 |
| Cuedc1 | 8.075469404 | 8.342420491 | 8.571136313 | -0.228715822 | -0.495666909 |
| Gldc | 6.694352671 | 7.160670706 | 7.190255406 | -0.0295847 | -0.495902734 |
| Galm | 6.802798175 | 6.966229582 | 7.298795998 | -0.332566417 | -0.495997824 |
| Aif1l | 10.12465857 | 10.50394029 | 10.6206758 | -0.116735513 | -0.496017238 |
| Ppp1r9a | 6.415761406 | 6.576516272 | 6.912314394 | -0.335798122 | -0.496552988 |
| Trim7 | 9.354801947 | 9.543245344 | 9.851684083 | -0.308438739 | -0.496882136 |
| Hmgn3 | 8.316796807 | 8.48509395 | 8.814003446 | -0.328909497 | -0.497206639 |
| Rspo3 | 5.622552119 | 5.904936493 | 6.120095922 | -0.215159429 | -0.497543803 |
| Calcrl | 8.455237635 | 8.602588871 | 8.953296197 | -0.350707326 | -0.498058562 |
| Col6a6 | 5.791424737 | 5.937432374 | 6.290293435 | -0.352861061 | -0.498868698 |
| Smarca2 | 10.37788536 | 10.5431773 | 10.87682887 | -0.333651563 | -0.49894351 |

| | | | | | |
|---|---|---|---|---|---|
| Zfp974 | 6.276689985 | 6.619505244 | 6.775836038 | -0.156330794 | -0.499146053 |
| Setd7 | 10.72864825 | 11.03335934 | 11.22780094 | -0.194441597 | -0.499152683 |
| Zfp707 | 6.956930307 | 7.17555173 | 7.456254309 | -0.280702579 | -0.499324003 |
| Sptlc3 | 9.84725947 | 10.08860035 | 10.34717773 | -0.258577387 | -0.499918264 |
| Bivm | 7.67897717 | 8.015413324 | 8.178902562 | -0.163489238 | -0.499925392 |
| Samd12 | 5.191363409 | 5.220492006 | 5.691410606 | -0.4709186 | -0.500047197 |
| Cdkn1b | 9.893149382 | 10.10504892 | 10.3932095 | -0.28816058 | -0.500060118 |
| Slc7a10 | 8.182544976 | 8.130302352 | 8.682961223 | -0.55265887 | -0.500416246 |
| Emcn | 6.885653854 | 7.077185118 | 7.386113664 | -0.308928546 | -0.500459809 |
| Cd1d1 | 9.419983678 | 9.222436904 | 9.920482046 | -0.698045142 | -0.500498368 |
| Stac3 | 8.496385088 | 8.673206824 | 8.997025953 | -0.323819129 | -0.500640865 |
| Armh4 | 6.127424001 | 6.436257248 | 6.628130748 | -0.1918735 | -0.500706747 |
| Lep | 10.86430779 | 10.61215146 | 11.36508755 | -0.752936087 | -0.50077976 |
| Vegfc | 5.718625717 | 5.950807566 | 6.219469748 | -0.268662182 | -0.500844031 |
| Lifr | 10.5702373 | 10.50613126 | 11.07126911 | -0.565137853 | -0.501031807 |
| Cbx7 | 8.655804466 | 9.114514221 | 9.157009463 | -0.042495242 | -0.501204997 |
| Kif13a | 10.06603621 | 10.31872839 | 10.56790911 | -0.249180723 | -0.501872907 |
| Fam189a2 | 8.991068551 | 9.309899931 | 9.492983801 | -0.18308387 | -0.50191525 |
| Mphosph9 | 7.235703925 | 7.392139674 | 7.737680789 | -0.345541115 | -0.501976864 |
| Foxo4 | 9.550572973 | 9.746183865 | 10.05263359 | -0.306449725 | -0.502060617 |
| Rapgefl1 | 11.37619251 | 11.97734186 | 11.87826245 | 0.099079413 | -0.502069937 |
| Tenm2 | 9.115851201 | 9.469605709 | 9.617997909 | -0.1483922 | -0.502146708 |
| Il22ra2 | 7.795222632 | 7.72566671 | 8.297527257 | -0.571860547 | -0.502304625 |
| Cyp3a13 | 6.698786774 | 7.160039389 | 7.201170194 | -0.041130805 | -0.50238342 |
| Nqo2 | 8.043790841 | 8.200712891 | 8.546283822 | -0.345570931 | -0.502492981 |
| Calcoco1 | 9.877600239 | 10.28310867 | 10.38013628 | -0.097027613 | -0.502536041 |
| Btbd3 | 10.35489258 | 10.75418453 | 10.85750302 | -0.103318487 | -0.502610433 |
| Cyp2f2 | 10.05559329 | 10.22924744 | 10.55839159 | -0.32914415 | -0.502798301 |
| Atp1b2 | 8.972262215 | 9.087395702 | 9.475356959 | -0.387961257 | -0.503094744 |
| Pik3c2g | 5.056890624 | 5.4281744 | 5.560003511 | -0.131829111 | -0.503112887 |
| Gli2 | 7.17996176 | 7.694482849 | 7.683363974 | 0.011118874 | -0.503402214 |
| Scml2 | 6.620149219 | 6.764722756 | 7.12373138 | -0.359008624 | -0.503582161 |

| | | | | | |
|---|---|---|---|---|---|
| Tspan12 | 7.311492876 | 7.304869476 | 7.815092098 | -0.510222622 | -0.503599222 |
| Arl4a | 10.51653739 | 10.73921855 | 11.0201401 | -0.280921547 | -0.50360271 |
| Gm94 | 11.49108705 | 11.65237504 | 11.99496991 | -0.34259487 | -0.50388286 |
| Itgbl1 | 8.799989261 | 8.958275444 | 9.303963297 | -0.345687853 | -0.503974036 |
| Fam171a1 | 7.527834051 | 7.675562701 | 8.03185722 | -0.356294519 | -0.504023169 |
| Srgap2 | 10.48577703 | 10.79937754 | 10.99012173 | -0.19074419 | -0.5043447 |
| Kifap3 | 8.837098807 | 8.998554444 | 9.341503539 | -0.342949095 | -0.504404732 |
| Hbp1 | 10.69999679 | 11.00268302 | 11.20441564 | -0.20173262 | -0.504418857 |
| Enpp2 | 10.50844114 | 10.42603446 | 11.01319218 | -0.58715772 | -0.504751047 |
| Slc9a9 | 8.263413207 | 8.230617755 | 8.768228674 | -0.53761092 | -0.504815468 |
| Tuft1 | 10.7754256 | 11.09595439 | 11.28030047 | -0.184346083 | -0.50487487 |
| Ms4a7 | 7.257576156 | 7.797358653 | 7.762687634 | 0.034671019 | -0.505111478 |
| Zfp870 | 6.932023158 | 7.186045513 | 7.437424746 | -0.251379233 | -0.505401588 |
| Chst3 | 6.373265007 | 6.750056278 | 6.878830245 | -0.128773967 | -0.505565239 |
| Neo1 | 9.955432465 | 10.25910922 | 10.46149535 | -0.202386133 | -0.506062888 |
| Tmem246 | 7.336414567 | 7.598235903 | 7.842508578 | -0.244272675 | -0.506094011 |
| Adora1 | 7.195617142 | 7.48572632 | 7.701861125 | -0.216134805 | -0.506243983 |
| Slc16a14 | 8.569848185 | 9.018397089 | 9.076286846 | -0.057889757 | -0.506438661 |
| Zfp618 | 4.038999269 | 4.11927995 | 4.545831127 | -0.426551177 | -0.506831858 |
| Plxna3 | 8.562967011 | 9.140352582 | 9.069814652 | 0.07053793 | -0.506847641 |
| Zfp354c | 5.966402861 | 6.23393224 | 6.473408172 | -0.239475932 | -0.507005311 |
| Zim1 | 5.421137688 | 5.505458557 | 5.928170368 | -0.422711811 | -0.50703268 |
| Vtn | 6.07151427 | 6.124618082 | 6.578826228 | -0.454208146 | -0.507311958 |
| Crispld1 | 5.432449994 | 5.575017415 | 5.939852186 | -0.364834771 | -0.507402192 |
| Ildr2 | 6.435112109 | 6.535059341 | 6.942862259 | -0.407802918 | -0.50775015 |
| Col1a1 | 12.80166683 | 13.47250329 | 13.30963749 | 0.162865793 | -0.507970667 |
| Arxes2 | 6.285175878 | 6.240976412 | 6.793225686 | -0.552249274 | -0.508049808 |
| Klhl13 | 6.083283761 | 6.004452314 | 6.591457918 | -0.587005604 | -0.508174157 |
| Scn1b | 9.704700353 | 9.932602672 | 10.21298119 | -0.280378514 | -0.508280833 |
| Six1 | 5.698504398 | 5.724690598 | 6.206792459 | -0.482101861 | -0.508288061 |
| Aknad1 | 7.248080976 | 7.521707988 | 7.756611801 | -0.234903814 | -0.508530825 |
| Casq2 | 6.490222181 | 6.479804869 | 6.999014969 | -0.5192101 | -0.508792789 |

| | | | | | |
|---|---|---|---|---|---|
| Abca12 | 11.9814833 | 12.37120283 | 12.49041656 | -0.11921373 | -0.508933263 |
| Arrdc2 | 7.097268672 | 7.485018577 | 7.606529714 | -0.121511138 | -0.509261042 |
| Nedd4 | 12.75522267 | 13.00506092 | 13.26452728 | -0.259466357 | -0.509304607 |
| Gjc3 | 5.359348724 | 5.480245361 | 5.868920847 | -0.388675486 | -0.509572123 |
| Npr1 | 8.239828336 | 8.560308616 | 8.749438087 | -0.189129471 | -0.509609751 |
| 4931406C07Rik | 8.242344166 | 8.374974738 | 8.75196189 | -0.376987152 | -0.509617724 |
| Kcnj2 | 9.135432502 | 9.263999724 | 9.645258877 | -0.381259153 | -0.509826374 |
| Hoxc4 | 6.556498801 | 6.810146955 | 7.066399833 | -0.256252878 | -0.509901032 |
| Fgfrl1 | 8.0196406 | 8.02875405 | 8.530358283 | -0.501604233 | -0.510717683 |
| Rcan2 | 6.996700638 | 6.98042202 | 7.508089848 | -0.527667828 | -0.51138921 |
| Myom2 | 11.34532141 | 11.23748004 | 11.85708816 | -0.61960812 | -0.51176675 |
| Acad12 | 6.935301789 | 7.204396546 | 7.447082782 | -0.242686236 | -0.511780993 |
| Cdnf | 6.546060394 | 6.647099314 | 7.058199938 | -0.411100624 | -0.512139543 |
| Adgrf4 | 10.71937055 | 10.89303442 | 11.23157928 | -0.338544863 | -0.512208733 |
| Kdelc2 | 7.899152779 | 7.981043229 | 8.411691626 | -0.430648396 | -0.512538847 |
| Lpar4 | 5.390892219 | 5.482177604 | 5.903438359 | -0.421260755 | -0.51254614 |
| Nrip2 | 4.294556231 | 4.442871982 | 4.807184451 | -0.36431247 | -0.51262822 |
| Mlxipl | 8.207669325 | 8.581838117 | 8.720337445 | -0.138499328 | -0.512668119 |
| Lrrcc1 | 7.940392255 | 8.143452689 | 8.453253836 | -0.309801147 | -0.512861581 |
| Pgm1 | 9.777437484 | 9.899411581 | 10.29074064 | -0.391329059 | -0.513303156 |
| Kif21a | 10.86886708 | 11.04740397 | 11.38248422 | -0.335080247 | -0.513617137 |
| Ank3 | 9.893414065 | 10.00884428 | 10.40725575 | -0.398411473 | -0.513841688 |
| Pygb | 10.21953695 | 10.49631601 | 10.73339151 | -0.237075507 | -0.51385456 |
| Scrn3 | 8.333738683 | 8.526262396 | 8.848034084 | -0.321771688 | -0.514295401 |
| Zfp260 | 9.332217045 | 9.636294975 | 9.847333352 | -0.211038378 | -0.515116307 |
| Slc6a14 | 9.60694253 | 9.955771979 | 10.12215523 | -0.166383247 | -0.515212696 |
| Kif3c | 7.657080194 | 8.045283044 | 8.172821696 | -0.127538652 | -0.515741502 |
| Gm20219 | 9.140530781 | 9.580877468 | 9.656504244 | -0.075626776 | -0.515973462 |
| Slc16a10 | 8.200969187 | 8.854593398 | 8.717359873 | 0.137233526 | -0.516390685 |
| Sytl1 | 9.834478756 | 10.33950211 | 10.35088389 | -0.01138178 | -0.516405138 |
| Mid2 | 9.493345003 | 9.759992805 | 10.00978834 | -0.249795535 | -0.516443337 |
| Dcaf6 | 9.601781885 | 9.761520787 | 10.11823207 | -0.356711278 | -0.51645018 |

| | | | | | |
|---|---|---|---|---|---|
| Rab38 | 8.937704807 | 9.287109384 | 9.454238902 | -0.167129518 | -0.516534095 |
| Col6a5 | 5.023117374 | 5.258945832 | 5.539776505 | -0.280830672 | -0.516659131 |
| Narf | 9.576453656 | 9.828576849 | 10.09329876 | -0.264721908 | -0.516845101 |
| Sdr9c7 | 10.08407317 | 10.29709039 | 10.60095271 | -0.303862323 | -0.516879542 |
| Ehd2 | 11.68701569 | 12.08227416 | 12.20411049 | -0.121836333 | -0.517094803 |
| Gm11992 | 8.502980923 | 8.926101255 | 9.020243934 | -0.094142679 | -0.517263011 |
| Mblac2 | 7.677756151 | 7.888203188 | 8.19553594 | -0.307332752 | -0.517779789 |
| Egf | 6.844583022 | 6.878930469 | 7.362535175 | -0.483604706 | -0.517952153 |
| Mrtfb | 9.386575036 | 9.722475919 | 9.904595281 | -0.182119362 | -0.518020245 |
| Cd36 | 12.36008614 | 12.11216506 | 12.87837548 | -0.76621042 | -0.518289337 |
| Dsg1a | 14.26677677 | 14.52599393 | 14.7851639 | -0.25916997 | -0.51838713 |
| Chek2 | 6.994915269 | 7.1540232 | 7.513714634 | -0.359691433 | -0.518799365 |
| Itm2a | 9.069171506 | 9.145523515 | 9.588033761 | -0.442510246 | -0.518862255 |
| Rbm24 | 7.579668648 | 7.993987988 | 8.098707748 | -0.10471976 | -0.5190391 |
| Lgr6 | 10.00100609 | 10.28235493 | 10.52019741 | -0.237842477 | -0.519191319 |
| Gm9843 | 4.700829605 | 7.781649159 | 5.220221675 | 2.561427484 | -0.51939207 |
| Ephx1 | 10.28952214 | 10.40652088 | 10.80903091 | -0.402510037 | -0.519508777 |
| Slc16a9 | 7.441046498 | 7.588538083 | 7.960677624 | -0.372139542 | -0.519631127 |
| Ttc23 | 7.590904801 | 7.837334539 | 8.110979327 | -0.273644788 | -0.520074526 |
| Mki67 | 10.79056343 | 11.13911062 | 11.31128297 | -0.172172357 | -0.520719547 |
| Afdn | 11.9843433 | 12.41265409 | 12.50558657 | -0.092932483 | -0.521243267 |
| Cyp2w1 | 5.103688183 | 5.708005674 | 5.625417862 | 0.082587812 | -0.521729679 |
| Col4a4 | 6.031197865 | 6.299714276 | 6.553283118 | -0.253568842 | -0.522085254 |
| Scara3 | 9.333746783 | 9.397862842 | 9.855868156 | -0.458005314 | -0.522121373 |
| Tppp3 | 9.433543662 | 9.5365346 | 9.955767458 | -0.419232858 | -0.522223796 |
| Rasgrp1 | 9.313876428 | 9.754326885 | 9.836525897 | -0.082199012 | -0.522649468 |
| Skint9 | 9.470140924 | 9.773751164 | 9.992803851 | -0.219052687 | -0.522662928 |
| B3galnt1 | 6.993649443 | 7.050272816 | 7.516350159 | -0.466077343 | -0.522700716 |
| Cd55 | 9.622112022 | 9.778828231 | 10.1450013 | -0.366173071 | -0.52288928 |
| Mpp2 | 6.430300692 | 7.062227878 | 6.953341114 | 0.108886764 | -0.523040421 |
| Synm | 11.00065646 | 11.0140555 | 11.52482903 | -0.510773527 | -0.524172573 |
| Scn4b | 9.84434778 | 9.712858761 | 10.3690161 | -0.656157342 | -0.524668323 |

| | | | | | |
|---|---|---|---|---|---|
| Tmod4 | 9.176906177 | 9.11575017 | 9.702342613 | -0.586592443 | -0.525436436 |
| Idh1 | 10.84106221 | 10.95311701 | 11.36673487 | -0.41361786 | -0.525672653 |
| Mylip | 9.151017508 | 9.440394989 | 9.676723316 | -0.236328327 | -0.525705808 |
| Bmp6 | 6.213547602 | 6.373347004 | 6.739626703 | -0.366279699 | -0.526079101 |
| Usp13 | 9.405405894 | 9.602449133 | 9.931680164 | -0.329231031 | -0.52627427 |
| Klf4 | 12.21550126 | 12.56605366 | 12.74228628 | -0.176232623 | -0.526785023 |
| Fam221a | 5.754704261 | 6.133971597 | 6.282327828 | -0.148356231 | -0.527623567 |
| Prlr | 9.646774141 | 9.93850362 | 10.17463481 | -0.236131187 | -0.527860665 |
| Rps6ka6 | 8.572842916 | 8.935615261 | 9.100718269 | -0.165103008 | -0.527875353 |
| Dyrk1b | 7.743254985 | 8.190823335 | 8.271188558 | -0.080365223 | -0.527933573 |
| Perm1 | 7.639051958 | 8.201521606 | 8.167123073 | 0.034398533 | -0.528071116 |
| Sdc2 | 9.104191723 | 9.165185332 | 9.6324931 | -0.467307768 | -0.528301377 |
| Aldh1a1 | 10.07299326 | 9.895954696 | 10.60186206 | -0.70590736 | -0.528868794 |
| Agfg2 | 8.183684516 | 8.583971488 | 8.71306348 | -0.129091992 | -0.529378964 |
| Acadm | 10.62075657 | 10.62849441 | 11.15040877 | -0.521914357 | -0.529652193 |
| Lars2 | 13.81889527 | 13.2784147 | 14.34860797 | -1.070193267 | -0.529712693 |
| Cldn8 | 6.230773692 | 6.818475456 | 6.761118722 | 0.057356734 | -0.530345031 |
| Krt78 | 11.32804058 | 11.76257475 | 11.85872606 | -0.09615131 | -0.530685487 |
| Mup19 | 8.031165782 | 8.450587301 | 8.561966244 | -0.111378943 | -0.530800463 |
| Kcna1 | 5.828417707 | 6.191481484 | 6.359249424 | -0.16776794 | -0.530831717 |
| Sntb1 | 9.152829576 | 9.341500524 | 9.683778602 | -0.342278078 | -0.530949026 |
| Map2 | 10.52795098 | 10.83233607 | 11.05968624 | -0.227350173 | -0.531735267 |
| Rbl2 | 9.517488616 | 9.83568117 | 10.04947122 | -0.213790054 | -0.531982608 |
| Kifc2 | 5.979764207 | 6.475562358 | 6.512238417 | -0.036676059 | -0.53247421 |
| Ctdspl | 9.81148698 | 10.19916469 | 10.34433331 | -0.145168627 | -0.532846334 |
| Kctd12b | 7.140347904 | 7.002947345 | 7.674472289 | -0.671524944 | -0.534124385 |
| Sorbs1 | 9.755740533 | 9.805369872 | 10.29008765 | -0.484717775 | -0.534347113 |
| Nlrp10 | 10.12070485 | 10.31299438 | 10.65515578 | -0.342161403 | -0.534450931 |
| Serpinb7 | 10.89005117 | 11.04050206 | 11.42466357 | -0.38416151 | -0.534612397 |
| Tjp3 | 8.754268329 | 9.363904241 | 9.289223332 | 0.074680909 | -0.534955003 |
| Fbxl20 | 9.660712259 | 10.06161351 | 10.19581511 | -0.134201608 | -0.535102855 |
| Ache | 7.714755551 | 8.271964661 | 8.250205964 | 0.021758697 | -0.535450413 |

| | | | | | |
|---|---|---|---|---|---|
| Esrp1 | 11.70898554 | 12.07326973 | 12.2445881 | -0.171318377 | -0.53560256 |
| Dgat2 | 12.00599918 | 12.06043198 | 12.54161512 | -0.48118314 | -0.53561594 |
| Kcnj11 | 7.277754815 | 7.56731999 | 7.81365721 | -0.24633722 | -0.535902394 |
| Cd300ld4 | 5.759538476 | 6.059176799 | 6.295597156 | -0.236420357 | -0.536058681 |
| Tmod1 | 9.473971272 | 9.731020001 | 10.01008454 | -0.279064541 | -0.536113269 |
| Mybpc2 | 12.77690128 | 13.075719 | 13.31303211 | -0.237313107 | -0.536130827 |
| Myh1 | 13.57168598 | 13.81792087 | 14.10823234 | -0.290311473 | -0.53654636 |
| Cd99l2 | 9.118117307 | 9.313422781 | 9.65599709 | -0.34257431 | -0.537879783 |
| Tbx15 | 10.05475332 | 9.940567893 | 10.5926903 | -0.652122404 | -0.537936981 |
| Rnf150 | 8.203770881 | 8.179967656 | 8.741912836 | -0.561945181 | -0.538141955 |
| Wee1 | 8.660327322 | 9.131212967 | 9.198593157 | -0.06738019 | -0.538265835 |
| Epm2a | 7.768598557 | 7.805454126 | 8.30723223 | -0.501778103 | -0.538633672 |
| Sik2 | 9.155168335 | 9.381417115 | 9.69447807 | -0.313060956 | -0.539309735 |
| Myo6 | 10.38593915 | 10.86665 | 10.92544425 | -0.058794257 | -0.5395051 |
| Cav1 | 12.62016761 | 12.63476523 | 13.16044197 | -0.525676733 | -0.54027436 |
| Clip3 | 6.888026077 | 7.43134067 | 7.428371587 | 0.002969083 | -0.540345511 |
| Pof1b | 12.54023325 | 12.82211107 | 13.08079324 | -0.25868217 | -0.54055999 |
| Shroom4 | 8.025945399 | 8.517009854 | 8.567301027 | -0.050291173 | -0.541355628 |
| Myl1 | 13.11241073 | 13.40436631 | 13.65394185 | -0.24957554 | -0.541531117 |
| Oas1e | 4.931311345 | 5.356472984 | 5.472991803 | -0.116518819 | -0.541680458 |
| Depp1 | 6.362061081 | 6.933010792 | 6.904389519 | 0.028621273 | -0.542328438 |
| Alox12 | 7.598970017 | 8.063954917 | 8.141508226 | -0.077553309 | -0.542538209 |
| Slc26a10 | 4.003036204 | 4.14083255 | 4.545990884 | -0.405158334 | -0.54295468 |
| Tgfb3 | 7.610113878 | 7.892119963 | 8.153221133 | -0.261101169 | -0.543107254 |
| Dixdc1 | 6.934381768 | 7.37663086 | 7.477827936 | -0.101197076 | -0.543446168 |
| Thra | 9.639850709 | 9.916020382 | 10.18367635 | -0.267655965 | -0.543825637 |
| Gjb6 | 6.891362616 | 7.164013607 | 7.435359826 | -0.271346218 | -0.543997209 |
| Sh3gl2 | 7.430270349 | 7.958294045 | 7.97497311 | -0.016679065 | -0.544702761 |
| Ttll7 | 8.018343196 | 8.055279328 | 8.563244698 | -0.507965371 | -0.544901502 |
| Irs3 | 4.861193311 | 5.03510874 | 5.406184112 | -0.371075372 | -0.544990801 |
| Clmn | 8.215678938 | 8.316066357 | 8.761966763 | -0.445900406 | -0.546287825 |
| Gm15519 | 4.002444371 | 4.282932938 | 4.54948434 | -0.266551402 | -0.547039969 |

| | | | | | |
|---|---|---|---|---|---|
| Gpsm2 | 9.636971562 | 10.0399636 | 10.18421438 | -0.144250787 | -0.547242822 |
| Slc9a3 | 6.097035235 | 6.81561842 | 6.644444258 | 0.171174162 | -0.547409023 |
| 4930578C19Rik | 4.915668072 | 5.110977995 | 5.463431009 | -0.352453014 | -0.547762936 |
| Btc | 8.894590375 | 9.333924317 | 9.44264769 | -0.108723374 | -0.548057315 |
| Aaed1 | 9.10181372 | 9.370986872 | 9.649881926 | -0.278895054 | -0.548068206 |
| Ago4 | 8.655946435 | 9.022836103 | 9.205222742 | -0.182386638 | -0.549276307 |
| Kif1b | 11.43981705 | 11.78286149 | 11.98922576 | -0.206364267 | -0.549408707 |
| Gstt2 | 5.646034075 | 5.887604409 | 6.19553125 | -0.307926841 | -0.549497175 |
| Srpx2 | 7.293252822 | 7.467148451 | 7.842768563 | -0.375620112 | -0.549515741 |
| Fmo5 | 8.358523414 | 8.571812853 | 8.908296308 | -0.336483455 | -0.549772894 |
| Fgf10 | 6.713630254 | 6.744840059 | 7.263851924 | -0.519011865 | -0.55022167 |
| Cbs | 8.217422828 | 8.808937065 | 8.767796643 | 0.041140422 | -0.550373815 |
| Tnik | 8.998675869 | 9.362816468 | 9.549123345 | -0.186306878 | -0.550447476 |
| Nrap | 10.96066562 | 12.08380404 | 11.51112273 | 0.572681307 | -0.55045711 |
| Ift46 | 8.344478484 | 8.784831939 | 8.895554938 | -0.110722999 | -0.551076454 |
| Dnm1 | 8.020025165 | 8.316792041 | 8.571453013 | -0.254660972 | -0.551427848 |
| Gulp1 | 7.177886103 | 7.374034389 | 7.729435765 | -0.355401376 | -0.551549662 |
| Slc44a3 | 5.775687852 | 5.919570353 | 6.327293414 | -0.407723061 | -0.551605562 |
| Slc22a23 | 9.447104799 | 9.861445238 | 9.99875999 | -0.137314752 | -0.551655191 |
| Fyco1 | 10.50077731 | 10.76163237 | 11.0525954 | -0.29096303 | -0.551818093 |
| Peli3 | 6.467803943 | 6.795782708 | 7.019718694 | -0.223935985 | -0.55191475 |
| Smyd1 | 9.553351025 | 9.941074293 | 10.10618681 | -0.165112518 | -0.552835786 |
| Lmod1 | 7.606227361 | 7.923382861 | 8.159182402 | -0.235799541 | -0.552955042 |
| Gm8116 | 6.244096399 | 6.387883665 | 6.797192625 | -0.40930896 | -0.553096226 |
| Nrn1 | 7.688346908 | 7.756123272 | 8.241541905 | -0.485418633 | -0.553194997 |
| Nudt12 | 7.062086161 | 7.208643348 | 7.615632808 | -0.40698946 | -0.553546647 |
| Nicn1 | 7.099888114 | 7.47137148 | 7.653513749 | -0.182142269 | -0.553625636 |
| Pak3 | 9.555026699 | 9.844106269 | 10.10942987 | -0.265323599 | -0.554403169 |
| Plpp7 | 6.666626258 | 6.887400691 | 7.22142393 | -0.334023238 | -0.554797672 |
| Kcnq5 | 6.262189902 | 6.431822917 | 6.817473211 | -0.385650294 | -0.555283309 |
| Hdac11 | 7.088794894 | 7.522902741 | 7.644321224 | -0.121418483 | -0.55552633 |
| Pigp | 8.407169467 | 8.683576331 | 8.963373679 | -0.279797348 | -0.556204212 |

| | | | | | |
|---|---|---|---|---|---|
| Mef2c | 9.982949492 | 9.871661303 | 10.53920818 | -0.667546877 | -0.556258688 |
| Lum | 12.04762163 | 12.40850242 | 12.60396455 | -0.195462127 | -0.556342923 |
| Emp2 | 12.65832325 | 13.01371827 | 13.21468891 | -0.200970637 | -0.556365663 |
| Pdp1 | 8.428155585 | 8.474868736 | 8.985414133 | -0.510545397 | -0.557258548 |
| Pik3r2 | 8.462830793 | 9.162806284 | 9.020520165 | 0.142286118 | -0.557689373 |
| Ptpn13 | 10.95624488 | 11.43912928 | 11.51435255 | -0.075223267 | -0.558107673 |
| Kcnc1 | 6.572528044 | 6.816506548 | 7.131939412 | -0.315432864 | -0.559411368 |
| Abcb4 | 7.606752149 | 7.833873945 | 8.166264544 | -0.3323906 | -0.559512396 |
| Fam84a | 10.7969632 | 11.34262186 | 11.35652526 | -0.0139034 | -0.55956206 |
| Skint1 | 8.064556491 | 8.458876688 | 8.624528864 | -0.165652176 | -0.559972372 |
| Arhgef28 | 8.109374315 | 8.470890509 | 8.670402196 | -0.199511688 | -0.561027881 |
| Cbx2 | 8.546847846 | 9.037632032 | 9.108372779 | -0.070740747 | -0.561524933 |
| Dsg2 | 8.895294577 | 9.304617261 | 9.456891679 | -0.152274418 | -0.561597102 |
| Rora | 11.91783371 | 12.15463177 | 12.47945405 | -0.324822287 | -0.561620343 |
| Pde3a | 6.463777253 | 6.65240161 | 7.025546933 | -0.373145323 | -0.56176968 |
| Clec3b | 9.363093356 | 9.518207914 | 9.924978294 | -0.406770381 | -0.561884938 |
| Kiz | 7.539688899 | 7.781067034 | 8.101852701 | -0.320785666 | -0.562163801 |
| Ehhadh | 7.542403943 | 7.943328953 | 8.104877165 | -0.161548212 | -0.562473222 |
| Akap6 | 6.886094934 | 7.068657766 | 7.448935987 | -0.380278221 | -0.562841053 |
| Cd34 | 11.46744396 | 11.56762629 | 12.03053843 | -0.462912137 | -0.563094467 |
| Scd2 | 11.39434763 | 11.72590002 | 11.9575892 | -0.23168918 | -0.563241567 |
| Nipal3 | 8.111407948 | 8.347901836 | 8.675182631 | -0.327280795 | -0.563774684 |
| Kcnj16 | 5.823223082 | 5.978179382 | 6.388210664 | -0.410031283 | -0.564987582 |
| Agpat2 | 9.940603901 | 9.920193367 | 10.50561692 | -0.585423553 | -0.565013019 |
| Ppara | 6.269872507 | 6.682592891 | 6.834887488 | -0.152294597 | -0.565014981 |
| Ypel4 | 5.284312763 | 5.585309545 | 5.850073707 | -0.264764162 | -0.565760944 |
| Wwp1 | 10.33339269 | 10.47179189 | 10.89940287 | -0.427610977 | -0.56601018 |
| Id4 | 9.272083787 | 9.52955838 | 9.838108801 | -0.308550421 | -0.566025014 |
| Plekhh2 | 7.222560936 | 7.637759624 | 7.7893231 | -0.151563476 | -0.566762164 |
| Tmtc1 | 8.760317111 | 8.808334236 | 9.327116888 | -0.518782652 | -0.566799777 |
| Foxp2 | 5.07247396 | 5.373345374 | 5.639435152 | -0.266089779 | -0.566961193 |
| Nudt7 | 7.699056513 | 7.678252994 | 8.266887889 | -0.588634895 | -0.567831376 |

| | | | | | |
|---|---|---|---|---|---|
| Rab11fip4 | 10.094919 | 10.38642877 | 10.66350466 | -0.277075893 | -0.56858566 |
| Sptb | 9.004802804 | 9.357475221 | 9.573434053 | -0.215958832 | -0.568631248 |
| Gstt3 | 7.797319573 | 8.068443366 | 8.366169928 | -0.297726562 | -0.568850355 |
| Pfkfb1 | 7.264837789 | 6.998129607 | 7.833689844 | -0.835560237 | -0.568852055 |
| Aspa | 6.392791019 | 6.577211811 | 6.962014603 | -0.384802793 | -0.569223584 |
| Epha1 | 10.28097464 | 10.78120676 | 10.85040119 | -0.069194437 | -0.56942655 |
| Gm49017 | 5.094744714 | 5.477981614 | 5.664617617 | -0.186636003 | -0.569872903 |
| Slc24a3 | 9.176302282 | 9.25825764 | 9.746595143 | -0.488337503 | -0.570292862 |
| Nadk2 | 7.804920251 | 8.071842024 | 8.375834767 | -0.303992743 | -0.570914516 |
| Sesn3 | 8.840557005 | 9.202915082 | 9.411484346 | -0.208569265 | -0.570927341 |
| Srpx | 8.609815393 | 8.794769264 | 9.181478124 | -0.38670886 | -0.571662731 |
| Col1a2 | 13.28969559 | 13.85161341 | 13.86177411 | -0.0101607 | -0.57207852 |
| Dnase1l1 | 8.797251262 | 9.095267825 | 9.369558186 | -0.274290361 | -0.572306924 |
| Speg | 8.35155724 | 9.076731355 | 8.924300353 | 0.152431002 | -0.572743113 |
| Fgf13 | 7.549878973 | 7.385205821 | 8.123057529 | -0.737851707 | -0.573178556 |
| Evc | 7.804868992 | 8.125081183 | 8.378415281 | -0.253334098 | -0.573546289 |
| Bace1 | 8.443572525 | 8.640203584 | 9.017352113 | -0.37714853 | -0.573779589 |
| H1f0 | 10.85604965 | 10.94781629 | 11.43020332 | -0.482387023 | -0.57415367 |
| Adam33 | 7.606125395 | 8.083267328 | 8.181335496 | -0.098068168 | -0.575210101 |
| Klhl23 | 7.146741919 | 7.402642063 | 7.7220689 | -0.319426837 | -0.575326981 |
| Syne1 | 9.658116113 | 10.06281824 | 10.23355105 | -0.170732817 | -0.57543494 |
| Ppfibp2 | 9.071659463 | 9.464139433 | 9.647690736 | -0.183551302 | -0.576031273 |
| Aifm2 | 8.374821471 | 8.658333765 | 8.953106874 | -0.294773108 | -0.578285402 |
| Crebl2 | 8.006423032 | 8.245969349 | 8.584833581 | -0.338864232 | -0.578410549 |
| Adcy9 | 7.210006921 | 7.511139216 | 7.788433199 | -0.277293983 | -0.578426279 |
| Arhgef25 | 7.481164354 | 7.944870746 | 8.059863529 | -0.114992782 | -0.578699175 |
| Tcaf1 | 8.625637504 | 8.665532354 | 9.204758329 | -0.539225975 | -0.579120826 |
| Osbpl6 | 6.436975866 | 6.410239752 | 7.016172398 | -0.605932646 | -0.579196532 |
| Map2k6 | 6.848582657 | 7.277897573 | 7.428397261 | -0.150499687 | -0.579814604 |
| Adamtsl3 | 6.392978863 | 6.528993264 | 6.972825023 | -0.443831759 | -0.579846161 |
| Fcgrt | 9.922697336 | 9.968484004 | 10.50315756 | -0.534673556 | -0.580460224 |
| Zc3h6 | 6.848852475 | 6.737265144 | 7.429422734 | -0.69215759 | -0.580570259 |

| | | | | | |
|---|---|---|---|---|---|
| Cped1 | 9.415402017 | 9.493256424 | 9.996088143 | -0.502831719 | -0.580686126 |
| Steap3 | 8.875040848 | 9.270181124 | 9.455757003 | -0.185575878 | -0.580716155 |
| Reep1 | 7.188795274 | 7.398239956 | 7.769621157 | -0.371381201 | -0.580825883 |
| Cap2 | 8.583447965 | 8.903167866 | 9.16540071 | -0.262232844 | -0.581952745 |
| Xlr3b | 6.391260307 | 5.782780438 | 6.97348391 | -1.190703472 | -0.582223603 |
| Cdk19 | 9.494715738 | 9.849206473 | 10.07761205 | -0.228405577 | -0.582896312 |
| Ulk2 | 9.745503065 | 9.965473321 | 10.3288777 | -0.363404375 | -0.583374632 |
| Tpbgl | 7.400729213 | 7.601229898 | 7.984153806 | -0.382923907 | -0.583424592 |
| Mertk | 8.036612085 | 8.354769878 | 8.620524678 | -0.2657548 | -0.583912593 |
| Ptger3 | 7.933205776 | 7.732678469 | 8.518193502 | -0.785515032 | -0.584987725 |
| Olfml1 | 7.109409239 | 7.22343622 | 7.694518777 | -0.471082557 | -0.585109538 |
| Hfe2 | 9.623377187 | 9.78292218 | 10.20895158 | -0.426029396 | -0.585574388 |
| Nynrin | 7.553984219 | 7.940235379 | 8.140154965 | -0.199919586 | -0.586170745 |
| Unc45b | 8.718422111 | 9.487423521 | 9.305690329 | 0.181733192 | -0.587268218 |
| Vldlr | 8.864529035 | 8.993689786 | 9.451991833 | -0.458302047 | -0.587462799 |
| Sec14l5 | 6.363662918 | 6.459393894 | 6.951552784 | -0.49215889 | -0.587889866 |
| Slc16a7 | 7.195909426 | 7.337193277 | 7.783982424 | -0.446789147 | -0.588072998 |
| Zfp266 | 10.7556625 | 11.12857723 | 11.34374336 | -0.215166137 | -0.588080867 |
| Il22ra1 | 8.040496318 | 8.526057824 | 8.628754326 | -0.102696502 | -0.588258008 |
| Shpk | 6.635107954 | 6.86019223 | 7.224046896 | -0.363854666 | -0.588938942 |
| Paqr3 | 7.073328708 | 7.733269315 | 7.662682613 | 0.070586703 | -0.589353905 |
| Snx21 | 7.986642019 | 8.450915642 | 8.576036883 | -0.125121242 | -0.589394865 |
| Cacna2d1 | 9.901321112 | 9.968343031 | 10.49133702 | -0.522993985 | -0.590015905 |
| Mcf2l | 8.482324979 | 8.957887362 | 9.07321386 | -0.115326498 | -0.590888881 |
| Sgsm2 | 8.587188424 | 9.043452096 | 9.178445926 | -0.13499383 | -0.591257502 |
| Smco3 | 5.407146426 | 5.913584134 | 5.998860916 | -0.085276782 | -0.59171449 |
| Asb14 | 5.593806009 | 5.746798565 | 6.185524306 | -0.438725741 | -0.591718296 |
| Cyp2j6 | 9.268819984 | 9.586714556 | 9.860681731 | -0.273967174 | -0.591861747 |
| Pgm2l1 | 7.913966237 | 8.083209957 | 8.505857129 | -0.422647172 | -0.591890892 |
| Usp35 | 6.259177352 | 6.746638335 | 6.851247987 | -0.104609652 | -0.592070635 |
| Maml2 | 7.997172443 | 8.356749131 | 8.589320674 | -0.232571543 | -0.592148231 |
| Slc6a4 | 7.855236772 | 8.020337203 | 8.447404268 | -0.427067065 | -0.592167496 |

| | | | | | |
|---|---|---|---|---|---|
| Hoxd9 | 6.250577743 | 6.378321743 | 6.842796538 | -0.464474795 | -0.592218795 |
| Nr3c2 | 6.117616435 | 6.491902262 | 6.71033729 | -0.218435028 | -0.592720855 |
| Mypn | 9.281139538 | 9.533687753 | 9.874667517 | -0.340979764 | -0.593527979 |
| Cbr2 | 9.927746228 | 9.890736703 | 10.52197891 | -0.63124221 | -0.594232685 |
| Phldb2 | 8.651027687 | 9.178343085 | 9.245352049 | -0.067008964 | -0.594324362 |
| Nr2f2 | 7.746533735 | 7.889481696 | 8.341470668 | -0.451988972 | -0.594936933 |
| 2310002L09Rik | 7.397294369 | 8.465810944 | 7.992708057 | 0.473102887 | -0.595413688 |
| Arhgef6 | 8.434767274 | 8.546782354 | 9.031105682 | -0.484323328 | -0.596338408 |
| Fndc9 | 9.787783565 | 10.25868411 | 10.38416133 | -0.12547722 | -0.596377762 |
| Esco2 | 6.822781253 | 7.204231939 | 7.419367736 | -0.215135797 | -0.596586483 |
| Camsap3 | 10.22344789 | 10.79264999 | 10.82015058 | -0.027500587 | -0.596702692 |
| Ripply3 | 6.104460628 | 6.616893554 | 6.701460018 | -0.084566464 | -0.59699939 |
| Dst | 13.2600249 | 13.72700471 | 13.85714673 | -0.130142017 | -0.597121823 |
| Klhl4 | 5.099247153 | 5.329261436 | 5.697352863 | -0.368091427 | -0.598105711 |
| Ptpn21 | 10.1320224 | 10.52979618 | 10.73179559 | -0.20199941 | -0.599773188 |
| Arhgef9 | 5.79785461 | 6.151298817 | 6.398409209 | -0.247110392 | -0.6005546 |
| Clec2g | 7.119021772 | 7.423485202 | 7.720008636 | -0.296523434 | -0.600986864 |
| Ace2 | 9.313753422 | 9.674412095 | 9.915200372 | -0.240788277 | -0.60144695 |
| Ablim3 | 8.216419143 | 8.362535221 | 8.818097959 | -0.455562738 | -0.601678816 |
| Arhgap19 | 8.230110751 | 8.667222258 | 8.832738375 | -0.165516117 | -0.602627624 |
| Nr1d2 | 10.89001181 | 11.03024705 | 11.49269757 | -0.462450523 | -0.60268576 |
| Atp1b1 | 10.41744597 | 10.43612999 | 11.02104387 | -0.58491388 | -0.603597897 |
| Ston1 | 7.787591803 | 8.388209149 | 8.391515634 | -0.003306485 | -0.603923831 |
| Npr2 | 8.387509513 | 8.539503638 | 8.992973208 | -0.45346957 | -0.605463695 |
| Ism1 | 8.588686403 | 8.78256724 | 9.194243909 | -0.411676668 | -0.605557505 |
| Add2 | 5.110354739 | 5.519280832 | 5.71591352 | -0.196632688 | -0.605558781 |
| Crebrf | 9.111171807 | 9.441774926 | 9.716811345 | -0.275036419 | -0.605639538 |
| Fap | 7.081391662 | 7.073519525 | 7.687680284 | -0.614160759 | -0.606288622 |
| Mybpc1 | 11.52274205 | 11.67827839 | 12.12912324 | -0.450844843 | -0.606381183 |
| Snx32 | 5.629626349 | 5.890753919 | 6.236484275 | -0.345730355 | -0.606857925 |
| Aoc3 | 10.58516462 | 10.4814271 | 11.19228046 | -0.710853357 | -0.60711584 |
| Calm5 | 8.598043875 | 8.736409816 | 9.205739979 | -0.469330163 | -0.607696104 |

| | | | | | |
|---|---|---|---|---|---|
| Chpt1 | 10.58256541 | 10.43866488 | 11.19054156 | -0.75187668 | -0.607976153 |
| Ankrd29 | 6.779672165 | 7.102814402 | 7.387941601 | -0.285127199 | -0.608269436 |
| Srr | 7.703154366 | 7.963306228 | 8.311585761 | -0.348279533 | -0.608431395 |
| Acsbg1 | 11.30753057 | 11.75130404 | 11.91604715 | -0.164743107 | -0.60851658 |
| Lzts1 | 6.538794818 | 7.33406097 | 7.147817949 | 0.186243022 | -0.609023131 |
| Tle2 | 6.249119885 | 6.6201168 | 6.85837906 | -0.23826226 | -0.609259174 |
| Gm2026 | 6.589128233 | 6.662136184 | 7.200163637 | -0.538027453 | -0.611035404 |
| Ehbp1 | 8.096267466 | 8.572165115 | 8.708689874 | -0.136524759 | -0.612422409 |
| Stx2 | 7.628822956 | 8.06393827 | 8.241277396 | -0.177339126 | -0.61245444 |
| Pdzd2 | 11.34115887 | 11.83229645 | 11.95413735 | -0.121840897 | -0.612978477 |
| Cfh | 8.974115458 | 8.768168483 | 9.587383385 | -0.819214902 | -0.613267927 |
| Wnt7b | 9.540904048 | 10.15975109 | 10.15461055 | 0.005140543 | -0.613706502 |
| Agmo | 7.201065267 | 7.323746119 | 7.814859101 | -0.491112982 | -0.613793834 |
| Gm9830 | 5.10682655 | 5.577263007 | 5.720926342 | -0.143663335 | -0.614099792 |
| Upb1 | 6.082363237 | 6.592892547 | 6.697635899 | -0.104743352 | -0.615272662 |
| Adcy5 | 7.632327303 | 8.466195561 | 8.24806493 | 0.218130632 | -0.615737627 |
| Kantr | 7.064680381 | 7.578799426 | 7.681418602 | -0.102619176 | -0.616738221 |
| Mrc2 | 8.906596848 | 9.553965313 | 9.523569199 | 0.030396114 | -0.616972351 |
| Gm49601 | 6.826798781 | 7.577031702 | 7.444326653 | 0.132705049 | -0.617527873 |
| Acsl1 | 12.03269152 | 12.23381872 | 12.65043935 | -0.416620633 | -0.617747837 |
| Lmod3 | 8.247981893 | 8.309055594 | 8.865856075 | -0.556800482 | -0.617874183 |
| Pkib | 8.752013523 | 9.078279866 | 9.36988924 | -0.291609374 | -0.617875716 |
| Adcyap1r1 | 7.307931311 | 7.383934457 | 7.925999029 | -0.542064572 | -0.618067719 |
| Ano7 | 7.380581145 | 7.986610908 | 7.998816853 | -0.012205944 | -0.618235708 |
| Zfp395 | 8.637375577 | 9.064560043 | 9.256333372 | -0.191773329 | -0.618957795 |
| Thrb | 8.296286091 | 8.796827297 | 8.915297104 | -0.118469807 | -0.619011013 |
| Ptprb | 9.056965827 | 9.154855112 | 9.676814273 | -0.521959161 | -0.619848446 |
| Mrip-ps | 5.24737327 | 5.385218628 | 5.867548895 | -0.482330267 | -0.620175625 |
| Mup10 | 6.872224917 | 7.34647739 | 7.494602476 | -0.148125086 | -0.622377559 |
| Sord | 9.243532294 | 9.506115198 | 9.866286159 | -0.360170962 | -0.622753866 |
| Slc4a3 | 6.205290342 | 6.591749103 | 6.828137702 | -0.236388599 | -0.62284736 |
| Wnk2 | 8.8317745 | 9.539778632 | 9.454811534 | 0.084967098 | -0.623037034 |

| | | | | | |
|---|---|---|---|---|---|
| Phldb1 | 9.972786195 | 10.33435019 | 10.5959982 | -0.261648013 | -0.623212009 |
| Fabp4 | 13.12813254 | 12.9759122 | 13.75163505 | -0.775722857 | -0.623502513 |
| Kank3 | 7.604738742 | 7.850317131 | 8.230245173 | -0.379928041 | -0.62550643 |
| Ptgfr | 9.070080707 | 9.312937831 | 9.695693464 | -0.382755632 | -0.625612757 |
| Psd3 | 8.350428253 | 8.669403103 | 8.976523363 | -0.30712026 | -0.62609511 |
| Slc2a13 | 6.700902967 | 6.775296912 | 7.327307322 | -0.55201041 | -0.626404355 |
| Fancd2 | 6.564017022 | 6.838643131 | 7.191544135 | -0.352901004 | -0.627527113 |
| Atp6v1c2 | 10.03275335 | 10.59562371 | 10.66056038 | -0.064936673 | -0.627807032 |
| Plin4 | 12.15957101 | 12.68650533 | 12.7874029 | -0.100897573 | -0.627831887 |
| Rev3l | 9.138658624 | 9.644864599 | 9.766550078 | -0.121685478 | -0.627891454 |
| Lmod2 | 9.166416027 | 9.838710391 | 9.794341045 | 0.044369346 | -0.627925018 |
| Ttc3 | 10.3904578 | 10.6684929 | 11.02030258 | -0.35180968 | -0.629844773 |
| Gstt1 | 6.993125288 | 7.116921456 | 7.623024678 | -0.506103222 | -0.629899389 |
| Cd300ld5 | 6.338728899 | 6.858021153 | 6.968735703 | -0.110714549 | -0.630006804 |
| Clca2 | 10.39382963 | 10.95940791 | 11.02444749 | -0.065039583 | -0.630617863 |
| Plekha5 | 9.690521674 | 10.04099441 | 10.32117436 | -0.28017995 | -0.630652689 |
| Gpihbp1 | 6.75329216 | 6.749126172 | 7.384463006 | -0.635336834 | -0.631170846 |
| Il1f5 | 11.06778248 | 11.4082077 | 11.69960677 | -0.291399067 | -0.63182429 |
| Tmem38a | 10.5996169 | 10.82299591 | 11.23189686 | -0.408900953 | -0.632279959 |
| Dzip1 | 7.609319268 | 8.154772956 | 8.242069124 | -0.087296168 | -0.632749856 |
| Mtss1l | 7.74623447 | 8.374549991 | 8.379322611 | -0.00477262 | -0.633088141 |
| Atp1a2 | 12.18061941 | 12.50299513 | 12.81495247 | -0.311957333 | -0.634333057 |
| Filip1 | 7.189607585 | 7.299782819 | 7.824375942 | -0.524593123 | -0.634768357 |
| Zfp667 | 7.439560696 | 7.747257489 | 8.07433777 | -0.327080281 | -0.634777074 |
| Trim55 | 6.62069142 | 7.747735437 | 7.256134794 | 0.491600643 | -0.635443374 |
| Itga11 | 6.001177828 | 6.234608696 | 6.636917509 | -0.402308813 | -0.635739681 |
| Tsc22d3 | 11.38497612 | 11.84738222 | 12.02081889 | -0.173436673 | -0.635842767 |
| Sh3d21 | 8.715750698 | 9.309799437 | 9.352791568 | -0.042992131 | -0.63704087 |
| Otop3 | 6.849488576 | 7.582895753 | 7.487526295 | 0.095369458 | -0.638037719 |
| Hs3st6 | 6.91813873 | 7.77770403 | 7.556538409 | 0.221165621 | -0.638399679 |
| Grhl3 | 9.808933848 | 10.41207671 | 10.44734829 | -0.035271577 | -0.638414439 |
| Lgr5 | 7.807698604 | 8.187326683 | 8.446260233 | -0.25893355 | -0.638561629 |

| | | | | | |
|---|---|---|---|---|---|
| Aplnr | 6.615300752 | 6.790011427 | 7.254647549 | -0.464636122 | -0.639346797 |
| Il12rb2 | 8.17094119 | 8.899890558 | 8.810523754 | 0.089366805 | -0.639582564 |
| Pde3b | 8.433054346 | 8.229958066 | 9.073163934 | -0.843205868 | -0.640109588 |
| 2310007B03Rik | 7.840625333 | 8.622975381 | 8.480743272 | 0.142232109 | -0.640117939 |
| Gprc5b | 8.870911319 | 8.883145527 | 9.511067838 | -0.62792231 | -0.640156519 |
| Ucp3 | 6.494000676 | 6.71166896 | 7.134224879 | -0.422555919 | -0.640224203 |
| Aldh6a1 | 10.49129284 | 10.52226333 | 11.1317917 | -0.60952837 | -0.64049886 |
| Bche | 8.950601555 | 8.827485522 | 9.591186756 | -0.763701234 | -0.640585201 |
| Gm12912 | 5.491251451 | 5.967596797 | 6.133889407 | -0.16629261 | -0.642637956 |
| Gan | 11.33124278 | 11.93893014 | 11.97552308 | -0.03659294 | -0.6442803 |
| Igsf9 | 9.108571208 | 9.691958629 | 9.753475484 | -0.061516855 | -0.644904276 |
| Nfia | 10.16380641 | 10.48986412 | 10.8087664 | -0.318902283 | -0.644959993 |
| L1cam | 8.86356652 | 9.489965074 | 9.509776017 | -0.019810943 | -0.646209497 |
| P2ry4 | 6.94122491 | 7.32479606 | 7.588524241 | -0.263728181 | -0.647299331 |
| Cdc25b | 9.443949415 | 9.982098172 | 10.09235669 | -0.110258518 | -0.648407275 |
| Chd6 | 9.465727053 | 9.898603311 | 10.11442699 | -0.215823676 | -0.648699934 |
| Thbs3 | 8.175754288 | 8.329275799 | 8.826318859 | -0.49704306 | -0.650564571 |
| Ahnak2 | 12.0522512 | 12.60998762 | 12.70347684 | -0.09348922 | -0.651225647 |
| Mpdz | 9.180694532 | 9.494548196 | 9.832261713 | -0.337713518 | -0.651567182 |
| Wdr47 | 9.12155858 | 9.533913761 | 9.773201136 | -0.239287375 | -0.651642556 |
| Mfap2 | 6.9076226 | 7.457219623 | 7.559383798 | -0.102164175 | -0.651761198 |
| Slc5a3 | 9.304498888 | 10.01827037 | 9.956263006 | 0.062007369 | -0.651764117 |
| Ecm2 | 8.622795471 | 8.765014796 | 9.274572323 | -0.509557527 | -0.651776853 |
| Agtr1a | 5.509202933 | 5.698128752 | 6.164301143 | -0.466172391 | -0.65509821 |
| Spice1 | 7.456545239 | 7.978439864 | 8.112144076 | -0.133704212 | -0.655598836 |
| Hmcn2 | 8.337674259 | 8.559901352 | 8.994115415 | -0.434214064 | -0.656441156 |
| Smco1 | 6.093717081 | 5.954902805 | 6.750238068 | -0.795335263 | -0.656520987 |
| Aph1b | 7.97519022 | 8.400105526 | 8.63178245 | -0.231676924 | -0.656592229 |
| Oxtr | 7.330701517 | 8.036468467 | 7.987554817 | 0.04891365 | -0.6568533 |
| Myh2 | 10.75247833 | 10.81476263 | 11.41112661 | -0.59636398 | -0.65864828 |
| Pank3 | 10.59944913 | 10.87996629 | 11.25814235 | -0.378176063 | -0.658693217 |
| Vegfd | 8.123195861 | 8.452708676 | 8.782199279 | -0.329490603 | -0.659003418 |

| | | | | | |
|---|---|---|---|---|---|
| Pgap1 | 9.382339512 | 9.938109411 | 10.04201457 | -0.103905163 | -0.659675062 |
| Wnt10b | 5.894531552 | 6.300036151 | 6.554233475 | -0.254197324 | -0.659701923 |
| Ankef1 | 6.638751122 | 6.769340433 | 7.298670755 | -0.529330322 | -0.659919633 |
| Olfml2a | 7.067693916 | 7.423902377 | 7.728081353 | -0.304178976 | -0.660387437 |
| Rasgrf2 | 6.8938644 | 7.11188603 | 7.555013 | -0.443126969 | -0.6611486 |
| Cdh19 | 7.025280124 | 7.167587354 | 7.686813334 | -0.51922598 | -0.661533209 |
| Tob1 | 10.07672637 | 10.50236909 | 10.73930809 | -0.236939 | -0.662581713 |
| Abhd14b | 8.103960371 | 8.349990958 | 8.767987351 | -0.417996394 | -0.664026981 |
| Igfbp3 | 11.36405788 | 11.32470866 | 12.0282113 | -0.70350264 | -0.66415342 |
| Col3a1 | 14.22743966 | 14.51801299 | 14.8921756 | -0.374162613 | -0.66473594 |
| Fam171b | 6.624391102 | 7.105992021 | 7.290107412 | -0.184115391 | -0.66571631 |
| Htra3 | 10.52750952 | 11.02479755 | 11.19361822 | -0.168820677 | -0.6661087 |
| Skint8 | 6.633318595 | 6.976825229 | 7.300754929 | -0.3239297 | -0.667436334 |
| Pnpla3 | 8.419088213 | 8.89680298 | 9.088472438 | -0.191669458 | -0.669384226 |
| Rtn4rl1 | 7.168326277 | 7.581201158 | 7.838174761 | -0.256973602 | -0.669848484 |
| C1qtnf7 | 6.976649557 | 7.209476867 | 7.646671872 | -0.437195006 | -0.670022316 |
| Rgmb | 10.23467724 | 10.91472361 | 10.90482011 | 0.0099035 | -0.670142863 |
| Ebf2 | 5.876491273 | 5.898332298 | 6.546881824 | -0.648549526 | -0.670390551 |
| Lca5 | 5.787828107 | 6.024759964 | 6.45850245 | -0.433742486 | -0.670674344 |
| Atp12a | 8.760262812 | 9.62740486 | 9.431151489 | 0.196253372 | -0.670888677 |
| Bmf | 7.586909756 | 7.940654209 | 8.2588058 | -0.318151591 | -0.671896044 |
| Pcdh18 | 6.540654309 | 6.66646842 | 7.214642987 | -0.548174566 | -0.673988678 |
| Zfp185 | 9.295842643 | 9.92059763 | 9.972401181 | -0.051803551 | -0.676558538 |
| Ccr4 | 7.634285471 | 8.290422203 | 8.310858109 | -0.020435906 | -0.676572638 |
| Cbx6 | 8.741023852 | 9.273360993 | 9.418516875 | -0.145155883 | -0.677493023 |
| Cyp27a1 | 8.255994435 | 8.526197129 | 8.933732024 | -0.407534895 | -0.67773759 |
| Ccng2 | 8.667512766 | 9.06421869 | 9.345261227 | -0.281042537 | -0.677748461 |
| Ppp1r12b | 10.07224551 | 10.26890371 | 10.75060451 | -0.481700807 | -0.678359 |
| Mrgprb2 | 6.97500994 | 7.5048775 | 7.653460454 | -0.148582954 | -0.678450514 |
| Itih5 | 10.06102743 | 9.91849099 | 10.73991158 | -0.821420593 | -0.678884151 |
| Sh3d19 | 11.06352941 | 11.55732715 | 11.74333492 | -0.186007777 | -0.679805517 |
| Cmbl | 7.562756962 | 7.645682033 | 8.242825596 | -0.597143563 | -0.680068633 |

| | | | | | |
|---|---|---|---|---|---|
| Abcb1a | 6.628900477 | 6.682349171 | 7.310752383 | -0.628403212 | -0.681851906 |
| Mlf1 | 8.828478085 | 8.988521354 | 9.510554181 | -0.522032827 | -0.682076096 |
| Camkk1 | 8.862656267 | 9.421371297 | 9.545154632 | -0.123783335 | -0.682498365 |
| Fam117a | 7.398624119 | 7.863574622 | 8.081855884 | -0.218281262 | -0.683231765 |
| Glb1l2 | 9.211966869 | 9.206647947 | 9.895485143 | -0.688837196 | -0.683518274 |
| Lama2 | 9.477621685 | 9.497518678 | 10.161421 | -0.663902322 | -0.683799315 |
| Nat8f3 | 5.651776563 | 6.030208185 | 6.336914181 | -0.306705996 | -0.685137618 |
| Fam13c | 7.00476535 | 7.495664826 | 7.691670176 | -0.19600535 | -0.686904826 |
| Exph5 | 8.721093158 | 9.174838089 | 9.409318679 | -0.23448059 | -0.688225521 |
| Spink7 | 4.649412699 | 4.701853268 | 5.33777612 | -0.635922851 | -0.688363421 |
| Cpt1a | 9.972747101 | 10.58062994 | 10.66129759 | -0.080667653 | -0.688550492 |
| Eepd1 | 8.735539324 | 8.78008515 | 9.424116679 | -0.644031529 | -0.688577355 |
| Hoxc6 | 6.568559984 | 7.007803346 | 7.258018373 | -0.250215028 | -0.689458389 |
| Pparg | 8.755712576 | 9.145252456 | 9.445361374 | -0.300108917 | -0.689648798 |
| Serpinb12 | 6.658641015 | 6.798310326 | 7.348748689 | -0.550438363 | -0.690107674 |
| Aox3 | 5.090104807 | 5.526012392 | 5.780723497 | -0.254711105 | -0.69061869 |
| Sucnr1 | 5.61816983 | 5.852493376 | 6.309404398 | -0.456911022 | -0.691234568 |
| Myzap | 7.865741831 | 8.287592363 | 8.5571497 | -0.269557337 | -0.691407869 |
| Ppargc1a | 6.128980892 | 7.402708188 | 6.821020104 | 0.581688084 | -0.692039212 |
| Adam22 | 4.974551914 | 5.167833527 | 5.667701359 | -0.499867832 | -0.693149445 |
| Slc9a3r2 | 7.575869154 | 8.164454411 | 8.269189713 | -0.104735302 | -0.693320559 |
| Hcar1 | 6.079404214 | 6.447458015 | 6.773014514 | -0.325556499 | -0.6936103 |
| Gpr1 | 4.803072999 | 5.107934712 | 5.497843551 | -0.389908838 | -0.694770552 |
| Lrrc39 | 6.664827199 | 6.872753152 | 7.360548793 | -0.487795641 | -0.695721594 |
| Myh4 | 15.60881469 | 16.04520061 | 16.30551191 | -0.2603113 | -0.696697223 |
| Fhl1 | 12.14210415 | 12.53005159 | 12.84068066 | -0.31062907 | -0.69857651 |
| Ramp2 | 7.589223333 | 7.809262257 | 8.287956812 | -0.478694555 | -0.69873348 |
| Clca3a1 | 8.425061818 | 9.017602412 | 9.12497057 | -0.107368157 | -0.699908751 |
| Map3k12 | 6.59460713 | 7.179071525 | 7.295237824 | -0.116166299 | -0.700630693 |
| Alpk3 | 8.034475395 | 8.47109425 | 8.735885641 | -0.264791391 | -0.701410246 |
| Acss3 | 7.294871749 | 7.169773598 | 7.997620407 | -0.827846809 | -0.702748659 |
| Ppp2r3a | 10.78329963 | 11.0834346 | 11.48690825 | -0.403473653 | -0.70360862 |

| | | | | | |
|---|---|---|---|---|---|
| Gdf10 | 5.932198229 | 6.098998821 | 6.63780813 | -0.538809309 | -0.705609901 |
| Zbtb16 | 9.338629241 | 9.806982478 | 10.04568371 | -0.238701229 | -0.707054466 |
| Tril | 6.675387682 | 7.087496336 | 7.383217178 | -0.295720842 | -0.707829496 |
| Pygo1 | 7.133055025 | 7.43338333 | 7.841163243 | -0.407779913 | -0.708108218 |
| Tanc1 | 11.59001639 | 12.08066792 | 12.30006229 | -0.219394367 | -0.710045893 |
| Coro6 | 7.715335078 | 7.861633257 | 8.425451125 | -0.563817868 | -0.710116047 |
| Fcrls | 8.191229859 | 8.097303427 | 8.902318284 | -0.805014857 | -0.711088425 |
| Phtf2 | 9.346397802 | 9.48066197 | 10.05886816 | -0.578206188 | -0.712470356 |
| Fat4 | 6.776631911 | 7.07068619 | 7.489631112 | -0.418944922 | -0.712999201 |
| Tfdp2 | 8.588239171 | 8.894825672 | 9.30273376 | -0.407908088 | -0.714494589 |
| Ank1 | 8.9212459 | 9.213236869 | 9.63644371 | -0.42320684 | -0.715197809 |
| Folr2 | 8.396937111 | 8.049928993 | 9.112302197 | -1.062373204 | -0.715365086 |
| Pcolce2 | 9.334653213 | 9.639409661 | 10.05058289 | -0.411173229 | -0.715929677 |
| Nrg4 | 7.517809135 | 7.598565772 | 8.233874532 | -0.63530876 | -0.716065398 |
| Timp4 | 8.373922372 | 8.244514628 | 9.090491547 | -0.845976919 | -0.716569175 |
| Phka1 | 9.881747078 | 9.904731485 | 10.59924414 | -0.694512659 | -0.717497065 |
| Col4a5 | 8.018598302 | 8.41760844 | 8.73614515 | -0.318536709 | -0.717546847 |
| Cys1 | 6.173272917 | 6.49166667 | 6.891272725 | -0.399606055 | -0.717999808 |
| Myrip | 5.242672112 | 5.640921346 | 5.962894086 | -0.32197274 | -0.720221974 |
| Zbtb20 | 10.76554164 | 11.11943442 | 11.48636682 | -0.366932393 | -0.720825177 |
| Ar | 9.405623175 | 9.747158917 | 10.12687431 | -0.37971539 | -0.721251132 |
| Cyp2d22 | 6.352250605 | 6.609724616 | 7.073713233 | -0.463988617 | -0.721462628 |
| Adcy6 | 9.382854899 | 9.931207918 | 10.10555902 | -0.174351102 | -0.722704121 |
| Lgals12 | 8.372631896 | 8.60925749 | 9.095589387 | -0.486331898 | -0.722957491 |
| Appl2 | 9.60506949 | 10.04786945 | 10.32833086 | -0.280461417 | -0.723261373 |
| Etv1 | 6.794472009 | 7.174961438 | 7.518522735 | -0.343561297 | -0.724050726 |
| Acbd4 | 8.64918039 | 9.27341204 | 9.373610188 | -0.100198148 | -0.724429798 |
| Sparcl1 | 10.15817557 | 10.22161743 | 10.88465126 | -0.663033837 | -0.72647569 |
| Asah2 | 10.04923979 | 10.46835043 | 10.77602845 | -0.307678023 | -0.726788656 |
| Myoc | 7.55211579 | 7.591351679 | 8.280655663 | -0.689303984 | -0.728539873 |
| Col5a1 | 9.93127869 | 10.3216825 | 10.66058673 | -0.338904223 | -0.729308037 |
| Dpep1 | 9.177151256 | 9.345486751 | 9.906739546 | -0.561252795 | -0.72958829 |

| | | | | | |
|---|---|---|---|---|---|
| Txndc16 | 8.380621862 | 8.651349398 | 9.111998148 | -0.46064875 | -0.731376286 |
| Ppm1l | 8.698171073 | 8.806842575 | 9.429734968 | -0.622892392 | -0.731563895 |
| Nuak1 | 9.122735049 | 9.818299175 | 9.8555491 | -0.037249925 | -0.732814051 |
| Gm15429 | 5.816980015 | 5.90375212 | 6.549996766 | -0.646244646 | -0.733016751 |
| 4930438A08Rik | 8.447729769 | 8.99057608 | 9.181858684 | -0.191282605 | -0.734128915 |
| Suox | 8.04377074 | 8.590972943 | 8.78000554 | -0.189032596 | -0.7362348 |
| Mill1 | 8.570163243 | 9.226346967 | 9.306573298 | -0.080226331 | -0.736410055 |
| Stard9 | 7.927346359 | 8.318770189 | 8.66486071 | -0.346090521 | -0.737514351 |
| Mfap3l | 9.949856626 | 10.50246452 | 10.68822499 | -0.18576047 | -0.738368367 |
| Castor2 | 8.553510752 | 8.976260449 | 9.291935106 | -0.315674657 | -0.738424354 |
| Agl | 10.45043871 | 10.47206806 | 11.18913021 | -0.717062157 | -0.738691506 |
| Cttnbp2 | 8.515645987 | 9.040233707 | 9.255059539 | -0.214825832 | -0.739413553 |
| Car3 | 14.3705345 | 14.60770392 | 15.10997655 | -0.502272623 | -0.73944205 |
| Adamts12 | 7.350638691 | 7.52024598 | 8.090338682 | -0.570092701 | -0.73969999 |
| Fcna | 6.193194356 | 5.864122145 | 6.933856223 | -1.069734078 | -0.740661867 |
| Dok4 | 7.897997165 | 8.731683807 | 8.638971975 | 0.092711832 | -0.740974809 |
| Prkag3 | 6.267375146 | 6.494642359 | 7.008717419 | -0.514075061 | -0.741342273 |
| Vmac | 7.226442805 | 7.603191408 | 7.969760411 | -0.366569003 | -0.743317606 |
| Cep85l | 8.816418565 | 8.541872149 | 9.560161885 | -1.018289736 | -0.74374332 |
| Nos1 | 7.655800798 | 8.095030021 | 8.400007748 | -0.304977727 | -0.74420695 |
| Jph1 | 8.698401448 | 8.805322798 | 9.444730942 | -0.639408144 | -0.746329494 |
| Sec16b | 6.189035434 | 6.611427644 | 6.936345045 | -0.324917401 | -0.747309611 |
| Ndrg4 | 7.372240313 | 8.052756924 | 8.11989925 | -0.067142326 | -0.747658937 |
| Cc2d2a | 7.891837734 | 8.298132012 | 8.639570873 | -0.341438861 | -0.747733139 |
| Reps2 | 6.787227717 | 6.883130923 | 7.53643798 | -0.653307057 | -0.749210263 |
| Igfbp5 | 12.0380718 | 12.23264354 | 12.78878044 | -0.556136897 | -0.750708633 |
| Ttc28 | 8.173307263 | 8.736450144 | 8.92525333 | -0.188803186 | -0.751946067 |
| Casp14 | 10.65655589 | 11.06356782 | 11.40873577 | -0.345167953 | -0.752179883 |
| Tmem233 | 7.742795043 | 7.954126533 | 8.497162326 | -0.543035793 | -0.754367282 |
| Ace | 9.210146012 | 9.228266437 | 9.965471901 | -0.737205464 | -0.75532589 |
| Srl | 11.19346167 | 11.50934399 | 11.94981552 | -0.440471523 | -0.756353847 |
| Adgra3 | 10.3822526 | 10.82179016 | 11.13909365 | -0.31730349 | -0.75684105 |

| | | | | | |
|---|---|---|---|---|---|
| Alox12e | 10.21726348 | 11.19833134 | 10.97560413 | 0.22272721 | -0.75834065 |
| Gm973 | 7.637404887 | 8.066596018 | 8.396452576 | -0.329856557 | -0.759047688 |
| Ryr1 | 12.0212745 | 12.47166723 | 12.78216031 | -0.310493087 | -0.760885813 |
| Lpl | 13.04512966 | 12.8629565 | 13.80823735 | -0.945280853 | -0.763107697 |
| Sfrp4 | 6.463266192 | 6.171469324 | 7.226889607 | -1.055420283 | -0.763623415 |
| Klhl38 | 4.468949651 | 4.675674846 | 5.234235687 | -0.558560841 | -0.765286036 |
| Slc22a4 | 8.435710259 | 9.155471223 | 9.202437199 | -0.046965976 | -0.76672694 |
| Pdpr | 8.676082089 | 9.03446102 | 9.444915238 | -0.410454218 | -0.768833149 |
| Glcci1 | 6.648526546 | 7.123709842 | 7.420347604 | -0.296637762 | -0.771821058 |
| Xirp2 | 12.2717128 | 13.24744431 | 13.04517115 | 0.20227316 | -0.77345835 |
| Abca8b | 7.590596787 | 7.889135821 | 8.364319275 | -0.475183454 | -0.773722489 |
| Col14a1 | 8.652348696 | 8.675988896 | 9.426779794 | -0.750790897 | -0.774431098 |
| Irs1 | 9.070612695 | 9.551762525 | 9.845752299 | -0.293989774 | -0.775139603 |
| Abcd2 | 9.765587134 | 9.527470343 | 10.5413764 | -1.013906053 | -0.775789263 |
| Sspn | 8.749941443 | 8.80759765 | 9.526270655 | -0.718673005 | -0.776329211 |
| Flg2 | 13.87075508 | 14.50489597 | 14.64833654 | -0.143440573 | -0.777581467 |
| Myct1 | 5.380542057 | 5.791188155 | 6.159006768 | -0.367818613 | -0.778464711 |
| Lrrc14b | 6.82653979 | 7.205551522 | 7.605200003 | -0.399648481 | -0.778660212 |
| Prkaa2 | 9.09506888 | 9.022624256 | 9.873780672 | -0.851156416 | -0.778711791 |
| Mn1 | 7.39168476 | 7.723072506 | 8.170924608 | -0.447852103 | -0.779239848 |
| Arhgef10 | 8.394935742 | 9.021457505 | 9.175848368 | -0.154390863 | -0.780912626 |
| Ogn | 9.487433396 | 9.414771863 | 10.26840541 | -0.853633547 | -0.780972014 |
| Tmod2 | 8.448263012 | 8.74096098 | 9.229432615 | -0.488471635 | -0.781169603 |
| Dpp4 | 8.757652899 | 9.014066967 | 9.542200721 | -0.528133754 | -0.784547822 |
| Neu2 | 8.124185002 | 8.274866003 | 8.909103967 | -0.634237964 | -0.784918965 |
| Ptpn14 | 10.44285955 | 11.1215133 | 11.22827144 | -0.106758133 | -0.785411887 |
| Cyp39a1 | 7.612695489 | 8.113537104 | 8.39820641 | -0.284669306 | -0.785510921 |
| Ccdc141 | 5.877363043 | 6.223866607 | 6.663083232 | -0.439216625 | -0.785720189 |
| Prkg2 | 5.079856255 | 5.22964612 | 5.868202459 | -0.638556339 | -0.788346204 |
| Sgcg | 8.349006154 | 8.417808026 | 9.138671712 | -0.720863686 | -0.789665558 |
| Ell3 | 7.412713183 | 7.74926645 | 8.202808717 | -0.453542267 | -0.790095535 |
| Zfp521 | 6.457214041 | 6.969845844 | 7.24848616 | -0.278640315 | -0.791272118 |

| | | | | | |
|---|---|---|---|---|---|
| Cdo1 | 10.2981347 | 10.01398037 | 11.09128552 | -1.077305153 | -0.793150821 |
| Cyp2j9 | 6.744016228 | 7.185993417 | 7.537290266 | -0.351296849 | -0.793274038 |
| Cidec | 11.08879969 | 11.2389831 | 11.88246271 | -0.643479613 | -0.79366302 |
| Hmcn1 | 8.42494817 | 8.712880803 | 9.218968265 | -0.506087462 | -0.794020096 |
| Pcmtd2 | 9.199933115 | 9.73710909 | 9.994160709 | -0.257051619 | -0.794227594 |
| Rgs5 | 9.907120209 | 10.01501824 | 10.7038748 | -0.688856559 | -0.796754591 |
| Capn3 | 7.699072875 | 7.940704977 | 8.49696917 | -0.556264192 | -0.797896294 |
| Mxd4 | 10.76338786 | 11.44296114 | 11.56145448 | -0.11849334 | -0.798066623 |
| Ophn1 | 8.165563851 | 8.594779913 | 8.963782133 | -0.36900222 | -0.798218282 |
| Rgs7bp | 6.143172255 | 6.439642934 | 6.94168799 | -0.502045056 | -0.798515736 |
| Cygb | 8.233197293 | 8.345797106 | 9.032387251 | -0.686590145 | -0.799189958 |
| Tmem119 | 6.826552936 | 7.421833039 | 7.625932045 | -0.204099006 | -0.799379109 |
| Klhl33 | 8.798498615 | 8.488015873 | 9.599928685 | -1.111912812 | -0.80143007 |
| Map1a | 7.343205391 | 7.969263959 | 8.145712531 | -0.176448572 | -0.80250714 |
| Gpr34 | 5.343862921 | 5.315877839 | 6.147264828 | -0.831386989 | -0.803401907 |
| Ldb2 | 6.692497406 | 7.082143119 | 7.498134079 | -0.41599096 | -0.805636673 |
| Slc27a1 | 9.898521335 | 10.3569014 | 10.70454737 | -0.347645967 | -0.806026035 |
| Angptl7 | 7.888383644 | 8.450399528 | 8.695907625 | -0.245508097 | -0.807523981 |
| Camk2a | 8.966927481 | 9.27371255 | 9.775837916 | -0.502125366 | -0.808910435 |
| Gucy1b1 | 5.891465159 | 6.231074319 | 6.700784091 | -0.469709772 | -0.809318931 |
| Mmp11 | 6.202300113 | 6.628198786 | 7.012019934 | -0.383821148 | -0.809719821 |
| Aff3 | 8.126007213 | 8.738978247 | 8.935766788 | -0.196788541 | -0.809759576 |
| Fsd2 | 8.329609094 | 8.507213744 | 9.139742763 | -0.632529019 | -0.81013367 |
| Flt3l | 7.855543591 | 8.50284228 | 8.665945507 | -0.163103227 | -0.810401915 |
| Islr | 7.015075838 | 7.018779714 | 7.826756919 | -0.807977205 | -0.811681081 |
| Abcc9 | 8.870431652 | 9.011926479 | 9.683986234 | -0.672059755 | -0.813554583 |
| Rasgrp3 | 6.521190504 | 6.807922171 | 7.336978102 | -0.529055931 | -0.815787598 |
| Pamr1 | 7.356866567 | 7.32448573 | 8.172982955 | -0.848497225 | -0.816116388 |
| Robo1 | 8.640131501 | 9.423422739 | 9.458452811 | -0.035030072 | -0.81832131 |
| Klf15 | 7.174523752 | 7.492056146 | 7.993703682 | -0.501647537 | -0.81917993 |
| Ddit4l | 7.976787629 | 8.18998769 | 8.796825309 | -0.606837619 | -0.82003768 |
| Jam2 | 8.225446069 | 8.595334371 | 9.047177338 | -0.451842967 | -0.821731269 |

| | | | | | |
|---|---|---|---|---|---|
| Slc2a12 | 8.466514314 | 9.213473503 | 9.293303373 | -0.07982987 | -0.826789058 |
| Ushbp1 | 6.395264681 | 6.795621178 | 7.223576925 | -0.427955747 | -0.828312244 |
| Sh3rf2 | 9.622909359 | 10.40656515 | 10.45200387 | -0.045438723 | -0.829094514 |
| H60c | 8.835387935 | 9.381772748 | 9.668025652 | -0.286252904 | -0.832637717 |
| Aqp7 | 6.316068339 | 6.332525835 | 7.152168834 | -0.819642999 | -0.836100495 |
| Sostdc1 | 8.877439896 | 9.418384691 | 9.71469725 | -0.296312559 | -0.837257354 |
| Arhgap24 | 8.569187158 | 9.139902974 | 9.408081029 | -0.268178054 | -0.838893871 |
| Flg | 12.20499423 | 12.96952331 | 13.04395462 | -0.074431313 | -0.83896039 |
| Scn4a | 9.238373943 | 9.3119154 | 10.07772043 | -0.765805027 | -0.839346485 |
| Fam198b | 8.018585878 | 8.621810977 | 8.862984753 | -0.241173776 | -0.844398875 |
| Aox1 | 7.527432744 | 7.69703122 | 8.374565022 | -0.677533802 | -0.847132278 |
| Rragd | 8.86499491 | 9.135480605 | 9.713293703 | -0.577813097 | -0.848298793 |
| Mme | 8.407694696 | 8.760320802 | 9.25975956 | -0.499438758 | -0.852064864 |
| Slc2a4 | 9.55201997 | 9.561569497 | 10.40622841 | -0.844658917 | -0.854208443 |
| Chdh | 9.291743201 | 9.996516361 | 10.14687531 | -0.150358946 | -0.855132105 |
| Bbox1 | 10.01434213 | 10.6293436 | 10.87176446 | -0.24242086 | -0.857422326 |
| Gxylt2 | 7.994704199 | 8.503961209 | 8.8523432 | -0.348381991 | -0.857639001 |
| Clca3a2 | 12.13179198 | 13.07676313 | 12.99170276 | 0.085060373 | -0.859910783 |
| Sult1a1 | 8.416631063 | 8.241372074 | 9.276847252 | -1.035475179 | -0.860216189 |
| Fzd4 | 10.51444145 | 10.21475295 | 11.37478967 | -1.160036723 | -0.86034822 |
| Heph | 8.458319408 | 8.989515399 | 9.320431736 | -0.330916336 | -0.862112328 |
| Il20rb | 10.93300277 | 11.59130835 | 11.79944951 | -0.208141163 | -0.866446743 |
| Mmd | 9.836801034 | 10.01544298 | 10.70390935 | -0.688466367 | -0.867108316 |
| Lrrn4cl | 7.205702496 | 7.250583473 | 8.078097148 | -0.827513675 | -0.872394652 |
| Pkia | 9.9461446 | 9.997778059 | 10.81934952 | -0.821571458 | -0.873204917 |
| Clec14a | 5.67872424 | 5.900820635 | 6.55854738 | -0.657726745 | -0.87982314 |
| Afap1l2 | 8.878062427 | 9.091430811 | 9.75954376 | -0.668112949 | -0.881481333 |
| Meox2 | 7.207988094 | 7.543198969 | 8.092320688 | -0.549121719 | -0.884332594 |
| Tef | 11.38835075 | 11.85684057 | 12.27320012 | -0.41635955 | -0.884849367 |
| Mt4 | 7.190673733 | 7.30295294 | 8.07570552 | -0.77275258 | -0.885031788 |
| Retnla | 9.691775306 | 9.468428676 | 10.57683309 | -1.108404417 | -0.885057787 |
| Six4 | 4.979962302 | 5.333925373 | 5.865222305 | -0.531296933 | -0.885260003 |

| | | | | | |
|---|---|---|---|---|---|
| Fmo1 | 8.071899458 | 8.2886566 | 8.957192684 | -0.668536084 | -0.885293226 |
| Palm | 8.380800487 | 8.993022276 | 9.2661246 | -0.273102324 | -0.885324112 |
| Aqp4 | 7.409285239 | 7.219648365 | 8.295016243 | -1.075367878 | -0.885731004 |
| Nrep | 9.182041164 | 9.465241235 | 10.06790945 | -0.602668216 | -0.885868287 |
| Sema6c | 6.760880699 | 7.232555943 | 7.64740596 | -0.414850016 | -0.886525261 |
| Neb | 12.44567651 | 12.85350369 | 13.3335497 | -0.480046017 | -0.88787319 |
| Stard8 | 8.130705697 | 8.563721027 | 9.018955457 | -0.45523443 | -0.88824976 |
| Adamts14 | 7.742924928 | 8.640488975 | 8.631429924 | 0.009059052 | -0.888504996 |
| Cd209g | 6.847804879 | 6.836290384 | 7.7369868 | -0.900696416 | -0.889181922 |
| Cpm | 9.209667219 | 9.981909969 | 10.09955441 | -0.117644446 | -0.889887195 |
| Tmeff2 | 5.964129286 | 6.216032967 | 6.85464771 | -0.638614743 | -0.890518425 |
| Slco2b1 | 9.009394792 | 9.510406826 | 9.900750077 | -0.390343251 | -0.891355284 |
| Selenbp1 | 8.380130541 | 8.488077264 | 9.271892994 | -0.78381573 | -0.891762453 |
| Dnase1l2 | 9.03518049 | 9.642117925 | 9.928774778 | -0.286656853 | -0.893594288 |
| Mrgprb11-ps | 8.182065341 | 8.497008523 | 9.076649693 | -0.57964117 | -0.894584353 |
| Dbp | 9.4532108 | 9.770092027 | 10.34791862 | -0.577826596 | -0.894707824 |
| Klhl24 | 9.37858469 | 9.804634112 | 10.27410066 | -0.469466544 | -0.895515967 |
| Adgrl4 | 7.054517948 | 7.392499938 | 7.953092395 | -0.560592457 | -0.898574448 |
| Scara5 | 9.784049873 | 10.01345373 | 10.68331892 | -0.669865192 | -0.899269044 |
| AI464131 | 8.225612632 | 8.577829916 | 9.126396603 | -0.548566687 | -0.900783971 |
| Dlg2 | 8.1499163 | 8.823092474 | 9.054077898 | -0.230985424 | -0.904161598 |
| Srcin1 | 8.993244709 | 10.21593994 | 9.898405923 | 0.317534016 | -0.905161214 |
| Pxdn | 9.451568897 | 9.78818483 | 10.35756851 | -0.569383683 | -0.905999616 |
| Skint4 | 8.555312988 | 9.153131284 | 9.461379874 | -0.30824859 | -0.906066886 |
| Zfhx4 | 6.798389026 | 6.781958735 | 7.712376625 | -0.93041789 | -0.913987599 |
| Ivl | 10.70378158 | 11.3224863 | 11.6183874 | -0.295901097 | -0.914605813 |
| Chp2 | 6.237607121 | 6.507073255 | 7.152861218 | -0.645787963 | -0.915254097 |
| Rpgrip1l | 6.755586874 | 7.707386123 | 7.672004162 | 0.035381961 | -0.916417288 |
| Myot | 10.22633462 | 10.32563589 | 11.14815033 | -0.822514441 | -0.921815711 |
| Plscr4 | 7.198011121 | 7.604321338 | 8.12044357 | -0.516122232 | -0.922432449 |
| Btnl9 | 5.480020611 | 5.74161345 | 6.405012283 | -0.663398832 | -0.924991671 |
| Foxn3 | 10.4484632 | 11.19160989 | 11.3737788 | -0.182168913 | -0.925315597 |

| | | | | | |
|---|---|---|---|---|---|
| Prkar2b | 8.780396122 | 8.787702213 | 9.714031903 | -0.92632969 | -0.933635781 |
| Adh1 | 8.53718373 | 8.663564301 | 9.472478793 | -0.808914492 | -0.935295063 |
| Cdon | 8.675935469 | 9.317880355 | 9.620860016 | -0.302979661 | -0.944924547 |
| Synpo2l | 7.694084457 | 8.505373868 | 8.639562173 | -0.134188305 | -0.945477716 |
| Npy1r | 5.338120914 | 5.58487027 | 6.286995292 | -0.702125022 | -0.948874378 |
| Cd300lg | 7.570803531 | 7.798009909 | 8.520064533 | -0.722054624 | -0.949261002 |
| Lpin1 | 9.650426413 | 9.784934558 | 10.60202465 | -0.817090089 | -0.951598233 |
| Ano5 | 8.358218721 | 8.371655075 | 9.311695151 | -0.940040077 | -0.95347643 |
| Sfrp2 | 5.561955471 | 5.568318929 | 6.516432615 | -0.948113686 | -0.954477144 |
| Abcc5 | 9.587515871 | 10.61539987 | 10.54449072 | 0.070909147 | -0.956974849 |
| Clcn1 | 7.552037913 | 7.855015431 | 8.513031369 | -0.658015938 | -0.960993455 |
| Galnt15 | 8.884958447 | 8.72523004 | 9.847200339 | -1.121970299 | -0.962241892 |
| Mettl7a1 | 9.340173337 | 9.98526985 | 10.30268655 | -0.3174167 | -0.962513213 |
| Cd248 | 8.862796152 | 9.52148665 | 9.828219528 | -0.306732878 | -0.965423376 |
| Lipe | 10.35649729 | 10.7580143 | 11.32459574 | -0.56658144 | -0.968098453 |
| Mxra8 | 8.809106974 | 8.847607676 | 9.779692248 | -0.932084572 | -0.970585274 |
| Sox6 | 8.786134079 | 9.264144227 | 9.758261304 | -0.494117077 | -0.972127225 |
| Thrsp | 9.899276025 | 9.074188001 | 10.87688603 | -1.802698025 | -0.977610001 |
| Cdkn2c | 8.726426331 | 8.675220352 | 9.704352682 | -1.02913233 | -0.977926351 |
| Sesn1 | 9.255784477 | 9.968226114 | 10.24089853 | -0.272672412 | -0.985114049 |
| Adrb3 | 7.542440414 | 8.054499629 | 8.53056724 | -0.47606761 | -0.988126826 |
| Fam180a | 6.76451031 | 7.430931914 | 7.757887383 | -0.326955468 | -0.993377072 |
| Txlnb | 9.316257614 | 9.660633085 | 10.31283047 | -0.652197385 | -0.996572856 |
| Kcna2 | 7.445050473 | 7.899247885 | 8.441974668 | -0.542726783 | -0.996924195 |
| Trp53inp1 | 9.382582543 | 10.06525125 | 10.37959415 | -0.314342902 | -0.997011611 |
| Pkhd1l1 | 5.012000228 | 5.477766084 | 6.010230102 | -0.532464018 | -0.998229874 |
| Lrrc30 | 4.717909567 | 5.079538431 | 5.720927883 | -0.641389451 | -1.003018316 |
| Fam214a | 8.917085911 | 9.416185167 | 9.920779454 | -0.504594287 | -1.003693543 |
| Frem2 | 7.504803542 | 8.519991177 | 8.511447329 | 0.008543848 | -1.006643787 |
| Asb15 | 6.088299739 | 6.050922808 | 7.095163894 | -1.044241086 | -1.006864155 |
| Klhl31 | 8.572621389 | 8.649010304 | 9.579773822 | -0.930763518 | -1.007152433 |
| Lims2 | 6.966487825 | 7.257807087 | 7.974299472 | -0.716492385 | -1.007811647 |

| | | | | | |
|---|---|---|---|---|---|
| Nova1 | 6.806984749 | 7.269777471 | 7.825263465 | -0.555485994 | -1.018278716 |
| Hlf | 8.692499577 | 9.202165814 | 9.710803556 | -0.508637741 | -1.018303978 |
| Irf4 | 7.279786853 | 7.938486513 | 8.299682014 | -0.361195501 | -1.019895161 |
| Hmgcs2 | 7.356313203 | 7.929615854 | 8.377607905 | -0.447992051 | -1.021294702 |
| Reck | 8.609510682 | 9.048547381 | 9.633686804 | -0.585139422 | -1.024176122 |
| Osr2 | 6.75936293 | 7.332385905 | 7.784411967 | -0.452026062 | -1.025049037 |
| Tek | 7.441154357 | 7.700116995 | 8.474690226 | -0.774573231 | -1.033535869 |
| Peg3 | 8.028554924 | 8.767462261 | 9.062392765 | -0.294930503 | -1.033837841 |
| Sema3d | 9.186260414 | 9.606868561 | 10.22091259 | -0.614044027 | -1.034652174 |
| Rps12l1 | 10.79067911 | 8.468640521 | 11.828029 | -3.359388476 | -1.037349885 |
| Amot | 7.857061065 | 8.589691578 | 8.895850238 | -0.306158659 | -1.038789173 |
| Selenop | 12.12441712 | 12.13873712 | 13.17089332 | -1.032156203 | -1.046476203 |
| Per3 | 9.171494396 | 9.754429708 | 10.21861699 | -0.464187285 | -1.047122597 |
| Abca6 | 5.880847051 | 6.140901453 | 6.933401553 | -0.7925001 | -1.052554502 |
| Apcdd1 | 10.85175324 | 11.69467065 | 11.91340139 | -0.218730733 | -1.061648143 |
| C130074G19Rik | 8.486039165 | 8.532604623 | 9.549778635 | -1.017174012 | -1.06373947 |
| Cmya5 | 11.48808443 | 11.79694518 | 12.55388953 | -0.75694435 | -1.0658051 |
| Tppp | 8.677119957 | 9.48762862 | 9.748265474 | -0.260636854 | -1.071145517 |
| Coro2b | 6.486950314 | 7.346353265 | 7.559224274 | -0.212871008 | -1.07227396 |
| Gm10273 | 5.173252999 | 4.380963026 | 6.248552634 | -1.867589609 | -1.075299636 |
| Fbxo40 | 8.381981551 | 9.180745271 | 9.458427047 | -0.277681777 | -1.076445496 |
| Mturn | 8.002647207 | 8.387351823 | 9.083798734 | -0.696446911 | -1.081151527 |
| Mycl | 8.601332535 | 9.688728501 | 9.690009302 | -0.001280801 | -1.088676767 |
| 9330159F19Rik | 5.658906998 | 6.084470182 | 6.766479388 | -0.682009206 | -1.10757239 |
| Pik3ip1 | 7.404143911 | 7.969277993 | 8.515907323 | -0.54662933 | -1.111763412 |
| Fmo2 | 8.167426264 | 8.357653674 | 9.295634566 | -0.937980892 | -1.128208303 |
| Gm6565 | 5.154336318 | 6.303312894 | 6.306207635 | -0.002894741 | -1.151871317 |
| Abca9 | 7.883701697 | 7.968497339 | 9.041157799 | -1.07266046 | -1.157456102 |
| Gm28661 | 13.36707661 | 13.94272878 | 14.55166953 | -0.60894075 | -1.184592917 |
| Inmt | 5.172436499 | 5.065047813 | 6.358165633 | -1.29311782 | -1.185729134 |
| Chst1 | 7.37923445 | 7.74313435 | 8.600756089 | -0.857621739 | -1.221521639 |
| Rptn | 8.795896426 | 9.263647996 | 10.03264511 | -0.768997111 | -1.236748681 |

| | | | | | |
|---|---|---|---|---|---|
| Cavin2 | 8.952128838 | 9.245506156 | 10.21204649 | -0.96654033 | -1.259917648 |
| Plin1 | 10.0059348 | 10.52576707 | 11.27020291 | -0.744435843 | -1.264268107 |
| C1qtnf9 | 5.969237734 | 6.335666004 | 7.249014796 | -0.913348792 | -1.279777062 |
| Trarg1 | 7.827162999 | 8.746763731 | 9.108817884 | -0.362054152 | -1.281654884 |
| A530016L24Rik | 6.927450063 | 6.980719933 | 8.212366612 | -1.231646679 | -1.284916549 |
| Fbxo32 | 8.182481119 | 8.763331723 | 9.471547002 | -0.708215279 | -1.289065883 |
| Fam13a | 7.740709088 | 8.16039835 | 9.032007924 | -0.871609574 | -1.291298836 |
| Gucy1a1 | 5.900713843 | 6.36078905 | 7.198387904 | -0.837598854 | -1.297674061 |
| Gm28437 | 12.36605991 | 13.13974346 | 13.68818551 | -0.548442053 | -1.322125607 |
| Nnat | 8.4060419 | 9.191342105 | 9.736760311 | -0.545418206 | -1.330718411 |
| Ppp1r3a | 8.604381067 | 8.448067665 | 9.956395221 | -1.508327556 | -1.352014154 |
| Skint7 | 6.763365461 | 7.751704221 | 8.127682898 | -0.375978676 | -1.364317437 |
| Aspn | 8.684636985 | 9.204057498 | 10.07075639 | -0.866698896 | -1.386119409 |
| 3425401B19Rik | 8.130329152 | 8.62358007 | 9.525477377 | -0.901897307 | -1.395148225 |
| Akr1c14 | 5.19948553 | 5.678396599 | 6.62613458 | -0.947737981 | -1.42664905 |
| Cd209f | 7.593889489 | 7.442870572 | 9.036558202 | -1.59368763 | -1.442668712 |
| Ccl21b | 6.098374994 | 7.632623441 | 7.547630154 | 0.084993287 | -1.44925516 |
| Lrrc17 | 6.538516721 | 7.176388271 | 8.007743215 | -0.831354944 | -1.469226494 |
| Angptl1 | 5.550826931 | 6.016354077 | 7.041417296 | -1.025063219 | -1.490590365 |
| Ppp1r3c | 9.220219238 | 10.09128098 | 10.74783179 | -0.65655081 | -1.527612555 |
| Cyp2e1 | 8.720711822 | 8.88811106 | 10.26378919 | -1.375678131 | -1.543077369 |
| Mstn | 8.312000266 | 8.592216024 | 9.862260225 | -1.270044201 | -1.550259959 |
| Acvr1c | 6.287888107 | 6.946665924 | 7.911652208 | -0.964986284 | -1.623764101 |
| Abca8a | 8.599094858 | 9.108393615 | 10.27731132 | -1.168917708 | -1.678216466 |
| Ky | 7.178543995 | 7.680934205 | 8.917020367 | -1.236086162 | -1.738476372 |
| Gm28438 | 7.655150254 | 7.545115395 | 9.436079763 | -1.890964368 | -1.780929509 |
| Mylk4 | 10.2500839 | 9.962204512 | 12.08574293 | -2.123538418 | -1.835659031 |
| B3galt2 | 6.842833527 | 7.564816551 | 8.739488309 | -1.174671758 | -1.896654782 |
| Pck1 | 7.484621318 | 9.382721695 | 10.28204865 | -0.899326955 | -2.797427332 |

| GeneSymbol | b-gluc._6h | Mann._6h | Sal._6h | Mann./Sal._6h | b-gluc./Sal._6h | b-gluc._24h | Mann._24h | Sal._24h | Mann./Sal._24h | b-gluc./Sal._24h |
|---|---|---|---|---|---|---|---|---|---|---|
| Slfn4 | 9.110213 | 10.0415 | 6.5021 | 3.539438212 | 2.608150412 | 9.334598 | 12.417689 | 6.98007 | 5.437623146 | 2.354531778 |
| Cxcl10 | 8.791419 | 11.09098 | 7.7933 | 3.297637555 | 0.998077289 | 8.2480676 | 12.465978 | 7.64122 | 4.824758946 | 0.606848419 |
| Isg15 | 9.282128 | 11.24273 | 8.2383 | 3.004434057 | 1.043830215 | 9.8700515 | 13.271299 | 8.66715 | 4.604149699 | 1.202902373 |
| Cxcl9 | 9.111431 | 10.06715 | 8.6201 | 1.447009468 | 0.491295053 | 8.3600515 | 12.525762 | 8.00359 | 4.522172286 | 0.356461499 |
| Ifit3b | 8.187147 | 10.70227 | 7.5239 | 3.178367107 | 0.663241694 | 8.8585529 | 12.282409 | 7.88603 | 4.396382573 | 0.97252667 |
| Ifit3 | 10.41197 | 12.83686 | 9.7476 | 3.089279245 | 0.664389422 | 11.193441 | 14.481172 | 10.166 | 4.315166307 | 1.027435007 |
| Il1rn | 8.615571 | 10.41242 | 6.3266 | 4.085818795 | 2.288967255 | 9.1441963 | 11.021309 | 6.71867 | 4.302641281 | 2.425528713 |
| Gbp10 | 7.879292 | 9.160876 | 7.2729 | 1.887980095 | 0.606395328 | 8.1903889 | 12.2162 | 7.95238 | 4.263819211 | 0.238007959 |
| Rsad2 | 9.053658 | 11.59597 | 8.0855 | 3.510475661 | 0.968164494 | 9.4803521 | 12.87201 | 8.61666 | 4.255350341 | 0.863692205 |
| Serpina3f | 8.503364 | 9.401199 | 8.2114 | 1.189793405 | 0.291958605 | 7.5477123 | 11.670299 | 7.51961 | 4.150691069 | 0.028104139 |
| Oasl2 | 9.56388 | 11.42547 | 8.9257 | 2.499765832 | 0.63817181 | 10.402487 | 13.451778 | 9.33885 | 4.112923294 | 1.063631958 |
| Oas2 | 9.320056 | 11.08857 | 8.58 | 2.508531897 | 0.740017105 | 10.493283 | 13.432385 | 9.40295 | 4.029434894 | 1.090332494 |
| Ifi204 | 8.998875 | 10.25596 | 7.4952 | 2.760712155 | 1.503628134 | 8.8780747 | 11.615905 | 7.59186 | 4.0240496 | 1.286219469 |
| Iigp1 | 9.63821 | 10.33421 | 9.0777 | 1.256476393 | 0.560478713 | 8.6644952 | 12.645867 | 8.73965 | 3.906216788 | -0.07515478 |
| Ifit1 | 9.530366 | 11.69941 | 8.7141 | 2.985307149 | 0.816260369 | 9.9489211 | 13.001189 | 9.0957 | 3.905484766 | 0.853217285 |
| Cxcl11 | 3.728582 | 5.476439 | 3.6546 | 1.821828289 | 0.073971885 | 3.9024379 | 7.46441 | 3.61482 | 3.849585417 | 0.287613301 |
| Irf7 | 11.15087 | 12.8282 | 10.625 | 2.203013368 | 0.525679848 | 12.083392 | 14.82607 | 10.981 | 3.845085996 | 1.102408038 |
| Oas3 | 10.03732 | 11.92694 | 9.3246 | 2.602311143 | 0.712695315 | 10.800437 | 13.6753 | 9.86373 | 3.811566198 | 0.93670291 |
| Gm12185 | 7.529636 | 8.629147 | 7.3469 | 1.282212149 | 0.182701732 | 7.2132283 | 10.94265 | 7.16881 | 3.773844123 | 0.044422121 |
| Mx1 | 9.735444 | 11.70181 | 9.1903 | 2.51148579 | 0.545123545 | 10.369895 | 13.613741 | 9.84501 | 3.768732823 | 0.524886972 |
| Usp18 | 9.422133 | 11.3811 | 8.8776 | 2.503519761 | 0.544549919 | 10.06853 | 13.073183 | 9.31218 | 3.760998355 | 0.756345007 |
| Trim30c | 6.994607 | 8.685957 | 6.3509 | 2.335105414 | 0.643755428 | 7.0068122 | 10.233854 | 6.48615 | 3.747708399 | 0.52066665 |
| Gbp11 | 4.941891 | 5.490122 | 4.696 | 0.794114575 | 0.245882909 | 5.0043796 | 8.5116326 | 4.78244 | 3.729193555 | 0.221940531 |
| Ifit2 | 10.05859 | 12.16666 | 9.7369 | 2.429753086 | 0.32167867 | 10.404847 | 13.691614 | 9.97178 | 3.719836913 | 0.433069401 |
| Gbp2 | 10.55197 | 11.43378 | 10.33 | 1.104048774 | 0.222235674 | 10.413123 | 13.995146 | 10.2959 | 3.699199964 | 0.11717681 |
| Isg20 | 8.48571 | 10.17639 | 7.9775 | 2.198861279 | 0.508181891 | 9.1924459 | 12.055549 | 8.36564 | 3.689908753 | 0.826805668 |
| Ms4a6d | 8.031897 | 9.168423 | 6.7727 | 2.395687568 | 1.259161319 | 8.7098986 | 10.499199 | 6.83534 | 3.663857934 | 1.874557837 |
| Mx2 | 7.780093 | 9.932173 | 7.4444 | 2.487740406 | 0.335660316 | 8.9060879 | 11.68589 | 8.03462 | 3.651266934 | 0.871464715 |
| Zbp1 | 10.97886 | 12.39641 | 10.477 | 1.919566926 | 0.50201768 | 11.448313 | 14.389496 | 10.749 | 3.640540202 | 0.699356291 |
| Gzmb | 7.842857 | 8.872527 | 7.1897 | 1.682791039 | 0.653120656 | 8.4463452 | 11.31197 | 7.69804 | 3.613929158 | 0.74830398 |
| Gm5431 | 6.958867 | 8.560708 | 6.6245 | 1.93618222 | 0.33434178 | 7.5803516 | 10.419521 | 6.82859 | 3.590933199 | 0.751763603 |

| | | | | | | | | | | |
|---|---|---|---|---|---|---|---|---|---|---|
| Fcgr1 | 7.311836 | 8.226444 | 6.1699 | 2.05658222 | 1.141974173 | 7.2458515 | 9.5749923 | 5.9877 | 3.587289459 | 1.258148647 |
| Gm12250 | 8.794423 | 10.07941 | 8.2282 | 1.851185425 | 0.566203634 | 8.9281833 | 11.886375 | 8.34891 | 3.537463567 | 0.579272318 |
| Tgtp1 | 10.62616 | 11.44614 | 10.393 | 1.053352962 | 0.23337071 | 10.382463 | 13.819686 | 10.3734 | 3.446250637 | 0.009027464 |
| Oasl1 | 9.224093 | 11.13989 | 8.7141 | 2.425751812 | 0.509956626 | 9.9491803 | 12.562852 | 9.12627 | 3.436582512 | 0.822910861 |
| Ifi211 | 8.75543 | 10.03046 | 7.896 | 2.134451516 | 0.859417647 | 8.5148142 | 11.420249 | 7.99008 | 3.430169901 | 0.524735229 |
| Oas1a | 8.918976 | 10.45559 | 8.3926 | 2.063031656 | 0.526413083 | 9.7882665 | 12.091894 | 8.6844 | 3.407491196 | 1.103863963 |
| Socs1 | 7.230421 | 7.58966 | 6.8052 | 0.784456014 | 0.42521712 | 7.0581157 | 9.9469922 | 6.54939 | 3.397600527 | 0.508724033 |
| Gbp6 | 10.83855 | 11.97856 | 10.427 | 1.551961156 | 0.411948656 | 10.53191 | 13.902281 | 10.5221 | 3.380189916 | 0.009819061 |
| Oas1g | 5.615756 | 6.817784 | 5.2182 | 1.599561031 | 0.397533607 | 6.4500999 | 8.8173518 | 5.44504 | 3.372307103 | 1.005055232 |
| Ifitm3 | 10.56652 | 11.15644 | 9.7283 | 1.428157871 | 0.838231345 | 10.698479 | 12.756931 | 9.44995 | 3.306978068 | 1.248525542 |
| Ifi44 | 7.322809 | 8.406291 | 6.9795 | 1.42683219 | 0.34335088 | 7.2014327 | 10.236591 | 6.96129 | 3.275303842 | 0.24014515 |
| Hk3 | 6.925041 | 7.941864 | 6.4379 | 1.503933591 | 0.487110401 | 7.058796 | 9.392259 | 6.1209 | 3.271359054 | 0.937895979 |
| Bst2 | 10.17322 | 11.07457 | 9.734 | 1.340593703 | 0.439247619 | 10.303076 | 12.927369 | 9.69835 | 3.22901401 | 0.604721876 |
| Cmpk2 | 9.519603 | 11.31526 | 8.8671 | 2.44812055 | 0.652462277 | 9.927438 | 12.33901 | 9.11716 | 3.221853177 | 0.810281548 |
| Ifng | 5.795687 | 6.544449 | 6.0163 | 0.528194562 | -0.220567743 | 5.3019055 | 8.9765175 | 5.7567 | 3.219813548 | -0.454798498 |
| Ly6i | 7.72312 | 7.459586 | 7.5732 | -0.113601338 | 0.149932476 | 7.805666 | 10.120967 | 6.92281 | 3.198158554 | 0.882857178 |
| Gm43302 | 4.011067 | 4.098216 | 3.5766 | 0.521614976 | 0.434466143 | 4.482374 | 7.8378874 | 4.68437 | 3.153520436 | -0.201992962 |
| Ifit1bl1 | 9.458973 | 11.10027 | 9.2967 | 1.803593734 | 0.162299855 | 9.9750333 | 12.733179 | 9.58207 | 3.151111046 | 0.392965772 |
| Phf11a | 7.265905 | 8.698101 | 7.2486 | 1.449497001 | 0.017300416 | 8.1836603 | 10.683678 | 7.53616 | 3.147518956 | 0.64750081 |
| Batf2 | 4.691127 | 5.375041 | 4.4974 | 0.877645347 | 0.193731361 | 4.7986919 | 7.6149671 | 4.47609 | 3.138881492 | 0.322606252 |
| Daxx | 10.54667 | 11.86434 | 10.369 | 1.495663814 | 0.177993818 | 10.969093 | 13.665329 | 10.5345 | 3.13081522 | 0.434579228 |
| Igtp | 10.84672 | 11.87328 | 10.557 | 1.315788022 | 0.289224922 | 10.869314 | 13.802346 | 10.6763 | 3.126053284 | 0.193021045 |
| Irgm1 | 10.87318 | 12.20225 | 10.6 | 1.602413826 | 0.273347304 | 11.17849 | 13.846007 | 10.7303 | 3.115704514 | 0.448187233 |
| Phf11d | 8.141232 | 9.315411 | 7.8729 | 1.442514412 | 0.26833486 | 8.4533174 | 11.088966 | 7.97902 | 3.10994412 | 0.474295552 |
| Rnf213 | 13.70513 | 14.98901 | 13.281 | 1.707846196 | 0.42396357 | 14.194123 | 16.807168 | 13.7095 | 3.097674804 | 0.484630338 |
| Ifi205 | 9.356084 | 10.3657 | 8.6349 | 1.730758053 | 0.721138547 | 9.1441864 | 11.569114 | 8.47463 | 3.094481581 | 0.669554007 |
| Parp12 | 8.355459 | 9.517538 | 8.0417 | 1.475812383 | 0.31373302 | 8.7450192 | 11.283546 | 8.18927 | 3.094272045 | 0.555745119 |
| Gm6545 | 8.978513 | 10.69329 | 8.4865 | 2.206803469 | 0.492029161 | 9.69166 | 12.010988 | 8.97174 | 3.039243734 | 0.719915576 |
| Serpina3g | 10.68667 | 11.51209 | 10.508 | 1.004077072 | 0.178659042 | 10.771455 | 13.309819 | 10.282 | 3.027770504 | 0.489406721 |
| Phf11b | 9.600621 | 11.07804 | 9.3639 | 1.714125606 | 0.236704204 | 10.185959 | 12.731658 | 9.71003 | 3.021629482 | 0.475930424 |
| Slfn5 | 12.3672 | 13.84733 | 11.702 | 2.145578384 | 0.665450554 | 12.461753 | 14.962666 | 11.9477 | 3.014989954 | 0.514077199 |
| Dhx58 | 9.446024 | 10.9297 | 9.0183 | 1.911369148 | 0.427688827 | 10.216757 | 12.392187 | 9.41675 | 2.975432489 | 0.800002369 |

| | | | | | | | | | | |
|---|---|---|---|---|---|---|---|---|---|---|
| Plac8 | 11.26964 | 12.28926 | 10.924 | 1.36550846 | 0.345892286 | 12.161691 | 13.989255 | 11.0227 | 2.966565194 | 1.139001272 |
| Tgtp2 | 9.721807 | 10.91978 | 9.3591 | 1.560655481 | 0.362679675 | 9.9669892 | 12.683127 | 9.72471 | 2.958419806 | 0.242282418 |
| Acod1 | 8.575921 | 8.712007 | 5.7599 | 2.952097294 | 2.816011331 | 7.9706429 | 9.4596333 | 6.51122 | 2.948416963 | 1.45942659 |
| Olfr56 | 6.027448 | 6.833928 | 5.6712 | 1.162746376 | 0.356266927 | 6.1355428 | 8.6732472 | 5.76184 | 2.911410613 | 0.373706166 |
| Herc6 | 10.09247 | 11.33083 | 9.7511 | 1.579700887 | 0.341338148 | 10.479331 | 12.995181 | 10.0878 | 2.907357418 | 0.391506864 |
| Nos2 | 5.141314 | 5.761823 | 4.9512 | 0.810596321 | 0.190086751 | 4.7423496 | 8.0173264 | 5.13099 | 2.88633254 | -0.388644189 |
| Klra2 | 6.487502 | 6.935312 | 5.2906 | 1.644755449 | 1.196945082 | 6.0639842 | 8.2815184 | 5.40785 | 2.873668167 | 0.656133925 |
| Rtp4 | 9.394775 | 10.83759 | 8.8497 | 1.987910924 | 0.545092548 | 9.9967342 | 12.078107 | 9.21352 | 2.864588643 | 0.783215482 |
| Chil3 | 9.355652 | 9.339788 | 6.8507 | 2.48909198 | 2.504956334 | 8.9413444 | 10.11745 | 7.27524 | 2.842204778 | 1.666099472 |
| Ifi47 | 11.3034 | 12.453 | 11.105 | 1.347791134 | 0.198189756 | 11.464731 | 13.906927 | 11.077 | 2.829877282 | 0.38768197 |
| Xaf1 | 9.296784 | 10.5717 | 8.9718 | 1.599846536 | 0.324934428 | 9.8672373 | 12.087814 | 9.25801 | 2.829804906 | 0.60922799 |
| Gbp7 | 10.98536 | 11.92062 | 10.702 | 1.218491552 | 0.283231586 | 10.773173 | 13.521804 | 10.6999 | 2.82194524 | 0.073314534 |
| Ly6a | 11.34987 | 12.04726 | 11.055 | 0.991898602 | 0.294512006 | 11.984788 | 13.891836 | 11.0739 | 2.817910464 | 0.910862106 |
| Fcgr4 | 5.651537 | 6.150231 | 5.0324 | 1.117802705 | 0.619108405 | 6.2255626 | 7.9486335 | 5.148 | 2.800632894 | 1.077561957 |
| Irgm2 | 9.773454 | 10.89182 | 9.5802 | 1.311652031 | 0.193285057 | 9.9815826 | 12.556101 | 9.76635 | 2.789746721 | 0.215228478 |
| Ifih1 | 9.404942 | 10.76738 | 8.9488 | 1.818597913 | 0.456155087 | 9.5400007 | 11.819419 | 9.03028 | 2.789140643 | 0.509722167 |
| Gbp4 | 11.28802 | 12.13961 | 11.163 | 0.977056868 | 0.125472634 | 11.497431 | 14.170296 | 11.3933 | 2.777024606 | 0.104159082 |
| Mlkl | 6.326817 | 6.973101 | 6.1184 | 0.85469941 | 0.208415316 | 6.1788074 | 8.6212908 | 5.85026 | 2.77102768 | 0.328544346 |
| Ccrl2 | 8.361856 | 9.641314 | 7.4728 | 2.168510042 | 0.889051266 | 9.4006914 | 10.553817 | 7.79348 | 2.760341453 | 1.607215491 |
| Ifi207 | 9.055898 | 9.610784 | 7.8251 | 1.785661642 | 1.230775519 | 8.4781181 | 10.520899 | 7.76836 | 2.752539512 | 0.709758833 |
| AW011738 | 7.007611 | 7.915651 | 6.4723 | 1.443390857 | 0.535350189 | 7.7375711 | 9.5483056 | 6.83846 | 2.709842884 | 0.899108363 |
| Ifi27l2a | 10.86194 | 11.84471 | 10.555 | 1.290139986 | 0.307368134 | 11.798078 | 13.462046 | 10.7766 | 2.685418174 | 1.021449659 |
| Calhm6 | 7.626611 | 8.349228 | 7.6624 | 0.686847995 | -0.035768287 | 7.8733569 | 10.181496 | 7.49638 | 2.685116508 | 0.376977251 |
| Stat2 | 11.20999 | 12.26189 | 10.939 | 1.322763916 | 0.270862742 | 11.760063 | 13.994711 | 11.3234 | 2.671296542 | 0.436648546 |
| I830127L07Rik | 5.122796 | 5.249207 | 4.1161 | 1.133077344 | 1.006667018 | 4.3449608 | 6.9020256 | 4.23309 | 2.668937859 | 0.111873069 |
| Gbp5 | 10.30272 | 11.18449 | 10.205 | 0.979564478 | 0.097788042 | 10.250605 | 12.854163 | 10.1956 | 2.658523128 | 0.054965716 |
| Ube2l6 | 8.845812 | 9.74642 | 8.8174 | 0.928981857 | 0.028374093 | 9.1861778 | 11.572407 | 8.92483 | 2.647573557 | 0.261344469 |
| Gm49339 | 8.342705 | 9.103271 | 6.5929 | 2.51036091 | 1.749795533 | 8.033603 | 9.2114318 | 6.58649 | 2.624937525 | 1.447108735 |
| Ly6c2 | 11.23093 | 11.84226 | 11.003 | 0.838846198 | 0.227508654 | 11.497056 | 13.692565 | 11.0757 | 2.616907774 | 0.421398579 |
| Pkib | 8.820497 | 9.77225 | 8.6697 | 1.102561749 | 0.150808686 | 9.0081877 | 11.299903 | 8.68746 | 2.612438809 | 0.320722992 |
| Ccl2 | 7.028895 | 7.994577 | 5.7828 | 2.211810522 | 1.246128731 | 5.55538 | 8.2500361 | 5.65742 | 2.592618563 | -0.102037527 |
| Xdh | 9.725249 | 9.972655 | 9.116 | 0.856684569 | 0.609278038 | 9.3555757 | 11.551251 | 8.97164 | 2.579609577 | 0.383934318 |

| | | | | | | | | | | |
|---|---|---|---|---|---|---|---|---|---|---|
| Tor3a | 10.04294 | 11.43898 | 9.8556 | 1.583340549 | 0.187304946 | 10.6855 | 12.586889 | 10.0162 | 2.570695026 | 0.669305542 |
| Lilrb4a | 9.028752 | 9.423174 | 7.5123 | 1.910870271 | 1.516447353 | 9.0534433 | 10.370858 | 7.80667 | 2.564183955 | 1.246769576 |
| Cfb | 11.01797 | 10.80238 | 10.754 | 0.048051716 | 0.263636826 | 10.331877 | 12.600355 | 10.0403 | 2.560022541 | 0.291544525 |
| Plau | 6.994316 | 8.402161 | 6.5369 | 1.865239606 | 0.457394405 | 7.0484401 | 8.8129984 | 6.25522 | 2.557779579 | 0.793221255 |
| Lilr4b | 8.789564 | 9.563892 | 7.1558 | 2.408055026 | 1.633727477 | 8.6437987 | 10.095274 | 7.53923 | 2.556040993 | 1.104565878 |
| Ifi213 | 11.61574 | 12.80021 | 11.253 | 1.547246114 | 0.362777602 | 12.368205 | 14.364867 | 11.829 | 2.535860722 | 0.539198916 |
| Trafd1 | 10.68301 | 11.68203 | 10.394 | 1.288420896 | 0.289398586 | 11.036847 | 13.286377 | 10.7575 | 2.528858494 | 0.279328259 |
| Inhba | 4.215241 | 5.138192 | 3.7567 | 1.381477143 | 0.458526734 | 4.3119501 | 6.5407476 | 4.02102 | 2.519723785 | 0.29092628 |
| Nt5c3 | 9.358054 | 10.26499 | 9.3712 | 0.893794929 | -0.013141009 | 9.4773548 | 11.887997 | 9.39054 | 2.497458622 | 0.08681614 |
| Ppa1 | 8.915115 | 9.410651 | 8.6106 | 0.800047617 | 0.304511829 | 9.028375 | 11.212515 | 8.73369 | 2.478825298 | 0.29468516 |
| Gpr141 | 6.807188 | 8.061503 | 6.3965 | 1.664971141 | 0.410655849 | 6.8230276 | 8.6969327 | 6.22806 | 2.468873554 | 0.594968429 |
| Gm4951 | 6.116807 | 6.629663 | 5.8852 | 0.744460789 | 0.231604525 | 5.3086053 | 8.1186531 | 5.65987 | 2.458785761 | -0.351262084 |
| Il18bp | 7.352004 | 7.436298 | 7.2509 | 0.185422532 | 0.101128138 | 7.1144821 | 9.2202135 | 6.76857 | 2.45164561 | 0.345914121 |
| Il12rb1 | 6.675762 | 6.996713 | 6.5977 | 0.399022161 | 0.078071192 | 6.5688185 | 9.0288369 | 6.59741 | 2.431427309 | -0.028591045 |
| Lgals3bp | 11.16193 | 12.0201 | 10.83 | 1.190156316 | 0.331989342 | 11.472364 | 13.334946 | 10.921 | 2.413994608 | 0.551412057 |
| Eppk1 | 5.595189 | 6.448636 | 5.1906 | 1.258075778 | 0.404628123 | 6.1189521 | 8.0151469 | 5.60684 | 2.408311614 | 0.512116777 |
| Clec4e | 7.937098 | 8.691998 | 5.6557 | 3.036342479 | 2.28144159 | 7.7289948 | 8.2898257 | 5.8856 | 2.404223051 | 1.84339212 |
| Il27 | 3.641005 | 4.081659 | 3.4362 | 0.6454708 | 0.204816676 | 3.5641531 | 5.7706677 | 3.37606 | 2.394612135 | 0.188097496 |
| AA467197 | 3.749229 | 4.218744 | 3.4805 | 0.738254899 | 0.26874023 | 3.893378 | 5.9881781 | 3.59506 | 2.393113579 | 0.298313418 |
| Gzma | 8.283356 | 8.673695 | 8.0405 | 0.633196403 | 0.242857299 | 8.1683044 | 10.485723 | 8.09751 | 2.38821555 | 0.070796454 |
| Phf11c | 8.803232 | 9.86661 | 8.5842 | 1.282431319 | 0.219052916 | 9.1761774 | 11.376309 | 8.98833 | 2.387981258 | 0.187849246 |
| Parp14 | 12.26298 | 13.43046 | 11.915 | 1.515287416 | 0.347813346 | 12.311909 | 14.466895 | 12.0818 | 2.385065128 | 0.230079053 |
| 1600014C10Rik | 10.11906 | 10.90289 | 9.9742 | 0.928715384 | 0.144885914 | 10.229084 | 12.301776 | 9.92252 | 2.379256288 | 0.306564527 |
| BC147527 | 8.88283 | 9.784304 | 8.7247 | 1.059638546 | 0.158164611 | 9.026506 | 11.178217 | 8.80104 | 2.377177217 | 0.225466258 |
| Upp1 | 7.181871 | 7.37104 | 6.7635 | 0.607520872 | 0.418352029 | 6.7495775 | 8.544793 | 6.17369 | 2.371107369 | 0.575891883 |
| Trim30a | 11.85009 | 13.00049 | 11.457 | 1.543055664 | 0.3926554 | 12.274703 | 14.049438 | 11.7042 | 2.345227062 | 0.57049162 |
| Cd300lf | 7.897564 | 8.127363 | 6.8541 | 1.273264732 | 1.043466112 | 8.0528663 | 9.3287107 | 6.99569 | 2.333017479 | 1.057173122 |
| Ms4a4c | 11.28925 | 12.34819 | 11.164 | 1.183995962 | 0.125059214 | 11.903123 | 13.720645 | 11.3955 | 2.325176064 | 0.507653881 |
| Nampt | 10.65712 | 11.43918 | 10.283 | 1.155689544 | 0.373624392 | 10.558619 | 12.743785 | 10.4231 | 2.320719006 | 0.135552188 |
| Aoah | 5.944224 | 6.548393 | 5.5746 | 0.97374699 | 0.369578434 | 6.1102107 | 7.6682977 | 5.36872 | 2.299578722 | 0.741491668 |
| Sdc3 | 10.67642 | 11.20807 | 10.165 | 1.042789372 | 0.51113422 | 10.641491 | 12.31823 | 10.0231 | 2.295129051 | 0.618390319 |
| Gbp3 | 9.825303 | 10.55582 | 9.7339 | 0.821910905 | 0.091389249 | 9.8117876 | 11.995392 | 9.7056 | 2.289795671 | 0.106191561 |

| | | | | | | | | | | |
|---|---|---|---|---|---|---|---|---|---|---|
| Trim30d | 10.14228 | 11.34335 | 9.8243 | 1.519081487 | 0.318011282 | 10.590208 | 12.327968 | 10.05 | 2.277920561 | 0.540160598 |
| Il1a | 5.260119 | 6.452361 | 4.4932 | 1.959148722 | 0.766905985 | 5.5638417 | 6.6389324 | 4.36538 | 2.273549264 | 1.198458607 |
| P2ry14 | 6.713455 | 6.86685 | 6.374 | 0.492805834 | 0.339410906 | 6.4383838 | 8.5287685 | 6.26137 | 2.267399885 | 0.177015179 |
| Parp9 | 10.79661 | 11.79795 | 10.519 | 1.278656084 | 0.27731042 | 10.884204 | 12.840262 | 10.5754 | 2.264867878 | 0.30881045 |
| Sct | 2.962069 | 3.38033 | 2.9673 | 0.413032711 | -0.005228083 | 3.0711735 | 5.2284598 | 2.97821 | 2.250251685 | 0.092965461 |
| Eif2ak2 | 9.903425 | 11.01905 | 9.4698 | 1.549269919 | 0.433640451 | 10.173816 | 12.196016 | 9.9517 | 2.244316416 | 0.222115614 |
| Tmem106a | 7.863025 | 8.357349 | 7.4021 | 0.955202454 | 0.460877988 | 8.1357565 | 9.5890584 | 7.35101 | 2.238052842 | 0.784750894 |
| Parp10 | 10.36381 | 11.22897 | 10.175 | 1.053844438 | 0.188692252 | 10.756676 | 12.63092 | 10.393 | 2.237874862 | 0.363630516 |
| Slamf8 | 7.559369 | 8.635777 | 7.4234 | 1.212408083 | 0.136000543 | 7.8487029 | 9.3605783 | 7.12374 | 2.236834957 | 0.724959501 |
| Ifi208 | 11.55601 | 12.79322 | 11.289 | 1.50391164 | 0.26670141 | 11.813481 | 13.736039 | 11.5053 | 2.230763344 | 0.308205509 |
| Gm18853 | 7.827207 | 8.719381 | 7.3667 | 1.352653089 | 0.46047994 | 7.5961745 | 9.7943279 | 7.566 | 2.228331559 | 0.030178242 |
| Ifi35 | 9.797667 | 10.72226 | 9.5504 | 1.171868322 | 0.247277663 | 10.228394 | 11.938199 | 9.71388 | 2.224316031 | 0.514511267 |
| Arhgef37 | 6.144547 | 6.839642 | 5.0439 | 1.795745285 | 1.10065009 | 6.2321706 | 7.3844151 | 5.1648 | 2.219614123 | 1.067369679 |
| Hcar2 | 8.051575 | 8.48425 | 6.2169 | 2.267391396 | 1.834716505 | 7.9248038 | 8.5488813 | 6.33059 | 2.218294769 | 1.59421729 |
| Stat1 | 13.18062 | 14.04244 | 12.975 | 1.06715221 | 0.205330522 | 13.422882 | 15.36764 | 13.1517 | 2.215966056 | 0.271207095 |
| Ifi214 | 9.791565 | 11.11651 | 9.6382 | 1.478287903 | 0.153338721 | 10.387908 | 12.176936 | 9.97077 | 2.206163035 | 0.417135535 |
| Fcgr3 | 8.23448 | 8.043877 | 7.0262 | 1.017688452 | 1.208291144 | 7.9196655 | 8.83253 | 6.65177 | 2.180763643 | 1.26789913 |
| Tnfaip2 | 9.251322 | 9.612166 | 8.1263 | 1.485899999 | 1.125056391 | 8.6864537 | 10.07332 | 7.89688 | 2.17643708 | 0.789570408 |
| Slfn8 | 10.42378 | 11.61971 | 10.139 | 1.480680517 | 0.284748835 | 10.672185 | 12.491956 | 10.3195 | 2.172479467 | 0.352708116 |
| Fcer1g | 9.27576 | 9.538184 | 8.7023 | 0.83589296 | 0.573468679 | 9.6306768 | 10.702685 | 8.53255 | 2.170136908 | 1.098128911 |
| Slfn1 | 8.918782 | 9.798145 | 8.655 | 1.143164015 | 0.263800592 | 8.974422 | 11.017169 | 8.84979 | 2.167374655 | 0.124627876 |
| Rilpl1 | 6.199025 | 7.107698 | 6.0672 | 1.040541081 | 0.131868768 | 6.0291736 | 7.9321877 | 5.78366 | 2.148524188 | 0.24551009 |
| Nts | 4.477729 | 4.780518 | 3.8575 | 0.922989022 | 0.620200867 | 4.1055037 | 5.9626286 | 3.82317 | 2.139458829 | 0.282334011 |
| Ptafr | 8.296001 | 9.330578 | 7.4819 | 1.848705555 | 0.814129324 | 8.7262707 | 9.7001153 | 7.56152 | 2.138594051 | 1.164749497 |
| Mov10 | 9.661551 | 10.62145 | 9.5269 | 1.094550463 | 0.134650036 | 9.8881307 | 11.840084 | 9.70572 | 2.134362103 | 0.18240853 |
| Irf1 | 11.66189 | 12.04306 | 11.54 | 0.503506258 | 0.122340932 | 11.500658 | 13.576822 | 11.4449 | 2.131893308 | 0.055728719 |
| Rasa4 | 9.066613 | 9.871069 | 9.1745 | 0.696543642 | -0.107912187 | 9.5658644 | 11.476041 | 9.34533 | 2.13071158 | 0.220534882 |
| Vcan | 8.217895 | 8.406244 | 5.2012 | 3.205058509 | 3.016710346 | 6.9898988 | 8.1492528 | 6.01974 | 2.129512353 | 0.970158406 |
| Gzmc | 6.139814 | 6.967394 | 6.2852 | 0.682228631 | -0.14535138 | 6.0849014 | 8.1956101 | 6.06665 | 2.128958457 | 0.018249775 |
| Ly6c1 | 9.095469 | 9.028646 | 9.033 | -0.004336924 | 0.062485881 | 8.3016177 | 10.736687 | 8.61298 | 2.123709898 | -0.311359278 |
| Cebpb | 7.734067 | 7.758937 | 6.7878 | 0.971174183 | 0.946303447 | 7.5526967 | 8.7656657 | 6.64355 | 2.122114718 | 0.909145731 |
| Ddx60 | 9.675614 | 10.38903 | 9.1851 | 1.203974249 | 0.490560139 | 9.5391191 | 11.531406 | 9.4185 | 2.112900701 | 0.120614238 |

| | | | | | | | | | | |
|---|---|---|---|---|---|---|---|---|---|---|
| Ccl24 | 3.964788 | 5.107037 | 3.6262 | 1.480883494 | 0.338634173 | 3.6121329 | 5.7317091 | 3.62844 | 2.103273377 | -0.01630276 |
| Tmem140 | 7.036088 | 7.962397 | 7.2855 | 0.676870157 | -0.249439322 | 7.4618221 | 9.3273034 | 7.22405 | 2.103254309 | 0.237772958 |
| Ffar2 | 6.050868 | 6.157104 | 5.9968 | 0.160274237 | 0.054038157 | 7.2999336 | 8.2445473 | 6.16232 | 2.082226361 | 1.137612695 |
| Samhd1 | 13.39586 | 13.91057 | 13.245 | 0.665336982 | 0.150627788 | 13.384886 | 15.344683 | 13.2721 | 2.072556536 | 0.112758984 |
| F10 | 6.235866 | 6.462217 | 4.5977 | 1.864469455 | 1.638119063 | 6.0456669 | 7.0030558 | 4.93718 | 2.065872036 | 1.10848315 |
| Tlr7 | 9.869578 | 11.02726 | 9.604 | 1.423291089 | 0.265612548 | 10.33851 | 11.819116 | 9.75325 | 2.065861375 | 0.585255875 |
| Mpeg1 | 12.3467 | 12.69669 | 12.078 | 0.618491196 | 0.268499532 | 11.8944 | 13.695519 | 11.6299 | 2.065576844 | 0.264457996 |
| Csf3r | 8.815585 | 9.219471 | 7.4187 | 1.800746613 | 1.396860056 | 9.4121135 | 9.3711753 | 7.30995 | 2.061227012 | 2.10216521 |
| Ms4a6c | 9.772208 | 10.00708 | 8.9748 | 1.032259494 | 0.797384197 | 9.7707148 | 11.029683 | 8.97624 | 2.053440628 | 0.79447215 |
| Clec5a | 6.396916 | 6.964899 | 5.3777 | 1.587222765 | 1.019239392 | 6.3022682 | 7.3329044 | 5.2833 | 2.049601534 | 1.018965352 |
| Gbp9 | 11.83168 | 12.65158 | 11.727 | 0.924280634 | 0.104382872 | 12.018507 | 13.993175 | 11.9445 | 2.04866423 | 0.073996302 |
| Dtx3l | 11.38554 | 12.33728 | 11.182 | 1.15482048 | 0.20307273 | 11.472955 | 13.301932 | 11.2592 | 2.04272449 | 0.213748039 |
| H2-T24 | 10.28556 | 11.33026 | 10.029 | 1.301060817 | 0.256356178 | 10.722052 | 12.287688 | 10.257 | 2.030718135 | 0.465081619 |
| Carmil1 | 8.231421 | 9.097776 | 8.3429 | 0.754892351 | -0.111462306 | 8.3013014 | 10.239083 | 8.21014 | 2.028942928 | 0.091161581 |
| Ccl4 | 7.522766 | 8.433009 | 6.7163 | 1.716669235 | 0.806426625 | 7.3947502 | 8.7204175 | 6.69204 | 2.028381829 | 0.702714529 |
| Apod | 5.744863 | 5.833858 | 5.3872 | 0.446696904 | 0.35770238 | 5.3012938 | 7.2904071 | 5.26221 | 2.028196592 | 0.03908333 |
| Pml | 12.102 | 13.02855 | 11.955 | 1.073494888 | 0.146945086 | 12.56521 | 14.225383 | 12.1981 | 2.02728885 | 0.367115535 |
| Gbp8 | 9.429955 | 9.888041 | 9.4244 | 0.46362795 | 0.005541562 | 9.2685161 | 11.651902 | 9.63133 | 2.020575606 | -0.362810043 |
| Trex1 | 8.815462 | 9.327596 | 8.5364 | 0.791192423 | 0.279058238 | 8.9600056 | 10.336391 | 8.31701 | 2.01938241 | 0.642996892 |
| Fgl2 | 9.706668 | 9.969266 | 9.2019 | 0.767337283 | 0.504738992 | 9.1662258 | 11.077885 | 9.06246 | 2.015430074 | 0.103770436 |
| Gm15056 | 3.385123 | 3.434901 | 3.1258 | 0.309099224 | 0.259321354 | 3.3000507 | 5.0606325 | 3.04774 | 2.012888956 | 0.252307118 |
| Gvin1 | 12.98238 | 13.89524 | 12.714 | 1.18099811 | 0.26814017 | 13.233899 | 14.868182 | 12.8583 | 2.009907464 | 0.375624694 |
| Gm18752 | 5.389238 | 6.210813 | 5.3692 | 0.841578085 | 0.020002691 | 5.9857982 | 7.6216274 | 5.61492 | 2.00670848 | 0.370879344 |
| Lgals3 | 9.388004 | 10.45103 | 8.7991 | 1.651954383 | 0.588928161 | 9.1273403 | 10.729818 | 8.72743 | 2.002383775 | 0.399906175 |
| Il1b | 10.66818 | 11.16479 | 8.3146 | 2.850173309 | 2.353560106 | 11.052129 | 10.638413 | 8.63957 | 1.998846012 | 2.412561581 |
| Gm6158 | 3.448631 | 3.929476 | 4.076 | -0.146507398 | -0.627352292 | 4.22429 | 5.2594427 | 3.2679 | 1.991546719 | 0.956394031 |
| Ifi206 | 11.69884 | 12.90397 | 11.461 | 1.443295946 | 0.238161866 | 11.99829 | 13.65952 | 11.6726 | 1.986937626 | 0.325707829 |
| Gm18852 | 6.159632 | 7.067464 | 6.0124 | 1.055080561 | 0.147248138 | 6.4204017 | 7.9994049 | 6.01313 | 1.986277136 | 0.407273966 |
| Wars | 10.14059 | 10.20781 | 10.052 | 0.156293378 | 0.08907804 | 10.215655 | 11.9739 | 9.99021 | 1.983687831 | 0.225442754 |
| Gm4070 | 11.99353 | 12.95837 | 11.661 | 1.297465652 | 0.33262397 | 12.147254 | 13.908386 | 11.9351 | 1.973280072 | 0.212148172 |
| Nlrc5 | 11.98064 | 12.56471 | 11.907 | 0.65732763 | 0.07325211 | 12.021964 | 14.091502 | 12.1192 | 1.972326244 | -0.097211597 |
| Cd274 | 10.40989 | 11.23062 | 10.341 | 0.889324674 | 0.06860039 | 10.623402 | 12.462318 | 10.4925 | 1.969792198 | 0.130876273 |

| | | | | | | | | | | |
|---|---|---|---|---|---|---|---|---|---|---|
| Clec7a | 9.167474 | 10.2023 | 7.7159 | 2.486356571 | 1.451526622 | 8.9318929 | 9.5291746 | 7.56408 | 1.965093585 | 1.367811837 |
| Tppp3 | 5.256832 | 5.553261 | 4.8841 | 0.66913844 | 0.372710219 | 5.5299861 | 6.8233491 | 4.86442 | 1.958924165 | 0.665561215 |
| Spon1 | 6.398792 | 6.441547 | 6.1913 | 0.250281304 | 0.207526432 | 8.0909683 | 8.1722431 | 6.21525 | 1.956995298 | 1.875720544 |
| Setdb2 | 9.211004 | 10.23126 | 9.1611 | 1.070116557 | 0.049863829 | 9.5122999 | 11.432047 | 9.47969 | 1.952354688 | 0.032608041 |
| Znfx1 | 11.02585 | 11.74409 | 10.842 | 0.902581352 | 0.184343398 | 10.992808 | 12.81243 | 10.8653 | 1.947125828 | 0.127504289 |
| Slc25a22 | 9.34646 | 10.36735 | 9.1652 | 1.202170682 | 0.181284878 | 9.9126534 | 11.502462 | 9.55581 | 1.946653394 | 0.356844656 |
| Oas1b | 7.677081 | 8.524527 | 7.5591 | 0.965423781 | 0.117977526 | 8.5057553 | 10.023123 | 8.08495 | 1.938176592 | 0.420808787 |
| Ifi209 | 12.66314 | 13.82853 | 12.507 | 1.321649704 | 0.156262284 | 13.123684 | 14.701416 | 12.7668 | 1.93464271 | 0.35691032 |
| Fpr2 | 5.909513 | 6.652972 | 4.6061 | 2.046869809 | 1.303411684 | 6.0437788 | 6.8635133 | 4.93722 | 1.926291503 | 1.10655699 |
| Hck | 9.995132 | 10.68925 | 9.6678 | 1.021475221 | 0.327355383 | 10.491418 | 11.614502 | 9.69046 | 1.924045752 | 0.800962473 |
| Casp4 | 8.656661 | 8.947943 | 8.2844 | 0.663572999 | 0.372290976 | 8.5634487 | 10.268544 | 8.34536 | 1.923182911 | 0.218088075 |
| Mefv | 7.220259 | 7.660393 | 6.279 | 1.381358451 | 0.941224136 | 7.4946369 | 8.289226 | 6.37033 | 1.918891327 | 1.124302203 |
| Il12rb2 | 6.758461 | 6.934793 | 6.5126 | 0.422148261 | 0.245815599 | 6.3037358 | 8.2179108 | 6.30454 | 1.913366404 | -0.000808634 |
| Gm6548 | 7.654068 | 8.564944 | 7.4057 | 1.159214538 | 0.248338691 | 8.0887972 | 9.6106615 | 7.7036 | 1.907065705 | 0.385201372 |
| Ccl7 | 6.97948 | 7.207377 | 5.5075 | 1.699859201 | 1.471962427 | 5.320262 | 7.1501816 | 5.2542 | 1.895980652 | 0.066061065 |
| A530032D15Rik | 7.085217 | 7.477379 | 6.8086 | 0.6687997 | 0.276637819 | 7.3049776 | 9.0888906 | 7.19366 | 1.895235207 | 0.111322217 |
| Il33 | 8.509193 | 7.973923 | 6.95 | 1.023966952 | 1.559236886 | 7.5897441 | 8.5917297 | 6.69786 | 1.893871924 | 0.891886311 |
| Nmral1 | 7.504962 | 8.455969 | 7.512 | 0.943967247 | -0.007040162 | 8.2736469 | 9.7780858 | 7.88665 | 1.89143523 | 0.386996304 |
| Tmem184b | 9.531632 | 10.49391 | 9.4448 | 1.049130943 | 0.086849738 | 9.9246949 | 11.374321 | 9.48657 | 1.887753869 | 0.438127329 |
| Gm8113 | 8.295564 | 9.063597 | 8.1857 | 0.877903429 | 0.109870556 | 9.2301182 | 10.636672 | 8.74925 | 1.887420524 | 0.480866505 |
| Hdc | 8.987004 | 8.35397 | 6.4035 | 1.950430461 | 2.583463911 | 7.8576935 | 8.711924 | 6.82834 | 1.883587327 | 1.029356778 |
| Tdrd7 | 9.063947 | 10.08408 | 9.0283 | 1.055786694 | 0.035656909 | 9.5647307 | 11.215071 | 9.35117 | 1.863898537 | 0.213558575 |
| Sp140 | 11.09261 | 11.54947 | 10.971 | 0.578645254 | 0.121788388 | 11.339946 | 13.035195 | 11.1782 | 1.857040956 | 0.161791927 |
| Arhgef10l | 6.432931 | 6.730479 | 5.978 | 0.752466162 | 0.454917521 | 7.1676068 | 7.7136887 | 5.86117 | 1.852514645 | 1.306432701 |
| Procr | 7.476178 | 7.889577 | 7.2628 | 0.626807228 | 0.213408245 | 6.8872594 | 8.6827276 | 6.83738 | 1.84534418 | 0.049875906 |
| Adap2 | 6.251078 | 6.726136 | 6.215 | 0.511133329 | 0.036075422 | 6.1050659 | 7.7293968 | 5.88779 | 1.84160429 | 0.217273388 |
| Trim30b | 9.183838 | 10.11109 | 8.9226 | 1.188457562 | 0.261208696 | 10.161403 | 11.335092 | 9.49783 | 1.837259785 | 0.663571145 |
| Rfc3 | 8.027264 | 8.47451 | 8.1178 | 0.35667887 | -0.090566549 | 8.2239409 | 9.9124187 | 8.08427 | 1.828144597 | 0.139666785 |
| Lacc1 | 6.813138 | 7.478105 | 6.7328 | 0.745255802 | 0.080288258 | 6.8672615 | 8.6325285 | 6.80662 | 1.825905244 | 0.060638224 |
| Gm15931 | 4.31881 | 4.754718 | 4.3082 | 0.446557347 | 0.010649291 | 5.6355165 | 6.2433675 | 4.418 | 1.825371239 | 1.217520232 |
| Slfn3 | 6.867837 | 7.459699 | 6.685 | 0.774731778 | 0.182869885 | 6.7711654 | 8.6623321 | 6.83824 | 1.824096531 | -0.067070239 |
| Tlr9 | 10.1515 | 10.64778 | 10.126 | 0.522242509 | 0.025964537 | 10.66831 | 11.927163 | 10.1033 | 1.82390916 | 0.565056191 |

| | | | | | | | | | | |
|---|---|---|---|---|---|---|---|---|---|---|
| Tapbp | 12.31405 | 12.92392 | 12.224 | 0.700283032 | 0.090415808 | 12.62776 | 13.966041 | 12.1527 | 1.813320838 | 0.475039889 |
| Pik3r6 | 5.673454 | 6.132013 | 5.8041 | 0.327956721 | -0.130601919 | 6.0502918 | 7.4421277 | 5.6387 | 1.803432285 | 0.411596402 |
| Asb13 | 8.878358 | 9.466158 | 8.7873 | 0.678849625 | 0.091049501 | 9.2921918 | 10.948094 | 9.14626 | 1.801838315 | 0.145935731 |
| Nmi | 9.785105 | 10.45315 | 9.6679 | 0.785244639 | 0.117203054 | 9.9293851 | 11.537054 | 9.73999 | 1.797067308 | 0.189398125 |
| Gm8995 | 12.90501 | 13.6609 | 12.791 | 0.870099818 | 0.114206472 | 13.162643 | 14.880786 | 13.0848 | 1.795975256 | 0.077831934 |
| Psmb10 | 10.35646 | 10.87491 | 10.289 | 0.585435025 | 0.066982523 | 10.73811 | 12.061477 | 10.2696 | 1.791912814 | 0.4685451 |
| Sgcb | 7.728335 | 7.985789 | 7.613 | 0.372749943 | 0.115296253 | 7.8385703 | 9.5329735 | 7.74996 | 1.783015078 | 0.08861189 |
| Hp | 8.238864 | 7.890069 | 6.0976 | 1.792440316 | 2.141235477 | 8.0299593 | 8.2810423 | 6.50283 | 1.778208523 | 1.527125485 |
| Prdx5 | 9.867607 | 10.04001 | 9.6985 | 0.341509032 | 0.169106604 | 10.209293 | 11.379845 | 9.60678 | 1.773063794 | 0.602512253 |
| Slc15a3 | 9.012287 | 9.679083 | 8.3404 | 1.338645395 | 0.671849301 | 9.1913639 | 10.17635 | 8.40347 | 1.77288315 | 0.787896947 |
| Gm9733 | 5.956522 | 6.151205 | 5.0599 | 1.091342603 | 0.896659351 | 6.698494 | 7.0337712 | 5.26693 | 1.766842084 | 1.431564866 |
| Ccl3 | 7.863739 | 8.666759 | 7.0152 | 1.651527792 | 0.848507461 | 7.5134231 | 8.475794 | 6.71661 | 1.759180448 | 0.796809504 |
| Pnp | 9.643605 | 10.18277 | 9.4548 | 0.728007843 | 0.188839106 | 9.9742739 | 11.174404 | 9.41975 | 1.754650836 | 0.554520268 |
| Ogfr | 10.62125 | 11.32734 | 10.421 | 0.906828 | 0.200741754 | 10.90779 | 12.281963 | 10.5275 | 1.754447898 | 0.380274965 |
| Cxcl2 | 9.145143 | 9.087063 | 6.8892 | 2.197836905 | 2.255916469 | 8.7035313 | 8.6181367 | 6.86511 | 1.753023342 | 1.838417861 |
| AC125149.3 | 7.497948 | 8.141387 | 7.4376 | 0.703806961 | 0.060368337 | 7.8591555 | 9.1647374 | 7.41175 | 1.752991264 | 0.447409312 |
| Lipg | 7.501091 | 7.093335 | 6.3667 | 0.726664387 | 1.134420752 | 6.2510842 | 7.486081 | 5.73664 | 1.749442483 | 0.514445699 |
| Sectm1a | 3.094764 | 2.990759 | 2.962 | 0.028754763 | 0.132759561 | 3.0489836 | 4.698681 | 2.95123 | 1.747452786 | 0.097755387 |
| Csprs | 7.964031 | 8.826365 | 7.7364 | 1.089983116 | 0.227649309 | 8.6165037 | 9.882638 | 8.14765 | 1.734986092 | 0.468851833 |
| Ifi203 | 12.79171 | 13.7141 | 12.647 | 1.067479258 | 0.14509352 | 13.026525 | 14.626686 | 12.8946 | 1.732115228 | 0.131953574 |
| Lst1 | 6.81073 | 7.199495 | 6.7712 | 0.428276537 | 0.039510908 | 7.7742833 | 8.4728634 | 6.74148 | 1.7313863 | 1.032806205 |
| Trim21 | 9.687558 | 10.38121 | 9.6417 | 0.73949851 | 0.045850829 | 9.8912963 | 11.501998 | 9.78302 | 1.71898007 | 0.108278412 |
| Tpst1 | 8.768247 | 9.593388 | 8.5855 | 1.007886005 | 0.182745452 | 9.1464018 | 10.370097 | 8.66326 | 1.706834869 | 0.483139847 |
| Slfn9 | 9.106005 | 9.940045 | 9.0026 | 0.937425348 | 0.103385125 | 9.2139766 | 10.849464 | 9.14946 | 1.700002418 | 0.064514665 |
| Tgm2 | 10.93297 | 10.78269 | 10.016 | 0.766871241 | 0.917148473 | 10.005994 | 11.23317 | 9.53332 | 1.69985284 | 0.472677619 |
| Ddx58 | 11.48638 | 12.4241 | 11.244 | 1.179705274 | 0.241985354 | 11.741425 | 13.199774 | 11.5012 | 1.69856356 | 0.240214474 |
| Cdkn1a | 9.114628 | 9.61806 | 8.5981 | 1.019945922 | 0.516514158 | 8.7826174 | 10.312377 | 8.61942 | 1.692957663 | 0.163198137 |
| Mmp14 | 8.64652 | 9.259027 | 8.1213 | 1.137771494 | 0.525264705 | 8.621098 | 9.4230142 | 7.73186 | 1.691152008 | 0.889235864 |
| Nlrp3 | 8.080672 | 8.39727 | 6.4099 | 1.98733728 | 1.670739141 | 7.8040746 | 8.0036928 | 6.31759 | 1.686100843 | 1.486482682 |
| Tap1 | 12.39695 | 12.86223 | 12.365 | 0.497135594 | 0.031855674 | 12.736292 | 14.133279 | 12.45 | 1.683263682 | 0.286275904 |
| H2-T22 | 11.71552 | 12.10574 | 11.65 | 0.455496718 | 0.06527213 | 11.899319 | 13.35896 | 11.6804 | 1.678536324 | 0.218896024 |
| Gng12 | 8.606325 | 8.60685 | 8.4563 | 0.150515632 | 0.149990448 | 8.0929637 | 9.7389441 | 8.06224 | 1.676708452 | 0.030727986 |

| | | | | | | | | | | |
|---|---|---|---|---|---|---|---|---|---|---|
| Gpnmb | 5.464755 | 6.894206 | 5.373 | 1.521230821 | 0.091779515 | 5.2158411 | 6.9136732 | 5.24066 | 1.673011929 | -0.024820191 |
| Parp11 | 9.097488 | 9.616643 | 9.1214 | 0.495283674 | -0.023870454 | 9.5256631 | 11.158455 | 9.48876 | 1.669697778 | 0.03690549 |
| Sirpb1c | 6.029383 | 6.577938 | 5.2067 | 1.371222558 | 0.822667406 | 6.9960203 | 7.269509 | 5.59992 | 1.669588463 | 1.396099718 |
| Mndal | 12.17806 | 12.93323 | 12.104 | 0.829149158 | 0.073974382 | 12.609972 | 14.098145 | 12.43 | 1.668149892 | 0.179976587 |
| Psmb9 | 11.04933 | 11.31226 | 11.004 | 0.308300836 | 0.04537774 | 11.361544 | 12.732367 | 11.0659 | 1.66645834 | 0.295634961 |
| Mitd1 | 8.86667 | 9.641238 | 8.6861 | 0.955187111 | 0.180618637 | 9.3038189 | 10.5797 | 8.92313 | 1.656571239 | 0.38068993 |
| Nupr1 | 6.213214 | 6.091383 | 5.9523 | 0.1391161 | 0.260947663 | 6.6839875 | 7.4712226 | 5.81656 | 1.654664173 | 0.867429062 |
| H2-T23 | 12.74171 | 13.15387 | 12.654 | 0.499728032 | 0.087561156 | 13.047566 | 14.333794 | 12.6846 | 1.649212264 | 0.362984499 |
| Ptgs2 | 9.574458 | 9.877371 | 7.7559 | 2.121507657 | 1.818594974 | 9.5317323 | 9.9855281 | 8.33692 | 1.648607013 | 1.194811214 |
| Zup1 | 9.964805 | 10.68432 | 9.9097 | 0.774604357 | 0.055087523 | 10.252613 | 11.762449 | 10.115 | 1.647435929 | 0.137600192 |
| Samd9l | 12.18071 | 13.04929 | 12.062 | 0.987262132 | 0.118687914 | 12.686868 | 14.099048 | 12.4572 | 1.641815014 | 0.229635234 |
| Aplnr | 5.685323 | 5.924683 | 5.6749 | 0.249798671 | 0.01043828 | 6.1614538 | 7.1529429 | 5.52563 | 1.627308432 | 0.635819364 |
| Lgals9 | 11.28061 | 12.07991 | 11.114 | 0.965589722 | 0.166288406 | 11.573596 | 12.829549 | 11.2054 | 1.624189708 | 0.368237081 |
| Cdh17 | 6.969228 | 7.118378 | 6.5426 | 0.575790088 | 0.426640023 | 7.170239 | 8.4434463 | 6.82138 | 1.622062944 | 0.348855715 |
| Mxd1 | 10.0041 | 10.41481 | 9.702 | 0.712806728 | 0.302104223 | 9.9595768 | 11.455585 | 9.83366 | 1.621927939 | 0.125919821 |
| Il3ra | 7.757159 | 7.588694 | 7.4389 | 0.149749644 | 0.318214883 | 7.8352619 | 8.8567287 | 7.23506 | 1.621664589 | 0.600197831 |
| Cd69 | 11.16921 | 11.60698 | 11.075 | 0.532138864 | 0.094375316 | 10.5735 | 12.433489 | 10.8154 | 1.618040476 | -0.2419481 |
| Slco3a1 | 9.249588 | 9.571518 | 9.1751 | 0.396452322 | 0.074521707 | 9.1747618 | 10.785665 | 9.1687 | 1.616960895 | 0.006058112 |
| Mfsd7a | 4.725753 | 4.838423 | 4.583 | 0.255374453 | 0.142705098 | 5.0922091 | 6.3547324 | 4.74065 | 1.614080636 | 0.351557314 |
| Cd86 | 9.703722 | 10.07682 | 9.5612 | 0.515589463 | 0.142487585 | 9.6327065 | 11.158303 | 9.54584 | 1.612465204 | 0.086868356 |
| Aif1 | 8.166814 | 8.155085 | 8.0136 | 0.141437707 | 0.153166806 | 8.7089723 | 9.4733533 | 7.86193 | 1.611420469 | 0.847039469 |
| Ccl12 | 6.890641 | 6.301475 | 5.0222 | 1.279321264 | 1.868487615 | 5.1393243 | 6.4925596 | 4.88574 | 1.606822896 | 0.253587604 |
| Plin2 | 8.542126 | 9.265373 | 8.3194 | 0.945950588 | 0.22270365 | 8.7607824 | 9.910194 | 8.30622 | 1.603972562 | 0.454561032 |
| Gm4841 | 5.051135 | 5.421863 | 5.0222 | 0.399689638 | 0.028961547 | 4.8209689 | 6.7008304 | 5.09725 | 1.60358342 | -0.276278026 |
| Uba7 | 11.46646 | 12.07802 | 11.524 | 0.553921384 | -0.057641638 | 12.298222 | 13.54521 | 11.9474 | 1.597834676 | 0.350846735 |
| Smpdl3b | 4.858739 | 5.121857 | 4.5713 | 0.550570239 | 0.287451839 | 4.6920736 | 6.228252 | 4.63562 | 1.592630302 | 0.056451949 |
| Igsf6 | 7.485045 | 8.344873 | 7.1699 | 1.174954856 | 0.315126769 | 8.4493418 | 8.7053374 | 7.11308 | 1.592259497 | 1.336263825 |
| Smchd1 | 12.1206 | 12.71166 | 12.043 | 0.668556432 | 0.07750278 | 12.390301 | 13.810455 | 12.2207 | 1.58975545 | 0.169601298 |
| Jaml | 9.823241 | 10.28578 | 9.6535 | 0.632293268 | 0.169750257 | 9.7164442 | 11.33824 | 9.75308 | 1.585159613 | -0.036635839 |
| Sirpb1b | 6.378429 | 6.598992 | 5.47 | 1.129016032 | 0.908452804 | 6.9655907 | 7.2515883 | 5.67073 | 1.580859061 | 1.294861494 |
| Mif | 9.661015 | 9.891559 | 9.5686 | 0.322964936 | 0.092420717 | 10.138122 | 11.043884 | 9.47059 | 1.573294873 | 0.667532907 |
| Nod2 | 7.069139 | 7.506083 | 6.6677 | 0.83835066 | 0.401406656 | 6.8807756 | 8.3789456 | 6.80583 | 1.57311606 | 0.074946107 |

| | | | | | | | | | | |
|---|---|---|---|---|---|---|---|---|---|---|
| Timp1 | 4.981001 | 5.591751 | 4.6408 | 0.950937073 | 0.340187132 | 5.0737568 | 5.9533406 | 4.38103 | 1.572307299 | 0.692723513 |
| Anxa4 | 8.943558 | 9.405729 | 8.9392 | 0.466532939 | 0.004362256 | 8.9737095 | 10.43601 | 8.86385 | 1.572158146 | 0.109857452 |
| Gm1966 | 12.45908 | 13.25294 | 12.242 | 1.01044447 | 0.216580796 | 12.719016 | 14.010891 | 12.439 | 1.571852352 | 0.279976659 |
| Capn5 | 7.954497 | 8.418117 | 8.014 | 0.404087325 | -0.059532496 | 8.0887009 | 9.4500854 | 7.8818 | 1.568289167 | 0.206904684 |
| Gm9843 | 8.398049 | 8.626317 | 7.133 | 1.493334222 | 1.265066398 | 8.0787898 | 8.7069367 | 7.13902 | 1.567919947 | 0.939773037 |
| Osm | 8.577909 | 8.739182 | 7.5999 | 1.139245175 | 0.977971845 | 8.535095 | 9.2574442 | 7.69205 | 1.565394575 | 0.843045323 |
| Serping1 | 9.059609 | 8.647232 | 8.6539 | -0.006699575 | 0.405678188 | 8.4030226 | 9.789694 | 8.22518 | 1.564515298 | 0.177843926 |
| Dram1 | 8.384017 | 8.337274 | 7.8277 | 0.509621107 | 0.556363688 | 7.8862849 | 8.8902729 | 7.32959 | 1.560680457 | 0.556692487 |
| Ccr5 | 8.744419 | 8.707309 | 7.8628 | 0.844519272 | 0.881628649 | 8.6460918 | 9.8283141 | 8.26779 | 1.560519408 | 0.378297077 |
| Lyve1 | 9.932811 | 9.572616 | 8.2985 | 1.274117971 | 1.634313433 | 8.624327 | 9.5495197 | 7.99621 | 1.553312886 | 0.628120155 |
| Tspo | 10.00394 | 10.34999 | 9.9041 | 0.4458553 | 0.099803171 | 10.433243 | 11.375225 | 9.82265 | 1.552579615 | 0.610597371 |
| Gm49342 | 8.345201 | 8.761678 | 8.1225 | 0.639133563 | 0.222656951 | 8.1909526 | 9.587939 | 8.03564 | 1.552294116 | 0.155307741 |
| Spint1 | 5.597247 | 5.74549 | 5.4053 | 0.340152003 | 0.19190814 | 6.3097227 | 7.0796549 | 5.53097 | 1.548684447 | 0.778752195 |
| Tlr3 | 8.144541 | 8.699491 | 8.082 | 0.617518365 | 0.06256776 | 8.0032201 | 9.5321252 | 7.98653 | 1.545599066 | 0.016693998 |
| Tnfsf10 | 9.678751 | 10.01481 | 9.5997 | 0.41513449 | 0.079073845 | 9.1710966 | 10.884902 | 9.34079 | 1.544115268 | -0.169690391 |
| Tnc | 9.374148 | 9.125865 | 8.0361 | 1.089723782 | 1.338006787 | 7.7785156 | 9.0514478 | 7.50866 | 1.542792203 | 0.269860048 |
| Gm21188 | 8.092069 | 8.523885 | 7.7765 | 0.747341265 | 0.315525107 | 8.2989611 | 9.2337273 | 7.69407 | 1.539659036 | 0.604892796 |
| Vdr | 5.520067 | 5.484768 | 5.0506 | 0.434120283 | 0.46941961 | 5.3286092 | 6.6109634 | 5.07136 | 1.539607424 | 0.257253199 |
| Sh3bp2 | 10.26734 | 10.88196 | 10.299 | 0.582495798 | -0.032126498 | 10.805279 | 11.927003 | 10.3879 | 1.539151592 | 0.417427781 |
| Pla2g16 | 9.207821 | 9.52453 | 9.2624 | 0.26216203 | -0.054547779 | 9.0875315 | 10.809151 | 9.27057 | 1.538576168 | -0.183043233 |
| Axl | 9.534339 | 9.976713 | 9.46 | 0.516753508 | 0.074379201 | 8.8017624 | 10.497829 | 8.95955 | 1.538278096 | -0.157788369 |
| Ms4a4a | 7.053405 | 7.536789 | 6.1969 | 1.339847789 | 0.856464243 | 7.5061151 | 7.7327633 | 6.1992 | 1.533563647 | 1.306915509 |
| Helz2 | 13.63026 | 14.52936 | 13.446 | 1.083095834 | 0.183998284 | 14.252629 | 15.389751 | 13.8578 | 1.531933168 | 0.394811272 |
| Hsh2d | 9.460187 | 10.24314 | 9.2291 | 1.014070256 | 0.231117117 | 9.8996341 | 10.916463 | 9.38703 | 1.529428983 | 0.512600053 |
| B430306N03Rik | 6.972418 | 7.608613 | 6.6619 | 0.946725432 | 0.310531123 | 7.2955395 | 8.3704923 | 6.84421 | 1.526284689 | 0.451331919 |
| Pla2g7 | 8.194954 | 8.706619 | 7.4406 | 1.26604847 | 0.754383477 | 9.1878657 | 8.8968354 | 7.37237 | 1.524465658 | 1.815495906 |
| Hmgn3 | 7.556872 | 7.810614 | 7.5789 | 0.231672203 | -0.022068841 | 7.4858258 | 9.0100596 | 7.48652 | 1.523536764 | -0.000697043 |
| Chmp4b | 11.16259 | 11.41599 | 10.923 | 0.493374478 | 0.239969702 | 11.192184 | 12.504536 | 10.9814 | 1.523170826 | 0.210818597 |
| Fanca | 7.68578 | 8.35799 | 7.7784 | 0.579551991 | -0.092658182 | 8.6118814 | 9.757845 | 8.24026 | 1.517582614 | 0.37161906 |
| Dnase1l3 | 9.76948 | 9.905041 | 9.6693 | 0.235731512 | 0.10016977 | 9.3832887 | 10.768595 | 9.2675 | 1.50109513 | 0.115788624 |
| Ccl6 | 8.273797 | 8.265253 | 6.7753 | 1.489905563 | 1.498449979 | 7.548779 | 8.1232557 | 6.62232 | 1.500931623 | 0.926454925 |
| Slit1 | 3.487068 | 3.915279 | 3.6227 | 0.292532156 | -0.135678437 | 4.0010777 | 4.8732885 | 3.37359 | 1.499697225 | 0.627486394 |

| | | | | | | | | | | |
|---|---|---|---|---|---|---|---|---|---|---|
| Cybb | 12.28892 | 13.14203 | 12.124 | 1.01850039 | 0.165391524 | 12.852627 | 13.730162 | 12.2326 | 1.497531968 | 0.619997373 |
| Mid1 | 8.413809 | 8.675853 | 8.2528 | 0.423022345 | 0.160979033 | 8.1080007 | 9.7698684 | 8.28171 | 1.488161578 | -0.173706137 |
| Slfn2 | 12.04777 | 12.90688 | 11.917 | 0.990252892 | 0.131149196 | 12.378044 | 13.50464 | 12.0204 | 1.484279928 | 0.357683056 |
| Slc7a8 | 6.191575 | 6.598351 | 5.7609 | 0.837451081 | 0.430674558 | 6.0767885 | 7.0541311 | 5.5706 | 1.483534119 | 0.506191553 |
| Ifi203-ps | 4.28848 | 4.691097 | 4.3396 | 0.351529029 | -0.051087834 | 4.2448842 | 5.8226936 | 4.34282 | 1.479869984 | -0.097939421 |
| Gpr171 | 9.88356 | 10.39986 | 9.6261 | 0.773720667 | 0.257424654 | 9.9797899 | 11.31399 | 9.83464 | 1.479348067 | 0.145148018 |
| Grn | 11.07392 | 11.19822 | 10.921 | 0.277388248 | 0.153088938 | 10.959833 | 12.109129 | 10.6381 | 1.470984794 | 0.321689561 |
| Pilra | 6.31282 | 6.95491 | 5.4314 | 1.523461735 | 0.881371363 | 7.0105036 | 7.0012709 | 5.53609 | 1.465179721 | 1.474412397 |
| Ncoa7 | 9.589069 | 10.20048 | 9.4448 | 0.755682584 | 0.144270711 | 9.3287794 | 10.877912 | 9.4171 | 1.460810737 | -0.088321628 |
| Fam241a | 7.625373 | 7.956205 | 7.5151 | 0.441108472 | 0.110277015 | 7.4045464 | 8.9170725 | 7.46597 | 1.451101778 | -0.061424326 |
| Psme1 | 12.15928 | 12.37095 | 12.068 | 0.303225216 | 0.091546138 | 12.268567 | 13.518492 | 12.0676 | 1.450913648 | 0.200989369 |
| Gm47283 | 9.415734 | 10.03373 | 9.5434 | 0.490288527 | -0.127706176 | 10.027048 | 11.097564 | 9.65333 | 1.444236919 | 0.373720533 |
| Adar | 11.94854 | 12.6661 | 11.795 | 0.871306 | 0.15375125 | 12.19941 | 13.396079 | 11.9519 | 1.444132444 | 0.247463708 |
| Tent4a | 8.704901 | 9.390168 | 8.4461 | 0.944035255 | 0.258768152 | 8.4724075 | 9.9064172 | 8.46453 | 1.441887316 | 0.007877531 |
| Scimp | 7.478311 | 7.813527 | 7.4092 | 0.404334928 | 0.06911852 | 7.6517229 | 8.673229 | 7.23308 | 1.440151301 | 0.418645178 |
| Pde7b | 5.759321 | 6.22115 | 5.6404 | 0.58078337 | 0.118954295 | 5.9332956 | 7.0512131 | 5.6119 | 1.439317462 | 0.321400028 |
| Gca | 6.846675 | 7.683538 | 6.7484 | 0.935161432 | 0.09829771 | 6.9655864 | 8.3823867 | 6.94315 | 1.439237843 | 0.022437455 |
| Gmppb | 8.280622 | 8.514848 | 8.2084 | 0.306480033 | 0.072254792 | 8.3647215 | 9.4975427 | 8.06426 | 1.433280691 | 0.300459542 |
| Tmem67 | 6.000636 | 6.121156 | 5.926 | 0.195127315 | 0.074607225 | 6.2749592 | 7.5137971 | 6.08711 | 1.426688692 | 0.187850791 |
| Naa20 | 8.462609 | 8.702672 | 8.4098 | 0.292887619 | 0.052824327 | 8.5470707 | 9.7910381 | 8.36737 | 1.423670156 | 0.179702789 |
| Tlr13 | 7.836637 | 7.965987 | 7.0546 | 0.911426474 | 0.78207703 | 7.676696 | 8.0991973 | 6.68076 | 1.418437616 | 0.995936258 |
| Csf1 | 9.451495 | 9.714831 | 8.9063 | 0.808542691 | 0.545206401 | 8.7827007 | 10.332394 | 8.92999 | 1.402404466 | -0.147288403 |
| Tor1aip2 | 10.56573 | 11.22344 | 10.497 | 0.726673366 | 0.068969492 | 10.597125 | 12.08441 | 10.6826 | 1.401819376 | -0.085465445 |
| Dll1 | 6.89875 | 7.148098 | 6.9639 | 0.184168499 | -0.065179859 | 6.8528825 | 7.8726187 | 6.47272 | 1.399897407 | 0.380161253 |
| Klrk1 | 8.786449 | 9.018409 | 8.7692 | 0.249160021 | 0.017199776 | 8.4662123 | 9.9190028 | 8.52123 | 1.397770059 | -0.055020433 |
| C130026I21Rik | 7.690789 | 8.142492 | 7.7965 | 0.346002392 | -0.105700825 | 8.1906962 | 9.4728982 | 8.08388 | 1.389018015 | 0.106816017 |
| Cd200r4 | 5.241035 | 5.548662 | 5.2151 | 0.333546736 | 0.025919773 | 5.2022398 | 6.6448092 | 5.26186 | 1.382948452 | -0.059620997 |
| Art2b | 8.319875 | 8.660536 | 8.298 | 0.362525399 | 0.0218648 | 8.2197839 | 9.750683 | 8.3682 | 1.382479455 | -0.148419651 |
| Psme2 | 11.4205 | 11.66863 | 11.338 | 0.330428276 | 0.082297596 | 11.57036 | 12.851699 | 11.4787 | 1.372956098 | 0.091617167 |
| Evi2a | 9.594489 | 9.934591 | 9.4483 | 0.486269414 | 0.146167802 | 9.716874 | 10.714708 | 9.34365 | 1.371054066 | 0.373219661 |
| Lrrc25 | 7.276155 | 7.538246 | 6.7866 | 0.75162544 | 0.48953473 | 7.6050201 | 8.0368779 | 6.66739 | 1.369487759 | 0.937629912 |
| Gipc3 | 5.051891 | 5.230621 | 4.8177 | 0.412908414 | 0.234179007 | 4.9108286 | 6.4279661 | 5.06251 | 1.365458883 | -0.151678702 |

| | | | | | | | | | | |
|---|---|---|---|---|---|---|---|---|---|---|
| Psmb8 | 12.06412 | 12.27965 | 11.973 | 0.30680898 | 0.091281798 | 12.235366 | 13.266259 | 11.9071 | 1.359187364 | 0.328294604 |
| Acot4 | 6.733078 | 7.128956 | 6.6803 | 0.448643347 | 0.052765452 | 6.8832623 | 8.2269486 | 6.86899 | 1.357962941 | 0.014276657 |
| H2-Q4 | 12.69854 | 12.77342 | 12.65 | 0.122954572 | 0.048067958 | 12.75766 | 14.069008 | 12.7126 | 1.356399274 | 0.045051155 |
| Bcl2l14 | 6.765463 | 6.894466 | 6.7871 | 0.107356854 | -0.021645779 | 6.8067267 | 8.055594 | 6.70135 | 1.354245911 | 0.105378696 |
| Rabepk | 8.021061 | 8.688668 | 7.9675 | 0.721208239 | 0.053601518 | 8.4074592 | 9.4067111 | 8.06076 | 1.345949045 | 0.34669719 |
| Abtb2 | 9.263967 | 9.298814 | 9.0099 | 0.288882321 | 0.254035291 | 9.0793339 | 10.337587 | 8.99552 | 1.342068529 | 0.083815754 |
| Rab44 | 5.120572 | 5.729509 | 4.8953 | 0.834167539 | 0.225229971 | 5.333747 | 6.0991269 | 4.75803 | 1.341092346 | 0.575712505 |
| Rnf114 | 11.44996 | 12.04834 | 11.379 | 0.668906028 | 0.070531218 | 11.696917 | 12.749815 | 11.4104 | 1.339384096 | 0.286486524 |
| Perm1 | 4.573692 | 4.992376 | 4.3442 | 0.648208998 | 0.229525037 | 4.8374796 | 5.9614412 | 4.62307 | 1.338375392 | 0.214413768 |
| Wfdc17 | 7.428543 | 7.850401 | 6.3844 | 1.465975111 | 1.044117195 | 7.7988537 | 7.9968635 | 6.65945 | 1.337417897 | 1.13940802 |
| Dusp28 | 6.043812 | 6.168911 | 5.8905 | 0.278429294 | 0.153330389 | 6.1303473 | 7.1961762 | 5.86149 | 1.334684626 | 0.268855718 |
| Slc16a3 | 7.603887 | 7.003394 | 6.1824 | 0.821024429 | 1.421517531 | 7.1963301 | 7.6600607 | 6.32579 | 1.334272628 | 0.870542024 |
| A530064D06Rik | 5.660583 | 6.288388 | 5.1759 | 1.112504975 | 0.484699108 | 5.8431038 | 6.7710354 | 5.43689 | 1.334146931 | 0.406215342 |
| Cmklr1 | 6.991774 | 7.557247 | 6.6627 | 0.894566143 | 0.329093175 | 7.0225064 | 7.8129723 | 6.48035 | 1.332619656 | 0.542153714 |
| Irf8 | 12.31493 | 12.47156 | 12.216 | 0.255989392 | 0.09936415 | 12.340261 | 13.314385 | 11.9828 | 1.331580498 | 0.357457166 |
| Ms4a4d | 5.919273 | 5.88746 | 5.5146 | 0.372844246 | 0.404656634 | 5.4919229 | 6.7058798 | 5.37562 | 1.330257919 | 0.116300962 |
| Fam177a | 7.479468 | 8.133761 | 7.017 | 1.116795327 | 0.462501919 | 7.1124792 | 8.2052375 | 6.88519 | 1.320046439 | 0.227288196 |
| Zcchc2 | 10.43786 | 11.22464 | 10.179 | 1.045179279 | 0.258395787 | 10.438906 | 11.771715 | 10.454 | 1.317755356 | -0.015054529 |
| Socs3 | 11.11054 | 10.58228 | 10.024 | 0.558533092 | 1.086796958 | 10.938465 | 11.15118 | 9.84048 | 1.310702518 | 1.097986889 |
| Tifab | 8.088477 | 8.252383 | 7.978 | 0.274379213 | 0.110473467 | 8.337013 | 9.1234764 | 7.81287 | 1.310608003 | 0.524144588 |
| Stx3 | 5.765685 | 6.036916 | 5.7471 | 0.289857403 | 0.018625643 | 5.3138605 | 6.7032807 | 5.39458 | 1.308705136 | -0.080715057 |
| Themis2 | 10.98871 | 11.30054 | 10.771 | 0.529343818 | 0.217514934 | 11.265076 | 12.047907 | 10.7406 | 1.307286324 | 0.524456207 |
| Max | 10.79334 | 11.09474 | 10.688 | 0.406245028 | 0.104845906 | 10.892094 | 12.024295 | 10.7192 | 1.305135994 | 0.172934934 |
| Cxcl13 | 9.464297 | 9.65374 | 8.3347 | 1.319048318 | 1.129604529 | 9.4060355 | 9.2991776 | 7.99518 | 1.303993722 | 1.41085167 |
| Olfr1372-ps1 | 7.459178 | 7.694065 | 7.2625 | 0.431594092 | 0.196707476 | 8.0974151 | 9.1485731 | 7.84619 | 1.302380034 | 0.251222037 |
| Pnpt1 | 9.679735 | 10.1698 | 9.3848 | 0.785006994 | 0.294944758 | 9.7449313 | 10.977638 | 9.67797 | 1.299669911 | 0.066963106 |
| Tarm1 | 5.726791 | 5.970455 | 4.6844 | 1.286042335 | 1.04237836 | 5.4628381 | 6.1081753 | 4.80871 | 1.2994688 | 0.654131587 |
| Ubr4 | 12.46181 | 12.76338 | 12.479 | 0.284691298 | -0.016882522 | 12.836416 | 14.024501 | 12.7259 | 1.298578364 | 0.110493862 |
| Cd14 | 8.987703 | 8.346896 | 6.1975 | 2.149433885 | 2.790240334 | 8.5078149 | 7.7436862 | 6.44576 | 1.297922978 | 2.062051655 |
| Gm12573 | 5.232594 | 5.794974 | 5.1374 | 0.657578057 | 0.095198285 | 5.8341986 | 6.8137321 | 5.51962 | 1.294115279 | 0.314581849 |
| Gm16464 | 5.290402 | 5.846596 | 5.2032 | 0.643417082 | 0.087223123 | 5.5412553 | 6.6511868 | 5.3611 | 1.290086069 | 0.180154575 |
| H2-T10 | 9.872238 | 10.17166 | 9.9687 | 0.202976088 | -0.096446982 | 10.279065 | 11.304385 | 10.0158 | 1.288556786 | 0.263236228 |

| | | | | | | | | | | |
|---|---|---|---|---|---|---|---|---|---|---|
| Dtx4 | 6.924733 | 7.399558 | 6.7926 | 0.60693695 | 0.132112004 | 7.3044665 | 8.248918 | 6.96075 | 1.288168344 | 0.34371689 |
| Mafk | 9.807864 | 10.22954 | 9.7041 | 0.525461485 | 0.103784024 | 9.7276754 | 10.886222 | 9.5984 | 1.287818399 | 0.129272258 |
| Tubb6 | 7.604943 | 7.546084 | 6.6092 | 0.936872903 | 0.995732108 | 7.1633564 | 7.7005981 | 6.4129 | 1.287703111 | 0.75046137 |
| Aida | 10.41814 | 11.00009 | 10.456 | 0.544283742 | -0.037668166 | 10.620768 | 11.805583 | 10.5183 | 1.287260598 | 0.102445471 |
| Klrc1 | 7.345773 | 7.476823 | 7.4067 | 0.070098718 | -0.060951455 | 7.1425328 | 8.4513964 | 7.16604 | 1.285351458 | -0.023512121 |
| B4galt5 | 10.24484 | 10.83717 | 9.8936 | 0.94357499 | 0.351240236 | 10.182383 | 11.477215 | 10.195 | 1.282168095 | -0.012663144 |
| Eomes | 8.346422 | 8.731231 | 8.5512 | 0.179992756 | -0.204817161 | 8.2465617 | 9.7192227 | 8.4401 | 1.279120446 | -0.193540592 |
| Pgd | 10.443 | 10.79416 | 10.33 | 0.46462775 | 0.113464096 | 10.64221 | 11.567262 | 10.2901 | 1.27712078 | 0.352068776 |
| Sp110 | 11.88543 | 12.29308 | 11.807 | 0.486433726 | 0.078790316 | 12.228643 | 13.222828 | 11.9461 | 1.27668221 | 0.282496697 |
| Gm6904 | 5.427319 | 5.930695 | 5.2396 | 0.691137492 | 0.187761351 | 5.9292247 | 6.8509429 | 5.57478 | 1.276166288 | 0.354448129 |
| Pik3ap1 | 11.91185 | 12.4311 | 11.808 | 0.623220828 | 0.103966502 | 12.336252 | 13.130864 | 11.8559 | 1.27498856 | 0.480376835 |
| Arf4 | 9.688498 | 10.02512 | 9.6523 | 0.372805205 | 0.036180393 | 9.7170353 | 10.849545 | 9.58028 | 1.269266421 | 0.136756932 |
| Ly86 | 10.8611 | 11.32011 | 10.849 | 0.471500504 | 0.012495294 | 11.396684 | 12.110862 | 10.842 | 1.26882782 | 0.554649953 |
| Mthfr | 8.588195 | 9.087005 | 8.7353 | 0.351672586 | -0.14713687 | 9.0330888 | 10.060873 | 8.79338 | 1.267493754 | 0.239709371 |
| Dnaja1 | 12.14178 | 12.81681 | 12.105 | 0.711880386 | 0.036848282 | 11.798705 | 13.393201 | 12.1279 | 1.265293808 | -0.32920231 |
| Ell3 | 8.166574 | 8.402063 | 8.1686 | 0.233416635 | -0.002072622 | 8.7782662 | 9.4413343 | 8.17614 | 1.265193687 | 0.602125561 |
| Treml2 | 10.70629 | 11.14899 | 10.581 | 0.567640034 | 0.124947462 | 10.914214 | 11.855755 | 10.5943 | 1.26148006 | 0.319938675 |
| Agrn | 8.740037 | 9.12692 | 8.4965 | 0.630437836 | 0.243554986 | 8.8823728 | 9.9286372 | 8.66732 | 1.261317135 | 0.215052704 |
| Ldlr | 9.647713 | 9.634433 | 9.1528 | 0.481659804 | 0.494939679 | 9.4141282 | 10.109673 | 8.8488 | 1.260872896 | 0.565328427 |
| Il6 | 5.270914 | 5.344049 | 4.4337 | 0.910397363 | 0.837262547 | 4.5378565 | 5.7879094 | 4.5292 | 1.25870824 | 0.008655328 |
| Cd68 | 9.04397 | 9.429023 | 8.6187 | 0.810283208 | 0.42523012 | 8.9499085 | 9.5175928 | 8.26047 | 1.257123096 | 0.689438846 |
| Zfp365 | 6.508479 | 6.953336 | 6.325 | 0.62829024 | 0.183433078 | 6.3876146 | 7.768384 | 6.51141 | 1.256972673 | -0.123796725 |
| Zxda | 4.143203 | 3.510132 | 3.7463 | -0.236157128 | 0.396913466 | 3.852778 | 4.8230031 | 3.56649 | 1.256517658 | 0.286292554 |
| H2-K2 | 9.407822 | 9.838101 | 9.542 | 0.296085272 | -0.1341945 | 9.9431053 | 10.997761 | 9.74266 | 1.255105329 | 0.200450038 |
| F830016B08Rik | 4.958537 | 5.362076 | 4.9926 | 0.36946183 | -0.034077107 | 4.7279163 | 6.2351175 | 4.98013 | 1.254986174 | -0.252215085 |
| Trim12c | 11.02236 | 11.66868 | 10.866 | 0.80315646 | 0.156840814 | 11.401061 | 12.320394 | 11.0655 | 1.254891524 | 0.335558974 |
| Cables1 | 4.84626 | 5.053267 | 4.7791 | 0.274168674 | 0.067160945 | 4.7305539 | 5.7760566 | 4.52267 | 1.253388052 | 0.207885384 |
| Scarf1 | 6.95187 | 6.87694 | 6.7868 | 0.090129116 | 0.165058616 | 6.3727986 | 7.3669653 | 6.11383 | 1.253138461 | 0.258971796 |
| Nectin4 | 5.169073 | 5.524343 | 5.1579 | 0.366402494 | 0.01113247 | 5.3398113 | 6.3278308 | 5.07533 | 1.252498207 | 0.264478718 |
| Slamf7 | 10.0079 | 10.40355 | 10.128 | 0.275058374 | -0.120587121 | 9.9496123 | 11.395917 | 10.1504 | 1.245538804 | -0.200765974 |
| 38412 | 8.543512 | 8.966139 | 8.4716 | 0.494547273 | 0.071920636 | 8.5715712 | 9.7417171 | 8.49861 | 1.243107972 | 0.072962132 |
| Sco1 | 7.71062 | 7.724379 | 7.6706 | 0.053752621 | 0.039994057 | 7.5107561 | 8.6862255 | 7.44348 | 1.242746342 | 0.067276896 |

| | | | | | | | | | | |
|---|---|---|---|---|---|---|---|---|---|---|
| Trim25 | 11.19207 | 11.89001 | 11.038 | 0.85216574 | 0.154229952 | 11.343727 | 12.51376 | 11.2715 | 1.24230472 | 0.072272302 |
| Fgf23 | 4.513005 | 4.637225 | 3.5002 | 1.137025689 | 1.012805298 | 4.0535692 | 4.8328272 | 3.59341 | 1.239418971 | 0.460161032 |
| Slpi | 7.153141 | 8.184876 | 6.4997 | 1.685152682 | 0.653417644 | 8.0823762 | 8.4149807 | 7.17635 | 1.238630178 | 0.906025728 |
| Rmi2 | 6.663413 | 6.672013 | 6.596 | 0.076056845 | 0.06745642 | 6.5453633 | 8.0804474 | 6.84269 | 1.237760576 | -0.297323544 |
| Trim34a | 10.70968 | 11.49454 | 10.732 | 0.76270162 | -0.02216221 | 11.242423 | 12.150577 | 10.9174 | 1.233214088 | 0.325059885 |
| Gm7609 | 6.499737 | 7.211812 | 6.3258 | 0.886059726 | 0.173984821 | 7.0993538 | 8.0056678 | 6.77794 | 1.227725059 | 0.321411038 |
| Hk2 | 8.215935 | 8.576493 | 7.8299 | 0.746605948 | 0.386048369 | 8.1822645 | 9.2249312 | 7.99774 | 1.227195268 | 0.184528608 |
| Myd88 | 10.1142 | 10.14346 | 9.6467 | 0.496810415 | 0.467549531 | 10.125946 | 10.83128 | 9.60708 | 1.224198065 | 0.518863747 |
| Selenow | 11.13605 | 11.56141 | 11.108 | 0.453800652 | 0.02843384 | 11.618934 | 12.347317 | 11.1236 | 1.223736838 | 0.49535346 |
| Trim14 | 8.318806 | 9.210299 | 8.4925 | 0.717782092 | -0.173710932 | 8.6389808 | 9.765872 | 8.54242 | 1.223448604 | 0.096557345 |
| Pilrb1 | 4.963681 | 5.178234 | 4.621 | 0.557264386 | 0.342710719 | 5.2929141 | 5.5510105 | 4.32819 | 1.222816686 | 0.964720262 |
| N4bp1 | 10.19003 | 10.66974 | 10.115 | 0.554523599 | 0.074812745 | 10.155128 | 11.361492 | 10.1449 | 1.216607096 | 0.010242527 |
| Cgas | 7.338127 | 7.834756 | 7.182 | 0.652734454 | 0.156105132 | 7.157033 | 8.5119799 | 7.29607 | 1.215904994 | -0.139041951 |
| H2-Q6 | 12.98841 | 13.12252 | 13.038 | 0.084787208 | -0.049321028 | 13.187201 | 14.338896 | 13.1233 | 1.215549072 | 0.06385443 |
| Pmepa1 | 7.247 | 7.596819 | 7.2309 | 0.365877006 | 0.016058936 | 6.9303036 | 8.3304039 | 7.11504 | 1.215368668 | -0.184731566 |
| Rundc3b | 7.483749 | 7.945879 | 7.4392 | 0.506661396 | 0.044530663 | 7.3824792 | 8.6804823 | 7.47003 | 1.210452702 | -0.087550372 |
| Atp10a | 8.987677 | 9.155552 | 8.872 | 0.283516505 | 0.115641441 | 8.6731493 | 10.190622 | 8.98141 | 1.209209369 | -0.308263309 |
| Ccnyl1 | 7.895433 | 8.299526 | 7.6214 | 0.678151021 | 0.27405886 | 7.8073214 | 9.0035174 | 7.79439 | 1.209128075 | 0.012932069 |
| Ifitm6 | 6.614016 | 6.23949 | 5.1043 | 1.135142632 | 1.509667764 | 6.0936647 | 6.5183885 | 5.31002 | 1.208369906 | 0.783646089 |
| Tapbpl | 10.2957 | 10.49723 | 10.277 | 0.220569242 | 0.019040196 | 10.417727 | 11.4786 | 10.271 | 1.207621024 | 0.146748064 |
| Tap2 | 11.73049 | 11.88478 | 11.672 | 0.212334404 | 0.058044524 | 11.873018 | 12.898675 | 11.6938 | 1.204872864 | 0.179216064 |
| Ctsc | 11.76044 | 11.99296 | 11.544 | 0.448748148 | 0.21622477 | 11.941226 | 12.721108 | 11.5163 | 1.204792622 | 0.424910501 |
| Ccr1 | 8.027168 | 7.681294 | 6.0537 | 1.627583135 | 1.973457111 | 7.2524475 | 7.389452 | 6.18622 | 1.203228634 | 1.066224151 |
| Serpinb9 | 10.52174 | 10.64898 | 10.42 | 0.228801422 | 0.101568256 | 9.8284086 | 11.25058 | 10.0482 | 1.202343371 | -0.219828158 |
| Usp25 | 11.65517 | 12.20457 | 11.534 | 0.670104878 | 0.120703468 | 11.611455 | 12.710244 | 11.5093 | 1.200898748 | 0.102110467 |
| Cnp | 11.7247 | 12.15322 | 11.686 | 0.46706242 | 0.038539228 | 11.936787 | 12.784616 | 11.5847 | 1.199880768 | 0.352051634 |
| Gm11331 | 5.214212 | 5.853884 | 4.7724 | 1.08146732 | 0.441795409 | 6.0234522 | 6.6998449 | 5.50333 | 1.196514305 | 0.52012158 |
| Lif | 5.893137 | 5.991208 | 5.6255 | 0.365673581 | 0.267602356 | 5.2626966 | 6.5676287 | 5.37254 | 1.195092206 | -0.109839819 |
| Lrp1 | 9.430858 | 9.14604 | 8.4807 | 0.665357065 | 0.950174485 | 9.3636495 | 9.5167538 | 8.32471 | 1.192047348 | 1.038942977 |
| Msrb1 | 8.978479 | 8.959357 | 8.4134 | 0.545955224 | 0.565076491 | 9.1703031 | 9.5275013 | 8.33677 | 1.190729129 | 0.833530858 |
| Mvb12a | 8.638589 | 8.898275 | 8.6007 | 0.297619209 | 0.037932307 | 8.8838977 | 9.7161137 | 8.52625 | 1.189858957 | 0.357643 |
| Clic4 | 11.99728 | 12.16253 | 11.606 | 0.556818598 | 0.391564276 | 11.703507 | 12.797568 | 11.6083 | 1.189314218 | 0.09525261 |

| | | | | | | | | | | |
|---|---|---|---|---|---|---|---|---|---|---|
| Mmp19 | 6.330567 | 6.984537 | 4.9597 | 2.024817011 | 1.370847596 | 5.743942 | 6.3111227 | 5.1219 | 1.189221332 | 0.622040648 |
| H2-K1 | 15.05553 | 15.2271 | 15.083 | 0.143685758 | -0.027881028 | 15.262512 | 16.253719 | 15.0689 | 1.184802874 | 0.193596065 |
| C2 | 7.72485 | 7.483834 | 7.5255 | -0.041662412 | 0.199354032 | 6.8166039 | 8.4217117 | 7.23804 | 1.183674391 | -0.421433371 |
| Vcpip1 | 10.92402 | 11.48529 | 10.877 | 0.60871702 | 0.047448006 | 10.924992 | 12.110952 | 10.9275 | 1.183476542 | -0.00248332 |
| Tfec | 5.336649 | 5.779675 | 4.8822 | 0.89751424 | 0.454488362 | 5.3432359 | 5.8836882 | 4.70134 | 1.18234652 | 0.641894256 |
| Scand1 | 6.291481 | 6.346774 | 6.2016 | 0.145158073 | 0.089865724 | 6.5835023 | 7.1288017 | 5.9476 | 1.181202798 | 0.635903385 |
| Car13 | 6.451624 | 7.023222 | 6.2966 | 0.726603834 | 0.155005968 | 6.4204726 | 7.2911757 | 6.11062 | 1.180559382 | 0.309856253 |
| St6galnac4 | 9.450847 | 9.754237 | 9.289 | 0.465194706 | 0.161804242 | 9.6275107 | 10.504179 | 9.32713 | 1.177045384 | 0.300376916 |
| Smagp | 7.447282 | 7.503579 | 7.5644 | -0.060830625 | -0.117127727 | 7.3990887 | 8.3307079 | 7.15557 | 1.175140072 | 0.243520778 |
| Zcchc18 | 8.236815 | 8.551459 | 8.1986 | 0.352865665 | 0.038221277 | 8.392463 | 9.177326 | 8.00257 | 1.17475393 | 0.389890908 |
| Gm48499 | 3.554934 | 3.648599 | 3.5105 | 0.138126247 | 0.044461283 | 4.3732304 | 4.9344072 | 3.76414 | 1.1702629 | 0.609086111 |
| AI837181 | 9.247449 | 9.680037 | 9.3588 | 0.321202978 | -0.111385107 | 9.5692505 | 10.372488 | 9.20369 | 1.168792966 | 0.365555625 |
| Emp1 | 8.697506 | 8.482476 | 7.559 | 0.923461284 | 1.138491103 | 7.6356852 | 8.6350925 | 7.46799 | 1.167101386 | 0.167694054 |
| Clec4n | 8.380151 | 8.462626 | 6.9845 | 1.47810036 | 1.395625411 | 8.2438621 | 7.7628662 | 6.59709 | 1.165771478 | 1.646767378 |
| Sowahc | 7.885766 | 7.956055 | 7.4902 | 0.465903613 | 0.395614047 | 7.344117 | 8.5467963 | 7.38137 | 1.165426312 | -0.037252964 |
| C3 | 12.82482 | 12.61386 | 12.152 | 0.461860248 | 0.672821216 | 11.895425 | 12.944224 | 11.7789 | 1.165351978 | 0.116552858 |
| Tyrobp | 10.07306 | 10.15011 | 9.7683 | 0.381821964 | 0.30477215 | 10.213378 | 10.604852 | 9.44023 | 1.164618336 | 0.773144835 |
| Stip1 | 11.0046 | 11.64357 | 11.024 | 0.61996772 | -0.0189997 | 11.015254 | 12.166807 | 11.0035 | 1.163332928 | 0.011779875 |
| Il15ra | 7.964605 | 8.12765 | 7.9709 | 0.156771216 | -0.006274028 | 7.7404178 | 9.2745068 | 8.11197 | 1.162534023 | -0.371554991 |
| Mvp | 10.51943 | 10.67993 | 10.262 | 0.417485304 | 0.256979269 | 10.627524 | 11.405745 | 10.2454 | 1.16035596 | 0.382135004 |
| Arid5a | 10.81456 | 10.99418 | 10.45 | 0.544513302 | 0.364896936 | 10.874446 | 11.878179 | 10.7189 | 1.159260322 | 0.155526581 |
| Serpinh1 | 8.516949 | 8.823507 | 8.133 | 0.690548667 | 0.383990667 | 8.8510131 | 8.8304692 | 7.67205 | 1.158417876 | 1.178961736 |
| Iqsec2 | 7.120104 | 7.035386 | 6.8237 | 0.211721222 | 0.296440154 | 6.6672969 | 7.5412764 | 6.39743 | 1.143847025 | 0.269867546 |
| Etv6 | 9.996659 | 10.04841 | 9.6743 | 0.374120154 | 0.322368365 | 9.8871033 | 10.8197 | 9.68281 | 1.136887152 | 0.204290844 |
| Rbl1 | 9.983921 | 10.3412 | 10.008 | 0.332945859 | -0.024336951 | 9.9747936 | 11.286744 | 10.1499 | 1.13680396 | -0.175146215 |
| A230050P20Rik | 8.363873 | 8.576349 | 8.4325 | 0.143891708 | -0.068585131 | 8.8386225 | 9.6650503 | 8.53032 | 1.1347276 | 0.308299837 |
| Pcgf5 | 9.543743 | 10.02584 | 9.5473 | 0.478512969 | -0.003582909 | 9.4470174 | 10.736674 | 9.60264 | 1.134033482 | -0.15562356 |
| H2-Q7 | 13.36642 | 13.43892 | 13.34 | 0.098764316 | 0.02626337 | 13.60324 | 14.58588 | 13.4521 | 1.133782042 | 0.151141264 |
| Csf2ra | 8.150701 | 8.077265 | 7.7884 | 0.288836408 | 0.362271693 | 8.3738756 | 9.0756205 | 7.94233 | 1.133291382 | 0.431546422 |
| Gramd3 | 10.35971 | 10.73566 | 10.372 | 0.363249746 | -0.012703878 | 10.41726 | 11.623544 | 10.4907 | 1.13280455 | -0.073479544 |
| AC132444.2 | 4.777095 | 5.074959 | 4.7018 | 0.373183908 | 0.075319653 | 4.9699769 | 5.92343 | 4.79317 | 1.130261475 | 0.176808396 |
| AC132444.4 | 4.777095 | 5.074959 | 4.7018 | 0.373183908 | 0.075319653 | 4.9699769 | 5.92343 | 4.79317 | 1.130261475 | 0.176808396 |

| | | | | | | | | | | |
|---|---|---|---|---|---|---|---|---|---|---|
| AC133103.6 | 4.777095 | 5.074959 | 4.7018 | 0.373183908 | 0.075319653 | 4.9699769 | 5.92343 | 4.79317 | 1.130261475 | 0.176808396 |
| I830077J02Rik | 7.257329 | 7.150356 | 7.0402 | 0.110124547 | 0.217097398 | 7.6639831 | 8.0589384 | 6.92877 | 1.130171706 | 0.735216431 |
| BC023105 | 3.307616 | 3.354379 | 3.2055 | 0.148852125 | 0.102089617 | 3.0167758 | 4.2404524 | 3.11265 | 1.127801145 | -0.095875526 |
| Bst1 | 8.19997 | 7.845655 | 7.9513 | -0.105647505 | 0.248667577 | 7.9051874 | 8.5580878 | 7.4306 | 1.127490183 | 0.474589822 |
| Cacybp | 9.95113 | 10.44435 | 9.9258 | 0.518513929 | 0.025290384 | 9.8852911 | 11.041337 | 9.9143 | 1.127037137 | -0.029008805 |
| Slc15a2 | 8.189105 | 7.315275 | 7.6368 | -0.321474915 | 0.55235523 | 6.9718584 | 8.5794625 | 7.45272 | 1.126744383 | -0.480859759 |
| Slc11a1 | 7.654741 | 7.694902 | 6.8992 | 0.79572386 | 0.75556243 | 7.5283242 | 7.5947786 | 6.46874 | 1.126040942 | 1.059586555 |
| Gpr35 | 8.471314 | 8.137359 | 7.5396 | 0.597728204 | 0.931683502 | 8.5132696 | 8.6556388 | 7.53192 | 1.123714577 | 0.981345426 |
| Sp100 | 13.29689 | 13.83442 | 13.137 | 0.697561996 | 0.160033912 | 13.580518 | 14.49231 | 13.3723 | 1.119962212 | 0.208170233 |
| Nr1h3 | 7.536127 | 8.36477 | 7.1493 | 1.215481398 | 0.386837567 | 7.5256699 | 7.9690273 | 6.84952 | 1.119510783 | 0.676153323 |
| Clec4a3 | 7.35018 | 7.551453 | 6.7976 | 0.753817613 | 0.552545189 | 8.4750356 | 8.0614752 | 6.9458 | 1.115673642 | 1.529234009 |
| Bbx | 10.01644 | 10.31659 | 9.9902 | 0.326387351 | 0.026227484 | 9.9680209 | 11.235422 | 10.1201 | 1.115321777 | -0.152079375 |
| Zbtb5 | 7.944703 | 8.215645 | 8.0837 | 0.13192065 | -0.13902186 | 8.0645139 | 9.0614311 | 7.94637 | 1.11505816 | 0.118140959 |
| Ctsz | 11.43898 | 11.70997 | 11.346 | 0.36431266 | 0.093324786 | 11.716547 | 12.29774 | 11.1848 | 1.112958686 | 0.531766181 |
| Dck | 9.90578 | 10.56718 | 9.9359 | 0.631261747 | -0.030134128 | 9.9821321 | 11.041936 | 9.93003 | 1.111901817 | 0.052097594 |
| Med12l | 5.506883 | 5.838085 | 5.5579 | 0.28016303 | -0.051039031 | 5.5175125 | 6.5916878 | 5.48 | 1.111686546 | 0.037511337 |
| Vps54 | 10.33558 | 10.57101 | 10.298 | 0.273140109 | 0.037703438 | 10.235032 | 11.596324 | 10.487 | 1.109317624 | -0.251974643 |
| Txn1 | 10.24063 | 10.29829 | 10.163 | 0.135027168 | 0.077367107 | 10.32264 | 11.298676 | 10.1908 | 1.107842785 | 0.131806552 |
| Keap1 | 10.06017 | 10.42822 | 9.939 | 0.489168476 | 0.121119751 | 10.090371 | 11.073165 | 9.966 | 1.107168176 | 0.124374337 |
| Tlr2 | 9.00063 | 9.046524 | 8.3173 | 0.729176294 | 0.683282814 | 8.6802328 | 9.0931776 | 7.98885 | 1.104331546 | 0.691386767 |
| Clec4d | 7.312192 | 7.027802 | 5.1312 | 1.896571607 | 2.180962014 | 6.2581439 | 6.2239604 | 5.12158 | 1.102382975 | 1.13656651 |
| Cacna1h | 6.293305 | 6.658354 | 6.7318 | -0.073430358 | -0.438479431 | 6.2546251 | 7.4813935 | 6.3791 | 1.1022967 | -0.124471768 |
| B2m | 15.09033 | 15.31983 | 15.085 | 0.234991792 | 0.005486306 | 15.274606 | 16.20229 | 15.1008 | 1.10144983 | 0.173765221 |
| Naa25 | 10.13433 | 10.41831 | 10.068 | 0.350634929 | 0.066655559 | 10.127323 | 11.160197 | 10.0608 | 1.099420208 | 0.066546992 |
| Gpr84 | 3.53338 | 4.60581 | 3.348 | 1.25780339 | 0.185373811 | 3.8901004 | 4.5086456 | 3.41 | 1.098645272 | 0.480100077 |
| Cysltr2 | 5.554541 | 5.643487 | 5.4776 | 0.165901858 | 0.076956086 | 5.5541312 | 6.7068933 | 5.60902 | 1.097868617 | -0.054893476 |
| Ctss | 12.90138 | 13.11196 | 12.762 | 0.349662382 | 0.139075556 | 13.213727 | 13.81194 | 12.715 | 1.096947344 | 0.498734916 |
| Il15 | 7.43175 | 7.601253 | 7.3267 | 0.274507192 | 0.105004565 | 6.8976722 | 8.1219419 | 7.03017 | 1.091774807 | -0.132494842 |
| Angpt2 | 7.717904 | 7.70833 | 6.9057 | 0.80263616 | 0.812209473 | 6.4116102 | 7.4871332 | 6.39561 | 1.091524366 | 0.01600132 |
| C5ar1 | 6.839998 | 6.807006 | 4.9736 | 1.833442768 | 1.866434443 | 6.5780128 | 6.1304047 | 5.04346 | 1.086947249 | 1.534555403 |
| Apol6 | 2.381341 | 2.492874 | 2.4146 | 0.078256872 | -0.033275918 | 2.3532495 | 3.5421079 | 2.45746 | 1.08464498 | -0.104213412 |
| Nceh1 | 9.228072 | 9.379355 | 9.2248 | 0.15455172 | 0.003268753 | 8.9903569 | 10.066989 | 8.98446 | 1.082527532 | 0.005895086 |

| | | | | | | | | | | |
|---|---|---|---|---|---|---|---|---|---|---|
| Ccnd2 | 12.20998 | 12.68475 | 12.059 | 0.62611237 | 0.151345302 | 12.086297 | 13.170114 | 12.0878 | 1.082274026 | -0.00154385 |
| Psme2b | 9.358003 | 9.960242 | 9.4538 | 0.506452212 | -0.095787225 | 9.6476339 | 10.629513 | 9.54869 | 1.080820755 | 0.098941333 |
| Vwa3b | 4.080729 | 4.117365 | 4.1671 | -0.04978363 | -0.086419642 | 3.8956794 | 4.9718046 | 3.89206 | 1.079741786 | 0.003616579 |
| Phc2 | 10.8039 | 11.0921 | 10.754 | 0.338185606 | 0.049981426 | 10.944775 | 11.762716 | 10.683 | 1.079714514 | 0.261773671 |
| Metrnl | 6.865317 | 6.7568 | 6.0069 | 0.749901446 | 0.858417901 | 6.6150134 | 6.9685738 | 5.89086 | 1.077710186 | 0.724149774 |
| Ahsa1 | 10.71579 | 11.09557 | 10.765 | 0.331002152 | -0.04877801 | 10.727116 | 11.756883 | 10.6795 | 1.07737011 | 0.047603317 |
| Serpine1 | 5.786109 | 5.773045 | 5.5407 | 0.232363936 | 0.245427547 | 5.2186582 | 6.338647 | 5.26129 | 1.077356068 | -0.042632741 |
| Ppm1k | 9.379306 | 10.31825 | 9.4652 | 0.853032149 | -0.085914615 | 9.5044823 | 10.631277 | 9.55476 | 1.076517407 | -0.050277192 |
| Plod3 | 8.97052 | 9.02938 | 8.6461 | 0.383291451 | 0.324430825 | 8.9742922 | 9.6606188 | 8.58507 | 1.075547905 | 0.38922129 |
| Gimap4 | 12.33715 | 12.4368 | 12.353 | 0.083880968 | -0.015770908 | 12.507402 | 13.433648 | 12.36 | 1.07369682 | 0.147450525 |
| Mmp3 | 5.004225 | 5.757523 | 4.441 | 1.316565601 | 0.563267197 | 5.6940919 | 5.6997699 | 4.63075 | 1.069022238 | 1.063344162 |
| Edn1 | 4.359217 | 4.685973 | 4.3934 | 0.292602284 | -0.034153903 | 4.1235593 | 5.342565 | 4.27393 | 1.06863145 | -0.150374277 |
| Igkv4-68 | 7.578734 | 7.987534 | 7.5454 | 0.442140843 | 0.033341003 | 7.9648085 | 8.3689411 | 7.30113 | 1.067813819 | 0.663681221 |
| Gm7582 | 4.232147 | 4.516048 | 4.1995 | 0.316512032 | 0.032611129 | 4.2347798 | 5.1195246 | 4.05251 | 1.06701754 | 0.182272756 |
| Chic1 | 6.372885 | 6.791575 | 6.3547 | 0.43684241 | 0.01815293 | 6.6948648 | 7.6336339 | 6.56891 | 1.064720583 | 0.125951431 |
| Serpini1 | 8.431033 | 9.135445 | 8.3929 | 0.742548317 | 0.038136066 | 8.8028113 | 9.4843059 | 8.42231 | 1.061992024 | 0.380497456 |
| Gm4294 | 6.695285 | 5.966676 | 7.3808 | -1.414166192 | -0.685557381 | 5.4538871 | 7.4416428 | 6.3798 | 1.061843 | -0.925912616 |
| Gm2564 | 7.519355 | 7.120892 | 6.464 | 0.656919615 | 1.055382616 | 6.8685329 | 7.1461431 | 6.0849 | 1.06123879 | 0.78362863 |
| Gm10053 | 9.2789 | 9.329555 | 9.0415 | 0.288097846 | 0.237442743 | 9.2611188 | 10.136997 | 9.07614 | 1.060851927 | 0.184974135 |
| Sfrp1 | 4.860069 | 4.468385 | 4.86 | -0.391624715 | 5.9553E-05 | 4.3565948 | 5.5673571 | 4.50967 | 1.057682313 | -0.153079933 |
| Baiap3 | 7.790938 | 7.907138 | 8.0276 | -0.120430289 | -0.236630231 | 8.3093389 | 8.9396659 | 7.8849 | 1.054765659 | 0.424438732 |
| Ly6k | 5.502199 | 5.451245 | 5.3969 | 0.054329478 | 0.105283658 | 5.4224208 | 6.2020622 | 5.14769 | 1.054375813 | 0.274734406 |
| P4ha1 | 9.066711 | 9.482728 | 9.0332 | 0.449541228 | 0.033523794 | 8.9801297 | 10.055274 | 9.00217 | 1.053108344 | -0.022036304 |
| Pnpla2 | 9.870639 | 10.09894 | 9.8284 | 0.270512587 | 0.042215991 | 10.125966 | 10.902236 | 9.85242 | 1.049819036 | 0.273549228 |
| Sap30 | 6.801534 | 7.360987 | 6.6641 | 0.696895075 | 0.137442154 | 6.8570599 | 7.7476592 | 6.69842 | 1.049237719 | 0.158638443 |
| Slamf9 | 6.367231 | 6.301065 | 6.2404 | 0.06068308 | 0.126848392 | 6.3368216 | 7.1138755 | 6.06558 | 1.048297615 | 0.271243659 |
| Slc25a28 | 9.356991 | 9.517711 | 9.4204 | 0.097280904 | -0.063439279 | 9.7127199 | 10.592657 | 9.54521 | 1.047449938 | 0.167513205 |
| 1500011B03Rik | 7.872922 | 8.304306 | 7.8504 | 0.453886946 | 0.022502609 | 8.2720528 | 9.0464954 | 7.99971 | 1.046789826 | 0.27234728 |
| Tmem33 | 9.540873 | 9.865183 | 9.4355 | 0.429660861 | 0.105351011 | 9.7376099 | 10.476841 | 9.4311 | 1.045744347 | 0.306513394 |
| Abcb1a | 8.540485 | 8.651983 | 8.5675 | 0.08452665 | -0.026971027 | 8.0918141 | 9.3427206 | 8.29731 | 1.045415137 | -0.205491355 |
| Ifitm2 | 9.696921 | 9.430637 | 8.7997 | 0.630942582 | 0.897226415 | 9.6299414 | 9.510913 | 8.46606 | 1.044856681 | 1.163885097 |
| Grina | 9.864338 | 9.953305 | 9.6703 | 0.282957908 | 0.193991697 | 10.121966 | 10.794609 | 9.75239 | 1.042218155 | 0.369575299 |

| | | | | | | | | | | |
|---|---|---|---|---|---|---|---|---|---|---|
| Ncr1 | 6.901056 | 7.250858 | 7.0741 | 0.17678561 | -0.173016043 | 6.6354542 | 7.884541 | 6.84252 | 1.042021673 | -0.207065114 |
| Lyz2 | 13.47515 | 13.51512 | 12.693 | 0.821818912 | 0.78184498 | 13.059033 | 13.151792 | 12.1153 | 1.036490802 | 0.943731207 |
| Hmox2 | 9.782483 | 10.16926 | 9.7618 | 0.407428589 | 0.020652589 | 9.9192347 | 10.717511 | 9.68131 | 1.036198331 | 0.23792227 |
| S1pr2 | 6.727615 | 7.264679 | 6.8535 | 0.411206236 | -0.125856894 | 6.99057 | 7.6317991 | 6.59721 | 1.034589176 | 0.39336012 |
| Cited2 | 8.959961 | 9.211489 | 9.1066 | 0.104908264 | -0.146619619 | 8.5989788 | 9.7257475 | 8.6912 | 1.034545842 | -0.092222866 |
| Col6a3 | 7.221891 | 8.084908 | 7.0782 | 1.006674417 | 0.143657988 | 7.624234 | 7.9412196 | 6.90956 | 1.031660357 | 0.714674722 |
| Igkv4-70 | 6.206576 | 7.380402 | 6.1952 | 1.185199831 | 0.011373427 | 6.5596798 | 7.1800985 | 6.1485 | 1.031600983 | 0.411182263 |
| Hspa5 | 13.42895 | 13.69074 | 13.227 | 0.463931492 | 0.202146826 | 13.048807 | 13.946398 | 12.9187 | 1.027673228 | 0.130081735 |
| Noc4l | 8.956198 | 9.177154 | 8.9516 | 0.225586501 | 0.004630257 | 9.3238733 | 10.091434 | 9.06413 | 1.027304081 | 0.25974345 |
| Atp6v0c | 11.17497 | 11.43235 | 10.984 | 0.448788298 | 0.191403174 | 11.409743 | 11.882829 | 10.8563 | 1.026564978 | 0.553478654 |
| Stoml1 | 9.772443 | 10.04306 | 9.8641 | 0.178920529 | -0.09169243 | 10.171265 | 10.957249 | 9.93077 | 1.026479149 | 0.240494251 |
| Psma7 | 10.55658 | 10.64302 | 10.504 | 0.13942453 | 0.052984308 | 10.79112 | 11.509063 | 10.4834 | 1.025663488 | 0.307719815 |
| Gda | 9.910708 | 9.508761 | 8.5276 | 0.981145041 | 1.383092063 | 8.6987557 | 9.0673656 | 8.04306 | 1.024304939 | 0.655694971 |
| Mgst1 | 6.53948 | 6.394163 | 5.9893 | 0.404870589 | 0.550187319 | 6.1015654 | 6.8105682 | 5.78755 | 1.023018382 | 0.314015528 |
| Kif9 | 5.861175 | 6.343537 | 5.9272 | 0.416319354 | -0.066042066 | 6.1549307 | 7.116944 | 6.09448 | 1.022468721 | 0.060455502 |
| Irf9 | 11.38126 | 11.96074 | 11.2 | 0.760709078 | 0.18122738 | 11.723442 | 12.340936 | 11.3186 | 1.022304596 | 0.404810303 |
| Sod3 | 6.381237 | 6.478675 | 6.2275 | 0.251136081 | 0.153697889 | 6.4170253 | 7.1582914 | 6.13983 | 1.018457988 | 0.277191849 |
| Tmbim6 | 12.69669 | 12.99899 | 12.624 | 0.374864536 | 0.07256126 | 12.909599 | 13.634557 | 12.6177 | 1.016858504 | 0.291901357 |
| Trmt61a | 8.117711 | 8.293492 | 7.7394 | 0.554072508 | 0.378291428 | 8.3654875 | 8.7186933 | 7.70199 | 1.016700673 | 0.663494847 |
| Ahsa2 | 8.644802 | 9.070879 | 8.6674 | 0.403476761 | -0.022600426 | 8.5660898 | 9.6728138 | 8.65628 | 1.016532777 | -0.090191273 |
| Cd47 | 12.2779 | 12.58655 | 12.217 | 0.369831172 | 0.061183644 | 12.296776 | 13.190443 | 12.1743 | 1.01613251 | 0.122466278 |
| Parp3 | 9.8275 | 10.02264 | 9.8294 | 0.193271924 | -0.001867397 | 10.179792 | 11.006095 | 9.99015 | 1.015942956 | 0.189640142 |
| Emilin2 | 8.586535 | 8.479116 | 6.8582 | 1.620935514 | 1.728354797 | 8.3310456 | 7.9264244 | 6.91176 | 1.014663997 | 1.419285169 |
| 9930111J21Rik2 | 12.51205 | 13.10676 | 12.531 | 0.575421924 | -0.019286376 | 12.785955 | 13.663194 | 12.6488 | 1.014391722 | 0.137153446 |
| Hspa8 | 15.4079 | 15.84243 | 15.395 | 0.447209884 | 0.012676726 | 15.121928 | 16.269751 | 15.2565 | 1.013250674 | -0.134572793 |
| Xcl1 | 7.20565 | 7.530644 | 7.2548 | 0.275814863 | -0.049180017 | 7.0172905 | 8.1743042 | 7.16154 | 1.012759441 | -0.144254249 |
| 1700030K09Rik | 7.989547 | 8.524269 | 8.2008 | 0.323455316 | -0.211266852 | 8.323661 | 9.1040337 | 8.09295 | 1.011086296 | 0.230713643 |
| Msr1 | 8.415167 | 9.036995 | 7.2926 | 1.744422508 | 1.122594527 | 8.1477936 | 8.3094384 | 7.30005 | 1.009391141 | 0.847746385 |
| Psma5 | 9.95816 | 10.01586 | 9.8748 | 0.141041106 | 0.083341126 | 9.9714501 | 10.821895 | 9.81504 | 1.006855925 | 0.156411529 |
| Ubald2 | 9.527826 | 9.821462 | 9.4627 | 0.35877057 | 0.065134424 | 9.5565356 | 10.426496 | 9.4204 | 1.006094283 | 0.136134035 |
| Nfkbib | 8.819009 | 9.032207 | 8.7395 | 0.292661594 | 0.079463625 | 8.9761307 | 9.6277944 | 8.62189 | 1.005903022 | 0.35423926 |
| Rcc2 | 12.04212 | 12.24778 | 12.032 | 0.215446726 | 0.009786662 | 12.28561 | 13.091218 | 12.086 | 1.005266706 | 0.199658975 |

| | | | | | | | | | | |
|---|---|---|---|---|---|---|---|---|---|---|
| Rbm43 | 9.117128 | 9.586409 | 9.1946 | 0.391853947 | -0.077426463 | 9.3152559 | 10.205159 | 9.20086 | 1.00429882 | 0.114395532 |
| Pla2g4a | 6.269193 | 6.385718 | 6.094 | 0.291690558 | 0.17516494 | 6.5225017 | 6.9222289 | 5.91831 | 1.003918604 | 0.604191398 |
| Pstpip2 | 5.578571 | 5.673572 | 5.33 | 0.343548932 | 0.248548244 | 5.2742228 | 6.0259565 | 5.02237 | 1.003585171 | 0.251851445 |
| Cers6 | 8.924619 | 9.367366 | 8.7856 | 0.581792631 | 0.139045524 | 9.0571015 | 9.9950287 | 8.99182 | 1.003212619 | 0.065285479 |
| Ccl8 | 8.009743 | 7.675287 | 7.3577 | 0.317559513 | 0.652015717 | 6.809671 | 7.8790001 | 6.87803 | 1.000974919 | -0.068354199 |
| Idnk | 9.467425 | 9.748543 | 9.4582 | 0.29033505 | 0.009217124 | 9.8103286 | 10.618003 | 9.61762 | 1.000384337 | 0.192709928 |
| Vwa5a | 9.878978 | 9.964775 | 9.6567 | 0.30806909 | 0.222272759 | 9.7260116 | 10.621137 | 9.62101 | 1.000128132 | 0.105003059 |
| Bcl2a1a | 7.040147 | 7.745801 | 7.1694 | 0.57641962 | -0.129234533 | 7.469213 | 8.137492 | 7.13892 | 0.998575776 | 0.330296736 |
| Apobec1 | 9.299161 | 9.635814 | 9.1554 | 0.480402033 | 0.143749479 | 9.6083686 | 10.21716 | 9.21872 | 0.998443057 | 0.389652083 |
| Tbx21 | 7.516847 | 7.820588 | 7.4632 | 0.357347153 | 0.053606394 | 7.4486519 | 8.5885582 | 7.59364 | 0.994913312 | -0.144993083 |
| Armcx6 | 5.804481 | 5.688867 | 5.8354 | -0.146533753 | -0.030919199 | 5.4836986 | 6.5042799 | 5.50985 | 0.994426878 | -0.026154378 |
| Cox17 | 8.554192 | 8.626428 | 8.3654 | 0.260992274 | 0.188755536 | 8.8340321 | 9.2760356 | 8.28313 | 0.992902301 | 0.550898837 |
| Fam111a | 11.57522 | 12.16938 | 11.565 | 0.604305436 | 0.010139758 | 11.954705 | 12.681809 | 11.6901 | 0.991726952 | 0.26462295 |
| Stra6l | 5.24986 | 6.071682 | 5.5067 | 0.565022278 | -0.256800051 | 5.4688223 | 6.6373607 | 5.64733 | 0.990034836 | -0.178503489 |
| Gm48774 | 5.526069 | 5.896599 | 5.357 | 0.539607404 | 0.169077984 | 5.790133 | 6.442811 | 5.45353 | 0.989279956 | 0.336601985 |
| Zfp513 | 8.992227 | 9.176839 | 8.9643 | 0.212491763 | 0.027879698 | 9.1808502 | 10.003583 | 9.01549 | 0.988091021 | 0.165358645 |
| Gadd45g | 8.14522 | 7.923536 | 7.9255 | -0.001929083 | 0.219754566 | 8.0155018 | 8.669438 | 7.6836 | 0.985835929 | 0.331899757 |
| Gm12216 | 5.761044 | 5.738029 | 5.5892 | 0.148787448 | 0.171803162 | 5.829235 | 6.7571126 | 5.77288 | 0.984235855 | 0.056358276 |
| Gm14654 | 4.799853 | 5.041512 | 4.683 | 0.358544482 | 0.116885289 | 5.0591897 | 5.9707868 | 4.98711 | 0.98367864 | 0.072081603 |
| Dop1b | 9.989294 | 10.14888 | 10.017 | 0.132335479 | -0.027252796 | 10.186719 | 11.102952 | 10.1201 | 0.982872868 | 0.066639713 |
| Hook2 | 7.127147 | 6.880743 | 6.8116 | 0.069109544 | 0.315513408 | 7.0951488 | 7.9774163 | 6.99518 | 0.982231714 | 0.099964163 |
| Klk1b26 | 6.012513 | 6.219716 | 6.8778 | -0.658100349 | -0.865302675 | 8.1591936 | 7.1376633 | 6.15726 | 0.980400117 | 2.001930427 |
| Unc93b1 | 12.22944 | 12.35617 | 12.112 | 0.244207308 | 0.117481472 | 12.783981 | 13.228111 | 12.2478 | 0.980325014 | 0.536194389 |
| Rab20 | 7.021865 | 7.200598 | 6.3749 | 0.825677296 | 0.646944123 | 6.645494 | 7.2686514 | 6.28939 | 0.979261734 | 0.356104302 |
| Aldh1b1 | 7.196248 | 7.609797 | 7.0459 | 0.563930929 | 0.150381494 | 7.2678394 | 7.9551544 | 6.97639 | 0.978761222 | 0.291446177 |
| Shisa5 | 14.48205 | 15.03401 | 14.542 | 0.492454142 | -0.059507274 | 14.974781 | 15.582232 | 14.6038 | 0.978445042 | 0.370994105 |
| Il18 | 6.560389 | 6.487061 | 6.4289 | 0.058162734 | 0.131491232 | 6.519917 | 7.209292 | 6.2317 | 0.977592371 | 0.288217382 |
| Il22 | 2.733928 | 2.866761 | 2.6648 | 0.201946566 | 0.069113352 | 2.9305482 | 3.7380554 | 2.76181 | 0.97624193 | 0.168734813 |
| Glipr2 | 10.32612 | 10.35653 | 10.207 | 0.149137432 | 0.118728078 | 10.295424 | 11.122534 | 10.149 | 0.973511242 | 0.146400718 |
| Cycs | 9.071951 | 9.219526 | 8.9988 | 0.220724608 | 0.073150036 | 9.2860417 | 10.034256 | 9.06603 | 0.968222684 | 0.220008627 |
| Gja4 | 5.463013 | 5.668713 | 5.5861 | 0.082651121 | -0.123048825 | 5.4954202 | 6.4168889 | 5.44958 | 0.967309839 | 0.04584113 |
| Thbs1 | 10.81769 | 10.46605 | 8.7412 | 1.724822424 | 2.07646066 | 9.5896564 | 9.7999676 | 8.83479 | 0.965182226 | 0.754870959 |

| | | | | | | | | | | |
|---|---|---|---|---|---|---|---|---|---|---|
| Ms4a8a | 4.856972 | 4.846351 | 4.5166 | 0.329783344 | 0.340404379 | 4.4523741 | 5.3647217 | 4.39956 | 0.965161447 | 0.052813908 |
| Csrp1 | 10.78136 | 10.92206 | 10.732 | 0.190306008 | 0.049605302 | 10.683246 | 11.556915 | 10.5924 | 0.964507894 | 0.090838923 |
| Gypc | 9.109374 | 9.484814 | 9.1836 | 0.301239931 | -0.074199987 | 9.057805 | 10.121805 | 9.15789 | 0.963916835 | -0.100083436 |
| Cd300a | 8.310733 | 8.656204 | 7.5401 | 1.116107369 | 0.770636304 | 8.644316 | 8.1899165 | 7.22623 | 0.963688842 | 1.418088426 |
| Rmdn3 | 8.356019 | 8.469018 | 8.2935 | 0.175478707 | 0.06247972 | 8.423595 | 9.4536984 | 8.49159 | 0.962110772 | -0.067992641 |
| Hap1 | 7.407342 | 7.446745 | 7.7074 | -0.260609315 | -0.300011732 | 7.0421561 | 8.0659684 | 7.10409 | 0.961876746 | -0.061935515 |
| Mfsd12 | 7.721068 | 8.11459 | 7.6806 | 0.433979096 | 0.040458079 | 7.7543317 | 8.4070347 | 7.44632 | 0.960710143 | 0.308007097 |
| Tor1aip1 | 11.64495 | 12.1275 | 11.557 | 0.570510076 | 0.087957688 | 11.706455 | 12.616563 | 11.6565 | 0.960042572 | 0.04993446 |
| Chordc1 | 10.33178 | 10.91492 | 10.375 | 0.540028408 | -0.04310815 | 10.108611 | 11.35214 | 10.3922 | 0.959933716 | -0.283595582 |
| Rpl36 | 12.23212 | 12.33798 | 12.109 | 0.228802714 | 0.122938698 | 12.52452 | 12.898746 | 11.9401 | 0.958669972 | 0.584443876 |
| Ccl19 | 8.182558 | 7.958933 | 7.3917 | 0.567217971 | 0.790843242 | 7.8195399 | 8.0623919 | 7.1044 | 0.957995311 | 0.715143279 |
| P2ry13 | 7.824033 | 7.454754 | 7.3355 | 0.119290245 | 0.488569599 | 7.328233 | 7.6739022 | 6.71712 | 0.956779847 | 0.611110601 |
| Gm45551 | 6.885423 | 6.9745 | 6.6592 | 0.315322177 | 0.226244492 | 6.8535203 | 7.3812614 | 6.42614 | 0.955118732 | 0.427377648 |
| Tmem51 | 5.704241 | 5.871684 | 5.4928 | 0.378865706 | 0.21142213 | 5.8409933 | 6.1874965 | 5.23246 | 0.955038016 | 0.608534805 |
| Ifitm1 | 7.448694 | 7.321644 | 6.38 | 0.941628652 | 1.068679312 | 8.0472636 | 7.2879578 | 6.33417 | 0.953789059 | 1.713094796 |
| D17Wsu92e | 11.28868 | 11.583 | 11.175 | 0.408313614 | 0.114002648 | 11.156794 | 12.123206 | 11.1706 | 0.952568692 | -0.013843302 |
| Atp6v0d2 | 3.775945 | 4.626565 | 3.7926 | 0.834007074 | -0.01661244 | 3.8580283 | 4.8198721 | 3.87008 | 0.949788328 | -0.012055468 |
| Vmn1r233 | 5.720565 | 6.073072 | 5.7023 | 0.370772209 | 0.018265517 | 5.9051577 | 6.6459417 | 5.69835 | 0.94759169 | 0.206807672 |
| Cstb | 8.72414 | 9.111409 | 8.6116 | 0.499845271 | 0.112576215 | 8.8682649 | 9.4972771 | 8.55083 | 0.946443718 | 0.317431479 |
| Aen | 9.055677 | 9.282216 | 8.9942 | 0.287983143 | 0.061443853 | 9.2389083 | 9.9459242 | 9.00087 | 0.945054723 | 0.238038844 |
| Ascc3 | 10.75888 | 11.08942 | 10.615 | 0.474038296 | 0.143499732 | 10.859667 | 11.711184 | 10.7668 | 0.944423532 | 0.092907062 |
| Apol7e | 9.558437 | 9.807929 | 9.6537 | 0.154232564 | -0.095259222 | 9.4286736 | 10.564518 | 9.62028 | 0.94423731 | -0.191607183 |
| Gm7666 | 5.228221 | 5.428418 | 5.4546 | -0.026214095 | -0.226411432 | 5.1559652 | 6.3650867 | 5.4262 | 0.938881787 | -0.270239708 |
| Ddx24 | 11.35797 | 11.6226 | 11.168 | 0.45448327 | 0.189860998 | 11.417864 | 12.139377 | 11.2035 | 0.935881934 | 0.21436881 |
| Gm15433 | 4.716809 | 4.98663 | 4.5252 | 0.461464803 | 0.191644344 | 4.9717938 | 5.6615577 | 4.72568 | 0.935874675 | 0.246110777 |
| Gpr65 | 9.28857 | 9.614101 | 9.1864 | 0.427707322 | 0.102175731 | 9.205348 | 10.067276 | 9.13317 | 0.934110305 | 0.072182091 |
| Gm14219 | 3.114323 | 3.102006 | 3.5029 | -0.400893785 | -0.388576666 | 2.8740562 | 4.0008463 | 3.06724 | 0.9336067 | -0.193183372 |
| Ighg1 | 7.609189 | 8.635172 | 7.9724 | 0.662753818 | -0.363229541 | 8.6225249 | 7.9660407 | 7.03275 | 0.93328644 | 1.58977062 |
| Svbp | 7.824324 | 7.991699 | 7.826 | 0.165694674 | -0.001680356 | 7.9461764 | 8.5494696 | 7.61764 | 0.931830408 | 0.328537212 |
| Ptges3 | 10.82706 | 11.15522 | 10.694 | 0.461437278 | 0.133269094 | 10.805703 | 11.671659 | 10.7401 | 0.931521872 | 0.065565181 |
| Tomm70a | 10.89 | 10.98718 | 10.83 | 0.157409196 | 0.060231806 | 10.831841 | 11.747319 | 10.818 | 0.92934783 | 0.013869455 |
| Atp13a1 | 10.73294 | 10.91096 | 10.749 | 0.161547228 | -0.016480574 | 11.094625 | 11.808341 | 10.8795 | 0.928854992 | 0.215138881 |

| | | | | | | | | | | |
|---|---|---|---|---|---|---|---|---|---|---|
| Mrpl30 | 9.785604 | 10.03323 | 9.7121 | 0.321154114 | 0.073531572 | 10.022615 | 10.72728 | 9.79901 | 0.928265103 | 0.223600616 |
| Nfkbie | 10.29905 | 10.60465 | 10.24 | 0.36509336 | 0.059489888 | 10.508306 | 11.213146 | 10.2861 | 0.927018682 | 0.22217852 |
| Zbtb32 | 5.545306 | 5.380996 | 5.7485 | -0.367506279 | -0.203196835 | 5.6705728 | 6.7899396 | 5.86353 | 0.926405022 | -0.192961768 |
| Ifit1bl2 | 5.398432 | 6.06371 | 5.22 | 0.843746063 | 0.178467977 | 5.6551526 | 6.3204042 | 5.3943 | 0.926100717 | 0.260849105 |
| Klre1 | 6.711454 | 6.804099 | 6.5177 | 0.286414533 | 0.193770152 | 6.5457148 | 7.485998 | 6.56278 | 0.923221222 | -0.017061989 |
| Col23a1 | 6.615438 | 7.208474 | 6.6171 | 0.591348632 | -0.001687437 | 6.8203802 | 7.7451358 | 6.82221 | 0.922922043 | -0.001833556 |
| H2afj | 7.895766 | 7.982661 | 7.7066 | 0.276087148 | 0.189192254 | 8.1964432 | 8.4427091 | 7.52157 | 0.921138162 | 0.674872283 |
| Gns | 11.42383 | 11.66644 | 11.399 | 0.267778234 | 0.025168224 | 11.465175 | 12.234204 | 11.3136 | 0.920576894 | 0.15154831 |
| Ch25h | 7.793752 | 7.632636 | 7.2818 | 0.350876541 | 0.511992544 | 6.1788753 | 7.3547026 | 6.43455 | 0.920153633 | -0.255673668 |
| Hspd1 | 11.8462 | 12.13979 | 11.757 | 0.383041048 | 0.089450702 | 11.720393 | 12.730676 | 11.8114 | 0.919244698 | -0.091037951 |
| Cars | 9.117522 | 9.063036 | 8.8513 | 0.211697843 | 0.266183486 | 9.5959602 | 10.318826 | 9.40001 | 0.918816242 | 0.195950069 |
| Htr7 | 5.490348 | 5.510778 | 4.9146 | 0.596171671 | 0.575741573 | 5.8315784 | 5.9249232 | 5.00624 | 0.91867952 | 0.825334699 |
| Prf1 | 9.23449 | 9.455631 | 9.1553 | 0.30031911 | 0.079178175 | 9.1091646 | 10.014118 | 9.09555 | 0.918568017 | 0.013614545 |
| Fcrl5 | 8.134884 | 8.013418 | 7.7464 | 0.267049586 | 0.388516356 | 8.2549186 | 8.7396656 | 7.82153 | 0.918140552 | 0.433393553 |
| Tbxas1 | 5.643226 | 5.460382 | 5.5757 | -0.115289416 | 0.067553638 | 5.5870346 | 6.3797344 | 5.46182 | 0.917912705 | 0.125212973 |
| Cd80 | 7.057005 | 7.175199 | 6.8064 | 0.368819434 | 0.250626079 | 7.008342 | 7.7662387 | 6.84847 | 0.917767147 | 0.159870481 |
| Fn1 | 10.04455 | 9.899498 | 8.5599 | 1.339641413 | 1.484694545 | 8.6860123 | 9.328778 | 8.41158 | 0.917200359 | 0.274434647 |
| Bcl3 | 10.01066 | 9.524383 | 9.1473 | 0.377124893 | 0.863404263 | 9.5502984 | 10.103993 | 9.18682 | 0.917173471 | 0.363479071 |
| Ramp3 | 6.282928 | 6.319789 | 6.1733 | 0.146448206 | 0.109586323 | 5.9393848 | 7.1537918 | 6.23738 | 0.916409128 | -0.297997838 |
| Ms4a4b | 12.442 | 12.69938 | 12.435 | 0.264048596 | 0.006662732 | 12.209301 | 13.360034 | 12.4437 | 0.916373448 | -0.234358997 |
| Lamp2 | 10.92011 | 10.86624 | 10.559 | 0.307688622 | 0.361552988 | 10.795614 | 11.343556 | 10.4278 | 0.915774688 | 0.367832038 |
| Tpgs1 | 6.672936 | 6.732071 | 6.6318 | 0.100233285 | 0.041099058 | 6.9894542 | 7.2450671 | 6.33075 | 0.914317119 | 0.658704233 |
| Il12b | 5.381947 | 5.738479 | 5.4181 | 0.320350554 | -0.036181907 | 5.1585138 | 6.2268528 | 5.31287 | 0.91398756 | -0.154351452 |
| Saa3 | 4.947578 | 5.696169 | 4.3356 | 1.360587411 | 0.611996282 | 6.6523707 | 5.5589579 | 4.64627 | 0.912688049 | 2.006100861 |
| Hspe1 | 11.17269 | 11.33702 | 11.17 | 0.167286858 | 0.002959594 | 11.269571 | 12.070013 | 11.1617 | 0.908277622 | 0.107835583 |
| Hsph1 | 11.266 | 11.77011 | 11.375 | 0.395595398 | -0.108506798 | 10.606921 | 12.420892 | 11.5155 | 0.90537949 | -0.908591545 |
| Znhit1 | 7.240948 | 7.291948 | 7.1839 | 0.108030548 | 0.057030367 | 7.4363637 | 8.0727181 | 7.1697 | 0.903020892 | 0.266666427 |
| Mucl2 | 8.320176 | 6.745441 | 7.3076 | -0.562207367 | 1.012527031 | 8.4348771 | 7.3869454 | 6.48497 | 0.901978717 | 1.949910484 |
| AC166110.3 | 4.010717 | 4.098752 | 3.6215 | 0.477265324 | 0.389230267 | 3.7112172 | 4.563793 | 3.66239 | 0.901406832 | 0.048831031 |
| Dbnl | 11.7284 | 11.92048 | 11.719 | 0.201781534 | 0.009696252 | 12.084419 | 12.617074 | 11.7161 | 0.901007522 | 0.368352877 |
| Insl6 | 6.158919 | 6.25699 | 6.0736 | 0.183353037 | 0.08528234 | 6.3675178 | 7.1367885 | 6.23601 | 0.900782929 | 0.131512208 |
| Tha1 | 7.060704 | 6.769593 | 6.628 | 0.141563193 | 0.432674005 | 7.1506769 | 7.4520551 | 6.55155 | 0.900503338 | 0.599125169 |

| | | | | | | | | | | |
|---|---|---|---|---|---|---|---|---|---|---|
| Col9a3 | 4.718868 | 4.93601 | 4.6763 | 0.259699394 | 0.04255773 | 4.9898726 | 5.7955315 | 4.89718 | 0.898356448 | 0.092697511 |
| Ubd | 10.34948 | 9.834513 | 10.102 | -0.267395692 | 0.247575094 | 8.4296555 | 9.698041 | 8.7998 | 0.898237528 | -0.370148014 |
| Gjb2 | 4.438291 | 4.801633 | 4.1666 | 0.635017138 | 0.271675351 | 4.5130724 | 5.0777377 | 4.18053 | 0.897206951 | 0.332541731 |
| Casp1 | 8.939542 | 9.130562 | 8.9158 | 0.214714054 | 0.02369449 | 9.2058388 | 9.8598699 | 8.96326 | 0.896608593 | 0.242577459 |
| Fas | 8.227091 | 8.586617 | 8.0406 | 0.546015641 | 0.186489967 | 7.9034171 | 8.9871041 | 8.09077 | 0.896330977 | -0.187356048 |
| Gbp2b | 4.960854 | 4.592378 | 5.2622 | -0.669812231 | -0.301336211 | 4.5762171 | 5.9131351 | 5.01706 | 0.896071476 | -0.440846564 |
| Smg7 | 11.3065 | 11.71833 | 11.26 | 0.458799648 | 0.04697007 | 11.342704 | 12.234702 | 11.3404 | 0.894273468 | 0.00227493 |
| Crlf2 | 8.327079 | 8.397409 | 8.1299 | 0.267515537 | 0.197185275 | 8.8667839 | 8.9784865 | 8.08435 | 0.894133656 | 0.782431063 |
| Pkig | 9.431252 | 9.641529 | 9.4056 | 0.235965131 | 0.025687825 | 9.6472819 | 10.027785 | 9.13476 | 0.893026876 | 0.512523979 |
| Zc3hav1 | 12.10704 | 12.47986 | 12.004 | 0.475978066 | 0.103162292 | 12.070236 | 12.978769 | 12.0879 | 0.890867938 | -0.017665335 |
| Lax1 | 11.17781 | 11.38877 | 11.192 | 0.196817134 | -0.014144592 | 11.500756 | 12.172972 | 11.2826 | 0.890413628 | 0.218197925 |
| Tmem221 | 5.566834 | 5.637602 | 5.4971 | 0.140462886 | 0.069695366 | 5.4542959 | 6.1882048 | 5.2996 | 0.888607321 | 0.154698435 |
| Spi1 | 9.683753 | 9.93759 | 9.5524 | 0.385158864 | 0.131322705 | 10.031125 | 10.389602 | 9.50374 | 0.885858984 | 0.527381659 |
| Gm10108 | 3.996862 | 3.804739 | 4.4143 | -0.6095133 | -0.417390092 | 4.0384598 | 4.7863681 | 3.90056 | 0.885809294 | 0.137901002 |
| Snx10 | 9.123412 | 9.281215 | 8.8591 | 0.422065387 | 0.264263181 | 9.0614615 | 9.8351276 | 8.95009 | 0.885037886 | 0.111371861 |
| Psma4 | 10.46094 | 10.51648 | 10.382 | 0.13446864 | 0.078929076 | 10.484608 | 11.253961 | 10.3702 | 0.88379956 | 0.11444704 |
| Dpp4 | 11.02101 | 11.28823 | 10.986 | 0.301957152 | 0.034734766 | 11.143313 | 12.09267 | 11.2096 | 0.883071162 | -0.066285823 |
| Rab29 | 7.810717 | 7.978845 | 7.7444 | 0.234478465 | 0.066350787 | 7.9992282 | 8.6202264 | 7.73759 | 0.882631869 | 0.261633713 |
| Larp1 | 11.94903 | 12.18984 | 11.889 | 0.300388066 | 0.059580164 | 11.967563 | 12.856239 | 11.9738 | 0.882453892 | -0.006221585 |
| Klk1b9 | 4.65734 | 5.47357 | 6.2177 | -0.744102904 | -1.560332892 | 7.2115765 | 6.3126964 | 5.4324 | 0.880300275 | 1.779180386 |
| Vars | 12.66899 | 12.87809 | 12.565 | 0.313079086 | 0.103971254 | 13.04652 | 13.5477 | 12.6681 | 0.879618718 | 0.378437786 |
| H2-D1 | 14.81223 | 14.85742 | 14.775 | 0.082570326 | 0.037376274 | 14.872498 | 15.610179 | 14.731 | 0.879187858 | 0.141506197 |
| Gm2666 | 3.964404 | 4.42071 | 4.0058 | 0.414947318 | -0.041359247 | 4.2837835 | 5.0172934 | 4.13901 | 0.87828212 | 0.144772153 |
| Omp | 3.669922 | 3.77966 | 3.5407 | 0.238967277 | 0.129229556 | 3.7858532 | 4.5611097 | 3.6863 | 0.874811462 | 0.099554998 |
| Nrg1 | 3.600162 | 3.539346 | 3.2599 | 0.279461916 | 0.340277645 | 3.5162227 | 4.1009975 | 3.22745 | 0.873545567 | 0.288770758 |
| H2-Q10 | 9.940572 | 10.0847 | 10.059 | 0.025847229 | -0.118280961 | 10.376934 | 11.250898 | 10.3777 | 0.873155348 | -0.000808552 |
| Igkv15-103 | 7.716412 | 7.760391 | 7.7031 | 0.057319003 | 0.013339509 | 7.9959834 | 8.229294 | 7.3583 | 0.870989375 | 0.637678745 |
| Mycbp2 | 12.8925 | 13.12079 | 12.948 | 0.172459772 | -0.05583312 | 12.90741 | 13.959726 | 13.0892 | 0.87055256 | -0.181763053 |
| Slc7a2 | 4.769539 | 4.826341 | 4.7239 | 0.102476031 | 0.045674827 | 4.4928498 | 5.3942701 | 4.52382 | 0.870450628 | -0.030969625 |
| Gm5960 | 1.665443 | 1.680741 | 1.6489 | 0.031852914 | 0.016554905 | 1.6736815 | 2.546737 | 1.67643 | 0.870309246 | -0.002746179 |
| Cct3 | 11.51287 | 11.70279 | 11.429 | 0.273844364 | 0.083926954 | 11.547221 | 12.287489 | 11.4176 | 0.869908128 | 0.129639409 |
| Casp8 | 10.87862 | 11.0872 | 10.796 | 0.290871976 | 0.082295274 | 11.008116 | 11.762975 | 10.8936 | 0.869398118 | 0.114539572 |

| | | | | | | | | | | |
|---|---|---|---|---|---|---|---|---|---|---|
| Gsdmd | 10.06915 | 10.28896 | 10.127 | 0.161760032 | -0.058044047 | 10.614228 | 11.230866 | 10.362 | 0.868899906 | 0.252261913 |
| Lpxn | 10.28832 | 10.66108 | 10.301 | 0.360325386 | -0.012430894 | 10.614544 | 11.228352 | 10.3602 | 0.868140906 | 0.254332899 |
| Msh3 | 8.991172 | 8.982097 | 8.8566 | 0.125455145 | 0.134529639 | 9.0311402 | 9.7675917 | 8.89977 | 0.867822969 | 0.131371404 |
| 9930111J21Rik1 | 10.86291 | 11.43357 | 10.808 | 0.625614654 | 0.054954408 | 11.024337 | 11.919245 | 11.052 | 0.867273816 | -0.027633652 |
| Cybc1 | 11.04144 | 11.46595 | 11.064 | 0.401973292 | -0.022540872 | 11.372402 | 12.041522 | 11.1751 | 0.866405778 | 0.19728525 |
| Gm47056 | 6.397855 | 6.800216 | 6.4898 | 0.310432135 | -0.091929251 | 6.7362851 | 7.5395783 | 6.67368 | 0.865897742 | 0.062604574 |
| Fkbp4 | 10.74561 | 11.13132 | 10.742 | 0.389072676 | 0.003363468 | 10.731127 | 11.474091 | 10.6087 | 0.865408352 | 0.122444527 |
| Ly6e | 14.63695 | 14.89596 | 14.596 | 0.299478004 | 0.040472322 | 15.002922 | 15.412505 | 14.5491 | 0.863441096 | 0.453857966 |
| Slc31a2 | 7.324576 | 7.432287 | 7.27 | 0.162288913 | 0.054578171 | 7.2406717 | 7.9605784 | 7.09752 | 0.863060904 | 0.143154162 |
| Rnf139 | 10.08063 | 10.36968 | 10.072 | 0.297930325 | 0.008882477 | 10.180688 | 10.926796 | 10.0639 | 0.862936503 | 0.116828049 |
| C3ar1 | 4.248866 | 4.918679 | 4.0804 | 0.838318863 | 0.168506515 | 4.4513538 | 4.7870122 | 3.92531 | 0.861700915 | 0.526042516 |
| 1110038F14Rik | 8.341951 | 8.379963 | 8.267 | 0.112979099 | 0.074967439 | 8.4663105 | 8.9853774 | 8.12379 | 0.861586047 | 0.3425191 |
| Cd177 | 6.307793 | 6.19288 | 5.0207 | 1.172199286 | 1.287112318 | 5.6873084 | 6.046204 | 5.18474 | 0.861465187 | 0.502569588 |
| Ankfy1 | 11.5425 | 12.08471 | 11.559 | 0.52529187 | -0.016919628 | 11.624065 | 12.448925 | 11.5878 | 0.86110604 | 0.036246574 |
| Col3a1 | 8.576989 | 8.576172 | 8.239 | 0.337207743 | 0.338024853 | 9.5488377 | 9.0211547 | 8.16021 | 0.860947519 | 1.388630558 |
| Laptm4a | 11.0228 | 11.0335 | 10.899 | 0.134520548 | 0.123812924 | 10.925989 | 11.574385 | 10.7151 | 0.859286226 | 0.210889971 |
| Ighv1-82 | 8.538128 | 8.612582 | 8.3749 | 0.237643009 | 0.16318888 | 8.6702293 | 9.194199 | 8.33695 | 0.857250335 | 0.333280623 |
| Zeb2 | 9.777196 | 10.11886 | 9.4003 | 0.718606977 | 0.37693949 | 9.7824844 | 10.221123 | 9.36395 | 0.857176792 | 0.418538492 |
| Col1a1 | 7.862628 | 8.048204 | 7.5697 | 0.478465709 | 0.292889677 | 8.3582516 | 8.1758309 | 7.31947 | 0.856358599 | 1.038779232 |
| Cmtr1 | 11.51753 | 11.92281 | 11.405 | 0.518221586 | 0.112937416 | 11.67961 | 12.383546 | 11.5284 | 0.855119394 | 0.151183373 |
| Arfgef1 | 11.77581 | 11.8569 | 11.758 | 0.09897642 | 0.017891986 | 11.920639 | 12.738729 | 11.8838 | 0.854939638 | 0.03684996 |
| Gsap | 8.786899 | 8.911713 | 8.7255 | 0.186168427 | 0.061354274 | 8.7638072 | 9.533232 | 8.67911 | 0.854125436 | 0.084700716 |
| Csf1r | 10.47751 | 10.50256 | 10.128 | 0.374246631 | 0.34918893 | 10.917175 | 10.489628 | 9.63765 | 0.851979432 | 1.279526477 |
| Lap3 | 8.895113 | 9.021909 | 8.7589 | 0.263003525 | 0.136208268 | 8.7972716 | 9.6434359 | 8.79212 | 0.851312605 | 0.005148286 |
| C1qb | 9.147461 | 8.976773 | 8.5973 | 0.379507142 | 0.550195504 | 8.4487314 | 8.9511268 | 8.10051 | 0.850618328 | 0.348222956 |
| Eif1a | 9.988532 | 9.91136 | 9.5911 | 0.320226933 | 0.397398749 | 9.6948551 | 10.38753 | 9.53705 | 0.85048221 | 0.157807107 |
| Mmp25 | 8.334892 | 8.604887 | 8.3696 | 0.235309321 | -0.034685073 | 7.9795216 | 9.2996035 | 8.44971 | 0.849894323 | -0.470187589 |
| Atf3 | 9.383547 | 9.238626 | 8.5499 | 0.688711803 | 0.833632828 | 8.4148154 | 9.412519 | 8.56326 | 0.84926127 | -0.148442383 |
| Ywhag | 10.80478 | 11.14932 | 10.67 | 0.479439704 | 0.134898672 | 10.851125 | 11.513007 | 10.6642 | 0.84879563 | 0.186913189 |
| Morc3 | 11.2387 | 11.54562 | 11.168 | 0.377989466 | 0.0710749 | 11.197773 | 12.093657 | 11.2456 | 0.848085872 | -0.047798424 |
| Serpinb9b | 5.274374 | 5.334288 | 5.4483 | -0.114032471 | -0.17394671 | 5.1878668 | 6.093399 | 5.2456 | 0.847800069 | -0.057732171 |
| Kat2a | 10.12273 | 10.54401 | 10.093 | 0.450950139 | 0.029665551 | 10.390845 | 10.958865 | 10.1115 | 0.847326388 | 0.279306124 |

| | | | | | | | | | | |
|---|---|---|---|---|---|---|---|---|---|---|
| Gpr18 | 10.43629 | 10.57575 | 10.382 | 0.194227592 | 0.05477352 | 10.673643 | 11.161986 | 10.3157 | 0.846286976 | 0.35794426 |
| Ms4a6b | 12.28363 | 12.48405 | 12.241 | 0.243129412 | 0.04271485 | 12.325451 | 13.20607 | 12.36 | 0.846103144 | -0.034516166 |
| Vps37b | 10.21898 | 10.60365 | 10.135 | 0.468632749 | 0.08396301 | 10.227587 | 11.310731 | 10.466 | 0.844712534 | -0.238431989 |
| C1ra | 9.148292 | 8.656964 | 8.8237 | -0.166738747 | 0.324589377 | 8.4125143 | 9.1989669 | 8.3558 | 0.843168754 | 0.056716172 |
| Rnf19b | 10.85615 | 11.09598 | 10.771 | 0.325449426 | 0.085617524 | 10.539957 | 11.544598 | 10.7015 | 0.843106374 | -0.161534899 |
| Platr17 | 4.92664 | 5.131424 | 5.0299 | 0.101554932 | -0.103229422 | 5.1386265 | 5.7815829 | 4.93969 | 0.841891671 | 0.198935351 |
| Gm5970 | 2.431704 | 2.543529 | 2.3777 | 0.165860531 | 0.054035134 | 2.4239006 | 3.1825674 | 2.34088 | 0.841684445 | 0.083017659 |
| Psma2 | 10.83316 | 10.91948 | 10.84 | 0.079887864 | -0.006437986 | 11.004 | 11.684624 | 10.8432 | 0.841467322 | 0.160842753 |
| Aldoa | 12.60094 | 12.65966 | 12.545 | 0.1149587 | 0.056242254 | 12.949706 | 13.393675 | 12.5525 | 0.841212736 | 0.397244042 |
| Rere | 11.55597 | 11.95629 | 11.66 | 0.295866198 | -0.104452994 | 11.798972 | 12.643781 | 11.8026 | 0.84119714 | -0.003611205 |
| Hsp90ab1 | 15.11642 | 15.31545 | 15.201 | 0.114867958 | -0.08416372 | 15.014949 | 15.859775 | 15.0188 | 0.840925814 | -0.0038992 |
| Tmem192 | 8.048195 | 8.13655 | 8.0437 | 0.092873506 | 0.004517986 | 8.1424857 | 8.8402809 | 8.00092 | 0.839362026 | 0.14156678 |
| Lhx2 | 3.775882 | 3.716271 | 3.98 | -0.263686261 | -0.204075976 | 3.5892356 | 4.5885129 | 3.74988 | 0.838633954 | -0.160643327 |
| Ndufb7 | 9.239805 | 9.228316 | 9.1517 | 0.076626608 | 0.08811588 | 9.5437008 | 9.8475066 | 9.01119 | 0.836318672 | 0.532512909 |
| Nod1 | 9.763752 | 10.03401 | 9.8504 | 0.183565654 | -0.086688323 | 9.8601676 | 10.676025 | 9.84043 | 0.835597762 | 0.019740199 |
| Irf2 | 11.38696 | 11.61595 | 11.353 | 0.263037298 | 0.034042336 | 11.4051 | 12.119624 | 11.285 | 0.834587374 | 0.12006334 |
| Hist2h2aa2 | 5.492071 | 5.632656 | 5.7608 | -0.128099472 | -0.268683943 | 5.7316635 | 6.1466079 | 5.31213 | 0.834481104 | 0.419536657 |
| Klk1b16 | 4.345376 | 4.932544 | 5.7987 | -0.866107031 | -1.45327519 | 6.7542557 | 5.976187 | 5.14325 | 0.832937592 | 1.611006312 |
| Sidt1 | 10.5483 | 10.97552 | 10.594 | 0.381579498 | -0.045641818 | 10.625602 | 11.564986 | 10.7323 | 0.83267721 | -0.106706021 |
| Plek | 12.13925 | 12.2268 | 11.673 | 0.553394982 | 0.465838222 | 12.018965 | 12.478919 | 11.6463 | 0.832572884 | 0.372618391 |
| Tcof1 | 11.82062 | 12.15017 | 11.753 | 0.397364388 | 0.067807958 | 12.042752 | 12.648772 | 11.8169 | 0.83191819 | 0.225897443 |
| Notch1 | 11.50322 | 11.33132 | 11.264 | 0.06696105 | 0.23885954 | 10.865921 | 12.159164 | 11.3289 | 0.830276408 | -0.462966152 |
| Ddx4 | 3.040202 | 3.18896 | 3.0033 | 0.185655665 | 0.036897681 | 3.0749474 | 3.9091272 | 3.07934 | 0.82979211 | -0.004387681 |
| Shmt2 | 10.36137 | 10.54125 | 10.314 | 0.227617674 | 0.047735618 | 11.033967 | 11.441242 | 10.6134 | 0.827819752 | 0.42054451 |
| Resf1 | 10.87655 | 11.21923 | 10.815 | 0.404577694 | 0.06189659 | 10.81681 | 11.678591 | 10.8514 | 0.827213722 | -0.034568247 |
| Tbkbp1 | 6.738749 | 6.629169 | 6.5456 | 0.083521132 | 0.193101163 | 6.1652445 | 6.8521612 | 6.02616 | 0.825998938 | 0.139082292 |
| Plscr3 | 8.135767 | 8.101382 | 8.0556 | 0.045813402 | 0.080197899 | 8.2122521 | 8.6984768 | 7.87276 | 0.825718853 | 0.339494183 |
| Lpl | 8.370469 | 9.617568 | 7.655 | 1.962519588 | 0.715419972 | 7.7473469 | 8.3009072 | 7.47541 | 0.825495473 | 0.271935149 |
| Pdk3 | 8.485587 | 8.818727 | 8.4327 | 0.385985788 | 0.052845636 | 8.9252959 | 9.3903152 | 8.56539 | 0.824922932 | 0.35990361 |
| Dnajc7 | 11.66516 | 11.95987 | 11.756 | 0.203989658 | -0.090727702 | 11.957026 | 12.804557 | 11.9799 | 0.824620238 | -0.022910815 |
| Nfil3 | 8.31249 | 8.228212 | 7.5909 | 0.637340457 | 0.721618389 | 8.4223882 | 8.7249363 | 7.90273 | 0.822204407 | 0.519656304 |
| Il10ra | 11.22062 | 11.61168 | 11.264 | 0.347889304 | -0.043166972 | 11.473073 | 12.102953 | 11.2824 | 0.820560742 | 0.190680444 |

| | | | | | | | | | | |
|---|---|---|---|---|---|---|---|---|---|---|
| Ptpro | 7.065216 | 7.497927 | 6.9785 | 0.519443629 | 0.086732825 | 7.266788 | 7.81724 | 6.99732 | 0.819916199 | 0.269464186 |
| H2-Q5 | 10.36901 | 10.48284 | 10.526 | -0.043289954 | -0.157119799 | 10.77638 | 11.792669 | 10.9735 | 0.81913106 | -0.197158167 |
| Tnfrsf14 | 8.309951 | 8.354756 | 8.2118 | 0.142941319 | 0.098136318 | 8.6374228 | 9.2648661 | 8.44667 | 0.818200521 | 0.19075723 |
| Dcp2 | 10.28869 | 10.34211 | 10.204 | 0.138408487 | 0.084991761 | 10.23276 | 11.110464 | 10.2937 | 0.816774272 | -0.060929071 |
| Ighg3 | 9.179931 | 9.558039 | 8.6862 | 0.871833441 | 0.493724994 | 8.747778 | 8.9447671 | 8.12813 | 0.816640172 | 0.619651049 |
| Stxbp3 | 9.701789 | 9.807524 | 9.6813 | 0.126190482 | 0.020455107 | 9.6135012 | 10.463666 | 9.6473 | 0.816366367 | -0.033797955 |
| Med28 | 9.415216 | 9.63444 | 9.4462 | 0.188263042 | -0.030960753 | 9.7667997 | 10.264872 | 9.44892 | 0.815951223 | 0.317878452 |
| BC035044 | 8.253258 | 8.71334 | 8.1253 | 0.588029092 | 0.127946863 | 8.1785329 | 8.8189401 | 8.00327 | 0.815673979 | 0.175266782 |
| Zfp455 | 6.212566 | 6.293819 | 6.2336 | 0.060181635 | -0.021071422 | 6.366579 | 6.9722489 | 6.15744 | 0.814809839 | 0.209139924 |
| Rsph3b | 8.917461 | 8.85496 | 9.0532 | -0.198224684 | -0.135723225 | 8.9339583 | 9.7119657 | 8.89744 | 0.814525522 | 0.036518192 |
| Tgfbi | 11.15047 | 10.97089 | 10.396 | 0.5745925 | 0.754174472 | 11.703562 | 10.920876 | 10.1069 | 0.813954137 | 1.59664005 |
| Map3k8 | 9.295682 | 9.487745 | 9.1058 | 0.38192405 | 0.18986018 | 9.3048924 | 10.074799 | 9.2618 | 0.812999315 | 0.043092537 |
| 5430427O19Rik | 8.327101 | 8.88428 | 8.3664 | 0.517897924 | -0.039281132 | 8.596961 | 9.234152 | 8.42224 | 0.811914877 | 0.174723853 |
| H2-M3 | 10.06721 | 10.19319 | 10.089 | 0.104673345 | -0.021303294 | 10.424826 | 10.992026 | 10.1804 | 0.811665692 | 0.24446568 |
| Kalrn | 5.96469 | 6.200659 | 5.9974 | 0.203287038 | -0.032681682 | 5.7760591 | 6.7277627 | 5.91652 | 0.81124227 | -0.140461366 |
| Clic5 | 5.909481 | 5.888874 | 5.9938 | -0.104899112 | -0.084291172 | 5.3859406 | 6.7184862 | 5.90777 | 0.810719443 | -0.521826152 |
| Abi3 | 9.834422 | 9.978625 | 9.7026 | 0.276006688 | 0.131802904 | 10.310359 | 10.488543 | 9.67978 | 0.808762953 | 0.630579063 |
| Tent5a | 10.06259 | 10.62729 | 10.155 | 0.472341143 | -0.09235987 | 10.093901 | 10.905937 | 10.1006 | 0.805345879 | -0.006690089 |
| Sod2 | 9.536426 | 9.772773 | 9.3567 | 0.416072833 | 0.179725252 | 9.4226557 | 10.118027 | 9.31327 | 0.804756915 | 0.109385624 |
| Tbrg1 | 10.31634 | 10.52492 | 10.328 | 0.19699447 | -0.011586972 | 10.440622 | 11.160021 | 10.3562 | 0.8038474 | 0.084448334 |
| Polr2l | 7.906479 | 8.0585 | 7.889 | 0.169497047 | 0.01747663 | 8.2677562 | 8.5065842 | 7.70322 | 0.803361995 | 0.56453399 |
| Sssca1 | 7.597155 | 7.79923 | 7.5198 | 0.279413074 | 0.077338645 | 7.9475759 | 8.1713408 | 7.3689 | 0.802441891 | 0.578676985 |
| Ptpn18 | 9.84117 | 9.827492 | 9.7471 | 0.080359087 | 0.094037077 | 9.9334743 | 10.351933 | 9.55183 | 0.800104746 | 0.381645653 |
| Rgs12 | 7.652353 | 7.908036 | 7.7604 | 0.147628785 | -0.108054143 | 7.4270198 | 8.5383554 | 7.73866 | 0.799692484 | -0.311643093 |
| Xpo1 | 11.774 | 11.86377 | 11.795 | 0.068759952 | -0.02100674 | 11.89055 | 12.702511 | 11.9029 | 0.799657304 | -0.012303683 |
| Hrh2 | 8.001932 | 8.273544 | 7.9654 | 0.308130484 | 0.03651913 | 8.3143706 | 8.7536708 | 7.95635 | 0.797318283 | 0.358018072 |
| Epsti1 | 12.0353 | 12.32374 | 12.075 | 0.24907563 | -0.03935644 | 11.965775 | 12.936532 | 12.1401 | 0.79644209 | -0.174314534 |
| Hk1 | 10.66914 | 10.78528 | 10.687 | 0.097816054 | -0.018317098 | 10.534038 | 11.375744 | 10.5795 | 0.79627576 | -0.045430458 |
| Gm21748 | 3.711816 | 3.853713 | 3.6295 | 0.224194106 | 0.082296728 | 3.730736 | 4.493147 | 3.6969 | 0.796251943 | 0.033840879 |
| Gm21860 | 3.711816 | 3.853713 | 3.6295 | 0.224194106 | 0.082296728 | 3.730736 | 4.493147 | 3.6969 | 0.796251943 | 0.033840879 |
| Psmb2 | 9.923463 | 9.986506 | 9.8607 | 0.125809952 | 0.062767566 | 10.165247 | 10.603006 | 9.80799 | 0.795018816 | 0.357259866 |
| Timm10 | 7.174693 | 7.402748 | 7.0358 | 0.366975945 | 0.138920598 | 7.4098406 | 7.91967 | 7.12504 | 0.794625227 | 0.28479585 |

| | | | | | | | | | | |
|---|---|---|---|---|---|---|---|---|---|---|
| Tmem243 | 10.05413 | 10.24447 | 10.033 | 0.211088119 | 0.020752011 | 10.285786 | 10.750571 | 9.95668 | 0.793895313 | 0.329110417 |
| Grwd1 | 8.753726 | 8.942613 | 8.661 | 0.281638658 | 0.09275205 | 8.9777959 | 9.3945299 | 8.6014 | 0.793133961 | 0.37639992 |
| Rnf126 | 8.65504 | 8.829598 | 8.6949 | 0.134729711 | -0.039828462 | 9.0011068 | 9.4011889 | 8.60913 | 0.792059476 | 0.391977378 |
| Nr1h2 | 10.06215 | 10.21819 | 10.169 | 0.04917738 | -0.106867942 | 10.52949 | 11.068856 | 10.2775 | 0.7913647 | 0.251998886 |
| Ogfrl1 | 9.788002 | 9.959762 | 9.7486 | 0.211201112 | 0.039440881 | 9.454043 | 10.443148 | 9.65294 | 0.79020548 | -0.198899238 |
| Psmg4 | 7.43477 | 7.501752 | 7.3639 | 0.137837037 | 0.070854671 | 7.5426502 | 8.1524784 | 7.36396 | 0.788521288 | 0.178693085 |
| Hmox1 | 9.043041 | 8.86997 | 8.1562 | 0.713736294 | 0.886807082 | 7.8029819 | 8.1888514 | 7.40044 | 0.788413983 | 0.402544435 |
| Ccdc86 | 8.57218 | 8.669363 | 8.3259 | 0.343478103 | 0.246294921 | 8.5593056 | 9.0610371 | 8.27384 | 0.787195818 | 0.285464381 |
| Fam208b | 10.95326 | 11.28413 | 10.87 | 0.414056712 | 0.083180934 | 11.168725 | 12.073028 | 11.2861 | 0.786962664 | -0.117340485 |
| Ackr2 | 6.56266 | 6.446641 | 6.3916 | 0.055028208 | 0.171047659 | 5.6510752 | 6.484531 | 5.69844 | 0.786088533 | -0.047367359 |
| Mrgbp | 9.183521 | 9.482107 | 9.2243 | 0.257761466 | -0.040824245 | 9.4810359 | 10.007842 | 9.22219 | 0.785651033 | 0.258844455 |
| Evi2 | 7.322405 | 7.594237 | 7.382 | 0.212241431 | -0.059589969 | 7.7277443 | 8.2198788 | 7.43594 | 0.783942581 | 0.291808152 |
| Hist3h2a | 7.571785 | 7.739465 | 7.6161 | 0.123386429 | -0.044293232 | 7.5814381 | 8.1490714 | 7.36514 | 0.783926828 | 0.216293469 |
| Scgb1b27 | 7.582453 | 6.25929 | 6.72 | -0.460704045 | 0.86245876 | 7.8417952 | 6.6314402 | 5.8481 | 0.783338915 | 1.993693885 |
| Npc2 | 12.33268 | 12.46777 | 12.329 | 0.138816678 | 0.003727994 | 12.29001 | 13.055726 | 12.2725 | 0.783180022 | 0.017464796 |
| Phf6 | 9.818436 | 10.02171 | 9.7862 | 0.235493691 | 0.03222062 | 9.8678445 | 10.63747 | 9.85442 | 0.783051432 | 0.013425573 |
| Amigo3 | 4.87758 | 5.146944 | 5.0988 | 0.048142451 | -0.22122144 | 5.2456808 | 5.6998013 | 4.9171 | 0.78269698 | 0.328576408 |
| Rsbn1l | 10.71638 | 10.81968 | 10.728 | 0.091224468 | -0.012074974 | 10.647355 | 11.536436 | 10.7544 | 0.782083856 | -0.106997047 |
| Taldo1 | 11.24136 | 11.48977 | 11.163 | 0.32722158 | 0.078811334 | 11.671862 | 11.933848 | 11.1535 | 0.780318272 | 0.518332092 |
| Tnfrsf1a | 9.788631 | 9.519323 | 9.2516 | 0.267754239 | 0.537062879 | 9.2420743 | 9.7560182 | 8.97621 | 0.779803285 | 0.26585943 |
| Gm43802 | 4.961635 | 5.359735 | 5.3936 | -0.033880416 | -0.431980096 | 4.9198987 | 6.045126 | 5.26533 | 0.779792299 | -0.345434996 |
| Ass1 | 10.48806 | 10.65013 | 10.504 | 0.145715238 | -0.016350376 | 10.508536 | 11.399445 | 10.62 | 0.779457598 | -0.111452309 |
| Stat3 | 12.47017 | 12.32749 | 11.974 | 0.353291656 | 0.495973976 | 12.156383 | 12.918191 | 12.139 | 0.779200286 | 0.017392723 |
| Trim12a | 11.10101 | 11.38193 | 11 | 0.381628262 | 0.100713322 | 11.272687 | 11.967567 | 11.19 | 0.7775702 | 0.082690434 |
| Kcnk5 | 8.11635 | 8.508617 | 8.1213 | 0.387339816 | -0.004926721 | 8.2535034 | 8.9881761 | 8.21069 | 0.77748401 | 0.042811251 |
| Scgb2b27 | 7.424529 | 6.110927 | 6.4746 | -0.363721384 | 0.94988043 | 7.6951218 | 6.5918894 | 5.81514 | 0.776749543 | 1.879981993 |
| Clec4a1 | 7.183859 | 7.532596 | 6.6991 | 0.833493755 | 0.484756415 | 8.0931268 | 7.492949 | 6.71628 | 0.776669407 | 1.376847158 |
| Uqcrq | 8.801694 | 8.877034 | 8.669 | 0.208061423 | 0.132721433 | 9.2440064 | 9.4610246 | 8.68524 | 0.775783784 | 0.558765514 |
| Elob | 9.557882 | 9.717219 | 9.5054 | 0.21184774 | 0.052510733 | 10.017665 | 10.209823 | 9.43471 | 0.775110216 | 0.582951739 |
| Litaf | 10.12152 | 9.999074 | 9.5692 | 0.429840848 | 0.552291757 | 9.9689868 | 10.422374 | 9.64727 | 0.775103388 | 0.321716097 |
| Ltbr | 7.577269 | 7.291253 | 6.9438 | 0.347437347 | 0.63345361 | 7.2721921 | 7.6911256 | 6.91654 | 0.774583046 | 0.355649545 |
| Srm | 9.090302 | 9.202131 | 8.9381 | 0.264008137 | 0.15217865 | 9.3404282 | 9.7329188 | 8.95997 | 0.772952086 | 0.380461426 |

| | | | | | | | | | | |
|---|---|---|---|---|---|---|---|---|---|---|
| Apobec3 | 12.50195 | 12.69856 | 12.492 | 0.20647909 | 0.009871382 | 12.826535 | 13.435052 | 12.6621 | 0.772931694 | 0.164414287 |
| Ccl9 | 8.751392 | 9.502402 | 7.2305 | 2.271918453 | 1.520908246 | 7.7754126 | 7.6101688 | 6.83906 | 0.771112761 | 0.936356545 |
| Card9 | 6.380637 | 6.262189 | 6.1006 | 0.161618438 | 0.280066128 | 6.7604252 | 7.0650282 | 6.29543 | 0.769594404 | 0.464991423 |
| Triobp | 9.795827 | 9.865517 | 9.7377 | 0.127785815 | 0.058094959 | 9.871641 | 10.47402 | 9.70509 | 0.768931971 | 0.166552606 |
| C1rl | 5.548156 | 5.028111 | 5.5159 | -0.487742289 | 0.032302814 | 5.4688608 | 5.6104329 | 4.8422 | 0.76823445 | 0.626662332 |
| Mgat1 | 10.9398 | 11.21657 | 10.902 | 0.315044226 | 0.038274108 | 11.215474 | 11.81606 | 11.048 | 0.76807235 | 0.167486615 |
| Cd40 | 9.964973 | 10.15387 | 9.9995 | 0.154379014 | -0.034520823 | 10.306273 | 10.946803 | 10.1797 | 0.767130456 | 0.126601068 |
| Pfkp | 11.07095 | 10.99861 | 10.92 | 0.079044278 | 0.151381474 | 10.942428 | 11.758224 | 10.9916 | 0.766602452 | -0.049194041 |
| Cst3 | 11.64608 | 11.77561 | 11.571 | 0.204515862 | 0.074982396 | 11.556753 | 12.072263 | 11.306 | 0.766254376 | 0.250743849 |
| Pttg1 | 8.416394 | 8.69943 | 8.4043 | 0.295147126 | 0.012110794 | 8.6030998 | 9.1221841 | 8.35664 | 0.765544037 | 0.246459737 |
| Skint8 | 5.53758 | 6.132281 | 5.6103 | 0.521999784 | -0.072701266 | 5.6923195 | 6.4549913 | 5.68985 | 0.765139293 | 0.002467509 |
| Emb | 10.81589 | 10.84839 | 10.7 | 0.148366756 | 0.115867412 | 10.470119 | 11.402645 | 10.6389 | 0.763731236 | -0.168794167 |
| Cabp4 | 3.940183 | 3.82024 | 3.8284 | -0.008193612 | 0.111749391 | 4.1699164 | 4.6316092 | 3.86829 | 0.76331542 | 0.301622652 |
| Uqcc2 | 7.81357 | 7.919405 | 7.8056 | 0.113797937 | 0.007962996 | 8.2586597 | 8.4526654 | 7.69059 | 0.762073873 | 0.568068207 |
| Rab5c | 11.49287 | 11.65812 | 11.363 | 0.29492943 | 0.129672506 | 11.711899 | 12.074287 | 11.3147 | 0.759588286 | 0.397201003 |
| Sell | 14.28861 | 14.47155 | 14.195 | 0.276683912 | 0.09374787 | 14.398274 | 14.952771 | 14.1937 | 0.759113944 | 0.204617186 |
| Eef1akmt4 | 6.801608 | 6.95306 | 6.6714 | 0.281657654 | 0.130205753 | 6.8864035 | 7.364123 | 6.60501 | 0.759113011 | 0.281393571 |
| Inhbb | 5.411718 | 5.561045 | 4.4401 | 1.120985296 | 0.97165907 | 5.0571426 | 5.5291525 | 4.77007 | 0.759079987 | 0.287070057 |
| Atp2b4 | 8.651348 | 8.691087 | 8.5533 | 0.137810576 | 0.098071915 | 8.8374868 | 9.3293737 | 8.57046 | 0.758908777 | 0.267021901 |
| Cish | 8.226162 | 8.481311 | 8.0474 | 0.4338703 | 0.178721856 | 7.8366347 | 8.9913545 | 8.2332 | 0.758153288 | -0.39656652 |
| Psmb5 | 9.456615 | 9.607222 | 9.3736 | 0.233573339 | 0.082966215 | 9.7807198 | 9.9953446 | 9.23739 | 0.757952163 | 0.543327371 |
| Mrps34 | 8.388614 | 8.413575 | 8.3251 | 0.088439622 | 0.063478626 | 8.4044172 | 8.9259894 | 8.16842 | 0.75757129 | 0.235999117 |
| Smox | 7.931036 | 7.957799 | 7.2701 | 0.68767282 | 0.660908927 | 7.7990397 | 8.1424374 | 7.38509 | 0.757342999 | 0.413945264 |
| Gtpbp2 | 10.70158 | 10.85245 | 10.785 | 0.067460287 | -0.083417173 | 11.211756 | 11.892646 | 11.1356 | 0.757084774 | 0.076195098 |
| Pdcd1lg2 | 7.468461 | 7.696618 | 7.4198 | 0.276836844 | 0.048680621 | 7.809566 | 8.2990165 | 7.54231 | 0.756709318 | 0.267258878 |
| Inpp1 | 9.56385 | 9.830033 | 9.551 | 0.279032513 | 0.012849197 | 9.4719813 | 10.383991 | 9.62919 | 0.754798598 | -0.157211108 |
| Manf | 9.865804 | 10.01753 | 9.6362 | 0.381337798 | 0.229616219 | 9.8700097 | 10.369084 | 9.61592 | 0.753164097 | 0.254089956 |
| Slc35e2 | 8.65112 | 8.959162 | 8.7763 | 0.182851026 | -0.125190938 | 8.7443183 | 9.5173932 | 8.76461 | 0.752781219 | -0.020293721 |
| Timm13 | 8.536473 | 8.642515 | 8.4815 | 0.160973398 | 0.054931832 | 8.9362814 | 9.1945405 | 8.44312 | 0.751417624 | 0.493158603 |
| Lpar6 | 8.568082 | 8.782379 | 8.7969 | -0.014564852 | -0.228862024 | 8.9125765 | 9.6449491 | 8.89355 | 0.751403917 | 0.019031314 |
| Mrps12 | 7.286114 | 7.262092 | 7.3129 | -0.050858075 | -0.026835532 | 7.5239752 | 7.7906528 | 7.03952 | 0.751135222 | 0.484457597 |
| Casz1 | 7.861109 | 8.103673 | 7.9778 | 0.125902053 | -0.116661937 | 7.9247571 | 8.6404149 | 7.88945 | 0.750963954 | 0.035306178 |

| | | | | | | | | | | |
|---|---|---|---|---|---|---|---|---|---|---|
| Lyn | 12.04202 | 12.27502 | 11.897 | 0.37784038 | 0.14484258 | 12.286974 | 12.614356 | 11.8636 | 0.750781522 | 0.423399701 |
| Gimap9 | 10.1692 | 10.36607 | 10.242 | 0.124271818 | -0.072594639 | 10.316793 | 10.961408 | 10.2111 | 0.750277256 | 0.105661984 |
| Herc3 | 10.552 | 10.947 | 10.454 | 0.49326219 | 0.098257542 | 10.483663 | 11.387018 | 10.6374 | 0.749576518 | -0.15377884 |
| 4931414P19Rik | 6.521799 | 6.53225 | 6.5578 | -0.02552226 | -0.035973003 | 6.7532019 | 7.1772527 | 6.42816 | 0.749097127 | 0.32504637 |
| Igkv4-53 | 7.166201 | 7.918394 | 7.2559 | 0.662537271 | -0.089655837 | 7.7292678 | 8.1666374 | 7.41849 | 0.748151238 | 0.310781659 |
| Gcnt2 | 7.933675 | 7.508914 | 6.9293 | 0.579663361 | 1.004424121 | 7.8004608 | 7.6583533 | 6.91106 | 0.747293475 | 0.889401012 |
| Prkcd | 11.29548 | 11.8363 | 11.4 | 0.435809442 | -0.105016334 | 11.679073 | 12.400259 | 11.6532 | 0.74708042 | 0.025894264 |
| Fes | 9.000451 | 8.946459 | 8.8733 | 0.073155995 | 0.127147788 | 9.2787829 | 9.4657055 | 8.72011 | 0.745592606 | 0.558670046 |
| Usb1 | 8.584217 | 8.731381 | 8.5586 | 0.172800134 | 0.025636176 | 8.9040349 | 9.4575293 | 8.71374 | 0.743784922 | 0.190290461 |
| Insl3 | 4.35579 | 4.125023 | 4.283 | -0.157953785 | 0.072813276 | 4.6412724 | 4.8913847 | 4.14819 | 0.743194837 | 0.493082523 |
| Ppan | 9.424727 | 9.594021 | 9.3467 | 0.247315276 | 0.078021289 | 9.8816156 | 10.359902 | 9.6173 | 0.742596664 | 0.26431072 |
| Ciao2b | 8.174803 | 8.396162 | 8.1773 | 0.218867597 | -0.002490768 | 8.6675506 | 8.9761364 | 8.23403 | 0.742106516 | 0.433520745 |
| Igkv10-96 | 9.980865 | 10.29631 | 10.175 | 0.121361059 | -0.19408879 | 10.305627 | 10.64893 | 9.90706 | 0.74187435 | 0.398571465 |
| Gimap7 | 9.329308 | 9.598914 | 9.3835 | 0.215406571 | -0.054198974 | 9.794696 | 10.274987 | 9.53324 | 0.741750762 | 0.261459397 |
| Mertk | 6.941991 | 7.034084 | 6.6887 | 0.345372636 | 0.253279881 | 6.193479 | 6.8646106 | 6.12311 | 0.741504024 | 0.070372461 |
| Atm | 10.78277 | 10.88905 | 10.85 | 0.039269116 | -0.067001996 | 10.710466 | 11.756176 | 11.0148 | 0.741353936 | -0.304356219 |
| Cxcr5 | 11.65601 | 11.91749 | 11.586 | 0.331451118 | 0.069967994 | 12.007538 | 12.299827 | 11.5601 | 0.73973038 | 0.447441666 |
| Rpl27 | 11.71105 | 11.8801 | 11.683 | 0.1971584 | 0.02811603 | 12.211844 | 12.317308 | 11.5777 | 0.739566874 | 0.634103384 |
| Pla1a | 3.896421 | 3.921064 | 3.7345 | 0.186524381 | 0.161881144 | 3.976595 | 4.6214381 | 3.88243 | 0.739012576 | 0.094169445 |
| Vcam1 | 9.899869 | 10.2715 | 9.6215 | 0.649984043 | 0.278348341 | 8.7283679 | 9.7102934 | 8.97214 | 0.738153582 | -0.24377193 |
| Mocos | 5.509489 | 5.658152 | 5.3294 | 0.328786365 | 0.180123777 | 5.2636149 | 5.9823441 | 5.24504 | 0.737301655 | 0.01857244 |
| Usf1 | 10.23281 | 10.55639 | 10.228 | 0.328099753 | 0.004523558 | 10.506533 | 11.016708 | 10.2805 | 0.736203724 | 0.226028561 |
| S1pr3 | 7.803091 | 7.735578 | 7.2167 | 0.51889912 | 0.58641164 | 7.3221797 | 7.8054103 | 7.06957 | 0.735840808 | 0.252610234 |
| Oas1c | 8.068099 | 8.661805 | 8.156 | 0.505837688 | -0.087868135 | 8.620586 | 9.2439722 | 8.51 | 0.733976027 | 0.110589774 |
| Tcirg1 | 9.99372 | 10.16931 | 10.115 | 0.054799744 | -0.120786746 | 10.692888 | 11.044893 | 10.3112 | 0.733737218 | 0.381731655 |
| Stx16 | 10.37635 | 10.36741 | 10.271 | 0.096457508 | 0.105389825 | 10.550604 | 11.184631 | 10.4537 | 0.730905894 | 0.096877974 |
| Sap18b | 9.457245 | 9.640291 | 9.48 | 0.160254887 | -0.022790813 | 9.8323408 | 10.033126 | 9.30331 | 0.729820362 | 0.529034637 |
| Olr1 | 3.910967 | 3.938532 | 3.5647 | 0.373845477 | 0.346280971 | 4.193936 | 4.3313965 | 3.60211 | 0.729286226 | 0.591825771 |
| Magohb | 6.683498 | 6.683435 | 6.4333 | 0.250152373 | 0.250215924 | 6.666547 | 7.3041254 | 6.57554 | 0.72858652 | 0.091008157 |
| Etnk1 | 12.59313 | 12.94022 | 12.547 | 0.39282961 | 0.04574007 | 12.72383 | 13.3494 | 12.622 | 0.727448674 | 0.101878292 |
| Ctrl | 4.184133 | 4.621258 | 4.3649 | 0.256359693 | -0.180765331 | 4.6001906 | 5.2360237 | 4.50898 | 0.727038952 | 0.091205855 |
| Mrpl54 | 8.148465 | 8.137535 | 8.0228 | 0.114756447 | 0.125686801 | 8.3929147 | 8.763072 | 8.03632 | 0.726755196 | 0.3565979 |

| | | | | | | | | | | |
|---|---|---|---|---|---|---|---|---|---|---|
| Exoc3l4 | 5.779353 | 5.811119 | 5.969 | -0.157854144 | -0.189620299 | 5.8048759 | 6.4424665 | 5.71602 | 0.726448887 | 0.088858344 |
| Tnf | 8.458291 | 8.50976 | 8.4891 | 0.02065044 | -0.03081871 | 8.4649505 | 9.1107381 | 8.38452 | 0.726216072 | 0.080428525 |
| Cyba | 11.2064 | 11.38531 | 11.179 | 0.206479926 | 0.027560886 | 11.623618 | 11.792105 | 11.066 | 0.72613173 | 0.557644989 |
| Fndc3a | 10.23719 | 10.49924 | 10.131 | 0.368283935 | 0.106231411 | 10.099843 | 10.962422 | 10.2369 | 0.725565683 | -0.137012935 |
| Alpk1 | 6.975526 | 6.815877 | 6.6112 | 0.204685574 | 0.364334287 | 6.4672403 | 7.1172554 | 6.3918 | 0.725457562 | 0.075442513 |
| S100a6 | 9.486653 | 9.421933 | 9.241 | 0.180967721 | 0.245687859 | 9.6194624 | 10.05818 | 9.33446 | 0.723716586 | 0.284999124 |
| Ubfd1 | 9.048782 | 9.504091 | 9.0762 | 0.427900367 | -0.027408638 | 9.2601056 | 9.99623 | 9.27317 | 0.723059684 | -0.013064687 |
| Mat2a | 10.33661 | 10.39007 | 10.337 | 0.053302526 | -0.000156736 | 10.052805 | 10.758563 | 10.0363 | 0.722277416 | 0.016519281 |
| Fdps | 8.776341 | 8.981955 | 8.7325 | 0.249424424 | 0.043810603 | 9.0121069 | 9.3664926 | 8.64457 | 0.721922818 | 0.367537087 |
| Igkv2-112 | 5.539566 | 5.879348 | 5.5101 | 0.369250707 | 0.029468333 | 5.6459451 | 6.1602154 | 5.43966 | 0.720553543 | 0.206283265 |
| Mtm1 | 8.102961 | 8.214867 | 8.0293 | 0.185586866 | 0.073680094 | 8.0956946 | 8.799085 | 8.07855 | 0.720532099 | 0.017141717 |
| 2310057J18Rik | 4.938356 | 5.072164 | 5.5863 | -0.514130433 | -0.647938541 | 6.573499 | 5.8063358 | 5.08593 | 0.720401029 | 1.48756422 |
| Klk1b4 | 4.522255 | 4.993609 | 5.572 | -0.578374654 | -1.049728725 | 6.7165986 | 5.8996894 | 5.18025 | 0.719436208 | 1.536345417 |
| Klk1b3 | 4.182568 | 4.663844 | 5.3401 | -0.676276456 | -1.157552664 | 6.3201937 | 5.5376912 | 4.81879 | 0.718900783 | 1.501403274 |
| Rpl10a | 12.8958 | 12.9471 | 12.847 | 0.100506848 | 0.049207846 | 13.455162 | 13.477755 | 12.7592 | 0.71855124 | 0.695958363 |
| Gpr33 | 3.57155 | 3.715735 | 3.583 | 0.132781655 | -0.011402755 | 3.5323609 | 4.226721 | 3.50823 | 0.718494907 | 0.024134855 |
| Clcn7 | 9.827907 | 9.989661 | 9.8453 | 0.144351488 | -0.017402464 | 10.157735 | 10.695843 | 9.97741 | 0.718432778 | 0.180325196 |
| Pnp2 | 3.58509 | 3.73757 | 3.6329 | 0.104714876 | -0.047765915 | 3.5780528 | 4.1990898 | 3.48067 | 0.718423788 | 0.097386744 |
| Ripk1 | 10.2322 | 10.27113 | 10.021 | 0.249639693 | 0.210706954 | 9.9182004 | 10.730702 | 10.013 | 0.717721166 | -0.094780552 |
| Rps27l | 8.260242 | 8.274669 | 8.135 | 0.139688266 | 0.125261269 | 8.3577739 | 8.7891014 | 8.07175 | 0.71735599 | 0.286028567 |
| Gm11427 | 9.607073 | 9.789814 | 9.5472 | 0.242659188 | 0.059918354 | 9.809224 | 10.378128 | 9.66107 | 0.717058261 | 0.1481546 |
| Tuba8 | 4.093875 | 4.234835 | 3.8718 | 0.363054194 | 0.222093334 | 3.9574191 | 4.5049754 | 3.78884 | 0.716138455 | 0.168582169 |
| Lipe | 8.942539 | 9.080271 | 8.8698 | 0.210469199 | 0.072736741 | 9.1660192 | 9.6990358 | 8.98293 | 0.716106925 | 0.183090286 |
| BC037034 | 9.680089 | 10.03511 | 9.72 | 0.315157903 | -0.039867387 | 10.160151 | 10.485221 | 9.76921 | 0.716013245 | 0.390942361 |
| Gm21762 | 5.890913 | 6.207773 | 6.0523 | 0.155492996 | -0.161366668 | 5.3513555 | 6.1367034 | 5.42131 | 0.715397845 | -0.069950108 |
| CT010433.1 | 4.391378 | 4.586606 | 4.2831 | 0.303556219 | 0.108328422 | 4.3259917 | 4.9193304 | 4.20429 | 0.715036824 | 0.121698191 |
| Med31 | 6.267364 | 6.389485 | 6.3101 | 0.079364197 | -0.042756365 | 6.4465035 | 6.7450769 | 6.03075 | 0.714323678 | 0.415750266 |
| Mrps28 | 7.227413 | 7.431474 | 7.1435 | 0.288018463 | 0.083957401 | 7.4185934 | 7.8553266 | 7.14113 | 0.714201173 | 0.277467952 |
| Tlr12 | 7.586204 | 7.823682 | 7.6367 | 0.186963181 | -0.050514567 | 7.5690055 | 8.2232241 | 7.50909 | 0.714134081 | 0.059915502 |
| Tnp2 | 2.487533 | 2.507656 | 2.4586 | 0.049027116 | 0.028904033 | 2.3946397 | 3.2096712 | 2.4961 | 0.713574471 | -0.101457001 |
| Tnfsf8 | 8.916104 | 8.972601 | 8.862 | 0.110592285 | 0.054095089 | 8.533419 | 9.5573358 | 8.84413 | 0.713207934 | -0.310708884 |
| Atp1a3 | 7.106253 | 7.135659 | 6.7607 | 0.374968328 | 0.345562559 | 6.7426838 | 7.3696671 | 6.65677 | 0.712894847 | 0.085911574 |

| | | | | | | | | | | |
|---|---|---|---|---|---|---|---|---|---|---|
| Sos1 | 9.767198 | 10.02009 | 9.7439 | 0.276236204 | 0.023347665 | 9.5166483 | 10.414695 | 9.70199 | 0.712705608 | -0.185341447 |
| Spata5 | 8.990978 | 9.138036 | 8.9777 | 0.160347312 | 0.013289959 | 9.056268 | 9.6120327 | 8.8998 | 0.712228521 | 0.15646381 |
| Traf5 | 9.247154 | 9.58315 | 9.3347 | 0.248434605 | -0.087561428 | 9.4500036 | 9.9630511 | 9.25225 | 0.710799799 | 0.197752319 |
| Hpse | 8.734026 | 8.730443 | 8.7189 | 0.011536321 | 0.01511948 | 9.0361012 | 9.2873363 | 8.57667 | 0.710664342 | 0.459429217 |
| Rcn1 | 8.508369 | 8.467218 | 8.3672 | 0.099979838 | 0.141130368 | 8.1964555 | 8.9061042 | 8.19569 | 0.71041561 | 0.000766967 |
| Bak1 | 9.695182 | 9.910461 | 9.7126 | 0.197851196 | -0.017427614 | 9.9487021 | 10.425267 | 9.71595 | 0.709314321 | 0.232749511 |
| Katna1 | 8.439299 | 8.573543 | 8.5012 | 0.072371408 | -0.061871775 | 8.5566484 | 9.2496126 | 8.54032 | 0.709290533 | 0.016326329 |
| Ptx3 | 3.015246 | 2.813995 | 2.5948 | 0.219167237 | 0.420418254 | 2.7901521 | 3.3362959 | 2.62778 | 0.708512241 | 0.162368433 |
| Dynll1 | 10.64918 | 11.05238 | 10.667 | 0.385515888 | -0.017677424 | 10.806393 | 11.357089 | 10.649 | 0.708123438 | 0.157427739 |
| Car2 | 7.458693 | 8.153311 | 7.5749 | 0.578386798 | -0.116230554 | 7.616677 | 8.2026977 | 7.49503 | 0.707667025 | 0.121646346 |
| Rgs1 | 9.900645 | 10.44264 | 9.8104 | 0.632228246 | 0.090236227 | 9.503054 | 10.607739 | 9.90087 | 0.706866231 | -0.397818623 |
| Icam1 | 11.62313 | 11.68024 | 11.386 | 0.293926638 | 0.23681519 | 11.172035 | 11.932093 | 11.2266 | 0.705518388 | -0.054539629 |
| Tstd1 | 4.379954 | 4.245037 | 4.2711 | -0.026100505 | 0.108816209 | 4.6454496 | 5.010042 | 4.30455 | 0.705496399 | 0.340903997 |
| Crem | 8.635917 | 8.629421 | 8.1939 | 0.435535208 | 0.442031323 | 8.2482684 | 9.4666727 | 8.76231 | 0.704367703 | -0.514036675 |
| Kcne4 | 6.135879 | 5.627205 | 5.2967 | 0.33047104 | 0.839144826 | 5.2290025 | 5.8264582 | 5.12252 | 0.703941572 | 0.106485865 |
| Ndufa12 | 8.551223 | 8.682524 | 8.5833 | 0.099198599 | -0.032102208 | 8.7331816 | 9.1741206 | 8.47187 | 0.70225279 | 0.261313858 |
| Nacc2 | 6.985132 | 7.029312 | 7.0655 | -0.03622316 | -0.080403483 | 6.8256236 | 7.6166615 | 6.91458 | 0.702085099 | -0.088952773 |
| Nme2 | 11.13751 | 11.15316 | 11.01 | 0.142934112 | 0.127284206 | 11.411978 | 11.7008 | 10.9993 | 0.701514754 | 0.412692961 |
| Als2 | 8.951745 | 9.105121 | 9.016 | 0.089167517 | -0.064208321 | 8.9069405 | 9.7344745 | 9.03314 | 0.701332239 | -0.126201806 |
| Ranbp1 | 8.708262 | 8.843773 | 8.6671 | 0.176673898 | 0.041163033 | 8.8389593 | 9.4183837 | 8.7178 | 0.700588344 | 0.121163979 |
| Gm12187 | 2.170835 | 2.307039 | 2.1491 | 0.157904323 | 0.021700582 | 2.2292027 | 2.8845963 | 2.18417 | 0.700422636 | 0.045029067 |
| Klk1 | 8.73256 | 7.021923 | 7.1852 | -0.16327688 | 1.547360156 | 7.0849395 | 7.0247608 | 6.32439 | 0.70036662 | 0.760545372 |
| Mcemp1 | 7.410327 | 7.339417 | 6.8782 | 0.461206573 | 0.532116571 | 7.385097 | 7.6253115 | 6.92509 | 0.70022387 | 0.46000936 |
| Pgam1 | 10.37108 | 10.34356 | 10.291 | 0.052307012 | 0.079829862 | 10.493861 | 10.932458 | 10.2345 | 0.697925048 | 0.25932773 |
| Rarres2 | 5.754158 | 5.407299 | 5.75 | -0.34272747 | 0.004131 | 5.3901422 | 6.0431527 | 5.34523 | 0.697920583 | 0.044909992 |
| Gm7367 | 4.604265 | 4.542043 | 4.3351 | 0.206930811 | 0.26915306 | 4.8902761 | 4.9357144 | 4.23846 | 0.697256438 | 0.651818173 |
| Gm8093 | 3.352998 | 3.262384 | 3.2325 | 0.029931418 | 0.120546121 | 3.2858104 | 3.8335114 | 3.13683 | 0.69668523 | 0.148984179 |
| C130050O18Rik | 4.524128 | 4.363086 | 4.1344 | 0.228720145 | 0.389762455 | 4.9272892 | 4.901927 | 4.20546 | 0.696471717 | 0.72183391 |
| Ccl11 | 5.739449 | 5.457445 | 5.0491 | 0.408314086 | 0.690318876 | 4.8431355 | 5.571065 | 4.87478 | 0.696283264 | -0.031646227 |
| Tmem160 | 7.430002 | 7.557067 | 7.3462 | 0.210915312 | 0.083850454 | 7.7534984 | 7.8936633 | 7.19752 | 0.696142338 | 0.555977454 |
| Ndufaf8 | 6.607694 | 6.728483 | 6.6634 | 0.06505783 | -0.055731329 | 6.7695331 | 7.2385226 | 6.54284 | 0.695679308 | 0.2266898 |
| Aars | 10.54964 | 10.55799 | 10.389 | 0.168651188 | 0.16029747 | 11.000902 | 11.410504 | 10.7149 | 0.695627044 | 0.286024428 |

| | | | | | | | | | | |
|---|---|---|---|---|---|---|---|---|---|---|
| Gm12253 | 3.363448 | 3.462451 | 3.2246 | 0.23788711 | 0.13888389 | 3.401129 | 3.9614506 | 3.26647 | 0.694980105 | 0.134658521 |
| C4b | 9.500937 | 9.07584 | 9.0163 | 0.059495105 | 0.484591554 | 8.6095217 | 9.4443665 | 8.74951 | 0.694857833 | -0.139986971 |
| Tmem50b | 8.957299 | 9.016889 | 8.8789 | 0.138026127 | 0.078436384 | 9.1600151 | 9.4967326 | 8.80287 | 0.693860398 | 0.357142876 |
| Fam102a | 11.42404 | 11.55338 | 11.366 | 0.187804834 | 0.05846552 | 11.29569 | 11.991123 | 11.2974 | 0.693694746 | -0.001738685 |
| Gad1-ps | 3.02672 | 3.272338 | 2.9119 | 0.360394551 | 0.114777207 | 3.3706345 | 3.5995274 | 2.90703 | 0.692498945 | 0.463606036 |
| Acer3 | 8.279259 | 8.404767 | 7.7628 | 0.641944389 | 0.516436679 | 8.165416 | 8.5154873 | 7.82321 | 0.692272845 | 0.342201517 |
| Sla | 11.68346 | 11.95924 | 11.691 | 0.268627254 | -0.007158508 | 11.585676 | 12.270302 | 11.5785 | 0.69179118 | 0.007165488 |
| Psmb4 | 10.46712 | 10.54738 | 10.489 | 0.058117904 | -0.022137978 | 10.642132 | 10.989052 | 10.2974 | 0.691681884 | 0.344761684 |
| Apex1 | 9.185392 | 9.235214 | 9.0578 | 0.177449336 | 0.127627127 | 9.4047305 | 9.7739349 | 9.08242 | 0.691519761 | 0.322315375 |
| Wdr62 | 6.595751 | 6.808411 | 6.8602 | -0.05175857 | -0.26441817 | 6.9317287 | 7.6225824 | 6.93135 | 0.691232829 | 0.000379134 |
| Tubb3 | 4.296599 | 4.404594 | 4.4228 | -0.01822511 | -0.126219835 | 4.2275203 | 4.888816 | 4.19762 | 0.691192679 | 0.029896975 |
| Ninj1 | 8.311226 | 8.285491 | 8.0786 | 0.206841489 | 0.232577281 | 7.9351221 | 8.3020289 | 7.61095 | 0.691079842 | 0.324173049 |
| Tpsab1 | 3.414453 | 3.813366 | 3.4956 | 0.317799047 | -0.081114649 | 3.4230395 | 4.0202419 | 3.32926 | 0.690981359 | 0.093778877 |
| Pdia3 | 12.58877 | 12.64046 | 12.442 | 0.197966952 | 0.146272018 | 12.47138 | 13.133396 | 12.4426 | 0.690792838 | 0.028776338 |
| Zdhhc14 | 6.967005 | 7.287918 | 6.8856 | 0.40235982 | 0.08144606 | 7.2056739 | 7.5793665 | 6.88882 | 0.690550787 | 0.316858121 |
| Tceanc2 | 9.174641 | 9.313135 | 9.108 | 0.205099695 | 0.066606043 | 9.2125251 | 9.6489237 | 8.95863 | 0.690288972 | 0.25389035 |
| Tifa | 8.711618 | 8.640977 | 8.5539 | 0.087092541 | 0.157732617 | 8.4150962 | 8.8972627 | 8.20819 | 0.689071471 | 0.206904925 |
| Ears2 | 8.253815 | 8.479071 | 8.1843 | 0.294751383 | 0.069495537 | 8.3219981 | 8.8272742 | 8.13895 | 0.688320086 | 0.183043988 |
| Klra8 | 5.726638 | 6.038224 | 5.8346 | 0.203595828 | -0.107990232 | 5.4621542 | 6.2283967 | 5.54051 | 0.687888628 | -0.078353862 |
| Stk19 | 8.379341 | 8.288206 | 8.2542 | 0.034040351 | 0.125174624 | 8.5366998 | 9.0301263 | 8.34228 | 0.687848257 | 0.194421783 |
| Itpr1 | 11.47586 | 11.61515 | 11.418 | 0.196752426 | 0.05746312 | 11.411494 | 12.234014 | 11.5462 | 0.687805636 | -0.134715287 |
| Ccnb1ip1 | 5.824071 | 5.316887 | 5.6811 | -0.364208986 | 0.142974976 | 6.6834429 | 6.2623248 | 5.57475 | 0.687578843 | 1.10869698 |
| Gm10478 | 5.166936 | 5.543348 | 5.2283 | 0.31505488 | -0.061356544 | 5.7093904 | 6.114145 | 5.42663 | 0.687519989 | 0.282765334 |
| Nudc | 10.09517 | 10.25994 | 10.08 | 0.179844394 | 0.015069728 | 10.278411 | 10.770981 | 10.0838 | 0.687228996 | 0.194658915 |
| Olfr1271 | 4.133595 | 4.25428 | 4.0883 | 0.165974105 | 0.045288731 | 4.1885137 | 4.9437868 | 4.25661 | 0.687177721 | -0.068095319 |
| Rhou | 5.01529 | 5.091716 | 4.5252 | 0.56653547 | 0.490109251 | 4.5529459 | 5.0755312 | 4.38836 | 0.687174149 | 0.164588841 |
| Tmem263 | 8.277267 | 8.456417 | 8.2415 | 0.214881894 | 0.03573204 | 8.1208362 | 8.8971873 | 8.21021 | 0.686977716 | -0.089373365 |
| Wdr43 | 10.94697 | 10.92017 | 10.751 | 0.168735212 | 0.195539212 | 11.012603 | 11.611517 | 10.925 | 0.686506234 | 0.087592225 |
| Syvn1 | 10.96066 | 11.27241 | 11.019 | 0.253716936 | -0.05802483 | 11.509926 | 11.777928 | 11.092 | 0.685910864 | 0.417909192 |
| Rplp1 | 14.41599 | 14.58189 | 14.355 | 0.227010326 | 0.061112126 | 14.799906 | 14.896738 | 14.2112 | 0.685558592 | 0.588726594 |
| Clec2d | 11.59596 | 11.83503 | 11.565 | 0.270367796 | 0.03129307 | 11.536095 | 12.23666 | 11.552 | 0.68469905 | -0.015865876 |
| Uqcr11 | 8.403859 | 8.354912 | 8.348 | 0.006896271 | 0.05584344 | 8.7384048 | 8.7964194 | 8.11235 | 0.684066104 | 0.626051504 |

| | | | | | | | | | | |
|---|---|---|---|---|---|---|---|---|---|---|
| Mybbp1a | 12.16695 | 12.1938 | 12.061 | 0.132369792 | 0.105522 | 12.360309 | 12.82516 | 12.1422 | 0.68292918 | 0.218078462 |
| Slc2a6 | 7.567442 | 7.944932 | 7.6057 | 0.339200736 | -0.038289276 | 7.8706618 | 8.3470424 | 7.66451 | 0.682535344 | 0.206154721 |
| Pomp | 10.07318 | 10.274 | 10.214 | 0.0601992 | -0.140619718 | 10.341147 | 10.795292 | 10.1128 | 0.682484606 | 0.228339378 |
| F13a1 | 9.293058 | 8.761003 | 7.2483 | 1.512660259 | 2.044714376 | 7.1776543 | 7.5641739 | 6.88269 | 0.681479683 | 0.294960075 |
| Sh3gl1 | 10.27803 | 10.48777 | 10.294 | 0.194084176 | -0.01565459 | 10.532594 | 10.962778 | 10.2815 | 0.681249582 | 0.251065681 |
| Pigr | 6.648975 | 6.571398 | 6.5584 | 0.012966762 | 0.090544317 | 6.8182968 | 7.3364537 | 6.65662 | 0.679832296 | 0.161675357 |
| Phlpp1 | 8.976563 | 9.302741 | 8.9849 | 0.317833837 | -0.008344931 | 8.8292207 | 9.5923326 | 8.91251 | 0.679820663 | -0.083291253 |
| H3f3b | 13.15887 | 12.9726 | 13.049 | -0.076297924 | 0.109981878 | 12.868196 | 13.328407 | 12.6491 | 0.67926135 | 0.219050469 |
| Taf7 | 7.989526 | 7.937746 | 7.9385 | -0.000706271 | 0.051073207 | 7.7734805 | 8.4033325 | 7.72466 | 0.678676525 | 0.048824575 |
| Pdpn | 4.637737 | 4.741302 | 4.0884 | 0.652929197 | 0.549364586 | 4.5232501 | 4.8556793 | 4.17771 | 0.677967803 | 0.345538547 |
| Efhd2 | 10.60296 | 10.69336 | 10.466 | 0.227408202 | 0.137014052 | 10.733221 | 11.024992 | 10.3472 | 0.677767934 | 0.385996696 |
| Coch | 5.23296 | 5.195717 | 5.7459 | -0.550189086 | -0.512946236 | 5.2412938 | 6.3809058 | 5.70385 | 0.677058494 | -0.462553498 |
| Anxa2 | 10.84873 | 11.03928 | 10.621 | 0.418478466 | 0.227927408 | 10.900257 | 11.342661 | 10.666 | 0.67665828 | 0.234254596 |
| Flt4 | 7.661705 | 7.559901 | 7.0418 | 0.518138439 | 0.619942259 | 6.7387152 | 7.4354728 | 6.75919 | 0.676286468 | -0.0204711 |
| Bola1 | 5.155416 | 4.977822 | 5.018 | -0.040205546 | 0.137388065 | 5.2671912 | 5.5395189 | 4.86401 | 0.675507347 | 0.40317969 |
| Zfp213 | 7.558988 | 7.711321 | 7.5619 | 0.149400565 | -0.002932476 | 7.5498126 | 8.1100634 | 7.43635 | 0.673709885 | 0.113459091 |
| Psma3 | 9.953159 | 10.13903 | 10.123 | 0.016045477 | -0.169822262 | 10.366556 | 10.775891 | 10.1037 | 0.672215707 | 0.262881489 |
| Tnfsf4 | 3.802363 | 3.886134 | 3.8764 | 0.009753561 | -0.074017506 | 3.848392 | 4.4306536 | 3.75845 | 0.672202927 | 0.089941385 |
| Gm2808 | 2.167598 | 2.142457 | 2.1209 | 0.02152524 | 0.046667093 | 2.1039489 | 2.7716306 | 2.09998 | 0.671649012 | 0.003967331 |
| Zfp281 | 9.833152 | 9.903012 | 9.6865 | 0.216527397 | 0.146667826 | 9.5640487 | 10.402001 | 9.73045 | 0.671547324 | -0.16640542 |
| Alkbh1 | 9.019039 | 9.16613 | 8.9317 | 0.234402768 | 0.08731215 | 9.2647375 | 9.7908122 | 9.12009 | 0.670723898 | 0.144649215 |
| Sirpa | 10.3813 | 10.61596 | 10.168 | 0.448114014 | 0.213453614 | 10.854663 | 10.518224 | 9.84816 | 0.670061036 | 1.006500099 |
| Atp5e | 10.53899 | 10.60103 | 10.499 | 0.101974886 | 0.039926636 | 10.976966 | 11.103288 | 10.4357 | 0.667626586 | 0.541304803 |
| Muc1 | 4.306304 | 4.141122 | 4.0954 | 0.045686018 | 0.210867785 | 4.0722465 | 4.5497414 | 3.88353 | 0.666211891 | 0.188716903 |
| Cd209c | 4.597538 | 4.49236 | 4.4711 | 0.021275325 | 0.126452539 | 4.6426333 | 5.1264784 | 4.46118 | 0.665295633 | 0.181450463 |
| Fbn1 | 6.935615 | 7.117618 | 6.7418 | 0.375796396 | 0.193793004 | 6.8617627 | 7.4070945 | 6.7422 | 0.66489345 | 0.119561701 |
| Gm45837 | 9.264452 | 8.945276 | 9.0357 | -0.090432312 | 0.228743655 | 9.5489657 | 10.101512 | 9.43733 | 0.664185639 | 0.111639675 |
| Itga5 | 8.492115 | 8.542278 | 8.3074 | 0.234898517 | 0.184736191 | 7.8513798 | 8.685193 | 8.02227 | 0.662927574 | -0.170885547 |
| Fasl | 5.335971 | 5.439393 | 5.3768 | 0.062614699 | -0.040806905 | 5.2125718 | 5.8974531 | 5.23454 | 0.66290847 | -0.021972798 |
| Gm15459 | 4.904935 | 5.375916 | 5.0284 | 0.347510855 | -0.123469888 | 5.0269671 | 5.8001393 | 5.1375 | 0.662635876 | -0.110536306 |
| Alas1 | 8.568624 | 8.926366 | 8.4907 | 0.435624171 | 0.077881415 | 8.8648818 | 9.1586645 | 8.49603 | 0.662630668 | 0.368847971 |
| Ccdc6 | 9.61472 | 9.808043 | 9.5949 | 0.213101538 | 0.019778999 | 9.6473654 | 10.441134 | 9.77858 | 0.662554463 | -0.13121405 |

| | | | | | | | | | | |
|---|---|---|---|---|---|---|---|---|---|---|
| Uqcr10 | 8.97028 | 8.972945 | 8.9421 | 0.030890648 | 0.028225361 | 9.2119478 | 9.4657949 | 8.80358 | 0.662212668 | 0.408365552 |
| Glrx | 8.704695 | 8.784184 | 8.6605 | 0.123641476 | 0.044152316 | 8.9064729 | 9.42493 | 8.76326 | 0.661667004 | 0.14320987 |
| Psmb6 | 10.144 | 10.25589 | 10.091 | 0.165025296 | 0.053127539 | 10.427661 | 10.723049 | 10.0619 | 0.661177912 | 0.365789187 |
| Mrpl23 | 6.122741 | 5.998346 | 5.9037 | 0.094696031 | 0.219090818 | 6.4500847 | 6.6350713 | 5.97395 | 0.661124837 | 0.47613831 |
| Ptprcap | 10.1284 | 10.30803 | 10.181 | 0.126856913 | -0.052770526 | 10.586447 | 10.683471 | 10.0224 | 0.661035444 | 0.564011221 |
| Txnl4a | 8.124938 | 8.493426 | 8.343 | 0.150376806 | -0.218110943 | 8.3765028 | 8.8959229 | 8.23596 | 0.659964743 | 0.140544624 |
| Lgmn | 10.79672 | 10.9811 | 10.483 | 0.498417512 | 0.314036992 | 11.052988 | 11.070862 | 10.4111 | 0.65978277 | 0.641907994 |
| Fnbp4 | 11.32825 | 11.60393 | 11.465 | 0.138646168 | -0.137038 | 11.852189 | 12.532968 | 11.8733 | 0.659652504 | -0.021125614 |
| Rbx1 | 9.779093 | 9.910567 | 9.7513 | 0.159279424 | 0.027805679 | 10.045822 | 10.424514 | 9.76515 | 0.65936103 | 0.280668381 |
| Zfp710 | 9.587643 | 9.901105 | 9.6315 | 0.269588158 | -0.043873374 | 9.8237799 | 10.324844 | 9.66648 | 0.658362302 | 0.157297825 |
| Wfdc12 | 4.852197 | 4.955949 | 5.4517 | -0.495765286 | -0.599517226 | 6.5548244 | 5.7225612 | 5.0657 | 0.656864038 | 1.489127245 |
| Rps13 | 13.31113 | 13.38777 | 13.315 | 0.07279222 | -0.003853848 | 13.456354 | 13.698815 | 13.0428 | 0.655996366 | 0.413534713 |
| Lman2l | 8.210015 | 8.460293 | 8.3172 | 0.143075951 | -0.107201475 | 8.2482481 | 8.9004906 | 8.2447 | 0.655786997 | 0.00354449 |
| Pfkl | 10.04749 | 10.07801 | 10.02 | 0.057655643 | 0.027140193 | 10.144445 | 10.668557 | 10.0133 | 0.655288359 | 0.131176139 |
| Runx3 | 10.63774 | 10.76955 | 10.522 | 0.247073568 | 0.115258328 | 10.736508 | 11.145757 | 10.4907 | 0.655098392 | 0.245849249 |
| Rab43 | 9.619672 | 9.406989 | 9.5009 | -0.093882342 | 0.118800939 | 9.4875265 | 9.9461455 | 9.29108 | 0.655063696 | 0.196444719 |
| Gm13835 | 9.128362 | 9.103696 | 8.9582 | 0.145465027 | 0.170130163 | 8.8963827 | 9.5886484 | 8.93393 | 0.654713666 | -0.037551969 |
| Sharpin | 9.986881 | 10.0637 | 10.064 | -0.000701017 | -0.077523047 | 10.402152 | 10.872226 | 10.2178 | 0.654433332 | 0.18435991 |
| Olfr318 | 5.343555 | 5.608423 | 5.3261 | 0.282293066 | 0.017425267 | 5.7081132 | 6.3829881 | 5.72865 | 0.65433384 | -0.020541023 |
| Myc | 11.17802 | 11.53264 | 11.002 | 0.530986528 | 0.17637445 | 10.629081 | 11.500211 | 10.8472 | 0.653016328 | -0.218113378 |
| Ddx39b | 12.31243 | 12.50196 | 12.346 | 0.155580132 | -0.03395313 | 12.676208 | 13.143408 | 12.4907 | 0.65271048 | 0.185510103 |
| Vmn2r59 | 5.097667 | 5.301842 | 5.1119 | 0.18995102 | -0.014224391 | 5.2466522 | 5.9626839 | 5.31068 | 0.652000501 | -0.064031198 |
| Klri2 | 7.004161 | 7.263898 | 6.9484 | 0.315452026 | 0.055714363 | 6.7692822 | 7.6032718 | 6.95159 | 0.651684121 | -0.182305405 |
| Itgax | 9.603717 | 9.777677 | 9.5318 | 0.245922404 | 0.071962509 | 10.089565 | 9.9695967 | 9.31849 | 0.651107241 | 0.771075174 |
| Slc27a3 | 5.633373 | 5.692787 | 5.5486 | 0.144228879 | 0.084814931 | 5.4871762 | 6.0891124 | 5.43943 | 0.649682405 | 0.047746209 |
| Fbxo34 | 8.212024 | 8.321345 | 8.1293 | 0.192018586 | 0.082697497 | 8.2146294 | 8.7711702 | 8.12185 | 0.649316066 | 0.092775251 |
| Kars | 10.91282 | 11.02188 | 10.896 | 0.125598968 | 0.016538904 | 11.020153 | 11.493491 | 10.8457 | 0.647784314 | 0.17444601 |
| Cul2 | 8.989307 | 8.981095 | 9.0171 | -0.036032571 | -0.027820376 | 9.0486322 | 9.5645041 | 8.91674 | 0.647764929 | 0.131892973 |
| Chchd2 | 10.77069 | 10.79663 | 10.716 | 0.080221418 | 0.054286536 | 10.822797 | 11.240733 | 10.593 | 0.647687606 | 0.229751787 |
| Odc1 | 11.11353 | 10.96453 | 10.942 | 0.02254603 | 0.17154355 | 10.657089 | 11.402891 | 10.7553 | 0.647614742 | -0.098186933 |
| Bcl2a1d | 8.700675 | 9.229422 | 8.7751 | 0.45437025 | -0.074376543 | 8.6999172 | 9.383475 | 8.7359 | 0.647570926 | -0.035986883 |
| Cenpj | 8.26696 | 8.39307 | 8.3997 | -0.006661965 | -0.132772774 | 8.1304847 | 8.8803548 | 8.23287 | 0.64748272 | -0.102387389 |

| | | | | | | | | | | |
|---|---|---|---|---|---|---|---|---|---|---|
| Rnf31 | 10.54301 | 10.71132 | 10.322 | 0.389149038 | 0.220845618 | 10.548131 | 11.156409 | 10.5091 | 0.64729923 | 0.039020859 |
| Zfp1 | 8.769939 | 8.925485 | 8.8373 | 0.088180744 | -0.067365536 | 8.4934193 | 9.1874815 | 8.54038 | 0.64709852 | -0.046963721 |
| Sfxn2 | 8.388819 | 8.708089 | 8.3668 | 0.341257154 | 0.021987053 | 8.6094187 | 9.0725806 | 8.42555 | 0.64703206 | 0.183870141 |
| Srxn1 | 5.960076 | 5.909326 | 5.4248 | 0.484563453 | 0.535313571 | 5.3631541 | 6.1898669 | 5.54339 | 0.646481394 | -0.180231454 |
| Gadd45gip1 | 7.989373 | 8.215046 | 8.0252 | 0.189871232 | -0.035801679 | 8.285126 | 8.5364134 | 7.89084 | 0.645576518 | 0.394289085 |
| Igkv4-57-1 | 6.471084 | 6.560166 | 6.7355 | -0.175359964 | -0.264442695 | 6.6981066 | 7.4043784 | 6.75941 | 0.644972162 | -0.061299632 |
| Myof | 8.270706 | 8.665261 | 8.3905 | 0.274774116 | -0.119780803 | 8.5222872 | 9.0267969 | 8.38254 | 0.644254171 | 0.139744499 |
| Fbxo4 | 7.630738 | 7.628234 | 7.6171 | 0.011147282 | 0.013650583 | 7.5793035 | 8.1138595 | 7.46961 | 0.644250173 | 0.109694107 |
| Slirp | 7.793933 | 7.83797 | 7.682 | 0.155942489 | 0.111905418 | 7.944431 | 8.4227688 | 7.77907 | 0.64369452 | 0.16535667 |
| Scfd1 | 9.585315 | 9.585952 | 9.5899 | -0.003942626 | -0.004579582 | 9.5826764 | 10.157505 | 9.51566 | 0.641846544 | 0.067017988 |
| Chac2 | 5.647088 | 5.807588 | 5.6291 | 0.178466773 | 0.017966829 | 5.7281329 | 6.2813448 | 5.64058 | 0.640768018 | 0.087556044 |
| Nkg7 | 10.24214 | 10.27434 | 10.193 | 0.081101626 | 0.048902755 | 10.392177 | 10.833074 | 10.1924 | 0.6406318 | 0.199734763 |
| Sertad3 | 7.264928 | 7.463986 | 7.3367 | 0.127302995 | -0.071754891 | 7.2719775 | 7.7873854 | 7.14699 | 0.640390528 | 0.124982583 |
| Mrpl27 | 7.390275 | 7.360213 | 7.3737 | -0.013471328 | 0.016591401 | 7.4307251 | 7.8692439 | 7.22932 | 0.639928729 | 0.20140999 |
| Fzd5 | 7.458899 | 7.406739 | 7.3802 | 0.026495281 | 0.078655535 | 7.056816 | 7.9805328 | 7.34102 | 0.639509444 | -0.284207348 |
| AC129217.1 | 4.356744 | 4.768923 | 4.448 | 0.320874034 | -0.091304693 | 4.7183574 | 5.0226765 | 4.38327 | 0.639402372 | 0.335083281 |
| Phip | 12.05786 | 12.36151 | 12.019 | 0.342469724 | 0.038820752 | 11.915081 | 12.820192 | 12.181 | 0.639177214 | -0.265933887 |
| Atf6b | 10.19813 | 10.27346 | 10.122 | 0.151503239 | 0.076170187 | 10.547203 | 10.708225 | 10.0693 | 0.63892581 | 0.477903955 |
| Eif5a | 12.72657 | 12.76303 | 12.636 | 0.127370612 | 0.09091262 | 12.854077 | 13.217667 | 12.5795 | 0.638211224 | 0.2746215 |
| Hspbp1 | 8.741038 | 8.677657 | 8.5926 | 0.085021694 | 0.148402827 | 8.7933504 | 9.2052055 | 8.56701 | 0.638197766 | 0.226342721 |
| Trim65 | 9.228736 | 9.689899 | 9.2932 | 0.396657071 | -0.064506581 | 9.4008925 | 9.9277278 | 9.28991 | 0.637818868 | 0.110983551 |
| Ifrd2 | 8.315796 | 8.389405 | 8.2855 | 0.103929848 | 0.030320398 | 8.6317333 | 9.0712683 | 8.43351 | 0.637755068 | 0.19822006 |
| Tnks1bp1 | 9.672354 | 9.711121 | 9.5852 | 0.125964665 | 0.087197965 | 9.5656198 | 10.088014 | 9.45031 | 0.637702922 | 0.115308233 |
| Tlr6 | 7.188759 | 7.084926 | 7.0652 | 0.019747859 | 0.123580732 | 7.2826724 | 7.7228666 | 7.08537 | 0.637492574 | 0.197298305 |
| Med7 | 8.011415 | 8.039198 | 7.8881 | 0.151122957 | 0.12333989 | 7.9555544 | 8.4843525 | 7.84696 | 0.63738825 | 0.108590149 |
| Mrpl38 | 8.654856 | 8.964117 | 8.6317 | 0.332405227 | 0.023143463 | 8.9154065 | 9.4354509 | 8.79861 | 0.636843315 | 0.116798895 |
| Rin2 | 7.645935 | 7.536088 | 7.2271 | 0.308987628 | 0.418834642 | 7.1848387 | 7.8181129 | 7.18139 | 0.636722256 | 0.003448067 |
| Pstpip1 | 10.44597 | 10.55921 | 10.472 | 0.087660258 | -0.025581504 | 10.6504 | 11.211934 | 10.5752 | 0.636693902 | 0.075160355 |
| Cox18 | 7.3762 | 7.504358 | 7.2833 | 0.221107429 | 0.092948958 | 7.7337143 | 8.0129193 | 7.37641 | 0.63650883 | 0.35730382 |
| Got1 | 8.926732 | 9.059664 | 8.8445 | 0.215134526 | 0.082203067 | 8.9826717 | 9.7624882 | 9.126 | 0.636490614 | -0.143325826 |
| Fbl | 11.32448 | 11.41763 | 11.256 | 0.161903608 | 0.068761506 | 11.46701 | 11.873024 | 11.2366 | 0.636437776 | 0.230424359 |
| B230307C23Rik | 7.067154 | 7.137947 | 6.9341 | 0.203834589 | 0.133041212 | 7.266412 | 7.6667817 | 7.03074 | 0.636044672 | 0.235675008 |

| | | | | | | | | | | |
|---|---|---|---|---|---|---|---|---|---|---|
| Asap3 | 5.605606 | 5.340044 | 5.2878 | 0.052291492 | 0.317853868 | 5.0270099 | 5.5749764 | 4.9396 | 0.635372826 | 0.087406312 |
| Jak3 | 11.52487 | 11.24074 | 11.055 | 0.18616383 | 0.470293066 | 11.865315 | 11.805582 | 11.1704 | 0.635183022 | 0.694916738 |
| Dtx1 | 10.45213 | 10.87836 | 10.486 | 0.392445326 | -0.033781266 | 11.321497 | 11.208834 | 10.5743 | 0.634535384 | 0.747198257 |
| Gnl3 | 10.13058 | 10.03215 | 10.018 | 0.013893441 | 0.112323498 | 10.234142 | 10.755103 | 10.1207 | 0.634388012 | 0.113427664 |
| Hbegf | 6.533232 | 6.563329 | 5.898 | 0.665333758 | 0.635236347 | 5.4664183 | 6.2140587 | 5.57991 | 0.634148874 | -0.113491538 |
| Klk1b21 | 4.166727 | 4.607885 | 5.2538 | -0.645920307 | -1.087078126 | 6.0220714 | 5.3460419 | 4.71237 | 0.633667932 | 1.309697475 |
| Mppe1 | 10.13619 | 10.27954 | 10.244 | 0.035834322 | -0.107511005 | 10.4375 | 10.916466 | 10.2835 | 0.632916006 | 0.153950302 |
| Pop7 | 7.354003 | 7.387195 | 7.3389 | 0.048288966 | 0.015097155 | 7.5825235 | 7.7082404 | 7.07547 | 0.632772527 | 0.507055629 |
| Pilrb2 | 4.637695 | 4.947852 | 4.3369 | 0.610920921 | 0.300763663 | 5.043256 | 4.9889056 | 4.35683 | 0.632076605 | 0.686427054 |
| Tmem170b | 8.582538 | 8.778761 | 8.555 | 0.223730145 | 0.027506621 | 8.5729492 | 9.2881606 | 8.65647 | 0.631688939 | -0.083522532 |
| St13 | 11.12351 | 11.44136 | 11.117 | 0.32441527 | 0.006556602 | 10.940212 | 11.716297 | 11.0849 | 0.631412834 | -0.144672321 |
| Pgs1 | 9.736148 | 9.75825 | 9.4841 | 0.274105719 | 0.252004037 | 9.8341588 | 10.197272 | 9.56586 | 0.631411974 | 0.268298493 |
| Lpar1 | 6.141932 | 5.826091 | 5.5273 | 0.298788288 | 0.614629444 | 6.0643907 | 6.0058389 | 5.37483 | 0.631008433 | 0.689560296 |
| Nop2 | 10.06659 | 10.15493 | 9.9381 | 0.216777803 | 0.128436912 | 10.121561 | 10.560744 | 9.93023 | 0.630510868 | 0.191328745 |
| Thap7 | 7.647152 | 7.716111 | 7.5892 | 0.126897703 | 0.057938922 | 8.0314574 | 8.1892394 | 7.5588 | 0.630434586 | 0.472652552 |
| Snrpb | 10.98802 | 11.12881 | 10.868 | 0.260820846 | 0.120036466 | 11.247478 | 11.468544 | 10.8389 | 0.629594916 | 0.408529888 |
| Syngr2 | 11.19434 | 11.28559 | 11.187 | 0.098290298 | 0.007040828 | 11.416657 | 11.692943 | 11.0635 | 0.629435246 | 0.353148813 |
| Irf2bp2 | 10.14894 | 10.22854 | 10.084 | 0.144452129 | 0.064853461 | 9.9604452 | 10.573189 | 9.94483 | 0.628357015 | 0.015613179 |
| Atox1 | 8.991641 | 9.162237 | 9.044 | 0.118244217 | -0.05235244 | 9.3698506 | 9.5474399 | 8.91958 | 0.627856781 | 0.450267482 |
| Tbc1d1 | 10.99425 | 11.33812 | 11.007 | 0.331372684 | -0.012490296 | 11.143422 | 11.675641 | 11.0478 | 0.627795898 | 0.095576881 |
| Klk1b11 | 4.880203 | 5.172832 | 5.805 | -0.632196898 | -0.924826368 | 6.8752569 | 5.8694211 | 5.24182 | 0.6276042 | 1.633440047 |
| Plet1 | 4.283602 | 3.964918 | 3.9756 | -0.010649002 | 0.308034757 | 4.4584519 | 4.4808949 | 3.85456 | 0.626333961 | 0.603890937 |
| Sh3pxd2b | 7.250059 | 7.384785 | 6.7378 | 0.647028804 | 0.51230268 | 6.7830496 | 7.1659265 | 6.54039 | 0.625539487 | 0.242662567 |
| Eif2s2 | 11.04917 | 11.06957 | 10.982 | 0.087756264 | 0.0673473 | 11.2339 | 11.710916 | 11.0862 | 0.624747622 | 0.14773091 |
| Map2k1 | 10.89887 | 11.20581 | 10.895 | 0.310426362 | 0.00349112 | 11.185132 | 11.57236 | 10.948 | 0.624346514 | 0.237118421 |
| Eif1ax | 9.913777 | 9.724693 | 9.6586 | 0.06611428 | 0.255197628 | 9.9182369 | 10.259555 | 9.63523 | 0.624324142 | 0.283006364 |
| Il2rb | 11.83485 | 11.98862 | 11.823 | 0.165915332 | 0.012149906 | 11.531698 | 12.344495 | 11.7207 | 0.623812958 | -0.188984323 |
| Cd38 | 10.93318 | 10.80681 | 10.666 | 0.141306148 | 0.267677322 | 11.28287 | 11.282088 | 10.6585 | 0.623557126 | 0.62433829 |
| Gm46223 | 12.3724 | 12.2729 | 12.44 | -0.16696525 | -0.067472646 | 12.402838 | 12.943618 | 12.3209 | 0.622723522 | 0.081944073 |
| Nub1 | 10.88076 | 11.05989 | 10.982 | 0.077874842 | -0.10125702 | 11.037542 | 11.602392 | 10.9807 | 0.621664304 | 0.056814102 |
| Sppl2a | 11.23632 | 11.43179 | 11.179 | 0.252559966 | 0.057096498 | 11.268361 | 11.771153 | 11.1496 | 0.621556354 | 0.118763937 |
| Adrm1 | 9.882406 | 9.983679 | 9.7763 | 0.20739623 | 0.106123792 | 10.067061 | 10.345068 | 9.72362 | 0.621445171 | 0.343437884 |

| | | | | | | | | | | |
|---|---|---|---|---|---|---|---|---|---|---|
| Rpl28 | 12.47171 | 12.64232 | 12.506 | 0.136199106 | -0.034412272 | 12.920148 | 13.046911 | 12.4259 | 0.621028446 | 0.494265373 |
| St3gal6 | 9.77675 | 9.86652 | 9.7754 | 0.091096138 | 0.001326904 | 9.7562117 | 10.254131 | 9.63353 | 0.620602859 | 0.122683217 |
| Bop1 | 10.06173 | 10.10433 | 9.9368 | 0.167556683 | 0.124947581 | 10.3729 | 10.57709 | 9.95758 | 0.619514778 | 0.415325179 |
| Blvrb | 8.819193 | 8.72275 | 8.7183 | 0.00443566 | 0.100878907 | 9.1386124 | 9.3462068 | 8.72694 | 0.6192672 | 0.411672828 |
| Slit2 | 3.860226 | 3.959773 | 3.835 | 0.124745766 | 0.025198673 | 4.1668258 | 4.4175271 | 3.79893 | 0.618596255 | 0.367895003 |
| Tnfrsf26 | 8.335028 | 8.754716 | 8.5417 | 0.213055625 | -0.20663284 | 8.6526389 | 9.2090082 | 8.59074 | 0.61826851 | 0.061899183 |
| Dpm3 | 7.169399 | 7.31142 | 7.0822 | 0.229239358 | 0.087218475 | 7.6965366 | 7.5655665 | 6.94765 | 0.61791439 | 0.748884453 |
| Zfp90 | 7.764516 | 7.947177 | 7.7475 | 0.199684769 | 0.017024216 | 7.8599912 | 8.36575 | 7.74847 | 0.617281792 | 0.111523065 |
| Prrc2a | 12.81107 | 12.87146 | 12.769 | 0.102261118 | 0.041867076 | 13.107918 | 13.491462 | 12.8743 | 0.617123604 | 0.233579501 |
| Ctsb | 12.42332 | 12.77486 | 12.174 | 0.60092738 | 0.24938979 | 12.059881 | 12.463947 | 11.847 | 0.616904046 | 0.212837132 |
| Ruvbl2 | 9.46426 | 9.571055 | 9.3925 | 0.17856855 | 0.07177373 | 9.7796428 | 10.028713 | 9.41213 | 0.616581613 | 0.36751175 |
| Rnf217 | 4.54299 | 4.401004 | 4.4581 | -0.05713412 | 0.084851774 | 4.2181347 | 4.8075655 | 4.19136 | 0.616205334 | 0.026774539 |
| Sphk2 | 9.143563 | 9.354337 | 9.1812 | 0.173152978 | -0.0376213 | 9.4092099 | 9.6342319 | 9.01892 | 0.615311752 | 0.390289739 |
| Nes | 5.48478 | 5.901949 | 5.0191 | 0.882809517 | 0.465640511 | 5.2404959 | 5.7199061 | 5.10486 | 0.615041474 | 0.135631312 |
| Capza2 | 11.75863 | 11.87927 | 11.684 | 0.195745942 | 0.075109824 | 11.704107 | 12.216129 | 11.6022 | 0.613917336 | 0.101895114 |
| Pirb | 10.2624 | 10.15941 | 10.027 | 0.132273478 | 0.235270622 | 10.871963 | 10.858949 | 10.2455 | 0.61344918 | 0.626463288 |
| Acot7 | 8.119127 | 8.209014 | 8.1436 | 0.065432582 | -0.024455165 | 8.3961277 | 8.8628628 | 8.25003 | 0.612835192 | 0.146100083 |
| Egln2 | 9.706293 | 9.726827 | 9.6048 | 0.122014098 | 0.101479877 | 10.007717 | 10.206764 | 9.59395 | 0.612814555 | 0.413767948 |
| Tpd52 | 10.55835 | 10.67815 | 10.336 | 0.34253533 | 0.222731922 | 10.946909 | 10.876566 | 10.2638 | 0.612721682 | 0.68306451 |
| Ndufc1 | 7.346438 | 7.373253 | 7.1478 | 0.22542818 | 0.198612568 | 7.744573 | 7.7647831 | 7.15235 | 0.612436955 | 0.59222682 |
| Klra7 | 6.706104 | 6.739509 | 6.8231 | -0.083582099 | -0.116986761 | 6.6369673 | 7.3491756 | 6.73675 | 0.612429626 | -0.099778644 |
| Rrp9 | 9.073492 | 9.322564 | 9.0643 | 0.258297889 | 0.009225796 | 9.2737609 | 9.7213225 | 9.1097 | 0.611619557 | 0.164057953 |
| Rnf123 | 10.82231 | 11.04762 | 10.966 | 0.081274894 | -0.144037346 | 11.312136 | 11.754045 | 11.1428 | 0.611287504 | 0.169378154 |
| Edem1 | 11.41942 | 11.60377 | 11.321 | 0.283087566 | 0.09873976 | 11.359777 | 12.045584 | 11.4345 | 0.611131936 | -0.074674184 |
| Rbm42 | 9.957277 | 10.02263 | 9.8461 | 0.17651899 | 0.11116463 | 10.228023 | 10.380805 | 9.76968 | 0.611124301 | 0.458342218 |
| Nup210 | 12.77893 | 12.85788 | 12.73 | 0.12824765 | 0.04929435 | 12.827764 | 13.504075 | 12.8931 | 0.611022906 | -0.065288297 |
| Zfp53 | 8.767834 | 8.980056 | 8.7911 | 0.188941945 | -0.023279623 | 8.5439749 | 9.2371568 | 8.62618 | 0.610972875 | -0.082209032 |
| Plxnb2 | 9.862249 | 9.648365 | 9.6749 | -0.026559307 | 0.187324852 | 9.7345895 | 9.9065648 | 9.296 | 0.610566645 | 0.438591377 |
| Tlnrd1 | 8.219777 | 8.129613 | 8.0627 | 0.066936395 | 0.157100184 | 8.1178826 | 8.4246557 | 7.81416 | 0.610494197 | 0.303721027 |
| Plat | 6.148904 | 6.384274 | 5.7384 | 0.645898308 | 0.410527638 | 5.7698535 | 6.5162448 | 5.90581 | 0.610430054 | -0.135961237 |
| Snrpc | 8.976328 | 9.025149 | 8.9908 | 0.034336445 | -0.014484294 | 9.2754643 | 9.4704254 | 8.86037 | 0.610057466 | 0.415096388 |
| Cope | 10.12417 | 10.24418 | 10.07 | 0.17462181 | 0.054612631 | 10.405936 | 10.618069 | 10.0092 | 0.608874966 | 0.396742092 |

| | | | | | | | | | | |
|---|---|---|---|---|---|---|---|---|---|---|
| Snrpd1 | 9.106913 | 9.064388 | 9.0166 | 0.047793075 | 0.090317711 | 9.1901932 | 9.6623267 | 9.05394 | 0.608381874 | 0.136248372 |
| Trib1 | 9.176808 | 8.975423 | 8.566 | 0.409378472 | 0.610763221 | 8.3273108 | 8.7818699 | 8.17386 | 0.608010238 | 0.153451111 |
| Gm21887 | 5.370106 | 5.54826 | 5.2763 | 0.271911309 | 0.093758083 | 5.5771997 | 6.1268933 | 5.51976 | 0.607131019 | 0.05743734 |
| Ndufab1 | 8.720432 | 8.77453 | 8.7056 | 0.068948281 | 0.01485091 | 8.9329463 | 9.3707827 | 8.7637 | 0.607077996 | 0.169241574 |
| Ssbp2 | 8.178873 | 8.221794 | 8.3221 | -0.100305783 | -0.143227365 | 7.7889215 | 8.6825384 | 8.0756 | 0.606935248 | -0.286681625 |
| Coro2a | 10.47401 | 10.69717 | 10.491 | 0.20605312 | -0.017111236 | 10.59854 | 11.071787 | 10.4651 | 0.606698352 | 0.13345128 |
| Ift22 | 6.8905 | 7.036847 | 7.0576 | -0.020740554 | -0.167088249 | 7.5158398 | 7.6503265 | 7.04383 | 0.606499241 | 0.472012594 |
| Mrpl14 | 7.221059 | 7.530073 | 7.3211 | 0.208999771 | -0.100014384 | 7.730489 | 7.8593366 | 7.25304 | 0.606293992 | 0.477446362 |
| Stard3 | 9.094869 | 9.280319 | 8.9598 | 0.320534471 | 0.135084676 | 9.3389447 | 9.7641586 | 9.15797 | 0.606184886 | 0.18097093 |
| Osbpl5 | 9.188677 | 9.576224 | 9.2109 | 0.365319004 | -0.022227254 | 9.4614762 | 9.795576 | 9.18952 | 0.606055385 | 0.271955534 |
| Rbms1 | 10.70897 | 10.82458 | 10.656 | 0.168323024 | 0.052712882 | 10.702118 | 11.237479 | 10.6314 | 0.606054168 | 0.070692886 |
| Igkv1-135 | 9.25215 | 9.60585 | 9.586 | 0.019822114 | -0.333878215 | 9.6742349 | 9.9489097 | 9.34316 | 0.605748033 | 0.331073248 |
| Tmem171 | 2.238776 | 2.406118 | 2.1877 | 0.218463542 | 0.051121981 | 2.2989835 | 2.8219456 | 2.21622 | 0.605728135 | 0.082766054 |
| Adap1 | 8.486254 | 8.647683 | 8.5903 | 0.057381781 | -0.104047297 | 9.1208534 | 9.3489218 | 8.74319 | 0.605727379 | 0.377658961 |
| Whamm | 8.837158 | 8.850797 | 8.7887 | 0.06214595 | 0.048507304 | 8.8303998 | 9.4232901 | 8.8182 | 0.60509225 | 0.012201949 |
| Rex1bd | 7.378525 | 7.309786 | 7.1909 | 0.118930364 | 0.18766913 | 7.5978595 | 7.6721654 | 7.06732 | 0.604845508 | 0.530539644 |
| Mrpl53 | 6.744091 | 6.779947 | 6.6947 | 0.085203316 | 0.049347387 | 6.8638392 | 7.0796678 | 6.47503 | 0.604636961 | 0.38880828 |
| Dedd2 | 10.06591 | 10.44412 | 10.116 | 0.3279805 | -0.050224235 | 10.168281 | 10.813158 | 10.2087 | 0.604467702 | -0.040409595 |
| 9530053A07Rik | 3.508268 | 3.587755 | 3.4987 | 0.089052701 | 0.009565346 | 3.6205755 | 4.1673299 | 3.56319 | 0.604141962 | 0.05738754 |
| Snx20 | 9.55696 | 9.853351 | 9.6107 | 0.24263155 | -0.053759963 | 9.9402381 | 10.227402 | 9.6237 | 0.603703642 | 0.316539296 |
| Itm2b | 12.9288 | 12.89913 | 12.929 | -0.030031488 | -0.00036036 | 12.822334 | 13.367701 | 12.764 | 0.603652948 | 0.058286337 |
| Apol11b | 4.178099 | 4.103165 | 4.0025 | 0.100646659 | 0.175580388 | 3.8584635 | 4.6459101 | 4.04232 | 0.603594999 | -0.1838516 |
| Il1r2 | 8.302287 | 8.128203 | 6.8827 | 1.245467088 | 1.419551227 | 8.3564839 | 7.6744285 | 7.07157 | 0.602860483 | 1.284915881 |
| Inka1 | 5.790183 | 6.14558 | 5.956 | 0.189600244 | -0.16579626 | 6.2366279 | 6.4910092 | 5.8884 | 0.602609342 | 0.348228069 |
| Cnot6l | 11.83077 | 12.33747 | 11.839 | 0.498905976 | -0.007791026 | 11.827838 | 12.462642 | 11.8603 | 0.602318222 | -0.03248544 |
| Rrbp1 | 11.2778 | 11.15045 | 11.154 | -0.003321834 | 0.12403186 | 11.233876 | 11.763107 | 11.1619 | 0.601193356 | 0.071961925 |
| Pdia4 | 11.35975 | 11.73262 | 11.302 | 0.430900896 | 0.05802855 | 11.552728 | 11.826457 | 11.2263 | 0.60014762 | 0.326418914 |
| Washc4 | 10.45333 | 10.878 | 10.464 | 0.414103962 | -0.010562944 | 10.389501 | 11.118675 | 10.5189 | 0.59978178 | -0.129391874 |
| Lgals8 | 10.20541 | 10.3968 | 10.023 | 0.373473891 | 0.182086212 | 10.290926 | 10.894491 | 10.2947 | 0.599771344 | -0.003793163 |
| Ptges3-ps | 8.49048 | 8.747998 | 8.2262 | 0.521799973 | 0.264282607 | 8.1659678 | 9.0698538 | 8.47012 | 0.599730614 | -0.304155433 |
| Chpf | 7.783556 | 7.776564 | 7.7411 | 0.035414817 | 0.042407236 | 7.7184029 | 7.9746466 | 7.37545 | 0.599195513 | 0.342951801 |
| Sdcbp | 11.89431 | 12.01236 | 11.732 | 0.28083444 | 0.162791712 | 11.816338 | 12.246968 | 11.6479 | 0.599059048 | 0.168429105 |

| | | | | | | | | | | |
|---|---|---|---|---|---|---|---|---|---|---|
| F7 | 1.735527 | 1.73597 | 1.6962 | 0.039743788 | 0.039300156 | 1.7432713 | 2.2673302 | 1.66838 | 0.598948793 | 0.074889924 |
| Igkv1-88 | 6.292518 | 6.41531 | 6.9002 | -0.484925153 | -0.607717197 | 6.4206751 | 6.5468826 | 5.94823 | 0.59864868 | 0.472441158 |
| Gpatch4 | 9.09657 | 9.176544 | 8.9136 | 0.262932717 | 0.182958593 | 9.1751145 | 9.6736132 | 9.07541 | 0.598201541 | 0.099702751 |
| Olfr46 | 5.194549 | 5.304253 | 5.2419 | 0.062315891 | -0.047388764 | 5.3320153 | 5.982341 | 5.38451 | 0.597826716 | -0.052498943 |
| Banp | 8.682082 | 8.809908 | 8.6405 | 0.169413522 | 0.041587984 | 8.700016 | 9.1181616 | 8.52034 | 0.597817505 | 0.179671939 |
| Tspan31 | 9.055915 | 9.141688 | 8.985 | 0.156709431 | 0.07093557 | 9.0123781 | 9.4765438 | 8.87883 | 0.597713826 | 0.133548093 |
| Tmco4 | 9.635508 | 9.550653 | 9.6235 | -0.072822456 | 0.012032498 | 9.6674415 | 10.093787 | 9.49664 | 0.597147077 | 0.170801433 |
| Llgl1 | 9.760968 | 9.87949 | 9.6604 | 0.219075955 | 0.10055428 | 9.8975074 | 10.186509 | 9.58946 | 0.597046594 | 0.308045334 |
| Nectin2 | 7.777463 | 7.622523 | 7.5153 | 0.10718847 | 0.26212826 | 7.0067406 | 7.3742524 | 6.77728 | 0.596973977 | 0.22946214 |
| Rplp2 | 13.61325 | 13.72183 | 13.544 | 0.178140718 | 0.069555424 | 13.804377 | 14.030462 | 13.4335 | 0.596953144 | 0.370868809 |
| Arhgap30 | 12.86722 | 13.03957 | 12.802 | 0.237171078 | 0.064818342 | 12.995646 | 13.473152 | 12.8767 | 0.596501024 | 0.118995524 |
| Trim26 | 10.92422 | 11.19592 | 10.879 | 0.316731504 | 0.045035198 | 11.119408 | 11.630337 | 11.0339 | 0.59644656 | 0.085517454 |
| C1s1 | 9.720532 | 9.157892 | 9.5596 | -0.401738802 | 0.160901491 | 8.5792161 | 9.3439652 | 8.74824 | 0.595721039 | -0.169028051 |
| Klk1b8 | 3.73684 | 4.554613 | 5.3907 | -0.836065732 | -1.653838647 | 6.332654 | 5.3863753 | 4.79073 | 0.59564185 | 1.54192054 |
| Olfr735 | 4.786317 | 5.097546 | 4.8658 | 0.231729858 | -0.079498474 | 5.1362989 | 5.8445504 | 5.2492 | 0.595347734 | -0.112903772 |
| Nufip1 | 8.587291 | 8.633231 | 8.5278 | 0.105405065 | 0.059464529 | 8.6472724 | 9.1876245 | 8.59271 | 0.594917591 | 0.054565477 |
| Sema7a | 9.643442 | 9.42003 | 9.4951 | -0.075110584 | 0.148301503 | 9.3859009 | 9.8296325 | 9.23487 | 0.59476064 | 0.151029043 |
| Plekhg4 | 2.280419 | 2.219618 | 2.2677 | -0.048039064 | 0.012762168 | 2.3164255 | 2.8608538 | 2.2662 | 0.594650196 | 0.050221944 |
| Ehd4 | 10.29503 | 10.48665 | 10.185 | 0.301212088 | 0.10959252 | 10.202877 | 10.756648 | 10.1626 | 0.594081686 | 0.040310463 |
| Psmd11 | 10.3054 | 10.36861 | 10.139 | 0.229617723 | 0.166400933 | 10.266217 | 10.741965 | 10.1481 | 0.5939045 | 0.118156486 |
| Snrpd2 | 9.661654 | 9.762176 | 9.6629 | 0.099239211 | -0.001282487 | 9.949926 | 10.210429 | 9.6169 | 0.593529539 | 0.333026157 |
| Tmtc4 | 6.977985 | 7.13989 | 6.9197 | 0.220144006 | 0.058239479 | 6.6548552 | 7.716421 | 7.12326 | 0.593160355 | -0.468405415 |
| Tgfb1i1 | 6.467639 | 6.719673 | 6.2555 | 0.464194811 | 0.212160897 | 6.5848603 | 6.8754675 | 6.28269 | 0.592780764 | 0.302173563 |
| Chchd1 | 8.079141 | 8.099773 | 8.0545 | 0.045275706 | 0.024643934 | 8.3758493 | 8.596791 | 8.00458 | 0.592214577 | 0.371272912 |
| Sec61b | 9.540376 | 9.503812 | 9.4101 | 0.093699266 | 0.130262379 | 9.6568588 | 9.8707709 | 9.27891 | 0.591864287 | 0.377952177 |
| Ecscr | 7.874639 | 7.691323 | 7.6646 | 0.026680503 | 0.209996562 | 7.0568432 | 7.5939904 | 7.00214 | 0.591851468 | 0.054704323 |
| Sub1 | 11.17034 | 11.20764 | 11.114 | 0.09359838 | 0.056295176 | 11.198424 | 11.64867 | 11.0571 | 0.591579734 | 0.141333536 |
| Mars | 10.06629 | 10.15958 | 9.974 | 0.185631732 | 0.092340135 | 10.663561 | 10.916777 | 10.3252 | 0.591560528 | 0.338344724 |
| Rfc5 | 8.260818 | 8.336785 | 8.4078 | -0.07103708 | -0.14700429 | 8.6313767 | 9.1709038 | 8.57992 | 0.590985079 | 0.051457996 |
| Naip5 | 7.774558 | 7.890828 | 7.7527 | 0.138138125 | 0.021867946 | 7.9064957 | 8.4863289 | 7.89542 | 0.590912058 | 0.01107886 |
| Cox6a1 | 9.786193 | 9.921441 | 9.7863 | 0.135181103 | -6.60704E-05 | 10.169309 | 10.315483 | 9.72541 | 0.590077703 | 0.443903431 |
| Mrpl12 | 9.05962 | 9.070319 | 9.0276 | 0.042742589 | 0.03204369 | 9.1308601 | 9.5034747 | 8.91342 | 0.59005235 | 0.217437731 |

| | | | | | | | | | | |
|---|---|---|---|---|---|---|---|---|---|---|
| Il2rg | 13.19377 | 13.30849 | 13.118 | 0.19019968 | 0.075472944 | 13.330061 | 13.842179 | 13.2528 | 0.58933872 | 0.077220131 |
| Cflar | 11.28775 | 11.46379 | 11.28 | 0.183451914 | 0.007411242 | 11.357968 | 11.933238 | 11.3441 | 0.589164912 | 0.013894435 |
| Slc39a1 | 9.844469 | 9.729517 | 9.6063 | 0.123247469 | 0.238199537 | 9.7261919 | 9.9365812 | 9.3477 | 0.588881837 | 0.378492486 |
| Gbgt1 | 2.934503 | 2.966489 | 3.1047 | -0.138195957 | -0.170181863 | 3.1163476 | 3.6004064 | 3.01166 | 0.588746377 | 0.104687568 |
| Tmem219 | 8.634544 | 8.674976 | 8.6366 | 0.038343771 | -0.002088518 | 8.8232544 | 8.9547632 | 8.36616 | 0.588604168 | 0.457095442 |
| Evl | 11.9945 | 12.2464 | 12.11 | 0.13609392 | -0.115807342 | 12.29281 | 12.755566 | 12.1673 | 0.588222866 | 0.125467208 |
| Sult2b1 | 3.580262 | 3.780127 | 3.5825 | 0.197649423 | -0.00221513 | 3.5996198 | 4.0571532 | 3.46987 | 0.587288019 | 0.129754665 |
| Timm22 | 8.062577 | 7.916532 | 7.8 | 0.116516179 | 0.262560573 | 8.0516959 | 8.396153 | 7.8094 | 0.586757626 | 0.242300503 |
| Rnf122 | 8.897625 | 8.95077 | 8.832 | 0.118726632 | 0.065582202 | 8.80366 | 9.2658172 | 8.67916 | 0.586661753 | 0.124504628 |
| Mrpl52 | 8.607767 | 8.538551 | 8.4474 | 0.091170434 | 0.160386734 | 8.8434251 | 8.9329977 | 8.34634 | 0.586659735 | 0.497087172 |
| Entpd1 | 9.447051 | 9.147453 | 9.0598 | 0.087673745 | 0.38727125 | 8.9531177 | 9.4727915 | 8.88615 | 0.586641955 | 0.066968089 |
| Napsa | 11.68563 | 11.86691 | 11.613 | 0.253565504 | 0.072281436 | 12.056266 | 12.139316 | 11.5532 | 0.586162204 | 0.50311275 |
| Fxr1 | 10.6671 | 10.83436 | 10.659 | 0.175686334 | 0.00842962 | 10.538833 | 11.261639 | 10.6763 | 0.585312616 | -0.137493432 |
| Gm2619 | 3.388889 | 3.618176 | 3.3633 | 0.254875682 | 0.025588522 | 3.518112 | 4.1010493 | 3.51622 | 0.584832041 | 0.001894726 |
| Gng5 | 9.623997 | 9.713724 | 9.592 | 0.12171574 | 0.03198901 | 9.7640364 | 10.075814 | 9.49145 | 0.584359723 | 0.272582253 |
| Pafah1b3 | 8.106859 | 8.039993 | 8.1684 | -0.128363229 | -0.061496857 | 8.2891856 | 8.5903642 | 8.00652 | 0.583843349 | 0.282664791 |
| Sirpb1a | 6.206356 | 6.379057 | 6.0685 | 0.310534629 | 0.137833561 | 7.22546 | 6.9031452 | 6.31931 | 0.583832842 | 0.906147612 |
| Klra4 | 6.233919 | 6.459288 | 6.1877 | 0.27155636 | 0.046187627 | 6.0083547 | 6.7625876 | 6.17903 | 0.583554787 | -0.170678137 |
| Ndufs7 | 8.723615 | 8.83064 | 8.7841 | 0.046554664 | -0.060470768 | 9.1894887 | 9.2901704 | 8.70701 | 0.583160801 | 0.482479101 |
| Rps4l | 5.637727 | 5.576226 | 5.5476 | 0.028657336 | 0.090157739 | 5.9305806 | 5.9478373 | 5.36485 | 0.582988852 | 0.565732145 |
| Pim1 | 11.29908 | 11.3223 | 10.823 | 0.499371402 | 0.476147002 | 10.973774 | 11.448172 | 10.8659 | 0.58226469 | 0.107866361 |
| Rps20 | 13.63717 | 13.70625 | 13.833 | -0.127152048 | -0.196227 | 13.786272 | 14.235089 | 13.6531 | 0.582034322 | 0.133217074 |
| Khnyn | 10.94245 | 11.17505 | 11.043 | 0.131796442 | -0.10079948 | 11.190665 | 11.725865 | 11.1441 | 0.581762136 | 0.046562117 |
| Gpr15 | 5.334073 | 5.430833 | 5.2362 | 0.19465989 | 0.097899974 | 5.1592345 | 5.8463088 | 5.26484 | 0.581465224 | -0.10560911 |
| Arrdc3 | 9.265213 | 9.13843 | 9.4071 | -0.268623333 | -0.141839643 | 8.8797333 | 9.5472924 | 8.96612 | 0.581173408 | -0.086385673 |
| Tnfrsf9 | 8.651847 | 8.893964 | 8.5197 | 0.374243286 | 0.132126672 | 8.5237258 | 9.1818245 | 8.60134 | 0.580483665 | -0.077615067 |
| Dhfr | 7.181153 | 7.187621 | 7.3297 | -0.142105563 | -0.148573848 | 7.2848791 | 8.0977043 | 7.51731 | 0.580393969 | -0.232431229 |
| Ndufs6 | 8.422643 | 8.445097 | 8.2992 | 0.145896697 | 0.123443172 | 8.6981081 | 8.8790382 | 8.29907 | 0.579966547 | 0.399036512 |
| Gngt2 | 9.161456 | 9.234867 | 9.0245 | 0.21036161 | 0.136950142 | 9.8481914 | 9.6830293 | 9.10361 | 0.579420378 | 0.744582452 |
| Sbno2 | 11.56288 | 11.29962 | 11.025 | 0.274810236 | 0.538072134 | 11.574433 | 11.744308 | 11.165 | 0.579283378 | 0.409409271 |
| Tinagl1 | 7.791653 | 7.617607 | 7.4897 | 0.127890047 | 0.301935865 | 7.1579999 | 7.7134696 | 7.13431 | 0.579156911 | 0.02368718 |
| Sdhb | 9.739518 | 9.700477 | 9.6638 | 0.036630737 | 0.075671819 | 9.8893124 | 10.153957 | 9.57485 | 0.579107528 | 0.314463062 |

| | | | | | | | | | | |
|---|---|---|---|---|---|---|---|---|---|---|
| Sem1 | 9.933385 | 9.909352 | 9.8117 | 0.097656075 | 0.121688153 | 10.018626 | 10.310488 | 9.73179 | 0.578698625 | 0.286836978 |
| Arrdc4 | 7.715379 | 7.789419 | 7.6875 | 0.101909202 | 0.027869191 | 7.0597936 | 7.9401512 | 7.36157 | 0.578583026 | -0.301774574 |
| Phyh | 9.425191 | 9.545162 | 9.3578 | 0.187407968 | 0.067437583 | 9.6337282 | 9.999219 | 9.42066 | 0.578560397 | 0.213069646 |
| Rhcg | 1.2138 | 1.24157 | 1.2124 | 0.029143355 | 0.001372699 | 1.2204256 | 1.8397329 | 1.26186 | 0.577877719 | -0.041429621 |
| Trir | 9.479057 | 9.614246 | 9.5496 | 0.064621687 | -0.070567705 | 9.7264948 | 10.050425 | 9.47264 | 0.577782427 | 0.253851795 |
| Isoc1 | 9.900661 | 10.0544 | 9.9026 | 0.151802022 | -0.001941164 | 9.9239745 | 10.446017 | 9.86946 | 0.576556568 | 0.054514259 |
| Gm6472 | 10.73562 | 11.22955 | 11.756 | -0.526828253 | -1.020763212 | 11.218234 | 11.589865 | 11.0134 | 0.576508157 | 0.204877711 |
| Igkv4-90 | 3.99222 | 4.601012 | 4.0316 | 0.569394079 | -0.039398344 | 4.2452646 | 4.5127145 | 3.93622 | 0.576499074 | 0.309049126 |
| Farsa | 9.80965 | 9.95313 | 9.8199 | 0.133182639 | -0.010296947 | 10.141654 | 10.443139 | 9.86672 | 0.576419424 | 0.274934232 |
| Azi2 | 10.50508 | 10.68013 | 10.458 | 0.221955758 | 0.046909328 | 10.487934 | 11.139378 | 10.563 | 0.576417558 | -0.075025628 |
| Srsf7 | 11.47308 | 11.68868 | 11.607 | 0.081737084 | -0.13385934 | 11.844591 | 12.331187 | 11.7553 | 0.575910898 | 0.089314359 |
| Ndufa13 | 9.647955 | 9.690987 | 9.6217 | 0.069271971 | 0.02624066 | 9.9166617 | 10.085144 | 9.50944 | 0.575704896 | 0.407222589 |
| Hat1 | 9.211737 | 9.363937 | 9.3065 | 0.057435978 | -0.094764465 | 9.3512399 | 9.9752855 | 9.3997 | 0.575583048 | -0.048462578 |
| Ap5z1 | 8.867099 | 9.118238 | 8.9245 | 0.193733617 | -0.057406212 | 9.3891597 | 9.6624488 | 9.08708 | 0.575364437 | 0.30207538 |
| Fbxw17 | 6.939891 | 7.041329 | 7.1341 | -0.092728787 | -0.194166519 | 6.964859 | 7.8334244 | 7.25823 | 0.575199139 | -0.293366227 |
| Cd63 | 9.531531 | 9.939119 | 9.4252 | 0.513881459 | 0.106293445 | 9.2832946 | 9.5711724 | 8.99617 | 0.575005965 | 0.287128173 |
| Igkv5-43 | 8.451944 | 8.838591 | 8.4361 | 0.402529774 | 0.015882712 | 8.6643173 | 9.0022927 | 8.42755 | 0.574747649 | 0.236772214 |
| Gm42715 | 9.415504 | 9.696247 | 9.5521 | 0.144172854 | -0.136569833 | 9.8295411 | 10.266374 | 9.69181 | 0.574561366 | 0.137728127 |
| Vill | 6.437204 | 6.303234 | 6.2554 | 0.04781271 | 0.181782659 | 6.9751965 | 6.8925061 | 6.31799 | 0.574515754 | 0.657206209 |
| Ppp1r35 | 6.854356 | 7.040834 | 6.9143 | 0.126579174 | -0.059899362 | 7.2422734 | 7.4054669 | 6.83111 | 0.574352776 | 0.411159295 |
| Nsd3 | 11.9485 | 12.18556 | 11.904 | 0.28126276 | 0.044198172 | 11.834183 | 12.557712 | 11.9835 | 0.5742275 | -0.149300775 |
| Nip7 | 9.157538 | 9.135714 | 9.0235 | 0.112223502 | 0.13404847 | 9.202998 | 9.5323517 | 8.95822 | 0.574126986 | 0.244773291 |
| Dph3 | 8.477552 | 8.427522 | 8.3838 | 0.0437026 | 0.093732757 | 8.7350827 | 8.822841 | 8.24893 | 0.573915843 | 0.486157526 |
| Smim3 | 6.790533 | 6.877934 | 6.5381 | 0.339801257 | 0.252400364 | 7.052783 | 7.007346 | 6.43378 | 0.573562252 | 0.618999239 |
| Trim56 | 11.78362 | 12.10379 | 11.699 | 0.404351202 | 0.084172548 | 11.78447 | 12.346349 | 11.7733 | 0.573039488 | 0.011160439 |
| Ighg2b | 9.932935 | 11.20188 | 9.8894 | 1.31249227 | 0.043545383 | 10.344809 | 11.314755 | 10.7418 | 0.572928738 | -0.397016584 |
| Slc39a2 | 3.830881 | 3.944523 | 3.917 | 0.027474732 | -0.086167078 | 4.1751954 | 4.6616627 | 4.08884 | 0.572824223 | 0.086356963 |
| Marcksl1 | 11.05779 | 11.30484 | 10.898 | 0.407338174 | 0.160294454 | 10.842123 | 11.244533 | 10.672 | 0.572553434 | 0.170143963 |
| Rps12 | 10.11744 | 10.07905 | 10.02 | 0.059008956 | 0.097396723 | 10.593331 | 10.62521 | 10.0532 | 0.572031539 | 0.540152525 |
| Rps12-ps3 | 14.03533 | 14.19906 | 14.028 | 0.17087794 | 0.00714643 | 14.35094 | 14.539954 | 13.9684 | 0.57156322 | 0.382549686 |
| Ighv3-1 | 5.765442 | 6.075402 | 6.162 | -0.086575639 | -0.396536229 | 5.7465401 | 6.3651362 | 5.79411 | 0.571026605 | -0.047569513 |
| Gm10177 | 9.419863 | 9.478046 | 9.3886 | 0.08943022 | 0.031246753 | 9.6111684 | 9.9401702 | 9.36989 | 0.570275618 | 0.241273853 |

| | | | | | | | | | | |
|---|---|---|---|---|---|---|---|---|---|---|
| Vmp1 | 9.539425 | 9.605695 | 9.4289 | 0.176757908 | 0.110487758 | 9.6029489 | 9.9504259 | 9.38056 | 0.569862297 | 0.222385327 |
| Gapdh | 13.31998 | 13.36654 | 13.402 | -0.035623898 | -0.082188954 | 13.658635 | 13.984353 | 13.4149 | 0.569441402 | 0.243723639 |
| Pou6f1 | 9.692453 | 9.618844 | 9.706 | -0.087167381 | -0.013558547 | 9.9080771 | 10.387743 | 9.81885 | 0.568894877 | 0.089229138 |
| Spo11 | 4.131882 | 4.123211 | 3.8334 | 0.289771764 | 0.2984422 | 4.3250031 | 4.7712012 | 4.20287 | 0.568333491 | 0.122135339 |
| Ctu2 | 8.463429 | 8.824461 | 8.5599 | 0.264549953 | -0.096482023 | 8.9286551 | 9.27043 | 8.70223 | 0.568203012 | 0.226428145 |
| Map2k2 | 10.13958 | 10.20675 | 10.173 | 0.03356033 | -0.033609967 | 10.435868 | 10.615922 | 10.0478 | 0.568112577 | 0.388058537 |
| CAAA01118383.1 | 7.448431 | 7.52157 | 7.4089 | 0.1126961 | 0.039557053 | 7.9494565 | 8.1030321 | 7.53493 | 0.568100353 | 0.414524696 |
| Vmn1r88 | 6.345022 | 6.610463 | 6.5377 | 0.072812572 | -0.192628025 | 6.626959 | 7.2054364 | 6.63816 | 0.567276945 | -0.011200438 |
| Hmga1 | 10.51354 | 10.60424 | 10.5 | 0.104356235 | 0.01365502 | 10.75455 | 11.116067 | 10.5491 | 0.566969262 | 0.205452109 |
| Fgr | 10.17699 | 10.15412 | 9.5988 | 0.555325936 | 0.578191524 | 10.776212 | 10.319926 | 9.7531 | 0.566826564 | 1.023112707 |
| Sf3b5 | 8.495529 | 8.462064 | 8.3849 | 0.077178346 | 0.110643018 | 8.8259212 | 8.943071 | 8.37682 | 0.566254239 | 0.449104365 |
| Armc5 | 8.405701 | 8.63378 | 8.3732 | 0.260598465 | 0.032519413 | 8.6909678 | 8.8319923 | 8.26692 | 0.56506795 | 0.424043412 |
| AC168977.1 | 5.141304 | 5.239758 | 5.007 | 0.232713816 | 0.134259372 | 5.4097982 | 5.9940665 | 5.42948 | 0.564584276 | -0.019684033 |
| Rab19 | 8.016267 | 8.179378 | 8.1346 | 0.044784349 | -0.118326247 | 8.2953775 | 8.6590391 | 8.09487 | 0.564168853 | 0.200507266 |
| Blnk | 11.00342 | 11.31827 | 11.085 | 0.233168326 | -0.081679506 | 11.564287 | 11.690952 | 11.127 | 0.563991576 | 0.437326364 |
| Gm48269 | 5.810161 | 5.963346 | 5.9991 | -0.035741671 | -0.188926143 | 5.8887859 | 6.7790981 | 6.21512 | 0.563981084 | -0.326331058 |
| Acsl5 | 11.08039 | 11.01937 | 11.007 | 0.012053292 | 0.07307296 | 11.102665 | 11.602466 | 11.0386 | 0.563832986 | 0.064032936 |
| Dctn3 | 8.606557 | 8.544837 | 8.3532 | 0.19163659 | 0.253357053 | 8.6255107 | 8.8745347 | 8.3112 | 0.563331608 | 0.314307566 |
| Timm10b | 7.832127 | 7.926189 | 7.8114 | 0.114812188 | 0.020749561 | 8.2279127 | 8.2585977 | 7.69549 | 0.563104194 | 0.532419206 |
| AC132444.1 | 3.992035 | 4.292419 | 4.0848 | 0.207591825 | -0.092791446 | 4.2521419 | 4.8792672 | 4.31656 | 0.562704484 | -0.064420831 |
| Sco2 | 7.124788 | 7.202966 | 7.0134 | 0.189572495 | 0.111394782 | 7.2779315 | 7.5090012 | 6.94701 | 0.561990083 | 0.330920363 |
| Zyx | 11.34057 | 11.32521 | 11.199 | 0.12581144 | 0.141169648 | 11.336226 | 11.606017 | 11.0442 | 0.561853418 | 0.292062156 |
| Tpi1 | 10.26617 | 10.07406 | 10.216 | -0.142417695 | 0.049696153 | 10.418661 | 10.794075 | 10.2327 | 0.56138965 | 0.185976113 |
| Gm38431 | 5.322549 | 5.654285 | 5.3451 | 0.309207686 | -0.022527698 | 5.9268719 | 6.0242597 | 5.46305 | 0.561213724 | 0.463825885 |
| Ptpn22 | 10.82579 | 10.95627 | 10.918 | 0.038650144 | -0.091835508 | 10.547014 | 11.351985 | 10.7908 | 0.56121087 | -0.243760649 |
| Snx2 | 12.34107 | 12.56106 | 12.314 | 0.246705578 | 0.026714956 | 12.662126 | 12.966299 | 12.4053 | 0.561007908 | 0.256834412 |
| CAAA01147332.1 | 6.383596 | 6.501562 | 6.2326 | 0.268938836 | 0.150972799 | 6.5251521 | 6.7328903 | 6.17255 | 0.560344781 | 0.352606562 |
| Dlgap4 | 9.972317 | 10.17759 | 10.137 | 0.040465547 | -0.164804081 | 10.134819 | 10.675908 | 10.1159 | 0.560023064 | 0.018934049 |
| Jmjd6 | 9.442485 | 9.593197 | 9.4677 | 0.125458193 | -0.025254593 | 9.5931663 | 10.029278 | 9.4694 | 0.559880814 | 0.123768915 |
| Cdk2ap2 | 9.113353 | 9.245113 | 9.0189 | 0.226169252 | 0.094408628 | 9.4590421 | 9.4987943 | 8.93924 | 0.559556234 | 0.519804038 |
| Mrps6 | 7.638875 | 7.756555 | 7.593 | 0.16355021 | 0.045870088 | 7.5892811 | 8.1737512 | 7.61422 | 0.559534893 | -0.024935202 |
| Igkv6-32 | 8.292965 | 8.699833 | 8.4667 | 0.23314157 | -0.173725772 | 8.7478997 | 8.8432641 | 8.28383 | 0.559436709 | 0.464072349 |

| | | | | | | | | | | |
|---|---|---|---|---|---|---|---|---|---|---|
| Rbbp8 | 8.73348 | 8.881167 | 8.6828 | 0.198412358 | 0.050725018 | 8.8392528 | 9.3173937 | 8.75817 | 0.559219559 | 0.081078605 |
| Wdfy1 | 8.496072 | 8.509396 | 8.5454 | -0.036023993 | -0.049347604 | 8.6880756 | 9.293173 | 8.73402 | 0.559148466 | -0.045948906 |
| Igkv6-14 | 6.470728 | 6.613227 | 6.3766 | 0.236636357 | 0.094137646 | 6.6261047 | 6.9300664 | 6.37188 | 0.558190956 | 0.25422927 |
| Rpp25l | 7.448341 | 7.575597 | 7.5058 | 0.069756412 | -0.057499593 | 7.7799427 | 7.8498838 | 7.29215 | 0.557733866 | 0.487792747 |
| Serpinb1a | 11.43743 | 11.38074 | 11.338 | 0.042656062 | 0.099347146 | 11.643875 | 11.674064 | 11.1163 | 0.557733434 | 0.527544911 |
| 9530068E07Rik | 10.37658 | 10.50155 | 10.337 | 0.164334476 | 0.039369764 | 10.448358 | 10.793953 | 10.2363 | 0.557690628 | 0.212095494 |
| Ccdc102a | 6.95941 | 6.971044 | 6.938 | 0.033074452 | 0.021440308 | 6.8638386 | 7.4508436 | 6.89363 | 0.557212786 | -0.029792218 |
| Nop56 | 11.21759 | 11.21654 | 11.294 | -0.07749849 | -0.076457038 | 11.417371 | 11.936048 | 11.379 | 0.557070096 | 0.038392732 |
| Slc37a2 | 8.492692 | 8.662727 | 8.4983 | 0.164443196 | -0.0055918 | 8.5293948 | 8.9463713 | 8.39028 | 0.556087566 | 0.139111127 |
| Nhp2 | 8.907388 | 8.987918 | 8.8654 | 0.122483362 | 0.041953616 | 9.1803514 | 9.4078358 | 8.85184 | 0.555999229 | 0.328514737 |
| Gfi1 | 7.243606 | 7.176795 | 7.1957 | -0.018903776 | 0.047907507 | 7.033791 | 7.7389136 | 7.18348 | 0.555428724 | -0.149693873 |
| Gm28043 | 8.892089 | 9.31081 | 8.9152 | 0.395633848 | -0.023087035 | 9.156298 | 9.8243554 | 9.26919 | 0.55516357 | -0.11289384 |
| Slc7a5 | 9.974805 | 9.767952 | 9.5883 | 0.179644327 | 0.386496608 | 10.279284 | 10.507483 | 9.95233 | 0.555156797 | 0.326957654 |
| Fstl1 | 7.990019 | 8.195615 | 7.933 | 0.262645813 | 0.057049882 | 7.6905783 | 8.1560651 | 7.60132 | 0.554745385 | 0.089258529 |
| Klk1b22 | 4.52037 | 4.892654 | 5.6676 | -0.774989628 | -1.147274074 | 6.559768 | 5.5996569 | 5.04501 | 0.554646627 | 1.514757759 |
| 4930518I15Rik | 3.99661 | 4.132669 | 4.0631 | 0.069546123 | -0.066513096 | 4.3490022 | 4.8213569 | 4.26677 | 0.554582413 | 0.082227732 |
| Mfsd5 | 8.818831 | 8.924438 | 8.7216 | 0.202812061 | 0.097205776 | 8.8786866 | 9.1019644 | 8.54763 | 0.554337701 | 0.331059897 |
| Usmg5 | 9.160486 | 9.174057 | 9.1191 | 0.054973265 | 0.041402644 | 9.3868753 | 9.6709582 | 9.11672 | 0.554241748 | 0.270158822 |
| Atg13 | 9.340027 | 9.545907 | 9.462 | 0.083875829 | -0.122004489 | 9.4239729 | 10.028895 | 9.47474 | 0.554152439 | -0.050769219 |
| Gtpbp6 | 8.570893 | 8.746431 | 8.6073 | 0.139124302 | -0.036413381 | 9.2668927 | 9.3258348 | 8.77173 | 0.554108742 | 0.495166645 |
| 2510039O18Rik | 9.763496 | 9.926048 | 9.7303 | 0.195740466 | 0.033189226 | 9.9569772 | 10.142749 | 9.58987 | 0.552877753 | 0.367106455 |
| Car6 | 7.174328 | 5.866785 | 6.3834 | -0.516617709 | 0.79092554 | 7.4729114 | 6.5501595 | 5.99742 | 0.552737378 | 1.475489274 |
| Col1a2 | 8.424575 | 8.392355 | 8.1911 | 0.201298868 | 0.233519266 | 8.6671845 | 8.5316174 | 7.9789 | 0.552719229 | 0.688286335 |
| Cox5a | 9.800007 | 9.806583 | 9.773 | 0.033552381 | 0.026976099 | 9.9063402 | 10.29763 | 9.74505 | 0.552583575 | 0.161294078 |
| Txndc17 | 9.455614 | 9.412305 | 9.3845 | 0.027821872 | 0.071130172 | 9.2965771 | 9.9034898 | 9.35114 | 0.552345714 | -0.054566947 |
| Creld2 | 8.666143 | 8.862135 | 8.5233 | 0.33885015 | 0.142858198 | 8.7593792 | 9.0623511 | 8.51007 | 0.55227929 | 0.249307398 |
| Klra9 | 5.289794 | 5.274812 | 5.2199 | 0.054874578 | 0.069856214 | 5.2192924 | 5.7671015 | 5.21501 | 0.55208693 | 0.004277808 |
| Kcnab1 | 3.244023 | 3.517683 | 3.2215 | 0.296221369 | 0.022561272 | 3.5230645 | 3.8751232 | 3.32308 | 0.552047422 | 0.199988664 |
| Mllt3 | 9.549006 | 9.744986 | 9.5991 | 0.14591949 | -0.050060214 | 9.2287782 | 10.175816 | 9.62388 | 0.551940848 | -0.395097435 |
| Psmb3 | 10.14636 | 10.3228 | 10.182 | 0.14087384 | -0.035564188 | 10.537728 | 10.716738 | 10.1656 | 0.55110685 | 0.372096966 |
| Psmg3 | 7.340425 | 7.351858 | 7.2517 | 0.10013478 | 0.088702373 | 7.5991103 | 7.7728938 | 7.22213 | 0.550765302 | 0.376981801 |
| Dph2 | 8.028675 | 8.296262 | 8.1317 | 0.164605259 | -0.10298136 | 8.6109237 | 8.8069836 | 8.25625 | 0.550737505 | 0.354677681 |

| | | | | | | | | | | |
|---|---|---|---|---|---|---|---|---|---|---|
| Tma16 | 8.170065 | 8.195022 | 8.1083 | 0.086741799 | 0.061784907 | 8.3364818 | 8.8463974 | 8.29587 | 0.5505241 | 0.040608546 |
| Tut4 | 11.65062 | 11.8997 | 11.702 | 0.19746372 | -0.051612986 | 11.703661 | 12.323151 | 11.7733 | 0.549842742 | -0.069647225 |
| Csf2rb | 10.9717 | 10.90992 | 10.536 | 0.374011292 | 0.435786444 | 10.516181 | 10.948778 | 10.3994 | 0.54940742 | 0.116810258 |
| Dus3l | 9.434208 | 9.511211 | 9.4638 | 0.047447794 | -0.029555371 | 10.027193 | 10.07263 | 9.52332 | 0.549312879 | 0.503876353 |
| 2310039H08Rik | 6.037832 | 5.813276 | 5.9458 | -0.132551321 | 0.092004837 | 6.2560224 | 6.4402225 | 5.89106 | 0.549166384 | 0.364966221 |
| Slc2a1 | 10.13083 | 9.995672 | 9.9306 | 0.065113041 | 0.200273439 | 9.9481614 | 10.25108 | 9.702 | 0.549081036 | 0.246162616 |
| Cct7 | 11.84915 | 11.96443 | 11.84 | 0.124733654 | 0.009455442 | 11.939072 | 12.425711 | 11.877 | 0.548723896 | 0.06208418 |
| Mrps36 | 7.7509 | 7.810702 | 7.7584 | 0.05229615 | -0.007506075 | 7.9716134 | 8.2688985 | 7.72027 | 0.54862498 | 0.251339836 |
| Crybg1 | 11.03878 | 11.29224 | 10.984 | 0.308590202 | 0.05512965 | 10.971685 | 11.507621 | 10.9597 | 0.54793561 | 0.011999943 |
| Psma1 | 10.4366 | 10.42149 | 10.401 | 0.020899144 | 0.036017826 | 10.683719 | 10.944133 | 10.3963 | 0.547842504 | 0.287429265 |
| Timm23 | 9.550913 | 9.659832 | 9.2471 | 0.412731379 | 0.303811862 | 9.6971705 | 9.9912365 | 9.44371 | 0.54752247 | 0.253456491 |
| Morf4l2 | 10.39386 | 10.42448 | 10.22 | 0.20411359 | 0.173491996 | 10.218778 | 10.809797 | 10.2623 | 0.547522212 | -0.043496422 |
| Pja1 | 9.449914 | 9.610937 | 9.4292 | 0.181711683 | 0.020688153 | 9.425608 | 9.8754425 | 9.32812 | 0.547320095 | 0.097485646 |
| Zfyve26 | 10.04818 | 10.25842 | 10.026 | 0.23264067 | 0.022398478 | 10.36477 | 10.944945 | 10.3976 | 0.547318529 | -0.032857242 |
| Cox5b | 9.710879 | 9.762263 | 9.735 | 0.027311321 | -0.024072858 | 10.114196 | 10.423148 | 9.87585 | 0.547301113 | 0.238349109 |
| Rbck1 | 11.1978 | 11.40492 | 11.229 | 0.175426002 | -0.031688784 | 11.560946 | 11.886857 | 11.3399 | 0.546963986 | 0.221053423 |
| Tex14 | 3.506001 | 3.653432 | 3.4649 | 0.188572312 | 0.041140917 | 3.6812328 | 4.144426 | 3.59773 | 0.546698989 | 0.083505769 |
| Psmc5 | 10.20383 | 10.28842 | 10.166 | 0.122382921 | 0.037787257 | 10.376658 | 10.702491 | 10.156 | 0.546490534 | 0.22065688 |
| Med11 | 8.516492 | 8.699509 | 8.5347 | 0.164824707 | -0.018192854 | 8.8427789 | 9.0714772 | 8.52511 | 0.546371381 | 0.317673083 |
| Focad | 8.774247 | 8.818283 | 8.7187 | 0.099572462 | 0.055536651 | 8.7993929 | 9.3582 | 8.81296 | 0.545241585 | -0.013565495 |
| Sdf2l1 | 8.461758 | 8.57346 | 8.2158 | 0.357628381 | 0.245927053 | 8.6312806 | 8.8364051 | 8.29135 | 0.545056153 | 0.339931635 |
| Apol9b | 6.029232 | 6.027189 | 6.1887 | -0.161551625 | -0.159509055 | 5.8756138 | 6.4798152 | 5.93522 | 0.544595216 | -0.059606168 |
| Tmem104 | 8.173728 | 8.26584 | 8.2453 | 0.020553739 | -0.071558346 | 8.0737119 | 8.4858446 | 7.94161 | 0.544239258 | 0.13210655 |
| Rhoc | 7.270151 | 7.474592 | 6.9418 | 0.53283422 | 0.328393443 | 6.9180616 | 7.482101 | 6.93855 | 0.543546615 | -0.020492727 |
| Cd164 | 12.44035 | 12.59869 | 12.413 | 0.185523068 | 0.027187554 | 12.340738 | 12.87685 | 12.334 | 0.542869 | 0.006757032 |
| Frmd4a | 8.794969 | 8.843762 | 8.6592 | 0.184514547 | 0.135721546 | 8.439076 | 9.1608008 | 8.6182 | 0.542601504 | -0.179123257 |
| Ptgir | 6.489179 | 6.504023 | 6.3308 | 0.173182013 | 0.158337304 | 6.2662127 | 6.67139 | 6.12885 | 0.54253959 | 0.137362246 |
| Suv39h1 | 9.075306 | 9.153548 | 9.1599 | -0.006381902 | -0.084623655 | 9.2060538 | 9.6844466 | 9.1421 | 0.542342129 | 0.063949359 |
| Cmc1 | 7.710703 | 7.482434 | 7.6918 | -0.209389712 | 0.018879235 | 7.7297338 | 8.2185837 | 7.6763 | 0.542282559 | 0.053432734 |
| Psmc4 | 10.24812 | 10.24841 | 10.175 | 0.073508708 | 0.073222752 | 10.433488 | 10.734551 | 10.1926 | 0.541934636 | 0.240871599 |
| Trmt112-ps2 | 6.936376 | 7.068564 | 6.8739 | 0.194679429 | 0.062490745 | 7.2437549 | 7.3357556 | 6.79389 | 0.541865261 | 0.449864493 |
| Slc25a12 | 9.898656 | 9.93524 | 9.9505 | -0.015239205 | -0.051823276 | 10.021619 | 10.403549 | 9.86183 | 0.541715107 | 0.159785331 |

| | | | | | | | | | | |
|---|---|---|---|---|---|---|---|---|---|---|
| Phgdh | 10.29158 | 10.46136 | 10.413 | 0.048494044 | -0.121278636 | 10.654836 | 11.040965 | 10.4995 | 0.541509816 | 0.155381012 |
| Gm12551 | 4.258521 | 4.382331 | 4.263 | 0.119347783 | -0.004461477 | 4.3099567 | 4.8152884 | 4.27379 | 0.541502783 | 0.036171065 |
| Trmt1l | 9.51431 | 9.793097 | 9.6361 | 0.157035764 | -0.121751231 | 9.6781852 | 10.342107 | 9.80103 | 0.541073538 | -0.122847999 |
| Akr1a1 | 11.63606 | 11.6765 | 11.623 | 0.053124492 | 0.012676656 | 11.67937 | 12.001384 | 11.4607 | 0.540671606 | 0.218658022 |
| Pprc1 | 10.20668 | 10.43134 | 10.138 | 0.293535261 | 0.068874489 | 10.628568 | 10.986007 | 10.4457 | 0.540315498 | 0.182876929 |
| Olfr624 | 7.317687 | 7.267027 | 7.2529 | 0.014112865 | 0.064772643 | 7.089686 | 7.8076572 | 7.26782 | 0.539841286 | -0.178129945 |
| Tlk2 | 10.04667 | 10.20937 | 9.9554 | 0.253924857 | 0.09122232 | 9.9107521 | 10.659085 | 10.1193 | 0.539797786 | -0.208535468 |
| Rrp15 | 8.610019 | 8.741114 | 8.4719 | 0.269179641 | 0.138085195 | 8.6261578 | 9.0884446 | 8.54931 | 0.539132942 | 0.076846098 |
| Trmt1 | 10.38151 | 10.36773 | 10.38 | -0.012214103 | 0.001561064 | 10.745973 | 11.106012 | 10.567 | 0.539060444 | 0.179021873 |
| Nkiras2 | 8.431195 | 8.573673 | 8.4065 | 0.167165489 | 0.024686767 | 8.5996909 | 8.847045 | 8.30843 | 0.538612418 | 0.291258289 |
| Kat2b | 9.393135 | 9.619956 | 9.411 | 0.208938721 | -0.017882746 | 9.3065569 | 10.107686 | 9.56917 | 0.538515203 | -0.262614381 |
| Mcmbp | 10.71287 | 11.0279 | 10.781 | 0.246644502 | -0.068385442 | 10.824085 | 11.445695 | 10.9074 | 0.53833975 | -0.083270059 |
| Ddx39 | 10.60315 | 10.68759 | 10.529 | 0.158286734 | 0.073847618 | 10.96066 | 11.217608 | 10.6795 | 0.538144124 | 0.281196102 |
| Mrpl39 | 8.422048 | 8.580507 | 8.4635 | 0.116958008 | -0.041501526 | 8.7830366 | 9.0210881 | 8.48301 | 0.538082867 | 0.30003134 |
| Bco2 | 4.953676 | 4.826948 | 4.9237 | -0.096774069 | 0.029953142 | 4.7848028 | 5.3154292 | 4.7785 | 0.53692926 | 0.006302886 |
| H13 | 11.03004 | 11.07519 | 10.957 | 0.118604548 | 0.073450182 | 10.946664 | 11.359603 | 10.8233 | 0.536319088 | 0.123380045 |
| Serpina10 | 6.538279 | 5.65465 | 5.5807 | 0.073919838 | 0.957548212 | 5.7847122 | 5.871996 | 5.3358 | 0.536196162 | 0.448912376 |
| Eif4e3 | 8.471954 | 8.528853 | 8.5159 | 0.012975466 | -0.04392412 | 8.4532687 | 9.0332172 | 8.49713 | 0.536088407 | -0.04386016 |
| Hax1 | 9.05901 | 9.132621 | 8.967 | 0.165611682 | 0.092000119 | 9.2859134 | 9.6183899 | 9.0823 | 0.536087663 | 0.20361114 |
| Jdp2 | 6.310601 | 6.312198 | 5.8321 | 0.480117569 | 0.478521337 | 6.2782587 | 6.3019936 | 5.76625 | 0.535739249 | 0.512004427 |
| Coa7 | 7.001735 | 7.153213 | 7.0029 | 0.150291489 | -0.001186724 | 7.1146592 | 7.3390637 | 6.80348 | 0.535586631 | 0.311182122 |
| C1qbp | 10.05595 | 10.16188 | 10.083 | 0.078429937 | -0.027501964 | 10.208939 | 10.685202 | 10.1498 | 0.535381892 | 0.059118818 |
| Taf4b | 8.789705 | 8.918596 | 8.6726 | 0.246020278 | 0.117129197 | 8.6899822 | 9.1629637 | 8.62823 | 0.53473636 | 0.06175486 |
| Cdk5r1 | 6.584127 | 6.896864 | 6.5089 | 0.387923451 | 0.07518595 | 6.8203522 | 7.0381954 | 6.50376 | 0.534434766 | 0.316591549 |
| Igkv8-28 | 6.923389 | 6.756664 | 7.3755 | -0.618872295 | -0.4521474 | 6.9581829 | 7.1275937 | 6.59354 | 0.534051673 | 0.3646409 |
| Rpl12 | 10.20505 | 10.14246 | 10.116 | 0.026305258 | 0.088894123 | 10.195094 | 10.607905 | 10.0741 | 0.53385522 | 0.121043365 |
| Fbxo39 | 2.367046 | 2.485744 | 2.3043 | 0.181492007 | 0.062793406 | 2.3612535 | 2.8708792 | 2.33779 | 0.533089589 | 0.023463889 |
| Raver1 | 10.98548 | 11.10202 | 11.005 | 0.097468376 | -0.019067478 | 11.154289 | 11.417345 | 10.8851 | 0.532261476 | 0.269204871 |
| Klk1b5 | 4.609894 | 4.403037 | 4.7794 | -0.376364237 | -0.169506664 | 5.8434557 | 5.1614803 | 4.62947 | 0.532013746 | 1.213989194 |
| Edf1 | 9.684185 | 9.862643 | 9.6874 | 0.175227202 | -0.003230463 | 9.955778 | 10.177825 | 9.64589 | 0.531934539 | 0.309887346 |
| Zfp800 | 9.677217 | 9.979312 | 9.6294 | 0.349958078 | 0.047863404 | 9.7630441 | 10.231613 | 9.69975 | 0.531862413 | 0.063293123 |
| Gm18342 | 3.314547 | 3.350063 | 3.1886 | 0.161482628 | 0.125966901 | 3.4101596 | 3.7828309 | 3.25128 | 0.531549451 | 0.158878118 |

| | | | | | | | | | | |
|---|---|---|---|---|---|---|---|---|---|---|
| Kti12 | 8.339379 | 8.617247 | 8.3882 | 0.229044082 | -0.048824218 | 8.5425586 | 8.7879221 | 8.25657 | 0.531350648 | 0.285987178 |
| Ccnd3 | 11.21921 | 11.28903 | 11.196 | 0.093338056 | 0.02351603 | 11.0217 | 11.557779 | 11.0267 | 0.531119976 | -0.004959138 |
| Tmsb10 | 13.83492 | 13.96425 | 13.866 | 0.098471742 | -0.030859942 | 13.948207 | 14.425035 | 13.894 | 0.531081056 | 0.054253067 |
| Gm3650 | 5.388429 | 5.321007 | 5.4827 | -0.161655896 | -0.094233307 | 5.5797666 | 5.9226151 | 5.39162 | 0.530992161 | 0.188143596 |
| Ndufa11 | 8.335181 | 8.335195 | 8.2975 | 0.037646717 | 0.037633052 | 8.6735141 | 8.7605472 | 8.23005 | 0.530501059 | 0.443467882 |
| Chpf2 | 9.444283 | 9.528069 | 9.3725 | 0.155540476 | 0.071755015 | 9.6353678 | 9.8519533 | 9.32162 | 0.530331633 | 0.313746164 |
| Uxt | 7.238842 | 7.17175 | 7.2154 | -0.043637322 | 0.023454119 | 7.3042182 | 7.6389688 | 7.10911 | 0.529863554 | 0.195113008 |
| Mosmo | 8.297333 | 8.443009 | 8.2619 | 0.181109617 | 0.035434048 | 8.330057 | 8.7949139 | 8.26506 | 0.529858751 | 0.065001887 |
| Prdx1 | 11.49849 | 11.64241 | 11.369 | 0.273131472 | 0.129204494 | 11.495147 | 11.891161 | 11.3616 | 0.529535426 | 0.133520685 |
| Dgat2 | 5.615542 | 6.066522 | 5.3573 | 0.709218737 | 0.258238446 | 5.5014006 | 5.9866678 | 5.45805 | 0.52861351 | 0.04334627 |
| Pmvk | 8.239935 | 8.315551 | 8.3019 | 0.013626645 | -0.061988656 | 8.5073209 | 8.8797342 | 8.35253 | 0.527199763 | 0.154786495 |
| Ldha | 12.4479 | 12.38125 | 12.261 | 0.120624776 | 0.18727676 | 12.490102 | 12.792669 | 12.2657 | 0.527003126 | 0.224436347 |
| Gm48981 | 4.219697 | 4.450532 | 4.215 | 0.235520374 | 0.004684807 | 4.4595141 | 4.817124 | 4.29031 | 0.52681466 | 0.169204758 |
| Igkv14-111 | 8.885256 | 9.399049 | 8.8263 | 0.572750871 | 0.058957666 | 9.030682 | 9.3639192 | 8.83765 | 0.526271228 | 0.193033963 |
| Mrps11 | 7.279654 | 7.331837 | 7.2605 | 0.071378477 | 0.019195361 | 7.4951195 | 7.7304501 | 7.20425 | 0.526196498 | 0.290865932 |
| Igkv6-23 | 8.973722 | 9.20069 | 8.6409 | 0.559783057 | 0.33281469 | 8.9356175 | 8.9541176 | 8.428 | 0.526120416 | 0.507620271 |
| Zfp628 | 8.29683 | 8.534788 | 8.2708 | 0.26397692 | 0.026018823 | 8.4477827 | 8.6429369 | 8.11695 | 0.525988847 | 0.330834718 |
| Dek | 11.6406 | 11.75495 | 11.652 | 0.103378454 | -0.01097795 | 11.585375 | 12.28366 | 11.7578 | 0.525815308 | -0.172469779 |
| Capn15 | 9.255232 | 9.430787 | 9.3409 | 0.089909926 | -0.085644504 | 9.560288 | 9.8508312 | 9.32513 | 0.525702559 | 0.235159362 |
| Mrpl2 | 8.076824 | 8.082525 | 8.1092 | -0.026643001 | -0.032344458 | 8.6932684 | 8.6549934 | 8.12945 | 0.525541344 | 0.563816385 |
| Stat4 | 10.6009 | 10.80419 | 10.666 | 0.137914026 | -0.06537587 | 10.554964 | 11.160231 | 10.6348 | 0.525443618 | -0.079824053 |
| Eif4ebp3 | 4.925049 | 4.767006 | 4.9288 | -0.161788991 | -0.003746103 | 4.739456 | 5.0807725 | 4.55544 | 0.525330611 | 0.184014112 |
| Phb | 10.01412 | 10.16477 | 9.9869 | 0.177883203 | 0.027232181 | 10.250644 | 10.486352 | 9.96103 | 0.525317543 | 0.28961014 |
| Tti2 | 8.339755 | 8.46478 | 8.3432 | 0.121622743 | -0.003401506 | 8.52882 | 8.8325518 | 8.30738 | 0.525174971 | 0.221443205 |
| Pold2 | 8.760344 | 8.815989 | 8.8288 | -0.01280498 | -0.068450623 | 9.2533533 | 9.2410766 | 8.71701 | 0.524071123 | 0.536347854 |
| Dnah17 | 5.719784 | 6.391962 | 5.8498 | 0.542120286 | -0.130057608 | 5.8880747 | 6.8959207 | 6.37209 | 0.52383 | -0.48401606 |
| Nhsl2 | 6.554669 | 6.56688 | 6.3825 | 0.184376098 | 0.172165485 | 6.2406311 | 6.769853 | 6.24624 | 0.52361267 | -0.005609202 |
| Penk | 6.23631 | 6.487523 | 5.9979 | 0.489595659 | 0.238383076 | 5.9691575 | 6.6296533 | 6.10672 | 0.522931849 | -0.137563861 |
| Rbm47 | 6.640899 | 6.471322 | 6.6331 | -0.16175565 | 0.00782123 | 6.7047321 | 6.7911163 | 6.2682 | 0.522915261 | 0.436531016 |
| Rsrp1 | 11.60818 | 11.70924 | 11.816 | -0.10663639 | -0.207694288 | 11.854863 | 12.374064 | 11.8515 | 0.52260629 | 0.003405501 |
| Med21 | 8.128871 | 8.30946 | 8.2385 | 0.070914186 | -0.109674721 | 8.3871493 | 8.6974608 | 8.17548 | 0.521983036 | 0.211671493 |
| Gfer | 7.733036 | 7.85145 | 7.8159 | 0.035502342 | -0.082912421 | 7.956189 | 8.1020067 | 7.58019 | 0.521815565 | 0.375997892 |

| | | | | | | | | | | |
|---|---|---|---|---|---|---|---|---|---|---|
| Pdcd10 | 10.19332 | 10.3529 | 10.196 | 0.15665846 | -0.002924716 | 10.334959 | 10.758188 | 10.2369 | 0.521246252 | 0.098017743 |
| Dnajb11 | 10.47197 | 10.54937 | 10.331 | 0.21797408 | 0.140570072 | 10.519902 | 10.89147 | 10.3708 | 0.520655832 | 0.149087382 |
| 2310009B15Rik | 5.611798 | 5.713745 | 5.7173 | -0.003535275 | -0.105482562 | 5.8214364 | 6.0817941 | 5.5613 | 0.520495148 | 0.260137449 |
| Slc6a9 | 4.735301 | 4.795598 | 4.424 | 0.371628349 | 0.311331032 | 4.7052928 | 5.1293589 | 4.60923 | 0.520132747 | 0.096066561 |
| Col18a1 | 7.338318 | 7.871872 | 7.2444 | 0.627456929 | 0.093903142 | 7.468497 | 7.4138725 | 6.89415 | 0.519720783 | 0.574345295 |
| Srgn | 12.56321 | 12.67598 | 12.4 | 0.276225462 | 0.163454384 | 12.581746 | 12.81101 | 12.292 | 0.519040592 | 0.28977649 |
| Ube2l3 | 10.81862 | 10.89972 | 10.827 | 0.072423264 | -0.008678762 | 10.850807 | 11.257537 | 10.739 | 0.518557812 | 0.111827183 |
| Peli1 | 11.45427 | 11.76585 | 11.528 | 0.237777496 | -0.0738046 | 11.192237 | 11.97657 | 11.4585 | 0.518068712 | -0.26626464 |
| Gm49356 | 2.68576 | 2.853787 | 2.6482 | 0.205635309 | 0.037608486 | 2.6867224 | 3.1066499 | 2.58873 | 0.51792003 | 0.097992593 |
| Lsm7 | 8.085359 | 8.226314 | 8.0806 | 0.145695745 | 0.004740696 | 8.5499826 | 8.631795 | 8.11394 | 0.517852686 | 0.436040281 |
| Pip | 6.628438 | 5.229891 | 5.8121 | -0.582186578 | 0.816360265 | 6.6998884 | 5.6551073 | 5.13729 | 0.517819855 | 1.562600975 |
| F2rl2 | 5.244791 | 5.168058 | 5.1208 | 0.04725221 | 0.12398463 | 4.9727944 | 5.5861794 | 5.06873 | 0.517449211 | -0.095935728 |
| Tomm40 | 9.134579 | 9.207197 | 9.0899 | 0.117264247 | 0.044646533 | 9.5026876 | 9.6700796 | 9.15288 | 0.517194842 | 0.349802798 |
| Itpk1 | 9.084733 | 9.185616 | 9.139 | 0.04662618 | -0.05425671 | 8.7865779 | 9.6205356 | 9.10345 | 0.517084303 | -0.316873398 |
| Ptpn23 | 9.629403 | 9.742218 | 9.5786 | 0.163640242 | 0.050825176 | 9.9319104 | 10.225616 | 9.70874 | 0.516879602 | 0.22317438 |
| Gm42742 | 8.886054 | 9.035476 | 8.8753 | 0.160133397 | 0.010711379 | 9.038453 | 9.2840146 | 8.76764 | 0.516374234 | 0.270812607 |
| Arhgap8 | 3.450489 | 3.434656 | 3.304 | 0.130653002 | 0.14648628 | 3.4911031 | 3.9029274 | 3.38678 | 0.516145639 | 0.104321328 |
| Wdr18 | 9.340049 | 9.421352 | 9.2302 | 0.191184244 | 0.109880831 | 9.7008189 | 9.9444898 | 9.42935 | 0.515141827 | 0.271470872 |
| Rars | 9.948614 | 10.08602 | 9.8549 | 0.231101557 | 0.09369788 | 9.9141034 | 10.473461 | 9.95873 | 0.51472689 | -0.044630719 |
| Akr1b8 | 3.376848 | 3.521951 | 3.2175 | 0.30443176 | 0.15932931 | 3.4516751 | 3.6607039 | 3.14664 | 0.514061508 | 0.305032675 |
| Commd5 | 6.991319 | 6.999953 | 7.0706 | -0.07059793 | -0.079232169 | 7.2729411 | 7.4053218 | 6.89186 | 0.513458843 | 0.381078115 |
| Casp2 | 10.18635 | 10.39146 | 10.302 | 0.089771124 | -0.115334092 | 10.448359 | 10.883146 | 10.3698 | 0.513395524 | 0.078607981 |
| Slc31a1 | 8.607085 | 8.557023 | 8.3806 | 0.176441532 | 0.226503083 | 8.6355625 | 8.9864977 | 8.47326 | 0.513237455 | 0.162302222 |
| Arg2 | 5.063832 | 5.253735 | 4.3609 | 0.892869808 | 0.702966282 | 5.0275978 | 4.8892565 | 4.37604 | 0.513221203 | 0.65156258 |
| Tep1 | 9.957601 | 9.924028 | 10.121 | -0.197127757 | -0.163555347 | 10.236283 | 10.607414 | 10.0948 | 0.512632859 | 0.141501721 |
| Chmp2a | 10.08786 | 10.12545 | 10.031 | 0.094016615 | 0.056426258 | 10.23918 | 10.562969 | 10.0504 | 0.512530794 | 0.188741048 |
| Atp6v1d | 9.797826 | 9.97364 | 9.9291 | 0.044588663 | -0.131225275 | 9.9159057 | 10.408597 | 9.89612 | 0.512477642 | 0.019786169 |
| Atp6v1f | 9.975088 | 10.0872 | 10.015 | 0.072365186 | -0.039749016 | 10.3471 | 10.402239 | 9.89005 | 0.512189382 | 0.457050275 |
| Pin1 | 8.492821 | 8.465402 | 8.3975 | 0.067933697 | 0.095352727 | 8.5475421 | 8.8777846 | 8.36682 | 0.510965838 | 0.180723292 |
| Pgf | 3.534711 | 3.600255 | 3.4408 | 0.159474227 | 0.093930014 | 3.5041234 | 3.8213172 | 3.31096 | 0.510359642 | 0.193165907 |
| Mrpl17 | 8.407714 | 8.243071 | 8.1714 | 0.071714731 | 0.236357618 | 8.3724542 | 8.6512059 | 8.14102 | 0.510190825 | 0.231439131 |
| Pus1 | 9.010225 | 9.211248 | 8.9793 | 0.231935646 | 0.030912995 | 9.2067756 | 9.5413448 | 9.03119 | 0.510150262 | 0.175581105 |

| | | | | | | | | | | |
|---|---|---|---|---|---|---|---|---|---|---|
| Eme2 | 9.172165 | 9.165294 | 9.0629 | 0.1023884 | 0.109259445 | 9.7895119 | 9.907998 | 9.39802 | 0.509973299 | 0.39148716 |
| Pim2 | 10.19738 | 9.855407 | 9.8532 | 0.002224877 | 0.344194278 | 9.9611018 | 10.406602 | 9.89691 | 0.509695997 | 0.064196174 |
| Gm2436 | 2.57256 | 2.831476 | 3.1414 | -0.309952296 | -0.568869093 | 3.4737407 | 3.1640643 | 2.65477 | 0.509298795 | 0.818975246 |
| Dynll2 | 9.783771 | 9.88501 | 9.8389 | 0.046147432 | -0.055092266 | 9.4313642 | 10.111985 | 9.60277 | 0.509217421 | -0.171403654 |
| 4932411N23Rik | 2.407416 | 2.495281 | 2.3721 | 0.123206827 | 0.035342236 | 2.4664142 | 2.9940536 | 2.48504 | 0.509010249 | -0.018629099 |
| Col20a1 | 5.533443 | 5.718805 | 5.5548 | 0.163987626 | -0.021373937 | 6.0365852 | 6.4092367 | 5.90024 | 0.508992977 | 0.136341481 |
| Rpl23a-ps3 | 5.585595 | 5.664437 | 5.5902 | 0.074261362 | -0.004580589 | 5.6531439 | 6.1220267 | 5.61341 | 0.508619622 | 0.039736757 |
| Ppp1r16b | 11.67822 | 11.71835 | 11.568 | 0.15006882 | 0.10994009 | 11.469757 | 12.051964 | 11.5434 | 0.508529552 | -0.073678066 |
| AW822252 | 7.227368 | 6.95461 | 7.6739 | -0.719290018 | -0.446532042 | 8.304028 | 7.7586501 | 7.25024 | 0.508414821 | 1.053792779 |
| Hmgb1-ps5 | 4.327023 | 4.368103 | 4.3486 | 0.019486763 | -0.021592634 | 4.273375 | 4.8801525 | 4.37198 | 0.508170078 | -0.098607475 |
| Mmp13 | 2.957021 | 4.776322 | 2.9011 | 1.875234979 | 0.055934433 | 2.9786128 | 3.4182076 | 2.9101 | 0.508107907 | 0.068513177 |
| Pfdn2 | 9.162972 | 9.306112 | 9.1495 | 0.1566379 | 0.013497785 | 9.3631573 | 9.7307148 | 9.22296 | 0.507754019 | 0.140196528 |
| Gm10800 | 1.767592 | 1.763863 | 1.7254 | 0.038461432 | 0.042190452 | 1.8639884 | 2.2652752 | 1.75754 | 0.507736282 | 0.106449495 |
| Wdr75 | 9.659593 | 9.903784 | 9.5327 | 0.371090721 | 0.126899567 | 9.820578 | 10.197925 | 9.69059 | 0.507331969 | 0.129985276 |
| Zbtb21 | 8.648303 | 8.915842 | 8.6489 | 0.266925938 | -0.000612563 | 8.4744261 | 9.1915117 | 8.6846 | 0.506913518 | -0.21017209 |
| Lrp8 | 6.850392 | 7.317347 | 6.9386 | 0.378718133 | -0.08823647 | 6.7461857 | 7.6316572 | 7.12532 | 0.506340913 | -0.379130583 |
| Gm4759 | 9.951484 | 10.10655 | 9.8541 | 0.252495243 | 0.097429287 | 10.06028 | 10.853168 | 10.3468 | 0.506340063 | -0.286548451 |
| Pisd | 9.681479 | 9.72172 | 9.5497 | 0.171995582 | 0.131754881 | 9.814035 | 10.068666 | 9.56236 | 0.506304968 | 0.251673936 |
| Lsg1 | 9.713569 | 9.806711 | 9.6065 | 0.200172232 | 0.107030215 | 9.6844107 | 10.058375 | 9.55211 | 0.506264058 | 0.132300032 |
| Zdhhc13 | 9.120483 | 9.198785 | 9.1633 | 0.035463495 | -0.042839017 | 9.2546373 | 9.5296739 | 9.02368 | 0.505991569 | 0.230954986 |
| Igkv17-121 | 8.551632 | 8.53417 | 8.3454 | 0.188748011 | 0.206209929 | 8.653875 | 8.7927264 | 8.2876 | 0.505122771 | 0.366271459 |
| Rspo1 | 2.632093 | 2.623931 | 2.6157 | 0.008263682 | 0.016425487 | 2.6293582 | 3.0688652 | 2.56385 | 0.505015998 | 0.065509038 |
| Eif6 | 10.01821 | 10.15978 | 9.9698 | 0.189976796 | 0.048411754 | 10.262214 | 10.454348 | 9.94959 | 0.504761571 | 0.312628029 |
| Ormdl2 | 7.867802 | 8.053795 | 7.8654 | 0.188423983 | 0.00243038 | 8.0475083 | 8.3590284 | 7.8545 | 0.504532735 | 0.193012583 |
| Txndc9 | 9.698167 | 9.847707 | 9.6721 | 0.17559156 | 0.026051502 | 9.7021645 | 10.117519 | 9.613 | 0.504518469 | 0.089164309 |
| Nolc1 | 10.76674 | 10.79843 | 10.692 | 0.106263102 | 0.074574398 | 10.885211 | 11.271532 | 10.7672 | 0.50429048 | 0.117969011 |
| Rnaset2a | 8.623645 | 8.746706 | 8.7321 | 0.014639026 | -0.108421787 | 8.7759738 | 9.0236533 | 8.51953 | 0.504119527 | 0.256440033 |
| Ppid | 9.633783 | 9.696847 | 9.6814 | 0.015403713 | -0.047660787 | 9.8273056 | 10.321734 | 9.81775 | 0.503985497 | 0.00955689 |
| Rufy3 | 8.23721 | 8.256766 | 8.155 | 0.101731443 | 0.082175326 | 8.3806676 | 8.6973662 | 8.19343 | 0.503937795 | 0.18723918 |
| Nsun4 | 8.127558 | 8.264685 | 8.223 | 0.041728198 | -0.09539856 | 8.3812166 | 8.7442304 | 8.24036 | 0.503868123 | 0.140854301 |
| Scgb2b26 | 6.762772 | 5.296915 | 5.6848 | -0.387900343 | 1.077957267 | 6.6034261 | 5.7318371 | 5.22822 | 0.503614837 | 1.375203868 |
| Adamts14 | 6.54418 | 7.066554 | 6.6593 | 0.407288383 | -0.115085545 | 6.4056096 | 7.2057796 | 6.70258 | 0.503197978 | -0.296972024 |

| | | | | | | | | | | |
|---|---|---|---|---|---|---|---|---|---|---|
| Ndufa7 | 9.023417 | 9.154902 | 9.0968 | 0.058090395 | -0.073395298 | 9.5829676 | 9.5515726 | 9.04856 | 0.503013591 | 0.534408624 |
| Sf3b4 | 9.89216 | 9.984139 | 9.8683 | 0.115860863 | 0.023881572 | 10.113335 | 10.290769 | 9.78781 | 0.502957372 | 0.325523216 |
| Ndufa2 | 9.522043 | 9.605963 | 9.4804 | 0.125568646 | 0.041648612 | 9.9055127 | 9.9350626 | 9.43239 | 0.502673589 | 0.473123742 |
| Nop16 | 8.436808 | 8.685219 | 8.3922 | 0.293068766 | 0.044657613 | 8.6753249 | 8.9301002 | 8.42757 | 0.502533669 | 0.247758374 |
| Atp5j | 9.848536 | 9.731976 | 9.8276 | -0.095670746 | 0.020888954 | 9.9135617 | 10.271193 | 9.76905 | 0.502144402 | 0.144513511 |
| Gm3052 | 4.659046 | 4.672134 | 4.5177 | 0.154434054 | 0.141346742 | 4.8366576 | 5.3541633 | 4.85208 | 0.502080841 | -0.015424889 |
| Pgk1 | 11.50392 | 11.5249 | 11.475 | 0.049692918 | 0.028710424 | 11.686663 | 11.94614 | 11.4441 | 0.50203901 | 0.242561609 |
| D8Ertd738e | 9.509395 | 9.571475 | 9.4557 | 0.11576348 | 0.053683091 | 9.8061481 | 9.9401527 | 9.43818 | 0.501976499 | 0.367971955 |
| Igkv19-93 | 9.151898 | 9.435155 | 9.6212 | -0.186001184 | -0.469258156 | 9.5337336 | 9.6243456 | 9.12244 | 0.501903622 | 0.411291653 |
| Igkv3-4 | 8.22412 | 8.637284 | 8.0323 | 0.604955984 | 0.191791612 | 8.450281 | 8.5749604 | 8.07411 | 0.500849429 | 0.376169965 |
| Sestd1 | 6.586654 | 6.813112 | 6.387 | 0.42614365 | 0.19968574 | 6.4080419 | 6.9046602 | 6.40412 | 0.500544663 | 0.003926312 |
| Rps2-ps10 | 7.064379 | 7.413646 | 7.1735 | 0.240098122 | -0.109169088 | 7.6122444 | 7.6311544 | 7.13062 | 0.500534485 | 0.481624524 |
| Herc4 | 10.78726 | 10.91412 | 10.972 | -0.057676086 | -0.184530724 | 10.912533 | 11.452463 | 10.9521 | 0.500315412 | -0.039614426 |
| Atp8b4 | 9.46157 | 9.645103 | 9.3243 | 0.320768454 | 0.137235535 | 9.301858 | 10.10736 | 9.60708 | 0.500276788 | -0.305224751 |
| Pik3ip1 | 10.4934 | 10.68916 | 10.479 | 0.210327166 | 0.014570828 | 10.115861 | 10.738136 | 10.2379 | 0.500258829 | -0.122015442 |
| Orai1 | 8.606589 | 8.804123 | 8.5736 | 0.230491085 | 0.032957127 | 8.713163 | 8.9265808 | 8.42633 | 0.500251064 | 0.286833349 |
| Galk1 | 7.834722 | 7.62085 | 7.6698 | -0.048970089 | 0.164901764 | 8.1064308 | 8.11408 | 7.61415 | 0.499934138 | 0.492284871 |
| Sirt6 | 7.425187 | 7.60348 | 7.4179 | 0.185533128 | 0.007240547 | 7.9986883 | 8.0538916 | 7.55413 | 0.499760527 | 0.444557256 |
| Tbx15 | 4.625592 | 4.450189 | 4.3738 | 0.076347233 | 0.251749996 | 4.1681313 | 4.542212 | 4.04278 | 0.499428125 | 0.12534748 |
| AC133103.1 | 4.359504 | 4.422576 | 4.2521 | 0.170460625 | 0.107388166 | 4.2585034 | 4.8351109 | 4.33593 | 0.499179839 | -0.077427695 |
| Otud5 | 11.0972 | 11.30615 | 11.157 | 0.149493302 | -0.059455556 | 11.356421 | 11.820583 | 11.3216 | 0.49901944 | 0.034858178 |
| Itgam | 9.679918 | 9.772961 | 8.7383 | 1.03463828 | 0.941595298 | 9.624275 | 9.0364796 | 8.53789 | 0.498593633 | 1.086389011 |
| Mrps18c | 8.022991 | 8.166828 | 8.105 | 0.061850156 | -0.081987131 | 8.2417523 | 8.5758789 | 8.07757 | 0.498304672 | 0.16417808 |
| Mrpl32 | 8.41233 | 8.611274 | 8.5096 | 0.101635841 | -0.097307485 | 8.5438561 | 9.0166968 | 8.51843 | 0.498266502 | 0.025425806 |
| Aimp2 | 8.409263 | 8.440884 | 8.2794 | 0.16147462 | 0.129853509 | 8.6325416 | 8.8566943 | 8.35846 | 0.498232328 | 0.274079582 |
| Ssbp1 | 8.742402 | 8.806581 | 8.7271 | 0.079493962 | 0.015315314 | 8.938333 | 9.3386065 | 8.84043 | 0.498172173 | 0.097898667 |
| Urm1 | 8.512179 | 8.600523 | 8.6431 | -0.042557419 | -0.1309015 | 8.9171204 | 9.015936 | 8.51834 | 0.497596919 | 0.398781304 |
| Zmiz2 | 11.84676 | 11.98583 | 11.899 | 0.087191584 | -0.051876818 | 12.257658 | 12.48482 | 11.9875 | 0.497336538 | 0.270174338 |
| Rrp12 | 9.479043 | 9.644956 | 9.3236 | 0.321386063 | 0.155473476 | 9.712592 | 9.9397368 | 9.44265 | 0.497085178 | 0.269940342 |
| Creb3 | 8.991014 | 8.996868 | 8.9701 | 0.026760734 | 0.020906156 | 9.0996705 | 9.4572058 | 8.96042 | 0.49678398 | 0.139248686 |
| Rrp7a | 9.694121 | 9.957041 | 9.8287 | 0.128300238 | -0.134619796 | 10.109553 | 10.459512 | 9.96284 | 0.496672776 | 0.146713647 |
| Heatr1 | 11.20466 | 11.3088 | 11.087 | 0.222123232 | 0.11798144 | 11.268793 | 11.71631 | 11.22 | 0.496298362 | 0.048781138 |

| | | | | | | | | | | |
|---|---|---|---|---|---|---|---|---|---|---|
| Rasgrp4 | 7.859801 | 7.918194 | 7.9177 | 0.000506075 | -0.057887027 | 8.4042066 | 8.3912641 | 7.89516 | 0.496103135 | 0.509045583 |
| A130010J15Rik | 7.063962 | 7.008876 | 7.0145 | -0.005607423 | 0.049478556 | 7.1454182 | 7.3854521 | 6.88951 | 0.495940282 | 0.255906394 |
| Clpb | 8.412395 | 8.618771 | 8.3509 | 0.267918266 | 0.061542384 | 8.4405784 | 8.7364687 | 8.2409 | 0.495565082 | 0.199674792 |
| Psmd3 | 10.47313 | 10.52986 | 10.417 | 0.113324322 | 0.056593158 | 10.70231 | 10.893499 | 10.3979 | 0.495550428 | 0.304361521 |
| Prrg4 | 6.072335 | 6.309212 | 6.1364 | 0.172809288 | -0.064068095 | 5.7174972 | 6.5643918 | 6.06997 | 0.494419678 | -0.352474939 |
| Tpst2 | 9.633107 | 9.701235 | 9.6663 | 0.034896516 | -0.033231388 | 9.6132783 | 10.202977 | 9.70867 | 0.494310736 | -0.09538749 |
| Ighv2-3 | 5.945661 | 6.076177 | 5.7879 | 0.288302075 | 0.15778644 | 5.8675723 | 6.1405182 | 5.64623 | 0.494284575 | 0.221338686 |
| Slc35b1 | 8.987296 | 8.968129 | 8.8652 | 0.102946508 | 0.122113999 | 9.2030585 | 9.4003034 | 8.9063 | 0.493999003 | 0.296754102 |
| Ipo4 | 9.706844 | 9.754507 | 9.4487 | 0.305793146 | 0.258130404 | 9.6158246 | 10.171437 | 9.67747 | 0.493968437 | -0.061643994 |
| Carhsp1 | 8.812435 | 8.853356 | 8.7255 | 0.127831845 | 0.086911181 | 8.6563232 | 9.0990989 | 8.60525 | 0.493846901 | 0.051071226 |
| Mettl1 | 8.463443 | 8.530621 | 8.3389 | 0.191707886 | 0.124529782 | 8.8504656 | 9.0283539 | 8.53459 | 0.493759782 | 0.315871471 |
| Fam84b | 8.940782 | 8.963137 | 8.9909 | -0.027769502 | -0.050123922 | 8.8038093 | 9.3330326 | 8.83936 | 0.493670559 | -0.035552748 |
| Igkc | 14.47074 | 14.97362 | 14.623 | 0.350202762 | -0.152680562 | 15.02838 | 15.112357 | 14.6192 | 0.493206338 | 0.409228831 |
| Mrpl43 | 8.556142 | 8.673942 | 8.6409 | 0.033061987 | -0.08473793 | 8.7906926 | 8.9696043 | 8.47658 | 0.49302233 | 0.314110605 |
| Dnpep | 8.887939 | 8.895073 | 8.9721 | -0.07707151 | -0.084205135 | 9.1934074 | 9.2268332 | 8.73386 | 0.492977811 | 0.459552042 |
| Gm10705 | 6.89445 | 6.947194 | 6.8278 | 0.119386157 | 0.066641939 | 6.9349783 | 7.3619859 | 6.86938 | 0.492607233 | 0.06559962 |
| Hif1a | 11.4373 | 11.42035 | 11.281 | 0.139267756 | 0.156221322 | 11.204737 | 11.786604 | 11.294 | 0.49257353 | -0.089293797 |
| Mrps7 | 8.9287 | 9.025553 | 8.896 | 0.129582553 | 0.032730189 | 9.0790129 | 9.4216409 | 8.92934 | 0.492300847 | 0.149672824 |
| Tbk1 | 10.14209 | 10.22894 | 10.118 | 0.111034213 | 0.024177556 | 10.195076 | 10.807359 | 10.3152 | 0.492191451 | -0.120091412 |
| Csrnp1 | 11.23382 | 11.30941 | 10.971 | 0.3381885 | 0.262599528 | 10.980828 | 11.307769 | 10.8156 | 0.492119404 | 0.165177908 |
| Alox5ap | 8.495706 | 8.344664 | 8.2086 | 0.136090609 | 0.287132475 | 9.0864419 | 8.7156499 | 8.22377 | 0.491876967 | 0.862669011 |
| Cacnb4 | 3.728181 | 3.769602 | 3.6447 | 0.124930484 | 0.083509589 | 3.5833869 | 4.2926809 | 3.80109 | 0.491593189 | -0.217700848 |
| Dtx2 | 8.797432 | 9.039285 | 8.8302 | 0.20904365 | -0.032809341 | 8.8979476 | 9.1401668 | 8.64859 | 0.491581481 | 0.249362255 |
| Treml4 | 5.03634 | 5.714584 | 5.1248 | 0.589758377 | -0.088485591 | 5.7918157 | 5.4468413 | 4.95534 | 0.491499214 | 0.836473579 |
| Ubl5 | 9.844983 | 9.795472 | 9.8144 | -0.018882536 | 0.030629095 | 10.077625 | 10.239606 | 9.74816 | 0.491447197 | 0.329466186 |
| Has2 | 4.965881 | 5.150216 | 4.5106 | 0.639660258 | 0.455324916 | 4.4301717 | 5.0600476 | 4.56866 | 0.491385144 | -0.13849079 |
| Tra2a | 10.47447 | 10.84761 | 10.504 | 0.343379502 | -0.029762122 | 10.640036 | 11.258721 | 10.7674 | 0.491364652 | -0.127320243 |
| Zfp219 | 7.832772 | 7.682106 | 7.821 | -0.138864227 | 0.011802331 | 7.5386378 | 8.1158867 | 7.62473 | 0.491153384 | -0.086095495 |
| Pomt2 | 6.915288 | 7.102676 | 6.8757 | 0.226953459 | 0.039564807 | 6.9851315 | 7.4915917 | 7.00075 | 0.490839964 | -0.015620236 |
| Eif5b | 11.38454 | 11.41127 | 11.272 | 0.139529546 | 0.112800728 | 11.258089 | 11.667525 | 11.177 | 0.490557544 | 0.081121575 |
| Cct8 | 11.77061 | 11.89176 | 11.703 | 0.188317948 | 0.067164578 | 11.798527 | 12.233665 | 11.7433 | 0.490341626 | 0.055203444 |
| Ubtd1 | 6.649159 | 7.009282 | 6.7057 | 0.303546859 | -0.056576359 | 6.9992729 | 7.1038184 | 6.61348 | 0.490336257 | 0.38579078 |

| | | | | | | | | | | |
|---|---|---|---|---|---|---|---|---|---|---|
| Acp2 | 8.81695 | 9.157162 | 8.6668 | 0.490319449 | 0.150107491 | 9.0797248 | 9.1063311 | 8.6161 | 0.490234175 | 0.463627886 |
| Loxl4 | 3.99623 | 4.005645 | 3.6328 | 0.372860552 | 0.363445806 | 3.9080517 | 4.0499857 | 3.55988 | 0.490102173 | 0.348168121 |
| Anpep | 8.671244 | 8.695641 | 8.5356 | 0.160072478 | 0.135674875 | 8.8573881 | 8.9927619 | 8.50276 | 0.490003701 | 0.354629947 |
| Stx12 | 8.833759 | 8.6785 | 8.5836 | 0.094867287 | 0.250126652 | 8.7575879 | 9.0238121 | 8.53404 | 0.489771443 | 0.223547286 |
| Irf5 | 11.0173 | 11.31837 | 11.048 | 0.26991783 | -0.031150636 | 11.312787 | 11.565923 | 11.0764 | 0.489545936 | 0.236410022 |
| Cdhr5 | 2.891667 | 2.923197 | 2.7916 | 0.131558225 | 0.100028651 | 2.7669268 | 3.2450827 | 2.75561 | 0.489469673 | 0.011313821 |
| Zfp653 | 7.949809 | 8.005215 | 7.9759 | 0.029284645 | -0.026121743 | 8.3209492 | 8.485666 | 7.9963 | 0.489367507 | 0.324650756 |
| Lmna | 7.910365 | 7.990308 | 7.4915 | 0.498842619 | 0.418899626 | 7.6181688 | 8.1261697 | 7.63681 | 0.48936279 | -0.018638096 |
| Batf | 8.548814 | 8.483194 | 8.2188 | 0.264433358 | 0.33005286 | 8.6006576 | 8.8491616 | 8.35995 | 0.489211406 | 0.240707433 |
| Pex14 | 8.33324 | 8.382277 | 8.3546 | 0.027685432 | -0.021351839 | 8.540402 | 8.8848315 | 8.39649 | 0.488344687 | 0.143915249 |
| Psmd7 | 9.806858 | 9.817399 | 9.7885 | 0.02892756 | 0.018386544 | 9.7779255 | 10.238464 | 9.75039 | 0.488074627 | 0.027536133 |
| 1600012H06Rik | 7.915324 | 7.928761 | 8.0324 | -0.10361304 | -0.117049957 | 8.1930758 | 8.3488379 | 7.8609 | 0.487941989 | 0.332179855 |
| Alg3 | 7.622724 | 7.845852 | 7.5486 | 0.29723422 | 0.074106496 | 7.9734098 | 8.1678401 | 7.68016 | 0.487682678 | 0.293252366 |
| Fam53c | 9.560865 | 9.866299 | 9.5009 | 0.365388891 | 0.059955075 | 9.6566183 | 10.037868 | 9.55046 | 0.487411199 | 0.106161045 |
| Tbl3 | 9.351564 | 9.490157 | 9.3501 | 0.140070588 | 0.001477726 | 9.68983 | 9.8751127 | 9.38775 | 0.487367655 | 0.302084908 |
| Gfra1 | 3.378113 | 3.82832 | 3.3793 | 0.449010724 | -0.001196335 | 3.4104101 | 3.7824852 | 3.29512 | 0.487366779 | 0.115291653 |
| Trib3 | 5.03325 | 5.161147 | 5.3978 | -0.236684801 | -0.364581143 | 5.6337924 | 5.8360744 | 5.34943 | 0.486643884 | 0.284361884 |
| Cass4 | 7.242539 | 6.768153 | 6.3367 | 0.431406333 | 0.905791807 | 6.6922703 | 6.8596288 | 6.37306 | 0.486564044 | 0.319205543 |
| Ly6d | 11.78108 | 12.17323 | 11.818 | 0.355082654 | -0.03706837 | 12.438574 | 12.221051 | 11.7349 | 0.486193808 | 0.703716982 |
| Med1 | 10.79811 | 11.08539 | 10.822 | 0.263875684 | -0.023401624 | 10.849749 | 11.433343 | 10.9475 | 0.485849256 | -0.097744347 |
| Ndufb2 | 7.609247 | 7.709562 | 7.602 | 0.107538526 | 0.007223713 | 7.924076 | 7.957732 | 7.47248 | 0.485256172 | 0.451600228 |
| Vmn1r197 | 4.828494 | 5.14899 | 4.8088 | 0.340181714 | 0.019685641 | 4.936447 | 5.5064983 | 5.02179 | 0.484711177 | -0.085340039 |
| Ppp1r14b | 8.561007 | 8.527912 | 8.5223 | 0.005657952 | 0.038752889 | 8.6275936 | 8.8563938 | 8.37177 | 0.484623493 | 0.255823325 |
| Cox15 | 8.990454 | 9.213593 | 9.0158 | 0.197837945 | -0.02530099 | 9.1741969 | 9.5545251 | 9.06993 | 0.484598806 | 0.104270574 |
| Safb | 11.15884 | 11.39916 | 11.255 | 0.1444047 | -0.095918494 | 11.325125 | 11.855187 | 11.3707 | 0.484507202 | -0.045554419 |
| Gimap1 | 10.95398 | 11.11138 | 11.019 | 0.092237922 | -0.065166098 | 11.222725 | 11.43292 | 10.9486 | 0.484361294 | 0.274166015 |
| Mea1 | 8.979135 | 9.106774 | 9.0552 | 0.051587407 | -0.076051745 | 9.2819203 | 9.4501627 | 8.96634 | 0.483826815 | 0.315584417 |
| Cndp2 | 10.26846 | 10.29364 | 10.048 | 0.245440602 | 0.220253488 | 10.282991 | 10.579818 | 10.096 | 0.483820767 | 0.186993821 |
| Sat1 | 11.39434 | 11.41795 | 11.337 | 0.080597922 | 0.05698808 | 11.254936 | 11.660266 | 11.1765 | 0.483803858 | 0.078474211 |
| Atp6v1g1 | 9.888216 | 9.929968 | 9.8665 | 0.063514783 | 0.02176275 | 10.035678 | 10.204065 | 9.7203 | 0.483765491 | 0.315378947 |
| Mrpl1 | 7.958349 | 8.056639 | 8.0432 | 0.013425355 | -0.084864777 | 8.0273148 | 8.5440604 | 8.06031 | 0.483754226 | -0.032991331 |
| Apba3 | 7.862673 | 8.112375 | 8.0257 | 0.086672545 | -0.163030106 | 8.2279371 | 8.5413822 | 8.05812 | 0.483263713 | 0.169818672 |

| | | | | | | | | | | |
|---|---|---|---|---|---|---|---|---|---|---|
| Dnah2 | 5.000793 | 5.134553 | 5.0399 | 0.094686415 | -0.039073818 | 5.3718839 | 5.303529 | 4.82035 | 0.483175984 | 0.551530839 |
| Ppp6r1 | 12.33307 | 12.50233 | 12.338 | 0.164556124 | -0.00471085 | 12.391139 | 12.781059 | 12.2983 | 0.482761596 | 0.092842199 |
| Adprm | 8.04113 | 8.155786 | 8.0809 | 0.074888969 | -0.039766633 | 8.30505 | 8.6545348 | 8.17198 | 0.482558539 | 0.133073717 |
| Cst7 | 8.974307 | 9.021087 | 8.9917 | 0.02941967 | -0.017360419 | 9.0392091 | 9.4479917 | 8.96598 | 0.48200776 | 0.073225122 |
| Alad | 7.980073 | 8.09245 | 7.9822 | 0.110227442 | -0.002149227 | 8.3811813 | 8.4722428 | 7.99088 | 0.481365706 | 0.390304195 |
| 1810037I17Rik | 8.865441 | 8.959635 | 8.8728 | 0.086804384 | -0.007389199 | 9.0994731 | 9.324093 | 8.84283 | 0.481267747 | 0.256647795 |
| Igkv12-98 | 6.103041 | 6.294718 | 6.2697 | 0.025001943 | -0.166675062 | 6.6717004 | 6.9957989 | 6.51464 | 0.481159232 | 0.157060734 |
| Pqlc2 | 7.218151 | 7.543079 | 7.2361 | 0.30693096 | -0.017996686 | 7.8628367 | 7.8621104 | 7.38105 | 0.481058107 | 0.481784346 |
| Acadl | 10.03931 | 10.05954 | 9.9889 | 0.070667559 | 0.050439646 | 9.9975371 | 10.465682 | 9.9847 | 0.480981021 | 0.012835912 |
| Mydgf | 8.69714 | 8.644624 | 8.5257 | 0.118903911 | 0.171419122 | 8.7117552 | 8.9507115 | 8.46994 | 0.480776216 | 0.241819916 |
| Prpf31 | 9.898536 | 9.961161 | 9.8433 | 0.117836527 | 0.055212341 | 10.119641 | 10.368355 | 9.88762 | 0.480738564 | 0.232024205 |
| Rab5if | 8.868255 | 8.989987 | 8.8664 | 0.123566311 | 0.001834411 | 8.9223242 | 9.2943995 | 8.81381 | 0.480589213 | 0.108513934 |
| Gnb4 | 8.370855 | 8.332716 | 8.4262 | -0.093529639 | -0.055390351 | 8.1206453 | 8.7912224 | 8.31067 | 0.480548443 | -0.190028642 |
| Lrp4 | 4.936844 | 5.140065 | 4.9866 | 0.153502505 | -0.049718842 | 4.9698907 | 5.3059856 | 4.82552 | 0.480464222 | 0.144369249 |
| Nuak2 | 9.761585 | 9.921339 | 9.7022 | 0.219179083 | 0.0594258 | 9.9465844 | 10.221446 | 9.74123 | 0.48021906 | 0.205357273 |
| Tmbim4 | 8.685287 | 8.629559 | 8.7028 | -0.073223024 | -0.017495108 | 8.7857412 | 9.1383394 | 8.6583 | 0.480038189 | 0.127439905 |
| Apaf1 | 9.976939 | 9.763674 | 9.7636 | 6.72802E-05 | 0.213331403 | 9.9065144 | 10.214205 | 9.73424 | 0.479964231 | 0.17227367 |
| Prpf38a | 8.920906 | 8.992479 | 8.8794 | 0.113030954 | 0.041457729 | 9.1336339 | 9.4620303 | 8.98213 | 0.479896107 | 0.151499693 |
| Ccdc124 | 8.463282 | 8.50752 | 8.317 | 0.190542157 | 0.146304135 | 8.6654734 | 8.8183795 | 8.33851 | 0.479872622 | 0.32696656 |
| Mrps22 | 7.229534 | 7.276597 | 7.1975 | 0.079072901 | 0.032010577 | 7.4272664 | 7.7495164 | 7.26971 | 0.479805266 | 0.157555285 |
| Eva1b | 6.455378 | 6.456835 | 6.4247 | 0.032139816 | 0.03068322 | 6.4062651 | 6.661416 | 6.18185 | 0.479564914 | 0.224414094 |
| Ube2f | 9.456846 | 9.549635 | 9.3685 | 0.181165767 | 0.088376998 | 9.5599171 | 9.7758859 | 9.2976 | 0.478288094 | 0.262319278 |
| Med29 | 7.201527 | 7.350822 | 7.1623 | 0.188531874 | 0.039236383 | 7.3511636 | 7.6345063 | 7.15626 | 0.478250616 | 0.194907865 |
| Lsm12 | 9.512299 | 9.546784 | 9.5077 | 0.039107487 | 0.004622551 | 9.7208826 | 9.9675604 | 9.48953 | 0.478028588 | 0.231350796 |
| Ran | 11.77316 | 11.82417 | 11.713 | 0.111067022 | 0.06005685 | 11.933173 | 12.189054 | 11.7112 | 0.477891466 | 0.222010555 |
| Eif4ebp1 | 7.213681 | 7.028095 | 7.1088 | -0.080729063 | 0.104857149 | 7.4773639 | 7.476828 | 6.99905 | 0.477774851 | 0.478310744 |
| B3gnt3 | 7.348655 | 7.100459 | 7.2522 | -0.151769618 | 0.096426622 | 6.861146 | 7.1901099 | 6.71237 | 0.477740175 | 0.148776274 |
| Ints5 | 9.082899 | 9.227415 | 9.0584 | 0.168973142 | 0.024457014 | 9.1780693 | 9.4226787 | 8.94551 | 0.477163891 | 0.23255451 |
| Trip6 | 6.67211 | 6.787357 | 6.5731 | 0.21430373 | 0.099057123 | 6.8843679 | 7.1864093 | 6.7093 | 0.477108069 | 0.175066598 |
| Fcf1 | 8.634143 | 8.808294 | 8.6587 | 0.149613328 | -0.024537973 | 8.9856036 | 9.2690771 | 8.79205 | 0.477023199 | 0.193549757 |
| Vegfa | 7.872501 | 7.713915 | 6.8608 | 0.853162 | 1.011747947 | 7.5021806 | 7.7511508 | 7.27428 | 0.476873771 | 0.227903615 |
| Adgrg7 | 1.491343 | 1.600537 | 1.4643 | 0.136222888 | 0.027028768 | 1.557495 | 1.9617073 | 1.48519 | 0.476519405 | 0.072307077 |

| | | | | | | | | | | |
|---|---|---|---|---|---|---|---|---|---|---|
| Pole4 | 9.851437 | 9.964227 | 9.8035 | 0.160700622 | 0.04791113 | 9.921619 | 10.300952 | 9.82462 | 0.476328167 | 0.096995252 |
| Cd53 | 13.31194 | 13.4226 | 13.201 | 0.221961684 | 0.111299278 | 13.492 | 13.701476 | 13.2256 | 0.475894126 | 0.266417731 |
| Fasn | 11.2935 | 11.31229 | 11.2 | 0.112028734 | 0.093231406 | 11.612307 | 11.834163 | 11.3587 | 0.475475818 | 0.253619247 |
| Padi4 | 4.570906 | 4.319412 | 4.179 | 0.14044507 | 0.391938849 | 4.29522 | 4.5710319 | 4.09557 | 0.475460549 | 0.199648654 |
| Manbal | 7.866243 | 7.934608 | 7.781 | 0.1535765 | 0.085210816 | 7.841796 | 8.1426806 | 7.66725 | 0.475426344 | 0.174541703 |
| 2010111I01Rik | 8.557447 | 8.592667 | 8.4702 | 0.122496655 | 0.08727734 | 8.601349 | 9.1212429 | 8.64586 | 0.475387225 | -0.044506647 |
| Slc12a9 | 8.473113 | 8.396413 | 8.3228 | 0.073652043 | 0.150352097 | 8.7164586 | 8.9752954 | 8.50007 | 0.475224818 | 0.216388085 |
| A430005L14Rik | 7.769968 | 7.835619 | 7.7908 | 0.044859155 | -0.020792068 | 7.9599495 | 8.1235357 | 7.64878 | 0.474759456 | 0.311173182 |
| Mrps18b | 8.80274 | 8.876151 | 8.7859 | 0.09028644 | 0.016876024 | 9.1653518 | 9.3731057 | 8.89859 | 0.474510972 | 0.266757024 |
| Dsc2 | 4.86059 | 4.835031 | 4.5401 | 0.294889826 | 0.320449067 | 5.3014329 | 4.9165925 | 4.44223 | 0.474361654 | 0.859202061 |
| Atp5k | 9.093305 | 9.093214 | 8.9612 | 0.132011164 | 0.132102338 | 9.2393493 | 9.5304423 | 9.05626 | 0.474183373 | 0.183090376 |
| Aire | 5.013329 | 5.084775 | 5.0618 | 0.02302401 | -0.04842211 | 5.3962011 | 5.8502817 | 5.37652 | 0.473761224 | 0.019680603 |
| Snrpf | 8.910717 | 8.802455 | 8.8189 | -0.016409557 | 0.091852105 | 8.9151755 | 9.2824267 | 8.80885 | 0.473579168 | 0.106327974 |
| Psma6 | 10.57186 | 10.56324 | 10.499 | 0.06418384 | 0.07280474 | 10.676691 | 10.990285 | 10.5171 | 0.473222518 | 0.159628925 |
| Aldh1l1 | 5.494603 | 4.951884 | 5.5327 | -0.580822442 | -0.038103092 | 4.9583705 | 5.6139829 | 5.14095 | 0.473030275 | -0.182582143 |
| Pi4k2a | 9.810076 | 9.972049 | 9.7954 | 0.17667876 | 0.014705547 | 9.8537641 | 10.286854 | 9.81393 | 0.472924872 | 0.039834497 |
| AC158774.1 | 3.77505 | 4.078596 | 3.8263 | 0.252247149 | -0.051299191 | 3.8488865 | 4.5378831 | 4.06541 | 0.472476964 | -0.216519592 |
| Sytl3 | 7.909848 | 8.050988 | 7.8802 | 0.170780236 | 0.029640136 | 7.5646266 | 8.2878436 | 7.81583 | 0.472015167 | -0.251201871 |
| Gm45799 | 6.823003 | 6.602353 | 6.5979 | 0.004464897 | 0.225115443 | 7.5129281 | 7.1010948 | 6.62929 | 0.471806155 | 0.883639465 |
| Pet100 | 7.912774 | 7.93568 | 8.006 | -0.07028108 | -0.093187039 | 8.3055125 | 8.4086227 | 7.93687 | 0.471754128 | 0.368643983 |
| Ptges | 4.038296 | 4.206434 | 3.9135 | 0.292935103 | 0.124796839 | 4.1289705 | 4.3968688 | 3.92522 | 0.471647924 | 0.203749633 |
| Ccn1 | 8.383585 | 8.185732 | 8.0972 | 0.088575721 | 0.286428993 | 7.3243559 | 7.8474161 | 7.37583 | 0.471583671 | -0.051476472 |
| Vmn1r184 | 3.913719 | 4.032295 | 3.9837 | 0.048586359 | -0.069989309 | 4.181136 | 4.6495315 | 4.17797 | 0.471565773 | 0.003170323 |
| Pfdn6 | 8.811774 | 8.841236 | 8.7436 | 0.09766277 | 0.068201479 | 9.0093482 | 9.3008409 | 8.82937 | 0.471472619 | 0.179979943 |
| Nos3 | 6.39043 | 6.547551 | 5.9772 | 0.570338935 | 0.413217417 | 5.9199064 | 6.2720367 | 5.80072 | 0.471319362 | 0.119189072 |
| Gdf15 | 4.090595 | 4.698747 | 3.748 | 0.950737784 | 0.342585609 | 3.7844138 | 4.1556054 | 3.68479 | 0.47081739 | 0.099625807 |
| Clip2 | 8.185076 | 8.23917 | 7.9909 | 0.248253949 | 0.194160525 | 8.0093871 | 8.6453697 | 8.17459 | 0.470782504 | -0.16520011 |
| Atp5g1 | 9.537715 | 9.737512 | 9.5956 | 0.14188417 | -0.057913252 | 9.9104509 | 10.015399 | 9.54491 | 0.470490861 | 0.365542461 |
| Tmem229b | 10.69056 | 10.94005 | 10.735 | 0.205133644 | -0.04435415 | 10.734732 | 11.078277 | 10.6085 | 0.469747722 | 0.126202795 |
| Cc2d1a | 9.227039 | 9.277759 | 9.2554 | 0.022345481 | -0.028374539 | 9.3600598 | 9.6004736 | 9.13139 | 0.469081829 | 0.228667965 |
| Nup153 | 11.84633 | 12.02003 | 11.832 | 0.18793504 | 0.0142351 | 11.83692 | 12.330149 | 11.8612 | 0.468960142 | -0.024268972 |
| Rab3ip | 8.70008 | 9.034572 | 8.7701 | 0.264425803 | -0.07006625 | 8.582199 | 9.2704295 | 8.80155 | 0.468878281 | -0.219352241 |

| | | | | | | | | | | |
|---|---|---|---|---|---|---|---|---|---|---|
| Rnf135 | 7.028769 | 7.123428 | 7.1681 | -0.04463701 | -0.139296555 | 7.312586 | 7.7790359 | 7.31028 | 0.468758724 | 0.002308781 |
| Luzp1 | 8.118431 | 8.084088 | 8.0498 | 0.034277998 | 0.068621233 | 7.4491671 | 8.4189329 | 7.95022 | 0.468708218 | -0.501057505 |
| Hsbp1 | 9.486168 | 9.47399 | 9.4804 | -0.006400065 | 0.005777774 | 9.664842 | 9.8769435 | 9.40825 | 0.468692885 | 0.256591378 |
| Igkv3-2 | 7.553965 | 8.191601 | 7.721 | 0.470603592 | -0.167032381 | 8.5449863 | 8.3488722 | 7.88024 | 0.468629284 | 0.664743345 |
| Trp53i11 | 11.22119 | 11.25141 | 11.069 | 0.18250799 | 0.152284554 | 11.424582 | 11.583679 | 11.1152 | 0.46845698 | 0.309360661 |
| Rab11a | 9.599103 | 9.469681 | 9.5395 | -0.069779912 | 0.059642312 | 9.3487608 | 9.7483872 | 9.27993 | 0.468453773 | 0.068827384 |
| Zfp593 | 7.715547 | 7.813868 | 7.6791 | 0.134799218 | 0.036478083 | 7.9276034 | 8.1250375 | 7.6568 | 0.468238197 | 0.270804051 |
| Tomm6 | 9.415093 | 9.511819 | 9.4005 | 0.111314561 | 0.014588631 | 9.787795 | 9.8087473 | 9.3406 | 0.468149896 | 0.447197619 |
| Lamtor5 | 8.887762 | 8.916739 | 8.8468 | 0.06989857 | 0.040921201 | 9.0184207 | 9.3613283 | 8.89357 | 0.467756109 | 0.124848574 |
| Nrros | 10.99432 | 11.34373 | 10.933 | 0.410235026 | 0.060820132 | 11.283996 | 11.442962 | 10.9753 | 0.467681048 | 0.308715095 |
| Nfix | 7.548842 | 7.64279 | 7.6009 | 0.041922149 | -0.052025586 | 7.2039765 | 7.7982235 | 7.33077 | 0.467450147 | -0.126796879 |
| Rsl24d1 | 9.700283 | 9.777821 | 9.609 | 0.168808161 | 0.091270819 | 9.818453 | 10.107267 | 9.63987 | 0.467399708 | 0.178585494 |
| Rexo4 | 9.800101 | 9.880894 | 9.8298 | 0.051086979 | -0.029706527 | 10.093305 | 10.380851 | 9.91348 | 0.467371018 | 0.1798248 |
| Rnasel | 8.996157 | 8.983515 | 9.0526 | -0.069125413 | -0.056483404 | 8.917401 | 9.4345073 | 8.96741 | 0.467102219 | -0.050004091 |
| Ncl | 13.76803 | 13.73786 | 13.731 | 0.006634572 | 0.03680808 | 13.651756 | 14.204563 | 13.7376 | 0.466997832 | -0.085809626 |
| Necap1 | 8.092524 | 8.259943 | 8.2193 | 0.040677168 | -0.126741476 | 8.3502308 | 8.6934807 | 8.22651 | 0.466968701 | 0.123718852 |
| Mrps18a | 8.24849 | 8.346659 | 8.2384 | 0.10827685 | 0.010107311 | 8.5471326 | 8.7509303 | 8.28398 | 0.466953355 | 0.263155741 |
| Ndufc2 | 8.608464 | 8.564273 | 8.5485 | 0.01575344 | 0.059945194 | 8.8378972 | 8.8995642 | 8.43293 | 0.466634008 | 0.404966995 |
| Fam207a | 8.938394 | 9.030133 | 8.8501 | 0.180052332 | 0.088312846 | 8.8632762 | 9.1837675 | 8.7173 | 0.466466868 | 0.145975594 |
| Chpt1 | 8.219705 | 8.071947 | 8.1993 | -0.127360113 | 0.020397754 | 8.3701164 | 8.5207812 | 8.05447 | 0.466307296 | 0.315642461 |
| Fkbpl | 5.472241 | 5.241269 | 5.2347 | 0.00656385 | 0.237536161 | 5.2984779 | 5.6231569 | 5.1569 | 0.466254296 | 0.141575286 |
| Yrdc | 8.600204 | 8.632462 | 8.5657 | 0.066776646 | 0.034517918 | 8.5712709 | 8.9452689 | 8.47917 | 0.466099964 | 0.092102017 |
| Cds1 | 4.824322 | 4.911948 | 4.6451 | 0.26683225 | 0.179206519 | 4.8256324 | 5.0679295 | 4.60197 | 0.465957278 | 0.223660164 |
| Atp8a2 | 5.429549 | 5.498718 | 5.2578 | 0.240920936 | 0.171751707 | 5.3724377 | 5.9528925 | 5.48694 | 0.46595481 | -0.114499992 |
| Emc8 | 8.112034 | 8.324833 | 8.3456 | -0.020720448 | -0.233520046 | 8.451301 | 8.6818072 | 8.21596 | 0.465845698 | 0.235339536 |
| Golt1b | 8.205964 | 8.31165 | 8.1085 | 0.20310015 | 0.097414287 | 8.0588701 | 8.4810266 | 8.01551 | 0.465514298 | 0.04335778 |
| Znhit2 | 6.675206 | 6.731811 | 6.6566 | 0.07519479 | 0.018589458 | 7.0358971 | 7.0172712 | 6.55177 | 0.465504976 | 0.484130822 |
| Ptpn1 | 11.65631 | 11.64842 | 11.4 | 0.248794968 | 0.256690236 | 11.506901 | 11.808863 | 11.3435 | 0.465374458 | 0.163412638 |
| Csf2 | 3.353751 | 3.87168 | 3.351 | 0.520700332 | 0.002771364 | 3.3070005 | 3.8349907 | 3.36964 | 0.465347828 | -0.062642383 |
| Rbks | 6.278737 | 6.197789 | 6.2033 | -0.005556793 | 0.075390918 | 6.5596262 | 6.6699563 | 6.20466 | 0.465293075 | 0.354962978 |
| Wrap73 | 7.957788 | 8.087967 | 8.1019 | -0.01390208 | -0.144081194 | 8.1858735 | 8.4000821 | 7.93529 | 0.464789658 | 0.250581087 |
| Gm47626 | 6.043564 | 6.020097 | 5.7336 | 0.286539022 | 0.310006487 | 6.0199656 | 6.6870816 | 6.22232 | 0.464760311 | -0.202355661 |

| | | | | | | | | | | |
|---|---|---|---|---|---|---|---|---|---|---|
| Sra1 | 9.087175 | 9.156627 | 9.0713 | 0.085308022 | 0.015856131 | 9.2982504 | 9.5000295 | 9.03534 | 0.464688412 | 0.262909316 |
| Ndufb9 | 8.956751 | 8.900297 | 8.8391 | 0.061244872 | 0.117698941 | 9.0432813 | 9.2729753 | 8.80887 | 0.464105097 | 0.234411169 |
| Mettl2 | 8.430488 | 8.435391 | 8.3359 | 0.099537209 | 0.094634003 | 8.8080731 | 8.9744258 | 8.51057 | 0.463858056 | 0.297505315 |
| Pno1 | 8.662326 | 8.689487 | 8.547 | 0.142441008 | 0.115280259 | 8.8150961 | 8.9893189 | 8.52573 | 0.463584263 | 0.289361418 |
| Cox7a2 | 9.58807 | 9.558637 | 9.5574 | 0.001207946 | 0.030641048 | 9.7841727 | 10.086193 | 9.62281 | 0.463386646 | 0.161365837 |
| Timm8a1 | 7.958631 | 7.879095 | 7.8075 | 0.071627523 | 0.151163151 | 8.0892545 | 8.2048807 | 7.74161 | 0.463270818 | 0.347644638 |
| Fads3 | 6.040229 | 6.084098 | 5.8802 | 0.203861581 | 0.159992811 | 5.6406936 | 5.7409956 | 5.27783 | 0.463170004 | 0.362868001 |
| Tcf4 | 10.77268 | 11.12335 | 10.802 | 0.320953044 | -0.029716382 | 10.938917 | 11.250746 | 10.7887 | 0.462081276 | 0.150252266 |
| Mapkapk2 | 11.11542 | 11.15598 | 10.902 | 0.25354766 | 0.2129853 | 11.083671 | 11.321356 | 10.8593 | 0.462040408 | 0.224355794 |
| Ndufs4 | 9.069308 | 9.148004 | 9.0244 | 0.123566973 | 0.044870977 | 9.2566419 | 9.6293175 | 9.16729 | 0.462031012 | 0.089355419 |
| Dvl1 | 9.703374 | 9.757318 | 9.6828 | 0.074511718 | 0.020568177 | 10.012461 | 10.236739 | 9.77483 | 0.461908056 | 0.237630542 |
| Filip1l | 11.247 | 11.63209 | 11.351 | 0.280918234 | -0.104168424 | 11.545346 | 11.860942 | 11.399 | 0.461896658 | 0.146300098 |
| Birc2 | 10.8744 | 10.87852 | 10.793 | 0.085290132 | 0.081170432 | 10.60197 | 11.284903 | 10.8231 | 0.461832036 | -0.221100253 |
| Slc44a5 | 3.443408 | 3.705135 | 3.5547 | 0.150476645 | -0.111250719 | 3.5710341 | 4.2332463 | 3.77145 | 0.46179327 | -0.200418985 |
| Pdcd2l | 7.712212 | 7.781722 | 7.7217 | 0.060041872 | -0.009468459 | 7.9841492 | 8.1865491 | 7.72482 | 0.461728616 | 0.259328696 |
| Tomm5 | 8.473126 | 8.449969 | 8.5225 | -0.072518982 | -0.049362062 | 8.7511288 | 8.8542232 | 8.39299 | 0.461229249 | 0.358134842 |
| Misp | 1.617824 | 1.640327 | 1.6245 | 0.015839013 | -0.006664126 | 1.5877159 | 2.0946065 | 1.63338 | 0.46122282 | -0.045667827 |
| Mlxip | 11.25155 | 11.46958 | 11.257 | 0.212552272 | -0.005478742 | 11.184351 | 11.901247 | 11.4401 | 0.461191274 | -0.25570392 |
| Spryd7 | 7.19583 | 7.100515 | 7.1007 | -0.000225041 | 0.09509023 | 7.1901115 | 7.4898999 | 7.02939 | 0.46051203 | 0.160723658 |
| Ndufaf4 | 8.15239 | 8.004815 | 8.0612 | -0.056375059 | 0.091199971 | 8.2218157 | 8.5238217 | 8.06336 | 0.460465826 | 0.158459836 |
| Gm8369 | 10.7784 | 10.93782 | 10.879 | 0.058350238 | -0.101067956 | 10.839945 | 11.184837 | 10.7244 | 0.460461482 | 0.115569998 |
| Mrps25 | 7.765662 | 8.082972 | 7.718 | 0.364930607 | 0.047620762 | 7.9180976 | 8.2165849 | 7.75616 | 0.460421722 | 0.161934408 |
| Gpatch3 | 6.837919 | 7.140129 | 7.2354 | -0.095268987 | -0.39747892 | 7.206957 | 7.5372255 | 7.07712 | 0.460106777 | 0.129838241 |
| Mad2l2 | 7.451055 | 7.637247 | 7.5421 | 0.095098363 | -0.091093754 | 7.904462 | 8.1208922 | 7.66097 | 0.459917473 | 0.243487252 |
| Minpp1 | 8.882353 | 9.027372 | 8.9569 | 0.07049439 | -0.074524312 | 8.7804043 | 9.3024903 | 8.84261 | 0.459883619 | -0.062202432 |
| Vav3 | 8.551025 | 8.477354 | 8.2416 | 0.235797598 | 0.309469171 | 8.2352791 | 8.8552575 | 8.39587 | 0.459386708 | -0.160591658 |
| Fmc1 | 6.578992 | 6.601019 | 6.4977 | 0.103344751 | 0.0813173 | 6.7894178 | 6.9771645 | 6.518 | 0.459166175 | 0.271419459 |
| Tmem128 | 9.2613 | 9.294932 | 9.302 | -0.007018838 | -0.040651048 | 9.4648832 | 9.8068954 | 9.34803 | 0.458866007 | 0.116853831 |
| Smug1 | 6.90749 | 6.802822 | 6.8369 | -0.034079121 | 0.070588849 | 7.1235316 | 7.3677131 | 6.90896 | 0.458755465 | 0.21457393 |
| Xbp1 | 10.80584 | 10.83206 | 10.512 | 0.320513984 | 0.294300506 | 10.68538 | 10.855811 | 10.3971 | 0.458683594 | 0.288252494 |
| Coa3 | 8.558667 | 8.694257 | 8.5556 | 0.138609685 | 0.003019121 | 8.9074323 | 8.9641037 | 8.50602 | 0.458088424 | 0.401417049 |
| Atp5d | 10.37241 | 10.48488 | 10.373 | 0.112048738 | -0.00042001 | 10.788751 | 10.775843 | 10.3178 | 0.458076635 | 0.470984378 |

| | | | | | | | | | | |
|---|---|---|---|---|---|---|---|---|---|---|
| Sec24d | 8.685436 | 8.665969 | 8.3168 | 0.349217139 | 0.368684285 | 8.091064 | 8.7829716 | 8.32542 | 0.45755405 | -0.234353572 |
| Shq1 | 7.08432 | 7.230778 | 7.0308 | 0.199997417 | 0.053538812 | 7.0818026 | 7.2801198 | 6.82271 | 0.457412805 | 0.259095661 |
| Tbcb | 8.563237 | 8.587358 | 8.6275 | -0.040122016 | -0.064243589 | 8.839541 | 8.913539 | 8.45616 | 0.457377853 | 0.383379847 |
| Ndufb6 | 8.202415 | 8.278077 | 8.1445 | 0.133602517 | 0.057940617 | 8.4856592 | 8.5666602 | 8.10978 | 0.456884227 | 0.375883277 |
| Thoc6 | 8.344937 | 8.384352 | 8.2877 | 0.096697252 | 0.057282194 | 8.723515 | 8.8686983 | 8.41212 | 0.45657608 | 0.311392838 |
| Mtx1 | 8.289256 | 8.316823 | 8.3405 | -0.023664285 | -0.051231761 | 8.6540808 | 8.8704692 | 8.4139 | 0.456568671 | 0.240180325 |
| Rpl29 | 13.17545 | 13.25145 | 13.182 | 0.069446854 | -0.00655992 | 13.387193 | 13.524245 | 13.0678 | 0.45647763 | 0.319425643 |
| Siglec1 | 9.797111 | 8.53845 | 9.417 | -0.878592882 | 0.380068398 | 8.406337 | 8.9568406 | 8.50083 | 0.456005863 | -0.094497761 |
| Gm5150 | 4.056411 | 4.246069 | 3.6544 | 0.591703155 | 0.402045262 | 4.8556516 | 4.2212615 | 3.76534 | 0.455921464 | 1.090311548 |
| D17H6S53E | 7.873853 | 7.922972 | 7.8712 | 0.051750654 | 0.00263102 | 7.9333823 | 8.1136349 | 7.65772 | 0.455915729 | 0.275663192 |
| Olfr986 | 6.641181 | 7.074455 | 6.8565 | 0.217957249 | -0.215316414 | 7.0250729 | 7.6831957 | 7.22746 | 0.455735531 | -0.202387256 |
| Igkv13-84 | 7.353324 | 7.713035 | 7.5177 | 0.195337935 | -0.16437339 | 8.0562948 | 7.9690583 | 7.51367 | 0.455392875 | 0.542629306 |
| Tubb2a | 8.194436 | 8.304251 | 8.3368 | -0.032501355 | -0.142317072 | 8.3829644 | 8.65026 | 8.1952 | 0.455055193 | 0.187759512 |
| Mrpl20 | 8.499805 | 8.387996 | 8.3628 | 0.025239825 | 0.137048598 | 8.4960576 | 8.702351 | 8.24763 | 0.45472441 | 0.24843099 |
| Tmem101 | 6.529647 | 6.73449 | 6.6466 | 0.087882362 | -0.116960106 | 6.8318074 | 6.9144702 | 6.45982 | 0.454649745 | 0.371986996 |
| Serinc3 | 14.3132 | 14.45075 | 14.325 | 0.125952906 | -0.011599064 | 14.24996 | 14.581852 | 14.1275 | 0.454380626 | 0.122488551 |
| Map3k6 | 6.139768 | 5.911283 | 5.482 | 0.429322239 | 0.657807713 | 5.2844943 | 5.5586056 | 5.10466 | 0.453946123 | 0.179834741 |
| Ccdc88b | 10.73762 | 10.79068 | 10.667 | 0.124166126 | 0.071105432 | 10.617276 | 11.15964 | 10.7058 | 0.453882846 | -0.088480795 |
| Ier5l | 5.049137 | 4.880924 | 4.8699 | 0.010986142 | 0.179198805 | 4.6913672 | 4.9501837 | 4.49672 | 0.453460258 | 0.194643746 |
| Smyd5 | 9.080496 | 9.252611 | 9.0968 | 0.155794652 | -0.016320266 | 9.4055035 | 9.7865497 | 9.33312 | 0.453434412 | 0.072388154 |
| Prol1 | 6.45922 | 5.242911 | 5.7726 | -0.529730019 | 0.686579063 | 6.6905887 | 5.6853605 | 5.23204 | 0.453320848 | 1.458548996 |
| Ntmt1 | 6.999692 | 6.968806 | 6.8466 | 0.122162861 | 0.153048151 | 7.2008295 | 7.3730942 | 6.91992 | 0.453173462 | 0.280908731 |
| Nacc1 | 10.57256 | 10.70396 | 10.549 | 0.15515205 | 0.023752554 | 10.646913 | 10.928333 | 10.4759 | 0.45244655 | 0.171026056 |
| Cdh11 | 5.733064 | 5.682423 | 5.2091 | 0.473334083 | 0.523974867 | 5.6774918 | 5.5051963 | 5.05297 | 0.45222931 | 0.624524807 |
| Rhog | 10.86583 | 11.08627 | 10.964 | 0.122729072 | -0.097707108 | 11.323292 | 11.414993 | 10.9629 | 0.452109048 | 0.36040767 |
| Psap | 14.4369 | 14.51456 | 14.315 | 0.19939902 | 0.121741806 | 14.381439 | 14.533325 | 14.0814 | 0.451911506 | 0.30002484 |
| Nsmaf | 10.88356 | 10.9811 | 10.995 | -0.014133264 | -0.111679478 | 11.047574 | 11.532972 | 11.0814 | 0.451557788 | -0.033839977 |
| Man2a1 | 11.3057 | 11.45677 | 11.174 | 0.282328668 | 0.13126426 | 11.240622 | 11.619641 | 11.1688 | 0.450857746 | 0.071838982 |
| Ebna1bp2 | 9.664255 | 9.693478 | 9.5336 | 0.159899285 | 0.130676567 | 9.7815508 | 10.096026 | 9.64517 | 0.450857009 | 0.136381758 |
| Tm2d3 | 7.523929 | 7.621345 | 7.4931 | 0.128240913 | 0.030824287 | 7.849616 | 7.946024 | 7.49523 | 0.450790816 | 0.354382887 |
| Ece1 | 10.45586 | 10.27868 | 10.303 | -0.02392956 | 0.153251997 | 10.074819 | 10.192887 | 9.74249 | 0.450396555 | 0.332329405 |
| Mzb1 | 9.85913 | 10.05615 | 9.8859 | 0.170219486 | -0.026795567 | 10.316586 | 10.260865 | 9.81076 | 0.45010445 | 0.505825236 |

| | | | | | | | | | | |
|---|---|---|---|---|---|---|---|---|---|---|
| Traf1 | 11.06283 | 11.29306 | 11.164 | 0.1290733 | -0.101163234 | 11.292888 | 11.8222 | 11.3722 | 0.44997856 | -0.079333357 |
| Pxn | 10.68014 | 10.77748 | 10.598 | 0.179930748 | 0.08258833 | 10.600351 | 11.037182 | 10.5872 | 0.449967678 | 0.013136348 |
| Fbxl19 | 7.947923 | 7.958374 | 7.9019 | 0.056500203 | 0.046049892 | 7.7645742 | 8.2054885 | 7.75599 | 0.449500943 | 0.008586666 |
| Milr1 | 7.851782 | 7.988973 | 7.8003 | 0.188720712 | 0.051529339 | 7.9646742 | 8.3022415 | 7.8531 | 0.449143851 | 0.111576553 |
| Rnh1 | 10.52615 | 10.45903 | 10.375 | 0.083986566 | 0.151114846 | 10.667144 | 10.805079 | 10.3568 | 0.44823049 | 0.310295983 |
| Nlrp1a | 4.522442 | 4.732114 | 4.181 | 0.551126874 | 0.34145538 | 4.7770701 | 4.9075049 | 4.45936 | 0.448143575 | 0.317708725 |
| Ccdc25 | 8.96819 | 9.059659 | 8.9535 | 0.106164362 | 0.014694903 | 8.969289 | 9.4127026 | 8.96466 | 0.448037678 | 0.004624138 |
| Rnpep | 9.409319 | 9.513896 | 9.4092 | 0.104729016 | 0.000152401 | 9.7218919 | 9.9480507 | 9.50018 | 0.447868869 | 0.22171003 |
| Pdlim2 | 9.739613 | 9.825433 | 9.8015 | 0.023963427 | -0.061856368 | 10.000112 | 10.146847 | 9.69907 | 0.447776293 | 0.301041637 |
| Mrpl46 | 7.811939 | 7.86743 | 7.8487 | 0.01872759 | -0.036763302 | 8.0241118 | 8.2412438 | 7.79348 | 0.4477657 | 0.230633665 |
| Aprt | 9.123183 | 9.28827 | 9.2777 | 0.010521051 | -0.154565734 | 9.5378026 | 9.6082085 | 9.16046 | 0.447747079 | 0.37734117 |
| Fpr1 | 4.320078 | 4.133732 | 3.435 | 0.698772006 | 0.885117436 | 3.8800919 | 4.0527967 | 3.60529 | 0.447503398 | 0.274798619 |
| Rrp36 | 8.312552 | 8.447625 | 8.3935 | 0.054107416 | -0.080965191 | 8.5889409 | 8.7061967 | 8.25914 | 0.447055033 | 0.329799243 |
| Pdcd2 | 8.343378 | 8.404669 | 8.2371 | 0.167607673 | 0.106315948 | 8.5035677 | 8.6762397 | 8.22955 | 0.446692964 | 0.274020982 |
| Csnk1e | 8.654971 | 8.903137 | 8.577 | 0.3261139 | 0.077947602 | 8.689886 | 9.0802743 | 8.63389 | 0.446384445 | 0.055996132 |
| Mtln | 6.323591 | 6.312743 | 6.26 | 0.052781793 | 0.063630082 | 6.579128 | 6.5136876 | 6.06739 | 0.44629351 | 0.511733869 |
| Exosc4 | 7.743611 | 7.687815 | 7.5971 | 0.09070399 | 0.146500147 | 7.9247083 | 8.0282701 | 7.58198 | 0.446292329 | 0.342730467 |
| Trem3 | 3.733992 | 4.063931 | 3.3735 | 0.690388986 | 0.360449855 | 3.9793959 | 3.8369452 | 3.39066 | 0.446285319 | 0.588736102 |
| Chchd4 | 7.558923 | 7.708902 | 7.4796 | 0.229274599 | 0.079296307 | 7.7158921 | 7.9500449 | 7.50386 | 0.446186552 | 0.212033788 |
| Plekhf2 | 9.888157 | 10.11068 | 9.8403 | 0.2704178 | 0.04789914 | 9.8756114 | 10.185954 | 9.73999 | 0.445961457 | 0.135618507 |
| Sertad2 | 9.71328 | 9.609456 | 9.5848 | 0.024700239 | 0.128523917 | 9.448632 | 9.8863477 | 9.4406 | 0.445745534 | 0.008029751 |
| Katnb1 | 8.269715 | 8.2627 | 8.3928 | -0.130105384 | -0.123089694 | 8.571673 | 8.7206299 | 8.27504 | 0.445588681 | 0.296631799 |
| Mrpl34 | 6.853916 | 6.992542 | 6.8465 | 0.146047754 | 0.007422259 | 6.9985461 | 7.1822203 | 6.73684 | 0.445384598 | 0.261710348 |
| Igkv8-27 | 7.59621 | 7.665954 | 7.4488 | 0.217121006 | 0.147377415 | 7.931687 | 7.9361595 | 7.491 | 0.445157954 | 0.440685435 |
| Znhit6 | 8.139699 | 8.242412 | 8.1061 | 0.1363125 | 0.033599501 | 8.2192595 | 8.6423075 | 8.19715 | 0.445155572 | 0.02210753 |
| Col5a3 | 3.933874 | 4.169663 | 3.9834 | 0.186276619 | -0.049512512 | 4.3404055 | 4.4345422 | 3.98987 | 0.444668333 | 0.3505316 |
| C1qc | 8.804896 | 8.327486 | 8.3901 | -0.062570769 | 0.414839927 | 7.8009691 | 8.2252273 | 7.78075 | 0.44447904 | 0.02022081 |
| Opa3 | 9.09602 | 9.212716 | 9.0652 | 0.147536938 | 0.030841201 | 9.0621523 | 9.4643712 | 9.01995 | 0.444421157 | 0.04220223 |
| Ufm1 | 9.332128 | 9.599324 | 9.1883 | 0.411066367 | 0.143870911 | 9.5181361 | 9.7782603 | 9.3339 | 0.444355838 | 0.184231617 |
| Chrm4 | 2.848338 | 2.839789 | 2.8412 | -0.001395309 | 0.007153999 | 2.8298901 | 3.2247372 | 2.78059 | 0.444150598 | 0.049303538 |
| Sertad1 | 9.017631 | 8.770487 | 8.8955 | -0.125010649 | 0.122133968 | 8.4308396 | 8.7865769 | 8.34259 | 0.443982508 | 0.08824527 |
| Gpr146 | 9.236861 | 8.99964 | 8.9341 | 0.065515965 | 0.302737256 | 8.8804737 | 9.2170883 | 8.77365 | 0.443435748 | 0.10682116 |

| | | | | | | | | | | |
|---|---|---|---|---|---|---|---|---|---|---|
| Usp15 | 10.4692 | 10.52992 | 10.492 | 0.037911612 | -0.022804826 | 10.389167 | 10.915435 | 10.4721 | 0.443286236 | -0.082982262 |
| Fastk | 9.208325 | 9.26691 | 9.275 | -0.008058461 | -0.066643095 | 9.7314442 | 9.8025905 | 9.35934 | 0.443251193 | 0.372104947 |
| Cox7c | 9.840492 | 9.847298 | 9.7861 | 0.061188978 | 0.054382556 | 10.128661 | 10.186634 | 9.74341 | 0.443222885 | 0.385250809 |
| Nutf2-ps1 | 8.798824 | 9.020025 | 8.8739 | 0.146105797 | -0.075094891 | 9.0039362 | 9.3106689 | 8.86781 | 0.442860023 | 0.136127296 |
| Bbc3 | 6.727024 | 6.942216 | 6.8816 | 0.060662475 | -0.154529776 | 6.3375015 | 6.9913673 | 6.54857 | 0.442798099 | -0.211067743 |
| Gale | 6.99216 | 6.934582 | 6.9297 | 0.004921348 | 0.062499355 | 7.3542734 | 7.4514098 | 7.00863 | 0.442780026 | 0.345643646 |
| Cox7b | 10.22676 | 10.12397 | 9.9772 | 0.146728567 | 0.249524134 | 10.452309 | 10.712394 | 10.2697 | 0.442665418 | 0.182580829 |
| Nit2 | 7.802259 | 7.945132 | 7.9204 | 0.024762628 | -0.118110089 | 7.8512853 | 8.3051029 | 7.86264 | 0.442461098 | -0.011356511 |
| Ube2n | 10.1577 | 10.30324 | 10.124 | 0.178931798 | 0.033391518 | 10.407757 | 10.653986 | 10.2119 | 0.44206945 | 0.195839773 |
| Cuta | 8.848664 | 8.860165 | 8.8971 | -0.036918004 | -0.048419739 | 9.2546687 | 9.2877251 | 8.84648 | 0.441243129 | 0.408186795 |
| Tmed5 | 10.99788 | 11.09402 | 10.812 | 0.28153508 | 0.185390284 | 10.911954 | 11.295345 | 10.8543 | 0.441022704 | 0.057631494 |
| Nfxl1 | 7.597216 | 7.686248 | 7.6424 | 0.04385448 | -0.045177361 | 7.44006 | 8.0123502 | 7.57195 | 0.440399104 | -0.131891139 |
| Ino80b | 7.911796 | 7.855555 | 7.7905 | 0.065022567 | 0.121262984 | 8.0017574 | 8.1389307 | 7.69854 | 0.440390819 | 0.303217539 |
| Scrib | 10.20582 | 10.32908 | 10.22 | 0.108934198 | -0.014327971 | 10.653773 | 10.786418 | 10.3462 | 0.440253556 | 0.30760856 |
| Stx11 | 8.75491 | 8.888959 | 8.4794 | 0.409585914 | 0.275537176 | 8.5055861 | 9.1556362 | 8.71539 | 0.440248014 | -0.209802095 |
| Pld3 | 10.57302 | 10.77951 | 10.539 | 0.240082224 | 0.033589078 | 10.594574 | 10.952211 | 10.5122 | 0.440002076 | 0.082365633 |
| Dnaja2 | 11.30903 | 11.39858 | 11.301 | 0.097215426 | 0.007672214 | 11.27348 | 11.689145 | 11.2501 | 0.439060396 | 0.023395466 |
| Ppp1r11 | 9.361463 | 9.529821 | 9.3999 | 0.129936328 | -0.03842135 | 9.6859816 | 9.8873814 | 9.44838 | 0.439004515 | 0.237604755 |
| Ddx54 | 11.17284 | 11.40117 | 11.139 | 0.261784884 | 0.033452898 | 11.637233 | 11.583048 | 11.1445 | 0.438595094 | 0.49277959 |
| Fkbp2 | 8.325348 | 8.394107 | 8.1906 | 0.203517697 | 0.134759544 | 8.5557204 | 8.6553532 | 8.21681 | 0.438542005 | 0.338909198 |
| Nubp2 | 9.205169 | 9.271321 | 9.2284 | 0.042903851 | -0.023248001 | 9.5949612 | 9.656751 | 9.21839 | 0.438363442 | 0.3765737 |
| Zfpl1 | 7.323069 | 7.270954 | 7.3349 | -0.063924917 | -0.011810383 | 7.5672197 | 7.6480867 | 7.20975 | 0.438336375 | 0.357469325 |
| Insig1 | 9.257637 | 9.054172 | 9.059 | -0.004841148 | 0.198623504 | 9.0883239 | 9.1491023 | 8.71115 | 0.437952702 | 0.377174245 |
| Clmp | 4.439455 | 4.575622 | 4.3435 | 0.232126078 | 0.095958588 | 4.4652382 | 4.789678 | 4.35173 | 0.437945592 | 0.113505818 |
| Tmed1 | 6.444756 | 6.455453 | 6.5301 | -0.074672686 | -0.085369587 | 6.518071 | 6.7659437 | 6.3289 | 0.437045075 | 0.189172423 |
| Gm20431 | 5.753325 | 5.767271 | 5.8197 | -0.052468717 | -0.066414574 | 6.3310441 | 6.0461748 | 5.60969 | 0.436487532 | 0.721356885 |
| Rnf149 | 9.171547 | 9.177342 | 8.6974 | 0.479894334 | 0.474099776 | 9.1989272 | 9.2955343 | 8.85913 | 0.436407225 | 0.339800103 |
| Ddit3 | 8.419673 | 8.28113 | 8.367 | -0.085861923 | 0.052680793 | 8.3561238 | 8.8414201 | 8.40509 | 0.436327902 | -0.048968463 |
| Arsi | 3.620969 | 3.779649 | 3.6043 | 0.175302465 | 0.016622113 | 4.1506774 | 4.0095579 | 3.57348 | 0.436081104 | 0.577200557 |
| Klhl18 | 9.315983 | 9.52262 | 9.2587 | 0.263968922 | 0.057331963 | 9.5624424 | 9.764982 | 9.3291 | 0.435879304 | 0.233339623 |
| Eef1e1 | 8.68734 | 8.896872 | 8.6105 | 0.286327166 | 0.076795668 | 8.9251792 | 9.0966882 | 8.66088 | 0.435811519 | 0.264302535 |
| Higd1a | 8.420381 | 8.358134 | 8.3431 | 0.01506474 | 0.077310913 | 8.4755223 | 8.7459407 | 8.31063 | 0.435315254 | 0.164896897 |

| | | | | | | | | | | |
|---|---|---|---|---|---|---|---|---|---|---|
| Amdhd2 | 8.011618 | 8.048226 | 8.0999 | -0.051688117 | -0.088296926 | 8.4955199 | 8.4397041 | 8.00446 | 0.435248158 | 0.491063957 |
| D13Ertd608e | 3.285437 | 3.530349 | 3.362 | 0.168367716 | -0.076543801 | 3.2493149 | 3.5901907 | 3.15504 | 0.435150215 | 0.094274434 |
| Vmn1r47 | 2.979428 | 2.921065 | 2.8861 | 0.03499075 | 0.093353543 | 3.0147136 | 3.389702 | 2.95471 | 0.4349954 | 0.060007057 |
| C1galt1c1 | 7.752608 | 7.798109 | 7.7004 | 0.097709002 | 0.052207923 | 7.7256544 | 8.1313785 | 7.69666 | 0.4347176 | 0.028993452 |
| Tbc1d22b | 9.397339 | 9.588603 | 9.462 | 0.126562684 | -0.064701617 | 9.3159368 | 9.8184388 | 9.38407 | 0.434369067 | -0.068132962 |
| Llph | 9.2505 | 9.227891 | 9.154 | 0.073907406 | 0.096516634 | 9.30836 | 9.683769 | 9.24953 | 0.434238856 | 0.058829914 |
| Dna2 | 7.096528 | 7.008468 | 7.0773 | -0.06881904 | 0.019241105 | 7.3023553 | 7.7105333 | 7.27633 | 0.43419843 | 0.026020382 |
| Mtfp1 | 4.6675 | 4.706034 | 4.5904 | 0.115628837 | 0.077094909 | 4.7285639 | 4.9075626 | 4.47339 | 0.434168257 | 0.25516953 |
| Pkm | 13.12071 | 13.12266 | 13.011 | 0.11125195 | 0.109308794 | 13.222236 | 13.481378 | 13.0479 | 0.433521392 | 0.174379142 |
| Bcl2a1b | 9.626024 | 10.1884 | 9.7413 | 0.447123788 | -0.115252026 | 9.7726256 | 10.202253 | 9.76875 | 0.433502288 | 0.003874563 |
| Klrc2 | 6.339135 | 6.330293 | 6.1772 | 0.153076352 | 0.161918487 | 6.151443 | 6.5816232 | 6.14913 | 0.432492861 | 0.002312703 |
| Rbm8a | 9.759783 | 9.750665 | 9.7213 | 0.029392133 | 0.038510681 | 9.7397303 | 10.140211 | 9.70776 | 0.432451152 | 0.031970543 |
| Krtcap2 | 9.104253 | 9.122998 | 9.1369 | -0.013930141 | -0.032674895 | 9.4023515 | 9.5239301 | 9.09158 | 0.4323485 | 0.310769848 |
| Rnpepl1 | 10.41361 | 10.51913 | 10.43 | 0.089429918 | -0.016091308 | 10.596312 | 10.881994 | 10.4498 | 0.43217501 | 0.146492562 |
| Szrd1 | 10.80357 | 11.00466 | 10.728 | 0.2767731 | 0.075676614 | 10.862754 | 11.188193 | 10.756 | 0.432149132 | 0.106710646 |
| Noc2l | 11.28116 | 11.42214 | 11.257 | 0.16517493 | 0.024186998 | 11.496326 | 11.790272 | 11.3583 | 0.431936824 | 0.137991049 |
| Cant1 | 10.4002 | 10.61363 | 10.379 | 0.234531644 | 0.021109034 | 10.535768 | 10.783091 | 10.3513 | 0.431818046 | 0.184494297 |
| Casp7 | 8.196385 | 8.183579 | 8.2361 | -0.052529044 | -0.039722684 | 8.2453518 | 8.7027623 | 8.2714 | 0.431361994 | -0.026048494 |
| Minos1 | 9.403683 | 9.391073 | 9.364 | 0.027060937 | 0.039670345 | 9.5414496 | 9.7811109 | 9.34992 | 0.431191222 | 0.191529966 |
| AC151990.1 | 1.088141 | 1.090892 | 1.0829 | 0.007944683 | 0.005193545 | 1.0794568 | 1.5255471 | 1.09451 | 0.431040312 | -0.015050017 |
| Erh | 9.930573 | 9.981454 | 9.8724 | 0.109083296 | 0.058202589 | 10.173786 | 10.399202 | 9.96823 | 0.430972094 | 0.205556084 |
| Gm4332 | 12.37431 | 12.46104 | 11.937 | 0.523704838 | 0.436976702 | 12.374673 | 12.798742 | 12.368 | 0.430730494 | 0.006661035 |
| Srsf2 | 11.81691 | 11.77229 | 11.82 | -0.047940318 | -0.003322792 | 12.104434 | 12.283666 | 11.853 | 0.430668186 | 0.251436074 |
| Ddt | 8.058324 | 8.157696 | 8.1223 | 0.035408207 | -0.063963063 | 8.4427809 | 8.4471809 | 8.01652 | 0.430665138 | 0.42626514 |
| Plpp2 | 5.899502 | 6.024456 | 5.8928 | 0.131696737 | 0.00674302 | 6.0062283 | 6.2094754 | 5.77898 | 0.430490499 | 0.227243458 |
| Gpt2 | 6.073526 | 5.931572 | 5.4037 | 0.527857673 | 0.669812288 | 5.6723396 | 5.9527848 | 5.52271 | 0.430079172 | 0.14963389 |
| Dnajc13 | 11.02477 | 11.13395 | 10.942 | 0.191898312 | 0.082716116 | 11.010647 | 11.570594 | 11.1407 | 0.429911424 | -0.130036221 |
| Rbm3 | 13.22379 | 12.9632 | 13.147 | -0.184066238 | 0.07652207 | 13.054974 | 13.327907 | 12.8981 | 0.429822578 | 0.156889021 |
| Stra6 | 5.733425 | 5.985966 | 5.3792 | 0.606783316 | 0.354242613 | 5.3983829 | 5.4463261 | 5.01652 | 0.429804888 | 0.381861715 |
| Slc39a7 | 10.73776 | 10.89461 | 10.706 | 0.18894821 | 0.032103218 | 10.865241 | 11.092665 | 10.6631 | 0.429611542 | 0.202187708 |
| Emilin1 | 7.283371 | 7.49684 | 7.1353 | 0.361531301 | 0.148062871 | 7.448133 | 7.3985791 | 6.96932 | 0.429257444 | 0.478811344 |
| Actb | 16.47357 | 16.64482 | 16.473 | 0.171338016 | 9.257E-05 | 16.809978 | 16.859825 | 16.4306 | 0.429219922 | 0.379372737 |

| | | | | | | | | | | |
|---|---|---|---|---|---|---|---|---|---|---|
| Chmp5 | 9.702806 | 9.925389 | 9.7009 | 0.224458144 | 0.001874406 | 9.8964473 | 10.176512 | 9.74733 | 0.429183985 | 0.149119215 |
| B3galt4 | 7.116689 | 7.14543 | 7.1799 | -0.034516411 | -0.063257831 | 7.3543921 | 7.4787324 | 7.04991 | 0.428819913 | 0.304479666 |
| Rfc2 | 9.947833 | 10.02419 | 9.9822 | 0.041955868 | -0.0343995 | 10.394511 | 10.540355 | 10.1117 | 0.428634954 | 0.282791411 |
| Gtf2f1 | 9.91342 | 9.963088 | 9.8384 | 0.124639805 | 0.074972444 | 10.100519 | 10.364898 | 9.93632 | 0.428578925 | 0.164200039 |
| Tsr1 | 10.27212 | 10.27655 | 10.096 | 0.180280646 | 0.175844704 | 10.410056 | 10.664673 | 10.2361 | 0.428554292 | 0.173937272 |
| Adgrg5 | 9.44581 | 9.467632 | 9.4352 | 0.03247661 | 0.010654647 | 9.6804726 | 10.059125 | 9.63062 | 0.428504468 | 0.049852249 |
| Hmgn2 | 11.47334 | 11.26585 | 11.453 | -0.186750642 | 0.020744668 | 11.627366 | 11.926926 | 11.4991 | 0.427834616 | 0.128273823 |
| Arpc1b | 12.64085 | 12.68029 | 12.529 | 0.151265122 | 0.111826988 | 12.952413 | 12.900425 | 12.4727 | 0.427705696 | 0.479693696 |
| Usp38 | 10.48005 | 10.67749 | 10.455 | 0.22292158 | 0.02547991 | 10.360123 | 10.859736 | 10.4323 | 0.427483386 | -0.072129404 |
| Ccz1 | 9.46424 | 9.458546 | 9.4532 | 0.005303862 | 0.0109983 | 9.7223575 | 10.085839 | 9.65846 | 0.427382803 | 0.063901536 |
| Nle1 | 8.474401 | 8.700465 | 8.5871 | 0.113407768 | -0.112655904 | 9.1708602 | 9.3037925 | 8.87653 | 0.427261487 | 0.294329174 |
| Ubb | 14.3524 | 14.35662 | 14.42 | -0.063808066 | -0.068029622 | 14.426939 | 14.738769 | 14.3117 | 0.427032022 | 0.115201748 |
| Tmem115 | 8.078871 | 8.287408 | 8.1483 | 0.139081556 | -0.069455588 | 8.2395902 | 8.3256699 | 7.89884 | 0.426833372 | 0.340753603 |
| Rab22a | 9.744366 | 9.86776 | 9.8129 | 0.054881501 | -0.068512053 | 9.6813337 | 10.208455 | 9.78189 | 0.426565906 | -0.100555525 |
| Smdt1 | 8.888085 | 8.737856 | 8.7686 | -0.030752311 | 0.119477248 | 9.0721196 | 9.1857331 | 8.75922 | 0.426510422 | 0.312896922 |
| Irf2bp1 | 8.589958 | 8.715561 | 8.6084 | 0.107119726 | -0.01848357 | 8.7302581 | 8.9492213 | 8.52276 | 0.426462797 | 0.207499623 |
| Rrs1 | 9.180047 | 9.20716 | 8.9984 | 0.208772614 | 0.181659134 | 9.2139101 | 9.4365658 | 9.01023 | 0.426332789 | 0.203677148 |
| Stx4a | 10.04109 | 10.05396 | 10.031 | 0.022785442 | 0.009912813 | 10.284857 | 10.582916 | 10.1567 | 0.426251628 | 0.128192211 |
| Drap1 | 10.25688 | 10.26307 | 10.284 | -0.021405011 | -0.027597913 | 10.507977 | 10.649789 | 10.2236 | 0.426191101 | 0.284378933 |
| Prm1 | 0.839667 | 0.874811 | 0.8371 | 0.037712953 | 0.002569288 | 0.8633455 | 1.2671421 | 0.84122 | 0.425922619 | 0.022125995 |
| Bax | 9.915186 | 9.982862 | 9.9521 | 0.030748599 | -0.036927153 | 10.172425 | 10.352868 | 9.92704 | 0.425826197 | 0.24538331 |
| Hcls1 | 12.48868 | 12.59992 | 12.427 | 0.173196018 | 0.061962696 | 12.75146 | 12.869933 | 12.4442 | 0.425754322 | 0.307281815 |
| Ruvbl1 | 9.471876 | 9.45575 | 9.3928 | 0.06297607 | 0.079102055 | 9.5780116 | 9.8416226 | 9.41597 | 0.425652285 | 0.162041296 |
| Aig1 | 7.226633 | 7.326716 | 7.201 | 0.12572074 | 0.02563758 | 7.5228429 | 7.7162023 | 7.29065 | 0.425553978 | 0.232194565 |
| Cct5 | 12.1491 | 12.27976 | 12.134 | 0.145627622 | 0.014968058 | 12.20922 | 12.582036 | 12.1566 | 0.425452476 | 0.052637254 |
| Psmb7 | 10.37025 | 10.36266 | 10.377 | -0.014405974 | -0.00681531 | 10.508867 | 10.714135 | 10.2887 | 0.425419016 | 0.220150827 |
| Elp5 | 8.711489 | 8.797421 | 8.7106 | 0.086778171 | 0.000845632 | 8.9006899 | 9.0530572 | 8.62767 | 0.425392073 | 0.273024799 |
| Hsp90aa1 | 13.46674 | 13.61708 | 13.549 | 0.068181702 | -0.082158386 | 13.102172 | 13.993291 | 13.5679 | 0.425349194 | -0.465769206 |
| Pdia6 | 11.16651 | 11.35447 | 11.018 | 0.336770612 | 0.148809234 | 11.434817 | 11.548629 | 11.1234 | 0.425181718 | 0.311369534 |
| Nfkb2 | 11.62183 | 11.84327 | 11.578 | 0.265619736 | 0.044176524 | 11.93094 | 12.045664 | 11.6205 | 0.425167246 | 0.310442918 |
| P2rx4 | 8.741379 | 8.766201 | 8.681 | 0.085160742 | 0.060339507 | 9.1253047 | 9.2224824 | 8.79757 | 0.424915566 | 0.327737894 |
| Clpp | 8.033802 | 8.095672 | 8.1158 | -0.020129059 | -0.081999085 | 8.4268851 | 8.3961899 | 7.97165 | 0.424536065 | 0.455231247 |

| | | | | | | | | | | |
|---|---|---|---|---|---|---|---|---|---|---|
| Rnf34 | 8.813336 | 9.077628 | 8.9067 | 0.170939484 | -0.09335226 | 9.0557167 | 9.3044471 | 8.88017 | 0.424280463 | 0.175550041 |
| Fam110a | 6.810559 | 6.87517 | 6.7994 | 0.075745931 | 0.011134518 | 6.8356689 | 7.2062918 | 6.78219 | 0.424099445 | 0.053476522 |
| Map1s | 9.027095 | 8.848812 | 8.7411 | 0.107697814 | 0.285980946 | 9.2052792 | 9.2779417 | 8.85386 | 0.424084304 | 0.35142173 |
| Raf1 | 11.02426 | 11.11142 | 11.05 | 0.061350566 | -0.025812558 | 11.254268 | 11.624382 | 11.2004 | 0.423983882 | 0.053869454 |
| Ngf | 4.139864 | 4.56515 | 5.1359 | -0.570716444 | -0.996001895 | 5.9087458 | 5.2370187 | 4.8137 | 0.423314862 | 1.095041917 |
| Bbip1 | 8.659299 | 8.740245 | 8.5358 | 0.204469188 | 0.123523366 | 8.9054577 | 8.9497376 | 8.52657 | 0.42316421 | 0.378884355 |
| Psmf1 | 9.294009 | 9.340568 | 9.3373 | 0.003250684 | -0.043308468 | 9.4863565 | 9.7921832 | 9.36921 | 0.422977754 | 0.117151021 |
| Nfkbil1 | 7.29975 | 7.200247 | 7.2413 | -0.041065891 | 0.058436737 | 7.2602752 | 7.4911817 | 7.06836 | 0.422821619 | 0.191915127 |
| Abhd17a | 9.205495 | 9.328459 | 9.2765 | 0.05191339 | -0.071050753 | 9.3933164 | 9.3791818 | 8.95663 | 0.422549946 | 0.436684522 |
| Prelid1 | 10.68174 | 10.74611 | 10.707 | 0.038989344 | -0.02537555 | 10.86082 | 11.060522 | 10.6382 | 0.422275396 | 0.222573262 |
| Ndufs3 | 8.465493 | 8.544803 | 8.4889 | 0.055952088 | -0.023357652 | 8.7479322 | 9.005437 | 8.58327 | 0.422169443 | 0.164664595 |
| Nlrp1b | 5.609993 | 5.377185 | 5.4691 | -0.091941982 | 0.140865707 | 5.6477143 | 6.0793944 | 5.65724 | 0.42215073 | -0.009529369 |
| Ppp1r15b | 11.25457 | 11.31213 | 11.224 | 0.087682562 | 0.030124478 | 11.279566 | 11.652409 | 11.2303 | 0.42212468 | 0.049281768 |
| Mdn1 | 11.73398 | 11.97101 | 11.804 | 0.16743625 | -0.069593338 | 11.835534 | 12.511222 | 12.0891 | 0.422081272 | -0.253606911 |
| Ssc5d | 3.678463 | 3.523126 | 3.4637 | 0.059412206 | 0.214749386 | 3.580289 | 3.8750043 | 3.45294 | 0.422062396 | 0.127347093 |
| Jpt1 | 11.11945 | 11.19469 | 11.106 | 0.08900384 | 0.013767238 | 11.25392 | 11.440913 | 11.0189 | 0.42204731 | 0.235054487 |
| Prr5l | 7.436026 | 7.20444 | 7.5054 | -0.300913413 | -0.069327688 | 7.1040816 | 7.5765102 | 7.15479 | 0.421718897 | -0.05070969 |
| Snrpa1 | 9.093203 | 9.067731 | 9.0193 | 0.048421386 | 0.073892738 | 9.3225663 | 9.5301926 | 9.10871 | 0.421482738 | 0.213856364 |
| Derl2 | 9.142752 | 9.23485 | 9.1656 | 0.069258405 | -0.022838977 | 9.2924384 | 9.5408474 | 9.11952 | 0.421326472 | 0.172917492 |
| Hhat | 6.051725 | 6.313746 | 6.227 | 0.086788492 | -0.175232411 | 6.2800224 | 6.6017694 | 6.18059 | 0.421182205 | 0.099435247 |
| Ankib1 | 10.43157 | 10.62699 | 10.488 | 0.139143932 | -0.056277572 | 10.351566 | 10.914395 | 10.4934 | 0.42103574 | -0.141793285 |
| Mcub | 6.496565 | 6.208529 | 6.2074 | 0.00110202 | 0.289137197 | 6.369451 | 6.4531556 | 6.03247 | 0.420681943 | 0.336977328 |
| Apof | 3.09086 | 3.234263 | 3.2352 | -0.000964229 | -0.144367024 | 3.2144353 | 3.6521408 | 3.23183 | 0.420315531 | -0.01739001 |
| Rpf2 | 8.498884 | 8.448286 | 8.4062 | 0.042118121 | 0.092716396 | 8.5269859 | 8.8801215 | 8.45981 | 0.420312485 | 0.067176842 |
| Stk39 | 7.93102 | 7.63355 | 7.7085 | -0.074936741 | 0.222533665 | 7.3795882 | 8.3040427 | 7.88381 | 0.420235733 | -0.504218798 |
| Gm13394 | 2.981374 | 2.974151 | 3.0343 | -0.060122959 | -0.052900132 | 3.1912694 | 3.2911524 | 2.87099 | 0.420164377 | 0.320281356 |
| Ccl20 | 4.20059 | 4.329943 | 3.7926 | 0.537392231 | 0.408039183 | 4.6630646 | 4.3374933 | 3.91761 | 0.419878517 | 0.745449801 |
| Asprv1 | 6.198119 | 6.293164 | 5.578 | 0.715116019 | 0.620070537 | 5.8447041 | 5.9358281 | 5.51598 | 0.419848322 | 0.328724365 |
| Cyb561d2 | 7.530193 | 7.554862 | 7.53 | 0.024872549 | 0.000203047 | 7.7200318 | 7.7758842 | 7.35611 | 0.41977412 | 0.363921755 |
| Sh3glb1 | 11.16725 | 11.27431 | 11.183 | 0.091370528 | -0.015686566 | 11.229007 | 11.583063 | 11.1634 | 0.419705788 | 0.06564998 |
| Ppp2r3d | 7.050639 | 7.239099 | 7.0749 | 0.164158483 | -0.024301518 | 7.6686786 | 7.7360834 | 7.3164 | 0.419686353 | 0.352281577 |
| Atp5g2 | 7.099395 | 7.080197 | 7.0415 | 0.038667802 | 0.057866261 | 7.480713 | 7.3691426 | 6.94955 | 0.419595196 | 0.531165583 |

| | | | | | | | | | | |
|---|---|---|---|---|---|---|---|---|---|---|
| Slc25a38 | 8.651271 | 8.784574 | 8.7674 | 0.017187455 | -0.116116174 | 8.8608568 | 9.1766478 | 8.75716 | 0.419488879 | 0.103697918 |
| Cdc34 | 9.067563 | 9.019179 | 9.0076 | 0.011590508 | 0.059975022 | 9.3361585 | 9.4380474 | 9.01856 | 0.419487732 | 0.317598821 |
| Zc3h3 | 8.885672 | 9.002173 | 8.9278 | 0.074327405 | -0.042174017 | 9.0180052 | 9.285754 | 8.86691 | 0.418844023 | 0.151095172 |
| Igkv16-104 | 8.277902 | 8.536572 | 8.0918 | 0.444808774 | 0.18613821 | 8.3133238 | 8.5920744 | 8.17327 | 0.418807576 | 0.140056966 |
| Pdcd5 | 8.537384 | 8.420264 | 8.3937 | 0.026526322 | 0.143646881 | 8.5682433 | 8.7355299 | 8.31675 | 0.418782716 | 0.251496095 |
| Pmf1 | 8.526324 | 8.646424 | 8.6019 | 0.044480922 | -0.075618653 | 8.911935 | 9.0885407 | 8.67009 | 0.418447431 | 0.241841706 |
| Snrpa | 9.151818 | 9.113239 | 9.0251 | 0.088143992 | 0.126722547 | 9.6669943 | 9.6707106 | 9.25227 | 0.418440705 | 0.414724444 |
| Ndufa3 | 9.412616 | 9.562767 | 9.4719 | 0.090864957 | -0.059286628 | 9.8675143 | 9.9229527 | 9.50467 | 0.418282285 | 0.362843864 |
| Nsmce4a | 10.16795 | 10.13348 | 10.084 | 0.049887867 | 0.084360348 | 10.202621 | 10.442848 | 10.0247 | 0.418164025 | 0.177937319 |
| Pcbp1 | 11.62852 | 11.64915 | 11.518 | 0.131054814 | 0.110426208 | 11.703714 | 11.926049 | 11.5089 | 0.417164788 | 0.194830355 |
| Nono | 12.40005 | 12.50445 | 12.429 | 0.075362612 | -0.029029462 | 12.524249 | 12.875281 | 12.4582 | 0.417107096 | 0.066075901 |
| St14 | 8.534214 | 8.527436 | 8.5217 | 0.005687063 | 0.012465494 | 8.7901082 | 8.8984947 | 8.48141 | 0.417089011 | 0.308702468 |
| Snw1 | 10.85687 | 10.92928 | 10.851 | 0.077831432 | 0.00542602 | 10.86184 | 11.248964 | 10.8319 | 0.41708548 | 0.029960921 |
| Emc4 | 8.388163 | 8.416273 | 8.3089 | 0.107355715 | 0.079245633 | 8.5837706 | 8.7852361 | 8.36815 | 0.417081385 | 0.215615834 |
| Olfr1307 | 6.545869 | 6.751596 | 6.6319 | 0.119663505 | -0.086063297 | 6.6847925 | 7.3209588 | 6.90404 | 0.416919364 | -0.219247013 |
| Wdr55 | 8.709592 | 8.831888 | 8.6076 | 0.224285484 | 0.101989127 | 8.9515658 | 9.1280324 | 8.71227 | 0.415767003 | 0.239300419 |
| Ap2s1 | 9.348741 | 9.242052 | 9.2891 | -0.046998154 | 0.059690393 | 9.2858484 | 9.5997792 | 9.18404 | 0.415738713 | 0.101807878 |
| Ddx18 | 10.3668 | 10.50973 | 10.333 | 0.176659486 | 0.033728166 | 10.621658 | 10.808312 | 10.3926 | 0.415730932 | 0.229076901 |
| Exosc5 | 8.982787 | 9.076963 | 8.9489 | 0.128104524 | 0.033928152 | 9.3862836 | 9.3061752 | 8.89051 | 0.415665369 | 0.49577373 |
| Acaa1a | 9.989571 | 10.04073 | 9.8981 | 0.142606907 | 0.091447953 | 10.383321 | 10.417063 | 10.0017 | 0.41540491 | 0.381662293 |
| Lsm6 | 9.356721 | 9.297933 | 9.1746 | 0.123298226 | 0.182086427 | 9.3534501 | 9.6535727 | 9.23835 | 0.415224994 | 0.115102352 |
| Trim34b | 5.203995 | 5.247875 | 5.0628 | 0.18502988 | 0.141149278 | 5.3288964 | 5.7702915 | 5.35525 | 0.415044352 | -0.026350757 |
| Ctla2b | 5.765552 | 5.671559 | 5.6493 | 0.022258571 | 0.116251523 | 5.5454493 | 5.8558764 | 5.44107 | 0.414808897 | 0.104381817 |
| Skap2 | 10.07351 | 9.772994 | 9.7893 | -0.016258163 | 0.284261874 | 10.111875 | 10.157794 | 9.7431 | 0.414693269 | 0.36877379 |
| Hist1h2ap | 6.095343 | 6.33624 | 6.2024 | 0.133831267 | -0.107065993 | 6.1326625 | 6.3308184 | 5.91648 | 0.414337346 | 0.216181422 |
| Adamts15 | 3.547029 | 3.478409 | 3.3218 | 0.156628237 | 0.225248705 | 3.3337884 | 3.5781154 | 3.16386 | 0.414253899 | 0.169926907 |
| Ccdc173 | 4.643469 | 4.64703 | 4.5131 | 0.133958443 | 0.130397381 | 4.550428 | 5.1536397 | 4.73976 | 0.413882599 | -0.189329093 |
| Ankrd49 | 7.570967 | 7.515687 | 7.5217 | -0.006028681 | 0.049251368 | 7.778289 | 7.9926767 | 7.5788 | 0.41387767 | 0.199489903 |
| Tmem173 | 11.13035 | 11.18971 | 11.105 | 0.084571716 | 0.025214142 | 11.290431 | 11.578005 | 11.1643 | 0.413717938 | 0.126143606 |
| Ankle2 | 10.61556 | 10.78193 | 10.728 | 0.05359936 | -0.112768136 | 10.684009 | 11.137013 | 10.7234 | 0.413581548 | -0.039422357 |
| Fiz1 | 8.838229 | 8.924726 | 8.7811 | 0.143588032 | 0.057091418 | 8.8664627 | 9.1179436 | 8.7047 | 0.413248594 | 0.161767606 |
| Nop10 | 9.35486 | 9.485824 | 9.4202 | 0.065657986 | -0.065305436 | 9.6408684 | 9.8185462 | 9.40547 | 0.413079638 | 0.235401884 |

| | | | | | | | | | | |
|---|---|---|---|---|---|---|---|---|---|---|
| Ccdc117 | 10.14864 | 10.59969 | 10.204 | 0.395402426 | -0.05565309 | 10.045197 | 10.655982 | 10.243 | 0.413009162 | -0.197775272 |
| Cnpy3 | 10.01647 | 10.05054 | 9.9817 | 0.068834677 | 0.034764501 | 10.348912 | 10.414134 | 10.0015 | 0.412631956 | 0.347409294 |
| Dpm1 | 7.574223 | 7.297841 | 7.5382 | -0.240391025 | 0.035991481 | 8.2154706 | 7.9872206 | 7.57474 | 0.412483758 | 0.640733814 |
| Igkv4-63 | 5.296979 | 5.473284 | 5.3442 | 0.129112108 | -0.047193534 | 5.5472176 | 5.7663171 | 5.35384 | 0.41247929 | 0.193379788 |
| Sfi1 | 9.5056 | 9.563251 | 9.6289 | -0.065662818 | -0.123313838 | 9.9714026 | 10.463704 | 10.0515 | 0.412184877 | -0.080116411 |
| Sqle | 8.052621 | 8.080884 | 7.881 | 0.199878295 | 0.171615947 | 8.0506907 | 8.0502393 | 7.63836 | 0.411879582 | 0.412330965 |
| Zkscan14 | 7.817277 | 8.038279 | 7.8831 | 0.155162881 | -0.065838798 | 7.7731052 | 8.0764881 | 7.66481 | 0.411682943 | 0.108300032 |
| Trappc4 | 8.739749 | 8.717764 | 8.7254 | -0.00759806 | 0.014386583 | 9.1122866 | 9.0882658 | 8.6766 | 0.411662051 | 0.43568283 |
| Hspb1 | 7.189328 | 7.17713 | 7.2685 | -0.09139202 | -0.079194159 | 6.6646081 | 7.267485 | 6.85597 | 0.411513264 | -0.191363559 |
| Cox14 | 8.346815 | 8.311706 | 8.2733 | 0.038407043 | 0.073516306 | 8.4958529 | 8.613541 | 8.20228 | 0.411265239 | 0.29357712 |
| Usp42 | 8.998542 | 9.109767 | 9.0629 | 0.046916126 | -0.064309042 | 9.0478466 | 9.4866714 | 9.07551 | 0.411156453 | -0.027668353 |
| Tlr8 | 5.622655 | 5.80022 | 4.9915 | 0.808765263 | 0.631200593 | 5.4061083 | 5.262758 | 4.85188 | 0.410877134 | 0.554227367 |
| Gm9797 | 6.017855 | 6.133588 | 5.8752 | 0.258388794 | 0.14265609 | 5.8594525 | 6.3397827 | 5.92895 | 0.410834983 | -0.069495223 |
| Igkv5-48 | 7.850422 | 7.981033 | 7.9066 | 0.074457065 | -0.056153754 | 8.4571087 | 8.2811832 | 7.87102 | 0.41015889 | 0.586084463 |
| Rps2 | 14.31421 | 14.43204 | 14.375 | 0.057254004 | -0.060574122 | 14.660647 | 14.782866 | 14.3729 | 0.409958714 | 0.287739978 |
| Utp20 | 10.38277 | 10.45647 | 10.359 | 0.097603452 | 0.023903978 | 10.542236 | 10.930127 | 10.5203 | 0.409861316 | 0.021970117 |
| Alkbh4 | 7.967504 | 8.173303 | 7.9709 | 0.202443462 | -0.003354954 | 7.877941 | 8.2631888 | 7.85346 | 0.409727921 | 0.024480138 |
| Rps26 | 13.02842 | 13.1147 | 13.097 | 0.017866112 | -0.068407016 | 13.312749 | 13.415857 | 13.0062 | 0.409614708 | 0.306506772 |
| Cxcl3 | 4.00764 | 3.597677 | 3.063 | 0.534668423 | 0.944632343 | 3.7136065 | 3.5555431 | 3.14596 | 0.40958079 | 0.567644193 |
| Wwox | 6.911465 | 6.915607 | 6.8307 | 0.084908103 | 0.080765608 | 6.9527993 | 7.3100697 | 6.90053 | 0.409537348 | 0.052266976 |
| 4931406B18Rik | 3.671306 | 3.725011 | 3.5652 | 0.159789723 | 0.106084677 | 3.6811669 | 4.0088972 | 3.5995 | 0.409396626 | 0.081666266 |
| Psmd4 | 10.68478 | 10.73856 | 10.706 | 0.032267274 | -0.021508518 | 10.901711 | 11.148069 | 10.7389 | 0.409138388 | 0.162780852 |
| Cavin3 | 6.36163 | 6.348748 | 6.1971 | 0.151692579 | 0.164574704 | 5.9611747 | 6.0881559 | 5.67904 | 0.409115675 | 0.282134488 |
| Gnl1 | 10.15161 | 10.2247 | 10.13 | 0.094272965 | 0.021181547 | 10.249869 | 10.447831 | 10.0388 | 0.409062599 | 0.211100467 |
| Plgrkt | 9.714876 | 9.900231 | 9.7426 | 0.157605681 | -0.0277495 | 10.019381 | 10.426933 | 10.0179 | 0.409060831 | 0.001508107 |
| Rhbdf2 | 9.788254 | 9.869273 | 9.6973 | 0.172014996 | 0.090995646 | 10.15286 | 10.193755 | 9.78473 | 0.409024397 | 0.368128711 |
| Sars2 | 7.905154 | 7.926227 | 7.848 | 0.078235627 | 0.057162269 | 8.3555909 | 8.3732046 | 7.96426 | 0.408947542 | 0.391333871 |
| Mrto4 | 9.096221 | 9.0682 | 9.0119 | 0.056340944 | 0.084362356 | 9.3519825 | 9.5714572 | 9.16254 | 0.40891675 | 0.189442025 |
| Dtwd1 | 6.526381 | 6.655036 | 6.4494 | 0.205681461 | 0.077025942 | 6.5514014 | 6.9532934 | 6.54446 | 0.408829467 | 0.0069375 |
| Uba1 | 12.60843 | 12.60936 | 12.55 | 0.059359404 | 0.058422252 | 12.743717 | 12.996155 | 12.5876 | 0.408529786 | 0.156091455 |
| Rps29 | 14.1527 | 14.35 | 14.142 | 0.207898826 | 0.010593774 | 14.52377 | 14.488278 | 14.0799 | 0.408405886 | 0.443897994 |
| Ythdf2 | 10.09518 | 10.26649 | 10.192 | 0.074882458 | -0.096430623 | 10.068686 | 10.468785 | 10.0606 | 0.408153128 | 0.008054149 |

| | | | | | | | | | | |
|---|---|---|---|---|---|---|---|---|---|---|
| Gm47933 | 5.015484 | 5.09505 | 4.9757 | 0.11934692 | 0.03978089 | 5.1551139 | 5.5287355 | 5.12083 | 0.407905699 | 0.034284131 |
| Igkv4-77 | 4.701535 | 4.443789 | 4.4104 | 0.033413823 | 0.291159922 | 4.4818064 | 4.7199005 | 4.31207 | 0.407829343 | 0.169735292 |
| Dtymk | 8.754303 | 8.654438 | 8.6744 | -0.019981689 | 0.079882562 | 8.9122867 | 9.12725 | 8.71948 | 0.407768412 | 0.192805116 |
| Pcolce | 7.100252 | 7.023377 | 7.0284 | -0.005052743 | 0.071822923 | 6.9992353 | 6.8133969 | 6.40571 | 0.407682004 | 0.593520338 |
| Gm48082 | 6.666313 | 6.813479 | 6.7553 | 0.058220641 | -0.088944989 | 6.8484679 | 7.5786 | 7.17092 | 0.407679606 | -0.322452445 |
| Mgat4a | 10.21951 | 10.27566 | 10.185 | 0.090196381 | 0.034046286 | 10.293002 | 10.699588 | 10.292 | 0.407569048 | 0.00098339 |
| Sf3a2 | 9.96149 | 10.09681 | 10.013 | 0.083439981 | -0.051876911 | 10.179806 | 10.397462 | 9.98992 | 0.407538487 | 0.189882611 |
| Gm11225 | 5.482013 | 5.365628 | 5.6416 | -0.275923609 | -0.159538511 | 6.3445407 | 5.9589897 | 5.55167 | 0.407319964 | 0.792870914 |
| Rpp21 | 7.654852 | 7.830142 | 7.7471 | 0.083034769 | -0.092255377 | 7.896511 | 8.0250418 | 7.61788 | 0.40716335 | 0.27863257 |
| Tuba1a | 10.64479 | 10.79173 | 10.72 | 0.072026824 | -0.074915492 | 10.723283 | 10.978865 | 10.5718 | 0.40711167 | 0.151529836 |
| Gsr | 9.169656 | 9.168905 | 8.6113 | 0.557610676 | 0.558361448 | 9.2527671 | 8.8464074 | 8.43932 | 0.407085915 | 0.813445595 |
| Ticam1 | 8.852929 | 9.000999 | 8.8777 | 0.123289307 | -0.024781199 | 9.0255482 | 9.2129854 | 8.80593 | 0.407053205 | 0.219615985 |
| Rpl37rt | 10.17516 | 10.49193 | 10.835 | -0.342920256 | -0.659690822 | 10.568931 | 10.708912 | 10.3019 | 0.406993761 | 0.267012039 |
| Coa5 | 10.97251 | 11.04467 | 10.984 | 0.060358122 | -0.01180804 | 11.042436 | 11.339979 | 10.9332 | 0.406777314 | 0.109233741 |
| Mrpl21 | 8.368954 | 8.400407 | 8.2847 | 0.115700365 | 0.084246659 | 8.6383365 | 8.7499777 | 8.34371 | 0.406266284 | 0.294625028 |
| Hspa9 | 11.86097 | 11.88844 | 11.723 | 0.165879288 | 0.138415962 | 11.875635 | 12.215315 | 11.8094 | 0.405874346 | 0.066195035 |
| AL731706.1 | 6.430837 | 6.431788 | 6.5062 | -0.074403778 | -0.075354246 | 6.6343255 | 6.7437222 | 6.33789 | 0.405833325 | 0.296436589 |
| Nop58 | 10.66177 | 10.74123 | 10.693 | 0.048118674 | -0.031345616 | 10.59297 | 11.136138 | 10.7303 | 0.405823442 | -0.137345127 |
| Gcn1l1 | 11.26967 | 11.37556 | 11.374 | 0.001197774 | -0.10468779 | 11.599489 | 11.77334 | 11.3677 | 0.405598488 | 0.231747653 |
| Psmc2 | 10.39422 | 10.40838 | 10.35 | 0.058450172 | 0.04428445 | 10.465858 | 10.80054 | 10.395 | 0.405587788 | 0.070905762 |
| Gm7592 | 3.640828 | 3.921307 | 3.5825 | 0.338795601 | 0.058316348 | 4.0232864 | 4.2580745 | 3.85255 | 0.40552554 | 0.170737416 |
| Zc3hav1l | 6.593944 | 6.854081 | 6.5691 | 0.285021681 | 0.024884001 | 6.5749953 | 7.1986209 | 6.79316 | 0.405464367 | -0.218161187 |
| Cd2ap | 11.43474 | 11.46554 | 11.35 | 0.116024326 | 0.08522334 | 11.39492 | 11.814099 | 11.4087 | 0.405391546 | -0.013787945 |
| Mbnl2 | 10.12041 | 10.2187 | 10.105 | 0.113610726 | 0.015322277 | 10.061423 | 10.475929 | 10.0706 | 0.405326627 | -0.009179453 |
| Arf5 | 10.14933 | 10.07911 | 10.104 | -0.024804021 | 0.045412501 | 10.206559 | 10.36886 | 9.96356 | 0.405301956 | 0.243001658 |
| Mrm3 | 6.578681 | 6.627278 | 6.4486 | 0.178692612 | 0.130095394 | 6.6684297 | 6.802113 | 6.39734 | 0.404774641 | 0.271091355 |
| Rpl41 | 13.89849 | 14.03588 | 14.008 | 0.027534382 | -0.10984787 | 14.368022 | 14.382654 | 13.9781 | 0.404533774 | 0.389902519 |
| Igkv6-17 | 8.508817 | 8.992687 | 8.5853 | 0.407435091 | -0.076434969 | 9.1349447 | 8.9902452 | 8.58584 | 0.404407981 | 0.549107504 |
| Cmtm6 | 11.52522 | 11.5664 | 11.444 | 0.122412296 | 0.08123464 | 11.510653 | 11.814968 | 11.4109 | 0.404068148 | 0.099753312 |
| Med15 | 10.4153 | 10.4978 | 10.353 | 0.144327532 | 0.061830434 | 10.561219 | 10.846946 | 10.443 | 0.403983334 | 0.118256269 |
| Gm8624 | 4.958205 | 5.193543 | 5.0859 | 0.107655316 | -0.127682105 | 5.499781 | 5.4254479 | 5.02151 | 0.40393783 | 0.478270932 |
| Mrps10 | 6.902737 | 6.784652 | 6.8769 | -0.092199422 | 0.025885313 | 6.9473621 | 7.2272649 | 6.82357 | 0.403696262 | 0.12379346 |

| | | | | | | | | | | |
|---|---|---|---|---|---|---|---|---|---|---|
| Vps13a | 10.91729 | 10.99607 | 11.046 | -0.050101758 | -0.128890014 | 10.875311 | 11.441844 | 11.0382 | 0.403647218 | -0.162885254 |
| Dcakd | 7.159592 | 7.305549 | 7.281 | 0.024559981 | -0.121397067 | 7.5174266 | 7.6695152 | 7.26604 | 0.403471559 | 0.25138296 |
| Stub1 | 9.592685 | 9.587184 | 9.5834 | 0.003832468 | 0.009333762 | 9.9384547 | 10.040497 | 9.63711 | 0.403386074 | 0.301344077 |
| Nutf2 | 6.537607 | 6.206986 | 6.1036 | 0.103372692 | 0.433993588 | 6.3494342 | 6.7660607 | 6.3628 | 0.403257657 | -0.013368866 |
| Enpp4 | 8.355025 | 8.457676 | 8.3755 | 0.082182435 | -0.020469181 | 7.8476383 | 8.4907387 | 8.08764 | 0.403098803 | -0.240001639 |
| Bysl | 8.745607 | 8.88433 | 8.5877 | 0.296580551 | 0.157857232 | 8.9533812 | 9.1610924 | 8.75816 | 0.40293647 | 0.195225191 |
| Fcgr2b | 10.55168 | 10.53719 | 10.375 | 0.162628086 | 0.177112958 | 11.097117 | 10.993334 | 10.5904 | 0.402911988 | 0.506694707 |
| Tkt | 11.7236 | 11.84318 | 11.727 | 0.116368816 | -0.00320983 | 12.04249 | 12.164101 | 11.7613 | 0.402798676 | 0.281187037 |
| Armc3 | 5.506622 | 5.460418 | 5.292 | 0.1684293 | 0.214633109 | 5.4409989 | 5.7790951 | 5.37649 | 0.402601686 | 0.064505469 |
| Tnn | 1.95295 | 1.897947 | 1.8916 | 0.006388606 | 0.061391592 | 2.0169783 | 2.3288259 | 1.92635 | 0.402479615 | 0.090631974 |
| Klhdc4 | 9.296763 | 9.335433 | 9.1586 | 0.176823238 | 0.138153612 | 9.5176109 | 9.8471011 | 9.44473 | 0.402368971 | 0.072878696 |
| Rps28 | 12.99178 | 13.18532 | 13.062 | 0.12320341 | -0.070331118 | 13.262224 | 13.322897 | 12.9205 | 0.402358624 | 0.341685961 |
| Fgd4 | 6.917398 | 6.879593 | 6.687 | 0.192612606 | 0.230418025 | 6.8791548 | 7.3578247 | 6.95548 | 0.402346364 | -0.07632353 |
| Rap2a | 8.358479 | 8.709132 | 8.2595 | 0.449599388 | 0.098946789 | 8.0588749 | 8.528136 | 8.12598 | 0.402159621 | -0.067101419 |
| Gm4202 | 2.787136 | 2.826518 | 3.4718 | -0.645304571 | -0.684686748 | 2.7668368 | 2.8779879 | 2.47588 | 0.402110606 | 0.290959552 |
| Mcl1 | 12.95521 | 12.89843 | 12.84 | 0.058507304 | 0.115292842 | 12.636011 | 12.989252 | 12.5872 | 0.402085796 | 0.048844644 |
| Ist1 | 10.97784 | 11.03619 | 11.026 | 0.01041401 | -0.04793551 | 11.146807 | 11.378257 | 10.9763 | 0.401924134 | 0.170474113 |
| Gm44502 | 1.62675 | 1.734665 | 1.6339 | 0.100747309 | -0.007168176 | 1.9548429 | 1.9714514 | 1.56961 | 0.401841947 | 0.385233465 |
| Gm5692 | 1.859563 | 1.94283 | 1.8583 | 0.08451675 | 0.001249725 | 1.8814683 | 2.2762195 | 1.87441 | 0.401805647 | 0.007054425 |
| Ighv5-16 | 6.638414 | 6.62792 | 6.6589 | -0.031006031 | -0.020511741 | 6.6718863 | 6.8866412 | 6.48487 | 0.40176723 | 0.187012343 |
| Piga | 8.162603 | 8.195618 | 8.1536 | 0.042027934 | 0.009013878 | 8.1444698 | 8.7036904 | 8.30204 | 0.401649174 | -0.157571388 |
| Tuba1b | 12.85037 | 12.88735 | 12.865 | 0.02218535 | -0.014794892 | 12.975815 | 13.291962 | 12.8903 | 0.401615772 | 0.08546868 |
| D1Ertd622e | 9.91763 | 10.12739 | 9.8322 | 0.295222705 | 0.085466781 | 9.8216731 | 10.099941 | 9.69849 | 0.401451996 | 0.123184261 |
| Zfp691 | 6.890937 | 6.985967 | 6.7269 | 0.259042582 | 0.164013023 | 7.2247127 | 7.207635 | 6.80622 | 0.401410635 | 0.418488316 |
| Atp5g3 | 10.67454 | 10.60617 | 10.639 | -0.032377024 | 0.035992676 | 10.708262 | 10.905429 | 10.5046 | 0.40083192 | 0.203665146 |
| Mob1b | 8.820637 | 9.035672 | 8.8947 | 0.141009436 | -0.074025209 | 8.7060891 | 9.3194924 | 8.91873 | 0.400759624 | -0.212643681 |
| Flt1 | 9.00995 | 8.606771 | 8.7901 | -0.183326416 | 0.219851734 | 7.9687734 | 8.997762 | 8.5972 | 0.400557633 | -0.628430986 |
| Gm8430 | 8.389989 | 8.260727 | 8.0494 | 0.211355594 | 0.340618103 | 7.8478315 | 8.5436818 | 8.14347 | 0.400212198 | -0.295638127 |
| Dimt1 | 7.698101 | 7.835995 | 7.6662 | 0.16979138 | 0.031897477 | 8.0152952 | 8.1964119 | 7.79635 | 0.400062917 | 0.218946141 |
| Gm48976 | 4.578471 | 4.575799 | 4.4916 | 0.084167039 | 0.086839029 | 4.5952205 | 4.9932337 | 4.59347 | 0.399766565 | 0.001753281 |
| Cdhr4 | 2.721624 | 2.649382 | 2.6757 | -0.02633898 | 0.045903765 | 2.668977 | 3.1407038 | 2.74097 | 0.399734274 | -0.071992533 |
| Impdh2 | 11.38353 | 11.40812 | 11.394 | 0.013649102 | -0.01093737 | 11.542639 | 11.772664 | 11.373 | 0.399674936 | 0.169649664 |

| | | | | | | | | | | |
|---|---|---|---|---|---|---|---|---|---|---|
| Gm19684 | 4.518972 | 4.728408 | 4.5359 | 0.192488591 | -0.016947112 | 4.7099702 | 4.9768387 | 4.57737 | 0.399468618 | 0.132600128 |
| Gpn2 | 7.404474 | 7.44249 | 7.3025 | 0.140031598 | 0.102015578 | 7.7183475 | 7.8457984 | 7.44646 | 0.399338459 | 0.271887501 |
| Cox6b1 | 10.424 | 10.43009 | 10.429 | 0.001025866 | -0.005063926 | 10.68854 | 10.821514 | 10.4226 | 0.398943702 | 0.265970143 |
| Nudt13 | 7.51012 | 7.811616 | 7.7087 | 0.102920913 | -0.198574554 | 7.6406579 | 8.1440863 | 7.74528 | 0.39880594 | -0.10462239 |
| Adamts4 | 6.508157 | 6.154461 | 5.3149 | 0.839600518 | 1.193296757 | 5.4969217 | 5.7010413 | 5.30225 | 0.398795637 | 0.194676024 |
| Them6 | 8.164985 | 8.274088 | 8.0927 | 0.18142711 | 0.072324425 | 8.3471271 | 8.4473179 | 8.04872 | 0.398596811 | 0.298405967 |
| 3110082I17Rik | 6.406882 | 6.218704 | 6.2591 | -0.040418286 | 0.147759526 | 6.5209624 | 6.6473739 | 6.24881 | 0.398565644 | 0.272154171 |
| Gm47406 | 5.900877 | 6.101225 | 5.9429 | 0.158366185 | -0.041982309 | 6.1170104 | 6.5245603 | 6.12603 | 0.398531058 | -0.009018844 |
| Txndc16 | 10.38257 | 10.55297 | 10.389 | 0.164042982 | -0.006353268 | 10.576876 | 10.985359 | 10.5871 | 0.398301352 | -0.01018255 |
| Slc25a1 | 8.282449 | 8.224133 | 8.3707 | -0.146528444 | -0.088212292 | 8.5342399 | 8.7106193 | 8.31235 | 0.398273049 | 0.221893671 |
| Adal | 7.317134 | 7.406768 | 7.3767 | 0.03008469 | -0.059548753 | 7.6168217 | 7.8063449 | 7.40809 | 0.398250154 | 0.208726884 |
| Cmtm3 | 8.032496 | 8.112826 | 7.9382 | 0.174674003 | 0.094344545 | 7.8032142 | 8.1582401 | 7.76031 | 0.397934765 | 0.042908788 |
| Smn1 | 8.338917 | 8.354161 | 8.3611 | -0.006941078 | -0.022184878 | 8.5465956 | 8.7447242 | 8.34682 | 0.397900703 | 0.199772106 |
| Utp23 | 8.254751 | 8.247431 | 8.0454 | 0.202004969 | 0.20932574 | 8.1839941 | 8.5625628 | 8.16501 | 0.39755538 | 0.018986715 |
| Romo1 | 7.402356 | 7.468683 | 7.303 | 0.165731867 | 0.099405598 | 7.7887871 | 7.5877256 | 7.19019 | 0.397533551 | 0.598594978 |
| Agfg1 | 11.12229 | 11.10523 | 10.964 | 0.140862602 | 0.15792084 | 10.978271 | 11.360966 | 10.9634 | 0.397528654 | 0.014833535 |
| Fis1 | 10.21204 | 10.26026 | 10.205 | 0.055303583 | 0.007079784 | 10.434249 | 10.461992 | 10.0647 | 0.397273977 | 0.369531271 |
| Tfg | 9.313282 | 9.309743 | 9.2726 | 0.037123859 | 0.040663532 | 9.1363236 | 9.4089858 | 9.01193 | 0.397051815 | 0.124389608 |
| Erp29 | 10.63566 | 10.74613 | 10.637 | 0.10866018 | -0.001805188 | 10.909975 | 10.987076 | 10.5903 | 0.396795516 | 0.319694793 |
| Ak4 | 3.253793 | 3.182603 | 3.2487 | -0.066079746 | 0.005110558 | 3.239543 | 3.7808278 | 3.38404 | 0.396788587 | -0.144496182 |
| Spry2 | 6.837309 | 6.784921 | 6.7275 | 0.057445814 | 0.10983414 | 6.3264592 | 6.6564188 | 6.25973 | 0.396686695 | 0.066727096 |
| Rangap1 | 10.8523 | 11.01358 | 10.866 | 0.147792474 | -0.013489076 | 10.935349 | 11.242126 | 10.8455 | 0.396620098 | 0.089843379 |
| Fkbp8 | 11.22974 | 11.21097 | 11.219 | -0.008023894 | 0.010743918 | 11.585359 | 11.661952 | 11.2656 | 0.396379268 | 0.319786212 |
| Gm48054 | 3.114273 | 3.106875 | 2.9742 | 0.132699196 | 0.140097514 | 3.0291486 | 3.4909316 | 3.09468 | 0.396248328 | -0.065534654 |
| Endod1 | 8.676463 | 8.902213 | 8.7918 | 0.11037156 | -0.115378062 | 8.6296635 | 9.1108063 | 8.71471 | 0.396100862 | -0.085041977 |
| Apool | 7.413377 | 7.504515 | 7.5549 | -0.05041832 | -0.141556759 | 7.7645158 | 8.0200027 | 7.6242 | 0.395803198 | 0.140316341 |
| Polr1d | 10.39422 | 10.43179 | 10.417 | 0.014531116 | -0.023036948 | 10.475301 | 10.760652 | 10.3652 | 0.395444578 | 0.11009337 |
| Cdk16 | 9.125858 | 9.282272 | 9.2829 | -0.000589815 | -0.157003821 | 9.5975275 | 9.8006979 | 9.40527 | 0.395425723 | 0.192255247 |
| Olfr862 | 2.334729 | 2.431399 | 2.3976 | 0.033842727 | -0.062827967 | 2.41643 | 2.8880186 | 2.49269 | 0.395330439 | -0.076258099 |
| Eif4h | 11.70906 | 11.74571 | 11.652 | 0.09356485 | 0.056918886 | 11.831373 | 12.078882 | 11.6836 | 0.395277014 | 0.147768512 |
| Rai14 | 8.22386 | 7.930255 | 7.7946 | 0.135610448 | 0.42921586 | 7.4028086 | 7.656969 | 7.26193 | 0.395039819 | 0.140879487 |
| Vamp8 | 8.888193 | 8.954085 | 8.953 | 0.001076377 | -0.064815782 | 9.0937575 | 9.3021791 | 8.90742 | 0.394758186 | 0.186336598 |

| | | | | | | | | | | |
|---|---|---|---|---|---|---|---|---|---|---|
| Pkdcc | 6.832386 | 6.71741 | 6.275 | 0.442375253 | 0.557351981 | 6.3845931 | 6.2676696 | 5.87298 | 0.394688711 | 0.511612235 |
| Arel1 | 10.46717 | 10.52252 | 10.435 | 0.0878835 | 0.032531994 | 10.496821 | 10.861633 | 10.4673 | 0.394293082 | 0.029480608 |
| Knop1 | 9.314151 | 9.325134 | 9.3257 | -0.000569439 | -0.011551766 | 9.3847715 | 9.5845086 | 9.19066 | 0.393848141 | 0.194110967 |
| Apol9a | 4.554272 | 4.70624 | 4.8015 | -0.095221588 | -0.247189577 | 4.2656709 | 5.0431738 | 4.64941 | 0.39376021 | -0.383742738 |
| Iars | 10.49358 | 10.46044 | 10.356 | 0.104157736 | 0.137296288 | 10.787367 | 10.993542 | 10.6006 | 0.392990538 | 0.18681568 |
| Dctpp1 | 7.048652 | 6.89718 | 7.0294 | -0.132258453 | 0.019213162 | 7.3546288 | 7.3831066 | 6.99031 | 0.392796156 | 0.364318391 |
| Foxred1 | 9.029535 | 8.988299 | 8.9341 | 0.054201066 | 0.095436802 | 9.3426978 | 9.5162376 | 9.12349 | 0.392747801 | 0.219207983 |
| Spag7 | 9.055053 | 9.12895 | 9.0383 | 0.090695412 | 0.016798477 | 9.1848211 | 9.4145904 | 9.02195 | 0.392642392 | 0.162873088 |
| Soat2 | 5.246093 | 5.084299 | 4.9358 | 0.148524865 | 0.310319306 | 5.3046284 | 5.3712316 | 4.97888 | 0.392348803 | 0.325745566 |
| Pgap2 | 9.16984 | 9.281815 | 9.294 | -0.012221095 | -0.124195695 | 9.2810943 | 9.722447 | 9.33022 | 0.392227787 | -0.04912494 |
| Tysnd1 | 8.005672 | 8.151285 | 7.8241 | 0.327201534 | 0.181589369 | 8.2111004 | 8.2320253 | 7.83984 | 0.392188698 | 0.371263716 |
| Hspa4 | 11.80637 | 11.78085 | 11.762 | 0.019037556 | 0.04455837 | 11.716798 | 12.16938 | 11.7774 | 0.392024254 | -0.06055747 |
| Igkv4-73 | 3.242366 | 3.400171 | 3.2504 | 0.149792648 | -0.008011706 | 3.356091 | 3.7090231 | 3.31728 | 0.391740687 | 0.038808635 |
| Snrpe | 10.08416 | 10.05235 | 10.086 | -0.033752259 | -0.001937425 | 10.227773 | 10.474454 | 10.0831 | 0.391338089 | 0.144657372 |
| Gm7536 | 13.47119 | 13.51272 | 13.507 | 0.005913174 | -0.035620878 | 13.66654 | 13.850089 | 13.4588 | 0.391250064 | 0.207700879 |
| Scarb2 | 10.27069 | 10.36954 | 10.217 | 0.152399624 | 0.053546872 | 10.147435 | 10.545358 | 10.1541 | 0.391239722 | -0.006683703 |
| Kdm5c | 11.68816 | 11.79872 | 11.524 | 0.274714442 | 0.164162336 | 11.690129 | 11.994774 | 11.6039 | 0.390921548 | 0.086276349 |
| Ost4 | 9.552297 | 9.568624 | 9.5173 | 0.051345199 | 0.035018177 | 9.9090921 | 9.9083551 | 9.51784 | 0.39051383 | 0.391250801 |
| Igkv17-127 | 8.783139 | 8.939798 | 8.7411 | 0.198724923 | 0.042066108 | 9.2486477 | 9.3149645 | 8.92479 | 0.390174768 | 0.323858057 |
| Zfp773 | 6.495275 | 6.657366 | 6.581 | 0.076382125 | -0.085708677 | 6.6398427 | 7.0006632 | 6.61052 | 0.390145266 | 0.029324722 |
| Tmem248 | 9.969764 | 9.972286 | 9.9277 | 0.044584955 | 0.042062358 | 9.955578 | 10.22916 | 9.83912 | 0.390044748 | 0.116462689 |
| Adamts9 | 7.93475 | 7.650868 | 6.6028 | 1.048090973 | 1.331972543 | 6.8453822 | 7.0617517 | 6.67195 | 0.389801552 | 0.173431971 |
| Plscr1 | 8.270728 | 7.743512 | 7.6437 | 0.099774058 | 0.626990762 | 7.8550818 | 7.9446429 | 7.55491 | 0.389730048 | 0.300169025 |
| Nfkbia | 12.20578 | 12.29028 | 12.093 | 0.197704738 | 0.113204862 | 11.951934 | 12.237554 | 11.8479 | 0.389682418 | 0.104062966 |
| Nedd8 | 9.990107 | 10.16014 | 10.026 | 0.134112825 | -0.035922281 | 10.26692 | 10.398313 | 10.0086 | 0.389669603 | 0.258276764 |
| Zfp952 | 8.194626 | 8.36204 | 8.335 | 0.027085178 | -0.140328684 | 8.4396177 | 8.806124 | 8.41692 | 0.389202774 | 0.022696505 |
| Aurkaip1 | 9.301523 | 9.356374 | 9.3236 | 0.032742814 | -0.022108315 | 9.6423155 | 9.7252784 | 9.33627 | 0.389008917 | 0.306046018 |
| Pelp1 | 9.384523 | 9.434456 | 9.393 | 0.041444983 | -0.008487538 | 9.4772048 | 9.6869475 | 9.29797 | 0.388981674 | 0.179239039 |
| Ftl1 | 13.78172 | 13.88495 | 13.771 | 0.113664454 | 0.010437724 | 13.866036 | 13.939556 | 13.5508 | 0.388771428 | 0.315251764 |
| Gm42417 | 3.038147 | 2.878303 | 2.8657 | 0.012563887 | 0.172407978 | 2.7251147 | 3.0241543 | 2.63553 | 0.388623657 | 0.089584061 |
| Grpel1 | 9.124322 | 9.217748 | 9.1185 | 0.099274934 | 0.005848637 | 9.2766807 | 9.5062316 | 9.11762 | 0.388613417 | 0.15906257 |
| Ddx41 | 10.45584 | 10.54798 | 10.464 | 0.083489236 | -0.008652696 | 10.835941 | 10.921642 | 10.5331 | 0.388540482 | 0.30283968 |

| | | | | | | | | | | |
|---|---|---|---|---|---|---|---|---|---|---|
| Ccdc137 | 8.257439 | 8.397137 | 8.3725 | 0.024630993 | -0.115067181 | 8.7328599 | 8.9786448 | 8.59011 | 0.388536427 | 0.142751471 |
| Ap5b1 | 7.879585 | 8.008236 | 7.9356 | 0.072670596 | -0.055981291 | 8.0783308 | 8.1528232 | 7.7643 | 0.388519449 | 0.314027035 |
| Rnaset2b | 11.28773 | 11.28921 | 11.296 | -0.006557154 | -0.008031386 | 11.264662 | 11.623982 | 11.2356 | 0.388413018 | 0.029092956 |
| Sec23b | 10.13812 | 10.29729 | 10.033 | 0.26460587 | 0.105431887 | 10.08684 | 10.445855 | 10.0577 | 0.388204656 | 0.029190151 |
| Srsf6 | 11.72407 | 11.72989 | 11.696 | 0.034274082 | 0.02846004 | 12.00604 | 12.286459 | 11.8983 | 0.38811104 | 0.107692395 |
| Cachd1 | 3.933957 | 3.963052 | 3.9109 | 0.052170606 | 0.023075407 | 3.627602 | 4.0868311 | 3.69884 | 0.387989473 | -0.071239609 |
| Rnf25 | 8.381192 | 8.579475 | 8.5154 | 0.06409098 | -0.134191831 | 8.7575183 | 9.0020886 | 8.61438 | 0.38770497 | 0.143134634 |
| Csnk2b | 10.45953 | 10.41714 | 10.442 | -0.024971478 | 0.017424906 | 10.599266 | 10.813 | 10.4255 | 0.387537778 | 0.173804097 |
| Ubxn4 | 10.3347 | 10.52236 | 10.334 | 0.188408639 | 0.00075491 | 10.432421 | 10.853247 | 10.4658 | 0.387402124 | -0.033423866 |
| Pi16 | 5.598535 | 5.373888 | 5.1737 | 0.200214438 | 0.424861057 | 5.6051068 | 5.2862138 | 4.89898 | 0.387231397 | 0.706124413 |
| Bspry | 4.920024 | 4.810811 | 4.8949 | -0.084048029 | 0.025165006 | 4.96969 | 5.3193508 | 4.93228 | 0.387073672 | 0.037412868 |
| Polr3e | 9.30573 | 9.318149 | 9.2296 | 0.088505308 | 0.076086919 | 9.4615007 | 9.6435503 | 9.25706 | 0.386492843 | 0.204443188 |
| Mrpl51 | 8.680428 | 8.691983 | 8.6882 | 0.003820132 | -0.007735424 | 8.7917569 | 8.9459524 | 8.55946 | 0.386489378 | 0.232293952 |
| Ndufa8 | 8.941673 | 8.987844 | 9 | -0.012107272 | -0.058278211 | 9.0177273 | 9.1631633 | 8.77684 | 0.386326813 | 0.240890855 |
| Naip2 | 8.582971 | 8.666336 | 8.5256 | 0.140763953 | 0.057399026 | 8.5357002 | 8.7574902 | 8.37126 | 0.386230726 | 0.16444068 |
| Polr2j | 8.306067 | 8.411174 | 8.323 | 0.088217188 | -0.016889666 | 8.5976755 | 8.7207696 | 8.33476 | 0.386012117 | 0.26291802 |
| Bag1 | 9.878553 | 9.931253 | 9.8445 | 0.08675651 | 0.034056112 | 9.9134589 | 10.170393 | 9.78448 | 0.385912991 | 0.128978563 |
| Iglv2 | 8.175221 | 8.189107 | 8.1679 | 0.021175975 | 0.007289734 | 8.1117578 | 8.3473069 | 7.96141 | 0.385895355 | 0.150346245 |
| Hsp90b1 | 13.02159 | 13.19482 | 12.877 | 0.317450738 | 0.144218058 | 13.032744 | 13.29475 | 12.9089 | 0.38585324 | 0.123847417 |
| Atp5l | 10.74548 | 10.75944 | 10.596 | 0.163374798 | 0.149415354 | 11.058951 | 11.018477 | 10.6328 | 0.385681114 | 0.426155233 |
| AC116766.1 | 3.934569 | 3.776781 | 4.0054 | -0.228607309 | -0.070819568 | 4.2158789 | 4.1488651 | 3.76324 | 0.385626001 | 0.452639804 |
| Ctnnb1 | 11.8177 | 11.9223 | 11.781 | 0.141430208 | 0.03682912 | 11.947812 | 12.227792 | 11.8422 | 0.385601546 | 0.105621808 |
| Nmt1 | 10.77405 | 10.74908 | 10.716 | 0.032909664 | 0.057880022 | 10.872162 | 11.126117 | 10.7407 | 0.385410438 | 0.131455626 |
| Saysd1 | 6.978973 | 7.047827 | 6.9811 | 0.066737031 | -0.002116647 | 7.2028894 | 7.3306429 | 6.94524 | 0.385401712 | 0.257648248 |
| Gm47611 | 4.929915 | 5.057889 | 5.1065 | -0.048640623 | -0.176614299 | 5.101022 | 5.5625 | 5.17734 | 0.385162731 | -0.076315327 |
| Ppdpf | 9.388825 | 9.567473 | 9.529 | 0.038442555 | -0.140205656 | 9.7720161 | 9.8274804 | 9.44276 | 0.384722953 | 0.329258618 |
| Tpd52l2 | 9.420376 | 9.36055 | 9.4098 | -0.04920891 | 0.010616883 | 9.5118481 | 9.6262557 | 9.24171 | 0.384544822 | 0.27013715 |
| Cldnd1 | 9.132662 | 9.242058 | 9.1091 | 0.132966396 | 0.023570745 | 9.0998951 | 9.5360145 | 9.15159 | 0.384425525 | -0.051693867 |
| Amigo2 | 7.898681 | 8.002913 | 7.8727 | 0.130201354 | 0.025969773 | 7.6730249 | 8.1137455 | 7.72969 | 0.384057154 | -0.056663402 |
| Isca2 | 8.142381 | 8.137695 | 8.1249 | 0.01280678 | 0.017492473 | 8.4844066 | 8.4527869 | 8.06879 | 0.383996439 | 0.415616088 |
| Rtl5 | 6.90695 | 6.478061 | 7.0313 | -0.553245557 | -0.12435685 | 6.9423441 | 7.3627824 | 6.97886 | 0.383920028 | -0.036518306 |
| Eif4a1 | 12.82528 | 12.77088 | 12.715 | 0.055892444 | 0.110296944 | 12.847438 | 13.148835 | 12.7649 | 0.383918334 | 0.082521848 |

| | | | | | | | | | | |
|---|---|---|---|---|---|---|---|---|---|---|
| Zdhhc12 | 6.770442 | 6.821642 | 6.8458 | -0.024202803 | -0.075402814 | 7.1174085 | 7.1399259 | 6.75609 | 0.383836377 | 0.361319002 |
| Tubd1 | 6.668134 | 6.822829 | 6.7518 | 0.071017227 | -0.083677535 | 6.7571098 | 6.9660477 | 6.58225 | 0.383797309 | 0.174859426 |
| Pold3 | 9.348535 | 9.38633 | 9.3508 | 0.035516006 | -0.002278315 | 9.4282163 | 9.972638 | 9.58896 | 0.383679831 | -0.160741852 |
| Nadk | 10.64211 | 10.81686 | 10.536 | 0.280977404 | 0.106232348 | 10.882508 | 10.944722 | 10.5612 | 0.383533164 | 0.32131975 |
| Polb | 8.783747 | 8.87627 | 8.7108 | 0.16546476 | 0.072942596 | 8.8289482 | 9.2197722 | 8.83628 | 0.383496948 | -0.007327108 |
| Anxa7 | 10.58036 | 10.64691 | 10.513 | 0.133901416 | 0.067347034 | 10.6221 | 10.877255 | 10.4938 | 0.383453458 | 0.128298231 |
| Igkj2 | 6.357237 | 6.825917 | 6.3425 | 0.483458114 | 0.01477754 | 6.6484304 | 6.7358622 | 6.35249 | 0.38337031 | 0.295938494 |
| Rbmxl1 | 10.21213 | 10.26382 | 10.229 | 0.034638134 | -0.017049424 | 10.339123 | 10.600325 | 10.2173 | 0.383018722 | 0.121816938 |
| Nfe2l2 | 10.46695 | 10.55523 | 10.236 | 0.319544468 | 0.231262434 | 10.327629 | 10.528216 | 10.1455 | 0.382714988 | 0.18212866 |
| Nol10 | 8.976907 | 8.991439 | 8.8027 | 0.188711875 | 0.174180271 | 9.0454983 | 9.3330298 | 8.95036 | 0.38266723 | 0.095135711 |
| Scn5a | 3.95447 | 3.743741 | 3.5884 | 0.155353664 | 0.366083177 | 3.5024245 | 3.8836927 | 3.50105 | 0.382647047 | 0.001378854 |
| Fndc5 | 4.088876 | 3.874524 | 3.9777 | -0.103201808 | 0.11115028 | 4.0717935 | 4.2689913 | 3.88647 | 0.382523736 | 0.18532592 |
| Polr2f | 8.152507 | 8.312636 | 8.2751 | 0.037515812 | -0.122613032 | 8.40536 | 8.5720288 | 8.1896 | 0.382431744 | 0.215762859 |
| Dync1h1 | 12.85849 | 12.90829 | 12.845 | 0.063583968 | 0.013788056 | 12.852295 | 13.31142 | 12.9291 | 0.382361548 | -0.076762676 |
| 37316 | 10.3139 | 10.30376 | 10.379 | -0.074973466 | -0.064832818 | 10.318573 | 10.564605 | 10.1823 | 0.38232063 | 0.136289074 |
| Igkv2-109 | 7.230775 | 7.431991 | 7.1409 | 0.291060921 | 0.089845306 | 7.5235945 | 7.5277402 | 7.14558 | 0.382164594 | 0.378018886 |
| Pelo | 7.395319 | 7.409168 | 7.3161 | 0.093029241 | 0.079180695 | 7.2841789 | 7.5841268 | 7.20197 | 0.382158997 | 0.082211049 |
| Panx1 | 9.284742 | 9.379246 | 9.3366 | 0.042666284 | -0.051837358 | 9.2891334 | 9.5266059 | 9.14484 | 0.381764818 | 0.144292335 |
| Coq4 | 5.913063 | 5.925948 | 5.8644 | 0.061537137 | 0.048652479 | 6.4288686 | 6.3882462 | 6.00665 | 0.381594767 | 0.422217218 |
| Bend6 | 3.872507 | 3.947836 | 3.8882 | 0.059631277 | -0.015698063 | 4.1075697 | 4.1746893 | 3.79322 | 0.381466077 | 0.314346424 |
| Gstt1 | 5.78998 | 5.365547 | 5.6484 | -0.282876392 | 0.14155656 | 5.4892228 | 5.7471505 | 5.36591 | 0.381244111 | 0.123316438 |
| Gtf3c6 | 9.059001 | 8.926592 | 8.9716 | -0.044976688 | 0.087432682 | 8.9478525 | 9.251963 | 8.87073 | 0.381233743 | 0.077123161 |
| Cog8 | 9.275272 | 9.57937 | 9.3786 | 0.200770219 | -0.103327606 | 9.4257329 | 9.6832266 | 9.30212 | 0.381107947 | 0.123614241 |
| Ankrd17 | 11.75154 | 11.92031 | 11.694 | 0.226031932 | 0.057260978 | 11.609671 | 12.214207 | 11.8333 | 0.380885656 | -0.223650381 |
| Zfand3 | 9.982519 | 10.06347 | 9.9491 | 0.114336986 | 0.033390678 | 9.9328807 | 10.251822 | 9.87097 | 0.380854556 | 0.061913714 |
| Rnasek | 9.165272 | 9.267493 | 9.2214 | 0.046105594 | -0.056115528 | 9.4984248 | 9.468802 | 9.08795 | 0.380850049 | 0.410472871 |
| Txnrd3 | 6.505147 | 6.361073 | 6.3859 | -0.02480794 | 0.119265775 | 6.4151962 | 6.4999651 | 6.11926 | 0.380707499 | 0.295938645 |
| Cnn3 | 10.80982 | 10.98414 | 10.761 | 0.223103246 | 0.048786136 | 11.034456 | 11.110378 | 10.7297 | 0.380637798 | 0.304715579 |
| Cluh | 10.5374 | 10.63232 | 10.544 | 0.087879826 | -0.007035328 | 10.76438 | 10.893073 | 10.5125 | 0.380622136 | 0.25192945 |
| Igkv12-44 | 9.233077 | 8.864763 | 9.1922 | -0.327437076 | 0.040877232 | 8.9818071 | 9.0047599 | 8.62419 | 0.380573008 | 0.357620219 |
| Trp53 | 11.19662 | 11.41874 | 11.288 | 0.130943986 | -0.091177724 | 11.488117 | 11.658556 | 11.2782 | 0.380362776 | 0.209923587 |
| Gm6034 | 1.930842 | 2.028056 | 1.9356 | 0.09249914 | -0.004715088 | 2.0240575 | 2.3195719 | 1.93923 | 0.380342304 | 0.084827932 |

| | | | | | | | | | | |
|---|---|---|---|---|---|---|---|---|---|---|
| Fhod1 | 9.239608 | 9.477337 | 9.3135 | 0.163861262 | -0.073867147 | 9.5613796 | 9.8561808 | 9.476 | 0.380183843 | 0.085382616 |
| Lysmd2 | 6.398836 | 6.335173 | 6.341 | -0.00585742 | 0.057805294 | 6.215352 | 6.5463606 | 6.16638 | 0.37997854 | 0.048969858 |
| Rpl36al | 11.55898 | 11.50766 | 11.454 | 0.053389046 | 0.10470516 | 11.655619 | 11.86398 | 11.4841 | 0.379909254 | 0.171547827 |
| Shmt1 | 7.855998 | 7.756482 | 7.8578 | -0.101289113 | -0.001773125 | 7.8297467 | 8.2963099 | 7.91641 | 0.379894971 | -0.086668286 |
| Mafb | 8.16763 | 7.24884 | 7.1551 | 0.093729148 | 1.012519257 | 7.3723715 | 6.9092726 | 6.52948 | 0.379789034 | 0.842887861 |
| Thap4 | 7.773858 | 7.961291 | 7.9253 | 0.035957533 | -0.151474773 | 7.7247772 | 8.0586928 | 7.67905 | 0.379642969 | 0.045727408 |
| Snapc2 | 7.643259 | 7.910519 | 7.7536 | 0.156966876 | -0.11029313 | 7.848773 | 7.9539085 | 7.57432 | 0.379592991 | 0.274457519 |
| Ghitm | 11.41195 | 11.55911 | 11.366 | 0.192798956 | 0.04563648 | 11.596056 | 11.836758 | 11.4573 | 0.379483176 | 0.138780509 |
| Card11 | 11.63817 | 11.78103 | 11.731 | 0.049969226 | -0.092888722 | 11.933138 | 12.169007 | 11.7896 | 0.379417104 | 0.143547769 |
| Tmem131 | 11.11194 | 11.21454 | 11.037 | 0.177103626 | 0.074497242 | 11.080142 | 11.566721 | 11.1874 | 0.379344112 | -0.107235346 |
| AC133103.2 | 3.011691 | 3.035218 | 2.9267 | 0.108523697 | 0.084997334 | 3.0229381 | 3.3264329 | 2.94717 | 0.379258711 | 0.075763905 |
| Aph1a | 10.11791 | 10.08606 | 10.099 | -0.013026305 | 0.018831397 | 10.142098 | 10.343533 | 9.96451 | 0.379022601 | 0.177587833 |
| Tsta3 | 8.957547 | 8.927311 | 8.9913 | -0.063959957 | -0.03372345 | 9.1126113 | 9.3519809 | 8.97301 | 0.378969737 | 0.1396002 |
| C1qtnf12 | 5.478103 | 5.430781 | 5.2273 | 0.203496606 | 0.25081783 | 5.6538351 | 5.6916485 | 5.3129 | 0.378748022 | 0.340934705 |
| Arl6ip4 | 8.325513 | 8.349987 | 8.4111 | -0.061127361 | -0.085600906 | 8.5669301 | 8.7369477 | 8.35835 | 0.378595577 | 0.208577933 |
| Ap1s3 | 9.851104 | 10.228 | 9.7892 | 0.438761705 | 0.061869772 | 10.139263 | 10.301462 | 9.9229 | 0.378560682 | 0.216361298 |
| Lfng | 10.3838 | 10.52869 | 10.411 | 0.117892248 | -0.027005804 | 10.500392 | 10.697238 | 10.3187 | 0.378521093 | 0.18167512 |
| Pde3b | 10.54706 | 10.75036 | 10.571 | 0.179327248 | -0.023976174 | 10.196419 | 10.960199 | 10.5819 | 0.378305574 | -0.385474473 |
| Zbtb17 | 8.956472 | 9.015764 | 8.9608 | 0.054973249 | -0.00431847 | 9.2107825 | 9.4665051 | 9.08833 | 0.378177295 | 0.122454711 |
| Lman2 | 10.97982 | 11.15685 | 10.95 | 0.206752552 | 0.029724508 | 11.228601 | 11.398973 | 11.0209 | 0.378105116 | 0.207733887 |
| Ndufa5 | 8.012538 | 8.045826 | 7.9476 | 0.098236546 | 0.06494868 | 8.1390764 | 8.3485364 | 7.97043 | 0.378104408 | 0.168644409 |
| Mrps23 | 8.339869 | 8.382089 | 8.3576 | 0.02445831 | -0.017761692 | 8.6005868 | 8.6481567 | 8.27017 | 0.377987848 | 0.330417874 |
| Atp5j2 | 9.908954 | 9.97483 | 9.9539 | 0.020933917 | -0.044942162 | 10.21083 | 10.356209 | 9.97851 | 0.377696545 | 0.232317721 |
| Ppp1r16a | 7.90915 | 7.995903 | 8.0282 | -0.032290152 | -0.119043245 | 8.2304862 | 8.472216 | 8.09455 | 0.377661271 | 0.135931481 |
| Clta | 11.09101 | 11.16778 | 11.114 | 0.054061076 | -0.022714664 | 11.252005 | 11.42083 | 11.0432 | 0.377652832 | 0.208827309 |
| Gng2 | 10.4841 | 10.51175 | 10.512 | 0.000224478 | -0.027427148 | 10.644928 | 10.876519 | 10.4989 | 0.377633158 | 0.146042705 |
| Pck2 | 9.512504 | 9.933302 | 9.7255 | 0.20782593 | -0.21297247 | 10.046393 | 10.019857 | 9.64232 | 0.377539561 | 0.404075686 |
| Anks1 | 9.061944 | 9.131271 | 8.9954 | 0.135894489 | 0.066567677 | 9.1883008 | 9.5823063 | 9.20523 | 0.37708035 | -0.01692513 |
| Tars | 10.02897 | 10.02222 | 9.9022 | 0.119976102 | 0.126723165 | 10.235546 | 10.614232 | 10.2372 | 0.377058917 | -0.001627259 |
| Rnaseh2a | 8.527526 | 8.509525 | 8.6149 | -0.105424058 | -0.087422551 | 8.6872822 | 8.9318964 | 8.55517 | 0.376729562 | 0.132115334 |
| Riox1 | 7.519321 | 7.643642 | 7.6545 | -0.010817973 | -0.135139097 | 7.5660405 | 7.8875288 | 7.51085 | 0.376674831 | 0.055186519 |
| Stk16 | 8.803373 | 8.790099 | 8.8022 | -0.012091409 | 0.00118198 | 9.1824195 | 9.3520972 | 8.97566 | 0.376439062 | 0.206761388 |

| | | | | | | | | | | |
|---|---|---|---|---|---|---|---|---|---|---|
| Map3k15 | 5.632323 | 5.591207 | 5.4286 | 0.1626555 | 0.203770929 | 5.6953269 | 5.949111 | 5.57273 | 0.376380964 | 0.122596878 |
| Cdc42ep2 | 6.554897 | 6.551485 | 6.2847 | 0.266767343 | 0.27017968 | 6.188993 | 6.3953144 | 6.01896 | 0.376351815 | 0.170030484 |
| Xpot | 9.724942 | 9.894462 | 9.6929 | 0.201571309 | 0.032051501 | 9.7170781 | 10.078083 | 9.70208 | 0.376000422 | 0.014995541 |
| Trim44 | 10.86231 | 10.90861 | 10.83 | 0.078478736 | 0.032186848 | 10.899404 | 11.36926 | 10.9933 | 0.375970662 | -0.093885314 |
| Mapk9 | 9.759084 | 9.766169 | 9.868 | -0.101851021 | -0.108935968 | 9.8231009 | 10.212281 | 9.83634 | 0.375936192 | -0.013243421 |
| Psenen | 10.01101 | 10.06117 | 10.016 | 0.044968547 | -0.005196924 | 10.331589 | 10.434644 | 10.0588 | 0.375813371 | 0.272758483 |
| Ddah1 | 5.040359 | 5.087806 | 4.7494 | 0.338417123 | 0.290969683 | 4.8429149 | 4.8863491 | 4.51056 | 0.375788889 | 0.33235464 |
| Cdc42ep4 | 7.780911 | 7.542037 | 7.4731 | 0.068935398 | 0.307809341 | 7.256331 | 7.5444692 | 7.16873 | 0.375742655 | 0.087604419 |
| Znrd1 | 8.756066 | 8.794886 | 8.7096 | 0.08526063 | 0.046440727 | 9.0306796 | 9.1329117 | 8.75729 | 0.375622887 | 0.2733907 |
| Tsr3 | 7.767731 | 7.839558 | 7.7252 | 0.114323202 | 0.042496709 | 7.9165299 | 8.2071367 | 7.83162 | 0.375519698 | 0.084912893 |
| Atp5o | 10.37716 | 10.43071 | 10.362 | 0.068333899 | 0.014785096 | 10.676082 | 10.810198 | 10.4347 | 0.375496402 | 0.241380485 |
| Igkv2-137 | 7.245818 | 7.507325 | 7.3545 | 0.152804265 | -0.108703546 | 7.651494 | 7.7030671 | 7.32769 | 0.375377794 | 0.323804677 |
| Tra2b | 11.66248 | 11.86611 | 11.654 | 0.212460346 | 0.008830966 | 11.755247 | 12.167066 | 11.7919 | 0.375144446 | -0.03667456 |
| Tg | 4.343695 | 4.477775 | 4.2331 | 0.244711325 | 0.110631066 | 4.2613135 | 4.6010315 | 4.22611 | 0.374917395 | 0.035199417 |
| Prmt1 | 10.60727 | 10.55646 | 10.548 | 0.008544826 | 0.059350354 | 10.758629 | 10.884813 | 10.51 | 0.374784608 | 0.248600322 |
| Proser3 | 6.320288 | 6.454742 | 6.3609 | 0.093828481 | -0.040626114 | 6.6628897 | 6.8928464 | 6.51807 | 0.374772924 | 0.144816224 |
| Klk1b1 | 3.200813 | 3.822007 | 4.3813 | -0.559301304 | -1.180495105 | 5.0622925 | 4.4227879 | 4.04808 | 0.374712 | 1.014216602 |
| Atp6v1b2 | 10.70426 | 11.01219 | 10.625 | 0.387526564 | 0.079600386 | 10.683583 | 10.951272 | 10.5766 | 0.374665434 | 0.106976321 |
| Ltbp1 | 6.339766 | 6.142404 | 6.1261 | 0.016337707 | 0.213699417 | 5.9707329 | 6.6768359 | 6.30227 | 0.374563834 | -0.331539077 |
| Arpp19 | 9.706369 | 9.838109 | 9.7635 | 0.074626177 | -0.057113466 | 9.9160195 | 10.122744 | 9.74829 | 0.374456255 | 0.167732101 |
| Lyar | 8.909506 | 8.916084 | 8.8614 | 0.054689366 | 0.048111929 | 8.9665351 | 9.3204639 | 8.94608 | 0.374385811 | 0.020457008 |
| Tnfaip8l3 | 2.538993 | 2.636863 | 2.5451 | 0.091810496 | -0.006059529 | 2.5373363 | 2.9212214 | 2.54686 | 0.374358522 | -0.009526502 |
| Ido1 | 2.35675 | 2.431857 | 2.4306 | 0.001263561 | -0.073843368 | 2.3490627 | 2.8226871 | 2.44841 | 0.374274918 | -0.099349451 |
| Vmn2r66 | 8.977444 | 8.955111 | 8.8961 | 0.059054043 | 0.081386854 | 8.8983569 | 9.1930564 | 8.8189 | 0.374151576 | 0.079452116 |
| Dnajc15 | 9.093747 | 9.211929 | 9.1107 | 0.101223208 | -0.016958461 | 9.2238429 | 9.5181678 | 9.14404 | 0.374126121 | 0.079801251 |
| A4galt | 6.472976 | 6.244793 | 6.2446 | 0.000221776 | 0.228404455 | 5.7305457 | 6.2165676 | 5.84255 | 0.374021634 | -0.112000218 |
| Ptms | 9.970996 | 9.888919 | 10.092 | -0.203271915 | -0.121194592 | 9.904228 | 10.31038 | 9.93659 | 0.373794083 | -0.032357802 |
| Adamts5 | 5.57997 | 5.57583 | 4.8757 | 0.700170392 | 0.704310609 | 5.5050486 | 5.2688864 | 4.89589 | 0.373000112 | 0.609162343 |
| Timm8b | 7.754495 | 7.734458 | 7.6401 | 0.094403702 | 0.114440751 | 7.9808515 | 7.96785 | 7.59506 | 0.3727889 | 0.385790344 |
| Olfr843 | 3.291349 | 3.283545 | 3.2193 | 0.064294566 | 0.07209878 | 3.3297008 | 3.6772977 | 3.30458 | 0.372719457 | 0.025122621 |
| Igkv9-120 | 8.334869 | 8.466054 | 8.2512 | 0.214870712 | 0.083685435 | 8.6352408 | 8.6731049 | 8.30041 | 0.372694687 | 0.334830547 |
| Dohh | 7.863933 | 7.894665 | 7.6638 | 0.230844036 | 0.200112236 | 7.8711797 | 7.9788967 | 7.60636 | 0.372532407 | 0.264815363 |

| | | | | | | | | | | |
|---|---|---|---|---|---|---|---|---|---|---|
| Sgta | 9.504049 | 9.662482 | 9.4503 | 0.21217012 | 0.053736616 | 9.8165263 | 9.8855895 | 9.51332 | 0.372268722 | 0.303205467 |
| Klra3 | 5.185721 | 5.253868 | 5.3199 | -0.066039974 | -0.134186747 | 5.2002795 | 5.544052 | 5.17194 | 0.372109587 | 0.028337021 |
| Gm47331 | 4.128908 | 4.324617 | 4.1795 | 0.145103135 | -0.050606444 | 4.2221013 | 4.6814605 | 4.30942 | 0.37203598 | -0.087323287 |
| Tmem14c | 9.526487 | 9.531824 | 9.3946 | 0.137218306 | 0.131880834 | 9.6984083 | 9.7076444 | 9.3357 | 0.371940194 | 0.362704145 |
| Ndufs8 | 9.478779 | 9.394643 | 9.3842 | 0.010464947 | 0.094600448 | 9.7696671 | 9.6825715 | 9.31071 | 0.371857293 | 0.458952918 |
| Rpusd1 | 6.471234 | 6.646373 | 6.5922 | 0.054198893 | -0.120940342 | 6.7913554 | 6.8587983 | 6.48694 | 0.371857269 | 0.304414341 |
| Fam117a | 9.52221 | 9.630571 | 9.5656 | 0.064951565 | -0.043409208 | 9.5768399 | 10.017945 | 9.64627 | 0.371671202 | -0.069433437 |
| Fabp5 | 10.02061 | 10.54732 | 9.7265 | 0.820788391 | 0.294079865 | 10.693196 | 10.182762 | 9.81112 | 0.371645485 | 0.882078947 |
| Slc3a2 | 10.97488 | 10.97743 | 10.958 | 0.019864616 | 0.017315472 | 11.022237 | 11.362428 | 10.9908 | 0.371587864 | 0.031397662 |
| Map7 | 9.057972 | 9.312627 | 9.1583 | 0.154311579 | -0.100343653 | 9.2611289 | 9.5496012 | 9.17808 | 0.371525939 | 0.083053655 |
| Xab2 | 10.58396 | 10.66993 | 10.576 | 0.093912672 | 0.007943408 | 10.897411 | 11.032704 | 10.6612 | 0.371496016 | 0.236203247 |
| Exosc1 | 8.424814 | 8.5043 | 8.48 | 0.024269392 | -0.055216237 | 8.6676748 | 8.8449433 | 8.47352 | 0.371424824 | 0.194156263 |
| Plagl2 | 9.431719 | 9.603143 | 9.4956 | 0.107528652 | -0.063896283 | 9.3765659 | 9.8503326 | 9.479 | 0.371331318 | -0.102435373 |
| Taf1a | 7.999588 | 7.944688 | 7.8526 | 0.092053223 | 0.146953317 | 8.1390831 | 8.3869774 | 8.01567 | 0.371303192 | 0.123408809 |
| Dis3l2 | 8.861792 | 8.99935 | 8.8849 | 0.114471891 | -0.02308624 | 8.9678689 | 9.2147867 | 8.84355 | 0.371231666 | 0.124313935 |
| Gatad2a | 11.11512 | 11.22196 | 11.022 | 0.200379788 | 0.093539002 | 10.94032 | 11.304374 | 10.9333 | 0.371099264 | 0.007045338 |
| Ppia | 13.48942 | 13.63156 | 13.502 | 0.129203272 | -0.01294163 | 13.711701 | 13.873043 | 13.5021 | 0.370951238 | 0.209609616 |
| Tcam1 | 3.313622 | 3.284815 | 3.2466 | 0.038220365 | 0.067027942 | 3.2002351 | 3.6571341 | 3.28627 | 0.370864022 | -0.086035012 |
| Enah | 6.328469 | 6.033774 | 6.0302 | 0.00360912 | 0.298303858 | 5.8437049 | 6.1128859 | 5.74208 | 0.370809906 | 0.101628938 |
| Dcaf15 | 9.973489 | 9.991656 | 10.007 | -0.014993433 | -0.033160335 | 10.302283 | 10.438652 | 10.068 | 0.370697392 | 0.234328442 |
| AI413582 | 7.931867 | 7.965369 | 8.0858 | -0.120394595 | -0.153896864 | 8.3153551 | 8.3297138 | 7.95905 | 0.370662196 | 0.356303528 |
| Rpl13 | 14.44328 | 14.48232 | 14.412 | 0.070532122 | 0.031494376 | 14.625367 | 14.726308 | 14.3557 | 0.370604116 | 0.269662919 |
| 1700123O20Rik | 8.520164 | 8.628838 | 8.623 | 0.005869258 | -0.102804698 | 8.6013875 | 8.8491078 | 8.47863 | 0.370475221 | 0.122754955 |
| Banf1 | 10.01339 | 10.09074 | 10.043 | 0.048094014 | -0.029261795 | 10.319918 | 10.429572 | 10.0593 | 0.370318727 | 0.260664533 |
| Adamts6 | 9.056258 | 9.267312 | 9.2226 | 0.044717779 | -0.166335701 | 9.0131987 | 9.6568098 | 9.28653 | 0.370275936 | -0.273335144 |
| Eif4a3 | 10.83322 | 10.80254 | 10.739 | 0.063735418 | 0.094418962 | 10.886437 | 11.070966 | 10.7009 | 0.37003582 | 0.185506552 |
| Igkv5-39 | 9.051575 | 9.736097 | 9.3016 | 0.434459221 | -0.250062403 | 8.9338577 | 8.8646081 | 8.49463 | 0.369979051 | 0.439228596 |
| Slc8b1 | 8.510542 | 8.64175 | 8.5372 | 0.104555491 | -0.026651815 | 8.5331189 | 8.7397099 | 8.36982 | 0.369887282 | 0.163296214 |
| Tnfsf12 | 5.866632 | 6.022605 | 5.8951 | 0.127479447 | -0.028494131 | 5.4642488 | 5.8737126 | 5.50387 | 0.369842614 | -0.039621178 |
| Dse | 8.98257 | 8.929901 | 8.8806 | 0.049297765 | 0.101966958 | 8.5301101 | 8.9398397 | 8.57022 | 0.36961896 | -0.040110598 |
| Mrpl44 | 8.048179 | 8.217828 | 8.1544 | 0.063450373 | -0.106198953 | 8.25392 | 8.4882877 | 8.11869 | 0.369596385 | 0.135228673 |
| Wsb2 | 9.202562 | 9.393078 | 9.1293 | 0.263730503 | 0.07321458 | 9.2417911 | 9.4829932 | 9.11343 | 0.369561467 | 0.128359331 |

| | | | | | | | | | | |
|---|---|---|---|---|---|---|---|---|---|---|
| Inpp5j | 4.042259 | 4.105111 | 4.2326 | -0.127473999 | -0.190325322 | 4.2740127 | 4.6253926 | 4.25594 | 0.369452681 | 0.018072818 |
| Dcbld1 | 6.21088 | 6.17135 | 6.0763 | 0.095047828 | 0.134578052 | 5.9197559 | 6.4573318 | 6.08819 | 0.369146333 | -0.168429526 |
| Ikbke | 10.69353 | 11.13326 | 10.861 | 0.272330922 | -0.167401318 | 10.716538 | 11.444889 | 11.076 | 0.368927742 | -0.359423128 |
| Fen1 | 8.942049 | 8.881143 | 8.9878 | -0.10669118 | -0.045784856 | 9.0965391 | 9.3919402 | 9.02302 | 0.368917503 | 0.073516372 |
| Pfdn1 | 8.624184 | 8.59086 | 8.5843 | 0.006511839 | 0.03983582 | 8.8198552 | 8.9039915 | 8.5354 | 0.368595263 | 0.28445892 |
| Anapc16 | 9.584111 | 9.637468 | 9.4853 | 0.152201226 | 0.098844948 | 9.6151716 | 9.7094995 | 9.34117 | 0.368328213 | 0.274000329 |
| Nudt19 | 7.889874 | 7.927097 | 7.7702 | 0.156913311 | 0.119691103 | 7.9252826 | 8.1762748 | 7.80796 | 0.368313907 | 0.1173217 |
| Mrpl41 | 7.694199 | 7.628697 | 7.7242 | -0.09546573 | -0.029964537 | 7.7986604 | 7.9741604 | 7.606 | 0.368155728 | 0.192655748 |
| Rnps1 | 10.97007 | 10.98951 | 10.981 | 0.008171176 | -0.01126798 | 11.176506 | 11.32543 | 10.9573 | 0.368117302 | 0.219193437 |
| Pole3 | 8.457466 | 8.448558 | 8.4702 | -0.021625179 | -0.0127169 | 8.6264367 | 8.8079247 | 8.43989 | 0.368039558 | 0.18655147 |
| Brd2 | 12.71695 | 12.64664 | 12.751 | -0.1038888 | -0.033575774 | 12.553691 | 12.905402 | 12.5376 | 0.367848136 | 0.016137316 |
| Mt2 | 7.261599 | 6.569418 | 5.5522 | 1.017171857 | 1.709353209 | 6.2060459 | 6.1911022 | 5.82338 | 0.367723679 | 0.382667457 |
| Tagap1 | 8.756285 | 8.923911 | 8.8971 | 0.026813065 | -0.140812693 | 9.2366002 | 9.4835069 | 9.11591 | 0.367601631 | 0.120694883 |
| Mrps21 | 8.410082 | 8.519423 | 8.4532 | 0.066263649 | -0.04307733 | 8.6507047 | 8.8445037 | 8.47706 | 0.367445809 | 0.173646812 |
| Tmem17 | 4.387592 | 4.471927 | 4.4399 | 0.032057818 | -0.052277052 | 4.6092895 | 4.8666467 | 4.49936 | 0.367285317 | 0.10992816 |
| Nop53 | 12.2848 | 12.28124 | 12.279 | 0.002223094 | 0.005783858 | 12.362368 | 12.495729 | 12.1287 | 0.367048524 | 0.233687525 |
| 1110008P14Rik | 7.152622 | 7.185242 | 7.1594 | 0.025861064 | -0.006758798 | 7.5705099 | 7.3117519 | 6.94471 | 0.36703826 | 0.625796255 |
| Gtf2f2 | 8.545056 | 8.593018 | 8.4528 | 0.14024478 | 0.09228265 | 8.5280092 | 8.8530558 | 8.48614 | 0.366914593 | 0.041868054 |
| Ccdc71 | 9.066531 | 9.081283 | 9.009 | 0.072280709 | 0.057529369 | 9.2946547 | 9.2745927 | 8.90769 | 0.366906348 | 0.38696837 |
| Polm | 9.371252 | 9.499534 | 9.3776 | 0.121983166 | -0.006298773 | 9.9138167 | 9.9150156 | 9.54812 | 0.366898601 | 0.365699735 |
| Nagpa | 8.140833 | 8.418576 | 8.3081 | 0.110481338 | -0.167262088 | 8.6715028 | 8.6906845 | 8.32398 | 0.366700121 | 0.347518428 |
| Psmd12 | 9.866029 | 9.915269 | 9.7626 | 0.152644909 | 0.103404251 | 9.8558981 | 10.194055 | 9.82748 | 0.366577687 | 0.028420536 |
| Ston2 | 5.300046 | 5.191481 | 5.0277 | 0.163782883 | 0.272347934 | 5.2249191 | 5.4857263 | 5.11933 | 0.366397761 | 0.105590625 |
| Fam98a | 8.774294 | 8.868115 | 8.7548 | 0.113299835 | 0.01947903 | 8.8253527 | 9.0773856 | 8.71137 | 0.366011657 | 0.11397868 |
| Tmem259 | 10.34795 | 10.42506 | 10.4 | 0.025446572 | -0.051655336 | 10.843294 | 10.858087 | 10.4922 | 0.365860036 | 0.351066757 |
| Gm49273 | 5.749499 | 5.445884 | 5.5454 | -0.099524352 | 0.204090842 | 5.3349544 | 5.898543 | 5.53295 | 0.36558802 | -0.198000622 |
| Esrra | 8.952409 | 9.154795 | 9.0271 | 0.12767822 | -0.074707177 | 9.4090053 | 9.4105564 | 9.04507 | 0.365486444 | 0.363935392 |
| Zfp263 | 9.982862 | 10.13654 | 9.9069 | 0.229630578 | 0.075947837 | 10.166611 | 10.332429 | 9.96716 | 0.365265332 | 0.199448029 |
| Gpr31b | 1.285178 | 1.269407 | 1.1989 | 0.070507015 | 0.08627714 | 1.2685178 | 1.5811414 | 1.21592 | 0.365220028 | 0.052596451 |
| Gabarap | 11.42261 | 11.43866 | 11.422 | 0.017105622 | 0.001053836 | 11.590499 | 11.740191 | 11.3751 | 0.365127934 | 0.215435922 |
| Ap3s2 | 9.244751 | 9.087892 | 9.166 | -0.078110269 | 0.078748534 | 9.155289 | 9.5620791 | 9.19721 | 0.364871758 | -0.041918341 |
| Ccnh | 8.250427 | 8.262708 | 8.2586 | 0.004062151 | -0.008219191 | 8.4449822 | 8.6138417 | 8.24929 | 0.364553781 | 0.195694332 |

| | | | | | | | | | | |
|---|---|---|---|---|---|---|---|---|---|---|
| Tnip3 | 4.890017 | 5.148561 | 4.52 | 0.62858713 | 0.370043149 | 4.7874952 | 4.9287677 | 4.5645 | 0.364267494 | 0.222995036 |
| Alkbh7 | 5.888801 | 6.24507 | 6.0136 | 0.23150978 | -0.124759296 | 6.2937427 | 6.3889823 | 6.02474 | 0.364240895 | 0.269001289 |
| H2-DMa | 11.68203 | 11.65632 | 11.77 | -0.113658622 | -0.087950608 | 12.187164 | 12.019848 | 11.6558 | 0.364047214 | 0.53136348 |
| Igkv3-7 | 7.189018 | 7.093447 | 6.7623 | 0.33118498 | 0.426756265 | 7.0078868 | 6.9902119 | 6.62634 | 0.363876407 | 0.381551245 |
| Mrpl57 | 7.28425 | 7.192172 | 7.29 | -0.097845568 | -0.005767027 | 7.4382299 | 7.4365576 | 7.0727 | 0.363857911 | 0.365530199 |
| Bloc1s4 | 6.59459 | 6.647126 | 6.5832 | 0.063924782 | 0.011388425 | 6.7086403 | 6.7398877 | 6.37614 | 0.363750224 | 0.332502775 |
| Txnl1 | 9.565747 | 9.643386 | 9.5291 | 0.114242338 | 0.036603296 | 9.625568 | 9.8955836 | 9.53193 | 0.363658156 | 0.093642622 |
| Aifm2 | 5.020052 | 4.832058 | 4.8798 | -0.047704441 | 0.140289572 | 5.0075587 | 5.3516187 | 4.98811 | 0.363513608 | 0.01945356 |
| Pigu | 8.62912 | 8.727813 | 8.6995 | 0.028323402 | -0.070369579 | 8.8454221 | 9.0231757 | 8.65974 | 0.363440119 | 0.185686565 |
| Sema3a | 6.963164 | 7.033883 | 6.0776 | 0.95627263 | 0.885553683 | 6.4717679 | 6.7449092 | 6.38147 | 0.363434206 | 0.090292908 |
| Necap2 | 10.27522 | 10.28748 | 10.26 | 0.027593428 | 0.015330416 | 10.618258 | 10.692048 | 10.3287 | 0.363372484 | 0.289582082 |
| Grip2 | 9.030066 | 8.753228 | 8.9248 | -0.171581218 | 0.105256698 | 10.009105 | 9.3538899 | 8.99091 | 0.362979975 | 1.018195405 |
| Sec61g | 7.02504 | 6.999661 | 6.8619 | 0.137735689 | 0.163114839 | 7.0667506 | 7.3418355 | 6.97908 | 0.36275813 | 0.0876732 |
| Gtpbp8 | 6.950618 | 6.998428 | 7.1071 | -0.108648825 | -0.156458768 | 7.0299316 | 7.3379922 | 6.97566 | 0.362333423 | 0.054272766 |
| Usp19 | 11.36013 | 11.52896 | 11.369 | 0.160089316 | -0.008740158 | 11.610819 | 11.883539 | 11.5213 | 0.362229554 | 0.0895089 |
| Fbxl12 | 9.428821 | 9.50261 | 9.4907 | 0.011941176 | -0.061847779 | 9.7175552 | 9.9673276 | 9.6051 | 0.362223119 | 0.112450729 |
| Bnip3 | 7.300684 | 7.178127 | 7.2408 | -0.062717995 | 0.059838492 | 7.2351721 | 7.5235176 | 7.16132 | 0.36219292 | 0.073847398 |
| Adat3 | 4.452112 | 4.548402 | 4.4358 | 0.112589229 | 0.01629978 | 4.4581395 | 4.6254867 | 4.26339 | 0.362093729 | 0.194746574 |
| Mthfd2 | 9.571935 | 9.408851 | 9.4521 | -0.043243487 | 0.119839835 | 9.768087 | 10.139605 | 9.77758 | 0.362023485 | -0.009495 |
| Gm8752 | 3.71029 | 3.573369 | 3.5857 | -0.012379131 | 0.124542061 | 3.2779154 | 3.70003 | 3.33808 | 0.361948571 | -0.060166013 |
| Stom | 8.61727 | 8.509317 | 8.5663 | -0.057022719 | 0.050930325 | 8.3631236 | 8.7335525 | 8.37183 | 0.361720633 | -0.008708316 |
| Cox16 | 8.44487 | 8.43284 | 8.3479 | 0.08489054 | 0.096920312 | 8.5690138 | 8.6321798 | 8.27088 | 0.361295533 | 0.298129507 |
| Gm48836 | 4.565619 | 4.431333 | 4.6344 | -0.203030073 | -0.068743534 | 4.20456 | 5.3644682 | 5.0033 | 0.361170971 | -0.79873727 |
| Zfp58 | 7.295654 | 7.512012 | 7.4407 | 0.071356494 | -0.145002163 | 7.7038308 | 7.7946422 | 7.43354 | 0.361103759 | 0.270292274 |
| Tapt1 | 10.41207 | 10.47261 | 10.558 | -0.085221264 | -0.145759202 | 10.484366 | 10.938668 | 10.5777 | 0.36091816 | -0.093383436 |
| Gm13502 | 4.797039 | 5.257777 | 4.9961 | 0.2617108 | -0.199026874 | 4.7903669 | 5.0430825 | 4.68217 | 0.360910092 | 0.108194446 |
| Arap1 | 11.73924 | 11.75193 | 11.59 | 0.162285032 | 0.149589034 | 11.96708 | 11.989001 | 11.6281 | 0.360906578 | 0.33898503 |
| Otub1 | 9.809646 | 9.758453 | 9.8028 | -0.044318444 | 0.00687374 | 10.047991 | 10.200732 | 9.83996 | 0.360772114 | 0.208031102 |
| Gm28048 | 10.4734 | 10.56459 | 10.439 | 0.126041472 | 0.034844272 | 10.352312 | 10.711762 | 10.3511 | 0.360662452 | 0.001212372 |
| Flad1 | 8.952905 | 9.025844 | 8.9878 | 0.038092616 | -0.034846314 | 9.3539648 | 9.3587512 | 8.99832 | 0.360429924 | 0.355643546 |
| Chtop | 11.6918 | 11.82784 | 11.669 | 0.158594952 | 0.02255706 | 11.838994 | 12.051919 | 11.6916 | 0.360362924 | 0.147437548 |
| Apeh | 10.39093 | 10.43681 | 10.464 | -0.026984438 | -0.07287088 | 10.766108 | 10.70604 | 10.3457 | 0.360341068 | 0.420409353 |

| | | | | | | | | | | |
|---|---|---|---|---|---|---|---|---|---|---|
| Gm6565 | 2.99145 | 3.153233 | 3.1914 | -0.038152339 | -0.199935294 | 2.8111156 | 3.116078 | 2.75576 | 0.360314661 | 0.05535227 |
| Cd248 | 3.548042 | 3.73247 | 3.5145 | 0.217984724 | 0.033556449 | 3.6296143 | 3.8390386 | 3.47876 | 0.360283451 | 0.150859153 |
| Furin | 9.713846 | 9.839412 | 9.7505 | 0.08895385 | -0.036612829 | 9.6049788 | 9.9014674 | 9.54143 | 0.36003513 | 0.063546511 |
| Khsrp | 10.86886 | 10.91635 | 10.827 | 0.089331772 | 0.041843116 | 11.083183 | 11.285701 | 10.9258 | 0.359926136 | 0.157407525 |
| Mapk7 | 8.47858 | 8.547017 | 8.5282 | 0.018845389 | -0.049590637 | 8.5859372 | 8.6727129 | 8.31285 | 0.359867106 | 0.27309136 |
| Ftsj3 | 10.80441 | 10.85297 | 10.769 | 0.084419388 | 0.035849744 | 10.932351 | 11.266614 | 10.9068 | 0.359805454 | 0.025542514 |
| Xrn2 | 11.83434 | 11.96589 | 11.828 | 0.137782168 | 0.00623022 | 11.902621 | 12.2673 | 11.9076 | 0.359674654 | -0.005004789 |
| Eif3g | 10.73057 | 10.72809 | 10.607 | 0.120719438 | 0.123196848 | 10.809633 | 11.015462 | 10.6564 | 0.359060276 | 0.153231455 |
| Tbxa2r | 7.808649 | 8.099787 | 7.9037 | 0.196117916 | -0.095020427 | 7.9571469 | 8.1271245 | 7.76829 | 0.358835683 | 0.18885808 |
| Ctu1 | 6.613935 | 6.850163 | 6.5988 | 0.251405724 | 0.015177729 | 6.5889134 | 6.7050661 | 6.34647 | 0.358592169 | 0.242439454 |
| Sac3d1 | 6.362838 | 6.467353 | 6.4368 | 0.030536738 | -0.073977897 | 6.4139486 | 6.5630781 | 6.20467 | 0.358409842 | 0.209280383 |
| Mrpl47 | 7.458036 | 7.626022 | 7.5533 | 0.072672339 | -0.095314251 | 7.865957 | 7.9634821 | 7.60508 | 0.35840047 | 0.260875306 |
| Cyp51 | 9.024428 | 9.08819 | 8.9588 | 0.129350575 | 0.065588065 | 9.1129067 | 9.1592915 | 8.80099 | 0.358304298 | 0.311919493 |
| Taf1d | 10.31684 | 10.25814 | 10.4 | -0.14226895 | -0.083578653 | 10.470502 | 10.811889 | 10.4536 | 0.358292428 | 0.01690567 |
| Prdx2 | 9.859673 | 9.855976 | 9.9155 | -0.059478702 | -0.055782247 | 10.009212 | 10.172377 | 9.81411 | 0.358270328 | 0.195105297 |
| Sag | 5.869529 | 5.725425 | 5.8054 | -0.079997679 | 0.064105862 | 5.8912303 | 6.339101 | 5.9809 | 0.358199291 | -0.089671377 |
| Tmub1 | 8.085894 | 8.279305 | 8.2347 | 0.044620394 | -0.148790359 | 8.4766068 | 8.4203157 | 8.06212 | 0.358191382 | 0.414482546 |
| Zrsr2 | 8.980736 | 8.983336 | 8.9154 | 0.067939797 | 0.065340036 | 8.9840079 | 9.282905 | 8.92487 | 0.358031927 | 0.059134754 |
| Bud31 | 9.490868 | 9.559064 | 9.4836 | 0.075461911 | 0.007266268 | 9.7094817 | 9.9350621 | 9.57706 | 0.358006586 | 0.132426209 |
| Ywhae | 11.85163 | 11.90314 | 11.726 | 0.17732322 | 0.125819582 | 11.870782 | 12.130613 | 11.7727 | 0.357868818 | 0.09803763 |
| Atp6v1e1 | 9.897879 | 10.01927 | 9.8911 | 0.128150621 | 0.006757069 | 9.9737963 | 10.09218 | 9.73435 | 0.357833318 | 0.239449964 |
| Tut7 | 11.7921 | 11.8055 | 11.702 | 0.10331675 | 0.089910188 | 11.61171 | 12.024069 | 11.6665 | 0.357616268 | -0.054742669 |
| Ino80 | 10.35515 | 10.51478 | 10.349 | 0.165574622 | 0.005942898 | 10.357739 | 10.80484 | 10.4473 | 0.357571996 | -0.089529651 |
| Phf5a | 8.909645 | 9.019238 | 9.0128 | 0.006441297 | -0.103151825 | 9.2387451 | 9.3815677 | 9.0241 | 0.357472542 | 0.21465 |
| Mrpl28 | 8.147645 | 8.191545 | 8.2161 | -0.024543729 | -0.068443841 | 8.434183 | 8.5218619 | 8.16446 | 0.357400439 | 0.269721516 |
| Olfr695 | 3.232561 | 3.219686 | 3.18 | 0.039703078 | 0.052578454 | 3.4112326 | 3.6471261 | 3.28989 | 0.357231775 | 0.121338298 |
| Ptma | 14.37764 | 14.35681 | 14.343 | 0.014271356 | 0.035103132 | 14.423175 | 14.673585 | 14.3164 | 0.35719471 | 0.106784502 |
| Ipo7 | 11.31358 | 11.46074 | 11.263 | 0.19822516 | 0.051067148 | 11.337857 | 11.667221 | 11.31 | 0.357177646 | 0.027813381 |
| Ociad1 | 11.31352 | 11.34857 | 11.247 | 0.10130689 | 0.066253026 | 11.398053 | 11.670504 | 11.3134 | 0.357094822 | 0.084643627 |
| Gemin4 | 7.996791 | 7.942099 | 8.1142 | -0.172150128 | -0.117458268 | 8.1690553 | 8.388535 | 8.03185 | 0.356681271 | 0.137201585 |
| Foxp4 | 9.231048 | 9.351242 | 9.2851 | 0.066157842 | -0.05403556 | 9.4259439 | 9.6226662 | 9.26604 | 0.35662278 | 0.159900506 |
| Fdx1l | 7.735433 | 7.851398 | 7.6837 | 0.167716011 | 0.051750174 | 7.8995133 | 8.0693474 | 7.71275 | 0.356600594 | 0.186766481 |

| | | | | | | | | | | |
|---|---|---|---|---|---|---|---|---|---|---|
| Cox8a | 10.1292 | 10.15838 | 10.125 | 0.033230073 | 0.004056609 | 10.429189 | 10.506139 | 10.1496 | 0.356547091 | 0.279597662 |
| Irf3 | 10.3121 | 10.40914 | 10.374 | 0.034811393 | -0.06223471 | 10.949404 | 10.97879 | 10.6224 | 0.356395844 | 0.327009447 |
| Rnase6 | 10.21709 | 10.35001 | 10.351 | -0.00109878 | -0.13401925 | 10.643197 | 10.655234 | 10.2989 | 0.356377362 | 0.344340017 |
| Rsc1a1 | 7.034963 | 7.341879 | 6.9477 | 0.39421956 | 0.087303766 | 7.2977778 | 7.232682 | 6.87679 | 0.355889791 | 0.42098557 |
| Gm44245 | 4.025851 | 4.165684 | 4.0116 | 0.154040332 | 0.014207526 | 4.5812658 | 4.5305068 | 4.17469 | 0.355813742 | 0.406572816 |
| Ube2e1 | 9.777511 | 9.984206 | 9.7916 | 0.192635125 | -0.01405975 | 9.8241713 | 10.119704 | 9.7639 | 0.355803466 | 0.060270892 |
| Asb6 | 8.912442 | 9.019675 | 8.8718 | 0.147902633 | 0.040669841 | 9.0782813 | 9.2831498 | 8.92752 | 0.35562914 | 0.150760708 |
| Lrrc59 | 10.09178 | 10.14445 | 9.9998 | 0.144607879 | 0.091928662 | 10.230339 | 10.344526 | 9.9889 | 0.355621576 | 0.241434726 |
| Itga2b | 7.904017 | 7.710741 | 7.4716 | 0.239169812 | 0.432445858 | 7.5958673 | 7.8134208 | 7.45783 | 0.355589708 | 0.138036156 |
| Nsun5 | 8.23909 | 8.419569 | 8.2067 | 0.212873121 | 0.032395032 | 8.7675786 | 8.8759036 | 8.52038 | 0.355525843 | 0.247200776 |
| Ube2q1 | 10.99703 | 11.12663 | 11.063 | 0.063131842 | -0.066465474 | 11.144366 | 11.349401 | 10.994 | 0.355393092 | 0.150357978 |
| Pak1ip1 | 9.889353 | 9.886343 | 9.841 | 0.04538334 | 0.048393757 | 10.01351 | 10.187966 | 9.83268 | 0.355283552 | 0.180827392 |
| Optn | 6.866694 | 6.827867 | 6.8125 | 0.015321831 | 0.054149546 | 6.80307 | 7.2038129 | 6.84863 | 0.355181701 | -0.045561228 |
| Tmem185b | 8.541528 | 8.65263 | 8.5642 | 0.088464627 | -0.022637156 | 8.7088086 | 8.8460591 | 8.49121 | 0.35485086 | 0.217600401 |
| Abhd11 | 8.374786 | 8.526876 | 8.3604 | 0.16648969 | 0.014399672 | 8.8189984 | 8.9217409 | 8.56694 | 0.354805748 | 0.252063304 |
| Bid | 8.992084 | 9.160975 | 9.0467 | 0.114250988 | -0.054640287 | 9.2374169 | 9.4180235 | 9.06323 | 0.354797502 | 0.174190898 |
| Clic1 | 12.1147 | 12.10694 | 12.092 | 0.014886224 | 0.022646338 | 12.245872 | 12.40405 | 12.0495 | 0.354507052 | 0.196329741 |
| Tmem199 | 7.306247 | 7.332779 | 7.3229 | 0.009864757 | -0.016667173 | 7.7880804 | 7.8449217 | 7.4906 | 0.354317315 | 0.297476058 |
| Ndufa1 | 8.737667 | 8.66291 | 8.5985 | 0.064450206 | 0.139207541 | 8.7015128 | 8.8607443 | 8.50644 | 0.354305588 | 0.195074104 |
| Rab32 | 8.265426 | 8.319436 | 8.1843 | 0.135185892 | 0.081176182 | 8.1832446 | 8.3118205 | 7.95762 | 0.354204975 | 0.225629092 |
| Dnajb12 | 8.943103 | 8.966986 | 8.9649 | 0.002054032 | -0.021828855 | 8.8806625 | 9.1469351 | 8.79287 | 0.354069259 | 0.087796659 |
| Ss18l1 | 6.84155 | 6.985991 | 6.974 | 0.011972514 | -0.132468606 | 6.9022904 | 7.4822109 | 7.12815 | 0.354063207 | -0.225857276 |
| BC003965 | 7.956351 | 7.932351 | 7.8109 | 0.121460591 | 0.145461418 | 8.2400223 | 8.2994156 | 7.94556 | 0.353850739 | 0.29445743 |
| Igkv4-92 | 4.588775 | 4.698744 | 4.5272 | 0.171573844 | 0.061604979 | 4.5664078 | 4.8153516 | 4.462 | 0.353347787 | 0.104404071 |
| Igkv12-41 | 8.101355 | 8.588507 | 7.8971 | 0.691368304 | 0.204215967 | 8.2521656 | 8.2461662 | 7.89329 | 0.352881054 | 0.358880414 |
| Ndufs5 | 8.175726 | 8.117038 | 8.0825 | 0.034499794 | 0.093187716 | 8.3747313 | 8.3608304 | 8.00803 | 0.352799459 | 0.366700313 |
| Ripk2 | 8.703703 | 8.846046 | 8.7272 | 0.118830883 | -0.023512053 | 8.5940288 | 9.0823053 | 8.72969 | 0.352614787 | -0.135661677 |
| Vps51 | 9.309775 | 9.339545 | 9.3981 | -0.058547764 | -0.088317834 | 9.6869849 | 9.8472652 | 9.49479 | 0.35247753 | 0.192197224 |
| Itpa | 8.478489 | 8.476914 | 8.4818 | -0.004915806 | -0.003340341 | 8.6491432 | 8.8546155 | 8.50216 | 0.352456519 | 0.14698424 |
| Map3k14 | 10.5266 | 10.62894 | 10.556 | 0.073121962 | -0.02921766 | 10.597554 | 10.821597 | 10.4692 | 0.352420494 | 0.128377675 |
| Pex16 | 7.833896 | 7.85372 | 7.9017 | -0.047993722 | -0.067818258 | 8.0156704 | 8.1276546 | 7.7755 | 0.352151242 | 0.240167054 |
| Cpsf4 | 8.770517 | 8.91346 | 8.8329 | 0.080549282 | -0.062393449 | 8.9560458 | 9.1513351 | 8.79957 | 0.351767829 | 0.156478507 |

| | | | | | | | | | | |
|---|---|---|---|---|---|---|---|---|---|---|
| Cct2 | 12.37368 | 12.44518 | 12.361 | 0.084201676 | 0.012701192 | 12.471069 | 12.741681 | 12.3904 | 0.351254744 | 0.080642326 |
| Kdelr2 | 8.867853 | 8.972225 | 8.8323 | 0.139958944 | 0.035586462 | 8.9946349 | 9.0626566 | 8.71144 | 0.351214182 | 0.283192502 |
| Mrpl55 | 6.983629 | 7.08935 | 7.1211 | -0.031713265 | -0.137434543 | 7.2693325 | 7.2576099 | 6.90645 | 0.351160491 | 0.36288311 |
| Gadd45b | 10.07273 | 10.07653 | 9.975 | 0.101521703 | 0.097723554 | 9.9532569 | 10.292641 | 9.94155 | 0.351090565 | 0.011706042 |
| Clip3 | 5.164716 | 4.978548 | 4.764 | 0.214545175 | 0.400713392 | 4.8570976 | 5.2199736 | 4.86898 | 0.350992244 | -0.011883835 |
| Plpp5 | 6.573102 | 6.641603 | 6.631 | 0.010638305 | -0.057862522 | 6.7668111 | 6.99105 | 6.64008 | 0.350973286 | 0.126734428 |
| Ythdf1 | 10.23118 | 10.51654 | 10.177 | 0.339879338 | 0.054523592 | 10.139884 | 10.569362 | 10.2184 | 0.350946554 | -0.078531696 |
| Serpinb10 | 4.668145 | 4.83196 | 4.3773 | 0.45466606 | 0.290851782 | 4.1564729 | 4.5276327 | 4.17671 | 0.35092188 | -0.020237968 |
| Ier3ip1 | 8.84846 | 9.147476 | 8.968 | 0.179453216 | -0.119562663 | 9.0970886 | 9.2008017 | 8.85007 | 0.350736277 | 0.247023125 |
| Dcaf1 | 10.24234 | 10.36795 | 10.273 | 0.095051038 | -0.030557126 | 10.217305 | 10.521649 | 10.171 | 0.350647154 | 0.046303076 |
| Eny2 | 8.847425 | 8.875824 | 8.7715 | 0.104308271 | 0.075908854 | 8.932843 | 9.1318264 | 8.78123 | 0.350592175 | 0.151608783 |
| Scpep1 | 10.22253 | 10.23512 | 10.254 | -0.018453628 | -0.031044828 | 10.277604 | 10.560842 | 10.2105 | 0.350326118 | 0.067087725 |
| Wdr4 | 8.790203 | 8.807055 | 8.7515 | 0.05553687 | 0.038684947 | 9.0461533 | 9.2002056 | 8.8501 | 0.350110562 | 0.196058206 |
| Scamp3 | 9.978287 | 10.1231 | 10.026 | 0.097082257 | -0.047726198 | 10.255323 | 10.323007 | 9.97297 | 0.350033598 | 0.282348921 |
| Rab10 | 11.26234 | 11.28358 | 11.179 | 0.104881732 | 0.083643006 | 11.168331 | 11.429211 | 11.0793 | 0.349904492 | 0.089025228 |
| Pex7 | 7.809331 | 7.899311 | 7.8498 | 0.049505774 | -0.0404741 | 7.9019307 | 8.1085931 | 7.75901 | 0.349580962 | 0.142918573 |
| Asna1 | 9.295977 | 9.388119 | 9.3151 | 0.073043589 | -0.019098877 | 9.6542209 | 9.6837039 | 9.33427 | 0.349431315 | 0.319948324 |
| Emg1 | 10.8303 | 10.89891 | 10.799 | 0.100057904 | 0.031452694 | 11.03149 | 11.15549 | 10.8062 | 0.349327902 | 0.225328102 |
| Mrpl36 | 8.051687 | 8.050473 | 8.0082 | 0.042223899 | 0.043437779 | 8.3453008 | 8.37189 | 8.02282 | 0.349072117 | 0.322482987 |
| Tespa1 | 9.982096 | 10.02322 | 9.9825 | 0.040733621 | -0.000391768 | 9.8530228 | 10.299758 | 9.95087 | 0.348887491 | -0.097847885 |
| Eapp | 9.412001 | 9.523454 | 9.3484 | 0.175045139 | 0.063592119 | 9.3730031 | 9.6471574 | 9.2984 | 0.348754743 | 0.074600445 |
| Psmb1 | 11.33273 | 11.33647 | 11.305 | 0.031608442 | 0.027864206 | 11.482743 | 11.65498 | 11.3064 | 0.348565746 | 0.176328227 |
| Esd | 10.00134 | 9.982724 | 9.9994 | -0.016663367 | 0.001950377 | 10.095204 | 10.284989 | 9.93646 | 0.348529447 | 0.158744585 |
| Baz1a | 11.84767 | 12.05912 | 11.735 | 0.324255588 | 0.112806248 | 11.864123 | 12.137031 | 11.7886 | 0.348462394 | 0.075554896 |
| Fjx1 | 3.057923 | 3.125195 | 2.7961 | 0.329129796 | 0.261857499 | 2.8024227 | 3.1224176 | 2.77425 | 0.348170545 | 0.028175568 |
| Mia2 | 11.07285 | 11.17357 | 11.086 | 0.087933652 | -0.012778656 | 11.073522 | 11.453597 | 11.1055 | 0.34812691 | -0.031947967 |
| Cd320 | 6.022083 | 6.174733 | 6.1265 | 0.04827722 | -0.104372649 | 6.2581722 | 6.4167551 | 6.06864 | 0.348119481 | 0.189536532 |
| Tph1 | 5.059266 | 4.969877 | 4.6658 | 0.304037944 | 0.393426996 | 4.7361725 | 4.7597926 | 4.41182 | 0.347976892 | 0.324356764 |
| Tcp1 | 11.96561 | 12.03231 | 11.886 | 0.146661 | 0.079957042 | 11.904899 | 12.269769 | 11.9219 | 0.347874296 | -0.016995202 |
| Arl3 | 7.701209 | 7.752846 | 7.672 | 0.080864579 | 0.029227156 | 7.7926072 | 8.0119217 | 7.66419 | 0.347728269 | 0.128413856 |
| Ndufv2 | 9.741721 | 9.693787 | 9.7246 | -0.030816515 | 0.017117107 | 9.891654 | 10.021283 | 9.67374 | 0.347543935 | 0.217915129 |
| Mul1 | 7.943346 | 8.052848 | 7.9325 | 0.120390076 | 0.010888131 | 7.9877726 | 8.2251942 | 7.87774 | 0.347455121 | 0.110033482 |

| | | | | | | | | | | |
|---|---|---|---|---|---|---|---|---|---|---|
| Ranbp2 | 12.38309 | 12.47688 | 12.296 | 0.180495484 | 0.086707588 | 12.18684 | 12.672612 | 12.3252 | 0.347384464 | -0.138387828 |
| Eif4a-ps4 | 11.06573 | 11.28842 | 11.085 | 0.20336565 | -0.019329628 | 11.259124 | 11.586816 | 11.2395 | 0.347322582 | 0.019631376 |
| Ranbp3 | 10.54412 | 10.62773 | 10.5 | 0.127520082 | 0.043916634 | 10.714161 | 10.960822 | 10.6137 | 0.347115048 | 0.100453838 |
| Mon1a | 7.98459 | 8.145754 | 8.0499 | 0.095825496 | -0.065338767 | 8.1195897 | 8.2791313 | 7.93205 | 0.347085073 | 0.187543457 |
| Atp5f1 | 11.05555 | 11.05821 | 11.106 | -0.047314276 | -0.049981608 | 11.092931 | 11.312361 | 10.9654 | 0.347008814 | 0.127577985 |
| Ube2k | 9.674826 | 9.728522 | 9.6293 | 0.099177865 | 0.045481761 | 9.6947612 | 9.8709191 | 9.52397 | 0.346952087 | 0.170794131 |
| Asf1a | 7.915732 | 8.104303 | 8.026 | 0.078326779 | -0.11024468 | 8.1120842 | 8.3729214 | 8.02615 | 0.346770337 | 0.08593308 |
| Hnrnph2 | 11.29946 | 11.28693 | 11.095 | 0.192208676 | 0.20473564 | 11.206351 | 11.471494 | 11.1248 | 0.346665888 | 0.081523527 |
| Gm13722 | 6.041216 | 5.933035 | 6.0211 | -0.088110981 | 0.020069588 | 5.9483571 | 6.5883994 | 6.2419 | 0.346501041 | -0.293541317 |
| Cited4 | 3.713476 | 3.798026 | 3.8465 | -0.048466462 | -0.133015796 | 3.5618476 | 4.1441974 | 3.79789 | 0.346307379 | -0.236042403 |
| Igkv10-94 | 7.037112 | 7.584275 | 7.1955 | 0.388747323 | -0.158416031 | 7.5501247 | 7.5850497 | 7.23879 | 0.346261817 | 0.311336803 |
| Sdk2 | 3.132756 | 3.309851 | 3.1487 | 0.16117537 | -0.01591991 | 3.2189355 | 3.5680446 | 3.22182 | 0.346226325 | -0.002882744 |
| Trp53inp1 | 11.28086 | 11.57104 | 11.393 | 0.177686088 | -0.11249035 | 11.060031 | 11.658623 | 11.3124 | 0.346201504 | -0.252390425 |
| Gal3st3 | 3.114552 | 3.158665 | 3.1224 | 0.036247055 | -0.007865337 | 3.1405446 | 3.4997775 | 3.15382 | 0.34595332 | -0.013279554 |
| Rbm15 | 9.89497 | 10.0402 | 9.9927 | 0.04748737 | -0.097743022 | 10.017716 | 10.462453 | 10.1167 | 0.345713892 | -0.099023121 |
| 2610001J05Rik | 8.773252 | 8.843386 | 8.6584 | 0.184950748 | 0.114816162 | 8.84846 | 8.9798232 | 8.63428 | 0.34553987 | 0.214176681 |
| BC005537 | 12.20388 | 12.4198 | 12.191 | 0.228631328 | 0.012711908 | 12.028841 | 12.520719 | 12.1752 | 0.345511534 | -0.146366678 |
| Fbxo6 | 7.870113 | 7.767752 | 7.9381 | -0.170358267 | -0.067997524 | 7.9962491 | 8.1146432 | 7.76914 | 0.345498781 | 0.227104705 |
| Nme1 | 9.627581 | 9.788275 | 9.6671 | 0.121188187 | -0.039505008 | 9.9683849 | 9.9356162 | 9.59036 | 0.345258473 | 0.378027157 |
| Ninj2 | 2.468861 | 2.471991 | 2.4682 | 0.003741611 | 0.000611981 | 2.449163 | 2.8482939 | 2.50315 | 0.345141911 | -0.053989066 |
| Cops6 | 9.821828 | 9.807251 | 9.7812 | 0.026005666 | 0.040582995 | 9.9463631 | 10.108499 | 9.76337 | 0.345124936 | 0.182988745 |
| Abcb11 | 0.805615 | 0.888638 | 0.832 | 0.056609621 | -0.026412664 | 0.9335678 | 1.1528562 | 0.8081 | 0.344759438 | 0.125471045 |
| Hikeshi | 8.097022 | 8.296537 | 8.1424 | 0.154168017 | -0.045347341 | 8.2920191 | 8.5229527 | 8.17819 | 0.344758143 | 0.113824536 |
| Il7r | 11.92764 | 12.14118 | 12.063 | 0.077759542 | -0.13577099 | 11.021142 | 11.903107 | 11.5586 | 0.34453768 | -0.537426805 |
| Psmd9 | 8.058651 | 8.15281 | 8.1819 | -0.029132537 | -0.123291424 | 8.2427839 | 8.4484082 | 8.10406 | 0.344350917 | 0.138726616 |
| Srsf3 | 12.34539 | 12.50478 | 12.415 | 0.08978512 | -0.069604516 | 12.486906 | 12.852416 | 12.5081 | 0.344266924 | -0.021243037 |
| Gm5483 | 5.295867 | 4.423323 | 3.7358 | 0.687546509 | 1.560091277 | 3.9428781 | 4.0364904 | 3.69229 | 0.344197669 | 0.250585372 |
| Gch1 | 8.225094 | 8.424139 | 8.1762 | 0.247916298 | 0.048871688 | 8.3039313 | 8.7177285 | 8.37363 | 0.344101563 | -0.06969572 |
| Tbcc | 6.955375 | 6.839579 | 6.7617 | 0.077843548 | 0.193639382 | 7.0444011 | 7.0425478 | 6.69876 | 0.343787218 | 0.345640516 |
| Igkv4-79 | 5.055454 | 5.272653 | 5.0468 | 0.225885906 | 0.008686871 | 5.5358759 | 5.4016987 | 5.05799 | 0.343711369 | 0.477888515 |
| Atf4 | 12.01862 | 11.93631 | 11.986 | -0.049919374 | 0.032389926 | 11.985652 | 12.240398 | 11.8967 | 0.343657286 | 0.088910804 |
| Thyn1 | 7.340966 | 7.373734 | 7.3454 | 0.028378227 | -0.004389249 | 7.420364 | 7.6291411 | 7.2858 | 0.343339369 | 0.134562274 |

| | | | | | | | | | | |
|---|---|---|---|---|---|---|---|---|---|---|
| Tma7 | 10.38222 | 10.47421 | 10.409 | 0.065426662 | -0.026570448 | 10.730589 | 10.748304 | 10.4051 | 0.343199466 | 0.325484574 |
| St6galnac6 | 6.907026 | 6.936355 | 6.7655 | 0.170868006 | 0.14153897 | 7.0757662 | 7.1593528 | 6.81635 | 0.343005397 | 0.259418798 |
| 0610037L13Rik | 8.894932 | 8.883402 | 8.808 | 0.07543408 | 0.086964969 | 9.1379136 | 9.2504081 | 8.90749 | 0.342921488 | 0.230426916 |
| Magoh | 9.298583 | 9.25534 | 9.2258 | 0.029545926 | 0.072788559 | 9.3785858 | 9.5541588 | 9.21128 | 0.342874025 | 0.167301058 |
| Epop | 6.064119 | 5.905791 | 5.8512 | 0.054621215 | 0.21294872 | 6.1044542 | 6.0439123 | 5.70115 | 0.342761262 | 0.403303183 |
| Gzmk | 3.00328 | 3.105931 | 3.0815 | 0.024396391 | -0.078254859 | 2.9366219 | 3.3338617 | 2.99111 | 0.342748938 | -0.054490884 |
| Tmem50a | 10.78772 | 10.94784 | 10.814 | 0.133959702 | -0.026159926 | 10.76126 | 11.004512 | 10.6619 | 0.342615286 | 0.099363742 |
| Pagr1a | 5.221825 | 5.527669 | 5.8372 | -0.309560174 | -0.615403819 | 5.7136588 | 5.9909938 | 5.64838 | 0.342609589 | 0.065274565 |
| Rps19bp1 | 6.901745 | 6.860018 | 6.8215 | 0.03850453 | 0.080231481 | 6.9319252 | 7.1285805 | 6.786 | 0.342576315 | 0.145921003 |
| Mak16 | 10.22277 | 10.26307 | 10.218 | 0.044583402 | 0.004284577 | 10.383767 | 10.592087 | 10.2496 | 0.342449494 | 0.134129512 |
| Col5a2 | 5.621309 | 5.973933 | 5.8932 | 0.080777698 | -0.271845762 | 6.1077929 | 5.8287655 | 5.48656 | 0.342205644 | 0.621233115 |
| Fam78b | 4.955597 | 5.122705 | 4.8073 | 0.315442734 | 0.148334715 | 4.6524827 | 4.8269386 | 4.48475 | 0.342188842 | 0.16773293 |
| Slco2a1 | 7.549136 | 7.976696 | 7.0079 | 0.968766859 | 0.541206131 | 7.4057107 | 7.0800431 | 6.73799 | 0.342058005 | 0.667725642 |
| Copz1 | 10.50548 | 10.60146 | 10.55 | 0.051558032 | -0.044422848 | 10.715343 | 10.820671 | 10.4786 | 0.342050834 | 0.236722678 |
| Snrpd3 | 10.25023 | 10.30389 | 10.194 | 0.110366674 | 0.05670127 | 10.230881 | 10.506484 | 10.1645 | 0.341948639 | 0.066345456 |
| Capn10 | 7.019146 | 6.97517 | 7.0981 | -0.122892411 | -0.078916377 | 7.4100704 | 7.6969212 | 7.35529 | 0.34162979 | 0.054778992 |
| Rpl27a | 12.51919 | 12.28542 | 12.32 | -0.03469468 | 0.19907502 | 12.465633 | 12.666813 | 12.3252 | 0.341568906 | 0.140389162 |
| Prmt6 | 8.136266 | 8.195822 | 8.1332 | 0.062668025 | 0.003112024 | 8.1361717 | 8.4147594 | 8.07341 | 0.341353597 | 0.062765913 |
| Scin | 5.23089 | 5.583632 | 5.2715 | 0.312085542 | -0.040656518 | 5.530403 | 5.843875 | 5.50264 | 0.341239667 | 0.027767675 |
| Gm48128 | 8.191177 | 8.429366 | 7.9144 | 0.514918779 | 0.276730136 | 8.0248043 | 8.8814663 | 8.54029 | 0.341178668 | -0.515483393 |
| Dhx34 | 9.060881 | 9.106205 | 9.0047 | 0.101484484 | 0.056159898 | 9.3657072 | 9.4453403 | 9.10425 | 0.341086092 | 0.261453 |
| Mcrip1 | 9.195294 | 9.300524 | 9.2353 | 0.065205275 | -0.040024106 | 9.4930959 | 9.4378908 | 9.097 | 0.34088868 | 0.396093765 |
| Exosc2 | 9.121647 | 9.060769 | 9.0423 | 0.018433596 | 0.079312013 | 9.309981 | 9.4420917 | 9.10147 | 0.340618534 | 0.208507808 |
| Snx3 | 10.42484 | 10.64847 | 10.543 | 0.10506589 | -0.118564836 | 10.672821 | 10.802413 | 10.4619 | 0.340558474 | 0.210965964 |
| Zfp768 | 5.937503 | 5.868645 | 5.9959 | -0.127239155 | -0.058381474 | 5.8431315 | 6.1303165 | 5.79004 | 0.340272428 | 0.053087355 |
| Anapc10 | 7.510763 | 7.508488 | 7.5537 | -0.045250451 | -0.042975691 | 7.5075829 | 7.7398797 | 7.39968 | 0.340202897 | 0.107906162 |
| Igkj3 | 6.372651 | 6.798623 | 6.3307 | 0.467957264 | 0.041985161 | 6.6178822 | 6.8312363 | 6.49113 | 0.340109747 | 0.126755652 |
| Alpl | 5.624221 | 5.810864 | 5.6353 | 0.175592283 | -0.011050671 | 5.6309545 | 5.9152832 | 5.57529 | 0.339990562 | 0.055661796 |
| Pmpcb | 9.281441 | 9.408864 | 9.3641 | 0.04473687 | -0.082685559 | 9.4534343 | 9.7397688 | 9.4 | 0.339767068 | 0.053432508 |
| Btnl2 | 6.629398 | 6.700564 | 6.5886 | 0.111939187 | 0.040773253 | 6.9640455 | 7.1676473 | 6.82804 | 0.339605415 | 0.136003626 |
| Dbi | 8.669217 | 8.511375 | 8.5944 | -0.083050055 | 0.074791706 | 8.7011456 | 8.8542869 | 8.51487 | 0.339416741 | 0.186275447 |
| Hyou1 | 11.35433 | 11.53692 | 11.211 | 0.32586876 | 0.143273406 | 11.47429 | 11.551688 | 11.2123 | 0.339342746 | 0.261944612 |

| | | | | | | | | | | |
|---|---|---|---|---|---|---|---|---|---|---|
| Zc3h10 | 9.133777 | 9.142164 | 9.2414 | -0.099275429 | -0.107662533 | 8.9366643 | 9.1183512 | 8.77911 | 0.33923998 | 0.157553048 |
| Rcc1l | 8.109199 | 7.972224 | 7.853 | 0.119224908 | 0.256199686 | 8.2625142 | 8.3076632 | 7.96846 | 0.339200388 | 0.294051355 |
| Cfap126 | 3.488467 | 3.428371 | 3.4507 | -0.022356779 | 0.037739009 | 3.4817558 | 3.6413871 | 3.30223 | 0.339160312 | 0.179528974 |
| Rpp14 | 8.199767 | 8.41201 | 8.3815 | 0.030491565 | -0.181751769 | 8.3494138 | 8.5672996 | 8.22826 | 0.339043945 | 0.121158088 |
| Mrpl24 | 10.4678 | 10.58023 | 10.498 | 0.08272913 | -0.029704676 | 10.691126 | 10.962889 | 10.6242 | 0.338737398 | 0.066973779 |
| Eepd1 | 6.269024 | 6.084806 | 6.3202 | -0.235406631 | -0.051188738 | 5.8269659 | 6.0815509 | 5.74284 | 0.338710836 | 0.084125843 |
| Zc3h11a | 11.68308 | 11.7264 | 11.705 | 0.021525866 | -0.021794808 | 11.686365 | 12.125613 | 11.7872 | 0.338436034 | -0.100812209 |
| Gm10175 | 11.18461 | 11.23326 | 11.238 | -0.004411346 | -0.053063122 | 11.468206 | 11.505904 | 11.1675 | 0.338400034 | 0.300701827 |
| Nfe2 | 4.16708 | 4.119316 | 3.8209 | 0.298462649 | 0.346226661 | 4.1113782 | 4.051827 | 3.71357 | 0.338259779 | 0.397811006 |
| Gpaa1 | 8.458157 | 8.436987 | 8.367 | 0.069971075 | 0.091140809 | 8.7724831 | 8.8237639 | 8.48563 | 0.338130376 | 0.286849588 |
| Sdf4 | 11.73569 | 11.67073 | 11.701 | -0.03047191 | 0.034491828 | 11.679261 | 11.912175 | 11.5743 | 0.337914956 | 0.105001272 |
| Ppp1r21 | 9.887381 | 10.11112 | 10.028 | 0.083591223 | -0.140151404 | 9.6506936 | 10.065151 | 9.72743 | 0.337724599 | -0.076732586 |
| Rps21 | 13.03005 | 13.1136 | 12.992 | 0.122050724 | 0.038505446 | 13.342663 | 13.267463 | 12.9298 | 0.337656358 | 0.412856654 |
| Spout1 | 8.692674 | 8.738939 | 8.633 | 0.1059121 | 0.059647266 | 8.9106756 | 9.0120462 | 8.67463 | 0.337417057 | 0.236046423 |
| Psen1 | 10.38039 | 10.53926 | 10.347 | 0.192246788 | 0.033380828 | 10.494804 | 10.691371 | 10.354 | 0.33733478 | 0.14076828 |
| Dpy19l1 | 9.705192 | 9.898051 | 9.7667 | 0.131366005 | -0.0614929 | 9.5694464 | 10.105874 | 9.76876 | 0.337118845 | -0.199309103 |
| Fubp1 | 11.33 | 11.48024 | 11.289 | 0.191126532 | 0.040878416 | 11.576962 | 11.801107 | 11.464 | 0.337101288 | 0.112956625 |
| Ube2a | 8.746835 | 8.64396 | 8.6634 | -0.019441735 | 0.08343292 | 8.7583015 | 8.9562543 | 8.6193 | 0.336953285 | 0.139000525 |
| Elmo2 | 9.879192 | 9.845691 | 9.8139 | 0.031838859 | 0.065340347 | 9.9005464 | 10.24498 | 9.90806 | 0.336919577 | -0.007513802 |
| Gtf2a2 | 9.238978 | 9.036553 | 9.1063 | -0.06975759 | 0.132668206 | 9.321619 | 9.5825955 | 9.24578 | 0.336811031 | 0.075834537 |
| Myo1f | 10.0725 | 10.17863 | 10.084 | 0.094394984 | -0.011741227 | 10.350463 | 10.451106 | 10.1144 | 0.336721739 | 0.236078274 |
| AC115631.1 | 9.326606 | 9.412906 | 9.3422 | 0.070676336 | -0.01562458 | 9.5057993 | 9.7248629 | 9.38825 | 0.33661781 | 0.117554185 |
| Ccdc153 | 5.66656 | 5.382068 | 5.6527 | -0.270653025 | 0.013838704 | 5.7172042 | 5.9087786 | 5.57227 | 0.336503755 | 0.144929326 |
| Serpina3i | 3.174456 | 3.301358 | 3.0143 | 0.287096682 | 0.160195278 | 3.0788787 | 3.4433922 | 3.10692 | 0.336477013 | -0.028036494 |
| Igkv4-55 | 7.985286 | 8.83205 | 7.8454 | 0.986692052 | 0.139928232 | 8.517486 | 8.4851551 | 8.14871 | 0.336441647 | 0.368772583 |
| Taf1c | 8.792848 | 9.081927 | 8.8583 | 0.22366889 | -0.065409518 | 9.1628833 | 9.2033277 | 8.86705 | 0.33627607 | 0.2958317 |
| Mrpl23-ps1 | 8.796396 | 8.732137 | 8.814 | -0.081890094 | -0.017630831 | 9.1360575 | 9.1253177 | 8.7891 | 0.33621775 | 0.34695759 |
| Bambi-ps1 | 6.434599 | 6.663939 | 6.7742 | -0.110283273 | -0.339623274 | 6.3728226 | 6.7801856 | 6.44435 | 0.33583471 | -0.07152829 |
| Ppp2r2a | 10.4875 | 10.54397 | 10.444 | 0.100230682 | 0.04375113 | 10.335763 | 10.762882 | 10.4271 | 0.33573205 | -0.091386342 |
| 9130401M01Rik | 7.057537 | 7.089156 | 6.9975 | 0.091693069 | 0.060073967 | 7.1500937 | 7.277002 | 6.94135 | 0.335654396 | 0.20874613 |
| Ftl1-ps1 | 5.623484 | 5.892132 | 5.5635 | 0.328585261 | 0.059937074 | 6.1120513 | 5.8943448 | 5.55872 | 0.335622551 | 0.553329071 |
| Itprip | 10.67969 | 10.81483 | 10.612 | 0.202848054 | 0.067708104 | 10.727543 | 11.018038 | 10.6825 | 0.33556946 | 0.045074382 |

| | | | | | | | | | | |
|---|---|---|---|---|---|---|---|---|---|---|
| Arl4a | 8.201181 | 8.142809 | 8.2393 | -0.096514157 | -0.038141631 | 7.8293199 | 8.3750299 | 8.03946 | 0.33556504 | -0.210144999 |
| Ybx3 | 9.627626 | 9.730298 | 9.6311 | 0.099193258 | -0.003478104 | 9.8140342 | 9.9739731 | 9.63851 | 0.335464471 | 0.175525554 |
| Spire2 | 2.108855 | 2.372989 | 2.1206 | 0.252340346 | -0.011793097 | 2.1115532 | 2.4269796 | 2.09155 | 0.335431116 | 0.020004727 |
| Ppp1r37 | 8.9842 | 9.158843 | 8.9306 | 0.228291323 | 0.053648424 | 9.2900339 | 9.4637014 | 9.12839 | 0.335309892 | 0.161642371 |
| Gm12240 | 4.790975 | 4.767191 | 4.7219 | 0.045325457 | 0.069108714 | 4.9853787 | 5.1675379 | 4.83226 | 0.335274629 | 0.15311542 |
| Mrpl4 | 9.122262 | 9.158075 | 9.0786 | 0.079465266 | 0.043651855 | 9.3491908 | 9.310708 | 8.97569 | 0.335018047 | 0.373500829 |
| Ttyh2 | 6.221549 | 6.656044 | 6.3352 | 0.320846279 | -0.113647972 | 6.1371583 | 6.4717108 | 6.13675 | 0.334963996 | 0.000411541 |
| Mdga1 | 2.846409 | 2.782513 | 2.8243 | -0.041738524 | 0.022157408 | 2.9849769 | 3.0589122 | 2.72415 | 0.334765076 | 0.260829717 |
| Cbfa2t3 | 9.485493 | 9.484262 | 9.3833 | 0.101001061 | 0.102231269 | 9.3131225 | 9.7282474 | 9.39354 | 0.33470966 | -0.080415275 |
| Vmn1r-ps144 | 5.231167 | 5.231121 | 5.344 | -0.112889372 | -0.112843273 | 5.4344421 | 5.9182185 | 5.58372 | 0.334496028 | -0.149280386 |
| Stoml2 | 8.902365 | 8.970234 | 8.8055 | 0.164783354 | 0.096914412 | 9.0283969 | 9.1152692 | 8.78097 | 0.334300563 | 0.247428269 |
| Zfp706 | 10.88577 | 10.99711 | 10.842 | 0.15538393 | 0.044041446 | 10.895094 | 11.13199 | 10.7978 | 0.334186026 | 0.097290516 |
| Chrac1 | 8.635062 | 8.712059 | 8.6211 | 0.090923319 | 0.013925817 | 8.8688703 | 8.9612973 | 8.62716 | 0.334133053 | 0.241706069 |
| Ddx56 | 9.432676 | 9.441248 | 9.3517 | 0.089498802 | 0.080926503 | 9.6135839 | 9.7979127 | 9.46383 | 0.334078014 | 0.149749192 |
| Vapa | 10.3216 | 10.41898 | 10.333 | 0.085631284 | -0.011747506 | 10.313432 | 10.567157 | 10.2333 | 0.333813542 | 0.080088645 |
| Tomm20 | 10.45139 | 10.47499 | 10.533 | -0.058356048 | -0.081961068 | 10.684651 | 10.863878 | 10.5301 | 0.333812736 | 0.154586023 |
| Tarbp2 | 8.547013 | 8.534222 | 8.6081 | -0.073841848 | -0.061051443 | 8.9318472 | 8.9965341 | 8.66274 | 0.333791567 | 0.26910469 |
| Atad3a | 9.448966 | 9.567058 | 9.4572 | 0.109889518 | -0.008201906 | 9.8393366 | 10.082907 | 9.74919 | 0.333719594 | 0.090148908 |
| Rnaseh1 | 7.88391 | 7.874795 | 7.7727 | 0.102135756 | 0.111250254 | 8.2133272 | 8.3002075 | 7.9666 | 0.333603978 | 0.246723663 |
| Rpl7a | 13.99017 | 14.06009 | 14.044 | 0.016280114 | -0.053635656 | 14.224759 | 14.367377 | 14.0338 | 0.33355852 | 0.190941247 |
| Sdhaf2 | 9.731579 | 9.809744 | 9.7111 | 0.098647056 | 0.020482424 | 9.926784 | 10.059235 | 9.72571 | 0.333528282 | 0.201077596 |
| Mrgpra2b | 5.248601 | 4.224064 | 3.5382 | 0.685839075 | 1.710376552 | 4.131355 | 4.0762747 | 3.7429 | 0.333371927 | 0.388452305 |
| Pou3f1 | 2.147948 | 2.296977 | 2.1272 | 0.169798908 | 0.020769548 | 2.1221788 | 2.4652045 | 2.13202 | 0.333187804 | -0.009837839 |
| Sipa1l1 | 10.89305 | 11.06139 | 10.886 | 0.175133582 | 0.00679551 | 10.795481 | 11.269144 | 10.936 | 0.33318748 | -0.140476086 |
| Txnl4b | 6.61877 | 6.685199 | 6.7541 | -0.068908413 | -0.135338283 | 6.8200408 | 7.0227475 | 6.68971 | 0.333032972 | 0.1303263 |
| Uchl5 | 8.991916 | 8.945175 | 8.9529 | -0.00774763 | 0.038992655 | 8.8822918 | 9.2962195 | 8.9633 | 0.332917658 | -0.081010032 |
| Rpl24 | 13.08907 | 13.10415 | 13.12 | -0.015889548 | -0.030969928 | 13.17463 | 13.405931 | 13.0731 | 0.3327949 | 0.101493947 |
| Ndufaf1 | 7.402411 | 7.500106 | 7.5719 | -0.071762824 | -0.169457438 | 7.6796215 | 7.7927722 | 7.46005 | 0.332721092 | 0.21957037 |
| Ccnd1 | 10.18979 | 10.14312 | 9.8552 | 0.287917338 | 0.33458804 | 9.5694594 | 9.8292976 | 9.49666 | 0.33264049 | 0.072802292 |
| Nol12 | 8.558785 | 8.654895 | 8.566 | 0.088874329 | -0.007236575 | 8.902901 | 8.9640066 | 8.63145 | 0.332558582 | 0.271452985 |
| Fgd2 | 10.45709 | 10.54892 | 10.42 | 0.128883376 | 0.037055863 | 11.105826 | 10.961706 | 10.6293 | 0.3324241 | 0.476544504 |
| Npffr1 | 4.149826 | 3.974059 | 4.2254 | -0.251304904 | -0.07553854 | 4.5098137 | 4.5113664 | 4.17901 | 0.332354924 | 0.330802233 |

| | | | | | | | | | | |
|---|---|---|---|---|---|---|---|---|---|---|
| Tmem167 | 9.773184 | 9.939148 | 9.7211 | 0.218056808 | 0.052092219 | 9.9018278 | 10.143489 | 9.81117 | 0.332315912 | 0.090654475 |
| Myl4 | 6.400427 | 6.529437 | 6.2555 | 0.273923269 | 0.144913549 | 6.9690856 | 6.7163152 | 6.38405 | 0.332270129 | 0.585040529 |
| Mrpl58 | 8.44258 | 8.430547 | 8.4429 | -0.012318937 | -0.000286156 | 8.6633771 | 8.7868204 | 8.4548 | 0.332018061 | 0.208574756 |
| Rbm14 | 9.333418 | 9.661593 | 9.3711 | 0.290504359 | -0.037670979 | 9.5893954 | 9.7527655 | 9.4208 | 0.331965183 | 0.168595093 |
| Gm48365 | 4.543203 | 4.615228 | 4.9476 | -0.332404417 | -0.404430264 | 4.9217423 | 5.0736478 | 4.74193 | 0.331719051 | 0.179813525 |
| Fbxl15 | 5.01283 | 5.125282 | 4.9474 | 0.17789218 | 0.065440366 | 5.2226276 | 5.2313616 | 4.89967 | 0.331689777 | 0.322955744 |
| Sdad1 | 10.48532 | 10.59428 | 10.384 | 0.209991368 | 0.101032704 | 10.502108 | 10.753421 | 10.4218 | 0.331668386 | 0.080355035 |
| Hsd3b7 | 7.549051 | 7.260649 | 7.2675 | -0.006802158 | 0.281599969 | 7.2031595 | 7.4220251 | 7.09056 | 0.331463156 | 0.112597615 |
| Gripap1 | 10.66486 | 10.73006 | 10.678 | 0.051651184 | -0.013551738 | 11.283174 | 11.435946 | 11.1048 | 0.331168468 | 0.178396297 |
| Jak2 | 11.52137 | 11.58368 | 11.474 | 0.109872258 | 0.047557204 | 11.075744 | 11.66956 | 11.3384 | 0.331144914 | -0.262670626 |
| Upf1 | 11.00703 | 11.11229 | 10.951 | 0.161594266 | 0.056336186 | 11.204839 | 11.402293 | 11.0712 | 0.331077396 | 0.133624171 |
| Trim11 | 9.441252 | 9.569454 | 9.4525 | 0.116920881 | -0.01128112 | 9.7840696 | 9.907048 | 9.57613 | 0.330915944 | 0.207937548 |
| Rhox8 | 2.690107 | 2.708548 | 2.7088 | -0.00023663 | -0.018677557 | 2.7064251 | 3.0647404 | 2.73396 | 0.330778101 | -0.027537196 |
| Cops9 | 8.765115 | 8.763097 | 8.7707 | -0.007578242 | -0.005560443 | 8.9675707 | 9.0429027 | 8.71215 | 0.330751327 | 0.255419329 |
| Cdt1 | 9.322169 | 9.561236 | 9.4709 | 0.090322138 | -0.148745468 | 9.6494175 | 9.7853325 | 9.45461 | 0.330723389 | 0.194808468 |
| Eif3b | 11.69317 | 11.74907 | 11.6 | 0.14927282 | 0.093368482 | 11.738678 | 12.006375 | 11.676 | 0.330415748 | 0.062718845 |
| Pop1 | 7.125557 | 7.295726 | 7.1839 | 0.11183745 | -0.058331203 | 7.2396397 | 7.4045966 | 7.07426 | 0.330338764 | 0.165381796 |
| Rtraf | 10.49735 | 10.44086 | 10.467 | -0.026595712 | 0.029897948 | 10.455739 | 10.821818 | 10.4919 | 0.32993769 | -0.036141338 |
| Fam174a | 7.885693 | 7.781084 | 7.7983 | -0.017176303 | 0.087432589 | 7.8652699 | 7.9891555 | 7.65934 | 0.329819825 | 0.205934169 |
| Rfx2 | 6.113715 | 6.033072 | 5.7508 | 0.282316737 | 0.362959534 | 5.750503 | 6.1551358 | 5.82534 | 0.329796106 | -0.074836757 |
| Dmtn | 4.451156 | 4.579794 | 4.5474 | 0.032390456 | -0.096247942 | 4.2207786 | 4.6269317 | 4.29714 | 0.329793941 | -0.076359189 |
| Zbtb48 | 8.175918 | 8.299112 | 8.1319 | 0.167201961 | 0.044008601 | 8.6736166 | 8.8221266 | 8.49234 | 0.329791124 | 0.181281136 |
| Cmip | 10.28783 | 10.37063 | 10.137 | 0.233981979 | 0.151177567 | 10.245333 | 10.443822 | 10.1147 | 0.329148651 | 0.130659215 |
| Slc28a2 | 11.14535 | 11.44895 | 11.29 | 0.159128494 | -0.14447018 | 11.664052 | 11.86887 | 11.5401 | 0.328734946 | 0.123917328 |
| Serf2 | 11.806 | 11.88393 | 11.826 | 0.058260642 | -0.019665162 | 12.100607 | 12.140649 | 11.812 | 0.328659502 | 0.288617358 |
| Ppp1r3f | 6.893478 | 6.868469 | 6.8943 | -0.025827035 | -0.0008179 | 6.8498598 | 7.1143259 | 6.78571 | 0.328620337 | 0.064154188 |
| Psmc1 | 10.19465 | 10.09867 | 10.113 | -0.014475752 | 0.081503027 | 10.228084 | 10.462654 | 10.1342 | 0.328499578 | 0.09392955 |
| Mphosph10 | 9.408049 | 9.426701 | 9.3482 | 0.078550588 | 0.059899039 | 9.3968438 | 9.7210357 | 9.39258 | 0.328453028 | 0.004261101 |
| Coro1b | 10.76502 | 10.76518 | 10.79 | -0.024691766 | -0.02485514 | 10.981934 | 11.141809 | 10.8134 | 0.32837244 | 0.168497174 |
| Mrpl11 | 8.427252 | 8.367555 | 8.3573 | 0.010218998 | 0.069916032 | 8.5483021 | 8.6593067 | 8.33096 | 0.328347043 | 0.217342449 |
| Igkv4-80 | 5.398441 | 5.696334 | 5.5979 | 0.098419815 | -0.199473891 | 6.5300879 | 5.8869654 | 5.55879 | 0.32817756 | 0.971300084 |
| Mrfap1 | 10.95658 | 11.00784 | 10.881 | 0.126453668 | 0.075199654 | 10.95264 | 11.148877 | 10.8207 | 0.328171136 | 0.131934221 |

| | | | | | | | | | | |
|---|---|---|---|---|---|---|---|---|---|---|
| Sumo1 | 10.43136 | 10.45165 | 10.425 | 0.026663342 | 0.00637932 | 10.383465 | 10.708126 | 10.38 | 0.328083582 | 0.003422464 |
| Polr2i | 7.099333 | 7.247152 | 7.2006 | 0.046515445 | -0.101303234 | 7.393062 | 7.3512168 | 7.02325 | 0.327965728 | 0.369810855 |
| Psmd6 | 9.392904 | 9.437771 | 9.3546 | 0.083156056 | 0.038289065 | 9.4725511 | 9.6967834 | 9.36883 | 0.32794898 | 0.103716752 |
| Pex10 | 6.85833 | 6.907482 | 6.8101 | 0.097406333 | 0.048254225 | 6.9031718 | 7.0173793 | 6.68954 | 0.327837406 | 0.213629912 |
| Spp1 | 5.780507 | 4.97961 | 4.6814 | 0.298183904 | 1.099081256 | 4.9606602 | 5.0503909 | 4.72276 | 0.327630925 | 0.237900258 |
| Exosc8 | 9.518119 | 9.589571 | 9.5386 | 0.050980138 | -0.0204719 | 9.7035523 | 10.093626 | 9.76605 | 0.327574506 | -0.062499141 |
| Ube2n-ps1 | 7.712066 | 7.834136 | 7.6611 | 0.172993317 | 0.050923213 | 7.7433498 | 7.9984725 | 7.67099 | 0.32747902 | 0.072356366 |
| Zfp830 | 7.828609 | 7.939064 | 7.9051 | 0.033918749 | -0.076536011 | 7.8166825 | 8.0472681 | 7.72031 | 0.326957753 | 0.096372184 |
| Selenot | 11.98645 | 12.1303 | 11.958 | 0.171844918 | 0.027992584 | 11.97706 | 12.266283 | 11.9394 | 0.326924168 | 0.037701049 |
| Rad9a | 8.713236 | 8.823163 | 8.7146 | 0.108526367 | -0.001400798 | 9.195389 | 9.2402807 | 8.9134 | 0.326877798 | 0.28198608 |
| Azin2 | 4.040195 | 4.119666 | 4.0881 | 0.031608579 | -0.047862048 | 4.1713946 | 4.475261 | 4.14843 | 0.326833157 | 0.022966801 |
| Klrb1c | 7.869309 | 8.025264 | 7.8617 | 0.163543481 | 0.007588841 | 7.4651626 | 7.8294671 | 7.50307 | 0.326393934 | -0.037910597 |
| Stx8 | 8.400217 | 8.480084 | 8.3942 | 0.085875781 | 0.006008061 | 8.3797866 | 8.7674033 | 8.4411 | 0.326298769 | -0.061317997 |
| Rarres1 | 2.104203 | 2.226367 | 2.0665 | 0.159912081 | 0.037748553 | 2.147156 | 2.3300916 | 2.0038 | 0.326293656 | 0.14335799 |
| Umps | 9.0064 | 9.141744 | 8.9674 | 0.174387652 | 0.039042935 | 9.1320895 | 9.3597785 | 9.03356 | 0.326218762 | 0.098529766 |
| Hint3 | 7.309802 | 7.360723 | 7.3184 | 0.042346328 | -0.008574874 | 7.402581 | 7.5067425 | 7.18076 | 0.325982717 | 0.221821185 |
| Cript | 8.880557 | 9.060279 | 9.0138 | 0.046491313 | -0.133230536 | 9.1411539 | 9.2873005 | 8.96143 | 0.325869461 | 0.179722892 |
| Eno1 | 12.75438 | 12.74689 | 12.752 | -0.00487104 | 0.00262271 | 12.997281 | 13.039102 | 12.7133 | 0.325838806 | 0.28401774 |
| Trmo | 7.899495 | 8.073323 | 7.9706 | 0.102694598 | -0.071133616 | 8.1476285 | 8.2922224 | 7.96656 | 0.325659773 | 0.181065852 |
| Mrpl19 | 8.714299 | 8.816068 | 8.6211 | 0.194988082 | 0.09321884 | 8.9542681 | 9.125656 | 8.80002 | 0.325638811 | 0.154250899 |
| Ackr3 | 4.970994 | 4.879962 | 4.486 | 0.39395164 | 0.484983816 | 4.4828389 | 4.8922344 | 4.56688 | 0.325358288 | -0.084037187 |
| Rab7 | 11.73804 | 11.80669 | 11.669 | 0.137607788 | 0.068949426 | 11.772254 | 11.924598 | 11.5993 | 0.325319386 | 0.17297529 |
| Ubqln1 | 11.36872 | 11.57435 | 11.327 | 0.2476496 | 0.042014474 | 11.453368 | 11.656789 | 11.3315 | 0.325284356 | 0.121863311 |
| Tomm7 | 9.757228 | 9.789673 | 9.7336 | 0.056079373 | 0.023634827 | 9.9108184 | 10.070326 | 9.74505 | 0.325278177 | 0.16577094 |
| Bud13 | 8.560478 | 8.786568 | 8.4846 | 0.301949966 | 0.075860633 | 8.6269443 | 8.9480466 | 8.62282 | 0.325223712 | 0.004121457 |
| Dhx37 | 9.254362 | 9.510098 | 9.3759 | 0.134235679 | -0.121500261 | 9.4548086 | 9.8731783 | 9.54807 | 0.325107497 | -0.093262118 |
| Atp6v0e | 9.921032 | 10.03519 | 9.9902 | 0.044989064 | -0.069172196 | 10.219694 | 10.254713 | 9.92978 | 0.324930924 | 0.289912009 |
| Nxt1 | 7.947301 | 7.945388 | 7.9157 | 0.029731071 | 0.031644375 | 8.0730072 | 8.123399 | 7.79849 | 0.324904513 | 0.274512684 |
| Uspl1 | 10.04835 | 10.20988 | 10.007 | 0.202390848 | 0.040860841 | 9.9123565 | 10.37102 | 10.0462 | 0.324835569 | -0.133827958 |
| Clp1 | 8.783886 | 8.718259 | 8.8105 | -0.092211897 | -0.026584896 | 8.7533717 | 8.9283539 | 8.60359 | 0.324768044 | 0.149785793 |
| Chtf8 | 11.19854 | 11.4294 | 11.168 | 0.261799442 | 0.030943962 | 11.23919 | 11.404794 | 11.0801 | 0.324714888 | 0.159110155 |
| Mrps24 | 7.921505 | 8.041483 | 7.9786 | 0.062923527 | -0.057054813 | 8.1522736 | 8.1789484 | 7.85439 | 0.324558275 | 0.297883459 |

| | | | | | | | | | | |
|---|---|---|---|---|---|---|---|---|---|---|
| Zc3h8 | 6.633142 | 6.766012 | 6.8261 | -0.060112738 | -0.19298241 | 6.7968931 | 7.0874577 | 6.76301 | 0.324451766 | 0.033887252 |
| Crebrf | 10.54252 | 10.72941 | 10.571 | 0.158241376 | -0.02864309 | 10.47506 | 10.950163 | 10.6257 | 0.324440766 | -0.150661503 |
| Csn1s2b | 2.73308 | 2.836514 | 2.7235 | 0.112969189 | 0.009535193 | 2.7178028 | 3.1544952 | 2.83008 | 0.324420157 | -0.112272215 |
| Taf10 | 9.981206 | 10.09733 | 10.042 | 0.05534777 | -0.060774893 | 10.462811 | 10.381757 | 10.0575 | 0.32427257 | 0.405327034 |
| Men1 | 8.448719 | 8.605077 | 8.5131 | 0.091998769 | -0.064358847 | 8.5328955 | 8.7339717 | 8.4098 | 0.324172477 | 0.123096289 |
| Eif3k | 11.31527 | 11.44434 | 11.398 | 0.046800386 | -0.082267086 | 11.624192 | 11.682506 | 11.3584 | 0.324103226 | 0.265789053 |
| Mbd3 | 9.89216 | 10.08643 | 9.9188 | 0.167642274 | -0.026630469 | 10.118816 | 10.261097 | 9.937 | 0.324094657 | 0.181813514 |
| Jun | 12.28226 | 11.9106 | 12.251 | -0.340815206 | 0.03084214 | 10.808868 | 11.485619 | 11.1615 | 0.324080422 | -0.352670008 |
| Gtpbp3 | 8.191982 | 8.252261 | 8.2131 | 0.039181531 | -0.021097381 | 8.5071834 | 8.5097951 | 8.18577 | 0.324023561 | 0.321411856 |
| Ccl5 | 11.40237 | 11.49973 | 11.451 | 0.049009248 | -0.048354644 | 11.061922 | 11.682661 | 11.3587 | 0.323940928 | -0.29679724 |
| Neil2 | 5.110043 | 5.286273 | 5.2146 | 0.071720192 | -0.104509415 | 5.4223188 | 5.5506058 | 5.22682 | 0.323786882 | 0.195499843 |
| Gm8979 | 4.366648 | 4.60281 | 4.387 | 0.215855986 | -0.020306316 | 4.4051917 | 4.7457052 | 4.42206 | 0.323646755 | -0.016866735 |
| En2 | 0.925453 | 0.972252 | 0.9154 | 0.05688308 | 0.010083235 | 0.909154 | 1.2583276 | 0.93479 | 0.3235371 | -0.025636439 |
| Pitpnm1 | 10.19759 | 10.35919 | 10.324 | 0.035326976 | -0.126274541 | 10.432934 | 10.720717 | 10.3973 | 0.323459352 | 0.035676247 |
| Pip4p1 | 9.461746 | 9.636552 | 9.6286 | 0.007966482 | -0.16683995 | 9.686148 | 9.8884574 | 9.56506 | 0.323402256 | 0.121092828 |
| Trappc5 | 9.037563 | 9.15304 | 9.0133 | 0.139715305 | 0.024238892 | 9.2669758 | 9.2552632 | 8.93188 | 0.323382347 | 0.335094899 |
| U2af1l4 | 7.611603 | 7.518743 | 7.6689 | -0.150158301 | -0.057299004 | 7.6639416 | 7.9288361 | 7.60548 | 0.323353311 | 0.058458814 |
| Dcpp3 | 3.158592 | 3.43784 | 2.5526 | 0.885204357 | 0.605955743 | 5.3259933 | 3.1591302 | 2.83589 | 0.323241033 | 2.490104131 |
| Nfkbiz | 10.9571 | 10.71254 | 10.621 | 0.091968026 | 0.336529694 | 10.430928 | 10.817226 | 10.4941 | 0.323162014 | -0.063136017 |
| Khdc3 | 2.928823 | 2.937028 | 2.8968 | 0.04024818 | 0.032043935 | 2.756207 | 3.2216939 | 2.89855 | 0.323138912 | -0.142347957 |
| 2410004B18Rik | 7.554678 | 7.475256 | 7.5709 | -0.095619787 | -0.016197998 | 7.6626992 | 7.8742936 | 7.55118 | 0.323110197 | 0.111515795 |
| Unc50 | 8.108039 | 8.274694 | 8.1533 | 0.121380579 | -0.045274332 | 8.2904483 | 8.4743081 | 8.15144 | 0.32287287 | 0.139013085 |
| Cct4 | 11.84187 | 11.95892 | 11.868 | 0.091228538 | -0.025818002 | 11.880888 | 12.233387 | 11.9106 | 0.322794204 | -0.029705533 |
| Dusp5 | 8.989603 | 8.917333 | 8.7366 | 0.180701583 | 0.252971613 | 8.5954549 | 9.1023722 | 8.77969 | 0.322677398 | -0.184239832 |
| F3 | 4.488745 | 4.635116 | 4.3241 | 0.310973541 | 0.164603031 | 4.2313394 | 4.5264702 | 4.20382 | 0.322652892 | 0.027522086 |
| Prdm2 | 11.36461 | 11.61428 | 11.434 | 0.17998113 | -0.069697578 | 11.533737 | 11.872438 | 11.5498 | 0.322591694 | -0.016109233 |
| Zmat5 | 7.95424 | 8.025274 | 7.9305 | 0.09472797 | 0.023694315 | 8.2286259 | 8.2156308 | 7.89306 | 0.322571531 | 0.335566607 |
| C1qa | 8.933341 | 8.458761 | 8.631 | -0.172221108 | 0.302359038 | 8.2612993 | 8.3311129 | 8.0086 | 0.322515348 | 0.252701791 |
| Naaa | 10.20635 | 9.870343 | 9.9713 | -0.100996842 | 0.235011225 | 10.056344 | 10.164948 | 9.84254 | 0.322408711 | 0.21380439 |
| Ssr2 | 9.791919 | 9.831984 | 9.6977 | 0.134328011 | 0.094263499 | 9.9437241 | 10.014963 | 9.69262 | 0.322341918 | 0.2511028 |
| Msmo1 | 8.480774 | 8.470335 | 8.3026 | 0.167691762 | 0.178130915 | 8.4819004 | 8.5822262 | 8.25997 | 0.322259461 | 0.221933688 |
| Cttnbp2nl | 8.227784 | 8.127185 | 8.186 | -0.058787875 | 0.041811156 | 7.9010237 | 8.3614098 | 8.03916 | 0.322253422 | -0.138132772 |

| | | | | | | | | | | |
|---|---|---|---|---|---|---|---|---|---|---|
| Evi2b | 11.01925 | 11.2281 | 11.095 | 0.133591504 | -0.07526481 | 11.502996 | 11.647016 | 11.3249 | 0.322138552 | 0.178118582 |
| Epb41l5 | 5.623553 | 5.415555 | 5.4396 | -0.024038089 | 0.183959779 | 5.4186165 | 5.9290629 | 5.60699 | 0.322072866 | -0.188373527 |
| B230303A05Rik | 1.25736 | 1.335195 | 1.1838 | 0.151411577 | 0.073576439 | 1.2504842 | 1.5529802 | 1.23099 | 0.321994715 | 0.019498728 |
| Gm49388 | 6.408166 | 6.646586 | 6.5405 | 0.106109474 | -0.132310363 | 6.6056568 | 6.4875674 | 6.16566 | 0.321909436 | 0.439998896 |
| Mss51 | 4.579445 | 4.727597 | 4.6709 | 0.056735654 | -0.091415981 | 4.7924783 | 5.2158341 | 4.89397 | 0.321860429 | -0.101495385 |
| Orai3 | 8.812031 | 8.93412 | 8.8695 | 0.064624774 | -0.057464436 | 9.1629181 | 9.1159404 | 8.79418 | 0.321762205 | 0.368739812 |
| Uvrag | 10.74087 | 10.97359 | 10.865 | 0.108588622 | -0.124135122 | 10.832161 | 11.173473 | 10.8517 | 0.321746708 | -0.019565166 |
| Slc1a3 | 6.007124 | 5.795487 | 5.6341 | 0.161366724 | 0.373004367 | 5.4069849 | 5.4978475 | 5.17618 | 0.321665973 | 0.230803455 |
| Gpbp1 | 11.69082 | 11.85819 | 11.691 | 0.167047392 | -0.000315976 | 11.51353 | 11.993353 | 11.6718 | 0.321571944 | -0.158250717 |
| Zdhhc22 | 3.261565 | 3.448531 | 3.2934 | 0.155113581 | -0.031852282 | 3.3635099 | 3.7582243 | 3.43667 | 0.321557211 | -0.073157178 |
| Dnajb6 | 10.76725 | 10.88456 | 10.677 | 0.207972544 | 0.090663632 | 10.778335 | 11.160985 | 10.8396 | 0.321406942 | -0.061242782 |
| Qtrt2 | 8.669177 | 8.811544 | 8.6389 | 0.172673992 | 0.030307109 | 8.7251571 | 9.1882202 | 8.86687 | 0.321352368 | -0.141710736 |
| Zbtb12 | 7.405068 | 7.497 | 7.3738 | 0.123205638 | 0.031273741 | 7.5087168 | 7.6656315 | 7.34434 | 0.321288353 | 0.164373635 |
| Polr1b | 9.362497 | 9.588566 | 9.4599 | 0.128644061 | -0.097424409 | 9.6720378 | 9.9415177 | 9.62045 | 0.321070251 | 0.051590378 |
| Thop1 | 7.447978 | 7.307467 | 7.5665 | -0.259066406 | -0.118554662 | 7.6639195 | 7.73751 | 7.41648 | 0.321027654 | 0.247437112 |
| Oxa1l | 9.918454 | 10.00567 | 9.9391 | 0.06658723 | -0.020628402 | 10.038922 | 10.256701 | 9.93586 | 0.320835974 | 0.103056971 |
| Slc25a5 | 12.51219 | 12.46442 | 12.43 | 0.034457328 | 0.08222037 | 12.61948 | 12.716071 | 12.3953 | 0.320806054 | 0.22421505 |
| Prcc | 9.515238 | 9.690644 | 9.6185 | 0.07211994 | -0.103286317 | 9.627676 | 9.8443977 | 9.52406 | 0.320339884 | 0.103618276 |
| Rab8a | 11.44559 | 11.55664 | 11.361 | 0.195475368 | 0.08442077 | 11.454696 | 11.722087 | 11.402 | 0.320126054 | 0.052735816 |
| Rpl38 | 12.98687 | 13.03314 | 13.021 | 0.012419348 | -0.033852442 | 13.154397 | 13.296774 | 12.9767 | 0.320025964 | 0.177649595 |
| Gm6210 | 2.866611 | 2.866139 | 2.8329 | 0.03326936 | 0.033741911 | 2.8512527 | 3.0670336 | 2.74711 | 0.319926714 | 0.104145738 |
| Terf1 | 8.491691 | 8.591552 | 8.5186 | 0.072987091 | -0.026873448 | 8.3529741 | 8.8231812 | 8.50335 | 0.319834269 | -0.150372821 |
| Gm14760 | 4.137649 | 4.259097 | 4.0764 | 0.18270411 | 0.061256742 | 4.4312087 | 4.308022 | 3.98821 | 0.319815816 | 0.443002465 |
| Mlf2 | 10.04755 | 10.15409 | 10.157 | -0.002484531 | -0.109016715 | 10.310793 | 10.320014 | 10.0004 | 0.319569185 | 0.310348368 |
| Atp6v0b | 9.814078 | 9.929483 | 9.8996 | 0.029914412 | -0.085490421 | 10.201196 | 10.218977 | 9.8995 | 0.319477314 | 0.301695682 |
| Sarnp | 10.04355 | 10.14849 | 10.039 | 0.109719575 | 0.004770665 | 10.117246 | 10.332169 | 10.0127 | 0.319445288 | 0.10452181 |
| Polr2e | 9.897523 | 10.04073 | 9.917 | 0.123777563 | -0.019430253 | 10.150503 | 10.179316 | 9.85992 | 0.319397728 | 0.290584682 |
| Cops7a | 8.980123 | 9.15501 | 9.0416 | 0.113456636 | -0.061430576 | 9.3266614 | 9.3796332 | 9.0603 | 0.319329061 | 0.266357205 |
| Llph-ps2 | 7.637264 | 7.808934 | 7.887 | -0.078023385 | -0.249693392 | 7.6957147 | 8.0167619 | 7.69777 | 0.318991693 | -0.002055516 |
| Abcb1b | 8.259065 | 8.28748 | 8.1361 | 0.15133036 | 0.12291512 | 7.9913773 | 8.5259091 | 8.20697 | 0.318936978 | -0.21559491 |
| Polr2h | 8.092389 | 8.131229 | 8.1469 | -0.015625319 | -0.054465836 | 8.3346986 | 8.4227731 | 8.1039 | 0.318876349 | 0.230801854 |
| Ndufb4c | 7.86167 | 7.915551 | 7.7625 | 0.153082578 | 0.099201354 | 7.9463677 | 8.0342779 | 7.71564 | 0.31863727 | 0.230727044 |

| | | | | | | | | | | |
|---|---|---|---|---|---|---|---|---|---|---|
| 4930453N24Rik | 9.522527 | 9.697422 | 9.6176 | 0.079795981 | -0.095099641 | 9.3666561 | 9.8982652 | 9.57973 | 0.318538107 | -0.213071027 |
| AB124611 | 10.7145 | 10.6428 | 10.704 | -0.061518936 | 0.010180496 | 10.997197 | 10.94905 | 10.6306 | 0.318474774 | 0.366622282 |
| Fabp3 | 1.059283 | 1.180912 | 1.0413 | 0.139626394 | 0.017997561 | 1.0468216 | 1.3481051 | 1.02971 | 0.318390843 | 0.017107318 |
| Snx17 | 10.40187 | 10.43946 | 10.387 | 0.052633734 | 0.015040918 | 10.485303 | 10.599846 | 10.2815 | 0.318308746 | 0.203765573 |
| Crebzf | 9.300375 | 9.366255 | 9.4017 | -0.035449101 | -0.101328819 | 9.6751437 | 9.8906133 | 9.5727 | 0.317908615 | 0.102439024 |
| Snf8 | 9.259516 | 9.208959 | 9.172 | 0.036931137 | 0.087487704 | 9.3409891 | 9.4987931 | 9.18102 | 0.317769202 | 0.159965187 |
| Ebag9 | 8.989268 | 8.833673 | 8.9238 | -0.090137074 | 0.065458239 | 8.7811554 | 9.131518 | 8.81381 | 0.31770515 | -0.032657452 |
| BC031181 | 9.273043 | 9.410247 | 9.283 | 0.127200476 | -0.01000331 | 9.4594994 | 9.57396 | 9.25631 | 0.317652492 | 0.203191902 |
| Dnlz | 7.785434 | 7.896355 | 7.8667 | 0.029632475 | -0.081288712 | 7.905144 | 8.0898797 | 7.77242 | 0.317463363 | 0.132727651 |
| Igkv1-122 | 5.224565 | 5.602699 | 5.2333 | 0.369401424 | -0.00873269 | 5.4178938 | 5.5279883 | 5.21065 | 0.317343093 | 0.207248597 |
| Nphp4 | 4.891039 | 5.053965 | 4.7545 | 0.299444593 | 0.136519398 | 5.211415 | 5.3640166 | 5.04675 | 0.31727046 | 0.164668901 |
| Lamtor3 | 8.778917 | 8.914963 | 8.7373 | 0.177684132 | 0.041638282 | 8.9468348 | 9.1300082 | 8.81306 | 0.316947442 | 0.133773971 |
| Abt1 | 7.989474 | 7.964619 | 7.944 | 0.020655493 | 0.045510381 | 7.921799 | 8.072815 | 7.75591 | 0.316902581 | 0.16588654 |
| Ostc | 9.475763 | 9.45627 | 9.4046 | 0.051667163 | 0.071160257 | 9.5526835 | 9.6208014 | 9.30405 | 0.316752691 | 0.248634747 |
| Sec13 | 10.08647 | 10.14548 | 10.031 | 0.114146972 | 0.055141275 | 10.404754 | 10.433514 | 10.1168 | 0.316697119 | 0.287937015 |
| Rps5 | 14.2649 | 14.38495 | 14.324 | 0.06075247 | -0.05929844 | 14.564849 | 14.573258 | 14.2566 | 0.31669453 | 0.308285385 |
| Lemd2 | 9.367728 | 9.494014 | 9.5004 | -0.006417599 | -0.132703765 | 9.6614952 | 9.8401036 | 9.52343 | 0.316677797 | 0.138069417 |
| Traf2 | 9.949395 | 10.12516 | 9.9509 | 0.174219085 | -0.001547766 | 10.071083 | 10.282969 | 9.96656 | 0.316412982 | 0.104526728 |
| Nr3c1 | 11.43322 | 11.71024 | 11.412 | 0.298687622 | 0.021662638 | 11.224853 | 11.683816 | 11.3675 | 0.316360028 | -0.142603022 |
| Efcab14 | 9.604574 | 9.744691 | 9.8153 | -0.070646787 | -0.210764284 | 9.524809 | 10.068728 | 9.75242 | 0.316305822 | -0.227612908 |
| Olfr77 | 6.512669 | 6.658396 | 6.6307 | 0.027705363 | -0.118021838 | 6.540268 | 6.7776238 | 6.46142 | 0.316207533 | 0.078851743 |
| Fastkd2 | 8.447145 | 8.516593 | 8.6266 | -0.110013864 | -0.179462442 | 8.824453 | 9.0234083 | 8.70739 | 0.316021596 | 0.117066312 |
| Timm9 | 7.765172 | 7.728111 | 7.6749 | 0.053196164 | 0.090257535 | 8.1112264 | 8.2929142 | 7.97707 | 0.315841195 | 0.134153414 |
| Abcf2 | 10.02914 | 10.08721 | 9.9969 | 0.090308851 | 0.032233994 | 10.128353 | 10.313941 | 9.9981 | 0.315837309 | 0.130249708 |
| Hsf1 | 9.624665 | 9.744116 | 9.6302 | 0.11391606 | -0.005534944 | 9.9582356 | 10.145381 | 9.82957 | 0.315806638 | 0.128661108 |
| Tefm | 6.576413 | 6.585999 | 6.5518 | 0.034234576 | 0.024648728 | 6.6502415 | 6.8090585 | 6.49333 | 0.315731717 | 0.156914712 |
| Paip1 | 9.63453 | 9.710851 | 9.5295 | 0.181368783 | 0.105047897 | 9.5015471 | 9.7933849 | 9.47789 | 0.315494343 | 0.023656533 |
| Nol9 | 9.998284 | 10.09322 | 9.9513 | 0.141882021 | 0.046944735 | 10.081 | 10.297943 | 9.98254 | 0.315405457 | 0.098462372 |
| Rtn4rl2 | 2.723394 | 2.578941 | 2.4482 | 0.130735583 | 0.27518823 | 2.6470035 | 2.7175142 | 2.40229 | 0.315219797 | 0.244709046 |
| Nifk | 9.520366 | 9.56919 | 9.3815 | 0.187645408 | 0.138821593 | 9.4388754 | 9.6841262 | 9.36892 | 0.315207021 | 0.069956227 |
| Gm48551 | 5.176332 | 5.37729 | 5.3113 | 0.065959767 | -0.134997681 | 5.2990767 | 5.4874053 | 5.17244 | 0.314968715 | 0.126640136 |
| Psmd14 | 10.36372 | 10.28957 | 10.251 | 0.039042936 | 0.113196852 | 10.347732 | 10.555487 | 10.2407 | 0.31478844 | 0.107033834 |

| | | | | | | | | | | |
|---|---|---|---|---|---|---|---|---|---|---|
| Atxn7l1 | 9.345026 | 9.577343 | 9.5404 | 0.036925976 | -0.195391137 | 9.5345443 | 9.9911497 | 9.67664 | 0.31451107 | -0.142094285 |
| Dynlt1b | 7.816834 | 7.960346 | 8.0462 | -0.085867274 | -0.229379428 | 8.2805401 | 8.2182204 | 7.90373 | 0.31448977 | 0.376809468 |
| Mitf | 5.846631 | 5.771623 | 5.6676 | 0.104026624 | 0.179034588 | 5.4524714 | 5.7039537 | 5.38959 | 0.314365172 | 0.062882842 |
| Nlrp1c-ps | 3.364005 | 3.314045 | 3.2981 | 0.015988013 | 0.0659488 | 3.4038386 | 3.6752765 | 3.36092 | 0.314353463 | 0.042915557 |
| Id1 | 5.155574 | 5.06509 | 4.9662 | 0.098840887 | 0.18932496 | 4.6603517 | 4.9904645 | 4.67618 | 0.314288168 | -0.015824609 |
| Nxpe4 | 6.317818 | 6.489737 | 6.2786 | 0.211179895 | 0.039261101 | 6.4975079 | 6.3688291 | 6.05458 | 0.314250661 | 0.442929501 |
| Cyc1 | 10.1064 | 10.04754 | 9.8983 | 0.149264315 | 0.208122657 | 10.203299 | 10.274626 | 9.96048 | 0.314142634 | 0.242816248 |
| Chl1 | 4.466684 | 4.247965 | 4.008 | 0.239982979 | 0.458701197 | 4.6529542 | 4.3589756 | 4.04497 | 0.314005387 | 0.607983975 |
| Lzts1 | 5.393429 | 5.518237 | 5.4055 | 0.112717167 | -0.01209069 | 5.344536 | 5.693998 | 5.38019 | 0.313810675 | -0.035651339 |
| Fblim1 | 5.605565 | 5.776825 | 5.4421 | 0.334766977 | 0.163506615 | 5.4190228 | 5.7382465 | 5.42446 | 0.313790858 | -0.005432871 |
| Ngrn | 7.849032 | 7.895551 | 7.9233 | -0.027740139 | -0.074259055 | 8.1282995 | 8.2293863 | 7.91569 | 0.313699498 | 0.212612715 |
| Eif2b5 | 9.868501 | 9.858832 | 9.8249 | 0.033956361 | 0.043625408 | 10.026687 | 10.17964 | 9.86599 | 0.31365041 | 0.160697497 |
| Fam162a | 7.917823 | 7.872088 | 7.8758 | -0.003724729 | 0.042010343 | 7.9691172 | 8.1164122 | 7.80285 | 0.313560576 | 0.166265617 |
| Oaz1 | 12.14444 | 12.19886 | 12.173 | 0.025843254 | -0.028578178 | 12.301498 | 12.382186 | 12.0687 | 0.313437196 | 0.232748687 |
| Mau2 | 11.15593 | 11.39777 | 11.238 | 0.159410448 | -0.082426982 | 11.47181 | 11.698415 | 11.385 | 0.313421828 | 0.086816486 |
| Rpf1 | 8.920162 | 8.91028 | 8.9175 | -0.007194713 | 0.002686722 | 9.0399628 | 9.2637456 | 8.95037 | 0.313379623 | 0.089596789 |
| Mdfic | 9.138433 | 9.170614 | 9.0947 | 0.075945679 | 0.04376461 | 8.8451812 | 9.2630973 | 8.95007 | 0.313023287 | -0.104892866 |
| Adarb2 | 4.124078 | 4.226256 | 4.0935 | 0.132713651 | 0.030534857 | 4.2290476 | 4.5482529 | 4.23531 | 0.312946247 | -0.006259093 |
| Atp8a1 | 10.81733 | 10.93969 | 10.783 | 0.15628976 | 0.033930442 | 10.907672 | 11.236825 | 10.924 | 0.312850388 | -0.016302642 |
| Cd27 | 10.17752 | 10.29804 | 10.287 | 0.01124815 | -0.109271487 | 10.252862 | 10.676978 | 10.3642 | 0.312757092 | -0.111359032 |
| Rpl14 | 13.64502 | 13.73403 | 13.686 | 0.048251772 | -0.040748612 | 13.80629 | 13.908216 | 13.5955 | 0.312686272 | 0.210760298 |
| Slc30a7 | 9.443188 | 9.448548 | 9.4305 | 0.018096151 | 0.012736385 | 9.3068975 | 9.7563887 | 9.44373 | 0.312659464 | -0.136831718 |
| Sptlc2 | 10.36189 | 10.32203 | 10.254 | 0.067736168 | 0.107587264 | 10.198537 | 10.381426 | 10.0688 | 0.312656123 | 0.129767181 |
| Lsm2 | 7.693083 | 7.911001 | 7.7553 | 0.155670508 | -0.06224819 | 8.0962002 | 8.2736396 | 7.96107 | 0.312567973 | 0.135128513 |
| Zdhhc16 | 8.232463 | 8.105273 | 8.24 | -0.134768411 | -0.007578627 | 8.4432528 | 8.5062805 | 8.19371 | 0.312567697 | 0.249540029 |
| Calm1 | 13.25303 | 13.27993 | 13.281 | -0.001182976 | -0.028081528 | 13.470388 | 13.59341 | 13.2808 | 0.31256349 | 0.189541063 |
| Igkv12-89 | 5.757056 | 6.542608 | 5.7944 | 0.748225755 | -0.037327041 | 5.9684028 | 5.9883446 | 5.67598 | 0.312362092 | 0.292420242 |
| Sys1 | 9.121989 | 9.1087 | 9.0895 | 0.01924865 | 0.032537039 | 9.3452864 | 9.294771 | 8.98243 | 0.31234039 | 0.362855855 |
| Atp6v1h | 9.87043 | 9.887871 | 9.7637 | 0.124165854 | 0.106725413 | 9.8556886 | 10.039908 | 9.7278 | 0.312110238 | 0.127891379 |
| Btf3 | 12.44685 | 12.46827 | 12.441 | 0.027295622 | 0.005874846 | 12.590097 | 12.738981 | 12.4269 | 0.312032826 | 0.16314858 |
| Ccdc189 | 3.107176 | 3.03619 | 3.0484 | -0.012244127 | 0.05874193 | 3.1231593 | 3.1631937 | 2.85125 | 0.311945931 | 0.271911453 |
| Naip6 | 7.110887 | 7.012214 | 7.1395 | -0.127298705 | -0.028625459 | 7.2301317 | 7.6173159 | 7.30552 | 0.311791629 | -0.075392558 |

| | | | | | | | | | | |
|---|---|---|---|---|---|---|---|---|---|---|
| Hic1 | 4.165593 | 3.945647 | 3.9766 | -0.031000302 | 0.188945621 | 4.0233849 | 4.0865157 | 3.77496 | 0.311559959 | 0.248429148 |
| Tm9sf4 | 9.80064 | 9.823181 | 9.6913 | 0.131899133 | 0.109358563 | 9.8561385 | 10.037328 | 9.72599 | 0.311335937 | 0.130146802 |
| Polr2c | 9.072344 | 9.087856 | 9.0295 | 0.058372516 | 0.042860698 | 9.2769196 | 9.3826284 | 9.07129 | 0.311335131 | 0.205626376 |
| Ltb | 12.22466 | 12.50821 | 12.301 | 0.20689243 | -0.076652118 | 12.589122 | 12.542542 | 12.2312 | 0.311315328 | 0.357895087 |
| Cisd2 | 9.630201 | 9.699831 | 9.6522 | 0.047629993 | -0.021999644 | 9.6358635 | 9.8507996 | 9.53957 | 0.311231756 | 0.096295616 |
| Gm42927 | 8.474517 | 8.797271 | 8.5736 | 0.223669379 | -0.099085044 | 8.6808489 | 9.5373374 | 9.22611 | 0.311228125 | -0.545260377 |
| Bahd1 | 10.05296 | 10.15103 | 10.076 | 0.074939693 | -0.023134582 | 10.082352 | 10.319097 | 10.008 | 0.311097487 | 0.074352243 |
| Phrf1 | 10.87244 | 11.04814 | 10.913 | 0.135608334 | -0.040096112 | 11.013612 | 11.212909 | 10.9019 | 0.311048644 | 0.111751661 |
| Vdac2 | 11.03409 | 11.03716 | 11.017 | 0.020465184 | 0.017394284 | 11.11019 | 11.357214 | 11.0462 | 0.311013616 | 0.063989868 |
| Ndufa4 | 9.851861 | 9.943713 | 9.9403 | 0.003401588 | -0.088449659 | 10.11587 | 10.219216 | 9.90821 | 0.311005151 | 0.207659215 |
| Sinhcaf | 10.13458 | 10.37326 | 10.177 | 0.196219674 | -0.042459631 | 10.183435 | 10.568584 | 10.2576 | 0.310994892 | -0.07415404 |
| Tox4 | 10.93828 | 11.08222 | 10.988 | 0.09436681 | -0.049569852 | 10.905388 | 11.257831 | 10.9469 | 0.310979056 | -0.041464267 |
| Lsm4 | 8.717406 | 8.749107 | 8.6022 | 0.14689655 | 0.115195822 | 9.0122211 | 9.039187 | 8.72836 | 0.310825461 | 0.283859519 |
| Mctp2 | 9.078955 | 9.334224 | 9.06 | 0.274253125 | 0.018984432 | 9.0027921 | 9.3673239 | 9.05654 | 0.310781867 | -0.053750007 |
| Rexo1 | 10.58295 | 10.59119 | 10.548 | 0.042864498 | 0.034621292 | 10.851702 | 10.959585 | 10.6489 | 0.310722658 | 0.202839483 |
| Uba52 | 11.46138 | 11.60265 | 11.512 | 0.090970992 | -0.050301512 | 11.681511 | 11.779057 | 11.4684 | 0.310634132 | 0.213087356 |
| Uhrf2 | 9.964458 | 10.20726 | 10.086 | 0.121667683 | -0.121132893 | 10.069143 | 10.566896 | 10.2564 | 0.310482844 | -0.187270079 |
| Tdrp | 8.861501 | 9.043369 | 9.1336 | -0.090263861 | -0.272131263 | 8.3945767 | 9.2096307 | 8.89923 | 0.310399676 | -0.504654362 |
| Pycard | 9.441946 | 9.485376 | 9.4883 | -0.002918384 | -0.046348117 | 9.6046597 | 9.8509556 | 9.54056 | 0.310392196 | 0.06409627 |
| Efemp2 | 5.203747 | 5.329295 | 5.2397 | 0.089589861 | -0.035958823 | 5.236475 | 5.3065009 | 4.99612 | 0.310379045 | 0.240353105 |
| Plod1 | 8.152784 | 7.990067 | 8.1315 | -0.141421847 | 0.021295441 | 7.7968582 | 8.0180716 | 7.7077 | 0.310369517 | 0.089156083 |
| Tmem151b | 4.365183 | 4.423908 | 4.2611 | 0.162853875 | 0.104129222 | 4.1871761 | 4.5286843 | 4.21843 | 0.310254757 | -0.031253368 |
| Gm17828 | 4.493338 | 4.838408 | 4.6838 | 0.154601602 | -0.190468471 | 4.7232217 | 5.0789219 | 4.76875 | 0.31016851 | -0.045531678 |
| Mapk1ip1l | 11.02011 | 11.1055 | 11.015 | 0.090225432 | 0.004837436 | 11.006711 | 11.315601 | 11.0055 | 0.310105186 | 0.001215498 |
| Rars2 | 8.140729 | 8.128893 | 8.1874 | -0.058532655 | -0.046696208 | 8.3395141 | 8.5614178 | 8.25142 | 0.3100026 | 0.088098888 |
| Selenos | 9.419242 | 9.350702 | 9.2469 | 0.103830041 | 0.172369379 | 9.3764227 | 9.5170478 | 9.20705 | 0.310001698 | 0.169376607 |
| Hmbs | 7.919689 | 7.786335 | 7.9301 | -0.143813035 | -0.01045885 | 8.0792978 | 8.0948059 | 7.78491 | 0.309895369 | 0.294387276 |
| Srsf4 | 10.16744 | 10.28932 | 10.186 | 0.103650259 | -0.018225399 | 10.265629 | 10.488295 | 10.1786 | 0.309744193 | 0.087078378 |
| Pym1 | 9.410698 | 9.564506 | 9.5066 | 0.057886163 | -0.095920955 | 9.7167537 | 9.8255469 | 9.5159 | 0.309643043 | 0.200849838 |
| Nob1 | 9.324411 | 9.509708 | 9.3756 | 0.134103816 | -0.051192502 | 9.56446 | 9.727323 | 9.4179 | 0.309421659 | 0.146558707 |
| Tut1 | 9.293748 | 9.43145 | 9.3464 | 0.085044405 | -0.052657859 | 9.5112847 | 9.720427 | 9.41102 | 0.309402644 | 0.100260298 |
| Arpc1a | 9.394988 | 9.353474 | 9.3338 | 0.019713271 | 0.061227918 | 9.4509052 | 9.5705958 | 9.26129 | 0.309310519 | 0.18961995 |

| | | | | | | | | | | |
|---|---|---|---|---|---|---|---|---|---|---|
| Nt5m | 6.87738 | 7.026028 | 6.8888 | 0.137236436 | -0.011411495 | 6.9149329 | 7.0854438 | 6.77628 | 0.30916103 | 0.138650147 |
| Swi5 | 10.02627 | 10.04816 | 10.009 | 0.039338715 | 0.017448243 | 10.094528 | 10.264766 | 9.95565 | 0.309111744 | 0.138873568 |
| Tmem119 | 4.46521 | 4.528256 | 4.2481 | 0.280189702 | 0.217143155 | 4.2174717 | 4.3787717 | 4.06972 | 0.309050445 | 0.147750477 |
| Tomm22 | 10.15808 | 10.23326 | 10.219 | 0.014145844 | -0.061029836 | 10.376816 | 10.486926 | 10.1779 | 0.30902804 | 0.198918264 |
| Ppp1r7 | 9.5677 | 9.828521 | 9.7666 | 0.061945217 | -0.198876429 | 9.704366 | 10.038206 | 9.72923 | 0.308977743 | -0.024861757 |
| Ggta1 | 10.40555 | 10.45056 | 10.264 | 0.187030258 | 0.142013188 | 10.316437 | 10.540732 | 10.2318 | 0.3089511 | 0.084655593 |
| Cep57l1 | 7.734485 | 7.82416 | 7.9111 | -0.08690933 | -0.176583873 | 7.7391929 | 8.2708438 | 7.96235 | 0.308493693 | -0.22315726 |
| Trmt10c | 8.681289 | 8.715687 | 8.6775 | 0.038141137 | 0.003742714 | 8.7482442 | 8.9416432 | 8.63327 | 0.308377793 | 0.114978792 |
| Ddx21 | 12.34356 | 12.39165 | 12.219 | 0.173119982 | 0.125029468 | 12.302505 | 12.575894 | 12.2676 | 0.308245248 | 0.034856024 |
| Rad23b | 11.19399 | 11.22018 | 11.231 | -0.010761284 | -0.03694833 | 11.248975 | 11.412232 | 11.1041 | 0.30809295 | 0.144835892 |
| Igkv14-100 | 7.160555 | 7.472051 | 7.1301 | 0.341971976 | 0.03047668 | 7.4430255 | 7.3951032 | 7.08706 | 0.308041094 | 0.355963382 |
| Nat14 | 4.577801 | 4.593503 | 4.4983 | 0.095178491 | 0.07947648 | 4.3566163 | 4.8455444 | 4.5377 | 0.307842937 | -0.181085189 |
| Ncam1 | 4.312082 | 4.340137 | 4.26 | 0.080098175 | 0.052043234 | 4.3872875 | 4.85084 | 4.54321 | 0.307632161 | -0.155920338 |
| Eif4e2 | 10.2271 | 10.30515 | 10.256 | 0.049638122 | -0.02840729 | 10.319149 | 10.628339 | 10.3208 | 0.307506418 | -0.001683929 |
| Mrpl16 | 8.252867 | 8.270071 | 8.2655 | 0.004556472 | -0.012647888 | 8.5047788 | 8.5414849 | 8.23449 | 0.306996483 | 0.270290445 |
| Ak7 | 2.831447 | 2.989735 | 2.8566 | 0.133126892 | -0.025160252 | 2.9217528 | 3.3116591 | 3.00491 | 0.306748702 | -0.083157636 |
| Ube2cbp | 5.673435 | 5.757361 | 5.6875 | 0.069845494 | -0.014080513 | 5.7059363 | 6.0180983 | 5.71143 | 0.306663889 | -0.005498085 |
| Gm4034 | 3.70589 | 3.690016 | 3.8591 | -0.169091594 | -0.15321689 | 4.1410573 | 4.1332024 | 3.8267 | 0.306500662 | 0.314355637 |
| Ager | 5.400295 | 5.637084 | 5.3809 | 0.256133927 | 0.019345349 | 5.555699 | 5.8358385 | 5.52961 | 0.306232037 | 0.026092555 |
| Nvl | 9.920647 | 9.922148 | 9.9506 | -0.028451106 | -0.029952323 | 10.268518 | 10.493031 | 10.1868 | 0.306183937 | 0.081670537 |
| Kctd19 | 2.714813 | 2.654624 | 2.767 | -0.112332764 | -0.052143413 | 2.6126364 | 2.9158348 | 2.60969 | 0.306142198 | 0.002943753 |
| Zcrb1 | 8.715727 | 8.626689 | 8.6807 | -0.053988568 | 0.035049977 | 8.7779018 | 9.0487961 | 8.74273 | 0.306065877 | 0.035171504 |
| Cyb5r4 | 10.19664 | 10.14914 | 10.16 | -0.010887896 | 0.036611848 | 10.123608 | 10.350483 | 10.0446 | 0.305843456 | 0.078968716 |
| Asns | 5.863584 | 5.614996 | 5.5296 | 0.085391526 | 0.333979527 | 5.873105 | 5.9134966 | 5.6077 | 0.305795271 | 0.265403639 |
| Cfap77 | 2.807056 | 2.868812 | 2.8151 | 0.053751881 | -0.008003598 | 2.9032327 | 3.1590068 | 2.85327 | 0.305741173 | 0.049967071 |
| Eif2b2 | 9.338828 | 9.406927 | 9.3195 | 0.08737746 | 0.01927848 | 9.6079988 | 9.6783196 | 9.37269 | 0.305624652 | 0.23530392 |
| Adsl | 9.837614 | 9.863862 | 9.7531 | 0.110753997 | 0.084505975 | 10.053569 | 10.18192 | 9.87641 | 0.305514431 | 0.177163389 |
| Parp4 | 10.9523 | 10.99004 | 10.93 | 0.060487844 | 0.02275217 | 10.954169 | 11.31734 | 11.0121 | 0.305261972 | -0.057909863 |
| Ighv1-78 | 6.875112 | 7.009574 | 6.6168 | 0.392739579 | 0.258277647 | 7.2692562 | 6.9648439 | 6.65963 | 0.305217734 | 0.60963002 |
| Hps6 | 7.239 | 7.253964 | 7.2625 | -0.00856906 | -0.023532599 | 7.333439 | 7.4110044 | 7.10588 | 0.305119416 | 0.227554017 |
| Hspb8 | 3.308177 | 3.3861 | 3.3371 | 0.048994903 | -0.028928344 | 3.5633557 | 3.6550912 | 3.35001 | 0.305078907 | 0.213343469 |
| Rsrc1 | 8.65993 | 8.816755 | 8.7809 | 0.035899004 | -0.120926138 | 8.7380857 | 9.0781248 | 8.77317 | 0.304955682 | -0.035083466 |

| | | | | | | | | | | |
|---|---|---|---|---|---|---|---|---|---|---|
| Rgl1 | 9.448784 | 9.352089 | 9.1903 | 0.161814309 | 0.2585089 | 9.2099205 | 9.260133 | 8.95525 | 0.304879269 | 0.254666771 |
| Ehmt2 | 11.14354 | 11.23676 | 11.211 | 0.025589208 | -0.067627908 | 11.333722 | 11.556579 | 11.2517 | 0.304832866 | 0.081976826 |
| Mfsd2a | 4.365836 | 4.352104 | 4.22 | 0.132102883 | 0.14583553 | 4.0456037 | 4.6252989 | 4.32058 | 0.304714784 | -0.274980352 |
| H2-T-ps | 7.280716 | 7.443834 | 7.2836 | 0.160252346 | -0.002866291 | 7.5857054 | 7.5785521 | 7.27425 | 0.304303764 | 0.311457076 |
| Tnip2 | 8.506851 | 8.263113 | 8.3254 | -0.062324919 | 0.181413654 | 8.3989912 | 8.6098911 | 8.30591 | 0.303980224 | 0.093080335 |
| Cpxm1 | 7.433093 | 7.230673 | 6.9142 | 0.316428971 | 0.518848879 | 6.8447196 | 6.7894955 | 6.4856 | 0.303899343 | 0.35912338 |
| Spryd4 | 6.777314 | 6.71317 | 6.6894 | 0.023804693 | 0.087949283 | 6.7722064 | 6.9603132 | 6.65647 | 0.30384785 | 0.11574106 |
| Aamp | 10.90475 | 10.94498 | 10.869 | 0.075579444 | 0.035344554 | 11.003225 | 11.070774 | 10.767 | 0.303817322 | 0.236268534 |
| Khdc4 | 10.29584 | 10.30789 | 10.277 | 0.031271307 | 0.019214451 | 10.535529 | 10.705503 | 10.4018 | 0.30374396 | 0.133769894 |
| Psmd13 | 10.0276 | 10.10207 | 10.063 | 0.038718723 | -0.035754217 | 10.25037 | 10.413155 | 10.1094 | 0.303705502 | 0.140919986 |
| Bri3 | 8.101081 | 8.209178 | 8.1393 | 0.069912887 | -0.038184718 | 8.3265499 | 8.4128962 | 8.10919 | 0.303703877 | 0.217357576 |
| Gpld1 | 3.323091 | 3.267904 | 3.1929 | 0.074992076 | 0.13017892 | 3.2055203 | 3.4390614 | 3.13537 | 0.303687728 | 0.070146639 |
| Cactin | 9.332731 | 9.459902 | 9.4868 | -0.026878291 | -0.154049224 | 9.7773733 | 9.8627857 | 9.5593 | 0.303480904 | 0.21806856 |
| Tmem11 | 8.127815 | 8.281202 | 8.1616 | 0.119636821 | -0.033749335 | 8.2403541 | 8.5416297 | 8.23825 | 0.303375762 | 0.002100186 |
| Slc1a4 | 5.358112 | 5.195355 | 5.0792 | 0.116105021 | 0.278862135 | 6.0425631 | 6.1216467 | 5.81829 | 0.303353756 | 0.224270195 |
| Gtf3c5 | 8.360808 | 8.33758 | 8.4191 | -0.081532131 | -0.058303939 | 8.5409681 | 8.7762514 | 8.47291 | 0.303337011 | 0.068053728 |
| Sf3b3 | 12.27854 | 12.36553 | 12.232 | 0.13395067 | 0.046965036 | 12.38859 | 12.568758 | 12.2656 | 0.303175926 | 0.123007893 |
| Ino80e | 9.408391 | 9.633466 | 9.3993 | 0.234170943 | 0.009095464 | 9.6413478 | 9.6565464 | 9.35355 | 0.302995943 | 0.287797361 |
| Map4k1 | 11.54435 | 11.54134 | 11.57 | -0.028521382 | -0.02550648 | 11.857187 | 11.878587 | 11.5757 | 0.302922744 | 0.281523094 |
| Arl6ip1 | 11.91687 | 12.00578 | 11.838 | 0.167544854 | 0.07862721 | 11.975531 | 12.196508 | 11.8936 | 0.302904554 | 0.081928051 |
| Igkv1-117 | 9.896909 | 10.03583 | 9.7224 | 0.313411304 | 0.174489373 | 10.081956 | 10.150648 | 9.84793 | 0.30271455 | 0.234023287 |
| Gm7899 | 3.238309 | 3.313357 | 3.1462 | 0.167192048 | 0.09214423 | 2.9884277 | 3.420914 | 3.11822 | 0.302689164 | -0.129797128 |
| Tnfrsf1b | 11.09726 | 11.33301 | 11.046 | 0.286970966 | 0.051214884 | 10.997279 | 11.480497 | 11.178 | 0.30250912 | -0.180709088 |
| Cxcl16 | 9.638565 | 9.823968 | 9.7233 | 0.100690873 | -0.084711891 | 9.3977559 | 10.207433 | 9.90497 | 0.302463354 | -0.507213951 |
| Armc8 | 9.9671 | 10.00286 | 9.8871 | 0.115756372 | 0.079994975 | 9.9638596 | 10.259936 | 9.95786 | 0.302079643 | 0.006002983 |
| Chst14 | 5.955613 | 5.840337 | 5.9831 | -0.142730453 | -0.027454236 | 6.1156134 | 6.0414674 | 5.73972 | 0.30174695 | 0.375892955 |
| Tnrc18 | 10.35768 | 10.48704 | 10.352 | 0.135230783 | 0.00586693 | 10.368832 | 10.660093 | 10.3584 | 0.301690618 | 0.010429814 |
| Zdhhc8 | 8.200887 | 8.17476 | 8.139 | 0.035800354 | 0.061926849 | 8.3062508 | 8.3249536 | 8.02399 | 0.300964631 | 0.282261799 |
| Ccnj | 7.620111 | 7.804643 | 7.8225 | -0.017881892 | -0.202414299 | 7.9565848 | 8.0063371 | 7.7054 | 0.300933091 | 0.251180824 |
| Maea | 10.39345 | 10.48269 | 10.429 | 0.053841196 | -0.035401856 | 10.532723 | 10.646689 | 10.3458 | 0.300888624 | 0.186921893 |
| Suclg1 | 9.724518 | 9.723927 | 9.6929 | 0.031036795 | 0.031627892 | 9.8232519 | 9.9401965 | 9.6395 | 0.300692922 | 0.183748321 |
| Psmc3 | 10.80365 | 10.87951 | 10.797 | 0.082500872 | 0.006638626 | 10.948967 | 11.128367 | 10.8281 | 0.300303904 | 0.12090333 |

| | | | | | | | | | | |
|---|---|---|---|---|---|---|---|---|---|---|
| Cpne1 | 10.97644 | 10.93996 | 10.872 | 0.06804018 | 0.104515096 | 10.976563 | 11.149479 | 10.8494 | 0.300036904 | 0.127121165 |
| Vrk1 | 9.827147 | 9.775074 | 9.769 | 0.006082971 | 0.05815531 | 9.8592614 | 10.135259 | 9.83523 | 0.300032699 | 0.024035332 |
| Uchl3 | 8.937616 | 8.940577 | 9.0727 | -0.132098296 | -0.135058626 | 9.0531728 | 9.1604751 | 8.86052 | 0.299954037 | 0.192651707 |
| Abr | 10.3709 | 10.47904 | 10.396 | 0.083188474 | -0.024958612 | 10.380271 | 10.704347 | 10.4046 | 0.299748272 | -0.024328044 |
| Trp53rkb | 5.836038 | 5.815786 | 5.9134 | -0.097619675 | -0.077368281 | 6.1122052 | 6.1027751 | 5.80305 | 0.299721962 | 0.309152036 |
| Wrap53 | 7.836734 | 7.866621 | 7.984 | -0.117350763 | -0.147237677 | 8.0242705 | 8.1734335 | 7.87372 | 0.299718205 | 0.150555232 |
| Zc3h7a | 10.97794 | 11.0556 | 11.043 | 0.012567874 | -0.06509708 | 11.253275 | 11.791263 | 11.4917 | 0.299519996 | -0.238468096 |
| Arf1 | 12.14953 | 12.22182 | 12.099 | 0.123268432 | 0.05097885 | 12.208787 | 12.30242 | 12.003 | 0.299433744 | 0.205800413 |
| Igkj5 | 7.240039 | 7.49149 | 7.1931 | 0.298384461 | 0.046933253 | 7.2731213 | 7.3933275 | 7.0939 | 0.29943182 | 0.17922558 |
| Psmc6 | 10.16245 | 10.13317 | 10.114 | 0.01897589 | 0.048257725 | 10.21026 | 10.512783 | 10.2135 | 0.299290325 | -0.00323355 |
| Tssc4 | 7.77642 | 7.895169 | 7.8842 | 0.010981203 | -0.107768003 | 8.1209783 | 8.0747359 | 7.77549 | 0.299244959 | 0.345487445 |
| Med20 | 8.603551 | 8.736188 | 8.672 | 0.064223983 | -0.068412796 | 8.7891067 | 9.0194565 | 8.7203 | 0.299158328 | 0.068808494 |
| Sar1a | 10.56012 | 10.56611 | 10.426 | 0.140378964 | 0.134381482 | 10.677351 | 10.766899 | 10.4678 | 0.299134872 | 0.209586892 |
| Srsf9 | 9.090536 | 9.115591 | 9.0704 | 0.045161298 | 0.020106984 | 9.1118271 | 9.2263009 | 8.9273 | 0.29899874 | 0.184524991 |
| Ncf2 | 10.80577 | 10.8563 | 10.621 | 0.234859256 | 0.18432705 | 11.153137 | 10.994394 | 10.6955 | 0.298915312 | 0.457658835 |
| Cr1l | 10.75586 | 10.79479 | 10.744 | 0.05046404 | 0.011529978 | 10.797957 | 10.979456 | 10.6807 | 0.298774936 | 0.117275743 |
| Nr2c2ap | 8.533869 | 8.620285 | 8.5456 | 0.074678545 | -0.011737356 | 8.8418614 | 8.8414785 | 8.54274 | 0.298733659 | 0.299116526 |
| Mavs | 9.218571 | 9.50084 | 9.245 | 0.255807001 | -0.026461701 | 9.2567624 | 9.4689571 | 9.17026 | 0.298697307 | 0.086502627 |
| Eif3m | 11.54633 | 11.53831 | 11.539 | -0.00019911 | 0.007821522 | 11.586047 | 11.809566 | 11.5109 | 0.298688244 | 0.075168755 |
| Lsm10 | 7.23508 | 7.298688 | 7.2236 | 0.075103578 | 0.0114961 | 7.4926258 | 7.4053202 | 7.1067 | 0.298620636 | 0.385926246 |
| Rwdd1 | 9.372848 | 9.349095 | 9.2666 | 0.082542882 | 0.106295794 | 9.5014199 | 9.5878707 | 9.28939 | 0.298483553 | 0.212032811 |
| Prpf40b | 7.456643 | 7.59885 | 7.5155 | 0.083383638 | -0.058823498 | 7.6320195 | 7.8987574 | 7.60029 | 0.298463966 | 0.031726045 |
| Nudt6 | 6.521738 | 6.766109 | 6.5969 | 0.169174762 | -0.075195558 | 6.9066615 | 6.9702254 | 6.67179 | 0.298437901 | 0.234874021 |
| Relt | 9.395481 | 9.517213 | 9.4261 | 0.091136229 | -0.03059653 | 9.6696844 | 9.7115008 | 9.41333 | 0.298171521 | 0.256355071 |
| Ndufa6 | 9.774652 | 9.595625 | 9.6735 | -0.077885597 | 0.101141724 | 9.8345372 | 9.9632358 | 9.66515 | 0.298088299 | 0.16938966 |
| Tm6sf2 | 3.167707 | 3.182799 | 3.0263 | 0.156458876 | 0.141366208 | 3.3359319 | 3.5604754 | 3.26271 | 0.297761071 | 0.073217627 |
| Eif3d | 11.44247 | 11.47228 | 11.424 | 0.048594176 | 0.018784124 | 11.593103 | 11.801051 | 11.5033 | 0.297709576 | 0.089762318 |
| Nfkb1 | 12.3115 | 12.46327 | 12.325 | 0.13836201 | -0.013408014 | 12.385427 | 12.62761 | 12.3299 | 0.297704164 | 0.055520667 |
| Cmss1 | 6.428636 | 6.468111 | 6.5241 | -0.056023897 | -0.095499375 | 6.5909882 | 6.8313817 | 6.53372 | 0.297666187 | 0.057272619 |
| Tradd | 8.154594 | 8.312058 | 8.1805 | 0.131562786 | -0.025900868 | 8.4780763 | 8.5670047 | 8.26943 | 0.297570958 | 0.208642544 |
| Gm27029 | 4.536091 | 4.538604 | 4.6658 | -0.127230032 | -0.129743247 | 4.6337758 | 4.8601951 | 4.56269 | 0.297507541 | 0.071088257 |
| Os9 | 11.4637 | 11.45276 | 11.451 | 0.00134834 | 0.012292902 | 11.594566 | 11.700341 | 11.4029 | 0.297455568 | 0.191681313 |

| | | | | | | | | | | |
|---|---|---|---|---|---|---|---|---|---|---|
| Igkv13-85 | 7.627369 | 7.594109 | 7.6697 | -0.075598766 | -0.042338584 | 7.8394664 | 7.7668891 | 7.46948 | 0.297407179 | 0.36998448 |
| Samd4b | 9.620469 | 9.623644 | 9.5151 | 0.108573799 | 0.105398254 | 9.5920302 | 9.6830039 | 9.38566 | 0.297348764 | 0.206375131 |
| Arhgdia | 12.96234 | 13.03386 | 12.955 | 0.079112 | 0.007586276 | 13.244004 | 13.208977 | 12.9118 | 0.297219478 | 0.332247226 |
| Spag1 | 5.503839 | 5.617166 | 5.4193 | 0.197873427 | 0.084546629 | 5.6642924 | 5.9721145 | 5.67499 | 0.297126337 | -0.010695794 |
| Inpp4a | 8.841006 | 8.77182 | 8.8386 | -0.06682939 | 0.002356324 | 8.7782331 | 9.2739166 | 8.97679 | 0.29712503 | -0.198558513 |
| Ube2v1 | 11.12426 | 11.1001 | 11.1 | -0.000160398 | 0.023998712 | 11.198181 | 11.407275 | 11.1105 | 0.296764828 | 0.087671228 |
| Gar1 | 9.006737 | 8.928332 | 8.8909 | 0.037453424 | 0.115858715 | 9.1041748 | 9.3325027 | 9.03576 | 0.296739181 | 0.068411247 |
| Mob3c | 8.889569 | 8.800111 | 8.8243 | -0.024213349 | 0.065245003 | 8.768496 | 9.0820188 | 8.78532 | 0.296698778 | -0.016824037 |
| Cnr2 | 11.03731 | 11.31112 | 11.08 | 0.230738512 | -0.04307575 | 11.476924 | 11.384612 | 11.088 | 0.296661284 | 0.388973423 |
| Ints9 | 9.296309 | 9.346716 | 9.3996 | -0.052903577 | -0.103310637 | 9.4490313 | 9.6709688 | 9.37446 | 0.296512596 | 0.074575159 |
| Mapkapk3 | 9.270497 | 9.173895 | 9.2824 | -0.10849199 | -0.011890484 | 9.1420222 | 9.4953768 | 9.19891 | 0.296463216 | -0.056891399 |
| Lag3 | 7.679413 | 7.50063 | 7.7573 | -0.256676807 | -0.077893757 | 7.1409252 | 7.7868647 | 7.49052 | 0.296342473 | -0.349597017 |
| Adprhl2 | 8.773186 | 8.923962 | 8.8131 | 0.110892948 | -0.039883775 | 9.1825584 | 9.1329045 | 8.83665 | 0.296250558 | 0.34590438 |
| Spcs2 | 10.92789 | 10.97218 | 10.885 | 0.087673914 | 0.043375936 | 11.014667 | 11.141867 | 10.8456 | 0.296239372 | 0.169039844 |
| Wdr74 | 8.92958 | 8.977483 | 8.9269 | 0.050596958 | 0.002694247 | 9.1878913 | 9.3527694 | 9.05669 | 0.296077868 | 0.131199784 |
| Tnfaip8l2 | 9.721729 | 9.750029 | 9.8179 | -0.067902702 | -0.096202315 | 10.079941 | 9.9741729 | 9.67823 | 0.295942218 | 0.401710712 |
| Tbc1d20 | 10.37737 | 10.5702 | 10.45 | 0.119697938 | -0.073132146 | 10.421098 | 10.643443 | 10.3475 | 0.29591204 | 0.073567419 |
| Ociad2 | 3.734574 | 3.764348 | 3.7695 | -0.005167468 | -0.03494207 | 3.9712096 | 4.0951612 | 3.79932 | 0.295838613 | 0.171886959 |
| Sephs2 | 8.878384 | 8.687758 | 8.7969 | -0.109139573 | 0.081485948 | 8.6990781 | 8.783675 | 8.48793 | 0.29574046 | 0.211143517 |
| Ssc4d | 3.943347 | 4.231412 | 3.9113 | 0.320064572 | 0.032000359 | 4.3273982 | 4.2625493 | 3.96688 | 0.295671958 | 0.360520837 |
| S100a11 | 10.6286 | 10.71471 | 10.597 | 0.117708418 | 0.031595994 | 10.921788 | 10.935122 | 10.6395 | 0.295671084 | 0.282337007 |
| Chchd6 | 5.357294 | 5.394035 | 5.3755 | 0.018542835 | -0.018198964 | 5.3867058 | 5.6338261 | 5.33817 | 0.295654412 | 0.048534112 |
| Ngdn | 9.428007 | 9.58446 | 9.4025 | 0.181931829 | 0.025478358 | 9.6152446 | 9.8512736 | 9.5557 | 0.295569615 | 0.059540543 |
| Lrrc41 | 8.983718 | 8.973614 | 8.883 | 0.090572114 | 0.100675578 | 8.8730624 | 8.9890215 | 8.69347 | 0.295554455 | 0.179595327 |
| Cpne3 | 10.33181 | 10.25664 | 10.146 | 0.110168262 | 0.18534219 | 10.140505 | 10.406314 | 10.1109 | 0.295463309 | 0.029654119 |
| Bclaf1 | 11.71384 | 11.83268 | 11.787 | 0.045678708 | -0.07317058 | 11.829995 | 12.240711 | 11.9453 | 0.295433904 | -0.115281311 |
| Nelfa | 8.979275 | 9.188546 | 9.0701 | 0.118434213 | -0.090837159 | 9.161523 | 9.2843319 | 8.98932 | 0.295013179 | 0.172204251 |
| Mtx2 | 8.151899 | 8.071903 | 8.1766 | -0.104703371 | -0.024707195 | 8.18555 | 8.3087965 | 8.01379 | 0.295005277 | 0.171758758 |
| Ube2g2 | 9.53021 | 9.603023 | 9.4347 | 0.168350253 | 0.095537724 | 9.6270092 | 9.7219546 | 9.42705 | 0.294905568 | 0.199960243 |
| Yars | 9.626679 | 9.516621 | 9.4886 | 0.027981052 | 0.138039081 | 9.9545176 | 10.218768 | 9.92405 | 0.294717053 | 0.0304668 |
| Ebp | 8.733696 | 8.840992 | 8.7326 | 0.108441609 | 0.001145439 | 9.0868832 | 9.0839591 | 8.78931 | 0.294644142 | 0.297568305 |
| Pars2 | 6.75475 | 6.858132 | 6.7785 | 0.079639714 | -0.02374187 | 6.9745273 | 6.9535141 | 6.65894 | 0.294569246 | 0.315582452 |

| | | | | | | | | | | |
|---|---|---|---|---|---|---|---|---|---|---|
| Apoc2 | 3.460838 | 3.396718 | 3.1703 | 0.226423807 | 0.290542947 | 3.5085157 | 3.448354 | 3.15381 | 0.294545362 | 0.354707044 |
| Cemip2 | 9.401191 | 9.063456 | 8.772 | 0.291504238 | 0.629239308 | 8.5554201 | 8.8219639 | 8.52745 | 0.294516613 | 0.027972862 |
| Mrps17 | 9.151881 | 9.150182 | 9.0543 | 0.095881207 | 0.09757955 | 9.3069089 | 9.2999124 | 9.00546 | 0.294455463 | 0.301451909 |
| Psmg1 | 7.608482 | 7.619101 | 7.5738 | 0.045337281 | 0.034718603 | 7.7120755 | 7.8198532 | 7.5254 | 0.294451121 | 0.186673405 |
| Fgfr1op | 8.76573 | 8.930852 | 9.0184 | -0.087533301 | -0.252655012 | 9.0527415 | 9.3517182 | 9.05735 | 0.294366201 | -0.004610543 |
| Igkj1 | 5.864188 | 6.107906 | 5.7091 | 0.39878971 | 0.15507213 | 6.0580454 | 6.0334772 | 5.73912 | 0.294354109 | 0.318922236 |
| Smarcb1 | 10.14815 | 10.19677 | 10.147 | 0.049313228 | 0.000698341 | 10.428665 | 10.416703 | 10.1225 | 0.294155296 | 0.306117945 |
| Grk5 | 8.119998 | 8.091868 | 7.7018 | 0.390068557 | 0.418198365 | 7.6898075 | 7.9751046 | 7.68122 | 0.293883232 | 0.008586208 |
| Med8 | 8.924329 | 8.873595 | 8.9179 | -0.044351275 | 0.006382144 | 9.071511 | 9.1498295 | 8.85612 | 0.293710931 | 0.215392394 |
| Luc7l | 10.24608 | 10.31294 | 10.149 | 0.163988762 | 0.09712545 | 10.211138 | 10.631459 | 10.3379 | 0.293582346 | -0.126738294 |
| Yipf6 | 8.947805 | 8.88332 | 8.923 | -0.0396437 | 0.024840831 | 9.019329 | 9.1008962 | 8.80754 | 0.293354158 | 0.211786957 |
| Bola2 | 9.009482 | 9.015017 | 9.0115 | 0.003540985 | -0.001993992 | 9.2751478 | 9.2154573 | 8.92213 | 0.293327998 | 0.353018489 |
| Zfp429 | 8.346009 | 8.347711 | 8.4855 | -0.137755068 | -0.139456921 | 8.3991026 | 8.7370898 | 8.44409 | 0.292996476 | -0.044990675 |
| Hist1h4m | 5.122166 | 5.051095 | 5.3125 | -0.261453839 | -0.190382812 | 5.6753781 | 5.5228346 | 5.22988 | 0.292958674 | 0.445502167 |
| Adora2b | 3.676966 | 3.766741 | 3.375 | 0.3917294 | 0.30195444 | 3.4033672 | 3.6469158 | 3.35405 | 0.292861963 | 0.049313288 |
| Hspa4l | 7.423674 | 7.420994 | 7.5032 | -0.082206742 | -0.079526827 | 7.0557399 | 7.7610823 | 7.46826 | 0.292825212 | -0.41251722 |
| Zfp207 | 11.25625 | 11.48881 | 11.339 | 0.150177278 | -0.082382 | 11.559293 | 11.728117 | 11.4353 | 0.292825116 | 0.124001083 |
| Rbm17 | 10.32756 | 10.35966 | 10.373 | -0.01328332 | -0.045383876 | 10.448938 | 10.726648 | 10.4339 | 0.292704118 | 0.014994187 |
| Rtcb | 11.60633 | 11.55966 | 11.628 | -0.068073414 | -0.021404184 | 11.620116 | 11.853312 | 11.5607 | 0.292611042 | 0.059415616 |
| Elof1 | 8.406225 | 8.416019 | 8.4507 | -0.034666879 | -0.044460231 | 8.6977438 | 8.6550332 | 8.36245 | 0.292580509 | 0.335291173 |
| Igkv9-124 | 7.799123 | 7.230284 | 7.2267 | 0.003552686 | 0.572392184 | 7.1264482 | 7.3291043 | 7.03655 | 0.29255125 | 0.089895208 |
| Gm21992 | 6.187558 | 6.429976 | 6.3237 | 0.106315231 | -0.136102654 | 6.4533464 | 6.6934029 | 6.40088 | 0.29252316 | 0.052466696 |
| Ube2d3 | 12.72337 | 12.84453 | 12.664 | 0.180493848 | 0.059331894 | 12.861916 | 13.052847 | 12.7604 | 0.292486784 | 0.101555355 |
| Ccdc184 | 4.202596 | 4.276579 | 4.0294 | 0.247158494 | 0.173176211 | 3.9805 | 4.298938 | 4.00675 | 0.292191832 | -0.026246127 |
| Slc25a44 | 8.929216 | 9.041198 | 8.9137 | 0.127530934 | 0.015548352 | 8.9746519 | 9.179547 | 8.88754 | 0.292011551 | 0.087116531 |
| Mrps30 | 8.669102 | 8.731396 | 8.7528 | -0.021361324 | -0.083654523 | 8.7824893 | 8.9778642 | 8.68596 | 0.291908515 | 0.096533575 |
| Ppp4c | 10.89652 | 11.01081 | 10.871 | 0.139345882 | 0.025058354 | 11.064382 | 11.29779 | 11.0059 | 0.291849796 | 0.058441914 |
| Rsl1d1 | 10.5421 | 10.61956 | 10.486 | 0.134044558 | 0.056579034 | 10.551192 | 10.856137 | 10.5644 | 0.291784816 | -0.013160452 |
| Pld4 | 9.949958 | 9.907238 | 9.988 | -0.080751014 | -0.038030853 | 10.453656 | 10.403773 | 10.1121 | 0.291660924 | 0.341543731 |
| Rps18 | 14.51189 | 14.62338 | 14.542 | 0.081457034 | -0.030030854 | 14.728773 | 14.825571 | 14.5339 | 0.291623284 | 0.194826044 |
| Bmp10 | 0.361983 | 0.361194 | 0.3491 | 0.012092155 | 0.012881319 | 0.3726165 | 0.6433478 | 0.35185 | 0.291499375 | 0.020768075 |
| Igkv1-132 | 3.442867 | 3.381392 | 3.3116 | 0.069753231 | 0.131228294 | 3.3516652 | 3.7057008 | 3.4142 | 0.291497234 | -0.062538379 |

| | | | | | | | | | | |
|---|---|---|---|---|---|---|---|---|---|---|
| Polr2k | 7.947661 | 7.887415 | 7.7643 | 0.123066468 | 0.183312717 | 8.0379819 | 8.1510076 | 7.8596 | 0.291411498 | 0.178385767 |
| Ppargc1b | 7.06254 | 7.162774 | 7.194 | -0.031179918 | -0.131413487 | 7.3314139 | 7.3431842 | 7.05182 | 0.291363413 | 0.279593096 |
| Dcaf13 | 9.147006 | 9.185874 | 9.1882 | -0.002277777 | -0.041146337 | 9.3075855 | 9.5128424 | 9.22151 | 0.291332612 | 0.086075692 |
| Dus4l | 6.98684 | 6.928522 | 7.0022 | -0.0736764 | -0.01535866 | 7.2655467 | 7.3519871 | 7.06087 | 0.291113199 | 0.20467283 |
| Ing4 | 8.86403 | 8.975234 | 8.9928 | -0.017579056 | -0.128783185 | 9.2862562 | 9.4524365 | 9.16148 | 0.290959859 | 0.12477956 |
| Mdk | 4.448216 | 4.611642 | 4.4788 | 0.132802231 | -0.030623494 | 4.7003515 | 4.7016721 | 4.41086 | 0.290814471 | 0.289493842 |
| Caml | 7.474618 | 7.678463 | 7.6704 | 0.008098271 | -0.195746486 | 7.6250078 | 7.971872 | 7.68112 | 0.290751439 | -0.056112734 |
| Gemin7 | 8.550577 | 8.713989 | 8.6065 | 0.107475501 | -0.055935855 | 8.8238513 | 8.8364466 | 8.54582 | 0.290624248 | 0.278028999 |
| Bcat1 | 5.71711 | 5.713947 | 5.7581 | -0.04418666 | -0.041024252 | 5.8627293 | 6.2105064 | 5.92003 | 0.290479035 | -0.057298054 |
| Pwp2 | 9.941456 | 10.07899 | 9.8234 | 0.25560155 | 0.118065991 | 10.100736 | 10.187766 | 9.89734 | 0.290422886 | 0.203392985 |
| Bcl6 | 10.25834 | 10.37812 | 10.242 | 0.135861198 | 0.016079248 | 10.16132 | 10.322713 | 10.0323 | 0.290404694 | 0.129012442 |
| Mrpl40 | 8.12532 | 8.044864 | 8.2083 | -0.163470303 | -0.083013746 | 8.3093545 | 8.3852661 | 8.09509 | 0.29017949 | 0.21426789 |
| Gm5148 | 7.442685 | 7.707367 | 7.6065 | 0.100891432 | -0.163789753 | 7.5858465 | 7.660156 | 7.37011 | 0.290041661 | 0.215732211 |
| Ercc2 | 9.337048 | 9.387444 | 9.4732 | -0.085774066 | -0.136170074 | 9.6218907 | 9.904193 | 9.61458 | 0.289609716 | 0.007307423 |
| Gm5831 | 3.384612 | 3.386942 | 3.3448 | 0.042094968 | 0.039765032 | 3.3876972 | 3.6909929 | 3.40157 | 0.289420574 | -0.013875145 |
| Mettl23 | 9.389947 | 9.435069 | 9.4435 | -0.008431359 | -0.053553625 | 9.7531095 | 9.773418 | 9.48412 | 0.289298326 | 0.268989834 |
| Set | 12.42768 | 12.40329 | 12.385 | 0.018409864 | 0.042803978 | 12.521963 | 12.629412 | 12.3402 | 0.289248936 | 0.181800171 |
| Alpk2 | 6.066678 | 6.175813 | 6.0616 | 0.114225464 | 0.005090283 | 6.3120547 | 6.5171249 | 6.22804 | 0.289088591 | 0.084018373 |
| Zfp36 | 13.37161 | 13.4165 | 13.189 | 0.227529392 | 0.18263711 | 12.876468 | 13.051912 | 12.7629 | 0.289039986 | 0.113595764 |
| Abca17 | 2.712103 | 2.506391 | 2.5887 | -0.082278707 | 0.123433545 | 2.6514838 | 2.8224408 | 2.53356 | 0.28888019 | 0.117923169 |
| Mtcp1 | 5.627106 | 5.622999 | 5.6522 | -0.02919587 | -0.025089336 | 6.072643 | 6.1823443 | 5.89366 | 0.288686596 | 0.178985239 |
| Uqcrb | 9.396949 | 9.412143 | 9.369 | 0.043130984 | 0.027936545 | 9.493885 | 9.6319651 | 9.34329 | 0.28867303 | 0.150592918 |
| Armc7 | 9.268559 | 9.242554 | 9.2896 | -0.047049764 | -0.02104501 | 9.1329084 | 9.3933664 | 9.10473 | 0.288637079 | 0.028179121 |
| Hnrnpa3 | 12.6763 | 12.57031 | 12.625 | -0.054486164 | 0.051511282 | 12.732747 | 12.981109 | 12.6925 | 0.288625848 | 0.040263485 |
| Abcb6 | 7.001206 | 6.959768 | 7.0312 | -0.071450765 | -0.030013345 | 7.2051358 | 7.1502835 | 6.86169 | 0.288594189 | 0.343446447 |
| Plekha2 | 12.25372 | 12.44467 | 12.302 | 0.143142478 | -0.047809242 | 12.286989 | 12.507468 | 12.219 | 0.288512414 | 0.068033457 |
| Gm20796 | 2.877957 | 2.921501 | 2.8967 | 0.024805262 | -0.018738723 | 2.8972456 | 3.1784675 | 2.8901 | 0.288370258 | 0.007148389 |
| Zfp791 | 4.632625 | 4.57328 | 4.6467 | -0.073463853 | -0.014118658 | 4.6949334 | 5.0855776 | 4.79725 | 0.288323548 | -0.102320677 |
| Selenom | 5.524746 | 5.550714 | 5.581 | -0.030266554 | -0.056235175 | 5.6238278 | 5.6623285 | 5.37431 | 0.288021061 | 0.24952036 |
| Timm50 | 8.575777 | 8.596792 | 8.5893 | 0.007527425 | -0.013487743 | 9.1136536 | 9.0510904 | 8.76319 | 0.287903431 | 0.350466684 |
| Usp10 | 9.594981 | 9.59291 | 9.5705 | 0.022382626 | 0.024453 | 9.7562187 | 9.9827794 | 9.6949 | 0.287881762 | 0.061321096 |
| Dlgap1 | 2.969123 | 3.093563 | 2.8858 | 0.20779225 | 0.083352322 | 3.1093391 | 3.3033454 | 3.01569 | 0.287659552 | 0.09365331 |

| | | | | | | | | | | |
|---|---|---|---|---|---|---|---|---|---|---|
| Srsf1 | 10.51608 | 10.57091 | 10.484 | 0.087254654 | 0.032424468 | 10.861343 | 10.98163 | 10.6941 | 0.287505352 | 0.167218776 |
| Sugt1 | 10.14699 | 10.16604 | 10.273 | -0.106658842 | -0.125714552 | 10.183742 | 10.491401 | 10.2039 | 0.287478874 | -0.020179528 |
| Ubl4a | 8.476507 | 8.501164 | 8.5148 | -0.013654827 | -0.0383117 | 8.8055913 | 8.8783101 | 8.59084 | 0.28747111 | 0.214752346 |
| Maged2 | 5.954124 | 5.961033 | 6.0764 | -0.1153444 | -0.12225397 | 5.9642779 | 6.0917017 | 5.80426 | 0.287437456 | 0.160013599 |
| Dnaja4 | 7.167984 | 7.445647 | 7.1378 | 0.307886073 | 0.030223359 | 6.8010191 | 7.439026 | 7.15159 | 0.287432966 | -0.350573966 |
| Tktl1 | 3.270127 | 3.416032 | 3.3611 | 0.054913107 | -0.090992306 | 3.2605177 | 3.6857717 | 3.39835 | 0.287420708 | -0.137833298 |
| Mrpl37 | 9.106318 | 9.193834 | 9.1219 | 0.071976854 | -0.015538662 | 9.4129145 | 9.5591379 | 9.27177 | 0.287368833 | 0.141145393 |
| Cipc | 9.329361 | 9.457744 | 9.2304 | 0.227325631 | 0.098942572 | 9.6081483 | 9.5838414 | 9.29656 | 0.287285569 | 0.311592471 |
| Tbrg4 | 9.78943 | 9.826775 | 9.7511 | 0.075639294 | 0.03829454 | 10.218251 | 10.289871 | 10.0026 | 0.287251232 | 0.215630914 |
| Gna13 | 11.81586 | 11.80043 | 11.785 | 0.015018658 | 0.030455632 | 11.584017 | 12.041163 | 11.754 | 0.287113672 | -0.170032075 |
| Pa2g4 | 11.38779 | 11.40295 | 11.335 | 0.068096468 | 0.052939306 | 11.49254 | 11.679536 | 11.3925 | 0.286999046 | 0.100003827 |
| Kansl3 | 10.29013 | 10.34928 | 10.308 | 0.041548948 | -0.017602616 | 10.195948 | 10.568426 | 10.2815 | 0.28693815 | -0.08554039 |
| Fbxl8 | 7.113587 | 7.22464 | 7.2868 | -0.062140913 | -0.173193494 | 7.2382804 | 7.3798209 | 7.09299 | 0.286834047 | 0.145293496 |
| Med18 | 6.285563 | 6.338144 | 6.202 | 0.136138082 | 0.083556512 | 6.4779917 | 6.586151 | 6.29939 | 0.286757917 | 0.178598637 |
| Cept1 | 10.17544 | 10.10557 | 10.108 | -0.002198526 | 0.067673886 | 10.022397 | 10.106348 | 9.81969 | 0.286655922 | 0.202704925 |
| Ighg2c | 10.77944 | 12.11821 | 9.9817 | 2.136537732 | 0.797771441 | 10.069414 | 11.865153 | 11.5785 | 0.286652102 | -1.509087595 |
| Dhrs7b | 7.687416 | 7.837356 | 7.7366 | 0.100804304 | -0.049135972 | 7.8171069 | 7.9747283 | 7.68814 | 0.286585405 | 0.12896395 |
| Mrpl15 | 9.059798 | 9.1653 | 9.0334 | 0.131881162 | 0.026378942 | 9.1749764 | 9.3517166 | 9.06557 | 0.286143748 | 0.109403508 |
| Mettl6 | 8.653961 | 8.679146 | 8.6891 | -0.009912162 | -0.035097501 | 8.629517 | 9.0048881 | 8.71899 | 0.285897458 | -0.089473638 |
| Tmem214 | 9.871535 | 9.919349 | 9.7895 | 0.129844879 | 0.082030955 | 9.9825368 | 9.9889601 | 9.7031 | 0.285863893 | 0.279440633 |
| Klra6 | 5.55245 | 5.428986 | 5.5567 | -0.127704239 | -0.004239742 | 5.5792501 | 5.848482 | 5.56262 | 0.285858744 | 0.016626853 |
| Dkc1 | 10.36033 | 10.47943 | 10.325 | 0.154453774 | 0.035353102 | 10.47363 | 10.73351 | 10.4477 | 0.285849302 | 0.02596928 |
| Lamtor1 | 9.432874 | 9.43707 | 9.5037 | -0.066603939 | -0.070800568 | 9.6599446 | 9.6452043 | 9.35976 | 0.285442442 | 0.300182688 |
| Ndufa9 | 9.877633 | 9.865418 | 9.8001 | 0.065333312 | 0.077548337 | 9.9194244 | 10.083626 | 9.7983 | 0.285327028 | 0.12112579 |
| Sf3a3 | 10.14086 | 10.18187 | 10.162 | 0.01970543 | -0.021300964 | 10.215806 | 10.442781 | 10.1577 | 0.285040066 | 0.058064998 |
| Ddx27 | 10.26695 | 10.34542 | 10.283 | 0.06261763 | -0.015850316 | 10.433258 | 10.550006 | 10.2651 | 0.284915204 | 0.168166881 |
| Fcrl1 | 11.41456 | 11.61957 | 11.445 | 0.174706354 | -0.030299676 | 12.056606 | 11.9431 | 11.6582 | 0.284901772 | 0.398408409 |
| Gm16976 | 2.216557 | 2.243017 | 2.165 | 0.077979968 | 0.05151924 | 2.2431175 | 2.5158229 | 2.23094 | 0.284883611 | 0.01217826 |
| Arl11 | 4.929922 | 4.934995 | 4.9244 | 0.01054507 | 0.00547265 | 5.3455388 | 5.0902966 | 4.80548 | 0.284819476 | 0.540061687 |
| Igkv4-50 | 5.629276 | 5.924009 | 5.7837 | 0.140273587 | -0.154459765 | 5.9975185 | 5.7757307 | 5.49099 | 0.284739037 | 0.506526787 |
| Fkrp | 8.868246 | 9.017117 | 8.882 | 0.135159984 | -0.013710814 | 8.9857088 | 9.151524 | 8.86681 | 0.284716106 | 0.118900965 |
| Fbxo42 | 9.376519 | 9.556588 | 9.3868 | 0.16983792 | -0.010230949 | 9.1974969 | 9.5752916 | 9.2909 | 0.284389018 | -0.093405648 |

| | | | | | | | | | | |
|---|---|---|---|---|---|---|---|---|---|---|
| Cyb561a3 | 11.24211 | 11.33705 | 11.331 | 0.006253018 | -0.088686828 | 11.622546 | 11.636659 | 11.3523 | 0.284346446 | 0.270233664 |
| Styk1 | 4.376842 | 4.269723 | 4.5169 | -0.24720094 | -0.140081852 | 4.4807655 | 4.8331041 | 4.54878 | 0.284326626 | -0.06801202 |
| Ccnl2 | 11.31898 | 11.5023 | 11.446 | 0.055840644 | -0.127473388 | 11.659001 | 11.986487 | 11.7022 | 0.28429738 | -0.043189148 |
| Raly | 11.62479 | 11.65114 | 11.604 | 0.046705374 | 0.020354874 | 11.61571 | 11.841191 | 11.5571 | 0.284059972 | 0.058578634 |
| Ascc2 | 9.698266 | 9.854904 | 9.7424 | 0.11254042 | -0.044097684 | 9.9450804 | 10.099805 | 9.81578 | 0.284025118 | 0.129300059 |
| Adprh | 9.916489 | 9.858931 | 9.7979 | 0.061019331 | 0.118577465 | 10.035798 | 9.9788323 | 9.69493 | 0.283904687 | 0.340870588 |
| Zfp598 | 10.00887 | 10.11488 | 10.033 | 0.081939676 | -0.024063273 | 10.31136 | 10.391789 | 10.108 | 0.283777994 | 0.203348931 |
| Ada | 6.56748 | 6.622752 | 6.7408 | -0.118074745 | -0.173347063 | 6.6338941 | 6.9420111 | 6.65833 | 0.283685324 | -0.024431709 |
| Bcs1l | 7.522952 | 7.562347 | 7.6141 | -0.051729306 | -0.091124333 | 7.8022214 | 7.9601355 | 7.67648 | 0.283655234 | 0.125741168 |
| Klhdc3 | 9.39116 | 9.532744 | 9.5058 | 0.026914765 | -0.114668953 | 9.6686716 | 9.7883157 | 9.5047 | 0.283614952 | 0.163970825 |
| Commd1 | 8.168124 | 8.24219 | 8.181 | 0.061182958 | -0.012882776 | 8.3448121 | 8.4835475 | 8.20007 | 0.283478571 | 0.144743209 |
| Aacs | 8.709553 | 8.902652 | 8.7674 | 0.135242094 | -0.057856225 | 9.2274525 | 8.9922829 | 8.70881 | 0.283470586 | 0.518640176 |
| Gfpt2 | 5.697369 | 5.766557 | 5.2921 | 0.47448747 | 0.405300137 | 5.1787387 | 5.5787344 | 5.29548 | 0.283257806 | -0.116737933 |
| Ank2 | 5.346289 | 5.281286 | 5.151 | 0.130304707 | 0.195307299 | 5.0778839 | 5.5730584 | 5.28985 | 0.283203907 | -0.211970606 |
| Rpl18a | 14.68677 | 14.80331 | 14.793 | 0.010670682 | -0.105867726 | 15.031278 | 14.99925 | 14.7161 | 0.283136532 | 0.315164536 |
| Smim12 | 7.742256 | 7.8154 | 7.7758 | 0.039579128 | -0.033565274 | 7.9179857 | 7.8773929 | 7.59431 | 0.28308735 | 0.323680126 |
| Ccdc12 | 9.704096 | 9.780138 | 9.6966 | 0.083546853 | 0.007504848 | 10.016502 | 10.075986 | 9.7929 | 0.28308362 | 0.223600051 |
| Ado | 7.335909 | 7.392047 | 7.3207 | 0.071374689 | 0.015236617 | 7.4192106 | 7.5056507 | 7.22262 | 0.283034466 | 0.196594412 |
| Gm8172 | 3.45114 | 3.482537 | 3.4812 | 0.001291136 | -0.030105761 | 3.6540541 | 3.6093653 | 3.32633 | 0.283031458 | 0.327720221 |
| Nelfe | 8.228083 | 8.206747 | 8.2037 | 0.003081042 | 0.024416663 | 8.3486327 | 8.494655 | 8.21169 | 0.282969453 | 0.136947196 |
| Snrnp27 | 8.328016 | 8.381009 | 8.2546 | 0.12645378 | 0.073461141 | 8.6507748 | 8.638269 | 8.35533 | 0.282935523 | 0.295441312 |
| Maff | 7.549826 | 7.191837 | 7.0587 | 0.133095471 | 0.491084968 | 6.9857672 | 7.2125062 | 6.92971 | 0.28279167 | 0.056052701 |
| Abracl | 10.12026 | 10.1463 | 10.14 | 0.006535194 | -0.019506038 | 10.157904 | 10.372295 | 10.0897 | 0.282575738 | 0.068183921 |
| Klrb1b | 7.072573 | 6.910649 | 6.5285 | 0.382145417 | 0.544069967 | 6.361275 | 7.1962767 | 6.91377 | 0.282503646 | -0.552498059 |
| Sema3f | 7.558471 | 7.359148 | 7.2248 | 0.134324896 | 0.333647581 | 6.8702786 | 7.0850756 | 6.80258 | 0.282495273 | 0.067698286 |
| Nars | 11.05179 | 11.06199 | 10.845 | 0.217052114 | 0.206855874 | 11.19888 | 11.359498 | 11.0771 | 0.282380458 | 0.121762419 |
| Metap2 | 10.50837 | 10.54381 | 10.516 | 0.027636246 | -0.007803716 | 10.368998 | 10.667176 | 10.3851 | 0.282043228 | -0.016135055 |
| Sh2d5 | 5.250777 | 5.320189 | 4.9155 | 0.404642171 | 0.335230033 | 5.0670128 | 5.4032314 | 5.12133 | 0.281902852 | -0.054315827 |
| Lypla2 | 10.32582 | 10.33131 | 10.33 | 0.001693455 | -0.003796912 | 10.626378 | 10.584335 | 10.3025 | 0.281871935 | 0.323915371 |
| Igkv9-129 | 4.70994 | 4.80155 | 4.5857 | 0.21589551 | 0.124286142 | 4.7124622 | 4.8264125 | 4.54469 | 0.281726444 | 0.167776217 |
| Mag | 4.214451 | 3.952383 | 4.0581 | -0.105723182 | 0.15634472 | 4.1211543 | 4.3069248 | 4.02554 | 0.281387806 | 0.095617298 |
| Tmem9 | 7.697283 | 7.631801 | 7.8571 | -0.225305824 | -0.159823517 | 7.7392812 | 7.9034613 | 7.62211 | 0.281348943 | 0.117168806 |

| | | | | | | | | | | |
|---|---|---|---|---|---|---|---|---|---|---|
| St3gal4 | 8.952446 | 8.675819 | 8.641 | 0.03478068 | 0.311408203 | 8.704125 | 8.8387163 | 8.55738 | 0.281336363 | 0.146745031 |
| Prkcsh | 10.28008 | 10.36893 | 10.333 | 0.036109631 | -0.052738953 | 10.534942 | 10.616998 | 10.3357 | 0.281334548 | 0.199278854 |
| Olfr822 | 2.088322 | 2.155153 | 2.0596 | 0.095567095 | 0.02873624 | 2.1137641 | 2.4255437 | 2.14425 | 0.28129793 | -0.030481667 |
| Snu13 | 10.3524 | 10.32932 | 10.324 | 0.005446296 | 0.028529362 | 10.419856 | 10.614106 | 10.333 | 0.281063196 | 0.086813049 |
| Wdr25 | 6.181924 | 6.179096 | 6.0006 | 0.178472404 | 0.181300266 | 6.3701835 | 6.4217805 | 6.14074 | 0.281045098 | 0.229448109 |
| Kmo | 10.43589 | 10.79378 | 10.472 | 0.321397266 | -0.036484948 | 10.897614 | 10.914678 | 10.6336 | 0.2810412 | 0.263976402 |
| Dennd1a | 9.288264 | 9.231107 | 9.3348 | -0.103736692 | -0.046579703 | 9.4751114 | 9.7076155 | 9.42671 | 0.280902854 | 0.048398729 |
| Rtn4 | 10.70882 | 10.77504 | 10.667 | 0.107605316 | 0.041385542 | 10.547661 | 10.929734 | 10.6489 | 0.280882224 | -0.101190812 |
| Tcf7l2 | 5.899096 | 5.823297 | 5.8251 | -0.001833409 | 0.073965551 | 5.5506462 | 6.0470583 | 5.76631 | 0.280743367 | -0.21566874 |
| Ppif | 8.83107 | 8.897827 | 8.9661 | -0.068315599 | -0.135073012 | 9.0540525 | 9.1476715 | 8.86695 | 0.280718276 | 0.187099259 |
| Lhx6 | 5.153786 | 4.953302 | 4.9511 | 0.002233468 | 0.202717376 | 4.6772531 | 4.9596036 | 4.67896 | 0.280647864 | -0.001702553 |
| Eloc | 9.508173 | 9.590528 | 9.523 | 0.067574581 | -0.01478057 | 9.6163012 | 9.9028769 | 9.62225 | 0.280629055 | -0.005946606 |
| Gm4707 | 4.389844 | 4.376337 | 4.6923 | -0.315918369 | -0.302411891 | 4.6276587 | 5.091747 | 4.81124 | 0.280510351 | -0.183577961 |
| Cd2 | 11.63714 | 11.70879 | 11.681 | 0.027295552 | -0.04436084 | 11.651484 | 11.902421 | 11.6221 | 0.28035734 | 0.029420508 |
| Gm16286 | 10.44962 | 10.47002 | 10.421 | 0.049433574 | 0.029037254 | 10.641096 | 10.787937 | 10.5076 | 0.280348006 | 0.133506535 |
| Aup1 | 10.73968 | 10.64661 | 10.712 | -0.065441456 | 0.027631864 | 11.111912 | 11.080777 | 10.8005 | 0.280326272 | 0.311461829 |
| Pih1d1 | 8.618223 | 8.61228 | 8.5651 | 0.047204861 | 0.053147936 | 8.908263 | 8.8457878 | 8.5655 | 0.280285295 | 0.342760558 |
| Fbxo7 | 8.953732 | 8.938526 | 8.8783 | 0.060242042 | 0.075448289 | 8.9691189 | 9.2720616 | 8.99183 | 0.280229457 | -0.02271324 |
| Ssh2 | 12.42335 | 12.53046 | 12.313 | 0.217073294 | 0.109965534 | 12.10532 | 12.557033 | 12.2768 | 0.280209126 | -0.1715043 |
| Rnf167 | 11.57218 | 11.63817 | 11.608 | 0.030001358 | -0.03598869 | 11.873614 | 12.011026 | 11.7311 | 0.279930958 | 0.14251939 |
| Prmt5 | 9.937132 | 10.01786 | 9.9059 | 0.11200272 | 0.03127802 | 9.9994802 | 10.346691 | 10.0668 | 0.279908819 | -0.067301657 |
| Clns1a | 9.649301 | 9.672313 | 9.6212 | 0.051077998 | 0.02806588 | 9.8248736 | 9.9213405 | 9.6417 | 0.279640937 | 0.183174063 |
| Ndufv3 | 9.107349 | 9.23448 | 9.1971 | 0.03735896 | -0.089771556 | 9.4524467 | 9.5302605 | 9.2507 | 0.27956315 | 0.201749349 |
| Maip1 | 7.742813 | 7.674196 | 7.5634 | 0.110750156 | 0.179366638 | 7.636814 | 7.9316389 | 7.65209 | 0.27954962 | -0.015275188 |
| Tmem70 | 8.240671 | 8.173657 | 8.1697 | 0.003982937 | 0.070996845 | 8.1621406 | 8.3286492 | 8.04916 | 0.279484508 | 0.112975845 |
| Rpain | 7.901228 | 7.980557 | 7.9447 | 0.035868278 | -0.043461409 | 8.1541027 | 8.3095267 | 8.03023 | 0.279298264 | 0.123874258 |
| Dnajc2 | 10.25658 | 10.22057 | 10.227 | -0.006845678 | 0.029165168 | 10.291574 | 10.5069 | 10.2276 | 0.279293666 | 0.063967138 |
| Xrn1 | 10.91895 | 11.08678 | 10.839 | 0.248164324 | 0.080334694 | 10.765544 | 11.310869 | 11.0317 | 0.27916392 | -0.266161518 |
| Map3k19 | 8.044459 | 8.355527 | 8.0884 | 0.267103427 | -0.043965287 | 8.7602932 | 8.8361241 | 8.55701 | 0.279111027 | 0.203280085 |
| Mllt1 | 9.462732 | 9.572355 | 9.4258 | 0.146528537 | 0.036905343 | 9.5604437 | 9.7192576 | 9.44025 | 0.279008152 | 0.120194201 |
| Npm1 | 13.67729 | 13.67355 | 13.71 | -0.036141604 | -0.032401156 | 13.729116 | 14.043083 | 13.7643 | 0.278779068 | -0.035187512 |
| Gm12338 | 6.929494 | 6.817824 | 6.8592 | -0.041422441 | 0.070247333 | 6.8096809 | 7.1540205 | 6.87567 | 0.278350057 | -0.065989564 |

| | | | | | | | | | | |
|---|---|---|---|---|---|---|---|---|---|---|
| Tmem59 | 10.59792 | 10.62166 | 10.633 | -0.011508016 | -0.035244672 | 10.627375 | 10.748295 | 10.4703 | 0.277946188 | 0.157026333 |
| Syne4 | 4.312814 | 4.39875 | 4.3913 | 0.007423309 | -0.078512788 | 4.3308567 | 4.6749927 | 4.39707 | 0.277927249 | -0.06620878 |
| Coq2 | 9.003284 | 9.155748 | 9.0409 | 0.114826585 | -0.037636811 | 9.4005134 | 9.5281817 | 9.25026 | 0.277920504 | 0.150252163 |
| Mettl17 | 8.670547 | 8.950857 | 8.9256 | 0.025294152 | -0.255016526 | 9.2120689 | 9.38471 | 9.10679 | 0.277919353 | 0.105278174 |
| Grcc10 | 9.959954 | 10.03838 | 10.017 | 0.021426469 | -0.05700345 | 10.389478 | 10.208024 | 9.93032 | 0.277706197 | 0.459159802 |
| Ugt1a7c | 7.347588 | 7.009258 | 6.8795 | 0.129771964 | 0.468102169 | 7.2859752 | 6.7035551 | 6.42585 | 0.277700314 | 0.860120405 |
| Pmpca | 9.965452 | 9.941953 | 9.8614 | 0.080519298 | 0.104018609 | 10.255752 | 10.285465 | 10.0078 | 0.27769092 | 0.247977572 |
| Sipa1l3 | 9.164143 | 9.270842 | 9.0537 | 0.217187829 | 0.110488637 | 9.3145456 | 9.3531792 | 9.07549 | 0.277687121 | 0.239053526 |
| Kpna3 | 10.28661 | 10.37736 | 10.253 | 0.124142402 | 0.03338923 | 10.118556 | 10.523361 | 10.2458 | 0.277589332 | -0.127215579 |
| Pdap1 | 9.881851 | 9.718636 | 9.8171 | -0.098492227 | 0.06472272 | 10.000148 | 10.082686 | 9.80517 | 0.277517313 | 0.194979518 |
| Exog | 7.332837 | 7.614184 | 7.451 | 0.163137932 | -0.11820961 | 7.3722815 | 7.7031705 | 7.4257 | 0.27747061 | -0.053418332 |
| Rpl22l1 | 11.25329 | 11.27585 | 11.276 | -0.000386078 | -0.022945786 | 11.323469 | 11.501881 | 11.2244 | 0.277447204 | 0.099035018 |
| Rundc1 | 8.929918 | 9.023916 | 8.9843 | 0.039649194 | -0.05434853 | 8.9956806 | 9.1871686 | 8.90983 | 0.277335248 | 0.085847184 |
| Rpl15 | 13.09101 | 13.13046 | 13.153 | -0.022783068 | -0.06223018 | 13.257676 | 13.384342 | 13.1071 | 0.277264272 | 0.150598573 |
| Nosip | 9.448535 | 9.500562 | 9.4645 | 0.036027258 | -0.015999533 | 9.6274623 | 9.8115303 | 9.53428 | 0.277251988 | 0.093183926 |
| Tmem86b | 6.797433 | 6.892193 | 6.8115 | 0.080734515 | -0.014026075 | 7.1735762 | 7.3703311 | 7.09314 | 0.277191228 | 0.080436339 |
| 1110059E24Rik | 9.36199 | 9.373323 | 9.2634 | 0.109877681 | 0.09854527 | 9.2518917 | 9.3848147 | 9.10769 | 0.277128784 | 0.144205818 |
| Cebpd | 6.284364 | 5.852625 | 5.9714 | -0.118750549 | 0.312988562 | 5.701345 | 5.5204054 | 5.24331 | 0.277093082 | 0.458032661 |
| Gm10020 | 6.111569 | 6.23905 | 6.3371 | -0.098002672 | -0.22548387 | 6.3185544 | 6.4581836 | 6.18114 | 0.277039065 | 0.137409852 |
| Sars | 11.0607 | 11.06844 | 11.037 | 0.031152788 | 0.023414284 | 11.358727 | 11.449725 | 11.1727 | 0.27702511 | 0.186026927 |
| Hamp | 3.290124 | 3.129471 | 3.1147 | 0.014771066 | 0.175423953 | 2.9722371 | 3.1675314 | 2.89077 | 0.276760239 | 0.081465939 |
| Rab39 | 6.43781 | 6.646221 | 6.5235 | 0.12269781 | -0.085713254 | 6.8179462 | 6.8774492 | 6.6007 | 0.276746204 | 0.217243148 |
| Fancf | 5.435076 | 5.371335 | 5.3387 | 0.032669639 | 0.096410653 | 5.3994088 | 5.500941 | 5.22422 | 0.276722919 | 0.175190801 |
| Pigm | 10.0538 | 10.07624 | 10.121 | -0.044971465 | -0.067409232 | 10.441245 | 10.631195 | 10.3545 | 0.276710154 | 0.086760271 |
| Gatd1 | 9.172509 | 9.264268 | 9.3176 | -0.053321542 | -0.145080955 | 9.508872 | 9.4876926 | 9.21106 | 0.276632463 | 0.297811884 |
| Gtf2ird2 | 7.983645 | 8.160489 | 8.0785 | 0.082013919 | -0.094829971 | 8.1084604 | 8.459727 | 8.18322 | 0.276511433 | -0.074755159 |
| Camk2d | 10.98575 | 10.94165 | 10.886 | 0.056085194 | 0.100183442 | 11.11324 | 11.143448 | 10.867 | 0.276402548 | 0.246194538 |
| Relb | 10.42865 | 10.57837 | 10.414 | 0.164630978 | 0.014915288 | 10.43224 | 10.777986 | 10.5019 | 0.276049352 | -0.069696748 |
| Scaf1 | 10.46363 | 10.40329 | 10.417 | -0.013390542 | 0.046954446 | 10.599468 | 10.8104 | 10.5344 | 0.276020722 | 0.065088685 |
| Cdkn1b | 11.13019 | 11.33495 | 11.311 | 0.024179096 | -0.18058045 | 11.098069 | 11.422473 | 11.1465 | 0.275995988 | -0.048408458 |
| Ptcd2 | 9.238191 | 9.28424 | 9.2162 | 0.068043586 | 0.021994582 | 9.3097531 | 9.5252542 | 9.24935 | 0.275906435 | 0.060405387 |
| Serbp1 | 12.71959 | 12.74742 | 12.684 | 0.063232114 | 0.035399294 | 12.607859 | 12.964659 | 12.6888 | 0.275880804 | -0.080918711 |

| | | | | | | | | | | |
|---|---|---|---|---|---|---|---|---|---|---|
| Tmem179b | 8.723827 | 8.681483 | 8.7051 | -0.023575207 | 0.01876816 | 9.0291552 | 9.0347035 | 8.75885 | 0.275855839 | 0.270307491 |
| Gm20219 | 4.702276 | 4.570294 | 4.6165 | -0.046212534 | 0.085768872 | 4.9757833 | 5.1825384 | 4.9067 | 0.275834033 | 0.069078897 |
| Gm11400 | 4.837946 | 5.025833 | 4.8698 | 0.156043853 | -0.031843833 | 5.0433888 | 5.2768731 | 5.00118 | 0.275691934 | 0.042207597 |
| Dxo | 8.150968 | 8.254673 | 8.269 | -0.014315803 | -0.118021256 | 8.6256178 | 8.6312835 | 8.35567 | 0.275608755 | 0.269943026 |
| D11Wsu47e | 7.456344 | 7.523816 | 7.4291 | 0.094727813 | 0.027256323 | 7.6163403 | 7.7065583 | 7.431 | 0.275557692 | 0.185339688 |
| Tsr2 | 8.379922 | 8.30357 | 8.4094 | -0.105807566 | -0.029455409 | 8.5711141 | 8.6308765 | 8.35538 | 0.275495878 | 0.215733403 |
| Gars | 10.47148 | 10.44931 | 10.429 | 0.020343334 | 0.042510042 | 10.581095 | 10.769801 | 10.4943 | 0.27547977 | 0.086773921 |
| Ube2d-ps | 7.139578 | 7.093315 | 7.0008 | 0.09254404 | 0.138806638 | 7.179927 | 7.33625 | 7.0608 | 0.275450789 | 0.119127771 |
| Gm7332 | 7.053679 | 7.085253 | 7.024 | 0.061282768 | 0.029709025 | 6.9529126 | 7.1738076 | 6.8984 | 0.27541049 | 0.054515474 |
| Sumo3 | 10.27431 | 10.22677 | 10.244 | -0.017709303 | 0.029831628 | 10.157067 | 10.481323 | 10.2061 | 0.275223336 | -0.049031797 |
| Ppt2 | 7.954362 | 8.046789 | 8.0414 | 0.005398703 | -0.087027794 | 8.2003151 | 8.1871766 | 7.91205 | 0.275123661 | 0.288262164 |
| Ggps1 | 9.740317 | 9.813801 | 9.6423 | 0.171550013 | 0.098065793 | 9.8566465 | 10.04948 | 9.77438 | 0.27509708 | 0.082263587 |
| Tmco6 | 8.071122 | 8.20506 | 8.1326 | 0.072489272 | -0.0614488 | 8.6313712 | 8.5719485 | 8.29686 | 0.275088306 | 0.334510973 |
| Xpo6 | 11.5499 | 11.61636 | 11.491 | 0.12564416 | 0.059185002 | 11.6447 | 11.900081 | 11.625 | 0.275047978 | 0.019667294 |
| Syt13 | 4.945647 | 5.275766 | 4.8641 | 0.411678352 | 0.081559006 | 4.7710274 | 5.1434961 | 4.86884 | 0.274655997 | -0.097812694 |
| Comp | 3.504381 | 3.445846 | 3.489 | -0.043186263 | 0.015349134 | 3.4916731 | 3.7375535 | 3.46294 | 0.274616856 | 0.028736442 |
| Trap1 | 10.50382 | 10.6525 | 10.54 | 0.112431622 | -0.036250194 | 10.741087 | 10.911402 | 10.6368 | 0.274589536 | 0.104274515 |
| Jtb | 8.647872 | 8.611955 | 8.5721 | 0.039843096 | 0.075760606 | 8.7788432 | 8.8929614 | 8.61838 | 0.274582001 | 0.160463799 |
| Ctdp1 | 9.687201 | 9.893211 | 9.7463 | 0.146903104 | -0.05910643 | 9.7421506 | 9.9727467 | 9.69854 | 0.274208674 | 0.043612535 |
| Otub2 | 5.877875 | 6.208682 | 6.0091 | 0.199608334 | -0.131198282 | 6.3678738 | 6.150214 | 5.87608 | 0.274134665 | 0.491794489 |
| Ttc9 | 2.798924 | 2.963603 | 2.7695 | 0.194106003 | 0.029427188 | 2.7383136 | 2.9389186 | 2.66491 | 0.274004799 | 0.073399799 |
| Arfrp1 | 9.925041 | 10.06211 | 9.9593 | 0.102805846 | -0.034262492 | 10.149749 | 10.317585 | 10.0437 | 0.273929272 | 0.106093352 |
| Pfn1 | 13.62504 | 13.66917 | 13.609 | 0.060132646 | 0.016002636 | 13.896749 | 13.854406 | 13.5805 | 0.27391072 | 0.316253938 |
| Polr1c | 9.457678 | 9.486426 | 9.5478 | -0.061323663 | -0.090071679 | 9.7017506 | 9.8092103 | 9.53535 | 0.273864028 | 0.166404368 |
| Pef1 | 8.748215 | 8.882142 | 8.7881 | 0.093998443 | -0.039927826 | 9.0310589 | 8.9913454 | 8.71756 | 0.273787021 | 0.313500536 |
| Sin3b | 10.59359 | 10.62424 | 10.522 | 0.10263354 | 0.071985634 | 10.661244 | 10.79538 | 10.5216 | 0.273786758 | 0.13965062 |
| Dnajc30 | 7.102827 | 7.024776 | 7.044 | -0.019241299 | 0.058810025 | 7.1781658 | 7.1468298 | 6.87306 | 0.273772296 | 0.305108305 |
| Renbp | 7.62238 | 7.567386 | 7.5909 | -0.023500639 | 0.031493066 | 7.7428704 | 7.6814784 | 7.40812 | 0.273353419 | 0.334745485 |
| Slc12a4 | 8.727079 | 8.768487 | 8.5316 | 0.236888155 | 0.195480632 | 8.4419619 | 8.8454036 | 8.57207 | 0.273336364 | -0.130105343 |
| Nqo2 | 7.397374 | 7.246096 | 7.392 | -0.145854788 | 0.005422742 | 7.24031 | 7.4977184 | 7.2244 | 0.273321395 | 0.015912965 |
| Atp5b | 13.22906 | 13.28872 | 13.272 | 0.017013246 | -0.042644588 | 13.505382 | 13.572944 | 13.2996 | 0.273314632 | 0.205753337 |
| Gopc | 10.03358 | 10.13426 | 10.017 | 0.116835321 | 0.016157051 | 9.9069729 | 10.251345 | 9.97804 | 0.273299947 | -0.071071844 |

| | | | | | | | | | | |
|---|---|---|---|---|---|---|---|---|---|---|
| Letmd1 | 9.400991 | 9.435193 | 9.4152 | 0.019977013 | -0.014224738 | 9.6672364 | 9.7089356 | 9.4358 | 0.273132952 | 0.231433775 |
| Fbxl14 | 9.901055 | 9.992426 | 9.9184 | 0.073984571 | -0.017386506 | 9.9043592 | 10.032734 | 9.75962 | 0.273113494 | 0.144738329 |
| Pigt | 10.29951 | 10.28336 | 10.209 | 0.074584874 | 0.090736514 | 10.498952 | 10.434276 | 10.1612 | 0.273046549 | 0.337722702 |
| Gnb2 | 11.68116 | 11.76102 | 11.587 | 0.174250804 | 0.094384004 | 11.857015 | 11.803191 | 11.5302 | 0.272968828 | 0.326792458 |
| Palld | 6.752734 | 6.814239 | 6.5648 | 0.249482329 | 0.187976396 | 6.5621078 | 6.6744777 | 6.40158 | 0.27290149 | 0.16053158 |
| Akirin1 | 10.02053 | 10.18763 | 10.009 | 0.178415606 | 0.011322661 | 10.063601 | 10.242632 | 9.96976 | 0.272872039 | 0.093841258 |
| Naa38 | 7.715941 | 7.83487 | 7.6912 | 0.14367298 | 0.024744441 | 7.9027687 | 7.9918971 | 7.71907 | 0.272825759 | 0.183697313 |
| Gata1 | 4.836356 | 4.762093 | 4.7478 | 0.014305046 | 0.088567898 | 4.5315446 | 4.8693876 | 4.59659 | 0.272799618 | -0.065043374 |
| Eef1akmt1 | 7.246074 | 7.152994 | 7.2316 | -0.078571933 | 0.01450809 | 7.3041998 | 7.4393197 | 7.16675 | 0.272573963 | 0.137454024 |
| Ssb | 11.34037 | 11.26547 | 11.309 | -0.043967308 | 0.030926776 | 11.234673 | 11.611497 | 11.3391 | 0.272432514 | -0.104391096 |
| Snrpb2 | 9.523001 | 9.572197 | 9.5105 | 0.061739983 | 0.012543852 | 9.6447587 | 9.9168853 | 9.64448 | 0.27240667 | 0.000280091 |
| Jund | 10.20606 | 9.875237 | 10.021 | -0.145360089 | 0.185463653 | 9.489407 | 9.6002529 | 9.32804 | 0.272208249 | 0.161362308 |
| Klk1b24 | 3.151205 | 3.574948 | 4.0198 | -0.444896188 | -0.868639061 | 4.654683 | 4.0742981 | 3.8022 | 0.272099568 | 0.852484492 |
| Rpl34 | 13.48599 | 13.48633 | 13.45 | 0.036813684 | 0.036478736 | 13.598376 | 13.696058 | 13.4241 | 0.272002376 | 0.174319794 |
| Olfr1120 | 4.532348 | 4.534562 | 4.4562 | 0.078352345 | 0.076138204 | 4.5444549 | 5.0331326 | 4.76116 | 0.271971396 | -0.216706278 |
| Bccip | 9.96389 | 10.00653 | 9.9508 | 0.055717823 | 0.013081517 | 10.149404 | 10.25826 | 9.98635 | 0.27190774 | 0.163051917 |
| C2cd3 | 10.78007 | 10.88818 | 10.827 | 0.061086268 | -0.047023172 | 10.78898 | 11.18722 | 10.9154 | 0.271866756 | -0.126373548 |
| Adssl1 | 6.928756 | 6.811987 | 6.7629 | 0.049109465 | 0.165878289 | 6.9015735 | 6.8062577 | 6.53441 | 0.271843878 | 0.367159623 |
| Fkbp10 | 6.539962 | 6.471573 | 6.3886 | 0.082963268 | 0.151352697 | 6.3439528 | 6.1774274 | 5.90561 | 0.271819398 | 0.438344718 |
| Snx1 | 10.64258 | 10.64435 | 10.62 | 0.02393625 | 0.022161708 | 10.821679 | 10.917565 | 10.6458 | 0.271746014 | 0.175859426 |
| Upf3a | 8.956256 | 9.096071 | 8.9787 | 0.117398458 | -0.022416799 | 8.9283395 | 9.281442 | 9.00985 | 0.27159569 | -0.08150681 |
| Ifrd1 | 9.76077 | 9.753608 | 9.5696 | 0.184017352 | 0.191179068 | 9.6007752 | 10.027635 | 9.75611 | 0.271526059 | -0.155333969 |
| Stk11 | 10.53773 | 10.67845 | 10.547 | 0.131706478 | -0.00901443 | 10.799511 | 10.876635 | 10.6054 | 0.271213378 | 0.194089072 |
| Ttc1 | 8.725097 | 8.932918 | 8.7611 | 0.171856883 | -0.035964168 | 8.7536173 | 9.0134662 | 8.74252 | 0.270945239 | 0.011096412 |
| Pusl1 | 7.200699 | 7.187822 | 7.1666 | 0.021243359 | 0.034121094 | 7.5435582 | 7.6229968 | 7.35207 | 0.270923793 | 0.191485159 |
| Rad51ap1 | 6.221347 | 6.253813 | 6.1059 | 0.147882375 | 0.115416312 | 6.1445274 | 6.6909578 | 6.42013 | 0.270828514 | -0.275601917 |
| Atad1 | 10.50521 | 10.50382 | 10.504 | -0.000327156 | 0.001065616 | 10.644328 | 10.725458 | 10.4547 | 0.270749114 | 0.18961917 |
| Dot1l | 10.11195 | 10.20483 | 10.107 | 0.097435619 | 0.004558989 | 10.317223 | 10.606592 | 10.336 | 0.27058375 | -0.018784719 |
| Pde8b | 2.889898 | 3.033662 | 3.0425 | -0.008852489 | -0.152617315 | 2.8916906 | 3.2749283 | 3.00442 | 0.27050771 | -0.112729919 |
| Lamp1 | 12.10664 | 12.0621 | 12.084 | -0.021458736 | 0.023079286 | 12.139619 | 12.2411 | 11.9706 | 0.270466548 | 0.168986363 |
| Rnmt | 9.156774 | 9.218762 | 9.1809 | 0.037874445 | -0.024113329 | 9.2380956 | 9.423489 | 9.15305 | 0.270434326 | 0.085040932 |
| Rnf4 | 10.79373 | 10.88639 | 10.793 | 0.093604184 | 0.000942806 | 10.9409 | 11.185702 | 10.9153 | 0.270372106 | 0.025569787 |

| | | | | | | | | | | |
|---|---|---|---|---|---|---|---|---|---|---|
| Synj1 | 10.00899 | 9.973695 | 9.9329 | 0.040793601 | 0.076088633 | 9.8443393 | 10.252925 | 9.98257 | 0.270352194 | -0.138233195 |
| Gm11847 | 5.305182 | 4.958769 | 5.0072 | -0.048423961 | 0.29798914 | 4.6247317 | 5.2403536 | 4.97026 | 0.270095948 | -0.345525937 |
| Sfmbt1 | 9.232528 | 9.433883 | 9.1921 | 0.241815067 | 0.040459546 | 9.0923612 | 9.4954865 | 9.22546 | 0.270026301 | -0.133098996 |
| Acsl1 | 8.358746 | 8.452444 | 8.2884 | 0.164086688 | 0.070388634 | 8.4085171 | 8.5324202 | 8.26256 | 0.269856321 | 0.145953153 |
| Rer1 | 10.35442 | 10.336 | 10.323 | 0.012537072 | 0.030956924 | 10.402189 | 10.507608 | 10.2378 | 0.26979705 | 0.164377805 |
| M6pr | 11.85655 | 11.95935 | 11.81 | 0.149105232 | 0.046311944 | 11.937468 | 12.090098 | 11.8204 | 0.26968875 | 0.117058116 |
| Cebpz | 10.69436 | 10.70515 | 10.629 | 0.07646331 | 0.06567955 | 10.67802 | 10.90196 | 10.6323 | 0.269624906 | 0.045684857 |
| Trpm3 | 4.584172 | 4.598493 | 4.6105 | -0.011993721 | -0.026314788 | 5.1325507 | 5.1239528 | 4.8544 | 0.269549507 | 0.278147401 |
| Tex2 | 9.261334 | 9.137257 | 9.1646 | -0.027384269 | 0.096693123 | 8.9149441 | 9.3023408 | 9.03301 | 0.269332001 | -0.118064667 |
| Gm45038 | 3.826063 | 3.618013 | 3.6686 | -0.050554201 | 0.157495348 | 3.7576771 | 4.0385115 | 3.76933 | 0.269183765 | -0.011650603 |
| Gon7 | 6.947318 | 7.066394 | 6.9961 | 0.070264748 | -0.048811642 | 7.2843441 | 7.3949958 | 7.12603 | 0.268962566 | 0.158310873 |
| Gm47191 | 3.488155 | 3.640698 | 3.4397 | 0.200961005 | 0.048417092 | 3.6304143 | 3.9429251 | 3.67421 | 0.268712947 | -0.043797859 |
| Vmn1r26 | 4.233167 | 4.303259 | 4.434 | -0.130721027 | -0.20081271 | 4.4352996 | 4.7778063 | 4.50914 | 0.268662277 | -0.073844448 |
| Rab1a | 10.54917 | 10.66713 | 10.46 | 0.206897606 | 0.08893702 | 10.424856 | 10.771964 | 10.5033 | 0.268661558 | -0.078446413 |
| Arl2 | 7.535036 | 7.551423 | 7.5729 | -0.021478728 | -0.037865683 | 7.8052924 | 7.7381288 | 7.46967 | 0.268460487 | 0.335624032 |
| Hist2h3c1 | 4.041056 | 4.032015 | 4.015 | 0.016999508 | 0.026040193 | 3.9397632 | 4.2864243 | 4.01809 | 0.268330995 | -0.078330156 |
| E030030I06Rik | 5.459145 | 5.598087 | 5.4903 | 0.107783325 | -0.031159219 | 5.6704856 | 5.6620259 | 5.39377 | 0.268256025 | 0.276715681 |
| Hars | 9.889132 | 9.943627 | 9.8186 | 0.125061023 | 0.070566544 | 9.969592 | 10.11426 | 9.84619 | 0.268067581 | 0.12339938 |
| Stat6 | 12.02301 | 12.20398 | 12.079 | 0.125356292 | -0.055619384 | 12.204548 | 12.427411 | 12.1594 | 0.26804833 | 0.045184985 |
| B9d2 | 7.6637 | 7.618956 | 7.6576 | -0.038680203 | 0.006064157 | 7.8842016 | 7.8342422 | 7.56634 | 0.267903431 | 0.317862821 |
| 1810009A15Rik | 8.504173 | 8.547179 | 8.4677 | 0.079432504 | 0.036426615 | 8.7734774 | 8.8433134 | 8.5757 | 0.267613382 | 0.197777385 |
| Klra1 | 6.259534 | 5.860056 | 6.1216 | -0.261538775 | 0.137939077 | 5.8675413 | 6.1679963 | 5.90044 | 0.267551775 | -0.032903254 |
| Vmn2r53 | 7.6555 | 7.706126 | 7.6848 | 0.021336149 | -0.029289973 | 7.9394441 | 8.684251 | 8.41681 | 0.267444415 | -0.477362454 |
| Zfp516 | 8.321535 | 8.511238 | 8.3197 | 0.191565712 | 0.001862948 | 8.2136769 | 8.5910396 | 8.32362 | 0.267418028 | -0.109944627 |
| Eif4e | 10.23058 | 10.3686 | 10.19 | 0.178114738 | 0.04009983 | 10.207133 | 10.498754 | 10.2314 | 0.267372806 | -0.02424781 |
| Atxn7 | 9.958204 | 9.99249 | 9.9839 | 0.00857965 | -0.025705841 | 9.8459826 | 10.31356 | 10.0463 | 0.267277032 | -0.200300121 |
| Pgam5 | 9.326891 | 9.370924 | 9.2074 | 0.163549532 | 0.119516224 | 9.3456957 | 9.5106387 | 9.24356 | 0.267082636 | 0.102139588 |
| Tgfb3 | 6.654672 | 6.511436 | 6.407 | 0.104459621 | 0.247695716 | 6.7176689 | 6.7737206 | 6.50673 | 0.266994507 | 0.210942767 |
| Snapc1 | 8.525817 | 8.444101 | 8.5685 | -0.124445135 | -0.04272933 | 8.7146596 | 8.8881673 | 8.62126 | 0.266903564 | 0.093395845 |
| Neu1 | 10.24526 | 10.23132 | 10.296 | -0.064474247 | -0.050537368 | 10.155062 | 10.390492 | 10.1237 | 0.266824644 | 0.0313944 |
| Calr | 13.17643 | 13.27716 | 13.055 | 0.221672798 | 0.120944128 | 13.337758 | 13.392454 | 13.1257 | 0.26673305 | 0.212037067 |
| Rpl32 | 14.27927 | 14.27755 | 14.282 | -0.004592092 | -0.00287787 | 14.309696 | 14.526785 | 14.2604 | 0.266396884 | 0.049308214 |

| | | | | | | | | | | |
|---|---|---|---|---|---|---|---|---|---|---|
| Rab11b | 11.02405 | 11.12654 | 11.102 | 0.02439389 | -0.07809611 | 11.286904 | 11.25593 | 10.9896 | 0.26635851 | 0.29733223 |
| Elavl1 | 10.56343 | 10.74133 | 10.582 | 0.15915967 | -0.018737624 | 10.672307 | 10.816487 | 10.5504 | 0.266107698 | 0.121927632 |
| Rps24 | 15.08774 | 15.10349 | 15.112 | -0.008095094 | -0.023841272 | 15.17664 | 15.319485 | 15.0534 | 0.266086704 | 0.123241994 |
| Rpl18 | 13.78873 | 13.85301 | 13.793 | 0.059776634 | -0.004500072 | 13.990578 | 14.022146 | 13.7561 | 0.266046912 | 0.234478718 |
| Prr15 | 3.440033 | 3.438574 | 3.4109 | 0.027666122 | 0.029125646 | 3.505973 | 3.6421058 | 3.37621 | 0.265892538 | 0.129759769 |
| 1110059G10Rik | 7.626468 | 7.469651 | 7.6212 | -0.151531754 | 0.005285734 | 7.8271427 | 7.8971654 | 7.6313 | 0.265870344 | 0.195847652 |
| Loxl2 | 5.368439 | 5.555956 | 5.2256 | 0.330348651 | 0.142831065 | 5.1779051 | 5.464244 | 5.19841 | 0.265836093 | -0.020502762 |
| Mrpl35 | 7.950976 | 7.981223 | 7.995 | -0.013785697 | -0.044031863 | 8.0625598 | 8.3246636 | 8.05901 | 0.265652685 | 0.003548861 |
| Lysmd1 | 7.712884 | 7.731598 | 7.6853 | 0.04624905 | 0.027534822 | 7.6066745 | 7.9007325 | 7.63512 | 0.265613636 | -0.028444367 |
| Wipf1 | 12.3782 | 12.5085 | 12.364 | 0.14423684 | 0.013932472 | 12.34743 | 12.55659 | 12.291 | 0.265542918 | 0.056382532 |
| Tvp23a | 2.722259 | 2.771452 | 2.7563 | 0.015161758 | -0.034031679 | 2.6735368 | 2.954823 | 2.68929 | 0.265529886 | -0.015756353 |
| Sft2d1 | 9.404426 | 9.228024 | 9.3388 | -0.110748636 | 0.065653023 | 9.6903621 | 9.6424463 | 9.37696 | 0.265481989 | 0.313397722 |
| Commd9 | 7.311912 | 7.395158 | 7.3781 | 0.017080282 | -0.066165597 | 7.628862 | 7.4930497 | 7.22765 | 0.265402689 | 0.401215014 |
| Tdp1 | 8.146723 | 8.296021 | 8.2989 | -0.002902754 | -0.15220127 | 8.3865178 | 8.4891161 | 8.22388 | 0.265238646 | 0.162640286 |
| Stx18 | 8.25545 | 8.260645 | 8.1582 | 0.102419722 | 0.097224183 | 8.2378341 | 8.483379 | 8.21815 | 0.265233675 | 0.019688868 |
| Ercc1 | 7.937237 | 7.837818 | 7.8805 | -0.042640585 | 0.056778153 | 8.1524619 | 8.1208552 | 7.85562 | 0.265232272 | 0.296838939 |
| Rps10 | 13.85126 | 13.96646 | 13.871 | 0.09549418 | -0.019701576 | 14.056534 | 14.075688 | 13.8105 | 0.265162982 | 0.246009053 |
| Gm45351 | 3.250238 | 3.236975 | 3.3581 | -0.121121739 | -0.107858569 | 3.2001036 | 3.4871969 | 3.22218 | 0.265015818 | -0.022077524 |
| Coa4 | 6.501319 | 6.504954 | 6.4613 | 0.043625909 | 0.039991462 | 6.7016202 | 6.7138839 | 6.44896 | 0.264926809 | 0.252663128 |
| Rps27 | 8.750875 | 8.630574 | 8.7195 | -0.088947405 | 0.031353926 | 9.0572233 | 9.0031472 | 8.73825 | 0.264892567 | 0.318968677 |
| Rps15 | 14.29379 | 14.4295 | 14.326 | 0.103255364 | -0.032453944 | 14.655381 | 14.585783 | 14.321 | 0.264740398 | 0.334339107 |
| Borcs6 | 7.682873 | 7.570723 | 7.6517 | -0.080978744 | 0.031171245 | 7.5950443 | 7.609924 | 7.34519 | 0.264733075 | 0.249853346 |
| Trmt112 | 9.333943 | 9.368803 | 9.3775 | -0.008743913 | -0.04360352 | 9.560077 | 9.5813386 | 9.31672 | 0.264614773 | 0.243353107 |
| Serpinb6b | 9.934485 | 10.06235 | 10.061 | 0.001368191 | -0.126492089 | 9.6929374 | 10.409136 | 10.1446 | 0.264582367 | -0.45161601 |
| Rbm26 | 10.43478 | 10.57378 | 10.427 | 0.146432676 | 0.007432348 | 10.456709 | 10.817773 | 10.5532 | 0.264574346 | -0.096489442 |
| Eif2s1 | 10.45976 | 10.46894 | 10.266 | 0.202951622 | 0.193766794 | 10.52342 | 10.777486 | 10.513 | 0.264530638 | 0.010464476 |
| Slc26a6 | 6.162345 | 6.242873 | 6.2347 | 0.008194734 | -0.072333867 | 6.2017835 | 6.350058 | 6.08563 | 0.264424832 | 0.116150378 |
| Trim8 | 9.963982 | 10.1491 | 10.029 | 0.120336595 | -0.064781973 | 10.040548 | 10.182669 | 9.91828 | 0.264387116 | 0.122266123 |
| Adgrd1 | 4.377115 | 4.282971 | 4.2865 | -0.003495674 | 0.090647789 | 4.3559216 | 4.5598469 | 4.29558 | 0.264270106 | 0.060344843 |
| Dync1i2 | 10.65057 | 10.67653 | 10.688 | -0.011704372 | -0.037673354 | 10.587599 | 10.858396 | 10.5942 | 0.264244436 | -0.006552694 |
| Slc25a53 | 8.571812 | 8.622374 | 8.4283 | 0.194086109 | 0.143523273 | 8.8605937 | 8.8286302 | 8.56442 | 0.264208842 | 0.296172356 |
| Igkj4 | 6.713276 | 6.967229 | 6.5072 | 0.460067834 | 0.206114562 | 6.8703402 | 6.9337729 | 6.66961 | 0.264160784 | 0.200728117 |

| | | | | | | | | | | |
|---|---|---|---|---|---|---|---|---|---|---|
| Vps13d | 11.59566 | 11.56853 | 11.556 | 0.012083564 | 0.03921225 | 11.413329 | 11.897819 | 11.6337 | 0.264095042 | -0.220395603 |
| Gm20390 | 10.3825 | 10.4808 | 10.228 | 0.252681339 | 0.154388335 | 10.624763 | 10.623755 | 10.3597 | 0.264059546 | 0.265067197 |
| E130309D02Rik | 8.728716 | 8.877229 | 8.6837 | 0.193577681 | 0.045064489 | 8.9068986 | 8.9053148 | 8.64135 | 0.263960609 | 0.265544412 |
| Clptm1l | 10.79915 | 10.86568 | 10.787 | 0.078445478 | 0.011921832 | 11.032189 | 11.16727 | 10.9033 | 0.263937968 | 0.128856635 |
| Chd9 | 12.80867 | 12.66506 | 12.825 | -0.160433754 | -0.016823254 | 12.812679 | 12.984513 | 12.7206 | 0.263936056 | 0.092102103 |
| Camta1 | 6.900378 | 6.773663 | 6.9006 | -0.126934778 | -0.000219927 | 7.0103565 | 6.9911476 | 6.72725 | 0.263896901 | 0.2831058 |
| Tnfaip6 | 4.225883 | 4.093283 | 3.6099 | 0.483366512 | 0.615966575 | 3.6850167 | 4.013506 | 3.74964 | 0.263867916 | -0.064621399 |
| Mcm9 | 8.501783 | 8.60254 | 8.5098 | 0.092732753 | -0.008024014 | 8.7707417 | 9.0207181 | 8.75691 | 0.263805879 | 0.013829441 |
| Akna | 12.69862 | 13.03316 | 12.794 | 0.239492406 | -0.095049306 | 12.635908 | 12.985828 | 12.722 | 0.263797818 | -0.086122946 |
| Adora3 | 5.06214 | 4.992439 | 4.9816 | 0.010868372 | 0.080569022 | 5.0970511 | 5.3223311 | 5.05875 | 0.263577158 | 0.038297154 |
| Usp29 | 3.695579 | 3.663632 | 3.6855 | -0.021887137 | 0.010058994 | 3.8960676 | 4.0807546 | 3.81726 | 0.263491347 | 0.078804352 |
| L3mbtl3 | 9.905744 | 10.13162 | 9.8991 | 0.232530075 | 0.006658248 | 9.9568427 | 10.181099 | 9.91811 | 0.262989447 | 0.038732727 |
| Mvd | 7.664663 | 7.905134 | 7.6628 | 0.242328857 | 0.00185727 | 7.7855616 | 7.832631 | 7.56969 | 0.262945158 | 0.215875735 |
| Wac | 11.24388 | 11.29206 | 11.192 | 0.100482504 | 0.052302934 | 11.071158 | 11.306047 | 11.0431 | 0.26293211 | 0.028042875 |
| Ccl19-ps1 | 3.622246 | 3.442976 | 3.3264 | 0.116567856 | 0.295837275 | 3.2611635 | 3.3990968 | 3.13617 | 0.262927775 | 0.124994443 |
| Olfr629 | 4.354423 | 4.530487 | 4.4473 | 0.083187861 | -0.092876171 | 4.5033761 | 4.7298441 | 4.46704 | 0.262808059 | 0.036340074 |
| Mrpl45 | 8.914639 | 8.974733 | 8.9386 | 0.036129088 | -0.023965346 | 9.1160128 | 9.272538 | 9.0098 | 0.262741086 | 0.106215869 |
| Idi1 | 9.472111 | 9.561426 | 9.3052 | 0.256229308 | 0.166913735 | 9.5018092 | 9.6204712 | 9.35775 | 0.262716972 | 0.144054914 |
| Map3k11 | 10.22654 | 10.26158 | 10.226 | 0.035430938 | 0.000389296 | 10.34594 | 10.398115 | 10.1354 | 0.262686369 | 0.21051098 |
| Fastkd5 | 7.73836 | 7.871856 | 7.7253 | 0.146528187 | 0.01303148 | 7.8997212 | 8.0442004 | 7.78152 | 0.262677522 | 0.118198318 |
| Rpl8 | 14.44528 | 14.51274 | 14.421 | 0.091758174 | 0.024298632 | 14.590093 | 14.595475 | 14.3329 | 0.262531472 | 0.257150221 |
| Igkv3-12 | 7.41441 | 7.494922 | 7.222 | 0.272900946 | 0.192388902 | 7.6432155 | 7.4955986 | 7.23313 | 0.262469271 | 0.410086189 |
| Sart1 | 10.88564 | 10.8793 | 10.887 | -0.007428548 | -0.001087634 | 11.154805 | 11.258988 | 10.9966 | 0.262396784 | 0.158213694 |
| Pcgf6 | 8.232866 | 8.220053 | 8.3183 | -0.098278793 | -0.085466406 | 8.4557823 | 8.6668648 | 8.40474 | 0.262124558 | 0.051042051 |
| Xylb | 4.343746 | 4.376487 | 4.3823 | -0.00583449 | -0.038575698 | 4.1879971 | 4.4052577 | 4.14316 | 0.262101072 | 0.044840438 |
| Pgm1 | 7.663005 | 7.56961 | 7.7591 | -0.189527942 | -0.096133599 | 7.7341041 | 7.9481629 | 7.68607 | 0.262087921 | 0.048029073 |
| Tmed9 | 10.9272 | 10.90695 | 10.839 | 0.068409368 | 0.088660188 | 10.945113 | 11.035713 | 10.7737 | 0.262042596 | 0.171442467 |
| Nat10 | 9.855436 | 9.700858 | 9.6523 | 0.048582552 | 0.203159685 | 10.023071 | 10.286798 | 10.0248 | 0.261973214 | -0.001753513 |
| Rpl37 | 12.87877 | 12.83652 | 12.71 | 0.126538882 | 0.168787964 | 12.91524 | 13.12697 | 12.865 | 0.261951692 | 0.050221426 |
| Ccnq | 7.701518 | 7.793114 | 7.673 | 0.12016327 | 0.028567389 | 7.8773031 | 7.8657344 | 7.60382 | 0.261918554 | 0.273487268 |
| Rxrb | 10.18031 | 10.29691 | 10.203 | 0.09422552 | -0.02237442 | 10.406553 | 10.449085 | 10.1873 | 0.26182826 | 0.219296649 |
| Gm11518 | 11.09387 | 11.18813 | 11.18 | 0.008389794 | -0.085870208 | 11.163802 | 11.444499 | 11.1828 | 0.261667278 | -0.019029872 |

| | | | | | | | | | | |
|---|---|---|---|---|---|---|---|---|---|---|
| Rpl27-ps3 | 11.09387 | 11.18813 | 11.18 | 0.008389794 | -0.085870208 | 11.163802 | 11.444499 | 11.1828 | 0.261667278 | -0.019029872 |
| Sdhd | 10.05013 | 10.0507 | 10.064 | -0.013786797 | -0.014363777 | 10.231945 | 10.279456 | 10.0179 | 0.261602953 | 0.21409199 |
| Dcun1d5 | 9.422162 | 9.435799 | 9.3686 | 0.067231692 | 0.053594262 | 9.5761826 | 9.8134384 | 9.55199 | 0.261447513 | 0.024191757 |
| Thbs3 | 4.790224 | 4.867297 | 4.7823 | 0.085041981 | 0.007969518 | 4.6965058 | 4.8731417 | 4.61179 | 0.261354428 | 0.084718512 |
| Fez2 | 8.121313 | 7.998089 | 7.957 | 0.041045235 | 0.164269292 | 7.6803349 | 7.9978322 | 7.73664 | 0.261192326 | -0.056304983 |
| Adck1 | 8.022366 | 8.099629 | 8.0773 | 0.022302896 | -0.05495986 | 8.1409014 | 8.331672 | 8.07057 | 0.261098842 | 0.070328261 |
| Traf3ip3 | 11.20296 | 11.31841 | 11.336 | -0.017823058 | -0.13326992 | 11.686259 | 11.803298 | 11.5422 | 0.26109876 | 0.144059801 |
| Gm6627 | 3.964274 | 4.074902 | 4.059 | 0.015925591 | -0.094702452 | 4.3247696 | 4.3398773 | 4.07878 | 0.261096597 | 0.245988942 |
| Rnf121 | 8.804944 | 8.909815 | 8.7785 | 0.131316715 | 0.026445434 | 8.8945933 | 8.9734905 | 8.71264 | 0.260853744 | 0.181956594 |
| Ip6k1 | 11.30808 | 11.54916 | 11.436 | 0.112778778 | -0.128308714 | 11.571352 | 11.827681 | 11.5668 | 0.260837086 | 0.004507797 |
| Pex11a | 4.065551 | 3.959553 | 3.9803 | -0.020707574 | 0.085290129 | 3.8318485 | 4.1083457 | 3.84761 | 0.260736006 | -0.015761182 |
| Zfp688 | 7.461982 | 7.67225 | 7.6477 | 0.024529798 | -0.18573892 | 7.9329071 | 7.8860313 | 7.62532 | 0.260715643 | 0.307591405 |
| Nlgn2 | 6.521834 | 6.382913 | 6.2671 | 0.115772851 | 0.254694388 | 6.2602391 | 6.5930498 | 6.33239 | 0.26066197 | -0.072148734 |
| Pank4 | 9.749342 | 9.834624 | 9.8669 | -0.032292463 | -0.117574932 | 10.157275 | 10.249702 | 9.98914 | 0.260564245 | 0.168137544 |
| Josd2 | 8.222384 | 8.342578 | 8.2454 | 0.09714996 | -0.023043777 | 8.8316713 | 8.7289515 | 8.46848 | 0.260466614 | 0.363186435 |
| Ahcy | 10.35573 | 10.44671 | 10.397 | 0.049747106 | -0.04124118 | 10.5651 | 10.661942 | 10.4016 | 0.260299742 | 0.163458102 |
| Pax6 | 3.064362 | 3.120741 | 3.0937 | 0.027040712 | -0.029337769 | 3.1900455 | 3.1823008 | 2.92201 | 0.260288338 | 0.26803306 |
| Mindy3 | 8.240003 | 8.256049 | 8.2658 | -0.009708228 | -0.025754527 | 8.2709937 | 8.6504642 | 8.3902 | 0.260265589 | -0.119204984 |
| Ankmy1 | 3.637628 | 3.680189 | 3.7147 | -0.034559476 | -0.077120424 | 3.6125485 | 3.8794458 | 3.61922 | 0.260220987 | -0.006676312 |
| Mrpl50 | 8.157644 | 8.147237 | 8.1377 | 0.009556766 | 0.019963782 | 8.2335399 | 8.3721592 | 8.11209 | 0.260073434 | 0.121454095 |
| Mocs1 | 8.247696 | 8.299744 | 8.2482 | 0.051590559 | -0.000457398 | 8.245399 | 8.4532182 | 8.19327 | 0.259946302 | 0.052127052 |
| Pdcd6 | 9.594123 | 9.648629 | 9.6289 | 0.01970309 | -0.034802473 | 9.74023 | 9.8269476 | 9.56706 | 0.259889915 | 0.173172392 |
| Bmyc | 7.38203 | 7.583277 | 7.4455 | 0.137738185 | -0.063508839 | 7.9245045 | 7.5466038 | 7.28694 | 0.259662189 | 0.637562817 |
| Isg20l2 | 9.912658 | 9.961734 | 9.9218 | 0.03998038 | -0.009095356 | 9.9014467 | 10.164616 | 9.90503 | 0.25958986 | -0.003579683 |
| Srp19 | 8.853677 | 8.812127 | 8.8175 | -0.005326544 | 0.03622339 | 8.9075361 | 9.1058591 | 8.84639 | 0.259471387 | 0.061148331 |
| Mgrn1 | 11.03291 | 11.06846 | 11.026 | 0.042194564 | 0.006647802 | 11.259396 | 11.263762 | 11.0044 | 0.259410992 | 0.255044125 |
| Zfp292 | 11.11049 | 11.2071 | 11.179 | 0.028364294 | -0.06824093 | 10.956986 | 11.464842 | 11.2055 | 0.259378304 | -0.248477866 |
| Gm21962 | 2.9551 | 2.987636 | 3.0122 | -0.024541944 | -0.057078672 | 2.997782 | 3.2564535 | 2.99718 | 0.259277915 | 0.000606403 |
| Plekha1 | 10.03043 | 10.09155 | 9.9679 | 0.123689927 | 0.062569776 | 10.009887 | 10.359755 | 10.1006 | 0.259166424 | -0.090701876 |
| Mis12 | 9.293511 | 9.284775 | 9.2689 | 0.015858713 | 0.024594645 | 9.146573 | 9.537188 | 9.27805 | 0.259138528 | -0.131476458 |
| Lcmt2 | 7.354295 | 7.148461 | 7.3023 | -0.153814013 | 0.052019582 | 7.0461491 | 7.1789453 | 6.91987 | 0.259072193 | 0.126275992 |
| Zswim1 | 7.204986 | 7.189059 | 7.3126 | -0.123538537 | -0.107611877 | 7.2975234 | 7.305303 | 7.04632 | 0.258978269 | 0.251198603 |

| | | | | | | | | | | |
|---|---|---|---|---|---|---|---|---|---|---|
| Tpm3 | 13.38808 | 13.29257 | 13.26 | 0.032178778 | 0.127694368 | 13.439524 | 13.57284 | 13.3139 | 0.258968444 | 0.12565256 |
| Tagln2 | 11.94083 | 12.01876 | 11.917 | 0.10163591 | 0.023708478 | 11.912598 | 12.149444 | 11.8906 | 0.258853044 | 0.022006529 |
| Tmtc2 | 5.873584 | 5.925328 | 6.028 | -0.102708208 | -0.154452396 | 5.539137 | 6.1786965 | 5.91984 | 0.258851901 | -0.380707625 |
| Mapk13 | 3.658959 | 3.553 | 3.5729 | -0.019945526 | 0.086013479 | 3.9131016 | 3.704464 | 3.44567 | 0.258797123 | 0.467434815 |
| Slc25a26 | 6.72284 | 6.42934 | 6.6085 | -0.179119209 | 0.114381414 | 6.6925976 | 6.84311 | 6.58432 | 0.258792069 | 0.10827967 |
| Mcoln3 | 7.248453 | 7.423036 | 7.3637 | 0.059328893 | -0.115253839 | 7.2902937 | 7.5892946 | 7.33056 | 0.25873202 | -0.040268901 |
| Sumo2 | 11.69841 | 11.64795 | 11.666 | -0.018406556 | 0.032054586 | 11.726655 | 11.871931 | 11.6133 | 0.258672476 | 0.113396697 |
| Erap1 | 11.64882 | 11.60004 | 11.54 | 0.059695098 | 0.10847628 | 11.445201 | 11.755443 | 11.4968 | 0.258662668 | -0.051579468 |
| Wdr36 | 9.861649 | 9.866132 | 9.828 | 0.038089043 | 0.033605885 | 9.917734 | 10.058035 | 9.79956 | 0.258471977 | 0.118170539 |
| Olfr539 | 5.781161 | 5.748935 | 5.6478 | 0.101173731 | 0.133400329 | 6.473763 | 6.3751778 | 6.11679 | 0.25838378 | 0.356968977 |
| Bcl9 | 9.540699 | 9.583774 | 9.4866 | 0.09720245 | 0.05412738 | 9.3771011 | 9.9579072 | 9.69958 | 0.258324003 | -0.322482178 |
| Cebpa | 7.355597 | 7.12623 | 6.9923 | 0.133966251 | 0.363333101 | 6.9555215 | 6.8838933 | 6.62564 | 0.25825199 | 0.329880142 |
| Pold1 | 10.35345 | 10.34923 | 10.394 | -0.044686319 | -0.040462408 | 10.82746 | 10.667343 | 10.4092 | 0.25818023 | 0.418296353 |
| Gm38356 | 5.614486 | 5.62223 | 5.7258 | -0.103598941 | -0.111343893 | 5.6761841 | 6.3375353 | 6.07936 | 0.258175086 | -0.403176101 |
| Dynlt1f | 8.535451 | 8.442228 | 8.5565 | -0.114271433 | -0.021048998 | 8.9570881 | 8.6784737 | 8.42043 | 0.258039868 | 0.53665426 |
| Fbxo46 | 8.519274 | 8.722562 | 8.5198 | 0.202739499 | -0.000548265 | 8.7453467 | 8.7415061 | 8.48352 | 0.257982742 | 0.261823342 |
| Cd9 | 9.386268 | 9.20466 | 9.1738 | 0.030888567 | 0.212496361 | 8.8751009 | 9.103387 | 8.84556 | 0.257824591 | 0.029538483 |
| Srprb | 9.220466 | 9.340605 | 9.0964 | 0.244252902 | 0.124113755 | 9.393776 | 9.5360678 | 9.27839 | 0.257680576 | 0.115388813 |
| Nudt1 | 6.573143 | 6.763226 | 6.6998 | 0.063466581 | -0.12661643 | 6.5972434 | 6.8354437 | 6.57794 | 0.257503756 | 0.019303431 |
| Pram1 | 5.248208 | 5.102287 | 5.0009 | 0.101397206 | 0.247317963 | 5.291799 | 5.1995063 | 4.94204 | 0.257467173 | 0.349759896 |
| Mcts1 | 8.983806 | 8.911382 | 8.8782 | 0.033138029 | 0.105562419 | 9.0974931 | 9.2844223 | 9.02697 | 0.257451866 | 0.070522723 |
| Cox4i1 | 11.71421 | 11.68788 | 11.642 | 0.045806532 | 0.07213431 | 11.829917 | 11.881283 | 11.6238 | 0.257451052 | 0.206084697 |
| Rbm10 | 10.45458 | 10.63345 | 10.529 | 0.104483384 | -0.074386726 | 10.645212 | 10.807734 | 10.5503 | 0.25739684 | 0.094874738 |
| Wdr83os | 9.120026 | 9.18061 | 9.1796 | 0.001012657 | -0.059572038 | 9.5493676 | 9.3859317 | 9.1288 | 0.257132598 | 0.420568494 |
| Lnx2 | 9.443431 | 9.48347 | 9.4648 | 0.018709504 | -0.021329958 | 9.2291215 | 9.4927607 | 9.23571 | 0.257055025 | -0.006584203 |
| Vrk3 | 9.985581 | 10.0257 | 10.051 | -0.0255349 | -0.065650262 | 10.194308 | 10.21109 | 9.95412 | 0.256970749 | 0.240188505 |
| Dapk2 | 4.972513 | 4.935965 | 4.8139 | 0.122071387 | 0.158618683 | 4.6647816 | 5.1663655 | 4.90941 | 0.256959222 | -0.244624672 |
| Gm2000 | 11.88002 | 11.94202 | 11.905 | 0.036976592 | -0.025028202 | 11.985618 | 12.13625 | 11.8793 | 0.25694016 | 0.106307512 |
| Yju2 | 7.92556 | 7.994143 | 7.8598 | 0.134376697 | 0.065794269 | 8.1419697 | 8.2473657 | 7.99046 | 0.256904004 | 0.151508002 |
| Psme3 | 11.06344 | 11.1686 | 11.098 | 0.070578154 | -0.034584142 | 11.022731 | 11.323412 | 11.0668 | 0.256614364 | -0.044066658 |
| Apol11a | 2.766097 | 2.621256 | 2.6774 | -0.056156887 | 0.088683993 | 2.566797 | 2.8364317 | 2.57982 | 0.256609431 | -0.013025269 |
| Kat8 | 8.70622 | 8.868926 | 8.7795 | 0.08944185 | -0.073263584 | 8.9655611 | 9.1444266 | 8.88783 | 0.25659639 | 0.077730902 |

| | | | | | | | | | | |
|---|---|---|---|---|---|---|---|---|---|---|
| Bcap31 | 10.6272 | 10.68502 | 10.617 | 0.06819734 | 0.010375408 | 10.835731 | 10.861498 | 10.6049 | 0.25657538 | 0.230808354 |
| Usp21 | 8.919591 | 9.05875 | 9.0444 | 0.014304763 | -0.124854108 | 9.2022118 | 9.3631202 | 9.1066 | 0.256524704 | 0.095616365 |
| Dapk3 | 8.80225 | 8.849932 | 8.6963 | 0.153663835 | 0.105982133 | 8.8745513 | 8.8592959 | 8.60282 | 0.256474559 | 0.271729892 |
| Acbd6 | 8.517625 | 8.457644 | 8.4665 | -0.008821684 | 0.051159196 | 8.5957735 | 8.6206471 | 8.36425 | 0.256398027 | 0.231524442 |
| Tmem208 | 8.328399 | 8.458343 | 8.3454 | 0.112907626 | -0.017036708 | 8.7433369 | 8.7454643 | 8.48927 | 0.256194624 | 0.254067239 |
| Wdr89 | 7.837393 | 7.820743 | 7.7744 | 0.046330673 | 0.062980721 | 8.0961658 | 8.0715724 | 7.81548 | 0.256088501 | 0.280681869 |
| Psmc3ip | 5.633611 | 5.708852 | 5.595 | 0.113895163 | 0.038653889 | 5.7450677 | 5.9559497 | 5.7 | 0.255951864 | 0.045069879 |
| H2-Q1 | 2.990058 | 2.858419 | 2.9185 | -0.06012653 | 0.071512703 | 2.871252 | 3.0558821 | 2.80008 | 0.255799899 | 0.071169844 |
| Kat6a | 12.27901 | 12.51765 | 12.224 | 0.29378991 | 0.055149706 | 12.114819 | 12.478661 | 12.223 | 0.255620074 | -0.108222169 |
| Cggbp1 | 11.15492 | 11.22374 | 11.155 | 0.068367672 | -0.000447606 | 11.050098 | 11.29651 | 11.041 | 0.255543448 | 0.009130903 |
| Epn1 | 11.00439 | 10.94493 | 10.965 | -0.020446396 | 0.039007106 | 11.214756 | 11.177219 | 10.9219 | 0.255342978 | 0.292879894 |
| Ighv9-4 | 5.360501 | 5.494654 | 5.2223 | 0.272330178 | 0.138176785 | 5.1357686 | 5.3503975 | 5.09506 | 0.25533832 | 0.040709419 |
| Lyrm1 | 6.434067 | 6.306446 | 6.3343 | -0.027836774 | 0.099784251 | 6.4190633 | 6.5544593 | 6.2992 | 0.255254869 | 0.119858838 |
| Dedd | 8.968913 | 9.084938 | 9.0207 | 0.064213202 | -0.051811723 | 8.8396235 | 9.261489 | 9.00634 | 0.255145418 | -0.166720098 |
| Emc10 | 10.21 | 10.20848 | 10.102 | 0.106964259 | 0.10848575 | 10.430853 | 10.305085 | 10.05 | 0.255122639 | 0.380889907 |
| Phb2 | 10.9398 | 10.92146 | 10.962 | -0.040904998 | -0.022566654 | 11.201033 | 11.201456 | 10.9465 | 0.25498525 | 0.254562274 |
| Gm29216 | 13.73261 | 13.7249 | 13.941 | -0.216472648 | -0.208762666 | 13.774742 | 14.063598 | 13.8087 | 0.254923824 | -0.03393158 |
| Smim27 | 6.169014 | 6.293549 | 6.2641 | 0.029423758 | -0.095111669 | 6.4460967 | 6.5461641 | 6.29126 | 0.254901398 | 0.154833944 |
| Elovl5 | 10.65458 | 10.81401 | 10.721 | 0.093056754 | -0.06636802 | 10.464942 | 10.893505 | 10.6387 | 0.254852742 | -0.173710631 |
| Pfkfb3 | 11.31038 | 11.56617 | 11.421 | 0.14477067 | -0.111018344 | 11.078285 | 11.683816 | 11.4292 | 0.254655676 | -0.350875424 |
| Scamp2 | 10.13846 | 10.19619 | 10.154 | 0.042015386 | -0.01571358 | 10.347188 | 10.539478 | 10.2848 | 0.254651384 | 0.062361298 |
| Elovl1 | 9.312796 | 9.346879 | 9.368 | -0.021094815 | -0.055177481 | 9.6637957 | 9.6228278 | 9.36819 | 0.254642449 | 0.295610375 |
| Gemin6 | 6.80287 | 6.864884 | 6.8025 | 0.062427747 | 0.000413488 | 6.7824014 | 7.0506977 | 6.79607 | 0.254630416 | -0.013665888 |
| Hmgn1 | 10.94482 | 11.01752 | 11.051 | -0.03339766 | -0.106097836 | 11.215285 | 11.284097 | 11.0295 | 0.25460278 | 0.185791141 |
| Immt | 10.71801 | 10.78894 | 10.675 | 0.113592486 | 0.042655992 | 10.689075 | 10.97236 | 10.7178 | 0.254598056 | -0.028686827 |
| Cchcr1 | 6.803189 | 6.964416 | 6.8413 | 0.123165619 | -0.038061426 | 6.8056196 | 7.2850122 | 7.0306 | 0.254410403 | -0.224982239 |
| Gm3200 | 2.255491 | 2.330075 | 2.2104 | 0.119716157 | 0.045131944 | 2.1978571 | 2.4458112 | 2.1915 | 0.254312489 | 0.006358382 |
| Zfp35 | 8.801225 | 8.799194 | 8.8014 | -0.002252467 | -0.000221645 | 8.8375833 | 9.0026236 | 8.74833 | 0.254288848 | 0.089248581 |
| Hint1 | 10.14692 | 10.09297 | 10.107 | -0.013720284 | 0.040229546 | 10.23192 | 10.275595 | 10.0214 | 0.25421581 | 0.210540655 |
| Prr36 | 3.129764 | 3.220422 | 3.1654 | 0.055026508 | -0.035631991 | 3.2144148 | 3.4934461 | 3.2393 | 0.254144487 | -0.02488677 |
| Tmem258 | 9.113613 | 9.152427 | 9.1071 | 0.045301537 | 0.006487367 | 9.3797877 | 9.4223986 | 9.16827 | 0.254127076 | 0.211516157 |
| Synpo | 7.516585 | 7.304102 | 7.0467 | 0.257423676 | 0.469905929 | 6.6660243 | 7.0863312 | 6.83228 | 0.254049489 | -0.166257405 |

| | | | | | | | | | | |
|---|---|---|---|---|---|---|---|---|---|---|
| Dbndd2 | 6.63005 | 6.666562 | 6.4948 | 0.171720446 | 0.135208489 | 6.755296 | 6.7470565 | 6.49311 | 0.253941624 | 0.262181183 |
| Mcat | 7.600831 | 7.786848 | 7.7277 | 0.059115781 | -0.126902009 | 7.9288239 | 7.9026469 | 7.64913 | 0.253520522 | 0.27969753 |
| Asb8 | 8.448985 | 8.369477 | 8.4871 | -0.117642295 | -0.038133982 | 8.5135012 | 8.4837976 | 8.23041 | 0.253387044 | 0.283090601 |
| Psmd1 | 10.69823 | 10.68602 | 10.64 | 0.045720718 | 0.057934422 | 10.696553 | 10.885195 | 10.6318 | 0.253383632 | 0.06474168 |
| Nudcd3 | 10.43995 | 10.31523 | 10.378 | -0.062337818 | 0.062383554 | 10.473208 | 10.655792 | 10.4024 | 0.253373878 | 0.070789 |
| Stambp | 7.808052 | 7.915805 | 7.9164 | -0.00059174 | -0.108344432 | 7.7101958 | 8.0772383 | 7.82393 | 0.253307047 | -0.113735484 |
| Wdr3 | 10.1808 | 10.31377 | 10.143 | 0.170536237 | 0.037559024 | 10.130436 | 10.456619 | 10.2034 | 0.253181348 | -0.073001997 |
| Cdkl5 | 6.567599 | 6.823285 | 6.636 | 0.187262161 | -0.0684238 | 6.6033561 | 6.8354127 | 6.58227 | 0.253140423 | 0.021083777 |
| Cdipt | 10.19561 | 10.23254 | 10.167 | 0.065968639 | 0.029039804 | 10.397771 | 10.447178 | 10.1941 | 0.2531188 | 0.203711385 |
| Ier5 | 11.69987 | 11.7162 | 11.717 | -0.000766478 | -0.01709493 | 11.322522 | 11.604111 | 11.351 | 0.253061854 | -0.028526542 |
| Creb3l1 | 4.874959 | 4.88052 | 4.7546 | 0.12591165 | 0.120350882 | 4.9452886 | 5.0017002 | 4.74903 | 0.252667653 | 0.196256077 |
| Frg1 | 9.470185 | 9.621632 | 9.4819 | 0.13970603 | -0.011741129 | 9.6315714 | 9.8094944 | 9.55714 | 0.252349972 | 0.074427032 |
| Hgh1 | 8.06435 | 8.113248 | 8.0157 | 0.097582897 | 0.048685157 | 8.3186556 | 8.4139899 | 8.16167 | 0.252323996 | 0.156989706 |
| Ssna1 | 8.254111 | 8.242761 | 8.3136 | -0.070827725 | -0.059477863 | 8.5634726 | 8.5008976 | 8.24864 | 0.25225282 | 0.314827854 |
| Rad23a | 10.11358 | 10.17988 | 10.138 | 0.041762854 | -0.024541317 | 10.425596 | 10.421753 | 10.1696 | 0.252202632 | 0.256045849 |
| Igkv6-13 | 5.956332 | 6.455274 | 6.1717 | 0.283578623 | -0.215362613 | 6.4387921 | 6.3557614 | 6.10361 | 0.252154643 | 0.335185373 |
| Tent5c | 9.989056 | 10.2369 | 10.067 | 0.169456317 | -0.078389483 | 9.4299803 | 10.190663 | 9.93855 | 0.252108705 | -0.508573703 |
| Rabggtb | 9.509037 | 9.583981 | 9.5517 | 0.032321064 | -0.042623228 | 9.7894819 | 9.9583127 | 9.70633 | 0.251982235 | 0.083151513 |
| Fap | 4.995614 | 5.04708 | 5.0185 | 0.028585687 | -0.022880989 | 5.0810269 | 5.2341401 | 4.98218 | 0.251960421 | 0.098847213 |
| Ccl21b | 4.440616 | 4.245165 | 3.9458 | 0.299319217 | 0.494771089 | 4.0508745 | 3.8593638 | 3.60803 | 0.251336834 | 0.442847459 |
| Taok2 | 10.89663 | 11.03995 | 10.875 | 0.165051966 | 0.021733352 | 11.135125 | 11.276808 | 11.0258 | 0.251051408 | 0.109368582 |
| Hist1h4d | 5.083474 | 5.055233 | 5.2689 | -0.213628185 | -0.185387103 | 5.6953663 | 5.5179477 | 5.26716 | 0.250790765 | 0.428209357 |
| Ndufaf5 | 7.076716 | 7.08673 | 7.2015 | -0.11479211 | -0.124806404 | 7.2912728 | 7.4526665 | 7.20254 | 0.250124124 | 0.088730413 |
| Hnrnpm | 12.04445 | 12.05408 | 12.053 | 0.000647532 | -0.008982356 | 11.993564 | 12.327452 | 12.0773 | 0.250117446 | -0.083770566 |
| Cops5 | 9.481497 | 9.498031 | 9.4202 | 0.077842474 | 0.061309084 | 9.6266053 | 9.768899 | 9.51905 | 0.249847176 | 0.107553422 |
| Cdk4 | 10.32693 | 10.32788 | 10.419 | -0.090910508 | -0.091863624 | 10.599857 | 10.601163 | 10.3513 | 0.24984131 | 0.248535519 |
| Pdss1 | 6.223004 | 6.040675 | 6.186 | -0.145285969 | 0.037043111 | 6.1854141 | 6.6254826 | 6.37566 | 0.249824221 | -0.190244267 |
| Rps23 | 13.39621 | 13.36448 | 13.415 | -0.050753664 | -0.019024446 | 13.528771 | 13.642147 | 13.3924 | 0.249790722 | 0.136415404 |
| Olfr635 | 6.03146 | 6.085299 | 6.1599 | -0.074647651 | -0.128486703 | 6.2621708 | 6.6500451 | 6.40041 | 0.249634787 | -0.138239485 |
| Ddx28 | 6.982661 | 7.074548 | 7.0949 | -0.020338061 | -0.1122249 | 7.1999382 | 7.2453663 | 6.99578 | 0.249582107 | 0.204154036 |
| Rpp38 | 6.309336 | 6.276915 | 6.2294 | 0.047542953 | 0.079963603 | 6.3564876 | 6.595315 | 6.34585 | 0.249460542 | 0.010633151 |
| Cpa6 | 4.228606 | 4.412093 | 4.3155 | 0.096638658 | -0.086848489 | 4.1695443 | 4.6532462 | 4.40382 | 0.249424669 | -0.234277288 |

| | | | | | | | | | | |
|---|---|---|---|---|---|---|---|---|---|---|
| Pitpna | 11.62395 | 11.72888 | 11.581 | 0.148091376 | 0.043160498 | 11.769807 | 11.827904 | 11.5785 | 0.24937692 | 0.191279616 |
| Trappc6a | 7.640521 | 7.574009 | 7.7131 | -0.139136807 | -0.07262509 | 7.9863485 | 7.8591882 | 7.60989 | 0.249294569 | 0.376454889 |
| Olfr76 | 3.038165 | 3.271203 | 3.1499 | 0.121341609 | -0.111696342 | 3.1508699 | 3.4689125 | 3.2198 | 0.249115753 | -0.068926894 |
| Gm9794 | 10.74032 | 10.56197 | 10.346 | 0.216124384 | 0.394475401 | 10.485497 | 10.6979 | 10.4489 | 0.248977117 | 0.036574027 |
| Pgls | 9.324096 | 9.360444 | 9.3254 | 0.035018525 | -0.001328903 | 9.5397577 | 9.3964509 | 9.14748 | 0.248966089 | 0.392272899 |
| Eif3j2 | 9.500277 | 9.630349 | 9.4329 | 0.197491219 | 0.067419975 | 9.4208648 | 9.7251781 | 9.47634 | 0.24884244 | -0.055470845 |
| Eif3j1 | 9.202133 | 9.232959 | 9.1214 | 0.111556832 | 0.080730675 | 9.1436744 | 9.3180371 | 9.0692 | 0.248833369 | 0.074470738 |
| Plod2 | 4.936454 | 4.761214 | 4.796 | -0.034738409 | 0.140502061 | 4.5759189 | 4.8701442 | 4.6214 | 0.248743733 | -0.045481587 |
| Ndufb5 | 9.483897 | 9.424447 | 9.4414 | -0.016971437 | 0.04247861 | 9.6561166 | 9.6809831 | 9.43228 | 0.248700945 | 0.223834473 |
| Gba | 9.951554 | 9.975663 | 9.9596 | 0.016023758 | -0.008084981 | 10.062146 | 10.183723 | 9.93505 | 0.248674865 | 0.127098117 |
| Olfr914 | 4.219114 | 4.315295 | 4.2944 | 0.020938579 | -0.075242897 | 4.250476 | 4.6113716 | 4.36293 | 0.248436607 | -0.112458956 |
| Ighv1-34 | 4.59319 | 4.3341 | 4.6317 | -0.297552892 | -0.038463465 | 4.6013031 | 4.664079 | 4.41571 | 0.248366211 | 0.185590349 |
| Hnrnpa2b1 | 13.70209 | 13.71496 | 13.608 | 0.106527862 | 0.093656384 | 13.781728 | 14.07689 | 13.8285 | 0.248353782 | -0.046808839 |
| Lima1 | 9.263811 | 8.938157 | 8.9392 | -0.001049819 | 0.324604426 | 8.6169279 | 8.9630969 | 8.71482 | 0.248273163 | -0.097895869 |
| Cdk9 | 10.65102 | 10.72653 | 10.676 | 0.050040908 | -0.025472558 | 10.820634 | 10.98215 | 10.7339 | 0.248228344 | 0.08671188 |
| Pdlim1 | 10.43721 | 10.31269 | 10.232 | 0.080527375 | 0.205044751 | 10.094389 | 10.253058 | 10.0048 | 0.248222192 | 0.089553258 |
| A630089N07Rik | 6.743688 | 7.057705 | 6.8816 | 0.176132744 | -0.137884957 | 6.9581649 | 7.3136451 | 7.06555 | 0.248095745 | -0.107384538 |
| Zfp169 | 7.797965 | 8.028005 | 7.7218 | 0.306218556 | 0.076178749 | 8.1209866 | 8.5050315 | 8.25716 | 0.247869888 | -0.136175022 |
| Rps6 | 14.71164 | 14.74956 | 14.644 | 0.105254084 | 0.067335356 | 14.809793 | 14.84715 | 14.5995 | 0.247674684 | 0.210317854 |
| Fggy | 4.833746 | 4.875097 | 4.92 | -0.044863238 | -0.086214174 | 4.6287346 | 4.8884822 | 4.64083 | 0.247654411 | -0.012093168 |
| Zmynd19 | 8.219602 | 8.339644 | 8.2809 | 0.058766144 | -0.061275355 | 8.2097151 | 8.5382731 | 8.29068 | 0.247595666 | -0.080962363 |
| Nudt9 | 8.716436 | 8.860867 | 8.8072 | 0.053692492 | -0.090738428 | 8.7623389 | 9.0348146 | 8.78729 | 0.247525768 | -0.024949993 |
| Ywhah | 11.09688 | 11.29632 | 11.099 | 0.197386204 | -0.002050758 | 11.11361 | 11.364398 | 11.1169 | 0.247497804 | -0.003289659 |
| Bdp1 | 10.62413 | 10.84698 | 10.659 | 0.187891878 | -0.034951534 | 10.596424 | 10.940974 | 10.6935 | 0.247463822 | -0.097086682 |
| Ndufb11 | 9.863996 | 9.789722 | 9.8724 | -0.082635877 | -0.008362554 | 9.9397199 | 9.9902207 | 9.74284 | 0.247380017 | 0.196879227 |
| Zcchc10 | 7.586394 | 7.692707 | 7.6863 | 0.006450788 | -0.099862569 | 7.6514345 | 7.8486907 | 7.60145 | 0.247240921 | 0.049984691 |
| Aar2 | 8.758836 | 8.840395 | 8.6962 | 0.144216059 | 0.062657034 | 8.8223469 | 8.8417677 | 8.59453 | 0.247238645 | 0.227817855 |
| Oaz1-ps | 8.135588 | 7.863668 | 7.9943 | -0.130657795 | 0.141262277 | 8.4548789 | 8.1158024 | 7.86861 | 0.247196473 | 0.58627294 |
| Rad1 | 7.620535 | 7.795165 | 7.8492 | -0.05400635 | -0.228637077 | 7.8948267 | 7.9876608 | 7.74054 | 0.247122649 | 0.154288593 |
| 2010005H15Rik | 2.193715 | 2.116797 | 2.104 | 0.012774839 | 0.089692681 | 2.0757752 | 2.3250225 | 2.07796 | 0.247063719 | -0.002183606 |
| Igkv10-95 | 4.431977 | 4.615159 | 4.6203 | -0.005120404 | -0.188302205 | 4.7249547 | 4.8382605 | 4.59133 | 0.246935444 | 0.133629606 |
| Des | 7.843828 | 7.910763 | 7.4042 | 0.506526874 | 0.439592544 | 7.3536715 | 7.0500977 | 6.80325 | 0.246851841 | 0.550425633 |

| | | | | | | | | | | |
|---|---|---|---|---|---|---|---|---|---|---|
| Mknk1 | 9.55097 | 9.465013 | 9.4767 | -0.011704217 | 0.074253282 | 9.6365054 | 9.6797425 | 9.4329 | 0.246847113 | 0.203610017 |
| Igkv4-58 | 5.533257 | 5.572338 | 5.6673 | -0.09496161 | -0.134043259 | 5.75957 | 5.7848951 | 5.53826 | 0.246636064 | 0.221310963 |
| Alkbh6 | 8.141941 | 8.219182 | 8.1973 | 0.021881509 | -0.055360039 | 8.3508836 | 8.519308 | 8.27277 | 0.2465416 | 0.078117282 |
| Gm44578 | 2.815233 | 2.742391 | 2.7602 | -0.017814 | 0.05502814 | 2.7493028 | 3.0042921 | 2.75779 | 0.246505469 | -0.0084838 |
| Ackr1 | 7.90016 | 7.648061 | 7.7488 | -0.100769667 | 0.151329547 | 6.5633489 | 6.9597032 | 6.71327 | 0.246432968 | -0.149921306 |
| Ccnl1 | 11.60284 | 11.70252 | 11.626 | 0.076113458 | -0.023566768 | 11.635975 | 12.101466 | 11.8551 | 0.246324602 | -0.219166428 |
| Mid1ip1 | 8.260304 | 8.150931 | 8.2483 | -0.097391137 | 0.011981689 | 8.4362125 | 8.3893632 | 8.14319 | 0.246177735 | 0.293026949 |
| Rpl11 | 13.78127 | 13.80343 | 13.779 | 0.024526798 | 0.002367494 | 13.883367 | 13.967159 | 13.721 | 0.246168384 | 0.16237613 |
| Ppie | 9.131205 | 9.102906 | 9.0699 | 0.033002744 | 0.061302095 | 9.4281616 | 9.3981112 | 9.15194 | 0.246167634 | 0.276218002 |
| Tm9sf1 | 9.80703 | 9.761603 | 9.7323 | 0.029335612 | 0.074762255 | 9.913423 | 9.9426267 | 9.69651 | 0.246121151 | 0.21691743 |
| Uqcrfs1 | 10.11956 | 10.04351 | 10.075 | -0.031685589 | 0.044371025 | 10.13887 | 10.243246 | 9.99731 | 0.245935746 | 0.141559785 |
| Polr1a | 10.61374 | 10.78099 | 10.592 | 0.188942856 | 0.021686658 | 10.612861 | 10.94351 | 10.6976 | 0.24588139 | -0.084767061 |
| Rnpc3 | 7.828477 | 7.828894 | 7.8191 | 0.0098207 | 0.00940363 | 8.0270541 | 8.1938516 | 7.94801 | 0.24583819 | 0.079040693 |
| Golga7 | 10.04871 | 10.12127 | 10.091 | 0.030104148 | -0.04245253 | 10.115335 | 10.268611 | 10.0228 | 0.245837593 | 0.092562067 |
| Fem1a | 10.07477 | 10.13661 | 10.041 | 0.096090542 | 0.034242641 | 10.162936 | 10.238628 | 9.99284 | 0.245783929 | 0.170091927 |
| Vmn2r89 | 4.001048 | 4.133165 | 4.0359 | 0.097218923 | -0.034898364 | 4.112785 | 4.449078 | 4.20333 | 0.245748347 | -0.090544683 |
| Msl3l2 | 5.375688 | 5.318272 | 5.3584 | -0.040109279 | 0.017307219 | 5.4181037 | 5.5721978 | 5.32651 | 0.24568799 | 0.091593929 |
| Ccdc97 | 9.993196 | 9.996594 | 10.035 | -0.038635219 | -0.042033095 | 10.156433 | 10.288035 | 10.0424 | 0.245609808 | 0.114007695 |
| Tnfrsf21 | 7.500942 | 7.270428 | 6.9797 | 0.290704878 | 0.521219086 | 7.3362986 | 7.0392113 | 6.79371 | 0.245504927 | 0.542592242 |
| Nfu1 | 8.81544 | 8.83824 | 8.8745 | -0.036235172 | -0.059035698 | 9.2443598 | 9.1641522 | 8.91865 | 0.245503266 | 0.325710848 |
| Cherp | 10.67465 | 10.7141 | 10.773 | -0.05861402 | -0.098055656 | 10.894923 | 10.965727 | 10.7203 | 0.24545388 | 0.174650197 |
| Psmd2 | 11.54942 | 11.59717 | 11.493 | 0.104493426 | 0.056745124 | 11.589237 | 11.764877 | 11.5195 | 0.245396158 | 0.069756495 |
| Zbtb7c | 3.746981 | 3.770197 | 3.5081 | 0.262126455 | 0.238910623 | 3.6795442 | 3.8072908 | 3.56191 | 0.245381126 | 0.117634585 |
| Dhh | 3.308022 | 3.250686 | 3.0643 | 0.186418742 | 0.243754604 | 3.0639764 | 3.3132638 | 3.06799 | 0.245271401 | -0.004016017 |
| Tubb4b | 10.82423 | 10.88221 | 10.858 | 0.023729382 | -0.034252572 | 10.885261 | 11.020396 | 10.7752 | 0.245197886 | 0.11006257 |
| Gm49391 | 7.225206 | 6.849046 | 6.8917 | -0.042694864 | 0.33346552 | 6.5238033 | 6.8438917 | 6.59875 | 0.245142037 | -0.074946394 |
| Mpp1 | 9.916595 | 9.988678 | 9.837 | 0.151633653 | 0.079550674 | 9.8634153 | 10.160997 | 9.91601 | 0.244987831 | -0.052593377 |
| Safb2 | 11.19348 | 11.30668 | 11.179 | 0.127277346 | 0.014074114 | 11.472622 | 11.776489 | 11.5316 | 0.244907982 | -0.05895922 |
| Chmp1a | 10.41264 | 10.4038 | 10.419 | -0.01540693 | -0.006565516 | 10.60563 | 10.546445 | 10.3016 | 0.244887984 | 0.304072828 |
| Comtd1 | 5.912544 | 6.103701 | 6.0327 | 0.07104116 | -0.120116047 | 6.2181594 | 6.1950304 | 5.95015 | 0.244877418 | 0.268006446 |
| Ndrg3 | 10.67614 | 10.62343 | 10.619 | 0.004285936 | 0.056993624 | 10.579085 | 10.775837 | 10.5311 | 0.244756918 | 0.04800548 |
| Exoc7 | 9.430314 | 9.249482 | 9.4131 | -0.163607473 | 0.017224726 | 9.3269559 | 9.5092957 | 9.26464 | 0.2446573 | 0.062317426 |

| | | | | | | | | | | |
|---|---|---|---|---|---|---|---|---|---|---|
| Bsg | 10.60164 | 10.57092 | 10.62 | -0.049415326 | -0.018691268 | 10.653064 | 10.732966 | 10.4884 | 0.244591012 | 0.164688826 |
| Naa15 | 11.11527 | 11.25165 | 11.203 | 0.049022794 | -0.087363868 | 11.005619 | 11.391486 | 11.1469 | 0.244561208 | -0.141305597 |
| Gm14542 | 1.151608 | 1.21452 | 1.1954 | 0.019092282 | -0.043819867 | 1.1606184 | 1.353904 | 1.10938 | 0.244527545 | 0.051241936 |
| Vps25 | 9.822484 | 9.787008 | 9.8243 | -0.03726813 | -0.001792436 | 10.123066 | 10.041696 | 9.79738 | 0.24431706 | 0.325686322 |
| Cdk11b | 11.07931 | 11.17927 | 11.101 | 0.078304414 | -0.021650426 | 11.220178 | 11.542567 | 11.2984 | 0.24420564 | -0.078183969 |
| Bloc1s1 | 8.972939 | 8.975706 | 8.9722 | 0.00353462 | 0.000768061 | 9.30868 | 9.2472433 | 9.00318 | 0.244064446 | 0.305501139 |
| Sdc4 | 10.75458 | 10.94205 | 10.568 | 0.374064166 | 0.186590292 | 10.904508 | 10.754384 | 10.5104 | 0.243947186 | 0.394071015 |
| Klrc3 | 2.585869 | 2.438542 | 2.4869 | -0.048350318 | 0.098976569 | 2.4608065 | 2.6119051 | 2.36808 | 0.24382124 | 0.092722651 |
| Rrn3 | 9.990637 | 10.06163 | 9.9472 | 0.114416841 | 0.043420345 | 9.9996861 | 10.148441 | 9.90477 | 0.243666536 | 0.094911509 |
| Gm47036 | 3.552076 | 3.49432 | 3.5046 | -0.010247876 | 0.047507801 | 3.850046 | 3.6789144 | 3.43552 | 0.243394632 | 0.414526232 |
| Prpf40a | 11.1522 | 11.15084 | 11.045 | 0.10582939 | 0.107197512 | 11.102988 | 11.324675 | 11.0814 | 0.243225494 | 0.021538419 |
| Zc3h12d | 9.6291 | 9.648172 | 9.5874 | 0.060761516 | 0.041689541 | 9.6369903 | 9.7871943 | 9.54403 | 0.243159959 | 0.092955928 |
| H2-Oa | 11.14992 | 11.11001 | 11.252 | -0.142261148 | -0.102350526 | 11.681218 | 11.422738 | 11.1797 | 0.243082224 | 0.501561462 |
| Vmn2r95 | 4.575216 | 4.655852 | 4.7616 | -0.105716386 | -0.186353316 | 4.7453645 | 5.1018074 | 4.8588 | 0.243008501 | -0.113434325 |
| Gm9762 | 7.535912 | 7.571848 | 7.7151 | -0.143298963 | -0.179235146 | 7.6832975 | 7.6310924 | 7.38819 | 0.242903971 | 0.295109003 |
| Bag5 | 8.685651 | 8.685577 | 8.7297 | -0.044077269 | -0.04400358 | 8.7044118 | 8.7661983 | 8.5234 | 0.242799998 | 0.181013528 |
| Commd3 | 9.210592 | 9.183339 | 9.3373 | -0.153943942 | -0.126690151 | 9.5147566 | 9.5997515 | 9.35705 | 0.242703183 | 0.157708295 |
| Cltb | 8.655408 | 8.672567 | 8.534 | 0.138593153 | 0.121433851 | 8.5676356 | 8.5809376 | 8.3383 | 0.242641671 | 0.229339679 |
| Usp31 | 8.827573 | 8.903864 | 8.6833 | 0.220568707 | 0.144278168 | 8.648873 | 9.1256768 | 8.88313 | 0.242545038 | -0.234258735 |
| Ddx51 | 8.937322 | 9.096729 | 9.0718 | 0.024896396 | -0.134510561 | 9.6234386 | 9.5723618 | 9.32984 | 0.242522313 | 0.293599091 |
| Selenok | 9.702814 | 9.749806 | 9.7005 | 0.049321678 | 0.002329343 | 9.6649183 | 9.8134668 | 9.57095 | 0.242516481 | 0.093967986 |
| Htra1 | 7.137391 | 7.218653 | 7.0283 | 0.190304745 | 0.109042761 | 6.725761 | 6.9481414 | 6.70567 | 0.242470734 | 0.020090359 |
| Myo1a | 2.740723 | 2.756227 | 2.7915 | -0.035285978 | -0.050789797 | 2.8552569 | 3.0321298 | 2.78978 | 0.242345484 | 0.06547259 |
| Rrnad1 | 8.590924 | 8.698866 | 8.6696 | 0.029315547 | -0.078627218 | 9.0563097 | 9.1784632 | 8.93625 | 0.242210601 | 0.120057086 |
| Tmem189 | 9.717093 | 9.947173 | 9.7377 | 0.209495995 | -0.020584373 | 10.009972 | 10.048482 | 9.80627 | 0.24220788 | 0.203697973 |
| Ms4a1 | 13.37003 | 13.6051 | 13.362 | 0.242621098 | 0.00755128 | 13.937329 | 13.703654 | 13.4616 | 0.242078448 | 0.475753811 |
| Sez6l2 | 5.979805 | 6.038924 | 5.802 | 0.236907127 | 0.177788742 | 5.9902215 | 6.1896486 | 5.9477 | 0.241948064 | 0.042521056 |
| Zbtb6 | 8.822899 | 8.897828 | 8.8889 | 0.008957991 | -0.065970726 | 9.0749938 | 9.3069844 | 9.06508 | 0.241904384 | 0.009913767 |
| St7 | 7.030205 | 6.953392 | 7.06 | -0.106590167 | -0.029777063 | 6.9191411 | 7.1547177 | 6.91283 | 0.241883583 | 0.006306941 |
| Glrx5 | 7.887627 | 7.77062 | 7.9042 | -0.13357568 | -0.016569124 | 7.9648647 | 7.9724899 | 7.73062 | 0.241867331 | 0.234242222 |
| Prmt7 | 9.108355 | 9.232896 | 9.1542 | 0.078744244 | -0.045796358 | 9.4087787 | 9.4290563 | 9.1874 | 0.24165173 | 0.221374049 |
| Gm12481 | 8.006196 | 8.373962 | 8.2916 | 0.082361999 | -0.285404235 | 8.8344701 | 8.5758769 | 8.33429 | 0.241587299 | 0.500180495 |

| | | | | | | | | | | |
|---|---|---|---|---|---|---|---|---|---|---|
| Rrp1b | 9.23486 | 9.108411 | 9.0514 | 0.057004685 | 0.183453248 | 9.1593176 | 9.4190685 | 9.17751 | 0.241558204 | -0.018192684 |
| Rpl35 | 12.64408 | 12.67723 | 12.726 | -0.049214802 | -0.082355762 | 12.835985 | 12.937403 | 12.6959 | 0.241530668 | 0.140111952 |
| Lars2 | 13.38395 | 13.70734 | 13.842 | -0.134716928 | -0.45810901 | 13.754612 | 14.190398 | 13.9489 | 0.241505954 | -0.19428054 |
| Gtf2b | 9.092237 | 9.17807 | 9.1841 | -0.006059076 | -0.091891634 | 9.1928696 | 9.329938 | 9.08852 | 0.241421671 | 0.104353321 |
| Gm9803 | 6.868667 | 6.858763 | 6.8326 | 0.026133256 | 0.036037211 | 6.8946624 | 7.2252134 | 6.9839 | 0.241310229 | -0.08924079 |
| Abhd8 | 9.102582 | 9.117617 | 9.1348 | -0.017218198 | -0.032252585 | 9.0086649 | 9.1024786 | 8.86126 | 0.241221687 | 0.147408015 |
| Tsacc | 4.772611 | 4.848539 | 4.794 | 0.054555183 | -0.021372043 | 5.1045837 | 5.1034273 | 4.86238 | 0.241051405 | 0.242207863 |
| Uqcc3 | 8.466696 | 8.468817 | 8.489 | -0.020191484 | -0.022312354 | 8.6168702 | 8.5638621 | 8.32286 | 0.241001995 | 0.294010118 |
| Med19 | 8.374557 | 8.449263 | 8.3145 | 0.134727917 | 0.060022186 | 8.4057753 | 8.5037754 | 8.26278 | 0.240994839 | 0.14299476 |
| Zfp933 | 7.242665 | 7.234464 | 7.2305 | 0.00393741 | 0.012138146 | 7.2895052 | 7.5700947 | 7.32911 | 0.240983604 | -0.039605972 |
| Rpl36-ps3 | 2.563835 | 2.506168 | 2.6013 | -0.095148161 | -0.037481236 | 2.6799148 | 2.7670429 | 2.52613 | 0.240915497 | 0.153787443 |
| Gfm1 | 9.294284 | 9.383953 | 9.2685 | 0.115490404 | 0.025820806 | 9.4386719 | 9.5750816 | 9.33419 | 0.240893147 | 0.104483467 |
| Utp18 | 9.602154 | 9.601236 | 9.527 | 0.074247135 | 0.075165408 | 9.6005165 | 9.7859043 | 9.54505 | 0.240858933 | 0.055471061 |
| Znrd1as | 5.061914 | 5.119541 | 5.0176 | 0.101926365 | 0.0442993 | 5.0811186 | 5.3087153 | 5.06798 | 0.240733246 | 0.013136557 |
| Rapgef2 | 9.519188 | 9.689469 | 9.4805 | 0.208973512 | 0.038692518 | 9.0681127 | 9.6892973 | 9.44879 | 0.240502412 | -0.380682166 |
| Cnih4 | 8.160232 | 8.116851 | 8.2771 | -0.160238035 | -0.116856705 | 8.2024201 | 8.3503644 | 8.10989 | 0.240473354 | 0.092529049 |
| Uhrf1bp1 | 7.049126 | 7.104594 | 7.0726 | 0.031947976 | -0.023519311 | 7.3551787 | 7.4687671 | 7.22839 | 0.240381308 | 0.126792971 |
| Fau | 13.71973 | 13.76542 | 13.84 | -0.074705672 | -0.12039101 | 14.025673 | 13.955904 | 13.7157 | 0.240233544 | 0.310002291 |
| Orc4 | 9.473966 | 9.595339 | 9.4585 | 0.136858515 | 0.015485129 | 9.4908827 | 9.9136391 | 9.67374 | 0.239896676 | -0.182859719 |
| Utp3 | 9.781978 | 9.793599 | 9.7669 | 0.026704187 | 0.015083445 | 9.768452 | 10.001926 | 9.76216 | 0.23976476 | 0.006290631 |
| Car4 | 2.954645 | 3.009441 | 2.8556 | 0.153822569 | 0.099026657 | 2.9104684 | 3.1500093 | 2.91025 | 0.239758814 | 0.000217891 |
| Atf7 | 10.3463 | 10.55105 | 10.426 | 0.125108784 | -0.079638432 | 10.669034 | 10.891676 | 10.652 | 0.239700904 | 0.017059771 |
| Efnb3 | 3.695821 | 3.778167 | 3.6968 | 0.081404251 | -0.000941667 | 3.7509855 | 4.019869 | 3.78029 | 0.239582505 | -0.029300992 |
| E2f4 | 10.13548 | 10.2545 | 10.201 | 0.053583984 | -0.065436454 | 10.414579 | 10.523483 | 10.2839 | 0.239566538 | 0.130662647 |
| Tpr | 12.61167 | 12.60195 | 12.7 | -0.097701126 | -0.087989902 | 12.670665 | 12.972774 | 12.7333 | 0.239508336 | -0.062600864 |
| Vmn1r213 | 2.372974 | 2.500274 | 2.328 | 0.172265154 | 0.044965026 | 2.4750281 | 2.5891185 | 2.34978 | 0.239339094 | 0.125248683 |
| Zdhhc7 | 9.025033 | 9.187902 | 9.0945 | 0.093421865 | -0.069447225 | 9.2388566 | 9.2132175 | 8.97395 | 0.239264869 | 0.264903892 |
| Hsdl2 | 8.864539 | 8.952378 | 8.8936 | 0.058777956 | -0.029060766 | 8.8609146 | 9.1446541 | 8.90549 | 0.239166747 | -0.04457274 |
| Gpsm2 | 7.122589 | 7.022345 | 6.9905 | 0.031883804 | 0.132127801 | 6.9770922 | 7.5460862 | 7.30706 | 0.239021305 | -0.329972651 |
| Sh2d1b2 | 2.437206 | 2.628839 | 2.5461 | 0.082748989 | -0.108883636 | 2.6074411 | 2.827096 | 2.58849 | 0.238610097 | 0.018955168 |
| Rpl6 | 14.58434 | 14.59036 | 14.609 | -0.01905142 | -0.025065594 | 14.736719 | 14.831173 | 14.5926 | 0.238593736 | 0.144139926 |
| Rab24 | 8.946141 | 8.792609 | 8.8224 | -0.029798399 | 0.123733426 | 9.2379932 | 9.2309161 | 8.99234 | 0.238575231 | 0.245652344 |

| | | | | | | | | | | |
|---|---|---|---|---|---|---|---|---|---|---|
| Ndufb8 | 9.077403 | 9.026503 | 9.0824 | -0.055905309 | -0.005004471 | 9.2558862 | 9.2465018 | 9.00794 | 0.238558082 | 0.247942464 |
| Dda1 | 9.384259 | 9.481179 | 9.3315 | 0.149674072 | 0.052754091 | 9.5103669 | 9.5813751 | 9.34285 | 0.238528261 | 0.167520125 |
| Cyp27a1 | 7.534168 | 7.679645 | 7.9039 | -0.224261816 | -0.369739118 | 8.2682264 | 8.3344687 | 8.09597 | 0.23850038 | 0.172258105 |
| Cse1l | 10.85384 | 10.95616 | 10.808 | 0.147929082 | 0.045616276 | 10.837202 | 11.107637 | 10.8691 | 0.238498954 | -0.031936587 |
| Canx | 12.45418 | 12.61518 | 12.396 | 0.218688506 | 0.057692532 | 12.485964 | 12.640002 | 12.4016 | 0.238363232 | 0.084325755 |
| Smim26 | 6.283608 | 6.319578 | 6.1564 | 0.163176957 | 0.127206871 | 6.4549566 | 6.5922567 | 6.3539 | 0.23835998 | 0.101059893 |
| Tnfsfm13 | 5.672338 | 5.412564 | 5.53 | -0.117465656 | 0.142307582 | 5.457468 | 5.3823267 | 5.14413 | 0.238192545 | 0.313333765 |
| Poldip3 | 10.92201 | 11.03875 | 10.917 | 0.121518566 | 0.00478228 | 11.056038 | 11.217771 | 10.9797 | 0.238032438 | 0.076298879 |
| B4galt3 | 9.067208 | 9.039655 | 8.9787 | 0.060989943 | 0.088542162 | 9.3764006 | 9.3245232 | 9.0865 | 0.238023779 | 0.28990116 |
| Trpc4ap | 11.49224 | 11.51505 | 11.504 | 0.011226576 | -0.011577228 | 11.517068 | 11.709505 | 11.4716 | 0.237903406 | 0.045466262 |
| Zbtb45 | 7.36405 | 7.510216 | 7.5726 | -0.062386375 | -0.2085527 | 7.5131168 | 7.740363 | 7.50255 | 0.237811188 | 0.010565044 |
| Adam15 | 8.749744 | 8.635308 | 8.4683 | 0.166988348 | 0.281424812 | 8.3267602 | 8.2952444 | 8.05752 | 0.237723967 | 0.26923981 |
| Ighv5-12 | 4.637775 | 5.784148 | 4.744 | 1.040165545 | -0.106207652 | 5.0195815 | 4.7939404 | 4.55639 | 0.237546607 | 0.463187648 |
| Gm15793 | 4.886724 | 5.077106 | 4.9818 | 0.095258865 | -0.095122998 | 5.173227 | 5.308 | 5.07054 | 0.237464413 | 0.102691363 |
| Ddx49 | 9.271367 | 9.299687 | 9.3487 | -0.048967805 | -0.077287614 | 9.6036384 | 9.7317263 | 9.49429 | 0.237434989 | 0.109347059 |
| Txn2 | 10.38713 | 10.49522 | 10.386 | 0.108872348 | 0.000786206 | 10.633737 | 10.645056 | 10.4076 | 0.237420266 | 0.226101064 |
| Fbrs | 11.04948 | 11.12703 | 11.064 | 0.062923138 | -0.014621276 | 11.149387 | 11.391342 | 11.1539 | 0.237405218 | -0.004549628 |
| Nfe2l1 | 11.21666 | 11.41777 | 11.299 | 0.118671114 | -0.08243493 | 11.270544 | 11.316779 | 11.0795 | 0.237272602 | 0.19103724 |
| Slc8a1 | 7.303099 | 7.134604 | 7.0145 | 0.120098092 | 0.288593662 | 7.7221226 | 7.1160389 | 6.87885 | 0.237190393 | 0.843274109 |
| Bloc1s6 | 9.039641 | 9.028632 | 8.9913 | 0.037314428 | 0.048322973 | 9.0610843 | 9.1635812 | 8.92644 | 0.23714001 | 0.134643202 |
| Mst1r | 2.832778 | 2.901798 | 2.7891 | 0.112744515 | 0.043724254 | 2.9614662 | 2.989651 | 2.75256 | 0.237088737 | 0.208903939 |
| Dad1 | 10.26124 | 10.33107 | 10.247 | 0.084075188 | 0.014244628 | 10.495407 | 10.494106 | 10.2572 | 0.236901517 | 0.238202415 |
| Entr1 | 10.02084 | 10.17574 | 10.098 | 0.078084615 | -0.076815637 | 10.343847 | 10.457082 | 10.2203 | 0.236771608 | 0.123536308 |
| Pum3 | 10.18387 | 10.23466 | 10.174 | 0.060934021 | 0.01014572 | 10.3885 | 10.558844 | 10.3221 | 0.236742346 | 0.066397803 |
| Ocstamp | 4.727792 | 4.70094 | 4.4675 | 0.233483245 | 0.260335679 | 4.8690343 | 4.7603356 | 4.52369 | 0.236649048 | 0.345347751 |
| Arpc3 | 11.8161 | 11.81608 | 11.738 | 0.077730824 | 0.07774891 | 11.963912 | 12.004047 | 11.7676 | 0.23648395 | 0.196349854 |
| Mterf1a | 6.571258 | 6.756432 | 6.6107 | 0.145718361 | -0.039455161 | 6.8484299 | 6.8742217 | 6.63782 | 0.236402401 | 0.210610548 |
| Hmgxb3 | 9.80135 | 9.910603 | 9.815 | 0.095607792 | -0.013644473 | 9.8147303 | 10.043319 | 9.80694 | 0.236381661 | 0.007793152 |
| Slc44a2 | 12.6756 | 12.68635 | 12.639 | 0.047485722 | 0.036730534 | 12.92199 | 12.942871 | 12.7065 | 0.236342766 | 0.215461659 |
| Tcp11l1 | 5.034082 | 5.195989 | 4.9222 | 0.273788095 | 0.111881479 | 4.9879425 | 5.1311565 | 4.89482 | 0.236333644 | 0.09311957 |
| Pinx1 | 7.705596 | 7.7914 | 7.6034 | 0.188013109 | 0.102209162 | 7.6829851 | 8.0027272 | 7.7664 | 0.236329302 | -0.083412853 |
| Pdcd1 | 7.27903 | 7.129135 | 7.1288 | 0.000294015 | 0.150188341 | 6.6498643 | 7.3345894 | 7.09832 | 0.236269579 | -0.448455495 |

| | | | | | | | | | | |
|---|---|---|---|---|---|---|---|---|---|---|
| Schip1 | 4.821563 | 4.798603 | 4.6633 | 0.135266541 | 0.158226529 | 4.3617854 | 4.6264018 | 4.39029 | 0.236114144 | -0.028502281 |
| Exo1 | 6.050943 | 6.080809 | 6.0775 | 0.003278766 | -0.026587788 | 6.0306885 | 6.5059138 | 6.26982 | 0.236095487 | -0.239129836 |
| Hgs | 9.629518 | 9.643256 | 9.5391 | 0.104178998 | 0.090440778 | 9.6309649 | 9.9005742 | 9.66479 | 0.235787944 | -0.033821378 |
| Igkv11-125 | 4.56146 | 4.679656 | 4.5198 | 0.15988091 | 0.041685169 | 4.5543526 | 4.7566412 | 4.52089 | 0.235748918 | 0.033460362 |
| Sart3 | 10.13366 | 10.34288 | 10.242 | 0.100809294 | -0.108405645 | 10.303258 | 10.562794 | 10.3271 | 0.235723582 | -0.023812143 |
| D230025D16Rik | 8.310397 | 8.322846 | 8.345 | -0.022157383 | -0.034606805 | 8.3052085 | 8.5272163 | 8.2915 | 0.235719929 | 0.013712176 |
| Scnm1 | 8.500647 | 8.456944 | 8.5016 | -0.044643604 | -0.000940679 | 8.8643814 | 8.9492198 | 8.71354 | 0.235679465 | 0.15084098 |
| Dennd4a | 11.22424 | 11.15234 | 11.079 | 0.073186494 | 0.145084504 | 11.306302 | 11.605007 | 11.3693 | 0.235657476 | -0.063047311 |
| Cox6c | 10.07199 | 9.992723 | 9.9629 | 0.029818534 | 0.109082911 | 10.069687 | 10.188409 | 9.95282 | 0.235588981 | 0.116866536 |
| Ttyh1 | 3.715833 | 3.965305 | 3.6653 | 0.300004338 | 0.050532406 | 3.9206081 | 3.8412109 | 3.60566 | 0.235546816 | 0.314944054 |
| Igkv4-72 | 7.714552 | 7.908428 | 7.6592 | 0.249267665 | 0.055392053 | 8.1284817 | 8.0174006 | 7.78188 | 0.235518385 | 0.346599481 |
| Dnajc17 | 7.573898 | 7.199613 | 7.4016 | -0.201963463 | 0.172321648 | 7.5340874 | 7.6165642 | 7.38112 | 0.235447129 | 0.152970368 |
| Amh | 2.827786 | 2.977695 | 2.9525 | 0.025156156 | -0.124753294 | 2.8957816 | 3.0949655 | 2.85954 | 0.235421554 | 0.036237619 |
| Spred1 | 9.001861 | 9.056367 | 8.8332 | 0.223122584 | 0.168617395 | 8.3199565 | 8.9282524 | 8.69307 | 0.23518611 | -0.373109816 |
| Tsc22d4 | 10.94538 | 11.08141 | 11.053 | 0.028907106 | -0.107118612 | 11.14776 | 11.185058 | 10.9499 | 0.235181586 | 0.197884163 |
| Yipf3 | 9.535517 | 9.688443 | 9.602 | 0.086486801 | -0.066438987 | 9.824359 | 9.8416463 | 9.6065 | 0.235148191 | 0.21786093 |
| Trim47 | 7.229975 | 6.769497 | 7.0171 | -0.247603694 | 0.212874085 | 6.542151 | 6.6159799 | 6.38083 | 0.235147228 | 0.161318301 |
| Ehbp1l1 | 10.41887 | 10.40922 | 10.341 | 0.068067388 | 0.07772131 | 10.649525 | 10.627622 | 10.3928 | 0.234831368 | 0.256734537 |
| Tufm | 10.05655 | 10.12598 | 10.1 | 0.026058603 | -0.043376414 | 10.135313 | 10.268471 | 10.0337 | 0.234802769 | 0.101644309 |
| P3h1 | 6.213705 | 6.239626 | 6.4429 | -0.203273866 | -0.229194625 | 6.2031573 | 6.3531836 | 6.11839 | 0.23479624 | 0.084769882 |
| Dnajc19 | 7.975083 | 7.784036 | 8.0056 | -0.221531686 | -0.030485024 | 8.100076 | 8.1236363 | 7.88889 | 0.23474938 | 0.211189098 |
| Gm14494 | 7.332159 | 7.402415 | 7.3361 | 0.066303076 | -0.003953491 | 7.4675004 | 7.6165718 | 7.38183 | 0.234741142 | 0.085669772 |
| Nckipsd | 7.211521 | 7.310926 | 7.124 | 0.186953692 | 0.087548452 | 7.3331725 | 7.5263525 | 7.29161 | 0.234741118 | 0.041561174 |
| Tbca | 9.454764 | 9.474139 | 9.4816 | -0.007423534 | -0.026798995 | 9.4621156 | 9.663877 | 9.42915 | 0.234722031 | 0.032960601 |
| Psmd5 | 9.34504 | 9.39462 | 9.3472 | 0.047414315 | -0.002165631 | 9.4588954 | 9.5637774 | 9.3294 | 0.234376115 | 0.129494126 |
| Gm48113 | 4.080871 | 3.73424 | 3.5368 | 0.197425437 | 0.544055966 | 3.7268081 | 3.8696884 | 3.63532 | 0.234372963 | 0.091492602 |
| Msh4 | 4.225803 | 4.250033 | 4.2129 | 0.037090793 | 0.012861011 | 4.3518851 | 4.6092389 | 4.37514 | 0.234099524 | -0.023254302 |
| Kif1a | 6.423973 | 6.10745 | 6.0588 | 0.048667054 | 0.365189235 | 5.8101779 | 6.2659965 | 6.03204 | 0.233960426 | -0.221858179 |
| Bag2 | 4.775101 | 4.783568 | 4.8322 | -0.048656086 | -0.057123362 | 4.6634166 | 5.056454 | 4.82259 | 0.233865906 | -0.159171487 |
| Dok1 | 9.200548 | 9.201966 | 9.2618 | -0.059848718 | -0.061265916 | 9.3310421 | 9.4105181 | 9.17674 | 0.233779216 | 0.154303204 |
| Tiparp | 9.677054 | 9.658 | 9.5228 | 0.135192501 | 0.154246515 | 9.5074775 | 9.7529225 | 9.51937 | 0.23355387 | -0.011891132 |
| Dmac1 | 8.808119 | 8.828411 | 8.7148 | 0.113563685 | 0.09327132 | 9.0188771 | 8.8654648 | 8.63195 | 0.23351304 | 0.386925323 |

| | | | | | | | | | | |
|---|---|---|---|---|---|---|---|---|---|---|
| Prmt3 | 9.573721 | 9.633813 | 9.5113 | 0.122550501 | 0.062458849 | 9.6371796 | 9.854166 | 9.62069 | 0.233480819 | 0.016494449 |
| Klra14-ps | 3.372229 | 3.332955 | 3.311 | 0.021907231 | 0.061181078 | 3.2543963 | 3.5261485 | 3.29269 | 0.233460778 | -0.038291479 |
| D16Ertd472e | 9.902216 | 9.99061 | 9.9813 | 0.009296063 | -0.079098208 | 9.8056304 | 10.195861 | 9.96241 | 0.233452074 | -0.156778162 |
| Sltm | 11.11896 | 11.16405 | 11.14 | 0.024529908 | -0.02055189 | 11.044852 | 11.447935 | 11.2146 | 0.233295616 | -0.169787305 |
| Dhx38 | 10.56062 | 10.57011 | 10.514 | 0.05582186 | 0.046330546 | 10.747196 | 10.812236 | 10.579 | 0.233280612 | 0.168240553 |
| Zfp446 | 7.157854 | 7.162029 | 7.2076 | -0.045561163 | -0.049736262 | 7.4227919 | 7.6209299 | 7.38765 | 0.233276394 | 0.035138346 |
| Ifnar2 | 10.24093 | 9.926377 | 10.208 | -0.281644382 | 0.032907646 | 9.9420725 | 10.160359 | 9.92709 | 0.23326695 | 0.014980853 |
| Ube2v2 | 9.11041 | 9.181212 | 9.0487 | 0.132557135 | 0.061754696 | 9.1536541 | 9.3107875 | 9.07759 | 0.233200375 | 0.076066932 |
| Ap1s1 | 8.375422 | 8.357261 | 8.4673 | -0.110086274 | -0.09192499 | 8.3766 | 8.5681609 | 8.33506 | 0.233100833 | 0.041540002 |
| Gm10282 | 6.388935 | 6.313468 | 6.3609 | -0.047412094 | 0.028054321 | 6.5456002 | 6.7423241 | 6.50923 | 0.23309093 | 0.036367082 |
| Cbwd1 | 7.30329 | 7.358743 | 7.2654 | 0.093355479 | 0.037902254 | 7.0648595 | 7.3839922 | 7.15098 | 0.233014899 | -0.086117793 |
| Tbl1x | 11.10566 | 11.17832 | 11.181 | -0.003152612 | -0.075809586 | 11.01713 | 11.418325 | 11.1853 | 0.233005708 | -0.168189366 |
| Disc1 | 4.789359 | 4.959696 | 4.8203 | 0.139414601 | -0.030921992 | 4.9264203 | 5.1027888 | 4.86985 | 0.23293745 | 0.056568942 |
| Arf6 | 11.2314 | 11.2887 | 11.219 | 0.069963938 | 0.012665856 | 11.217937 | 11.303634 | 11.0707 | 0.232921434 | 0.147224981 |
| Dcaf11 | 10.74366 | 10.94246 | 10.783 | 0.15938743 | -0.039418532 | 10.954541 | 11.042571 | 10.8097 | 0.232822544 | 0.144793229 |
| 2310022A10Rik | 8.909671 | 9.115808 | 9.0742 | 0.041614588 | -0.164522153 | 9.1296085 | 9.2696268 | 9.03683 | 0.23279497 | 0.092776651 |
| Dok3 | 10.91855 | 10.97334 | 10.954 | 0.019337792 | -0.035451084 | 11.654353 | 11.298457 | 11.0658 | 0.232669038 | 0.588565028 |
| Ptp4a1 | 10.81143 | 10.63894 | 10.65 | -0.010724342 | 0.161764408 | 10.576685 | 10.709921 | 10.4773 | 0.232622376 | 0.099386959 |
| AC168977.2 | 2.012366 | 2.054273 | 1.9656 | 0.088718635 | 0.046811113 | 1.9342572 | 2.2627089 | 2.03012 | 0.232589792 | -0.095861928 |
| Gm8233 | 2.727544 | 2.723956 | 2.6814 | 0.042588338 | 0.046176205 | 2.6297126 | 2.9327153 | 2.70026 | 0.232456265 | -0.070546424 |
| Hmgcr | 10.20762 | 10.26087 | 10.227 | 0.033971838 | -0.019283988 | 10.351088 | 10.429281 | 10.197 | 0.232247316 | 0.15405477 |
| Nudcd1 | 8.673855 | 8.74683 | 8.5405 | 0.206319486 | 0.133344523 | 8.7047373 | 8.7891457 | 8.5569 | 0.232241372 | 0.147832992 |
| Cntf | 4.244766 | 4.324134 | 4.3753 | -0.051119474 | -0.130487242 | 4.3485741 | 4.6206629 | 4.38848 | 0.232178147 | -0.039910668 |
| Clec4a2 | 6.197664 | 6.574993 | 5.8161 | 0.758919831 | 0.381591421 | 7.2480151 | 6.2945068 | 6.06241 | 0.232094638 | 1.185602864 |
| Wdr83 | 7.830064 | 7.841682 | 7.8806 | -0.038870312 | -0.050489168 | 8.2418491 | 8.2374629 | 8.00544 | 0.232024539 | 0.236410748 |
| Ube2m | 9.660856 | 9.687459 | 9.5868 | 0.100663547 | 0.074059968 | 9.7567845 | 9.8724896 | 9.64081 | 0.231675325 | 0.115970289 |
| Vps28 | 10.44811 | 10.41953 | 10.446 | -0.026452984 | 0.002131292 | 10.693831 | 10.6862 | 10.4545 | 0.231663308 | 0.239294423 |
| Prkrip1 | 7.912798 | 8.061259 | 7.9065 | 0.154753801 | 0.006292154 | 8.1506532 | 8.3268672 | 8.09531 | 0.231559105 | 0.055345176 |
| Ik | 11.25399 | 11.28385 | 11.264 | 0.019394548 | -0.01046293 | 11.291121 | 11.500799 | 11.2694 | 0.231412064 | 0.021734109 |
| Hist1h4n | 4.589682 | 4.684235 | 4.6902 | -0.005917453 | -0.100469886 | 4.7423713 | 4.8894471 | 4.6581 | 0.231348088 | 0.084272307 |
| Noc3l | 9.19464 | 9.353907 | 9.1941 | 0.159764215 | 0.000497404 | 9.2991392 | 9.4616308 | 9.2303 | 0.231330064 | 0.068838487 |
| Nprl2 | 8.289272 | 8.266164 | 8.3094 | -0.043262526 | -0.020154886 | 8.5721091 | 8.6265509 | 8.39531 | 0.231239437 | 0.176797612 |

| | | | | | | | | | | |
|---|---|---|---|---|---|---|---|---|---|---|
| Ntrk1 | 4.338694 | 3.918112 | 3.9571 | -0.038942171 | 0.381639123 | 3.8347609 | 4.0240852 | 3.79292 | 0.231166156 | 0.041841887 |
| Ssr4 | 9.936309 | 9.976723 | 9.9197 | 0.05705351 | 0.016639993 | 10.119119 | 10.144758 | 9.91374 | 0.231022424 | 0.205382884 |
| Naa80 | 7.351087 | 7.446835 | 7.3537 | 0.093089634 | -0.002657618 | 7.8781842 | 7.7677375 | 7.53697 | 0.230769074 | 0.341215836 |
| Rtf2 | 10.49037 | 10.52521 | 10.435 | 0.090046106 | 0.055202518 | 10.557118 | 10.65725 | 10.4266 | 0.230697214 | 0.130565391 |
| Gpr108 | 9.646226 | 9.685762 | 9.7089 | -0.023111004 | -0.062646262 | 9.8322385 | 9.9331006 | 9.70251 | 0.230590071 | 0.129727979 |
| Rsbn1 | 10.32212 | 10.45084 | 10.347 | 0.103612502 | -0.02510104 | 10.225037 | 10.519346 | 10.2888 | 0.230565776 | -0.063743913 |
| Zfp385a | 8.04296 | 8.15192 | 8.0932 | 0.058696931 | -0.050262817 | 8.4691399 | 8.2526966 | 8.02236 | 0.230339951 | 0.446783311 |
| Npsr1 | 1.119874 | 1.289814 | 1.1876 | 0.102240647 | -0.067699644 | 1.1290148 | 1.3384439 | 1.10813 | 0.230313961 | 0.020884872 |
| Pbx2 | 9.976709 | 10.16331 | 10.112 | 0.051063232 | -0.135536612 | 10.300709 | 10.349693 | 10.1194 | 0.230295713 | 0.18131201 |
| Fam133b | 9.32348 | 9.343398 | 9.2983 | 0.045091831 | 0.025174192 | 9.4557701 | 9.6582731 | 9.42799 | 0.230284938 | 0.027782003 |
| Tor2a | 9.014674 | 8.794922 | 8.8421 | -0.047208486 | 0.172543078 | 9.1181715 | 8.9783751 | 8.74813 | 0.230247331 | 0.37004374 |
| Slc37a1 | 9.225383 | 9.326761 | 9.2799 | 0.046873046 | -0.05450541 | 9.392607 | 9.5329934 | 9.30279 | 0.23020221 | 0.089815844 |
| Rpl10 | 13.45095 | 13.28348 | 13.335 | -0.051155624 | 0.116311892 | 13.461833 | 13.414105 | 13.1841 | 0.230008026 | 0.277735329 |
| Ankrd66 | 0.839882 | 0.912469 | 0.8301 | 0.0824109 | 0.009823083 | 0.8769368 | 1.0539488 | 0.82406 | 0.229886957 | 0.052874947 |
| Spred3 | 4.75071 | 4.792143 | 4.8971 | -0.104999818 | -0.146432876 | 4.9773003 | 4.9734356 | 4.74367 | 0.229762749 | 0.233627516 |
| Acrbp | 6.843859 | 7.155668 | 7.1299 | 0.025782513 | -0.28602627 | 7.6492305 | 7.7922264 | 7.56248 | 0.229745738 | 0.086749878 |
| Ctdnep1 | 10.46945 | 10.49768 | 10.472 | 0.025882008 | -0.002344354 | 10.520594 | 10.553974 | 10.3247 | 0.22929993 | 0.195920367 |
| Zbtb7a | 10.75924 | 10.75973 | 10.714 | 0.046052212 | 0.045563628 | 10.837157 | 10.797742 | 10.5685 | 0.229193586 | 0.268608378 |
| Dcaf5 | 10.55644 | 10.59468 | 10.619 | -0.024373512 | -0.062619426 | 10.597308 | 10.734653 | 10.5055 | 0.229115434 | 0.091771358 |
| N4bp2l2 | 10.60279 | 10.69187 | 10.643 | 0.049131034 | -0.03994988 | 10.722462 | 10.902986 | 10.6739 | 0.22910768 | 0.04858341 |
| Bfar | 9.947282 | 10.0424 | 10.019 | 0.023003474 | -0.072118264 | 10.173709 | 10.364917 | 10.1359 | 0.22905746 | 0.03784948 |
| Elf1 | 12.45204 | 12.61925 | 12.477 | 0.14262808 | -0.024580496 | 12.379224 | 12.662916 | 12.4339 | 0.22900432 | -0.054687695 |
| Nptn | 10.29398 | 10.34638 | 10.344 | 0.00269243 | -0.049707298 | 10.325907 | 10.483762 | 10.2548 | 0.228976226 | 0.071121447 |
| Akap2 | 10.4824 | 10.39411 | 10.3 | 0.094045572 | 0.1823419 | 9.9662846 | 10.525193 | 10.2963 | 0.228922548 | -0.329985548 |
| Clasrp | 9.244892 | 9.326946 | 9.4305 | -0.103597431 | -0.185651706 | 9.8289194 | 9.9342927 | 9.70539 | 0.22890685 | 0.123533565 |
| Khdc1c | 1.732635 | 1.794479 | 1.8603 | -0.065796292 | -0.127640243 | 1.7506524 | 1.9673092 | 1.73841 | 0.228901413 | 0.012244629 |
| Tnfsf13 | 5.136373 | 4.791319 | 4.8898 | -0.098478976 | 0.246575214 | 4.9742945 | 4.9228109 | 4.69393 | 0.228883469 | 0.280367039 |
| Gm4366 | 2.62964 | 2.686406 | 2.8483 | -0.16191601 | -0.218681971 | 2.9594011 | 2.8783611 | 2.64964 | 0.228719854 | 0.309759844 |
| Fam151b | 4.909303 | 4.977268 | 4.9795 | -0.002240721 | -0.070205907 | 5.0242492 | 5.2042719 | 4.97572 | 0.22854765 | 0.048524995 |
| Col16a1 | 4.860835 | 4.798551 | 4.6 | 0.198551784 | 0.260836131 | 4.7764389 | 4.8849912 | 4.65652 | 0.228469812 | 0.119917567 |
| Fbxl6 | 9.225348 | 9.328124 | 9.3773 | -0.049142356 | -0.151918679 | 9.6995292 | 9.5698689 | 9.3416 | 0.228269959 | 0.357930223 |
| Mbd6 | 10.17809 | 10.23476 | 10.193 | 0.041901522 | -0.01476465 | 10.407743 | 10.562732 | 10.3345 | 0.228233092 | 0.073243458 |

| | | | | | | | | | | |
|---|---|---|---|---|---|---|---|---|---|---|
| Supv3l1 | 8.594087 | 8.541036 | 8.5752 | -0.034119493 | 0.018930921 | 8.6550353 | 8.8231022 | 8.5949 | 0.228200923 | 0.060134005 |
| Ddx31 | 8.637624 | 8.660558 | 8.6708 | -0.010270107 | -0.033204296 | 8.5506268 | 8.8472109 | 8.61919 | 0.228018499 | -0.068565545 |
| Brix1 | 9.353118 | 9.25386 | 9.3995 | -0.145681778 | -0.046424402 | 9.525746 | 9.5924353 | 9.36443 | 0.228000498 | 0.16131122 |
| Zbtb7b | 10.09813 | 10.27015 | 10.141 | 0.128927755 | -0.043097987 | 10.054465 | 10.293112 | 10.0654 | 0.227737579 | -0.010909154 |
| Tacstd2 | 5.631912 | 5.713424 | 5.6466 | 0.06684772 | -0.014663898 | 5.7219756 | 5.7745944 | 5.54692 | 0.227671071 | 0.175052308 |
| Hacd4 | 5.643542 | 5.360758 | 5.3614 | -0.000660196 | 0.28212378 | 6.0653221 | 5.3390409 | 5.11141 | 0.227635866 | 0.953917058 |
| Fuca1 | 10.04265 | 9.954583 | 10.067 | -0.11235873 | -0.024291845 | 10.205073 | 10.403794 | 10.1762 | 0.227632399 | 0.028910563 |
| Tex261 | 10.02306 | 10.09224 | 10.129 | -0.036325496 | -0.1055096 | 10.332622 | 10.249391 | 10.0218 | 0.227617839 | 0.310848316 |
| Plekhj1 | 9.028156 | 9.147482 | 9.1913 | -0.043851416 | -0.163178068 | 9.4142722 | 9.397626 | 9.17014 | 0.227486278 | 0.244132506 |
| Slc25a33 | 6.273139 | 6.235607 | 6.1851 | 0.050515784 | 0.088048217 | 6.3241251 | 6.28377 | 6.05642 | 0.227347606 | 0.267702705 |
| 9130208D14Rik | 3.474362 | 3.456395 | 3.3319 | 0.124500148 | 0.142467629 | 3.3327951 | 3.4697926 | 3.24248 | 0.227307972 | 0.090310435 |
| Mier3 | 9.111412 | 9.260053 | 9.0168 | 0.243283684 | 0.09464306 | 8.9111355 | 9.3534341 | 9.12617 | 0.227266163 | -0.215032415 |
| Gm47476 | 5.23146 | 5.256492 | 5.3302 | -0.07366949 | -0.0987016 | 5.0962703 | 5.6204054 | 5.39324 | 0.227165133 | -0.296969943 |
| Tcea1 | 11.25077 | 11.31719 | 11.243 | 0.07433698 | 0.007917728 | 11.268364 | 11.408024 | 11.1809 | 0.22715145 | 0.087491612 |
| Ppme1 | 9.235873 | 9.118305 | 9.0241 | 0.094215771 | 0.21178369 | 9.0565815 | 9.2349071 | 9.00788 | 0.227028869 | 0.048703271 |
| Mmadhc | 8.892767 | 8.853157 | 8.809 | 0.044109817 | 0.083720245 | 8.7560505 | 8.9436133 | 8.71671 | 0.22690157 | 0.039338753 |
| Gtdc1 | 8.347836 | 8.446004 | 8.3426 | 0.103394557 | 0.005226282 | 8.2454939 | 8.4309982 | 8.2041 | 0.22689605 | 0.041391802 |
| Vti1b | 9.289048 | 9.311346 | 9.2329 | 0.078493062 | 0.05619483 | 9.3085247 | 9.4258294 | 9.1991 | 0.226724739 | 0.109419987 |
| Pgam2 | 2.797633 | 2.852452 | 2.8176 | 0.034896775 | -0.01992185 | 2.7086805 | 2.9506326 | 2.72408 | 0.226552192 | -0.01539988 |
| Ptpn2 | 9.859608 | 9.820782 | 9.8114 | 0.009355572 | 0.04818178 | 9.7008889 | 9.9699572 | 9.74365 | 0.226309459 | -0.042758852 |
| Igkv4-86 | 7.270826 | 7.085294 | 6.9241 | 0.161193498 | 0.346725487 | 7.1989121 | 7.0316473 | 6.80534 | 0.22630516 | 0.393569889 |
| Agpat1 | 9.753598 | 9.911743 | 9.7847 | 0.127049369 | -0.031095459 | 10.167205 | 10.054373 | 9.82808 | 0.226293729 | 0.339126345 |
| Ubxn1 | 10.30403 | 10.34285 | 10.368 | -0.025314054 | -0.06413083 | 10.503004 | 10.522629 | 10.2965 | 0.226160994 | 0.206535613 |
| Psmd10 | 7.526967 | 7.58707 | 7.5841 | 0.002965393 | -0.057138377 | 7.6282454 | 7.7769024 | 7.55107 | 0.225835329 | 0.077178326 |
| 3110001I22Rik | 6.870765 | 6.898017 | 6.905 | -0.007028843 | -0.034280473 | 7.051199 | 7.3537549 | 7.12802 | 0.225731946 | -0.076823912 |
| Zfp987 | 4.60742 | 4.622474 | 4.4864 | 0.136099942 | 0.121045853 | 4.5576843 | 4.6816657 | 4.45594 | 0.225723763 | 0.101742408 |
| Rnf138 | 9.626291 | 9.594556 | 9.6586 | -0.06403363 | -0.032297907 | 9.4394707 | 9.7487452 | 9.52316 | 0.225582417 | -0.083692169 |
| Exosc3 | 8.729967 | 8.752273 | 8.6757 | 0.076527195 | 0.0542218 | 8.7219656 | 8.8774144 | 8.65191 | 0.225508953 | 0.07006019 |
| Asah1 | 9.980068 | 9.771334 | 9.8971 | -0.12573634 | 0.08299717 | 9.8253159 | 9.7997964 | 9.57444 | 0.225355109 | 0.250874604 |
| Pcna | 11.34373 | 11.23374 | 11.35 | -0.115882546 | -0.005885828 | 11.140481 | 11.505714 | 11.2806 | 0.225137016 | -0.140096303 |
| Kat5 | 9.313632 | 9.240739 | 9.4898 | -0.24909168 | -0.176199236 | 9.6246905 | 9.566173 | 9.34109 | 0.225086938 | 0.283604387 |
| Puf60 | 11.20037 | 11.24405 | 11.198 | 0.046180834 | 0.002496734 | 11.378156 | 11.407205 | 11.1825 | 0.224663958 | 0.195615403 |

| | | | | | | | | | | |
|---|---|---|---|---|---|---|---|---|---|---|
| Fzd1 | 6.200482 | 6.028334 | 5.8406 | 0.187712323 | 0.359860784 | 5.8057204 | 5.8029991 | 5.57844 | 0.224563279 | 0.227284566 |
| Fryl | 11.44505 | 11.52859 | 11.512 | 0.01638386 | -0.06715742 | 11.282851 | 11.938652 | 11.7142 | 0.224469582 | -0.431331368 |
| Zfp39 | 7.814357 | 7.851161 | 7.9099 | -0.058692114 | -0.095495475 | 7.6130663 | 7.9866982 | 7.76225 | 0.224446005 | -0.149185867 |
| Gm11808 | 13.95511 | 14.03729 | 13.978 | 0.059663598 | -0.022520302 | 14.24458 | 14.177667 | 13.9534 | 0.224272422 | 0.291185456 |
| 1600002K03Rik | 6.821087 | 7.01581 | 6.8171 | 0.198713927 | 0.003990754 | 7.0056894 | 7.176431 | 6.95219 | 0.224241852 | 0.053500281 |
| Crlf3 | 11.54952 | 11.87479 | 11.667 | 0.207571964 | -0.117689138 | 11.572991 | 11.855223 | 11.6311 | 0.224129192 | -0.058103643 |
| Cdkn2a | 3.178599 | 3.206564 | 3.2202 | -0.013595128 | -0.041559678 | 3.2120717 | 3.3733667 | 3.14928 | 0.224088311 | 0.062793314 |
| Insrr | 3.795183 | 3.572316 | 3.7656 | -0.19332365 | 0.029543349 | 3.5271562 | 3.8730562 | 3.6491 | 0.223952616 | -0.121947385 |
| Col11a2 | 7.123309 | 7.389554 | 7.2755 | 0.114097958 | -0.152147401 | 7.1954541 | 7.5836242 | 7.35975 | 0.223877471 | -0.164292667 |
| Actr2 | 13.4797 | 13.55418 | 13.412 | 0.142405612 | 0.067925816 | 13.455083 | 13.610281 | 13.3865 | 0.223803494 | 0.068605896 |
| Apol10b | 6.518712 | 6.477922 | 6.5844 | -0.106501077 | -0.065710718 | 5.9164451 | 6.7291151 | 6.50532 | 0.223795796 | -0.588874157 |
| Cwc15 | 10.07376 | 10.18468 | 10.148 | 0.037121924 | -0.073800144 | 10.207216 | 10.429263 | 10.2055 | 0.223791154 | 0.001743791 |
| Galns | 8.779732 | 8.722916 | 8.8254 | -0.102448289 | -0.045632927 | 8.9485936 | 8.9548948 | 8.73123 | 0.223666176 | 0.217364991 |
| Eif3c | 12.42801 | 12.3869 | 12.317 | 0.069413014 | 0.110520842 | 12.391928 | 12.539015 | 12.3154 | 0.223609202 | 0.076522258 |
| Id2 | 10.24318 | 10.22349 | 10.214 | 0.009192128 | 0.028882518 | 9.7204233 | 10.20301 | 9.97946 | 0.223551771 | -0.2590354 |
| Zfp574 | 8.303784 | 8.424647 | 8.4227 | 0.001916528 | -0.118947084 | 8.3224484 | 8.4499225 | 8.22638 | 0.223541246 | 0.09606723 |
| Arpc5l | 10.88153 | 10.90288 | 10.905 | -0.001644106 | -0.022989498 | 11.106447 | 11.066145 | 10.8426 | 0.223517882 | 0.263819444 |
| Hdgfl2 | 10.08378 | 10.14555 | 10.121 | 0.024862725 | -0.036908962 | 10.358638 | 10.533338 | 10.3099 | 0.223447438 | 0.048747835 |
| Rpsa-ps10 | 4.63239 | 4.434295 | 4.6211 | -0.186803383 | 0.011291485 | 4.6589995 | 4.5331249 | 4.3098 | 0.223325395 | 0.349199995 |
| Tsc22d3 | 12.54223 | 12.73604 | 12.565 | 0.171491464 | -0.022324512 | 12.280368 | 12.534089 | 12.3109 | 0.22314519 | -0.030575582 |
| Alg8 | 7.683272 | 7.791769 | 7.6993 | 0.092517048 | -0.01598019 | 8.0281293 | 8.2839934 | 8.0609 | 0.223089991 | -0.032774068 |
| Igkv3-10 | 7.066789 | 7.210163 | 6.9219 | 0.288227783 | 0.144854139 | 7.3415114 | 7.3772823 | 7.15431 | 0.222975778 | 0.187204852 |
| Tcerg1 | 10.72853 | 10.74408 | 10.707 | 0.036756542 | 0.021204881 | 10.628356 | 11.038692 | 10.8158 | 0.222907704 | -0.187428368 |
| Mcm6 | 10.73327 | 10.64004 | 10.766 | -0.12582884 | -0.032592312 | 10.630294 | 11.105448 | 10.8825 | 0.222906368 | -0.252248171 |
| Slc35a4 | 10.28185 | 10.34021 | 9.9559 | 0.384332006 | 0.325968384 | 10.219252 | 10.376211 | 10.1535 | 0.222703959 | 0.065745023 |
| Gm1123 | 2.279551 | 2.28194 | 2.2576 | 0.024299436 | 0.021910301 | 2.208918 | 2.4124007 | 2.18979 | 0.222610141 | 0.019127413 |
| Spsb2 | 5.637583 | 5.680958 | 5.7743 | -0.093387251 | -0.136762456 | 5.8082963 | 5.8068485 | 5.58429 | 0.222562747 | 0.224010577 |
| Parva | 7.323334 | 7.19942 | 7.1792 | 0.020232354 | 0.144146104 | 7.0035445 | 6.9996786 | 6.77718 | 0.222498754 | 0.226364653 |
| Commd2 | 8.496129 | 8.491035 | 8.5275 | -0.036420421 | -0.031326262 | 8.6722426 | 8.6838727 | 8.46156 | 0.222314231 | 0.210684201 |
| Txlna | 10.49818 | 10.51067 | 10.454 | 0.056422252 | 0.043938888 | 10.356433 | 10.583696 | 10.3614 | 0.222269816 | -0.004993202 |
| Arhgef10 | 7.645633 | 7.030475 | 7.6025 | -0.572029939 | 0.043128395 | 7.3352445 | 7.5448251 | 7.32259 | 0.222230767 | 0.012650077 |
| Uckl1 | 9.207633 | 9.21426 | 9.2666 | -0.052378221 | -0.059005484 | 9.9512436 | 9.8737398 | 9.65158 | 0.222161119 | 0.29966493 |

| | | | | | | | | | | |
|---|---|---|---|---|---|---|---|---|---|---|
| Dctn5 | 10.00846 | 10.08999 | 10.034 | 0.05605961 | -0.025474055 | 10.187783 | 10.28997 | 10.0679 | 0.22210312 | 0.119916067 |
| Plppr4 | 4.142949 | 4.205344 | 4.1554 | 0.049939202 | -0.012455902 | 4.0293895 | 4.2878376 | 4.06578 | 0.222057618 | -0.036390468 |
| Sparc | 9.982827 | 10.09467 | 9.9226 | 0.172051136 | 0.060210545 | 10.177703 | 9.7399236 | 9.51789 | 0.22203735 | 0.659816747 |
| Lsm3 | 8.542912 | 8.563328 | 8.566 | -0.002679289 | -0.023094726 | 8.759622 | 8.8318778 | 8.60985 | 0.222030592 | 0.149774852 |
| Rpl19 | 14.60755 | 14.70358 | 14.638 | 0.065888218 | -0.030143026 | 14.824915 | 14.842484 | 14.6205 | 0.221955542 | 0.204386371 |
| Repin1 | 6.10649 | 6.073716 | 6.0958 | -0.022078132 | 0.010695092 | 6.3687832 | 6.4188418 | 6.19694 | 0.221900286 | 0.171841659 |
| Slc7a6os | 8.954754 | 8.913501 | 8.9664 | -0.052864931 | -0.011611827 | 9.2498507 | 9.3664148 | 9.14473 | 0.221681595 | 0.105117508 |
| Ap4s1 | 7.970142 | 7.868901 | 7.8501 | 0.018818009 | 0.120059022 | 7.9885142 | 8.0665224 | 7.84496 | 0.221557882 | 0.143549665 |
| Cops8 | 9.427159 | 9.494214 | 9.5285 | -0.03428224 | -0.101337141 | 9.7009758 | 9.7901017 | 9.56856 | 0.221546666 | 0.132420775 |
| Utp15 | 9.597503 | 9.689734 | 9.6145 | 0.075263494 | -0.01696726 | 9.6642495 | 9.8456669 | 9.62423 | 0.2214408 | 0.040023405 |
| Olfr1380 | 2.000911 | 2.017383 | 2.0378 | -0.020433152 | -0.036905874 | 2.0950097 | 2.2328165 | 2.01144 | 0.221381083 | 0.083574306 |
| Ube2i | 11.66055 | 11.69929 | 11.637 | 0.062743506 | 0.023998798 | 11.763579 | 11.912966 | 11.6916 | 0.221377794 | 0.07199085 |
| Vmn1r81 | 5.388497 | 5.380678 | 5.4848 | -0.104138201 | -0.096319061 | 5.6962108 | 5.9247026 | 5.7035 | 0.221201627 | -0.007290122 |
| Gm21056 | 3.063572 | 3.061687 | 3.1236 | -0.061925927 | -0.060040248 | 3.1126941 | 3.4043652 | 3.18323 | 0.221134025 | -0.070537061 |
| Jagn1 | 7.807863 | 7.889849 | 7.8424 | 0.047462584 | -0.03452344 | 8.1223397 | 8.0997103 | 7.87863 | 0.221083488 | 0.243712912 |
| Rbm34 | 8.941628 | 8.982064 | 8.9942 | -0.012108975 | -0.052544864 | 8.9145362 | 9.1330727 | 8.91199 | 0.221081469 | 0.00254499 |
| Rps10-ps1 | 3.959254 | 4.082118 | 3.9636 | 0.11853467 | -0.004329388 | 4.0859586 | 4.0653016 | 3.84428 | 0.221021217 | 0.241678167 |
| Hpcal1 | 9.493397 | 9.634262 | 9.5306 | 0.10363801 | -0.037227347 | 9.496498 | 9.6040732 | 9.38318 | 0.220893514 | 0.113318331 |
| Znhit3 | 6.976714 | 6.85462 | 6.7247 | 0.129953121 | 0.252047208 | 6.9779031 | 7.1883345 | 6.9675 | 0.22083907 | 0.0104076 |
| Gm5176 | 2.509491 | 2.586581 | 2.5557 | 0.030869503 | -0.046220747 | 2.6239068 | 2.7314561 | 2.51073 | 0.220722903 | 0.113173581 |
| Dgcr2 | 10.37643 | 10.50111 | 10.412 | 0.088641492 | -0.03603676 | 10.420729 | 10.563158 | 10.3426 | 0.22060534 | 0.078176194 |
| Telo2 | 8.893857 | 8.866451 | 8.8893 | -0.022885563 | 0.004519846 | 9.072489 | 9.2621967 | 9.04164 | 0.220558823 | 0.030851084 |
| Ndufb3 | 8.592692 | 8.68873 | 8.5915 | 0.097212139 | 0.001174201 | 8.8824515 | 8.8543519 | 8.63385 | 0.220502066 | 0.248601676 |
| Cd3eap | 7.882986 | 7.834911 | 7.9277 | -0.092800466 | -0.044725644 | 7.9992839 | 8.0960533 | 7.87558 | 0.22047355 | 0.123704067 |
| Lyl1 | 10.08381 | 10.00514 | 10.034 | -0.028666587 | 0.050006191 | 10.626976 | 10.187196 | 9.96689 | 0.220306095 | 0.660085302 |
| Hopx | 8.928531 | 9.03195 | 8.7788 | 0.253153061 | 0.149734538 | 9.0636914 | 9.077284 | 8.85721 | 0.22007216 | 0.206479541 |
| Slc9a1 | 9.938962 | 10.10313 | 9.9769 | 0.126272198 | -0.037894878 | 9.8838836 | 10.219771 | 9.99984 | 0.219927298 | -0.115960179 |
| Zfp637 | 6.867251 | 7.006571 | 6.9098 | 0.096818587 | -0.042501321 | 6.9132762 | 6.9320036 | 6.71249 | 0.219517391 | 0.200790043 |
| Timp3 | 10.00878 | 9.578633 | 9.7431 | -0.164421618 | 0.265727046 | 8.5527436 | 9.4715144 | 9.25202 | 0.21949797 | -0.699272859 |
| Apba1 | 5.735085 | 5.726703 | 5.738 | -0.011315173 | -0.002933149 | 5.7683173 | 5.9172639 | 5.69781 | 0.219450765 | 0.070504076 |
| Tepsin | 8.154997 | 8.295836 | 8.219 | 0.076880459 | -0.063958889 | 8.5739919 | 8.634677 | 8.41528 | 0.219395092 | 0.15871001 |
| 2810004N23Rik | 8.872457 | 8.872012 | 8.8524 | 0.019588081 | 0.02003279 | 9.0150023 | 9.0354578 | 8.81629 | 0.219163122 | 0.198707627 |

| | | | | | | | | | | |
|---|---|---|---|---|---|---|---|---|---|---|
| Arhgap17 | 11.08981 | 11.17481 | 11.206 | -0.03085051 | -0.115853616 | 11.48101 | 11.557295 | 11.3383 | 0.218992622 | 0.142708179 |
| Rcl1 | 8.688545 | 8.751854 | 8.5906 | 0.161275481 | 0.097966913 | 8.6818348 | 8.9109495 | 8.69202 | 0.218933236 | -0.010181459 |
| Ankrd37 | 6.761665 | 6.74071 | 7.0026 | -0.261852796 | -0.240897556 | 7.4651597 | 7.2622132 | 7.04391 | 0.218301936 | 0.42124845 |
| Zkscan1 | 8.916871 | 9.035852 | 8.9895 | 0.046401427 | -0.072578935 | 8.8396192 | 9.230787 | 9.01252 | 0.218270224 | -0.172897567 |
| Obsl1 | 3.617196 | 3.705607 | 3.6404 | 0.065211739 | -0.023199725 | 3.6047616 | 3.6784777 | 3.46021 | 0.218266918 | 0.144550822 |
| Fastkd1 | 8.169049 | 8.292445 | 8.1865 | 0.10598907 | -0.017407338 | 8.1502335 | 8.3799645 | 8.16186 | 0.218101544 | -0.011629423 |
| Nme6 | 6.285456 | 6.19331 | 6.0923 | 0.101034178 | 0.193180014 | 6.2104316 | 6.3723488 | 6.15426 | 0.21809346 | 0.056176262 |
| Ube2j2 | 9.362508 | 9.469479 | 9.3732 | 0.096271159 | -0.010700082 | 9.5060532 | 9.7052788 | 9.48729 | 0.217987299 | 0.018761769 |
| Tmed4 | 8.860077 | 8.910063 | 8.9303 | -0.020279847 | -0.07026612 | 9.0659654 | 9.018682 | 8.80078 | 0.217900709 | 0.265184119 |
| Ring1 | 8.776427 | 8.839292 | 8.8044 | 0.034865035 | -0.027999701 | 9.1772992 | 9.1242211 | 8.90644 | 0.21777875 | 0.270856875 |
| Tnfsf18 | 2.396714 | 2.659617 | 2.3103 | 0.349300267 | 0.086396785 | 2.4143385 | 2.5761815 | 2.35848 | 0.217698761 | 0.055855722 |
| Lmbrd1 | 10.93835 | 11.06251 | 10.893 | 0.169507424 | 0.045354992 | 11.224539 | 11.104822 | 10.8872 | 0.217646812 | 0.337363855 |
| Desi1 | 9.181114 | 9.297265 | 9.1715 | 0.125743423 | 0.009592407 | 9.146691 | 9.3459749 | 9.12838 | 0.217592065 | 0.018308238 |
| 3830406C13Rik | 8.216819 | 8.060628 | 8.2393 | -0.178646471 | -0.022454927 | 8.4552149 | 8.6867797 | 8.46924 | 0.217538536 | -0.014026289 |
| Ighv1-84 | 3.797143 | 4.043797 | 3.7468 | 0.296988706 | 0.050335067 | 3.9460627 | 4.0223376 | 3.80486 | 0.217482333 | 0.141207403 |
| Kdm8 | 6.926527 | 6.941458 | 6.9651 | -0.023666999 | -0.03859774 | 7.3981617 | 7.2108584 | 6.99363 | 0.217230815 | 0.404534144 |
| Sugp1 | 9.818934 | 9.836845 | 9.839 | -0.002130653 | -0.020042133 | 9.9968782 | 10.169768 | 9.95266 | 0.217110148 | 0.044220262 |
| Gm29721 | 0.923903 | 0.857941 | 0.8973 | -0.039398588 | 0.026563333 | 0.8237123 | 1.0386998 | 0.8216 | 0.217102186 | 0.002114723 |
| Ybey | 6.106724 | 6.087229 | 6.2249 | -0.137676455 | -0.11818152 | 6.2851112 | 6.4596311 | 6.24261 | 0.217017018 | 0.042497204 |
| Atp5h | 10.96007 | 10.90543 | 10.908 | -0.002633646 | 0.05200542 | 11.067455 | 11.099351 | 10.8825 | 0.216810092 | 0.184914144 |
| Slc19a1 | 8.188269 | 8.338016 | 8.1379 | 0.20008506 | 0.050338302 | 8.5831565 | 8.4464477 | 8.2297 | 0.216752629 | 0.35346143 |
| Fgf9 | 2.666513 | 2.650032 | 2.6642 | -0.014136124 | 0.002344954 | 2.7064654 | 2.9346375 | 2.71789 | 0.21674543 | -0.011426625 |
| Tpm3-rs7 | 4.589534 | 4.650011 | 4.5489 | 0.101126335 | 0.040649901 | 4.8111323 | 4.7472991 | 4.53073 | 0.216568904 | 0.280402077 |
| Ccdc9 | 10.10448 | 10.07402 | 10.06 | 0.013909231 | 0.044374213 | 10.319039 | 10.600551 | 10.3842 | 0.216399938 | -0.065111955 |
| Ighv1-71 | 5.832957 | 5.906245 | 5.7115 | 0.194700577 | 0.121412027 | 6.3265882 | 6.038528 | 5.82218 | 0.21635006 | 0.504410237 |
| Emc6 | 8.583332 | 8.698859 | 8.4481 | 0.250715566 | 0.135188376 | 8.711147 | 8.7485973 | 8.53228 | 0.216321782 | 0.178871471 |
| Slc7a11 | 7.69972 | 7.788576 | 6.9861 | 0.80248987 | 0.713634077 | 7.1916055 | 7.4880139 | 7.27182 | 0.216188938 | -0.080219514 |
| Polr3b | 10.24242 | 10.39277 | 10.244 | 0.148981812 | -0.001363406 | 10.268796 | 10.414322 | 10.1982 | 0.216162788 | 0.070636443 |
| Dusp16 | 8.812091 | 8.791421 | 8.585 | 0.206434052 | 0.227103944 | 8.7524268 | 8.8923794 | 8.67633 | 0.216052986 | 0.07610036 |
| Chid1 | 8.087371 | 8.116341 | 8.0334 | 0.082893219 | 0.053923456 | 7.9656366 | 8.1564182 | 7.94059 | 0.215825928 | 0.025044367 |
| Mrps27 | 8.366856 | 8.452455 | 8.4135 | 0.038914456 | -0.046684524 | 8.6367063 | 8.6340541 | 8.41832 | 0.215734395 | 0.218386573 |
| Psd | 3.439786 | 3.528896 | 3.5128 | 0.016050864 | -0.07305913 | 3.5562721 | 3.6806394 | 3.46497 | 0.215671795 | 0.091304492 |

| | | | | | | | | | | |
|---|---|---|---|---|---|---|---|---|---|---|
| Il23a | 2.731446 | 2.957915 | 2.7093 | 0.248622836 | 0.022154333 | 2.7837291 | 2.8369143 | 2.62125 | 0.21566073 | 0.162475498 |
| Kif19a | 5.956281 | 5.708981 | 5.6498 | 0.059135468 | 0.306434679 | 5.9812634 | 5.7726545 | 5.55709 | 0.215565806 | 0.424174716 |
| Fermt3 | 12.06373 | 12.08752 | 12.007 | 0.08051354 | 0.056721444 | 12.371223 | 12.222739 | 12.0073 | 0.215477196 | 0.363961477 |
| A3galt2 | 4.044425 | 4.155251 | 4.022 | 0.133237777 | 0.022411966 | 4.0376127 | 4.3238193 | 4.10834 | 0.215476035 | -0.070730582 |
| Sh3bp5l | 8.233132 | 8.332856 | 8.3183 | 0.014555701 | -0.085167996 | 8.501172 | 8.592063 | 8.37665 | 0.21541095 | 0.124520002 |
| Gps1 | 9.93509 | 9.976825 | 9.9679 | 0.008949065 | -0.032785762 | 10.32918 | 10.247272 | 10.032 | 0.215295383 | 0.297203328 |
| Paqr5 | 3.913364 | 3.935897 | 3.9615 | -0.025627211 | -0.048160098 | 3.7886789 | 3.9192082 | 3.70396 | 0.215253035 | 0.08472373 |
| Tmem97 | 6.644307 | 6.614317 | 6.8473 | -0.233031557 | -0.203041064 | 6.9896433 | 6.9017721 | 6.68652 | 0.215251705 | 0.303122927 |
| Med9 | 7.116255 | 7.074119 | 7.1468 | -0.072692497 | -0.030556277 | 7.2843091 | 7.281724 | 7.06648 | 0.21524758 | 0.217832619 |
| Aatf | 9.036196 | 9.002893 | 9.1123 | -0.109430808 | -0.076128564 | 9.0810931 | 9.3135924 | 9.09845 | 0.215146049 | -0.017353182 |
| Aip | 9.243063 | 9.383462 | 9.249 | 0.134461896 | -0.005936741 | 9.5451237 | 9.5066528 | 9.29155 | 0.215106457 | 0.253577345 |
| Pbdc1 | 9.071335 | 9.153212 | 9.0393 | 0.113873489 | 0.031996326 | 9.2919468 | 9.316406 | 9.10136 | 0.215043168 | 0.190583928 |
| Rpa3 | 7.283101 | 7.324277 | 7.2849 | 0.039355722 | -0.00182042 | 7.4608441 | 7.5308798 | 7.3161 | 0.214779915 | 0.144744249 |
| Tmem129 | 8.164646 | 8.357718 | 8.265 | 0.092670932 | -0.100401377 | 8.5078649 | 8.5127791 | 8.29807 | 0.214713346 | 0.209799221 |
| Hdgf | 11.18075 | 11.16662 | 11.053 | 0.113715518 | 0.127850354 | 11.180536 | 11.297684 | 11.083 | 0.21467874 | 0.097531413 |
| Reep4 | 8.236892 | 8.428149 | 8.4671 | -0.038948806 | -0.23020605 | 8.713316 | 8.8631318 | 8.64869 | 0.214443428 | 0.064627629 |
| Ndufaf3 | 7.2112 | 7.220627 | 7.3384 | -0.117800422 | -0.127227627 | 7.693216 | 7.5940674 | 7.37965 | 0.214417593 | 0.313566164 |
| Gm44715 | 4.58666 | 4.712954 | 4.6425 | 0.070493708 | -0.05580021 | 4.601618 | 5.1607152 | 4.94656 | 0.214159362 | -0.344937797 |
| Oxsr1 | 9.920893 | 9.962242 | 9.8504 | 0.111817826 | 0.070468771 | 9.7019572 | 10.053593 | 9.83953 | 0.214066281 | -0.137569222 |
| Utp4 | 9.459121 | 9.395281 | 9.4127 | -0.017443217 | 0.046396646 | 9.420918 | 9.6135598 | 9.3995 | 0.214058802 | 0.021416989 |
| 1810024B03Rik | 4.442501 | 4.514385 | 4.3313 | 0.183106251 | 0.111222042 | 4.5137676 | 4.7466685 | 4.53271 | 0.213955146 | -0.018945696 |
| Ssr3 | 11.6335 | 11.62721 | 11.5 | 0.127603054 | 0.133897522 | 11.452882 | 11.642583 | 11.4288 | 0.213819324 | 0.024117789 |
| Gm10131 | 7.254751 | 7.373026 | 7.3729 | 0.0001645 | -0.118110289 | 7.2141158 | 7.2576938 | 7.04393 | 0.213766444 | 0.170188446 |
| Btbd2 | 8.713626 | 8.730783 | 8.7699 | -0.039138727 | -0.056294872 | 8.9510307 | 8.8880933 | 8.67438 | 0.213709778 | 0.276647127 |
| Bad | 7.063839 | 6.999987 | 7.0068 | -0.006857028 | 0.056995533 | 7.3279702 | 7.1608268 | 6.94712 | 0.213706614 | 0.380849985 |
| Washc3 | 8.661991 | 8.705498 | 8.7993 | -0.093766357 | -0.137273324 | 8.7486332 | 8.9816243 | 8.76795 | 0.213675008 | -0.019316078 |
| Hypk | 7.835325 | 7.603773 | 7.5702 | 0.033543669 | 0.265094794 | 8.0192815 | 8.0352607 | 7.8216 | 0.213665178 | 0.197685991 |
| Zfp689 | 6.543953 | 6.448737 | 6.5258 | -0.077013083 | 0.018202218 | 6.5986803 | 6.474064 | 6.26065 | 0.213418433 | 0.3380347 |
| Mfsd2b | 3.881479 | 3.871006 | 3.8884 | -0.017385928 | -0.006913549 | 3.9325446 | 4.3135433 | 4.10033 | 0.213216026 | -0.167782676 |
| Cops7b | 8.204256 | 8.234913 | 8.1698 | 0.065119002 | 0.034461701 | 8.485603 | 8.4809018 | 8.26773 | 0.213169674 | 0.217870829 |
| Cnot8 | 9.986796 | 10.11062 | 10.081 | 0.029505738 | -0.094320599 | 10.17068 | 10.291549 | 10.0785 | 0.213038639 | 0.092170371 |
| Vps37c | 8.298256 | 8.228477 | 8.186 | 0.042504432 | 0.112283163 | 7.9769229 | 8.1931094 | 7.9801 | 0.213007547 | -0.00317889 |

| | | | | | | | | | | |
|---|---|---|---|---|---|---|---|---|---|---|
| Fam136a | 8.129083 | 8.139322 | 8.1513 | -0.012013496 | -0.022253132 | 8.1625962 | 8.3119353 | 8.09904 | 0.21289745 | 0.063558305 |
| Kcmf1 | 9.877915 | 9.864108 | 9.8779 | -0.01382685 | -1.99558E-05 | 9.7232286 | 10.047649 | 9.83484 | 0.212804749 | -0.111615288 |
| Galnt2 | 7.651447 | 7.471826 | 7.5205 | -0.048627766 | 0.130992485 | 7.464726 | 7.2498376 | 7.03707 | 0.212770001 | 0.42765849 |
| Leng1 | 7.557301 | 7.734791 | 7.6801 | 0.05466763 | -0.122822446 | 7.5613238 | 7.7861621 | 7.57341 | 0.212754757 | -0.01208352 |
| Coq5 | 8.230639 | 8.269729 | 8.2828 | -0.013100738 | -0.052190748 | 8.3545603 | 8.5070254 | 8.29467 | 0.212353697 | 0.059888585 |
| Tusc2 | 8.135528 | 8.057127 | 8.1574 | -0.100279551 | -0.021878833 | 8.2232348 | 8.2341138 | 8.02178 | 0.212336164 | 0.201457231 |
| Ampd2 | 9.190971 | 9.168992 | 9.2413 | -0.072274665 | -0.050294942 | 9.4163774 | 9.542107 | 9.32981 | 0.212292124 | 0.086562518 |
| Alyref | 9.427103 | 9.493901 | 9.5249 | -0.031020285 | -0.097817524 | 9.6454584 | 9.7451584 | 9.53293 | 0.212226599 | 0.11252662 |
| Gnl2 | 9.906874 | 9.885667 | 9.9009 | -0.015204603 | 0.006002612 | 10.075796 | 10.250945 | 10.0388 | 0.212143386 | 0.036993768 |
| Gins4 | 8.118196 | 8.283441 | 8.2658 | 0.017606977 | -0.147638113 | 8.2836644 | 8.4733262 | 8.26119 | 0.212138042 | 0.022476224 |
| Tnfrsf11b | 2.86445 | 2.939193 | 2.6536 | 0.285558892 | 0.210815776 | 2.7595841 | 2.9176788 | 2.70557 | 0.212104315 | 0.054009644 |
| Ptrh2 | 8.001435 | 7.898523 | 7.944 | -0.045500685 | 0.057410985 | 8.2044842 | 8.1886607 | 7.97662 | 0.212036049 | 0.227859572 |
| Coasy | 8.248306 | 8.140929 | 8.283 | -0.142036978 | -0.034660285 | 8.6662124 | 8.4764172 | 8.2646 | 0.211813565 | 0.401608787 |
| Dhx16 | 10.35438 | 10.29858 | 10.397 | -0.098270785 | -0.042471462 | 10.778847 | 10.871749 | 10.66 | 0.21174284 | 0.118841646 |
| Dnajc12 | 5.514447 | 5.842766 | 5.6681 | 0.174623594 | -0.153695058 | 5.6665279 | 5.8833243 | 5.67168 | 0.21164183 | -0.00515464 |
| Cbx1 | 9.991655 | 10.12705 | 10.046 | 0.081171041 | -0.05422389 | 10.075803 | 10.276809 | 10.0652 | 0.211633776 | 0.010628049 |
| Ptk2 | 8.247405 | 8.364036 | 8.253 | 0.1110387 | -0.005591737 | 8.181159 | 8.3284346 | 8.11682 | 0.211614802 | 0.064339257 |
| Pknox1 | 9.864365 | 10.01694 | 9.91 | 0.106949839 | -0.045622783 | 9.7151833 | 9.9839627 | 9.77254 | 0.211425607 | -0.057353818 |
| Atf5 | 7.282351 | 7.408215 | 7.1657 | 0.242516097 | 0.116651814 | 7.2679834 | 7.1955557 | 6.98418 | 0.211372222 | 0.283799859 |
| Masp2 | 2.10631 | 2.173683 | 2.108 | 0.065633746 | -0.001739818 | 2.3213279 | 2.3074058 | 2.09604 | 0.21137055 | 0.225292669 |
| Rps7 | 14.29804 | 14.39511 | 14.269 | 0.1262995 | 0.029228148 | 14.518877 | 14.454106 | 14.2428 | 0.211354756 | 0.27612618 |
| BC024978 | 7.644287 | 7.791337 | 7.5082 | 0.283132651 | 0.136082192 | 7.9410783 | 8.0430784 | 7.83178 | 0.211293698 | 0.109293668 |
| G0s2 | 6.570733 | 6.043926 | 5.7788 | 0.265112408 | 0.791919377 | 6.2605919 | 5.8332544 | 5.62218 | 0.211078795 | 0.638416268 |
| Ppp1r12c | 10.53785 | 10.43297 | 10.576 | -0.142952264 | -0.038072459 | 10.836205 | 10.965972 | 10.7549 | 0.211042984 | 0.081275889 |
| Utrn | 11.78591 | 11.99212 | 11.863 | 0.12941456 | -0.076797286 | 11.429602 | 12.213851 | 12.0028 | 0.2110143 | -0.573234859 |
| Sash3 | 12.20274 | 12.40585 | 12.26 | 0.145983782 | -0.057122632 | 12.515554 | 12.479343 | 12.2684 | 0.210943692 | 0.247154996 |
| Spcs1 | 9.597875 | 9.577789 | 9.5821 | -0.004304575 | 0.015781572 | 9.5976704 | 9.7269758 | 9.51604 | 0.210938032 | 0.081632584 |
| Msn | 14.53535 | 14.64408 | 14.483 | 0.161414836 | 0.052685584 | 14.651732 | 14.743935 | 14.533 | 0.21088587 | 0.11868293 |
| Col4a2 | 9.906025 | 10.03121 | 9.4037 | 0.627500382 | 0.502319134 | 9.8666092 | 9.4955055 | 9.28463 | 0.210878809 | 0.581982563 |
| Zfyve21 | 4.771141 | 4.932505 | 4.9578 | -0.025311992 | -0.186675335 | 5.0126274 | 4.8200961 | 4.60923 | 0.210861803 | 0.403393114 |
| Commd4 | 9.236846 | 9.28357 | 9.2775 | 0.006021276 | -0.040702272 | 9.6325032 | 9.5785074 | 9.36766 | 0.21085227 | 0.264848041 |
| Rpl9 | 14.50243 | 14.48919 | 14.483 | 0.00633802 | 0.019584242 | 14.583564 | 14.652152 | 14.4414 | 0.210717464 | 0.142129131 |

| | | | | | | | | | | |
|---|---|---|---|---|---|---|---|---|---|---|
| Cd300lb | 5.948916 | 5.889553 | 5.2087 | 0.680887953 | 0.740251579 | 5.8963176 | 5.4549746 | 5.24431 | 0.210660473 | 0.652003466 |
| Alg1 | 8.673052 | 8.805014 | 8.5953 | 0.209726634 | 0.077764991 | 8.9308422 | 8.9320196 | 8.72137 | 0.210644805 | 0.209467411 |
| Bap1 | 10.39811 | 10.45441 | 10.379 | 0.0756441 | 0.019338162 | 10.576351 | 10.614029 | 10.4034 | 0.210640124 | 0.172962818 |
| Nasp | 9.941225 | 9.924077 | 10.004 | -0.079764185 | -0.062616328 | 9.9070274 | 10.317108 | 10.1066 | 0.210510996 | -0.199569592 |
| Ppih | 8.021433 | 8.04996 | 8.0535 | -0.003495861 | -0.032023058 | 8.3190346 | 8.3883852 | 8.17791 | 0.210478123 | 0.141127471 |
| P2rx7 | 7.793379 | 7.724035 | 7.9737 | -0.249666431 | -0.180321865 | 7.7089392 | 7.8067762 | 7.59631 | 0.210463757 | 0.112626739 |
| Letm1 | 10.12819 | 10.25931 | 10.212 | 0.046839294 | -0.084280623 | 10.338163 | 10.435319 | 10.2249 | 0.210371296 | 0.113215522 |
| Wnt6 | 2.124623 | 2.11789 | 2.1387 | -0.020797182 | -0.014064232 | 2.3440351 | 2.3071813 | 2.09686 | 0.210323277 | 0.247177095 |
| Pom121 | 10.47212 | 10.6517 | 10.512 | 0.139255382 | -0.040324182 | 10.562456 | 10.66107 | 10.4509 | 0.210186894 | 0.111573553 |
| Derl1 | 10.63275 | 10.6357 | 10.582 | 0.053232146 | 0.050284698 | 10.721198 | 10.720379 | 10.5104 | 0.20998415 | 0.210803208 |
| Etf1 | 11.26414 | 11.17997 | 11.161 | 0.018649958 | 0.102823164 | 11.104507 | 11.279903 | 11.0699 | 0.209977104 | 0.034581615 |
| Btbd1 | 9.864863 | 9.857478 | 9.8755 | -0.018026006 | -0.010640853 | 9.8874211 | 10.040563 | 9.83059 | 0.209969625 | 0.056828128 |
| Itk | 12.05005 | 12.23245 | 12.162 | 0.070166776 | -0.112236842 | 11.753089 | 12.419258 | 12.2093 | 0.209937102 | -0.456232518 |
| Gm12504 | 2.8345 | 2.658222 | 2.5664 | 0.091847227 | 0.268125306 | 2.5171007 | 2.5819844 | 2.37217 | 0.20981174 | 0.144928039 |
| Got2 | 10.84056 | 10.81659 | 10.902 | -0.085439866 | -0.061462836 | 10.978754 | 11.067095 | 10.8574 | 0.209707438 | 0.12136689 |
| Gm3226 | 2.95344 | 2.90031 | 2.9685 | -0.068195772 | -0.015065163 | 2.9495932 | 3.0455138 | 2.83591 | 0.209599634 | 0.113679034 |
| Tgs1 | 9.69794 | 9.919905 | 9.753 | 0.166954779 | -0.055010412 | 9.6545711 | 9.9689939 | 9.75961 | 0.209383945 | -0.105038876 |
| Ccnk | 9.994664 | 10.04848 | 10.079 | -0.030874037 | -0.084692331 | 10.234214 | 10.298166 | 10.0889 | 0.209253797 | 0.145301312 |
| Bub3 | 10.25312 | 10.42922 | 10.276 | 0.15313941 | -0.02295741 | 10.354318 | 10.455572 | 10.2464 | 0.209165876 | 0.107911147 |
| Cd2bp2 | 10.62846 | 10.5844 | 10.645 | -0.060451606 | -0.016399018 | 10.668797 | 10.834421 | 10.6253 | 0.20916107 | 0.043536491 |
| Hs3st1 | 7.856199 | 7.891065 | 7.5196 | 0.371490576 | 0.336624935 | 7.9316311 | 7.7206919 | 7.51156 | 0.209130291 | 0.420069547 |
| G6pdx | 9.740158 | 9.68072 | 9.6168 | 0.063907043 | 0.12334508 | 9.8767396 | 9.8672511 | 9.65813 | 0.209122923 | 0.218611414 |
| Eif4g1 | 12.71791 | 12.76839 | 12.653 | 0.115550666 | 0.065068556 | 12.794488 | 12.934477 | 12.7254 | 0.209053898 | 0.069064742 |
| Jakmip2 | 2.883681 | 2.739148 | 2.8799 | -0.140791098 | 0.003742038 | 2.9635386 | 3.1282139 | 2.91924 | 0.208978034 | 0.04430274 |
| Park7 | 10.28739 | 10.28866 | 10.27 | 0.018885316 | 0.017616699 | 10.519514 | 10.502968 | 10.294 | 0.208972198 | 0.225517753 |
| Mocs2 | 8.160635 | 8.118483 | 8.0935 | 0.02502861 | 0.067181507 | 8.1266611 | 8.299952 | 8.09102 | 0.20893649 | 0.035645599 |
| Igkv4-57 | 7.407511 | 7.588861 | 7.3211 | 0.267759897 | 0.086410359 | 8.0738518 | 7.8669415 | 7.65807 | 0.20887189 | 0.41578218 |
| CR974586.5 | 3.883048 | 3.603364 | 3.4088 | 0.194528149 | 0.474212547 | 3.6597023 | 3.5708765 | 3.36208 | 0.20879212 | 0.297617942 |
| Ccdc22 | 8.851292 | 8.861832 | 8.8659 | -0.004041671 | -0.014581402 | 8.8597803 | 8.9564305 | 8.74769 | 0.208745257 | 0.112095028 |
| Fibp | 9.080167 | 9.000839 | 9.1287 | -0.127895798 | -0.048567595 | 9.3670798 | 9.3554604 | 9.14676 | 0.20870486 | 0.220324223 |
| Magi3 | 7.884838 | 7.913978 | 7.9079 | 0.006095457 | -0.023045049 | 7.8707346 | 8.1243418 | 7.91566 | 0.208677526 | -0.044929646 |
| Twf1 | 8.875533 | 8.740751 | 8.7446 | -0.00385875 | 0.13092357 | 8.4800497 | 8.7103946 | 8.50173 | 0.208666378 | -0.021678489 |

| | | | | | | | | | | |
|---|---|---|---|---|---|---|---|---|---|---|
| Rpl39 | 13.39935 | 13.37529 | 13.4 | -0.024914438 | -0.000851836 | 13.461772 | 13.574452 | 13.3658 | 0.20862256 | 0.095942115 |
| Ncstn | 10.69254 | 10.7239 | 10.704 | 0.01988603 | -0.011468718 | 10.98248 | 10.979659 | 10.7711 | 0.208527788 | 0.211348067 |
| Rfesd | 7.211285 | 7.212316 | 7.378 | -0.16563979 | -0.166670352 | 7.363251 | 7.519821 | 7.31133 | 0.208489175 | 0.05191919 |
| Ly75 | 10.42843 | 10.43285 | 10.375 | 0.057384618 | 0.05296576 | 10.204533 | 10.621186 | 10.4127 | 0.20843647 | -0.208216676 |
| Sec61a1 | 11.35364 | 11.41707 | 11.264 | 0.153337366 | 0.089912766 | 11.401789 | 11.512455 | 11.3041 | 0.208321378 | 0.097655317 |
| Ywhaz | 13.48101 | 13.47641 | 13.394 | 0.08272154 | 0.087317638 | 13.419932 | 13.577461 | 13.3693 | 0.20813515 | 0.050606256 |
| Alg11 | 7.94387 | 8.135457 | 8.1908 | -0.0553004 | -0.246887237 | 8.3588852 | 8.3219772 | 8.11413 | 0.207851426 | 0.244759339 |
| Cyb5b | 10.54306 | 10.56802 | 10.522 | 0.04640063 | 0.021446662 | 10.572778 | 10.719803 | 10.5121 | 0.207657398 | 0.060632145 |
| Sik3 | 10.03038 | 10.11776 | 10.095 | 0.022450891 | -0.064928873 | 9.915282 | 10.257831 | 10.0503 | 0.207571138 | -0.134977875 |
| Prr7 | 5.367953 | 5.2794 | 5.1885 | 0.090888004 | 0.179440827 | 5.4545099 | 5.2940077 | 5.08661 | 0.207393207 | 0.367895451 |
| Hras | 8.159365 | 8.217979 | 8.1287 | 0.089232279 | 0.030617626 | 8.2163059 | 8.215898 | 8.00852 | 0.207380299 | 0.20778819 |
| Wsb1 | 10.73496 | 10.60852 | 10.7 | -0.091698048 | 0.034738336 | 10.633642 | 10.917897 | 10.711 | 0.20694265 | -0.077311948 |
| Pebp1 | 10.32493 | 10.38942 | 10.326 | 0.063101816 | -0.001390888 | 10.511769 | 10.468249 | 10.2615 | 0.206731276 | 0.250251571 |
| Fbxw9 | 5.949417 | 5.975075 | 6.1164 | -0.141304332 | -0.166962439 | 6.3433853 | 6.3091903 | 6.10248 | 0.206711718 | 0.24090673 |
| Chmp3 | 9.978622 | 9.98623 | 10.047 | -0.06060479 | -0.068212674 | 10.132918 | 10.134755 | 9.92806 | 0.206690372 | 0.204853364 |
| Gm48493 | 7.065416 | 7.160721 | 7.4301 | -0.269364295 | -0.36466966 | 6.8063865 | 7.9676989 | 7.76104 | 0.206660286 | -0.954652163 |
| Trim28 | 11.73883 | 11.87784 | 11.837 | 0.040579194 | -0.098432744 | 11.948547 | 12.084854 | 11.8782 | 0.206632448 | 0.070324762 |
| Nsun2 | 11.20262 | 11.26509 | 11.129 | 0.136427144 | 0.0739581 | 11.305073 | 11.442869 | 11.2364 | 0.206510848 | 0.068714874 |
| Ptov1 | 7.2565 | 7.217684 | 7.3819 | -0.164237398 | -0.125420496 | 6.8939536 | 7.2517193 | 7.04529 | 0.206427063 | -0.151338717 |
| Ddx47 | 10.48565 | 10.65692 | 10.583 | 0.073572786 | -0.097701368 | 10.720557 | 10.786977 | 10.5807 | 0.206283864 | 0.139864617 |
| Selenoh | 8.514155 | 8.435406 | 8.5468 | -0.1113884 | -0.032639086 | 8.7353033 | 8.6988196 | 8.49264 | 0.206182424 | 0.242666107 |
| Trappc3 | 9.207374 | 9.243056 | 9.2049 | 0.038172172 | 0.002489692 | 9.2974656 | 9.3376421 | 9.13159 | 0.206056437 | 0.165879869 |
| Pdcd11 | 10.78891 | 10.88222 | 10.752 | 0.129775668 | 0.036463076 | 10.807171 | 11.009912 | 10.8039 | 0.206041854 | 0.003300891 |
| Slc36a1 | 8.298892 | 8.268853 | 8.3045 | -0.035623456 | -0.005584653 | 8.3847508 | 8.614226 | 8.40819 | 0.206040808 | -0.023434406 |
| Foxk1 | 11.08332 | 11.27751 | 11.125 | 0.152753796 | -0.04143567 | 11.002276 | 11.3171 | 11.1113 | 0.20582661 | -0.108997549 |
| Prr29 | 1.745297 | 1.764958 | 1.678 | 0.086937262 | 0.067276629 | 1.7601303 | 1.8746193 | 1.669 | 0.2056237 | 0.091134723 |
| Gm9726 | 2.199247 | 2.466983 | 2.1559 | 0.311061171 | 0.043325011 | 2.1623908 | 2.3284629 | 2.12296 | 0.205504523 | 0.039432476 |
| Dnttip2 | 10.27928 | 10.29389 | 10.271 | 0.022789226 | 0.008181458 | 10.248641 | 10.399804 | 10.1944 | 0.205436028 | 0.054272648 |
| Scyl2 | 9.882277 | 10.05459 | 9.9059 | 0.148706503 | -0.023605449 | 9.7556677 | 10.216497 | 10.0113 | 0.205244311 | -0.25558525 |
| Med6 | 7.439196 | 7.527589 | 7.393 | 0.134565305 | 0.046172253 | 7.6899244 | 7.7939217 | 7.58873 | 0.20518774 | 0.101190446 |
| Ssr1 | 11.10628 | 11.03205 | 10.906 | 0.125605412 | 0.199839064 | 10.777549 | 10.95278 | 10.7476 | 0.205176532 | 0.029945546 |
| Strap | 10.55008 | 10.53007 | 10.524 | 0.005999832 | 0.026014246 | 10.522118 | 10.654596 | 10.4495 | 0.205106652 | 0.072628937 |

| | | | | | | | | | | |
|---|---|---|---|---|---|---|---|---|---|---|
| Fbxo8 | 8.319984 | 8.292894 | 8.4026 | -0.109696947 | -0.082607261 | 8.1450883 | 8.4663533 | 8.26125 | 0.205099423 | -0.116165619 |
| Usp39 | 8.629355 | 8.681608 | 8.5536 | 0.128034169 | 0.075780963 | 8.6711917 | 8.7849662 | 8.57994 | 0.205026536 | 0.091252052 |
| Elp1 | 9.925705 | 9.955942 | 9.9798 | -0.023893945 | -0.054130094 | 10.057657 | 10.175276 | 9.97031 | 0.204967066 | 0.087348405 |
| Tmem68 | 8.769932 | 8.78545 | 8.7962 | -0.010775981 | -0.026293056 | 8.8761992 | 9.0530063 | 8.84811 | 0.204899328 | 0.028092286 |
| Diexf | 9.234228 | 9.518703 | 9.2223 | 0.296448224 | 0.011973833 | 9.2789607 | 9.6323797 | 9.42751 | 0.204873129 | -0.148545885 |
| Klk1b27 | 4.496669 | 4.865792 | 5.5898 | -0.724042053 | -1.093164643 | 5.7351331 | 5.1229164 | 4.91809 | 0.204822833 | 0.817039516 |
| Bglap3 | 4.658436 | 4.660787 | 4.9672 | -0.306432951 | -0.308784402 | 5.6412372 | 5.0514849 | 4.8467 | 0.204789265 | 0.794541586 |
| Gm10060 | 4.131114 | 4.255286 | 4.0119 | 0.243384252 | 0.119211799 | 4.2119998 | 4.4146834 | 4.2099 | 0.204780247 | 0.002096596 |
| Cdc42se1 | 11.63204 | 11.75907 | 11.668 | 0.0906673 | -0.036367458 | 11.795311 | 11.792536 | 11.5881 | 0.204418688 | 0.207194259 |
| Pdf | 6.326352 | 6.307556 | 6.2971 | 0.010476055 | 0.029272342 | 6.3621022 | 6.4208134 | 6.21649 | 0.204319214 | 0.145607967 |
| Casp3 | 9.474793 | 9.435412 | 9.622 | -0.186549393 | -0.147168747 | 9.3920671 | 9.9315673 | 9.72731 | 0.204262045 | -0.33523818 |
| Rpl17-ps10 | 3.43485 | 3.273079 | 3.3204 | -0.047271586 | 0.11449887 | 3.2020114 | 3.3726362 | 3.16845 | 0.204187071 | 0.033562348 |
| Sfpq | 12.34203 | 12.54859 | 12.377 | 0.17171692 | -0.034837548 | 12.204301 | 12.563096 | 12.359 | 0.20406475 | -0.154730373 |
| Glmp | 10.2637 | 10.38338 | 10.29 | 0.093807608 | -0.025874132 | 10.540732 | 10.512415 | 10.3084 | 0.204062164 | 0.23237909 |
| Ccn2 | 5.71582 | 5.841183 | 5.7638 | 0.077384822 | -0.047978105 | 5.6765075 | 5.7498365 | 5.54605 | 0.203787925 | 0.130458921 |
| Fam49b | 11.95553 | 11.9051 | 11.869 | 0.036105258 | 0.086536498 | 12.104193 | 12.054515 | 11.8507 | 0.203769398 | 0.253447482 |
| Lrrc73 | 2.500106 | 2.508299 | 2.4196 | 0.088663365 | 0.08047058 | 2.4815382 | 2.6315891 | 2.42818 | 0.203407513 | 0.053356548 |
| Eif1ad | 9.54455 | 9.595946 | 9.5981 | -0.00220093 | -0.053596961 | 9.7285191 | 9.7812929 | 9.57789 | 0.203404155 | 0.150630385 |
| Gm3854 | 4.515528 | 4.479418 | 4.4618 | 0.017664709 | 0.053774544 | 4.5473669 | 4.9650641 | 4.76169 | 0.203373172 | -0.214323972 |
| Gm28041 | 3.638042 | 3.634946 | 3.5071 | 0.127843622 | 0.130940233 | 3.7583588 | 3.7194532 | 3.51609 | 0.203360979 | 0.242266562 |
| Igkv12-46 | 8.512127 | 8.924271 | 8.2846 | 0.639643617 | 0.22750038 | 9.2456449 | 8.9435778 | 8.74025 | 0.203323243 | 0.505390327 |
| Cd8a | 12.01371 | 12.02324 | 12.043 | -0.019302484 | -0.028831222 | 11.822798 | 12.246937 | 12.0438 | 0.203164648 | -0.220974538 |
| Atg4d | 8.746729 | 8.82476 | 8.7795 | 0.045302954 | -0.032727789 | 9.0602502 | 8.9973424 | 8.79418 | 0.203158031 | 0.266065845 |
| Cog4 | 9.179121 | 9.230212 | 9.2548 | -0.024574535 | -0.075664942 | 9.2888631 | 9.5876959 | 9.38458 | 0.203114055 | -0.095718746 |
| Ube2b | 10.63518 | 10.71122 | 10.616 | 0.095363516 | 0.019324002 | 10.688315 | 10.880694 | 10.6776 | 0.203109962 | 0.010730229 |
| Arfgap3 | 7.91691 | 7.763235 | 7.8454 | -0.082169325 | 0.07150553 | 7.6227925 | 7.9873068 | 7.78421 | 0.203096823 | -0.161417444 |
| Olfr775 | 3.654514 | 3.673481 | 3.5428 | 0.13066805 | 0.111700395 | 3.547162 | 3.8910041 | 3.68793 | 0.203073117 | -0.140768943 |
| Vps29 | 9.119955 | 9.156381 | 9.1405 | 0.015861351 | -0.020564067 | 9.2768167 | 9.3385913 | 9.13554 | 0.203050391 | 0.141275729 |
| Prelid3b | 9.152977 | 9.115372 | 9.0612 | 0.054149572 | 0.09175375 | 9.1286528 | 9.3441072 | 9.14108 | 0.203025692 | -0.012428672 |
| Mocs3 | 6.473898 | 6.411885 | 6.5381 | -0.12623851 | -0.064225076 | 6.6389251 | 6.6336579 | 6.43104 | 0.202616661 | 0.20788381 |
| Dnajc11 | 9.82146 | 9.989103 | 9.8518 | 0.137333255 | -0.030310242 | 9.8793445 | 10.211125 | 10.0086 | 0.202553371 | -0.129227005 |
| Isy1 | 10.13708 | 10.13367 | 10.089 | 0.044215305 | 0.047631037 | 10.110319 | 10.380305 | 10.1778 | 0.202552586 | -0.067433131 |

| | | | | | | | | | | |
|---|---|---|---|---|---|---|---|---|---|---|
| Tlr4 | 8.323984 | 8.342677 | 8.2702 | 0.072491404 | 0.053799129 | 8.3508073 | 8.2929401 | 8.09055 | 0.202386952 | 0.260254116 |
| Ube2z | 10.4841 | 10.50936 | 10.486 | 0.023617272 | -0.001642584 | 10.39749 | 10.644606 | 10.4423 | 0.202350278 | -0.044766118 |
| Pcdh10 | 2.605079 | 2.67865 | 2.5508 | 0.1278728 | 0.054302158 | 2.5551617 | 2.7994884 | 2.59714 | 0.202343726 | -0.041982937 |
| Tspan14 | 10.43097 | 10.5806 | 10.458 | 0.122812924 | -0.026817426 | 10.481591 | 10.596714 | 10.3944 | 0.20229859 | 0.087175354 |
| Zfp397 | 9.270064 | 9.315433 | 9.2166 | 0.098839636 | 0.053470671 | 9.3976837 | 9.5329355 | 9.33088 | 0.202051601 | 0.066799811 |
| Rps16 | 13.69662 | 13.67545 | 13.735 | -0.059983902 | -0.038814602 | 13.892093 | 13.864231 | 13.6623 | 0.201980908 | 0.229843381 |
| Srpx | 5.370467 | 5.338516 | 5.2176 | 0.120871526 | 0.152822827 | 5.2189466 | 5.1393243 | 4.93735 | 0.201978839 | 0.281601079 |
| Pdgfrl | 1.603734 | 1.588745 | 1.5714 | 0.017391686 | 0.032380916 | 1.6301647 | 1.7605067 | 1.55854 | 0.201969023 | 0.071627059 |
| Eif1 | 12.83566 | 12.86122 | 12.809 | 0.052178758 | 0.026621724 | 12.770203 | 12.97008 | 12.7682 | 0.201883994 | 0.002007058 |
| Ip6k3 | 4.363245 | 4.496081 | 4.4289 | 0.067206554 | -0.065629621 | 4.6420891 | 4.737761 | 4.53598 | 0.201784088 | 0.106112198 |
| Ptcd1 | 8.740114 | 8.95726 | 8.8026 | 0.154705929 | -0.062439756 | 8.8794245 | 8.942307 | 8.74055 | 0.201760694 | 0.13887818 |
| Taf8 | 9.550033 | 9.76941 | 9.6464 | 0.122977773 | -0.096398852 | 9.9621634 | 10.101311 | 9.89973 | 0.20157615 | 0.062428576 |
| Zfp945 | 8.231458 | 8.043287 | 8.2485 | -0.205226798 | -0.017055275 | 8.3632201 | 8.5982674 | 8.39687 | 0.201394 | -0.033653306 |
| Scly | 9.0853 | 8.86403 | 8.8903 | -0.026262253 | 0.195007766 | 8.9878353 | 9.1338201 | 8.93254 | 0.201284163 | 0.055299415 |
| Slc40a1 | 5.942121 | 6.697032 | 5.7504 | 0.946644858 | 0.191733252 | 5.4804901 | 5.1538302 | 4.9526 | 0.201234038 | 0.527893917 |
| Ciao1 | 9.393256 | 9.456289 | 9.345 | 0.111287264 | 0.048254317 | 9.5370913 | 9.6202689 | 9.41911 | 0.201159964 | 0.117982427 |
| Mettl13 | 8.45291 | 8.57853 | 8.4422 | 0.136310314 | 0.010690937 | 8.7346817 | 8.7685308 | 8.56741 | 0.201117155 | 0.16726811 |
| Gm9522 | 3.265006 | 3.165456 | 3.2802 | -0.114700446 | -0.015150821 | 3.095484 | 3.2058803 | 3.00478 | 0.201102174 | 0.090705849 |
| Cstf2t | 9.722799 | 9.870678 | 9.7599 | 0.11082794 | -0.037051459 | 9.8313846 | 9.8813794 | 9.68034 | 0.201039503 | 0.151044719 |
| Naca | 13.13458 | 13.11836 | 13.152 | -0.033310038 | -0.01708232 | 13.197797 | 13.352189 | 13.1512 | 0.201030944 | 0.046638633 |
| Gm48270 | 5.101928 | 5.342402 | 5.2668 | 0.075596699 | -0.164877635 | 5.1420542 | 5.674116 | 5.47311 | 0.201006491 | -0.331055287 |
| Tpm4 | 11.90996 | 12.03446 | 11.802 | 0.23250804 | 0.108006578 | 12.039053 | 12.078732 | 11.8778 | 0.200956148 | 0.16127689 |
| 1110065P20Rik | 5.830865 | 5.690074 | 5.728 | -0.037890659 | 0.102899509 | 5.6935684 | 5.7845117 | 5.5836 | 0.200908927 | 0.109965559 |
| Nubp1 | 9.637125 | 9.657597 | 9.6287 | 0.028928353 | 0.008455955 | 9.9147866 | 9.857504 | 9.65661 | 0.200892564 | 0.2581752 |
| Tbcd | 10.05152 | 9.900047 | 9.9916 | -0.091565122 | 0.059908586 | 9.952018 | 10.02307 | 9.82222 | 0.200849648 | 0.129797897 |
| Psme4 | 10.67927 | 10.82331 | 10.698 | 0.12541187 | -0.018624534 | 10.649696 | 11.013312 | 10.8126 | 0.200746474 | -0.162870241 |
| Ttc39b | 10.28291 | 10.1011 | 10.142 | -0.040763126 | 0.14104753 | 9.9963204 | 10.520592 | 10.3198 | 0.200745832 | -0.323525562 |
| U2af1 | 10.36526 | 10.40207 | 10.34 | 0.062540418 | 0.025722516 | 10.428632 | 10.694844 | 10.4942 | 0.200606232 | -0.065605753 |
| Rpl5 | 9.497714 | 9.487855 | 9.47 | 0.017864819 | 0.027723952 | 9.7797611 | 9.859512 | 9.65895 | 0.200564274 | 0.120813458 |
| Brk1 | 10.74218 | 10.81841 | 10.764 | 0.053996578 | -0.022232664 | 10.865408 | 10.898024 | 10.6975 | 0.200478628 | 0.167862799 |
| Pycrl | 7.371169 | 7.433797 | 7.4988 | -0.064964861 | -0.127593233 | 7.7789916 | 7.6557031 | 7.45523 | 0.200469137 | 0.323757635 |
| Rbm3-ps | 2.374669 | 2.372165 | 2.3426 | 0.029591347 | 0.032095249 | 2.3028966 | 2.4607625 | 2.26033 | 0.200429947 | 0.042564081 |

| | | | | | | | | | | |
|---|---|---|---|---|---|---|---|---|---|---|
| Dnajc8 | 10.86642 | 10.81708 | 10.818 | -0.000800508 | 0.048542434 | 10.881921 | 11.009258 | 10.8089 | 0.200383104 | 0.073046349 |
| C1d | 8.598441 | 8.546915 | 8.5331 | 0.013820778 | 0.065347678 | 8.6617614 | 8.7303867 | 8.53003 | 0.200355682 | 0.131730403 |
| Npff | 5.070405 | 5.156046 | 5.0875 | 0.068577438 | -0.017063121 | 5.290088 | 5.672249 | 5.47193 | 0.20031484 | -0.18184612 |
| Mybpc2 | 6.383415 | 6.298163 | 6.2672 | 0.030963888 | 0.116215921 | 6.5852477 | 6.5318275 | 6.33156 | 0.200266217 | 0.253686409 |
| Cpox | 8.563462 | 8.670974 | 8.539 | 0.131983044 | 0.02447177 | 8.5741812 | 8.8457411 | 8.6455 | 0.200242592 | -0.071317375 |
| Olfr1160 | 3.215437 | 3.173282 | 3.1924 | -0.01912712 | 0.023028239 | 3.2154053 | 3.4283971 | 3.22818 | 0.200216807 | -0.012774951 |
| App | 11.43716 | 11.17349 | 11.265 | -0.091233508 | 0.172427812 | 10.737691 | 10.981498 | 10.7813 | 0.200192064 | -0.043614265 |
| Amn1 | 7.202628 | 7.243038 | 7.1031 | 0.139977147 | 0.099566686 | 7.2010397 | 7.3801224 | 7.17998 | 0.200144304 | 0.021061606 |
| Riok3 | 10.7947 | 10.862 | 10.781 | 0.08112368 | 0.013821366 | 10.741776 | 11.006472 | 10.8067 | 0.199798332 | -0.064897611 |
| Lrrc47 | 10.25388 | 10.3905 | 10.261 | 0.129077686 | -0.00754344 | 10.261894 | 10.455027 | 10.2554 | 0.19967531 | 0.006542469 |
| Map7d1 | 9.891937 | 9.937446 | 9.7695 | 0.167991605 | 0.122482256 | 9.8921363 | 10.07756 | 9.8779 | 0.199664267 | 0.014240224 |
| 2700062C07Rik | 6.928599 | 6.75452 | 6.9531 | -0.198609398 | -0.024530827 | 7.1204348 | 7.2307142 | 7.03111 | 0.199603854 | 0.089324495 |
| Casp12 | 5.824078 | 5.398621 | 5.6235 | -0.224919663 | 0.200536984 | 5.1254974 | 5.5501074 | 5.35051 | 0.199601841 | -0.225008098 |
| Spata2l | 6.275648 | 6.539765 | 6.2587 | 0.281022504 | 0.016905427 | 6.4304092 | 6.5773947 | 6.37783 | 0.19956845 | 0.052582881 |
| Olfr893 | 2.749225 | 2.838795 | 2.7618 | 0.076948232 | -0.012622174 | 2.8684674 | 3.0915554 | 2.89214 | 0.199414535 | -0.023673464 |
| Olfr552 | 5.150143 | 5.279629 | 5.1348 | 0.144787264 | 0.015300887 | 5.21676 | 5.5423792 | 5.3431 | 0.199280328 | -0.126338857 |
| Nol8 | 10.02811 | 10.14638 | 9.9568 | 0.189590888 | 0.071320199 | 10.020581 | 10.174658 | 9.97547 | 0.199183243 | 0.045106303 |
| Plaa | 10.47023 | 10.45145 | 10.454 | -0.003012762 | 0.015761568 | 10.362031 | 10.628799 | 10.4297 | 0.199102842 | -0.067665577 |
| Phactr1 | 4.333348 | 3.927313 | 4.6469 | -0.719606413 | -0.313571252 | 3.9259673 | 4.2057226 | 4.00667 | 0.199050899 | -0.080704353 |
| Chmp7 | 9.315096 | 9.385409 | 9.3614 | 0.024006884 | -0.04630535 | 9.4467784 | 9.5560278 | 9.35699 | 0.199033363 | 0.089783974 |
| Atp5mpl | 9.451845 | 9.500241 | 9.2608 | 0.239395353 | 0.190998774 | 9.6551783 | 9.6479006 | 9.44889 | 0.199013131 | 0.20629088 |
| Gm10358 | 0.000175 | 0.000296 | 0.066 | -0.065656283 | -0.065776932 | 0.0024237 | 0.1997624 | 0.00104 | 0.198721014 | 0.001382292 |
| Srp9 | 9.64776 | 9.612061 | 9.6512 | -0.039107919 | -0.00340823 | 9.5237386 | 9.7229724 | 9.52445 | 0.198522431 | -0.000711366 |
| Med27 | 7.979776 | 8.046325 | 8.1094 | -0.063101617 | -0.129651322 | 8.1634957 | 8.2004413 | 8.00211 | 0.198334419 | 0.161388781 |
| Gm8773 | 2.452374 | 2.541824 | 2.4993 | 0.042555737 | -0.046894357 | 2.4585665 | 2.7015635 | 2.50325 | 0.198312057 | -0.044684949 |
| 5730480H06Rik | 7.101887 | 6.993154 | 7.2349 | -0.241775756 | -0.133043273 | 7.3167545 | 7.3407465 | 7.14268 | 0.198065747 | 0.174073702 |
| Pik3r5 | 10.60569 | 10.64645 | 10.542 | 0.10441518 | 0.063657388 | 10.430939 | 10.712884 | 10.5149 | 0.198033572 | -0.083911208 |
| Dbr1 | 8.461251 | 8.575508 | 8.5013 | 0.074233526 | -0.040023481 | 8.3905648 | 8.6002194 | 8.40228 | 0.19793721 | -0.011717349 |
| 4833439L19Rik | 9.484626 | 9.536136 | 9.5284 | 0.00771547 | -0.043793933 | 9.6281017 | 9.7110414 | 9.51311 | 0.197934813 | 0.114995182 |
| Gab2 | 7.761078 | 7.677988 | 7.4579 | 0.220081276 | 0.303171723 | 7.6780571 | 7.5088455 | 7.31108 | 0.197767412 | 0.366979024 |
| Gm15542 | 6.274504 | 6.248202 | 6.215 | 0.03317496 | 0.059476996 | 5.5139908 | 6.5639081 | 6.36618 | 0.197723878 | -0.852193447 |
| Dgcr8 | 8.585512 | 8.642488 | 8.7206 | -0.078139126 | -0.135114693 | 8.5102944 | 8.6559688 | 8.45835 | 0.19762219 | 0.051947786 |

| | | | | | | | | | | |
|---|---|---|---|---|---|---|---|---|---|---|
| Trem2 | 3.073329 | 3.195007 | 2.8626 | 0.332426426 | 0.21074835 | 2.95238 | 3.0077921 | 2.81023 | 0.197563753 | 0.142151651 |
| Tor1a | 8.682559 | 8.5899 | 8.6637 | -0.073797497 | 0.018861315 | 8.9755933 | 8.9699821 | 8.77248 | 0.197500964 | 0.203112232 |
| 4930426L09Rik | 3.807227 | 3.679001 | 3.8762 | -0.197167735 | -0.068940956 | 3.8938486 | 3.9862953 | 3.78897 | 0.19732576 | 0.104878994 |
| Psmg2 | 8.395083 | 8.455037 | 8.4476 | 0.007482719 | -0.052471183 | 8.6624313 | 8.7009087 | 8.50359 | 0.197317912 | 0.158840498 |
| Hcfc1 | 11.80837 | 11.89405 | 11.815 | 0.078986806 | -0.006698178 | 11.969767 | 12.051721 | 11.8544 | 0.197288012 | 0.115333906 |
| Cryab | 4.008589 | 4.026682 | 4.0178 | 0.008855164 | -0.009238429 | 4.0238502 | 4.0764067 | 3.87913 | 0.197277569 | 0.144721102 |
| Kdr | 9.643581 | 9.44845 | 9.4555 | -0.007012793 | 0.188118617 | 8.9896647 | 9.4785607 | 9.2813 | 0.197263106 | -0.291632799 |
| Dynlrb1 | 9.541959 | 9.648158 | 9.671 | -0.022803727 | -0.129002342 | 9.8741471 | 9.8411387 | 9.64393 | 0.197210337 | 0.230218719 |
| Vac14 | 9.863629 | 9.97358 | 9.9595 | 0.014040131 | -0.095910775 | 10.135926 | 10.141375 | 9.94429 | 0.197081533 | 0.19163248 |
| Icosl | 12.28744 | 12.44053 | 12.346 | 0.09498673 | -0.05810308 | 12.763879 | 12.682355 | 12.4854 | 0.196973144 | 0.278497311 |
| 3110053B16Rik | 4.199795 | 4.135867 | 4.0729 | 0.063013553 | 0.126941821 | 4.0484156 | 4.1588509 | 3.96198 | 0.196873504 | 0.086438173 |
| Ttc27 | 8.969981 | 9.011683 | 8.8648 | 0.146847708 | 0.105146082 | 8.9512311 | 9.2295557 | 9.03286 | 0.196695222 | -0.081629386 |
| Mgat2 | 9.950107 | 9.827932 | 9.8509 | -0.022958975 | 0.099215988 | 9.7614648 | 9.8531005 | 9.6565 | 0.196598813 | 0.104963079 |
| LTO1 | 8.069264 | 8.102332 | 8.1462 | -0.043873254 | -0.076940777 | 8.330646 | 8.417294 | 8.22075 | 0.196542896 | 0.10989487 |
| Hsd17b12 | 9.539339 | 9.498878 | 9.402 | 0.096896776 | 0.137358136 | 9.7563886 | 9.6082064 | 9.41175 | 0.196452642 | 0.344634855 |
| Mettl22 | 7.033389 | 6.80337 | 7.0557 | -0.252351657 | -0.022332718 | 7.3404417 | 7.2351344 | 7.03869 | 0.196441028 | 0.301748319 |
| Lpin2 | 9.167859 | 9.403757 | 9.3618 | 0.041981951 | -0.193915901 | 9.3759362 | 9.8044985 | 9.60813 | 0.196369245 | -0.232193074 |
| Utp14a | 10.32959 | 10.37427 | 10.271 | 0.103077176 | 0.058400534 | 10.408939 | 10.5524 | 10.3562 | 0.196226938 | 0.052765865 |
| Aimp1 | 9.648342 | 9.655943 | 9.679 | -0.02304608 | -0.030647469 | 9.7704285 | 9.8535974 | 9.65739 | 0.196203118 | 0.113034275 |
| Rpl36-ps12 | 3.079243 | 3.111963 | 3.095 | 0.016999167 | -0.015721277 | 3.217254 | 3.1997339 | 3.00362 | 0.196109675 | 0.21362978 |
| Dnajb2 | 8.424649 | 8.517601 | 8.4744 | 0.043214671 | -0.049737571 | 8.5567578 | 8.6893068 | 8.49342 | 0.19588277 | 0.063333745 |
| B3gat3 | 9.120091 | 9.228306 | 9.1498 | 0.078522001 | -0.029693235 | 9.5030176 | 9.368816 | 9.17311 | 0.195706807 | 0.329908454 |
| AC169509.1 | 4.18149 | 4.002304 | 3.9995 | 0.002757956 | 0.181943667 | 4.0678719 | 4.3339941 | 4.13836 | 0.195629477 | -0.070492758 |
| Arglu1 | 10.51159 | 10.5968 | 10.502 | 0.094399607 | 0.009186828 | 10.888972 | 11.024921 | 10.8293 | 0.195611888 | 0.059662921 |
| 2410015M20Rik | 9.287658 | 9.299821 | 9.3421 | -0.042232383 | -0.054395983 | 9.5124158 | 9.4930102 | 9.29753 | 0.195482388 | 0.214887996 |
| Gm10134 | 2.566444 | 2.623004 | 2.5717 | 0.051327101 | -0.00523348 | 2.5545412 | 2.7862645 | 2.5908 | 0.195466291 | -0.03625696 |
| Pnkp | 9.595129 | 9.640828 | 9.7075 | -0.066719588 | -0.112418585 | 9.9941949 | 9.8589573 | 9.66357 | 0.195384076 | 0.33062161 |
| Vmn2r79 | 5.558824 | 5.648222 | 5.6014 | 0.046850541 | -0.042547246 | 5.5278259 | 6.0846872 | 5.88936 | 0.19533175 | -0.361529536 |
| Map1lc3b | 10.71629 | 10.74136 | 10.755 | -0.013329166 | -0.038397526 | 10.770319 | 10.831282 | 10.636 | 0.195319846 | 0.134356511 |
| Gtf2h4 | 7.671306 | 7.748429 | 7.6811 | 0.067292608 | -0.009829912 | 8.0710871 | 8.017721 | 7.82241 | 0.195309521 | 0.248675618 |
| Gm7027 | 2.788945 | 2.786063 | 2.8523 | -0.066262701 | -0.063380367 | 2.7911756 | 3.0253895 | 2.83012 | 0.195265879 | -0.038947993 |
| Pop5 | 8.429817 | 8.623883 | 8.512 | 0.111834371 | -0.082231478 | 8.6832356 | 8.7749094 | 8.57966 | 0.195246869 | 0.103573049 |

| | | | | | | | | | | |
|---|---|---|---|---|---|---|---|---|---|---|
| Tbc1d30 | 7.039703 | 7.196067 | 7.281 | -0.084949907 | -0.241314701 | 6.656232 | 7.053742 | 6.85852 | 0.195220848 | -0.202289199 |
| Vps11 | 10.42585 | 10.44738 | 10.398 | 0.048898234 | 0.02737065 | 10.610243 | 10.67955 | 10.4844 | 0.19519693 | 0.125889515 |
| Skiv2l | 10.87986 | 10.85643 | 10.9 | -0.043505514 | -0.020084736 | 11.278333 | 11.279954 | 11.0848 | 0.195190716 | 0.193570093 |
| AC122306.1 | 2.528311 | 2.623958 | 2.4639 | 0.160085349 | 0.064437822 | 2.6152662 | 2.7145274 | 2.51949 | 0.195037695 | 0.095776425 |
| Pkd1l3 | 5.710117 | 5.633752 | 5.5503 | 0.08349181 | 0.159856421 | 5.7862887 | 6.1470561 | 5.95229 | 0.194763155 | -0.166004299 |
| Hnrnpa0 | 11.39687 | 11.30821 | 11.318 | -0.010199864 | 0.078461394 | 11.25573 | 11.341723 | 11.147 | 0.194746654 | 0.108753399 |
| Ighv9-3 | 8.518352 | 8.858089 | 8.4707 | 0.387401098 | 0.047664706 | 8.6486771 | 8.9476656 | 8.75293 | 0.194732498 | -0.104256009 |
| Rps27a | 14.02217 | 14.03182 | 14.037 | -0.004996104 | -0.014643154 | 14.094887 | 14.193735 | 13.9991 | 0.194684066 | 0.095835695 |
| Adam1a | 4.814065 | 4.830066 | 4.9485 | -0.118421161 | -0.134422001 | 4.9292422 | 5.0363763 | 4.84176 | 0.194618003 | 0.087483834 |
| Nup88 | 10.25569 | 10.19387 | 10.267 | -0.072749785 | -0.010930488 | 10.507667 | 10.589453 | 10.395 | 0.194476956 | 0.112690539 |
| Csnk2a2 | 10.01732 | 10.10989 | 10.08 | 0.029723049 | -0.062847022 | 9.9885588 | 10.233201 | 10.0388 | 0.1944246 | -0.050218033 |
| Wdr77 | 9.511872 | 9.635042 | 9.4859 | 0.149125165 | 0.025955258 | 9.7026007 | 9.7086357 | 9.51423 | 0.194409993 | 0.188374943 |
| Bin3 | 9.446388 | 9.341435 | 9.4915 | -0.150111229 | -0.045157608 | 9.576835 | 9.5981657 | 9.40389 | 0.194277609 | 0.172946958 |
| Gm5619 | 3.807082 | 3.879885 | 3.7797 | 0.100147322 | 0.027344824 | 3.7716666 | 4.0135214 | 3.81932 | 0.194197061 | -0.04765769 |
| Gm28285 | 5.678204 | 5.710484 | 5.7858 | -0.075293024 | -0.107572487 | 5.7976366 | 5.9158702 | 5.72172 | 0.194150836 | 0.075917148 |
| Rbm6 | 10.64245 | 10.72113 | 10.763 | -0.04213815 | -0.120822316 | 10.82195 | 11.083551 | 10.8895 | 0.194070874 | -0.067530767 |
| Eps15l1 | 10.44093 | 10.56146 | 10.535 | 0.026731724 | -0.093802298 | 10.630664 | 10.834288 | 10.6402 | 0.194062648 | -0.009561858 |
| Uqcrc1 | 9.872116 | 9.875396 | 9.8574 | 0.01804383 | 0.014763768 | 10.027056 | 9.9780819 | 9.78404 | 0.194040372 | 0.243014352 |
| Wars2 | 7.948068 | 7.891747 | 7.9858 | -0.094088558 | -0.037767605 | 8.0261467 | 8.1246562 | 7.93071 | 0.19394362 | 0.095434065 |
| Atpif1 | 8.515313 | 8.496343 | 8.6035 | -0.107161901 | -0.088191264 | 8.6896471 | 8.6534571 | 8.45955 | 0.193911574 | 0.230101622 |
| Sympk | 11.21972 | 11.35436 | 11.268 | 0.086478314 | -0.048168248 | 11.552916 | 11.578873 | 11.3852 | 0.193627462 | 0.167670801 |
| Tbc1d8 | 9.277707 | 9.32195 | 9.3169 | 0.005011234 | -0.039231159 | 9.0752381 | 9.2956759 | 9.10214 | 0.193531153 | -0.026906654 |
| Gm21761 | 1.697582 | 1.71484 | 1.6353 | 0.079555028 | 0.062296525 | 1.7314 | 1.8735869 | 1.68012 | 0.193471736 | 0.051284788 |
| Klra13-ps | 6.129832 | 6.219506 | 6.1977 | 0.021782322 | -0.067891342 | 6.3498842 | 6.6801046 | 6.4867 | 0.193404062 | -0.136816369 |
| Gm8355 | 1.962204 | 1.900782 | 1.9482 | -0.047402553 | 0.014019955 | 1.8809266 | 2.1040647 | 1.91066 | 0.193402423 | -0.029735628 |
| Acot8 | 7.431316 | 7.495999 | 7.5066 | -0.010556912 | -0.075239454 | 7.8623287 | 7.8248149 | 7.63144 | 0.193375726 | 0.230889505 |
| Praf2 | 7.620997 | 7.596278 | 7.6485 | -0.052202639 | -0.027483484 | 7.6459585 | 7.664642 | 7.47135 | 0.193294632 | 0.174611206 |
| Crcp | 8.485741 | 8.542575 | 8.5183 | 0.02432248 | -0.032512238 | 8.5263271 | 8.7100413 | 8.51693 | 0.193108548 | 0.009394376 |
| Htra4 | 2.18652 | 2.285105 | 2.1725 | 0.112596931 | 0.014012104 | 2.1229441 | 2.3648791 | 2.17179 | 0.193093417 | -0.048841586 |
| AA986860 | 3.210325 | 3.225682 | 3.3109 | -0.085174717 | -0.1005316 | 3.1789974 | 3.389954 | 3.19687 | 0.193082876 | -0.017873771 |
| Prpsap1 | 9.424147 | 9.596747 | 9.6199 | -0.023201595 | -0.195802238 | 9.5489336 | 9.8421346 | 9.64908 | 0.193056876 | -0.100144183 |
| Tmem147 | 8.242564 | 8.255498 | 8.1853 | 0.070218433 | 0.057284228 | 8.4988628 | 8.3650347 | 8.1722 | 0.192834977 | 0.326663062 |

| | | | | | | | | | | |
|---|---|---|---|---|---|---|---|---|---|---|
| Enkd1 | 5.096225 | 5.100906 | 4.9453 | 0.155608147 | 0.150926698 | 5.2673084 | 5.3624727 | 5.16968 | 0.192792107 | 0.097627804 |
| Aars2 | 7.762184 | 7.897299 | 7.902 | -0.004743053 | -0.139857591 | 8.2151971 | 8.2705738 | 8.07795 | 0.192622368 | 0.137245627 |
| Hspa1b | 13.81423 | 13.61506 | 13.886 | -0.27116788 | -0.072002224 | 12.948011 | 13.403558 | 13.211 | 0.19259764 | -0.26294924 |
| Kpna4 | 11.7868 | 11.99958 | 11.779 | 0.22079202 | 0.008011198 | 11.766734 | 11.974636 | 11.7821 | 0.192503232 | -0.015398642 |
| Gp6 | 2.477278 | 2.491029 | 2.5128 | -0.021749534 | -0.035500551 | 2.4616162 | 2.7918241 | 2.59939 | 0.192431336 | -0.137776584 |
| Tfe3 | 9.435832 | 9.461316 | 9.318 | 0.143266765 | 0.117783234 | 9.4515006 | 9.4799763 | 9.28757 | 0.192410321 | 0.163934585 |
| Lnpep | 12.44121 | 12.55482 | 12.329 | 0.22579277 | 0.112184432 | 12.105757 | 12.646177 | 12.4539 | 0.192300924 | -0.348119056 |
| Ccdc115 | 8.913812 | 8.728651 | 8.7908 | -0.062112178 | 0.123048819 | 8.9119175 | 8.8547016 | 8.66242 | 0.192282957 | 0.249498897 |
| Tmem79 | 4.247409 | 4.221273 | 4.281 | -0.059717781 | -0.033582657 | 4.6150935 | 4.4321497 | 4.24011 | 0.192040014 | 0.374983763 |
| Chchd10 | 8.40175 | 8.342241 | 8.3751 | -0.032908636 | 0.026599681 | 8.7142851 | 8.4248815 | 8.23287 | 0.192011236 | 0.481414904 |
| Hip1r | 11.38708 | 11.56314 | 11.489 | 0.07391672 | -0.102135918 | 11.829522 | 11.810028 | 11.618 | 0.191987864 | 0.211481458 |
| Mtf1 | 9.35684 | 9.260289 | 9.4244 | -0.164108055 | -0.067557704 | 9.1828641 | 9.432661 | 9.2407 | 0.191960196 | -0.057836721 |
| Hsf2bp | 2.306857 | 2.436152 | 2.2974 | 0.138730351 | 0.009435353 | 2.2941778 | 2.4683008 | 2.27637 | 0.191935794 | 0.017812816 |
| Nsg2 | 11.08209 | 11.17111 | 11.139 | 0.032236818 | -0.056791676 | 10.707998 | 11.222085 | 11.0302 | 0.191924826 | -0.32216216 |
| Bloc1s2 | 8.952982 | 9.025683 | 9.119 | -0.09330593 | -0.166007579 | 9.3173662 | 9.3766644 | 9.18481 | 0.19185206 | 0.132553875 |
| Ralb | 8.766105 | 8.571407 | 8.5516 | 0.019763395 | 0.214461035 | 8.3758973 | 8.4262042 | 8.23436 | 0.191843317 | 0.141536461 |
| Rheb | 9.367314 | 9.410929 | 9.2489 | 0.162053565 | 0.118439261 | 9.4037283 | 9.4490716 | 9.25733 | 0.191744253 | 0.146401 |
| Slbp | 9.946619 | 9.963016 | 10.013 | -0.050208501 | -0.066605321 | 10.014014 | 10.246669 | 10.0549 | 0.191727962 | -0.0409268 |
| Gm29094 | 4.819341 | 5.254509 | 5.0625 | 0.192011222 | -0.243156965 | 4.6188049 | 4.9548507 | 4.76312 | 0.191726923 | -0.144318875 |
| Itfg2 | 8.413174 | 8.623419 | 8.3612 | 0.262179008 | 0.051933595 | 8.6770227 | 8.7366303 | 8.54495 | 0.191683834 | 0.132076235 |
| Dirc2 | 8.447708 | 8.488783 | 8.4747 | 0.014074431 | -0.027000307 | 8.1096008 | 8.3407431 | 8.1491 | 0.191646364 | -0.039495967 |
| Akr1b3 | 11.03335 | 11.14697 | 11.054 | 0.092929128 | -0.020686506 | 11.249905 | 11.238301 | 11.0467 | 0.191605198 | 0.203209756 |
| Traf3ip1 | 7.153584 | 7.284175 | 7.0725 | 0.211711103 | 0.081120298 | 7.278645 | 7.416661 | 7.22508 | 0.191579757 | 0.0535637 |
| Caprin1 | 12.71856 | 12.83442 | 12.666 | 0.168414524 | 0.052553878 | 12.617166 | 12.893515 | 12.7019 | 0.191575266 | -0.08477408 |
| Ctps | 9.196466 | 9.199957 | 9.0962 | 0.103750956 | 0.100259226 | 9.1752053 | 9.2904032 | 9.09886 | 0.191544839 | 0.076346973 |
| Mettl21a | 6.7039 | 6.611157 | 6.7798 | -0.168644044 | -0.075901333 | 6.8184951 | 6.7309344 | 6.53944 | 0.191491038 | 0.279051678 |
| Lpcat2 | 7.701491 | 7.589699 | 7.4214 | 0.1683004 | 0.280092471 | 7.7693574 | 7.5966499 | 7.40523 | 0.191416493 | 0.364124002 |
| Loxl1 | 2.612155 | 2.801689 | 2.5947 | 0.206967387 | 0.017433773 | 2.7085006 | 2.768344 | 2.57696 | 0.191388019 | 0.131544604 |
| Ift20 | 8.042312 | 8.071193 | 7.9755 | 0.09573968 | 0.066859088 | 8.1598783 | 8.041087 | 7.84972 | 0.191366725 | 0.310158058 |
| Steap2 | 2.697267 | 2.697968 | 2.7228 | -0.02478693 | -0.025487909 | 2.7186379 | 2.785097 | 2.59375 | 0.191351306 | 0.124892168 |
| Tsku | 5.067956 | 4.884093 | 4.9676 | -0.08354357 | 0.100318965 | 4.7040352 | 4.7690058 | 4.57769 | 0.191315978 | 0.126345365 |
| Mrpl22 | 7.60451 | 7.834584 | 7.6108 | 0.2237672 | -0.006306451 | 7.8351065 | 7.8263156 | 7.63503 | 0.191290487 | 0.200081399 |

| | | | | | | | | | | |
|---|---|---|---|---|---|---|---|---|---|---|
| Pcbp2 | 12.93813 | 12.9065 | 12.95 | -0.043898444 | -0.012268458 | 13.002643 | 13.162031 | 12.9708 | 0.191270744 | 0.031882354 |
| Hoxb4 | 5.777343 | 5.662087 | 5.7365 | -0.074440423 | 0.040815279 | 5.5019309 | 5.6865787 | 5.49531 | 0.191264514 | 0.006616719 |
| Zfp131 | 9.68751 | 9.874123 | 9.7508 | 0.123342854 | -0.063270348 | 9.709352 | 10.003165 | 9.81191 | 0.191257148 | -0.102555917 |
| Gm37601 | 3.418655 | 3.458434 | 3.294 | 0.164446802 | 0.124667897 | 3.5413222 | 3.7043279 | 3.51314 | 0.191189307 | 0.028183598 |
| Gm12751 | 4.513914 | 4.437316 | 4.6463 | -0.208999075 | -0.132401319 | 5.1310899 | 4.7962408 | 4.60514 | 0.191099145 | 0.525948235 |
| Cbln2 | 1.367261 | 1.395058 | 1.3736 | 0.021506666 | -0.006290837 | 1.3670322 | 1.5516493 | 1.36059 | 0.191063023 | 0.006445959 |
| Cdc42bpb | 8.6368 | 8.514495 | 8.4248 | 0.089671409 | 0.21197707 | 8.0376274 | 8.5553794 | 8.36434 | 0.191037772 | -0.326714209 |
| Trim27 | 10.04195 | 10.10323 | 10.118 | -0.015127061 | -0.076405193 | 10.138907 | 10.265822 | 10.0748 | 0.191031529 | 0.064116894 |
| Dpm2 | 8.586583 | 8.520754 | 8.6415 | -0.120775204 | -0.05494622 | 8.9536175 | 8.8772786 | 8.68627 | 0.191005317 | 0.267344179 |
| Kctd6 | 6.900628 | 7.036455 | 6.8699 | 0.166580912 | 0.030754581 | 6.851856 | 7.0043107 | 6.81333 | 0.190985028 | 0.03853031 |
| Rpl26 | 14.06169 | 14.02884 | 14.004 | 0.024768202 | 0.05761673 | 14.131847 | 14.211452 | 14.0205 | 0.190965232 | 0.111359614 |
| BC005624 | 9.744673 | 9.722922 | 9.6868 | 0.036148488 | 0.057900337 | 9.893689 | 9.9797417 | 9.78879 | 0.190949985 | 0.104897256 |
| Polrmt | 8.877075 | 9.026695 | 8.9858 | 0.040868492 | -0.108751724 | 9.2166849 | 9.4305292 | 9.23959 | 0.190937145 | -0.022907066 |
| Sgf29 | 8.360264 | 8.326151 | 8.3108 | 0.015398935 | 0.049511658 | 8.5625366 | 8.5987149 | 8.40781 | 0.190909144 | 0.154730799 |
| Nbl1 | 4.829926 | 4.722596 | 4.8881 | -0.165535822 | -0.058205838 | 4.5556818 | 4.9069143 | 4.71606 | 0.190853132 | -0.160379322 |
| Mtrex | 10.39719 | 10.53315 | 10.473 | 0.059826338 | -0.076130081 | 10.518252 | 10.669082 | 10.4782 | 0.19084673 | 0.040016832 |
| Iqcb1 | 7.62281 | 7.739164 | 7.905 | -0.165882795 | -0.282236845 | 7.6332559 | 8.0840585 | 7.89358 | 0.190482148 | -0.260320435 |
| Gm10123 | 12.35062 | 12.20604 | 12.234 | -0.028443924 | 0.116130698 | 12.252894 | 12.359552 | 12.1692 | 0.190346482 | 0.0836883 |
| Akr7a5 | 7.341599 | 7.483042 | 7.2981 | 0.18493098 | 0.043487582 | 7.5433906 | 7.4510501 | 7.26079 | 0.190256561 | 0.282596996 |
| Slc38a4 | 5.657275 | 5.28473 | 5.3055 | -0.020817668 | 0.35172755 | 5.0990368 | 5.3427188 | 5.15265 | 0.19006691 | -0.053615098 |
| Serpina3n | 8.525105 | 8.56101 | 7.6258 | 0.935182377 | 0.899276944 | 8.1699512 | 7.5143975 | 7.32452 | 0.189874076 | 0.845427783 |
| Ddit4 | 10.91153 | 10.87071 | 10.867 | 0.004051074 | 0.044874542 | 10.66569 | 10.753814 | 10.564 | 0.189803014 | 0.101678974 |
| Drg1 | 9.876698 | 9.854662 | 9.9316 | -0.076932176 | -0.054896697 | 9.8991421 | 9.9852026 | 9.79541 | 0.18979199 | 0.103731552 |
| 4932438A13Rik | 12.16506 | 12.18113 | 12.179 | 0.00257061 | -0.013500986 | 11.894523 | 12.494436 | 12.3047 | 0.189752176 | -0.410160935 |
| Wdr46 | 9.455495 | 9.583229 | 9.401 | 0.182234388 | 0.054500235 | 9.6075145 | 9.6446007 | 9.45519 | 0.189415302 | 0.152329133 |
| Utp11 | 9.031266 | 9.101064 | 9.1054 | -0.004314224 | -0.074111424 | 9.1362643 | 9.23244 | 9.04307 | 0.189372244 | 0.093196573 |
| Gm15421 | 3.611866 | 3.555208 | 3.6375 | -0.082340531 | -0.02568317 | 3.6260879 | 3.7288714 | 3.53953 | 0.189340312 | 0.086556742 |
| Nsmce2 | 8.068131 | 8.052787 | 8.1323 | -0.07950611 | -0.064161564 | 8.1897317 | 8.2638282 | 8.07462 | 0.189205579 | 0.115109092 |
| Uros | 7.752912 | 7.837492 | 7.7798 | 0.057675744 | -0.026903876 | 8.062534 | 8.138481 | 7.94934 | 0.189139781 | 0.113192781 |
| Brd4 | 11.74396 | 11.82026 | 11.731 | 0.08893237 | 0.012638228 | 11.739175 | 11.9967 | 11.8077 | 0.189026662 | -0.068498349 |
| Snd1 | 11.27961 | 11.38615 | 11.236 | 0.150197208 | 0.043653818 | 11.32825 | 11.50588 | 11.3169 | 0.188984834 | 0.011354714 |
| Gm32536 | 2.617774 | 2.657218 | 2.7007 | -0.043451437 | -0.082896183 | 2.5848723 | 2.7867908 | 2.59782 | 0.188968218 | -0.012950299 |

| | | | | | | | | | | |
|---|---|---|---|---|---|---|---|---|---|---|
| Satb1 | 12.55056 | 12.57181 | 12.458 | 0.113540584 | 0.092293132 | 12.211816 | 12.637341 | 12.4485 | 0.18888128 | -0.236643591 |
| Gm5617 | 4.864171 | 4.693169 | 4.789 | -0.095871214 | 0.075131302 | 4.9993371 | 4.9290254 | 4.74019 | 0.188832039 | 0.259143787 |
| Hnrnpab | 12.13521 | 12.08301 | 12.028 | 0.05522492 | 0.107433962 | 11.984206 | 12.224484 | 12.0358 | 0.188651614 | -0.051626604 |
| Nr2f6 | 6.121103 | 6.049667 | 6.2745 | -0.224845699 | -0.153409743 | 6.1744851 | 6.1042913 | 5.91577 | 0.18851799 | 0.258711752 |
| Fbxw7 | 8.915785 | 8.976592 | 8.8858 | 0.090750705 | 0.029944028 | 8.7640877 | 9.1897374 | 9.0013 | 0.188432427 | -0.237217215 |
| Mtch2 | 8.646837 | 8.653792 | 8.6146 | 0.039163822 | 0.032209366 | 8.7391587 | 8.9014725 | 8.7131 | 0.188367679 | 0.026053831 |
| Utp6 | 10.00088 | 10.07935 | 9.9655 | 0.113815577 | 0.035352792 | 9.9689495 | 10.177572 | 9.98929 | 0.188286752 | -0.020335581 |
| Nfam1 | 9.717077 | 9.865321 | 9.6276 | 0.237708626 | 0.089465167 | 10.089593 | 9.6212392 | 9.43311 | 0.188131881 | 0.656485703 |
| Slc45a4 | 8.292139 | 8.597868 | 8.4354 | 0.162455282 | -0.143274564 | 8.402161 | 8.537428 | 8.34946 | 0.187972344 | 0.052705368 |
| Leng9 | 7.170817 | 7.135548 | 7.1784 | -0.042855524 | -0.007586613 | 7.4890806 | 7.4061754 | 7.21822 | 0.187958866 | 0.270864055 |
| Prkx | 10.11131 | 10.13628 | 10.03 | 0.106023932 | 0.081056577 | 9.9482724 | 10.166322 | 9.97841 | 0.187908714 | -0.030141086 |
| Tspan2 | 7.396526 | 7.332261 | 7.4321 | -0.099812126 | -0.035547578 | 7.5662865 | 7.8341921 | 7.6466 | 0.187597 | -0.080308671 |
| Gm6863 | 4.463358 | 4.525275 | 4.4208 | 0.104500288 | 0.042583197 | 4.703258 | 4.7915497 | 4.60396 | 0.187584788 | 0.09929307 |
| BC049715 | 4.216223 | 4.330451 | 4.3582 | -0.027787005 | -0.142015396 | 4.4088965 | 4.5563729 | 4.36891 | 0.187467708 | 0.03999126 |
| Twnk | 8.161883 | 8.1678 | 8.1647 | 0.003147466 | -0.002768989 | 8.3234208 | 8.3795982 | 8.19219 | 0.187411129 | 0.131233719 |
| Zfp746 | 9.730863 | 9.874734 | 9.8124 | 0.062381187 | -0.081489613 | 9.6254918 | 9.7532726 | 9.56591 | 0.187360016 | 0.059579195 |
| Cyp11a1 | 4.158417 | 4.086965 | 3.9581 | 0.128866238 | 0.200318075 | 3.7621817 | 4.1074758 | 3.92012 | 0.187354052 | -0.157940016 |
| Vmn1r218 | 5.469788 | 5.375395 | 5.6191 | -0.243673308 | -0.149279735 | 5.4935104 | 5.8886555 | 5.70143 | 0.187228705 | -0.207916393 |
| Rpl31 | 13.06676 | 13.10818 | 13.214 | -0.105748714 | -0.147175032 | 13.256535 | 13.297172 | 13.1099 | 0.187223022 | 0.146586686 |
| Mien1 | 8.736349 | 8.801039 | 8.7075 | 0.09352966 | 0.028840051 | 8.9673244 | 8.9803879 | 8.79317 | 0.187214957 | 0.174151522 |
| Cbx5 | 10.25368 | 10.34152 | 10.309 | 0.03223432 | -0.055604575 | 10.308286 | 10.652556 | 10.4654 | 0.187170346 | -0.157099376 |
| Cep170b | 6.450579 | 6.357102 | 6.6416 | -0.284499503 | -0.191022274 | 6.1960621 | 6.583944 | 6.39685 | 0.187098787 | -0.200783064 |
| Gemin5 | 9.717252 | 9.907233 | 9.6598 | 0.247442919 | 0.057461446 | 9.6687678 | 9.9064323 | 9.71934 | 0.187091052 | -0.050573501 |
| Lrp10 | 11.82 | 11.74348 | 11.651 | 0.092091282 | 0.168611394 | 11.960166 | 11.934715 | 11.7478 | 0.186909758 | 0.212360454 |
| Gm9512 | 3.71474 | 3.704312 | 3.7852 | -0.080905562 | -0.070478141 | 3.6982242 | 3.8098039 | 3.62294 | 0.186861727 | 0.075282029 |
| Gm20662 | 1.984264 | 1.95082 | 2.0678 | -0.116951239 | -0.083507871 | 2.0693067 | 2.2050519 | 2.01822 | 0.186834412 | 0.05108917 |
| Igkv8-21 | 7.499796 | 7.76105 | 7.4146 | 0.346417379 | 0.085162961 | 7.7829593 | 7.7830255 | 7.59639 | 0.186634802 | 0.186568555 |
| Gm29013 | 2.181457 | 2.19334 | 2.4312 | -0.237896084 | -0.249779043 | 2.2395279 | 2.2968661 | 2.11039 | 0.186471359 | 0.129133177 |
| Rpl35a | 13.61002 | 13.65254 | 13.629 | 0.023095702 | -0.01942383 | 13.74398 | 13.780398 | 13.594 | 0.18640943 | 0.149991659 |
| Abhd16a | 9.536553 | 9.52338 | 9.4759 | 0.047465961 | 0.060639001 | 9.7910855 | 9.7922745 | 9.60589 | 0.186381556 | 0.185192488 |
| Zmat2 | 10.21793 | 10.25743 | 10.215 | 0.042880476 | 0.003381322 | 10.429487 | 10.538309 | 10.3519 | 0.186369614 | 0.077547279 |
| BC048403 | 7.531498 | 7.503536 | 7.4789 | 0.024669675 | 0.052631169 | 7.5713822 | 7.6086094 | 7.42232 | 0.186292833 | 0.149065646 |

| | | | | | | | | | | |
|---|---|---|---|---|---|---|---|---|---|---|
| Rpl23a | 14.34344 | 14.39205 | 14.368 | 0.023589948 | -0.025020074 | 14.478308 | 14.569467 | 14.3833 | 0.186194032 | 0.095035077 |
| Psat1 | 9.291521 | 9.23511 | 9.2983 | -0.063219813 | -0.006808865 | 9.6280674 | 9.8236821 | 9.63752 | 0.186159092 | -0.00945567 |
| Dr1 | 10.46646 | 10.51387 | 10.467 | 0.046724596 | -0.000675754 | 10.486307 | 10.531669 | 10.3455 | 0.18613007 | 0.140767753 |
| 2410002F23Rik | 10.16845 | 10.09068 | 10.149 | -0.05849627 | 0.019271439 | 10.460716 | 10.556978 | 10.3709 | 0.186092724 | 0.089829996 |
| Susd6 | 11.28083 | 11.42236 | 11.155 | 0.26713876 | 0.12560956 | 11.176552 | 11.329578 | 11.1436 | 0.185958082 | 0.032932789 |
| Bex6 | 3.939609 | 3.816146 | 3.8857 | -0.06959065 | 0.053872901 | 4.2432351 | 4.051091 | 3.86514 | 0.18594625 | 0.378090324 |
| Tnfrsf23 | 5.874348 | 5.948749 | 5.8238 | 0.124976772 | 0.050575169 | 6.0274103 | 5.7613341 | 5.57558 | 0.185749621 | 0.451825875 |
| Clk3 | 10.7966 | 10.67568 | 10.793 | -0.117019056 | 0.00390118 | 10.787847 | 11.021219 | 10.8355 | 0.185734028 | -0.047638446 |
| Nipa2 | 9.300362 | 9.358093 | 9.1976 | 0.160542726 | 0.10281192 | 9.2705846 | 9.4828717 | 9.29718 | 0.185694482 | -0.026592594 |
| Mrpl33 | 8.976116 | 8.939176 | 8.8918 | 0.047342101 | 0.084282494 | 9.1095767 | 9.0763973 | 8.89107 | 0.185331806 | 0.218511202 |
| Cubn | 2.991478 | 2.901685 | 2.9711 | -0.069451309 | 0.02034133 | 2.7693692 | 3.0840839 | 2.89898 | 0.185108075 | -0.129606616 |
| Luc7l3 | 10.5322 | 10.72916 | 10.632 | 0.096791924 | -0.100170462 | 10.981534 | 11.178156 | 10.9931 | 0.185007292 | -0.011614439 |
| Cox11 | 6.742939 | 6.691847 | 6.7356 | -0.043763144 | 0.007328876 | 6.6416368 | 6.9219691 | 6.73703 | 0.18493661 | -0.095395616 |
| Gm10548 | 5.099214 | 5.285511 | 5.62 | -0.334449572 | -0.520746313 | 5.3943193 | 5.4746475 | 5.2898 | 0.184845864 | 0.104517696 |
| Papola | 11.82478 | 11.86 | 11.786 | 0.073680526 | 0.038455602 | 11.754838 | 11.995346 | 11.8105 | 0.184839618 | -0.055668772 |
| Lins1 | 7.541057 | 7.556846 | 7.4341 | 0.122762991 | 0.106973459 | 7.5487849 | 7.6765291 | 7.49169 | 0.184836869 | 0.057092731 |
| Zfyve27 | 9.295034 | 9.427328 | 9.3642 | 0.063147924 | -0.069146132 | 9.7388862 | 9.7730302 | 9.5882 | 0.184832821 | 0.150688852 |
| Lss | 8.494021 | 8.429272 | 8.3169 | 0.112325212 | 0.177074596 | 8.5054471 | 8.5738087 | 8.38904 | 0.184766734 | 0.116405153 |
| Zap70 | 10.35709 | 10.30979 | 10.333 | -0.02302181 | 0.024276904 | 10.206843 | 10.497024 | 10.3123 | 0.18472284 | -0.105457818 |
| Snx15 | 7.736554 | 7.76545 | 7.7815 | -0.016087229 | -0.044983291 | 7.8466582 | 7.8984591 | 7.71388 | 0.184574098 | 0.132773207 |
| Cep104 | 8.789252 | 8.683711 | 8.7382 | -0.054476797 | 0.051063288 | 8.84353 | 8.9203081 | 8.73576 | 0.184551694 | 0.107773606 |
| Nup54 | 9.319751 | 9.304866 | 9.1985 | 0.106391373 | 0.121276747 | 9.2263171 | 9.5332904 | 9.34899 | 0.184303717 | -0.122669518 |
| Rnf26 | 8.473084 | 8.454953 | 8.4781 | -0.023172087 | -0.005041383 | 8.5655056 | 8.627941 | 8.44373 | 0.184209266 | 0.121773897 |
| Med17 | 9.482696 | 9.632694 | 9.4969 | 0.135752412 | -0.014246138 | 9.6661727 | 9.7355966 | 9.55139 | 0.184208529 | 0.114784663 |
| Ighv7-1 | 4.712037 | 4.713096 | 4.643 | 0.070103404 | 0.069044948 | 4.6181851 | 4.7147611 | 4.53059 | 0.18417007 | 0.087594153 |
| Ggct | 8.190093 | 8.262833 | 8.1994 | 0.063475865 | -0.009264854 | 8.2107943 | 8.2872007 | 8.10305 | 0.184154617 | 0.107748221 |
| Cd300c2 | 6.581418 | 6.495072 | 6.5272 | -0.03211927 | 0.054226676 | 7.3774419 | 6.6540126 | 6.47 | 0.18401694 | 0.907446202 |
| Mrpl13 | 8.737426 | 8.756754 | 8.7614 | -0.004670168 | -0.023998051 | 9.0113001 | 8.9880008 | 8.80406 | 0.183938542 | 0.20723781 |
| Spats2l | 4.870473 | 4.861806 | 4.6935 | 0.168321173 | 0.176988498 | 4.4587264 | 4.8207047 | 4.63686 | 0.183846205 | -0.178132097 |
| Polr2d | 8.102505 | 8.066707 | 8.0888 | -0.022114008 | 0.013684115 | 8.2714617 | 8.1916769 | 8.00789 | 0.183787622 | 0.263572475 |
| Anxa5 | 10.44725 | 10.53517 | 10.384 | 0.150841824 | 0.062914888 | 10.419398 | 10.52055 | 10.3375 | 0.183061504 | 0.081909566 |
| Rplp0 | 15.35189 | 15.45491 | 15.389 | 0.066201946 | -0.036826836 | 15.554396 | 15.53402 | 15.3512 | 0.182836702 | 0.203212654 |

| | | | | | | | | | | |
|---|---|---|---|---|---|---|---|---|---|---|
| Top1mt | 6.80484 | 6.855551 | 6.9574 | -0.101897129 | -0.152608742 | 7.1397482 | 7.018335 | 6.83553 | 0.182800258 | 0.304213388 |
| Dgat1 | 8.251225 | 8.176542 | 8.1224 | 0.054108463 | 0.128790685 | 8.5862318 | 8.767254 | 8.5846 | 0.18265497 | 0.001632723 |
| Ddx23 | 11.06048 | 11.11122 | 11.066 | 0.045343288 | -0.005391324 | 11.195475 | 11.301245 | 11.1186 | 0.18263688 | 0.076866692 |
| Tigd2 | 9.271863 | 9.276987 | 9.2698 | 0.007184872 | 0.002061155 | 9.2650126 | 9.368557 | 9.18597 | 0.182585838 | 0.079041423 |
| Ncf1 | 11.28354 | 11.2772 | 11.307 | -0.0293289 | -0.02299142 | 11.517108 | 11.523222 | 11.3407 | 0.182512168 | 0.17639886 |
| Dmkn | 2.64683 | 2.565703 | 2.199 | 0.366727698 | 0.447855239 | 2.2890867 | 2.4014197 | 2.21894 | 0.182476776 | 0.070143809 |
| Brf2 | 6.98644 | 7.036102 | 7.015 | 0.021081571 | -0.028581083 | 7.2149791 | 7.1457925 | 6.96335 | 0.18244709 | 0.251633697 |
| Lrpap1 | 9.223876 | 9.15708 | 9.3046 | -0.147518376 | -0.080721902 | 9.3993185 | 9.3942513 | 9.21182 | 0.182430119 | 0.187497326 |
| Mlst8 | 8.589042 | 8.739167 | 8.7766 | -0.037431055 | -0.187555602 | 8.9630383 | 9.001351 | 8.81894 | 0.182410692 | 0.144097958 |
| Lgi1 | 2.13926 | 2.178683 | 1.9722 | 0.206488787 | 0.167065664 | 2.0349947 | 2.2798271 | 2.09743 | 0.182394745 | -0.062437598 |
| Ywhaq | 11.36069 | 11.37434 | 11.314 | 0.060756038 | 0.047109756 | 11.330735 | 11.406186 | 11.2238 | 0.18238936 | 0.106937931 |
| Kansl2 | 10.195 | 10.28957 | 10.248 | 0.041731322 | -0.05284212 | 10.219616 | 10.349171 | 10.1668 | 0.1823327 | 0.052777347 |
| Hspbap1 | 8.3426 | 8.449149 | 8.3375 | 0.111628458 | 0.005078842 | 8.4073904 | 8.5024424 | 8.32014 | 0.182306874 | 0.087254847 |
| Ppp1r3d | 3.810859 | 3.668001 | 3.4453 | 0.222686146 | 0.365543893 | 3.5605741 | 3.5584297 | 3.3764 | 0.182034469 | 0.184178934 |
| Gm14137 | 3.349386 | 3.360211 | 3.4703 | -0.1100533 | -0.120878464 | 3.2789586 | 3.4772805 | 3.29529 | 0.181989414 | -0.016332484 |
| D030056L22Rik | 7.865793 | 7.920168 | 7.9114 | 0.008769155 | -0.045605851 | 7.9929343 | 8.0751452 | 7.89324 | 0.181905463 | 0.099694604 |
| Slc38a7 | 8.189285 | 8.26589 | 8.2968 | -0.030952095 | -0.107556617 | 8.3127017 | 8.4041429 | 8.22257 | 0.18157783 | 0.090136635 |
| Atp1b3 | 12.33205 | 12.41249 | 12.349 | 0.063384486 | -0.017047756 | 12.018925 | 12.359136 | 12.1777 | 0.181463198 | -0.158748634 |
| Rras2 | 9.937383 | 10.06389 | 9.9702 | 0.093724508 | -0.032784143 | 9.9336654 | 10.074993 | 9.89357 | 0.181427002 | 0.040098985 |
| Igkv1-133 | 5.400096 | 5.488219 | 5.5375 | -0.04930948 | -0.13743232 | 5.3782804 | 5.5765412 | 5.39527 | 0.181273135 | -0.016987633 |
| Mri1 | 8.605516 | 8.771599 | 8.6031 | 0.168547412 | 0.002465006 | 8.9916337 | 8.8374181 | 8.65616 | 0.181254477 | 0.335470086 |
| Nup98 | 11.35734 | 11.38829 | 11.246 | 0.142462638 | 0.111506488 | 11.185268 | 11.429388 | 11.2482 | 0.181221118 | -0.062898877 |
| Tfam | 9.963298 | 9.939533 | 10.009 | -0.069127944 | -0.045363283 | 10.032371 | 10.230159 | 10.049 | 0.181145342 | -0.016642604 |
| Mthfsl | 8.162017 | 8.142463 | 8.1472 | -0.004708567 | 0.014845786 | 8.4260972 | 8.3996962 | 8.21868 | 0.181012129 | 0.207413153 |
| Cxcr1 | 2.629361 | 2.605614 | 2.5152 | 0.090451388 | 0.114199008 | 2.6896937 | 2.7185634 | 2.53758 | 0.18097933 | 0.152109623 |
| Sec22b | 9.329965 | 9.300261 | 9.2121 | 0.088160549 | 0.11786492 | 9.1566165 | 9.3874625 | 9.20651 | 0.180951495 | -0.049894479 |
| Sypl | 10.83118 | 10.86666 | 10.799 | 0.067391088 | 0.03191758 | 10.82014 | 10.837135 | 10.6562 | 0.180891544 | 0.163896212 |
| Pdlim4 | 7.140142 | 7.272421 | 7.1944 | 0.078069517 | -0.054209208 | 7.2536178 | 7.1683251 | 6.98751 | 0.180811139 | 0.266103882 |
| Arhgap27 | 10.66657 | 10.65198 | 10.762 | -0.109841426 | -0.095250614 | 10.892553 | 10.975986 | 10.7953 | 0.180735294 | 0.097302155 |
| Bcl7c | 8.077186 | 8.066846 | 8.2936 | -0.226732849 | -0.216392635 | 8.2722927 | 8.2559721 | 8.07539 | 0.180577929 | 0.196898524 |
| Nox4 | 3.629735 | 3.519744 | 3.7043 | -0.184526655 | -0.074536437 | 3.7386627 | 3.8477196 | 3.66717 | 0.180554297 | 0.071497409 |
| Rab2a | 9.966855 | 9.895857 | 9.9393 | -0.043438851 | 0.027559681 | 9.8006975 | 9.9276458 | 9.74716 | 0.180490082 | 0.053541833 |

| | | | | | | | | | | |
|---|---|---|---|---|---|---|---|---|---|---|
| Trmt11 | 7.869233 | 7.822269 | 7.8119 | 0.010351568 | 0.057315631 | 7.9458283 | 7.9733628 | 7.79288 | 0.180484046 | 0.152949557 |
| Rps26-ps1 | 2.995299 | 2.843361 | 2.9886 | -0.14519295 | 0.006744524 | 2.8975424 | 3.0342269 | 2.85375 | 0.180479705 | 0.043795221 |
| Pou2af1 | 11.94844 | 12.0996 | 11.908 | 0.191644578 | 0.040483784 | 12.313159 | 12.041174 | 11.8608 | 0.180420364 | 0.452405936 |
| Sphk1 | 5.05909 | 4.977668 | 4.8858 | 0.091907479 | 0.17332949 | 4.4262661 | 4.9342084 | 4.7538 | 0.180405838 | -0.327536425 |
| Tsx | 1.270292 | 1.291627 | 1.2679 | 0.023735414 | 0.002400632 | 1.3460764 | 1.4618269 | 1.28154 | 0.180282851 | 0.064532313 |
| Gpank1 | 8.813131 | 8.929371 | 8.874 | 0.055393831 | -0.060846454 | 8.889696 | 9.0079302 | 8.82776 | 0.1801652 | 0.061931034 |
| Gm20721 | 4.518186 | 4.532481 | 4.619 | -0.086498654 | -0.100793777 | 4.5830135 | 4.6031128 | 4.42299 | 0.180123273 | 0.16002398 |
| Ppig | 10.75277 | 10.69392 | 10.712 | -0.017837702 | 0.041011098 | 10.679706 | 10.918049 | 10.738 | 0.18009572 | -0.058247848 |
| Mrpl10 | 9.319284 | 9.35688 | 9.3187 | 0.038205671 | 0.00060942 | 9.5240157 | 9.4929702 | 9.31298 | 0.179992575 | 0.211038052 |
| Gtf2h5 | 8.485426 | 8.482536 | 8.4671 | 0.015432974 | 0.018323789 | 8.6618845 | 8.6779807 | 8.49802 | 0.179961104 | 0.163864894 |
| Gm13092 | 7.37838 | 7.533134 | 7.4601 | 0.073030551 | -0.081723736 | 7.4360002 | 7.7697131 | 7.5898 | 0.179912357 | -0.153800606 |
| Aldh16a1 | 9.306994 | 9.223218 | 9.3991 | -0.175865574 | -0.092089783 | 9.6644656 | 9.5866028 | 9.40683 | 0.179773549 | 0.257636422 |
| Pi4kb | 10.3062 | 10.43883 | 10.332 | 0.10657325 | -0.026054674 | 10.333357 | 10.452592 | 10.2729 | 0.179729122 | 0.060494203 |
| Vmn1r65 | 2.322115 | 2.370525 | 2.3525 | 0.018006664 | -0.030403086 | 2.3126837 | 2.575162 | 2.3956 | 0.179566162 | -0.082912085 |
| Cep57 | 9.660694 | 9.744282 | 9.6955 | 0.048813629 | -0.034774345 | 9.7428179 | 9.9927334 | 9.81319 | 0.179540786 | -0.070374735 |
| Mid1-ps1 | 1.832661 | 1.849699 | 1.8237 | 0.025985983 | 0.008948422 | 1.8631011 | 1.9886208 | 1.80916 | 0.179460621 | 0.05394086 |
| Esf1 | 9.480063 | 9.479837 | 9.407 | 0.072880311 | 0.07310635 | 9.3403342 | 9.621814 | 9.44246 | 0.179356921 | -0.102122878 |
| Rgs19 | 10.42402 | 10.37901 | 10.393 | -0.013893758 | 0.031114868 | 10.717636 | 10.555427 | 10.3761 | 0.17930051 | 0.341509422 |
| Duoxa2 | 1.700142 | 1.636531 | 1.6431 | -0.006579219 | 0.057032066 | 1.7258969 | 1.8076069 | 1.62831 | 0.179292285 | 0.097582334 |
| Dctn4 | 9.74632 | 9.687862 | 9.6286 | 0.059300131 | 0.117757574 | 9.7206461 | 9.7851252 | 9.60583 | 0.179290543 | 0.114811388 |
| Tns3 | 9.474741 | 9.449312 | 9.337 | 0.112279186 | 0.137708153 | 9.264619 | 9.3124753 | 9.13332 | 0.179150777 | 0.131294504 |
| Gnpnat1 | 8.3875 | 8.403064 | 8.3838 | 0.019233746 | 0.003670499 | 8.2934942 | 8.5519753 | 8.37284 | 0.179138576 | -0.079342568 |
| Il1f9 | 5.17419 | 5.746069 | 4.5585 | 1.187588688 | 0.615709383 | 5.4696771 | 4.9901231 | 4.811 | 0.179125605 | 0.658679679 |
| Llgl2 | 8.123621 | 8.05345 | 8.1322 | -0.078735013 | -0.008564066 | 8.354582 | 8.1002506 | 7.92129 | 0.178959133 | 0.433290481 |
| Hcn3 | 6.000705 | 6.129489 | 5.9285 | 0.200991677 | 0.072207862 | 6.1141462 | 6.1802721 | 6.00148 | 0.178795223 | 0.112669308 |
| Sema4d | 12.03248 | 12.08487 | 11.947 | 0.137776436 | 0.085386404 | 12.199412 | 12.162089 | 11.9833 | 0.178753486 | 0.216076208 |
| Ythdc1 | 11.46218 | 11.61998 | 11.498 | 0.122083796 | -0.03571332 | 11.437957 | 11.769265 | 11.5905 | 0.178750524 | -0.152557733 |
| Vmn2r31 | 3.069696 | 3.045315 | 3.1227 | -0.077343144 | -0.052962318 | 3.1703495 | 3.3961314 | 3.21742 | 0.178709943 | -0.04707199 |
| Gm47685 | 3.126842 | 3.095526 | 3.1192 | -0.023627296 | 0.007689122 | 3.0127791 | 3.3007769 | 3.12212 | 0.178655166 | -0.109342561 |
| Seh1l | 10.12861 | 10.08314 | 10.059 | 0.024283257 | 0.069748924 | 10.164207 | 10.263062 | 10.0844 | 0.178633902 | 0.079778919 |
| Cacna1s | 5.206462 | 5.384745 | 5.241 | 0.143774213 | -0.034508644 | 5.3735171 | 5.4707668 | 5.2922 | 0.178566086 | 0.081316374 |
| Bet1l | 8.504864 | 8.479299 | 8.5053 | -0.025961216 | -0.000396144 | 8.6930873 | 8.6278289 | 8.44935 | 0.178477599 | 0.243735934 |

| | | | | | | | | | | |
|---|---|---|---|---|---|---|---|---|---|---|
| Krit1 | 9.934477 | 9.970639 | 10.018 | -0.047448048 | -0.083610428 | 10.033584 | 10.388422 | 10.2101 | 0.178314264 | -0.176524328 |
| Ncor2 | 10.94402 | 11.06239 | 11.03 | 0.032089908 | -0.08628127 | 11.032155 | 11.163171 | 10.9851 | 0.178065536 | 0.047049191 |
| Sod1 | 11.03446 | 10.84613 | 10.97 | -0.123965456 | 0.064365804 | 11.00472 | 11.086893 | 10.9088 | 0.178052312 | 0.095879106 |
| Slc25a45 | 9.186178 | 9.335633 | 9.3039 | 0.031728697 | -0.117726706 | 9.5028632 | 9.4764121 | 9.29843 | 0.177986345 | 0.204437414 |
| Gpi1 | 12.66964 | 12.60213 | 12.67 | -0.067913416 | -0.000412248 | 12.695316 | 12.810387 | 12.6325 | 0.177893096 | 0.062822042 |
| Fam118b | 8.26089 | 8.422728 | 8.3002 | 0.122487824 | -0.039349513 | 8.2827071 | 8.402292 | 8.2244 | 0.177886962 | 0.058302074 |
| Pam16 | 5.770216 | 5.730964 | 5.6078 | 0.123153575 | 0.162406178 | 6.2319971 | 6.0287 | 5.85102 | 0.17767682 | 0.380973905 |
| Zswim7 | 5.31199 | 5.230173 | 5.3022 | -0.072042457 | 0.00977399 | 5.4258196 | 5.3889039 | 5.21133 | 0.177578725 | 0.214494389 |
| Igkv6-15 | 9.160371 | 9.561705 | 9.1277 | 0.434053645 | 0.032720218 | 9.6215325 | 9.5587986 | 9.38124 | 0.177562139 | 0.240295987 |
| Vmn1r202 | 3.357552 | 3.264633 | 3.2647 | -9.27014E-05 | 0.092826658 | 3.3674931 | 3.520885 | 3.34333 | 0.177551603 | 0.024159679 |
| Itgb1bp1 | 8.583011 | 8.582073 | 8.5885 | -0.006435168 | -0.00549702 | 8.5245429 | 8.5782194 | 8.40071 | 0.177511787 | 0.123835299 |
| Cd160 | 7.743214 | 7.839112 | 7.7974 | 0.041695785 | -0.05420209 | 7.751135 | 8.0277716 | 7.85037 | 0.177399251 | -0.099237375 |
| Mbip | 7.499138 | 7.423142 | 7.5473 | -0.124163622 | -0.048168395 | 7.5876888 | 7.680809 | 7.50353 | 0.177277802 | 0.084157505 |
| Tpra1 | 8.684505 | 8.861647 | 8.8466 | 0.015053545 | -0.162088458 | 8.9619105 | 8.9741374 | 8.7969 | 0.177235975 | 0.165009047 |
| Pik3r4 | 9.919184 | 10.1514 | 9.9848 | 0.166576933 | -0.065644206 | 10.371076 | 10.255334 | 10.0782 | 0.177101768 | 0.292843832 |
| Trub2 | 8.200578 | 8.544856 | 8.375 | 0.169899226 | -0.174378016 | 8.791469 | 8.658685 | 8.48165 | 0.177032826 | 0.309816835 |
| Selenon | 7.865355 | 7.463003 | 7.2498 | 0.213196487 | 0.615548952 | 7.613356 | 7.2202432 | 7.04322 | 0.177023401 | 0.570136142 |
| Eef1d | 11.62995 | 11.6977 | 11.668 | 0.029639692 | -0.038113882 | 11.833677 | 11.901777 | 11.7249 | 0.176873778 | 0.108773121 |
| Mrps2 | 8.589559 | 8.606324 | 8.5562 | 0.050164473 | 0.033399448 | 8.7010295 | 8.7070117 | 8.53016 | 0.176855223 | 0.170873023 |
| Dpf2 | 10.24602 | 10.38111 | 10.278 | 0.103050686 | -0.032043055 | 10.483144 | 10.540026 | 10.3632 | 0.176820556 | 0.11993858 |
| Mmp12 | 6.794247 | 8.564441 | 6.2742 | 2.290247508 | 0.520053812 | 6.6162453 | 6.3484803 | 6.17179 | 0.17669039 | 0.444455387 |
| Cog1 | 9.345697 | 9.605513 | 9.5265 | 0.079036504 | -0.180779349 | 9.6184468 | 9.7785823 | 9.60198 | 0.176606512 | 0.016470967 |
| Senp1 | 10.46377 | 10.59106 | 10.419 | 0.171945908 | 0.044662318 | 10.499753 | 10.827099 | 10.6507 | 0.176410646 | -0.150935139 |
| Rs1 | 1.873047 | 1.909551 | 1.895 | 0.014517042 | -0.021986729 | 1.7979081 | 2.0623821 | 1.886 | 0.17638655 | -0.088087456 |
| Pmm2 | 9.780611 | 9.691248 | 9.6544 | 0.036829922 | 0.126193049 | 9.7051509 | 10.008413 | 9.83203 | 0.176384703 | -0.12687762 |
| Palb2 | 7.095745 | 7.150412 | 7.0601 | 0.090330276 | 0.035662838 | 7.1936784 | 7.3485533 | 7.17232 | 0.176234672 | 0.021359818 |
| Zfp729a | 11.92263 | 11.83097 | 11.799 | 0.032025452 | 0.12368619 | 11.819795 | 11.866423 | 11.6903 | 0.176090514 | 0.129462282 |
| Polr1e | 8.281032 | 8.394541 | 8.2414 | 0.1531289 | 0.039620556 | 8.5023102 | 8.486912 | 8.31091 | 0.175997671 | 0.191395866 |
| Clk2 | 10.11618 | 10.20872 | 10.214 | -0.005603706 | -0.098140056 | 10.408358 | 10.57719 | 10.4012 | 0.175980327 | 0.007148709 |
| Nme3 | 7.054339 | 7.069077 | 7.0108 | 0.058306725 | 0.043568236 | 7.3537202 | 7.1937878 | 7.01793 | 0.17585667 | 0.335789076 |
| Rpl7l1 | 10.25418 | 10.40282 | 10.304 | 0.098633266 | -0.050005824 | 10.4188 | 10.555977 | 10.3802 | 0.17581347 | 0.038636348 |
| Nus1 | 10.04084 | 10.00322 | 10.004 | -0.000658591 | 0.036965369 | 9.9276644 | 10.095284 | 9.91952 | 0.175764293 | 0.00814505 |

| | | | | | | | | | | |
|---|---|---|---|---|---|---|---|---|---|---|
| Taf13 | 8.096648 | 8.18283 | 8.1408 | 0.042021701 | -0.04416056 | 8.0391372 | 8.299598 | 8.12395 | 0.175643399 | -0.084817389 |
| Stxbp2 | 10.58058 | 10.58346 | 10.594 | -0.01024589 | -0.013126314 | 11.113864 | 11.009734 | 10.8341 | 0.175636182 | 0.279765837 |
| Ppp1r1b | 2.549797 | 2.254925 | 2.3805 | -0.125619185 | 0.169252622 | 2.398707 | 2.6039108 | 2.42836 | 0.175555583 | -0.029648139 |
| Spns1 | 9.142672 | 9.430811 | 9.2885 | 0.142336925 | -0.145801993 | 9.6551402 | 9.5773253 | 9.40183 | 0.175492798 | 0.253307741 |
| Guca1b | 3.785475 | 3.795612 | 3.6996 | 0.096041017 | 0.085904085 | 3.9506013 | 4.0677253 | 3.8923 | 0.175429707 | 0.058305739 |
| Gm9844 | 2.140822 | 2.129851 | 2.1618 | -0.031992251 | -0.021021709 | 2.178075 | 2.1785629 | 2.00315 | 0.175415442 | 0.174927555 |
| Ppargc1a | 5.140587 | 5.107833 | 5.0405 | 0.067314263 | 0.100068049 | 5.0343292 | 5.3040954 | 5.12869 | 0.175408021 | -0.094358223 |
| Ly9 | 11.63227 | 11.74412 | 11.711 | 0.03335824 | -0.078492162 | 11.752641 | 11.851193 | 11.6759 | 0.175325756 | 0.076774583 |
| Gm5884 | 4.142338 | 4.007973 | 4.0563 | -0.048373152 | 0.085992399 | 4.1192667 | 3.9869314 | 3.81162 | 0.17531235 | 0.307647709 |
| Tgfb2 | 4.088398 | 4.013299 | 4.3563 | -0.34295467 | -0.267854912 | 4.1557118 | 4.3602447 | 4.18497 | 0.17527735 | -0.029255501 |
| Fam84a | 2.999262 | 2.637106 | 2.695 | -0.057870191 | 0.304285845 | 2.7710343 | 2.8868277 | 2.71158 | 0.175251184 | 0.059457773 |
| Gabarapl2 | 9.881085 | 9.953055 | 9.9566 | -0.00355487 | -0.075524572 | 9.9406035 | 9.9626887 | 9.78746 | 0.175227107 | 0.153141889 |
| Gm37348 | 2.215422 | 2.300177 | 2.2178 | 0.082338425 | -0.002416738 | 2.1941614 | 2.3414122 | 2.16621 | 0.175203981 | 0.027953166 |
| AI597479 | 8.000369 | 8.057929 | 8.0582 | -0.000224565 | -0.057784392 | 8.1237387 | 8.3161161 | 8.14098 | 0.175134836 | -0.017242542 |
| 9430025C20Rik | 2.652149 | 2.643633 | 2.6653 | -0.021622395 | -0.013106878 | 2.7214004 | 2.8940274 | 2.71893 | 0.17509815 | 0.002471174 |
| Coq8b | 8.326349 | 8.275486 | 8.2393 | 0.036219991 | 0.087083052 | 8.591099 | 8.4628383 | 8.28779 | 0.175052634 | 0.303313356 |
| Ildr1 | 6.802901 | 6.882087 | 6.839 | 0.043046549 | -0.036139374 | 6.5180397 | 7.0309955 | 6.85607 | 0.174925494 | -0.338030342 |
| Alkbh5 | 10.92747 | 11.00974 | 10.939 | 0.071151786 | -0.011117608 | 10.77618 | 11.006202 | 10.8313 | 0.174883876 | -0.055138009 |
| Usp4 | 11.21779 | 11.29182 | 11.207 | 0.085094166 | 0.011066072 | 11.227733 | 11.387116 | 11.2122 | 0.174874004 | 0.015490671 |
| Fuk | 7.580755 | 7.872414 | 7.6416 | 0.230786945 | -0.060872229 | 7.8183096 | 7.9303968 | 7.75555 | 0.174849725 | 0.062762516 |
| 0610030E20Rik | 8.809887 | 8.900523 | 8.8117 | 0.088857533 | -0.00177874 | 9.1374199 | 9.1388012 | 8.96398 | 0.174821342 | 0.173440062 |
| C1qtnf6 | 6.508225 | 6.71814 | 6.8036 | -0.085459307 | -0.29537391 | 6.5028424 | 6.5395478 | 6.36473 | 0.174814865 | 0.138109486 |
| Txndc12 | 8.871231 | 8.822897 | 8.9075 | -0.084558202 | -0.036224176 | 8.9844987 | 9.0137 | 8.83897 | 0.174734763 | 0.14553338 |
| Gm10146 | 1.394882 | 1.788193 | 1.4116 | 0.376552969 | -0.016758744 | 1.496736 | 1.6999981 | 1.52528 | 0.174714824 | -0.028547369 |
| Pla2r1 | 4.539661 | 4.103334 | 4.3222 | -0.218859807 | 0.217467039 | 3.9829534 | 4.3529697 | 4.17857 | 0.174400569 | -0.195615749 |
| Hif1an | 9.675761 | 9.711133 | 9.6839 | 0.027195986 | -0.008176671 | 9.7617453 | 9.9179666 | 9.74359 | 0.174377044 | 0.01815568 |
| Gm43738 | 1.767407 | 1.890477 | 1.8292 | 0.06129496 | -0.061774817 | 1.8330446 | 2.0827309 | 1.90867 | 0.174057656 | -0.075628628 |
| Nmd3 | 9.506623 | 9.42191 | 9.499 | -0.077129219 | 0.007583525 | 9.6195126 | 9.6863737 | 9.51239 | 0.173980544 | 0.107119461 |
| Ssu72 | 9.730718 | 9.603362 | 9.6779 | -0.074494911 | 0.052861581 | 9.9420663 | 9.8537153 | 9.67977 | 0.173946842 | 0.262297861 |
| Ndufaf2 | 6.244083 | 6.283739 | 6.3327 | -0.048991725 | -0.088648009 | 6.307846 | 6.3628748 | 6.18897 | 0.173904854 | 0.118876046 |
| Ighv14-2 | 7.19792 | 7.374399 | 7.2479 | 0.126451681 | -0.050027221 | 7.4541831 | 7.3454788 | 7.17158 | 0.173894408 | 0.282598723 |
| Crls1 | 7.19698 | 7.297782 | 7.2215 | 0.076237129 | -0.024565192 | 7.2498623 | 7.4801937 | 7.30638 | 0.173817136 | -0.056514307 |

| | | | | | | | | | | |
|---|---|---|---|---|---|---|---|---|---|---|
| Csmd3 | 2.258839 | 2.288273 | 2.2321 | 0.056191332 | 0.026756554 | 2.1873599 | 2.4650099 | 2.29124 | 0.17376739 | -0.103882584 |
| Zfp296 | 7.307324 | 7.338531 | 7.4175 | -0.078968382 | -0.110176012 | 7.3746866 | 7.3628107 | 7.18907 | 0.173745585 | 0.185621487 |
| Sh2b3 | 11.28503 | 11.36703 | 11.302 | 0.065260292 | -0.016737776 | 11.540294 | 11.453839 | 11.2802 | 0.173677894 | 0.260133076 |
| Btg3 | 7.943423 | 8.004406 | 8.0636 | -0.059238564 | -0.120220956 | 7.8362885 | 8.1401684 | 7.9665 | 0.173669973 | -0.130209934 |
| Strc | 2.040339 | 2.11396 | 2.1546 | -0.040628407 | -0.114249879 | 2.0635554 | 2.3307706 | 2.1572 | 0.173567504 | -0.093647677 |
| Mfsd11 | 8.214933 | 8.397694 | 8.2919 | 0.105830999 | -0.076930019 | 8.1752248 | 8.2599589 | 8.0864 | 0.173562017 | 0.088827914 |
| Klhl33 | 3.781814 | 3.820814 | 3.6974 | 0.123374387 | 0.084374688 | 3.7477765 | 4.0054288 | 3.83189 | 0.173539949 | -0.084112324 |
| Ergic1 | 9.480682 | 9.485208 | 9.4255 | 0.059680346 | 0.055154414 | 9.4173799 | 9.5681601 | 9.39483 | 0.173334341 | 0.022554154 |
| Rhbg | 0.307861 | 0.300017 | 0.3017 | -0.001682833 | 0.006160572 | 0.3823462 | 0.4822953 | 0.30897 | 0.173324168 | 0.073375048 |
| Mbd2 | 11.00407 | 11.04152 | 11.005 | 0.036784112 | -0.000658962 | 11.011925 | 11.136943 | 10.9638 | 0.173172628 | 0.048154843 |
| Ung | 7.388353 | 7.419242 | 7.4293 | -0.010023956 | -0.040912718 | 7.4315033 | 7.6221384 | 7.44902 | 0.173115641 | -0.017519536 |
| Nfatc1 | 11.4675 | 11.52038 | 11.495 | 0.025359806 | -0.027521992 | 11.357415 | 11.666221 | 11.4932 | 0.172972776 | -0.135833802 |
| Tmem161a | 8.50295 | 8.629794 | 8.4435 | 0.186339584 | 0.059495357 | 8.8512923 | 8.837204 | 8.6644 | 0.172803037 | 0.186891405 |
| Pdzd11 | 7.65427 | 7.582625 | 7.7776 | -0.194972396 | -0.123327477 | 7.9666018 | 7.8725074 | 7.69972 | 0.172785605 | 0.266879966 |
| Gm20604 | 6.352079 | 6.2023 | 6.3437 | -0.141402251 | 0.008377148 | 6.3792509 | 6.323638 | 6.15095 | 0.172689684 | 0.228302635 |
| Nop14 | 9.951285 | 9.921684 | 9.8329 | 0.088751146 | 0.118352062 | 10.024044 | 10.188674 | 10.0161 | 0.172555428 | 0.007925665 |
| Cep85l | 8.41758 | 8.602398 | 8.6537 | -0.051305855 | -0.236123887 | 8.3266222 | 8.6726162 | 8.50016 | 0.172458857 | -0.173535211 |
| Actr5 | 8.551242 | 8.471896 | 8.5074 | -0.035549875 | 0.043796503 | 8.7035348 | 8.693112 | 8.52084 | 0.172274941 | 0.182697717 |
| Gid8 | 9.672348 | 9.731467 | 9.7019 | 0.029542413 | -0.029577024 | 9.778095 | 9.817756 | 9.64549 | 0.172263527 | 0.132602566 |
| Srp68 | 10.33987 | 10.4166 | 10.337 | 0.079369808 | 0.002631282 | 10.465689 | 10.481301 | 10.3093 | 0.172011304 | 0.156399057 |
| Rae1 | 9.917703 | 9.938424 | 9.925 | 0.01339273 | -0.00732822 | 10.096632 | 10.287613 | 10.1157 | 0.171878479 | -0.019102757 |
| Gm12715 | 2.403365 | 2.500793 | 2.3774 | 0.123366469 | 0.025937868 | 2.5513987 | 2.6204613 | 2.44871 | 0.171753951 | 0.102691356 |
| Nsfl1c | 10.14455 | 10.18299 | 10.16 | 0.022534052 | -0.015898666 | 10.340246 | 10.42099 | 10.2494 | 0.171620618 | 0.090876478 |
| Myo16 | 3.847776 | 3.591362 | 3.5046 | 0.086742479 | 0.343156565 | 3.5804423 | 3.6654679 | 3.4939 | 0.171565891 | 0.086540324 |
| BC028528 | 7.94068 | 7.728093 | 7.8901 | -0.162022078 | 0.05056564 | 8.0729744 | 8.0222806 | 7.85072 | 0.171564882 | 0.222258672 |
| Mrpl18 | 9.088508 | 9.078873 | 9.1536 | -0.074757097 | -0.065122127 | 9.110194 | 9.2864845 | 9.11494 | 0.171546539 | -0.004743973 |
| Serpinb2 | 2.460442 | 2.675839 | 2.5579 | 0.117917167 | -0.097479282 | 2.2902101 | 2.4607216 | 2.2892 | 0.17152248 | 0.001010915 |
| Eif3i | 11.24257 | 11.28971 | 11.255 | 0.03449383 | -0.01265221 | 11.455425 | 11.384459 | 11.213 | 0.17146371 | 0.24242942 |
| Tmsb15l | 5.632273 | 5.778076 | 5.8606 | -0.082504778 | -0.228307726 | 5.5578572 | 5.6328723 | 5.46142 | 0.171454405 | 0.096439361 |
| Rps11 | 14.15645 | 14.18951 | 14.232 | -0.042080464 | -0.075135636 | 14.337255 | 14.389494 | 14.2181 | 0.171384254 | 0.119145883 |
| Cdkl3 | 5.810247 | 5.574591 | 5.794 | -0.219376156 | 0.016279994 | 5.5979542 | 5.932034 | 5.7608 | 0.171236091 | -0.162843703 |
| Bcl2l11 | 9.574634 | 9.772895 | 9.5744 | 0.198512901 | 0.000252094 | 9.3732329 | 9.7102097 | 9.53901 | 0.171199853 | -0.165776912 |

| | | | | | | | | | | |
|---|---|---|---|---|---|---|---|---|---|---|
| Vmn1r172 | 2.638698 | 2.608457 | 2.5674 | 0.041097281 | 0.071337902 | 2.6625579 | 2.8927481 | 2.72159 | 0.171161641 | -0.059028469 |
| Samd15 | 3.651884 | 3.710472 | 3.6606 | 0.049830049 | -0.008758917 | 3.7796407 | 3.9920683 | 3.82096 | 0.171108201 | -0.041319399 |
| Atp6v0d1 | 9.911834 | 9.985568 | 9.8745 | 0.111112593 | 0.037378776 | 10.111528 | 10.041585 | 9.87062 | 0.170965086 | 0.240908676 |
| Slc38a10 | 10.73282 | 10.77411 | 10.721 | 0.05330732 | 0.012017294 | 10.872149 | 10.827221 | 10.6563 | 0.170890594 | 0.215818575 |
| Nol6 | 10.50764 | 10.65157 | 10.521 | 0.130149018 | -0.013775108 | 10.79067 | 10.777695 | 10.6069 | 0.170790168 | 0.183765743 |
| Cdh1 | 6.578532 | 6.346213 | 6.0406 | 0.305565609 | 0.537884276 | 6.3118121 | 6.108114 | 5.93749 | 0.170624618 | 0.374322725 |
| Ssbp4 | 9.044841 | 8.721488 | 8.8652 | -0.143751684 | 0.179600712 | 9.2039624 | 8.8867184 | 8.71615 | 0.17057149 | 0.487815557 |
| Vipas39 | 9.517821 | 9.52727 | 9.4787 | 0.048574184 | 0.039125211 | 9.5714825 | 9.7681631 | 9.59765 | 0.170517017 | -0.026163491 |
| Il1rapl2 | 2.294339 | 2.35359 | 2.2497 | 0.103861297 | 0.044610229 | 2.259837 | 2.4521882 | 2.28172 | 0.170468586 | -0.02188268 |
| Ubac1 | 8.614215 | 8.516328 | 8.546 | -0.029686041 | 0.068201061 | 8.4978305 | 8.7784992 | 8.60805 | 0.170454081 | -0.110214615 |
| Zfpm1 | 8.74866 | 8.580415 | 8.8824 | -0.302020629 | -0.133775608 | 8.8090765 | 8.8665624 | 8.69611 | 0.170452436 | 0.112966626 |
| Mtif3 | 8.350102 | 8.317654 | 8.3236 | -0.005912989 | 0.026535102 | 8.4370611 | 8.4812612 | 8.31099 | 0.170269406 | 0.126069278 |
| 4930539E08Rik | 4.703307 | 4.711287 | 4.8388 | -0.127525176 | -0.135505384 | 4.678635 | 4.9481031 | 4.77795 | 0.170148327 | -0.099319754 |
| Lime1 | 9.569418 | 9.547825 | 9.6943 | -0.146469568 | -0.124876255 | 9.7540241 | 9.9078831 | 9.73778 | 0.170103455 | 0.016244426 |
| Sf3a1 | 10.96423 | 11.03436 | 10.955 | 0.079340374 | 0.009214696 | 11.112354 | 11.181789 | 11.0117 | 0.170082388 | 0.100647447 |
| Aftph | 11.01465 | 11.07054 | 11.047 | 0.023345186 | -0.032543954 | 10.970913 | 11.230204 | 11.0602 | 0.16997083 | -0.089319229 |
| Xntrpc | 6.578922 | 6.858554 | 6.7144 | 0.144137734 | -0.135493563 | 6.9795708 | 6.9751383 | 6.80525 | 0.169885735 | 0.174318192 |
| Snx6 | 10.00795 | 10.00051 | 9.9453 | 0.05523389 | 0.062675247 | 9.9824773 | 10.14418 | 9.9743 | 0.169877008 | 0.008174002 |
| Patz1 | 9.478158 | 9.725855 | 9.6204 | 0.105428608 | -0.142268637 | 9.5485994 | 9.6618201 | 9.49216 | 0.1696621 | 0.056441399 |
| 2200002D01Rik | 5.031438 | 5.166392 | 5.0699 | 0.096488155 | -0.038465738 | 4.8736184 | 5.0126977 | 4.84309 | 0.169611129 | 0.030531857 |
| Fam3b | 1.927392 | 1.935445 | 1.9189 | 0.016544708 | 0.008491276 | 1.9692055 | 2.0452762 | 1.87567 | 0.169606467 | 0.093535783 |
| Adgrb1 | 1.374022 | 1.28938 | 1.2481 | 0.041288733 | 0.125930574 | 1.2836764 | 1.4170703 | 1.24748 | 0.169594421 | 0.036200519 |
| Rpn2 | 11.64715 | 11.70242 | 11.631 | 0.071693582 | 0.01641839 | 11.858675 | 11.828955 | 11.6594 | 0.169584446 | 0.199304539 |
| Galnt18 | 6.495635 | 6.520252 | 6.0987 | 0.421586688 | 0.396970109 | 6.4140526 | 6.2212474 | 6.05199 | 0.16925374 | 0.36205896 |
| Wdr91 | 9.294151 | 9.556058 | 9.4741 | 0.08192391 | -0.179983153 | 9.327343 | 9.5374141 | 9.36845 | 0.16896429 | -0.041106811 |
| Csf2rb2 | 10.78096 | 10.38914 | 10.515 | -0.125893818 | 0.265927876 | 9.8565396 | 10.268542 | 10.0996 | 0.168956639 | -0.243045811 |
| Ess2 | 9.030764 | 9.025994 | 9.1002 | -0.074248663 | -0.069479172 | 9.2621986 | 9.3043259 | 9.1355 | 0.168821791 | 0.126694525 |
| Get4 | 9.036229 | 9.015921 | 9.0847 | -0.068808482 | -0.048500022 | 9.3046637 | 9.2240882 | 9.05528 | 0.168805753 | 0.249381184 |
| Thap3 | 7.972799 | 7.954371 | 8.0673 | -0.112881789 | -0.094454346 | 8.2230059 | 8.2742233 | 8.10543 | 0.168797831 | 0.117580388 |
| Ppat | 9.461643 | 9.441425 | 9.3999 | 0.041553944 | 0.061772226 | 9.3051968 | 9.5316326 | 9.36285 | 0.168785353 | -0.057650527 |
| Tomm34 | 10.29009 | 10.42868 | 10.374 | 0.054305058 | -0.084283542 | 10.559946 | 10.590292 | 10.4216 | 0.168727494 | 0.13838122 |
| Hemk1 | 6.71403 | 6.599859 | 6.8112 | -0.211309583 | -0.097137947 | 7.2152847 | 7.3443515 | 7.17576 | 0.168587362 | 0.039520541 |

| | | | | | | | | | | |
|---|---|---|---|---|---|---|---|---|---|---|
| Gm43587 | 4.717948 | 4.76187 | 4.7974 | -0.0355087 | -0.079430387 | 4.6411026 | 4.9109736 | 4.74246 | 0.168513277 | -0.101357642 |
| Prmt9 | 8.725778 | 8.906489 | 8.8061 | 0.100349682 | -0.080360898 | 8.7763589 | 9.0117996 | 8.84336 | 0.168441991 | -0.066998713 |
| Rps9 | 14.90873 | 14.96379 | 14.872 | 0.091906868 | 0.036847566 | 15.023241 | 15.003251 | 14.8348 | 0.16844072 | 0.188430974 |
| Sesn1 | 10.2471 | 10.50657 | 10.362 | 0.144263238 | -0.115210185 | 10.257869 | 10.432522 | 10.2641 | 0.168393419 | -0.00625927 |
| Mrps9 | 9.131683 | 9.144165 | 9.1561 | -0.011937307 | -0.024419616 | 9.2345148 | 9.4368791 | 9.26852 | 0.168363137 | -0.034001126 |
| Tsg101 | 9.312865 | 9.340996 | 9.344 | -0.003029531 | -0.031160877 | 9.4949995 | 9.4771319 | 9.30882 | 0.168315379 | 0.186183033 |
| Ubiad1 | 7.848418 | 7.935524 | 7.8273 | 0.108254384 | 0.021148261 | 8.0666945 | 7.9814775 | 7.81316 | 0.168314637 | 0.253531696 |
| U2af2 | 12.19828 | 12.28733 | 12.186 | 0.100933356 | 0.01188777 | 12.439937 | 12.47518 | 12.3069 | 0.168290968 | 0.1330475 |
| Lrp11 | 3.1901 | 3.278909 | 3.1943 | 0.084648378 | -0.004160703 | 3.2852231 | 3.3013993 | 3.13362 | 0.167781305 | 0.151605144 |
| Rps19 | 14.42431 | 14.49298 | 14.407 | 0.086468734 | 0.017807708 | 14.576722 | 14.619951 | 14.4522 | 0.167716312 | 0.124486324 |
| Rfx1 | 9.551796 | 9.644369 | 9.5621 | 0.082301774 | -0.010271012 | 9.7187709 | 9.7594002 | 9.59178 | 0.16762514 | 0.126995829 |
| G3bp2 | 11.76423 | 11.76108 | 11.711 | 0.050128034 | 0.053281688 | 11.711103 | 11.85601 | 11.6884 | 0.167572836 | 0.022665497 |
| Ccdc85b | 6.946706 | 6.823432 | 6.9519 | -0.128459487 | -0.005184852 | 7.0206033 | 6.873077 | 6.70551 | 0.167571239 | 0.315097495 |
| Chd5 | 2.022503 | 2.081162 | 1.9612 | 0.119951881 | 0.061292616 | 2.0593102 | 2.1191856 | 1.95168 | 0.167503676 | 0.107628267 |
| Capg | 11.28152 | 11.37079 | 11.259 | 0.111580036 | 0.022316526 | 11.650508 | 11.49923 | 11.3318 | 0.167423632 | 0.31870224 |
| Glt8d1 | 6.863106 | 6.959091 | 6.8425 | 0.116615859 | 0.020630764 | 7.1316402 | 7.0783474 | 6.91093 | 0.167418988 | 0.220711808 |
| Nphs1 | 5.04765 | 5.144142 | 4.9889 | 0.155216376 | 0.05872521 | 5.1376627 | 5.1499282 | 4.98258 | 0.167351773 | 0.155086321 |
| Cpne2 | 8.044313 | 8.035733 | 8.075 | -0.039260875 | -0.030680569 | 7.9358371 | 8.2591552 | 8.09185 | 0.1673084 | -0.156009717 |
| 6030445D17Rik | 1.363609 | 1.403302 | 1.3652 | 0.038142249 | -0.001549894 | 1.3837316 | 1.5274346 | 1.36013 | 0.167307726 | 0.023604705 |
| Tcrg-C4 | 6.222977 | 6.404521 | 6.2374 | 0.167090937 | -0.014453196 | 5.9589704 | 6.4169891 | 6.24969 | 0.167295183 | -0.290723512 |
| H2afy | 11.81011 | 11.78216 | 11.842 | -0.060084864 | -0.032142332 | 11.962635 | 12.049904 | 11.8827 | 0.167182004 | 0.079913156 |
| Slc30a1 | 7.766566 | 7.962565 | 7.6068 | 0.355745164 | 0.159745896 | 7.4471339 | 7.7589805 | 7.59191 | 0.167071394 | -0.144775214 |
| Exosc10 | 10.65958 | 10.6527 | 10.641 | 0.011862054 | 0.018744682 | 10.866415 | 10.898439 | 10.7315 | 0.16695685 | 0.134933579 |
| Vil1 | 1.539499 | 1.492107 | 1.3885 | 0.103607662 | 0.150999753 | 1.4442194 | 1.5660693 | 1.39912 | 0.166951799 | 0.045101877 |
| 1700066B19Rik | 4.485371 | 4.264977 | 4.4168 | -0.151823283 | 0.068570903 | 4.3631545 | 4.4951271 | 4.32822 | 0.166907074 | 0.034934494 |
| Nxf1 | 11.79394 | 11.87822 | 11.936 | -0.057982872 | -0.142268178 | 12.134249 | 12.299308 | 12.1324 | 0.166890358 | 0.001831395 |
| Trnt1 | 9.671568 | 9.735319 | 9.6459 | 0.089408917 | 0.025657726 | 9.6561924 | 9.8651128 | 9.69839 | 0.16672722 | -0.042193153 |
| Slc29a3 | 9.712957 | 9.895641 | 9.782 | 0.113624366 | -0.069059631 | 9.7894493 | 9.910237 | 9.74361 | 0.166630052 | 0.045842419 |
| Rela | 10.89851 | 10.92592 | 10.888 | 0.037734836 | 0.01033407 | 11.070623 | 11.060463 | 10.8939 | 0.166544342 | 0.176704102 |
| Tubb2b | 7.298832 | 7.532605 | 7.6255 | -0.092906376 | -0.32667874 | 7.5733615 | 7.8571534 | 7.69061 | 0.166540349 | -0.117251576 |
| Basp1 | 8.766603 | 8.939134 | 8.7668 | 0.172289227 | -0.000242194 | 8.5652922 | 8.9617189 | 8.79526 | 0.166460603 | -0.229966073 |
| Dnaja3 | 9.576557 | 9.698658 | 9.525 | 0.173663055 | 0.051561495 | 9.6242333 | 9.6874373 | 9.52098 | 0.166456103 | 0.103252056 |

| | | | | | | | | | | |
|---|---|---|---|---|---|---|---|---|---|---|
| Gm16301 | 0.614823 | 0.656262 | 0.6139 | 0.042402509 | 0.000963496 | 0.6140725 | 0.7851991 | 0.61877 | 0.166427794 | -0.004698772 |
| Olfr1264 | 2.236615 | 2.260243 | 2.2149 | 0.045301781 | 0.021673989 | 2.2459214 | 2.3945357 | 2.22819 | 0.166346304 | 0.017731974 |
| Slc9a3r1 | 10.47904 | 10.64822 | 10.582 | 0.06582721 | -0.103349154 | 10.599627 | 10.688267 | 10.5219 | 0.166336814 | 0.077696691 |
| Brms1 | 9.262952 | 9.169773 | 9.1537 | 0.016048913 | 0.10922825 | 9.3670387 | 9.4047774 | 9.23853 | 0.166249984 | 0.128511276 |
| 1190007I07Rik | 5.815287 | 5.720982 | 5.9847 | -0.263673284 | -0.169369091 | 6.1398633 | 6.1230546 | 5.95688 | 0.16617814 | 0.182986827 |
| Proser1 | 8.825209 | 8.937035 | 8.9177 | 0.019361502 | -0.092464887 | 9.1485859 | 9.2658266 | 9.09966 | 0.166163333 | 0.048922635 |
| Lsm8 | 8.612092 | 8.561845 | 8.627 | -0.065200341 | -0.01495328 | 8.6526034 | 8.7688978 | 8.60274 | 0.166162446 | 0.049868051 |
| Abca12 | 1.553482 | 1.51037 | 1.4693 | 0.041022803 | 0.084134152 | 1.5094074 | 1.6640813 | 1.49797 | 0.166108468 | 0.011434583 |
| Tagap | 11.2262 | 11.38795 | 11.31 | 0.078243338 | -0.083502884 | 10.709387 | 11.212374 | 11.0463 | 0.16608169 | -0.336905686 |
| Adrb3 | 0.716546 | 0.683696 | 0.671 | 0.012730794 | 0.045581405 | 0.7505016 | 0.8494818 | 0.68343 | 0.166054557 | 0.067074337 |
| Cnot4 | 9.54668 | 9.699227 | 9.6207 | 0.078509965 | -0.074036647 | 9.5621421 | 9.7936805 | 9.62766 | 0.166019316 | -0.065519071 |
| Rab5a | 9.947499 | 9.912424 | 9.8606 | 0.051848147 | 0.08692282 | 9.6635902 | 9.9537206 | 9.78774 | 0.165981308 | -0.124149099 |
| Tmem203 | 6.433663 | 6.329941 | 6.3332 | -0.003259293 | 0.100462139 | 6.6346351 | 6.5257744 | 6.35981 | 0.165961172 | 0.274821832 |
| Tmed10 | 11.40044 | 11.32067 | 11.305 | 0.015590896 | 0.095356552 | 11.309903 | 11.409483 | 11.2436 | 0.165921006 | 0.066341098 |
| Gm10382 | 2.623293 | 2.616755 | 2.6763 | -0.059505765 | -0.052967675 | 2.636356 | 2.7344014 | 2.56855 | 0.16585125 | 0.067805835 |
| Pak4 | 7.269605 | 7.328605 | 7.3414 | -0.012816889 | -0.071817227 | 7.2884863 | 7.3759292 | 7.21019 | 0.165743381 | 0.078300422 |
| Dnal1 | 4.176523 | 4.181317 | 4.2198 | -0.038435936 | -0.043229836 | 4.101042 | 4.4811202 | 4.31543 | 0.165694472 | -0.21438368 |
| Tpcn2 | 7.983987 | 8.067726 | 8.0724 | -0.004691171 | -0.08843063 | 8.2683524 | 8.3630433 | 8.19738 | 0.165666473 | 0.070975534 |
| 2610002M06Rik | 8.972184 | 8.922315 | 8.8448 | 0.07755973 | 0.127428801 | 8.6762478 | 9.0379851 | 8.87234 | 0.165646745 | -0.196090569 |
| Ptrh1 | 5.021289 | 5.038421 | 4.958 | 0.080418065 | 0.063286518 | 5.188957 | 5.066352 | 4.90075 | 0.165605412 | 0.288210336 |
| Atic | 10.80378 | 10.87073 | 10.741 | 0.12945087 | 0.062494908 | 10.905473 | 10.987876 | 10.8223 | 0.16560154 | 0.083199314 |
| Gm29668 | 4.222828 | 4.075221 | 4.0631 | 0.012133442 | 0.159740341 | 4.1742388 | 4.4159444 | 4.25058 | 0.165363158 | -0.076342402 |
| Eya4 | 3.296756 | 3.238949 | 3.2969 | -0.057981307 | -0.000173327 | 3.293063 | 3.3967545 | 3.23143 | 0.16532575 | 0.06163422 |
| S1pr4 | 9.928653 | 10.14626 | 10.081 | 0.065039244 | -0.152564468 | 10.287515 | 10.156839 | 9.99156 | 0.165279801 | 0.295955968 |
| Mief1 | 9.424647 | 9.488746 | 9.4185 | 0.070268599 | 0.006169228 | 9.5309659 | 9.5921584 | 9.4269 | 0.165254358 | 0.10406187 |
| Cmc2 | 6.60835 | 6.619453 | 6.7793 | -0.15984168 | -0.170945353 | 6.7869662 | 6.8981162 | 6.73288 | 0.165232834 | 0.054082818 |
| Olfr397 | 1.840462 | 1.849477 | 1.8069 | 0.042533389 | 0.033518299 | 1.845839 | 2.094528 | 1.92939 | 0.165140868 | -0.083548132 |
| Wnt4 | 4.335004 | 4.251955 | 4.3266 | -0.074663963 | 0.00838511 | 4.4453723 | 4.4377508 | 4.27268 | 0.16507475 | 0.172696202 |
| Myrfl | 0.780201 | 0.759049 | 0.7721 | -0.013019291 | 0.008131881 | 0.8111587 | 0.9895675 | 0.82463 | 0.16493613 | -0.013472647 |
| Dph7 | 8.290001 | 8.423715 | 8.4046 | 0.019146647 | -0.114567822 | 8.5487841 | 8.547835 | 8.38295 | 0.164880627 | 0.165829689 |
| Arl6ip5 | 11.34106 | 11.28564 | 11.276 | 0.0092645 | 0.064688806 | 11.236053 | 11.32059 | 11.1559 | 0.16473105 | 0.08019376 |
| Rmc1 | 9.422864 | 9.547965 | 9.4621 | 0.085850912 | -0.03924968 | 9.7073259 | 9.762181 | 9.59749 | 0.164686143 | 0.109831033 |

| | | | | | | | | | | |
|---|---|---|---|---|---|---|---|---|---|---|
| Elk4 | 11.16862 | 11.30844 | 11.228 | 0.08007913 | -0.059742204 | 11.324362 | 11.670277 | 11.5056 | 0.16463112 | -0.181283656 |
| Zfat | 7.203201 | 7.329662 | 7.2905 | 0.039149675 | -0.087311789 | 7.079886 | 7.3319761 | 7.16767 | 0.164304847 | -0.087785262 |
| Gm614 | 6.639572 | 6.446767 | 6.6307 | -0.183952624 | 0.008851658 | 6.5804175 | 6.7768649 | 6.61257 | 0.164296451 | -0.032150938 |
| Lmnb2 | 8.321658 | 8.414159 | 8.3038 | 0.110383388 | 0.017882173 | 8.6379762 | 8.6145452 | 8.45027 | 0.164274657 | 0.187705664 |
| Prkcq | 11.1682 | 11.19122 | 11.232 | -0.040913342 | -0.063931678 | 10.864953 | 11.359249 | 11.195 | 0.164274236 | -0.330022445 |
| Patj | 9.423534 | 9.555676 | 9.4747 | 0.080946863 | -0.051195012 | 9.3894163 | 9.8332045 | 9.66922 | 0.163981087 | -0.279807119 |
| Abca4 | 2.526479 | 2.365048 | 2.5268 | -0.161771525 | -0.000340968 | 2.4016809 | 2.6244839 | 2.46053 | 0.163958248 | -0.058844828 |
| Ptrhd1 | 6.776615 | 6.700776 | 6.7006 | 0.000161717 | 0.07600153 | 6.9482929 | 6.9098594 | 6.7459 | 0.163954471 | 0.202388054 |
| Narf | 9.153818 | 9.397675 | 9.3158 | 0.081924284 | -0.161933214 | 9.3914352 | 9.4860846 | 9.32215 | 0.163938981 | 0.069289557 |
| Isoc2a | 8.454555 | 8.544185 | 8.3798 | 0.164433261 | 0.074803693 | 8.5557968 | 8.4461192 | 8.28219 | 0.16393068 | 0.273608231 |
| Prr14 | 10.53251 | 10.54039 | 10.517 | 0.02363572 | 0.01575708 | 10.693958 | 10.776975 | 10.6131 | 0.163891578 | 0.080874711 |
| Ccdc32 | 7.855335 | 7.79427 | 7.8328 | -0.038564417 | 0.022501219 | 7.7920133 | 7.8532943 | 7.68943 | 0.1638636 | 0.102582571 |
| Abhd17b | 10.23658 | 10.31439 | 10.241 | 0.073413538 | -0.004396557 | 10.186775 | 10.214242 | 10.0504 | 0.163800056 | 0.136332366 |
| P4ha2 | 4.73624 | 4.783068 | 4.6417 | 0.141393807 | 0.094566007 | 4.502623 | 4.6304682 | 4.46676 | 0.163710997 | 0.035865763 |
| Prr13 | 10.32021 | 10.26622 | 10.364 | -0.097436721 | -0.043441228 | 10.576539 | 10.499489 | 10.3358 | 0.163660268 | 0.240711012 |
| Gm8991 | 2.613649 | 2.522232 | 2.6032 | -0.081010091 | 0.010406755 | 2.5624013 | 2.6766937 | 2.51311 | 0.16358751 | 0.049295046 |
| Cntrl | 11.67687 | 11.78383 | 11.735 | 0.048542566 | -0.058416682 | 11.735087 | 12.119144 | 11.9558 | 0.163389846 | -0.220667758 |
| 0610009B22Rik | 5.835366 | 5.865883 | 5.8507 | 0.015223016 | -0.01529419 | 5.9155261 | 5.9964366 | 5.8331 | 0.163339312 | 0.082428765 |
| Vdac3 | 10.00061 | 10.052 | 10.033 | 0.018745754 | -0.032645176 | 10.089 | 10.154919 | 9.9916 | 0.163320628 | 0.097401889 |
| Scnn1a | 3.572834 | 3.493299 | 3.3549 | 0.138358735 | 0.217894207 | 3.4665189 | 3.5379725 | 3.37474 | 0.163227551 | 0.091773919 |
| Tmem177 | 6.550769 | 6.651934 | 6.5526 | 0.099358223 | -0.001806261 | 6.7007625 | 6.6545534 | 6.49136 | 0.163190755 | 0.209399835 |
| Mterf4 | 7.05601 | 7.222791 | 7.0912 | 0.131598501 | -0.035182073 | 7.3377244 | 7.2381754 | 7.07504 | 0.16313923 | 0.262688304 |
| Rpp30 | 7.463927 | 7.392493 | 7.4297 | -0.037158144 | 0.0342754 | 7.5476274 | 7.6800093 | 7.5169 | 0.163108367 | 0.030726526 |
| Dazap2 | 12.73068 | 12.66031 | 12.743 | -0.083001098 | -0.012628724 | 12.83862 | 12.815537 | 12.6526 | 0.162904134 | 0.185987114 |
| Pygo2 | 9.249068 | 9.336696 | 9.3793 | -0.042590755 | -0.130218729 | 9.4550506 | 9.4261602 | 9.26326 | 0.162904087 | 0.191794577 |
| Mbd1 | 10.05069 | 10.20541 | 10.092 | 0.113131567 | -0.041590655 | 10.16192 | 10.351919 | 10.189 | 0.162876566 | -0.027122261 |
| Api5 | 11.35534 | 11.4184 | 11.326 | 0.091898596 | 0.028845428 | 11.324149 | 11.45568 | 11.293 | 0.162717856 | 0.031186362 |
| Fam174b | 7.421212 | 7.253447 | 7.0022 | 0.251227936 | 0.41899265 | 6.9335785 | 7.1158589 | 6.95332 | 0.162541122 | -0.019739209 |
| Arpc2 | 12.81255 | 12.77694 | 12.762 | 0.01444789 | 0.05006507 | 12.892722 | 12.876344 | 12.7138 | 0.16250571 | 0.178884118 |
| Agtrap | 8.638959 | 8.478903 | 8.6244 | -0.145506729 | 0.014549504 | 8.6128181 | 8.855943 | 8.69347 | 0.162474643 | -0.080650286 |
| Cgrrf1 | 6.749316 | 6.77204 | 6.7431 | 0.028891707 | 0.00616773 | 6.9490664 | 6.9102131 | 6.74779 | 0.162419094 | 0.201272357 |
| Cfap45 | 4.235413 | 4.321943 | 4.2898 | 0.032117437 | -0.054412137 | 4.229917 | 4.4917837 | 4.32941 | 0.16237678 | -0.099489903 |

| | | | | | | | | | | |
|---|---|---|---|---|---|---|---|---|---|---|
| Atp6v1c1 | 8.845971 | 8.763668 | 8.7301 | 0.033592793 | 0.115895401 | 8.7913284 | 8.777876 | 8.61561 | 0.162264555 | 0.175716972 |
| Gm8818 | 5.605718 | 5.426288 | 5.5251 | -0.098769874 | 0.080659486 | 5.335266 | 5.539968 | 5.37779 | 0.162182424 | -0.042519651 |
| Nipsnap3b | 8.854691 | 8.768081 | 8.7459 | 0.022185808 | 0.10879559 | 8.9151993 | 8.9790751 | 8.81695 | 0.162126865 | 0.098251053 |
| Prpf4 | 9.453411 | 9.583289 | 9.5182 | 0.065104324 | -0.064773604 | 9.6236732 | 9.7093195 | 9.5473 | 0.162018279 | 0.076371929 |
| Mrpl42 | 8.82766 | 8.738385 | 8.8558 | -0.117376313 | -0.028101118 | 8.9647404 | 8.9955097 | 8.83352 | 0.161989994 | 0.131220739 |
| Lsm5 | 8.364106 | 8.285086 | 8.3207 | -0.035614366 | 0.043405991 | 8.5270991 | 8.493694 | 8.33171 | 0.161984934 | 0.195389984 |
| Lamc2 | 5.631488 | 5.739037 | 5.3018 | 0.437201827 | 0.329652805 | 5.2485991 | 5.2176313 | 5.05575 | 0.161884777 | 0.192852507 |
| Fhit | 4.637246 | 4.587958 | 4.5156 | 0.072347318 | 0.121635935 | 4.3704916 | 4.478335 | 4.3165 | 0.161839767 | 0.053996375 |
| Igfbp2 | 2.839007 | 3.037454 | 2.6965 | 0.340942769 | 0.142495693 | 3.1831169 | 2.8813284 | 2.71957 | 0.161753809 | 0.463542297 |
| Ndrg1 | 9.955374 | 9.613512 | 9.7283 | -0.114808142 | 0.22705323 | 9.1651272 | 9.569724 | 9.40804 | 0.161681367 | -0.242915384 |
| Zfp991 | 7.651753 | 7.623309 | 7.6516 | -0.028328186 | 0.000115578 | 7.5784547 | 7.8514637 | 7.68982 | 0.161638691 | -0.111370323 |
| Arpc5 | 12.2936 | 12.3026 | 12.202 | 0.100325448 | 0.09132151 | 12.3763 | 12.313825 | 12.1523 | 0.161546474 | 0.224020906 |
| Txlng | 9.175347 | 9.152106 | 9.046 | 0.106132774 | 0.129373477 | 8.8447953 | 9.2394425 | 9.07792 | 0.161526472 | -0.233120777 |
| Duox2 | 2.840394 | 2.851109 | 2.6941 | 0.156976679 | 0.146262017 | 2.8069835 | 2.9279692 | 2.7665 | 0.16147166 | 0.040485926 |
| Rabif | 8.895135 | 8.955457 | 8.8488 | 0.106668543 | 0.046347146 | 9.0066009 | 9.0864616 | 8.92512 | 0.161342948 | 0.081482191 |
| Uck1 | 8.971292 | 9.103741 | 9.0729 | 0.030794894 | -0.101653976 | 9.3274975 | 9.2769317 | 9.11571 | 0.161222192 | 0.211787988 |
| Babam2 | 9.164064 | 9.291886 | 9.0523 | 0.239546975 | 0.111724839 | 9.2041721 | 9.3682125 | 9.20709 | 0.161122163 | -0.002918249 |
| Ints13 | 9.723665 | 9.868802 | 9.7384 | 0.130388523 | -0.014748021 | 9.6748106 | 10.031834 | 9.87074 | 0.161092674 | -0.195931089 |
| Vmn2r120 | 6.772629 | 6.843836 | 6.7778 | 0.066057418 | -0.005149534 | 7.1513215 | 7.4070681 | 7.24598 | 0.161089486 | -0.094657124 |
| Rps14 | 14.20944 | 14.33481 | 14.295 | 0.039631728 | -0.085733656 | 14.460258 | 14.375723 | 14.2147 | 0.161070638 | 0.245605269 |
| Rpl28-ps1 | 12.52419 | 12.58281 | 12.562 | 0.021021776 | -0.037596836 | 12.621464 | 12.768756 | 12.6078 | 0.160926614 | 0.013633736 |
| Ighv1-75 | 6.702475 | 6.690182 | 6.8184 | -0.128248948 | -0.115956055 | 7.1427551 | 7.2736044 | 7.11268 | 0.16092589 | 0.030076536 |
| Ddost | 10.83492 | 10.85342 | 10.846 | 0.00752495 | -0.01097269 | 11.079811 | 10.965144 | 10.8042 | 0.160921368 | 0.275587862 |
| Cyfip1 | 9.916356 | 9.935045 | 9.7467 | 0.188373328 | 0.169684213 | 9.9248757 | 9.9257734 | 9.76492 | 0.160856613 | 0.159958931 |
| Spopl | 9.090297 | 8.906134 | 8.8912 | 0.014981883 | 0.199144316 | 8.9453016 | 9.1995027 | 9.03866 | 0.160843757 | -0.093357298 |
| Pnpo | 8.530039 | 8.424606 | 8.4112 | 0.013455031 | 0.118888639 | 8.5891778 | 8.5544136 | 8.39369 | 0.160718715 | 0.195482848 |
| Dnah7b | 2.716354 | 2.715821 | 2.6851 | 0.03072289 | 0.031255757 | 2.558758 | 2.8304988 | 2.66982 | 0.16067553 | -0.111065284 |
| Rps2-ps13 | 3.451298 | 3.372635 | 3.5613 | -0.188633168 | -0.109970374 | 3.4374947 | 3.5222293 | 3.36156 | 0.16067012 | 0.075935566 |
| Gstt3 | 5.712843 | 5.520962 | 5.6828 | -0.161837287 | 0.030043638 | 5.788129 | 5.7426914 | 5.58205 | 0.160640771 | 0.206078295 |
| Cytip | 12.89768 | 12.9219 | 12.685 | 0.237202962 | 0.212975476 | 12.728099 | 12.877874 | 12.7173 | 0.16058379 | 0.010809228 |
| Phldb3 | 7.782571 | 7.831891 | 7.7872 | 0.044646073 | -0.004674682 | 7.8638198 | 7.8892412 | 7.72874 | 0.16050495 | 0.135083527 |
| Gtf3a | 8.814087 | 8.903014 | 8.8624 | 0.040566443 | -0.048360435 | 9.1044506 | 9.0331024 | 8.87273 | 0.160370824 | 0.231719058 |

| | | | | | | | | | | |
|---|---|---|---|---|---|---|---|---|---|---|
| Aebp1 | 8.079906 | 7.948457 | 7.8729 | 0.07558013 | 0.207028679 | 7.435019 | 7.3742309 | 7.21389 | 0.160337809 | 0.221125836 |
| Ighv9-1 | 6.507936 | 6.413596 | 6.1797 | 0.233931954 | 0.328271562 | 6.1806553 | 6.2863556 | 6.12605 | 0.160304172 | 0.054603898 |
| Usp9x | 11.90359 | 11.98857 | 11.804 | 0.184268202 | 0.099290742 | 11.684063 | 11.940249 | 11.78 | 0.16029631 | -0.095889293 |
| Mettl3 | 9.216714 | 9.34182 | 9.2856 | 0.056192929 | -0.068912804 | 9.7461569 | 9.814164 | 9.65393 | 0.160237524 | 0.092230415 |
| 0610012G03Rik | 7.056873 | 6.962302 | 6.9842 | -0.021891775 | 0.072678874 | 7.1209666 | 7.1280169 | 6.96781 | 0.160208593 | 0.1531583 |
| Prpf38b | 11.88658 | 11.91942 | 11.896 | 0.023597578 | -0.009244228 | 11.9528 | 12.247944 | 12.0878 | 0.160183816 | -0.134959261 |
| Vps72 | 9.262461 | 9.382062 | 9.3193 | 0.062755875 | -0.056845273 | 9.4092624 | 9.5335268 | 9.37336 | 0.160164234 | 0.035899812 |
| Hip1 | 8.968953 | 9.034006 | 8.8588 | 0.175234408 | 0.110181927 | 8.6106569 | 8.7058004 | 8.54566 | 0.160139469 | 0.064995988 |
| Mecp2 | 10.71335 | 10.74407 | 10.595 | 0.1488599 | 0.118144956 | 10.686871 | 10.840232 | 10.6802 | 0.160077626 | 0.006717081 |
| Snrnp40 | 8.782184 | 8.735338 | 8.7657 | -0.030313318 | 0.016532229 | 8.9051058 | 8.9361893 | 8.77613 | 0.160064206 | 0.128980697 |
| Lrrc51 | 3.526889 | 3.468789 | 3.5709 | -0.102130665 | -0.04403123 | 3.6743519 | 3.8169868 | 3.65706 | 0.15992762 | 0.017292728 |
| Hadhb | 10.43298 | 10.33813 | 10.448 | -0.110000002 | -0.015147116 | 10.422379 | 10.638933 | 10.479 | 0.159901854 | -0.05665141 |
| Ppp6c | 10.28045 | 10.29595 | 10.204 | 0.092165654 | 0.076665384 | 10.198871 | 10.347966 | 10.1881 | 0.159891876 | 0.010796994 |
| Smim20 | 7.940037 | 8.031063 | 7.9525 | 0.078583531 | -0.012442709 | 8.2165979 | 8.2961448 | 8.13627 | 0.159879011 | 0.080332181 |
| Fth1 | 13.64765 | 13.84118 | 13.688 | 0.15296293 | -0.040570018 | 13.818192 | 13.777752 | 13.6179 | 0.159875372 | 0.20031556 |
| Cib1 | 9.259036 | 9.236451 | 9.269 | -0.032576455 | -0.009991896 | 9.576518 | 9.4423343 | 9.28252 | 0.159817503 | 0.294001187 |
| Sf3b2 | 12.46336 | 12.47992 | 12.464 | 0.01606239 | -0.000497548 | 12.638089 | 12.656841 | 12.4971 | 0.159749178 | 0.140997367 |
| Vav1 | 11.74849 | 11.77188 | 11.671 | 0.10039381 | 0.0770028 | 11.768002 | 11.827493 | 11.6677 | 0.15974828 | 0.100257303 |
| Dars | 10.66132 | 10.66433 | 10.61 | 0.05410994 | 0.05109764 | 10.696811 | 10.753466 | 10.5937 | 0.159739928 | 0.103085639 |
| 4921524J17Rik | 8.009667 | 8.183872 | 7.9814 | 0.202471405 | 0.028265923 | 7.9638933 | 8.2269816 | 8.0673 | 0.159677749 | -0.103410553 |
| Rpl17-ps3 | 2.952741 | 2.89597 | 3.1895 | -0.293576644 | -0.236806004 | 3.0868164 | 3.1485639 | 2.98901 | 0.159549648 | 0.097802096 |
| Fbxl18 | 7.596144 | 7.726674 | 7.6114 | 0.115271723 | -0.015258365 | 7.8886422 | 7.9723902 | 7.81294 | 0.159453398 | 0.075705437 |
| Pex26 | 7.577861 | 8.088358 | 7.6914 | 0.396971273 | -0.113525475 | 7.6353704 | 7.8089647 | 7.64951 | 0.159450057 | -0.01414423 |
| Rpl13a | 15.17732 | 15.26549 | 15.224 | 0.040998154 | -0.047167366 | 15.375463 | 15.329548 | 15.1701 | 0.15944089 | 0.205355769 |
| Rpl30-ps9 | 2.655253 | 2.469213 | 2.5784 | -0.109170255 | 0.076870269 | 2.702368 | 2.7481849 | 2.58884 | 0.159343526 | 0.11352656 |
| Polr2m | 11.01054 | 11.03392 | 11.035 | -0.001539168 | -0.02491194 | 11.072106 | 11.148099 | 10.9888 | 0.159274756 | 0.083281824 |
| Celf2 | 12.2254 | 12.3898 | 12.279 | 0.110974368 | -0.053422628 | 12.357438 | 12.48497 | 12.3258 | 0.159142278 | 0.031610072 |
| Gm48130 | 1.784536 | 1.754395 | 1.7523 | 0.002140194 | 0.03228034 | 1.7114423 | 1.9258694 | 1.76677 | 0.159103798 | -0.055323255 |
| Copb2 | 11.24573 | 11.23597 | 11.17 | 0.065739614 | 0.075500974 | 11.259093 | 11.304053 | 11.145 | 0.159051576 | 0.114091432 |
| Thumpd3 | 9.743825 | 9.716005 | 9.6422 | 0.073853469 | 0.101672891 | 9.877973 | 9.8871169 | 9.72816 | 0.158960241 | 0.14981634 |
| Bgn | 9.147081 | 8.973864 | 9.0029 | -0.029014928 | 0.14420167 | 8.6839333 | 8.7116569 | 8.5527 | 0.158953334 | 0.131229712 |
| Zfp335 | 9.942633 | 10.10656 | 9.9428 | 0.163777756 | -0.000149037 | 10.090052 | 10.130024 | 9.97109 | 0.158930803 | 0.118958573 |

| | | | | | | | | | | |
|---|---|---|---|---|---|---|---|---|---|---|
| Adgrf2 | 1.212896 | 1.255213 | 1.2682 | -0.012979254 | -0.055296378 | 1.245482 | 1.4255511 | 1.26666 | 0.158893206 | -0.021175918 |
| Rps16-ps2 | 13.31311 | 13.51958 | 13.414 | 0.105985808 | -0.100485956 | 13.670601 | 13.558776 | 13.3999 | 0.158843922 | 0.270669415 |
| Irgc1 | 5.618556 | 5.435938 | 5.5594 | -0.123471735 | 0.05914629 | 6.0989647 | 5.737265 | 5.57843 | 0.158835812 | 0.520535543 |
| Ssrp1 | 12.10126 | 12.02345 | 12.015 | 0.008629248 | 0.086440488 | 12.131961 | 12.287929 | 12.1291 | 0.158786922 | 0.002819462 |
| Klc2 | 8.221419 | 8.169473 | 8.1867 | -0.017198524 | 0.034747653 | 8.2697956 | 8.4582774 | 8.29953 | 0.158745532 | -0.029736193 |
| Arl8b | 10.04703 | 10.13647 | 10.002 | 0.134096267 | 0.044657906 | 10.037353 | 10.13596 | 9.97738 | 0.158575663 | 0.059968827 |
| Wfdc3 | 1.73608 | 1.764564 | 1.7948 | -0.030246189 | -0.058729878 | 1.7518415 | 1.9655583 | 1.80709 | 0.158469448 | -0.055247309 |
| Zfp622 | 9.827528 | 9.896042 | 9.953 | -0.056997492 | -0.125510891 | 9.8631831 | 10.025586 | 9.86713 | 0.158459075 | -0.003943472 |
| Gm10130 | 3.996409 | 4.047127 | 3.9151 | 0.131999869 | 0.0812817 | 3.9826494 | 3.9893623 | 3.83099 | 0.158371045 | 0.151658185 |
| Rbbp6 | 11.15941 | 11.24044 | 11.192 | 0.048896852 | -0.03213978 | 11.141864 | 11.524433 | 11.3661 | 0.158320014 | -0.224248623 |
| Birc6 | 12.66615 | 12.70091 | 12.72 | -0.018623794 | -0.053381944 | 12.59092 | 12.943474 | 12.7853 | 0.158222746 | -0.194331218 |
| Fbrsl1 | 9.421097 | 9.502992 | 9.4362 | 0.066801465 | -0.015093364 | 9.6505364 | 9.7306582 | 9.57256 | 0.158097189 | 0.077975314 |
| Cfl1 | 14.17486 | 14.19998 | 14.169 | 0.031179152 | 0.006054764 | 14.49144 | 14.35186 | 14.1938 | 0.158067382 | 0.297647134 |
| Srrt | 11.63026 | 11.7092 | 11.712 | -0.002501022 | -0.081433604 | 12.045552 | 12.135564 | 11.9775 | 0.158019154 | 0.06800714 |
| Pnisr | 10.50131 | 10.63178 | 10.641 | -0.008962827 | -0.139433817 | 11.297587 | 11.470667 | 11.3127 | 0.157959588 | -0.015121166 |
| Gm21750 | 3.808521 | 3.958616 | 3.9717 | -0.013125456 | -0.163219839 | 3.7706478 | 4.0637302 | 3.90594 | 0.157793091 | -0.135289313 |
| Maf1 | 10.49685 | 10.58044 | 10.635 | -0.054868248 | -0.138453974 | 10.694496 | 10.741274 | 10.5837 | 0.15761738 | 0.110839488 |
| Zfyve1 | 9.142435 | 9.227858 | 9.1682 | 0.059631917 | -0.025790646 | 8.8723879 | 9.2955374 | 9.13837 | 0.157168689 | -0.26598084 |
| Wdr38 | 3.268715 | 3.38382 | 3.3319 | 0.051959525 | -0.06314561 | 3.572279 | 3.5812061 | 3.42406 | 0.15714713 | 0.148220036 |
| Ccnt2 | 10.36284 | 10.48507 | 10.387 | 0.097603579 | -0.024624257 | 10.510275 | 10.855862 | 10.6987 | 0.1571451 | -0.188442434 |
| Irak2 | 9.924004 | 10.1819 | 9.9947 | 0.187182392 | -0.070714016 | 10.153846 | 10.26208 | 10.1053 | 0.15676866 | 0.048535138 |
| Pik3c2g | 0.932229 | 1.031344 | 0.9179 | 0.113466355 | 0.01435103 | 0.9440121 | 1.1058993 | 0.94915 | 0.156744417 | -0.005142732 |
| Plekhh1 | 4.185126 | 4.457624 | 4.3452 | 0.11244234 | -0.160055915 | 4.2746181 | 4.3343069 | 4.1776 | 0.156710852 | 0.097022006 |
| Dclre1c | 8.765 | 8.749352 | 8.7002 | 0.049191458 | 0.064839061 | 8.9474742 | 9.1931359 | 9.03643 | 0.15670762 | -0.088954089 |
| Serp1 | 12.94483 | 13.00197 | 12.943 | 0.059021312 | 0.001879192 | 13.201056 | 13.058218 | 12.9017 | 0.156555204 | 0.299393475 |
| Dalrd3 | 8.760339 | 8.749566 | 8.7543 | -0.004708934 | 0.006064178 | 9.5955617 | 9.391039 | 9.23453 | 0.156508632 | 0.361031353 |
| Elac2 | 9.291522 | 9.295348 | 9.2745 | 0.020805703 | 0.016979884 | 9.4395144 | 9.5736883 | 9.41718 | 0.156506805 | 0.022332853 |
| Hps1 | 9.088019 | 9.228295 | 9.1967 | 0.031617948 | -0.108658775 | 9.6221057 | 9.4096994 | 9.25321 | 0.156485928 | 0.368892258 |
| Trim6 | 3.380258 | 3.388779 | 3.3708 | 0.017958002 | 0.009436654 | 3.582287 | 3.6944027 | 3.53794 | 0.156459195 | 0.044343525 |
| Dpep2 | 4.906231 | 4.821049 | 4.6733 | 0.147708445 | 0.232890505 | 5.3016971 | 4.8763002 | 4.71996 | 0.156343967 | 0.581740836 |
| Zdhhc21 | 9.501532 | 9.492192 | 9.4645 | 0.02773145 | 0.03707177 | 9.5097414 | 9.7973654 | 9.64102 | 0.15634134 | -0.1312827 |
| Ogfod2 | 7.534781 | 7.564414 | 7.6166 | -0.052193874 | -0.081826031 | 7.7537896 | 7.7225987 | 7.5663 | 0.156294993 | 0.187485832 |

| | | | | | | | | | | |
|---|---|---|---|---|---|---|---|---|---|---|
| Polr3d | 8.175521 | 8.198571 | 8.1261 | 0.072473801 | 0.049424017 | 8.4922945 | 8.4125558 | 8.2563 | 0.156257703 | 0.235996449 |
| Slc4a2 | 10.04916 | 9.974463 | 10.049 | -0.074692519 | 4.6958E-06 | 10.147069 | 10.182074 | 10.0258 | 0.156233858 | 0.121229015 |
| Srek1ip1 | 7.825516 | 7.727311 | 7.9383 | -0.210998416 | -0.112793575 | 7.9420353 | 8.0119119 | 7.85575 | 0.156162934 | 0.086286366 |
| Pdp2 | 8.251963 | 8.352564 | 8.3653 | -0.012735841 | -0.113337426 | 8.301524 | 8.4741242 | 8.318 | 0.156125474 | -0.016474753 |
| Lpin3 | 4.38602 | 4.439962 | 4.2979 | 0.142034564 | 0.088092488 | 3.9724459 | 4.2909061 | 4.13482 | 0.156086004 | -0.162374277 |
| Lrmda | 3.592567 | 3.460571 | 3.6251 | -0.164498759 | -0.032502407 | 3.5692617 | 3.6570457 | 3.50097 | 0.156079011 | 0.068294949 |
| 2610507B11Rik | 12.06152 | 12.01837 | 12.117 | -0.098272842 | -0.05512444 | 12.048344 | 12.305633 | 12.1497 | 0.155883984 | -0.101404657 |
| Cemip | 2.943842 | 2.952451 | 2.9351 | 0.01739065 | 0.008781138 | 2.8928863 | 3.0177056 | 2.86183 | 0.15587181 | 0.031052428 |
| Mfsd10 | 8.672739 | 8.646661 | 8.6113 | 0.035385847 | 0.061463356 | 9.0677754 | 8.6615005 | 8.50563 | 0.15587033 | 0.562145178 |
| Celf4 | 4.527585 | 4.513138 | 4.4777 | 0.035479811 | 0.049926716 | 4.6279828 | 4.5459804 | 4.39017 | 0.155809365 | 0.237811738 |
| Cuedc2 | 8.967976 | 8.829431 | 8.9508 | -0.121340826 | 0.017204426 | 9.221048 | 8.9742767 | 8.81849 | 0.155784573 | 0.402555873 |
| Apbb3 | 8.475912 | 8.543193 | 8.646 | -0.102773887 | -0.170055456 | 8.94966 | 9.198563 | 9.04301 | 0.155552057 | -0.09335091 |
| Calhm5 | 2.801618 | 2.780063 | 2.738 | 0.042101222 | 0.063655478 | 2.628882 | 2.8658038 | 2.71027 | 0.155531563 | -0.081390205 |
| Lama1 | 0.514428 | 0.55151 | 0.4888 | 0.062700226 | 0.025618521 | 0.504331 | 0.6918917 | 0.5364 | 0.155490771 | -0.03206986 |
| Cln8 | 7.453509 | 7.457833 | 7.2485 | 0.209311819 | 0.204987916 | 7.2611705 | 7.4280944 | 7.27262 | 0.15547621 | -0.011447699 |
| Wdr61 | 8.831831 | 8.867903 | 8.874 | -0.006116082 | -0.042187865 | 9.0955266 | 9.0320593 | 8.87669 | 0.15537098 | 0.218838264 |
| Slc29a2 | 5.488679 | 5.81257 | 5.441 | 0.371528581 | 0.047637093 | 5.7618832 | 5.7782939 | 5.62298 | 0.155309717 | 0.138899024 |
| Tmem38b | 7.474336 | 7.442585 | 7.3964 | 0.046191936 | 0.077942862 | 7.5584459 | 7.513367 | 7.35808 | 0.155288191 | 0.200367001 |
| Ubac2 | 10.201 | 10.22565 | 10.198 | 0.027377071 | 0.00273181 | 10.235942 | 10.331836 | 10.1766 | 0.155196287 | 0.059302706 |
| Fbxo33 | 9.066929 | 9.204502 | 9.2725 | -0.067981819 | -0.205555517 | 9.1496663 | 9.4589408 | 9.30375 | 0.155188428 | -0.154085985 |
| Pax9 | 1.658701 | 1.495383 | 1.5743 | -0.078966472 | 0.084351652 | 1.5819838 | 1.7024326 | 1.54728 | 0.155157226 | 0.034708468 |
| Cxcr2 | 7.805507 | 7.56162 | 6.6963 | 0.865283077 | 1.109170658 | 7.3509183 | 6.6247814 | 6.46969 | 0.155095116 | 0.88123196 |
| Gm10335 | 4.033997 | 4.019986 | 4.0892 | -0.069178477 | -0.055167006 | 3.9630762 | 4.1320241 | 3.97694 | 0.155084153 | -0.01386368 |
| Mfsd4b5 | 2.526175 | 2.511395 | 2.5058 | 0.005629023 | 0.020409007 | 2.5155158 | 2.7028341 | 2.54775 | 0.15508104 | -0.032237248 |
| Tmem127 | 10.62471 | 10.72784 | 10.65 | 0.077781818 | -0.025347666 | 10.730201 | 10.724067 | 10.569 | 0.155066646 | 0.161199968 |
| Erlec1 | 9.12991 | 8.943412 | 9.1505 | -0.207121936 | -0.020623694 | 9.1038277 | 9.2213375 | 9.06637 | 0.154966649 | 0.037456888 |
| Gem | 10.19832 | 9.774752 | 10.039 | -0.263865401 | 0.159702286 | 9.2051239 | 9.7942555 | 9.63931 | 0.154946745 | -0.434184821 |
| Rpp40 | 6.985293 | 6.888028 | 6.8904 | -0.002342581 | 0.094922261 | 6.9766334 | 7.109078 | 6.95432 | 0.154759093 | 0.022314467 |
| Gm3755 | 2.92896 | 2.95099 | 3.0181 | -0.067120572 | -0.089150666 | 3.0664381 | 3.2657667 | 3.11106 | 0.154702567 | -0.044626068 |
| Flnc | 3.156833 | 3.304345 | 2.965 | 0.339366158 | 0.191853594 | 2.9227022 | 3.098935 | 2.94427 | 0.154666471 | -0.021566325 |
| Sav1 | 10.07862 | 10.05751 | 10.099 | -0.041899334 | -0.020791724 | 9.9245157 | 10.130667 | 9.97605 | 0.154620433 | -0.051530649 |
| Gm5844 | 2.811927 | 2.808219 | 2.8344 | -0.026139096 | -0.022430329 | 2.7773189 | 2.9887931 | 2.83418 | 0.154617767 | -0.056856466 |

| | | | | | | | | | | |
|---|---|---|---|---|---|---|---|---|---|---|
| Gmps | 10.06959 | 10.03879 | 10.056 | -0.016938554 | 0.01386282 | 9.9743816 | 10.156367 | 10.0019 | 0.154478018 | -0.027507311 |
| Pdp1 | 7.578597 | 7.567362 | 7.5591 | 0.00827865 | 0.019513696 | 7.4188498 | 7.6303506 | 7.47602 | 0.154330741 | -0.057170072 |
| Olfr352 | 3.074669 | 3.280823 | 3.3495 | -0.068653673 | -0.27480747 | 3.1110978 | 3.4253738 | 3.27108 | 0.154297788 | -0.159978226 |
| Gm14434 | 5.43947 | 5.974889 | 6.1679 | -0.193032131 | -0.728451936 | 6.1320088 | 6.04335 | 5.88917 | 0.154184915 | 0.242843689 |
| Olfr1487 | 2.792776 | 2.894004 | 2.7552 | 0.138799029 | 0.037570145 | 2.9359552 | 2.9684523 | 2.81427 | 0.154178685 | 0.121681575 |
| Olfr607 | 5.104539 | 4.971735 | 5.0567 | -0.084922363 | 0.04788163 | 4.9743163 | 5.5212486 | 5.36707 | 0.154174826 | -0.392757457 |
| Vmn1r86 | 3.057752 | 3.151632 | 3.12 | 0.031642802 | -0.062237862 | 3.1752708 | 3.1920814 | 3.03793 | 0.154155391 | 0.137344806 |
| Cdk5rap3 | 9.667088 | 9.500305 | 9.697 | -0.196714596 | -0.029931653 | 9.8662989 | 9.8858867 | 9.73186 | 0.154021758 | 0.134433894 |
| Igkv12-38 | 4.825755 | 4.750442 | 4.6951 | 0.05530266 | 0.130614932 | 4.9417161 | 4.8574368 | 4.70342 | 0.154018303 | 0.238297601 |
| Coq6 | 7.736779 | 7.974587 | 7.8122 | 0.162376254 | -0.075431814 | 8.1379097 | 8.1732543 | 8.01927 | 0.153987968 | 0.118643334 |
| Igkv9-123 | 4.253595 | 4.226879 | 4.1492 | 0.077690323 | 0.10440622 | 4.3262814 | 4.3053885 | 4.15141 | 0.153980519 | 0.174873396 |
| Ints11 | 9.384566 | 9.562123 | 9.6063 | -0.044132398 | -0.22168876 | 9.7536478 | 9.7542956 | 9.60033 | 0.15396368 | 0.153315828 |
| Sulf1 | 8.762325 | 8.483179 | 8.6396 | -0.156383934 | 0.122762444 | 8.078158 | 7.9621767 | 7.80829 | 0.15388224 | 0.269863511 |
| Slc44a3 | 1.63069 | 1.356292 | 1.3439 | 0.012367148 | 0.286765092 | 1.4562328 | 1.480988 | 1.32717 | 0.15382169 | 0.129066516 |
| Gm49051 | 0.760487 | 0.720288 | 0.7398 | -0.019472999 | 0.02072635 | 0.7797572 | 0.8417864 | 0.68799 | 0.153800607 | 0.091771324 |
| Tmem253 | 1.504974 | 1.511675 | 1.5022 | 0.009457096 | 0.002755716 | 1.5255486 | 1.5796291 | 1.42583 | 0.153799144 | 0.099718669 |
| Igkv5-37 | 3.154736 | 3.169751 | 3.3175 | -0.147699395 | -0.162714723 | 3.3342853 | 3.3207865 | 3.16704 | 0.153742402 | 0.167241255 |
| Irak4 | 9.063414 | 8.905281 | 8.9056 | -0.000282266 | 0.157851089 | 9.1924316 | 9.0409685 | 8.8873 | 0.153665582 | 0.305128703 |
| Rtf1 | 10.82553 | 10.85694 | 10.852 | 0.004890976 | -0.026512158 | 10.775765 | 11.003157 | 10.8496 | 0.153532802 | -0.073859933 |
| Emd | 9.827532 | 9.850644 | 9.8569 | -0.006241347 | -0.029353936 | 10.003182 | 9.9754289 | 9.82201 | 0.15342244 | 0.18117568 |
| Tamm41 | 7.190349 | 7.144068 | 7.3395 | -0.195428443 | -0.149147624 | 7.3464258 | 7.507819 | 7.35444 | 0.153380231 | -0.008012986 |
| Hes1 | 8.0707 | 7.892748 | 7.8391 | 0.053620935 | 0.23157266 | 8.161392 | 7.8286467 | 7.67534 | 0.153302838 | 0.486048154 |
| Gm4925 | 3.79197 | 3.261404 | 3.3467 | -0.085282349 | 0.44528405 | 3.2921542 | 3.408455 | 3.25521 | 0.15324057 | 0.036939816 |
| Dhx30 | 10.70109 | 10.87586 | 10.74 | 0.135895422 | -0.038868578 | 10.881879 | 11.091973 | 10.9388 | 0.153197794 | -0.056895538 |
| 4932414N04Rik | 2.551844 | 2.659976 | 2.4928 | 0.167149348 | 0.059017595 | 2.7069309 | 2.8025172 | 2.64935 | 0.153165762 | 0.057579404 |
| Rrp1 | 11.41028 | 11.38066 | 11.404 | -0.023573116 | 0.00604434 | 11.543759 | 11.576171 | 11.4231 | 0.153096384 | 0.12068435 |
| Ahnak2 | 2.436151 | 2.590877 | 2.515 | 0.07589054 | -0.078835987 | 2.4266975 | 2.6307892 | 2.47774 | 0.15304444 | -0.051047252 |
| Pcca | 8.413089 | 8.345377 | 8.3684 | -0.023005889 | 0.044706265 | 8.3974471 | 8.5451685 | 8.39214 | 0.153024277 | 0.005302822 |
| Zc3h15 | 10.73895 | 10.83136 | 10.724 | 0.107592406 | 0.015185144 | 10.783015 | 11.03004 | 10.877 | 0.153011088 | -0.094013491 |
| Vmn1r236 | 3.597072 | 3.461687 | 3.4524 | 0.009318038 | 0.144703047 | 3.4701761 | 3.7962938 | 3.64339 | 0.152904803 | -0.173212953 |
| Ccdc59 | 9.144596 | 9.099733 | 9.1322 | -0.032486676 | 0.012376351 | 9.2548563 | 9.3763683 | 9.22351 | 0.152859426 | 0.031347421 |
| Smarca4 | 11.56217 | 11.62037 | 11.564 | 0.056653036 | -0.001544422 | 11.644641 | 11.766535 | 11.6137 | 0.152824694 | 0.030930201 |

| | | | | | | | | | | |
|---|---|---|---|---|---|---|---|---|---|---|
| Iah1 | 8.204928 | 8.314772 | 8.3026 | 0.01212493 | -0.097719536 | 8.5075911 | 8.3229898 | 8.17018 | 0.152809867 | 0.33741115 |
| Foxs1 | 3.001069 | 3.113905 | 2.9451 | 0.168848272 | 0.056013085 | 2.8557674 | 3.0009436 | 2.84814 | 0.15280569 | 0.007629509 |
| Spata2 | 8.94933 | 9.076661 | 8.9832 | 0.09347399 | -0.033856717 | 8.9967266 | 9.1652002 | 9.01243 | 0.152773603 | -0.015700064 |
| Pus7 | 8.899195 | 8.944505 | 8.7027 | 0.241802298 | 0.196492133 | 8.8928038 | 9.1640363 | 9.01129 | 0.152743215 | -0.118489246 |
| Zfp687 | 9.258011 | 9.239324 | 9.2526 | -0.013270693 | 0.005416452 | 9.3257716 | 9.4772235 | 9.32455 | 0.152673484 | 0.001221507 |
| Sp2 | 9.895115 | 9.993958 | 9.942 | 0.051935912 | -0.046906784 | 10.163395 | 10.021362 | 9.86871 | 0.152650849 | 0.294684223 |
| Fam53a | 9.236737 | 9.290859 | 9.275 | 0.01581955 | -0.038302317 | 9.2338171 | 9.3164734 | 9.16389 | 0.152586919 | 0.06993067 |
| Ighv1-50 | 7.201456 | 7.834967 | 7.1004 | 0.734595807 | 0.101084313 | 7.4135652 | 7.2806853 | 7.12813 | 0.152551085 | 0.285430972 |
| Gm14496 | 5.175569 | 5.153524 | 5.1454 | 0.008098945 | 0.030144144 | 5.2167556 | 5.2430086 | 5.09056 | 0.152446518 | 0.126193489 |
| P2ry10 | 11.73123 | 11.90688 | 11.665 | 0.24186962 | 0.066220222 | 11.677789 | 11.889796 | 11.7375 | 0.152341854 | -0.059664829 |
| Tnxb | 6.204523 | 6.09722 | 5.8042 | 0.293018895 | 0.40032162 | 5.476879 | 5.581002 | 5.42875 | 0.152256092 | 0.048133099 |
| Slc27a4 | 8.380184 | 8.52294 | 8.4091 | 0.113854272 | -0.028901878 | 8.4107403 | 8.3674346 | 8.21521 | 0.152226917 | 0.195532619 |
| Rab34 | 4.119179 | 3.999271 | 4.0386 | -0.039289769 | 0.080618458 | 3.8999511 | 4.0294123 | 3.87719 | 0.152221037 | 0.022759815 |
| Washc1 | 9.27604 | 9.284632 | 9.4219 | -0.137266626 | -0.145858564 | 9.5676453 | 9.633006 | 9.48079 | 0.152219058 | 0.086858364 |
| Gspt1 | 10.87624 | 10.97188 | 10.843 | 0.128786954 | 0.033142044 | 10.890842 | 11.118233 | 10.9663 | 0.151921592 | -0.075469602 |
| Gm47356 | 5.06142 | 5.254305 | 4.9562 | 0.298105336 | 0.10522093 | 5.2178517 | 5.5558118 | 5.40396 | 0.15185326 | -0.186106802 |
| Gm7816 | 2.48037 | 2.471188 | 2.3207 | 0.15048921 | 0.159671096 | 2.2627802 | 2.5742053 | 2.42238 | 0.151828927 | -0.159596174 |
| Gga3 | 10.28544 | 10.39369 | 10.246 | 0.147294093 | 0.039044382 | 10.595333 | 10.563062 | 10.4113 | 0.151804884 | 0.184076176 |
| 4931428L18Rik | 4.149011 | 4.069671 | 4.1249 | -0.055223528 | 0.02411661 | 4.2575863 | 4.3763118 | 4.22462 | 0.15169656 | 0.032971049 |
| Sipa1 | 12.3955 | 12.47257 | 12.459 | 0.013849878 | -0.063220468 | 12.949927 | 12.74954 | 12.5979 | 0.151677876 | 0.35206475 |
| Tmem69 | 7.039035 | 6.983969 | 7.0581 | -0.074087217 | -0.019021315 | 7.1587025 | 7.2061104 | 7.05445 | 0.151658053 | 0.104250187 |
| Imp4 | 10.229 | 10.25204 | 10.189 | 0.063186995 | 0.040147571 | 10.462775 | 10.322534 | 10.1712 | 0.151337432 | 0.291578214 |
| Erg28 | 8.508388 | 8.507296 | 8.4597 | 0.047546967 | 0.048639146 | 8.7343519 | 8.7287044 | 8.5774 | 0.151302036 | 0.156949569 |
| Cnbp | 12.8972 | 12.86087 | 12.832 | 0.029092522 | 0.065417678 | 12.906922 | 12.979289 | 12.8281 | 0.151187494 | 0.078820556 |
| Supt4a | 9.955715 | 10.11559 | 10.078 | 0.038044099 | -0.121833553 | 10.156157 | 10.241383 | 10.0903 | 0.151101251 | 0.065875267 |
| Cdca4 | 9.692219 | 9.711975 | 9.7593 | -0.047310713 | -0.06706625 | 9.8208991 | 9.94367 | 9.79257 | 0.151096683 | 0.028325825 |
| Aste1 | 7.694583 | 7.582235 | 7.6418 | -0.059519873 | 0.052828606 | 7.7610291 | 7.7606343 | 7.60955 | 0.151079393 | 0.15147421 |
| Cwc27 | 8.431616 | 8.395818 | 8.5152 | -0.119357538 | -0.083558978 | 8.5733351 | 8.6932911 | 8.54226 | 0.151027357 | 0.031071348 |
| Psmd8 | 10.46721 | 10.51576 | 10.464 | 0.05217288 | 0.003621178 | 10.668673 | 10.67521 | 10.5242 | 0.151011576 | 0.144474206 |
| Lyrm2 | 7.283841 | 7.333804 | 7.4393 | -0.105488938 | -0.155451201 | 7.5180865 | 7.4884296 | 7.33743 | 0.151003605 | 0.180660507 |
| Smg8 | 8.615643 | 8.737858 | 8.6 | 0.137820564 | 0.015605677 | 8.6018683 | 8.6555757 | 8.50462 | 0.150955196 | 0.097247789 |
| Arl1 | 9.964232 | 9.835947 | 9.8555 | -0.019512354 | 0.108772196 | 9.8036027 | 9.9685823 | 9.81772 | 0.150862133 | -0.014117554 |

| | | | | | | | | | | |
|---|---|---|---|---|---|---|---|---|---|---|
| Setdb1 | 10.11751 | 10.16994 | 10.151 | 0.019101684 | -0.033321253 | 10.121428 | 10.338167 | 10.1873 | 0.150850103 | -0.065888798 |
| Vmn1r208 | 1.62283 | 1.594327 | 1.6641 | -0.069724325 | -0.041221113 | 1.7167366 | 1.8475424 | 1.69672 | 0.150823502 | 0.020017723 |
| Gm5417 | 1.663869 | 1.561611 | 1.5111 | 0.050523716 | 0.152782235 | 1.658539 | 1.726939 | 1.57613 | 0.150804054 | 0.082404045 |
| Gpr176 | 4.178587 | 4.297162 | 3.9835 | 0.313695222 | 0.195120125 | 4.2378925 | 4.0356736 | 3.88489 | 0.150780247 | 0.352999167 |
| Vta1 | 9.043785 | 9.067153 | 9.059 | 0.008169244 | -0.015198816 | 9.1375064 | 9.2336455 | 9.08292 | 0.150724596 | 0.054585464 |
| Ppil2 | 10.50315 | 10.48457 | 10.54 | -0.055762912 | -0.03717424 | 10.579823 | 10.784012 | 10.6334 | 0.150562226 | -0.053627261 |
| Mepce | 10.31736 | 10.41241 | 10.395 | 0.017741126 | -0.07730856 | 10.36814 | 10.510702 | 10.3602 | 0.15048249 | 0.007920451 |
| Rce1 | 8.370496 | 8.364995 | 8.362 | 0.003017715 | 0.008519339 | 8.700465 | 8.5924933 | 8.44207 | 0.150425369 | 0.258397011 |
| Tmem126a | 7.739587 | 7.736841 | 7.8291 | -0.092297943 | -0.089551653 | 7.9857979 | 7.9269119 | 7.77675 | 0.150166121 | 0.209052062 |
| Shkbp1 | 9.472386 | 9.562683 | 9.5239 | 0.038788306 | -0.051508445 | 9.8209923 | 9.7988955 | 9.64881 | 0.150087203 | 0.172183946 |
| Stxbp1 | 7.246318 | 7.16773 | 7.2953 | -0.127606279 | -0.04901835 | 6.6876614 | 7.3321905 | 7.18228 | 0.149909664 | -0.49461943 |
| Alg5 | 8.789978 | 8.657035 | 8.6831 | -0.026040698 | 0.106902231 | 8.7882741 | 8.8613752 | 8.71153 | 0.149848008 | 0.076746904 |
| Arhgap26 | 10.42702 | 10.54145 | 10.406 | 0.135061358 | 0.020628036 | 10.550021 | 10.789227 | 10.6394 | 0.149783234 | -0.089422876 |
| Ncf4 | 10.45995 | 10.48075 | 10.516 | -0.035414042 | -0.056209118 | 10.945287 | 10.825787 | 10.676 | 0.149783228 | 0.269283814 |
| Tulp1 | 2.936617 | 2.859155 | 2.9504 | -0.091212609 | -0.013750768 | 2.9550496 | 3.0517085 | 2.90204 | 0.14966925 | 0.0530104 |
| Gm47628 | 3.988965 | 3.967016 | 4.0449 | -0.077861272 | -0.055912572 | 3.9495296 | 4.2677712 | 4.11836 | 0.149415817 | -0.16882579 |
| Git1 | 9.763088 | 9.712829 | 9.7343 | -0.021490295 | 0.028769044 | 9.663418 | 9.8921201 | 9.74282 | 0.149303043 | -0.079399099 |
| Ttc32 | 6.182992 | 6.211008 | 6.1565 | 0.054487836 | 0.026471259 | 6.1155944 | 6.1611332 | 6.01189 | 0.149245765 | 0.103707022 |
| Gm10155 | 3.420625 | 3.487648 | 3.5298 | -0.042151406 | -0.109174756 | 3.381881 | 3.5639953 | 3.41485 | 0.149146317 | -0.032968016 |
| Rbm19 | 9.235727 | 9.349513 | 9.2253 | 0.124187 | 0.010400883 | 9.3295153 | 9.4904466 | 9.3413 | 0.149143328 | -0.011787952 |
| Atl2 | 9.292643 | 9.506672 | 9.3385 | 0.168163457 | -0.04586548 | 9.3053868 | 9.5961226 | 9.44699 | 0.149133673 | -0.141602184 |
| Srrd | 7.042036 | 6.942854 | 7.0102 | -0.067380716 | 0.031800597 | 7.2009051 | 7.3033826 | 7.15425 | 0.14913217 | 0.046654629 |
| Naf1 | 7.833657 | 7.789054 | 7.6423 | 0.146745078 | 0.191348361 | 7.5121356 | 7.8427941 | 7.69368 | 0.14911786 | -0.181540723 |
| Gpat2 | 2.264577 | 2.174183 | 2.1851 | -0.01089393 | 0.07949974 | 2.4164399 | 2.4897042 | 2.34063 | 0.14907787 | 0.075813648 |
| Pctp | 7.377585 | 7.247405 | 7.3009 | -0.053449408 | 0.076730102 | 7.3672433 | 7.2704595 | 7.12139 | 0.149069901 | 0.245853659 |
| Pyroxd2 | 4.900432 | 4.80316 | 4.9553 | -0.152164553 | -0.054892148 | 4.8567758 | 5.2059981 | 5.05695 | 0.149053 | -0.200169269 |
| Cnot10 | 9.998942 | 10.07521 | 10.031 | 0.044518009 | -0.031746947 | 10.152164 | 10.271862 | 10.1228 | 0.149023974 | 0.029326158 |
| Nlrp5 | 2.277362 | 2.231374 | 2.2324 | -0.000989196 | 0.044999037 | 2.2498098 | 2.4182677 | 2.26935 | 0.148916638 | -0.019541345 |
| Mif4gd | 11.01106 | 10.90848 | 10.888 | 0.02001907 | 0.122600848 | 11.583269 | 11.190158 | 11.0413 | 0.148840948 | 0.541951279 |
| Uimc1 | 10.33857 | 10.38434 | 10.374 | 0.010195966 | -0.03557515 | 10.422285 | 10.55819 | 10.4094 | 0.148804108 | 0.012899792 |
| Lsm14a | 11.47737 | 11.57633 | 11.497 | 0.079792578 | -0.019158942 | 11.31113 | 11.545151 | 11.3963 | 0.148802674 | -0.085218654 |
| Igkv6-20 | 7.224945 | 7.250588 | 7.316 | -0.06542222 | -0.091065159 | 7.4019735 | 7.2594821 | 7.11079 | 0.148695368 | 0.291186786 |

| | | | | | | | | | | |
|---|---|---|---|---|---|---|---|---|---|---|
| 1700109H08Rik | 5.766177 | 5.814174 | 5.9432 | -0.129046566 | -0.177043186 | 5.7747696 | 5.9890139 | 5.84043 | 0.148580704 | -0.065663595 |
| Rpl9-ps7 | 2.91754 | 2.879913 | 2.8067 | 0.073181286 | 0.110808098 | 3.0417918 | 2.9011803 | 2.75264 | 0.148544056 | 0.289155627 |
| Ergic2 | 9.758013 | 9.811898 | 9.7742 | 0.037665688 | -0.016219143 | 9.7090018 | 10.015261 | 9.86675 | 0.148508783 | -0.157750106 |
| Hmg20b | 8.599599 | 8.49602 | 8.6661 | -0.170045564 | -0.066466268 | 8.9046179 | 8.9452753 | 8.79714 | 0.148137569 | 0.10748017 |
| Rbm25 | 12.03002 | 12.01282 | 12.044 | -0.030775164 | -0.0135827 | 12.170422 | 12.376317 | 12.2282 | 0.148135956 | -0.057759308 |
| Unc93a | 1.121919 | 1.172945 | 1.1628 | 0.010132334 | -0.040893178 | 1.1017041 | 1.2915439 | 1.14345 | 0.148090915 | -0.041748852 |
| Ccdc51 | 7.157756 | 7.183247 | 7.1922 | -0.008931555 | -0.0344231 | 7.3395931 | 7.3531226 | 7.20517 | 0.147950545 | 0.134421033 |
| Mrpl3 | 9.409364 | 9.347114 | 9.2724 | 0.074685171 | 0.136934385 | 9.5608695 | 9.5430513 | 9.39517 | 0.147879561 | 0.165697698 |
| Pofut2 | 8.829957 | 8.888416 | 8.6853 | 0.203162418 | 0.144703005 | 8.9664982 | 8.9120337 | 8.76421 | 0.147822115 | 0.202286535 |
| Itm2c | 10.89424 | 10.86511 | 10.91 | -0.044649764 | -0.015520588 | 10.699427 | 10.860042 | 10.7123 | 0.147788758 | -0.012826835 |
| Gm48040 | 3.153462 | 3.265763 | 3.2797 | -0.013918499 | -0.126219038 | 3.1538044 | 3.4307896 | 3.28301 | 0.147775996 | -0.129209156 |
| Lrrc45 | 8.757645 | 8.993901 | 8.9099 | 0.083952555 | -0.152304078 | 9.2646887 | 9.3102479 | 9.1625 | 0.14774634 | 0.102187144 |
| Ecpas | 10.73291 | 10.77365 | 10.852 | -0.078353046 | -0.11909726 | 10.703189 | 11.032372 | 10.8847 | 0.147683074 | -0.181500262 |
| Sec24b | 10.67998 | 10.80681 | 10.645 | 0.161506564 | 0.034674078 | 10.611459 | 10.845887 | 10.6983 | 0.147607474 | -0.086820307 |
| Gm12543 | 2.859937 | 2.812464 | 2.7919 | 0.020588151 | 0.068061498 | 2.7997326 | 3.0944142 | 2.94687 | 0.147540564 | -0.147141019 |
| Ppp5c | 10.0617 | 10.076 | 10.082 | -0.005901501 | -0.020206043 | 10.562538 | 10.403512 | 10.2561 | 0.147379598 | 0.306405859 |
| Cdk18 | 3.593603 | 3.724872 | 3.6551 | 0.069721943 | -0.061546562 | 3.6023563 | 3.5906309 | 3.44328 | 0.147355643 | 0.15908099 |
| Dcaf7 | 11.65459 | 11.82354 | 11.643 | 0.180741104 | 0.011791932 | 11.697405 | 11.771388 | 11.6241 | 0.147333016 | 0.073350063 |
| Rab21 | 10.94504 | 11.10634 | 11.028 | 0.078828596 | -0.082469812 | 10.923043 | 11.067157 | 10.9199 | 0.147211208 | 0.003097442 |
| Ccdc127 | 8.938624 | 9.074606 | 8.9467 | 0.127885004 | -0.008097118 | 8.8457334 | 9.0404525 | 8.89333 | 0.147121264 | -0.047597849 |
| Med25 | 9.310787 | 9.37483 | 9.2948 | 0.080015863 | 0.015972709 | 9.5968119 | 9.5099585 | 9.36286 | 0.14710208 | 0.23395546 |
| Gla | 8.322277 | 8.452703 | 8.1561 | 0.296645839 | 0.166220456 | 8.1765984 | 8.4391451 | 8.29206 | 0.147082457 | -0.115464264 |
| Mtmr3 | 11.54177 | 11.67109 | 11.609 | 0.061836828 | -0.067488744 | 11.567081 | 11.717041 | 11.5701 | 0.146947556 | -0.003012315 |
| Ufc1 | 9.284718 | 9.102293 | 9.1834 | -0.081079195 | 0.101346513 | 9.3104377 | 9.298573 | 9.15164 | 0.146928599 | 0.158793352 |
| Camkk2 | 9.300933 | 9.517777 | 9.2323 | 0.285481367 | 0.068637789 | 9.3951981 | 9.3588673 | 9.21209 | 0.146774733 | 0.18310553 |
| Bod1 | 7.160469 | 7.146065 | 7.1383 | 0.007767509 | 0.022171449 | 7.0477474 | 7.1702024 | 7.02343 | 0.146773395 | 0.024318424 |
| Ubxn2b | 8.391608 | 8.502684 | 8.3862 | 0.116459488 | 0.005382727 | 8.4469058 | 8.6230575 | 8.47636 | 0.146700456 | -0.02945119 |
| Gm17084 | 4.735264 | 4.836688 | 4.8452 | -0.008473371 | -0.109896649 | 4.7636512 | 5.0426568 | 4.89597 | 0.146683056 | -0.132322486 |
| Vmn2r7 | 1.648362 | 1.711117 | 1.657 | 0.054067614 | -0.008687275 | 1.6295652 | 1.754952 | 1.60846 | 0.146489695 | 0.021102904 |
| Twf2 | 10.38473 | 10.43334 | 10.421 | 0.012350304 | -0.036255488 | 10.566992 | 10.511915 | 10.3655 | 0.14643745 | 0.201513792 |
| Dyrk4 | 2.796764 | 2.856711 | 2.7502 | 0.10656091 | 0.046613781 | 2.9356127 | 2.9790135 | 2.83267 | 0.146346691 | 0.102945864 |
| Slc9a2 | 2.179565 | 2.156545 | 2.1805 | -0.023974851 | -0.000954316 | 2.2422994 | 2.3824043 | 2.2361 | 0.146308616 | 0.006203806 |

| | | | | | | | | | | |
|---|---|---|---|---|---|---|---|---|---|---|
| Gnptg | 8.964238 | 8.980201 | 9.0017 | -0.021487484 | -0.037449767 | 9.2063207 | 9.2591719 | 9.11286 | 0.146308382 | 0.093457248 |
| Gm20683 | 5.607301 | 5.637995 | 5.6074 | 0.030602998 | -9.05052E-05 | 5.9865716 | 5.9767948 | 5.83062 | 0.146177289 | 0.155954127 |
| Mrpl9 | 9.597643 | 9.623714 | 9.6594 | -0.035655749 | -0.061726333 | 9.7569378 | 9.8953616 | 9.74922 | 0.146136856 | 0.007713102 |
| Rpl21-ps8 | 3.210209 | 3.347791 | 3.3221 | 0.025662013 | -0.111919776 | 3.2338692 | 3.4394989 | 3.29336 | 0.146134262 | -0.059495446 |
| L3mbtl4 | 2.435234 | 2.452107 | 2.4435 | 0.008579642 | -0.008293449 | 2.4508835 | 2.5897514 | 2.44366 | 0.14609231 | 0.00722443 |
| Timm44 | 9.563707 | 9.603424 | 9.5196 | 0.083775799 | 0.044059374 | 9.7896387 | 9.8866639 | 9.74068 | 0.145988305 | 0.048963142 |
| Ighv2-2 | 7.307845 | 7.619997 | 7.4058 | 0.214212357 | -0.097939517 | 7.6307094 | 7.4792962 | 7.33333 | 0.145963062 | 0.297376208 |
| Tsfm | 7.425028 | 7.535655 | 7.3545 | 0.18110865 | 0.07048235 | 7.5877474 | 7.7485959 | 7.60263 | 0.145961338 | -0.014887125 |
| Supt3 | 7.40248 | 7.362991 | 7.4183 | -0.055357167 | -0.01586844 | 7.3565431 | 7.4852391 | 7.33928 | 0.145955301 | 0.017259285 |
| Heatr5b | 9.729918 | 9.888458 | 9.7755 | 0.112994432 | -0.045545388 | 9.7313608 | 9.9424118 | 9.79647 | 0.145939758 | -0.065111294 |
| Vmn2r93 | 1.664624 | 1.712427 | 1.6703 | 0.042164332 | -0.005638309 | 1.6802629 | 1.8580244 | 1.7122 | 0.14582293 | -0.031938557 |
| Ilkap | 9.82536 | 9.774325 | 9.8051 | -0.030782574 | 0.020252277 | 9.988706 | 9.9989021 | 9.85309 | 0.145811277 | 0.135615158 |
| Slc2a4rg-ps | 8.155116 | 8.347271 | 8.3306 | 0.016652799 | -0.175502987 | 8.9353084 | 9.058398 | 8.91259 | 0.145803109 | 0.022713468 |
| Abraxas1 | 7.401018 | 7.532881 | 7.5292 | 0.003727138 | -0.128135954 | 7.6447311 | 7.7798353 | 7.63404 | 0.145795234 | 0.010691 |
| Gm10031 | 3.00271 | 3.129884 | 3.1191 | 0.010740066 | -0.116434635 | 2.8812679 | 3.287553 | 3.14178 | 0.145776636 | -0.26050841 |
| Il17rc | 3.9697 | 4.098526 | 3.9823 | 0.116238257 | -0.01258772 | 4.3830275 | 3.9702349 | 3.82457 | 0.145662271 | 0.558454922 |
| Cdc26 | 9.25277 | 9.282746 | 9.2805 | 0.002246325 | -0.02772994 | 9.3855902 | 9.4137899 | 9.26826 | 0.145534671 | 0.11733501 |
| Prdm16 | 5.64183 | 6.115087 | 5.897 | 0.218127777 | -0.255129022 | 5.4085655 | 6.11227 | 5.9668 | 0.145465288 | -0.55823918 |
| Gm48841 | 4.88104 | 4.854605 | 5.0095 | -0.154899284 | -0.1284644 | 5.1699246 | 5.3052093 | 5.15977 | 0.145441334 | 0.010156633 |
| Rpl37a | 13.81552 | 13.90421 | 13.873 | 0.03072184 | -0.057965978 | 14.112948 | 13.990232 | 13.8448 | 0.14543819 | 0.268154015 |
| Naa10 | 9.49236 | 9.45419 | 9.4975 | -0.043307183 | -0.005136509 | 9.7119088 | 9.6905392 | 9.54512 | 0.145421149 | 0.166790701 |
| Prelid2 | 4.972942 | 4.854356 | 4.8142 | 0.040140953 | 0.158726184 | 4.7799944 | 4.9858871 | 4.84048 | 0.14541154 | -0.060481137 |
| Glrp1 | 1.056621 | 1.074428 | 1.0296 | 0.044868883 | 0.027062587 | 1.0902183 | 1.2061833 | 1.06093 | 0.145249734 | 0.029284772 |
| Yjefn3 | 0.979779 | 0.972023 | 0.9457 | 0.026346563 | 0.034102875 | 1.0382303 | 1.1104145 | 0.96518 | 0.145236774 | 0.073052578 |
| Ppp1r13b | 10.26678 | 10.33143 | 10.344 | -0.012769486 | -0.07742429 | 10.011321 | 10.268624 | 10.1234 | 0.145226984 | -0.112076333 |
| Taf5l | 9.950944 | 9.866571 | 9.888 | -0.021449208 | 0.062923401 | 9.8750849 | 9.9950457 | 9.84985 | 0.145199992 | 0.025239184 |
| Gm10275 | 14.81821 | 14.73717 | 14.816 | -0.078682798 | 0.002356326 | 14.849087 | 14.907144 | 14.7621 | 0.145036968 | 0.086980298 |
| Stx7 | 10.17114 | 10.1308 | 10.141 | -0.009716186 | 0.03062422 | 10.242755 | 10.173988 | 10.029 | 0.145029079 | 0.21379651 |
| Hspa1a | 13.2503 | 12.9143 | 13.225 | -0.310518594 | 0.025482296 | 12.341371 | 12.845364 | 12.7003 | 0.145022914 | -0.358970257 |
| Gng4 | 2.42318 | 2.606473 | 2.5357 | 0.070809363 | -0.112483934 | 2.5368214 | 2.7191238 | 2.57412 | 0.1449989 | -0.037303535 |
| Arsj | 1.060701 | 1.105127 | 1.0326 | 0.072534345 | 0.028108862 | 1.0936651 | 1.2138769 | 1.0689 | 0.144978819 | 0.02476694 |
| Olfm1 | 8.042287 | 7.545168 | 7.4783 | 0.066856735 | 0.563976174 | 7.8251905 | 7.7070075 | 7.56204 | 0.144969247 | 0.263152243 |

| | | | | | | | | | | |
|---|---|---|---|---|---|---|---|---|---|---|
| Mcpt2 | 0.947267 | 1.023664 | 0.9312 | 0.092480571 | 0.016083841 | 1.1190252 | 1.0693733 | 0.92447 | 0.144907821 | 0.194559751 |
| Bdh1 | 8.563218 | 8.525543 | 8.5307 | -0.005135971 | 0.032539384 | 8.4141317 | 8.5120559 | 8.36715 | 0.144901506 | 0.046977256 |
| Gm9696 | 1.450752 | 1.492042 | 1.4706 | 0.021409471 | -0.01988042 | 1.4448209 | 1.5827969 | 1.43796 | 0.144837543 | 0.006861497 |
| Gtf2h2 | 8.504522 | 8.521794 | 8.5251 | -0.003264693 | -0.020536528 | 8.7203689 | 8.8325802 | 8.68776 | 0.144818272 | 0.03260704 |
| Zfp787 | 7.82963 | 7.667579 | 7.8544 | -0.186773699 | -0.024722919 | 7.8855779 | 7.86081 | 7.71603 | 0.144784354 | 0.169552255 |
| B3galt6 | 6.856979 | 6.935504 | 6.7527 | 0.182836204 | 0.104311059 | 6.8583793 | 6.9009114 | 6.7562 | 0.144715956 | 0.102183925 |
| Olfr1031 | 1.72501 | 1.845861 | 1.6969 | 0.148933647 | 0.028082628 | 1.7712088 | 1.9329258 | 1.78839 | 0.144536069 | -0.017180845 |
| Rpl23 | 14.54139 | 14.51625 | 14.536 | -0.019267562 | 0.005873034 | 14.590447 | 14.706301 | 14.5619 | 0.144393876 | 0.028539633 |
| Snrnp35 | 6.755963 | 6.580634 | 6.4431 | 0.137536037 | 0.312864666 | 6.7263218 | 6.7642966 | 6.61994 | 0.144357885 | 0.106383132 |
| Gm6611 | 3.021172 | 2.984043 | 3.0365 | -0.052452072 | -0.01532321 | 3.0070485 | 3.1782043 | 3.03395 | 0.144255786 | -0.026900007 |
| Wdr44 | 9.387754 | 9.46793 | 9.4037 | 0.064261525 | -0.015914384 | 9.3715432 | 9.5196009 | 9.37537 | 0.144228125 | -0.003829541 |
| Edem2 | 9.532326 | 9.562485 | 9.5157 | 0.046792971 | 0.016634817 | 9.6564377 | 9.720947 | 9.57682 | 0.144122093 | 0.079612764 |
| Zbtb26 | 6.91338 | 7.147663 | 7.0567 | 0.090980616 | -0.143301918 | 7.6514887 | 7.5246596 | 7.38061 | 0.144049959 | 0.270879057 |
| 1810022K09Rik | 8.486329 | 8.697563 | 8.3127 | 0.38485471 | 0.173620686 | 8.715188 | 8.8923084 | 8.7483 | 0.144011509 | -0.033108937 |
| Arid5b | 9.876547 | 9.711296 | 9.7045 | 0.006783868 | 0.172035342 | 9.5659272 | 9.9646607 | 9.82065 | 0.144011209 | -0.254722346 |
| Dcun1d2 | 8.495869 | 8.674375 | 8.5582 | 0.116210371 | -0.06229493 | 8.4977367 | 8.6730733 | 8.52919 | 0.143887429 | -0.031449264 |
| Rab13 | 5.088566 | 4.803912 | 5.128 | -0.324063045 | -0.039409094 | 4.8354071 | 5.0012849 | 4.85742 | 0.143862165 | -0.022015648 |
| Tcrg-V1 | 3.420522 | 3.511842 | 3.4801 | 0.031760338 | -0.059560261 | 3.4917068 | 3.5173496 | 3.37353 | 0.143815224 | 0.11817249 |
| Gm12070 | 4.650327 | 4.684965 | 4.8973 | -0.212310417 | -0.246948086 | 4.9694622 | 4.8150426 | 4.67127 | 0.143771702 | 0.298191314 |
| Gm13868 | 1.410476 | 1.405303 | 1.4588 | -0.053475501 | -0.04830243 | 1.3883719 | 1.5441918 | 1.40048 | 0.143711728 | -0.012108241 |
| Cbx4 | 9.956847 | 10.05418 | 10.003 | 0.051271898 | -0.046059229 | 9.780208 | 10.106818 | 9.96312 | 0.143697505 | -0.182912605 |
| Zfp524 | 7.504492 | 7.549859 | 7.4753 | 0.074509744 | 0.029143364 | 7.6685614 | 7.5867367 | 7.44308 | 0.143652816 | 0.225477497 |
| Gm7143 | 4.981949 | 5.11503 | 5.2152 | -0.100128161 | -0.233209542 | 5.2508436 | 5.2626632 | 5.11907 | 0.14359493 | 0.131775246 |
| Pira2 | 8.020083 | 8.095963 | 7.9982 | 0.09774677 | 0.021867109 | 8.6498885 | 8.1915001 | 8.04797 | 0.143526425 | 0.601914836 |
| Zfp993 | 5.896695 | 5.817911 | 5.8891 | -0.071216751 | 0.007567222 | 5.7660467 | 6.0412707 | 5.89778 | 0.14349355 | -0.131730441 |
| Ovgp1 | 6.576064 | 6.346904 | 6.6679 | -0.320982934 | -0.091822796 | 6.8647559 | 6.9250702 | 6.78167 | 0.143395815 | 0.083081473 |
| Cmas | 9.22919 | 9.192767 | 9.2043 | -0.011526462 | 0.024896978 | 9.0788729 | 9.2866102 | 9.14322 | 0.143389526 | -0.064347847 |
| Aldh1a2 | 5.73831 | 6.149639 | 5.9146 | 0.234997144 | -0.176331237 | 5.6574222 | 5.9447145 | 5.8014 | 0.1433153 | -0.143977041 |
| Hist1h4b | 3.663667 | 3.798515 | 3.7774 | 0.021159882 | -0.113688542 | 3.967778 | 3.8940325 | 3.75076 | 0.143270983 | 0.217016534 |
| Siah1a | 8.098203 | 8.204604 | 8.2599 | -0.055266833 | -0.161667929 | 8.0326157 | 8.2740084 | 8.13077 | 0.143242148 | -0.098150544 |
| Chuk | 10.39235 | 10.35652 | 10.368 | -0.011022666 | 0.024805826 | 10.4284 | 10.723568 | 10.5804 | 0.14314246 | -0.152025011 |
| Hspa14 | 9.728822 | 9.712367 | 9.7493 | -0.036916688 | -0.020462436 | 9.9434911 | 9.9859895 | 9.8429 | 0.143088876 | 0.100590426 |

| | | | | | | | | | | |
|---|---|---|---|---|---|---|---|---|---|---|
| Prodh | 4.842231 | 4.660225 | 4.7053 | -0.045063771 | 0.136942619 | 4.5888085 | 4.8668338 | 4.72398 | 0.14285191 | -0.135173367 |
| Ube2d1 | 9.332218 | 9.512342 | 9.4053 | 0.107034552 | -0.073089347 | 9.6150385 | 9.5674437 | 9.42466 | 0.142788001 | 0.190382748 |
| Rab8b | 11.27745 | 11.4064 | 11.232 | 0.174287208 | 0.04533781 | 10.981124 | 11.371501 | 11.2287 | 0.142783238 | -0.247593953 |
| Bin1 | 11.42402 | 11.55954 | 11.463 | 0.096217738 | -0.039301752 | 11.801801 | 11.670036 | 11.5273 | 0.142715956 | 0.274480969 |
| Coa6 | 6.885388 | 6.648088 | 6.7123 | -0.064209301 | 0.173090285 | 6.947348 | 6.8868895 | 6.74455 | 0.142338624 | 0.202797113 |
| Plekhm2 | 8.904125 | 9.135407 | 9.1202 | 0.015164441 | -0.216116949 | 9.0000124 | 9.1794667 | 9.03719 | 0.142274995 | -0.037179337 |
| Erlin1 | 8.738005 | 8.674898 | 8.612 | 0.062906804 | 0.126013824 | 8.7193459 | 8.6473519 | 8.50509 | 0.142261573 | 0.214255591 |
| Yars2 | 7.735277 | 7.71605 | 7.6067 | 0.109303174 | 0.128529896 | 7.8066104 | 7.9683273 | 7.82614 | 0.142184657 | -0.01953217 |
| Tmem256 | 7.39735 | 7.258934 | 7.2155 | 0.043407348 | 0.181823512 | 7.5717995 | 7.3587136 | 7.2166 | 0.142111247 | 0.355197087 |
| Map3k10 | 7.207703 | 7.305338 | 7.1625 | 0.142886593 | 0.045251181 | 7.3494545 | 7.1602725 | 7.01825 | 0.142017554 | 0.331199641 |
| Eya3 | 9.826403 | 9.897456 | 9.9474 | -0.049934191 | -0.120986982 | 9.8391727 | 10.059553 | 9.91754 | 0.142012694 | -0.07836784 |
| Myg1 | 8.770275 | 8.85071 | 8.8075 | 0.043181929 | -0.037252308 | 9.1768991 | 9.1097715 | 8.96779 | 0.141977928 | 0.209105496 |
| Wdr70 | 8.798035 | 8.864769 | 8.7642 | 0.100566068 | 0.033832651 | 8.8950516 | 8.9651946 | 8.8233 | 0.141893045 | 0.07175 |
| Rnf220 | 10.31164 | 10.47482 | 10.349 | 0.125640694 | -0.037545874 | 10.535096 | 10.534626 | 10.3928 | 0.141808318 | 0.142278581 |
| Zcchc9 | 8.484319 | 8.524292 | 8.477 | 0.047293361 | 0.007320469 | 8.6694677 | 8.7174623 | 8.57575 | 0.14171681 | 0.093722281 |
| Ppp2r5b | 6.730675 | 6.820424 | 6.785 | 0.035442205 | -0.054306678 | 6.9112849 | 6.8227768 | 6.68116 | 0.141620694 | 0.230128778 |
| Metap1 | 9.453225 | 9.641347 | 9.5099 | 0.131457884 | -0.056663787 | 9.5998038 | 9.756534 | 9.61498 | 0.141556513 | -0.015173725 |
| Il17f | 2.240625 | 2.127597 | 2.2037 | -0.076080005 | 0.036948153 | 2.2462272 | 2.374696 | 2.23314 | 0.141551259 | 0.013082453 |
| Glis3 | 4.497133 | 4.399041 | 4.3346 | 0.06442852 | 0.162520306 | 4.3062592 | 4.5263868 | 4.38487 | 0.141516948 | -0.078610588 |
| Mpi | 7.556033 | 7.582983 | 7.638 | -0.055043674 | -0.081993643 | 7.6263464 | 7.7344602 | 7.59298 | 0.141476183 | 0.03336247 |
| Gm48670 | 3.02922 | 2.939701 | 2.9366 | 0.003147734 | 0.092666241 | 2.9414083 | 3.1668446 | 3.02551 | 0.141336061 | -0.084100254 |
| Gm20075 | 4.553654 | 4.324172 | 4.4883 | -0.164130429 | 0.065351351 | 3.9101756 | 4.5216647 | 4.38035 | 0.141314198 | -0.470174881 |
| Rps6kb2 | 7.903587 | 7.851427 | 7.7335 | 0.117902142 | 0.170062379 | 8.0985453 | 8.2173146 | 8.07605 | 0.141260985 | 0.022491743 |
| Vsx2 | 3.172347 | 3.19771 | 3.2746 | -0.076894023 | -0.102256556 | 3.4174181 | 3.4569078 | 3.31577 | 0.141137634 | 0.101647859 |
| Urb1 | 10.00492 | 10.12482 | 9.9104 | 0.214430957 | 0.094525491 | 10.027289 | 10.161649 | 10.0205 | 0.141128505 | 0.006768605 |
| Uqcc1 | 8.695735 | 8.912403 | 8.7064 | 0.206041534 | -0.010626499 | 8.8976966 | 8.9028922 | 8.76179 | 0.141100947 | 0.135905321 |
| 1810030O07Rik | 9.746467 | 9.813532 | 9.7794 | 0.034151386 | -0.032913696 | 9.798317 | 9.8963523 | 9.75525 | 0.14110089 | 0.043065543 |
| Nkrf | 7.796366 | 7.808071 | 7.653 | 0.155069119 | 0.143364507 | 7.6822659 | 7.8348865 | 7.69383 | 0.141052911 | -0.011567699 |
| Lonp1 | 10.11194 | 10.15823 | 10.096 | 0.061875492 | 0.01558601 | 10.267874 | 10.340668 | 10.1997 | 0.140970814 | 0.068177139 |
| Rtp3 | 2.628237 | 2.587038 | 2.5916 | -0.004530815 | 0.036667767 | 2.6969042 | 2.8084053 | 2.66746 | 0.140944153 | 0.02944305 |
| Ciao2a | 9.670904 | 9.540428 | 9.5474 | -0.006930858 | 0.1235447 | 9.6851616 | 9.67113 | 9.5302 | 0.140932712 | 0.154964349 |
| Spop | 11.4437 | 11.54058 | 11.451 | 0.089640272 | -0.007237214 | 11.361248 | 11.551635 | 11.4107 | 0.140920954 | -0.049466412 |

| | | | | | | | | | | |
|---|---|---|---|---|---|---|---|---|---|---|
| Nmnat2 | 5.022809 | 5.016814 | 5.0038 | 0.013059856 | 0.019054466 | 4.9137282 | 5.010379 | 4.8695 | 0.140884016 | 0.044233151 |
| Tmem201 | 8.272835 | 8.443333 | 8.3295 | 0.113853689 | -0.056643422 | 8.4634208 | 8.5565515 | 8.41572 | 0.140827464 | 0.047696752 |
| Unk | 9.211049 | 9.13403 | 9.1552 | -0.021180355 | 0.055837997 | 9.6739209 | 9.6831368 | 9.5424 | 0.140735644 | 0.131519715 |
| Nkap | 9.046438 | 9.016463 | 9.1122 | -0.095764823 | -0.065789883 | 9.0747368 | 9.2051647 | 9.06447 | 0.140698938 | 0.010271045 |
| Olfr755-ps1 | 2.217969 | 2.152511 | 2.2394 | -0.086901148 | -0.021443131 | 2.3027074 | 2.3806098 | 2.24001 | 0.140601169 | 0.062698806 |
| Gm42957 | 2.517656 | 2.678274 | 2.5885 | 0.089801403 | -0.070816847 | 2.5595134 | 2.6837223 | 2.54319 | 0.140533431 | 0.016324569 |
| Pdhb | 10.22922 | 10.2527 | 10.168 | 0.084290025 | 0.060809123 | 10.313752 | 10.356883 | 10.2165 | 0.140398766 | 0.097268131 |
| Ensa | 10.83288 | 10.82032 | 10.777 | 0.043377388 | 0.055946532 | 10.881503 | 10.937064 | 10.7969 | 0.140144558 | 0.084583071 |
| Dpp3 | 10.22591 | 10.27749 | 10.28 | -0.002756482 | -0.05433527 | 10.471367 | 10.441942 | 10.3019 | 0.140024536 | 0.16944975 |
| Tspyl1 | 10.49789 | 10.48782 | 10.496 | -0.008039482 | 0.002029794 | 10.324191 | 10.434456 | 10.2945 | 0.139938286 | 0.029673913 |
| Thbs4 | 3.007142 | 3.182768 | 3.0852 | 0.097612694 | -0.0780137 | 3.2051925 | 3.3327054 | 3.19281 | 0.139892412 | 0.012379529 |
| Lpo | 4.322514 | 3.527849 | 3.8329 | -0.305020913 | 0.4896434 | 4.2995054 | 3.9285334 | 3.78865 | 0.139886231 | 0.510858287 |
| Gorab | 7.084611 | 6.990971 | 7.1118 | -0.120791778 | -0.027152045 | 6.9868113 | 7.0893247 | 6.94952 | 0.13980895 | 0.037295551 |
| Ccl25 | 5.52635 | 5.459928 | 5.3506 | 0.109345662 | 0.17576767 | 5.4966304 | 5.6460996 | 5.50629 | 0.139807605 | -0.009661591 |
| Cnep1r1 | 8.632376 | 8.609889 | 8.668 | -0.058112515 | -0.035625849 | 8.6528015 | 8.7221048 | 8.58236 | 0.139744257 | 0.070440912 |
| Pigx | 8.051207 | 8.127201 | 8.1682 | -0.040975346 | -0.116970177 | 8.167357 | 8.2692424 | 8.12956 | 0.139679186 | 0.037793828 |
| Slc6a21 | 1.695393 | 1.708189 | 1.7019 | 0.006242808 | -0.006553173 | 1.7188741 | 1.8611373 | 1.72158 | 0.139557639 | -0.002705584 |
| Lhcgr | 3.847104 | 3.850721 | 3.7882 | 0.062558693 | 0.058941247 | 3.7660139 | 4.0181753 | 3.87867 | 0.139501643 | -0.112659734 |
| Pus7l | 7.85578 | 7.803465 | 7.7819 | 0.021554833 | 0.073869096 | 8.0308232 | 7.8633634 | 7.72389 | 0.139471604 | 0.306931385 |
| Cpeb1 | 2.731217 | 2.754798 | 2.6935 | 0.061309649 | 0.037729282 | 2.7366461 | 2.8196367 | 2.68027 | 0.13937016 | 0.05637955 |
| Glrx2 | 8.24744 | 8.122154 | 8.2281 | -0.105924457 | 0.019361536 | 8.3557572 | 8.3276525 | 8.18835 | 0.139306024 | 0.167410692 |
| Wbp4 | 9.457834 | 9.526081 | 9.5615 | -0.035399949 | -0.103647384 | 9.4521108 | 9.7687095 | 9.62944 | 0.139267262 | -0.177331364 |
| Slc1a2 | 4.490829 | 4.475059 | 4.5311 | -0.055999521 | -0.040230009 | 4.6885974 | 4.5566608 | 4.41741 | 0.13925226 | 0.271188909 |
| Gm6218 | 2.279436 | 2.276033 | 2.2323 | 0.043714413 | 0.047117214 | 2.293133 | 2.332527 | 2.19328 | 0.139251926 | 0.099857894 |
| Arfgap2 | 10.52983 | 10.58138 | 10.603 | -0.02133527 | -0.072887824 | 10.835517 | 10.856529 | 10.7173 | 0.139229818 | 0.118217217 |
| Rab28 | 8.135283 | 8.083549 | 8.1176 | -0.034011125 | 0.0177232 | 8.2601815 | 8.2276474 | 8.08848 | 0.139164013 | 0.171698082 |
| Csnk1d | 11.29126 | 11.16625 | 11.192 | -0.026009574 | 0.09899958 | 11.124409 | 11.301418 | 11.1623 | 0.13907803 | -0.037931062 |
| D630033A02Rik | 2.641332 | 2.652272 | 2.717 | -0.064692717 | -0.075632401 | 2.6050294 | 2.7840786 | 2.64504 | 0.139040518 | -0.040008733 |
| Asb3 | 8.497663 | 8.587693 | 8.5319 | 0.055819255 | -0.034210874 | 8.4342936 | 8.7021489 | 8.56314 | 0.139012805 | -0.128842485 |
| Bri3bp | 9.27579 | 9.270341 | 9.372 | -0.101630244 | -0.096181131 | 9.358619 | 9.2821561 | 9.14319 | 0.13896912 | 0.215431956 |
| Nif3l1 | 7.766698 | 7.729992 | 7.8392 | -0.10921182 | -0.072505816 | 7.8206095 | 7.9209332 | 7.782 | 0.138932864 | 0.038609131 |
| Gm14667 | 3.718037 | 3.61834 | 3.8115 | -0.193132675 | -0.093434755 | 4.0056801 | 3.9064057 | 3.76756 | 0.138843841 | 0.238118207 |

| | | | | | | | | | | |
|---|---|---|---|---|---|---|---|---|---|---|
| Parg | 9.981315 | 10.06263 | 9.9096 | 0.15302678 | 0.071707308 | 9.9503222 | 10.057578 | 9.91883 | 0.138748882 | 0.031493187 |
| Anapc13 | 8.127135 | 8.174588 | 8.1227 | 0.051911629 | 0.004457872 | 8.4118217 | 8.3326012 | 8.19393 | 0.138666917 | 0.217887379 |
| Olfr907 | 3.183654 | 3.180904 | 3.2118 | -0.030919536 | -0.028169543 | 3.372506 | 3.3338698 | 3.19534 | 0.138530826 | 0.177166973 |
| Aldh18a1 | 9.864732 | 9.735835 | 9.807 | -0.071175909 | 0.057721753 | 10.188829 | 10.266855 | 10.1285 | 0.138382552 | 0.060356586 |
| Tsn | 10.68491 | 10.69095 | 10.705 | -0.014274296 | -0.020310124 | 10.704658 | 10.759501 | 10.6213 | 0.138197668 | 0.083355411 |
| Pced1a | 8.01536 | 8.09343 | 8.0468 | 0.04661918 | -0.031450733 | 8.4329479 | 8.4210855 | 8.28297 | 0.138119685 | 0.149982131 |
| Ddo | 2.90991 | 2.840475 | 2.7412 | 0.099230427 | 0.168665441 | 2.9502639 | 3.0013697 | 2.86339 | 0.137975571 | 0.086869839 |
| Rnf115 | 10.53513 | 10.59686 | 10.6 | -0.002875638 | -0.064600668 | 10.368506 | 10.697879 | 10.5599 | 0.13797241 | -0.19140111 |
| Gm8186 | 7.675417 | 7.748653 | 7.6553 | 0.09334828 | 0.020112357 | 7.6760227 | 7.8040155 | 7.66636 | 0.137655098 | 0.009662233 |
| Rras | 7.64915 | 7.605681 | 7.7607 | -0.154997661 | -0.111527939 | 7.7612764 | 7.726678 | 7.58907 | 0.137610081 | 0.172208552 |
| Nudt3 | 10.17063 | 10.23018 | 10.181 | 0.04919187 | -0.010359947 | 10.10505 | 10.151458 | 10.0139 | 0.137508851 | 0.091101366 |
| Gm10036 | 4.749747 | 4.67026 | 4.6319 | 0.038332246 | 0.117819282 | 4.7308124 | 4.7348234 | 4.59735 | 0.13747348 | 0.133462408 |
| Txk | 9.830477 | 9.728944 | 9.7207 | 0.008264214 | 0.109797164 | 9.5849234 | 9.9755248 | 9.83812 | 0.137404551 | -0.253196779 |
| Mtap | 9.64977 | 9.651307 | 9.5907 | 0.060598671 | 0.0590623 | 9.6990149 | 9.7884079 | 9.65104 | 0.137363368 | 0.047970371 |
| Gm8798 | 2.535032 | 2.514821 | 2.5587 | -0.043904382 | -0.023693251 | 2.5542594 | 2.6810562 | 2.5437 | 0.137360736 | 0.010563967 |
| Pin4 | 8.00295 | 7.96713 | 7.9925 | -0.025371 | 0.010449475 | 8.0003209 | 8.140183 | 8.00283 | 0.137356549 | -0.00250548 |
| Dhdds | 9.101251 | 9.098302 | 9.004 | 0.094255621 | 0.097204102 | 9.1762966 | 9.1203354 | 8.98315 | 0.137182055 | 0.193143196 |
| Emc7 | 9.350819 | 9.290711 | 9.2992 | -0.008470931 | 0.051636948 | 9.3645634 | 9.4148688 | 9.2777 | 0.137172125 | 0.08686677 |
| Stbd1 | 4.746045 | 4.769587 | 4.7736 | -0.004058809 | -0.027600774 | 4.5697342 | 4.7447851 | 4.60764 | 0.137141137 | -0.037909749 |
| Flywch1 | 8.785935 | 8.858132 | 8.8514 | 0.006727089 | -0.065469532 | 8.7211121 | 8.8415519 | 8.70446 | 0.137092127 | 0.016652307 |
| Olfr1369-ps1 | 1.891078 | 1.889094 | 1.8568 | 0.032337368 | 0.034321242 | 1.9635789 | 2.0414448 | 1.9046 | 0.136848545 | 0.058982706 |
| Rsl1 | 7.89201 | 7.997065 | 7.931 | 0.06611344 | -0.038940995 | 8.2059531 | 8.1889923 | 8.05215 | 0.136843576 | 0.153804323 |
| Vmn1r75 | 1.913358 | 1.928061 | 1.9596 | -0.031543101 | -0.046245748 | 1.9421101 | 2.1517919 | 2.01501 | 0.136786069 | -0.072895653 |
| Src | 7.363653 | 7.519222 | 7.267 | 0.252216329 | 0.096647927 | 7.0535775 | 7.3066052 | 7.16994 | 0.13666562 | -0.116361993 |
| Rack1 | 14.93597 | 14.92685 | 14.968 | -0.041574 | -0.03245299 | 15.128296 | 15.079343 | 14.9427 | 0.136646908 | 0.185599706 |
| Napa | 10.34643 | 10.29349 | 10.378 | -0.084640484 | -0.031702488 | 10.582059 | 10.614141 | 10.4775 | 0.136635376 | 0.104553177 |
| Dnajc24 | 6.958404 | 6.98479 | 6.7616 | 0.22318277 | 0.196796647 | 6.8816343 | 6.936327 | 6.79979 | 0.136541951 | 0.081849268 |
| Gm5786 | 2.548671 | 2.640949 | 2.5971 | 0.043816625 | -0.048461471 | 2.664389 | 2.672933 | 2.5364 | 0.136529827 | 0.127985743 |
| Ighv1-52 | 6.840047 | 7.00866 | 6.8979 | 0.110770809 | -0.05784264 | 7.2293611 | 7.0257168 | 6.88919 | 0.136529591 | 0.340173824 |
| Rpl17-ps5 | 2.328325 | 2.578407 | 2.2641 | 0.314281683 | 0.064200095 | 2.197179 | 2.3841932 | 2.24767 | 0.136518283 | -0.050495906 |
| 2310011J03Rik | 8.736123 | 8.786404 | 8.7113 | 0.075129602 | 0.024848583 | 8.8753784 | 8.7797294 | 8.64325 | 0.136480889 | 0.232129919 |
| Adpgk | 10.38926 | 10.51479 | 10.312 | 0.20307463 | 0.077546042 | 10.680164 | 10.55534 | 10.4189 | 0.136477464 | 0.261301884 |

| | | | | | | | | | | |
|---|---|---|---|---|---|---|---|---|---|---|
| Eps8l2 | 3.807535 | 3.676874 | 3.8931 | -0.216262823 | -0.085601545 | 3.9223922 | 3.9415829 | 3.80511 | 0.136474713 | 0.117283986 |
| Sc5d | 8.905303 | 8.807928 | 8.9196 | -0.111703091 | -0.014328198 | 8.8939825 | 8.8149223 | 8.67849 | 0.136432897 | 0.215493139 |
| Zcchc3 | 4.187449 | 4.279799 | 4.2791 | 0.00066932 | -0.091680939 | 4.2700988 | 4.3018612 | 4.16548 | 0.136385558 | 0.104623196 |
| Cd19 | 13.17125 | 13.32518 | 13.282 | 0.043510786 | -0.110418134 | 13.981847 | 13.732061 | 13.5957 | 0.136343328 | 0.386129774 |
| Scfd2 | 8.092338 | 8.133107 | 8.0205 | 0.112570243 | 0.071800479 | 8.0222942 | 8.1715843 | 8.03527 | 0.136312686 | -0.012977388 |
| Trmt2a | 8.952942 | 8.991151 | 8.9657 | 0.02546642 | -0.012742562 | 9.1224208 | 9.2288204 | 9.09266 | 0.136161454 | 0.029761812 |
| Ostm1 | 9.527428 | 9.388899 | 9.5262 | -0.137260241 | 0.001269153 | 9.4871039 | 9.5088802 | 9.37275 | 0.136129048 | 0.114352676 |
| Gm47932 | 3.540744 | 3.638079 | 3.5372 | 0.100892244 | 0.003557547 | 3.6203671 | 3.859398 | 3.72332 | 0.136075137 | -0.102955771 |
| Rps23-ps1 | 13.77604 | 13.74444 | 13.673 | 0.071700972 | 0.10329564 | 13.746587 | 13.874696 | 13.7387 | 0.13602964 | 0.007920984 |
| Pcmt1 | 8.79035 | 8.778514 | 8.6474 | 0.131152918 | 0.142988993 | 8.8328381 | 8.7762937 | 8.64034 | 0.135949327 | 0.192493682 |
| Bcl7b | 9.555234 | 9.567872 | 9.5868 | -0.018913531 | -0.031551974 | 9.7643222 | 9.7210157 | 9.58508 | 0.135936474 | 0.179242917 |
| Gm5141 | 5.924761 | 5.990033 | 5.9793 | 0.010770014 | -0.054502182 | 5.9667467 | 6.1636005 | 6.0277 | 0.135899669 | -0.060954192 |
| B4galt4 | 6.523408 | 6.30394 | 6.5496 | -0.245674125 | -0.026206209 | 6.1767466 | 6.2239058 | 6.08801 | 0.135891657 | 0.088732431 |
| Gm21983 | 0.773909 | 0.833391 | 0.7715 | 0.061875001 | 0.002392364 | 0.8833908 | 0.994584 | 0.8587 | 0.135887003 | 0.024693804 |
| Tpp2 | 10.89975 | 10.94305 | 10.828 | 0.115445378 | 0.072136672 | 10.846747 | 11.004568 | 10.8687 | 0.135835392 | -0.021985478 |
| Gm48990 | 2.232522 | 2.17692 | 2.1499 | 0.027043473 | 0.082645276 | 2.3300275 | 2.468978 | 2.3332 | 0.135780782 | -0.003169645 |
| Efcab11 | 3.742352 | 3.868048 | 3.9203 | -0.052283992 | -0.177980326 | 3.8464352 | 3.9418628 | 3.80617 | 0.135696608 | 0.040268922 |
| Wipf2 | 9.835758 | 9.826767 | 9.7646 | 0.062144766 | 0.071135486 | 9.5956162 | 9.8784496 | 9.74283 | 0.135617905 | -0.147215432 |
| Ergic3 | 10.04173 | 9.909603 | 10.047 | -0.137544044 | -0.005417271 | 10.168518 | 10.098999 | 9.9634 | 0.135595591 | 0.205114761 |
| Exosc7 | 8.645594 | 8.587797 | 8.4929 | 0.094925375 | 0.152722163 | 8.7809888 | 8.7964646 | 8.66091 | 0.135555892 | 0.120080076 |
| Iscu | 9.289853 | 9.28648 | 9.3303 | -0.043825473 | -0.040452447 | 9.3710137 | 9.4241399 | 9.28876 | 0.135382068 | 0.082255833 |
| Rpl36a | 13.16598 | 13.10962 | 13.104 | 0.006018506 | 0.06238651 | 13.168182 | 13.23795 | 13.1027 | 0.135200232 | 0.065432252 |
| Hdhd5 | 9.427625 | 9.568205 | 9.5587 | 0.009483529 | -0.131095641 | 9.6619642 | 9.8084725 | 9.67329 | 0.135182289 | -0.011326043 |
| Hus1 | 8.321745 | 8.486564 | 8.3474 | 0.139149902 | -0.025668414 | 8.4068081 | 8.4748215 | 8.33976 | 0.135057717 | 0.067044353 |
| Olfr1156 | 3.143861 | 3.367871 | 3.3472 | 0.020631261 | -0.203378374 | 3.3338164 | 3.4702268 | 3.3353 | 0.13492347 | -0.00148691 |
| Dynlt1c | 6.170868 | 6.067596 | 6.3347 | -0.267110833 | -0.163838788 | 6.5967633 | 6.8431956 | 6.70834 | 0.134851641 | -0.111580633 |
| Slc22a13 | 0.801701 | 0.832658 | 0.8492 | -0.016567847 | -0.047525437 | 0.8792819 | 0.954453 | 0.81962 | 0.134832397 | 0.059661391 |
| Cnot3 | 10.61579 | 10.62748 | 10.576 | 0.051103746 | 0.039417594 | 10.7158 | 10.798148 | 10.6633 | 0.134809422 | 0.052461268 |
| Ttc39c | 6.54337 | 6.412824 | 6.3279 | 0.084898489 | 0.215444302 | 6.2214332 | 6.7923988 | 6.65771 | 0.134692274 | -0.43627334 |
| Vmn1r13 | 5.258896 | 5.100965 | 5.4965 | -0.395557439 | -0.23762646 | 5.4960348 | 5.4633292 | 5.32865 | 0.134675901 | 0.167381545 |
| Bpnt1 | 8.121421 | 8.498767 | 8.2162 | 0.282517957 | -0.094827635 | 8.0260058 | 8.2211575 | 8.08651 | 0.13464613 | -0.060505554 |
| Rps15a-ps7 | 2.06102 | 1.941206 | 2.0263 | -0.085092375 | 0.034721694 | 1.9934558 | 2.0630296 | 1.92844 | 0.134585323 | 0.065011442 |

| | | | | | | | | | | |
|---|---|---|---|---|---|---|---|---|---|---|
| Rabep2 | 9.868817 | 9.887455 | 9.922 | -0.034555104 | -0.053193651 | 10.277732 | 10.117081 | 9.9825 | 0.134579143 | 0.295230911 |
| Gm7842 | 0.238211 | 0.25651 | 0.2175 | 0.039036998 | 0.020738555 | 0.3487816 | 0.3544508 | 0.21987 | 0.134578551 | 0.128909359 |
| Cyld | 11.62936 | 11.66509 | 11.64 | 0.025214988 | -0.010511694 | 11.459296 | 11.75776 | 11.6232 | 0.134555108 | -0.163908412 |
| Dmap1 | 8.173787 | 8.240012 | 8.2547 | -0.014729432 | -0.080954878 | 8.4048192 | 8.4325111 | 8.29796 | 0.134554628 | 0.106862695 |
| Gm6254 | 3.85963 | 3.863136 | 3.8519 | 0.011208691 | 0.007702642 | 3.9234566 | 3.9531946 | 3.81866 | 0.134534919 | 0.104796906 |
| Igkv3-1 | 5.464465 | 5.713427 | 5.5597 | 0.153733749 | -0.095228788 | 5.800618 | 5.8227534 | 5.68823 | 0.134523965 | 0.11238854 |
| Olfr620 | 2.319451 | 2.323216 | 2.3615 | -0.038249629 | -0.042014575 | 2.3423235 | 2.3856165 | 2.25111 | 0.134510531 | 0.091217528 |
| Vps16 | 10.64858 | 10.75745 | 10.704 | 0.053091958 | -0.055785652 | 11.091176 | 11.177448 | 11.043 | 0.134468022 | 0.048195634 |
| Cwf19l2 | 9.451511 | 9.434319 | 9.4505 | -0.016193858 | 0.000997684 | 9.3586361 | 9.6629737 | 9.52855 | 0.134428337 | -0.169909258 |
| Lamp3 | 5.665247 | 5.959716 | 5.8478 | 0.111930028 | -0.182538912 | 6.0187575 | 6.06999 | 5.93562 | 0.13436544 | 0.083133014 |
| Nubpl | 6.555355 | 6.534294 | 6.4353 | 0.098956129 | 0.120016815 | 6.6402972 | 6.66292 | 6.52858 | 0.134336197 | 0.111713369 |
| Gm43213 | 3.706608 | 3.802726 | 3.642 | 0.16069946 | 0.064581514 | 3.7090154 | 4.0078665 | 3.87358 | 0.134289003 | -0.16456213 |
| Ints8 | 9.833075 | 10.01568 | 9.8212 | 0.194485677 | 0.011879181 | 9.7759561 | 10.049229 | 9.91514 | 0.134088063 | -0.139185054 |
| Dap3 | 9.871548 | 9.944425 | 9.9039 | 0.040552705 | -0.032325042 | 9.9717564 | 10.154595 | 10.0205 | 0.134059865 | -0.048779149 |
| Pgpep1 | 7.26706 | 7.499906 | 7.254 | 0.245948871 | 0.013102465 | 7.4962829 | 7.1744436 | 7.04041 | 0.134031231 | 0.455870481 |
| Gm5157 | 3.078679 | 3.122734 | 3.1237 | -0.001014686 | -0.045069561 | 3.0981044 | 3.3192209 | 3.18525 | 0.133966113 | -0.087150359 |
| Chil1 | 4.095671 | 4.494719 | 3.6646 | 0.830106384 | 0.431057912 | 4.3247287 | 3.9099262 | 3.7761 | 0.133826938 | 0.548629405 |
| Srfbp1 | 8.186936 | 8.400022 | 8.1603 | 0.239755533 | 0.026669332 | 8.2742696 | 8.5462731 | 8.41266 | 0.133608653 | -0.138394801 |
| Chchd5 | 7.221395 | 7.087192 | 7.2971 | -0.209921452 | -0.075717928 | 7.4270049 | 7.3708225 | 7.23726 | 0.133560849 | 0.189743173 |
| Casc1 | 3.012481 | 3.133495 | 3.0756 | 0.057863316 | -0.063150119 | 3.0865629 | 3.3032356 | 3.16974 | 0.133499255 | -0.083173497 |
| Gm48443 | 6.7465 | 6.890198 | 6.8712 | 0.019035709 | -0.124662002 | 7.0765039 | 7.3653459 | 7.23189 | 0.13345107 | -0.155390925 |
| Xrcc1 | 9.174397 | 9.281026 | 9.3418 | -0.060818877 | -0.167448283 | 9.676559 | 9.6296658 | 9.49635 | 0.133315696 | 0.180208888 |
| Brip1 | 6.775326 | 6.503348 | 6.584 | -0.080602764 | 0.191375273 | 6.2475753 | 6.9526293 | 6.81935 | 0.133274668 | -0.571779318 |
| Srebf2 | 12.20277 | 12.23648 | 12.139 | 0.097189812 | 0.063473888 | 12.479406 | 12.18982 | 12.0566 | 0.133263828 | 0.422849464 |
| Eif4a2 | 13.28958 | 13.39429 | 13.353 | 0.04085816 | -0.063847824 | 13.403913 | 13.548713 | 13.4155 | 0.133181606 | -0.011619136 |
| Slc7a3 | 3.015519 | 3.066922 | 2.9958 | 0.071149769 | 0.019746222 | 3.1777205 | 3.3578468 | 3.22476 | 0.133090691 | -0.047035561 |
| Gm3448 | 3.886617 | 3.978987 | 3.8127 | 0.166277985 | 0.073908788 | 4.0497007 | 4.2373894 | 4.1043 | 0.13308892 | -0.05459973 |
| Hnrnpdl | 11.99698 | 11.93278 | 12.02 | -0.08683038 | -0.022630284 | 12.272176 | 12.375158 | 12.2421 | 0.13302148 | 0.030040103 |
| Mettl5 | 7.209851 | 7.215981 | 7.3009 | -0.08491299 | -0.091042274 | 7.4664282 | 7.5351418 | 7.40226 | 0.132882115 | 0.064168511 |
| Ube2g1 | 9.716215 | 9.789208 | 9.7033 | 0.085947187 | 0.012954766 | 9.5068401 | 9.6662903 | 9.5335 | 0.13278874 | -0.026661559 |
| Fam19a3 | 3.533061 | 3.558861 | 3.5247 | 0.034133647 | 0.008333992 | 3.5978667 | 3.6785828 | 3.5459 | 0.13268145 | 0.051965265 |
| Mcm2 | 10.32109 | 10.26329 | 10.312 | -0.049026123 | 0.008770568 | 10.507113 | 10.556616 | 10.424 | 0.132576046 | 0.083072921 |

| | | | | | | | | | | |
|---|---|---|---|---|---|---|---|---|---|---|
| Numbl | 5.6744 | 5.569097 | 5.5518 | 0.017282289 | 0.12258613 | 5.8399798 | 5.5636407 | 5.4311 | 0.132538517 | 0.408877599 |
| Wnk1 | 13.1826 | 13.26536 | 13.208 | 0.057776512 | -0.02498324 | 13.032565 | 13.445165 | 13.3126 | 0.132534696 | -0.280065268 |
| Ifngr1 | 11.1123 | 10.98318 | 11.013 | -0.029528942 | 0.099597092 | 11.046094 | 11.101503 | 10.969 | 0.132497504 | 0.077088927 |
| Gm15753 | 1.501752 | 1.596147 | 1.5233 | 0.072799166 | -0.021595431 | 1.5421103 | 1.7060673 | 1.57365 | 0.132418901 | -0.031538107 |
| Vps33a | 9.881117 | 9.978211 | 9.9272 | 0.051001343 | -0.04609252 | 10.011709 | 10.128731 | 9.9965 | 0.132230333 | 0.01520803 |
| Prps1 | 9.50809 | 9.433131 | 9.4107 | 0.022445677 | 0.097404694 | 9.4984399 | 9.5251613 | 9.39301 | 0.132152682 | 0.105431267 |
| Ak2 | 10.27412 | 10.33566 | 10.311 | 0.02512365 | -0.03641739 | 10.517728 | 10.454049 | 10.3222 | 0.131887152 | 0.195566134 |
| Zgpat | 10.6543 | 10.84981 | 10.763 | 0.08666798 | -0.10883775 | 10.902517 | 11.022597 | 10.8907 | 0.131853484 | 0.011773557 |
| Itpka | 2.041579 | 2.136128 | 1.9815 | 0.154665049 | 0.060116397 | 1.9780433 | 2.1801512 | 2.04831 | 0.131843467 | -0.07026446 |
| Srp54b | 7.987765 | 8.143066 | 8.1461 | -0.003030329 | -0.158330999 | 8.5971321 | 8.2591768 | 8.12743 | 0.131743119 | 0.469698413 |
| Etfa | 9.756035 | 9.672117 | 9.7511 | -0.078946287 | 0.004971707 | 9.7883411 | 9.8732908 | 9.74165 | 0.131642143 | 0.046692496 |
| Nup62 | 9.641319 | 9.729225 | 9.7214 | 0.007854857 | -0.080050952 | 9.7064828 | 9.8169763 | 9.68535 | 0.131630864 | 0.021137326 |
| Bean1 | 2.550775 | 2.621549 | 2.5392 | 0.0823122 | 0.011538563 | 2.6354674 | 2.7092982 | 2.57774 | 0.131555285 | 0.057724509 |
| Mterf3 | 8.973794 | 9.089705 | 8.9941 | 0.095633214 | -0.020277595 | 9.2743682 | 9.3622237 | 9.23073 | 0.13149605 | 0.043640526 |
| Tgif1 | 10.22916 | 10.5729 | 10.272 | 0.300882748 | -0.042853736 | 10.125749 | 10.585285 | 10.4539 | 0.131415934 | -0.328120149 |
| Plbd2 | 9.647912 | 9.568455 | 9.5264 | 0.042092502 | 0.121549181 | 9.7333712 | 9.558134 | 9.42673 | 0.131403387 | 0.306640593 |
| Mrps15 | 8.364448 | 8.423721 | 8.4317 | -0.007963361 | -0.06723645 | 8.582218 | 8.6133853 | 8.482 | 0.131388569 | 0.100221185 |
| Usp12 | 10.32305 | 10.4542 | 10.405 | 0.049586074 | -0.081567046 | 10.028019 | 10.400224 | 10.269 | 0.131250681 | -0.240954072 |
| Rbm28 | 9.869637 | 9.945888 | 9.8868 | 0.059122028 | -0.017128678 | 10.172997 | 10.344855 | 10.2136 | 0.131216359 | -0.0406411 |
| Dtd2 | 6.218704 | 6.081316 | 6.2583 | -0.17700972 | -0.03962226 | 6.3530473 | 6.1703864 | 6.0392 | 0.131185747 | 0.313846579 |
| Ciapin1 | 8.927107 | 8.888713 | 8.8815 | 0.007167095 | 0.045560564 | 9.1575882 | 9.1681056 | 9.03698 | 0.131124285 | 0.120606867 |
| Zfp444 | 7.490858 | 7.750166 | 7.7231 | 0.027086141 | -0.232222236 | 7.771302 | 7.8806605 | 7.7496 | 0.131063739 | 0.021705192 |
| Uqcrc2 | 11.20509 | 11.19732 | 11.172 | 0.025615676 | 0.033382298 | 11.279861 | 11.364855 | 11.2338 | 0.131043164 | 0.046048854 |
| Slc6a19 | 7.22831 | 7.477648 | 7.523 | -0.045312801 | -0.294651619 | 7.0079074 | 7.4411593 | 7.31012 | 0.131034541 | -0.302217333 |
| Trnau1ap | 8.705205 | 8.707143 | 8.717 | -0.009866423 | -0.011803806 | 8.9018829 | 8.8426903 | 8.71171 | 0.130982369 | 0.19017493 |
| Elp6 | 6.924193 | 6.954065 | 6.9362 | 0.017820594 | -0.012051409 | 7.1977965 | 7.1367991 | 7.00594 | 0.130854631 | 0.191852018 |
| Cstf1 | 9.103693 | 9.325946 | 9.3072 | 0.018745938 | -0.203507068 | 9.2057493 | 9.3233125 | 9.19246 | 0.130853782 | 0.013290591 |
| Medag | 4.385523 | 4.48161 | 4.191 | 0.290659394 | 0.194572243 | 4.2559524 | 4.3033388 | 4.17249 | 0.130844346 | 0.083457942 |
| Prr3 | 8.410212 | 8.566587 | 8.4367 | 0.129878494 | -0.026496685 | 8.704438 | 8.797623 | 8.66678 | 0.130843736 | 0.037658687 |
| Rara | 9.949858 | 9.992297 | 9.848 | 0.144314313 | 0.101874976 | 9.9880124 | 9.9704808 | 9.83965 | 0.130831697 | 0.148363271 |
| Uhrf1bp1l | 10.5545 | 10.65416 | 10.515 | 0.138851684 | 0.03919617 | 10.525943 | 10.761011 | 10.6302 | 0.130823606 | -0.10424433 |
| Commd7 | 9.385574 | 9.41844 | 9.4324 | -0.013941572 | -0.046807138 | 9.5493092 | 9.4279955 | 9.29719 | 0.130805411 | 0.252119141 |

| | | | | | | | | | | |
|---|---|---|---|---|---|---|---|---|---|---|
| Cmpk1 | 10.46744 | 10.31106 | 10.402 | -0.091354334 | 0.065019966 | 10.343939 | 10.411616 | 10.2809 | 0.130745431 | 0.063068356 |
| Gins1 | 5.850892 | 5.706627 | 5.7876 | -0.080963104 | 0.06330153 | 5.9053027 | 6.0887318 | 5.95799 | 0.130744609 | -0.052684503 |
| Rpl31-ps9 | 1.779016 | 1.688901 | 1.7165 | -0.027549766 | 0.062565149 | 1.6470663 | 1.7663995 | 1.63619 | 0.130212983 | 0.010879811 |
| Adam9 | 9.408345 | 9.274374 | 9.0996 | 0.174749317 | 0.308719914 | 8.9600602 | 9.0555607 | 8.92537 | 0.130185981 | 0.0346855 |
| Med4 | 8.852197 | 8.905457 | 8.8192 | 0.08625072 | 0.032991024 | 8.8920626 | 8.9804953 | 8.85043 | 0.130066478 | 0.04163373 |
| Atxn7l3b | 10.62542 | 10.64499 | 10.633 | 0.012482346 | -0.007087434 | 10.700814 | 10.659962 | 10.5299 | 0.130048158 | 0.170900337 |
| Olfr90 | 6.112928 | 6.233037 | 6.1922 | 0.040866247 | -0.079242388 | 6.4261382 | 6.6707065 | 6.54074 | 0.129962286 | -0.114606011 |
| Gm11960 | 4.27555 | 4.209593 | 4.2467 | -0.037061145 | 0.028895511 | 4.2248051 | 4.5012337 | 4.3714 | 0.129835924 | -0.146592719 |
| C1rb | 2.071312 | 1.908011 | 1.9983 | -0.090247111 | 0.073054521 | 2.0250994 | 1.9871709 | 1.85742 | 0.129752384 | 0.16768088 |
| Rnf113a1 | 5.73242 | 5.55873 | 5.5804 | -0.02167858 | 0.152011689 | 5.669837 | 5.6949738 | 5.56526 | 0.1297182 | 0.104581434 |
| Surf2 | 9.191901 | 9.174761 | 9.1971 | -0.022363146 | -0.005223933 | 9.2975432 | 9.3455041 | 9.21581 | 0.129689665 | 0.081728721 |
| Rpl21 | 14.13132 | 14.13408 | 14.139 | -0.004618086 | -0.007376764 | 14.129684 | 14.232536 | 14.1029 | 0.12963092 | 0.026779004 |
| Ehd1 | 9.857281 | 9.861446 | 9.6703 | 0.19113391 | 0.186968368 | 9.7659968 | 9.6872872 | 9.55766 | 0.1296251 | 0.208334673 |
| Dph1 | 8.349486 | 8.458417 | 8.3937 | 0.06473705 | -0.04419444 | 8.7723961 | 8.5868588 | 8.45724 | 0.12961829 | 0.315155538 |
| Fam219a | 7.241544 | 7.304113 | 7.2924 | 0.011755801 | -0.050812748 | 7.3705732 | 7.4149093 | 7.28535 | 0.129558299 | 0.085222144 |
| Gm8000 | 1.852947 | 1.896031 | 1.8033 | 0.092778568 | 0.049694709 | 1.9246767 | 1.9923287 | 1.86282 | 0.129503756 | 0.061851725 |
| Pkn3 | 6.562823 | 6.418129 | 6.5905 | -0.172347477 | -0.027653385 | 6.2776473 | 6.4590476 | 6.32957 | 0.129477386 | -0.051922843 |
| Adgrf3 | 1.3675 | 1.226421 | 1.3456 | -0.119179525 | 0.021899225 | 1.2512898 | 1.3545808 | 1.22511 | 0.129472629 | 0.026181707 |
| Klhl32 | 3.542835 | 3.546144 | 3.6382 | -0.092095234 | -0.095404311 | 3.7509358 | 3.696864 | 3.56743 | 0.129431964 | 0.183503735 |
| Ak6 | 9.66308 | 9.699483 | 9.7399 | -0.040394906 | -0.076797537 | 9.8606958 | 9.848353 | 9.71894 | 0.129413747 | 0.141756575 |
| Phkg2 | 9.652502 | 9.688537 | 9.8 | -0.111435133 | -0.147470495 | 9.9848259 | 10.068701 | 9.93929 | 0.129412401 | 0.045537572 |
| Zscan22 | 7.439605 | 7.58893 | 7.5175 | 0.071427377 | -0.077898061 | 7.5948022 | 7.674433 | 7.54503 | 0.129403973 | 0.049773186 |
| Gm12033 | 0.271799 | 0.271912 | 0.3933 | -0.121406933 | -0.121519863 | 0.3248565 | 0.4020563 | 0.27265 | 0.129403538 | 0.052203769 |
| Arpc4 | 11.68893 | 11.62081 | 11.562 | 0.058374022 | 0.126492682 | 11.779517 | 11.653374 | 11.5241 | 0.129305484 | 0.255449049 |
| Ppfia1 | 10.15231 | 10.143 | 10.096 | 0.046838392 | 0.056152704 | 10.123602 | 10.289917 | 10.1606 | 0.129291101 | -0.037023883 |
| Tmub2 | 9.06028 | 9.161779 | 9.1747 | -0.012942033 | -0.1144411 | 9.4362237 | 9.3136049 | 9.1844 | 0.129201932 | 0.251820724 |
| Faap100 | 8.268146 | 8.281231 | 8.1747 | 0.106527443 | 0.093443123 | 8.5738108 | 8.3966035 | 8.26742 | 0.129182525 | 0.306389885 |
| Tbc1d22a | 9.144062 | 9.075927 | 9.1909 | -0.114971592 | -0.046835984 | 9.1811348 | 9.1766654 | 9.0475 | 0.129169633 | 0.133639061 |
| Zfp738 | 8.250903 | 8.421739 | 8.301 | 0.120752959 | -0.050083475 | 8.1762336 | 8.4536459 | 8.32454 | 0.129105221 | -0.148307049 |
| Cenpt | 7.595164 | 7.721596 | 7.6746 | 0.047018417 | -0.079412803 | 7.7980606 | 7.9548306 | 7.82576 | 0.129068392 | -0.027701611 |
| Midn | 11.08655 | 11.06926 | 10.945 | 0.12470123 | 0.141984122 | 11.175268 | 11.028107 | 10.8991 | 0.128967784 | 0.276128715 |
| Gart | 10.69765 | 10.74547 | 10.684 | 0.061288348 | 0.013463258 | 10.832534 | 10.918616 | 10.7899 | 0.12872038 | 0.042638087 |

| | | | | | | | | | | |
|---|---|---|---|---|---|---|---|---|---|---|
| Eef1g | 13.96909 | 14.02055 | 13.979 | 0.041746566 | -0.009712614 | 14.129234 | 14.138048 | 14.0095 | 0.128554198 | 0.119740251 |
| Tbp | 8.906185 | 8.998439 | 8.9864 | 0.012058167 | -0.080196268 | 9.1370129 | 9.1981165 | 9.0696 | 0.128517219 | 0.067413617 |
| Olfr652 | 1.605168 | 1.661664 | 1.6535 | 0.008188984 | -0.048306844 | 1.6545646 | 1.7165236 | 1.58809 | 0.128435409 | 0.066476386 |
| Gm49325 | 5.009605 | 5.109116 | 4.2932 | 0.815924508 | 0.716413705 | 5.3231548 | 5.2423302 | 5.11391 | 0.128420784 | 0.209245415 |
| Csnk1g2 | 10.86398 | 10.80321 | 10.905 | -0.102279742 | -0.041511386 | 11.01012 | 10.930086 | 10.8017 | 0.128375952 | 0.208409763 |
| Trip12 | 12.8004 | 12.87057 | 12.765 | 0.106037144 | 0.035864868 | 12.696968 | 12.967477 | 12.8391 | 0.128358816 | -0.142150305 |
| Flrt3 | 7.179189 | 6.928407 | 6.4691 | 0.459304296 | 0.71008714 | 6.3156804 | 6.5168254 | 6.38851 | 0.128320202 | -0.072824807 |
| Zfp260 | 9.619823 | 9.748608 | 9.6598 | 0.088776626 | -0.040008528 | 9.7866451 | 9.9064492 | 9.77814 | 0.128307112 | 0.008503043 |
| Rabggta | 9.353968 | 9.433294 | 9.3605 | 0.072800901 | -0.006524796 | 9.8768302 | 9.7650607 | 9.63683 | 0.128230775 | 0.240000315 |
| Gm11249 | 3.019124 | 3.093012 | 3.0535 | 0.039540341 | -0.03434755 | 3.0748127 | 3.2139536 | 3.08596 | 0.127991321 | -0.011149529 |
| Tbc1d2 | 4.348896 | 4.189073 | 4.3336 | -0.144551914 | 0.015271122 | 4.3993989 | 4.5311099 | 4.40314 | 0.127973084 | -0.00373795 |
| Dppa2 | 1.340522 | 1.337113 | 1.4075 | -0.070383498 | -0.066974003 | 1.3755118 | 1.4770256 | 1.34937 | 0.127658997 | 0.026145205 |
| Rpgrip1l | 6.136125 | 6.66241 | 6.2746 | 0.387771361 | -0.138513137 | 6.278576 | 6.7923684 | 6.66474 | 0.127630672 | -0.386161695 |
| Atp5c1 | 11.39125 | 11.31259 | 11.383 | -0.070264156 | 0.008390038 | 11.507705 | 11.486378 | 11.3588 | 0.12754259 | 0.148869232 |
| Zfp456 | 7.928099 | 8.0374 | 7.9471 | 0.090279301 | -0.019022413 | 8.188576 | 8.1997077 | 8.0722 | 0.127512024 | 0.116380281 |
| Dnd1 | 4.136945 | 4.240929 | 4.2199 | 0.021059936 | -0.082924599 | 4.4189649 | 4.4477044 | 4.3202 | 0.127500451 | 0.098761011 |
| Boll | 4.388258 | 4.419492 | 4.4609 | -0.041420306 | -0.072654374 | 4.0328675 | 4.568564 | 4.44121 | 0.127353253 | -0.408343206 |
| Fam32a | 10.52788 | 10.61471 | 10.576 | 0.038825272 | -0.048000526 | 10.651166 | 10.668197 | 10.5409 | 0.127329672 | 0.110298896 |
| Grin1 | 2.334999 | 2.333373 | 2.2249 | 0.108501077 | 0.110126915 | 2.223287 | 2.3729617 | 2.24567 | 0.127293296 | -0.022381407 |
| Tnfrsf25 | 7.618961 | 7.794199 | 7.8175 | -0.023261812 | -0.198500203 | 7.7805524 | 7.92536 | 7.79812 | 0.127235371 | -0.017572218 |
| Cpne8 | 6.851531 | 6.650846 | 6.933 | -0.28218039 | -0.081495368 | 6.6690676 | 6.6897454 | 6.56266 | 0.127082869 | 0.106405096 |
| Ubqln4 | 9.934091 | 9.948215 | 9.8632 | 0.085055732 | 0.07093107 | 9.9266789 | 10.024367 | 9.89753 | 0.126838579 | 0.029150751 |
| Gm48641 | 4.916796 | 4.852891 | 5.073 | -0.220114827 | -0.156210351 | 4.7297034 | 4.9227408 | 4.79592 | 0.126820404 | -0.066217036 |
| Marveld2 | 7.556337 | 7.666561 | 7.648 | 0.018543276 | -0.091681263 | 7.6432131 | 7.7778908 | 7.6511 | 0.126791951 | -0.00788575 |
| Upk1a | 1.950027 | 1.931814 | 1.9968 | -0.065034669 | -0.046821337 | 2.0002557 | 2.1113044 | 1.98457 | 0.126732398 | 0.015683655 |
| Eif2b3 | 7.982507 | 7.960076 | 7.9116 | 0.048488015 | 0.070919262 | 7.9850626 | 8.0620858 | 7.93544 | 0.126647437 | 0.049624196 |
| Syf2 | 10.47627 | 10.54339 | 10.55 | -0.00654943 | -0.073674586 | 10.637163 | 10.665218 | 10.5386 | 0.126646706 | 0.098591877 |
| Zfp51 | 8.77894 | 8.894657 | 8.8633 | 0.031340397 | -0.084376563 | 8.6805009 | 8.917752 | 8.79111 | 0.126638889 | -0.110612157 |
| Nsrp1 | 9.193376 | 9.151433 | 9.1256 | 0.025829716 | 0.067772586 | 9.1297157 | 9.2844229 | 9.15779 | 0.126630622 | -0.028076592 |
| Gm6637 | 3.763733 | 3.733385 | 3.8357 | -0.102364925 | -0.072016716 | 3.8117869 | 4.0006828 | 3.87412 | 0.126560833 | -0.062335072 |
| Tmem179 | 1.633885 | 1.624302 | 1.6702 | -0.045909749 | -0.036326587 | 1.7945997 | 1.783982 | 1.65743 | 0.126556783 | 0.137174422 |
| Ptpn6 | 13.31385 | 13.27331 | 13.266 | 0.007330258 | 0.047877488 | 13.734818 | 13.574206 | 13.4477 | 0.12653001 | 0.287141756 |

| | | | | | | | | | | |
|---|---|---|---|---|---|---|---|---|---|---|
| Ptprn | 2.605533 | 2.794562 | 2.4956 | 0.298953451 | 0.109924266 | 2.5568038 | 2.5949936 | 2.46849 | 0.126502106 | 0.088312293 |
| Srp14 | 9.782127 | 9.803484 | 9.7658 | 0.037718923 | 0.01636191 | 9.9381314 | 9.849095 | 9.72261 | 0.12648201 | 0.215518404 |
| Exoc8 | 8.16795 | 8.298645 | 8.1594 | 0.139214833 | 0.008519975 | 8.2053777 | 8.2888789 | 8.16244 | 0.126441031 | 0.042939857 |
| Ankrd12 | 10.55542 | 10.5488 | 10.786 | -0.237051512 | -0.230428426 | 10.579284 | 10.817857 | 10.6914 | 0.126436738 | -0.112135883 |
| Rnf40 | 11.00747 | 11.00578 | 10.985 | 0.020867418 | 0.022556808 | 11.139115 | 11.212248 | 11.0858 | 0.126424114 | 0.053290696 |
| Samd3 | 7.1099 | 7.118243 | 7.0827 | 0.035520216 | 0.027177167 | 6.9364544 | 7.3612531 | 7.23498 | 0.126275184 | -0.298523494 |
| Nudt21 | 9.479911 | 9.641127 | 9.5177 | 0.123437088 | -0.037779295 | 9.8254078 | 9.8316515 | 9.70541 | 0.126240899 | 0.11999723 |
| Metrn | 7.083431 | 7.113272 | 7.1574 | -0.044113021 | -0.073953282 | 7.1082968 | 7.0815689 | 6.95544 | 0.126131222 | 0.152859167 |
| Gm8281 | 1.668054 | 1.643593 | 1.6238 | 0.019779883 | 0.044241321 | 1.679019 | 1.7331918 | 1.6071 | 0.126095849 | 0.071923094 |
| Rusc2 | 5.536287 | 5.629483 | 5.4599 | 0.169632668 | 0.076436855 | 5.3256522 | 5.1633209 | 5.03723 | 0.126088921 | 0.288420174 |
| Gm5541 | 2.085201 | 2.213882 | 2.0413 | 0.172595304 | 0.043914337 | 2.0631834 | 2.1776367 | 2.05168 | 0.125956172 | 0.011502824 |
| Pdxdc1 | 9.52923 | 9.287838 | 9.4209 | -0.133080298 | 0.108312374 | 9.6507954 | 9.5248731 | 9.39892 | 0.125948389 | 0.251870732 |
| Rps15a-ps3 | 2.434819 | 2.517024 | 2.4641 | 0.05290013 | -0.029304738 | 2.5955754 | 2.6280569 | 2.50215 | 0.12590567 | 0.09342417 |
| Zfp874a | 7.566431 | 7.944097 | 7.7349 | 0.209197586 | -0.168468916 | 7.7395601 | 7.7963306 | 7.67057 | 0.125761775 | 0.068991243 |
| Pias1 | 10.40496 | 10.49768 | 10.455 | 0.042762602 | -0.049956616 | 10.293194 | 10.558767 | 10.433 | 0.125741682 | -0.139831421 |
| Imp3 | 9.221508 | 9.091069 | 9.1774 | -0.086313031 | 0.044126739 | 9.2338948 | 9.156594 | 9.0309 | 0.125689286 | 0.202989994 |
| Me3 | 4.138776 | 3.992586 | 4.1529 | -0.160332119 | -0.014141856 | 3.9506876 | 4.0474152 | 3.92175 | 0.12566098 | 0.028933407 |
| Igkv4-62 | 2.386289 | 2.51705 | 2.4137 | 0.103313955 | -0.027447734 | 2.4798701 | 2.5452568 | 2.4197 | 0.125553935 | 0.060167286 |
| Olfr729 | 3.521693 | 3.414565 | 3.594 | -0.179412167 | -0.072283678 | 3.950561 | 3.6600472 | 3.53452 | 0.125530086 | 0.416043947 |
| Camkk1 | 7.138078 | 6.607646 | 6.6535 | -0.045885794 | 0.48454615 | 6.9801377 | 6.6661747 | 6.54067 | 0.125504745 | 0.439467743 |
| Nploc4 | 10.1507 | 10.17699 | 10.157 | 0.019973564 | -0.006321682 | 10.091958 | 10.282186 | 10.1567 | 0.125476042 | -0.064751136 |
| Nom1 | 9.564768 | 9.792209 | 9.5595 | 0.232691114 | 0.005249696 | 9.478564 | 9.7892995 | 9.66395 | 0.125353343 | -0.185382104 |
| Morf4l1 | 12.27897 | 12.26206 | 12.301 | -0.038934522 | -0.022022272 | 12.394532 | 12.400606 | 12.2754 | 0.125246684 | 0.119172993 |
| Gm29288 | 4.021476 | 4.169675 | 4.0326 | 0.137098573 | -0.011100301 | 4.1297205 | 4.3322116 | 4.20703 | 0.125183272 | -0.077307873 |
| Rap1gap2 | 8.108354 | 7.895992 | 8.0354 | -0.139429094 | 0.072933077 | 8.0951946 | 7.976012 | 7.85084 | 0.125173225 | 0.244355812 |
| Sar1b | 8.942216 | 8.967998 | 8.9565 | 0.011494245 | -0.014287552 | 9.0470819 | 9.0899739 | 8.96485 | 0.125119571 | 0.082227527 |
| Wfs1 | 6.449587 | 6.297402 | 6.3885 | -0.091121365 | 0.061063569 | 6.4687255 | 6.4354862 | 6.31042 | 0.125061954 | 0.158301312 |
| Enpp6 | 2.550426 | 2.406202 | 2.4161 | -0.009899921 | 0.134324141 | 2.52338 | 2.5456358 | 2.42061 | 0.125028461 | 0.102772727 |
| Secisbp2 | 9.436681 | 9.475714 | 9.3668 | 0.108888247 | 0.069854347 | 9.6520116 | 9.6974275 | 9.57243 | 0.124995251 | 0.079579352 |
| Gm13147 | 2.10628 | 2.075562 | 2.0443 | 0.031220935 | 0.061939733 | 2.2984626 | 2.2197373 | 2.09479 | 0.124949695 | 0.20367506 |
| Atxn7l2 | 7.66973 | 7.591759 | 7.637 | -0.045274389 | 0.032696189 | 7.7018124 | 7.8916466 | 7.76677 | 0.124881604 | -0.064952674 |
| Kctd11 | 6.249674 | 6.139579 | 6.0796 | 0.059952024 | 0.170046369 | 6.1907021 | 6.198802 | 6.07398 | 0.124821021 | 0.116721142 |

| | | | | | | | | | | |
|---|---|---|---|---|---|---|---|---|---|---|
| Rnf14 | 9.556987 | 9.52257 | 9.6292 | -0.106599613 | -0.072183156 | 9.8076806 | 9.8254469 | 9.70063 | 0.124814237 | 0.107048001 |
| Gm12537 | 0.410989 | 0.282393 | 0.3185 | -0.036124067 | 0.092471923 | 0.2844893 | 0.4078699 | 0.28312 | 0.124751742 | 0.0013711 |
| Kcp | 1.022524 | 1.066911 | 0.9992 | 0.067664025 | 0.023276693 | 1.0038156 | 1.0845669 | 0.95986 | 0.124706733 | 0.043955437 |
| 4933408B17Rik | 3.642815 | 3.54187 | 3.6519 | -0.11005242 | -0.009107986 | 3.6894866 | 3.8193368 | 3.69474 | 0.12459511 | -0.005255087 |
| Exosc9 | 8.896581 | 8.692226 | 8.8045 | -0.112254177 | 0.092100094 | 8.9622153 | 8.9239964 | 8.79941 | 0.124581882 | 0.162800745 |
| Ctbp1 | 11.49984 | 11.51221 | 11.533 | -0.020412044 | -0.03277741 | 11.609124 | 11.455318 | 11.3308 | 0.124544626 | 0.278350163 |
| Eipr1 | 8.291834 | 8.441188 | 8.2302 | 0.21094092 | 0.06158694 | 8.3781627 | 8.5977461 | 8.47323 | 0.124520436 | -0.095062925 |
| Tpt1 | 15.67671 | 15.66221 | 15.682 | -0.019517676 | -0.005015672 | 15.655098 | 15.786313 | 15.6618 | 0.124518484 | -0.006696868 |
| Lrfn2 | 0.527197 | 0.520714 | 0.4703 | 0.050420999 | 0.056903798 | 0.4663397 | 0.5934066 | 0.46902 | 0.124385943 | -0.002680963 |
| Ighv13-2 | 4.934375 | 4.956248 | 4.9418 | 0.014488517 | -0.007384098 | 5.1578013 | 5.0539181 | 4.92967 | 0.124245655 | 0.228128841 |
| Morn3 | 0.446277 | 0.450717 | 0.4593 | -0.008533802 | -0.012973999 | 0.4524451 | 0.6017766 | 0.47759 | 0.124183229 | -0.025148274 |
| Trit1 | 8.765704 | 8.799013 | 8.742 | 0.057009971 | 0.023701011 | 8.8597826 | 8.8702682 | 8.74609 | 0.124175352 | 0.113689746 |
| Sec62 | 11.09771 | 11.2023 | 11.043 | 0.15975613 | 0.05517184 | 11.049802 | 11.275425 | 11.1513 | 0.124161278 | -0.101461786 |
| Gimap5 | 10.41551 | 10.2097 | 10.321 | -0.111659394 | 0.094154768 | 10.357706 | 10.350174 | 10.226 | 0.124155258 | 0.131687217 |
| Exoc6 | 9.850608 | 9.836905 | 9.8385 | -0.001553372 | 0.012149796 | 9.851088 | 9.8862596 | 9.76217 | 0.124089633 | 0.088918039 |
| Desi2 | 8.272519 | 8.240679 | 8.2294 | 0.011244133 | 0.043083858 | 8.0570437 | 8.2859018 | 8.16184 | 0.124065236 | -0.104792945 |
| Fdxr | 8.15717 | 8.31473 | 8.1933 | 0.121456344 | -0.036103013 | 8.5306945 | 8.4983542 | 8.37429 | 0.124063282 | 0.15640359 |
| Qtrt1 | 8.016895 | 8.108254 | 8.0884 | 0.019815608 | -0.07154303 | 8.3675923 | 8.5452439 | 8.4212 | 0.124045651 | -0.053605962 |
| Gm15694 | 1.905476 | 1.936091 | 1.9384 | -0.002317571 | -0.032932079 | 1.8677405 | 2.0143345 | 1.89033 | 0.12400637 | -0.022587639 |
| Tmem159 | 5.636123 | 5.472667 | 5.7859 | -0.313234527 | -0.149778479 | 5.3402337 | 5.5424565 | 5.4185 | 0.123953781 | -0.078268929 |
| Gm16147 | 1.176236 | 1.190246 | 1.1671 | 0.023100732 | 0.009090355 | 1.1511399 | 1.3079528 | 1.18414 | 0.123814198 | -0.032998696 |
| Gm10719 | 1.040352 | 1.033122 | 1.0608 | -0.027657055 | -0.02042739 | 1.0772693 | 1.2613002 | 1.13754 | 0.12375668 | -0.060274216 |
| Sox9 | 1.982691 | 1.951461 | 1.9035 | 0.047981786 | 0.079211438 | 1.9668502 | 2.0010851 | 1.87734 | 0.123741885 | 0.089506944 |
| Rps4x | 15.20639 | 15.21569 | 15.214 | 0.002032856 | -0.00726661 | 15.300976 | 15.319893 | 15.1962 | 0.123681514 | 0.104763871 |
| Padi3 | 4.133496 | 4.127421 | 4.1242 | 0.003262246 | 0.009337412 | 4.2319035 | 4.2367575 | 4.11308 | 0.123680919 | 0.118826914 |
| Dph5 | 8.927895 | 8.763279 | 8.87 | -0.106723496 | 0.057892489 | 9.0084113 | 9.0198967 | 8.89622 | 0.123675396 | 0.112189963 |
| Gm3272 | 2.837686 | 2.779299 | 2.8255 | -0.046242541 | 0.012144923 | 3.0485789 | 2.9122443 | 2.78862 | 0.123628305 | 0.259962893 |
| 2300009A05Rik | 5.928669 | 5.862419 | 5.9589 | -0.096528953 | -0.030279028 | 6.0840162 | 6.1621064 | 6.03849 | 0.123612541 | 0.04552227 |
| Ltv1 | 9.474212 | 9.490431 | 9.359 | 0.131396599 | 0.115178175 | 9.5732432 | 9.6342999 | 9.51075 | 0.123552354 | 0.062495635 |
| Zfp330 | 9.415014 | 9.371003 | 9.3162 | 0.054768699 | 0.098779423 | 9.4424424 | 9.5789037 | 9.45538 | 0.123519709 | -0.012941646 |
| Gatm | 5.976401 | 5.637011 | 5.4939 | 0.143113576 | 0.482502934 | 5.8667644 | 5.7575724 | 5.63412 | 0.123456769 | 0.232648741 |
| Gm8909 | 0.882885 | 0.843265 | 0.8402 | 0.003069905 | 0.042689769 | 0.8012577 | 0.9308709 | 0.80749 | 0.123384213 | -0.00622892 |

| | | | | | | | | | | |
|---|---|---|---|---|---|---|---|---|---|---|
| Mrps33 | 8.680458 | 8.630599 | 8.5632 | 0.067355855 | 0.117215649 | 8.792838 | 8.8079381 | 8.68457 | 0.123369523 | 0.108269485 |
| Hmgcs1 | 9.768432 | 9.847378 | 9.7017 | 0.145629122 | 0.066682852 | 9.7726788 | 9.7230758 | 9.59971 | 0.123366592 | 0.172969536 |
| Ttf1 | 9.635912 | 9.678926 | 9.6275 | 0.05139683 | 0.008382609 | 9.707814 | 9.9287578 | 9.8054 | 0.123357923 | -0.097585865 |
| Serpinb13 | 0.572043 | 0.6194 | 0.5661 | 0.05332188 | 0.005964981 | 0.6015949 | 0.6948973 | 0.57163 | 0.123270316 | 0.029967887 |
| Igkv4-91 | 7.544534 | 7.662398 | 7.7567 | -0.094274829 | -0.212138788 | 7.8726113 | 7.5813864 | 7.45812 | 0.123265499 | 0.414490376 |
| Fcrlb | 2.360542 | 2.380514 | 2.4642 | -0.083721953 | -0.103693175 | 2.473425 | 2.5075989 | 2.38436 | 0.123237354 | 0.08906345 |
| Klra5 | 2.703781 | 2.697732 | 2.71 | -0.012302999 | -0.006254625 | 2.6997482 | 2.765631 | 2.64245 | 0.123177305 | 0.057294473 |
| Snrnp70 | 11.88415 | 11.96659 | 11.905 | 0.061993234 | -0.020451684 | 12.181114 | 12.334146 | 12.2111 | 0.123018288 | -0.030014079 |
| Cfl2 | 7.448009 | 7.25964 | 7.3006 | -0.040929267 | 0.147439109 | 7.185665 | 7.4036122 | 7.28061 | 0.123004401 | -0.094942767 |
| Xpnpep1 | 9.25331 | 9.281196 | 9.2691 | 0.012087023 | -0.015798769 | 9.4386203 | 9.3220256 | 9.19919 | 0.122832773 | 0.239427528 |
| St8sia3 | 2.17309 | 2.24103 | 2.1242 | 0.11681171 | 0.04887186 | 2.1633878 | 2.351548 | 2.22885 | 0.122698039 | -0.065462196 |
| Fnip2 | 7.448405 | 7.365204 | 7.0769 | 0.2883158 | 0.371516596 | 7.2062895 | 7.263194 | 7.14055 | 0.122647477 | 0.065742984 |
| Gstp1 | 9.397332 | 9.347111 | 9.3955 | -0.048360042 | 0.001861119 | 9.6307251 | 9.3941161 | 9.27148 | 0.122639722 | 0.359248672 |
| Tbpl1 | 9.161742 | 9.307967 | 9.2372 | 0.070753202 | -0.075472095 | 9.3087881 | 9.4834849 | 9.36092 | 0.122563373 | -0.052133351 |
| Ighv1-55 | 7.907891 | 8.374614 | 7.9266 | 0.448030752 | -0.01869229 | 8.2455424 | 8.0704807 | 7.94795 | 0.122527549 | 0.297589258 |
| Dnah14 | 0.909679 | 0.89557 | 1.0024 | -0.106796422 | -0.092687752 | 1.0203825 | 1.0093862 | 0.88693 | 0.122459129 | 0.133455378 |
| Gm12896 | 0.915143 | 0.958234 | 0.9867 | -0.028516167 | -0.071606659 | 0.9778211 | 1.1083848 | 0.98594 | 0.122447797 | -0.008115939 |
| Pex6 | 9.451668 | 9.492798 | 9.4907 | 0.002073836 | -0.039056313 | 9.9044643 | 9.7310819 | 9.60875 | 0.122332858 | 0.29571523 |
| AC165327.2 | 3.094848 | 3.16563 | 3.0608 | 0.104797251 | 0.034015583 | 3.2503673 | 3.1499511 | 3.02766 | 0.122290523 | 0.222706762 |
| Adat1 | 6.918415 | 7.150226 | 7.0226 | 0.127584473 | -0.104226491 | 7.3458588 | 7.38218 | 7.25992 | 0.122264448 | 0.085943187 |
| Pheta1 | 5.19883 | 5.226408 | 5.2759 | -0.04949713 | -0.077074913 | 5.476128 | 5.2821415 | 5.16004 | 0.122100884 | 0.316087391 |
| Anp32b | 11.97822 | 11.88941 | 12.036 | -0.146460366 | -0.057653972 | 12.026032 | 12.136164 | 12.0141 | 0.122078092 | 0.011946991 |
| Pcsk7 | 9.836876 | 9.875039 | 9.9035 | -0.028510359 | -0.066673533 | 9.8890295 | 10.162318 | 10.0403 | 0.121969777 | -0.151319199 |
| Ppp1r13l | 3.431306 | 3.380953 | 3.4296 | -0.048659241 | 0.001693585 | 3.5595826 | 3.5827421 | 3.4608 | 0.121940927 | 0.09878139 |
| 3110040N11Rik | 7.680095 | 7.631345 | 7.5414 | 0.089983767 | 0.138734135 | 7.7543633 | 7.7885605 | 7.66673 | 0.121832654 | 0.087635383 |
| Mthfd1 | 9.95252 | 9.935173 | 9.867 | 0.068181575 | 0.085528354 | 10.134837 | 10.104184 | 9.98236 | 0.121825702 | 0.152478617 |
| Nat9 | 7.098537 | 7.046505 | 7.0872 | -0.040707644 | 0.011324702 | 7.5291132 | 7.3672268 | 7.24541 | 0.121813647 | 0.283700096 |
| Gm4691 | 2.177712 | 2.239466 | 2.2023 | 0.037154253 | -0.024600221 | 2.2686026 | 2.3572717 | 2.23548 | 0.121794186 | 0.033124995 |
| Zfp668 | 7.774908 | 7.715301 | 7.6684 | 0.046896183 | 0.106503088 | 7.8925725 | 7.9319489 | 7.8102 | 0.121747904 | 0.08237155 |
| Wdr82 | 11.33721 | 11.48747 | 11.388 | 0.09990449 | -0.050352892 | 11.2329 | 11.432441 | 11.3107 | 0.12174492 | -0.077795725 |
| Olfr1303 | 0.94663 | 0.914005 | 0.9609 | -0.04689869 | -0.014273581 | 0.9539529 | 1.0569799 | 0.93525 | 0.121730836 | 0.018703767 |
| Il21r | 12.18691 | 12.07193 | 12.057 | 0.014916542 | 0.129901374 | 12.261503 | 12.225129 | 12.1034 | 0.121707798 | 0.15808086 |

| | | | | | | | | | | |
|---|---|---|---|---|---|---|---|---|---|---|
| Col25a1 | 2.11071 | 2.23783 | 2.1797 | 0.058114795 | -0.069004622 | 2.1632826 | 2.2629161 | 2.14126 | 0.121652551 | 0.022019092 |
| Gm38393 | 2.105921 | 2.158891 | 2.0798 | 0.079088664 | 0.026119064 | 2.2235537 | 2.1984262 | 2.07698 | 0.121441805 | 0.146569291 |
| Ranbp6 | 8.593702 | 8.697189 | 8.6362 | 0.060940832 | -0.042545564 | 8.8680054 | 8.8517059 | 8.73027 | 0.1214344 | 0.137733908 |
| Selplg | 12.13832 | 12.16698 | 12.205 | -0.037826298 | -0.066493216 | 12.15235 | 12.239897 | 12.1185 | 0.121428496 | 0.033881096 |
| Nelfcd | 9.337068 | 9.120645 | 9.3118 | -0.191106115 | 0.025316545 | 9.6153605 | 9.5148912 | 9.39348 | 0.121416128 | 0.221885435 |
| Apon | 1.640204 | 1.672089 | 1.5956 | 0.076466121 | 0.044580675 | 1.6464974 | 1.7659219 | 1.64457 | 0.121353545 | 0.001928982 |
| Tm2d1 | 7.553993 | 7.663882 | 7.6908 | -0.026916379 | -0.136806001 | 7.6603616 | 7.6761268 | 7.55478 | 0.121348826 | 0.10558369 |
| Ap2m1 | 11.72512 | 11.71067 | 11.679 | 0.03184209 | 0.046295916 | 11.843086 | 11.786937 | 11.6656 | 0.1213331 | 0.177482017 |
| Gm12276 | 2.650514 | 2.612279 | 2.5948 | 0.017497613 | 0.055731692 | 2.5969729 | 2.7121207 | 2.59079 | 0.121332144 | 0.006184375 |
| Mpc2 | 8.147659 | 8.091609 | 8.1443 | -0.052699445 | 0.003349825 | 8.2581246 | 8.1783839 | 8.05711 | 0.12126929 | 0.201009996 |
| Wdsub1 | 6.230172 | 6.415275 | 6.4603 | -0.045024586 | -0.230127017 | 6.4234551 | 6.5051112 | 6.38385 | 0.121257005 | 0.03960088 |
| Brpf1 | 10.56055 | 10.59548 | 10.589 | 0.006900058 | -0.028030554 | 10.784453 | 10.77678 | 10.6555 | 0.121232738 | 0.128905855 |
| Trappc2l | 8.021816 | 7.767086 | 7.9505 | -0.183428093 | 0.071301513 | 8.2967765 | 8.1042044 | 7.98298 | 0.121222352 | 0.313794426 |
| Echdc1 | 7.726152 | 7.949264 | 7.6798 | 0.269469174 | 0.046356861 | 7.7656361 | 7.8110274 | 7.68987 | 0.121153819 | 0.075762566 |
| Lipt1 | 5.248801 | 5.369678 | 5.2401 | 0.129622823 | 0.008745501 | 5.4512182 | 5.5195025 | 5.39839 | 0.121108278 | 0.052824012 |
| Trappc9 | 9.283984 | 9.390504 | 9.323 | 0.067471239 | -0.039048478 | 9.4074847 | 9.3817586 | 9.26068 | 0.121075229 | 0.146801272 |
| Timmdc1 | 7.724143 | 7.765308 | 7.6922 | 0.073127018 | 0.031962046 | 7.8377252 | 7.9406013 | 7.81962 | 0.120983827 | 0.018107712 |
| Ccn4 | 2.943508 | 2.845217 | 2.8687 | -0.023485861 | 0.074804574 | 2.8734609 | 2.9804744 | 2.8595 | 0.120976873 | 0.013963308 |
| Ogfod3 | 6.701023 | 6.597828 | 6.6983 | -0.100487243 | 0.002707351 | 6.6879047 | 6.731522 | 6.61057 | 0.1209476 | 0.077330288 |
| Kremen2 | 3.556719 | 3.476649 | 3.6006 | -0.123947089 | -0.043877599 | 3.6014688 | 3.6387057 | 3.51778 | 0.120921371 | 0.083684563 |
| Abhd2 | 11.39805 | 11.48509 | 11.397 | 0.088234474 | 0.001196084 | 11.25916 | 11.492836 | 11.372 | 0.120837624 | -0.112837745 |
| Fmnl2 | 8.778498 | 8.801245 | 8.7065 | 0.094725491 | 0.071977936 | 7.9841669 | 8.7213834 | 8.60065 | 0.120728522 | -0.616487941 |
| Fbn2 | 3.46573 | 3.196526 | 3.4269 | -0.230343911 | 0.038859828 | 3.1058598 | 3.2795329 | 3.1589 | 0.120629352 | -0.053043783 |
| Skp2 | 6.048582 | 5.978293 | 6.1573 | -0.179019909 | -0.108731539 | 6.1636976 | 6.343753 | 6.22315 | 0.120602902 | -0.059452472 |
| Fem1c | 9.932463 | 10.01178 | 9.8574 | 0.154379763 | 0.075058116 | 9.776272 | 9.9249934 | 9.80441 | 0.120585424 | -0.028135977 |
| Gm14749 | 1.204359 | 1.267622 | 1.2101 | 0.057510857 | -0.005752087 | 1.2275046 | 1.3018214 | 1.18139 | 0.120433491 | 0.046116626 |
| Meis2 | 5.494136 | 5.189459 | 5.1619 | 0.027513482 | 0.332190091 | 4.7504476 | 5.1530301 | 5.03262 | 0.120415074 | -0.282167521 |
| Ufd1 | 9.549463 | 9.467814 | 9.5344 | -0.066539509 | 0.01510961 | 9.6294258 | 9.7166283 | 9.59625 | 0.120383139 | 0.033180679 |
| BC035947 | 3.001738 | 2.966212 | 2.9887 | -0.022527452 | 0.012999114 | 3.0130128 | 3.0595138 | 2.9392 | 0.120311702 | 0.073810642 |
| Slc39a6 | 9.114612 | 9.125014 | 9.1446 | -0.019571994 | -0.029973664 | 9.0654404 | 9.1107706 | 8.99046 | 0.12030807 | 0.074977833 |
| Paqr3 | 5.04288 | 5.090373 | 4.8783 | 0.212074584 | 0.164581679 | 4.8571096 | 5.1667836 | 5.04652 | 0.120260081 | -0.189413885 |
| Vcpkmt | 6.841513 | 6.876942 | 6.8458 | 0.031100196 | -0.004328563 | 7.0048758 | 7.0994305 | 6.97924 | 0.120191914 | 0.025637171 |

| | | | | | | | | | | |
|---|---|---|---|---|---|---|---|---|---|---|
| Rab12 | 8.083881 | 8.100734 | 8.1337 | -0.032916593 | -0.049769083 | 8.0330252 | 8.2867552 | 8.16668 | 0.120077388 | -0.133652586 |
| Dusp11 | 11.54795 | 11.75147 | 11.697 | 0.054775024 | -0.14874881 | 11.709459 | 11.979861 | 11.8598 | 0.120076414 | -0.150324983 |
| Zkscan6 | 8.724988 | 8.767087 | 8.755 | 0.012052444 | -0.030046261 | 8.7940471 | 8.8677634 | 8.74773 | 0.120030173 | 0.046313874 |
| Gm5740 | 1.404703 | 1.455323 | 1.4053 | 0.050058759 | -0.000561758 | 1.4145885 | 1.5293193 | 1.40935 | 0.119967535 | 0.005236726 |
| Dclk2 | 9.031462 | 8.7791 | 9.0245 | -0.245387975 | 0.006974183 | 9.3776731 | 9.4143435 | 9.29453 | 0.119816226 | 0.083145919 |
| Hirip3 | 8.443428 | 8.54405 | 8.3871 | 0.156900762 | 0.056278486 | 8.4718044 | 8.8758977 | 8.75615 | 0.11974761 | -0.28434571 |
| Sdhaf4 | 6.814944 | 6.808133 | 6.7858 | 0.022311272 | 0.029122451 | 6.9116886 | 6.9231169 | 6.80348 | 0.119637764 | 0.108209395 |
| Sec14l4 | 0.544303 | 0.550441 | 0.6398 | -0.089317215 | -0.095455572 | 0.753536 | 0.7016441 | 0.58207 | 0.119573067 | 0.171464984 |
| Gm45855 | 3.889098 | 3.988084 | 4.1017 | -0.113608597 | -0.21259456 | 3.9309198 | 4.0544637 | 3.93504 | 0.119421855 | -0.004122062 |
| Gfra3 | 0.623999 | 0.657034 | 0.606 | 0.051010062 | 0.017975064 | 0.5904766 | 0.7099232 | 0.59061 | 0.119318049 | -0.00012855 |
| Pes1 | 10.63424 | 10.72023 | 10.611 | 0.10904606 | 0.02306342 | 10.880681 | 10.848002 | 10.7287 | 0.119289338 | 0.151968472 |
| Zc3h18 | 10.6121 | 10.6851 | 10.639 | 0.046347868 | -0.02665079 | 10.724069 | 10.794134 | 10.6749 | 0.11926363 | 0.049198639 |
| Rpn1 | 10.79085 | 10.85033 | 10.728 | 0.122283386 | 0.062801336 | 11.021443 | 10.908201 | 10.7889 | 0.11925116 | 0.232493059 |
| Pigo | 9.331684 | 9.500219 | 9.3874 | 0.112812457 | -0.055722059 | 9.8056442 | 9.7008097 | 9.58156 | 0.119244738 | 0.22407927 |
| Ercc3 | 9.755773 | 9.798443 | 9.7459 | 0.052534576 | 0.009863966 | 9.8067358 | 9.8510102 | 9.73181 | 0.119199033 | 0.074924661 |
| Zfp488 | 6.313096 | 6.274915 | 6.2041 | 0.070773947 | 0.108955587 | 6.3377869 | 6.7896398 | 6.67047 | 0.119168205 | -0.332684737 |
| Ppp1r2 | 10.05196 | 9.918523 | 9.9372 | -0.01866442 | 0.114772847 | 9.8848157 | 9.885141 | 9.76598 | 0.11916413 | 0.118838886 |
| Chmp6 | 8.383762 | 8.286008 | 8.3687 | -0.082686303 | 0.01506812 | 8.4190926 | 8.3344038 | 8.21531 | 0.119093761 | 0.203782555 |
| Gmppa | 8.916993 | 8.934874 | 8.9541 | -0.019202072 | -0.037082724 | 9.293589 | 9.1949709 | 9.07593 | 0.119040946 | 0.217659024 |
| Lct | 2.68401 | 2.679831 | 2.715 | -0.035216508 | -0.031037292 | 2.6780589 | 2.8078347 | 2.68884 | 0.118992775 | -0.010783043 |
| Coro1a | 14.45195 | 14.4985 | 14.452 | 0.046972482 | 0.000417072 | 14.870248 | 14.556599 | 14.4376 | 0.118967732 | 0.4326163 |
| Grhl3 | 0.641502 | 0.695223 | 0.6408 | 0.054404796 | 0.000684002 | 0.6081284 | 0.7688154 | 0.64985 | 0.118961249 | -0.041725741 |
| Sf3b6 | 9.78004 | 9.941939 | 9.8497 | 0.092239364 | -0.069658932 | 9.8807431 | 10.008312 | 9.88939 | 0.118924655 | -0.008643787 |
| Rab9 | 8.352238 | 8.450355 | 8.4873 | -0.036939836 | -0.135056947 | 8.3449036 | 8.7139912 | 8.5951 | 0.118894551 | -0.250193077 |
| Sirt7 | 10.4071 | 10.51129 | 10.488 | 0.02317322 | -0.081020512 | 10.834506 | 10.757711 | 10.6389 | 0.11877473 | 0.195570217 |
| Nlrp9b | 2.589168 | 2.545653 | 2.5691 | -0.023421678 | 0.020093676 | 2.6697502 | 2.7410084 | 2.62227 | 0.118733866 | 0.04747566 |
| Mdh2 | 11.25977 | 11.23971 | 11.207 | 0.032427918 | 0.052481894 | 11.425506 | 11.373817 | 11.2551 | 0.11871916 | 0.170408504 |
| Klhl6 | 11.35064 | 11.34541 | 11.347 | -0.002043866 | 0.003191512 | 11.240863 | 11.325674 | 11.207 | 0.11871844 | 0.033908213 |
| Gm12097 | 0.773123 | 0.734895 | 0.7634 | -0.028491208 | 0.009737308 | 0.7489405 | 0.8718272 | 0.75314 | 0.118684947 | -0.00420171 |
| Bcdin3d | 7.261049 | 7.198737 | 7.297 | -0.098235705 | -0.03592347 | 7.4482181 | 7.2687115 | 7.15008 | 0.118631688 | 0.298138227 |
| Gm45471 | 1.371982 | 1.437444 | 1.3759 | 0.061523468 | -0.003937959 | 1.3608058 | 1.4925474 | 1.37397 | 0.118579946 | -0.013161709 |
| Olfr433 | 3.553412 | 3.644904 | 3.5634 | 0.081495845 | -0.009996955 | 3.7107816 | 3.8019114 | 3.68336 | 0.118548932 | 0.027419136 |

| | | | | | | | | | | |
|---|---|---|---|---|---|---|---|---|---|---|
| Pex12 | 7.306326 | 7.249482 | 7.2904 | -0.040892509 | 0.015951249 | 7.507458 | 7.4222692 | 7.30382 | 0.118447403 | 0.203636121 |
| Gm45665 | 2.813736 | 2.813591 | 2.88 | -0.066456437 | -0.066310952 | 3.0099671 | 3.1437257 | 3.02529 | 0.118435994 | -0.015322535 |
| Plrg1 | 9.382265 | 9.422189 | 9.4166 | 0.005630283 | -0.034293323 | 9.566529 | 9.6463031 | 9.52787 | 0.11843364 | 0.038659506 |
| Dhx33 | 9.597564 | 9.620082 | 9.5544 | 0.065686371 | 0.043168041 | 9.4112856 | 9.79901 | 9.68066 | 0.118354635 | -0.269369696 |
| Ptprj | 10.94842 | 11.13233 | 10.903 | 0.229080174 | 0.04516919 | 11.190778 | 10.942234 | 10.824 | 0.118247226 | 0.366790971 |
| Ccne1 | 6.781405 | 6.852267 | 7.0006 | -0.148362943 | -0.219224325 | 6.9626565 | 7.1401674 | 7.02195 | 0.11821723 | -0.059293662 |
| Cers2 | 10.83044 | 10.83308 | 10.774 | 0.059547058 | 0.056912232 | 11.056487 | 10.97443 | 10.8562 | 0.118214078 | 0.200270739 |
| Gna11 | 8.586997 | 8.574068 | 8.5982 | -0.024163801 | -0.011234329 | 8.7197637 | 8.7559866 | 8.63781 | 0.118172589 | 0.081949654 |
| Gm10447 | 3.600594 | 3.66025 | 3.5597 | 0.100544808 | 0.040888065 | 3.7469073 | 3.9568145 | 3.83865 | 0.118161121 | -0.091746031 |
| Miga2 | 7.386508 | 7.670779 | 7.5059 | 0.164896117 | -0.119375266 | 7.5922335 | 7.7063234 | 7.58821 | 0.118112639 | 0.004022807 |
| Zc3h14 | 10.31884 | 10.36055 | 10.41 | -0.04923713 | -0.090950692 | 10.389935 | 10.526818 | 10.4087 | 0.118095028 | -0.018787436 |
| Pola2 | 9.406786 | 9.313402 | 9.4308 | -0.117421067 | -0.024036963 | 9.5530771 | 9.6833279 | 9.5653 | 0.118032901 | -0.012217915 |
| Olfr1349 | 1.898589 | 1.764391 | 1.7749 | -0.010499486 | 0.123698527 | 1.7911028 | 1.92362 | 1.80568 | 0.117935933 | -0.014581245 |
| Lipt2 | 4.160693 | 4.030877 | 4.2317 | -0.200848848 | -0.07103258 | 4.2426845 | 4.244979 | 4.12705 | 0.117930562 | 0.115636059 |
| Idh3g | 10.66352 | 10.66855 | 10.689 | -0.020940462 | -0.02596883 | 10.929726 | 10.842167 | 10.7243 | 0.117891742 | 0.205450016 |
| Cryzl1 | 8.437275 | 8.410267 | 8.428 | -0.017776188 | 0.009232692 | 8.5053969 | 8.6078997 | 8.49008 | 0.117823081 | 0.015320339 |
| Son | 13.50839 | 13.52155 | 13.522 | -0.00015474 | -0.013315164 | 13.550011 | 13.813008 | 13.6954 | 0.117639748 | -0.145356824 |
| Mtfmt | 7.958258 | 8.001245 | 7.9842 | 0.017061372 | -0.025925475 | 8.1988659 | 8.4029453 | 8.28537 | 0.117579232 | -0.086500182 |
| Taf12 | 9.038488 | 9.085082 | 8.9724 | 0.112659093 | 0.066065299 | 9.0582659 | 9.064468 | 8.94691 | 0.117556742 | 0.11135461 |
| Cops2 | 10.49888 | 10.44534 | 10.294 | 0.151809268 | 0.205351686 | 10.289545 | 10.407344 | 10.2898 | 0.117502492 | -0.000296545 |
| Ctsw | 10.66809 | 10.65098 | 10.644 | 0.007184272 | 0.024286556 | 10.716086 | 10.932796 | 10.8153 | 0.117502162 | -0.09920801 |
| Actl6a | 9.655007 | 9.736963 | 9.6177 | 0.119257853 | 0.037301397 | 9.8365781 | 9.8215831 | 9.70408 | 0.117499793 | 0.132494835 |
| Itgad | 3.739441 | 3.988996 | 3.8516 | 0.137423845 | -0.112131063 | 3.6601528 | 3.827008 | 3.70965 | 0.117361731 | -0.049493499 |
| Gas8 | 6.15561 | 5.978968 | 5.986 | -0.007066455 | 0.169575847 | 5.9382381 | 6.1053367 | 5.98798 | 0.117354757 | -0.049743877 |
| 1700007E05Rik | 2.30456 | 2.349789 | 2.3621 | -0.012265546 | -0.057493639 | 2.3490594 | 2.5015508 | 2.38424 | 0.117312055 | -0.035179342 |
| AU022252 | 8.097049 | 8.418789 | 8.2145 | 0.204244147 | -0.117496196 | 8.2852777 | 8.1322832 | 8.015 | 0.117286156 | 0.270280688 |
| Usp16 | 10.04387 | 10.25386 | 10.022 | 0.232219387 | 0.022229361 | 10.235582 | 10.359544 | 10.2423 | 0.117265257 | -0.006697044 |
| Tmem42 | 6.98358 | 6.940461 | 7.0388 | -0.09831507 | -0.055196182 | 7.1454081 | 7.1791913 | 7.06193 | 0.117257251 | 0.083474056 |
| Stard5 | 8.324031 | 8.464594 | 8.4673 | -0.002700388 | -0.143263686 | 8.7549698 | 8.5604647 | 8.44323 | 0.117236153 | 0.311741202 |
| Chmp1b | 9.882184 | 9.904597 | 9.9187 | -0.014123097 | -0.036535562 | 9.8350512 | 9.9437368 | 9.82651 | 0.117229881 | 0.008544338 |
| Ss18l2 | 6.416624 | 6.418619 | 6.4311 | -0.012453117 | -0.014448818 | 6.5747611 | 6.5918887 | 6.47468 | 0.117209275 | 0.10008161 |
| Sdcbp2 | 5.263784 | 5.219244 | 5.2758 | -0.056535207 | -0.011995099 | 5.3440652 | 5.5743276 | 5.45714 | 0.117184898 | -0.113077507 |

| | | | | | | | | | | |
|---|---|---|---|---|---|---|---|---|---|---|
| Golga1 | 8.297315 | 8.044666 | 8.1081 | -0.063448052 | 0.18920172 | 8.5596979 | 8.5624649 | 8.44543 | 0.117029928 | 0.11426292 |
| Fbxw13 | 2.299227 | 2.30499 | 2.2611 | 0.043878121 | 0.038115694 | 2.2583819 | 2.4500877 | 2.33307 | 0.117013173 | -0.074692634 |
| Epc1 | 10.34761 | 10.47143 | 10.411 | 0.060120182 | -0.063693512 | 10.340538 | 10.471159 | 10.3542 | 0.11699941 | -0.013620817 |
| Larp4 | 10.89321 | 10.93241 | 10.814 | 0.118066108 | 0.078862144 | 10.635535 | 10.903976 | 10.787 | 0.116978106 | -0.151463649 |
| Igkv1-110 | 9.540646 | 9.706405 | 9.6386 | 0.067766375 | -0.097992451 | 9.7252177 | 9.796138 | 9.67917 | 0.11697261 | 0.046052315 |
| Nanp | 6.865077 | 6.854799 | 7.1268 | -0.272046001 | -0.261768103 | 6.7958726 | 6.8978771 | 6.78094 | 0.116933866 | 0.014929328 |
| Dsg1b | 1.740932 | 1.906809 | 1.7826 | 0.124221359 | -0.041655665 | 1.7842619 | 1.8567816 | 1.73989 | 0.116896227 | 0.044376542 |
| Tnnc1 | 2.063139 | 2.101857 | 2.0344 | 0.067462158 | 0.028743565 | 2.1220568 | 2.1496909 | 2.03283 | 0.116857514 | 0.089223391 |
| Preb | 10.25854 | 10.23456 | 10.238 | -0.003319907 | 0.02065963 | 10.513702 | 10.510945 | 10.3941 | 0.1168546 | 0.119611068 |
| Cd70 | 1.630747 | 1.629449 | 1.586 | 0.043425609 | 0.044723088 | 1.5704954 | 1.7157698 | 1.59895 | 0.116818024 | -0.028456322 |
| Vps26a | 9.875492 | 9.91966 | 9.8448 | 0.074840552 | 0.030672308 | 9.7988508 | 9.8036646 | 9.68688 | 0.116784232 | 0.111970449 |
| Ccdc170 | 2.768316 | 2.861524 | 2.7401 | 0.121419791 | 0.028211086 | 2.6792569 | 2.8838913 | 2.76716 | 0.116735304 | -0.087899102 |
| Ceacam19 | 3.163721 | 3.162029 | 3.1221 | 0.03997229 | 0.041663756 | 3.194149 | 3.22658 | 3.10986 | 0.116718434 | 0.084287401 |
| 4933427D14Rik | 8.09215 | 7.937448 | 7.7899 | 0.147574339 | 0.302276709 | 8.1166813 | 8.0163844 | 7.89979 | 0.116593929 | 0.216890828 |
| Gyg | 9.783647 | 9.807402 | 9.9134 | -0.106029192 | -0.129783972 | 9.9939356 | 10.095363 | 9.97887 | 0.116490997 | 0.015063453 |
| Siah1b | 5.17859 | 5.146257 | 5.3272 | -0.180912218 | -0.148579028 | 5.1643225 | 5.3990627 | 5.28262 | 0.116445341 | -0.118294788 |
| Abhd10 | 7.521819 | 7.468946 | 7.4056 | 0.063319142 | 0.11619154 | 7.5097239 | 7.6265996 | 7.51022 | 0.116377603 | -0.000498059 |
| Tagln3 | 1.791934 | 1.770809 | 1.7995 | -0.028736753 | -0.007611844 | 1.7742683 | 1.8832009 | 1.76687 | 0.116334767 | 0.007402148 |
| Gps2 | 9.786833 | 9.91536 | 9.8497 | 0.065666332 | -0.062861269 | 10.165087 | 10.125766 | 10.0095 | 0.116278243 | 0.155598941 |
| Adamtsl5 | 2.510367 | 2.617947 | 2.6275 | -0.009509933 | -0.117090089 | 2.5857344 | 2.6918214 | 2.57561 | 0.116212318 | 0.01012532 |
| Pnpla1 | 1.968774 | 1.993391 | 2.0249 | -0.031480426 | -0.056097399 | 2.0458077 | 2.1073016 | 1.99111 | 0.116189478 | 0.054695558 |
| Leprotl1 | 10.51753 | 10.58648 | 10.59 | -0.003410126 | -0.07235829 | 10.287439 | 10.650476 | 10.5344 | 0.116112518 | -0.246924065 |
| Cdk5rap1 | 7.304909 | 7.563852 | 7.6112 | -0.047339801 | -0.306283247 | 7.6156764 | 7.6155469 | 7.49948 | 0.116069632 | 0.116199124 |
| A030005L19Rik | 0.609077 | 0.62325 | 0.6152 | 0.008092275 | -0.006080484 | 0.6504308 | 0.7328582 | 0.61682 | 0.116040559 | 0.03361313 |
| Phospho2 | 8.536673 | 8.637452 | 8.5778 | 0.059681333 | -0.041097611 | 8.5331231 | 8.5342282 | 8.41822 | 0.116011078 | 0.114905966 |
| Atxn7l3 | 10.78997 | 10.88615 | 10.83 | 0.056021166 | -0.040158914 | 10.86925 | 10.88794 | 10.772 | 0.115904684 | 0.097215478 |
| Cntn3 | 2.043719 | 1.991988 | 2.1208 | -0.128859911 | -0.077128649 | 2.1482854 | 2.145403 | 2.02956 | 0.115845937 | 0.11872832 |
| Rnaseh2b | 7.76472 | 7.670335 | 7.7991 | -0.128730594 | -0.034345811 | 7.738931 | 8.015907 | 7.90007 | 0.115838552 | -0.161137387 |
| Mroh8 | 1.937418 | 1.887332 | 1.9118 | -0.02451242 | 0.025573081 | 1.8802577 | 1.985309 | 1.86948 | 0.115824893 | 0.010773582 |
| Miip | 9.388248 | 9.519794 | 9.4983 | 0.021500372 | -0.11004551 | 9.9909227 | 9.8699486 | 9.75419 | 0.115762533 | 0.236736585 |
| Adora2a | 8.649785 | 8.797252 | 8.7288 | 0.068456529 | -0.079009799 | 8.4981371 | 8.8621592 | 8.7464 | 0.115761933 | -0.248260164 |
| Mta3 | 10.08048 | 10.15882 | 10.152 | 0.006497099 | -0.071841561 | 10.147153 | 10.185935 | 10.0702 | 0.115722618 | 0.07694055 |

| | | | | | | | | | | |
|---|---|---|---|---|---|---|---|---|---|---|
| Fbxw14 | 0.934583 | 0.964416 | 0.9447 | 0.019670534 | -0.010162951 | 0.975595 | 1.0591034 | 0.94344 | 0.115664252 | 0.03215592 |
| Plekha3 | 9.424172 | 9.489138 | 9.4624 | 0.026760146 | -0.038206379 | 9.3599986 | 9.4666401 | 9.351 | 0.115640792 | 0.008999315 |
| Bet1 | 7.958084 | 7.922505 | 7.893 | 0.029513048 | 0.065092623 | 7.9410181 | 7.9256579 | 7.81004 | 0.115618978 | 0.130979128 |
| Phex | 2.862096 | 2.871255 | 2.9446 | -0.073347677 | -0.082507292 | 2.9182696 | 3.0662288 | 2.95072 | 0.115507757 | -0.032451522 |
| Olfr420 | 0.981949 | 1.05101 | 0.9747 | 0.076274266 | 0.007213303 | 0.9562968 | 1.1158097 | 1.00036 | 0.115448186 | -0.044064789 |
| Rasgef1c | 0.802669 | 0.84222 | 0.772 | 0.070244565 | 0.030694191 | 0.8031836 | 0.8894441 | 0.77408 | 0.1153607 | 0.029100165 |
| Dhrs9 | 4.846602 | 4.407425 | 4.0463 | 0.361084868 | 0.800261618 | 4.2434582 | 4.1816846 | 4.06637 | 0.115316368 | 0.177090042 |
| Vcp | 12.47752 | 12.47275 | 12.434 | 0.039113488 | 0.043877636 | 12.423826 | 12.530965 | 12.4157 | 0.115287172 | 0.008148044 |
| Rps25 | 13.8498 | 13.86822 | 13.879 | -0.010639486 | -0.029059172 | 13.935093 | 13.931172 | 13.816 | 0.115189184 | 0.119109321 |
| Wbp1l | 10.37043 | 10.35609 | 10.319 | 0.037490014 | 0.051829542 | 10.163696 | 10.361993 | 10.2468 | 0.115176866 | -0.083120172 |
| Ezr | 12.73669 | 12.80923 | 12.7 | 0.10954971 | 0.037010916 | 12.980265 | 12.904356 | 12.7892 | 0.115150158 | 0.191059371 |
| Ddrgk1 | 9.760922 | 9.728664 | 9.7447 | -0.016028407 | 0.016230063 | 9.9488756 | 9.9913326 | 9.87638 | 0.114955513 | 0.072498467 |
| Gabrg3 | 2.13399 | 2.153635 | 2.138 | 0.015680514 | -0.003964876 | 2.1485011 | 2.1573421 | 2.04246 | 0.114885883 | 0.106044832 |
| Hscb | 7.913005 | 7.761471 | 7.9364 | -0.174964217 | -0.023430332 | 7.869415 | 8.018551 | 7.90379 | 0.114762567 | -0.034373421 |
| Gm3531 | 2.534879 | 2.512693 | 2.5737 | -0.061027415 | -0.038841198 | 2.5005657 | 2.5635677 | 2.449 | 0.114569646 | 0.051567727 |
| Zfp26 | 9.544677 | 9.579829 | 9.6387 | -0.058905415 | -0.094056922 | 9.3557654 | 9.7055528 | 9.591 | 0.114550369 | -0.235237037 |
| Ints14 | 9.383359 | 9.41018 | 9.3644 | 0.045819316 | 0.018998479 | 9.443418 | 9.4679247 | 9.35338 | 0.114547032 | 0.090040298 |
| Mogs | 10.07844 | 10.0954 | 10.08 | 0.015504251 | -0.001450954 | 10.213299 | 10.190519 | 10.076 | 0.114543901 | 0.137324599 |
| Hprt | 10.44988 | 10.34765 | 10.384 | -0.03642183 | 0.06581187 | 10.448118 | 10.484636 | 10.3702 | 0.114477074 | 0.077959529 |
| Eral1 | 8.431482 | 8.498081 | 8.5429 | -0.044845664 | -0.1114449 | 8.8531348 | 8.7967177 | 8.68231 | 0.114403088 | 0.170820158 |
| Smarce1 | 10.89963 | 10.89605 | 10.865 | 0.030675982 | 0.034253284 | 10.795734 | 10.986105 | 10.8718 | 0.114308856 | -0.076062666 |
| Mb21d2 | 4.748181 | 4.468219 | 4.4147 | 0.053492124 | 0.333454409 | 4.1922174 | 4.4475843 | 4.33329 | 0.114293743 | -0.141073172 |
| Vmn2r117 | 2.967662 | 2.857869 | 2.9702 | -0.112362504 | -0.002569869 | 3.0330548 | 3.1653869 | 3.05113 | 0.114260605 | -0.018071436 |
| Mcfd2 | 9.008795 | 8.993604 | 8.8674 | 0.126195609 | 0.141386859 | 8.9973049 | 8.8864335 | 8.77221 | 0.114223995 | 0.225095434 |
| Mrps14 | 8.888996 | 8.843145 | 8.8227 | 0.020407441 | 0.066258377 | 9.0610303 | 8.9962701 | 8.88208 | 0.114190401 | 0.17895057 |
| Ltb4r1 | 5.035298 | 4.928967 | 4.728 | 0.201009013 | 0.307339501 | 5.0183865 | 4.8295706 | 4.71541 | 0.114164688 | 0.302980679 |
| Gm47325 | 3.170367 | 3.204326 | 3.2368 | -0.032490147 | -0.066448961 | 3.1516887 | 3.4217041 | 3.30755 | 0.114156269 | -0.155859108 |
| Abce1 | 11.33473 | 11.37539 | 11.277 | 0.098643956 | 0.057979778 | 11.411032 | 11.429746 | 11.3156 | 0.11413893 | 0.095425253 |
| Ttc30a1 | 5.703298 | 5.768988 | 5.7402 | 0.028812521 | -0.036877285 | 5.9306716 | 5.7981856 | 5.68406 | 0.114120725 | 0.246606729 |
| Lars | 10.55215 | 10.21513 | 10.438 | -0.222586164 | 0.114427812 | 10.726891 | 10.795851 | 10.6818 | 0.114057644 | 0.045097324 |
| Rac3 | 2.72521 | 2.743894 | 2.7801 | -0.036221014 | -0.054905069 | 2.7249168 | 2.7801901 | 2.66617 | 0.114015675 | 0.058742382 |
| Olfr713 | 3.130392 | 3.061242 | 3.1305 | -0.069224465 | -7.47094E-05 | 3.2570502 | 3.2592224 | 3.14536 | 0.113857542 | 0.111685256 |

| | | | | | | | | | | |
|---|---|---|---|---|---|---|---|---|---|---|
| Gm15807 | 3.952902 | 3.975098 | 3.9231 | 0.051995021 | 0.029798268 | 4.0324661 | 4.2146 | 4.10081 | 0.113789771 | -0.068344055 |
| Malsu1 | 7.052856 | 7.003923 | 6.9275 | 0.076422373 | 0.125354623 | 7.0258948 | 7.0766538 | 6.96288 | 0.11377095 | 0.063012006 |
| Klhdc7a | 3.448532 | 3.26987 | 3.3572 | -0.087358053 | 0.091303099 | 3.537909 | 3.4897583 | 3.37605 | 0.113709526 | 0.161860187 |
| Gm12208 | 0.491005 | 0.55084 | 0.5085 | 0.042379235 | -0.017456694 | 0.5603646 | 0.6134949 | 0.49982 | 0.113675147 | 0.060544801 |
| Cd79a | 13.78347 | 13.90154 | 13.862 | 0.039901254 | -0.078169714 | 14.496149 | 13.970177 | 13.8566 | 0.113571028 | 0.639542965 |
| Aoc1 | 0.754316 | 0.750041 | 0.755 | -0.004923478 | -0.000648795 | 0.863565 | 0.8258785 | 0.71239 | 0.113484047 | 0.151170518 |
| Gm14326 | 6.655704 | 6.416288 | 6.6186 | -0.202306372 | 0.037109487 | 6.4256076 | 6.5044907 | 6.39108 | 0.113415391 | 0.034532286 |
| Xiap | 10.96091 | 11.1446 | 10.993 | 0.151369184 | -0.03232358 | 10.806586 | 11.101604 | 10.9882 | 0.11338992 | -0.181628031 |
| Coq7 | 6.111508 | 6.00636 | 6.1342 | -0.127862341 | -0.022715166 | 6.2915082 | 6.1928263 | 6.0796 | 0.113228182 | 0.211910107 |
| Igkv4-74 | 5.895681 | 6.823884 | 6.0228 | 0.801130682 | -0.127072503 | 6.8354686 | 6.6232967 | 6.51011 | 0.113184603 | 0.32535649 |
| Sugp2 | 9.59477 | 9.479765 | 9.5372 | -0.057421505 | 0.05758421 | 9.9389382 | 9.8774394 | 9.76434 | 0.113101352 | 0.174600195 |
| Hmcn2 | 4.058959 | 4.183623 | 4.1201 | 0.063529974 | -0.061134662 | 4.0136326 | 4.2358298 | 4.12279 | 0.113040505 | -0.109156677 |
| Ranbp3l | 3.187488 | 3.160647 | 3.1957 | -0.035056723 | -0.008215623 | 3.2699454 | 3.310701 | 3.1978 | 0.112897836 | 0.072142257 |
| Gm10293 | 1.529192 | 1.503085 | 1.516 | -0.012913007 | 0.013194367 | 1.5996006 | 1.5986441 | 1.4858 | 0.112845995 | 0.113802549 |
| Gls | 11.4379 | 11.57129 | 11.445 | 0.1258348 | -0.007546504 | 11.402452 | 11.6654 | 11.5526 | 0.112828128 | -0.15011984 |
| Apex2 | 7.624713 | 7.620055 | 7.5462 | 0.073894589 | 0.078552757 | 7.6772289 | 7.7879977 | 7.67518 | 0.112818345 | 0.0020495 |
| Ndst2 | 9.115085 | 9.160466 | 9.0618 | 0.098672203 | 0.053291333 | 9.4024571 | 9.3169357 | 9.20415 | 0.112781747 | 0.198303196 |
| Gm8213 | 2.873032 | 2.8342 | 2.8034 | 0.030792531 | 0.069624115 | 2.8937924 | 2.9285217 | 2.81575 | 0.112774087 | 0.078044712 |
| Wdr24 | 8.929741 | 8.957784 | 8.9706 | -0.012790215 | -0.04083335 | 9.412005 | 9.2274653 | 9.1147 | 0.112767536 | 0.297307239 |
| Gm9958 | 3.445733 | 3.45581 | 3.503 | -0.047232546 | -0.057310079 | 3.6890188 | 3.6117735 | 3.49904 | 0.112736645 | 0.189981939 |
| Crtc2 | 10.69511 | 10.80728 | 10.819 | -0.01143322 | -0.123598988 | 10.976728 | 11.043452 | 10.9308 | 0.1126581 | 0.04593423 |
| Usp8 | 10.49501 | 10.49693 | 10.508 | -0.011061762 | -0.012983246 | 10.587847 | 10.700358 | 10.5877 | 0.112657512 | 0.000146173 |
| Zfp940 | 6.353471 | 6.448308 | 6.6391 | -0.190806117 | -0.285643128 | 6.5061177 | 6.5430652 | 6.43046 | 0.112608879 | 0.075661383 |
| Gimap3 | 12.32369 | 12.43524 | 12.431 | 0.004586358 | -0.106963252 | 12.464494 | 12.558522 | 12.4459 | 0.11260802 | 0.018580331 |
| Lrch1 | 10.48268 | 10.38665 | 10.434 | -0.04690526 | 0.049123348 | 10.330376 | 10.721636 | 10.6092 | 0.112401426 | -0.278858695 |
| Spa17 | 3.11449 | 2.922 | 3.0349 | -0.112868579 | 0.079620969 | 3.1089418 | 3.0769731 | 2.96467 | 0.112304758 | 0.144273512 |
| Rpl31-ps14 | 1.696586 | 1.659502 | 1.6349 | 0.024625453 | 0.061710144 | 1.7404866 | 1.7432855 | 1.63098 | 0.112303674 | 0.109504764 |
| Olfr1152 | 3.561039 | 3.639531 | 3.5103 | 0.12927878 | 0.05078684 | 3.54404 | 3.5844057 | 3.4722 | 0.112207598 | 0.071841954 |
| Card19 | 7.765338 | 7.692043 | 7.6658 | 0.02619601 | 0.099491151 | 7.9460471 | 7.6101151 | 7.49801 | 0.112103775 | 0.448035742 |
| Zfp408 | 8.606664 | 8.564383 | 8.4698 | 0.094590651 | 0.136871115 | 8.6904238 | 8.7562647 | 8.64417 | 0.112091672 | 0.046250763 |
| Gpat4 | 10.1299 | 10.13556 | 10.037 | 0.098560796 | 0.092905498 | 10.071859 | 10.194928 | 10.0829 | 0.112050039 | -0.011018828 |
| Clcc1 | 8.94469 | 8.965209 | 8.9648 | 0.00038356 | -0.020135905 | 9.0823954 | 9.1276988 | 9.01565 | 0.112045712 | 0.06674232 |

| | | | | | | | | | | |
|---|---|---|---|---|---|---|---|---|---|---|
| Hoxd4 | 2.684 | 2.612079 | 2.6231 | -0.010993503 | 0.060926973 | 2.5319945 | 2.7860231 | 2.67401 | 0.112012805 | -0.142015798 |
| Cd200r3 | 4.128194 | 4.170936 | 4.1929 | -0.021995065 | -0.064737712 | 4.0525313 | 4.1900328 | 4.07806 | 0.11197317 | -0.025528339 |
| Tmem223 | 7.475309 | 7.520322 | 7.5056 | 0.014690837 | -0.030322287 | 7.8406017 | 7.6176836 | 7.50578 | 0.111902878 | 0.33482101 |
| Polr3k | 8.67767 | 8.703988 | 8.7314 | -0.027395321 | -0.053713416 | 8.8141074 | 8.762838 | 8.65097 | 0.111866772 | 0.163136129 |
| Parvb | 7.268273 | 6.864463 | 6.9092 | -0.04470707 | 0.359102675 | 6.7173669 | 6.6239275 | 6.51206 | 0.111865569 | 0.205304897 |
| Gm8520 | 0.444783 | 0.433275 | 0.4416 | -0.008313371 | 0.003194623 | 0.4896015 | 0.5269491 | 0.4151 | 0.111847202 | 0.074499638 |
| Rps8 | 14.58255 | 14.60294 | 14.588 | 0.014883246 | -0.005505976 | 14.716444 | 14.666148 | 14.5543 | 0.111813694 | 0.16210998 |
| 2700097O09Rik | 6.795863 | 6.848901 | 6.78 | 0.068890911 | 0.015853283 | 6.8194017 | 7.0663662 | 6.95456 | 0.111805443 | -0.135159089 |
| Phf10 | 8.69334 | 8.554026 | 8.5771 | -0.023093929 | 0.116221042 | 8.6064513 | 8.6374298 | 8.52563 | 0.111800089 | 0.080821654 |
| Tmem87a | 10.22236 | 10.19833 | 10.209 | -0.010482639 | 0.013546909 | 10.24409 | 10.403808 | 10.292 | 0.111794358 | -0.047924202 |
| Myrf | 3.944386 | 4.104735 | 3.9714 | 0.133351003 | -0.026997957 | 4.0396296 | 4.1698971 | 4.05811 | 0.11178819 | -0.018479325 |
| Wdr12 | 8.979343 | 9.087184 | 8.9484 | 0.138750121 | 0.030909507 | 9.076217 | 9.3010528 | 9.18928 | 0.111777769 | -0.113057986 |
| Myh8 | 1.622049 | 1.64902 | 1.6547 | -0.005695088 | -0.032666623 | 1.6202519 | 1.7319768 | 1.62034 | 0.111634859 | -8.9993E-05 |
| Naa50 | 10.80219 | 10.92937 | 10.785 | 0.144772436 | 0.01759601 | 10.794993 | 10.882373 | 10.7709 | 0.111457848 | 0.024077214 |
| Gm17555 | 4.129912 | 4.283426 | 4.2888 | -0.005369251 | -0.158883202 | 4.347384 | 4.5161865 | 4.40477 | 0.11141722 | -0.057385263 |
| Gm8730 | 3.579308 | 3.538471 | 3.6744 | -0.135897347 | -0.095061163 | 3.4457644 | 3.5581221 | 3.44673 | 0.111394158 | -0.000963557 |
| Akt1s1 | 7.49016 | 7.273102 | 7.602 | -0.328876366 | -0.111818968 | 7.680937 | 7.599545 | 7.48828 | 0.11126578 | 0.192657802 |
| Gm47678 | 4.389504 | 4.439722 | 4.4288 | 0.010917343 | -0.03930134 | 4.4586885 | 4.7596209 | 4.64843 | 0.111189425 | -0.189742932 |
| Stab1 | 9.120957 | 9.04819 | 8.3561 | 0.692099465 | 0.764866131 | 8.467836 | 8.6614562 | 8.55036 | 0.111091786 | -0.082528411 |
| Smndc1 | 10.15884 | 10.11832 | 10.085 | 0.032932916 | 0.073454292 | 10.08709 | 10.233678 | 10.1229 | 0.110792035 | -0.035796486 |
| Dock11 | 12.31366 | 12.4103 | 12.36 | 0.050678496 | -0.045952632 | 12.249675 | 12.425491 | 12.3148 | 0.110737168 | -0.065078323 |
| Rpl22 | 12.72786 | 12.70117 | 12.755 | -0.05361332 | -0.026922274 | 12.892902 | 12.886278 | 12.7756 | 0.110691596 | 0.117315055 |
| AY074887 | 1.607439 | 1.654738 | 1.6316 | 0.023112211 | -0.024186997 | 1.6217413 | 1.7408265 | 1.63018 | 0.1106513 | -0.008433896 |
| Pnrc2 | 11.21153 | 11.28999 | 11.277 | 0.013282122 | -0.06517785 | 11.346928 | 11.362463 | 11.2519 | 0.110607626 | 0.095072841 |
| Vmn2r60 | 4.93063 | 4.86573 | 4.7842 | 0.081508316 | 0.146408009 | 4.9779233 | 5.1515055 | 5.04095 | 0.110552873 | -0.063029391 |
| Ighv1-14 | 2.962518 | 3.06233 | 2.8116 | 0.250760397 | 0.150948169 | 2.9986629 | 2.9058222 | 2.79529 | 0.110535517 | 0.203376184 |
| Pknox2 | 3.845657 | 3.847379 | 3.7478 | 0.099557731 | 0.097836226 | 3.8685978 | 3.9501912 | 3.83971 | 0.11048404 | 0.028890697 |
| Pde6h | 3.272993 | 3.201734 | 3.2568 | -0.055026591 | 0.016232277 | 3.4418421 | 3.4499796 | 3.33952 | 0.110456897 | 0.102319435 |
| Plcb3 | 8.124173 | 8.222313 | 8.111 | 0.111291389 | 0.013151376 | 8.2685227 | 8.260051 | 8.14961 | 0.110440868 | 0.118912573 |
| Gm47102 | 6.205365 | 6.25921 | 6.3825 | -0.123274302 | -0.177118609 | 6.0468873 | 6.9568565 | 6.84646 | 0.110395988 | -0.799573189 |
| Rpsa-ps11 | 3.367363 | 3.508783 | 3.4797 | 0.029114915 | -0.112305557 | 3.4124948 | 3.5673649 | 3.45714 | 0.110228782 | -0.044641338 |
| Rbm7 | 10.24469 | 10.26276 | 10.297 | -0.034695018 | -0.052765248 | 10.282554 | 10.423436 | 10.3132 | 0.110216908 | -0.030665484 |

| | | | | | | | | | | |
|---|---|---|---|---|---|---|---|---|---|---|
| Vps18 | 8.989314 | 8.956714 | 8.9395 | 0.017190387 | 0.049790332 | 8.9758315 | 8.917696 | 8.80748 | 0.110216385 | 0.168351928 |
| Tango2 | 7.20612 | 7.132622 | 7.1292 | 0.003392324 | 0.076889387 | 7.2798567 | 7.4562184 | 7.34605 | 0.110170265 | -0.066191429 |
| Vnn3 | 3.314164 | 3.372003 | 3.1754 | 0.196629755 | 0.138790805 | 3.2678006 | 3.200043 | 3.0899 | 0.110141888 | 0.177899455 |
| Gm7488 | 8.288342 | 8.267664 | 8.2805 | -0.012862425 | 0.007815731 | 8.1775835 | 8.4051604 | 8.29509 | 0.110073742 | -0.117503157 |
| Cpsf6 | 10.90283 | 11.05302 | 10.927 | 0.126015342 | -0.024176942 | 11.001002 | 11.072683 | 10.9627 | 0.109977068 | 0.038296724 |
| Vmn2r108 | 4.378395 | 4.40746 | 4.343 | 0.064501168 | 0.03543663 | 4.5881321 | 4.7118285 | 4.60193 | 0.109896849 | -0.013799574 |
| Gm10718 | 1.225738 | 1.258295 | 1.2419 | 0.016378978 | -0.016177617 | 1.2911235 | 1.4110254 | 1.30133 | 0.10969078 | -0.01021105 |
| Surf4 | 11.49126 | 11.49272 | 11.426 | 0.066470378 | 0.065006746 | 11.503519 | 11.457933 | 11.3483 | 0.109628336 | 0.155214302 |
| Idi2 | 2.064827 | 2.205105 | 2.0742 | 0.130860653 | -0.009417766 | 1.9921133 | 2.1058386 | 1.99624 | 0.109603203 | -0.004122004 |
| Sun2 | 12.97601 | 12.9207 | 12.994 | -0.073648166 | -0.018339712 | 13.036399 | 13.093033 | 12.9835 | 0.109521228 | 0.052887111 |
| Arhgap45 | 13.93449 | 14.09164 | 14.016 | 0.075636636 | -0.08151244 | 14.247219 | 14.254894 | 14.1454 | 0.109518406 | 0.101842993 |
| Klhl9 | 9.814964 | 9.940308 | 9.9084 | 0.031882962 | -0.09346064 | 9.7114855 | 9.8992258 | 9.78972 | 0.109507055 | -0.07823319 |
| Gm3558 | 1.832868 | 1.73397 | 1.8231 | -0.089146504 | 0.009752139 | 1.6694167 | 1.9083768 | 1.79909 | 0.10928806 | -0.129672058 |
| Gm12792 | 1.719619 | 1.730506 | 1.7682 | -0.037733497 | -0.048620829 | 1.839664 | 1.8689861 | 1.75977 | 0.109219254 | 0.07989713 |
| Dtnbp1 | 8.248434 | 8.265066 | 8.359 | -0.09395931 | -0.110591525 | 8.3222455 | 8.4123897 | 8.3032 | 0.109185474 | 0.019041264 |
| Ftsj1 | 7.599075 | 7.652269 | 7.6232 | 0.029039931 | -0.024153769 | 7.7594711 | 7.6399304 | 7.53087 | 0.10905545 | 0.228596133 |
| Samm50 | 10.43222 | 10.45732 | 10.429 | 0.028674812 | 0.003573434 | 10.639184 | 10.622454 | 10.5135 | 0.108968094 | 0.125697572 |
| Gnat2 | 1.303364 | 1.321321 | 1.408 | -0.086722626 | -0.104679531 | 1.3312344 | 1.4137763 | 1.30492 | 0.108856588 | 0.026314658 |
| Syap1 | 9.309104 | 9.29322 | 9.2268 | 0.06644407 | 0.082328185 | 9.2666899 | 9.3275457 | 9.2187 | 0.108849071 | 0.047993284 |
| Tstd2 | 9.218166 | 9.280914 | 9.3348 | -0.053895421 | -0.116643229 | 9.469797 | 9.4348125 | 9.32609 | 0.108727483 | 0.143711969 |
| Kbtbd11 | 12.14828 | 11.93692 | 12.001 | -0.063894972 | 0.147462062 | 11.612779 | 11.910962 | 11.8022 | 0.108725166 | -0.189457639 |
| Lcat | 3.41893 | 3.3162 | 3.3463 | -0.030112335 | 0.072617945 | 3.4718864 | 3.5585367 | 3.44982 | 0.108714327 | 0.022064052 |
| Gm5835 | 2.62491 | 2.460183 | 2.5175 | -0.057288478 | 0.107437918 | 2.4992245 | 2.6045185 | 2.49585 | 0.108672943 | 0.003378954 |
| Eef2kmt | 8.270604 | 8.295805 | 8.2964 | -0.000629528 | -0.025830304 | 8.5777027 | 8.4857546 | 8.37709 | 0.1086617 | 0.200609798 |
| Dusp12 | 7.862053 | 7.651153 | 7.6555 | -0.004300841 | 0.2065992 | 7.8770113 | 7.7497466 | 7.64109 | 0.108659502 | 0.235924168 |
| Rpsa-ps4 | 1.342328 | 1.309942 | 1.3674 | -0.057440619 | -0.025054458 | 1.4272232 | 1.4208429 | 1.31223 | 0.108613995 | 0.114994282 |
| Eif1b | 8.954418 | 9.005959 | 8.8726 | 0.133349621 | 0.081808795 | 9.0556622 | 9.0078516 | 8.89927 | 0.108582576 | 0.156393178 |
| Gm10563 | 1.710259 | 1.664353 | 1.7087 | -0.044322499 | 0.001583675 | 1.7438619 | 1.8446336 | 1.73608 | 0.108554216 | 0.007782551 |
| Mup5 | 3.067432 | 2.396211 | 2.5506 | -0.154411776 | 0.516809089 | 2.5681393 | 2.7177475 | 2.60923 | 0.108521353 | -0.04108689 |
| Ighv5-4 | 7.839415 | 7.350615 | 8.1941 | -0.843453064 | -0.354653839 | 7.7075597 | 7.4219817 | 7.31352 | 0.108463501 | 0.394041468 |
| Sgcg | 4.626568 | 4.680702 | 4.7958 | -0.115120416 | -0.169254066 | 4.7333095 | 5.0394323 | 4.93097 | 0.108458988 | -0.197663865 |
| Bend4 | 8.19173 | 8.219083 | 8.21 | 0.009094825 | -0.018257657 | 7.7152073 | 8.2077833 | 8.09937 | 0.108416361 | -0.384159661 |

| | | | | | | | | | | |
|---|---|---|---|---|---|---|---|---|---|---|
| Mier2 | 8.415123 | 8.475858 | 8.4643 | 0.011544417 | -0.049190028 | 8.6489706 | 8.7412465 | 8.63294 | 0.108311162 | 0.016035263 |
| Fzr1 | 9.202342 | 9.261236 | 9.2507 | 0.010544568 | -0.048348753 | 9.3985764 | 9.4570521 | 9.34879 | 0.10826503 | 0.049789301 |
| Cc2d2b | 2.994722 | 3.093049 | 3.028 | 0.065022195 | -0.033304457 | 3.1115604 | 3.1419833 | 3.03382 | 0.108160032 | 0.077737096 |
| Chkb | 9.561832 | 9.545218 | 9.7245 | -0.179321218 | -0.162706559 | 10.094146 | 10.024586 | 9.91653 | 0.10805438 | 0.177614949 |
| Vmn1r220 | 2.467022 | 2.458105 | 2.5029 | -0.04477071 | -0.035853758 | 2.4905888 | 2.619181 | 2.51113 | 0.108053913 | -0.020538267 |
| Pdk1 | 11.17994 | 11.07862 | 11.303 | -0.224874268 | -0.123561212 | 10.811635 | 11.24525 | 11.1372 | 0.108044472 | -0.325570036 |
| AC107664.2 | 0.384087 | 0.40284 | 0.4276 | -0.024770615 | -0.043523032 | 0.4234906 | 0.5330955 | 0.4251 | 0.107999108 | -0.001605776 |
| Trbj1-5 | 4.859689 | 4.938727 | 4.8353 | 0.10338247 | 0.024344712 | 4.8421071 | 5.0002136 | 4.89225 | 0.107967069 | -0.050139431 |
| Bach1 | 9.868218 | 9.864494 | 9.6138 | 0.250697463 | 0.254421086 | 9.5508012 | 9.6493474 | 9.54138 | 0.107962816 | 0.009416549 |
| Snrpg | 10.23514 | 10.28916 | 10.334 | -0.045088145 | -0.099116151 | 10.548964 | 10.455815 | 10.348 | 0.107842082 | 0.200990995 |
| Parp8 | 9.815416 | 9.944146 | 9.9849 | -0.040754766 | -0.1694843 | 9.921885 | 10.145894 | 10.0381 | 0.107823966 | -0.116185346 |
| Gm26509 | 1.204863 | 1.204955 | 1.1852 | 0.0197208 | 0.019629767 | 1.2874475 | 1.3423603 | 1.23457 | 0.107791799 | 0.052878998 |
| AC133103.5 | 0.286277 | 0.231084 | 0.2715 | -0.040383716 | 0.014808915 | 0.2330863 | 0.3767944 | 0.26905 | 0.107741825 | -0.035966247 |
| Apoo | 6.328945 | 6.145723 | 6.3536 | -0.207906753 | -0.024684849 | 6.3713131 | 6.3747609 | 6.26704 | 0.107717617 | 0.104269872 |
| D430042O09Rik | 8.217377 | 8.210612 | 8.3928 | -0.182214138 | -0.175449847 | 8.6207144 | 8.5598957 | 8.45219 | 0.107707942 | 0.168526646 |
| Cox19 | 7.423652 | 7.365378 | 7.3961 | -0.030684546 | 0.027589154 | 7.474263 | 7.5396674 | 7.432 | 0.107667036 | 0.042262593 |
| Zfp595 | 7.934509 | 8.029606 | 7.8922 | 0.137364085 | 0.04226711 | 8.0473479 | 8.1771742 | 8.06952 | 0.107653029 | -0.022173254 |
| Gpr85 | 2.706604 | 2.659766 | 2.682 | -0.022263513 | 0.024574481 | 2.5527461 | 2.6288853 | 2.52126 | 0.107620775 | 0.031481545 |
| Olfr1415 | 1.482197 | 1.530565 | 1.4749 | 0.055655743 | 0.007286967 | 1.4990299 | 1.5424599 | 1.43492 | 0.107539606 | 0.064109674 |
| Adh5 | 10.33819 | 10.30621 | 10.345 | -0.038575236 | -0.006590286 | 10.520705 | 10.492117 | 10.3846 | 0.107505874 | 0.136094233 |
| Ubxn6 | 9.540703 | 9.538317 | 9.4869 | 0.051403102 | 0.053789484 | 9.733958 | 9.6177641 | 9.51027 | 0.107495421 | 0.223689279 |
| 4930431P19Rik | 3.902252 | 3.840221 | 3.8529 | -0.012675677 | 0.04935521 | 3.9826684 | 3.9357657 | 3.82827 | 0.107495357 | 0.154398061 |
| Sufu | 9.768134 | 9.889722 | 9.7964 | 0.093356548 | -0.028231238 | 9.8119611 | 9.8804646 | 9.77311 | 0.107358405 | 0.038854933 |
| Dpp7 | 8.125188 | 8.118565 | 8.1676 | -0.04907666 | -0.042453502 | 8.5433768 | 8.4904803 | 8.38313 | 0.107352484 | 0.160249035 |
| Rom1 | 6.208795 | 6.053263 | 6.2167 | -0.163392695 | -0.007860629 | 6.2285515 | 6.1911205 | 6.08382 | 0.107304948 | 0.144735969 |
| Rac1 | 12.02542 | 11.96781 | 11.996 | -0.02839543 | 0.029214284 | 11.9658 | 11.998515 | 11.8914 | 0.10707328 | 0.074358247 |
| Adck5 | 7.65928 | 7.647642 | 7.7267 | -0.079060496 | -0.067421904 | 8.0369168 | 8.0339772 | 7.92691 | 0.107068942 | 0.110008581 |
| Vmn1r198 | 3.41894 | 3.209619 | 3.4038 | -0.194133371 | 0.015188185 | 3.2744416 | 3.4740563 | 3.36703 | 0.107024283 | -0.092590461 |
| Tcrg-C2 | 6.933345 | 6.531246 | 6.8567 | -0.325496502 | 0.0766021 | 6.7155191 | 6.8929952 | 6.78601 | 0.106981928 | -0.070494186 |
| Atp6ap2 | 10.28954 | 10.28267 | 10.275 | 0.007245372 | 0.014113318 | 10.252839 | 10.233121 | 10.1262 | 0.10692906 | 0.126647258 |
| Slc25a39 | 9.119442 | 8.99014 | 9.1655 | -0.175322126 | -0.046020036 | 9.3278756 | 9.1693529 | 9.06246 | 0.106892106 | 0.265414731 |
| Lamtor4 | 8.476185 | 8.405999 | 8.451 | -0.044980047 | 0.025206115 | 8.7452245 | 8.4744456 | 8.36756 | 0.106883247 | 0.377662181 |

| | | | | | | | | | | |
|---|---|---|---|---|---|---|---|---|---|---|
| Nlgn3 | 4.124103 | 4.142784 | 4.1007 | 0.042067025 | 0.023385536 | 4.0407158 | 4.2619401 | 4.15513 | 0.10681111 | -0.114413237 |
| Klhl30 | 1.573477 | 1.619658 | 1.6306 | -0.010934651 | -0.057116061 | 1.6327824 | 1.7524065 | 1.64571 | 0.106696865 | -0.012927236 |
| Col8a2 | 1.837286 | 1.878737 | 1.744 | 0.134688023 | 0.093237228 | 1.702233 | 1.8950678 | 1.78845 | 0.106620782 | -0.086213988 |
| Ighv7-4 | 3.140578 | 3.122865 | 3.1154 | 0.007498866 | 0.025211452 | 3.2323098 | 3.1946359 | 3.08803 | 0.10660697 | 0.144280858 |
| Ndufb10 | 9.491954 | 9.42161 | 9.4128 | 0.008798151 | 0.079142836 | 9.5408194 | 9.5396794 | 9.43308 | 0.106599059 | 0.107739134 |
| Nptx2 | 2.597352 | 2.768846 | 2.7845 | -0.015696803 | -0.187190914 | 2.6751688 | 2.7318505 | 2.62525 | 0.106597392 | 0.049915624 |
| Cxcl5 | 1.128203 | 1.085396 | 1.0521 | 0.033304254 | 0.076111556 | 1.0910778 | 1.1923248 | 1.08574 | 0.10658151 | 0.005334537 |
| Ccdc90b | 7.269127 | 7.171665 | 7.3358 | -0.164165472 | -0.066704077 | 7.0818085 | 7.2992541 | 7.1927 | 0.106550902 | -0.110894706 |
| Cep68 | 9.955816 | 10.14885 | 10.107 | 0.042197762 | -0.150838101 | 10.002753 | 10.113249 | 10.0068 | 0.106428925 | -0.004066844 |
| Fam208a | 10.59988 | 10.65226 | 10.621 | 0.030769346 | -0.021609066 | 10.50846 | 10.730044 | 10.6238 | 0.10625226 | -0.115332396 |
| Tacr1 | 2.961225 | 2.791676 | 2.974 | -0.182293223 | -0.012743939 | 2.8859078 | 2.9981581 | 2.89195 | 0.106207215 | -0.006043127 |
| Gnptab | 10.4282 | 10.56555 | 10.446 | 0.119653976 | -0.017698442 | 10.393526 | 10.616855 | 10.5107 | 0.10618174 | -0.11714698 |
| Btf3l4 | 9.436226 | 9.394412 | 9.4716 | -0.077207828 | -0.035393499 | 9.6095456 | 9.5467417 | 9.44057 | 0.106173205 | 0.168977061 |
| Nol11 | 9.756303 | 9.777659 | 9.7049 | 0.072795046 | 0.051439421 | 9.7548399 | 9.9229705 | 9.81692 | 0.10604897 | -0.062081624 |
| Gm20458 | 1.761985 | 1.84338 | 1.8706 | -0.027223646 | -0.108618079 | 1.9537464 | 1.8864909 | 1.78046 | 0.106026795 | 0.173282254 |
| Spon2 | 1.396197 | 1.443135 | 1.3796 | 0.06349426 | 0.016556142 | 1.459114 | 1.5024929 | 1.39647 | 0.106019179 | 0.062640289 |
| Ppib | 11.68595 | 11.59405 | 11.615 | -0.020691754 | 0.071206754 | 11.760127 | 11.694686 | 11.5887 | 0.105961164 | 0.171402596 |
| Lims1 | 10.85516 | 10.99694 | 10.917 | 0.080235292 | -0.061544202 | 10.865826 | 11.076698 | 10.9708 | 0.105928748 | -0.10494283 |
| Gm7331 | 5.775638 | 6.048614 | 6.0817 | -0.033067047 | -0.306043163 | 6.4280495 | 6.3044042 | 6.19855 | 0.105852288 | 0.229497595 |
| Kcnh4 | 4.108946 | 4.096144 | 3.9668 | 0.129376349 | 0.142178706 | 4.0589568 | 4.1485887 | 4.04284 | 0.105748388 | 0.016116432 |
| Trrap | 11.65198 | 11.70792 | 11.702 | 0.006042574 | -0.049889304 | 11.682135 | 12.004072 | 11.8984 | 0.105630906 | -0.216306212 |
| Kctd21 | 5.636141 | 5.712846 | 5.7048 | 0.008016896 | -0.068688025 | 5.604308 | 5.7265725 | 5.62107 | 0.10550138 | -0.016763164 |
| Olfr678 | 1.462098 | 1.444179 | 1.4409 | 0.003297714 | 0.021217191 | 1.502384 | 1.5307121 | 1.42528 | 0.10543411 | 0.077105991 |
| Cox6a2 | 3.918301 | 3.900268 | 3.9206 | -0.020336213 | -0.002303163 | 3.6880457 | 3.7739538 | 3.66864 | 0.105314329 | 0.019406243 |
| Vbp1 | 9.886646 | 9.926527 | 9.961 | -0.034482439 | -0.074363216 | 9.972611 | 10.050623 | 9.94533 | 0.105293021 | 0.027281327 |
| Rab1b | 10.67807 | 10.5108 | 10.586 | -0.075301524 | 0.091967104 | 10.791626 | 10.618999 | 10.5138 | 0.105244344 | 0.277871378 |
| Batf3 | 6.212548 | 6.31017 | 6.2724 | 0.037751656 | -0.059870444 | 6.3302021 | 6.5894517 | 6.48423 | 0.105224616 | -0.154024991 |
| Gm10662 | 1.077641 | 1.077939 | 1.1773 | -0.099331572 | -0.099629021 | 1.2807457 | 1.2979792 | 1.19278 | 0.105203819 | 0.087970312 |
| Gm5891 | 1.077641 | 1.077939 | 1.1773 | -0.099331572 | -0.099629021 | 1.2807457 | 1.2979792 | 1.19278 | 0.105203819 | 0.087970312 |
| Hrk | 1.171495 | 1.144044 | 1.1794 | -0.035389241 | -0.007938934 | 1.1092569 | 1.2174472 | 1.11231 | 0.105139423 | -0.00305081 |
| Vps4a | 9.454011 | 9.58946 | 9.4721 | 0.117408687 | -0.018040379 | 9.5727419 | 9.6064149 | 9.50157 | 0.104847643 | 0.071174625 |
| Trbj1-7 | 3.524338 | 3.667949 | 3.6697 | -0.001791812 | -0.145402388 | 3.5363139 | 3.6875592 | 3.58273 | 0.104825231 | -0.046420056 |

| | | | | | | | | | | |
|---|---|---|---|---|---|---|---|---|---|---|
| AC159289.2 | 0.492856 | 0.455602 | 0.5339 | -0.078256955 | -0.041002852 | 0.4469426 | 0.5560883 | 0.45134 | 0.104744763 | -0.004400942 |
| Timm8a2 | 1.115994 | 1.180043 | 1.1579 | 0.022159604 | -0.041890097 | 1.1540902 | 1.2004709 | 1.09577 | 0.104704584 | 0.058323889 |
| Mapk6 | 10.19202 | 10.11035 | 10.021 | 0.089020053 | 0.170693999 | 9.8633415 | 10.152234 | 10.0476 | 0.104633885 | -0.184258661 |
| 4931440P22Rik | 4.494095 | 4.634783 | 4.4085 | 0.226294414 | 0.085606684 | 4.5634434 | 4.7397122 | 4.63509 | 0.104625141 | -0.07164367 |
| Thap11 | 8.307028 | 8.296009 | 8.4032 | -0.107225795 | -0.096207032 | 8.3008911 | 8.3402296 | 8.23573 | 0.104495422 | 0.065156983 |
| Zfp672 | 8.247001 | 8.140915 | 8.2656 | -0.124657879 | -0.018572537 | 8.3774112 | 8.2510881 | 8.14668 | 0.104412959 | 0.23073604 |
| Ccnjl | 2.777393 | 2.811106 | 2.7354 | 0.075701328 | 0.041987735 | 2.6466176 | 2.8920207 | 2.78764 | 0.104376055 | -0.141027064 |
| Prdm1 | 7.929333 | 7.78103 | 7.4177 | 0.363347027 | 0.511650844 | 7.3227153 | 7.7204272 | 7.61607 | 0.104356604 | -0.293355344 |
| Gm4152 | 1.860931 | 1.796954 | 1.9124 | -0.115474528 | -0.051497068 | 1.928966 | 1.9493858 | 1.84519 | 0.104199053 | 0.083779242 |
| Mapkapk5 | 9.258486 | 9.23565 | 9.2976 | -0.061973151 | -0.039136641 | 9.2564941 | 9.4634053 | 9.35921 | 0.104194665 | -0.10271654 |
| Il20rb | 4.965333 | 4.955527 | 4.9346 | 0.020963217 | 0.030769317 | 4.9047738 | 4.923146 | 4.81902 | 0.104129168 | 0.085756994 |
| Cutc | 7.596763 | 7.828589 | 7.6545 | 0.174080217 | -0.057745773 | 7.775985 | 7.7320922 | 7.62797 | 0.104120466 | 0.148013294 |
| Dnajb1 | 11.43733 | 11.48181 | 11.541 | -0.05933658 | -0.103814554 | 10.714925 | 11.293332 | 11.1892 | 0.104097914 | -0.474308654 |
| Rpl30 | 13.98195 | 13.98816 | 13.961 | 0.027628822 | 0.021412194 | 14.015794 | 14.094174 | 13.9901 | 0.104058428 | 0.025677963 |
| L1td1 | 0.705983 | 0.670547 | 0.6544 | 0.01613433 | 0.05157079 | 0.7045404 | 0.7772737 | 0.67325 | 0.10402017 | 0.031286901 |
| Ovca2 | 8.420012 | 8.462661 | 8.3617 | 0.100916421 | 0.058267237 | 8.724903 | 8.5784278 | 8.47448 | 0.103952163 | 0.250427297 |
| Pcnx3 | 10.89942 | 11.03377 | 10.989 | 0.044685416 | -0.089666256 | 11.391684 | 11.441627 | 11.3377 | 0.10393699 | 0.053994178 |
| Art3 | 4.478236 | 4.387642 | 4.596 | -0.208353474 | -0.117760034 | 4.1024631 | 4.6231579 | 4.5193 | 0.103862036 | -0.416832705 |
| Palm | 6.352918 | 6.71816 | 6.7189 | -0.00076444 | -0.36600615 | 6.4879963 | 6.5789203 | 6.47509 | 0.103825506 | 0.012901502 |
| Pfdn5 | 11.65558 | 11.65386 | 11.714 | -0.059833186 | -0.058104214 | 11.761516 | 11.773569 | 11.6698 | 0.103757512 | 0.09170443 |
| Dars2 | 7.981153 | 7.915863 | 7.9798 | -0.063932819 | 0.001356324 | 7.9841714 | 8.122973 | 8.01928 | 0.103694159 | -0.035107475 |
| Gsk3a | 10.7653 | 10.81497 | 10.804 | 0.011080252 | -0.038585098 | 10.978067 | 10.860915 | 10.7572 | 0.10368565 | 0.220838328 |
| Gm20661 | 1.331334 | 1.329655 | 1.3106 | 0.019051829 | 0.02073126 | 1.3124502 | 1.4803088 | 1.37668 | 0.103630179 | -0.06422842 |
| Nup43 | 6.572743 | 6.649181 | 6.5639 | 0.085310819 | 0.008873319 | 6.671586 | 6.8764806 | 6.77286 | 0.103620236 | -0.101274341 |
| Sppl2b | 8.44864 | 8.493091 | 8.4346 | 0.058446951 | 0.013996545 | 8.7103624 | 8.6531956 | 8.54964 | 0.103553043 | 0.160719905 |
| Rpl31-ps8 | 12.2924 | 12.31371 | 12.258 | 0.055762806 | 0.034448616 | 12.244421 | 12.409194 | 12.3058 | 0.103383662 | -0.061389005 |
| Fam50a | 9.472214 | 9.533363 | 9.4752 | 0.058173261 | -0.002975935 | 9.6365969 | 9.6043827 | 9.5011 | 0.103280379 | 0.135494636 |
| Yipf5 | 9.093549 | 9.145394 | 8.9847 | 0.160665112 | 0.108820157 | 9.1902263 | 9.1732258 | 9.07 | 0.103227247 | 0.120227736 |
| Osr1 | 3.260219 | 2.915476 | 3.0813 | -0.165859455 | 0.178883563 | 2.8388637 | 3.0153076 | 2.9121 | 0.103209743 | -0.073234236 |
| Ffar1 | 9.677124 | 9.699825 | 9.5898 | 0.109978985 | 0.087278317 | 10.226263 | 9.9823769 | 9.87917 | 0.103209527 | 0.347095767 |
| Mrgprf | 2.5313 | 2.72503 | 2.5916 | 0.133477317 | -0.060253275 | 2.7095666 | 2.5336626 | 2.43047 | 0.103194602 | 0.279098612 |
| Egfl8 | 4.13358 | 4.263957 | 4.3745 | -0.110561363 | -0.240938218 | 4.3480706 | 4.3619431 | 4.25886 | 0.103084601 | 0.089212063 |

| | | | | | | | | | | |
|---|---|---|---|---|---|---|---|---|---|---|
| Efhc2 | 1.154228 | 1.102278 | 1.1307 | -0.028457072 | 0.023493519 | 1.076643 | 1.2016036 | 1.09852 | 0.103080241 | -0.021880289 |
| Cyth1 | 11.88467 | 12.02789 | 11.747 | 0.28063938 | 0.137418704 | 11.616091 | 11.864268 | 11.7612 | 0.10303192 | -0.145144668 |
| Zfp948 | 6.178038 | 6.195684 | 6.0861 | 0.10963229 | 0.091986862 | 6.0027611 | 6.092175 | 5.98916 | 0.10301671 | 0.013602868 |
| Ipo5 | 11.25468 | 11.19245 | 11.189 | 0.003392588 | 0.065615884 | 11.310458 | 11.319368 | 11.2164 | 0.102946178 | 0.094036861 |
| Rnf17 | 3.000256 | 2.971122 | 2.8829 | 0.088251075 | 0.117384949 | 2.952243 | 3.136914 | 3.03398 | 0.102932014 | -0.081738934 |
| Gpm6b | 7.886958 | 7.548891 | 7.8464 | -0.297477092 | 0.040590283 | 7.1961262 | 7.4697782 | 7.36685 | 0.102926913 | -0.170725048 |
| Nsmce3 | 8.044839 | 8.022772 | 8.106 | -0.083270387 | -0.061203375 | 7.9430016 | 8.0722451 | 7.96938 | 0.102865071 | -0.026378409 |
| Htr1b | 2.165015 | 2.215115 | 2.2001 | 0.014973914 | -0.035126215 | 2.2815128 | 2.3355283 | 2.23268 | 0.102852601 | 0.048837175 |
| Rapgefl1 | 3.421116 | 3.394792 | 3.467 | -0.072185391 | -0.045860666 | 3.3799871 | 3.4556436 | 3.3528 | 0.102841328 | 0.027184837 |
| Rpl3 | 15.03606 | 15.02041 | 15.051 | -0.030184862 | -0.014539508 | 15.123459 | 15.10269 | 14.9999 | 0.102829984 | 0.123598525 |
| Gm20056 | 0.932743 | 1.209673 | 0.8976 | 0.312116402 | 0.035186641 | 0.9173843 | 1.00381 | 0.90116 | 0.10264968 | 0.016223923 |
| Gorasp2 | 10.36068 | 10.45301 | 10.403 | 0.050268322 | -0.04206179 | 10.460866 | 10.421867 | 10.3192 | 0.10263497 | 0.141634503 |
| Polr2g | 9.58118 | 9.636306 | 9.6289 | 0.007359156 | -0.047766025 | 9.8919478 | 9.8692015 | 9.76658 | 0.102623657 | 0.125369948 |
| Dpp9 | 10.1459 | 10.24811 | 10.123 | 0.125525713 | 0.023314989 | 10.320597 | 10.206007 | 10.1034 | 0.102562539 | 0.217152624 |
| Zfp697 | 2.944626 | 2.845272 | 2.8167 | 0.028560846 | 0.127915222 | 2.890739 | 2.9612306 | 2.85867 | 0.102560951 | 0.032069281 |
| Gpx1 | 11.067 | 11.09376 | 11.09 | 0.00377804 | -0.022982392 | 11.395591 | 10.973693 | 10.8712 | 0.102533566 | 0.52443196 |
| Eprs | 11.40986 | 11.34519 | 11.309 | 0.036177252 | 0.100838784 | 11.434003 | 11.476789 | 11.3744 | 0.102430384 | 0.059643709 |
| Ccs | 8.169796 | 8.212425 | 8.2445 | -0.032112784 | -0.074742028 | 8.1866266 | 8.277402 | 8.17498 | 0.102417361 | 0.01164199 |
| Ube2s | 10.02934 | 10.08642 | 10.019 | 0.067539927 | 0.010452336 | 10.011028 | 10.177343 | 10.0749 | 0.102412772 | -0.063901609 |
| Met | 7.416217 | 7.440189 | 7.3147 | 0.125451986 | 0.101480617 | 6.8525268 | 6.8173689 | 6.71496 | 0.10240607 | 0.137564058 |
| Fkbp1a | 10.33806 | 10.45503 | 10.344 | 0.11053556 | -0.006440216 | 10.458433 | 10.434971 | 10.3326 | 0.102401474 | 0.125863645 |
| Dzip3 | 8.015012 | 8.209579 | 7.8735 | 0.336062972 | 0.141496614 | 8.2740233 | 8.4193219 | 8.31694 | 0.10238399 | -0.042914647 |
| Polr3a | 9.436388 | 9.51811 | 9.4957 | 0.022410988 | -0.059311362 | 9.6473239 | 9.8000057 | 9.69763 | 0.102370712 | -0.050311019 |
| Pidd1 | 7.269686 | 7.284021 | 7.3533 | -0.069325548 | -0.083660468 | 7.6033308 | 7.680975 | 7.57861 | 0.102362163 | 0.024717897 |
| Nkapd1 | 9.177161 | 9.075211 | 9.0521 | 0.023068316 | 0.125018111 | 9.0852709 | 9.1388479 | 9.03651 | 0.102342549 | 0.048765548 |
| Gm17823 | 1.034362 | 1.064801 | 1.0817 | -0.016894009 | -0.047332964 | 1.0148867 | 1.1496783 | 1.04734 | 0.102338047 | -0.032453524 |
| Glrx3 | 10.09168 | 10.02834 | 10.084 | -0.055391897 | 0.007943749 | 10.273739 | 10.189389 | 10.0872 | 0.102195212 | 0.186545997 |
| Aptx | 8.213614 | 8.168704 | 8.2181 | -0.049388934 | -0.004478491 | 8.4117314 | 8.3488031 | 8.24671 | 0.102092501 | 0.165020769 |
| Rbm27 | 10.59689 | 10.68462 | 10.56 | 0.124211454 | 0.036486504 | 10.467066 | 10.635261 | 10.5332 | 0.102053082 | -0.066141525 |
| 5430437J10Rik | 3.883357 | 4.043372 | 3.9389 | 0.104456929 | -0.055558449 | 3.9313675 | 4.0326011 | 3.93057 | 0.102030948 | 0.000797314 |
| Gm28551 | 2.42772 | 2.377284 | 2.5346 | -0.157355722 | -0.106918945 | 2.5540187 | 2.5547626 | 2.45277 | 0.101987693 | 0.10124383 |
| Pcyt2 | 9.417895 | 9.37126 | 9.4483 | -0.077027757 | -0.03039286 | 9.8184886 | 9.7350579 | 9.63308 | 0.101980931 | 0.185411597 |

| | | | | | | | | | | |
|---|---|---|---|---|---|---|---|---|---|---|
| Csn3 | 1.736342 | 1.695465 | 1.6571 | 0.03838594 | 0.079262435 | 1.7057524 | 1.8059823 | 1.70401 | 0.10197254 | 0.001742562 |
| Adamts17 | 3.291237 | 3.229476 | 3.273 | -0.04349201 | 0.018269765 | 3.2907814 | 3.3430658 | 3.24111 | 0.101950961 | 0.049666514 |
| Plac9a | 1.975081 | 2.060614 | 2.0753 | -0.014648857 | -0.100181421 | 2.0217331 | 2.0032419 | 1.90134 | 0.101898313 | 0.120389511 |
| A630010A05Rik | 3.18869 | 3.121383 | 3.197 | -0.075594835 | -0.008288065 | 3.4246938 | 3.3230584 | 3.22118 | 0.101880899 | 0.203516317 |
| Ybx1 | 12.43104 | 12.40468 | 12.445 | -0.040631972 | -0.014271694 | 12.572551 | 12.590962 | 12.4891 | 0.101869698 | 0.083458515 |
| Wfdc2 | 1.449095 | 1.277362 | 1.3216 | -0.044236386 | 0.127496045 | 1.3076402 | 1.4228999 | 1.32112 | 0.10177993 | -0.013479768 |
| Fam173b | 7.766659 | 7.968232 | 7.7545 | 0.21373152 | 0.012158691 | 7.9448591 | 7.9441925 | 7.84248 | 0.101716551 | 0.102383143 |
| Srd5a3 | 8.684951 | 8.776954 | 8.8045 | -0.027574272 | -0.119577161 | 8.9057545 | 8.8020847 | 8.70038 | 0.101701369 | 0.205371096 |
| Cep350 | 11.39221 | 11.64317 | 11.515 | 0.128434328 | -0.122524808 | 11.209969 | 11.641379 | 11.5398 | 0.101570834 | -0.329839313 |
| 1700030J22Rik | 2.460869 | 2.457826 | 2.3728 | 0.085075576 | 0.08811912 | 2.503295 | 2.5354133 | 2.43384 | 0.101569016 | 0.069450668 |
| Muc13 | 3.431183 | 3.388038 | 3.4077 | -0.019613178 | 0.023531765 | 3.4132385 | 3.3333465 | 3.23178 | 0.101568399 | 0.181460442 |
| Epm2aip1 | 9.411188 | 9.412869 | 9.5173 | -0.104463499 | -0.106144443 | 9.9855314 | 9.9119937 | 9.81045 | 0.101547677 | 0.175085346 |
| Gm12254 | 0.518243 | 0.533748 | 0.509 | 0.02472044 | 0.009215554 | 0.5291594 | 0.5611978 | 0.45975 | 0.101451615 | 0.069413285 |
| Rpusd3 | 5.948261 | 5.881449 | 6.0097 | -0.128247942 | -0.061435872 | 6.3858197 | 5.9429113 | 5.84148 | 0.101432287 | 0.544340704 |
| Megf6 | 2.850246 | 3.041952 | 2.8377 | 0.204294815 | 0.012588064 | 2.8540496 | 2.7712241 | 2.66979 | 0.101431952 | 0.184257458 |
| Rps2-ps6 | 4.664825 | 4.687778 | 4.777 | -0.089193035 | -0.112145298 | 4.8075942 | 4.9043226 | 4.8029 | 0.101423733 | 0.004695399 |
| Plekho1 | 10.43583 | 10.52873 | 10.498 | 0.030580214 | -0.062314508 | 10.738328 | 10.560145 | 10.4588 | 0.10137711 | 0.279559853 |
| Gadl1 | 3.166919 | 3.153874 | 3.1218 | 0.032122417 | 0.045167353 | 3.0799194 | 3.1664939 | 3.06515 | 0.10134333 | 0.014768825 |
| Cpsf2 | 10.61069 | 10.75644 | 10.609 | 0.147314756 | 0.001564522 | 10.59606 | 10.670764 | 10.5694 | 0.10134169 | 0.02663786 |
| Igkv4-56 | 1.348211 | 1.354189 | 1.4127 | -0.058462181 | -0.064440441 | 1.3795105 | 1.4282863 | 1.32704 | 0.101248544 | 0.052472736 |
| Arl8a | 9.549896 | 9.66056 | 9.5875 | 0.073035593 | -0.037627853 | 9.7460282 | 9.6861049 | 9.58488 | 0.101225021 | 0.161148338 |
| Exo5 | 7.719736 | 7.605369 | 7.85 | -0.244636356 | -0.130269199 | 7.8639016 | 7.8115047 | 7.7103 | 0.101206904 | 0.153603785 |
| Utf1 | 2.582179 | 2.608732 | 2.6134 | -0.004712045 | -0.031264883 | 2.5908576 | 2.7828267 | 2.68167 | 0.101158249 | -0.09081086 |
| Gm17374 | 1.984607 | 1.954579 | 2.0381 | -0.083531211 | -0.053503074 | 1.9488233 | 2.1129846 | 2.01184 | 0.10114278 | -0.063018462 |
| Col5a1 | 6.070548 | 6.447415 | 6.0513 | 0.396126635 | 0.019259533 | 6.4907512 | 5.8936202 | 5.79249 | 0.101130366 | 0.698261326 |
| Ugcg | 11.47038 | 11.44957 | 11.415 | 0.034364766 | 0.055180774 | 11.21565 | 11.477562 | 11.3765 | 0.101046876 | -0.160865291 |
| Mxra8 | 5.617798 | 5.552351 | 5.7665 | -0.214100802 | -0.148654309 | 5.3052246 | 5.5719591 | 5.47097 | 0.100987482 | -0.165746994 |
| Ccdc106 | 0.989139 | 0.963502 | 1.0395 | -0.07601073 | -0.050373791 | 1.0205178 | 1.1061956 | 1.00522 | 0.100975745 | 0.015297948 |
| Trappc1 | 8.29887 | 8.17537 | 8.2839 | -0.108558552 | 0.014941185 | 8.4988418 | 8.3848107 | 8.28385 | 0.100958642 | 0.214989715 |
| Ilf2 | 10.20949 | 10.19257 | 10.245 | -0.051995064 | -0.035071002 | 10.395406 | 10.457931 | 10.357 | 0.100909808 | 0.038385366 |
| Gm20257 | 5.353372 | 5.338428 | 5.2296 | 0.108783885 | 0.123727689 | 5.9251198 | 5.7636815 | 5.66278 | 0.100901611 | 0.262339974 |
| Arhgap39 | 8.0691 | 8.056975 | 8.0324 | 0.024535267 | 0.036660562 | 7.9618644 | 8.4610594 | 8.36033 | 0.100733614 | -0.398461435 |

| | | | | | | | | | | |
|---|---|---|---|---|---|---|---|---|---|---|
| Trav15n-1 | 0.860632 | 0.855849 | 0.832 | 0.023886372 | 0.028669154 | 0.857221 | 0.9347314 | 0.834 | 0.100733141 | 0.02322269 |
| Cetn2 | 9.175083 | 9.026058 | 9.1271 | -0.101090292 | 0.047935293 | 9.1929804 | 9.1267629 | 9.02605 | 0.100717743 | 0.166935285 |
| Pdyn | 0.376088 | 0.276215 | 0.3291 | -0.052856187 | 0.047016923 | 0.2981605 | 0.3833848 | 0.28268 | 0.100704132 | 0.01547979 |
| Abcb9 | 7.601088 | 7.514736 | 7.6703 | -0.155587978 | -0.069236723 | 7.8224796 | 7.8212201 | 7.72053 | 0.100690051 | 0.101949526 |
| Siah2 | 8.324757 | 8.257815 | 8.2931 | -0.035279201 | 0.031663125 | 7.9574267 | 8.3068769 | 8.20625 | 0.100625018 | -0.248825258 |
| Bicdl2 | 1.518768 | 1.483089 | 1.5601 | -0.077053193 | -0.041374486 | 1.4938942 | 1.6853758 | 1.5848 | 0.100576734 | -0.090904902 |
| Slc39a3 | 8.157439 | 8.265507 | 8.254 | 0.011544473 | -0.096523628 | 8.4285315 | 8.4038753 | 8.30335 | 0.100527238 | 0.12518348 |
| Olfml2b | 4.372836 | 4.360146 | 4.2958 | 0.064375209 | 0.077065506 | 4.2892472 | 4.2368064 | 4.13633 | 0.100476556 | 0.152917435 |
| Gm10226 | 3.078868 | 3.070749 | 3.1627 | -0.09197968 | -0.08386037 | 3.1373437 | 3.1547284 | 3.05433 | 0.100396344 | 0.083011665 |
| Tnfrsf13c | 11.11278 | 11.05625 | 11.128 | -0.071488934 | -0.014955226 | 11.497137 | 11.210116 | 11.1098 | 0.100276274 | 0.387297674 |
| Stk40 | 10.0996 | 10.19153 | 9.8911 | 0.300394127 | 0.208461984 | 10.08944 | 10.027706 | 9.92751 | 0.100199838 | 0.161933393 |
| Ing2 | 7.88599 | 7.656733 | 7.9351 | -0.278381855 | -0.049124667 | 7.8368794 | 7.8654447 | 7.76529 | 0.100151933 | 0.071586641 |
| Zbtb2 | 9.233446 | 9.375435 | 9.2706 | 0.104825428 | -0.03716335 | 9.0292588 | 9.2130432 | 9.11296 | 0.100081235 | -0.083703195 |
| Naprt | 5.393669 | 5.24895 | 5.3184 | -0.069407392 | 0.07531132 | 5.5327454 | 5.3542455 | 5.25419 | 0.100051147 | 0.278551063 |
| Inip | 9.240951 | 9.38323 | 9.2442 | 0.139025845 | -0.003253465 | 9.4244979 | 9.4367321 | 9.33674 | 0.0999922 | 0.087758028 |
| Pcnp | 10.39175 | 10.40397 | 10.343 | 0.06075738 | 0.048536344 | 10.392148 | 10.620705 | 10.5207 | 0.09998311 | -0.128573604 |
| Piwil2 | 4.743012 | 4.634441 | 4.6003 | 0.034117166 | 0.142687881 | 4.8560238 | 4.7794176 | 4.67946 | 0.099955479 | 0.176561704 |
| Klhl42 | 8.614831 | 8.839326 | 8.7363 | 0.102995345 | -0.121498779 | 9.0036127 | 8.9949079 | 8.89496 | 0.099951033 | 0.108655834 |
| Ormdl3 | 9.133361 | 9.21442 | 9.0821 | 0.132354091 | 0.051294214 | 9.036934 | 9.0584979 | 8.95858 | 0.099919715 | 0.078355849 |
| Mob1a | 12.48208 | 12.61315 | 12.471 | 0.142273882 | 0.01120371 | 12.517973 | 12.510763 | 12.4109 | 0.099882456 | 0.107092068 |
| Tor1b | 9.738072 | 9.784321 | 9.8146 | -0.030298055 | -0.076546769 | 9.9445468 | 10.018165 | 9.91842 | 0.099742851 | 0.026124511 |
| Gm12416 | 3.070906 | 3.080872 | 3.118 | -0.037126232 | -0.047092899 | 3.1500026 | 3.2006209 | 3.10089 | 0.099726259 | 0.049107908 |
| Thap8 | 1.988583 | 1.914085 | 1.9804 | -0.066343879 | 0.008153708 | 1.994326 | 2.0413657 | 1.94173 | 0.099637919 | 0.052598284 |
| Prokr1 | 1.59236 | 1.557674 | 1.5609 | -0.003266289 | 0.031419713 | 1.52849 | 1.6570776 | 1.55752 | 0.099558146 | -0.029029432 |
| Rraga | 9.780109 | 9.674828 | 9.7362 | -0.061340255 | 0.043940642 | 9.7840227 | 9.6517362 | 9.55227 | 0.099465022 | 0.23175153 |
| Rnf44 | 11.0412 | 11.15513 | 11.022 | 0.13321716 | 0.019294612 | 11.247049 | 11.282903 | 11.1835 | 0.099432824 | 0.063579124 |
| Inafm1 | 7.621046 | 7.489058 | 7.5028 | -0.013771706 | 0.118216724 | 7.7257735 | 7.6722079 | 7.57278 | 0.099429195 | 0.152994757 |
| 2310033P09Rik | 8.070408 | 8.078268 | 8.103 | -0.024685479 | -0.032544865 | 8.2515157 | 8.1848032 | 8.0854 | 0.0994078 | 0.166120265 |
| Igkv4-69 | 4.818437 | 5.075258 | 5.12 | -0.044771727 | -0.301592693 | 5.2241401 | 5.3858869 | 5.2865 | 0.099385385 | -0.062361491 |
| Glyr1 | 11.43131 | 11.51612 | 11.486 | 0.029871642 | -0.05494136 | 11.475234 | 11.509959 | 11.4106 | 0.099363404 | 0.06463864 |
| Sec24c | 11.74096 | 11.77768 | 11.749 | 0.029090172 | -0.007633692 | 11.885332 | 11.933128 | 11.8338 | 0.099339824 | 0.051543592 |
| Nat2 | 7.540905 | 7.542756 | 7.5471 | -0.004297603 | -0.006148733 | 7.8303794 | 7.8029373 | 7.7037 | 0.099233966 | 0.126676132 |

| | | | | | | | | | | |
|---|---|---|---|---|---|---|---|---|---|---|
| Igkv3-9 | 1.803734 | 1.890235 | 1.8235 | 0.06673027 | -0.019770944 | 1.9475802 | 1.847938 | 1.74878 | 0.099153479 | 0.198795674 |
| D5Ertd579e | 9.354127 | 9.359324 | 9.3038 | 0.055538096 | 0.050340859 | 9.3641959 | 9.3865645 | 9.28748 | 0.099082522 | 0.076713882 |
| Nras | 10.88138 | 10.99833 | 10.907 | 0.090908896 | -0.026039806 | 10.910245 | 10.996768 | 10.8977 | 0.099078686 | 0.012556541 |
| Gm6335 | 0.779034 | 0.888968 | 0.7917 | 0.097271606 | -0.012663106 | 0.7877094 | 0.8780045 | 0.77893 | 0.099072515 | 0.008777422 |
| Gm14295 | 5.691916 | 5.671993 | 5.6162 | 0.055755614 | 0.075677982 | 5.6585077 | 5.613928 | 5.51486 | 0.099071671 | 0.143651281 |
| Ppp1r8 | 9.449224 | 9.535459 | 9.5066 | 0.028891561 | -0.057342987 | 9.4540868 | 9.570653 | 9.47174 | 0.09891235 | -0.017653822 |
| Psmb11 | 3.756616 | 3.660742 | 3.6849 | -0.02418009 | 0.071694388 | 3.7688498 | 3.866046 | 3.76714 | 0.098909915 | 0.001713771 |
| Armcx3 | 8.075962 | 8.098393 | 7.978 | 0.120408351 | 0.09797737 | 7.959993 | 8.0858374 | 7.98699 | 0.09884761 | -0.02699675 |
| Sema3b | 4.932425 | 4.559162 | 4.5359 | 0.023311052 | 0.396574244 | 4.5315164 | 4.6023629 | 4.50367 | 0.098692511 | 0.027846011 |
| Chst11 | 8.523699 | 8.178418 | 8.0608 | 0.117663392 | 0.46294395 | 8.0558113 | 7.9915845 | 7.89294 | 0.098641475 | 0.162868299 |
| Olfr1124 | 2.289709 | 2.26253 | 2.302 | -0.039519699 | -0.012341201 | 2.3036981 | 2.3902434 | 2.29164 | 0.098607117 | 0.01206176 |
| Slc25a3 | 12.46139 | 12.47016 | 12.502 | -0.031806504 | -0.040580458 | 12.587237 | 12.632604 | 12.534 | 0.09857889 | 0.053211236 |
| Gm17131 | 6.425896 | 6.54318 | 6.3695 | 0.173676039 | 0.05639193 | 6.6294831 | 7.0713393 | 6.97281 | 0.098528928 | -0.343327268 |
| Gm49383 | 5.469109 | 5.386895 | 5.5748 | -0.187874886 | -0.105661589 | 5.5254511 | 5.5460788 | 5.44769 | 0.098390183 | 0.077762453 |
| Strn4 | 10.28354 | 10.46932 | 10.395 | 0.074109588 | -0.111670188 | 10.611078 | 10.637872 | 10.5395 | 0.098386046 | 0.071591528 |
| Naaladl2 | 1.974891 | 1.948597 | 1.9452 | 0.003405395 | 0.029698709 | 1.9527094 | 1.9800779 | 1.8817 | 0.098375394 | 0.071006934 |
| Gm37108 | 4.814559 | 4.899824 | 4.9834 | -0.083576321 | -0.168840972 | 5.0407292 | 5.1027689 | 5.00445 | 0.098315656 | 0.03627589 |
| Lhx5 | 1.438231 | 1.415747 | 1.3727 | 0.043091167 | 0.065574555 | 1.4032828 | 1.5233095 | 1.42505 | 0.098259553 | -0.021767141 |
| Gm49327 | 3.388218 | 3.223885 | 3.4113 | -0.187433736 | -0.023100135 | 3.199828 | 3.3534615 | 3.25523 | 0.098236115 | -0.055397407 |
| Tymp | 1.614085 | 1.542809 | 1.6351 | -0.0922744 | -0.020998463 | 1.5626818 | 1.6552814 | 1.55705 | 0.098228982 | 0.005629347 |
| Gm3788 | 2.602921 | 2.590428 | 2.7868 | -0.196403083 | -0.18391078 | 2.8657006 | 2.8842158 | 2.78607 | 0.098141233 | 0.079626002 |
| Cdk15 | 2.273835 | 2.309069 | 2.3451 | -0.036073515 | -0.071306622 | 2.2289379 | 2.4237227 | 2.3256 | 0.098121571 | -0.09666314 |
| Hist1h4h | 4.430985 | 4.319642 | 4.3893 | -0.069635344 | 0.041708581 | 4.5660078 | 4.5303325 | 4.43228 | 0.09805292 | 0.133728156 |
| Setd7 | 10.19084 | 10.21788 | 10.191 | 0.02649939 | -0.000540545 | 10.002272 | 10.378387 | 10.2804 | 0.098000054 | -0.278115237 |
| Rab15 | 2.129963 | 2.014616 | 2.0615 | -0.046887602 | 0.068460088 | 2.163548 | 2.1333652 | 2.03538 | 0.097984688 | 0.128167486 |
| Maob | 3.741916 | 3.84639 | 3.7885 | 0.057912874 | -0.046560751 | 3.4859375 | 3.7588502 | 3.66088 | 0.097970016 | -0.174942628 |
| Tfcp2l1 | 5.656944 | 4.862924 | 5.3644 | -0.501458789 | 0.292560443 | 5.8177887 | 5.2992248 | 5.20127 | 0.097959372 | 0.61652321 |
| Dennd4b | 11.52217 | 11.57129 | 11.469 | 0.102310388 | 0.053188414 | 11.874527 | 11.741061 | 11.6432 | 0.09782216 | 0.231288392 |
| Chrnb2 | 6.049816 | 5.964058 | 6.1547 | -0.190667615 | -0.104908749 | 6.0514863 | 6.3858475 | 6.28807 | 0.097775686 | -0.236585491 |
| 1110004F10Rik | 10.07035 | 10.09701 | 10.064 | 0.032530835 | 0.005878464 | 10.138444 | 10.149507 | 10.0518 | 0.097721291 | 0.086658257 |
| 1700037C18Rik | 5.979332 | 5.826459 | 5.9033 | -0.076824623 | 0.076048265 | 6.178268 | 6.011841 | 5.91415 | 0.097691326 | 0.26411839 |
| Klrd1 | 8.95296 | 8.937915 | 9.0813 | -0.143335853 | -0.128290559 | 8.9065464 | 9.167385 | 9.06974 | 0.097640643 | -0.163197962 |

| | | | | | | | | | | |
|---|---|---|---|---|---|---|---|---|---|---|
| Cabyr | 1.822878 | 1.832733 | 1.8006 | 0.032131052 | 0.022275394 | 1.8747788 | 1.9378208 | 1.8402 | 0.097622347 | 0.034580356 |
| Gm12188 | 1.206909 | 1.178589 | 1.2179 | -0.039283199 | -0.010963556 | 1.1599268 | 1.328084 | 1.23048 | 0.097607721 | -0.070549498 |
| Dll3 | 0.759148 | 0.704974 | 0.7852 | -0.080250131 | -0.026075982 | 0.7367544 | 0.8408761 | 0.74328 | 0.097596589 | -0.006525072 |
| Nsmce1 | 9.574379 | 9.39727 | 9.4605 | -0.063218636 | 0.113889881 | 9.6714622 | 9.5869784 | 9.48944 | 0.097534535 | 0.182018376 |
| Taf15 | 11.32651 | 11.29052 | 11.31 | -0.019938004 | 0.016051812 | 11.329774 | 11.575766 | 11.4783 | 0.097467242 | -0.148525237 |
| Ak9 | 3.296158 | 3.344541 | 3.3689 | -0.024337328 | -0.072720584 | 3.3964951 | 3.6476851 | 3.55025 | 0.097436062 | -0.153753907 |
| G3bp1 | 12.06886 | 12.10228 | 12.011 | 0.091465964 | 0.058047148 | 12.152668 | 12.167057 | 12.0696 | 0.097415888 | 0.083026311 |
| Arfgap1 | 9.817253 | 9.859366 | 9.8816 | -0.022197014 | -0.064310496 | 9.8004556 | 9.8822943 | 9.7849 | 0.097396579 | 0.015557922 |
| Btnl4 | 1.643941 | 1.650213 | 1.6292 | 0.02104318 | 0.014770828 | 1.7173773 | 1.7994591 | 1.70207 | 0.097389555 | 0.015307679 |
| Gm16470 | 0.638004 | 0.629377 | 0.5946 | 0.034799648 | 0.043427337 | 0.6082635 | 0.6829063 | 0.58559 | 0.097315148 | 0.022672349 |
| Sbds | 9.700218 | 9.669032 | 9.6956 | -0.026578503 | 0.004607695 | 9.9376231 | 9.9601147 | 9.86296 | 0.097153028 | 0.074661378 |
| Gm15539 | 2.460473 | 2.431049 | 2.3913 | 0.039705112 | 0.06912867 | 2.4702397 | 2.5456302 | 2.44858 | 0.097051926 | 0.021661465 |
| Atp5a1 | 12.11596 | 12.1029 | 12.1 | 0.003139062 | 0.016198124 | 12.205533 | 12.176858 | 12.0798 | 0.097013346 | 0.125687847 |
| Usp47 | 10.87036 | 10.84777 | 10.863 | -0.015160248 | 0.00743725 | 10.660423 | 10.936308 | 10.8393 | 0.0969942 | -0.178890582 |
| Smg9 | 9.299834 | 9.345408 | 9.3189 | 0.026516674 | -0.019057601 | 9.6208404 | 9.6309364 | 9.53398 | 0.096957898 | 0.086861928 |
| Rnaseh2c | 7.231604 | 7.483671 | 7.527 | -0.043300069 | -0.295367465 | 7.7631474 | 7.4090088 | 7.31212 | 0.09689375 | 0.451032357 |
| Rpl36a-ps3 | 1.143346 | 1.065387 | 1.0718 | -0.00640051 | 0.071558054 | 1.070563 | 1.1918427 | 1.09496 | 0.096883683 | -0.024396042 |
| Olfr64 | 1.439213 | 1.488108 | 1.4129 | 0.075169031 | 0.026273889 | 1.4201924 | 1.5616608 | 1.4648 | 0.096859423 | -0.044608993 |
| Tead4 | 3.891741 | 3.673209 | 3.6983 | -0.025084518 | 0.193447964 | 3.5258656 | 3.712723 | 3.61601 | 0.09671268 | -0.09014467 |
| Nek11 | 2.404267 | 2.391142 | 2.3438 | 0.047391006 | 0.060516791 | 2.4559836 | 2.5327817 | 2.43618 | 0.096604778 | 0.019806736 |
| Rtca | 8.487816 | 8.454282 | 8.5324 | -0.078076521 | -0.044543037 | 8.5779359 | 8.6666326 | 8.57003 | 0.096598722 | 0.007902029 |
| Tmem138 | 6.777507 | 6.876559 | 6.8751 | 0.00146276 | -0.097589769 | 7.1504321 | 7.0779615 | 6.98138 | 0.096583337 | 0.169053965 |
| Gm8349 | 2.035128 | 2.081454 | 2.0348 | 0.046667701 | 0.00034139 | 2.116027 | 2.1758322 | 2.07944 | 0.096391756 | 0.036586475 |
| Gm6682 | 2.306093 | 2.311368 | 2.2251 | 0.08631462 | 0.081039578 | 2.2922127 | 2.2795869 | 2.18321 | 0.096378378 | 0.109004155 |
| Gm49721 | 2.351738 | 2.557913 | 2.4735 | 0.084390987 | -0.121784449 | 2.3338093 | 2.6814088 | 2.58504 | 0.096367775 | -0.251231748 |
| Gm8839 | 0.977994 | 1.039492 | 1.0182 | 0.021265063 | -0.040232263 | 0.9992045 | 1.1578152 | 1.06147 | 0.096347439 | -0.062263238 |
| Hist1h3c | 2.228474 | 2.100999 | 2.2175 | -0.116456796 | 0.011018463 | 2.2885352 | 2.205423 | 2.1091 | 0.096321862 | 0.179434092 |
| Dcun1d1 | 9.905575 | 10.04855 | 9.9056 | 0.142981956 | 2.4622E-06 | 9.7417803 | 9.862296 | 9.76598 | 0.096319654 | -0.024196118 |
| Olfr904 | 2.341183 | 2.308981 | 2.4288 | -0.1198556 | -0.087654185 | 2.3851851 | 2.4406715 | 2.34437 | 0.096296627 | 0.040810151 |
| Zfp780b | 8.167625 | 8.280913 | 8.2691 | 0.011859688 | -0.101427641 | 8.1709316 | 8.3648783 | 8.26859 | 0.096284823 | -0.097661878 |
| Bcorl1 | 7.525158 | 7.80373 | 7.8507 | -0.046976785 | -0.325548376 | 6.9539182 | 7.2510593 | 7.1548 | 0.096263553 | -0.200877581 |
| Cyb5r3 | 9.645541 | 9.517187 | 9.6886 | -0.171401613 | -0.043047701 | 9.734819 | 9.5853542 | 9.4891 | 0.096255741 | 0.245720591 |

| | | | | | | | | | | |
|---|---|---|---|---|---|---|---|---|---|---|
| Lrpprc | 10.42726 | 10.41925 | 10.382 | 0.037408458 | 0.045425202 | 10.564493 | 10.581496 | 10.4853 | 0.096240864 | 0.079238013 |
| Tmprss4 | 1.438417 | 1.386822 | 1.472 | -0.085189837 | -0.03359472 | 1.4162023 | 1.5713007 | 1.47515 | 0.096149009 | -0.058949341 |
| Ncdn | 9.584457 | 9.64069 | 9.5408 | 0.099916293 | 0.043682406 | 9.7461488 | 9.5647419 | 9.46868 | 0.096063472 | 0.277470423 |
| Homer1 | 8.043052 | 8.01243 | 7.7541 | 0.25830161 | 0.288923334 | 7.5610843 | 7.9065956 | 7.81054 | 0.096053915 | -0.249457394 |
| Slc17a8 | 2.376247 | 2.488225 | 2.3602 | 0.128025114 | 0.016047549 | 2.4005177 | 2.5005139 | 2.40448 | 0.096036478 | -0.003959638 |
| Rnf168 | 8.312622 | 8.398766 | 8.3473 | 0.051418638 | -0.034725696 | 8.2678746 | 8.4161542 | 8.32013 | 0.096025511 | -0.052254092 |
| Alkbh2 | 5.958987 | 6.168046 | 6.1378 | 0.030265364 | -0.1787938 | 6.1727404 | 6.1612103 | 6.06528 | 0.095926374 | 0.107456448 |
| Gm9442 | 1.797506 | 1.696413 | 1.6754 | 0.021057272 | 0.122150703 | 1.7386297 | 1.7819786 | 1.68606 | 0.095918395 | 0.052569494 |
| Gm21988 | 6.228177 | 6.270498 | 6.216 | 0.05447578 | 0.012154393 | 6.2785517 | 6.0794029 | 5.98365 | 0.09575203 | 0.294900848 |
| Gm6736 | 0.642115 | 0.66606 | 0.6404 | 0.025681448 | 0.00173655 | 0.6959523 | 0.6895812 | 0.59388 | 0.095700812 | 0.102071922 |
| Has1 | 1.195194 | 1.134413 | 1.0292 | 0.105182459 | 0.165964017 | 1.0414415 | 1.0975025 | 1.0018 | 0.095699292 | 0.039638358 |
| Apln | 4.444215 | 4.470404 | 4.1444 | 0.326053079 | 0.299863659 | 4.3006811 | 4.3652699 | 4.26962 | 0.09565478 | 0.031065964 |
| Msra | 8.080113 | 8.15776 | 8.0719 | 0.085820135 | 0.008172825 | 8.1266084 | 8.1596349 | 8.06423 | 0.095404281 | 0.062377792 |
| Gm47201 | 0.924887 | 0.930226 | 0.932 | -0.001739828 | -0.007078714 | 0.9373411 | 0.9577422 | 0.86235 | 0.095387619 | 0.074986516 |
| Smu1 | 10.75779 | 10.67002 | 10.716 | -0.04572588 | 0.042044666 | 10.789466 | 10.7737 | 10.6784 | 0.095281168 | 0.111046988 |
| Atp8b2 | 11.04176 | 10.89544 | 10.785 | 0.110133094 | 0.256446908 | 10.853209 | 11.105108 | 11.0098 | 0.095259914 | -0.156639456 |
| Pde6d | 7.846095 | 7.929508 | 7.8702 | 0.059309608 | -0.024102805 | 8.0438219 | 8.0112607 | 7.91616 | 0.095103485 | 0.127664707 |
| Ipcef1 | 11.14224 | 11.25322 | 11.256 | -0.00305899 | -0.114044772 | 10.810038 | 11.278476 | 11.1834 | 0.095073156 | -0.373364843 |
| Rpl23a-ps2 | 1.129799 | 1.133842 | 1.1461 | -0.012306049 | -0.016349229 | 1.1403772 | 1.2413721 | 1.14633 | 0.095041368 | -0.005953522 |
| Ccdc116 | 5.124371 | 5.403858 | 5.2588 | 0.145018162 | -0.134469041 | 5.418575 | 5.4699861 | 5.37498 | 0.095008121 | 0.043597028 |
| Ehd3 | 10.73753 | 10.63423 | 10.612 | 0.022287256 | 0.125583596 | 10.163985 | 10.48619 | 10.3912 | 0.094975606 | -0.227228936 |
| Itpripl2 | 8.295981 | 8.394183 | 8.2006 | 0.193601245 | 0.095398595 | 7.9432382 | 8.0901885 | 7.99524 | 0.094948522 | -0.052001799 |
| Ighv1-49 | 3.731048 | 3.609592 | 3.6335 | -0.023896787 | 0.097559133 | 3.6590294 | 3.8039433 | 3.70905 | 0.094891875 | -0.050021995 |
| Lamtor2 | 9.164263 | 9.108442 | 9.1526 | -0.044141802 | 0.011678973 | 9.3335144 | 9.2146656 | 9.11981 | 0.094851086 | 0.213699896 |
| Zfp384 | 10.25738 | 10.32847 | 10.292 | 0.036415202 | -0.03466998 | 10.327666 | 10.390811 | 10.296 | 0.094798038 | 0.031652497 |
| Ankrd40 | 10.14199 | 10.18628 | 10.218 | -0.031690017 | -0.075972454 | 10.210077 | 10.311549 | 10.2168 | 0.094760298 | -0.00671169 |
| Gm14150 | 1.872016 | 1.860826 | 1.8903 | -0.029435043 | -0.018245274 | 1.9065006 | 1.9423131 | 1.84757 | 0.094744883 | 0.058932392 |
| Cabin1 | 11.11373 | 11.20978 | 11.064 | 0.145776294 | 0.049721496 | 11.253118 | 11.320703 | 11.2261 | 0.094557178 | 0.026972124 |
| Zfp949 | 7.406517 | 7.601912 | 7.5167 | 0.085172258 | -0.110223307 | 7.206388 | 7.5896953 | 7.49516 | 0.094533604 | -0.288773726 |
| Crym | 1.707644 | 1.709775 | 1.6992 | 0.010595313 | 0.008464754 | 1.7592438 | 1.7226589 | 1.62823 | 0.094433313 | 0.131018256 |
| Gm14633 | 2.586896 | 2.623194 | 2.6275 | -0.004330968 | -0.040628686 | 2.728241 | 2.7490543 | 2.65467 | 0.094387169 | 0.073573882 |
| Olfr718-ps1 | 4.808805 | 4.941905 | 4.9392 | 0.0027461 | -0.130353618 | 5.1112316 | 5.099577 | 5.0052 | 0.094372258 | 0.106026909 |

| | | | | | | | | | | |
|---|---|---|---|---|---|---|---|---|---|---|
| E4f1 | 9.396295 | 9.495547 | 9.4509 | 0.044678961 | -0.054572675 | 9.6925217 | 9.6901612 | 9.59581 | 0.09435547 | 0.096715937 |
| Rabac1 | 10.01339 | 10.12845 | 10.082 | 0.046513925 | -0.068554561 | 10.344744 | 10.049726 | 9.9554 | 0.094326686 | 0.389343856 |
| Adamts1 | 9.696423 | 9.193102 | 8.6049 | 0.58819966 | 1.091521117 | 7.7771219 | 8.282815 | 8.18851 | 0.094308289 | -0.411384845 |
| Olfr1246 | 2.89094 | 2.9442 | 2.8405 | 0.10369305 | 0.050433335 | 2.8745046 | 2.9950296 | 2.90074 | 0.094289704 | -0.026235245 |
| Babam1 | 9.823182 | 9.865866 | 9.7986 | 0.067229579 | 0.02454606 | 10.131996 | 10.015619 | 9.92134 | 0.094277431 | 0.210654427 |
| Xpo5 | 10.3145 | 10.33701 | 10.349 | -0.012239624 | -0.03474695 | 10.364823 | 10.420124 | 10.3259 | 0.094174924 | 0.038874628 |
| Meaf6 | 8.61276 | 8.71885 | 8.6788 | 0.040082567 | -0.066006815 | 8.9677378 | 8.9497756 | 8.85561 | 0.094162048 | 0.112124307 |
| Smarcc1 | 10.18888 | 10.16566 | 10.205 | -0.039746239 | -0.016528576 | 10.197236 | 10.326253 | 10.2321 | 0.094109332 | -0.034907398 |
| Zfp618 | 2.81943 | 2.866693 | 2.8412 | 0.02551625 | -0.021746415 | 2.7484661 | 2.9943842 | 2.90033 | 0.094054039 | -0.151864076 |
| Rpl19-ps12 | 2.047603 | 2.087989 | 2.0479 | 0.040114129 | -0.000271574 | 2.058739 | 2.0891406 | 1.99509 | 0.094045987 | 0.063644439 |
| Naxe | 8.231245 | 8.19149 | 8.2377 | -0.046233024 | -0.006477234 | 8.3434577 | 8.1880695 | 8.09406 | 0.094009364 | 0.249397587 |
| Spsb3 | 8.833163 | 8.901995 | 8.9615 | -0.059514113 | -0.128345679 | 9.0746815 | 9.0644753 | 8.97047 | 0.094006748 | 0.104212964 |
| Rpl27-ps1 | 3.094093 | 3.186893 | 3.0129 | 0.174039291 | 0.081239985 | 2.9603697 | 3.1084058 | 3.01445 | 0.093959393 | -0.054076706 |
| Fars2 | 7.668106 | 7.827123 | 7.8498 | -0.022666018 | -0.18168242 | 8.0602991 | 8.078593 | 7.98474 | 0.093855476 | 0.075561574 |
| Dmc1 | 0.833187 | 0.772377 | 0.812 | -0.039624831 | 0.02118514 | 0.7780233 | 0.8656031 | 0.77177 | 0.093836905 | 0.006257078 |
| Gemin8 | 6.431831 | 6.466313 | 6.4623 | 0.003968657 | -0.030513706 | 6.361086 | 6.5872421 | 6.49341 | 0.093836657 | -0.132319439 |
| Purg | 5.034289 | 5.061943 | 5.0981 | -0.036139232 | -0.063793928 | 5.126667 | 5.1707458 | 5.07693 | 0.093819022 | 0.049740275 |
| Gm7030 | 2.457604 | 2.38515 | 2.475 | -0.089863754 | -0.017409276 | 2.4283948 | 2.552002 | 2.4582 | 0.093802663 | -0.029804616 |
| Ggh | 7.793957 | 7.615849 | 7.6558 | -0.039959178 | 0.138148531 | 7.6801813 | 7.7140732 | 7.62028 | 0.093793069 | 0.059901163 |
| Dnajc3 | 11.1006 | 11.14804 | 11.085 | 0.063326202 | 0.015889248 | 10.86587 | 11.050487 | 10.9567 | 0.093770506 | -0.090846447 |
| Ttc12 | 3.399038 | 3.365057 | 3.3568 | 0.008214585 | 0.042195453 | 3.4025952 | 3.4756822 | 3.38195 | 0.093734108 | 0.020647106 |
| Gm5822 | 1.792261 | 1.723985 | 1.7328 | -0.008847101 | 0.059429624 | 1.7582348 | 1.7361223 | 1.64259 | 0.093530856 | 0.115643263 |
| Zbtb3 | 5.02348 | 5.235304 | 5.31 | -0.07469452 | -0.286518281 | 5.3503233 | 5.2119965 | 5.11848 | 0.093513495 | 0.231840226 |
| Taf6 | 9.027922 | 9.19259 | 9.16 | 0.032575294 | -0.132092442 | 9.2220976 | 9.3825958 | 9.2891 | 0.093495126 | -0.067003064 |
| Gm19220 | 8.182868 | 8.142056 | 8.1631 | -0.021057639 | 0.01975413 | 8.7029668 | 8.5739526 | 8.48048 | 0.093475107 | 0.222489291 |
| Gtpbp4 | 10.81391 | 10.73432 | 10.637 | 0.097179818 | 0.176761762 | 10.749357 | 10.881745 | 10.7883 | 0.093449006 | -0.038939481 |
| Gm4977 | 3.727535 | 3.512827 | 3.4185 | 0.094313454 | 0.309020793 | 3.4107795 | 3.4535674 | 3.36013 | 0.093438857 | 0.050650911 |
| Sobp | 1.919556 | 1.965168 | 1.9011 | 0.064043629 | 0.018431698 | 1.9230457 | 1.995417 | 1.9021 | 0.0933203 | 0.020948982 |
| Itgal | 12.91164 | 12.99601 | 12.881 | 0.114822048 | 0.03045163 | 12.935777 | 13.051757 | 12.9585 | 0.093274572 | -0.02270535 |
| Rita1 | 5.70492 | 5.906324 | 5.6102 | 0.296121422 | 0.094717567 | 5.8858785 | 5.9362187 | 5.84296 | 0.093254702 | 0.042914468 |
| Trav6d-5 | 2.61815 | 2.654019 | 2.594 | 0.060034998 | 0.024165971 | 2.6024595 | 2.672705 | 2.57951 | 0.093194096 | 0.022948657 |
| Zdhhc4 | 8.728212 | 8.695036 | 8.7945 | -0.099460469 | -0.06628461 | 8.9288943 | 8.9800593 | 8.88687 | 0.093192612 | 0.042027544 |

| | | | | | | | | | | |
|---|---|---|---|---|---|---|---|---|---|---|
| Myl12a | 11.48236 | 11.45526 | 11.422 | 0.032880416 | 0.059983126 | 11.599807 | 11.4942 | 11.4011 | 0.09308077 | 0.198687846 |
| Gpa33 | 1.284187 | 1.183953 | 1.2427 | -0.058704604 | 0.041529477 | 1.1277831 | 1.257511 | 1.16446 | 0.093050584 | -0.036677334 |
| Rpl17 | 14.72057 | 14.71468 | 14.715 | -0.000398768 | 0.005485488 | 14.78862 | 14.809906 | 14.7169 | 0.09298829 | 0.071702862 |
| Gpr89 | 8.472849 | 8.573341 | 8.4573 | 0.116036057 | 0.015543427 | 8.4948449 | 8.629002 | 8.53602 | 0.092978934 | -0.041178171 |
| Ankrd54 | 8.439829 | 8.574301 | 8.581 | -0.006733867 | -0.141205093 | 8.7861642 | 8.699154 | 8.60622 | 0.09293329 | 0.179943505 |
| Siglece | 5.591604 | 5.658604 | 4.7443 | 0.914269445 | 0.847269972 | 5.6896833 | 4.9121662 | 4.81927 | 0.092892348 | 0.870409422 |
| Gm12657 | 1.198769 | 1.216533 | 1.2147 | 0.001836279 | -0.015928155 | 1.2434071 | 1.3008531 | 1.20811 | 0.092747604 | 0.035301577 |
| Tars2 | 9.805798 | 9.859667 | 9.833 | 0.026626079 | -0.027242621 | 10.052386 | 10.143108 | 10.0504 | 0.092720712 | 0.001998443 |
| Gm8738 | 2.784908 | 2.819912 | 2.7642 | 0.05572468 | 0.020720214 | 2.7529041 | 2.8760339 | 2.78336 | 0.092672546 | -0.030457299 |
| Gm18113 | 2.903956 | 2.851766 | 2.787 | 0.064778708 | 0.116968633 | 2.9014499 | 2.8838366 | 2.79119 | 0.092645723 | 0.110259002 |
| Poldip2 | 8.748016 | 8.600817 | 8.7842 | -0.183339419 | -0.036140439 | 8.9114765 | 8.8444746 | 8.75185 | 0.092627392 | 0.159629268 |
| Rps2-ps7 | 1.685599 | 1.735026 | 1.726 | 0.009008714 | -0.040418358 | 1.7616089 | 1.7968305 | 1.70423 | 0.092604033 | 0.057382453 |
| Hist2h2ac | 4.500174 | 4.492833 | 4.536 | -0.043172368 | -0.035830773 | 4.6831558 | 4.6595408 | 4.567 | 0.092545381 | 0.116160375 |
| Anp32a | 12.0514 | 12.01235 | 12.099 | -0.086322052 | -0.047266494 | 11.967583 | 12.178551 | 12.086 | 0.092529284 | -0.118438403 |
| Hist2h3c2 | 2.559216 | 2.507323 | 2.5142 | -0.006894465 | 0.044998634 | 2.5447746 | 2.6171377 | 2.52467 | 0.092464014 | 0.020100954 |
| Naxd | 7.862942 | 7.847618 | 7.8261 | 0.021494037 | 0.036817974 | 8.2314038 | 8.2200305 | 8.12758 | 0.092445649 | 0.103819024 |
| Gm6807 | 2.98345 | 3.082413 | 2.9639 | 0.118477308 | 0.019514441 | 3.0019163 | 3.0978751 | 3.00553 | 0.09234776 | -0.00361111 |
| Syne2 | 9.313171 | 9.293592 | 9.3546 | -0.061057804 | -0.041479114 | 9.2883568 | 9.5022226 | 9.40989 | 0.092335499 | -0.121530289 |
| Bag3 | 7.667907 | 7.614615 | 7.5353 | 0.079338877 | 0.132630655 | 7.0116313 | 7.4721755 | 7.37991 | 0.09226485 | -0.368279302 |
| Xrcc3 | 5.550738 | 5.746288 | 5.9167 | -0.170389198 | -0.365939441 | 6.1103066 | 6.101955 | 6.0097 | 0.092259005 | 0.100610586 |
| Cisd1 | 6.779375 | 6.849269 | 6.9219 | -0.072629507 | -0.14252353 | 6.829204 | 7.0265485 | 6.93432 | 0.092226062 | -0.105118368 |
| Prx | 4.669584 | 4.545372 | 4.6314 | -0.085989165 | 0.038223104 | 4.531509 | 4.5661157 | 4.4739 | 0.092216756 | 0.057610117 |
| Pex11g | 4.813513 | 4.637346 | 4.7259 | -0.088550097 | 0.087616563 | 5.07487 | 4.8328589 | 4.74068 | 0.092183847 | 0.334194973 |
| Tdgf1 | 3.785223 | 3.712201 | 3.8012 | -0.089038008 | -0.016016529 | 3.7767574 | 3.8830441 | 3.79087 | 0.092178619 | -0.014108119 |
| Mall | 3.82201 | 3.747801 | 3.7226 | 0.025219134 | 0.099428188 | 3.7309887 | 3.7621173 | 3.67006 | 0.092060479 | 0.060931883 |
| Cyb5rl | 7.176986 | 7.482611 | 7.3394 | 0.143231979 | -0.162393704 | 7.7060459 | 7.6112508 | 7.51924 | 0.092006186 | 0.186801267 |
| Trdj1 | 2.256775 | 2.253341 | 2.2737 | -0.020356803 | -0.0169227 | 2.178227 | 2.3327738 | 2.24086 | 0.091915144 | -0.062631682 |
| Gm18997 | 2.66713 | 2.615813 | 2.6328 | -0.01695082 | 0.034366587 | 2.7354297 | 2.7371599 | 2.64527 | 0.091887851 | 0.090157624 |
| Sptlc1 | 9.438068 | 9.469253 | 9.5316 | -0.062357293 | -0.093542383 | 9.6386828 | 9.532882 | 9.44099 | 0.09188701 | 0.197687846 |
| Hcfc1r1 | 8.184387 | 8.210268 | 8.3389 | -0.128616724 | -0.154498301 | 8.524613 | 8.2711642 | 8.1793 | 0.091867466 | 0.345316237 |
| Cdc42ep5 | 2.297918 | 2.219483 | 2.2079 | 0.011594667 | 0.090030173 | 2.1193179 | 2.2407331 | 2.14887 | 0.091866046 | -0.029549075 |
| Hs3st6 | 1.948646 | 1.847093 | 1.943 | -0.095934254 | 0.005618976 | 1.8346983 | 1.9341007 | 1.84225 | 0.091850375 | -0.007552054 |

| | | | | | | | | | | |
|---|---|---|---|---|---|---|---|---|---|---|
| Crk | 10.40734 | 10.49895 | 10.34 | 0.158510104 | 0.066895102 | 10.327614 | 10.431306 | 10.3395 | 0.091818632 | -0.011873488 |
| Efna5 | 3.155899 | 3.256576 | 3.3122 | -0.055619479 | -0.156296851 | 3.2746252 | 3.2194077 | 3.12775 | 0.091654511 | 0.146872007 |
| Scamp4 | 9.595843 | 9.634035 | 9.5529 | 0.081094815 | 0.042903283 | 9.6752539 | 9.6383533 | 9.54672 | 0.091634891 | 0.128535501 |
| Grm1 | 2.876581 | 2.982899 | 2.9787 | 0.004210423 | -0.10210786 | 3.0293071 | 3.0420654 | 2.95046 | 0.091605234 | 0.078846991 |
| Gm5898 | 1.669941 | 1.736846 | 1.6944 | 0.042401062 | -0.024503547 | 1.6968 | 1.6599329 | 1.56852 | 0.091415284 | 0.128282358 |
| Hist1h2ag | 1.852544 | 1.751531 | 1.8131 | -0.061562392 | 0.039450529 | 1.7478682 | 1.8177399 | 1.72639 | 0.091348019 | 0.021476306 |
| Unc5cl | 5.969402 | 5.959615 | 5.9327 | 0.026870603 | 0.036657864 | 5.8971603 | 5.9602899 | 5.86898 | 0.091307514 | 0.028177952 |
| Rpsa-ps9 | 4.640879 | 4.731495 | 4.618 | 0.11347734 | 0.022861597 | 4.8283925 | 4.8046646 | 4.71336 | 0.091306473 | 0.115034382 |
| Hormad2 | 1.443709 | 1.424963 | 1.4835 | -0.058497483 | -0.039751368 | 1.4096361 | 1.4963867 | 1.40508 | 0.091304184 | 0.004553576 |
| Tardbp | 11.65425 | 11.78407 | 11.646 | 0.138198372 | 0.008383046 | 11.931862 | 12.012476 | 11.9212 | 0.091299606 | 0.010685549 |
| Rnf181 | 8.661438 | 8.657224 | 8.758 | -0.100732438 | -0.096518453 | 8.9041967 | 8.839866 | 8.74858 | 0.091290642 | 0.155621288 |
| Olfr173 | 2.346044 | 2.35184 | 2.2011 | 0.150753918 | 0.144958407 | 2.2996212 | 2.4621225 | 2.37084 | 0.091278281 | -0.07122301 |
| Gm11362 | 0.119114 | 0.133677 | 0.0866 | 0.047069114 | 0.032505876 | 0.0799227 | 0.1574285 | 0.06617 | 0.091255674 | 0.013749929 |
| Olfr1122 | 2.311592 | 2.277239 | 2.3624 | -0.085161363 | -0.050808003 | 2.3315603 | 2.4278559 | 2.33661 | 0.091250517 | -0.005045122 |
| Pus3 | 8.216728 | 8.308839 | 8.2231 | 0.08578668 | -0.006324303 | 8.356917 | 8.3552751 | 8.2641 | 0.091171251 | 0.092813193 |
| Lrrc63 | 1.903004 | 1.979311 | 1.9193 | 0.05998507 | -0.016321994 | 2.0161198 | 2.0375879 | 1.94643 | 0.091155132 | 0.069687026 |
| Caln1 | 2.199247 | 2.236007 | 2.2177 | 0.018347698 | -0.018412403 | 2.2062918 | 2.2774884 | 2.18639 | 0.091096429 | 0.019899818 |
| Gm47759 | 0.981393 | 0.94893 | 0.9994 | -0.050504557 | -0.018041639 | 0.9831302 | 1.0552688 | 0.96417 | 0.091095344 | 0.018956725 |
| Mat2b | 10.60546 | 10.64291 | 10.644 | -0.001092104 | -0.038541288 | 10.6607 | 10.696498 | 10.6054 | 0.091048622 | 0.055250146 |
| Ccdc162 | 4.322865 | 4.478342 | 4.3158 | 0.162569497 | 0.007092578 | 4.4772747 | 4.5076102 | 4.41666 | 0.090945319 | 0.060609792 |
| Dennd2a | 5.339081 | 5.294443 | 5.3856 | -0.091199598 | -0.046561863 | 5.2204622 | 5.5562302 | 5.46532 | 0.090905771 | -0.244862232 |
| Dnajc16 | 8.573112 | 8.478345 | 8.5385 | -0.060156387 | 0.03461123 | 8.7569087 | 8.6490451 | 8.55824 | 0.090800656 | 0.19866428 |
| Matn4 | 2.432449 | 2.413544 | 2.3308 | 0.082742515 | 0.101647662 | 2.4219749 | 2.5507783 | 2.46003 | 0.090745612 | -0.038057741 |
| Olfr1245 | 2.202497 | 2.179874 | 2.197 | -0.017107668 | 0.00551532 | 2.2653516 | 2.3938641 | 2.30317 | 0.090698224 | -0.037814235 |
| Ighv1-53 | 7.778777 | 7.493672 | 7.0965 | 0.397182131 | 0.682287503 | 7.5765506 | 7.4167749 | 7.32611 | 0.09066814 | 0.250443784 |
| Zbtb9 | 8.817104 | 8.952513 | 8.9295 | 0.022984855 | -0.112424188 | 8.7738524 | 8.9282445 | 8.83762 | 0.090623021 | -0.063769028 |
| Ubtf | 11.59701 | 11.67532 | 11.608 | 0.066958728 | -0.011350576 | 11.674653 | 11.699571 | 11.609 | 0.090591234 | 0.065673175 |
| Prr12 | 10.08644 | 10.2322 | 10.251 | -0.019241493 | -0.164994944 | 10.098355 | 10.165476 | 10.075 | 0.090435132 | 0.023314044 |
| Olfr976 | 1.182169 | 1.250482 | 1.2139 | 0.03659452 | -0.031717826 | 1.3625343 | 1.316892 | 1.22654 | 0.090351154 | 0.135993511 |
| Gm48501 | 5.383657 | 5.34887 | 5.4027 | -0.053866265 | -0.019079162 | 5.5168756 | 5.7638297 | 5.67349 | 0.09033614 | -0.156617994 |
| Cyb561d1 | 9.196607 | 9.292307 | 9.184 | 0.108295803 | 0.012595877 | 9.3740202 | 9.2719409 | 9.18161 | 0.090334411 | 0.192413733 |
| Rcan1 | 8.084293 | 7.701976 | 7.6818 | 0.020144857 | 0.402462014 | 7.5535378 | 7.6441073 | 7.55384 | 0.090262811 | -0.000306678 |

| | | | | | | | | | | |
|---|---|---|---|---|---|---|---|---|---|---|
| Crygn | 1.697367 | 1.484965 | 1.637 | -0.152040555 | 0.060361778 | 1.5306378 | 1.6125152 | 1.52228 | 0.090232615 | 0.00835515 |
| Cxxc1 | 10.52481 | 10.58971 | 10.54 | 0.050134314 | -0.014764558 | 10.88945 | 10.872154 | 10.7821 | 0.090082636 | 0.107378589 |
| Zfp91 | 11.27689 | 11.34446 | 11.288 | 0.056001428 | -0.011566404 | 11.173799 | 11.361909 | 11.2718 | 0.090077452 | -0.098033079 |
| Tnfrsf12a | 6.058464 | 5.808642 | 5.8735 | -0.064861222 | 0.184961132 | 5.600633 | 5.8088014 | 5.71875 | 0.090053222 | -0.118115172 |
| P4hb | 12.10969 | 12.04639 | 12.048 | -0.00132444 | 0.061978144 | 12.08688 | 11.969289 | 11.8792 | 0.090044992 | 0.207635869 |
| Fam83e | 1.413065 | 1.378178 | 1.4461 | -0.06793579 | -0.033048801 | 1.4236639 | 1.4901989 | 1.40016 | 0.090041231 | 0.023506222 |
| Vmn1r73 | 1.535724 | 1.472855 | 1.6043 | -0.13149434 | -0.068625709 | 1.5413112 | 1.6131842 | 1.52321 | 0.089974215 | 0.018101237 |
| Adamts19 | 0.919697 | 0.918616 | 0.9618 | -0.043204956 | -0.042123671 | 0.9439852 | 1.0286195 | 0.9387 | 0.089922518 | 0.005288242 |
| Picalm | 12.27661 | 12.07544 | 11.995 | 0.080028278 | 0.28119713 | 12.002384 | 12.09706 | 12.0071 | 0.089913906 | -0.004762066 |
| Gm9887 | 1.183387 | 1.15507 | 1.0746 | 0.080508628 | 0.108825451 | 1.0892631 | 1.1720455 | 1.08215 | 0.089894932 | 0.007112552 |
| Poll | 8.170954 | 8.158912 | 8.2351 | -0.076141338 | -0.064099714 | 8.3771797 | 8.3086905 | 8.219 | 0.089685751 | 0.158174924 |
| Gpbp1l1 | 10.97559 | 10.9875 | 11.044 | -0.056561424 | -0.068472752 | 11.026049 | 11.251212 | 11.1616 | 0.0896355 | -0.135527576 |
| Slc6a20b | 2.656669 | 2.642927 | 2.6504 | -0.007440764 | 0.006301351 | 2.6882809 | 2.8396411 | 2.75009 | 0.089556055 | -0.061804068 |
| Mttp | 5.233705 | 5.072224 | 5.241 | -0.168790307 | -0.007309955 | 5.266237 | 5.4376875 | 5.34815 | 0.089542259 | -0.081908168 |
| Scg3 | 3.099635 | 3.038417 | 3.0607 | -0.022305958 | 0.038911781 | 2.9412809 | 3.0018137 | 2.91228 | 0.089537451 | 0.029004662 |
| Gm10408 | 1.616466 | 1.766661 | 1.686 | 0.08066393 | -0.069531677 | 1.8526504 | 1.6599973 | 1.5705 | 0.089498077 | 0.28215118 |
| Gm12922 | 2.04322 | 2.116773 | 2.1476 | -0.030796442 | -0.104349683 | 2.1436421 | 2.2299166 | 2.14043 | 0.089488144 | 0.003213653 |
| Pdrg1 | 9.558664 | 9.668453 | 9.5804 | 0.08801899 | -0.021769768 | 9.7268985 | 9.7083503 | 9.61887 | 0.089482695 | 0.108030873 |
| Gm17241 | 2.786723 | 2.703729 | 2.7552 | -0.0514685 | 0.031525657 | 2.8633431 | 2.8100167 | 2.72056 | 0.089456764 | 0.142783228 |
| Anxa9 | 4.171661 | 4.103312 | 4.1 | 0.003359844 | 0.07170872 | 4.3123454 | 4.2395718 | 4.15024 | 0.089328954 | 0.162102547 |
| Snx8 | 10.11025 | 10.22752 | 10.054 | 0.173325394 | 0.056049947 | 10.478321 | 10.333094 | 10.2438 | 0.089324156 | 0.234551417 |
| Got2-ps1 | 0.916726 | 0.858865 | 0.8707 | -0.011839692 | 0.046021259 | 0.9147833 | 0.9886851 | 0.89939 | 0.089297351 | 0.015395489 |
| Trem1 | 6.842011 | 6.272583 | 5.2023 | 1.070327011 | 1.639755688 | 6.1421031 | 5.2550503 | 5.16583 | 0.08921723 | 0.976270032 |
| Opcml | 2.294139 | 2.196545 | 2.2048 | -0.008285379 | 0.089308609 | 2.3150417 | 2.3095773 | 2.2204 | 0.089173611 | 0.094637997 |
| Slc17a6 | -0.430746 | -0.37322 | -0.4319 | 0.058726473 | 0.001200467 | -0.4225631 | -0.312243 | -0.4014 | 0.089171659 | -0.021148373 |
| Slc17a7 | 2.313967 | 2.343322 | 2.109 | 0.234279954 | 0.204924768 | 2.060353 | 2.2397017 | 2.15055 | 0.089152522 | -0.090196148 |
| Vmn1r68 | 4.1289 | 4.193804 | 4.0848 | 0.108998754 | 0.04409567 | 4.217424 | 4.5566251 | 4.46753 | 0.089094177 | -0.250106899 |
| Pitpnb | 9.772694 | 9.729064 | 9.68 | 0.049105214 | 0.092735805 | 9.6318331 | 9.807582 | 9.71849 | 0.089090282 | -0.086658651 |
| Prpf3 | 9.338855 | 9.33584 | 9.4194 | -0.083584259 | -0.080568757 | 9.6086014 | 9.6991738 | 9.61013 | 0.089045086 | -0.00152739 |
| Bud23 | 8.920006 | 8.920348 | 8.8326 | 0.087741742 | 0.087399777 | 9.2362496 | 9.1423653 | 9.05334 | 0.089020484 | 0.182904794 |
| Vmn1r-ps14 | 2.072583 | 2.146759 | 2.1579 | -0.011107378 | -0.085283847 | 2.1857168 | 2.1976597 | 2.10865 | 0.089010948 | 0.077068096 |
| Ctbp2 | 6.570377 | 6.497249 | 6.5539 | -0.056649067 | 0.016479089 | 6.2311598 | 6.5085465 | 6.41958 | 0.08896786 | -0.188418863 |

| | | | | | | | | | | |
|---|---|---|---|---|---|---|---|---|---|---|
| Ywhab | 11.84404 | 11.94913 | 11.852 | 0.096822778 | -0.008261482 | 11.842437 | 11.87076 | 11.7818 | 0.08892463 | 0.060602082 |
| Cldnd2 | 2.780249 | 2.827823 | 2.7901 | 0.037686885 | -0.009886388 | 2.9010871 | 2.936382 | 2.84748 | 0.088902668 | 0.053607736 |
| Fam173a | 8.868273 | 8.829146 | 8.9594 | -0.130232956 | -0.091106437 | 9.1717503 | 8.9165208 | 8.82763 | 0.088890606 | 0.344120081 |
| Ogg1 | 7.211245 | 7.146997 | 7.2348 | -0.087811438 | -0.023563185 | 7.5419425 | 7.3140956 | 7.22522 | 0.088870757 | 0.316717657 |
| Wdr5 | 9.884785 | 9.917943 | 9.8327 | 0.085253084 | 0.052095434 | 9.9841118 | 9.9931697 | 9.90438 | 0.088793296 | 0.079735475 |
| Ppp1r3b | 6.284152 | 6.063955 | 6.0222 | 0.041795264 | 0.261992319 | 6.0203127 | 6.0692801 | 5.9806 | 0.088679352 | 0.039711949 |
| Sgk3 | 9.870846 | 10.11275 | 9.8434 | 0.269375457 | 0.027467445 | 9.9062529 | 9.930116 | 9.84145 | 0.088665123 | 0.064801964 |
| Gm10698 | 8.356896 | 8.328134 | 8.0604 | 0.267723024 | 0.296485209 | 8.0261528 | 8.0420686 | 7.9534 | 0.088664957 | 0.072749157 |
| Akt3 | 10.36131 | 10.29525 | 10.383 | -0.087373228 | -0.021314732 | 10.275344 | 10.4992 | 10.4106 | 0.088583486 | -0.135272896 |
| Olfr1109 | 1.790811 | 1.82012 | 1.8213 | -0.001166983 | -0.030476503 | 1.9127633 | 1.9564749 | 1.86791 | 0.088566555 | 0.044854942 |
| Rbbp5 | 9.975898 | 10.03273 | 9.9837 | 0.049014086 | -0.007817607 | 10.083229 | 10.182773 | 10.0943 | 0.088503688 | -0.011040807 |
| Dus1l | 9.324407 | 9.271572 | 9.3447 | -0.073164981 | -0.020330686 | 9.6509144 | 9.5687903 | 9.4803 | 0.088486431 | 0.170610547 |
| Rps6-ps1 | 2.026032 | 1.967584 | 1.957 | 0.010563781 | 0.069011439 | 1.9526004 | 2.0773812 | 1.98893 | 0.08845234 | -0.036328514 |
| Igkv4-78 | 2.902528 | 2.94699 | 3.0111 | -0.064128012 | -0.108589764 | 3.0350887 | 3.0639344 | 2.97549 | 0.08844325 | 0.059597527 |
| Myadm | 9.542637 | 9.509987 | 9.4273 | 0.08272945 | 0.115379029 | 9.6151857 | 9.427032 | 9.3386 | 0.088436016 | 0.27658976 |
| Npnt | 4.496158 | 4.872469 | 4.4385 | 0.433950313 | 0.057638761 | 4.4222963 | 4.5680044 | 4.47966 | 0.088348531 | -0.057359517 |
| Tigd3 | 4.063538 | 4.089502 | 4.1614 | -0.071914247 | -0.097878113 | 4.0602584 | 4.2316998 | 4.14339 | 0.088313006 | -0.083128396 |
| Olfr906 | 2.926544 | 2.82586 | 2.9449 | -0.119032812 | -0.018348604 | 3.1688756 | 2.9828736 | 2.89457 | 0.088305835 | 0.274307817 |
| Gm47968 | 4.183567 | 4.213033 | 4.2358 | -0.022785943 | -0.052251825 | 4.0411684 | 4.3114655 | 4.22325 | 0.088219699 | -0.182077393 |
| Arf2 | 8.671401 | 8.493012 | 8.5589 | -0.06590044 | 0.112488615 | 8.554467 | 8.5115245 | 8.42338 | 0.088145907 | 0.131088358 |
| Tsen54 | 8.260078 | 8.258272 | 8.231 | 0.027288938 | 0.029094316 | 8.5134173 | 8.3372201 | 8.24914 | 0.088078614 | 0.264275867 |
| Rnf10 | 11.15623 | 11.23904 | 11.194 | 0.045295878 | -0.037519046 | 11.225905 | 11.313964 | 11.2261 | 0.087844862 | -0.000214949 |
| Usp40 | 7.954587 | 7.856587 | 7.9257 | -0.069067852 | 0.02893272 | 7.905042 | 8.065575 | 7.97773 | 0.087844119 | -0.072688951 |
| Phyhip | 1.67549 | 1.663113 | 1.4569 | 0.206232252 | 0.218609168 | 1.5248898 | 1.5976356 | 1.50981 | 0.08782566 | 0.015079902 |
| Vmn2r67 | 3.616223 | 3.571878 | 3.6658 | -0.093875872 | -0.049530541 | 3.6988598 | 3.8546362 | 3.76682 | 0.087819082 | -0.067957359 |
| Acta1 | 0.990601 | 0.90368 | 0.9332 | -0.029537832 | 0.05738367 | 0.9379107 | 0.9704419 | 0.88273 | 0.087711881 | 0.055180708 |
| Igkv4-54 | 2.652028 | 2.76787 | 2.6782 | 0.089624084 | -0.026217629 | 2.8611023 | 2.8533958 | 2.76571 | 0.087685018 | 0.095391485 |
| Gm5819 | 1.669892 | 1.635024 | 1.6198 | 0.015198111 | 0.050066922 | 1.5733447 | 1.7623386 | 1.67468 | 0.087659424 | -0.101334495 |
| Gm2961 | 2.547939 | 2.522468 | 2.6822 | -0.159687262 | -0.134215975 | 2.7131452 | 2.6304474 | 2.54284 | 0.087611186 | 0.17030899 |
| Mon1b | 9.438915 | 9.55059 | 9.4451 | 0.10547132 | -0.00620411 | 9.4660241 | 9.5152131 | 9.42763 | 0.087587244 | 0.038398237 |
| Pdzd2 | 7.183526 | 7.177159 | 7.2384 | -0.061231462 | -0.054863774 | 6.6581456 | 7.3492522 | 7.26169 | 0.087566462 | -0.603540086 |
| Olfr24 | 1.097971 | 1.082032 | 1.1023 | -0.020291177 | -0.004351576 | 1.215487 | 1.157914 | 1.07037 | 0.087548234 | 0.145121168 |

| | | | | | | | | | | |
|---|---|---|---|---|---|---|---|---|---|---|
| Frzb | 1.165037 | 1.517492 | 1.1332 | 0.384317119 | 0.031861807 | 1.1796932 | 1.2542206 | 1.16667 | 0.087547833 | 0.013020443 |
| Btbd3 | 7.174829 | 6.920125 | 6.9061 | 0.01399356 | 0.268697559 | 6.3886793 | 6.5102656 | 6.42277 | 0.087499544 | -0.0340868 |
| Pgpep1l | 4.630737 | 4.689042 | 4.6038 | 0.085218129 | 0.026913471 | 4.3552079 | 4.5895048 | 4.50203 | 0.087476452 | -0.146820462 |
| Limd1 | 11.44562 | 11.55034 | 11.474 | 0.07641759 | -0.028307676 | 11.556105 | 11.540197 | 11.4528 | 0.087410542 | 0.103318387 |
| Rif1 | 10.90796 | 10.98211 | 10.879 | 0.102759948 | 0.028609122 | 10.823232 | 11.083799 | 10.9964 | 0.087353962 | -0.173212461 |
| Apopt1 | 8.212785 | 8.179024 | 8.1196 | 0.059417987 | 0.093178084 | 8.3210358 | 8.1804233 | 8.09313 | 0.087294803 | 0.227907328 |
| Mafg | 8.680869 | 8.595202 | 8.7583 | -0.163093253 | -0.077425786 | 8.9013706 | 9.0177715 | 8.93049 | 0.087284103 | -0.029116808 |
| Fam129b | 8.481119 | 8.392793 | 7.9791 | 0.413665539 | 0.501991129 | 7.8782928 | 7.9074338 | 7.82015 | 0.087279139 | 0.058138137 |
| Igkv6-25 | 7.554501 | 7.94849 | 7.7843 | 0.164152726 | -0.229835954 | 8.1352756 | 7.9614447 | 7.87419 | 0.08725218 | 0.261083135 |
| Hps3 | 10.42484 | 10.49702 | 10.457 | 0.040053734 | -0.032132023 | 10.622828 | 10.604635 | 10.5175 | 0.08710453 | 0.105297397 |
| Majin | 1.555751 | 1.559543 | 1.581 | -0.021503913 | -0.025296043 | 1.5671679 | 1.6494369 | 1.56237 | 0.087063172 | 0.004794238 |
| Cryba4 | 1.673851 | 1.622596 | 1.6809 | -0.058293847 | -0.007038671 | 1.6360704 | 1.6786661 | 1.59161 | 0.087056684 | 0.044460981 |
| Bdkrb1 | 0.841249 | 0.848546 | 0.8168 | 0.031793863 | 0.02449715 | 0.8863018 | 0.9918466 | 0.90482 | 0.087027208 | -0.018517521 |
| Olfr1393 | 1.203149 | 1.247988 | 1.1729 | 0.07509391 | 0.030254504 | 1.2967889 | 1.3322768 | 1.24527 | 0.087005912 | 0.051518022 |
| Rbfox3 | 0.974266 | 0.901885 | 0.9321 | -0.030167052 | 0.042214821 | 0.8255619 | 0.9159046 | 0.82896 | 0.086946527 | -0.003396122 |
| Akt2 | 11.08681 | 10.9723 | 10.968 | 0.004576308 | 0.11907905 | 11.086431 | 11.123881 | 11.0369 | 0.086939682 | 0.049488978 |
| Rhoh | 11.59519 | 11.6842 | 11.609 | 0.075269308 | -0.01374023 | 11.318766 | 11.429683 | 11.3428 | 0.086881568 | -0.0240355 |
| Acy1 | 7.314663 | 7.102633 | 7.2421 | -0.13945339 | 0.072576642 | 7.4563057 | 7.286283 | 7.1994 | 0.086879929 | 0.256902629 |
| Gm29257 | 2.974665 | 2.911672 | 2.9087 | 0.003000793 | 0.065993096 | 3.2257991 | 3.1176806 | 3.03082 | 0.086855626 | 0.194974122 |
| Nlgn1 | 3.589247 | 3.526664 | 3.615 | -0.088322473 | -0.025739443 | 3.9425014 | 3.7883665 | 3.70152 | 0.086850067 | 0.240985055 |
| Gm5905 | 2.074211 | 2.089458 | 2.0677 | 0.021748906 | 0.006501701 | 2.1746741 | 2.1297883 | 2.04301 | 0.086782823 | 0.131668645 |
| Ero1lb | 10.9636 | 11.03803 | 10.933 | 0.104553156 | 0.030124128 | 11.140362 | 11.068202 | 10.9817 | 0.086533994 | 0.158693822 |
| Gm7730 | 2.105693 | 2.069211 | 2.0776 | -0.008424865 | 0.028057552 | 2.0664853 | 2.153682 | 2.06717 | 0.086512095 | -0.000684579 |
| Tent2 | 10.00984 | 10.03244 | 10.101 | -0.068957485 | -0.091552747 | 10.015103 | 10.231769 | 10.1453 | 0.086483723 | -0.130182189 |
| Ptp4a3 | 11.76087 | 11.79805 | 11.842 | -0.044073208 | -0.081258478 | 12.299994 | 11.853371 | 11.7669 | 0.08646442 | 0.53308757 |
| Tmem9b | 9.089096 | 9.141967 | 9.1065 | 0.03542644 | -0.017445043 | 9.0221419 | 9.08437 | 8.99793 | 0.086435077 | 0.024207063 |
| Gm20517 | 3.709824 | 3.770933 | 3.6427 | 0.128231158 | 0.067122598 | 3.772268 | 3.851479 | 3.76505 | 0.0864294 | 0.007218433 |
| Tyw3 | 5.958926 | 5.735238 | 5.5126 | 0.222610278 | 0.446298227 | 5.8431254 | 5.8718735 | 5.78561 | 0.086267164 | 0.057519068 |
| Gm7232 | 1.890996 | 1.854931 | 1.7897 | 0.065185166 | 0.101250204 | 1.8500541 | 1.9158292 | 1.82957 | 0.086261594 | 0.020486416 |
| Aatk | 4.675268 | 4.356628 | 4.3437 | 0.012917653 | 0.331557968 | 4.5068076 | 4.3007787 | 4.21461 | 0.0861697 | 0.292198667 |
| AI429214 | 3.247889 | 3.068585 | 3.1065 | -0.037946072 | 0.141358602 | 3.0843064 | 3.0732884 | 2.98713 | 0.086156039 | 0.097174081 |
| Gpn1 | 8.962227 | 8.982497 | 8.956 | 0.026518602 | 0.006247974 | 9.2548276 | 9.1684123 | 9.08244 | 0.08597518 | 0.172390507 |

| | | | | | | | | | | |
|---|---|---|---|---|---|---|---|---|---|---|
| Rpusd4 | 8.783503 | 8.920835 | 8.8466 | 0.074253625 | -0.063078081 | 9.0696896 | 8.9850296 | 8.8991 | 0.08592885 | 0.170588878 |
| Gm15427 | 4.857286 | 4.922991 | 4.8301 | 0.092923459 | 0.027217949 | 4.944256 | 5.0064126 | 4.92051 | 0.085904796 | 0.02374813 |
| Gldn | 0.732015 | 0.727835 | 0.7773 | -0.049436951 | -0.0452564 | 0.7023889 | 0.8099769 | 0.7241 | 0.085872582 | -0.021715392 |
| Ppp4r1l-ps | 7.938932 | 8.134634 | 8.2074 | -0.072742555 | -0.2684448 | 8.321759 | 8.3777938 | 8.29197 | 0.085819604 | 0.029784766 |
| Igkv4-75 | 1.46568 | 1.520563 | 1.3997 | 0.120846182 | 0.06596264 | 1.4631974 | 1.5494922 | 1.46371 | 0.085785917 | -0.000508866 |
| H2-Bl | 0.92644 | 0.841965 | 0.8532 | -0.011255736 | 0.073219532 | 0.8534167 | 0.959647 | 0.87389 | 0.085759455 | -0.020470907 |
| Rbms2 | 9.102163 | 9.090305 | 8.9635 | 0.126813654 | 0.138672017 | 9.2566824 | 9.1737745 | 9.08811 | 0.08566686 | 0.168574769 |
| Hspa12b | 6.052345 | 6.052481 | 5.813 | 0.239438071 | 0.239302125 | 5.7442362 | 5.7716005 | 5.68606 | 0.085538714 | 0.058174397 |
| Srrm2 | 13.49942 | 13.52837 | 13.541 | -0.012136912 | -0.0410868 | 13.854351 | 14.01142 | 13.9259 | 0.085529556 | -0.071539888 |
| Scyl1 | 9.808137 | 9.746583 | 9.7335 | 0.013043518 | 0.07459685 | 10.089331 | 9.9151442 | 9.82967 | 0.085470113 | 0.259656541 |
| 3110009E18Rik | 5.451459 | 5.49171 | 5.4035 | 0.088236283 | 0.047985688 | 5.6575193 | 5.6194227 | 5.53402 | 0.08540608 | 0.123502659 |
| Traf7 | 10.79908 | 10.83995 | 10.733 | 0.106484584 | 0.065614796 | 11.093176 | 11.064163 | 10.9788 | 0.085368752 | 0.114381623 |
| Zfand2a | 8.760767 | 8.622754 | 8.6774 | -0.054636007 | 0.083376654 | 8.377575 | 8.7140048 | 8.62865 | 0.085355346 | -0.251074456 |
| Gmpr2 | 8.324163 | 8.352313 | 8.3299 | 0.022368935 | -0.005780232 | 8.4448153 | 8.502329 | 8.417 | 0.085325047 | 0.027811365 |
| Pum1 | 9.862119 | 10.07248 | 9.9358 | 0.136719375 | -0.073642094 | 9.762756 | 9.9593106 | 9.87412 | 0.085194868 | -0.111359706 |
| Trp53rka | 8.285576 | 8.460074 | 8.4073 | 0.052801741 | -0.121695874 | 8.6054875 | 8.5550716 | 8.46999 | 0.085077276 | 0.135493203 |
| Rps15a | 14.37962 | 14.42902 | 14.391 | 0.037564984 | -0.011835192 | 14.431399 | 14.467101 | 14.3821 | 0.085021678 | 0.049319639 |
| Fem1b | 9.859601 | 9.86136 | 9.8624 | -0.001066394 | -0.002825214 | 9.7815621 | 9.9426566 | 9.85766 | 0.084996925 | -0.076097487 |
| Taf2 | 9.32712 | 9.371172 | 9.3832 | -0.011994461 | -0.056045723 | 9.607256 | 9.6193793 | 9.53447 | 0.084914199 | 0.072790899 |
| Hsd11b2 | 1.755213 | 1.884317 | 1.7584 | 0.125952995 | -0.003151089 | 1.8266754 | 1.7965355 | 1.71165 | 0.084882822 | 0.115022753 |
| D630023F18Rik | 0.600163 | 0.631295 | 0.6849 | -0.053641954 | -0.084773853 | 0.670529 | 0.7487128 | 0.66388 | 0.084836856 | 0.006652995 |
| Cops3 | 9.361464 | 9.280967 | 9.3607 | -0.079767286 | 0.000729486 | 9.5698956 | 9.6125062 | 9.52768 | 0.084827784 | 0.042217201 |
| Vav2 | 10.13846 | 10.23867 | 10.259 | -0.02034202 | -0.120548316 | 10.524458 | 10.45958 | 10.3748 | 0.084790578 | 0.149669239 |
| Ppox | 9.179041 | 9.200893 | 9.3063 | -0.105403725 | -0.127255473 | 9.6994331 | 9.5389804 | 9.4542 | 0.084775535 | 0.245228271 |
| Grik4 | 2.431764 | 2.433931 | 2.4509 | -0.016928421 | -0.019095612 | 2.4325799 | 2.5168956 | 2.43217 | 0.084723469 | 0.00040779 |
| Lzts3 | 5.05555 | 4.91575 | 4.9431 | -0.0273355 | 0.112464188 | 4.8666355 | 5.0890537 | 5.00434 | 0.084712589 | -0.137705575 |
| Olfr344 | 5.128738 | 5.09837 | 4.9925 | 0.105911833 | 0.136280421 | 5.3285206 | 5.546412 | 5.46176 | 0.084656477 | -0.133234872 |
| Zfp536 | 1.44911 | 1.552278 | 1.453 | 0.099323088 | -0.003845037 | 1.475396 | 1.5240342 | 1.43944 | 0.084591571 | 0.035953359 |
| Dync2h1 | 7.769695 | 7.992129 | 7.7991 | 0.19303385 | -0.029400196 | 7.5523303 | 7.908823 | 7.82427 | 0.084554164 | -0.271938476 |
| Rbmx2 | 7.471256 | 7.420324 | 7.4884 | -0.068050675 | -0.01711845 | 7.614024 | 7.6512617 | 7.56673 | 0.084536146 | 0.04729853 |
| Hes6 | 8.282014 | 8.207096 | 8.3992 | -0.192057911 | -0.117139499 | 8.5262958 | 8.2958616 | 8.21146 | 0.084398954 | 0.314833195 |
| Oaf | 4.896773 | 5.056647 | 4.9231 | 0.133524906 | -0.026349257 | 5.0740177 | 4.9757843 | 4.89141 | 0.084373621 | 0.182606994 |

| | | | | | | | | | | |
|---|---|---|---|---|---|---|---|---|---|---|
| Nudt16 | 7.771881 | 7.625775 | 7.7246 | -0.098872232 | 0.047233212 | 7.749132 | 7.8125974 | 7.72822 | 0.084373281 | 0.02090791 |
| Fam71b | 4.828188 | 5.055261 | 5.0732 | -0.017975959 | -0.245048613 | 4.7705726 | 5.1867133 | 5.10249 | 0.084223169 | -0.331917485 |
| AC163399.1 | 3.353934 | 3.331293 | 3.3542 | -0.022942963 | -0.000301349 | 3.3539624 | 3.4018134 | 3.3177 | 0.084110472 | 0.036259559 |
| Hibadh | 9.586898 | 9.538356 | 9.606 | -0.06761127 | -0.01906904 | 9.5467854 | 9.5820453 | 9.49804 | 0.084003647 | 0.048743767 |
| Rev1 | 9.046137 | 9.148508 | 9.1677 | -0.019239659 | -0.121610236 | 9.1720977 | 9.1929887 | 9.10904 | 0.083951088 | 0.063060085 |
| Olfm4 | 3.02157 | 2.849028 | 2.7248 | 0.124268663 | 0.296810722 | 2.8000673 | 2.756453 | 2.67258 | 0.083873226 | 0.127487481 |
| Ccdc71l | 8.728162 | 8.892168 | 8.7346 | 0.157588506 | -0.006417729 | 8.7474225 | 8.7131131 | 8.62925 | 0.083861852 | 0.118171267 |
| Zcchc8 | 9.698128 | 9.742827 | 9.7791 | -0.036301702 | -0.081000019 | 9.7870465 | 9.8886523 | 9.8048 | 0.083851401 | -0.017754414 |
| Cfdp1 | 9.749533 | 9.802775 | 9.7266 | 0.076204547 | 0.022962767 | 9.7383821 | 9.81861 | 9.73491 | 0.083699338 | 0.003471457 |
| 9930022D16Rik | 2.363303 | 2.370941 | 2.3502 | 0.020722462 | 0.013084594 | 2.4304904 | 2.4427296 | 2.35904 | 0.083690807 | 0.071451554 |
| AC116841.1 | 1.867041 | 1.81743 | 1.8566 | -0.039179107 | 0.010431916 | 1.7503586 | 1.8155521 | 1.73195 | 0.083606209 | 0.018412713 |
| Gm5845 | 0.644166 | 0.538492 | 0.5754 | -0.036860008 | 0.068814239 | 0.5377261 | 0.5940026 | 0.51044 | 0.083562873 | 0.027286382 |
| Ap3b2 | 2.879011 | 2.895682 | 2.9398 | -0.04415571 | -0.060827078 | 2.9055734 | 3.0928875 | 3.00934 | 0.083548603 | -0.103765447 |
| Akip1 | 6.766667 | 6.736466 | 6.867 | -0.13051302 | -0.100312381 | 6.9367335 | 6.9625565 | 6.87903 | 0.083527145 | 0.057704118 |
| Rbbp9 | 7.087607 | 7.066212 | 7.141 | -0.074750138 | -0.053355284 | 6.9427156 | 7.0737561 | 6.99026 | 0.083499965 | -0.04754059 |
| Glod4 | 9.359584 | 9.267352 | 9.3139 | -0.04652953 | 0.045702503 | 9.1449219 | 9.3108126 | 9.22737 | 0.08344615 | -0.082444556 |
| Gm49205 | 0.444032 | 0.504201 | 0.4577 | 0.046515623 | -0.013653497 | 0.4621663 | 0.5551368 | 0.47172 | 0.083413664 | -0.009556744 |
| Parl | 9.176373 | 8.975677 | 9.0353 | -0.059591076 | 0.141104443 | 9.3093227 | 9.2303561 | 9.147 | 0.083356824 | 0.162323372 |
| Oxsm | 7.674594 | 7.583663 | 7.6393 | -0.055599215 | 0.035332163 | 7.6326507 | 7.6441895 | 7.56086 | 0.083332754 | 0.07179397 |
| Sh2d1a | 7.899336 | 8.061288 | 7.9953 | 0.065992199 | -0.095959792 | 7.8980941 | 8.1970405 | 8.11371 | 0.08333258 | -0.2156138 |
| Dhx40 | 10.02545 | 10.17165 | 10.056 | 0.115863294 | -0.030337741 | 10.109344 | 10.107587 | 10.0243 | 0.083298455 | 0.085054644 |
| Timm17a | 8.891221 | 8.875555 | 8.7143 | 0.161207599 | 0.176873883 | 8.8704956 | 8.9353552 | 8.85206 | 0.083295893 | 0.018436383 |
| Fundc1 | 8.996316 | 9.007795 | 8.9749 | 0.032925418 | 0.021447309 | 8.8391323 | 8.9587097 | 8.87543 | 0.083280767 | -0.036296706 |
| Crnkl1 | 9.741019 | 9.786776 | 9.7422 | 0.044621592 | -0.001136376 | 9.7891972 | 9.8959223 | 9.81267 | 0.083255176 | -0.02346999 |
| Gm28530 | 3.434399 | 3.40183 | 3.4063 | -0.004435823 | 0.028133058 | 3.4296317 | 3.4430938 | 3.35985 | 0.083241999 | 0.069779817 |
| Fam57a | 5.915544 | 6.02103 | 5.9327 | 0.08835127 | -0.017134101 | 5.9423599 | 5.9907248 | 5.90751 | 0.083219169 | 0.034854286 |
| Gm8797 | 4.865262 | 4.921821 | 4.7754 | 0.146457624 | 0.089899073 | 4.7437533 | 4.9815675 | 4.89836 | 0.08320661 | -0.154607642 |
| Scrn1 | 2.437422 | 2.587101 | 2.6066 | -0.01949956 | -0.169179457 | 2.3348169 | 2.4772724 | 2.39407 | 0.083201924 | -0.059253563 |
| Mettl4 | 8.816302 | 8.883316 | 8.8282 | 0.055126885 | -0.011887071 | 8.8742992 | 8.9096586 | 8.82649 | 0.083168115 | 0.047808765 |
| Tfap4 | 7.795313 | 7.762052 | 7.7573 | 0.004732309 | 0.037993389 | 7.8734524 | 7.9339103 | 7.85084 | 0.083066539 | 0.022608709 |
| Dmp1 | 1.049473 | 1.063825 | 1.02 | 0.043776118 | 0.029423562 | 1.0581079 | 1.1187606 | 1.03585 | 0.082905944 | 0.022253299 |
| Sap30bp | 9.733222 | 9.712491 | 9.7299 | -0.017447556 | 0.003283427 | 9.717524 | 9.7799656 | 9.69708 | 0.082889513 | 0.020447913 |

| | | | | | | | | | | |
|---|---|---|---|---|---|---|---|---|---|---|
| Hist1h2bj | 3.364813 | 3.321972 | 3.2808 | 0.041219199 | 0.084060385 | 3.4823692 | 3.4256914 | 3.34286 | 0.082835671 | 0.139513461 |
| Lrrk1 | 10.25696 | 10.26656 | 10.279 | -0.012878936 | -0.022481758 | 10.469913 | 10.578081 | 10.4953 | 0.082751466 | -0.025415756 |
| Ror2 | 1.062064 | 1.10705 | 1.0908 | 0.016284153 | -0.028701198 | 1.1803071 | 1.1019561 | 1.01923 | 0.082722497 | 0.161073572 |
| Laptm5 | 14.33561 | 14.42949 | 14.356 | 0.073489044 | -0.02039315 | 14.571219 | 14.573748 | 14.4911 | 0.082661212 | 0.080133055 |
| Galnt3 | 4.058561 | 3.93556 | 3.829 | 0.106570296 | 0.229571066 | 3.7814598 | 3.926893 | 3.84426 | 0.082630861 | -0.06280237 |
| Islr | 4.32915 | 4.443654 | 4.4326 | 0.011079307 | -0.103425359 | 4.3193508 | 4.3287299 | 4.24611 | 0.082622887 | 0.073243784 |
| Gm13436 | 1.603772 | 1.590501 | 1.5469 | 0.043599786 | 0.05687036 | 1.5838663 | 1.6052706 | 1.52267 | 0.082603248 | 0.061198984 |
| Narfl | 8.648024 | 8.590857 | 8.5703 | 0.020557917 | 0.077725367 | 8.856234 | 8.7714918 | 8.68889 | 0.082602888 | 0.167345101 |
| Gm15712 | 0.914501 | 0.912022 | 0.9056 | 0.006449559 | 0.008928623 | 0.9160353 | 1.0461546 | 0.96356 | 0.082596182 | -0.047523127 |
| Hes7 | 3.202822 | 3.203914 | 3.337 | -0.133052411 | -0.134144861 | 3.3079094 | 3.4370195 | 3.35443 | 0.082588808 | -0.046521279 |
| Eps8l1 | 8.936739 | 8.567866 | 9.2197 | -0.651838382 | -0.282965042 | 8.1945776 | 8.7246894 | 8.64211 | 0.082575079 | -0.447536679 |
| Pum2 | 12.4769 | 12.52958 | 12.475 | 0.054354806 | 0.001675298 | 12.305195 | 12.521723 | 12.4392 | 0.082563686 | -0.133964663 |
| Lgr6 | 3.246707 | 3.291217 | 3.1962 | 0.095006762 | 0.050496193 | 3.2549963 | 3.3957451 | 3.3132 | 0.0825453 | -0.058203493 |
| Ndn | 3.859392 | 3.845779 | 3.7453 | 0.100453299 | 0.114065731 | 3.6173545 | 3.7531374 | 3.67063 | 0.082511834 | -0.053271022 |
| Yae1d1 | 8.061497 | 8.317573 | 8.0288 | 0.288736223 | 0.032660572 | 8.1924756 | 8.2421799 | 8.15968 | 0.082500286 | 0.032795977 |
| St6galnac1 | 0.93574 | 0.935697 | 0.9464 | -0.010661108 | -0.010618583 | 0.9759626 | 1.0329784 | 0.95052 | 0.082458478 | 0.025442654 |
| Lmf2 | 9.433172 | 9.518909 | 9.4496 | 0.069287341 | -0.01644905 | 9.6571109 | 9.5719771 | 9.48953 | 0.082448814 | 0.167582613 |
| Sh2d4b | 4.113152 | 4.047837 | 4.101 | -0.05320938 | 0.012104872 | 4.3114555 | 4.2130622 | 4.13062 | 0.082441079 | 0.180834379 |
| Tinf2 | 9.667683 | 9.59536 | 9.5381 | 0.057286756 | 0.129609247 | 9.8214622 | 9.8001256 | 9.71769 | 0.082432403 | 0.103768958 |
| Gm8502 | 1.633519 | 1.680718 | 1.6359 | 0.04478053 | -0.002418261 | 1.7020147 | 1.6967438 | 1.61435 | 0.082394206 | 0.087665091 |
| Hnrnpc | 12.30192 | 12.36646 | 12.257 | 0.109876262 | 0.045336252 | 12.479782 | 12.50788 | 12.4256 | 0.082314768 | 0.054216418 |
| Lekr1 | 3.227852 | 3.314802 | 3.2091 | 0.105748224 | 0.018798197 | 3.3052757 | 3.3705455 | 3.28834 | 0.082207766 | 0.016937964 |
| Zfp358 | 7.180363 | 7.054487 | 7.2867 | -0.232216144 | -0.106340817 | 7.1380731 | 7.2302762 | 7.14807 | 0.082203411 | -0.009999671 |
| Hspd1-ps3 | 3.001454 | 3.022726 | 3.0567 | -0.033925108 | -0.055196963 | 2.9552816 | 3.0195744 | 2.93738 | 0.082197046 | 0.017904241 |
| Asb11 | 0.66967 | 0.726307 | 0.6877 | 0.038655604 | -0.01798052 | 0.6757033 | 0.7724646 | 0.69029 | 0.082170009 | -0.014591322 |
| Itgb6 | 1.985791 | 1.968705 | 2.0437 | -0.075026286 | -0.057940209 | 2.188387 | 2.2087852 | 2.12664 | 0.082140374 | 0.061742232 |
| Msto1 | 8.055576 | 7.995446 | 8.1206 | -0.125191074 | -0.065061181 | 8.2510589 | 8.1589064 | 8.07677 | 0.082131786 | 0.174284278 |
| AC154486.2 | 0.081905 | 0.105596 | 0.0833 | 0.022319403 | -0.001371296 | 0.0876543 | 0.1915903 | 0.10947 | 0.082124404 | -0.021811538 |
| Ilvbl | 9.020404 | 8.893483 | 9.0262 | -0.132748227 | -0.005827351 | 9.3656175 | 9.1579159 | 9.07581 | 0.082106129 | 0.289807765 |
| Ctsa | 11.44436 | 11.37151 | 11.361 | 0.01095932 | 0.083807238 | 11.962887 | 11.470059 | 11.388 | 0.082073866 | 0.574902326 |
| Zfand2b | 8.176866 | 8.085821 | 8.2205 | -0.134636548 | -0.043590952 | 8.46883 | 8.2589895 | 8.17699 | 0.082001532 | 0.291842007 |
| Hps4 | 8.198781 | 8.277812 | 8.2783 | -0.000470221 | -0.079501633 | 8.5025776 | 8.4358584 | 8.35388 | 0.081975178 | 0.148694403 |

| | | | | | | | | | | |
|---|---|---|---|---|---|---|---|---|---|---|
| Tfip11 | 9.807024 | 9.74169 | 9.7849 | -0.043236558 | 0.022096642 | 9.8331814 | 9.8934102 | 9.81149 | 0.081920237 | 0.021691464 |
| Emc3 | 9.6074 | 9.495929 | 9.548 | -0.052109994 | 0.059360348 | 9.601173 | 9.6211878 | 9.5393 | 0.081885519 | 0.061870686 |
| Siglecf | 2.870073 | 2.724852 | 2.7237 | 0.001194907 | 0.146416097 | 2.8563012 | 2.6513775 | 2.56956 | 0.081813419 | 0.286737065 |
| Sap30l | 7.523121 | 7.547394 | 7.5412 | 0.006219243 | -0.0180541 | 7.468436 | 7.6478166 | 7.56601 | 0.081806754 | -0.097573801 |
| Nck2 | 8.223445 | 8.241352 | 8.2473 | -0.005992435 | -0.023899589 | 7.8786093 | 8.2433944 | 8.16159 | 0.081802132 | -0.282983029 |
| Galnt9 | 4.807004 | 4.890886 | 4.8437 | 0.047139869 | -0.03674269 | 4.6069592 | 4.844674 | 4.76289 | 0.081784463 | -0.155930329 |
| Gm15760 | 3.702182 | 3.665842 | 3.6904 | -0.024522729 | 0.011817046 | 3.995496 | 4.0232457 | 3.94153 | 0.081713531 | 0.053963799 |
| Gm4149 | 3.218096 | 3.30317 | 3.378 | -0.074781561 | -0.159855423 | 3.2465628 | 3.2768682 | 3.19516 | 0.081712868 | 0.051407475 |
| Ppp2ca | 11.61928 | 11.64587 | 11.541 | 0.104799598 | 0.078210198 | 11.529037 | 11.656287 | 11.5746 | 0.081641734 | -0.045608894 |
| Gpatch1 | 8.148366 | 8.256784 | 8.2474 | 0.009433915 | -0.098984222 | 8.1827137 | 8.227998 | 8.14645 | 0.081546788 | 0.036262499 |
| Gm11969 | 1.307126 | 1.250686 | 1.2753 | -0.024568659 | 0.031871491 | 1.3239059 | 1.3809143 | 1.29945 | 0.081468918 | 0.024460561 |
| Olfr117 | 2.542482 | 2.548467 | 2.6255 | -0.077057627 | -0.083043308 | 2.5570056 | 2.6478131 | 2.56638 | 0.081433525 | -0.009373974 |
| Olfr1202 | 2.320449 | 2.274783 | 2.2837 | -0.008877346 | 0.036789053 | 2.2605005 | 2.5100651 | 2.42866 | 0.08140932 | -0.168155288 |
| Gm5896 | 1.860821 | 1.881493 | 1.8468 | 0.034725377 | 0.014053131 | 1.8415292 | 1.9166163 | 1.83527 | 0.081345187 | 0.006258129 |
| Atp2a2 | 11.20148 | 11.19727 | 11.043 | 0.1546926 | 0.158904166 | 11.040608 | 11.140498 | 11.0592 | 0.081266716 | -0.01862295 |
| Zwint | 9.381156 | 9.396409 | 9.4389 | -0.042532388 | -0.057785552 | 9.3633484 | 9.4821521 | 9.40096 | 0.081187636 | -0.037616068 |
| Urb2 | 9.099925 | 9.177366 | 9.0426 | 0.134756999 | 0.057316023 | 9.0741543 | 9.1800627 | 9.09891 | 0.081151191 | -0.024757181 |
| Olfr1425 | 0.960856 | 0.906063 | 0.9455 | -0.039432716 | 0.015360074 | 1.0328233 | 1.0165238 | 0.93539 | 0.081134233 | 0.097433811 |
| Ctcfl | 1.777428 | 1.689563 | 1.7875 | -0.097931115 | -0.010066248 | 1.7617489 | 1.8461524 | 1.76507 | 0.081084618 | -0.003318898 |
| Gm5075 | 3.558046 | 3.47018 | 3.5117 | -0.041490944 | 0.046375527 | 3.6923556 | 3.5546588 | 3.47361 | 0.081050025 | 0.218746798 |
| Brcc3 | 9.240839 | 9.255584 | 9.2293 | 0.026292937 | 0.011548198 | 9.2905158 | 9.4284519 | 9.34742 | 0.081031471 | -0.056904706 |
| Glis1 | 1.415804 | 1.472597 | 1.4209 | 0.051712145 | -0.005080003 | 1.599142 | 1.4780362 | 1.39717 | 0.080866194 | 0.20197196 |
| Slc38a6 | 9.18925 | 9.309422 | 9.3081 | 0.001371527 | -0.118800297 | 9.4390292 | 9.7910301 | 9.71019 | 0.080842543 | -0.271158342 |
| Orc6 | 7.896796 | 7.729378 | 7.9102 | -0.180799361 | -0.013382124 | 7.961387 | 8.1188475 | 8.03803 | 0.080813669 | -0.076646816 |
| Acd | 9.149656 | 9.205887 | 9.2977 | -0.091807733 | -0.148038752 | 9.5203627 | 9.3466805 | 9.26589 | 0.08079334 | 0.254475569 |
| Olfr571 | 1.587851 | 1.656432 | 1.6405 | 0.01594574 | -0.052635116 | 1.5939364 | 1.6555355 | 1.57478 | 0.080752324 | 0.019153178 |
| Vmn1r216 | 4.082494 | 4.120357 | 3.9775 | 0.142843598 | 0.104980147 | 4.1166863 | 4.4079638 | 4.32733 | 0.080634222 | -0.210643345 |
| 1700025G04Rik | 8.175076 | 7.899036 | 8.0702 | -0.171119788 | 0.104920186 | 7.887168 | 8.1405146 | 8.05989 | 0.080622071 | -0.172724539 |
| Gm3248 | 0.761196 | 0.686364 | 0.701 | -0.014635335 | 0.060196179 | 0.6490265 | 0.7384752 | 0.65786 | 0.08061409 | -0.008834555 |
| Wdr53 | 6.99136 | 7.036161 | 6.9499 | 0.086224376 | 0.041423304 | 6.9290096 | 7.0608395 | 6.98028 | 0.08056015 | -0.051269679 |
| Gm15823 | 1.65874 | 1.664689 | 1.6597 | 0.004961854 | -0.000987195 | 1.7324716 | 1.7479321 | 1.66753 | 0.080402347 | 0.064941883 |
| Fam3c | 10.50714 | 10.49551 | 10.575 | -0.079369014 | -0.067747344 | 10.463375 | 10.624593 | 10.5444 | 0.080196574 | -0.081021244 |

| | | | | | | | | | | |
|---|---|---|---|---|---|---|---|---|---|---|
| Upf2 | 10.44003 | 10.43854 | 10.457 | -0.018696938 | -0.017198462 | 10.284711 | 10.510171 | 10.43 | 0.080156568 | -0.145303779 |
| AC121135.2 | 2.433171 | 2.44859 | 2.4089 | 0.039649563 | 0.02423003 | 2.5821638 | 2.5573389 | 2.4772 | 0.080140741 | 0.104965651 |
| Mdfi | 2.997308 | 2.875421 | 2.927 | -0.051627866 | 0.070259225 | 2.8865687 | 2.9574793 | 2.87738 | 0.080094698 | 0.00918414 |
| Eif5 | 12.58961 | 12.62679 | 12.662 | -0.03524956 | -0.072436888 | 12.473972 | 12.665825 | 12.5857 | 0.080093314 | -0.11175994 |
| Farsb | 10.07531 | 10.05335 | 10.032 | 0.021311098 | 0.043272886 | 10.196038 | 10.145915 | 10.0658 | 0.080087914 | 0.13021076 |
| Fkbp15 | 10.77728 | 10.77625 | 10.814 | -0.0375301 | -0.036503556 | 10.927731 | 10.948695 | 10.8686 | 0.080048494 | 0.05908441 |
| AC133509.1 | 3.617217 | 3.630594 | 3.7797 | -0.149147435 | -0.1625243 | 3.6881822 | 3.7584539 | 3.67843 | 0.080023703 | 0.009751957 |
| St6galnac5 | 2.102264 | 2.029097 | 2.0452 | -0.016097873 | 0.05706877 | 1.9564274 | 2.0361523 | 1.95617 | 0.079986228 | 0.00026137 |
| Eif4enif1 | 10.3648 | 10.44967 | 10.507 | -0.057018786 | -0.141895318 | 10.528264 | 10.664728 | 10.5848 | 0.079957036 | -0.056507176 |
| Rbm22 | 10.50243 | 10.47695 | 10.511 | -0.033845986 | -0.008371184 | 10.674889 | 10.666191 | 10.5862 | 0.079948314 | 0.088645912 |
| Iba57 | 7.240328 | 7.361632 | 7.3624 | -0.000719487 | -0.122023211 | 7.3405741 | 7.2933559 | 7.21356 | 0.079796809 | 0.127015039 |
| Sfmbt2 | 6.946315 | 6.926929 | 7.0981 | -0.171130006 | -0.151744504 | 6.7155989 | 7.2438435 | 7.16405 | 0.07979642 | -0.448448121 |
| Eftud2 | 11.0857 | 11.14015 | 11.108 | 0.032594118 | -0.021857018 | 11.286326 | 11.353949 | 11.2742 | 0.079795526 | 0.012171987 |
| Zdhhc1 | 6.17649 | 6.076215 | 6.1407 | -0.064479767 | 0.035794996 | 6.1521962 | 6.2527566 | 6.17298 | 0.079781418 | -0.020779064 |
| Gm7986 | 2.033129 | 2.036578 | 2.019 | 0.017577488 | 0.014128362 | 2.045234 | 2.0877934 | 2.00814 | 0.079654566 | 0.037095145 |
| Olfr873 | 0.706827 | 0.719396 | 0.7372 | -0.017783763 | -0.030352573 | 0.777467 | 0.8106435 | 0.73103 | 0.079615361 | 0.046438869 |
| Atg12 | 9.549184 | 9.605688 | 9.5992 | 0.006497119 | -0.050006606 | 9.5314897 | 9.6068311 | 9.52723 | 0.079602546 | 0.004261111 |
| Pde6a | 2.591861 | 2.589611 | 2.7084 | -0.118815158 | -0.116565928 | 2.6894186 | 2.7536046 | 2.67416 | 0.079444353 | 0.015258348 |
| Necab3 | 1.739372 | 1.835116 | 1.7462 | 0.088916608 | -0.006827098 | 1.8111776 | 1.9084605 | 1.82903 | 0.079429186 | -0.017853734 |
| Haspin | 6.474693 | 6.444714 | 6.4818 | -0.037071223 | -0.007091911 | 6.5386589 | 6.6278232 | 6.5484 | 0.079422024 | -0.009742342 |
| Krt18 | 3.693648 | 3.241002 | 3.5201 | -0.279102785 | 0.173544081 | 3.9704703 | 3.7501643 | 3.67088 | 0.079284908 | 0.299590959 |
| Gpd1l | 10.71712 | 10.84818 | 10.796 | 0.05265684 | -0.07840261 | 10.847494 | 10.761298 | 10.682 | 0.07926936 | 0.165464439 |
| Senp2 | 10.54378 | 10.61248 | 10.511 | 0.101322094 | 0.032619346 | 10.486124 | 10.646432 | 10.5673 | 0.079152984 | -0.081154306 |
| Olfr481 | 1.110391 | 1.158138 | 1.1274 | 0.030773657 | -0.016973595 | 1.1999587 | 1.2515195 | 1.1724 | 0.07911543 | 0.027554645 |
| Olfr1459 | 5.281807 | 5.182646 | 5.0457 | 0.136904886 | 0.236065875 | 5.4120892 | 5.4990744 | 5.41996 | 0.079113917 | -0.007871348 |
| Gm4858 | 1.56495 | 1.722505 | 1.613 | 0.109516535 | -0.048037815 | 1.5790232 | 1.6434676 | 1.5644 | 0.079063663 | 0.014619341 |
| Taz | 9.675296 | 9.746666 | 9.7157 | 0.031010363 | -0.040359258 | 10.19643 | 9.9876035 | 9.90859 | 0.079013641 | 0.28784003 |
| Slc50a1 | 10.07855 | 10.02092 | 10.124 | -0.103086252 | -0.045461226 | 10.354171 | 10.246125 | 10.1672 | 0.078965236 | 0.187011196 |
| Rinl | 10.79816 | 10.80756 | 10.94 | -0.13211805 | -0.141515084 | 10.805141 | 11.059439 | 10.9806 | 0.078868088 | -0.175430005 |
| Vapb | 9.796522 | 9.777885 | 9.8368 | -0.058870207 | -0.040233385 | 9.6052598 | 9.6610662 | 9.58234 | 0.078726087 | 0.022919704 |
| Mark3 | 10.02447 | 10.10799 | 10.027 | 0.080789727 | -0.002728044 | 10.07871 | 10.24974 | 10.1711 | 0.078631119 | -0.092398093 |
| Rps18-ps3 | 1.580023 | 1.572895 | 1.5576 | 0.015273548 | 0.022401019 | 1.5864056 | 1.6600178 | 1.58142 | 0.078596982 | 0.004984807 |

| | | | | | | | | | | |
|---|---|---|---|---|---|---|---|---|---|---|
| Mad2l1bp | 8.257607 | 8.138547 | 8.3645 | -0.225997291 | -0.106936863 | 8.3469065 | 8.3081425 | 8.22956 | 0.078580102 | 0.117344138 |
| Olfr1346 | 1.195356 | 1.157436 | 1.2196 | -0.062213263 | -0.024292944 | 1.2081405 | 1.2897031 | 1.21113 | 0.078573342 | -0.002989278 |
| Pwwp2b | 4.28786 | 4.313348 | 4.3432 | -0.029810168 | -0.055297517 | 4.3153226 | 4.3692188 | 4.29065 | 0.078568849 | 0.02467264 |
| Hspb6 | 6.019067 | 5.779429 | 6.0044 | -0.224991233 | 0.014646618 | 6.1113251 | 6.0481827 | 5.96963 | 0.07855214 | 0.141694522 |
| Vmn1r77 | 1.156406 | 1.207311 | 1.211 | -0.003689575 | -0.054594806 | 1.194508 | 1.2508797 | 1.17238 | 0.078496574 | 0.022124867 |
| Wdyhv1 | 6.705783 | 6.830553 | 6.8035 | 0.027012098 | -0.097757416 | 6.8501731 | 6.9491548 | 6.87072 | 0.078430204 | -0.020551502 |
| Qars | 11.10309 | 10.91771 | 11.052 | -0.133837826 | 0.051543682 | 11.214215 | 11.22939 | 11.151 | 0.078364952 | 0.063189942 |
| Ms4a7 | 4.973708 | 5.389792 | 5.0198 | 0.369984999 | -0.046099472 | 5.0978159 | 4.7950243 | 4.71676 | 0.078266836 | 0.381058439 |
| Tnfrsf18 | 9.082477 | 9.047868 | 8.9214 | 0.126458387 | 0.161067776 | 8.9692072 | 9.05014 | 8.97191 | 0.078229767 | -0.002703045 |
| Madd | 11.25297 | 11.45547 | 11.406 | 0.049487736 | -0.153012914 | 11.18588 | 11.464864 | 11.3867 | 0.07818634 | -0.20079822 |
| Zfp553 | 8.058917 | 8.182112 | 8.1348 | 0.047288949 | -0.075906242 | 8.1108508 | 8.109427 | 8.03126 | 0.078165522 | 0.079589326 |
| Usp17la | 3.282842 | 3.20391 | 3.2556 | -0.051648412 | 0.027283592 | 3.3774755 | 3.4228621 | 3.34477 | 0.078089159 | 0.032702587 |
| Pde10a | 3.854909 | 3.801663 | 3.882 | -0.080353738 | -0.027107422 | 3.8745559 | 3.6708371 | 3.59275 | 0.078085745 | 0.281804623 |
| Tubgcp3 | 9.961721 | 9.95072 | 9.9199 | 0.030805159 | 0.041806651 | 10.219318 | 10.265343 | 10.1873 | 0.078026839 | 0.032002497 |
| Aven | 8.465643 | 8.444415 | 8.3267 | 0.117710515 | 0.138938377 | 8.1705956 | 8.4687932 | 8.39084 | 0.077957712 | -0.220239822 |
| Gjb1 | 1.768336 | 1.510449 | 1.5957 | -0.085200867 | 0.172686409 | 1.675704 | 1.6190456 | 1.54118 | 0.077866582 | 0.134524961 |
| Gm8454 | 1.626746 | 1.568371 | 1.5936 | -0.025182156 | 0.033192879 | 1.59403 | 1.6748443 | 1.59702 | 0.077823034 | -0.002991204 |
| Rpia | 9.584635 | 9.624595 | 9.6424 | -0.017827182 | -0.057787271 | 9.6326384 | 9.6086638 | 9.53086 | 0.077808096 | 0.101782714 |
| Dpysl5 | 2.397833 | 2.363759 | 2.4186 | -0.054825846 | -0.020751843 | 2.3732002 | 2.4420101 | 2.36422 | 0.077789756 | 0.00897982 |
| Arl9 | 2.131833 | 2.058919 | 2.1156 | -0.056688674 | 0.016225468 | 2.2755062 | 2.2709114 | 2.19318 | 0.077734246 | 0.082329017 |
| Hcn2 | 2.249807 | 2.319747 | 2.2663 | 0.053447882 | -0.016492689 | 2.3841142 | 2.3994872 | 2.32185 | 0.077632309 | 0.062259374 |
| Med10 | 9.230282 | 9.188745 | 9.2727 | -0.083991367 | -0.042454675 | 9.3647798 | 9.3514635 | 9.27384 | 0.07762021 | 0.090936514 |
| Gm4045 | 1.727942 | 1.721306 | 1.6911 | 0.030177866 | 0.036813219 | 1.7599973 | 1.7511911 | 1.67362 | 0.077570116 | 0.08637627 |
| Rpe | 9.367118 | 9.460787 | 9.3716 | 0.089211588 | -0.004457203 | 9.2934199 | 9.4902873 | 9.41278 | 0.077505052 | -0.119362382 |
| Zfyve9 | 6.741693 | 6.628435 | 6.2813 | 0.34710922 | 0.460366645 | 6.3020979 | 6.2340314 | 6.15661 | 0.077417227 | 0.145483763 |
| Gm9625 | 5.389001 | 5.258758 | 5.2651 | -0.006302927 | 0.123939154 | 5.2127126 | 5.3225838 | 5.24521 | 0.07737474 | -0.032496493 |
| Calcoco1 | 9.450513 | 9.48542 | 9.5968 | -0.111387648 | -0.146293964 | 9.5499434 | 9.5788775 | 9.50153 | 0.077347832 | 0.048413745 |
| Rgmb | 7.644503 | 7.767334 | 7.8141 | -0.046807927 | -0.169638817 | 7.4114445 | 7.7170301 | 7.63977 | 0.077256024 | -0.228329578 |
| Alox15 | 3.293791 | 3.076138 | 3.0811 | -0.004968926 | 0.212684043 | 2.9979695 | 3.0776814 | 3.00055 | 0.077129778 | -0.002582149 |
| Itga3 | 7.142902 | 6.857192 | 6.7171 | 0.140073314 | 0.425782781 | 6.4346893 | 6.4788951 | 6.40181 | 0.077084089 | 0.0328783 |
| Gm13002 | 3.293796 | 3.38072 | 3.3632 | 0.017527799 | -0.069395393 | 3.5019915 | 3.5379011 | 3.46094 | 0.076960242 | 0.041050695 |
| Anxa11 | 10.6824 | 10.63372 | 10.634 | -0.000197584 | 0.048487952 | 10.805864 | 10.621056 | 10.5442 | 0.076897638 | 0.261705812 |

| | | | | | | | | | | |
|---|---|---|---|---|---|---|---|---|---|---|
| Vmn2r3 | 3.633917 | 3.657299 | 3.7173 | -0.060046648 | -0.083428783 | 3.676905 | 3.9176436 | 3.84079 | 0.076856372 | -0.16388227 |
| Olfr753-ps1 | 1.224973 | 1.159969 | 1.2252 | -0.065194024 | -0.000190364 | 1.1240704 | 1.2204589 | 1.14364 | 0.076819887 | -0.019568632 |
| Efl1 | 9.078123 | 9.08208 | 9.0229 | 0.059157953 | 0.055201114 | 9.02346 | 9.1230264 | 9.04633 | 0.076696795 | -0.022869644 |
| Lck | 12.45494 | 12.46203 | 12.488 | -0.025715948 | -0.0328071 | 12.281483 | 12.598599 | 12.5219 | 0.076664742 | -0.240451218 |
| Gm15878 | 1.113602 | 1.101919 | 1.1409 | -0.038983334 | -0.027299884 | 1.1203682 | 1.201444 | 1.12482 | 0.076627404 | -0.004448403 |
| Tm2d2 | 7.832052 | 7.663441 | 7.8439 | -0.180411472 | -0.011800743 | 7.8899594 | 7.6776044 | 7.60102 | 0.07658873 | 0.288943729 |
| Olfr341 | 1.887965 | 1.908631 | 1.8995 | 0.009164744 | -0.011500865 | 1.9575685 | 1.9742627 | 1.89769 | 0.076573286 | 0.059879086 |
| Hspa2 | 6.740957 | 6.41275 | 6.7053 | -0.292577346 | 0.035629353 | 6.3188892 | 6.4599309 | 6.38336 | 0.076569578 | -0.064472075 |
| Lxn | 6.502569 | 6.728511 | 6.6136 | 0.114867757 | -0.111074662 | 6.5956915 | 6.6199374 | 6.54343 | 0.076503296 | 0.052257377 |
| Gm16026 | 4.821202 | 4.827973 | 4.8447 | -0.016754341 | -0.023525468 | 4.9176108 | 5.0905044 | 5.01401 | 0.076499131 | -0.096394475 |
| Zfp511 | 8.565309 | 8.590762 | 8.667 | -0.076189516 | -0.101642759 | 9.076989 | 8.9576565 | 8.88116 | 0.076491927 | 0.195824454 |
| Cd46 | 6.301778 | 6.305892 | 6.0499 | 0.256033464 | 0.251920037 | 6.3271087 | 6.6893239 | 6.61289 | 0.076434101 | -0.285781086 |
| Gm9670 | 2.011379 | 1.974914 | 2.0123 | -0.037420638 | -0.000955435 | 2.0916555 | 2.0875317 | 2.0112 | 0.076328705 | 0.080452471 |
| Mrps16 | 8.956542 | 8.958101 | 8.9606 | -0.002496191 | -0.004055724 | 9.1016785 | 9.0687619 | 8.99243 | 0.076327795 | 0.109244415 |
| Nsa2 | 12.21267 | 12.16062 | 12.204 | -0.04368073 | 0.008367532 | 12.205986 | 12.275608 | 12.1993 | 0.076313806 | 0.006691435 |
| Rgsl1 | 1.974448 | 2.032012 | 2.0381 | -0.006070528 | -0.06363426 | 2.028636 | 2.0605218 | 1.98424 | 0.076283815 | 0.04439808 |
| Vmn1r72 | 3.167387 | 3.223563 | 3.3098 | -0.086266008 | -0.142441615 | 3.230122 | 3.5092151 | 3.43297 | 0.076248759 | -0.202844419 |
| Prss2 | 1.839637 | 1.804647 | 1.8048 | -0.000165234 | 0.034824278 | 1.7662039 | 1.8729945 | 1.79678 | 0.076210222 | -0.030580347 |
| Ighv1-80 | 8.169875 | 8.742794 | 7.795 | 0.947798032 | 0.374878502 | 8.4188736 | 7.9747495 | 7.89858 | 0.076167276 | 0.52029135 |
| Cebpe | 1.345959 | 1.34053 | 1.3538 | -0.013260637 | -0.007831512 | 1.4106001 | 1.4317866 | 1.35563 | 0.076155136 | 0.054968639 |
| Spaca6 | 2.885658 | 2.713328 | 2.8026 | -0.089240221 | 0.08308958 | 2.8293628 | 2.8363248 | 2.76019 | 0.076135727 | 0.069173738 |
| Nol7 | 10.21275 | 10.2071 | 10.212 | -0.005110849 | 0.000529761 | 10.195963 | 10.301112 | 10.2251 | 0.076003792 | -0.029145268 |
| AC114990.1 | 0.502234 | 0.479555 | 0.4221 | 0.057462831 | 0.080142215 | 0.4526438 | 0.5189805 | 0.44299 | 0.075994129 | 0.009657373 |
| Ccdc58 | 7.266322 | 7.129048 | 7.3593 | -0.230277839 | -0.093003809 | 7.4936446 | 7.5686514 | 7.49268 | 0.075970244 | 0.000963414 |
| Gm14659 | 1.12423 | 1.08629 | 1.1045 | -0.018203203 | 0.01973633 | 1.1418234 | 1.2082337 | 1.13245 | 0.075779643 | 0.009369314 |
| Cdk7 | 8.807533 | 8.748294 | 8.6901 | 0.058146083 | 0.117385706 | 8.7705266 | 8.9946938 | 8.91898 | 0.075715555 | -0.148451701 |
| Olfr509 | 4.622332 | 4.89856 | 4.7053 | 0.193297183 | -0.082930963 | 4.8271549 | 5.2144517 | 5.13878 | 0.075675246 | -0.31162163 |
| Lhfpl4 | 3.575718 | 3.609655 | 3.5863 | 0.023395437 | -0.010541704 | 3.588625 | 3.7795586 | 3.70395 | 0.075608041 | -0.115325506 |
| Zkscan17 | 9.703989 | 9.657737 | 9.7034 | -0.045687703 | 0.000564205 | 9.6998766 | 9.7032285 | 9.62763 | 0.075594461 | 0.072242536 |
| Rem1 | 3.958207 | 3.867499 | 3.9858 | -0.118252697 | -0.027543874 | 3.9800622 | 4.0506174 | 3.97507 | 0.075546954 | 0.004991812 |
| Slf2 | 10.4104 | 10.4166 | 10.446 | -0.02975825 | -0.035961204 | 10.314593 | 10.51204 | 10.4365 | 0.075529222 | -0.121917967 |
| Pkn1 | 12.27763 | 12.29252 | 12.342 | -0.049644706 | -0.064529742 | 12.492022 | 12.502995 | 12.4276 | 0.075380854 | 0.064407739 |

| | | | | | | | | | | |
|---|---|---|---|---|---|---|---|---|---|---|
| Gcm1 | 1.39297 | 1.344655 | 1.4015 | -0.056834627 | -0.008519529 | 1.4076465 | 1.422063 | 1.34671 | 0.075348205 | 0.060931714 |
| Gm6983 | 1.7444 | 1.791948 | 1.7574 | 0.034554527 | -0.012994003 | 1.7202026 | 1.8290071 | 1.75367 | 0.075340409 | -0.033464109 |
| Tmpo | 11.48546 | 11.60139 | 11.612 | -0.010488544 | -0.12642516 | 11.637992 | 11.755485 | 11.6802 | 0.075331988 | -0.042161675 |
| Cyp27b1 | 2.104657 | 2.379219 | 2.0878 | 0.291416068 | 0.016854313 | 2.1753889 | 2.2852068 | 2.20989 | 0.075319252 | -0.034498652 |
| Kbtbd4 | 7.464506 | 7.500559 | 7.5208 | -0.020240687 | -0.056294203 | 7.5989262 | 7.6168955 | 7.5417 | 0.075196036 | 0.057226705 |
| Msi2 | 10.24556 | 10.39895 | 10.287 | 0.111762702 | -0.041623106 | 10.067549 | 10.35268 | 10.2775 | 0.075195288 | -0.209935567 |
| Srpr | 10.93369 | 10.76922 | 10.7 | 0.06965289 | 0.234121998 | 10.782705 | 10.909876 | 10.8347 | 0.075141402 | -0.052029255 |
| Sdhc | 9.708054 | 9.789022 | 9.7264 | 0.062585254 | -0.018383423 | 10.085787 | 9.8675685 | 9.79263 | 0.074939911 | 0.293158742 |
| Ino80d | 10.69845 | 10.74705 | 10.836 | -0.08872129 | -0.13732909 | 10.687512 | 11.085005 | 11.0101 | 0.07493918 | -0.322553507 |
| Vmn2r27 | 1.824503 | 1.822826 | 1.855 | -0.032220072 | -0.030543071 | 1.8410715 | 1.9047042 | 1.82978 | 0.074925557 | 0.011292834 |
| Dera | 8.781934 | 8.712849 | 8.7462 | -0.033313208 | 0.03577118 | 8.9210248 | 8.9104588 | 8.83559 | 0.074864125 | 0.085430151 |
| Ttll12 | 8.446498 | 8.564789 | 8.4569 | 0.107865061 | -0.01042602 | 8.4259175 | 8.5375255 | 8.46266 | 0.074861392 | -0.036746639 |
| Slc41a1 | 9.62543 | 9.555282 | 9.4797 | 0.075549228 | 0.145696589 | 9.5697077 | 9.6118412 | 9.53698 | 0.074860075 | 0.032726633 |
| Tmem198 | 3.916204 | 3.884278 | 3.9602 | -0.075938444 | -0.044012743 | 4.0745479 | 3.9775742 | 3.90277 | 0.074804936 | 0.171778595 |
| Sucla2 | 9.326709 | 9.266409 | 9.2703 | -0.003873274 | 0.056427425 | 9.1749938 | 9.2514854 | 9.17668 | 0.074801075 | -0.001690546 |
| Prpf4b | 11.47282 | 11.49918 | 11.449 | 0.050623574 | 0.024263514 | 11.584426 | 11.672713 | 11.5979 | 0.074774848 | -0.013512259 |
| Olfr1356 | 1.874403 | 1.892086 | 1.8654 | 0.026696781 | 0.009013912 | 1.8946377 | 2.0073531 | 1.93264 | 0.074715325 | -0.038000062 |
| Pigq | 9.77099 | 9.792053 | 9.8653 | -0.073229437 | -0.094291778 | 10.104089 | 10.038581 | 9.96388 | 0.074702885 | 0.140210943 |
| Vmn2r113 | 2.535156 | 2.519472 | 2.4996 | 0.019852844 | 0.035536379 | 2.6093675 | 2.6833288 | 2.60864 | 0.07468792 | 0.000726541 |
| Pkmyt1 | 8.535488 | 8.520218 | 8.6868 | -0.166556337 | -0.151286164 | 8.9135985 | 8.9001699 | 8.82549 | 0.074683813 | 0.08811239 |
| Hnrnpd | 11.90816 | 11.76938 | 11.868 | -0.098977454 | 0.039802708 | 11.90275 | 12.06307 | 11.9885 | 0.074601198 | -0.085719074 |
| Abca7 | 11.05694 | 11.14631 | 11.17 | -0.023781232 | -0.113152852 | 11.375384 | 11.372933 | 11.2984 | 0.074527148 | 0.076978807 |
| Scd3 | 3.846584 | 3.901265 | 3.9111 | -0.009851591 | -0.06453251 | 3.8823423 | 3.9231273 | 3.8487 | 0.074422718 | 0.033637754 |
| Junb | 12.56582 | 12.37989 | 12.409 | -0.02951171 | 0.156421742 | 11.684909 | 11.816647 | 11.7422 | 0.074410772 | -0.057327709 |
| Vma21 | 9.01372 | 8.96488 | 8.9108 | 0.054078749 | 0.102918232 | 8.991669 | 9.0154274 | 8.94105 | 0.074382294 | 0.05062388 |
| Shisa4 | 3.426147 | 3.36752 | 3.4401 | -0.072592025 | -0.013965381 | 3.4741225 | 3.5020095 | 3.4277 | 0.074313641 | 0.046426662 |
| Uba5 | 9.288248 | 9.301798 | 9.2142 | 0.087644207 | 0.074094389 | 9.2027844 | 9.3287507 | 9.25444 | 0.074306213 | -0.051660139 |
| Sft2d3 | 5.884545 | 5.833676 | 6.0353 | -0.201658106 | -0.150789099 | 5.9625688 | 6.1835104 | 6.10921 | 0.074303125 | -0.146638444 |
| Trav7d-3 | 3.497777 | 3.43892 | 3.5229 | -0.083960607 | -0.025103716 | 3.3523739 | 3.5287419 | 3.45445 | 0.074290477 | -0.102077485 |
| Nudt5 | 9.113775 | 9.1593 | 9.1205 | 0.038760797 | -0.006764276 | 9.2302839 | 9.2264779 | 9.1522 | 0.074273284 | 0.078079296 |
| Mcur1 | 9.678696 | 9.773593 | 9.7541 | 0.019510689 | -0.075385693 | 9.6122562 | 9.7765935 | 9.70233 | 0.074263738 | -0.090073605 |
| Hist1h3f | 2.903142 | 2.801391 | 2.8631 | -0.061720288 | 0.04003109 | 2.8136605 | 2.9040841 | 2.82984 | 0.07424005 | -0.01618347 |

| | | | | | | | | | | |
|---|---|---|---|---|---|---|---|---|---|---|
| Mybpc3 | 3.19197 | 3.104736 | 3.0457 | 0.059020282 | 0.146254554 | 3.1710386 | 3.2908462 | 3.21662 | 0.07422165 | -0.045585958 |
| Fndc8 | 1.172086 | 1.203974 | 1.1589 | 0.045050356 | 0.01316301 | 1.1347165 | 1.1600042 | 1.0858 | 0.07420003 | 0.04891234 |
| Dnal4 | 6.828444 | 6.741107 | 6.8362 | -0.095094698 | -0.007757518 | 6.9466879 | 6.9545845 | 6.88041 | 0.074174251 | 0.066277636 |
| Cpsf7 | 11.21802 | 11.30671 | 11.293 | 0.014019486 | -0.074673196 | 11.508724 | 11.596652 | 11.5225 | 0.074166448 | -0.01376121 |
| Tpbg | 1.900617 | 1.824723 | 1.8499 | -0.025190138 | 0.05070318 | 1.8722267 | 1.8271584 | 1.753 | 0.074156321 | 0.119224647 |
| Ighv2-9-1 | 6.148186 | 6.1411 | 6.3048 | -0.163717256 | -0.156631926 | 6.5445346 | 6.2400417 | 6.1659 | 0.07414651 | 0.378639429 |
| Tbc1d10a | 9.072464 | 9.17549 | 9.0267 | 0.148796767 | 0.045770095 | 9.1371563 | 9.1771321 | 9.10319 | 0.073942689 | 0.033966899 |
| Brd3 | 9.725227 | 9.888144 | 9.8543 | 0.033810682 | -0.129106331 | 9.8186133 | 9.9298917 | 9.85604 | 0.073852703 | -0.037425644 |
| Pde5a | 6.00566 | 5.802468 | 5.9412 | -0.138745763 | 0.064446461 | 5.9595538 | 5.9163026 | 5.84246 | 0.07384104 | 0.117092205 |
| Supt16 | 11.17604 | 11.22329 | 11.189 | 0.0339738 | -0.013275598 | 11.221026 | 11.320022 | 11.2462 | 0.073819462 | -0.02517691 |
| Ccn5 | 2.06473 | 2.24328 | 2.0169 | 0.226422218 | 0.04787274 | 2.1489383 | 2.1044536 | 2.03064 | 0.073811635 | 0.118296381 |
| Cd151 | 8.340016 | 8.327202 | 8.3451 | -0.017860984 | -0.005047196 | 8.3038461 | 8.1220026 | 8.0482 | 0.07380178 | 0.25564533 |
| Dnmt3l | 0.63541 | 0.700106 | 0.6651 | 0.035012188 | -0.02968402 | 0.6689992 | 0.7324155 | 0.65862 | 0.073793921 | 0.010377666 |
| Gm42857 | 5.593352 | 5.608357 | 5.9505 | -0.342193137 | -0.357198208 | 5.6347815 | 6.0061067 | 5.93233 | 0.073771786 | -0.297553432 |
| Zfp943 | 8.783348 | 8.892295 | 8.8609 | 0.031394789 | -0.077552544 | 8.9832779 | 8.9758736 | 8.90211 | 0.073763013 | 0.08116736 |
| Tnfsf13b | 5.216395 | 5.065068 | 4.8981 | 0.167001698 | 0.318328962 | 4.8846432 | 4.8033326 | 4.72958 | 0.073751979 | 0.155062548 |
| Smap1 | 9.717202 | 9.742288 | 9.7843 | -0.041997045 | -0.067082419 | 9.5414759 | 9.6084534 | 9.53478 | 0.073675958 | 0.006698481 |
| S100a14 | 0.714313 | 0.737429 | 0.778 | -0.040545957 | -0.063661676 | 0.7215243 | 0.7700917 | 0.69642 | 0.073668415 | 0.025101053 |
| Arf3 | 11.74016 | 11.84399 | 11.784 | 0.060313436 | -0.043521792 | 11.751156 | 11.791875 | 11.7182 | 0.073649428 | 0.032930577 |
| Sec11a | 10.10159 | 10.15368 | 10.075 | 0.078360621 | 0.026279146 | 10.01057 | 10.198262 | 10.1247 | 0.073604631 | -0.114087496 |
| Atpaf2 | 8.438347 | 8.508409 | 8.4292 | 0.079244354 | 0.009183074 | 8.6767896 | 8.6710291 | 8.59743 | 0.073596206 | 0.079356628 |
| Abcg4 | 1.543142 | 1.440278 | 1.4403 | -6.31524E-05 | 0.102800536 | 1.5617591 | 1.5771566 | 1.50359 | 0.07356175 | 0.058164228 |
| Ubc | 14.23679 | 14.14987 | 14.296 | -0.145709668 | -0.058789478 | 13.660349 | 13.954205 | 13.8807 | 0.073518098 | -0.220337682 |
| Gm15710 | 3.014403 | 3.146071 | 3.1685 | -0.022391081 | -0.154059862 | 3.1730336 | 3.2715569 | 3.19813 | 0.073427383 | -0.025095862 |
| Casp9 | 8.951339 | 8.890761 | 9.0105 | -0.119762452 | -0.059185241 | 9.1549922 | 9.0101707 | 8.93681 | 0.073361784 | 0.21818327 |
| Arrdc1 | 9.891428 | 9.874301 | 9.9241 | -0.049816267 | -0.03268995 | 10.410057 | 10.210081 | 10.1368 | 0.07331848 | 0.273294083 |
| Cenpi | 6.221116 | 6.155895 | 6.2959 | -0.139967962 | -0.07474753 | 6.3880575 | 6.6037595 | 6.53052 | 0.073244056 | -0.142457954 |
| Gm49355 | 1.596243 | 1.597049 | 1.5934 | 0.003683993 | 0.002878408 | 1.6344486 | 1.6854132 | 1.61217 | 0.073243973 | 0.022279344 |
| D10Wsu102e | 11.42019 | 11.50286 | 11.437 | 0.066200124 | -0.016474692 | 11.580119 | 11.901412 | 11.8282 | 0.073171098 | -0.248121229 |
| Pqbp1 | 10.28023 | 10.25267 | 10.316 | -0.06339392 | -0.035840914 | 10.438853 | 10.32456 | 10.2516 | 0.073008536 | 0.187301621 |
| Dram2 | 9.192936 | 8.898045 | 9.0559 | -0.157824014 | 0.137066367 | 9.0149535 | 8.7675337 | 8.69453 | 0.072998854 | 0.320418676 |
| Tnni1 | 2.54771 | 2.540788 | 2.4879 | 0.052865281 | 0.059787427 | 2.5023752 | 2.5632939 | 2.4903 | 0.072994241 | 0.012075557 |

| | | | | | | | | | | |
|---|---|---|---|---|---|---|---|---|---|---|
| Cd247 | 11.0203 | 11.10909 | 11.102 | 0.006774572 | -0.082008714 | 10.842205 | 11.170729 | 11.0978 | 0.072953232 | -0.255570139 |
| Xlr4e-ps | 1.620399 | 1.611665 | 1.5917 | 0.019994528 | 0.028728792 | 1.5200832 | 1.6395392 | 1.56661 | 0.072929103 | -0.046526887 |
| 4933430I17Rik | 3.001942 | 3.060517 | 3.0199 | 0.040654713 | -0.017921048 | 3.0998598 | 3.1111439 | 3.03825 | 0.072891912 | 0.061607777 |
| Aldh3b1 | 7.58988 | 7.329253 | 7.4398 | -0.110502029 | 0.150125054 | 7.9276776 | 7.5296562 | 7.45682 | 0.072832455 | 0.470853875 |
| Mkrn1 | 10.31444 | 10.32576 | 10.255 | 0.070404654 | 0.05907869 | 10.299236 | 10.403245 | 10.3304 | 0.07280863 | -0.031200425 |
| Phf23 | 10.32839 | 10.20764 | 10.31 | -0.10251072 | 0.018237286 | 10.167651 | 10.276172 | 10.2034 | 0.072764453 | -0.035756987 |
| Lgr5 | 3.390552 | 3.448909 | 3.5075 | -0.058578551 | -0.116935665 | 3.5207517 | 3.5993054 | 3.5266 | 0.072701841 | -0.005851841 |
| Zdhhc6 | 8.866165 | 8.83966 | 8.9023 | -0.062676823 | -0.036172125 | 8.906751 | 9.0329596 | 8.96026 | 0.072700431 | -0.053508218 |
| Nufip2 | 10.4875 | 10.68912 | 10.496 | 0.192629436 | -0.008997229 | 10.566195 | 10.815222 | 10.7425 | 0.072681666 | -0.176345388 |
| Mcrs1 | 9.611692 | 9.561713 | 9.6706 | -0.108882288 | -0.058903652 | 9.9788447 | 9.7559133 | 9.68323 | 0.072681311 | 0.295612712 |
| Ift172 | 8.862544 | 9.056353 | 8.9302 | 0.126135015 | -0.06767404 | 9.2464992 | 9.2796741 | 9.20703 | 0.072648053 | 0.039473133 |
| Usp37 | 9.54108 | 9.60844 | 9.4488 | 0.159619326 | 0.092258885 | 9.4303183 | 9.6004515 | 9.52785 | 0.072599352 | -0.09753383 |
| Gm10110 | 3.296208 | 3.342488 | 3.2353 | 0.107206777 | 0.060927381 | 3.3689817 | 3.4159999 | 3.34344 | 0.072558029 | 0.025539843 |
| Slc6a2 | 2.00513 | 2.078188 | 2.0005 | 0.077655547 | 0.004597105 | 2.0045751 | 2.1332306 | 2.0607 | 0.072527069 | -0.056128435 |
| Gtf3c2 | 10.6474 | 10.7234 | 10.811 | -0.088024462 | -0.164023044 | 10.936351 | 11.066634 | 10.9943 | 0.072290846 | -0.057992329 |
| Vps53 | 9.613578 | 9.503601 | 9.5438 | -0.040234121 | 0.069743294 | 9.6809912 | 9.6179143 | 9.54566 | 0.072255575 | 0.135332495 |
| Rpl3-ps2 | 1.880097 | 1.877518 | 1.9006 | -0.023102472 | -0.020524008 | 1.898407 | 2.001245 | 1.9291 | 0.07214388 | -0.030694087 |
| Ror1 | 3.959512 | 3.879126 | 3.7672 | 0.111881516 | 0.192267012 | 3.6413166 | 3.716762 | 3.64462 | 0.072143267 | -0.003302105 |
| Dpt | 5.887249 | 5.657772 | 5.6493 | 0.008513639 | 0.237990017 | 5.3006862 | 5.2969332 | 5.22482 | 0.072113804 | 0.075866775 |
| Gm10029 | 2.48276 | 2.418507 | 2.5377 | -0.119144598 | -0.054891958 | 2.6037402 | 2.4811411 | 2.40903 | 0.072113124 | 0.194712184 |
| 1700028J19Rik | 3.016134 | 2.99469 | 3.0351 | -0.04036299 | -0.018918485 | 2.9849995 | 3.0443454 | 2.97226 | 0.072082572 | 0.012736701 |
| Atp23 | 6.784721 | 6.644353 | 6.7955 | -0.151169657 | -0.01080217 | 6.9766931 | 6.8206411 | 6.74857 | 0.072073317 | 0.228125349 |
| Apom | 1.274533 | 1.253738 | 1.2921 | -0.038385606 | -0.017590644 | 1.3202274 | 1.2996983 | 1.22765 | 0.072045798 | 0.092574908 |
| Olfr237-ps1 | 2.55441 | 2.431436 | 2.514 | -0.082551014 | 0.040423464 | 2.5029797 | 2.6174047 | 2.54545 | 0.071957124 | -0.042467883 |
| Card14 | 1.127578 | 1.151747 | 1.1422 | 0.009564866 | -0.01460431 | 1.1066229 | 1.2148971 | 1.14295 | 0.071951804 | -0.036322455 |
| Vmn2r23 | 3.39264 | 3.657414 | 3.6148 | 0.042657789 | -0.222116087 | 3.5927467 | 3.7377188 | 3.66577 | 0.071951745 | -0.073020317 |
| Ptk2b | 12.69745 | 12.8323 | 12.755 | 0.077026842 | -0.057827408 | 13.033328 | 12.917879 | 12.8459 | 0.071929692 | 0.187378901 |
| Acsm2 | 2.314931 | 2.367148 | 2.3831 | -0.0159979 | -0.068214085 | 2.3878373 | 2.4753674 | 2.40348 | 0.071890384 | -0.01563976 |
| Capzb | 12.42629 | 12.38834 | 12.354 | 0.034721658 | 0.072675576 | 12.552598 | 12.419283 | 12.3474 | 0.071875364 | 0.205190075 |
| Pop4 | 7.839633 | 7.713693 | 7.8922 | -0.178516454 | -0.052576334 | 8.0136477 | 7.9261043 | 7.85437 | 0.071730716 | 0.159274056 |
| Atat1 | 5.792289 | 5.941078 | 5.9392 | 0.001895284 | -0.146893546 | 6.0826938 | 6.3264516 | 6.25478 | 0.071670924 | -0.172086798 |
| Amd2 | 2.854777 | 2.855438 | 2.8951 | -0.039653629 | -0.04031445 | 2.8947359 | 2.8791233 | 2.80751 | 0.07161222 | 0.087224812 |

| | | | | | | | | | | |
|---|---|---|---|---|---|---|---|---|---|---|
| Uqcrh | 11.26282 | 11.29722 | 11.312 | -0.014800872 | -0.049198624 | 11.328403 | 11.336655 | 11.2651 | 0.071566044 | 0.063314674 |
| Reg2 | 4.297572 | 4.435645 | 4.3403 | 0.095376214 | -0.042697208 | 4.3030888 | 4.6557224 | 4.58418 | 0.071538135 | -0.281095408 |
| Rbm38 | 10.42533 | 10.6479 | 10.558 | 0.089474287 | -0.133095938 | 10.738485 | 10.6008 | 10.5293 | 0.0714701 | 0.209155065 |
| Faf2 | 10.29247 | 10.23819 | 10.145 | 0.093659232 | 0.147942074 | 10.140447 | 10.216555 | 10.1451 | 0.071421804 | -0.004686486 |
| Sfn | 8.787054 | 8.809021 | 8.8341 | -0.02510525 | -0.047071796 | 8.9618986 | 8.7712859 | 8.69987 | 0.071416106 | 0.26202883 |
| Prrc2c | 13.00843 | 13.16041 | 13.02 | 0.14050068 | -0.011478454 | 13.023528 | 13.287646 | 13.2163 | 0.07136748 | -0.192750047 |
| Npc1l1 | 1.626627 | 1.662711 | 1.6858 | -0.023099483 | -0.059183234 | 1.6773094 | 1.7909113 | 1.71955 | 0.071357403 | -0.042244444 |
| Pcdhgb5 | 3.698295 | 3.517461 | 3.5522 | -0.034696523 | 0.146137283 | 3.568282 | 3.737732 | 3.66658 | 0.071152818 | -0.098297243 |
| Gm16089 | 1.052427 | 1.024264 | 1.0705 | -0.046227607 | -0.018065362 | 1.0602911 | 1.0633033 | 0.9922 | 0.071103589 | 0.068091318 |
| Dock3 | 1.485288 | 1.511259 | 1.2905 | 0.220738006 | 0.194767471 | 1.2132387 | 1.3104478 | 1.23936 | 0.071091876 | -0.026117152 |
| Nacad | 0.952706 | 0.848996 | 0.8819 | -0.032917051 | 0.070793039 | 0.8285029 | 0.9034528 | 0.83241 | 0.071041208 | -0.003908738 |
| Zic5 | 1.131729 | 1.152276 | 1.1168 | 0.035457748 | 0.014910141 | 1.1534322 | 1.1921042 | 1.12107 | 0.071032819 | 0.032360773 |
| Gm10163 | 11.82787 | 11.74778 | 11.845 | -0.096767856 | -0.016685902 | 11.729012 | 11.853027 | 11.782 | 0.071032224 | -0.052982879 |
| AL603804.1 | 3.40037 | 3.476818 | 3.4845 | -0.007669895 | -0.084118128 | 3.4072072 | 3.4972692 | 3.42628 | 0.070990582 | -0.019071501 |
| Atg5 | 9.177378 | 9.238476 | 9.1997 | 0.038744926 | -0.022353486 | 9.2563984 | 9.2096935 | 9.1387 | 0.070990001 | 0.11769492 |
| Gnaq | 8.562965 | 8.62792 | 8.6645 | -0.036595114 | -0.101550643 | 8.6438323 | 8.6576273 | 8.58666 | 0.070964194 | 0.057169158 |
| B4galt2 | 3.002444 | 2.967356 | 2.8878 | 0.079574087 | 0.114662231 | 2.8587057 | 2.9576801 | 2.88675 | 0.07092714 | -0.028047218 |
| Clca2 | 1.179514 | 1.251608 | 1.1899 | 0.061743072 | -0.010350793 | 1.2451722 | 1.2593594 | 1.18844 | 0.070916829 | 0.056729604 |
| Axdnd1 | 3.845524 | 3.979519 | 3.9288 | 0.050763367 | -0.083231963 | 4.0342503 | 4.0725532 | 4.00166 | 0.070893421 | 0.032590523 |
| Unc119b | 10.01636 | 10.00914 | 9.986 | 0.023176696 | 0.030395444 | 10.054892 | 10.022616 | 9.95175 | 0.070861419 | 0.103137215 |
| Map3k5 | 9.96185 | 9.935448 | 9.9158 | 0.01968406 | 0.046085768 | 9.8869306 | 10.102291 | 10.0315 | 0.07080666 | -0.14455336 |
| Klrb1a | 4.091955 | 4.132514 | 4.2022 | -0.069709508 | -0.110268341 | 3.8816688 | 4.1082445 | 4.03749 | 0.070755707 | -0.155820001 |
| Sec11c | 10.19159 | 10.19337 | 10.187 | 0.006657009 | 0.004880382 | 10.282821 | 10.305076 | 10.2344 | 0.070698486 | 0.04844353 |
| Hoxb3 | 3.929283 | 3.950068 | 3.9681 | -0.017990773 | -0.038775906 | 4.0088462 | 3.9711851 | 3.90052 | 0.070663487 | 0.108324645 |
| Mgarp | 2.291014 | 2.267076 | 2.2325 | 0.03460392 | 0.058542033 | 2.2409844 | 2.4244955 | 2.35383 | 0.070660657 | -0.112850495 |
| Vmn1r183 | 4.424751 | 4.381839 | 4.4468 | -0.064946627 | -0.022034398 | 4.4996675 | 4.6887707 | 4.6182 | 0.070573765 | -0.118529373 |
| Cd79b | 13.06382 | 13.20089 | 13.209 | -0.00845358 | -0.145523204 | 13.650121 | 13.208171 | 13.1376 | 0.070563684 | 0.512513794 |
| Tmem242 | 8.615795 | 8.540564 | 8.6919 | -0.151296746 | -0.076066272 | 8.6708929 | 8.6947409 | 8.62418 | 0.070559688 | 0.046711671 |
| Ccdc78 | 2.801525 | 2.784114 | 2.8231 | -0.039024072 | -0.02161303 | 2.9605678 | 3.0037796 | 2.93323 | 0.070546311 | 0.027334501 |
| Slc25a47 | 4.116303 | 4.001527 | 4.0604 | -0.05884319 | 0.055932791 | 4.126737 | 4.1481684 | 4.07764 | 0.070526418 | 0.049094975 |
| Gm13255 | 0.689257 | 0.660169 | 0.6752 | -0.014996554 | 0.014091796 | 0.7157186 | 0.7695405 | 0.69906 | 0.070477477 | 0.016655527 |
| Vmn2r115 | 5.003875 | 5.011368 | 5.1688 | -0.157445036 | -0.164937455 | 5.0656823 | 5.4525669 | 5.38213 | 0.07043641 | -0.31644814 |

| | | | | | | | | | | |
|---|---|---|---|---|---|---|---|---|---|---|
| Gm8451 | 2.315279 | 2.239085 | 2.2474 | -0.008341076 | 0.06785279 | 2.217608 | 2.2927324 | 2.22231 | 0.070418031 | -0.004706301 |
| Gm6030 | 3.567263 | 3.580347 | 3.633 | -0.052612882 | -0.065696705 | 3.6444793 | 3.7317634 | 3.66138 | 0.070380204 | -0.016903929 |
| Tctex1d2 | 5.887852 | 5.86863 | 5.8989 | -0.030229399 | -0.011007928 | 6.0881936 | 5.990551 | 5.92018 | 0.070373264 | 0.16801589 |
| Ighv8-5 | 4.238661 | 4.340667 | 4.348 | -0.007341763 | -0.109348258 | 4.5486599 | 4.4178224 | 4.3475 | 0.070326104 | 0.201163624 |
| Gm28044 | 1.94282 | 1.998196 | 1.9192 | 0.079009047 | 0.023633113 | 1.9562256 | 2.0795204 | 2.0092 | 0.070325069 | -0.052969796 |
| Gm49111 | 1.017875 | 1.004254 | 1.0193 | -0.015057392 | -0.001436214 | 1.0299516 | 1.0864077 | 1.01627 | 0.07014104 | 0.013684933 |
| Synj2bp | 9.980187 | 10.0527 | 9.931 | 0.121671941 | 0.049156803 | 9.9083329 | 9.9725865 | 9.90248 | 0.070101754 | 0.005848146 |
| Cd8b1 | 11.7535 | 11.81192 | 11.858 | -0.046138302 | -0.104559226 | 11.657611 | 11.889893 | 11.8198 | 0.070101204 | -0.162180749 |
| Dlst | 11.05184 | 11.10732 | 11.094 | 0.013537394 | -0.041942098 | 11.140319 | 11.1958 | 11.1258 | 0.070029758 | 0.014549239 |
| Tusc1 | 3.606623 | 3.682931 | 3.6249 | 0.058043937 | -0.018264394 | 3.5152934 | 3.5239051 | 3.4539 | 0.07000971 | 0.061398045 |
| Olfr39 | 1.055917 | 1.064761 | 1.1362 | -0.071461449 | -0.08030477 | 1.1085305 | 1.1562559 | 1.08625 | 0.070003954 | 0.022278523 |
| Eqtn | 1.83041 | 1.825004 | 1.8624 | -0.037375859 | -0.031969068 | 1.8624627 | 1.9229308 | 1.85304 | 0.069893404 | 0.00942522 |
| Aco2 | 11.46887 | 11.47866 | 11.441 | 0.037703572 | 0.027914234 | 11.677194 | 11.542819 | 11.4729 | 0.06989001 | 0.204264461 |
| Padi1 | 2.966753 | 3.149812 | 3.2077 | -0.057850021 | -0.240909028 | 3.0097958 | 3.1891303 | 3.11925 | 0.069884491 | -0.109449974 |
| Trbv21 | 0.815849 | 0.83098 | 0.8564 | -0.025434901 | -0.040565193 | 0.8581183 | 0.9066642 | 0.83679 | 0.069873634 | 0.021327744 |
| Gm20784 | 3.391459 | 3.267396 | 3.271 | -0.003592862 | 0.120469686 | 3.0846705 | 3.2102687 | 3.14044 | 0.069831933 | -0.055766294 |
| Zfp992 | 6.717686 | 6.624592 | 6.718 | -0.09337906 | -0.000285245 | 6.6107207 | 6.7479411 | 6.67814 | 0.069805085 | -0.067415316 |
| Zcchc17 | 8.050659 | 7.89771 | 7.9872 | -0.089441853 | 0.063507675 | 7.944358 | 7.9570105 | 7.88724 | 0.069766032 | 0.057113516 |
| Tfpt | 7.631733 | 7.704973 | 7.7678 | -0.06286883 | -0.136109089 | 7.8697865 | 7.8248811 | 7.75538 | 0.069505781 | 0.114411242 |
| Suds3 | 10.20025 | 10.35535 | 10.271 | 0.083888216 | -0.071205524 | 10.147954 | 10.357539 | 10.288 | 0.069497062 | -0.140088137 |
| Lrrtm2 | 0.702832 | 0.699981 | 0.7187 | -0.018694054 | -0.015842984 | 0.7345038 | 0.7644338 | 0.69499 | 0.069448761 | 0.039518682 |
| Stn1 | 8.259557 | 8.199082 | 8.3073 | -0.108241212 | -0.04776636 | 8.4813311 | 8.3008009 | 8.23141 | 0.069394579 | 0.249924794 |
| Guca1a | 1.833833 | 1.850364 | 1.8377 | 0.012675789 | -0.003855043 | 1.8983653 | 1.904327 | 1.83504 | 0.069291185 | 0.063329425 |
| Aoc2 | 7.004395 | 7.126864 | 7.1105 | 0.016398767 | -0.10606978 | 7.4659354 | 7.5008554 | 7.43162 | 0.069233051 | 0.034313079 |
| Dhps | 8.754288 | 8.76164 | 8.8086 | -0.046990117 | -0.05434206 | 9.1524148 | 9.051861 | 8.98268 | 0.069181626 | 0.16973542 |
| Chek2 | 6.530792 | 6.456955 | 6.7413 | -0.284370441 | -0.210532918 | 6.8037103 | 6.7096862 | 6.64053 | 0.0691605 | 0.163184609 |
| Camk1 | 6.487216 | 6.074992 | 6.2993 | -0.224271461 | 0.187952403 | 6.3787274 | 5.9850351 | 5.91591 | 0.069128542 | 0.462820751 |
| Uap1l1 | 8.532734 | 8.651893 | 8.4634 | 0.188489691 | 0.06933085 | 9.0946383 | 8.5951061 | 8.526 | 0.069108319 | 0.568640539 |
| Gm13464 | 2.349112 | 2.391417 | 2.3605 | 0.030921144 | -0.011383668 | 2.4789292 | 2.3430759 | 2.27398 | 0.069096272 | 0.204949548 |
| Gak | 11.41456 | 11.44147 | 11.343 | 0.098928358 | 0.072019814 | 11.445854 | 11.541262 | 11.4722 | 0.069061744 | -0.026345728 |
| Ola1 | 9.847524 | 9.86009 | 9.8604 | -0.000315842 | -0.012881908 | 9.852609 | 9.8988941 | 9.82984 | 0.069051127 | 0.022765979 |
| Rhox4d | 2.64131 | 2.599203 | 2.7546 | -0.15537065 | -0.113263888 | 2.827337 | 2.786091 | 2.71706 | 0.069029537 | 0.110275604 |

| | | | | | | | | | | |
|---|---|---|---|---|---|---|---|---|---|---|
| Cnrip1 | 6.418975 | 6.516451 | 6.4093 | 0.107145027 | 0.009669799 | 6.7503575 | 6.51313 | 6.44413 | 0.068996462 | 0.306223993 |
| Toe1 | 8.611823 | 8.591003 | 8.6121 | -0.021127156 | -0.000307948 | 8.8474309 | 8.7338306 | 8.66484 | 0.068985884 | 0.182586253 |
| Topors | 9.608549 | 9.675062 | 9.6604 | 0.014702191 | -0.051811346 | 9.3975588 | 9.663145 | 9.59416 | 0.068983979 | -0.1966022 |
| Efna2 | 4.37875 | 4.422436 | 4.523 | -0.100577919 | -0.144264464 | 4.397141 | 4.515476 | 4.44653 | 0.068947919 | -0.049387116 |
| Dcn | 9.613386 | 9.053303 | 9.4238 | -0.370478026 | 0.189605338 | 8.5372299 | 8.9369727 | 8.86804 | 0.068933418 | -0.330809378 |
| Nudt16l1 | 9.039801 | 9.130323 | 9.0766 | 0.05370642 | -0.036815309 | 9.1627318 | 9.140602 | 9.0717 | 0.068903911 | 0.091033785 |
| Gm43960 | 4.204381 | 4.043184 | 4.1814 | -0.138217376 | 0.022979742 | 4.1261261 | 4.2981664 | 4.22927 | 0.068901078 | -0.103139255 |
| Tm4sf19 | 1.490004 | 1.934555 | 1.4541 | 0.480433509 | 0.035882843 | 1.4693254 | 1.5133474 | 1.44449 | 0.068853512 | 0.02483155 |
| Abcf1 | 11.10377 | 11.30669 | 11.183 | 0.12382697 | -0.079098298 | 11.149316 | 11.413485 | 11.3447 | 0.068784396 | -0.19538446 |
| Saal1 | 8.426594 | 8.401034 | 8.5087 | -0.107700424 | -0.082140893 | 8.5987113 | 8.5512124 | 8.48243 | 0.068783612 | 0.116282558 |
| Pcdhga5 | 3.387128 | 3.304454 | 3.3803 | -0.075874461 | 0.006799305 | 3.4409334 | 3.3459026 | 3.27722 | 0.068686517 | 0.163717335 |
| Amer2 | 1.596547 | 1.579371 | 1.5811 | -0.001679747 | 0.015496454 | 1.6859492 | 1.5702222 | 1.50154 | 0.068680769 | 0.184407702 |
| Smarcd2 | 11.2271 | 11.27763 | 11.196 | 0.081869018 | 0.03134293 | 11.397226 | 11.286265 | 11.2176 | 0.068641892 | 0.179603008 |
| Gm45140 | 9.728317 | 10.00781 | 9.7198 | 0.287960878 | 0.00847003 | 9.7038071 | 9.8127766 | 9.7442 | 0.068574077 | -0.040395449 |
| Gm49028 | 1.665222 | 1.732424 | 1.6664 | 0.065988822 | -0.001213216 | 1.7049584 | 1.7832825 | 1.71471 | 0.068573087 | -0.009751 |
| Cald1 | 7.870012 | 8.1405 | 7.584 | 0.556527027 | 0.286039333 | 7.7323104 | 7.4515732 | 7.383 | 0.06856846 | 0.34930558 |
| Anapc15 | 8.201442 | 8.087415 | 8.3193 | -0.231846383 | -0.117820202 | 8.4441965 | 8.3411596 | 8.27263 | 0.068529252 | 0.171566158 |
| Lmtk3 | 2.735935 | 2.820492 | 2.9291 | -0.108648099 | -0.193204663 | 2.7684899 | 2.7976327 | 2.72916 | 0.06846977 | 0.039326889 |
| 0610010K14Rik | 8.655279 | 8.559192 | 8.6863 | -0.1270684 | -0.030981301 | 8.9130673 | 8.7711291 | 8.70271 | 0.068423881 | 0.210362134 |
| Prkar1a | 12.5012 | 12.5217 | 12.419 | 0.102778768 | 0.082286362 | 12.449539 | 12.496816 | 12.4284 | 0.068386024 | 0.021109043 |
| Ighv1-72 | 8.036838 | 8.264912 | 8.1496 | 0.115270291 | -0.112803871 | 8.2761559 | 8.0599493 | 7.99168 | 0.068267831 | 0.284474365 |
| Tshz3 | 4.870753 | 4.692693 | 4.7502 | -0.05748052 | 0.120579114 | 4.4076963 | 4.7717693 | 4.7036 | 0.068172924 | -0.295900126 |
| Gm46209 | 4.101267 | 4.056473 | 4.087 | -0.03050647 | 0.014287698 | 4.1116829 | 4.1843797 | 4.11623 | 0.068147304 | -0.004549534 |
| Cux2 | 3.021149 | 2.969848 | 3.1014 | -0.13151521 | -0.080214412 | 2.8311805 | 3.0523281 | 2.98419 | 0.068134721 | -0.153012864 |
| Olfr1161 | 0.523722 | 0.480162 | 0.4591 | 0.021018468 | 0.06457894 | 0.5184852 | 0.6012485 | 0.53318 | 0.068071745 | -0.014691605 |
| Dio2 | 1.920901 | 1.920303 | 1.9725 | -0.052192968 | -0.051594811 | 1.8734555 | 2.031668 | 1.96363 | 0.068040874 | -0.090171645 |
| Tmem151a | 2.081409 | 2.105929 | 2.1828 | -0.076860791 | -0.101380475 | 2.1277058 | 2.1825285 | 2.11453 | 0.068000877 | 0.013178243 |
| 4930404N11Rik | 2.716072 | 2.781473 | 2.6817 | 0.099808238 | 0.034407192 | 2.8543259 | 2.8332153 | 2.76524 | 0.067978102 | 0.089088752 |
| Msantd2 | 8.01819 | 8.246412 | 8.161 | 0.085458675 | -0.142763362 | 8.1472766 | 8.4866841 | 8.4188 | 0.067881086 | -0.271526465 |
| Fgfr1op2 | 10.90609 | 10.85437 | 10.912 | -0.058128332 | -0.006413164 | 10.961383 | 10.996606 | 10.9288 | 0.067826748 | 0.032603445 |
| Timm29 | 9.141318 | 9.149162 | 9.1408 | 0.008375359 | 0.000532231 | 9.094418 | 9.0965961 | 9.02878 | 0.067813646 | 0.065635632 |
| Rap1b | 12.75009 | 12.73468 | 12.689 | 0.045519694 | 0.060928004 | 12.677106 | 12.727901 | 12.6601 | 0.067812636 | 0.017017846 |

| | | | | | | | | | | |
|---|---|---|---|---|---|---|---|---|---|---|
| Nomo1 | 10.14283 | 10.17193 | 10.111 | 0.060833746 | 0.031736886 | 10.209074 | 10.213411 | 10.1456 | 0.067762156 | 0.063425003 |
| Akap8 | 11.89503 | 11.94112 | 11.967 | -0.025499458 | -0.071590502 | 12.031949 | 12.186189 | 12.1184 | 0.067753536 | -0.08648674 |
| Rpl17-ps4 | 2.062698 | 2.119796 | 2.119 | 0.000752203 | -0.056346273 | 2.0593735 | 2.1836918 | 2.11606 | 0.067636079 | -0.056682318 |
| Dgcr6 | 5.111445 | 5.066691 | 5.1096 | -0.042868974 | 0.001884996 | 5.0996653 | 5.1430745 | 5.07548 | 0.067595434 | 0.024186221 |
| Dcun1d3 | 7.924666 | 7.961239 | 7.9063 | 0.054950469 | 0.018377219 | 7.7189338 | 7.8863392 | 7.81876 | 0.067577227 | -0.099828087 |
| Atp8b1 | 6.820256 | 6.39424 | 6.6303 | -0.236088359 | 0.189927365 | 5.870697 | 5.9546419 | 5.88707 | 0.067574405 | -0.016370534 |
| Etaa1 | 8.211636 | 8.452932 | 8.4302 | 0.02276837 | -0.21852744 | 8.3014136 | 8.4655112 | 8.398 | 0.06751149 | -0.096586113 |
| Mphosph6 | 7.598445 | 7.500687 | 7.5598 | -0.059129791 | 0.038628249 | 7.5948171 | 7.6252298 | 7.55784 | 0.067391748 | 0.036979006 |
| Trav6d-4 | 2.947413 | 2.965406 | 2.8277 | 0.137684111 | 0.119691744 | 2.8488758 | 2.9377617 | 2.87039 | 0.06737022 | -0.021515678 |
| Galk2 | 8.553787 | 8.536438 | 8.4776 | 0.058811146 | 0.076160416 | 8.5563365 | 8.5903487 | 8.52301 | 0.067339848 | 0.033327711 |
| 1110012L19Rik | 7.193686 | 7.148781 | 7.2716 | -0.122788179 | -0.077883066 | 7.1588865 | 7.2322876 | 7.16498 | 0.067307543 | -0.0060936 |
| Med24 | 10.02344 | 10.04868 | 10.142 | -0.093037511 | -0.11827341 | 10.399169 | 10.265056 | 10.1978 | 0.067298134 | 0.201411466 |
| Psg22 | 2.413169 | 2.452003 | 2.4675 | -0.015508816 | -0.054343296 | 2.430047 | 2.5170572 | 2.44978 | 0.067273091 | -0.019737109 |
| Nup35 | 8.077141 | 8.159042 | 8.1643 | -0.005249316 | -0.08714987 | 8.248138 | 8.2871921 | 8.21993 | 0.067265047 | 0.02821096 |
| Gtf2e2 | 8.15858 | 8.335257 | 8.1997 | 0.135552283 | -0.04112464 | 8.2963261 | 8.3296705 | 8.26249 | 0.067180646 | 0.033836281 |
| Mark4 | 8.966026 | 8.86322 | 8.8257 | 0.037474947 | 0.140281353 | 8.6136305 | 8.8341397 | 8.76699 | 0.067152884 | -0.153356284 |
| Fxyd3 | 2.302684 | 2.105097 | 2.1741 | -0.06901971 | 0.128566601 | 2.352878 | 2.2153438 | 2.1482 | 0.067146158 | 0.204680362 |
| Gm7324 | 0.746439 | 0.823446 | 0.7709 | 0.052594351 | -0.024413096 | 0.7782607 | 0.7989329 | 0.73181 | 0.067120175 | 0.046447958 |
| A2ml1 | 2.763213 | 3.023864 | 2.6284 | 0.395442302 | 0.134791948 | 3.8603164 | 2.924686 | 2.85768 | 0.067009115 | 1.002639605 |
| Cdyl2 | 7.588284 | 7.680998 | 7.7377 | -0.056744347 | -0.14945891 | 7.6519861 | 7.7044631 | 7.63749 | 0.066971342 | 0.014494339 |
| Mlc1 | 2.256694 | 2.255528 | 2.0818 | 0.173772907 | 0.174938967 | 2.1160161 | 2.1747546 | 2.1078 | 0.066952728 | 0.008214261 |
| Zfp65 | 8.31297 | 8.241704 | 8.3131 | -0.071375579 | -0.000110342 | 8.2878166 | 8.3716443 | 8.30477 | 0.066869397 | -0.016958281 |
| Gm21994 | 6.842718 | 6.902707 | 6.8537 | 0.048963054 | -0.011025298 | 6.7670752 | 6.7445563 | 6.67774 | 0.066811416 | 0.089330333 |
| Vmn1r179 | 2.817509 | 2.683738 | 2.7433 | -0.059587519 | 0.074183512 | 2.8029726 | 2.8331202 | 2.76631 | 0.066810435 | 0.036662825 |
| Smarcd3 | 3.128121 | 3.149557 | 3.1206 | 0.02894628 | 0.007509913 | 3.0728768 | 3.1361667 | 3.0694 | 0.066771323 | 0.003481381 |
| Cacna1c | 4.37919 | 4.695628 | 4.2861 | 0.409558732 | 0.09312123 | 4.4412702 | 4.3595794 | 4.29282 | 0.066756625 | 0.148447411 |
| Atg3 | 9.785617 | 9.739468 | 9.747 | -0.007494531 | 0.038653934 | 9.796109 | 9.7784484 | 9.71169 | 0.066754103 | 0.084414633 |
| Dolk | 6.469709 | 6.474184 | 6.3846 | 0.089549302 | 0.085074557 | 6.8062011 | 6.5743552 | 6.50761 | 0.066749439 | 0.298595301 |
| Vamp3 | 9.42415 | 9.28113 | 9.4003 | -0.119195938 | 0.023823226 | 9.3572019 | 9.2366414 | 9.16989 | 0.066746468 | 0.187306978 |
| Fam161b | 4.033163 | 4.061319 | 4.0056 | 0.05568366 | 0.027527314 | 4.1955365 | 4.1509809 | 4.08429 | 0.06669211 | 0.11124768 |
| Olfr1555-ps1 | 2.561559 | 2.612272 | 2.6473 | -0.03505552 | -0.08576835 | 2.6088255 | 2.7604355 | 2.69377 | 0.066665865 | -0.084944147 |
| Mcm5 | 10.34794 | 10.26126 | 10.376 | -0.114244133 | -0.02756575 | 10.558961 | 10.662524 | 10.5959 | 0.066645744 | -0.036916358 |

| | | | | | | | | | | |
|---|---|---|---|---|---|---|---|---|---|---|
| Zpr1 | 9.807081 | 9.960938 | 9.8577 | 0.103276128 | -0.05058085 | 10.049555 | 10.151866 | 10.0853 | 0.066560163 | -0.035750674 |
| Zdhhc5 | 10.64175 | 10.70168 | 10.587 | 0.114951074 | 0.055022966 | 10.528573 | 10.571267 | 10.5047 | 0.06654833 | 0.02385437 |
| Folh1 | 1.777775 | 1.711658 | 1.8298 | -0.118133838 | -0.052017008 | 1.7284715 | 1.8642598 | 1.79773 | 0.066529967 | -0.069258405 |
| Stam2 | 9.053499 | 8.934439 | 8.9164 | 0.018066639 | 0.13712696 | 8.7340003 | 8.9748476 | 8.90832 | 0.066526889 | -0.174320356 |
| Plek2 | 1.996409 | 1.95311 | 1.9445 | 0.008571144 | 0.051870088 | 2.038462 | 2.0907999 | 2.02434 | 0.066455639 | 0.014117721 |
| Zbtb8os | 8.697235 | 8.671496 | 8.6775 | -0.00597991 | 0.01975946 | 8.7580397 | 8.8313894 | 8.76494 | 0.06644882 | -0.006900823 |
| Dyrk1b | 7.003868 | 7.117386 | 7.1223 | -0.004930401 | -0.118449043 | 6.9883859 | 7.0897542 | 7.02333 | 0.066422805 | -0.034945548 |
| Kdelr3 | 3.745598 | 3.755782 | 3.7368 | 0.018985919 | 0.008801132 | 3.8137794 | 3.6626111 | 3.59622 | 0.066386141 | 0.217554437 |
| Mcm10 | 6.482388 | 6.418272 | 6.6306 | -0.21231543 | -0.14819891 | 6.4463981 | 6.9694754 | 6.90311 | 0.066365732 | -0.456711648 |
| Vmn1r224 | 1.500236 | 1.510817 | 1.4874 | 0.023419681 | 0.012839347 | 1.5059707 | 1.6105421 | 1.5442 | 0.066346482 | -0.038224982 |
| F12 | 2.78823 | 2.910847 | 2.8118 | 0.099031976 | -0.023585681 | 3.017307 | 2.9001305 | 2.83382 | 0.066309701 | 0.183486165 |
| Gm49274 | 3.401409 | 3.435806 | 3.491 | -0.055164274 | -0.089560942 | 3.4390665 | 3.6365492 | 3.57025 | 0.066295706 | -0.131186934 |
| Rsad1 | 7.125301 | 7.123431 | 7.0348 | 0.088629086 | 0.090499063 | 7.3721881 | 7.196 | 7.12971 | 0.066285895 | 0.242474035 |
| Tmem130 | 1.229695 | 1.199366 | 1.1692 | 0.03012613 | 0.060454861 | 1.1874238 | 1.2232218 | 1.15694 | 0.06627813 | 0.030480112 |
| Cnksr1 | 1.090428 | 1.156167 | 1.1033 | 0.052825787 | -0.012913171 | 1.1143384 | 1.2408988 | 1.17464 | 0.066260463 | -0.06029995 |
| Skp1a | 10.84427 | 10.80463 | 10.7 | 0.104695048 | 0.144340708 | 10.428111 | 10.785944 | 10.7197 | 0.06625582 | -0.291576562 |
| Aldh8a1 | 2.541149 | 2.466802 | 2.4619 | 0.004911479 | 0.079258215 | 2.446209 | 2.500755 | 2.43451 | 0.066243523 | 0.011697443 |
| Homer2 | 4.535287 | 4.056005 | 4.2135 | -0.157458084 | 0.321824361 | 4.4964392 | 4.3188401 | 4.25262 | 0.066224661 | 0.243823751 |
| Zfp438 | 6.489083 | 6.481413 | 6.4738 | 0.007611849 | 0.015281809 | 6.4286658 | 6.4972693 | 6.43117 | 0.066094414 | -0.002509144 |
| Gm6344 | -0.139365 | -0.13926 | -0.0155 | -0.123771084 | -0.123872095 | -0.0831078 | -0.072583 | -0.1387 | 0.066083506 | 0.055558608 |
| Iqsec1 | 11.53213 | 11.52079 | 11.504 | 0.016712842 | 0.028057016 | 11.631268 | 11.603906 | 11.5378 | 0.066072464 | 0.093434822 |
| Mcrip2 | 4.596953 | 4.562241 | 4.7056 | -0.143368633 | -0.108656821 | 4.652711 | 4.5986073 | 4.53258 | 0.066029916 | 0.120133565 |
| F8a | 6.795957 | 6.777077 | 6.707 | 0.070092721 | 0.088973277 | 6.8414044 | 6.6418752 | 6.57585 | 0.06602294 | 0.265552102 |
| Flot1 | 9.296397 | 8.813368 | 8.8222 | -0.008785404 | 0.474244057 | 9.2877034 | 8.8845053 | 8.81849 | 0.066016821 | 0.469214933 |
| Fat3 | 1.057357 | 1.055446 | 1.0719 | -0.016411939 | -0.014500535 | 1.0062192 | 1.0336902 | 0.96773 | 0.065961275 | 0.038490318 |
| AA413626 | 3.298707 | 3.271076 | 3.326 | -0.054890663 | -0.027259348 | 3.3967013 | 3.4960832 | 3.4302 | 0.065879895 | -0.033502057 |
| Slc7a13 | 1.815738 | 1.789492 | 1.8972 | -0.107660414 | -0.081414943 | 1.9386026 | 1.8896365 | 1.82376 | 0.065874947 | 0.114841111 |
| Polr3h | 7.63889 | 7.418928 | 7.6125 | -0.193524083 | 0.026438591 | 7.8127958 | 7.5884193 | 7.52255 | 0.065865249 | 0.290241715 |
| Ppp3cc | 10.48601 | 10.44819 | 10.509 | -0.061245326 | -0.02342457 | 10.557944 | 10.598946 | 10.5331 | 0.065865172 | 0.024863034 |
| Dgkk | 2.859429 | 2.924604 | 2.8402 | 0.084441163 | 0.019266139 | 2.8411406 | 2.9636374 | 2.8978 | 0.065832451 | -0.056664314 |
| Tnfaip1 | 9.156044 | 9.041675 | 9.0666 | -0.024955292 | 0.089413422 | 9.0922975 | 9.0764435 | 9.01062 | 0.065825136 | 0.081679128 |
| Pts | 8.726185 | 8.561122 | 8.7307 | -0.169535099 | -0.004471554 | 8.9238107 | 8.8128967 | 8.74711 | 0.065782403 | 0.176696413 |

| | | | | | | | | | | |
|---|---|---|---|---|---|---|---|---|---|---|
| Mmachc | 7.001157 | 7.172376 | 7.0431 | 0.129322304 | -0.041896846 | 7.2612451 | 7.1108407 | 7.04511 | 0.065729858 | 0.216134304 |
| Ncaph2 | 10.61496 | 10.58695 | 10.67 | -0.082629542 | -0.054618184 | 10.975264 | 10.927326 | 10.8616 | 0.065679672 | 0.113617999 |
| Fxr2 | 10.08945 | 10.1515 | 10.228 | -0.076219993 | -0.138265148 | 10.327448 | 10.34768 | 10.282 | 0.06566792 | 0.045436325 |
| Tbx1 | 4.080848 | 3.938845 | 3.7507 | 0.188134015 | 0.330136521 | 3.6618528 | 3.8601655 | 3.79454 | 0.065622621 | -0.132690142 |
| Sdhaf1 | 8.016299 | 8.033541 | 8.146 | -0.112420539 | -0.129662972 | 8.0455043 | 8.0668873 | 8.00135 | 0.065539144 | 0.044156183 |
| Ccdc136 | 2.080407 | 2.015715 | 2.1539 | -0.138167863 | -0.073475934 | 2.1925741 | 2.1252951 | 2.05978 | 0.065518957 | 0.132797964 |
| Cdc20b | 0.247465 | 0.283782 | 0.2684 | 0.015371841 | -0.02094534 | 0.2534229 | 0.3374942 | 0.272 | 0.065491154 | -0.018580137 |
| Glb1l2 | 2.406967 | 2.375571 | 2.3767 | -0.001104109 | 0.030292038 | 2.3530495 | 2.3903006 | 2.32482 | 0.065481253 | 0.028230184 |
| Ccdc166 | 6.263116 | 6.13672 | 6.295 | -0.158309565 | -0.031913786 | 6.2913698 | 6.3195923 | 6.25413 | 0.06546528 | 0.03724278 |
| Hist1h4a | 1.859418 | 1.780068 | 1.9226 | -0.14255299 | -0.063202639 | 1.8901028 | 1.9044853 | 1.83907 | 0.065414815 | 0.051032353 |
| Fam114a2 | 9.912096 | 9.971534 | 9.9813 | -0.009755682 | -0.06919311 | 9.9852166 | 10.089147 | 10.0238 | 0.065354126 | -0.038576598 |
| Cltc | 12.71728 | 12.68508 | 12.607 | 0.077995468 | 0.110193686 | 12.680288 | 12.685138 | 12.6198 | 0.06533545 | 0.060484719 |
| Ap3s1-ps2 | 0.447576 | 0.398985 | 0.4358 | -0.036844094 | 0.011746806 | 0.4338758 | 0.4451044 | 0.37978 | 0.065320579 | 0.054091982 |
| Cdca7 | 8.556519 | 8.466643 | 8.5502 | -0.083599817 | 0.006275984 | 8.6437696 | 8.7335371 | 8.66822 | 0.065317658 | -0.024449892 |
| Kcnk10 | 3.679466 | 3.549192 | 3.7531 | -0.203911349 | -0.073637572 | 3.6649265 | 3.6608898 | 3.59568 | 0.065208127 | 0.069244809 |
| Tmc1 | 2.252984 | 2.292376 | 2.2504 | 0.041967208 | 0.002575226 | 2.3626634 | 2.3793138 | 2.31425 | 0.065063009 | 0.048412607 |
| mt-Co3 | 15.86995 | 16.02072 | 16 | 0.021074508 | -0.129694728 | 16.122503 | 16.03062 | 15.9656 | 0.065046658 | 0.156929427 |
| Erv3 | 1.225826 | 1.289498 | 1.2817 | 0.007774198 | -0.055897285 | 1.2376813 | 1.2752537 | 1.21031 | 0.064941587 | 0.02736919 |
| Tmem246 | 3.356215 | 3.393884 | 3.3701 | 0.023797972 | -0.013870463 | 3.2764518 | 3.3879241 | 3.32304 | 0.06487961 | -0.046592718 |
| Olfr459 | 0.413325 | 0.424402 | 0.3935 | 0.030858448 | 0.019781565 | 0.4404836 | 0.4865686 | 0.42172 | 0.064853031 | 0.018768068 |
| Ucp2 | 13.25281 | 13.2521 | 13.239 | 0.013169108 | 0.013878578 | 13.383675 | 13.157744 | 13.093 | 0.064755536 | 0.290686043 |
| Nrl | 1.828964 | 1.89299 | 1.8251 | 0.067896233 | 0.003869832 | 1.9124742 | 1.9908908 | 1.92614 | 0.064754292 | -0.01366234 |
| Kcnj8 | 6.036112 | 5.705322 | 5.733 | -0.027658258 | 0.303131899 | 5.6721902 | 5.5111934 | 5.44646 | 0.064729728 | 0.225726556 |
| Ighd | 14.27919 | 14.33546 | 14.338 | -0.00237468 | -0.05863996 | 14.871835 | 14.462841 | 14.3982 | 0.064684428 | 0.473678352 |
| Csmd2 | 2.388272 | 2.343871 | 2.3653 | -0.021403405 | 0.022997051 | 2.4003565 | 2.4117018 | 2.3471 | 0.064606147 | 0.053260814 |
| Ints12 | 8.520688 | 8.587474 | 8.5889 | -0.001395735 | -0.068182125 | 8.4359226 | 8.6197326 | 8.55513 | 0.064604614 | -0.119205448 |
| Pllp | 1.413894 | 1.493486 | 1.5177 | -0.024233543 | -0.103825604 | 1.5391492 | 1.5131247 | 1.44855 | 0.064577131 | 0.090601566 |
| Marf1 | 11.16845 | 11.35317 | 11.277 | 0.07638831 | -0.108332918 | 11.345487 | 11.584124 | 11.5196 | 0.06455898 | -0.174078377 |
| Hdac3 | 9.822374 | 9.733816 | 9.8945 | -0.160696228 | -0.072138211 | 9.9489522 | 9.9449329 | 9.88054 | 0.064388455 | 0.068407776 |
| Vmn1r191 | 3.009328 | 2.913867 | 2.9871 | -0.073223816 | 0.02223643 | 3.0622877 | 3.0773614 | 3.01298 | 0.064382145 | 0.049308417 |
| Mier1 | 10.99195 | 11.08646 | 11.147 | -0.060329154 | -0.15483284 | 10.908509 | 11.196596 | 11.1322 | 0.064346832 | -0.223740004 |
| Tst | 4.203776 | 4.333726 | 4.2502 | 0.083501828 | -0.046447583 | 4.2641437 | 4.2949492 | 4.2306 | 0.064345969 | 0.033540446 |

| | | | | | | | | | | |
|---|---|---|---|---|---|---|---|---|---|---|
| Pygm | 5.40438 | 5.232378 | 5.4223 | -0.189872702 | -0.017871238 | 5.427768 | 5.5123505 | 5.44802 | 0.064329457 | -0.020253061 |
| Gm49396 | 8.100256 | 8.191216 | 8.3444 | -0.153145248 | -0.244105513 | 8.8083301 | 8.8423699 | 8.77805 | 0.064323195 | 0.030283358 |
| Trbj2-5 | 4.315696 | 4.665884 | 4.5723 | 0.093627363 | -0.256560336 | 4.770126 | 4.6737756 | 4.60947 | 0.064304298 | 0.16065478 |
| Ccdc130 | 7.82366 | 7.96626 | 7.8622 | 0.104013917 | -0.038585302 | 8.1628068 | 8.0020348 | 7.93774 | 0.064291391 | 0.225063343 |
| Fbxo44 | 4.15503 | 4.117287 | 4.2722 | -0.154944124 | -0.117201861 | 4.2423009 | 4.1894163 | 4.12521 | 0.064201967 | 0.117086601 |
| Tial1 | 10.17535 | 10.2354 | 10.268 | -0.032773156 | -0.092822913 | 10.326421 | 10.452323 | 10.3882 | 0.064166996 | -0.061734736 |
| Gm8326 | 1.421419 | 1.418472 | 1.4299 | -0.011406072 | -0.008459048 | 1.3931499 | 1.4038161 | 1.33969 | 0.06412598 | 0.05345972 |
| Hmces | 8.492366 | 8.607586 | 8.5849 | 0.022715547 | -0.092504204 | 8.7410956 | 8.7142553 | 8.65022 | 0.064036003 | 0.090876306 |
| Acyp2 | 3.409601 | 3.479174 | 3.5223 | -0.043164158 | -0.112737323 | 3.4499244 | 3.5094907 | 3.44547 | 0.064022958 | 0.004456674 |
| Vmn1r227 | 5.665895 | 5.491626 | 5.4591 | 0.032509939 | 0.206778858 | 5.5326608 | 5.7763136 | 5.71232 | 0.063997777 | -0.179655048 |
| Rgl2 | 10.6062 | 10.88751 | 10.758 | 0.130001592 | -0.151301632 | 11.193389 | 11.017636 | 10.9537 | 0.06398547 | 0.239738599 |
| Hist2h3b | 2.706982 | 2.608562 | 2.6123 | -0.003715898 | 0.094704726 | 2.702724 | 2.7037051 | 2.63973 | 0.063970283 | 0.062989131 |
| Zbtb1 | 10.06163 | 10.11948 | 10.153 | -0.033201658 | -0.091046284 | 10.159455 | 10.368059 | 10.3041 | 0.0639522 | -0.144651697 |
| Inf2 | 9.122808 | 9.180274 | 9.1438 | 0.036467087 | -0.020999407 | 9.0797648 | 9.1663929 | 9.1025 | 0.063895569 | -0.022732478 |
| Slc25a19 | 9.643067 | 9.569831 | 9.512 | 0.0578222 | 0.131057835 | 10.084974 | 9.8531591 | 9.78927 | 0.063886398 | 0.295701374 |
| Gm49322 | 6.625656 | 7.088971 | 7.0309 | 0.058031194 | -0.405283995 | 7.0298677 | 7.1588496 | 7.09511 | 0.063741696 | -0.065240219 |
| Serpinb1b | 2.749402 | 2.577063 | 2.6862 | -0.109128143 | 0.063211388 | 2.6243257 | 2.7996409 | 2.73595 | 0.063694745 | -0.111620458 |
| Esr1 | 6.26406 | 6.560196 | 6.4299 | 0.130319307 | -0.165816592 | 6.5027972 | 6.6178359 | 6.55423 | 0.063601817 | -0.05143686 |
| Rps11-ps1 | 2.655539 | 2.680869 | 2.7433 | -0.062457428 | -0.087787547 | 2.6899852 | 2.8079409 | 2.74436 | 0.063582883 | -0.054372836 |
| Triap1 | 8.216825 | 8.174804 | 8.2138 | -0.03895943 | 0.003061011 | 8.4455201 | 8.3247468 | 8.26117 | 0.063576992 | 0.184350359 |
| Haus6 | 9.115393 | 9.195809 | 9.1248 | 0.070979902 | -0.009436168 | 9.0473646 | 9.2205701 | 9.15703 | 0.063536795 | -0.109668717 |
| Mettl16 | 9.567554 | 9.73296 | 9.5746 | 0.158390659 | -0.007015424 | 9.5650355 | 9.6774353 | 9.61405 | 0.063384831 | -0.049014956 |
| Gm16199 | 5.41476 | 5.496064 | 5.4462 | 0.049901298 | -0.031403085 | 5.3644614 | 5.4457864 | 5.38242 | 0.063368689 | -0.017956323 |
| Arid4a | 10.81693 | 10.92602 | 10.89 | 0.036264618 | -0.072824646 | 10.590916 | 10.918401 | 10.8551 | 0.063326546 | -0.264158933 |
| Actr1a | 10.76858 | 10.87896 | 10.802 | 0.076983652 | -0.033390826 | 10.889258 | 10.843356 | 10.7801 | 0.06327652 | 0.109179167 |
| Hmgb1-rs16 | 1.563043 | 1.589516 | 1.5831 | 0.006390743 | -0.020081717 | 1.5635484 | 1.62479 | 1.56156 | 0.063226199 | 0.001984634 |
| Olfr102 | 1.82442 | 1.791108 | 1.8137 | -0.022564545 | 0.010748056 | 1.7371742 | 1.8198774 | 1.75666 | 0.063219572 | -0.019483641 |
| Ciz1 | 10.32681 | 10.28945 | 10.41 | -0.120831743 | -0.083477252 | 10.631096 | 10.690473 | 10.6273 | 0.063199236 | 0.003822321 |
| Gm19000 | 2.949904 | 2.935486 | 2.9581 | -0.022612895 | -0.008194459 | 2.9849116 | 3.1297177 | 3.06653 | 0.063189788 | -0.081616349 |
| Minar1 | 2.072203 | 2.087633 | 2.0597 | 0.027886541 | 0.01245647 | 1.9944528 | 2.1663907 | 2.10323 | 0.06316224 | -0.108775707 |
| Gm19016 | 0.900944 | 0.87908 | 0.8919 | -0.012830745 | 0.00903279 | 0.9589118 | 0.9461544 | 0.8831 | 0.063058223 | 0.0758156 |
| Zfp52 | 8.67402 | 8.832072 | 8.7765 | 0.055609958 | -0.102442629 | 8.5118597 | 8.6349496 | 8.57191 | 0.063038613 | -0.060051333 |

| | | | | | | | | | | |
|---|---|---|---|---|---|---|---|---|---|---|
| Gm48808 | 0.234739 | 0.198023 | 0.2317 | -0.033641134 | 0.003075032 | 0.2627259 | 0.2795563 | 0.21665 | 0.062901865 | 0.046071447 |
| Rpl17-ps7 | 3.239595 | 3.16146 | 3.1964 | -0.034905081 | 0.043230204 | 3.2427469 | 3.3319839 | 3.26909 | 0.06289609 | -0.026340891 |
| Ppp1r12a | 11.62895 | 11.67317 | 11.613 | 0.059808744 | 0.01558787 | 11.42478 | 11.745891 | 11.6831 | 0.062823978 | -0.258286397 |
| 1700030C10Rik | 1.468344 | 1.452217 | 1.5879 | -0.135646788 | -0.119519491 | 1.5016176 | 1.5197515 | 1.45694 | 0.062812554 | 0.044678619 |
| Igkv4-51 | 4.358313 | 4.392746 | 4.2187 | 0.174020814 | 0.139587893 | 4.3597329 | 4.1640467 | 4.10125 | 0.062797335 | 0.258483502 |
| Afap1l2 | 4.086225 | 4.137088 | 4.0207 | 0.116374334 | 0.065510887 | 4.2097297 | 4.1434644 | 4.08071 | 0.062758122 | 0.129023436 |
| Ighv1-79 | 0.15499 | 0.132174 | 0.1567 | -0.024523397 | -0.001707104 | 0.1372539 | 0.1961403 | 0.13344 | 0.062696731 | 0.003810364 |
| Sri | 11.06385 | 11.04731 | 11.019 | 0.028015408 | 0.044553282 | 11.12104 | 11.071924 | 11.0092 | 0.062691918 | 0.111807434 |
| Gm6311 | 0.543809 | 0.499919 | 0.539 | -0.039096087 | 0.004793526 | 0.5700499 | 0.5973999 | 0.53483 | 0.062573071 | 0.035223093 |
| Fam221b | 4.780667 | 4.717194 | 4.8027 | -0.085468996 | -0.021995529 | 4.7326564 | 4.7624261 | 4.69986 | 0.06256612 | 0.032796343 |
| Eif2b4 | 8.923327 | 8.903989 | 8.9525 | -0.048480077 | -0.029142402 | 9.2709967 | 9.0776654 | 9.01514 | 0.062527635 | 0.255858964 |
| Vcp-rs | 2.725425 | 2.723586 | 2.9713 | -0.247668366 | -0.24582926 | 2.6095423 | 2.8535103 | 2.791 | 0.062507438 | -0.181460586 |
| Gsdmc4 | 1.938024 | 1.939823 | 1.8908 | 0.049009885 | 0.047211605 | 1.9335394 | 1.9966227 | 1.93413 | 0.062495421 | -0.000587893 |
| Tmem43 | 9.68014 | 9.651685 | 9.674 | -0.022275255 | 0.006179125 | 9.632766 | 9.6023865 | 9.53992 | 0.062464775 | 0.092844285 |
| Gm8125 | 1.570225 | 1.513196 | 1.583 | -0.069769726 | -0.012741024 | 1.569976 | 1.6391007 | 1.57664 | 0.062458153 | -0.006666563 |
| Sulf2 | 8.412813 | 8.352938 | 8.6283 | -0.275319422 | -0.215444227 | 8.4765121 | 8.4386836 | 8.37629 | 0.062391939 | 0.100220403 |
| Cfap46 | 2.239973 | 2.162946 | 2.1774 | -0.014482007 | 0.062544812 | 2.2162031 | 2.3343061 | 2.27194 | 0.062363898 | -0.055739117 |
| Igbp1 | 10.20831 | 10.19914 | 10.346 | -0.146390284 | -0.137222582 | 10.259001 | 10.304037 | 10.2417 | 0.062306042 | 0.017270144 |
| Hoxb8 | 2.194937 | 2.113383 | 2.1048 | 0.008583725 | 0.090138588 | 1.9864225 | 2.0639147 | 2.00166 | 0.062254491 | -0.015237682 |
| Hivep1 | 10.51135 | 10.65585 | 10.472 | 0.183998408 | 0.039498332 | 10.454563 | 10.873139 | 10.8111 | 0.06205864 | -0.356517118 |
| Pik3c3 | 9.892559 | 9.921985 | 9.9811 | -0.059097554 | -0.088524042 | 9.8640968 | 9.9709458 | 9.90895 | 0.061996583 | -0.044852445 |
| Tent4b | 10.20342 | 10.14256 | 10.167 | -0.024451875 | 0.036409022 | 9.9683061 | 10.163445 | 10.1015 | 0.061991915 | -0.133147431 |
| Arv1 | 7.002863 | 7.10657 | 7.0965 | 0.01008467 | -0.093622082 | 7.3347359 | 7.3390633 | 7.27708 | 0.061980622 | 0.057653211 |
| Ccdc57 | 3.229048 | 3.120598 | 3.1276 | -0.006956338 | 0.101493732 | 3.0795323 | 3.2069139 | 3.14494 | 0.06196925 | -0.065412354 |
| Gm9027 | 3.349895 | 3.474662 | 3.2763 | 0.198336324 | 0.073569655 | 3.3977669 | 3.4246105 | 3.36265 | 0.061957905 | 0.035114337 |
| Gm10269 | 1.453449 | 1.43722 | 1.4579 | -0.020679157 | -0.004450385 | 1.3813838 | 1.46579 | 1.40385 | 0.0619401 | -0.022466153 |
| Dact2 | 1.808561 | 1.808233 | 1.8979 | -0.089626179 | -0.08929781 | 1.7997835 | 1.8318394 | 1.76997 | 0.061873622 | 0.029817732 |
| CT010583.1 | 0.845401 | 0.88247 | 0.858 | 0.024501355 | -0.012567109 | 0.8609984 | 0.9005409 | 0.83867 | 0.061868242 | 0.022325727 |
| Gm11839 | 2.125533 | 2.07622 | 2.1039 | -0.027711038 | 0.021602323 | 2.2121349 | 2.2030042 | 2.14114 | 0.061866069 | 0.070996857 |
| Iglj3 | 3.279829 | 3.285291 | 3.259 | 0.026321659 | 0.020859287 | 3.3484309 | 3.2512364 | 3.18938 | 0.061859409 | 0.159053933 |
| Zic1 | 1.943077 | 1.965282 | 1.9693 | -0.003969699 | -0.026175024 | 2.0068373 | 2.0495124 | 1.98766 | 0.061852178 | 0.019177069 |
| Gm13680 | 3.066578 | 3.113513 | 3.0899 | 0.023652209 | -0.02328288 | 3.0291081 | 3.1114045 | 3.04957 | 0.061832077 | -0.020464318 |

| | | | | | | | | | | |
|---|---|---|---|---|---|---|---|---|---|---|
| Cfap70 | 0.922135 | 0.895174 | 0.8652 | 0.029985274 | 0.056946438 | 0.9636197 | 0.9539628 | 0.89217 | 0.061792913 | 0.071449805 |
| Gm5616 | 2.847218 | 2.947293 | 3.1951 | -0.247767865 | -0.347843503 | 3.0487833 | 2.8941533 | 2.83237 | 0.061786091 | 0.21641613 |
| Ppp1r9b | 11.56033 | 11.54183 | 11.51 | 0.031927216 | 0.050429902 | 11.391186 | 11.426513 | 11.3647 | 0.061774792 | 0.026447008 |
| Gm9892 | 2.759495 | 3.124412 | 2.7519 | 0.372497853 | 0.007581227 | 2.7051739 | 3.0878105 | 3.02608 | 0.061725718 | -0.32091094 |
| Srsf5 | 12.51909 | 12.03263 | 12.453 | -0.4199264 | 0.06653756 | 12.440316 | 12.381044 | 12.3193 | 0.061715494 | 0.120986874 |
| Rps27rt | 14.6486 | 14.69622 | 14.671 | 0.025076192 | -0.0225426 | 14.835786 | 14.705263 | 14.6436 | 0.06169731 | 0.192220805 |
| Ubxn7 | 10.66996 | 10.8914 | 10.764 | 0.127566192 | -0.09387439 | 10.594859 | 10.882191 | 10.8205 | 0.061654794 | -0.225677177 |
| Zfp276 | 8.901149 | 9.142885 | 9.0455 | 0.097390932 | -0.144344821 | 9.2197747 | 9.339793 | 9.27815 | 0.061641194 | -0.05837718 |
| Ston1 | 5.839303 | 5.806984 | 5.9485 | -0.141501979 | -0.109182086 | 5.5528419 | 5.5976575 | 5.53603 | 0.061628401 | 0.016812831 |
| 8030462N17Rik | 8.726351 | 8.754058 | 8.6708 | 0.08330375 | 0.055596912 | 8.5499802 | 8.6822664 | 8.62065 | 0.061620934 | -0.070665211 |
| Snap25 | 3.079714 | 3.101031 | 2.6986 | 0.402398207 | 0.381081666 | 2.8578125 | 2.9005321 | 2.83891 | 0.061618823 | 0.018899208 |
| Gm13216 | 1.547918 | 1.469315 | 1.4567 | 0.012597251 | 0.091200289 | 1.5398893 | 1.514613 | 1.45308 | 0.061535704 | 0.086811992 |
| Tomm40l | 8.597425 | 8.54382 | 8.6899 | -0.146094324 | -0.092489034 | 8.9114075 | 8.7778067 | 8.71629 | 0.061517252 | 0.195118125 |
| Rab3c | 2.99874 | 2.933074 | 2.9027 | 0.030328352 | 0.095995017 | 2.9024863 | 3.02127 | 2.95978 | 0.061492118 | -0.057291562 |
| Vps52 | 10.21426 | 10.33036 | 10.299 | 0.031422932 | -0.084676421 | 10.45459 | 10.437675 | 10.3763 | 0.06141914 | 0.078334461 |
| Gm47265 | 1.388043 | 1.319289 | 1.3518 | -0.032543632 | 0.036209749 | 1.3629647 | 1.4340157 | 1.37261 | 0.061403378 | -0.00964765 |
| Rab27b | 4.924265 | 4.699346 | 4.7756 | -0.076293034 | 0.148625543 | 4.5797262 | 4.6615295 | 4.60015 | 0.061376627 | -0.020426677 |
| Gm48557 | 2.197226 | 2.234961 | 2.2428 | -0.007802211 | -0.045537361 | 2.2078515 | 2.2823053 | 2.22106 | 0.061248605 | -0.013205182 |
| Gigyf1 | 10.66909 | 10.76958 | 10.693 | 0.076179626 | -0.024309877 | 10.916752 | 10.951256 | 10.89 | 0.06124835 | 0.026743425 |
| Gm14336 | 1.277678 | 1.233462 | 1.2029 | 0.0305263 | 0.074742153 | 1.1904656 | 1.2463959 | 1.18515 | 0.061244847 | 0.005314487 |
| Sgsm3 | 6.819813 | 6.897564 | 7.0433 | -0.145695368 | -0.223446206 | 7.0670076 | 7.1610999 | 7.09986 | 0.061244315 | -0.032847894 |
| Cacna2d3 | 2.990519 | 3.004789 | 2.9343 | 0.070454296 | 0.056184197 | 2.8794997 | 2.9968508 | 2.93562 | 0.061226537 | -0.056124597 |
| Olfr1467 | 1.235637 | 1.272756 | 1.2258 | 0.046949728 | 0.009831083 | 1.2337039 | 1.36629 | 1.30513 | 0.061159287 | -0.071426738 |
| Zfp809 | 8.26688 | 8.64227 | 8.4512 | 0.191047313 | -0.1843427 | 8.471452 | 8.5411435 | 8.47999 | 0.061149597 | -0.008541962 |
| Ptpre | 8.978263 | 9.026823 | 8.9405 | 0.08633838 | 0.037778152 | 9.1117724 | 8.88571 | 8.82458 | 0.061125129 | 0.287187472 |
| Rbm45 | 8.848998 | 8.647377 | 8.7877 | -0.140340517 | 0.061280947 | 9.0249056 | 8.9050999 | 8.84398 | 0.061119873 | 0.180925502 |
| Slco1a5 | 1.269219 | 1.30277 | 1.3306 | -0.02786076 | -0.061411962 | 1.3258646 | 1.2706551 | 1.20956 | 0.061097051 | 0.116306477 |
| Kxd1 | 9.650435 | 9.597054 | 9.6762 | -0.079150813 | -0.025769652 | 9.8285836 | 9.8250479 | 9.76397 | 0.061081011 | 0.064616691 |
| Gm7061 | 3.148856 | 3.174899 | 3.1873 | -0.012353696 | -0.038396657 | 3.2807564 | 3.2780538 | 3.21706 | 0.060994052 | 0.063696592 |
| Dapp1 | 10.16023 | 10.33788 | 10.226 | 0.111784933 | -0.06586224 | 10.158994 | 10.38505 | 10.3241 | 0.060910122 | -0.16514588 |
| Cep131 | 6.689648 | 6.836832 | 6.759 | 0.077845658 | -0.069337638 | 7.1111074 | 6.7400183 | 6.67913 | 0.06089145 | 0.431980483 |
| Oser1 | 9.390972 | 9.431458 | 9.5134 | -0.081934861 | -0.122420573 | 9.3811103 | 9.5454045 | 9.48452 | 0.060885532 | -0.103408699 |

| | | | | | | | | | | |
|---|---|---|---|---|---|---|---|---|---|---|
| Nop9 | 9.128697 | 9.119254 | 9.0969 | 0.02238797 | 0.031831732 | 9.3829543 | 9.2589483 | 9.19807 | 0.060875587 | 0.184881516 |
| Pcdhga8 | 1.872916 | 1.886108 | 1.9221 | -0.036019097 | -0.049211183 | 1.9304849 | 1.902777 | 1.84198 | 0.060794205 | 0.08850208 |
| Tgm4 | 2.709738 | 2.654096 | 2.6584 | -0.004338814 | 0.051303033 | 2.6636166 | 2.7977779 | 2.737 | 0.060778233 | -0.073383081 |
| Nrxn1 | 4.040958 | 3.876029 | 3.9496 | -0.07358099 | 0.091347608 | 3.81318 | 3.8699697 | 3.80926 | 0.060705516 | 0.003915781 |
| Gpc1 | 7.562153 | 7.533574 | 7.6107 | -0.077152406 | -0.048572891 | 7.2861417 | 7.5663619 | 7.50566 | 0.060702465 | -0.21951779 |
| Dvl3 | 9.012067 | 9.096825 | 9.0132 | 0.083663175 | -0.001095212 | 9.1966495 | 9.2611724 | 9.20052 | 0.060650566 | -0.003872372 |
| Dync1li1 | 9.786313 | 9.818802 | 9.7692 | 0.049571878 | 0.017083274 | 9.6879162 | 9.7676314 | 9.70707 | 0.06056402 | -0.019151205 |
| Osgep | 9.054891 | 9.158739 | 9.0643 | 0.094410782 | -0.009437305 | 9.2492562 | 9.1440001 | 9.08344 | 0.060555153 | 0.165811331 |
| Klkb1 | 2.386806 | 2.334242 | 2.4354 | -0.101150969 | -0.048587126 | 2.4166557 | 2.3919736 | 2.33142 | 0.060548954 | 0.085231061 |
| Gm48590 | 0.429406 | 0.464932 | 0.4169 | 0.048070549 | 0.012544779 | 0.433728 | 0.5218168 | 0.46132 | 0.060501255 | -0.027587533 |
| Smap2 | 12.16231 | 12.34095 | 12.202 | 0.13865585 | -0.039983042 | 12.337632 | 12.245359 | 12.1849 | 0.060490092 | 0.152762832 |
| Slc35b4 | 9.204726 | 9.372375 | 9.2622 | 0.110180195 | -0.057468701 | 9.2796078 | 9.2761268 | 9.21564 | 0.060483584 | 0.063964536 |
| Parp2 | 9.493242 | 9.492138 | 9.6057 | -0.113573903 | -0.112470012 | 9.7530727 | 9.7807935 | 9.72033 | 0.060466353 | 0.032745548 |
| Fam98b | 9.252535 | 9.238144 | 9.2983 | -0.060196256 | -0.045805078 | 9.3121365 | 9.2824974 | 9.22205 | 0.060443974 | 0.090083083 |
| Khdrbs2 | 1.65242 | 1.667625 | 1.8014 | -0.13377705 | -0.148981135 | 1.656949 | 1.7856393 | 1.72522 | 0.060418432 | -0.068271937 |
| Rab4b | 9.875534 | 9.812483 | 9.8127 | -0.000232516 | 0.062818873 | 10.322232 | 9.9786643 | 9.91825 | 0.060411644 | 0.403979761 |
| Zfp428 | 3.162629 | 3.181352 | 3.093 | 0.088328834 | 0.069605842 | 3.2660336 | 3.1443804 | 3.08398 | 0.060398605 | 0.182051804 |
| Rps24-ps3 | 1.554655 | 1.562996 | 1.5734 | -0.010377713 | -0.018719127 | 1.5402819 | 1.5912497 | 1.53086 | 0.060389706 | 0.009421898 |
| Gm14398 | 1.50698 | 1.552662 | 1.5587 | -0.006008016 | -0.051690213 | 1.5981776 | 1.5572837 | 1.4969 | 0.060388063 | 0.101282016 |
| Vmn1r214 | 1.425596 | 1.330709 | 1.3268 | 0.003905238 | 0.098792415 | 1.3381928 | 1.4333159 | 1.37294 | 0.060373307 | -0.034749758 |
| Pick1 | 7.796106 | 7.739535 | 7.8315 | -0.091990004 | -0.035418482 | 8.215498 | 8.0946986 | 8.03437 | 0.060328425 | 0.181127766 |
| Polr3c | 10.18295 | 10.20312 | 10.284 | -0.080668258 | -0.100836216 | 10.318962 | 10.420183 | 10.3599 | 0.060302098 | -0.040918852 |
| Gm18911 | 3.336802 | 3.296311 | 3.214 | 0.082271669 | 0.122763214 | 3.2099285 | 3.3922506 | 3.33195 | 0.0603001 | -0.122021997 |
| Vmn2r-ps130 | 1.091154 | 1.083623 | 1.1215 | -0.037925499 | -0.030394495 | 1.142005 | 1.1843791 | 1.1241 | 0.060274105 | 0.017900058 |
| Prkd2 | 11.87933 | 12.08803 | 11.953 | 0.134661266 | -0.074031738 | 12.124668 | 12.086476 | 12.0263 | 0.060172308 | 0.098364599 |
| Gm5805 | 2.301748 | 2.009304 | 2.0241 | -0.014749168 | 0.277695576 | 2.0066439 | 2.0601569 | 2.00015 | 0.060007856 | 0.006494907 |
| AC122453.1 | 0.728822 | 0.739031 | 0.7534 | -0.014378229 | -0.024587097 | 0.7699096 | 0.7690761 | 0.70918 | 0.059895863 | 0.06072939 |
| Cbln3 | 2.363827 | 2.316899 | 2.3109 | 0.006028735 | 0.052956314 | 2.3855192 | 2.408563 | 2.3487 | 0.059858288 | 0.036814482 |
| Sycp2 | 2.34834 | 2.090499 | 2.0467 | 0.043831213 | 0.301672274 | 2.128683 | 2.1065154 | 2.04667 | 0.059846708 | 0.082014279 |
| Ndufv1 | 10.03484 | 9.966519 | 10.027 | -0.060425634 | 0.007892003 | 10.281865 | 10.128485 | 10.0686 | 0.059844507 | 0.213224855 |
| Tmem167b | 9.798043 | 9.726444 | 9.7416 | -0.015194868 | 0.056404073 | 9.6379459 | 9.6803149 | 9.62049 | 0.05982766 | 0.017458618 |
| Gm11449 | 1.243969 | 1.207618 | 1.2458 | -0.038204209 | -0.001853859 | 1.2377408 | 1.275045 | 1.21533 | 0.059717521 | 0.022413322 |

| | | | | | | | | | | |
|---|---|---|---|---|---|---|---|---|---|---|
| Gm12013 | 6.096694 | 6.040583 | 6.2235 | -0.182936029 | -0.126825507 | 5.8022668 | 6.1544223 | 6.09475 | 0.059673213 | -0.292482327 |
| Retn | 1.209571 | 1.18942 | 1.2096 | -0.020151344 | -6.092E-07 | 1.2031861 | 1.1886736 | 1.12902 | 0.059656888 | 0.074169432 |
| Gm18492 | 1.766659 | 1.681392 | 1.6809 | 0.000508173 | 0.085774925 | 1.7551459 | 1.7833913 | 1.72375 | 0.059639429 | 0.031394008 |
| Usp50 | 2.934492 | 3.004136 | 2.9486 | 0.055561241 | -0.014082788 | 3.1008991 | 2.9729904 | 2.91337 | 0.05961689 | 0.187525509 |
| Zfp319 | 8.730777 | 8.838781 | 8.7809 | 0.057889455 | -0.050114914 | 8.6938929 | 8.7732272 | 8.71367 | 0.059558016 | -0.019776349 |
| Fcnb | 1.120475 | 1.125864 | 1.0734 | 0.052510621 | 0.047121651 | 1.1222323 | 1.1862104 | 1.12668 | 0.059535197 | -0.00444292 |
| Pdcl3 | 8.935917 | 8.972649 | 8.9911 | -0.018425636 | -0.05515687 | 8.8490967 | 8.9592158 | 8.89971 | 0.059501683 | -0.050617387 |
| Trappc13 | 9.001862 | 9.001148 | 8.964 | 0.037131888 | 0.037845756 | 8.8562697 | 8.9968397 | 8.93737 | 0.059474276 | -0.081095704 |
| Kif3c | 7.845919 | 7.920581 | 7.8453 | 0.075256125 | 0.000594358 | 7.6138671 | 7.817243 | 7.75784 | 0.059398817 | -0.143977146 |
| Olfr166 | 3.184869 | 3.08778 | 3.1259 | -0.038107473 | 0.058981247 | 3.0906242 | 3.2596178 | 3.20031 | 0.05930387 | -0.109689821 |
| Gm5766 | 1.676151 | 1.599952 | 1.6599 | -0.059927776 | 0.016271137 | 1.7911635 | 1.7275222 | 1.66824 | 0.059286569 | 0.122927774 |
| Bpifa2 | 4.001262 | 2.279121 | 2.1416 | 0.137483481 | 1.859624911 | 3.6726657 | 2.2321942 | 2.17294 | 0.059252588 | 1.499724128 |
| Slc22a13b-ps | 2.203908 | 2.234872 | 2.305 | -0.070094455 | -0.10105883 | 2.2634833 | 2.3082516 | 2.24906 | 0.059195454 | 0.014427161 |
| Pycr2 | 8.782095 | 8.729115 | 8.6945 | 0.034576677 | 0.087556639 | 9.0988156 | 8.8404062 | 8.78124 | 0.059162366 | 0.317571849 |
| Thap1 | 7.205763 | 7.205624 | 7.279 | -0.073385832 | -0.073247279 | 7.3004405 | 7.3897456 | 7.33063 | 0.059112898 | -0.0301922 |
| 37500 | 10.786 | 10.76708 | 10.672 | 0.094627816 | 0.113543404 | 10.745395 | 10.898162 | 10.8391 | 0.059016878 | -0.093750399 |
| Rap1a | 11.2676 | 11.27194 | 11.234 | 0.0379411 | 0.03360068 | 11.19404 | 11.263425 | 11.2044 | 0.058977924 | -0.010407622 |
| Rab3gap1 | 10.79466 | 10.8563 | 10.78 | 0.07679406 | 0.015152334 | 10.800692 | 10.939652 | 10.8807 | 0.058946436 | -0.080014254 |
| Hist1h2bh | 3.363557 | 3.33977 | 3.3649 | -0.025087724 | -0.001300833 | 3.5327516 | 3.4171154 | 3.35821 | 0.058903857 | 0.174540089 |
| Epb41l4a | 2.217516 | 2.135248 | 2.1229 | 0.01232362 | 0.094591128 | 2.0926887 | 2.2075294 | 2.14865 | 0.058879762 | -0.055960904 |
| Pcnt | 10.65847 | 10.44462 | 10.683 | -0.238153507 | -0.024302464 | 10.738533 | 10.828086 | 10.7692 | 0.058878554 | -0.030674577 |
| Pcsk2 | 3.22451 | 3.112996 | 3.1009 | 0.012054244 | 0.123568178 | 3.0593723 | 3.2751309 | 3.21625 | 0.058877063 | -0.156881495 |
| Rpsa-ps1 | 3.000941 | 3.259068 | 3.1659 | 0.093190715 | -0.164936535 | 3.2408605 | 3.1688675 | 3.10999 | 0.05887602 | 0.130869025 |
| Gm7511 | 0.647916 | 0.705737 | 0.6325 | 0.073283855 | 0.015462835 | 0.6673953 | 0.6958075 | 0.63696 | 0.05885055 | 0.03043831 |
| Map1a | 4.117301 | 4.327163 | 4.1421 | 0.185086793 | -0.02477442 | 3.9350941 | 4.1856735 | 4.12685 | 0.058818734 | -0.19176065 |
| Olfr596 | 1.519474 | 1.505714 | 1.4793 | 0.026401686 | 0.040161229 | 1.5141641 | 1.5897262 | 1.53092 | 0.058801694 | -0.01676034 |
| Gm12174 | 2.033016 | 2.052915 | 2.1321 | -0.079143266 | -0.099042268 | 2.0575218 | 2.0791183 | 2.02032 | 0.058797521 | 0.037201043 |
| Vmn1r46 | 6.40746 | 6.375548 | 6.4138 | -0.038205676 | -0.006294008 | 6.2949366 | 6.7590138 | 6.70022 | 0.058796563 | -0.405280696 |
| Myl12b | 12.77552 | 12.70074 | 12.677 | 0.023544074 | 0.098318662 | 12.890907 | 12.720575 | 12.6618 | 0.058781156 | 0.229112983 |
| Senp5 | 9.824769 | 9.842863 | 9.7601 | 0.082750515 | 0.064656203 | 9.6473416 | 9.7695943 | 9.71087 | 0.058726241 | -0.063526485 |
| Sqstm1 | 11.67993 | 11.7229 | 11.751 | -0.028137414 | -0.071108498 | 11.630402 | 11.763461 | 11.7048 | 0.058664002 | -0.074394635 |
| Dazl | 0.600531 | 0.591467 | 0.5791 | 0.012342581 | 0.02140647 | 0.5801321 | 0.6796769 | 0.62111 | 0.058562158 | -0.040982613 |

| | | | | | | | | | | |
|---|---|---|---|---|---|---|---|---|---|---|
| Olfr853 | 1.385483 | 1.406982 | 1.4128 | -0.005859461 | -0.027358174 | 1.4296739 | 1.4453562 | 1.38681 | 0.058547939 | 0.042865674 |
| Rhov | 1.764222 | 1.63331 | 1.5858 | 0.047529108 | 0.178441377 | 1.8838488 | 1.6831485 | 1.62468 | 0.058464429 | 0.259164754 |
| Gm48360 | 0.787331 | 0.767872 | 0.7064 | 0.061475413 | 0.080933854 | 0.7670444 | 0.7864378 | 0.72799 | 0.058447094 | 0.039053689 |
| Slc35c2 | 9.657413 | 9.707723 | 9.6548 | 0.052885038 | 0.002574285 | 10.102231 | 9.7109181 | 9.65257 | 0.05834328 | 0.44965569 |
| Ubap2l | 11.9586 | 11.91118 | 11.892 | 0.019107002 | 0.066532096 | 11.758616 | 11.959212 | 11.9009 | 0.058276124 | -0.142319131 |
| Gm43215 | 0.580002 | 0.609359 | 0.5996 | 0.009775788 | -0.019580456 | 0.5874281 | 0.6649755 | 0.6067 | 0.0582732 | -0.019274233 |
| S1pr5 | 3.715931 | 3.868724 | 3.8398 | 0.028885991 | -0.123906878 | 3.8767578 | 3.8770457 | 3.8188 | 0.058248936 | 0.057961059 |
| Tmem126b | 8.118945 | 8.196179 | 8.132 | 0.064196781 | -0.013037214 | 8.2117281 | 8.269007 | 8.21081 | 0.058200563 | 0.000921713 |
| Slco4c1 | 1.225233 | 1.159825 | 1.1505 | 0.009365307 | 0.074773732 | 1.2299891 | 1.1908847 | 1.13282 | 0.058068839 | 0.097173279 |
| Atg4b | 9.686236 | 9.766234 | 9.6702 | 0.096064124 | 0.016066745 | 9.6187898 | 9.8269711 | 9.76897 | 0.058001186 | -0.150180121 |
| Apba2 | 1.518771 | 1.419534 | 1.4633 | -0.043741636 | 0.055495278 | 1.4043198 | 1.5256102 | 1.46768 | 0.057934645 | -0.063355712 |
| 9130230L23Rik | 3.198231 | 3.129808 | 3.3431 | -0.213271959 | -0.144848819 | 3.1026056 | 3.2978375 | 3.2399 | 0.057934351 | -0.137297555 |
| Olfr190 | 1.95007 | 1.929375 | 1.9084 | 0.020951991 | 0.041647649 | 1.9650518 | 2.0676472 | 2.00972 | 0.057931758 | -0.044663593 |
| AC133103.7 | 1.104523 | 0.950618 | 0.9696 | -0.0189868 | 0.134918109 | 0.9903914 | 0.9614039 | 0.90348 | 0.057923705 | 0.086911263 |
| Thumpd1 | 9.933699 | 9.882193 | 9.8868 | -0.004594279 | 0.046912401 | 9.9231658 | 9.9895699 | 9.93169 | 0.057880563 | -0.008523569 |
| Zfp105 | 3.50621 | 3.507638 | 3.4034 | 0.104192503 | 0.102764612 | 3.3067887 | 3.3623473 | 3.3045 | 0.057843215 | 0.002284679 |
| Gm6680 | 1.333415 | 1.332886 | 1.3675 | -0.034659241 | -0.034130623 | 1.3207292 | 1.423709 | 1.3659 | 0.057808317 | -0.045171432 |
| Rpl7 | 14.72235 | 14.72996 | 14.734 | -0.003813922 | -0.011427762 | 14.776456 | 14.791378 | 14.7336 | 0.05775289 | 0.042830673 |
| Plcd3 | 3.825718 | 3.718057 | 3.8503 | -0.132224126 | -0.024563307 | 3.7373711 | 3.7725052 | 3.71484 | 0.05766874 | 0.022534592 |
| Tmem260 | 9.008356 | 9.043932 | 9.0777 | -0.033778454 | -0.069354296 | 8.9600923 | 9.1274883 | 9.06982 | 0.057667079 | -0.109728928 |
| Stfa2l1 | 3.581536 | 2.955527 | 2.726 | 0.2295157 | 0.855525535 | 2.9684835 | 2.7895544 | 2.73189 | 0.057664023 | 0.236593153 |
| Ube2o | 8.995528 | 9.098396 | 9.0678 | 0.030569593 | -0.072298277 | 9.0134454 | 8.9984095 | 8.94076 | 0.057645843 | 0.072681691 |
| Cystm1 | 3.994796 | 3.791454 | 3.9133 | -0.121870007 | 0.081472809 | 4.2963995 | 4.2406593 | 4.18306 | 0.057603262 | 0.113343477 |
| Gm11474 | 1.60618 | 1.66503 | 1.576 | 0.089000202 | 0.030149876 | 1.6172818 | 1.6584141 | 1.60084 | 0.057577587 | 0.016445272 |
| Ighv1-62-3 | 2.688474 | 2.696867 | 2.6102 | 0.086630885 | 0.078237869 | 2.6176274 | 2.6134895 | 2.55598 | 0.057511622 | 0.061649468 |
| Gm16589 | 0.994091 | 1.001611 | 0.9808 | 0.020762954 | 0.013242364 | 1.101508 | 1.0539541 | 0.99646 | 0.057490358 | 0.105044296 |
| Nabp1 | 10.29421 | 10.19377 | 10.188 | 0.005949976 | 0.10638936 | 10.164212 | 10.408071 | 10.3507 | 0.057407128 | -0.186451397 |
| Idh3a | 9.803426 | 9.759558 | 9.7221 | 0.037415925 | 0.081283197 | 9.8818024 | 9.800843 | 9.74347 | 0.057371518 | 0.138330871 |
| Tmem183a | 9.629523 | 9.546412 | 9.5837 | -0.037330841 | 0.045780324 | 9.733164 | 9.6646159 | 9.60725 | 0.057369548 | 0.125917616 |
| U2surp | 11.03057 | 10.9995 | 11.008 | -0.008108966 | 0.022959354 | 10.99547 | 11.112507 | 11.0552 | 0.057341334 | -0.059695424 |
| Btnl10 | 2.183941 | 2.092083 | 2.1188 | -0.026725266 | 0.065133552 | 2.1589967 | 2.2110865 | 2.15375 | 0.057337139 | 0.005247375 |
| Chst8 | 1.99441 | 2.099437 | 2.0324 | 0.067021042 | -0.03800647 | 2.0128392 | 2.0111384 | 1.95383 | 0.05730422 | 0.059005048 |

| | | | | | | | | | | |
|---|---|---|---|---|---|---|---|---|---|---|
| Gm14648 | 1.815479 | 1.89867 | 1.8642 | 0.03446412 | -0.04872693 | 2.3019687 | 1.8858752 | 1.82862 | 0.057250512 | 0.473344042 |
| Mpst | 8.381367 | 8.299256 | 8.3186 | -0.019303411 | 0.062807364 | 8.4298117 | 8.2969001 | 8.23966 | 0.057244733 | 0.190156346 |
| Mtmr14 | 8.934594 | 9.054504 | 9.0572 | -0.002665457 | -0.122575439 | 8.9540275 | 9.1617686 | 9.10457 | 0.057196767 | -0.150544424 |
| Usp5 | 10.53663 | 10.58915 | 10.587 | 0.002303084 | -0.050225872 | 10.882111 | 10.806948 | 10.7498 | 0.057175846 | 0.132339206 |
| Gm44958 | 1.018193 | 1.021377 | 0.9859 | 0.035471764 | 0.032287675 | 1.0168021 | 1.0442514 | 0.98711 | 0.057141387 | 0.029692076 |
| Mrps26 | 8.454198 | 8.368398 | 8.4171 | -0.048683603 | 0.037116756 | 8.5856008 | 8.5024006 | 8.44527 | 0.05712646 | 0.140326744 |
| Vmn1r168 | 2.88165 | 2.94346 | 3.0084 | -0.064918003 | -0.126728144 | 3.0135113 | 3.0538224 | 2.99673 | 0.057095542 | 0.016784393 |
| Slc30a6 | 7.588922 | 7.758772 | 7.6581 | 0.10063587 | -0.069214548 | 7.7826917 | 7.8427608 | 7.78574 | 0.057023356 | -0.003045737 |
| AC126807.1 | 1.2108 | 1.118302 | 1.1534 | -0.035132969 | 0.057365293 | 1.0771237 | 1.204689 | 1.14768 | 0.057004498 | -0.070560851 |
| Pnn | 10.86101 | 10.90106 | 10.895 | 0.005895086 | -0.03415646 | 11.102296 | 11.254597 | 11.1976 | 0.056958118 | -0.09534321 |
| AC126438.1 | 1.487682 | 1.47248 | 1.4386 | 0.033910399 | 0.049111854 | 1.5289109 | 1.528158 | 1.4712 | 0.056957908 | 0.057710824 |
| Olfr95 | 2.082155 | 1.922962 | 1.9637 | -0.04078413 | 0.118409016 | 1.9219553 | 2.0701753 | 2.01323 | 0.056947577 | -0.091272425 |
| Trbv22 | 0.747574 | 0.782178 | 0.7507 | 0.031494465 | -0.003110353 | 0.6981121 | 0.7783651 | 0.72145 | 0.056910115 | -0.023342805 |
| Nup214 | 10.72679 | 10.72495 | 10.7 | 0.02503628 | 0.026871444 | 10.828468 | 11.007619 | 10.9508 | 0.056846456 | -0.122304953 |
| Slc23a3 | 2.195309 | 2.268305 | 2.357 | -0.088684794 | -0.161680751 | 2.3520166 | 2.3032562 | 2.24645 | 0.056806002 | 0.105566354 |
| Icam2 | 9.731689 | 9.917362 | 9.8182 | 0.099129571 | -0.086542909 | 10.084268 | 9.9687225 | 9.91194 | 0.056778508 | 0.172323643 |
| Ptges3l | 4.213757 | 4.186246 | 4.0193 | 0.16691607 | 0.194427504 | 4.4774055 | 4.4002285 | 4.34346 | 0.056768159 | 0.133945121 |
| Eaf2 | 4.958002 | 5.191495 | 4.9786 | 0.212847717 | -0.020645152 | 4.8688079 | 5.1721776 | 5.11545 | 0.056728675 | -0.246641002 |
| Gm45268 | 3.93146 | 3.96662 | 3.8284 | 0.138183094 | 0.103022752 | 3.8713747 | 4.0953963 | 4.03876 | 0.056631819 | -0.167389736 |
| Igkv4-60 | 3.06315 | 3.027981 | 2.9792 | 0.04882649 | 0.083996088 | 3.1132495 | 3.1016694 | 3.04505 | 0.056615897 | 0.068195998 |
| Lgi2 | 3.250183 | 3.113517 | 2.8788 | 0.234760015 | 0.371425944 | 2.9968462 | 2.8582996 | 2.80173 | 0.05656936 | 0.195115985 |
| Fxyd1 | 3.355381 | 3.315155 | 3.3319 | -0.01679016 | 0.023435402 | 3.110127 | 3.1826407 | 3.1261 | 0.056544678 | -0.015969067 |
| Gpsm3 | 10.73026 | 10.70537 | 10.771 | -0.065809724 | -0.040921818 | 10.955945 | 10.757067 | 10.7005 | 0.056540036 | 0.255417096 |
| Pigv | 8.538595 | 8.814313 | 8.5955 | 0.218832524 | -0.056885034 | 8.7551982 | 8.762597 | 8.70612 | 0.056478244 | 0.049079474 |
| Nab2 | 9.744047 | 9.866063 | 9.7262 | 0.139879522 | 0.017863539 | 9.8448919 | 9.8528884 | 9.79642 | 0.056464719 | 0.048468253 |
| Shf | 5.505964 | 5.392164 | 5.4815 | -0.089357315 | 0.02444197 | 5.3779314 | 5.416184 | 5.35983 | 0.056354815 | 0.018102236 |
| Macc1 | 2.194939 | 2.252178 | 2.2403 | 0.011886381 | -0.045353032 | 2.2922884 | 2.3235225 | 2.26717 | 0.056351363 | 0.025117268 |
| Gm4735 | 2.29072 | 2.401818 | 2.4296 | -0.027826794 | -0.138924457 | 2.2484204 | 2.3539486 | 2.29763 | 0.056319488 | -0.049208701 |
| Map3k21 | 0.887304 | 0.768003 | 0.8839 | -0.115847655 | 0.003453147 | 0.8166676 | 0.8579831 | 0.80169 | 0.056288743 | 0.014973161 |
| Gm13194 | 1.715855 | 1.481479 | 1.4257 | 0.05578109 | 0.290157024 | 1.5199881 | 1.5393631 | 1.48312 | 0.05624032 | 0.036865314 |
| Olfr447 | 1.77183 | 1.744262 | 1.748 | -0.00377146 | 0.023796972 | 1.6613999 | 1.8527767 | 1.79655 | 0.056222431 | -0.135154383 |
| Jph2 | 2.363967 | 2.48436 | 2.3363 | 0.148101929 | 0.027708188 | 2.4555346 | 2.4554356 | 2.39922 | 0.056215013 | 0.056314019 |

| | | | | | | | | | | |
|---|---|---|---|---|---|---|---|---|---|---|
| Foxj1 | 2.655459 | 2.711623 | 2.6379 | 0.073750838 | 0.017586862 | 2.5910255 | 2.7300575 | 2.67398 | 0.056081926 | -0.082950041 |
| Bnip2 | 10.20091 | 10.21383 | 10.201 | 0.013131692 | 0.00021212 | 10.247764 | 10.290288 | 10.2342 | 0.056079114 | 0.013555043 |
| Gjd4 | 1.582497 | 1.551898 | 1.5811 | -0.029161479 | 0.001437052 | 1.6020103 | 1.6941496 | 1.63813 | 0.056015835 | -0.036123434 |
| Tctex1d4 | 2.229093 | 2.119416 | 2.1487 | -0.029314942 | 0.080362545 | 2.2151325 | 2.2187628 | 2.16275 | 0.056010055 | 0.052379783 |
| Gm42669 | 6.894461 | 6.979913 | 7.0385 | -0.058574297 | -0.144026153 | 7.1854226 | 7.4321942 | 7.37621 | 0.055985877 | -0.190785714 |
| Gabrb1 | 2.147482 | 2.194839 | 2.1655 | 0.0293074 | -0.018049627 | 2.2414408 | 2.2375825 | 2.18162 | 0.055964939 | 0.059823192 |
| Mobp | 2.069265 | 2.211484 | 1.9445 | 0.266983137 | 0.124764105 | 1.9488554 | 2.1292047 | 2.07325 | 0.055951044 | -0.124398233 |
| Tm4sf1 | 9.074885 | 8.544687 | 8.5084 | 0.036246639 | 0.566443999 | 7.72841 | 8.0460791 | 7.99014 | 0.055942074 | -0.261727001 |
| Gm16181 | 1.843893 | 1.874487 | 1.8597 | 0.014808499 | -0.015785604 | 1.8398847 | 1.8847941 | 1.82895 | 0.05583976 | 0.01093036 |
| Olfr740 | 1.708037 | 1.669373 | 1.67 | -0.000593148 | 0.038070546 | 1.7105157 | 1.7131162 | 1.6573 | 0.055811801 | 0.053211293 |
| Nkapl | 0.854277 | 0.82551 | 0.926 | -0.100511649 | -0.071745298 | 0.8496103 | 0.8971059 | 0.84132 | 0.055789968 | 0.008294343 |
| Gm9104 | 0.744716 | 0.728262 | 0.7467 | -0.018424381 | -0.00196949 | 0.6956596 | 0.7251911 | 0.6694 | 0.055786346 | 0.026254817 |
| Prrc1 | 9.33844 | 9.337392 | 9.1733 | 0.164126021 | 0.165173994 | 9.7900488 | 9.3897248 | 9.33398 | 0.055748156 | 0.456072161 |
| Pak1 | 7.807702 | 7.64863 | 7.9869 | -0.338255942 | -0.179183192 | 7.8001328 | 7.8439729 | 7.78827 | 0.055697999 | 0.011857879 |
| Lalba | 1.731791 | 1.743305 | 1.7476 | -0.004312325 | -0.015826682 | 1.727952 | 1.7274132 | 1.67173 | 0.055682252 | 0.056221 |
| Prdx3 | 9.971048 | 9.995655 | 10.024 | -0.028817374 | -0.053424984 | 10.114339 | 10.007643 | 9.95198 | 0.055664787 | 0.162361352 |
| Gm45492 | 1.027628 | 1.056972 | 1.0038 | 0.053177803 | 0.023833948 | 1.0209432 | 1.0649891 | 1.00935 | 0.055636839 | 0.011590924 |
| Olfr204 | 3.035884 | 3.04456 | 3.114 | -0.069424735 | -0.078100684 | 3.1444914 | 3.1822486 | 3.12668 | 0.055573046 | 0.01781585 |
| Cptp | 7.189252 | 7.304067 | 7.3697 | -0.065666664 | -0.18048181 | 7.2710808 | 7.3204293 | 7.26488 | 0.055549269 | 0.006200767 |
| Mtg2 | 8.436569 | 8.513405 | 8.4873 | 0.026112456 | -0.050723982 | 8.792765 | 8.6891514 | 8.63377 | 0.055384575 | 0.158998175 |
| BC055324 | 6.790405 | 6.618893 | 6.8033 | -0.184455134 | -0.012943291 | 6.9920112 | 6.9155933 | 6.86021 | 0.055382668 | 0.131800485 |
| Nrf1 | 9.502235 | 9.64617 | 9.5642 | 0.081973062 | -0.061961508 | 9.5175723 | 9.6335701 | 9.5782 | 0.055374248 | -0.060623506 |
| Olfr575 | 2.716746 | 2.701692 | 2.7812 | -0.079508044 | -0.064454311 | 2.7169242 | 2.8495641 | 2.79426 | 0.055306541 | -0.077333323 |
| Fam169b | 10.41517 | 10.62245 | 10.536 | 0.08625893 | -0.12101754 | 10.562082 | 10.758299 | 10.703 | 0.055305146 | -0.140912575 |
| Klhl31 | 2.249852 | 2.197084 | 2.2808 | -0.08369384 | -0.030926159 | 2.2508022 | 2.2794165 | 2.22425 | 0.055166894 | 0.026552649 |
| Olfr109 | 2.228134 | 2.164904 | 2.222 | -0.05710175 | 0.006128277 | 2.2615185 | 2.233336 | 2.17818 | 0.055153921 | 0.083336429 |
| Vrk2 | 8.978486 | 8.945969 | 8.8837 | 0.062309118 | 0.094825907 | 9.0272862 | 8.940629 | 8.88549 | 0.055141006 | 0.14179824 |
| Olfr1009 | 0.461429 | 0.454853 | 0.4478 | 0.007087405 | 0.013663214 | 0.4252943 | 0.5146377 | 0.45953 | 0.055107432 | -0.034235901 |
| Gm48663 | 1.553296 | 1.52134 | 1.5703 | -0.048929322 | -0.016973329 | 1.6008114 | 1.6248241 | 1.56972 | 0.055103364 | 0.031090631 |
| Gnb1 | 12.14138 | 12.18258 | 12.156 | 0.02615854 | -0.01504407 | 12.24107 | 12.206216 | 12.1511 | 0.055094504 | 0.089948373 |
| Stk35 | 8.119881 | 8.153494 | 8.2078 | -0.054309771 | -0.087922904 | 8.0619915 | 8.2590585 | 8.20398 | 0.055082879 | -0.141984187 |
| Zmynd10 | 3.318267 | 3.407271 | 3.3021 | 0.105200062 | 0.016196195 | 3.5158302 | 3.623223 | 3.56823 | 0.054997649 | -0.052395125 |

| | | | | | | | | | | |
|---|---|---|---|---|---|---|---|---|---|---|
| Lancl2 | 7.264106 | 7.399674 | 7.3739 | 0.02578969 | -0.10977803 | 7.3965658 | 7.4164496 | 7.36146 | 0.054990649 | 0.035106819 |
| 4930562C15Rik | 1.73464 | 1.757132 | 1.7525 | 0.004603466 | -0.017888447 | 1.7918989 | 1.821713 | 1.76673 | 0.054978388 | 0.025164296 |
| Mpz | 3.47071 | 3.298974 | 3.2547 | 0.044242469 | 0.215978758 | 3.1784031 | 3.232559 | 3.17758 | 0.054974567 | 0.000818589 |
| Slc10a7 | 8.746884 | 8.663088 | 8.752 | -0.088880108 | -0.005084097 | 8.7272102 | 8.8570604 | 8.80219 | 0.054865769 | -0.074984483 |
| Pls1 | 3.541803 | 3.601842 | 3.6242 | -0.022392206 | -0.08243112 | 3.3984318 | 3.6063633 | 3.55154 | 0.054824286 | -0.153107157 |
| Adamtsl4 | 5.871314 | 5.83891 | 6.0004 | -0.161506695 | -0.129102668 | 5.8911838 | 6.08466 | 6.02984 | 0.0548194 | -0.138656871 |
| Gm20481 | 4.470183 | 4.490051 | 4.7591 | -0.269002224 | -0.288869698 | 4.3089151 | 4.3299494 | 4.27515 | 0.054802383 | 0.033768087 |
| Arih1 | 11.46163 | 11.58193 | 11.423 | 0.159067498 | 0.038770142 | 11.263618 | 11.537343 | 11.4826 | 0.054786818 | -0.218938717 |
| Rasl10b | 1.490599 | 1.472261 | 1.408 | 0.064211447 | 0.082549728 | 1.4744594 | 1.4315548 | 1.37679 | 0.054762436 | 0.097666984 |
| Gatad1 | 8.795054 | 8.890825 | 8.8177 | 0.073133566 | -0.022637834 | 8.6425359 | 8.776503 | 8.72176 | 0.054739226 | -0.07922787 |
| Zfp988 | 1.525749 | 1.606037 | 1.5229 | 0.083103271 | 0.002816098 | 1.5399063 | 1.5354384 | 1.4807 | 0.054734116 | 0.059202009 |
| Plekhm1 | 10.43376 | 10.54548 | 10.362 | 0.183487528 | 0.071761234 | 10.605925 | 10.416694 | 10.362 | 0.054714166 | 0.243944745 |
| Olfr157 | 1.177779 | 1.193052 | 1.2257 | -0.032612131 | -0.047884196 | 1.1906822 | 1.2543471 | 1.19964 | 0.054704402 | -0.008960459 |
| Serpinb8 | 5.428588 | 6.016608 | 5.2811 | 0.735549774 | 0.147529681 | 5.1783219 | 4.9899116 | 4.93521 | 0.054697024 | 0.243107336 |
| Gm12693 | 2.219161 | 2.19463 | 2.526 | -0.331398776 | -0.306866899 | 2.0875528 | 2.3704032 | 2.31572 | 0.054681261 | -0.228169126 |
| Tada3 | 8.493585 | 8.567938 | 8.5735 | -0.005519183 | -0.079872276 | 8.8525387 | 8.7318226 | 8.67719 | 0.054629945 | 0.17534605 |
| Spg21 | 9.922022 | 9.880824 | 9.8541 | 0.026697131 | 0.067895963 | 9.9575777 | 9.8318458 | 9.77722 | 0.054622398 | 0.180354296 |
| Rlim | 10.85317 | 10.78033 | 10.779 | 0.000876384 | 0.073715556 | 10.806534 | 10.949507 | 10.8949 | 0.054611394 | -0.08836149 |
| Kctd3 | 8.547971 | 8.656014 | 8.5264 | 0.129653884 | 0.021611687 | 8.8976478 | 8.8924862 | 8.8379 | 0.054585642 | 0.059747202 |
| Sdccag8 | 8.531124 | 8.461617 | 8.5878 | -0.126163176 | -0.056656216 | 8.6505211 | 8.7312757 | 8.67675 | 0.054523585 | -0.026230973 |
| 4933412A08Rik | 1.726389 | 1.798152 | 1.7119 | 0.086217241 | 0.014453796 | 1.7035428 | 1.7947299 | 1.74021 | 0.054515163 | -0.036671996 |
| Ttll9 | 1.856542 | 1.868143 | 1.8526 | 0.01558383 | 0.003983068 | 1.7844261 | 1.938764 | 1.88427 | 0.054496176 | -0.09984179 |
| Acin1 | 12.37602 | 12.42809 | 12.42 | 0.007670446 | -0.044395346 | 12.54465 | 12.703743 | 12.6493 | 0.054458854 | -0.1046343 |
| Aggf1 | 9.406753 | 9.37143 | 9.4573 | -0.085859777 | -0.05053692 | 9.2811273 | 9.4699417 | 9.41548 | 0.054456997 | -0.134357331 |
| Olfr414 | 2.348135 | 2.410588 | 2.3954 | 0.015216381 | -0.04723653 | 2.4442511 | 2.4939257 | 2.43949 | 0.054436908 | 0.004762289 |
| Casp8ap2 | 9.732552 | 9.799511 | 9.791 | 0.008492617 | -0.058466481 | 9.4934713 | 9.8007896 | 9.74635 | 0.05443478 | -0.252883557 |
| Supt5 | 12.03395 | 12.1075 | 12.105 | 0.002606972 | -0.070940062 | 12.202888 | 12.206217 | 12.1518 | 0.054434704 | 0.051105555 |
| Olfr1196 | 2.25874 | 2.25866 | 2.3265 | -0.067830263 | -0.067750543 | 2.3273325 | 2.374498 | 2.32008 | 0.054421194 | 0.007255684 |
| Ptgis | 1.123785 | 1.208314 | 1.1275 | 0.080777103 | -0.003751456 | 1.0680588 | 1.133213 | 1.07882 | 0.05438815 | -0.01076608 |
| Dnttip1 | 8.782053 | 8.818456 | 8.7698 | 0.048693303 | 0.012290402 | 8.9437047 | 8.9641312 | 8.90976 | 0.054373764 | 0.03394726 |
| Gm9973 | 5.561641 | 5.490175 | 5.434 | 0.05617766 | 0.127643598 | 5.5343333 | 5.6105627 | 5.55622 | 0.054344646 | -0.02188478 |
| Prps1l3 | 9.347082 | 9.477889 | 9.3181 | 0.159761172 | 0.028953778 | 9.3465165 | 9.4366813 | 9.3824 | 0.054279603 | -0.035885132 |

| | | | | | | | | | | |
|---|---|---|---|---|---|---|---|---|---|---|
| Mtpn | 12.04789 | 12.09784 | 12.037 | 0.060606914 | 0.010665746 | 12.127495 | 12.048953 | 11.9947 | 0.054265752 | 0.132808002 |
| Slc3a1 | 3.803368 | 3.826712 | 3.7274 | 0.09928921 | 0.075945236 | 3.8064821 | 3.8713551 | 3.81712 | 0.054231947 | -0.010641065 |
| Rnf113a2 | 7.918594 | 7.826686 | 7.7737 | 0.053022303 | 0.144930812 | 7.989812 | 7.9128404 | 7.85861 | 0.054227169 | 0.131198747 |
| Il11 | 1.73436 | 1.795634 | 1.7934 | 0.002201373 | -0.059072864 | 1.7853245 | 1.7865247 | 1.73231 | 0.054217857 | 0.053017683 |
| AC126944.1 | 1.459914 | 1.415907 | 1.4521 | -0.036178344 | 0.007828898 | 1.4875087 | 1.5849307 | 1.53079 | 0.054143106 | -0.043278835 |
| Olfr1192-ps1 | 4.266569 | 4.281924 | 4.3392 | -0.05725463 | -0.072609556 | 4.4146188 | 4.6065843 | 4.55253 | 0.054053497 | -0.137912056 |
| Tmem273 | 1.618345 | 1.620346 | 1.6012 | 0.019106705 | 0.017106173 | 1.7593879 | 1.6247874 | 1.57078 | 0.054003472 | 0.188603987 |
| Xndc1 | 8.700477 | 8.802929 | 8.9461 | -0.143211086 | -0.24566358 | 9.1244078 | 9.1477505 | 9.09377 | 0.053979794 | 0.030637173 |
| Gm4462 | 2.596541 | 2.581276 | 2.6188 | -0.037525462 | -0.022260176 | 2.6583133 | 2.6890661 | 2.6351 | 0.053966831 | 0.023214015 |
| Adipor1 | 10.94498 | 10.96157 | 10.984 | -0.022911918 | -0.039506712 | 11.111979 | 10.943275 | 10.8893 | 0.053943792 | 0.222648092 |
| Ccbe1 | 3.530585 | 3.680197 | 3.5947 | 0.085493907 | -0.064118341 | 3.3912243 | 3.5813976 | 3.52748 | 0.053918896 | -0.136254445 |
| Cox10 | 8.563452 | 8.566262 | 8.5409 | 0.025394198 | 0.022583898 | 8.6103403 | 8.5329579 | 8.47909 | 0.053866894 | 0.131249282 |
| Ddx43 | 4.298511 | 4.162686 | 4.1396 | 0.023083408 | 0.158908483 | 4.1133694 | 4.3639936 | 4.31022 | 0.05377778 | -0.196846468 |
| Gm11343 | 2.089627 | 2.018712 | 2.1112 | -0.092482848 | -0.021567663 | 2.1426585 | 2.1250035 | 2.07127 | 0.053729186 | 0.07138414 |
| Six1 | 2.304115 | 2.284624 | 2.2366 | 0.048020932 | 0.067511465 | 2.3920869 | 2.3406923 | 2.28697 | 0.053725333 | 0.10511993 |
| Rnf185 | 8.760696 | 8.738988 | 8.7305 | 0.008491913 | 0.030199797 | 8.6784229 | 8.6523427 | 8.59866 | 0.053685911 | 0.079766137 |
| Gm11361 | 2.178728 | 2.270279 | 2.2734 | -0.003140994 | -0.094691326 | 2.2505459 | 2.2822148 | 2.22858 | 0.053637695 | 0.021968762 |
| Tmem222 | 8.695352 | 8.627818 | 8.7472 | -0.119418995 | -0.051884984 | 9.0792122 | 8.8451008 | 8.79146 | 0.05363591 | 0.287747386 |
| Gm13199 | 1.133926 | 1.136176 | 1.1324 | 0.003783566 | 0.001533749 | 1.2232593 | 1.1862516 | 1.13263 | 0.053624566 | 0.090632327 |
| Lrrc4 | 1.633101 | 1.569447 | 1.6424 | -0.072911677 | -0.009257651 | 1.5340926 | 1.6470636 | 1.59346 | 0.053600949 | -0.059370147 |
| Brd7 | 10.15934 | 10.12189 | 10.165 | -0.0428638 | -0.005423658 | 10.188648 | 10.229661 | 10.1761 | 0.053560202 | 0.012547313 |
| Iglj3p | 2.954235 | 3.183549 | 2.97 | 0.213571828 | -0.0157425 | 2.9972494 | 3.1073321 | 3.05379 | 0.053538389 | -0.056544288 |
| Rpl7a-ps3 | 2.48175 | 2.426006 | 2.4318 | -0.005817883 | 0.049925838 | 2.4735585 | 2.427979 | 2.37454 | 0.053439548 | 0.099019048 |
| Olfr313 | 1.741265 | 1.77177 | 1.8485 | -0.076769797 | -0.107275332 | 1.8980382 | 1.9160875 | 1.86276 | 0.053328409 | 0.035279135 |
| Olfr720 | 1.541676 | 1.448102 | 1.4339 | 0.014218176 | 0.107791997 | 1.4901012 | 1.5577433 | 1.50443 | 0.053308339 | -0.01433382 |
| Fcmr | 12.95203 | 13.13295 | 13.036 | 0.096983642 | -0.08393014 | 13.46441 | 13.157798 | 13.1046 | 0.05320867 | 0.359821203 |
| Ffar3 | 2.93834 | 2.931996 | 2.9226 | 0.009408996 | 0.015752976 | 3.0788977 | 3.1584256 | 3.10524 | 0.053180679 | -0.026347227 |
| Chgb | 0.943265 | 0.890133 | 0.8438 | 0.046284828 | 0.099416828 | 0.8473907 | 0.9184954 | 0.86533 | 0.053165701 | -0.017938926 |
| Myh3 | 1.076505 | 1.081175 | 1.1446 | -0.063419567 | -0.068089695 | 1.0827233 | 1.1398416 | 1.08668 | 0.05316325 | -0.003954965 |
| Cd1d2 | 2.617509 | 2.635181 | 2.6653 | -0.030142037 | -0.047814262 | 2.6921468 | 2.716241 | 2.66314 | 0.053097672 | 0.029003529 |
| Cldn7 | 1.179132 | 1.196679 | 1.2032 | -0.006513497 | -0.024060756 | 1.36563 | 1.2809614 | 1.22788 | 0.053081119 | 0.137749727 |
| Hbs1l | 9.859706 | 9.906273 | 9.917 | -0.010735095 | -0.057302833 | 9.9993011 | 10.162647 | 10.1096 | 0.052998264 | -0.110347283 |

| | | | | | | | | | | |
|---|---|---|---|---|---|---|---|---|---|---|
| Gm2033 | 0.951015 | 0.878568 | 0.8984 | -0.019799101 | 0.052647785 | 0.8956704 | 0.9814603 | 0.92848 | 0.05297745 | -0.032812511 |
| Ctnnbl1 | 9.675194 | 9.685005 | 9.6967 | -0.011728604 | -0.021539245 | 9.7327985 | 9.7586704 | 9.70578 | 0.052890922 | 0.027019009 |
| Trak1 | 10.48234 | 10.59152 | 10.446 | 0.145562514 | 0.036383798 | 10.446531 | 10.529835 | 10.477 | 0.052814674 | -0.030490109 |
| Jpt2 | 8.948424 | 8.962272 | 9.089 | -0.12670976 | -0.140557896 | 9.0888495 | 8.9647922 | 8.91199 | 0.052805794 | 0.176863136 |
| Zc3h13 | 10.43438 | 10.50765 | 10.447 | 0.060794856 | -0.01247655 | 10.382961 | 10.570865 | 10.5181 | 0.052774394 | -0.135128881 |
| Rnf7 | 8.827568 | 8.854645 | 8.828 | 0.026691842 | -0.000385692 | 8.8537407 | 8.8013991 | 8.74864 | 0.052763832 | 0.105105434 |
| Mtdh | 11.64724 | 11.59792 | 11.696 | -0.098377852 | -0.04906261 | 11.624921 | 11.685491 | 11.6328 | 0.05273744 | -0.007833216 |
| Gucy2c | 2.261655 | 2.219276 | 2.3008 | -0.081546587 | -0.039167344 | 2.2295232 | 2.257839 | 2.20511 | 0.052733968 | 0.024418186 |
| Scel | 1.852784 | 1.937918 | 1.8759 | 0.061998984 | -0.023135812 | 1.8605378 | 1.9235391 | 1.87086 | 0.052680639 | -0.010320666 |
| Hnrnph3 | 8.913251 | 8.912776 | 8.9564 | -0.043621098 | -0.043145402 | 9.0791492 | 9.4145362 | 9.36186 | 0.05267761 | -0.282709314 |
| Gm6377 | 9.393548 | 9.447798 | 9.198 | 0.249781004 | 0.195531258 | 8.9735775 | 9.0156538 | 8.96298 | 0.052670214 | 0.010593953 |
| Smim11 | 7.627224 | 7.638624 | 7.6752 | -0.036546978 | -0.047947467 | 7.7658659 | 7.7620697 | 7.7094 | 0.052667155 | 0.056463268 |
| Kctd2 | 8.872064 | 8.894769 | 8.9613 | -0.066525132 | -0.089229817 | 9.1654169 | 9.0238603 | 8.97119 | 0.052665647 | 0.1942223 |
| Cldn4 | 3.020914 | 2.773651 | 2.9829 | -0.209236964 | 0.038026558 | 2.9645856 | 2.9416615 | 2.889 | 0.05265649 | 0.075580633 |
| Mipep | 8.026761 | 8.071468 | 7.9669 | 0.104614619 | 0.059907494 | 8.2099559 | 8.2119126 | 8.15935 | 0.05256508 | 0.050608343 |
| Gm17893 | 2.540536 | 2.541858 | 2.5739 | -0.032009598 | -0.033332009 | 2.5920829 | 2.6581021 | 2.60557 | 0.052535441 | -0.013483731 |
| H2-DMb2 | 12.78379 | 12.84378 | 12.852 | -0.008501964 | -0.06849618 | 13.426811 | 12.922508 | 12.87 | 0.05252706 | 0.556830265 |
| Use1 | 11.03265 | 11.05416 | 11.058 | -0.00400419 | -0.025505966 | 11.179164 | 11.008583 | 10.9561 | 0.052520944 | 0.223101791 |
| Hist1h3a | 3.307937 | 3.263722 | 3.2795 | -0.015767757 | 0.028447042 | 3.3436441 | 3.3518741 | 3.29937 | 0.052506995 | 0.044277022 |
| Trabd | 10.10352 | 10.0548 | 10.077 | -0.022163168 | 0.026557899 | 10.23123 | 10.096497 | 10.044 | 0.052497351 | 0.187230571 |
| Pik3cd | 12.9101 | 12.83837 | 12.793 | 0.045531838 | 0.117266254 | 12.929882 | 12.889657 | 12.8372 | 0.052475888 | 0.092700844 |
| Rpl22-ps1 | 1.242283 | 1.258047 | 1.2735 | -0.01549738 | -0.031261909 | 1.3287273 | 1.3638776 | 1.31146 | 0.05242025 | 0.017269973 |
| Gm11989 | 1.31576 | 1.28218 | 1.2835 | -0.001312104 | 0.032267835 | 1.2974957 | 1.3180619 | 1.26567 | 0.052389861 | 0.03182367 |
| Ggnbp1 | 5.814708 | 5.938627 | 5.86 | 0.0785948 | -0.045324364 | 6.2684594 | 6.0884454 | 6.03613 | 0.052317146 | 0.232331208 |
| Med12 | 11.88777 | 11.94642 | 11.896 | 0.050679762 | -0.007977268 | 12.148841 | 12.231818 | 12.1795 | 0.052271012 | -0.030705286 |
| E2f3 | 7.701334 | 7.61879 | 7.6876 | -0.068817835 | 0.013725718 | 7.6476678 | 7.75392 | 7.70172 | 0.052202754 | -0.054049462 |
| Tsc2 | 10.30688 | 10.17933 | 10.279 | -0.09922259 | 0.028326231 | 10.446789 | 10.490821 | 10.4387 | 0.05211063 | 0.008078347 |
| Lrrk2 | 11.42919 | 11.5438 | 11.52 | 0.024284762 | -0.09032149 | 11.893126 | 11.764052 | 11.712 | 0.052083672 | 0.181157391 |
| Chic2 | 9.207983 | 9.14449 | 9.0954 | 0.049044645 | 0.112537702 | 9.0490218 | 9.1086205 | 9.05654 | 0.052079776 | -0.007518918 |
| Abhd13 | 9.219939 | 9.299576 | 9.2228 | 0.07673129 | -0.002905705 | 9.2612511 | 9.2637068 | 9.21166 | 0.052047441 | 0.049591717 |
| Npm3 | 9.765941 | 9.788311 | 9.7993 | -0.010943473 | -0.033314032 | 9.9070311 | 9.758323 | 9.70632 | 0.051998657 | 0.200706789 |
| Stk32b | 0.80968 | 0.844793 | 0.8161 | 0.028720233 | -0.006392602 | 0.85959 | 0.9024951 | 0.85051 | 0.051981911 | 0.009076905 |

| | | | | | | | | | | |
|---|---|---|---|---|---|---|---|---|---|---|
| Stk25 | 10.1427 | 10.16928 | 10.194 | -0.024946972 | -0.051532986 | 10.323427 | 10.385675 | 10.3338 | 0.051880645 | -0.010367247 |
| Mrpl49 | 8.752123 | 8.772288 | 8.7192 | 0.05306191 | 0.032897334 | 8.9307463 | 8.809904 | 8.7581 | 0.051807705 | 0.172649993 |
| Arl6ip6 | 8.629611 | 8.678631 | 8.6307 | 0.047890682 | -0.001128764 | 8.5955499 | 8.6887003 | 8.63689 | 0.051806127 | -0.041344313 |
| Aspscr1 | 9.535904 | 9.503364 | 9.575 | -0.071672095 | -0.039131789 | 9.8017734 | 9.790231 | 9.73843 | 0.05179753 | 0.063339874 |
| Pcdh15 | 6.21305 | 6.104692 | 5.6865 | 0.418193621 | 0.526551837 | 5.7421424 | 5.7477919 | 5.69617 | 0.051622211 | 0.045972702 |
| Riox2 | 9.583847 | 9.637755 | 9.6907 | -0.052990199 | -0.106898954 | 9.8010299 | 9.7975294 | 9.74591 | 0.051620921 | 0.055121456 |
| Acot6 | 5.123817 | 4.952761 | 5.115 | -0.16220739 | 0.008847894 | 5.1703826 | 5.2159015 | 5.16446 | 0.051437854 | 0.005918977 |
| Vmn2r26 | 2.647357 | 2.520001 | 2.5659 | -0.045912354 | 0.081443623 | 2.5576659 | 2.6353683 | 2.58396 | 0.051410242 | -0.026292181 |
| Arhgef17 | 5.682503 | 5.626566 | 5.501 | 0.125528743 | 0.181464868 | 5.2502083 | 5.2741513 | 5.22275 | 0.051398382 | 0.027455377 |
| Grin2a | 1.201874 | 1.39131 | 1.1933 | 0.198020693 | 0.008583892 | 1.2546627 | 1.2296354 | 1.17824 | 0.051391179 | 0.076418481 |
| Rnls | 3.22302 | 3.164043 | 3.1503 | 0.01377339 | 0.072750168 | 3.1534889 | 3.1578507 | 3.10659 | 0.051258687 | 0.046896923 |
| Amdhd1 | 1.337596 | 1.421345 | 1.3339 | 0.087426129 | 0.003676776 | 1.3327279 | 1.4405036 | 1.38928 | 0.05122647 | -0.056549259 |
| Zkscan3 | 9.882902 | 9.935928 | 9.9031 | 0.032855805 | -0.020170187 | 10.034016 | 10.23713 | 10.1859 | 0.051205138 | -0.151909076 |
| Tubb5 | 13.25096 | 13.27222 | 13.371 | -0.098447338 | -0.11970339 | 13.412767 | 13.447124 | 13.3959 | 0.051204386 | 0.016847076 |
| Eif3a | 12.89624 | 12.88809 | 12.884 | 0.003828074 | 0.011975386 | 12.822104 | 12.993761 | 12.9426 | 0.051186654 | -0.120470363 |
| Gm14327 | 3.990733 | 3.779578 | 4.0659 | -0.286307639 | -0.075152429 | 3.9718689 | 3.9430584 | 3.89196 | 0.051093964 | 0.079904484 |
| Rhod | 3.982281 | 3.916711 | 3.9715 | -0.054835809 | 0.010733877 | 3.8013207 | 3.8959595 | 3.84489 | 0.051066941 | -0.043571838 |
| Det1 | 8.566257 | 8.524604 | 8.4762 | 0.048426088 | 0.090078607 | 8.6724421 | 8.5526023 | 8.50156 | 0.05104633 | 0.170886141 |
| Dcdc2a | 1.89733 | 1.846394 | 1.9049 | -0.058514334 | -0.007577637 | 1.861241 | 1.9389412 | 1.88792 | 0.051019294 | -0.026680873 |
| Trbj2-7 | 6.183425 | 6.47139 | 6.2439 | 0.227467167 | -0.06049775 | 6.1640062 | 6.2112113 | 6.16021 | 0.051000637 | 0.003795561 |
| Uchl1 | 3.380401 | 3.366697 | 3.2884 | 0.07828868 | 0.091992726 | 3.2589761 | 3.4333684 | 3.38243 | 0.050939629 | -0.123452607 |
| Rasl12 | 4.981727 | 5.094786 | 5.1444 | -0.049575924 | -0.162634434 | 5.0633912 | 5.0458746 | 4.99495 | 0.050922918 | 0.068439523 |
| Zbtb42 | 6.215791 | 6.191449 | 6.3254 | -0.133907825 | -0.109565483 | 6.4975925 | 6.3355353 | 6.28462 | 0.050916012 | 0.212973156 |
| Lca5l | 2.09774 | 2.134304 | 2.1493 | -0.014975595 | -0.051539405 | 2.1140938 | 2.1867071 | 2.13581 | 0.050893955 | -0.021719381 |
| Srsf10 | 10.98747 | 11.06085 | 10.922 | 0.1385729 | 0.06519776 | 11.120784 | 11.152533 | 11.1017 | 0.0508282 | 0.019079539 |
| Olfr1277 | 1.823885 | 1.820454 | 1.882 | -0.061512755 | -0.058082173 | 1.8256452 | 1.8880408 | 1.83722 | 0.050825166 | -0.011570424 |
| Scarf2 | 3.756335 | 3.652696 | 3.662 | -0.009290522 | 0.094348287 | 3.5444304 | 3.4879723 | 3.43715 | 0.050822494 | 0.107280591 |
| Pkn2 | 10.25279 | 10.28883 | 10.221 | 0.068063022 | 0.032024712 | 10.127771 | 10.364935 | 10.3143 | 0.050682166 | -0.186481961 |
| Olfr905 | 2.222667 | 2.260727 | 2.1321 | 0.128587256 | 0.090527348 | 2.2857984 | 2.3009516 | 2.25031 | 0.050639402 | 0.035486216 |
| Zbtb49 | 7.559902 | 7.605933 | 7.5484 | 0.057498668 | 0.011467716 | 7.9672093 | 7.8169318 | 7.76635 | 0.050581598 | 0.200859014 |
| Syna | 1.22757 | 1.206439 | 1.2311 | -0.024656948 | -0.003526086 | 1.2221152 | 1.2528715 | 1.20229 | 0.05057923 | 0.019822935 |
| Tgfbr3l | 4.851805 | 4.997456 | 4.8854 | 0.112096461 | -0.033554578 | 5.2245133 | 5.2600901 | 5.20951 | 0.050576555 | 0.014999739 |

| | | | | | | | | | | |
|---|---|---|---|---|---|---|---|---|---|---|
| Atxn2l | 12.07465 | 12.01739 | 12.067 | -0.049869352 | 0.007390584 | 12.196855 | 12.249632 | 12.1991 | 0.050548108 | -0.002229206 |
| Gm10097 | 0.723139 | 0.752721 | 0.6849 | 0.067845585 | 0.038263082 | 0.805081 | 0.7830438 | 0.73252 | 0.050526405 | 0.072563609 |
| Gls2 | 1.656219 | 1.614449 | 1.6753 | -0.060821638 | -0.019052081 | 1.6262685 | 1.6682211 | 1.61771 | 0.050508114 | 0.008555562 |
| Gm15276 | 0.680109 | 0.679842 | 0.6769 | 0.002937381 | 0.003204521 | 0.7212526 | 0.7354696 | 0.685 | 0.050473924 | 0.036257008 |
| Vmn1r57 | 0.957805 | 0.994492 | 1.0152 | -0.020666577 | -0.057353323 | 1.0789322 | 1.0920224 | 1.04159 | 0.050436391 | 0.037346107 |
| Camk2n1 | 5.400432 | 5.429853 | 5.322 | 0.107852125 | 0.078431585 | 4.8010096 | 5.1567442 | 5.10638 | 0.050368609 | -0.305366039 |
| Eno1b | 1.608957 | 1.693438 | 1.712 | -0.018568329 | -0.103049133 | 1.7893102 | 1.8144638 | 1.76411 | 0.0503495 | 0.025195934 |
| Haus3 | 8.60431 | 8.639871 | 8.6559 | -0.016009745 | -0.051570891 | 8.385329 | 8.7491956 | 8.69885 | 0.05034731 | -0.313519237 |
| Inpp5b | 10.29363 | 10.36262 | 10.368 | -0.00542531 | -0.07441408 | 10.244197 | 10.517218 | 10.4669 | 0.050322694 | -0.222698405 |
| Meig1 | 0.405964 | 0.392222 | 0.3922 | 1.27164E-05 | 0.013754521 | 0.3784907 | 0.4436999 | 0.39339 | 0.050306539 | -0.014902603 |
| Rex2 | 0.062239 | 0.056546 | 0.0674 | -0.010806902 | -0.005113205 | 0.1550295 | 0.199751 | 0.14945 | 0.05030279 | 0.005581253 |
| Ubap1l | 1.442206 | 1.40679 | 1.4496 | -0.042811716 | -0.007396131 | 1.5025585 | 1.4697949 | 1.41952 | 0.050275673 | 0.083039295 |
| Slc15a5 | 0.530163 | 0.518734 | 0.4982 | 0.020576947 | 0.032005872 | 0.5320165 | 0.5695907 | 0.51932 | 0.050270668 | 0.012696528 |
| Zfp579 | 6.536193 | 6.584744 | 6.7109 | -0.12618018 | -0.174731513 | 6.942326 | 6.9176542 | 6.8674 | 0.050258469 | 0.074930229 |
| Pla2g4c | 2.575722 | 2.627594 | 2.5206 | 0.106974538 | 0.055103063 | 2.5552566 | 2.6393779 | 2.58916 | 0.050213044 | -0.033908185 |
| Yif1a | 7.960663 | 7.875064 | 7.9644 | -0.089308725 | -0.003709243 | 8.0648226 | 7.8383503 | 7.78825 | 0.050101354 | 0.276573695 |
| Ift88 | 6.528222 | 6.491281 | 6.6371 | -0.145774045 | -0.10883286 | 6.7440564 | 6.6803939 | 6.63036 | 0.050032408 | 0.113694877 |
| Adamts2 | 5.34438 | 5.472264 | 5.4942 | -0.021944898 | -0.149829022 | 5.4549566 | 5.4979229 | 5.44791 | 0.050011162 | 0.007044893 |
| Gm20521 | 0.797226 | 0.684345 | 0.6294 | 0.054899065 | 0.167779975 | 0.7061711 | 0.7146098 | 0.66462 | 0.049989491 | 0.041550819 |
| Myl10 | 2.299578 | 2.290361 | 2.2563 | 0.03405231 | 0.043269603 | 2.2089578 | 2.2747316 | 2.22476 | 0.049973525 | -0.015800349 |
| Olfr6 | 2.527215 | 2.547073 | 2.5325 | 0.014533699 | -0.00532511 | 2.5767395 | 2.6374526 | 2.58753 | 0.04992707 | -0.010786051 |
| Spg11 | 10.01086 | 10.1516 | 10.05 | 0.101717403 | -0.03902604 | 10.074072 | 10.200218 | 10.1503 | 0.049887522 | -0.076258023 |
| Vezt | 8.033698 | 7.900244 | 7.9881 | -0.087824923 | 0.045628776 | 7.9407452 | 8.0848073 | 8.03498 | 0.049822892 | -0.094239157 |
| Htatip2 | 7.784468 | 7.647717 | 7.8951 | -0.24734942 | -0.110598534 | 7.7750584 | 7.8785199 | 7.82876 | 0.049756993 | -0.053704596 |
| Tpt1-ps5 | 1.510935 | 1.468384 | 1.5147 | -0.046315357 | -0.003764584 | 1.5215586 | 1.5178482 | 1.4681 | 0.049752862 | 0.053463282 |
| Gm17782 | 0.964974 | 1.018734 | 0.9118 | 0.10694651 | 0.053186823 | 0.9132528 | 1.0179244 | 0.96817 | 0.049750956 | -0.054920683 |
| Fgfrl1 | 5.070458 | 5.090579 | 5.1934 | -0.102824428 | -0.122944959 | 5.1260931 | 4.9670326 | 4.91731 | 0.049718804 | 0.208779261 |
| Sec63 | 10.57257 | 10.47914 | 10.532 | -0.053257512 | 0.04017529 | 10.485953 | 10.642711 | 10.593 | 0.049690248 | -0.10706807 |
| Denr | 9.642577 | 9.520167 | 9.5734 | -0.053270475 | 0.069139164 | 9.6814079 | 9.7611119 | 9.7115 | 0.049616184 | -0.030087781 |
| Nrip1 | 9.491876 | 9.641444 | 9.5122 | 0.129211282 | -0.020355931 | 9.0130442 | 9.4000955 | 9.35048 | 0.049613076 | -0.337438174 |
| Mpdu1 | 9.06741 | 9.019792 | 9.0451 | -0.025279101 | 0.022338517 | 9.2402726 | 9.101459 | 9.0519 | 0.04955864 | 0.188372249 |
| Spdl1 | 6.548433 | 6.652098 | 6.7478 | -0.095725432 | -0.199391197 | 6.4703075 | 6.8222164 | 6.77266 | 0.04955736 | -0.302351544 |

| | | | | | | | | | | |
|---|---|---|---|---|---|---|---|---|---|---|
| Lsamp | 2.273223 | 2.333389 | 2.1493 | 0.18409498 | 0.123929506 | 2.245823 | 2.194108 | 2.14458 | 0.049524979 | 0.101239961 |
| Nrbf2 | 7.634072 | 7.660164 | 7.7919 | -0.131779147 | -0.157871036 | 7.6831371 | 7.7386782 | 7.68917 | 0.049511852 | -0.006029262 |
| Celf1 | 10.57657 | 10.7166 | 10.643 | 0.073113504 | -0.06691039 | 10.717557 | 10.885838 | 10.8364 | 0.04941802 | -0.118863585 |
| Otud3 | 6.42929 | 6.701118 | 6.5192 | 0.181905519 | -0.089922077 | 6.3781231 | 6.4677736 | 6.41838 | 0.049395 | -0.040255537 |
| Fxyd5 | 11.92338 | 11.88486 | 11.89 | -0.00540547 | 0.03311639 | 11.981553 | 11.999619 | 11.9503 | 0.049365806 | 0.03129975 |
| Mab21l3 | 6.233484 | 6.41142 | 6.0599 | 0.351529037 | 0.173593593 | 5.6065792 | 6.0084564 | 5.95912 | 0.049333823 | -0.352543464 |
| Ttc9c | 9.764915 | 9.978751 | 9.8723 | 0.106416717 | -0.107419865 | 9.9443694 | 10.058814 | 10.0095 | 0.049290833 | -0.06515396 |
| Smg5 | 10.91274 | 10.95312 | 10.942 | 0.011076332 | -0.029302758 | 11.100021 | 11.241365 | 11.1921 | 0.049257 | -0.092086826 |
| Gm15720 | 0.236961 | 0.224587 | 0.2455 | -0.020867187 | -0.008492989 | 0.2080272 | 0.2654708 | 0.21633 | 0.04913985 | -0.008303667 |
| Sec31a | 11.19137 | 11.16356 | 11.116 | 0.047729324 | 0.075545286 | 11.261458 | 11.195673 | 11.1466 | 0.049040934 | 0.114825283 |
| Fgf6 | 1.005806 | 0.987903 | 0.9718 | 0.016080062 | 0.033982826 | 0.9948336 | 1.0592513 | 1.01021 | 0.049037093 | -0.015380577 |
| Gm49387 | 0.707321 | 0.654074 | 0.7117 | -0.057593548 | -0.00434595 | 0.8020353 | 0.7763221 | 0.72731 | 0.049016907 | 0.074730065 |
| A730061H03Rik | 1.696901 | 1.682257 | 1.7309 | -0.048637053 | -0.033992611 | 1.7666463 | 1.749099 | 1.7001 | 0.048994359 | 0.066541668 |
| Aldh3b3 | 0.857194 | 0.874094 | 0.8598 | 0.014339775 | -0.002559774 | 0.837361 | 0.950012 | 0.90106 | 0.048956877 | -0.063694113 |
| Fbxw2 | 10.1413 | 10.22718 | 10.139 | 0.088259002 | 0.002373022 | 10.17523 | 10.181217 | 10.1323 | 0.048956703 | 0.042969163 |
| Shc2 | 0.904717 | 0.845638 | 0.8702 | -0.024536412 | 0.034543366 | 0.9487495 | 0.8907771 | 0.84183 | 0.048946133 | 0.106918539 |
| Olfr1205 | 3.015356 | 2.96077 | 3.0663 | -0.105482001 | -0.050896118 | 3.0455048 | 3.190322 | 3.14138 | 0.048940446 | -0.095876726 |
| Gm6477 | 1.88164 | 1.778328 | 1.8103 | -0.031935019 | 0.071376462 | 1.7841385 | 1.8647894 | 1.81585 | 0.048937308 | -0.031713598 |
| Tmem262 | 0.212893 | 0.210753 | 0.2191 | -0.008303426 | -0.006163885 | 0.2260262 | 0.2921412 | 0.24324 | 0.048903849 | -0.017211145 |
| Eed | 9.929334 | 9.989852 | 9.9536 | 0.036268631 | -0.024248628 | 9.9334643 | 10.268444 | 10.2196 | 0.0488073 | -0.286172125 |
| Ccl27a | 4.524558 | 4.351919 | 4.4725 | -0.120626972 | 0.052011878 | 4.4850588 | 4.3402459 | 4.29144 | 0.048806719 | 0.193619659 |
| Iglc4 | 2.864872 | 2.896585 | 2.9612 | -0.064601019 | -0.096314494 | 2.8836293 | 2.9398656 | 2.89109 | 0.048779905 | -0.007456439 |
| Gm10343 | 0.976858 | 0.982248 | 0.9518 | 0.03042576 | 0.02503549 | 0.9561869 | 1.0088477 | 0.96007 | 0.048777941 | -0.003882814 |
| Omg | 1.004351 | 0.959585 | 0.9434 | 0.016169102 | 0.060934864 | 0.917513 | 0.9695983 | 0.92088 | 0.048714741 | -0.003370497 |
| Cfhr1 | 0.424397 | 0.39235 | 0.3921 | 0.000297493 | 0.032343669 | 0.3922413 | 0.4789761 | 0.43034 | 0.048634463 | -0.038100377 |
| Ddx20 | 8.81057 | 8.839179 | 8.8138 | 0.025417761 | -0.003190872 | 8.8222812 | 8.9170584 | 8.86844 | 0.048618953 | -0.04615826 |
| Pmp22 | 6.723242 | 6.567975 | 6.3088 | 0.259189315 | 0.414455965 | 5.9452496 | 5.8505445 | 5.80196 | 0.048580339 | 0.143285423 |
| Hnrnpa1 | 13.70388 | 13.6287 | 13.737 | -0.10785109 | -0.032670716 | 13.796747 | 13.833852 | 13.7853 | 0.048557342 | 0.011451413 |
| B3galnt1 | 4.034408 | 3.906304 | 3.8669 | 0.039387704 | 0.167491408 | 3.6633867 | 3.7216588 | 3.67311 | 0.048544816 | -0.009727323 |
| Edc4 | 10.66925 | 10.70407 | 10.71 | -0.006158244 | -0.04097001 | 10.957476 | 10.862466 | 10.814 | 0.048501908 | 0.14351166 |
| Oas1e | 0.367686 | 0.35731 | 0.3381 | 0.019242658 | 0.029618089 | 0.400458 | 0.4209642 | 0.37256 | 0.048407872 | 0.02790166 |
| Ttc30a2 | 0.588556 | 0.579137 | 0.6055 | -0.026325004 | -0.016906031 | 0.5625445 | 0.6333068 | 0.5849 | 0.048406643 | -0.022355734 |

| | | | | | | | | | | |
|---|---|---|---|---|---|---|---|---|---|---|
| Synb | 1.775189 | 1.750155 | 1.7688 | -0.018613316 | 0.006420858 | 1.7551795 | 1.824355 | 1.77603 | 0.048328009 | -0.020847464 |
| Eif3h | 12.32845 | 12.32246 | 12.363 | -0.04087444 | -0.034881234 | 12.390005 | 12.379307 | 12.331 | 0.048324264 | 0.059022635 |
| Med16 | 9.105457 | 8.794356 | 9.0319 | -0.237557442 | 0.073544301 | 9.1997698 | 9.0139399 | 8.96563 | 0.048314411 | 0.23414428 |
| 2700099C18Rik | 5.32229 | 5.184044 | 5.3682 | -0.184119214 | -0.045872401 | 5.4336033 | 5.6249251 | 5.57676 | 0.048168482 | -0.143153368 |
| Ift122 | 5.961289 | 5.643548 | 6.0099 | -0.366336329 | -0.048595674 | 5.8334001 | 5.9591762 | 5.91101 | 0.048166056 | -0.077610023 |
| Pde12 | 7.968332 | 7.889168 | 7.8 | 0.089182749 | 0.168346884 | 7.7546713 | 7.7699278 | 7.72177 | 0.048155547 | 0.032898988 |
| Bzw2 | 10.23969 | 10.27467 | 10.188 | 0.08713035 | 0.052153998 | 10.211776 | 10.252984 | 10.2049 | 0.048129912 | 0.006921979 |
| Mrgpra2a | 2.154791 | 1.674963 | 1.6605 | 0.014497138 | 0.49432582 | 1.6299923 | 1.6562561 | 1.60832 | 0.047931419 | 0.021667596 |
| Slc35g2 | 2.24006 | 2.125553 | 2.1575 | -0.031972238 | 0.082534327 | 2.1560133 | 2.1741439 | 2.12628 | 0.047867015 | 0.029736389 |
| Dsg1a | 1.691311 | 1.745494 | 1.7255 | 0.02004411 | -0.034138748 | 1.6621421 | 1.7132213 | 1.66537 | 0.047854169 | -0.003224978 |
| Suclg2 | 9.69473 | 9.625761 | 9.6315 | -0.005752154 | 0.06321649 | 9.5319914 | 9.6637251 | 9.61589 | 0.047837452 | -0.083896208 |
| Ms4a14 | 1.887986 | 1.987468 | 1.8642 | 0.123272431 | 0.023790378 | 1.8514736 | 1.8778709 | 1.83005 | 0.047819878 | 0.02142255 |
| Il16 | 12.07835 | 12.37907 | 12.246 | 0.132602042 | -0.168111366 | 12.290035 | 12.293786 | 12.246 | 0.047750346 | 0.043999474 |
| Fam89a | 4.180764 | 4.206631 | 4.241 | -0.034337495 | -0.060204251 | 4.1027201 | 4.195126 | 4.14741 | 0.047716183 | -0.044689673 |
| Atg2b | 10.1651 | 10.2479 | 10.165 | 0.082768299 | -3.2588E-05 | 10.147105 | 10.281482 | 10.2338 | 0.047716028 | -0.086661693 |
| Kpnb1 | 12.64411 | 12.58329 | 12.566 | 0.016905346 | 0.07772283 | 12.6083 | 12.719114 | 12.6714 | 0.047685722 | -0.063128902 |
| Gm20489 | 1.637468 | 1.576605 | 1.6024 | -0.025811828 | 0.035051345 | 1.6511547 | 1.7680868 | 1.72041 | 0.047681404 | -0.069250675 |
| Grk3 | 8.766247 | 8.958104 | 8.9657 | -0.007580079 | -0.199437157 | 8.96983 | 9.2877802 | 9.2401 | 0.047680632 | -0.270269576 |
| Trav13d-4 | 1.610927 | 1.555103 | 1.6266 | -0.071530357 | -0.015706171 | 1.5932713 | 1.6455727 | 1.59791 | 0.047666579 | -0.004634812 |
| Cyp4x1 | 2.163184 | 2.06561 | 2.1095 | -0.043915735 | 0.053657788 | 2.0445152 | 2.1951485 | 2.14752 | 0.047624722 | -0.103008593 |
| Bnip1 | 8.552754 | 8.608614 | 8.5997 | 0.008904623 | -0.046954994 | 8.9043292 | 8.7060001 | 8.65839 | 0.047614046 | 0.245943124 |
| Apoa2 | 1.150576 | 1.093213 | 1.1772 | -0.083951296 | -0.026587967 | 1.1363482 | 1.1688417 | 1.12123 | 0.047610886 | 0.015117388 |
| Myl6 | 12.68362 | 12.73035 | 12.718 | 0.01234547 | -0.034386702 | 12.833969 | 12.728539 | 12.6809 | 0.04759897 | 0.153028586 |
| Gm16570 | 0.040663 | -0.02149 | 0.0039 | -0.025403787 | 0.036751556 | 0.0187768 | 0.0572209 | 0.00965 | 0.047567709 | 0.009123546 |
| Mcm7 | 10.30713 | 10.16144 | 10.375 | -0.213591563 | -0.067905984 | 10.264751 | 10.48379 | 10.4362 | 0.04756278 | -0.171475738 |
| Vps45 | 7.778299 | 7.699437 | 7.8295 | -0.130052784 | -0.051191105 | 8.0353555 | 7.8447362 | 7.79719 | 0.047548168 | 0.238167454 |
| Oard1 | 8.067933 | 7.984169 | 8.1006 | -0.116407987 | -0.032643845 | 8.1763373 | 8.0717119 | 8.0242 | 0.047514352 | 0.152139784 |
| Gm49395 | 1.594386 | 1.49401 | 1.5967 | -0.102647107 | -0.002270189 | 1.4695385 | 1.5919113 | 1.5444 | 0.047508891 | -0.074863885 |
| Calml4 | 2.223759 | 2.225019 | 2.2551 | -0.03007649 | -0.031336728 | 2.2554359 | 2.155266 | 2.1078 | 0.047467624 | 0.147637595 |
| Lair1 | 8.218579 | 8.130964 | 8.1151 | 0.01587412 | 0.1034883 | 8.318737 | 7.712827 | 7.66541 | 0.04741746 | 0.653327374 |
| Fitm2 | 6.526829 | 6.764106 | 6.6439 | 0.120252609 | -0.117024045 | 6.6433925 | 6.6505051 | 6.60321 | 0.047297718 | 0.040185119 |
| Trgv2 | 5.173839 | 5.122984 | 5.07 | 0.052993092 | 0.103848267 | 4.9605508 | 5.1602724 | 5.11299 | 0.047277759 | -0.152443777 |

| | | | | | | | | | | |
|---|---|---|---|---|---|---|---|---|---|---|
| Kri1 | 10.56751 | 10.41753 | 10.559 | -0.14152053 | 0.008461498 | 10.753351 | 10.819002 | 10.7717 | 0.04726194 | -0.018389211 |
| AC154369.1 | 1.991922 | 1.953313 | 2.0412 | -0.087857808 | -0.049248918 | 2.0011263 | 2.1060132 | 2.0588 | 0.047209935 | -0.057676955 |
| Gm8249 | 1.097907 | 1.042894 | 1.0281 | 0.014772109 | 0.069785382 | 1.0516788 | 1.1167992 | 1.06975 | 0.047044736 | -0.018075666 |
| Dhrs4 | 7.824258 | 7.770925 | 7.8282 | -0.057256438 | -0.003924032 | 8.0871966 | 7.8493118 | 7.80228 | 0.047027463 | 0.284912224 |
| Vmn2r100 | 3.176594 | 3.031941 | 3.1661 | -0.134168067 | 0.010484315 | 3.2514748 | 3.2766828 | 3.22966 | 0.047018476 | 0.021810421 |
| Tdh | 2.142004 | 2.014888 | 2.0783 | -0.063461202 | 0.06365542 | 2.0118401 | 2.0901877 | 2.04324 | 0.046952371 | -0.031395267 |
| Bcl2l12 | 7.744748 | 8.002223 | 7.8563 | 0.145889895 | -0.111585499 | 8.3881695 | 7.9541884 | 7.90724 | 0.046947654 | 0.480928708 |
| Igkv3-3 | 2.17707 | 2.265736 | 2.124 | 0.141738467 | 0.053072734 | 2.1775207 | 2.1582627 | 2.11132 | 0.046938784 | 0.066196829 |
| Mycbp | 6.422469 | 6.505387 | 6.5099 | -0.004541068 | -0.087459013 | 6.4785218 | 6.6761689 | 6.62923 | 0.046938412 | -0.150708651 |
| Gm16505 | 2.090248 | 2.125141 | 2.2 | -0.074853853 | -0.109746971 | 2.1540592 | 2.1512093 | 2.10428 | 0.046932532 | 0.049782493 |
| Taok3 | 11.04036 | 11.13079 | 10.934 | 0.197009886 | 0.106582214 | 11.04857 | 11.088842 | 11.0419 | 0.046920788 | 0.006648751 |
| Crhbp | 0.445039 | 0.418403 | 0.5102 | -0.091836911 | -0.065200683 | 0.5559112 | 0.4827177 | 0.43585 | 0.046868847 | 0.120062376 |
| Apbb1ip | 11.35352 | 11.48532 | 11.409 | 0.076463946 | -0.055341284 | 11.363131 | 11.366499 | 11.3197 | 0.046844146 | 0.04347574 |
| Gm12941 | 1.981061 | 2.04075 | 1.9463 | 0.0944814 | 0.034791811 | 2.08711 | 2.1180004 | 2.07117 | 0.046827734 | 0.015937301 |
| Gm3919 | 1.473329 | 1.401625 | 1.4416 | -0.039950888 | 0.031753985 | 1.4534825 | 1.4781203 | 1.43137 | 0.046753527 | 0.02211577 |
| Cic | 11.83299 | 11.75299 | 11.708 | 0.045021662 | 0.125022144 | 11.847211 | 11.869549 | 11.8228 | 0.046734404 | 0.024396163 |
| Lrrc14b | 0.502935 | 0.523172 | 0.5333 | -0.010104392 | -0.030341338 | 0.5691314 | 0.6048246 | 0.55812 | 0.046699873 | 0.011006634 |
| Yif1b | 8.092381 | 7.962912 | 8.0543 | -0.091368295 | 0.038100735 | 8.3048334 | 8.0384918 | 7.99188 | 0.046613405 | 0.312954977 |
| Olfr1310 | 3.675144 | 3.636964 | 3.7306 | -0.093663858 | -0.055484405 | 3.7866573 | 3.8558503 | 3.80927 | 0.046576604 | -0.022616464 |
| Cks2 | 7.258362 | 7.23197 | 7.3865 | -0.154502101 | -0.12811048 | 7.3341095 | 7.5165194 | 7.46996 | 0.046557428 | -0.135852456 |
| Oprm1 | 1.759684 | 1.737267 | 1.8273 | -0.090076264 | -0.067659636 | 1.7461438 | 1.8033704 | 1.75685 | 0.046521471 | -0.010705135 |
| Ets2 | 10.13798 | 10.20414 | 9.8037 | 0.400462983 | 0.334300398 | 9.8909701 | 9.9190817 | 9.87257 | 0.046508706 | 0.018397086 |
| Cnppd1 | 10.59288 | 10.5183 | 10.601 | -0.082739088 | -0.008162984 | 10.758956 | 10.527053 | 10.4806 | 0.046481252 | 0.278384923 |
| Gm7701 | 2.275613 | 2.254555 | 2.3229 | -0.068352447 | -0.047293951 | 2.3441003 | 2.4526204 | 2.40615 | 0.046474712 | -0.062045455 |
| Eef1b2 | 13.63889 | 13.65329 | 13.678 | -0.024783372 | -0.03918799 | 13.668199 | 13.71203 | 13.6656 | 0.046464038 | 0.002632818 |
| Igkv8-16 | 5.597539 | 6.03825 | 5.9026 | 0.13567663 | -0.305034361 | 5.9738732 | 5.8636891 | 5.81733 | 0.046358163 | 0.156542261 |
| Ints1 | 10.66304 | 10.71249 | 10.761 | -0.048307654 | -0.097754744 | 10.936912 | 10.910509 | 10.8642 | 0.046314452 | 0.072717309 |
| Dbx2 | 1.609603 | 1.293421 | 1.2861 | 0.007281613 | 0.323462715 | 1.2874119 | 1.3756175 | 1.32931 | 0.046309527 | -0.041896123 |
| Rpl13-ps6 | 1.487344 | 1.402893 | 1.5045 | -0.101601811 | -0.01715084 | 1.7055776 | 1.469302 | 1.42302 | 0.046278208 | 0.282553834 |
| Gm21769 | 1.432705 | 1.532799 | 1.4959 | 0.036922012 | -0.063172448 | 1.4986285 | 1.500512 | 1.45428 | 0.046236112 | 0.044352592 |
| Armh2 | 0.863976 | 0.915874 | 0.8559 | 0.059992354 | 0.008094178 | 0.9074793 | 0.8978604 | 0.85167 | 0.046192387 | 0.055811298 |
| Gm10721 | 0.412527 | 0.331859 | 0.3673 | -0.035480282 | 0.045187937 | 0.3428999 | 0.4413171 | 0.39515 | 0.046166998 | -0.052250207 |

| | | | | | | | | | | |
|---|---|---|---|---|---|---|---|---|---|---|
| Bmper | 3.182484 | 3.297143 | 3.2754 | 0.021762368 | -0.092895776 | 3.3291371 | 3.2564047 | 3.21024 | 0.04616594 | 0.118898256 |
| Zfp119b | 6.1809 | 6.143676 | 6.2034 | -0.059705951 | -0.022481499 | 6.1885372 | 6.1997415 | 6.15365 | 0.046087228 | 0.034882938 |
| Gm49141 | 0.968956 | 1.004854 | 1.0078 | -0.002988541 | -0.038886717 | 0.9974689 | 1.0692467 | 1.02319 | 0.046060815 | -0.025716936 |
| Rragc | 9.956341 | 9.900555 | 9.8738 | 0.026732897 | 0.0825191 | 9.7641983 | 9.9522307 | 9.90619 | 0.046036451 | -0.141995993 |
| Copb1 | 11.14781 | 11.17281 | 11.148 | 0.024761532 | -0.000243978 | 11.257124 | 11.196544 | 11.1505 | 0.046030884 | 0.106610277 |
| Islr2 | 3.167543 | 3.064626 | 3.1528 | -0.088197461 | 0.014719597 | 3.0021649 | 3.1162255 | 3.07024 | 0.045982854 | -0.068077746 |
| Lat | 11.50592 | 11.54207 | 11.55 | -0.007554374 | -0.043701204 | 11.395629 | 11.543352 | 11.4974 | 0.04592858 | -0.101794933 |
| Gusb | 9.994138 | 10.03899 | 9.9642 | 0.074769709 | 0.029918799 | 9.9832059 | 9.9239381 | 9.87807 | 0.045872919 | 0.105140805 |
| Pak6 | 3.169764 | 3.192667 | 3.3103 | -0.117607827 | -0.140510903 | 3.2840481 | 3.3161231 | 3.27027 | 0.045848312 | 0.013773323 |
| Zfp747 | 7.928939 | 7.79944 | 7.7849 | 0.014531262 | 0.144030559 | 7.7617928 | 7.7955186 | 7.74971 | 0.045809012 | 0.012083199 |
| Olfr57 | 3.70673 | 3.555388 | 3.6537 | -0.098347264 | 0.052994924 | 3.6672703 | 3.8101348 | 3.76434 | 0.045790725 | -0.097073769 |
| Gm15667 | 0.914183 | 0.990889 | 0.9417 | 0.049187106 | -0.027518844 | 0.8873536 | 0.9992699 | 0.9535 | 0.045771317 | -0.066145032 |
| Gm6153 | 1.624969 | 1.662236 | 1.6507 | 0.011530684 | -0.025735948 | 1.6484476 | 1.6462842 | 1.60055 | 0.045739131 | 0.047902439 |
| Fbxo31 | 8.298734 | 8.33631 | 8.4164 | -0.080114783 | -0.11769146 | 8.507542 | 8.3908902 | 8.34517 | 0.04572508 | 0.162376945 |
| Tenm1 | 3.20338 | 3.237942 | 3.2498 | -0.01181395 | -0.046375218 | 3.2418429 | 3.430639 | 3.38496 | 0.045683898 | -0.143112156 |
| Shoc2 | 10.15036 | 10.35095 | 10.242 | 0.109320694 | -0.09126725 | 10.021698 | 10.158378 | 10.1127 | 0.045646255 | -0.091034452 |
| Olfr1420 | 0.397921 | 0.393583 | 0.392 | 0.001586372 | 0.005924662 | 0.4059357 | 0.4661327 | 0.4205 | 0.045631503 | -0.014565477 |
| Vmn2r90 | 3.812278 | 3.836795 | 3.823 | 0.013828138 | -0.010688719 | 3.8370363 | 3.9754849 | 3.92987 | 0.045614876 | -0.092833711 |
| Igkv8-18 | 3.718261 | 3.857666 | 3.7638 | 0.093872461 | -0.045532613 | 4.2743982 | 3.7767161 | 3.73111 | 0.045607309 | 0.543289401 |
| Zfyve19 | 8.178805 | 8.119899 | 8.3039 | -0.18399083 | -0.125084102 | 8.374186 | 8.3553747 | 8.30988 | 0.045497349 | 0.064308663 |
| Crbn | 9.882245 | 9.905944 | 10.034 | -0.128473415 | -0.152172224 | 9.8787222 | 9.9714208 | 9.92594 | 0.045481725 | -0.047216844 |
| Olfr1330 | 2.527178 | 2.517208 | 2.4478 | 0.069381767 | 0.07935165 | 2.5766413 | 2.6269891 | 2.58156 | 0.045428044 | -0.004919793 |
| Calr3 | 2.860791 | 2.908288 | 2.9276 | -0.019346746 | -0.066843756 | 3.077497 | 3.0180265 | 2.97266 | 0.045362796 | 0.104833276 |
| Mpc1 | 9.307123 | 9.25026 | 9.2879 | -0.037611063 | 0.019252052 | 9.4309803 | 9.2998604 | 9.25453 | 0.045327165 | 0.176447069 |
| Nudt22 | 6.36323 | 6.060824 | 6.207 | -0.146148839 | 0.156257862 | 6.3064262 | 6.3566601 | 6.31134 | 0.045321137 | -0.004912773 |
| Gm49686 | 1.009627 | 1.004607 | 0.9877 | 0.016931256 | 0.021951898 | 1.0101314 | 1.0350344 | 0.98971 | 0.045319762 | 0.020416825 |
| Lrrfip2 | 8.387124 | 8.406887 | 8.4408 | -0.03391329 | -0.05367657 | 8.3144291 | 8.5828666 | 8.53756 | 0.045306401 | -0.223131153 |
| Anapc11 | 8.525404 | 8.574087 | 8.5477 | 0.026382697 | -0.022299653 | 8.7944768 | 8.6203493 | 8.57505 | 0.045302445 | 0.21942994 |
| Pramel4 | 1.215113 | 1.212122 | 1.2133 | -0.00114563 | 0.001845827 | 1.2221603 | 1.338958 | 1.29369 | 0.045263627 | -0.071534051 |
| Ncln | 10.22068 | 10.30612 | 10.217 | 0.088688033 | 0.003244233 | 10.600046 | 10.545064 | 10.4998 | 0.045262134 | 0.100243983 |
| AC090881.3 | 1.592667 | 1.706586 | 1.6135 | 0.093088526 | -0.020830483 | 1.5842648 | 1.6341943 | 1.58895 | 0.045245334 | -0.004684129 |
| Gm6767 | 2.752522 | 2.708771 | 2.7536 | -0.044798879 | -0.001047774 | 2.7467895 | 2.7839378 | 2.73872 | 0.045219887 | 0.008071557 |

| | | | | | | | | | | |
|---|---|---|---|---|---|---|---|---|---|---|
| Clrn1 | 2.049679 | 2.07975 | 2.1127 | -0.032938075 | -0.063008236 | 2.1098344 | 2.1934663 | 2.14826 | 0.045204591 | -0.038427325 |
| Vegfc | 4.781593 | 4.944851 | 4.6334 | 0.311402503 | 0.148144558 | 4.7244757 | 4.5672813 | 4.5221 | 0.045177957 | 0.2023724 |
| Gm20716 | 11.1006 | 11.17274 | 11.156 | 0.01672203 | -0.055413044 | 11.069004 | 11.197411 | 11.1523 | 0.045120782 | -0.083286262 |
| Zfp72 | 5.457404 | 5.581357 | 5.6367 | -0.055325223 | -0.179278828 | 5.7412265 | 5.5396538 | 5.49456 | 0.045090292 | 0.246663021 |
| Vps33b | 8.916584 | 8.901331 | 8.9484 | -0.047073079 | -0.031819701 | 9.1258672 | 9.147733 | 9.10274 | 0.044992918 | 0.023127095 |
| Adamts20 | 2.81505 | 2.644052 | 2.5904 | 0.053632477 | 0.224630385 | 2.7009668 | 2.5390677 | 2.49408 | 0.044988793 | 0.206887878 |
| Sgpl1 | 10.9307 | 10.99617 | 10.798 | 0.198073244 | 0.132600898 | 10.899536 | 10.958143 | 10.9132 | 0.044901222 | -0.01370624 |
| Cib3 | 2.42953 | 2.291177 | 2.432 | -0.140789561 | -0.00243624 | 2.2784568 | 2.4308773 | 2.386 | 0.044877647 | -0.107542916 |
| Eif3l | 11.24916 | 11.26781 | 11.316 | -0.048280014 | -0.066928462 | 11.422143 | 11.362512 | 11.3177 | 0.044855982 | 0.104486433 |
| Cend1 | 0.873822 | 0.85418 | 0.8113 | 0.042925233 | 0.062566884 | 0.8647639 | 0.8591223 | 0.81435 | 0.044774483 | 0.050416038 |
| Idh3b | 10.61591 | 10.58567 | 10.59 | -0.004059854 | 0.02617497 | 10.853405 | 10.71515 | 10.6704 | 0.04474534 | 0.1830006 |
| Trbj2-6 | 4.510619 | 4.847882 | 4.555 | 0.292912321 | -0.044350456 | 4.6470979 | 4.7471992 | 4.70247 | 0.044724472 | -0.055376847 |
| Gm47630 | 1.88589 | 1.896839 | 1.9443 | -0.047467886 | -0.058417481 | 1.8604805 | 2.0477045 | 2.00304 | 0.044661909 | -0.142562075 |
| Olfr676 | 1.602935 | 1.498294 | 1.6068 | -0.108527569 | -0.003886415 | 1.5706316 | 1.6051228 | 1.56049 | 0.044631637 | 0.010140374 |
| Hlx | 6.736638 | 6.846457 | 6.7738 | 0.07262887 | -0.037190547 | 6.7610654 | 6.5586119 | 6.51399 | 0.044624554 | 0.247078053 |
| Ganab | 11.92293 | 11.96253 | 11.944 | 0.01892201 | -0.02068496 | 12.080978 | 12.064608 | 12.02 | 0.044566562 | 0.06093724 |
| mt-Atp6 | 15.28387 | 15.51148 | 15.472 | 0.039381606 | -0.188231676 | 15.49618 | 15.476454 | 15.432 | 0.04449201 | 0.064217589 |
| Immp1l | 8.331027 | 8.375675 | 8.3609 | 0.014803168 | -0.029844612 | 8.4857732 | 8.3683873 | 8.32392 | 0.044466818 | 0.161852742 |
| Ppara | 1.206029 | 1.184749 | 1.3375 | -0.152730443 | -0.131450501 | 1.2444028 | 1.3094092 | 1.26501 | 0.044397955 | -0.020608408 |
| Prrt1 | 5.706368 | 5.729353 | 5.6775 | 0.051842747 | 0.028857896 | 5.6440292 | 5.7982725 | 5.75388 | 0.044391638 | -0.109851665 |
| Aasdhppt | 8.132319 | 8.187299 | 8.1917 | -0.004415163 | -0.059395169 | 8.1405036 | 8.1951438 | 8.15079 | 0.044357793 | -0.010282425 |
| Gm49288 | 1.553154 | 1.593716 | 1.5913 | 0.002379021 | -0.038183599 | 1.5702565 | 1.6774271 | 1.63311 | 0.044321684 | -0.062848938 |
| 1700001O22Rik | 2.228675 | 2.210693 | 2.2801 | -0.069386765 | -0.051405033 | 2.327987 | 2.3907087 | 2.34653 | 0.044178218 | -0.018543557 |
| Kcnn2 | 1.096784 | 0.984654 | 1.0012 | -0.016552168 | 0.095577319 | 0.9898815 | 1.0333713 | 0.98921 | 0.044166067 | 0.000676283 |
| Bicdl1 | 8.960898 | 8.938003 | 8.9834 | -0.045391564 | -0.022497152 | 8.5206734 | 8.9102936 | 8.86615 | 0.044140533 | -0.345479671 |
| Ptch2 | 1.456662 | 1.454923 | 1.4812 | -0.026250489 | -0.024511883 | 1.5607146 | 1.4526491 | 1.40852 | 0.044131307 | 0.152196732 |
| Uba3 | 9.292119 | 9.362351 | 9.3244 | 0.037935137 | -0.03229738 | 9.2261169 | 9.3872321 | 9.34316 | 0.04406751 | -0.117047754 |
| Snx25 | 9.227838 | 9.338903 | 9.2134 | 0.125458738 | 0.014393256 | 9.1667682 | 9.2068295 | 9.16277 | 0.044055312 | 0.003994092 |
| Gm12407 | 1.781701 | 1.731848 | 1.7047 | 0.027122647 | 0.076975639 | 1.6606073 | 1.7452231 | 1.70118 | 0.044042738 | -0.040573102 |
| Dkk1 | 1.03349 | 1.028733 | 1.0231 | 0.005641883 | 0.010399076 | 1.0331954 | 1.1230642 | 1.07905 | 0.04401455 | -0.045854257 |
| Gm43712 | 1.745638 | 1.769285 | 1.7564 | 0.012911335 | -0.010735565 | 1.7138132 | 1.8655603 | 1.8216 | 0.043956774 | -0.107790421 |
| Prpf19 | 11.00036 | 10.99795 | 10.947 | 0.051263602 | 0.05367884 | 11.166991 | 11.082997 | 11.0391 | 0.043911124 | 0.1279047 |

| | | | | | | | | | | |
|---|---|---|---|---|---|---|---|---|---|---|
| Poc5 | 8.381397 | 8.327502 | 8.4413 | -0.113815702 | -0.059920211 | 8.3538407 | 8.347564 | 8.30373 | 0.043836429 | 0.050113059 |
| Rnf146 | 9.839465 | 9.985833 | 9.8824 | 0.103410632 | -0.042957254 | 9.8424407 | 9.9573882 | 9.91361 | 0.043782623 | -0.071164868 |
| Foxc1 | 3.272219 | 3.036147 | 3.2243 | -0.188166044 | 0.047905902 | 2.9018494 | 3.0132988 | 2.96953 | 0.04376806 | -0.067681272 |
| Sult2a7 | 0.969252 | 0.973726 | 0.951 | 0.02272573 | 0.018251715 | 0.9760362 | 0.9743304 | 0.93061 | 0.043724704 | 0.045430441 |
| Vmn1r203 | 4.977874 | 5.179022 | 4.9556 | 0.223421422 | 0.022273391 | 4.9495269 | 5.1523254 | 5.10861 | 0.04372023 | -0.159078238 |
| Olfr243 | 3.809066 | 3.850108 | 3.8912 | -0.041097562 | -0.082139599 | 3.9369509 | 4.0371091 | 3.99343 | 0.043674942 | -0.056483248 |
| Gm13498 | 1.621977 | 1.520013 | 1.5066 | 0.013392684 | 0.115356024 | 1.6211545 | 1.5919372 | 1.54828 | 0.043655256 | 0.072872563 |
| 9530050K03Rik | 3.386011 | 3.448698 | 3.3974 | 0.051305769 | -0.011380916 | 3.4340379 | 3.5547044 | 3.51111 | 0.043598531 | -0.077067967 |
| Gm8956 | 1.666224 | 1.669775 | 1.7077 | -0.037938576 | -0.041490104 | 1.6535855 | 1.759974 | 1.71643 | 0.043546803 | -0.062841689 |
| Dyrk2 | 10.13268 | 10.19982 | 10.26 | -0.060254298 | -0.127393005 | 10.047322 | 10.32174 | 10.2782 | 0.04354643 | -0.230871512 |
| Ift74 | 6.179991 | 6.086585 | 6.1696 | -0.083044707 | 0.010361191 | 6.2780703 | 6.0547294 | 6.01121 | 0.043518427 | 0.266859358 |
| Memo1 | 8.739721 | 8.69924 | 8.6967 | 0.002537554 | 0.043019291 | 8.60116 | 8.7046352 | 8.66115 | 0.04348345 | -0.059991761 |
| Rragd | 6.534918 | 6.063138 | 6.2192 | -0.156109 | 0.315670261 | 5.9544323 | 6.3809915 | 6.33752 | 0.043466681 | -0.383092529 |
| Hmgb1-ps3 | 3.849505 | 3.967301 | 3.9934 | -0.026099339 | -0.143894799 | 3.8637663 | 3.8962644 | 3.8528 | 0.043464205 | 0.010966052 |
| Rnf41 | 9.547695 | 9.540161 | 9.4627 | 0.07747141 | 0.085005365 | 9.5872327 | 9.4454153 | 9.40197 | 0.043449918 | 0.185267256 |
| Gm5424 | 1.088577 | 1.109354 | 1.1302 | -0.020860239 | -0.041637562 | 1.1846885 | 1.146599 | 1.10322 | 0.04338155 | 0.08147102 |
| Vasp | 12.41527 | 12.41152 | 12.33 | 0.08130445 | 0.085055152 | 12.680407 | 12.444367 | 12.401 | 0.043344666 | 0.279384996 |
| Cndp1 | 2.007819 | 1.976266 | 1.9968 | -0.020519433 | 0.011033752 | 1.995853 | 2.0570742 | 2.01373 | 0.043340774 | -0.017880457 |
| Pld2 | 7.042136 | 7.056845 | 7.1845 | -0.127622267 | -0.142331159 | 7.3215586 | 7.4044433 | 7.36118 | 0.04326481 | -0.039619915 |
| Gm45647 | 0.418201 | 0.439363 | 0.4878 | -0.048397181 | -0.069559219 | 0.4529082 | 0.5085161 | 0.46529 | 0.043228646 | -0.012379234 |
| Grsf1 | 9.860353 | 9.808711 | 9.801 | 0.007728802 | 0.059371045 | 9.6574127 | 9.7399323 | 9.69671 | 0.043227161 | -0.039292475 |
| Klra17 | 4.961066 | 4.961739 | 4.9117 | 0.050073526 | 0.049401247 | 4.5105993 | 4.6300347 | 4.58683 | 0.043206783 | -0.076228598 |
| Calhm4 | 0.986279 | 1.063065 | 0.9409 | 0.122147788 | 0.045362338 | 0.9814141 | 0.9945831 | 0.9514 | 0.043179879 | 0.030010947 |
| Gm47773 | 1.230625 | 1.208256 | 1.2405 | -0.032244542 | -0.00987628 | 1.2097842 | 1.3348562 | 1.29169 | 0.043162853 | -0.081909132 |
| Trim17 | 1.568413 | 1.568408 | 1.5571 | 0.01131138 | 0.011316211 | 1.6480166 | 1.5809985 | 1.53788 | 0.043121417 | 0.11013946 |
| Fdft1 | 8.499146 | 8.511105 | 8.4092 | 0.101946134 | 0.089986957 | 8.4564948 | 8.4307678 | 8.38767 | 0.043100646 | 0.068827656 |
| Myo19 | 7.248167 | 7.079303 | 7.0729 | 0.00641247 | 0.175275692 | 7.3206764 | 7.3896135 | 7.34652 | 0.043095826 | -0.025841316 |
| Olfr1457 | 0.489317 | 0.557444 | 0.5126 | 0.044879314 | -0.023247454 | 0.5905128 | 0.5430879 | 0.5 | 0.043091837 | 0.090516764 |
| Rpl10-ps5 | 5.646205 | 5.807556 | 5.752 | 0.055567905 | -0.105782702 | 5.7216745 | 5.8213023 | 5.77826 | 0.043043287 | -0.056584464 |
| Cebpzos | 6.641356 | 6.576745 | 6.765 | -0.188302523 | -0.123690577 | 6.7259699 | 6.5926664 | 6.54967 | 0.042997503 | 0.176300981 |
| Xpnpep2 | 2.368276 | 2.392776 | 2.3445 | 0.048257979 | 0.023758249 | 2.3023354 | 2.4646512 | 2.42169 | 0.042960995 | -0.119354759 |
| Tram1l1 | 0.892162 | 0.874674 | 0.9719 | -0.097198375 | -0.079710158 | 0.8771925 | 0.9223453 | 0.87947 | 0.04287641 | -0.002276358 |

| | | | | | | | | | | |
|---|---|---|---|---|---|---|---|---|---|---|
| Chchd3 | 9.907667 | 9.915113 | 9.9106 | 0.004554775 | -0.002890863 | 10.148815 | 10.077651 | 10.0348 | 0.042823425 | 0.113987314 |
| Dpp6 | 3.111247 | 3.005356 | 2.8993 | 0.106087569 | 0.211977973 | 3.0872282 | 3.1031456 | 3.06033 | 0.042811089 | 0.026893699 |
| Zfp735 | 0.91848 | 0.883952 | 0.8862 | -0.002225025 | 0.032303151 | 0.87514 | 0.9391607 | 0.89637 | 0.04279132 | -0.021229323 |
| Gm14270 | 0.209834 | 0.171599 | 0.2469 | -0.075303972 | -0.037068668 | 0.1788269 | 0.2283474 | 0.18569 | 0.04265454 | -0.006865934 |
| Ptbp1 | 12.53134 | 12.49408 | 12.56 | -0.066000528 | -0.028741904 | 12.690363 | 12.611808 | 12.5692 | 0.042643594 | 0.121198586 |
| Olfr767 | 3.946025 | 3.988616 | 3.8871 | 0.101539286 | 0.058948942 | 3.9789926 | 4.2347265 | 4.1921 | 0.042631228 | -0.21310263 |
| Ambra1 | 10.43564 | 10.43884 | 10.459 | -0.020604524 | -0.02379758 | 10.36933 | 10.520829 | 10.4782 | 0.042623864 | -0.108875471 |
| Gm44984 | 1.995959 | 2.048406 | 2.0331 | 0.015330977 | -0.037116088 | 2.1354842 | 2.0712747 | 2.02876 | 0.042510689 | 0.106720158 |
| Riok2 | 9.063784 | 8.953059 | 8.9591 | -0.006054099 | 0.104671396 | 8.886481 | 8.9722006 | 8.92972 | 0.042481366 | -0.04323825 |
| Gm14226 | 2.035745 | 2.031745 | 2.0059 | 0.025820134 | 0.029820015 | 2.0287842 | 2.1227448 | 2.08027 | 0.042469907 | -0.051490736 |
| Fam222a | 4.766907 | 4.724569 | 4.7684 | -0.043872037 | -0.001533634 | 4.6672405 | 4.5868738 | 4.54445 | 0.042424139 | 0.122790784 |
| Gm10109 | 0.909052 | 0.897264 | 1.0081 | -0.110849292 | -0.099061536 | 0.8720188 | 0.9602596 | 0.91785 | 0.042409374 | -0.045831408 |
| Gpr26 | 2.103894 | 2.147637 | 2.0416 | 0.106072606 | 0.062329778 | 2.0932723 | 2.1029014 | 2.0605 | 0.042400864 | 0.032771817 |
| Tnfrsf4 | 8.674367 | 8.740387 | 8.6828 | 0.057575759 | -0.008444265 | 8.7737711 | 8.9100628 | 8.86774 | 0.042320363 | -0.093971425 |
| Skint4 | 2.025109 | 2.088768 | 2.0471 | 0.04165993 | -0.02199978 | 2.0412589 | 2.0908382 | 2.04852 | 0.042319668 | -0.00725959 |
| Ugp2 | 9.629286 | 9.579544 | 9.6106 | -0.031093101 | 0.018649321 | 9.6020257 | 9.5689224 | 9.52661 | 0.042310798 | 0.075414154 |
| Abcb10 | 8.013128 | 8.064657 | 8.0873 | -0.022639621 | -0.074169158 | 8.283217 | 8.0497004 | 8.00747 | 0.042231359 | 0.27574801 |
| Crb2 | 1.227178 | 1.169935 | 1.1808 | -0.010856825 | 0.046386104 | 1.2431276 | 1.2292229 | 1.18701 | 0.042217617 | 0.056122368 |
| Dld | 10.15967 | 10.11398 | 10.033 | 0.080491151 | 0.126174666 | 10.018911 | 10.192665 | 10.1505 | 0.042199974 | -0.131553165 |
| Celsr1 | 9.498413 | 9.261513 | 9.3749 | -0.113343843 | 0.123557059 | 9.0632677 | 9.3354492 | 9.29328 | 0.042171879 | -0.230009579 |
| 2010109I03Rik | 5.728434 | 5.315713 | 5.3873 | -0.071578389 | 0.3411428 | 6.028384 | 5.7571935 | 5.71506 | 0.042137578 | 0.313328076 |
| 1700037H04Rik | 7.588892 | 7.462852 | 7.5004 | -0.037561129 | 0.088479318 | 7.5608619 | 7.3443848 | 7.30228 | 0.042104593 | 0.258581764 |
| Zim1 | 1.398555 | 1.381473 | 1.461 | -0.079568723 | -0.062487009 | 1.4881194 | 1.4988012 | 1.45674 | 0.042061346 | 0.031379549 |
| Ankzf1 | 8.915013 | 8.829902 | 8.9477 | -0.117798478 | -0.032688077 | 9.2440464 | 9.2927096 | 9.25068 | 0.042027831 | -0.006635298 |
| Stim2 | 9.553416 | 9.630778 | 9.556 | 0.074794577 | -0.002567507 | 9.3535738 | 9.469331 | 9.42735 | 0.041983573 | -0.07377364 |
| Fcamr | 3.009623 | 2.992701 | 2.7696 | 0.223099319 | 0.24002119 | 2.7531096 | 2.7794258 | 2.73746 | 0.041968148 | 0.015651966 |
| Agpat5 | 8.092876 | 7.898722 | 8.0241 | -0.125391352 | 0.068761963 | 7.7156016 | 7.8875419 | 7.8456 | 0.041936951 | -0.130003353 |
| Rrad | 8.378742 | 8.325346 | 8.2618 | 0.063499009 | 0.116895263 | 8.2569158 | 8.3437335 | 8.3019 | 0.041832647 | -0.044985112 |
| Mc5r | 0.676832 | 0.683769 | 0.7172 | -0.033383825 | -0.040320862 | 0.6824698 | 0.7391022 | 0.69731 | 0.041796655 | -0.014835774 |
| Rps12-ps4 | 3.284395 | 3.203353 | 3.1337 | 0.069628024 | 0.150670627 | 3.2530047 | 3.2870828 | 3.24529 | 0.041790441 | 0.007712304 |
| Zfp985 | 2.27991 | 2.378015 | 2.4229 | -0.044889157 | -0.142993416 | 2.3055662 | 2.4004933 | 2.35883 | 0.041666309 | -0.053260734 |
| Hist2h2bb | 4.310416 | 4.268287 | 4.5095 | -0.241192692 | -0.199063838 | 4.4953105 | 4.5186125 | 4.47695 | 0.041658618 | 0.018356622 |

| | | | | | | | | | | |
|---|---|---|---|---|---|---|---|---|---|---|
| Slc43a3 | 8.723918 | 8.342423 | 8.4855 | -0.143035921 | 0.238459483 | 8.1147421 | 8.277205 | 8.2356 | 0.041601748 | -0.120861235 |
| Rap2c | 10.74003 | 10.80967 | 10.692 | 0.11780978 | 0.048164742 | 10.627227 | 10.628212 | 10.5866 | 0.04157462 | 0.040589765 |
| Gm14173 | 0.328833 | 0.323678 | 0.3138 | 0.00987717 | 0.015031919 | 0.3001706 | 0.320035 | 0.27848 | 0.041555624 | 0.02169123 |
| Ccdc73 | 3.098192 | 3.006307 | 3.1621 | -0.155752421 | -0.063867434 | 3.1383678 | 3.0920846 | 3.05053 | 0.041549815 | 0.087833047 |
| Slc35e1 | 10.51362 | 10.49064 | 10.433 | 0.057774404 | 0.080751496 | 10.418979 | 10.422278 | 10.3808 | 0.041517312 | 0.038218511 |
| Gm15842 | 1.783209 | 1.6931 | 1.786 | -0.092875137 | -0.002766745 | 1.7503105 | 1.7958233 | 1.75434 | 0.041484946 | -0.004027911 |
| Actr1b | 9.890941 | 9.9392 | 9.9315 | 0.007713183 | -0.040546331 | 10.347998 | 10.258117 | 10.2166 | 0.04148395 | 0.131365661 |
| Ydjc | 5.547149 | 5.448949 | 5.5085 | -0.059569613 | 0.038630771 | 5.7073083 | 5.5234252 | 5.48198 | 0.041445235 | 0.225328337 |
| Kctd1 | 6.767607 | 6.707743 | 6.6859 | 0.021838646 | 0.081702056 | 6.7654028 | 6.6034923 | 6.56208 | 0.041407603 | 0.20331814 |
| Mfap2 | 2.407308 | 2.398983 | 2.2432 | 0.155770001 | 0.164094347 | 2.2977178 | 2.2879949 | 2.24661 | 0.041385393 | 0.051108274 |
| 1700001J03Rik | 1.080147 | 1.075854 | 1.112 | -0.036174682 | -0.031882211 | 1.1174207 | 1.1306699 | 1.08929 | 0.041380341 | 0.028131176 |
| Foxc2 | 2.258084 | 2.216506 | 2.2219 | -0.005407544 | 0.036170636 | 2.2326546 | 2.2278908 | 2.18654 | 0.041355459 | 0.046119217 |
| Xirp1 | 1.161429 | 1.247151 | 1.2392 | 0.007944977 | -0.077776505 | 1.2016618 | 1.2461502 | 1.2048 | 0.041354713 | -0.003133746 |
| Phf8 | 10.26125 | 10.36076 | 10.308 | 0.052368492 | -0.047140904 | 10.35066 | 10.317308 | 10.276 | 0.041316248 | 0.074668917 |
| Ubtfl1 | 2.455951 | 2.49626 | 2.5541 | -0.057792405 | -0.098102155 | 2.525476 | 2.6437716 | 2.60253 | 0.041245221 | -0.077050406 |
| Gpr135 | 1.560862 | 1.709004 | 1.6611 | 0.047897483 | -0.100244741 | 1.658547 | 1.6819184 | 1.64067 | 0.041244929 | 0.017873516 |
| Gm15066 | 1.28314 | 1.259344 | 1.3009 | -0.041546925 | -0.017751267 | 1.3255259 | 1.3418333 | 1.30069 | 0.041139986 | 0.024832602 |
| Vmn1r221 | 1.381003 | 1.39578 | 1.3888 | 0.00701076 | -0.007766325 | 1.3742694 | 1.4409928 | 1.39988 | 0.041115022 | -0.025608371 |
| Gm2986 | 0.94375 | 0.895335 | 0.9282 | -0.032913754 | 0.015501155 | 0.9800589 | 0.9745654 | 0.93348 | 0.041084036 | 0.046577525 |
| Faim | 8.023895 | 7.822504 | 7.911 | -0.088538492 | 0.112852782 | 7.8202998 | 7.9170507 | 7.87604 | 0.041011124 | -0.055739773 |
| Smcr8 | 10.77241 | 10.85475 | 10.807 | 0.048206456 | -0.034136434 | 10.778633 | 10.979983 | 10.939 | 0.040962102 | -0.160388269 |
| Obox8 | 2.313237 | 2.39979 | 2.3341 | 0.065734258 | -0.020818416 | 2.3829994 | 2.3869889 | 2.34606 | 0.040928687 | 0.036939152 |
| AC154779.4 | 0.61069 | 0.599475 | 0.6151 | -0.015596577 | -0.004381537 | 0.6099022 | 0.6070831 | 0.56617 | 0.04091095 | 0.043730023 |
| Trip10 | 8.049712 | 7.780827 | 7.8198 | -0.038941983 | 0.229943208 | 7.5981454 | 7.6874477 | 7.64654 | 0.040905868 | -0.048396515 |
| Gm9521 | 1.561804 | 1.525737 | 1.5286 | -0.002835707 | 0.03323207 | 1.5448115 | 1.5868954 | 1.54604 | 0.040858488 | -0.001225451 |
| Gm4832 | 0.798209 | 0.722919 | 0.7357 | -0.012812651 | 0.062477483 | 0.7246419 | 0.7516663 | 0.71082 | 0.040847456 | 0.01382302 |
| Gm12346 | 3.231286 | 3.190723 | 3.2872 | -0.096518822 | -0.055955971 | 3.0426343 | 3.2980334 | 3.25725 | 0.040780319 | -0.214618778 |
| Hoxa1 | 1.784012 | 1.723598 | 1.7878 | -0.064185456 | -0.003771308 | 1.786049 | 1.7643698 | 1.7237 | 0.04067008 | 0.062349244 |
| Gm11231 | 1.540559 | 1.524642 | 1.485 | 0.039667822 | 0.055585039 | 1.4792962 | 1.5776805 | 1.53702 | 0.040658871 | -0.057725388 |
| Tcap | 0.612897 | 0.6246 | 0.6773 | -0.052681902 | -0.06438486 | 0.6285145 | 0.6648166 | 0.62416 | 0.040654554 | 0.004352415 |
| L3mbtl2 | 9.463429 | 9.45434 | 9.5066 | -0.052279071 | -0.043190012 | 9.5786714 | 9.511253 | 9.4706 | 0.040651732 | 0.108070156 |
| Mlf1 | 2.561034 | 2.540796 | 2.5298 | 0.011043132 | 0.031281653 | 2.5792827 | 2.5558725 | 2.51522 | 0.040648125 | 0.064058348 |

| | | | | | | | | | | |
|---|---|---|---|---|---|---|---|---|---|---|
| Nox1 | 4.074904 | 4.105042 | 4.1779 | -0.072901139 | -0.103039456 | 3.9451417 | 4.3518752 | 4.31126 | 0.040614843 | -0.366118703 |
| Hmgb1-ps2 | 1.46842 | 1.395131 | 1.4148 | -0.019695377 | 0.05359404 | 1.4820161 | 1.4516576 | 1.41109 | 0.040572558 | 0.070931131 |
| Ppm1g | 11.28612 | 11.23335 | 11.293 | -0.059632936 | -0.00686683 | 11.311567 | 11.306587 | 11.2661 | 0.040526178 | 0.045505671 |
| Ppp1ca | 11.49709 | 11.51966 | 11.53 | -0.009865196 | -0.032442858 | 11.735692 | 11.553623 | 11.5131 | 0.04052012 | 0.222588728 |
| Gm42814 | 2.251895 | 2.287152 | 2.3022 | -0.015019616 | -0.05027615 | 2.2873805 | 2.2476669 | 2.20722 | 0.040450549 | 0.080164141 |
| Vmn2r10 | 1.361077 | 1.363598 | 1.3113 | 0.052313083 | 0.049791678 | 1.3486361 | 1.3865127 | 1.3461 | 0.040412991 | 0.002536418 |
| Trav2 | 2.720779 | 2.672863 | 2.6437 | 0.029200803 | 0.077117027 | 2.6002651 | 2.6650352 | 2.62484 | 0.040194487 | -0.024575557 |
| Vmn1r234 | 2.396283 | 2.45321 | 2.4057 | 0.047493773 | -0.009433579 | 2.4505531 | 2.5183133 | 2.47819 | 0.040125635 | -0.027634537 |
| 4930430A15Rik | 0.15053 | 0.141243 | 0.1288 | 0.012412836 | 0.021700187 | 0.106672 | 0.1955312 | 0.15541 | 0.040117036 | -0.048742171 |
| Htr2a | 1.313218 | 1.331563 | 1.3709 | -0.039385328 | -0.057731044 | 1.2874199 | 1.3196732 | 1.27957 | 0.040104091 | 0.007850761 |
| Ndufb4 | 6.455532 | 6.496648 | 6.5334 | -0.036792165 | -0.077908424 | 6.8301202 | 6.7422796 | 6.70219 | 0.040091652 | 0.127932269 |
| Pxmp4 | 7.703022 | 7.619066 | 7.8639 | -0.24482923 | -0.160873094 | 7.8800533 | 7.6802408 | 7.6402 | 0.040045379 | 0.239857926 |
| Hes5 | 2.61295 | 2.653897 | 2.5525 | 0.101444045 | 0.060496512 | 2.7152821 | 2.7156364 | 2.67563 | 0.040005332 | 0.039651055 |
| Gm15304 | 1.312863 | 1.304026 | 1.2683 | 0.035719085 | 0.044556007 | 1.291954 | 1.3306542 | 1.29073 | 0.039927292 | 0.001227097 |
| Gm47842 | 2.993859 | 3.046797 | 3.1464 | -0.09955813 | -0.152496337 | 3.0339205 | 3.1345943 | 3.0947 | 0.039895984 | -0.06077786 |
| Prorsd1 | 7.418207 | 7.428748 | 7.396 | 0.032766476 | 0.022224864 | 7.4987798 | 7.4414622 | 7.40159 | 0.03986921 | 0.097186848 |
| Lypd5 | 0.524294 | 0.515908 | 0.6039 | -0.088004455 | -0.079618571 | 0.5420831 | 0.5702434 | 0.53039 | 0.039856649 | 0.011696443 |
| Gm1141 | 0.959789 | 0.89799 | 0.958 | -0.06004406 | 0.001754301 | 0.9522403 | 0.9444127 | 0.90459 | 0.039822503 | 0.047650059 |
| Slx1b | 7.264923 | 7.020856 | 7.2398 | -0.218991006 | 0.025076257 | 7.4584624 | 7.2985184 | 7.25872 | 0.039794496 | 0.1997385 |
| Dcst1 | 2.778438 | 2.776647 | 2.8469 | -0.070277327 | -0.068485574 | 2.6824652 | 2.757078 | 2.71729 | 0.039790913 | -0.034821839 |
| Mamdc4 | 5.596194 | 5.793897 | 5.67 | 0.123883011 | -0.073819958 | 5.5697886 | 5.7025432 | 5.66276 | 0.039782954 | -0.092971611 |
| Grin3a | 0.905272 | 0.884704 | 0.8799 | 0.004800552 | 0.025368318 | 0.8417232 | 0.8925231 | 0.85276 | 0.039761515 | -0.011038448 |
| Gdf3 | 2.809385 | 2.907853 | 2.8203 | 0.08757623 | -0.01089162 | 3.1778863 | 2.8252023 | 2.78546 | 0.039743614 | 0.392427637 |
| Vgll3 | 0.741959 | 0.754525 | 0.7918 | -0.037321464 | -0.049887523 | 0.8132209 | 0.7952201 | 0.75549 | 0.039727916 | 0.057728654 |
| Bricd5 | 0.468154 | 0.441774 | 0.4292 | 0.012621917 | 0.039002559 | 0.4249344 | 0.4684305 | 0.42877 | 0.039661163 | -0.00383486 |
| Spock1 | 2.16241 | 2.147207 | 2.0636 | 0.083563706 | 0.098766844 | 2.0366223 | 2.1273985 | 2.0878 | 0.039598355 | -0.051177778 |
| 2210016F16Rik | 8.8845 | 8.73546 | 8.797 | -0.061569286 | 0.087470943 | 8.9019627 | 8.8658401 | 8.82624 | 0.039597299 | 0.075719966 |
| Gm38247 | 1.366359 | 1.415931 | 1.3771 | 0.03886234 | -0.010709558 | 1.402583 | 1.4770582 | 1.4375 | 0.039560217 | -0.034915012 |
| Trmt10a | 7.253932 | 7.110193 | 7.2973 | -0.187139096 | -0.043399839 | 7.3930542 | 7.4572491 | 7.4177 | 0.039550201 | -0.024644707 |
| Msantd3 | 2.294303 | 2.408069 | 2.2115 | 0.196547714 | 0.082781685 | 2.3743103 | 2.3078338 | 2.26831 | 0.039522813 | 0.105999378 |
| Itprid1 | 1.69389 | 1.517212 | 1.7067 | -0.189512726 | -0.012834567 | 1.8352724 | 1.6599648 | 1.62048 | 0.039487786 | 0.214795336 |
| Hist1h3i | 2.545641 | 2.545462 | 2.6439 | -0.098390001 | -0.098211348 | 2.6488086 | 2.5705392 | 2.53107 | 0.039470674 | 0.11774003 |

| | | | | | | | | | | |
|---|---|---|---|---|---|---|---|---|---|---|
| Gli3 | 3.183503 | 3.317419 | 3.2923 | 0.025128678 | -0.108787432 | 3.1535373 | 3.167086 | 3.12765 | 0.03943645 | 0.025887783 |
| Gm49309 | 0.514288 | 0.480604 | 0.5129 | -0.032274783 | 0.001408303 | 0.5184885 | 0.5496191 | 0.51019 | 0.03942841 | 0.008297809 |
| Psmb7-ps2 | 1.291237 | 1.325595 | 1.2904 | 0.035181174 | 0.000823233 | 1.3010029 | 1.3363416 | 1.29693 | 0.039408341 | 0.004069609 |
| Cyp2b19 | 0.540482 | 0.536235 | 0.6142 | -0.077983393 | -0.073735879 | 0.6358941 | 0.6116136 | 0.57221 | 0.039398998 | 0.063679469 |
| Gm14416 | 2.233414 | 2.08742 | 2.2093 | -0.121864681 | 0.02412953 | 2.0127713 | 2.2764358 | 2.23707 | 0.039363856 | -0.224300661 |
| Gpc4 | 3.493454 | 3.442023 | 3.454 | -0.012021469 | 0.039410202 | 3.2390266 | 3.351785 | 3.31243 | 0.0393553 | -0.073403101 |
| Mecr | 7.764623 | 7.775592 | 7.7569 | 0.018675154 | 0.00770632 | 8.0308174 | 8.0922348 | 8.0529 | 0.039336733 | -0.022080735 |
| Taf9 | 5.843701 | 5.846843 | 5.8519 | -0.005065348 | -0.008207015 | 5.9065191 | 5.8985963 | 5.8593 | 0.039300604 | 0.047223352 |
| Phactr3 | 1.740295 | 1.721442 | 1.7502 | -0.028717481 | -0.009864165 | 1.6795559 | 1.8261947 | 1.78693 | 0.039261387 | -0.107377349 |
| Bhlhe22 | 0.647205 | 0.564827 | 0.5538 | 0.011025696 | 0.093404369 | 0.5418201 | 0.5849479 | 0.5457 | 0.039252645 | -0.003875146 |
| Gm5403 | 0.541434 | 0.516325 | 0.4686 | 0.047744577 | 0.072853423 | 0.5556743 | 0.5447772 | 0.50553 | 0.039250787 | 0.050147911 |
| Cyp19a1 | 1.445478 | 1.51354 | 1.4796 | 0.033923561 | -0.034138076 | 1.5306006 | 1.4958809 | 1.45665 | 0.039234339 | 0.073954027 |
| Cenpw | 5.734195 | 5.631659 | 5.6441 | -0.012456473 | 0.090080376 | 5.7480981 | 5.8239012 | 5.7847 | 0.039200745 | -0.036602348 |
| Gm18584 | -0.248953 | -0.23126 | -0.215 | -0.016220388 | -0.033909761 | -0.2438818 | -0.200979 | -0.2402 | 0.039199059 | -0.003703466 |
| Gpr107 | 9.942463 | 9.864999 | 9.8274 | 0.037619809 | 0.115084011 | 9.9172517 | 9.857764 | 9.81857 | 0.039194079 | 0.098681793 |
| Cinp | 7.953437 | 8.023151 | 7.9592 | 0.063928894 | -0.005784634 | 8.1900418 | 8.2124288 | 8.17324 | 0.039193777 | 0.016806743 |
| Hist1h3h | 0.616841 | 0.68582 | 0.6606 | 0.025218558 | -0.043760717 | 0.5773793 | 0.6485773 | 0.60939 | 0.039188387 | -0.03200961 |
| Vmn1r201 | 3.879417 | 3.899683 | 3.7762 | 0.123496704 | 0.103230154 | 3.7874273 | 3.9405084 | 3.90138 | 0.039127895 | -0.11395321 |
| Igkv2-116 | 1.631408 | 1.662755 | 1.7016 | -0.038890675 | -0.07023732 | 1.7180607 | 1.6713496 | 1.63223 | 0.03911915 | 0.085830244 |
| Cadm4 | 3.211246 | 3.181997 | 3.1178 | 0.064164486 | 0.093412834 | 3.2898004 | 3.046729 | 3.00763 | 0.039102636 | 0.282174065 |
| Ighv1-4 | 5.311516 | 5.370813 | 5.5619 | -0.191112783 | -0.250410196 | 5.3912719 | 5.32659 | 5.2875 | 0.039087551 | 0.103769432 |
| Pate2 | 1.245764 | 1.245223 | 1.2526 | -0.00740685 | -0.006866521 | 1.3805142 | 1.3125528 | 1.27347 | 0.039080403 | 0.107041778 |
| Vmn1r185 | 2.093638 | 2.110902 | 2.164 | -0.053096154 | -0.070359605 | 2.1558323 | 2.2236111 | 2.18454 | 0.039071946 | -0.028706886 |
| Gm2420 | 1.476031 | 1.499762 | 1.4751 | 0.024652726 | 0.000921829 | 1.4619631 | 1.5043117 | 1.46535 | 0.038965606 | -0.003382989 |
| Gm4613 | 1.394568 | 1.479368 | 1.5064 | -0.026987012 | -0.111787255 | 1.5341348 | 1.5388127 | 1.49985 | 0.038960047 | 0.034282143 |
| Fam189b | 9.69058 | 9.775693 | 9.7536 | 0.022114628 | -0.062998424 | 9.8084875 | 9.727431 | 9.68854 | 0.038890576 | 0.119947098 |
| Zscan25 | 8.221387 | 8.354077 | 8.1803 | 0.173778755 | 0.041088247 | 8.2723283 | 8.2343691 | 8.19549 | 0.038882683 | 0.076841911 |
| Otud1 | 8.953462 | 8.852382 | 9.0663 | -0.213901551 | -0.112822045 | 8.5449022 | 8.5605858 | 8.52173 | 0.038855607 | 0.023171989 |
| Gm15538 | 2.817534 | 2.851435 | 2.891 | -0.039542806 | -0.073444187 | 2.9267036 | 2.9951786 | 2.95633 | 0.038845229 | -0.029629773 |
| Adamts8 | 1.503324 | 1.476499 | 1.3426 | 0.133926748 | 0.160751171 | 1.4597645 | 1.3700509 | 1.33123 | 0.038825687 | 0.128539286 |
| Fst | 3.646589 | 3.597387 | 3.3498 | 0.247602412 | 0.296804683 | 3.273139 | 3.416511 | 3.3777 | 0.038815775 | -0.104556285 |
| Ppil1 | 8.611311 | 8.74254 | 8.7366 | 0.005919823 | -0.12530912 | 8.9339481 | 8.8191177 | 8.78034 | 0.038780093 | 0.153610465 |

| | | | | | | | | | | |
|---|---|---|---|---|---|---|---|---|---|---|
| Gm47495 | 0.705119 | 0.768233 | 0.6774 | 0.090825833 | 0.027712513 | 0.6800267 | 0.7446288 | 0.70586 | 0.038770374 | -0.02583172 |
| Olfr1268-ps1 | 3.234467 | 3.231498 | 3.1748 | 0.056717579 | 0.05968675 | 3.2423368 | 3.3868775 | 3.34815 | 0.038730822 | -0.105809878 |
| Ccdc187 | 2.958135 | 2.938845 | 2.9417 | -0.002845876 | 0.016443832 | 3.1211069 | 3.0163222 | 2.9776 | 0.038726691 | 0.143511347 |
| Camkv | 1.121945 | 1.125373 | 1.0389 | 0.086492945 | 0.083064987 | 1.0380101 | 1.0848361 | 1.04615 | 0.038686884 | -0.008139181 |
| Olfr1419 | 1.528685 | 1.526127 | 1.4871 | 0.039037413 | 0.041595656 | 1.5514358 | 1.6168391 | 1.57821 | 0.038627669 | -0.026775604 |
| Traf6 | 10.04698 | 10.20824 | 10.092 | 0.116432673 | -0.04483261 | 9.8909207 | 10.1368 | 10.0982 | 0.038621888 | -0.207257235 |
| Tmem181c-ps | 2.957951 | 2.955626 | 2.8905 | 0.065125589 | 0.067450529 | 2.9422969 | 2.9965688 | 2.958 | 0.038570473 | -0.015701427 |
| Serpina1c | 0.704464 | 0.729356 | 0.7846 | -0.05525511 | -0.080147337 | 0.7343574 | 0.7441244 | 0.70557 | 0.038556022 | 0.028789002 |
| Nnmt | 1.757244 | 1.810164 | 1.7214 | 0.088754425 | 0.035834237 | 1.7566405 | 1.7513931 | 1.71285 | 0.038541334 | 0.043788782 |
| Gm5697 | 3.206643 | 3.166171 | 3.2396 | -0.073456456 | -0.032985282 | 3.1889511 | 3.2951309 | 3.25665 | 0.038484393 | -0.067695418 |
| Ighv1-81 | 7.226221 | 7.523117 | 7.295 | 0.228127491 | -0.068768037 | 7.6014391 | 7.4593533 | 7.42093 | 0.038424625 | 0.18051047 |
| Faxc | 2.190989 | 2.149899 | 2.1395 | 0.01037892 | 0.051469427 | 2.0626069 | 2.1611817 | 2.1228 | 0.038377526 | -0.060197279 |
| Gm11425 | 4.015642 | 4.095125 | 4.0491 | 0.046061119 | -0.033421915 | 3.952999 | 4.1419565 | 4.1036 | 0.038351732 | -0.15060573 |
| Zbtb34 | 7.592087 | 7.732543 | 7.8635 | -0.130951505 | -0.271408306 | 7.7824906 | 7.8431118 | 7.8048 | 0.038313179 | -0.022308043 |
| Ttc33 | 8.734353 | 8.794793 | 8.6995 | 0.09526345 | 0.034823363 | 8.6020887 | 8.6244853 | 8.58619 | 0.03830006 | 0.015903465 |
| Scaf4 | 10.0875 | 10.05503 | 9.9863 | 0.068765439 | 0.101236117 | 10.336252 | 10.350431 | 10.3122 | 0.03819205 | 0.024013395 |
| Gm5292 | 0.560893 | 0.53681 | 0.5693 | -0.032447519 | -0.008364804 | 0.6175372 | 0.6482067 | 0.61003 | 0.038178368 | 0.007508935 |
| Eci1 | 8.098499 | 8.129825 | 8.2183 | -0.088520043 | -0.119845822 | 8.4129085 | 8.2098314 | 8.17169 | 0.038146134 | 0.241223288 |
| 9330182L06Rik | 1.389115 | 1.412585 | 1.407 | 0.005623083 | -0.017847126 | 1.5800292 | 1.4544354 | 1.4163 | 0.038139236 | 0.16373299 |
| Gm3591 | 0.681638 | 0.661118 | 0.65 | 0.011131194 | 0.031651268 | 0.689576 | 0.6592187 | 0.62111 | 0.03811301 | 0.068470292 |
| Olfr1431 | 0.863745 | 0.832438 | 0.8538 | -0.021363243 | 0.009943764 | 0.8125152 | 0.8923694 | 0.85426 | 0.038109064 | -0.041745114 |
| Gm14176 | 0.718471 | 0.744074 | 0.7503 | -0.006226879 | -0.031830026 | 0.7410907 | 0.684942 | 0.64684 | 0.038104912 | 0.094253544 |
| Arhgef1 | 13.86687 | 13.95995 | 13.952 | 0.007706084 | -0.085370608 | 14.27877 | 14.201375 | 14.1633 | 0.038044042 | 0.115439503 |
| Ribc1 | 3.724705 | 3.682222 | 3.7817 | -0.099458338 | -0.056975035 | 3.8420269 | 3.7860622 | 3.74804 | 0.038023041 | 0.093987734 |
| Mok | 1.753583 | 1.807267 | 1.8152 | -0.007913092 | -0.06159627 | 1.8034249 | 1.7973222 | 1.75931 | 0.03801612 | 0.044118796 |
| Erp44 | 10.14965 | 10.21599 | 10.155 | 0.061410698 | -0.004935098 | 10.317221 | 10.317734 | 10.2797 | 0.037997348 | 0.037484196 |
| Olfr1295 | 0.597885 | 0.596898 | 0.6352 | -0.038297733 | -0.037310024 | 0.5809608 | 0.6306179 | 0.59271 | 0.037909303 | -0.011747797 |
| Olfr1278 | 2.701331 | 2.610569 | 2.7644 | -0.153875581 | -0.063113679 | 2.6524204 | 2.6995817 | 2.66183 | 0.037751879 | -0.009409454 |
| Eef1a2 | 1.646903 | 1.722217 | 1.5652 | 0.157051652 | 0.081737575 | 1.5755331 | 1.6408266 | 1.6031 | 0.037727446 | -0.027566073 |
| Pigs | 10.08661 | 10.06621 | 10.094 | -0.028188244 | -0.007785186 | 10.298411 | 10.0603 | 10.0226 | 0.037724827 | 0.275835223 |
| Gm11478 | 2.789224 | 2.769045 | 2.8012 | -0.03215544 | -0.011976833 | 2.8037681 | 2.7870688 | 2.74935 | 0.037720117 | 0.054419393 |
| Myh10 | 5.594344 | 5.7041 | 5.6364 | 0.06768864 | -0.042068208 | 5.5378319 | 5.6857654 | 5.64809 | 0.037679092 | -0.110254321 |

| | | | | | | | | | | |
|---|---|---|---|---|---|---|---|---|---|---|
| Trbj1-1 | 5.640381 | 5.800725 | 5.5689 | 0.231817799 | 0.071473476 | 5.2323784 | 5.7069019 | 5.66929 | 0.037613535 | -0.436909933 |
| Wdr46-ps | 1.471683 | 1.45074 | 1.4017 | 0.048997504 | 0.069940351 | 1.4179969 | 1.4836588 | 1.44609 | 0.037570724 | -0.028091129 |
| Map10 | 5.434337 | 5.249208 | 5.371 | -0.121758328 | 0.063371579 | 5.4316338 | 5.3239368 | 5.28637 | 0.037570011 | 0.145266965 |
| Rpl6l | 2.038301 | 1.99042 | 1.9204 | 0.070031256 | 0.117912475 | 2.0537068 | 2.0221789 | 1.98462 | 0.037561083 | 0.069088978 |
| N6amt1 | 7.919488 | 7.810307 | 7.8401 | -0.029799193 | 0.079382332 | 7.9870539 | 7.9164833 | 7.87898 | 0.037505631 | 0.108076215 |
| Pigf | 6.415544 | 6.547364 | 6.5111 | 0.036221734 | -0.095598036 | 6.524216 | 6.5411159 | 6.50361 | 0.037504147 | 0.020604249 |
| Copg1 | 11.24235 | 11.26226 | 11.268 | -0.00562294 | -0.025530902 | 11.448801 | 11.4502 | 11.4128 | 0.03742699 | 0.036028122 |
| Gm7862 | 1.845103 | 1.812477 | 1.8014 | 0.011112554 | 0.043739258 | 1.8865965 | 1.8325781 | 1.79517 | 0.037412533 | 0.091430882 |
| Gm15772 | 2.987604 | 3.019483 | 3.0873 | -0.067821477 | -0.099700289 | 3.1527053 | 3.1998914 | 3.16255 | 0.037344101 | -0.009842069 |
| Aldh3b2 | 1.06157 | 1.035442 | 0.9959 | 0.039562165 | 0.065690314 | 1.1552018 | 1.0862859 | 1.04899 | 0.037293822 | 0.10620966 |
| Gtf2e1 | 8.808898 | 8.8529 | 8.8044 | 0.048461402 | 0.004459625 | 8.7362601 | 8.7796333 | 8.74234 | 0.03729168 | -0.006081533 |
| Olfr912 | 2.660774 | 2.679795 | 2.6513 | 0.028464443 | 0.009443064 | 2.7641538 | 2.7250656 | 2.68786 | 0.037202435 | 0.076290602 |
| Serf1 | 3.32904 | 3.41612 | 3.5305 | -0.114420528 | -0.201500422 | 3.4353622 | 3.4821848 | 3.44504 | 0.037146015 | -0.009676575 |
| Gm43218 | 2.764985 | 2.824012 | 2.7181 | 0.105950351 | 0.046922913 | 2.8328857 | 2.7829804 | 2.74585 | 0.037130349 | 0.08703572 |
| Cbr4 | 6.886388 | 6.897483 | 7.083 | -0.185505192 | -0.196599418 | 6.9214726 | 6.8962724 | 6.85917 | 0.037100836 | 0.062300961 |
| Tom1 | 8.226333 | 8.222429 | 8.2076 | 0.014799001 | 0.018703088 | 8.368577 | 8.2075636 | 8.1705 | 0.03706341 | 0.198076801 |
| Aass | 1.429029 | 1.394796 | 1.5496 | -0.154815408 | -0.120582644 | 1.6353665 | 1.4238325 | 1.38677 | 0.037062375 | 0.248596442 |
| Gm11407 | 1.422525 | 1.480343 | 1.5059 | -0.02559163 | -0.083409659 | 1.3933921 | 1.4202846 | 1.38323 | 0.037053656 | 0.01016122 |
| Baiap2l2 | 1.201428 | 1.200235 | 1.2804 | -0.080115803 | -0.078923009 | 1.1552268 | 1.2501195 | 1.21307 | 0.037053611 | -0.057839066 |
| Ccr3 | 2.98248 | 3.036904 | 3.0141 | 0.022845692 | -0.031578096 | 2.9886576 | 3.0684785 | 3.03149 | 0.036993099 | -0.042827765 |
| AC147227.1 | 2.661594 | 2.794218 | 2.7399 | 0.054293013 | -0.078331525 | 2.6953504 | 2.7336775 | 2.69673 | 0.036950631 | -0.001376457 |
| Klhl26 | 7.483719 | 7.416291 | 7.3983 | 0.018020098 | 0.08544838 | 7.439036 | 7.5616799 | 7.52474 | 0.036943104 | -0.085700735 |
| Pcdhb18 | 2.20588 | 2.108046 | 2.0452 | 0.062821132 | 0.160655092 | 2.0746176 | 2.0531723 | 2.01627 | 0.036907079 | 0.058352379 |
| Slc6a5 | 1.456239 | 1.498286 | 1.4903 | 0.008010386 | -0.03403692 | 1.5255959 | 1.6483315 | 1.61143 | 0.036899618 | -0.085835944 |
| Aipl1 | 2.231425 | 2.177323 | 2.2596 | -0.082271391 | -0.028169329 | 2.2799826 | 2.261177 | 2.22434 | 0.036838431 | 0.05564403 |
| Atr | 10.26489 | 10.24785 | 10.314 | -0.066374739 | -0.049332479 | 10.261993 | 10.461905 | 10.4251 | 0.036829974 | -0.163082539 |
| Gm7206 | 1.940258 | 2.058552 | 2.0791 | -0.020574629 | -0.138869256 | 2.1819762 | 2.0237951 | 1.98699 | 0.036801942 | 0.194983055 |
| Pik3c2a | 10.1002 | 10.12812 | 10.092 | 0.035816809 | 0.007893537 | 9.8476885 | 10.095094 | 10.0583 | 0.036768897 | -0.210636898 |
| Dis3 | 9.193672 | 9.214735 | 9.2139 | 0.000841742 | -0.020221647 | 9.2202526 | 9.4184834 | 9.38177 | 0.036718035 | -0.161512712 |
| Commd6 | 8.695556 | 8.688823 | 8.7102 | -0.021402984 | -0.014670459 | 8.9709162 | 8.7643267 | 8.7277 | 0.036625941 | 0.243215499 |
| Togaram1 | 9.782527 | 9.811124 | 9.8257 | -0.014567659 | -0.043164472 | 9.8033492 | 9.8859187 | 9.8493 | 0.036620303 | -0.045949194 |
| Pank3 | 10.44323 | 10.51252 | 10.409 | 0.103266566 | 0.03397804 | 10.376093 | 10.384147 | 10.3476 | 0.036579534 | 0.028524893 |

| | | | | | | | | | | |
|---|---|---|---|---|---|---|---|---|---|---|
| Chsy1 | 10.32637 | 10.40531 | 10.306 | 0.099225742 | 0.020287532 | 10.276611 | 10.230633 | 10.1941 | 0.03657077 | 0.082548245 |
| Gm8848 | 1.779531 | 1.810338 | 1.7035 | 0.10682368 | 0.07601671 | 1.8968702 | 1.7598303 | 1.72327 | 0.036562435 | 0.173602319 |
| Rnf5 | 8.544938 | 8.604007 | 8.5895 | 0.014504764 | -0.044564587 | 8.7699731 | 8.6024728 | 8.56593 | 0.036544284 | 0.204044574 |
| Ppp2cb | 9.452221 | 9.348654 | 9.4491 | -0.10041491 | 0.003152541 | 9.3633359 | 9.3633794 | 9.32685 | 0.036525904 | 0.036482412 |
| Gm4342 | 0.901216 | 0.93925 | 0.9152 | 0.024028001 | -0.01400543 | 0.9788438 | 0.9341239 | 0.89761 | 0.036516641 | 0.08123655 |
| Cpsf1 | 10.91038 | 11.0127 | 10.928 | 0.084483288 | -0.017836354 | 11.419069 | 11.431819 | 11.3954 | 0.03646853 | 0.023719027 |
| Gm42420 | 0.484885 | 0.586152 | 0.5847 | 0.001431581 | -0.099836355 | 0.4875054 | 0.5730507 | 0.53662 | 0.036426226 | -0.049119098 |
| Ablim1 | 13.66377 | 13.65847 | 13.629 | 0.028973612 | 0.034282388 | 13.656966 | 13.721106 | 13.6847 | 0.036419194 | -0.02772081 |
| Gm36128 | 3.161786 | 3.183974 | 3.1925 | -0.008533054 | -0.030720682 | 3.1095874 | 3.2686999 | 3.23229 | 0.036413383 | -0.122699109 |
| Brms1l | 7.811633 | 7.666482 | 7.7028 | -0.03633102 | 0.108819607 | 7.8765301 | 7.8911238 | 7.85491 | 0.036218777 | 0.021625046 |
| Dnajc25 | 6.972783 | 6.893862 | 6.9201 | -0.026223107 | 0.052698018 | 6.7925712 | 6.7272823 | 6.69111 | 0.036169822 | 0.101458767 |
| Olfr1414 | 0.799506 | 0.773109 | 0.7954 | -0.022324897 | 0.004071458 | 0.7863875 | 0.8552926 | 0.81915 | 0.036141107 | -0.032763988 |
| Gm6055 | 4.067124 | 4.292754 | 4.2899 | 0.002838136 | -0.222791691 | 3.9413552 | 4.2079978 | 4.17187 | 0.036123662 | -0.23051899 |
| Bex2 | 1.623334 | 1.5597 | 1.6038 | -0.044104509 | 0.019529239 | 1.5586573 | 1.5995752 | 1.56347 | 0.036102486 | -0.004815403 |
| Ighv1-56 | 2.035823 | 2.134095 | 2.0981 | 0.03596121 | -0.062310444 | 2.0848979 | 2.1358854 | 2.09979 | 0.036100029 | -0.014887459 |
| Dnajc21 | 9.106618 | 9.111176 | 9.1028 | 0.008374229 | 0.003815568 | 8.8910211 | 9.1350795 | 9.09899 | 0.036088968 | -0.207969511 |
| Ighv3-3 | 3.447823 | 3.526544 | 3.4551 | 0.071415919 | -0.007304829 | 3.5827437 | 3.5093587 | 3.47333 | 0.036033372 | 0.109418346 |
| Pde11a | 5.543717 | 5.358492 | 5.4968 | -0.138265611 | 0.046959362 | 5.4109018 | 5.5343317 | 5.49833 | 0.036004817 | -0.087425041 |
| Olfr1342 | 0.633958 | 0.635326 | 0.7204 | -0.085028076 | -0.086396414 | 0.6273343 | 0.6481389 | 0.61223 | 0.035911949 | 0.015107287 |
| Bcas2 | 9.806451 | 9.800163 | 9.8045 | -0.004344223 | 0.001944536 | 9.8661171 | 9.8052482 | 9.76934 | 0.035904561 | 0.096773509 |
| Gnat1 | 2.21584 | 2.218956 | 2.236 | -0.017088264 | -0.02020397 | 2.1791878 | 2.2993983 | 2.26355 | 0.035846995 | -0.08436356 |
| Olfr239 | 1.732516 | 1.682363 | 1.7334 | -0.051049385 | -0.000896751 | 1.7402618 | 1.7379941 | 1.70216 | 0.035835512 | 0.038103267 |
| Speer6-ps1 | 3.147281 | 3.146304 | 3.1692 | -0.02293025 | -0.021952954 | 3.1262923 | 3.278936 | 3.24312 | 0.03581736 | -0.116826386 |
| Sema4c | 7.680676 | 7.22088 | 7.2388 | -0.017937102 | 0.441858437 | 6.7225242 | 7.0508603 | 7.01505 | 0.035807615 | -0.292528472 |
| Gm8543 | 1.052233 | 1.03782 | 1.0577 | -0.019862093 | -0.005448923 | 1.0580123 | 1.059899 | 1.02411 | 0.035791489 | 0.033904751 |
| Hoxd3 | 0.314851 | 0.365513 | 0.3553 | 0.010209957 | -0.040451779 | 0.2906192 | 0.3645899 | 0.32881 | 0.035776443 | -0.038194211 |
| Rbm33 | 11.4718 | 11.55529 | 11.495 | 0.06046306 | -0.023019084 | 11.344231 | 11.686275 | 11.6505 | 0.035750912 | -0.30629338 |
| Gm18959 | 0.534012 | 0.508075 | 0.515 | -0.006949696 | 0.018987988 | 0.5283675 | 0.5759806 | 0.54031 | 0.035672827 | -0.011940354 |
| Wdr33 | 10.69977 | 10.69951 | 10.623 | 0.076179582 | 0.076447072 | 10.664596 | 10.757256 | 10.7217 | 0.035603782 | -0.057057095 |
| Gm6988 | 3.255036 | 3.180288 | 3.3015 | -0.121178735 | -0.046430142 | 3.4164938 | 3.3098403 | 3.27426 | 0.03557845 | 0.142231979 |
| Lin37 | 8.037697 | 8.056688 | 8.21 | -0.15331364 | -0.172304648 | 8.3701967 | 8.269244 | 8.23368 | 0.035567649 | 0.136520397 |
| Gm5415 | 0.620503 | 0.645767 | 0.6512 | -0.005449314 | -0.030713408 | 0.6226403 | 0.6828793 | 0.64737 | 0.035512074 | -0.02472698 |

| | | | | | | | | | | |
|---|---|---|---|---|---|---|---|---|---|---|
| Olfr1048 | 1.424478 | 1.376737 | 1.351 | 0.025771485 | 0.073511854 | 1.4040049 | 1.3799652 | 1.34458 | 0.035389772 | 0.059429521 |
| Ropn1l | 1.922963 | 1.866917 | 1.8964 | -0.029499797 | 0.02654667 | 1.8908236 | 1.8752989 | 1.83994 | 0.03535551 | 0.050880227 |
| Cebpg | 9.70129 | 9.744426 | 9.6025 | 0.141932111 | 0.098795642 | 9.9090231 | 9.8831334 | 9.84779 | 0.035342501 | 0.061232212 |
| Zfp457 | 3.303372 | 3.261184 | 3.3011 | -0.039877288 | 0.002311275 | 3.3561746 | 3.3833177 | 3.348 | 0.035315311 | 0.00817218 |
| Olfr273 | 1.332916 | 1.298464 | 1.3461 | -0.047593734 | -0.013141852 | 1.2621707 | 1.3484523 | 1.31316 | 0.0352948 | -0.050986826 |
| 9330159F19Rik | 2.815107 | 2.752535 | 2.7225 | 0.030034832 | 0.092606647 | 2.7540797 | 2.7853092 | 2.75004 | 0.035269299 | 0.004039749 |
| Thnsl2 | 3.973897 | 3.774091 | 3.9483 | -0.174162793 | 0.025642452 | 3.7266349 | 3.7367705 | 3.70151 | 0.035260075 | 0.025124517 |
| Abcb8 | 8.095224 | 7.987198 | 8.1165 | -0.129333325 | -0.021307291 | 8.4562456 | 8.1949251 | 8.15969 | 0.035234693 | 0.296555209 |
| Olfr129 | 0.787815 | 0.807099 | 0.7886 | 0.018519179 | -0.000764127 | 0.7790881 | 0.8718727 | 0.8367 | 0.035174003 | -0.05761056 |
| Pacsin2 | 10.0321 | 10.08981 | 9.9697 | 0.120100182 | 0.062390805 | 10.052911 | 10.088366 | 10.0532 | 0.035153042 | -0.000302666 |
| Ddx10 | 10.08289 | 10.12102 | 10.052 | 0.069140899 | 0.031013765 | 9.9481879 | 10.079994 | 10.0449 | 0.035140458 | -0.09666564 |
| Gm17509 | 3.070597 | 3.148197 | 3.0973 | 0.050898678 | -0.02670131 | 3.1862331 | 3.1787102 | 3.14357 | 0.035136206 | 0.042659078 |
| Gm45713 | 5.011976 | 4.837585 | 5.0714 | -0.233794113 | -0.059403176 | 5.4424827 | 4.8788059 | 4.8437 | 0.035101194 | 0.598777934 |
| Tmem120a | 7.294225 | 7.322136 | 7.3514 | -0.029271795 | -0.057183662 | 7.3830704 | 7.3283191 | 7.29323 | 0.035089193 | 0.089840412 |
| Zfp324 | 6.88382 | 6.888083 | 6.9578 | -0.069697908 | -0.073961614 | 7.0466309 | 6.9936112 | 6.95853 | 0.035079194 | 0.088098883 |
| Synpo2l | 2.481842 | 2.47111 | 2.4613 | 0.009785291 | 0.020516933 | 2.4511218 | 2.5499838 | 2.51492 | 0.035062913 | -0.063799042 |
| Mmab | 7.216641 | 7.084253 | 7.2121 | -0.12789025 | 0.004498035 | 7.4482104 | 7.3034087 | 7.26841 | 0.03499872 | 0.179800378 |
| Sec23ip | 10.25675 | 10.3387 | 10.284 | 0.054596224 | -0.02736188 | 10.16442 | 10.252283 | 10.2175 | 0.0348301 | -0.053033098 |
| Tada2a | 6.660255 | 6.671795 | 6.8449 | -0.173110955 | -0.184651427 | 6.6756465 | 6.7594825 | 6.72469 | 0.034792626 | -0.049043369 |
| Nudcd2 | 8.25901 | 8.086554 | 8.2483 | -0.161754574 | 0.010701283 | 8.0823799 | 8.2313341 | 8.1966 | 0.034732247 | -0.11422192 |
| Rnf215 | 8.288164 | 8.223799 | 8.3162 | -0.09236961 | -0.028004912 | 8.5777971 | 8.4874953 | 8.45277 | 0.034722114 | 0.12502388 |
| Gm14429 | 0.126494 | 0.075661 | 0.0961 | -0.020464453 | 0.030369428 | 0.0703375 | 0.0957296 | 0.06103 | 0.034704076 | 0.009311999 |
| Olfr73 | 0.88266 | 0.862468 | 0.8684 | -0.005911351 | 0.014280024 | 0.8967622 | 0.9190612 | 0.8844 | 0.034661042 | 0.012362006 |
| Vasn | 5.053539 | 5.126235 | 4.9286 | 0.197665857 | 0.124970534 | 4.8630593 | 4.8507561 | 4.81615 | 0.034606111 | 0.046909361 |
| Tm7sf2 | 6.238991 | 6.175729 | 6.3485 | -0.172727293 | -0.109464984 | 6.564912 | 6.3267476 | 6.29217 | 0.034580171 | 0.272744489 |
| Sephs1 | 9.075754 | 9.127589 | 9.0353 | 0.092286899 | 0.040452086 | 9.0308681 | 9.2247764 | 9.1902 | 0.034575078 | -0.159333148 |
| Vmn2r28 | 1.160654 | 1.127017 | 1.1571 | -0.030090779 | 0.003545919 | 1.1260197 | 1.2136036 | 1.17904 | 0.034568322 | -0.053015563 |
| Olfr510 | 4.661773 | 4.832079 | 4.6174 | 0.214702804 | 0.04439699 | 4.6127355 | 4.6393402 | 4.60484 | 0.034504672 | 0.007899998 |
| Ndufaf6 | 6.052681 | 5.911924 | 5.9056 | 0.006360044 | 0.147117779 | 6.2922623 | 6.1145592 | 6.08014 | 0.034417682 | 0.212120847 |
| Sh2d1b1 | 4.537366 | 4.739415 | 4.5572 | 0.182257471 | -0.019791232 | 4.8387376 | 4.7307142 | 4.69631 | 0.03440005 | 0.142423397 |
| Nemp1 | 8.869166 | 8.796878 | 8.878 | -0.081168035 | -0.008880082 | 9.0279117 | 9.0580373 | 9.02372 | 0.034315444 | 0.004189873 |
| Gm4869 | 1.144382 | 1.108715 | 1.1024 | 0.006347036 | 0.042013939 | 1.1000665 | 1.185078 | 1.15081 | 0.034266069 | -0.050745508 |

| | | | | | | | | | | |
|---|---|---|---|---|---|---|---|---|---|---|
| Limd2 | 13.05665 | 13.08282 | 13.137 | -0.054093068 | -0.08026638 | 13.188237 | 13.106065 | 13.0718 | 0.034248872 | 0.116420199 |
| Rpl13-ps3 | 1.097321 | 1.080751 | 1.1222 | -0.041481291 | -0.024911659 | 1.1159042 | 1.1158712 | 1.08163 | 0.034239098 | 0.034272128 |
| Ppp2r2d | 9.015402 | 9.007116 | 9.0253 | -0.018189368 | -0.009902589 | 8.8613633 | 9.0645472 | 9.03038 | 0.034167459 | -0.169016414 |
| Gga1 | 10.20015 | 10.18443 | 10.218 | -0.033556892 | -0.017840798 | 10.409247 | 10.371003 | 10.3368 | 0.034161982 | 0.072405362 |
| Rufy2 | 8.700145 | 8.588543 | 8.7502 | -0.161647354 | -0.050045153 | 8.8930891 | 9.0830431 | 9.04889 | 0.034156777 | -0.155797285 |
| Trmt10b | 7.30629 | 7.29727 | 7.1739 | 0.123384851 | 0.132404869 | 7.2927099 | 7.226664 | 7.1926 | 0.034068693 | 0.100114574 |
| Slc36a3 | 3.277198 | 3.248152 | 3.2906 | -0.042403343 | -0.013356893 | 3.34581 | 3.295223 | 3.26134 | 0.033884982 | 0.084471974 |
| Csmd1 | 5.528037 | 5.188545 | 5.1702 | 0.018361146 | 0.357853773 | 4.9562175 | 5.0404205 | 5.00654 | 0.033881113 | -0.050321878 |
| Actr10 | 9.969346 | 10.01323 | 9.9725 | 0.040742636 | -0.003139903 | 9.9123546 | 10.002725 | 9.96889 | 0.033837982 | -0.0565324 |
| Klhl41 | 1.230581 | 1.220076 | 1.3048 | -0.084700994 | -0.074195913 | 1.2218968 | 1.2618624 | 1.22804 | 0.033825553 | -0.006140026 |
| Naa16 | 8.885396 | 8.995067 | 8.9442 | 0.050861353 | -0.058809798 | 8.7819351 | 9.0817242 | 9.04799 | 0.033734699 | -0.266054355 |
| Noa1 | 8.497891 | 8.615716 | 8.5494 | 0.066271523 | -0.051553827 | 8.6130373 | 8.518867 | 8.48514 | 0.033722412 | 0.127892722 |
| Scp2-ps2 | 3.686425 | 3.730226 | 3.6304 | 0.099814211 | 0.056013316 | 3.589253 | 3.7334668 | 3.69979 | 0.033675257 | -0.110538564 |
| A830005F24Rik | 2.478332 | 2.50399 | 2.5381 | -0.034102574 | -0.059760389 | 2.4252233 | 2.5564482 | 2.52279 | 0.033653668 | -0.097571261 |
| Abca13 | 4.031247 | 4.185823 | 3.7152 | 0.470576825 | 0.316000744 | 3.7732305 | 3.9203965 | 3.88681 | 0.033590602 | -0.113575383 |
| Gm16060 | 2.668543 | 2.756423 | 2.8217 | -0.065256382 | -0.153135991 | 2.7459846 | 2.7290822 | 2.69549 | 0.033587857 | 0.050490324 |
| Olfr291 | 1.067533 | 1.026108 | 1.0529 | -0.026837856 | 0.014586938 | 1.0345235 | 1.0805271 | 1.04703 | 0.033494758 | -0.012508824 |
| Asl | 8.284798 | 8.349123 | 8.5073 | -0.158160377 | -0.222485725 | 8.4138748 | 8.4429303 | 8.40948 | 0.033450475 | 0.004394997 |
| Lhx4 | 0.239959 | 0.269236 | 0.2406 | 0.028639502 | -0.000636703 | 0.2429968 | 0.2777404 | 0.24432 | 0.033421551 | -0.001322045 |
| Piezo1 | 10.86856 | 10.84095 | 10.813 | 0.027948782 | 0.055563254 | 10.784696 | 10.911932 | 10.8785 | 0.033414096 | -0.093821747 |
| Adrb2 | 9.611518 | 9.459633 | 9.7556 | -0.295981616 | -0.144097411 | 9.2891687 | 9.2966042 | 9.26321 | 0.033389342 | 0.025953774 |
| Tmem39b | 7.584945 | 7.643627 | 7.6685 | -0.02486596 | -0.083548662 | 7.935142 | 7.8391772 | 7.80583 | 0.033347882 | 0.12931272 |
| Dnase1l2 | 6.027125 | 6.197954 | 6.1689 | 0.029023976 | -0.141804439 | 6.6744228 | 6.560186 | 6.52689 | 0.033296089 | 0.147532913 |
| Asb5 | 0.939946 | 0.967825 | 0.9479 | 0.019960846 | -0.007918426 | 0.9603662 | 1.0269138 | 0.99366 | 0.033256631 | -0.033290969 |
| Grin2b | 1.958746 | 1.959591 | 1.8656 | 0.094032188 | 0.093187074 | 1.8463953 | 1.9507579 | 1.91756 | 0.033201232 | -0.071161359 |
| Chchd7 | 7.232519 | 7.046482 | 7.1391 | -0.092570365 | 0.09346654 | 7.3837659 | 7.3130485 | 7.27994 | 0.033110417 | 0.103827843 |
| Ddx55 | 8.96216 | 9.021054 | 9.0609 | -0.039865906 | -0.098759912 | 9.279651 | 9.2512895 | 9.21821 | 0.033079237 | 0.061440768 |
| Gm5475 | 0.644306 | 0.661997 | 0.6433 | 0.018736371 | 0.001045134 | 0.655202 | 0.699237 | 0.66619 | 0.033047101 | -0.01098788 |
| Ddx52 | 10.0589 | 9.984344 | 10.102 | -0.117523918 | -0.042970552 | 10.19321 | 10.271271 | 10.2383 | 0.032924416 | -0.045136346 |
| AC157214.1 | 1.180816 | 1.128871 | 1.199 | -0.070114898 | -0.018169491 | 1.1416707 | 1.1767331 | 1.14388 | 0.032848847 | -0.00221354 |
| Gm49392 | 4.376144 | 4.363085 | 4.3784 | -0.015353383 | -0.002294558 | 3.8765602 | 4.2389477 | 4.20611 | 0.032837483 | -0.329550051 |
| Brf1 | 9.982066 | 10.05337 | 10.045 | 0.00876057 | -0.062541538 | 10.240205 | 10.338729 | 10.3059 | 0.032812554 | -0.06571208 |

| | | | | | | | | | | |
|---|---|---|---|---|---|---|---|---|---|---|
| Klrg2 | 2.38585 | 2.320231 | 2.4416 | -0.12132257 | -0.055703497 | 2.3273167 | 2.3996209 | 2.36683 | 0.032793704 | -0.039510529 |
| Tgif2 | 8.418872 | 8.251242 | 8.2547 | -0.003478099 | 0.164152664 | 8.2519217 | 8.5446368 | 8.51196 | 0.032677995 | -0.260037084 |
| Paip2 | 11.16424 | 11.2512 | 11.226 | 0.024958994 | -0.062002472 | 11.263904 | 11.272303 | 11.2396 | 0.032677748 | 0.024278444 |
| Fzd2 | 1.234847 | 1.296185 | 1.2282 | 0.068029903 | 0.006691934 | 1.2980158 | 1.21185 | 1.1792 | 0.032645788 | 0.118811624 |
| Gm20767 | -0.129595 | -0.10646 | -0.0843 | -0.022150282 | -0.045285385 | -0.135193 | -0.052232 | -0.0849 | 0.032644035 | -0.05031656 |
| Faim2 | 3.206319 | 3.080272 | 3.0263 | 0.053930756 | 0.179978122 | 3.1644422 | 3.144346 | 3.11174 | 0.032603531 | 0.052699746 |
| Jade2 | 10.92872 | 11.01147 | 10.96 | 0.051015092 | -0.031734524 | 10.71408 | 10.854396 | 10.8218 | 0.032599036 | -0.107717319 |
| Mcmdc2 | 3.765957 | 3.996088 | 3.9422 | 0.053886504 | -0.176244577 | 3.8171188 | 4.1102345 | 4.07767 | 0.032560148 | -0.260555565 |
| Il13ra1 | 9.159419 | 8.878732 | 8.7849 | 0.09385324 | 0.374540047 | 8.5694962 | 8.3912212 | 8.35866 | 0.032557353 | 0.21083231 |
| Camk2b | 6.807503 | 6.748505 | 6.9431 | -0.194566853 | -0.135569145 | 6.6485242 | 6.8738452 | 6.84134 | 0.032505748 | -0.192815275 |
| Slc7a6 | 9.556288 | 9.725676 | 9.568 | 0.157725337 | -0.011661902 | 9.4872309 | 9.6719221 | 9.63947 | 0.032451661 | -0.15223947 |
| Dagla | 2.986044 | 2.853264 | 2.7239 | 0.129343537 | 0.262122785 | 2.8329749 | 2.7831273 | 2.75069 | 0.032434309 | 0.082281828 |
| Gm3222 | 0.721271 | 0.727812 | 0.6968 | 0.030996584 | 0.024455525 | 0.7192793 | 0.7270541 | 0.69463 | 0.032426857 | 0.0246521 |
| Gm10268 | -0.041966 | -0.04649 | -0.0485 | 0.001976429 | 0.006504373 | -0.0361945 | -0.049298 | -0.0817 | 0.032410923 | 0.045514234 |
| Igkv14-130 | 4.492657 | 4.549637 | 4.4508 | 0.098871044 | 0.041890763 | 4.4903736 | 4.4928732 | 4.46046 | 0.032409146 | 0.029909552 |
| Cpne9 | 2.097245 | 2.066222 | 2.0983 | -0.032061034 | -0.00103809 | 2.1816903 | 2.2214008 | 2.18901 | 0.032393857 | -0.007316671 |
| Cwh43 | 0.980444 | 0.911973 | 0.9716 | -0.059589292 | 0.008881294 | 1.0700943 | 1.0393919 | 1.00701 | 0.032382265 | 0.063084598 |
| Gm14681 | 1.51355 | 1.371325 | 1.3466 | 0.024716277 | 0.166942049 | 1.3396263 | 1.4033981 | 1.37106 | 0.032335839 | -0.031435938 |
| Fam110b | 3.917932 | 3.830523 | 3.9238 | -0.093292132 | -0.005882618 | 3.885209 | 3.8677406 | 3.83543 | 0.032313588 | 0.049782032 |
| Pgp | 8.326375 | 8.243914 | 8.3037 | -0.059802543 | 0.022657909 | 8.3706499 | 8.1500048 | 8.1177 | 0.0323063 | 0.252951356 |
| Gm10704 | 1.678241 | 1.628857 | 1.7301 | -0.101200016 | -0.051816223 | 1.6310813 | 1.6799741 | 1.64767 | 0.032306261 | -0.016586446 |
| D6Wsu163e | 8.36213 | 8.348925 | 8.3154 | 0.033545146 | 0.046749999 | 8.575751 | 8.4584192 | 8.42616 | 0.032257379 | 0.149589202 |
| Pnkd | 8.113851 | 8.10467 | 8.1045 | 0.000194488 | 0.009375216 | 8.150996 | 7.9900738 | 7.95783 | 0.032241924 | 0.193164144 |
| Adgb | 2.426252 | 2.462863 | 2.4313 | 0.031532526 | -0.005078514 | 2.3391585 | 2.4882311 | 2.45604 | 0.032193429 | -0.116879236 |
| Ank1 | 4.003928 | 3.924484 | 3.8832 | 0.04126849 | 0.120711962 | 4.0296694 | 4.0613861 | 4.02925 | 0.032135732 | 0.000418998 |
| Gnmt | 2.997298 | 3.023825 | 2.9067 | 0.117159877 | 0.090632883 | 3.0462857 | 3.051689 | 3.01957 | 0.032121419 | 0.026718132 |
| Gm15666 | 1.130033 | 1.087138 | 1.2044 | -0.117287876 | -0.074392445 | 1.1236817 | 1.177594 | 1.14548 | 0.03210993 | -0.021802426 |
| Klra10 | 2.25105 | 2.230498 | 2.2107 | 0.019825981 | 0.040377178 | 2.2400937 | 2.3285243 | 2.29643 | 0.032091288 | -0.056339273 |
| Gm40020 | 3.074541 | 2.965265 | 3.004 | -0.038757404 | 0.070518253 | 3.0053233 | 3.0882375 | 3.05615 | 0.032084615 | -0.050829526 |
| Fam120a | 11.76451 | 11.77841 | 11.794 | -0.015379686 | -0.029276302 | 11.721173 | 11.773573 | 11.7415 | 0.032074996 | -0.020325184 |
| Frmd4b | 8.439303 | 7.837294 | 7.9925 | -0.155194243 | 0.446814351 | 7.6736853 | 7.8912041 | 7.85915 | 0.032053159 | -0.18546568 |
| Snip1 | 8.907807 | 8.916732 | 8.983 | -0.066309349 | -0.075234563 | 8.9489985 | 8.9640478 | 8.93204 | 0.032003629 | 0.016954281 |

| | | | | | | | | | | |
|---|---|---|---|---|---|---|---|---|---|---|
| Atp6v1g3 | 1.023689 | 0.936705 | 0.9676 | -0.030903646 | 0.056080763 | 0.9677235 | 1.0343312 | 1.00237 | 0.031965723 | -0.034641979 |
| Nr6a1 | 5.247258 | 5.497908 | 5.395 | 0.102883328 | -0.147766835 | 5.4438137 | 5.4893384 | 5.45741 | 0.031927187 | -0.013597442 |
| Gm8108 | 2.357267 | 2.411002 | 2.3944 | 0.016620197 | -0.037114735 | 2.2091037 | 2.3746886 | 2.34283 | 0.031858045 | -0.133726824 |
| Ankrd16 | 8.927645 | 8.97172 | 9.009 | -0.037233028 | -0.08130877 | 9.633431 | 9.4532167 | 9.42136 | 0.031857331 | 0.212071606 |
| Ighv1-83 | 1.518292 | 1.50025 | 1.4397 | 0.060501502 | 0.078542923 | 1.4769188 | 1.4850344 | 1.45332 | 0.031711338 | 0.023595775 |
| Igfals | 1.304577 | 1.335697 | 1.3256 | 0.010121083 | -0.020999487 | 1.4208123 | 1.3024698 | 1.27087 | 0.031604607 | 0.149947066 |
| Xpnpep3 | 8.528971 | 8.557959 | 8.6313 | -0.073291969 | -0.102279566 | 8.6374146 | 8.7420655 | 8.71049 | 0.031572411 | -0.073078496 |
| Garem1 | 1.256542 | 1.311537 | 1.2535 | 0.058078284 | 0.003082949 | 1.279868 | 1.2607888 | 1.22927 | 0.031521385 | 0.050600604 |
| Gm42421 | 2.565951 | 2.500173 | 2.5598 | -0.059614094 | 0.0061636 | 2.5277859 | 2.6447622 | 2.61345 | 0.031307924 | -0.085668336 |
| Gm10912 | 0.676236 | 0.64936 | 0.6932 | -0.043835429 | -0.016959012 | 0.7622064 | 0.7127506 | 0.68145 | 0.031302909 | 0.080758663 |
| Olfr887 | 2.600215 | 2.598661 | 2.5904 | 0.008250082 | 0.009804498 | 2.6098601 | 2.7456004 | 2.71435 | 0.031247598 | -0.104492671 |
| Gm6264 | 4.660272 | 4.695168 | 4.647 | 0.048141406 | 0.013245721 | 4.8598905 | 5.0521574 | 5.02095 | 0.031208025 | -0.16105884 |
| Olfr457 | 1.054138 | 1.009124 | 1.0505 | -0.041371258 | 0.00364252 | 1.0699304 | 1.099626 | 1.06843 | 0.031194222 | 0.00149864 |
| Mup6 | 2.711407 | 2.77753 | 2.7782 | -0.00071887 | -0.066842334 | 2.7851291 | 2.8554543 | 2.82426 | 0.031191944 | -0.039133268 |
| Tex264 | 8.817048 | 8.775569 | 8.8756 | -0.100051787 | -0.058572759 | 8.8421214 | 8.8227804 | 8.79162 | 0.031156197 | 0.050497204 |
| E130208F15Rik | 4.040766 | 3.894614 | 4.0415 | -0.146871471 | -0.000720192 | 3.7849194 | 4.0135493 | 3.9824 | 0.03114872 | -0.197481145 |
| Gpr153 | 2.401712 | 2.550192 | 2.3207 | 0.229507025 | 0.081027144 | 2.5761799 | 2.3654565 | 2.33437 | 0.031089154 | 0.241812631 |
| Fgfbp3 | 4.587596 | 4.506882 | 4.498 | 0.008889544 | 0.089603857 | 4.5423638 | 4.517354 | 4.48635 | 0.031004356 | 0.056014151 |
| Lrfn3 | 1.854536 | 1.909956 | 1.8904 | 0.019585667 | -0.035835105 | 1.8236963 | 1.8257219 | 1.79474 | 0.030978667 | 0.028953014 |
| Cd300c | 4.79946 | 4.604673 | 4.9012 | -0.296494139 | -0.101706873 | 4.3822903 | 4.6078571 | 4.5769 | 0.030959797 | -0.194607006 |
| Anapc5 | 11.75584 | 11.79177 | 11.813 | -0.021290908 | -0.057219078 | 11.977572 | 11.96388 | 11.9329 | 0.03093042 | 0.044622767 |
| Gm4032 | 2.591535 | 2.543958 | 2.6507 | -0.106706438 | -0.059129127 | 2.8115786 | 2.6394477 | 2.60852 | 0.030924298 | 0.203055183 |
| Gm49486 | 1.877257 | 1.765311 | 1.8188 | -0.053450742 | 0.058494877 | 1.8091293 | 1.7832346 | 1.75231 | 0.030922422 | 0.056817146 |
| Tram1 | 11.55559 | 11.53757 | 11.574 | -0.036870396 | -0.018853266 | 11.594635 | 11.475903 | 11.445 | 0.03091045 | 0.149642254 |
| Klri1 | 5.19566 | 5.173129 | 5.2108 | -0.037695826 | -0.015165237 | 4.9721228 | 5.1740298 | 5.14316 | 0.030867817 | -0.171039221 |
| Pnma2 | 1.429757 | 1.476182 | 1.4553 | 0.020836339 | -0.025589461 | 1.5099324 | 1.506046 | 1.47519 | 0.030851282 | 0.034737698 |
| Cdo1 | 3.877649 | 3.936932 | 3.8041 | 0.132837705 | 0.073554871 | 3.9215641 | 4.0375275 | 4.00673 | 0.030798365 | -0.085165 |
| Thoc5 | 9.674751 | 9.74571 | 9.6744 | 0.071272023 | 0.000312566 | 9.8816393 | 9.799096 | 9.76831 | 0.030782908 | 0.113326221 |
| Tmem161b | 8.725886 | 8.719977 | 8.6693 | 0.050635824 | 0.056544773 | 8.6719309 | 8.7066956 | 8.67593 | 0.030768681 | -0.00399602 |
| Actr6 | 7.999036 | 8.046922 | 8.0253 | 0.021619377 | -0.026266055 | 8.0005165 | 8.1293013 | 8.09856 | 0.030737478 | -0.098047295 |
| Nelfb | 10.58796 | 10.71717 | 10.699 | 0.018583934 | -0.110631142 | 10.839944 | 10.791954 | 10.7613 | 0.030658604 | 0.078648983 |
| H2-DMb1 | 10.85114 | 10.86658 | 10.933 | -0.066547942 | -0.081988424 | 11.620522 | 11.256351 | 11.2257 | 0.030649488 | 0.394820621 |

| | | | | | | | | | | |
|---|---|---|---|---|---|---|---|---|---|---|
| Igkv11-106 | 1.148785 | 1.178761 | 1.2041 | -0.025345313 | -0.055321701 | 1.2669321 | 1.2209653 | 1.19033 | 0.030638294 | 0.076605169 |
| Gskip | 8.131889 | 8.172261 | 8.2215 | -0.049281888 | -0.089654303 | 8.1803548 | 8.2674402 | 8.2368 | 0.030635314 | -0.05645006 |
| Gm15191 | 0.692168 | 0.703883 | 0.7119 | -0.00803181 | -0.019746142 | 0.779426 | 0.7123908 | 0.68178 | 0.030612918 | 0.097648144 |
| Brat1 | 8.908946 | 8.907908 | 8.9602 | -0.052319983 | -0.051281691 | 9.1781507 | 8.985641 | 8.95505 | 0.03058979 | 0.223099513 |
| Ube3b | 10.49963 | 10.60466 | 10.618 | -0.013624938 | -0.11865629 | 10.608767 | 10.598777 | 10.5682 | 0.030560228 | 0.04055021 |
| Gm44216 | 0.97659 | 0.966468 | 0.9958 | -0.029306161 | -0.019183473 | 0.9969256 | 0.992101 | 0.9616 | 0.030504429 | 0.035329052 |
| Arhgef33 | 1.662788 | 1.633702 | 1.6841 | -0.050433708 | -0.02134782 | 1.6599474 | 1.6850803 | 1.65462 | 0.030464619 | 0.005331783 |
| Gpr1 | 1.340482 | 1.395468 | 1.4104 | -0.014972557 | -0.069958399 | 1.3104133 | 1.3269797 | 1.29656 | 0.030422319 | 0.013855962 |
| Vmn2r4 | 3.799143 | 3.825714 | 3.8052 | 0.020523999 | -0.006046902 | 3.8195329 | 3.9746254 | 3.94426 | 0.030364285 | -0.124728248 |
| Zfp57 | 3.175653 | 3.125097 | 3.1432 | -0.01808236 | 0.032473732 | 3.109009 | 3.0395669 | 3.00923 | 0.030336443 | 0.099778554 |
| Gabrd | 2.772115 | 2.725068 | 2.6837 | 0.041323352 | 0.088370372 | 2.6962279 | 2.7660937 | 2.73585 | 0.030243209 | -0.039622601 |
| Ren1 | 3.541659 | 5.763907 | 6.432 | -0.668094221 | -2.890342583 | 7.8118415 | 4.1049288 | 4.07474 | 0.030188213 | 3.737100916 |
| Serpinf2 | 2.634901 | 2.585982 | 2.7109 | -0.124902206 | -0.07598268 | 2.7809513 | 2.7428878 | 2.7127 | 0.030186512 | 0.068249954 |
| Nxpe5 | 4.010316 | 3.539043 | 3.4245 | 0.114500109 | 0.585773184 | 4.1172199 | 3.4988771 | 3.46872 | 0.030160164 | 0.648503005 |
| Tjap1 | 8.40424 | 8.402929 | 8.3186 | 0.084299282 | 0.085610033 | 8.4250256 | 8.5230214 | 8.49288 | 0.030137328 | -0.067858518 |
| Slc25a14 | 6.00547 | 6.060968 | 6.0866 | -0.025612169 | -0.081110192 | 5.9762672 | 6.1454333 | 6.1153 | 0.030133767 | -0.139032349 |
| Gm19001 | 0.695727 | 0.705519 | 0.7192 | -0.013650952 | -0.023442995 | 0.7209654 | 0.741786 | 0.7117 | 0.030088695 | 0.009268086 |
| Zfp202 | 6.53109 | 6.504899 | 6.4361 | 0.068789754 | 0.094980148 | 6.4496836 | 6.4557668 | 6.42571 | 0.030059731 | 0.023976543 |
| Leo1 | 8.478111 | 8.559137 | 8.3902 | 0.168891668 | 0.087866205 | 8.41529 | 8.5231849 | 8.49316 | 0.030023119 | -0.077871846 |
| Olfr1341 | 2.191304 | 2.152665 | 2.1219 | 0.030780667 | 0.069420304 | 2.1787111 | 2.2344221 | 2.20441 | 0.030016247 | -0.025694794 |
| Vmn1r90 | 3.448526 | 3.40276 | 3.3277 | 0.075052934 | 0.120818779 | 3.3980331 | 3.5037687 | 3.47377 | 0.029995372 | -0.075740159 |
| Hist1h2bq | 3.784364 | 3.600407 | 3.711 | -0.110622695 | 0.073334165 | 3.6912508 | 3.8035473 | 3.77362 | 0.029930015 | -0.082366427 |
| Hipk2 | 8.950484 | 8.766384 | 8.7189 | 0.0474753 | 0.231575316 | 8.5489643 | 9.0256476 | 8.99581 | 0.029837997 | -0.446845284 |
| Abi3bp | 6.674801 | 6.308141 | 6.4821 | -0.173927955 | 0.19273126 | 6.218574 | 6.3231974 | 6.29337 | 0.029827144 | -0.074796231 |
| Setd2 | 11.74934 | 11.90161 | 11.879 | 0.022732164 | -0.129542564 | 11.807695 | 12.051639 | 12.0219 | 0.029777774 | -0.214165985 |
| Tceal3 | 2.378864 | 2.372924 | 2.2872 | 0.085727039 | 0.091667118 | 2.3716687 | 2.4186299 | 2.38888 | 0.029748474 | -0.017212738 |
| Nrn1l | 1.167077 | 1.20133 | 1.1909 | 0.010477981 | -0.023774795 | 1.2889212 | 1.2866413 | 1.25693 | 0.029715725 | 0.03199572 |
| Fzd7 | 5.890401 | 5.872346 | 5.6375 | 0.234849317 | 0.252904425 | 5.5809935 | 5.6542546 | 5.62458 | 0.029678075 | -0.043582997 |
| Hmgn2-ps1 | 1.606107 | 1.587412 | 1.6074 | -0.020024737 | -0.001329728 | 1.6501915 | 1.7555067 | 1.72585 | 0.029654994 | -0.075660162 |
| Kcnj15 | 6.367558 | 6.118274 | 5.9495 | 0.168775159 | 0.418059219 | 5.6000549 | 5.6008972 | 5.57126 | 0.029632795 | 0.028790572 |
| 9130409I23Rik | 0.815555 | 0.735673 | 0.7648 | -0.029119922 | 0.050762036 | 0.7499271 | 0.8664807 | 0.83686 | 0.029617909 | -0.086935751 |
| Sema3e | 0.915154 | 0.90371 | 0.9484 | -0.044711259 | -0.033267129 | 0.8616528 | 0.9140797 | 0.88452 | 0.029556491 | -0.022870391 |

| | | | | | | | | | | |
|---|---|---|---|---|---|---|---|---|---|---|
| Gm48862 | 1.724164 | 1.727866 | 1.6968 | 0.031057766 | 0.027355607 | 1.7544025 | 1.840774 | 1.81122 | 0.029552719 | -0.056818793 |
| Trappc6b | 9.334121 | 9.348455 | 9.3858 | -0.037362438 | -0.05169611 | 9.3530506 | 9.4345011 | 9.40499 | 0.029512924 | -0.051937543 |
| Inpp5k | 10.23653 | 10.22367 | 10.345 | -0.120964148 | -0.10810404 | 10.427458 | 10.394319 | 10.3648 | 0.029503318 | 0.062642834 |
| Ptcd3 | 9.807135 | 9.902976 | 9.891 | 0.011955548 | -0.083884628 | 10.064533 | 10.183315 | 10.1538 | 0.029493291 | -0.089288 |
| Mapk10 | 2.952444 | 2.972013 | 2.819 | 0.153006775 | 0.133437328 | 2.7485149 | 2.8558457 | 2.82638 | 0.029464681 | -0.077866198 |
| Gm7476 | 0.916196 | 0.915489 | 0.9675 | -0.052003278 | -0.05129668 | 0.9700172 | 0.9623561 | 0.9329 | 0.029456998 | 0.037118025 |
| Ppcs | 7.811234 | 7.749889 | 7.7858 | -0.035906869 | 0.025438695 | 8.0832204 | 7.839616 | 7.81017 | 0.029449327 | 0.273053711 |
| AC122119.2 | 1.435319 | 1.401712 | 1.3746 | 0.02714576 | 0.060753417 | 1.4191166 | 1.4600285 | 1.43059 | 0.029441011 | -0.011470924 |
| Gm17999 | 0.623944 | 0.586028 | 0.5989 | -0.012863956 | 0.025051636 | 0.6456437 | 0.6288466 | 0.59943 | 0.029414773 | 0.046211883 |
| Olfr578 | 0.67837 | 0.658596 | 0.6682 | -0.009633289 | 0.010140604 | 0.6738237 | 0.7153143 | 0.68597 | 0.029348937 | -0.012141653 |
| Zfp493 | 5.334337 | 5.459447 | 5.2724 | 0.187084031 | 0.061973187 | 5.2555453 | 5.442961 | 5.41362 | 0.029339056 | -0.158076634 |
| Ighv8-6 | 2.196345 | 2.153634 | 2.1945 | -0.040885854 | 0.001825291 | 2.1413393 | 2.1035161 | 2.0742 | 0.029318281 | 0.067141483 |
| Apip | 7.166604 | 7.133841 | 7.1817 | -0.047906374 | -0.015142814 | 7.3648481 | 7.2344423 | 7.20513 | 0.02931611 | 0.159721873 |
| Gm19196 | 3.062151 | 2.950644 | 3.0258 | -0.07514238 | 0.036364511 | 2.9814842 | 3.0259895 | 2.99667 | 0.029316072 | -0.015189217 |
| Rbpms | 7.54571 | 7.407153 | 7.6017 | -0.194571433 | -0.056014198 | 7.2563592 | 7.2602212 | 7.23092 | 0.029305469 | 0.025443536 |
| Clec2l | 1.255254 | 1.338851 | 1.3044 | 0.034404259 | -0.049192528 | 1.3316276 | 1.3243584 | 1.29508 | 0.029277457 | 0.036546683 |
| Zfp865 | 9.344422 | 9.12156 | 9.3198 | -0.198254266 | 0.024607618 | 9.4467673 | 9.480944 | 9.45169 | 0.029252821 | -0.004923923 |
| Wdpcp | 4.656247 | 4.958044 | 4.8388 | 0.119250864 | -0.182546195 | 4.5906725 | 4.6107332 | 4.58149 | 0.029238741 | 0.009178109 |
| Hacd1 | 3.163051 | 3.129618 | 3.0139 | 0.115705215 | 0.149138247 | 3.197763 | 3.1168719 | 3.08764 | 0.029233485 | 0.110124648 |
| Spata31d1a | 1.751541 | 1.773886 | 1.6897 | 0.084232579 | 0.06188768 | 1.6350644 | 1.7362852 | 1.70709 | 0.029193578 | -0.072027214 |
| Kcnma1 | 3.926522 | 3.823535 | 3.7902 | 0.033327288 | 0.136313825 | 4.0754983 | 3.9125742 | 3.88338 | 0.029190938 | 0.192115077 |
| Unc80 | 1.519195 | 1.405973 | 1.507 | -0.10103217 | 0.01218986 | 1.3239346 | 1.4441024 | 1.41493 | 0.029174503 | -0.090993298 |
| Gclc | 9.483924 | 9.703082 | 9.5289 | 0.174222817 | -0.044935338 | 9.4715037 | 9.6025874 | 9.57342 | 0.029165818 | -0.1019178 |
| Gm47853 | 2.655354 | 2.670583 | 2.7667 | -0.096138693 | -0.11136726 | 2.7040149 | 2.7580359 | 2.72888 | 0.029154503 | -0.024866465 |
| Gm20594 | 11.58697 | 11.44165 | 11.613 | -0.171372118 | -0.026060372 | 10.896452 | 11.365976 | 11.3369 | 0.02908166 | -0.440442693 |
| Mtrr | 8.436072 | 8.508341 | 8.481 | 0.027336307 | -0.04493253 | 8.3930737 | 8.5263514 | 8.49732 | 0.029035492 | -0.10424228 |
| Awat2 | 3.079413 | 3.120502 | 3.0605 | 0.059975861 | 0.018886476 | 3.0857808 | 3.1713143 | 3.14232 | 0.028992516 | -0.056540985 |
| Nt5c | 9.454047 | 9.421164 | 9.4627 | -0.041515608 | -0.008633215 | 9.7047357 | 9.5104256 | 9.48143 | 0.0289915 | 0.223301623 |
| Olfr1256 | 1.070127 | 1.056244 | 1.0543 | 0.001935031 | 0.015818062 | 1.1059705 | 1.1266591 | 1.09767 | 0.028984788 | 0.008296212 |
| Lats1 | 9.105895 | 9.134946 | 9.0726 | 0.062317483 | 0.033266752 | 9.0777005 | 9.2103996 | 9.18146 | 0.028937688 | -0.103761368 |
| Arg1 | 1.948428 | 2.006988 | 1.9813 | 0.025706288 | -0.032853714 | 2.0468666 | 2.001621 | 1.97271 | 0.028911396 | 0.074157043 |
| Gm8337 | 2.512473 | 2.545101 | 2.5254 | 0.019742724 | -0.012885737 | 2.432478 | 2.5620815 | 2.53318 | 0.028901606 | -0.100701904 |

| | | | | | | | | | | |
|---|---|---|---|---|---|---|---|---|---|---|
| Itih4 | 3.142738 | 3.011303 | 2.9515 | 0.059771235 | 0.191206085 | 2.9077189 | 2.9932354 | 2.96439 | 0.028843069 | -0.056673416 |
| Gm3365 | 4.176119 | 4.224532 | 4.1449 | 0.079657055 | 0.031243856 | 4.3301883 | 4.3161228 | 4.28729 | 0.028832701 | 0.042898221 |
| Ahcyl1 | 10.63433 | 10.61526 | 10.551 | 0.064053366 | 0.083127078 | 10.697024 | 10.73742 | 10.7086 | 0.02882005 | -0.011575069 |
| Phax | 8.80965 | 8.828719 | 8.8331 | -0.004388908 | -0.023457086 | 8.6837098 | 8.8325656 | 8.80378 | 0.028781621 | -0.120074111 |
| Synpr | 1.505486 | 1.496564 | 1.4405 | 0.056069555 | 0.064991415 | 1.449104 | 1.5241969 | 1.49542 | 0.028778618 | -0.046314301 |
| Rpap3-ps2 | 1.988685 | 2.020446 | 1.9955 | 0.024972582 | -0.006788683 | 1.9690535 | 2.1011754 | 2.07254 | 0.028632883 | -0.10348898 |
| Rfxap | 7.355994 | 7.587018 | 7.4755 | 0.111491753 | -0.119532723 | 7.4709862 | 7.5372717 | 7.50867 | 0.028598057 | -0.037687496 |
| Zfp108 | 2.595655 | 2.539779 | 2.6402 | -0.100422985 | -0.044546548 | 2.6446466 | 2.5837461 | 2.55519 | 0.028556331 | 0.089456861 |
| Krt10 | 3.903195 | 3.87985 | 4.0227 | -0.142851898 | -0.119507111 | 4.0337049 | 3.9498817 | 3.92136 | 0.028525425 | 0.112348631 |
| Garnl3 | 4.431618 | 4.421613 | 4.95 | -0.528430637 | -0.518424707 | 4.3765188 | 4.59551 | 4.56699 | 0.028521572 | -0.19046958 |
| Atp10b | 4.086125 | 4.336599 | 4.2148 | 0.121762618 | -0.128710508 | 4.3598633 | 4.4176385 | 4.38916 | 0.028477999 | -0.029297219 |
| Ctla2a | 9.837719 | 9.492731 | 9.8255 | -0.332720837 | 0.012267757 | 8.7032619 | 9.0805628 | 9.05212 | 0.028446075 | -0.348854817 |
| Gm5093 | 0.934911 | 0.957761 | 0.916 | 0.041741844 | 0.018892072 | 0.9251043 | 0.9320658 | 0.90362 | 0.0284447 | 0.021483234 |
| Gm21897 | 5.55107 | 5.36315 | 5.1828 | 0.180338912 | 0.368258762 | 5.4663461 | 5.8236921 | 5.79527 | 0.028425124 | -0.328920827 |
| Arhgap15 | 11.22632 | 11.19159 | 11.269 | -0.077161978 | -0.042428222 | 11.239369 | 11.322184 | 11.2939 | 0.028279886 | -0.05453516 |
| Gm38392 | 0.611481 | 0.64988 | 0.5767 | 0.073190504 | 0.03479139 | 0.5972371 | 0.6392059 | 0.61094 | 0.028268565 | -0.013700222 |
| Gm14324 | 3.437942 | 3.506411 | 3.573 | -0.066613685 | -0.135082861 | 3.4596711 | 3.5186518 | 3.4904 | 0.028247562 | -0.030733233 |
| Qrich1 | 10.84019 | 10.86661 | 10.888 | -0.021629344 | -0.048047964 | 10.803809 | 10.917127 | 10.8889 | 0.028236442 | -0.085081158 |
| Oxld1 | 5.229804 | 5.08298 | 5.1602 | -0.077243428 | 0.069581426 | 5.3709978 | 5.1699559 | 5.14173 | 0.028226427 | 0.229268339 |
| Msln | 1.22648 | 1.173577 | 1.2374 | -0.063786959 | -0.010884499 | 1.2606097 | 1.3144423 | 1.28623 | 0.028214617 | -0.025617999 |
| Zmat4 | 0.854192 | 1.098947 | 0.906 | 0.192908726 | -0.051846889 | 0.8212944 | 0.8781971 | 0.84999 | 0.028208714 | -0.028694014 |
| Boc | 3.337458 | 3.358263 | 3.3979 | -0.039643303 | -0.060448095 | 3.3154299 | 3.3128314 | 3.28465 | 0.028183201 | 0.030781714 |
| Gm34248 | 0.927982 | 0.913248 | 0.9401 | -0.026880146 | -0.012146955 | 0.9452145 | 0.9991218 | 0.97104 | 0.028084626 | -0.025822671 |
| Irak1bp1 | 3.676356 | 3.609207 | 3.5462 | 0.063008961 | 0.130157964 | 3.6934081 | 3.805531 | 3.77747 | 0.028063508 | -0.084059333 |
| Katnbl1 | 7.811926 | 7.839038 | 7.88 | -0.040955633 | -0.068067867 | 7.8483607 | 7.9510926 | 7.92303 | 0.028059709 | -0.074672132 |
| Frmpd3 | 1.142126 | 1.133777 | 1.1398 | -0.006015919 | 0.002332894 | 1.0968907 | 1.1809256 | 1.153 | 0.027923187 | -0.05611168 |
| Ptpa | 10.89856 | 10.90005 | 10.87 | 0.030543782 | 0.029052382 | 11.033384 | 10.888094 | 10.8602 | 0.027915212 | 0.173205126 |
| Grb2 | 12.10211 | 12.0839 | 12.01 | 0.074134862 | 0.092337502 | 12.089507 | 11.981931 | 11.9541 | 0.027872168 | 0.135447591 |
| Trav13d-3 | 2.329706 | 2.307705 | 2.2918 | 0.015897208 | 0.037897775 | 2.3457466 | 2.394018 | 2.36616 | 0.027856742 | -0.020414591 |
| Prop1 | 0.632067 | 0.599551 | 0.6569 | -0.057321781 | -0.024805735 | 0.6393025 | 0.6869862 | 0.65914 | 0.027848202 | -0.019835497 |
| Ppp4r2 | 11.3207 | 11.49168 | 11.278 | 0.213181544 | 0.04220593 | 10.941801 | 11.291266 | 11.2635 | 0.027794954 | -0.321669959 |
| Olfr1423 | 3.09882 | 2.961352 | 2.9919 | -0.030507984 | 0.10695959 | 2.9933774 | 3.1627625 | 3.13502 | 0.027747527 | -0.141637601 |

| | | | | | | | | | | |
|---|---|---|---|---|---|---|---|---|---|---|
| Dlx6 | 2.617485 | 2.621941 | 2.5999 | 0.022012078 | 0.017555614 | 2.7718305 | 2.6919054 | 2.66425 | 0.027654305 | 0.107579428 |
| Kbtbd2 | 10.0594 | 10.12632 | 10.043 | 0.083570291 | 0.016652032 | 9.7563955 | 10.080594 | 10.053 | 0.027633116 | -0.296564938 |
| Hpca | 1.755035 | 1.71557 | 1.6824 | 0.033196149 | 0.072660385 | 1.6714085 | 1.7191735 | 1.69156 | 0.027617739 | -0.020147273 |
| Sema3c | 3.659244 | 3.600135 | 3.8528 | -0.252668727 | -0.193560245 | 3.702506 | 3.6250496 | 3.59746 | 0.027592373 | 0.105048767 |
| Haus7 | 7.756732 | 7.689311 | 7.7822 | -0.092863209 | -0.025441385 | 8.0526647 | 7.8974915 | 7.86994 | 0.027551984 | 0.182725169 |
| Gm2574 | 0.605013 | 0.614787 | 0.5546 | 0.060193529 | 0.050418884 | 0.6187678 | 0.59995 | 0.5724 | 0.02754557 | 0.046363463 |
| Gm5138 | 0.605013 | 0.614787 | 0.5546 | 0.060193529 | 0.050418884 | 0.6187678 | 0.59995 | 0.5724 | 0.02754557 | 0.046363463 |
| Gm13226 | 1.015068 | 1.027214 | 1.0023 | 0.024923103 | 0.012777101 | 1.0033324 | 1.0252935 | 0.99781 | 0.027480788 | 0.005519683 |
| Cxcl14 | 1.541184 | 1.500036 | 1.4666 | 0.033450425 | 0.074598719 | 1.5443698 | 1.5488985 | 1.52143 | 0.027469059 | 0.022940404 |
| Nit1 | 8.766864 | 8.716556 | 8.7543 | -0.037771731 | 0.012535826 | 8.819778 | 8.8297435 | 8.80228 | 0.027465341 | 0.017499797 |
| Rorb | 1.893349 | 1.711116 | 1.7848 | -0.073639884 | 0.108592617 | 1.8398641 | 1.8624646 | 1.83503 | 0.027436462 | 0.004835899 |
| Serpina3h | 1.581995 | 1.581365 | 1.4922 | 0.089116857 | 0.08974697 | 1.5296495 | 1.53542 | 1.50804 | 0.027375461 | 0.021604908 |
| Gm6610 | 0.998604 | 1.057594 | 1.0234 | 0.034179 | -0.02481132 | 1.016882 | 1.099082 | 1.07171 | 0.027374028 | -0.054826039 |
| Rerg | 1.68055 | 1.69014 | 1.6956 | -0.00545439 | -0.015043423 | 1.7367048 | 1.6650899 | 1.63772 | 0.027370625 | 0.098985448 |
| Gm7867 | 1.368236 | 1.397097 | 1.3537 | 0.043431091 | 0.014569927 | 1.4097749 | 1.368999 | 1.34165 | 0.027347045 | 0.068122891 |
| Zkscan2 | 2.803245 | 2.818615 | 2.7554 | 0.063194942 | 0.04782526 | 2.7777797 | 2.9291544 | 2.90181 | 0.027341952 | -0.124032776 |
| Smco1 | 0.46685 | 0.484335 | 0.4906 | -0.00622311 | -0.023707497 | 0.4907777 | 0.5598061 | 0.53248 | 0.027321621 | -0.041706761 |
| Hmbox1 | 9.444714 | 9.492251 | 9.443 | 0.049215347 | 0.001678623 | 9.4391232 | 9.6770864 | 9.64977 | 0.02732032 | -0.2106429 |
| Glrb | 1.786659 | 1.837045 | 1.7773 | 0.059708605 | 0.009322713 | 1.7929292 | 1.8657762 | 1.83846 | 0.02731651 | -0.045530413 |
| Gm8873 | 1.490967 | 1.491498 | 1.5331 | -0.041593207 | -0.042123871 | 1.5755466 | 1.526018 | 1.49874 | 0.027280492 | 0.076809089 |
| Olfr392 | 2.255961 | 2.240248 | 2.2923 | -0.052062514 | -0.03634975 | 2.2966766 | 2.355581 | 2.32832 | 0.027259866 | -0.031644544 |
| Nt5e | 8.506438 | 8.251926 | 8.4766 | -0.224640721 | 0.029871271 | 8.0666024 | 8.5972137 | 8.56999 | 0.027225835 | -0.503385452 |
| Olfr75-ps1 | 2.418926 | 2.563199 | 2.5263 | 0.036910216 | -0.107362945 | 2.5194557 | 2.6073926 | 2.58018 | 0.027210024 | -0.060726937 |
| N4bp2l1 | 9.206653 | 9.379989 | 9.4174 | -0.037402272 | -0.210738167 | 9.2457356 | 9.5438396 | 9.51663 | 0.027205895 | -0.270898155 |
| Mt3 | 1.57712 | 1.564658 | 1.5619 | 0.002763092 | 0.015224719 | 1.5008046 | 1.5181378 | 1.49094 | 0.027202568 | 0.009869405 |
| Gldc | 0.918667 | 0.849372 | 0.8749 | -0.025533904 | 0.043761467 | 0.8843033 | 0.8910159 | 0.86383 | 0.027181556 | 0.020468962 |
| Lrrfip1 | 11.86425 | 11.96426 | 11.855 | 0.10933322 | 0.009315706 | 11.800553 | 11.996813 | 11.9696 | 0.027180004 | -0.169079803 |
| Arntl2 | 5.115418 | 5.046725 | 5.0752 | -0.028431425 | 0.040261536 | 5.3026885 | 5.2897909 | 5.26262 | 0.027175371 | 0.040072991 |
| Gm48541 | 4.989612 | 5.090083 | 5.2028 | -0.112687407 | -0.213157834 | 5.2377669 | 5.363881 | 5.33673 | 0.027154863 | -0.098959267 |
| Nefm | 0.613978 | 0.572804 | 0.4991 | 0.073665247 | 0.114838841 | 0.5114072 | 0.5560616 | 0.52891 | 0.027154229 | -0.017500106 |
| Adipor2 | 10.83151 | 10.89821 | 10.794 | 0.104427974 | 0.03772873 | 10.685258 | 10.776519 | 10.7494 | 0.02712234 | -0.064138243 |
| Spock3 | 2.556979 | 2.419798 | 2.4255 | -0.005695695 | 0.131484713 | 2.2563589 | 2.4004032 | 2.37329 | 0.027109387 | -0.116934895 |

| | | | | | | | | | | |
|---|---|---|---|---|---|---|---|---|---|---|
| Morn1 | 4.507848 | 4.613277 | 4.5604 | 0.052924334 | -0.052504365 | 4.6856913 | 4.5753406 | 4.54829 | 0.027047026 | 0.137397747 |
| Nab1 | 10.51028 | 10.56501 | 10.401 | 0.164179638 | 0.109455834 | 10.329757 | 10.429454 | 10.4025 | 0.02695112 | -0.072746066 |
| Slc26a3 | 3.464642 | 3.476061 | 3.4895 | -0.013436329 | -0.024855178 | 3.5252461 | 3.5553305 | 3.52841 | 0.026918596 | -0.003165787 |
| Olfr898 | 2.938645 | 2.993874 | 2.9221 | 0.071748163 | 0.016519329 | 3.0345889 | 3.1135181 | 3.08661 | 0.026904159 | -0.052024957 |
| C330022B21Rik | 1.447935 | 1.371293 | 1.4046 | -0.033309285 | 0.043332249 | 1.3246228 | 1.3721017 | 1.3452 | 0.026898934 | -0.02058001 |
| Olfr981 | 3.01836 | 3.002088 | 3.0376 | -0.03549784 | -0.019225654 | 3.0458069 | 3.2095366 | 3.18268 | 0.0268602 | -0.136869461 |
| Rpl21-ps10 | 1.164729 | 1.190868 | 1.1972 | -0.006360444 | -0.032499931 | 1.220711 | 1.2175121 | 1.19067 | 0.026844393 | 0.03004335 |
| Eif3f | 12.65465 | 12.66093 | 12.719 | -0.057893296 | -0.064168802 | 12.740022 | 12.686332 | 12.6595 | 0.026815454 | 0.080505219 |
| Tmem184a | 1.02313 | 0.977836 | 0.9823 | -0.004427309 | 0.040867229 | 1.2736907 | 1.1130726 | 1.08629 | 0.026784759 | 0.187402909 |
| Fam76b | 9.958881 | 9.935645 | 9.9874 | -0.051738508 | -0.028503014 | 9.9456647 | 10.054304 | 10.0275 | 0.026765564 | -0.081873435 |
| Amd-ps3 | -0.525985 | -0.55094 | -0.5676 | 0.016645168 | 0.041599904 | -0.5861644 | -0.560476 | -0.5872 | 0.026759475 | 0.001070863 |
| Rpsa | 15.60237 | 15.63245 | 15.618 | 0.014317782 | -0.015761152 | 15.741852 | 15.65149 | 15.6248 | 0.026729746 | 0.117091572 |
| Zfp641 | 1.641279 | 1.63648 | 1.6365 | -3.78164E-05 | 0.004761091 | 1.5606787 | 1.6430959 | 1.61639 | 0.026705583 | -0.055711689 |
| Ighv1-5 | 6.378391 | 6.535123 | 6.6984 | -0.163260242 | -0.319992681 | 6.6920487 | 6.267168 | 6.24047 | 0.026696332 | 0.451577031 |
| Zscan21 | 7.9209 | 7.897652 | 7.956 | -0.058338034 | -0.035089714 | 7.7946557 | 7.8811587 | 7.85451 | 0.026650905 | -0.059852083 |
| Rnf157 | 9.008798 | 8.906254 | 8.9119 | -0.005638139 | 0.096905869 | 8.8508878 | 8.7940918 | 8.76755 | 0.026542873 | 0.083338805 |
| Fscn1 | 11.74539 | 11.82493 | 11.881 | -0.056259298 | -0.135803038 | 11.666623 | 11.886637 | 11.8601 | 0.026495858 | -0.19351864 |
| Thsd7b | 2.505758 | 2.500231 | 2.4784 | 0.021869848 | 0.027397182 | 2.4546329 | 2.616329 | 2.58987 | 0.026454171 | -0.135241944 |
| Gm11889 | 0.576238 | 0.566313 | 0.5097 | 0.05659354 | 0.066518629 | 0.5433216 | 0.5356197 | 0.50918 | 0.026437826 | 0.034139709 |
| Mcm3 | 10.28459 | 10.21018 | 10.414 | -0.204142394 | -0.129734016 | 10.543738 | 10.585195 | 10.5588 | 0.026405476 | -0.015051923 |
| Afg3l2 | 10.05622 | 10.00776 | 10.017 | -0.009212796 | 0.039243852 | 10.136283 | 10.091765 | 10.0654 | 0.026389763 | 0.070908517 |
| Olfr1150-ps1 | 1.83941 | 1.903895 | 1.82 | 0.083917777 | 0.019432796 | 1.863913 | 1.9130971 | 1.88672 | 0.026373782 | -0.022810305 |
| Zfp772 | 6.335533 | 6.281087 | 6.3951 | -0.114038244 | -0.059591957 | 6.4571758 | 6.3183543 | 6.29205 | 0.026305528 | 0.165127126 |
| Gm43467 | 0.454387 | 0.461153 | 0.4163 | 0.044841651 | 0.038075091 | 0.4305959 | 0.4719009 | 0.44565 | 0.026253606 | -0.015051397 |
| Trim9 | 1.514451 | 1.46019 | 1.3891 | 0.071043273 | 0.125304722 | 1.4497299 | 1.4317661 | 1.40551 | 0.026252842 | 0.044216692 |
| Gm6198 | 0.462807 | 0.443091 | 0.4903 | -0.047257017 | -0.027541284 | 0.4222267 | 0.4646977 | 0.43845 | 0.026245199 | -0.016225778 |
| Rnf187 | 10.75521 | 10.78357 | 10.779 | 0.00465894 | -0.023693354 | 10.909542 | 10.716557 | 10.6903 | 0.026237358 | 0.219222624 |
| Olfr1284 | 2.064741 | 2.034555 | 2.1379 | -0.103294577 | -0.073109192 | 2.0161756 | 2.1262151 | 2.1 | 0.02621563 | -0.083823853 |
| Esp8 | 0.738671 | 0.674834 | 0.8032 | -0.12837258 | -0.064535506 | 0.8078042 | 0.7544503 | 0.72828 | 0.02617359 | 0.079527437 |
| Mlycd | 7.994104 | 8.012786 | 8.1086 | -0.095773595 | -0.114455625 | 8.3365941 | 8.1031312 | 8.07698 | 0.026149016 | 0.259611923 |
| Gm6710 | 5.102638 | 5.120762 | 5.3218 | -0.201051692 | -0.219175852 | 5.1414827 | 5.065548 | 5.03941 | 0.026134562 | 0.102069235 |
| Olfr1118 | 2.233471 | 2.210834 | 2.1437 | 0.067129627 | 0.089766757 | 2.1652804 | 2.2868384 | 2.26072 | 0.026115631 | -0.095442415 |

| | | | | | | | | | | |
|---|---|---|---|---|---|---|---|---|---|---|
| Traj58 | 0.703148 | 0.687351 | 0.7222 | -0.034812643 | -0.019015855 | 0.7255997 | 0.7123177 | 0.6863 | 0.026018998 | 0.039301033 |
| Gm6129 | 0.64892 | 0.622081 | 0.6873 | -0.065239965 | -0.038400185 | 0.7629172 | 0.6598286 | 0.63388 | 0.025947411 | 0.129036056 |
| Ighmbp2 | 8.217464 | 8.349635 | 8.2162 | 0.133480869 | 0.001309945 | 8.4231242 | 8.3005772 | 8.27469 | 0.025889585 | 0.148436526 |
| Gm6981 | 1.150527 | 1.133285 | 1.1649 | -0.031567539 | -0.014326059 | 1.2531831 | 1.1799355 | 1.15406 | 0.025873337 | 0.099120838 |
| Gm5789 | 0.196223 | 0.198343 | 0.1908 | 0.007545265 | 0.005425062 | 0.215873 | 0.2544262 | 0.2286 | 0.025823227 | -0.012729993 |
| Atg16l1 | 9.517682 | 9.594824 | 9.5506 | 0.044220308 | -0.032922452 | 9.6873132 | 9.859673 | 9.8339 | 0.02577354 | -0.146586292 |
| Gm4204 | 1.291429 | 1.314928 | 1.3744 | -0.059473163 | -0.082971695 | 1.3895962 | 1.3008173 | 1.27513 | 0.02569208 | 0.11447093 |
| Crim1 | 9.854988 | 9.753613 | 9.6556 | 0.098005664 | 0.199380571 | 9.1712857 | 9.4126502 | 9.38698 | 0.025666682 | -0.215697787 |
| Plch2 | 1.170735 | 1.097269 | 1.2297 | -0.132473218 | -0.059006476 | 1.0858048 | 1.1630152 | 1.13739 | 0.025621576 | -0.051588821 |
| Pole2 | 7.461524 | 7.295463 | 7.4255 | -0.130001226 | 0.036059601 | 7.4190106 | 7.5458778 | 7.52026 | 0.025617524 | -0.101249653 |
| Odaph | 0.738478 | 0.706675 | 0.7191 | -0.012474478 | 0.019329302 | 0.8597314 | 0.7101695 | 0.68457 | 0.025596379 | 0.175158254 |
| Als2cr12 | 1.596211 | 1.649616 | 1.6659 | -0.01624533 | -0.069650549 | 1.6316161 | 1.6325916 | 1.60703 | 0.025566321 | 0.024590821 |
| Slc24a2 | 1.318037 | 1.448834 | 1.3334 | 0.115461487 | -0.015335115 | 1.2652991 | 1.2700035 | 1.2445 | 0.02550727 | 0.020802809 |
| Crisp1 | 3.157164 | 2.892315 | 3.5166 | -0.624300469 | -0.359451629 | 3.818833 | 3.4437596 | 3.41826 | 0.025504327 | 0.400577709 |
| Gm38699 | 3.315335 | 3.238516 | 3.3696 | -0.131129843 | -0.054311371 | 3.1643238 | 3.3909757 | 3.3655 | 0.025478873 | -0.201173115 |
| Adcy10 | 0.948814 | 0.935393 | 0.9629 | -0.027478271 | -0.014057492 | 0.9335223 | 1.0261492 | 1.00069 | 0.025461167 | -0.067165648 |
| Gm3417 | 1.281735 | 1.201135 | 1.2323 | -0.031135961 | 0.049464099 | 1.2465414 | 1.2485155 | 1.22312 | 0.025395617 | 0.023421472 |
| Rpl13-ps1 | 2.119961 | 2.198484 | 2.1807 | 0.017765984 | -0.060756192 | 2.185522 | 2.1220113 | 2.09662 | 0.025393739 | 0.088904432 |
| Impg1 | 2.021459 | 1.988837 | 1.9812 | 0.007662304 | 0.040283651 | 1.958349 | 2.0331731 | 2.00785 | 0.025325317 | -0.049498719 |
| Trim72 | 0.637826 | 0.637427 | 0.6426 | -0.005165858 | -0.004766308 | 0.653772 | 0.686497 | 0.66117 | 0.025325208 | -0.007399716 |
| Surf1 | 8.535284 | 8.362607 | 8.5956 | -0.232959371 | -0.060282314 | 8.6291002 | 8.6797851 | 8.65449 | 0.025290607 | -0.025394302 |
| Ighv1-63 | 4.717541 | 4.781002 | 4.8505 | -0.069516639 | -0.132976665 | 4.8976123 | 4.6991908 | 4.67403 | 0.025164518 | 0.223585996 |
| Nup85 | 9.567619 | 9.639158 | 9.6054 | 0.033781984 | -0.037756773 | 9.8697018 | 9.9004846 | 9.87533 | 0.025149641 | -0.005633196 |
| Zwilch | 6.685315 | 6.452178 | 6.6208 | -0.168628142 | 0.06450899 | 6.8182514 | 6.8942512 | 6.86914 | 0.025113492 | -0.050886355 |
| Trav8d-2 | 3.157618 | 3.135518 | 3.1215 | 0.013977889 | 0.036077619 | 2.9645285 | 3.1224457 | 3.09734 | 0.025102762 | -0.132814396 |
| Snhg11 | 6.065066 | 6.191072 | 6.4126 | -0.221565816 | -0.347572068 | 6.4377107 | 6.6156758 | 6.59064 | 0.025039209 | -0.15292595 |
| Cyth2 | 9.132314 | 9.211434 | 9.2634 | -0.051997215 | -0.131117588 | 9.3660717 | 9.3173313 | 9.29233 | 0.024999891 | 0.073740301 |
| Ykt6 | 9.897838 | 9.928489 | 9.8926 | 0.035859078 | 0.005208819 | 9.9930877 | 9.9558643 | 9.93091 | 0.02495007 | 0.0621735 |
| Elp2 | 10.50362 | 10.51774 | 10.517 | 0.001211286 | -0.012906922 | 10.567771 | 10.611009 | 10.5861 | 0.024919178 | -0.018319148 |
| Gm3565 | 0.199409 | 0.233839 | 0.2013 | 0.032543062 | -0.001886562 | 0.1910928 | 0.2203452 | 0.19546 | 0.024887391 | -0.004364956 |
| Gm1604a | 0.424128 | 0.380736 | 0.4266 | -0.045889587 | -0.002497273 | 0.3915326 | 0.4445139 | 0.41967 | 0.02484312 | -0.028138122 |
| Ptgfr | 0.506793 | 0.422106 | 0.5127 | -0.090616898 | -0.005929378 | 0.4414965 | 0.4707133 | 0.4459 | 0.024815186 | -0.004401541 |

| | | | | | | | | | | |
|---|---|---|---|---|---|---|---|---|---|---|
| Il20ra | 0.848693 | 0.86406 | 0.941 | -0.076935893 | -0.092303405 | 0.8427718 | 0.8737541 | 0.84896 | 0.024797448 | -0.006184801 |
| Vmn2r63 | 4.05496 | 3.997783 | 4.0642 | -0.066434531 | -0.009258164 | 4.0897594 | 4.2509225 | 4.22613 | 0.024791613 | -0.136371506 |
| Fkbp1b | 3.612312 | 3.343969 | 3.501 | -0.157035815 | 0.111307649 | 3.4621062 | 3.4160122 | 3.39128 | 0.024735862 | 0.070829792 |
| Lman1 | 9.73841 | 9.68327 | 9.5678 | 0.115423544 | 0.170564178 | 9.5465053 | 9.5145132 | 9.48978 | 0.024731685 | 0.056723721 |
| Faap24 | 7.326799 | 7.266933 | 7.3915 | -0.124605042 | -0.064739307 | 7.5869179 | 7.4399014 | 7.41522 | 0.024682752 | 0.171699287 |
| Fbxw5 | 9.576549 | 9.520814 | 9.5608 | -0.039965411 | 0.015769658 | 9.8651522 | 9.7821581 | 9.75748 | 0.024679764 | 0.10767391 |
| Tmem209 | 9.739654 | 9.826245 | 9.7699 | 0.056389724 | -0.030201868 | 9.8189294 | 9.8881301 | 9.86352 | 0.024612859 | -0.04458785 |
| Pou5f2 | 2.028759 | 2.007386 | 2.1003 | -0.09286521 | -0.071492234 | 2.0595611 | 2.1761969 | 2.15159 | 0.024608209 | -0.092027648 |
| Gdf9 | 1.396654 | 1.358683 | 1.3779 | -0.019260507 | 0.018711228 | 1.3868698 | 1.4017478 | 1.37714 | 0.02460752 | 0.00972947 |
| Nudt14 | 7.230351 | 7.049873 | 7.2213 | -0.171467419 | 0.009011088 | 7.3422126 | 7.1510443 | 7.12645 | 0.024590135 | 0.215758431 |
| Guk1 | 8.172723 | 8.042137 | 8.177 | -0.134896343 | -0.004309653 | 8.3590722 | 8.1560697 | 8.1315 | 0.024571306 | 0.227573751 |
| Gabrb3 | 0.854402 | 0.857609 | 0.8041 | 0.053506528 | 0.050299171 | 0.7797921 | 0.8572131 | 0.83266 | 0.024549478 | -0.052871582 |
| Dgkh | 8.078352 | 8.052285 | 7.9521 | 0.100163647 | 0.126229925 | 7.7257286 | 8.1254255 | 8.10088 | 0.024549347 | -0.375147488 |
| Morn2 | 3.98309 | 3.870053 | 3.8518 | 0.01824494 | 0.131281321 | 3.8967968 | 3.9337349 | 3.9092 | 0.024530185 | -0.012407916 |
| Sigmar1 | 8.303003 | 8.292741 | 8.2715 | 0.021269442 | 0.031531684 | 8.4192693 | 8.2397603 | 8.21526 | 0.02449783 | 0.204006864 |
| Tmem53 | 1.540158 | 1.523759 | 1.5083 | 0.015415296 | 0.031813751 | 1.5070363 | 1.4859545 | 1.46161 | 0.024343719 | 0.045425562 |
| Tspan11 | 3.356704 | 3.379316 | 3.3071 | 0.072264341 | 0.049651727 | 3.3269146 | 3.3031747 | 3.27886 | 0.024318544 | 0.048058437 |
| Gm10364 | 0.66912 | 0.649861 | 0.6324 | 0.017448395 | 0.036707761 | 0.6706545 | 0.6420511 | 0.61776 | 0.024286184 | 0.05288956 |
| Srd5a1 | 3.03283 | 3.095893 | 3.1135 | -0.017638512 | -0.080701145 | 3.1666914 | 3.0431463 | 3.01887 | 0.024274255 | 0.147819338 |
| Gm11957 | 0.584825 | 0.626766 | 0.6102 | 0.016543499 | -0.025397444 | 0.6052831 | 0.6086796 | 0.58445 | 0.024227844 | 0.020831326 |
| Sgpp2 | 1.593558 | 1.58163 | 1.5776 | 0.004053792 | 0.015981371 | 1.5398843 | 1.6257664 | 1.60159 | 0.024177172 | -0.061704885 |
| Blzf1 | 8.086962 | 7.860154 | 7.993 | -0.132860803 | 0.093947042 | 8.0936351 | 7.8987554 | 7.8746 | 0.024150428 | 0.219030132 |
| Trgj4 | 1.321268 | 1.260039 | 1.2375 | 0.022514425 | 0.083743693 | 1.2729677 | 1.2749096 | 1.25077 | 0.024139307 | 0.022197401 |
| Ina | 0.472788 | 0.461099 | 0.3778 | 0.083270327 | 0.094959212 | 0.3858684 | 0.439066 | 0.41493 | 0.024136797 | -0.0290608 |
| C1qtnf3 | 0.491139 | 0.449803 | 0.4626 | -0.012819893 | 0.028516038 | 0.4698153 | 0.5031649 | 0.47905 | 0.024112834 | -0.009236729 |
| Gm14735 | 3.887343 | 4.000042 | 4.0378 | -0.037732345 | -0.150431155 | 3.922552 | 4.1613535 | 4.13728 | 0.024073978 | -0.214727534 |
| Etnppl | 2.199384 | 2.227201 | 2.184 | 0.043208985 | 0.015392102 | 2.1365391 | 2.1908436 | 2.16677 | 0.024070889 | -0.030233647 |
| Apoc1 | 2.584055 | 2.435378 | 2.6001 | -0.164721134 | -0.016043975 | 2.4864905 | 2.4247031 | 2.40067 | 0.02403135 | 0.085818765 |
| Mdp1 | 9.686671 | 9.804489 | 9.7393 | 0.065154841 | -0.052663515 | 10.037522 | 9.890625 | 9.86662 | 0.024006988 | 0.170903881 |
| Mc1r | 1.30662 | 1.271339 | 1.3157 | -0.044338216 | -0.009056478 | 1.3843423 | 1.3642307 | 1.34025 | 0.023978763 | 0.044090447 |
| Gm17911 | 0.189209 | 0.159235 | 0.1854 | -0.026130979 | 0.003842849 | 0.1850645 | 0.2098609 | 0.18589 | 0.023968313 | -0.000828039 |
| Mfsd14a | 10.11431 | 10.23936 | 10.129 | 0.110457407 | -0.014586744 | 10.120647 | 10.211948 | 10.188 | 0.02394637 | -0.067355344 |

| | | | | | | | | | | |
|---|---|---|---|---|---|---|---|---|---|---|
| Eif2a | 10.16299 | 10.03258 | 10.02 | 0.012241059 | 0.142651433 | 9.9434075 | 10.151778 | 10.1278 | 0.023937634 | -0.184432777 |
| Olfr654 | 3.565154 | 3.51696 | 3.572 | -0.055062241 | -0.00686876 | 3.5651377 | 3.7620036 | 3.73813 | 0.023875774 | -0.172990109 |
| Gm9812 | 0.745208 | 0.70383 | 0.7371 | -0.033222165 | 0.008156159 | 0.6952683 | 0.6932411 | 0.66941 | 0.023834041 | 0.025861183 |
| Olfr794 | 1.344689 | 1.371758 | 1.3511 | 0.020686856 | -0.006382019 | 1.4554315 | 1.4125936 | 1.3888 | 0.023798062 | 0.066635924 |
| Csnk1a1 | 12.16841 | 12.15546 | 12.126 | 0.029190918 | 0.042138792 | 12.015742 | 12.106771 | 12.083 | 0.023723452 | -0.067305503 |
| Afg3l1 | 9.591848 | 9.444406 | 9.6712 | -0.226834872 | -0.07939324 | 9.8608767 | 9.6799561 | 9.6563 | 0.023655401 | 0.204575928 |
| Gdap1 | 1.667448 | 1.601429 | 1.6324 | -0.030953696 | 0.035065475 | 1.5946536 | 1.5563066 | 1.53266 | 0.02365031 | 0.061997341 |
| Vmn2r114 | 5.82815 | 5.856281 | 5.8584 | -0.00209497 | -0.030226103 | 5.8252562 | 6.2733358 | 6.24969 | 0.02364766 | -0.424431963 |
| Hoxa4 | 1.956041 | 1.906921 | 1.9317 | -0.024735123 | 0.024384685 | 1.8973675 | 1.9118814 | 1.88829 | 0.023587082 | 0.009073143 |
| Tpsg1 | 1.189693 | 1.237909 | 1.2245 | 0.013413091 | -0.034802872 | 1.1472623 | 1.1972449 | 1.17367 | 0.02357443 | -0.026408142 |
| Gm9722 | -0.20663 | -0.21527 | -0.1906 | -0.024671281 | -0.016032143 | -0.2074441 | -0.176228 | -0.1997 | 0.023516386 | -0.007699536 |
| Gm13341 | 2.772024 | 2.871468 | 2.8996 | -0.028103066 | -0.127547549 | 2.906789 | 2.8916219 | 2.86811 | 0.023514578 | 0.038681671 |
| Ceacam15 | 2.184931 | 2.200492 | 2.1318 | 0.068648349 | 0.053087203 | 2.1903085 | 2.2823704 | 2.25887 | 0.023505186 | -0.06855679 |
| Mcam | 8.146939 | 7.948888 | 7.6289 | 0.32000314 | 0.51805373 | 7.636589 | 7.3791582 | 7.35567 | 0.023487556 | 0.280918334 |
| Gm11549 | 1.330859 | 1.309843 | 1.3114 | -0.001543791 | 0.019471602 | 1.2764944 | 1.3516508 | 1.3282 | 0.023452868 | -0.051703484 |
| Trmt6 | 9.24477 | 9.06958 | 9.2693 | -0.199703141 | -0.02451327 | 9.3863427 | 9.3581 | 9.33467 | 0.023429775 | 0.051672431 |
| Mest | 2.917132 | 2.977864 | 2.8343 | 0.143600054 | 0.082868257 | 3.0183479 | 3.05698 | 3.03355 | 0.023427788 | -0.015204346 |
| Myh1 | 1.520378 | 1.492618 | 1.4559 | 0.036767799 | 0.064527402 | 1.4872778 | 1.5273379 | 1.50394 | 0.023400318 | -0.016659769 |
| Gm21742 | 1.333426 | 1.294308 | 1.2818 | 0.01252289 | 0.051640189 | 1.2214155 | 1.2739325 | 1.25065 | 0.023280792 | -0.029236189 |
| Rpl31-ps11 | 1.73036 | 1.685066 | 1.6784 | 0.006703447 | 0.051997793 | 1.6869456 | 1.7242404 | 1.701 | 0.023241387 | -0.014053378 |
| Sfr1 | 10.22289 | 10.12642 | 10.241 | -0.115068386 | -0.018595394 | 10.129151 | 10.138843 | 10.1156 | 0.023232404 | 0.013540201 |
| Rps15a-ps4 | 3.872557 | 3.924536 | 4.048 | -0.12351303 | -0.175491964 | 4.1003548 | 4.0249416 | 4.00173 | 0.023208576 | 0.09862173 |
| Zfp874b | 7.621172 | 7.851832 | 7.5727 | 0.27912721 | 0.048467438 | 7.4488178 | 7.7857445 | 7.7626 | 0.023145775 | -0.313780932 |
| Vmn2r75 | 1.223298 | 1.238668 | 1.2519 | -0.013243214 | -0.028612933 | 1.2313051 | 1.2772865 | 1.25418 | 0.023103207 | -0.022878115 |
| Gm8225 | 0.554743 | 0.609517 | 0.5782 | 0.031280936 | -0.023492988 | 0.5745468 | 0.576177 | 0.55308 | 0.023092612 | 0.021462317 |
| Sh2d2a | 10.61958 | 10.596 | 10.689 | -0.09289367 | -0.069311176 | 10.613737 | 10.748176 | 10.7251 | 0.023084238 | -0.111355251 |
| Vmn2r18 | 0.898803 | 0.86319 | 0.8446 | 0.018552441 | 0.05416547 | 0.821828 | 0.9132515 | 0.89017 | 0.023083541 | -0.068339951 |
| Car5a | 2.548243 | 2.551719 | 2.5085 | 0.043257142 | 0.039780888 | 2.5504963 | 2.624425 | 2.60135 | 0.023074457 | -0.050854272 |
| Ecsit | 8.315583 | 8.238511 | 8.2668 | -0.028243089 | 0.048828356 | 8.5264698 | 8.2770614 | 8.25399 | 0.023072494 | 0.272480899 |
| Slc2a5 | 0.995718 | 0.829198 | 0.8526 | -0.023434196 | 0.1430852 | 0.8401178 | 0.823155 | 0.8001 | 0.023058254 | 0.040021037 |
| Gm10247 | 3.144409 | 3.236009 | 3.1801 | 0.055956995 | -0.035643704 | 3.1128722 | 3.2344239 | 3.21138 | 0.023039644 | -0.098512038 |
| Ercc6l2 | 9.24142 | 9.18766 | 9.3578 | -0.170144519 | -0.116384127 | 9.2268916 | 9.3532109 | 9.33018 | 0.023033053 | -0.103286235 |

| | | | | | | | | | | |
|---|---|---|---|---|---|---|---|---|---|---|
| Gm10340 | 0.472467 | 0.429934 | 0.4609 | -0.030986835 | 0.011546131 | 0.3980189 | 0.444902 | 0.42188 | 0.023017498 | -0.023865548 |
| Shisal1 | 2.461469 | 2.410305 | 2.4698 | -0.059452922 | -0.00828847 | 2.4147662 | 2.4561889 | 2.43319 | 0.023002531 | -0.018420184 |
| Mis18a | 8.125613 | 8.002323 | 8.0751 | -0.072732878 | 0.050557914 | 8.0124835 | 8.0563368 | 8.03334 | 0.02299222 | -0.020861062 |
| 2210016L21Rik | 8.468329 | 8.23649 | 8.4462 | -0.209684654 | 0.022154433 | 8.70926 | 8.3439914 | 8.32104 | 0.022948026 | 0.388216566 |
| Gm5637 | 1.737247 | 1.792241 | 1.7581 | 0.034106278 | -0.020887201 | 1.7120955 | 1.7491227 | 1.72618 | 0.022940773 | -0.014086468 |
| Wdr13 | 8.475833 | 8.361227 | 8.5305 | -0.169317177 | -0.054710941 | 9.0278102 | 8.8029642 | 8.78006 | 0.022903229 | 0.247749207 |
| Vinac1 | 0.406789 | 0.416032 | 0.4504 | -0.034409309 | -0.043652892 | 0.4551348 | 0.4378269 | 0.41494 | 0.022884387 | 0.040192239 |
| Fam71e1 | 3.640581 | 3.480718 | 3.7103 | -0.229622072 | -0.069759725 | 3.869114 | 3.6184468 | 3.59564 | 0.022805131 | 0.273472359 |
| Zfp629 | 7.4462 | 7.247129 | 7.3492 | -0.10207957 | 0.096991302 | 7.2201162 | 7.3745832 | 7.35185 | 0.022732984 | -0.131733948 |
| Atp5l2-ps | 0.324775 | 0.304773 | 0.3235 | -0.018755195 | 0.001246599 | 0.3079907 | 0.3236672 | 0.30094 | 0.022726413 | 0.007049852 |
| Gm13423 | 2.91287 | 2.862271 | 2.9273 | -0.06500992 | -0.014410426 | 2.9599638 | 2.959044 | 2.93632 | 0.022724913 | 0.023644669 |
| Erc2 | 1.087468 | 1.070433 | 1.0408 | 0.029593782 | 0.046628553 | 1.0171883 | 1.0935969 | 1.07091 | 0.022686865 | -0.053721827 |
| Syce2 | 5.8265 | 5.615686 | 5.8663 | -0.250631991 | -0.03981734 | 5.8106733 | 5.9904754 | 5.96781 | 0.022663921 | -0.157138228 |
| Lrfn1 | 3.517402 | 3.339262 | 3.3004 | 0.038860001 | 0.216999261 | 3.3038247 | 3.3001042 | 3.27744 | 0.022663527 | 0.026384029 |
| Rps12-ps26 | 1.670216 | 1.637038 | 1.5983 | 0.038734714 | 0.071912514 | 1.6398773 | 1.6132289 | 1.59058 | 0.022648741 | 0.049297218 |
| Hist2h2aa1 | 5.013009 | 4.982269 | 4.9832 | -0.000928733 | 0.029810852 | 5.6800578 | 5.232481 | 5.20986 | 0.022624337 | 0.470201192 |
| Ier2 | 10.57038 | 10.36455 | 10.474 | -0.109158376 | 0.096673171 | 9.8068502 | 9.7490972 | 9.72652 | 0.022574342 | 0.080327337 |
| Gm5244 | 1.385292 | 1.3645 | 1.376 | -0.011504495 | 0.009286894 | 1.3349743 | 1.3923943 | 1.36983 | 0.022567554 | -0.03485241 |
| Trim43b | 4.237215 | 4.389797 | 4.5 | -0.110188285 | -0.26277021 | 4.6070892 | 4.6093971 | 4.58687 | 0.022526131 | 0.020218255 |
| Gm4247 | 0.820712 | 0.826962 | 0.788 | 0.038946315 | 0.032696021 | 0.8863867 | 0.8670213 | 0.84456 | 0.022463598 | 0.041828958 |
| Kif1b | 10.51122 | 10.42296 | 10.346 | 0.077434872 | 0.165696646 | 10.197528 | 10.481226 | 10.4588 | 0.022452194 | -0.261245812 |
| 1700028B04Rik | 0.696117 | 0.644109 | 0.6503 | -0.006230224 | 0.04577773 | 0.7104168 | 0.7309709 | 0.70853 | 0.022439746 | 0.001885655 |
| Gm11701 | 0.170667 | 0.127743 | 0.1451 | -0.017341782 | 0.0255823 | 0.1372653 | 0.1804364 | 0.15808 | 0.022356172 | -0.020814953 |
| AC121967.1 | 1.168049 | 1.17094 | 1.226 | -0.05509495 | -0.057986238 | 1.2900994 | 1.3006763 | 1.27832 | 0.022352498 | 0.01177568 |
| Arid2 | 10.27395 | 10.39164 | 10.27 | 0.121683591 | 0.003991557 | 10.044347 | 10.3495 | 10.3272 | 0.022313786 | -0.282839242 |
| Gm17383 | 0.594915 | 0.605644 | 0.6135 | -0.007853803 | -0.018582923 | 0.593382 | 0.6556072 | 0.63331 | 0.022296319 | -0.039928841 |
| Ltbp3 | 5.84014 | 5.360835 | 5.3411 | 0.019685652 | 0.498991235 | 5.2235173 | 5.3856126 | 5.36334 | 0.022276401 | -0.139818882 |
| Nnat | 1.575645 | 1.62992 | 1.5404 | 0.089470706 | 0.035195199 | 1.5983507 | 1.5773668 | 1.55514 | 0.022222226 | 0.043206084 |
| Asphd2 | 0.846399 | 0.781018 | 0.7515 | 0.029513315 | 0.094894883 | 0.7144382 | 0.7661499 | 0.74411 | 0.022042405 | -0.029669273 |
| Becn1 | 11.06855 | 11.05343 | 10.996 | 0.057634836 | 0.072752362 | 11.035902 | 11.106772 | 11.0847 | 0.022030864 | -0.048839247 |
| Chp1 | 10.66748 | 10.7336 | 10.712 | 0.021763872 | -0.044359468 | 10.679751 | 10.671221 | 10.6492 | 0.02200843 | 0.030539101 |
| Mkln1 | 10.49713 | 10.50702 | 10.469 | 0.038203356 | 0.028306308 | 10.337342 | 10.460698 | 10.4387 | 0.021983724 | -0.101372428 |

| | | | | | | | | | | |
|---|---|---|---|---|---|---|---|---|---|---|
| Olfr1535 | 0.700912 | 0.648772 | 0.75 | -0.101230432 | -0.049091153 | 0.7100147 | 0.6838845 | 0.66197 | 0.021910308 | 0.048040491 |
| 9430038I01Rik | 6.785751 | 6.782406 | 6.8259 | -0.043497112 | -0.040151358 | 6.8836573 | 6.807655 | 6.78582 | 0.021830403 | 0.097832718 |
| Hnrnpf | 13.88602 | 13.89958 | 13.825 | 0.07451571 | 0.060952088 | 13.837029 | 13.85992 | 13.8381 | 0.021797746 | -0.001093068 |
| Olfr418 | 2.444742 | 2.447972 | 2.3734 | 0.074549356 | 0.071319493 | 2.5203042 | 2.502414 | 2.48062 | 0.021793475 | 0.039683725 |
| Thoc3 | 8.850974 | 8.858715 | 8.8569 | 0.001797925 | -0.005942766 | 8.969259 | 8.9186246 | 8.89685 | 0.021773411 | 0.072407861 |
| Nktr | 11.22384 | 11.20919 | 11.224 | -0.014433516 | 0.000216862 | 11.465901 | 11.616771 | 11.595 | 0.021759352 | -0.129111012 |
| Ikzf4 | 7.268582 | 7.289036 | 7.5309 | -0.241848573 | -0.262302446 | 7.1804128 | 7.5180279 | 7.49632 | 0.021711293 | -0.315903818 |
| Gm12091 | 3.153167 | 3.133865 | 3.1435 | -0.009654633 | 0.009647274 | 3.0665004 | 3.2759641 | 3.25427 | 0.021696162 | -0.187767503 |
| Ddx3y | 1.156914 | 1.308212 | 1.1965 | 0.111718369 | -0.039579866 | 1.2544852 | 1.2647692 | 1.24314 | 0.021628068 | 0.011344112 |
| A130006I12Rik | 1.592443 | 1.618898 | 1.5989 | 0.019991458 | -0.006463552 | 1.64798 | 1.642185 | 1.62057 | 0.02161022 | 0.027405219 |
| Mob4 | 9.596176 | 9.59076 | 9.5433 | 0.047441755 | 0.052858017 | 9.5517879 | 9.6997957 | 9.6782 | 0.021596445 | -0.12641135 |
| Gatc | 7.314232 | 7.327159 | 7.3922 | -0.06504852 | -0.07797626 | 7.4898184 | 7.3105824 | 7.28903 | 0.02155595 | 0.200791951 |
| Olfr566 | 0.535821 | 0.54287 | 0.5673 | -0.024436048 | -0.031484938 | 0.6027442 | 0.6024651 | 0.58095 | 0.021510362 | 0.021789403 |
| Hist1h4j | 1.846374 | 1.838387 | 1.8805 | -0.042116446 | -0.034129253 | 1.9124864 | 1.8074699 | 1.78597 | 0.021496205 | 0.126512658 |
| Zglp1 | 1.562544 | 1.523403 | 1.4912 | 0.032163257 | 0.071303576 | 1.6379433 | 1.6082812 | 1.58699 | 0.021296084 | 0.050958168 |
| Tesmin | 1.611948 | 1.594362 | 1.6262 | -0.031816501 | -0.01423053 | 1.6432095 | 1.5656616 | 1.54438 | 0.021283497 | 0.098831357 |
| Ighv1-43 | 1.83225 | 1.786516 | 1.8367 | -0.050166803 | -0.004431953 | 1.7684287 | 1.831317 | 1.81003 | 0.021282098 | -0.041606182 |
| Tesk1 | 9.642358 | 9.78809 | 9.6656 | 0.122509261 | -0.023222548 | 9.7692721 | 9.6439506 | 9.62267 | 0.021281689 | 0.146603174 |
| Zdhhc19 | 2.200448 | 2.127748 | 2.0639 | 0.063843421 | 0.136543158 | 2.1504867 | 2.1875777 | 2.1663 | 0.021274492 | -0.015816555 |
| Kcnd1 | 4.983695 | 5.006786 | 4.9648 | 0.042032577 | 0.018941147 | 5.1569003 | 5.2598584 | 5.23864 | 0.021220201 | -0.08173792 |
| Rest | 9.965242 | 10.04641 | 9.9804 | 0.065987755 | -0.015183988 | 10.004059 | 10.015292 | 9.99413 | 0.021165904 | 0.009932769 |
| Zfp976 | 3.844619 | 3.742284 | 3.8875 | -0.145167229 | -0.042832386 | 3.7930418 | 3.8200357 | 3.79889 | 0.021141362 | -0.005852606 |
| Ep400 | 12.23618 | 12.21377 | 12.259 | -0.045631456 | -0.023217138 | 12.223306 | 12.456616 | 12.4355 | 0.021130718 | -0.212179557 |
| Olfr851 | 3.155182 | 3.054339 | 2.9923 | 0.062011584 | 0.1628551 | 2.964478 | 3.2000186 | 3.17901 | 0.021008975 | -0.214531616 |
| Gm9626 | 0.833368 | 0.859516 | 0.8344 | 0.025074496 | -0.001073115 | 0.8730148 | 0.8689606 | 0.84796 | 0.02100011 | 0.025054248 |
| Rps19-ps4 | 2.291149 | 2.370949 | 2.3033 | 0.067616809 | -0.012182691 | 2.4143381 | 2.3812306 | 2.36024 | 0.020987142 | 0.054094668 |
| Fn3krp | 6.724809 | 6.734652 | 6.667 | 0.067677336 | 0.057833754 | 6.6747658 | 6.8306983 | 6.80978 | 0.020921737 | -0.135010737 |
| Gm6061 | 1.957152 | 1.915945 | 1.9908 | -0.074883645 | -0.033676672 | 2.012112 | 2.0529218 | 2.03201 | 0.020914301 | -0.019895569 |
| Hist1h2ai | 2.548178 | 2.535414 | 2.5366 | -0.001214576 | 0.011549655 | 2.5903606 | 2.5039998 | 2.48309 | 0.020908218 | 0.107269047 |
| Tuba4a | 10.38007 | 10.40383 | 10.522 | -0.118272292 | -0.142036456 | 10.429801 | 10.343786 | 10.3229 | 0.020902946 | 0.106917977 |
| Rps10-ps2 | 1.163507 | 1.19297 | 1.2929 | -0.099917225 | -0.129380144 | 1.243012 | 1.3109862 | 1.29012 | 0.020863322 | -0.047110854 |
| Gm20768 | 0.59743 | 0.599553 | 0.6204 | -0.020820631 | -0.022943589 | 0.5912394 | 0.6093878 | 0.58853 | 0.020855314 | 0.002706878 |

| | | | | | | | | | | |
|---|---|---|---|---|---|---|---|---|---|---|
| Klf16 | 8.836426 | 8.747897 | 8.8232 | -0.075265208 | 0.013263449 | 8.7326637 | 8.7889696 | 8.76818 | 0.020787836 | -0.035518056 |
| Ankrd61 | 0.748185 | 0.766106 | 0.7312 | 0.034888429 | 0.016966509 | 0.7351815 | 0.8027662 | 0.78198 | 0.020781424 | -0.046803276 |
| Gm4724 | 6.477236 | 6.275454 | 6.1806 | 0.094843759 | 0.296625866 | 6.3906745 | 6.5320478 | 6.51138 | 0.020663431 | -0.120709958 |
| Coprs | 3.926569 | 3.851164 | 3.8901 | -0.038960892 | 0.036444351 | 3.716864 | 3.8285829 | 3.80793 | 0.020654753 | -0.091064176 |
| Fastkd3 | 7.539262 | 7.747641 | 7.6216 | 0.125996548 | -0.082381759 | 7.7058645 | 7.8599947 | 7.83936 | 0.020639561 | -0.133490662 |
| Gm10362 | 1.143615 | 1.189315 | 1.1794 | 0.009953744 | -0.035745393 | 1.2821025 | 1.2376846 | 1.21705 | 0.020630563 | 0.06504847 |
| Ltb4r2 | 2.10954 | 2.041539 | 2.1136 | -0.072066375 | -0.004064772 | 2.0916412 | 2.0711937 | 2.05058 | 0.020616895 | 0.041064418 |
| Gm11824 | 0.903084 | 0.898297 | 0.9344 | -0.03606449 | -0.031277398 | 0.9376747 | 0.9350849 | 0.91448 | 0.020602811 | 0.023192694 |
| Gm12917 | 1.733576 | 1.73439 | 1.7079 | 0.026495554 | 0.025682119 | 1.8506773 | 1.7989816 | 1.77839 | 0.020590577 | 0.072286206 |
| Ank3 | 4.37633 | 4.39204 | 4.3057 | 0.08636209 | 0.070652434 | 4.4795413 | 4.6428246 | 4.62226 | 0.0205653 | -0.14271801 |
| Gm9157 | 1.009777 | 0.974703 | 0.9813 | -0.006641663 | 0.028433032 | 1.0563776 | 0.9949472 | 0.97441 | 0.020539763 | 0.081970155 |
| Sntg2 | 1.092867 | 1.013531 | 1.0526 | -0.039096302 | 0.040239443 | 1.0835737 | 1.0103268 | 0.9898 | 0.020531685 | 0.09377853 |
| Mtcl1 | 2.904548 | 2.899151 | 2.9878 | -0.088648076 | -0.083251014 | 2.9412541 | 2.8420244 | 2.82149 | 0.020529982 | 0.119759648 |
| Wdr41 | 8.749 | 8.769708 | 8.6707 | 0.098967793 | 0.078260454 | 8.8692899 | 8.8385684 | 8.81804 | 0.020526055 | 0.051247629 |
| Olfr889 | 2.382022 | 2.380801 | 2.4495 | -0.068650678 | -0.067429429 | 2.3559634 | 2.4569597 | 2.43645 | 0.020511335 | -0.080484924 |
| Smarcd1 | 9.397267 | 9.158213 | 9.3208 | -0.162604307 | 0.076449524 | 9.4772678 | 9.4363858 | 9.4159 | 0.020488579 | 0.06137057 |
| AC157903.1 | 0.478763 | 0.541734 | 0.5539 | -0.012197065 | -0.075167177 | 0.5144246 | 0.5082891 | 0.48781 | 0.020476581 | 0.026612062 |
| Tdg | 9.248579 | 9.229733 | 9.2593 | -0.029580794 | -0.010735084 | 9.1210335 | 9.3195829 | 9.29912 | 0.020457909 | -0.17809149 |
| Pcdhb22 | 3.406739 | 3.129128 | 3.1181 | 0.010980628 | 0.288591004 | 3.0825901 | 3.0842447 | 3.06379 | 0.020457837 | 0.018803218 |
| Gm10916 | 1.912305 | 2.090138 | 2.0519 | 0.038253034 | -0.139579234 | 2.0929685 | 2.0780159 | 2.05757 | 0.020450729 | 0.035403302 |
| Lrtm1 | 1.003198 | 1.015276 | 1.0188 | -0.003569773 | -0.015647464 | 1.0231181 | 1.0975567 | 1.07712 | 0.020437952 | -0.054000669 |
| Dcps | 9.13202 | 9.231693 | 9.2067 | 0.025019297 | -0.074653172 | 9.2518307 | 9.1748701 | 9.15447 | 0.020395313 | 0.097355933 |
| Dcpp1 | 3.45349 | 3.148484 | 2.5151 | 0.633347482 | 0.938353235 | 5.0267462 | 2.6313588 | 2.61098 | 0.020381438 | 2.415768889 |
| Mamstr | 4.071413 | 3.92042 | 4.0802 | -0.159824536 | -0.00883105 | 3.7771496 | 3.9017544 | 3.88137 | 0.020381022 | -0.104223696 |
| Rps13-ps4 | 4.315149 | 4.420536 | 4.3162 | 0.104300149 | -0.00108691 | 4.5224654 | 4.4545782 | 4.43428 | 0.020296035 | 0.088183253 |
| Gm9616 | 2.379503 | 2.312662 | 2.3663 | -0.053588549 | 0.013253146 | 2.282119 | 2.3489393 | 2.32868 | 0.020261676 | -0.046558611 |
| Dlx1 | 1.187565 | 1.222441 | 1.1724 | 0.050002162 | 0.015126683 | 1.1963143 | 1.280638 | 1.26039 | 0.020244007 | -0.064079706 |
| Ttll4 | 9.796365 | 9.843528 | 9.7771 | 0.066386531 | 0.019222869 | 9.6867556 | 9.9317241 | 9.91156 | 0.020162458 | -0.224806041 |
| Podxl2 | 1.460927 | 1.41884 | 1.4198 | -0.00092314 | 0.041164108 | 1.4318083 | 1.4688434 | 1.44868 | 0.020159558 | -0.01687551 |
| Rbmx | 9.703757 | 9.603297 | 9.6819 | -0.078568539 | 0.021892248 | 10.034115 | 9.934605 | 9.91448 | 0.020121161 | 0.119631233 |
| Vmn2r-ps104 | 2.982323 | 3.058875 | 3.1427 | -0.083811757 | -0.160364327 | 3.0759106 | 3.0617005 | 3.04159 | 0.020106233 | 0.034316362 |
| Vmn1r55 | 5.663097 | 5.690997 | 5.7162 | -0.025183942 | -0.053083699 | 5.6909561 | 6.0644959 | 6.0444 | 0.020098173 | -0.353441693 |

| | | | | | | | | | | |
|---|---|---|---|---|---|---|---|---|---|---|
| Eif2s3y | 0.519931 | 0.465384 | 0.5015 | -0.036077767 | 0.018470003 | 0.6341927 | 0.4893836 | 0.46932 | 0.020060091 | 0.164869205 |
| Pfn4 | 1.451609 | 1.38961 | 1.4202 | -0.030581327 | 0.031416972 | 1.3532445 | 1.4437793 | 1.42373 | 0.02005341 | -0.070481419 |
| Pla2g5 | 2.404641 | 2.256605 | 2.2521 | 0.004472556 | 0.15250853 | 2.3042516 | 2.3419399 | 2.32191 | 0.020025569 | -0.017662704 |
| Pcdhb21 | 1.178753 | 1.104137 | 1.1453 | -0.041197941 | 0.033418121 | 1.0781499 | 1.0827983 | 1.06279 | 0.020005135 | 0.015356776 |
| Ddn | 2.292321 | 2.169472 | 2.1056 | 0.063859297 | 0.186708307 | 2.1754826 | 2.1998579 | 2.17992 | 0.019942302 | -0.00443294 |
| Gm12411 | 1.987512 | 1.936438 | 1.9224 | 0.014078042 | 0.065151814 | 1.9873494 | 1.9620967 | 1.94219 | 0.019907012 | 0.045159761 |
| Ntng2 | 7.33828 | 7.289932 | 7.3936 | -0.103623555 | -0.055275603 | 7.4610134 | 7.5601554 | 7.54031 | 0.019841368 | -0.079300677 |
| Rps7-ps2 | 0.149924 | 0.123359 | 0.1639 | -0.040522339 | -0.013957608 | 0.1019255 | 0.1615373 | 0.14171 | 0.019827489 | -0.039784276 |
| Grtp1 | 0.677418 | 0.658391 | 0.6932 | -0.034859163 | -0.015832003 | 0.6203343 | 0.7104725 | 0.69065 | 0.019825602 | -0.070312661 |
| Tbc1d23 | 9.057258 | 8.919255 | 9.1479 | -0.228655021 | -0.090652031 | 9.2640731 | 9.3600784 | 9.34026 | 0.019814109 | -0.076191264 |
| Thrap3 | 12.54811 | 12.52325 | 12.529 | -0.006018474 | 0.018838866 | 12.449726 | 12.716371 | 12.6966 | 0.019788732 | -0.246856516 |
| Slu7 | 10.16863 | 10.1883 | 10.249 | -0.061027844 | -0.080694831 | 10.193207 | 10.25733 | 10.2375 | 0.0197832 | -0.044338878 |
| Nova1 | 4.192303 | 4.14875 | 4.2892 | -0.140492852 | -0.096939921 | 4.446974 | 4.3162395 | 4.29647 | 0.019774313 | 0.150508829 |
| Gm49027 | 1.259761 | 1.198768 | 1.293 | -0.094273683 | -0.033281029 | 1.2606029 | 1.2401063 | 1.22037 | 0.019734813 | 0.040231405 |
| Gm10481 | 1.695122 | 1.71285 | 1.6537 | 0.059184292 | 0.041455682 | 1.7055651 | 1.7170065 | 1.69729 | 0.019721476 | 0.008280032 |
| Zfp54 | 6.375166 | 6.418071 | 6.3616 | 0.056500355 | 0.013595692 | 6.3410707 | 6.4367135 | 6.41699 | 0.019719992 | -0.075922773 |
| Il5 | 0.870533 | 0.861498 | 0.7435 | 0.11795479 | 0.126989288 | 0.7428787 | 0.741144 | 0.72145 | 0.019692868 | 0.02142762 |
| Yipf1 | 9.111194 | 9.08863 | 9.116 | -0.02732634 | -0.00476276 | 9.2978314 | 9.2084021 | 9.18871 | 0.019691392 | 0.109120685 |
| Gm5539 | 0.519377 | 0.499279 | 0.5094 | -0.010133689 | 0.009964474 | 0.5377071 | 0.5361416 | 0.51646 | 0.019680236 | 0.021245765 |
| Hdlbp | 11.79054 | 11.77116 | 11.678 | 0.092725348 | 0.112104748 | 11.648696 | 11.727728 | 11.7081 | 0.019643282 | -0.059389093 |
| Olfr1286 | 1.938574 | 2.028642 | 2.0129 | 0.015759441 | -0.074308136 | 1.985717 | 1.9954194 | 1.97579 | 0.019629804 | 0.009927476 |
| Lgals7 | 1.089588 | 1.134372 | 1.1865 | -0.052121049 | -0.096904641 | 1.1985399 | 1.1429529 | 1.12336 | 0.019595227 | 0.075182262 |
| Slc25a43 | 2.793482 | 2.748297 | 2.846 | -0.097717241 | -0.052532427 | 2.8452679 | 2.8814349 | 2.86191 | 0.019521214 | -0.016645772 |
| Ap2a2 | 10.34382 | 10.26119 | 10.269 | -0.007698622 | 0.074925712 | 10.277492 | 10.141259 | 10.1218 | 0.019507306 | 0.155740327 |
| Rala | 9.455923 | 9.518838 | 9.428 | 0.0907971 | 0.027882739 | 9.371882 | 9.387215 | 9.36776 | 0.019455761 | 0.004122759 |
| Hexdc | 8.440028 | 8.42995 | 8.6046 | -0.174675437 | -0.164596953 | 8.8020475 | 8.7648125 | 8.74536 | 0.019452638 | 0.05668766 |
| Acat2 | 7.981039 | 8.190776 | 8.0526 | 0.138173151 | -0.071563871 | 8.092585 | 8.0474237 | 8.02798 | 0.01944129 | 0.06460256 |
| Hdx | 2.29534 | 2.125729 | 2.2077 | -0.082016043 | 0.087594797 | 2.0877279 | 2.1326731 | 2.11327 | 0.01940269 | -0.025542504 |
| Gabpb1 | 9.385195 | 9.465968 | 9.4496 | 0.016379524 | -0.064393436 | 9.4600271 | 9.4477668 | 9.4284 | 0.019368122 | 0.031628439 |
| Tsga10ip | 1.264608 | 1.216446 | 1.2263 | -0.009867401 | 0.038294023 | 1.2783928 | 1.2107901 | 1.19143 | 0.019356597 | 0.086959326 |
| Epx | 0.547655 | 0.524896 | 0.5494 | -0.024545952 | -0.001787725 | 0.52265 | 0.4832722 | 0.46401 | 0.019265091 | 0.05864294 |
| Snph | 1.399368 | 1.388401 | 1.32 | 0.068391378 | 0.079358522 | 1.4013134 | 1.3614444 | 1.3423 | 0.019145967 | 0.059014907 |

| | | | | | | | | | | |
|---|---|---|---|---|---|---|---|---|---|---|
| Plk3 | 8.637593 | 8.336049 | 8.301 | 0.035095973 | 0.336639887 | 8.3424781 | 8.2186047 | 8.19952 | 0.019088645 | 0.142962123 |
| Gm13136 | 1.091954 | 1.053774 | 1.0748 | -0.020977884 | 0.017202166 | 1.0715196 | 1.1374552 | 1.11842 | 0.019037273 | -0.04689841 |
| Zfp160 | 8.518406 | 8.603376 | 8.5681 | 0.035291886 | -0.049678103 | 8.6616888 | 8.7945248 | 8.7755 | 0.019029383 | -0.113806633 |
| Olfr1274-ps | 1.388108 | 1.423408 | 1.4013 | 0.022116228 | -0.013183858 | 1.3946147 | 1.4852673 | 1.46628 | 0.018991644 | -0.071660969 |
| Flii | 11.61997 | 11.54633 | 11.608 | -0.061408914 | 0.012231588 | 11.814934 | 11.735105 | 11.7161 | 0.01897996 | 0.09880849 |
| Tgm3 | 2.356521 | 2.258466 | 2.3446 | -0.086114011 | 0.011941304 | 2.305419 | 2.3517095 | 2.33274 | 0.018971146 | -0.027319337 |
| Gipc2 | 2.049857 | 1.998993 | 2.019 | -0.019994698 | 0.030869853 | 1.9847675 | 2.0209752 | 2.00202 | 0.018959244 | -0.017248462 |
| Mrgprx2 | 2.009394 | 2.013277 | 2.1789 | -0.165588698 | -0.169472312 | 1.9694751 | 1.9866188 | 1.96769 | 0.018933384 | 0.001789722 |
| Adam4 | 3.54046 | 3.633613 | 3.5372 | 0.096402285 | 0.003249017 | 3.6007265 | 3.7095844 | 3.69069 | 0.018897154 | -0.089960755 |
| Ppl | 4.548631 | 4.546507 | 4.4832 | 0.063352023 | 0.065476144 | 4.2352597 | 4.2913074 | 4.27245 | 0.018857443 | -0.03719029 |
| Lmo1 | 2.885012 | 2.896576 | 2.9088 | -0.012242008 | -0.02380592 | 2.7047252 | 2.8161558 | 2.79734 | 0.018813869 | -0.092616765 |
| Onecut2 | 2.09621 | 2.069711 | 2.0735 | -0.003764168 | 0.022735141 | 2.0873792 | 2.0848651 | 2.06605 | 0.018813793 | 0.021327911 |
| Grhpr | 7.600576 | 7.589271 | 7.6211 | -0.03179939 | -0.020494911 | 7.9831433 | 7.4904546 | 7.47164 | 0.018810974 | 0.511499676 |
| Tmbim1 | 7.965741 | 7.752446 | 7.9171 | -0.16464135 | 0.048653715 | 7.8264639 | 7.5932535 | 7.57447 | 0.018784283 | 0.2519947 |
| Olfr1438-ps1 | 2.85527 | 2.773128 | 2.8857 | -0.112601802 | -0.030459464 | 2.8626672 | 3.0044709 | 2.9857 | 0.018772015 | -0.123031617 |
| Emc1 | 9.429832 | 9.55553 | 9.5123 | 0.043247974 | -0.082449593 | 9.8156198 | 9.6451162 | 9.62638 | 0.018737439 | 0.189241046 |
| Fzd9 | 0.791253 | 0.782605 | 0.7789 | 0.003679199 | 0.012326846 | 0.7670619 | 0.7743832 | 0.75565 | 0.018735813 | 0.011414509 |
| Ostf1 | 11.03974 | 11.08623 | 11 | 0.086350286 | 0.039853748 | 11.029125 | 11.052735 | 11.034 | 0.01871416 | -0.004896313 |
| Senp3 | 10.02224 | 10.07697 | 10.132 | -0.055254397 | -0.109988001 | 10.310338 | 10.425311 | 10.4067 | 0.018638914 | -0.096333891 |
| Zbed3 | 6.838216 | 6.807243 | 6.9683 | -0.161048419 | -0.130075062 | 7.2141317 | 6.9910362 | 6.97243 | 0.018610437 | 0.241705963 |
| Fam185a | 6.724865 | 6.83053 | 6.6146 | 0.215901197 | 0.110236901 | 6.9071146 | 6.9168895 | 6.89828 | 0.018607173 | 0.008832272 |
| Dmbt1 | 2.759843 | 2.726375 | 2.8017 | -0.07533204 | -0.041863501 | 2.5863498 | 2.7068067 | 2.6882 | 0.01860182 | -0.101855059 |
| P2ry2 | 4.415361 | 4.438767 | 4.0136 | 0.425185051 | 0.401779815 | 4.462525 | 3.9570964 | 3.93854 | 0.018555068 | 0.523983653 |
| AF067061 | 1.687424 | 1.646148 | 1.642 | 0.004177211 | 0.045452993 | 1.6371859 | 1.6690261 | 1.65054 | 0.018490249 | -0.013350036 |
| Gm12366 | 2.430286 | 2.473074 | 2.3605 | 0.112533838 | 0.06974664 | 2.5033416 | 2.4917607 | 2.47328 | 0.018481412 | 0.030062376 |
| Rabl2 | 4.084129 | 4.017225 | 3.9104 | 0.106851506 | 0.173755608 | 4.2280993 | 3.9887695 | 3.97034 | 0.01842624 | 0.257756045 |
| Sh2b1 | 10.31193 | 10.29727 | 10.341 | -0.043543698 | -0.028882802 | 10.719406 | 10.638063 | 10.6197 | 0.018364696 | 0.0997075 |
| Tmem170 | 6.939506 | 7.000313 | 6.9313 | 0.068986915 | 0.008179568 | 6.7944017 | 6.8753537 | 6.85704 | 0.018311723 | -0.062640236 |
| 4933434E20Rik | 10.11677 | 10.2549 | 10.139 | 0.116347566 | -0.021776116 | 10.360246 | 10.327037 | 10.3087 | 0.018310126 | 0.051518994 |
| Ppp1r26 | 3.001787 | 2.835762 | 2.9784 | -0.142668217 | 0.023356631 | 2.7701765 | 2.8814556 | 2.86317 | 0.018286486 | -0.092992542 |
| Fcho1 | 10.73295 | 10.72791 | 10.769 | -0.040874602 | -0.035832928 | 11.078483 | 10.947508 | 10.9292 | 0.01825978 | 0.149235148 |
| S100a1 | 5.631948 | 5.654651 | 5.8362 | -0.181582886 | -0.204286294 | 6.0439549 | 5.7253414 | 5.70709 | 0.018252153 | 0.336865621 |

| | | | | | | | | | | |
|---|---|---|---|---|---|---|---|---|---|---|
| Gm10128 | -0.169418 | -0.16911 | -0.1444 | -0.024668675 | -0.024981273 | -0.1998331 | -0.156876 | -0.1751 | 0.018246141 | -0.02471077 |
| 6030458C11Rik | 8.749817 | 8.771213 | 8.7388 | 0.032364356 | 0.010968694 | 8.902039 | 8.9065201 | 8.88833 | 0.01818558 | 0.013704517 |
| Adh6b | 1.776196 | 1.709223 | 1.6518 | 0.057447474 | 0.124420696 | 1.6516597 | 1.6998512 | 1.68167 | 0.018176891 | -0.030014545 |
| Dach2 | 2.132167 | 1.970543 | 2.0417 | -0.071114872 | 0.090508705 | 2.018274 | 2.0480369 | 2.02989 | 0.018142587 | -0.011620341 |
| Usp27x | 4.536053 | 4.406247 | 4.4758 | -0.069539702 | 0.060266627 | 4.1255693 | 4.3529284 | 4.3348 | 0.018125933 | -0.209233117 |
| Pdgfb | 7.239738 | 7.368195 | 7.2211 | 0.147104538 | 0.018648239 | 7.0935467 | 7.1724099 | 7.15434 | 0.018071669 | -0.060791572 |
| Pde6g | 2.384151 | 2.458159 | 2.4016 | 0.056551438 | -0.017456011 | 2.3722753 | 2.5504706 | 2.53244 | 0.018031086 | -0.160164152 |
| Slc25a48 | 0.614912 | 0.550943 | 0.6128 | -0.061849658 | 0.002119038 | 0.5774861 | 0.5853319 | 0.56731 | 0.018018169 | 0.010172284 |
| Timm21 | 7.974196 | 8.072288 | 8.0352 | 0.037100386 | -0.060991826 | 8.0744865 | 8.0099552 | 7.99196 | 0.017994863 | 0.082526253 |
| Olfr448 | 1.743903 | 1.773508 | 1.7076 | 0.065955316 | 0.036349948 | 1.8268209 | 1.8247276 | 1.80675 | 0.017973766 | 0.020067084 |
| Pkd1l2 | 1.977816 | 1.940424 | 1.8819 | 0.058483102 | 0.095875294 | 1.9146399 | 1.9160887 | 1.89813 | 0.017959829 | 0.016511107 |
| Olfr802 | 2.402813 | 2.385535 | 2.3671 | 0.018464377 | 0.035741857 | 2.4160246 | 2.4815346 | 2.46366 | 0.017873233 | -0.047636783 |
| Osbpl9 | 11.25046 | 11.18358 | 11.302 | -0.118844414 | -0.051970284 | 11.28783 | 11.282996 | 11.2652 | 0.017843804 | 0.022677184 |
| Spata48 | 2.860609 | 2.873079 | 2.9002 | -0.027087534 | -0.039557524 | 3.0548123 | 2.8814534 | 2.86364 | 0.017817793 | 0.191176622 |
| Trim59 | 10.30388 | 10.47417 | 10.538 | -0.064036426 | -0.234324116 | 10.3727 | 10.406799 | 10.389 | 0.017808958 | -0.016290412 |
| Stx19 | 0.538749 | 0.552669 | 0.5708 | -0.018084298 | -0.032003641 | 0.5750203 | 0.6280045 | 0.61024 | 0.017760758 | -0.035223452 |
| Gm48349 | 2.585028 | 2.607263 | 2.6037 | 0.003557451 | -0.018677677 | 2.5859703 | 2.6390548 | 2.62133 | 0.017725831 | -0.035358621 |
| Ndst4 | 3.700503 | 3.606382 | 3.7045 | -0.098074063 | -0.003953581 | 3.7073611 | 3.6621977 | 3.6445 | 0.017694372 | 0.062857769 |
| Slc9b1 | 0.852689 | 0.856345 | 0.9142 | -0.057879253 | -0.061535289 | 0.8769744 | 0.9304672 | 0.91278 | 0.017687372 | -0.035805436 |
| Rnd3 | 6.825647 | 6.858385 | 6.7342 | 0.124143774 | 0.09140627 | 6.3743791 | 6.3540478 | 6.33637 | 0.017676119 | 0.038007391 |
| Cfap20 | 8.229833 | 8.06778 | 8.151 | -0.083191036 | 0.078862001 | 8.1724511 | 8.2628481 | 8.24521 | 0.017636175 | -0.072760762 |
| Ncan | 2.883917 | 2.906105 | 2.8649 | 0.041217877 | 0.019029053 | 2.8795816 | 2.8926422 | 2.87505 | 0.017588026 | 0.004527488 |
| Nme7 | 6.816922 | 6.75991 | 6.7827 | -0.022808399 | 0.034203477 | 6.7077526 | 6.9870373 | 6.96949 | 0.0175493 | -0.261735391 |
| 2810459M11Rik | 2.310642 | 2.221102 | 2.2812 | -0.060123255 | 0.029416336 | 2.3163268 | 2.2623983 | 2.24486 | 0.017541029 | 0.071469551 |
| Snx27 | 9.827996 | 10.02928 | 9.8486 | 0.180720196 | -0.020564128 | 9.8756684 | 9.9782049 | 9.96067 | 0.017535717 | -0.085000849 |
| Tbx5 | 1.750241 | 1.822278 | 1.8434 | -0.02115782 | -0.093195427 | 1.8226224 | 1.8635229 | 1.84599 | 0.017533632 | -0.023366891 |
| Dennd6b | 8.581967 | 8.698212 | 8.8417 | -0.143534532 | -0.259779451 | 9.5215584 | 9.2604805 | 9.24308 | 0.01739999 | 0.278477892 |
| Chd1 | 11.14501 | 11.06733 | 11.061 | 0.006712228 | 0.084399518 | 10.720317 | 11.028775 | 11.0114 | 0.017371612 | -0.291085496 |
| Traj49 | 1.803175 | 1.69563 | 1.7515 | -0.055873474 | 0.051671013 | 1.7593619 | 1.76612 | 1.74876 | 0.017357415 | 0.010599366 |
| Akirin2 | 9.097161 | 9.251751 | 9.092 | 0.159793087 | 0.005202647 | 9.0676596 | 9.1841137 | 9.16678 | 0.017338313 | -0.099115773 |
| Wiz | 9.566798 | 9.504296 | 9.5104 | -0.006061959 | 0.056439841 | 9.4966225 | 9.5415562 | 9.52423 | 0.017331113 | -0.027602523 |
| Srp72 | 11.22078 | 11.21071 | 11.209 | 0.001941344 | 0.012012982 | 11.242491 | 11.223104 | 11.2058 | 0.017294498 | 0.036681925 |

| | | | | | | | | | | |
|---|---|---|---|---|---|---|---|---|---|---|
| Gm15920 | 1.092244 | 1.090485 | 1.1331 | -0.042616497 | -0.040856903 | 1.1228902 | 1.1527093 | 1.13543 | 0.017276117 | -0.01254294 |
| Lrch4 | 11.015 | 11.11749 | 11.11 | 0.00747401 | -0.095020076 | 11.419912 | 11.203729 | 11.1865 | 0.017258954 | 0.233441607 |
| Gm6161 | 4.440992 | 4.657257 | 4.5619 | 0.095402558 | -0.120862752 | 4.3195796 | 4.5376389 | 4.52041 | 0.0172288 | -0.200830514 |
| Wfikkn1 | 3.140429 | 2.995389 | 3.0668 | -0.071403093 | 0.073636961 | 3.0192862 | 3.0414565 | 3.02432 | 0.017136368 | -0.005033921 |
| Olfr479 | 0.943276 | 1.005922 | 0.9839 | 0.022070477 | -0.040575949 | 0.9838109 | 0.9845399 | 0.9674 | 0.017135031 | 0.016406049 |
| Unc13c | 0.824933 | 0.865634 | 0.8097 | 0.055908415 | 0.015207672 | 0.8532091 | 0.838402 | 0.82129 | 0.017111657 | 0.031918671 |
| Krr1 | 9.68415 | 9.832966 | 9.6753 | 0.15765702 | 0.00884054 | 9.6720283 | 9.7170087 | 9.69991 | 0.017095143 | -0.027885272 |
| Clec2i | 9.974911 | 9.931596 | 10.08 | -0.148218973 | -0.104904774 | 10.074241 | 10.136378 | 10.1194 | 0.016929804 | -0.045206996 |
| Tmem59l | 0.762843 | 0.823427 | 0.7351 | 0.088282118 | 0.027698273 | 0.740793 | 0.7740704 | 0.75721 | 0.016858128 | -0.016419252 |
| Ccdc84 | 7.987229 | 8.023921 | 8.2547 | -0.230747133 | -0.267438658 | 8.6623636 | 8.5428111 | 8.52597 | 0.016837176 | 0.136389654 |
| Lpcat2b | 1.767201 | 1.805162 | 1.7823 | 0.022865675 | -0.015096121 | 1.7319443 | 1.8221631 | 1.80534 | 0.016819618 | -0.073399199 |
| Zfp617 | 7.805049 | 7.802533 | 7.9975 | -0.194949221 | -0.192434112 | 7.9966934 | 7.8509939 | 7.83418 | 0.016813289 | 0.162512731 |
| Wwc1 | 2.78295 | 2.465451 | 2.587 | -0.121510206 | 0.195989077 | 2.7447252 | 2.5623133 | 2.54555 | 0.016764231 | 0.199176164 |
| Bnc2 | 3.107098 | 3.073029 | 3.06 | 0.013046863 | 0.047115977 | 3.2587887 | 3.0874196 | 3.07081 | 0.0166047 | 0.18797383 |
| Gm15039 | 1.028882 | 1.019302 | 1.0089 | 0.010392139 | 0.01997221 | 1.0804539 | 1.0655297 | 1.04902 | 0.016509845 | 0.03143401 |
| Olfr1316 | 1.981369 | 1.972158 | 2.0323 | -0.060142393 | -0.050932158 | 2.0182389 | 2.0826843 | 2.06625 | 0.016433527 | -0.048011851 |
| Mesd | 9.587889 | 9.516255 | 9.5562 | -0.039924505 | 0.031709193 | 9.4180262 | 9.5029406 | 9.48652 | 0.016425034 | -0.068489381 |
| Lmf1 | 7.676 | 7.714578 | 7.7225 | -0.00796896 | -0.046547023 | 7.8679176 | 7.8172826 | 7.80089 | 0.016392008 | 0.06702699 |
| Igfbp6 | 3.037681 | 3.07704 | 2.9106 | 0.16639266 | 0.12703328 | 3.4706425 | 2.9055188 | 2.88913 | 0.016388029 | 0.581511704 |
| Dpysl2 | 8.820748 | 8.824662 | 8.8469 | -0.022233863 | -0.026147883 | 8.8614105 | 8.8524633 | 8.83608 | 0.016382506 | 0.025329747 |
| Trmt5 | 7.444532 | 7.368243 | 7.2942 | 0.074078282 | 0.150367379 | 7.396834 | 7.576113 | 7.55973 | 0.016381343 | -0.162897609 |
| Zfp575 | 0.810951 | 0.79709 | 0.7753 | 0.021795333 | 0.035656129 | 0.8114927 | 0.8021654 | 0.78581 | 0.016354623 | 0.025681913 |
| Adtrp | 2.196796 | 2.232391 | 2.1348 | 0.097549205 | 0.061954693 | 2.1845514 | 2.1859405 | 2.16959 | 0.016351043 | 0.014961912 |
| Gm48909 | 1.030692 | 0.966845 | 1.0101 | -0.043274986 | 0.020572024 | 0.9902238 | 1.0073426 | 0.99102 | 0.016318959 | -0.000799834 |
| Rab17 | 0.872917 | 0.771576 | 0.8674 | -0.095790902 | 0.005550211 | 0.861582 | 0.8705501 | 0.85424 | 0.016314038 | 0.007345869 |
| Rbm39 | 12.63162 | 12.71788 | 12.686 | 0.031958012 | -0.05430248 | 12.712976 | 12.92341 | 12.9071 | 0.016278332 | -0.194155753 |
| Dhx15 | 12.25025 | 12.32278 | 12.239 | 0.083491794 | 0.010961464 | 12.301455 | 12.353315 | 12.3371 | 0.01624966 | -0.035610196 |
| Surf6 | 9.86162 | 9.817157 | 9.7248 | 0.092391211 | 0.136853989 | 9.9048219 | 9.8986245 | 9.88239 | 0.016232551 | 0.022429969 |
| Gm13245 | 0.415315 | 0.395196 | 0.3967 | -0.001526818 | 0.018593018 | 0.4627608 | 0.4363932 | 0.42018 | 0.016212541 | 0.042580202 |
| Gm6180 | 1.343943 | 1.377672 | 1.3584 | 0.019298046 | -0.01443099 | 1.3835551 | 1.374378 | 1.35818 | 0.016194127 | 0.025371255 |
| Olfr975 | 2.8804 | 2.886149 | 2.9015 | -0.015331894 | -0.021080943 | 2.9594995 | 3.0047027 | 2.98853 | 0.016173228 | -0.029029975 |
| Hoxb6 | 3.172433 | 3.067108 | 3.2164 | -0.149263349 | -0.043938238 | 2.8158753 | 2.9247278 | 2.90859 | 0.01613579 | -0.092716749 |

| | | | | | | | | | | |
|---|---|---|---|---|---|---|---|---|---|---|
| Ebi3 | 7.191022 | 7.265173 | 7.3302 | -0.065007623 | -0.139158338 | 7.2681201 | 7.3110235 | 7.29492 | 0.016107134 | -0.026796277 |
| Sez6 | 2.035569 | 2.001415 | 1.9233 | 0.078106721 | 0.112260525 | 1.9758299 | 1.9687435 | 1.95265 | 0.016090604 | 0.02317699 |
| Clvs1 | 0.509132 | 0.449558 | 0.5425 | -0.092964191 | -0.033390818 | 0.5186833 | 0.4733624 | 0.4573 | 0.016063911 | 0.061384828 |
| Gm7645 | 2.160232 | 2.148132 | 2.1982 | -0.050018837 | -0.037919478 | 2.1336295 | 2.1448789 | 2.12882 | 0.016059587 | 0.00481011 |
| Hspb7 | 1.497193 | 1.563064 | 1.468 | 0.095099012 | 0.029228451 | 1.4025054 | 1.5606686 | 1.54461 | 0.016054078 | -0.142109062 |
| Gm13323 | 3.566601 | 3.541077 | 3.5279 | 0.013127822 | 0.038651926 | 3.6224911 | 3.6882022 | 3.67216 | 0.016044916 | -0.049666224 |
| Abcf3 | 9.953478 | 9.950497 | 9.9705 | -0.019954473 | -0.016973033 | 10.057277 | 10.045078 | 10.0291 | 0.015998977 | 0.028197777 |
| Csnk2a1 | 11.47554 | 11.49635 | 11.5 | -0.003784394 | -0.024593788 | 11.437609 | 11.514618 | 11.4987 | 0.015947412 | -0.061062131 |
| Olfr1463 | 0.708185 | 0.7127 | 0.726 | -0.013291535 | -0.017806044 | 0.667542 | 0.7538273 | 0.73788 | 0.015946066 | -0.070339175 |
| Olfml1 | 2.812996 | 2.698439 | 2.778 | -0.07960548 | 0.034950959 | 2.7140458 | 2.7025903 | 2.68665 | 0.015945185 | 0.027400705 |
| Wdr86 | 4.974233 | 4.821453 | 4.7846 | 0.036823682 | 0.189604204 | 4.7280004 | 4.7770888 | 4.76116 | 0.015929226 | -0.033159113 |
| Tfb1m | 6.97386 | 7.112882 | 7.0591 | 0.053776273 | -0.085245212 | 7.1852979 | 7.1295447 | 7.11362 | 0.015928959 | 0.071682118 |
| Traj52 | 0.397368 | 0.398725 | 0.4255 | -0.026727285 | -0.028084003 | 0.4165937 | 0.4813866 | 0.46546 | 0.01592801 | -0.048864878 |
| Stx1b | 4.239572 | 4.362895 | 4.1333 | 0.22963862 | 0.10631559 | 4.1234845 | 4.2935958 | 4.2777 | 0.015895017 | -0.154216298 |
| Gm8557 | 0.650237 | 0.746192 | 0.6713 | 0.074938779 | -0.021016391 | 0.6835291 | 0.6868108 | 0.67093 | 0.015885569 | 0.012603861 |
| Olfr788 | 0.498224 | 0.496061 | 0.4874 | 0.008648882 | 0.0108124 | 0.5162553 | 0.5333808 | 0.51752 | 0.015865618 | -0.001259832 |
| Tada1 | 9.073273 | 9.127102 | 9.0926 | 0.03453319 | -0.019296412 | 9.176894 | 9.1010532 | 9.08525 | 0.01580733 | 0.091648092 |
| Fam184a | 2.937692 | 2.809921 | 2.7677 | 0.042262649 | 0.170033481 | 2.6432519 | 2.7876646 | 2.77187 | 0.015795056 | -0.128617711 |
| Tnip1 | 10.90286 | 11.02189 | 10.962 | 0.060075318 | -0.058953608 | 10.752214 | 10.917858 | 10.9021 | 0.01578389 | -0.149859666 |
| Msl2 | 11.47548 | 11.54638 | 11.488 | 0.058415572 | -0.012484132 | 11.363543 | 11.505959 | 11.4902 | 0.01577007 | -0.126646469 |
| Gm8556 | 1.802296 | 1.788312 | 1.8232 | -0.034876145 | -0.0208925 | 1.8101846 | 1.8844776 | 1.86872 | 0.015761035 | -0.058532005 |
| Gm6397 | 1.170918 | 1.251411 | 1.2008 | 0.050563942 | -0.029929095 | 1.1723988 | 1.2022367 | 1.1865 | 0.015733871 | -0.014104086 |
| H60b | 2.880266 | 2.966905 | 2.8553 | 0.111592122 | 0.024952564 | 2.7543918 | 2.9335657 | 2.91788 | 0.015689355 | -0.163484588 |
| Gm5791 | 1.897596 | 1.961167 | 1.9352 | 0.026000334 | -0.03757041 | 1.9565041 | 1.9985478 | 1.98286 | 0.015683377 | -0.026360357 |
| Rims4 | 0.850229 | 0.909617 | 0.8399 | 0.069686291 | 0.010298861 | 0.8405917 | 0.8769908 | 0.86131 | 0.015680528 | -0.02071858 |
| Acbd3 | 10.15662 | 10.25237 | 10.16 | 0.092572034 | -0.003180914 | 10.026602 | 10.197324 | 10.1817 | 0.015655036 | -0.155066307 |
| AC148013.1 | 2.473294 | 2.498698 | 2.5375 | -0.038828175 | -0.064231836 | 2.4248417 | 2.5495098 | 2.53386 | 0.015652471 | -0.1090156 |
| Zfp87 | 8.751496 | 8.708254 | 8.758 | -0.049795126 | -0.006552682 | 8.8457607 | 8.6072751 | 8.59169 | 0.015587488 | 0.254073102 |
| Prkag3 | 0.59886 | 0.575601 | 0.5807 | -0.005085097 | 0.018174252 | 0.589611 | 0.6168878 | 0.60132 | 0.015571802 | -0.011705045 |
| Ftcd | 1.73677 | 1.752765 | 1.7229 | 0.029911561 | 0.013916082 | 1.7786279 | 1.7752586 | 1.75974 | 0.015520737 | 0.018890013 |
| Gm20900 | 2.972422 | 3.013855 | 2.8953 | 0.118580838 | 0.077147645 | 3.1300197 | 3.0011051 | 2.9856 | 0.015508549 | 0.144423161 |
| Hsd17b13 | 0.993313 | 1.010313 | 0.98 | 0.030292837 | 0.013292498 | 1.0080218 | 1.0364956 | 1.021 | 0.01549737 | -0.012976402 |

| | | | | | | | | | | |
|---|---|---|---|---|---|---|---|---|---|---|
| Ube2t | 6.004336 | 6.127347 | 6.1749 | -0.047528388 | -0.170539773 | 6.1579816 | 6.3703035 | 6.35481 | 0.015497123 | -0.196824786 |
| Ewsr1 | 12.57815 | 12.55387 | 12.541 | 0.012924596 | 0.037207238 | 12.691789 | 12.851356 | 12.8359 | 0.01549365 | -0.144073235 |
| Umad1 | 7.928344 | 7.859946 | 7.8915 | -0.031528711 | 0.036869718 | 7.8992073 | 7.8976618 | 7.88222 | 0.015440523 | 0.016986055 |
| Fip1l1 | 10.74326 | 10.83026 | 10.791 | 0.039391274 | -0.047613874 | 10.822585 | 10.843861 | 10.8284 | 0.015420576 | -0.005854904 |
| Snapc3 | 8.010254 | 8.084081 | 8.0598 | 0.024242766 | -0.049584576 | 8.3435657 | 8.30231 | 8.2869 | 0.015410312 | 0.056666007 |
| Gm15645 | 1.922274 | 1.973887 | 1.9606 | 0.013250541 | -0.038363337 | 2.0416907 | 1.9641052 | 1.9487 | 0.015409673 | 0.092995231 |
| Greb1l | 0.918494 | 0.898316 | 0.8918 | 0.00655098 | 0.026728711 | 0.8444143 | 0.9073269 | 0.89196 | 0.015367899 | -0.047544649 |
| Gm6039 | 0.039341 | 0.026702 | 0.0664 | -0.03973581 | -0.027096714 | -0.0071753 | 0.0358786 | 0.02052 | 0.015356442 | -0.027697399 |
| Ighv8-4 | 1.669461 | 1.766108 | 1.5741 | 0.192054115 | 0.095406451 | 1.7204705 | 1.5709119 | 1.5556 | 0.015310471 | 0.16486903 |
| Gm3953 | 1.921267 | 1.914651 | 1.9372 | -0.022528411 | -0.015912266 | 2.0052918 | 1.9497579 | 1.93449 | 0.015266685 | 0.070800579 |
| Fa2h | 1.574613 | 1.557567 | 1.6694 | -0.111855024 | -0.094808973 | 1.78334 | 1.6986184 | 1.68336 | 0.015257868 | 0.099979469 |
| Exd1 | 3.83009 | 3.8678 | 3.7888 | 0.079010211 | 0.041300281 | 4.2114582 | 3.9183533 | 3.90313 | 0.015219686 | 0.308324595 |
| Amotl2 | 6.137535 | 6.174259 | 5.9612 | 0.213045723 | 0.176321559 | 5.8043949 | 5.9998816 | 5.98468 | 0.01520231 | -0.180284356 |
| Olfr1129 | 1.281285 | 1.271146 | 1.2907 | -0.019568993 | -0.009429233 | 1.3060593 | 1.3314438 | 1.31626 | 0.015182897 | -0.010201646 |
| Olfr1410 | 1.802184 | 1.752721 | 1.7529 | -0.000186711 | 0.049276836 | 1.8251677 | 1.8709018 | 1.85572 | 0.015180768 | -0.030553356 |
| Dnase2b | 0.785133 | 0.778249 | 0.7848 | -0.006592624 | 0.000291561 | 0.7663721 | 0.7717813 | 0.75664 | 0.01514313 | 0.00973397 |
| Svopl | 1.234539 | 1.168094 | 1.2945 | -0.126381421 | -0.059936673 | 1.1850986 | 1.2297429 | 1.21463 | 0.015111745 | -0.029532602 |
| Smtnl2 | 1.989458 | 2.124568 | 2.0037 | 0.12085258 | -0.014257353 | 2.0602286 | 2.0357233 | 2.02064 | 0.015086744 | 0.039592029 |
| Hsf5 | 0.311652 | 0.26994 | 0.3052 | -0.035273786 | 0.006438217 | 0.3011048 | 0.2674712 | 0.25249 | 0.014980406 | 0.048613996 |
| Gm9770 | 0.474167 | 0.450135 | 0.4898 | -0.039618302 | -0.015586441 | 0.4429755 | 0.4830706 | 0.46809 | 0.014980048 | -0.025115065 |
| Znf41-ps | 8.169588 | 8.159197 | 8.2506 | -0.091390382 | -0.080999448 | 8.1492818 | 8.176282 | 8.16134 | 0.014945668 | -0.012054532 |
| Smgc | 2.705384 | 2.028474 | 1.9167 | 0.111731409 | 0.788641881 | 2.419445 | 1.9904444 | 1.97551 | 0.014939229 | 0.443939818 |
| Cnga3 | 3.804236 | 3.795067 | 3.7046 | 0.090425568 | 0.099594838 | 3.8743188 | 3.9336552 | 3.91874 | 0.014914237 | -0.044422192 |
| Vmn2r56 | 3.557884 | 3.549559 | 3.5383 | 0.011255691 | 0.019580253 | 3.6367021 | 3.7718371 | 3.75694 | 0.014900732 | -0.120234271 |
| Gm13135 | 0.552973 | 0.558176 | 0.5685 | -0.010285598 | -0.015488248 | 0.564636 | 0.6001608 | 0.5853 | 0.014862737 | -0.02066201 |
| Dctd | 6.233855 | 6.157543 | 6.0209 | 0.136638538 | 0.212949876 | 6.2368845 | 6.2669331 | 6.2521 | 0.014833712 | -0.015214884 |
| Wdr45b | 9.100893 | 9.169308 | 9.1252 | 0.044091128 | -0.024324098 | 9.1436719 | 9.1022478 | 9.08755 | 0.01469441 | 0.056118498 |
| Gm7299 | 1.379325 | 1.377119 | 1.4149 | -0.037737058 | -0.03553066 | 1.3618875 | 1.3840936 | 1.36945 | 0.014646603 | -0.007559492 |
| Gm11510 | -0.00173 | -0.02933 | 0.0152 | -0.044505394 | -0.016902788 | -0.0316709 | 0.0155624 | 0.00093 | 0.014633553 | -0.032599781 |
| Hnrnpu | 12.37673 | 12.33034 | 12.342 | -0.011447636 | 0.034937652 | 12.397376 | 12.503134 | 12.4885 | 0.014584948 | -0.091172552 |
| Pdcd6ip | 11.77943 | 11.7739 | 11.716 | 0.058094882 | 0.063620572 | 11.603198 | 11.749641 | 11.7351 | 0.014508994 | -0.131933822 |
| Ddhd1 | 9.189682 | 9.41854 | 9.2918 | 0.126701557 | -0.102155774 | 9.1423707 | 9.5109032 | 9.49651 | 0.014388289 | -0.354144245 |

| | | | | | | | | | | |
|---|---|---|---|---|---|---|---|---|---|---|
| Lsmem1 | 3.20812 | 3.150569 | 3.2281 | -0.077501866 | -0.019951144 | 3.160282 | 3.2118332 | 3.19751 | 0.014323542 | -0.037227687 |
| Smg1 | 13.29025 | 13.34227 | 13.237 | 0.105430146 | 0.053412696 | 13.170622 | 13.525877 | 13.5116 | 0.014269474 | -0.340985596 |
| Gm10357 | 1.461256 | 1.366495 | 1.3831 | -0.016631101 | 0.07812925 | 1.4305133 | 1.4114108 | 1.39717 | 0.014245778 | 0.033348255 |
| Gm2606 | 1.192367 | 1.262998 | 1.1198 | 0.143165843 | 0.072534139 | 1.4945013 | 1.1802579 | 1.16612 | 0.01413986 | 0.32838323 |
| Wdr54 | 1.480484 | 1.405562 | 1.4345 | -0.028895032 | 0.046027317 | 1.4823498 | 1.5300551 | 1.51593 | 0.014126135 | -0.033579123 |
| Hspg2 | 9.862342 | 9.587239 | 9.7179 | -0.130663991 | 0.144439036 | 9.2090492 | 9.2296384 | 9.21551 | 0.014125226 | -0.006463924 |
| Vmn1r50 | 2.785959 | 2.808913 | 2.8535 | -0.044597863 | -0.067552496 | 2.9363728 | 2.9260885 | 2.91199 | 0.014102647 | 0.02438696 |
| Rabl3 | 7.237218 | 7.148658 | 7.1821 | -0.033437304 | 0.055122825 | 7.5590112 | 7.3549617 | 7.34087 | 0.014094511 | 0.218144044 |
| Prrx2 | 0.914268 | 0.874166 | 0.881 | -0.006839431 | 0.033263363 | 0.8662139 | 0.8528005 | 0.83872 | 0.014077579 | 0.027491006 |
| Gpr12 | 0.395532 | 0.364391 | 0.3903 | -0.025913485 | 0.005227183 | 0.4490107 | 0.4125899 | 0.39855 | 0.014037884 | 0.0504587 |
| E2f1 | 6.878723 | 6.755294 | 7.0775 | -0.322176682 | -0.19874688 | 7.0834748 | 7.1221102 | 7.10808 | 0.014032568 | -0.024602827 |
| Grik3 | 1.937959 | 1.931941 | 1.8905 | 0.041439475 | 0.04745701 | 1.9420597 | 1.9168905 | 1.90292 | 0.013965973 | 0.039135152 |
| Gm10775 | 1.127926 | 1.130144 | 1.1099 | 0.020195985 | 0.017978027 | 1.0939496 | 1.1302418 | 1.11633 | 0.013907668 | -0.022384541 |
| Trim52 | 1.489654 | 1.493971 | 1.4644 | 0.029605012 | 0.025287871 | 1.4148304 | 1.47491 | 1.46101 | 0.013899753 | -0.046179861 |
| Pou1f1 | 0.641727 | 0.646138 | 0.7166 | -0.070416933 | -0.074827743 | 0.6840231 | 0.7147475 | 0.70091 | 0.013840676 | -0.016883744 |
| Hoxb1 | 2.25677 | 2.299513 | 2.3418 | -0.042309865 | -0.085052677 | 2.4284905 | 2.3638158 | 2.34999 | 0.013827877 | 0.078502546 |
| Smpd4 | 9.19672 | 9.180184 | 9.2547 | -0.074528391 | -0.057992034 | 9.5241402 | 9.4105358 | 9.39672 | 0.013820648 | 0.127425023 |
| Olfr1318 | 1.873877 | 1.904597 | 1.8159 | 0.088660519 | 0.057939922 | 1.8833061 | 1.9560465 | 1.94223 | 0.013815926 | -0.058924471 |
| Kcng4 | 1.365745 | 1.271617 | 1.3598 | -0.088188556 | 0.005939184 | 1.2475085 | 1.2438547 | 1.2301 | 0.013751422 | 0.017405259 |
| Clnk | 4.442814 | 4.441672 | 4.6108 | -0.169116279 | -0.167974678 | 4.1988246 | 4.4548848 | 4.44113 | 0.013751332 | -0.242308878 |
| Adgrg6 | 6.788043 | 6.672428 | 6.592 | 0.08045917 | 0.196074654 | 6.4424947 | 6.7049154 | 6.69118 | 0.013740418 | -0.24868029 |
| Spin1 | 10.95728 | 11.03607 | 11.01 | 0.026005588 | -0.052780842 | 10.871331 | 10.961048 | 10.9474 | 0.013640974 | -0.076076831 |
| Zfp148-ps1 | 0.003987 | -0.00585 | -0.0031 | -0.002716184 | 0.007119712 | 0.0130452 | 0.0164969 | 0.00291 | 0.013585279 | 0.010133601 |
| Vmn2r30 | 1.986938 | 1.937166 | 1.9608 | -0.023584754 | 0.026187848 | 1.9984372 | 1.9688252 | 1.95524 | 0.013581407 | 0.043193433 |
| Rhox4b | 0.707633 | 0.792936 | 0.7759 | 0.017000841 | -0.068301725 | 0.6795765 | 0.7685104 | 0.75493 | 0.01357879 | -0.075355094 |
| Zfp990 | 2.074143 | 2.029651 | 2.0161 | 0.013554013 | 0.058046819 | 2.0877042 | 2.0950706 | 2.08152 | 0.013549826 | 0.006183426 |
| Gm6939 | 1.24925 | 1.233801 | 1.3178 | -0.084045527 | -0.068596835 | 1.2671797 | 1.2724035 | 1.25888 | 0.013520678 | 0.008296884 |
| Olfr769 | 4.793884 | 4.875126 | 4.9446 | -0.069500006 | -0.150741359 | 4.99896 | 5.1416129 | 5.12814 | 0.0134712 | -0.129181647 |
| Olfr1444 | 1.180065 | 1.220176 | 1.2102 | 0.009937696 | -0.030173476 | 1.2816285 | 1.2732543 | 1.25979 | 0.013466929 | 0.021841139 |
| Olfr1220 | 2.432989 | 2.4311 | 2.4772 | -0.0460667 | -0.044177728 | 2.3942784 | 2.5067724 | 2.49332 | 0.013455293 | -0.099038775 |
| Rab35 | 10.37999 | 10.44083 | 10.386 | 0.05483921 | -0.005996762 | 10.346308 | 10.320446 | 10.307 | 0.013432534 | 0.039294998 |
| Vmn2r29 | 1.76657 | 1.708558 | 1.6372 | 0.071399728 | 0.129412006 | 1.7454888 | 1.8054836 | 1.79206 | 0.013420191 | -0.046574681 |

| | | | | | | | | | | |
|---|---|---|---|---|---|---|---|---|---|---|
| Stx2 | 8.349657 | 8.362825 | 8.3566 | 0.00625582 | -0.006911886 | 8.1221787 | 8.2622653 | 8.24886 | 0.013406602 | -0.126680091 |
| Anks3 | 8.910923 | 8.979192 | 9.072 | -0.092848045 | -0.161116343 | 9.4289378 | 9.3385714 | 9.32517 | 0.013401589 | 0.10376799 |
| Cyfip2 | 12.83147 | 12.82454 | 12.824 | 0.000448782 | 0.007380114 | 12.940817 | 12.879358 | 12.866 | 0.013396076 | 0.074854381 |
| Napb | 4.228516 | 4.303154 | 4.1914 | 0.111793444 | 0.037155322 | 4.0881627 | 4.1496275 | 4.13625 | 0.013380063 | -0.048084705 |
| Cdh26 | 0.695185 | 0.620626 | 0.6407 | -0.020091183 | 0.054467555 | 0.6052662 | 0.6348046 | 0.62144 | 0.013361668 | -0.016176725 |
| Rbm12b2 | 8.898678 | 9.12286 | 8.9354 | 0.187460287 | -0.036722008 | 9.2800136 | 9.2372464 | 9.2239 | 0.01334884 | 0.056116042 |
| Gm12895 | 0.415938 | 0.385289 | 0.4413 | -0.056038578 | -0.025389759 | 0.4827736 | 0.4061829 | 0.39283 | 0.013347985 | 0.089938722 |
| Gm8464 | 1.043218 | 1.111025 | 1.1056 | 0.005468591 | -0.06233804 | 1.092467 | 1.0881133 | 1.07478 | 0.013335643 | 0.017689294 |
| Gm4750 | 1.238727 | 1.309958 | 1.3141 | -0.004109905 | -0.07534052 | 1.300771 | 1.2858059 | 1.27247 | 0.013334868 | 0.02829996 |
| Olfr338 | 3.593589 | 3.731663 | 3.7242 | 0.007475601 | -0.130598095 | 3.7819079 | 3.8121224 | 3.7988 | 0.013318515 | -0.016895935 |
| Ighm | 15.67506 | 15.85211 | 15.703 | 0.14892144 | -0.02813224 | 15.86359 | 15.661757 | 15.6485 | 0.013303724 | 0.215136914 |
| Prss34 | 0.631044 | 0.560677 | 0.6651 | -0.104446511 | -0.034079719 | 0.5456391 | 0.5759511 | 0.56265 | 0.01330006 | -0.017011905 |
| Etv3 | 11.37921 | 11.51275 | 11.351 | 0.162229782 | 0.02868756 | 11.164421 | 11.434494 | 11.4212 | 0.01325613 | -0.256816424 |
| Tiprl | 9.369934 | 9.327714 | 9.2827 | 0.045038387 | 0.087257817 | 9.3507735 | 9.4334154 | 9.42018 | 0.013238661 | -0.069403326 |
| Gm10025 | 2.625569 | 2.678922 | 2.7364 | -0.057473803 | -0.110826795 | 2.6991887 | 2.7326702 | 2.71946 | 0.013211052 | -0.020270515 |
| Gosr2 | 11.02847 | 10.99498 | 10.931 | 0.06415507 | 0.097643196 | 10.973731 | 10.926523 | 10.9133 | 0.01320667 | 0.060413986 |
| Dcstamp | 2.64568 | 2.91869 | 2.6553 | 0.263361702 | -0.009648054 | 2.6244472 | 2.6941684 | 2.68097 | 0.013194844 | -0.05652629 |
| Wdr31 | 5.190951 | 5.228404 | 5.179 | 0.049395566 | 0.011942128 | 5.1529721 | 5.2102656 | 5.19709 | 0.013178959 | -0.04411456 |
| Sorcs1 | 1.889136 | 1.90531 | 1.8691 | 0.036208044 | 0.02003403 | 1.874407 | 1.8928883 | 1.87974 | 0.01315059 | -0.005330725 |
| 1700028K03Rik | 1.552967 | 1.455796 | 1.4669 | -0.011065353 | 0.086105994 | 1.5788191 | 1.4751155 | 1.4621 | 0.013018496 | 0.116722014 |
| Fam89b | 8.079111 | 8.130257 | 8.2144 | -0.08412269 | -0.135268373 | 8.3707002 | 8.0637532 | 8.05074 | 0.013016411 | 0.319963426 |
| Armc12 | 0.402689 | 0.435343 | 0.4375 | -0.00216242 | -0.034815947 | 0.4755499 | 0.4809795 | 0.468 | 0.012978161 | 0.007548531 |
| Cym | 1.355057 | 1.296661 | 1.3494 | -0.052780348 | 0.005615939 | 1.2835003 | 1.2914485 | 1.27848 | 0.012971795 | 0.005023628 |
| Gpt | 6.078629 | 6.02481 | 6.1789 | -0.154102139 | -0.100283081 | 6.372721 | 6.2005693 | 6.18761 | 0.012956975 | 0.185108608 |
| Hal | 2.32372 | 2.257341 | 2.1817 | 0.075646683 | 0.1420255 | 2.3047679 | 2.2529521 | 2.24 | 0.01294849 | 0.064764355 |
| Dusp3 | 8.789987 | 8.538852 | 8.4387 | 0.100143029 | 0.351278578 | 8.4263791 | 8.1761109 | 8.16329 | 0.01282443 | 0.26309267 |
| Plch1 | 1.827016 | 1.871382 | 1.7764 | 0.094945303 | 0.050579903 | 1.8111549 | 1.7066854 | 1.69389 | 0.012795979 | 0.117265468 |
| Nyap2 | 0.655208 | 0.659639 | 0.6185 | 0.04111071 | 0.036679838 | 0.6433055 | 0.6848909 | 0.67219 | 0.012697159 | -0.028888154 |
| Rab25 | 2.142935 | 2.16334 | 2.1733 | -0.009925421 | -0.030330188 | 2.2757373 | 2.1501462 | 2.13746 | 0.01268709 | 0.138278267 |
| Acmsd | 0.673137 | 0.655485 | 0.6952 | -0.039742495 | -0.022090387 | 0.6696453 | 0.6829722 | 0.67032 | 0.012652541 | -0.000674386 |
| Gm48417 | 5.118873 | 5.514885 | 5.2498 | 0.265060769 | -0.13095075 | 4.9660908 | 5.9089173 | 5.89627 | 0.012644396 | -0.930182069 |
| Gm10132 | 3.302037 | 3.435691 | 3.2153 | 0.220365843 | 0.086711586 | 3.0193698 | 3.3076008 | 3.29496 | 0.012642236 | -0.275588737 |

| | | | | | | | | | | |
|---|---|---|---|---|---|---|---|---|---|---|
| Cnih2 | 2.425648 | 2.347294 | 2.2461 | 0.101185408 | 0.179539154 | 2.3135209 | 2.3463832 | 2.33375 | 0.012633418 | -0.020228909 |
| Gm14269 | 0.656478 | 0.692401 | 0.661 | 0.031425833 | -0.004497444 | 0.6939508 | 0.7472648 | 0.73466 | 0.012600797 | -0.040713214 |
| Gm12816 | 2.689091 | 2.800117 | 2.7136 | 0.086537781 | -0.024487692 | 2.7640049 | 2.7465058 | 2.73391 | 0.012597604 | 0.03009673 |
| Gm15454 | 0.952953 | 0.973277 | 0.9781 | -0.004843762 | -0.025167868 | 1.0120518 | 0.9003243 | 0.88775 | 0.012569324 | 0.124296822 |
| Lrrc7 | 2.11758 | 2.082647 | 2.0923 | -0.009643854 | 0.025289554 | 2.116912 | 2.1618797 | 2.14934 | 0.012537194 | -0.03243055 |
| Vmn2r1 | 2.687589 | 2.58127 | 2.5313 | 0.049997538 | 0.156316649 | 2.560056 | 2.595295 | 2.58282 | 0.012477052 | -0.022761949 |
| Gm49527 | 0.510461 | 0.61227 | 0.5694 | 0.042855398 | -0.058952958 | 0.4919933 | 0.5537457 | 0.54127 | 0.01247542 | -0.049277053 |
| 1700029I15Rik | 3.902791 | 3.949757 | 4.1164 | -0.166606108 | -0.213572156 | 4.0623638 | 4.0774741 | 4.06506 | 0.012413908 | -0.002696393 |
| Olfr177 | 0.933374 | 0.968605 | 0.9392 | 0.029359544 | -0.005871782 | 0.9573949 | 0.958442 | 0.94604 | 0.012406346 | 0.011359294 |
| Stac3 | 4.469962 | 4.327189 | 4.5493 | -0.222069321 | -0.079296229 | 4.765391 | 4.7267088 | 4.71434 | 0.012368009 | 0.051050144 |
| Champ1 | 9.169962 | 9.171845 | 9.2149 | -0.043030907 | -0.044913656 | 9.1587721 | 9.1418619 | 9.12953 | 0.012336402 | 0.029246644 |
| Cdkn2d | 9.696994 | 9.258516 | 9.3507 | -0.092233802 | 0.346244225 | 9.4513365 | 9.152949 | 9.14066 | 0.01228544 | 0.310672953 |
| Trav13-4-dv7 | 4.367619 | 4.397234 | 4.3658 | 0.031410751 | 0.001794859 | 4.2822228 | 4.4407251 | 4.42844 | 0.01228466 | -0.146217641 |
| Rpl18a-ps1 | 0.254063 | 0.26905 | 0.2696 | -0.000518424 | -0.015505457 | 0.2820641 | 0.307966 | 0.29574 | 0.012224538 | -0.013677337 |
| Gm48930 | 1.289446 | 1.266808 | 1.3101 | -0.043321133 | -0.020683664 | 1.4365551 | 1.3258723 | 1.31365 | 0.012222944 | 0.122905749 |
| Ddi2 | 10.77089 | 10.7154 | 10.651 | 0.064773732 | 0.120263822 | 10.657961 | 10.741074 | 10.7289 | 0.012222124 | -0.070890895 |
| Tsen15 | 6.924087 | 6.857619 | 6.9297 | -0.072081456 | -0.005613313 | 6.9689073 | 6.9734982 | 6.9613 | 0.012193409 | 0.007602447 |
| Ankef1 | 3.41432 | 3.558095 | 3.5346 | 0.023463888 | -0.120311148 | 3.4873743 | 3.6536765 | 3.6415 | 0.012171507 | -0.154130654 |
| Gm17137 | 0.339919 | 0.362921 | 0.3683 | -0.005339694 | -0.028341749 | 0.3446705 | 0.3715431 | 0.3594 | 0.012144032 | -0.014728578 |
| Xlr4b | 8.624337 | 8.607722 | 8.6974 | -0.089658485 | -0.073043482 | 8.6546405 | 8.6590192 | 8.6469 | 0.012117532 | 0.007738838 |
| Oosp1 | 2.973211 | 3.139585 | 3.1028 | 0.036770215 | -0.129604087 | 3.1763587 | 3.0329257 | 3.02081 | 0.012113233 | 0.155546251 |
| Gm12231 | 0.197172 | 0.177386 | 0.218 | -0.040626262 | -0.020840507 | 0.2037912 | 0.2010532 | 0.18903 | 0.012022599 | 0.01476057 |
| Cdkl4 | 3.864979 | 3.922681 | 3.9668 | -0.044075107 | -0.101777472 | 3.9296047 | 3.9616378 | 3.94968 | 0.011954201 | -0.020078895 |
| Gm49336 | 10.4253 | 10.40124 | 10.322 | 0.079006784 | 0.103065992 | 10.460139 | 10.431387 | 10.4195 | 0.01192758 | 0.040680369 |
| Rlf | 10.21598 | 10.17306 | 10.239 | -0.066106843 | -0.023188823 | 10.00058 | 10.181642 | 10.1697 | 0.011919759 | -0.169142339 |
| Ap1g1 | 10.80734 | 10.79023 | 10.813 | -0.022885414 | -0.00577591 | 10.745942 | 10.88697 | 10.8751 | 0.011892706 | -0.129135458 |
| Piwil4 | 2.535961 | 2.480233 | 2.4862 | -0.005938383 | 0.049790556 | 2.5081484 | 2.4788152 | 2.46695 | 0.011866995 | 0.04120018 |
| Pglyrp2 | 10.01981 | 10.03968 | 9.9845 | 0.055199278 | 0.035333724 | 10.114031 | 10.009112 | 9.99725 | 0.011857632 | 0.11677668 |
| 9130019O22Rik | 6.921301 | 6.823995 | 6.9875 | -0.163536154 | -0.066230431 | 6.9894871 | 6.9255421 | 6.91376 | 0.011782788 | 0.075727775 |
| Bco1 | 1.233453 | 1.218315 | 1.276 | -0.057689545 | -0.042551768 | 1.2875452 | 1.314347 | 1.30259 | 0.011755437 | -0.015046346 |
| Naip3-ps1 | 1.170034 | 1.160683 | 1.217 | -0.056359736 | -0.047009042 | 1.1580964 | 1.1756857 | 1.16394 | 0.011747775 | -0.005841475 |
| Ppt1 | 10.93702 | 10.65791 | 10.865 | -0.207147184 | 0.0719675 | 10.993452 | 10.816875 | 10.8052 | 0.011684576 | 0.188261657 |

| | | | | | | | | | | |
|---|---|---|---|---|---|---|---|---|---|---|
| Gm16216 | 1.668872 | 1.648225 | 1.6608 | -0.012618526 | 0.008028204 | 1.660423 | 1.7097618 | 1.69814 | 0.01162139 | -0.037717413 |
| Gm3468 | 3.385313 | 3.44787 | 3.4836 | -0.03572734 | -0.098284195 | 3.3945678 | 3.519544 | 3.50793 | 0.011617073 | -0.11335914 |
| Rpl7-ps7 | 2.212951 | 2.334566 | 2.2769 | 0.057666848 | -0.063948046 | 2.409243 | 2.3396894 | 2.32808 | 0.011609226 | 0.081162782 |
| Gm9385 | 1.580515 | 1.498157 | 1.5516 | -0.053477064 | 0.028881138 | 1.428369 | 1.4897122 | 1.47814 | 0.011575835 | -0.049767388 |
| Smarcal1 | 9.246497 | 9.232649 | 9.2364 | -0.003797725 | 0.010049625 | 9.4509409 | 9.3109575 | 9.29938 | 0.011574982 | 0.151558337 |
| Crybb3 | 1.594308 | 1.559303 | 1.601 | -0.041693136 | -0.006687234 | 1.5676511 | 1.588108 | 1.57654 | 0.011570209 | -0.008886676 |
| Sumf2 | 7.657496 | 7.665966 | 7.6553 | 0.010657913 | 0.002187918 | 7.7358843 | 7.6107563 | 7.59928 | 0.011472927 | 0.136600943 |
| Ighj1 | 6.466092 | 6.769319 | 6.7365 | 0.032806107 | -0.27042049 | 6.7695989 | 6.5949563 | 6.58349 | 0.011466112 | 0.186108759 |
| Hpn | 3.43203 | 3.621494 | 3.3994 | 0.222120311 | 0.032656404 | 3.6749246 | 3.4941517 | 3.4827 | 0.011452094 | 0.192224941 |
| Pdzd3 | 0.670233 | 0.660155 | 0.5882 | 0.071923981 | 0.082001127 | 0.5992452 | 0.6037664 | 0.59234 | 0.011422699 | 0.006901521 |
| Ndor1 | 9.809853 | 9.984145 | 10.032 | -0.047726815 | -0.222019455 | 10.422325 | 10.365691 | 10.3543 | 0.011422332 | 0.068055909 |
| Gm9025 | 0.797897 | 0.799341 | 0.8694 | -0.070050056 | -0.071493969 | 0.7990072 | 0.7750766 | 0.76368 | 0.011392329 | 0.035322936 |
| H2-T3 | 0.412327 | 0.352128 | 0.3404 | 0.011683162 | 0.071882524 | 0.3529909 | 0.3294223 | 0.31806 | 0.01136463 | 0.034933231 |
| Rab27a | 10.09332 | 10.05218 | 10.045 | 0.007556376 | 0.0486999 | 9.6219901 | 9.8130754 | 9.80172 | 0.011351461 | -0.179733897 |
| Gm8258 | 0.416412 | 0.458285 | 0.4622 | -0.003918458 | -0.045791166 | 0.3899536 | 0.4570964 | 0.44578 | 0.01132122 | -0.055821509 |
| Gm6853 | 1.577144 | 1.585534 | 1.5506 | 0.034949737 | 0.026560403 | 1.5759821 | 1.651098 | 1.63986 | 0.011242464 | -0.063873492 |
| Gm20763 | 1.241135 | 1.229772 | 1.2199 | 0.009841562 | 0.021204217 | 1.1903964 | 1.2455632 | 1.2344 | 0.011164728 | -0.044002088 |
| Stmn3 | 0.891717 | 0.951319 | 0.8397 | 0.111574924 | 0.051973435 | 0.927847 | 0.8720115 | 0.86089 | 0.011121483 | 0.066956989 |
| Gm29036 | 2.406821 | 2.449205 | 2.4382 | 0.011031699 | -0.031352809 | 2.442859 | 2.4265124 | 2.41541 | 0.011106484 | 0.027453162 |
| Trim29 | 0.683138 | 0.627987 | 0.6441 | -0.016064739 | 0.039087094 | 0.7238885 | 0.6529357 | 0.64183 | 0.011104407 | 0.082057186 |
| Hspa1l | 1.884742 | 1.848253 | 1.8777 | -0.029417696 | 0.007071852 | 1.7869968 | 1.8357552 | 1.82465 | 0.011102719 | -0.037655718 |
| Otud4 | 11.29668 | 11.32502 | 11.249 | 0.075647292 | 0.047314136 | 11.267162 | 11.418462 | 11.4074 | 0.011063096 | -0.140237346 |
| Zfp978 | -0.075246 | -0.00084 | -0.0382 | 0.037366831 | -0.037042449 | -0.0847079 | -0.065939 | -0.077 | 0.01104045 | -0.007728742 |
| 2610042L04Rik | 0.489914 | 0.492755 | 0.5004 | -0.007631249 | -0.010472236 | 0.4725979 | 0.5610391 | 0.55001 | 0.011025554 | -0.077415691 |
| Gm10224 | 2.23181 | 2.188915 | 2.1494 | 0.039501028 | 0.082396011 | 2.2099498 | 2.1500942 | 2.13912 | 0.010971025 | 0.070826614 |
| Spx | 1.929485 | 2.014733 | 1.9969 | 0.017818346 | -0.067429039 | 2.0231433 | 2.1188607 | 2.1079 | 0.010958075 | -0.084759251 |
| Ccdc150 | 1.251692 | 1.215877 | 1.3061 | -0.09023742 | -0.054422837 | 1.1741648 | 1.2253525 | 1.21441 | 0.010940864 | -0.040246827 |
| Ooep | 0.723792 | 0.70946 | 0.6914 | 0.018101164 | 0.032433627 | 0.7186587 | 0.7116649 | 0.70079 | 0.010870156 | 0.017864034 |
| Tigar | 7.878498 | 7.997421 | 7.6852 | 0.312199986 | 0.193277084 | 7.9986424 | 7.9664243 | 7.95558 | 0.010843781 | 0.043061866 |
| Traf4 | 9.165894 | 9.373986 | 9.1403 | 0.233667086 | 0.025574149 | 9.4569045 | 9.3177264 | 9.3069 | 0.010828774 | 0.150006887 |
| Gm5566 | 0.395499 | 0.371259 | 0.3581 | 0.013111728 | 0.03735213 | 0.3568221 | 0.4244657 | 0.41367 | 0.010797037 | -0.056846516 |
| Olfr1381 | 1.015023 | 0.998273 | 1.074 | -0.075737387 | -0.058987003 | 0.9944256 | 1.0656244 | 1.05484 | 0.010782147 | -0.0604167 |

| | | | | | | | | | | |
|---|---|---|---|---|---|---|---|---|---|---|
| Hykk | 5.550436 | 5.113675 | 5.3143 | -0.200613844 | 0.236147818 | 5.5124355 | 5.3655361 | 5.35477 | 0.010770147 | 0.157669518 |
| Gm42785 | 5.985502 | 5.913045 | 6.0938 | -0.180723408 | -0.108265927 | 6.0226007 | 6.2815196 | 6.27084 | 0.010681679 | -0.248237171 |
| Zbtb8a | 6.455096 | 6.377056 | 6.5426 | -0.165540476 | -0.087500323 | 6.6278688 | 6.5567932 | 6.54616 | 0.010633123 | 0.081708718 |
| Gm45162 | 0.162117 | 0.213609 | 0.2169 | -0.003255066 | -0.054747353 | 0.1911207 | 0.1646144 | 0.15404 | 0.010571804 | 0.037078147 |
| Uba1y | 1.706416 | 1.650313 | 1.6959 | -0.045588455 | 0.010514394 | 1.6988355 | 1.7698511 | 1.7593 | 0.010547257 | -0.06046826 |
| Ttc8 | 4.695849 | 4.567887 | 4.5918 | -0.023894889 | 0.104067018 | 4.7300626 | 4.5383506 | 4.5278 | 0.010546391 | 0.202258431 |
| Cdkal1 | 8.419418 | 8.559334 | 8.5395 | 0.019849833 | -0.12006581 | 8.6408426 | 8.6472303 | 8.63678 | 0.010452408 | 0.004064721 |
| Cask | 7.111546 | 7.204226 | 6.7091 | 0.495119608 | 0.40243957 | 6.7211925 | 6.891083 | 6.88069 | 0.010389141 | -0.15950132 |
| Slitrk2 | 2.728819 | 2.678491 | 2.6561 | 0.022347587 | 0.072675545 | 2.7338197 | 2.7322412 | 2.7219 | 0.01033678 | 0.011915344 |
| Gm7600 | 3.424271 | 3.524245 | 3.4675 | 0.056772869 | -0.043201311 | 3.5023262 | 3.5194305 | 3.50912 | 0.010307805 | -0.006796529 |
| Slc4a3 | 3.312577 | 3.286181 | 3.2702 | 0.015972866 | 0.042368515 | 3.220767 | 3.2821348 | 3.27183 | 0.010305493 | -0.051062308 |
| Epha7 | 1.134256 | 1.149429 | 1.1099 | 0.039506657 | 0.024333569 | 1.0649921 | 1.174017 | 1.16374 | 0.010280351 | -0.098744552 |
| Kbtbd7 | 8.432215 | 8.463789 | 8.5405 | -0.076661621 | -0.108235065 | 8.3747589 | 8.2657882 | 8.25553 | 0.010259392 | 0.119230038 |
| Neurl1a | 2.059484 | 2.06572 | 1.986 | 0.079681595 | 0.073446336 | 2.0312574 | 2.0234585 | 2.01326 | 0.010197942 | 0.017996863 |
| Tpt1-ps6 | 2.040714 | 2.103882 | 2.0437 | 0.060139817 | -0.003028058 | 2.0944739 | 2.1259574 | 2.11579 | 0.01017212 | -0.021311376 |
| Olfr188-ps1 | 1.369077 | 1.362952 | 1.3791 | -0.016121244 | -0.009995977 | 1.4321966 | 1.4368616 | 1.42671 | 0.01015594 | 0.005490965 |
| Nipal2 | 5.479962 | 5.400648 | 5.57 | -0.169328483 | -0.090014164 | 5.5747746 | 5.7574399 | 5.74732 | 0.010123583 | -0.172541788 |
| Mettl25 | 6.890839 | 6.767114 | 6.9038 | -0.1367298 | -0.013004884 | 6.9779239 | 6.8883536 | 6.87824 | 0.010116855 | 0.099687105 |
| Lamc1 | 9.945078 | 9.682456 | 9.6637 | 0.018769693 | 0.281391638 | 9.2099591 | 9.4509589 | 9.44085 | 0.010112823 | -0.23088698 |
| Crkl | 10.799 | 10.94063 | 10.836 | 0.104316882 | -0.037314548 | 10.75186 | 10.785477 | 10.7754 | 0.010096932 | -0.023519983 |
| Gm6363 | 1.320929 | 1.313407 | 1.2702 | 0.043240655 | 0.05076289 | 1.2879755 | 1.3148656 | 1.30478 | 0.010082158 | -0.016807987 |
| Gm16415 | 1.954774 | 2.053871 | 2.0185 | 0.03535184 | -0.063744911 | 1.9727463 | 2.0197278 | 2.00966 | 0.010070178 | -0.036911363 |
| 5530401A14Rik | 1.595238 | 1.620069 | 1.6313 | -0.011201836 | -0.036032416 | 1.6662153 | 1.6550926 | 1.64503 | 0.010060269 | 0.021182963 |
| Agap3 | 9.326958 | 9.368477 | 9.3574 | 0.011060381 | -0.030458252 | 9.1632903 | 9.2163349 | 9.20628 | 0.010057888 | -0.042986692 |
| Gm8319 | 0.746784 | 0.720104 | 0.7402 | -0.020144332 | 0.006535709 | 0.7318824 | 0.7405588 | 0.7305 | 0.010057666 | 0.00138128 |
| Gad2 | 1.60347 | 1.656722 | 1.5499 | 0.106842421 | 0.053590651 | 1.6993083 | 1.5745935 | 1.56459 | 0.010004686 | 0.134719522 |
| Gpx4 | 10.24574 | 10.11536 | 10.215 | -0.100135927 | 0.030245268 | 10.323159 | 10.045305 | 10.0353 | 0.010003893 | 0.287857471 |
| Cd200r2 | 0.453193 | 0.381262 | 0.4127 | -0.03142205 | 0.040509316 | 0.4534301 | 0.4271321 | 0.41715 | 0.009980894 | 0.036278932 |
| Ptprd | 3.888886 | 3.645106 | 3.7944 | -0.149319094 | 0.09446134 | 3.9201767 | 3.8380138 | 3.82804 | 0.009977064 | 0.092139955 |
| Zbtb39 | 8.055791 | 8.069957 | 8.0484 | 0.021536598 | 0.007369724 | 8.1437002 | 7.9528683 | 7.94294 | 0.009930538 | 0.20076245 |
| Golph3 | 10.61541 | 10.67156 | 10.582 | 0.08961262 | 0.033461076 | 10.474634 | 10.526537 | 10.5166 | 0.009888802 | -0.042014213 |
| Klhdc8a | 2.053438 | 2.077205 | 2.1405 | -0.063335192 | -0.087102377 | 2.0923018 | 1.9986762 | 1.98882 | 0.009856256 | 0.103481834 |

| | | | | | | | | | | |
|---|---|---|---|---|---|---|---|---|---|---|
| Gm3851 | 0.409912 | 0.423899 | 0.433 | -0.009068996 | -0.023055594 | 0.4295972 | 0.4110735 | 0.40126 | 0.009815405 | 0.028339122 |
| Gm9877 | 1.031204 | 1.096932 | 1.0651 | 0.031821071 | -0.033907349 | 1.1399267 | 1.0966851 | 1.08689 | 0.009797971 | 0.053039599 |
| Gm21319 | 2.996537 | 2.955849 | 3.0181 | -0.062272662 | -0.021585406 | 3.05768 | 3.1229488 | 3.11318 | 0.009773588 | -0.055495209 |
| Fam71a | 0.76226 | 0.696599 | 0.6417 | 0.054864324 | 0.120525666 | 0.6511359 | 0.6298696 | 0.62012 | 0.009752221 | 0.031018505 |
| Hs3st4 | 0.599633 | 0.588706 | 0.614 | -0.025323606 | -0.014396878 | 0.6472553 | 0.6111258 | 0.60139 | 0.009733882 | 0.045863337 |
| Lipo3 | 7.676273 | 7.663187 | 7.703 | -0.039768235 | -0.02668198 | 7.6663898 | 7.6961157 | 7.6864 | 0.009716274 | -0.020009626 |
| Tmc5 | 1.049726 | 1.15691 | 1.1479 | 0.009047894 | -0.09813684 | 1.0307844 | 1.120483 | 1.11078 | 0.009706682 | -0.079991875 |
| Gfpt1 | 10.53598 | 10.4746 | 10.549 | -0.074876042 | -0.013497184 | 10.305322 | 10.542369 | 10.5327 | 0.009687362 | -0.227359481 |
| Gm15739 | 0.836825 | 0.811959 | 0.8319 | -0.019972775 | 0.004892489 | 0.8335079 | 0.8960027 | 0.88632 | 0.009681922 | -0.052812863 |
| Ttc25 | 3.623592 | 3.656163 | 3.7291 | -0.072949897 | -0.105521621 | 3.5762287 | 3.7229539 | 3.71332 | 0.009633883 | -0.137091371 |
| Patl1 | 10.13161 | 10.19944 | 10.125 | 0.074361794 | 0.006534245 | 10.13616 | 10.221198 | 10.2116 | 0.009607748 | -0.075430255 |
| Fmr1nb | -0.653373 | -0.74136 | -0.7172 | -0.024163869 | 0.063822183 | -0.7284979 | -0.73158 | -0.7412 | 0.009588694 | 0.012670763 |
| Hist1h4k | 0.368378 | 0.337055 | 0.3465 | -0.009415529 | 0.021907498 | 0.4226047 | 0.4193067 | 0.40976 | 0.009541908 | 0.012839896 |
| Ggt6 | 2.835226 | 2.647937 | 2.9158 | -0.267821305 | -0.080531704 | 2.6358067 | 2.5631559 | 2.55363 | 0.009527195 | 0.08217802 |
| Slc2a10 | 3.407173 | 3.492059 | 3.4048 | 0.08725624 | 0.002369572 | 3.2382107 | 3.33302 | 3.32353 | 0.009485637 | -0.085323688 |
| Slc35a2 | 8.198333 | 8.358819 | 8.2067 | 0.152090152 | -0.008395749 | 8.6604261 | 8.2033501 | 8.19388 | 0.009471586 | 0.466547655 |
| AI464131 | 1.607125 | 1.59921 | 1.7256 | -0.126418577 | -0.118503874 | 1.7200249 | 1.596944 | 1.5875 | 0.009448823 | 0.132529685 |
| Gsto2 | 1.440487 | 1.420429 | 1.4129 | 0.007542808 | 0.027600819 | 1.4869783 | 1.4668401 | 1.4574 | 0.009439457 | 0.029577654 |
| Rasl2-9 | 2.038228 | 1.927453 | 1.986 | -0.058575603 | 0.052199464 | 2.0521363 | 1.9896984 | 1.98033 | 0.009371538 | 0.071809499 |
| Dcx | 1.607202 | 1.534845 | 1.5517 | -0.0168056 | 0.055551044 | 1.6594635 | 1.626155 | 1.6168 | 0.009352757 | 0.042661282 |
| Ighv1-67 | 4.122254 | 4.121519 | 4.1247 | -0.003133195 | -0.002397628 | 4.2668988 | 4.2380669 | 4.22873 | 0.009336808 | 0.038168792 |
| Adam17 | 10.70124 | 10.84141 | 10.725 | 0.116436046 | -0.023735958 | 10.701723 | 10.758062 | 10.7488 | 0.009293984 | -0.047044821 |
| Slc7a14 | 2.714446 | 2.766126 | 2.7919 | -0.025802626 | -0.077482855 | 2.7269207 | 2.876688 | 2.86741 | 0.009282078 | -0.140485198 |
| Spata19 | 2.262996 | 2.292109 | 2.3366 | -0.044462365 | -0.073575428 | 2.291467 | 2.3349961 | 2.32573 | 0.00926738 | -0.034261632 |
| Plekha4 | 4.877774 | 4.999952 | 4.713 | 0.286942649 | 0.164764529 | 4.8733954 | 5.0389051 | 5.02966 | 0.009247604 | -0.156262048 |
| Zfp639 | 9.266616 | 9.204887 | 9.248 | -0.043131577 | 0.01859754 | 9.3569037 | 9.4144447 | 9.40525 | 0.009199205 | -0.048341843 |
| Hist1h4f | 1.271661 | 1.299259 | 1.3479 | -0.048676048 | -0.076274108 | 1.3356358 | 1.3088896 | 1.29972 | 0.009172969 | 0.035919194 |
| Gm10167 | 1.598514 | 1.541338 | 1.6171 | -0.075777874 | -0.018601844 | 1.6270771 | 1.5966787 | 1.58753 | 0.009150976 | 0.039549348 |
| Vstm5 | 0.740131 | 0.718254 | 0.7482 | -0.02989613 | -0.008019667 | 0.7815228 | 0.7207746 | 0.71164 | 0.009137736 | 0.06988599 |
| Gm13081 | 4.195604 | 4.210817 | 4.0853 | 0.125476833 | 0.110264375 | 4.5396615 | 4.4455309 | 4.4364 | 0.009131249 | 0.103261787 |
| Gm5977 | -0.403863 | -0.38772 | -0.3564 | -0.031332052 | -0.047477508 | -0.3525225 | -0.384359 | -0.3934 | 0.009082017 | 0.040918428 |
| Aqp5 | 2.880524 | 2.46949 | 2.6384 | -0.168901261 | 0.242133505 | 2.8667172 | 2.6273857 | 2.61831 | 0.009077367 | 0.248408845 |

| | | | | | | | | | | |
|---|---|---|---|---|---|---|---|---|---|---|
| Usp14 | 9.752336 | 9.758306 | 9.5858 | 0.172524391 | 0.166554535 | 9.6201179 | 9.7969628 | 9.78794 | 0.009018781 | -0.167826053 |
| Lnx1 | 0.781517 | 0.70652 | 0.7187 | -0.012213884 | 0.062783014 | 0.7520458 | 0.748042 | 0.73906 | 0.008983555 | 0.012987333 |
| Olfr642 | 1.340281 | 1.27021 | 1.2566 | 0.013576085 | 0.083647247 | 1.2506858 | 1.3341119 | 1.32514 | 0.008976789 | -0.074449307 |
| Hpcal4 | 2.19639 | 2.221871 | 1.9957 | 0.226205424 | 0.200723779 | 2.0391379 | 2.0809997 | 2.07216 | 0.00884407 | -0.033017741 |
| Gm5513 | 1.144689 | 0.949945 | 0.9399 | 0.010091887 | 0.204835985 | 1.028479 | 1.0025875 | 0.99381 | 0.008773901 | 0.03466543 |
| Traj36 | 0.113949 | 0.125516 | 0.1006 | 0.024883865 | 0.013316242 | 0.0549731 | 0.0838298 | 0.0751 | 0.008734644 | -0.020122022 |
| Zmym3 | 9.446045 | 9.434759 | 9.5162 | -0.081486971 | -0.070200046 | 9.723882 | 9.5789105 | 9.57023 | 0.008676996 | 0.153648537 |
| Rhbdl1 | 4.599541 | 4.621802 | 4.6876 | -0.065837895 | -0.088098804 | 4.7319426 | 4.6961688 | 4.68752 | 0.008646821 | 0.044420649 |
| Syce1l | 0.598681 | 0.577062 | 0.5899 | -0.012823173 | 0.008795227 | 0.5942661 | 0.6043465 | 0.59574 | 0.008603438 | -0.001476935 |
| Slc1a1 | 1.494525 | 1.527512 | 1.5064 | 0.0211255 | -0.011862089 | 1.3947472 | 1.4892781 | 1.48068 | 0.008598113 | -0.085932714 |
| Pcsk4 | 5.068209 | 4.892298 | 5.0773 | -0.185013712 | -0.009103051 | 5.2639815 | 5.0119434 | 5.00335 | 0.008596858 | 0.260634953 |
| 4930519P11Rik | 0.366903 | 0.391813 | 0.3769 | 0.014896964 | -0.010013214 | 0.4073172 | 0.4182872 | 0.4097 | 0.008588016 | -0.002381965 |
| Aurkc | 1.000782 | 0.943181 | 0.9897 | -0.046486041 | 0.011114289 | 1.0132031 | 0.9757996 | 0.96725 | 0.008554305 | 0.045957723 |
| Plcd1 | 6.001182 | 5.777673 | 5.8358 | -0.058167175 | 0.165342005 | 5.7840335 | 5.7257857 | 5.71723 | 0.008552906 | 0.066800694 |
| Gm8493 | 2.008722 | 2.027541 | 1.9853 | 0.042192027 | 0.023373609 | 2.0701047 | 2.1238663 | 2.11535 | 0.008521009 | -0.04524053 |
| Gm17334 | 2.141303 | 2.19409 | 2.1501 | 0.043972877 | -0.008813828 | 2.1489203 | 2.224814 | 2.21632 | 0.008497324 | -0.06739636 |
| Igkv14-126-1 | 1.228608 | 1.232374 | 1.2064 | 0.026009664 | 0.022243278 | 1.3367369 | 1.310486 | 1.30204 | 0.008443191 | 0.034694026 |
| Evc2 | 4.210866 | 4.310393 | 4.2472 | 0.063225997 | -0.036301091 | 3.8245372 | 3.9825175 | 3.9741 | 0.008421815 | -0.149558482 |
| 3110070M22Rik | 2.648003 | 2.585092 | 2.647 | -0.061884186 | 0.001027011 | 2.630467 | 2.6496953 | 2.64133 | 0.008365383 | -0.010862941 |
| Kirrel | 5.820762 | 5.97983 | 5.6587 | 0.321110579 | 0.162042577 | 5.6177254 | 5.4540147 | 5.44566 | 0.008352734 | 0.17206347 |
| Olfr466 | 1.897721 | 1.831729 | 1.9346 | -0.102915474 | -0.036923329 | 1.9151929 | 1.9332283 | 1.92492 | 0.008310387 | -0.009724935 |
| Myt1 | 1.177914 | 1.133859 | 1.1503 | -0.016480408 | 0.027574786 | 1.0872236 | 1.1782875 | 1.17002 | 0.008271933 | -0.082791913 |
| Stc2 | 5.570571 | 5.363441 | 5.2767 | 0.086699949 | 0.293829544 | 4.9726215 | 5.070088 | 5.06185 | 0.008238362 | -0.089228182 |
| Chrna2 | 1.755691 | 1.738472 | 1.7684 | -0.029926276 | -0.012706557 | 1.7776761 | 1.8383954 | 1.83016 | 0.008235437 | -0.052483786 |
| Il1rap | 8.333918 | 8.348075 | 8.031 | 0.317040326 | 0.302882674 | 8.1942465 | 8.0052237 | 7.997 | 0.008226917 | 0.197249813 |
| Gm7638 | 0.912072 | 0.93113 | 0.9735 | -0.042358491 | -0.06141614 | 0.9158871 | 0.9586777 | 0.95047 | 0.008207758 | -0.034582891 |
| Celsr3 | 1.800839 | 1.635788 | 1.4964 | 0.139353532 | 0.304403702 | 1.7647997 | 1.6427815 | 1.63458 | 0.008203735 | 0.13022192 |
| Cntnap5a | 1.602008 | 1.57958 | 1.6329 | -0.053290347 | -0.030862309 | 1.6574056 | 1.6582191 | 1.65004 | 0.008181144 | 0.007367697 |
| Mro | 0.642936 | 0.661397 | 0.6146 | 0.046755615 | 0.028294901 | 0.6132718 | 0.6646475 | 0.65648 | 0.008164938 | -0.043210744 |
| Tmigd1 | 0.67239 | 0.563998 | 0.632 | -0.068018673 | 0.040373483 | 0.5852718 | 0.5983301 | 0.5902 | 0.00812887 | -0.004929414 |
| Kcnf1 | 1.138414 | 1.079289 | 1.0951 | -0.01585298 | 0.043271544 | 1.0471014 | 1.051272 | 1.04315 | 0.008126702 | 0.003956114 |
| Mllt6 | 12.14711 | 12.17046 | 12.111 | 0.059498828 | 0.036147192 | 12.142019 | 12.164703 | 12.1566 | 0.008100482 | -0.014583096 |

| | | | | | | | | | | |
|---|---|---|---|---|---|---|---|---|---|---|
| Olfr901 | 1.4468 | 1.35112 | 1.342 | 0.0091473 | 0.104827826 | 1.3579314 | 1.4187337 | 1.41071 | 0.008026806 | -0.052775458 |
| Otof | 1.602218 | 1.619006 | 1.611 | 0.007974573 | -0.008813016 | 1.5790281 | 1.5784685 | 1.57046 | 0.008011291 | 0.008570931 |
| Lrp2bp | 0.637293 | 0.579666 | 0.6366 | -0.056915843 | 0.000711126 | 0.7482387 | 0.6239573 | 0.61598 | 0.007977668 | 0.132259103 |
| Wnt9a | 0.89313 | 0.83628 | 0.8391 | -0.002778137 | 0.054072147 | 0.8099456 | 0.8658177 | 0.85786 | 0.007954062 | -0.047918021 |
| Cdca7l | 8.746159 | 8.605012 | 8.7765 | -0.171439136 | -0.030292257 | 8.4333515 | 8.7739927 | 8.76608 | 0.007908611 | -0.332732638 |
| Pnoc | 1.241044 | 1.211789 | 1.2213 | -0.009514186 | 0.019740559 | 1.1914488 | 1.2499766 | 1.24208 | 0.007897879 | -0.050629908 |
| Uba2 | 10.84363 | 10.85112 | 10.868 | -0.01720781 | -0.024696692 | 10.930762 | 10.911104 | 10.9032 | 0.007879806 | 0.027537058 |
| Spta1 | 0.449775 | 0.529708 | 0.4717 | 0.057961947 | -0.021970295 | 0.4164841 | 0.4440473 | 0.43621 | 0.007834583 | -0.019728642 |
| Gm48552 | 4.05849 | 4.007003 | 4.1327 | -0.125665044 | -0.074177382 | 4.1849866 | 4.1408586 | 4.13303 | 0.007832671 | 0.051960602 |
| Inhbe | 3.05013 | 2.958564 | 3.0043 | -0.045760342 | 0.045805394 | 3.0910519 | 3.1147678 | 3.10696 | 0.007811279 | -0.015904638 |
| Adra2a | 4.227752 | 3.97449 | 4.0746 | -0.100111078 | 0.153151536 | 3.9840872 | 3.8147013 | 3.80691 | 0.007792903 | 0.177178859 |
| Rnf180 | 3.568198 | 3.633542 | 3.7734 | -0.139865926 | -0.205209729 | 3.786719 | 3.6420999 | 3.63435 | 0.007745151 | 0.152364229 |
| Igf2bp2 | 4.888582 | 5.096151 | 4.7788 | 0.317361984 | 0.109792591 | 5.0440562 | 5.0660821 | 5.05834 | 0.007742609 | -0.014283309 |
| Lrfn4 | 3.003024 | 3.086408 | 3.031 | 0.055420755 | -0.027963858 | 3.0324738 | 2.9418295 | 2.93409 | 0.007734979 | 0.098379272 |
| Gm13305 | 0.22248 | 0.222328 | 0.2826 | -0.060285688 | -0.060133014 | 0.2102112 | 0.2297722 | 0.22204 | 0.007734187 | -0.011826772 |
| Copz2 | 4.169081 | 4.046278 | 4.2373 | -0.190997971 | -0.0681945 | 4.1704536 | 4.1123067 | 4.10457 | 0.007732909 | 0.065879758 |
| Yeats4 | 9.361096 | 9.235704 | 9.3664 | -0.130674483 | -0.005282469 | 9.3067546 | 9.2650177 | 9.25729 | 0.007723872 | 0.049460695 |
| Zfp566 | 6.315212 | 6.668668 | 6.6211 | 0.047553361 | -0.305903349 | 6.5161382 | 6.469703 | 6.46199 | 0.007710615 | 0.054145796 |
| Pfdn4 | 7.410906 | 7.578781 | 7.4465 | 0.132280766 | -0.035594283 | 7.4579286 | 7.5273093 | 7.5196 | 0.007708781 | -0.061671919 |
| Igkv12-40 | 2.208104 | 2.270812 | 2.2123 | 0.05855334 | -0.004154978 | 2.1765878 | 2.1736167 | 2.16597 | 0.007647173 | 0.01061825 |
| Ddi1 | 0.477974 | 0.442151 | 0.4243 | 0.017856981 | 0.053679802 | 0.4971742 | 0.4720648 | 0.46443 | 0.007633284 | 0.032742706 |
| Megf11 | 3.308192 | 3.067832 | 3.485 | -0.417168083 | -0.176808615 | 3.0649798 | 3.1377867 | 3.13018 | 0.007610353 | -0.065196604 |
| Igdcc3 | 0.609117 | 0.613864 | 0.6191 | -0.005193063 | -0.009940171 | 0.5400453 | 0.5424627 | 0.53488 | 0.007579363 | 0.005161981 |
| Nova2 | 4.608747 | 4.557198 | 4.5699 | -0.012719209 | 0.038829715 | 4.1364613 | 4.2530436 | 4.2455 | 0.007546037 | -0.109036305 |
| Vmn1r49 | 2.121653 | 2.133497 | 2.1175 | 0.015958118 | 0.00411385 | 2.0797175 | 2.1574284 | 2.14995 | 0.007480896 | -0.070230064 |
| Slc30a5 | 10.45125 | 10.42359 | 10.481 | -0.057426234 | -0.029763598 | 10.514176 | 10.440716 | 10.4333 | 0.007398904 | 0.080858491 |
| Vmn2r111 | 1.301288 | 1.371122 | 1.3961 | -0.024994516 | -0.09482786 | 1.3484593 | 1.4019598 | 1.39461 | 0.00735037 | -0.046150121 |
| Slc25a2 | 2.220096 | 2.20784 | 2.2139 | -0.006099051 | 0.00615721 | 2.1714303 | 2.2996704 | 2.29236 | 0.007308767 | -0.120931396 |
| Tmem168 | 10.05265 | 10.22175 | 10.099 | 0.123246567 | -0.045849841 | 10.104314 | 10.209006 | 10.2017 | 0.007289765 | -0.097401693 |
| Rad52 | 8.172514 | 8.198323 | 8.269 | -0.070713071 | -0.096521876 | 8.5736298 | 8.456515 | 8.44929 | 0.007226096 | 0.124340892 |
| Gm4734 | 1.309737 | 1.385905 | 1.3615 | 0.024415886 | -0.051752254 | 1.3603209 | 1.3893219 | 1.38212 | 0.007199981 | -0.02180097 |
| Tnfrsf22 | 8.162795 | 8.408324 | 8.4523 | -0.043945106 | -0.289474043 | 8.2958099 | 8.3113565 | 8.30421 | 0.007144868 | -0.00840171 |

| | | | | | | | | | | |
|---|---|---|---|---|---|---|---|---|---|---|
| Fxn | 6.55144 | 6.668764 | 6.6217 | 0.047024833 | -0.070299704 | 6.7512911 | 6.8014913 | 6.79436 | 0.007126429 | -0.04307371 |
| Olfr668 | 1.018467 | 0.979407 | 1.0077 | -0.028271192 | 0.01078871 | 1.033801 | 0.9993691 | 0.99232 | 0.007044557 | 0.041476441 |
| Trim69 | 3.12029 | 3.107946 | 3.186 | -0.078058911 | -0.06571548 | 3.1848774 | 3.2519068 | 3.24488 | 0.007029848 | -0.059999524 |
| Gjc3 | 2.891485 | 2.820211 | 2.8976 | -0.077405443 | -0.006131249 | 2.9536104 | 3.0076125 | 3.00059 | 0.007024789 | -0.046977315 |
| Pcdhga4 | 2.977068 | 2.891224 | 2.9402 | -0.048983109 | 0.036861141 | 2.8175781 | 2.8011211 | 2.7941 | 0.007019833 | 0.023476805 |
| Vmn2r8 | 1.380702 | 1.41753 | 1.4492 | -0.031701276 | -0.068529054 | 1.381026 | 1.4398876 | 1.4329 | 0.006988134 | -0.051873465 |
| Gm14917 | 1.041557 | 1.040656 | 1.0655 | -0.024869745 | -0.023968756 | 1.0100919 | 1.1295102 | 1.12255 | 0.00695758 | -0.112460674 |
| Scn11a | -0.063858 | -0.05636 | -0.0454 | -0.010909251 | -0.01841158 | -0.0781755 | -0.095078 | -0.102 | 0.006941812 | 0.02384404 |
| Ttc37 | 9.725703 | 9.787046 | 9.7204 | 0.066690065 | 0.005346921 | 9.7671281 | 9.7666235 | 9.75972 | 0.006904488 | 0.007409103 |
| Hydin | 1.094543 | 1.156124 | 1.1281 | 0.028058317 | -0.033522003 | 1.1888165 | 1.1641949 | 1.15736 | 0.006839682 | 0.03146126 |
| Aarsd1 | 8.505418 | 8.527053 | 8.4989 | 0.028111751 | 0.006476066 | 8.8013413 | 8.6155241 | 8.60878 | 0.006742007 | 0.192559226 |
| Tmem125 | 0.031731 | 0.0089 | 0.0426 | -0.033747088 | -0.01091518 | 0.0163916 | 0.0364945 | 0.02976 | 0.006736438 | -0.013366465 |
| Naip3 | 0.764023 | 0.769627 | 0.7871 | -0.017436162 | -0.023040012 | 0.7158313 | 0.7350438 | 0.72833 | 0.006715705 | -0.012496762 |
| Ralbp1 | 10.32731 | 10.35431 | 10.374 | -0.019334028 | -0.046329252 | 10.219922 | 10.251334 | 10.2446 | 0.00669189 | -0.024719832 |
| Aff1 | 11.43344 | 11.56457 | 11.456 | 0.108611624 | -0.02251843 | 11.275438 | 11.462508 | 11.4558 | 0.006663234 | -0.180407258 |
| Paics | 10.96319 | 10.89159 | 10.951 | -0.058934512 | 0.012666626 | 11.004627 | 10.959039 | 10.9524 | 0.006593992 | 0.052181322 |
| Gm44845 | 1.86679 | 1.786337 | 1.7752 | 0.011153987 | 0.091607252 | 1.7358223 | 1.848072 | 1.84148 | 0.006592312 | -0.105657481 |
| Tgm5 | 1.52641 | 1.551717 | 1.5112 | 0.04050704 | 0.015200887 | 1.5159466 | 1.5473213 | 1.54073 | 0.006590698 | -0.024784009 |
| Eri3 | 8.273591 | 8.132422 | 8.198 | -0.065538554 | 0.075630367 | 8.4793857 | 8.371543 | 8.36497 | 0.006569132 | 0.114411907 |
| Gm27186 | -0.045867 | -0.036 | -0.0122 | -0.023848726 | -0.0337166 | -0.0533953 | -0.01455 | -0.0211 | 0.006541816 | -0.032303576 |
| Bhmt | 2.037936 | 2.033124 | 2.0668 | -0.033695836 | -0.02888365 | 2.0575304 | 2.0462161 | 2.03968 | 0.006538572 | 0.017852813 |
| Scube1 | 2.194662 | 2.218374 | 2.1843 | 0.034089128 | 0.010377376 | 2.1771093 | 2.2507975 | 2.24427 | 0.006532335 | -0.067155909 |
| Vmn2r-ps117 | 1.154218 | 1.126196 | 1.1837 | -0.057476159 | -0.029454851 | 1.1532879 | 1.1716872 | 1.16527 | 0.006413163 | -0.011986158 |
| Gm44954 | 0.921861 | 0.963519 | 0.8846 | 0.078951601 | 0.037293372 | 0.9583906 | 0.9446622 | 0.93829 | 0.006376231 | 0.020104661 |
| Alyref2 | 6.599641 | 6.517578 | 6.6819 | -0.164294994 | -0.082231473 | 6.8337665 | 6.7491003 | 6.74275 | 0.006353157 | 0.091019433 |
| Rab40b | 0.879531 | 0.86203 | 0.9401 | -0.078104996 | -0.060603755 | 0.8770757 | 0.8682728 | 0.86192 | 0.0063517 | 0.015154596 |
| Cnot11 | 8.849965 | 8.854733 | 8.9176 | -0.062840792 | -0.06760868 | 8.8015948 | 8.8596603 | 8.85333 | 0.00633276 | -0.05173274 |
| Abcc2 | 2.218076 | 2.245757 | 2.2191 | 0.026651298 | -0.001029298 | 2.3049807 | 2.2814782 | 2.27515 | 0.006330691 | 0.02983313 |
| Gm26727 | 0.351775 | 0.333102 | 0.35 | -0.016932154 | 0.001740916 | 0.3760456 | 0.4052745 | 0.39896 | 0.006318713 | -0.022910231 |
| Rgs17 | 3.296665 | 3.124006 | 3.3426 | -0.218574233 | -0.045915804 | 3.1481738 | 3.2555951 | 3.24931 | 0.00628724 | -0.101134147 |
| Rpl17-ps9 | 4.349177 | 4.341487 | 4.4892 | -0.147685035 | -0.139995446 | 4.2899796 | 4.4169484 | 4.41066 | 0.006285782 | -0.120683019 |
| Fancg | 8.110592 | 8.070586 | 8.1466 | -0.076024612 | -0.036018182 | 8.2835066 | 8.3100485 | 8.30376 | 0.006284302 | -0.020257549 |

| | | | | | | | | | | |
|---|---|---|---|---|---|---|---|---|---|---|
| Mzt2 | 6.07292 | 5.991842 | 6.0781 | -0.086208995 | -0.005130764 | 6.196005 | 6.0043939 | 5.99819 | 0.006203641 | 0.197814766 |
| Clec2g | 1.168206 | 1.109279 | 1.1685 | -0.059198833 | -0.000271762 | 1.1776638 | 1.1050818 | 1.09892 | 0.006162631 | 0.078744604 |
| Zfp748 | 8.337289 | 8.535958 | 8.6014 | -0.065451265 | -0.264120221 | 8.5681253 | 8.5611117 | 8.555 | 0.00610742 | 0.013121007 |
| Gm29394 | 2.627077 | 2.632696 | 2.7863 | -0.153649939 | -0.159268847 | 2.6924533 | 2.6959405 | 2.68984 | 0.006099076 | 0.002611873 |
| Gm14469 | 1.522662 | 1.538162 | 1.5184 | 0.019765648 | 0.004266497 | 1.5355856 | 1.6160263 | 1.60993 | 0.006095375 | -0.074345346 |
| Ccno | 2.541075 | 2.453046 | 2.6429 | -0.189849709 | -0.101820634 | 2.4319764 | 2.5920636 | 2.58605 | 0.006013321 | -0.154073939 |
| Nell2 | 0.961378 | 0.880816 | 0.8215 | 0.05930459 | 0.139866628 | 0.7834178 | 0.8534831 | 0.84749 | 0.005992807 | -0.064072517 |
| Eloa | 10.51954 | 10.60084 | 10.594 | 0.007258098 | -0.074041516 | 10.605909 | 10.607344 | 10.6014 | 0.00598447 | 0.004549561 |
| Sgtb | 2.441558 | 2.404723 | 2.6 | -0.195261971 | -0.158426795 | 2.3648788 | 2.3491216 | 2.34316 | 0.005956904 | 0.021714091 |
| Scn1a | 1.059552 | 1.048384 | 1.0258 | 0.02259618 | 0.033764596 | 1.0338185 | 1.0435174 | 1.0376 | 0.005921153 | -0.003777762 |
| Gm12106 | 2.099978 | 2.042138 | 2.0987 | -0.056560177 | 0.00127988 | 2.1397781 | 2.095122 | 2.08923 | 0.005896584 | 0.050552683 |
| Cenpv | 4.807224 | 5.013529 | 4.8871 | 0.126379048 | -0.079925366 | 4.9208021 | 4.9752678 | 4.96938 | 0.005888619 | -0.048577018 |
| Frmpd1 | 0.838458 | 0.733566 | 0.7146 | 0.018997156 | 0.12388932 | 0.6945816 | 0.705363 | 0.69948 | 0.005880506 | -0.004900825 |
| Hadha | 11.20289 | 11.17639 | 11.188 | -0.011306934 | 0.015188284 | 11.166269 | 11.168888 | 11.1631 | 0.005800192 | 0.003181101 |
| Olfr1432 | 2.852163 | 2.805993 | 2.8404 | -0.034433872 | 0.011736857 | 2.9423155 | 2.9258609 | 2.92009 | 0.005769613 | 0.022224193 |
| Tmed7 | 10.92894 | 10.87661 | 10.782 | 0.09479186 | 0.14712092 | 10.699563 | 10.685053 | 10.6793 | 0.00575628 | 0.020266193 |
| Swsap1 | 6.324975 | 6.460402 | 6.4248 | 0.035634273 | -0.09979206 | 6.5570018 | 6.2717589 | 6.26602 | 0.005739588 | 0.290982461 |
| Eef2 | 16.10314 | 16.10187 | 16.127 | -0.025371648 | -0.024096508 | 16.141381 | 16.009121 | 16.0034 | 0.005722042 | 0.137981835 |
| Gm2423 | -0.20054 | -0.16085 | -0.1862 | 0.025335946 | -0.014351964 | -0.1060149 | -0.155366 | -0.161 | 0.005656835 | 0.055007505 |
| A630001G21Rik | 10.06452 | 10.11525 | 9.9931 | 0.122193196 | 0.071459538 | 10.461673 | 10.473292 | 10.4677 | 0.005623742 | -0.00599514 |
| Gm20695 | 2.610764 | 2.625804 | 2.6449 | -0.019090506 | -0.034131134 | 2.4698922 | 2.6658945 | 2.66028 | 0.005612682 | -0.190389687 |
| Tgoln1 | 11.8232 | 11.8457 | 11.79 | 0.056189302 | 0.033691516 | 11.763736 | 11.752585 | 11.7471 | 0.005488264 | 0.016638907 |
| Mettl18 | 6.683863 | 6.657622 | 6.6924 | -0.034809325 | -0.008568048 | 6.68005 | 6.6292221 | 6.62374 | 0.005484008 | 0.056311905 |
| D630003M21Rik | 1.192228 | 1.181824 | 1.1912 | -0.009355536 | 0.001048416 | 1.1654404 | 1.1885101 | 1.1832 | 0.005311599 | -0.017758093 |
| Rnf6 | 9.674743 | 9.597925 | 9.7351 | -0.137127391 | -0.060308916 | 9.6911919 | 9.7208793 | 9.71561 | 0.005272195 | -0.024415211 |
| Relch | 10.07278 | 10.12196 | 10.219 | -0.097138299 | -0.146320894 | 10.187898 | 10.334644 | 10.3294 | 0.005232352 | -0.14151441 |
| Rap1gap | 6.180729 | 5.729539 | 5.9786 | -0.249053054 | 0.202136878 | 5.7752491 | 5.8960673 | 5.89093 | 0.005139904 | -0.115678347 |
| Vmn2r82 | 0.556533 | 0.548085 | 0.5377 | 0.010364545 | 0.018812083 | 0.5239803 | 0.5398418 | 0.53474 | 0.005097581 | -0.010763888 |
| Dgkb | 1.294858 | 1.221707 | 1.2904 | -0.06867102 | 0.004480256 | 1.1852928 | 1.3063138 | 1.30122 | 0.005096092 | -0.115924982 |
| Senp6 | 11.02165 | 11.07187 | 11.038 | 0.034192364 | -0.016034864 | 10.988118 | 11.095057 | 11.09 | 0.005080914 | -0.101857872 |
| Dcp1a | 10.21199 | 10.24611 | 10.19 | 0.056414624 | 0.022286152 | 10.2337 | 10.223912 | 10.2188 | 0.005075512 | 0.014863636 |
| Pdzrn3 | 2.483046 | 2.533171 | 2.607 | -0.073873779 | -0.123998142 | 2.4180539 | 2.4773658 | 2.47232 | 0.005041684 | -0.054270189 |

| | | | | | | | | | | |
|---|---|---|---|---|---|---|---|---|---|---|
| 2310061I04Rik | 7.693482 | 7.861383 | 7.8566 | 0.004785957 | -0.16311522 | 8.1916029 | 8.0180037 | 8.01302 | 0.004986729 | 0.17858587 |
| Flrt1 | 0.485202 | 0.47256 | 0.523 | -0.050431357 | -0.037789672 | 0.5106901 | 0.4970787 | 0.49211 | 0.004965648 | 0.018576977 |
| AI987944 | 8.332745 | 8.414661 | 8.4722 | -0.057517403 | -0.139433559 | 8.5005048 | 8.4122935 | 8.40735 | 0.004943582 | 0.093154831 |
| Cyp2r1 | 5.622613 | 5.506133 | 5.5722 | -0.066028428 | 0.050450705 | 5.7955749 | 5.6202045 | 5.61528 | 0.004926738 | 0.180297141 |
| Tmod4 | 4.083069 | 4.150902 | 4.0479 | 0.102982142 | 0.035148533 | 4.2129747 | 4.2792245 | 4.27433 | 0.004893382 | -0.061356432 |
| Colec11 | 3.320096 | 3.138107 | 3.4368 | -0.298718151 | -0.116728812 | 3.0694538 | 3.2149754 | 3.21014 | 0.004838889 | -0.140682638 |
| A930035D04Rik | 1.363894 | 1.318123 | 1.3608 | -0.042704932 | 0.003065833 | 1.4377279 | 1.3622806 | 1.35746 | 0.004821401 | 0.080268774 |
| Ifnz | 0.31365 | 0.258363 | 0.2142 | 0.044202734 | 0.09949041 | 0.2429921 | 0.2726784 | 0.26793 | 0.004752916 | -0.024933326 |
| Mmp28 | 3.55247 | 3.644691 | 3.8165 | -0.171771171 | -0.263992027 | 3.4837439 | 3.57701 | 3.57226 | 0.004751283 | -0.088514834 |
| Gm13285 | 1.985234 | 2.000929 | 2.0528 | -0.051872037 | -0.06756651 | 1.9277003 | 1.9275494 | 1.92282 | 0.004726731 | 0.004877628 |
| Rsph9 | 2.393817 | 2.405104 | 2.4427 | -0.037640916 | -0.048928144 | 2.6101233 | 2.485339 | 2.48061 | 0.004724167 | 0.129508375 |
| Syt6 | 3.226671 | 3.120219 | 3.0459 | 0.074334275 | 0.180786242 | 2.9603032 | 3.0198287 | 3.01511 | 0.004718425 | -0.054807036 |
| Zswim4 | 8.709372 | 8.547732 | 8.551 | -0.003312236 | 0.158328098 | 8.7472013 | 8.5735232 | 8.56884 | 0.004683726 | 0.178361841 |
| Galntl6 | 1.409879 | 1.332541 | 1.3638 | -0.031268641 | 0.046069718 | 1.371774 | 1.4785694 | 1.47393 | 0.004634723 | -0.10216068 |
| Gm8850 | 1.005536 | 0.987553 | 0.9763 | 0.011267815 | 0.029251392 | 0.9226691 | 1.0001787 | 0.99555 | 0.004623986 | -0.072885624 |
| Trmt13 | 8.034286 | 7.890428 | 8.0375 | -0.147028553 | -0.00317018 | 8.3346121 | 8.3382075 | 8.3336 | 0.004610094 | 0.001014656 |
| AC122861.1 | 2.254926 | 2.251348 | 2.3053 | -0.0539742 | -0.050396771 | 2.21796 | 2.377925 | 2.37332 | 0.00460477 | -0.155360245 |
| Ang | 2.421058 | 2.256854 | 2.3683 | -0.111405939 | 0.052798174 | 2.3951876 | 2.3044313 | 2.29985 | 0.004582181 | 0.095338495 |
| Pde4c | 3.479224 | 3.573578 | 3.5861 | -0.012529596 | -0.106883706 | 3.6781565 | 3.6926946 | 3.68821 | 0.00448916 | -0.010048984 |
| Mpp3 | 3.912632 | 3.928592 | 3.9251 | 0.003521594 | -0.012438002 | 3.8869792 | 4.0163279 | 4.01188 | 0.00445037 | -0.124898315 |
| Il4ra | 12.54452 | 12.41675 | 12.444 | -0.027407762 | 0.100368672 | 12.584178 | 12.486804 | 12.4824 | 0.004437902 | 0.101812076 |
| Gm13762 | 1.18005 | 1.203232 | 1.2311 | -0.027832617 | -0.051014837 | 1.2147915 | 1.1951454 | 1.19077 | 0.004378535 | 0.024024678 |
| Hcn1 | 0.546961 | 0.516276 | 0.5035 | 0.012766827 | 0.043452285 | 0.4657425 | 0.5118646 | 0.50752 | 0.004342943 | -0.041779168 |
| Gm9194 | 1.559223 | 1.551377 | 1.5416 | 0.009765075 | 0.017611537 | 1.5802887 | 1.5904038 | 1.58613 | 0.004270533 | -0.005844613 |
| Olfr484 | 1.904788 | 1.915355 | 1.8302 | 0.085129581 | 0.074562852 | 1.8531811 | 1.9584986 | 1.95424 | 0.004260073 | -0.101057371 |
| Fmn2 | 0.990192 | 1.047894 | 0.8712 | 0.17673612 | 0.11903431 | 0.7992832 | 0.8271499 | 0.82289 | 0.004255405 | -0.023611307 |
| Zfp811 | 2.509316 | 2.598553 | 2.5822 | 0.016401727 | -0.072835201 | 2.5147259 | 2.6769479 | 2.67272 | 0.004227958 | -0.157994087 |
| Bend5 | 8.69123 | 8.654048 | 8.7254 | -0.071322189 | -0.034140079 | 8.8118444 | 8.4729579 | 8.46883 | 0.004125715 | 0.343012149 |
| Rpl34-ps1 | 0.540752 | 0.38649 | 0.4165 | -0.029982828 | 0.124278913 | 0.4546263 | 0.4160751 | 0.41196 | 0.00411499 | 0.042666279 |
| Hus1b | 1.549988 | 1.534915 | 1.5733 | -0.038340411 | -0.023267422 | 1.5980814 | 1.6274388 | 1.62334 | 0.004101406 | -0.025255958 |
| Rpl19-ps11 | 3.410973 | 3.176955 | 3.2306 | -0.053624914 | 0.180392883 | 3.2437804 | 3.3773589 | 3.37329 | 0.004069881 | -0.129508576 |
| Gm37988 | 1.667114 | 1.722239 | 1.5958 | 0.126479011 | 0.071354463 | 1.7202828 | 1.698117 | 1.69407 | 0.004050469 | 0.026216241 |

| | | | | | | | | | | |
|---|---|---|---|---|---|---|---|---|---|---|
| Mkrn2os | 1.069584 | 1.084862 | 1.1583 | -0.073455466 | -0.088733833 | 1.02668 | 1.0970249 | 1.093 | 0.004027842 | -0.066317039 |
| Gm28374 | 0.342775 | 0.330281 | 0.3295 | 0.000785138 | 0.013279953 | 0.359592 | 0.3625539 | 0.35854 | 0.004014133 | 0.001052274 |
| Blvra | 8.029093 | 8.135205 | 8.0346 | 0.100589198 | -0.005522774 | 8.2032107 | 7.9936726 | 7.98971 | 0.003962234 | 0.213500407 |
| Kcna5 | 1.306485 | 1.282616 | 1.3045 | -0.021840046 | 0.0020282 | 1.2339454 | 1.2613121 | 1.25738 | 0.003936269 | -0.023430416 |
| Mal | 0.867505 | 0.792118 | 0.7403 | 0.051828278 | 0.127215447 | 0.7570495 | 0.7464864 | 0.74256 | 0.003926636 | 0.014489776 |
| Gm14049 | 0.525954 | 0.499554 | 0.5386 | -0.039035863 | -0.012635206 | 0.46505 | 0.5344264 | 0.5306 | 0.003827398 | -0.065549029 |
| Baz1b | 11.91612 | 11.92919 | 11.921 | 0.008087532 | -0.004983494 | 11.853493 | 11.992741 | 11.9889 | 0.003815874 | -0.135431699 |
| Olfr30 | 0.753883 | 0.725915 | 0.7037 | 0.022205121 | 0.050173553 | 0.7159039 | 0.7634063 | 0.75961 | 0.003798799 | -0.043703577 |
| Stk32a | 1.577107 | 1.510202 | 1.5233 | -0.01312242 | 0.053782749 | 1.5932529 | 1.605401 | 1.60162 | 0.003785252 | -0.008362787 |
| Gm7432 | 1.593736 | 1.616285 | 1.7132 | -0.096876503 | -0.119425375 | 1.6105503 | 1.6718927 | 1.66814 | 0.003757666 | -0.057584711 |
| Cfap61 | 2.249059 | 2.213166 | 2.1792 | 0.033939264 | 0.069831936 | 2.2694775 | 2.2412453 | 2.2375 | 0.003742664 | 0.031974899 |
| Atp11b | 11.82817 | 11.774 | 11.847 | -0.073415618 | -0.019246734 | 11.52702 | 11.865527 | 11.8618 | 0.00374048 | -0.334766554 |
| Vmn1r59 | 0.130269 | 0.158895 | 0.13 | 0.028943547 | 0.000317982 | 0.1030278 | 0.1401352 | 0.13649 | 0.003647316 | -0.033460128 |
| Rab26 | 3.550296 | 3.572896 | 3.5596 | 0.013324386 | -0.009276379 | 3.6446455 | 3.6795194 | 3.67591 | 0.003612094 | -0.031261815 |
| Gm42791 | 1.949703 | 1.993139 | 2.1002 | -0.107081341 | -0.1505168 | 1.9244733 | 1.93737 | 1.93379 | 0.003579843 | -0.00931682 |
| Pura | 9.981691 | 10.03529 | 9.9876 | 0.047708038 | -0.005892365 | 10.062119 | 10.116437 | 10.1129 | 0.003545592 | -0.050772381 |
| Tma7-ps | -0.156869 | -0.14268 | -0.1436 | 0.000962354 | -0.013223625 | -0.1575483 | -0.124404 | -0.1279 | 0.003494156 | -0.029650652 |
| 9430015G10Rik | 8.674617 | 8.820819 | 8.7875 | 0.033284424 | -0.112917392 | 8.9981408 | 9.0465345 | 9.04307 | 0.003465153 | -0.044928549 |
| Elfn1 | 0.623226 | 0.596062 | 0.5821 | 0.013981899 | 0.041145796 | 0.6594937 | 0.5943114 | 0.59085 | 0.003460348 | 0.068642647 |
| Trnp1 | 3.923722 | 3.912892 | 3.9244 | -0.011506669 | -0.000676438 | 3.6514798 | 3.628029 | 3.62462 | 0.003407638 | 0.026858493 |
| Mboat4 | 0.819792 | 0.859175 | 0.827 | 0.032223314 | -0.007159097 | 0.8235876 | 0.7878303 | 0.7845 | 0.003330568 | 0.039087829 |
| Gm21811 | 7.152803 | 7.194801 | 7.0092 | 0.185574653 | 0.143576884 | 7.2707846 | 7.2766711 | 7.2734 | 0.003275894 | -0.00261061 |
| AC131185.2 | 0.881848 | 0.839441 | 0.8392 | 0.00028995 | 0.042696967 | 0.8263309 | 0.8427506 | 0.8395 | 0.003250421 | -0.013169332 |
| Dact3 | 3.438451 | 3.381547 | 3.3183 | 0.063223852 | 0.12012709 | 3.3954539 | 3.2979809 | 3.29479 | 0.003191488 | 0.100664487 |
| Gm8623 | 2.227952 | 2.204433 | 2.1425 | 0.061972184 | 0.08549109 | 2.4055906 | 2.2434339 | 2.24025 | 0.00318343 | 0.165340144 |
| BC030499 | 3.74842 | 3.607913 | 3.6848 | -0.076850242 | 0.063656867 | 3.8492133 | 3.805508 | 3.80232 | 0.003183022 | 0.046888382 |
| Rasef | 2.268375 | 2.172508 | 2.2078 | -0.03525897 | 0.060608054 | 2.4172022 | 2.1646599 | 2.16153 | 0.003125026 | 0.255667358 |
| Uty | 2.783762 | 2.900455 | 2.8593 | 0.041184616 | -0.075508741 | 2.8989095 | 3.1276105 | 3.12449 | 0.003123277 | -0.225577695 |
| Gbx2 | 0.751522 | 0.73776 | 0.7155 | 0.022269065 | 0.036031779 | 0.8001249 | 0.709059 | 0.70594 | 0.003120719 | 0.094186602 |
| Olfr867 | 0.340708 | 0.366101 | 0.3881 | -0.021976875 | -0.047369867 | 0.3596296 | 0.3927216 | 0.38961 | 0.003107596 | -0.029984433 |
| Trav14n-3 | 5.166808 | 5.020862 | 5.1272 | -0.106381688 | 0.039563537 | 4.7980148 | 4.9200294 | 4.91695 | 0.003076553 | -0.118938068 |
| Ube4b | 10.36211 | 10.24696 | 10.416 | -0.168561806 | -0.053410424 | 10.448334 | 10.378205 | 10.3752 | 0.003051512 | 0.073180507 |

| | | | | | | | | | | |
|---|---|---|---|---|---|---|---|---|---|---|
| Kctd10 | 10.4709 | 10.57989 | 10.524 | 0.055536962 | -0.053453328 | 10.435665 | 10.353871 | 10.3508 | 0.003042198 | 0.084836433 |
| Axin1 | 10.82979 | 10.86086 | 10.824 | 0.036743586 | 0.005665474 | 10.736191 | 10.722227 | 10.7192 | 0.003007608 | 0.016972143 |
| Rpl17-ps8 | 3.089829 | 2.991603 | 3.0936 | -0.102015666 | -0.00379005 | 3.05144 | 3.0212207 | 3.01823 | 0.002988011 | 0.033207329 |
| Tspan4 | 7.603614 | 7.393922 | 7.4886 | -0.094673691 | 0.115018811 | 6.8909979 | 6.8781154 | 6.87515 | 0.002965661 | 0.015848202 |
| Acnat1 | 2.549337 | 2.560037 | 2.5248 | 0.035211385 | 0.024511422 | 2.4473316 | 2.5472012 | 2.54428 | 0.002919556 | -0.096950043 |
| Trav9d-4 | 0.914493 | 0.887701 | 0.9445 | -0.056824173 | -0.030032401 | 0.8586183 | 0.7909716 | 0.78813 | 0.002839472 | 0.070486172 |
| Ttc4 | 9.02545 | 9.132809 | 9.0288 | 0.103960718 | -0.003398428 | 9.3881694 | 9.3048723 | 9.30205 | 0.002819691 | 0.086116824 |
| Runx1t1 | 3.284363 | 3.257108 | 3.3594 | -0.102285984 | -0.075031115 | 3.2802689 | 3.2193823 | 3.21657 | 0.002814545 | 0.06370114 |
| Gtpbp1 | 10.9888 | 11.15988 | 11.164 | -0.004267192 | -0.17535301 | 11.091661 | 11.156853 | 11.154 | 0.002804268 | -0.062388235 |
| 4930519G04Rik | 1.385033 | 1.413162 | 1.4497 | -0.036493443 | -0.064622222 | 1.3974673 | 1.4547761 | 1.45198 | 0.002793844 | -0.05451501 |
| E430018J23Rik | 6.69855 | 6.544281 | 6.7752 | -0.23088357 | -0.076614289 | 6.8695967 | 6.7791612 | 6.77642 | 0.002741384 | 0.093176855 |
| Ugt1a6b | 0.535349 | 0.492821 | 0.4237 | 0.06916353 | 0.111691697 | 0.6337019 | 0.4178305 | 0.4151 | 0.002733553 | 0.218605029 |
| Insyn1 | 1.089411 | 1.140637 | 1.058 | 0.082623238 | 0.031396984 | 1.1108453 | 1.0543986 | 1.05171 | 0.002685986 | 0.059132691 |
| Trav17 | 3.957827 | 3.83818 | 3.922 | -0.083833378 | 0.035814348 | 3.7101755 | 3.8530395 | 3.85037 | 0.002669675 | -0.140194385 |
| Serpinb1c | 0.916982 | 0.993433 | 0.9528 | 0.040653796 | -0.035797563 | 0.9015126 | 0.9835006 | 0.98084 | 0.002660457 | -0.079327623 |
| Alx4 | 1.364312 | 1.363265 | 1.4188 | -0.05548726 | -0.054441048 | 1.3920337 | 1.4822915 | 1.47964 | 0.002655586 | -0.087602198 |
| Mrap2 | 1.578521 | 1.596901 | 1.5699 | 0.027045795 | 0.008665485 | 1.6043389 | 1.6211725 | 1.61855 | 0.002624747 | -0.014208782 |
| Asph | 7.127596 | 7.03408 | 7.0241 | 0.010007326 | 0.103523909 | 6.755629 | 6.7594456 | 6.75685 | 0.00259754 | -0.001219054 |
| Ighv3-4 | 2.965191 | 3.022011 | 2.8483 | 0.17366972 | 0.11684974 | 2.9170358 | 2.7902389 | 2.78769 | 0.002545073 | 0.129341884 |
| Gtf3c4 | 9.003579 | 8.980658 | 8.8648 | 0.115892317 | 0.138813296 | 8.794964 | 8.8434824 | 8.84095 | 0.002533186 | -0.045985164 |
| Olfr403 | 0.720772 | 0.698588 | 0.7436 | -0.044965919 | -0.022782431 | 0.7211824 | 0.7048805 | 0.70236 | 0.002519477 | 0.018821399 |
| Pip5k1a | 9.824865 | 9.883099 | 9.8782 | 0.004936437 | -0.053297306 | 9.7771405 | 9.807402 | 9.80488 | 0.002517424 | -0.027744133 |
| Hist1h2bk | 2.435465 | 2.38988 | 2.4901 | -0.100171145 | -0.054585754 | 2.6422962 | 2.4715132 | 2.469 | 0.002513813 | 0.173296836 |
| Gm9480 | 2.807098 | 2.715543 | 2.7772 | -0.061706486 | 0.029847857 | 2.6474205 | 2.7142423 | 2.71173 | 0.00250959 | -0.064312228 |
| Ccdc89 | 0.402387 | 0.414901 | 0.3961 | 0.018790161 | 0.006276645 | 0.3835843 | 0.4632416 | 0.46075 | 0.002487042 | -0.077170249 |
| Zswim8 | 10.93779 | 10.87442 | 10.98 | -0.105795178 | -0.042426798 | 11.191821 | 11.296156 | 11.2937 | 0.002478484 | -0.101857025 |
| Syncrip | 11.76624 | 11.74069 | 11.698 | 0.043176972 | 0.06873211 | 11.658655 | 11.764978 | 11.7625 | 0.002451752 | -0.103871051 |
| Ifitm5 | 1.247776 | 1.240218 | 1.2629 | -0.022710733 | -0.01515344 | 1.3616021 | 1.2374569 | 1.23505 | 0.002403522 | 0.126548631 |
| Nsdhl | 7.942613 | 7.886183 | 7.9622 | -0.075980487 | -0.019549844 | 7.9525196 | 7.9406241 | 7.93824 | 0.00238542 | 0.014280859 |
| Olfr545 | 1.435022 | 1.470167 | 1.446 | 0.024192919 | -0.010952111 | 1.4966685 | 1.5141928 | 1.51192 | 0.002270297 | -0.015253986 |
| Tbce | 9.045389 | 8.978252 | 9.1966 | -0.218304706 | -0.151168139 | 9.5641103 | 9.5784994 | 9.57632 | 0.002180562 | -0.012208526 |
| Fkbp11 | 4.918407 | 4.97729 | 4.916 | 0.061320501 | 0.002437098 | 5.1289262 | 4.7586191 | 4.75653 | 0.002087221 | 0.372394341 |

| | | | | | | | | | | |
|---|---|---|---|---|---|---|---|---|---|---|
| Gm2287 | 0.839248 | 0.81549 | 0.8765 | -0.060988301 | -0.037230055 | 0.8210089 | 0.8171309 | 0.81505 | 0.002079365 | 0.005957325 |
| Gm5564 | -0.119627 | -0.1342 | -0.1177 | -0.01645221 | -0.001883211 | -0.0740729 | -0.135078 | -0.1372 | 0.002079059 | 0.063083943 |
| 5730455P16Rik | 8.153569 | 7.820362 | 8.3514 | -0.531023188 | -0.197815657 | 8.4528708 | 8.2665673 | 8.26451 | 0.002061779 | 0.188365299 |
| Gm47605 | 0.869543 | 0.940557 | 0.8316 | 0.108942959 | 0.03792908 | 0.8214027 | 0.907243 | 0.90519 | 0.002048672 | -0.083791654 |
| Olfr1340 | 0.644466 | 0.622588 | 0.6313 | -0.008719258 | 0.013158243 | 0.6015779 | 0.6717538 | 0.66971 | 0.002044899 | -0.068130997 |
| Gm6169 | 1.941115 | 1.968392 | 2.1161 | -0.147728576 | -0.175006353 | 2.0370653 | 1.9354074 | 1.93338 | 0.002028149 | 0.103686021 |
| Mapk8ip1 | 4.436601 | 4.703839 | 4.6215 | 0.082300041 | -0.184938204 | 4.6460747 | 4.432517 | 4.4305 | 0.002020037 | 0.215577764 |
| Phf12 | 10.89774 | 10.95869 | 11.044 | -0.08505912 | -0.146002694 | 11.019595 | 11.110048 | 11.1081 | 0.001995618 | -0.088458324 |
| Gm266 | 1.25756 | 1.23641 | 1.2362 | 0.000197723 | 0.021347985 | 1.2182362 | 1.2897273 | 1.28775 | 0.001974009 | -0.069517089 |
| Rpap3 | 8.275632 | 8.243681 | 8.2573 | -0.013639575 | 0.01831105 | 8.4391011 | 8.3379434 | 8.33599 | 0.001952971 | 0.10311059 |
| Rslcan18 | 4.575167 | 4.60097 | 4.4105 | 0.19048716 | 0.164683843 | 4.5751122 | 4.7361027 | 4.73415 | 0.00194969 | -0.15904081 |
| Mrpl48 | 8.822759 | 8.768908 | 8.8372 | -0.068265762 | -0.01441449 | 8.9076763 | 8.8745295 | 8.87261 | 0.001918806 | 0.035065571 |
| Gm49530 | 1.206491 | 1.163466 | 1.1693 | -0.005847133 | 0.037178117 | 1.1725259 | 1.1944488 | 1.19254 | 0.001913401 | -0.020009458 |
| Rbm48 | 7.78376 | 7.967601 | 7.9719 | -0.004319195 | -0.188160182 | 7.9309897 | 8.0004468 | 7.99853 | 0.001912138 | -0.067545031 |
| Sat2 | 0.785721 | 0.754106 | 0.7917 | -0.037603232 | -0.005988765 | 0.7603856 | 0.728976 | 0.72712 | 0.001857655 | 0.033267316 |
| Gm12846 | 0.547745 | 0.49079 | 0.5351 | -0.044343346 | 0.012612292 | 0.5012902 | 0.5090027 | 0.50716 | 0.001841975 | -0.00587053 |
| Tle4 | 10.32644 | 10.42514 | 10.366 | 0.058843504 | -0.039850554 | 10.213709 | 10.308773 | 10.307 | 0.00176766 | -0.09329608 |
| Gm11127 | 0.286353 | 0.292241 | 0.272 | 0.020269553 | 0.014382059 | 0.3293862 | 0.2679994 | 0.26625 | 0.001747798 | 0.063134606 |
| Shisa7 | 2.114848 | 2.049077 | 2.0717 | -0.022636506 | 0.043133979 | 1.9687333 | 1.9275803 | 1.92584 | 0.001741215 | 0.04289424 |
| Sh2d6 | 0.807529 | 0.814097 | 0.8293 | -0.015245205 | -0.021813271 | 0.8385038 | 0.8041438 | 0.80241 | 0.001737819 | 0.03609785 |
| Esp18 | -0.034761 | -0.15399 | -0.1153 | -0.038710679 | 0.08051887 | -0.0408612 | -0.151545 | -0.1532 | 0.001703454 | 0.112387456 |
| Trpc6 | 2.129253 | 2.164153 | 2.1818 | -0.017624253 | -0.05252466 | 2.168387 | 2.1625065 | 2.16084 | 0.001661837 | 0.007542344 |
| Ighv1-70 | 2.063262 | 2.093812 | 2.1504 | -0.056632045 | -0.08718274 | 2.2378147 | 2.1717756 | 2.17015 | 0.001621508 | 0.067660606 |
| Gm10778 | 6.816172 | 6.876921 | 7.0063 | -0.129360758 | -0.190109416 | 6.7170134 | 6.7203253 | 6.71871 | 0.001611251 | -0.001700581 |
| Gcc2 | 8.729175 | 8.770663 | 8.8847 | -0.114000693 | -0.155488815 | 8.8112338 | 8.8738385 | 8.87225 | 0.001587272 | -0.06101736 |
| Aox3 | 1.209326 | 1.109737 | 1.2102 | -0.100431988 | -0.000843294 | 1.147221 | 1.1654611 | 1.16394 | 0.001521058 | -0.016719054 |
| Ugt1a1 | -0.035606 | -0.05683 | -0.1289 | 0.072017806 | 0.093244578 | -0.0366828 | -0.125372 | -0.1269 | 0.001504324 | 0.090193693 |
| Rps8-ps4 | 2.470632 | 2.426083 | 2.4642 | -0.038164626 | 0.006384237 | 2.428817 | 2.4513445 | 2.44987 | 0.001477874 | -0.021049635 |
| Rpl30-ps8 | 1.646642 | 1.534315 | 1.6209 | -0.086583534 | 0.025743455 | 1.6736986 | 1.5478545 | 1.54638 | 0.001476358 | 0.127320386 |
| Gm11928 | -0.275717 | -0.27735 | -0.2671 | -0.010260105 | -0.008629311 | -0.2966709 | -0.2949 | -0.2964 | 0.00147443 | -0.000296669 |
| Pik3cb | 8.259452 | 8.313708 | 8.1928 | 0.120861476 | 0.066605593 | 8.193157 | 8.0487771 | 8.04734 | 0.001433538 | 0.145813417 |
| Ncoa3 | 11.84143 | 11.96702 | 11.853 | 0.114128732 | -0.011459302 | 11.860702 | 12.019933 | 12.0185 | 0.00142994 | -0.157801103 |

| | | | | | | | | | | |
|---|---|---|---|---|---|---|---|---|---|---|
| Gm10985 | 1.171733 | 1.249 | 1.2123 | 0.036714209 | -0.040552801 | 1.2283547 | 1.2231442 | 1.2218 | 0.001343761 | 0.006554254 |
| Sirt2 | 9.91491 | 9.855579 | 9.9561 | -0.100554961 | -0.041223836 | 9.997781 | 9.8885956 | 9.88729 | 0.001308571 | 0.110494007 |
| Cacfd1 | 9.280636 | 9.215638 | 9.2594 | -0.04380141 | 0.021197169 | 9.4956735 | 9.2442516 | 9.24295 | 0.001306027 | 0.252727942 |
| Acsl4 | 9.990304 | 9.980232 | 10.013 | -0.032635796 | -0.022563532 | 9.7922846 | 9.9023987 | 9.90112 | 0.001274961 | -0.108839124 |
| Fstl3 | 3.498763 | 3.509249 | 3.5292 | -0.019937725 | -0.030424174 | 3.5529466 | 3.4189458 | 3.41774 | 0.001202571 | 0.135203377 |
| Sult2a8 | 0.371514 | 0.390948 | 0.4335 | -0.042574764 | -0.062008993 | 0.3664741 | 0.4352102 | 0.43402 | 0.001190486 | -0.067545652 |
| Gm16223 | 0.356441 | 0.366117 | 0.3348 | 0.031268912 | 0.021592756 | 0.3418188 | 0.3899405 | 0.38876 | 0.001180778 | -0.046940864 |
| Pced1b | 10.03768 | 10.16013 | 10.075 | 0.085530158 | -0.036919016 | 10.152977 | 10.111385 | 10.1102 | 0.001173344 | 0.042765804 |
| Zfp984 | 9.463984 | 9.563652 | 9.6122 | -0.048530925 | -0.148198883 | 9.7273975 | 9.7025854 | 9.70142 | 0.00116974 | 0.025981839 |
| Ap2a1 | 10.08708 | 9.968104 | 10.077 | -0.109236255 | 0.009737809 | 10.105314 | 10.11477 | 10.1136 | 0.001149989 | -0.008305901 |
| Tbc1d10b | 9.921572 | 10.08372 | 9.9225 | 0.161174681 | -0.000971839 | 9.7854542 | 10.101559 | 10.1004 | 0.001136415 | -0.314968532 |
| H2-Ab1 | 14.76289 | 14.69452 | 14.759 | -0.064625502 | 0.003746396 | 15.250667 | 14.713523 | 14.7124 | 0.001126422 | 0.538269472 |
| Rfx5 | 10.22776 | 10.33642 | 10.282 | 0.05475171 | -0.05390983 | 10.553295 | 10.521976 | 10.5209 | 0.001115594 | 0.032434662 |
| Gcc1 | 9.375258 | 9.163337 | 9.0652 | 0.098162079 | 0.310082395 | 9.1262844 | 9.1037029 | 9.10262 | 0.001082378 | 0.0236638 |
| Klhl35 | 2.261797 | 2.175699 | 2.2332 | -0.057549412 | 0.028548673 | 2.2750948 | 2.2518674 | 2.25079 | 0.0010811 | 0.024308543 |
| Olfr830 | 0.897382 | 0.912341 | 0.9546 | -0.042306037 | -0.057264502 | 0.9592817 | 0.9617926 | 0.96073 | 0.001058628 | -0.001452283 |
| Tmem40 | 1.224821 | 1.214221 | 1.2218 | -0.007593756 | 0.003006495 | 1.2236328 | 1.2108336 | 1.20978 | 0.001056488 | 0.013855679 |
| Edem3 | 9.875047 | 9.861506 | 9.8408 | 0.020752995 | 0.034293918 | 9.4948397 | 9.7029413 | 9.70189 | 0.001050083 | -0.207051477 |
| H3f3c | 1.465166 | 1.369881 | 1.5114 | -0.141507064 | -0.046222208 | 1.4129359 | 1.371006 | 1.36997 | 0.001036906 | 0.042966743 |
| Gm47355 | 4.170353 | 4.176345 | 4.1517 | 0.024688897 | 0.018696759 | 4.1270495 | 4.4542157 | 4.45319 | 0.001026692 | -0.326139534 |
| Cfp | 9.397022 | 9.16444 | 9.0136 | 0.150817314 | 0.383399461 | 9.4933386 | 8.7210586 | 8.72006 | 0.000999638 | 0.773279615 |
| Cblb | 11.00743 | 11.097 | 11.066 | 0.03078769 | -0.058776448 | 10.798147 | 11.099861 | 11.0989 | 0.000992826 | -0.300721044 |
| Cfap43 | 1.829581 | 1.847737 | 1.8504 | -0.002662607 | -0.020818989 | 1.7518284 | 1.8914549 | 1.89051 | 0.00094045 | -0.138686105 |
| Oxr1 | 10.133 | 10.1555 | 10.152 | 0.003808872 | -0.018697674 | 10.091912 | 10.00874 | 10.0078 | 0.00093567 | 0.084107595 |
| Bcl2l15 | 1.849267 | 1.764774 | 1.79 | -0.025218237 | 0.059274188 | 1.8043231 | 1.7918662 | 1.79099 | 0.000877523 | 0.013334454 |
| Tnpo3 | 10.65769 | 10.59036 | 10.684 | -0.093883128 | -0.026547318 | 10.654602 | 10.689646 | 10.6888 | 0.000869998 | -0.034174446 |
| Setd3 | 10.40314 | 10.35244 | 10.403 | -0.050946528 | -0.000247648 | 10.357336 | 10.430143 | 10.4293 | 0.000863764 | -0.071942791 |
| Srrm3 | 1.825747 | 1.854574 | 1.8193 | 0.035290283 | 0.006463415 | 1.8557117 | 1.8069829 | 1.80614 | 0.000845506 | 0.049574352 |
| Gm43498 | 3.896158 | 3.950172 | 3.8252 | 0.124954537 | 0.07094036 | 3.8416969 | 4.051889 | 4.05105 | 0.000836536 | -0.209355527 |
| Tasp1 | 8.04341 | 7.90522 | 7.9918 | -0.086562077 | 0.051628324 | 7.8333275 | 7.9396308 | 7.93883 | 0.000799468 | -0.105503854 |
| Lypd6 | 0.539258 | 0.538795 | 0.6465 | -0.107657272 | -0.107193656 | 0.5645702 | 0.5689582 | 0.56816 | 0.000798983 | -0.003589051 |
| Rgs20 | 1.147878 | 1.107008 | 1.1182 | -0.011217882 | 0.029651658 | 1.0751311 | 1.13547 | 1.13468 | 0.00079498 | -0.05954398 |

| | | | | | | | | | | |
|---|---|---|---|---|---|---|---|---|---|---|
| Atp1a1 | 12.65311 | 12.72074 | 12.546 | 0.174489562 | 0.106857848 | 12.59617 | 12.523353 | 12.5226 | 0.000767718 | 0.073584874 |
| Gm19353 | 0.316038 | 0.30942 | 0.3179 | -0.008441429 | -0.001823705 | 0.3910393 | 0.320049 | 0.31932 | 0.000730083 | 0.071720416 |
| Ap5s1 | 7.000307 | 6.948352 | 7.0013 | -0.05293169 | -0.00097656 | 7.0057205 | 6.9206464 | 6.91993 | 0.000714675 | 0.085788871 |
| Olfr133 | 1.826238 | 1.789785 | 1.785 | 0.004795051 | 0.041247836 | 1.9213988 | 1.8997197 | 1.89905 | 0.000665785 | 0.022344849 |
| Soat1 | 10.11542 | 10.31923 | 9.946 | 0.37318209 | 0.169374173 | 10.177588 | 10.042693 | 10.042 | 0.000660797 | 0.135555509 |
| Gm17655 | 6.250763 | 6.194021 | 6.1526 | 0.04138506 | 0.098127732 | 6.208301 | 6.4290456 | 6.42838 | 0.000660655 | -0.220083861 |
| Tmem132e | 2.920977 | 2.990615 | 2.6757 | 0.314870089 | 0.245231905 | 2.7527151 | 2.8487037 | 2.84804 | 0.000658679 | -0.095329932 |
| Hddc2 | 7.775336 | 7.57859 | 7.8158 | -0.237163615 | -0.040417048 | 7.9217032 | 7.7598537 | 7.75922 | 0.000630025 | 0.162479503 |
| Ttyh3 | 11.20166 | 11.09022 | 11.107 | -0.016678256 | 0.094753794 | 11.304255 | 11.099331 | 11.0987 | 0.000614606 | 0.205538132 |
| Galnt13 | 0.538904 | 0.536653 | 0.5463 | -0.009619905 | -0.00736894 | 0.5306789 | 0.5586924 | 0.55808 | 0.000608127 | -0.027405392 |
| Tmed10-ps | 1.522964 | 1.464053 | 1.5649 | -0.100878184 | -0.041967057 | 1.6025992 | 1.4918687 | 1.49128 | 0.000586716 | 0.111317209 |
| Gnaz | 3.451621 | 3.478764 | 3.4892 | -0.010459441 | -0.037602601 | 3.5970313 | 3.4588889 | 3.45831 | 0.000581067 | 0.138723499 |
| Gm7420 | 0.853011 | 0.906943 | 0.9087 | -0.001736664 | -0.05566881 | 0.8556671 | 0.8762251 | 0.87566 | 0.00056304 | -0.019994927 |
| Vmn1r188 | 3.226794 | 3.193065 | 3.1966 | -0.003518243 | 0.030210594 | 3.177237 | 3.2836097 | 3.28307 | 0.000544543 | -0.105828132 |
| BC005561 | 7.51575 | 7.614106 | 7.5358 | 0.078282406 | -0.020073027 | 7.4729922 | 7.6196256 | 7.61909 | 0.000537009 | -0.146096434 |
| Ighv5-17 | 7.541598 | 7.636706 | 7.648 | -0.01134189 | -0.106450805 | 7.8829432 | 7.5592935 | 7.55882 | 0.000472616 | 0.324122316 |
| Cpz | 0.968189 | 0.868316 | 0.9089 | -0.040622486 | 0.059250382 | 0.8460178 | 0.8338915 | 0.83351 | 0.000380088 | 0.012506377 |
| Trav4-2 | 2.694625 | 2.605831 | 2.654 | -0.048123046 | 0.040670663 | 2.6605131 | 2.5981295 | 2.59786 | 0.000267141 | 0.062650775 |
| Foxo3 | 10.33602 | 10.51085 | 10.401 | 0.11012726 | -0.064696086 | 10.294222 | 10.290969 | 10.2907 | 0.000259052 | 0.003511552 |
| Pkp4 | 8.715996 | 8.647044 | 8.6749 | -0.027897865 | 0.04105475 | 8.6205028 | 8.6640638 | 8.66385 | 0.000211409 | -0.043349616 |
| Kctd5 | 8.714418 | 8.713907 | 8.6386 | 0.075266007 | 0.075777107 | 8.6930406 | 8.6855526 | 8.68535 | 0.000206841 | 0.007694828 |
| Gm11687 | 0.890992 | 0.8757 | 0.9077 | -0.031988208 | -0.016696441 | 0.8915292 | 0.9160553 | 0.91586 | 0.000197981 | -0.024328128 |
| Gm29761 | 0.566813 | 0.568751 | 0.6227 | -0.053965748 | -0.055903534 | 0.5791991 | 0.6039556 | 0.6038 | 0.000151801 | -0.024604698 |
| Cacna1f | 1.529524 | 1.474768 | 1.5447 | -0.069914722 | -0.015158629 | 1.5133318 | 1.5644451 | 1.56435 | 9.73028E-05 | -0.05101603 |
| Olfr1471 | 0.423551 | 0.398774 | 0.4327 | -0.033971051 | -0.009194449 | 0.5135617 | 0.4113219 | 0.41124 | 8.6704E-05 | 0.102326495 |
| Mlph | 4.498603 | 4.456469 | 4.6749 | -0.218413311 | -0.176279462 | 4.7026305 | 4.5277849 | 4.52773 | 5.62412E-05 | 0.17490187 |
| Pla2g4e | 1.952285 | 1.940327 | 1.9033 | 0.037022659 | 0.048980835 | 1.9612507 | 1.9727347 | 1.9727 | 3.79724E-05 | -0.011446101 |
| Gm11620 | 0.464962 | 0.494604 | 0.4256 | 0.06896386 | 0.039321472 | 0.4787916 | 0.4835765 | 0.48356 | 1.43634E-05 | -0.00477055 |
| BC049352 | 3.186708 | 3.155996 | 3.2556 | -0.099571057 | -0.068858718 | 3.1873767 | 3.2490885 | 3.24908 | 1.10902E-05 | -0.061700694 |
| Chil6 | 0.610241 | 0.598418 | 0.5789 | 0.019478856 | 0.03130137 | 0.5928101 | 0.5711452 | 0.57117 | -2.5941E-05 | 0.02163894 |
| Tmem238 | 4.522331 | 4.423969 | 4.4887 | -0.064722586 | 0.033639308 | 4.5381594 | 4.3013339 | 4.30137 | -4.03734E-05 | 0.236785089 |
| C330007P06Rik | 9.434881 | 9.412034 | 9.4549 | -0.042833762 | -0.019986976 | 9.4970699 | 9.4996371 | 9.49968 | -4.52278E-05 | -0.002612486 |

| | | | | | | | | | | |
|---|---|---|---|---|---|---|---|---|---|---|
| Fstl4 | 1.802613 | 1.838468 | 1.8702 | -0.031756165 | -0.067611248 | 1.7605888 | 1.8312864 | 1.83137 | -8.41414E-05 | -0.070781705 |
| Rnf166 | 9.487101 | 9.514638 | 9.529 | -0.014400286 | -0.041937398 | 9.6156288 | 9.4654319 | 9.46558 | -0.000150684 | 0.150046214 |
| Katnal2 | 0.716702 | 0.714138 | 0.674 | 0.040091256 | 0.042655416 | 0.6645956 | 0.6771939 | 0.67735 | -0.000152755 | -0.012751072 |
| Olfr1157 | 1.550303 | 1.485903 | 1.5044 | -0.018460389 | 0.045940187 | 1.5485141 | 1.5444499 | 1.54465 | -0.00020253 | 0.003861698 |
| Trav13n-3 | 2.013054 | 1.847045 | 1.9078 | -0.060739687 | 0.105269559 | 1.8492254 | 1.8688334 | 1.86905 | -0.000219052 | -0.019827085 |
| Gm2805 | 0.435382 | 0.375936 | 0.3854 | -0.009496932 | 0.049948831 | 0.4116736 | 0.3926752 | 0.3929 | -0.000226766 | 0.018771601 |
| Vmn1r27 | 1.289398 | 1.271924 | 1.3214 | -0.049524371 | -0.03205042 | 1.2783495 | 1.289836 | 1.29008 | -0.000245061 | -0.011731617 |
| Sqor | 6.630757 | 6.601811 | 6.4762 | 0.125590933 | 0.154536962 | 6.3994961 | 6.2028391 | 6.20316 | -0.000324128 | 0.19633283 |
| Rps25-ps1 | 1.706805 | 1.715999 | 1.7578 | -0.041761485 | -0.050955805 | 1.7224179 | 1.7353418 | 1.7357 | -0.000356726 | -0.013280665 |
| Ddr2 | 7.571317 | 7.284679 | 7.1273 | 0.157358752 | 0.443996747 | 6.6130266 | 6.6063434 | 6.60675 | -0.000405953 | 0.006277277 |
| Prdm15 | 8.860687 | 9.059696 | 8.9211 | 0.138570663 | -0.060437719 | 8.8146105 | 9.0519581 | 9.05237 | -0.000410003 | -0.237757587 |
| Prdm11 | 8.867863 | 9.084831 | 8.8177 | 0.267121908 | 0.050154521 | 8.9951092 | 8.9058237 | 8.90624 | -0.000420122 | 0.088865374 |
| Gdap2 | 8.607361 | 8.672266 | 8.6422 | 0.030069793 | -0.034835293 | 8.5539289 | 8.5765896 | 8.57714 | -0.000554331 | -0.02321502 |
| Olfr1406 | 0.642701 | 0.592956 | 0.5846 | 0.008384555 | 0.058129823 | 0.5596438 | 0.6517241 | 0.65228 | -0.000559569 | -0.092639833 |
| Npdc1 | 7.023613 | 6.89469 | 6.7362 | 0.158504946 | 0.287427421 | 6.6416026 | 6.4795749 | 6.48015 | -0.000575123 | 0.161452488 |
| Usp44 | 1.539203 | 1.519796 | 1.5814 | -0.061566573 | -0.042160052 | 1.512625 | 1.5229416 | 1.52353 | -0.000585428 | -0.010902023 |
| Gm18860 | 0.021133 | 0.015038 | 0.0437 | -0.028621687 | -0.022527045 | 0.0452776 | 0.0668147 | 0.06748 | -0.000666504 | -0.022203572 |
| Esp6 | -0.132903 | -0.14677 | -0.1423 | -0.004439388 | 0.009430083 | -0.1612361 | -0.094634 | -0.094 | -0.00067841 | -0.06728099 |
| 4930578G10Rik | 0.538025 | 0.49867 | 0.504 | -0.005332673 | 0.034022386 | 0.4696268 | 0.4653131 | 0.46601 | -0.000695808 | 0.003617871 |
| Ubr2 | 11.09468 | 11.19708 | 11.187 | 0.009780146 | -0.092617548 | 11.161768 | 11.36279 | 11.3635 | -0.000712186 | -0.201734805 |
| Rps27a-ps1 | 1.619693 | 1.557142 | 1.5711 | -0.013914678 | 0.048636196 | 1.6012568 | 1.5717125 | 1.57242 | -0.000712251 | 0.028832064 |
| Rps3 | 15.09379 | 15.0704 | 15.025 | 0.045263106 | 0.068654984 | 15.045716 | 15.007714 | 15.0084 | -0.00072605 | 0.037275965 |
| Hdac1 | 11.36182 | 11.32716 | 11.408 | -0.080940708 | -0.046280938 | 11.559959 | 11.422708 | 11.4234 | -0.000731112 | 0.136519588 |
| Clcn6 | 8.927972 | 8.97023 | 9.0262 | -0.055979058 | -0.098237516 | 9.0412707 | 9.0992351 | 9.09997 | -0.000731491 | -0.058695878 |
| Plb1 | 1.598806 | 1.652176 | 1.593 | 0.059181711 | 0.005811351 | 1.5471543 | 1.5269587 | 1.52774 | -0.000785171 | 0.019410445 |
| Gimap6 | 13.12739 | 13.1021 | 13.176 | -0.074026312 | -0.048740596 | 13.109461 | 13.08083 | 13.0816 | -0.000790166 | 0.027840479 |
| Nxph3 | 1.217603 | 1.182383 | 1.34 | -0.15757415 | -0.122353982 | 1.139182 | 1.2247725 | 1.22561 | -0.000834545 | -0.086425022 |
| Cdiptos | 3.304538 | 3.188259 | 3.1778 | 0.010500765 | 0.126779736 | 3.0786404 | 3.1700683 | 3.17091 | -0.000844375 | -0.092272322 |
| 1190005I06Rik | 1.824231 | 1.797196 | 1.7624 | 0.03478524 | 0.061819596 | 1.8189653 | 1.8932166 | 1.89407 | -0.000854282 | -0.075105574 |
| Cdpf1 | 7.14885 | 7.233533 | 7.2403 | -0.006762687 | -0.091445702 | 7.3682221 | 7.2595677 | 7.26043 | -0.000857813 | 0.107796564 |
| Gm20637 | 1.581159 | 1.589912 | 1.6101 | -0.020201233 | -0.028954176 | 1.5788549 | 1.5692565 | 1.57012 | -0.000859786 | 0.008738623 |
| Ncoa5 | 9.787278 | 9.833004 | 9.8581 | -0.025133807 | -0.070860426 | 9.916095 | 9.9043595 | 9.90523 | -0.000868715 | 0.010866715 |

| | | | | | | | | | | |
|---|---|---|---|---|---|---|---|---|---|---|
| Olfr1033 | 12.33244 | 12.22924 | 12.054 | 0.175155272 | 0.278352872 | 12.112292 | 12.024493 | 12.0254 | -0.000873522 | 0.086925399 |
| Teddm2 | 3.252969 | 3.310186 | 3.3398 | -0.029589889 | -0.086806506 | 3.2984051 | 3.3997868 | 3.40067 | -0.000882066 | -0.102263842 |
| Clptm1 | 10.77217 | 10.76901 | 10.766 | 0.003060182 | 0.006221338 | 10.880317 | 10.792284 | 10.7932 | -0.000884018 | 0.087148554 |
| Zcchc4 | 8.280889 | 8.286274 | 8.3791 | -0.092816184 | -0.098201766 | 8.4977861 | 8.384734 | 8.38563 | -0.000897649 | 0.112154439 |
| Trav8-1 | 4.139457 | 4.029766 | 4.0175 | 0.012256268 | 0.121947468 | 3.9132825 | 4.0016469 | 4.00256 | -0.000909435 | -0.089273779 |
| Nradd | 4.927493 | 4.844624 | 4.8922 | -0.047561767 | 0.035307206 | 5.0772799 | 4.948991 | 4.94992 | -0.000928729 | 0.127360098 |
| Olfr588-ps1 | 0.69299 | 0.681402 | 0.7171 | -0.035667036 | -0.024078863 | 0.6776348 | 0.7765804 | 0.77752 | -0.000943769 | -0.099889329 |
| Ighv1-37 | 2.272004 | 2.227317 | 2.2274 | -9.69516E-05 | 0.044590693 | 2.2562207 | 2.1697758 | 2.17072 | -0.000945167 | 0.085499778 |
| Olfr394 | 1.092151 | 1.105807 | 1.1065 | -0.000698162 | -0.01435502 | 1.0920832 | 1.1468761 | 1.14785 | -0.000978352 | -0.055771296 |
| Dmxl2 | 6.391825 | 6.238746 | 5.8013 | 0.437454489 | 0.590533217 | 6.1867703 | 6.0062907 | 6.00729 | -0.000998835 | 0.179480756 |
| Pycr1 | 5.052034 | 5.015921 | 4.827 | 0.188912765 | 0.225025024 | 5.3067613 | 5.3317576 | 5.33276 | -0.001005428 | -0.026001772 |
| Tc2n | 4.522254 | 4.230658 | 4.3417 | -0.111073212 | 0.180523139 | 4.1573194 | 4.2451542 | 4.24621 | -0.001056791 | -0.088891572 |
| Atp6v1fnb | 0.738233 | 0.742072 | 0.7859 | -0.043832632 | -0.047671778 | 0.7761021 | 0.7678172 | 0.76889 | -0.001071437 | 0.007213455 |
| Fbxw11 | 10.12914 | 10.20682 | 10.069 | 0.137332973 | 0.059650565 | 9.8478992 | 10.057531 | 10.0586 | -0.001110352 | -0.210742466 |
| Gm14537 | 1.650925 | 1.62423 | 1.6113 | 0.012962403 | 0.039657291 | 1.6605613 | 1.6822691 | 1.6834 | -0.001128043 | -0.022835895 |
| Vmn1r38 | 1.537051 | 1.453872 | 1.5631 | -0.10921392 | -0.026034524 | 1.4905637 | 1.5182825 | 1.51941 | -0.001129983 | -0.028848727 |
| Afg1l | 6.826042 | 6.842031 | 6.8235 | 0.018511078 | 0.002521747 | 6.761918 | 6.8989442 | 6.90008 | -0.001137555 | -0.138163698 |
| Trav12d-1 | 2.307434 | 2.386974 | 2.3547 | 0.032235049 | -0.04730509 | 2.3073054 | 2.3263368 | 2.32748 | -0.001144874 | -0.020176222 |
| Trav12n-1 | 2.307434 | 2.386974 | 2.3547 | 0.032235049 | -0.04730509 | 2.3073054 | 2.3263368 | 2.32748 | -0.001144874 | -0.020176222 |
| Osbpl8 | 10.61535 | 10.64776 | 10.64 | 0.00776608 | -0.02464744 | 10.442499 | 10.60481 | 10.606 | -0.001145112 | -0.163456634 |
| Henmt1 | 2.586708 | 2.589791 | 2.5152 | 0.074560844 | 0.071477785 | 2.5342081 | 2.5497545 | 2.55094 | -0.001182066 | -0.016728477 |
| Rictor | 10.83967 | 10.87777 | 10.848 | 0.029782094 | -0.008321192 | 10.715214 | 10.94757 | 10.9488 | -0.001217328 | -0.233572489 |
| Olfr967 | 1.358472 | 1.367294 | 1.421 | -0.053679989 | -0.062502627 | 1.4106108 | 1.3980421 | 1.39926 | -0.001221699 | 0.011346915 |
| Cd276 | 4.91115 | 5.267002 | 4.9081 | 0.358920658 | 0.003068414 | 5.1936035 | 4.9674225 | 4.96868 | -0.001256969 | 0.224924053 |
| Adora1 | 1.649965 | 1.621175 | 1.4841 | 0.137122806 | 0.165912725 | 1.5381368 | 1.5357388 | 1.53701 | -0.001268809 | 0.001129155 |
| Btnl7-ps | 1.977312 | 2.029937 | 1.9852 | 0.044760672 | -0.007864362 | 1.9476178 | 2.0564589 | 2.05773 | -0.00127141 | -0.110112445 |
| Olfr600 | 3.588193 | 3.638641 | 3.5235 | 0.115120368 | 0.064672533 | 3.6078122 | 3.6440658 | 3.64534 | -0.001274711 | -0.037528289 |
| Pdcd7 | 8.491577 | 8.612995 | 8.521 | 0.091977348 | -0.029441028 | 8.5137758 | 8.5211121 | 8.52243 | -0.001320791 | -0.008657072 |
| Gm12291 | 1.189696 | 1.011098 | 1.1417 | -0.130571635 | 0.048026355 | 0.9834535 | 1.0163481 | 1.01768 | -0.001332231 | -0.034226816 |
| 4930474N05Rik | 0.409178 | 0.436073 | 0.4571 | -0.021001174 | -0.047896102 | 0.3896038 | 0.4197294 | 0.42107 | -0.001339932 | -0.031465569 |
| Ighv3-2 | 3.10213 | 3.096949 | 3.0176 | 0.0793485 | 0.084528526 | 2.934514 | 3.0469177 | 3.04827 | -0.001352499 | -0.113756182 |
| Igkv1-99 | 5.587961 | 5.859424 | 5.2461 | 0.613307193 | 0.341843697 | 5.5620842 | 5.5271464 | 5.52851 | -0.001365446 | 0.033572445 |

| | | | | | | | | | | |
|---|---|---|---|---|---|---|---|---|---|---|
| Tmigd3 | 0.47866 | 0.445592 | 0.4822 | -0.036644787 | -0.003576355 | 0.397941 | 0.4377524 | 0.43913 | -0.001382458 | -0.041193819 |
| Prpf39 | 10.58179 | 10.51947 | 10.597 | -0.077216059 | -0.014897498 | 10.616837 | 10.851926 | 10.8534 | -0.001471392 | -0.236559754 |
| Crisp2 | 0.764179 | 0.813278 | 0.8107 | 0.002545974 | -0.046553427 | 0.8496967 | 0.8321787 | 0.83366 | -0.001482657 | 0.016035318 |
| Ighv5-15 | 4.076902 | 4.66405 | 4.2462 | 0.417799899 | -0.169347742 | 4.2195168 | 4.1597127 | 4.16122 | -0.001504823 | 0.058299251 |
| 37681 | 6.454864 | 6.428926 | 6.4955 | -0.066553849 | -0.040615843 | 6.2592571 | 6.3411063 | 6.34263 | -0.001521929 | -0.083371103 |
| Ighv3-5 | 5.085116 | 5.684884 | 5.572 | 0.112891615 | -0.486876164 | 5.1423757 | 5.189141 | 5.19077 | -0.001631635 | -0.048396934 |
| Eif2ak1 | 10.19103 | 10.199 | 10.213 | -0.014368122 | -0.022339664 | 10.222106 | 10.184272 | 10.1859 | -0.0016438 | 0.036190608 |
| Smarca5-ps | 2.226064 | 2.165474 | 2.2007 | -0.035208616 | 0.025380858 | 2.1752994 | 2.2857749 | 2.28742 | -0.001646491 | -0.112122066 |
| Slc25a17 | 9.025871 | 9.10894 | 9.1905 | -0.081602823 | -0.16467215 | 9.2142726 | 9.0793057 | 9.08099 | -0.00168724 | 0.133279725 |
| Hpdl | 0.802886 | 0.810591 | 0.8901 | -0.079510473 | -0.087215564 | 0.8601161 | 0.8222176 | 0.82392 | -0.001698313 | 0.036200189 |
| Ppil3 | 8.440614 | 8.431067 | 8.413 | 0.018064581 | 0.027611566 | 8.5618768 | 8.5393688 | 8.54109 | -0.001721594 | 0.020786426 |
| Ccin | 1.173538 | 1.158677 | 1.1923 | -0.033613242 | -0.018753103 | 1.1378577 | 1.1686639 | 1.17039 | -0.001722675 | -0.032528903 |
| Rps17 | 13.35348 | 13.35751 | 13.377 | -0.019498248 | -0.023528072 | 13.400259 | 13.370885 | 13.3726 | -0.001752932 | 0.027620367 |
| Tex38 | 1.417839 | 1.456512 | 1.4328 | 0.023719943 | -0.014953139 | 1.4255505 | 1.4074512 | 1.40927 | -0.001820042 | 0.016279268 |
| Gm6278 | 0.538398 | 0.513446 | 0.5208 | -0.007383445 | 0.017568187 | 0.552537 | 0.5637214 | 0.56557 | -0.001847529 | -0.013031955 |
| Bicd2 | 9.799609 | 9.875771 | 9.8455 | 0.03025721 | -0.045905058 | 9.6867613 | 9.7946621 | 9.79657 | -0.001912272 | -0.109813076 |
| Gfod2 | 6.357712 | 6.628536 | 6.47 | 0.158498229 | -0.11232542 | 6.2898247 | 6.3262357 | 6.32816 | -0.001928815 | -0.038339825 |
| Wnt16 | 1.509887 | 1.580847 | 1.5169 | 0.063913545 | -0.007046557 | 1.5762209 | 1.5696581 | 1.57162 | -0.001964315 | 0.004598526 |
| Agpat3 | 10.71839 | 10.76781 | 10.714 | 0.053751526 | 0.004327468 | 10.919708 | 10.751235 | 10.7532 | -0.001989064 | 0.166483654 |
| Pcdhb11 | 1.319256 | 1.26566 | 1.2898 | -0.02418466 | 0.029411859 | 1.3271569 | 1.2665316 | 1.26852 | -0.001990049 | 0.058635256 |
| Olfr1039 | 2.34116 | 2.277803 | 2.3157 | -0.037883105 | 0.025473965 | 2.2443122 | 2.4741202 | 2.47612 | -0.001996268 | -0.231804251 |
| Ighv1-47 | 5.283083 | 5.333724 | 5.2919 | 0.041823957 | -0.008816849 | 5.485739 | 5.2943243 | 5.29633 | -0.0020101 | 0.189404604 |
| Gli2 | 2.669921 | 2.965766 | 2.7407 | 0.22510655 | -0.070738653 | 2.7473642 | 2.5982477 | 2.60026 | -0.002011222 | 0.147105223 |
| Cstf3 | 8.765085 | 8.703571 | 8.8959 | -0.192354693 | -0.130840786 | 9.0736784 | 9.1082789 | 9.11029 | -0.00201149 | -0.036611981 |
| Gm9894 | 2.181249 | 2.150118 | 2.1736 | -0.023437952 | 0.007692613 | 2.1307358 | 2.1521492 | 2.1542 | -0.002053774 | -0.023467145 |
| Gm48401 | 3.902148 | 4.163563 | 4.0178 | 0.145757013 | -0.11565794 | 4.0918762 | 4.2322834 | 4.23434 | -0.00205541 | -0.142462637 |
| Matr3-ps2 | 0.731098 | 0.61692 | 0.651 | -0.034114152 | 0.080063991 | 0.6373743 | 0.6662144 | 0.66832 | -0.002109549 | -0.03094971 |
| Qsox2 | 7.905312 | 7.825852 | 7.8041 | 0.02172756 | 0.101187158 | 7.9532154 | 7.7456291 | 7.74775 | -0.0021259 | 0.205460442 |
| Kank3 | 6.337214 | 6.423844 | 6.4733 | -0.04944964 | -0.136079202 | 6.2803712 | 6.2034074 | 6.20556 | -0.002149437 | 0.074814396 |
| Tlcd2 | 5.358878 | 5.060918 | 5.1902 | -0.129249688 | 0.168710324 | 5.1877063 | 5.0125999 | 5.01479 | -0.002187101 | 0.172919327 |
| Olfr1293-ps | 1.868719 | 1.81483 | 1.7797 | 0.035110244 | 0.088999087 | 1.7596077 | 1.7904715 | 1.79267 | -0.002203025 | -0.033066826 |
| Chrne | 1.945516 | 1.969905 | 2.0503 | -0.080344927 | -0.104734056 | 2.0534715 | 1.9851323 | 1.98735 | -0.002221363 | 0.066117893 |

| | | | | | | | | | | |
|---|---|---|---|---|---|---|---|---|---|---|
| Fam104a | 9.076603 | 9.112986 | 9.0575 | 0.055480197 | 0.01909712 | 9.1670527 | 8.962446 | 8.96467 | -0.002226071 | 0.202380648 |
| Tti1 | 9.410016 | 9.494101 | 9.431 | 0.063077296 | -0.02100747 | 9.5062828 | 9.5159738 | 9.51823 | -0.002252072 | -0.011942989 |
| Sfrp2 | 2.315702 | 2.496958 | 2.3597 | 0.13722274 | -0.044032987 | 2.300118 | 2.2237411 | 2.22602 | -0.002274394 | 0.074102479 |
| Sv2b | 2.063561 | 2.026775 | 2.037 | -0.010182476 | 0.026603218 | 1.989508 | 2.0680221 | 2.07034 | -0.002315205 | -0.080829284 |
| Crebbp | 11.05676 | 11.19958 | 11.128 | 0.071179436 | -0.071641248 | 10.74565 | 11.168398 | 11.1707 | -0.002338114 | -0.425086554 |
| Gpkow | 9.906407 | 10.04624 | 9.8609 | 0.185370687 | 0.045540327 | 10.064362 | 10.022083 | 10.0245 | -0.002368474 | 0.039910406 |
| Gm5915 | 2.004492 | 1.953875 | 1.8625 | 0.091364028 | 0.141981185 | 1.9240841 | 2.015728 | 2.0181 | -0.00237335 | -0.094017223 |
| Cacng7 | 2.297947 | 2.277308 | 2.2423 | 0.035031107 | 0.055670011 | 2.2165817 | 2.184075 | 2.18651 | -0.002436107 | 0.030070599 |
| Nrsn1 | 0.684515 | 0.754451 | 0.6394 | 0.115040234 | 0.045105177 | 0.6382109 | 0.6768346 | 0.67927 | -0.0024385 | -0.041062255 |
| Gm8615 | 1.151505 | 1.110628 | 1.1612 | -0.050606905 | -0.009730346 | 1.1414544 | 1.2300989 | 1.23263 | -0.002526645 | -0.091171121 |
| Tnni3 | 4.416202 | 4.328476 | 4.5894 | -0.260919777 | -0.173193381 | 4.5833255 | 4.7498039 | 4.75235 | -0.002542734 | -0.169021086 |
| Fancl | 7.86182 | 7.850264 | 7.8849 | -0.034647941 | -0.023091974 | 8.0001172 | 7.9210916 | 7.92364 | -0.002546465 | 0.076479162 |
| Gm5931 | 1.658436 | 1.523504 | 1.6482 | -0.124672717 | 0.01025952 | 1.8238137 | 1.4779795 | 1.48053 | -0.002551719 | 0.34328247 |
| Fam92a | 6.808168 | 6.457678 | 6.563 | -0.105274787 | 0.245215237 | 6.4750169 | 6.7164385 | 6.71899 | -0.00255237 | -0.243973976 |
| Fcor | 1.945878 | 1.900928 | 1.9683 | -0.067392296 | -0.022441453 | 2.0099062 | 1.9233442 | 1.92591 | -0.002567375 | 0.083994554 |
| Pou5f1 | 0.684613 | 0.70424 | 0.6837 | 0.020570354 | 0.000943937 | 0.735994 | 0.6917294 | 0.69433 | -0.002597109 | 0.041667555 |
| Pla2g3 | 1.861964 | 1.868423 | 1.9023 | -0.033857223 | -0.040316174 | 1.8552287 | 1.9124057 | 1.91501 | -0.00260715 | -0.059784117 |
| Ano7 | 0.836846 | 0.83337 | 0.8185 | 0.014849114 | 0.018324887 | 0.9810545 | 0.9165656 | 0.91919 | -0.002622006 | 0.061866827 |
| Gm46136 | 4.607673 | 4.561874 | 4.6127 | -0.050818049 | -0.005018757 | 4.4215719 | 4.7734323 | 4.77608 | -0.002645588 | -0.354506064 |
| Mmp24 | 1.038107 | 1.003432 | 0.9881 | 0.015301179 | 0.049977069 | 1.1358644 | 0.964649 | 0.96735 | -0.002696416 | 0.168518938 |
| Gm20589 | 3.254957 | 3.095581 | 3.2539 | -0.15834132 | 0.001034098 | 3.2332087 | 3.2735057 | 3.27621 | -0.002702621 | -0.042999645 |
| Trdc | 8.494906 | 8.547667 | 8.4993 | 0.048393137 | -0.004368522 | 8.1459919 | 8.3810417 | 8.38375 | -0.002709085 | -0.237758931 |
| Acads | 8.604028 | 8.60053 | 8.6564 | -0.055849234 | -0.052351249 | 8.8523689 | 8.5575557 | 8.56027 | -0.002711325 | 0.292101836 |
| Zfand5 | 10.64448 | 10.48302 | 10.575 | -0.092442772 | 0.069015256 | 10.372358 | 10.427495 | 10.4302 | -0.002747818 | -0.057884111 |
| Aqp4 | 2.301214 | 2.30792 | 2.252 | 0.055960805 | 0.049254791 | 2.2871033 | 2.3101483 | 2.3129 | -0.002752528 | -0.025797585 |
| Rps6kb1 | 10.03541 | 10.07111 | 9.9668 | 0.104306732 | 0.068604742 | 9.9192956 | 10.133568 | 10.1364 | -0.002801241 | -0.217073425 |
| Fam72a | 4.533448 | 4.674385 | 4.5516 | 0.122739991 | -0.018197168 | 4.4920995 | 4.7600384 | 4.76291 | -0.002874979 | -0.270813912 |
| Ccdc68 | 0.317025 | 0.280714 | 0.4379 | -0.157187868 | -0.120877346 | 0.3607233 | 0.2695556 | 0.27243 | -0.002876758 | 0.088290983 |
| Rps6ka4 | 9.169609 | 8.983356 | 9.1135 | -0.130130983 | 0.056122219 | 9.3449333 | 9.0645517 | 9.06743 | -0.002881234 | 0.277500314 |
| Mcm8 | 6.302168 | 6.401026 | 6.535 | -0.133950566 | -0.232808054 | 6.4436816 | 6.5699476 | 6.57287 | -0.002919742 | -0.129185772 |
| Fxyd7 | 1.613069 | 1.540134 | 1.5612 | -0.021079045 | 0.051855938 | 1.5448998 | 1.4946422 | 1.49759 | -0.002943669 | 0.047313946 |
| Mff | 9.893516 | 9.872883 | 9.9824 | -0.10947944 | -0.088845845 | 9.9523028 | 9.8837023 | 9.88667 | -0.00297128 | 0.065629146 |

| | | | | | | | | | | |
|---|---|---|---|---|---|---|---|---|---|---|
| Gm6433 | 0.354479 | 0.415727 | 0.3364 | 0.079297769 | 0.018049813 | 0.3945685 | 0.3690717 | 0.37205 | -0.002976059 | 0.022520774 |
| Olfr1254 | 2.520488 | 2.481302 | 2.499 | -0.017690514 | 0.021495755 | 2.6469941 | 2.6084977 | 2.61148 | -0.002977375 | 0.035519048 |
| Olfr750 | 0.694594 | 0.725472 | 0.6892 | 0.036258866 | 0.005380203 | 0.7217296 | 0.740902 | 0.74389 | -0.002989936 | -0.02216234 |
| Zfp941 | 2.157559 | 2.167699 | 2.1375 | 0.030234335 | 0.020094135 | 2.0971018 | 2.1679486 | 2.17099 | -0.003044201 | -0.073891033 |
| Nsun7 | 3.41416 | 3.458897 | 3.5049 | -0.045974993 | -0.090711858 | 3.5078781 | 3.5132922 | 3.51638 | -0.003090879 | -0.008505036 |
| 1700001C19Rik | 0.274717 | 0.245833 | 0.2986 | -0.052770204 | -0.023886773 | 0.2303923 | 0.2676966 | 0.27083 | -0.003136324 | -0.040440586 |
| Lrrc40 | 8.392221 | 8.242337 | 8.4752 | -0.232821492 | -0.082938286 | 8.4500172 | 8.4575808 | 8.46077 | -0.003187778 | -0.010751447 |
| Gm13864 | 2.918697 | 3.022873 | 2.9672 | 0.055710338 | -0.048465679 | 3.0342166 | 3.0137618 | 3.01699 | -0.003227869 | 0.017226903 |
| Prrg2 | 5.832753 | 5.785784 | 5.7431 | 0.042681673 | 0.089651183 | 5.8871786 | 5.7860288 | 5.78926 | -0.003229599 | 0.097920189 |
| Reep3 | 9.876045 | 9.947697 | 9.9049 | 0.042799973 | -0.028851166 | 9.6658072 | 9.8622473 | 9.86552 | -0.003276747 | -0.199716857 |
| Nsd2 | 9.779425 | 9.589816 | 9.921 | -0.331231515 | -0.141623273 | 9.7540017 | 9.9008223 | 9.90411 | -0.003288664 | -0.150109216 |
| Rab14 | 11.0236 | 11.04285 | 11.04 | 0.00306784 | -0.016191168 | 11.096463 | 11.139066 | 11.1424 | -0.003289282 | -0.045892728 |
| Olfr1130 | 2.760147 | 2.734931 | 2.7692 | -0.034305394 | -0.009090187 | 2.7902591 | 2.7951296 | 2.79844 | -0.003312597 | -0.00818316 |
| AC166779.2 | 3.962531 | 4.061558 | 4.0316 | 0.02998981 | -0.069036968 | 3.9817855 | 4.050847 | 4.05417 | -0.003318238 | -0.072379719 |
| Tprn | 7.619189 | 7.464896 | 7.6444 | -0.179523279 | -0.02523026 | 7.3034293 | 7.4338581 | 7.43726 | -0.003400185 | -0.133829004 |
| Oaz3 | 2.227213 | 2.229986 | 2.3949 | -0.164900314 | -0.167673751 | 2.2783087 | 2.3110934 | 2.3146 | -0.003504428 | -0.036289132 |
| Fam187b | 0.808172 | 0.827683 | 0.8415 | -0.013768858 | -0.033279651 | 0.9018098 | 0.8434029 | 0.84693 | -0.003530726 | 0.054876195 |
| Nlrp12 | 2.338421 | 2.471309 | 2.2171 | 0.254224235 | 0.121335635 | 2.4744658 | 2.2007461 | 2.2043 | -0.003552743 | 0.270166945 |
| Amhr2 | 2.529742 | 2.485679 | 2.4961 | -0.010430181 | 0.033633408 | 2.5906909 | 2.5223586 | 2.52592 | -0.0035582 | 0.064774084 |
| Rbbp4 | 11.74484 | 11.70851 | 11.759 | -0.050593794 | -0.014265922 | 11.700965 | 11.693563 | 11.6972 | -0.003587768 | 0.003814192 |
| Gm44821 | 3.61077 | 3.577562 | 3.5669 | 0.010652285 | 0.043860701 | 3.5640992 | 3.6288237 | 3.63242 | -0.00360059 | -0.068325144 |
| Rph3a | 2.430071 | 2.525789 | 2.4474 | 0.078339705 | -0.017378911 | 2.4015888 | 2.4798537 | 2.48355 | -0.003699642 | -0.081964584 |
| Hhipl2 | 0.687589 | 0.802355 | 0.6361 | 0.166289465 | 0.051523356 | 0.5830587 | 0.5965199 | 0.60028 | -0.003763647 | -0.017224839 |
| Aox2 | 1.048323 | 1.09545 | 1.0572 | 0.038296189 | -0.008830147 | 1.1627475 | 1.0795828 | 1.08337 | -0.003783851 | 0.079380858 |
| Rnft1 | 8.10798 | 8.085259 | 8.1869 | -0.101686619 | -0.078965769 | 7.9248124 | 7.8706959 | 7.87449 | -0.003791124 | 0.050325366 |
| Gm47803 | 0.458874 | 0.433084 | 0.4513 | -0.018208521 | 0.007581424 | 0.4633582 | 0.4135797 | 0.41739 | -0.003813228 | 0.04596528 |
| Mapk4 | 0.980623 | 1.024507 | 1.0542 | -0.029694279 | -0.073578436 | 0.9567707 | 1.0502441 | 1.05406 | -0.003820216 | -0.09729364 |
| Ighv14-1 | 5.546828 | 5.727806 | 5.4693 | 0.258544279 | 0.077566314 | 5.6909748 | 5.4183412 | 5.42216 | -0.00382211 | 0.268811518 |
| Dusp7 | 9.471063 | 9.596649 | 9.5559 | 0.040742191 | -0.084843964 | 9.7152335 | 9.4127717 | 9.41662 | -0.00384659 | 0.298615237 |
| Gde1 | 8.379217 | 8.450847 | 8.4017 | 0.049152585 | -0.022476993 | 8.4054171 | 8.4412181 | 8.44513 | -0.003907119 | -0.039708058 |
| Gm6483 | 4.607716 | 4.638022 | 4.7046 | -0.066587738 | -0.096893278 | 4.7370551 | 4.77576 | 4.77968 | -0.003922953 | -0.042627907 |
| Rpl31-ps4 | 0.611955 | 0.549948 | 0.5364 | 0.013531331 | 0.075538042 | 0.5957995 | 0.5227283 | 0.52666 | -0.003933926 | 0.069137219 |

| | | | | | | | | | | |
|---|---|---|---|---|---|---|---|---|---|---|
| Taf3 | 9.237625 | 9.290139 | 9.3065 | -0.016382175 | -0.068896845 | 9.3111061 | 9.2591384 | 9.26311 | -0.003974168 | 0.047993609 |
| Dcpp2 | 2.531082 | 2.986003 | 2.243 | 0.743022021 | 0.288100822 | 4.482988 | 2.459329 | 2.46336 | -0.004033751 | 2.019625273 |
| Crb1 | 2.818847 | 2.777874 | 2.7685 | 0.009419876 | 0.050392941 | 3.01686 | 2.9770738 | 2.98114 | -0.004061806 | 0.035724343 |
| Gm18289 | 0.280762 | 0.273543 | 0.2449 | 0.028640263 | 0.035859047 | 0.2808575 | 0.2715756 | 0.27568 | -0.004106918 | 0.005175002 |
| Col27a1 | 7.522787 | 7.653307 | 7.3659 | 0.28741074 | 0.156891279 | 7.4751353 | 7.5649385 | 7.56907 | -0.004134719 | -0.093937863 |
| Pcdhb10 | 1.293023 | 1.21759 | 1.2292 | -0.011606618 | 0.063826385 | 1.2278402 | 1.1912969 | 1.19544 | -0.004140097 | 0.032403204 |
| Asphd1 | 1.144475 | 1.129622 | 1.1308 | -0.001218037 | 0.013635654 | 1.1094106 | 1.1303442 | 1.13449 | -0.004144663 | -0.025078227 |
| Zfp947 | 3.397125 | 3.407642 | 3.5795 | -0.171835723 | -0.18235325 | 3.4630433 | 3.4417522 | 3.44594 | -0.004188699 | 0.017102449 |
| Olfr1250 | 2.50444 | 2.613813 | 2.5604 | 0.053381382 | -0.055992449 | 2.5613201 | 2.6840627 | 2.6883 | -0.004241499 | -0.126984076 |
| Tmem121 | 0.789474 | 0.799546 | 0.765 | 0.034570189 | 0.024497475 | 0.8691112 | 0.747831 | 0.75216 | -0.004331468 | 0.116948791 |
| Ddx1 | 9.969763 | 9.74598 | 9.7761 | -0.030077797 | 0.193705472 | 9.7777827 | 9.8655777 | 9.86991 | -0.004334525 | -0.092129538 |
| Cdc5l | 10.88774 | 11.03623 | 10.963 | 0.072876192 | -0.075613948 | 10.935803 | 10.966943 | 10.9713 | -0.004336598 | -0.035476119 |
| 3300002I08Rik | 4.837572 | 4.955677 | 4.8826 | 0.073121988 | -0.044982614 | 4.764663 | 4.810251 | 4.81461 | -0.004361874 | -0.049949802 |
| Aldh1a3 | 3.275467 | 3.312685 | 3.2327 | 0.080030457 | 0.042812908 | 3.1570657 | 3.2196547 | 3.22403 | -0.004375783 | -0.066964807 |
| Nfs1 | 8.862479 | 8.85999 | 8.8626 | -0.002574148 | -8.5843E-05 | 9.1189775 | 8.9408305 | 8.94521 | -0.00437825 | 0.173768739 |
| Vmn1r226 | 0.872172 | 0.869387 | 0.927 | -0.057611207 | -0.054826855 | 0.8545237 | 0.9076531 | 0.91204 | -0.004390885 | -0.057520288 |
| Foxf2 | 0.222581 | 0.26487 | 0.2614 | 0.003444827 | -0.038843628 | 0.2952079 | 0.2460229 | 0.25043 | -0.004403547 | 0.04478148 |
| 1700008O03Rik | 0.943644 | 0.877228 | 0.8283 | 0.048943325 | 0.115358768 | 0.8986106 | 0.9183976 | 0.92282 | -0.004425891 | -0.024212903 |
| Nptx1 | 2.70985 | 2.64731 | 2.6382 | 0.009083663 | 0.071623844 | 2.4777413 | 2.5105671 | 2.51501 | -0.004445087 | -0.037270885 |
| Paxbp1 | 10.09297 | 10.14224 | 10.145 | -0.003159044 | -0.052427294 | 10.23734 | 10.483249 | 10.4878 | -0.00451649 | -0.250426202 |
| Castor1 | 6.878311 | 6.568986 | 6.8413 | -0.272358358 | 0.036966379 | 6.8955013 | 6.8423491 | 6.84693 | -0.004578723 | 0.048573534 |
| Chrnb3 | 0.865089 | 0.820856 | 0.9025 | -0.081664149 | -0.037431733 | 0.8573307 | 0.9291542 | 0.93374 | -0.004582797 | -0.076406332 |
| Nol3 | 4.485026 | 4.409596 | 4.303 | 0.106620361 | 0.182050136 | 4.2028117 | 4.1202032 | 4.1248 | -0.004592528 | 0.078015901 |
| Trp63 | 1.667633 | 1.681272 | 1.7545 | -0.073208404 | -0.086847815 | 1.6246522 | 1.6399965 | 1.6446 | -0.004599253 | -0.01994353 |
| Gm30646 | 0.561962 | 0.55265 | 0.5213 | 0.031363875 | 0.040675805 | 0.4615394 | 0.4817408 | 0.48635 | -0.004611938 | -0.024813412 |
| Arhgap25 | 11.27253 | 11.45299 | 11.333 | 0.120400492 | -0.060060326 | 11.484234 | 11.401676 | 11.4063 | -0.00463838 | 0.077919076 |
| Ten1 | 7.782961 | 7.642109 | 7.8308 | -0.188649547 | -0.0477976 | 7.9952773 | 7.770583 | 7.77522 | -0.004639794 | 0.22005453 |
| Cul4a | 9.511639 | 9.472649 | 9.5315 | -0.058878228 | -0.019888418 | 9.494426 | 9.5763114 | 9.58097 | -0.004661058 | -0.086546512 |
| Aes | 11.9698 | 11.95418 | 12.055 | -0.100517696 | -0.084893974 | 12.189921 | 11.946704 | 11.9514 | -0.004680842 | 0.238536423 |
| Lztr1 | 9.319543 | 9.292022 | 9.529 | -0.236935044 | -0.209414537 | 9.5952535 | 9.5008146 | 9.50551 | -0.004691726 | 0.089747236 |
| 1700061G19Rik | 2.755782 | 2.755857 | 2.7081 | 0.047716459 | 0.047641514 | 2.7576955 | 2.6231507 | 2.62785 | -0.004700455 | 0.129844337 |
| Ypel5 | 9.644544 | 9.586114 | 9.6256 | -0.039501029 | 0.018928674 | 9.6688376 | 9.7336586 | 9.73836 | -0.00470193 | -0.069522937 |

| | | | | | | | | | | |
|---|---|---|---|---|---|---|---|---|---|---|
| Zfp92 | 2.335836 | 2.3764 | 2.3797 | -0.003296769 | -0.043861142 | 2.3204721 | 2.4342008 | 2.43891 | -0.004704456 | -0.118433148 |
| Caps2 | 1.57165 | 1.561471 | 1.6069 | -0.045455087 | -0.035276301 | 1.6024609 | 1.6076422 | 1.61239 | -0.004750432 | -0.009931767 |
| Clec16a | 8.788529 | 8.885803 | 8.9547 | -0.068890925 | -0.166164123 | 8.7876043 | 8.9111063 | 8.91592 | -0.004817174 | -0.128319149 |
| Gm10093 | 2.232878 | 2.310152 | 2.2098 | 0.10037919 | 0.023105235 | 2.2643281 | 2.3903155 | 2.39515 | -0.004829802 | -0.130817226 |
| Zfp759 | 6.062003 | 6.139834 | 6.2096 | -0.069800056 | -0.147631868 | 5.9806878 | 6.2314468 | 6.23629 | -0.004841092 | -0.255600001 |
| Gm11517 | 1.292818 | 1.364354 | 1.3746 | -0.010212397 | -0.081747821 | 1.3259699 | 1.3586087 | 1.36351 | -0.004906207 | -0.037544986 |
| Gm15547 | 1.874255 | 1.883554 | 1.8774 | 0.006179776 | -0.003119454 | 1.8461923 | 1.9343804 | 1.93933 | -0.004952685 | -0.093140783 |
| Gm18294 | 2.76794 | 2.76241 | 2.7681 | -0.005692283 | -0.000162268 | 2.7063286 | 2.8116412 | 2.8166 | -0.004956102 | -0.110268682 |
| Vpreb1 | 2.257148 | 2.317133 | 2.2321 | 0.085055742 | 0.025070218 | 2.2368063 | 2.2479563 | 2.25294 | -0.004980815 | -0.016130762 |
| Gm10291 | 1.431848 | 1.485343 | 1.4682 | 0.017179747 | -0.036315043 | 1.4869192 | 1.4421982 | 1.44718 | -0.004981407 | 0.039739609 |
| Olfr705 | 3.052407 | 3.05355 | 3.1155 | -0.061915579 | -0.063058558 | 3.0701856 | 3.1347246 | 3.13971 | -0.004984397 | -0.069523408 |
| Olfr1437 | 0.852957 | 0.827958 | 0.8617 | -0.033760315 | -0.008760916 | 0.8084959 | 0.8933524 | 0.89834 | -0.004987578 | -0.089844085 |
| Prpf6 | 10.81022 | 10.7392 | 10.803 | -0.063362496 | 0.00765676 | 10.773848 | 10.765242 | 10.7703 | -0.00503083 | 0.00357526 |
| Lmbr1l | 9.860654 | 9.908573 | 9.9719 | -0.06334683 | -0.111265863 | 10.210419 | 10.183935 | 10.189 | -0.00504335 | 0.02144012 |
| Dgki | 1.174445 | 1.072751 | 1.1571 | -0.08437327 | 0.017320854 | 1.0986739 | 1.0816642 | 1.08671 | -0.005050729 | 0.011958913 |
| Arhgap10 | 7.382983 | 7.428858 | 7.3739 | 0.055006449 | 0.009131696 | 7.1702352 | 7.4205634 | 7.42562 | -0.00505616 | -0.255384357 |
| Plxnb3 | 1.890493 | 1.817302 | 1.8918 | -0.074543742 | -0.001353298 | 1.9602939 | 1.9059096 | 1.91097 | -0.005063073 | 0.049321253 |
| Larp7 | 9.784314 | 9.672753 | 9.7203 | -0.047520728 | 0.064040788 | 9.8049082 | 9.783723 | 9.7888 | -0.005075065 | 0.016110206 |
| Gm11639 | 0.61204 | 0.646051 | 0.6145 | 0.031559527 | -0.002451922 | 0.5875838 | 0.652227 | 0.65732 | -0.005091973 | -0.06973523 |
| Fam221a | 4.475071 | 4.700851 | 4.6961 | 0.004789804 | -0.22098971 | 4.7654696 | 4.9644773 | 4.9696 | -0.005117892 | -0.204125519 |
| Olfr1062 | 1.100499 | 1.028805 | 1.0507 | -0.021891989 | 0.049801867 | 1.0952044 | 1.1234642 | 1.12859 | -0.005124671 | -0.033384511 |
| Gm9077 | 0.388354 | 0.415182 | 0.4176 | -0.00245927 | -0.029287015 | 0.3761329 | 0.3946912 | 0.39982 | -0.005128193 | -0.023686569 |
| Drg2 | 9.117066 | 9.144233 | 9.0538 | 0.090385637 | 0.063218632 | 9.4070093 | 9.2931684 | 9.29838 | -0.005207098 | 0.108633793 |
| Faah | 9.866714 | 10.02068 | 10.116 | -0.095328764 | -0.249291051 | 9.8494877 | 10.236157 | 10.2415 | -0.005329251 | -0.39199833 |
| Tmem87b | 8.845472 | 8.879343 | 8.8946 | -0.01528613 | -0.049156438 | 8.9655104 | 9.0506155 | 9.05595 | -0.005338567 | -0.090443605 |
| Kdf1 | 0.893859 | 0.897093 | 0.9415 | -0.04437309 | -0.047606768 | 0.9122823 | 0.9445959 | 0.94996 | -0.005359289 | -0.03767291 |
| Bzw1 | 12.50188 | 12.48191 | 12.439 | 0.042523346 | 0.062489786 | 12.437967 | 12.399231 | 12.4046 | -0.005364518 | 0.033371426 |
| Zfp449 | 3.459448 | 3.462637 | 3.6263 | -0.163681492 | -0.166871063 | 3.5071868 | 3.4221996 | 3.42768 | -0.005480847 | 0.07950639 |
| Rpl30-ps10 | 1.088964 | 1.200936 | 0.977 | 0.223887542 | 0.111916063 | 1.0863587 | 1.02859 | 1.03413 | -0.005536693 | 0.052232085 |
| Trgj1 | 0.432961 | 0.397221 | 0.4689 | -0.071667567 | -0.035928357 | 0.4239144 | 0.4269822 | 0.43252 | -0.005536978 | -0.008604764 |
| Tctn3 | 6.821147 | 6.811703 | 6.8219 | -0.010154646 | -0.000710747 | 7.068563 | 6.8393521 | 6.8449 | -0.005551039 | 0.223659851 |
| Gm29719 | 0.539339 | 0.5011 | 0.4919 | 0.009190192 | 0.047429641 | 0.4511308 | 0.5080691 | 0.51364 | -0.005570644 | -0.062508936 |

| | | | | | | | | | | |
|---|---|---|---|---|---|---|---|---|---|---|
| Alg12 | 8.130222 | 8.26331 | 8.1914 | 0.071907461 | -0.061180622 | 8.5892275 | 8.4260626 | 8.43166 | -0.005592878 | 0.157572037 |
| Tbx19 | 1.022767 | 1.075052 | 1.0675 | 0.007505998 | -0.044778814 | 1.0732545 | 1.0576208 | 1.06323 | -0.005605796 | 0.010027847 |
| Cd22 | 13.63768 | 13.61912 | 13.642 | -0.02239394 | -0.003834888 | 14.229433 | 13.821683 | 13.8273 | -0.005633368 | 0.402115899 |
| Gm9396 | 0.936172 | 0.894345 | 0.933 | -0.038639725 | 0.003187599 | 0.8942601 | 0.8631617 | 0.8688 | -0.005638762 | 0.025459632 |
| Cd37 | 13.23504 | 13.27095 | 13.263 | 0.007995378 | -0.027914012 | 13.808762 | 13.357541 | 13.3632 | -0.005671216 | 0.445549279 |
| Top1 | 11.32331 | 11.3096 | 11.227 | 0.08288023 | 0.096592944 | 11.182705 | 11.263347 | 11.269 | -0.005696414 | -0.086338207 |
| Olfr1105 | 0.007472 | 0.000718 | 0.0157 | -0.014971145 | -0.008217054 | 0.0150284 | 0.0599832 | 0.06568 | -0.005699067 | -0.050653911 |
| Agk | 8.31362 | 8.520429 | 8.5126 | 0.007828496 | -0.19897975 | 8.5132259 | 8.4152803 | 8.42099 | -0.005706491 | 0.092239095 |
| Mrgprc2-ps | 3.486939 | 3.522587 | 3.3275 | 0.195126709 | 0.159479071 | 3.5641202 | 3.685785 | 3.69149 | -0.005709841 | -0.127374644 |
| Pbld2 | 3.377244 | 3.250859 | 3.4618 | -0.210916132 | -0.084531811 | 3.5064041 | 3.3535579 | 3.35928 | -0.005725493 | 0.147120717 |
| Lbhd1 | 6.811698 | 6.28549 | 7.0998 | -0.814270225 | -0.288062272 | 7.0191308 | 6.9134045 | 6.91914 | -0.005734999 | 0.099991361 |
| Gm48254 | 1.458866 | 1.413947 | 1.472 | -0.058102292 | -0.013183768 | 1.4238331 | 1.4500214 | 1.45581 | -0.005785633 | -0.031973946 |
| Bves | 2.097818 | 2.062359 | 2.0797 | -0.017336814 | 0.018121951 | 2.0508161 | 2.0645767 | 2.07038 | -0.005800373 | -0.019561032 |
| Fam8a1 | 8.689891 | 8.744155 | 8.7234 | 0.020719166 | -0.033544825 | 8.5287042 | 8.6224703 | 8.62827 | -0.005802632 | -0.099568772 |
| Lrrtm4 | 1.40468 | 1.385778 | 1.4026 | -0.016825758 | 0.002075891 | 1.4373371 | 1.4548808 | 1.46069 | -0.005805501 | -0.023349138 |
| Gm47951 | 4.080448 | 4.098727 | 4.067 | 0.031687197 | 0.013408766 | 3.933459 | 4.2100209 | 4.21584 | -0.005820175 | -0.282382091 |
| Gstp-ps | 1.784607 | 1.843178 | 1.8124 | 0.030753304 | -0.027817139 | 1.8383429 | 1.8270608 | 1.83292 | -0.005859475 | 0.0054227 |
| Pomc | 2.673707 | 2.769157 | 2.7485 | 0.020663025 | -0.074787187 | 2.841962 | 2.7788437 | 2.78474 | -0.005894729 | 0.057223584 |
| Mfsd3 | 6.106384 | 6.17567 | 6.2113 | -0.035669644 | -0.104955968 | 6.2936594 | 6.1645563 | 6.17047 | -0.005911648 | 0.123191452 |
| Gm5846 | 0.910264 | 0.899278 | 0.9205 | -0.021226286 | -0.010240599 | 0.93821 | 0.8641408 | 0.87007 | -0.005929303 | 0.06813981 |
| Morc1 | 1.173487 | 1.158619 | 1.2223 | -0.063672338 | -0.04880419 | 1.1790358 | 1.1842212 | 1.19016 | -0.005937168 | -0.011122503 |
| Fignl1 | 7.242065 | 6.932031 | 7.2138 | -0.281817496 | 0.028216412 | 7.0226402 | 7.2677013 | 7.27368 | -0.005973743 | -0.251034889 |
| Sfxn1 | 9.597212 | 9.572036 | 9.6793 | -0.10730311 | -0.082127513 | 9.755991 | 9.6531232 | 9.65913 | -0.006008794 | 0.09685894 |
| Traj33 | 0.873226 | 0.864091 | 0.8201 | 0.04395015 | 0.053084719 | 0.8417149 | 0.8310029 | 0.83702 | -0.006020039 | 0.004691982 |
| Ocel1 | 7.181172 | 7.222079 | 7.3933 | -0.171232191 | -0.212139517 | 7.3872246 | 7.5464537 | 7.55249 | -0.006038789 | -0.165267917 |
| Zfp410 | 9.149608 | 9.254206 | 9.1891 | 0.065081153 | -0.039517169 | 9.2851751 | 9.3698384 | 9.37589 | -0.006051033 | -0.090714269 |
| Ttc13 | 9.793314 | 9.755579 | 9.9125 | -0.156919426 | -0.119184544 | 9.8408168 | 9.8904543 | 9.89651 | -0.006059521 | -0.055696946 |
| Mtor | 10.23945 | 10.05055 | 10.132 | -0.081398953 | 0.107505564 | 10.063733 | 10.206934 | 10.213 | -0.006095196 | -0.149296408 |
| Nfyc | 9.635976 | 9.645333 | 9.7074 | -0.06206982 | -0.071426556 | 9.8154937 | 9.767603 | 9.77371 | -0.00610865 | 0.041782036 |
| Gm49337 | 1.488375 | 1.31582 | 1.3837 | -0.067896715 | 0.104658091 | 1.3724092 | 1.3374615 | 1.34361 | -0.006149108 | 0.028798597 |
| Fcgbp | 2.167778 | 2.047052 | 2.1018 | -0.054711612 | 0.066014118 | 2.1227256 | 2.0601834 | 2.06634 | -0.006158556 | 0.056383655 |
| Fahd1 | 5.356049 | 5.228356 | 5.5007 | -0.272388511 | -0.14469553 | 5.5834939 | 5.333517 | 5.33968 | -0.00616717 | 0.243809749 |

| | | | | | | | | | | |
|---|---|---|---|---|---|---|---|---|---|---|
| 1700001P01Rik | 0.923918 | 0.890686 | 0.9036 | -0.012953962 | 0.020278149 | 0.8968169 | 0.8975135 | 0.90369 | -0.006178692 | -0.006875289 |
| Prmt2 | 7.344498 | 7.453203 | 7.4162 | 0.03696886 | -0.071736433 | 7.4152474 | 7.4263662 | 7.43258 | -0.00621506 | -0.017333841 |
| Egf | 6.832881 | 7.32984 | 8.2147 | -0.884825539 | -1.381784257 | 9.158596 | 7.8774207 | 7.88365 | -0.006225355 | 1.27494999 |
| Gm49376 | 3.225654 | 3.262693 | 3.2924 | -0.029679976 | -0.066719598 | 3.2938037 | 3.2786178 | 3.2849 | -0.006283771 | 0.008902092 |
| Rbbp7 | 11.4248 | 11.30508 | 11.407 | -0.102192102 | 0.017526566 | 11.444148 | 11.440013 | 11.4463 | -0.006287808 | -0.002153691 |
| Nxf7 | 0.82945 | 0.797099 | 0.837 | -0.039874477 | -0.007523592 | 0.8206958 | 0.8314499 | 0.83782 | -0.006369748 | -0.017123889 |
| Mapre1 | 11.60431 | 11.58229 | 11.479 | 0.103359886 | 0.125381078 | 11.409183 | 11.599566 | 11.6059 | -0.00637025 | -0.196752991 |
| P4ha3 | 1.126422 | 1.100484 | 1.1589 | -0.058405987 | -0.032467766 | 1.1421076 | 1.2070256 | 1.21341 | -0.006387328 | -0.071305364 |
| Zfp638 | 11.34454 | 11.40238 | 11.293 | 0.109869304 | 0.052026904 | 11.396829 | 11.436849 | 11.4433 | -0.00641459 | -0.046434655 |
| Dcdc2c | 1.514532 | 1.527324 | 1.5239 | 0.003392381 | -0.009399333 | 1.4399918 | 1.5809802 | 1.5874 | -0.006415274 | -0.147403671 |
| Ntrk2 | 5.031247 | 4.926414 | 5.0288 | -0.102368033 | 0.00246471 | 4.6406729 | 4.8183904 | 4.82483 | -0.006437446 | -0.184154968 |
| Sptbn1 | 13.08459 | 13.0573 | 13.084 | -0.026350566 | 0.000936938 | 12.809525 | 13.080534 | 13.087 | -0.006461162 | -0.277470229 |
| Ramac | 9.579264 | 9.473296 | 9.5737 | -0.100395859 | 0.005572124 | 9.5463597 | 9.5255471 | 9.53202 | -0.006476609 | 0.014335958 |
| Gm14388 | 1.505067 | 1.554587 | 1.5871 | -0.032468763 | -0.081989085 | 1.6115743 | 1.6164503 | 1.62295 | -0.006501817 | -0.011377798 |
| Mdh1 | 11.18374 | 11.17926 | 11.263 | -0.084125084 | -0.07964055 | 11.385315 | 11.216661 | 11.2232 | -0.006506978 | 0.162146877 |
| Gm9320 | 3.22725 | 3.317104 | 3.2314 | 0.085686826 | -0.004166796 | 3.3051123 | 3.3317213 | 3.33826 | -0.006542951 | -0.033151947 |
| Pmaip1 | 9.586033 | 9.62804 | 9.3596 | 0.268464272 | 0.226457111 | 9.1319417 | 9.2982546 | 9.30483 | -0.00657478 | -0.172887681 |
| Rps19-ps1 | 1.341435 | 1.365141 | 1.3142 | 0.050918768 | 0.027212776 | 1.3603386 | 1.3178019 | 1.3244 | -0.006598057 | 0.035938633 |
| Rps19-ps2 | 1.341435 | 1.365141 | 1.3142 | 0.050918768 | 0.027212776 | 1.3603386 | 1.3178019 | 1.3244 | -0.006598057 | 0.035938633 |
| Rufy4 | 3.645426 | 3.580585 | 3.6436 | -0.062978795 | 0.001862489 | 3.6387018 | 3.5238235 | 3.53045 | -0.00662223 | 0.108256072 |
| Gm7808 | 1.496492 | 1.533386 | 1.5205 | 0.012869303 | -0.024024648 | 1.5431113 | 1.4825652 | 1.48923 | -0.006667911 | 0.053878132 |
| Esrp1 | 2.021566 | 1.741577 | 1.9502 | -0.208639318 | 0.071349784 | 2.010886 | 1.8784141 | 1.88522 | -0.006805432 | 0.125666407 |
| Olfr597 | 0.408511 | 0.384331 | 0.387 | -0.002648901 | 0.021531064 | 0.3611106 | 0.3463322 | 0.35315 | -0.006817803 | 0.00796061 |
| Vmn2r14 | 0.083444 | 0.089088 | 0.103 | -0.013874098 | -0.019518115 | 0.065422 | 0.0861194 | 0.09295 | -0.006829493 | -0.027526827 |
| Snrnp48 | 10.03649 | 10.04483 | 10.179 | -0.134221502 | -0.142562678 | 10.289844 | 10.399419 | 10.4063 | -0.006844469 | -0.116419645 |
| Dpep1 | 4.133033 | 3.91058 | 4.2042 | -0.293656207 | -0.071203252 | 3.8139718 | 3.8854282 | 3.89228 | -0.006854457 | -0.07831081 |
| Grem2 | 1.69961 | 1.521081 | 1.416 | 0.105041174 | 0.283570423 | 1.4452444 | 1.4481089 | 1.45499 | -0.006880193 | -0.009744722 |
| Gm5558 | 0.545367 | 0.545349 | 0.5184 | 0.026953676 | 0.026971313 | 0.5792203 | 0.5235503 | 0.53043 | -0.006880799 | 0.048789159 |
| Ccdc63 | 4.335448 | 3.917738 | 4.1615 | -0.243796923 | 0.173912838 | 4.1408722 | 4.0584645 | 4.06538 | -0.00691892 | 0.075488783 |
| Pemt | 4.1094 | 4.089709 | 4.1542 | -0.064521876 | -0.044830672 | 4.1667669 | 4.048939 | 4.05591 | -0.006968213 | 0.110859675 |
| Actr8 | 9.014601 | 9.002773 | 8.9826 | 0.020199134 | 0.032027002 | 9.1896173 | 9.0487645 | 9.05579 | -0.007025379 | 0.133827397 |
| Gpr157 | 6.666189 | 6.620178 | 6.8342 | -0.213987238 | -0.167975852 | 6.4707342 | 6.7650595 | 6.77217 | -0.007111886 | -0.301437129 |

| | | | | | | | | | | |
|---|---|---|---|---|---|---|---|---|---|---|
| Fbxo2 | 0.650528 | 0.616247 | 0.6438 | -0.027535925 | 0.006745088 | 0.6067855 | 0.5718887 | 0.57905 | -0.007164848 | 0.027731996 |
| Scnn1b | 1.413077 | 1.316384 | 1.3603 | -0.043915288 | 0.052777124 | 1.4173836 | 1.3415836 | 1.34875 | -0.00716589 | 0.068634115 |
| Rtn3 | 10.9143 | 10.89655 | 10.928 | -0.031705154 | -0.013954492 | 10.875048 | 10.849845 | 10.8571 | -0.007235982 | 0.01796725 |
| Gm10045 | 1.246088 | 1.21896 | 1.2512 | -0.032235761 | -0.005107869 | 1.1692625 | 1.1968548 | 1.20409 | -0.007236512 | -0.034828719 |
| Klhdc9 | 1.4566 | 1.424689 | 1.4584 | -0.033753173 | -0.001841922 | 1.4019065 | 1.458053 | 1.46534 | -0.00728792 | -0.063434366 |
| Slc35c1 | 9.830662 | 9.943655 | 9.868 | 0.075691825 | -0.037301347 | 10.157615 | 9.9243501 | 9.93165 | -0.007297113 | 0.225967797 |
| Stt3a | 11.28568 | 11.19773 | 11 | 0.198171548 | 0.286130924 | 11.092341 | 11.074525 | 11.0818 | -0.00731381 | 0.010501733 |
| Rnf223 | 2.464356 | 2.442266 | 2.4849 | -0.042668427 | -0.020577916 | 2.4103995 | 2.5602049 | 2.56752 | -0.007317445 | -0.157122823 |
| Sms | 10.08936 | 10.04444 | 10.179 | -0.134442419 | -0.089522113 | 10.111826 | 10.173923 | 10.1812 | -0.0073203 | -0.069417025 |
| Patl2 | 0.692401 | 0.623349 | 0.6518 | -0.028405623 | 0.040645664 | 0.6361365 | 0.6713824 | 0.67873 | -0.007351883 | -0.042597808 |
| Olfr1508 | 0.234861 | 0.139293 | 0.124 | 0.015282505 | 0.110850929 | 0.1000198 | 0.1207068 | 0.12806 | -0.007352871 | -0.028039901 |
| Gm9118 | 1.345154 | 1.393156 | 1.3639 | 0.029262537 | -0.018739606 | 1.3483648 | 1.3175978 | 1.325 | -0.007404251 | 0.023362753 |
| Por | 9.729587 | 9.693916 | 9.4545 | 0.239424137 | 0.275095848 | 9.5601354 | 9.5042007 | 9.51162 | -0.007418174 | 0.048516484 |
| Zfp729b | 8.704624 | 8.72618 | 8.6969 | 0.029261655 | 0.007705555 | 8.6927964 | 8.6897291 | 8.69716 | -0.007427208 | -0.004359933 |
| Xkr8 | 6.245182 | 5.82092 | 6.0505 | -0.229591495 | 0.194669997 | 6.0532459 | 5.9792921 | 5.98678 | -0.007492865 | 0.066460997 |
| Lsr | 6.258159 | 6.180984 | 6.2346 | -0.053619967 | 0.023554647 | 6.5155542 | 6.246353 | 6.25387 | -0.007517145 | 0.26168405 |
| Vmn1r207-ps | 1.636271 | 1.611851 | 1.5606 | 0.051292787 | 0.07571289 | 1.6456811 | 1.6904509 | 1.69797 | -0.007518316 | -0.052288023 |
| Olfr287 | 1.367182 | 1.393807 | 1.3768 | 0.017030085 | -0.009595311 | 1.4956578 | 1.3963737 | 1.40396 | -0.007590738 | 0.091693406 |
| Ttc14 | 9.851842 | 9.757889 | 9.8613 | -0.103439046 | -0.009486563 | 9.959737 | 10.089188 | 10.0968 | -0.007617962 | -0.137068737 |
| Fadd | 7.918713 | 7.766286 | 8.0394 | -0.27310902 | -0.120682017 | 8.0858241 | 7.9540034 | 7.96163 | -0.007629772 | 0.12419089 |
| Gm4756 | 1.75077 | 1.773327 | 1.7809 | -0.007527851 | -0.030084522 | 1.891076 | 1.8034637 | 1.81112 | -0.00765176 | 0.079960564 |
| Pim3 | 9.696954 | 9.466556 | 9.4887 | -0.022101958 | 0.208295969 | 9.109981 | 8.9826575 | 8.99033 | -0.007670638 | 0.119652935 |
| Gm12248 | 0.589114 | 0.613308 | 0.6147 | -0.001357449 | -0.025552167 | 0.6007622 | 0.5970473 | 0.60473 | -0.007679861 | -0.003964957 |
| Gm5859 | 1.774469 | 1.709303 | 1.7823 | -0.073021337 | -0.007855576 | 1.5609337 | 1.5851215 | 1.59281 | -0.007691886 | -0.031879632 |
| Ighv1-20 | 4.099991 | 4.089049 | 3.9176 | 0.17141669 | 0.182357994 | 4.3420748 | 3.992292 | 4.00004 | -0.007747442 | 0.342035345 |
| Gm3791 | 1.887542 | 1.965575 | 1.9911 | -0.025560648 | -0.103593681 | 1.9553198 | 2.0441734 | 2.05195 | -0.007779983 | -0.096633552 |
| Tnpo1 | 11.84573 | 11.91354 | 11.864 | 0.049316014 | -0.018495184 | 11.861832 | 11.971991 | 11.9798 | -0.007845128 | -0.118003426 |
| Magef1 | 6.110247 | 5.989402 | 6.2723 | -0.282917679 | -0.162073106 | 6.2719661 | 6.2268068 | 6.23466 | -0.007856535 | 0.037302835 |
| Enox1 | 0.993008 | 1.007658 | 1.0513 | -0.043610499 | -0.05826106 | 0.9952927 | 0.9371739 | 0.94506 | -0.007885507 | 0.05023325 |
| Efs | 2.109047 | 2.038686 | 2.1274 | -0.088710084 | -0.018348987 | 2.2562352 | 2.049457 | 2.05735 | -0.007888373 | 0.198889801 |
| Heyl | 2.991509 | 3.289557 | 3.0271 | 0.262447308 | -0.035600735 | 3.0950326 | 3.0362292 | 3.04415 | -0.007924885 | 0.050878563 |
| Ttr | 1.143032 | 1.138323 | 0.9917 | 0.146654788 | 0.151363786 | 1.0453336 | 0.9487056 | 0.95683 | -0.008121489 | 0.088506476 |

| | | | | | | | | | | |
|---|---|---|---|---|---|---|---|---|---|---|
| Snx5 | 12.58782 | 12.68514 | 12.734 | -0.048432788 | -0.145760614 | 12.825055 | 12.890786 | 12.8989 | -0.00814986 | -0.073880608 |
| Morc2a | 10.23524 | 10.11779 | 10.113 | 0.004946536 | 0.122389284 | 10.24255 | 10.272066 | 10.2803 | -0.008189327 | -0.037705016 |
| Atp9b | 10.42703 | 10.40706 | 10.48 | -0.073390374 | -0.05342017 | 10.598767 | 10.637031 | 10.6452 | -0.008199082 | -0.046463261 |
| Cd82 | 11.73467 | 11.8584 | 11.726 | 0.132475748 | 0.008738494 | 11.756795 | 11.652121 | 11.6604 | -0.008321744 | 0.096352781 |
| Tyk2 | 10.74983 | 10.83262 | 10.787 | 0.045375322 | -0.037417276 | 10.88981 | 10.894421 | 10.9028 | -0.008330674 | -0.012941566 |
| Dctn1 | 10.74839 | 10.74487 | 10.74 | 0.005156132 | 0.008677136 | 10.959884 | 10.807773 | 10.8161 | -0.008357816 | 0.143752684 |
| Colec12 | 7.967423 | 8.531846 | 7.8392 | 0.692624645 | 0.128201857 | 7.2932543 | 7.3177127 | 7.32608 | -0.008363294 | -0.03282167 |
| Mrps31 | 7.896348 | 7.752642 | 7.8326 | -0.079969621 | 0.063736906 | 7.9111262 | 7.8933376 | 7.9017 | -0.00836368 | 0.009424928 |
| Olfr646 | 2.074105 | 2.0797 | 2.0911 | -0.011358374 | -0.016953598 | 2.0935771 | 2.1455271 | 2.15397 | -0.008439419 | -0.06038939 |
| Fbxw4 | 8.333401 | 8.25018 | 8.384 | -0.133848615 | -0.0506277 | 8.6657378 | 8.7071382 | 8.71562 | -0.008484255 | -0.049884596 |
| Bms1 | 10.92047 | 11.02309 | 10.937 | 0.08645874 | -0.016155294 | 10.955383 | 10.983134 | 10.9916 | -0.008489066 | -0.036239695 |
| Il17a | 1.879795 | 1.817143 | 1.816 | 0.001134782 | 0.063787455 | 1.8019316 | 1.8551567 | 1.86368 | -0.008519412 | -0.061744516 |
| Cwf19l1 | 8.352792 | 8.357994 | 8.3409 | 0.017084775 | 0.011882693 | 8.4775661 | 8.3985962 | 8.40712 | -0.008519441 | 0.07045045 |
| Avpi1 | 6.828004 | 6.677858 | 6.8799 | -0.20207347 | -0.051927365 | 6.9685968 | 6.665896 | 6.67446 | -0.008559643 | 0.294141149 |
| Asb18 | 1.54728 | 1.594183 | 1.5755 | 0.018666828 | -0.028235544 | 1.6020104 | 1.6189661 | 1.62756 | -0.008590968 | -0.025546608 |
| Paf1 | 10.10329 | 10.1862 | 10.194 | -0.007461529 | -0.09037362 | 10.128955 | 10.310432 | 10.3191 | -0.008622539 | -0.190098785 |
| Prkn | 3.077046 | 3.057807 | 3.1492 | -0.091359601 | -0.072120377 | 3.0408756 | 3.1405097 | 3.14918 | -0.008666616 | -0.108300746 |
| C030006K11Rik | 7.359175 | 7.147858 | 7.3999 | -0.252080429 | -0.04076353 | 7.6602986 | 7.2459105 | 7.25459 | -0.008680353 | 0.40570774 |
| Cdh3 | 2.52064 | 2.455071 | 2.581 | -0.125923644 | -0.060354591 | 2.4542318 | 2.4789189 | 2.48762 | -0.00869967 | -0.033386756 |
| Gpatch11 | 8.063647 | 8.12719 | 8.1898 | -0.062639699 | -0.126182761 | 8.3133562 | 8.2780318 | 8.28678 | -0.008749229 | 0.026575178 |
| Ahctf1 | 11.2279 | 11.14388 | 11.164 | -0.0196274 | 0.064389786 | 11.185123 | 11.205982 | 11.2148 | -0.008819116 | -0.029678118 |
| Nup205 | 10.59496 | 10.66585 | 10.751 | -0.084976742 | -0.155867662 | 10.809263 | 10.828937 | 10.8378 | -0.008840282 | -0.028514126 |
| Pex11b | 8.326322 | 8.185595 | 8.3384 | -0.15285084 | -0.012124283 | 8.4238312 | 8.1970445 | 8.2059 | -0.00885481 | 0.217931893 |
| Akr1c19 | 1.213422 | 1.080556 | 1.2446 | -0.164073201 | -0.031206441 | 1.1877691 | 1.1281716 | 1.13704 | -0.008873143 | 0.050724342 |
| Dnajb13 | 4.770192 | 4.618304 | 4.7395 | -0.121205867 | 0.030682062 | 4.8488731 | 4.837711 | 4.8466 | -0.008889153 | 0.002272869 |
| Pou3f2 | 0.876194 | 0.910593 | 0.9347 | -0.024091409 | -0.058490351 | 0.8807983 | 0.9664239 | 0.97534 | -0.008920331 | -0.094546003 |
| Impdh2-ps | 0.971761 | 0.922172 | 0.8555 | 0.0667146 | 0.116303211 | 1.0012177 | 0.9290003 | 0.93795 | -0.008945314 | 0.063272125 |
| Mob2 | 8.890217 | 9.024952 | 9.0875 | -0.062583766 | -0.197319107 | 8.8939039 | 8.9700569 | 8.979 | -0.008945767 | -0.085098736 |
| Gm17018 | 7.653006 | 7.557459 | 7.5814 | -0.023955144 | 0.07159157 | 7.9418765 | 7.6635422 | 7.67253 | -0.00898683 | 0.269347484 |
| Cracr2b | 5.808945 | 5.300549 | 5.8005 | -0.499948121 | 0.008448038 | 5.0094027 | 5.1580085 | 5.16701 | -0.009000773 | -0.157606531 |
| Osbpl7 | 9.110164 | 9.133886 | 9.2095 | -0.075614138 | -0.099336701 | 9.4399955 | 9.3928322 | 9.4019 | -0.009069199 | 0.038094168 |
| Rtl8b | 6.561724 | 6.406085 | 6.4447 | -0.038616013 | 0.117022967 | 6.4234789 | 6.2702942 | 6.27937 | -0.009072535 | 0.144112122 |

| | | | | | | | | | | |
|---|---|---|---|---|---|---|---|---|---|---|
| Atp7b | 3.281298 | 3.243382 | 3.2152 | 0.028203736 | 0.066120513 | 3.3390262 | 3.3484541 | 3.35754 | -0.009083753 | -0.018511703 |
| Enoph1 | 7.931578 | 7.877796 | 7.9489 | -0.071077886 | -0.017295191 | 8.0322556 | 7.9436347 | 7.95278 | -0.009149317 | 0.079471645 |
| Capza1 | 12.72416 | 12.7801 | 12.63 | 0.149672958 | 0.093726694 | 12.743195 | 12.668259 | 12.6774 | -0.009175512 | 0.065760025 |
| Cnot2 | 10.79355 | 10.89106 | 10.838 | 0.052988264 | -0.044516006 | 10.767682 | 10.858044 | 10.8672 | -0.009183764 | -0.099546213 |
| Tax1bp1 | 11.92009 | 11.90567 | 11.906 | -3.0208E-05 | 0.014385324 | 11.894795 | 11.883207 | 11.8924 | -0.009200596 | 0.002386533 |
| Gm16487 | 0.930026 | 0.98034 | 0.9517 | 0.028641796 | -0.021671931 | 0.9790538 | 1.0107289 | 1.01994 | -0.009209649 | -0.040884758 |
| Gm48515 | 0.664694 | 0.68891 | 0.7095 | -0.020597472 | -0.044813455 | 0.6915775 | 0.7095386 | 0.71877 | -0.009231379 | -0.027192478 |
| Gm47974 | 2.828196 | 2.731766 | 2.8189 | -0.087092655 | 0.009338066 | 2.7536101 | 2.8451647 | 2.85441 | -0.00924581 | -0.100800374 |
| Rpl10-ps3 | 13.51683 | 13.66949 | 13.71 | -0.04055421 | -0.193223634 | 13.839128 | 13.775419 | 13.7848 | -0.009331838 | 0.054376572 |
| Slc28a3 | 1.245301 | 1.260598 | 1.3216 | -0.06101356 | -0.076310634 | 1.374113 | 1.3223951 | 1.33174 | -0.009344411 | 0.042373463 |
| AC123853.3 | 0.84751 | 0.802148 | 0.8156 | -0.013459285 | 0.031902523 | 0.8347421 | 0.8634116 | 0.87278 | -0.009366028 | -0.038035515 |
| Spr-ps1 | 1.797774 | 1.694975 | 1.7113 | -0.016299212 | 0.086499936 | 1.7347033 | 1.8013905 | 1.81076 | -0.009374286 | -0.076061413 |
| Thoc7 | 8.997489 | 8.994619 | 8.9999 | -0.00524471 | -0.002375085 | 9.0890837 | 9.1144675 | 9.12386 | -0.009387656 | -0.034771391 |
| Olfr1085 | 1.563088 | 1.514508 | 1.5306 | -0.016107086 | 0.03247342 | 1.6111964 | 1.584302 | 1.59369 | -0.009390843 | 0.017503629 |
| Dhcr24 | 8.183582 | 8.205429 | 8.1061 | 0.099320282 | 0.077473649 | 8.2831073 | 7.9385595 | 7.94802 | -0.009463041 | 0.335084709 |
| Rdh11 | 8.811588 | 8.761432 | 8.8135 | -0.052018825 | -0.00186222 | 8.8469405 | 8.7337055 | 8.74328 | -0.009578506 | 0.103656455 |
| Gm14914 | 2.341253 | 2.301522 | 2.2825 | 0.019047498 | 0.05877849 | 2.2483456 | 2.3768523 | 2.38644 | -0.00958805 | -0.138094762 |
| Ywhaq-ps3 | 2.521132 | 2.451555 | 2.4737 | -0.022098473 | 0.047478639 | 2.4565857 | 2.6751384 | 2.68478 | -0.009646188 | -0.228198903 |
| Ak5 | 1.093608 | 1.066634 | 1.003 | 0.063626845 | 0.090600809 | 0.9881462 | 1.0576971 | 1.06739 | -0.009690103 | -0.079241059 |
| Sirt3 | 7.187341 | 6.951595 | 7.1984 | -0.24681569 | -0.011069913 | 7.2208056 | 7.1645909 | 7.17429 | -0.009700302 | 0.046514349 |
| Has3 | 1.931221 | 1.958125 | 2.0586 | -0.100426503 | -0.127330373 | 1.9202787 | 2.001135 | 2.01084 | -0.009707371 | -0.090563629 |
| Cntfr | 3.937306 | 3.76267 | 3.7676 | -0.00489885 | 0.169737222 | 3.7411051 | 3.3911221 | 3.40083 | -0.009711359 | 0.340271666 |
| Gm8960 | 1.311025 | 1.186612 | 1.2105 | -0.023931609 | 0.10048166 | 1.2278567 | 1.2102425 | 1.21998 | -0.009740424 | 0.007873785 |
| Rasal3 | 12.26196 | 12.28918 | 12.41 | -0.120899408 | -0.148114488 | 12.670539 | 12.649776 | 12.6595 | -0.009752612 | 0.01101001 |
| Cdc37 | 11.60467 | 11.64183 | 11.62 | 0.022208634 | -0.0149456 | 11.783879 | 11.614684 | 11.6245 | -0.009805174 | 0.159390405 |
| Gm17657 | 0.645447 | 0.612538 | 0.674 | -0.061483009 | -0.028573458 | 0.7046184 | 0.6631579 | 0.673 | -0.009844187 | 0.031616278 |
| Nid1 | 9.185162 | 9.352464 | 8.9066 | 0.445844587 | 0.278542293 | 8.8091331 | 8.7941987 | 8.80408 | -0.009880647 | 0.00505373 |
| Ppp1r32 | 4.023861 | 4.110348 | 4.073 | 0.037333669 | -0.049153551 | 4.2671632 | 4.432465 | 4.44236 | -0.009893839 | -0.175195685 |
| Slc30a9 | 9.620421 | 9.784521 | 9.6743 | 0.110186026 | -0.053913892 | 9.5751251 | 9.6968307 | 9.70676 | -0.009928029 | -0.131633555 |
| Eid2 | 4.525168 | 4.552816 | 4.4412 | 0.111580678 | 0.083932135 | 4.3838935 | 4.4122908 | 4.42223 | -0.009934807 | -0.038332122 |
| Gm16409 | 3.717446 | 3.46072 | 3.4749 | -0.01418674 | 0.242538854 | 3.3961036 | 3.4537327 | 3.46367 | -0.009935172 | -0.067564278 |
| Gm5643 | 1.368518 | 1.197037 | 1.2118 | -0.014740193 | 0.156740514 | 1.1755917 | 1.2429819 | 1.25295 | -0.009964934 | -0.077355154 |

| | | | | | | | | | | |
|---|---|---|---|---|---|---|---|---|---|---|
| Iqsec3 | 1.734649 | 1.761201 | 1.694 | 0.067237532 | 0.040685888 | 1.721076 | 1.7138575 | 1.72382 | -0.00996682 | -0.002748326 |
| Naif1 | 7.174364 | 7.323892 | 7.2636 | 0.060302361 | -0.089226052 | 7.198735 | 7.1871041 | 7.19713 | -0.010028326 | 0.001602549 |
| Fosl1 | 1.988177 | 1.843674 | 1.8002 | 0.043513232 | 0.188016677 | 1.8149081 | 1.9165826 | 1.92662 | -0.010039175 | -0.111713756 |
| Gm9042 | 2.288181 | 2.268944 | 2.2655 | 0.003395687 | 0.022632623 | 2.2613251 | 2.3667208 | 2.37689 | -0.010167308 | -0.115563014 |
| Kif28 | 0.591037 | 0.567251 | 0.5794 | -0.012150497 | 0.011635046 | 0.6049752 | 0.5886624 | 0.59888 | -0.010219422 | 0.006093348 |
| Hoxc4 | 3.354552 | 3.136955 | 3.2701 | -0.133146244 | 0.084450145 | 3.1605038 | 3.169141 | 3.17936 | -0.010220728 | -0.018857895 |
| Tnr | 2.097961 | 2.078835 | 2.0737 | 0.005135291 | 0.024261744 | 2.1358246 | 2.1215385 | 2.13176 | -0.010222212 | 0.004063873 |
| Borcs7 | 7.053609 | 6.836995 | 6.9743 | -0.137283437 | 0.079330069 | 7.1266866 | 6.9576618 | 6.96798 | -0.010320536 | 0.158704252 |
| Tmem240 | 2.63228 | 2.731055 | 2.7162 | 0.014858414 | -0.083916855 | 2.4966423 | 2.5830633 | 2.59339 | -0.010323147 | -0.096744157 |
| Slc52a2 | 8.405104 | 8.262274 | 8.3302 | -0.067880632 | 0.074949479 | 8.5585921 | 8.364719 | 8.37513 | -0.010408151 | 0.18346498 |
| Flcn | 9.85133 | 9.916991 | 9.9421 | -0.025108972 | -0.090769898 | 10.021271 | 9.971329 | 9.98174 | -0.010410014 | 0.039531483 |
| Fkbp3 | 10.42734 | 10.29447 | 10.466 | -0.171741152 | -0.038873724 | 10.5106 | 10.426867 | 10.4373 | -0.010424248 | 0.073308293 |
| Vmn2r5 | 2.234893 | 2.175646 | 2.1867 | -0.011058683 | 0.048188044 | 2.2369513 | 2.235488 | 2.24592 | -0.010429569 | -0.008966236 |
| Irs3 | 0.951975 | 0.971305 | 0.9929 | -0.021581362 | -0.040911666 | 0.9732658 | 0.9729929 | 0.98344 | -0.01044785 | -0.010174953 |
| Ulk4 | 2.332157 | 2.598087 | 2.4902 | 0.107906757 | -0.158023187 | 2.3746188 | 2.4585262 | 2.46899 | -0.010464767 | -0.094372134 |
| Uba52-ps | 0.968462 | 0.942658 | 0.9328 | 0.009812469 | 0.035616595 | 0.9776341 | 0.9158702 | 0.92635 | -0.01047982 | 0.051284061 |
| Tex45 | 0.399707 | 0.381593 | 0.376 | 0.00554383 | 0.023658306 | 0.4164032 | 0.3562684 | 0.36675 | -0.010481904 | 0.049652885 |
| Myom2 | 3.335109 | 3.232824 | 3.3599 | -0.127066162 | -0.024780679 | 3.272021 | 3.3724767 | 3.38299 | -0.010514595 | -0.110970314 |
| Foxr1 | 2.491422 | 2.47948 | 2.4837 | -0.004212187 | 0.007730261 | 2.4947097 | 2.4270754 | 2.43759 | -0.010516298 | 0.057117975 |
| Baiap2l1 | 1.706516 | 1.791102 | 1.7417 | 0.049360233 | -0.035225657 | 1.8068716 | 1.6990107 | 1.70953 | -0.01051754 | 0.097343363 |
| Gm3219 | 6.157644 | 6.012936 | 6.0775 | -0.064612546 | 0.080094951 | 6.155085 | 6.0027412 | 6.0133 | -0.010562768 | 0.141781034 |
| Klf2 | 11.2375 | 11.13313 | 11.332 | -0.198499418 | -0.09412635 | 10.720423 | 10.668884 | 10.6795 | -0.010580004 | 0.040959094 |
| Tmem63b | 7.878806 | 7.856366 | 7.9448 | -0.08843519 | -0.065995844 | 8.1082581 | 8.167433 | 8.17807 | -0.010641189 | -0.069816076 |
| Gm11027 | 1.510098 | 1.512416 | 1.5136 | -0.001226197 | -0.003544252 | 1.6087739 | 1.6236344 | 1.63431 | -0.010675987 | -0.025536517 |
| Pros1 | 8.149924 | 7.805893 | 7.8111 | -0.005167912 | 0.338863806 | 7.2876929 | 7.3673454 | 7.37803 | -0.010679905 | -0.090332382 |
| Pet117 | 0.737918 | 0.756359 | 0.7995 | -0.043109177 | -0.061549688 | 0.6698267 | 0.7172795 | 0.72797 | -0.010689609 | -0.058142369 |
| Plekhs1 | 2.519038 | 2.562862 | 2.6163 | -0.053467681 | -0.097291757 | 2.478457 | 2.6614188 | 2.67213 | -0.01071235 | -0.193674095 |
| Mgat4c | 2.020873 | 1.984083 | 2.0114 | -0.02735389 | 0.009436017 | 1.9963331 | 2.063493 | 2.07423 | -0.010739433 | -0.077899299 |
| Nefl | 0.73391 | 0.719987 | 0.5978 | 0.122232971 | 0.136155846 | 0.6208753 | 0.6332376 | 0.64403 | -0.010792937 | -0.023155179 |
| Fsbp | 3.323002 | 3.035749 | 3.1741 | -0.13832069 | 0.148931875 | 3.1231133 | 3.2033711 | 3.21417 | -0.010802349 | -0.091060213 |
| Bpifb1 | 0.719977 | 0.616809 | 0.6488 | -0.031970152 | 0.071198643 | 0.6454757 | 0.6275789 | 0.63839 | -0.010815511 | 0.007081333 |
| Kcnq2 | 1.541638 | 1.622465 | 1.5453 | 0.077134116 | -0.003693251 | 1.4801253 | 1.4984232 | 1.50934 | -0.01091682 | -0.029214685 |

| | | | | | | | | | | |
|---|---|---|---|---|---|---|---|---|---|---|
| Ap1m1 | 10.90288 | 10.94567 | 10.925 | 0.020360432 | -0.022428736 | 11.214595 | 10.975874 | 10.9868 | -0.010920028 | 0.227800976 |
| Unc13a | 4.317297 | 4.419514 | 4.3146 | 0.104909358 | 0.002691924 | 4.4743218 | 4.334432 | 4.34537 | -0.01093723 | 0.128952629 |
| Olfr780 | 1.1698 | 1.170535 | 1.1704 | 0.000104401 | -0.000630574 | 1.1871871 | 1.2658437 | 1.27679 | -0.010943219 | -0.089599828 |
| Mpnd | 9.863954 | 9.668443 | 9.8 | -0.131531939 | 0.063979417 | 10.062086 | 9.8832057 | 9.89415 | -0.010946493 | 0.167933498 |
| B4galt7 | 9.056833 | 9.064206 | 9.0843 | -0.020050246 | -0.027422916 | 9.1750514 | 8.9919031 | 9.00289 | -0.010987006 | 0.172161265 |
| Olfr1265 | 2.228903 | 2.280789 | 2.2263 | 0.054521469 | 0.0026354 | 2.3104949 | 2.2851531 | 2.29615 | -0.010992057 | 0.014349826 |
| Gm16459 | 1.390723 | 1.212661 | 1.2495 | -0.036847745 | 0.141214135 | 1.1830074 | 1.2381222 | 1.24912 | -0.010998027 | -0.06611275 |
| Olfr801 | 1.623683 | 1.606824 | 1.6336 | -0.026772715 | -0.00991409 | 1.5932418 | 1.6661321 | 1.67718 | -0.01104964 | -0.083939939 |
| Reep6 | 2.08801 | 2.05541 | 2.0865 | -0.031069954 | 0.001529911 | 2.0225758 | 2.0931425 | 2.1042 | -0.011057808 | -0.081624537 |
| Zfa-ps | 1.037512 | 1.037387 | 0.9956 | 0.041805658 | 0.041930968 | 0.9719692 | 1.0193258 | 1.03041 | -0.011081522 | -0.058438126 |
| Gm13237 | -0.040507 | -0.08305 | -0.0095 | -0.073517977 | -0.03097172 | -0.0815936 | -0.039833 | -0.0287 | -0.011098952 | -0.052859791 |
| Greb1 | 0.567927 | 0.617569 | 0.8211 | -0.203533447 | -0.25317599 | 0.6344577 | 0.5560479 | 0.56719 | -0.011139753 | 0.067270079 |
| Vps36 | 9.731733 | 9.720977 | 9.7319 | -0.010957055 | -0.000200357 | 9.7709743 | 9.7371605 | 9.74847 | -0.011313462 | 0.022500352 |
| B230359F08Rik | 2.135153 | 2.114544 | 2.1801 | -0.065521862 | -0.044912683 | 2.1345698 | 2.1085 | 2.11989 | -0.011388633 | 0.014681182 |
| Elf5 | 3.021475 | 2.867981 | 3.0834 | -0.215435217 | -0.061941814 | 3.1886101 | 3.0646069 | 3.07601 | -0.011404103 | 0.112599095 |
| Olfr1029 | 0.231849 | 0.271634 | 0.2686 | 0.003013986 | -0.036771137 | 0.2393271 | 0.2711726 | 0.28259 | -0.011419925 | -0.043265453 |
| F830045P16Rik | 0.620054 | 0.654923 | 0.6194 | 0.035549318 | 0.000680938 | 0.7021881 | 0.5609484 | 0.57242 | -0.011471687 | 0.129767998 |
| Crtap | 7.015498 | 6.967554 | 7.06 | -0.092470901 | -0.044526409 | 7.1228396 | 6.8300633 | 6.84161 | -0.011547088 | 0.281229274 |
| Mfap3 | 9.332431 | 9.313321 | 9.4036 | -0.090285259 | -0.071175258 | 9.7222992 | 9.556907 | 9.56849 | -0.01158117 | 0.153811014 |
| Tpt1-ps3 | 14.15307 | 14.26176 | 14.165 | 0.096733754 | -0.01194963 | 14.31562 | 14.239448 | 14.2511 | -0.011607408 | 0.06456519 |
| Skint10 | 0.379334 | 0.288798 | 0.4328 | -0.143956215 | -0.053420718 | 0.301825 | 0.2853937 | 0.297 | -0.011608056 | 0.004823289 |
| Vwce | 3.303493 | 3.281969 | 3.3392 | -0.057182673 | -0.03565848 | 3.340021 | 3.4315963 | 3.44325 | -0.011655711 | -0.103230995 |
| Igkv6-29 | 1.827619 | 1.766167 | 1.8704 | -0.104272918 | -0.042821693 | 1.8593197 | 1.8295316 | 1.84126 | -0.011724797 | 0.018063281 |
| Zfp989 | 1.626183 | 1.607309 | 1.5735 | 0.033769353 | 0.052642493 | 1.567787 | 1.6538941 | 1.66562 | -0.011725957 | -0.097833008 |
| Dnaaf3 | 6.182909 | 5.949037 | 6.2123 | -0.263306822 | -0.029435104 | 6.2468678 | 6.4179388 | 6.42969 | -0.01174867 | -0.182819622 |
| Klc1 | 9.525402 | 9.47423 | 9.3293 | 0.14496395 | 0.196136583 | 9.6089463 | 9.6056317 | 9.61739 | -0.011761968 | -0.008447342 |
| Kcnip3 | 5.234266 | 5.185181 | 5.2844 | -0.099171808 | -0.050087082 | 5.327008 | 5.1515653 | 5.16334 | -0.011770082 | 0.163672617 |
| Fam160b2 | 9.330822 | 9.350704 | 9.4358 | -0.085098801 | -0.104980706 | 9.5692061 | 9.4651998 | 9.47697 | -0.011772275 | 0.092234088 |
| Gm2099 | 1.813162 | 1.811545 | 1.7959 | 0.015689979 | 0.017307151 | 1.8019537 | 1.8971161 | 1.9089 | -0.011783698 | -0.106946119 |
| Bfsp1 | 1.768329 | 1.857211 | 1.8312 | 0.026049252 | -0.062833414 | 1.9457079 | 1.8410528 | 1.85287 | -0.01181414 | 0.092840978 |
| Mroh6 | 0.077533 | 0.054995 | 0.0511 | 0.003844942 | 0.026383598 | 0.0940408 | 0.0741513 | 0.08597 | -0.011816074 | 0.008073495 |
| Gm2792 | 3.689637 | 3.721605 | 3.765 | -0.043375255 | -0.075342778 | 3.9192388 | 3.8673009 | 3.87915 | -0.011844443 | 0.04009339 |

| | | | | | | | | | | |
|---|---|---|---|---|---|---|---|---|---|---|
| Hnrnpk | 13.4656 | 13.49007 | 13.463 | 0.026855336 | 0.002383504 | 13.495718 | 13.483156 | 13.495 | -0.011874508 | 0.000687543 |
| Nccrp1 | 3.055217 | 3.053463 | 3.1141 | -0.060669337 | -0.058914714 | 3.0474995 | 2.9277864 | 2.93969 | -0.011908064 | 0.107805079 |
| Trav9d-2 | 0.845682 | 0.901238 | 0.8471 | 0.054102841 | -0.001453334 | 0.8689571 | 0.8752928 | 0.88721 | -0.011918659 | -0.01825442 |
| Trav9n-2 | 0.845682 | 0.901238 | 0.8471 | 0.054102841 | -0.001453334 | 0.8689571 | 0.8752928 | 0.88721 | -0.011918659 | -0.01825442 |
| Gm5709 | 0.717559 | 0.699446 | 0.6958 | 0.003614456 | 0.02172715 | 0.6611812 | 0.6642948 | 0.67638 | -0.012086277 | -0.015199903 |
| Olfr145 | 0.574363 | 0.538559 | 0.5848 | -0.046213485 | -0.010408761 | 0.5973815 | 0.5459196 | 0.55804 | -0.012124033 | 0.039337866 |
| Stag2 | 12.19726 | 12.24759 | 12.185 | 0.06222339 | 0.011896834 | 12.072353 | 12.115043 | 12.1273 | -0.012212092 | -0.054901937 |
| Gm49321 | 3.274798 | 3.368928 | 3.2397 | 0.129186812 | 0.035057021 | 3.5120027 | 3.2857119 | 3.29794 | -0.012229255 | 0.214061501 |
| Satb2 | 2.141379 | 2.124157 | 2.1666 | -0.042458355 | -0.025236432 | 2.1035105 | 2.1888046 | 2.20109 | -0.012289675 | -0.097583734 |
| Gm26920 | 3.425154 | 3.049066 | 3.3666 | -0.317514735 | 0.058573836 | 3.2917233 | 3.2321771 | 3.24457 | -0.01239266 | 0.047153539 |
| Psip1 | 10.97241 | 10.98813 | 11.013 | -0.02536497 | -0.041085816 | 10.945061 | 11.070521 | 11.0829 | -0.012419106 | -0.137879038 |
| Olfr465-ps1 | 0.318365 | 0.296678 | 0.3232 | -0.026484455 | -0.004798002 | 0.2934993 | 0.3426686 | 0.35514 | -0.01246905 | -0.061638414 |
| Rbfox2 | 5.859198 | 5.670066 | 5.8259 | -0.155821446 | 0.033310988 | 5.8328579 | 5.3570413 | 5.36953 | -0.012492334 | 0.463324336 |
| Ephb2 | 4.245978 | 4.352553 | 4.1007 | 0.251846022 | 0.145270862 | 4.1726797 | 4.1531171 | 4.16562 | -0.012498593 | 0.007064045 |
| Gm13230 | 0.971071 | 0.986111 | 0.9915 | -0.005385047 | -0.020424688 | 0.9708198 | 0.9752274 | 0.98773 | -0.012503918 | -0.016911479 |
| Olfr1321 | 2.269468 | 2.271716 | 2.3111 | -0.039350017 | -0.041598399 | 2.4018396 | 2.3426105 | 2.35517 | -0.012559895 | 0.04666923 |
| Snapc4 | 9.498026 | 9.520544 | 9.5878 | -0.067226304 | -0.089743768 | 9.9961997 | 9.9457442 | 9.95838 | -0.012635747 | 0.037819737 |
| 38596 | 4.054744 | 4.212319 | 3.937 | 0.275302493 | 0.117726715 | 3.8231625 | 3.9187674 | 3.93148 | -0.012707783 | -0.108312665 |
| Cox7a1 | 1.584496 | 1.588037 | 1.6214 | -0.033404439 | -0.036945544 | 1.6779563 | 1.6121067 | 1.62485 | -0.012746078 | 0.053103603 |
| Slc2a4 | 2.494172 | 2.331856 | 2.5254 | -0.193537671 | -0.031221922 | 2.3499438 | 2.3294281 | 2.34222 | -0.012794671 | 0.007721078 |
| Gm2531 | 1.723695 | 1.685515 | 1.6691 | 0.016390749 | 0.054570563 | 1.7675269 | 1.7614158 | 1.77422 | -0.012807953 | -0.006696828 |
| Ppp2r5c | 11.51464 | 11.46123 | 11.558 | -0.096820914 | -0.04340964 | 11.545006 | 11.466184 | 11.479 | -0.0128607 | 0.065961561 |
| Zscan12 | 6.719553 | 6.543842 | 6.5616 | -0.017761796 | 0.157949136 | 6.4196901 | 6.6583673 | 6.67123 | -0.012865057 | -0.251542276 |
| Mybl2 | 6.852016 | 6.432261 | 6.9099 | -0.47759846 | -0.057844018 | 6.8346862 | 7.0662763 | 7.0792 | -0.012925616 | -0.24451574 |
| P2ry12 | 5.785089 | 5.471032 | 5.6192 | -0.148133226 | 0.165923694 | 5.9023619 | 5.5385543 | 5.55152 | -0.01296662 | 0.350840941 |
| Zfp821 | 7.623411 | 7.614006 | 7.7709 | -0.156862204 | -0.147457039 | 7.6764853 | 7.5985468 | 7.61155 | -0.01300353 | 0.064934901 |
| Palm3 | 0.888798 | 0.889278 | 0.8877 | 0.001528439 | 0.001049045 | 0.9248223 | 0.8603918 | 0.87344 | -0.013053023 | 0.0513775 |
| AC169506.2 | 1.536145 | 1.536938 | 1.5251 | 0.011854203 | 0.011061251 | 1.6455311 | 1.6232243 | 1.63629 | -0.013069563 | 0.009237243 |
| Triml2 | 0.589311 | 0.577964 | 0.5745 | 0.003507631 | 0.014854705 | 0.5711117 | 0.5993795 | 0.61253 | -0.013149002 | -0.041416757 |
| Trim43c | 0.275462 | 0.290987 | 0.2403 | 0.050657527 | 0.035132272 | 0.2449725 | 0.2347842 | 0.24795 | -0.013169883 | -0.002981582 |
| Ccdc174 | 8.836255 | 8.795456 | 8.9184 | -0.122978454 | -0.082179741 | 8.6889743 | 8.8953737 | 8.90865 | -0.013271594 | -0.219670992 |
| Ighv8-8 | 8.36183 | 8.499871 | 8.3436 | 0.156297949 | 0.0182562 | 8.6657504 | 8.5021107 | 8.51541 | -0.013299141 | 0.150340547 |

| | | | | | | | | | | |
|---|---|---|---|---|---|---|---|---|---|---|
| Pabpc5 | 0.672475 | 0.638558 | 0.6233 | 0.015234041 | 0.049151263 | 0.7028917 | 0.6378619 | 0.65117 | -0.013304831 | 0.051724988 |
| Vmn1r74 | 1.033256 | 1.029966 | 1.0493 | -0.0193587 | -0.016069277 | 1.1180208 | 1.1035835 | 1.11689 | -0.013307461 | 0.001129826 |
| Gm15515 | 2.56898 | 2.538457 | 2.4905 | 0.047940288 | 0.078463761 | 2.7849613 | 2.6365782 | 2.64993 | -0.013348833 | 0.135034241 |
| Isyna1 | 9.638608 | 9.564529 | 9.6139 | -0.049379483 | 0.024700163 | 9.8972765 | 9.5791706 | 9.59253 | -0.013355049 | 0.30475078 |
| Szt2 | 10.58493 | 10.64471 | 10.582 | 0.063127402 | 0.003342906 | 10.764875 | 10.939031 | 10.9524 | -0.013396114 | -0.187552247 |
| Pdxk | 9.209102 | 9.310523 | 9.1911 | 0.119450066 | 0.018029248 | 9.2176982 | 9.1427545 | 9.15616 | -0.013404449 | 0.061539228 |
| Chodl | 1.053874 | 0.993747 | 1.0236 | -0.029899505 | 0.030227268 | 1.1991656 | 1.0640914 | 1.07768 | -0.01358739 | 0.121486805 |
| Hist1h3d | 2.275374 | 2.328982 | 2.3909 | -0.061869042 | -0.11547727 | 2.3814228 | 2.3439984 | 2.3576 | -0.013606444 | 0.023817945 |
| Olfr307 | 1.219783 | 1.189812 | 1.2084 | -0.018630493 | 0.011340615 | 1.2692059 | 1.2542579 | 1.26788 | -0.013624103 | 0.001323803 |
| Gm20167 | 0.718928 | 0.646218 | 0.6763 | -0.030063332 | 0.042646379 | 0.6951397 | 0.6609441 | 0.67461 | -0.013661052 | 0.02053456 |
| Igkv20-101-2 | 1.339795 | 1.388414 | 1.4352 | -0.046801388 | -0.095420359 | 1.4058518 | 1.3562173 | 1.36989 | -0.013670865 | 0.035963673 |
| P2rx5 | 3.871792 | 3.942019 | 3.9561 | -0.014088263 | -0.084314539 | 3.9759608 | 3.9167696 | 3.93049 | -0.013721345 | 0.045469825 |
| Adgre4 | 6.880601 | 7.817685 | 7.2299 | 0.587790413 | -0.349293228 | 8.0979708 | 7.0834089 | 7.09713 | -0.013723367 | 1.000838516 |
| Uchl4 | 1.392238 | 1.353984 | 1.3775 | -0.023565825 | 0.014688173 | 1.3943264 | 1.4239488 | 1.43768 | -0.013734322 | -0.043356725 |
| Spef2 | 5.581455 | 5.388752 | 5.5195 | -0.130714396 | 0.061988312 | 5.6570658 | 5.5954619 | 5.6092 | -0.013739152 | 0.047864736 |
| Gnrh1 | 2.579434 | 2.564657 | 2.5868 | -0.022166833 | -0.007390063 | 2.6088723 | 2.6537655 | 2.66751 | -0.013748211 | -0.058641376 |
| Gm19246 | 0.923274 | 0.944571 | 0.9055 | 0.039025165 | 0.017727929 | 0.9661489 | 0.9532603 | 0.96704 | -0.013777491 | -0.00088882 |
| Timm17b | 8.601274 | 8.554589 | 8.571 | -0.016379919 | 0.030304734 | 8.5532193 | 8.4992891 | 8.51307 | -0.013780752 | 0.040149453 |
| Hoxc5 | 0.619053 | 0.684686 | 0.6131 | 0.071586674 | 0.005953751 | 0.7137686 | 0.6557243 | 0.66951 | -0.013784448 | 0.04425981 |
| Olfr1416 | 2.736414 | 2.722239 | 2.8077 | -0.085472979 | -0.071297833 | 2.8577152 | 2.9028899 | 2.91668 | -0.013793637 | -0.058968341 |
| Cap1 | 13.3018 | 13.29414 | 13.227 | 0.067594576 | 0.07526015 | 13.429075 | 13.260133 | 13.274 | -0.013834198 | 0.155107707 |
| Lcn2 | 6.770554 | 6.323884 | 5.6952 | 0.628731461 | 1.075401768 | 6.8145385 | 5.9570086 | 5.97085 | -0.013845917 | 0.84368393 |
| Ralgds | 9.587873 | 9.529841 | 9.4953 | 0.03454557 | 0.092577895 | 9.3473209 | 9.4040212 | 9.41788 | -0.013863422 | -0.070563762 |
| Pitrm1 | 9.919385 | 9.933378 | 9.9353 | -0.001896107 | -0.01588956 | 10.062123 | 10.044205 | 10.0581 | -0.013877792 | 0.00404028 |
| Syt14 | 1.268922 | 1.287445 | 1.3269 | -0.0394877 | -0.058010753 | 1.1990904 | 1.298872 | 1.31281 | -0.013934943 | -0.113716457 |
| Gm10327 | 1.593467 | 1.668674 | 1.5599 | 0.108764089 | 0.033556313 | 1.5405838 | 1.6144293 | 1.62838 | -0.013948511 | -0.087793991 |
| Enkur | 0.855275 | 0.870997 | 0.9343 | -0.063330109 | -0.079052519 | 0.8625973 | 0.8931347 | 0.90709 | -0.013957112 | -0.044494568 |
| Olfr1238 | 4.469948 | 4.489618 | 4.5375 | -0.047853579 | -0.0675239 | 4.6413883 | 4.7596881 | 4.77371 | -0.01402093 | -0.132320786 |
| Pcdh20 | 0.09402 | 0.164732 | 0.1317 | 0.033081317 | -0.037631054 | 0.0868189 | 0.0738841 | 0.08791 | -0.014025704 | -0.00109089 |
| Traj27 | 2.963057 | 2.946006 | 2.9278 | 0.018162531 | 0.035213001 | 2.8014736 | 2.9381501 | 2.95224 | -0.014092155 | -0.150768588 |
| Asgr2 | 1.900378 | 1.777602 | 1.7931 | -0.015451989 | 0.107324107 | 1.7533347 | 1.7757468 | 1.78985 | -0.014107584 | -0.03651963 |
| Zfp273 | 6.550512 | 6.696117 | 6.7023 | -0.006149182 | -0.151753535 | 6.4818922 | 6.4780771 | 6.49219 | -0.014115245 | -0.010300147 |

| | | | | | | | | | | |
|---|---|---|---|---|---|---|---|---|---|---|
| Itgav | 9.344129 | 9.384119 | 9.4068 | -0.022719332 | -0.062709104 | 9.4173648 | 9.3941919 | 9.40833 | -0.014133287 | 0.009039623 |
| Gm10762 | 4.401592 | 4.43849 | 4.3862 | 0.052292018 | 0.015394645 | 4.6352668 | 4.3370485 | 4.35122 | -0.014167886 | 0.284050478 |
| Vmn1r89 | 0.807648 | 0.773907 | 0.8049 | -0.031022823 | 0.002717386 | 0.8179345 | 0.8257221 | 0.8399 | -0.014175438 | -0.021962964 |
| Gm15696 | 3.083857 | 3.040095 | 3.0109 | 0.029231194 | 0.072992703 | 3.1757521 | 3.1453564 | 3.15956 | -0.014200464 | 0.016195205 |
| Vmn2r101 | 0.345603 | 0.374933 | 0.3935 | -0.018532456 | -0.04786199 | 0.3831306 | 0.4004797 | 0.4147 | -0.014219841 | -0.031568958 |
| Tmprss11g | 1.731884 | 1.71162 | 1.6544 | 0.057232253 | 0.077495783 | 1.6851468 | 1.7204652 | 1.73469 | -0.014224731 | -0.049543111 |
| Olfr1385 | 6.719421 | 6.763834 | 6.8178 | -0.053944901 | -0.098356929 | 6.8477418 | 7.0313549 | 7.04565 | -0.014293126 | -0.197906165 |
| Fbxl16 | 2.16116 | 2.178983 | 1.9893 | 0.189690331 | 0.17186722 | 2.0429072 | 1.9824429 | 1.99676 | -0.014317408 | 0.046146883 |
| Cdh10 | 2.098098 | 2.000358 | 2.0456 | -0.04524004 | 0.052499866 | 1.9703228 | 2.0189432 | 2.03329 | -0.014342512 | -0.062962882 |
| Olfr398 | 2.485434 | 2.558804 | 2.5094 | 0.049374543 | -0.02399517 | 2.4873758 | 2.5859371 | 2.60031 | -0.014372542 | -0.112933927 |
| Srcin1 | 1.872785 | 1.768805 | 1.8278 | -0.059011609 | 0.044968599 | 1.7712947 | 1.7825753 | 1.797 | -0.014427847 | -0.025708482 |
| Gstz1 | 7.333325 | 7.267954 | 7.4467 | -0.178753666 | -0.113383148 | 7.6201933 | 7.6140454 | 7.62848 | -0.014438869 | -0.008290952 |
| Rfc4 | 7.538166 | 7.550274 | 7.5957 | -0.045377849 | -0.057485202 | 7.732149 | 7.6271224 | 7.64157 | -0.014446416 | 0.090580167 |
| Zfp277 | 9.153317 | 9.171026 | 9.2539 | -0.082825625 | -0.100534445 | 9.2470769 | 9.1795733 | 9.19408 | -0.014508637 | 0.052994908 |
| Lrp12 | 6.32752 | 6.720447 | 6.4317 | 0.288706075 | -0.104220867 | 5.9693017 | 6.2136327 | 6.22814 | -0.014510275 | -0.258841311 |
| Gm5621 | 1.094821 | 1.107984 | 1.1187 | -0.010706581 | -0.023869206 | 1.1545124 | 1.1048649 | 1.11944 | -0.014579708 | 0.035067826 |
| Psph | 7.601881 | 7.677794 | 7.829 | -0.151157823 | -0.22707045 | 8.0172757 | 7.7931482 | 7.80784 | -0.014691327 | 0.209436142 |
| Gm11084 | 0.149423 | 0.186044 | 0.1623 | 0.023770617 | -0.012850272 | 0.1506713 | 0.1482713 | 0.16297 | -0.014694281 | -0.012294269 |
| Cd200 | 10.20763 | 10.12516 | 10.12 | 0.005284379 | 0.087755953 | 9.8797278 | 9.8656089 | 9.88033 | -0.014725886 | -0.000606941 |
| Olfr1454 | 0.707414 | 0.744856 | 0.7431 | 0.001794157 | -0.03564806 | 0.7635395 | 0.7768598 | 0.79159 | -0.014733096 | -0.028053397 |
| Olfr488 | 0.88628 | 0.936251 | 0.9068 | 0.029471098 | -0.020500251 | 0.9157857 | 0.9085941 | 0.9234 | -0.014803728 | -0.007612145 |
| Sdf2 | 9.386617 | 9.457118 | 9.4453 | 0.011847862 | -0.058652723 | 9.7720742 | 9.5708182 | 9.58565 | -0.014827409 | 0.186428641 |
| Plekho2 | 11.17092 | 11.13266 | 10.927 | 0.205815464 | 0.244077446 | 11.170329 | 10.956662 | 10.9715 | -0.014834108 | 0.198832622 |
| Mef2b | 5.025082 | 5.068222 | 4.9428 | 0.125399331 | 0.082258983 | 5.1958965 | 4.968861 | 4.98373 | -0.014867585 | 0.212167958 |
| Dhx9 | 12.45858 | 12.46586 | 12.429 | 0.036763812 | 0.029493342 | 12.389423 | 12.452487 | 12.4674 | -0.014879758 | -0.077943213 |
| Cacnb2 | 3.027892 | 2.965886 | 3.005 | -0.039101495 | 0.022904513 | 3.0107361 | 3.0292353 | 3.04413 | -0.014897945 | -0.033397134 |
| Nfatc4 | 5.802916 | 5.765661 | 5.9191 | -0.153487985 | -0.116233307 | 5.5812538 | 5.5868284 | 5.60175 | -0.014925082 | -0.020499598 |
| Gm48216 | 5.108996 | 5.048205 | 5.2052 | -0.156991196 | -0.096199946 | 4.9547804 | 5.4478056 | 5.46275 | -0.014945797 | -0.507971081 |
| Olfr983 | 1.398713 | 1.392967 | 1.4615 | -0.068492242 | -0.062746417 | 1.3896679 | 1.4529453 | 1.46794 | -0.014995209 | -0.07827267 |
| Tmem251 | 8.109493 | 8.080554 | 8.1243 | -0.043739689 | -0.014800669 | 8.2045046 | 8.0729516 | 8.08803 | -0.01507691 | 0.116476132 |
| Arhgap11a | 8.960233 | 9.088716 | 9.1121 | -0.023364294 | -0.151846596 | 8.9406516 | 9.1404939 | 9.15561 | -0.015111566 | -0.21495393 |
| Fgf16 | 1.286793 | 1.336126 | 1.3458 | -0.009632634 | -0.058965716 | 1.3664337 | 1.3443299 | 1.35945 | -0.015118671 | 0.006985103 |

| | | | | | | | | | | |
|---|---|---|---|---|---|---|---|---|---|---|
| Cnot9 | 9.620078 | 9.504098 | 9.5909 | -0.086834803 | 0.029144699 | 9.6371082 | 9.5436398 | 9.55877 | -0.015127763 | 0.078340561 |
| AC163277.1 | 1.4621 | 1.423463 | 1.4427 | -0.019265177 | 0.019372521 | 1.4951011 | 1.5115429 | 1.52669 | -0.015142779 | -0.031584502 |
| Orm3 | 0.048199 | -0.03437 | 0.0194 | -0.053754231 | 0.028810678 | -0.0516779 | -0.044665 | -0.0295 | -0.015157962 | -0.02217109 |
| Rab5b | 10.63825 | 10.68191 | 10.671 | 0.011126834 | -0.032529702 | 10.712581 | 10.508684 | 10.5238 | -0.015159166 | 0.188737391 |
| H2afz | 11.33156 | 11.15756 | 11.329 | -0.171266564 | 0.002728374 | 11.209874 | 11.285911 | 11.3011 | -0.015213576 | -0.091249865 |
| Mib2 | 8.807155 | 8.949337 | 8.957 | -0.007709469 | -0.149891187 | 9.1411675 | 9.0070405 | 9.02228 | -0.015242806 | 0.11888425 |
| Cacul1 | 10.35925 | 10.3666 | 10.273 | 0.093913768 | 0.086559502 | 10.21039 | 10.280974 | 10.2962 | -0.015242848 | -0.085826624 |
| Trbj2-3 | 5.446857 | 5.513245 | 5.5242 | -0.010975666 | -0.07736304 | 5.2409522 | 5.4841599 | 5.49942 | -0.015259606 | -0.258467307 |
| Gm28230 | 1.595115 | 1.548789 | 1.7009 | -0.152119399 | -0.105793336 | 1.5253906 | 1.558184 | 1.57347 | -0.015285444 | -0.048078842 |
| Ighv1-7 | 7.000078 | 7.293507 | 7.0046 | 0.288925923 | -0.004502854 | 7.5117705 | 7.0789968 | 7.09429 | -0.015293346 | 0.417480323 |
| Zfp786 | 1.315918 | 1.202808 | 1.3535 | -0.150693412 | -0.037583921 | 1.3688241 | 1.3090682 | 1.32438 | -0.015310704 | 0.044445167 |
| Igkv8-19 | 7.086249 | 7.595459 | 7.3738 | 0.221688227 | -0.287521584 | 7.569174 | 7.4705966 | 7.48597 | -0.015369016 | 0.083208378 |
| Gpr37 | 5.481235 | 5.451688 | 5.6209 | -0.16924497 | -0.13969834 | 5.6891413 | 5.8565501 | 5.87192 | -0.01537074 | -0.182779498 |
| Pcdhga2 | 2.704938 | 2.604493 | 2.6007 | 0.003807386 | 0.104252026 | 2.564536 | 2.5366006 | 2.55199 | -0.015389192 | 0.012546272 |
| Pyurf | 6.798842 | 6.860838 | 6.7469 | 0.113935843 | 0.051940212 | 6.9856133 | 6.7908868 | 6.8063 | -0.015410594 | 0.179315936 |
| Slc6a18 | 1.128982 | 1.128273 | 1.1482 | -0.019964436 | -0.019255565 | 1.0778123 | 1.1160338 | 1.13146 | -0.015422666 | -0.053644193 |
| Hexim1 | 10.32716 | 10.12802 | 10.477 | -0.348589724 | -0.149447032 | 9.5994617 | 9.7787759 | 9.79422 | -0.015448025 | -0.194762202 |
| Foxn2 | 10.50919 | 10.58697 | 10.481 | 0.105807896 | 0.02802576 | 10.144791 | 10.420352 | 10.4358 | -0.015474792 | -0.291036166 |
| Olfr384 | 1.620059 | 1.644022 | 1.5513 | 0.092769435 | 0.068805906 | 1.7447917 | 1.7121562 | 1.72766 | -0.015505185 | 0.017130326 |
| Traj22 | 2.983622 | 2.882925 | 2.8553 | 0.027627889 | 0.128324878 | 2.7999541 | 2.9095957 | 2.92513 | -0.015531721 | -0.125173352 |
| Ggnbp2 | 10.71688 | 10.63326 | 10.726 | -0.09289249 | -0.009270112 | 10.642814 | 10.812363 | 10.828 | -0.015610724 | -0.185159647 |
| Gm2897 | 2.048554 | 1.959882 | 1.9948 | -0.034894973 | 0.053776461 | 1.8923072 | 2.0131004 | 2.02876 | -0.01566415 | -0.136457355 |
| Zfp345 | 2.24944 | 2.299884 | 2.324 | -0.024075538 | -0.074519243 | 2.4314512 | 2.3461203 | 2.36184 | -0.015715049 | 0.069615863 |
| Acat3 | 1.857475 | 1.868373 | 1.847 | 0.021421227 | 0.010523331 | 2.0957618 | 1.9655324 | 1.98126 | -0.015725006 | 0.114504429 |
| Tfr2 | 2.078256 | 2.180481 | 2.2339 | -0.053411867 | -0.155637336 | 2.1216307 | 2.0978819 | 2.11363 | -0.015749309 | 0.007999427 |
| Pik3ca | 10.39795 | 10.14347 | 10.323 | -0.179638306 | 0.074847874 | 10.316558 | 10.268534 | 10.2843 | -0.015801929 | 0.032221723 |
| Zbtb11 | 10.14455 | 10.28678 | 10.149 | 0.137353199 | -0.004878871 | 10.023604 | 10.191098 | 10.2069 | -0.015803755 | -0.183298151 |
| Plin4 | 2.214117 | 2.270607 | 2.2455 | 0.025116346 | -0.031373502 | 2.0456214 | 2.1295915 | 2.14543 | -0.015835753 | -0.099805921 |
| Gm19035 | 1.235207 | 1.232831 | 1.2551 | -0.022223831 | -0.019847866 | 1.2722763 | 1.2743776 | 1.29023 | -0.015852435 | -0.017953749 |
| Fut9 | 0.158508 | 0.139242 | 0.1056 | 0.033621812 | 0.052888524 | 0.1056129 | 0.102433 | 0.11829 | -0.01585284 | -0.012672957 |
| Vmn1r29 | 0.361762 | 0.373638 | 0.3663 | 0.007325355 | -0.004550495 | 0.399534 | 0.4028404 | 0.41885 | -0.016010873 | -0.019317218 |
| Paqr9 | 1.057932 | 0.9688 | 1.0147 | -0.045921788 | 0.04321049 | 0.9338625 | 0.9207928 | 0.9368 | -0.016011523 | -0.002941811 |

| | | | | | | | | | | |
|---|---|---|---|---|---|---|---|---|---|---|
| Gpr45 | 2.618999 | 2.495365 | 2.5613 | -0.065929367 | 0.057705103 | 2.6264472 | 2.6081753 | 2.62419 | -0.016012456 | 0.002259493 |
| Gm14830 | 1.601572 | 1.537184 | 1.5737 | -0.036516602 | 0.027871289 | 1.5840969 | 1.5837816 | 1.5998 | -0.016013929 | -0.015698552 |
| Dusp23 | 6.399048 | 6.271796 | 6.5899 | -0.318136304 | -0.190884615 | 6.592925 | 6.3685334 | 6.38455 | -0.01602017 | 0.208371446 |
| Gm16580 | 3.390823 | 3.428524 | 3.497 | -0.068515314 | -0.106216114 | 3.5732641 | 3.5492003 | 3.56525 | -0.01604788 | 0.008015885 |
| Bloc1s3 | 7.139761 | 7.063622 | 7.2045 | -0.14087997 | -0.064740889 | 7.1302709 | 7.0790673 | 7.09513 | -0.016060099 | 0.035143436 |
| Smim15 | 9.816378 | 9.823126 | 9.8592 | -0.036038263 | -0.042786182 | 9.8168592 | 9.7664479 | 9.78257 | -0.016125867 | 0.034285438 |
| Gins3 | 7.11067 | 7.10144 | 7.0987 | 0.002752265 | 0.011981917 | 7.082014 | 7.3415414 | 7.3577 | -0.016154689 | -0.275682099 |
| Guca2b | 0.525014 | 0.540738 | 0.5087 | 0.032005056 | 0.016280746 | 0.5942244 | 0.4825331 | 0.4987 | -0.016164164 | 0.095527137 |
| Vmn1r85 | 4.995377 | 5.207369 | 5.0831 | 0.12426988 | -0.087721944 | 5.173973 | 5.1476804 | 5.16386 | -0.01618091 | 0.010111727 |
| Rnf219 | 7.733095 | 7.767129 | 7.735 | 0.032142725 | -0.001891782 | 7.8591353 | 7.813449 | 7.82967 | -0.016225951 | 0.029460364 |
| Plaur | 10.33965 | 10.54426 | 10.153 | 0.391670601 | 0.187066813 | 10.183111 | 10.179826 | 10.1961 | -0.016244325 | -0.012959226 |
| Prss53 | 2.496369 | 2.614112 | 2.5363 | 0.07779442 | -0.039948257 | 2.6461695 | 2.4916031 | 2.50802 | -0.01641204 | 0.138154323 |
| Ints3 | 10.30183 | 10.45153 | 10.338 | 0.113686656 | -0.036008774 | 10.390888 | 10.384959 | 10.4014 | -0.016427004 | -0.010498121 |
| Olfr1233 | 2.440303 | 2.382549 | 2.3841 | -0.001511718 | 0.056242282 | 2.3268102 | 2.4298092 | 2.44626 | -0.016455111 | -0.119454147 |
| Gm11263 | 1.907043 | 1.715927 | 1.7196 | -0.003688532 | 0.187427971 | 1.7164795 | 1.6971209 | 1.71359 | -0.016468363 | 0.002890235 |
| Olfr107 | 2.636247 | 2.686909 | 2.7756 | -0.088705598 | -0.139367565 | 2.645138 | 2.6891873 | 2.70567 | -0.01648523 | -0.060534491 |
| Usp7 | 12.00323 | 11.96046 | 11.957 | 0.003060168 | 0.0458315 | 11.862211 | 12.102253 | 12.1189 | -0.01661313 | -0.256654743 |
| Vmn1r173 | 1.911725 | 1.959888 | 1.9236 | 0.036289029 | -0.011874248 | 1.9910072 | 1.9356321 | 1.95225 | -0.016615608 | 0.038759514 |
| Clba1 | 1.79715 | 1.831327 | 1.7725 | 0.058871581 | 0.024694162 | 1.7937928 | 1.8321422 | 1.84879 | -0.016645018 | -0.054994475 |
| Smoc2 | 2.098192 | 2.061175 | 2.0082 | 0.052966454 | 0.089983626 | 2.0802813 | 2.0460436 | 2.0627 | -0.016652187 | 0.017585508 |
| Cxcl17 | 1.508271 | 1.420025 | 1.4193 | 0.000735665 | 0.088981755 | 1.5477258 | 1.4352158 | 1.45188 | -0.01666035 | 0.095849683 |
| Atp11a | 9.300629 | 9.11709 | 9.1378 | -0.020682901 | 0.162855591 | 9.1419312 | 9.473328 | 9.49001 | -0.016682674 | -0.348079417 |
| Rab3b | 0.496669 | 0.486422 | 0.4663 | 0.020142727 | 0.030389701 | 0.4556247 | 0.4802283 | 0.49698 | -0.016748122 | -0.04135174 |
| Grifin | 2.221912 | 2.20842 | 2.2265 | -0.018129559 | -0.004637574 | 2.2955968 | 2.2351724 | 2.25197 | -0.016799489 | 0.043624862 |
| Efhb | 1.103584 | 1.045318 | 1.0345 | 0.010775472 | 0.069041085 | 1.1212699 | 1.0489088 | 1.06584 | -0.016929894 | 0.055431291 |
| Sla2 | 8.806198 | 8.885974 | 8.8225 | 0.063437158 | -0.01633903 | 8.6374481 | 8.8484596 | 8.86543 | -0.01697074 | -0.227982246 |
| Gm5173 | 1.091089 | 1.047458 | 1.0663 | -0.018834829 | 0.024795935 | 1.0986807 | 1.1053827 | 1.12242 | -0.017036837 | -0.023738829 |
| Chmp2b | 9.258447 | 9.273661 | 9.2284 | 0.045210684 | 0.029996818 | 9.1627645 | 9.101577 | 9.11862 | -0.01704768 | 0.044139886 |
| Gm5518 | 0.452811 | 0.473116 | 0.483 | -0.009841534 | -0.030146307 | 0.4828854 | 0.4774601 | 0.49453 | -0.017071619 | -0.011646317 |
| Avpr1a | 2.127115 | 2.008157 | 1.9915 | 0.016623085 | 0.13558207 | 2.0061916 | 1.9786211 | 1.99573 | -0.017110682 | 0.010459795 |
| Sgpp1 | 10.18199 | 10.20621 | 10.234 | -0.027779212 | -0.051997442 | 9.9715935 | 10.038826 | 10.0559 | -0.017123237 | -0.084355608 |
| Uso1 | 10.79977 | 10.82205 | 10.764 | 0.058431062 | 0.036159198 | 10.665904 | 10.75861 | 10.7757 | -0.017136424 | -0.109843186 |

| | | | | | | | | | | |
|---|---|---|---|---|---|---|---|---|---|---|
| Zscan29 | 9.400256 | 9.486138 | 9.4588 | 0.027372819 | -0.05850898 | 9.2570786 | 9.3895093 | 9.40667 | -0.017158483 | -0.149589127 |
| Trbc1 | 11.58492 | 11.59875 | 11.62 | -0.020912046 | -0.034743672 | 11.48937 | 11.602196 | 11.6194 | -0.017234132 | -0.130060659 |
| Gcnt3 | 1.355806 | 1.345533 | 1.3379 | 0.007611966 | 0.017884396 | 1.3512313 | 1.3895881 | 1.40683 | -0.017246287 | -0.055603145 |
| Ubtd2 | 5.263294 | 5.260395 | 5.2536 | 0.006836885 | 0.009735685 | 5.1147246 | 5.1720848 | 5.18934 | -0.017257938 | -0.074618127 |
| Gm14730 | 2.717783 | 2.809538 | 2.7115 | 0.098043321 | 0.006288267 | 2.7372069 | 2.7821395 | 2.79941 | -0.017268928 | -0.062201599 |
| Olfm2 | 1.200629 | 1.28056 | 1.2838 | -0.003251101 | -0.083182572 | 1.1846271 | 1.251418 | 1.26869 | -0.017274929 | -0.084065779 |
| Olfr714 | 1.9624 | 2.020096 | 2.0336 | -0.01349478 | -0.071191189 | 2.0627996 | 2.0731794 | 2.09051 | -0.017331129 | -0.02771084 |
| Fermt1 | 1.112337 | 1.147266 | 1.1402 | 0.007050166 | -0.027879109 | 1.2787978 | 1.147697 | 1.16505 | -0.017351049 | 0.113749718 |
| Gm12689 | 0.338526 | 0.381597 | 0.3893 | -0.007663517 | -0.050734319 | 0.4067661 | 0.3664843 | 0.38384 | -0.017353162 | 0.022928585 |
| Mcoln2 | 8.83324 | 8.215165 | 8.6255 | -0.410293794 | 0.207781402 | 8.8224629 | 8.795775 | 8.81316 | -0.01738268 | 0.009305145 |
| Rc3h1 | 10.67571 | 10.79717 | 10.758 | 0.039417632 | -0.082044756 | 10.549287 | 10.819392 | 10.8368 | -0.017390602 | -0.287496124 |
| Gm10330 | 2.262334 | 2.249404 | 2.2645 | -0.015108405 | -0.002178538 | 2.2550789 | 2.2249072 | 2.24232 | -0.017408635 | 0.012762991 |
| Ccdc191 | 7.316922 | 7.098895 | 7.2592 | -0.160311114 | 0.057716301 | 7.0656514 | 7.3827412 | 7.40016 | -0.017414031 | -0.33450387 |
| Nlrx1 | 7.000741 | 6.839986 | 6.8381 | 0.00188785 | 0.162643057 | 7.1352455 | 6.7529749 | 6.77039 | -0.017416392 | 0.364854205 |
| Gm10051 | 1.287063 | 1.26065 | 1.305 | -0.044369848 | -0.017957203 | 1.3360056 | 1.2788006 | 1.29622 | -0.017420223 | 0.039784728 |
| Slc6a1 | 2.189759 | 2.194821 | 2.1689 | 0.02593499 | 0.020873192 | 2.0738347 | 2.1639133 | 2.18134 | -0.01742694 | -0.107505478 |
| Bbs7 | 4.041299 | 4.087985 | 4.1542 | -0.066257397 | -0.112943084 | 4.0757705 | 4.0188634 | 4.03634 | -0.01748153 | 0.039425598 |
| Rtl4 | 0.306038 | 0.326166 | 0.3648 | -0.038600393 | -0.058728023 | 0.328802 | 0.3305101 | 0.34803 | -0.017520029 | -0.019228069 |
| Krt8 | 3.008314 | 2.869238 | 2.9436 | -0.074360819 | 0.064715023 | 3.2259757 | 2.9621018 | 2.97968 | -0.017579679 | 0.24629428 |
| Glp1r | 1.031831 | 1.043026 | 1.0652 | -0.0221549 | -0.033349241 | 1.0140455 | 1.0091116 | 1.0267 | -0.017585871 | -0.012651963 |
| Ighv6-5 | 0.817051 | 0.89053 | 0.8107 | 0.079792066 | 0.006312468 | 0.8218114 | 0.8240483 | 0.8417 | -0.01765255 | -0.01988944 |
| Myadml2 | 0.872053 | 0.877435 | 0.9228 | -0.045322624 | -0.050705382 | 0.9048677 | 0.8860629 | 0.90373 | -0.017671194 | 0.001133575 |
| Pecr | 5.480115 | 5.479431 | 5.6309 | -0.151438563 | -0.150754331 | 5.7893694 | 5.5584674 | 5.57616 | -0.017695644 | 0.213206393 |
| Dao | 2.672775 | 2.692474 | 2.7451 | -0.052668206 | -0.07236705 | 2.7421605 | 2.7107756 | 2.72848 | -0.017708763 | 0.013676173 |
| Sptbn4 | 2.636861 | 2.652064 | 2.6506 | 0.001435264 | -0.013767634 | 2.6917139 | 2.6756727 | 2.69338 | -0.017710491 | -0.001669247 |
| Rnf165 | 3.550858 | 3.712044 | 3.5181 | 0.193935888 | 0.032750055 | 3.5284952 | 3.5561778 | 3.57392 | -0.017737739 | -0.045420389 |
| Tekt2 | 0.739608 | 0.76619 | 0.7651 | 0.001098796 | -0.025483117 | 0.7496247 | 0.7616142 | 0.77936 | -0.017749102 | -0.029738588 |
| Rpl9-ps4 | 1.503289 | 1.422751 | 1.4779 | -0.055148815 | 0.025388692 | 1.4259962 | 1.4989953 | 1.51681 | -0.01781615 | -0.090815252 |
| Srebf1 | 10.78427 | 10.55471 | 10.727 | -0.17236613 | 0.057193468 | 10.841905 | 10.708055 | 10.7259 | -0.01784768 | 0.116002372 |
| Capn6 | 1.113656 | 1.184427 | 1.1674 | 0.017023816 | -0.053747006 | 1.1527468 | 1.1926881 | 1.21059 | -0.017898743 | -0.057840084 |
| Amn | 2.06095 | 2.055912 | 2.0242 | 0.03168739 | 0.03672463 | 2.1824291 | 2.0867334 | 2.10467 | -0.017939459 | 0.077756288 |
| Tshb | 2.633648 | 2.613709 | 2.7007 | -0.086970937 | -0.067032114 | 2.6705863 | 2.6829877 | 2.70094 | -0.017947353 | -0.030348755 |

| | | | | | | | | | | |
|---|---|---|---|---|---|---|---|---|---|---|
| Grrp1 | 6.138333 | 5.919716 | 5.9866 | -0.066900904 | 0.151715617 | 5.3103458 | 5.4545227 | 5.47247 | -0.017951437 | -0.162128312 |
| AC110913.1 | 0.845782 | 0.840187 | 0.8614 | -0.021261297 | -0.015666848 | 0.8794131 | 0.8188081 | 0.83679 | -0.017978505 | 0.042626562 |
| Dnajc5 | 11.42693 | 11.55576 | 11.478 | 0.077860888 | -0.05096679 | 11.429809 | 11.466389 | 11.4844 | -0.017993584 | -0.054573223 |
| Mtpap | 9.239721 | 9.319584 | 9.3097 | 0.009842716 | -0.070020251 | 9.2845663 | 9.3221664 | 9.34016 | -0.017994115 | -0.055594262 |
| Rassf1 | 9.28697 | 9.158756 | 9.2306 | -0.071831709 | 0.056382271 | 9.2269319 | 9.2263411 | 9.24438 | -0.018034441 | -0.017443618 |
| Trav13-3 | 0.734567 | 0.705972 | 0.7052 | 0.000777742 | 0.029372546 | 0.7684297 | 0.7161611 | 0.7342 | -0.01804304 | 0.034225592 |
| Abhd12 | 9.305615 | 9.298529 | 9.3882 | -0.089658221 | -0.082572513 | 9.616838 | 9.2920691 | 9.31012 | -0.018052453 | 0.306716441 |
| Zfc3h1 | 9.587743 | 9.320223 | 9.6806 | -0.36033641 | -0.092816443 | 10.079462 | 10.226135 | 10.2442 | -0.018053791 | -0.164727059 |
| Diablo | 9.457854 | 9.453711 | 9.4032 | 0.050539855 | 0.054682847 | 9.4769869 | 9.3979419 | 9.41602 | -0.018076161 | 0.060968799 |
| Hist1h2ab | 0.347545 | 0.319656 | 0.3585 | -0.038819456 | -0.010930802 | 0.3167864 | 0.3581123 | 0.3762 | -0.018084323 | -0.059410159 |
| Agbl4 | 1.031128 | 1.005568 | 1.0263 | -0.020751252 | 0.004809173 | 1.0292294 | 1.0321675 | 1.05026 | -0.018089758 | -0.021027815 |
| Kcns3 | 1.599097 | 1.510354 | 1.5919 | -0.081582599 | 0.007160072 | 1.5710277 | 1.5173138 | 1.53543 | -0.01811741 | 0.035596496 |
| Rps19-ps6 | 2.511775 | 2.582704 | 2.4966 | 0.086127137 | 0.015198684 | 2.520919 | 2.5263005 | 2.54445 | -0.018147466 | -0.023529027 |
| Lrrc8e | 2.936618 | 2.864964 | 3.0177 | -0.152764649 | -0.081109794 | 2.9788455 | 3.0116997 | 3.02986 | -0.018158752 | -0.05101292 |
| Slc45a1 | 0.952168 | 1.017923 | 0.926 | 0.091922214 | 0.026166716 | 0.9593744 | 0.9644997 | 0.98267 | -0.018172991 | -0.023298251 |
| Gm8206 | 2.477485 | 2.059712 | 1.9528 | 0.10687037 | 0.524644201 | 1.9711805 | 2.0457054 | 2.06389 | -0.018184752 | -0.092709647 |
| Gm26712 | 2.588039 | 2.677453 | 2.5307 | 0.146752645 | 0.057338112 | 2.5695038 | 2.5835328 | 2.60177 | -0.018233263 | -0.032262316 |
| Gm44511 | 2.808847 | 2.667141 | 2.7241 | -0.056967306 | 0.084737834 | 2.5434674 | 2.5915502 | 2.60979 | -0.018239299 | -0.066322114 |
| Apobec2 | 2.659811 | 2.648268 | 2.6439 | 0.004375044 | 0.01591783 | 2.6652505 | 2.7073494 | 2.72559 | -0.018244649 | -0.06034353 |
| Micall2 | 3.447521 | 3.441227 | 3.4192 | 0.022055243 | 0.028348609 | 3.4361827 | 3.4696036 | 3.48786 | -0.01825569 | -0.051676628 |
| Duxbl2 | 0.280227 | 0.335908 | 0.2341 | 0.1018474 | 0.046166412 | 0.2124557 | 0.2347602 | 0.25302 | -0.018255778 | -0.040560262 |
| Med13 | 12.01604 | 12.18584 | 12.082 | 0.104302704 | -0.065498972 | 11.798823 | 12.008719 | 12.027 | -0.018313978 | -0.228210017 |
| Lpp | 9.778001 | 9.886196 | 9.8118 | 0.074413333 | -0.033781003 | 9.8414307 | 10.063481 | 10.0819 | -0.018453391 | -0.240503551 |
| Gm48786 | 4.481641 | 4.512011 | 4.5653 | -0.053261633 | -0.083631838 | 4.4905294 | 4.6598007 | 4.67828 | -0.018474412 | -0.187745686 |
| Trbc2 | 12.37491 | 12.43307 | 12.492 | -0.058941132 | -0.117097772 | 12.332307 | 12.435075 | 12.4536 | -0.018476022 | -0.121244514 |
| Zfp141 | 8.147038 | 8.310042 | 8.2976 | 0.012473458 | -0.150530852 | 8.1851694 | 8.3015189 | 8.32001 | -0.018487754 | -0.134837223 |
| Gm2225 | 2.836911 | 2.719523 | 2.8031 | -0.083558717 | 0.033828431 | 2.7136792 | 2.8363131 | 2.85484 | -0.018524181 | -0.141158114 |
| Nek9 | 11.35052 | 11.34998 | 11.37 | -0.019713792 | -0.019173026 | 11.400205 | 11.366188 | 11.3847 | -0.018548286 | 0.015468343 |
| Ret | 5.040474 | 4.953136 | 4.9186 | 0.034560701 | 0.12189851 | 4.7426219 | 4.5776944 | 4.59625 | -0.018553784 | 0.146373656 |
| Vmn2r105 | 1.411938 | 1.370226 | 1.4321 | -0.061848548 | -0.020136667 | 1.3869224 | 1.4234444 | 1.442 | -0.018560266 | -0.05508229 |
| 6820431F20Rik | 11.89126 | 11.8777 | 11.935 | -0.057565672 | -0.04400765 | 12.08042 | 12.164613 | 12.1832 | -0.018566374 | -0.102759052 |
| Phf1 | 10.85163 | 10.87841 | 10.978 | -0.099859414 | -0.1266395 | 11.194707 | 10.93467 | 10.9532 | -0.018569388 | 0.241467531 |

| | | | | | | | | | | |
|---|---|---|---|---|---|---|---|---|---|---|
| Slc38a11 | 2.211822 | 2.149468 | 2.2248 | -0.075312119 | -0.012958046 | 2.1824442 | 2.1887937 | 2.20742 | -0.018622144 | -0.024971688 |
| Rplp0-ps1 | 1.512721 | 1.596909 | 1.5747 | 0.022250741 | -0.061937398 | 1.5476049 | 1.5455502 | 1.56423 | -0.018679844 | -0.016625212 |
| Tfap2c | 1.096303 | 1.008404 | 1.0169 | -0.008533285 | 0.079366657 | 1.0729095 | 0.9931815 | 1.01187 | -0.018687634 | 0.061040356 |
| Ly6g6e | 0.754441 | 0.692815 | 0.7307 | -0.037869495 | 0.023756001 | 0.7029321 | 0.6412472 | 0.65997 | -0.018724184 | 0.042960693 |
| Rims3 | 5.386394 | 5.486935 | 5.4038 | 0.083087122 | -0.017453722 | 5.4292854 | 5.5861425 | 5.6049 | -0.018755363 | -0.175612502 |
| Vat1l | 1.52674 | 1.472856 | 1.4843 | -0.011491516 | 0.042392692 | 1.4338272 | 1.4930748 | 1.51184 | -0.018763179 | -0.078010769 |
| Fh1 | 9.782953 | 9.828564 | 9.9003 | -0.071776357 | -0.117387674 | 10.094202 | 9.8490944 | 9.8679 | -0.018805887 | 0.226301313 |
| Spaar | 3.627069 | 3.58595 | 3.7148 | -0.128819905 | -0.087701037 | 3.6785582 | 3.504199 | 3.52304 | -0.018837698 | 0.155521517 |
| Pi4k2b | 6.730542 | 6.664963 | 6.8021 | -0.137172724 | -0.071593178 | 6.504525 | 6.7111548 | 6.73003 | -0.018877565 | -0.225507355 |
| Gm8290 | 0.967624 | 0.967465 | 1.0102 | -0.04274726 | -0.042588297 | 0.9303027 | 1.014911 | 1.03382 | -0.018913043 | -0.103521306 |
| Gramd4 | 11.13856 | 11.15649 | 11.034 | 0.122985448 | 0.105060362 | 11.17509 | 11.069512 | 11.0885 | -0.01897064 | 0.086607459 |
| Gm45692 | 0.222993 | 0.221464 | 0.27 | -0.048517157 | -0.046987992 | 0.1707088 | 0.1908169 | 0.20979 | -0.018971995 | -0.039080108 |
| Gm5559 | 1.454927 | 1.410348 | 1.4624 | -0.052034702 | -0.007456202 | 1.4425346 | 1.4142856 | 1.43326 | -0.018973046 | 0.009275917 |
| Gpx7 | 3.841101 | 3.782242 | 3.9122 | -0.129960977 | -0.071101757 | 3.8645796 | 3.5999429 | 3.61895 | -0.019005339 | 0.245631339 |
| Pus10 | 8.781378 | 8.690403 | 8.8637 | -0.173286982 | -0.082311962 | 8.6935267 | 8.8634098 | 8.88242 | -0.01901155 | -0.18889465 |
| Magee2 | 0.953929 | 1.038586 | 1.0021 | 0.036441437 | -0.048215312 | 0.9448976 | 0.9906417 | 1.00967 | -0.019023304 | -0.064767364 |
| Gm9057 | 0.398632 | 0.437 | 0.3966 | 0.040368575 | 0.002001007 | 0.4075382 | 0.4298832 | 0.44891 | -0.019026046 | -0.041371046 |
| Mcm3ap | 10.54653 | 10.49432 | 10.609 | -0.11487753 | -0.0626608 | 10.801589 | 10.718542 | 10.7376 | -0.01908486 | 0.063961834 |
| Olfr890 | 1.386986 | 1.388791 | 1.4051 | -0.016353551 | -0.018158443 | 1.4187678 | 1.4886768 | 1.50777 | -0.019089027 | -0.088998048 |
| Clip4 | 2.116278 | 2.13209 | 2.1414 | -0.009330264 | -0.025142146 | 2.1278887 | 2.1720136 | 2.19117 | -0.019154577 | -0.063279488 |
| Slc16a13 | 6.064017 | 6.052594 | 6.2474 | -0.194770148 | -0.183347525 | 5.9073032 | 5.9930159 | 6.01218 | -0.019163069 | -0.104875751 |
| Upp2 | 1.444774 | 1.403599 | 1.463 | -0.059420065 | -0.018244835 | 1.4173386 | 1.3582442 | 1.37747 | -0.019229452 | 0.039865003 |
| P2rx6 | 0.740773 | 0.762882 | 0.7591 | 0.003791249 | -0.018316939 | 0.7609723 | 0.79854 | 0.8178 | -0.019263198 | -0.056830891 |
| Olfr613 | 4.303879 | 4.274045 | 4.2554 | 0.018632771 | 0.048467173 | 4.2632494 | 4.4328626 | 4.4522 | -0.019333714 | -0.188946991 |
| Lsm1 | 8.022013 | 7.968048 | 7.8409 | 0.127173942 | 0.181138275 | 8.1242525 | 7.9789594 | 7.99832 | -0.019362867 | 0.125930221 |
| Oprl1 | 1.771165 | 1.738201 | 1.8069 | -0.06874724 | -0.035783591 | 1.7622616 | 1.7564061 | 1.77578 | -0.019377285 | -0.013521836 |
| Kcna4 | 0.922037 | 0.9437 | 0.9759 | -0.032198447 | -0.053861323 | 1.002095 | 1.0360342 | 1.05543 | -0.01939245 | -0.053331632 |
| Tfap2a | 2.343231 | 2.276411 | 2.3454 | -0.069003818 | -0.002183324 | 2.3216126 | 2.3742773 | 2.39369 | -0.019415814 | -0.072080547 |
| Cttn | 7.296904 | 7.314508 | 7.047 | 0.267473666 | 0.249869596 | 6.8415905 | 6.7787329 | 6.79816 | -0.01942779 | 0.043429835 |
| Gm49673 | 2.929171 | 2.798974 | 2.7946 | 0.004327219 | 0.134524233 | 3.0191122 | 3.1598691 | 3.17931 | -0.019444874 | -0.160201766 |
| Tomt | 1.204072 | 1.234751 | 1.1778 | 0.05694943 | 0.026270921 | 1.2242819 | 1.174158 | 1.19361 | -0.019451377 | 0.030672527 |
| Rgs7 | 2.47502 | 2.458041 | 2.4791 | -0.021085898 | -0.004106713 | 2.4648586 | 2.501884 | 2.52134 | -0.019459998 | -0.056485438 |

| | | | | | | | | | | |
|---|---|---|---|---|---|---|---|---|---|---|
| Ankrd34c | 0.781765 | 0.791286 | 0.7701 | 0.021150378 | 0.011628942 | 0.8539537 | 0.7915145 | 0.81099 | -0.019477606 | 0.042961638 |
| Gpr4 | 5.734006 | 5.494917 | 5.5136 | -0.018709278 | 0.220379978 | 5.356809 | 5.2030736 | 5.22262 | -0.019547197 | 0.134188233 |
| Rps19-ps3 | 3.151166 | 3.130693 | 3.2449 | -0.114207718 | -0.093734463 | 3.2147944 | 3.225171 | 3.24474 | -0.019573504 | -0.029950079 |
| Olfr381 | 1.779991 | 1.799382 | 1.7788 | 0.020576147 | 0.001185808 | 1.7753818 | 1.8753192 | 1.89493 | -0.019610977 | -0.119548394 |
| Fhad1 | 0.982284 | 0.96911 | 1.0411 | -0.072020373 | -0.058846351 | 0.9900598 | 1.0281587 | 1.04779 | -0.019629902 | -0.05772886 |
| Otulinl | 10.19332 | 10.23208 | 10.3 | -0.067494024 | -0.106252772 | 10.328056 | 10.246674 | 10.2663 | -0.019646918 | 0.061734722 |
| Pds5b | 10.66802 | 10.71159 | 10.706 | 0.00557067 | -0.037998102 | 10.532185 | 10.730018 | 10.7497 | -0.019650656 | -0.217483378 |
| Prmt8 | 0.922905 | 0.977467 | 0.9733 | 0.004136078 | -0.050425539 | 0.953783 | 0.9390182 | 0.95869 | -0.019672943 | -0.004908223 |
| Tmem56 | 1.161444 | 0.870084 | 0.9885 | -0.118398761 | 0.172960762 | 0.9225511 | 0.8768143 | 0.89653 | -0.019713461 | 0.026023286 |
| Gpx8 | 3.245606 | 3.220484 | 3.2184 | 0.002083035 | 0.02720444 | 3.2737331 | 3.1136966 | 3.13349 | -0.019791766 | 0.140244739 |
| Gm6345 | 0.993338 | 0.984069 | 1.01 | -0.025977177 | -0.016708481 | 0.9171972 | 0.9998699 | 1.01969 | -0.019824971 | -0.102497666 |
| Far2 | 3.241061 | 3.256187 | 3.2308 | 0.025389177 | 0.010262741 | 3.1845926 | 3.2532611 | 3.27314 | -0.019874273 | -0.088542752 |
| Gm14057 | 2.469338 | 2.517062 | 2.4578 | 0.059247278 | 0.011523776 | 2.5428427 | 2.5062426 | 2.52629 | -0.020050989 | 0.016549132 |
| Grm8 | 2.687972 | 2.619147 | 2.7019 | -0.08274817 | -0.013923323 | 2.4507837 | 2.4855213 | 2.50557 | -0.020053212 | -0.054790884 |
| Bcan | 1.338561 | 1.335867 | 1.241 | 0.094886146 | 0.097580161 | 1.2190191 | 1.2211365 | 1.24121 | -0.020074806 | -0.02219222 |
| Gm14165 | 0.586988 | 0.621611 | 0.5991 | 0.022539193 | -0.01208372 | 0.626095 | 0.5857336 | 0.60584 | -0.02010369 | 0.020257659 |
| Rag2 | 1.559995 | 1.618092 | 1.5982 | 0.019867585 | -0.03822912 | 1.6725466 | 1.602212 | 1.62233 | -0.020116715 | 0.050217892 |
| Hnmt | 2.30544 | 2.202494 | 2.1835 | 0.019010475 | 0.121956799 | 2.3004741 | 2.1376474 | 2.15778 | -0.020134798 | 0.142691942 |
| Gm14554 | 0.751454 | 0.737126 | 0.7051 | 0.032024752 | 0.046352431 | 0.6950154 | 0.732243 | 0.75245 | -0.02020648 | -0.057434069 |
| Ccdc96 | 2.286802 | 2.335374 | 2.3524 | -0.016997815 | -0.06556965 | 2.3337883 | 2.3403097 | 2.36055 | -0.020241835 | -0.026763213 |
| Zfp418 | 5.70571 | 5.885644 | 5.8184 | 0.067248328 | -0.112685827 | 5.7886321 | 5.751639 | 5.7719 | -0.020261807 | 0.016731307 |
| Mesp2 | 0.435589 | 0.407743 | 0.402 | 0.005697384 | 0.033543826 | 0.3699296 | 0.4059697 | 0.42628 | -0.020308143 | -0.056348291 |
| Gm12286 | 2.582482 | 2.615904 | 2.7259 | -0.110015536 | -0.1434369 | 2.6291906 | 2.6959123 | 2.71624 | -0.020329894 | -0.087051633 |
| Prkar1b | 4.444694 | 4.418323 | 4.579 | -0.160712517 | -0.13434097 | 4.4701192 | 4.5068187 | 4.52715 | -0.020331727 | -0.057031198 |
| Npas3 | 0.729353 | 0.688195 | 0.8452 | -0.156987468 | -0.115829719 | 0.6807321 | 0.6634623 | 0.6838 | -0.020334638 | -0.003064841 |
| Pm20d2 | 1.114325 | 1.03122 | 1.0862 | -0.054942536 | 0.02816318 | 1.0540794 | 1.0230706 | 1.04342 | -0.020348399 | 0.010660387 |
| Gm49057 | 1.20186 | 1.18533 | 1.1853 | 5.35998E-05 | 0.016584309 | 1.2007261 | 1.275766 | 1.29613 | -0.020363016 | -0.095402943 |
| Olfr732 | 1.185113 | 1.183714 | 1.2144 | -0.030647562 | -0.029248501 | 1.1820452 | 1.2365368 | 1.25692 | -0.020380103 | -0.07487169 |
| Alg9 | 8.141542 | 8.227249 | 8.3173 | -0.090026867 | -0.17573414 | 8.4047903 | 8.373552 | 8.39395 | -0.020395042 | 0.010843324 |
| Gm12902 | 2.078447 | 2.125992 | 2.117 | 0.008952963 | -0.038591797 | 2.1948646 | 2.1215713 | 2.14197 | -0.020400104 | 0.052893234 |
| Tbc1d17 | 9.414523 | 9.476328 | 9.6547 | -0.178390004 | -0.240195181 | 9.8122559 | 9.7424225 | 9.76288 | -0.020457935 | 0.049375468 |
| Chst9 | 0.817372 | 0.754678 | 0.7997 | -0.045070217 | 0.017624066 | 1.0139508 | 0.7899422 | 0.81042 | -0.020480969 | 0.203527634 |

| | | | | | | | | | | |
|---|---|---|---|---|---|---|---|---|---|---|
| Rxfp1 | 2.763624 | 2.696476 | 2.6944 | 0.002056128 | 0.069203502 | 2.8077956 | 2.7821952 | 2.80268 | -0.020484965 | 0.005115415 |
| Nckap5 | 4.886344 | 4.497241 | 4.613 | -0.115754287 | 0.273348297 | 4.3794735 | 4.6972103 | 4.7177 | -0.020491471 | -0.33822826 |
| Gm11032 | 0.771269 | 0.813177 | 0.7807 | 0.032501065 | -0.009407032 | 0.7760933 | 0.7791353 | 0.79967 | -0.020530739 | -0.023572707 |
| Mark2 | 11.65627 | 11.71289 | 11.74 | -0.02675711 | -0.083373102 | 11.626873 | 11.725825 | 11.7464 | -0.02055108 | -0.119502332 |
| Icmt | 9.757365 | 9.971735 | 9.7691 | 0.202617006 | -0.011752817 | 9.8776097 | 9.7786151 | 9.79917 | -0.020551932 | 0.07844269 |
| Col4a6 | 1.324131 | 1.251459 | 1.3048 | -0.053338706 | 0.019333793 | 1.2017127 | 1.2628854 | 1.28349 | -0.020602528 | -0.081775262 |
| Olfr250 | 2.687 | 2.728304 | 2.6246 | 0.103743126 | 0.062439027 | 2.6944573 | 2.7285603 | 2.74917 | -0.020604812 | -0.05470774 |
| Mycbpap | 0.585477 | 0.599495 | 0.5723 | 0.027180143 | 0.013162619 | 0.5506849 | 0.5678122 | 0.58848 | -0.020665066 | -0.037792425 |
| Pip5k1c | 11.0485 | 11.14213 | 11.115 | 0.026826014 | -0.066807926 | 11.30967 | 11.033297 | 11.054 | -0.020712108 | 0.255661861 |
| Nudt7 | 6.877206 | 6.80653 | 7.0589 | -0.252409199 | -0.181732744 | 6.909632 | 6.839029 | 6.85974 | -0.020715478 | 0.04988759 |
| Cenpk | 4.706859 | 4.518463 | 4.7074 | -0.188969826 | -0.000574087 | 4.58223 | 4.7650179 | 4.78576 | -0.020738066 | -0.203526039 |
| Col22a1 | 3.070367 | 3.025876 | 2.9353 | 0.090550705 | 0.135041588 | 2.8262648 | 3.0481302 | 3.06889 | -0.020756676 | -0.242622137 |
| Hoxb7 | 3.273377 | 3.087564 | 3.0936 | -0.006013897 | 0.179799665 | 2.8984631 | 2.8890593 | 2.90982 | -0.020762611 | -0.011358812 |
| Iffo1 | 8.843566 | 8.828676 | 8.7866 | 0.042117362 | 0.05700805 | 8.9673769 | 9.0027877 | 9.02355 | -0.020764929 | -0.056175698 |
| Olfr671 | 1.735883 | 1.69501 | 1.7161 | -0.021055465 | 0.019818028 | 1.7479826 | 1.7575979 | 1.77838 | -0.020778482 | -0.030393801 |
| Gm16027 | 1.431962 | 1.441474 | 1.4059 | 0.035590704 | 0.026078976 | 1.4829955 | 1.4087812 | 1.42956 | -0.020780859 | 0.053433412 |
| Rps6kc1 | 7.788569 | 7.80328 | 7.8573 | -0.054009522 | -0.068720865 | 7.6630773 | 7.6291303 | 7.64994 | -0.020812581 | 0.013134434 |
| Med26 | 8.095062 | 8.25216 | 8.1703 | 0.081887652 | -0.07521003 | 8.0984872 | 8.08192 | 8.10277 | -0.020845656 | -0.004278543 |
| Kmt2d | 12.53362 | 12.62372 | 12.573 | 0.0506184 | -0.039485232 | 12.466413 | 12.998198 | 13.019 | -0.020850474 | -0.552634894 |
| Gm5903 | -0.055498 | -0.09251 | -0.091 | -0.001554426 | 0.03545947 | -0.0886237 | -0.084061 | -0.0632 | -0.020876785 | -0.025439532 |
| Chrm2 | 0.924733 | 0.81671 | 0.8732 | -0.05649164 | 0.051531422 | 0.8820593 | 0.8473554 | 0.86825 | -0.020894987 | 0.013808849 |
| Gm7222 | 0.246969 | 0.256161 | 0.2415 | 0.01463987 | 0.005447697 | 0.2231018 | 0.2346669 | 0.25556 | -0.020895036 | -0.032460061 |
| Dsc3 | 6.693113 | 6.319363 | 6.1187 | 0.200650491 | 0.574400268 | 6.5140155 | 6.0720223 | 6.09294 | -0.020920046 | 0.42107315 |
| Olfr295 | 1.193107 | 1.177795 | 1.1576 | 0.020219028 | 0.035530556 | 1.1631736 | 1.1688001 | 1.18973 | -0.020925195 | -0.026551643 |
| Pdlim7 | 8.192726 | 8.2631 | 8.164 | 0.099145057 | 0.028771539 | 8.4278434 | 8.0190418 | 8.04001 | -0.020963499 | 0.387838156 |
| Mknk2 | 12.44118 | 12.51753 | 12.5 | 0.017261474 | -0.059087918 | 12.569121 | 12.316136 | 12.3371 | -0.020966186 | 0.232018184 |
| Rab18 | 10.34601 | 10.19966 | 10.179 | 0.020530212 | 0.166880492 | 9.8837053 | 9.9862542 | 10.0072 | -0.020984653 | -0.123533542 |
| Pcdhb12 | 0.986328 | 0.959535 | 0.978 | -0.018422739 | 0.008370083 | 0.958621 | 0.9483013 | 0.96929 | -0.020985246 | -0.010665558 |
| Rpl5-ps2 | 3.162123 | 3.208956 | 3.1861 | 0.022902788 | -0.023930064 | 3.1685887 | 3.1648212 | 3.18583 | -0.021008018 | -0.017240513 |
| Ube2d2a | 11.37872 | 11.41372 | 11.443 | -0.029558258 | -0.064566128 | 11.335877 | 11.386565 | 11.4076 | -0.021015474 | -0.071702916 |
| Pcsk1 | 4.114005 | 4.315523 | 4.1718 | 0.143681327 | -0.057836405 | 4.2625998 | 4.2084095 | 4.22943 | -0.021024557 | 0.03316576 |
| Gja5 | 4.075098 | 4.239404 | 4.1376 | 0.101776198 | -0.062529779 | 3.9973617 | 4.0185828 | 4.03963 | -0.021042919 | -0.04226404 |

| | | | | | | | | | | |
|---|---|---|---|---|---|---|---|---|---|---|
| Trbv23 | 3.96728 | 3.965791 | 3.9684 | -0.002604017 | -0.001114813 | 3.8966394 | 3.9070155 | 3.92809 | -0.021070622 | -0.031446731 |
| Tcea2 | 6.903358 | 6.793688 | 6.9331 | -0.139452961 | -0.029783694 | 7.3059329 | 7.0785356 | 7.09961 | -0.021070702 | 0.206326595 |
| Syngr1 | 2.986588 | 2.943915 | 2.9851 | -0.041155039 | 0.001518335 | 2.7325956 | 2.7717607 | 2.79287 | -0.021107664 | -0.060272738 |
| Eif4ebp2 | 12.00777 | 12.08517 | 12.096 | -0.011020212 | -0.08842547 | 11.912746 | 11.910253 | 11.9314 | -0.021152992 | -0.018660877 |
| Vmn1r80 | 2.455499 | 2.423066 | 2.4525 | -0.029385171 | 0.003047905 | 2.4633594 | 2.5459437 | 2.56711 | -0.021166353 | -0.103750602 |
| Pid1 | 7.217169 | 7.426395 | 7.0838 | 0.342574352 | 0.133349094 | 7.0789213 | 6.9230642 | 6.94425 | -0.021184299 | 0.134672785 |
| Ighv1-13 | 0.742346 | 0.772346 | 0.7485 | 0.023890379 | -0.006110234 | 0.7523627 | 0.7197515 | 0.74096 | -0.021208075 | 0.011403147 |
| Olfr1191-ps1 | 3.507866 | 3.47515 | 3.5197 | -0.044594004 | -0.01187806 | 3.5207234 | 3.5884913 | 3.6097 | -0.021208918 | -0.088976755 |
| Nkx6-1 | 2.163416 | 2.092356 | 2.1237 | -0.03132627 | 0.039733709 | 2.1751038 | 2.0899511 | 2.1112 | -0.021252126 | 0.063900528 |
| Mvk | 6.907944 | 6.937994 | 6.9211 | 0.016854291 | -0.013196485 | 7.1502887 | 6.8826055 | 6.90387 | -0.021260511 | 0.246422626 |
| Frrs1l | 2.642826 | 2.617477 | 2.5381 | 0.079400454 | 0.104749235 | 2.5985914 | 2.5982446 | 2.61953 | -0.021288994 | -0.020942135 |
| Ankub1 | 2.59401 | 2.660983 | 2.6083 | 0.05265194 | -0.014321284 | 2.6391904 | 2.6220543 | 2.64335 | -0.021300338 | -0.004164197 |
| Serpinb6a | 9.114145 | 8.90114 | 9.1872 | -0.28608856 | -0.073082748 | 8.6833925 | 8.7575694 | 8.77889 | -0.02132305 | -0.095500035 |
| Cdv3 | 11.33759 | 11.46285 | 11.372 | 0.091291056 | -0.033971918 | 11.399809 | 11.374092 | 11.3954 | -0.021330046 | 0.004386853 |
| Zfand6 | 10.35485 | 10.37699 | 10.402 | -0.024854632 | -0.046997134 | 10.343546 | 10.436576 | 10.4579 | -0.021351308 | -0.114381196 |
| Camta2 | 10.10631 | 10.18282 | 10.148 | 0.035077434 | -0.041429572 | 10.287106 | 10.255386 | 10.2767 | -0.021352524 | 0.01036724 |
| Gabbr2 | 1.241401 | 1.277379 | 1.2033 | 0.074039101 | 0.038061068 | 1.1782497 | 1.1455222 | 1.16689 | -0.021364934 | 0.011362641 |
| Rnf222 | 0.853681 | 0.817245 | 0.8277 | -0.010449249 | 0.02598679 | 0.8405969 | 0.8346449 | 0.85601 | -0.021368467 | -0.015416515 |
| Wdr59 | 8.6255 | 8.62995 | 8.5659 | 0.064029934 | 0.05957935 | 8.4046303 | 8.5076924 | 8.52913 | -0.021440596 | -0.124502637 |
| Gm13623 | 0.472372 | 0.512681 | 0.5175 | -0.004796304 | -0.045105577 | 0.475073 | 0.4772004 | 0.49868 | -0.021475301 | -0.023602693 |
| Map9 | 4.164095 | 4.192422 | 4.1199 | 0.072475577 | 0.044148535 | 3.9349507 | 4.1473884 | 4.16891 | -0.021524307 | -0.233962009 |
| Ppp1r1a | 1.56267 | 1.52326 | 1.5028 | 0.020444573 | 0.059854684 | 1.4567294 | 1.390599 | 1.41213 | -0.021534984 | 0.044595373 |
| Crp | 0.643419 | 0.618265 | 0.6409 | -0.022592331 | 0.002561211 | 0.647722 | 0.5862979 | 0.60786 | -0.021560932 | 0.039863216 |
| Rps15a-ps5 | 0.989353 | 0.994145 | 0.9729 | 0.021243847 | 0.016451514 | 1.0372586 | 0.9778736 | 0.99944 | -0.021569375 | 0.037815594 |
| Rpl10-ps6 | 1.913269 | 1.941741 | 1.9477 | -0.005943521 | -0.034415896 | 1.9857688 | 1.9312872 | 1.95286 | -0.021569998 | 0.032911554 |
| Tm4sf20 | 0.426261 | 0.433083 | 0.3662 | 0.066906063 | 0.060084599 | 0.4642458 | 0.3651237 | 0.38675 | -0.021621525 | 0.077500527 |
| Clcnkb | 0.933222 | 0.816264 | 0.8625 | -0.046245275 | 0.070712289 | 1.0497677 | 0.8764923 | 0.89816 | -0.021665978 | 0.151609361 |
| Gpn3 | 8.480831 | 8.456821 | 8.4639 | -0.007044285 | 0.016965176 | 8.8254543 | 8.5391902 | 8.56087 | -0.021678174 | 0.264585919 |
| Smim7 | 9.503012 | 9.36023 | 9.4313 | -0.071085071 | 0.071695949 | 9.4904336 | 9.3614065 | 9.38309 | -0.021686561 | 0.107340526 |
| Wbp11 | 10.67284 | 10.68198 | 10.617 | 0.06450785 | 0.055372654 | 10.596744 | 10.641732 | 10.6634 | -0.021694362 | -0.066682456 |
| Sema5b | 1.624656 | 1.502565 | 1.516 | -0.013466882 | 0.108623962 | 1.5127711 | 1.4015707 | 1.42327 | -0.021696399 | 0.089504006 |
| Olfr1352 | 1.642801 | 1.588858 | 1.5989 | -0.010064828 | 0.043878242 | 1.5965184 | 1.61359 | 1.6353 | -0.021705105 | -0.038776698 |

| | | | | | | | | | | |
|---|---|---|---|---|---|---|---|---|---|---|
| Drp2 | 2.455438 | 2.411339 | 2.4379 | -0.026547983 | 0.017551241 | 2.5363045 | 2.4121501 | 2.43386 | -0.02171034 | 0.102444032 |
| Btc | 3.67306 | 3.707455 | 3.5114 | 0.196075432 | 0.161680565 | 3.5310059 | 3.6876794 | 3.70941 | -0.021733858 | -0.178407341 |
| Pon2 | 9.200429 | 9.146979 | 9.1074 | 0.039544577 | 0.092994581 | 9.110292 | 9.0690499 | 9.09086 | -0.021811269 | 0.019430833 |
| Qrsl1 | 8.326349 | 8.34293 | 8.3423 | 0.000653397 | -0.015927856 | 8.6712674 | 8.5436728 | 8.5655 | -0.021825512 | 0.105769025 |
| 1810013L24Rik | 10.13187 | 10.13117 | 10.092 | 0.039436199 | 0.040133059 | 9.9961565 | 10.182789 | 10.2046 | -0.021826992 | -0.208459232 |
| Trp53i13 | 7.478637 | 7.238426 | 7.3013 | -0.062914876 | 0.177296227 | 7.4808693 | 7.3342712 | 7.3561 | -0.021831612 | 0.124766526 |
| Oscar | 1.651766 | 1.738892 | 1.7004 | 0.03845134 | -0.048674156 | 1.7625725 | 1.7814675 | 1.80336 | -0.021892893 | -0.040787938 |
| Espnl | 0.522309 | 0.545529 | 0.515 | 0.030513287 | 0.007292912 | 0.5407936 | 0.4924955 | 0.5144 | -0.021904894 | 0.02639322 |
| Mpp6 | 8.683428 | 8.787184 | 8.8371 | -0.049959162 | -0.153715116 | 8.5486083 | 8.803354 | 8.82528 | -0.021922146 | -0.276667839 |
| Bola3 | 7.559051 | 7.552728 | 7.5639 | -0.011203235 | -0.004880158 | 7.8825905 | 7.6502627 | 7.67219 | -0.02192245 | 0.210405338 |
| Hagh | 7.595208 | 7.58892 | 7.6117 | -0.022815131 | -0.016527752 | 7.8387375 | 7.6330402 | 7.65499 | -0.021950734 | 0.183746602 |
| Tsen34 | 7.53995 | 7.562656 | 7.6234 | -0.060744734 | -0.083450295 | 7.723858 | 7.4966976 | 7.51866 | -0.021959316 | 0.205201044 |
| Nat8f2 | 1.702905 | 1.589409 | 1.628 | -0.038578347 | 0.074918108 | 1.734161 | 1.6525549 | 1.67452 | -0.021960421 | 0.059645656 |
| Gm17970 | 2.664969 | 2.694352 | 2.7277 | -0.033313977 | -0.062696981 | 2.7406174 | 2.7841926 | 2.80617 | -0.021975435 | -0.065550649 |
| Slc38a8 | 3.029611 | 3.040227 | 3.0271 | 0.013150252 | 0.002534478 | 3.1139669 | 3.024367 | 3.04636 | -0.021991884 | 0.067608021 |
| Apobr | 8.675425 | 8.684713 | 8.4829 | 0.201855221 | 0.192567578 | 8.6008181 | 8.4917439 | 8.51385 | -0.022106531 | 0.086967598 |
| Rybp | 10.03961 | 10.17172 | 9.9988 | 0.172906573 | 0.040787596 | 9.951382 | 9.9362702 | 9.9584 | -0.022127961 | -0.007016155 |
| Mapkap1 | 9.317769 | 9.3903 | 9.3753 | 0.015038373 | -0.057491886 | 9.2581537 | 9.342342 | 9.36447 | -0.022130497 | -0.10631883 |
| Fam192a | 9.652861 | 9.559691 | 9.5916 | -0.031953253 | 0.061216902 | 9.5030876 | 9.6285413 | 9.65069 | -0.02214401 | -0.147597679 |
| Tex11 | 0.137534 | 0.062274 | 0.0987 | -0.036460881 | 0.03879927 | 0.1371228 | 0.0789509 | 0.10127 | -0.02232203 | 0.035849926 |
| Ube2w | 8.971395 | 9.007623 | 8.9207 | 0.086933548 | 0.050705473 | 8.9455843 | 8.9031015 | 8.92547 | -0.022364515 | 0.020118275 |
| Gm47145 | 1.715436 | 1.729666 | 1.7492 | -0.01949512 | -0.033724832 | 1.7488516 | 1.7723941 | 1.79479 | -0.022397677 | -0.045940134 |
| Slc27a5 | 1.305568 | 1.326909 | 1.3502 | -0.023294783 | -0.044635499 | 1.4007387 | 1.3849969 | 1.40751 | -0.022512079 | -0.006770275 |
| Tmem234 | 10.43102 | 10.26871 | 10.47 | -0.201006828 | -0.03869153 | 10.6013 | 10.458434 | 10.481 | -0.022561776 | 0.120304185 |
| Snap91 | 2.984499 | 2.926819 | 2.8168 | 0.110043558 | 0.167723104 | 2.8941147 | 2.8674627 | 2.89007 | -0.02260526 | 0.004046754 |
| Fam172a | 10.10317 | 10.0416 | 10.027 | 0.014417183 | 0.075989799 | 9.9457977 | 10.072127 | 10.0947 | -0.022605657 | -0.148935241 |
| Gm13303 | 0.330245 | 0.292466 | 0.2575 | 0.034939526 | 0.072717656 | 0.2642466 | 0.2706132 | 0.29322 | -0.022611318 | -0.028977845 |
| Zfhx3 | 5.33012 | 5.328212 | 5.2745 | 0.053734649 | 0.055642654 | 4.9608829 | 5.3245288 | 5.34714 | -0.02261516 | -0.386261075 |
| Rhob | 10.18297 | 9.801616 | 10.24 | -0.438821407 | -0.057471684 | 8.7107515 | 8.9793681 | 9.002 | -0.022635531 | -0.291252149 |
| Gm48494 | 1.166368 | 1.28511 | 1.1588 | 0.1263584 | 0.0076167 | 1.1394534 | 1.2212217 | 1.24395 | -0.022729547 | -0.104497911 |
| Gm14740 | 2.419145 | 2.447666 | 2.4078 | 0.039905263 | 0.011384182 | 2.3697964 | 2.4605334 | 2.48328 | -0.022743207 | -0.113480179 |
| Olfr1276 | 0.924233 | 0.96586 | 0.9643 | 0.001581494 | -0.040045978 | 0.9104082 | 0.9513964 | 0.97415 | -0.022749672 | -0.063737878 |

| | | | | | | | | | | |
|---|---|---|---|---|---|---|---|---|---|---|
| Mcpt-ps1 | 1.336368 | 1.391864 | 1.2857 | 0.10613654 | 0.050639977 | 1.3401098 | 1.2683748 | 1.29114 | -0.022768503 | 0.04896651 |
| Adra2b | 1.988295 | 1.956129 | 1.9751 | -0.01895185 | 0.013213694 | 1.9508401 | 2.0110974 | 2.03387 | -0.02277296 | -0.083030269 |
| Gm48592 | 1.673038 | 1.65105 | 1.6905 | -0.039490532 | -0.01750244 | 1.7554483 | 1.7615742 | 1.78454 | -0.02296675 | -0.029092598 |
| Gm43247 | 0.891729 | 0.924566 | 0.9226 | 0.001988373 | -0.030848998 | 0.9230112 | 0.8943952 | 0.91742 | -0.023022515 | 0.005593497 |
| B3gnt5 | 11.47204 | 11.38676 | 11.46 | -0.073030714 | 0.012244516 | 11.611282 | 11.426833 | 11.4499 | -0.02302822 | 0.161421164 |
| Cfap97 | 9.318842 | 9.396606 | 9.4502 | -0.053640804 | -0.131404634 | 9.4744606 | 9.3460871 | 9.36914 | -0.023056359 | 0.105317131 |
| Gm12229 | 1.212022 | 1.188017 | 1.2211 | -0.033082393 | -0.009077257 | 1.1842601 | 1.241415 | 1.26449 | -0.023075471 | -0.080230335 |
| Clic3 | 1.84059 | 1.894568 | 1.8859 | 0.008658124 | -0.045320297 | 1.9871686 | 1.8210864 | 1.84417 | -0.023081381 | 0.143000787 |
| Zbed4 | 9.583789 | 9.696513 | 9.5764 | 0.120112649 | 0.007389227 | 9.5786512 | 9.5855414 | 9.60864 | -0.023100213 | -0.029990477 |
| Gm18225 | 1.030809 | 1.001952 | 1.0584 | -0.056476845 | -0.027619758 | 1.054855 | 1.0476148 | 1.07072 | -0.023101339 | -0.015861148 |
| Gm13292 | 0.718246 | 0.670577 | 0.7471 | -0.076558046 | -0.028889366 | 0.6998706 | 0.6568873 | 0.68 | -0.023114444 | 0.019868803 |
| Twistnb | 9.438616 | 9.458212 | 9.4616 | -0.003429602 | -0.023025433 | 9.4428133 | 9.4554513 | 9.47863 | -0.023174317 | -0.0358123 |
| Amph | 2.811496 | 2.894427 | 2.8276 | 0.06683653 | -0.016094855 | 2.6957952 | 2.7016968 | 2.72493 | -0.023232978 | -0.029134619 |
| Cfap54 | 1.720594 | 1.759906 | 1.6899 | 0.0699762 | 0.030664363 | 1.8148061 | 1.7349911 | 1.75826 | -0.02327157 | 0.056543367 |
| Nbn | 8.807976 | 8.672985 | 8.7032 | -0.030260731 | 0.1047297 | 8.5572796 | 8.7204108 | 8.74368 | -0.023273182 | -0.186404326 |
| Cyhr1 | 9.972622 | 10.03282 | 10.042 | -0.009513437 | -0.069707853 | 10.051267 | 10.002239 | 10.0255 | -0.023304182 | 0.02572313 |
| Olfr727 | 1.047862 | 1.032664 | 1.0647 | -0.032026959 | -0.016829088 | 1.0105288 | 1.0458906 | 1.06923 | -0.023336649 | -0.058698453 |
| Fgf13 | 7.911028 | 7.94193 | 8.0914 | -0.149488315 | -0.180390268 | 7.7997035 | 7.7829465 | 7.80629 | -0.023345119 | -0.00658807 |
| Pex19 | 9.28506 | 9.23586 | 9.271 | -0.035097537 | 0.014103096 | 9.3964783 | 9.2293405 | 9.25269 | -0.023345561 | 0.143792237 |
| Trbv12-1 | 7.245822 | 7.195157 | 7.2962 | -0.10108379 | -0.050418876 | 6.959146 | 7.2393943 | 7.26275 | -0.023360008 | -0.303608324 |
| Olfr93 | 0.157731 | 0.155628 | 0.1808 | -0.025184961 | -0.023081663 | 0.1338105 | 0.1222845 | 0.14565 | -0.023363199 | -0.011837255 |
| Tmem150a | 3.671496 | 3.771834 | 3.6945 | 0.077358675 | -0.022979293 | 3.6057485 | 3.5367184 | 3.5601 | -0.023385354 | 0.045644793 |
| 4930516K23Rik | 2.621704 | 2.640767 | 2.69 | -0.049208908 | -0.068272209 | 2.6148299 | 2.7145466 | 2.73798 | -0.023435749 | -0.123152476 |
| Mfap5 | 2.438582 | 2.501258 | 2.6979 | -0.196651102 | -0.259327263 | 2.5123356 | 2.5394994 | 2.56294 | -0.023436212 | -0.050600062 |
| Mob3a | 11.37959 | 11.51874 | 11.427 | 0.09157498 | -0.04757728 | 11.417624 | 11.376312 | 11.3998 | -0.023444776 | 0.017867616 |
| Impg2 | 5.786697 | 5.894092 | 6.0928 | -0.198669527 | -0.306063742 | 5.8026683 | 6.0305372 | 6.05399 | -0.023451124 | -0.25131997 |
| Pyroxd1 | 8.239315 | 8.030313 | 8.0107 | 0.019647496 | 0.228649487 | 8.1326187 | 8.21498 | 8.23857 | -0.023586869 | -0.10594816 |
| Chrna1 | 1.768933 | 1.710641 | 1.7231 | -0.012446429 | 0.045845364 | 1.7436597 | 1.7470501 | 1.77067 | -0.023619587 | -0.027009995 |
| Muc5ac | 0.45779 | 0.48633 | 0.4757 | 0.010649792 | -0.017890694 | 0.4147345 | 0.4306575 | 0.45433 | -0.023668663 | -0.039591644 |
| Gm17981 | 1.373472 | 1.370712 | 1.395 | -0.02429823 | -0.021538251 | 1.4156317 | 1.389806 | 1.41351 | -0.023707966 | 0.002117748 |
| Ighv8-7 | 1.94982 | 1.960268 | 1.9922 | -0.031896551 | -0.042343653 | 1.9563168 | 1.9595017 | 1.98322 | -0.02371838 | -0.0269033 |
| Dcaf12l1 | 1.76476 | 1.74446 | 1.7678 | -0.023376682 | -0.003077564 | 1.7183572 | 1.7201575 | 1.74388 | -0.023719913 | -0.025520208 |

| | | | | | | | | | | |
|---|---|---|---|---|---|---|---|---|---|---|
| Slc9a4 | 0.826245 | 0.710068 | 0.822 | -0.111972115 | 0.004204916 | 0.7385692 | 0.7627654 | 0.7865 | -0.023736178 | -0.047932341 |
| Kcnj10 | 4.432527 | 4.367278 | 4.2312 | 0.136037257 | 0.201287032 | 4.1169384 | 3.9804829 | 4.00423 | -0.023748358 | 0.112707179 |
| Olfr1247 | 4.931616 | 5.005261 | 5.104 | -0.098732631 | -0.172377501 | 4.8051068 | 5.3875951 | 5.41136 | -0.023763147 | -0.606251447 |
| Gm3629 | 0.803099 | 0.912425 | 0.8113 | 0.101145855 | -0.008179843 | 0.7994016 | 0.8199723 | 0.84375 | -0.023773209 | -0.044343935 |
| Avil | 3.271683 | 3.310281 | 3.3437 | -0.033376794 | -0.071975266 | 3.1833203 | 3.42994 | 3.45372 | -0.023775941 | -0.270395612 |
| Slc35f6 | 8.244395 | 8.177257 | 8.0243 | 0.152996038 | 0.220134728 | 8.0884962 | 8.0913698 | 8.11527 | -0.023903814 | -0.026777473 |
| Kdm4dl | 0.776578 | 0.729747 | 0.7872 | -0.057444069 | -0.010612509 | 0.7075457 | 0.7516002 | 0.77551 | -0.023912472 | -0.067967058 |
| Olfr1489 | 3.240426 | 3.24795 | 3.3181 | -0.070191575 | -0.077715481 | 3.2545284 | 3.4316208 | 3.45556 | -0.023939292 | -0.2010317 |
| Noxa1 | 0.912673 | 0.861653 | 0.8805 | -0.018886118 | 0.032133677 | 0.8144004 | 0.8657562 | 0.88972 | -0.023960241 | -0.075316002 |
| Gm36262 | 0.454108 | 0.428972 | 0.4322 | -0.003237837 | 0.021898896 | 0.4450775 | 0.4383578 | 0.46234 | -0.023982867 | -0.017263172 |
| Fyb2 | 2.171297 | 2.214189 | 2.331 | -0.116797525 | -0.159690205 | 2.1185156 | 2.1307026 | 2.1547 | -0.023996263 | -0.036183182 |
| Gm10157 | 1.375024 | 1.454197 | 1.5418 | -0.087605033 | -0.16677826 | 1.4612248 | 1.412856 | 1.43688 | -0.024019373 | 0.024349435 |
| Gm8670 | 2.016498 | 2.105803 | 1.9296 | 0.176232932 | 0.086927721 | 2.0314378 | 2.0705642 | 2.09461 | -0.024044115 | -0.063170551 |
| Olfr774 | 2.338929 | 2.320303 | 2.3271 | -0.006815987 | 0.01180995 | 2.3347729 | 2.3436833 | 2.36773 | -0.024051277 | -0.03296167 |
| Tmem165 | 8.827076 | 8.920689 | 8.9696 | -0.048953203 | -0.142566579 | 8.8634297 | 9.0269492 | 9.05103 | -0.024076485 | -0.187595921 |
| Atp13a2 | 10.06323 | 10.08332 | 10.219 | -0.135984803 | -0.156072638 | 10.474344 | 10.205766 | 10.2299 | -0.024097813 | 0.244480204 |
| Pip4k2c | 9.948255 | 9.959096 | 9.9873 | -0.02823471 | -0.039076558 | 10.045716 | 9.950225 | 9.97434 | -0.024116229 | 0.071374353 |
| Gm29689 | 1.427634 | 1.416111 | 1.4672 | -0.051094636 | -0.039572385 | 1.5287858 | 1.4403156 | 1.46447 | -0.024156171 | 0.064313957 |
| Gm7451 | 1.54104 | 1.487584 | 1.56 | -0.072377631 | -0.018921286 | 1.5157984 | 1.5344573 | 1.55862 | -0.024161724 | -0.042820566 |
| Fyb | 11.61408 | 11.70616 | 11.686 | 0.020584598 | -0.07149462 | 11.366776 | 11.698934 | 11.7231 | -0.024183324 | -0.356340995 |
| Iglon5 | 2.217616 | 2.169397 | 2.2331 | -0.063685007 | -0.015465997 | 2.2456619 | 2.2154996 | 2.23974 | -0.024236258 | 0.005926061 |
| Gemin2 | 7.473049 | 7.568772 | 7.5726 | -0.00387756 | -0.099600895 | 7.6745259 | 7.6351666 | 7.65944 | -0.024269471 | 0.015089848 |
| Fntb | 8.276534 | 8.226587 | 8.2642 | -0.037608232 | 0.012338973 | 8.0917361 | 8.1907362 | 8.21507 | -0.024329138 | -0.123329245 |
| Vmn1r61 | 2.29575 | 2.355218 | 2.3937 | -0.038440363 | -0.097908245 | 2.4382108 | 2.4308308 | 2.45518 | -0.024345465 | -0.016965482 |
| Hoxa6 | 0.844618 | 0.757731 | 0.8257 | -0.067969811 | 0.018917078 | 0.7941784 | 0.7608317 | 0.78519 | -0.024356403 | 0.00899035 |
| Fgf14 | 0.573505 | 0.49503 | 0.5525 | -0.05751582 | 0.020958694 | 0.5750729 | 0.509449 | 0.53389 | -0.024443192 | 0.041180668 |
| Gm15481 | 1.007752 | 1.025469 | 1.0579 | -0.032458817 | -0.050176154 | 1.0572953 | 1.0168742 | 1.04132 | -0.024444985 | 0.015976152 |
| Ephx4 | 2.032818 | 2.04622 | 2.0284 | 0.017773495 | 0.004371706 | 1.9660025 | 1.9592938 | 1.98375 | -0.024454411 | -0.017745756 |
| Aff2 | 2.746333 | 2.773359 | 2.8792 | -0.105819122 | -0.132844674 | 2.8747306 | 2.8111988 | 2.8357 | -0.024505064 | 0.039026699 |
| Wrnip1 | 9.589043 | 9.719116 | 9.7071 | 0.011992488 | -0.118080319 | 9.7177916 | 9.7240468 | 9.74856 | -0.024515011 | -0.030770272 |
| Pgghg | 8.594021 | 8.574609 | 8.595 | -0.020427602 | -0.001015931 | 8.8979511 | 8.6628362 | 8.68735 | -0.024515777 | 0.21059911 |
| Hist1h2bf | 1.743845 | 1.859508 | 1.8542 | 0.005340225 | -0.110322552 | 1.7852306 | 1.78063 | 1.80518 | -0.024545997 | -0.019945475 |

| | | | | | | | | | | |
|---|---|---|---|---|---|---|---|---|---|---|
| Snx18 | 9.605144 | 9.596544 | 9.3907 | 0.205804814 | 0.214404977 | 9.3545246 | 9.4557673 | 9.48037 | -0.024604122 | -0.125846799 |
| Gm18990 | 1.28442 | 1.238857 | 1.3112 | -0.072320137 | -0.026757016 | 1.2539117 | 1.3315507 | 1.35619 | -0.024641906 | -0.102280857 |
| Pld5 | 3.051994 | 2.850189 | 3.0686 | -0.218460289 | -0.016655764 | 2.8887332 | 3.0394629 | 3.06411 | -0.024647808 | -0.175377445 |
| Kmt2a | 12.36485 | 12.49904 | 12.461 | 0.037695668 | -0.096495858 | 12.200092 | 12.695697 | 12.7204 | -0.02465471 | -0.520259557 |
| Shc4 | 2.755748 | 2.650124 | 2.6642 | -0.014057003 | 0.091567277 | 2.6855683 | 2.7321754 | 2.75685 | -0.024671668 | -0.071278815 |
| Spats1 | 2.571148 | 2.494095 | 2.5603 | -0.066228631 | 0.010823676 | 2.6557262 | 2.559817 | 2.5845 | -0.024686682 | 0.071222531 |
| Gpr137 | 7.470968 | 7.27463 | 7.324 | -0.049414797 | 0.146922869 | 7.7165874 | 7.4327873 | 7.45747 | -0.024687322 | 0.259112747 |
| Slc1a5 | 11.32129 | 11.27954 | 11.406 | -0.126034366 | -0.084279562 | 11.731 | 11.570913 | 11.5956 | -0.02470542 | 0.13538105 |
| Trav16n | 5.373464 | 5.364695 | 5.48 | -0.115292082 | -0.106522577 | 5.3923646 | 5.3502947 | 5.37502 | -0.024729438 | 0.017340471 |
| Ak8 | 3.722334 | 3.344823 | 3.6068 | -0.261957859 | 0.115552774 | 3.4062226 | 3.3636524 | 3.38848 | -0.024825475 | 0.017744786 |
| Tectb | 0.677333 | 0.678758 | 0.761 | -0.08219446 | -0.083618786 | 0.6707307 | 0.62366 | 0.64859 | -0.024931983 | 0.022138789 |
| Abhd3 | 1.209325 | 1.102348 | 1.196 | -0.093644416 | 0.013332145 | 1.1111755 | 1.1248608 | 1.14983 | -0.024969085 | -0.038654399 |
| Gm11389 | 2.446294 | 2.402266 | 2.4177 | -0.01542092 | 0.028606638 | 2.4585045 | 2.4107553 | 2.43573 | -0.024973051 | 0.022776184 |
| Rps15a-ps6 | 2.894628 | 2.89701 | 2.9878 | -0.090803093 | -0.093184666 | 2.8644925 | 2.9157865 | 2.94078 | -0.024992156 | -0.076286077 |
| Olfr1322 | 2.139622 | 2.153664 | 2.2024 | -0.048731379 | -0.062774193 | 2.1701094 | 2.2042494 | 2.22925 | -0.025003556 | -0.059143622 |
| Cd300e | 3.321059 | 3.288034 | 3.447 | -0.15894545 | -0.12592068 | 3.5728299 | 3.2959571 | 3.32099 | -0.025033308 | 0.251839499 |
| Wrn | 9.961193 | 10.02195 | 10.045 | -0.023233247 | -0.08398574 | 9.830067 | 10.06079 | 10.0858 | -0.025047066 | -0.255769791 |
| Olfr461 | 0.547773 | 0.559396 | 0.5536 | 0.005754548 | -0.005868288 | 0.5666435 | 0.5710338 | 0.59609 | -0.025060992 | -0.029451281 |
| Olfr760-ps1 | 0.307729 | 0.265676 | 0.2913 | -0.025672966 | 0.016380345 | 0.2436398 | 0.2674245 | 0.29251 | -0.025087695 | -0.048872396 |
| Suz12 | 11.33005 | 11.16895 | 11.258 | -0.088551362 | 0.072547648 | 10.822952 | 11.083113 | 11.1083 | -0.0251678 | -0.285328408 |
| Gm14124 | 0.872291 | 0.837473 | 0.8936 | -0.056154633 | -0.021336764 | 0.8082502 | 0.837514 | 0.86271 | -0.025193041 | -0.054456797 |
| Olfr1106 | 0.68402 | 0.71299 | 0.6943 | 0.018658223 | -0.010311587 | 0.7211999 | 0.7187557 | 0.74405 | -0.025297712 | -0.022853479 |
| Gm10406 | 1.741328 | 1.839084 | 1.8826 | -0.043547066 | -0.14130369 | 1.8294366 | 1.7576159 | 1.78296 | -0.025346094 | 0.046474568 |
| Gm2427 | 2.350647 | 2.33393 | 2.2394 | 0.094510916 | 0.111227372 | 2.2490055 | 2.3195486 | 2.34492 | -0.025370238 | -0.09591338 |
| Tmem91 | 1.600288 | 1.629721 | 1.5723 | 0.057437359 | 0.028004165 | 1.4619869 | 1.4622412 | 1.48763 | -0.025384299 | -0.025638606 |
| Gm49594 | -0.033376 | -0.04516 | 0.0019 | -0.047097067 | -0.035316066 | -0.0039418 | -0.045153 | -0.0197 | -0.025435304 | 0.015776083 |
| Fam71f2 | 2.178797 | 2.171799 | 2.1538 | 0.017971949 | 0.024970555 | 2.1791519 | 2.2078049 | 2.23327 | -0.025464857 | -0.054117925 |
| Prim1 | 8.047453 | 7.92073 | 8.1576 | -0.23682686 | -0.110103795 | 8.1338135 | 8.1759898 | 8.20148 | -0.025489446 | -0.067665739 |
| Nucb2 | 8.664175 | 8.380911 | 8.5208 | -0.139912114 | 0.143351887 | 8.339771 | 8.3489093 | 8.37442 | -0.025510558 | -0.034648837 |
| Slc52a3 | 6.001915 | 5.87482 | 5.932 | -0.057144448 | 0.069950343 | 5.7669384 | 5.8255497 | 5.85106 | -0.025514964 | -0.084126304 |
| Thg1l | 7.726016 | 7.569279 | 7.6917 | -0.122401947 | 0.034334773 | 7.8953188 | 7.8529553 | 7.87852 | -0.025560019 | 0.016803445 |
| Rassf9 | 3.672428 | 3.587622 | 3.6684 | -0.080786823 | 0.004019044 | 3.3262476 | 3.4857799 | 3.51134 | -0.025562327 | -0.185094629 |

| | | | | | | | | | | |
|---|---|---|---|---|---|---|---|---|---|---|
| Inafm2 | 7.840453 | 7.803922 | 7.8993 | -0.095379103 | -0.058847542 | 7.661149 | 7.509841 | 7.53541 | -0.025572218 | 0.125735795 |
| Gria4 | 1.044801 | 1.03466 | 0.9878 | 0.046873184 | 0.057014785 | 0.9682898 | 0.9763388 | 1.00194 | -0.025605993 | -0.033654969 |
| Wdr34 | 5.007519 | 5.211448 | 5.1572 | 0.05421143 | -0.149717578 | 5.3776585 | 5.2663407 | 5.29197 | -0.025624468 | 0.085693304 |
| Gm12247 | 0.881025 | 0.894873 | 0.9074 | -0.01257066 | -0.026419302 | 0.8943631 | 0.8600117 | 0.88565 | -0.025641717 | 0.008709651 |
| Hpf1 | 8.837041 | 8.736339 | 8.8598 | -0.123413025 | -0.022711706 | 9.0124214 | 9.0451933 | 9.07085 | -0.025654166 | -0.05842611 |
| Map2k3 | 9.262168 | 9.202577 | 9.0958 | 0.10679272 | 0.166384053 | 9.1868213 | 9.1138436 | 9.13951 | -0.025666103 | 0.047311545 |
| Gm19040 | 1.530179 | 1.624027 | 1.5412 | 0.082843026 | -0.011004811 | 1.5518468 | 1.5921749 | 1.61788 | -0.025705526 | -0.066033629 |
| Vwa5b2 | 0.323707 | 0.306289 | 0.3712 | -0.064928017 | -0.047510785 | 0.3091484 | 0.3279717 | 0.35372 | -0.02575146 | -0.044574692 |
| Usp1 | 10.43358 | 10.39837 | 10.418 | -0.019301448 | 0.015914844 | 10.311593 | 10.346484 | 10.3722 | -0.025759974 | -0.060651482 |
| Chrna5 | 1.115439 | 1.167098 | 1.1152 | 0.051919889 | 0.000260916 | 1.0918396 | 1.1300042 | 1.15578 | -0.025780662 | -0.06394533 |
| Ncbp2 | 8.555173 | 8.612511 | 8.5463 | 0.066252868 | 0.008914868 | 8.9004508 | 8.6983052 | 8.7241 | -0.02579205 | 0.176353572 |
| F11r | 8.460366 | 8.302509 | 8.3078 | -0.005270828 | 0.152585864 | 8.0530694 | 8.0319518 | 8.05777 | -0.025819929 | -0.004702299 |
| Clec1b | 3.750716 | 3.733067 | 3.8199 | -0.08686398 | -0.069214553 | 3.4627353 | 3.496875 | 3.5227 | -0.025821215 | -0.059960867 |
| Gsto1 | 8.304414 | 8.414169 | 8.5735 | -0.159329866 | -0.269085229 | 8.8221618 | 8.4190765 | 8.44493 | -0.025853843 | 0.37723144 |
| Cd1d1 | 8.655721 | 8.785422 | 8.6779 | 0.107551366 | -0.022149771 | 9.013087 | 8.8319855 | 8.85787 | -0.025888227 | 0.155213237 |
| Zranb2 | 9.964234 | 10.09822 | 10.068 | 0.029809655 | -0.104179082 | 10.235109 | 10.171482 | 10.1974 | -0.025889969 | 0.037737076 |
| Churc1 | 8.783245 | 8.675027 | 8.7898 | -0.114812639 | -0.006594878 | 8.9558097 | 8.7651692 | 8.79118 | -0.026012456 | 0.164628058 |
| Rps8-ps1 | 2.1475 | 2.130903 | 2.0869 | 0.044044275 | 0.060641212 | 2.1675287 | 2.1787053 | 2.20472 | -0.026014902 | -0.037191556 |
| Porcn | 2.624016 | 2.507107 | 2.7018 | -0.194725499 | -0.077816582 | 2.670411 | 2.4984776 | 2.52452 | -0.02603815 | 0.145895273 |
| Syt4 | 0.581843 | 0.628813 | 0.5655 | 0.063288714 | 0.016318365 | 0.5135285 | 0.5123293 | 0.53839 | -0.02605624 | -0.02485705 |
| Lypd2 | 1.164263 | 1.11161 | 1.1618 | -0.0501622 | 0.002490512 | 1.0359914 | 1.0342059 | 1.0603 | -0.026093276 | -0.024307781 |
| Gm5641 | 13.39403 | 13.25462 | 13.179 | 0.075162848 | 0.214574172 | 13.206534 | 13.283446 | 13.3096 | -0.02614922 | -0.103061216 |
| Vmn2r-ps54 | 2.474806 | 2.471736 | 2.4217 | 0.050006234 | 0.053076095 | 2.4828751 | 2.5434126 | 2.5696 | -0.026186786 | -0.086724299 |
| Thoc1 | 9.459059 | 9.471453 | 9.4573 | 0.014115742 | 0.001721282 | 9.7592172 | 9.7918057 | 9.81801 | -0.026208478 | -0.058796982 |
| Gm8226 | 3.032389 | 2.831754 | 3.1705 | -0.338779654 | -0.138144146 | 3.0325417 | 3.0649665 | 3.0912 | -0.026233469 | -0.058658307 |
| 9030624G23Rik | 2.81982 | 2.714068 | 2.9689 | -0.254821123 | -0.149068289 | 2.7734855 | 2.7628564 | 2.78909 | -0.02623419 | -0.015605075 |
| Npm2 | 0.398565 | 0.418211 | 0.4378 | -0.019573394 | -0.03921952 | 0.4077633 | 0.4304419 | 0.45671 | -0.02627046 | -0.048949112 |
| Irak1 | 10.97349 | 11.01325 | 11.024 | -0.010801204 | -0.05056075 | 11.291717 | 11.209151 | 11.2355 | -0.026337198 | 0.056229151 |
| Sec16a | 11.58782 | 11.54905 | 11.55 | -0.000488722 | 0.038282928 | 11.605284 | 11.613532 | 11.6399 | -0.026339832 | -0.03458705 |
| Amz1 | 6.844745 | 6.778137 | 6.7433 | 0.034794849 | 0.101402364 | 7.0454722 | 6.7437918 | 6.77018 | -0.026387851 | 0.275292619 |
| Zfp981 | 0.216258 | 0.214072 | 0.2578 | -0.04375145 | -0.041565004 | 0.2024868 | 0.1986114 | 0.22501 | -0.026401191 | -0.022525769 |
| Nmrk2 | 2.289249 | 2.320417 | 2.3302 | -0.009747666 | -0.040915567 | 2.2786911 | 2.3634433 | 2.38986 | -0.026412096 | -0.111164222 |

| | | | | | | | | | | |
|---|---|---|---|---|---|---|---|---|---|---|
| Tgm1 | 6.454544 | 6.72297 | 6.1902 | 0.53274965 | 0.264323946 | 5.7007189 | 5.7983039 | 5.82472 | -0.026414383 | -0.123999454 |
| Serinc4 | 2.346416 | 2.32249 | 2.3533 | -0.0307823 | -0.006856803 | 2.2141703 | 2.3028666 | 2.3293 | -0.026429127 | -0.115125487 |
| Cd3e | 11.51315 | 11.51488 | 11.547 | -0.03165632 | -0.033381758 | 11.386591 | 11.561197 | 11.5877 | -0.026506222 | -0.201111791 |
| Arl14ep | 8.397927 | 8.280657 | 8.2554 | 0.025302047 | 0.142572482 | 8.3095563 | 8.2625373 | 8.28905 | -0.026516961 | 0.020501969 |
| Herpud2 | 9.837752 | 9.978866 | 9.9227 | 0.056132579 | -0.084981482 | 9.8644383 | 9.8690035 | 9.89554 | -0.026539563 | -0.0311047 |
| Enho | 1.7919 | 1.669919 | 1.7521 | -0.082140994 | 0.03984002 | 1.7563715 | 1.7380941 | 1.76464 | -0.026547371 | -0.008269977 |
| Zfp354a | 3.232174 | 3.301883 | 3.1938 | 0.108033807 | 0.038325613 | 3.1160798 | 3.2221972 | 3.24876 | -0.026561613 | -0.132679018 |
| Ednra | 4.087297 | 3.935921 | 4.1264 | -0.190497711 | -0.039121319 | 3.6966765 | 3.7819637 | 3.80859 | -0.026630316 | -0.11191749 |
| Olfr569 | 1.054747 | 1.119408 | 1.1244 | -0.004943622 | -0.069604178 | 1.1204839 | 1.1624381 | 1.18915 | -0.026710009 | -0.068664294 |
| Olfr628 | 2.317323 | 2.25713 | 2.3008 | -0.043655561 | 0.01653704 | 2.3582587 | 2.3158751 | 2.34259 | -0.026711101 | 0.015672496 |
| Eif3e | 12.45372 | 12.37285 | 12.485 | -0.112397572 | -0.031523942 | 12.388718 | 12.407331 | 12.4341 | -0.026759988 | -0.045373059 |
| Slf1 | 8.544891 | 8.435445 | 8.5064 | -0.070971695 | 0.038474404 | 8.5417867 | 8.631814 | 8.65859 | -0.026775325 | -0.116802682 |
| Vmn2r13 | 2.216169 | 2.192607 | 2.2341 | -0.041493148 | -0.017930843 | 2.2464703 | 2.2920581 | 2.31891 | -0.026856075 | -0.072443937 |
| Kcnk7 | 2.479897 | 2.46754 | 2.4768 | -0.009298742 | 0.003057847 | 2.5286868 | 2.5046319 | 2.53149 | -0.026857772 | -0.002802872 |
| Olfr1116 | 4.645061 | 4.549486 | 4.5444 | 0.005068059 | 0.100643297 | 4.5093852 | 4.7006995 | 4.72758 | -0.026885115 | -0.218199354 |
| Dhrs11 | 7.096232 | 7.003174 | 7.1688 | -0.165641726 | -0.072583441 | 7.3624474 | 7.0475377 | 7.07443 | -0.026892187 | 0.288017586 |
| Rftn1 | 10.88481 | 10.96853 | 10.95 | 0.018964852 | -0.064756238 | 10.879543 | 10.887658 | 10.9146 | -0.026901174 | -0.035016629 |
| Mapre3 | 6.011599 | 6.152258 | 6.2613 | -0.109009238 | -0.249668015 | 6.0407643 | 6.1487254 | 6.17563 | -0.026902703 | -0.134863856 |
| Fam186b | 1.163062 | 1.168924 | 1.2117 | -0.04280798 | -0.048669346 | 1.2431054 | 1.1227095 | 1.1497 | -0.026988488 | 0.093407441 |
| Ly6g | 1.07907 | 1.033136 | 0.9736 | 0.059488182 | 0.105421653 | 1.0109634 | 0.96201 | 0.98902 | -0.027007387 | 0.021945995 |
| B330016D10Rik | 5.036047 | 4.833712 | 4.8823 | -0.048540837 | 0.153794169 | 5.1165269 | 4.9151194 | 4.94217 | -0.027051875 | 0.174355561 |
| Rilp | 2.694133 | 2.579122 | 2.6154 | -0.036302679 | 0.078708861 | 2.5951249 | 2.6352092 | 2.66226 | -0.02705474 | -0.067139104 |
| Gm14682 | 0.46203 | 0.507227 | 0.4752 | 0.031982667 | -0.013214108 | 0.4846417 | 0.4578863 | 0.48496 | -0.027069645 | -0.000314254 |
| Siva1 | 6.550904 | 6.558694 | 6.7401 | -0.181418295 | -0.189208539 | 6.8554907 | 6.752301 | 6.77938 | -0.027082523 | 0.076107231 |
| Pttg1ip | 9.933472 | 9.989009 | 9.9924 | -0.003344745 | -0.058881938 | 9.8375466 | 9.7644245 | 9.79151 | -0.027086539 | 0.046035559 |
| Fbxo36 | 3.459026 | 3.330142 | 3.4859 | -0.155796092 | -0.026911179 | 3.2839352 | 3.3349364 | 3.3621 | -0.027159058 | -0.078160278 |
| Olfr373 | 2.946593 | 2.960059 | 2.9054 | 0.054645494 | 0.041179491 | 2.9139672 | 2.879576 | 2.90677 | -0.027195301 | 0.007195918 |
| Copg2 | 9.695171 | 9.750149 | 9.7716 | -0.021491846 | -0.076469614 | 9.876497 | 9.7326685 | 9.75992 | -0.027256245 | 0.11657216 |
| Gm10499 | 3.2015 | 3.18904 | 3.2709 | -0.081883276 | -0.069423145 | 3.2724868 | 3.2247902 | 3.25206 | -0.027266893 | 0.020429679 |
| Nudt12 | 4.042088 | 4.007288 | 4.0486 | -0.041269004 | -0.006468834 | 3.9929076 | 4.0191696 | 4.04649 | -0.02732087 | -0.05358288 |
| Gm29666 | 5.162903 | 5.110706 | 5.2346 | -0.1239059 | -0.071708953 | 5.4602488 | 5.4354256 | 5.46276 | -0.027330871 | -0.002507746 |
| Ntm | 1.603227 | 1.5245 | 1.625 | -0.100524235 | -0.021796403 | 1.5586406 | 1.4882226 | 1.51559 | -0.027367763 | 0.04305025 |

| | | | | | | | | | | |
|---|---|---|---|---|---|---|---|---|---|---|
| Trav4-4-dv10 | 4.897643 | 4.781415 | 4.7811 | 0.000318608 | 0.116546738 | 4.5094042 | 4.6439871 | 4.67137 | -0.027385004 | -0.161967925 |
| Ap3d1 | 11.20679 | 11.28401 | 11.305 | -0.02092336 | -0.098142888 | 11.429925 | 11.441107 | 11.4685 | -0.027399832 | -0.038581464 |
| Epn3 | 1.277576 | 1.037471 | 1.1464 | -0.108902309 | 0.131201977 | 1.2082065 | 1.0385564 | 1.06596 | -0.02740819 | 0.142241846 |
| Gm5314 | 0.29572 | 0.311129 | 0.3072 | 0.003961525 | -0.011448242 | 0.3356835 | 0.3100832 | 0.33749 | -0.027408597 | -0.001808272 |
| Wfikkn2 | 5.237804 | 5.17574 | 5.2337 | -0.057966251 | 0.004097654 | 5.0445706 | 5.102763 | 5.13017 | -0.027408607 | -0.08560092 |
| Senp8 | 5.826939 | 5.953675 | 5.9727 | -0.018975903 | -0.145711663 | 6.1275204 | 5.9495111 | 5.97699 | -0.027476578 | 0.150532714 |
| Klc3 | 4.834703 | 5.029313 | 4.9589 | 0.070370951 | -0.124239441 | 5.0308529 | 4.7939765 | 4.82148 | -0.027501042 | 0.209375373 |
| Gm9800 | -0.010652 | 0.039884 | -0.0116 | 0.051450595 | 0.000914241 | -0.0356258 | 0.0137243 | 0.04123 | -0.027503198 | -0.076853317 |
| Ripk3 | 9.27845 | 9.278792 | 9.3473 | -0.068518481 | -0.068860385 | 9.5774287 | 9.5402142 | 9.56773 | -0.027512376 | 0.009702125 |
| Serp2 | 3.979824 | 4.17073 | 4.0531 | 0.117597997 | -0.073308182 | 4.1157369 | 3.9848259 | 4.01234 | -0.027512654 | 0.103398319 |
| Cdc45 | 6.96098 | 6.65732 | 6.8829 | -0.225582108 | 0.078078563 | 6.9276758 | 7.158391 | 7.18591 | -0.027520011 | -0.258235223 |
| Gm48119 | 6.462796 | 6.613322 | 6.4127 | 0.200597214 | 0.050071425 | 6.6683881 | 6.9265937 | 6.95413 | -0.027538022 | -0.285743681 |
| Gm44212 | 1.260454 | 1.196517 | 1.2299 | -0.03342885 | 0.030507898 | 1.1772161 | 1.2064457 | 1.23401 | -0.027564737 | -0.056794364 |
| Hmgb1-ps6 | 1.113234 | 0.951752 | 0.9404 | 0.011319876 | 0.172801744 | 0.9205109 | 0.9607206 | 0.98831 | -0.027584971 | -0.067794662 |
| Prss30 | 4.547772 | 4.468831 | 4.5575 | -0.088650516 | -0.00970992 | 4.5324835 | 4.3702247 | 4.39784 | -0.027615919 | 0.13464289 |
| Skor1 | 0.778763 | 0.829378 | 0.7992 | 0.030201956 | -0.020413168 | 0.7653534 | 0.7609503 | 0.78862 | -0.027668666 | -0.023265548 |
| Spag8 | 0.77523 | 0.791607 | 0.8024 | -0.010821693 | -0.027198601 | 0.8039349 | 0.7955071 | 0.82322 | -0.02770998 | -0.01928216 |
| Igkv4-81 | 2.943349 | 2.973891 | 3.0714 | -0.097537852 | -0.128079731 | 2.9643124 | 2.9374548 | 2.96518 | -0.027728977 | -0.000871414 |
| Nap1l5 | 1.033319 | 1.054765 | 1.0314 | 0.023324461 | 0.001878404 | 0.9958574 | 0.9537573 | 0.9815 | -0.027743209 | 0.014356934 |
| Rnf128 | 3.38164 | 3.341419 | 3.4356 | -0.094205879 | -0.053984778 | 3.4464462 | 3.4287975 | 3.45654 | -0.02774328 | -0.010094576 |
| Ckm | 2.618771 | 2.540552 | 2.542 | -0.001482121 | 0.07673657 | 2.5196729 | 2.5903349 | 2.61809 | -0.027759941 | -0.098422001 |
| Gm21960 | 1.852176 | 1.771335 | 1.7727 | -0.001410719 | 0.079430674 | 1.8685967 | 1.8142688 | 1.84209 | -0.027822559 | 0.026505404 |
| Zc3h7b | 10.42938 | 10.54713 | 10.492 | 0.055065716 | -0.06268582 | 10.527037 | 10.523676 | 10.5515 | -0.027827474 | -0.024466185 |
| Nipal4 | 1.819564 | 1.82371 | 1.8406 | -0.016896489 | -0.021042116 | 1.8815692 | 1.7767265 | 1.80456 | -0.027832542 | 0.077010109 |
| Slc25a46 | 9.515762 | 9.536567 | 9.5569 | -0.020322273 | -0.04112679 | 9.424476 | 9.4756699 | 9.50353 | -0.027860639 | -0.079054503 |
| Dhx8 | 10.62054 | 10.7622 | 10.61 | 0.1518285 | 0.010168216 | 10.714514 | 10.62663 | 10.6545 | -0.027905778 | 0.059978214 |
| Gna14 | 2.174724 | 2.293619 | 2.195 | 0.098573031 | -0.020322058 | 2.1464339 | 2.1114222 | 2.13933 | -0.027908912 | 0.007102795 |
| Rexo2 | 9.448293 | 9.353434 | 9.4336 | -0.080177431 | 0.014682124 | 9.539542 | 9.4036108 | 9.43153 | -0.027920014 | 0.108011237 |
| Ighv4-2 | 2.780978 | 2.833974 | 2.6949 | 0.139092289 | 0.086096101 | 2.9265628 | 2.7986619 | 2.82661 | -0.027948829 | 0.099952091 |
| Tmem202 | 1.956483 | 1.944998 | 1.9344 | 0.010626573 | 0.022111913 | 2.0695299 | 1.9851514 | 2.01311 | -0.027955906 | 0.05642257 |
| Vmn2r32 | 0.604887 | 0.621237 | 0.5655 | 0.05571408 | 0.039364317 | 0.574892 | 0.6155483 | 0.64351 | -0.027966051 | -0.068622383 |
| Zranb3 | 6.931104 | 6.951883 | 7.0065 | -0.054613873 | -0.075392516 | 7.1251988 | 7.0888904 | 7.11687 | -0.027975366 | 0.008333005 |

| | | | | | | | | | | |
|---|---|---|---|---|---|---|---|---|---|---|
| Aloxe3 | 2.068727 | 2.183023 | 2.1467 | 0.03632546 | -0.077970087 | 2.1903605 | 2.0875388 | 2.11553 | -0.0279898 | 0.074831945 |
| Kcnk6 | 7.988236 | 8.178424 | 8.0149 | 0.163569734 | -0.026617808 | 7.7735181 | 7.9242507 | 7.95229 | -0.028034719 | -0.178767314 |
| Vmn1r1 | 2.450695 | 2.360821 | 2.4073 | -0.046448799 | 0.043425655 | 2.3737231 | 2.4125384 | 2.44058 | -0.0280375 | -0.066852764 |
| Pigz | 2.897266 | 2.903295 | 2.9498 | -0.046528963 | -0.052557149 | 3.0810873 | 2.8664425 | 2.89451 | -0.02806699 | 0.186577859 |
| Rpl19-ps1 | 2.437398 | 2.210356 | 2.2647 | -0.054386147 | 0.172655852 | 2.1787412 | 2.2486779 | 2.27675 | -0.028071033 | -0.098007767 |
| Zc3h4 | 11.10246 | 11.1431 | 11.137 | 0.006448952 | -0.034190308 | 10.892292 | 10.95111 | 10.9792 | -0.028083444 | -0.086901096 |
| AC125149.2 | 0.668053 | 0.619027 | 0.6177 | 0.001285228 | 0.050310871 | 0.6265331 | 0.6166601 | 0.64479 | -0.02813005 | -0.018257058 |
| Olfr633 | 1.060372 | 1.003142 | 1.0796 | -0.076479762 | -0.019249689 | 1.050051 | 1.0520116 | 1.08015 | -0.028139145 | -0.030099805 |
| Akap6 | 3.738997 | 3.859318 | 3.744 | 0.115349031 | -0.004972051 | 3.715817 | 3.704328 | 3.73248 | -0.028151504 | -0.016662461 |
| Pias2 | 9.07269 | 8.992358 | 9.0423 | -0.049916077 | 0.030415 | 8.9714722 | 9.0056564 | 9.03384 | -0.028183588 | -0.062367757 |
| Extl2 | 6.442562 | 6.453513 | 6.3905 | 0.063055216 | 0.052104353 | 6.2386401 | 6.1711118 | 6.19932 | -0.028203894 | 0.039324364 |
| Atp6v1a | 10.41802 | 10.32882 | 10.288 | 0.04127399 | 0.130470612 | 10.356516 | 10.118856 | 10.1471 | -0.028239258 | 0.20942099 |
| Olig3 | 0.84681 | 0.837515 | 0.8474 | -0.009926319 | -0.000631816 | 0.8417366 | 0.818597 | 0.84685 | -0.028248585 | -0.005109007 |
| Esco1 | 10.72586 | 10.78577 | 10.725 | 0.060687708 | 0.000772042 | 10.747311 | 10.674926 | 10.7032 | -0.028288252 | 0.044097146 |
| BC004004 | 9.64484 | 9.480555 | 9.5807 | -0.100100192 | 0.064184571 | 9.9020851 | 9.5059364 | 9.53424 | -0.028301878 | 0.367846781 |
| Olfr362 | 0.771619 | 0.764573 | 0.7809 | -0.016339467 | -0.009294202 | 0.7574173 | 0.7866736 | 0.81498 | -0.028307049 | -0.057563338 |
| Gm43743 | 0.220323 | 0.190284 | 0.1887 | 0.001599725 | 0.031638767 | 0.1820508 | 0.183413 | 0.21176 | -0.02834583 | -0.029708031 |
| Cldn12 | 5.157919 | 5.013729 | 5.1246 | -0.110825969 | 0.033363416 | 4.9229837 | 4.90369 | 4.93208 | -0.028386387 | -0.009092711 |
| Adipoq | 1.248747 | 1.264825 | 1.2604 | 0.004411734 | -0.011666748 | 1.2421205 | 1.2081836 | 1.23667 | -0.028491123 | 0.005445761 |
| Mt1 | 7.586414 | 7.267129 | 6.5642 | 0.702949929 | 1.022234838 | 6.9738306 | 6.6209462 | 6.64944 | -0.028496839 | 0.324387593 |
| Kcnn4 | 9.88382 | 9.92657 | 10.017 | -0.090159965 | -0.132909979 | 9.8668467 | 9.9437677 | 9.97227 | -0.02849734 | -0.105418427 |
| Mlec | 11.67975 | 11.69029 | 11.565 | 0.125688132 | 0.115149224 | 11.612559 | 11.66791 | 11.6964 | -0.028507624 | -0.083858767 |
| Bmi1 | 10.39242 | 10.41139 | 10.278 | 0.132987112 | 0.114017876 | 10.368787 | 10.398579 | 10.4271 | -0.028510212 | -0.058302633 |
| Sgcd | 1.629814 | 1.627323 | 1.6516 | -0.024242953 | -0.02175222 | 1.5332048 | 1.5017272 | 1.53029 | -0.028567636 | 0.002909931 |
| Hspa13 | 8.789241 | 8.734038 | 8.696 | 0.03799376 | 0.093197153 | 8.4455841 | 8.4158582 | 8.44444 | -0.028582533 | 0.001143381 |
| Igkv4-71 | 2.556923 | 2.700633 | 2.549 | 0.151638191 | 0.007927391 | 2.7518845 | 2.6303697 | 2.65896 | -0.028594223 | 0.092920638 |
| Fam122a | 6.907039 | 6.916114 | 6.8341 | 0.081985858 | 0.072911278 | 6.7805201 | 6.7529283 | 6.78155 | -0.02862649 | -0.001034667 |
| Msantd1 | 4.041476 | 4.146332 | 4.0463 | 0.100008666 | -0.004846837 | 4.1545025 | 4.2549934 | 4.28362 | -0.028631017 | -0.129121883 |
| Gm10037 | 1.911558 | 1.870232 | 1.8942 | -0.023967419 | 0.017358004 | 1.9316685 | 1.8913369 | 1.91997 | -0.028633484 | 0.011698146 |
| Nek10 | 2.393427 | 2.301979 | 2.2882 | 0.013807982 | 0.10525601 | 2.2381076 | 2.3451878 | 2.37383 | -0.02864502 | -0.135725185 |
| Cspp1 | 8.742065 | 8.811804 | 8.6547 | 0.157096003 | 0.087356939 | 8.6810443 | 8.9663303 | 8.99498 | -0.028652699 | -0.313938692 |
| Lrrn2 | 2.074603 | 2.05551 | 1.944 | 0.11148017 | 0.130573363 | 1.9078984 | 1.8731582 | 1.90185 | -0.028687088 | 0.006053027 |

| | | | | | | | | | | |
|---|---|---|---|---|---|---|---|---|---|---|
| Tk2 | 10.01509 | 9.903799 | 10.126 | -0.222040862 | -0.110749043 | 10.380982 | 10.116427 | 10.1452 | -0.028730733 | 0.235824011 |
| Slc5a8 | 1.018187 | 0.912111 | 0.936 | -0.023896762 | 0.082178496 | 1.1096638 | 0.8835181 | 0.9123 | -0.028786441 | 0.197359248 |
| Gm49286 | 2.23815 | 2.274169 | 2.3071 | -0.032915478 | -0.068934272 | 2.2906683 | 2.27677 | 2.30559 | -0.028819529 | -0.014921289 |
| Rpl30-ps1 | 0.168349 | 0.202497 | 0.1969 | 0.005604952 | -0.028543236 | 0.1570921 | 0.1286011 | 0.15744 | -0.028841865 | -0.000350842 |
| Ecm1 | 8.017113 | 7.909116 | 7.9065 | 0.00264705 | 0.110644297 | 7.8740366 | 7.9448567 | 7.97371 | -0.028858315 | -0.099678338 |
| Cmtr2 | 7.331247 | 7.25632 | 7.2606 | -0.004230668 | 0.070696403 | 7.2366875 | 7.2395099 | 7.2685 | -0.028993529 | -0.031815964 |
| C330021F23Rik | 1.697819 | 1.711589 | 1.7241 | -0.012551876 | -0.026322026 | 1.7163656 | 1.726612 | 1.75565 | -0.029037189 | -0.039283564 |
| Mms19 | 10.44027 | 10.30293 | 10.52 | -0.217360104 | -0.08001971 | 10.648586 | 10.654969 | 10.6841 | -0.029087778 | -0.035471041 |
| Cenpc1 | 8.426487 | 8.371479 | 8.4577 | -0.086195348 | -0.031186958 | 8.5094723 | 8.466158 | 8.49525 | -0.029094204 | 0.014220105 |
| G6pc3 | 8.613839 | 8.65222 | 8.6967 | -0.044506629 | -0.082887021 | 9.1161465 | 8.8654433 | 8.89454 | -0.029096756 | 0.221606468 |
| Gprasp2 | 5.357261 | 5.262658 | 5.2487 | 0.013954395 | 0.108557886 | 5.1428008 | 5.4453586 | 5.47448 | -0.029123705 | -0.331681436 |
| Hey2 | 1.515954 | 1.52268 | 1.5779 | -0.055217327 | -0.061943162 | 1.52613 | 1.5278683 | 1.557 | -0.029127949 | -0.03086624 |
| Nfatc2ip | 8.986853 | 9.187718 | 8.995 | 0.192679175 | -0.008185316 | 9.1580286 | 9.1341807 | 9.16332 | -0.029137824 | -0.005289892 |
| Icam5 | 3.56922 | 3.518137 | 3.6741 | -0.156005017 | -0.10492211 | 3.5320483 | 3.4934909 | 3.52263 | -0.029139187 | 0.009418177 |
| Gm48850 | 3.021036 | 2.923701 | 3.0838 | -0.160066316 | -0.062731698 | 3.1279641 | 3.2729569 | 3.30213 | -0.029170008 | -0.174162821 |
| Ppp1r1c | 1.510454 | 1.523805 | 1.5055 | 0.018306433 | 0.004955883 | 1.5498166 | 1.5270828 | 1.55626 | -0.029176105 | -0.006442348 |
| Zbtb44 | 10.02937 | 10.18145 | 10.06 | 0.121559512 | -0.030513157 | 9.8940594 | 10.045946 | 10.0751 | -0.029182925 | -0.181069565 |
| Htra3 | 4.25172 | 3.895735 | 4.0952 | -0.199485597 | 0.156500223 | 4.0329769 | 3.9294577 | 3.95868 | -0.029224466 | 0.074294733 |
| 2010109A12Rik | 2.951233 | 2.92972 | 2.9771 | -0.047378111 | -0.025865441 | 3.0590845 | 2.9597278 | 2.98896 | -0.029228311 | 0.070128399 |
| Ighv1-2 | 4.414259 | 4.623597 | 4.4895 | 0.134139145 | -0.075199069 | 4.5847201 | 4.3783044 | 4.40754 | -0.029238621 | 0.177177139 |
| Prss8 | 1.560893 | 1.372171 | 1.3618 | 0.010401448 | 0.199123532 | 1.4848245 | 1.4187597 | 1.44803 | -0.029270082 | 0.036794653 |
| Gm49059 | 0.834238 | 0.851107 | 0.8278 | 0.023294233 | 0.006425079 | 0.8548632 | 0.7934244 | 0.82271 | -0.029281214 | 0.03215759 |
| A2m | 1.635772 | 1.569354 | 1.6235 | -0.054188517 | 0.012229493 | 1.4974616 | 1.5209224 | 1.55021 | -0.029285021 | -0.052745818 |
| Nrxn2 | 2.970721 | 2.988916 | 2.8484 | 0.140508493 | 0.122313808 | 2.8779298 | 2.8117481 | 2.84106 | -0.029314096 | 0.036867608 |
| Gm13302 | 0.393962 | 0.293996 | 0.3668 | -0.07279057 | 0.027174778 | 0.2706741 | 0.2710389 | 0.30036 | -0.029325038 | -0.029689832 |
| Tssk6 | 1.031017 | 0.960303 | 0.9869 | -0.026612439 | 0.044102444 | 0.9933459 | 0.9529269 | 0.98233 | -0.029399087 | 0.011019912 |
| Ccm2 | 11.08137 | 11.2702 | 11.231 | 0.039217718 | -0.149604686 | 11.292194 | 11.198742 | 11.2282 | -0.029413386 | 0.064038186 |
| Ube2ql1 | 2.984114 | 2.986746 | 2.9462 | 0.040575746 | 0.037944043 | 2.9454143 | 3.0752752 | 3.1047 | -0.029420632 | -0.159281583 |
| Ptcra | 0.96195 | 0.923706 | 0.9609 | -0.03721499 | 0.001028832 | 1.0048974 | 0.9691828 | 0.99861 | -0.029425848 | 0.006288703 |
| Gm42718 | 3.136829 | 3.167033 | 3.1711 | -0.004074037 | -0.034277655 | 3.1558182 | 3.2708513 | 3.30031 | -0.029453859 | -0.144486997 |
| Tcf25 | 12.19542 | 12.19413 | 12.199 | -0.005118408 | -0.00382623 | 12.23969 | 12.221944 | 12.2514 | -0.029456354 | -0.011710552 |
| Vmn2r70 | 0.647804 | 0.640857 | 0.6093 | 0.031530797 | 0.038478094 | 0.6015983 | 0.6838202 | 0.71328 | -0.029458205 | -0.111680086 |

| | | | | | | | | | | |
|---|---|---|---|---|---|---|---|---|---|---|
| Gm9985 | 2.390398 | 2.380421 | 2.3424 | 0.038029758 | 0.04800722 | 2.5178187 | 2.43133 | 2.46084 | -0.029507136 | 0.056981569 |
| Rhebl1 | 6.376446 | 6.312448 | 6.5456 | -0.233138907 | -0.169140672 | 6.6577798 | 6.6172018 | 6.64672 | -0.029513772 | 0.011064199 |
| Gm12080 | 2.217782 | 2.149309 | 2.1471 | 0.002230896 | 0.070703171 | 2.112782 | 2.178074 | 2.2076 | -0.029526466 | -0.094818412 |
| Rabl6 | 9.900532 | 9.803013 | 9.8962 | -0.093224511 | 0.004293936 | 10.074491 | 10.080971 | 10.1106 | -0.029613563 | -0.036093271 |
| Trav3-3 | 5.55702 | 5.517431 | 5.5483 | -0.030825468 | 0.008763238 | 5.1969136 | 5.2640534 | 5.29368 | -0.029624689 | -0.09676452 |
| Rpl19-ps6 | 1.389416 | 1.370628 | 1.4026 | -0.031978364 | -0.013189863 | 1.4410257 | 1.3012606 | 1.33089 | -0.029629249 | 0.110135845 |
| Gm49496 | 1.905511 | 1.714029 | 1.8168 | -0.102803217 | 0.088679181 | 1.6389673 | 1.740549 | 1.7702 | -0.02965471 | -0.131236424 |
| Cdk2ap1 | 10.11272 | 10.06659 | 10.136 | -0.069227973 | -0.023090725 | 10.215051 | 10.02973 | 10.0594 | -0.029674722 | 0.155646131 |
| Arhgef19 | 0.773568 | 0.733651 | 0.6644 | 0.069213266 | 0.10913082 | 0.8794731 | 0.7014233 | 0.7311 | -0.029675858 | 0.148373918 |
| Mpped2 | 0.804344 | 0.826056 | 0.8208 | 0.00523014 | -0.016481354 | 0.836937 | 0.8273696 | 0.85705 | -0.029679454 | -0.020112052 |
| Rnf214 | 8.68585 | 8.708678 | 8.6807 | 0.027981224 | 0.005152578 | 8.6507084 | 8.7971639 | 8.82685 | -0.029686878 | -0.176142359 |
| Tmem196 | 0.490709 | 0.534983 | 0.4951 | 0.039885361 | -0.004389136 | 0.4823115 | 0.4879844 | 0.51768 | -0.029694019 | -0.035366896 |
| Gm10600 | 2.717472 | 2.525436 | 2.6661 | -0.140703025 | 0.051333192 | 2.555327 | 2.504503 | 2.53423 | -0.029730557 | 0.021093491 |
| Rtn4r | 1.68058 | 1.622229 | 1.7026 | -0.08039309 | -0.022041424 | 1.640193 | 1.5622815 | 1.59202 | -0.029741426 | 0.048170067 |
| Stard3nl | 7.957767 | 7.885869 | 7.9816 | -0.09574622 | -0.023848905 | 7.9735035 | 7.9443816 | 7.97415 | -0.029767161 | -0.00064521 |
| Entpd3 | 3.391489 | 3.012559 | 2.9597 | 0.052812358 | 0.43174201 | 3.9945564 | 3.0758073 | 3.10558 | -0.029774378 | 0.888974649 |
| Stfa3 | 1.162147 | 1.04568 | 1.0526 | -0.006943893 | 0.109523075 | 1.0532707 | 1.0133689 | 1.04317 | -0.029800287 | 0.010101539 |
| Plcl2 | 11.00607 | 10.96885 | 11.041 | -0.072598572 | -0.035378268 | 10.927357 | 10.990975 | 11.0208 | -0.02980069 | -0.093419244 |
| Lzts2 | 5.81546 | 5.589242 | 5.7553 | -0.16610662 | 0.060110977 | 5.4176356 | 5.3953768 | 5.42518 | -0.029801622 | -0.007542845 |
| Gm18733 | 0.800301 | 0.777541 | 0.7585 | 0.019085387 | 0.041845151 | 0.803352 | 0.7890855 | 0.8189 | -0.029811384 | -0.015544968 |
| Rarb | 1.775516 | 1.702462 | 1.7305 | -0.028045454 | 0.045008373 | 1.601476 | 1.660274 | 1.6901 | -0.029828051 | -0.088626128 |
| Aaas | 9.231978 | 9.176522 | 9.2949 | -0.118370402 | -0.062914339 | 9.4498815 | 9.4485612 | 9.4784 | -0.029835038 | -0.028514727 |
| Ercc8 | 7.683687 | 7.751216 | 7.4587 | 0.292512177 | 0.2249836 | 7.8199241 | 7.7280563 | 7.7579 | -0.02983934 | 0.062028443 |
| Gm45871 | 7.120369 | 7.141051 | 7.3921 | -0.250999769 | -0.271681088 | 7.3329479 | 7.2261508 | 7.25606 | -0.029904402 | 0.07689266 |
| Rnf207 | 1.534321 | 1.467152 | 1.5178 | -0.050617832 | 0.016551298 | 1.5472003 | 1.5258802 | 1.55582 | -0.029934916 | -0.00861475 |
| Olfr477 | 1.09953 | 1.155172 | 1.1481 | 0.007070529 | -0.048572068 | 1.1948892 | 1.1971404 | 1.22711 | -0.029965327 | -0.032216498 |
| Angptl6 | 4.204088 | 4.161998 | 4.2898 | -0.127801174 | -0.085711104 | 4.2025667 | 4.2852216 | 4.31527 | -0.030046601 | -0.112701501 |
| Nid2 | 6.39456 | 5.908288 | 6.2449 | -0.336642625 | 0.149629521 | 5.9080344 | 5.9922264 | 6.02229 | -0.030066351 | -0.114258296 |
| Gm7860 | 1.860488 | 1.918411 | 1.9329 | -0.014455231 | -0.072378622 | 1.9559329 | 1.9024172 | 1.93249 | -0.030072477 | 0.023443136 |
| BC085271 | 0.285448 | 0.215063 | 0.2282 | -0.013185832 | 0.057198851 | 0.2036898 | 0.2265154 | 0.2566 | -0.030080585 | -0.052906154 |
| Nepro | 8.397855 | 8.448946 | 8.2381 | 0.210841115 | 0.159750335 | 8.2750063 | 8.3476993 | 8.37779 | -0.030095659 | -0.102788697 |
| Gm6189 | 2.149759 | 2.138443 | 2.1103 | 0.028187725 | 0.03950387 | 2.137589 | 2.1075896 | 2.13772 | -0.030127816 | -0.000128359 |

| | | | | | | | | | | |
|---|---|---|---|---|---|---|---|---|---|---|
| Taco1 | 7.584225 | 7.50255 | 7.5386 | -0.036036894 | 0.045637907 | 7.7881421 | 7.5955122 | 7.62568 | -0.030167145 | 0.162462679 |
| Cadm2 | 2.149604 | 2.064108 | 1.8618 | 0.202352911 | 0.287848869 | 2.0154905 | 1.8954192 | 1.92559 | -0.030170665 | 0.089900619 |
| Gm6793 | 1.829503 | 1.752397 | 1.8348 | -0.082401517 | -0.005295641 | 1.8152341 | 1.8007414 | 1.83092 | -0.030183051 | -0.015690363 |
| Gm15950 | 0.802352 | 0.77837 | 0.7812 | -0.002786007 | 0.021196097 | 0.8239097 | 0.78409 | 0.8143 | -0.030210619 | 0.009609081 |
| Olfr13 | 1.372016 | 1.375917 | 1.4235 | -0.047625821 | -0.051526735 | 1.4319809 | 1.4879718 | 1.51819 | -0.030215411 | -0.086206298 |
| Slc4a11 | 6.077958 | 5.732711 | 6.015 | -0.282310042 | 0.062936791 | 5.9573734 | 6.0602722 | 6.0905 | -0.030230212 | -0.133129007 |
| Gm14418 | 2.555492 | 2.543715 | 2.7119 | -0.168167491 | -0.156390442 | 2.4960582 | 2.4398307 | 2.47006 | -0.030234187 | 0.025993231 |
| Ninl | 4.029452 | 3.994379 | 3.8514 | 0.143021397 | 0.178094165 | 3.8435335 | 4.0062053 | 4.0365 | -0.030289814 | -0.192961607 |
| Spag16 | 0.631638 | 0.62781 | 0.6008 | 0.027051129 | 0.030879655 | 0.597257 | 0.6334727 | 0.6638 | -0.030331178 | -0.066546848 |
| Gabre | 1.738622 | 1.743797 | 1.8354 | -0.091606328 | -0.096781056 | 1.7352677 | 1.8488605 | 1.87924 | -0.030382265 | -0.143975094 |
| Tceal8 | 5.983132 | 5.715777 | 5.8221 | -0.106306837 | 0.161047677 | 5.491208 | 5.4422191 | 5.47263 | -0.030409514 | 0.018579404 |
| Gm45880 | 1.28037 | 1.230047 | 1.2574 | -0.027326255 | 0.022996748 | 1.2492652 | 1.2170196 | 1.24748 | -0.030460051 | 0.001785484 |
| Gm4118 | 1.037816 | 0.9599 | 0.9527 | 0.007194766 | 0.085110738 | 1.0241782 | 0.9969597 | 1.02742 | -0.030464331 | -0.003245889 |
| Gkn3 | 0.42124 | 0.346512 | 0.4932 | -0.146646923 | -0.071918072 | 0.3488276 | 0.3769715 | 0.40748 | -0.030513383 | -0.05865731 |
| Cdan1 | 9.352832 | 9.420837 | 9.3656 | 0.055190786 | -0.01281436 | 9.2694774 | 9.3928562 | 9.42338 | -0.030522398 | -0.153901185 |
| Tdg-ps | -0.122224 | -0.14164 | -0.1682 | 0.026564273 | 0.045984183 | -0.1763772 | -0.14414 | -0.1136 | -0.030542278 | -0.062779618 |
| Rd3 | 2.001504 | 1.960418 | 2.0638 | -0.103346261 | -0.062259669 | 1.9064927 | 1.9862374 | 2.01678 | -0.030544618 | -0.110289342 |
| Exoc3 | 10.15072 | 10.13614 | 10.154 | -0.017407152 | -0.002826306 | 10.317904 | 10.215695 | 10.2462 | -0.03054685 | 0.071662812 |
| K230010J24Rik | 0.779526 | 0.758037 | 0.8012 | -0.043160892 | -0.02167125 | 0.8351407 | 0.7446708 | 0.77522 | -0.030547617 | 0.059922311 |
| Pcdhb17 | 4.007877 | 3.661686 | 3.7381 | -0.076380864 | 0.26981026 | 3.6996909 | 3.652655 | 3.68321 | -0.030552704 | 0.0164832 |
| Mdm2 | 10.1549 | 10.2837 | 10.232 | 0.05121798 | -0.077576707 | 10.046949 | 10.260307 | 10.2909 | -0.030563444 | -0.243921857 |
| Nxpe3 | 10.33774 | 10.37146 | 10.41 | -0.038454942 | -0.072170682 | 10.394753 | 10.336478 | 10.367 | -0.030567856 | 0.02770751 |
| Me1 | 5.090729 | 5.023304 | 4.9366 | 0.086727212 | 0.15415216 | 4.71017 | 4.5562219 | 4.58681 | -0.030592418 | 0.123355767 |
| Actg-ps1 | 2.127859 | 2.001265 | 2.0352 | -0.033978027 | 0.092615657 | 2.0013718 | 1.97293 | 2.00353 | -0.030597213 | -0.002155408 |
| Zbtb8b | 1.1215 | 1.114373 | 1.1471 | -0.032698439 | -0.025571627 | 1.2397356 | 1.1938679 | 1.22447 | -0.030606127 | 0.015261592 |
| Fbxw20 | 1.885025 | 1.931295 | 1.8741 | 0.057198524 | 0.010928974 | 1.9225616 | 1.900139 | 1.93075 | -0.03060688 | -0.008184286 |
| Slc37a3 | 8.875964 | 8.764759 | 8.7893 | -0.024550071 | 0.086654869 | 8.8808787 | 8.8862113 | 8.91686 | -0.030646079 | -0.0359787 |
| Serpina3c | 1.260756 | 1.240032 | 1.2284 | 0.011604007 | 0.032328078 | 1.1203346 | 1.1167342 | 1.14739 | -0.03065475 | -0.027054285 |
| Hist1h2bp | 1.477886 | 1.41452 | 1.4345 | -0.019961717 | 0.043404262 | 1.4250853 | 1.4094161 | 1.44019 | -0.030771997 | -0.015102799 |
| Tns2 | 7.500895 | 7.065622 | 7.0896 | -0.023983144 | 0.411289705 | 6.7770814 | 6.9354359 | 6.96622 | -0.030786231 | -0.189140695 |
| Gm4761 | 0.6557 | 0.670704 | 0.6977 | -0.027010015 | -0.042013628 | 0.7229855 | 0.6331619 | 0.66403 | -0.030872592 | 0.05895101 |
| Lrriq3 | 1.748048 | 1.750716 | 1.7942 | -0.043510145 | -0.046177704 | 1.800192 | 1.7695116 | 1.80042 | -0.030910381 | -0.000230006 |

| | | | | | | | | | | |
|---|---|---|---|---|---|---|---|---|---|---|
| Fancc | 8.093545 | 8.150196 | 8.0811 | 0.069077372 | 0.012425619 | 8.3138808 | 8.2651332 | 8.29606 | -0.030921915 | 0.017825709 |
| Olfr715 | 1.376564 | 1.428172 | 1.4085 | 0.019662748 | -0.031945665 | 1.4234162 | 1.4144984 | 1.44544 | -0.030940739 | -0.022022903 |
| Fkbp14 | 3.563934 | 3.595593 | 3.6276 | -0.031968674 | -0.063627756 | 3.4646301 | 3.4229799 | 3.45392 | -0.030944902 | 0.01070525 |
| Gm13736 | 6.095492 | 6.130988 | 6.0749 | 0.056090602 | 0.020594094 | 6.2991858 | 5.8494988 | 5.88047 | -0.030971788 | 0.418715191 |
| Scg2 | 0.93577 | 0.988704 | 0.8609 | 0.127757378 | 0.074823835 | 0.9254134 | 0.9001488 | 0.93113 | -0.030985706 | -0.005721172 |
| Lin7a | 1.568463 | 1.488042 | 1.5324 | -0.044314924 | 0.036105449 | 1.5374555 | 1.4718251 | 1.50285 | -0.031023513 | 0.034606874 |
| Pigl | 7.169311 | 7.167851 | 7.0695 | 0.098376875 | 0.099837101 | 7.3938289 | 7.3165573 | 7.34758 | -0.031025731 | 0.046245881 |
| Gm17949 | 0.279959 | 0.288364 | 0.3003 | -0.011895685 | -0.020301014 | 0.3087756 | 0.2752742 | 0.3064 | -0.031120935 | 0.002380474 |
| Lrrc58 | 9.472 | 9.39346 | 9.3811 | 0.012341567 | 0.090882339 | 9.109596 | 9.3296447 | 9.36077 | -0.031129493 | -0.251178181 |
| Tyw5 | 6.929443 | 6.817655 | 6.6404 | 0.177304567 | 0.2890927 | 6.9782403 | 7.1053651 | 7.13655 | -0.031181914 | -0.1583067 |
| Gm45925 | 0.701853 | 0.67903 | 0.6833 | -0.004309416 | 0.01851433 | 0.7007268 | 0.7299308 | 0.76112 | -0.03118626 | -0.060390314 |
| Gm9974 | 2.434565 | 2.34956 | 2.4454 | -0.095864905 | -0.01086002 | 2.6437 | 2.4615799 | 2.49279 | -0.031212169 | 0.150907926 |
| Olfr1090 | 0.797497 | 0.78914 | 0.7798 | 0.009307804 | 0.017664319 | 0.7516344 | 0.8130884 | 0.84432 | -0.031227273 | -0.092681292 |
| Gm5205 | 1.222182 | 1.146607 | 1.1252 | 0.021363298 | 0.096937803 | 1.1285651 | 1.1657566 | 1.19703 | -0.03127756 | -0.068469076 |
| 2410131K14Rik | 8.412178 | 8.543852 | 8.509 | 0.034863593 | -0.096810442 | 8.5146453 | 8.4555951 | 8.48688 | -0.031288316 | 0.0277619 |
| Ybx2 | 0.048096 | -0.04217 | -0.0259 | -0.01624246 | 0.074021267 | 0.0428798 | -0.048304 | -0.0169 | -0.031367993 | 0.059815545 |
| Gm7446 | 3.400946 | 3.34884 | 3.3168 | 0.032015578 | 0.08412176 | 3.3430283 | 3.3533121 | 3.38472 | -0.031407722 | -0.041691537 |
| Irx2 | 0.038023 | -0.09485 | -0.0757 | -0.019196089 | 0.113675811 | -0.015843 | -0.069594 | -0.0381 | -0.031458499 | 0.022292466 |
| Gm6542 | 2.375446 | 2.305727 | 2.2472 | 0.058483317 | 0.128202204 | 2.4050228 | 2.3525366 | 2.38401 | -0.031475536 | 0.021010669 |
| AW209491 | 6.70516 | 6.795555 | 6.7726 | 0.022920967 | -0.067474328 | 6.8426304 | 6.812244 | 6.84373 | -0.031483154 | -0.001096823 |
| Pou3f3 | 1.296512 | 1.328292 | 1.3164 | 0.011845548 | -0.019934926 | 1.2811478 | 1.3497827 | 1.38129 | -0.031506839 | -0.100141795 |
| Cnr1 | 2.338559 | 2.320184 | 2.3408 | -0.020607038 | -0.002231271 | 2.2771956 | 2.2020687 | 2.23359 | -0.03152279 | 0.04360417 |
| Olfr943 | 0.271773 | 0.256164 | 0.2851 | -0.028963951 | -0.013354914 | 0.2562239 | 0.3016073 | 0.33313 | -0.031524651 | -0.076908006 |
| Gm19098 | 1.997398 | 1.93597 | 1.9756 | -0.039614276 | 0.021814481 | 1.9148591 | 2.0334687 | 2.06505 | -0.03157763 | -0.150187255 |
| Gm6286 | 1.920621 | 1.980525 | 1.9386 | 0.041889717 | -0.018014855 | 1.9394108 | 1.8952839 | 1.9269 | -0.031614513 | 0.012512403 |
| Krt83 | 2.498133 | 2.451152 | 2.417 | 0.034131998 | 0.081112374 | 2.3217315 | 2.2882228 | 2.31988 | -0.031660876 | 0.001847823 |
| BC064078 | 6.508066 | 6.354317 | 6.5844 | -0.230070873 | -0.076321625 | 6.6069126 | 6.5383721 | 6.57013 | -0.031753357 | 0.036787176 |
| Tmem39a | 8.896144 | 9.006695 | 8.99 | 0.016675002 | -0.093876097 | 9.0776087 | 9.1282339 | 9.16005 | -0.031812513 | -0.082437723 |
| Gm6563 | 4.967546 | 4.392487 | 4.5301 | -0.137650511 | 0.437408191 | 4.5789208 | 4.2797595 | 4.31159 | -0.031832129 | 0.26732913 |
| Gm15500 | 15.23939 | 15.2318 | 15.243 | -0.011048274 | -0.003459766 | 15.269372 | 15.193957 | 15.2259 | -0.0319323 | 0.04348276 |
| Hist2h2ab | 1.971589 | 1.998353 | 1.9652 | 0.033167179 | 0.006403274 | 2.0168725 | 1.9856904 | 2.01764 | -0.031949226 | -0.000767081 |
| Tmprss11a | 4.34103 | 4.152261 | 4.3118 | -0.159584973 | 0.029184407 | 4.3896084 | 4.3649701 | 4.39693 | -0.031963672 | -0.007325388 |

| | | | | | | | | | | |
|---|---|---|---|---|---|---|---|---|---|---|
| Rpl18-ps1 | 0.771874 | 0.814538 | 0.7743 | 0.040279112 | -0.002385321 | 0.7638151 | 0.7622426 | 0.79421 | -0.031963938 | -0.030391452 |
| Cdh22 | 3.213787 | 3.231346 | 3.1193 | 0.111998054 | 0.094439689 | 3.0390283 | 3.0489152 | 3.08089 | -0.031973351 | -0.041860267 |
| Ighv1-22 | 7.153481 | 7.016834 | 6.8606 | 0.156277565 | 0.292924542 | 7.0690039 | 6.8298281 | 6.8618 | -0.031976247 | 0.207199562 |
| Ikbip | 6.321942 | 6.109803 | 6.3411 | -0.23132231 | -0.019182968 | 6.1819045 | 6.1040967 | 6.13613 | -0.032037297 | 0.045770569 |
| Gjc1 | 4.718909 | 4.715337 | 4.5859 | 0.129469121 | 0.133040811 | 4.2984366 | 4.3621733 | 4.39424 | -0.032062243 | -0.095798916 |
| Serpine2 | 5.676853 | 5.834133 | 5.7797 | 0.054474548 | -0.10280533 | 5.4807093 | 5.7007069 | 5.73281 | -0.032104046 | -0.252101648 |
| Pappa2 | 3.229393 | 3.407293 | 3.5646 | -0.157342396 | -0.335242344 | 3.4503181 | 3.2116023 | 3.24375 | -0.032151577 | 0.20656423 |
| Psma8 | 1.898493 | 1.975285 | 1.893 | 0.082295314 | 0.00550337 | 1.9587693 | 1.8562192 | 1.88844 | -0.032221661 | 0.070328427 |
| Cd3d | 11.49274 | 11.48007 | 11.526 | -0.046260488 | -0.033589464 | 11.344373 | 11.49555 | 11.5278 | -0.032277122 | -0.183453866 |
| Erich3 | 1.22939 | 1.248853 | 1.2021 | 0.046747482 | 0.027284401 | 1.1466769 | 1.1792401 | 1.21153 | -0.032291903 | -0.064855112 |
| Olfr328 | 1.686055 | 1.782011 | 1.6637 | 0.118263895 | 0.022308257 | 1.6708893 | 1.7082271 | 1.74052 | -0.032293378 | -0.069631208 |
| Pcolce2 | 2.265821 | 2.23949 | 2.3489 | -0.109448834 | -0.083117717 | 2.196232 | 2.2257138 | 2.25801 | -0.032299126 | -0.061780935 |
| Colca2 | 1.42864 | 1.375056 | 1.6605 | -0.285418517 | -0.2318344 | 1.4245318 | 1.3915565 | 1.42388 | -0.032318802 | 0.000656537 |
| Bard1 | 6.848198 | 6.817704 | 6.9223 | -0.104577841 | -0.07408455 | 6.9559278 | 7.0579084 | 7.09033 | -0.032418647 | -0.13439921 |
| Iws1 | 10.46915 | 10.44143 | 10.503 | -0.062030096 | -0.034310225 | 10.392899 | 10.559054 | 10.5915 | -0.03246571 | -0.198620538 |
| Ccdc85c | 5.618497 | 5.542786 | 5.7169 | -0.174065905 | -0.098355681 | 5.3237651 | 5.4886678 | 5.52116 | -0.032489191 | -0.197391898 |
| Olfr691 | 1.920411 | 1.862292 | 1.9255 | -0.063224852 | -0.005105761 | 1.9156347 | 1.8779409 | 1.91045 | -0.032508254 | 0.005185615 |
| Sbk2 | 1.653812 | 1.566994 | 1.6393 | -0.072341951 | 0.014475757 | 1.5116483 | 1.5316668 | 1.56423 | -0.032560207 | -0.052578677 |
| Proser2 | 5.855439 | 5.635052 | 5.9907 | -0.355639992 | -0.135252997 | 5.3604218 | 5.1821605 | 5.21472 | -0.032561006 | 0.145700281 |
| C2cd4c | 1.130981 | 1.150605 | 1.1637 | -0.013066205 | -0.032689836 | 1.2190648 | 1.1920875 | 1.22466 | -0.032568421 | -0.005591164 |
| Nrip3 | 2.421978 | 2.21245 | 2.1338 | 0.078639661 | 0.288167515 | 2.1667181 | 2.2335623 | 2.26614 | -0.032574126 | -0.099418304 |
| Lama2 | 3.108815 | 3.060256 | 2.9347 | 0.125563095 | 0.174122448 | 3.1255508 | 2.8815455 | 2.91418 | -0.032636544 | 0.211368673 |
| Klhdc10 | 8.854735 | 8.907093 | 8.8443 | 0.062835578 | 0.010478223 | 8.7462422 | 8.7849803 | 8.81764 | -0.032658212 | -0.07139635 |
| Gm9130 | 2.887076 | 2.944198 | 2.9815 | -0.037295088 | -0.0944161 | 2.8785076 | 2.9079247 | 2.94064 | -0.032714296 | -0.062131317 |
| Vmn2r99 | 3.173201 | 3.144929 | 3.1694 | -0.024425702 | 0.003846184 | 3.2389323 | 3.3223704 | 3.3551 | -0.032731583 | -0.116169678 |
| F5 | 5.06049 | 4.813916 | 4.5444 | 0.269533679 | 0.516108207 | 4.0991683 | 4.2895687 | 4.3223 | -0.032736042 | -0.223136415 |
| Bclaf3 | 8.552702 | 8.404747 | 8.5408 | -0.136092115 | 0.011862855 | 8.579706 | 8.725613 | 8.75835 | -0.032741054 | -0.17864803 |
| Ppef2 | 3.959183 | 3.967732 | 3.8501 | 0.117660323 | 0.109111162 | 3.9588628 | 4.04107 | 4.07391 | -0.03284008 | -0.115047217 |
| Mrap | 1.060137 | 0.988364 | 1.0428 | -0.054397018 | 0.017375871 | 0.962943 | 0.9451623 | 0.97802 | -0.032861085 | -0.015080415 |
| Gm6211 | 5.025212 | 5.030681 | 5.0435 | -0.012833661 | -0.018302273 | 4.942632 | 5.1704771 | 5.20336 | -0.032880068 | -0.260725209 |
| Olfr462 | 1.381966 | 1.344785 | 1.3145 | 0.030329163 | 0.067510549 | 1.404564 | 1.3311701 | 1.36405 | -0.032882337 | 0.040511534 |
| Gja8 | 2.418434 | 2.447002 | 2.38 | 0.066996933 | 0.038429142 | 2.3540009 | 2.4099643 | 2.44286 | -0.032890978 | -0.08885443 |

| | | | | | | | | | | |
|---|---|---|---|---|---|---|---|---|---|---|
| Rhobtb2 | 10.71209 | 10.75001 | 10.734 | 0.015724734 | -0.022191632 | 10.80898 | 10.705763 | 10.7387 | -0.032907784 | 0.070309314 |
| Matn1 | 1.081163 | 1.05247 | 1.1426 | -0.090126433 | -0.061433241 | 1.0355744 | 1.099601 | 1.13251 | -0.032908838 | -0.096935466 |
| Ggn | 2.631601 | 2.795397 | 2.6894 | 0.10604601 | -0.057750709 | 2.7921453 | 2.6614443 | 2.69438 | -0.032933665 | 0.097767323 |
| Gpx2-ps1 | 0.48248 | 0.525685 | 0.4684 | 0.057250712 | 0.014046363 | 0.5228829 | 0.4986332 | 0.53158 | -0.032947134 | -0.008697447 |
| Olfr1 | 0.30459 | 0.279644 | 0.2874 | -0.007760078 | 0.017185537 | 0.2842717 | 0.2873258 | 0.32031 | -0.032983411 | -0.036037555 |
| S100a7a | 0.3152 | 0.295241 | 0.3081 | -0.012847842 | 0.007110952 | 0.3385625 | 0.2783503 | 0.31139 | -0.033038383 | 0.027173866 |
| Gtf2a1 | 10.12978 | 10.09027 | 10.117 | -0.026901558 | 0.012608713 | 9.9513237 | 10.162054 | 10.1951 | -0.033051853 | -0.243782261 |
| Elavl3 | 1.551739 | 1.682846 | 1.5712 | 0.11166987 | -0.019437179 | 1.5246337 | 1.5651096 | 1.59831 | -0.033196106 | -0.073671969 |
| Mfsd13b | 1.247729 | 1.188467 | 1.4669 | -0.278452112 | -0.219190689 | 1.3294276 | 1.2457548 | 1.27904 | -0.033280699 | 0.050392041 |
| Nupl1 | 1.076056 | 1.079981 | 1.0961 | -0.016117551 | -0.020042921 | 1.0587291 | 1.0859005 | 1.11918 | -0.033281089 | -0.060452448 |
| Rassf10 | 1.539124 | 1.529092 | 1.6124 | -0.083298109 | -0.073266191 | 1.5395673 | 1.4982594 | 1.53158 | -0.033316818 | 0.00799111 |
| Hnrnph1 | 12.62782 | 12.56213 | 12.672 | -0.110212126 | -0.044526686 | 12.689047 | 12.812411 | 12.8458 | -0.033338808 | -0.156702719 |
| Pwp1 | 9.390194 | 9.385178 | 9.3296 | 0.055531989 | 0.060547197 | 9.4795831 | 9.3760518 | 9.40941 | -0.033356756 | 0.070174567 |
| Adam32 | 0.802057 | 0.82993 | 0.7686 | 0.061283698 | 0.033410616 | 0.7586749 | 0.8240584 | 0.85752 | -0.033465946 | -0.098849402 |
| Gm14569 | 2.059336 | 2.078185 | 2.1341 | -0.055867137 | -0.074716124 | 2.1373513 | 2.184455 | 2.21794 | -0.033488648 | -0.080592361 |
| Olfr1348 | 1.598601 | 1.65519 | 1.5646 | 0.090564926 | 0.0339762 | 1.5654425 | 1.60146 | 1.63496 | -0.03349869 | -0.069516163 |
| Mzf1 | 3.457399 | 3.351542 | 3.444 | -0.09248529 | 0.013371924 | 3.4401583 | 3.3721024 | 3.40563 | -0.033524925 | 0.034530932 |
| CR974586.1 | 2.045575 | 1.994111 | 2.0868 | -0.092648544 | -0.041183954 | 1.8913201 | 1.8900641 | 1.9236 | -0.033536906 | -0.032280963 |
| Acp1 | 9.176321 | 9.011091 | 9.02 | -0.008881757 | 0.15634888 | 9.1220131 | 9.0582508 | 9.09181 | -0.033556304 | 0.030205998 |
| Vars2 | 9.028293 | 8.980891 | 9.004 | -0.023111171 | 0.024290871 | 9.4881837 | 9.2928316 | 9.32639 | -0.033558244 | 0.161793902 |
| Eef1akmt2 | 7.644857 | 7.482537 | 7.5017 | -0.01914309 | 0.14317695 | 7.5839341 | 7.5019434 | 7.53554 | -0.033593285 | 0.048397339 |
| Car8 | 1.798997 | 1.808939 | 1.8244 | -0.015457399 | -0.025399714 | 1.908967 | 1.8410556 | 1.87468 | -0.033620697 | 0.034290705 |
| Shld1 | 7.130939 | 7.285386 | 7.275 | 0.010390909 | -0.144055677 | 7.3193245 | 7.1188061 | 7.15243 | -0.033621289 | 0.16689712 |
| D10Jhu81e | 9.090244 | 8.978687 | 9.0091 | -0.03038951 | 0.081167095 | 9.1899813 | 8.922508 | 8.95613 | -0.033621422 | 0.233851841 |
| 6030408B16Rik | 2.881936 | 2.845824 | 2.8563 | -0.010501597 | 0.025610181 | 2.8154898 | 2.9439804 | 2.97762 | -0.033638007 | -0.162128607 |
| Cnot1 | 12.53161 | 12.50393 | 12.575 | -0.071317018 | -0.043636786 | 12.488317 | 12.634653 | 12.6684 | -0.033729272 | -0.180065929 |
| Gmeb2 | 9.665194 | 9.737844 | 9.7536 | -0.01574476 | -0.088394865 | 9.7922058 | 9.8602122 | 9.89396 | -0.033744226 | -0.101750628 |
| Gm8566 | 0.279996 | 0.296697 | 0.3026 | -0.00591146 | -0.022611988 | 0.2676923 | 0.2668451 | 0.3006 | -0.033758863 | -0.032911693 |
| Gm15185 | 1.280435 | 1.207149 | 1.1814 | 0.025748339 | 0.099034262 | 1.2999321 | 1.2393373 | 1.27313 | -0.03379534 | 0.026799428 |
| Vsig2 | 3.382891 | 3.303812 | 3.3382 | -0.034433457 | 0.044644973 | 3.2469019 | 3.2162743 | 3.25009 | -0.033811165 | -0.00318364 |
| Vopp1 | 9.138875 | 9.204366 | 9.1329 | 0.071483657 | 0.005992375 | 8.9900385 | 9.1164502 | 9.15027 | -0.03381925 | -0.16023086 |
| Olfr20 | 1.431835 | 1.424231 | 1.4251 | -0.000845444 | 0.006758321 | 1.3376612 | 1.4712826 | 1.50511 | -0.033822472 | -0.167443811 |

| | | | | | | | | | | |
|---|---|---|---|---|---|---|---|---|---|---|
| Srsf11 | 11.84925 | 11.85818 | 11.869 | -0.010693898 | -0.019621566 | 11.950992 | 11.991526 | 12.0254 | -0.033906774 | -0.074441161 |
| Gpr165 | 1.286389 | 1.247206 | 1.2619 | -0.01466337 | 0.024519625 | 1.3150804 | 1.2148121 | 1.24882 | -0.034009113 | 0.066259179 |
| Olfr1537 | 0.447125 | 0.357766 | 0.347 | 0.010788602 | 0.10014773 | 0.3535148 | 0.360745 | 0.39481 | -0.034069423 | -0.041299689 |
| Hars2 | 9.333651 | 9.400979 | 9.3322 | 0.068792566 | 0.001463969 | 9.5766699 | 9.5300798 | 9.56419 | -0.034112435 | 0.01247765 |
| Ccnb2-ps | 1.29675 | 1.330426 | 1.3472 | -0.016742257 | -0.050417995 | 1.3652907 | 1.3506929 | 1.38481 | -0.034120245 | -0.019522363 |
| Plekhf1 | 4.016448 | 3.910539 | 3.9794 | -0.068899417 | 0.037010137 | 3.9610792 | 4.0099789 | 4.04412 | -0.034140895 | -0.08304057 |
| Fbxw19 | 2.400309 | 2.370351 | 2.4254 | -0.055013636 | -0.025055483 | 2.3827011 | 2.472629 | 2.50686 | -0.034235866 | -0.124163773 |
| Sfrp4 | 2.292034 | 2.304471 | 2.3027 | 0.001739349 | -0.010698013 | 2.4253633 | 2.4458422 | 2.48012 | -0.034281649 | -0.054760509 |
| Trav13n-4 | 3.473769 | 3.604183 | 3.572 | 0.032186059 | -0.09822788 | 3.4175889 | 3.4353234 | 3.46961 | -0.034289012 | -0.052023503 |
| Vmn2r69 | 3.224078 | 3.25816 | 3.2596 | -0.001461242 | -0.035543896 | 3.4234324 | 3.3300526 | 3.36436 | -0.034304494 | 0.059075273 |
| Celf6 | 0.328798 | 0.261027 | 0.2759 | -0.014839046 | 0.052932341 | 0.2461685 | 0.281475 | 0.31581 | -0.034337653 | -0.069644083 |
| Gm5879 | 1.936417 | 1.976923 | 1.9957 | -0.018780468 | -0.059286961 | 1.91609 | 1.9537748 | 1.98811 | -0.034338982 | -0.072023834 |
| Trav9d-1 | 1.151902 | 1.216464 | 1.1631 | 0.053398352 | -0.011163983 | 1.0820769 | 1.1645843 | 1.19896 | -0.034379007 | -0.116886426 |
| Gm6044 | 0.757691 | 0.766329 | 0.7353 | 0.031015017 | 0.022377182 | 0.7383683 | 0.7467267 | 0.78114 | -0.034416538 | -0.042774974 |
| Rnd2 | 2.91605 | 3.095066 | 2.9311 | 0.16395144 | -0.015064592 | 2.8425956 | 2.8112584 | 2.84571 | -0.034447866 | -0.003110593 |
| C730034F03Rik | 4.820571 | 4.686422 | 4.8692 | -0.182735681 | -0.048587025 | 4.7555716 | 4.7920311 | 4.82648 | -0.034452791 | -0.070912368 |
| Gm527 | 4.29681 | 4.299773 | 4.3894 | -0.089584277 | -0.092547847 | 4.2150267 | 4.172603 | 4.20707 | -0.034469849 | 0.007953901 |
| Gm6392 | 2.520575 | 2.577023 | 2.5745 | 0.002556743 | -0.05389054 | 2.5999132 | 2.5058406 | 2.54032 | -0.03447876 | 0.059593822 |
| Gm38098 | 0.440128 | 0.418878 | 0.4806 | -0.061686535 | -0.040436184 | 0.4429899 | 0.446602 | 0.48109 | -0.034492196 | -0.038104265 |
| Tbc1d13 | 10.04378 | 10.03771 | 9.9163 | 0.121447219 | 0.127518117 | 10.047738 | 10.05993 | 10.0944 | -0.034504774 | -0.046697028 |
| Ghrl | 1.252136 | 1.206099 | 1.2532 | -0.047087847 | -0.001050843 | 1.2933652 | 1.1633695 | 1.19793 | -0.034562103 | 0.095433609 |
| Gulo | 1.057824 | 0.982651 | 1.1417 | -0.159044739 | -0.083871911 | 0.9634862 | 0.9451318 | 0.97973 | -0.034601892 | -0.01624756 |
| Vmn2r107 | 0.603117 | 0.604012 | 0.6209 | -0.016871205 | -0.01776638 | 0.645616 | 0.6282723 | 0.66289 | -0.034621439 | -0.017277728 |
| Cluap1 | 8.264449 | 8.220581 | 8.1936 | 0.027013794 | 0.070882239 | 8.4414833 | 8.2877341 | 8.32236 | -0.034628797 | 0.119120438 |
| Nckap1l | 12.80266 | 12.85539 | 12.827 | 0.028821 | -0.023909686 | 13.079722 | 12.874307 | 12.9089 | -0.034630124 | 0.170785025 |
| C2cd4b | 0.548222 | 0.484152 | 0.623 | -0.138884075 | -0.074814337 | 0.532071 | 0.5118061 | 0.54649 | -0.034683899 | -0.01441899 |
| Pex2 | 8.298564 | 8.246534 | 8.4664 | -0.219896943 | -0.167867065 | 8.5036213 | 8.2671959 | 8.30191 | -0.034718677 | 0.201706701 |
| Cbx3-ps7 | -0.106478 | -0.13454 | -0.1598 | 0.025267832 | 0.053325734 | -0.0225588 | -0.156659 | -0.1219 | -0.034738784 | 0.099361655 |
| Specc1 | 6.940195 | 7.085409 | 7.0533 | 0.032090568 | -0.113123945 | 7.2000377 | 7.2726146 | 7.30739 | -0.034771189 | -0.107348099 |
| Lgals12 | 0.34471 | 0.309172 | 0.3402 | -0.031015314 | 0.004523357 | 0.3338369 | 0.3100131 | 0.34482 | -0.034807943 | -0.010984149 |
| Apc-ps1 | 1.74024 | 1.779671 | 1.8216 | -0.041910982 | -0.081341455 | 1.7873762 | 1.7918216 | 1.82663 | -0.03480971 | -0.039255177 |
| Rwdd4a | 8.412323 | 8.484973 | 8.3973 | 0.087706847 | 0.015056785 | 8.5655714 | 8.3734331 | 8.40825 | -0.034813805 | 0.157324512 |

| | | | | | | | | | | |
|---|---|---|---|---|---|---|---|---|---|---|
| Hdhd2 | 8.341488 | 8.374999 | 8.3 | 0.07497307 | 0.041461734 | 8.5504526 | 8.3772731 | 8.4121 | -0.034822719 | 0.138356766 |
| Efr3a | 11.27348 | 11.21106 | 11.208 | 0.002674122 | 0.065094534 | 11.095879 | 11.2039 | 11.2387 | -0.034843334 | -0.142864398 |
| Tcte3 | 1.055324 | 1.024881 | 1.063 | -0.038101934 | -0.00765918 | 1.1305923 | 1.0441523 | 1.079 | -0.034850319 | 0.051589773 |
| Vmn2r73 | 3.221596 | 3.242649 | 3.3055 | -0.062831679 | -0.083883951 | 3.1865923 | 3.3808897 | 3.41576 | -0.034874434 | -0.22917188 |
| Dusp18 | 3.639669 | 3.563746 | 3.5793 | -0.015578191 | 0.060344264 | 3.3741877 | 3.4088723 | 3.44375 | -0.034875284 | -0.069559875 |
| Nup133 | 9.318749 | 9.434325 | 9.319 | 0.115329151 | -0.000247228 | 9.5388254 | 9.3840701 | 9.419 | -0.034926771 | 0.119828496 |
| Grasp | 6.658314 | 6.383997 | 6.5596 | -0.175640086 | 0.098676394 | 6.2138069 | 6.369009 | 6.40397 | -0.034962281 | -0.190164375 |
| Fam167b | 4.857444 | 4.697399 | 4.8282 | -0.130832925 | 0.029211556 | 4.6296949 | 4.4879461 | 4.52292 | -0.034978839 | 0.106769944 |
| Scube2 | 0.791798 | 0.757857 | 0.8432 | -0.085362197 | -0.051420327 | 0.7735495 | 0.7077938 | 0.74279 | -0.034992195 | 0.03076343 |
| 1700092M07Rik | 1.500367 | 1.465554 | 1.4893 | -0.023704621 | 0.011109238 | 1.4624696 | 1.5010689 | 1.53609 | -0.035020284 | -0.07361961 |
| Gm5460 | 3.684736 | 3.668435 | 3.7314 | -0.062928364 | -0.04662716 | 3.7852494 | 3.7177742 | 3.75281 | -0.035035807 | 0.03243934 |
| Fam241b | 3.670982 | 3.699473 | 3.6985 | 0.000980297 | -0.027510536 | 3.7847326 | 3.7084115 | 3.74346 | -0.035050635 | 0.041270428 |
| Arpp21 | 2.430987 | 2.428342 | 2.4017 | 0.026625554 | 0.029270341 | 2.4295663 | 2.309106 | 2.34422 | -0.035110414 | 0.085349887 |
| Sh3bgr | 2.406972 | 2.328521 | 2.2984 | 0.030156695 | 0.108607247 | 2.3015989 | 2.2472654 | 2.28239 | -0.035127517 | 0.019206013 |
| Tmem45b | 1.413499 | 1.423463 | 1.4071 | 0.016388 | 0.006423607 | 1.3212449 | 1.3543987 | 1.38955 | -0.035149461 | -0.068303263 |
| Gm5182 | 3.103201 | 3.143668 | 3.044 | 0.099683725 | 0.059216497 | 3.0431401 | 3.0904259 | 3.12558 | -0.035154718 | -0.082440439 |
| Tmem37 | 3.537306 | 3.423839 | 3.6264 | -0.202577574 | -0.089110836 | 3.5830013 | 3.5141579 | 3.54937 | -0.03521685 | 0.033626532 |
| Olfr675 | 1.9007 | 1.984609 | 1.9534 | 0.031206878 | -0.052702515 | 2.0158789 | 1.9294619 | 1.96469 | -0.035225846 | 0.051191158 |
| Ncald | 5.221729 | 4.853524 | 5.1499 | -0.296363439 | 0.071841157 | 5.0175502 | 4.805702 | 4.84095 | -0.035245231 | 0.176602929 |
| Rps3a1 | 15.14557 | 15.1657 | 15.164 | 0.001769412 | -0.018352044 | 15.217786 | 15.112212 | 15.1475 | -0.03525434 | 0.070319822 |
| Ano4 | 2.545421 | 2.493015 | 2.5069 | -0.013874542 | 0.038530974 | 2.6260884 | 2.5674418 | 2.60273 | -0.035291654 | 0.023354931 |
| Tab2 | 11.49471 | 11.56256 | 11.539 | 0.023489564 | -0.04436498 | 11.434874 | 11.500818 | 11.5362 | -0.035337564 | -0.101281462 |
| Pax3 | 0.730671 | 0.732939 | 0.7884 | -0.055484905 | -0.057752877 | 0.7667687 | 0.7493751 | 0.78473 | -0.035355893 | -0.017962225 |
| Gm11336 | 1.813996 | 1.803151 | 1.8261 | -0.022992217 | -0.012146346 | 1.8638221 | 1.7566607 | 1.79207 | -0.035409757 | 0.071751636 |
| Serpina9 | 1.728979 | 1.777376 | 1.861 | -0.083600486 | -0.131997559 | 1.7433114 | 1.7224657 | 1.75795 | -0.035487197 | -0.014641579 |
| Kpna2 | 9.434097 | 9.45352 | 9.5326 | -0.079065011 | -0.098487622 | 9.6168217 | 9.6417903 | 9.67729 | -0.035498001 | -0.060466654 |
| Gm27149 | 0.353364 | 0.402951 | 0.3533 | 0.049652608 | 6.51036E-05 | 0.4443499 | 0.3784299 | 0.41394 | -0.035511546 | 0.030408504 |
| Clec12b | 0.727282 | 0.723353 | 0.7149 | 0.008453745 | 0.01238224 | 0.7332679 | 0.6774461 | 0.71296 | -0.035514719 | 0.020307092 |
| Gnas | 13.17048 | 13.12642 | 13.205 | -0.078424438 | -0.034365204 | 13.291371 | 13.138244 | 13.1738 | -0.035578804 | 0.117547625 |
| Dffa | 9.149648 | 9.148742 | 9.188 | -0.039208569 | -0.038302703 | 9.2832947 | 9.0640467 | 9.09963 | -0.035583941 | 0.183664087 |
| 1700021F05Rik | 6.561081 | 6.380153 | 6.5954 | -0.215265985 | -0.034337277 | 6.5674356 | 6.5342987 | 6.56989 | -0.035593448 | -0.002456494 |
| Gm30285 | 1.781875 | 1.754535 | 1.8071 | -0.052519956 | -0.025179518 | 1.8473213 | 1.862108 | 1.89773 | -0.035617114 | -0.050403855 |

| | | | | | | | | | | |
|---|---|---|---|---|---|---|---|---|---|---|
| Fam49a | 10.21812 | 10.46702 | 10.3 | 0.16712353 | -0.081776258 | 10.168259 | 10.275698 | 10.3113 | -0.035631334 | -0.143069365 |
| Arhgap1 | 10.58092 | 10.59276 | 10.658 | -0.065264878 | -0.077112142 | 10.732114 | 10.566524 | 10.6022 | -0.0356375 | 0.12995202 |
| Pde1b | 9.029433 | 8.944016 | 9.1599 | -0.215925231 | -0.130508001 | 9.3241763 | 9.3095418 | 9.34523 | -0.035684942 | -0.021050485 |
| Pi4ka | 11.40533 | 11.47427 | 11.491 | -0.016649384 | -0.0855888 | 11.588384 | 11.534904 | 11.5706 | -0.035698536 | 0.017782235 |
| Edil3 | 1.939183 | 1.905955 | 1.869 | 0.036999709 | 0.070227064 | 1.9590815 | 1.9468582 | 1.98264 | -0.035779944 | -0.023556616 |
| Slc6a12 | 2.311585 | 2.144482 | 2.5008 | -0.356288016 | -0.189185687 | 2.2144423 | 2.1625079 | 2.19835 | -0.035844639 | 0.016089753 |
| Pm20d1 | 1.793141 | 1.684221 | 1.8163 | -0.132070172 | -0.023150034 | 1.7180348 | 1.7251198 | 1.76097 | -0.035854361 | -0.042939426 |
| Gm10073 | 3.970779 | 3.889467 | 4.1604 | -0.270912557 | -0.189600371 | 3.9865037 | 3.9383872 | 3.97425 | -0.03586312 | 0.01225345 |
| Gm43380 | 2.055042 | 2.013532 | 2.056 | -0.042511411 | -0.001002082 | 2.01518 | 2.1244296 | 2.1603 | -0.035875375 | -0.145124977 |
| Gmfg | 11.001 | 11.01568 | 11.035 | -0.018970814 | -0.03365207 | 11.120358 | 10.989708 | 11.0256 | -0.035886614 | 0.094763373 |
| Gm49430 | 0.51579 | 0.524756 | 0.4752 | 0.049590679 | 0.040624385 | 0.5820295 | 0.4788053 | 0.51469 | -0.035887113 | 0.067337112 |
| Sertad4 | 1.872573 | 1.807426 | 1.9637 | -0.156250963 | -0.091103315 | 1.9540589 | 1.7815427 | 1.81743 | -0.035890044 | 0.136626176 |
| 7530416G11Rik | 0.757169 | 0.711724 | 0.8003 | -0.088528442 | -0.043083949 | 0.7322461 | 0.7606079 | 0.79652 | -0.035908554 | -0.064270422 |
| Gabra3 | 2.432887 | 2.477995 | 2.3808 | 0.097171513 | 0.052063118 | 2.39555 | 2.3565453 | 2.39247 | -0.03592408 | 0.003080608 |
| Adhfe1 | 2.553796 | 2.429526 | 2.4332 | -0.003629365 | 0.120640508 | 2.4089478 | 2.4397434 | 2.47575 | -0.036007241 | -0.066802909 |
| Ppp4r3a | 11.31076 | 11.33256 | 11.324 | 0.00899638 | -0.012803442 | 11.187485 | 11.363518 | 11.3995 | -0.036022416 | -0.212054932 |
| Krt17 | 0.721869 | 0.700488 | 0.7304 | -0.029928142 | -0.008547271 | 0.7596552 | 0.7249591 | 0.76101 | -0.036046359 | -0.001350261 |
| Fmr1 | 10.19416 | 10.25909 | 10.217 | 0.04162787 | -0.023299924 | 10.198978 | 10.330551 | 10.3667 | -0.036117902 | -0.167691413 |
| Slc13a1 | 2.441694 | 2.502148 | 2.428 | 0.074103763 | 0.01364937 | 2.5207674 | 2.4913658 | 2.5275 | -0.036129555 | -0.006727985 |
| Ptpru | 1.331474 | 1.267742 | 1.3005 | -0.03275562 | 0.03097565 | 1.2441973 | 1.2654444 | 1.30158 | -0.036136773 | -0.057383835 |
| Dleu7 | 1.697516 | 1.670231 | 1.6308 | 0.039452562 | 0.066737472 | 1.6503579 | 1.6534398 | 1.68959 | -0.036153128 | -0.039235032 |
| Ppp2r3c | 8.510371 | 8.445021 | 8.6615 | -0.216487259 | -0.151137783 | 8.7191945 | 8.7393918 | 8.77564 | -0.036247692 | -0.056445028 |
| Tax1bp3 | 7.807656 | 7.798173 | 7.8282 | -0.029996669 | -0.020513419 | 7.9201039 | 7.649048 | 7.6853 | -0.036249404 | 0.23480643 |
| Col24a1 | 0.69495 | 0.661586 | 0.6925 | -0.030934988 | 0.00242894 | 0.6534759 | 0.6799735 | 0.71623 | -0.036255401 | -0.062753026 |
| Ss18 | 10.68786 | 10.69932 | 10.65 | 0.049163622 | 0.037706754 | 10.647003 | 10.680665 | 10.717 | -0.036292004 | -0.06995359 |
| Mum1l1 | 1.26399 | 1.396074 | 1.2337 | 0.162330845 | 0.030247344 | 1.1891913 | 1.1946737 | 1.23101 | -0.036337216 | -0.041819604 |
| Mucl1 | 1.363456 | 1.318003 | 1.3465 | -0.028530052 | 0.016922545 | 1.2401642 | 1.2909415 | 1.32728 | -0.03634232 | -0.087119611 |
| Dcaf12l2 | 1.067302 | 1.131443 | 1.0875 | 0.043962908 | -0.020177616 | 1.040325 | 1.0513404 | 1.0877 | -0.036359282 | -0.047374646 |
| Chfr | 9.980908 | 9.989969 | 9.9423 | 0.047643896 | 0.038583228 | 9.985029 | 9.9969821 | 10.0334 | -0.036401848 | -0.048354878 |
| Vmn1r223 | 0.317991 | 0.324741 | 0.3088 | 0.015893302 | 0.009143559 | 0.3282613 | 0.3414014 | 0.37784 | -0.036435578 | -0.049575601 |
| Gtf2i | 11.41159 | 11.4068 | 11.57 | -0.163361282 | -0.158576042 | 11.205526 | 11.346465 | 11.3829 | -0.036447308 | -0.177385862 |
| Oxnad1 | 7.178107 | 7.144134 | 7.4006 | -0.256486646 | -0.222513642 | 7.3161145 | 7.4076452 | 7.44409 | -0.036447362 | -0.127977987 |

| | | | | | | | | | | |
|---|---|---|---|---|---|---|---|---|---|---|
| 1700067K01Rik | 0.567203 | 0.595691 | 0.6327 | -0.036969248 | -0.065456883 | 0.5965296 | 0.5812534 | 0.61775 | -0.036491825 | -0.021215568 |
| Hdhd3 | 2.126119 | 2.125238 | 2.0981 | 0.02714379 | 0.028024281 | 2.172327 | 2.0917351 | 2.12824 | -0.036508673 | 0.044083199 |
| Caap1 | 7.846442 | 7.689437 | 7.8993 | -0.209850113 | -0.052844425 | 7.8252967 | 7.7934091 | 7.82992 | -0.036509026 | -0.004621409 |
| Lrrc18 | 1.361181 | 1.284018 | 1.3781 | -0.094072839 | -0.016909674 | 1.2536572 | 1.2432811 | 1.2798 | -0.036517014 | -0.026140918 |
| Lpar4 | 2.913884 | 2.849718 | 2.9097 | -0.059936838 | 0.004229172 | 2.868962 | 2.8693054 | 2.90584 | -0.036533199 | -0.036876538 |
| Trpv4 | 5.07373 | 4.886804 | 4.8451 | 0.04168058 | 0.228606798 | 4.8740374 | 4.765828 | 4.80243 | -0.036605689 | 0.071603742 |
| Rpl31-ps10 | 1.061568 | 1.075997 | 1.0767 | -0.000669979 | -0.015099201 | 1.0544442 | 1.0638765 | 1.10054 | -0.03666537 | -0.046097611 |
| Efemp1 | 2.178528 | 2.124061 | 2.251 | -0.126920267 | -0.072454084 | 2.2448355 | 2.1345191 | 2.17122 | -0.036700165 | 0.073616284 |
| Fubp3 | 8.381956 | 8.360542 | 8.3801 | -0.019601862 | 0.001811416 | 8.2829299 | 8.4811949 | 8.51793 | -0.036738893 | -0.235003864 |
| Il10 | 4.217729 | 3.893657 | 4.1239 | -0.230249894 | 0.093822495 | 3.8096065 | 3.9047785 | 3.94155 | -0.03677398 | -0.131945921 |
| Taf5 | 7.686537 | 7.621594 | 7.6952 | -0.073635287 | -0.008692705 | 7.6381915 | 7.6976558 | 7.73445 | -0.0367938 | -0.096258113 |
| Atg4a | 7.504485 | 7.434965 | 7.4974 | -0.062435857 | 0.007084399 | 7.4365088 | 7.5151003 | 7.55191 | -0.036808693 | -0.115400201 |
| Best1 | 1.94365 | 1.909048 | 1.92 | -0.010931203 | 0.023670518 | 1.8864536 | 1.9616056 | 1.99844 | -0.036830931 | -0.111982952 |
| Tmem178b | 1.966201 | 1.917787 | 1.8965 | 0.021336678 | 0.069750421 | 1.8752296 | 1.9358589 | 1.97269 | -0.036833751 | -0.097463065 |
| Ect2l | 2.897514 | 2.860293 | 2.9318 | -0.071499437 | -0.034278183 | 2.8836283 | 2.8185353 | 2.85538 | -0.036841681 | 0.02825133 |
| Adgra3 | 4.799265 | 4.615765 | 4.6967 | -0.080973791 | 0.102526207 | 4.5776488 | 4.5963341 | 4.63325 | -0.03691868 | -0.055604005 |
| Olfr1226 | 1.57529 | 1.516054 | 1.6009 | -0.084811519 | -0.02557579 | 1.5555764 | 1.5341734 | 1.57111 | -0.036938077 | -0.01553504 |
| Olfr1309 | 4.098607 | 4.102226 | 4.0948 | 0.007377348 | 0.003757456 | 4.1163863 | 4.2930938 | 4.33004 | -0.036949628 | -0.213657069 |
| Sh3bp1 | 10.59802 | 10.50014 | 10.57 | -0.069600194 | 0.02827335 | 10.765229 | 10.61741 | 10.6544 | -0.036953096 | 0.110865138 |
| Olfr212 | 0.681576 | 0.606353 | 0.676 | -0.069682648 | 0.005539795 | 0.6603259 | 0.6126975 | 0.64967 | -0.036973204 | 0.010655221 |
| Tnfsf15 | 5.396384 | 5.340975 | 5.4994 | -0.158420159 | -0.103011769 | 4.5393947 | 5.1476157 | 5.18463 | -0.037017331 | -0.645238338 |
| Tmprss11d | 4.269885 | 4.201814 | 4.3102 | -0.108342846 | -0.040272658 | 4.3235325 | 4.3018195 | 4.33887 | -0.037048971 | -0.015335966 |
| Plpp7 | 1.703226 | 1.718993 | 1.7056 | 0.013407361 | -0.002359141 | 1.7351934 | 1.7201965 | 1.75727 | -0.03707568 | -0.022078771 |
| Zfp346 | 8.793617 | 8.826881 | 8.825 | 0.001877318 | -0.031386416 | 9.0198678 | 8.8877746 | 8.92487 | -0.037098321 | 0.094994861 |
| Mfap3l | 2.148222 | 2.12693 | 2.1313 | -0.004361646 | 0.016930198 | 2.2830384 | 2.0536773 | 2.09081 | -0.03713682 | 0.192224221 |
| Speer5-ps1 | 3.303224 | 3.426405 | 3.2802 | 0.146185137 | 0.023004184 | 3.3750917 | 3.4322527 | 3.46939 | -0.037140603 | -0.094301658 |
| Olfr715b | 2.791919 | 2.700778 | 2.7127 | -0.011954498 | 0.079186552 | 2.7626894 | 2.7534482 | 2.7906 | -0.037148109 | -0.027906906 |
| Gm609 | 1.46123 | 1.454438 | 1.451 | 0.003414888 | 0.01020641 | 1.49308 | 1.4751299 | 1.51229 | -0.03716055 | -0.019210421 |
| Gabra2 | 2.863325 | 2.809891 | 2.7821 | 0.027763388 | 0.081197929 | 2.9129652 | 2.8213105 | 2.8585 | -0.037188872 | 0.054465839 |
| Olfr1336 | 5.226183 | 5.386583 | 5.272 | 0.114613737 | -0.045786213 | 5.4037417 | 5.4965901 | 5.53381 | -0.037216271 | -0.130064667 |
| Naalad2 | 4.53132 | 4.077629 | 4.4441 | -0.366460189 | 0.087230334 | 4.2622944 | 4.0420618 | 4.07928 | -0.037217849 | 0.18301484 |
| Gm6501 | 2.255875 | 2.134205 | 2.1891 | -0.054889856 | 0.066780046 | 2.3336725 | 2.2286593 | 2.26589 | -0.03722627 | 0.067786939 |

| | | | | | | | | | | |
|---|---|---|---|---|---|---|---|---|---|---|
| Zfp866 | 9.067683 | 9.215273 | 9.0769 | 0.138375102 | -0.009214914 | 9.1734282 | 9.1544374 | 9.19166 | -0.037226627 | -0.018235904 |
| Cenph | 5.303487 | 5.15477 | 5.4097 | -0.254925351 | -0.106208541 | 5.3490269 | 5.5679884 | 5.60526 | -0.037276417 | -0.256237972 |
| Dguok | 8.932856 | 8.935502 | 9.0312 | -0.095670904 | -0.098316368 | 9.2024376 | 9.0381357 | 9.07552 | -0.037380701 | 0.126921212 |
| Gm14267 | 2.182594 | 2.098155 | 2.0389 | 0.059249211 | 0.143687794 | 2.0593282 | 2.1467565 | 2.18415 | -0.037388578 | -0.124816874 |
| Cpsf3 | 10.92431 | 10.96293 | 10.967 | -0.004139308 | -0.042767 | 11.192967 | 11.048164 | 11.0856 | -0.037388912 | 0.10741353 |
| Ubp1 | 10.72877 | 10.74964 | 10.854 | -0.103895572 | -0.124768274 | 10.990274 | 10.925879 | 10.9633 | -0.037401136 | 0.026993493 |
| Lhfpl3 | 0.720483 | 0.726962 | 0.7266 | 0.000362345 | -0.00611638 | 0.7194377 | 0.6888898 | 0.72629 | -0.037405042 | -0.006857189 |
| Zfp212 | 8.60838 | 8.712008 | 8.6622 | 0.049793707 | -0.053834122 | 8.4872324 | 8.4844831 | 8.52193 | -0.037447379 | -0.034698092 |
| Lrrc6 | 1.956885 | 1.872439 | 1.8552 | 0.017255979 | 0.101701674 | 1.8583955 | 1.8017295 | 1.83918 | -0.037452776 | 0.019213301 |
| Cdr2l | 5.573999 | 5.346062 | 5.3673 | -0.021214492 | 0.206721918 | 5.1488672 | 5.0347258 | 5.07219 | -0.037467416 | 0.076673958 |
| Tmem131l | 11.57232 | 11.63486 | 11.705 | -0.069967428 | -0.132509354 | 11.973193 | 11.852583 | 11.8901 | -0.037552422 | 0.083057284 |
| Gm49384 | 5.495867 | 5.568567 | 5.6418 | -0.073183452 | -0.145883366 | 5.2988769 | 5.4898537 | 5.52744 | -0.037588869 | -0.228565669 |
| Arl5a | 10.65675 | 10.79159 | 10.77 | 0.02188772 | -0.112957906 | 10.552029 | 10.800651 | 10.8383 | -0.037637668 | -0.286259468 |
| Gm15682 | 0.877194 | 0.849832 | 0.9015 | -0.051660702 | -0.024298981 | 0.8445384 | 0.854729 | 0.89239 | -0.03765754 | -0.047848141 |
| P2rx1 | 3.379889 | 3.305563 | 3.2955 | 0.010094161 | 0.084420877 | 3.3677933 | 3.3523314 | 3.39003 | -0.037703453 | -0.022241527 |
| Wnt2 | 0.657704 | 0.716108 | 0.6776 | 0.038511969 | -0.01989192 | 0.6288282 | 0.6124791 | 0.65031 | -0.037833124 | -0.021484018 |
| Hgf | 4.152825 | 4.022261 | 4.0828 | -0.06058283 | 0.06998047 | 3.8450176 | 3.8957285 | 3.93359 | -0.037857322 | -0.088568193 |
| Olfr622 | 0.280754 | 0.254994 | 0.2567 | -0.001696087 | 0.024063862 | 0.2511542 | 0.2760943 | 0.31399 | -0.037892467 | -0.062832515 |
| Lin9 | 7.948888 | 7.996932 | 8.1346 | -0.137641098 | -0.185684392 | 7.8701498 | 8.0149754 | 8.05288 | -0.037901982 | -0.182727622 |
| Kctd18 | 8.393005 | 8.260482 | 8.4825 | -0.222004721 | -0.089482075 | 8.4401438 | 8.5435595 | 8.58147 | -0.037907757 | -0.141323388 |
| Gm5108 | 1.064328 | 1.015829 | 1.0789 | -0.063104877 | -0.014605945 | 1.1384158 | 1.118079 | 1.156 | -0.037917585 | -0.017580811 |
| Marveld3 | 2.598672 | 2.36668 | 2.512 | -0.145306954 | 0.086684895 | 2.4621544 | 2.3756542 | 2.41357 | -0.037920621 | 0.048579587 |
| Kcnmb1 | 3.141497 | 3.206647 | 3.1184 | 0.088234655 | 0.023084116 | 3.076668 | 3.1238223 | 3.16179 | -0.037971657 | -0.085125939 |
| 9130213A22Rik | 1.607069 | 1.559143 | 1.5753 | -0.016165529 | 0.03176083 | 1.6642527 | 1.6746508 | 1.71269 | -0.038038071 | -0.048436189 |
| Eid2b | 5.205589 | 5.120049 | 5.2755 | -0.155491777 | -0.069952336 | 5.0169789 | 5.1267524 | 5.16482 | -0.038063558 | -0.147837135 |
| Gm16363 | 0.511042 | 0.475968 | 0.5207 | -0.044710236 | -0.009636249 | 0.5227315 | 0.4992102 | 0.53733 | -0.038124243 | -0.014603004 |
| Gm14438 | 1.259292 | 1.277959 | 1.2907 | -0.01271082 | -0.031378038 | 1.2570959 | 1.3116807 | 1.34982 | -0.038135685 | -0.092720449 |
| Rps8-ps2 | 1.259292 | 1.277959 | 1.2907 | -0.01271082 | -0.031378038 | 1.2570959 | 1.3116807 | 1.34982 | -0.038135685 | -0.092720449 |
| Slc35a5 | 8.616571 | 8.619011 | 8.5676 | 0.051415363 | 0.048975484 | 8.7054381 | 8.5481946 | 8.58635 | -0.038156653 | 0.119086861 |
| Ighv1-36 | 4.572805 | 4.853563 | 4.5698 | 0.283793317 | 0.003035475 | 4.7880483 | 4.5018589 | 4.54002 | -0.038158664 | 0.248030756 |
| Gm5917 | 1.866475 | 1.802596 | 1.8254 | -0.022815204 | 0.041063935 | 1.7376116 | 1.8388871 | 1.87706 | -0.038169959 | -0.139445492 |
| Bbs10 | 3.819195 | 3.84517 | 3.893 | -0.047812156 | -0.073786916 | 3.8434603 | 3.8520496 | 3.89033 | -0.038280325 | -0.046869572 |

| | | | | | | | | | | |
|---|---|---|---|---|---|---|---|---|---|---|
| Rundc3a | 5.055997 | 5.134088 | 4.8797 | 0.254349321 | 0.176258259 | 4.809615 | 4.866843 | 4.90515 | -0.038311118 | -0.095539151 |
| Gm20878 | 2.227956 | 2.107391 | 2.2596 | -0.152249955 | -0.031685388 | 2.1396084 | 2.1672418 | 2.20565 | -0.038411158 | -0.066044502 |
| Ubox5 | 7.54463 | 7.407641 | 7.5434 | -0.135784436 | 0.001205223 | 7.5393734 | 7.4492144 | 7.48764 | -0.038422441 | 0.051736571 |
| Zfp472 | 8.272327 | 8.36634 | 8.4419 | -0.075533945 | -0.169547037 | 8.1951293 | 8.304072 | 8.3425 | -0.038432714 | -0.14737544 |
| Zfp334 | 2.813521 | 2.645499 | 2.7654 | -0.119944544 | 0.048077438 | 2.5747953 | 2.6241622 | 2.66262 | -0.0384568 | -0.087823697 |
| Srpk1 | 11.38276 | 11.46798 | 11.419 | 0.048693034 | -0.036522044 | 11.074788 | 11.436822 | 11.4753 | -0.03849227 | -0.400526846 |
| Olfr542-ps1 | 0.034981 | -0.00396 | 0.005 | -0.008981738 | 0.029963656 | 0.0406676 | 0.0297872 | 0.06831 | -0.038523617 | -0.027643192 |
| Nkpd1 | 2.119701 | 1.982479 | 2.1879 | -0.205425804 | -0.068203849 | 2.0244124 | 1.9391175 | 1.97767 | -0.03855175 | 0.046743189 |
| Ric1 | 9.812467 | 9.917737 | 9.8312 | 0.086493936 | -0.018775849 | 9.7630284 | 9.9327069 | 9.9713 | -0.038594134 | -0.20827262 |
| Ldb3 | 1.186992 | 1.304876 | 1.2781 | 0.02677428 | -0.091110051 | 1.1794356 | 1.2050975 | 1.24374 | -0.038647199 | -0.06430915 |
| Vmn1r17 | 2.911075 | 3.033071 | 3.0529 | -0.019816813 | -0.14181265 | 2.9941668 | 3.0559259 | 3.09462 | -0.038692548 | -0.100451621 |
| Gigyf2 | 10.58686 | 10.71521 | 10.666 | 0.048940084 | -0.079410378 | 10.664235 | 10.675553 | 10.7143 | -0.038697354 | -0.050014653 |
| Krt7 | 4.03389 | 4.012899 | 4.0211 | -0.008249453 | 0.012741413 | 4.1463831 | 4.1181688 | 4.15687 | -0.03869957 | -0.010485241 |
| Prkaa1 | 9.932639 | 9.971521 | 9.9695 | 0.002005406 | -0.036876596 | 9.7806392 | 9.7784127 | 9.81712 | -0.03870761 | -0.036481106 |
| Nhlrc2 | 9.573285 | 9.567308 | 9.5241 | 0.043166846 | 0.049143591 | 9.6805923 | 9.6664682 | 9.70518 | -0.038707926 | -0.024583869 |
| Nup50 | 10.2431 | 10.34909 | 10.326 | 0.022770276 | -0.08321715 | 10.164596 | 10.297808 | 10.3365 | -0.038741528 | -0.171953617 |
| C1qtnf5 | 2.261077 | 2.230217 | 2.2125 | 0.017760513 | 0.048619872 | 2.3194907 | 2.1207674 | 2.15955 | -0.038778375 | 0.159944864 |
| Selenof | 11.02757 | 10.93294 | 11.017 | -0.084092962 | 0.01054053 | 11.130117 | 10.962661 | 11.0015 | -0.038857966 | 0.128597613 |
| Gm6641 | 2.578271 | 2.546382 | 2.5401 | 0.006251741 | 0.03814047 | 2.4952581 | 2.6607934 | 2.69968 | -0.038887553 | -0.2044228 |
| Rpsa-ps12 | 0.991355 | 0.958852 | 1.0004 | -0.041581277 | -0.009077515 | 0.957148 | 0.9898265 | 1.02882 | -0.038997051 | -0.071675526 |
| Tmem30a | 11.17821 | 11.15724 | 11.188 | -0.03065342 | -0.009683802 | 11.023827 | 11.095446 | 11.1344 | -0.03899791 | -0.110617233 |
| Gm12463 | 1.507048 | 1.488473 | 1.5017 | -0.013229637 | 0.005344996 | 1.6083796 | 1.5201567 | 1.55921 | -0.039058143 | 0.04916476 |
| Ighv2-7 | 0.382556 | 0.28031 | 0.2887 | -0.008423556 | 0.093821979 | 0.417282 | 0.2766803 | 0.31576 | -0.039075167 | 0.101526537 |
| Rasl10a | 2.323178 | 2.205757 | 2.4018 | -0.196036251 | -0.078615054 | 2.2910685 | 2.1989443 | 2.23805 | -0.039107899 | 0.053016342 |
| Olfr137 | 3.212618 | 3.066983 | 3.1579 | -0.090964911 | 0.054670373 | 3.0815832 | 3.3338427 | 3.37296 | -0.039117834 | -0.291377275 |
| Lef1 | 11.12915 | 11.21403 | 11.243 | -0.028657776 | -0.113538188 | 10.945575 | 11.160504 | 11.1996 | -0.039143132 | -0.25407242 |
| Cys1 | 1.742951 | 1.663027 | 1.8742 | -0.211197691 | -0.131273233 | 1.6032682 | 1.636669 | 1.67583 | -0.039159424 | -0.072560239 |
| Foxq1 | 1.147851 | 1.036948 | 1.1633 | -0.126361338 | -0.015458814 | 1.1086171 | 1.0360751 | 1.07531 | -0.039236618 | 0.033305418 |
| Mansc4 | 1.16534 | 1.10499 | 1.1599 | -0.054959505 | 0.005390382 | 1.1071985 | 1.111007 | 1.15025 | -0.039239598 | -0.043048117 |
| Rbm20 | 1.261819 | 1.225789 | 1.2578 | -0.031967148 | 0.004062816 | 1.1640685 | 1.1666227 | 1.20601 | -0.03938378 | -0.041938041 |
| Creld1 | 7.075979 | 7.105059 | 6.9927 | 0.112312406 | 0.083232787 | 7.3114692 | 7.25331 | 7.29271 | -0.039395861 | 0.018763277 |
| Stau1 | 9.972754 | 10.11595 | 10.059 | 0.056921508 | -0.086279605 | 10.046863 | 10.211985 | 10.2514 | -0.039429298 | -0.204550993 |

| | | | | | | | | | | |
|---|---|---|---|---|---|---|---|---|---|---|
| Gm10309 | 2.390993 | 2.270175 | 2.2771 | -0.006969971 | 0.113847783 | 2.1312429 | 2.1086853 | 2.14813 | -0.039442933 | -0.016885333 |
| Fbxo38 | 10.53131 | 10.62247 | 10.549 | 0.073860894 | -0.017290364 | 10.505344 | 10.534586 | 10.574 | -0.039445536 | -0.068687791 |
| Gm8399 | 3.491901 | 3.385777 | 3.3691 | 0.01672043 | 0.122844331 | 3.5752756 | 3.4499252 | 3.48942 | -0.039490045 | 0.085860371 |
| Stx5a | 10.16067 | 10.18201 | 10.248 | -0.065894008 | -0.087233769 | 10.284847 | 10.304844 | 10.3444 | -0.039519884 | -0.059517704 |
| Car10 | 1.340544 | 1.362417 | 1.3279 | 0.034485286 | 0.012613007 | 1.3375192 | 1.3949282 | 1.43445 | -0.039520324 | -0.096929344 |
| Gm12569 | 1.479726 | 1.509564 | 1.4844 | 0.025205551 | -0.004631739 | 1.5557165 | 1.4838918 | 1.52343 | -0.039539838 | 0.032284891 |
| Nalcn | 1.206905 | 1.266091 | 1.179 | 0.087099448 | 0.027913429 | 1.2018433 | 1.2551695 | 1.29477 | -0.039598948 | -0.092925096 |
| Zfp341 | 6.841627 | 6.956265 | 6.8843 | 0.072003086 | -0.042634656 | 7.0079508 | 6.8900468 | 6.92966 | -0.039608538 | 0.07829548 |
| Unc45a | 10.0619 | 10.10339 | 9.9982 | 0.105148854 | 0.063660051 | 10.173548 | 9.9898986 | 10.0295 | -0.039620229 | 0.144029442 |
| Cpt2 | 7.500607 | 7.35809 | 7.4302 | -0.07212573 | 0.0703918 | 7.6293776 | 7.3557841 | 7.39544 | -0.039659759 | 0.233933815 |
| Bcl6b | 5.359216 | 5.062371 | 4.9693 | 0.093068075 | 0.389913569 | 4.9077259 | 4.9739831 | 5.01369 | -0.039706414 | -0.105963618 |
| Asb2 | 6.410146 | 6.310144 | 6.1633 | 0.146828748 | 0.246830698 | 6.000509 | 6.2269681 | 6.26669 | -0.039720702 | -0.266179845 |
| Rbm18 | 9.302449 | 9.199638 | 9.2624 | -0.062722992 | 0.040088222 | 9.1579511 | 9.1007877 | 9.1406 | -0.039809496 | 0.017353901 |
| Ripk4 | 1.798552 | 1.707469 | 1.7937 | -0.086192394 | 0.004890869 | 1.8395688 | 1.7614782 | 1.80132 | -0.039836955 | 0.038253683 |
| Arl4c | 11.77589 | 11.5864 | 11.669 | -0.082412064 | 0.107074524 | 11.316772 | 11.392737 | 11.4326 | -0.039840094 | -0.11580516 |
| Socs7 | 9.458471 | 9.603942 | 9.4766 | 0.12737349 | -0.018097217 | 9.2297326 | 9.4078079 | 9.44767 | -0.039862577 | -0.217937842 |
| Serinc2 | 1.24664 | 1.150826 | 1.2283 | -0.077523774 | 0.018290112 | 1.2191676 | 1.1760319 | 1.21591 | -0.039881241 | 0.00325452 |
| Mia | 1.308145 | 1.23418 | 1.2831 | -0.048921347 | 0.0250434 | 1.3804425 | 1.2806732 | 1.32056 | -0.039886405 | 0.059882864 |
| Pank2 | 9.856818 | 9.913479 | 9.8996 | 0.013910102 | -0.04275127 | 9.7296996 | 9.8038676 | 9.8438 | -0.039933779 | -0.11410179 |
| Gabra1 | 0.5791 | 0.653248 | 0.556 | 0.09721937 | 0.023071263 | 0.4966727 | 0.5324872 | 0.57244 | -0.039952415 | -0.07576692 |
| Slc38a3 | 0.888311 | 0.893674 | 0.8681 | 0.025529623 | 0.020166455 | 0.8653218 | 0.8856117 | 0.92558 | -0.039965176 | -0.060255018 |
| Igf2bp1 | 2.018081 | 2.004981 | 1.9909 | 0.014119295 | 0.0272198 | 2.0268978 | 2.0679374 | 2.1079 | -0.039966134 | -0.081005705 |
| Ncam2 | 1.681992 | 1.652242 | 1.6917 | -0.039464717 | -0.009715118 | 1.659262 | 1.7355393 | 1.77558 | -0.040038316 | -0.116315642 |
| Mgst2 | 8.467018 | 8.421717 | 8.6829 | -0.26122696 | -0.215925923 | 8.4853228 | 8.7314972 | 8.77155 | -0.040054504 | -0.286228901 |
| Maz | 11.13199 | 11.17095 | 11.223 | -0.051946404 | -0.090904166 | 11.386328 | 11.287419 | 11.3275 | -0.040080078 | 0.058828896 |
| Fbxl2 | 3.213582 | 3.256536 | 3.3834 | -0.126845011 | -0.169799689 | 3.3253003 | 3.4569239 | 3.49701 | -0.040081925 | -0.171705503 |
| Dyrk1a | 11.24397 | 11.2707 | 11.306 | -0.034997802 | -0.061733126 | 11.017788 | 11.147582 | 11.1877 | -0.040115604 | -0.169909523 |
| Slc25a34 | 0.775839 | 0.661126 | 0.7113 | -0.050189431 | 0.064523679 | 0.6984877 | 0.6561429 | 0.69626 | -0.040118143 | 0.002226595 |
| Slc5a1 | 0.197559 | 0.105184 | 0.1769 | -0.07171249 | 0.020662394 | 0.1970665 | 0.1077804 | 0.14796 | -0.040180636 | 0.049105495 |
| Nfx1 | 11.03724 | 11.07201 | 11.096 | -0.023785764 | -0.058554592 | 11.1818 | 11.213241 | 11.2534 | -0.040194078 | -0.071634794 |
| Cntd1 | 3.041559 | 3.023826 | 3.0472 | -0.023365763 | -0.005632446 | 3.0926512 | 3.1373051 | 3.1775 | -0.040199571 | -0.084853384 |
| Olfr1183 | 3.140554 | 3.128575 | 2.9824 | 0.146182109 | 0.158160399 | 3.1764613 | 3.162864 | 3.20308 | -0.040217949 | -0.026620708 |

| | | | | | | | | | | |
|---|---|---|---|---|---|---|---|---|---|---|
| Sf3b1 | 13.77774 | 13.80572 | 13.809 | -0.003493566 | -0.031471938 | 13.850195 | 13.864795 | 13.905 | -0.040250094 | -0.054849758 |
| Nectin3 | 5.185822 | 4.892924 | 5.0462 | -0.153311869 | 0.139586404 | 4.7263011 | 4.91136 | 4.95163 | -0.040273172 | -0.225332125 |
| Olfr446 | 3.423467 | 3.343641 | 3.4254 | -0.081719512 | -0.001893213 | 3.4472779 | 3.5315372 | 3.57183 | -0.040288445 | -0.12454777 |
| Uox | 0.983216 | 0.941482 | 0.9474 | -0.005894903 | 0.035838965 | 0.9087886 | 0.9144362 | 0.95476 | -0.040324097 | -0.045971698 |
| Pip5kl1 | 0.776227 | 0.767401 | 0.8027 | -0.035251436 | -0.026425549 | 0.8282907 | 0.7889023 | 0.82923 | -0.040324146 | -0.000935751 |
| Prdx6 | 10.50053 | 10.47663 | 10.542 | -0.065506962 | -0.041608062 | 10.45059 | 10.427043 | 10.4674 | -0.040326546 | -0.016780357 |
| Kpna1 | 10.57973 | 10.6998 | 10.567 | 0.132667812 | 0.012597882 | 10.457798 | 10.519115 | 10.5595 | -0.04035335 | -0.101670221 |
| Serinc1 | 11.02298 | 11.02045 | 10.987 | 0.033279016 | 0.035809726 | 10.947076 | 10.935241 | 10.9756 | -0.04035389 | -0.028518317 |
| Tro | 0.60427 | 0.574475 | 0.5085 | 0.065959629 | 0.095754272 | 0.6666688 | 0.4909023 | 0.53126 | -0.040360413 | 0.13540611 |
| Gm49373 | 2.283763 | 2.307151 | 2.3456 | -0.038416457 | -0.061804914 | 2.4721661 | 2.4916488 | 2.53204 | -0.040386828 | -0.059869575 |
| Plxna1 | 8.251019 | 8.55746 | 7.7365 | 0.820934202 | 0.514493458 | 7.269344 | 7.6410045 | 7.6814 | -0.040392629 | -0.412053067 |
| G530012D18Rik | 0.514238 | 0.563785 | 0.5071 | 0.056732697 | 0.007185357 | 0.5884142 | 0.5115842 | 0.552 | -0.040412922 | 0.036417051 |
| Capn1 | 11.16026 | 11.11183 | 11.166 | -0.054588684 | -0.006159938 | 11.020738 | 11.056922 | 11.0974 | -0.040437948 | -0.076621557 |
| Gm48676 | 0.324141 | 0.325569 | 0.3442 | -0.018616439 | -0.020044548 | 0.3177535 | 0.3521732 | 0.39266 | -0.040481869 | -0.074901594 |
| Bicra | 10.11326 | 10.1975 | 10.21 | -0.01260201 | -0.096838748 | 10.213118 | 10.240788 | 10.2813 | -0.04048258 | -0.068153064 |
| Aqr | 11.15615 | 11.15587 | 11.091 | 0.065143112 | 0.06543193 | 11.025603 | 11.035756 | 11.0763 | -0.04052002 | -0.050672852 |
| Gm973 | 1.649266 | 1.58952 | 1.5611 | 0.028384083 | 0.088130381 | 1.6133454 | 1.6275299 | 1.66809 | -0.04055773 | -0.054742254 |
| Ppp6r3 | 11.11679 | 11.05748 | 11.111 | -0.053621594 | 0.005682296 | 10.940951 | 11.089977 | 11.1305 | -0.040559114 | -0.189585857 |
| Plekhb2 | 9.984767 | 10.03734 | 10.129 | -0.091492072 | -0.144060097 | 9.9711199 | 10.058652 | 10.0992 | -0.040589638 | -0.128121348 |
| Ankrd9 | 4.73505 | 4.781252 | 4.8128 | -0.03156638 | -0.077768114 | 4.9583292 | 4.7688989 | 4.8095 | -0.040605162 | 0.148825181 |
| Snx30 | 10.78039 | 10.97466 | 10.902 | 0.072462218 | -0.121802188 | 10.874169 | 10.825453 | 10.8661 | -0.040615124 | 0.008100828 |
| Olfr825 | 1.216383 | 1.196282 | 1.2091 | -0.012835469 | 0.007265594 | 1.1784458 | 1.2229136 | 1.26356 | -0.040646384 | -0.085114206 |
| Ighv1-85 | 5.658041 | 5.357855 | 5.364 | -0.006126594 | 0.294059211 | 5.7120147 | 5.433982 | 5.47467 | -0.040687944 | 0.237344762 |
| Cldn8 | 0.668637 | 0.578106 | 0.6356 | -0.057462285 | 0.033068591 | 0.6575382 | 0.5610514 | 0.60176 | -0.040709341 | 0.055777416 |
| Gpr173 | 0.967733 | 1.140933 | 1.0853 | 0.055626567 | -0.117573448 | 1.0134051 | 0.932191 | 0.97292 | -0.040733176 | 0.040480907 |
| Zbtb33 | 9.942537 | 10.05932 | 9.9714 | 0.087925149 | -0.028857517 | 10.055481 | 9.9324131 | 9.97315 | -0.040735454 | 0.082332237 |
| Slc9a5 | 5.427149 | 5.628178 | 5.5676 | 0.060569183 | -0.140459693 | 5.8001658 | 5.6427729 | 5.68352 | -0.040743764 | 0.116649146 |
| Susd5 | 0.703853 | 0.704435 | 0.7738 | -0.069376347 | -0.069958903 | 0.7047474 | 0.7042333 | 0.74506 | -0.040829413 | -0.040315316 |
| Gm14435 | 4.023098 | 3.950833 | 3.8578 | 0.093035899 | 0.16530026 | 3.8295597 | 3.8655058 | 3.90637 | -0.040868182 | -0.076814272 |
| Adgrb3 | 2.140416 | 2.189473 | 2.1381 | 0.051422935 | 0.002365883 | 2.1531105 | 2.2226319 | 2.26352 | -0.040891872 | -0.110413277 |
| Zfp879 | 2.351238 | 2.354299 | 2.2491 | 0.105154624 | 0.102093689 | 2.4192453 | 2.3233023 | 2.36421 | -0.040904742 | 0.055038285 |
| Setd1b | 10.55737 | 10.59482 | 10.553 | 0.04141205 | 0.003959068 | 10.442202 | 10.524709 | 10.5657 | -0.040985764 | -0.123493062 |

| | | | | | | | | | | |
|---|---|---|---|---|---|---|---|---|---|---|
| Gm10420 | 3.213169 | 3.190624 | 3.157 | 0.033609551 | 0.056154339 | 3.115756 | 3.1368107 | 3.1779 | -0.041087251 | -0.062142007 |
| Ttpal | 10.92534 | 11.09486 | 10.977 | 0.118112334 | -0.0514069 | 11.21051 | 11.026101 | 11.0672 | -0.041087748 | 0.143321592 |
| Olfr1280 | 2.30787 | 2.325062 | 2.3401 | -0.015027208 | -0.032219564 | 2.3452705 | 2.3596136 | 2.40077 | -0.041156076 | -0.05549917 |
| Gm12712 | 0.576345 | 0.592025 | 0.6245 | -0.032479728 | -0.04815983 | 0.6183642 | 0.6342445 | 0.67546 | -0.041214011 | -0.057094265 |
| Edc3 | 9.72997 | 9.783388 | 9.6929 | 0.090531437 | 0.037113079 | 9.6366236 | 9.5613833 | 9.60261 | -0.041230591 | 0.034009701 |
| Runx2 | 8.185176 | 8.113689 | 8.3245 | -0.210806482 | -0.139319401 | 8.0700272 | 8.0837037 | 8.12494 | -0.041236315 | -0.054912808 |
| Wdr73 | 8.47075 | 8.499458 | 8.4554 | 0.044080285 | 0.015372366 | 8.5264726 | 8.4932789 | 8.53452 | -0.041243871 | -0.008050183 |
| Vmn1r200 | 1.412265 | 1.42956 | 1.3706 | 0.058961121 | 0.04166591 | 1.387573 | 1.3983744 | 1.43966 | -0.041286493 | -0.052087892 |
| Gm10297 | 2.726699 | 2.640022 | 2.7056 | -0.065548924 | 0.02112792 | 2.4590525 | 2.6248415 | 2.66617 | -0.041332037 | -0.207121043 |
| Gm3912 | 0.410635 | 0.384609 | 0.3671 | 0.017504887 | 0.043531146 | 0.4469341 | 0.4120412 | 0.45338 | -0.041341777 | -0.006448829 |
| Mylk4 | 3.230154 | 3.153796 | 3.1969 | -0.043116791 | 0.033241427 | 3.0918951 | 3.21899 | 3.26035 | -0.04135923 | -0.168454148 |
| Gm13653 | 1.229728 | 1.29272 | 1.3221 | -0.029382577 | -0.092374828 | 1.2724232 | 1.2766613 | 1.31804 | -0.041374081 | -0.045612205 |
| Arih2 | 9.979149 | 10.07772 | 9.9566 | 0.121133265 | 0.022563726 | 9.9640269 | 10.072268 | 10.1137 | -0.041384915 | -0.149626306 |
| Gm13574 | 0.553899 | 0.486864 | 0.4722 | 0.014643422 | 0.08167823 | 0.4961195 | 0.49867 | 0.54007 | -0.041396149 | -0.04394665 |
| Nln | 8.263826 | 8.266246 | 8.149 | 0.11724062 | 0.114820951 | 8.2266203 | 8.2295036 | 8.27093 | -0.041422097 | -0.044305403 |
| Epha10 | 2.510152 | 2.49836 | 2.5413 | -0.042948635 | -0.031156445 | 2.5406751 | 2.5839565 | 2.62539 | -0.041431386 | -0.084712793 |
| Gm5124 | 3.020133 | 3.118355 | 3.0154 | 0.102939144 | 0.004717102 | 3.0338818 | 3.0129506 | 3.05453 | -0.041574498 | -0.020643301 |
| Capns1 | 11.36721 | 11.28947 | 11.379 | -0.089919736 | -0.012182886 | 11.379978 | 11.238466 | 11.2801 | -0.041592624 | 0.099918651 |
| Traj37 | 2.804103 | 2.794202 | 2.6746 | 0.119589215 | 0.12949091 | 2.67142 | 2.6434758 | 2.68511 | -0.041632671 | -0.013688434 |
| Fhod3 | 1.059915 | 1.069483 | 1.0648 | 0.004710431 | -0.004857482 | 1.0955651 | 1.0653931 | 1.10703 | -0.041636913 | -0.011464914 |
| 2310022B05Rik | 7.533949 | 7.424447 | 7.4519 | -0.027422061 | 0.082080067 | 7.3345145 | 7.1939664 | 7.23565 | -0.041684574 | 0.09886356 |
| Gm7045 | 2.543709 | 2.490033 | 2.6097 | -0.119683823 | -0.066007782 | 2.5164463 | 2.5390105 | 2.58074 | -0.041729029 | -0.06429319 |
| Vmn2r20 | 3.395842 | 3.442063 | 3.4306 | 0.011501232 | -0.034719045 | 3.4011903 | 3.5877312 | 3.62952 | -0.04178824 | -0.228329073 |
| Slc39a11 | 8.128929 | 8.095057 | 8.1337 | -0.038637382 | -0.004765346 | 8.1059925 | 7.8630436 | 7.90484 | -0.041798794 | 0.201150154 |
| Atf6 | 10.69587 | 10.58641 | 10.532 | 0.054080846 | 0.163544528 | 10.704647 | 10.517783 | 10.5596 | -0.041819636 | 0.145043812 |
| Mrgprb2 | 1.878876 | 1.880336 | 1.9256 | -0.045260614 | -0.046721418 | 1.8044526 | 1.8076971 | 1.84954 | -0.041839938 | -0.045084429 |
| Gm48035 | 2.821103 | 2.707955 | 2.7009 | 0.007089139 | 0.120237078 | 2.7141189 | 2.6892531 | 2.73114 | -0.041885348 | -0.01701961 |
| Gm7517 | 1.587915 | 1.540196 | 1.5113 | 0.028942086 | 0.076660628 | 1.5683825 | 1.456499 | 1.49841 | -0.041913015 | 0.069970493 |
| Tulp3 | 7.479472 | 7.496629 | 7.4809 | 0.015744951 | -0.001412496 | 7.4327438 | 7.5036898 | 7.54561 | -0.041923499 | -0.1128695 |
| Gm16588 | 2.702744 | 2.501822 | 2.6654 | -0.163616197 | 0.037306164 | 2.5390753 | 2.6905049 | 2.73244 | -0.041938174 | -0.193367767 |
| Foxk2 | 9.772391 | 9.989444 | 9.8868 | 0.102623535 | -0.114429602 | 9.7558062 | 9.9617188 | 10.0037 | -0.041950322 | -0.247862949 |
| Epha1 | 1.705428 | 1.68091 | 1.8178 | -0.136900454 | -0.112381725 | 1.7505704 | 1.6725284 | 1.71449 | -0.041962067 | 0.036079893 |

| | | | | | | | | | | |
|---|---|---|---|---|---|---|---|---|---|---|
| Olfr1175-ps | 3.830186 | 3.76498 | 3.7549 | 0.010121416 | 0.075327095 | 3.7274522 | 3.8903188 | 3.93231 | -0.041987742 | -0.204854347 |
| Prpsap2 | 9.351483 | 9.369156 | 9.3656 | 0.003602031 | -0.014071333 | 9.4299405 | 9.3629669 | 9.40506 | -0.042096269 | 0.024877331 |
| Brd8 | 10.7218 | 10.73699 | 10.857 | -0.120248188 | -0.135440254 | 10.87627 | 10.867044 | 10.9091 | -0.04209798 | -0.032871906 |
| Rpa2 | 8.605776 | 8.550223 | 8.8588 | -0.308540614 | -0.252987574 | 8.9297938 | 8.6914906 | 8.7336 | -0.042106872 | 0.196196358 |
| Prdm6 | -0.020142 | 0.010127 | -0.0088 | 0.018882665 | -0.011386495 | -0.0013688 | -0.033582 | 0.00854 | -0.042118385 | -0.009905599 |
| Hapln4 | 3.042645 | 3.023393 | 3.0605 | -0.037081504 | -0.017829415 | 3.1173596 | 3.0467667 | 3.08889 | -0.042125919 | 0.028467011 |
| Cul4b | 9.881685 | 9.830159 | 9.8879 | -0.057709969 | -0.006183296 | 9.763365 | 9.7781881 | 9.82032 | -0.042132416 | -0.056955528 |
| Gm12394 | 0.003656 | -0.02849 | 0.0279 | -0.056379759 | -0.024235897 | -0.0538262 | -0.053556 | -0.0114 | -0.04213379 | -0.042403813 |
| Top3a | 8.745271 | 8.77697 | 8.7798 | -0.002784856 | -0.034483959 | 8.813192 | 8.8489666 | 8.89111 | -0.0421436 | -0.077918126 |
| Gm12435 | 1.695509 | 1.650974 | 1.6303 | 0.020720303 | 0.065255256 | 1.574117 | 1.6872963 | 1.72945 | -0.042156263 | -0.155335597 |
| Dnah12 | 2.035463 | 2.090423 | 2.0558 | 0.034619358 | -0.020340788 | 2.0992234 | 2.0381531 | 2.08035 | -0.042197087 | 0.01887326 |
| Slc16a1 | 9.3389 | 9.24371 | 9.3154 | -0.071665718 | 0.023524442 | 9.2321294 | 9.2829307 | 9.32514 | -0.042207937 | -0.093009203 |
| 1810010D01Rik | 1.138775 | 1.136906 | 1.1245 | 0.012378534 | 0.014247853 | 1.1980303 | 1.2013093 | 1.24353 | -0.042215927 | -0.045494926 |
| Ddx50 | 10.60795 | 10.52974 | 10.605 | -0.075523636 | 0.002686344 | 10.490052 | 10.536921 | 10.5791 | -0.042223096 | -0.089091541 |
| Tmod1 | 2.50838 | 2.587847 | 2.5854 | 0.002449511 | -0.077017008 | 2.6484354 | 2.5751534 | 2.61738 | -0.042228004 | 0.03105403 |
| Gm14086 | 1.830092 | 1.747961 | 1.7532 | -0.005218246 | 0.076911865 | 1.8640991 | 1.8277423 | 1.87002 | -0.042276695 | -0.005919954 |
| 1700088E04Rik | 2.68294 | 2.698233 | 2.7664 | -0.068206741 | -0.083499024 | 2.7965251 | 2.6856207 | 2.72796 | -0.042341883 | 0.068562475 |
| Rab36 | 2.825111 | 2.607458 | 2.6271 | -0.019690541 | 0.197961997 | 2.79942 | 2.7224302 | 2.76478 | -0.042352696 | 0.034637096 |
| Gm15013 | 1.38078 | 1.444656 | 1.473 | -0.028341291 | -0.092216935 | 1.4724903 | 1.4088673 | 1.45123 | -0.042366452 | 0.02125652 |
| Gm2398 | 0.598764 | 0.603788 | 0.6013 | 0.00247109 | -0.002552912 | 0.6156089 | 0.6223598 | 0.66475 | -0.042388496 | -0.049139428 |
| Ankrd1 | 2.199292 | 2.211408 | 2.2146 | -0.003145891 | -0.015261673 | 2.2340191 | 2.2060142 | 2.24841 | -0.042396877 | -0.014391961 |
| Smad2 | 10.01024 | 10.02371 | 10.045 | -0.021422206 | -0.034894306 | 9.8887365 | 9.853768 | 9.8962 | -0.042429173 | -0.007460649 |
| Cdh24 | 6.833921 | 7.02188 | 6.9666 | 0.055273706 | -0.132685441 | 6.9575459 | 6.9214563 | 6.96389 | -0.042434335 | -0.006344692 |
| Olfr645 | 0.497163 | 0.495786 | 0.4775 | 0.018300364 | 0.019677395 | 0.4488587 | 0.4746496 | 0.51709 | -0.042441165 | -0.068232016 |
| Atxn1l | 9.769615 | 9.629692 | 9.5707 | 0.058996922 | 0.198920206 | 9.6756433 | 9.5321104 | 9.57455 | -0.042442892 | 0.101090018 |
| Hbb-bh1 | 0.117278 | 0.069898 | 0.1092 | -0.039320212 | 0.008060251 | 0.1652087 | 0.0915416 | 0.13401 | -0.042468896 | 0.031198239 |
| Fam107a | 4.388231 | 4.345341 | 4.5298 | -0.184469722 | -0.141579451 | 3.9531182 | 4.0781212 | 4.1206 | -0.042482431 | -0.167485413 |
| Sh3yl1 | 3.562692 | 3.6585 | 3.647 | 0.011508032 | -0.084300288 | 3.7423109 | 3.7270608 | 3.76954 | -0.042483278 | -0.027233193 |
| Gm5129 | 2.715467 | 2.626597 | 2.6433 | -0.016670275 | 0.072200402 | 2.6747472 | 2.6513409 | 2.69384 | -0.04249866 | -0.019092383 |
| Traj43 | 1.086782 | 1.132631 | 1.1622 | -0.029616366 | -0.075465066 | 1.069762 | 1.0510732 | 1.09361 | -0.042535355 | -0.023846614 |
| Bmp8b | 1.681765 | 1.70747 | 1.6833 | 0.024160339 | -0.001544892 | 1.6630378 | 1.6542654 | 1.69682 | -0.042557634 | -0.033785241 |
| Kctd13 | 7.852191 | 7.648535 | 7.8477 | -0.199198592 | 0.004457375 | 8.095794 | 7.9420748 | 7.98463 | -0.042557736 | 0.111161439 |

| | | | | | | | | | | |
|---|---|---|---|---|---|---|---|---|---|---|
| Traj11 | 2.317895 | 2.278457 | 2.4067 | -0.128206639 | -0.088768551 | 2.2690034 | 2.3150688 | 2.35763 | -0.042561208 | -0.088626681 |
| Traj38 | 0.484641 | 0.467295 | 0.5015 | -0.03421229 | -0.016866289 | 0.4447893 | 0.4895338 | 0.53215 | -0.042616103 | -0.087360642 |
| Rmi1 | 9.519091 | 9.545401 | 9.5773 | -0.031944222 | -0.058254111 | 9.7053549 | 9.5617185 | 9.60435 | -0.04263445 | 0.101001917 |
| Ascc1 | 8.658841 | 8.627482 | 8.6887 | -0.06126116 | -0.029902029 | 8.6327042 | 8.567745 | 8.61039 | -0.042648435 | 0.022310744 |
| Creb3l4 | 1.750288 | 1.610149 | 1.6923 | -0.082152839 | 0.057986646 | 1.7684441 | 1.5879587 | 1.63062 | -0.042663993 | 0.137821335 |
| Ttc36 | 1.574087 | 1.493423 | 1.6056 | -0.112139384 | -0.03147506 | 1.4765868 | 1.5428719 | 1.5856 | -0.042728072 | -0.109013188 |
| Wnk4 | 2.834922 | 3.045488 | 2.84 | 0.205444591 | -0.005121114 | 2.968462 | 2.8333673 | 2.8761 | -0.042729383 | 0.092365254 |
| Cds2 | 11.57407 | 11.61136 | 11.578 | 0.033702626 | -0.00358096 | 11.624699 | 11.664727 | 11.7075 | -0.04281191 | -0.082840189 |
| Asic2 | 0.975877 | 0.994416 | 1.0143 | -0.019906244 | -0.038444846 | 0.912614 | 0.941097 | 0.98393 | -0.042835141 | -0.071318114 |
| 1700024J04Rik | 1.560326 | 1.616461 | 1.6724 | -0.055972429 | -0.112107362 | 1.689771 | 1.6492006 | 1.69216 | -0.042960752 | -0.002390304 |
| Ndufa10 | 9.992637 | 10.09602 | 10.118 | -0.021881047 | -0.125266151 | 10.202396 | 10.079426 | 10.1224 | -0.042997224 | 0.079972313 |
| Chac1 | 1.945901 | 1.845891 | 1.824 | 0.021929333 | 0.121939002 | 1.8146541 | 1.8367784 | 1.87979 | -0.043012203 | -0.065136502 |
| Sppl3 | 9.513828 | 9.640174 | 9.6218 | 0.01841018 | -0.107936191 | 9.4856599 | 9.6357798 | 9.6788 | -0.043024019 | -0.193143981 |
| Rangrf | 6.067917 | 5.994828 | 6.0222 | -0.027387212 | 0.045701369 | 6.1299285 | 6.0798232 | 6.12286 | -0.043038419 | 0.007066853 |
| Usp49 | 7.316095 | 7.174722 | 7.402 | -0.227318898 | -0.085945881 | 7.3419615 | 7.3452395 | 7.38828 | -0.043041791 | -0.046319759 |
| Tex22 | 1.49474 | 1.543792 | 1.5694 | -0.025646709 | -0.074698911 | 1.5669845 | 1.5800327 | 1.62309 | -0.043055296 | -0.056103457 |
| Slc22a19 | 1.41345 | 1.414939 | 1.4478 | -0.032835901 | -0.034324938 | 1.3859034 | 1.5097462 | 1.55282 | -0.04307593 | -0.166918693 |
| A630095N17Rik | 0.710528 | 0.678633 | 0.7107 | -0.032086739 | -0.000191718 | 0.7148431 | 0.6736857 | 0.71683 | -0.043145234 | -0.001987876 |
| Rdh9 | 1.704498 | 1.587604 | 1.5696 | 0.018039108 | 0.134932975 | 1.6777712 | 1.6815226 | 1.72467 | -0.04314608 | -0.046897495 |
| Adamts13 | 7.506165 | 7.369506 | 7.6045 | -0.234963666 | -0.098305108 | 7.2949212 | 7.2821817 | 7.32535 | -0.043172872 | -0.030433317 |
| Prph | 2.21849 | 2.21305 | 2.2592 | -0.04616446 | -0.040724171 | 2.0801809 | 2.1684434 | 2.21165 | -0.043211454 | -0.131473953 |
| Gm12439 | 0.540658 | 0.530854 | 0.5544 | -0.023550535 | -0.013746386 | 0.5255756 | 0.532083 | 0.5753 | -0.043218965 | -0.049726359 |
| Vmn2r-ps62 | 0.117534 | 0.111738 | 0.1245 | -0.012779245 | -0.006983606 | 0.090181 | 0.0926743 | 0.13594 | -0.043261376 | -0.045754644 |
| Gm13608 | 1.246392 | 1.216861 | 1.2512 | -0.034350705 | -0.004820201 | 1.230018 | 1.2129965 | 1.25627 | -0.043277017 | -0.026255547 |
| Pnpla6 | 9.221794 | 9.243907 | 9.3751 | -0.13123508 | -0.153348947 | 9.5205973 | 9.6174728 | 9.66078 | -0.043305786 | -0.140181288 |
| Mtmr4 | 10.42198 | 10.55155 | 10.495 | 0.056892484 | -0.072679656 | 10.37732 | 10.368151 | 10.4115 | -0.043330942 | -0.034161855 |
| Hebp2 | 3.119593 | 3.127011 | 3.1866 | -0.059611013 | -0.067028675 | 3.2690329 | 3.1815148 | 3.22491 | -0.043397115 | 0.044120995 |
| Rspo3 | 1.366691 | 1.366375 | 1.4148 | -0.04837555 | -0.048059624 | 1.3314091 | 1.3562697 | 1.39967 | -0.043401325 | -0.068261917 |
| Syt3 | 1.114243 | 1.073219 | 1.1086 | -0.035407024 | 0.0056174 | 1.0426489 | 1.0588255 | 1.10224 | -0.043409641 | -0.059586186 |
| Vmn1r39 | 1.293275 | 1.279263 | 1.2874 | -0.008174459 | 0.005837297 | 1.3344948 | 1.3030491 | 1.34648 | -0.043428534 | -0.011982853 |
| Wdr19 | 6.372822 | 6.137895 | 6.5325 | -0.394590431 | -0.159663624 | 6.3991711 | 6.3818854 | 6.42532 | -0.043438334 | -0.026152633 |
| Cox20 | 6.956131 | 6.978606 | 7.0125 | -0.033858326 | -0.056332699 | 7.0322173 | 7.0322444 | 7.07571 | -0.043464056 | -0.043491087 |

| | | | | | | | | | | |
|---|---|---|---|---|---|---|---|---|---|---|
| Traj14 | 1.97617 | 1.969847 | 1.8424 | 0.127409679 | 0.13373343 | 1.8936808 | 1.9074764 | 1.95099 | -0.043513825 | -0.057309393 |
| Igsf11 | 3.395835 | 3.542551 | 3.6412 | -0.098686582 | -0.245402314 | 3.4789172 | 3.5360141 | 3.57954 | -0.043522411 | -0.100619286 |
| Gm12403 | 1.587655 | 1.60598 | 1.6405 | -0.034496825 | -0.052822303 | 1.6011508 | 1.6824346 | 1.72596 | -0.043529657 | -0.124813538 |
| Crabp2 | 0.859828 | 0.840327 | 0.8107 | 0.02967546 | 0.049177093 | 0.8417075 | 0.8103866 | 0.85393 | -0.04354775 | -0.012226817 |
| Shank2 | 1.833413 | 1.790161 | 1.8831 | -0.092902959 | -0.049650926 | 1.9159682 | 1.7796142 | 1.82317 | -0.043555696 | 0.09279831 |
| Glyctk | 7.464442 | 7.596374 | 7.3838 | 0.212525812 | 0.080593715 | 7.6764578 | 7.4898446 | 7.53346 | -0.043615786 | 0.142997407 |
| Lhx8 | 2.585234 | 2.587756 | 2.7133 | -0.125514908 | -0.128036959 | 2.6218179 | 2.7027488 | 2.74642 | -0.043667423 | -0.124598314 |
| Ficd | 7.198306 | 7.075384 | 7.1076 | -0.032230513 | 0.090690672 | 7.0591959 | 6.9897095 | 7.03339 | -0.043682139 | 0.025804226 |
| Rab33a | 0.999475 | 1.059262 | 1.0516 | 0.007661515 | -0.052126175 | 1.0798586 | 1.0458018 | 1.08955 | -0.043746563 | -0.009689834 |
| Oip5 | 4.181633 | 4.080605 | 4.284 | -0.203431454 | -0.10240359 | 4.2892999 | 4.345132 | 4.3889 | -0.043763123 | -0.09959518 |
| Frk | 2.144938 | 2.035447 | 2.0625 | -0.027101297 | 0.082390541 | 1.9996299 | 1.9451918 | 1.98897 | -0.043774228 | 0.010663941 |
| Pir | 2.566336 | 2.510862 | 2.6601 | -0.149201715 | -0.093727969 | 2.5705816 | 2.5747342 | 2.61851 | -0.04378079 | -0.047933353 |
| Dcst2 | 1.756758 | 1.686932 | 1.7991 | -0.112201667 | -0.042376094 | 1.6624998 | 1.6492252 | 1.69301 | -0.043786272 | -0.030511696 |
| Vmn1r60 | 3.397773 | 3.304305 | 3.2838 | 0.020487824 | 0.113955362 | 3.4197608 | 3.4100719 | 3.45387 | -0.043795123 | -0.034106176 |
| Ttpa | 1.549888 | 1.458923 | 1.5708 | -0.111857136 | -0.020892851 | 1.3781493 | 1.3784953 | 1.4223 | -0.043802462 | -0.044148413 |
| Sel1l3 | 2.971774 | 2.881504 | 2.9095 | -0.028000479 | 0.062269498 | 3.1603428 | 2.8098023 | 2.85363 | -0.043824588 | 0.306715912 |
| Olig2 | 0.987153 | 0.940169 | 0.9365 | 0.003661256 | 0.05064529 | 0.8676189 | 0.8900735 | 0.93393 | -0.043852195 | -0.066306856 |
| Olfr920 | 4.732315 | 4.558539 | 4.5073 | 0.051269983 | 0.225045983 | 4.589358 | 4.6345779 | 4.67845 | -0.043875401 | -0.089095279 |
| Lbhd2 | 1.095122 | 1.033657 | 1.0324 | 0.001213695 | 0.062678555 | 0.981918 | 1.0352007 | 1.07908 | -0.043880717 | -0.097163414 |
| Vmn2r78 | 1.747998 | 1.708131 | 1.696 | 0.012110329 | 0.051977078 | 1.7678433 | 1.8035461 | 1.84743 | -0.043885103 | -0.079587942 |
| Retreg3 | 11.3291 | 11.44235 | 11.414 | 0.028459516 | -0.08478952 | 11.500759 | 11.355155 | 11.399 | -0.043891602 | 0.101711913 |
| Tmx1 | 10.27631 | 10.27712 | 10.311 | -0.034237704 | -0.035056202 | 10.190291 | 10.189935 | 10.2339 | -0.0439392 | -0.043582825 |
| Fam209 | 2.446326 | 2.410842 | 2.4121 | -0.001242562 | 0.034241299 | 2.5331614 | 2.4365481 | 2.48052 | -0.043973322 | 0.052639956 |
| Cog3 | 9.745538 | 9.796064 | 9.798 | -0.001965363 | -0.052491664 | 9.648205 | 9.7316029 | 9.77558 | -0.043973588 | -0.127371469 |
| Epb42 | 1.765749 | 1.702015 | 1.7482 | -0.046150869 | 0.01758241 | 1.804832 | 1.7381107 | 1.78212 | -0.044013002 | 0.022708265 |
| Calb1 | 2.504136 | 2.705734 | 2.6075 | 0.098232539 | -0.103365609 | 2.641508 | 2.6093325 | 2.65336 | -0.044027772 | -0.0118523 |
| 6430531B16Rik | 1.333993 | 1.30013 | 1.3495 | -0.04937219 | -0.015509018 | 1.3297871 | 1.2934812 | 1.33754 | -0.044056879 | -0.007750942 |
| Ighv4-1 | 7.773225 | 7.796609 | 7.5647 | 0.23192213 | 0.208538198 | 7.7828085 | 7.3668647 | 7.41093 | -0.044064681 | 0.371879103 |
| Pip4k2a | 11.47808 | 11.44149 | 11.428 | 0.013694652 | 0.050288382 | 11.296304 | 11.376181 | 11.4203 | -0.044079074 | -0.123955898 |
| Gm13339 | 3.01069 | 3.077683 | 3.1243 | -0.046642977 | -0.113636136 | 3.1496894 | 3.118763 | 3.16286 | -0.044093666 | -0.013167297 |
| Tmem30b | 1.876697 | 1.655652 | 1.7209 | -0.06524207 | 0.155802737 | 1.8353713 | 1.7639765 | 1.80827 | -0.044289951 | 0.027104871 |
| Bmp4 | 3.354105 | 3.397373 | 3.6113 | -0.213946375 | -0.257214421 | 3.2521078 | 3.178824 | 3.22312 | -0.044299031 | 0.028984768 |

| | | | | | | | | | | |
|---|---|---|---|---|---|---|---|---|---|---|
| Hira | 9.77916 | 9.847955 | 9.835 | 0.01292275 | -0.055872588 | 9.9194209 | 9.8562455 | 9.9006 | -0.044357845 | 0.018817539 |
| Mtx3 | 7.676595 | 7.580261 | 7.7269 | -0.146656799 | -0.050323058 | 8.0832873 | 7.9741549 | 8.01856 | -0.044408183 | 0.06472418 |
| Gm14411 | 2.219802 | 2.234968 | 2.1903 | 0.04465111 | 0.029484872 | 2.1312424 | 2.1402605 | 2.18468 | -0.0444243 | -0.053442371 |
| Kctd7 | 6.07273 | 5.933713 | 6.1076 | -0.173861887 | -0.034845149 | 6.0480199 | 5.9052169 | 5.94965 | -0.044429712 | 0.09837325 |
| Tpm2 | 5.789273 | 6.073826 | 5.9669 | 0.106957433 | -0.177595828 | 6.1068247 | 5.6990439 | 5.74347 | -0.044430918 | 0.363349914 |
| Gm20537 | 0.622977 | 0.647064 | 0.6017 | 0.0453649 | 0.021277726 | 0.6836358 | 0.6286528 | 0.67311 | -0.044461487 | 0.010521462 |
| Zkscan4 | 1.629999 | 1.625565 | 1.6938 | -0.068186601 | -0.063752876 | 1.5173679 | 1.5523623 | 1.59686 | -0.044499465 | -0.07949379 |
| Vmn1r230 | 1.50087 | 1.523759 | 1.5543 | -0.030574331 | -0.053463585 | 1.4494301 | 1.5460536 | 1.59055 | -0.044500019 | -0.141123535 |
| Smyd2 | 7.540473 | 7.476818 | 7.4622 | 0.014588665 | 0.078243382 | 7.3630755 | 7.3860826 | 7.43059 | -0.044509667 | -0.067516737 |
| Apol8 | 2.344937 | 2.271337 | 2.4307 | -0.159406232 | -0.085805568 | 2.1078487 | 2.1270095 | 2.17152 | -0.044513721 | -0.063674444 |
| Pcdhb4 | 1.893593 | 1.841973 | 1.8921 | -0.050086288 | 0.001534455 | 1.8085616 | 1.7891919 | 1.83371 | -0.04451932 | -0.025149593 |
| 6430548M08Rik | 6.813975 | 6.75149 | 6.7739 | -0.022451136 | 0.040034704 | 6.5695159 | 6.3397573 | 6.38433 | -0.04457042 | 0.185188258 |
| 4931406P16Rik | 9.725586 | 9.596915 | 9.734 | -0.137106246 | -0.00843522 | 9.3121518 | 9.4851051 | 9.52973 | -0.04462861 | -0.217581857 |
| Gm42543 | 1.299032 | 1.370593 | 1.3389 | 0.031688808 | -0.039871671 | 1.3301517 | 1.3016715 | 1.34636 | -0.04468924 | -0.016208968 |
| Olfr531 | 0.588803 | 0.625032 | 0.623 | 0.002078647 | -0.034150969 | 0.6280123 | 0.587016 | 0.63171 | -0.044692184 | -0.003695867 |
| Eva1c | 3.307289 | 3.249713 | 3.1406 | 0.109158494 | 0.166735227 | 3.2862936 | 3.1515066 | 3.19621 | -0.044707622 | 0.090079383 |
| Rpl4 | 15.46044 | 15.44058 | 15.461 | -0.020307376 | -0.000454832 | 15.465163 | 15.417962 | 15.4627 | -0.044744344 | 0.002457236 |
| Mypop | 3.981279 | 4.052763 | 3.9656 | 0.087135687 | 0.015651981 | 3.9888523 | 3.9066672 | 3.95143 | -0.044758028 | 0.037427044 |
| Gm7729 | 2.594156 | 2.56856 | 2.5877 | -0.019155332 | 0.006439851 | 2.5995924 | 2.6302505 | 2.67505 | -0.044802748 | -0.075460835 |
| Pnldc1 | 3.034738 | 3.028865 | 3.1762 | -0.147304632 | -0.141431045 | 3.0496789 | 3.1650523 | 3.20991 | -0.044861163 | -0.160234582 |
| Glb1l | 6.449373 | 6.463529 | 6.6248 | -0.161317441 | -0.175474248 | 6.6252977 | 6.4924247 | 6.53729 | -0.04486961 | 0.088003399 |
| Plcg2 | 11.9352 | 11.98814 | 11.94 | 0.048392736 | -0.004541018 | 12.254912 | 11.904427 | 11.9493 | -0.044885164 | 0.305599851 |
| Trav1 | 3.247137 | 3.282435 | 3.3302 | -0.047759175 | -0.083057357 | 3.0837224 | 3.1853436 | 3.23023 | -0.0448852 | -0.14650638 |
| Avpr2 | 1.034807 | 0.998579 | 1.0346 | -0.035977204 | 0.000250063 | 1.0531524 | 1.0017326 | 1.04664 | -0.044908575 | 0.006511209 |
| Wnt3 | 2.4636 | 2.545757 | 2.4684 | 0.07736278 | -0.004793993 | 2.3832842 | 2.3387031 | 2.38369 | -0.044986686 | -0.000405498 |
| Ccdc163 | 7.518926 | 7.382414 | 7.5588 | -0.176356569 | -0.039844942 | 7.8518623 | 7.7772199 | 7.82224 | -0.045019496 | 0.029622838 |
| Gpr161 | 1.812131 | 1.887031 | 1.8463 | 0.040765276 | -0.034134924 | 1.7861298 | 1.7025128 | 1.74756 | -0.045042381 | 0.038574574 |
| Sbf1 | 11.79512 | 11.92534 | 12.017 | -0.091671768 | -0.221894548 | 12.04853 | 12.084679 | 12.1298 | -0.045106582 | -0.081255595 |
| Ntn5 | 1.379704 | 1.503089 | 1.3847 | 0.118404053 | -0.004980545 | 1.3495599 | 1.4090757 | 1.45419 | -0.045119164 | -0.104634993 |
| Ucp3 | 1.050487 | 1.008137 | 1.0599 | -0.051715085 | -0.009365047 | 1.0327491 | 1.0188344 | 1.06399 | -0.045150831 | -0.031236141 |
| Trim67 | 2.36674 | 2.372915 | 2.388 | -0.015061864 | -0.021236754 | 2.4361549 | 2.4465172 | 2.49167 | -0.045152515 | -0.055514802 |
| Rpl36a-ps1 | 2.541822 | 2.753879 | 2.5547 | 0.19920103 | -0.012855787 | 2.6752775 | 2.6057663 | 2.65092 | -0.04515502 | 0.024356194 |

| | | | | | | | | | | |
|---|---|---|---|---|---|---|---|---|---|---|
| Dclk3 | 0.557258 | 0.570395 | 0.4996 | 0.070746576 | 0.057609092 | 0.4942391 | 0.488089 | 0.53328 | -0.045188327 | -0.039038216 |
| Gm11703 | 1.916542 | 1.773138 | 1.8221 | -0.048914022 | 0.094490047 | 1.8209421 | 1.7719744 | 1.81739 | -0.045416454 | 0.003551263 |
| Igkv1-108 | 0.815137 | 0.779743 | 0.8125 | -0.032741595 | 0.002652807 | 0.8574041 | 0.796168 | 0.84159 | -0.045417831 | 0.015818308 |
| Ldhal6b | 2.097189 | 2.130459 | 2.0662 | 0.064292127 | 0.031022655 | 2.1607368 | 2.1613053 | 2.20676 | -0.045455588 | -0.046024123 |
| Zfp280d | 10.03345 | 9.973939 | 10.094 | -0.120279184 | -0.060772731 | 9.8490131 | 10.104593 | 10.1501 | -0.045462064 | -0.301041903 |
| Ctsk | 4.454086 | 4.431661 | 4.4075 | 0.024150861 | 0.04657555 | 4.3659035 | 4.3073389 | 4.35282 | -0.045483359 | 0.013081206 |
| Elf3 | 0.52579 | 0.531233 | 0.5634 | -0.032208686 | -0.037651715 | 0.6848072 | 0.5841778 | 0.62971 | -0.045536227 | 0.055093124 |
| Gm47632 | 3.379304 | 3.641711 | 3.541 | 0.100686283 | -0.161721522 | 3.5712064 | 3.7661791 | 3.81173 | -0.04555213 | -0.240524828 |
| Zfp94 | 6.883608 | 6.868713 | 7.0383 | -0.16958024 | -0.154686178 | 7.142228 | 7.109086 | 7.15468 | -0.045595805 | -0.012453783 |
| Lbx2 | 1.125853 | 1.11812 | 1.1421 | -0.023993735 | -0.016261162 | 1.1833493 | 1.108582 | 1.15421 | -0.045625605 | 0.029141687 |
| Stoml3 | 2.111986 | 2.128295 | 2.1408 | -0.012536998 | -0.028845807 | 2.1956831 | 2.2536336 | 2.29926 | -0.045627588 | -0.103578091 |
| Tent5b | 0.966905 | 1.009746 | 0.9822 | 0.027550042 | -0.015290745 | 0.986362 | 0.9339294 | 0.97956 | -0.045632494 | 0.006800081 |
| Ctns | 8.28923 | 8.207768 | 8.3253 | -0.117527147 | -0.036065318 | 8.4177058 | 8.3082497 | 8.3539 | -0.045651766 | 0.063804312 |
| Vmn1r43 | 1.725095 | 1.693041 | 1.7124 | -0.019318718 | 0.012735258 | 1.7055331 | 1.7489742 | 1.79468 | -0.045708266 | -0.089149377 |
| Syt12 | 2.688462 | 2.772359 | 2.7233 | 0.04902119 | -0.034875423 | 2.7029295 | 2.6542035 | 2.69997 | -0.045762798 | 0.002963244 |
| Olfr506 | 1.483369 | 1.531216 | 1.5295 | 0.001707639 | -0.046139042 | 1.4746761 | 1.5573165 | 1.6031 | -0.045781484 | -0.128421917 |
| Ube3a | 11.12557 | 11.12279 | 11.06 | 0.062736294 | 0.065509874 | 11.117253 | 11.077816 | 11.1236 | -0.045781774 | -0.00634521 |
| Lrrc14 | 7.969474 | 8.021404 | 8.0961 | -0.074672185 | -0.126602434 | 8.1662296 | 8.035489 | 8.08127 | -0.045784349 | 0.084956246 |
| Cdc16 | 9.33552 | 9.246803 | 9.3251 | -0.078336551 | 0.010379772 | 9.364826 | 9.3162587 | 9.36204 | -0.045785746 | 0.002781477 |
| Gm16204 | 1.985766 | 2.026803 | 2.0768 | -0.049989064 | -0.091026908 | 2.1073666 | 1.985382 | 2.0312 | -0.045814247 | 0.07617033 |
| Gm3667 | 2.429677 | 2.355279 | 2.4154 | -0.060082019 | 0.014316449 | 2.3092872 | 2.3527412 | 2.39857 | -0.045823793 | -0.089277869 |
| 9930104L06Rik | 8.397705 | 8.405549 | 8.2452 | 0.160339133 | 0.152494432 | 8.4943669 | 8.4540228 | 8.49986 | -0.045838014 | -0.005493919 |
| 1500009C09Rik | 0.604186 | 0.641596 | 0.6145 | 0.027121281 | -0.010289207 | 0.6104134 | 0.5965596 | 0.6424 | -0.045840445 | -0.031986642 |
| Klhl28 | 9.224427 | 9.333988 | 9.3036 | 0.030343309 | -0.079217397 | 9.3648952 | 9.2030171 | 9.2489 | -0.045880684 | 0.115997468 |
| Supt6 | 11.75002 | 11.7442 | 11.719 | 0.025515726 | 0.031330496 | 11.736321 | 11.741074 | 11.787 | -0.045884508 | -0.050636865 |
| 6430571L13Rik | 3.198517 | 3.263219 | 3.2748 | -0.011608088 | -0.076310304 | 3.3908298 | 3.3176536 | 3.36356 | -0.045905594 | 0.027270648 |
| Cog2 | 9.396079 | 9.348049 | 9.4146 | -0.066529446 | -0.018499131 | 9.5185246 | 9.4822277 | 9.52814 | -0.045910628 | -0.00961379 |
| Cops4 | 9.781499 | 9.730933 | 9.8061 | -0.075166557 | -0.02460067 | 9.7237973 | 9.6758049 | 9.72172 | -0.045912415 | 0.002079996 |
| Gm9903 | 1.737436 | 1.646957 | 1.7672 | -0.120269352 | -0.029790303 | 1.7644872 | 1.694968 | 1.7409 | -0.045927537 | 0.023591651 |
| Hoxc6 | 2.026631 | 2.099621 | 2.1605 | -0.060841628 | -0.133831382 | 2.0443279 | 2.03062 | 2.07655 | -0.045928712 | -0.032220792 |
| Plcd4 | 2.632006 | 2.585027 | 2.6353 | -0.050302073 | -0.003322335 | 2.5226508 | 2.6237745 | 2.66973 | -0.045960298 | -0.147084014 |
| Pcid2 | 9.325844 | 9.328653 | 9.2459 | 0.082723863 | 0.079915137 | 9.2488578 | 9.2135653 | 9.25955 | -0.045989126 | -0.010696625 |

| | | | | | | | | | | |
|---|---|---|---|---|---|---|---|---|---|---|
| Gm4922 | 3.149338 | 3.149126 | 3.1524 | -0.003262973 | -0.003051093 | 3.3162019 | 3.1919938 | 3.23798 | -0.045989941 | 0.078218104 |
| Morf4l1-ps1 | 8.921558 | 8.76394 | 8.8384 | -0.074412612 | 0.083205418 | 8.5922901 | 8.8700775 | 8.91609 | -0.046017262 | -0.323804625 |
| Dhrs3 | 5.878739 | 5.964082 | 6.0357 | -0.071602637 | -0.156944949 | 5.9467222 | 5.9656871 | 6.01177 | -0.046086814 | -0.065051697 |
| Cs | 11.80748 | 11.76683 | 11.839 | -0.072216352 | -0.031569868 | 11.969684 | 11.766309 | 11.8124 | -0.046091308 | 0.157284246 |
| Tchp | 7.940661 | 8.040694 | 8.1401 | -0.09945247 | -0.199485287 | 8.3303155 | 8.313742 | 8.35984 | -0.04610101 | -0.029527505 |
| Nxnl1 | 0.017871 | 0.033757 | 0.031 | 0.002753893 | -0.013132358 | -0.0084715 | -0.00235 | 0.04376 | -0.046104853 | -0.052226704 |
| Lpcat3 | 8.74688 | 8.791921 | 8.7415 | 0.050382647 | 0.005341831 | 8.7882648 | 8.8420214 | 8.88816 | -0.046139473 | -0.099896011 |
| A330044P14Rik | 3.621463 | 3.590771 | 3.6365 | -0.045700825 | -0.015009365 | 3.560905 | 3.6626019 | 3.70875 | -0.04614806 | -0.147844936 |
| Rps6-ps2 | 0.625463 | 0.638351 | 0.6489 | -0.010569757 | -0.023458126 | 0.5907307 | 0.6366724 | 0.68287 | -0.046197812 | -0.092139506 |
| H1fx | 1.796875 | 1.78924 | 1.8065 | -0.01724546 | -0.009611103 | 1.7426652 | 1.7849251 | 1.83117 | -0.046249541 | -0.088509496 |
| Gm43028 | 1.796595 | 1.883609 | 1.7378 | 0.14576122 | 0.058747304 | 1.8108767 | 1.7702012 | 1.81646 | -0.046254499 | -0.005579037 |
| Mthfd1l | 10.16071 | 10.20818 | 10.135 | 0.072865801 | 0.025401642 | 10.200413 | 10.159065 | 10.2053 | -0.046275708 | -0.004927461 |
| Vmn1r76 | 5.193993 | 5.145902 | 5.1423 | 0.003611677 | 0.051702469 | 5.1148622 | 5.3422855 | 5.38857 | -0.046287468 | -0.273710783 |
| Gm12879 | 2.266989 | 2.156902 | 2.3252 | -0.168326273 | -0.05823948 | 2.1891636 | 2.2109731 | 2.25727 | -0.046293082 | -0.068102548 |
| Fam222b | 7.378111 | 7.286096 | 7.3432 | -0.057140089 | 0.034874248 | 7.2720391 | 7.2624202 | 7.30874 | -0.046323624 | -0.036704708 |
| Anapc7 | 9.33133 | 9.487062 | 9.3932 | 0.093887034 | -0.061844955 | 9.5069343 | 9.4595745 | 9.50592 | -0.046343249 | 0.001016473 |
| Egfr | 5.668759 | 5.57164 | 5.3682 | 0.203425287 | 0.30054396 | 4.9924175 | 5.2165125 | 5.2629 | -0.0463911 | -0.270486078 |
| Ms4a12 | 2.290251 | 2.230489 | 2.2898 | -0.059324728 | 0.000437991 | 2.2975247 | 2.2846276 | 2.33104 | -0.046416504 | -0.033519446 |
| Catsperg2 | 1.620709 | 1.58842 | 1.5728 | 0.015658154 | 0.04794714 | 1.5569377 | 1.6052608 | 1.6517 | -0.046443021 | -0.094766155 |
| Ikbkb | 11.8132 | 11.8126 | 11.839 | -0.026620698 | -0.0260193 | 11.996447 | 11.927535 | 11.974 | -0.04645469 | 0.022457289 |
| Ift43 | 4.539882 | 4.34177 | 4.5425 | -0.20077263 | -0.002660955 | 4.378115 | 4.3231118 | 4.36957 | -0.046462077 | 0.008541113 |
| Trav10d | 3.219318 | 3.324715 | 3.24 | 0.084731061 | -0.020666017 | 3.1700915 | 3.2012578 | 3.24773 | -0.046473628 | -0.077639928 |
| Trav4d-3 | 2.624765 | 2.567752 | 2.6214 | -0.053696409 | 0.003316424 | 2.5184251 | 2.523529 | 2.57004 | -0.046515725 | -0.05161963 |
| Trav4n-3 | 2.624765 | 2.567752 | 2.6214 | -0.053696409 | 0.003316424 | 2.5184251 | 2.523529 | 2.57004 | -0.046515725 | -0.05161963 |
| Kcnk9 | 0.11468 | 0.133867 | 0.1048 | 0.029082135 | 0.009896081 | 0.093865 | 0.0901311 | 0.13668 | -0.046551273 | -0.042817426 |
| Fbxo48 | 4.953495 | 4.640315 | 4.87 | -0.22965311 | 0.083526545 | 4.8489393 | 4.8584887 | 4.90508 | -0.046593625 | -0.056142985 |
| Gm19046 | 2.487144 | 2.488765 | 2.4359 | 0.052821738 | 0.051201516 | 2.495848 | 2.4653473 | 2.51195 | -0.046605465 | -0.016104734 |
| Mllt11 | 6.693368 | 6.84076 | 6.6676 | 0.173149977 | 0.025757956 | 6.9668412 | 6.8942276 | 6.94085 | -0.046625295 | 0.025988287 |
| Gm48991 | 1.063704 | 1.010745 | 1.0329 | -0.022141893 | 0.030816901 | 1.0622243 | 0.9870777 | 1.03381 | -0.046730346 | 0.028416206 |
| Gm13083 | 0.562671 | 0.59106 | 0.5946 | -0.003587179 | -0.031976919 | 0.5639999 | 0.5844184 | 0.63121 | -0.046789465 | -0.067207925 |
| Syce1 | 0.714226 | 0.733513 | 0.7405 | -0.00698775 | -0.026274574 | 0.7406451 | 0.7417724 | 0.78856 | -0.046791164 | -0.047918455 |
| Cbfa2t2 | 9.045563 | 9.057498 | 9.2131 | -0.155594259 | -0.167529359 | 8.9707546 | 9.0911595 | 9.13796 | -0.046795866 | -0.167200815 |

| | | | | | | | | | | |
|---|---|---|---|---|---|---|---|---|---|---|
| Hist1h2br | 1.835867 | 1.69727 | 1.9178 | -0.2205572 | -0.081960436 | 1.957378 | 1.8579932 | 1.90479 | -0.0467998 | 0.052585021 |
| Prss35 | 1.398677 | 1.375024 | 1.4494 | -0.074413239 | -0.05076005 | 1.380768 | 1.4124911 | 1.45932 | -0.046827678 | -0.078550708 |
| Trav9-4 | 4.530138 | 4.300899 | 4.3238 | -0.022946435 | 0.206292209 | 4.2897342 | 4.3064544 | 4.3533 | -0.046848818 | -0.063569009 |
| Atp13a4 | 0.744542 | 0.717219 | 0.7769 | -0.059661148 | -0.032337392 | 0.7393535 | 0.7072603 | 0.75413 | -0.046871738 | -0.014778521 |
| Gm17771 | 1.69575 | 1.693779 | 1.7055 | -0.011715962 | -0.009745517 | 1.5865131 | 1.6192439 | 1.66615 | -0.046902126 | -0.079632935 |
| Ptges2 | 7.149026 | 7.073472 | 7.2979 | -0.224454012 | -0.148900001 | 7.4303559 | 7.1002669 | 7.14719 | -0.046926319 | 0.283162698 |
| Tmem30c | 2.970748 | 2.943832 | 2.9735 | -0.029716406 | -0.002800239 | 3.0336055 | 3.0591472 | 3.10611 | -0.046958657 | -0.072500433 |
| Gm6654 | 0.927083 | 0.973017 | 0.9747 | -0.001634853 | -0.047569019 | 1.0194644 | 0.9536492 | 1.00062 | -0.046972446 | 0.018842775 |
| Mafa | 1.679432 | 1.709713 | 1.7827 | -0.072968676 | -0.103250406 | 1.6717497 | 1.6974852 | 1.74451 | -0.047024502 | -0.072760002 |
| Stambpl1 | 9.23857 | 9.241416 | 9.3024 | -0.061018542 | -0.063864453 | 9.1143255 | 9.2404214 | 9.28745 | -0.047025389 | -0.173121277 |
| Gm3550 | 2.149038 | 2.078638 | 2.2075 | -0.128825977 | -0.058426297 | 2.1694345 | 2.1386315 | 2.18567 | -0.047034957 | -0.016231901 |
| Snx12 | 9.28365 | 9.223522 | 9.2822 | -0.058725458 | 0.001402956 | 9.3122229 | 9.2275498 | 9.2746 | -0.047055196 | 0.037617887 |
| Gm10638 | 0.990494 | 0.960432 | 1.0181 | -0.057646723 | -0.027584296 | 1.0160234 | 0.978724 | 1.02579 | -0.047064883 | -0.009765461 |
| Ilf3 | 10.94739 | 10.98043 | 10.994 | -0.013087328 | -0.04612698 | 11.005574 | 11.196479 | 11.2436 | -0.0470816 | -0.237986582 |
| Olfr704 | 3.807844 | 3.764155 | 3.8378 | -0.073618769 | -0.029929782 | 3.7529901 | 3.7309176 | 3.77801 | -0.047090236 | -0.025017688 |
| Ankrd34a | 0.207646 | 0.156887 | 0.1481 | 0.008752019 | 0.059511253 | 0.2292452 | 0.1601054 | 0.2072 | -0.047094223 | 0.022045594 |
| T2 | 2.021701 | 1.97585 | 1.9263 | 0.049525201 | 0.095376234 | 1.9915457 | 2.0152804 | 2.06238 | -0.047097317 | -0.070832085 |
| Ift27 | 7.714771 | 7.602309 | 7.7877 | -0.185436855 | -0.072974504 | 7.9919393 | 7.7954981 | 7.84261 | -0.047109654 | 0.149331509 |
| Gm5171 | 0.608376 | 0.573242 | 0.5564 | 0.016801669 | 0.0519356 | 0.5745002 | 0.5744104 | 0.62153 | -0.047120675 | -0.047030894 |
| Rbm8a2 | 1.13915 | 1.109412 | 1.1706 | -0.061173106 | -0.031434495 | 1.2189816 | 1.1558264 | 1.20297 | -0.047139657 | 0.016015503 |
| Chtf18 | 6.927991 | 6.78622 | 7.0914 | -0.305169959 | -0.163398239 | 7.3579977 | 7.2564796 | 7.30363 | -0.047147825 | 0.054370248 |
| Agrp | 2.198255 | 2.097908 | 2.2243 | -0.126415675 | -0.026069127 | 2.2595411 | 2.1738982 | 2.22107 | -0.04717012 | 0.038472844 |
| Trmt44 | 7.164294 | 7.223406 | 7.1844 | 0.039052663 | -0.020059545 | 7.3183532 | 7.3467087 | 7.39392 | -0.047206819 | -0.075562324 |
| Vax2 | 1.427986 | 1.361696 | 1.4323 | -0.07056065 | -0.004270406 | 1.3412752 | 1.4006325 | 1.44785 | -0.047222198 | -0.106579528 |
| R3hdm1 | 10.50842 | 10.35725 | 10.457 | -0.099596596 | 0.051572582 | 10.346988 | 10.442987 | 10.4902 | -0.047244452 | -0.143243669 |
| Olfr798 | 1.319588 | 1.316307 | 1.374 | -0.057717428 | -0.054436309 | 1.441729 | 1.3303004 | 1.37755 | -0.04725363 | 0.064174995 |
| Nup93 | 9.530821 | 9.500535 | 9.5097 | -0.009130871 | 0.021155194 | 9.7303862 | 9.6228279 | 9.6701 | -0.047273145 | 0.060285234 |
| Vmn1r180 | 1.635184 | 1.595237 | 1.6242 | -0.029011986 | 0.01093517 | 1.7559074 | 1.6797412 | 1.72714 | -0.047397867 | 0.02876832 |
| H2-Eb1 | 14.81055 | 14.73884 | 14.787 | -0.04865038 | 0.02306175 | 15.262709 | 14.77707 | 14.8245 | -0.047409534 | 0.438229709 |
| Gcnt1 | 9.967723 | 9.868583 | 9.9951 | -0.126559888 | -0.027419486 | 9.7157672 | 9.6581958 | 9.70563 | -0.0474353 | 0.010136121 |
| Sil1 | 8.350508 | 8.086354 | 8.2496 | -0.163271442 | 0.100882113 | 8.3250303 | 8.2030442 | 8.25052 | -0.047474651 | 0.074511507 |
| Heatr3 | 9.217526 | 9.151385 | 9.1649 | -0.013506337 | 0.052635531 | 9.0614501 | 9.1421803 | 9.18969 | -0.047509473 | -0.128239645 |

| | | | | | | | | | | |
|---|---|---|---|---|---|---|---|---|---|---|
| Gm4994 | 0.629084 | 0.664378 | 0.6367 | 0.027689831 | -0.007604689 | 0.5896495 | 0.6260972 | 0.67367 | -0.047576095 | -0.084023743 |
| Pou4f1 | 2.221906 | 2.172586 | 2.2278 | -0.055234055 | -0.005913438 | 2.1826661 | 2.167896 | 2.2155 | -0.04760656 | -0.032836447 |
| Gm5869 | 2.789286 | 2.870328 | 2.7466 | 0.12376526 | 0.042722373 | 2.9047292 | 2.8438475 | 2.89146 | -0.047614614 | 0.013267022 |
| Cntn2 | 1.497405 | 1.610107 | 1.4574 | 0.152731872 | 0.040029358 | 1.3498884 | 1.4227533 | 1.47038 | -0.047628252 | -0.12049313 |
| Gm43720 | 0.971981 | 0.918272 | 0.9203 | -0.002014868 | 0.051693953 | 0.9079182 | 0.8950958 | 0.94273 | -0.047633789 | -0.034811349 |
| Echdc2 | 1.841665 | 1.882876 | 1.8207 | 0.062204462 | 0.020993439 | 1.9303618 | 1.8344964 | 1.88216 | -0.047659377 | 0.048205953 |
| Sesn2 | 7.47744 | 7.459977 | 7.4425 | 0.01751524 | 0.03497861 | 7.6145218 | 7.4233313 | 7.47101 | -0.047677615 | 0.143512817 |
| Stx1a | 6.796018 | 6.85452 | 6.9805 | -0.125986984 | -0.184489031 | 6.7900075 | 6.8714687 | 6.91918 | -0.047709211 | -0.129170397 |
| Syn2 | 3.521905 | 3.446179 | 3.4762 | -0.030016754 | 0.045709723 | 3.0649838 | 3.4009194 | 3.44864 | -0.047720197 | -0.383655868 |
| Stkld1 | 2.019685 | 2.023094 | 2.1538 | -0.130745941 | -0.134154887 | 2.0239177 | 2.0831488 | 2.13087 | -0.047720229 | -0.106951291 |
| Pcdhgb8 | 1.650162 | 1.631793 | 1.5988 | 0.033035064 | 0.051403907 | 1.5671157 | 1.5575142 | 1.60527 | -0.04775788 | -0.038156401 |
| Styxl1 | 1.793139 | 1.783158 | 1.7851 | -0.001942182 | 0.0080385 | 1.8184486 | 1.7321322 | 1.77992 | -0.047785712 | 0.038530654 |
| Top3b | 9.991766 | 10.11593 | 10.087 | 0.029052416 | -0.095111527 | 10.403506 | 10.280133 | 10.3279 | -0.04781394 | 0.075558579 |
| Ethe1 | 7.009481 | 7.000118 | 7.0625 | -0.062372367 | -0.053009361 | 7.1321005 | 6.9014744 | 6.94932 | -0.047844666 | 0.182781438 |
| Slc38a2 | 12.14749 | 12.2067 | 12.253 | -0.046726028 | -0.105934242 | 12.075982 | 12.271405 | 12.3193 | -0.047853916 | -0.243276928 |
| 1700020D05Rik | 3.094574 | 3.090039 | 3.0754 | 0.014667624 | 0.019202943 | 3.3464285 | 3.1186827 | 3.1666 | -0.047913954 | 0.179831867 |
| Ppm1n | 0.688456 | 0.70228 | 0.6935 | 0.008771917 | -0.005052815 | 0.7650278 | 0.6837079 | 0.73166 | -0.0479524 | 0.033367555 |
| Vmn2r65 | 1.975808 | 1.960155 | 2.0404 | -0.080226661 | -0.064573751 | 2.1385271 | 2.0832783 | 2.13127 | -0.047990386 | 0.007258414 |
| Hoxa5 | 3.50054 | 3.402976 | 3.5457 | -0.142753288 | -0.045189408 | 3.2814148 | 3.2389774 | 3.28699 | -0.048016541 | -0.005579183 |
| Frat1 | 7.116883 | 7.024086 | 7.1716 | -0.147471858 | -0.054675006 | 6.8625219 | 6.8308534 | 6.87889 | -0.048032261 | -0.016363716 |
| Gm45380 | 2.430183 | 2.347968 | 2.4192 | -0.071208283 | 0.011006958 | 2.3724103 | 2.398233 | 2.44629 | -0.048056326 | -0.07387907 |
| Rhpn1 | 1.579043 | 1.643351 | 1.6713 | -0.027914682 | -0.092223044 | 1.6447509 | 1.591818 | 1.6399 | -0.048078682 | 0.004854219 |
| Gucy1a2 | 2.527464 | 2.448653 | 2.462 | -0.013312257 | 0.065498749 | 2.3584758 | 2.4191517 | 2.46724 | -0.048085226 | -0.108761126 |
| Pirt | 1.621875 | 1.598029 | 1.6149 | -0.01689433 | 0.006951582 | 1.5987614 | 1.6543983 | 1.7025 | -0.048101301 | -0.103738195 |
| Rimkla | 1.505115 | 1.434616 | 1.4048 | 0.029801163 | 0.100300418 | 1.4261732 | 1.3831593 | 1.43129 | -0.048130403 | -0.005116505 |
| Arsa | 8.918943 | 8.842673 | 8.9437 | -0.101042394 | -0.024772773 | 9.021297 | 8.5082878 | 8.55648 | -0.048195902 | 0.464813278 |
| C4bp | 3.720732 | 3.822369 | 3.7073 | 0.11505154 | 0.013414659 | 3.9014312 | 3.9064481 | 3.95465 | -0.048198445 | -0.053215347 |
| Krt23 | 1.936976 | 1.650044 | 1.7979 | -0.147886081 | 0.139045233 | 1.9045362 | 1.6920313 | 1.74025 | -0.048218202 | 0.164286687 |
| Npr3 | 2.330606 | 2.333007 | 2.3314 | 0.001597001 | -0.000803603 | 2.2721118 | 2.2116024 | 2.25982 | -0.048219844 | 0.012289584 |
| Man1a2 | 10.25852 | 10.40565 | 10.224 | 0.181464959 | 0.034334908 | 10.049803 | 10.207042 | 10.2553 | -0.04822816 | -0.205467763 |
| Tceanc | 6.939718 | 6.815292 | 6.8916 | -0.076265585 | 0.048161168 | 7.1334805 | 7.0223415 | 7.07057 | -0.048232354 | 0.06290664 |
| Olfr918 | 1.525885 | 1.537362 | 1.5394 | -0.002052384 | -0.013529088 | 1.6037718 | 1.5467517 | 1.59502 | -0.048270054 | 0.008750072 |

| | | | | | | | | | | |
|---|---|---|---|---|---|---|---|---|---|---|
| Amy2a5 | 0.995063 | 1.011111 | 1.0518 | -0.040719086 | -0.056766316 | 1.0035104 | 1.0265447 | 1.07483 | -0.048286733 | -0.071321036 |
| Gm6296 | 2.117055 | 2.025114 | 2.0049 | 0.02021844 | 0.112159421 | 2.061612 | 2.039107 | 2.0874 | -0.048288685 | -0.025783609 |
| Nae1 | 9.589304 | 9.603211 | 9.588 | 0.0151944 | 0.001286821 | 9.694228 | 9.7294784 | 9.77777 | -0.048294328 | -0.083544714 |
| Slc2a9 | 7.838482 | 7.762066 | 8.0278 | -0.265778593 | -0.189362085 | 7.8528386 | 7.8066186 | 7.85494 | -0.048318747 | -0.002098827 |
| 0610040J01Rik | 3.642329 | 3.354149 | 3.262 | 0.092128405 | 0.380308483 | 4.0019042 | 3.4349921 | 3.48333 | -0.048336968 | 0.518575061 |
| Btn2a2 | 1.821096 | 1.715497 | 1.7799 | -0.064385189 | 0.041213301 | 1.8561585 | 1.7171279 | 1.7655 | -0.04837251 | 0.09065808 |
| Gm17809 | 3.371379 | 3.277194 | 3.2595 | 0.017727241 | 0.111912087 | 3.3668692 | 3.3933011 | 3.44171 | -0.048409097 | -0.074841049 |
| Gpr27 | 1.285276 | 1.35834 | 1.3415 | 0.016799067 | -0.05626508 | 1.3068614 | 1.2083956 | 1.25681 | -0.048410409 | 0.050055352 |
| Rps3a2 | 2.798227 | 2.923788 | 2.9327 | -0.008862695 | -0.134423776 | 2.8201429 | 2.9274986 | 2.97592 | -0.048419297 | -0.155775053 |
| Tuba1c | 10.17459 | 9.93095 | 10.07 | -0.138663325 | 0.104976387 | 9.9001454 | 9.8265051 | 9.87494 | -0.048430364 | 0.025209945 |
| Nudt18 | 7.85357 | 7.769685 | 7.8494 | -0.079725029 | 0.004159485 | 7.8551089 | 7.7043453 | 7.75278 | -0.048430799 | 0.102332714 |
| Dock2 | 13.36405 | 13.42771 | 13.385 | 0.04312478 | -0.020533876 | 13.39325 | 13.427856 | 13.4763 | -0.048469222 | -0.083075221 |
| Mtfr2 | 5.03626 | 5.008318 | 5.2229 | -0.214600134 | -0.186658764 | 4.9910059 | 5.1649487 | 5.21344 | -0.048487068 | -0.222429942 |
| Il18rap | 9.129229 | 8.857052 | 8.9738 | -0.116748434 | 0.155428511 | 8.8696471 | 9.0922511 | 9.14075 | -0.048502374 | -0.271106312 |
| Kctd14 | 8.017267 | 8.213925 | 8.1288 | 0.085080606 | -0.111576619 | 8.2166209 | 8.0554676 | 8.10398 | -0.048512192 | 0.112641158 |
| BC049762 | 0.656618 | 0.71899 | 0.6852 | 0.033818756 | -0.028553086 | 0.6705229 | 0.646756 | 0.69528 | -0.048528591 | -0.024761695 |
| Zfp959 | 7.687302 | 7.7213 | 7.6953 | 0.025996986 | -0.008000524 | 7.7900901 | 7.7340853 | 7.78262 | -0.048538211 | 0.007466549 |
| Gm21974 | 0.124976 | 0.125073 | 0.1935 | -0.06847469 | -0.068571427 | 0.1449849 | 0.1891224 | 0.23775 | -0.048628352 | -0.092765801 |
| Tmem144 | 3.361752 | 3.299799 | 3.497 | -0.197247095 | -0.135294418 | 3.2849213 | 3.3247569 | 3.3734 | -0.048641847 | -0.088477397 |
| Skil | 10.90963 | 10.94592 | 10.86 | 0.085454692 | 0.04915937 | 10.581303 | 10.838408 | 10.8871 | -0.048662352 | -0.305767003 |
| Gm13408 | 3.453501 | 3.053814 | 3.1321 | -0.078255441 | 0.321431067 | 3.2159812 | 3.187572 | 3.23627 | -0.04869325 | -0.020284084 |
| Tspan3 | 9.59269 | 9.509598 | 9.5643 | -0.054746755 | 0.028344957 | 8.9185156 | 9.2340156 | 9.28272 | -0.048707266 | -0.364207198 |
| Olfr1260 | 2.172434 | 2.196726 | 2.1606 | 0.036140327 | 0.011847742 | 2.2809253 | 2.1592191 | 2.20793 | -0.048708049 | 0.072998212 |
| Myo9b | 11.58276 | 11.64689 | 11.741 | -0.094000294 | -0.158130736 | 11.707748 | 11.750961 | 11.7997 | -0.048742208 | -0.091955175 |
| Gm13238 | 0.865063 | 0.889828 | 0.8869 | 0.002911757 | -0.021853827 | 0.855838 | 0.8816916 | 0.93043 | -0.048743381 | -0.074596905 |
| Hinfp | 9.125142 | 9.114151 | 9.1445 | -0.030348671 | -0.019357579 | 9.3810786 | 9.276362 | 9.32515 | -0.048785866 | 0.055930676 |
| Gm28557 | 5.593906 | 5.69631 | 5.8547 | -0.158367295 | -0.260771565 | 5.7446192 | 5.8337789 | 5.88262 | -0.048840607 | -0.138000335 |
| Cntnap4 | 1.46368 | 1.384227 | 1.3804 | 0.003855371 | 0.083308571 | 1.429928 | 1.4148643 | 1.46374 | -0.048873307 | -0.033809533 |
| Traj48 | 1.555999 | 1.541112 | 1.5303 | 0.010794454 | 0.025681471 | 1.5836538 | 1.5021963 | 1.55108 | -0.048885068 | 0.032572392 |
| Akr1b10 | 8.123987 | 8.136386 | 8.2133 | -0.076961179 | -0.089360383 | 8.510176 | 8.4686204 | 8.51753 | -0.048905276 | -0.007349639 |
| Iglj1 | 3.185426 | 3.284064 | 3.1085 | 0.175592804 | 0.076955205 | 3.1002126 | 3.0351432 | 3.08406 | -0.048916814 | 0.016152565 |
| Gm3005 | 0.719992 | 0.686993 | 0.8013 | -0.114262233 | -0.081263362 | 0.7764213 | 0.713212 | 0.76216 | -0.048943149 | 0.014266142 |

| | | | | | | | | | | |
|---|---|---|---|---|---|---|---|---|---|---|
| Gstp2 | 3.357304 | 3.240139 | 3.4324 | -0.19227314 | -0.075108386 | 3.6998333 | 3.5424626 | 3.59158 | -0.049122194 | 0.108248574 |
| Hint2 | 6.964815 | 6.77324 | 7.0324 | -0.259175802 | -0.067600279 | 7.1939966 | 6.8920524 | 6.94121 | -0.049154255 | 0.252789881 |
| Olfr1344 | 0.298373 | 0.26882 | 0.3179 | -0.049071762 | -0.019518495 | 0.2962529 | 0.3329434 | 0.38212 | -0.04917637 | -0.085866865 |
| Pdxk-ps | 1.399283 | 1.446437 | 1.419 | 0.027437205 | -0.019717405 | 1.4571355 | 1.3980003 | 1.44722 | -0.049216877 | 0.009918328 |
| H2-M5 | 0.664323 | 0.590581 | 0.7139 | -0.123279439 | -0.049537585 | 0.6115703 | 0.6148902 | 0.66423 | -0.049340513 | -0.052660396 |
| Nagk | 7.694055 | 7.451926 | 7.7797 | -0.327738363 | -0.085608899 | 7.6756348 | 7.7194938 | 7.76886 | -0.049366549 | -0.09322552 |
| Igll1 | 0.35713 | 0.354138 | 0.4701 | -0.115974991 | -0.112983388 | 0.3474828 | 0.3709246 | 0.42029 | -0.049368448 | -0.072810276 |
| Dlg2 | 4.695991 | 4.636047 | 4.508 | 0.128016545 | 0.187959941 | 4.323856 | 4.2534007 | 4.30277 | -0.049370242 | 0.021085105 |
| Trim3 | 7.770328 | 7.692088 | 7.7772 | -0.085144117 | -0.006904634 | 7.9670835 | 7.7434282 | 7.7928 | -0.049373792 | 0.174281486 |
| Jph1 | 2.442562 | 2.41895 | 2.4607 | -0.041791311 | -0.018179541 | 2.2654884 | 2.1944102 | 2.24379 | -0.049377447 | 0.02170078 |
| Rpl9-ps6 | 2.956608 | 3.072677 | 2.9608 | 0.111850936 | -0.004217859 | 3.0663401 | 2.9528316 | 3.00222 | -0.049385019 | 0.064123469 |
| Vps35l | 10.10473 | 10.17943 | 10.194 | -0.014608128 | -0.089305128 | 10.294915 | 10.12184 | 10.1712 | -0.0493902 | 0.123685268 |
| Pappa | 1.485629 | 1.455471 | 1.3432 | 0.112302201 | 0.14246019 | 1.3202853 | 1.3253465 | 1.37474 | -0.049396337 | -0.054457489 |
| Mnt | 9.520482 | 9.462715 | 9.4537 | 0.008966248 | 0.06673307 | 9.7214246 | 9.5439672 | 9.5934 | -0.049436093 | 0.12802131 |
| Tekt5 | 0.809797 | 0.784559 | 0.7566 | 0.027980321 | 0.053219051 | 0.7517241 | 0.7292753 | 0.77872 | -0.049446087 | -0.026997256 |
| Slit3 | 5.579033 | 5.569915 | 5.3788 | 0.191102521 | 0.200221037 | 5.166763 | 4.9701477 | 5.0196 | -0.04945551 | 0.147159788 |
| Nos1ap | 3.735218 | 3.59193 | 3.6804 | -0.088468216 | 0.054820317 | 3.6066918 | 3.5082237 | 3.55771 | -0.049488699 | 0.048979389 |
| Gm2163 | 1.891903 | 1.81008 | 1.8251 | -0.014983555 | 0.066839543 | 1.7476093 | 1.7021304 | 1.75162 | -0.049492782 | -0.004013795 |
| Plscr2 | 5.92624 | 5.405388 | 5.8881 | -0.482739505 | 0.03811189 | 5.0513772 | 5.3797604 | 5.42927 | -0.049511751 | -0.377894957 |
| Slc8a2 | 2.726262 | 2.667119 | 2.8148 | -0.147721248 | -0.088577402 | 2.6478347 | 2.5583999 | 2.60792 | -0.049518256 | 0.039916484 |
| Ccdc92 | 6.366641 | 6.403024 | 6.1208 | 0.282234411 | 0.24585209 | 6.0537306 | 6.1795043 | 6.22903 | -0.049529901 | -0.175303574 |
| Lyrm4 | 6.697362 | 6.63296 | 6.6146 | 0.018383194 | 0.082785313 | 6.5885177 | 6.5596703 | 6.60924 | -0.049573088 | -0.020725693 |
| Gatb | 7.555014 | 7.617219 | 7.5133 | 0.103921903 | 0.041717275 | 7.8312177 | 7.5016071 | 7.55124 | -0.049636561 | 0.279974032 |
| Gm49776 | -0.227285 | -0.24485 | -0.2585 | 0.013602491 | 0.031166235 | -0.2865113 | -0.286114 | -0.2364 | -0.049674055 | -0.050071267 |
| Lbp | 6.564557 | 6.350553 | 6.4562 | -0.105638493 | 0.108365301 | 6.0353735 | 5.9998193 | 6.04954 | -0.04971782 | -0.014163562 |
| Plekhb1 | 1.904927 | 1.887232 | 1.7985 | 0.088716266 | 0.1064112 | 1.8099878 | 1.8126661 | 1.86244 | -0.049769153 | -0.052447521 |
| Ido2 | 1.017643 | 0.993047 | 1.0533 | -0.06028769 | -0.035690796 | 0.9443142 | 0.976652 | 1.02646 | -0.049810508 | -0.082148346 |
| Slc16a4 | 1.21415 | 1.17898 | 1.2299 | -0.050877034 | -0.015706773 | 1.163409 | 1.153824 | 1.20364 | -0.049816674 | -0.04023174 |
| Bloc1s2-ps | 0.429429 | 0.325593 | 0.3238 | 0.001782513 | 0.105618267 | 0.5289417 | 0.3284528 | 0.37833 | -0.049877013 | 0.150611933 |
| Gm2974 | 0.799626 | 0.814197 | 0.7264 | 0.087842661 | 0.073271215 | 0.7288919 | 0.7208748 | 0.77077 | -0.04989941 | -0.041882252 |
| Trappc11 | 10.08659 | 10.06431 | 10.103 | -0.038729979 | -0.016444427 | 10.116838 | 10.043684 | 10.0936 | -0.049931454 | 0.023222632 |
| Cyp4a31 | 0.690212 | 0.686537 | 0.6572 | 0.029322712 | 0.032997051 | 0.6518184 | 0.6392961 | 0.68925 | -0.049950987 | -0.037428725 |

| | | | | | | | | | | |
|---|---|---|---|---|---|---|---|---|---|---|
| Tnnt3 | 4.904623 | 4.78021 | 4.9486 | -0.168397039 | -0.043983963 | 5.115673 | 5.0802206 | 5.13017 | -0.049953642 | -0.014501275 |
| AC148174.1 | 0.863917 | 0.896951 | 0.8973 | -0.000336946 | -0.033371866 | 0.9049973 | 0.9085947 | 0.95857 | -0.049974057 | -0.053571521 |
| Kcnj14 | 1.592547 | 1.611162 | 1.6817 | -0.070562678 | -0.089177979 | 1.6255013 | 1.5771419 | 1.62724 | -0.050093159 | -0.001733749 |
| Gm12166 | 1.757303 | 1.752597 | 1.8702 | -0.117618865 | -0.112912502 | 1.8455777 | 1.7454293 | 1.79553 | -0.050099719 | 0.050048646 |
| Mrps5 | 9.335026 | 9.344093 | 9.3401 | 0.003993034 | -0.005074073 | 9.3126696 | 9.3130334 | 9.36314 | -0.05010747 | -0.050471298 |
| Gm10038 | 2.388961 | 2.439868 | 2.4082 | 0.031647839 | -0.019259727 | 2.5831705 | 2.4547738 | 2.5049 | -0.050123989 | 0.078272692 |
| Lage3 | 7.563707 | 7.484049 | 7.633 | -0.148944382 | -0.069286176 | 7.841314 | 7.5564306 | 7.60656 | -0.050127206 | 0.234756255 |
| Cd74 | 16.63856 | 16.71876 | 16.672 | 0.046590998 | -0.033607336 | 17.247595 | 16.68351 | 16.7336 | -0.050128198 | 0.513956115 |
| Rngtt | 9.374245 | 9.393465 | 9.3567 | 0.036767978 | 0.017547553 | 9.1917056 | 9.3519405 | 9.4021 | -0.050159385 | -0.210394354 |
| BC022960 | 3.379947 | 3.353755 | 3.3973 | -0.043572976 | -0.017380962 | 3.5232571 | 3.3950793 | 3.44532 | -0.05024553 | 0.077932247 |
| Atoh8 | 3.260133 | 3.029442 | 3.2465 | -0.21701944 | 0.013671329 | 2.8629337 | 2.9731366 | 3.02338 | -0.050245953 | -0.160448887 |
| Gm13033 | 1.977667 | 1.860148 | 1.9113 | -0.051201192 | 0.066318014 | 1.9066503 | 1.8613219 | 1.91161 | -0.050292467 | -0.004964036 |
| Erp27 | 5.397641 | 5.379458 | 5.5983 | -0.21886492 | -0.200681778 | 5.941299 | 5.6305623 | 5.68086 | -0.05029546 | 0.260441261 |
| Foxj3 | 10.12924 | 10.13337 | 10.155 | -0.021536081 | -0.025668888 | 9.9139143 | 10.136114 | 10.1864 | -0.050301958 | -0.272502016 |
| Gm7589 | 1.363102 | 1.39166 | 1.3944 | -0.002763291 | -0.031321353 | 1.4825675 | 1.3719858 | 1.42229 | -0.050302195 | 0.060279578 |
| Cpt1b | 5.776595 | 5.806904 | 5.7901 | 0.016819918 | -0.013488514 | 6.1896715 | 6.149715 | 6.20008 | -0.050360424 | -0.010403852 |
| Gm18188 | 0.260424 | 0.238966 | 0.2656 | -0.026597298 | -0.00513907 | 0.2344744 | 0.2571798 | 0.30758 | -0.050403434 | -0.073108867 |
| Padi2 | 9.719128 | 9.77089 | 9.6946 | 0.076283172 | 0.02452147 | 9.5087244 | 9.4823547 | 9.53277 | -0.050412907 | -0.024043164 |
| Mmp16 | 2.249904 | 2.159535 | 2.1693 | -0.009791271 | 0.080577497 | 2.1510609 | 2.1333303 | 2.18376 | -0.050429603 | -0.032698923 |
| BC024139 | 1.37272 | 1.331885 | 1.3035 | 0.028356669 | 0.069191364 | 1.4024187 | 1.3388287 | 1.38937 | -0.050543708 | 0.013046315 |
| Vmn1r51 | 0.940089 | 0.849847 | 0.8752 | -0.025323124 | 0.064919118 | 0.9014911 | 0.9224542 | 0.97303 | -0.050571842 | -0.071534979 |
| Fam219b | 8.640102 | 8.649721 | 8.6594 | -0.009663327 | -0.01928153 | 8.8440297 | 8.5514583 | 8.60205 | -0.05058792 | 0.241983411 |
| Cacng2 | 1.359091 | 1.382712 | 1.333 | 0.04966457 | 0.026044193 | 1.3516541 | 1.3573028 | 1.4079 | -0.050595064 | -0.0562437 |
| Lrrc8d | 9.156288 | 9.073944 | 9.1632 | -0.089242928 | -0.006898557 | 9.2191111 | 9.0977782 | 9.14839 | -0.050612321 | 0.070720561 |
| Ptbp3 | 13.17365 | 13.20593 | 13.139 | 0.066718124 | 0.034432948 | 13.133133 | 13.092993 | 13.1436 | -0.050637924 | -0.010497466 |
| Olfr1328 | 1.09446 | 1.132685 | 1.1234 | 0.009279015 | -0.028946515 | 1.1676021 | 1.1005162 | 1.15117 | -0.05064889 | 0.01643709 |
| Gm6012 | 0.95567 | 0.951432 | 1.0286 | -0.077176782 | -0.072938729 | 0.9637714 | 0.9910343 | 1.04169 | -0.050659849 | -0.077922697 |
| Pcdhb13 | 2.611742 | 2.602363 | 2.6194 | -0.017025463 | -0.007646652 | 2.4427688 | 2.458407 | 2.50913 | -0.050718283 | -0.066356451 |
| Ccdc36 | 3.65767 | 3.566831 | 3.7945 | -0.227705315 | -0.136865772 | 3.6994182 | 3.7088368 | 3.75958 | -0.050739541 | -0.060158184 |
| Zfp712 | 7.383044 | 7.735298 | 7.4906 | 0.244686662 | -0.107567306 | 7.4767773 | 7.4493396 | 7.50011 | -0.050772315 | -0.023334613 |
| Ak1 | 2.578972 | 2.575328 | 2.5811 | -0.005762302 | -0.002118412 | 2.5496547 | 2.5063601 | 2.55716 | -0.050801759 | -0.00750716 |
| Gm28438 | 5.17929 | 3.19473 | 4.1166 | -0.921825752 | 1.062734475 | 4.0610887 | 3.8391189 | 3.88995 | -0.050834087 | 0.171135753 |

| | | | | | | | | | | |
|---|---|---|---|---|---|---|---|---|---|---|
| Ighv5-1 | 0.932823 | 0.959849 | 0.8961 | 0.063741855 | 0.036716246 | 0.9131908 | 0.8429114 | 0.89375 | -0.050842913 | 0.01943654 |
| Mta1 | 10.21613 | 10.23977 | 10.239 | 0.000468832 | -0.023169835 | 10.20886 | 10.167967 | 10.2188 | -0.050865532 | -0.009972618 |
| Vmn2r92 | 3.05745 | 3.003912 | 3.1686 | -0.164707359 | -0.111168523 | 3.1914404 | 3.1339471 | 3.18482 | -0.05087089 | 0.006622446 |
| Agbl2 | 1.643392 | 1.618046 | 1.6767 | -0.058680022 | -0.033333948 | 1.6656773 | 1.6971395 | 1.74802 | -0.050875813 | -0.082338026 |
| Glt1d1 | 2.454074 | 2.378186 | 2.442 | -0.063819201 | 0.012068415 | 2.4102217 | 2.4448685 | 2.49577 | -0.050902446 | -0.085549237 |
| Lmo4 | 9.227374 | 9.224406 | 9.2838 | -0.059421416 | -0.056453218 | 9.1618358 | 9.1464735 | 9.1976 | -0.051125551 | -0.035763267 |
| Gm5812 | -0.023542 | 0.009405 | -0.0251 | 0.034542147 | 0.001594815 | -0.0546067 | -0.034116 | 0.01701 | -0.051129504 | -0.071620484 |
| Slc23a4 | 3.138245 | 3.044165 | 3.1616 | -0.117397489 | -0.023317015 | 3.1489314 | 3.1114232 | 3.16256 | -0.051141064 | -0.013632868 |
| Lpcat4 | 8.887265 | 8.778186 | 9.0199 | -0.241759416 | -0.132680706 | 8.8623534 | 8.7566702 | 8.80795 | -0.051283159 | 0.054400024 |
| Olfr742 | 0.849486 | 0.897585 | 0.8732 | 0.024342921 | -0.023756187 | 0.8385576 | 0.8573996 | 0.90876 | -0.051360134 | -0.070202133 |
| Gm44524 | 0.640591 | 0.654498 | 0.6982 | -0.043716413 | -0.057624342 | 0.7116304 | 0.6605701 | 0.71193 | -0.051362015 | -0.00030166 |
| Cdh8 | 1.584358 | 1.574308 | 1.657 | -0.082707627 | -0.072657858 | 1.4534404 | 1.4922099 | 1.54358 | -0.051367043 | -0.090136555 |
| Nipa1 | 4.247676 | 4.367646 | 4.277 | 0.090663832 | -0.029305929 | 4.0496264 | 4.1678913 | 4.21926 | -0.051372514 | -0.169637368 |
| Gm49328 | 2.043164 | 1.96898 | 1.955 | 0.014006629 | 0.088190641 | 1.8814538 | 1.9158965 | 1.96729 | -0.051388767 | -0.085831512 |
| Ascl3 | 1.608891 | 1.519639 | 1.443 | 0.076662846 | 0.165914797 | 1.5122049 | 1.5909115 | 1.64235 | -0.051437943 | -0.130144532 |
| Gm7803 | 1.027893 | 1.0716 | 1.0549 | 0.016687262 | -0.027019856 | 1.0039935 | 1.0056614 | 1.05713 | -0.051464154 | -0.05313202 |
| E330034G19Rik | 1.424876 | 1.391635 | 1.4063 | -0.014710285 | 0.018530662 | 1.400118 | 1.4153212 | 1.46682 | -0.051494837 | -0.06669804 |
| Ddah2 | 5.94128 | 5.781697 | 5.8909 | -0.109186564 | 0.050396826 | 5.7944704 | 5.5917909 | 5.64329 | -0.051495376 | 0.151184059 |
| Fbxo43 | 0.993019 | 0.95637 | 0.9934 | -0.037011643 | -0.000362738 | 0.9439286 | 0.9546371 | 1.00621 | -0.05157072 | -0.062279261 |
| Kdm2b | 10.1309 | 10.31922 | 10.214 | 0.105403927 | -0.082912234 | 10.037191 | 10.171886 | 10.2235 | -0.051595795 | -0.186291246 |
| Arhgef2 | 11.21953 | 11.2843 | 11.283 | 0.00170555 | -0.063067246 | 11.307848 | 11.31152 | 11.3632 | -0.051645238 | -0.055318071 |
| Prss36 | 4.400193 | 4.429079 | 4.5653 | -0.136171769 | -0.165057676 | 4.6675177 | 4.5813865 | 4.63305 | -0.051664525 | 0.034466671 |
| Igf2 | 4.558571 | 4.58853 | 4.5408 | 0.047713889 | 0.017755486 | 4.1915385 | 4.3467557 | 4.39843 | -0.051679179 | -0.206896394 |
| Mcoln1 | 9.279093 | 9.239059 | 9.3228 | -0.083773616 | -0.043740219 | 9.7823529 | 9.664885 | 9.71659 | -0.0517042 | 0.065763623 |
| Peg12 | 1.762527 | 1.715628 | 1.728 | -0.012414061 | 0.034484634 | 1.725891 | 1.6320408 | 1.68376 | -0.051720725 | 0.042129549 |
| Rtkn | 3.133467 | 3.153415 | 3.143 | 0.010380989 | -0.009567174 | 3.0112669 | 3.099278 | 3.15106 | -0.05178564 | -0.139796799 |
| Rpl7a-ps7 | 0.326383 | 0.302459 | 0.2857 | 0.016767992 | 0.040692488 | 0.3155562 | 0.3063203 | 0.35815 | -0.051831608 | -0.042595719 |
| Gm16477 | 1.371042 | 1.372683 | 1.3827 | -0.010016457 | -0.011657748 | 1.3739888 | 1.3214241 | 1.37326 | -0.051832123 | 0.000732597 |
| Gm5380 | 2.569225 | 2.476164 | 2.5777 | -0.101490426 | -0.008430004 | 2.6032766 | 2.5227767 | 2.57465 | -0.051874135 | 0.028625809 |
| Usp45 | 9.579574 | 9.652207 | 9.6369 | 0.015274368 | -0.05735862 | 9.4557101 | 9.5554281 | 9.60732 | -0.051896584 | -0.151614646 |
| Pdgfrb | 7.535454 | 7.676004 | 7.0581 | 0.617888214 | 0.477338556 | 6.7425654 | 6.3208264 | 6.37273 | -0.051907449 | 0.369831601 |
| Odf3b | 0.341007 | 0.324042 | 0.4021 | -0.078084575 | -0.061119356 | 0.3617628 | 0.3341363 | 0.38616 | -0.052025726 | -0.024399217 |

| | | | | | | | | | | |
|---|---|---|---|---|---|---|---|---|---|---|
| Slitrk4 | 1.570435 | 1.52859 | 1.525 | 0.003574806 | 0.045420502 | 1.6080641 | 1.5807721 | 1.63284 | -0.052069127 | -0.02477721 |
| Tsga13 | 1.255542 | 1.242972 | 1.331 | -0.088020248 | -0.075450101 | 1.3343514 | 1.2917928 | 1.34389 | -0.052096115 | -0.00953758 |
| Slc35f1 | 1.686983 | 1.717649 | 1.6865 | 0.031138813 | 0.000472985 | 1.6418613 | 1.6920105 | 1.74411 | -0.052102929 | -0.102252086 |
| Fam216a | 6.515528 | 6.419744 | 6.5796 | -0.159903183 | -0.064119024 | 6.470736 | 6.3931035 | 6.44522 | -0.052118987 | 0.025513452 |
| Gm43951 | 2.999016 | 3.056181 | 2.9998 | 0.056349333 | -0.000815008 | 3.1188199 | 2.9683145 | 3.02045 | -0.052136711 | 0.09836861 |
| Rem2 | 1.391542 | 1.328345 | 1.3776 | -0.049216651 | 0.013979597 | 1.29302 | 1.3349013 | 1.38705 | -0.052148371 | -0.094029653 |
| Sox11 | 1.953456 | 1.900404 | 1.854 | 0.046370756 | 0.099423681 | 1.8535085 | 1.801755 | 1.85396 | -0.052209512 | -0.000456019 |
| Tdrd5 | 1.837339 | 1.847439 | 1.9686 | -0.121130257 | -0.131229873 | 1.9065051 | 1.8441626 | 1.89637 | -0.052209591 | 0.010132901 |
| Gm11874 | 1.586262 | 1.544177 | 1.541 | 0.003175613 | 0.045260237 | 1.516937 | 1.4832068 | 1.53543 | -0.052223583 | -0.018493326 |
| Trpc1 | 2.733052 | 2.683841 | 2.7702 | -0.086405647 | -0.037194214 | 2.5202769 | 2.580421 | 2.63266 | -0.052237807 | -0.112381909 |
| Trim63 | 0.435845 | 0.439969 | 0.4268 | 0.013207662 | 0.009083467 | 0.3733702 | 0.397296 | 0.44963 | -0.052336537 | -0.076262252 |
| Med30 | 8.120926 | 8.028212 | 8.1566 | -0.128342163 | -0.035628462 | 8.2719981 | 8.0506337 | 8.10301 | -0.052373519 | 0.168990803 |
| Dchs2 | 0.761951 | 0.776765 | 0.793 | -0.016247868 | -0.031062377 | 0.7246733 | 0.6985845 | 0.75099 | -0.052408288 | -0.026319436 |
| Sh3bp4 | 8.066385 | 8.038984 | 8.2259 | -0.186939252 | -0.159538787 | 7.6677364 | 7.934473 | 7.98692 | -0.052449577 | -0.319186172 |
| Lrrc74b | 1.177663 | 1.149352 | 1.1818 | -0.03241723 | -0.004106696 | 1.1172568 | 1.1139477 | 1.16642 | -0.052476937 | -0.049167814 |
| Pcdhb5 | 2.201804 | 2.128672 | 2.1736 | -0.044970762 | 0.028161038 | 2.0925138 | 2.0697077 | 2.12221 | -0.052498587 | -0.029692471 |
| Olfr1305 | 1.461012 | 1.571143 | 1.4972 | 0.073960406 | -0.036170127 | 1.5644089 | 1.5155828 | 1.56812 | -0.052534175 | -0.003708058 |
| Mcpt8 | 0.819684 | 0.7697 | 0.7327 | 0.037046332 | 0.08703059 | 0.8014229 | 0.7366179 | 0.78916 | -0.052543761 | 0.012261218 |
| Gm6421 | 0.94864 | 0.975892 | 0.9151 | 0.060743924 | 0.033492649 | 0.9345919 | 0.9346474 | 0.98722 | -0.052574283 | -0.052629783 |
| Kbtbd12 | 1.582102 | 1.634138 | 1.469 | 0.165138031 | 0.113102309 | 1.4670417 | 1.4241047 | 1.47669 | -0.052586738 | -0.009649687 |
| Gm5586 | 2.431354 | 2.414377 | 2.5053 | -0.090886542 | -0.073909946 | 2.4716741 | 2.2605155 | 2.31311 | -0.052595029 | 0.158563535 |
| Efcc1 | 2.264888 | 2.166853 | 2.2632 | -0.096381863 | 0.001652806 | 2.0560928 | 2.0499452 | 2.10255 | -0.052601085 | -0.046453446 |
| Ppard | 8.925103 | 8.848382 | 8.8067 | 0.041695547 | 0.118416378 | 8.7833492 | 8.6924982 | 8.74512 | -0.052623978 | 0.038227004 |
| Elp3 | 9.857066 | 9.834201 | 9.8723 | -0.038055274 | -0.015190199 | 10.011599 | 9.8755736 | 9.9282 | -0.052625378 | 0.083400371 |
| Ilk | 10.43091 | 10.4433 | 10.553 | -0.110151702 | -0.122542176 | 10.505754 | 10.403438 | 10.4561 | -0.052636824 | 0.049679965 |
| Ubap1 | 9.675159 | 9.658092 | 9.6384 | 0.019697986 | 0.036764588 | 9.5123256 | 9.6368303 | 9.68948 | -0.052645138 | -0.177149839 |
| Rep15 | 1.391493 | 1.333935 | 1.3314 | 0.002503832 | 0.060061264 | 1.3434189 | 1.2970976 | 1.34975 | -0.05264961 | -0.006328264 |
| Izumo1 | 2.278172 | 2.201879 | 2.2386 | -0.03669885 | 0.039594195 | 2.0794215 | 2.2211208 | 2.27385 | -0.052728563 | -0.194427844 |
| Gm8050 | 0.883541 | 0.883968 | 0.8875 | -0.003521367 | -0.003948781 | 0.9345293 | 0.880013 | 0.93276 | -0.052745302 | 0.001770981 |
| Gm49181 | 2.526915 | 2.674065 | 2.667 | 0.007026657 | -0.140122702 | 2.7378133 | 2.7117895 | 2.7646 | -0.052806694 | -0.026782904 |
| Pthlh | 2.772475 | 2.814282 | 2.7271 | 0.087218045 | 0.045411479 | 2.7111965 | 2.6819877 | 2.73483 | -0.052840968 | -0.023632099 |
| Rprm | 0.60803 | 0.576593 | 0.6219 | -0.04533276 | -0.01389571 | 0.5614309 | 0.5353352 | 0.58818 | -0.052847307 | -0.02675167 |

| | | | | | | | | | | |
|---|---|---|---|---|---|---|---|---|---|---|
| Ndel1 | 9.519475 | 9.61869 | 9.3002 | 0.318521524 | 0.219305662 | 9.2970481 | 9.3874318 | 9.44032 | -0.052883275 | -0.143267033 |
| Olfr160 | 1.761318 | 1.711799 | 1.6974 | 0.014413261 | 0.063932223 | 1.8142956 | 1.7319108 | 1.78488 | -0.052973321 | 0.029411442 |
| 1700020L24Rik | 2.02273 | 2.075333 | 2.0746 | 0.000709576 | -0.051892723 | 2.1481048 | 2.0359976 | 2.08899 | -0.05299584 | 0.059111389 |
| Dicer1 | 10.82728 | 10.80473 | 10.855 | -0.05016183 | -0.027616258 | 10.61067 | 10.684018 | 10.737 | -0.053029376 | -0.126376631 |
| Coro6 | 1.391397 | 1.444838 | 1.3999 | 0.044946531 | -0.008494516 | 1.4562643 | 1.4294263 | 1.48246 | -0.053033512 | -0.026195437 |
| Gm14139 | 1.711162 | 1.814663 | 1.8022 | 0.012417622 | -0.091083084 | 1.826441 | 1.7724075 | 1.82551 | -0.053106627 | 0.000926826 |
| Gm11879 | 2.030563 | 1.982557 | 2.0624 | -0.079836429 | -0.031829792 | 2.0749687 | 2.0216965 | 2.07481 | -0.053109182 | 0.000163023 |
| Gm10509 | 5.02611 | 4.9433 | 5.0399 | -0.096562229 | -0.013751722 | 5.0597932 | 4.9721102 | 5.02524 | -0.053127254 | 0.034555768 |
| Tspear | 3.10254 | 2.980249 | 3.1016 | -0.121362366 | 0.000928592 | 3.2342843 | 3.0850708 | 3.13822 | -0.053147904 | 0.096065612 |
| Suv39h2 | 5.325509 | 5.221326 | 5.3604 | -0.139036659 | -0.034853853 | 5.3537275 | 5.3816216 | 5.4348 | -0.053175957 | -0.081069994 |
| Trav13n-1 | 3.078145 | 3.162885 | 3.2082 | -0.045315393 | -0.130055353 | 3.0444171 | 3.0915276 | 3.14473 | -0.053197652 | -0.100308217 |
| Jph4 | 1.872982 | 1.997475 | 1.8308 | 0.166642979 | 0.042149757 | 1.8460909 | 1.8889014 | 1.94215 | -0.053246029 | -0.096056592 |
| Prelid3a | 2.720568 | 2.752874 | 2.8036 | -0.050761973 | -0.083067955 | 2.8652141 | 2.7510339 | 2.80429 | -0.053260924 | 0.060919253 |
| Atp4a | 1.054505 | 1.027639 | 1.1467 | -0.119031475 | -0.092165391 | 1.1688397 | 1.100296 | 1.15361 | -0.053310497 | 0.015233108 |
| C1s2 | 1.443191 | 1.44038 | 1.4086 | 0.031825311 | 0.034636146 | 1.3408853 | 1.3513478 | 1.40471 | -0.053358317 | -0.063820788 |
| Gm14853 | 0.40948 | 0.435244 | 0.4474 | -0.012138832 | -0.0379033 | 0.4540081 | 0.4382424 | 0.49161 | -0.053364987 | -0.037599203 |
| C87436 | 8.577291 | 8.56457 | 8.7787 | -0.21413073 | -0.201410476 | 8.7162025 | 8.633876 | 8.68726 | -0.053379411 | 0.028947061 |
| Rpl39-ps | 2.750926 | 2.755526 | 2.7872 | -0.031670748 | -0.036269994 | 2.7262596 | 2.7050927 | 2.75854 | -0.053446241 | -0.032279352 |
| Gm13785 | 2.500113 | 2.480455 | 2.5364 | -0.055924845 | -0.036266388 | 2.5096639 | 2.4844541 | 2.53792 | -0.053468287 | -0.028258537 |
| Lbr | 11.89339 | 11.96914 | 11.89 | 0.07962416 | 0.003872116 | 11.929313 | 11.868256 | 11.9217 | -0.053475652 | 0.007581376 |
| Fam83h | 3.485551 | 3.503835 | 3.4227 | 0.081139893 | 0.062855681 | 3.5889965 | 3.5076856 | 3.56124 | -0.053552786 | 0.027758175 |
| Btbd16 | 2.919489 | 2.922837 | 2.931 | -0.00817897 | -0.011527594 | 2.9484531 | 2.9447668 | 2.99834 | -0.053568555 | -0.049882165 |
| Actg1 | 15.19192 | 15.13831 | 15.118 | 0.019922214 | 0.073531974 | 15.331229 | 15.058598 | 15.1122 | -0.053572312 | 0.219059313 |
| Gm2387 | 2.883707 | 2.781309 | 2.8349 | -0.053550731 | 0.048847141 | 2.8695509 | 2.789728 | 2.84334 | -0.053611402 | 0.026211554 |
| Gm8989 | 2.136625 | 2.163411 | 2.2068 | -0.043377862 | -0.070164237 | 2.3338339 | 2.1456404 | 2.19926 | -0.053618956 | 0.134574527 |
| Yme1l1 | 11.3901 | 11.44317 | 11.439 | 0.004240938 | -0.04882832 | 11.514128 | 11.45686 | 11.5106 | -0.053714906 | 0.003552627 |
| Casc3 | 10.48988 | 10.57501 | 10.541 | 0.033968408 | -0.051165844 | 10.59221 | 10.579609 | 10.6334 | -0.05374593 | -0.041144492 |
| Iqcc | 7.499088 | 7.611366 | 7.7401 | -0.128723756 | -0.241002518 | 7.8780883 | 7.6380739 | 7.69185 | -0.053772596 | 0.186241776 |
| Mpv17l2 | 6.373383 | 6.309091 | 6.5007 | -0.191630288 | -0.127337976 | 6.7041878 | 6.4286121 | 6.4824 | -0.05379154 | 0.221784151 |
| Pgbd1 | 3.12696 | 3.179684 | 3.1602 | 0.019527477 | -0.033196847 | 3.1250419 | 3.1721177 | 3.22594 | -0.053817908 | -0.100893672 |
| Klhl21 | 8.617213 | 8.738807 | 8.6565 | 0.082263608 | -0.039330525 | 8.7363702 | 8.5888844 | 8.64272 | -0.053833616 | 0.093652268 |
| Socs4 | 10.08666 | 10.11301 | 10.109 | 0.004024042 | -0.022321617 | 10.199134 | 10.232497 | 10.2864 | -0.053876421 | -0.087239504 |

| | | | | | | | | | | |
|---|---|---|---|---|---|---|---|---|---|---|
| Cklf | 8.503761 | 8.505754 | 8.4776 | 0.028121332 | 0.026128248 | 8.9085553 | 8.6354633 | 8.68936 | -0.053896921 | 0.219195095 |
| Fam169a | 1.069392 | 1.105372 | 1.0706 | 0.034758891 | -0.001221313 | 1.0613602 | 1.037444 | 1.09136 | -0.053920794 | -0.030004656 |
| Ecd | 9.737845 | 9.863445 | 9.7751 | 0.088339868 | -0.037260465 | 10.009689 | 9.7838475 | 9.83782 | -0.053970007 | 0.171871844 |
| Nutm1 | 0.731474 | 0.705735 | 0.773 | -0.067285201 | -0.041546307 | 0.6770524 | 0.6590519 | 0.71318 | -0.05412341 | -0.036122873 |
| Grik2 | 2.739933 | 2.682954 | 2.8415 | -0.158555087 | -0.1015765 | 2.8279198 | 2.8590089 | 2.91326 | -0.054251384 | -0.085340501 |
| Cdk19 | 10.78981 | 10.76294 | 10.882 | -0.119190858 | -0.092320552 | 10.590653 | 10.661059 | 10.7153 | -0.05425411 | -0.124660164 |
| Rit1 | 8.419411 | 8.363452 | 8.3616 | 0.001896808 | 0.057855821 | 8.4322029 | 8.1962638 | 8.25063 | -0.054366398 | 0.181572735 |
| Fbxo11 | 10.67621 | 10.64821 | 10.692 | -0.04394513 | -0.015945642 | 10.380868 | 10.702399 | 10.7568 | -0.05436956 | -0.375900934 |
| Mcm4 | 10.68156 | 10.484 | 10.608 | -0.124463174 | 0.07309358 | 10.678966 | 10.638923 | 10.6933 | -0.05440797 | -0.014365508 |
| Cmc4 | 6.200698 | 5.900534 | 6.0195 | -0.118965234 | 0.181198267 | 6.2596599 | 6.1040849 | 6.15853 | -0.054440128 | 0.101134866 |
| Ctf2 | 2.0508 | 2.140833 | 2.0647 | 0.07617881 | -0.013854589 | 2.0516092 | 2.1403655 | 2.19482 | -0.054450809 | -0.143207176 |
| Mdm4-ps | 2.965657 | 2.916136 | 3.0294 | -0.113238672 | -0.063718075 | 2.9306599 | 3.0189786 | 3.07344 | -0.054460355 | -0.142779052 |
| Lactbl1 | 1.901031 | 1.894703 | 1.8632 | 0.031461258 | 0.037789238 | 1.9624069 | 1.8675716 | 1.92206 | -0.05448706 | 0.040348298 |
| Bpifb5 | 6.297587 | 6.337589 | 6.3693 | -0.03166539 | -0.071666673 | 5.675749 | 5.4931552 | 5.54768 | -0.054525369 | 0.128068494 |
| Gm2011 | 4.195419 | 4.248231 | 4.3481 | -0.099874753 | -0.152687262 | 4.3614457 | 4.3182617 | 4.37281 | -0.054550051 | -0.011366068 |
| Trav7-3 | 4.605669 | 4.661386 | 4.7684 | -0.107027134 | -0.162744185 | 4.6307131 | 4.6576258 | 4.71219 | -0.054568264 | -0.081480908 |
| Setd6 | 7.947816 | 7.786877 | 7.9174 | -0.130537073 | 0.030401277 | 8.0687058 | 8.0172443 | 8.07188 | -0.054630851 | -0.00316942 |
| Ighv11-1 | 1.363716 | 1.347332 | 1.3266 | 0.020691507 | 0.037074674 | 1.3246908 | 1.2954463 | 1.35008 | -0.05463785 | -0.025393394 |
| Hexa | 10.25608 | 10.12755 | 10.276 | -0.14854274 | -0.02001931 | 10.432359 | 10.143272 | 10.1979 | -0.054671256 | 0.234415805 |
| Dscc1 | 5.293608 | 5.007643 | 5.3405 | -0.332885166 | -0.046919601 | 5.4271808 | 5.3969653 | 5.45167 | -0.054702313 | -0.024486802 |
| Gm45717 | 1.38302 | 1.459542 | 1.3535 | 0.106003165 | 0.029480951 | 1.4249112 | 1.4177175 | 1.47243 | -0.054711113 | -0.047517421 |
| Adamts3 | 2.799563 | 2.779549 | 2.7222 | 0.057359303 | 0.077373286 | 2.7037953 | 2.7234118 | 2.77818 | -0.054769464 | -0.074385944 |
| Gm11346 | 9.404405 | 9.406 | 9.3678 | 0.03823257 | 0.036637121 | 9.6615378 | 9.5180758 | 9.57292 | -0.054847462 | 0.088614544 |
| Pcdhb15 | 1.890697 | 1.801942 | 1.8551 | -0.05319577 | 0.035559096 | 1.7801709 | 1.739998 | 1.79487 | -0.054871724 | -0.014698815 |
| Hacd2 | 7.897014 | 7.983292 | 7.7766 | 0.20666746 | 0.120390186 | 7.7638418 | 7.8944544 | 7.9494 | -0.054948949 | -0.185561452 |
| Ubl7 | 9.642192 | 9.606447 | 9.7447 | -0.138270151 | -0.102525377 | 9.7961197 | 9.6219373 | 9.67693 | -0.054992813 | 0.119189617 |
| Gm15549 | 1.077851 | 1.085318 | 1.1054 | -0.020091249 | -0.027557459 | 1.1502498 | 1.0826915 | 1.1377 | -0.055010532 | 0.012547783 |
| Frmd8 | 10.77036 | 10.96039 | 10.78 | 0.180395264 | -0.0096379 | 10.685245 | 10.615383 | 10.6704 | -0.05501926 | 0.014842801 |
| Slain1 | 9.365472 | 9.49224 | 9.4694 | 0.022814778 | -0.103953192 | 9.3817016 | 9.3512216 | 9.40628 | -0.055055029 | -0.02457504 |
| Olfr118 | 1.640469 | 1.519727 | 1.5718 | -0.052101043 | 0.068640947 | 1.5423227 | 1.5356708 | 1.59075 | -0.055081191 | -0.048429307 |
| Dnajb3 | 1.043915 | 1.045971 | 1.0168 | 0.029215291 | 0.027159629 | 1.0274681 | 1.0239175 | 1.07901 | -0.055087984 | -0.051537409 |
| Adgrg4 | 1.562096 | 1.560254 | 1.611 | -0.050768261 | -0.048925929 | 1.5909438 | 1.5972182 | 1.65231 | -0.055088402 | -0.061362776 |

| | | | | | | | | | | |
|---|---|---|---|---|---|---|---|---|---|---|
| Cdadc1 | 8.147086 | 8.058592 | 8.1772 | -0.118606536 | -0.030113041 | 8.0609959 | 8.0181175 | 8.07321 | -0.05509729 | -0.012218833 |
| Acly | 12.13104 | 12.19236 | 12.117 | 0.075405952 | 0.014084192 | 12.132678 | 12.03517 | 12.0903 | -0.055100944 | 0.04240722 |
| Gm3839 | 2.55656 | 2.653742 | 2.708 | -0.054252855 | -0.151435091 | 2.5264009 | 2.6326361 | 2.68779 | -0.055149799 | -0.161384947 |
| Gm8116 | 6.254931 | 6.35899 | 6.3707 | -0.011746066 | -0.115805207 | 6.2850861 | 6.0867209 | 6.14187 | -0.05515099 | 0.143214226 |
| Gltp | 10.79531 | 10.7542 | 10.858 | -0.104120048 | -0.06301064 | 10.824223 | 10.660626 | 10.7158 | -0.055195304 | 0.108402179 |
| Stac | 0.907991 | 0.933804 | 0.9765 | -0.042680394 | -0.068493033 | 0.9929052 | 0.9402199 | 0.99545 | -0.055232956 | -0.002547666 |
| Myo1c | 11.82476 | 11.92345 | 11.875 | 0.04832688 | -0.050369422 | 12.214041 | 11.964844 | 12.0201 | -0.055288432 | 0.193908582 |
| Cox6b2 | 3.037692 | 3.144877 | 3.1604 | -0.015541122 | -0.122726113 | 3.1460661 | 3.0144061 | 3.0697 | -0.055296891 | 0.076363139 |
| Kcnj3 | 1.241914 | 1.245327 | 1.2299 | 0.015444031 | 0.012030857 | 1.207005 | 1.1545981 | 1.20991 | -0.055312642 | -0.002905799 |
| Brsk1 | 3.094039 | 3.111 | 3.0787 | 0.032250625 | 0.015289736 | 3.0106951 | 3.0159345 | 3.07135 | -0.055414595 | -0.060654045 |
| Ighv1-12 | 5.866256 | 5.820304 | 5.768 | 0.052329366 | 0.098281562 | 6.0869888 | 5.9350131 | 5.99048 | -0.055466477 | 0.096509244 |
| Gm17849 | 0.55636 | 0.535928 | 0.5754 | -0.039429773 | -0.018997541 | 0.557459 | 0.5501458 | 0.60567 | -0.055520764 | -0.048207491 |
| Taf6l | 7.539949 | 7.513674 | 7.6026 | -0.08896777 | -0.062692468 | 7.5272673 | 7.6046086 | 7.66017 | -0.055559594 | -0.13290089 |
| Gm5239 | 2.268074 | 2.256147 | 2.1882 | 0.067933272 | 0.079860108 | 2.2174577 | 2.2153477 | 2.27092 | -0.055574471 | -0.053464507 |
| Necab1 | 1.434877 | 1.414722 | 1.4038 | 0.010884295 | 0.031038934 | 1.4144985 | 1.4320786 | 1.48766 | -0.055577701 | -0.073157814 |
| Fth-ps2 | 2.445543 | 2.459576 | 2.4154 | 0.044158797 | 0.030126607 | 2.4597741 | 2.4441653 | 2.49974 | -0.055578736 | -0.039969965 |
| Dlgap3 | 3.256177 | 3.198499 | 3.298 | -0.099491067 | -0.041812791 | 3.3034804 | 3.2732307 | 3.32881 | -0.055578922 | -0.025329248 |
| Vmn2r22 | 1.73011 | 1.743767 | 1.6736 | 0.070136265 | 0.056480026 | 1.7169917 | 1.7276313 | 1.78324 | -0.055609025 | -0.066248575 |
| Gm13212 | 8.743993 | 8.761306 | 8.9505 | -0.189179982 | -0.206493358 | 8.9111346 | 8.866814 | 8.92244 | -0.055626368 | -0.011305741 |
| Tdp2 | 8.778631 | 8.782726 | 8.8863 | -0.103538289 | -0.107633739 | 8.9414729 | 8.947164 | 9.00283 | -0.055666375 | -0.061357443 |
| Lypd1 | 1.611975 | 1.490347 | 1.5455 | -0.055137677 | 0.066490394 | 1.4161276 | 1.5002329 | 1.55591 | -0.0556778 | -0.139783132 |
| Myh7 | 4.810454 | 4.974083 | 4.8359 | 0.138147168 | -0.025481946 | 5.2551737 | 5.0951003 | 5.15082 | -0.055723413 | 0.104349984 |
| Gm14440 | 4.673776 | 3.735694 | 3.9802 | -0.244555355 | 0.693526477 | 3.872009 | 4.087628 | 4.14345 | -0.055823516 | -0.271442555 |
| Terf2ip | 8.803554 | 8.862745 | 8.792 | 0.070746133 | 0.011554681 | 8.6404497 | 8.6863632 | 8.7422 | -0.055836198 | -0.101749689 |
| Dnah10 | 3.151733 | 3.045355 | 3.0402 | 0.005137697 | 0.111515512 | 3.0419586 | 2.990384 | 3.04628 | -0.055899865 | -0.004325324 |
| Hist1h2bm | 1.57957 | 1.641192 | 1.6158 | 0.025362327 | -0.036260354 | 1.5889729 | 1.6011451 | 1.65706 | -0.055910854 | -0.068083133 |
| Syngap1 | 5.207169 | 5.257714 | 5.2196 | 0.038143372 | -0.012402248 | 4.9297655 | 5.0063053 | 5.06224 | -0.055938152 | -0.132477951 |
| Bcr | 9.116088 | 8.928111 | 9.0859 | -0.157757527 | 0.030218883 | 8.6877506 | 8.9918024 | 9.04775 | -0.055945963 | -0.359997817 |
| Olfr120 | 1.034359 | 1.033851 | 1.0312 | 0.002664432 | 0.003172728 | 1.0914405 | 1.0388517 | 1.09484 | -0.055985159 | -0.003396368 |
| Ighv2-9 | 5.753759 | 5.890786 | 5.3438 | 0.546950881 | 0.409923476 | 6.0239299 | 5.4973581 | 5.55335 | -0.055991952 | 0.470579861 |
| 4930447C04Rik | 0.578502 | 0.552078 | 0.5751 | -0.022990271 | 0.003433627 | 0.6137311 | 0.5438059 | 0.59982 | -0.056016643 | 0.013908577 |
| Atp10d | 10.36095 | 10.33045 | 10.407 | -0.076133494 | -0.045629092 | 10.346305 | 10.34739 | 10.4034 | -0.056033138 | -0.057117715 |

| | | | | | | | | | | |
|---|---|---|---|---|---|---|---|---|---|---|
| Zfp84 | 8.771766 | 8.942047 | 8.8381 | 0.103977307 | -0.066303712 | 8.8088935 | 8.8308646 | 8.88691 | -0.056049843 | -0.078020881 |
| Hsd17b11 | 9.549392 | 9.629137 | 9.6598 | -0.030686401 | -0.110431161 | 9.6470249 | 9.4099999 | 9.46608 | -0.0560821 | 0.180942893 |
| 4930444P10Rik | 0.929786 | 0.949485 | 0.9086 | 0.040884958 | 0.021186372 | 0.9203649 | 0.9385673 | 0.99467 | -0.056097802 | -0.074300252 |
| H2afy2 | 2.29368 | 2.28723 | 2.2785 | 0.008735681 | 0.015186489 | 2.2937873 | 2.213429 | 2.26954 | -0.056109109 | 0.024249167 |
| Abhd6 | 7.269618 | 7.399541 | 7.3348 | 0.06477995 | -0.065143127 | 7.19303 | 7.263015 | 7.31914 | -0.056126462 | -0.126111399 |
| Gm4767 | 4.808003 | 4.879706 | 4.9306 | -0.050847641 | -0.122550326 | 4.8141787 | 4.6404846 | 4.69662 | -0.056131027 | 0.117563126 |
| Kir3dl1 | 0.596881 | 0.656083 | 0.6156 | 0.040453127 | -0.01874923 | 0.575554 | 0.5417741 | 0.598 | -0.056224804 | -0.022444921 |
| Wif1 | 0.702447 | 0.647581 | 0.7507 | -0.103115648 | -0.048249555 | 0.6324504 | 0.6264527 | 0.68271 | -0.056253954 | -0.050256185 |
| Gp1ba | 0.898012 | 0.920685 | 0.915 | 0.005707618 | -0.016964973 | 0.8276424 | 0.8329271 | 0.88922 | -0.056289916 | -0.061574585 |
| Capns2 | 1.076637 | 1.139475 | 1.0909 | 0.04855767 | -0.014280678 | 1.13826 | 1.1242488 | 1.18054 | -0.056294106 | -0.042282888 |
| Galt | 8.290598 | 8.201716 | 8.5151 | -0.313429457 | -0.224547285 | 8.471094 | 8.2830168 | 8.33932 | -0.05630693 | 0.131770221 |
| Mslnl | 0.185585 | 0.174723 | 0.204 | -0.029319791 | -0.018457851 | 0.2455076 | 0.1588078 | 0.21522 | -0.05641132 | 0.030288406 |
| Gsdma2 | 2.663745 | 2.552962 | 2.6383 | -0.085309717 | 0.025473013 | 2.5627861 | 2.6551342 | 2.71155 | -0.056416697 | -0.14876481 |
| Gm13244 | 0.264719 | 0.280159 | 0.2887 | -0.008586088 | -0.024026071 | 0.2533954 | 0.2957575 | 0.35219 | -0.056431922 | -0.098793963 |
| Sbsn | 5.544273 | 5.5163 | 5.5969 | -0.080644083 | -0.052670843 | 5.4252196 | 5.5049138 | 5.56138 | -0.056462474 | -0.136156679 |
| Agt | 2.315794 | 1.925401 | 1.9504 | -0.024961069 | 0.365432037 | 2.1290177 | 1.8720301 | 1.9286 | -0.05656732 | 0.200420338 |
| Nars2 | 7.463964 | 7.62632 | 7.4565 | 0.169801009 | 0.007444487 | 7.1848856 | 7.493259 | 7.54983 | -0.05656792 | -0.364941348 |
| Slc9a8 | 9.993253 | 9.869292 | 9.9803 | -0.110962106 | 0.012998771 | 10.369643 | 10.241486 | 10.2981 | -0.056568543 | 0.071589122 |
| Olfr731 | 4.260691 | 4.270784 | 4.3076 | -0.036791778 | -0.046884294 | 4.4686477 | 4.3258386 | 4.38247 | -0.056635705 | 0.086173322 |
| Traj13 | 2.302787 | 2.386198 | 2.2919 | 0.094301953 | 0.010891118 | 2.3005839 | 2.3001755 | 2.35681 | -0.056635754 | -0.056227383 |
| Fut4 | 6.717505 | 6.875221 | 6.7901 | 0.085150301 | -0.0725659 | 6.4320006 | 6.4168742 | 6.47352 | -0.056646771 | -0.041520299 |
| Zfp322a | 8.70833 | 8.733364 | 8.6173 | 0.116108216 | 0.091074062 | 8.6770059 | 8.5182288 | 8.57489 | -0.05666272 | 0.102114351 |
| Maf | 9.405365 | 9.219511 | 9.332 | -0.112475136 | 0.073378586 | 8.7642886 | 9.2129591 | 9.26966 | -0.056704276 | -0.505374788 |
| Gin1 | 6.628953 | 6.667231 | 6.8654 | -0.198154922 | -0.236432833 | 6.9485439 | 6.8676805 | 6.92439 | -0.056709984 | 0.024153435 |
| Atp2a1 | 3.536145 | 3.709633 | 3.9404 | -0.230808099 | -0.404295705 | 3.6360064 | 3.5946208 | 3.65137 | -0.056748295 | -0.015362747 |
| Gm6630 | 1.039057 | 1.037041 | 1.0303 | 0.006735095 | 0.00875085 | 1.0296454 | 1.0547513 | 1.11151 | -0.056754081 | -0.081859961 |
| Shd | 0.52302 | 0.529578 | 0.5662 | -0.036624171 | -0.043182077 | 0.535266 | 0.5099481 | 0.5667 | -0.05675578 | -0.031437901 |
| Elmod3 | 7.86159 | 7.868301 | 7.9589 | -0.09062926 | -0.097340883 | 7.8985781 | 7.7372348 | 7.79402 | -0.056785575 | 0.104557731 |
| Pramef8 | 7.938834 | 7.921388 | 7.879 | 0.042354459 | 0.059800509 | 7.8892435 | 7.8882894 | 7.94508 | -0.056791883 | -0.055837786 |
| Cd52 | 13.10538 | 13.02518 | 13.081 | -0.056120344 | 0.024089022 | 13.343876 | 13.075653 | 13.1325 | -0.056806402 | 0.211416516 |
| Gm13577 | 1.409657 | 1.422849 | 1.3994 | 0.023472874 | 0.010280485 | 1.3635029 | 1.3644518 | 1.42128 | -0.056829913 | -0.057778796 |
| Foxd2 | 3.571799 | 3.541512 | 3.5509 | -0.009410219 | 0.020877017 | 3.5442005 | 3.4164164 | 3.47325 | -0.0568352 | 0.070948879 |

| | | | | | | | | | | |
|---|---|---|---|---|---|---|---|---|---|---|
| Nthl1 | 5.040791 | 5.060376 | 5.0206 | 0.039753681 | 0.020168454 | 5.1060452 | 4.9997329 | 5.05663 | -0.056894434 | 0.049417892 |
| Glra3 | 0.487221 | 0.480601 | 0.4971 | -0.016515575 | -0.009896168 | 0.4513366 | 0.4758916 | 0.5328 | -0.056906689 | -0.081461671 |
| Siglecg | 11.34909 | 11.43044 | 11.482 | -0.051897516 | -0.133242084 | 12.173579 | 11.668284 | 11.7252 | -0.05692272 | 0.448372848 |
| Gm12978 | 0.882409 | 0.827523 | 0.821 | 0.006541676 | 0.061427499 | 1.0097469 | 0.7905047 | 0.84743 | -0.056924214 | 0.162318 |
| Zscan30 | 2.406874 | 2.437214 | 2.3848 | 0.052372928 | 0.022033045 | 2.4633432 | 2.3633724 | 2.42034 | -0.05697174 | 0.042999036 |
| Rpsa-ps2 | 4.72515 | 5.00644 | 5.0003 | 0.006132652 | -0.275158175 | 4.9364386 | 5.0179628 | 5.07495 | -0.056987761 | -0.138512015 |
| Gm38317 | 0.587341 | 0.642423 | 0.5546 | 0.087796595 | 0.032714361 | 0.5639875 | 0.5284288 | 0.58543 | -0.057005048 | -0.021446362 |
| Uri1 | 9.755739 | 9.66764 | 9.7727 | -0.105085739 | -0.016986817 | 9.6331019 | 9.7227987 | 9.77985 | -0.057055176 | -0.146751985 |
| Mill2 | 5.160202 | 4.843889 | 5.2338 | -0.389920765 | -0.073607212 | 4.6010798 | 4.8058494 | 4.86297 | -0.057125339 | -0.261895005 |
| Nostrin | 7.226579 | 7.060996 | 7.2542 | -0.193155994 | -0.027572708 | 6.8686367 | 7.121703 | 7.17886 | -0.057154966 | -0.310221193 |
| Hist1h2ak | 1.519239 | 1.47677 | 1.5868 | -0.110041255 | -0.06757252 | 1.6201747 | 1.50759 | 1.56476 | -0.057174301 | 0.055410433 |
| Vmn1r211 | 1.562238 | 1.582405 | 1.5226 | 0.059843852 | 0.039676658 | 1.5801488 | 1.6619676 | 1.71922 | -0.057248499 | -0.139067337 |
| P2rx3 | 2.645677 | 2.597247 | 2.547 | 0.050223632 | 0.098653577 | 2.5608718 | 2.6146473 | 2.67191 | -0.057263801 | -0.111039317 |
| Fam160a2 | 8.923228 | 9.00508 | 8.9853 | 0.019818397 | -0.062033979 | 9.186475 | 9.1083188 | 9.16559 | -0.057273304 | 0.020882914 |
| Gm10012 | 1.579036 | 1.547791 | 1.5556 | -0.007780029 | 0.023465054 | 1.6232078 | 1.5719278 | 1.62922 | -0.057289029 | -0.006009024 |
| Foxh1 | 2.108612 | 2.186473 | 2.1876 | -0.001176343 | -0.079037145 | 2.1555087 | 2.0533006 | 2.11061 | -0.057309327 | 0.044898783 |
| Traj40 | 1.539785 | 1.4839 | 1.489 | -0.005127143 | 0.050757707 | 1.4290233 | 1.4260522 | 1.48339 | -0.057339627 | -0.054368506 |
| Zfp661 | 6.143292 | 6.278144 | 6.3628 | -0.084642065 | -0.219494223 | 6.2877319 | 6.3350865 | 6.39245 | -0.05736253 | -0.104717112 |
| Vmn2r116 | 3.116764 | 3.230079 | 3.1223 | 0.107754403 | -0.005560385 | 3.1920283 | 3.3606463 | 3.41801 | -0.057366257 | -0.22598431 |
| Rnf208 | 1.458858 | 1.535075 | 1.4924 | 0.042641957 | -0.033575274 | 1.4515733 | 1.4065314 | 1.46393 | -0.057395489 | -0.012353635 |
| F2rl1 | 8.552144 | 8.687148 | 8.6357 | 0.051417291 | -0.083586599 | 8.3776634 | 8.5379637 | 8.59545 | -0.057486311 | -0.217786582 |
| Lrat | 2.620367 | 2.651689 | 2.5307 | 0.120987613 | 0.089665505 | 2.6620193 | 2.4538307 | 2.51134 | -0.057509806 | 0.150678817 |
| Galnt17 | 3.686485 | 3.723581 | 3.5939 | 0.129660273 | 0.09256414 | 3.5428101 | 3.5108259 | 3.56836 | -0.057530942 | -0.025546739 |
| Defb42 | 2.390266 | 2.404231 | 2.3716 | 0.032667919 | 0.018703331 | 2.2941328 | 2.2906507 | 2.34829 | -0.057643323 | -0.054161197 |
| Gm525 | 1.463975 | 1.359209 | 1.4261 | -0.066875387 | 0.03789038 | 1.3228005 | 1.2675441 | 1.32519 | -0.057649174 | -0.002392779 |
| Cnpy2 | 8.094568 | 8.016998 | 8.1683 | -0.151336178 | -0.073765579 | 8.1814078 | 8.0372601 | 8.09493 | -0.057668834 | 0.086478801 |
| Nat8f3 | 1.463529 | 1.399353 | 1.5115 | -0.112135217 | -0.047959391 | 1.415323 | 1.3709296 | 1.42865 | -0.057721004 | -0.013327615 |
| Fam205a1 | 0.818157 | 0.892337 | 0.8957 | -0.003380167 | -0.077559508 | 0.8202128 | 0.774509 | 0.8323 | -0.057787349 | -0.01208355 |
| Ccr7 | 12.22311 | 12.29661 | 12.279 | 0.017254112 | -0.05624764 | 12.009967 | 12.21 | 12.2678 | -0.057790128 | -0.257822903 |
| Gm9083 | 0.749723 | 0.762742 | 0.7698 | -0.007081588 | -0.020100432 | 0.802922 | 0.7328957 | 0.79069 | -0.057798963 | 0.012227251 |
| Ip6k2 | 8.098212 | 8.036956 | 8.2087 | -0.171711812 | -0.110455639 | 7.9854343 | 8.1277517 | 8.18556 | -0.057806374 | -0.200123787 |
| Cry2 | 8.066827 | 8.038611 | 8.0164 | 0.022191404 | 0.050407247 | 8.0903039 | 8.0368085 | 8.09462 | -0.057812748 | -0.004317329 |

| | | | | | | | | | | |
|---|---|---|---|---|---|---|---|---|---|---|
| Tm9sf3 | 12.09417 | 11.99727 | 12.019 | -0.021583076 | 0.07531811 | 11.895943 | 11.849261 | 11.9071 | -0.057823504 | -0.011141003 |
| Mtbp | 8.040641 | 7.840283 | 8.0221 | -0.181772722 | 0.018585717 | 8.28686 | 8.2658811 | 8.32375 | -0.057872989 | -0.036894042 |
| Hpx | 1.000525 | 0.981009 | 0.9482 | 0.032839296 | 0.052355969 | 1.0136502 | 0.9422219 | 1.00016 | -0.057935664 | 0.013492697 |
| Ntn3 | 1.12147 | 1.038353 | 1.0438 | -0.005398994 | 0.07771754 | 0.9973788 | 0.9373548 | 0.99529 | -0.057938214 | 0.002085824 |
| Dph6 | 8.354741 | 8.421769 | 8.2318 | 0.190009707 | 0.122981704 | 8.5554428 | 8.4248541 | 8.4828 | -0.057940955 | 0.072647659 |
| Ndfip2 | 9.012263 | 8.869561 | 8.9045 | -0.034896492 | 0.107805189 | 8.8226682 | 8.7593353 | 8.81729 | -0.057951124 | 0.005381833 |
| Cdh16 | 0.537601 | 0.538382 | 0.6075 | -0.069077255 | -0.069857709 | 0.5754179 | 0.5451682 | 0.60316 | -0.057987665 | -0.027738031 |
| Ceacam2 | 2.556168 | 2.548001 | 2.491 | 0.056987261 | 0.06515417 | 2.6208122 | 2.4812842 | 2.5393 | -0.058013834 | 0.081514198 |
| Gm7964 | 2.132217 | 2.21616 | 2.2262 | -0.010044749 | -0.093987822 | 2.2331487 | 2.1710645 | 2.22914 | -0.058077818 | 0.004006434 |
| Gm48364 | 2.979829 | 2.7659 | 2.9835 | -0.217611879 | -0.003683052 | 2.9161841 | 2.9423977 | 3.00051 | -0.058114583 | -0.084328233 |
| Vsnl1 | 1.959841 | 1.962115 | 1.9275 | 0.03459315 | 0.03232008 | 1.7941529 | 1.8307686 | 1.8889 | -0.058126879 | -0.094742602 |
| Pcif1 | 10.81697 | 10.72451 | 10.779 | -0.054666334 | 0.03779402 | 11.063012 | 10.774937 | 10.8331 | -0.058165226 | 0.229909587 |
| Saraf | 11.83744 | 11.93031 | 11.944 | -0.013214166 | -0.106083998 | 11.59031 | 11.775319 | 11.8335 | -0.058227942 | -0.243237583 |
| Gm45902 | 6.638937 | 6.580259 | 6.5369 | 0.043409069 | 0.102087059 | 6.6439174 | 6.72139 | 6.77964 | -0.058247359 | -0.135719898 |
| Nxpe2 | 9.158527 | 9.24175 | 9.0439 | 0.197883468 | 0.114660447 | 9.0507997 | 8.9500205 | 9.0083 | -0.058281078 | 0.042498083 |
| Olfr922 | 0.634624 | 0.63599 | 0.6398 | -0.003821671 | -0.005187633 | 0.7245714 | 0.6445524 | 0.70289 | -0.058338136 | 0.021680898 |
| Wfdc1 | 0.909678 | 0.891902 | 0.8657 | 0.026157581 | 0.043933101 | 0.8571759 | 0.836246 | 0.89459 | -0.058340123 | -0.037410216 |
| Pclo | 2.039395 | 1.949296 | 1.8515 | 0.097811478 | 0.18790976 | 1.9303338 | 1.8876018 | 1.946 | -0.058396413 | -0.01566439 |
| Trav13-2 | 4.060564 | 4.094948 | 4.1334 | -0.038482165 | -0.072866079 | 4.0806261 | 4.05928 | 4.11771 | -0.058434598 | -0.037088571 |
| Vmn1r181 | 1.009854 | 0.98097 | 1.0202 | -0.0392708 | -0.010386471 | 1.0274363 | 1.0383833 | 1.09684 | -0.058460353 | -0.069407349 |
| Col7a1 | 4.41732 | 4.68328 | 4.675 | 0.00828305 | -0.257676444 | 4.9148441 | 4.8756173 | 4.9341 | -0.058482146 | -0.019255272 |
| Gas2l1 | 6.222998 | 6.363351 | 6.2301 | 0.133257072 | -0.007095384 | 6.1514451 | 6.0825164 | 6.14103 | -0.058513612 | 0.010415121 |
| Tecta | 1.023411 | 1.09108 | 1.0831 | 0.007958476 | -0.059710246 | 1.0889457 | 1.0870067 | 1.14552 | -0.05851555 | -0.056576474 |
| Igkv11-114 | 1.423573 | 1.414731 | 1.4297 | -0.014996936 | -0.006155523 | 1.3896997 | 1.3680637 | 1.42658 | -0.058516868 | -0.036880863 |
| Ephb3 | 2.302893 | 2.346137 | 2.4212 | -0.075083366 | -0.118327626 | 2.3662708 | 2.189172 | 2.24777 | -0.058598065 | 0.118500701 |
| Pxk | 11.71687 | 11.70331 | 11.759 | -0.055309202 | -0.041744472 | 12.018971 | 11.772395 | 11.831 | -0.058618424 | 0.187957129 |
| Brap | 10.3459 | 10.35364 | 10.336 | 0.017457664 | 0.009715358 | 10.347053 | 10.286842 | 10.3455 | -0.058697776 | 0.001513149 |
| Igkv7-33 | 2.471365 | 2.465211 | 2.5385 | -0.073253297 | -0.067098856 | 2.6271224 | 2.582902 | 2.64167 | -0.058763153 | -0.014542753 |
| Elmo3 | 6.648733 | 6.568807 | 6.6951 | -0.126325629 | -0.046398805 | 6.6605996 | 6.7760872 | 6.83492 | -0.058830297 | -0.174317872 |
| Dcbld2 | 5.431341 | 5.710191 | 5.6372 | 0.072949787 | -0.205899761 | 5.6535814 | 5.8934423 | 5.95229 | -0.058844693 | -0.298705572 |
| AC113953.1 | 1.818176 | 1.797041 | 1.8072 | -0.010163143 | 0.010972263 | 1.6990262 | 1.723929 | 1.78279 | -0.058865935 | -0.083768702 |
| Art5 | 2.747537 | 2.44375 | 2.5596 | -0.115894582 | 0.187892394 | 2.5712625 | 2.5250011 | 2.58387 | -0.058869073 | -0.012607694 |

| | | | | | | | | | | |
|---|---|---|---|---|---|---|---|---|---|---|
| Clint1 | 11.63043 | 11.64295 | 11.631 | 0.012312376 | -0.000209298 | 11.581492 | 11.565534 | 11.6244 | -0.058869252 | -0.042910598 |
| Gas7 | 8.815095 | 8.776464 | 8.6347 | 0.141793695 | 0.180424264 | 7.5904324 | 8.1102021 | 8.16912 | -0.058913288 | -0.578683025 |
| Lrif1 | 9.445049 | 9.431349 | 9.4567 | -0.025304077 | -0.011604525 | 9.6097634 | 9.5805008 | 9.63944 | -0.058938008 | -0.029675424 |
| Opn3 | 4.065384 | 3.807578 | 3.8341 | -0.02651651 | 0.231289362 | 4.0406916 | 3.8297207 | 3.88867 | -0.058946973 | 0.15202394 |
| Rac2 | 13.52926 | 13.57587 | 13.658 | -0.08216671 | -0.12877232 | 13.796579 | 13.520524 | 13.5795 | -0.058959986 | 0.217094443 |
| Tmem82 | 3.2513 | 3.349101 | 3.2717 | 0.077365484 | -0.020435669 | 3.2961982 | 3.2345727 | 3.29362 | -0.059047268 | 0.002578292 |
| Triml1 | 2.920426 | 2.888883 | 2.9716 | -0.082692396 | -0.051148689 | 2.9546496 | 2.9302053 | 2.98926 | -0.059059595 | -0.034615274 |
| Gm16253 | 0.332002 | 0.327053 | 0.3608 | -0.033727947 | -0.028778689 | 0.3670589 | 0.3218864 | 0.38096 | -0.059075629 | -0.01390305 |
| Rgl3 | 3.797675 | 3.784699 | 3.8942 | -0.109487803 | -0.096511247 | 3.7198783 | 3.666141 | 3.72523 | -0.059087642 | -0.005350305 |
| Ccng1 | 10.92454 | 10.91122 | 10.98 | -0.068975266 | -0.055649846 | 10.782449 | 10.864292 | 10.9234 | -0.05909144 | -0.140934755 |
| Eif4g2 | 14.10076 | 14.14135 | 14.081 | 0.060030726 | 0.019443902 | 14.10855 | 14.12485 | 14.1839 | -0.059095218 | -0.075395136 |
| Zbed5 | 4.550436 | 4.459661 | 4.5572 | -0.097548119 | -0.006772465 | 4.6001667 | 4.5419326 | 4.60104 | -0.059109033 | -0.000874938 |
| Lrrc27 | 3.136995 | 3.007421 | 3.0949 | -0.087474551 | 0.04209962 | 3.0116339 | 3.0050811 | 3.06419 | -0.059111166 | -0.05255828 |
| Adcy8 | 1.241249 | 1.215024 | 1.3022 | -0.08715775 | -0.060932723 | 1.3640096 | 1.2080795 | 1.2672 | -0.05912534 | 0.096804731 |
| Agpat2 | 7.412377 | 7.361736 | 7.5021 | -0.14033119 | -0.089690038 | 7.5015281 | 7.2928929 | 7.35206 | -0.059164524 | 0.149470667 |
| Barx2 | 1.213126 | 1.093761 | 1.1727 | -0.078959821 | 0.040404833 | 1.1451045 | 1.1665336 | 1.22574 | -0.059210631 | -0.080639802 |
| Zfp937 | 6.748253 | 6.816568 | 6.7141 | 0.102458372 | 0.034142617 | 6.6648915 | 6.5783372 | 6.63758 | -0.059239564 | 0.027314763 |
| Gm48416 | 7.13191 | 7.164414 | 7.3437 | -0.179301436 | -0.211805435 | 7.1314949 | 7.5469315 | 7.60625 | -0.059319952 | -0.474756535 |
| Nap1l4 | 11.2488 | 11.26582 | 11.202 | 0.064178268 | 0.047153622 | 11.249209 | 11.251623 | 11.311 | -0.059359628 | -0.061772718 |
| Sh3rf3 | 2.213607 | 2.043514 | 1.9123 | 0.131240713 | 0.301333942 | 1.9294758 | 1.9064348 | 1.96581 | -0.059379435 | -0.036338432 |
| Nt5dc2 | 5.302985 | 5.036979 | 5.2781 | -0.241093028 | 0.024913052 | 5.1630493 | 5.0324972 | 5.09188 | -0.059383044 | 0.071169063 |
| Tigd5 | 3.646917 | 3.677763 | 3.7647 | -0.086967633 | -0.117813007 | 3.6950538 | 3.6758272 | 3.73523 | -0.059398778 | -0.040172158 |
| Gm10490 | 2.460141 | 2.510752 | 2.5055 | 0.005216267 | -0.045394754 | 2.4655257 | 2.5499734 | 2.60943 | -0.059453737 | -0.143901425 |
| Map1b | 6.353404 | 6.192 | 6.134 | 0.05798586 | 0.219389879 | 5.79996 | 6.005968 | 6.06543 | -0.059466741 | -0.265474753 |
| Dgkg | 4.042154 | 3.837388 | 3.6899 | 0.147514965 | 0.352281491 | 3.6407086 | 3.7086056 | 3.76814 | -0.059534147 | -0.127431116 |
| Rgma | 2.403274 | 2.496845 | 2.4407 | 0.056116692 | -0.037454398 | 2.5363339 | 2.4354518 | 2.495 | -0.059550234 | 0.04133191 |
| Gpr75 | 1.083224 | 1.139328 | 1.1409 | -0.001581972 | -0.057685867 | 1.1293094 | 1.112613 | 1.17217 | -0.059560464 | -0.042864013 |
| Gm21378 | 2.797178 | 2.719242 | 2.8852 | -0.165929765 | -0.087993857 | 2.6416007 | 2.5681283 | 2.62771 | -0.059579904 | 0.013892532 |
| Kcnh7 | 1.70023 | 1.72952 | 1.7504 | -0.020850118 | -0.050140016 | 1.7891175 | 1.7377949 | 1.79744 | -0.059646782 | -0.008324164 |
| Sox8 | 1.909978 | 1.878371 | 1.8379 | 0.040512054 | 0.072119338 | 1.8410159 | 1.7360818 | 1.79574 | -0.059659902 | 0.045274154 |
| Slfn10-ps | 6.804584 | 6.745688 | 6.8013 | -0.055565228 | 0.003331153 | 6.5328382 | 6.812703 | 6.87239 | -0.059691339 | -0.339556147 |
| Gm42915 | 0.700731 | 0.613055 | 0.6476 | -0.034542485 | 0.05313296 | 0.6309173 | 0.6591352 | 0.71883 | -0.059696436 | -0.087914287 |

| | | | | | | | | | | |
|---|---|---|---|---|---|---|---|---|---|---|
| Zfp775 | 6.709017 | 6.781464 | 6.9229 | -0.141395045 | -0.213841588 | 6.6119618 | 6.4005824 | 6.46029 | -0.059710592 | 0.151668849 |
| Plcxd1 | 3.872478 | 3.97323 | 3.964 | 0.009267237 | -0.091485405 | 3.8436781 | 3.9084472 | 3.96819 | -0.059743505 | -0.124512536 |
| Mtmr2 | 9.942213 | 9.840528 | 9.8591 | -0.018598567 | 0.083086852 | 9.8476769 | 9.8265986 | 9.88636 | -0.059764361 | -0.038686051 |
| Mta2 | 12.12713 | 12.06492 | 12.12 | -0.054623918 | 0.007576958 | 12.132982 | 12.037016 | 12.0968 | -0.05977411 | 0.036192404 |
| Grb14 | 1.361671 | 1.387707 | 1.3594 | 0.028355059 | 0.002319827 | 1.3738845 | 1.2936329 | 1.35341 | -0.059778348 | 0.020473222 |
| Slc6a15 | 0.554004 | 0.581737 | 0.5251 | 0.056591272 | 0.028857546 | 0.6368275 | 0.5435519 | 0.60335 | -0.059801634 | 0.033473889 |
| Rcor2 | 4.450574 | 4.594424 | 4.5995 | -0.005104877 | -0.148954465 | 4.4079004 | 4.5475324 | 4.60735 | -0.05982206 | -0.199454062 |
| 1700057G04Rik | 0.945674 | 0.91672 | 0.9108 | 0.005921049 | 0.034875004 | 0.9134463 | 0.9154445 | 0.97531 | -0.059866782 | -0.061864939 |
| Prkab2 | 7.442565 | 7.535124 | 7.4542 | 0.080959618 | -0.011599865 | 7.8555253 | 7.8114409 | 7.87131 | -0.059867929 | -0.015783573 |
| Olfr1217 | 1.385865 | 1.374879 | 1.3817 | -0.00677459 | 0.004211386 | 1.4217148 | 1.3912659 | 1.45116 | -0.059896863 | -0.029448031 |
| Nol4 | 1.239856 | 1.217284 | 1.2325 | -0.015208068 | 0.007364528 | 1.2389263 | 1.2291768 | 1.28908 | -0.059901705 | -0.050152122 |
| Wtap | 11.14094 | 11.06295 | 11.032 | 0.03087212 | 0.108861676 | 10.970097 | 11.016818 | 11.0767 | -0.059931902 | -0.106652487 |
| Itpa-ps1 | 0.662619 | 0.689349 | 0.711 | -0.021693198 | -0.048423251 | 0.6950665 | 0.710496 | 0.77044 | -0.059941559 | -0.075370978 |
| Xrcc5 | 9.016518 | 9.008569 | 9.0905 | -0.081965745 | -0.074016852 | 9.06056 | 9.0327621 | 9.0927 | -0.059942605 | -0.03214477 |
| Gm21798 | 0.555194 | 0.547251 | 0.6333 | -0.086064389 | -0.078121736 | 0.5791194 | 0.5732308 | 0.63319 | -0.059957776 | -0.054069191 |
| Gm4617 | 2.910264 | 2.450377 | 2.6162 | -0.165779793 | 0.294107105 | 2.4633057 | 2.4115137 | 2.47148 | -0.059965079 | -0.00817305 |
| Pitx1 | 0.631294 | 0.516337 | 0.5142 | 0.002097755 | 0.117054752 | 0.6195167 | 0.5338128 | 0.59379 | -0.059973687 | 0.025730231 |
| Prokr2 | 3.054637 | 3.176389 | 3.2344 | -0.058016808 | -0.17976806 | 3.0459785 | 3.1641824 | 3.22417 | -0.059989381 | -0.178193227 |
| Rabgef1 | 9.280683 | 9.222874 | 9.2663 | -0.043384361 | 0.014424694 | 9.2408114 | 9.2058024 | 9.26584 | -0.060040821 | -0.025031822 |
| Iqck | 2.205195 | 2.122418 | 2.1331 | -0.01068845 | 0.072088794 | 2.2229637 | 2.1010143 | 2.16109 | -0.060071529 | 0.061877902 |
| Ppp1r14c | 2.22503 | 2.17758 | 2.0944 | 0.083146633 | 0.130595979 | 2.0917052 | 2.1041935 | 2.16434 | -0.060147743 | -0.072636063 |
| Vgll4 | 9.581749 | 9.787865 | 9.6666 | 0.121270784 | -0.084845484 | 9.4962251 | 9.5391714 | 9.59936 | -0.060191179 | -0.103137442 |
| Olfr31 | 3.580034 | 3.643452 | 3.6663 | -0.02288254 | -0.086300556 | 3.6415349 | 3.7175911 | 3.77779 | -0.060201885 | -0.13625802 |
| Catsperb | 0.508284 | 0.512748 | 0.5523 | -0.039598302 | -0.044062277 | 0.4695037 | 0.5096733 | 0.56992 | -0.060244171 | -0.10041384 |
| Cdkn2b | 5.007339 | 5.079875 | 5.0267 | 0.0532041 | -0.019331319 | 4.8762332 | 4.9431016 | 5.00336 | -0.060259832 | -0.127128286 |
| Pcdhb7 | 2.462861 | 2.296889 | 2.3261 | -0.029181332 | 0.136790238 | 2.3994776 | 2.2603904 | 2.32066 | -0.060266921 | 0.078820336 |
| Vmn1r12 | 2.56966 | 2.508484 | 2.5376 | -0.029082968 | 0.032093215 | 2.4731837 | 2.5704704 | 2.63075 | -0.060280079 | -0.157566784 |
| Oaz2 | 9.00638 | 8.840253 | 8.9594 | -0.119149222 | 0.046976971 | 8.8629156 | 8.6859115 | 8.74625 | -0.060333793 | 0.116670354 |
| Otog | 1.631508 | 1.585974 | 1.6717 | -0.085717747 | -0.040183672 | 1.6922797 | 1.6689771 | 1.72937 | -0.060397758 | -0.037095233 |
| Nrbp1 | 11.00351 | 10.95555 | 11.022 | -0.066528384 | -0.018567646 | 11.108613 | 11.09404 | 11.1545 | -0.060445 | -0.045871472 |
| Olfr1378 | 1.477504 | 1.386809 | 1.455 | -0.06819006 | 0.022505468 | 1.4523392 | 1.4362569 | 1.49672 | -0.060460438 | -0.044378134 |
| Gm17402 | 2.321485 | 2.245128 | 2.2208 | 0.024346771 | 0.100704321 | 2.2415334 | 2.2069362 | 2.26742 | -0.060484649 | -0.025887514 |

| | | | | | | | | | | |
|---|---|---|---|---|---|---|---|---|---|---|
| Sec1 | 1.936617 | 2.010712 | 1.8983 | 0.112434814 | 0.038340024 | 1.9179181 | 1.8848194 | 1.94531 | -0.06049448 | -0.027395709 |
| Gm18537 | 4.927315 | 5.054593 | 5.0556 | -0.000978734 | -0.128255972 | 5.4113588 | 5.321079 | 5.38163 | -0.060546464 | 0.029733284 |
| Ncbp1 | 10.5591 | 10.57425 | 10.573 | 0.000993276 | -0.01415928 | 10.500825 | 10.507075 | 10.5677 | -0.060626602 | -0.066876671 |
| Eif2s3x | 12.26694 | 12.25091 | 12.28 | -0.028609712 | -0.012579868 | 12.308149 | 12.207842 | 12.2685 | -0.060691158 | 0.039615395 |
| Rell2 | 5.650469 | 5.582481 | 5.7633 | -0.180780327 | -0.112792025 | 5.878652 | 5.8213104 | 5.88204 | -0.060725115 | -0.003383561 |
| Gm44911 | 1.875382 | 1.760451 | 1.8662 | -0.105763276 | 0.009167787 | 1.9991535 | 1.8367775 | 1.89757 | -0.060797272 | 0.101578778 |
| Kifc3 | 6.440101 | 6.225557 | 6.0987 | 0.126836695 | 0.34138101 | 5.9623298 | 5.9565999 | 6.01743 | -0.060833227 | -0.055103316 |
| Gm10774 | 1.958269 | 2.042336 | 1.9066 | 0.13575918 | 0.051692138 | 1.8957233 | 1.8968152 | 1.95771 | -0.060890308 | -0.061982187 |
| A930033H14Rik | 1.524519 | 1.464158 | 1.4648 | -0.000634275 | 0.059726414 | 1.4553345 | 1.413769 | 1.47466 | -0.06089038 | -0.019324903 |
| Endog | 5.79681 | 5.818742 | 5.8698 | -0.051012038 | -0.072944547 | 5.9690407 | 5.7222371 | 5.78315 | -0.060911588 | 0.185892007 |
| Agap2 | 10.3998 | 10.57809 | 10.434 | 0.144222516 | -0.034065825 | 10.381672 | 10.263335 | 10.3243 | -0.060936569 | 0.057400524 |
| Gm48141 | 0.689831 | 0.735949 | 0.7063 | 0.02969709 | -0.016421698 | 0.7617204 | 0.6998414 | 0.7608 | -0.060961873 | 0.000917174 |
| Vmn2r102 | 4.068585 | 3.973688 | 3.883 | 0.090660003 | 0.185556612 | 3.9122674 | 4.1287211 | 4.18976 | -0.06104005 | -0.277493731 |
| Elf4 | 11.96462 | 12.02689 | 11.95 | 0.07682439 | 0.014554664 | 11.856375 | 11.942681 | 12.0038 | -0.061069672 | -0.147375443 |
| Ppwd1 | 8.14385 | 8.080807 | 8.2188 | -0.137948561 | -0.074905812 | 8.2097795 | 8.1920333 | 8.25314 | -0.06111044 | -0.043364243 |
| Scd4 | 4.221766 | 4.021251 | 4.1575 | -0.13627977 | 0.064235277 | 4.1224429 | 4.1344694 | 4.1956 | -0.061129759 | -0.073156174 |
| Clec4b2 | 0.331726 | 0.285293 | 0.3095 | -0.024221339 | 0.02221136 | 0.2863242 | 0.2515097 | 0.31272 | -0.061205995 | -0.026391498 |
| Bhlha15 | 5.634016 | 5.269316 | 5.4641 | -0.194778141 | 0.169921306 | 5.2416114 | 5.106891 | 5.16812 | -0.0612264 | 0.073494009 |
| Mfsd4a | 5.124879 | 5.051331 | 5.2252 | -0.173911514 | -0.100364009 | 5.2248825 | 5.1065975 | 5.16784 | -0.061244171 | 0.057040756 |
| Tmem237 | 5.849267 | 5.840334 | 5.6589 | 0.181482589 | 0.190415425 | 5.7568085 | 5.8248921 | 5.88614 | -0.061251837 | -0.129335418 |
| Gimap8 | 12.14162 | 12.22189 | 12.185 | 0.0371867 | -0.043089188 | 12.264156 | 12.222595 | 12.2839 | -0.061256304 | -0.019695366 |
| Gm9918 | 1.025576 | 1.058606 | 1.045 | 0.01356761 | -0.019462471 | 1.0720642 | 1.0405488 | 1.10182 | -0.061268378 | -0.02975298 |
| Ckmt1 | 2.316671 | 2.211557 | 2.2603 | -0.048751914 | 0.056361502 | 2.329998 | 2.2065118 | 2.26779 | -0.061281017 | 0.062205142 |
| Bahcc1 | 5.934182 | 6.06907 | 6.0254 | 0.043696153 | -0.091191541 | 5.5340755 | 6.021276 | 6.08264 | -0.061368657 | -0.548569213 |
| Gapvd1 | 11.14859 | 11.21517 | 11.235 | -0.020112322 | -0.086688524 | 11.117326 | 11.171675 | 11.233 | -0.061371236 | -0.115720244 |
| Cfap97d2 | 3.641202 | 3.562528 | 3.6034 | -0.040886276 | 0.037788118 | 3.8242913 | 3.6044706 | 3.66586 | -0.061386757 | 0.158433954 |
| Gm6087 | 0.02876 | 0.081181 | 0.0727 | 0.008500952 | -0.043919983 | 0.0915761 | 0.0440569 | 0.10546 | -0.061407063 | -0.013887879 |
| Akr1cl | 2.291501 | 2.304021 | 2.2754 | 0.02866458 | 0.016144945 | 2.2679059 | 2.220435 | 2.28188 | -0.061449615 | -0.013978764 |
| Rrh | 1.103014 | 1.086798 | 1.131 | -0.04421245 | -0.027996194 | 1.0919303 | 1.0959307 | 1.15739 | -0.061460018 | -0.065460453 |
| Cacng4 | 2.818218 | 2.977204 | 2.6932 | 0.283964734 | 0.124977987 | 2.7209766 | 2.6754273 | 2.73692 | -0.061492286 | -0.015942976 |
| Gm48789 | 0.461796 | 0.425891 | 0.5017 | -0.075784111 | -0.039878941 | 0.4585467 | 0.4457108 | 0.50721 | -0.061502317 | -0.048666456 |
| Ighv9-2 | 5.488505 | 5.539403 | 5.3759 | 0.163511662 | 0.112613415 | 5.6282256 | 5.2198907 | 5.28143 | -0.061534353 | 0.346800566 |

| | | | | | | | | | | |
|---|---|---|---|---|---|---|---|---|---|---|
| Eef1a1 | 17.81099 | 17.82543 | 17.864 | -0.038605002 | -0.053037226 | 17.897792 | 17.765855 | 17.8274 | -0.061545424 | 0.070392255 |
| AW549877 | 10.47908 | 10.25148 | 10.474 | -0.222064906 | 0.005534276 | 10.551485 | 10.555625 | 10.6172 | -0.061551186 | -0.065690756 |
| Slc5a4b | 0.27601 | 0.274707 | 0.2678 | 0.006870196 | 0.008173616 | 0.2893689 | 0.244837 | 0.30641 | -0.061572205 | -0.017040314 |
| Ctdsp1 | 11.06193 | 11.05159 | 11.131 | -0.079154742 | -0.0688174 | 11.166986 | 10.883323 | 10.9449 | -0.061583116 | 0.222079598 |
| Gm17229 | 1.585013 | 1.57018 | 1.6435 | -0.073363558 | -0.058530106 | 1.7094291 | 1.5876356 | 1.64925 | -0.061611842 | 0.060181615 |
| Nrcam | 3.013057 | 2.874203 | 2.9017 | -0.027536615 | 0.111316828 | 2.8104519 | 2.8310867 | 2.89273 | -0.06164443 | -0.082279235 |
| Npb | 1.285789 | 1.248896 | 1.3372 | -0.088319391 | -0.051426011 | 1.3017333 | 1.2866396 | 1.34833 | -0.061691147 | -0.046597359 |
| Tnfsf9 | 5.597888 | 5.409358 | 5.3466 | 0.062744112 | 0.251274122 | 5.2249006 | 5.5045024 | 5.5662 | -0.061702116 | -0.341303956 |
| Olfr432 | 1.752215 | 1.712583 | 1.7777 | -0.065120079 | -0.025487779 | 1.8466563 | 1.7583514 | 1.82006 | -0.061706302 | 0.026598612 |
| Kiss1r | 2.403051 | 2.309219 | 2.3532 | -0.044006624 | 0.049825337 | 2.3156672 | 2.3219067 | 2.38364 | -0.061737758 | -0.067977249 |
| Slc2a2 | 0.119104 | 0.103107 | 0.1264 | -0.02329568 | -0.007298401 | 0.090707 | 0.1027305 | 0.16448 | -0.061747062 | -0.07377048 |
| Olfr167 | 1.185589 | 1.174106 | 1.2054 | -0.031341584 | -0.019858553 | 1.1669321 | 1.1794505 | 1.24121 | -0.061757739 | -0.074276124 |
| Chad | 0.354918 | 0.342542 | 0.3517 | -0.009163228 | 0.003212406 | 0.4616813 | 0.3223415 | 0.38417 | -0.061830542 | 0.077509254 |
| Cd300ld2 | 1.102339 | 0.97541 | 1.1435 | -0.168139581 | -0.041210211 | 0.9784403 | 0.9878731 | 1.04972 | -0.061846714 | -0.0712795 |
| Gm13456 | 4.273768 | 4.357482 | 4.4335 | -0.07597283 | -0.159686837 | 4.3858816 | 4.2894282 | 4.35131 | -0.061878865 | 0.034574538 |
| Rbm5 | 12.04566 | 12.10427 | 12.069 | 0.035510734 | -0.023100432 | 12.166693 | 12.226548 | 12.2885 | -0.061909526 | -0.121764654 |
| Dpy30 | 8.266083 | 8.307452 | 8.4136 | -0.106153398 | -0.147522635 | 8.5462856 | 8.3072673 | 8.36924 | -0.061969544 | 0.177048757 |
| Olfr1440 | 2.295485 | 2.303577 | 2.2617 | 0.041908988 | 0.033817108 | 2.3781102 | 2.3456723 | 2.40766 | -0.061984481 | -0.029546601 |
| Nt5c2 | 9.501244 | 9.506118 | 9.5808 | -0.074726406 | -0.079600054 | 9.562787 | 9.6183814 | 9.68037 | -0.061990383 | -0.117584856 |
| Tcf24 | 0.399397 | 0.350092 | 0.4117 | -0.061642096 | -0.012336945 | 0.361577 | 0.3418378 | 0.4039 | -0.062058954 | -0.042319726 |
| Olfr393 | 3.039446 | 2.980225 | 2.9702 | 0.010071734 | 0.069293321 | 3.0147681 | 3.1049042 | 3.16699 | -0.06208776 | -0.152223859 |
| Meiob | 0.775252 | 0.704666 | 0.7869 | -0.082259357 | -0.011673469 | 0.7193265 | 0.7096386 | 0.77174 | -0.062104552 | -0.052416722 |
| Zfp583 | 4.333927 | 4.345803 | 4.3824 | -0.03654807 | -0.048424773 | 4.4185773 | 4.3568321 | 4.41896 | -0.062125862 | -0.000380615 |
| Ankrd39 | 7.516637 | 7.387367 | 7.239 | 0.148357972 | 0.277628145 | 7.514706 | 7.3849717 | 7.4471 | -0.062126577 | 0.067607754 |
| Itga4 | 12.72706 | 12.89628 | 12.777 | 0.119397942 | -0.049826024 | 12.625269 | 12.73669 | 12.7988 | -0.062139126 | -0.173560281 |
| Ighv8-9 | 0.474012 | 0.50345 | 0.5127 | -0.009206334 | -0.03864446 | 0.4466385 | 0.4630765 | 0.52531 | -0.062237777 | -0.078675825 |
| Cnot7 | 9.900373 | 9.847362 | 9.9022 | -0.054834076 | -0.001823564 | 9.8521553 | 9.819424 | 9.88166 | -0.062238583 | -0.029507311 |
| Adnp2 | 9.208407 | 9.215134 | 9.1827 | 0.032454255 | 0.025727418 | 9.0867483 | 9.0748286 | 9.1371 | -0.062267194 | -0.050347527 |
| Mpp4 | 2.45053 | 2.445419 | 2.4711 | -0.02568477 | -0.020573578 | 2.531178 | 2.4668455 | 2.52918 | -0.062335487 | 0.001997039 |
| Sos2 | 9.522134 | 9.51084 | 9.6354 | -0.124544217 | -0.113250939 | 9.4905872 | 9.632942 | 9.69532 | -0.06237571 | -0.204730509 |
| Trav19 | 4.318254 | 4.242076 | 4.2444 | -0.002281365 | 0.073896101 | 4.0984617 | 4.2680087 | 4.33043 | -0.06242232 | -0.2319693 |
| Pabpc4 | 10.63847 | 10.55703 | 10.673 | -0.115487312 | -0.03404885 | 10.847115 | 10.691544 | 10.754 | -0.062426684 | 0.093144324 |

| | | | | | | | | | | |
|---|---|---|---|---|---|---|---|---|---|---|
| Vps35 | 10.78698 | 10.80947 | 10.806 | 0.003416852 | -0.019075256 | 10.829358 | 10.684265 | 10.7467 | -0.062441236 | 0.082651031 |
| 1110051M20Rik | 5.930195 | 5.892376 | 6.0853 | -0.192927587 | -0.155108657 | 5.962441 | 6.0899165 | 6.15237 | -0.062453946 | -0.189929369 |
| Gm20479 | 0.630357 | 0.690596 | 0.5994 | 0.091218985 | 0.030980648 | 0.6115222 | 0.5571998 | 0.61967 | -0.062471083 | -0.008148709 |
| Cwc22 | 9.945439 | 9.852174 | 9.9738 | -0.121647711 | -0.028383262 | 9.8744089 | 9.8949425 | 9.95749 | -0.06255102 | -0.083084579 |
| Vmn1r174 | 1.65818 | 1.680021 | 1.6915 | -0.011480598 | -0.033321064 | 1.8189322 | 1.6993576 | 1.76192 | -0.062562337 | 0.057012337 |
| Ppp1r2-ps4 | 0.647664 | 0.687593 | 0.6142 | 0.07342095 | 0.03349135 | 0.6435545 | 0.6536355 | 0.71625 | -0.062617827 | -0.072698848 |
| Unc5d | 3.427194 | 3.432194 | 3.5018 | -0.069593387 | -0.074593168 | 3.4790834 | 3.4212152 | 3.48387 | -0.062654335 | -0.004786092 |
| Rassf5 | 11.91454 | 11.98187 | 11.932 | 0.04998839 | -0.017337032 | 11.925922 | 11.822281 | 11.8849 | -0.06265468 | 0.040985698 |
| Gm5628 | 0.816845 | 0.816683 | 0.7796 | 0.037072918 | 0.03723512 | 0.7822642 | 0.7691511 | 0.83186 | -0.062713112 | -0.049599998 |
| Smim9 | 0.309939 | 0.273286 | 0.2433 | 0.030014337 | 0.066666796 | 0.2234254 | 0.2446289 | 0.30737 | -0.06273619 | -0.083939688 |
| Olfr1402 | 2.121534 | 2.127581 | 2.0968 | 0.030770871 | 0.02472364 | 2.1689085 | 2.1340684 | 2.19682 | -0.062749544 | -0.027909468 |
| Gm10571 | 1.979998 | 1.860507 | 1.9092 | -0.048735355 | 0.070755193 | 1.8490996 | 1.9186229 | 1.98141 | -0.062791823 | -0.132315105 |
| Vamp7 | 8.305976 | 8.295654 | 8.3018 | -0.006117543 | 0.004204439 | 8.2543803 | 8.127547 | 8.19035 | -0.062803517 | 0.064029823 |
| Ppp4r4 | 2.554428 | 2.512306 | 2.4401 | 0.072198726 | 0.114321423 | 2.5148379 | 2.4605095 | 2.52336 | -0.06285418 | -0.008525832 |
| Gm3696 | 3.77296 | 3.7735 | 3.7908 | -0.017342673 | -0.01788273 | 3.5864381 | 3.6093526 | 3.6723 | -0.062946186 | -0.085860752 |
| Ppfia3 | 1.480866 | 1.458844 | 1.451 | 0.007834262 | 0.02985677 | 1.4183234 | 1.4055802 | 1.4686 | -0.063023921 | -0.050280752 |
| Mex3a | 6.912455 | 6.8363 | 6.6578 | 0.178546708 | 0.254701099 | 6.530431 | 6.642065 | 6.70513 | -0.063064855 | -0.174698837 |
| Ivns1abp | 11.86988 | 11.9106 | 11.912 | -0.001399378 | -0.04212479 | 11.903931 | 11.892916 | 11.956 | -0.063070268 | -0.052055512 |
| Disp2 | 2.057989 | 2.130526 | 2.0059 | 0.124613229 | 0.052076193 | 2.0245433 | 2.0082057 | 2.07128 | -0.063073793 | -0.046736262 |
| Eid3 | 2.305639 | 2.346629 | 2.2974 | 0.049238173 | 0.008248407 | 2.3758216 | 2.3602237 | 2.42338 | -0.063159474 | -0.047561551 |
| Txndc15 | 8.597712 | 8.538963 | 8.6208 | -0.081851774 | -0.023103196 | 8.6116015 | 8.5124935 | 8.57572 | -0.063226012 | 0.035881998 |
| Tgds | 8.027322 | 8.07068 | 7.9972 | 0.073526163 | 0.030168406 | 8.0759742 | 8.0618727 | 8.12519 | -0.063317961 | -0.049216455 |
| Nup188 | 10.62895 | 10.54492 | 10.608 | -0.062963968 | 0.021059972 | 10.682616 | 10.586978 | 10.6504 | -0.06337934 | 0.032258683 |
| Gm11843 | 0.536315 | 0.560922 | 0.5352 | 0.02568768 | 0.001080373 | 0.570309 | 0.5253553 | 0.58874 | -0.063388296 | -0.018434621 |
| Zfp961 | 8.183014 | 8.287084 | 8.2147 | 0.072341611 | -0.031728203 | 8.343014 | 8.2315374 | 8.29498 | -0.063441404 | 0.048035155 |
| Gm10557 | 6.413115 | 6.690141 | 6.6395 | 0.050604149 | -0.22642171 | 6.678067 | 6.9174829 | 6.98093 | -0.063450826 | -0.302866705 |
| Mgst3 | 4.957546 | 4.89728 | 5.006 | -0.108679535 | -0.048413475 | 5.2341371 | 4.6880933 | 4.75155 | -0.063460511 | 0.482583351 |
| Hist3h2bb-ps | 2.259696 | 2.220903 | 2.2275 | -0.006627768 | 0.032165048 | 2.3175581 | 2.2256152 | 2.28908 | -0.063468274 | 0.028474658 |
| Vmn1r193 | 3.717364 | 3.640556 | 3.6338 | 0.006801374 | 0.083609371 | 3.801054 | 3.7639604 | 3.82747 | -0.063510243 | -0.02641656 |
| Stag3 | 4.061157 | 4.045131 | 4.2764 | -0.231237169 | -0.21521121 | 4.469558 | 4.265884 | 4.32942 | -0.063531375 | 0.140142625 |
| Myl6b | 2.979237 | 2.942331 | 3.0603 | -0.117969112 | -0.081063455 | 3.0224822 | 2.9913863 | 3.05492 | -0.063538116 | -0.032442208 |
| Lemd1 | 2.456075 | 2.347489 | 2.2858 | 0.061680615 | 0.170266674 | 2.3005113 | 2.3404403 | 2.40404 | -0.063596554 | -0.103525575 |

| | | | | | | | | | | |
|---|---|---|---|---|---|---|---|---|---|---|
| Xrcc6 | 9.254505 | 9.081054 | 9.2373 | -0.156217642 | 0.017232972 | 9.2784038 | 9.2142639 | 9.27795 | -0.063688832 | 0.000451052 |
| Gm18029 | 0.387451 | 0.325317 | 0.3181 | 0.00717938 | 0.069312833 | 0.3226969 | 0.3447732 | 0.40847 | -0.063691976 | -0.085768257 |
| Amy1 | 4.884668 | 3.735074 | 3.782 | -0.046922927 | 1.102671203 | 3.765518 | 3.7756219 | 3.83932 | -0.063697491 | -0.073801366 |
| Gm6831 | 0.255157 | 0.243479 | 0.297 | -0.053495518 | -0.041818031 | 0.2097956 | 0.2206963 | 0.28452 | -0.063827828 | -0.074728538 |
| Gm13981 | 2.01392 | 2.047365 | 2.0529 | -0.005486064 | -0.03893113 | 2.0420286 | 2.0376764 | 2.10153 | -0.063853176 | -0.059500987 |
| Zfp24 | 9.712044 | 9.595501 | 9.6354 | -0.03990922 | 0.076633821 | 9.4427359 | 9.4773295 | 9.5412 | -0.063870866 | -0.098464427 |
| Hnrnpa1l2-ps2 | 3.118889 | 3.177285 | 3.1287 | 0.048571195 | -0.00982478 | 3.1865547 | 3.1076543 | 3.17159 | -0.06393418 | 0.01496621 |
| Tecpr2 | 7.598024 | 7.757308 | 7.7624 | -0.005064375 | -0.164348026 | 7.6597908 | 7.7439225 | 7.80789 | -0.063969878 | -0.148101577 |
| Exoc2 | 10.29797 | 10.2913 | 10.301 | -0.009708142 | -0.003036404 | 10.279682 | 10.266083 | 10.3301 | -0.063986084 | -0.050387171 |
| Prom2 | 2.886006 | 2.385548 | 2.621 | -0.235476984 | 0.264981145 | 3.0374494 | 2.5988123 | 2.66289 | -0.064077265 | 0.374559822 |
| Atg2a | 9.171742 | 9.20427 | 9.2165 | -0.012208742 | -0.044736017 | 9.4189018 | 9.219116 | 9.28324 | -0.064122215 | 0.135663595 |
| Stat5a | 10.22304 | 10.19024 | 10.156 | 0.034149307 | 0.066944069 | 10.282088 | 10.137678 | 10.2018 | -0.064138752 | 0.080270837 |
| Ppfia2 | 1.417658 | 1.395231 | 1.3857 | 0.009516779 | 0.031944087 | 1.3494224 | 1.355765 | 1.41996 | -0.064190197 | -0.070532798 |
| Fam57b | 1.101774 | 1.16433 | 1.1569 | 0.007467483 | -0.055088499 | 1.1548556 | 1.1522373 | 1.21644 | -0.064197801 | -0.06157951 |
| Ranbp9 | 10.49014 | 10.52815 | 10.569 | -0.041027822 | -0.079043282 | 10.28358 | 10.471256 | 10.5355 | -0.06427229 | -0.251948666 |
| Cyren | 6.314322 | 6.585611 | 6.4255 | 0.160063569 | -0.111225371 | 6.9105989 | 6.5267795 | 6.59107 | -0.064288422 | 0.319531032 |
| mt-Co2 | 15.79507 | 15.83054 | 15.944 | -0.1134642 | -0.148931024 | 15.733079 | 15.805755 | 15.8701 | -0.064296954 | -0.136973494 |
| Trpc3 | -0.018272 | -0.02195 | -0.0006 | -0.0213894 | -0.017709597 | -0.0625822 | -0.015206 | 0.0491 | -0.064310525 | -0.111686807 |
| Gm17455 | 1.987748 | 1.898674 | 1.9548 | -0.056090389 | 0.032983483 | 2.0517049 | 1.9081132 | 1.97244 | -0.064322721 | 0.079268948 |
| Fcho2 | 9.963737 | 9.863887 | 10.012 | -0.148141403 | -0.04829194 | 9.6928874 | 9.9568184 | 10.0211 | -0.064328083 | -0.328259156 |
| Mmp8 | 5.974218 | 5.507057 | 4.6191 | 0.887971093 | 1.35513215 | 4.9100714 | 4.6087556 | 4.67311 | -0.064356783 | 0.236958996 |
| Thrb | 5.264992 | 5.136912 | 4.7864 | 0.35048869 | 0.478568402 | 5.1955509 | 5.2109158 | 5.27531 | -0.064397841 | -0.079762756 |
| Slc35b3 | 8.916657 | 8.874513 | 9.0072 | -0.132714851 | -0.090571044 | 8.9487823 | 8.9847694 | 9.04919 | -0.064424345 | -0.100411406 |
| Ccl17 | 5.691867 | 6.25346 | 5.9754 | 0.27808484 | -0.283507876 | 5.5878139 | 5.4874448 | 5.55188 | -0.06443618 | 0.035933008 |
| Ccar1 | 10.91594 | 10.97274 | 10.921 | 0.051999786 | -0.00479194 | 10.785265 | 10.872094 | 10.9365 | -0.064444974 | -0.151273583 |
| Kdelr1 | 9.927966 | 9.920282 | 9.98 | -0.059737429 | -0.052053323 | 10.031124 | 9.8296885 | 9.89417 | -0.064478363 | 0.136957248 |
| Fam181b | 0.736171 | 0.653281 | 0.6789 | -0.025623473 | 0.057267224 | 0.6872541 | 0.6252153 | 0.6897 | -0.064483836 | -0.002444997 |
| D6Ertd527e | 2.982053 | 2.993055 | 3.0168 | -0.0237615 | -0.034763533 | 2.9811724 | 2.9679616 | 3.03256 | -0.064593528 | -0.05138274 |
| Gm12568 | 2.151272 | 2.216355 | 2.1697 | 0.04665463 | -0.018428039 | 2.1905408 | 2.2012328 | 2.26584 | -0.064605467 | -0.075297484 |
| Xlr5b | 0.696393 | 0.66397 | 0.6999 | -0.035975221 | -0.003552259 | 0.5920553 | 0.592528 | 0.65719 | -0.064659961 | -0.065132715 |
| Gm5526 | 0.676952 | 0.628005 | 0.6454 | -0.017408787 | 0.031538257 | 0.5951765 | 0.6179676 | 0.68263 | -0.064665194 | -0.087456261 |
| Trav6d-3 | 1.511578 | 1.491555 | 1.5781 | -0.086569568 | -0.066546031 | 1.4614209 | 1.4707824 | 1.53548 | -0.0646989 | -0.074060346 |

| | | | | | | | | | | |
|---|---|---|---|---|---|---|---|---|---|---|
| Dnajc27 | 8.580941 | 8.550201 | 8.488 | 0.062245601 | 0.09298557 | 8.6435188 | 8.5685914 | 8.6333 | -0.064709953 | 0.010217406 |
| Lrrc24 | 3.261578 | 3.086634 | 3.0636 | 0.023037559 | 0.197981462 | 3.0882753 | 3.1656371 | 3.23047 | -0.064828404 | -0.142190202 |
| Gm18407 | 2.203155 | 2.217944 | 2.1832 | 0.034723795 | 0.019934596 | 2.21353 | 2.1417606 | 2.20661 | -0.064849433 | 0.006920002 |
| Traj35 | 0.227168 | 0.218683 | 0.2527 | -0.034058074 | -0.025573011 | 0.238992 | 0.2193917 | 0.28425 | -0.064854033 | -0.045253724 |
| Nr2f2 | 7.440324 | 7.060842 | 6.9286 | 0.132230863 | 0.511712984 | 6.2915427 | 6.3714118 | 6.43635 | -0.064940294 | -0.144809331 |
| Fbxl5 | 10.49458 | 10.39812 | 10.42 | -0.02208718 | 0.074372174 | 10.308183 | 10.347396 | 10.4124 | -0.064995334 | -0.104208077 |
| Ablim2 | 4.047524 | 3.869262 | 4.1711 | -0.301841426 | -0.123579251 | 3.6923201 | 3.789964 | 3.85515 | -0.065181704 | -0.162825658 |
| Nr2f1 | 4.361172 | 4.147582 | 4.329 | -0.18136854 | 0.032222077 | 4.3064833 | 4.1671796 | 4.23238 | -0.065203132 | 0.074100521 |
| Gm3002 | 0.731162 | 0.704825 | 0.6829 | 0.021906425 | 0.048243171 | 0.7303626 | 0.7364436 | 0.80167 | -0.065222069 | -0.071303018 |
| Mthfs | 7.736621 | 7.758315 | 7.7848 | -0.026495585 | -0.048188995 | 7.9647076 | 7.9137264 | 7.97896 | -0.065231424 | -0.014250138 |
| Slc35f3 | 0.44448 | 0.467982 | 0.4156 | 0.052375198 | 0.028872921 | 0.4327831 | 0.3948288 | 0.46006 | -0.06523232 | -0.027278068 |
| Ppfia4 | 7.326694 | 7.05296 | 7.273 | -0.220008563 | 0.053725892 | 7.4223028 | 6.9333547 | 6.99863 | -0.06527504 | 0.423673013 |
| Otud7a | 1.88338 | 1.760743 | 1.8664 | -0.105680677 | 0.016955961 | 1.7804195 | 1.8864665 | 1.95175 | -0.065285566 | -0.171332569 |
| Cfap73 | 1.603981 | 1.505532 | 1.4624 | 0.043179156 | 0.141628158 | 1.4028359 | 1.3986435 | 1.46395 | -0.065301968 | -0.061109547 |
| Eif2b1 | 8.771506 | 8.656032 | 8.6747 | -0.018641888 | 0.096832017 | 8.9003665 | 8.6857695 | 8.7511 | -0.065327797 | 0.14926924 |
| Stmn1-rs1 | -0.068709 | -0.06845 | -0.0655 | -0.002957743 | -0.003213412 | -0.0502262 | -0.074551 | -0.0092 | -0.065352328 | -0.041027267 |
| 1700016D06Rik | 0.601337 | 0.583556 | 0.6032 | -0.019635038 | -0.001854159 | 0.6220481 | 0.6005947 | 0.66597 | -0.06537276 | -0.043919355 |
| Acp5 | 11.09792 | 11.33534 | 11.182 | 0.153032162 | -0.084387196 | 11.232699 | 10.901758 | 10.9671 | -0.065372938 | 0.265567905 |
| Zgrf1 | 7.403809 | 7.299523 | 7.4015 | -0.102024416 | 0.002261087 | 7.5864361 | 7.5519772 | 7.61738 | -0.065400273 | -0.030941334 |
| Vmn1r16 | 1.648274 | 1.645631 | 1.6374 | 0.008194298 | 0.010837539 | 1.6593938 | 1.6701126 | 1.73553 | -0.065420976 | -0.076139791 |
| Fam171b | 3.483483 | 3.086364 | 3.1605 | -0.074102373 | 0.323016709 | 3.1919019 | 3.1025102 | 3.16794 | -0.065428959 | 0.023962727 |
| 1700012B07Rik | 0.826131 | 0.846749 | 0.8735 | -0.026755926 | -0.047373347 | 0.8454523 | 0.8062103 | 0.87166 | -0.065451606 | -0.026209668 |
| Krt14 | 0.766512 | 0.71947 | 0.7209 | -0.001448448 | 0.045593419 | 0.8643867 | 0.7617434 | 0.82722 | -0.065478085 | 0.037165229 |
| Tmem200a | 1.315105 | 1.292644 | 1.3625 | -0.069873561 | -0.04741282 | 1.3701065 | 1.3001395 | 1.36565 | -0.065505672 | 0.004461302 |
| Gm49377 | 0.85642 | 0.860875 | 0.8993 | -0.03840289 | -0.042857025 | 0.8278291 | 0.8419354 | 0.90751 | -0.065575794 | -0.079682012 |
| Ttc7 | 11.27002 | 11.23485 | 11.334 | -0.099610584 | -0.064441688 | 11.336996 | 11.304466 | 11.3701 | -0.06559555 | -0.033065481 |
| Rps23-ps2 | 2.055383 | 2.05465 | 2.0457 | 0.008994271 | 0.009727965 | 2.0800657 | 1.9858107 | 2.05142 | -0.065612677 | 0.028642318 |
| Olfr814 | 2.27065 | 2.294928 | 2.3963 | -0.101323759 | -0.12560199 | 2.2634081 | 2.2916168 | 2.35726 | -0.065641509 | -0.093850187 |
| Pcdhb9 | 2.782241 | 2.73349 | 2.853 | -0.119539842 | -0.070788208 | 2.74658 | 2.6322399 | 2.69789 | -0.065654643 | 0.048685466 |
| Syn1 | 2.462947 | 2.422828 | 2.3851 | 0.037747785 | 0.07786674 | 2.393995 | 2.3040077 | 2.36967 | -0.065658879 | 0.024328444 |
| Gm12789 | 1.301772 | 1.264949 | 1.1995 | 0.065477259 | 0.102300049 | 1.2876059 | 1.1952926 | 1.261 | -0.065703088 | 0.02661016 |
| Tom1l2 | 8.935544 | 8.824977 | 8.9483 | -0.123274683 | -0.012708401 | 8.9171746 | 9.0749637 | 9.14069 | -0.065727084 | -0.223516175 |

| | | | | | | | | | | |
|---|---|---|---|---|---|---|---|---|---|---|
| Cutal | 0.960502 | 0.961889 | 1.0101 | -0.048167247 | -0.049554423 | 0.8827137 | 0.8915437 | 0.95732 | -0.065775535 | -0.074605501 |
| Zfp938 | 6.166169 | 5.995202 | 6.1587 | -0.163544718 | 0.007421889 | 6.2583655 | 6.1374071 | 6.20319 | -0.065782425 | 0.055175938 |
| Gm18190 | 3.032339 | 3.04662 | 3.0593 | -0.012693063 | -0.026974694 | 3.2406918 | 3.0716134 | 3.1374 | -0.065790122 | 0.103288279 |
| Dzip1 | 8.608843 | 8.65444 | 8.8478 | -0.193350888 | -0.238948149 | 8.5029901 | 8.8340005 | 8.89985 | -0.065845644 | -0.396856094 |
| Gm29126 | 0.730512 | 0.786961 | 0.7334 | 0.053567733 | -0.002881535 | 0.7371358 | 0.7488954 | 0.81475 | -0.065857254 | -0.077616838 |
| Gm12424 | 3.272787 | 3.385065 | 3.3603 | 0.024745546 | -0.087532469 | 3.3565077 | 3.3851493 | 3.45102 | -0.065872389 | -0.094514019 |
| Ttc5 | 9.801654 | 9.803094 | 9.8721 | -0.069024168 | -0.070463493 | 9.9890604 | 9.8408251 | 9.90672 | -0.065890043 | 0.082345243 |
| Olfr366 | 4.678778 | 4.681674 | 4.7189 | -0.037247206 | -0.040143284 | 4.9002525 | 4.9553327 | 5.02125 | -0.065917115 | -0.120997345 |
| Slc22a18 | 3.119225 | 3.054723 | 3.0943 | -0.039583081 | 0.024918279 | 3.0048145 | 3.0043226 | 3.07029 | -0.065970432 | -0.065478503 |
| Srcap | 12.11911 | 12.11844 | 12.005 | 0.113154796 | 0.113823506 | 12.003053 | 12.195684 | 12.2617 | -0.065971032 | -0.258601658 |
| Ccdc169 | 1.560423 | 1.598854 | 1.5365 | 0.062357934 | 0.02392753 | 1.6105398 | 1.6219928 | 1.68798 | -0.065987783 | -0.077440841 |
| 4932438H23Rik | 1.520985 | 1.624537 | 1.5835 | 0.041038335 | -0.062513575 | 1.5467709 | 1.5006649 | 1.56674 | -0.0660757 | -0.019969729 |
| Prr14l | 10.75984 | 10.585 | 10.657 | -0.071811946 | 0.10302999 | 10.475692 | 10.585452 | 10.6515 | -0.06607809 | -0.175838687 |
| Veph1 | 3.412753 | 3.278611 | 3.2842 | -0.005577531 | 0.128564275 | 3.383662 | 3.3775053 | 3.4436 | -0.066097813 | -0.059941139 |
| Ift57 | 6.486145 | 6.2821 | 6.2827 | -0.000611237 | 0.203433488 | 6.4319701 | 6.2773298 | 6.34347 | -0.066138839 | 0.088501533 |
| Kif17 | 2.332071 | 2.337583 | 2.3578 | -0.020214892 | -0.025726827 | 2.2766479 | 2.3483729 | 2.41451 | -0.066140732 | -0.137865784 |
| Nhlrc3 | 9.091758 | 8.901247 | 9.0813 | -0.180079839 | 0.010431774 | 9.2020187 | 9.0188258 | 9.08498 | -0.066151992 | 0.11704088 |
| Gm14401 | 1.240923 | 1.217224 | 1.2667 | -0.049440662 | -0.02574241 | 1.2874654 | 1.2294431 | 1.29564 | -0.066194052 | -0.008171756 |
| Per2 | 8.089151 | 7.859794 | 7.8998 | -0.040035032 | 0.189322166 | 7.9839691 | 7.9009447 | 7.96715 | -0.066203729 | 0.016820713 |
| Gm8325 | 6.698609 | 6.678042 | 6.6289 | 0.049112777 | 0.069680599 | 6.4508778 | 6.4861142 | 6.55241 | -0.066296717 | -0.101533103 |
| Gm48208 | 2.324669 | 2.252109 | 2.2802 | -0.028047887 | 0.044512576 | 2.2323724 | 2.2719875 | 2.33831 | -0.066319017 | -0.105934172 |
| Rsph1 | 2.383283 | 2.281966 | 2.3296 | -0.047676411 | 0.053640776 | 2.4091792 | 2.2971621 | 2.3635 | -0.066338276 | 0.045678893 |
| Smad9 | 0.399241 | 0.34978 | 0.3747 | -0.02490101 | 0.024560288 | 0.3366827 | 0.3115827 | 0.37794 | -0.066353093 | -0.041253033 |
| Pepd | 9.127714 | 9.161817 | 9.0648 | 0.097049052 | 0.062945941 | 9.0986606 | 8.995636 | 9.06205 | -0.06641827 | 0.036606307 |
| Klf15 | 2.656304 | 2.539297 | 2.6406 | -0.101351237 | 0.015655648 | 2.4692455 | 2.3978897 | 2.46433 | -0.066438529 | 0.004917339 |
| Gm15784 | 1.925599 | 1.897655 | 1.8987 | -0.00102966 | 0.026914033 | 1.9915904 | 1.983381 | 2.04986 | -0.066477258 | -0.058267899 |
| Casd1 | 9.318538 | 9.341219 | 9.4198 | -0.078604189 | -0.101285406 | 9.4309483 | 9.1951802 | 9.26167 | -0.066492581 | 0.169275524 |
| Dop1a | 10.16017 | 10.30884 | 10.281 | 0.027570014 | -0.121093391 | 10.108244 | 10.233842 | 10.3004 | -0.066547508 | -0.192145701 |
| Traj42 | 1.685097 | 1.726503 | 1.704 | 0.022513054 | -0.018893678 | 1.6969954 | 1.6603207 | 1.72695 | -0.066626338 | -0.029951629 |
| Ppp3cb | 11.1119 | 11.09537 | 11.123 | -0.028102672 | -0.01157184 | 11.096014 | 11.141098 | 11.2078 | -0.06666215 | -0.111745994 |
| Gm49037 | 0.725709 | 0.822609 | 0.7232 | 0.099429381 | 0.002528664 | 0.8396965 | 0.8348956 | 0.90162 | -0.066729307 | -0.061928447 |
| Gria2 | 2.346346 | 2.409605 | 2.4307 | -0.021051236 | -0.084310286 | 2.4655841 | 2.3956899 | 2.46244 | -0.066750137 | 0.003144084 |

| | | | | | | | | | | |
|---|---|---|---|---|---|---|---|---|---|---|
| Pdhx | 7.86715 | 7.812058 | 7.9021 | -0.090010489 | -0.034918656 | 8.0011853 | 7.8481529 | 7.91495 | -0.066797455 | 0.086235006 |
| Olfr60 | 1.147666 | 1.092284 | 1.1227 | -0.030465516 | 0.024916599 | 1.1650193 | 1.1007744 | 1.16764 | -0.066863153 | -0.002618257 |
| Ogdhl | 2.681457 | 2.546329 | 2.5928 | -0.046471018 | 0.088657161 | 2.6119879 | 2.5457735 | 2.61265 | -0.066878432 | -0.000664072 |
| Trav3-1 | 2.927681 | 3.048361 | 3.0313 | 0.017103753 | -0.103576145 | 2.8890362 | 2.9108422 | 2.97778 | -0.066936875 | -0.088742866 |
| Ccr2 | 10.23512 | 9.711042 | 9.5965 | 0.114590277 | 0.638670114 | 10.308358 | 9.5277116 | 9.59465 | -0.066941596 | 0.713704782 |
| Gm17420 | 3.869931 | 3.94308 | 3.9508 | -0.007692729 | -0.080841729 | 3.9216446 | 4.0786054 | 4.14557 | -0.066961956 | -0.223922709 |
| Agbl1 | 1.25618 | 1.266979 | 1.2475 | 0.019517658 | 0.008718226 | 1.2446205 | 1.2367029 | 1.30369 | -0.066982386 | -0.05906475 |
| Ctnnbip1 | 5.822885 | 5.459082 | 5.8251 | -0.366010883 | -0.002207827 | 5.6404446 | 5.5239493 | 5.59095 | -0.066999732 | 0.04949558 |
| Slc22a3 | 4.208487 | 4.013743 | 4.1502 | -0.136407494 | 0.058336961 | 4.1001992 | 3.9694115 | 4.03647 | -0.067053819 | 0.063733809 |
| Adrb1 | 2.254914 | 2.183383 | 2.2384 | -0.054972605 | 0.016558505 | 2.095041 | 2.1919022 | 2.25896 | -0.06705736 | -0.163918618 |
| Gm8189 | 3.4142 | 3.67015 | 3.5313 | 0.138867454 | -0.11708232 | 3.5646941 | 3.6518318 | 3.71893 | -0.067099541 | -0.15423722 |
| Mgat3 | 3.437602 | 3.382818 | 3.3286 | 0.054190656 | 0.108974518 | 3.1340219 | 3.0948515 | 3.16198 | -0.067132932 | -0.027962475 |
| Smim4 | 6.656399 | 6.537796 | 6.8355 | -0.297666099 | -0.17906401 | 6.93791 | 6.7158242 | 6.78297 | -0.067144461 | 0.154941352 |
| Olfr1312 | 2.60466 | 2.53256 | 2.6336 | -0.100997025 | -0.028897302 | 2.6046834 | 2.5629054 | 2.63006 | -0.06715184 | -0.025373829 |
| Olfr51 | 0.864416 | 0.853232 | 0.8932 | -0.03993261 | -0.028748663 | 0.8781615 | 0.8736281 | 0.9408 | -0.067168897 | -0.062635534 |
| Rpl19-ps4 | 1.386628 | 1.412033 | 1.3622 | 0.049848076 | 0.024443014 | 1.3334545 | 1.3412469 | 1.40842 | -0.067174892 | -0.074967235 |
| Msgn1 | 1.415678 | 1.487865 | 1.4795 | 0.008325184 | -0.063862498 | 1.4844581 | 1.3955764 | 1.46276 | -0.067180021 | 0.021701701 |
| I0C0044D17Rik | 1.175917 | 1.141134 | 1.1426 | -0.001426093 | 0.033357063 | 1.1495332 | 1.1653386 | 1.23255 | -0.06720822 | -0.083013634 |
| Pifo | 2.304639 | 2.232979 | 2.3492 | -0.116244028 | -0.044584497 | 2.414242 | 2.2934762 | 2.36069 | -0.067210132 | 0.053555657 |
| Cpeb3 | 3.401436 | 3.415538 | 3.3905 | 0.025055788 | 0.010954037 | 3.492475 | 3.2627831 | 3.33001 | -0.067223485 | 0.162468457 |
| Iglv3 | 8.599879 | 8.472054 | 8.57 | -0.097926736 | 0.029898547 | 8.8130055 | 8.4865483 | 8.55388 | -0.067332703 | 0.25912447 |
| Efna4 | 1.299697 | 1.262577 | 1.31 | -0.04746283 | -0.010342592 | 1.2693528 | 1.1946113 | 1.26212 | -0.067508567 | 0.007232914 |
| Usp36 | 10.40426 | 10.53235 | 10.361 | 0.171317148 | 0.043226496 | 10.59134 | 10.545269 | 10.6128 | -0.067536348 | -0.02146572 |
| Ngef | 1.244217 | 1.204081 | 1.184 | 0.020032696 | 0.060168502 | 1.154581 | 1.1602386 | 1.22789 | -0.067648196 | -0.073305769 |
| Mal2 | 1.097182 | 1.099437 | 1.0997 | -0.000225826 | -0.002480518 | 1.1213957 | 1.0715161 | 1.13917 | -0.067658439 | -0.017778811 |
| 38961 | 12.01427 | 11.95678 | 12.029 | -0.072498046 | -0.015006096 | 12.031067 | 11.969452 | 12.0372 | -0.067712104 | -0.006098005 |
| Spata13 | 11.17774 | 11.24668 | 11.247 | -0.00011752 | -0.069050062 | 10.875873 | 11.002088 | 11.0698 | -0.067714866 | -0.193929385 |
| Olfr846 | 2.625969 | 2.62712 | 2.6807 | -0.053559287 | -0.054710889 | 2.7319887 | 2.7322744 | 2.80001 | -0.067736655 | -0.068022354 |
| A630073D07Rik | 3.68289 | 3.464142 | 3.4257 | 0.038404841 | 0.257152169 | 3.6161276 | 3.5994464 | 3.66719 | -0.067740304 | -0.051059089 |
| Adss | 10.64176 | 10.49128 | 10.611 | -0.119695952 | 0.030788252 | 10.599622 | 10.616176 | 10.6839 | -0.06774905 | -0.084303048 |
| Rpl19-ps3 | 1.482499 | 1.516669 | 1.4704 | 0.0462242 | 0.012054121 | 1.4472281 | 1.4720167 | 1.53979 | -0.067776004 | -0.092564684 |
| Bcl9l | 10.88767 | 11.11151 | 11.022 | 0.089794508 | -0.13405255 | 10.838234 | 10.756065 | 10.824 | -0.067940228 | 0.01422793 |

| | | | | | | | | | | |
|---|---|---|---|---|---|---|---|---|---|---|
| Rpl28-ps2 | 0.823449 | 0.804666 | 0.779 | 0.025657956 | 0.044440537 | 0.7751168 | 0.7624431 | 0.83043 | -0.067985423 | -0.055311671 |
| Gm15009 | 4.059142 | 4.221883 | 4.0657 | 0.156188373 | -0.00655203 | 4.2929129 | 4.2035207 | 4.27153 | -0.068014065 | 0.021378201 |
| Gm20422 | 3.355501 | 3.427752 | 3.4163 | 0.011403015 | -0.060847938 | 3.594185 | 3.5092033 | 3.57724 | -0.068037315 | 0.016944431 |
| Cdk10 | 8.328219 | 8.375065 | 8.4785 | -0.103441028 | -0.150286943 | 8.7436038 | 8.5866243 | 8.65471 | -0.068084279 | 0.088895277 |
| Inca1 | 2.8721 | 2.871994 | 2.8947 | -0.022728355 | -0.022622721 | 2.9406833 | 2.8629118 | 2.93102 | -0.068104524 | 0.009666936 |
| Slc16a6 | 9.50542 | 9.195339 | 9.4303 | -0.234967383 | 0.075113741 | 9.8557352 | 9.6629341 | 9.73104 | -0.068109504 | 0.124691642 |
| Trappc8 | 10.73301 | 10.81727 | 10.756 | 0.06101039 | -0.023244656 | 10.66492 | 10.700295 | 10.7684 | -0.068141964 | -0.103517396 |
| Itgb2l | 2.252622 | 2.19894 | 2.105 | 0.093954134 | 0.147635396 | 2.2179483 | 2.1022994 | 2.17048 | -0.068182389 | 0.047466429 |
| Olfr1396 | 1.327312 | 1.303639 | 1.3315 | -0.027846357 | -0.004173897 | 1.2847279 | 1.3665697 | 1.43477 | -0.06820213 | -0.150043919 |
| Ints6 | 10.15646 | 10.22681 | 9.8674 | 0.359397533 | 0.289053499 | 9.9005462 | 9.9603171 | 10.0286 | -0.068239863 | -0.128010746 |
| Hmgn5 | 8.855519 | 8.842898 | 8.9156 | -0.072700699 | -0.060079648 | 8.723082 | 8.8407506 | 8.90902 | -0.068271526 | -0.185940135 |
| Ntf3 | 1.5678 | 1.544497 | 1.4752 | 0.069278299 | 0.092580645 | 1.4614076 | 1.4164261 | 1.4847 | -0.068271793 | -0.023290282 |
| Ccndbp1 | 9.898627 | 9.849145 | 9.9289 | -0.079798779 | -0.03031678 | 10.080381 | 9.8780172 | 9.94629 | -0.068274747 | 0.134089466 |
| Leprot | 9.509109 | 9.279408 | 9.4912 | -0.211785708 | 0.01791582 | 9.3391335 | 9.2313908 | 9.29967 | -0.068283802 | 0.039458895 |
| Mtrf1l | 7.368863 | 7.425027 | 7.5567 | -0.131672294 | -0.187835591 | 7.5564167 | 7.5446404 | 7.61293 | -0.06829434 | -0.05651805 |
| Wdr20rt | 2.31075 | 2.245707 | 2.3046 | -0.058873441 | 0.006169709 | 2.2396101 | 2.330936 | 2.39932 | -0.068386748 | -0.159712676 |
| Msl3 | 9.070382 | 9.023249 | 9.1285 | -0.105203664 | -0.058070168 | 9.1237527 | 8.9657478 | 9.03415 | -0.068400074 | 0.08960481 |
| Olfr982 | 1.879661 | 1.875091 | 1.842 | 0.033138171 | 0.037707344 | 1.8602616 | 1.8153742 | 1.88384 | -0.068464878 | -0.023577511 |
| Vmn2r16 | 1.334913 | 1.34448 | 1.328 | 0.016518804 | 0.006951802 | 1.3641803 | 1.3663584 | 1.43484 | -0.068478283 | -0.070656318 |
| Slc45a3 | 5.432726 | 5.913514 | 5.5745 | 0.339051802 | -0.141736582 | 5.0311382 | 5.088671 | 5.15718 | -0.068513171 | -0.126045983 |
| Gm43941 | 1.175638 | 1.17204 | 1.179 | -0.006962754 | -0.003364147 | 1.2727336 | 1.2104965 | 1.27909 | -0.068597887 | -0.006360833 |
| Cmbl | 1.386087 | 1.404358 | 1.4921 | -0.087736855 | -0.106007897 | 1.3612653 | 1.3223696 | 1.39098 | -0.068609981 | -0.029714232 |
| Armc1 | 9.392275 | 9.470684 | 9.3566 | 0.114045828 | 0.035636213 | 9.3926792 | 9.2237652 | 9.29243 | -0.068662167 | 0.100251886 |
| Teddm3 | 1.835635 | 1.757991 | 1.8538 | -0.095783952 | -0.018139519 | 1.8726341 | 1.7338079 | 1.80252 | -0.068709928 | 0.07011634 |
| Pih1d2 | 2.232613 | 2.30183 | 2.2312 | 0.07059854 | 0.001381225 | 2.2701537 | 2.293284 | 2.36201 | -0.068723566 | -0.091853863 |
| 9330182O14Rik | 1.800951 | 1.795473 | 1.7928 | 0.002635578 | 0.008114501 | 1.8878209 | 1.8280661 | 1.89687 | -0.068803311 | -0.009048526 |
| Ajm1 | 2.370268 | 2.344448 | 2.4531 | -0.108624005 | -0.082804293 | 2.4039327 | 2.2718917 | 2.34078 | -0.068887223 | 0.063153788 |
| Snx19 | 9.899978 | 9.757573 | 9.8661 | -0.108512039 | 0.033893234 | 9.8977446 | 9.7523302 | 9.82127 | -0.068940539 | 0.076473844 |
| Trbj1-3 | 5.225001 | 5.335044 | 5.3103 | 0.0247011 | -0.085342575 | 5.0760712 | 5.3169756 | 5.38594 | -0.068966939 | -0.309871365 |
| Cep95 | 8.307509 | 8.339011 | 8.3111 | 0.027957428 | -0.003544735 | 8.4685911 | 8.5831067 | 8.6521 | -0.068988568 | -0.183504138 |
| Swt1 | 9.042383 | 9.273199 | 9.2589 | 0.014338614 | -0.216477153 | 9.0579189 | 9.3290795 | 9.39809 | -0.069014066 | -0.340174664 |
| 4930550C14Rik | 1.780719 | 1.897903 | 1.8704 | 0.027458838 | -0.089724934 | 1.7739232 | 1.7244681 | 1.79349 | -0.069021422 | -0.0195663 |

| | | | | | | | | | | |
|---|---|---|---|---|---|---|---|---|---|---|
| Col26a1 | 4.416492 | 4.457214 | 4.3723 | 0.084864284 | 0.044142648 | 4.6896994 | 4.3185455 | 4.38769 | -0.069145354 | 0.302008562 |
| 2610021A01Rik | 8.28107 | 8.186205 | 8.3666 | -0.180367152 | -0.08550186 | 8.1313793 | 8.1989347 | 8.26809 | -0.06915656 | -0.136711974 |
| Fbxo28 | 9.42714 | 9.518734 | 9.4305 | 0.088272982 | -0.003320612 | 9.1832807 | 9.3956569 | 9.46495 | -0.069289241 | -0.281665475 |
| Sec14l1 | 9.488886 | 9.313642 | 9.3371 | -0.023497806 | 0.151745928 | 9.2878433 | 9.1322899 | 9.20168 | -0.069390587 | 0.0861628 |
| Stk32c | 4.569996 | 4.492462 | 4.5788 | -0.086338894 | -0.008804391 | 4.3835819 | 4.3005704 | 4.36996 | -0.069391297 | 0.013620289 |
| Mras | 5.37701 | 5.456388 | 5.3129 | 0.143451143 | 0.064073847 | 4.9452666 | 5.200284 | 5.26973 | -0.06944118 | -0.324458563 |
| Olfr709-ps1 | 2.44545 | 2.438708 | 2.5447 | -0.105955415 | -0.099213443 | 2.5232664 | 2.5262644 | 2.59573 | -0.069469847 | -0.072467936 |
| Acad9 | 9.262738 | 9.32484 | 9.293 | 0.031869068 | -0.030233364 | 9.5148883 | 9.5736878 | 9.64317 | -0.069484654 | -0.128284238 |
| Hoxc9 | 1.379999 | 1.35605 | 1.3692 | -0.01314077 | 0.010808526 | 1.4141796 | 1.3947753 | 1.46429 | -0.069516742 | -0.050112367 |
| Grpr | 1.609364 | 1.657117 | 1.5806 | 0.076563241 | 0.028810423 | 1.5791178 | 1.5314041 | 1.60096 | -0.069551862 | -0.021838138 |
| Gm47967 | 2.501027 | 2.548301 | 2.5185 | 0.029774722 | -0.017499477 | 2.5730281 | 2.5180977 | 2.58768 | -0.069580996 | -0.014650577 |
| Cfd | 0.234676 | 0.216745 | 0.2766 | -0.059817811 | -0.041886918 | 0.204265 | 0.2153462 | 0.28494 | -0.06959023 | -0.080671371 |
| Hsdl1 | 9.953268 | 10.1103 | 10.065 | 0.045308099 | -0.111723289 | 9.9822062 | 9.8968486 | 9.96644 | -0.069592724 | 0.015764861 |
| Olfr1100 | 1.091229 | 1.058559 | 1.0889 | -0.030351242 | 0.002318601 | 1.033512 | 1.0626587 | 1.13226 | -0.069602095 | -0.098748809 |
| Stk36 | 1.828825 | 1.801843 | 1.7974 | 0.004448298 | 0.031429356 | 1.8674835 | 1.7165113 | 1.78615 | -0.069634191 | 0.08133793 |
| Ahi1 | 7.369215 | 7.368812 | 7.3417 | 0.027088208 | 0.027491117 | 7.2609311 | 7.2292289 | 7.29899 | -0.06975658 | -0.038054375 |
| Dffb | 7.955903 | 7.972218 | 7.9966 | -0.02434978 | -0.04066491 | 8.067929 | 7.9650243 | 8.03483 | -0.069808895 | 0.03309576 |
| Gli1 | 3.043037 | 3.220812 | 3.0682 | 0.152633892 | -0.025141105 | 3.0076981 | 2.9694327 | 3.03926 | -0.06983154 | -0.031566159 |
| Gm7633 | 1.824203 | 1.828065 | 1.8977 | -0.069633975 | -0.07349641 | 1.8330942 | 1.783757 | 1.85361 | -0.069852419 | -0.020515239 |
| Bag6 | 11.54052 | 11.52047 | 11.582 | -0.061281638 | -0.041228392 | 11.672485 | 11.543522 | 11.6134 | -0.069865414 | 0.059097146 |
| Olfr1269 | 3.49897 | 3.506855 | 3.417 | 0.089879942 | 0.081995375 | 3.5335655 | 3.565239 | 3.63512 | -0.069878989 | -0.101552431 |
| Tmprss3 | 2.794155 | 2.74863 | 2.8567 | -0.108111283 | -0.062586572 | 2.761229 | 2.775107 | 2.84502 | -0.069917078 | -0.083795111 |
| Ogdh | 12.03665 | 12.06322 | 12.032 | 0.031358696 | 0.0047852 | 12.182582 | 11.971855 | 12.0418 | -0.069941122 | 0.140785873 |
| Ighv5-6 | 5.913046 | 6.029474 | 6.1462 | -0.116693345 | -0.233121036 | 6.0169318 | 5.747358 | 5.81735 | -0.069988999 | 0.199584756 |
| Fam131b | 0.990586 | 0.979592 | 0.9611 | 0.018462736 | 0.029457106 | 0.9502408 | 0.9095508 | 0.97956 | -0.070007371 | -0.029317436 |
| Gm43552 | 0.226117 | 0.154142 | 0.1564 | -0.002229931 | 0.069745281 | 0.1145881 | 0.1149556 | 0.18498 | -0.070025711 | -0.0703932 |
| Brinp2 | 2.624061 | 2.46757 | 2.526 | -0.058425619 | 0.09806521 | 2.5239599 | 2.443062 | 2.51309 | -0.070029826 | 0.010868126 |
| Nlk | 9.712483 | 9.616481 | 9.6282 | -0.011688547 | 0.084314011 | 9.7582382 | 9.5563982 | 9.62643 | -0.070031828 | 0.131808212 |
| Retreg2 | 10.21324 | 10.12758 | 10.241 | -0.113020578 | -0.027357232 | 10.310121 | 10.094425 | 10.1645 | -0.070050013 | 0.145646295 |
| Gm12058 | 1.202251 | 1.251084 | 1.2216 | 0.029488006 | -0.019344667 | 1.2695172 | 1.203458 | 1.27355 | -0.070088564 | -0.004029346 |
| Six6 | 1.316592 | 1.31774 | 1.365 | -0.047251245 | -0.048400064 | 1.2714945 | 1.3363611 | 1.40645 | -0.070088874 | -0.134955487 |
| Vmn1r9 | 2.476688 | 2.455962 | 2.5341 | -0.078118499 | -0.057393108 | 2.476323 | 2.53953 | 2.60965 | -0.07011921 | -0.133326177 |

| | | | | | | | | | | |
|---|---|---|---|---|---|---|---|---|---|---|
| Cspg5 | 3.128185 | 3.062109 | 3.0329 | 0.029222949 | 0.095298667 | 3.0815078 | 3.0510325 | 3.12118 | -0.070150216 | -0.039674948 |
| Cacng8 | 0.947687 | 0.960952 | 0.9568 | 0.00413259 | -0.009132328 | 0.9214312 | 0.9164413 | 0.98662 | -0.070180347 | -0.065190393 |
| Poglut1 | 9.501333 | 9.580411 | 9.571 | 0.009370081 | -0.069708239 | 9.5231936 | 9.6420134 | 9.71221 | -0.070191891 | -0.189011715 |
| Sorbs3 | 7.33581 | 7.065466 | 7.2903 | -0.224852722 | 0.045492185 | 6.6710588 | 6.8441104 | 6.91436 | -0.070250814 | -0.243302373 |
| Vit | 2.628785 | 2.529076 | 2.6278 | -0.09871686 | 0.000991978 | 2.4586611 | 2.5121175 | 2.58237 | -0.070255478 | -0.1237119 |
| Osbpl10 | 3.154931 | 2.834761 | 2.886 | -0.051235159 | 0.268934322 | 2.8904524 | 2.7017004 | 2.772 | -0.070303563 | 0.118448407 |
| Fundc2 | 9.398118 | 9.32296 | 9.4854 | -0.162477534 | -0.087320398 | 9.3440121 | 9.3110841 | 9.38142 | -0.070334627 | -0.037406652 |
| Traj19 | 2.144969 | 2.17971 | 2.1027 | 0.076977233 | 0.042236658 | 2.0604522 | 2.1134996 | 2.18385 | -0.07034583 | -0.123393214 |
| Sbno1 | 11.76964 | 11.77612 | 11.715 | 0.060984098 | 0.054505822 | 11.690852 | 11.784023 | 11.8544 | -0.070387574 | -0.163558663 |
| Ttc30b | 4.351493 | 4.041988 | 4.3302 | -0.288216453 | 0.021288024 | 4.1097795 | 4.0747004 | 4.1451 | -0.070400814 | -0.035321657 |
| Srp54c | 10.34815 | 10.22982 | 10.265 | -0.035275396 | 0.083061574 | 10.267146 | 10.223717 | 10.2942 | -0.070445948 | -0.027016984 |
| Olfr323 | 1.553861 | 1.554643 | 1.6192 | -0.064539282 | -0.065321096 | 1.568731 | 1.5720568 | 1.64251 | -0.070455689 | -0.073781489 |
| Olfr754-ps1 | 1.670691 | 1.650626 | 1.658 | -0.007330835 | 0.012734143 | 1.8667614 | 1.6880395 | 1.75856 | -0.070520928 | 0.108201004 |
| Prr16 | 3.188633 | 3.131733 | 3.0395 | 0.092275279 | 0.149175327 | 3.069598 | 3.0036061 | 3.07423 | -0.070619551 | -0.004627664 |
| Sap25 | 8.547285 | 8.77476 | 8.8188 | -0.044004767 | -0.271479815 | 9.466953 | 9.038675 | 9.10936 | -0.070681572 | 0.357596375 |
| Gm10054 | 1.617745 | 1.611028 | 1.6075 | 0.003504329 | 0.010220962 | 1.7271426 | 1.599361 | 1.67006 | -0.070701033 | 0.057080497 |
| Zfp868 | 8.712691 | 8.805459 | 8.8873 | -0.081794745 | -0.174563185 | 8.7810608 | 8.7799854 | 8.85071 | -0.07072813 | -0.069652774 |
| Ermap | 4.358412 | 4.196751 | 4.5169 | -0.320105047 | -0.158444537 | 4.1693925 | 4.3042194 | 4.37495 | -0.070729644 | -0.205556585 |
| Tmem117 | 0.910103 | 0.836507 | 0.8739 | -0.037421155 | 0.036174872 | 0.8593124 | 0.8418935 | 0.91264 | -0.070742984 | -0.053324088 |
| Gpatch2 | 8.354435 | 8.334202 | 8.3397 | -0.005537157 | 0.014695285 | 8.3827111 | 8.2889646 | 8.35971 | -0.070743358 | 0.023003204 |
| Art2a-ps | 3.997451 | 3.885839 | 3.8961 | -0.01027564 | 0.101336409 | 3.7342861 | 3.7606684 | 3.83142 | -0.070753425 | -0.09713569 |
| Stard13 | 5.022841 | 4.686811 | 4.8196 | -0.132769672 | 0.203260239 | 4.4035041 | 4.3694689 | 4.44027 | -0.070797374 | -0.036762122 |
| Snx7 | 5.194965 | 5.006766 | 5.1528 | -0.146058545 | 0.042141182 | 4.9801666 | 4.8672957 | 4.93815 | -0.070853624 | 0.042017273 |
| Spcs3 | 10.72828 | 10.67206 | 10.681 | -0.008718406 | 0.04749571 | 10.635519 | 10.479704 | 10.5506 | -0.070918456 | 0.084896372 |
| Traj24 | 2.135152 | 2.094347 | 2.2057 | -0.111320862 | -0.070515675 | 2.0659987 | 2.1044723 | 2.17546 | -0.070983829 | -0.109457425 |
| Sox10 | 0.868152 | 0.821173 | 0.8221 | -0.000958602 | 0.046020892 | 0.8377541 | 0.7903306 | 0.86132 | -0.070986159 | -0.023562729 |
| Taf11 | 8.212998 | 8.148853 | 8.291 | -0.142157908 | -0.078012262 | 8.2839751 | 8.2914351 | 8.36243 | -0.07099182 | -0.078451812 |
| Scoc | 7.889736 | 7.876552 | 7.8134 | 0.063173206 | 0.076357348 | 7.8349783 | 7.6931952 | 7.76424 | -0.071042668 | 0.070740428 |
| Rint1 | 8.616304 | 8.763556 | 8.6472 | 0.116349338 | -0.030902559 | 8.6012491 | 8.6648408 | 8.73588 | -0.071043133 | -0.134634847 |
| Klhl10 | 2.055627 | 1.982524 | 2.1139 | -0.131331945 | -0.058228684 | 2.048448 | 2.0003097 | 2.07146 | -0.071146867 | -0.023008619 |
| Rpl39l | 2.727643 | 2.627222 | 2.6635 | -0.03629923 | 0.06412115 | 2.6729829 | 2.735764 | 2.80692 | -0.071157887 | -0.133938973 |
| Fam205a3 | 1.736129 | 1.691223 | 1.6636 | 0.027658718 | 0.072564993 | 1.598437 | 1.52443 | 1.59562 | -0.071185574 | 0.002821473 |

| | | | | | | | | | | |
|---|---|---|---|---|---|---|---|---|---|---|
| P2ry4 | 1.847596 | 1.811014 | 1.8276 | -0.016628916 | 0.019953393 | 1.8962777 | 1.8370275 | 1.90823 | -0.071198138 | -0.011947952 |
| Efcab12 | 1.559433 | 1.53248 | 1.581 | -0.048545693 | -0.021592202 | 1.5291201 | 1.5048924 | 1.57616 | -0.071272179 | -0.047044465 |
| Wdr49 | 1.898285 | 1.892837 | 1.9315 | -0.038666494 | -0.033218316 | 1.8983512 | 1.8804542 | 1.95175 | -0.0712952 | -0.053398257 |
| Gm10974 | 0.520035 | 0.469503 | 0.4971 | -0.027597149 | 0.022934211 | 0.4900955 | 0.509894 | 0.5812 | -0.071306418 | -0.091104858 |
| Kcne3 | 2.937419 | 2.885973 | 2.9412 | -0.055232826 | -0.003786231 | 2.9652276 | 2.8786261 | 2.95 | -0.071376852 | 0.015224654 |
| Vmn2r25 | 2.998395 | 2.90852 | 3.005 | -0.096465914 | -0.006590388 | 2.9693319 | 3.0675396 | 3.13892 | -0.071381356 | -0.16958907 |
| Tmem107 | 4.178311 | 4.190936 | 4.4274 | -0.23642052 | -0.249045502 | 4.4061055 | 4.2143541 | 4.28574 | -0.071390409 | 0.120360953 |
| Atg7 | 8.547086 | 8.441476 | 8.3758 | 0.065632634 | 0.171242341 | 8.4538721 | 8.2202579 | 8.29177 | -0.071514477 | 0.162099708 |
| Pagr1b | 3.108478 | 2.876749 | 3.1711 | -0.294392669 | -0.062663752 | 2.9659632 | 3.106335 | 3.17786 | -0.071521675 | -0.211893487 |
| Kmt2b | 11.4243 | 11.40579 | 11.505 | -0.099466848 | -0.080951168 | 11.585402 | 11.622872 | 11.6944 | -0.07155793 | -0.109028133 |
| Ppp2r1a | 11.53452 | 11.55239 | 11.58 | -0.027738426 | -0.04561036 | 11.76162 | 11.524819 | 11.5964 | -0.071567938 | 0.16523276 |
| Car15 | 5.44806 | 5.628091 | 5.3983 | 0.229827807 | 0.049796445 | 5.6796305 | 5.7694246 | 5.841 | -0.071575763 | -0.161369898 |
| Gm1848 | 0.58892 | 0.586684 | 0.5633 | 0.023420086 | 0.02565703 | 0.5491083 | 0.5636692 | 0.63529 | -0.071624783 | -0.086185598 |
| Nhlrc4 | 0.560351 | 0.569824 | 0.5571 | 0.012754353 | 0.003280582 | 0.630936 | 0.551498 | 0.62313 | -0.07163693 | 0.007801142 |
| Capn12 | 2.241383 | 2.150679 | 2.2669 | -0.116172148 | -0.025468911 | 2.1764059 | 2.1835413 | 2.25522 | -0.071676637 | -0.07881201 |
| Ufsp1 | 3.865922 | 3.662794 | 3.8629 | -0.200076722 | 0.003051102 | 3.9317 | 3.7945467 | 3.86624 | -0.071688381 | 0.065464981 |
| Gm13304 | 0.645174 | 0.76861 | 0.7446 | 0.023972033 | -0.09946407 | 0.8118116 | 0.6495682 | 0.72131 | -0.071737912 | 0.090505493 |
| Ndufs1 | 10.38105 | 10.32531 | 10.348 | -0.022639214 | 0.03310744 | 10.473524 | 10.382005 | 10.4537 | -0.071742458 | 0.01977637 |
| Tcte1 | 1.586769 | 1.605153 | 1.6306 | -0.02543329 | -0.043817113 | 1.6489509 | 1.6122507 | 1.68401 | -0.07175577 | -0.03505558 |
| Smim17 | 1.82846 | 1.736976 | 1.8771 | -0.140135939 | -0.048651795 | 1.7955253 | 1.7252793 | 1.79705 | -0.071767062 | -0.001521055 |
| Olfr716 | 1.96795 | 2.015141 | 2.032 | -0.016903158 | -0.064093903 | 2.0092565 | 1.9879063 | 2.05974 | -0.071833034 | -0.050482748 |
| Gm11545 | 1.237246 | 1.265338 | 1.3055 | -0.040137376 | -0.068229016 | 1.2488845 | 1.2271251 | 1.29898 | -0.071857105 | -0.050097742 |
| Gmnn | 7.080673 | 6.950871 | 7.2332 | -0.282320656 | -0.152518696 | 7.219741 | 7.2871532 | 7.35902 | -0.071865904 | -0.139278085 |
| Epha3 | 3.001674 | 3.030397 | 3.0765 | -0.046151158 | -0.074873516 | 3.0076213 | 2.9182451 | 2.99013 | -0.071882228 | 0.017493986 |
| Numa1 | 12.46641 | 12.49498 | 12.536 | -0.04079149 | -0.069365666 | 12.399892 | 12.448727 | 12.5206 | -0.071886122 | -0.120721019 |
| Ldb1 | 11.86429 | 11.91991 | 11.969 | -0.04868956 | -0.104311364 | 12.154135 | 11.951415 | 12.0234 | -0.071956912 | 0.130763426 |
| Slc5a5 | 1.038072 | 1.11199 | 1.0437 | 0.068290841 | -0.005626702 | 1.1629735 | 1.0314058 | 1.10338 | -0.071972915 | 0.059594741 |
| Adam28 | 2.198055 | 2.122009 | 2.2439 | -0.121912656 | -0.045866085 | 2.2611493 | 2.1679043 | 2.23989 | -0.071982705 | 0.0212623 |
| Tmem141 | 4.574984 | 4.559768 | 4.6679 | -0.108172856 | -0.092956662 | 4.4028069 | 4.3270859 | 4.39911 | -0.072024541 | 0.003696436 |
| Cela2a | 1.39026 | 1.443601 | 1.4311 | 0.012476141 | -0.040864353 | 1.4491083 | 1.4152989 | 1.48735 | -0.072051487 | -0.03824209 |
| Tmem230 | 8.652433 | 8.578059 | 8.735 | -0.156944793 | -0.082570753 | 8.8362419 | 8.6544765 | 8.72659 | -0.072113793 | 0.109651614 |
| 4933407L21Rik | 1.191256 | 1.165096 | 1.2488 | -0.08374747 | -0.057587161 | 1.1891246 | 1.2476059 | 1.31975 | -0.072148291 | -0.130629619 |

| | | | | | | | | | | |
|---|---|---|---|---|---|---|---|---|---|---|
| Mre11a | 9.509941 | 9.540584 | 9.5801 | -0.039499413 | -0.070141784 | 9.4751534 | 9.4666132 | 9.53881 | -0.072195416 | -0.063655288 |
| Oacyl | 1.15606 | 1.186304 | 1.1581 | 0.028237636 | -0.00200676 | 1.1902306 | 1.118824 | 1.19103 | -0.072203925 | -0.000797321 |
| Usp3 | 10.38905 | 10.36355 | 10.492 | -0.128182558 | -0.102685508 | 10.208759 | 10.412728 | 10.4849 | -0.072206218 | -0.276174915 |
| Reep5 | 10.20325 | 10.06424 | 10.26 | -0.195914225 | -0.056901875 | 10.183225 | 10.155941 | 10.2283 | -0.072334296 | -0.045049714 |
| Tmem265 | 6.67754 | 6.547579 | 6.6724 | -0.124837577 | 0.005123409 | 6.7461211 | 6.5871317 | 6.65947 | -0.072335935 | 0.086653425 |
| Gramd1a | 12.22849 | 12.3639 | 12.37 | -0.00639578 | -0.141812614 | 12.451711 | 12.457952 | 12.5304 | -0.072409606 | -0.078650446 |
| Olfr372 | -0.136214 | -0.13797 | -0.1378 | -0.000126611 | 0.001629348 | -0.1616721 | -0.163488 | -0.0911 | -0.072420272 | -0.070604388 |
| Cry1 | 8.63291 | 8.670466 | 8.6441 | 0.026382005 | -0.011174105 | 8.4266882 | 8.483133 | 8.55557 | -0.07243788 | -0.128882696 |
| Cacna2d1 | 3.964948 | 4.120777 | 3.7526 | 0.368173841 | 0.212344872 | 3.9493388 | 3.7302883 | 3.80279 | -0.072505347 | 0.146545189 |
| Cnga1 | 5.539231 | 5.725873 | 5.6179 | 0.108015203 | -0.078626672 | 5.3398405 | 5.5389552 | 5.61151 | -0.072553315 | -0.271668003 |
| Dytn | 0.676175 | 0.722723 | 0.6402 | 0.082563358 | 0.036015641 | 0.674787 | 0.6537777 | 0.72636 | -0.072582421 | -0.051573064 |
| Trbj2-4 | 5.231897 | 5.399413 | 5.2626 | 0.136811952 | -0.03070368 | 5.0934962 | 5.245056 | 5.31768 | -0.072626926 | -0.224186688 |
| Mief2 | 7.187516 | 7.096178 | 7.252 | -0.155820593 | -0.064482347 | 7.4801057 | 7.2227594 | 7.2954 | -0.072636855 | 0.184709451 |
| Dusp26 | 1.078577 | 0.970429 | 1.0607 | -0.090296236 | 0.017851761 | 1.0366548 | 0.959003 | 1.03175 | -0.072746984 | 0.004904843 |
| Klrb1f | 6.411654 | 6.385021 | 6.4382 | -0.053150426 | -0.026517224 | 6.2984792 | 6.2761228 | 6.34887 | -0.072751992 | -0.050395659 |
| Smoc1 | 3.915012 | 3.826221 | 3.8939 | -0.06763764 | 0.02115362 | 3.835056 | 3.8327217 | 3.90551 | -0.072788925 | -0.070454643 |
| Gm10409 | 1.75232 | 1.702317 | 1.8012 | -0.098884964 | -0.048882294 | 1.6414694 | 1.6622999 | 1.7351 | -0.072799197 | -0.093629695 |
| Gm3020 | 1.75232 | 1.702317 | 1.8012 | -0.098884964 | -0.048882294 | 1.6414694 | 1.6622999 | 1.7351 | -0.072799197 | -0.093629695 |
| 1700029H14Rik | 1.05047 | 1.044826 | 1.0933 | -0.048452937 | -0.042809306 | 1.0412952 | 1.0192715 | 1.09208 | -0.072806531 | -0.05078281 |
| AC124512.1 | 2.856629 | 2.847949 | 2.7544 | 0.093543436 | 0.102223862 | 2.8989778 | 2.8004383 | 2.87325 | -0.072807173 | 0.02573237 |
| Chd2 | 12.09194 | 12.14558 | 12.206 | -0.060898156 | -0.11453731 | 11.906137 | 12.180743 | 12.2536 | -0.072822702 | -0.347428439 |
| Dzank1 | 5.057226 | 5.217465 | 5.097 | 0.120499048 | -0.039740615 | 5.2918327 | 5.3240093 | 5.39685 | -0.072838783 | -0.105015339 |
| Ces2d-ps | 0.143788 | 0.133777 | 0.1789 | -0.045073819 | -0.03506315 | 0.111939 | 0.0993938 | 0.17232 | -0.072925895 | -0.060380724 |
| Csde1 | 12.87451 | 12.89985 | 12.872 | 0.027917036 | 0.002571852 | 12.795286 | 12.818364 | 12.8913 | -0.072926772 | -0.096004065 |
| Olfr224 | 0.737998 | 0.735962 | 0.7689 | -0.032986439 | -0.03095074 | 0.74308 | 0.7159365 | 0.78888 | -0.072946872 | -0.045803323 |
| C1ql2 | 1.048269 | 0.932876 | 0.9792 | -0.046351232 | 0.069040934 | 0.8817091 | 0.8728906 | 0.94584 | -0.072950108 | -0.064131639 |
| Gm49196 | 1.340923 | 1.423589 | 1.4812 | -0.057565199 | -0.140231399 | 1.3812118 | 1.3981568 | 1.47125 | -0.073096238 | -0.090041292 |
| Trav7n-6 | 4.40929 | 4.44221 | 4.4175 | 0.024737747 | -0.008182147 | 4.3329586 | 4.4152189 | 4.48833 | -0.073107416 | -0.155367716 |
| Pcdhga9 | 2.476611 | 2.429288 | 2.4885 | -0.059215951 | -0.011892437 | 2.4010217 | 2.3323171 | 2.40547 | -0.073148005 | -0.004443485 |
| Chd7 | 11.00798 | 11.08788 | 10.919 | 0.169365924 | 0.089467368 | 11.081836 | 11.145613 | 11.2188 | -0.073156406 | -0.136932688 |
| Olfr525 | 1.203118 | 1.184151 | 1.1696 | 0.014579974 | 0.033547165 | 1.1463214 | 1.1968413 | 1.27002 | -0.07317381 | -0.12369373 |
| Trav6n-5 | 1.108863 | 1.133654 | 1.1428 | -0.009132506 | -0.033922857 | 1.1407729 | 1.1347303 | 1.20791 | -0.073181781 | -0.067139192 |

| | | | | | | | | | | |
|---|---|---|---|---|---|---|---|---|---|---|
| Fam83g | 5.516997 | 5.464059 | 5.5306 | -0.066583258 | -0.013645247 | 5.6216925 | 5.4549649 | 5.52816 | -0.073194321 | 0.09353328 |
| Dnajc4 | 7.153653 | 7.135452 | 7.298 | -0.162556017 | -0.144354315 | 7.5822012 | 7.4067574 | 7.47996 | -0.07319777 | 0.102246004 |
| Lrig3 | 3.971709 | 3.926671 | 4.0873 | -0.160663336 | -0.115624851 | 3.9464318 | 3.9194573 | 3.99268 | -0.073218346 | -0.046243894 |
| Sms-ps | 7.998083 | 8.272317 | 8.1946 | 0.077690642 | -0.196543778 | 8.0500679 | 8.0445945 | 8.11781 | -0.073218688 | -0.067745224 |
| Cyp2u1 | 2.465618 | 2.458089 | 2.533 | -0.074931349 | -0.067401984 | 2.5650658 | 2.4781299 | 2.55141 | -0.073277772 | 0.013658157 |
| Reg3g | 3.475188 | 3.665082 | 3.4311 | 0.23401292 | 0.044119298 | 3.2665308 | 3.1763262 | 3.24963 | -0.073304467 | 0.016900148 |
| Prickle4 | 1.798736 | 1.775257 | 1.8906 | -0.115320584 | -0.091842269 | 1.8259271 | 1.7939077 | 1.86723 | -0.073325384 | -0.041305996 |
| Tbc1d25 | 7.737917 | 7.801329 | 7.8144 | -0.013098065 | -0.076510269 | 7.8319797 | 7.7291221 | 7.80252 | -0.07340162 | 0.029455984 |
| Gm28362 | -0.539987 | -0.52812 | -0.4914 | -0.036739564 | -0.048610893 | -0.5262312 | -0.537187 | -0.4638 | -0.073426672 | -0.062470558 |
| Grin2c | 2.253451 | 2.399252 | 2.332 | 0.06721642 | -0.078584401 | 2.3622363 | 2.2806363 | 2.35412 | -0.073479981 | 0.008120013 |
| Kcnt1 | 1.209024 | 1.235012 | 1.2911 | -0.056051041 | -0.082039211 | 1.1870637 | 1.1859816 | 1.25946 | -0.073482465 | -0.072400377 |
| Olfr1262 | 1.641172 | 1.571843 | 1.5851 | -0.013242574 | 0.056086178 | 1.6178867 | 1.6205582 | 1.69408 | -0.073517684 | -0.076189133 |
| Comt | 7.803359 | 7.647822 | 7.8132 | -0.165418639 | -0.009880928 | 7.8252199 | 7.5209944 | 7.59453 | -0.073536845 | 0.230688706 |
| Dmac2 | 9.813044 | 9.908645 | 10.026 | -0.117093649 | -0.212694509 | 10.475354 | 10.114622 | 10.1882 | -0.073549103 | 0.287182559 |
| Opn1sw | 2.311511 | 2.415429 | 2.3143 | 0.101153933 | -0.002763685 | 2.4374589 | 2.4476165 | 2.52119 | -0.073577854 | -0.083735408 |
| Thap2 | 8.244356 | 8.413603 | 8.2807 | 0.132937848 | -0.036309715 | 8.4716061 | 8.3642442 | 8.43783 | -0.073582545 | 0.033779371 |
| Etv4 | 1.396306 | 1.364612 | 1.4113 | -0.046723478 | -0.015029273 | 1.319991 | 1.2834064 | 1.35703 | -0.073621257 | -0.037036682 |
| Rbm11 | 2.720087 | 2.708563 | 2.7498 | -0.041255529 | -0.029731654 | 2.8089316 | 2.7595423 | 2.83319 | -0.07364447 | -0.024255143 |
| Bmp3 | 2.861602 | 2.85331 | 2.8327 | 0.020597253 | 0.028889797 | 2.9546537 | 2.7455395 | 2.8192 | -0.073659287 | 0.135454909 |
| Peg10 | 3.167706 | 3.278047 | 3.1513 | 0.12671562 | 0.016374902 | 3.2219179 | 3.1378429 | 3.21156 | -0.073716477 | 0.010358573 |
| Zswim5 | 1.957788 | 1.894189 | 1.9166 | -0.022436144 | 0.041162679 | 1.8272635 | 1.838492 | 1.91221 | -0.073721841 | -0.084950304 |
| Cenpq | 8.37115 | 8.380004 | 8.3177 | 0.062262781 | 0.053408452 | 8.553042 | 8.3925877 | 8.46632 | -0.073737289 | 0.086717035 |
| Slc30a3 | 0.841329 | 0.819613 | 0.9566 | -0.136964512 | -0.115248662 | 0.8013616 | 0.8280728 | 0.90186 | -0.073786402 | -0.100497629 |
| Aebp2 | 11.13279 | 11.19447 | 11.097 | 0.097811294 | 0.036130344 | 10.816973 | 11.088965 | 11.1628 | -0.073790498 | -0.345782512 |
| Btk | 11.0334 | 11.11819 | 11.049 | 0.069674642 | -0.015115892 | 11.390094 | 11.103234 | 11.1771 | -0.07386811 | 0.212991263 |
| Jchain | 11.11131 | 11.80747 | 10.978 | 0.829460792 | 0.133295552 | 10.509997 | 10.874164 | 10.9481 | -0.07391075 | -0.438077677 |
| Gm8696 | 4.054054 | 3.727006 | 4.1363 | -0.409304577 | -0.082257399 | 3.6241831 | 3.6992525 | 3.77317 | -0.073917277 | -0.148986685 |
| Zfp385c | 1.338788 | 1.386688 | 1.3305 | 0.056211994 | 0.008311838 | 1.3977688 | 1.2573655 | 1.3313 | -0.073930833 | 0.066472439 |
| Amacr | 6.087284 | 5.985722 | 6.0589 | -0.073168751 | 0.028393626 | 6.1281235 | 6.0982163 | 6.17221 | -0.073996123 | -0.04408894 |
| Acot9 | 9.186868 | 9.062482 | 9.1215 | -0.059014594 | 0.065371286 | 9.0699862 | 9.0094114 | 9.08344 | -0.074032118 | -0.013457383 |
| Cby1 | 6.45099 | 6.368855 | 6.5021 | -0.133265056 | -0.051129696 | 6.5954245 | 6.3488063 | 6.42286 | -0.074049837 | 0.172568414 |
| Vsir | 11.48516 | 11.27652 | 11.318 | -0.041832802 | 0.166807832 | 11.248143 | 11.082063 | 11.1561 | -0.074083734 | 0.091996909 |

| | | | | | | | | | | |
|---|---|---|---|---|---|---|---|---|---|---|
| Borcs8 | 8.538842 | 8.528727 | 8.5379 | -0.009160577 | 0.000954548 | 8.8729293 | 8.5940546 | 8.66818 | -0.074121879 | 0.204752863 |
| Fut1 | 1.212391 | 1.280512 | 1.2367 | 0.043833364 | -0.024287568 | 1.3148866 | 1.2013632 | 1.27551 | -0.074142861 | 0.039380592 |
| Gm21969 | 1.578829 | 1.575946 | 1.5714 | 0.004553036 | 0.007436273 | 1.5460851 | 1.5368498 | 1.61101 | -0.074155242 | -0.064919914 |
| Olfr1392 | 2.719775 | 2.688455 | 2.7501 | -0.061595637 | -0.030275696 | 2.755015 | 2.7485685 | 2.82278 | -0.074208532 | -0.067761986 |
| Hsd3b2 | 5.17567 | 5.104274 | 5.1497 | -0.045430779 | 0.025964878 | 5.0710502 | 5.401809 | 5.47605 | -0.074245243 | -0.405004012 |
| Trappc2 | 7.580454 | 7.522348 | 7.5041 | 0.018264203 | 0.076369748 | 7.6245338 | 7.485116 | 7.55937 | -0.074254317 | 0.065163468 |
| Fam120b | 9.935863 | 10.05039 | 9.9967 | 0.053662437 | -0.060861245 | 10.12796 | 10.103169 | 10.1775 | -0.074288568 | -0.049497865 |
| Ces2f | 4.203754 | 4.148775 | 4.3672 | -0.21843323 | -0.16345372 | 4.4489183 | 4.3724945 | 4.44681 | -0.074318744 | 0.002105092 |
| Nsmf | 7.731858 | 7.589299 | 7.845 | -0.255704275 | -0.113144391 | 8.0480501 | 7.8480577 | 7.92238 | -0.074322691 | 0.125669771 |
| Ksr1 | 9.069425 | 8.94657 | 9.0831 | -0.136553279 | -0.013698051 | 8.7643016 | 8.8708638 | 8.94526 | -0.07439875 | -0.180960984 |
| Samd4 | 3.89774 | 3.681258 | 3.6641 | 0.017184144 | 0.233666342 | 3.6575957 | 3.613277 | 3.68768 | -0.074407581 | -0.030088792 |
| Gm18180 | 0.589513 | 0.521625 | 0.5128 | 0.008852648 | 0.076740623 | 0.5394202 | 0.5107969 | 0.58521 | -0.074415968 | -0.045792688 |
| Psg16 | 7.674506 | 7.703297 | 7.6352 | 0.068138505 | 0.039346862 | 7.6198108 | 7.8646474 | 7.9391 | -0.074448665 | -0.319285324 |
| Uck2 | 9.678958 | 9.457648 | 9.4595 | -0.001825502 | 0.219485197 | 9.5240505 | 9.4172748 | 9.49173 | -0.074452106 | 0.032323592 |
| Rhpn2 | 3.976264 | 3.892891 | 4.2175 | -0.324639604 | -0.241266703 | 3.9972894 | 4.0870846 | 4.16154 | -0.074459739 | -0.164254991 |
| Gk | 7.701633 | 7.463047 | 7.1486 | 0.314440163 | 0.553026526 | 7.4594779 | 7.3145995 | 7.38913 | -0.074525964 | 0.070352442 |
| Dnajc22 | 1.509379 | 1.501894 | 1.5308 | -0.028858902 | -0.021373935 | 1.4809782 | 1.4973412 | 1.57191 | -0.074569311 | -0.090932357 |
| Zfp85 | 7.123165 | 6.998805 | 7.1998 | -0.200986874 | -0.076627117 | 7.1018506 | 7.0545899 | 7.12917 | -0.074576384 | -0.027315706 |
| Hnf1b | 1.52725 | 1.439892 | 1.4947 | -0.054764628 | 0.032594239 | 1.5116397 | 1.4299701 | 1.50458 | -0.074611513 | 0.007058122 |
| Iqcd | 1.872812 | 1.865045 | 1.8215 | 0.043564211 | 0.051331245 | 1.8859771 | 1.8114837 | 1.88615 | -0.074665004 | -0.000171609 |
| Psg19 | 1.437155 | 1.49487 | 1.5546 | -0.059692037 | -0.117407165 | 1.5050322 | 1.5314743 | 1.60616 | -0.074687442 | -0.101129578 |
| Cd244a | 7.629081 | 7.208684 | 7.0299 | 0.178823638 | 0.599221305 | 7.4894535 | 6.9662062 | 7.04093 | -0.07472587 | 0.448521487 |
| Fam163b | 0.882055 | 0.869273 | 0.8233 | 0.04593189 | 0.058713465 | 0.8015805 | 0.7968587 | 0.87159 | -0.074726646 | -0.070004838 |
| 4932415D10Rik | 0.199506 | 0.204079 | 0.2194 | -0.015315111 | -0.019888574 | 0.2397638 | 0.1794532 | 0.25421 | -0.074755143 | -0.014444566 |
| Efhd1 | 2.027923 | 1.983746 | 2.0012 | -0.01744968 | 0.026726447 | 1.9930949 | 1.9799513 | 2.05471 | -0.074759658 | -0.061616051 |
| Gm45778 | 3.865975 | 3.763247 | 4.0026 | -0.239317425 | -0.13658924 | 3.7633716 | 3.7870709 | 3.86189 | -0.074819358 | -0.098518635 |
| Hfm1 | 0.81081 | 0.761602 | 0.7268 | 0.034824377 | 0.084032419 | 0.7939055 | 0.7507228 | 0.82558 | -0.074856135 | -0.031673499 |
| Usp32 | 10.05621 | 9.939406 | 9.9565 | -0.017072336 | 0.099733124 | 9.8018138 | 9.953739 | 10.0286 | -0.074877329 | -0.226802525 |
| Rtl9 | 0.176821 | 0.124693 | 0.1626 | -0.03789697 | 0.014230854 | 0.1229315 | 0.1161514 | 0.19105 | -0.074903535 | -0.068123431 |
| Bloc1s5 | 7.375412 | 7.189919 | 7.4514 | -0.261450873 | -0.075957655 | 7.2191868 | 7.1933697 | 7.26828 | -0.074911269 | -0.049094164 |
| Zfp275 | 9.382448 | 9.529676 | 9.4591 | 0.070534287 | -0.076693368 | 9.5052486 | 9.46202 | 9.53702 | -0.074995715 | -0.03176707 |
| Pkp3 | 9.260844 | 9.115569 | 9.3378 | -0.222191443 | -0.07691632 | 9.3469434 | 9.0577324 | 9.13276 | -0.075031448 | 0.214179642 |

| | | | | | | | | | | |
|---|---|---|---|---|---|---|---|---|---|---|
| Olfr267 | 0.489461 | 0.489492 | 0.5075 | -0.018042979 | -0.018074058 | 0.5342461 | 0.4887917 | 0.56383 | -0.07503808 | -0.029583636 |
| Tfcp2 | 7.677064 | 7.687444 | 7.7299 | -0.042406735 | -0.052787225 | 7.882477 | 7.8490535 | 7.92409 | -0.07504101 | -0.041617514 |
| 39873 | 5.858051 | 5.943248 | 6.0642 | -0.120964212 | -0.206161775 | 5.6998537 | 5.7535059 | 5.82855 | -0.075044225 | -0.128696394 |
| Las1l | 10.16257 | 10.22238 | 10.176 | 0.046144802 | -0.013668264 | 10.334855 | 10.182924 | 10.258 | -0.075045232 | 0.076885868 |
| Gm18517 | 3.733529 | 3.781875 | 3.987 | -0.205099471 | -0.253445568 | 3.9503234 | 3.7844826 | 3.85964 | -0.075154552 | 0.090686165 |
| Ptk6 | 3.038054 | 3.137006 | 3.1486 | -0.011601605 | -0.110553205 | 3.1293672 | 3.1419615 | 3.21712 | -0.075158162 | -0.087752543 |
| Prr15l | 1.243544 | 1.171417 | 1.3049 | -0.133511738 | -0.061384507 | 1.3068198 | 1.2423288 | 1.3175 | -0.075171774 | -0.010680711 |
| Gm11344 | 1.397658 | 1.374264 | 1.3815 | -0.007197493 | 0.016196573 | 1.4926898 | 1.3342461 | 1.40945 | -0.075205984 | 0.083237689 |
| Casp16-ps | 0.841111 | 0.862578 | 0.8448 | 0.017729001 | -0.00373794 | 0.9202241 | 0.8944455 | 0.96966 | -0.075217843 | -0.049439221 |
| Rps13-ps1 | 2.267102 | 2.595845 | 2.0915 | 0.504315828 | 0.175572605 | 2.2109302 | 1.9801606 | 2.05539 | -0.075224932 | 0.155544661 |
| Kazald1 | 1.935263 | 1.935948 | 2.0117 | -0.075741273 | -0.076426064 | 1.9251814 | 1.9452889 | 2.02052 | -0.075234375 | -0.095341875 |
| Spata33 | 0.727391 | 0.660681 | 0.7826 | -0.12191523 | -0.055204839 | 0.715382 | 0.6865501 | 0.76179 | -0.075235915 | -0.046404008 |
| Avl9 | 10.33961 | 10.38476 | 10.35 | 0.034584152 | -0.010570154 | 10.20056 | 10.282112 | 10.3574 | -0.07526588 | -0.156817356 |
| Kif24 | 6.06049 | 6.27996 | 6.2019 | 0.078023574 | -0.141447153 | 6.1737123 | 6.4445956 | 6.51991 | -0.075317234 | -0.346200566 |
| Draxin | 2.967209 | 2.869285 | 2.9516 | -0.082328438 | 0.015595554 | 2.9354706 | 2.8700634 | 2.94539 | -0.075328494 | -0.009921351 |
| Gsg1 | 0.892299 | 0.865951 | 0.9522 | -0.08625634 | -0.059908724 | 0.9027885 | 0.8293255 | 0.90466 | -0.075334715 | -0.001871728 |
| Angptl3 | 0.612217 | 0.589737 | 0.5511 | 0.038612009 | 0.061091645 | 0.6646551 | 0.5565733 | 0.63192 | -0.075351251 | 0.032730488 |
| Cwc25 | 8.972818 | 8.834349 | 8.9314 | -0.097036788 | 0.041432323 | 9.0090379 | 8.9116411 | 8.987 | -0.075358772 | 0.02203806 |
| Gm20547 | 4.518691 | 4.257755 | 4.2189 | 0.038864753 | 0.299800302 | 4.6138266 | 4.3129095 | 4.38833 | -0.075415596 | 0.225501561 |
| Nhlrc1 | 3.22792 | 3.222532 | 3.1875 | 0.035077076 | 0.040464715 | 3.1373462 | 3.1083865 | 3.18381 | -0.07542795 | -0.046468275 |
| Lamc3 | 1.066568 | 0.857321 | 0.9436 | -0.086303532 | 0.122943358 | 0.9223082 | 0.9017992 | 0.97725 | -0.075446029 | -0.054937006 |
| Olfr329-ps | 0.667887 | 0.672392 | 0.6523 | 0.020093584 | 0.015588435 | 0.6560918 | 0.6414091 | 0.71688 | -0.075468796 | -0.060786102 |
| 5330417C22Rik | 2.789335 | 2.567384 | 2.6326 | -0.065237361 | 0.156713511 | 2.6775071 | 2.5657655 | 2.6413 | -0.07553415 | 0.036207431 |
| Gm17906 | 2.326485 | 2.263289 | 2.3332 | -0.069913503 | -0.006717273 | 2.250966 | 2.2939521 | 2.36963 | -0.07568113 | -0.118667287 |
| Kdm1a | 10.43073 | 10.47613 | 10.418 | 0.057647084 | 0.01224579 | 10.319599 | 10.393607 | 10.4693 | -0.075728114 | -0.149736834 |
| Zfp217 | 10.81905 | 10.83708 | 10.821 | 0.016135674 | -0.001888806 | 10.611748 | 10.624705 | 10.7005 | -0.07574583 | -0.088702162 |
| Ccdc126 | 6.687229 | 6.724175 | 6.6823 | 0.04184977 | 0.004904029 | 6.6270259 | 6.6091339 | 6.68495 | -0.075812742 | -0.057920721 |
| Traj5 | 1.89285 | 1.841768 | 1.8148 | 0.026949778 | 0.078032365 | 1.8368861 | 1.8338567 | 1.90974 | -0.075881175 | -0.072851754 |
| Plac9b | 4.176729 | 4.027753 | 4.0779 | -0.05010934 | 0.098866661 | 4.1222817 | 3.8167532 | 3.89268 | -0.075930015 | 0.229598401 |
| Gm49368 | 2.248821 | 2.39936 | 2.0854 | 0.313976748 | 0.163438107 | 2.2931756 | 2.1091932 | 2.18513 | -0.075934489 | 0.108047952 |
| Gm21970 | 3.07515 | 3.228488 | 3.1643 | 0.064229359 | -0.089107677 | 2.9209783 | 3.0567877 | 3.13275 | -0.075960439 | -0.211769794 |
| Fam189a1 | 1.429782 | 1.505052 | 1.7249 | -0.219849601 | -0.295119386 | 1.4017203 | 1.3975656 | 1.47354 | -0.075971393 | -0.071816675 |

| | | | | | | | | | | |
|---|---|---|---|---|---|---|---|---|---|---|
| Slc30a2 | 2.253943 | 2.191802 | 2.1712 | 0.020624656 | 0.082765943 | 2.2058824 | 2.1661215 | 2.24212 | -0.075995542 | -0.036234728 |
| Ddit4l | 1.789927 | 1.695462 | 1.9242 | -0.228775057 | -0.134309252 | 1.7554042 | 1.7790689 | 1.85507 | -0.075996904 | -0.099661692 |
| Oplah | 5.898346 | 5.763379 | 5.9715 | -0.208103924 | -0.073136236 | 5.7803077 | 5.7836278 | 5.8597 | -0.076067744 | -0.079387848 |
| Tslp | 2.269381 | 2.171483 | 2.3195 | -0.14801964 | -0.050121637 | 2.143443 | 2.0986408 | 2.17472 | -0.076075977 | -0.031273804 |
| Tmem71 | 10.16993 | 10.07714 | 10.165 | -0.087490779 | 0.005296335 | 10.099182 | 10.061378 | 10.1376 | -0.076185008 | -0.038380479 |
| Morn4 | 2.406043 | 2.364162 | 2.3959 | -0.031739043 | 0.010142796 | 2.3879458 | 2.2573543 | 2.33355 | -0.076192709 | 0.054398836 |
| Gm2237 | 6.052729 | 5.940551 | 6.0489 | -0.108387372 | 0.003791389 | 6.0208183 | 6.0467691 | 6.12298 | -0.076205947 | -0.102156708 |
| Olfr1204 | 1.48871 | 1.478096 | 1.4835 | -0.005408687 | 0.005205409 | 1.4715693 | 1.4950112 | 1.57123 | -0.076214688 | -0.099656589 |
| Pcdh18 | 2.801158 | 2.8497 | 2.9097 | -0.060027607 | -0.108569802 | 2.8398843 | 2.6667598 | 2.74299 | -0.076232794 | 0.096891613 |
| Ppp1r36 | 1.535735 | 1.453684 | 1.4138 | 0.039885736 | 0.121937408 | 1.5157856 | 1.4186083 | 1.49485 | -0.076242542 | 0.020934739 |
| Tirap | 7.751905 | 7.612374 | 7.6944 | -0.082055378 | 0.057475229 | 7.7729666 | 7.6585819 | 7.73487 | -0.076285893 | 0.03809879 |
| 1810018F18Rik | 0.799584 | 0.812132 | 0.7525 | 0.059679515 | 0.047132463 | 0.762258 | 0.7729131 | 0.84927 | -0.076361026 | -0.087016133 |
| Abhd17c | 8.478415 | 8.34329 | 8.3968 | -0.053488483 | 0.081636049 | 8.1117419 | 8.0858767 | 8.16232 | -0.076443856 | -0.050578647 |
| Gm12057 | 2.537386 | 2.435399 | 2.561 | -0.125641163 | -0.023653941 | 2.5791898 | 2.555667 | 2.63214 | -0.076472272 | -0.052949497 |
| Traj26 | 2.559406 | 2.542122 | 2.631 | -0.08888298 | -0.071598638 | 2.4589668 | 2.5097172 | 2.58621 | -0.076494956 | -0.127245359 |
| Trbj2-1 | 5.676444 | 5.658242 | 5.5541 | 0.104132877 | 0.122335535 | 5.3919424 | 5.5666188 | 5.64312 | -0.076497628 | -0.251174033 |
| Elmod2 | 8.192549 | 8.306188 | 8.3283 | -0.022070426 | -0.135709188 | 8.3118144 | 8.2909167 | 8.36749 | -0.076573212 | -0.05567549 |
| Hoxb5 | 1.961707 | 1.943826 | 1.9107 | 0.033124241 | 0.051005519 | 1.8300736 | 1.8607462 | 1.93739 | -0.076648719 | -0.107321277 |
| Pla2g15 | 8.330688 | 8.22115 | 8.1561 | 0.065039727 | 0.174577189 | 8.4614402 | 7.9167825 | 7.99343 | -0.076651415 | 0.468006348 |
| Etfb | 8.792679 | 8.743057 | 8.7934 | -0.050336868 | -0.000715105 | 9.0765126 | 8.7366428 | 8.81329 | -0.076651645 | 0.263218182 |
| Bcl10 | 10.10284 | 10.15314 | 10.059 | 0.093801664 | 0.043506384 | 10.193826 | 9.9394276 | 10.0161 | -0.076679791 | 0.177718157 |
| Tldc1 | 6.733912 | 6.663416 | 6.9311 | -0.267701091 | -0.19720474 | 6.9146745 | 6.8391838 | 6.91589 | -0.076705895 | -0.001215225 |
| Smarca1 | 1.872385 | 1.779348 | 1.8635 | -0.084176922 | 0.008859734 | 1.7473729 | 1.7028675 | 1.77968 | -0.076817167 | -0.032311682 |
| Mblac1 | 3.200453 | 3.084866 | 3.1331 | -0.048281277 | 0.067306491 | 3.1708503 | 3.0528147 | 3.12963 | -0.076818985 | 0.041216675 |
| Gm43518 | 2.161469 | 2.236945 | 2.2254 | 0.011539332 | -0.063937149 | 2.1407423 | 2.2379148 | 2.31475 | -0.076833727 | -0.174006237 |
| Tmem41a | 5.537696 | 5.57168 | 5.6248 | -0.053079438 | -0.087063052 | 5.5507126 | 5.4632545 | 5.54013 | -0.07687608 | 0.010582011 |
| Gm32742 | 3.139862 | 3.174119 | 3.2299 | -0.055770984 | -0.090027458 | 3.038024 | 3.1164448 | 3.19333 | -0.076888902 | -0.155309751 |
| Olfr49 | 3.215377 | 3.237447 | 3.239 | -0.001526861 | -0.023596966 | 3.2347018 | 3.3048684 | 3.38179 | -0.0769209 | -0.147087503 |
| Gm16044 | 2.44995 | 2.373422 | 2.3825 | -0.009101755 | 0.067426648 | 2.5022026 | 2.4021366 | 2.47911 | -0.076971062 | 0.023094884 |
| Hpse2 | 1.29715 | 1.302618 | 1.2902 | 0.012440607 | 0.00697301 | 1.364214 | 1.2520478 | 1.32905 | -0.077002316 | 0.035163936 |
| CT033749.1 | 1.16116 | 1.184068 | 1.236 | -0.051918019 | -0.074825182 | 1.3032027 | 1.1894357 | 1.26652 | -0.077085793 | 0.036681186 |
| Gm15937 | 1.909203 | 1.752413 | 1.8028 | -0.05042822 | 0.106361786 | 1.738732 | 1.759869 | 1.83698 | -0.077111838 | -0.098248819 |

| | | | | | | | | | | |
|---|---|---|---|---|---|---|---|---|---|---|
| Rimbp3 | 0.532508 | 0.541022 | 0.609 | -0.067984274 | -0.076497731 | 0.5519788 | 0.5621035 | 0.63926 | -0.077155514 | -0.087280253 |
| Carm1 | 9.824794 | 9.759734 | 9.8337 | -0.073995462 | -0.0089346 | 9.9097542 | 9.745946 | 9.82319 | -0.077241706 | 0.08656642 |
| Ighv1-23 | 1.561245 | 1.593741 | 1.6172 | -0.023481562 | -0.055977841 | 1.5799612 | 1.5455672 | 1.62284 | -0.077275919 | -0.042881895 |
| Gm15452 | 0.524969 | 0.361136 | 0.3402 | 0.020980713 | 0.184813319 | 0.3987574 | 0.3450306 | 0.42235 | -0.077319424 | -0.023592633 |
| Hoxa2 | 1.912478 | 1.772308 | 1.8917 | -0.119354488 | 0.020815221 | 1.7141018 | 1.7344123 | 1.81174 | -0.07733014 | -0.097640638 |
| Rhox4f | 1.32852 | 1.313268 | 1.3076 | 0.005618512 | 0.020870068 | 1.3505308 | 1.26871 | 1.34608 | -0.07737342 | 0.004447406 |
| Gm13597 | 4.149806 | 4.120533 | 4.1354 | -0.014899834 | 0.014373648 | 4.0933271 | 4.0828945 | 4.16029 | -0.077391624 | -0.066959013 |
| Dntt | 6.181354 | 6.190333 | 6.2612 | -0.070857939 | -0.079837469 | 5.8880119 | 5.9345631 | 6.01201 | -0.07744723 | -0.123998352 |
| Gm3500 | 1.278331 | 1.145703 | 1.3138 | -0.168069248 | -0.035441837 | 1.2358553 | 1.2363631 | 1.31383 | -0.07747041 | -0.077978181 |
| Palm2 | -0.162356 | -0.18283 | -0.1 | -0.082831803 | -0.062354321 | -0.1596809 | -0.18843 | -0.1109 | -0.07754021 | -0.048791565 |
| Bche | 1.887445 | 1.835261 | 1.9482 | -0.112979123 | -0.060794845 | 1.7302552 | 1.7096495 | 1.78722 | -0.077569677 | -0.056963977 |
| Olfr641 | 1.290636 | 1.311605 | 1.3264 | -0.014751688 | -0.035721091 | 1.2948165 | 1.3381179 | 1.41578 | -0.077658541 | -0.120960008 |
| Zfp119a | 6.452688 | 6.350953 | 6.5217 | -0.170709852 | -0.06897528 | 6.5084686 | 6.4517695 | 6.52947 | -0.077701039 | -0.021001964 |
| Snap23 | 9.826318 | 9.767402 | 9.6916 | 0.075768497 | 0.134684792 | 9.6291763 | 9.5515813 | 9.6294 | -0.0778233 | -0.00022838 |
| Fer | 5.421942 | 5.173632 | 5.2338 | -0.060201622 | 0.188108381 | 5.1203462 | 5.093612 | 5.17144 | -0.077832205 | -0.051097991 |
| Vmn2r84 | 3.06565 | 3.026715 | 3.1486 | -0.121909523 | -0.082974634 | 2.9954741 | 3.210032 | 3.28793 | -0.07789813 | -0.292455998 |
| Fbxo15 | 2.332599 | 2.333658 | 2.2852 | 0.048408157 | 0.047349997 | 2.2872221 | 2.3136038 | 2.3915 | -0.077899622 | -0.10428127 |
| Gucy2g | 0.431537 | 0.383359 | 0.4549 | -0.071521211 | -0.023342308 | 0.3809351 | 0.3719579 | 0.44988 | -0.077919974 | -0.068942789 |
| Hmgcl | 8.827807 | 8.720115 | 8.8116 | -0.09150858 | 0.016183204 | 9.097453 | 8.8195888 | 8.89756 | -0.077974088 | 0.19989007 |
| Trav6d-7 | 4.259082 | 4.126447 | 4.1559 | -0.029430002 | 0.103204777 | 4.0919299 | 4.0790878 | 4.15708 | -0.077991031 | -0.065148975 |
| Trav6n-7 | 4.259082 | 4.126447 | 4.1559 | -0.029430002 | 0.103204777 | 4.0919299 | 4.0790878 | 4.15708 | -0.077991031 | -0.065148975 |
| Gm9755 | 1.382229 | 1.380967 | 1.407 | -0.0260756 | -0.02481395 | 1.4827467 | 1.4527437 | 1.53077 | -0.078025604 | -0.048022578 |
| Gm15697 | 2.18544 | 2.156409 | 2.1348 | 0.021575442 | 0.050605976 | 2.178492 | 2.2284419 | 2.3065 | -0.078057821 | -0.128007735 |
| Hoxd10 | 2.459659 | 2.407867 | 2.5386 | -0.130767452 | -0.078975078 | 2.3051024 | 2.3437897 | 2.42187 | -0.078077354 | -0.116764706 |
| Arl16 | 7.280516 | 7.175539 | 7.2753 | -0.099719283 | 0.005257762 | 7.4582385 | 7.2108654 | 7.28895 | -0.078085468 | 0.169287691 |
| Trim66 | 1.985796 | 1.984425 | 2.0413 | -0.056876565 | -0.055505635 | 1.9476398 | 2.0058185 | 2.08392 | -0.078098716 | -0.13627737 |
| Eps15 | 11.42946 | 11.41606 | 11.48 | -0.063855126 | -0.050463478 | 11.440738 | 11.351733 | 11.4299 | -0.078183766 | 0.010821548 |
| Ubash3a | 9.855655 | 9.927761 | 9.8882 | 0.03957623 | -0.032529843 | 9.5957827 | 9.790401 | 9.86865 | -0.078251374 | -0.272869726 |
| Klf8 | 6.313579 | 6.227095 | 6.3678 | -0.140663563 | -0.054179354 | 6.3948475 | 6.3662617 | 6.44455 | -0.07829166 | -0.04970587 |
| Zfp189 | 6.578578 | 6.660705 | 6.7422 | -0.081528445 | -0.163655673 | 6.3783683 | 6.5501817 | 6.62848 | -0.078297413 | -0.250110875 |
| Nisch | 11.54332 | 11.51821 | 11.633 | -0.114591298 | -0.089484424 | 11.862117 | 11.90961 | 11.9879 | -0.078302298 | -0.125795303 |
| Smim5 | 5.073842 | 4.946557 | 4.9999 | -0.053390768 | 0.073893886 | 4.6489001 | 4.680254 | 4.75858 | -0.078328872 | -0.109682784 |

| | | | | | | | | | | |
|---|---|---|---|---|---|---|---|---|---|---|
| Rbm41 | 8.17631 | 8.097101 | 8.0362 | 0.060888533 | 0.140097371 | 8.1237255 | 8.0229621 | 8.10129 | -0.078330722 | 0.022432619 |
| Prrt4 | 1.143788 | 1.081157 | 1.1319 | -0.050725432 | 0.011905681 | 1.1405034 | 1.0453739 | 1.12371 | -0.078332319 | 0.016797248 |
| Alpk3 | 4.110454 | 4.033375 | 3.996 | 0.037330466 | 0.114410096 | 3.9742549 | 3.9602596 | 4.03863 | -0.078370972 | -0.064375647 |
| 9130008F23Rik | 3.897142 | 3.599384 | 3.7385 | -0.139145415 | 0.158612371 | 3.7638835 | 3.6940853 | 3.77248 | -0.078394318 | -0.008596112 |
| Setmar | 5.250986 | 5.073819 | 5.1722 | -0.098420635 | 0.078745722 | 5.1369698 | 5.0796908 | 5.15813 | -0.07843647 | -0.021157485 |
| Ankra2 | 7.993986 | 8.010243 | 7.9668 | 0.043400941 | 0.027144572 | 8.1497599 | 8.1069316 | 8.18542 | -0.078487553 | -0.035659242 |
| Tcf15 | 0.406508 | 0.383107 | 0.4414 | -0.058257943 | -0.034856673 | 0.4000684 | 0.3677266 | 0.44622 | -0.07849258 | -0.04615085 |
| Ankrd24 | 5.318616 | 5.2043 | 5.4499 | -0.245579148 | -0.13126293 | 5.3883318 | 5.4331292 | 5.5117 | -0.078574511 | -0.123371918 |
| Eda | 2.263387 | 2.209096 | 2.1834 | 0.02565488 | 0.07994553 | 2.3276017 | 2.1432175 | 2.22186 | -0.078641053 | 0.105743151 |
| Polr2b | 10.88212 | 10.79606 | 10.748 | 0.048517342 | 0.13458399 | 10.767178 | 10.584637 | 10.6633 | -0.078642242 | 0.103898957 |
| Fibcd1 | 3.369044 | 3.407128 | 3.3893 | 0.017854825 | -0.020228553 | 3.4170163 | 3.2912743 | 3.36993 | -0.078658151 | 0.047083829 |
| Olfr954 | 1.171652 | 1.157236 | 1.0996 | 0.057648065 | 0.072064354 | 1.1235432 | 1.123871 | 1.20253 | -0.078663972 | -0.078991771 |
| Vps50 | 8.891167 | 8.86908 | 8.8562 | 0.012842712 | 0.034929692 | 8.7327829 | 8.8708483 | 8.94955 | -0.078702637 | -0.216768115 |
| Lrrd1 | 1.03816 | 1.049008 | 1.0707 | -0.021649262 | -0.032497039 | 1.1163355 | 1.021725 | 1.10045 | -0.078724456 | 0.015886014 |
| Ppil4 | 10.313 | 10.33657 | 10.366 | -0.029296654 | -0.052868138 | 10.254264 | 10.373203 | 10.4519 | -0.078728978 | -0.197668424 |
| Il2ra | 9.529572 | 9.453465 | 9.6791 | -0.225639926 | -0.149532911 | 9.5912026 | 9.7241564 | 9.80291 | -0.078758582 | -0.211712339 |
| R3hdm4 | 10.4845 | 10.32241 | 10.453 | -0.130973996 | 0.031117658 | 10.617883 | 10.297971 | 10.3768 | -0.078808554 | 0.241103485 |
| Gm38394 | 9.984094 | 9.991543 | 10.198 | -0.206736561 | -0.214185855 | 10.148482 | 10.311535 | 10.3904 | -0.078856739 | -0.241910552 |
| Clk4 | 10.45695 | 10.56994 | 10.634 | -0.06445021 | -0.177444496 | 10.765464 | 10.849782 | 10.9287 | -0.078905416 | -0.16322269 |
| Cxcr3 | 8.588775 | 8.403442 | 8.7336 | -0.330157768 | -0.144824586 | 8.4913793 | 8.5462932 | 8.62523 | -0.078937922 | -0.133851818 |
| Gprin2 | 2.120208 | 2.055478 | 2.0594 | -0.003897203 | 0.060832538 | 2.1475531 | 2.0590486 | 2.13811 | -0.079063623 | 0.009440919 |
| Tmem269 | 1.415193 | 1.397734 | 1.4226 | -0.024828064 | -0.007369641 | 1.4224681 | 1.4208929 | 1.50001 | -0.079115664 | -0.077540442 |
| Fnta | 8.821195 | 8.67097 | 8.8699 | -0.19895741 | -0.048732518 | 8.7539957 | 8.7319928 | 8.81114 | -0.079147596 | -0.057144636 |
| Amigo1 | 6.055944 | 6.115609 | 6.0132 | 0.102366305 | 0.042701334 | 5.8918703 | 5.9150152 | 5.99417 | -0.079150383 | -0.102295242 |
| Olfr839-ps1 | 5.498973 | 5.599868 | 5.6416 | -0.041763041 | -0.142658795 | 5.807853 | 6.0764719 | 6.15563 | -0.079162469 | -0.347781377 |
| Rbm4b | 8.237538 | 8.437082 | 8.3116 | 0.125523635 | -0.074020776 | 8.2158612 | 8.3425166 | 8.42174 | -0.079223328 | -0.205878775 |
| Tmprss2 | 2.084309 | 1.968104 | 2.0432 | -0.075107581 | 0.041097491 | 2.3398841 | 2.0921627 | 2.1714 | -0.079234295 | 0.168487056 |
| Gm9855 | 1.988706 | 1.842196 | 2.0284 | -0.186158814 | -0.039649016 | 1.9849349 | 1.92929 | 2.00852 | -0.07923483 | -0.023589877 |
| Aldh1a7 | 0.501258 | 0.351865 | 0.3266 | 0.025225001 | 0.174618567 | 0.4228052 | 0.3322887 | 0.41162 | -0.07932729 | 0.011189143 |
| Olfr558 | 2.212561 | 2.232534 | 2.2857 | -0.053131219 | -0.073104784 | 2.1264255 | 2.1043012 | 2.18366 | -0.079357952 | -0.057233579 |
| Arhgef25 | 4.376081 | 4.372832 | 4.5932 | -0.220376129 | -0.217127538 | 4.3370816 | 4.2888686 | 4.36823 | -0.079359597 | -0.031146595 |
| Fam45a | 8.666485 | 8.554433 | 8.6377 | -0.083262377 | 0.028789448 | 8.7821856 | 8.5892984 | 8.66867 | -0.079372363 | 0.113514911 |

| | | | | | | | | | | |
|---|---|---|---|---|---|---|---|---|---|---|
| Prkd3 | 10.69848 | 10.7578 | 10.73 | 0.02733341 | -0.031989548 | 10.443637 | 10.616634 | 10.696 | -0.07938423 | -0.252380862 |
| Olfml3 | 5.656065 | 5.286424 | 5.4016 | -0.115216917 | 0.254424466 | 5.252746 | 5.1304613 | 5.20985 | -0.079387378 | 0.042897273 |
| Cyp4f17 | 5.447734 | 5.420676 | 5.5575 | -0.136777015 | -0.109719111 | 5.4690113 | 5.447436 | 5.52683 | -0.079393613 | -0.057818291 |
| Gm49359 | 8.148746 | 8.281719 | 8.2352 | 0.046515663 | -0.086457735 | 8.3049553 | 8.2957639 | 8.37517 | -0.079408992 | -0.070217533 |
| Lta | 8.798298 | 8.65802 | 8.7103 | -0.052264969 | 0.088012857 | 8.9890932 | 8.7407843 | 8.82021 | -0.079425943 | 0.168882931 |
| Ythdc2 | 9.107007 | 9.073373 | 9.1599 | -0.086559049 | -0.052925557 | 9.0283436 | 9.1269754 | 9.20646 | -0.079485676 | -0.178117543 |
| Olfr1023 | 1.406128 | 1.39546 | 1.3954 | 9.13588E-05 | 0.010759048 | 1.3581954 | 1.3755606 | 1.45506 | -0.079500385 | -0.096865572 |
| Cda | 2.698749 | 2.570859 | 2.502 | 0.068875036 | 0.196765496 | 2.4353164 | 2.3230304 | 2.40259 | -0.079562496 | 0.032723483 |
| Scn2a | 1.639565 | 1.705271 | 1.6657 | 0.039563557 | -0.026142374 | 1.6300399 | 1.6065855 | 1.68619 | -0.079607755 | -0.056153347 |
| Shroom1 | 2.663697 | 2.497348 | 2.7397 | -0.242323255 | -0.075974707 | 2.6242481 | 2.5240655 | 2.60368 | -0.079618621 | 0.020563975 |
| Gm17082 | 2.311048 | 2.242362 | 2.2985 | -0.056099907 | 0.012586456 | 2.2856866 | 2.3301395 | 2.40986 | -0.079724128 | -0.12417704 |
| Rsrc2 | 11.04944 | 11.10255 | 11.171 | -0.06833746 | -0.121445188 | 11.142897 | 11.177097 | 11.2569 | -0.07982132 | -0.114021076 |
| Cdc37l1 | 9.797723 | 9.652497 | 9.7608 | -0.10831384 | 0.03691165 | 9.9614464 | 9.8821693 | 9.96199 | -0.079822242 | -0.000545144 |
| Lat2 | 10.22175 | 10.0666 | 10.32 | -0.253057781 | -0.097915059 | 10.698369 | 10.293687 | 10.3735 | -0.079847605 | 0.324834586 |
| AC125059.1 | 1.650941 | 1.610489 | 1.6489 | -0.038371234 | 0.00208105 | 1.6594418 | 1.6254886 | 1.70534 | -0.079849416 | -0.045896188 |
| Ttc19 | 7.077814 | 6.990789 | 6.9677 | 0.02306929 | 0.110094921 | 6.8014825 | 6.9132047 | 6.99306 | -0.079855186 | -0.191577375 |
| Tmem116 | 2.637709 | 2.564145 | 2.5462 | 0.017944736 | 0.09150831 | 2.6625378 | 2.522504 | 2.60237 | -0.079862906 | 0.060170815 |
| Trav7d-5 | 2.701886 | 2.661079 | 2.7893 | -0.128218913 | -0.087411598 | 2.6674766 | 2.6146498 | 2.69463 | -0.079976558 | -0.027149739 |
| Trav7n-5 | 2.701886 | 2.661079 | 2.7893 | -0.128218913 | -0.087411598 | 2.6674766 | 2.6146498 | 2.69463 | -0.079976558 | -0.027149739 |
| Chga | 2.484174 | 2.544129 | 2.4393 | 0.104850574 | 0.044895802 | 2.3905693 | 2.4523105 | 2.53236 | -0.080053917 | -0.141795181 |
| Liph | 2.876683 | 2.891529 | 2.8596 | 0.031956739 | 0.017111348 | 2.7765732 | 2.7346001 | 2.81469 | -0.080091124 | -0.038118069 |
| Vmn2r86 | 1.911915 | 1.932085 | 1.9023 | 0.029760626 | 0.009590689 | 1.9454725 | 1.9203859 | 2.00048 | -0.080096311 | -0.055009766 |
| Trbj1-4 | 5.347401 | 5.315197 | 5.1138 | 0.201368973 | 0.233573381 | 5.1122894 | 5.1402204 | 5.22032 | -0.080102723 | -0.108033676 |
| Ppp1r14a | 2.967757 | 2.85659 | 2.9749 | -0.118271609 | -0.007104619 | 3.0358065 | 2.9323561 | 3.01247 | -0.080115756 | 0.023334722 |
| Ap4b1 | 8.91564 | 8.932819 | 9.006 | -0.073153296 | -0.090332489 | 9.1800658 | 8.952071 | 9.03222 | -0.080148028 | 0.147846817 |
| Srpx2 | 2.844077 | 2.439161 | 2.6643 | -0.225165877 | 0.179750246 | 2.4740729 | 2.4347365 | 2.51491 | -0.080169576 | -0.040833215 |
| Tmem60 | 7.330278 | 7.38741 | 7.3595 | 0.027955977 | -0.02917558 | 7.5919977 | 7.2387004 | 7.31889 | -0.080190307 | 0.273106927 |
| Gm19085 | 2.293925 | 2.346397 | 2.2468 | 0.099639172 | 0.047166915 | 2.229804 | 2.2613116 | 2.34153 | -0.080216925 | -0.111724557 |
| Ccdc7b | 1.56236 | 1.522293 | 1.4908 | 0.031456139 | 0.071523042 | 1.4730909 | 1.5414384 | 1.62172 | -0.080281805 | -0.148629283 |
| Trav13d-2 | 3.39755 | 3.319067 | 3.3637 | -0.044630862 | 0.033852017 | 3.353766 | 3.3092786 | 3.3896 | -0.080317413 | -0.035830016 |
| Trav13n-2 | 3.39755 | 3.319067 | 3.3637 | -0.044630862 | 0.033852017 | 3.353766 | 3.3092786 | 3.3896 | -0.080317413 | -0.035830016 |
| Slc15a4 | 10.34981 | 10.27942 | 10.306 | -0.02643181 | 0.04395831 | 10.534274 | 10.271366 | 10.3517 | -0.080335529 | 0.182572634 |

| | | | | | | | | | | |
|---|---|---|---|---|---|---|---|---|---|---|
| Itgb1bp2 | 2.757296 | 2.765836 | 2.7913 | -0.025462364 | -0.034002215 | 2.8408013 | 2.7949303 | 2.87529 | -0.080359374 | -0.034488404 |
| 1110032F04Rik | 5.554812 | 5.742257 | 5.7867 | -0.044404386 | -0.231849861 | 5.3836546 | 5.434642 | 5.51502 | -0.08038166 | -0.131369051 |
| Mast3 | 10.26161 | 10.29007 | 10.403 | -0.112862376 | -0.141330024 | 10.579033 | 10.290873 | 10.3713 | -0.080397996 | 0.207762425 |
| Gpr63 | 1.901977 | 2.048887 | 1.9008 | 0.148083587 | 0.001173573 | 1.9113172 | 1.8561868 | 1.93662 | -0.08042836 | -0.025297939 |
| Ccdc114 | 5.464269 | 5.371503 | 5.476 | -0.104518726 | -0.011752965 | 5.8435102 | 5.568337 | 5.64881 | -0.080470449 | 0.194702785 |
| Tmem220 | 4.04571 | 3.993716 | 4.0864 | -0.09272813 | -0.040733752 | 4.1209259 | 3.9814822 | 4.06198 | -0.0804962 | 0.058947517 |
| Scyl3 | 9.735258 | 9.685225 | 9.7357 | -0.050451073 | -0.000417259 | 9.800356 | 9.7117858 | 9.79228 | -0.080496692 | 0.008073478 |
| Gm13547 | 1.640945 | 1.654042 | 1.6921 | -0.038023451 | -0.05112054 | 1.6467167 | 1.6518296 | 1.73234 | -0.080509957 | -0.085622837 |
| Akap10 | 9.95481 | 10.0236 | 9.9857 | 0.037862472 | -0.030931271 | 9.9834662 | 10.015331 | 10.0958 | -0.080510092 | -0.112374892 |
| Celf5 | 1.373056 | 1.433891 | 1.3756 | 0.058284191 | -0.002550189 | 1.3096081 | 1.358414 | 1.43894 | -0.080521061 | -0.129326915 |
| Gm48418 | 4.061981 | 4.08528 | 4.0447 | 0.040538424 | 0.01723949 | 3.9189213 | 4.1068928 | 4.18742 | -0.080529078 | -0.268500551 |
| Gm7285 | 3.908866 | 3.852572 | 3.9367 | -0.08410682 | -0.027812542 | 4.0187841 | 3.8964664 | 3.97701 | -0.080541321 | 0.041776318 |
| Eef1akmt3 | 2.101731 | 2.088111 | 2.1503 | -0.062181923 | -0.048562145 | 2.1578653 | 2.0675318 | 2.14809 | -0.080554984 | 0.00977845 |
| Krcc1 | 9.970697 | 9.957201 | 9.9758 | -0.018569068 | -0.005073828 | 9.8771876 | 9.9265281 | 10.0071 | -0.080616996 | -0.129957552 |
| Esrp2 | 1.996186 | 2.061738 | 1.9744 | 0.087383465 | 0.021831053 | 1.9833323 | 2.0662418 | 2.14688 | -0.080639657 | -0.163549117 |
| Pabpc4l | 1.500546 | 1.43366 | 1.5374 | -0.103704012 | -0.036817943 | 1.3954806 | 1.3675252 | 1.44817 | -0.080648702 | -0.052693299 |
| Gm43106 | 1.333884 | 1.319769 | 1.314 | 0.005740624 | 0.019855777 | 1.3032336 | 1.2780452 | 1.35873 | -0.080684793 | -0.055496465 |
| Serpinb7 | 3.959333 | 4.022519 | 4.0482 | -0.025647401 | -0.088833129 | 4.0930236 | 3.9204831 | 4.00121 | -0.080724329 | 0.091816178 |
| Impa1 | 10.017 | 10.03579 | 9.9647 | 0.071109017 | 0.052314734 | 10.002913 | 9.9243027 | 10.005 | -0.080737832 | -0.002127538 |
| Shisa2 | 2.820089 | 2.659698 | 2.7816 | -0.121938784 | 0.038452079 | 2.6141157 | 2.6178403 | 2.6986 | -0.080761502 | -0.084486096 |
| Hmgb1-ps8 | 6.449682 | 6.41706 | 6.3313 | 0.08576122 | 0.118383747 | 6.961282 | 6.9335808 | 7.01447 | -0.08089008 | -0.053188874 |
| Btbd19 | 7.990739 | 7.864113 | 7.8573 | 0.006848644 | 0.133474926 | 8.2351783 | 7.8550301 | 7.936 | -0.080965715 | 0.299182495 |
| Dync2li1 | 4.330948 | 4.186328 | 4.3419 | -0.155528951 | -0.010909794 | 4.1036589 | 4.0943511 | 4.17544 | -0.081085465 | -0.071777691 |
| Sfswap | 10.2968 | 10.13545 | 10.269 | -0.133484501 | 0.027867711 | 10.268691 | 10.343225 | 10.4243 | -0.081091416 | -0.155624842 |
| H2afx | 8.437492 | 8.390825 | 8.5298 | -0.138925638 | -0.09225859 | 8.4213298 | 8.3225252 | 8.40362 | -0.08109266 | 0.017711892 |
| Wnt5b | 3.109598 | 3.189904 | 3.2088 | -0.018873512 | -0.099179973 | 3.0650472 | 3.0420631 | 3.1232 | -0.081135307 | -0.058151202 |
| Zan | 2.570551 | 2.446557 | 2.4383 | 0.008225393 | 0.132219252 | 2.5493625 | 2.5729615 | 2.6541 | -0.081138539 | -0.104737582 |
| Dync1li2 | 9.616943 | 9.626779 | 9.6925 | -0.065717734 | -0.07555388 | 9.6177081 | 9.6955871 | 9.77674 | -0.081153374 | -0.159032365 |
| Crip1 | 10.95035 | 10.98222 | 11.021 | -0.039195982 | -0.0710614 | 11.120453 | 10.873314 | 10.9545 | -0.081166674 | 0.165972049 |
| Fam43a | 10.17679 | 10.05903 | 10.215 | -0.156094045 | -0.038324278 | 9.6685131 | 9.4636564 | 9.54487 | -0.081215864 | 0.123640871 |
| Zfp592 | 11.45097 | 11.70437 | 11.537 | 0.16717082 | -0.086230206 | 11.454116 | 11.434171 | 11.5154 | -0.081218432 | -0.061273937 |
| Tnpo2 | 10.6554 | 10.72169 | 10.613 | 0.108327832 | 0.042036638 | 10.705792 | 10.62244 | 10.7037 | -0.081264154 | 0.002088028 |

| | | | | | | | | | | |
|---|---|---|---|---|---|---|---|---|---|---|
| Pfas | 9.403121 | 9.41778 | 9.4111 | 0.006635386 | -0.0080235 | 9.8301466 | 9.6471306 | 9.72841 | -0.081277367 | 0.101738656 |
| Fam19a2 | 2.082241 | 2.027161 | 2.0381 | -0.01093088 | 0.044149359 | 1.9136793 | 1.9355281 | 2.01695 | -0.081417923 | -0.103266732 |
| Tenm2 | 0.904377 | 0.905032 | 0.8736 | 0.031425342 | 0.03077051 | 0.9088934 | 0.8884428 | 0.96988 | -0.0814383 | -0.060987667 |
| Cenpm | 5.0417 | 4.879252 | 4.9456 | -0.066310917 | 0.096137191 | 5.352981 | 5.0326036 | 5.11406 | -0.081454725 | 0.238922687 |
| Igsf23 | 6.408177 | 6.44304 | 6.6232 | -0.18016573 | -0.21502879 | 6.2372031 | 6.3778518 | 6.45931 | -0.08145657 | -0.222105267 |
| Rpl3-ps1 | 3.409979 | 3.516537 | 3.4018 | 0.114706363 | 0.008147855 | 3.214994 | 3.3678917 | 3.44935 | -0.081462921 | -0.234360657 |
| Akr1c14 | 3.589218 | 3.460286 | 3.6553 | -0.195005643 | -0.066074401 | 3.3650655 | 3.5093986 | 3.59089 | -0.081490549 | -0.225823713 |
| Dlec1 | 2.024336 | 1.90634 | 1.9478 | -0.041438726 | 0.076556825 | 1.8923001 | 1.9288129 | 2.01035 | -0.081539846 | -0.118052712 |
| Mast1 | 3.608168 | 3.604049 | 3.6933 | -0.089236961 | -0.08511874 | 3.6492479 | 3.5451323 | 3.62669 | -0.081553734 | 0.022561868 |
| Slc25a20 | 8.537142 | 8.520388 | 8.6509 | -0.130499364 | -0.113744896 | 8.7387929 | 8.6747585 | 8.75632 | -0.081563253 | -0.017528895 |
| Ccar2 | 10.70348 | 10.63162 | 10.776 | -0.144414784 | -0.072551036 | 11.043161 | 10.788327 | 10.8699 | -0.08157189 | 0.173262344 |
| Gm10443 | 10.53327 | 10.3112 | 9.2532 | 1.05795866 | 1.280022903 | 10.331265 | 10.567043 | 10.6486 | -0.081589004 | -0.31736668 |
| Gm48691 | 0.417254 | 0.416881 | 0.4329 | -0.016003075 | -0.015630005 | 0.4336496 | 0.4279193 | 0.50953 | -0.081614744 | -0.075884473 |
| Ttc21a | 1.369875 | 1.344471 | 1.4066 | -0.062171606 | -0.036767719 | 1.3509648 | 1.4326121 | 1.51426 | -0.081643701 | -0.163291034 |
| Wapl | 11.41502 | 11.51552 | 11.438 | 0.077229484 | -0.023275176 | 11.230368 | 11.361226 | 11.4429 | -0.081646966 | -0.212505293 |
| Pcsk6 | 2.051395 | 1.874494 | 1.9989 | -0.124394623 | 0.052506406 | 1.8640803 | 1.8835331 | 1.96521 | -0.081681572 | -0.101134334 |
| Ptdss2 | 7.342782 | 7.460983 | 7.6812 | -0.220231286 | -0.338432927 | 7.4780448 | 7.5122074 | 7.59393 | -0.081727591 | -0.115890133 |
| Smarca5 | 11.66903 | 11.63817 | 11.656 | -0.017606022 | 0.013254598 | 11.509526 | 11.571166 | 11.653 | -0.08178595 | -0.143426444 |
| Mnd1 | 3.310432 | 3.18679 | 3.3039 | -0.117077283 | 0.006565045 | 3.1658072 | 3.1665309 | 3.24832 | -0.081787141 | -0.082510897 |
| Cfap300 | 3.009799 | 2.95764 | 3.0408 | -0.083126181 | -0.030966436 | 2.9708925 | 2.9529122 | 3.03472 | -0.081809271 | -0.063828966 |
| Slc26a4 | 0.83417 | 1.047497 | 0.7717 | 0.27583382 | 0.062506816 | 0.8446211 | 0.8089797 | 0.89088 | -0.081897619 | -0.046256189 |
| Vsig10l | 4.630117 | 4.453584 | 4.6431 | -0.189509118 | -0.012975658 | 4.5646087 | 4.4887582 | 4.57078 | -0.082022134 | -0.00617165 |
| Ppp1r18 | 12.12565 | 12.16896 | 12.169 | -0.000299954 | -0.043607288 | 12.227289 | 11.984135 | 12.0662 | -0.082029022 | 0.161124475 |
| Riiad1 | 2.002984 | 2.003909 | 1.986 | 0.017949772 | 0.017024741 | 1.9491122 | 2.0245428 | 2.1066 | -0.082054491 | -0.157485108 |
| Ptprt | 2.640096 | 2.412413 | 2.457 | -0.044616084 | 0.183066582 | 2.4756559 | 2.3857411 | 2.46781 | -0.082068007 | 0.007846849 |
| Hoxc8 | 2.364597 | 2.396771 | 2.3105 | 0.086222176 | 0.054048288 | 2.4027131 | 2.2379914 | 2.32009 | -0.082096351 | 0.082625282 |
| Dusp22 | 7.928179 | 7.963005 | 7.9534 | 0.009592011 | -0.025233891 | 8.2412662 | 7.9517581 | 8.0339 | -0.082140925 | 0.207367194 |
| F2 | 2.537839 | 2.369612 | 2.6355 | -0.265900775 | -0.097673879 | 2.3493572 | 2.3184442 | 2.4006 | -0.082160105 | -0.051247107 |
| Rp9 | 9.193311 | 9.253201 | 9.3884 | -0.135247391 | -0.195137824 | 9.4040075 | 9.217139 | 9.2994 | -0.082262356 | 0.10460619 |
| Plcb1 | 6.049798 | 6.063359 | 5.624 | 0.439367078 | 0.425805795 | 5.4387138 | 5.5032914 | 5.58556 | -0.082272446 | -0.146850052 |
| Hist1h4i | 5.902499 | 5.946041 | 6.1059 | -0.159827423 | -0.203369055 | 6.0753312 | 5.9810944 | 6.06339 | -0.082297458 | 0.011939324 |
| Hyal3 | 2.746237 | 2.647445 | 2.7958 | -0.148352401 | -0.049560705 | 2.7052017 | 2.735097 | 2.81742 | -0.082327729 | -0.112223024 |

| | | | | | | | | | | |
|---|---|---|---|---|---|---|---|---|---|---|
| Slc23a1 | 3.742444 | 3.778594 | 3.783 | -0.00437246 | -0.040523004 | 3.9161568 | 3.8859488 | 3.96828 | -0.082331972 | -0.052123999 |
| Smim8 | 7.456073 | 7.266016 | 7.3978 | -0.13178294 | 0.058273834 | 7.4329951 | 7.4237649 | 7.50611 | -0.082349848 | -0.073119667 |
| Trdv2-2 | 4.3157 | 4.333083 | 4.3875 | -0.054412213 | -0.0717946 | 4.11041 | 4.1580397 | 4.24041 | -0.082366159 | -0.129995795 |
| Gm6063 | 2.336199 | 2.245143 | 2.3094 | -0.064219644 | 0.026836298 | 2.2699601 | 2.2513607 | 2.33373 | -0.082374303 | -0.063774828 |
| Ighv1-61 | 5.793727 | 5.66618 | 5.6867 | -0.020545465 | 0.107001773 | 5.8827547 | 5.4879836 | 5.57038 | -0.082397504 | 0.312373543 |
| Gmeb1 | 9.262847 | 9.334153 | 9.2517 | 0.082481814 | 0.011176009 | 9.2565285 | 9.2893093 | 9.37171 | -0.082399352 | -0.115180142 |
| Txnrd1 | 10.42127 | 10.3968 | 10.196 | 0.201103936 | 0.225576054 | 10.197873 | 10.109865 | 10.1923 | -0.082410005 | 0.005597436 |
| Kin | 8.26162 | 8.2797 | 8.2651 | 0.014568639 | -0.003511411 | 8.3277686 | 8.4148506 | 8.49729 | -0.082434797 | -0.169516768 |
| Nkx1-2 | 2.077511 | 2.037396 | 1.9988 | 0.038575089 | 0.078690611 | 2.1488019 | 2.0346715 | 2.11712 | -0.082449096 | 0.031681317 |
| Zfp459 | 5.291209 | 5.301893 | 5.1402 | 0.161664708 | 0.150980154 | 5.3838274 | 5.3941799 | 5.47666 | -0.082479816 | -0.092832295 |
| Obox3 | 2.59154 | 2.603219 | 2.5674 | 0.035798345 | 0.024118995 | 2.6807118 | 2.6472428 | 2.72972 | -0.082479872 | -0.049010837 |
| Pou6f2 | 0.687933 | 0.572633 | 0.6072 | -0.034581556 | 0.080718486 | 0.5876939 | 0.5523408 | 0.63485 | -0.082504188 | -0.047151128 |
| Zzz3 | 10.30875 | 10.41738 | 10.401 | 0.015981864 | -0.092652799 | 10.256385 | 10.371265 | 10.4538 | -0.082525608 | -0.197404908 |
| Abraxas2 | 9.332842 | 9.393405 | 9.3452 | 0.048160108 | -0.012402173 | 9.1727989 | 9.2691644 | 9.35174 | -0.082573494 | -0.178939077 |
| Trdv1 | 3.393525 | 3.455674 | 3.505 | -0.049333759 | -0.111483078 | 3.4997902 | 3.4037963 | 3.48642 | -0.082627531 | 0.013366423 |
| Ddx46 | 11.54665 | 11.53547 | 11.577 | -0.041898868 | -0.030712878 | 11.678274 | 11.55555 | 11.6382 | -0.082627874 | 0.040095432 |
| Ccdc134 | 9.114236 | 9.199976 | 9.2215 | -0.021557474 | -0.107297403 | 9.5467405 | 9.2286502 | 9.31128 | -0.082633324 | 0.235456895 |
| Chek1 | 6.121556 | 6.023829 | 6.2515 | -0.227628075 | -0.129900996 | 6.1458715 | 6.3205579 | 6.40335 | -0.082793488 | -0.257479844 |
| Eci2 | 9.320139 | 9.179807 | 9.313 | -0.133218782 | 0.007113434 | 9.2689678 | 9.2220864 | 9.30489 | -0.082807106 | -0.035925719 |
| Qprt | 6.482042 | 6.514422 | 6.5306 | -0.016165747 | -0.048545461 | 6.2010553 | 6.4804703 | 6.56332 | -0.082849438 | -0.362264364 |
| Tnrc6a | 11.35705 | 11.40287 | 11.471 | -0.067902768 | -0.11372587 | 11.392402 | 11.58307 | 11.666 | -0.082967108 | -0.273635211 |
| Bcl2l13 | 9.94719 | 9.83204 | 9.8156 | 0.016488814 | 0.131639507 | 9.9505772 | 9.7305234 | 9.81351 | -0.082985035 | 0.137068775 |
| Gpx2 | 1.221577 | 1.206253 | 1.1666 | 0.039629521 | 0.054953429 | 1.2308921 | 1.1369312 | 1.21995 | -0.083016888 | 0.010943975 |
| Gm3488 | 2.882558 | 2.837187 | 2.7828 | 0.054383887 | 0.09975551 | 2.8470008 | 2.7890254 | 2.87207 | -0.083040357 | -0.025064936 |
| Vps9d1 | 8.631008 | 8.361436 | 8.5435 | -0.182096085 | 0.087475404 | 8.7539077 | 8.7719155 | 8.85498 | -0.083067646 | -0.101075466 |
| Rpap1 | 9.56723 | 9.621319 | 9.6381 | -0.016752503 | -0.070841984 | 9.7983481 | 9.692474 | 9.77556 | -0.083081404 | 0.022792611 |
| Zfp758 | 8.944259 | 8.992208 | 8.9784 | 0.013844003 | -0.034105248 | 9.0708597 | 8.9454392 | 9.02852 | -0.083083552 | 0.042337018 |
| Rab7b | 7.883091 | 7.728035 | 7.6431 | 0.08497415 | 0.240029634 | 7.3567593 | 7.1648689 | 7.24799 | -0.083118412 | 0.108772033 |
| Neto2 | 6.309446 | 6.170791 | 6.2844 | -0.113653181 | 0.025001674 | 6.0463512 | 6.1954695 | 6.27861 | -0.083136781 | -0.232255051 |
| Rhbdl2 | 2.955124 | 2.909361 | 2.9539 | -0.044530618 | 0.001232426 | 2.9465138 | 2.8569222 | 2.94011 | -0.083184846 | 0.006406686 |
| Appbp2 | 10.15589 | 10.12475 | 10.134 | -0.008829024 | 0.022310282 | 10.006866 | 10.032519 | 10.1157 | -0.083202446 | -0.108855371 |
| Gm37240 | 2.928633 | 3.248864 | 3.2307 | 0.018208965 | -0.302022756 | 3.0059436 | 3.043625 | 3.12686 | -0.083235199 | -0.1209166 |

| | | | | | | | | | | |
|---|---|---|---|---|---|---|---|---|---|---|
| Frs2 | 9.199027 | 9.257945 | 9.2645 | -0.006522475 | -0.06544039 | 9.0777993 | 9.1227921 | 9.20603 | -0.083242824 | -0.128235642 |
| Tatdn2 | 10.87918 | 10.86967 | 10.83 | 0.039743768 | 0.049251248 | 10.820916 | 10.83904 | 10.9224 | -0.08331606 | -0.101440621 |
| Nfyb | 9.389179 | 9.325048 | 9.2902 | 0.034826784 | 0.09895731 | 9.3301757 | 9.2206636 | 9.304 | -0.083334716 | 0.026177461 |
| Nkain3 | 2.047588 | 2.09514 | 2.074 | 0.021109152 | -0.026442852 | 2.032174 | 2.0792203 | 2.16261 | -0.083387049 | -0.130433373 |
| Gpr156 | 1.272436 | 1.286205 | 1.2587 | 0.027554187 | 0.013785047 | 1.2681373 | 1.206496 | 1.29 | -0.08350259 | -0.021861313 |
| Spata24 | 5.442771 | 5.230882 | 5.5237 | -0.292859368 | -0.080970296 | 5.6218939 | 5.4092157 | 5.4928 | -0.083580267 | 0.129097999 |
| Btg1 | 14.29132 | 14.31581 | 14.18 | 0.135388532 | 0.110892408 | 13.814123 | 13.870188 | 13.9538 | -0.083635302 | -0.139700995 |
| Trbv26 | 6.786903 | 6.612379 | 6.7383 | -0.125943727 | 0.048579821 | 6.284027 | 6.5243151 | 6.60797 | -0.083654722 | -0.323942869 |
| Ppp2r1b | 9.607621 | 9.656216 | 9.6622 | -0.005970157 | -0.054565443 | 9.5965228 | 9.6237495 | 9.70746 | -0.083706563 | -0.110933198 |
| 2610024D14Rik | 0.593165 | 0.582994 | 0.5911 | -0.008080019 | 0.002091125 | 0.5901751 | 0.6014785 | 0.6852 | -0.083717319 | -0.095020735 |
| Gab3 | 7.314215 | 7.369577 | 7.4866 | -0.117068374 | -0.172430031 | 7.4005133 | 7.4649653 | 7.5487 | -0.083732606 | -0.148184586 |
| Vmn2r112 | 4.172768 | 4.239784 | 4.2209 | 0.018907918 | -0.048107755 | 4.0686109 | 4.223775 | 4.30756 | -0.083789975 | -0.238954081 |
| Ddx19a | 10.07182 | 10.15014 | 10.067 | 0.083200862 | 0.004875575 | 10.26287 | 10.03702 | 10.1209 | -0.083897331 | 0.141952338 |
| Fam220a | 8.584212 | 8.576346 | 8.4615 | 0.114864851 | 0.122730576 | 8.5431027 | 8.4347382 | 8.51864 | -0.083900139 | 0.024464314 |
| Gsta3 | 3.047255 | 2.982995 | 3.0302 | -0.047239513 | 0.017020285 | 3.1138707 | 2.9721894 | 3.05624 | -0.084055387 | 0.057625943 |
| Rnf39 | 0.916542 | 0.956959 | 1.03 | -0.073066317 | -0.113483567 | 0.893305 | 0.8771646 | 0.96125 | -0.084088942 | -0.067948543 |
| Gm10136 | 1.443937 | 1.424106 | 1.4343 | -0.010166924 | 0.009663482 | 1.4889692 | 1.3892758 | 1.47337 | -0.084096638 | 0.015596712 |
| Map3k13 | 6.285272 | 6.265794 | 6.2361 | 0.029717463 | 0.049196116 | 6.294985 | 6.2119807 | 6.29613 | -0.084145265 | -0.001140997 |
| Gsdma | 0.853978 | 0.774673 | 0.7902 | -0.01553481 | 0.063770331 | 0.7645342 | 0.7941462 | 0.87832 | -0.084172932 | -0.113784966 |
| Ccdc60 | 2.309965 | 2.308523 | 2.37 | -0.061478211 | -0.060036339 | 2.3471307 | 2.3115926 | 2.39586 | -0.084271046 | -0.048732886 |
| Enthd1 | 2.08522 | 1.987927 | 2.0943 | -0.106420442 | -0.009128268 | 2.0737282 | 1.9603491 | 2.04466 | -0.08431586 | 0.029063226 |
| Rhbdl3 | 4.685864 | 4.517792 | 4.7273 | -0.209491136 | -0.04141894 | 4.5499359 | 4.3935189 | 4.47786 | -0.084345029 | 0.07207199 |
| Opalin | 2.508415 | 2.461961 | 2.5043 | -0.04235053 | 0.004103506 | 2.4681056 | 2.5450745 | 2.62947 | -0.084399227 | -0.161368208 |
| Kcnh6 | 1.216098 | 1.270666 | 1.2446 | 0.026035386 | -0.028533362 | 1.2222446 | 1.1738293 | 1.25824 | -0.084408271 | -0.035993002 |
| Ube4a | 10.53344 | 10.6339 | 10.546 | 0.087467694 | -0.012985324 | 10.548582 | 10.596512 | 10.681 | -0.084446592 | -0.132376005 |
| Nudt2 | 6.241423 | 6.102489 | 6.1096 | -0.007143802 | 0.131790349 | 6.2878803 | 6.0688494 | 6.15333 | -0.084482707 | 0.134548241 |
| Psg23 | 4.393816 | 4.206798 | 4.3089 | -0.102146943 | 0.084871127 | 4.3130842 | 4.1500009 | 4.23451 | -0.084511824 | 0.078571536 |
| Adra1a | 2.960252 | 2.868895 | 2.96 | -0.09109942 | 0.000257997 | 3.0929706 | 2.9350194 | 3.01959 | -0.08456915 | 0.073382008 |
| Tmem200b | 4.179477 | 4.118762 | 4.2538 | -0.135007648 | -0.074292584 | 4.1193196 | 4.0216597 | 4.10629 | -0.084625427 | 0.013034485 |
| Cabcoco1 | 2.817525 | 2.751028 | 2.789 | -0.037984469 | 0.028512361 | 2.7651677 | 2.7181313 | 2.80279 | -0.084654016 | -0.037617634 |
| AC154766.1 | 4.603538 | 4.483818 | 4.5673 | -0.083440867 | 0.036278312 | 4.3174053 | 4.4239418 | 4.50861 | -0.084670812 | -0.191207321 |
| Atp13a5 | 1.214405 | 1.179025 | 1.2072 | -0.02818832 | 0.007191995 | 1.4583034 | 1.1954125 | 1.28014 | -0.084731511 | 0.178159333 |

| | | | | | | | | | | |
|---|---|---|---|---|---|---|---|---|---|---|
| Gtse1 | 6.456418 | 6.481119 | 6.5743 | -0.093215636 | -0.117916385 | 6.5731029 | 6.6454917 | 6.73029 | -0.084794373 | -0.157183161 |
| Stxbp5l | 1.626781 | 1.565562 | 1.6888 | -0.123266529 | -0.062047833 | 1.6419203 | 1.6226754 | 1.70747 | -0.084794419 | -0.065549517 |
| Dcdc2b | 3.886965 | 3.921081 | 3.839 | 0.082100028 | 0.047983983 | 4.1831691 | 4.1285205 | 4.2134 | -0.084882767 | -0.030234214 |
| Col8a1 | 2.805726 | 2.833705 | 2.8506 | -0.016845931 | -0.044825254 | 2.7782737 | 2.7047664 | 2.78966 | -0.084893585 | -0.011386344 |
| Pot1b | 7.968385 | 8.048297 | 8.1136 | -0.065270255 | -0.145181801 | 8.3632073 | 8.2440086 | 8.32898 | -0.084972535 | 0.034226211 |
| AC132444.6 | 3.986432 | 4.078706 | 4.0257 | 0.053029537 | -0.039244041 | 3.9902931 | 4.2289151 | 4.31389 | -0.084972934 | -0.323594924 |
| Gpc3 | 3.064596 | 3.055933 | 3.0028 | 0.053101768 | 0.061765126 | 2.9118433 | 2.8946994 | 2.97977 | -0.085075147 | -0.067931189 |
| Ttc26 | 2.806153 | 2.628324 | 2.7171 | -0.088799926 | 0.08902915 | 2.6930495 | 2.715606 | 2.80071 | -0.085106449 | -0.107662943 |
| Ccdc120 | 2.89508 | 2.756991 | 2.8591 | -0.102122058 | 0.035967177 | 2.9183938 | 2.8813983 | 2.96653 | -0.085127085 | -0.0481315 |
| Lipo2 | 0.788003 | 0.785551 | 0.8091 | -0.023514913 | -0.021063171 | 0.7803529 | 0.7563956 | 0.84153 | -0.085133977 | -0.061176652 |
| Abcc10 | 7.941835 | 7.866426 | 7.8563 | 0.010098015 | 0.08550655 | 8.151534 | 8.048046 | 8.13321 | -0.085165625 | 0.018322406 |
| Prodh2 | 2.519764 | 2.526772 | 2.5615 | -0.034759188 | -0.041766297 | 2.495194 | 2.6528259 | 2.73801 | -0.085180851 | -0.242812812 |
| Sec14l2 | 3.93876 | 3.806869 | 4.0738 | -0.266961385 | -0.135070515 | 3.8665669 | 3.7373277 | 3.82255 | -0.085223968 | 0.04401517 |
| Trav15d-1-dv6d-1 | 3.488753 | 3.38105 | 3.5192 | -0.138172724 | -0.030470224 | 3.4948252 | 3.2776237 | 3.36291 | -0.085285556 | 0.131915956 |
| Gm37102 | 3.709463 | 3.818189 | 3.7409 | 0.077271134 | -0.031455307 | 3.7430889 | 3.8441871 | 3.92949 | -0.085300734 | -0.186398963 |
| Blk | 11.31012 | 11.29921 | 11.375 | -0.0760444 | -0.065132666 | 11.862228 | 11.446497 | 11.5318 | -0.085334316 | 0.33039594 |
| Rtn2 | 3.558184 | 3.480148 | 3.6811 | -0.200937836 | -0.12290247 | 3.6001294 | 3.6218593 | 3.70723 | -0.085372101 | -0.10710198 |
| Traj16 | 2.493773 | 2.375168 | 2.4041 | -0.028897668 | 0.089707253 | 2.2828151 | 2.3956016 | 2.481 | -0.085398646 | -0.19818522 |
| Plag1 | 4.239179 | 4.27636 | 4.469 | -0.192613435 | -0.229793734 | 4.2909356 | 4.3953565 | 4.48083 | -0.085471544 | -0.189892496 |
| Gm15186 | 1.415703 | 1.399888 | 1.4378 | -0.037946269 | -0.022131272 | 1.4245581 | 1.4178748 | 1.50339 | -0.085518003 | -0.078834646 |
| Gm48177 | 2.674847 | 2.638071 | 2.6011 | 0.037005179 | 0.073780877 | 2.6601635 | 2.6085155 | 2.69406 | -0.085547963 | -0.033899965 |
| E2f5 | 7.310148 | 7.210968 | 7.3244 | -0.113439494 | -0.014259154 | 7.1886149 | 7.2169632 | 7.30252 | -0.085556067 | -0.113904368 |
| Pibf1 | 7.976366 | 7.899627 | 8.0088 | -0.10915765 | -0.032418871 | 8.1763255 | 8.0658245 | 8.15149 | -0.085667509 | 0.024833489 |
| Kndc1 | 3.769939 | 3.784555 | 3.7983 | -0.013766004 | -0.028381495 | 3.7969641 | 3.8364311 | 3.92214 | -0.085707009 | -0.125174013 |
| Car11 | 2.97508 | 3.175315 | 3.0473 | 0.128019331 | -0.072215782 | 2.910682 | 3.015343 | 3.10108 | -0.085739114 | -0.190400094 |
| Tubb1 | 1.167653 | 1.142484 | 1.31 | -0.167533598 | -0.142364775 | 1.1235289 | 1.146542 | 1.2323 | -0.085762747 | -0.108775923 |
| Ndrg4 | 4.594805 | 4.599294 | 4.561 | 0.038334924 | 0.033846118 | 4.4246268 | 4.5111878 | 4.59699 | -0.085801516 | -0.172362469 |
| Rilpl2 | 9.213515 | 9.112262 | 9.079 | 0.033258665 | 0.134511796 | 8.9662741 | 9.0826349 | 9.16848 | -0.085846921 | -0.202207685 |
| Mpdz | 5.623699 | 5.661095 | 5.7608 | -0.099656495 | -0.137052824 | 5.1733856 | 5.2616236 | 5.34754 | -0.085921324 | -0.174159287 |
| Gm3194 | 2.441794 | 2.416749 | 2.501 | -0.084254539 | -0.059209312 | 2.4034492 | 2.4408007 | 2.52674 | -0.085935217 | -0.12328674 |
| Trav12d-2 | 4.568964 | 4.493167 | 4.5027 | -0.009510132 | 0.066286419 | 4.5770169 | 4.3833936 | 4.46934 | -0.085951114 | 0.107672214 |
| Trav12n-2 | 4.568964 | 4.493167 | 4.5027 | -0.009510132 | 0.066286419 | 4.5770169 | 4.3833936 | 4.46934 | -0.085951114 | 0.107672214 |

| | | | | | | | | | | |
|---|---|---|---|---|---|---|---|---|---|---|
| Cdc42 | 13.20924 | 13.23609 | 13.223 | 0.013070658 | -0.01377843 | 13.370406 | 13.146149 | 13.2321 | -0.085959876 | 0.13829704 |
| Slc11a2 | 9.129297 | 8.929425 | 9.1236 | -0.194130591 | 0.005740995 | 8.8252607 | 8.8527519 | 8.93877 | -0.086020357 | -0.113511605 |
| Cdc25c | 4.849202 | 4.765715 | 4.7636 | 0.002111135 | 0.085598115 | 4.9927687 | 5.0402712 | 5.12637 | -0.086093909 | -0.133596436 |
| Mmp2 | 5.118709 | 5.022992 | 5.0276 | -0.004572159 | 0.091145223 | 4.8083716 | 4.6980767 | 4.78417 | -0.086096043 | 0.02419885 |
| Cpxm2 | 2.223552 | 2.120434 | 2.2072 | -0.086759969 | 0.016358007 | 2.1393454 | 2.0264598 | 2.11257 | -0.086106839 | 0.026778781 |
| Gm45808 | 0.123273 | 0.085832 | 0.1536 | -0.06771913 | -0.030278308 | 0.1579155 | 0.0879604 | 0.17409 | -0.086130235 | -0.016175115 |
| Slamf6 | 10.84096 | 10.98733 | 11.007 | -0.019874842 | -0.16623799 | 11.154957 | 11.06732 | 11.1535 | -0.086133054 | 0.001503867 |
| Elobl | 2.656227 | 2.518367 | 2.7667 | -0.248360724 | -0.110500698 | 2.6096253 | 2.5436302 | 2.6298 | -0.086172097 | -0.020176976 |
| Ppp1r3e | 6.07824 | 6.075438 | 6.1758 | -0.100330698 | -0.097529363 | 5.9326597 | 5.8335256 | 5.9197 | -0.086174192 | 0.012959985 |
| Gm16503 | 3.007161 | 2.889423 | 2.9618 | -0.072334705 | 0.045403391 | 3.0174287 | 2.9775767 | 3.06386 | -0.086285507 | -0.046433428 |
| Gm47659 | 1.62058 | 1.589821 | 1.6552 | -0.065405475 | -0.0346461 | 1.6009207 | 1.6521053 | 1.73841 | -0.086304021 | -0.137488633 |
| Tmem255b | 2.838546 | 2.550716 | 2.8015 | -0.250825868 | 0.037004497 | 2.5698028 | 2.5931056 | 2.67947 | -0.086365338 | -0.109668115 |
| Pax7 | 0.692026 | 0.655989 | 0.6796 | -0.023605094 | 0.012432122 | 0.7682482 | 0.6396816 | 0.7261 | -0.086421511 | 0.042145048 |
| Klf6 | 12.12178 | 11.94826 | 12.111 | -0.163109584 | 0.010416286 | 11.396491 | 11.724382 | 11.8108 | -0.086429656 | -0.414320633 |
| Clpx | 8.894904 | 8.98605 | 8.9289 | 0.057190156 | -0.03395532 | 8.8568425 | 8.8880843 | 8.97457 | -0.086487509 | -0.117729218 |
| Trav3-4 | 2.428617 | 2.383875 | 2.4485 | -0.06465874 | -0.019916978 | 2.3225014 | 2.3134419 | 2.39995 | -0.086512338 | -0.077452918 |
| Neurod4 | 2.428304 | 2.516942 | 2.5032 | 0.01370637 | -0.074931212 | 2.430026 | 2.3799228 | 2.46649 | -0.086565713 | -0.036462564 |
| Tmeff2 | 2.30376 | 2.447009 | 2.3631 | 0.083951295 | -0.059297642 | 2.4192223 | 2.3820156 | 2.46858 | -0.086568832 | -0.049362097 |
| Tfpi2 | 4.072848 | 4.048166 | 3.9679 | 0.080270283 | 0.104952244 | 3.7022985 | 3.6371497 | 3.72373 | -0.086575726 | -0.021426923 |
| Cyp8b1 | 0.810639 | 0.73843 | 0.7463 | -0.007855452 | 0.06435406 | 0.7334913 | 0.6844915 | 0.77111 | -0.086623339 | -0.037623508 |
| Mgam | 2.676261 | 2.467393 | 2.3052 | 0.16214629 | 0.371014382 | 2.4746528 | 2.3941075 | 2.48074 | -0.086635553 | -0.006090268 |
| Amer1 | 7.589763 | 7.642754 | 7.6332 | 0.009509591 | -0.043481855 | 7.5611037 | 7.6325162 | 7.71922 | -0.086699891 | -0.158112454 |
| Gm6802 | 2.868351 | 2.856849 | 2.938 | -0.081172269 | -0.069670091 | 2.9660147 | 2.9440708 | 3.03079 | -0.086719068 | -0.064775129 |
| Anxa1 | 9.234112 | 8.987116 | 8.9304 | 0.056682388 | 0.303678254 | 8.9877631 | 8.6479086 | 8.73467 | -0.08676039 | 0.253094099 |
| Nudt8 | 6.758437 | 6.578908 | 6.5979 | -0.01897584 | 0.160553595 | 7.0017531 | 6.458941 | 6.54572 | -0.086779768 | 0.45603237 |
| S100pbp | 9.33217 | 9.406088 | 9.3981 | 0.007975284 | -0.065942406 | 9.3514112 | 9.4079932 | 9.49484 | -0.086843806 | -0.14342574 |
| Kat14 | 9.441896 | 9.548995 | 9.468 | 0.08098793 | -0.026110786 | 9.6172021 | 9.4291097 | 9.51601 | -0.086904824 | 0.101187551 |
| Ighv14-3 | 6.543911 | 6.624508 | 6.9128 | -0.288312434 | -0.368908604 | 7.4372342 | 6.5387319 | 6.62566 | -0.086926479 | 0.811575833 |
| 4930548H24Rik | 2.552312 | 2.598935 | 2.664 | -0.065081386 | -0.111704171 | 2.6018935 | 2.5410478 | 2.62799 | -0.086942977 | -0.026097312 |
| Lrrc39 | 1.393794 | 1.423243 | 1.3741 | 0.049191896 | 0.019742805 | 1.4389656 | 1.3324417 | 1.41939 | -0.086946458 | 0.019577432 |
| Lrrc2 | 2.428492 | 2.468911 | 2.448 | 0.02089104 | -0.019528453 | 2.5184041 | 2.3691652 | 2.45617 | -0.087004865 | 0.062234107 |
| Ighv8-11 | 1.437053 | 1.44139 | 1.4447 | -0.003317066 | -0.007653715 | 1.5140066 | 1.3917997 | 1.47884 | -0.087039616 | 0.035167222 |

| | | | | | | | | | | |
|---|---|---|---|---|---|---|---|---|---|---|
| Slc25a11 | 10.26682 | 10.23563 | 10.303 | -0.066906462 | -0.03572266 | 10.607949 | 10.238823 | 10.3259 | -0.087045632 | 0.282080679 |
| Gm16479 | 1.188979 | 1.203729 | 1.1707 | 0.03301465 | 0.018264391 | 1.2158611 | 1.1760717 | 1.26314 | -0.087071942 | -0.04728254 |
| Cts8-ps | 0.320971 | 0.283193 | 0.3364 | -0.053232492 | -0.015454585 | 0.3063718 | 0.2861993 | 0.3733 | -0.087104716 | -0.066932214 |
| Cenpx | 8.103901 | 8.123951 | 8.2553 | -0.131329813 | -0.151379819 | 8.4742143 | 8.154825 | 8.24195 | -0.087120817 | 0.232268523 |
| Blmh | 9.672187 | 9.591531 | 9.6418 | -0.050264105 | 0.030391701 | 9.6452977 | 9.6365435 | 9.7237 | -0.087156376 | -0.07840222 |
| Gm7102 | 0.937374 | 0.901159 | 0.9418 | -0.040636872 | -0.004422611 | 0.8978388 | 0.9136182 | 1.00079 | -0.087174304 | -0.102953732 |
| Zfp300 | 4.823275 | 4.796627 | 4.8017 | -0.005073732 | 0.021574668 | 4.9256911 | 4.762232 | 4.84941 | -0.087178388 | 0.0762807 |
| Yy2 | 6.865152 | 6.894628 | 6.7266 | 0.16805756 | 0.138581249 | 6.5332934 | 6.8524898 | 6.93975 | -0.08725849 | -0.406454834 |
| Supt7l | 8.235457 | 8.242086 | 8.3968 | -0.15470514 | -0.161333943 | 8.3298151 | 8.286056 | 8.37336 | -0.087300527 | -0.043541437 |
| Tmco1 | 9.302119 | 9.177189 | 9.2578 | -0.080619041 | 0.044310807 | 9.3381858 | 9.1466144 | 9.23393 | -0.08731508 | 0.104256391 |
| Rab42 | 1.501197 | 1.436925 | 1.4794 | -0.042483674 | 0.021787771 | 1.4927676 | 1.4052112 | 1.49255 | -0.087334713 | 0.000221649 |
| Lsm14b | 9.473027 | 9.584983 | 9.5497 | 0.035328496 | -0.07662678 | 9.4365289 | 9.4464136 | 9.5338 | -0.087383325 | -0.097268069 |
| Hs6st1 | 9.369328 | 9.500596 | 9.3852 | 0.115381784 | -0.01588679 | 9.2124932 | 9.1857281 | 9.27314 | -0.087409825 | -0.060644665 |
| Cdk5 | 8.320753 | 8.373028 | 8.4464 | -0.073371969 | -0.125647491 | 8.7687656 | 8.595475 | 8.68289 | -0.087413876 | 0.085876706 |
| Aqp9 | 5.700902 | 5.65837 | 5.3651 | 0.293298543 | 0.335831153 | 5.5770516 | 5.3420619 | 5.42948 | -0.08741821 | 0.147571476 |
| Trhr | 1.292197 | 1.27053 | 1.2675 | 0.003062819 | 0.024730316 | 1.241672 | 1.2659609 | 1.35347 | -0.087505169 | -0.111794068 |
| Trav15d-2-dv6d-2 | 2.83441 | 2.8037 | 2.8317 | -0.028044988 | 0.002664938 | 2.8177525 | 2.7071409 | 2.79466 | -0.08751746 | 0.023094101 |
| Trav15n-2 | 2.83441 | 2.8037 | 2.8317 | -0.028044988 | 0.002664938 | 2.8177525 | 2.7071409 | 2.79466 | -0.08751746 | 0.023094101 |
| Lilra5 | 2.145342 | 2.131134 | 2.12 | 0.01117478 | 0.025382707 | 2.2283593 | 2.0574645 | 2.14505 | -0.08758291 | 0.083311955 |
| Poli | 8.883999 | 8.796306 | 8.9153 | -0.119000706 | -0.031307862 | 9.0120355 | 8.9365035 | 9.02436 | -0.087859771 | -0.012327807 |
| Adck2 | 8.787713 | 8.91482 | 9.0019 | -0.087037385 | -0.214143911 | 9.007497 | 8.9123172 | 9.00018 | -0.087867197 | 0.007312548 |
| Olfr523 | 3.522393 | 3.511523 | 3.5601 | -0.048597438 | -0.037727545 | 3.5592322 | 3.5359155 | 3.6238 | -0.087880868 | -0.064564085 |
| Scap | 10.33737 | 10.41494 | 10.337 | 0.078191346 | 0.000623576 | 10.532796 | 10.362116 | 10.4501 | -0.087992278 | 0.082686864 |
| Traj29 | 0.906634 | 0.907947 | 0.9235 | -0.015538118 | -0.016851296 | 0.8725947 | 0.894049 | 0.98205 | -0.087998093 | -0.109452423 |
| Zdhhc3 | 10.32446 | 10.29038 | 10.297 | -0.006731802 | 0.027346386 | 10.296589 | 10.180894 | 10.2689 | -0.08800486 | 0.027690611 |
| Gm14686 | 2.308262 | 2.346638 | 2.324 | 0.022631934 | -0.015744634 | 2.1872774 | 2.3284305 | 2.41644 | -0.08800533 | -0.229158432 |
| Impa2 | 7.184309 | 7.220898 | 7.1685 | 0.052352447 | 0.015763714 | 7.239681 | 7.352935 | 7.44094 | -0.088008062 | -0.201262036 |
| Gm42878 | 4.170431 | 4.410418 | 4.1916 | 0.218783357 | -0.021203934 | 4.5088553 | 4.6381114 | 4.72613 | -0.088022457 | -0.217278617 |
| Scn3b | 3.98077 | 3.814964 | 3.9403 | -0.125340769 | 0.040465661 | 3.7050659 | 3.6970495 | 3.78513 | -0.088081929 | -0.08006554 |
| Topbp1 | 10.84839 | 10.73066 | 10.94 | -0.208883464 | -0.091151228 | 10.989193 | 11.029641 | 11.1177 | -0.088100048 | -0.128547885 |
| Gm18042 | 1.116372 | 1.106004 | 1.1625 | -0.056513574 | -0.04614515 | 1.1607474 | 1.1472208 | 1.23533 | -0.088105986 | -0.074579381 |
| Stk24 | 11.44927 | 11.45964 | 11.464 | -0.0045386 | -0.01490129 | 11.198522 | 11.198752 | 11.2869 | -0.088127314 | -0.08835734 |

| | | | | | | | | | | |
|---|---|---|---|---|---|---|---|---|---|---|
| Ginm1 | 7.704743 | 7.574548 | 7.6173 | -0.042792066 | 0.087402366 | 7.5759859 | 7.485761 | 7.57391 | -0.088152319 | 0.002072613 |
| Med22 | 8.725248 | 8.701088 | 8.7269 | -0.025787694 | -0.001627553 | 8.8266251 | 8.5905589 | 8.67875 | -0.088195352 | 0.147870905 |
| Gm10120 | 0.940461 | 0.863346 | 1.0102 | -0.146880492 | -0.069765269 | 0.827125 | 0.8277012 | 0.91591 | -0.088205465 | -0.08878164 |
| Psg25 | 1.627341 | 1.546659 | 1.576 | -0.029373122 | 0.051309092 | 1.5233575 | 1.5403515 | 1.6286 | -0.088244996 | -0.105238972 |
| Hist1h2bl | 1.959758 | 2.057104 | 2.0444 | 0.012749486 | -0.084596083 | 2.0270547 | 1.9851131 | 2.07345 | -0.088341874 | -0.046400278 |
| Sel1l | 10.54875 | 10.53567 | 10.583 | -0.04761504 | -0.034540178 | 10.513612 | 10.523866 | 10.6122 | -0.088349708 | -0.098603612 |
| Ldah | 9.598237 | 9.683768 | 9.5183 | 0.165442342 | 0.079911304 | 9.7717879 | 9.6141946 | 9.7026 | -0.088404684 | 0.069188533 |
| Gm10040 | 4.046785 | 3.937457 | 4.1675 | -0.230030522 | -0.120702958 | 4.122217 | 4.0769663 | 4.16538 | -0.088412966 | -0.043162203 |
| Samd13 | 1.857969 | 1.85174 | 1.7794 | 0.072313283 | 0.078541956 | 1.766723 | 1.7822542 | 1.87069 | -0.08844025 | -0.103971422 |
| Sntg1 | 1.4646 | 1.560093 | 1.5189 | 0.041145769 | -0.054347411 | 1.5418215 | 1.4468769 | 1.53532 | -0.088441775 | 0.006502822 |
| 6430550D23Rik | 3.577824 | 3.508659 | 3.6237 | -0.115052589 | -0.045887093 | 3.7565805 | 3.5635317 | 3.65197 | -0.088443164 | 0.104605664 |
| Gm45208 | 2.624298 | 2.721822 | 2.6813 | 0.040571421 | -0.05695257 | 2.7477474 | 2.6768207 | 2.76531 | -0.088489525 | -0.017562793 |
| Tmem268 | 9.132252 | 8.982663 | 8.9496 | 0.033022043 | 0.182610914 | 9.239047 | 8.9406927 | 9.0292 | -0.088504538 | 0.209849731 |
| Pbk | 5.53035 | 5.289821 | 5.5894 | -0.299589395 | -0.059060437 | 5.4672785 | 5.6638616 | 5.75241 | -0.088549282 | -0.285132368 |
| CT030661.1 | 8.792547 | 8.687073 | 8.8018 | -0.114710211 | -0.009236728 | 8.7020427 | 8.7526238 | 8.84118 | -0.088552051 | -0.139133214 |
| Pcsk5 | 3.890485 | 3.68277 | 3.8113 | -0.128514742 | 0.079200033 | 3.7062543 | 3.699661 | 3.78822 | -0.088562414 | -0.081969128 |
| Fam135b | 2.890979 | 2.829265 | 2.7511 | 0.07815335 | 0.139868057 | 2.8695464 | 2.8576721 | 2.94628 | -0.088603467 | -0.076729202 |
| C8a | 2.192859 | 2.201722 | 2.2354 | -0.033706429 | -0.042569535 | 2.1863207 | 2.1557559 | 2.24442 | -0.088661174 | -0.058096388 |
| F2r | 9.478363 | 9.211227 | 9.4061 | -0.194824098 | 0.072311318 | 8.8841358 | 9.0484953 | 9.13717 | -0.088672411 | -0.253031921 |
| Emp2 | 6.562093 | 6.256387 | 6.5013 | -0.244946854 | 0.060759503 | 6.2152672 | 5.975752 | 6.06446 | -0.088706172 | 0.150809025 |
| Nudt15 | 7.815963 | 7.974711 | 7.9748 | -0.000120671 | -0.158868384 | 7.9838884 | 7.8176124 | 7.90635 | -0.088741001 | 0.077535074 |
| Slc7a1 | 10.62444 | 10.53777 | 10.507 | 0.030875416 | 0.117546258 | 10.75921 | 10.687604 | 10.7764 | -0.08877988 | -0.017173593 |
| Maats1 | 3.407953 | 3.365632 | 3.3889 | -0.023269562 | 0.019051097 | 3.425562 | 3.3488279 | 3.43764 | -0.088815951 | -0.012081857 |
| Zfp111 | 5.256038 | 5.304766 | 5.4291 | -0.124366107 | -0.173093751 | 5.18077 | 5.201437 | 5.29026 | -0.088827147 | -0.1094941 |
| Dctn6 | 9.156432 | 9.018375 | 9.1404 | -0.121987621 | 0.016069235 | 9.0758263 | 9.0361257 | 9.12496 | -0.088831429 | -0.049130755 |
| Arl6 | 5.364179 | 5.157179 | 5.2901 | -0.132898895 | 0.074100952 | 5.160502 | 5.2012046 | 5.29007 | -0.08886689 | -0.129569418 |
| Cdip1 | 9.411346 | 9.377052 | 9.4898 | -0.112715838 | -0.078421718 | 9.5747993 | 9.344231 | 9.4332 | -0.088966337 | 0.14160202 |
| Flywch2 | 1.339087 | 1.320589 | 1.3321 | -0.011499131 | 0.0069993 | 1.2676624 | 1.2374374 | 1.32647 | -0.089034515 | -0.058809503 |
| Cdc123 | 9.48476 | 9.574452 | 9.5668 | 0.007695914 | -0.081996489 | 9.6204442 | 9.6300532 | 9.71913 | -0.089075406 | -0.098684441 |
| 2210418O10Rik | 6.193266 | 6.079826 | 6.0593 | 0.020476834 | 0.133916924 | 6.066616 | 6.0246808 | 6.11377 | -0.089091574 | -0.047156309 |
| Gm47726 | 1.041368 | 1.039762 | 0.992 | 0.047739144 | 0.049344507 | 1.082419 | 1.0235589 | 1.11265 | -0.089092423 | -0.030232265 |
| Ddx59 | 7.378663 | 7.187501 | 7.4328 | -0.245325905 | -0.054164175 | 7.4087578 | 7.2357337 | 7.32484 | -0.08910285 | 0.083921301 |

| | | | | | | | | | | |
|---|---|---|---|---|---|---|---|---|---|---|
| Vmn2r94 | 2.019801 | 2.017538 | 2.0218 | -0.004266879 | -0.002004158 | 2.153698 | 2.057641 | 2.14681 | -0.089167704 | 0.006889291 |
| Snrnp200 | 12.46002 | 12.44463 | 12.494 | -0.049481566 | -0.034096778 | 12.514869 | 12.473065 | 12.5622 | -0.08917594 | -0.04737138 |
| Zfp968 | 4.791395 | 4.891253 | 4.8425 | 0.048784517 | -0.051073066 | 4.855466 | 4.7703689 | 4.85956 | -0.089186992 | -0.004089814 |
| Cmtm1 | 1.735391 | 1.74408 | 1.6564 | 0.087660691 | 0.078971531 | 1.6900963 | 1.6054011 | 1.69459 | -0.089189527 | -0.004494276 |
| Traf3ip2 | 9.421572 | 9.40094 | 9.5863 | -0.185390007 | -0.164757886 | 9.7675956 | 9.612072 | 9.70129 | -0.089213936 | 0.066309661 |
| Saxo2 | 1.67707 | 1.607932 | 1.6416 | -0.033656546 | 0.035482153 | 1.6610386 | 1.5825882 | 1.67185 | -0.089256967 | -0.010806588 |
| Dctn2 | 10.5022 | 10.4177 | 10.543 | -0.125041322 | -0.040545776 | 10.69768 | 10.485761 | 10.575 | -0.089263194 | 0.12265611 |
| 4930469K13Rik | 1.820993 | 1.712471 | 1.7968 | -0.084342307 | 0.024179602 | 1.8139904 | 1.7405085 | 1.82978 | -0.089267298 | -0.015785458 |
| Csk | 12.64679 | 12.73619 | 12.71 | 0.026229706 | -0.063168616 | 12.830384 | 12.605957 | 12.6952 | -0.089286034 | 0.135141 |
| Fbxl13 | 1.156961 | 1.147079 | 1.2133 | -0.066266369 | -0.056384575 | 1.1277385 | 1.1093115 | 1.19861 | -0.089294697 | -0.070867645 |
| Bpifc | 2.770619 | 2.751996 | 2.7242 | 0.027839658 | 0.046463172 | 2.7415142 | 2.7220225 | 2.81134 | -0.089315416 | -0.069823676 |
| Nr1d2 | 9.782082 | 9.607464 | 9.8921 | -0.284636958 | -0.110019237 | 9.5692382 | 9.7900166 | 9.87934 | -0.089322073 | -0.310100496 |
| Gm4824 | 3.151853 | 3.207599 | 3.1713 | 0.036308875 | -0.019437784 | 3.2913881 | 3.0763394 | 3.16568 | -0.089344222 | 0.125704435 |
| Mark1 | 3.653575 | 3.653871 | 3.4997 | 0.154144082 | 0.15384836 | 3.4040297 | 3.275936 | 3.36532 | -0.089381716 | 0.038711922 |
| Hyal1 | 5.941426 | 5.832704 | 5.9862 | -0.153453546 | -0.04473169 | 6.0306788 | 5.927316 | 6.01671 | -0.089390446 | 0.0139723 |
| 1700056E22Rik | 4.19484 | 4.030989 | 4.2119 | -0.180933706 | -0.017082547 | 4.1355524 | 3.9911032 | 4.08053 | -0.089423258 | 0.055026015 |
| Wdr92 | 8.137338 | 8.12006 | 8.0978 | 0.022214362 | 0.039492122 | 8.1585016 | 7.958697 | 8.04812 | -0.089424143 | 0.110380444 |
| Rln3 | 3.751879 | 3.62499 | 3.6333 | -0.008302742 | 0.118586122 | 3.6323179 | 3.4985871 | 3.58802 | -0.089437519 | 0.044293326 |
| Krtcap3 | 5.950313 | 6.092933 | 6.1349 | -0.041974066 | -0.18459419 | 6.2739226 | 6.2525758 | 6.34201 | -0.089437879 | -0.068091101 |
| Mfap1b | 10.78185 | 10.78582 | 10.796 | -0.010527354 | -0.01449538 | 10.754746 | 10.692941 | 10.7824 | -0.089447494 | -0.027642328 |
| Tank | 10.37534 | 10.47303 | 10.424 | 0.048921738 | -0.048774792 | 10.253982 | 10.355051 | 10.4445 | -0.089449492 | -0.190518427 |
| Ighv15-2 | 4.829166 | 4.723626 | 4.6318 | 0.091835913 | 0.19737561 | 5.0573093 | 4.7521444 | 4.84164 | -0.089495226 | 0.215669677 |
| Ripply3 | 4.707845 | 4.452688 | 4.734 | -0.2813368 | -0.02617984 | 4.4138266 | 4.4883444 | 4.57786 | -0.089519889 | -0.164037714 |
| Flot2 | 9.363634 | 9.174591 | 9.2619 | -0.087343983 | 0.101699049 | 9.2987721 | 9.0799575 | 9.16956 | -0.089599791 | 0.129214743 |
| Fam171a2 | 3.065743 | 2.949855 | 2.8365 | 0.113380318 | 0.229268751 | 2.7194801 | 2.6545572 | 2.74416 | -0.08960368 | -0.024680707 |
| Acap1 | 12.32809 | 12.37227 | 12.434 | -0.061325892 | -0.10551028 | 12.912881 | 12.653468 | 12.7431 | -0.089633656 | 0.169778768 |
| Srgap1 | 4.494011 | 4.170387 | 4.3192 | -0.148819761 | 0.174803701 | 4.2058633 | 4.0998319 | 4.1895 | -0.089670812 | 0.016360589 |
| Tm9sf2 | 10.92203 | 10.86552 | 10.945 | -0.079004796 | -0.022493776 | 10.894025 | 10.814384 | 10.9041 | -0.089754184 | -0.010113259 |
| Tubgcp4 | 9.555995 | 9.645495 | 9.6398 | 0.005707009 | -0.083793245 | 9.8590882 | 9.9099341 | 9.99969 | -0.089757836 | -0.140603741 |
| Ell | 8.894623 | 8.829593 | 8.8277 | 0.001887213 | 0.066917217 | 8.8906315 | 8.7210631 | 8.81082 | -0.089761875 | 0.07980654 |
| Agpat4 | 7.867898 | 7.639676 | 7.6986 | -0.058922823 | 0.169298305 | 7.7275127 | 7.6795024 | 7.76933 | -0.089827888 | -0.0418176 |
| Trim41 | 8.910232 | 8.999928 | 8.9981 | 0.001837203 | -0.087858859 | 9.0901987 | 9.0302806 | 9.12012 | -0.089842035 | -0.029923956 |

| | | | | | | | | | | |
|---|---|---|---|---|---|---|---|---|---|---|
| Cyp4f41-ps | 3.222854 | 3.20356 | 3.2428 | -0.039199041 | -0.019905079 | 3.4402431 | 3.1918446 | 3.28169 | -0.089845353 | 0.158553125 |
| Sult4a1 | 1.752786 | 1.713806 | 1.629 | 0.084810388 | 0.123790683 | 1.6111713 | 1.6214618 | 1.71138 | -0.089915485 | -0.100205975 |
| Mtmr6 | 10.36989 | 10.38224 | 10.369 | 0.01305489 | 0.000707946 | 10.243274 | 10.267692 | 10.3576 | -0.08992717 | -0.114344854 |
| Dmrt2 | 1.513417 | 1.473421 | 1.6446 | -0.17115276 | -0.131155908 | 1.4603881 | 1.4285727 | 1.51854 | -0.089971034 | -0.058155613 |
| Nucks1 | 11.0483 | 10.96986 | 11.02 | -0.050458198 | 0.02797926 | 11.097063 | 11.018959 | 11.1089 | -0.089978454 | -0.011874435 |
| Rph3al | 6.728715 | 6.824038 | 6.7864 | 0.037634773 | -0.057687865 | 6.8621725 | 6.6312289 | 6.72122 | -0.089994309 | 0.140949321 |
| Kbtbd6 | 3.36277 | 3.223186 | 3.3561 | -0.132867229 | 0.006717025 | 3.2652954 | 3.3358435 | 3.42586 | -0.090016486 | -0.160564581 |
| D630045J12Rik | 3.72869 | 3.796533 | 3.8286 | -0.032036601 | -0.09988032 | 3.8645796 | 3.758096 | 3.84814 | -0.090045993 | 0.016437607 |
| Tspan13 | 10.75581 | 10.64263 | 10.692 | -0.049158684 | 0.064024406 | 10.757679 | 10.488943 | 10.579 | -0.09009057 | 0.178645276 |
| Kcnip4 | 1.181345 | 1.197981 | 1.1517 | 0.046259006 | 0.029623304 | 1.1696457 | 1.1846678 | 1.27482 | -0.090147921 | -0.105170008 |
| Rbm4 | 8.85222 | 8.9008 | 8.7598 | 0.14095143 | 0.092371706 | 9.0885642 | 8.9082058 | 8.99835 | -0.090148049 | 0.090210385 |
| Otos | 1.882323 | 1.809491 | 1.8924 | -0.082916132 | -0.010084371 | 1.9380745 | 1.877104 | 1.96727 | -0.09016584 | -0.029195387 |
| AC122117.1 | 1.659643 | 1.632525 | 1.7156 | -0.083038868 | -0.055921119 | 1.6739611 | 1.7104612 | 1.80067 | -0.090208829 | -0.126708915 |
| Eif2d | 9.206648 | 9.226694 | 9.2726 | -0.045884522 | -0.065930811 | 9.5140238 | 9.2637872 | 9.35406 | -0.09027233 | 0.159964263 |
| Lonrf2 | 1.449127 | 1.449958 | 1.4143 | 0.035706696 | 0.034875819 | 1.3916153 | 1.3983292 | 1.48866 | -0.090331314 | -0.097045287 |
| Slc10a2 | 2.96222 | 2.894823 | 2.9206 | -0.025737469 | 0.041660179 | 2.9857228 | 2.9161592 | 3.0065 | -0.090344544 | -0.020780982 |
| Ap3m2 | 9.490094 | 9.627908 | 9.4979 | 0.130000657 | -0.007813064 | 9.6246339 | 9.5854356 | 9.67579 | -0.090351847 | -0.051153477 |
| Vim | 13.14139 | 12.96935 | 13.139 | -0.169626062 | 0.00241775 | 12.986171 | 13.007181 | 13.0976 | -0.090427308 | -0.111437289 |
| Mfn2 | 10.08296 | 9.953825 | 10.058 | -0.104135705 | 0.025003619 | 10.085836 | 9.9050697 | 9.99566 | -0.090588743 | 0.090177514 |
| Gm17841 | 0.37107 | 0.349445 | 0.4043 | -0.054883114 | -0.033258241 | 0.356429 | 0.3726839 | 0.46331 | -0.090621857 | -0.106876764 |
| Ammecr1l | 9.366877 | 9.42724 | 9.38 | 0.047215481 | -0.01314778 | 9.1979311 | 9.2362631 | 9.32694 | -0.090676769 | -0.129008718 |
| 2510002D24Rik | 7.815484 | 7.765082 | 7.8875 | -0.122449415 | -0.072048069 | 8.0887833 | 7.9418541 | 8.03261 | -0.090752334 | 0.056176858 |
| Eva1a | 3.352235 | 3.156601 | 3.1109 | 0.045668438 | 0.241302226 | 3.1093795 | 2.9903534 | 3.08113 | -0.090774356 | 0.028251675 |
| Btbd10 | 8.306456 | 8.354047 | 8.2851 | 0.06892874 | 0.021338504 | 8.1219225 | 8.0930695 | 8.18385 | -0.090776431 | -0.061923463 |
| Paip2b | 9.680364 | 9.689164 | 9.7897 | -0.100564784 | -0.109364475 | 9.7969954 | 9.6393455 | 9.73014 | -0.090790308 | 0.066859672 |
| 4933427I04Rik | 1.73126 | 1.761731 | 1.8009 | -0.039156863 | -0.069628391 | 1.8183504 | 1.7856083 | 1.8764 | -0.090793512 | -0.058051486 |
| Rai2 | 2.776692 | 2.670298 | 2.8055 | -0.135241184 | -0.02884665 | 2.6031354 | 2.5807341 | 2.67154 | -0.090804789 | -0.068403545 |
| Mapk1 | 11.80884 | 11.83228 | 11.791 | 0.041249118 | 0.017805634 | 11.825967 | 11.73084 | 11.8219 | -0.091013394 | 0.004113144 |
| Mreg | 8.992817 | 9.174085 | 9.1097 | 0.064418674 | -0.116849428 | 8.7690758 | 9.1634595 | 9.2546 | -0.091139014 | -0.485522674 |
| Klhl22 | 7.940784 | 7.869153 | 8.1021 | -0.232925098 | -0.16129402 | 7.9501262 | 7.8507947 | 7.942 | -0.091208503 | 0.008122945 |
| Kpna6 | 10.04332 | 10.06656 | 9.9979 | 0.068657936 | 0.045414293 | 9.9967971 | 10.00918 | 10.1005 | -0.091284308 | -0.103667023 |
| Onecut3 | 1.271667 | 1.221826 | 1.2115 | 0.010299115 | 0.060140601 | 1.2872432 | 1.2047079 | 1.296 | -0.091295403 | -0.008760087 |

| | | | | | | | | | | |
|---|---|---|---|---|---|---|---|---|---|---|
| Kirrel3 | 2.947027 | 2.797071 | 3.0187 | -0.221618406 | -0.07166278 | 2.777461 | 2.6093237 | 2.70078 | -0.091457155 | 0.076680055 |
| Epb41l4b | 4.274461 | 3.892013 | 4.1711 | -0.279099087 | 0.103349602 | 3.9332162 | 4.0521938 | 4.14367 | -0.091478831 | -0.210456434 |
| Rgs8 | 1.872645 | 1.750024 | 1.6305 | 0.119518079 | 0.242139136 | 1.6544428 | 1.6221029 | 1.71362 | -0.091519981 | -0.059180081 |
| C87414 | 3.398396 | 3.449267 | 3.4382 | 0.011031481 | -0.039839554 | 3.378214 | 3.4572128 | 3.54873 | -0.09152036 | -0.170519216 |
| Nabp2 | 9.128161 | 9.095963 | 9.2432 | -0.147244822 | -0.115046977 | 9.3772751 | 9.0825456 | 9.17408 | -0.091529958 | 0.203199596 |
| Smim19 | 7.722781 | 7.660905 | 7.6951 | -0.034167527 | 0.02770923 | 7.6830236 | 7.5776077 | 7.66919 | -0.091584193 | 0.013831742 |
| Trav7n-4 | 2.437401 | 2.41083 | 2.3826 | 0.028244075 | 0.054815061 | 2.4561678 | 2.359957 | 2.4516 | -0.091647531 | 0.004563227 |
| Hapln3 | 1.47307 | 1.287993 | 1.4106 | -0.122579954 | 0.062497107 | 1.4104064 | 1.2898182 | 1.38148 | -0.091659144 | 0.028929012 |
| Wdr78 | 5.759304 | 5.778256 | 5.753 | 0.025246625 | 0.006294922 | 5.8978292 | 5.9107587 | 6.00245 | -0.091689823 | -0.104619351 |
| Fsip1 | 3.409734 | 3.399515 | 3.454 | -0.054447651 | -0.044228724 | 3.6389207 | 3.4364681 | 3.52816 | -0.0916931 | 0.110759501 |
| Olfr2 | 3.544276 | 3.668335 | 3.6275 | 0.040789501 | -0.0832695 | 3.7966909 | 3.8158593 | 3.90757 | -0.091713002 | -0.110881408 |
| Trav21-dv12 | 4.110856 | 4.09911 | 4.1327 | -0.0335636 | -0.021817144 | 3.9416097 | 3.9744224 | 4.06615 | -0.091726398 | -0.124539125 |
| Samd5 | 2.952169 | 3.149337 | 3.0584 | 0.090982138 | -0.106186097 | 2.9893878 | 2.9812851 | 3.07305 | -0.091760901 | -0.083658195 |
| Fpgs | 8.178795 | 8.178268 | 8.1053 | 0.072991872 | 0.073519649 | 8.4192197 | 8.1224437 | 8.21423 | -0.091784196 | 0.204991863 |
| Hcar1 | 1.657177 | 1.595631 | 1.6516 | -0.055961048 | 0.005584558 | 1.5913157 | 1.5169381 | 1.60873 | -0.091792043 | -0.017414503 |
| Camk2n2 | 2.04359 | 2.054812 | 1.9943 | 0.060549232 | 0.049327048 | 1.929601 | 1.9386099 | 2.03042 | -0.091805221 | -0.100814117 |
| Znrf1 | 10.42167 | 10.42291 | 10.532 | -0.109307477 | -0.110538431 | 10.239867 | 10.337348 | 10.4292 | -0.091816856 | -0.189297845 |
| Zfp654 | 9.605805 | 9.618174 | 9.7096 | -0.091450092 | -0.103818572 | 9.3916092 | 9.5102672 | 9.60215 | -0.091877955 | -0.210535944 |
| Tmem41b | 8.583138 | 8.415809 | 8.456 | -0.040185168 | 0.127143663 | 8.3746972 | 8.4407142 | 8.5326 | -0.09188627 | -0.157903344 |
| Gm8983 | 1.90113 | 1.790217 | 1.8689 | -0.078683219 | 0.032230283 | 1.8277718 | 1.7673586 | 1.85926 | -0.091898396 | -0.031485126 |
| Tmem92 | 1.249942 | 1.242153 | 1.2322 | 0.009986309 | 0.017775171 | 1.2307269 | 1.1912249 | 1.28315 | -0.091921683 | -0.052419714 |
| Auh | 8.495563 | 8.441041 | 8.5518 | -0.110742566 | -0.056220495 | 8.6736638 | 8.5841541 | 8.67623 | -0.092079196 | -0.002569521 |
| Twist1 | 1.119715 | 1.12339 | 1.1262 | -0.00284081 | -0.006515274 | 1.0994304 | 1.0598582 | 1.15204 | -0.092177635 | -0.052605504 |
| Pan3 | 10.86728 | 11.05228 | 11.02 | 0.032561462 | -0.152440486 | 10.863915 | 11.093302 | 11.1855 | -0.092217094 | -0.321604738 |
| Mkx | 0.708597 | 0.73521 | 0.6802 | 0.055033433 | 0.02842047 | 0.6955251 | 0.6902937 | 0.78259 | -0.092293667 | -0.087062252 |
| Gas1 | 2.581104 | 2.640191 | 2.7062 | -0.065978746 | -0.12506558 | 2.692641 | 2.4856398 | 2.57794 | -0.092303909 | 0.114697363 |
| Pafah1b1 | 11.89543 | 11.84125 | 11.759 | 0.082225202 | 0.136411002 | 11.599201 | 11.707055 | 11.7994 | -0.092306494 | -0.200160562 |
| Myo15b | 1.270665 | 1.241798 | 1.2248 | 0.016957337 | 0.045824191 | 1.2144733 | 1.216032 | 1.30835 | -0.092313828 | -0.093872525 |
| Gm12960 | 2.040557 | 1.963344 | 1.9417 | 0.021645014 | 0.09885847 | 1.9756284 | 1.911068 | 2.0034 | -0.092330303 | -0.027769816 |
| Rrm1 | 10.16872 | 10.09568 | 10.268 | -0.172049223 | -0.099012954 | 10.237301 | 10.353975 | 10.4463 | -0.09236756 | -0.209042033 |
| 4430402I18Rik | 1.843642 | 1.606825 | 1.8435 | -0.23665351 | 0.000162828 | 1.6513407 | 1.6427133 | 1.73515 | -0.092434682 | -0.083807238 |
| Fopnl | 9.031124 | 8.86988 | 9.003 | -0.133164466 | 0.028079507 | 9.0778339 | 8.8349287 | 8.92741 | -0.092478748 | 0.150426462 |

| | | | | | | | | | | |
|---|---|---|---|---|---|---|---|---|---|---|
| Tnfrsf8 | 5.982566 | 6.049689 | 6.1556 | -0.105871936 | -0.172995258 | 6.2804503 | 6.426159 | 6.51868 | -0.092522547 | -0.238231309 |
| Rad51b | 4.387621 | 4.400164 | 4.4837 | -0.083569362 | -0.096112042 | 4.4568017 | 4.5762105 | 4.66875 | -0.092541602 | -0.211950416 |
| Trav3d-3 | 4.948434 | 4.90637 | 4.9874 | -0.08099437 | -0.038931003 | 4.8369385 | 4.8498976 | 4.94249 | -0.092589312 | -0.105548394 |
| Trav3n-3 | 4.948434 | 4.90637 | 4.9874 | -0.08099437 | -0.038931003 | 4.8369385 | 4.8498976 | 4.94249 | -0.092589312 | -0.105548394 |
| Cdkn1c | 3.565598 | 3.614492 | 3.6328 | -0.018328707 | -0.067222277 | 3.546151 | 3.5627061 | 3.65532 | -0.092616663 | -0.109171786 |
| Stard6 | 4.141837 | 3.992571 | 3.8956 | 0.096992218 | 0.246258697 | 3.9575641 | 3.922657 | 4.01528 | -0.092623492 | -0.057716376 |
| Zbtb22 | 9.06998 | 9.148472 | 9.1232 | 0.025284212 | -0.053208005 | 9.1915106 | 8.8242721 | 8.91692 | -0.092651234 | 0.274587313 |
| Epdr1 | 3.186042 | 3.147644 | 3.2085 | -0.060852066 | -0.022453764 | 3.2871551 | 3.0773183 | 3.16997 | -0.092653228 | 0.117183523 |
| B4galnt1 | 12.62932 | 12.67583 | 12.731 | -0.05498733 | -0.101499414 | 12.863645 | 12.582305 | 12.675 | -0.092671996 | 0.188667745 |
| Kdm5d | 5.55262 | 5.697497 | 5.5239 | 0.173567314 | 0.028689808 | 5.6608115 | 5.6422052 | 5.73489 | -0.092680823 | -0.074074514 |
| Rmdn2 | 4.777273 | 4.854233 | 4.6211 | 0.23316604 | 0.156206509 | 4.5954079 | 4.6483616 | 4.74108 | -0.092716563 | -0.145670244 |
| Dbf4 | 8.509138 | 8.521567 | 8.6188 | -0.097188752 | -0.109616984 | 8.3055393 | 8.5339549 | 8.62669 | -0.092737018 | -0.32115269 |
| Snx9 | 10.31191 | 10.33064 | 10.297 | 0.03342754 | 0.014698242 | 10.495865 | 10.185167 | 10.2779 | -0.092756572 | 0.217941275 |
| Tab1 | 8.62456 | 8.307783 | 8.7141 | -0.406295957 | -0.089518795 | 8.7585317 | 8.5062394 | 8.59901 | -0.092768194 | 0.159524084 |
| Ubr5 | 11.76048 | 11.7345 | 11.854 | -0.119373656 | -0.093393348 | 11.762248 | 11.909211 | 12.002 | -0.09277167 | -0.239733927 |
| Tcp11 | 3.03654 | 3.0639 | 3.0217 | 0.042198328 | 0.014838148 | 2.9378442 | 3.0287417 | 3.12157 | -0.092824819 | -0.183722273 |
| Rpl15-ps2 | 1.183002 | 1.13849 | 1.1049 | 0.033566282 | 0.078078057 | 1.1845842 | 1.1098195 | 1.20267 | -0.092852827 | -0.018088123 |
| Gm14849 | 2.455812 | 2.558361 | 2.5118 | 0.046570198 | -0.055978443 | 2.5121264 | 2.5225787 | 2.61545 | -0.092874713 | -0.103327011 |
| Wdr20 | 8.572393 | 8.684749 | 8.5355 | 0.149277963 | 0.036921551 | 8.5793174 | 8.6288698 | 8.72176 | -0.092894917 | -0.142447326 |
| Ighv10-1 | 6.919446 | 7.344853 | 6.9256 | 0.419286251 | -0.006120918 | 7.1686129 | 6.7340411 | 6.82702 | -0.092977856 | 0.341593976 |
| Gm49450 | 5.41303 | 5.095828 | 5.4229 | -0.327026596 | -0.009824831 | 5.7519454 | 5.3372658 | 5.43024 | -0.09297836 | 0.321701243 |
| Ift52 | 9.093652 | 9.044966 | 9.034 | 0.010926117 | 0.059612329 | 9.191901 | 9.0699048 | 9.16292 | -0.093012013 | 0.028984221 |
| Tspan17 | 5.664374 | 5.581561 | 5.6199 | -0.038312333 | 0.0445001 | 5.8477759 | 5.646338 | 5.73938 | -0.093042391 | 0.108395436 |
| Olfr536 | 2.537626 | 2.498656 | 2.5407 | -0.04203877 | -0.003068429 | 2.5937638 | 2.5869 | 2.67998 | -0.093081915 | -0.08621809 |
| Kat7 | 10.7866 | 10.70912 | 10.777 | -0.068117006 | 0.009359514 | 10.588437 | 10.615487 | 10.7086 | -0.0930952 | -0.120145702 |
| Pigw | 7.1582 | 7.191737 | 7.1724 | 0.019296377 | -0.014240724 | 7.4665074 | 7.1973014 | 7.29049 | -0.09319057 | 0.176015396 |
| Btbd7 | 10.11234 | 10.18637 | 10.108 | 0.078507932 | 0.004479924 | 9.8366897 | 10.01687 | 10.1101 | -0.093230067 | -0.273410551 |
| 4930590J08Rik | 2.69962 | 2.609958 | 2.7786 | -0.168644873 | -0.078983369 | 2.6638044 | 2.6400418 | 2.73328 | -0.093237349 | -0.069474727 |
| Chrna9 | 1.299956 | 1.273437 | 1.3285 | -0.055102527 | -0.028583518 | 1.314911 | 1.2800478 | 1.3733 | -0.093254361 | -0.05839119 |
| Ighv1-59 | 5.939654 | 5.934278 | 5.8236 | 0.110647308 | 0.116023328 | 6.2576734 | 5.8097583 | 5.90311 | -0.093348422 | 0.354566627 |
| Gm14586 | 1.052768 | 0.997122 | 1.0726 | -0.075470012 | -0.019823349 | 1.067525 | 1.0531884 | 1.14655 | -0.093362087 | -0.079025574 |
| Klhl2 | 8.745028 | 8.611376 | 8.635 | -0.023598212 | 0.110053507 | 8.4959843 | 8.403074 | 8.49644 | -0.093366275 | -0.000455992 |

| | | | | | | | | | | |
|---|---|---|---|---|---|---|---|---|---|---|
| Gm3883 | 1.460743 | 1.513725 | 1.5012 | 0.012547161 | -0.040435121 | 1.3807419 | 1.3396986 | 1.43319 | -0.093487525 | -0.052444164 |
| Hist1h2ae | 2.650264 | 2.553699 | 2.647 | -0.093264371 | 0.003300972 | 2.6557561 | 2.5489738 | 2.64248 | -0.093505483 | 0.01327676 |
| Azin1 | 10.98169 | 10.99173 | 10.901 | 0.090468282 | 0.080430278 | 10.728862 | 10.887623 | 10.9812 | -0.093546848 | -0.252307742 |
| Klrb1 | 1.7457 | 1.582762 | 1.7298 | -0.146993867 | 0.015944563 | 1.6271861 | 1.6304975 | 1.72412 | -0.093625122 | -0.096936594 |
| S100a4 | 10.64596 | 10.72891 | 10.691 | 0.0378657 | -0.045086282 | 10.905 | 10.582545 | 10.6762 | -0.093628686 | 0.228825837 |
| Far1 | 10.96497 | 11.01423 | 10.985 | 0.029561136 | -0.01969234 | 11.022367 | 11.075432 | 11.1691 | -0.09363759 | -0.146702345 |
| Clcf1 | 8.607836 | 8.849689 | 8.6777 | 0.171973099 | -0.069880712 | 9.1598491 | 8.9685962 | 9.06225 | -0.09365818 | 0.097594735 |
| Cep126 | 2.847044 | 2.911897 | 2.9789 | -0.066988172 | -0.131841307 | 2.8131328 | 2.616207 | 2.70989 | -0.0936785 | 0.103247288 |
| Sdhaf3 | 6.492882 | 6.324579 | 6.4537 | -0.129159255 | 0.039143801 | 6.4940069 | 6.3975399 | 6.49127 | -0.093731753 | 0.002735211 |
| Slc4a1ap | 9.403986 | 9.469112 | 9.5719 | -0.102748858 | -0.16787502 | 9.5809593 | 9.4891811 | 9.58294 | -0.093758123 | -0.001979868 |
| 1700093K21Rik | 2.37138 | 2.320736 | 2.3277 | -0.006996438 | 0.043647955 | 2.2290709 | 2.2496516 | 2.34351 | -0.09385582 | -0.114436538 |
| Gm5538 | 2.125114 | 2.082229 | 2.1084 | -0.026199245 | 0.016685926 | 2.1462145 | 2.0817097 | 2.17559 | -0.093880666 | -0.029375868 |
| Dennd4c | 9.91014 | 9.986215 | 9.9325 | 0.053759747 | -0.022314645 | 9.5331658 | 9.6939173 | 9.78784 | -0.093920799 | -0.254672268 |
| Gm15320 | 4.515249 | 4.285084 | 4.3381 | -0.052986609 | 0.177178779 | 4.2912646 | 4.3512834 | 4.44523 | -0.093951142 | -0.153969935 |
| Cltrn | 1.679897 | 1.787019 | 1.6718 | 0.115196753 | 0.008074149 | 1.6146072 | 1.6380823 | 1.73211 | -0.094026729 | -0.117501845 |
| Dido1 | 10.90895 | 10.81893 | 10.812 | 0.007092468 | 0.097114364 | 10.961141 | 10.964412 | 11.0584 | -0.094028304 | -0.097299542 |
| Lgr4 | 4.417867 | 4.409023 | 4.3296 | 0.079446925 | 0.088290858 | 4.1940912 | 4.2425037 | 4.33656 | -0.094059142 | -0.142471709 |
| Podn | 3.071524 | 3.051094 | 3.1448 | -0.093682502 | -0.07325218 | 3.028241 | 2.9636752 | 3.05784 | -0.094160382 | -0.029594582 |
| Gkap1 | 5.236161 | 5.243683 | 5.2743 | -0.030622253 | -0.038144405 | 5.0360535 | 5.0265371 | 5.12076 | -0.094227807 | -0.084711436 |
| Zfp354b | 1.943712 | 1.949717 | 1.9469 | 0.00283779 | -0.003166957 | 1.9551433 | 1.8957981 | 1.99003 | -0.094229019 | -0.034883852 |
| Mroh2a | 1.921609 | 2.09599 | 1.8901 | 0.205925569 | 0.031544829 | 1.9237518 | 1.8623829 | 1.95663 | -0.094246591 | -0.032877686 |
| Sema4a | 11.14844 | 10.85948 | 11.04 | -0.180587118 | 0.108365706 | 10.880548 | 10.982804 | 11.0771 | -0.09427387 | -0.196529803 |
| Tmed2 | 11.28899 | 11.24079 | 11.299 | -0.058410528 | -0.010206658 | 11.198736 | 11.070345 | 11.1646 | -0.094285954 | 0.034105448 |
| Olfr1564 | 2.223275 | 2.174366 | 2.183 | -0.008629762 | 0.040278679 | 2.2558149 | 2.2111129 | 2.3054 | -0.094289775 | -0.049587784 |
| Krt20 | 1.415383 | 1.436968 | 1.4011 | 0.035861308 | 0.014276624 | 1.3330697 | 1.2989624 | 1.39327 | -0.094303362 | -0.060196074 |
| Olfr322 | 1.282596 | 1.258525 | 1.2753 | -0.016744942 | 0.007325998 | 1.2980912 | 1.341263 | 1.43563 | -0.094371398 | -0.137543206 |
| Dhx32 | 8.200029 | 8.034773 | 8.1987 | -0.163970794 | 0.001285416 | 8.0334725 | 7.9735803 | 8.06797 | -0.094385125 | -0.034492975 |
| Zp1 | 2.162513 | 2.126986 | 2.0654 | 0.061616957 | 0.09714398 | 2.0982491 | 2.0410802 | 2.13548 | -0.094398017 | -0.03722905 |
| Gm20605 | 9.913575 | 9.987652 | 10.116 | -0.12819837 | -0.202275633 | 10.627344 | 10.364062 | 10.4585 | -0.094403222 | 0.168878854 |
| Setd4 | 7.338176 | 7.192196 | 7.3449 | -0.152655371 | -0.006674731 | 7.7217216 | 7.3168286 | 7.41123 | -0.094405688 | 0.310487335 |
| Kif20b | 8.249911 | 8.1007 | 8.381 | -0.280343784 | -0.131132552 | 8.1645836 | 8.3171979 | 8.41171 | -0.09451607 | -0.24713036 |
| Alkbh3 | 8.260713 | 8.064794 | 8.2534 | -0.188600039 | 0.007319145 | 8.4383783 | 8.3409602 | 8.43554 | -0.094581256 | 0.002836891 |

| | | | | | | | | | | |
|---|---|---|---|---|---|---|---|---|---|---|
| Pde4d | 8.358325 | 8.453528 | 8.3826 | 0.070892096 | -0.024310222 | 8.1355184 | 8.4313137 | 8.52594 | -0.094625688 | -0.390420949 |
| Car3 | 2.48873 | 2.531219 | 2.3556 | 0.175570006 | 0.133081249 | 2.3157952 | 2.3401388 | 2.43482 | -0.094677974 | -0.119021514 |
| Rnf145 | 11.30441 | 11.24734 | 11.329 | -0.082035736 | -0.024967718 | 11.481195 | 11.377013 | 11.4717 | -0.094685838 | 0.009496466 |
| Terb1 | 5.526437 | 5.453317 | 5.3598 | 0.09350807 | 0.16662794 | 6.083429 | 5.5791827 | 5.67387 | -0.094686376 | 0.409559922 |
| Fbf1 | 8.423761 | 8.405643 | 8.6673 | -0.261675132 | -0.243557118 | 8.6916214 | 8.5600316 | 8.65472 | -0.094687666 | 0.03690211 |
| Pramel6 | 0.285842 | 0.286092 | 0.3006 | -0.014462346 | -0.014711878 | 0.3259683 | 0.283324 | 0.37805 | -0.094727528 | -0.052083226 |
| Mapk8ip2 | 1.357555 | 1.474954 | 1.424 | 0.050997425 | -0.066401967 | 1.3479543 | 1.3114216 | 1.40616 | -0.094734985 | -0.058202282 |
| Mlip | 1.612954 | 1.598969 | 1.6077 | -0.008702368 | 0.005282265 | 1.6384033 | 1.5236469 | 1.61838 | -0.09473668 | 0.020019758 |
| Slc23a2 | 9.925221 | 9.988185 | 9.8834 | 0.104787988 | 0.041824095 | 9.7746383 | 9.9294018 | 10.0242 | -0.09484564 | -0.249609129 |
| Gan | 8.230035 | 8.200549 | 8.2642 | -0.063684917 | -0.03419822 | 8.4932536 | 8.5093306 | 8.60429 | -0.094959437 | -0.111036417 |
| Wipi2 | 9.901061 | 10.03325 | 10.06 | -0.026272371 | -0.158463247 | 10.042243 | 10.035602 | 10.1306 | -0.094968956 | -0.088327963 |
| Myocd | 1.353623 | 1.402267 | 1.4316 | -0.029285651 | -0.077930037 | 1.3654774 | 1.3382844 | 1.43333 | -0.095044653 | -0.067851714 |
| Bcl11a | 11.28777 | 11.26895 | 11.175 | 0.094406 | 0.113226958 | 11.318053 | 11.009098 | 11.1042 | -0.095066664 | 0.213888721 |
| Fgl1 | 2.295926 | 2.319456 | 2.4004 | -0.080979167 | -0.104509885 | 2.3883393 | 2.3301713 | 2.42531 | -0.095136812 | -0.036968769 |
| Etfrf1 | 7.257324 | 7.030368 | 7.1901 | -0.159751753 | 0.067203973 | 7.2977844 | 7.2448852 | 7.34007 | -0.095185617 | -0.042286451 |
| Derl3 | 5.541908 | 5.816264 | 5.5528 | 0.263485062 | -0.010871742 | 5.7053252 | 5.6137871 | 5.70898 | -0.095197142 | -0.003659066 |
| Fam20b | 9.904928 | 9.953005 | 9.9442 | 0.008837766 | -0.03923914 | 9.7314889 | 9.7456349 | 9.84084 | -0.095204983 | -0.109350934 |
| Cybrd1 | 4.637313 | 4.443698 | 4.7074 | -0.263730904 | -0.070115772 | 4.6814655 | 4.5966423 | 4.69193 | -0.095291226 | -0.010468044 |
| Cnih1 | 9.179175 | 9.20024 | 9.2408 | -0.040543475 | -0.061607572 | 9.2190007 | 9.0396887 | 9.13499 | -0.095298267 | 0.084013702 |
| Hsd17b10 | 9.224301 | 9.189017 | 9.2397 | -0.050730638 | -0.015447011 | 9.3139525 | 9.073544 | 9.16887 | -0.095323017 | 0.145085528 |
| Ube2e2 | 5.3896 | 5.364736 | 5.4438 | -0.079023911 | -0.054159942 | 5.2322793 | 5.2684268 | 5.36377 | -0.095343968 | -0.131491474 |
| Coil | 7.670318 | 7.774785 | 7.7562 | 0.018594188 | -0.085873115 | 7.9821989 | 7.7526565 | 7.84806 | -0.095402471 | 0.134139986 |
| Cadps | 1.926388 | 1.931723 | 2.1053 | -0.173561195 | -0.178896554 | 1.8819568 | 1.972676 | 2.06813 | -0.095455918 | -0.186175094 |
| Qk | 11.20621 | 11.2648 | 11.287 | -0.022519362 | -0.081118132 | 11.164706 | 10.921083 | 11.0166 | -0.095479718 | 0.148142979 |
| Gm7972 | 1.789839 | 1.777304 | 1.8605 | -0.083184157 | -0.070648682 | 1.7682293 | 1.7656206 | 1.86114 | -0.09551717 | -0.092908459 |
| Vmn1r18 | 4.795855 | 4.74054 | 4.78 | -0.039457428 | 0.015857217 | 4.900642 | 4.9490803 | 5.04462 | -0.095539473 | -0.143977797 |
| F9 | 4.051864 | 4.060535 | 3.9361 | 0.124482414 | 0.115810726 | 4.1136493 | 4.0000818 | 4.09568 | -0.095593836 | 0.017973656 |
| H2-M9 | 1.57752 | 1.723181 | 1.8134 | -0.090254116 | -0.235915148 | 1.5267902 | 1.5338578 | 1.62946 | -0.095600087 | -0.102667709 |
| Socs6 | 8.59681 | 8.627175 | 8.646 | -0.018843125 | -0.049207623 | 8.4404374 | 8.5498362 | 8.64548 | -0.095642702 | -0.205041441 |
| Arcn1 | 11.57798 | 11.5213 | 11.443 | 0.078491724 | 0.13517258 | 11.353339 | 11.297062 | 11.3928 | -0.095700026 | -0.03942233 |
| Gm18186 | 3.837464 | 3.856294 | 3.9751 | -0.118852867 | -0.137682366 | 3.9897462 | 3.8883775 | 3.98412 | -0.095746641 | 0.005622101 |
| Olfr281 | 1.220468 | 1.157274 | 1.1826 | -0.025294635 | 0.037899293 | 1.1472926 | 1.1865833 | 1.28241 | -0.095823889 | -0.135114583 |

| | | | | | | | | | | |
|---|---|---|---|---|---|---|---|---|---|---|
| Plce1 | 5.123515 | 4.914277 | 5.0876 | -0.173357076 | 0.035880175 | 4.9309441 | 4.7873781 | 4.8832 | -0.095825628 | 0.047740335 |
| Copa | 12.3736 | 12.36196 | 12.386 | -0.024322434 | -0.012681802 | 12.346879 | 12.261035 | 12.3569 | -0.095825884 | -0.009982133 |
| Gm11353 | 1.884345 | 2.010179 | 1.9637 | 0.046506611 | -0.079327302 | 1.9523657 | 1.9542015 | 2.05003 | -0.09583134 | -0.097667123 |
| Tead2 | 5.069673 | 5.235548 | 5.1787 | 0.056892311 | -0.108982734 | 5.0858799 | 5.0630266 | 5.15887 | -0.095842959 | -0.072989691 |
| Sgce | 4.513437 | 4.319113 | 4.3875 | -0.068405843 | 0.125918274 | 4.3302151 | 4.0935846 | 4.18945 | -0.09586935 | 0.140761126 |
| Wwp2 | 10.49344 | 10.51198 | 10.488 | 0.023528192 | 0.004991984 | 10.736137 | 10.553261 | 10.6492 | -0.0959509 | 0.086925108 |
| Elp4 | 7.203234 | 6.845006 | 7.3964 | -0.551379974 | -0.193152167 | 7.1285195 | 6.9942932 | 7.09026 | -0.095969079 | 0.038257166 |
| Gm8274 | 1.103325 | 1.144607 | 1.0821 | 0.062534098 | 0.021252364 | 1.135356 | 1.1239659 | 1.21994 | -0.09597119 | -0.084581081 |
| Clec18a | 1.340909 | 1.300939 | 1.3524 | -0.051472093 | -0.011502545 | 1.2738709 | 1.3260996 | 1.42209 | -0.09599448 | -0.148223175 |
| Gm6480 | 1.858873 | 1.845835 | 1.837 | 0.008849755 | 0.021887268 | 1.8132714 | 1.8635493 | 1.95955 | -0.095997174 | -0.146275005 |
| Hrct1 | 1.888622 | 1.94222 | 1.9665 | -0.024234812 | -0.07783269 | 1.7920744 | 1.7874484 | 1.8835 | -0.096054548 | -0.091428483 |
| Traj39 | 1.600113 | 1.567797 | 1.6471 | -0.079325124 | -0.047009731 | 1.6608865 | 1.5242689 | 1.62033 | -0.096059227 | 0.040558411 |
| Six4 | 3.44992 | 3.351117 | 3.4571 | -0.105959707 | -0.007156274 | 3.4037078 | 3.344858 | 3.44093 | -0.096074493 | -0.037224718 |
| Dtna | 2.340323 | 2.269537 | 2.2766 | -0.007072273 | 0.063713981 | 2.2195459 | 2.23177 | 2.32789 | -0.096118527 | -0.108342667 |
| Ccn3 | 4.834576 | 4.67098 | 4.8457 | -0.174698909 | -0.011102859 | 4.5671789 | 4.3503846 | 4.44659 | -0.096201562 | 0.120592752 |
| Tjp3 | 7.206713 | 7.088486 | 7.3703 | -0.281804704 | -0.163577584 | 7.4560933 | 7.3417177 | 7.43793 | -0.096214078 | 0.01816153 |
| Pgap3 | 7.266371 | 7.321616 | 7.2043 | 0.117332645 | 0.062088352 | 7.4439598 | 7.2618421 | 7.35806 | -0.096222596 | 0.085895077 |
| Dnm1l | 10.61029 | 10.67968 | 10.661 | 0.01857054 | -0.050825834 | 10.596544 | 10.729934 | 10.8262 | -0.096276888 | -0.22966714 |
| Gm11743 | 1.740006 | 1.731337 | 1.7169 | 0.014443806 | 0.023113207 | 1.8567993 | 1.7352079 | 1.83154 | -0.096335425 | 0.025255962 |
| Tmem229a | 0.836925 | 0.786446 | 0.8938 | -0.107369911 | -0.056891412 | 0.7708176 | 0.7840131 | 0.88042 | -0.096407307 | -0.109602805 |
| Pkp1 | 2.487143 | 2.469008 | 2.5896 | -0.120569234 | -0.102433557 | 2.4680523 | 2.4835558 | 2.58 | -0.096440228 | -0.111943657 |
| Cyp2e1 | 1.630933 | 1.54676 | 1.6568 | -0.110054303 | -0.025881553 | 1.4958235 | 1.5059642 | 1.60241 | -0.096448638 | -0.106589294 |
| Chst10 | 9.045964 | 9.285086 | 9.0381 | 0.246977149 | 0.007856023 | 9.2835404 | 9.0674401 | 9.1639 | -0.096464177 | 0.119636152 |
| Ppp4r3b | 11.53893 | 11.66503 | 11.558 | 0.106614482 | -0.019486184 | 11.507807 | 11.585151 | 11.6816 | -0.096472316 | -0.173816057 |
| Vmn1r42 | 1.664107 | 1.603898 | 1.6012 | 0.002746854 | 0.06295636 | 1.7466387 | 1.6286243 | 1.72512 | -0.096496151 | 0.021518264 |
| Hsd17b7 | 7.473748 | 7.695232 | 7.3653 | 0.329891219 | 0.108406798 | 7.7786009 | 7.3595143 | 7.45605 | -0.096540143 | 0.322546384 |
| Zfp68 | 9.694189 | 9.716428 | 9.7212 | -0.004803786 | -0.027043352 | 9.7643067 | 9.6214321 | 9.71801 | -0.096580814 | 0.046293851 |
| Aldh3a1 | 2.420382 | 2.336576 | 2.2859 | 0.05069271 | 0.134498448 | 2.3516174 | 2.2129332 | 2.30957 | -0.096634235 | 0.042049912 |
| Gm12527 | 0.345236 | 0.341263 | 0.3188 | 0.022434571 | 0.026407899 | 0.3503794 | 0.3037352 | 0.40045 | -0.096711639 | -0.050067467 |
| Snca | 2.505228 | 2.31505 | 2.4776 | -0.162557835 | 0.027619259 | 2.3968867 | 2.277469 | 2.37419 | -0.096723155 | 0.022694554 |
| Ust | 6.980427 | 6.743617 | 6.8897 | -0.146113053 | 0.090697739 | 6.7050997 | 6.7835446 | 6.88028 | -0.096733425 | -0.175178299 |
| Gnb1l | 6.54428 | 6.574077 | 6.5653 | 0.00874853 | -0.021048116 | 6.8124917 | 6.4812815 | 6.57802 | -0.096734779 | 0.23447542 |

| | | | | | | | | | | |
|---|---|---|---|---|---|---|---|---|---|---|
| Nck1 | 9.606073 | 9.692592 | 9.682 | 0.010633186 | -0.075885976 | 9.4060686 | 9.4636282 | 9.56037 | -0.096739092 | -0.154298755 |
| Rdh14 | 7.457964 | 7.5042 | 7.5024 | 0.001828642 | -0.04440713 | 7.5639316 | 7.4374114 | 7.53418 | -0.096768394 | 0.02975186 |
| Smad4 | 10.41773 | 10.46293 | 10.521 | -0.058430726 | -0.103629492 | 10.374858 | 10.373381 | 10.4702 | -0.09680921 | -0.095331455 |
| Mkrn2 | 8.706719 | 8.621532 | 8.8414 | -0.219889971 | -0.134702651 | 9.009649 | 8.7513904 | 8.84822 | -0.096828379 | 0.161430301 |
| Exph5 | 2.53568 | 2.478939 | 2.5561 | -0.077147248 | -0.020405808 | 2.5161487 | 2.4316927 | 2.52863 | -0.096942042 | -0.012486 |
| Prkce | 9.553295 | 9.594437 | 9.5422 | 0.052245004 | 0.011103119 | 9.4620414 | 9.3730026 | 9.47001 | -0.097002772 | -0.007963908 |
| Aifm1 | 8.970711 | 8.884783 | 8.9966 | -0.111831995 | -0.025904006 | 9.0254835 | 8.9280703 | 9.02509 | -0.097018905 | 0.000394242 |
| D130007C19Rik | 2.13443 | 2.079598 | 2.0949 | -0.015310982 | 0.039521142 | 2.2090237 | 2.1066438 | 2.20367 | -0.097022819 | 0.005357018 |
| Armt1 | 9.020301 | 8.922627 | 9.024 | -0.101369815 | -0.003695836 | 9.1602032 | 8.9858631 | 9.08294 | -0.097077667 | 0.077262413 |
| Raet1e | 4.350742 | 4.229083 | 4.1778 | 0.051252418 | 0.172910905 | 4.0011991 | 4.0182993 | 4.1154 | -0.097105641 | -0.114205847 |
| Ldhd | 5.296998 | 5.170173 | 5.2902 | -0.120038308 | 0.006787284 | 5.1607931 | 5.1393642 | 5.23649 | -0.097125556 | -0.075696709 |
| Mmp7 | 2.049232 | 2.020258 | 2.079 | -0.058767107 | -0.029793133 | 2.1409517 | 2.0414838 | 2.13867 | -0.097182249 | 0.002285621 |
| Cbs | 3.526829 | 3.541682 | 3.473 | 0.068691364 | 0.05383848 | 3.521842 | 3.4709098 | 3.5681 | -0.09718582 | -0.046253631 |
| Khdrbs3 | 3.813626 | 3.73698 | 3.6134 | 0.123587306 | 0.200233158 | 3.6815849 | 3.6326433 | 3.72983 | -0.097189295 | -0.048247731 |
| Ranbp10 | 9.944221 | 10.00301 | 10.114 | -0.111224765 | -0.170009283 | 9.8872837 | 9.9262431 | 10.0235 | -0.097274394 | -0.136233789 |
| Olfr1123 | 1.383467 | 1.424892 | 1.4297 | -0.004797114 | -0.046223021 | 1.4428821 | 1.3996686 | 1.49696 | -0.097290322 | -0.05407681 |
| Tcte2 | 0.586189 | 0.595609 | 0.6424 | -0.046803323 | -0.056223415 | 0.6510695 | 0.5906038 | 0.6879 | -0.097297628 | -0.036831926 |
| Tmed3 | 8.927273 | 8.738542 | 8.9101 | -0.17157257 | 0.017158388 | 8.8131615 | 8.6035168 | 8.70082 | -0.097299071 | 0.112345644 |
| 1810055G02Rik | 5.81984 | 5.768523 | 5.9236 | -0.155061082 | -0.103744061 | 6.0532879 | 5.8258352 | 5.92315 | -0.097310695 | 0.13014193 |
| Fhl2 | 3.886418 | 3.814153 | 4.0168 | -0.202601572 | -0.130337149 | 3.8281096 | 3.827549 | 3.92487 | -0.097324586 | -0.09676399 |
| Rps6ka6 | 1.429629 | 1.315219 | 1.3347 | -0.019462742 | 0.094946784 | 1.2872374 | 1.2218687 | 1.3192 | -0.097330698 | -0.031961982 |
| Vegfd | 2.504169 | 2.447832 | 2.4765 | -0.028670163 | 0.027666141 | 2.3152287 | 2.3525766 | 2.44993 | -0.097352865 | -0.134700677 |
| Snta1 | 4.007442 | 3.952209 | 4.1263 | -0.174090374 | -0.118857149 | 4.2868512 | 3.7950657 | 3.89255 | -0.097486754 | 0.394298696 |
| S100a13 | 8.599751 | 8.38999 | 8.4942 | -0.104168798 | 0.105591468 | 8.4491 | 8.4197348 | 8.51733 | -0.097598138 | -0.068232929 |
| Cyp1a1 | 1.103289 | 1.06068 | 1.22 | -0.159275752 | -0.116666484 | 1.0748043 | 1.055196 | 1.15282 | -0.097625084 | -0.078016809 |
| 2410024N13Rik | 1.642175 | 1.644916 | 1.6153 | 0.029595975 | 0.026854777 | 1.5391099 | 1.6544933 | 1.75217 | -0.09767451 | -0.213057972 |
| Vti1a | 9.144965 | 9.164931 | 9.1005 | 0.06440327 | 0.044436934 | 9.219529 | 9.061015 | 9.15875 | -0.097731797 | 0.060782216 |
| Tmem267 | 6.823047 | 7.021197 | 6.6975 | 0.323674809 | 0.125524989 | 6.8401006 | 6.665589 | 6.76349 | -0.097896242 | 0.076615363 |
| Cdk3-ps | 3.512844 | 3.604591 | 3.5877 | 0.016913603 | -0.074833442 | 3.7364362 | 3.6429294 | 3.74091 | -0.097975757 | -0.004469013 |
| Bckdha | 9.019501 | 8.952461 | 9.0508 | -0.09831049 | -0.031271006 | 9.4648156 | 8.9063496 | 9.00434 | -0.09798601 | 0.460479994 |
| Mei4 | 3.004547 | 2.949087 | 3.0134 | -0.064339506 | -0.008880062 | 2.9514086 | 2.9942638 | 3.09228 | -0.098019407 | -0.140874557 |
| Slc25a13 | 6.276844 | 6.03327 | 6.0932 | -0.059965275 | 0.183609314 | 6.0742624 | 6.0452728 | 6.1433 | -0.098025974 | -0.069036382 |

| | | | | | | | | | | |
|---|---|---|---|---|---|---|---|---|---|---|
| Doc2b | 3.504931 | 3.264589 | 3.7053 | -0.440738318 | -0.200396327 | 3.1206508 | 2.9639051 | 3.06199 | -0.098084784 | 0.05866089 |
| Gle1 | 9.755129 | 9.79902 | 9.9075 | -0.108471175 | -0.152362263 | 9.9562236 | 9.929368 | 10.0275 | -0.098102463 | -0.071246842 |
| Rdh16 | 1.964868 | 1.984424 | 1.9827 | 0.001764823 | -0.017790831 | 2.0667201 | 1.9790738 | 2.07737 | -0.098296413 | -0.010650098 |
| Clvs2 | 1.323274 | 1.323494 | 1.2771 | 0.046439217 | 0.046218949 | 1.3408686 | 1.2762466 | 1.37455 | -0.098300738 | -0.033678727 |
| Iglc3 | 10.73197 | 10.74594 | 10.933 | -0.187229316 | -0.201198134 | 11.353444 | 10.81079 | 10.9091 | -0.098313284 | 0.444341188 |
| Nfrkb | 10.679 | 10.71493 | 10.711 | 0.004386696 | -0.03154335 | 10.748176 | 10.741621 | 10.84 | -0.098330226 | -0.0917755 |
| Slc35d3 | 1.595123 | 1.610633 | 1.4993 | 0.111342678 | 0.095833105 | 1.5022745 | 1.4442057 | 1.54254 | -0.098337286 | -0.04026846 |
| Fndc1 | 2.417134 | 2.587494 | 2.6101 | -0.022580414 | -0.192940842 | 2.513069 | 2.4845565 | 2.5829 | -0.098339722 | -0.069827257 |
| Samsn1 | 10.69363 | 10.53306 | 10.574 | -0.040945336 | 0.119619504 | 10.461198 | 10.513514 | 10.6119 | -0.098361476 | -0.150677096 |
| Mef2d | 11.60171 | 11.66922 | 11.672 | -0.003084048 | -0.070596432 | 11.68255 | 11.658264 | 11.7567 | -0.098461886 | -0.074175435 |
| Zfp784 | 5.226693 | 5.14662 | 5.2876 | -0.140972628 | -0.06090009 | 5.2289274 | 5.0947047 | 5.19319 | -0.09848645 | 0.035736266 |
| Aim2 | 9.69247 | 9.709513 | 9.8222 | -0.112713238 | -0.129756278 | 9.8888277 | 9.9054722 | 10.004 | -0.098551504 | -0.115196003 |
| Hs6st2 | 2.81473 | 2.693568 | 2.6174 | 0.076212256 | 0.197374109 | 2.5629413 | 2.5250223 | 2.62359 | -0.098568164 | -0.060649181 |
| Eng | 10.29722 | 10.02338 | 10.325 | -0.301263316 | -0.027423377 | 9.6877538 | 9.6104395 | 9.70902 | -0.098577093 | -0.021262802 |
| Ccdc155 | 1.858795 | 1.877502 | 1.9021 | -0.024599798 | -0.043307066 | 1.9466567 | 1.8538801 | 1.95248 | -0.098600221 | -0.005823641 |
| Fam124a | 4.699313 | 4.521347 | 4.6396 | -0.118259219 | 0.059707207 | 4.3867044 | 4.4859495 | 4.58456 | -0.098612409 | -0.197857525 |
| Sprn | 2.017455 | 2.050443 | 1.9716 | 0.078820903 | 0.045832424 | 2.0205749 | 1.8422237 | 1.94086 | -0.09863593 | 0.079715187 |
| Olfr1443 | 1.44899 | 1.470732 | 1.483 | -0.012268176 | -0.034009788 | 1.4795487 | 1.4506832 | 1.54941 | -0.098723946 | -0.06985842 |
| Npbwr1 | 1.043793 | 1.04052 | 1.0764 | -0.035878792 | -0.032605939 | 1.078111 | 1.0601699 | 1.15896 | -0.098792032 | -0.080850994 |
| Aagab | 9.55436 | 9.466572 | 9.5209 | -0.054304674 | 0.033483597 | 9.5302581 | 9.4470942 | 9.54591 | -0.098813858 | -0.015649946 |
| Star | 5.498884 | 5.114533 | 5.4387 | -0.324142456 | 0.060209378 | 5.620992 | 5.3805192 | 5.47936 | -0.098842311 | 0.141630561 |
| Gm5069 | 2.553121 | 2.476073 | 2.5158 | -0.039732714 | 0.037315347 | 2.5634058 | 2.4969993 | 2.59586 | -0.098857391 | -0.032450916 |
| Mybpc1 | 2.143211 | 2.078778 | 2.2013 | -0.122522526 | -0.05808905 | 2.0669268 | 2.107667 | 2.20662 | -0.09895079 | -0.13969105 |
| Fzd8 | 4.044765 | 4.027737 | 4.0769 | -0.049145946 | -0.032117454 | 3.8103618 | 3.729524 | 3.82852 | -0.098992607 | -0.018154785 |
| Kdm4d | 3.270214 | 3.21391 | 3.2767 | -0.062760468 | -0.006457148 | 3.1854635 | 3.1701604 | 3.26918 | -0.099017177 | -0.083714005 |
| Ptger1 | 4.076611 | 4.158811 | 4.1326 | 0.026202676 | -0.055997459 | 4.1364345 | 4.1039793 | 4.20304 | -0.099062425 | -0.066607261 |
| Mmp10 | 0.831722 | 0.779181 | 0.7857 | -0.006492894 | 0.046048422 | 0.8554288 | 0.7904934 | 0.88958 | -0.099087493 | -0.034152071 |
| Urod | 8.696703 | 8.670602 | 8.7335 | -0.062924763 | -0.036824057 | 8.983463 | 8.7751313 | 8.87424 | -0.099108235 | 0.109223509 |
| Zc3hc1 | 8.651346 | 8.634214 | 8.5418 | 0.092403003 | 0.109535036 | 8.9005356 | 8.6849146 | 8.78403 | -0.099113401 | 0.116507547 |
| Prtg | 2.211242 | 1.987084 | 2.134 | -0.14688935 | 0.077268755 | 1.9855088 | 1.9728235 | 2.07194 | -0.099118079 | -0.086432846 |
| H2al1m | 1.491194 | 1.524601 | 1.5535 | -0.028900135 | -0.062307724 | 1.5653669 | 1.4901613 | 1.58932 | -0.09916003 | -0.023954457 |
| Leng8 | 10.50394 | 10.2655 | 10.61 | -0.344749429 | -0.106314389 | 11.303285 | 11.010001 | 11.1093 | -0.099250594 | 0.194032888 |

| | | | | | | | | | | |
|---|---|---|---|---|---|---|---|---|---|---|
| Gh | 3.411235 | 3.36854 | 3.3684 | 0.000119273 | 0.042814305 | 3.6125096 | 3.3165619 | 3.41597 | -0.099406648 | 0.196540991 |
| Klf10 | 8.2334 | 7.913134 | 8.2424 | -0.329249898 | -0.008983523 | 8.159238 | 8.0250294 | 8.12445 | -0.099418351 | 0.034790232 |
| Gm13534 | 1.072303 | 1.043679 | 0.9782 | 0.065485873 | 0.094109676 | 1.1065607 | 0.9700616 | 1.06956 | -0.099498988 | 0.037000174 |
| Olfr1261 | 2.621556 | 2.599444 | 2.6917 | -0.092294259 | -0.070181855 | 2.6225034 | 2.5144351 | 2.61397 | -0.099533712 | 0.008534581 |
| Zfp825 | 7.542127 | 7.514956 | 7.674 | -0.159051236 | -0.13188005 | 7.664 | 7.582535 | 7.6821 | -0.099560308 | -0.018095271 |
| Rhbdd2 | 8.105914 | 8.192266 | 8.2187 | -0.026394529 | -0.112746722 | 8.5311436 | 8.3917636 | 8.49136 | -0.09959541 | 0.039784553 |
| Zfp503 | 4.75645 | 4.513319 | 4.7169 | -0.203592111 | 0.039539415 | 4.3326811 | 4.3578654 | 4.45752 | -0.099658583 | -0.124842857 |
| Mfsd13a | 5.383306 | 5.165636 | 5.4111 | -0.245487899 | -0.027818569 | 5.2172291 | 4.9686635 | 5.06833 | -0.099663887 | 0.148901634 |
| C9orf72 | 7.713284 | 7.735598 | 7.699 | 0.03655048 | 0.014236141 | 7.9382399 | 7.8579941 | 7.95779 | -0.099799128 | -0.019553296 |
| Slc46a1 | 3.310227 | 3.33097 | 3.1353 | 0.195630769 | 0.174888426 | 3.3754961 | 3.1851682 | 3.28503 | -0.099863169 | 0.090464698 |
| Ctxn1 | 2.882181 | 2.933026 | 2.9189 | 0.014170858 | -0.036673953 | 2.9743332 | 2.8111017 | 2.91101 | -0.09990413 | 0.063327279 |
| Trbv24 | 3.217725 | 3.317634 | 3.2261 | 0.091499347 | -0.008410358 | 3.1950077 | 3.225893 | 3.32581 | -0.09992062 | -0.13080597 |
| Abhd14a | 6.071392 | 6.130835 | 6.0924 | 0.038479566 | -0.020963063 | 6.2587941 | 5.9544258 | 6.05447 | -0.100044655 | 0.204323624 |
| Fam206a | 8.988066 | 8.909323 | 8.9697 | -0.060340104 | 0.018403081 | 9.0396496 | 9.0321906 | 9.13228 | -0.100091346 | -0.092632336 |
| Gm45105 | 1.903841 | 1.91673 | 1.9502 | -0.033468521 | -0.046357769 | 2.0199688 | 1.9132231 | 2.01333 | -0.100109858 | 0.006635881 |
| Hhip | 2.865789 | 3.240694 | 2.9547 | 0.285959878 | -0.088944879 | 2.8252747 | 2.7030025 | 2.80314 | -0.100136868 | 0.022135399 |
| Zfp972 | 3.984562 | 3.80888 | 4.0309 | -0.222037518 | -0.046355206 | 3.9978936 | 3.9967648 | 4.09705 | -0.100286522 | -0.099157795 |
| 1810046K07Rik | 2.346506 | 2.309477 | 2.4782 | -0.168707985 | -0.131678519 | 2.2709328 | 2.3177706 | 2.41806 | -0.100292441 | -0.147130267 |
| Yap1 | 7.226212 | 6.979187 | 7.0169 | -0.037676618 | 0.209348741 | 6.3498183 | 6.6904232 | 6.79073 | -0.100309745 | -0.440914649 |
| Togaram2 | 1.413322 | 1.428313 | 1.4535 | -0.025211106 | -0.040202786 | 1.4131859 | 1.3650551 | 1.46544 | -0.100381544 | -0.052250664 |
| Gon4l | 11.05485 | 11.1756 | 11.117 | 0.058335624 | -0.0624133 | 11.065285 | 11.123308 | 11.2237 | -0.100415528 | -0.158437945 |
| Zfp423 | 5.918232 | 5.897783 | 5.8437 | 0.054087666 | 0.07453684 | 5.2206301 | 5.0576913 | 5.15818 | -0.10048823 | 0.062450561 |
| Dnah1 | 3.126296 | 3.103789 | 3.0529 | 0.050859761 | 0.073366824 | 2.9717935 | 2.9731427 | 3.07366 | -0.10051683 | -0.101866064 |
| Cdk13 | 11.60801 | 11.57487 | 11.598 | -0.022933406 | 0.01021281 | 11.442248 | 11.563358 | 11.6639 | -0.100532022 | -0.221641738 |
| Vmn1r58 | 4.212191 | 4.153988 | 4.1998 | -0.045818627 | 0.01238443 | 4.0521217 | 4.1414479 | 4.24199 | -0.100537926 | -0.189864097 |
| Ankrd46 | 8.815749 | 8.68451 | 8.7519 | -0.067409108 | 0.063830447 | 8.8705104 | 8.654002 | 8.75458 | -0.100581336 | 0.115927058 |
| Hmga2 | 1.157863 | 1.159028 | 1.1521 | 0.006950848 | 0.00578637 | 1.1686784 | 1.1763601 | 1.27696 | -0.100602962 | -0.108284649 |
| Zbtb43 | 8.666944 | 8.76775 | 8.8269 | -0.059174192 | -0.159979653 | 8.8878499 | 8.8075667 | 8.90818 | -0.100614672 | -0.020331472 |
| Gm16558 | 1.141599 | 1.093043 | 1.1125 | -0.019497171 | 0.029058184 | 1.1580276 | 1.102719 | 1.20336 | -0.100639259 | -0.045330644 |
| Hsd17b1 | 3.246827 | 3.29767 | 3.2704 | 0.027248966 | -0.023593932 | 3.3440272 | 3.2399608 | 3.34062 | -0.100659705 | 0.003406677 |
| Cd302 | 6.653392 | 6.11086 | 6.2697 | -0.158877254 | 0.383654267 | 6.4236126 | 5.8563133 | 5.95712 | -0.100805243 | 0.466494122 |
| Myo18b | 3.540366 | 3.526752 | 3.5244 | 0.002363983 | 0.015978339 | 3.3229674 | 3.4194507 | 3.52029 | -0.100843825 | -0.197327102 |

| | | | | | | | | | | |
|---|---|---|---|---|---|---|---|---|---|---|
| Smco3 | 1.269351 | 1.312752 | 1.2865 | 0.026219728 | -0.017181827 | 1.3412378 | 1.2105549 | 1.3114 | -0.100846639 | 0.029836261 |
| Ccdc77 | 7.896853 | 8.008472 | 7.8703 | 0.138206486 | 0.026588134 | 7.8808372 | 8.0283617 | 8.12926 | -0.100900401 | -0.248424904 |
| B230118H07Rik | 4.547017 | 4.330972 | 4.5685 | -0.237493151 | -0.021448671 | 4.4139843 | 4.2714656 | 4.37237 | -0.100906761 | 0.041611957 |
| Vldlr | 5.392487 | 5.062427 | 5.3291 | -0.266624646 | 0.063435482 | 4.8193067 | 4.8073842 | 4.9083 | -0.100914968 | -0.08899249 |
| Ndufs2 | 10.67943 | 10.60115 | 10.694 | -0.093284638 | -0.015008364 | 10.825008 | 10.680759 | 10.7817 | -0.100945778 | 0.043302865 |
| Unc5a | 5.243759 | 5.136117 | 5.2006 | -0.064444992 | 0.043196609 | 5.1232102 | 5.0469629 | 5.14796 | -0.101001936 | -0.024754652 |
| Socs2 | 8.778583 | 8.77206 | 9.0468 | -0.274777219 | -0.268253595 | 8.6994073 | 8.9870953 | 9.08811 | -0.10101042 | -0.388698466 |
| Kcnq4 | 1.972021 | 1.984717 | 2.1546 | -0.169913834 | -0.182609506 | 1.9467126 | 1.9251669 | 2.02618 | -0.101013008 | -0.079467298 |
| Trmt12 | 7.101331 | 6.989035 | 7.1567 | -0.167686493 | -0.055391175 | 7.4356963 | 7.134814 | 7.23595 | -0.101132321 | 0.19974999 |
| Psd4 | 12.08426 | 12.1902 | 12.222 | -0.031488254 | -0.137428362 | 12.213256 | 12.089129 | 12.1903 | -0.101138072 | 0.022988322 |
| Slc35e4 | 5.08925 | 5.007178 | 4.8409 | 0.16632818 | 0.248399403 | 4.8829772 | 4.6480799 | 4.74927 | -0.101193286 | 0.133703946 |
| Ap1g2 | 10.42892 | 10.45553 | 10.511 | -0.055498489 | -0.082105449 | 10.799884 | 10.614652 | 10.7159 | -0.10123877 | 0.083993342 |
| Gm44041 | 1.743259 | 1.705872 | 1.7166 | -0.010730358 | 0.026656764 | 1.6951448 | 1.6547269 | 1.756 | -0.101268294 | -0.060850382 |
| Evpl | 2.370077 | 2.277796 | 2.4178 | -0.140032838 | -0.0477518 | 2.2334725 | 2.2830479 | 2.3844 | -0.101348041 | -0.150923432 |
| Traj30 | 2.183801 | 2.194066 | 2.1682 | 0.025840205 | 0.015576101 | 2.2216835 | 2.1260452 | 2.22741 | -0.101362234 | -0.005723909 |
| Olfr1167 | 1.813403 | 1.831985 | 1.8985 | -0.066534278 | -0.085115539 | 1.8480062 | 1.8223689 | 1.92385 | -0.101477877 | -0.075840547 |
| Pygo1 | 2.903936 | 2.857972 | 2.896 | -0.038042965 | 0.007920697 | 2.8505468 | 2.8533403 | 2.95482 | -0.101481556 | -0.104275091 |
| Aqp11 | 3.869654 | 3.633325 | 3.7102 | -0.076899219 | 0.159430194 | 3.705334 | 3.7479017 | 3.84939 | -0.101491832 | -0.144059522 |
| Pcnx | 9.815953 | 9.649902 | 10.002 | -0.352561546 | -0.186510301 | 9.7074693 | 9.6055906 | 9.70728 | -0.10169132 | 0.000187375 |
| Gm14387 | 2.270972 | 2.217033 | 2.2582 | -0.041214055 | 0.01272472 | 2.2908504 | 2.1640288 | 2.26572 | -0.1016956 | 0.025126025 |
| Sec61a2 | 6.573051 | 6.68999 | 6.568 | 0.121967817 | 0.005028609 | 6.7700918 | 6.7829774 | 6.88474 | -0.101764056 | -0.114649717 |
| Urgcp | 9.02738 | 8.846695 | 8.962 | -0.115298942 | 0.065385974 | 8.7101981 | 8.7341066 | 8.8359 | -0.101790714 | -0.125699277 |
| Cyp4v3 | 8.291448 | 8.406918 | 8.4419 | -0.035029738 | -0.150499607 | 8.4457267 | 8.2504333 | 8.35225 | -0.101812431 | 0.093481028 |
| Ighv1-42 | 6.873682 | 6.747632 | 6.6901 | 0.05752649 | 0.183576461 | 6.7509956 | 6.3228731 | 6.4247 | -0.101828914 | 0.326293584 |
| Tcf3 | 10.79144 | 10.82872 | 10.859 | -0.030137394 | -0.067417916 | 11.005628 | 10.688676 | 10.7907 | -0.102009782 | 0.214942074 |
| Olfr395 | 1.407237 | 1.395815 | 1.3379 | 0.057946175 | 0.069368722 | 1.3829306 | 1.3851129 | 1.48715 | -0.102033021 | -0.104215411 |
| Tnks | 10.13576 | 10.13316 | 10.076 | 0.057097055 | 0.059694161 | 10.103549 | 10.186027 | 10.2881 | -0.102034402 | -0.184512437 |
| Orc5 | 8.89671 | 8.911019 | 8.8875 | 0.023555925 | 0.009247384 | 8.901301 | 8.798647 | 8.90072 | -0.102068122 | 0.000585882 |
| Kif3b | 9.850987 | 9.915017 | 9.9011 | 0.013965549 | -0.050064839 | 9.8448892 | 9.8120867 | 9.91425 | -0.102162911 | -0.069360426 |
| Gm14085 | 8.925199 | 9.217562 | 9.1951 | 0.022508577 | -0.269854372 | 8.8446849 | 9.2034062 | 9.30565 | -0.102245588 | -0.46096688 |
| Rapgef6 | 12.88249 | 12.72944 | 12.77 | -0.04051493 | 0.112539522 | 12.498823 | 12.63531 | 12.7376 | -0.102247206 | -0.238733909 |
| Fxyd2 | 5.799724 | 5.440277 | 5.6213 | -0.181070198 | 0.178377442 | 5.2450131 | 5.3930706 | 5.49537 | -0.102296086 | -0.250353508 |

| | | | | | | | | | | |
|---|---|---|---|---|---|---|---|---|---|---|
| Cdhr3 | 1.398145 | 1.507851 | 1.4274 | 0.080457473 | -0.029248539 | 1.3667955 | 1.3760699 | 1.47837 | -0.102300926 | -0.111575394 |
| Ttc41 | 4.417666 | 4.53478 | 4.5606 | -0.025772887 | -0.142887167 | 4.5538943 | 4.5013966 | 4.60372 | -0.102320622 | -0.049822955 |
| Brd3os | 1.607069 | 1.566558 | 1.5496 | 0.016912826 | 0.057423883 | 1.5972683 | 1.5157294 | 1.61807 | -0.102340479 | -0.020801568 |
| Tmc8 | 10.0973 | 10.20889 | 10.168 | 0.040502403 | -0.07108757 | 10.402281 | 10.204794 | 10.3072 | -0.102359994 | 0.095127307 |
| Lrrc49 | 4.948274 | 4.615217 | 4.7824 | -0.167178003 | 0.165878923 | 4.6527096 | 4.5865883 | 4.68897 | -0.102384081 | -0.036262693 |
| Tead3 | 3.745878 | 3.615257 | 3.5839 | 0.031357106 | 0.16197756 | 3.5272594 | 3.4780694 | 3.58046 | -0.102387903 | -0.053197907 |
| Grin2d | 3.797466 | 3.680661 | 3.7836 | -0.102944841 | 0.013860274 | 3.6420056 | 3.6000685 | 3.70246 | -0.102394788 | -0.060457685 |
| Rassf7 | 6.305001 | 6.379082 | 6.5995 | -0.220441092 | -0.294522328 | 6.7798461 | 6.3543217 | 6.45672 | -0.102402735 | 0.323121641 |
| Itga10 | 5.698802 | 6.006175 | 5.9672 | 0.038960503 | -0.268412298 | 6.118982 | 6.1632477 | 6.26566 | -0.102411839 | -0.146677593 |
| 1700016K19Rik | 2.256083 | 2.273736 | 2.2001 | 0.073619329 | 0.055966379 | 2.4063108 | 2.1098484 | 2.21227 | -0.102418477 | 0.194043936 |
| Ptdss1 | 9.66573 | 9.60286 | 9.6179 | -0.015062694 | 0.047807052 | 9.551822 | 9.5872971 | 9.68973 | -0.102428235 | -0.137903325 |
| Spr | 8.02305 | 7.990391 | 7.9129 | 0.07753696 | 0.110195308 | 7.7901599 | 7.7969719 | 7.89942 | -0.10245116 | -0.109263137 |
| Nfya | 9.495306 | 9.634003 | 9.4822 | 0.151843435 | 0.013146681 | 9.4733836 | 9.4922656 | 9.59475 | -0.10248905 | -0.121371009 |
| Wdr45 | 8.381172 | 8.381954 | 8.4849 | -0.10295113 | -0.103733556 | 8.5541699 | 8.5766471 | 8.67929 | -0.102640444 | -0.125117644 |
| Ufsp2 | 8.420502 | 8.262804 | 8.4413 | -0.178532423 | -0.020834541 | 8.6363905 | 8.3517194 | 8.45438 | -0.102655759 | 0.182015334 |
| Nfe2l3 | 3.292834 | 3.265809 | 3.3089 | -0.043086275 | -0.016061182 | 3.2017477 | 3.326156 | 3.42884 | -0.102685416 | -0.227093789 |
| Golga4 | 10.6564 | 10.70361 | 10.688 | 0.015646798 | -0.031562894 | 10.715903 | 10.841879 | 10.9446 | -0.102718384 | -0.228694016 |
| Fam229b | 2.64516 | 2.579799 | 2.761 | -0.181229071 | -0.115867752 | 2.5789087 | 2.5610568 | 2.66384 | -0.102782909 | -0.084931024 |
| Zkscan5 | 7.179923 | 7.149122 | 7.2627 | -0.113578485 | -0.082777115 | 7.1993437 | 7.1425229 | 7.24531 | -0.10278374 | -0.045962975 |
| Gm4540 | 0.678881 | 0.627065 | 0.6873 | -0.060187921 | -0.008372596 | 0.7728679 | 0.6552994 | 0.75809 | -0.102792827 | 0.014775637 |
| Srp54a | 10.93757 | 10.72991 | 10.76 | -0.030440992 | 0.177217176 | 10.634883 | 10.641431 | 10.7443 | -0.10287359 | -0.10942143 |
| Scara3 | 6.092855 | 5.922507 | 5.8204 | 0.102068285 | 0.272416981 | 5.7744413 | 5.4881051 | 5.59101 | -0.102903505 | 0.183432692 |
| Samd8 | 8.765247 | 8.74813 | 8.7139 | 0.034195117 | 0.051311867 | 8.6220729 | 8.6661046 | 8.76903 | -0.10292734 | -0.146959057 |
| Blcap | 7.918582 | 7.88092 | 7.8528 | 0.028141683 | 0.065804252 | 8.0071245 | 7.8863874 | 7.98936 | -0.102976967 | 0.017760137 |
| Epb41 | 11.58314 | 11.62018 | 11.539 | 0.08074249 | 0.043708402 | 11.405995 | 11.380736 | 11.4838 | -0.103033104 | -0.077774159 |
| Zg16 | 2.171885 | 2.125546 | 2.1964 | -0.070855283 | -0.024517109 | 2.1494738 | 2.1317698 | 2.23485 | -0.103081574 | -0.085377493 |
| Mlh1 | 8.504135 | 8.564988 | 8.7069 | -0.141920323 | -0.202772958 | 9.0234393 | 8.8388478 | 8.94195 | -0.103103045 | 0.081488501 |
| Fn3k | 1.477905 | 1.385182 | 1.4528 | -0.067655741 | 0.025066799 | 1.4177093 | 1.3846121 | 1.4878 | -0.103192521 | -0.070095252 |
| Nup155 | 9.581749 | 9.633339 | 9.6326 | 0.000690709 | -0.050898394 | 9.8120433 | 9.7232202 | 9.82647 | -0.103245934 | -0.014422818 |
| Nxpe1-ps | 6.073187 | 6.059594 | 6.1619 | -0.102282101 | -0.088689011 | 6.364333 | 6.1446557 | 6.24792 | -0.103263965 | 0.116413333 |
| Jmjd8 | 8.272653 | 8.233392 | 8.2394 | -0.005996331 | 0.033264633 | 8.5172798 | 8.326667 | 8.42999 | -0.103318928 | 0.087293843 |
| Gstcd | 8.467236 | 8.443753 | 8.4839 | -0.040130228 | -0.016647213 | 8.6590355 | 8.4056792 | 8.50903 | -0.103348583 | 0.150007692 |

| | | | | | | | | | | |
|---|---|---|---|---|---|---|---|---|---|---|
| Riok1 | 9.650198 | 9.619631 | 9.6042 | 0.0154486 | 0.046014931 | 9.7538711 | 9.7918457 | 9.89521 | -0.103362279 | -0.141336901 |
| Nxnl2 | 2.223597 | 2.163765 | 2.1359 | 0.027836593 | 0.087668322 | 2.0616212 | 2.0593184 | 2.16273 | -0.103415749 | -0.101112951 |
| Kdsr | 8.390239 | 8.221571 | 8.4082 | -0.186608934 | -0.017941003 | 8.6500845 | 8.415221 | 8.5187 | -0.103482095 | 0.131381444 |
| Teddm1b | 1.369856 | 1.33581 | 1.3874 | -0.051625663 | -0.017580005 | 1.3452065 | 1.3199448 | 1.42348 | -0.103533071 | -0.078271352 |
| Pld1 | 6.442506 | 6.0737 | 6.3383 | -0.264587697 | 0.104219263 | 6.2746628 | 6.1736417 | 6.27721 | -0.103569149 | -0.002548085 |
| Gm49331 | 3.676907 | 3.610346 | 3.7035 | -0.093163614 | -0.026602396 | 3.6351289 | 3.6407779 | 3.74443 | -0.103649452 | -0.109298451 |
| Cspg4 | 2.880587 | 2.97871 | 2.9709 | 0.007827937 | -0.090295195 | 3.0971926 | 2.8767045 | 2.98038 | -0.10367465 | 0.116813424 |
| Prss57 | 1.581223 | 1.578978 | 1.6112 | -0.032261287 | -0.030016466 | 1.6183147 | 1.577577 | 1.68129 | -0.103711394 | -0.062973727 |
| Ap3b1 | 10.73941 | 10.72015 | 10.806 | -0.085856506 | -0.066603924 | 10.775452 | 10.674269 | 10.778 | -0.103740594 | -0.002557867 |
| D3Ertd254e | 7.139229 | 7.245358 | 7.2337 | 0.011681213 | -0.094447756 | 6.905337 | 7.0810101 | 7.18475 | -0.10374396 | -0.279417061 |
| B9d1 | 3.26227 | 3.318364 | 3.3744 | -0.056006134 | -0.112099848 | 3.3187489 | 3.3340746 | 3.43794 | -0.103865098 | -0.119190805 |
| Degs1 | 10.02684 | 10.03214 | 10.047 | -0.015354552 | -0.020651613 | 9.9430078 | 9.7516339 | 9.85553 | -0.103891216 | 0.087482706 |
| Slc13a3 | 3.058446 | 3.173408 | 3.0683 | 0.105133725 | -0.009828378 | 3.6927491 | 3.1302311 | 3.2342 | -0.103965946 | 0.458552065 |
| Tom1l1 | 2.84089 | 2.647727 | 2.812 | -0.164318356 | 0.028844721 | 2.6903353 | 2.6060788 | 2.71005 | -0.103972537 | -0.019715992 |
| Rubcnl | 10.32646 | 10.26879 | 10.424 | -0.155288816 | -0.097615616 | 10.721226 | 10.367899 | 10.4719 | -0.10400965 | 0.249317556 |
| Abhd12b | 0.321847 | 0.332376 | 0.282 | 0.050403843 | 0.039874897 | 0.2927572 | 0.2804653 | 0.38456 | -0.104096926 | -0.091805035 |
| Ipo13 | 9.36053 | 9.200049 | 9.3244 | -0.124370573 | 0.036110811 | 9.4763307 | 9.3324236 | 9.43661 | -0.104185248 | 0.039721819 |
| Pcdhga7 | 2.790275 | 2.737655 | 2.7471 | -0.009472031 | 0.04314778 | 2.8097469 | 2.6256435 | 2.72983 | -0.104189503 | 0.079913895 |
| Ube2r2 | 10.51422 | 10.39226 | 10.526 | -0.13329263 | -0.01133567 | 10.423162 | 10.372254 | 10.4764 | -0.10419555 | -0.05328765 |
| Zfp652 | 10.68764 | 10.72681 | 10.754 | -0.027382982 | -0.066547748 | 10.65429 | 10.542546 | 10.6468 | -0.104259492 | 0.007484536 |
| Cdkn2aip | 9.021813 | 8.936178 | 9.1198 | -0.183623526 | -0.09798878 | 8.9027236 | 8.8799004 | 8.98417 | -0.104272202 | -0.081449075 |
| Sucnr1 | 3.090065 | 3.087241 | 2.9752 | 0.11208779 | 0.114911671 | 3.1185183 | 2.8954605 | 2.99973 | -0.104273756 | 0.118784001 |
| Mkks | 7.415761 | 7.120398 | 7.1688 | -0.048439099 | 0.246923994 | 6.8398447 | 7.3239099 | 7.4282 | -0.104288408 | -0.588353617 |
| Aunip | 3.674012 | 3.444865 | 3.7023 | -0.257470976 | -0.028324162 | 3.7468267 | 3.704911 | 3.80925 | -0.104339036 | -0.062423409 |
| Ccdc40 | 3.622147 | 3.61623 | 3.7094 | -0.093177912 | -0.087260987 | 3.7811577 | 3.6286241 | 3.73297 | -0.104346946 | 0.048186587 |
| Mboat2 | 1.427513 | 1.462397 | 1.4214 | 0.040963477 | 0.00607892 | 1.4551319 | 1.3726347 | 1.47707 | -0.104433132 | -0.021935849 |
| Olfr524 | 1.285888 | 1.133502 | 1.205 | -0.071484803 | 0.08090082 | 1.1736822 | 1.1275475 | 1.23205 | -0.104501425 | -0.058366724 |
| Hist1h1e | 5.807325 | 5.630419 | 5.8245 | -0.194051534 | -0.017145428 | 6.1884462 | 5.88436 | 5.9889 | -0.104535372 | 0.199550827 |
| Erich5 | 1.942047 | 1.925285 | 1.8607 | 0.064563546 | 0.081325644 | 1.7884547 | 1.842357 | 1.9469 | -0.104543749 | -0.15844611 |
| Sh2d3c | 10.53469 | 10.68013 | 10.644 | 0.036026526 | -0.109413956 | 10.6249 | 10.430017 | 10.5346 | -0.104597958 | 0.090285233 |
| Nap1l3 | 2.887559 | 2.882449 | 3.0972 | -0.214732769 | -0.209622929 | 2.8535241 | 2.8779396 | 2.98263 | -0.104689018 | -0.129104484 |
| Trav8n-2 | 4.335972 | 4.329736 | 4.2227 | 0.107060879 | 0.11329698 | 4.1497746 | 4.126572 | 4.23149 | -0.104916474 | -0.081713922 |

| | | | | | | | | | | |
|---|---|---|---|---|---|---|---|---|---|---|
| Atad2 | 8.899534 | 8.840258 | 9.0153 | -0.175088937 | -0.115812129 | 8.8687851 | 8.9473064 | 9.05228 | -0.104975694 | -0.183496976 |
| Dok6 | 2.891572 | 2.878307 | 2.8453 | 0.033043014 | 0.046308597 | 2.8675481 | 2.8751401 | 2.98013 | -0.104985429 | -0.11257744 |
| Trappc10 | 10.59877 | 10.64229 | 10.66 | -0.0176222 | -0.06113249 | 10.612721 | 10.529676 | 10.6347 | -0.105007958 | -0.021962963 |
| Ubald1 | 9.789249 | 9.828041 | 9.8036 | 0.024473029 | -0.014319203 | 10.047813 | 9.6618848 | 9.7669 | -0.105017382 | 0.280910647 |
| 2810006K23Rik | 7.219478 | 7.247264 | 7.369 | -0.121747518 | -0.149533779 | 7.5000027 | 7.2940786 | 7.39914 | -0.105064263 | 0.100859844 |
| Ly96 | 6.29338 | 6.262395 | 6.4297 | -0.167291178 | -0.136306054 | 6.1848461 | 6.0861238 | 6.19121 | -0.10508699 | -0.006364617 |
| Gm6733 | 3.035461 | 3.005458 | 2.9916 | 0.013907119 | 0.043909607 | 3.1733766 | 3.1326781 | 3.23777 | -0.105092049 | -0.064393564 |
| Rhot1 | 9.530382 | 9.582298 | 9.5659 | 0.016444053 | -0.035471509 | 9.5281124 | 9.4877586 | 9.59291 | -0.105146937 | -0.064793104 |
| Kcnk3 | 2.73774 | 2.75714 | 2.931 | -0.173898317 | -0.193297751 | 2.8260187 | 2.7340103 | 2.83917 | -0.105157261 | -0.01314888 |
| Castor2 | 8.381733 | 8.552765 | 8.5991 | -0.046303742 | -0.217335826 | 8.553937 | 8.5608933 | 8.66611 | -0.105217387 | -0.112173644 |
| Fanci | 7.480796 | 7.364779 | 7.3646 | 0.000201413 | 0.116218764 | 7.4595367 | 7.7600694 | 7.86537 | -0.10529768 | -0.405830393 |
| Cfap53 | 2.564009 | 2.491141 | 2.4793 | 0.011853015 | 0.084720335 | 2.4322437 | 2.4462936 | 2.55163 | -0.105332383 | -0.119382259 |
| Cdc40 | 10.3464 | 10.34378 | 10.378 | -0.033744316 | -0.0311178 | 10.469775 | 10.333702 | 10.439 | -0.10534489 | 0.030727297 |
| Tmem88b | 2.758213 | 2.762072 | 2.7942 | -0.032106239 | -0.035965317 | 2.786327 | 2.7806009 | 2.88596 | -0.10536154 | -0.09963539 |
| Pex1 | 8.001359 | 7.738443 | 8.1202 | -0.381735245 | -0.118819134 | 8.1978882 | 8.2383194 | 8.34373 | -0.105406537 | -0.14583773 |
| Gata4 | 2.63484 | 2.535101 | 2.6257 | -0.09060419 | 0.009134806 | 2.4778614 | 2.3870064 | 2.49242 | -0.105412778 | -0.014557749 |
| Zfp422 | 8.800385 | 8.759764 | 8.751 | 0.008800571 | 0.049420992 | 8.8446496 | 8.67979 | 8.78528 | -0.105494701 | 0.059364922 |
| Vmn1r-ps47 | 2.17858 | 2.167341 | 2.2663 | -0.099002902 | -0.0877646 | 2.1769426 | 2.2139296 | 2.31943 | -0.105497455 | -0.142484505 |
| Gm48260 | 3.55419 | 3.639867 | 3.5362 | 0.103621277 | 0.017944515 | 3.7725161 | 3.7367796 | 3.8423 | -0.105522716 | -0.069786211 |
| Morc2b | 2.179951 | 2.202399 | 2.2303 | -0.027887625 | -0.050335995 | 2.2672838 | 2.2255844 | 2.33112 | -0.105537172 | -0.063837808 |
| Magix | 1.632156 | 1.509484 | 1.6231 | -0.113616901 | 0.009055698 | 1.5013765 | 1.5109779 | 1.61652 | -0.105544126 | -0.115145435 |
| Mgam2-ps | 0.893937 | 0.884854 | 0.8855 | -0.000685214 | 0.008398044 | 0.8262849 | 0.8239597 | 0.92963 | -0.105670619 | -0.103345476 |
| Cntnap2 | 1.098593 | 1.084318 | 1.0357 | 0.048602656 | 0.062877299 | 1.0262176 | 1.0110859 | 1.11676 | -0.105675161 | -0.090543416 |
| Slc46a3 | 8.653887 | 8.645347 | 8.9353 | -0.289998095 | -0.281457974 | 8.927125 | 8.8629078 | 8.9686 | -0.105694298 | -0.041477061 |
| Pth1r | 1.735596 | 1.730948 | 1.7383 | -0.007390288 | -0.002742377 | 1.7218712 | 1.6552944 | 1.76103 | -0.105740125 | -0.039163354 |
| Apoc4 | 3.63926 | 3.44919 | 3.4585 | -0.009277394 | 0.180791698 | 3.4497999 | 3.5103873 | 3.61613 | -0.105745833 | -0.166333258 |
| Gpr21 | 2.674249 | 2.770844 | 2.7452 | 0.02564473 | -0.070950641 | 2.6848232 | 2.7245565 | 2.83042 | -0.105863554 | -0.145596849 |
| Nup210l | 5.414777 | 5.523368 | 5.4 | 0.123356248 | 0.014764869 | 5.8521617 | 5.6363051 | 5.7422 | -0.105892569 | 0.109964015 |
| Olfr380 | 3.447444 | 3.402922 | 3.424 | -0.021120962 | 0.023401076 | 3.3658871 | 3.4666512 | 3.57258 | -0.105925426 | -0.2066895 |
| Khdrbs1 | 12.13292 | 11.9567 | 12.106 | -0.148999994 | 0.027221572 | 12.192185 | 12.009406 | 12.1154 | -0.105999 | 0.076780521 |
| Ica1l | 2.46707 | 2.458737 | 2.4493 | 0.009429783 | 0.017762338 | 2.3196809 | 2.3916463 | 2.49765 | -0.106003863 | -0.177969269 |
| Simc1 | 8.146996 | 8.233822 | 8.2438 | -0.010013247 | -0.096839489 | 8.2524763 | 8.2595011 | 8.36562 | -0.106117759 | -0.113142536 |

| | | | | | | | | | | |
|---|---|---|---|---|---|---|---|---|---|---|
| Grid1 | 1.333037 | 1.360353 | 1.3631 | -0.002757629 | -0.030074543 | 1.3520715 | 1.3475653 | 1.4537 | -0.106136358 | -0.101630114 |
| Nphp1 | 4.626275 | 4.376729 | 4.4673 | -0.090593929 | 0.158951349 | 4.3792282 | 4.3782916 | 4.48446 | -0.106163678 | -0.105227052 |
| Olfr11 | 1.762793 | 1.689059 | 1.6985 | -0.009420967 | 0.064313049 | 1.6778633 | 1.6105157 | 1.71669 | -0.106170586 | -0.038822968 |
| Tmem186 | 8.461065 | 8.42184 | 8.4327 | -0.010909037 | 0.028315669 | 8.5082181 | 8.3608707 | 8.46708 | -0.106205187 | 0.041142256 |
| Unkl | 9.203286 | 9.42281 | 9.4789 | -0.056094747 | -0.275619198 | 9.4003042 | 9.3870758 | 9.49345 | -0.106377745 | -0.093149432 |
| Trav7-1 | 2.97617 | 3.039262 | 3.0465 | -0.007220027 | -0.070311731 | 2.9769723 | 2.9068018 | 3.01321 | -0.106405745 | -0.036235319 |
| Trav8-2 | 2.908693 | 2.839486 | 2.8699 | -0.030425478 | 0.038782173 | 2.962411 | 2.8415817 | 2.94804 | -0.106461667 | 0.014367646 |
| Adamtsl1 | 3.522866 | 3.233328 | 3.33 | -0.096702241 | 0.19283651 | 3.2727363 | 3.338246 | 3.44479 | -0.106539985 | -0.172049724 |
| Brd1 | 10.78604 | 10.82121 | 10.746 | 0.074956058 | 0.039791746 | 10.556445 | 10.631683 | 10.7383 | -0.106620752 | -0.181859193 |
| Wfdc18 | 3.720544 | 3.469959 | 3.5974 | -0.127471445 | 0.123113553 | 3.9558352 | 3.3992705 | 3.50593 | -0.106661451 | 0.449903192 |
| Agps | 10.45432 | 10.60905 | 10.407 | 0.201640046 | 0.046900878 | 10.298681 | 10.298579 | 10.4053 | -0.106688285 | -0.106585996 |
| Gucy2f | 1.032149 | 0.97326 | 0.9694 | 0.003861287 | 0.062750477 | 0.9955015 | 0.9450303 | 1.05177 | -0.106738206 | -0.056267005 |
| Pcyt1a | 8.70032 | 8.831294 | 8.7493 | 0.081976013 | -0.048998258 | 8.6852615 | 8.5916182 | 8.69839 | -0.106772321 | -0.013128942 |
| Trarg1 | 3.53688 | 3.518986 | 3.6551 | -0.136128448 | -0.118234489 | 3.5835063 | 3.4481402 | 3.555 | -0.106863987 | 0.028502121 |
| E2f6 | 7.842332 | 7.84261 | 7.7807 | 0.061915169 | 0.061636871 | 7.8194167 | 7.8023294 | 7.90922 | -0.106889562 | -0.089802199 |
| AI661453 | 3.391115 | 3.352168 | 3.433 | -0.080808116 | -0.041861096 | 3.3912562 | 3.2190785 | 3.326 | -0.106924234 | 0.0652535 |
| Pwwp2a | 8.956474 | 9.052656 | 8.8735 | 0.179113447 | 0.082931813 | 8.9498157 | 9.0209208 | 9.12785 | -0.106927481 | -0.178032648 |
| Lrrc4c | 2.784426 | 2.587851 | 2.6006 | -0.012750625 | 0.183823792 | 2.5324973 | 2.291617 | 2.39858 | -0.106965199 | 0.133915082 |
| Nup62cl | 0.229414 | 0.157545 | 0.2132 | -0.055681761 | 0.016187016 | 0.1852313 | 0.1538404 | 0.26084 | -0.106995748 | -0.075604804 |
| Gm10709 | 2.477474 | 2.392502 | 2.4254 | -0.032858412 | 0.052114069 | 2.3890292 | 2.4078532 | 2.51489 | -0.107031968 | -0.125855949 |
| Igkv4-59 | 7.744991 | 8.777802 | 7.7395 | 1.038298609 | 0.005487655 | 8.3964205 | 8.3785056 | 8.48567 | -0.107166334 | -0.089251413 |
| Micu2 | 10.11106 | 10.01505 | 10.128 | -0.112564682 | -0.016550769 | 10.130274 | 10.043661 | 10.1508 | -0.107176677 | -0.020563779 |
| Gm12191 | 3.155064 | 3.227218 | 3.4377 | -0.210468062 | -0.282621451 | 3.1483799 | 3.1880925 | 3.29531 | -0.107217123 | -0.146929681 |
| Cul7 | 7.991461 | 7.889181 | 8.0962 | -0.207032115 | -0.104751838 | 7.9645657 | 7.6539737 | 7.76122 | -0.107245088 | 0.203346977 |
| Dnm2 | 11.82988 | 11.86517 | 11.891 | -0.025793966 | -0.061084678 | 11.934497 | 11.869457 | 11.9767 | -0.1072708 | -0.042230162 |
| Adam8 | 9.475896 | 9.495073 | 9.0924 | 0.402703047 | 0.383525887 | 8.7389052 | 9.205046 | 9.31235 | -0.107309009 | -0.573449767 |
| Gm49358 | 3.763159 | 3.734126 | 3.8561 | -0.12192854 | -0.092895346 | 3.3631577 | 3.6866389 | 3.79395 | -0.107309937 | -0.430791131 |
| Samd11 | 2.837666 | 2.781486 | 2.7949 | -0.013461264 | 0.042719548 | 2.8232244 | 2.7936487 | 2.90097 | -0.107318077 | -0.077742339 |
| Zfp873 | 6.453549 | 6.544477 | 6.5057 | 0.038783938 | -0.052143469 | 6.5775222 | 6.5046175 | 6.61206 | -0.107439799 | -0.034535121 |
| Tpd52l1 | 1.723804 | 1.67753 | 1.6677 | 0.00980098 | 0.056075853 | 1.6367861 | 1.5272946 | 1.63475 | -0.107459844 | 0.002031684 |
| Lcorl | 8.502661 | 8.565194 | 8.616 | -0.050829437 | -0.113362572 | 8.2986009 | 8.5385272 | 8.64599 | -0.107463281 | -0.347389507 |
| Olfr9 | 1.951174 | 1.88055 | 1.893 | -0.012401547 | 0.058222438 | 2.0023496 | 1.8783817 | 1.98586 | -0.107479053 | 0.01648887 |

| | | | | | | | | | | |
|---|---|---|---|---|---|---|---|---|---|---|
| Pparg | 4.582229 | 4.496126 | 4.5276 | -0.031445588 | 0.054656985 | 4.2062234 | 4.0568266 | 4.16432 | -0.107490924 | 0.041905847 |
| Slc33a1 | 8.599799 | 8.321646 | 8.4984 | -0.176721771 | 0.101431135 | 8.3913085 | 8.3986053 | 8.5061 | -0.10749093 | -0.114787683 |
| Orc2 | 9.240998 | 9.3045 | 9.2021 | 0.102371433 | 0.038869087 | 9.1934694 | 9.1508653 | 9.25837 | -0.107509313 | -0.06490526 |
| Gm3411 | 1.124615 | 1.078174 | 1.1097 | -0.031532766 | 0.014908238 | 1.0280467 | 1.089803 | 1.19732 | -0.107517853 | -0.169274123 |
| Exoc1 | 9.430841 | 9.408603 | 9.6637 | -0.2551419 | -0.232903642 | 9.914443 | 9.8296168 | 9.93716 | -0.107538629 | -0.022712449 |
| Olfr1198 | 1.923046 | 1.962899 | 1.9577 | 0.005164325 | -0.034688585 | 2.0109991 | 1.9306926 | 2.03836 | -0.107664032 | -0.027357476 |
| Gm48132 | 6.952093 | 6.975859 | 6.6774 | 0.298427483 | 0.274661438 | 7.3617144 | 7.343023 | 7.45077 | -0.107751647 | -0.089060251 |
| Sumf1 | 8.99595 | 8.986191 | 8.9604 | 0.025781716 | 0.035540868 | 9.0336161 | 8.846264 | 8.95403 | -0.107766261 | 0.079585818 |
| Sox6 | 1.243794 | 1.146249 | 1.2352 | -0.088926546 | 0.008618323 | 1.1367896 | 1.105546 | 1.21334 | -0.107791038 | -0.076547454 |
| Anxa6 | 12.78292 | 12.79014 | 12.808 | -0.01817911 | -0.02539944 | 12.72969 | 12.723125 | 12.831 | -0.10782743 | -0.10126239 |
| Rrp8 | 8.85665 | 9.008634 | 8.9106 | 0.098076995 | -0.053907168 | 9.0845763 | 8.8645852 | 8.97241 | -0.107829698 | 0.112161352 |
| Rnf2 | 10.40321 | 10.4347 | 10.471 | -0.036433076 | -0.06792614 | 10.410839 | 10.240565 | 10.3485 | -0.107904174 | 0.062370038 |
| Lpar2 | 8.226751 | 8.249594 | 8.3622 | -0.112624084 | -0.135466947 | 8.5747208 | 8.1969366 | 8.30485 | -0.107911446 | 0.269872671 |
| Cdkn2aipnl | 9.951149 | 9.996058 | 10.017 | -0.020748132 | -0.065657133 | 9.9487161 | 9.7630813 | 9.87115 | -0.108069363 | 0.077565395 |
| Pias4 | 8.80545 | 8.85149 | 8.8865 | -0.035037635 | -0.0810774 | 9.2751563 | 8.9020639 | 9.01014 | -0.108077471 | 0.265014884 |
| Vmn1r229 | 5.631138 | 5.509387 | 5.495 | 0.014388247 | 0.136138755 | 5.4505703 | 5.747356 | 5.85566 | -0.108299077 | -0.405084758 |
| Vamp4 | 9.11638 | 9.167804 | 9.2027 | -0.034856689 | -0.086280377 | 9.1155197 | 9.0120714 | 9.12038 | -0.108312816 | -0.004864536 |
| Snx16 | 6.363292 | 6.335563 | 6.3245 | 0.011110173 | 0.038839412 | 6.2876655 | 6.1932297 | 6.30161 | -0.108378074 | -0.013942265 |
| Malt1 | 12.04727 | 12.1462 | 12.071 | 0.075207154 | -0.023717772 | 11.974929 | 12.05066 | 12.1591 | -0.108488636 | -0.184220056 |
| Snx24 | 3.687048 | 3.475792 | 3.5374 | -0.061602332 | 0.149653856 | 3.459773 | 3.3288246 | 3.43735 | -0.108529288 | 0.022419115 |
| Rnf169 | 9.484809 | 9.622128 | 9.6049 | 0.0171793 | -0.120139978 | 9.3745258 | 9.5854326 | 9.69396 | -0.108529354 | -0.3194362 |
| Rc3h2 | 10.09477 | 10.02027 | 10.134 | -0.114097777 | -0.03959612 | 10.015114 | 10.212783 | 10.3213 | -0.108542932 | -0.30621193 |
| Man2c1 | 9.99102 | 10.04216 | 10.079 | -0.036717482 | -0.087853637 | 10.43587 | 10.220703 | 10.3293 | -0.108610035 | 0.106557202 |
| Plpp1 | 6.450779 | 6.408432 | 6.1637 | 0.244775537 | 0.287122533 | 6.2236571 | 5.9260185 | 6.03467 | -0.108648895 | 0.188989671 |
| Gas2 | 4.832367 | 4.831791 | 4.8849 | -0.053153874 | -0.052578362 | 4.9623675 | 4.6975471 | 4.80627 | -0.108726194 | 0.156094238 |
| Pikfyve | 10.33935 | 10.39768 | 10.35 | 0.047943253 | -0.010377964 | 10.372557 | 10.36014 | 10.4689 | -0.108763804 | -0.096346579 |
| Rexo5 | 7.160091 | 6.924632 | 7.3694 | -0.444743228 | -0.209284157 | 7.0598335 | 7.263425 | 7.3722 | -0.108777835 | -0.312369403 |
| Ighv5-2 | 5.063914 | 4.945322 | 4.9892 | -0.043854668 | 0.074737033 | 5.1926461 | 5.120355 | 5.22915 | -0.10879819 | -0.036507064 |
| Fam180a | 2.648452 | 2.581323 | 2.7833 | -0.20196273 | -0.134834393 | 2.324459 | 2.3020829 | 2.41088 | -0.108801538 | -0.086425412 |
| Traj28 | 2.131286 | 2.136811 | 2.0582 | 0.07857397 | 0.073049151 | 2.0678211 | 2.074788 | 2.18359 | -0.108806591 | -0.115773466 |
| Dpf3 | 2.217111 | 2.156489 | 2.1224 | 0.034061912 | 0.094683987 | 2.2313709 | 2.1799472 | 2.28883 | -0.108881635 | -0.057457984 |
| Tmem266 | 1.628856 | 1.606802 | 1.6419 | -0.035064251 | -0.01301032 | 1.5964326 | 1.5448373 | 1.65381 | -0.108968938 | -0.05737372 |

| | | | | | | | | | | |
|---|---|---|---|---|---|---|---|---|---|---|
| Acvr1c | 1.891926 | 1.995142 | 1.9409 | 0.054282185 | -0.048934566 | 1.8745286 | 1.9187299 | 2.02778 | -0.109053774 | -0.153255087 |
| Col6a4 | 6.471554 | 6.37969 | 6.4095 | -0.029837483 | 0.062026635 | 5.8492234 | 5.7734219 | 5.88249 | -0.109070857 | -0.033269357 |
| Megf8 | 7.020823 | 6.754472 | 7.0434 | -0.288964619 | -0.022613628 | 6.6259283 | 6.7609012 | 6.87003 | -0.109131711 | -0.244104607 |
| Apol7b | 3.64439 | 3.626722 | 3.6858 | -0.059036515 | -0.041368665 | 3.634286 | 3.5471668 | 3.65637 | -0.109199216 | -0.022079988 |
| Amfr | 11.1865 | 11.21892 | 11.234 | -0.014602854 | -0.047027774 | 11.209568 | 11.000946 | 11.1102 | -0.109211592 | 0.099410655 |
| Wdr81 | 9.941557 | 9.884077 | 10.034 | -0.149680865 | -0.092201221 | 10.261582 | 9.8515182 | 9.96086 | -0.109339117 | 0.300724522 |
| Npepps | 10.20014 | 10.26683 | 10.247 | 0.01949476 | -0.047193552 | 10.079753 | 10.186339 | 10.2957 | -0.109344127 | -0.215929323 |
| Slc22a2 | 1.760567 | 1.749645 | 1.869 | -0.119373781 | -0.108451839 | 1.7182084 | 1.7900587 | 1.89947 | -0.109408945 | -0.1812592 |
| Zfp146 | 9.365028 | 9.358764 | 9.396 | -0.037254145 | -0.030989619 | 9.5101237 | 9.4000218 | 9.50945 | -0.109425376 | 0.000676484 |
| Ppef1 | 1.328082 | 1.318781 | 1.2874 | 0.031399788 | 0.04070147 | 1.4004282 | 1.3132345 | 1.42272 | -0.109483732 | -0.02229011 |
| Lin52 | 7.897237 | 8.044205 | 8.0162 | 0.027979031 | -0.118989086 | 7.8519182 | 7.9026823 | 8.01218 | -0.109494122 | -0.160258258 |
| Cdk14 | 6.862077 | 6.915054 | 6.8267 | 0.088380467 | 0.03540397 | 6.5806599 | 6.734287 | 6.84402 | -0.10973731 | -0.263364397 |
| Spaca9 | 2.847916 | 2.837526 | 2.9675 | -0.129960588 | -0.119570051 | 3.0024558 | 2.8673798 | 2.97722 | -0.109838764 | 0.025237159 |
| Dixdc1 | 3.109246 | 3.194566 | 3.2464 | -0.051798531 | -0.137118532 | 3.2572943 | 2.9865413 | 3.09642 | -0.109880079 | 0.16087297 |
| Tmem205 | 4.839768 | 4.603855 | 4.7705 | -0.166632183 | 0.069280493 | 4.5922856 | 4.5405169 | 4.65049 | -0.109971872 | -0.058203143 |
| Gm4779 | 1.596111 | 1.650333 | 1.5536 | 0.096736449 | 0.042515199 | 1.7547035 | 1.569508 | 1.67953 | -0.110024367 | 0.075171063 |
| Rtbdn | 1.814306 | 1.824685 | 1.8738 | -0.04912773 | -0.059506707 | 1.904279 | 1.8310271 | 1.9411 | -0.110070549 | -0.036818622 |
| Vps4b | 10.82909 | 10.78289 | 10.848 | -0.065141202 | -0.018945684 | 10.747276 | 10.740558 | 10.8507 | -0.110113402 | -0.103395895 |
| Ccdc159 | 1.71252 | 1.675785 | 1.6403 | 0.035442012 | 0.072176914 | 1.5990977 | 1.5529848 | 1.66311 | -0.110125356 | -0.064012517 |
| Ice2 | 8.848308 | 8.992776 | 9.1505 | -0.157711485 | -0.30217978 | 9.1727793 | 9.1895861 | 9.29972 | -0.110131885 | -0.12693867 |
| Pik3r2 | 7.150897 | 6.957201 | 7.2048 | -0.247555139 | -0.053859052 | 7.3193384 | 7.1007011 | 7.21084 | -0.110142377 | 0.108494992 |
| Serpinc1 | 4.981358 | 5.094577 | 5.0361 | 0.058510398 | -0.054708239 | 4.7660407 | 4.8634879 | 4.97365 | -0.110160518 | -0.207607701 |
| Mgat4b | 7.534665 | 7.431658 | 7.4781 | -0.046467573 | 0.056539853 | 7.2541999 | 7.3325749 | 7.44274 | -0.110165166 | -0.188540185 |
| Cnnm4 | 8.287294 | 8.360732 | 8.3343 | 0.026453572 | -0.046983898 | 8.131904 | 8.1164979 | 8.22667 | -0.110171509 | -0.094765408 |
| Aasdh | 7.761666 | 7.582702 | 7.9029 | -0.320204317 | -0.141240091 | 7.969985 | 8.0115539 | 8.12175 | -0.110192936 | -0.151761816 |
| Kcnh1 | 4.182106 | 3.906028 | 4.0996 | -0.193585462 | 0.0824922 | 3.6353476 | 3.4046663 | 3.51488 | -0.110216213 | 0.120465061 |
| Tmod3 | 11.98497 | 12.04492 | 12.044 | 0.000831964 | -0.059124332 | 12.093075 | 11.942809 | 12.0531 | -0.110243012 | 0.040023627 |
| Mmp17 | 4.404323 | 4.475334 | 4.3536 | 0.12178308 | 0.05077232 | 4.3405064 | 4.2801593 | 4.39042 | -0.11026001 | -0.049912899 |
| Cercam | 5.642756 | 5.747054 | 5.5783 | 0.168758919 | 0.064461127 | 5.6003978 | 5.6652463 | 5.77553 | -0.110284585 | -0.175133065 |
| Zhx1 | 9.179576 | 9.142704 | 9.2095 | -0.066805753 | -0.029933627 | 9.1660288 | 9.0390793 | 9.14942 | -0.110343195 | 0.016606344 |
| Gm10221 | 3.949693 | 3.366761 | 5.0544 | -1.687679661 | -1.104747787 | 3.6036965 | 3.7618105 | 3.87216 | -0.110352628 | -0.268466592 |
| Lmcd1 | 4.359657 | 4.093904 | 4.1515 | -0.057577458 | 0.208175299 | 3.9517424 | 4.0247126 | 4.13514 | -0.110431589 | -0.183401788 |

| | | | | | | | | | | |
|---|---|---|---|---|---|---|---|---|---|---|
| Olfr1484 | 2.454779 | 2.478364 | 2.4792 | -0.000867116 | -0.024451849 | 2.4610072 | 2.4599155 | 2.57036 | -0.110443054 | -0.109351375 |
| Htra2 | 9.109708 | 9.135981 | 9.3112 | -0.175245077 | -0.20151822 | 9.4318161 | 9.2450828 | 9.35553 | -0.110447894 | 0.076285422 |
| Mgat5 | 9.383101 | 9.460386 | 9.3607 | 0.099716202 | 0.022431225 | 9.3357876 | 9.2992601 | 9.40972 | -0.110462835 | -0.073935313 |
| Phc1 | 8.890787 | 8.784515 | 8.9046 | -0.12003632 | -0.013764344 | 8.86911 | 8.6776372 | 8.78815 | -0.110510716 | 0.080962059 |
| Gm3336 | 0.911726 | 0.892255 | 0.9404 | -0.048164478 | -0.028693782 | 0.9881134 | 0.8717058 | 0.98225 | -0.110544223 | 0.005863402 |
| Asb4 | 3.422496 | 3.394723 | 3.3728 | 0.021888021 | 0.049660212 | 3.1682842 | 3.1456618 | 3.25621 | -0.110544905 | -0.087922463 |
| Limk1 | 8.897079 | 8.839712 | 8.9446 | -0.104903989 | -0.047537814 | 8.8713436 | 8.5821156 | 8.69268 | -0.110564205 | 0.178663735 |
| Fhl4 | 2.277659 | 2.280763 | 2.2961 | -0.015318126 | -0.01842216 | 2.3191494 | 2.2802201 | 2.39079 | -0.110567647 | -0.071638286 |
| Kcnj5 | 2.018913 | 1.980288 | 2.0043 | -0.024010225 | 0.0146144 | 2.1165845 | 2.0111074 | 2.12169 | -0.110582538 | -0.00510545 |
| Traj31 | 2.657774 | 2.693899 | 2.6112 | 0.082654869 | 0.046529275 | 2.5886821 | 2.5636855 | 2.67436 | -0.110676086 | -0.085679469 |
| Astn2 | 3.130714 | 3.056029 | 3.1026 | -0.046585584 | 0.028098941 | 2.903064 | 2.9201044 | 3.03088 | -0.110778733 | -0.127819113 |
| Ing5 | 9.660928 | 9.57655 | 9.5833 | -0.006715134 | 0.077662476 | 9.6493024 | 9.5637031 | 9.67452 | -0.110821716 | -0.02522246 |
| Gpatch8 | 10.34295 | 10.28364 | 10.373 | -0.089844458 | -0.030538296 | 10.29634 | 10.328563 | 10.4395 | -0.11093388 | -0.14315669 |
| Ap3s1 | 8.871833 | 8.753031 | 8.7975 | -0.044466578 | 0.074335673 | 8.5186713 | 8.5655814 | 8.67652 | -0.110941717 | -0.157851844 |
| Chrdl1 | 6.065536 | 5.826601 | 6.0144 | -0.187756818 | 0.051177778 | 5.3607983 | 5.5268397 | 5.63782 | -0.110983672 | -0.27702506 |
| 1700019D03Rik | 3.968687 | 3.897726 | 3.9723 | -0.074538277 | -0.003578117 | 3.7430746 | 3.8114169 | 3.92259 | -0.111168376 | -0.179510706 |
| Ctsd | 12.04904 | 12.16731 | 12.011 | 0.155956122 | 0.037689838 | 11.884184 | 11.720727 | 11.8319 | -0.111209112 | 0.052248132 |
| Cul1 | 10.75347 | 10.66511 | 10.793 | -0.128164486 | -0.039808424 | 10.688544 | 10.705443 | 10.8167 | -0.111223022 | -0.128122206 |
| Gal | 1.698512 | 1.767087 | 1.7654 | 0.001674099 | -0.066900763 | 1.7820124 | 1.7050818 | 1.81632 | -0.11123956 | -0.034309 |
| Gdi2 | 12.62983 | 12.58121 | 12.611 | -0.029780266 | 0.018840406 | 12.626977 | 12.492158 | 12.6034 | -0.111281028 | 0.023537398 |
| Igip | 7.639786 | 7.733594 | 7.6906 | 0.04304237 | -0.050765639 | 7.8643204 | 7.8163364 | 7.92763 | -0.111295209 | -0.063311179 |
| Nbas | 9.555828 | 9.609191 | 9.4723 | 0.136855821 | 0.083493043 | 9.6728722 | 9.5216472 | 9.63296 | -0.111310727 | 0.039914347 |
| Sytl4 | 3.293686 | 3.208348 | 3.3466 | -0.138203826 | -0.052865406 | 3.3011229 | 3.2029358 | 3.31428 | -0.111347077 | -0.01315998 |
| C1galt1 | 10.96261 | 11.09503 | 11.047 | 0.04822794 | -0.084196946 | 10.946168 | 10.805351 | 10.9167 | -0.111376566 | 0.029439957 |
| Srek1 | 9.897584 | 9.818166 | 9.9808 | -0.162660249 | -0.083242465 | 10.01882 | 10.003785 | 10.1152 | -0.111390522 | -0.096355382 |
| Mroh7 | 0.60073 | 0.567715 | 0.6275 | -0.059808157 | -0.026793529 | 0.6241073 | 0.5576226 | 0.66903 | -0.11140764 | -0.044922951 |
| Elf2 | 10.76645 | 10.70855 | 10.77 | -0.06179107 | -0.003892528 | 10.596914 | 10.579536 | 10.691 | -0.111448466 | -0.094070341 |
| Acad8 | 7.229012 | 7.170415 | 7.3106 | -0.140139234 | -0.081542644 | 7.1539965 | 7.3178558 | 7.42932 | -0.111461556 | -0.275320859 |
| Ankhd1 | 11.6492 | 11.5133 | 11.524 | -0.011054944 | 0.124848844 | 11.479196 | 11.45581 | 11.5674 | -0.11154958 | -0.088163377 |
| Psen2 | 9.802796 | 9.79325 | 9.8548 | -0.061572745 | -0.052026231 | 10.077325 | 9.8550145 | 9.96661 | -0.111590877 | 0.110719221 |
| Mylpf | 5.863674 | 5.954563 | 6.0034 | -0.048886754 | -0.139775808 | 6.0507867 | 5.8537485 | 5.96535 | -0.11160367 | 0.085434569 |
| Lrp3 | 3.579802 | 3.567533 | 3.4788 | 0.088742014 | 0.101011042 | 3.361327 | 3.2184432 | 3.33007 | -0.111631115 | 0.031252751 |

| | | | | | | | | | | |
|---|---|---|---|---|---|---|---|---|---|---|
| Lrrc26 | 1.905508 | 1.698233 | 1.7785 | -0.080261227 | 0.127014189 | 1.8587622 | 1.7802577 | 1.8919 | -0.111644409 | -0.03313991 |
| Cldn13 | 2.642315 | 2.426309 | 2.742 | -0.315723374 | -0.099716831 | 2.4086121 | 2.3727794 | 2.4845 | -0.11172548 | -0.075892776 |
| Samd1 | 8.605834 | 8.726657 | 8.7906 | -0.063906369 | -0.184728639 | 8.823321 | 8.6614244 | 8.77316 | -0.111732326 | 0.050164248 |
| Vps26b | 9.602038 | 9.598447 | 9.6203 | -0.021844542 | -0.018254316 | 9.2685441 | 9.4496952 | 9.56154 | -0.111847114 | -0.292998206 |
| Kl | 4.248329 | 4.084037 | 4.2039 | -0.119882714 | 0.044408655 | 4.1118914 | 4.2513599 | 4.36322 | -0.111860759 | -0.251329253 |
| Rps3a3 | 3.74982 | 3.411504 | 3.4634 | -0.051882512 | 0.28643274 | 3.5832554 | 3.4486463 | 3.56056 | -0.111914803 | 0.02269434 |
| Oat | 10.62064 | 10.49549 | 10.503 | -0.007535476 | 0.117615344 | 10.616483 | 10.350163 | 10.4621 | -0.111927258 | 0.154392606 |
| Ntng1 | 1.901143 | 1.733409 | 1.7994 | -0.066007735 | 0.101726823 | 1.7773777 | 1.6469507 | 1.75904 | -0.112092866 | 0.018334134 |
| Trav6-2 | 3.253827 | 3.242043 | 3.2544 | -0.012311116 | -0.000527113 | 3.1212909 | 3.1442645 | 3.25642 | -0.112157916 | -0.135131557 |
| Gm9830 | 5.344524 | 5.770806 | 6.0887 | -0.317943461 | -0.744224984 | 5.4767945 | 5.6038124 | 5.71599 | -0.11217511 | -0.239192988 |
| Gm28053 | 7.289958 | 7.235304 | 7.4023 | -0.167033026 | -0.11237911 | 7.3951174 | 7.3960417 | 7.50826 | -0.112214759 | -0.113139043 |
| Zfp935 | 1.621044 | 1.388298 | 1.3819 | 0.006389838 | 0.239135948 | 1.792655 | 1.4969021 | 1.60924 | -0.112338968 | 0.183413921 |
| Chd1l | 9.302781 | 9.342245 | 9.423 | -0.08077942 | -0.120243411 | 9.4271263 | 9.4058188 | 9.51823 | -0.112407548 | -0.091100036 |
| Rtl10 | 4.746789 | 4.808702 | 4.971 | -0.162305586 | -0.224218096 | 4.9110872 | 4.7601163 | 4.87254 | -0.112425049 | 0.038545856 |
| Mink1 | 10.75568 | 10.76742 | 10.842 | -0.074391606 | -0.086133284 | 10.787218 | 10.844204 | 10.9566 | -0.112426534 | -0.169413236 |
| Tnfrsf17 | 2.241759 | 2.256565 | 2.2826 | -0.026051913 | -0.040857691 | 2.1439895 | 2.1438805 | 2.25649 | -0.112607531 | -0.112498544 |
| Gata5 | 0.92854 | 0.911932 | 0.8986 | 0.013381329 | 0.029989039 | 0.9861745 | 0.9176089 | 1.03022 | -0.112611311 | -0.044045752 |
| Sass6 | 7.617441 | 7.599998 | 7.6736 | -0.073573763 | -0.056130618 | 7.5840908 | 7.7739372 | 7.8866 | -0.112663378 | -0.302509784 |
| Vps37a | 9.313671 | 9.234825 | 9.2368 | -0.001982035 | 0.076864228 | 9.1254052 | 9.0110366 | 9.12374 | -0.112704114 | 0.001664456 |
| Tcf20 | 11.23666 | 11.26525 | 11.299 | -0.033487102 | -0.062075822 | 11.170211 | 11.194203 | 11.307 | -0.112777388 | -0.136769131 |
| Txndc11 | 9.894155 | 9.898069 | 9.8395 | 0.058590745 | 0.054676987 | 9.8737295 | 9.7648339 | 9.87763 | -0.112793207 | -0.003897593 |
| Mpv17 | 8.12845 | 8.142049 | 8.2229 | -0.080872701 | -0.094472307 | 8.4105767 | 8.1163672 | 8.22917 | -0.112804866 | 0.18140461 |
| Gchfr | 2.539213 | 2.297554 | 2.4792 | -0.181672933 | 0.059986201 | 2.2336508 | 2.1970202 | 2.30992 | -0.11289502 | -0.076264388 |
| Trbv29 | 7.461646 | 7.330504 | 7.4145 | -0.083956717 | 0.047185169 | 7.1582085 | 7.2885931 | 7.40151 | -0.112918017 | -0.243302611 |
| Slc10a3 | 8.232447 | 8.179122 | 8.1686 | 0.010488708 | 0.063813811 | 8.2902513 | 7.9964255 | 8.10937 | -0.112947683 | 0.180878156 |
| C330018D20Rik | 7.485799 | 7.45544 | 7.4923 | -0.036822592 | -0.006463894 | 7.7584347 | 7.4356379 | 7.5486 | -0.112957668 | 0.209839086 |
| Itch | 11.08574 | 11.1409 | 11.141 | 0.000321494 | -0.054832242 | 10.939051 | 10.897881 | 11.0109 | -0.113029216 | -0.07185848 |
| Mon2 | 11.07558 | 11.12016 | 11.062 | 0.058046206 | 0.013459306 | 10.997572 | 11.043533 | 11.1566 | -0.113066536 | -0.159027879 |
| Ammecr1 | 7.618465 | 7.64956 | 7.6833 | -0.033773789 | -0.064868906 | 7.6160449 | 7.6038107 | 7.71688 | -0.113071932 | -0.100837767 |
| Gm12112 | 1.748391 | 1.784918 | 1.9517 | -0.1667408 | -0.20326764 | 1.7686697 | 1.8840815 | 1.9972 | -0.11311964 | -0.22853143 |
| Olfr111 | 2.697242 | 2.688437 | 2.6843 | 0.004132885 | 0.012938237 | 2.80148 | 2.6918295 | 2.80495 | -0.113122578 | -0.003472073 |
| Ogt | 11.58488 | 11.74704 | 11.676 | 0.071129196 | -0.091029352 | 12.11532 | 12.044843 | 12.158 | -0.11316974 | -0.04269264 |

| | | | | | | | | | | |
|---|---|---|---|---|---|---|---|---|---|---|
| Sin3a | 11.03086 | 10.99906 | 11.028 | -0.029312344 | 0.002490398 | 10.845657 | 10.933874 | 11.0471 | -0.113179686 | -0.201396575 |
| Gstm7 | 1.392732 | 1.342474 | 1.4183 | -0.07585347 | -0.025595144 | 1.3757157 | 1.2970731 | 1.41028 | -0.113202514 | -0.034559949 |
| Pln | 1.067738 | 1.113057 | 1.1136 | -0.000554075 | -0.045873448 | 1.0741798 | 1.0370478 | 1.1503 | -0.113247664 | -0.076115641 |
| Adam5 | 2.160337 | 2.016267 | 2.1805 | -0.164198771 | -0.020129667 | 1.9088881 | 1.924266 | 2.03752 | -0.113252888 | -0.128630804 |
| Ly6h | 4.075956 | 3.929397 | 4.045 | -0.115598017 | 0.030960553 | 4.0889243 | 4.0703405 | 4.18364 | -0.113303459 | -0.094719658 |
| Ctdsp2 | 11.22626 | 11.31196 | 11.417 | -0.104676096 | -0.190373048 | 11.183329 | 11.149514 | 11.2628 | -0.113326278 | -0.079510766 |
| Gm18194 | 4.368191 | 4.351201 | 4.3935 | -0.04229178 | -0.025301747 | 4.4165324 | 4.3836571 | 4.49714 | -0.113478969 | -0.080603678 |
| Samd14 | 5.417756 | 5.071334 | 5.2971 | -0.225720818 | 0.120701346 | 5.0434974 | 4.8858092 | 4.99929 | -0.113483146 | 0.04420501 |
| Vmn2r88 | 3.191647 | 3.173665 | 3.081 | 0.092617342 | 0.110599601 | 3.1542472 | 3.1309813 | 3.24448 | -0.11349953 | -0.090233642 |
| Arfip2 | 8.739165 | 8.70315 | 8.7664 | -0.06329402 | -0.027278713 | 9.0555066 | 8.7428294 | 8.85635 | -0.113518882 | 0.199158332 |
| Olfr1441 | 1.033638 | 1.027398 | 1.0482 | -0.020844155 | -0.0146039 | 1.0434681 | 1.0034188 | 1.11698 | -0.113564372 | -0.073515027 |
| Gm7902 | 2.359258 | 2.45298 | 2.4627 | -0.009708365 | -0.103429718 | 2.4471279 | 2.5083347 | 2.62194 | -0.113603091 | -0.174809846 |
| Spic | 6.86185 | 6.509705 | 6.4655 | 0.044174796 | 0.396319102 | 6.5185888 | 6.1930569 | 6.30669 | -0.113628619 | 0.21190328 |
| BC107364 | 1.404642 | 1.331861 | 1.4337 | -0.101805228 | -0.029024107 | 1.4084433 | 1.3188053 | 1.43244 | -0.113638585 | -0.024000494 |
| Thumpd2 | 7.377996 | 7.292338 | 7.3798 | -0.087467174 | -0.001808968 | 7.4491102 | 7.2977835 | 7.41145 | -0.113661684 | 0.037665063 |
| Gm3608 | 6.470354 | 6.591413 | 6.3744 | 0.217062492 | 0.096003632 | 6.509468 | 6.3216981 | 6.43538 | -0.113685718 | 0.074084189 |
| Ppp1cc | 11.5681 | 11.48573 | 11.602 | -0.116592186 | -0.034220596 | 11.543007 | 11.493134 | 11.6069 | -0.113736682 | -0.06386336 |
| Klhl23 | 3.115703 | 2.890635 | 2.9199 | -0.029293848 | 0.195773453 | 3.1020704 | 2.7525906 | 2.86636 | -0.113767098 | 0.235712787 |
| Tmem132b | 1.576049 | 1.632817 | 1.5151 | 0.117703789 | 0.060935183 | 1.4771294 | 1.5696046 | 1.6834 | -0.113794015 | -0.206269303 |
| Fgf7 | 2.632899 | 2.567658 | 2.6116 | -0.043938474 | 0.021302742 | 2.4725555 | 2.439418 | 2.55325 | -0.113828816 | -0.080691323 |
| Cgn | 1.886361 | 1.772191 | 1.8352 | -0.063003732 | 0.051166917 | 1.7436625 | 1.722932 | 1.83682 | -0.113889135 | -0.093158694 |
| CAAA01141682.1 | 4.124793 | 4.221302 | 4.2651 | -0.043762015 | -0.140271048 | 4.3084113 | 4.1370705 | 4.25099 | -0.113915699 | 0.057425141 |
| Gm2308 | -0.129918 | -0.11998 | -0.1089 | -0.011093493 | -0.021033543 | -0.1474921 | -0.147246 | -0.0332 | -0.114071575 | -0.114317657 |
| Cep295 | 9.730482 | 9.810565 | 9.7713 | 0.039293683 | -0.040789516 | 9.8188044 | 9.8227058 | 9.93682 | -0.114109215 | -0.118010684 |
| Zkscan16 | 1.073451 | 1.110912 | 1.1026 | 0.008307217 | -0.02915365 | 1.1195419 | 1.0609809 | 1.17514 | -0.114157348 | -0.055596387 |
| Nkd2 | 3.0824 | 3.014209 | 3.0017 | 0.01247615 | 0.080667727 | 2.8581746 | 2.7941111 | 2.90845 | -0.114334283 | -0.050270781 |
| Slc24a3 | 3.107109 | 2.977252 | 3.0138 | -0.036520959 | 0.093336304 | 2.8757215 | 2.954502 | 3.06884 | -0.114340081 | -0.19312061 |
| Inpp5a | 7.832016 | 7.677867 | 8.0474 | -0.369534482 | -0.215384998 | 7.7983754 | 7.7288359 | 7.8432 | -0.11436483 | -0.044825332 |
| Trav15-2-dv6-2 | 2.700508 | 2.74175 | 2.7174 | 0.024309536 | -0.016932659 | 2.632693 | 2.643621 | 2.75799 | -0.114370074 | -0.125298021 |
| Trim39 | 9.711557 | 9.750718 | 9.7649 | -0.014149771 | -0.053311366 | 9.9944982 | 9.9843433 | 10.0988 | -0.114410469 | -0.104255552 |
| Traj20 | 2.006904 | 1.932308 | 1.9598 | -0.027474335 | 0.047122259 | 1.9175027 | 1.9263113 | 2.04087 | -0.114559349 | -0.123367871 |
| Ppp2r5e | 10.08632 | 10.08207 | 10.111 | -0.029151943 | -0.024903052 | 9.9649752 | 9.9060471 | 10.0207 | -0.114622199 | -0.055694089 |

| | | | | | | | | | | |
|---|---|---|---|---|---|---|---|---|---|---|
| Il4 | 2.210294 | 2.183632 | 2.1747 | 0.008893721 | 0.035555135 | 2.05511 | 2.0733813 | 2.18803 | -0.114653008 | -0.132924313 |
| Hoxb2 | 2.901086 | 2.860372 | 2.8347 | 0.025666162 | 0.066380993 | 2.568851 | 2.6444082 | 2.7591 | -0.114691449 | -0.190248582 |
| Zcchc12 | 2.962663 | 2.874044 | 3.0295 | -0.155410461 | -0.066791795 | 2.9137795 | 2.8580301 | 2.97278 | -0.114746026 | -0.058996583 |
| Olfr648 | 4.892226 | 4.825695 | 4.8766 | -0.05094894 | 0.015582299 | 4.8887625 | 5.0503475 | 5.16515 | -0.11480596 | -0.276390917 |
| Klf5 | 2.815102 | 2.728551 | 2.8078 | -0.079280883 | 0.007269663 | 2.6308318 | 2.6856431 | 2.80047 | -0.114831055 | -0.169642279 |
| Olfr66 | 3.310662 | 3.317556 | 3.2804 | 0.03711236 | 0.030218317 | 3.3631022 | 3.4407316 | 3.55558 | -0.114848099 | -0.192477479 |
| Pard6a | 6.184241 | 6.088174 | 6.2276 | -0.139471943 | -0.04340512 | 6.2756171 | 6.1624837 | 6.27738 | -0.114895862 | -0.001762488 |
| Usp43 | 1.11735 | 1.067783 | 1.1007 | -0.03288688 | 0.016680001 | 1.0432644 | 1.0339539 | 1.14886 | -0.114910227 | -0.105599788 |
| Rgs6 | 1.809823 | 1.826113 | 1.9278 | -0.101663987 | -0.117954076 | 1.8029832 | 1.8111361 | 1.92617 | -0.11503158 | -0.123184498 |
| Tmx2 | 9.590133 | 9.602717 | 9.6595 | -0.056818529 | -0.069402288 | 9.8548847 | 9.6185867 | 9.73366 | -0.115077626 | 0.121220434 |
| Spryd3 | 9.368065 | 9.17606 | 9.1883 | -0.012289057 | 0.179716112 | 9.3085037 | 9.0727767 | 9.18789 | -0.11511564 | 0.120611377 |
| Pigyl | 6.314122 | 6.186344 | 6.2956 | -0.10922161 | 0.018555695 | 6.466047 | 6.1207766 | 6.236 | -0.115223197 | 0.230047179 |
| Ighv1-39 | 6.741083 | 7.006103 | 6.7171 | 0.288963371 | 0.023942495 | 7.04441 | 6.8626417 | 6.97787 | -0.115224425 | 0.066543913 |
| Cldn3 | 1.224087 | 1.0964 | 1.1788 | -0.082400186 | 0.045286291 | 1.3410319 | 1.1529145 | 1.26814 | -0.115230132 | 0.072887283 |
| Vmn1r87 | 4.683952 | 4.612565 | 4.6899 | -0.077345997 | -0.005958436 | 4.7702331 | 4.8374791 | 4.95275 | -0.115269709 | -0.182515755 |
| Arhgef16 | 1.479791 | 1.359177 | 1.3485 | 0.010717778 | 0.131332198 | 1.3604076 | 1.3829326 | 1.49821 | -0.115274232 | -0.137799202 |
| Kif5b | 11.66123 | 11.63265 | 11.566 | 0.066917602 | 0.095494914 | 11.464894 | 11.532558 | 11.6478 | -0.115276748 | -0.182940489 |
| Rapsn | 2.191693 | 2.185852 | 2.2422 | -0.056372504 | -0.050530855 | 2.1165188 | 2.113588 | 2.22886 | -0.115276779 | -0.112346019 |
| Gm13192 | 1.153469 | 1.091569 | 1.0856 | 0.005968667 | 0.067869014 | 1.0541904 | 1.0464147 | 1.16186 | -0.115446904 | -0.107671206 |
| Fgfr1 | 7.006599 | 6.546365 | 6.9355 | -0.389137134 | 0.071096696 | 6.362824 | 6.5008855 | 6.61635 | -0.115460629 | -0.253522189 |
| Carmil2 | 11.23277 | 11.28658 | 11.313 | -0.02656043 | -0.080368328 | 11.661749 | 11.387194 | 11.5027 | -0.115460718 | 0.15909416 |
| Gosr1 | 10.00146 | 9.788022 | 9.9697 | -0.181687717 | 0.031751259 | 9.5717373 | 9.7438971 | 9.85948 | -0.115586031 | -0.287745838 |
| Zw10 | 9.48415 | 9.586835 | 9.6022 | -0.015355336 | -0.118041164 | 9.6364185 | 9.5778378 | 9.69345 | -0.11561658 | -0.057035952 |
| Phospho1 | 4.161749 | 3.827814 | 3.8306 | -0.002736453 | 0.331198537 | 4.1448935 | 3.4857368 | 3.60136 | -0.115621145 | 0.543535559 |
| Zhx2 | 9.846223 | 10.02071 | 9.8858 | 0.13495171 | -0.039535154 | 9.8346916 | 9.6637529 | 9.77942 | -0.115662308 | 0.05527639 |
| Trim62 | 6.611362 | 6.69131 | 6.7269 | -0.03555432 | -0.115501995 | 6.5492294 | 6.421642 | 6.53734 | -0.115700572 | 0.011886791 |
| Nans | 9.564226 | 9.497083 | 9.6298 | -0.132684617 | -0.065541281 | 9.6034513 | 9.4789867 | 9.59472 | -0.115730839 | 0.008733799 |
| Large2 | 2.415065 | 2.38212 | 2.3501 | 0.032009717 | 0.064954379 | 2.5592436 | 2.370606 | 2.48635 | -0.115743674 | 0.072893968 |
| Gm43540 | 5.054624 | 4.812201 | 5.0591 | -0.24689729 | -0.004474482 | 5.0400397 | 5.2408791 | 5.35672 | -0.115845425 | -0.316684826 |
| Cop1 | 9.922585 | 9.919232 | 9.9264 | -0.007127144 | -0.003774204 | 9.8420082 | 9.852311 | 9.96819 | -0.115876851 | -0.126179631 |
| Washc2 | 10.8245 | 10.83232 | 10.82 | 0.012689808 | 0.004870328 | 10.897001 | 10.824989 | 10.9409 | -0.115886222 | -0.043874882 |
| Reep2 | 3.483545 | 3.478287 | 3.6505 | -0.172208754 | -0.166950921 | 3.5792382 | 3.4609239 | 3.57684 | -0.115916763 | 0.00239759 |

| | | | | | | | | | | |
|---|---|---|---|---|---|---|---|---|---|---|
| Golga2 | 10.3383 | 10.19128 | 10.319 | -0.128093679 | 0.01891936 | 10.361179 | 10.157326 | 10.2733 | -0.115947424 | 0.087905089 |
| Klf3 | 10.87294 | 11.02588 | 10.954 | 0.07167919 | -0.081259546 | 10.886696 | 10.777447 | 10.8934 | -0.115948204 | -0.006699323 |
| Pphln1 | 9.551083 | 9.480744 | 9.524 | -0.043261398 | 0.027077408 | 9.5892323 | 9.4323751 | 9.54851 | -0.11613294 | 0.040724283 |
| Sprtn | 9.441963 | 9.496934 | 9.482 | 0.014893652 | -0.04007707 | 9.5971012 | 9.4414297 | 9.55761 | -0.11618267 | 0.0394889 |
| Eefsec | 8.787808 | 8.835113 | 8.8393 | -0.004205659 | -0.051511326 | 8.997878 | 8.8632801 | 8.97951 | -0.116234103 | 0.018363884 |
| Sap130 | 10.16741 | 10.14701 | 10.182 | -0.035405279 | -0.0150087 | 10.105724 | 10.075148 | 10.1914 | -0.116235888 | -0.08566083 |
| Traj18 | 2.562562 | 2.581106 | 2.5631 | 0.017990241 | -0.000553914 | 2.4754917 | 2.5401605 | 2.6564 | -0.11623845 | -0.180907242 |
| Dpagt1 | 8.098517 | 7.934368 | 8.0497 | -0.115310074 | 0.04883953 | 8.0524229 | 7.9475131 | 8.06376 | -0.116246654 | -0.011336831 |
| Tmx3 | 10.85034 | 10.72377 | 10.761 | -0.03770684 | 0.088869594 | 10.610968 | 10.629238 | 10.7455 | -0.116249954 | -0.134520299 |
| Osbp2 | 1.314755 | 1.345921 | 1.3837 | -0.037827454 | -0.068993441 | 1.2794804 | 1.28568 | 1.402 | -0.116317012 | -0.122516656 |
| Dtl | 7.192531 | 6.989523 | 7.2931 | -0.303624136 | -0.100616109 | 7.0978858 | 7.496074 | 7.61241 | -0.116335894 | -0.514524154 |
| 8030474K03Rik | 0.072284 | 0.080982 | 0.0638 | 0.017230514 | 0.008532537 | 0.2162318 | 0.0675163 | 0.18386 | -0.116345743 | 0.032369814 |
| Adcy1 | 3.001305 | 3.044313 | 3.0813 | -0.037009396 | -0.080017163 | 2.9842003 | 2.9395637 | 3.05591 | -0.116347619 | -0.071711047 |
| Osbp | 10.33944 | 10.42119 | 10.37 | 0.051590384 | -0.030165604 | 10.247452 | 10.169961 | 10.2863 | -0.116385356 | -0.038894346 |
| Tcrg-V5 | 1.101115 | 1.053263 | 1.0858 | -0.032510603 | 0.015341896 | 1.0283716 | 1.0006959 | 1.11714 | -0.116441135 | -0.088765454 |
| Phkg1 | 4.964253 | 4.913564 | 5.0264 | -0.112794325 | -0.062105599 | 5.0217113 | 4.9804245 | 5.09692 | -0.116491888 | -0.075205113 |
| Naa40 | 9.629242 | 9.63847 | 9.6794 | -0.040966526 | -0.050194346 | 9.7943166 | 9.5788872 | 9.69539 | -0.116501373 | 0.098928029 |
| Tlcd1 | 5.291308 | 5.324051 | 5.3135 | 0.010591883 | -0.022151434 | 5.2165707 | 5.3720293 | 5.48854 | -0.116508967 | -0.271967527 |
| Gm16574 | 1.58554 | 1.51921 | 1.5858 | -0.06662831 | -0.000298238 | 1.5313423 | 1.5643321 | 1.68087 | -0.116540065 | -0.149529939 |
| Ndufaf7 | 7.802194 | 7.750821 | 7.8417 | -0.090886151 | -0.039513216 | 8.0065264 | 7.7748379 | 7.89146 | -0.116624753 | 0.115063769 |
| Ric8b | 8.293555 | 8.331683 | 8.3412 | -0.009549122 | -0.047677464 | 8.1602754 | 8.1992827 | 8.31593 | -0.116648175 | -0.155655415 |
| Sh3gl2 | 1.848406 | 1.769583 | 1.7732 | -0.003607016 | 0.075216776 | 1.7183627 | 1.7003082 | 1.81705 | -0.116736951 | -0.098682436 |
| Vash2 | 3.267213 | 3.071874 | 3.2647 | -0.19285489 | 0.002484294 | 3.1263392 | 3.0374059 | 3.15422 | -0.116810923 | -0.027877579 |
| Dok2 | 8.790888 | 8.92181 | 8.7995 | 0.122324805 | -0.008597163 | 8.768558 | 8.7073477 | 8.82418 | -0.116829866 | -0.055619538 |
| Fam83f | 4.587681 | 4.760163 | 4.7065 | 0.053621665 | -0.1188606 | 4.6834221 | 4.4781253 | 4.59502 | -0.116894944 | 0.088401818 |
| Ighv1-11 | 4.377522 | 4.747476 | 4.6523 | 0.095137023 | -0.274816726 | 4.7370859 | 4.2957081 | 4.41265 | -0.116938396 | 0.324439442 |
| Rnf227 | 4.878402 | 4.913084 | 4.8925 | 0.020555393 | -0.014126456 | 4.8921604 | 4.810406 | 4.92739 | -0.116979781 | -0.035225377 |
| Usp6nl | 10.07761 | 10.14976 | 10.057 | 0.093157154 | 0.02101288 | 10.057018 | 9.9424317 | 10.0595 | -0.1170486 | -0.00246229 |
| Slc39a9 | 9.806386 | 9.75188 | 9.7484 | 0.003478558 | 0.057984138 | 9.7986382 | 9.6934997 | 9.81061 | -0.117107726 | -0.011969226 |
| Ypel3 | 9.985476 | 10.09621 | 10.168 | -0.071305421 | -0.182036637 | 10.2941 | 10.013795 | 10.1309 | -0.117127749 | 0.163177556 |
| Mzt1 | 9.209779 | 9.11749 | 9.1684 | -0.050898175 | 0.041391094 | 9.104167 | 9.0270859 | 9.14428 | -0.117191832 | -0.040110689 |
| Hook3 | 9.870616 | 9.756265 | 9.8569 | -0.100603376 | 0.013747462 | 9.6681904 | 9.7108008 | 9.828 | -0.117197962 | -0.15980833 |

| | | | | | | | | | | |
|---|---|---|---|---|---|---|---|---|---|---|
| Tmem45a | 1.426099 | 1.445585 | 1.3477 | 0.097886687 | 0.078401004 | 1.4048253 | 1.3774263 | 1.49472 | -0.117296827 | -0.089897757 |
| Rbpms2 | 3.256325 | 3.348333 | 3.0597 | 0.288677503 | 0.196670003 | 3.089728 | 3.0533479 | 3.17075 | -0.117405873 | -0.081025758 |
| Tprgl | 10.68977 | 10.77248 | 10.809 | -0.036211946 | -0.118927176 | 10.857806 | 10.579399 | 10.6968 | -0.117407104 | 0.160999965 |
| Sdha | 12.03777 | 11.98685 | 11.988 | -0.001319646 | 0.049597384 | 12.016076 | 11.852664 | 11.9702 | -0.117519836 | 0.045892247 |
| Acsl3 | 7.843705 | 7.857591 | 7.8232 | 0.034364648 | 0.020477799 | 7.7369583 | 7.7696845 | 7.88723 | -0.117541618 | -0.150267825 |
| Gne | 9.335928 | 9.161636 | 9.2871 | -0.125466194 | 0.04882532 | 9.3190062 | 9.1197915 | 9.23735 | -0.117554476 | 0.081660218 |
| Zc3h12b | 4.387817 | 4.405067 | 4.5889 | -0.183855042 | -0.201105319 | 4.49273 | 4.3242707 | 4.44186 | -0.117589292 | 0.050870045 |
| Car9 | 3.758522 | 3.697696 | 3.6887 | 0.008989636 | 0.069816594 | 3.809787 | 3.6487456 | 3.76634 | -0.117598772 | 0.04344263 |
| Trav5-1 | 3.376925 | 3.480728 | 3.475 | 0.005750193 | -0.098052707 | 3.3024634 | 3.3871341 | 3.50475 | -0.117612483 | -0.202283189 |
| Lman1l | 3.090069 | 3.034949 | 3.1146 | -0.079635086 | -0.024515905 | 3.2985582 | 3.0184183 | 3.13607 | -0.117652388 | 0.162487476 |
| Cdh5 | 10.13538 | 9.791414 | 9.9509 | -0.159529276 | 0.184434249 | 9.1981232 | 9.2651264 | 9.38284 | -0.117718435 | -0.184721669 |
| Myh15 | 1.505058 | 1.482026 | 1.6473 | -0.165314815 | -0.142282784 | 1.4292395 | 1.3954661 | 1.51324 | -0.117770585 | -0.083997202 |
| Gm3379 | 3.141064 | 2.95131 | 2.9416 | 0.009698249 | 0.199452231 | 2.9651245 | 2.9592758 | 3.07706 | -0.117786076 | -0.11193743 |
| Cth | 2.089494 | 1.901234 | 1.9563 | -0.05508063 | 0.133179263 | 2.0013759 | 1.881281 | 1.9991 | -0.117814381 | 0.002280515 |
| Tbck | 9.151704 | 9.197974 | 9.1236 | 0.074330862 | 0.028061623 | 9.1356717 | 9.0107991 | 9.12861 | -0.117815395 | 0.007057108 |
| Golph3l | 9.314597 | 9.368671 | 9.4614 | -0.092727737 | -0.146801547 | 9.2857055 | 9.2835914 | 9.40146 | -0.117871481 | -0.115757377 |
| Ddx6 | 13.39009 | 13.36901 | 13.305 | 0.063905114 | 0.084980746 | 13.222424 | 13.212238 | 13.3301 | -0.117877002 | -0.107691355 |
| Pdzk1ip1 | 2.831311 | 2.904278 | 2.8731 | 0.031150969 | -0.041816079 | 2.7991923 | 2.7717327 | 2.88961 | -0.11788099 | -0.09042148 |
| Ppm1a | 10.19108 | 10.11783 | 10.244 | -0.125861415 | -0.052605806 | 10.115179 | 10.00425 | 10.1222 | -0.117915992 | -0.006987115 |
| Rhno1 | 8.806304 | 8.869863 | 8.9329 | -0.063077703 | -0.126636496 | 8.9304336 | 8.8767983 | 8.99486 | -0.118060373 | -0.06442517 |
| Lcmt1 | 7.529889 | 7.534724 | 7.4961 | 0.038636921 | 0.033802004 | 7.5903658 | 7.4375334 | 7.55563 | -0.118099069 | 0.034733303 |
| Ccdc141 | 4.825103 | 4.989924 | 4.9498 | 0.040135786 | -0.124685774 | 4.8243874 | 4.9360654 | 5.05418 | -0.118113658 | -0.229791715 |
| Sox5 | 4.339962 | 4.305786 | 4.4544 | -0.148589161 | -0.114413604 | 4.4596053 | 4.2441958 | 4.36235 | -0.118154145 | 0.09725541 |
| Acp6 | 8.334312 | 8.528249 | 8.5328 | -0.00457729 | -0.198514245 | 8.5730804 | 8.582221 | 8.7004 | -0.118179166 | -0.127319727 |
| Nr3c2 | 5.133004 | 4.896217 | 5.3188 | -0.422610233 | -0.185823362 | 5.0969342 | 4.9179072 | 5.0361 | -0.118188138 | 0.060838816 |
| Ccdc138 | 5.54676 | 5.531887 | 5.7345 | -0.202607644 | -0.18773462 | 5.5764093 | 5.5926502 | 5.7109 | -0.118246972 | -0.134487877 |
| Btbd9 | 9.459136 | 9.37964 | 9.3979 | -0.018251971 | 0.061244493 | 9.2727234 | 9.268965 | 9.38726 | -0.11829922 | -0.114540787 |
| Gm20696 | 3.872845 | 3.825443 | 3.8696 | -0.044136626 | 0.003265335 | 3.86698 | 3.889206 | 4.00752 | -0.118310506 | -0.140536544 |
| Nectin1 | 7.0827 | 7.169768 | 7.2894 | -0.119627242 | -0.20669477 | 6.9453298 | 6.9472793 | 7.06564 | -0.118364807 | -0.120314304 |
| Rgs14 | 10.47479 | 10.61184 | 10.595 | 0.016473342 | -0.120572938 | 10.829628 | 10.56051 | 10.6789 | -0.11841782 | 0.150700415 |
| Gm11737 | 2.206764 | 2.092486 | 2.1937 | -0.101255579 | 0.013022675 | 2.1549241 | 2.1973668 | 2.31584 | -0.118471751 | -0.160914468 |
| Zfp677 | 7.480713 | 7.370352 | 7.633 | -0.262664532 | -0.152304033 | 7.4538313 | 7.489797 | 7.6083 | -0.118503712 | -0.154469462 |

| | | | | | | | | | | |
|---|---|---|---|---|---|---|---|---|---|---|
| Vmn2r24 | 4.827563 | 4.874964 | 4.9071 | -0.032101427 | -0.079502373 | 4.8695375 | 4.9881985 | 5.10673 | -0.118531119 | -0.23719217 |
| Gm35315 | 5.687987 | 5.446655 | 5.4597 | -0.01299767 | 0.22833459 | 6.0190402 | 5.5831352 | 5.70167 | -0.118532284 | 0.31737279 |
| Gm2115 | 4.104138 | 4.149005 | 4.1783 | -0.029271932 | -0.074138902 | 3.9238969 | 4.060012 | 4.17865 | -0.118638394 | -0.254753556 |
| Ptprz1 | 1.217432 | 1.153783 | 1.1642 | -0.010428366 | 0.053220451 | 1.1096265 | 1.1328142 | 1.25147 | -0.118651806 | -0.141839536 |
| Kptn | 7.770563 | 7.841181 | 7.8737 | -0.032472499 | -0.103090609 | 8.2082015 | 7.8957423 | 8.01441 | -0.118667043 | 0.193792169 |
| Chrm1 | 2.115123 | 2.256319 | 2.1251 | 0.131241254 | -0.00995496 | 2.2425658 | 2.1225907 | 2.24132 | -0.118730711 | 0.001244431 |
| Zfp414 | 7.855414 | 7.622005 | 7.9438 | -0.321825538 | -0.088416741 | 8.2773618 | 7.9305584 | 8.04934 | -0.118777954 | 0.228025425 |
| Lemd3 | 9.289424 | 9.321176 | 9.2551 | 0.066080526 | 0.03432853 | 9.1143104 | 9.1718074 | 9.29077 | -0.118963822 | -0.176460799 |
| Cenpb | 10.25046 | 10.04582 | 10.199 | -0.153419808 | 0.05121359 | 10.339729 | 9.9057749 | 10.0247 | -0.118968534 | 0.314985564 |
| Prkaa2 | 2.460245 | 2.296599 | 2.2728 | 0.023756723 | 0.187402769 | 2.4101051 | 2.3434605 | 2.46246 | -0.118995763 | -0.052351093 |
| Olfr1297 | 1.783314 | 1.786203 | 1.7494 | 0.036772232 | 0.033883872 | 1.6799319 | 1.7258723 | 1.84491 | -0.119033909 | -0.164974224 |
| Gm5373 | 3.255751 | 3.197101 | 3.3458 | -0.148660916 | -0.090011664 | 3.2573354 | 3.2668811 | 3.38593 | -0.119049938 | -0.128595651 |
| Cdk5rap2 | 9.910585 | 9.984965 | 9.9821 | 0.002833653 | -0.071546417 | 10.026157 | 9.9526506 | 10.0717 | -0.119089175 | -0.045582628 |
| AC154486.3 | 2.487544 | 2.458682 | 2.5239 | -0.065218661 | -0.036356474 | 2.6247437 | 2.5022696 | 2.62137 | -0.119098745 | 0.003375433 |
| Cep295nl | 1.805503 | 1.815794 | 1.8745 | -0.058735514 | -0.069027123 | 1.8063269 | 1.7520984 | 1.8712 | -0.119104821 | -0.064876359 |
| Trav12n-3 | 5.413796 | 5.343879 | 5.4537 | -0.109800301 | -0.039882776 | 5.2615098 | 5.3393155 | 5.45842 | -0.119104855 | -0.196910618 |
| Gm49461 | 2.938843 | 2.851647 | 2.8006 | 0.05099787 | 0.138193321 | 3.1171767 | 2.8889881 | 3.00811 | -0.119126201 | 0.109062455 |
| Gse1 | 10.26653 | 10.34606 | 10.308 | 0.037795394 | -0.041732351 | 10.33975 | 10.276825 | 10.396 | -0.119188027 | -0.05626318 |
| Papolg | 9.543464 | 9.679106 | 9.6619 | 0.017207271 | -0.118434301 | 9.7328266 | 9.6148597 | 9.73408 | -0.119218701 | -0.001251795 |
| Paqr6 | 2.853487 | 2.77166 | 2.8511 | -0.079477257 | 0.002349189 | 2.8193037 | 2.8917932 | 3.01103 | -0.11923865 | -0.19172812 |
| Chn1 | 3.558742 | 3.533911 | 3.4548 | 0.079126059 | 0.103957716 | 3.3161277 | 3.3607791 | 3.48004 | -0.119264128 | -0.163915511 |
| Clec2h | 1.036721 | 1.016121 | 1.0047 | 0.01142307 | 0.032022849 | 1.007537 | 0.994593 | 1.11386 | -0.119267556 | -0.106323601 |
| Arid4b | 11.23237 | 11.23088 | 11.275 | -0.044590408 | -0.04310279 | 11.097983 | 11.159461 | 11.2787 | -0.119286912 | -0.180764679 |
| Susd4 | 3.251062 | 3.362078 | 3.3068 | 0.055234358 | -0.055781754 | 3.2432581 | 3.1833728 | 3.30266 | -0.119289875 | -0.059404496 |
| Snap29 | 10.29131 | 10.36781 | 10.26 | 0.10762826 | 0.031126134 | 10.597084 | 10.212055 | 10.3314 | -0.119321082 | 0.265707622 |
| Zfp607b | 4.801241 | 4.995404 | 4.858 | 0.137357292 | -0.05680644 | 4.9661325 | 4.9072051 | 5.02659 | -0.119386133 | -0.060458745 |
| H2-Q2 | 2.805338 | 2.833082 | 3.0239 | -0.190807381 | -0.21855079 | 2.8933534 | 2.8823214 | 3.00172 | -0.119394734 | -0.108362747 |
| Cplx1 | 1.819316 | 1.869546 | 1.6347 | 0.234893468 | 0.184663743 | 1.7239972 | 1.7029681 | 1.82242 | -0.119453825 | -0.0984247 |
| Xlr5a | 0.515703 | 0.452279 | 0.5003 | -0.048053172 | 0.015371018 | 0.4705187 | 0.4409612 | 0.56054 | -0.119576752 | -0.09001918 |
| Tns4 | 4.848196 | 4.951148 | 4.9096 | 0.041564832 | -0.061388056 | 5.1553546 | 4.7699081 | 4.8895 | -0.119587385 | 0.26585914 |
| Galnt14 | 3.332472 | 3.102675 | 3.18 | -0.077312312 | 0.152485031 | 3.2014696 | 3.0990421 | 3.21871 | -0.11966396 | -0.017236419 |
| Gm8170 | 1.158789 | 1.110588 | 1.1384 | -0.027830206 | 0.020369985 | 1.1961954 | 1.1041813 | 1.22389 | -0.11970753 | -0.027693477 |

| | | | | | | | | | | |
|---|---|---|---|---|---|---|---|---|---|---|
| 9330161L09Rik | 2.826206 | 2.771613 | 2.786 | -0.014427901 | 0.040165293 | 2.8372488 | 2.6931357 | 2.81291 | -0.119777355 | 0.024335757 |
| Traj3 | 2.926625 | 2.864278 | 2.9343 | -0.070002786 | -0.007655987 | 2.8101569 | 2.7711271 | 2.89091 | -0.119780154 | -0.080750426 |
| Nme9 | 1.564051 | 1.541239 | 1.6596 | -0.118369722 | -0.095557478 | 1.5554167 | 1.6034907 | 1.7233 | -0.119809924 | -0.167883983 |
| Gm6505 | 2.742631 | 2.870294 | 2.93 | -0.059655754 | -0.187318949 | 2.7684306 | 2.5667852 | 2.68667 | -0.119888423 | 0.081756959 |
| Gm340 | 7.869452 | 7.917931 | 7.8043 | 0.113585336 | 0.065106333 | 7.8674403 | 7.9773989 | 8.09729 | -0.119894474 | -0.229853047 |
| Nanog | 1.820148 | 1.767509 | 1.8112 | -0.04369561 | 0.008944128 | 1.9327939 | 1.7953464 | 1.91525 | -0.119900824 | 0.017546614 |
| Tcea3 | 4.176204 | 4.050662 | 4.1828 | -0.13209223 | -0.006550757 | 4.4016408 | 4.0565355 | 4.17655 | -0.12001658 | 0.225088707 |
| Slc25a15 | 7.630548 | 7.566649 | 7.6109 | -0.044255497 | 0.019643722 | 7.5599598 | 7.2938391 | 7.4139 | -0.12006284 | 0.14605792 |
| Ap2b1 | 12.0448 | 12.05962 | 12 | 0.059546854 | 0.044724164 | 12.094617 | 11.904251 | 12.0243 | -0.120081912 | 0.070283601 |
| Eml4 | 11.27129 | 11.28114 | 11.316 | -0.035169814 | -0.045014146 | 11.303464 | 11.221435 | 11.3415 | -0.120087544 | -0.0380586 |
| Tspan8 | 2.166701 | 1.99677 | 1.8993 | 0.097478329 | 0.267409742 | 2.017727 | 1.9231801 | 2.04332 | -0.12013626 | -0.025589349 |
| Dpp8 | 11.00968 | 10.9743 | 10.997 | -0.023098998 | 0.012275426 | 11.008827 | 10.943666 | 11.0639 | -0.120197544 | -0.055036053 |
| Ttc6 | 2.027059 | 2.116108 | 2.0677 | 0.048364027 | -0.040685144 | 2.0456044 | 1.916081 | 2.03634 | -0.120261264 | 0.009262081 |
| B3galt1 | 2.50844 | 2.384481 | 2.5025 | -0.118007764 | 0.005951765 | 2.3578783 | 2.2224884 | 2.34279 | -0.120296825 | 0.015093051 |
| Acox3 | 10.41231 | 10.21502 | 10.286 | -0.070485446 | 0.126803314 | 10.876855 | 10.437553 | 10.5579 | -0.120334904 | 0.318966762 |
| Fam240a | 3.490427 | 3.415344 | 3.5287 | -0.113404575 | -0.038321359 | 3.5182777 | 3.4007371 | 3.52111 | -0.120367944 | -0.002827379 |
| Tnfsf14 | 7.151706 | 7.300926 | 7.2408 | 0.0600793 | -0.089140925 | 7.0827165 | 7.0319383 | 7.15234 | -0.120403199 | -0.069625024 |
| Aqp7 | 2.717621 | 2.640156 | 2.7011 | -0.060907082 | 0.016558108 | 2.7241301 | 2.7634499 | 2.88388 | -0.120435019 | -0.159754817 |
| Tacc3 | 8.472824 | 8.320354 | 8.5237 | -0.203329829 | -0.050859719 | 8.5542628 | 8.6497436 | 8.77018 | -0.120440908 | -0.215921642 |
| Cct6b | 2.298282 | 2.291301 | 2.3319 | -0.040617457 | -0.033636878 | 2.3326616 | 2.2568639 | 2.37744 | -0.120574789 | -0.044777149 |
| Rhbdd3 | 5.940682 | 5.940555 | 6.0754 | -0.134844164 | -0.134717091 | 6.1838896 | 6.1324831 | 6.25312 | -0.120637005 | -0.069230484 |
| Pacsin1 | 9.549141 | 9.496876 | 9.6111 | -0.114272372 | -0.062007239 | 9.5445712 | 9.5108752 | 9.63152 | -0.120640819 | -0.086944803 |
| Gm43070 | 1.307911 | 1.226729 | 1.2849 | -0.058148966 | 0.023033245 | 1.3022254 | 1.2363119 | 1.35702 | -0.120708703 | -0.054795203 |
| Begain | 4.86541 | 5.040388 | 4.9179 | 0.122458477 | -0.0525198 | 5.0999727 | 5.0108692 | 5.13179 | -0.120923528 | -0.031820072 |
| Zfp114 | 2.99882 | 2.961122 | 2.9753 | -0.014210461 | 0.023488193 | 2.9160758 | 2.8664842 | 2.98745 | -0.120967295 | -0.071375635 |
| Dapk1 | 7.33944 | 7.453648 | 7.3265 | 0.127121558 | 0.012913462 | 6.6854559 | 6.9536225 | 7.07463 | -0.1210112 | -0.389177817 |
| Olfr1383 | 2.204425 | 2.110798 | 2.1642 | -0.053396227 | 0.04023083 | 2.2696806 | 2.1457719 | 2.26681 | -0.121040483 | 0.002868178 |
| Trav9n-4 | 5.868204 | 5.595376 | 5.8473 | -0.251884295 | 0.020943622 | 5.5525266 | 5.7224323 | 5.84352 | -0.121090086 | -0.290995823 |
| Runx1 | 10.09601 | 10.05158 | 10.147 | -0.095851923 | -0.051424138 | 9.9083503 | 10.097163 | 10.2184 | -0.121196401 | -0.310008794 |
| Sirt4 | 5.767707 | 5.746271 | 5.7085 | 0.037751106 | 0.059186311 | 6.2034611 | 5.8048571 | 5.92606 | -0.121202056 | 0.277401941 |
| Trav6-6 | 4.482 | 4.411294 | 4.44 | -0.028723133 | 0.041981946 | 4.3165087 | 4.2925106 | 4.41374 | -0.121232663 | -0.097234479 |
| Trav6d-6 | 4.482 | 4.411294 | 4.44 | -0.028723133 | 0.041981946 | 4.3165087 | 4.2925106 | 4.41374 | -0.121232663 | -0.097234479 |

| | | | | | | | | | | |
|---|---|---|---|---|---|---|---|---|---|---|
| Gcsh | 7.434517 | 7.067029 | 7.1012 | -0.034162588 | 0.333324949 | 7.1282335 | 7.1263039 | 7.24757 | -0.121264621 | -0.119335049 |
| 2610318N02Rik | 5.179227 | 5.196177 | 5.2541 | -0.057944877 | -0.074894892 | 5.4300945 | 5.3275491 | 5.44893 | -0.121380639 | -0.018835148 |
| Cbll1 | 8.998307 | 9.139525 | 8.9903 | 0.149272425 | 0.00805379 | 9.0209242 | 8.9510461 | 9.07258 | -0.121537461 | -0.051659402 |
| Gm2253 | 1.436112 | 1.444245 | 1.4251 | 0.019119782 | 0.01098619 | 1.4651329 | 1.4220312 | 1.54377 | -0.121737264 | -0.078635535 |
| Nwd1 | 2.328545 | 2.450763 | 2.4426 | 0.008137746 | -0.114079632 | 2.3896716 | 2.4604999 | 2.58225 | -0.121751165 | -0.192579452 |
| Plekha6 | 6.492959 | 6.656883 | 6.6806 | -0.023748143 | -0.187672367 | 6.3068959 | 6.3421766 | 6.46395 | -0.121775718 | -0.157056429 |
| Ireb2 | 10.62279 | 10.63793 | 10.559 | 0.07907655 | 0.063935912 | 10.64086 | 10.577251 | 10.6991 | -0.121871868 | -0.058263285 |
| Lias | 8.40527 | 8.343234 | 8.4158 | -0.07252456 | -0.010487819 | 8.5420165 | 8.4126436 | 8.53466 | -0.122016199 | 0.007356717 |
| Man2b1 | 11.65725 | 11.55195 | 11.702 | -0.149591416 | -0.04429093 | 11.856229 | 11.530403 | 11.6525 | -0.12206595 | 0.203759727 |
| Ei24 | 9.672805 | 9.625701 | 9.6568 | -0.03114546 | 0.015958167 | 9.6308395 | 9.4774129 | 9.59956 | -0.122145812 | 0.031280861 |
| Adamtsl3 | 2.426816 | 2.186924 | 2.2781 | -0.091181602 | 0.148710333 | 2.0398637 | 2.029281 | 2.15144 | -0.12215703 | -0.111574322 |
| Olfr161 | 4.364695 | 4.400003 | 4.3281 | 0.071895634 | 0.036587894 | 4.5296328 | 4.5274164 | 4.64967 | -0.122249648 | -0.120033264 |
| Slco1a6 | 1.47418 | 1.473406 | 1.527 | -0.053606623 | -0.052831992 | 1.4883918 | 1.4802649 | 1.60258 | -0.122310522 | -0.11418367 |
| Deaf1 | 8.68956 | 8.583199 | 8.7221 | -0.13885496 | -0.032493961 | 8.6537704 | 8.6427464 | 8.76509 | -0.12234016 | -0.111316197 |
| Tmem150b | 3.725465 | 3.453984 | 3.6854 | -0.23136811 | 0.04011258 | 3.5881528 | 3.4225883 | 3.54495 | -0.122356683 | 0.04320774 |
| Lrtm2 | 3.250284 | 3.220581 | 3.2051 | 0.015482105 | 0.045184115 | 3.2431792 | 3.0988627 | 3.22124 | -0.122377645 | 0.021938837 |
| Vmn1r206 | 2.162192 | 2.149573 | 2.1124 | 0.037166198 | 0.049784887 | 2.117305 | 2.2079015 | 2.33032 | -0.122420835 | -0.213017332 |
| Gm19236 | 1.278635 | 1.27373 | 1.269 | 0.004686384 | 0.009591663 | 1.3274919 | 1.2501716 | 1.37261 | -0.122443217 | -0.045122924 |
| Marcks | 10.56557 | 10.67331 | 10.43 | 0.243352184 | 0.135611594 | 10.310378 | 10.228568 | 10.3511 | -0.12251335 | -0.040703989 |
| Eif4g3 | 11.80529 | 11.8416 | 11.794 | 0.04758153 | 0.011267872 | 11.873762 | 11.808707 | 11.9313 | -0.122581176 | -0.057525828 |
| Ceacam16 | 5.530456 | 5.596762 | 5.6148 | -0.018061674 | -0.084367334 | 6.2559278 | 5.7175645 | 5.8402 | -0.12263428 | 0.415729077 |
| Yod1 | 8.46541 | 8.549231 | 8.5678 | -0.018577066 | -0.102397657 | 8.6560556 | 8.6745433 | 8.79722 | -0.122681341 | -0.141169064 |
| Ighv1-74 | 7.222992 | 6.881058 | 7.2195 | -0.338469966 | 0.003464603 | 7.5051947 | 6.3125191 | 6.43532 | -0.122803871 | 1.069871723 |
| Nf1 | 9.164666 | 9.164093 | 9.1328 | 0.031290439 | 0.031862975 | 9.2230069 | 9.1331681 | 9.25602 | -0.122849917 | -0.033011051 |
| Kansl1 | 11.34425 | 11.42846 | 11.44 | -0.011149234 | -0.09536839 | 11.22656 | 11.24811 | 11.3711 | -0.122958132 | -0.144508326 |
| Fndc11 | 1.539742 | 1.54589 | 1.5949 | -0.049017619 | -0.055165481 | 1.6498666 | 1.4781918 | 1.60117 | -0.122980231 | 0.048694626 |
| Caskin1 | 1.792312 | 1.76209 | 1.7397 | 0.022399651 | 0.052621565 | 1.6977789 | 1.7209734 | 1.84404 | -0.123067861 | -0.146262384 |
| Dnase2a | 9.142736 | 9.031701 | 9.1428 | -0.111125234 | -9.02804E-05 | 9.3561186 | 8.9403052 | 9.06339 | -0.123085465 | 0.292727891 |
| Nxf2 | 2.07091 | 1.942264 | 2.0224 | -0.080097524 | 0.048547741 | 2.0367483 | 2.0028136 | 2.1259 | -0.123090473 | -0.089155752 |
| Man1b1 | 10.31092 | 10.38034 | 10.404 | -0.024021434 | -0.093435036 | 10.347654 | 10.350301 | 10.4734 | -0.12311558 | -0.125761742 |
| Osmr | 8.418276 | 7.838003 | 7.8679 | -0.029902005 | 0.550371078 | 6.9333027 | 7.2072958 | 7.33043 | -0.123129753 | -0.39712279 |
| Fos | 12.46746 | 12.00707 | 12.304 | -0.297139088 | 0.163253752 | 11.069343 | 11.155201 | 11.2784 | -0.123198952 | -0.209057244 |

| | | | | | | | | | | |
|---|---|---|---|---|---|---|---|---|---|---|
| Col6a1 | 7.586458 | 7.659602 | 7.4922 | 0.16735847 | 0.094215171 | 7.2478565 | 6.790854 | 6.91412 | -0.123264227 | 0.333738256 |
| Usf2 | 10.58896 | 10.55396 | 10.66 | -0.105543442 | -0.070542936 | 10.59569 | 10.492855 | 10.6162 | -0.123326188 | -0.020490671 |
| Gm3298 | 1.633382 | 1.719368 | 1.7834 | -0.064002806 | -0.149987902 | 1.7884026 | 1.693265 | 1.81661 | -0.123345493 | -0.028207902 |
| Ap1ar | 9.432139 | 9.395444 | 9.4043 | -0.00883583 | 0.027859447 | 9.3534743 | 9.3268314 | 9.45019 | -0.123359151 | -0.096716315 |
| Gspt2 | 3.45362 | 3.328556 | 3.5138 | -0.185267869 | -0.060203935 | 3.1487037 | 3.0879503 | 3.21133 | -0.12337491 | -0.062621534 |
| Vmn2r6 | 2.854851 | 2.83287 | 2.9199 | -0.087040879 | -0.065059853 | 2.8956452 | 2.8833657 | 3.00677 | -0.123401806 | -0.111122348 |
| Msrb2 | 6.276824 | 6.44305 | 6.4081 | 0.034918748 | -0.131307429 | 6.3718145 | 6.2927375 | 6.41616 | -0.123422514 | -0.044345442 |
| Orc3 | 9.903333 | 9.866786 | 9.9311 | -0.064272613 | -0.027726083 | 9.9411552 | 9.9843961 | 10.1078 | -0.123440953 | -0.166681806 |
| Ccpg1os | 5.193346 | 5.265636 | 5.1265 | 0.139121331 | 0.066830782 | 5.2870862 | 5.2251947 | 5.34869 | -0.123497193 | -0.061605708 |
| Naip1 | 4.095376 | 3.904538 | 4.1095 | -0.204958902 | -0.014120623 | 3.7912616 | 3.7887302 | 3.91226 | -0.123531994 | -0.12100058 |
| Spin4 | 2.758635 | 2.694224 | 2.7592 | -0.064935939 | -0.000524908 | 2.6010419 | 2.5815534 | 2.70509 | -0.12353327 | -0.104044811 |
| Trav16d-dv11 | 3.587029 | 3.41964 | 3.4819 | -0.06221966 | 0.10516851 | 3.3870162 | 3.42085 | 3.5446 | -0.123750124 | -0.157583941 |
| Atf1 | 10.06852 | 10.01726 | 10.009 | 0.008043476 | 0.059303028 | 9.9330031 | 9.8965082 | 10.0203 | -0.123820471 | -0.087325523 |
| Srrm1 | 11.73598 | 11.6887 | 11.694 | -0.005332244 | 0.041956314 | 11.627012 | 11.751157 | 11.875 | -0.123857838 | -0.248003177 |
| Gm13241 | 1.533564 | 1.517372 | 1.5519 | -0.034552045 | -0.018359424 | 1.5032407 | 1.5471333 | 1.67112 | -0.123989533 | -0.167882188 |
| Ankrd42 | 2.326674 | 2.19531 | 2.3085 | -0.113236413 | 0.018128218 | 2.1963624 | 2.163482 | 2.28748 | -0.124002876 | -0.091122513 |
| 1700010I14Rik | 5.130172 | 5.160261 | 5.2414 | -0.081183398 | -0.111272391 | 5.170793 | 5.1307059 | 5.25472 | -0.12401157 | -0.083924523 |
| Hoxa9 | 1.411954 | 1.425915 | 1.5731 | -0.147186414 | -0.161146866 | 1.3796547 | 1.3304069 | 1.45443 | -0.124020255 | -0.07477249 |
| Gnal | 4.822654 | 4.844667 | 5.0618 | -0.217152417 | -0.239164983 | 4.8707311 | 4.7852374 | 4.90926 | -0.124023944 | -0.038530188 |
| Cul5 | 9.699208 | 9.642275 | 9.7126 | -0.070347693 | -0.013414066 | 9.5458653 | 9.6455527 | 9.76965 | -0.124095076 | -0.223782397 |
| Map2k5 | 8.417568 | 8.403426 | 8.5043 | -0.100907287 | -0.086764436 | 8.4716783 | 8.3083672 | 8.43255 | -0.124184426 | 0.039126675 |
| Gm9531 | 9.58318 | 9.7804 | 9.3613 | 0.419076071 | 0.221856058 | 9.482345 | 9.7956611 | 9.91985 | -0.124188742 | -0.437504831 |
| Bmp5 | 1.65083 | 1.64194 | 1.6927 | -0.050790722 | -0.04190011 | 1.6220148 | 1.5939585 | 1.71816 | -0.124197253 | -0.096140998 |
| Cyp2j13 | 2.851813 | 2.912952 | 2.8522 | 0.060784771 | -0.000353844 | 2.9518049 | 2.8708436 | 2.99506 | -0.124215833 | -0.043254531 |
| Prtn3 | 2.703192 | 2.538983 | 2.5814 | -0.042418691 | 0.121789972 | 2.7696587 | 2.6160423 | 2.74026 | -0.124219639 | 0.029396724 |
| Ublcp1 | 10.10558 | 10.09644 | 10.024 | 0.072383064 | 0.081525019 | 10.646074 | 10.093997 | 10.2182 | -0.124232945 | 0.427843445 |
| Gm5620 | 2.575607 | 2.597151 | 2.5951 | 0.002086262 | -0.01945761 | 2.5851916 | 2.5517575 | 2.67605 | -0.124290872 | -0.090856783 |
| Herpud1 | 10.61489 | 10.56789 | 10.651 | -0.083194402 | -0.036194098 | 10.51697 | 10.406456 | 10.5308 | -0.124316006 | -0.013802331 |
| Cyp2j9 | 2.99106 | 2.714563 | 2.9381 | -0.223581923 | 0.052915403 | 2.6816042 | 2.5849229 | 2.70927 | -0.124344633 | -0.027663415 |
| Lgi4 | 3.674989 | 3.568531 | 3.8838 | -0.315276291 | -0.208818153 | 3.6705695 | 3.5579436 | 3.68234 | -0.124398908 | -0.011773047 |
| Hsd17b14 | 2.362062 | 2.4068 | 2.5434 | -0.13655953 | -0.18129761 | 2.4643693 | 2.4240844 | 2.54849 | -0.124403698 | -0.084118806 |
| Lysmd3 | 9.232074 | 9.260488 | 9.2689 | -0.008410896 | -0.03682489 | 9.1469565 | 8.9776777 | 9.10212 | -0.124439371 | 0.044839403 |

| | | | | | | | | | | |
|---|---|---|---|---|---|---|---|---|---|---|
| Fbxo27 | 3.457463 | 3.377274 | 3.4327 | -0.055385467 | 0.024804035 | 3.5237667 | 3.3612407 | 3.48572 | -0.124476933 | 0.038049043 |
| Zfp655 | 9.655073 | 9.64626 | 9.6395 | 0.006773272 | 0.015586876 | 9.62118 | 9.5695733 | 9.69405 | -0.124479268 | -0.072872603 |
| Lrrc55 | 1.449222 | 1.476661 | 1.4857 | -0.009072327 | -0.036511983 | 1.4992667 | 1.3953373 | 1.51994 | -0.124605374 | -0.020675945 |
| Hnrnpul2 | 12.44508 | 12.41184 | 12.457 | -0.04557283 | -0.012330456 | 12.335821 | 12.289664 | 12.4143 | -0.124614964 | -0.078457849 |
| Igdcc4 | 3.836171 | 3.905785 | 3.9511 | -0.045292325 | -0.114906069 | 4.1071053 | 3.9493044 | 4.07393 | -0.124629138 | 0.033171768 |
| Dcp1b | 8.563239 | 8.484128 | 8.7055 | -0.221328153 | -0.142217433 | 8.7603312 | 8.6096288 | 8.73426 | -0.124630419 | 0.026071959 |
| Dcun1d4 | 7.412212 | 7.151568 | 7.4509 | -0.299381914 | -0.038737572 | 6.8381504 | 7.0712919 | 7.19595 | -0.124662426 | -0.357803997 |
| Ska1 | 4.499348 | 4.287727 | 4.4965 | -0.208754914 | 0.002865972 | 4.4472737 | 4.4785655 | 4.60327 | -0.124707845 | -0.155999645 |
| Rhoa | 13.10546 | 13.05232 | 13.047 | 0.005812236 | 0.058949226 | 13.155726 | 12.916249 | 13.041 | -0.124718786 | 0.114757632 |
| Gm19203 | 2.273069 | 2.225688 | 2.2424 | -0.016759797 | 0.030621393 | 2.2570737 | 2.2453251 | 2.37009 | -0.124762408 | -0.113013774 |
| Lysmd4 | 6.641918 | 6.531259 | 6.5771 | -0.045849626 | 0.064809371 | 6.7040145 | 6.589106 | 6.7139 | -0.124791804 | -0.009883222 |
| Gm9544 | 2.750744 | 2.794879 | 2.7581 | 0.036807492 | -0.007327688 | 2.8542537 | 2.7962523 | 2.92106 | -0.124810322 | -0.066808839 |
| Map4 | 11.29216 | 11.3702 | 11.433 | -0.06263426 | -0.14067382 | 11.377367 | 11.381226 | 11.5061 | -0.12487868 | -0.128737406 |
| Zfp711 | 2.377176 | 2.419306 | 2.3691 | 0.050166102 | 0.008036418 | 2.3107557 | 2.2425985 | 2.36749 | -0.124890252 | -0.05673304 |
| Zfp626 | 8.351612 | 8.387951 | 8.4169 | -0.028960688 | -0.065298976 | 8.4895554 | 8.4323736 | 8.55729 | -0.124918598 | -0.067736779 |
| Lefty1 | 3.852607 | 3.644143 | 3.7832 | -0.139065798 | 0.069397525 | 3.7357938 | 3.6232908 | 3.74825 | -0.124958312 | -0.012455352 |
| Prom1 | 3.063425 | 3.072955 | 3.1639 | -0.090984567 | -0.100513885 | 3.2657528 | 3.0871017 | 3.21208 | -0.124981698 | 0.053669376 |
| Ubr7 | 9.776657 | 9.793352 | 9.8021 | -0.008749797 | -0.025444697 | 9.8200433 | 9.6551447 | 9.78015 | -0.125008289 | 0.039890303 |
| Prdx4 | 7.732537 | 7.484011 | 7.6529 | -0.168852805 | 0.079673131 | 7.7430785 | 7.5152712 | 7.64033 | -0.125058983 | 0.102748299 |
| Gm19195 | 0.916831 | 0.869608 | 0.8716 | -0.001957466 | 0.045265656 | 0.8714122 | 0.8706111 | 0.99567 | -0.125063626 | -0.124262511 |
| Snn | 11.5028 | 11.26496 | 11.529 | -0.263576018 | -0.025730548 | 11.729947 | 11.22878 | 11.3539 | -0.125076246 | 0.37609128 |
| Selenoo | 8.872511 | 8.903249 | 8.9862 | -0.082991469 | -0.113730206 | 9.0709098 | 8.8415071 | 8.96662 | -0.125115831 | 0.10428691 |
| Gm37515 | 5.441638 | 5.600793 | 5.4869 | 0.113845626 | -0.045308951 | 5.5035867 | 5.6533222 | 5.7785 | -0.125174999 | -0.274910514 |
| Wnk2 | 2.835686 | 2.674743 | 2.794 | -0.11928862 | 0.04165499 | 2.7522795 | 2.7721239 | 2.89732 | -0.125201038 | -0.145045408 |
| Fyn | 11.50775 | 11.48681 | 11.537 | -0.050049176 | -0.029100166 | 11.215026 | 11.401389 | 11.5266 | -0.125255022 | -0.311617341 |
| Kif7 | 4.628188 | 4.525672 | 4.517 | 0.008707474 | 0.111222853 | 4.4632838 | 4.5180461 | 4.64335 | -0.125306421 | -0.180068659 |
| Abi1 | 11.58775 | 11.59262 | 11.593 | -0.00034193 | -0.00521041 | 11.607611 | 11.487288 | 11.6127 | -0.125366094 | -0.005043218 |
| Sugct | 3.404645 | 3.306433 | 3.3311 | -0.024648536 | 0.073563058 | 3.3044524 | 3.4782149 | 3.60359 | -0.125373226 | -0.299135791 |
| Olfr829 | 1.820634 | 1.802112 | 1.8011 | 0.00102312 | 0.019545395 | 1.8231639 | 1.8181454 | 1.94355 | -0.125402422 | -0.120383916 |
| Tmem38a | 4.506078 | 4.459778 | 4.5867 | -0.126898576 | -0.080598084 | 4.5901323 | 4.2798892 | 4.40533 | -0.125441453 | 0.184801626 |
| Parp6 | 9.522015 | 9.514945 | 9.6252 | -0.11021506 | -0.103145681 | 9.6942788 | 9.5775813 | 9.70316 | -0.125579617 | -0.008882141 |
| Clec11a | 3.0848 | 2.916447 | 3.0076 | -0.091138718 | 0.077214022 | 3.3342347 | 2.9290471 | 3.05473 | -0.125679326 | 0.27950826 |

| | | | | | | | | | | |
|---|---|---|---|---|---|---|---|---|---|---|
| Lurap1l | 2.549377 | 2.538241 | 2.6944 | -0.15617532 | -0.145039448 | 2.480496 | 2.4402362 | 2.56599 | -0.125749432 | -0.085489675 |
| Lhfpl1 | 1.84688 | 1.747087 | 1.8098 | -0.062707634 | 0.037084894 | 1.9393092 | 1.7651915 | 1.89098 | -0.125785273 | 0.048332423 |
| Syde2 | 4.08865 | 3.98228 | 4.1014 | -0.119125382 | -0.012756043 | 3.8245998 | 3.9041321 | 4.02994 | -0.125802933 | -0.205335196 |
| Arhgef5 | 6.623022 | 6.196812 | 6.6244 | -0.427567305 | -0.001357142 | 5.9883753 | 6.0755694 | 6.20141 | -0.12584269 | -0.213036751 |
| Aldh1l2 | 3.857228 | 3.775894 | 3.8299 | -0.054013754 | 0.027320281 | 3.8528862 | 4.0040864 | 4.13008 | -0.125990181 | -0.277190418 |
| Ccr9 | 9.251502 | 9.199221 | 9.2996 | -0.100426654 | -0.048146036 | 8.723988 | 8.9165101 | 9.04253 | -0.126024827 | -0.318546854 |
| Igkv3-5 | 7.889033 | 7.776666 | 7.4201 | 0.356598049 | 0.468964684 | 7.6572182 | 7.8101627 | 7.93628 | -0.126116 | -0.279060515 |
| Gm11870 | 3.638476 | 3.567902 | 3.5936 | -0.025662521 | 0.044911623 | 3.5762934 | 3.5084604 | 3.63461 | -0.126152388 | -0.058319333 |
| Tfdp1 | 9.453938 | 9.388688 | 9.5462 | -0.157492932 | -0.092243315 | 9.3388556 | 9.4928534 | 9.61914 | -0.126282986 | -0.280280767 |
| Trav5d-4 | 3.308083 | 3.247925 | 3.2887 | -0.040772105 | 0.019385956 | 3.2317166 | 3.1513668 | 3.27767 | -0.126304437 | -0.045954682 |
| Trav5n-4 | 3.308083 | 3.247925 | 3.2887 | -0.040772105 | 0.019385956 | 3.2317166 | 3.1513668 | 3.27767 | -0.126304437 | -0.045954682 |
| Egflam | 1.664974 | 1.688694 | 1.6819 | 0.006775154 | -0.016944816 | 1.6417118 | 1.5310658 | 1.6574 | -0.126338726 | -0.015692689 |
| Cd163 | 3.244978 | 3.049062 | 3.1762 | -0.127106874 | 0.068809025 | 2.9936675 | 3.0416873 | 3.16805 | -0.126361527 | -0.174381396 |
| Sf1 | 11.32309 | 11.30583 | 11.319 | -0.012842118 | 0.004419268 | 11.430586 | 11.428383 | 11.5548 | -0.126375394 | -0.124172053 |
| Mroh1 | 9.962624 | 9.861835 | 10.102 | -0.240511787 | -0.139723591 | 10.155075 | 9.9746326 | 10.101 | -0.126411474 | 0.054031121 |
| Cabp1 | 2.520453 | 2.31931 | 2.406 | -0.08668735 | 0.114455688 | 2.402143 | 2.3713739 | 2.49779 | -0.126415855 | -0.0956468 |
| Aplf | 6.679718 | 6.659327 | 6.8629 | -0.203529132 | -0.183138836 | 6.6845912 | 6.6134784 | 6.73991 | -0.126436374 | -0.055323525 |
| Gm14296 | 6.422311 | 6.335399 | 6.5115 | -0.17611213 | -0.089200341 | 6.3127573 | 6.3566313 | 6.48309 | -0.126459152 | -0.170333093 |
| Clec9a | 6.830017 | 6.677362 | 6.9371 | -0.259689695 | -0.107034319 | 6.302904 | 6.6773716 | 6.8039 | -0.12652509 | -0.500992766 |
| Arc | 8.043189 | 8.001652 | 8.1742 | -0.17259298 | -0.131056315 | 7.6267715 | 7.9330073 | 8.05965 | -0.126637915 | -0.432873649 |
| Tm4sf5 | 4.689559 | 4.587423 | 4.7908 | -0.203411013 | -0.101275098 | 4.7585235 | 4.5791201 | 4.70576 | -0.126639049 | 0.052764323 |
| Sv2a | 5.355072 | 5.152986 | 5.2706 | -0.117601297 | 0.084485123 | 5.4494211 | 5.1815542 | 5.30819 | -0.126639455 | 0.141227474 |
| Tmem164 | 9.672481 | 9.61962 | 9.7027 | -0.083100835 | -0.030239421 | 9.6022952 | 9.5483429 | 9.67506 | -0.126714588 | -0.072762367 |
| Plekhh2 | 3.630598 | 3.57675 | 3.6297 | -0.052963125 | 0.00088416 | 3.6389381 | 3.4966109 | 3.62341 | -0.126794873 | 0.01553229 |
| Catsper2 | 5.239597 | 5.04837 | 5.2833 | -0.234951689 | -0.043724514 | 5.3157025 | 5.3056782 | 5.43258 | -0.126898719 | -0.116874421 |
| Rnf20 | 11.00252 | 11.05031 | 10.976 | 0.074485174 | 0.026697956 | 11.029497 | 10.991303 | 11.1182 | -0.126940586 | -0.088747111 |
| Pheta2 | 3.728119 | 3.709763 | 3.7157 | -0.0059478 | 0.012408015 | 3.77709 | 3.5775926 | 3.70463 | -0.127041437 | 0.072455973 |
| Gm4787 | 2.5469 | 2.526385 | 2.5924 | -0.065992309 | -0.045476694 | 2.5413712 | 2.5556468 | 2.68271 | -0.127067803 | -0.141343353 |
| Vwa7 | 4.686308 | 4.701027 | 4.7695 | -0.068450502 | -0.083169294 | 4.9390661 | 4.7794262 | 4.90658 | -0.127154555 | 0.032485351 |
| Ubxn2a | 9.145292 | 9.199505 | 9.2317 | -0.032199238 | -0.086412231 | 8.955517 | 9.0313147 | 9.15861 | -0.127294236 | -0.203091923 |
| Egln1 | 9.351207 | 9.293451 | 9.413 | -0.119526477 | -0.06177045 | 9.166697 | 9.0716573 | 9.19899 | -0.1273283 | -0.032288667 |
| Pradc1 | 6.645904 | 6.744246 | 6.6357 | 0.108508732 | 0.010166842 | 6.7519535 | 6.7034468 | 6.83079 | -0.127341404 | -0.078834751 |

| | | | | | | | | | | |
|---|---|---|---|---|---|---|---|---|---|---|
| Traj17 | 2.831392 | 2.848133 | 2.857 | -0.008825975 | -0.025567467 | 2.7587306 | 2.764983 | 2.89236 | -0.127373529 | -0.133625964 |
| Ermn | 1.88277 | 1.766051 | 1.7673 | -0.001228082 | 0.115491002 | 1.8410208 | 1.7689097 | 1.89636 | -0.127445817 | -0.055334699 |
| Gm48793 | 3.304959 | 3.455932 | 3.4303 | 0.02562183 | -0.125350634 | 3.4499851 | 3.4047008 | 3.53216 | -0.127457624 | -0.082173312 |
| Actr3 | 14.24053 | 14.20096 | 14.13 | 0.070562786 | 0.110133652 | 14.150872 | 13.944424 | 14.072 | -0.127563282 | 0.078884527 |
| Odr4 | 9.017313 | 8.924348 | 9.0869 | -0.162578254 | -0.069613329 | 9.0178898 | 8.9688238 | 9.09639 | -0.127563554 | -0.078497577 |
| Tmsb4x | 15.58164 | 15.50849 | 15.56 | -0.0516519 | 0.021502302 | 15.612851 | 15.425805 | 15.5534 | -0.127610688 | 0.059434491 |
| Snapin | 8.891194 | 8.659014 | 8.9341 | -0.275126941 | -0.042946216 | 9.0511121 | 8.7395365 | 8.86715 | -0.127611541 | 0.183964097 |
| Hrh4 | 1.539807 | 1.547399 | 1.7532 | -0.205784263 | -0.213375915 | 1.5510782 | 1.4987209 | 1.62638 | -0.12765485 | -0.075297544 |
| Npas2 | 2.398532 | 2.367991 | 2.4824 | -0.114437083 | -0.083896122 | 2.4372022 | 2.2705874 | 2.39828 | -0.127690529 | 0.03892427 |
| Cd209g | 1.807614 | 1.523346 | 1.7418 | -0.218442338 | 0.065826308 | 1.5415106 | 1.5146242 | 1.64233 | -0.127708296 | -0.100821859 |
| Pbld1 | 1.630325 | 1.653214 | 1.7882 | -0.135034558 | -0.157923968 | 1.6477805 | 1.6282932 | 1.75608 | -0.127791561 | -0.108304313 |
| Acot1 | 2.775192 | 2.713405 | 2.7363 | -0.022922955 | 0.038863242 | 2.8065279 | 2.5888432 | 2.71665 | -0.127804822 | 0.089879814 |
| Fuom | 4.559474 | 4.457746 | 4.5181 | -0.0603532 | 0.041375067 | 4.6880922 | 4.3629247 | 4.49073 | -0.127807659 | 0.197359828 |
| Tbx20 | 1.696261 | 1.667657 | 1.7444 | -0.076742969 | -0.048138856 | 1.6994469 | 1.7466671 | 1.87449 | -0.127818357 | -0.175038576 |
| Mrc2 | 4.133417 | 4.253833 | 3.9481 | 0.305746896 | 0.185330472 | 4.4120758 | 3.956159 | 4.08398 | -0.127823866 | 0.328092909 |
| Olfr12 | 4.951241 | 4.770896 | 4.8979 | -0.127022717 | 0.053322214 | 4.8468416 | 4.8509949 | 4.97886 | -0.127865845 | -0.132019088 |
| Ntn1 | 5.186932 | 5.082668 | 5.0428 | 0.039910393 | 0.144174473 | 4.8465417 | 4.8530889 | 4.98098 | -0.127891066 | -0.134438265 |
| Olfr78 | 1.846892 | 1.764846 | 1.7881 | -0.023253944 | 0.058792601 | 1.7978404 | 1.8001654 | 1.92813 | -0.127968556 | -0.13029354 |
| Acer1 | 2.56889 | 2.516088 | 2.5425 | -0.026448263 | 0.026354108 | 2.5734049 | 2.5481771 | 2.67616 | -0.12798079 | -0.102752905 |
| Lhpp | 6.334972 | 6.318427 | 6.3099 | 0.008541428 | 0.025086863 | 6.3989376 | 6.1802633 | 6.30828 | -0.128020743 | 0.09065364 |
| Pdgfa | 5.240108 | 5.415193 | 5.1933 | 0.221850995 | 0.046766566 | 5.0267741 | 4.9481231 | 5.07615 | -0.128022766 | -0.049371794 |
| Sgms2 | 4.145286 | 3.672077 | 3.2489 | 0.423204776 | 0.896413948 | 3.4662089 | 3.6458406 | 3.77386 | -0.128023511 | -0.307655216 |
| Nupr1l | 2.8997 | 2.803288 | 2.8687 | -0.065433294 | 0.030978554 | 2.8428847 | 2.7687521 | 2.89678 | -0.128025698 | -0.053893157 |
| Kcnn1 | 2.424437 | 2.492669 | 2.3716 | 0.121065298 | 0.052833393 | 2.4223268 | 2.3199632 | 2.44799 | -0.128027166 | -0.025663563 |
| D630039A03Rik | 3.353528 | 3.229292 | 3.4136 | -0.184258566 | -0.060023357 | 3.3780018 | 3.1683764 | 3.29641 | -0.128034653 | 0.081590731 |
| Trdmt1 | 7.693234 | 7.33178 | 7.5056 | -0.173793363 | 0.187661077 | 7.8850751 | 7.5938515 | 7.72189 | -0.128035495 | 0.163188126 |
| Hnrnpul1 | 12.25524 | 12.19286 | 12.257 | -0.06442944 | -0.002047862 | 12.245303 | 12.110944 | 12.2391 | -0.128145846 | 0.00621354 |
| Gnpat | 10.21662 | 10.21152 | 10.295 | -0.08386181 | -0.078754786 | 10.577619 | 10.370247 | 10.4984 | -0.128162986 | 0.079209156 |
| Gpcpd1 | 10.94493 | 10.7851 | 10.965 | -0.179625008 | -0.019800524 | 10.943657 | 10.888392 | 11.0166 | -0.128168652 | -0.072903387 |
| Icam4 | 2.070718 | 1.943015 | 2.0215 | -0.078438175 | 0.049264582 | 1.9567399 | 1.9315512 | 2.05974 | -0.128187054 | -0.102998374 |
| Pkp2 | 2.673303 | 2.515266 | 2.5792 | -0.063941328 | 0.094095813 | 2.6182152 | 2.5334782 | 2.66167 | -0.128194117 | -0.043457076 |
| Ighv8-2 | 4.137664 | 4.39704 | 4.2129 | 0.184118041 | -0.075258071 | 4.2619725 | 4.1437663 | 4.27201 | -0.128245786 | -0.010039522 |

| | | | | | | | | | | |
|---|---|---|---|---|---|---|---|---|---|---|
| Arsb | 9.684469 | 9.488633 | 9.6844 | -0.195737878 | 9.80472E-05 | 9.3316306 | 9.4791817 | 9.60748 | -0.128298208 | -0.275849255 |
| Lrrc56 | 5.625719 | 5.668803 | 5.7631 | -0.094284962 | -0.13736809 | 5.9328415 | 5.6056489 | 5.73398 | -0.128326531 | 0.198866082 |
| Zfp30 | 4.089249 | 4.059012 | 4.1186 | -0.059612495 | -0.029376316 | 4.1575994 | 4.1419159 | 4.27028 | -0.128360393 | -0.112676923 |
| Fam217a | 1.839375 | 1.790052 | 1.8836 | -0.093538645 | -0.044215598 | 1.7937576 | 1.7094977 | 1.83798 | -0.128480494 | -0.044220652 |
| Lonrf3 | 3.382601 | 3.359896 | 3.3591 | 0.000828099 | 0.023532427 | 3.0713187 | 3.0836035 | 3.21214 | -0.128535303 | -0.140820089 |
| Rsg1 | 3.781708 | 3.892718 | 3.9019 | -0.009177153 | -0.120187177 | 4.0333449 | 3.8512085 | 3.97988 | -0.128673815 | 0.053462572 |
| Efcab8 | 2.698804 | 2.709617 | 2.7943 | -0.084661869 | -0.095475098 | 2.9175437 | 2.6359193 | 2.76467 | -0.128750118 | 0.152874223 |
| Crx | 2.687446 | 2.71009 | 2.6941 | 0.015953967 | -0.006690413 | 2.6539297 | 2.6509848 | 2.77978 | -0.128795789 | -0.125850933 |
| Sft2d2 | 10.87319 | 10.851 | 10.922 | -0.071068308 | -0.048884992 | 10.72259 | 10.657739 | 10.7865 | -0.128801376 | -0.063950782 |
| Cenpo | 8.025497 | 7.982463 | 8.1329 | -0.15043161 | -0.107397473 | 8.0121814 | 7.9963223 | 8.12516 | -0.128835087 | -0.112976018 |
| Arfip1 | 9.715822 | 9.626471 | 9.6659 | -0.039398519 | 0.049953117 | 9.5938138 | 9.4248832 | 9.55375 | -0.128867279 | 0.040063248 |
| Tbc1d24 | 6.063579 | 6.030126 | 6.1463 | -0.116204889 | -0.082751627 | 5.8640642 | 5.7826721 | 5.9116 | -0.128926517 | -0.047534499 |
| Upb1 | 5.620524 | 5.290697 | 5.7146 | -0.423930087 | -0.094103012 | 5.2340349 | 5.3413326 | 5.47027 | -0.128941376 | -0.236239065 |
| Daam1 | 7.939508 | 7.832606 | 7.9732 | -0.14054981 | -0.033647256 | 7.5002603 | 7.6876396 | 7.81673 | -0.129094938 | -0.316474157 |
| Tango6 | 8.02124 | 7.996023 | 7.9735 | 0.022530315 | 0.047747623 | 8.0657043 | 7.9850587 | 8.11418 | -0.129116888 | -0.048471286 |
| Ryr3 | 4.829562 | 4.852446 | 4.6893 | 0.163189547 | 0.140305822 | 4.8872084 | 4.7988736 | 4.92806 | -0.1291913 | -0.040856475 |
| Pear1 | 9.475871 | 9.433218 | 9.4955 | -0.062285578 | -0.019632426 | 9.6299421 | 9.2746476 | 9.40386 | -0.129215502 | 0.226079018 |
| Wdr37 | 9.899167 | 9.916645 | 9.9075 | 0.009135565 | -0.00834207 | 9.7168819 | 9.8866409 | 10.0161 | -0.129427239 | -0.299186222 |
| Map2k7 | 10.50229 | 10.41219 | 10.432 | -0.020034816 | 0.070066464 | 10.586771 | 10.457153 | 10.5866 | -0.129427772 | 0.000190114 |
| Emsy | 10.31582 | 10.37793 | 10.388 | -0.009818264 | -0.071924942 | 10.247162 | 10.352815 | 10.4823 | -0.129438568 | -0.235092118 |
| Mn1 | 3.288828 | 3.286476 | 3.3106 | -0.024083199 | -0.021731102 | 3.0942099 | 3.0765102 | 3.206 | -0.129494409 | -0.111794713 |
| Coro1c | 10.31842 | 10.21906 | 10.288 | -0.069064166 | 0.030290882 | 9.9484259 | 9.9578039 | 10.0874 | -0.129552259 | -0.138930188 |
| Ap3m1 | 9.852541 | 9.728013 | 9.6164 | 0.111610268 | 0.236138098 | 9.6328049 | 9.6215768 | 9.75114 | -0.129558852 | -0.118330819 |
| Pnma1 | 2.736043 | 2.695279 | 2.864 | -0.168770439 | -0.128006216 | 2.6574727 | 2.5854007 | 2.71503 | -0.129632275 | -0.05756035 |
| Bcar3 | 10.25942 | 10.34757 | 10.333 | 0.014883904 | -0.073260128 | 10.599651 | 10.358607 | 10.4883 | -0.129674528 | 0.111369192 |
| Anks6 | 3.743656 | 3.547187 | 3.6951 | -0.14786737 | 0.048601902 | 3.4489796 | 3.4246098 | 3.55437 | -0.129764727 | -0.105394883 |
| Plxna2 | 7.405488 | 7.147742 | 6.9549 | 0.192837658 | 0.450583155 | 6.2430866 | 6.5327167 | 6.66248 | -0.1297663 | -0.419396413 |
| Smt3h2-ps4 | 0.060871 | 0.096264 | 0.1168 | -0.020487026 | -0.055879314 | 0.1005184 | 0.0631925 | 0.19299 | -0.129792599 | -0.092466623 |
| Ighv1-38 | 2.790804 | 2.776214 | 2.8091 | -0.032846051 | -0.018256345 | 2.7745917 | 2.6367029 | 2.76655 | -0.129842567 | 0.008046218 |
| Cln3 | 9.060571 | 9.180709 | 9.2135 | -0.03276172 | -0.152899964 | 9.4354318 | 9.2784129 | 9.40826 | -0.129850434 | 0.027168413 |
| Dynlt1-ps1 | 4.806378 | 4.728085 | 4.388 | 0.340089361 | 0.418382245 | 4.2979777 | 4.6290496 | 4.759 | -0.129952218 | -0.461024098 |
| Slc7a7 | 7.65929 | 7.604422 | 7.3797 | 0.224682707 | 0.279551098 | 7.486222 | 7.2651254 | 7.39527 | -0.1301457 | 0.090950956 |

| | | | | | | | | | | |
|---|---|---|---|---|---|---|---|---|---|---|
| Ptprc | 15.16522 | 15.1326 | 15.122 | 0.010377798 | 0.04300022 | 15.187311 | 15.017083 | 15.1473 | -0.130177678 | 0.040050671 |
| Inppl1 | 8.34138 | 8.173476 | 8.5202 | -0.346771786 | -0.178867024 | 8.3447856 | 8.2423198 | 8.37254 | -0.130220079 | -0.027754354 |
| Vhl | 9.010701 | 8.90415 | 9.2072 | -0.303095435 | -0.196545256 | 8.9387281 | 8.9163292 | 9.04658 | -0.130253302 | -0.10785443 |
| Gm40466 | 2.883801 | 2.769878 | 2.9072 | -0.137281514 | -0.023358574 | 2.8288559 | 2.8092457 | 2.93951 | -0.130268222 | -0.110658033 |
| Olfr896-ps1 | 1.606828 | 1.467353 | 1.5997 | -0.132369688 | 0.007105063 | 1.4502141 | 1.4652773 | 1.59558 | -0.13030676 | -0.145369945 |
| Rcor1 | 10.48786 | 10.49206 | 10.524 | -0.031776046 | -0.035973836 | 10.236177 | 10.339493 | 10.4698 | -0.13030813 | -0.233624379 |
| Lyrm7 | 5.820491 | 6.013798 | 6.0695 | -0.055716047 | -0.249022233 | 6.1867046 | 6.09696 | 6.22733 | -0.130371925 | -0.040627303 |
| Gm2004 | 1.710468 | 1.553465 | 1.5868 | -0.033313919 | 0.123689662 | 1.6641191 | 1.5800874 | 1.71054 | -0.130451047 | -0.046419367 |
| Bod1l | 11.40964 | 11.42466 | 11.48 | -0.055378212 | -0.070394094 | 11.275486 | 11.389512 | 11.5201 | -0.130603894 | -0.244629561 |
| Npepl1 | 10.04289 | 10.00639 | 10.109 | -0.102722228 | -0.066221315 | 10.28626 | 10.069982 | 10.2007 | -0.130738967 | 0.085539255 |
| Aldh1a1 | 2.598174 | 2.62702 | 2.6575 | -0.030445495 | -0.059291329 | 2.4672512 | 2.4679362 | 2.59883 | -0.130889415 | -0.131574454 |
| Reep1 | 5.186268 | 4.86378 | 4.626 | 0.23775245 | 0.560240437 | 4.5172008 | 4.4925187 | 4.62347 | -0.130947089 | -0.106264953 |
| Pgk2 | 1.243405 | 1.21146 | 1.3135 | -0.102014412 | -0.070068652 | 1.3108531 | 1.1770326 | 1.30804 | -0.131006906 | 0.002813572 |
| Gm5511 | 5.961393 | 5.808845 | 5.9033 | -0.094437321 | 0.058110984 | 5.9701171 | 5.7890627 | 5.92009 | -0.131024132 | 0.050030264 |
| Gm14335 | 2.945266 | 2.966567 | 2.9794 | -0.012863972 | -0.034164885 | 3.0621564 | 3.10836 | 3.23939 | -0.131026255 | -0.177229776 |
| Clstn3 | 2.312936 | 2.295893 | 2.3032 | -0.007312737 | 0.009730561 | 2.1918741 | 2.2032035 | 2.33432 | -0.131119927 | -0.142449347 |
| Gm18716 | 4.103744 | 4.06816 | 4.211 | -0.142869922 | -0.107285341 | 4.0248943 | 4.1308907 | 4.26205 | -0.131157859 | -0.237154231 |
| Gm5590 | 2.900567 | 2.961509 | 2.9115 | 0.050030917 | -0.010911472 | 2.966722 | 2.8622974 | 2.99352 | -0.131227193 | -0.026802605 |
| Hdgfl3 | 8.309731 | 8.161593 | 8.2442 | -0.082593216 | 0.065545149 | 7.8340113 | 8.0433322 | 8.1746 | -0.131263566 | -0.340584465 |
| Zfp9 | 4.699797 | 4.501637 | 4.6094 | -0.10779252 | 0.090367298 | 4.3310585 | 4.2731359 | 4.40446 | -0.131327345 | -0.073404829 |
| R3hcc1l | 9.378079 | 9.40727 | 9.3129 | 0.094398588 | 0.065207279 | 9.1062372 | 9.3794278 | 9.51078 | -0.131349614 | -0.404540205 |
| Zfp383 | 6.66164 | 6.687355 | 6.6278 | 0.059552136 | 0.03383786 | 6.598102 | 6.4641705 | 6.59556 | -0.13138738 | 0.002544053 |
| Tmem100 | 3.522286 | 3.578165 | 3.5359 | 0.042312321 | -0.013566665 | 3.4945793 | 3.4578717 | 3.58927 | -0.13140152 | -0.094693882 |
| Ighv2-6-8 | 3.359035 | 3.306378 | 3.3854 | -0.079028559 | -0.026371687 | 3.3626691 | 3.2410672 | 3.37248 | -0.131409392 | -0.009807552 |
| Fsd1l | 3.722029 | 3.963861 | 4.0689 | -0.105045488 | -0.346877657 | 3.7693642 | 3.6971701 | 3.82861 | -0.131443996 | -0.059249942 |
| Gm4791 | 1.194233 | 1.184118 | 1.2344 | -0.050305045 | -0.040190846 | 1.1762373 | 1.2030747 | 1.33456 | -0.131484713 | -0.158322176 |
| Zfp619 | 6.482056 | 6.627904 | 6.5112 | 0.116669442 | -0.029178305 | 6.5221756 | 6.4398934 | 6.5714 | -0.131504741 | -0.049222494 |
| Zfp958 | 7.490296 | 7.52675 | 7.5927 | -0.0659036 | -0.102357248 | 7.415185 | 7.5340315 | 7.66556 | -0.131523965 | -0.250370415 |
| Erf | 8.814377 | 8.658052 | 8.7145 | -0.056489641 | 0.099835932 | 8.4161287 | 8.4671567 | 8.59871 | -0.131548427 | -0.182576425 |
| Taok1 | 10.91993 | 10.92364 | 10.881 | 0.042522212 | 0.038804214 | 10.659533 | 10.863154 | 10.9948 | -0.131622226 | -0.335243266 |
| Slc13a4 | 1.672766 | 1.687716 | 1.6708 | 0.016876187 | 0.001926178 | 1.6466465 | 1.7286947 | 1.86036 | -0.131667826 | -0.213716029 |
| Pcdh19 | 4.829753 | 4.71296 | 4.6825 | 0.030434331 | 0.147226635 | 4.2113799 | 4.4934218 | 4.62513 | -0.131712267 | -0.413754161 |

| | | | | | | | | | | |
|---|---|---|---|---|---|---|---|---|---|---|
| L3hypdh | 2.616726 | 2.601038 | 2.7229 | -0.121821652 | -0.106133592 | 2.5459211 | 2.4373268 | 2.56908 | -0.131750113 | -0.023155815 |
| Rimbp2 | 4.176844 | 4.248464 | 4.0613 | 0.18717788 | 0.115557826 | 4.3401389 | 4.2109124 | 4.34271 | -0.131795017 | -0.002568573 |
| Fam189a2 | 4.710871 | 4.61728 | 5.031 | -0.413677976 | -0.320087882 | 4.6275276 | 4.5802392 | 4.71208 | -0.131836062 | -0.084547576 |
| Cerkl | 5.341139 | 5.39053 | 5.3111 | 0.079458365 | 0.030066729 | 5.6013897 | 5.4653273 | 5.59723 | -0.131903414 | 0.004158983 |
| Lmntd1 | 2.738911 | 2.673003 | 2.66 | 0.012995446 | 0.07890288 | 2.8687798 | 2.6135727 | 2.74551 | -0.131936268 | 0.123270777 |
| Cpt1c | 4.406687 | 4.327069 | 4.515 | -0.187908687 | -0.108290938 | 4.2990909 | 4.4046786 | 4.53662 | -0.131936896 | -0.23752461 |
| Asnsd1 | 10.28074 | 10.27341 | 10.362 | -0.088654196 | -0.08132276 | 10.404904 | 10.184128 | 10.3161 | -0.131988552 | 0.08878745 |
| Ric3 | 3.455931 | 3.271352 | 3.482 | -0.210628787 | -0.026049521 | 3.2501744 | 3.2238359 | 3.35585 | -0.13201486 | -0.105676349 |
| Peli3 | 3.036581 | 3.019019 | 3.0762 | -0.057173222 | -0.039611336 | 3.0214186 | 2.8712155 | 3.00324 | -0.132024137 | 0.018178899 |
| Cpped1 | 7.202744 | 7.14796 | 7.1999 | -0.051972922 | 0.002811508 | 6.9896785 | 6.9021524 | 7.03426 | -0.132110659 | -0.044584523 |
| Duxbl1 | 1.767185 | 1.710689 | 1.781 | -0.070302444 | -0.013806644 | 1.6569146 | 1.6618894 | 1.794 | -0.13211343 | -0.137088316 |
| Kmt5a | 9.596854 | 9.652999 | 9.5893 | 0.063710134 | 0.007564625 | 9.5302993 | 9.5566198 | 9.68876 | -0.132141315 | -0.158461845 |
| Nrp2 | 9.598789 | 9.626223 | 9.5569 | 0.06927294 | 0.041839645 | 8.9818306 | 9.1447756 | 9.27698 | -0.132203309 | -0.29514835 |
| Cdc25a | 8.506495 | 8.535232 | 8.5827 | -0.047489945 | -0.076226333 | 8.478906 | 8.5550366 | 8.68725 | -0.132211956 | -0.208342575 |
| Pdlim5 | 10.37708 | 10.35326 | 10.286 | 0.06758535 | 0.091407718 | 10.187103 | 10.123347 | 10.2556 | -0.132264562 | -0.06850882 |
| Armc6 | 6.951817 | 7.221021 | 7.04 | 0.181061209 | -0.088142151 | 7.4310381 | 7.1133916 | 7.24572 | -0.132328646 | 0.18531781 |
| Rbm12b1 | 7.695416 | 8.064275 | 7.9053 | 0.159022934 | -0.20983543 | 8.0378606 | 8.0995901 | 8.23193 | -0.13233962 | -0.194069157 |
| Letm2 | 7.762937 | 7.840832 | 7.8917 | -0.05083668 | -0.128732077 | 8.1618079 | 7.9329456 | 8.06529 | -0.132340506 | 0.096521742 |
| Harbi1 | 7.066732 | 7.017495 | 7.264 | -0.246521945 | -0.197284734 | 7.1982489 | 7.1686889 | 7.30108 | -0.132393768 | -0.102833846 |
| Larp6 | 3.199955 | 3.175294 | 3.1496 | 0.025662517 | 0.050323006 | 2.9621853 | 2.8708413 | 3.00331 | -0.132471925 | -0.041127951 |
| Shox2 | 2.267028 | 2.271119 | 2.2836 | -0.012448339 | -0.016539163 | 2.3633937 | 2.1987833 | 2.33127 | -0.132490891 | 0.03211948 |
| Arap2 | 11.00951 | 11.12828 | 10.934 | 0.193796306 | 0.07502589 | 10.764599 | 10.738046 | 10.8706 | -0.132568404 | -0.106014594 |
| Gm13769 | 2.309273 | 2.349565 | 2.2666 | 0.08299782 | 0.04270678 | 2.2193017 | 2.2501366 | 2.38274 | -0.132598969 | -0.163433875 |
| Nptxr | 4.480488 | 4.42471 | 4.4088 | 0.015889918 | 0.071667215 | 4.2835328 | 4.2997421 | 4.43234 | -0.132599343 | -0.148808702 |
| Olfr984 | 2.695183 | 2.719728 | 2.7036 | 0.016125387 | -0.00841919 | 2.75355 | 2.5954151 | 2.72812 | -0.132701582 | 0.025433268 |
| Gphn | 8.33709 | 8.398436 | 8.4521 | -0.053621329 | -0.114967333 | 8.0110451 | 8.2768235 | 8.40953 | -0.13270526 | -0.398483658 |
| Gm49320 | 1.981391 | 1.87193 | 2.018 | -0.146099468 | -0.036638449 | 1.9892848 | 1.9485651 | 2.08137 | -0.13280305 | -0.092083361 |
| Pipox | 2.183997 | 2.039618 | 2.1388 | -0.099151263 | 0.045227738 | 2.034008 | 1.9421251 | 2.07493 | -0.132804512 | -0.040921544 |
| Clca3a2 | 2.667136 | 2.549288 | 2.5799 | -0.030634563 | 0.087213541 | 2.5671507 | 2.5016865 | 2.6345 | -0.132813645 | -0.067349481 |
| Ighv1-62-2 | 5.208858 | 5.494886 | 5.4271 | 0.067776258 | -0.218251712 | 5.4031991 | 5.2845958 | 5.4175 | -0.132900767 | -0.014297478 |
| Syde1 | 6.573639 | 6.616512 | 6.5739 | 0.042631812 | -0.000241629 | 6.4651069 | 6.2554154 | 6.3884 | -0.132987485 | 0.076704007 |
| Vmn1r83 | 3.047274 | 3.059743 | 3.1214 | -0.061635333 | -0.074104777 | 3.1337044 | 3.0676658 | 3.20093 | -0.133261728 | -0.067223104 |

| | | | | | | | | | | |
|---|---|---|---|---|---|---|---|---|---|---|
| Wdr48 | 10.05518 | 9.947729 | 10.125 | -0.177672687 | -0.070222804 | 10.158207 | 9.9947091 | 10.128 | -0.13326236 | 0.030236035 |
| Hspb11 | 7.119979 | 7.099436 | 7.2774 | -0.177918215 | -0.157374943 | 7.1120824 | 7.0181738 | 7.15144 | -0.133265799 | -0.039357163 |
| Il13 | 2.392221 | 2.398268 | 2.3321 | 0.066152696 | 0.060105845 | 2.2471639 | 2.2542609 | 2.38755 | -0.133293686 | -0.140390729 |
| Lyplal1 | 1.904775 | 1.98017 | 2.1096 | -0.129464495 | -0.204860081 | 1.9490083 | 1.8803051 | 2.01369 | -0.133380674 | -0.064677487 |
| Tecr | 10.47377 | 10.43502 | 10.585 | -0.149502802 | -0.1107526 | 10.75921 | 10.460727 | 10.5941 | -0.133402698 | 0.16508004 |
| Klf13 | 12.16801 | 12.11989 | 12.104 | 0.015951376 | 0.064074326 | 11.863308 | 11.814759 | 11.9482 | -0.133411668 | -0.084862639 |
| Alg13 | 7.031577 | 6.842245 | 7.0939 | -0.251697585 | -0.062365572 | 6.9525883 | 7.1263162 | 7.25973 | -0.133417282 | -0.307145269 |
| Hhipl1 | 1.766242 | 1.806944 | 1.7733 | 0.033627026 | -0.007074603 | 1.7673247 | 1.6526332 | 1.78605 | -0.133419984 | -0.018728486 |
| Arhgdib | 13.48282 | 13.55679 | 13.568 | -0.011242616 | -0.08521167 | 13.753064 | 13.398569 | 13.5321 | -0.13351076 | 0.220983371 |
| Hist1h2bc | 5.892813 | 5.773318 | 5.8758 | -0.102457121 | 0.017038088 | 6.0336055 | 5.8930161 | 6.02657 | -0.133558289 | 0.007031204 |
| Gramd1c | 5.886634 | 5.842661 | 5.9714 | -0.128759612 | -0.084786492 | 5.744385 | 5.6764236 | 5.81008 | -0.133658458 | -0.06569708 |
| Arid1a | 12.85946 | 12.97919 | 12.921 | 0.058282184 | -0.061445278 | 12.724914 | 12.790497 | 12.9242 | -0.133700432 | -0.199283879 |
| Fyco1 | 9.901577 | 9.887143 | 9.9418 | -0.054692627 | -0.040258654 | 10.198973 | 10.207358 | 10.3412 | -0.133794813 | -0.142179986 |
| Fdx1 | 6.634591 | 6.546172 | 6.6692 | -0.122982447 | -0.034563462 | 6.6207316 | 6.5921821 | 6.72602 | -0.133841329 | -0.105291816 |
| Acot2 | 7.918084 | 7.85473 | 7.9835 | -0.128763039 | -0.065408804 | 8.0068994 | 7.7878602 | 7.92179 | -0.133929394 | 0.08510982 |
| Kif5c | 4.924495 | 5.650039 | 5.3514 | 0.298628011 | -0.426916311 | 4.9053363 | 5.2174637 | 5.3515 | -0.13404115 | -0.446168535 |
| Atg4c | 6.618919 | 6.584464 | 6.7427 | -0.158200117 | -0.12374438 | 6.8393733 | 6.5294944 | 6.66356 | -0.134064453 | 0.175814394 |
| Stk3 | 7.574276 | 7.351252 | 7.3655 | -0.01423629 | 0.20878757 | 7.1553624 | 7.230602 | 7.36482 | -0.134219753 | -0.209459318 |
| Ccdc144b | 4.046097 | 3.887089 | 4.0908 | -0.203695849 | -0.044687512 | 3.9763998 | 4.1136928 | 4.24794 | -0.134242457 | -0.271535415 |
| Cacng3 | 0.930282 | 0.876784 | 0.9546 | -0.077774452 | -0.024276015 | 0.8374418 | 0.8677233 | 1.002 | -0.134277984 | -0.164559451 |
| Nin | 11.13076 | 11.22961 | 11.139 | 0.090261068 | -0.008583682 | 10.951211 | 11.033522 | 11.1678 | -0.1343074 | -0.216618061 |
| Gpd1 | 3.432642 | 3.330524 | 3.3777 | -0.047185364 | 0.054933089 | 3.6001804 | 3.2561998 | 3.39055 | -0.134353237 | 0.209627336 |
| Ddias | 5.697852 | 5.580139 | 5.6776 | -0.097479026 | 0.020234747 | 5.7406439 | 5.7835324 | 5.91791 | -0.134380433 | -0.177268939 |
| Araf | 10.68727 | 10.67682 | 10.709 | -0.03182287 | -0.021372568 | 10.986044 | 10.797844 | 10.9322 | -0.134402082 | 0.053798052 |
| Smarcad1 | 9.312485 | 9.370181 | 9.3035 | 0.066711059 | 0.009015346 | 9.3650033 | 9.2805766 | 9.41503 | -0.134452921 | -0.050026178 |
| Pomgnt1 | 8.676624 | 8.603592 | 8.6932 | -0.089592232 | -0.016559558 | 8.9551734 | 8.6281374 | 8.76259 | -0.134453055 | 0.192582968 |
| Dele1 | 9.13547 | 9.12661 | 9.1246 | 0.001998571 | 0.010858927 | 9.450153 | 9.148954 | 9.28347 | -0.13451106 | 0.166687918 |
| Gm16353 | 0.862112 | 0.847798 | 0.7945 | 0.053293256 | 0.067607138 | 0.8291045 | 0.7985784 | 0.93315 | -0.134567667 | -0.104041584 |
| Col4a1 | 10.56473 | 10.5957 | 10.41 | 0.185302081 | 0.154326135 | 10.453763 | 10.081627 | 10.2163 | -0.134683936 | 0.237452268 |
| Atp2b2 | 3.197123 | 3.067763 | 3.086 | -0.018239917 | 0.111120186 | 2.8598984 | 2.9130927 | 3.04778 | -0.13468642 | -0.187880752 |
| Trbj1-6 | 5.023885 | 5.118411 | 5.1622 | -0.04381807 | -0.138343936 | 5.1326064 | 5.0354268 | 5.17012 | -0.134692554 | -0.037512881 |
| Igkv8-34 | 2.919016 | 2.967233 | 3.016 | -0.048754935 | -0.096972353 | 3.04877 | 2.8935385 | 3.02826 | -0.134718737 | 0.020512767 |

| | | | | | | | | | | |
|---|---|---|---|---|---|---|---|---|---|---|
| Fkbp5 | 10.47639 | 10.51249 | 10.36 | 0.152557884 | 0.116453984 | 9.7214341 | 9.849905 | 9.98465 | -0.134741346 | -0.263212208 |
| Iqch | 2.293354 | 2.321364 | 2.3585 | -0.037124997 | -0.06513571 | 2.2605801 | 2.2804167 | 2.41517 | -0.134749729 | -0.154586378 |
| Ptger2 | 5.552051 | 5.385909 | 5.6836 | -0.297681176 | -0.131538872 | 5.42178 | 5.4759264 | 5.61069 | -0.134767588 | -0.188914016 |
| Rcc1 | 8.604528 | 8.608024 | 8.5756 | 0.032392787 | 0.028897609 | 8.7486528 | 8.5926421 | 8.72769 | -0.135049351 | 0.020961352 |
| Vipr2 | 1.707371 | 1.711866 | 1.8275 | -0.115658441 | -0.120153093 | 1.6045157 | 1.6132666 | 1.74836 | -0.135089329 | -0.143840173 |
| Kank1 | 7.027973 | 6.23025 | 6.6766 | -0.446366651 | 0.351355736 | 6.0156946 | 6.0173159 | 6.15241 | -0.135091217 | -0.136712515 |
| Zfp644 | 9.438872 | 9.500839 | 9.5037 | -0.002885681 | -0.064852924 | 9.4706025 | 9.4467646 | 9.58186 | -0.135098474 | -0.111260552 |
| Olfr916 | 1.665793 | 1.76888 | 1.7358 | 0.033036969 | -0.070050269 | 1.6840941 | 1.6325785 | 1.76768 | -0.135100858 | -0.083585262 |
| Hectd3 | 9.840703 | 9.806152 | 9.9416 | -0.13541025 | -0.100858905 | 10.175314 | 9.8547353 | 9.98992 | -0.135185099 | 0.185393317 |
| Mrm1 | 7.76664 | 7.93961 | 7.8828 | 0.056808051 | -0.116161969 | 8.3337137 | 8.1204596 | 8.25574 | -0.135281058 | 0.077973089 |
| Angpt1 | 4.058256 | 4.039605 | 4.0994 | -0.05976351 | -0.041112915 | 3.8960113 | 3.8642263 | 3.99951 | -0.135286675 | -0.103501666 |
| Adra1b | 2.449137 | 2.328675 | 2.5059 | -0.177216889 | -0.056754118 | 2.4529561 | 2.2524646 | 2.38775 | -0.135287363 | 0.065204169 |
| Kcnab3 | 3.271433 | 3.244919 | 3.2977 | -0.052781278 | -0.0262674 | 3.414844 | 3.1883285 | 3.32373 | -0.13539701 | 0.091118473 |
| Gm49626 | 2.988837 | 2.986981 | 3.0989 | -0.111957621 | -0.110101374 | 3.1099485 | 3.1014105 | 3.23685 | -0.135443181 | -0.126905238 |
| Ppp1cb | 11.82927 | 11.93369 | 11.845 | 0.089066268 | -0.015360072 | 11.764721 | 11.686271 | 11.8218 | -0.135547618 | -0.057097168 |
| H2-Aa | 15.34709 | 15.27172 | 15.39 | -0.118103668 | -0.042729384 | 15.857251 | 15.250784 | 15.3863 | -0.135560962 | 0.470906293 |
| Gm10643 | 4.279323 | 4.159516 | 4.3146 | -0.155090887 | -0.035283893 | 4.3198674 | 4.2484031 | 4.38407 | -0.135670635 | -0.064206278 |
| Olfr639 | 5.720123 | 5.656071 | 5.7066 | -0.050481596 | 0.013571272 | 5.7485016 | 6.0389618 | 6.17465 | -0.13568391 | -0.426144061 |
| 3830403N18Rik | 3.856547 | 3.834241 | 3.9098 | -0.075587429 | -0.053282172 | 3.9971331 | 3.8803439 | 4.01603 | -0.135685035 | -0.018895843 |
| Tmem35a | 2.370535 | 2.439021 | 2.3745 | 0.064545603 | -0.003940769 | 2.3743932 | 2.3292202 | 2.46491 | -0.135690699 | -0.090517639 |
| Glo1 | 9.404501 | 9.39603 | 9.51 | -0.113928983 | -0.105458348 | 9.4620972 | 9.2219146 | 9.35764 | -0.135727438 | 0.104455207 |
| Olfr667 | 2.694037 | 2.616903 | 2.6847 | -0.067786603 | 0.009346767 | 2.6407758 | 2.6623224 | 2.79807 | -0.135749241 | -0.157295841 |
| Sergef | 7.86712 | 7.78588 | 7.8221 | -0.036248361 | 0.044991968 | 8.0095273 | 7.6287764 | 7.76457 | -0.135798149 | 0.244952825 |
| Otulin | 9.923051 | 9.876376 | 9.9691 | -0.092677905 | -0.046002626 | 10.06822 | 9.7599617 | 9.89582 | -0.135858755 | 0.172399688 |
| AL732309.1 | 7.04486 | 6.900404 | 6.9206 | -0.020158066 | 0.124297118 | 7.3076972 | 6.9074582 | 7.04339 | -0.135927405 | 0.264311537 |
| Acot13 | 7.915774 | 7.622643 | 7.8634 | -0.240762836 | 0.052367822 | 7.9997284 | 7.7293129 | 7.86527 | -0.135953935 | 0.134461568 |
| Krt80 | 6.647275 | 6.30811 | 6.3823 | -0.074161171 | 0.265004035 | 6.8369719 | 6.1488533 | 6.28485 | -0.13600058 | 0.552117986 |
| Crispld1 | 1.50462 | 1.579003 | 1.5881 | -0.009053079 | -0.083435849 | 1.4542045 | 1.42605 | 1.56219 | -0.136140031 | -0.107985574 |
| Ighv5-9 | 5.383911 | 5.521794 | 5.2634 | 0.25836446 | 0.120481247 | 5.6754814 | 5.28955 | 5.42574 | -0.136188965 | 0.249742415 |
| Btaf1 | 10.61116 | 10.61175 | 10.578 | 0.034224872 | 0.033631468 | 10.383394 | 10.493386 | 10.6298 | -0.136376298 | -0.246368172 |
| Dhx36 | 10.45041 | 10.47897 | 10.497 | -0.017897698 | -0.046455654 | 10.54634 | 10.475214 | 10.6116 | -0.136376954 | -0.06525015 |
| Cep83 | 8.837011 | 8.653301 | 8.8329 | -0.179635509 | 0.004074983 | 8.7658612 | 8.852547 | 8.98897 | -0.136424791 | -0.223110557 |

| | | | | | | | | | | |
|---|---|---|---|---|---|---|---|---|---|---|
| Ino80c | 9.537473 | 9.462643 | 9.4657 | -0.003014091 | 0.071815438 | 9.7296657 | 9.3078469 | 9.44432 | -0.136474858 | 0.28534398 |
| Limk2 | 10.09231 | 10.14735 | 10.182 | -0.034738643 | -0.089778626 | 10.295253 | 10.195276 | 10.3318 | -0.1364794 | -0.036503022 |
| Erich1 | 7.363141 | 7.323394 | 7.2959 | 0.027489535 | 0.06723636 | 7.1412997 | 7.1798144 | 7.31633 | -0.136517483 | -0.175032184 |
| Dclre1b | 8.361908 | 8.38958 | 8.3661 | 0.023465998 | -0.004206273 | 8.3445311 | 8.1211462 | 8.25768 | -0.136528909 | 0.086855991 |
| Kctd17 | 7.234404 | 7.058405 | 7.0316 | 0.026788123 | 0.202787482 | 6.9499395 | 6.7804982 | 6.91704 | -0.136545623 | 0.032895687 |
| Mrps35 | 8.623777 | 8.588011 | 8.5881 | -0.000104857 | 0.035661267 | 8.5342425 | 8.4508075 | 8.58738 | -0.136570374 | -0.053135312 |
| Was | 11.38154 | 11.37752 | 11.388 | -0.010918066 | -0.006895214 | 11.607152 | 11.248557 | 11.3852 | -0.136658444 | 0.221936155 |
| Nrde2 | 8.930019 | 8.696079 | 8.9543 | -0.258192588 | -0.024253544 | 9.0815391 | 8.8895975 | 9.02638 | -0.136779577 | 0.055162001 |
| Terf2 | 9.722366 | 9.628437 | 9.7695 | -0.141047852 | -0.04711834 | 9.8523707 | 9.7470662 | 9.88391 | -0.136844907 | -0.031540379 |
| Ces2b | 2.963805 | 2.774591 | 2.9761 | -0.201459568 | -0.012245481 | 2.8089383 | 2.7292265 | 2.86621 | -0.136984375 | -0.057272621 |
| Cep63 | 9.248122 | 9.327256 | 9.2802 | 0.047103113 | -0.03203097 | 9.3409834 | 9.3548732 | 9.49192 | -0.137045944 | -0.150935753 |
| Tbc1d2b | 9.74815 | 9.706483 | 9.724 | -0.017492258 | 0.024174654 | 9.3293895 | 9.4111348 | 9.54821 | -0.137074036 | -0.218819254 |
| Cryaa | 2.774963 | 2.814408 | 2.8334 | -0.018961655 | -0.058406728 | 2.9578008 | 2.8066858 | 2.94381 | -0.137122494 | 0.01399248 |
| 2010300C02Rik | 4.445319 | 4.281718 | 4.2576 | 0.024116182 | 0.187717321 | 4.1317851 | 4.2323234 | 4.3695 | -0.137181097 | -0.237719397 |
| Gm9780 | 3.150961 | 3.081544 | 3.1969 | -0.115360949 | -0.045944209 | 2.9917927 | 2.914213 | 3.05141 | -0.137195935 | -0.059616197 |
| Rabgap1 | 9.962673 | 9.908421 | 9.8474 | 0.061033003 | 0.115285367 | 9.8399147 | 9.7626688 | 9.90007 | -0.137405559 | -0.060159677 |
| Gm8203 | 7.927227 | 8.187907 | 7.9259 | 0.26198941 | 0.001309848 | 7.9827726 | 8.1216664 | 8.25912 | -0.13744871 | -0.276342449 |
| Rpusd2 | 6.612043 | 6.48575 | 6.6595 | -0.173731911 | -0.047438436 | 6.7256705 | 6.4875676 | 6.62502 | -0.137455993 | 0.100646927 |
| Zfp184 | 3.113479 | 3.244854 | 3.1744 | 0.070405696 | -0.060969275 | 3.238182 | 3.0590593 | 3.19656 | -0.137500283 | 0.041622386 |
| Tbc1d7 | 5.608201 | 5.568456 | 5.7487 | -0.180212843 | -0.140467956 | 5.934381 | 5.635101 | 5.77276 | -0.137660095 | 0.161619964 |
| Yeats2 | 9.476469 | 9.461735 | 9.5599 | -0.098208593 | -0.083474628 | 9.3121602 | 9.6279817 | 9.7657 | -0.137713926 | -0.45353543 |
| Gm18588 | 0.57731 | 0.58037 | 0.54 | 0.040338482 | 0.037277756 | 0.623164 | 0.52784 | 0.66559 | -0.137750244 | -0.042426204 |
| Maml1 | 9.925425 | 9.998131 | 9.9103 | 0.087872713 | 0.015167564 | 9.7748332 | 9.6764285 | 9.8142 | -0.137768224 | -0.039363583 |
| Pafah1b2 | 10.13748 | 10.22607 | 10.169 | 0.05722638 | -0.03136725 | 10.038005 | 10.108054 | 10.2459 | -0.137806049 | -0.207855335 |
| Armc2 | 3.26825 | 3.473068 | 3.4859 | -0.012840384 | -0.217658658 | 3.4837009 | 3.3061777 | 3.44402 | -0.137842539 | 0.039680577 |
| Ncoa6 | 10.49777 | 10.40982 | 10.496 | -0.08620357 | 0.00174534 | 10.426157 | 10.352756 | 10.4906 | -0.137874196 | -0.064473156 |
| Gzmm | 5.443609 | 5.285762 | 5.6498 | -0.363991863 | -0.206144943 | 5.5849618 | 5.4010505 | 5.53894 | -0.137886323 | 0.04602497 |
| Grm5 | 3.852167 | 3.769909 | 3.7866 | -0.016722998 | 0.065535223 | 3.9172739 | 3.8424382 | 3.98043 | -0.137991494 | -0.063155794 |
| Olfr1178 | 3.18984 | 3.254889 | 3.3236 | -0.068704457 | -0.133752784 | 3.3374161 | 3.2570639 | 3.39512 | -0.138056688 | -0.057704486 |
| Nemp2 | 10.06375 | 9.934445 | 10.121 | -0.186730918 | -0.0574232 | 10.337024 | 10.135541 | 10.2736 | -0.13807691 | 0.063405843 |
| Tmem158 | 4.731507 | 4.739599 | 4.8482 | -0.10861809 | -0.11670964 | 4.3960756 | 4.5719883 | 4.71023 | -0.138245431 | -0.314158172 |
| Map7d3 | 3.018066 | 2.976653 | 3.1284 | -0.151772558 | -0.110359389 | 2.9116181 | 2.8547739 | 2.99305 | -0.138277468 | -0.081433272 |

| | | | | | | | | | | |
|---|---|---|---|---|---|---|---|---|---|---|
| Impad1 | 9.718111 | 9.635589 | 9.7092 | -0.073630919 | 0.008891174 | 9.413915 | 9.4691186 | 9.60748 | -0.13836128 | -0.193564889 |
| Stard8 | 6.825372 | 6.431793 | 6.7296 | -0.29783535 | 0.095743564 | 6.4469127 | 6.5367451 | 6.67515 | -0.138408834 | -0.2282412 |
| Gpc6 | 3.506867 | 3.185758 | 3.3093 | -0.123554302 | 0.197554148 | 3.0072502 | 3.0001915 | 3.13861 | -0.138420322 | -0.131361573 |
| 3000002C10Rik | 5.383019 | 5.249661 | 5.3585 | -0.108817087 | 0.024540505 | 5.88426 | 5.5405838 | 5.6791 | -0.138518186 | 0.205157918 |
| Strn3 | 9.302901 | 9.206518 | 9.2216 | -0.015089601 | 0.08129333 | 8.9721975 | 9.0162029 | 9.15477 | -0.138569106 | -0.182574475 |
| Zeb1 | 10.17668 | 10.25556 | 10.262 | -0.00610107 | -0.084987626 | 10.061607 | 10.197932 | 10.3365 | -0.13858227 | -0.274907835 |
| Eri1 | 9.965096 | 10.09048 | 9.9785 | 0.111969341 | -0.0134159 | 9.645076 | 9.846231 | 9.98506 | -0.138830867 | -0.339985816 |
| Fam193b | 10.46386 | 10.62923 | 10.641 | -0.011401293 | -0.1767661 | 10.907067 | 10.760524 | 10.8994 | -0.138849797 | 0.007692912 |
| Skida1 | 2.452394 | 2.610827 | 2.5177 | 0.093169697 | -0.065263731 | 2.4908404 | 2.4349109 | 2.57379 | -0.138879684 | -0.082950171 |
| Traj15 | 3.103109 | 3.119131 | 3.1303 | -0.011147362 | -0.027168972 | 2.9433463 | 2.9245883 | 3.06348 | -0.138891413 | -0.12013339 |
| Trav10 | 5.099836 | 5.147997 | 5.2463 | -0.098292592 | -0.146453607 | 5.0051707 | 5.1401896 | 5.27908 | -0.138893249 | -0.273912193 |
| Trav13-1 | 5.103827 | 5.153611 | 5.2626 | -0.108945437 | -0.15872951 | 5.0131814 | 4.9497407 | 5.08867 | -0.138927448 | -0.075486757 |
| Trp53bp1 | 10.2528 | 10.1911 | 10.264 | -0.0728009 | -0.011103288 | 10.25664 | 10.288076 | 10.4271 | -0.138978802 | -0.170415256 |
| Mboat1 | 7.273571 | 7.291199 | 7.3364 | -0.045183555 | -0.062811418 | 7.3053245 | 7.4015471 | 7.54062 | -0.139076375 | -0.23529906 |
| Tmem88 | 5.711506 | 5.229064 | 5.4712 | -0.242133263 | 0.240308969 | 5.2969209 | 4.9612214 | 5.10031 | -0.13908732 | 0.196612151 |
| Gcm2 | 2.528051 | 2.481473 | 2.628 | -0.146540477 | -0.099962624 | 2.6610242 | 2.4511565 | 2.59025 | -0.139092411 | 0.070775297 |
| Myo5c | 4.022237 | 3.602235 | 3.8645 | -0.262310334 | 0.157691864 | 3.8973169 | 3.7900945 | 3.92928 | -0.139187906 | -0.03196544 |
| B3gnt9 | 5.005423 | 5.082752 | 4.964 | 0.11871757 | 0.041387925 | 4.9733477 | 4.8304694 | 4.96967 | -0.139203905 | 0.003674415 |
| Arl5c | 10.0119 | 10.25474 | 10.236 | 0.0182869 | -0.224554634 | 10.131123 | 10.200061 | 10.3393 | -0.13928883 | -0.208226094 |
| mt-Co1 | 17.44295 | 17.35249 | 17.468 | -0.115992184 | -0.0255354 | 17.269192 | 17.337181 | 17.4765 | -0.139321602 | -0.207310581 |
| Atxn2 | 9.068006 | 9.039233 | 8.9274 | 0.111849082 | 0.1406224 | 9.0092256 | 8.9161958 | 9.05555 | -0.139355396 | -0.046325633 |
| Tmf1 | 10.3968 | 10.37282 | 10.421 | -0.048546192 | -0.024562706 | 10.372995 | 10.339157 | 10.4785 | -0.139362946 | -0.105525046 |
| Nup37 | 6.964851 | 6.943185 | 7.1019 | -0.158760916 | -0.137095442 | 7.1779395 | 7.0855905 | 7.22496 | -0.139371355 | -0.047022362 |
| Ypel1 | 3.840791 | 3.817823 | 3.9911 | -0.173238312 | -0.150270686 | 3.752538 | 3.7791382 | 3.91858 | -0.139439091 | -0.166039325 |
| Ankrd27 | 9.576201 | 9.469337 | 9.5312 | -0.061814935 | 0.045049162 | 9.4571716 | 9.4853394 | 9.62479 | -0.139451861 | -0.167619613 |
| Olfr1028 | 4.089606 | 4.104125 | 4.1759 | -0.071774539 | -0.086293081 | 4.250757 | 4.2832851 | 4.42274 | -0.139457166 | -0.171985263 |
| Gm43191 | 1.700516 | 1.581447 | 1.5443 | 0.037161784 | 0.156231637 | 1.4693592 | 1.5185407 | 1.65802 | -0.139476202 | -0.188657629 |
| Tollip | 10.11772 | 10.12938 | 10.11 | 0.019276554 | 0.007615508 | 10.214287 | 10.086116 | 10.2257 | -0.139540248 | -0.011369479 |
| Rragb | 6.28007 | 6.469963 | 6.4779 | -0.007927776 | -0.197820832 | 6.7275645 | 6.345633 | 6.48543 | -0.139796139 | 0.242135391 |
| Yy1 | 11.07545 | 11.14513 | 11.139 | 0.005971078 | -0.0637042 | 10.966473 | 11.001069 | 11.141 | -0.139901474 | -0.17449746 |
| Cdk12 | 10.9947 | 11.04211 | 11.005 | 0.037483302 | -0.009930984 | 11.08718 | 11.032086 | 11.172 | -0.139959324 | -0.084865792 |
| Rtl8a | 6.439978 | 6.321398 | 6.4846 | -0.163225348 | -0.044644725 | 6.3860108 | 6.1506896 | 6.29065 | -0.139962708 | 0.095358515 |

| | | | | | | | | | | |
|---|---|---|---|---|---|---|---|---|---|---|
| Mcee | 6.323239 | 6.352662 | 6.4818 | -0.129118987 | -0.158541582 | 6.5227853 | 6.3009017 | 6.44089 | -0.139991073 | 0.081892446 |
| Trav6n-6 | 4.138247 | 4.164804 | 4.1645 | 0.000321998 | -0.026235023 | 4.0447569 | 4.0219539 | 4.16203 | -0.140075112 | -0.117272073 |
| Tbx2 | 3.05643 | 3.039914 | 3.0745 | -0.034602298 | -0.018086818 | 2.9611206 | 2.9249303 | 3.06502 | -0.140091383 | -0.103901002 |
| Zfp707 | 7.550547 | 7.462395 | 7.5478 | -0.085434923 | 0.002716757 | 7.5198478 | 7.5096907 | 7.64981 | -0.140123144 | -0.129966118 |
| Cldn5 | 7.048676 | 6.841243 | 6.7125 | 0.128708901 | 0.336141565 | 6.086553 | 6.3832798 | 6.52341 | -0.140133813 | -0.43686066 |
| Zfp523 | 7.366553 | 7.229998 | 7.4992 | -0.269247864 | -0.132693383 | 7.5256024 | 7.4433505 | 7.58353 | -0.1401771 | -0.057925194 |
| Spib | 12.11001 | 12.24146 | 12.135 | 0.106172336 | -0.02527959 | 12.583977 | 11.879244 | 12.0194 | -0.140188602 | 0.564544841 |
| Slc4a9 | 2.022755 | 2.025832 | 2.0782 | -0.052344583 | -0.055421148 | 2.0027252 | 1.9312258 | 2.07156 | -0.140337659 | -0.06883828 |
| Cdh6 | 2.56354 | 2.494315 | 2.499 | -0.004656003 | 0.064568566 | 2.4154235 | 2.4207311 | 2.56115 | -0.140419644 | -0.145727253 |
| Tmppe | 7.577803 | 7.620103 | 7.5107 | 0.109358684 | 0.067059165 | 7.8091875 | 7.5664115 | 7.70686 | -0.140449806 | 0.102326222 |
| Uhrf1 | 9.19257 | 9.075302 | 9.3174 | -0.242097952 | -0.124830587 | 9.391147 | 9.5038922 | 9.64435 | -0.140459519 | -0.253204805 |
| Adgrv1 | 5.397228 | 5.272188 | 5.2563 | 0.015884542 | 0.140924557 | 5.6593591 | 5.6042042 | 5.7447 | -0.14050075 | -0.085345794 |
| Ythdf3 | 11.1251 | 11.08229 | 11.117 | -0.034405012 | 0.008411358 | 11.040691 | 10.976462 | 11.117 | -0.140506982 | -0.076278515 |
| Wasf3 | 3.159614 | 3.014286 | 3.1907 | -0.176389118 | -0.031061233 | 2.9905267 | 2.8763289 | 3.01689 | -0.140564783 | -0.026366889 |
| Pfn2 | 5.81109 | 5.718717 | 5.8397 | -0.121003586 | -0.028630226 | 5.6009531 | 5.5036379 | 5.64421 | -0.140570893 | -0.043255659 |
| Pax5 | 13.26136 | 13.27268 | 13.289 | -0.016752414 | -0.02807806 | 13.615598 | 13.27386 | 13.4145 | -0.140625138 | 0.201112841 |
| Mamdc2 | 1.732875 | 1.72267 | 1.8493 | -0.12666511 | -0.116459782 | 1.6748651 | 1.6932013 | 1.83393 | -0.140732506 | -0.159068662 |
| Alg2 | 8.565004 | 8.531187 | 8.5542 | -0.022969439 | 0.010848064 | 8.7216351 | 8.5292671 | 8.67 | -0.140737226 | 0.051630785 |
| Traj34 | 1.952225 | 1.961713 | 1.9291 | 0.032606665 | 0.023118681 | 1.8946549 | 1.8574168 | 1.99821 | -0.140797217 | -0.103559113 |
| Dcaf17 | 8.569631 | 8.500761 | 8.7081 | -0.207365716 | -0.138495631 | 8.8510629 | 8.5521999 | 8.69312 | -0.140918853 | 0.157944187 |
| Gcnt4 | 3.955307 | 3.997691 | 3.9862 | 0.011523335 | -0.030860834 | 3.9103746 | 3.9096826 | 4.05071 | -0.14102827 | -0.1403363 |
| Mfap1a | 10.41657 | 10.56347 | 10.46 | 0.103825928 | -0.043073232 | 10.489825 | 10.340192 | 10.4813 | -0.141122088 | 0.008510886 |
| Clhc1 | 1.838365 | 1.754035 | 1.8729 | -0.118818275 | -0.034488506 | 1.8293719 | 1.7684064 | 1.90959 | -0.141179999 | -0.080214504 |
| Ebf2 | 2.207977 | 2.331483 | 2.3161 | 0.015399292 | -0.108107026 | 2.1657041 | 2.1561093 | 2.29744 | -0.141333473 | -0.131738718 |
| Gm35339 | 6.730958 | 6.990532 | 6.7818 | 0.208744436 | -0.050829964 | 7.4351085 | 7.2963445 | 7.43773 | -0.141380557 | -0.00261653 |
| Ankrd11 | 12.42573 | 12.33327 | 12.444 | -0.110286386 | -0.017829396 | 12.449677 | 12.432309 | 12.5737 | -0.141394308 | -0.124026846 |
| Ptger3 | 2.194075 | 2.257403 | 2.2171 | 0.040257152 | -0.023070524 | 2.1882159 | 2.0705663 | 2.21205 | -0.141480895 | -0.02383127 |
| Gm36298 | 5.016023 | 4.885895 | 5.0192 | -0.133308631 | -0.003180705 | 4.9996623 | 4.8628675 | 5.00442 | -0.141556772 | -0.004761919 |
| Yipf4 | 9.122646 | 9.330578 | 9.2103 | 0.120238295 | -0.087693752 | 9.184909 | 9.1346139 | 9.2762 | -0.141587965 | -0.091292833 |
| Bcl2l1 | 8.883296 | 8.9249 | 8.768 | 0.156902471 | 0.115299409 | 8.4698544 | 8.5701149 | 8.71174 | -0.141620718 | -0.241881269 |
| Sgsm2 | 7.325597 | 7.153252 | 7.1417 | 0.011571201 | 0.183916197 | 7.1954058 | 7.0316977 | 7.1734 | -0.141700981 | 0.022007163 |
| Scp2 | 11.05508 | 11.10163 | 11.088 | 0.013860646 | -0.032689502 | 11.101974 | 10.890109 | 11.0318 | -0.141706504 | 0.070158311 |

| | | | | | | | | | | |
|---|---|---|---|---|---|---|---|---|---|---|
| Ncapd3 | 9.328036 | 9.372472 | 9.4116 | -0.039144171 | -0.083580599 | 9.5170211 | 9.5128374 | 9.65464 | -0.141803243 | -0.137619589 |
| Brdt | 5.048427 | 4.97414 | 5.0427 | -0.068529946 | 0.005757176 | 4.9201271 | 4.857515 | 4.99934 | -0.141823948 | -0.079211889 |
| Hoxb9 | 3.036446 | 2.759715 | 2.8849 | -0.125189371 | 0.151541918 | 2.7180188 | 2.6573842 | 2.79921 | -0.141829888 | -0.081195253 |
| Cad | 10.55832 | 10.76236 | 10.651 | 0.111068668 | -0.092973918 | 10.811631 | 10.645272 | 10.7871 | -0.14185052 | 0.024508736 |
| Ankrd13c | 10.13195 | 10.20519 | 10.203 | 0.001894333 | -0.071346793 | 9.888418 | 9.9855697 | 10.1274 | -0.141854529 | -0.239006245 |
| Rfx3 | 8.363333 | 8.436088 | 8.4901 | -0.054010222 | -0.126765679 | 8.3018317 | 8.4226873 | 8.56468 | -0.14199056 | -0.262846183 |
| Tex10 | 9.76839 | 9.826494 | 9.7977 | 0.028779469 | -0.029324454 | 9.758285 | 9.7227417 | 9.86474 | -0.142001145 | -0.10645786 |
| Cenps | 5.732661 | 5.614015 | 5.8483 | -0.234318029 | -0.115672194 | 6.0151299 | 5.7790784 | 5.92109 | -0.142013857 | 0.094037652 |
| Trdv5 | 3.852802 | 3.76773 | 3.9301 | -0.162402305 | -0.077330553 | 3.8578435 | 3.8275299 | 3.96962 | -0.142089463 | -0.111775863 |
| Snupn | 7.166112 | 6.958544 | 7.1218 | -0.163230149 | 0.044337995 | 7.1162391 | 6.9514583 | 7.0936 | -0.142137575 | 0.022643219 |
| Trbv17 | 6.789029 | 6.806531 | 6.7791 | 0.027425124 | 0.009923379 | 6.4860273 | 6.696402 | 6.83861 | -0.142206402 | -0.352581112 |
| Traj25 | 1.507102 | 1.471034 | 1.5123 | -0.041283785 | -0.005215876 | 1.4641892 | 1.4874078 | 1.62967 | -0.14225898 | -0.165477614 |
| Gnao1 | 3.853139 | 3.658897 | 3.7112 | -0.052269223 | 0.141973398 | 3.3388749 | 3.416966 | 3.55923 | -0.14226345 | -0.220354612 |
| Nupl2 | 6.859922 | 6.630406 | 6.8894 | -0.259043894 | -0.029527924 | 6.935686 | 6.7907759 | 6.93309 | -0.142310731 | 0.002599348 |
| Dnajb5 | 8.483013 | 8.388738 | 8.4173 | -0.028579056 | 0.065695635 | 8.3559649 | 8.1831129 | 8.32553 | -0.142413179 | 0.030438809 |
| Gm6451 | 4.17775 | 4.206421 | 4.2636 | -0.057204647 | -0.085875752 | 4.1247334 | 4.0134707 | 4.1559 | -0.14242612 | -0.031163382 |
| Ercc6l | 7.191044 | 7.100431 | 7.3186 | -0.21814311 | -0.127529833 | 7.0605626 | 7.2363663 | 7.37897 | -0.142601024 | -0.318404765 |
| Olfr63 | 5.417319 | 5.473179 | 5.6366 | -0.163441787 | -0.219301399 | 5.5161367 | 5.6541741 | 5.79681 | -0.142634188 | -0.28067168 |
| Dyrk3 | 4.286175 | 4.250281 | 4.2829 | -0.032615725 | 0.003278583 | 4.1396483 | 4.0713366 | 4.21403 | -0.142691126 | -0.074379379 |
| Pcdhgb2 | 4.302515 | 4.194201 | 4.154 | 0.040183277 | 0.148497031 | 4.0800664 | 3.9271024 | 4.06984 | -0.142742496 | 0.010221488 |
| St7l | 8.255043 | 8.436397 | 8.4105 | 0.025931557 | -0.155422432 | 8.1158394 | 8.2387757 | 8.38156 | -0.142786634 | -0.265722913 |
| Rab6a | 10.30261 | 10.25399 | 10.264 | -0.010035082 | 0.038585332 | 10.248513 | 10.041517 | 10.1844 | -0.142880637 | 0.064115576 |
| Kctd15 | 3.771201 | 3.358266 | 3.4381 | -0.07985372 | 0.333081944 | 3.3109049 | 3.3714917 | 3.51437 | -0.142881991 | -0.203468801 |
| Pcdhb14 | 4.109672 | 3.983028 | 3.9937 | -0.010627112 | 0.116016947 | 3.8805531 | 3.7624855 | 3.90537 | -0.142884719 | -0.024817196 |
| Gm10702 | 1.746489 | 1.659312 | 1.7548 | -0.095485686 | -0.008308414 | 1.7719643 | 1.6417966 | 1.78474 | -0.142942859 | -0.012775189 |
| Gm44126 | 4.802846 | 4.541133 | 4.7854 | -0.244227367 | 0.017485029 | 4.6813479 | 4.5654813 | 4.70843 | -0.14295127 | -0.027084642 |
| Lmo7 | 5.071301 | 4.890457 | 5.1564 | -0.265980134 | -0.085136205 | 4.7587689 | 4.7672048 | 4.91019 | -0.142986285 | -0.151422204 |
| Fhl5 | 1.972595 | 1.893483 | 2.0849 | -0.191427683 | -0.112315316 | 1.8306868 | 1.8291775 | 1.97224 | -0.143065164 | -0.141555885 |
| Apol7a | 1.148938 | 1.111893 | 1.0883 | 0.023638352 | 0.060683188 | 1.1655501 | 1.1040467 | 1.24719 | -0.143148152 | -0.081644802 |
| Gm48800 | 2.836714 | 2.789668 | 2.829 | -0.039350039 | 0.007696091 | 2.9627761 | 2.8844918 | 3.02766 | -0.143164593 | -0.064880268 |
| Pdzk1 | 1.767239 | 1.676205 | 1.74 | -0.063762561 | 0.027271693 | 1.6634568 | 1.6315682 | 1.77476 | -0.143195807 | -0.111307201 |
| Hesx1 | 1.036943 | 0.961827 | 0.9672 | -0.005370842 | 0.06974598 | 1.025133 | 0.9510718 | 1.09428 | -0.143205254 | -0.069144026 |

| | | | | | | | | | | |
|---|---|---|---|---|---|---|---|---|---|---|
| Rbsn | 9.125123 | 9.043075 | 9.1058 | -0.062683027 | 0.019364073 | 9.0458935 | 8.9081619 | 9.05147 | -0.143305334 | -0.00557381 |
| Acss3 | 3.858116 | 3.793516 | 3.922 | -0.128485134 | -0.063884455 | 3.9478552 | 3.8951525 | 4.03849 | -0.143341187 | -0.09063855 |
| Entpd7 | 7.386612 | 7.546859 | 7.338 | 0.208833039 | 0.048586612 | 7.5056818 | 7.3476704 | 7.49102 | -0.143353395 | 0.014657999 |
| Ttll13 | 2.247135 | 2.151332 | 2.1865 | -0.035127965 | 0.060674581 | 2.2389906 | 2.2033234 | 2.34674 | -0.143415607 | -0.107748384 |
| Rbm15b | 9.766747 | 9.817488 | 9.8178 | -0.000347859 | -0.051088465 | 9.4130075 | 9.6090359 | 9.75257 | -0.143536435 | -0.339564919 |
| Shc1 | 10.72283 | 10.71227 | 10.725 | -0.012943568 | -0.002381074 | 10.709043 | 10.507141 | 10.6507 | -0.143559486 | 0.058341689 |
| Ctif | 5.876057 | 5.65788 | 5.8261 | -0.168222198 | 0.049955174 | 5.4767917 | 5.4916824 | 5.63525 | -0.143572337 | -0.158463005 |
| Nek6 | 8.179249 | 8.253455 | 8.1773 | 0.076148238 | 0.001942271 | 7.8756792 | 7.9024579 | 8.04612 | -0.143663757 | -0.170442422 |
| Nap1l1 | 12.74472 | 12.66999 | 12.802 | -0.13245426 | -0.057719448 | 12.853087 | 12.736728 | 12.8805 | -0.143729632 | -0.027370156 |
| Slc39a14 | 8.886301 | 8.679966 | 8.7743 | -0.094299586 | 0.11203585 | 8.6351319 | 8.5165611 | 8.6603 | -0.143741224 | -0.025170372 |
| P2ry6 | 6.751515 | 6.28511 | 6.3224 | -0.037332412 | 0.429072775 | 6.5884101 | 5.9692816 | 6.11306 | -0.14377992 | 0.475348576 |
| Zfp110 | 10.03625 | 10.03435 | 10.054 | -0.019420509 | -0.017521044 | 10.063721 | 9.8899188 | 10.0338 | -0.143876618 | 0.029926053 |
| Rhot2 | 9.967822 | 9.989976 | 10.088 | -0.097806255 | -0.119960634 | 10.183623 | 10.058651 | 10.2026 | -0.143899934 | -0.018927565 |
| Vmn2r124 | 4.297482 | 4.487904 | 4.3765 | 0.11145243 | -0.078970454 | 4.4360047 | 4.502916 | 4.64683 | -0.143914476 | -0.210825784 |
| Cyp2j6 | 4.942399 | 4.63189 | 4.9275 | -0.295627949 | 0.014881397 | 4.4854617 | 4.4505666 | 4.59453 | -0.143961131 | -0.10906602 |
| Gtf2h1 | 9.730638 | 9.639717 | 9.7318 | -0.092054991 | -0.001133818 | 9.5456007 | 9.6564359 | 9.80043 | -0.143990561 | -0.254825761 |
| Jmjd4 | 8.275944 | 8.352589 | 8.2481 | 0.10453283 | 0.027888048 | 8.3161274 | 8.2533711 | 8.39737 | -0.143997017 | -0.081240732 |
| Ahcyl2 | 10.26108 | 10.5094 | 10.17 | 0.33939169 | 0.09106586 | 10.034821 | 10.190469 | 10.3345 | -0.14399809 | -0.299645884 |
| Zmym1 | 8.07876 | 8.131849 | 8.1148 | 0.017057693 | -0.036031834 | 7.9451418 | 8.0511988 | 8.1954 | -0.14419659 | -0.250253605 |
| Gm18811 | 2.100022 | 2.02789 | 2.0596 | -0.031707141 | 0.04042489 | 1.9988093 | 1.9832814 | 2.12753 | -0.14424969 | -0.128721768 |
| Sostdc1 | 3.596539 | 3.424984 | 3.4481 | -0.023068032 | 0.148486812 | 3.3967508 | 3.2338774 | 3.37813 | -0.144255627 | 0.018617734 |
| Id3 | 9.348495 | 9.066082 | 9.3636 | -0.297546313 | -0.015133508 | 8.9442794 | 8.7338712 | 8.87814 | -0.14426575 | 0.066142459 |
| Naa60 | 10.57919 | 10.48164 | 10.49 | -0.007871064 | 0.08967905 | 10.649962 | 10.407561 | 10.5519 | -0.144371468 | 0.098029736 |
| Ddx17 | 12.94182 | 12.95389 | 12.864 | 0.090350862 | 0.078277508 | 13.285588 | 13.04309 | 13.1876 | -0.144519602 | 0.097978526 |
| Ppm1e | 5.930733 | 5.849395 | 5.841 | 0.008370886 | 0.089709433 | 5.7815929 | 5.8641992 | 6.00883 | -0.144631785 | -0.227238009 |
| Jag2 | 6.672274 | 6.718536 | 6.5399 | 0.178658729 | 0.132396505 | 6.2287483 | 6.4404803 | 6.58512 | -0.144641306 | -0.356373316 |
| Sptan1 | 12.63628 | 12.62462 | 12.716 | -0.091586228 | -0.079925088 | 12.589377 | 12.546894 | 12.6916 | -0.144657858 | -0.102174547 |
| Cldn10 | 5.386204 | 5.302298 | 5.2977 | 0.004634732 | 0.088540833 | 5.4603045 | 5.1609287 | 5.30564 | -0.144711847 | 0.154663922 |
| Slc7a4 | 3.41599 | 3.16286 | 3.3309 | -0.168018603 | 0.08511181 | 3.1484671 | 3.1375932 | 3.28234 | -0.144745196 | -0.13387129 |
| Dnmt3a | 10.19604 | 10.38085 | 10.228 | 0.153108078 | -0.031703706 | 10.147244 | 10.257658 | 10.4024 | -0.144752504 | -0.255166064 |
| Opa1 | 10.10194 | 9.979464 | 10.081 | -0.101417014 | 0.021056501 | 10.059963 | 9.9515694 | 10.0965 | -0.144889474 | -0.036495893 |
| Rad54l | 6.673818 | 6.331544 | 6.6855 | -0.353959849 | -0.011685562 | 6.6222201 | 6.7243073 | 6.86923 | -0.144926438 | -0.24701363 |

| | | | | | | | | | | |
|---|---|---|---|---|---|---|---|---|---|---|
| Olfr354 | 3.253425 | 3.2049 | 3.2695 | -0.064623849 | -0.016099698 | 3.4977053 | 3.2631911 | 3.40818 | -0.144991692 | 0.089522549 |
| Il31ra | 3.31917 | 3.36227 | 3.2524 | 0.109901474 | 0.066801198 | 3.3029961 | 3.2005493 | 3.34557 | -0.145019627 | -0.042572813 |
| Phxr2 | 1.773018 | 1.780154 | 1.7573 | 0.022813446 | 0.015677778 | 1.8177144 | 1.6539165 | 1.79896 | -0.145047086 | 0.018750851 |
| Ago2 | 11.90042 | 11.90905 | 11.852 | 0.05748139 | 0.04885553 | 11.744593 | 11.805881 | 11.951 | -0.145112202 | -0.206400021 |
| Cc2d1b | 10.65879 | 10.49418 | 10.631 | -0.13669809 | 0.02791531 | 10.828993 | 10.722247 | 10.8676 | -0.145395716 | -0.038649265 |
| Rims2 | 1.474459 | 1.478832 | 1.4282 | 0.05066799 | 0.046295338 | 1.406663 | 1.3250645 | 1.47048 | -0.145414082 | -0.063815528 |
| Gm14322 | 2.05544 | 2.011905 | 1.9512 | 0.060688719 | 0.104224288 | 1.9836722 | 1.8840187 | 2.02947 | -0.145448912 | -0.04579538 |
| Rnf11 | 9.103521 | 9.098295 | 9.0789 | 0.019398761 | 0.024624337 | 8.8259647 | 8.734921 | 8.88045 | -0.145533799 | -0.05449008 |
| Ccdc47 | 10.56728 | 10.53199 | 10.609 | -0.07736968 | -0.042082762 | 10.472848 | 10.438692 | 10.5843 | -0.145606572 | -0.11145087 |
| Zfp36l1 | 13.25155 | 13.21938 | 13.107 | 0.112701098 | 0.144871394 | 12.63997 | 12.76681 | 12.9124 | -0.145621352 | -0.272460887 |
| Zfp106 | 11.81851 | 11.75282 | 11.803 | -0.050368808 | 0.015324124 | 11.715264 | 11.660749 | 11.8064 | -0.145631952 | -0.091116795 |
| St3gal2 | 8.341732 | 8.364683 | 8.3536 | 0.01110965 | -0.011841341 | 8.1079641 | 8.1055835 | 8.25122 | -0.14563544 | -0.143254818 |
| Cnn1 | 2.659276 | 2.672164 | 2.7626 | -0.090432119 | -0.103320367 | 2.665423 | 2.5416523 | 2.68731 | -0.145655313 | -0.021884644 |
| Micall1 | 10.05597 | 10.15195 | 10.1 | 0.052426796 | -0.043556294 | 10.071965 | 9.905937 | 10.0516 | -0.145676783 | 0.020351601 |
| Map3k7cl | 1.644438 | 1.630399 | 1.6864 | -0.056047776 | -0.042008623 | 1.6365298 | 1.5700443 | 1.71577 | -0.145724129 | -0.079238595 |
| Gm5316 | 2.508642 | 2.532125 | 2.542 | -0.009833324 | -0.033316719 | 2.4951213 | 2.6212024 | 2.76702 | -0.145816614 | -0.271897756 |
| Adamts18 | 2.882616 | 2.824156 | 3.03 | -0.205882008 | -0.147421353 | 2.951173 | 2.8631389 | 3.009 | -0.145860648 | -0.057826507 |
| Plxnd1 | 9.259974 | 9.67347 | 9.6116 | 0.061866358 | -0.35162912 | 9.4016876 | 9.4106878 | 9.55656 | -0.145867699 | -0.154867875 |
| Vezf1 | 9.786929 | 9.811552 | 9.8578 | -0.046234594 | -0.070858097 | 9.7172513 | 9.6204152 | 9.76629 | -0.145876431 | -0.04904037 |
| Ccnc | 8.622408 | 8.735648 | 8.7327 | 0.002962726 | -0.110276966 | 8.855793 | 8.7058231 | 8.8517 | -0.145878878 | 0.004090986 |
| Rsph10b | 4.418715 | 4.086634 | 4.2291 | -0.142439189 | 0.189641872 | 4.3514511 | 4.1326266 | 4.27854 | -0.145912955 | 0.072911558 |
| Fuca2 | 8.975822 | 8.895508 | 8.9158 | -0.020324298 | 0.059989607 | 8.9988746 | 8.6672618 | 8.81321 | -0.14594915 | 0.185663647 |
| D430041D05Rik | 2.419506 | 2.424274 | 2.3871 | 0.037147958 | 0.032380629 | 2.2905287 | 2.366495 | 2.51248 | -0.145988968 | -0.221955301 |
| Lrrc10b | 1.316215 | 1.305565 | 1.3792 | -0.073617391 | -0.062967238 | 1.3219159 | 1.2545842 | 1.40059 | -0.146004835 | -0.078673155 |
| Bcap29 | 9.385154 | 9.124392 | 9.3804 | -0.256037513 | 0.004724623 | 9.4448825 | 9.2827448 | 9.42877 | -0.146025922 | 0.016111782 |
| S100a3 | 2.563875 | 2.620082 | 2.729 | -0.108879821 | -0.165086435 | 2.5640177 | 2.4968069 | 2.64286 | -0.146057172 | -0.078846413 |
| Tead1 | 6.044697 | 5.969081 | 5.5688 | 0.400323731 | 0.475939404 | 5.7615336 | 5.4315877 | 5.57772 | -0.146135101 | 0.183810813 |
| Hsf2 | 7.985379 | 8.088482 | 8.1031 | -0.014660262 | -0.117763351 | 7.6522458 | 7.9952795 | 8.14144 | -0.14616186 | -0.489195571 |
| Mfsd1 | 10.23647 | 10.23921 | 10.168 | 0.071645673 | 0.068902092 | 10.15371 | 9.9710839 | 10.1173 | -0.146245703 | 0.036380149 |
| Esr2 | 4.127181 | 4.050581 | 4.1741 | -0.123548863 | -0.046949498 | 4.1666657 | 4.0467552 | 4.19305 | -0.146293227 | -0.026382697 |
| Ppp2r2c | 2.658991 | 2.637125 | 2.5839 | 0.053193426 | 0.075059589 | 2.6379454 | 2.5811525 | 2.72758 | -0.146429236 | -0.089636317 |
| Ccdc50 | 11.77077 | 11.92804 | 11.799 | 0.128668704 | -0.028602892 | 11.902967 | 11.777963 | 11.9244 | -0.146460438 | -0.021455598 |

| | | | | | | | | | | |
|---|---|---|---|---|---|---|---|---|---|---|
| Sdk1 | 3.599918 | 3.263437 | 3.5203 | -0.256849783 | 0.079631596 | 3.6306593 | 3.1876863 | 3.33424 | -0.146550343 | 0.296422663 |
| Vmn1r20 | 3.860847 | 3.739285 | 3.7316 | 0.007650892 | 0.129212472 | 3.8107231 | 3.7844211 | 3.93102 | -0.146601902 | -0.120299918 |
| Pcdhgc5 | 2.133388 | 2.171322 | 2.1646 | 0.006728585 | -0.031204891 | 2.1213806 | 2.1755869 | 2.32219 | -0.146602727 | -0.200809001 |
| Pdxp | 6.68879 | 6.915232 | 6.8342 | 0.081074105 | -0.145367459 | 6.7965806 | 6.6641012 | 6.81072 | -0.146614016 | -0.014134561 |
| Gm13340 | 6.495609 | 6.637233 | 6.7696 | -0.132389321 | -0.274013665 | 6.5818059 | 6.6572209 | 6.80387 | -0.14665188 | -0.222066943 |
| Slc5a11 | 1.674452 | 1.683176 | 1.7717 | -0.088494563 | -0.097219097 | 1.7059464 | 1.5492152 | 1.69595 | -0.146732271 | 0.009998949 |
| Pnck | 3.23807 | 3.19626 | 3.2292 | -0.032921078 | 0.008889053 | 3.040275 | 3.0214902 | 3.16824 | -0.14674675 | -0.127961981 |
| Rab9b | 1.82606 | 1.766572 | 1.7556 | 0.010925016 | 0.070413073 | 1.7225514 | 1.7340523 | 1.88086 | -0.146810602 | -0.158311587 |
| Wdr66 | 5.308053 | 5.52149 | 5.5171 | 0.004342024 | -0.209094936 | 5.63067 | 5.6931413 | 5.83996 | -0.146814894 | -0.209286122 |
| Ankrd29 | 4.036661 | 3.758695 | 3.9691 | -0.210388333 | 0.067577611 | 3.8353191 | 3.7567813 | 3.90361 | -0.146832459 | -0.068294723 |
| Tmc6 | 10.7368 | 10.73337 | 10.865 | -0.13141416 | -0.127981716 | 11.089305 | 10.841606 | 10.9885 | -0.146846062 | 0.100853528 |
| Kif3a | 6.287979 | 6.296852 | 6.329 | -0.032149726 | -0.0410229 | 6.1988256 | 6.0454053 | 6.19233 | -0.146929586 | 0.006490751 |
| Wfdc21 | 2.899409 | 2.846801 | 2.5567 | 0.290136745 | 0.342744867 | 2.7383775 | 2.4899178 | 2.63686 | -0.14694255 | 0.101517101 |
| Rpl15-ps3 | 11.92135 | 12.18352 | 12.165 | 0.018333352 | -0.243841862 | 12.369973 | 12.057938 | 12.205 | -0.147065238 | 0.164968939 |
| Ing3 | 9.286924 | 9.424349 | 9.3366 | 0.087708108 | -0.049716073 | 9.3152718 | 9.1140382 | 9.26113 | -0.147086955 | 0.054146671 |
| Trbv30 | 3.695494 | 3.730434 | 3.7098 | 0.020651461 | -0.014289032 | 3.5887263 | 3.6560281 | 3.80317 | -0.147139085 | -0.214440897 |
| Paxx | 8.161512 | 7.95126 | 8.2582 | -0.306933612 | -0.096681957 | 8.4567436 | 8.1370947 | 8.28424 | -0.147146777 | 0.172502084 |
| Thrsp | 3.102658 | 3.035846 | 3.1033 | -0.067428154 | -0.000616129 | 2.8015929 | 2.8520762 | 2.99923 | -0.147153575 | -0.197636917 |
| Cyp2b10 | 1.595141 | 1.688514 | 1.8402 | -0.15166993 | -0.245042714 | 1.7984102 | 1.7340081 | 1.88119 | -0.147183754 | -0.082781674 |
| Rubcn | 10.86105 | 11.085 | 11.046 | 0.038866632 | -0.185087668 | 11.254667 | 10.967065 | 11.1143 | -0.14718648 | 0.140415336 |
| Frs3 | 3.898199 | 3.805831 | 3.9041 | -0.09823955 | -0.005871686 | 3.7836395 | 3.7861052 | 3.93345 | -0.147341418 | -0.149807175 |
| Spink10 | 3.320164 | 3.410343 | 3.31 | 0.100367273 | 0.010188275 | 3.5180823 | 3.2689098 | 3.41627 | -0.1473573 | 0.101815231 |
| Slc10a6 | 5.953242 | 5.651579 | 5.6742 | -0.022664023 | 0.278999154 | 4.9164503 | 4.9102189 | 5.05761 | -0.147393359 | -0.141161938 |
| Map1lc3a | 6.176838 | 5.972946 | 6.2057 | -0.232720202 | -0.028828503 | 6.2722591 | 5.8592796 | 6.00672 | -0.147439467 | 0.265539983 |
| Col6a2 | 7.130356 | 7.305498 | 7.1084 | 0.197059754 | 0.02191757 | 7.0793677 | 6.489023 | 6.63657 | -0.147544957 | 0.442799686 |
| Oit3 | 1.081188 | 1.133684 | 1.1446 | -0.010927472 | -0.063423056 | 1.2102906 | 1.1065293 | 1.25413 | -0.147602744 | -0.043841374 |
| AC170998.1 | 3.74582 | 3.805479 | 3.8359 | -0.030426578 | -0.090086343 | 3.8591676 | 3.7546613 | 3.90231 | -0.14764816 | -0.043141827 |
| Dennd1b | 10.69131 | 10.68833 | 10.654 | 0.034336898 | 0.03731149 | 10.672902 | 10.607723 | 10.7554 | -0.147707352 | -0.082528484 |
| Ippk | 7.544334 | 7.425135 | 7.525 | -0.099830872 | 0.019367748 | 7.3249742 | 7.2544345 | 7.40219 | -0.147755636 | -0.077215908 |
| Kcng3 | 3.046036 | 2.892208 | 2.9497 | -0.057462509 | 0.096364934 | 2.7150245 | 2.6176102 | 2.76538 | -0.147767897 | -0.050353599 |
| Doc2g | 5.628753 | 5.822979 | 5.5969 | 0.226108305 | 0.031882463 | 6.2156218 | 5.7348985 | 5.88279 | -0.147891364 | 0.332831986 |
| Mrm2 | 6.241196 | 6.195882 | 6.2489 | -0.052975381 | -0.007661963 | 6.3846793 | 6.1124704 | 6.2604 | -0.147930103 | 0.124278806 |

| | | | | | | | | | | |
|---|---|---|---|---|---|---|---|---|---|---|
| Echdc3 | 3.313274 | 3.300426 | 3.2772 | 0.023188729 | 0.036037257 | 3.3445403 | 3.1775943 | 3.32553 | -0.147939127 | 0.019006881 |
| Lingo1 | 3.298844 | 3.244627 | 3.1269 | 0.11774917 | 0.171965951 | 3.1956959 | 3.0001252 | 3.14807 | -0.147942538 | 0.047628199 |
| Bsdc1 | 9.892798 | 10.10478 | 10.014 | 0.090388767 | -0.121593731 | 9.9873386 | 9.9745514 | 10.1225 | -0.147954882 | -0.135167705 |
| Vmn1r6 | 6.118551 | 6.193132 | 6.1661 | 0.027030735 | -0.047551172 | 5.877777 | 6.1450046 | 6.29297 | -0.14796161 | -0.415189243 |
| AC132481.2 | 1.076692 | 1.105112 | 1.0556 | 0.049538195 | 0.021118736 | 1.0472075 | 1.0162788 | 1.1644 | -0.14812332 | -0.117194645 |
| Bckdhb | 7.979988 | 7.941243 | 8.0717 | -0.130491094 | -0.0917463 | 7.8918009 | 7.8024405 | 7.95057 | -0.148133974 | -0.058773551 |
| Olfr1044 | 2.727279 | 2.757596 | 2.7255 | 0.032086026 | 0.001768296 | 2.7631479 | 2.903399 | 3.05158 | -0.148179224 | -0.288430318 |
| Wdcp | 7.070108 | 7.003905 | 7.1271 | -0.123224344 | -0.057021719 | 6.9921449 | 6.9512053 | 7.09944 | -0.148230482 | -0.107290845 |
| Rwdd2b | 7.357151 | 7.331971 | 7.3672 | -0.035267775 | -0.010087635 | 7.4195088 | 7.2811982 | 7.42955 | -0.148347095 | -0.010036491 |
| Zfhx4 | 3.683994 | 3.460875 | 3.4669 | -0.00604789 | 0.217071488 | 3.4542146 | 3.3645252 | 3.513 | -0.148478232 | -0.058788805 |
| AW146154 | 7.005375 | 7.093874 | 6.9618 | 0.132094637 | 0.043595746 | 7.2575934 | 7.0778503 | 7.22636 | -0.14850784 | 0.031235311 |
| Gm12183 | 2.381533 | 2.385274 | 2.3327 | 0.052544927 | 0.048804163 | 2.3131809 | 2.2610075 | 2.40958 | -0.148573998 | -0.096400603 |
| Rbfa | 8.973082 | 8.865849 | 9.0486 | -0.182742231 | -0.075509081 | 9.0339873 | 8.9143367 | 9.06312 | -0.148786503 | -0.029135844 |
| Dhdh | 7.147676 | 6.957849 | 7.0791 | -0.121216416 | 0.068610682 | 7.1763004 | 7.08243 | 7.23124 | -0.148812814 | -0.054942436 |
| Olfr656 | 2.232639 | 2.197534 | 2.1476 | 0.049903044 | 0.08500839 | 2.2413969 | 2.1883369 | 2.33717 | -0.148833392 | -0.095773392 |
| A830018L16Rik | 2.739087 | 2.671369 | 2.6926 | -0.021270488 | 0.046447486 | 2.8340109 | 2.6986013 | 2.84746 | -0.148855824 | -0.013446167 |
| Stxbp6 | 5.940536 | 5.654627 | 5.5324 | 0.122197641 | 0.408106826 | 5.6667508 | 5.4329163 | 5.5818 | -0.148885672 | 0.084948824 |
| Gucy2e | 2.496414 | 2.571179 | 2.5556 | 0.01554128 | -0.059223809 | 2.4683001 | 2.4900177 | 2.63903 | -0.149009768 | -0.170727447 |
| Slc6a20a | 3.066589 | 2.986409 | 3.1056 | -0.119194031 | -0.039014223 | 2.9639967 | 2.9466229 | 3.09565 | -0.14902833 | -0.131654527 |
| Olfr734 | 2.559085 | 2.555965 | 2.5843 | -0.028287185 | -0.025167444 | 2.4746649 | 2.4605649 | 2.6096 | -0.149035251 | -0.134935217 |
| Espn | 5.339928 | 4.981488 | 5.288 | -0.306479044 | 0.051960847 | 5.3715984 | 5.2366285 | 5.38571 | -0.149077613 | -0.014107688 |
| 1700102P08Rik | 4.383578 | 4.42473 | 4.3329 | 0.091840237 | 0.05068868 | 4.4414879 | 4.2867295 | 4.43581 | -0.149078023 | 0.005680322 |
| B3glct | 6.994804 | 6.812609 | 6.963 | -0.150432561 | 0.031762048 | 6.8590973 | 6.7727827 | 6.92189 | -0.149112174 | -0.062797584 |
| Sh3bgrl | 11.09337 | 10.99228 | 11.044 | -0.051879396 | 0.049216042 | 10.997524 | 10.800861 | 10.95 | -0.14913605 | 0.047526719 |
| Atmin | 9.38409 | 9.440828 | 9.4805 | -0.039677061 | -0.096415084 | 9.2968118 | 9.2066818 | 9.35584 | -0.14916233 | -0.059032302 |
| Syt1 | 4.691027 | 4.489272 | 4.4127 | 0.076602064 | 0.278356911 | 4.0602651 | 4.0917666 | 4.24095 | -0.149180373 | -0.18068191 |
| Zc3h12a | 9.302951 | 9.257199 | 9.423 | -0.165819891 | -0.120068498 | 9.5914336 | 9.3069553 | 9.45627 | -0.149319136 | 0.135159213 |
| Ighv7-3 | 5.816545 | 5.989243 | 5.8464 | 0.142860368 | -0.029837469 | 5.8621305 | 5.8055361 | 5.95486 | -0.149328472 | -0.092734053 |
| Drc3 | 2.576369 | 2.517992 | 2.6727 | -0.154712489 | -0.096335087 | 2.5306626 | 2.4720139 | 2.62136 | -0.14934112 | -0.090692417 |
| Ccdc149 | 4.195524 | 4.128626 | 4.1437 | -0.015089541 | 0.051809168 | 3.7822581 | 3.7497484 | 3.89909 | -0.149342079 | -0.116832332 |
| Gnai2 | 13.26966 | 13.19161 | 13.263 | -0.071817084 | 0.00623116 | 13.418072 | 13.041698 | 13.1911 | -0.149374148 | 0.226999491 |
| Emid1 | 7.773337 | 7.691686 | 7.6906 | 0.001094829 | 0.082745915 | 8.1129783 | 7.6504214 | 7.79984 | -0.149422606 | 0.313134204 |

| | | | | | | | | | | |
|---|---|---|---|---|---|---|---|---|---|---|
| Txndc5 | 10.8518 | 10.79396 | 10.842 | -0.048276146 | 0.00956802 | 11.007752 | 10.62726 | 10.7767 | -0.149450862 | 0.231041591 |
| Trav15-1-dv6-1 | 3.402881 | 3.414162 | 3.4248 | -0.010618165 | -0.021898899 | 3.3299192 | 3.2851355 | 3.43464 | -0.149508209 | -0.104724455 |
| Asah2 | 8.446415 | 8.263037 | 8.4017 | -0.138636999 | 0.044740513 | 7.8107888 | 8.1144769 | 8.26401 | -0.149534675 | -0.45322286 |
| C8g | 2.4746 | 2.550773 | 2.5766 | -0.025779124 | -0.10195177 | 2.6168533 | 2.4811731 | 2.63072 | -0.149542348 | -0.013862216 |
| Tmem18 | 7.923609 | 7.817927 | 7.9446 | -0.126655236 | -0.020973126 | 7.877434 | 7.8007092 | 7.95037 | -0.149659772 | -0.072934941 |
| Ankrd52 | 10.39541 | 10.3405 | 10.433 | -0.092915736 | -0.03800665 | 10.479776 | 10.476997 | 10.6267 | -0.149676524 | -0.146897609 |
| Klhl20 | 9.482355 | 9.537285 | 9.5362 | 0.001088295 | -0.053841791 | 9.7296439 | 9.4793773 | 9.62909 | -0.149708606 | 0.100558014 |
| Cmtm7 | 9.888874 | 9.907739 | 10.009 | -0.101466967 | -0.120332417 | 10.200723 | 9.873402 | 10.0232 | -0.149763966 | 0.177556627 |
| Ifnar1 | 10.75244 | 10.76346 | 10.811 | -0.047544476 | -0.05856654 | 10.771707 | 10.736031 | 10.8859 | -0.149829028 | -0.114152412 |
| Gabpa | 10.45835 | 10.47331 | 10.391 | 0.081941778 | 0.06697693 | 10.276923 | 10.323254 | 10.4731 | -0.149861482 | -0.19619233 |
| Zfr | 10.76607 | 10.75712 | 10.671 | 0.085864838 | 0.094820738 | 10.704368 | 10.748879 | 10.8988 | -0.149880352 | -0.194390774 |
| Arhgap6 | 6.680882 | 6.823044 | 6.7129 | 0.110182815 | -0.031979126 | 6.9640501 | 6.6128737 | 6.76287 | -0.149997374 | 0.20117897 |
| Gm6649 | 2.904859 | 2.857234 | 2.8629 | -0.005706415 | 0.041918128 | 2.9925514 | 2.8085508 | 2.95857 | -0.150020183 | 0.03398038 |
| Zfr2 | 4.895645 | 4.793186 | 4.9537 | -0.160506094 | -0.058046378 | 4.8948803 | 4.8336678 | 4.98384 | -0.150167246 | -0.088954768 |
| Dolpp1 | 7.819491 | 7.751783 | 7.8128 | -0.060993677 | 0.00671435 | 7.9933386 | 7.7472376 | 7.89746 | -0.150217953 | 0.095883051 |
| Mplkip | 8.257499 | 8.053806 | 8.1688 | -0.115006795 | 0.088686383 | 8.3379685 | 8.086541 | 8.23681 | -0.150273818 | 0.101153716 |
| Zfp13 | 2.661651 | 2.5031 | 2.7604 | -0.25727595 | -0.098725405 | 2.6806691 | 2.4079525 | 2.55824 | -0.150284302 | 0.122432391 |
| 4930523C07Rik | 11.25566 | 11.29867 | 11.533 | -0.234743398 | -0.277752644 | 11.23136 | 11.359685 | 11.51 | -0.150338764 | -0.278663592 |
| Itga11 | 1.882287 | 1.845612 | 1.9061 | -0.060462881 | -0.023788015 | 1.8339411 | 1.7734189 | 1.9238 | -0.15038575 | -0.089863546 |
| Esyt2 | 11.55181 | 11.62392 | 11.661 | -0.036641396 | -0.108744936 | 11.362667 | 11.498958 | 11.6494 | -0.1504073 | -0.286697574 |
| 4930503L19Rik | 8.315147 | 8.333079 | 8.413 | -0.079941497 | -0.09787352 | 8.5748564 | 8.371443 | 8.52195 | -0.150502139 | 0.052911266 |
| Zfx | 10.27767 | 10.36328 | 10.346 | 0.017313665 | -0.068296427 | 10.070377 | 10.172265 | 10.3228 | -0.150507965 | -0.252396004 |
| Tmem252 | 6.931474 | 6.820025 | 6.6995 | 0.120544239 | 0.231993383 | 6.0205243 | 5.9786882 | 6.12934 | -0.150649916 | -0.108813801 |
| Rnf170 | 8.030661 | 7.88518 | 8.0685 | -0.183328334 | -0.037846844 | 8.076222 | 7.8370998 | 7.98776 | -0.150657729 | 0.088464496 |
| Elk1 | 8.061742 | 8.012134 | 7.9292 | 0.08290753 | 0.132515637 | 7.9989037 | 7.75073 | 7.90143 | -0.150696217 | 0.097477538 |
| Capsl | 2.870847 | 2.920183 | 2.9622 | -0.042060397 | -0.09139626 | 2.9841545 | 2.8763249 | 3.02706 | -0.150734695 | -0.042905078 |
| Wdr1 | 12.60266 | 12.63033 | 12.581 | 0.048907768 | 0.0212379 | 12.734669 | 12.450446 | 12.6012 | -0.150772398 | 0.13345037 |
| Lig3 | 9.717762 | 9.86175 | 9.6412 | 0.220555875 | 0.07656794 | 9.6677099 | 9.5847381 | 9.73554 | -0.150798139 | -0.067826359 |
| Mtg1 | 7.64724 | 7.476656 | 7.6746 | -0.197989496 | -0.027405393 | 7.9538648 | 7.5991264 | 7.74998 | -0.150851766 | 0.203886624 |
| Lix1 | 2.280598 | 2.252837 | 2.385 | -0.132128648 | -0.104367518 | 2.2871966 | 2.2196398 | 2.37062 | -0.150983215 | -0.083426431 |
| Smyd1 | 3.634726 | 3.631934 | 3.7689 | -0.136935439 | -0.134143249 | 3.6094804 | 3.5433301 | 3.69432 | -0.150993098 | -0.08484282 |
| Abcg1 | 11.12449 | 11.13841 | 11.154 | -0.015639962 | -0.029567716 | 10.949076 | 11.201295 | 11.3523 | -0.151053422 | -0.403272125 |

| | | | | | | | | | | |
|---|---|---|---|---|---|---|---|---|---|---|
| Dnajc9 | 10.33675 | 10.21653 | 10.345 | -0.128511464 | -0.008284014 | 10.46264 | 10.24984 | 10.4009 | -0.151065322 | 0.061735108 |
| Scaf8 | 11.214 | 11.24443 | 11.223 | 0.021588548 | -0.008844016 | 11.12385 | 11.08643 | 11.2375 | -0.151073578 | -0.113653863 |
| Strada | 9.17197 | 9.090263 | 9.1743 | -0.083996914 | -0.002289121 | 9.3807665 | 9.2065579 | 9.35767 | -0.151107182 | 0.023101422 |
| Dzip1l | 4.226746 | 4.219247 | 4.1933 | 0.025977568 | 0.033475987 | 4.2222287 | 4.2166245 | 4.36776 | -0.151133462 | -0.145529249 |
| Itgb2 | 12.40627 | 12.47253 | 12.401 | 0.071850706 | 0.005596274 | 12.407589 | 12.20987 | 12.3612 | -0.151342336 | 0.04637652 |
| Prdm5 | 3.57056 | 3.531504 | 3.6227 | -0.091161482 | -0.05210507 | 3.4765971 | 3.4042696 | 3.55572 | -0.1514492 | -0.079121624 |
| Trpm5 | 1.762997 | 1.712537 | 1.6785 | 0.034061749 | 0.084522138 | 1.6589343 | 1.6197909 | 1.77125 | -0.151458768 | -0.112315323 |
| Zfp60 | 8.113666 | 8.235149 | 8.134 | 0.101126333 | -0.020357174 | 8.2798762 | 7.9141906 | 8.06576 | -0.151564685 | 0.214120921 |
| Olfr1221 | 4.12652 | 4.065589 | 4.1303 | -0.064715029 | -0.003784197 | 4.1926123 | 4.202123 | 4.35381 | -0.151686387 | -0.161197109 |
| Gbe1 | 8.200341 | 8.107628 | 8.2219 | -0.11424676 | -0.02153293 | 8.2505978 | 8.037805 | 8.18958 | -0.151779822 | 0.061012936 |
| Gm5466 | 2.541352 | 2.640493 | 2.6226 | 0.01785968 | -0.081280644 | 2.7623408 | 2.6054782 | 2.75726 | -0.151780895 | 0.00508168 |
| Rnf111 | 10.46393 | 10.54785 | 10.555 | -0.007179014 | -0.09109969 | 10.630098 | 10.550982 | 10.7028 | -0.151787982 | -0.072672059 |
| Nucb1 | 11.24871 | 11.14828 | 11.178 | -0.029476082 | 0.07095699 | 11.053546 | 10.950846 | 11.1027 | -0.151845014 | -0.04914538 |
| Tpsb2 | 7.222488 | 7.401276 | 7.2264 | 0.174848232 | -0.003939508 | 6.7879622 | 6.7973261 | 6.94941 | -0.152084563 | -0.161448483 |
| Card6 | 11.04747 | 11.2848 | 11.129 | 0.15573563 | -0.081592706 | 11.030875 | 10.951234 | 11.1033 | -0.152112402 | -0.072471229 |
| Nxt2 | 6.947901 | 6.985342 | 6.9373 | 0.047998541 | 0.010557383 | 7.0910254 | 6.8702534 | 7.02237 | -0.152112886 | 0.068659116 |
| Rmdn1 | 6.17575 | 5.941837 | 6.0895 | -0.147693935 | 0.086218576 | 6.2483536 | 5.9308005 | 6.08292 | -0.152118824 | 0.16543422 |
| Nexn | 3.528429 | 3.3507 | 3.3358 | 0.014943127 | 0.192671721 | 3.4282689 | 3.2474275 | 3.39963 | -0.152202561 | 0.028638795 |
| Parm1 | 6.829498 | 6.69144 | 6.6847 | 0.006690543 | 0.144748015 | 6.450589 | 6.1275791 | 6.27982 | -0.152238277 | 0.170771686 |
| Il10rb | 10.35833 | 10.31858 | 10.407 | -0.088038752 | -0.048284318 | 10.44932 | 10.138004 | 10.2903 | -0.15225592 | 0.159059776 |
| Grk2 | 13.1194 | 13.23324 | 13.213 | 0.020442832 | -0.093391048 | 13.433768 | 13.153991 | 13.3063 | -0.152358354 | 0.127418864 |
| Zfp64 | 8.108857 | 8.102351 | 8.1647 | -0.062306648 | -0.055800964 | 8.2152953 | 8.0029951 | 8.15545 | -0.152455985 | 0.059844147 |
| Zyg11b | 10.83234 | 10.85644 | 10.86 | -0.003340912 | -0.02744708 | 10.558784 | 10.529222 | 10.6817 | -0.152490688 | -0.122928828 |
| Acy3 | 4.515585 | 4.064261 | 4.4833 | -0.419062675 | 0.032261936 | 4.1108462 | 3.8958121 | 4.04839 | -0.152573613 | 0.062460525 |
| Nmt2 | 9.177188 | 9.255317 | 9.3567 | -0.101429632 | -0.179558615 | 9.0943774 | 9.2044388 | 9.35706 | -0.152619586 | -0.262680969 |
| Sync | 2.503063 | 2.554359 | 2.567 | -0.01263512 | -0.063931285 | 2.4165639 | 2.3951091 | 2.54775 | -0.152640058 | -0.131185329 |
| Gm2260 | 7.105691 | 7.113794 | 7.1977 | -0.083896816 | -0.092000048 | 7.2833436 | 7.0092354 | 7.16202 | -0.152780054 | 0.121328195 |
| Spata31d1b | 4.339387 | 4.529839 | 4.4314 | 0.098478308 | -0.091973023 | 4.2592954 | 4.4734294 | 4.62622 | -0.152792565 | -0.366926606 |
| Capn7 | 9.994479 | 9.923403 | 9.9977 | -0.07434628 | -0.003269912 | 9.8790206 | 9.9583694 | 10.1112 | -0.152811045 | -0.232159873 |
| Dhx35 | 8.890956 | 8.794639 | 8.867 | -0.072400558 | 0.023916757 | 9.0204485 | 8.9921026 | 9.14506 | -0.152954689 | -0.124608765 |
| Rsph3a | 6.372864 | 6.336089 | 6.5149 | -0.17880989 | -0.14203548 | 6.4890544 | 6.2613137 | 6.41429 | -0.152975177 | 0.074765512 |
| Atp8b5 | 2.013194 | 2.031736 | 2.1061 | -0.074340462 | -0.092882824 | 1.9405533 | 1.939815 | 2.09281 | -0.152992948 | -0.152254637 |

| | | | | | | | | | | |
|---|---|---|---|---|---|---|---|---|---|---|
| Olfr351 | 1.995629 | 2.001765 | 1.991 | 0.010800626 | 0.004664755 | 1.9835347 | 1.9780662 | 2.13111 | -0.153040787 | -0.147572332 |
| Smurf1 | 9.573344 | 9.566379 | 9.6925 | -0.126090173 | -0.119125108 | 9.2300971 | 9.4032981 | 9.55634 | -0.153043046 | -0.326244045 |
| Nsg1 | 4.503249 | 4.325574 | 4.2598 | 0.06577832 | 0.243453393 | 4.2181762 | 3.9276727 | 4.08072 | -0.153050612 | 0.137452863 |
| Dlat | 9.111961 | 9.001626 | 9.0003 | 0.001309515 | 0.111643972 | 9.0539057 | 8.9622835 | 9.1154 | -0.153121439 | -0.061499331 |
| Ern1 | 8.980371 | 8.762086 | 8.8408 | -0.078684762 | 0.139599773 | 8.6732564 | 8.6952272 | 8.84842 | -0.153189969 | -0.175160746 |
| Poc1a | 6.138965 | 6.146701 | 6.0997 | 0.047041743 | 0.039305597 | 6.2706424 | 6.2054279 | 6.3587 | -0.153268091 | -0.088053526 |
| Virma | 10.88692 | 10.9273 | 10.936 | -0.009084638 | -0.049465258 | 10.793338 | 10.72828 | 10.8816 | -0.153287242 | -0.088228787 |
| Rab38 | 3.861256 | 3.835433 | 4.1876 | -0.352156204 | -0.32633374 | 4.0921278 | 3.7863008 | 3.93961 | -0.153310743 | 0.152516209 |
| Uxs1 | 8.046821 | 7.866263 | 7.9032 | -0.036913758 | 0.143644487 | 7.7512236 | 7.7143919 | 7.86771 | -0.153320254 | -0.116488603 |
| Gm48391 | 3.545225 | 3.479157 | 3.5534 | -0.074248343 | -0.008180922 | 3.6126935 | 3.5533499 | 3.70673 | -0.153379153 | -0.094035542 |
| Gm21833 | 3.566191 | 3.546042 | 3.6451 | -0.099050929 | -0.07890253 | 3.6769322 | 3.6788316 | 3.83223 | -0.153402101 | -0.155301495 |
| Nek4 | 7.286488 | 7.153968 | 7.3312 | -0.17720009 | -0.044680134 | 7.3946604 | 7.383138 | 7.53671 | -0.153568193 | -0.142045815 |
| Pmel | 2.530111 | 2.589596 | 2.612 | -0.022401067 | -0.081886018 | 2.6112653 | 2.4955083 | 2.64911 | -0.153599372 | -0.037842377 |
| Gpr82 | 2.6077 | 2.379236 | 2.4372 | -0.057967506 | 0.170496529 | 2.582961 | 2.402681 | 2.55638 | -0.153702677 | 0.026577285 |
| Pcdhb16 | 3.769427 | 3.49524 | 3.5093 | -0.014052589 | 0.260134347 | 3.4363197 | 3.24177 | 3.39552 | -0.153754017 | 0.040795672 |
| Dazap1 | 10.23486 | 10.22036 | 10.292 | -0.07166232 | -0.057157992 | 10.46558 | 10.301667 | 10.4554 | -0.153757163 | 0.010155441 |
| Ric8a | 10.47456 | 10.41122 | 10.504 | -0.092390338 | -0.029046764 | 10.759126 | 10.388607 | 10.5424 | -0.153820528 | 0.216698307 |
| Trav7-6 | 3.459662 | 3.542708 | 3.573 | -0.030290941 | -0.113337334 | 3.5569238 | 3.5246145 | 3.67851 | -0.153899615 | -0.121590347 |
| Nup160 | 10.39377 | 10.25337 | 10.411 | -0.157693492 | -0.017289334 | 10.504769 | 10.292107 | 10.446 | -0.15391654 | 0.058744566 |
| Dym | 8.90174 | 8.869503 | 8.9569 | -0.08740097 | -0.055163933 | 8.9857161 | 8.7906337 | 8.94458 | -0.153942195 | 0.041140233 |
| Dok4 | 6.05381 | 5.843937 | 6.0461 | -0.202157078 | 0.007715523 | 6.0282893 | 5.7568156 | 5.91077 | -0.153955229 | 0.117518521 |
| Ccdc62 | 6.477507 | 6.525219 | 6.6194 | -0.094206547 | -0.14191879 | 6.6829388 | 6.4217046 | 6.57569 | -0.153983662 | 0.107250603 |
| Ddr1 | 7.442637 | 7.198757 | 7.305 | -0.106282817 | 0.137596968 | 7.4185769 | 6.8531954 | 7.00719 | -0.153989955 | 0.41139152 |
| Ptpn12 | 9.708131 | 9.666433 | 9.4328 | 0.233652841 | 0.275350752 | 8.9763216 | 9.1859721 | 9.34008 | -0.154111496 | -0.363762062 |
| Rell1 | 9.163244 | 9.096925 | 8.9914 | 0.105509074 | 0.17182848 | 8.748601 | 8.8038511 | 8.95823 | -0.154378202 | -0.209628347 |
| Gm48498 | 5.742099 | 5.680793 | 5.6857 | -0.004891524 | 0.056414021 | 5.7726878 | 5.82183 | 5.97621 | -0.15438017 | -0.203522368 |
| Pde9a | 3.347161 | 3.227417 | 3.4318 | -0.204392641 | -0.084648053 | 3.2579081 | 3.0519772 | 3.20651 | -0.154535124 | 0.051395707 |
| Phykpl | 7.822932 | 7.660759 | 7.798 | -0.137290202 | 0.024882777 | 8.1363579 | 7.6987624 | 7.85341 | -0.154646598 | 0.28294885 |
| Slc12a5 | 3.207743 | 3.256685 | 3.0203 | 0.23637785 | 0.187435459 | 3.0565333 | 3.0085602 | 3.16324 | -0.154679552 | -0.106706486 |
| Wdfy4 | 13.00933 | 13.09363 | 13.062 | 0.031510542 | -0.052780962 | 13.324436 | 12.935445 | 13.0901 | -0.154691732 | 0.234298993 |
| Pkd2l2 | 3.598378 | 3.937814 | 3.771 | 0.166838145 | -0.172596997 | 3.8897562 | 3.576509 | 3.7312 | -0.154691914 | 0.158555323 |
| Zfp143 | 9.557761 | 9.542216 | 9.5336 | 0.008654599 | 0.024199297 | 9.5739802 | 9.4194203 | 9.57413 | -0.154706619 | -0.000146691 |

| | | | | | | | | | | |
|---|---|---|---|---|---|---|---|---|---|---|
| 2810039B14Rik | 4.438738 | 4.529605 | 4.5893 | -0.059650249 | -0.150516775 | 4.7466347 | 4.5792321 | 4.73401 | -0.154775296 | 0.012627264 |
| Zfp708 | 5.51866 | 5.335567 | 5.4017 | -0.066149588 | 0.116943112 | 5.3403121 | 5.279578 | 5.43442 | -0.154839259 | -0.094105213 |
| Hgsnat | 8.738288 | 8.80564 | 8.7669 | 0.038774203 | -0.028578067 | 8.6600329 | 8.4924252 | 8.64739 | -0.154968493 | 0.01263923 |
| Gm28035 | 4.893674 | 5.099393 | 5.3954 | -0.295993649 | -0.501713163 | 5.6750212 | 5.3949194 | 5.54989 | -0.154974925 | 0.125126865 |
| Cep41 | 7.481696 | 7.556006 | 7.6811 | -0.125086816 | -0.19939656 | 7.7611681 | 7.6670573 | 7.82213 | -0.155069679 | -0.060958914 |
| Cetn3 | 9.62078 | 9.416377 | 9.6703 | -0.253937432 | -0.049534758 | 9.622034 | 9.4807263 | 9.6358 | -0.155075166 | -0.013767449 |
| Cfap298 | 7.271062 | 7.29756 | 7.2187 | 0.078863694 | 0.052366001 | 7.2917311 | 7.2555172 | 7.41072 | -0.155200399 | -0.118986522 |
| Hapln1 | 2.616835 | 2.672449 | 2.6383 | 0.034129871 | -0.021484703 | 2.4629514 | 2.5325316 | 2.68773 | -0.155202437 | -0.224782673 |
| Fndc7 | 5.571735 | 5.106676 | 5.6366 | -0.529877209 | -0.064817788 | 5.4113279 | 5.1521412 | 5.30737 | -0.155229709 | 0.10395694 |
| Trav4-3 | 4.835409 | 4.697356 | 4.9333 | -0.235942007 | -0.097889201 | 4.4916981 | 4.6721208 | 4.82736 | -0.155240995 | -0.335663718 |
| CT010429.1 | 1.983355 | 1.888395 | 1.9404 | -0.051988214 | 0.042972135 | 2.1005505 | 1.8355366 | 1.99098 | -0.155445943 | 0.109567967 |
| Gm45739 | 2.677727 | 2.626041 | 2.6649 | -0.038836339 | 0.012849504 | 2.6512801 | 2.5953779 | 2.7509 | -0.15552393 | -0.09962167 |
| Arhgef28 | 4.964793 | 4.618368 | 4.9747 | -0.356344461 | -0.009919302 | 4.6462613 | 4.4340378 | 4.58965 | -0.155609396 | 0.056614048 |
| Anapc4 | 9.602565 | 9.532948 | 9.5847 | -0.051765634 | 0.017851473 | 9.4569776 | 9.59933 | 9.75495 | -0.155617113 | -0.297969509 |
| Npas4 | 3.423659 | 3.250427 | 3.572 | -0.321540775 | -0.148309085 | 3.4054145 | 3.211886 | 3.36753 | -0.155645105 | 0.037883321 |
| Npr1 | 7.831348 | 7.370101 | 7.6289 | -0.258763649 | 0.202483516 | 6.6339429 | 6.8497293 | 7.00538 | -0.155648848 | -0.371435254 |
| Zfp771 | 5.204649 | 5.396451 | 5.2819 | 0.114511989 | -0.077290424 | 5.6887168 | 5.4371288 | 5.59279 | -0.155665099 | 0.095922931 |
| Plin1 | 1.89238 | 2.004018 | 1.8932 | 0.110793102 | -0.000844201 | 2.0920157 | 1.9086775 | 2.06437 | -0.155689824 | 0.027648381 |
| Ighv6-3 | 7.758139 | 7.715435 | 7.5237 | 0.1917037 | 0.234407922 | 7.3716767 | 7.369321 | 7.52501 | -0.155693319 | -0.153337669 |
| Gm10642 | 4.102657 | 3.963576 | 4.142 | -0.178412991 | -0.039331865 | 4.1265526 | 4.0097434 | 4.16546 | -0.155714375 | -0.038905193 |
| Strn | 10.19676 | 10.17043 | 10.216 | -0.045264622 | -0.018938754 | 9.8062198 | 9.9022974 | 10.0581 | -0.155764159 | -0.251841743 |
| Ndnf | 6.290748 | 6.356456 | 6.1713 | 0.185163221 | 0.119454887 | 5.8448985 | 6.1037971 | 6.25969 | -0.155890309 | -0.414788919 |
| Helz | 10.39558 | 10.48257 | 10.528 | -0.045634286 | -0.132624076 | 10.397259 | 10.59148 | 10.7474 | -0.155913354 | -0.350134863 |
| Ddx42 | 11.06457 | 10.99568 | 11.11 | -0.114693882 | -0.045810052 | 11.017623 | 11.052136 | 11.2082 | -0.156031886 | -0.190544033 |
| Fam198a | 2.29941 | 2.2805 | 2.2621 | 0.018399688 | 0.037310437 | 2.3743893 | 2.1872167 | 2.34337 | -0.156154736 | 0.031017948 |
| Fam227a | 5.132342 | 5.173894 | 5.1284 | 0.045507459 | 0.003955339 | 4.9925364 | 5.2499482 | 5.40617 | -0.156220626 | -0.413632422 |
| Ddx11 | 8.537712 | 8.734305 | 8.7193 | 0.014969836 | -0.181622963 | 8.9639611 | 8.8507021 | 9.00696 | -0.156259222 | -0.043000254 |
| Kif5a | 4.311361 | 4.308097 | 4.2502 | 0.057916698 | 0.06118023 | 4.1999552 | 4.0886451 | 4.24495 | -0.156303 | -0.044992906 |
| Dnah8 | 9.629552 | 9.359803 | 9.5692 | -0.209347422 | 0.06040162 | 9.6309015 | 9.7504003 | 9.90671 | -0.156313148 | -0.27581195 |
| Tmem47 | 5.948156 | 5.902321 | 5.961 | -0.058706545 | -0.012872007 | 5.5108387 | 5.3902646 | 5.54666 | -0.156393216 | -0.035819139 |
| Tmem81 | 7.363745 | 7.352403 | 7.4377 | -0.085282449 | -0.073940103 | 7.4874565 | 7.4067411 | 7.56314 | -0.156396623 | -0.07568117 |
| Pptc7 | 10.13652 | 10.2468 | 10.128 | 0.11886917 | 0.008589229 | 10.064199 | 9.8868251 | 10.0433 | -0.156438464 | 0.020935648 |

| | | | | | | | | | | |
|---|---|---|---|---|---|---|---|---|---|---|
| Mustn1 | 5.951727 | 5.735033 | 5.8407 | -0.105640641 | 0.11105415 | 5.5924556 | 5.4357242 | 5.59219 | -0.156470285 | 0.000261103 |
| Edrf1 | 8.906879 | 9.199947 | 9.0901 | 0.109891102 | -0.183177448 | 8.9995718 | 9.0824154 | 9.23891 | -0.156496799 | -0.239340434 |
| Tmem121b | 2.228477 | 2.170699 | 2.1831 | -0.012415375 | 0.04536258 | 2.1909024 | 2.0441303 | 2.20065 | -0.156522759 | -0.009750609 |
| Fam98c | 8.71098 | 8.657839 | 8.7783 | -0.120490947 | -0.06735009 | 8.9206308 | 8.6512311 | 8.80779 | -0.156561239 | 0.112838455 |
| Rcan2 | 2.700831 | 2.806641 | 2.8983 | -0.091660471 | -0.197470081 | 2.6859698 | 2.6385014 | 2.79512 | -0.156616136 | -0.109147738 |
| Vmn1r217 | 3.790192 | 3.799393 | 3.8866 | -0.087254099 | -0.096454693 | 3.881032 | 3.9230752 | 4.07972 | -0.156645822 | -0.198688955 |
| Adgrg2 | 3.37335 | 3.325546 | 3.4083 | -0.082795721 | -0.03499135 | 3.1160108 | 3.1483644 | 3.30508 | -0.156712775 | -0.1890664 |
| Il2 | 3.115266 | 2.862971 | 3.0045 | -0.14155059 | 0.110744108 | 2.8588729 | 2.8291083 | 2.98587 | -0.156764431 | -0.126999822 |
| Mtmr11 | 5.048818 | 5.007906 | 4.8067 | 0.201172848 | 0.242085018 | 4.8252716 | 4.6036866 | 4.76048 | -0.156796699 | 0.064788319 |
| Dcaf8 | 11.25303 | 11.19831 | 11.271 | -0.07282055 | -0.018100332 | 11.289753 | 11.15633 | 11.3131 | -0.156808512 | -0.023385773 |
| Rin1 | 4.440908 | 4.410646 | 4.6503 | -0.239691771 | -0.209430321 | 4.386825 | 4.3418112 | 4.49868 | -0.156867606 | -0.111853812 |
| Ctsh | 11.45946 | 11.42137 | 11.539 | -0.117729864 | -0.079637554 | 11.906661 | 11.488252 | 11.6452 | -0.156945474 | 0.261463538 |
| Kmt5c | 8.803044 | 8.734336 | 8.8143 | -0.079984569 | -0.011276513 | 9.1823208 | 8.9971624 | 9.15411 | -0.156945691 | 0.028212665 |
| Trav7-5 | 4.307232 | 4.296641 | 4.2118 | 0.08482196 | 0.095412969 | 4.2565964 | 4.2062068 | 4.36317 | -0.156959655 | -0.106569965 |
| 1300017J02Rik | 3.031722 | 2.892034 | 3.1078 | -0.215783774 | -0.076096272 | 2.8582562 | 2.7978501 | 2.95483 | -0.156984113 | -0.096577997 |
| Csf3 | 4.334997 | 3.600581 | 3.7644 | -0.163774104 | 0.570642251 | 3.9255462 | 3.7136476 | 3.87072 | -0.157069084 | 0.054829547 |
| Hp1bp3 | 11.99563 | 11.94922 | 12.01 | -0.060790194 | -0.014378216 | 11.921086 | 11.913299 | 12.0704 | -0.157147682 | -0.149361172 |
| Angptl4 | 5.947963 | 6.225228 | 5.746 | 0.479197331 | 0.201931924 | 6.3479488 | 6.2479627 | 6.40523 | -0.157271214 | -0.05728512 |
| Chd8 | 12.1899 | 12.10861 | 12.184 | -0.07491091 | 0.006381432 | 12.175239 | 12.16184 | 12.3191 | -0.157308658 | -0.143909742 |
| Fam129c | 8.5404 | 8.708107 | 8.8835 | -0.175342833 | -0.343049913 | 8.9265072 | 8.6941754 | 8.8516 | -0.15742317 | 0.074908667 |
| Smad6 | 3.209143 | 2.993058 | 3.2459 | -0.252851223 | -0.036765873 | 3.0856233 | 2.9477059 | 3.10518 | -0.157477299 | -0.019559888 |
| Cd3g | 11.88482 | 11.88386 | 11.925 | -0.040947856 | -0.039992452 | 11.711111 | 11.844938 | 12.0025 | -0.157586064 | -0.291413075 |
| Gipc1 | 8.181663 | 8.138848 | 8.2567 | -0.117871131 | -0.075056352 | 8.2488412 | 8.0603964 | 8.21805 | -0.157656203 | 0.03078857 |
| 38047 | 3.108599 | 2.971772 | 3.1836 | -0.211860021 | -0.075032591 | 2.8767926 | 2.8036145 | 2.9615 | -0.157888792 | -0.0847107 |
| Gm17205 | 3.309765 | 3.329501 | 3.2994 | 0.030070252 | 0.010333661 | 3.4454845 | 3.3721232 | 3.53002 | -0.157897077 | -0.084535771 |
| Pole | 8.45445 | 8.331578 | 8.6517 | -0.320148211 | -0.197276379 | 8.5176116 | 8.7081908 | 8.86609 | -0.157898782 | -0.348478028 |
| Zfp286 | 4.26259 | 4.258736 | 4.4622 | -0.203424269 | -0.199570015 | 4.406346 | 4.3595042 | 4.51743 | -0.157921289 | -0.111079528 |
| Wbp2 | 10.54153 | 10.51952 | 10.633 | -0.11316672 | -0.091157816 | 10.630292 | 10.436214 | 10.5942 | -0.157949854 | 0.036128091 |
| Mycn | 3.529473 | 3.443506 | 3.7116 | -0.268114192 | -0.182147316 | 3.3062436 | 3.278026 | 3.43598 | -0.15795616 | -0.129738658 |
| Rusc1 | 9.044863 | 9.071515 | 9.1832 | -0.111718453 | -0.138370784 | 9.3672589 | 9.0315152 | 9.18947 | -0.157958527 | 0.177785203 |
| Pcdhb20 | 2.778964 | 2.706342 | 2.8663 | -0.159981293 | -0.08735932 | 2.6603059 | 2.5484715 | 2.70643 | -0.157963275 | -0.046128937 |
| Sult5a1 | 3.584127 | 3.456439 | 3.7654 | -0.309008135 | -0.181319587 | 3.337413 | 3.4832338 | 3.64124 | -0.158002784 | -0.303823578 |

| | | | | | | | | | | |
|---|---|---|---|---|---|---|---|---|---|---|
| Adgrl3 | 4.04699 | 3.740187 | 3.9197 | -0.179475945 | 0.127327643 | 3.8356393 | 3.8083214 | 3.96637 | -0.158051509 | -0.130733621 |
| Prg2 | 4.72429 | 4.981719 | 4.7502 | 0.231534696 | -0.025894758 | 4.4849028 | 4.5785162 | 4.73657 | -0.158053576 | -0.251666962 |
| Eaf1 | 9.765284 | 9.762214 | 9.7368 | 0.025376789 | 0.028446953 | 9.7559611 | 9.6883598 | 9.84654 | -0.158176037 | -0.090574714 |
| Zfp871 | 10.79651 | 10.87443 | 10.939 | -0.06447243 | -0.142387312 | 10.868451 | 10.94481 | 11.103 | -0.158236026 | -0.234594512 |
| Ccdc43 | 8.726857 | 8.571231 | 8.584 | -0.012751353 | 0.142873953 | 8.5784135 | 8.4706113 | 8.62897 | -0.158361766 | -0.050559588 |
| Bbs5 | 3.470953 | 3.359005 | 3.311 | 0.048006482 | 0.159954369 | 3.4918073 | 3.3279991 | 3.4864 | -0.158404418 | 0.005403712 |
| Olfr45 | 4.351082 | 4.297348 | 4.4548 | -0.157416694 | -0.103681877 | 4.479474 | 4.4120541 | 4.57047 | -0.158420285 | -0.091000397 |
| Gnai3 | 11.64885 | 11.59147 | 11.587 | 0.00450617 | 0.061887652 | 11.450495 | 11.346969 | 11.5054 | -0.158476124 | -0.054949665 |
| Atf7ip | 11.66218 | 11.78663 | 11.686 | 0.100376342 | -0.024074064 | 11.462403 | 11.660655 | 11.8192 | -0.15853318 | -0.356785341 |
| Prr18 | 1.629191 | 1.55789 | 1.616 | -0.058077827 | 0.013223726 | 1.6285259 | 1.5674095 | 1.726 | -0.158593484 | -0.097477048 |
| Fut11 | 8.082087 | 8.032687 | 8.187 | -0.154317611 | -0.104917701 | 8.1046418 | 7.833475 | 7.9921 | -0.158625892 | 0.112540894 |
| Raet1d | 3.330562 | 3.258411 | 3.2865 | -0.028108619 | 0.044042997 | 3.2727203 | 3.2174865 | 3.37613 | -0.158643315 | -0.10340951 |
| Traj32 | 2.609797 | 2.536186 | 2.4809 | 0.055280181 | 0.128891323 | 2.4507228 | 2.4509289 | 2.60964 | -0.158711276 | -0.158917471 |
| Pcmtd2 | 9.908939 | 10.0542 | 10.026 | 0.02854945 | -0.116714123 | 10.121305 | 9.9175792 | 10.0763 | -0.158767389 | 0.044958523 |
| Tia1 | 10.04768 | 9.973986 | 10.042 | -0.068293659 | 0.005401977 | 10.304326 | 10.103119 | 10.2619 | -0.158776049 | 0.042431428 |
| Il34 | 3.50685 | 3.556953 | 3.4865 | 0.070450063 | 0.020347404 | 3.315533 | 3.23971 | 3.39855 | -0.158835258 | -0.083012283 |
| Phf13 | 7.687831 | 7.611872 | 7.7255 | -0.113637892 | -0.037678917 | 7.5312536 | 7.6088615 | 7.76771 | -0.158847471 | -0.236455363 |
| Dst | 8.70464 | 8.293973 | 8.5501 | -0.25614688 | 0.154519409 | 7.923703 | 8.2474514 | 8.40637 | -0.158922737 | -0.482671067 |
| Gm8822 | 4.249196 | 4.287307 | 4.245 | 0.04228225 | 0.004170917 | 4.2358182 | 4.1512364 | 4.31016 | -0.158923121 | -0.074341331 |
| Cdh23 | 3.355809 | 3.424733 | 3.4375 | -0.012748131 | -0.081672862 | 3.2737639 | 3.3184255 | 3.47735 | -0.158927792 | -0.203589435 |
| Ccl1 | 2.413854 | 2.284319 | 2.4004 | -0.116104861 | 0.013430567 | 2.1800683 | 2.2231582 | 2.38212 | -0.158960484 | -0.202050383 |
| Pcdhb19 | 1.987653 | 1.943619 | 1.9862 | -0.042574375 | 0.00146025 | 1.9188218 | 1.7908454 | 1.94982 | -0.158976321 | -0.030999886 |
| Myo1e | 11.08571 | 11.12947 | 11.057 | 0.072902468 | 0.029151308 | 11.334299 | 10.969811 | 11.1288 | -0.15898843 | 0.205499166 |
| Hist1h4c | 5.292009 | 5.301996 | 5.48 | -0.178003153 | -0.187989954 | 5.66212 | 5.26132 | 5.42045 | -0.159130626 | 0.241669325 |
| Rttn | 8.854165 | 8.726677 | 8.9294 | -0.20275092 | -0.075262639 | 8.9656648 | 8.9958582 | 9.15507 | -0.159215529 | -0.189408872 |
| Pecam1 | 12.59503 | 12.3366 | 12.619 | -0.282626458 | -0.02418999 | 12.239481 | 12.111731 | 12.271 | -0.159285696 | -0.031535701 |
| Gmfb | 10.64304 | 10.62519 | 10.662 | -0.036682104 | -0.018827684 | 10.840148 | 10.560713 | 10.7201 | -0.1593398 | 0.120095457 |
| Dnajc14 | 10.6904 | 10.64185 | 10.701 | -0.059315868 | -0.010760174 | 10.659955 | 10.486108 | 10.6455 | -0.159346678 | 0.014500062 |
| Tac4 | 1.72956 | 1.672617 | 1.6904 | -0.017823934 | 0.039118272 | 1.7134308 | 1.5873623 | 1.74674 | -0.159380835 | -0.033312336 |
| Zfp970 | 6.637071 | 6.532155 | 6.5494 | -0.017212667 | 0.087703981 | 6.6510573 | 6.5459723 | 6.70542 | -0.159445928 | -0.05436098 |
| Rad51c | 6.06635 | 6.24176 | 6.22 | 0.021782738 | -0.153627143 | 6.1075818 | 6.0796295 | 6.2391 | -0.15947029 | -0.131518018 |
| Pnpla7 | 10.18062 | 10.21638 | 10.353 | -0.136609687 | -0.172369338 | 10.578303 | 10.327225 | 10.4867 | -0.159497752 | 0.091580988 |

| | | | | | | | | | | |
|---|---|---|---|---|---|---|---|---|---|---|
| Fam217b | 5.313593 | 5.122633 | 4.9568 | 0.165826722 | 0.356786076 | 5.1116678 | 4.5918222 | 4.75138 | -0.159555624 | 0.360289979 |
| Egr2 | 7.537836 | 7.502253 | 7.5033 | -0.001010023 | 0.034572189 | 6.7214397 | 6.8102132 | 6.9698 | -0.159583567 | -0.248357075 |
| Zfp973 | 6.238376 | 6.225285 | 6.2405 | -0.015175589 | -0.00208412 | 6.1970602 | 6.1694225 | 6.32909 | -0.159666529 | -0.132028832 |
| Mbtd1 | 9.759852 | 9.888371 | 9.7483 | 0.140088418 | 0.011569229 | 9.6429844 | 9.6095476 | 9.76923 | -0.159682298 | -0.126245465 |
| Pomgnt2 | 2.908885 | 2.906309 | 2.8806 | 0.025710297 | 0.028286304 | 2.7997367 | 2.7242506 | 2.88397 | -0.159722908 | -0.084236834 |
| Strip1 | 10.12547 | 10.1616 | 10.165 | -0.003308856 | -0.039429385 | 10.2553 | 10.096862 | 10.2567 | -0.15986796 | -0.001430038 |
| Ppm1f | 9.779794 | 9.741386 | 9.7454 | -0.004012724 | 0.034395418 | 9.6577859 | 9.4793576 | 9.63935 | -0.159989272 | 0.018438962 |
| Pak2 | 12.33516 | 12.39967 | 12.319 | 0.08063424 | 0.01612406 | 12.227033 | 12.11325 | 12.2732 | -0.159990802 | -0.046207949 |
| Ndc1 | 7.961285 | 7.915899 | 7.9631 | -0.047223266 | -0.001837232 | 7.9300274 | 7.8003627 | 7.96044 | -0.160073927 | -0.030409273 |
| Pan2 | 10.35218 | 10.39175 | 10.394 | -0.002557184 | -0.042122259 | 10.689868 | 10.50858 | 10.6687 | -0.1601544 | 0.021134495 |
| Tes | 10.68135 | 10.76124 | 10.747 | 0.013750522 | -0.066136924 | 10.548568 | 10.526033 | 10.6863 | -0.160292188 | -0.137757251 |
| Gm3055 | 6.53664 | 6.489414 | 6.1939 | 0.295499949 | 0.342725883 | 6.1999874 | 6.2301583 | 6.39046 | -0.160301244 | -0.19047217 |
| Sema4g | 3.730941 | 3.731141 | 3.7676 | -0.036484624 | -0.036684241 | 3.9048272 | 3.7146726 | 3.87508 | -0.160405645 | 0.029748931 |
| Oma1 | 8.089865 | 8.207588 | 8.2171 | -0.009531259 | -0.127254688 | 8.2392757 | 8.0753719 | 8.23581 | -0.160437719 | 0.003466043 |
| Paxip1 | 9.41998 | 9.317041 | 9.3064 | 0.010650751 | 0.113589826 | 9.4837966 | 9.4367819 | 9.59722 | -0.160442758 | -0.113428075 |
| Peli2 | 5.155051 | 5.451314 | 5.4199 | 0.031404758 | -0.264858887 | 5.4362495 | 5.0054408 | 5.16598 | -0.160538118 | 0.270270626 |
| Entpd6 | 9.360512 | 9.384146 | 9.3662 | 0.01790573 | -0.005727749 | 9.6227321 | 9.3058151 | 9.46636 | -0.16054826 | 0.156368708 |
| Metap1d | 8.497864 | 8.493348 | 8.514 | -0.02069815 | -0.016181924 | 8.9913292 | 8.774047 | 8.93464 | -0.160592638 | 0.056689502 |
| Lztfl1 | 8.186985 | 8.322594 | 8.1622 | 0.160425995 | 0.02481664 | 8.4444291 | 8.296202 | 8.45684 | -0.160635746 | -0.012408676 |
| Hccs | 8.736206 | 8.683595 | 8.7056 | -0.021968873 | 0.030642498 | 8.8404157 | 8.6194581 | 8.78011 | -0.160652753 | 0.06030485 |
| Gas2l3 | 6.229398 | 5.936637 | 5.8755 | 0.061089579 | 0.353850961 | 5.6084227 | 5.7885627 | 5.94923 | -0.160662551 | -0.340802571 |
| Gzf1 | 8.244177 | 8.2826 | 8.319 | -0.036420592 | -0.074844308 | 8.1614196 | 8.0144016 | 8.17515 | -0.160746076 | -0.01372813 |
| Ccnt1 | 10.72048 | 10.92383 | 10.798 | 0.125914984 | -0.077441122 | 10.603006 | 10.742388 | 10.9032 | -0.160793412 | -0.300175569 |
| Arsg | 6.259209 | 5.902335 | 6.0405 | -0.138213288 | 0.218660456 | 5.9970649 | 6.0830081 | 6.24397 | -0.160962248 | -0.246905415 |
| Osbpl11 | 9.342963 | 9.40826 | 9.3878 | 0.020456969 | -0.04483933 | 9.1176349 | 9.0972826 | 9.2583 | -0.161017608 | -0.1406653 |
| Atp6ap1 | 11.35243 | 11.44501 | 11.306 | 0.138922342 | 0.046341994 | 11.787938 | 11.201717 | 11.3628 | -0.161105576 | 0.425115192 |
| Gm11991 | 3.4276 | 3.486439 | 3.3567 | 0.129788764 | 0.070950233 | 3.438996 | 3.3482933 | 3.50942 | -0.161128388 | -0.070425647 |
| Tfeb | 9.632202 | 9.490053 | 9.5909 | -0.100838312 | 0.041310704 | 9.7639162 | 9.5138267 | 9.67512 | -0.161294279 | 0.08879517 |
| Prr5 | 9.036198 | 9.123978 | 9.101 | 0.022982765 | -0.064797328 | 9.4735005 | 8.8021009 | 8.96342 | -0.161322308 | 0.510077255 |
| Sigirr | 9.674718 | 9.61776 | 9.7322 | -0.114434103 | -0.057475819 | 9.9106613 | 9.6161488 | 9.77755 | -0.161401394 | 0.133111058 |
| Itprid2 | 9.76235 | 9.73256 | 9.7375 | -0.004918748 | 0.024870467 | 9.6300521 | 9.4954727 | 9.65692 | -0.161452056 | -0.026872667 |
| Tubgcp6 | 10.32424 | 10.32926 | 10.29 | 0.03940003 | 0.034373958 | 10.419344 | 10.261353 | 10.4228 | -0.161474944 | -0.00348338 |

| | | | | | | | | | | |
|---|---|---|---|---|---|---|---|---|---|---|
| Vmn2r97 | 6.87383 | 6.799374 | 6.8193 | -0.019901794 | 0.054554451 | 7.2015375 | 7.0544075 | 7.21589 | -0.161479767 | -0.014349675 |
| Cpne7 | 3.825232 | 4.011417 | 4.0966 | -0.08520948 | -0.271394454 | 3.7966536 | 3.8441634 | 4.00565 | -0.161488432 | -0.2089982 |
| Trmt9b | 2.111642 | 2.046294 | 2.103 | -0.0567529 | 0.008594698 | 1.9968378 | 1.9567435 | 2.11826 | -0.161513487 | -0.121419148 |
| Olfr784 | 5.026926 | 5.097463 | 5.049 | 0.048457992 | -0.022078538 | 5.0600141 | 5.1738887 | 5.33558 | -0.161686977 | -0.275561547 |
| Lrrn4 | 2.69238 | 2.612369 | 2.7272 | -0.114859093 | -0.034848556 | 2.5754884 | 2.5361836 | 2.6979 | -0.161718857 | -0.122414027 |
| 4930526A20Rik | 2.267265 | 2.205894 | 2.1595 | 0.046425558 | 0.107796496 | 2.2541624 | 2.238048 | 2.39982 | -0.161775564 | -0.145661162 |
| Ddx5 | 15.15444 | 15.06688 | 15.093 | -0.026399168 | 0.061168166 | 14.981175 | 14.897934 | 15.0599 | -0.161925554 | -0.078684566 |
| Tcta | 7.058988 | 6.817404 | 7.0138 | -0.196376855 | 0.045207088 | 7.2434959 | 6.8375957 | 6.99953 | -0.161930739 | 0.243969385 |
| Naa35 | 9.168046 | 9.131273 | 9.1858 | -0.054529531 | -0.017757102 | 9.1020872 | 9.0719422 | 9.2339 | -0.161958615 | -0.131813626 |
| Ndst1 | 10.88279 | 10.73834 | 10.816 | -0.07790324 | 0.066545242 | 10.903678 | 10.499199 | 10.6612 | -0.161996564 | 0.242483156 |
| Nbr1 | 10.61301 | 10.76111 | 10.725 | 0.036592398 | -0.11150931 | 10.732409 | 10.701584 | 10.8636 | -0.16204858 | -0.131223457 |
| Trpm6 | 4.313187 | 4.39261 | 4.329 | 0.063581341 | -0.015841831 | 4.1411801 | 4.3127637 | 4.47483 | -0.162069528 | -0.333653088 |
| Atp9a | 6.830179 | 6.774706 | 6.7343 | 0.040444565 | 0.095918084 | 6.5747878 | 6.5511551 | 6.71332 | -0.162160695 | -0.138528016 |
| Phldb1 | 6.80187 | 6.84511 | 6.8255 | 0.019653791 | -0.023586278 | 6.5803148 | 6.7253565 | 6.88755 | -0.162195591 | -0.30723734 |
| Eps8l3 | 0.603781 | 0.613791 | 0.5557 | 0.058063652 | 0.048054387 | 0.5723871 | 0.528423 | 0.69068 | -0.162260501 | -0.118296467 |
| Lrrc15 | 2.74651 | 2.637247 | 2.7494 | -0.112109968 | -0.002847326 | 2.8252181 | 2.7372952 | 2.89967 | -0.162377424 | -0.074454536 |
| Efhc1 | 2.495837 | 2.430257 | 2.6274 | -0.197123226 | -0.131542371 | 2.4742554 | 2.3756159 | 2.538 | -0.162383796 | -0.063744304 |
| Gm13357 | 1.655531 | 1.570545 | 1.6838 | -0.113261454 | -0.028275346 | 1.6330503 | 1.6253581 | 1.78781 | -0.162451888 | -0.154759645 |
| Scml4 | 10.78332 | 10.91349 | 10.835 | 0.078494388 | -0.05167799 | 10.78121 | 10.767013 | 10.9295 | -0.16252029 | -0.148323564 |
| Cbr1 | 8.41653 | 8.385837 | 8.464 | -0.078137209 | -0.047444137 | 8.4383537 | 8.1664783 | 8.32902 | -0.162541282 | 0.109334073 |
| 3425401B19Rik | 2.59498 | 2.594006 | 2.5466 | 0.047413405 | 0.048387556 | 2.4986567 | 2.4368507 | 2.59957 | -0.162720578 | -0.10091456 |
| Fam20c | 5.895149 | 5.994556 | 5.2008 | 0.793755416 | 0.694348109 | 5.0853738 | 4.9420748 | 5.10481 | -0.162737529 | -0.019438516 |
| Prob1 | 3.943702 | 4.142074 | 3.9934 | 0.148653294 | -0.049718426 | 4.3515596 | 3.9583679 | 4.12112 | -0.16274936 | 0.230442344 |
| Prlr | 3.679714 | 3.349129 | 3.4166 | -0.067432526 | 0.263152572 | 3.5651609 | 3.3638394 | 3.52664 | -0.162804153 | 0.038517334 |
| Sec31b | 6.116429 | 5.988573 | 6.1774 | -0.188820359 | -0.060963887 | 6.1496459 | 6.0838746 | 6.24675 | -0.162874033 | -0.097102736 |
| Kcna7 | 5.027316 | 4.988802 | 5.0362 | -0.047377327 | -0.008863486 | 5.0005308 | 4.9173148 | 5.08028 | -0.162968487 | -0.079752493 |
| Srpk3 | 9.806358 | 9.853245 | 9.9162 | -0.06295704 | -0.109843647 | 10.937755 | 10.268613 | 10.4316 | -0.162968878 | 0.506172983 |
| Gm21182 | 1.971636 | 1.934926 | 1.9252 | 0.009774935 | 0.046484754 | 2.0693328 | 2.0117594 | 2.17498 | -0.163220751 | -0.105647377 |
| Mtif2 | 8.937096 | 8.702944 | 8.8979 | -0.1949224 | 0.039229801 | 8.9066076 | 8.7793436 | 8.94259 | -0.16324163 | -0.035977626 |
| Trav9-1 | 5.381379 | 5.363037 | 5.4387 | -0.075656312 | -0.057314304 | 5.2237341 | 5.349092 | 5.51238 | -0.163292816 | -0.288650759 |
| Ccdc33 | 2.684312 | 2.612131 | 2.5696 | 0.042562393 | 0.114744029 | 2.6032446 | 2.3815268 | 2.54485 | -0.163321016 | 0.058396779 |
| Klrg1 | 5.293431 | 5.337305 | 5.4278 | -0.090485963 | -0.13436075 | 5.4844317 | 5.7666512 | 5.93002 | -0.163367891 | -0.445587418 |

| | | | | | | | | | | |
|---|---|---|---|---|---|---|---|---|---|---|
| Kdm3a | 10.52259 | 10.59249 | 10.672 | -0.0792884 | -0.149184123 | 10.600489 | 10.632194 | 10.7956 | -0.163422208 | -0.195127171 |
| Ighv1-58 | 5.187359 | 5.081504 | 5.2536 | -0.172136467 | -0.066280953 | 5.2463882 | 4.9429081 | 5.10635 | -0.163442584 | 0.140037529 |
| Sdr42e1 | 5.635953 | 5.669533 | 5.6408 | 0.028739084 | -0.004840216 | 5.7533209 | 5.514644 | 5.67813 | -0.16348666 | 0.075190229 |
| Gm14548 | 6.92178 | 6.946939 | 6.6027 | 0.34424359 | 0.319084554 | 7.2945826 | 6.6871167 | 6.85066 | -0.163540571 | 0.443925307 |
| Cxxc4 | 2.239643 | 2.171686 | 2.1719 | -0.000234555 | 0.067722153 | 2.2283081 | 2.0476742 | 2.21141 | -0.16373151 | 0.016902388 |
| Spg7 | 9.938465 | 9.838337 | 10.027 | -0.189150178 | -0.089021965 | 10.068215 | 9.9719801 | 10.1357 | -0.163763994 | -0.067529168 |
| Ofd1 | 8.586016 | 8.496998 | 8.6989 | -0.201874731 | -0.112855985 | 8.6410616 | 8.5421774 | 8.70597 | -0.163797208 | -0.06491306 |
| Hdac8 | 7.279622 | 7.241819 | 7.3087 | -0.066844146 | -0.029040728 | 7.4191353 | 7.1396937 | 7.30352 | -0.163823745 | 0.115617897 |
| Cnnm3 | 8.620573 | 8.721985 | 8.6984 | 0.02356771 | -0.077845129 | 8.8245339 | 8.6465823 | 8.81042 | -0.163837713 | 0.014113905 |
| Hgd | 3.079424 | 2.942454 | 3.1771 | -0.234645235 | -0.09767475 | 2.855507 | 2.8944348 | 3.05833 | -0.16389371 | -0.202821538 |
| Akt1 | 10.84079 | 10.64166 | 10.754 | -0.112744952 | 0.086388978 | 10.77052 | 10.524765 | 10.6887 | -0.163899404 | 0.08185571 |
| Nmnat1 | 6.32083 | 6.305817 | 6.4288 | -0.122998823 | -0.107985844 | 6.4829473 | 6.2730187 | 6.43694 | -0.163921478 | 0.046007049 |
| Zfp120 | 9.00145 | 8.977329 | 8.9545 | 0.022867769 | 0.04698891 | 8.9612872 | 8.8328018 | 8.99674 | -0.163933683 | -0.035448291 |
| Prl8a1 | 5.793254 | 5.864364 | 5.8577 | 0.006619443 | -0.064491224 | 6.0483908 | 5.9428617 | 6.10684 | -0.163982749 | -0.058453677 |
| Nup107 | 9.58143 | 9.519213 | 9.5202 | -0.000989066 | 0.061227719 | 9.552449 | 9.5888133 | 9.75282 | -0.164010248 | -0.200374558 |
| Unc5b | 6.233405 | 6.268797 | 6.0758 | 0.192991155 | 0.15759939 | 5.8220326 | 5.7227669 | 5.88679 | -0.164025878 | -0.064760187 |
| Mrrf | 8.232654 | 8.073997 | 8.0313 | 0.04274677 | 0.201403308 | 7.9339467 | 7.8596485 | 8.02387 | -0.164223655 | -0.089925523 |
| Suco | 10.39507 | 10.05315 | 10.257 | -0.20387064 | 0.138051351 | 9.9360167 | 10.121069 | 10.2853 | -0.164245155 | -0.349297751 |
| Mgme1 | 6.969625 | 6.972189 | 6.9474 | 0.024791782 | 0.022228066 | 6.9002717 | 6.8053728 | 6.96964 | -0.164266264 | -0.069367416 |
| Vmn2r11 | 2.379595 | 2.376759 | 2.4581 | -0.081376145 | -0.078540014 | 2.3125536 | 2.416737 | 2.58105 | -0.164311099 | -0.268494433 |
| Glt8d2 | 3.186653 | 3.2631 | 3.3254 | -0.06232518 | -0.138773014 | 3.1217035 | 3.1452541 | 3.3096 | -0.164344159 | -0.187894733 |
| Dnaic2 | 1.631559 | 1.567891 | 1.6039 | -0.036017525 | 0.027650488 | 1.6684269 | 1.5553706 | 1.71975 | -0.164382015 | -0.051325736 |
| Gm21981 | 1.707878 | 1.74038 | 1.6883 | 0.052114685 | 0.019612079 | 1.7073586 | 1.6260031 | 1.79054 | -0.164532152 | -0.083176588 |
| Cbx8 | 7.189874 | 7.24932 | 7.2421 | 0.00725052 | -0.052195436 | 7.3801252 | 7.0056134 | 7.1702 | -0.164591474 | 0.209920336 |
| Cldn34c1 | 5.269643 | 5.074202 | 5.0799 | -0.005669911 | 0.189771076 | 5.1207473 | 4.927377 | 5.09215 | -0.164769172 | 0.028601112 |
| Trbj1-2 | 5.362939 | 5.431768 | 5.4118 | 0.019962336 | -0.048866609 | 5.2109742 | 5.2858952 | 5.4507 | -0.16480148 | -0.239722508 |
| Hist4h4 | 5.263213 | 5.140407 | 5.4418 | -0.301438219 | -0.178632525 | 5.1589028 | 5.1507557 | 5.31578 | -0.165022906 | -0.156875875 |
| Il17ra | 11.69764 | 11.81284 | 11.71 | 0.102783736 | -0.012416414 | 11.652226 | 11.499748 | 11.6648 | -0.1650457 | -0.012568485 |
| Nr4a3 | 9.43494 | 9.497132 | 9.1478 | 0.349344087 | 0.287152216 | 8.897526 | 9.5064292 | 9.67148 | -0.165047798 | -0.773950916 |
| Vps8 | 9.296418 | 9.128915 | 9.3664 | -0.237509752 | -0.070006644 | 9.4811939 | 9.0751368 | 9.2402 | -0.165061112 | 0.240995923 |
| Olfr152 | 4.402957 | 4.395909 | 4.3742 | 0.021690117 | 0.02873814 | 4.4608472 | 4.5535291 | 4.71859 | -0.165065371 | -0.257747304 |
| Traj7 | 2.933232 | 2.988781 | 3.0117 | -0.022930888 | -0.078479909 | 2.9504471 | 2.8804837 | 3.04559 | -0.16511049 | -0.095147106 |

| | | | | | | | | | | |
|---|---|---|---|---|---|---|---|---|---|---|
| H2afv | 8.781631 | 8.774701 | 8.9956 | -0.220892871 | -0.213962944 | 8.8600658 | 8.5364958 | 8.70163 | -0.165136046 | 0.158433927 |
| Gm3512 | 4.374103 | 4.463128 | 4.6495 | -0.18641815 | -0.275442947 | 4.4897477 | 4.3318864 | 4.49708 | -0.165198306 | -0.007337008 |
| Ermp1 | 9.713761 | 9.711405 | 9.6783 | 0.033102189 | 0.035457474 | 9.7735016 | 9.5603229 | 9.72557 | -0.165249861 | 0.047928858 |
| Gm19241 | 2.14579 | 2.060304 | 2.0867 | -0.026417756 | 0.059068519 | 2.0776933 | 2.0444679 | 2.20977 | -0.165306118 | -0.132080781 |
| Atg101 | 8.102373 | 8.064495 | 8.1805 | -0.115971458 | -0.078093148 | 8.1043201 | 7.9697917 | 8.13511 | -0.165319923 | -0.030791528 |
| Zfp112 | 4.239865 | 4.138403 | 4.299 | -0.16061483 | -0.059153621 | 4.2102152 | 4.0905976 | 4.25597 | -0.165369211 | -0.045751545 |
| Kcna6 | 2.742877 | 2.731107 | 2.7513 | -0.020166223 | -0.008396435 | 2.6189526 | 2.600102 | 2.76556 | -0.165459113 | -0.146608471 |
| Galc | 6.846026 | 6.757935 | 6.969 | -0.211114774 | -0.123024127 | 6.8381073 | 6.5935491 | 6.75903 | -0.165484976 | 0.079073251 |
| Tmem181b-ps | 7.202781 | 7.395983 | 7.3012 | 0.094828263 | -0.098373744 | 7.3731971 | 7.2835901 | 7.44909 | -0.165503461 | -0.075896454 |
| Ighv12-3 | 5.579175 | 5.616906 | 5.6175 | -0.000572323 | -0.038303229 | 5.8905918 | 5.3412883 | 5.50686 | -0.165573698 | 0.38372979 |
| Zbtb25 | 8.79852 | 8.79682 | 8.7919 | 0.004933084 | 0.006633795 | 8.7089999 | 8.7740552 | 8.93978 | -0.165724842 | -0.230780229 |
| Trav12-1 | 4.119428 | 4.038349 | 4.0021 | 0.036219079 | 0.117298503 | 4.0311618 | 4.0197792 | 4.1856 | -0.165819081 | -0.154436432 |
| Gm15382 | 3.793432 | 3.832947 | 3.8596 | -0.026656125 | -0.066171437 | 4.0592532 | 3.9269971 | 4.09282 | -0.165822461 | -0.033566433 |
| Gfm2 | 9.308815 | 9.384752 | 9.3645 | 0.020284863 | -0.055651888 | 9.4223166 | 9.3726166 | 9.53847 | -0.165851219 | -0.116151245 |
| Cd59b | 3.765523 | 3.688332 | 3.7324 | -0.044105466 | 0.033086013 | 3.7083224 | 3.7216657 | 3.88762 | -0.165956691 | -0.179299954 |
| B4galt1 | 10.92179 | 10.92051 | 11.004 | -0.083750552 | -0.082473966 | 10.756299 | 10.762716 | 10.9287 | -0.1659697 | -0.172386966 |
| Timd2 | 2.73719 | 2.786165 | 2.713 | 0.073147223 | 0.024171443 | 2.7299051 | 2.636741 | 2.80274 | -0.165997805 | -0.072833725 |
| Vps26c | 9.377596 | 9.300127 | 9.4413 | -0.1411505 | -0.063681514 | 9.3440292 | 9.2987945 | 9.46485 | -0.166054664 | -0.120819889 |
| Atp6v0e2 | 3.367332 | 3.369691 | 3.2966 | 0.073134643 | 0.070776406 | 3.2412739 | 3.1542155 | 3.32031 | -0.166091961 | -0.079033553 |
| Jmjd7 | 7.104907 | 6.896372 | 7.0748 | -0.178396939 | 0.030138024 | 7.4205171 | 6.8716987 | 7.03787 | -0.166174284 | 0.382644191 |
| Zfp819 | 1.614342 | 1.498983 | 1.6196 | -0.120602057 | -0.005243707 | 1.5535091 | 1.478645 | 1.64488 | -0.16623954 | -0.091375459 |
| Camsap3 | 3.491377 | 3.277185 | 3.6516 | -0.374399323 | -0.160207438 | 3.416499 | 3.3314069 | 3.49766 | -0.166251673 | -0.081159535 |
| Gngt1 | 4.18562 | 4.000331 | 4.2563 | -0.255970726 | -0.070681176 | 4.0555468 | 4.1615537 | 4.32783 | -0.166273359 | -0.272280207 |
| Pvrig | 4.092207 | 3.838442 | 4.1326 | -0.294174015 | -0.040409356 | 3.9614478 | 3.9516853 | 4.11798 | -0.166297743 | -0.156535225 |
| Slc2a12 | 3.02931 | 2.824553 | 3.2174 | -0.392821218 | -0.188063965 | 2.9065801 | 2.6944924 | 2.86084 | -0.166343368 | 0.045744285 |
| 1700017B05Rik | 9.432476 | 9.349004 | 9.4179 | -0.06887707 | 0.014594412 | 9.2698922 | 9.2457206 | 9.41223 | -0.166508628 | -0.142337013 |
| Gpr88 | 2.944099 | 2.659538 | 2.7911 | -0.131581487 | 0.152979275 | 2.6746511 | 2.551124 | 2.71771 | -0.16658136 | -0.043054334 |
| Ddx19b | 9.61281 | 9.424013 | 9.5107 | -0.08668693 | 0.102109469 | 9.6064739 | 9.411387 | 9.57802 | -0.166637037 | 0.028449884 |
| Mcu | 8.066518 | 8.024402 | 8.1686 | -0.144237505 | -0.102121914 | 8.0634097 | 7.9411585 | 8.10782 | -0.166657892 | -0.044406714 |
| Fchsd1 | 8.296942 | 8.12693 | 8.3673 | -0.24038013 | -0.070367852 | 8.174841 | 8.2138413 | 8.38056 | -0.166715266 | -0.205715581 |
| Gm47103 | 4.110833 | 4.259072 | 4.1377 | 0.121345985 | -0.026893294 | 4.1608917 | 4.2627793 | 4.42953 | -0.166751562 | -0.268639151 |
| Mllt10 | 10.65179 | 10.46629 | 10.66 | -0.19352927 | -0.008022216 | 10.496825 | 10.446524 | 10.6134 | -0.166831112 | -0.116529672 |

| | | | | | | | | | | |
|---|---|---|---|---|---|---|---|---|---|---|
| Prr22 | 3.508188 | 3.515837 | 3.587 | -0.07114417 | -0.078793034 | 3.7527309 | 3.5334728 | 3.70031 | -0.166835315 | 0.052422813 |
| Emc2 | 9.273165 | 9.164471 | 9.2905 | -0.126038035 | -0.017344055 | 9.2457561 | 9.0349175 | 9.20181 | -0.166895389 | 0.043943124 |
| Pygb | 10.47846 | 10.50649 | 10.522 | -0.015006496 | -0.043040656 | 10.426833 | 10.266986 | 10.434 | -0.166995158 | -0.007148316 |
| Golga5 | 9.845804 | 9.774523 | 9.9221 | -0.147590767 | -0.076309958 | 9.8712485 | 9.7495108 | 9.91656 | -0.167047423 | -0.045309643 |
| Negr1 | 2.275231 | 2.315909 | 2.3073 | 0.008560008 | -0.0321182 | 2.0915604 | 2.1279732 | 2.2951 | -0.167122209 | -0.203535073 |
| Ptpn11 | 10.7873 | 10.80501 | 10.771 | 0.033947472 | 0.016229704 | 10.679854 | 10.576707 | 10.7438 | -0.167141956 | -0.063995248 |
| Cmtm8 | 3.580482 | 3.491778 | 3.7317 | -0.239967302 | -0.15126293 | 3.5414351 | 3.3855976 | 3.55277 | -0.167170396 | -0.011332924 |
| Ace | 9.120835 | 8.71283 | 9.1555 | -0.442702073 | -0.034696533 | 8.492818 | 8.228305 | 8.3956 | -0.167295863 | 0.097217107 |
| Traj23 | 2.925546 | 2.882004 | 2.9106 | -0.028594827 | 0.014948094 | 2.8731022 | 2.7244735 | 2.89179 | -0.167312338 | -0.018683628 |
| Gm17195 | 3.085116 | 3.166855 | 3.1145 | 0.052334028 | -0.029405254 | 3.1515634 | 3.1068771 | 3.27421 | -0.167328493 | -0.122642213 |
| Unc13d | 10.95393 | 10.92024 | 11.008 | -0.08728065 | -0.05358554 | 11.289849 | 10.963317 | 11.1307 | -0.16733411 | 0.1591982 |
| Cyb561 | 6.220928 | 5.680752 | 5.9419 | -0.261184023 | 0.278991743 | 5.5690332 | 5.4728751 | 5.64022 | -0.16734551 | -0.071187487 |
| D330045A20Rik | 1.20298 | 1.232708 | 1.2763 | -0.043614981 | -0.073342824 | 1.2018377 | 1.2277055 | 1.39506 | -0.16735547 | -0.193223232 |
| Polr3f | 9.257755 | 9.29366 | 9.25 | 0.043687492 | 0.007782674 | 9.3489347 | 9.1634504 | 9.33086 | -0.1674057 | 0.018078568 |
| Ccdc181 | 6.780806 | 6.612758 | 6.7798 | -0.167083698 | 0.000963776 | 6.8348702 | 6.5390822 | 6.70671 | -0.167623528 | 0.12816442 |
| Gstm5 | 3.764907 | 3.54137 | 3.4986 | 0.04280993 | 0.266347099 | 3.4507582 | 3.2953886 | 3.46302 | -0.167632147 | -0.012262625 |
| Cep162 | 8.623425 | 8.837208 | 8.7213 | 0.115922683 | -0.097860575 | 8.5381489 | 8.7475283 | 8.91517 | -0.167638679 | -0.377018015 |
| Efcab5 | 2.888241 | 3.05974 | 2.9504 | 0.109314489 | -0.062184306 | 2.9521659 | 2.9492899 | 3.11693 | -0.167639293 | -0.164763375 |
| Pnrc1 | 11.62357 | 11.51834 | 11.703 | -0.184323486 | -0.079090108 | 11.076806 | 11.211622 | 11.3793 | -0.167671676 | -0.302487093 |
| Zfp563 | 6.442977 | 6.587653 | 6.6797 | -0.09201273 | -0.236688568 | 6.329293 | 6.4289434 | 6.59675 | -0.167807413 | -0.267457769 |
| Rbm24 | 1.728986 | 1.784972 | 1.8431 | -0.058109781 | -0.114095252 | 1.8633053 | 1.6963722 | 1.8643 | -0.167928828 | -0.000995673 |
| Dhodh | 8.211683 | 8.306589 | 8.238 | 0.068605789 | -0.026300709 | 8.6825204 | 8.2733645 | 8.44143 | -0.168065936 | 0.241089952 |
| Smim10l2a | 2.528083 | 2.571625 | 2.6593 | -0.087625793 | -0.131168332 | 2.4829566 | 2.3878671 | 2.55594 | -0.168077434 | -0.072987898 |
| Thoc2 | 11.19423 | 11.10424 | 11.087 | 0.017332308 | 0.107324012 | 11.053577 | 11.155622 | 11.3238 | -0.168222204 | -0.270267293 |
| Selenbp1 | 6.207197 | 5.71388 | 6.1564 | -0.442505659 | 0.050811201 | 5.7544218 | 5.7727118 | 5.94095 | -0.168241199 | -0.186531215 |
| Trpt1 | 5.265489 | 5.077811 | 5.1115 | -0.033702253 | 0.153976123 | 5.3541287 | 5.2216742 | 5.38996 | -0.168283747 | -0.035829209 |
| Akap12 | 9.342615 | 9.114414 | 8.9915 | 0.122872181 | 0.351073217 | 8.8537987 | 8.8070116 | 8.97534 | -0.168327585 | -0.121540512 |
| Slc18a1 | 4.339617 | 4.210461 | 4.5024 | -0.291893621 | -0.162737584 | 4.1850892 | 4.1413618 | 4.30971 | -0.168343279 | -0.12461585 |
| Larp4b | 10.70386 | 10.73987 | 10.741 | -0.001572004 | -0.037586286 | 10.673578 | 10.660321 | 10.8287 | -0.168351478 | -0.155094447 |
| Khk | 7.422807 | 7.271309 | 7.6005 | -0.329236933 | -0.177738912 | 7.5012609 | 7.1920776 | 7.36044 | -0.168358631 | 0.140824634 |
| Tcrg-V6 | 3.236562 | 3.226355 | 3.3003 | -0.073962133 | -0.063755909 | 3.1185386 | 3.0270425 | 3.1955 | -0.16845281 | -0.076956773 |
| Pign | 9.342418 | 9.47702 | 9.3378 | 0.139218091 | 0.0046159 | 9.4880123 | 9.4753585 | 9.64385 | -0.168493607 | -0.155839778 |

| | | | | | | | | | | |
|---|---|---|---|---|---|---|---|---|---|---|
| Mfsd14b | 10.06577 | 10.02255 | 9.9889 | 0.033614746 | 0.076839757 | 10.081627 | 9.9588978 | 10.1274 | -0.168543519 | -0.045814494 |
| Gm6170 | 4.859543 | 4.940357 | 4.8769 | 0.063450031 | -0.01736407 | 4.7912681 | 4.8339468 | 5.00256 | -0.16861095 | -0.211289632 |
| Pcgf2 | 5.260858 | 5.015102 | 5.2242 | -0.209064228 | 0.036691895 | 4.9046127 | 5.2438273 | 5.41246 | -0.168637199 | -0.507851882 |
| Map2 | 5.309628 | 5.211351 | 5.3279 | -0.116563785 | -0.018286238 | 5.172814 | 5.0947036 | 5.26334 | -0.168637619 | -0.090527169 |
| Olfr472 | 5.183964 | 5.207151 | 5.1696 | 0.03751583 | 0.014329064 | 5.3779374 | 5.3019472 | 5.47059 | -0.168645272 | -0.092655105 |
| Sp4 | 10.55317 | 10.54827 | 10.538 | 0.010146382 | 0.015051952 | 10.583327 | 10.504355 | 10.6731 | -0.16871766 | -0.089745933 |
| Ky | 1.477278 | 1.658023 | 1.4084 | 0.24962645 | 0.068880917 | 1.559647 | 1.4217256 | 1.59044 | -0.168718947 | -0.030797545 |
| Tmem184c | 8.412795 | 8.436651 | 8.5515 | -0.114888788 | -0.138745499 | 8.5842749 | 8.4740539 | 8.64281 | -0.168757746 | -0.058536714 |
| Ech1 | 9.599061 | 9.526457 | 9.6377 | -0.111193053 | -0.03858953 | 9.7980838 | 9.4240554 | 9.59287 | -0.168818228 | 0.205210173 |
| Bbs12 | 5.3763 | 5.262195 | 5.5348 | -0.27261837 | -0.158514051 | 5.4603117 | 5.4465015 | 5.61546 | -0.168956788 | -0.155146575 |
| Cir1 | 9.154146 | 9.116465 | 9.2463 | -0.129788304 | -0.09210661 | 9.2258947 | 9.0698824 | 9.23887 | -0.168982618 | -0.012970351 |
| Zpbp | 5.892657 | 5.939398 | 5.9897 | -0.050296714 | -0.097037876 | 5.8910744 | 5.7817487 | 5.95075 | -0.168997988 | -0.059672368 |
| Kyat3 | 5.884115 | 5.853522 | 5.8893 | -0.035774725 | -0.005182159 | 6.0704743 | 5.8510885 | 6.02019 | -0.169096513 | 0.050289303 |
| Cfap36 | 7.620364 | 7.620692 | 7.6251 | -0.004436775 | -0.004764829 | 7.6726058 | 7.5128803 | 7.68201 | -0.169129034 | -0.009403534 |
| Lrrc42 | 8.535248 | 8.653036 | 8.5828 | 0.070283215 | -0.047504429 | 8.7023647 | 8.4958515 | 8.66504 | -0.169193192 | 0.037319982 |
| Ephx3 | 3.551515 | 3.493018 | 3.4333 | 0.05975967 | 0.118255859 | 3.385944 | 3.1928997 | 3.36222 | -0.169317205 | 0.023727157 |
| Ocln | 2.882022 | 2.80942 | 2.8718 | -0.062334794 | 0.010267499 | 2.9458743 | 2.9523089 | 3.12171 | -0.169403295 | -0.175837879 |
| Myh13 | 2.055747 | 2.120614 | 2.2145 | -0.093925865 | -0.15879225 | 1.997105 | 1.9298795 | 2.09944 | -0.169559444 | -0.102333948 |
| Heatr6 | 9.597051 | 9.646386 | 9.6749 | -0.028524968 | -0.077859822 | 9.7922789 | 9.6621497 | 9.83179 | -0.169640188 | -0.039511012 |
| Nsl1 | 6.079216 | 5.79406 | 6.1292 | -0.335189645 | -0.050033726 | 6.0028954 | 6.1849573 | 6.35463 | -0.169673902 | -0.351735823 |
| Trbv16 | 7.847801 | 7.93651 | 7.9921 | -0.055634669 | -0.144343218 | 7.7177096 | 7.7498085 | 7.91949 | -0.169683152 | -0.201782032 |
| 1810062G17Rik | 3.014694 | 2.943783 | 2.9307 | 0.013041908 | 0.083952981 | 3.0102521 | 2.804289 | 2.97404 | -0.169754214 | 0.036208896 |
| Itgb3bp | 6.004441 | 5.934625 | 6.0248 | -0.090143116 | -0.020327916 | 6.1312031 | 5.8445028 | 6.01428 | -0.169777062 | 0.116923149 |
| Pold4 | 9.328359 | 9.307621 | 9.4014 | -0.093752371 | -0.073014532 | 9.8250101 | 9.3192147 | 9.48901 | -0.16979157 | 0.336003737 |
| Cep85 | 8.918032 | 8.866385 | 9.0631 | -0.196668522 | -0.145021768 | 8.8771191 | 8.9230055 | 9.09285 | -0.169848564 | -0.21573491 |
| Ndrg2 | 6.62031 | 6.778423 | 7.0874 | -0.308989187 | -0.467102949 | 6.5690821 | 6.6416532 | 6.81157 | -0.169913615 | -0.24248472 |
| B3galt2 | 3.280752 | 2.979838 | 3.0959 | -0.116013643 | 0.184900545 | 3.0011748 | 2.9384719 | 3.10843 | -0.169959991 | -0.107257142 |
| Olfr689 | 2.845328 | 2.817427 | 2.8869 | -0.069506171 | -0.041605203 | 2.9206403 | 2.88408 | 3.05404 | -0.169961543 | -0.133401227 |
| Havcr2 | 7.244249 | 7.059679 | 7.1969 | -0.137177886 | 0.047391625 | 7.0672414 | 7.0282713 | 7.19826 | -0.16998941 | -0.131019322 |
| Magt1 | 10.37561 | 10.45636 | 10.335 | 0.121760066 | 0.041004306 | 10.394022 | 10.182685 | 10.3527 | -0.1700218 | 0.041314719 |
| Ephb1 | 2.182732 | 2.158702 | 2.3103 | -0.151630662 | -0.127600201 | 2.2075123 | 2.0230567 | 2.19312 | -0.170058391 | 0.014397177 |
| Apmap | 9.070283 | 9.025276 | 9.0077 | 0.017559836 | 0.062567085 | 9.119184 | 8.8647466 | 9.03491 | -0.170161159 | 0.084276263 |

| | | | | | | | | | | |
|---|---|---|---|---|---|---|---|---|---|---|
| Ranbp17 | 3.224371 | 3.157749 | 3.2457 | -0.087999544 | -0.021377081 | 3.1539223 | 3.0553425 | 3.22562 | -0.170279028 | -0.071699239 |
| Angel1 | 7.819505 | 7.758489 | 7.8913 | -0.132829996 | -0.071814844 | 7.9934812 | 7.7293741 | 7.89974 | -0.170370335 | 0.093736819 |
| Higd2a | 9.057208 | 9.014129 | 9.1413 | -0.127154545 | -0.084075083 | 9.2324643 | 8.8722655 | 9.04264 | -0.17037777 | 0.189821011 |
| Akap9 | 10.76278 | 10.78218 | 10.889 | -0.10654984 | -0.125947586 | 10.695271 | 10.900249 | 11.0708 | -0.170597208 | -0.375575759 |
| Slc22a4 | 3.893727 | 3.809293 | 3.7311 | 0.078159806 | 0.162593381 | 3.8413791 | 3.5230907 | 3.69372 | -0.170625921 | 0.147662487 |
| Vstm4 | 2.94492 | 2.933981 | 3.0449 | -0.110952398 | -0.100013715 | 2.8830259 | 2.726987 | 2.89763 | -0.170646598 | -0.014607742 |
| Gm6551 | 2.390424 | 2.366589 | 2.3462 | 0.020366562 | 0.044200938 | 2.3602936 | 2.2898268 | 2.46049 | -0.170660466 | -0.100193596 |
| Tlk1 | 11.07156 | 11.04178 | 11.182 | -0.14056625 | -0.110785738 | 10.82946 | 10.91225 | 11.0829 | -0.170665534 | -0.253455206 |
| Tex15 | 3.717106 | 3.769155 | 3.8357 | -0.066496671 | -0.118545908 | 3.809665 | 3.6325371 | 3.80339 | -0.170854184 | 0.006273706 |
| Txnip | 13.03881 | 13.10759 | 13.237 | -0.129817516 | -0.198607402 | 12.86719 | 12.891517 | 13.0624 | -0.170861724 | -0.195188547 |
| Col13a1 | 3.053914 | 2.975261 | 2.9754 | -0.000140534 | 0.078513044 | 2.8332894 | 2.8566446 | 3.02772 | -0.171071843 | -0.194427086 |
| Trav9d-3 | 4.889529 | 4.790408 | 4.888 | -0.09756039 | 0.001559868 | 4.7152641 | 4.7579424 | 4.92905 | -0.171112072 | -0.213790319 |
| Trav9n-3 | 4.889529 | 4.790408 | 4.888 | -0.09756039 | 0.001559868 | 4.7152641 | 4.7579424 | 4.92905 | -0.171112072 | -0.213790319 |
| Pdcl | 9.298965 | 9.217516 | 9.3124 | -0.094917069 | -0.013467886 | 9.0099537 | 8.9820733 | 9.15321 | -0.171138113 | -0.143257649 |
| Phlda1 | 7.724851 | 7.345988 | 7.9106 | -0.564645441 | -0.185782562 | 6.8313803 | 7.2336481 | 7.40479 | -0.171146182 | -0.573413978 |
| Slc6a4 | 4.29027 | 4.489121 | 4.4743 | 0.014795955 | -0.184055046 | 4.1902044 | 4.2184182 | 4.38957 | -0.171148117 | -0.199361936 |
| Pdgfc | 4.15135 | 3.849693 | 3.8301 | 0.019583689 | 0.321240511 | 3.4330456 | 3.5353949 | 3.70657 | -0.171171557 | -0.273520875 |
| Cpa2 | 1.4212 | 1.386502 | 1.4084 | -0.021885225 | 0.012812449 | 1.365797 | 1.3318427 | 1.50306 | -0.171221339 | -0.137267008 |
| Fhdc1 | 3.971268 | 4.122838 | 4.1203 | 0.002524564 | -0.14904579 | 3.9779162 | 4.1016677 | 4.27291 | -0.171240384 | -0.29499188 |
| Svop | 2.056767 | 2.033536 | 2.0415 | -0.007965415 | 0.015265759 | 2.0584706 | 2.0014733 | 2.17272 | -0.171242503 | -0.114245165 |
| Abcd1 | 10.27373 | 10.2805 | 10.35 | -0.069239564 | -0.076002406 | 10.553857 | 10.179452 | 10.3507 | -0.171270864 | 0.203134233 |
| Twsg1 | 9.313486 | 9.354704 | 9.3654 | -0.010701751 | -0.051920066 | 9.2299835 | 9.1784357 | 9.34975 | -0.171311811 | -0.119764037 |
| Rft1 | 8.090615 | 7.955255 | 8.1284 | -0.17314822 | -0.037787942 | 8.4161768 | 8.1255682 | 8.29691 | -0.171337701 | 0.119270927 |
| Ikbkg | 10.1577 | 10.26315 | 10.195 | 0.068490109 | -0.036959067 | 10.314762 | 10.012487 | 10.1839 | -0.171371778 | 0.130903801 |
| Ssx2ip | 6.232454 | 6.216534 | 6.2782 | -0.061676649 | -0.045757053 | 6.2462091 | 5.9770783 | 6.14849 | -0.171409824 | 0.097721028 |
| Csdc2 | 3.600523 | 3.556459 | 3.5056 | 0.050864722 | 0.094928579 | 3.5995393 | 3.6332198 | 3.80463 | -0.171410455 | -0.205090946 |
| Mcc | 4.517616 | 4.555522 | 4.7148 | -0.15932213 | -0.197228686 | 4.657285 | 4.5746853 | 4.7461 | -0.17141711 | -0.088817402 |
| Dct | 4.128419 | 4.006227 | 4.0878 | -0.081557387 | 0.040634855 | 3.858421 | 3.9690694 | 4.1405 | -0.171428829 | -0.282077208 |
| Diaph1 | 12.21403 | 12.26354 | 12.197 | 0.066076 | 0.016564632 | 12.215394 | 12.004991 | 12.1764 | -0.171450302 | 0.038952409 |
| Ctnna1 | 10.66295 | 10.54751 | 10.601 | -0.053304102 | 0.062134778 | 10.286847 | 10.358886 | 10.5305 | -0.171570058 | -0.243608846 |
| Naga | 9.508775 | 9.291397 | 9.429 | -0.137624365 | 0.079753905 | 9.543478 | 9.2722038 | 9.44381 | -0.171604419 | 0.099669766 |
| Cd300ld | 7.644768 | 7.405112 | 6.9772 | 0.427959232 | 0.667615322 | 7.8292606 | 6.5516787 | 6.72336 | -0.171677303 | 1.105904623 |

| | | | | | | | | | | |
|---|---|---|---|---|---|---|---|---|---|---|
| Tbl2 | 8.710236 | 8.688746 | 8.565 | 0.123737449 | 0.145227429 | 8.7746833 | 8.6040113 | 8.77575 | -0.17173418 | -0.001062241 |
| Gm19065 | 1.392423 | 1.335368 | 1.3846 | -0.049245013 | 0.007810048 | 1.3380458 | 1.2911408 | 1.46296 | -0.171817424 | -0.124912466 |
| Xlr4c | 6.933563 | 6.966832 | 7.1405 | -0.173677562 | -0.206946959 | 6.7980123 | 7.0817783 | 7.2536 | -0.171822004 | -0.455588015 |
| Pcf11 | 11.12446 | 11.11873 | 11.154 | -0.035204904 | -0.029471526 | 11.042077 | 11.089551 | 11.2614 | -0.171880126 | -0.219354304 |
| Vamp5 | 6.282344 | 6.169559 | 6.4465 | -0.27693079 | -0.16414634 | 6.3230645 | 6.0149298 | 6.18682 | -0.171894265 | 0.136240445 |
| Gm2962 | 3.786253 | 3.72143 | 3.7848 | -0.063402171 | 0.001420785 | 3.9426101 | 3.7788282 | 3.95072 | -0.171894311 | -0.008112369 |
| Tbc1d10c | 10.80374 | 10.83627 | 10.92 | -0.083334406 | -0.115861506 | 11.303016 | 10.849766 | 11.0217 | -0.1719499 | 0.281299709 |
| Fads6 | 6.05395 | 5.876504 | 5.7785 | 0.098045677 | 0.27549121 | 5.9345885 | 5.6435851 | 5.81572 | -0.172131132 | 0.118872235 |
| Nrap | 2.941935 | 2.666515 | 2.9638 | -0.297256899 | -0.021836913 | 2.7387516 | 2.7025777 | 2.87477 | -0.172190396 | -0.136016579 |
| Ktn1 | 10.52361 | 10.37688 | 10.572 | -0.195068094 | -0.048331772 | 10.582994 | 10.603344 | 10.7756 | -0.172243748 | -0.192594027 |
| Rab4a | 4.956218 | 4.579337 | 4.758 | -0.178674842 | 0.198206873 | 4.5480649 | 4.5679191 | 4.74045 | -0.172528003 | -0.192382273 |
| Traj9 | 2.989055 | 2.84144 | 2.9402 | -0.098718401 | 0.048896215 | 2.8188245 | 2.8847367 | 3.05736 | -0.172619053 | -0.238531335 |
| Trav14-3 | 4.635487 | 4.573776 | 4.5851 | -0.011311949 | 0.050399382 | 4.596846 | 4.4469002 | 4.61954 | -0.172636661 | -0.022690787 |
| Dock7 | 6.755342 | 6.774112 | 6.7114 | 0.062721669 | 0.043951451 | 6.7179592 | 6.6067151 | 6.7794 | -0.172683811 | -0.061439747 |
| Ccny | 10.63162 | 10.61282 | 10.672 | -0.059142858 | -0.040338562 | 10.466903 | 10.308874 | 10.4817 | -0.172785934 | -0.014756733 |
| Tssk4 | 4.016241 | 4.249633 | 4.2567 | -0.007060599 | -0.240452775 | 4.336362 | 4.3288003 | 4.50168 | -0.172877943 | -0.165316306 |
| Pcdhga12 | 3.553407 | 3.538313 | 3.5539 | -0.015616341 | -0.000522702 | 3.4258122 | 3.3892021 | 3.56226 | -0.173054429 | -0.136444339 |
| Lzic | 8.539488 | 8.510251 | 8.4174 | 0.09281959 | 0.122057061 | 8.4345255 | 8.4624913 | 8.63562 | -0.173124978 | -0.201090714 |
| Plppr3 | 2.818568 | 2.667948 | 2.659 | 0.008923346 | 0.159544137 | 2.6379493 | 2.5407309 | 2.71395 | -0.173218107 | -0.075999703 |
| Pi15 | 4.922774 | 4.683584 | 4.7189 | -0.035281385 | 0.203908609 | 4.6773771 | 4.5692994 | 4.74253 | -0.173225643 | -0.065147958 |
| Ccdc171 | 7.203892 | 7.377606 | 7.2869 | 0.090704417 | -0.083009719 | 7.3973529 | 7.4396821 | 7.61291 | -0.173232507 | -0.215561707 |
| Fras1 | 1.683016 | 1.570683 | 1.6794 | -0.108686274 | 0.003647133 | 1.7047306 | 1.5387496 | 1.71202 | -0.173270657 | -0.007289685 |
| Fgf12 | 1.79652 | 1.759781 | 1.7306 | 0.02916723 | 0.065906173 | 1.7444417 | 1.758485 | 1.93185 | -0.173364818 | -0.187408124 |
| Traj4 | 2.730584 | 2.684918 | 2.8009 | -0.115941264 | -0.070275147 | 2.5751409 | 2.6204766 | 2.794 | -0.173522855 | -0.218858645 |
| Angel2 | 10.2801 | 10.14006 | 10.187 | -0.047083667 | 0.092955206 | 10.289132 | 10.190884 | 10.3645 | -0.173573792 | -0.075325205 |
| Zfp704 | 5.6435 | 5.383592 | 5.6623 | -0.278723859 | -0.018815648 | 5.2319734 | 5.3672846 | 5.54087 | -0.17358934 | -0.308900533 |
| Vdac1 | 10.42246 | 10.33577 | 10.37 | -0.034398144 | 0.052287316 | 10.298451 | 10.200271 | 10.374 | -0.173707848 | -0.075527461 |
| Gm8899 | 2.866382 | 2.901042 | 2.9184 | -0.017312063 | -0.051972295 | 2.9486526 | 2.8882282 | 3.06195 | -0.173719473 | -0.113295084 |
| Epha5 | 4.264185 | 4.291558 | 4.2426 | 0.048936147 | 0.021562571 | 4.3517385 | 4.2833856 | 4.45718 | -0.173793497 | -0.105440514 |
| Prex1 | 12.12808 | 12.20751 | 12.186 | 0.02118304 | -0.058241874 | 12.139436 | 11.987442 | 12.1613 | -0.17382397 | -0.021829528 |
| Hgfac | 2.842361 | 2.897967 | 2.9971 | -0.099146313 | -0.154751687 | 2.881492 | 2.7724035 | 2.94628 | -0.173874922 | -0.064786434 |
| Tex30 | 8.138255 | 8.094942 | 8.2041 | -0.109118393 | -0.065804905 | 8.0928433 | 8.0038989 | 8.17795 | -0.174049587 | -0.085105173 |

| | | | | | | | | | | |
|---|---|---|---|---|---|---|---|---|---|---|
| Fgd3 | 11.10003 | 11.28655 | 11.21 | 0.076061718 | -0.110461842 | 11.316724 | 11.039246 | 11.2133 | -0.174067176 | 0.103410444 |
| Slc25a51 | 9.751058 | 9.873816 | 9.8459 | 0.027882075 | -0.094876181 | 9.8622953 | 9.6761676 | 9.85028 | -0.174117356 | 0.012010312 |
| Mindy4 | 3.920322 | 3.792731 | 3.8699 | -0.077199583 | 0.05039159 | 3.7164332 | 3.593149 | 3.76727 | -0.174119146 | -0.050834919 |
| Macrod1 | 6.501141 | 6.436021 | 6.5441 | -0.10807169 | -0.042950875 | 6.6267155 | 6.3137962 | 6.48795 | -0.174153342 | 0.138765931 |
| Trav6-4 | 3.460148 | 3.408273 | 3.4524 | -0.044110588 | 0.007764482 | 3.4602817 | 3.3336999 | 3.50789 | -0.174192448 | -0.047610646 |
| Ulbp1 | 6.650822 | 6.617105 | 6.8154 | -0.198300475 | -0.164583378 | 6.2648498 | 6.6964813 | 6.87088 | -0.174399938 | -0.606031458 |
| Ubxn11 | 8.044749 | 8.0487 | 8.1263 | -0.077599034 | -0.081550387 | 8.5579558 | 8.3279843 | 8.5024 | -0.17442007 | 0.055551459 |
| Slc6a8 | 4.818454 | 4.967041 | 4.8209 | 0.146166054 | -0.002421535 | 4.5532442 | 4.5769845 | 4.75144 | -0.174453721 | -0.198194027 |
| Reln | 5.941915 | 5.428829 | 5.5616 | -0.132771334 | 0.380315495 | 5.1777608 | 5.1081777 | 5.28267 | -0.174487873 | -0.104904783 |
| Ighv1-64 | 8.23365 | 8.186492 | 8.1815 | 0.004984585 | 0.052142475 | 8.4047867 | 7.8413653 | 8.01592 | -0.174550212 | 0.388871202 |
| Zdhhc24 | 6.819068 | 6.775642 | 6.9325 | -0.156824196 | -0.113398172 | 7.1490295 | 6.8065888 | 6.98128 | -0.174686697 | 0.167753915 |
| Fam214b | 7.993435 | 7.64852 | 8.0612 | -0.412690651 | -0.067775718 | 8.0415715 | 7.7407369 | 7.91549 | -0.174757935 | 0.12607657 |
| Vmn2r57 | 3.05261 | 2.999223 | 3.0787 | -0.079510289 | -0.026122925 | 3.1491301 | 2.8807393 | 3.05558 | -0.174840271 | 0.093550453 |
| Gm3383 | 3.115113 | 3.029876 | 3.2073 | -0.177434373 | -0.092197476 | 2.9488722 | 3.0646283 | 3.23948 | -0.174847767 | -0.290603897 |
| Gprasp1 | 9.790782 | 9.712106 | 9.7803 | -0.068220983 | 0.010454515 | 9.6520194 | 9.6638417 | 9.83873 | -0.174889722 | -0.186712049 |
| Cttnbp2 | 4.020125 | 3.899104 | 3.9744 | -0.075278969 | 0.045741456 | 3.6455549 | 3.8643363 | 4.03924 | -0.174899087 | -0.393680428 |
| Lypla1 | 10.35085 | 10.32961 | 10.396 | -0.066475898 | -0.045238952 | 10.384715 | 10.1757 | 10.3507 | -0.174997084 | 0.034017979 |
| Rab11fip1 | 9.044292 | 9.134542 | 9.0262 | 0.1083108 | 0.018061261 | 8.9988338 | 8.932276 | 9.10728 | -0.175000637 | -0.108442825 |
| Adcyap1r1 | 5.418901 | 5.199921 | 5.5055 | -0.30559524 | -0.086615564 | 5.0603694 | 4.7129971 | 4.88805 | -0.175056738 | 0.1723155 |
| Spata7 | 4.494953 | 4.350965 | 4.4453 | -0.094344679 | 0.049643424 | 4.3407376 | 4.3005208 | 4.47565 | -0.175129856 | -0.13491305 |
| Dock10 | 12.84293 | 12.92486 | 12.903 | 0.02187147 | -0.06005827 | 12.942257 | 12.931468 | 13.1067 | -0.175236266 | -0.164447487 |
| Tmc7 | 3.19487 | 3.071942 | 3.2293 | -0.157349519 | -0.034421637 | 3.3058153 | 3.0284099 | 3.20367 | -0.175255607 | 0.102149817 |
| Rad17 | 9.143024 | 9.18371 | 9.2443 | -0.060619052 | -0.101305344 | 9.1313083 | 9.046392 | 9.22171 | -0.175317213 | -0.090400857 |
| Ccdc28a | 6.912375 | 6.788065 | 6.943 | -0.154962851 | -0.030652995 | 7.0915064 | 6.9172199 | 7.09263 | -0.175411544 | -0.001125006 |
| Otud6b | 8.890239 | 8.818938 | 8.8861 | -0.067178216 | 0.004122459 | 8.8599997 | 8.660306 | 8.83572 | -0.175417487 | 0.024276194 |
| Tet2 | 9.704027 | 9.648851 | 9.6798 | -0.030910661 | 0.024265458 | 9.6746103 | 9.9451631 | 10.1206 | -0.175476787 | -0.446029551 |
| Cpeb4 | 9.843797 | 9.891354 | 9.7818 | 0.109544019 | 0.06198758 | 9.5999593 | 9.7136202 | 9.88913 | -0.175509796 | -0.289170635 |
| Scaf11 | 12.02349 | 12.02023 | 12.013 | 0.0077256 | 0.010989544 | 11.960224 | 11.860847 | 12.0364 | -0.17553786 | -0.076160596 |
| Srms | 4.51686 | 4.569799 | 4.4878 | 0.082034216 | 0.029095162 | 4.4816863 | 4.3786553 | 4.55422 | -0.175561658 | -0.072530726 |
| Ano8 | 7.809977 | 7.876802 | 7.894 | -0.017191963 | -0.084016906 | 8.0156862 | 7.7379252 | 7.91359 | -0.175665919 | 0.102095137 |
| Trbj2-2 | 4.293998 | 4.34752 | 4.2898 | 0.057733045 | 0.004211036 | 4.1706963 | 4.2265673 | 4.4023 | -0.175728512 | -0.231599496 |
| St18 | 1.722677 | 1.546247 | 1.633 | -0.086728993 | 0.089701386 | 1.5008498 | 1.4883364 | 1.66409 | -0.175751757 | -0.163238324 |

| | | | | | | | | | | |
|---|---|---|---|---|---|---|---|---|---|---|
| Zfp532 | 4.812738 | 4.727612 | 4.9774 | -0.249812478 | -0.164686706 | 4.5337387 | 4.3701944 | 4.54596 | -0.175761459 | -0.012217173 |
| Ipo9 | 10.41406 | 10.42344 | 10.428 | -0.004846782 | -0.014225264 | 10.4051 | 10.408293 | 10.5841 | -0.175772224 | -0.178966048 |
| Trappc12 | 10.03185 | 9.939847 | 10.013 | -0.072689496 | 0.019317906 | 10.092826 | 9.9352952 | 10.1113 | -0.175985509 | -0.018454768 |
| Chst12 | 8.300597 | 8.396577 | 8.5242 | -0.12766359 | -0.223643166 | 8.5595308 | 8.1880044 | 8.36405 | -0.176042411 | 0.195483959 |
| Rasl11b | 5.402073 | 5.140889 | 5.7602 | -0.619327959 | -0.358143991 | 4.9162159 | 4.9575004 | 5.13358 | -0.176082682 | -0.217367232 |
| Dusp27 | 3.619853 | 3.442685 | 3.8192 | -0.376554048 | -0.199386782 | 3.1827724 | 3.1168379 | 3.29293 | -0.176087748 | -0.110153243 |
| Gm42672 | 2.197729 | 2.1661 | 2.2591 | -0.092958949 | -0.061329282 | 2.2315958 | 2.1599599 | 2.33605 | -0.176094253 | -0.104458295 |
| Hnrnpl | 12.49587 | 12.45372 | 12.628 | -0.174773636 | -0.132626622 | 12.695312 | 12.543922 | 12.7201 | -0.176136366 | -0.024746531 |
| Auts2 | 4.460889 | 4.392541 | 4.9969 | -0.604328419 | -0.535979743 | 4.363751 | 4.5534546 | 4.72966 | -0.176208296 | -0.365911881 |
| Aox4 | 2.42389 | 2.235235 | 2.3137 | -0.078428054 | 0.110226773 | 2.4138039 | 2.2467564 | 2.42301 | -0.176257834 | -0.009210257 |
| Galnt11 | 9.166895 | 8.958024 | 9.1459 | -0.187907251 | 0.020962789 | 9.2085768 | 9.0477986 | 9.2241 | -0.176302674 | -0.015524481 |
| Gm14305 | 6.493717 | 6.43328 | 6.6716 | -0.238369181 | -0.177932216 | 6.5291357 | 6.1922026 | 6.36856 | -0.17636099 | 0.160572058 |
| Zfp692 | 7.662294 | 7.764945 | 7.8984 | -0.133435185 | -0.236086166 | 8.3654515 | 7.9214598 | 8.09785 | -0.176388516 | 0.267603187 |
| Adk | 8.281591 | 8.179975 | 8.2141 | -0.034174492 | 0.067441667 | 8.1127707 | 8.052622 | 8.22902 | -0.176398241 | -0.116249582 |
| Cdkl2 | 4.48438 | 4.295427 | 4.4348 | -0.139326537 | 0.049625895 | 4.3695089 | 4.3179087 | 4.49442 | -0.17651446 | -0.124914339 |
| Pabpc1 | 14.72362 | 14.72292 | 14.844 | -0.120909104 | -0.120210242 | 14.766819 | 14.602105 | 14.7787 | -0.176562416 | -0.011848007 |
| Pgrmc2 | 9.143563 | 8.942258 | 9.1172 | -0.174962549 | 0.026343284 | 8.8983522 | 8.7874703 | 8.96403 | -0.176563389 | -0.065681457 |
| Amd1 | 10.30282 | 10.26122 | 10.345 | -0.08391517 | -0.042311822 | 10.258975 | 10.227725 | 10.4044 | -0.176704682 | -0.145454719 |
| Gnb5 | 4.868077 | 4.542259 | 4.8191 | -0.276800493 | 0.049018073 | 4.5902205 | 4.5746102 | 4.75138 | -0.176771848 | -0.161161582 |
| Zfp420 | 7.027825 | 7.170899 | 7.0746 | 0.096327216 | -0.046746549 | 7.2142221 | 7.2094694 | 7.38639 | -0.176920356 | -0.172167686 |
| Frg2f1 | 5.265359 | 5.261332 | 5.3865 | -0.125191973 | -0.121165508 | 5.4352674 | 5.289611 | 5.46654 | -0.176924852 | -0.031268442 |
| Tmem175 | 8.131645 | 7.906452 | 8.0513 | -0.144829585 | 0.080363252 | 8.2755309 | 8.0443534 | 8.22132 | -0.176962288 | 0.054215233 |
| Rps6-ps4 | 9.511464 | 9.306222 | 9.6482 | -0.34198434 | -0.136742799 | 9.0116616 | 9.646758 | 9.82378 | -0.177026065 | -0.812122459 |
| Mtus2 | 4.474161 | 4.658086 | 4.4114 | 0.24667642 | 0.06275134 | 4.6535884 | 4.4753853 | 4.65243 | -0.177048939 | 0.001154204 |
| Olfr279 | 4.119901 | 4.153252 | 4.0987 | 0.05454389 | 0.021193672 | 4.1974318 | 4.1132795 | 4.29034 | -0.177058577 | -0.092906293 |
| Pdss2 | 7.074822 | 7.068552 | 7.0666 | 0.001949058 | 0.008219147 | 7.0172023 | 6.9353734 | 7.11244 | -0.177061765 | -0.095232937 |
| Gm48366 | 3.368083 | 3.320686 | 3.3973 | -0.076608207 | -0.029211187 | 3.3467156 | 3.3469168 | 3.52406 | -0.177148221 | -0.177349438 |
| Zfp236 | 10.50383 | 10.47512 | 10.44 | 0.034847436 | 0.063549162 | 10.472588 | 10.355947 | 10.5331 | -0.177150094 | -0.06050881 |
| Sybu | 2.128868 | 2.148467 | 2.1763 | -0.027822027 | -0.047421122 | 2.1487192 | 1.9990277 | 2.17622 | -0.177188667 | -0.027497154 |
| Mpg | 7.778938 | 7.759446 | 7.9135 | -0.154068123 | -0.134575672 | 8.1744149 | 7.8259177 | 8.00326 | -0.177340962 | 0.171156166 |
| Mapk1ip1 | 8.110679 | 8.109437 | 8.1613 | -0.051898046 | -0.050655511 | 8.5142112 | 8.2626267 | 8.44012 | -0.177493164 | 0.074091394 |
| Srbd1 | 8.761276 | 8.789947 | 8.8092 | -0.019279281 | -0.047950167 | 8.8767325 | 8.7134913 | 8.891 | -0.177506845 | -0.014265691 |

| | | | | | | | | | | |
|---|---|---|---|---|---|---|---|---|---|---|
| Neurl1b | 5.653286 | 5.614082 | 5.7652 | -0.151155328 | -0.111951017 | 5.6371886 | 5.7434554 | 5.92097 | -0.177509993 | -0.283776765 |
| Rgr | 2.053794 | 1.862087 | 2.0987 | -0.23658691 | -0.044879776 | 1.8118613 | 1.8153707 | 1.99289 | -0.177521411 | -0.181030884 |
| B4galnt4 | 4.455877 | 4.402575 | 4.4884 | -0.085834476 | -0.032532605 | 4.6958373 | 4.3135268 | 4.49108 | -0.177555594 | 0.20475487 |
| Grpel2 | 8.950108 | 8.955569 | 8.8998 | 0.05576913 | 0.050307779 | 9.0334844 | 8.9055257 | 9.0831 | -0.177576209 | -0.049617546 |
| Cdc14a | 8.635626 | 8.629565 | 8.6118 | 0.017789024 | 0.023849582 | 8.3414053 | 8.4363749 | 8.61407 | -0.177698748 | -0.272668345 |
| Ryr1 | 6.064868 | 5.884329 | 6.2126 | -0.328278935 | -0.147739285 | 6.1046678 | 5.8932365 | 6.07106 | -0.177822454 | 0.033608905 |
| Zfp930 | 7.578887 | 7.528697 | 7.5661 | -0.037394072 | 0.012795501 | 7.8414716 | 7.6886783 | 7.86652 | -0.177837473 | -0.025044112 |
| Hacd3 | 9.398848 | 9.513312 | 9.4185 | 0.094837934 | -0.019626022 | 9.4922936 | 9.4149262 | 9.59276 | -0.17783763 | -0.100470191 |
| Prkg1 | 3.406397 | 3.373067 | 3.3847 | -0.011670572 | 0.021658965 | 3.2647318 | 3.2671182 | 3.44507 | -0.177954349 | -0.180340788 |
| Trib2 | 10.33526 | 10.32168 | 10.459 | -0.137291156 | -0.123707492 | 10.149727 | 10.068205 | 10.2462 | -0.178003543 | -0.096481878 |
| Zfp369 | 8.519196 | 8.760876 | 8.6461 | 0.114793301 | -0.126886633 | 8.7533814 | 8.7396145 | 8.91763 | -0.178010689 | -0.164243801 |
| Cep192 | 10.54381 | 10.44018 | 10.57 | -0.129735334 | -0.026107758 | 10.444534 | 10.507995 | 10.686 | -0.178011106 | -0.241471505 |
| Tyr | 2.198476 | 2.086849 | 2.2116 | -0.124780221 | -0.013153624 | 2.0261522 | 2.0873939 | 2.26547 | -0.178074838 | -0.239316539 |
| Asic1 | 3.056382 | 2.7412 | 3.0385 | -0.297277714 | 0.01790391 | 2.6924281 | 2.606204 | 2.78431 | -0.178106972 | -0.091882896 |
| Ptgr1 | 4.916545 | 4.68484 | 4.78 | -0.095165696 | 0.136539162 | 4.8515615 | 4.5669542 | 4.74513 | -0.178171606 | 0.10643567 |
| Acvr2a | 7.415397 | 7.204109 | 7.2008 | 0.003287933 | 0.214576401 | 6.8720911 | 6.9059718 | 7.08415 | -0.178179938 | -0.212060671 |
| Cbfb | 11.39905 | 11.42183 | 11.417 | 0.0050073 | -0.01777531 | 11.275533 | 11.195725 | 11.3739 | -0.178200056 | -0.09839219 |
| Rgs10 | 9.835458 | 9.853576 | 10.096 | -0.242704871 | -0.260823041 | 9.8026108 | 9.7732539 | 9.95169 | -0.178433963 | -0.149077121 |
| Syk | 12.73097 | 12.9446 | 12.83 | 0.114820788 | -0.098807112 | 13.034566 | 12.660326 | 12.8389 | -0.17861383 | 0.195626198 |
| Mutyh | 7.142546 | 7.143589 | 7.2099 | -0.066346199 | -0.067389767 | 7.6064681 | 7.2010402 | 7.37967 | -0.178631414 | 0.226796485 |
| Tceal1 | 3.477586 | 3.246686 | 3.441 | -0.194298808 | 0.036601927 | 3.2333318 | 3.1555903 | 3.33439 | -0.178802633 | -0.101061135 |
| Gm47147 | 4.463953 | 4.391714 | 4.3964 | -0.004649871 | 0.067588542 | 4.2401956 | 4.4529723 | 4.63185 | -0.178877071 | -0.391653801 |
| Clec12a | 8.045429 | 8.03165 | 8.066 | -0.03434572 | -0.020567023 | 8.217296 | 7.7871199 | 7.96607 | -0.178946704 | 0.251229334 |
| Htt | 11.08332 | 11.14087 | 11.072 | 0.06857476 | 0.011018504 | 11.242261 | 11.22209 | 11.4011 | -0.17901095 | -0.158839417 |
| Spty2d1 | 10.41953 | 10.34371 | 10.348 | -0.004396182 | 0.071424818 | 10.132788 | 10.200334 | 10.3794 | -0.17902096 | -0.246566032 |
| Mageh1 | 4.934956 | 4.648026 | 4.912 | -0.263980774 | 0.022949274 | 4.6374578 | 4.4193779 | 4.59842 | -0.179037492 | 0.039042404 |
| Pon3 | 5.963273 | 5.836045 | 5.9097 | -0.073609742 | 0.053618112 | 5.8477104 | 5.7762938 | 5.95537 | -0.179074497 | -0.107657973 |
| Ccl28 | 3.586992 | 3.192872 | 3.2454 | -0.052522796 | 0.341597474 | 3.5144306 | 3.2506784 | 3.42976 | -0.179086052 | 0.084666181 |
| Map4k3 | 7.928615 | 7.71383 | 7.8811 | -0.167265529 | 0.047519576 | 7.5566274 | 7.6505298 | 7.82967 | -0.179140822 | -0.273043239 |
| Zfp869 | 9.421232 | 9.352159 | 9.4382 | -0.086089534 | -0.017016522 | 9.3564884 | 9.1925902 | 9.37176 | -0.179173799 | -0.015275555 |
| 4931428F04Rik | 6.731584 | 6.568962 | 6.8381 | -0.269165164 | -0.106543325 | 6.8179784 | 6.8064466 | 6.98571 | -0.179263799 | -0.167732001 |
| Zxdb | 9.136755 | 9.20059 | 9.2044 | -0.003805574 | -0.067641079 | 8.9688321 | 8.9251639 | 9.10444 | -0.179273747 | -0.135605567 |

| | | | | | | | | | | |
|---|---|---|---|---|---|---|---|---|---|---|
| Trim35 | 10.44843 | 10.54168 | 10.493 | 0.048426634 | -0.044825222 | 10.657521 | 10.231029 | 10.4104 | -0.179324272 | 0.247168124 |
| Cog5 | 10.28908 | 10.25929 | 10.32 | -0.060671678 | -0.03088609 | 10.461123 | 10.321154 | 10.5005 | -0.179331394 | -0.039362837 |
| Myo15 | 3.079169 | 2.925881 | 3.115 | -0.189159543 | -0.035871802 | 3.1115453 | 3.027502 | 3.20691 | -0.179408529 | -0.095365287 |
| Cage1 | 4.726711 | 4.688315 | 4.8179 | -0.129569505 | -0.091174327 | 4.7714893 | 4.4619663 | 4.64148 | -0.179515274 | 0.130007752 |
| Lrr1 | 3.355105 | 3.517855 | 3.3696 | 0.148212757 | -0.014537692 | 3.5401581 | 3.4336514 | 3.61334 | -0.179685061 | -0.073178367 |
| Irs1 | 3.361218 | 3.426812 | 3.3766 | 0.050199141 | -0.015395733 | 3.3106534 | 3.1811086 | 3.361 | -0.179894344 | -0.050349513 |
| Ces1d | 2.322043 | 1.994929 | 1.9199 | 0.075023015 | 0.402136685 | 1.9583314 | 1.8943716 | 2.07434 | -0.179963728 | -0.116003981 |
| Zfp651 | 4.34045 | 4.223917 | 4.1902 | 0.033716731 | 0.150249652 | 4.0729689 | 3.9030095 | 4.08299 | -0.179983762 | -0.010024295 |
| Cog7 | 8.174124 | 7.963305 | 8.128 | -0.164712802 | 0.04610628 | 8.1596155 | 8.0028281 | 8.18285 | -0.18001846 | -0.023231017 |
| Prg4 | 4.943348 | 4.935445 | 4.7761 | 0.159360513 | 0.167263346 | 5.0268327 | 4.7572903 | 4.93737 | -0.180083111 | 0.089459341 |
| Adgra2 | 6.378891 | 6.202594 | 6.4207 | -0.218076207 | -0.041779198 | 6.2907523 | 5.8733922 | 6.05348 | -0.180092513 | 0.237267601 |
| Trp73 | 1.706088 | 1.691712 | 1.7723 | -0.080635371 | -0.066258951 | 1.699204 | 1.6130667 | 1.79318 | -0.180108896 | -0.093971546 |
| Zfp180 | 9.523562 | 9.517078 | 9.5776 | -0.060500101 | -0.054015298 | 9.431981 | 9.3448099 | 9.52494 | -0.18012555 | -0.092954481 |
| Slc36a2 | 3.175303 | 3.093658 | 3.0949 | -0.001209189 | 0.080435642 | 3.1101484 | 2.8145242 | 2.99465 | -0.180130358 | 0.115493817 |
| Gm14421 | 4.007346 | 3.982967 | 4.0608 | -0.077851604 | -0.053473242 | 4.1457034 | 4.0152114 | 4.19536 | -0.180146654 | -0.04965468 |
| Atp6v0a2 | 10.8278 | 10.67236 | 10.799 | -0.127138676 | 0.028308038 | 10.605631 | 10.575655 | 10.7558 | -0.180177976 | -0.150201796 |
| Rgs9bp | 3.082334 | 3.050977 | 3.0756 | -0.024593756 | 0.006763913 | 3.0511732 | 3.1011119 | 3.28135 | -0.180242462 | -0.23018114 |
| Scamp5 | 4.490861 | 4.105629 | 4.4661 | -0.360506307 | 0.024725831 | 4.1382009 | 4.0410296 | 4.22135 | -0.180324839 | -0.083153621 |
| Cyth4 | 11.50571 | 11.4509 | 11.477 | -0.02636763 | 0.028444802 | 11.623413 | 11.356002 | 11.5364 | -0.180395522 | 0.087016153 |
| Zdhhc15 | 7.876282 | 8.005737 | 8.0178 | -0.012021381 | -0.141477181 | 7.6002226 | 7.7193732 | 7.89978 | -0.180403498 | -0.299554132 |
| Pxmp2 | 4.486229 | 4.298013 | 4.627 | -0.328989909 | -0.140773343 | 4.2011915 | 4.2892466 | 4.46975 | -0.180501441 | -0.26855647 |
| 4632433K11Rik | 2.440102 | 2.356245 | 2.4835 | -0.127232271 | -0.043374749 | 2.2927692 | 2.3150827 | 2.49559 | -0.180503406 | -0.202816888 |
| Fbxo5 | 8.030381 | 7.639766 | 7.8797 | -0.239946234 | 0.150668526 | 7.9439115 | 7.909367 | 8.09002 | -0.180657082 | -0.14611266 |
| Pgm5 | 4.295855 | 4.32364 | 4.4959 | -0.17229026 | -0.200075569 | 4.1609776 | 4.1200129 | 4.30071 | -0.180697417 | -0.139732771 |
| Lhfp | 5.937643 | 5.959014 | 5.814 | 0.144994153 | 0.123622866 | 5.501997 | 5.3786705 | 5.55957 | -0.180901694 | -0.057575118 |
| Gm3362 | 4.469892 | 4.714539 | 4.4303 | 0.284234456 | 0.039587697 | 4.618515 | 4.4681651 | 4.64911 | -0.180940327 | -0.030590404 |
| Tigit | 6.994226 | 6.832272 | 7.0701 | -0.237826988 | -0.075873822 | 6.8257994 | 7.0762641 | 7.25725 | -0.18098924 | -0.431453934 |
| Gbf1 | 10.91169 | 10.78462 | 10.795 | -0.010486114 | 0.116585498 | 10.921221 | 10.838051 | 11.0191 | -0.181005288 | -0.097835349 |
| Hic2 | 6.096588 | 5.993246 | 6.0133 | -0.020086633 | 0.083255942 | 6.1020861 | 5.9020471 | 6.08312 | -0.181070848 | 0.01896812 |
| Slc38a9 | 9.138988 | 9.074737 | 9.2262 | -0.151468133 | -0.087217376 | 9.2218703 | 9.2407784 | 9.42194 | -0.181166018 | -0.200074052 |
| Pggt1b | 9.990997 | 9.832088 | 9.9117 | -0.079567888 | 0.079341015 | 9.9394908 | 9.8208048 | 10.002 | -0.181173899 | -0.062487893 |
| Nuggc | 4.384175 | 4.824094 | 4.4544 | 0.369694108 | -0.070225009 | 4.5913078 | 4.5619755 | 4.74326 | -0.181282462 | -0.151950159 |

| | | | | | | | | | | |
|---|---|---|---|---|---|---|---|---|---|---|
| Skap1 | 10.96389 | 10.90922 | 10.97 | -0.061169382 | -0.00650633 | 10.774069 | 10.829942 | 11.0113 | -0.18134204 | -0.237215561 |
| Tal1 | 5.517734 | 5.148885 | 5.5205 | -0.371609121 | -0.002760487 | 4.676746 | 4.7609566 | 4.9423 | -0.18134243 | -0.265553044 |
| Tedc2 | 6.607953 | 6.53768 | 6.5809 | -0.043219063 | 0.027054437 | 6.8987495 | 6.5853949 | 6.76697 | -0.181578203 | 0.131776348 |
| Plxnc1 | 10.77958 | 10.68972 | 10.762 | -0.072147376 | 0.017714532 | 10.9307 | 10.752745 | 10.9343 | -0.181589238 | -0.003634779 |
| Pabpn1 | 10.40902 | 10.32215 | 10.501 | -0.179024694 | -0.09214848 | 10.741008 | 10.475313 | 10.657 | -0.181639312 | 0.084055387 |
| Pcdhgb1 | 3.517509 | 3.386643 | 3.4963 | -0.109661947 | 0.02120428 | 3.3420255 | 3.201005 | 3.38271 | -0.181707219 | -0.040686684 |
| Fam76a | 8.989602 | 8.78399 | 9.0586 | -0.274588668 | -0.068976131 | 8.9145688 | 8.6259887 | 8.80785 | -0.181859461 | 0.106720649 |
| Apcdd1 | 6.08505 | 6.225455 | 6.085 | 0.140425707 | 2.09296E-05 | 5.8628224 | 5.3330927 | 5.51495 | -0.181861387 | 0.347868322 |
| Hdac5 | 9.770042 | 9.674492 | 9.8477 | -0.173244565 | -0.077694991 | 10.100948 | 9.6806702 | 9.86259 | -0.18192212 | 0.23835521 |
| Zfp407 | 9.33748 | 9.440545 | 9.3338 | 0.106757849 | 0.003692572 | 9.4367946 | 9.2607033 | 9.44267 | -0.181963084 | -0.005871844 |
| Mbnl1 | 13.78679 | 13.81914 | 13.8 | 0.018830548 | -0.013522316 | 13.739539 | 13.672817 | 13.8548 | -0.182016164 | -0.115294226 |
| Gm5421 | 5.897523 | 5.436144 | 5.6841 | -0.247953119 | 0.213426733 | 6.0384894 | 5.7000431 | 5.88211 | -0.182064072 | 0.156382219 |
| Kif21a | 4.865111 | 4.794149 | 4.6426 | 0.151571135 | 0.222532696 | 4.462516 | 4.4516919 | 4.63378 | -0.182085865 | -0.171261759 |
| Gm17132 | 6.35567 | 6.231871 | 6.4443 | -0.212411851 | -0.088613479 | 6.3245986 | 6.6702476 | 6.85236 | -0.182117257 | -0.527766223 |
| Tipin | 8.363442 | 8.107707 | 8.3299 | -0.222166666 | 0.033569093 | 8.3493778 | 8.0797975 | 8.26193 | -0.18212783 | 0.087452463 |
| Iglc2 | 11.64862 | 11.60209 | 11.799 | -0.196993676 | -0.150465398 | 12.101888 | 11.551488 | 11.7337 | -0.182199854 | 0.368199842 |
| Zfp142 | 9.530521 | 9.671563 | 9.4221 | 0.249474706 | 0.108432791 | 9.4693667 | 9.4333412 | 9.61556 | -0.182222761 | -0.146197223 |
| Iqcg | 3.233453 | 3.405226 | 3.3583 | 0.046890803 | -0.124882258 | 3.1560116 | 3.0275346 | 3.20997 | -0.182432551 | -0.053955553 |
| Trpm2 | 7.271294 | 7.101732 | 7.0945 | 0.007228281 | 0.17678944 | 6.6667422 | 6.4572734 | 6.63984 | -0.18257047 | 0.026898327 |
| AI182371 | 4.542287 | 4.311058 | 4.418 | -0.106984035 | 0.124244863 | 4.3444516 | 4.3704664 | 4.55308 | -0.182617393 | -0.208632194 |
| Slc5a3 | 7.836486 | 8.010297 | 7.906 | 0.104321752 | -0.069488868 | 7.5852598 | 8.1560339 | 8.33876 | -0.182724784 | -0.753498812 |
| Dmwd | 6.917791 | 6.619666 | 6.6396 | -0.019977963 | 0.278147138 | 6.991176 | 6.5143365 | 6.69712 | -0.182787725 | 0.294051741 |
| Sh3bgrl3 | 12.47025 | 12.42202 | 12.501 | -0.079167708 | -0.030936226 | 12.820619 | 12.247365 | 12.4302 | -0.182816666 | 0.390437425 |
| Neurl4 | 10.21091 | 10.37685 | 10.457 | -0.080371886 | -0.246307059 | 10.538258 | 10.39085 | 10.5738 | -0.182971622 | -0.035563378 |
| Mettl14 | 8.909186 | 9.0034 | 8.9533 | 0.050104733 | -0.044108941 | 8.8782749 | 8.7078861 | 8.89094 | -0.183054059 | -0.012665219 |
| Fabp1 | 1.99296 | 1.946309 | 1.8906 | 0.055682402 | 0.102333442 | 1.8431988 | 1.7989921 | 1.98208 | -0.18308995 | -0.138883251 |
| Gata6 | 4.084248 | 4.063174 | 4.1318 | -0.06860046 | -0.047526834 | 3.784444 | 3.8718668 | 4.05497 | -0.183100295 | -0.270523051 |
| Nemf | 10.5797 | 10.49646 | 10.476 | 0.020249174 | 0.103492983 | 10.530494 | 10.493314 | 10.6764 | -0.183111176 | -0.145930863 |
| Rbp7 | 2.298674 | 2.312511 | 2.2724 | 0.040098879 | 0.026261546 | 2.343759 | 2.1398949 | 2.32302 | -0.183124745 | 0.020739384 |
| Lpar3 | 3.916091 | 4.12838 | 4.0774 | 0.050980632 | -0.161308608 | 3.8983641 | 3.7929105 | 3.97614 | -0.183232811 | -0.077779256 |
| Slc17a9 | 8.842176 | 8.959272 | 9.0114 | -0.052162521 | -0.169258048 | 9.2055678 | 8.8602487 | 9.04359 | -0.18333685 | 0.161982215 |
| Gm16385 | 4.263468 | 4.246427 | 4.2867 | -0.040231768 | -0.023190762 | 4.348384 | 4.3093289 | 4.49276 | -0.183435156 | -0.144380105 |

| | | | | | | | | | | |
|---|---|---|---|---|---|---|---|---|---|---|
| Prnp | 8.906484 | 8.755525 | 8.9353 | -0.179787521 | -0.028828011 | 8.3293508 | 8.4949164 | 8.67837 | -0.183458046 | -0.349023627 |
| Trav6-1 | 3.351989 | 3.27261 | 3.2733 | -0.000721704 | 0.078657344 | 3.1876763 | 3.195832 | 3.37942 | -0.183585419 | -0.191741108 |
| Cep44 | 7.636063 | 7.606064 | 7.7129 | -0.106860689 | -0.076862411 | 7.8211002 | 7.6820794 | 7.86568 | -0.183597482 | -0.044576774 |
| Eif4b | 13.22592 | 13.15895 | 13.303 | -0.143827966 | -0.076863402 | 13.214159 | 13.067356 | 13.251 | -0.18366924 | -0.03686645 |
| Dnaaf2 | 6.479851 | 6.25143 | 6.4046 | -0.153155814 | 0.075265716 | 6.5193917 | 6.4034083 | 6.58708 | -0.183674024 | -0.067690644 |
| Cftr | 1.729232 | 1.698156 | 1.7178 | -0.019599254 | 0.011476926 | 1.8522721 | 1.755606 | 1.93929 | -0.183686707 | -0.087020581 |
| Tubg1 | 8.298828 | 8.298977 | 8.4 | -0.101011875 | -0.101160545 | 8.5847053 | 8.3495824 | 8.53335 | -0.183762985 | 0.051359904 |
| Zmym6 | 8.803021 | 8.998823 | 9.0468 | -0.047950675 | -0.24375333 | 9.0893602 | 8.9843799 | 9.16815 | -0.183773588 | -0.078793232 |
| Laptm4b | 7.975318 | 7.930313 | 8.0882 | -0.157877515 | -0.112872215 | 7.3706537 | 7.5284104 | 7.71243 | -0.184016096 | -0.34177279 |
| Tmem25 | 2.451537 | 2.368264 | 2.4994 | -0.131087865 | -0.047814452 | 2.5089901 | 2.3825126 | 2.56659 | -0.184073571 | -0.05759608 |
| Tpmt | 4.759423 | 4.692666 | 4.8165 | -0.123807452 | -0.057050513 | 4.9358375 | 4.7896243 | 4.97378 | -0.184151167 | -0.037937933 |
| Clk1 | 11.98426 | 12.03707 | 12.097 | -0.060395948 | -0.113208504 | 12.242262 | 12.314509 | 12.4987 | -0.184161266 | -0.256408835 |
| Pabpc1l | 3.632543 | 3.623518 | 3.6617 | -0.03820636 | -0.029181195 | 3.7419579 | 3.4380433 | 3.62221 | -0.184165611 | 0.119749028 |
| Fam78a | 10.94742 | 11.17768 | 11.12 | 0.057561118 | -0.17269618 | 10.781083 | 10.863248 | 11.0474 | -0.18419237 | -0.266357893 |
| Phlda3 | 4.558791 | 4.624979 | 4.6566 | -0.031651111 | -0.097839631 | 4.4451415 | 4.3682111 | 4.55243 | -0.18421763 | -0.107287286 |
| Arhgap12 | 9.106226 | 9.031513 | 9.1216 | -0.090047363 | -0.015334183 | 9.0327057 | 8.9780029 | 9.16224 | -0.184237215 | -0.129534355 |
| Ighj4 | 6.776838 | 7.371603 | 7.2099 | 0.161745785 | -0.433019235 | 7.4689946 | 7.1490503 | 7.33331 | -0.184263067 | 0.13568121 |
| Sptbn2 | 3.692269 | 3.704751 | 4.0367 | -0.331949683 | -0.344431126 | 3.8065024 | 3.5117358 | 3.69602 | -0.184282819 | 0.110483764 |
| Stk4 | 12.7823 | 12.84433 | 12.823 | 0.021428924 | -0.04059904 | 12.586503 | 12.596635 | 12.781 | -0.184324428 | -0.194456575 |
| Sh3d21 | 4.259955 | 4.227267 | 4.3751 | -0.147841514 | -0.115153391 | 4.2543027 | 4.194424 | 4.37876 | -0.18433231 | -0.124453676 |
| Atraid | 8.836503 | 8.761451 | 8.9592 | -0.19776424 | -0.122711953 | 8.9309876 | 8.6151843 | 8.79979 | -0.184605364 | 0.131197895 |
| Plppr2 | 3.903377 | 3.782536 | 3.9483 | -0.165798687 | -0.044958068 | 3.8348094 | 3.6011814 | 3.78585 | -0.184669925 | 0.048958064 |
| Trav11 | 3.29629 | 3.193584 | 3.3299 | -0.136297465 | -0.033591511 | 3.1702839 | 3.0101362 | 3.19494 | -0.184805653 | -0.024658017 |
| Grk1 | 5.694675 | 5.641879 | 5.744 | -0.10212161 | -0.049325425 | 5.7563088 | 5.7137987 | 5.89866 | -0.184857496 | -0.142347363 |
| Kazn | 4.381581 | 4.251074 | 4.0876 | 0.163435255 | 0.29394243 | 3.9150733 | 3.9054282 | 4.09055 | -0.185123548 | -0.175478445 |
| Hyls1 | 7.177875 | 7.232897 | 7.2418 | -0.008942424 | -0.063963903 | 7.2469591 | 7.1865722 | 7.37174 | -0.185168425 | -0.124781514 |
| Atp6v1g2 | 4.907462 | 4.761094 | 4.917 | -0.155931257 | -0.009563511 | 4.8625613 | 4.7430192 | 4.92824 | -0.185215974 | -0.065673971 |
| Tet3 | 10.80197 | 10.87563 | 10.876 | -0.000648152 | -0.07430992 | 10.722618 | 10.810515 | 10.9959 | -0.185350044 | -0.273247352 |
| Cblc | 2.104414 | 2.050525 | 1.9891 | 0.061381436 | 0.115270301 | 1.9215755 | 1.9588616 | 2.14423 | -0.185370926 | -0.22265706 |
| Nrg4 | 2.930639 | 2.851952 | 2.9133 | -0.061329637 | 0.017356723 | 2.9877445 | 2.8455657 | 3.03098 | -0.185412803 | -0.043234 |
| Gm9493 | 4.259571 | 4.149114 | 4.3147 | -0.165608924 | -0.055152044 | 4.3258123 | 4.1461766 | 4.33159 | -0.185415381 | -0.005779656 |
| Vamp2 | 9.911671 | 9.804446 | 10.053 | -0.248814582 | -0.141589958 | 9.7588899 | 9.6619537 | 9.84738 | -0.185429168 | -0.088492979 |

| | | | | | | | | | | |
|---|---|---|---|---|---|---|---|---|---|---|
| Ntan1 | 9.201118 | 9.084727 | 9.2678 | -0.183033212 | -0.066642437 | 9.4875524 | 9.156783 | 9.34233 | -0.185545654 | 0.145223767 |
| Mprip | 11.16157 | 11.15123 | 11.201 | -0.049893968 | -0.03955871 | 11.09479 | 10.994068 | 11.1796 | -0.185573478 | -0.084850895 |
| Traj6 | 3.080664 | 3.050481 | 3.0887 | -0.038192699 | -0.008009531 | 3.0602443 | 2.9197594 | 3.10534 | -0.185577544 | -0.045092589 |
| Rp2 | 9.814218 | 9.678045 | 9.718 | -0.039968324 | 0.096205284 | 9.8314035 | 9.478876 | 9.66448 | -0.185600493 | 0.166927036 |
| Ccdc15 | 6.798278 | 6.686928 | 6.7953 | -0.108420223 | 0.002928889 | 6.9467604 | 6.7224601 | 6.90807 | -0.185606611 | 0.038693734 |
| Trav7d-2 | 5.57257 | 5.499896 | 5.8403 | -0.340431306 | -0.267757173 | 4.7249851 | 5.3645977 | 5.55022 | -0.185619111 | -0.825231779 |
| Lonp2 | 10.38062 | 10.37082 | 10.439 | -0.06793588 | -0.05813499 | 10.322081 | 10.225534 | 10.4113 | -0.185732084 | -0.089184467 |
| Prkab1 | 9.557487 | 9.456798 | 9.5929 | -0.136065575 | -0.035376324 | 9.6519496 | 9.4837987 | 9.66953 | -0.185735128 | -0.017584272 |
| Fbxo24 | 1.944584 | 1.920226 | 1.9624 | -0.042215463 | -0.01785744 | 1.9218875 | 1.8439698 | 2.02993 | -0.185962877 | -0.108045119 |
| Mysm1 | 10.01339 | 9.98152 | 9.9366 | 0.044951465 | 0.076816348 | 9.965294 | 9.9482115 | 10.1342 | -0.186007595 | -0.168925138 |
| Mtmr9 | 8.980102 | 8.841824 | 8.9234 | -0.081570181 | 0.056707909 | 8.9859527 | 8.8027487 | 8.98877 | -0.186024483 | -0.002820458 |
| Igf2bp3 | 5.09623 | 4.958783 | 4.9892 | -0.030409747 | 0.107037443 | 5.1077429 | 4.826212 | 5.01231 | -0.186102265 | 0.095428646 |
| Shisa9 | 1.253066 | 1.188962 | 1.264 | -0.075018483 | -0.010914154 | 1.1922447 | 1.1497474 | 1.33618 | -0.186428044 | -0.143930698 |
| Strip2 | 7.958817 | 8.093509 | 8.1909 | -0.097367237 | -0.232059617 | 7.6989121 | 8.1406637 | 8.32712 | -0.186451929 | -0.628203539 |
| Zc3h12c | 8.044775 | 8.190456 | 8.1281 | 0.062371847 | -0.083309199 | 7.543035 | 7.9945183 | 8.18106 | -0.186542113 | -0.638025486 |
| Gm15801 | 2.425057 | 2.524263 | 2.4247 | 0.099594714 | 0.000389037 | 2.3161014 | 2.2922186 | 2.47882 | -0.186597071 | -0.162714279 |
| Gm8783 | 2.425057 | 2.524263 | 2.4247 | 0.099594714 | 0.000389037 | 2.3161014 | 2.2922186 | 2.47882 | -0.186597071 | -0.162714279 |
| Loxl3 | 5.916151 | 6.050423 | 5.9656 | 0.084816166 | -0.049455485 | 5.8984199 | 6.0944773 | 6.28111 | -0.186637417 | -0.382694864 |
| Aplp1 | 5.359145 | 5.266654 | 5.2912 | -0.024595822 | 0.067895676 | 4.9664493 | 4.9602994 | 5.14695 | -0.186650776 | -0.180500843 |
| Qdpr | 9.295599 | 9.253882 | 9.3783 | -0.124461468 | -0.082744741 | 9.4654502 | 9.0790515 | 9.26576 | -0.186704619 | 0.199694132 |
| Rnf141 | 8.857789 | 8.643201 | 8.705 | -0.061816339 | 0.152771342 | 8.7472426 | 8.6852674 | 8.87201 | -0.186744146 | -0.124768906 |
| Fbxl22 | 3.932849 | 4.048684 | 3.8838 | 0.164875961 | 0.049040877 | 3.9839912 | 3.9230054 | 4.10983 | -0.186828332 | -0.125842614 |
| Mcts2 | 5.257363 | 5.282776 | 5.3369 | -0.054125018 | -0.079537427 | 5.1672934 | 5.1943819 | 5.38137 | -0.18698416 | -0.214072686 |
| Fbxo9 | 8.798075 | 8.855753 | 8.8274 | 0.028349958 | -0.029328857 | 9.0487024 | 8.8214562 | 9.00871 | -0.187256261 | 0.039989953 |
| Layn | 3.721792 | 3.447158 | 3.6373 | -0.190168593 | 0.084466298 | 3.4596657 | 3.3406552 | 3.52794 | -0.187286393 | -0.068275828 |
| Prrx1 | 6.342275 | 6.361899 | 6.0507 | 0.311246 | 0.291621916 | 6.1003101 | 5.7315842 | 5.91888 | -0.187294483 | 0.181431434 |
| Tcrg-V4 | 4.760421 | 4.535225 | 4.7257 | -0.190452307 | 0.034742868 | 4.5786319 | 4.5677294 | 4.75503 | -0.187297527 | -0.176395059 |
| Cd72 | 11.58377 | 11.63582 | 11.582 | 0.053997572 | 0.0019441 | 12.027864 | 11.577464 | 11.7648 | -0.187303786 | 0.263096957 |
| Cela1 | 2.357845 | 2.303322 | 2.4166 | -0.113300846 | -0.058778146 | 2.3734977 | 2.3125789 | 2.49993 | -0.187352619 | -0.126433768 |
| Hunk | 4.064115 | 3.831665 | 4.0165 | -0.184825361 | 0.047624594 | 3.9517439 | 3.8775576 | 4.065 | -0.187440249 | -0.113254013 |
| Cib2 | 4.729344 | 4.574073 | 4.684 | -0.109884979 | 0.045385904 | 4.6666982 | 4.4474306 | 4.63489 | -0.187454797 | 0.031812807 |
| Matr3 | 12.2944 | 12.23102 | 12.241 | -0.010203984 | 0.053178644 | 12.215381 | 12.114864 | 12.3024 | -0.18748694 | -0.086969981 |

| | | | | | | | | | | |
|---|---|---|---|---|---|---|---|---|---|---|
| Srf | 9.696808 | 9.628554 | 9.6686 | -0.040082754 | 0.028171087 | 10.051029 | 9.4854475 | 9.67305 | -0.187605759 | 0.377975971 |
| Thnsl1 | 6.262651 | 6.219735 | 6.1925 | 0.02722647 | 0.07014272 | 6.2785994 | 6.0379949 | 6.22576 | -0.187761626 | 0.052842872 |
| Gss | 7.147239 | 7.173707 | 7.1689 | 0.004757379 | -0.021711135 | 7.1497007 | 6.8947393 | 7.08261 | -0.187866391 | 0.067094995 |
| Zfp558 | 3.872918 | 3.879091 | 3.7998 | 0.07929245 | 0.073120225 | 3.7934466 | 3.6579202 | 3.84581 | -0.187885256 | -0.052358917 |
| Cep152 | 8.63255 | 8.677549 | 8.6741 | 0.003482281 | -0.041517337 | 8.6037442 | 8.6411083 | 8.82903 | -0.187925504 | -0.225289621 |
| Mrtfa | 10.86433 | 10.98941 | 10.996 | -0.006360604 | -0.13143839 | 10.945986 | 10.829779 | 11.0177 | -0.187934824 | -0.071727993 |
| Zfp287 | 7.783659 | 7.724919 | 7.9484 | -0.223467683 | -0.164728167 | 7.8117785 | 7.8666386 | 8.05469 | -0.188053916 | -0.242914002 |
| Tceal9 | 8.437671 | 8.264904 | 8.4815 | -0.216632197 | -0.043865366 | 8.2809326 | 8.0889474 | 8.27703 | -0.188079454 | 0.003905703 |
| Rsf1 | 10.60169 | 10.6753 | 10.733 | -0.057307612 | -0.130919784 | 10.505452 | 10.590044 | 10.7782 | -0.188148088 | -0.272740004 |
| Rcbtb1 | 10.00084 | 10.12909 | 10.038 | 0.090908733 | -0.037341062 | 9.9228773 | 9.8705007 | 10.0587 | -0.188182647 | -0.135806075 |
| Jrk | 6.606806 | 6.411293 | 6.5783 | -0.166956819 | 0.028555577 | 6.8014912 | 6.5834795 | 6.7717 | -0.188218306 | 0.029793412 |
| Bcl2 | 11.35941 | 11.15391 | 11.38 | -0.226415806 | -0.02091958 | 11.214856 | 11.157568 | 11.3458 | -0.188230628 | -0.130942796 |
| AI467606 | 10.50144 | 10.52844 | 10.627 | -0.098301488 | -0.125300816 | 10.666241 | 10.339039 | 10.5273 | -0.18825311 | 0.138948879 |
| Tspyl2 | 8.967847 | 8.879285 | 9.0548 | -0.175522703 | -0.086960712 | 9.0708639 | 8.9842029 | 9.1725 | -0.188296654 | -0.101635677 |
| Rock2 | 10.76745 | 10.75088 | 10.726 | 0.024617336 | 0.041190662 | 10.37763 | 10.525177 | 10.7135 | -0.188322416 | -0.335869238 |
| Rfwd3 | 10.37382 | 10.39693 | 10.465 | -0.068043614 | -0.091161078 | 10.330543 | 10.30384 | 10.4922 | -0.188361682 | -0.161658303 |
| Lrrc1 | 7.329179 | 7.520743 | 7.5267 | -0.005951015 | -0.197514504 | 7.2251648 | 7.2299913 | 7.41846 | -0.188464182 | -0.193290689 |
| Brpf3 | 9.621983 | 9.624899 | 9.6946 | -0.069713585 | -0.072629502 | 9.3410617 | 9.4266684 | 9.61515 | -0.188480699 | -0.274087414 |
| Cab39 | 11.66171 | 11.64993 | 11.656 | -0.006066496 | 0.005710112 | 11.472296 | 11.459149 | 11.6476 | -0.18849665 | -0.175349891 |
| Baz2a | 11.91299 | 11.95867 | 12.022 | -0.063641164 | -0.1093228 | 11.917753 | 11.899633 | 12.0881 | -0.188511524 | -0.170391612 |
| Pde1a | 3.28088 | 3.339927 | 3.3734 | -0.033458769 | -0.092505469 | 3.0621618 | 3.1288003 | 3.31733 | -0.188525518 | -0.255164001 |
| Zfp953 | 7.819593 | 7.946445 | 7.9931 | -0.04667645 | -0.173528363 | 7.8415193 | 7.7599207 | 7.94877 | -0.188851759 | -0.107253186 |
| Wdtc1 | 9.461187 | 9.48384 | 9.4642 | 0.019615249 | -0.003037972 | 9.315958 | 9.2004229 | 9.38934 | -0.188917225 | -0.073382187 |
| Gm26965 | 0.999121 | 0.970959 | 0.9341 | 0.036819304 | 0.064981092 | 1.0264601 | 0.9493628 | 1.13857 | -0.189203356 | -0.112106054 |
| Clcn3 | 10.22095 | 10.26673 | 10.289 | -0.022077704 | -0.067847836 | 10.258701 | 10.170723 | 10.3601 | -0.189365488 | -0.101387562 |
| Abca8b | 3.567071 | 3.439162 | 3.5771 | -0.137918344 | -0.010008919 | 3.4451669 | 3.3834617 | 3.57284 | -0.189374599 | -0.127669413 |
| Ksr2 | 6.270658 | 6.394721 | 6.3593 | 0.035464755 | -0.088597546 | 6.0415762 | 6.2312009 | 6.42062 | -0.189421273 | -0.379045992 |
| Olfr149 | 2.481781 | 2.40783 | 2.4219 | -0.014068289 | 0.059882408 | 2.5289217 | 2.3553485 | 2.54478 | -0.189426627 | -0.015853416 |
| Lcor | 8.659344 | 8.612993 | 8.6314 | -0.018413235 | 0.027937926 | 8.5097711 | 8.6863093 | 8.87588 | -0.189571774 | -0.366109908 |
| Zfp971 | 6.785753 | 6.659844 | 6.6186 | 0.041247384 | 0.167156722 | 6.7243958 | 6.5753429 | 6.76493 | -0.189589853 | -0.04053693 |
| Larp1b | 8.297932 | 8.235236 | 8.1311 | 0.104143617 | 0.166839264 | 8.0447092 | 8.0231068 | 8.21277 | -0.189664725 | -0.168062257 |
| Gm21596 | 4.535899 | 4.344558 | 4.3669 | -0.022302946 | 0.169038531 | 4.3251191 | 4.1245539 | 4.31423 | -0.189671307 | 0.010893845 |

| | | | | | | | | | | |
|---|---|---|---|---|---|---|---|---|---|---|
| Zbtb24 | 9.144143 | 9.138664 | 9.1337 | 0.00494155 | 0.010420386 | 9.2086608 | 9.0117632 | 9.20144 | -0.189674475 | 0.007223179 |
| Maneal | 3.795254 | 3.876055 | 3.8732 | 0.002849171 | -0.077951864 | 3.7501547 | 3.6774331 | 3.86713 | -0.189693091 | -0.116971457 |
| Abhd5 | 7.048173 | 7.210428 | 6.9761 | 0.23433148 | 0.072076499 | 7.1147674 | 6.9723436 | 7.16209 | -0.189749585 | -0.047325725 |
| Inpp5e | 8.993261 | 8.953455 | 9.0631 | -0.109653654 | -0.069846755 | 9.2161732 | 9.0241338 | 9.21391 | -0.189780605 | 0.002258813 |
| Zcchc7 | 10.73381 | 10.57788 | 10.684 | -0.106351198 | 0.049585472 | 10.789706 | 10.662072 | 10.8519 | -0.189817678 | -0.062184554 |
| Hipk1 | 12.26091 | 12.28572 | 12.297 | -0.011533156 | -0.036342228 | 12.083386 | 12.024424 | 12.2143 | -0.18984245 | -0.13088073 |
| Sae1 | 10.19162 | 10.21028 | 10.238 | -0.028122564 | -0.046779328 | 10.118328 | 10.01281 | 10.2027 | -0.189843195 | -0.084325036 |
| Snx11 | 8.637101 | 8.558745 | 8.7151 | -0.156346127 | -0.077989466 | 8.9506416 | 8.8524541 | 9.04247 | -0.190013688 | -0.091826266 |
| Dusp1 | 11.88833 | 11.75813 | 11.738 | 0.020121512 | 0.15031773 | 10.957886 | 10.861629 | 11.0517 | -0.190031192 | -0.093774319 |
| 9030025P20Rik | 7.500776 | 7.345552 | 7.5191 | -0.173499685 | -0.018274881 | 7.8770807 | 7.6986342 | 7.88874 | -0.190101547 | -0.011655034 |
| Vmn2r76 | 6.116655 | 6.223114 | 6.1129 | 0.110177186 | 0.003718712 | 6.2490866 | 6.4265389 | 6.6167 | -0.190165683 | -0.367618065 |
| Mapt | 2.656847 | 2.71883 | 2.7368 | -0.017965548 | -0.079948744 | 2.9520065 | 2.6646372 | 2.85483 | -0.190192862 | 0.097176384 |
| Ercc6 | 9.55393 | 9.484095 | 9.4353 | 0.04879255 | 0.118627548 | 9.3414525 | 9.4176238 | 9.60784 | -0.190217036 | -0.266388317 |
| Pcdhga3 | 3.269359 | 3.051198 | 3.1656 | -0.1144334 | 0.103727602 | 2.9512342 | 2.867357 | 3.05758 | -0.190224919 | -0.106347697 |
| Dcaf10 | 8.839088 | 8.936137 | 8.8423 | 0.093849568 | -0.003199759 | 8.7384003 | 8.6347458 | 8.82497 | -0.190227939 | -0.086573435 |
| Macrod2 | 3.523939 | 3.560268 | 3.5637 | -0.003409085 | -0.039738078 | 3.41171 | 3.3149016 | 3.50527 | -0.190372853 | -0.093564518 |
| Ormdl1 | 9.106042 | 9.179094 | 9.1555 | 0.023639285 | -0.049412704 | 9.1791983 | 8.8879788 | 9.07836 | -0.190380121 | 0.100839381 |
| Zfp442 | 5.842235 | 5.783945 | 5.9546 | -0.170673848 | -0.112383972 | 5.8128384 | 5.7542201 | 5.94483 | -0.190611112 | -0.131992817 |
| Lum | 4.505512 | 4.49527 | 4.5723 | -0.077068142 | -0.06682631 | 4.4615198 | 4.4141907 | 4.60481 | -0.190617469 | -0.143288297 |
| Fbxo47 | 4.343474 | 4.220416 | 4.4452 | -0.224819312 | -0.101761425 | 4.4531077 | 4.2031329 | 4.39375 | -0.190619722 | 0.059355069 |
| Agtr1a | 3.312705 | 3.072076 | 3.2358 | -0.163691443 | 0.076937123 | 3.1083716 | 2.850714 | 3.04137 | -0.190654763 | 0.067002894 |
| Vmn1r209 | 2.500892 | 2.461891 | 2.4774 | -0.015497641 | 0.023503326 | 2.5055025 | 2.490135 | 2.68082 | -0.190689258 | -0.175321776 |
| Zscan26 | 10.40243 | 10.40718 | 10.421 | -0.013738617 | -0.018481528 | 10.572179 | 10.459514 | 10.6503 | -0.19078233 | -0.078117827 |
| Hdac10 | 9.899168 | 9.957665 | 10.06 | -0.10192158 | -0.160418398 | 10.700223 | 10.087054 | 10.2779 | -0.190835504 | 0.422332836 |
| Aspa | 3.178618 | 3.011749 | 3.0634 | -0.05167037 | 0.115199239 | 2.8404157 | 2.9351912 | 3.12607 | -0.190881247 | -0.285656739 |
| Zfp37 | 3.934432 | 3.899636 | 3.8771 | 0.022559118 | 0.057355323 | 3.8564082 | 3.6973846 | 3.88829 | -0.190907016 | -0.031883452 |
| Ubl3 | 10.83641 | 10.96151 | 10.835 | 0.126092352 | 0.000996178 | 10.678854 | 10.560847 | 10.7518 | -0.190979534 | -0.072971813 |
| Moap1 | 6.012692 | 5.902888 | 5.7525 | 0.150403634 | 0.260206837 | 5.8660598 | 5.7793445 | 5.97041 | -0.191061941 | -0.104346566 |
| Smad3 | 9.940209 | 9.862314 | 9.8661 | -0.00381133 | 0.074083575 | 9.5234235 | 9.5689792 | 9.76004 | -0.19106195 | -0.236617641 |
| Secisbp2l | 10.46246 | 10.43472 | 10.499 | -0.06475425 | -0.03701653 | 10.298282 | 10.342563 | 10.5337 | -0.191182654 | -0.235463098 |
| Mdm4 | 12.15102 | 12.19738 | 12.15 | 0.046953162 | 0.000598754 | 12.254912 | 12.38271 | 12.5741 | -0.191348174 | -0.319145983 |
| Rai1 | 9.808929 | 9.672356 | 9.7416 | -0.069202171 | 0.067371118 | 9.699749 | 9.5686873 | 9.76006 | -0.191369397 | -0.060307771 |

| | | | | | | | | | | |
|---|---|---|---|---|---|---|---|---|---|---|
| Btg2 | 12.30883 | 12.3398 | 12.284 | 0.056045618 | 0.025075268 | 11.955712 | 11.844261 | 12.0357 | -0.19148002 | -0.080029358 |
| Trbv15 | 7.750961 | 7.810459 | 7.9003 | -0.089838485 | -0.149336219 | 7.5156713 | 7.6008043 | 7.7924 | -0.191597851 | -0.276730886 |
| Tspan15 | 6.758901 | 6.172331 | 6.6059 | -0.433606047 | 0.152963474 | 5.9396313 | 5.8553941 | 6.04702 | -0.191626896 | -0.107389693 |
| Xpa | 7.881862 | 7.651036 | 7.8338 | -0.182744352 | 0.048081941 | 7.7591492 | 7.7764358 | 7.96807 | -0.191629935 | -0.20891655 |
| Slc5a6 | 7.535644 | 7.286038 | 7.5603 | -0.274310483 | -0.024705031 | 7.5833279 | 7.4196969 | 7.61145 | -0.191758139 | -0.028127128 |
| Xpc | 9.086649 | 9.119813 | 9.2257 | -0.105899092 | -0.139063722 | 9.0848174 | 8.897838 | 9.08968 | -0.191843873 | -0.004864505 |
| Nfasc | 3.986797 | 3.97381 | 3.6555 | 0.318295927 | 0.331283215 | 3.9056334 | 3.5887603 | 3.78072 | -0.191960801 | 0.124912368 |
| Tbx6 | 5.856527 | 6.220792 | 6.0603 | 0.160520293 | -0.203745199 | 6.5118792 | 6.3122266 | 6.5042 | -0.191971817 | 0.007680756 |
| Efcab2 | 6.937982 | 6.899294 | 6.7283 | 0.17104 | 0.209727903 | 6.9428098 | 6.7677556 | 6.95973 | -0.191974474 | -0.01692024 |
| Rfng | 7.99973 | 7.998588 | 8.1421 | -0.143525243 | -0.142383777 | 8.133848 | 7.8663905 | 8.05845 | -0.192057939 | 0.075399557 |
| Epha2 | 6.635357 | 6.370367 | 6.3113 | 0.059028463 | 0.324018292 | 6.1152024 | 6.1523972 | 6.34454 | -0.192144382 | -0.229339134 |
| Serpina1d | 2.597875 | 2.511344 | 2.611 | -0.099650705 | -0.0131195 | 2.3848479 | 2.3788278 | 2.57102 | -0.192192959 | -0.186172815 |
| Gm14698 | 7.484516 | 7.630331 | 7.6409 | -0.010538135 | -0.156352981 | 7.4139195 | 7.003248 | 7.1955 | -0.192250762 | 0.218420723 |
| Mef2a | 10.62586 | 10.59618 | 10.651 | -0.054459008 | -0.024782568 | 10.338122 | 10.3454 | 10.5377 | -0.192263916 | -0.199542182 |
| Esyt3 | 1.457807 | 1.505241 | 1.5581 | -0.052839016 | -0.100272341 | 1.4802895 | 1.4073519 | 1.59965 | -0.192297459 | -0.119359838 |
| Fzd10 | 3.451492 | 3.399563 | 3.4527 | -0.053135337 | -0.001206761 | 3.2478452 | 3.1562113 | 3.3486 | -0.192389594 | -0.100755686 |
| Setd1a | 10.68439 | 10.66361 | 10.749 | -0.085441242 | -0.064656044 | 10.756933 | 10.62771 | 10.8203 | -0.19254748 | -0.063324666 |
| Mtch1 | 10.04767 | 9.903378 | 10.081 | -0.178071371 | -0.033775691 | 10.317377 | 9.8506176 | 10.0433 | -0.192648711 | 0.274110399 |
| Vmn1r-ps15 | 2.796563 | 2.916333 | 2.8554 | 0.060923664 | -0.058845695 | 2.8609401 | 2.8299283 | 3.02264 | -0.1927089 | -0.161697143 |
| Abat | 3.147397 | 3.071072 | 3.1015 | -0.030401269 | 0.045923522 | 2.9852299 | 2.8793022 | 3.07205 | -0.192750582 | -0.0868229 |
| Ints7 | 10.4123 | 10.47423 | 10.436 | 0.037737424 | -0.024191898 | 10.576236 | 10.326556 | 10.5194 | -0.192825438 | 0.056854282 |
| Zfp862-ps | 7.430547 | 7.356069 | 7.3878 | -0.03177798 | 0.042700414 | 7.1357221 | 7.1522062 | 7.34508 | -0.192869033 | -0.209353105 |
| Msl1 | 11.06602 | 11.12667 | 11.15 | -0.02381578 | -0.084467282 | 10.970072 | 10.90593 | 11.099 | -0.193031376 | -0.128889383 |
| Gm18562 | 2.486472 | 2.588704 | 2.5016 | 0.087054415 | -0.015178148 | 2.4513033 | 2.4546009 | 2.64775 | -0.193150612 | -0.196448185 |
| Gm10288 | 9.25076 | 10.06407 | 8.992 | 1.072108191 | 0.25879365 | 9.0076977 | 10.258998 | 10.4522 | -0.193205398 | -1.444505554 |
| Epb41l2 | 11.07405 | 11.16305 | 11.126 | 0.036770414 | -0.052227466 | 11.195564 | 11.049214 | 11.2425 | -0.193315504 | -0.046965558 |
| Col28a1 | 2.568424 | 2.562431 | 2.601 | -0.038531293 | -0.032538232 | 2.5041669 | 2.3971052 | 2.59055 | -0.193443206 | -0.086381542 |
| Gm37568 | 3.476395 | 3.304228 | 3.4927 | -0.188444509 | -0.016278019 | 3.4491883 | 3.3607671 | 3.55424 | -0.19347017 | -0.105048935 |
| Tatdn1 | 7.176254 | 6.973909 | 7.1774 | -0.20348848 | -0.001143082 | 7.3407724 | 7.0093661 | 7.20289 | -0.193520997 | 0.13788529 |
| Ccser1 | 4.599224 | 4.310988 | 4.5566 | -0.245644157 | 0.042591936 | 4.5591238 | 4.3487971 | 4.54232 | -0.193527715 | 0.016799029 |
| Erbin | 12.27032 | 12.27276 | 12.263 | 0.009708636 | 0.007265558 | 11.997719 | 11.982262 | 12.1758 | -0.193560274 | -0.178103674 |
| Tmem254a | 5.08212 | 5.428196 | 5.3473 | 0.080931308 | -0.265145346 | 5.2246509 | 5.2155755 | 5.40914 | -0.193562374 | -0.184486984 |

| | | | | | | | | | | |
|---|---|---|---|---|---|---|---|---|---|---|
| Gm8251 | 3.375676 | 3.316948 | 3.3826 | -0.065689004 | -0.006960152 | 3.570244 | 3.3749493 | 3.56866 | -0.193708228 | 0.001586477 |
| Pex5 | 9.312587 | 9.248107 | 9.2916 | -0.043480284 | 0.020999715 | 9.3449831 | 9.2185348 | 9.41239 | -0.193854593 | -0.067406307 |
| Snrpn | 4.09634 | 4.095223 | 4.0376 | 0.05764929 | 0.058766651 | 3.9137109 | 3.7989561 | 3.99282 | -0.193861664 | -0.079106875 |
| Pkia | 4.468844 | 4.12146 | 4.4395 | -0.318080273 | 0.029303329 | 4.0835628 | 3.9978035 | 4.19181 | -0.194005011 | -0.108245734 |
| Dgkq | 8.831324 | 8.835019 | 9.0184 | -0.183359655 | -0.187054564 | 9.243227 | 9.0422219 | 9.23626 | -0.194033329 | 0.006971827 |
| Glt28d2 | 1.983794 | 1.996957 | 1.9567 | 0.040257106 | 0.027094197 | 1.9537311 | 1.9067755 | 2.10105 | -0.194275958 | -0.147320414 |
| Zfp251 | 8.407576 | 8.444014 | 8.4056 | 0.038445704 | 0.002007681 | 8.3348152 | 8.1357383 | 8.3301 | -0.194360065 | 0.004716821 |
| 1700029J07Rik | 5.493887 | 5.517359 | 5.4292 | 0.088203608 | 0.064731346 | 5.7626248 | 5.591078 | 5.78553 | -0.194455108 | -0.022908303 |
| Hnrnpll | 8.768034 | 8.520627 | 8.6342 | -0.113558961 | 0.133847907 | 8.4289819 | 8.5064435 | 8.701 | -0.194560993 | -0.27202262 |
| BC034090 | 2.737377 | 2.76852 | 2.7161 | 0.052445124 | 0.02130223 | 2.6655019 | 2.6780098 | 2.87278 | -0.194766425 | -0.207274312 |
| Rad9b | 6.266744 | 6.26718 | 6.3122 | -0.045030203 | -0.045465972 | 6.6281868 | 6.2856385 | 6.48048 | -0.194838816 | 0.147709489 |
| Pla2g12a | 6.844243 | 6.782081 | 6.696 | 0.086083496 | 0.148245612 | 6.849316 | 6.5949033 | 6.78979 | -0.194885884 | 0.059526852 |
| Cul9 | 8.208427 | 8.263289 | 8.478 | -0.214701754 | -0.269563805 | 8.2590229 | 8.2206941 | 8.4157 | -0.195004254 | -0.15667544 |
| Tln2 | 5.492191 | 5.186918 | 5.3945 | -0.207607783 | 0.097665699 | 5.0707633 | 5.0991782 | 5.2942 | -0.195019015 | -0.223433927 |
| Iglv1 | 9.595544 | 9.864667 | 9.6136 | 0.251062566 | -0.018060299 | 9.8920489 | 9.8280273 | 10.0231 | -0.195112983 | -0.131091444 |
| Pcdhga10 | 4.350689 | 4.175815 | 4.1405 | 0.035307679 | 0.210181435 | 3.9651286 | 3.9537694 | 4.14888 | -0.195115356 | -0.183756221 |
| Itgb3 | 9.951468 | 9.851966 | 9.8623 | -0.010345811 | 0.089156297 | 9.8162035 | 9.6413876 | 9.83675 | -0.195364999 | -0.020549074 |
| Usp48 | 10.98807 | 11.13183 | 11.103 | 0.028519672 | -0.115239508 | 11.085669 | 11.132034 | 11.3274 | -0.19537402 | -0.241740006 |
| Ubn2 | 10.20344 | 10.33425 | 10.354 | -0.019719638 | -0.150526434 | 10.323533 | 10.492746 | 10.6882 | -0.195419059 | -0.364631948 |
| Arntl | 8.408503 | 8.513053 | 8.5411 | -0.028049691 | -0.132600366 | 8.4770273 | 8.308177 | 8.50367 | -0.195490672 | -0.026640409 |
| B130006D01Rik | 3.536164 | 3.534415 | 3.4822 | 0.052263747 | 0.054013127 | 3.5020885 | 3.5155402 | 3.71103 | -0.195491719 | -0.208943434 |
| Id4 | 3.232647 | 3.227267 | 3.3897 | -0.162480541 | -0.157100088 | 3.1311932 | 2.9263299 | 3.12185 | -0.195520353 | 0.00934289 |
| Tbc1d32 | 6.256919 | 6.271158 | 6.305 | -0.033841126 | -0.048079817 | 6.3074331 | 6.3248675 | 6.52044 | -0.195568877 | -0.213003238 |
| Zswim3 | 6.047075 | 5.95671 | 6.0998 | -0.143131537 | -0.052765617 | 6.3097242 | 6.0692579 | 6.26488 | -0.195620824 | 0.044845512 |
| Phf3 | 11.32024 | 11.39492 | 11.282 | 0.113138556 | 0.038460346 | 11.06409 | 11.224119 | 11.4197 | -0.19562189 | -0.355651205 |
| Usp35 | 6.798358 | 6.662959 | 6.9615 | -0.298558379 | -0.163158883 | 6.8272032 | 7.0298495 | 7.22548 | -0.195626595 | -0.398272862 |
| Erbb2 | 4.690407 | 4.605005 | 4.5519 | 0.053140666 | 0.13854277 | 4.5206891 | 4.4262107 | 4.62185 | -0.195635247 | -0.10115686 |
| Scml2 | 2.118082 | 2.107336 | 2.0847 | 0.022586481 | 0.033331617 | 2.1250436 | 2.0182676 | 2.21397 | -0.195705513 | -0.0889295 |
| R3hdm2 | 10.69707 | 10.56541 | 10.693 | -0.127434758 | 0.004228222 | 10.703858 | 10.532888 | 10.7287 | -0.195772328 | -0.024802222 |
| Cnn2 | 13.02792 | 13.02171 | 13.005 | 0.016722606 | 0.022939152 | 13.181301 | 12.716218 | 12.912 | -0.195786954 | 0.269296512 |
| Poln | 3.970239 | 3.974611 | 4.0149 | -0.040302654 | -0.044674362 | 3.8661526 | 3.9570252 | 4.15284 | -0.195811764 | -0.286684321 |
| Fbxo17 | 5.552072 | 5.543908 | 5.5808 | -0.036863617 | -0.028699602 | 5.2879943 | 5.4234529 | 5.61934 | -0.195889478 | -0.331348021 |

| | | | | | | | | | | |
|---|---|---|---|---|---|---|---|---|---|---|
| Prpf18 | 9.938664 | 9.974492 | 9.8747 | 0.099764089 | 0.063936659 | 9.8808445 | 9.7883908 | 9.98434 | -0.195946991 | -0.103493301 |
| Thap6 | 7.380991 | 7.317476 | 7.4788 | -0.161298015 | -0.09778273 | 7.3867126 | 7.1617214 | 7.35771 | -0.195986212 | 0.029004921 |
| Commd10 | 8.132632 | 8.049021 | 8.2143 | -0.165273642 | -0.081662678 | 8.1556311 | 7.9340103 | 8.13002 | -0.196011313 | 0.025609454 |
| Ctrc | 3.589835 | 3.661161 | 3.5976 | 0.063538965 | -0.007787183 | 3.6941726 | 3.5060734 | 3.70209 | -0.1960181 | -0.007918819 |
| Npl | 5.293131 | 5.048686 | 5.1755 | -0.126768777 | 0.117676511 | 5.2959125 | 4.8340841 | 5.03013 | -0.19604426 | 0.265784155 |
| Cdca5 | 6.393223 | 6.116228 | 6.4648 | -0.34852447 | -0.071529464 | 6.2782912 | 6.3318341 | 6.52793 | -0.196094555 | -0.249637417 |
| Gm15501 | 5.844311 | 5.809501 | 5.9535 | -0.143972845 | -0.1091629 | 5.6472138 | 5.7787985 | 5.97498 | -0.196183416 | -0.327768105 |
| Olfr1505 | 3.040335 | 3.188852 | 3.2921 | -0.103290919 | -0.251807914 | 3.1548324 | 3.1226622 | 3.31888 | -0.196215165 | -0.164045013 |
| Gm10033 | 7.017959 | 7.015655 | 7.0404 | -0.024756763 | -0.02245305 | 7.076265 | 6.9012666 | 7.0975 | -0.196232202 | -0.021233878 |
| Als2cl | 11.56272 | 11.83881 | 11.752 | 0.08658434 | -0.189505432 | 11.563632 | 11.690135 | 11.8866 | -0.196474212 | -0.322976665 |
| Cacna2d2 | 4.991768 | 5.00028 | 5.2494 | -0.24914629 | -0.257658052 | 5.1287614 | 4.8830435 | 5.07965 | -0.196606059 | 0.049111824 |
| Cstf2 | 10.48368 | 10.53131 | 10.41 | 0.121681894 | 0.074051938 | 10.499577 | 10.27026 | 10.4669 | -0.196615768 | 0.032701172 |
| Hmgb1 | 12.11091 | 12.01942 | 12.124 | -0.104111012 | -0.012624972 | 12.010236 | 11.855963 | 12.0526 | -0.196634912 | -0.042361709 |
| Usp34 | 11.9678 | 11.83917 | 12.034 | -0.194999944 | -0.066371138 | 11.802983 | 11.907255 | 12.1039 | -0.196641018 | -0.300913468 |
| Rnase12 | 2.472407 | 2.327887 | 2.4208 | -0.092926099 | 0.051593232 | 2.5125149 | 2.2487775 | 2.44542 | -0.196645187 | 0.067092236 |
| Armc10 | 9.949728 | 9.878021 | 10.028 | -0.149763815 | -0.078057242 | 10.102648 | 9.7610399 | 9.95773 | -0.196688605 | 0.144920017 |
| Dnajb4 | 8.985419 | 8.846984 | 8.9857 | -0.138706494 | -0.000270858 | 8.3169113 | 8.5625031 | 8.75926 | -0.196756651 | -0.442348381 |
| Med14 | 10.25309 | 10.18739 | 10.253 | -0.066036922 | -0.000337818 | 10.075184 | 10.081283 | 10.2781 | -0.196768128 | -0.202866872 |
| Flvcr1 | 8.814163 | 8.666847 | 8.6194 | 0.047493955 | 0.194809742 | 8.6115278 | 8.4280914 | 8.62487 | -0.19677579 | -0.013339368 |
| Scaper | 9.083331 | 9.207213 | 9.1307 | 0.076491977 | -0.047389949 | 9.0237501 | 8.8929279 | 9.08986 | -0.196927162 | -0.066104902 |
| AW551984 | 3.07144 | 3.108373 | 2.8077 | 0.300638017 | 0.263705309 | 2.8270849 | 2.8289556 | 3.02597 | -0.19701473 | -0.198885434 |
| Helb | 9.306215 | 9.31816 | 9.4438 | -0.125630333 | -0.137575369 | 9.3079101 | 9.2306578 | 9.42768 | -0.197017977 | -0.119765681 |
| Rspry1 | 9.548572 | 9.581419 | 9.5588 | 0.022633218 | -0.010214328 | 9.3892844 | 9.3458974 | 9.543 | -0.19709784 | -0.153710764 |
| Rad51 | 6.924609 | 6.815304 | 7.0815 | -0.266220503 | -0.156915708 | 7.0235464 | 7.2277076 | 7.42482 | -0.197113044 | -0.401274237 |
| Mgmt | 4.800312 | 4.563276 | 4.7146 | -0.151327446 | 0.085708518 | 4.7450116 | 4.4914751 | 4.68859 | -0.197119351 | 0.056417168 |
| Flrt2 | 4.425212 | 4.469438 | 4.4351 | 0.034360941 | -0.009864664 | 4.1758917 | 4.1767614 | 4.37397 | -0.197213262 | -0.198082997 |
| Arsk | 7.832815 | 7.8314 | 7.9636 | -0.132245178 | -0.130830077 | 7.9478587 | 7.7162239 | 7.91344 | -0.197216519 | 0.034418255 |
| Gm5940 | 1.344804 | 1.226945 | 1.2891 | -0.062142037 | 0.055716759 | 1.2955358 | 1.1943111 | 1.39155 | -0.197237588 | -0.096012919 |
| Tmc4 | 6.839673 | 6.698286 | 6.6491 | 0.049153375 | 0.190539799 | 6.7458098 | 6.7433698 | 6.94065 | -0.197284672 | -0.194844657 |
| 9930021J03Rik | 9.671891 | 9.711298 | 9.715 | -0.003711875 | -0.043119359 | 9.3745538 | 9.3969894 | 9.59431 | -0.197323986 | -0.219759623 |
| Eml6 | 5.908841 | 5.748611 | 6.1666 | -0.417958808 | -0.257729148 | 5.5700819 | 5.6879063 | 5.8854 | -0.197496135 | -0.315320548 |
| Atl3 | 11.36079 | 11.36372 | 11.387 | -0.02340162 | -0.026324884 | 11.223778 | 11.156179 | 11.3537 | -0.197500352 | -0.129900871 |

| | | | | | | | | | | |
|---|---|---|---|---|---|---|---|---|---|---|
| Polk | 7.793198 | 8.018167 | 7.827 | 0.191153887 | -0.033814691 | 7.8171652 | 7.8611781 | 8.05884 | -0.19766137 | -0.241674252 |
| Gm20708 | 2.962902 | 2.93057 | 3.3138 | -0.383245123 | -0.35091368 | 3.1729069 | 3.1491762 | 3.34687 | -0.197693956 | -0.173963276 |
| Trav14-2 | 4.993257 | 4.946048 | 4.9694 | -0.023341086 | 0.023867863 | 4.8339076 | 4.9270879 | 5.12485 | -0.197763643 | -0.290943913 |
| Gm14308 | 4.663621 | 4.469639 | 5.0361 | -0.56645466 | -0.372472393 | 4.7868212 | 4.451463 | 4.64928 | -0.197820126 | 0.13753809 |
| Mad1l1 | 8.342259 | 8.30962 | 8.3331 | -0.023441853 | 0.009196792 | 8.4168119 | 8.1795068 | 8.37742 | -0.197908533 | 0.039396548 |
| Gdf10 | 4.895158 | 5.311026 | 5.124 | 0.187067996 | -0.228799874 | 4.4628036 | 4.6369467 | 4.83488 | -0.197934825 | -0.372077933 |
| Kdm2a | 11.9278 | 12.08503 | 11.999 | 0.086299812 | -0.070932856 | 12.170592 | 11.911549 | 12.1096 | -0.19802476 | 0.061018115 |
| Igkv2-107 | 2.112783 | 2.149582 | 2.0722 | 0.077407292 | 0.04060765 | 2.1348984 | 2.0097021 | 2.20774 | -0.198040437 | -0.072844169 |
| Smg6 | 10.14302 | 10.0414 | 10.099 | -0.058026401 | 0.043590052 | 10.16406 | 10.039629 | 10.2377 | -0.198053024 | -0.073622005 |
| 5830417I10Rik | 10.36536 | 10.40944 | 10.384 | 0.025739168 | -0.018341788 | 10.381978 | 10.285993 | 10.4841 | -0.198121078 | -0.102135821 |
| Uba6 | 9.235492 | 9.21563 | 9.2753 | -0.059624402 | -0.039762138 | 9.1775541 | 9.0595414 | 9.25769 | -0.198152152 | -0.080139427 |
| Lmo2 | 10.57513 | 10.46263 | 10.562 | -0.09903101 | 0.01347355 | 10.922971 | 10.397653 | 10.5958 | -0.198196038 | 0.327121719 |
| Idua | 9.075752 | 9.067743 | 9.1286 | -0.060877243 | -0.052868009 | 9.529924 | 9.1807944 | 9.37912 | -0.198324921 | 0.150804664 |
| Tspan10 | 4.187807 | 4.044353 | 4.2403 | -0.195912595 | -0.052458047 | 3.5126752 | 3.5350893 | 3.73351 | -0.198418617 | -0.22083265 |
| Heatr5a | 9.166138 | 9.210635 | 9.1348 | 0.075791293 | 0.03129397 | 9.0710085 | 8.9761014 | 9.17457 | -0.198466924 | -0.103559842 |
| Zfp580 | 4.129209 | 4.083209 | 4.0434 | 0.039846353 | 0.085845717 | 4.1897853 | 3.9327019 | 4.13126 | -0.198553867 | 0.058529554 |
| Nkiras1 | 7.230873 | 7.153994 | 7.276 | -0.122012492 | -0.045133037 | 7.3613902 | 7.1802722 | 7.37889 | -0.198622682 | -0.017504683 |
| Dcxr | 6.401185 | 6.230004 | 6.4464 | -0.216368613 | -0.045187345 | 6.731895 | 6.1449776 | 6.34368 | -0.198701901 | 0.388215529 |
| Brox | 10.14782 | 10.19769 | 10.215 | -0.017730611 | -0.067601015 | 10.255625 | 10.245905 | 10.4448 | -0.198896597 | -0.189176865 |
| Jak1 | 13.95005 | 13.92012 | 13.899 | 0.020711692 | 0.050641422 | 13.797372 | 13.61258 | 13.8115 | -0.198939856 | -0.014148154 |
| Pgm2l1 | 8.703811 | 8.768666 | 8.7301 | 0.038587669 | -0.026267375 | 8.4419004 | 8.5812711 | 8.78039 | -0.19911569 | -0.338486391 |
| Stard4 | 9.199853 | 9.184435 | 9.2341 | -0.049673039 | -0.034254779 | 9.0792916 | 8.8357425 | 9.03493 | -0.199183538 | 0.04436552 |
| Arxes2 | 4.40206 | 4.234052 | 4.2789 | -0.044830097 | 0.123177497 | 4.1827225 | 4.0486712 | 4.24788 | -0.199212236 | -0.065160904 |
| Kcnb1 | 6.335702 | 6.38824 | 6.5972 | -0.208923639 | -0.261462397 | 6.608014 | 6.1593266 | 6.35859 | -0.199258499 | 0.24942893 |
| Thap12 | 10.37283 | 10.27114 | 10.335 | -0.063422328 | 0.038268802 | 10.233044 | 10.070203 | 10.2695 | -0.199341414 | -0.036501142 |
| Tprkb | 8.551778 | 8.521263 | 8.5262 | -0.004981606 | 0.025533576 | 8.4942159 | 8.4770058 | 8.67641 | -0.199402288 | -0.182192189 |
| Stard9 | 6.6809 | 6.561623 | 6.7608 | -0.199144926 | -0.079867757 | 6.695168 | 6.6114163 | 6.81085 | -0.199428779 | -0.115677034 |
| Ighj2 | 6.531701 | 7.319496 | 6.7496 | 0.569849295 | -0.217945741 | 6.7993027 | 6.6643144 | 6.86381 | -0.199500573 | -0.064512244 |
| Tmem185a | 8.577634 | 8.517635 | 8.5836 | -0.065946855 | -0.005947766 | 8.5316995 | 8.2420383 | 8.44158 | -0.199536975 | 0.090124249 |
| Sema6a | 6.026137 | 5.712668 | 5.6759 | 0.036739169 | 0.35020805 | 4.9983839 | 5.3025963 | 5.50225 | -0.199657695 | -0.503870103 |
| Gm6430 | 2.536003 | 2.674915 | 2.6026 | 0.072353025 | -0.066558566 | 2.5475138 | 2.5493842 | 2.74911 | -0.199724979 | -0.201595357 |
| Ighv2-4 | 2.349319 | 2.440204 | 2.3763 | 0.063889079 | -0.026995788 | 2.4184981 | 2.244106 | 2.44398 | -0.199873038 | -0.025480885 |

| | | | | | | | | | | |
|---|---|---|---|---|---|---|---|---|---|---|
| Eea1 | 9.630158 | 9.523622 | 9.6008 | -0.07722185 | 0.029313939 | 9.2482617 | 9.3198827 | 9.51976 | -0.199875311 | -0.271496318 |
| Gsk3b | 10.30657 | 10.38071 | 10.269 | 0.111788594 | 0.037649546 | 9.9852861 | 10.075574 | 10.2755 | -0.1999667 | -0.290254493 |
| Fzd3 | 2.814016 | 2.949365 | 2.8452 | 0.104142384 | -0.031206684 | 2.73208 | 2.7328112 | 2.9328 | -0.199992353 | -0.200723541 |
| Tle6 | 3.35164 | 3.313317 | 3.3896 | -0.07632141 | -0.037997896 | 3.3198511 | 3.2750802 | 3.47514 | -0.200060371 | -0.155289511 |
| Sp1 | 11.82352 | 11.80586 | 11.801 | 0.005289502 | 0.022951362 | 11.697382 | 11.626889 | 11.827 | -0.200130476 | -0.129638103 |
| Nrd1 | 11.67426 | 11.5986 | 11.607 | -0.008468114 | 0.067188522 | 11.673835 | 11.667122 | 11.8674 | -0.200243178 | -0.193530638 |
| 39142 | 11.12096 | 11.03132 | 11.128 | -0.09636881 | -0.006735998 | 10.977158 | 10.958322 | 11.1587 | -0.20033072 | -0.18149472 |
| Prdm4 | 9.282785 | 9.350563 | 9.3213 | 0.029223648 | -0.038554291 | 9.3290767 | 9.2095641 | 9.40993 | -0.200363418 | -0.080850832 |
| Traj2 | 3.006061 | 3.052307 | 3.0941 | -0.041799579 | -0.0880447 | 2.8827413 | 2.9287211 | 3.12909 | -0.200366133 | -0.246345997 |
| Acad12 | 6.529559 | 6.455716 | 6.8433 | -0.387589303 | -0.313746606 | 6.5933635 | 6.4271128 | 6.62751 | -0.200398165 | -0.034147436 |
| Zfp467 | 7.293043 | 7.24747 | 7.4307 | -0.183275737 | -0.137702754 | 7.2954189 | 7.1030845 | 7.3035 | -0.200416317 | -0.008081903 |
| Ppm1h | 10.158 | 10.08483 | 10.126 | -0.041342514 | 0.031827248 | 9.8550936 | 9.9714335 | 10.172 | -0.200553352 | -0.316893204 |
| AC124502.3 | 4.136402 | 4.199422 | 4.0652 | 0.134235061 | 0.071214709 | 4.549827 | 4.1539586 | 4.35465 | -0.200687453 | 0.195180868 |
| Zranb1 | 9.988907 | 9.84176 | 9.9673 | -0.125579366 | 0.021568051 | 9.8334391 | 9.7953087 | 9.99602 | -0.200706964 | -0.162576583 |
| Akap17b | 6.593005 | 6.397297 | 6.6197 | -0.222386363 | -0.026678023 | 6.4776667 | 6.4416729 | 6.6424 | -0.200728879 | -0.1647351 |
| Ica1 | 5.865148 | 5.450612 | 5.8903 | -0.439666156 | -0.025130752 | 5.4581923 | 5.1353909 | 5.33615 | -0.200759597 | 0.122041725 |
| Fcrla | 11.23601 | 11.38402 | 11.379 | 0.004526986 | -0.143482298 | 11.645993 | 11.071791 | 11.2726 | -0.200781692 | 0.373419916 |
| Rnf24 | 6.76403 | 6.646653 | 6.6539 | -0.007292429 | 0.110084479 | 6.6683765 | 6.5891935 | 6.79002 | -0.200828789 | -0.121645718 |
| Atn1 | 10.15716 | 10.16187 | 10.026 | 0.135541482 | 0.130836346 | 9.9045788 | 9.8435028 | 10.0444 | -0.200859459 | -0.139783519 |
| Dact1 | 5.49262 | 5.368334 | 5.4032 | -0.034882581 | 0.089403984 | 4.7014601 | 4.8991099 | 5.10004 | -0.200927405 | -0.398577217 |
| Epc2 | 10.1986 | 10.40797 | 10.321 | 0.086479322 | -0.122883945 | 9.9960927 | 9.9327475 | 10.1338 | -0.201049471 | -0.137704313 |
| Zcwpw1 | 7.942619 | 7.876955 | 8.0893 | -0.212309987 | -0.146645755 | 8.0134426 | 7.9787619 | 8.17984 | -0.201082975 | -0.166402305 |
| Cgref1 | 1.907844 | 1.974912 | 1.9454 | 0.029475112 | -0.037593719 | 2.0277229 | 1.8388963 | 2.03998 | -0.201085078 | -0.012258518 |
| Parvg | 11.3615 | 11.32038 | 11.34 | -0.02003289 | 0.021084522 | 11.583242 | 11.202725 | 11.4038 | -0.201106714 | 0.179410514 |
| Prg3 | 3.573941 | 3.454371 | 3.5039 | -0.049552976 | 0.070016494 | 3.2340605 | 3.1522825 | 3.35342 | -0.201133869 | -0.119355873 |
| Cab39l | 9.866338 | 9.870993 | 9.865 | 0.005951955 | 0.001296378 | 9.9227728 | 9.7673246 | 9.9685 | -0.201171599 | -0.045723406 |
| Tbx18 | 3.233169 | 3.320518 | 3.2498 | 0.070677379 | -0.016671761 | 3.1040613 | 3.1037102 | 3.30492 | -0.201206293 | -0.200855181 |
| Hexim2 | 4.421436 | 4.413529 | 4.6272 | -0.213656106 | -0.205749459 | 4.4831638 | 4.5055955 | 4.70683 | -0.201237438 | -0.223669097 |
| Slc7a10 | 2.868103 | 2.854263 | 2.9524 | -0.098132672 | -0.084292838 | 2.9518493 | 2.7963658 | 2.99762 | -0.201255872 | -0.04577237 |
| Pfkfb4 | 8.249351 | 8.234939 | 8.2411 | -0.006199988 | 0.008211975 | 8.7261495 | 8.1801918 | 8.38148 | -0.201286205 | 0.344671484 |
| Ptk7 | 4.997151 | 5.098817 | 4.885 | 0.213823334 | 0.112157345 | 4.8731935 | 4.5874044 | 4.78871 | -0.201307043 | 0.084482123 |
| Ccdc9b | 4.55751 | 4.398312 | 4.432 | -0.033725096 | 0.125472675 | 4.296148 | 4.1313163 | 4.33263 | -0.201310006 | -0.036478288 |

| | | | | | | | | | | |
|---|---|---|---|---|---|---|---|---|---|---|
| Rdh10 | 8.219064 | 7.908643 | 8.3068 | -0.398203016 | -0.08778208 | 7.4729163 | 7.5166796 | 7.71813 | -0.201448274 | -0.24521156 |
| Fbxl21 | 2.69401 | 2.555935 | 2.6958 | -0.139834263 | -0.001759619 | 2.6940361 | 2.5311972 | 2.73273 | -0.201529298 | -0.038690451 |
| Immp2l | 5.85776 | 5.838531 | 5.8192 | 0.019292782 | 0.038522378 | 6.1808425 | 5.8900086 | 6.09161 | -0.201598996 | 0.089234935 |
| Bcas1 | 3.755226 | 3.71508 | 3.8857 | -0.170570002 | -0.130424661 | 3.9538385 | 3.5696245 | 3.77124 | -0.201613707 | 0.18260027 |
| Tmsb15b1 | 5.036901 | 5.001881 | 5.1232 | -0.121273781 | -0.08625436 | 5.0291359 | 4.854814 | 5.05643 | -0.201619733 | -0.027297835 |
| Cdc6 | 6.529908 | 6.233238 | 6.6477 | -0.414443791 | -0.117773048 | 6.71763 | 6.7682899 | 6.97009 | -0.201805028 | -0.25246492 |
| Calhm2 | 9.212418 | 9.461651 | 9.3317 | 0.129987063 | -0.11924554 | 9.7107623 | 9.1072087 | 9.3091 | -0.201889812 | 0.401663757 |
| Fance | 7.865548 | 7.773711 | 7.8999 | -0.126234793 | -0.034397792 | 8.0637366 | 7.7944765 | 7.99641 | -0.201936157 | 0.067323907 |
| Spats2 | 5.947787 | 5.612559 | 5.6082 | 0.004353911 | 0.339582477 | 5.5471868 | 5.1437617 | 5.34586 | -0.202094338 | 0.20133069 |
| Lynx1 | 9.364378 | 9.345357 | 9.2937 | 0.051639854 | 0.070660462 | 9.6986765 | 9.15094 | 9.3531 | -0.202157124 | 0.345579395 |
| Rnf216 | 10.33921 | 10.30236 | 10.339 | -0.036924854 | -7.6936E-05 | 10.291358 | 10.08866 | 10.2908 | -0.20216473 | 0.000532408 |
| Cbx3 | 11.47166 | 11.42561 | 11.471 | -0.045486246 | 0.000562118 | 11.358497 | 11.269514 | 11.4717 | -0.202193692 | -0.113210493 |
| Olfr1195 | 3.555594 | 3.504355 | 3.4664 | 0.037922493 | 0.089161428 | 3.5295769 | 3.4463052 | 3.64855 | -0.202240972 | -0.118969307 |
| Ndufa4l2 | 2.116688 | 2.06874 | 2.1926 | -0.123878959 | -0.075930875 | 2.041628 | 1.9552015 | 2.15744 | -0.202242167 | -0.115815751 |
| Hbp1 | 10.0749 | 10.15704 | 10.184 | -0.027169677 | -0.109311083 | 10.097711 | 10.07695 | 10.2792 | -0.202258772 | -0.181497948 |
| Kif2a | 9.938249 | 9.943298 | 9.8688 | 0.074512899 | 0.069463373 | 9.5124147 | 9.6085478 | 9.81081 | -0.202262147 | -0.298395267 |
| Plec | 11.83587 | 11.89131 | 11.787 | 0.104113948 | 0.048673686 | 11.58404 | 11.64258 | 11.8449 | -0.202294918 | -0.260834439 |
| Mapk8 | 9.201823 | 9.120586 | 9.0943 | 0.026331743 | 0.107568823 | 8.8978882 | 8.8681761 | 9.07053 | -0.202354816 | -0.172642691 |
| Washc5 | 9.958467 | 9.882929 | 9.9567 | -0.073761894 | 0.001776828 | 10.016854 | 9.7586282 | 9.96119 | -0.202558074 | 0.05566775 |
| Hist1h2bn | 2.980302 | 2.882371 | 2.9319 | -0.049515386 | 0.048415279 | 3.0534593 | 2.8329905 | 3.03557 | -0.202578603 | 0.017890251 |
| Gm20498 | 6.397764 | 6.614673 | 6.433 | 0.181651213 | -0.035257176 | 6.2146476 | 6.2501873 | 6.45278 | -0.202596354 | -0.238135985 |
| Efcab7 | 3.866722 | 3.839272 | 3.9191 | -0.079868091 | -0.052418751 | 3.9658642 | 3.8539378 | 4.05658 | -0.202643792 | -0.090717426 |
| Lrwd1 | 9.252805 | 9.39375 | 9.3244 | 0.069350344 | -0.071594068 | 9.5170952 | 9.2076962 | 9.41039 | -0.202694533 | 0.106704423 |
| Dynlt3 | 9.475558 | 9.311051 | 9.4149 | -0.103871314 | 0.06063635 | 9.5591113 | 9.2510079 | 9.45374 | -0.202734874 | 0.105368517 |
| Gm3636 | 5.681839 | 5.715495 | 5.7919 | -0.076356217 | -0.110011884 | 5.5038264 | 5.6435707 | 5.84635 | -0.202779923 | -0.342524227 |
| Tmem136 | 4.715529 | 4.620533 | 4.931 | -0.310460122 | -0.215463766 | 4.7711738 | 4.7963761 | 4.9992 | -0.202826069 | -0.228028369 |
| Wnt10b | 2.969755 | 2.821518 | 2.9429 | -0.12137208 | 0.026864806 | 2.8842549 | 2.7544055 | 2.95725 | -0.202847087 | -0.072997648 |
| Cep112 | 3.145692 | 3.139979 | 3.1877 | -0.047695417 | -0.041982224 | 3.048495 | 3.0174219 | 3.22032 | -0.202894041 | -0.171820933 |
| Tenm4 | 5.472122 | 5.392695 | 5.7563 | -0.363629991 | -0.284202837 | 5.6293636 | 5.0759004 | 5.2788 | -0.202901615 | 0.350561587 |
| Ubap2 | 10.28864 | 10.19896 | 10.133 | 0.066310667 | 0.155984918 | 10.044011 | 10.148719 | 10.3517 | -0.202974171 | -0.307682123 |
| Ankrd13a | 11.53837 | 11.5423 | 11.559 | -0.016906428 | -0.020835864 | 11.524404 | 11.320949 | 11.5239 | -0.20298814 | 0.000466801 |
| Bend7 | 2.989051 | 2.809488 | 2.9537 | -0.144237231 | 0.03532594 | 2.7901384 | 2.6269759 | 2.82999 | -0.203012262 | -0.039849799 |

| | | | | | | | | | | |
|---|---|---|---|---|---|---|---|---|---|---|
| Eif4e1b | 2.840066 | 2.920648 | 2.8272 | 0.093485312 | 0.012903995 | 2.8413097 | 2.8498508 | 3.053 | -0.203153705 | -0.211694759 |
| Alox12 | 3.779018 | 3.698843 | 4.0019 | -0.303071133 | -0.222895838 | 3.7895497 | 3.7852484 | 3.98845 | -0.203199778 | -0.198898483 |
| Prepl | 8.406986 | 8.520628 | 8.3788 | 0.141802064 | 0.02816009 | 8.6265943 | 8.4628382 | 8.66614 | -0.203304019 | -0.039547927 |
| Mterf1b | 4.907227 | 4.902926 | 5.1519 | -0.248992033 | -0.244691349 | 4.9827896 | 4.8897828 | 5.09327 | -0.203487788 | -0.110481012 |
| Slc26a8 | 2.89712 | 2.933924 | 2.8619 | 0.072057325 | 0.035252864 | 2.7962834 | 2.7219968 | 2.92554 | -0.203545864 | -0.129259349 |
| mt-Nd3 | 12.51108 | 12.40527 | 12.572 | -0.16678961 | -0.060976868 | 12.502519 | 12.308603 | 12.5122 | -0.203555978 | -0.009639176 |
| Dcaf12 | 9.980913 | 10.17422 | 10.095 | 0.078967253 | -0.1143379 | 10.152467 | 9.9401204 | 10.1437 | -0.203565607 | 0.008780648 |
| Zpbp2 | 4.875277 | 4.723904 | 4.7995 | -0.075603885 | 0.075768893 | 4.9962018 | 4.7027979 | 4.90652 | -0.203726249 | 0.089677674 |
| Zfp954 | 8.03723 | 8.083618 | 8.1408 | -0.057149567 | -0.103537975 | 8.2172429 | 8.0574536 | 8.26136 | -0.203908895 | -0.044119642 |
| Supt20 | 11.06919 | 11.10252 | 11.165 | -0.0629654 | -0.096291924 | 11.300101 | 11.152405 | 11.3564 | -0.203965492 | -0.056269577 |
| Stag1 | 10.64774 | 10.61396 | 10.683 | -0.069307704 | -0.035535064 | 10.487773 | 10.411494 | 10.6155 | -0.203981514 | -0.127703106 |
| Tmem154 | 9.182198 | 9.099919 | 9.3267 | -0.226755638 | -0.144477412 | 9.1250839 | 9.1146342 | 9.31866 | -0.204030714 | -0.193581011 |
| Tubb4a | 4.858719 | 4.793967 | 4.7253 | 0.068649411 | 0.13340172 | 4.6174194 | 4.5552267 | 4.75927 | -0.20404158 | -0.141848857 |
| Srr | 7.012017 | 6.618318 | 6.8307 | -0.21238926 | 0.181309193 | 7.0675338 | 6.7338418 | 6.93792 | -0.204082702 | 0.12960932 |
| Masp1 | 4.870038 | 4.877868 | 4.9024 | -0.024581635 | -0.032411524 | 5.0262989 | 4.8109807 | 5.01507 | -0.204090225 | 0.011228032 |
| Ache | 3.436885 | 3.184844 | 3.2574 | -0.072539387 | 0.179501742 | 3.1671249 | 3.0421659 | 3.24629 | -0.204126639 | -0.07916763 |
| Map2k4 | 9.714147 | 9.619515 | 9.7817 | -0.162160311 | -0.067528973 | 9.494306 | 9.5231026 | 9.72735 | -0.204250547 | -0.233047184 |
| 4930430F08Rik | 6.529876 | 6.650991 | 6.6255 | 0.025455574 | -0.095659781 | 6.5661522 | 6.3502437 | 6.55458 | -0.20433498 | 0.011573551 |
| Gm13509 | 1.119743 | 1.233231 | 1.2743 | -0.041026917 | -0.154515681 | 1.1239195 | 1.1525161 | 1.35687 | -0.204354936 | -0.232951544 |
| Gata2 | 6.199353 | 6.078485 | 6.1254 | -0.046865486 | 0.074002894 | 5.4030804 | 5.3224285 | 5.52684 | -0.204408654 | -0.123756753 |
| Cyp2ab1 | 3.403105 | 3.462191 | 3.5838 | -0.121586726 | -0.180673527 | 3.4264748 | 3.3117673 | 3.51621 | -0.204444829 | -0.089737386 |
| Golgb1 | 10.48935 | 10.27409 | 10.52 | -0.246038773 | -0.030778064 | 10.185594 | 10.565131 | 10.7696 | -0.204458046 | -0.583995201 |
| Erlin2 | 9.018959 | 9.287193 | 9.2685 | 0.018683493 | -0.249550107 | 9.2836633 | 8.9283584 | 9.13283 | -0.204469989 | 0.150834916 |
| Zdhhc18 | 11.77513 | 11.68861 | 11.886 | -0.196894792 | -0.110370112 | 11.930196 | 11.65894 | 11.8634 | -0.204482906 | 0.066773588 |
| Cbx7 | 11.11916 | 11.19543 | 11.259 | -0.063453512 | -0.13972228 | 11.58389 | 11.131342 | 11.3359 | -0.204590542 | 0.247956981 |
| Nol4l | 5.641219 | 5.921094 | 5.9671 | -0.045960943 | -0.32583498 | 5.6937597 | 5.5214123 | 5.72606 | -0.204647226 | -0.032299793 |
| Gnai1 | 3.276699 | 3.268682 | 3.2841 | -0.015423659 | -0.007406955 | 3.0811147 | 3.0061926 | 3.2109 | -0.204711311 | -0.129789213 |
| Zswim9 | 6.590319 | 6.432955 | 6.6598 | -0.226876091 | -0.069511774 | 6.6998506 | 6.6308724 | 6.83564 | -0.204769247 | -0.135791063 |
| Ccm2l | 4.829133 | 4.591917 | 4.7907 | -0.198738767 | 0.038476623 | 4.5726649 | 4.3417585 | 4.54665 | -0.204894565 | 0.026011858 |
| Slc20a2 | 7.401422 | 7.474216 | 7.403 | 0.071257079 | -0.00153615 | 7.5921998 | 7.4261507 | 7.63107 | -0.204915608 | -0.038866523 |
| Trav11d | 2.991926 | 2.814504 | 2.8424 | -0.027856885 | 0.149565934 | 2.8892468 | 2.8470563 | 3.052 | -0.204941862 | -0.162751311 |
| Glb1 | 9.283896 | 9.174793 | 9.3496 | -0.174784003 | -0.065680574 | 9.2128865 | 9.019456 | 9.2244 | -0.204945268 | -0.011514714 |

| | | | | | | | | | | |
|---|---|---|---|---|---|---|---|---|---|---|
| Ube3c | 10.06145 | 10.02456 | 10.003 | 0.021573713 | 0.058466957 | 9.8868983 | 9.814622 | 10.0196 | -0.205002524 | -0.132726218 |
| Rxrg | 3.266809 | 3.110154 | 3.2844 | -0.174230939 | -0.017575875 | 3.0715661 | 2.8923544 | 3.09743 | -0.205079164 | -0.025867459 |
| Ighv2-6 | 4.933086 | 5.049162 | 4.8981 | 0.151087199 | 0.035010632 | 5.0016741 | 4.9182729 | 5.12336 | -0.205087623 | -0.121686446 |
| Siah3 | 3.336313 | 3.308677 | 3.4672 | -0.158501719 | -0.130865751 | 3.2911124 | 3.278791 | 3.4839 | -0.205107306 | -0.192785961 |
| Atrip | 8.563301 | 8.561978 | 8.5135 | 0.048453161 | 0.04977533 | 8.6739902 | 8.4367324 | 8.64201 | -0.20527877 | 0.031979011 |
| Sh3tc2 | 3.59563 | 3.404567 | 3.5495 | -0.144970468 | 0.046092344 | 3.3521184 | 3.2319917 | 3.43735 | -0.205360419 | -0.085233653 |
| Fam167a | 8.333133 | 8.288884 | 8.3286 | -0.039686557 | 0.004562748 | 8.7128681 | 8.3928471 | 8.59834 | -0.20549608 | 0.114524933 |
| Ccdc28b | 7.307535 | 7.103744 | 7.3928 | -0.289024657 | -0.085233711 | 7.4348973 | 7.164629 | 7.37016 | -0.205526299 | 0.064742015 |
| BC029722 | 8.641876 | 8.613898 | 8.6136 | 0.000289679 | 0.028267948 | 8.8310151 | 8.4031613 | 8.60874 | -0.205581042 | 0.222272812 |
| Pamr1 | 3.565459 | 3.412352 | 3.4966 | -0.084245554 | 0.068861382 | 3.2405372 | 3.1829386 | 3.38854 | -0.20560597 | -0.148007404 |
| Txnrd2 | 7.055364 | 6.988735 | 7.0531 | -0.064351931 | 0.002277411 | 7.1836284 | 6.9444657 | 7.1501 | -0.205638076 | 0.033524651 |
| Gm12397 | 5.260965 | 5.553741 | 5.2576 | 0.296186669 | 0.003411175 | 5.1723567 | 5.1048073 | 5.31078 | -0.205967751 | -0.138418361 |
| Cxxc5 | 9.335319 | 9.447501 | 9.3849 | 0.06257216 | -0.049610159 | 9.4024434 | 8.9611218 | 9.16712 | -0.205994066 | 0.235327487 |
| Trbv1 | 7.661954 | 7.751548 | 7.7437 | 0.007802671 | -0.081791203 | 7.4809074 | 7.4464299 | 7.65252 | -0.206088404 | -0.17161086 |
| Myo6 | 8.165487 | 7.683037 | 8.0947 | -0.411707772 | 0.070742358 | 7.7321082 | 7.6127767 | 7.81888 | -0.206102673 | -0.086771115 |
| Rasgrf1 | 3.294301 | 3.220548 | 3.1391 | 0.081410978 | 0.155163494 | 3.1928397 | 3.1921972 | 3.39836 | -0.20616104 | -0.205518586 |
| Hdac2 | 10.02603 | 10.01541 | 10.009 | 0.00626536 | 0.016886813 | 9.8786238 | 9.8978688 | 10.1042 | -0.206291493 | -0.225536504 |
| Stt3b | 11.54634 | 11.46868 | 11.507 | -0.037959762 | 0.039706898 | 11.551437 | 11.219123 | 11.4255 | -0.206336034 | 0.125978461 |
| Mtrf1 | 5.870616 | 5.741175 | 6.0113 | -0.270127284 | -0.140686424 | 6.0439984 | 5.7221443 | 5.92851 | -0.206363498 | 0.115490629 |
| Mapk8ip3 | 10.78526 | 10.92739 | 10.914 | 0.013863646 | -0.128260599 | 11.246157 | 11.189316 | 11.3957 | -0.206368976 | -0.149527219 |
| Snx21 | 4.873617 | 4.424828 | 4.6152 | -0.190376421 | 0.258413002 | 4.3801832 | 4.3603926 | 4.56678 | -0.206383047 | -0.186592492 |
| Smad5 | 9.552183 | 9.566087 | 9.5544 | 0.011681712 | -0.00222227 | 9.3101515 | 9.3645944 | 9.57102 | -0.20642821 | -0.260871105 |
| Taf1b | 9.065415 | 9.027779 | 9.1854 | -0.157669564 | -0.120033351 | 9.2563833 | 9.0435514 | 9.25001 | -0.206461868 | 0.006370071 |
| Adamts7 | 4.197567 | 4.083566 | 4.1149 | -0.031323228 | 0.082678001 | 4.1191071 | 3.8332619 | 4.03998 | -0.206715215 | 0.079130034 |
| Faf1 | 9.712716 | 9.714859 | 9.7059 | 0.00898467 | 0.006841714 | 9.6629751 | 9.5487635 | 9.75556 | -0.206797431 | -0.092585831 |
| Zfp820 | 3.791955 | 3.709264 | 3.8773 | -0.168031924 | -0.085340689 | 3.6466004 | 3.4755223 | 3.68235 | -0.206829235 | -0.035751069 |
| Lrig2 | 9.529717 | 9.56897 | 9.5859 | -0.016913568 | -0.056166736 | 9.3950965 | 9.4429292 | 9.64976 | -0.206829581 | -0.25466226 |
| Rgp1 | 10.16463 | 10.11382 | 10.179 | -0.064825524 | -0.014011258 | 10.616711 | 10.24413 | 10.451 | -0.20690937 | 0.165671326 |
| Nqo1 | 3.556033 | 3.6175 | 3.6527 | -0.03516562 | -0.096632397 | 3.6182977 | 3.454697 | 3.66167 | -0.206973748 | -0.043373077 |
| Itpr2 | 11.64713 | 11.60474 | 11.619 | -0.0140307 | 0.028358866 | 11.578172 | 11.599267 | 11.8064 | -0.207139824 | -0.228235502 |
| Atad2b | 10.27637 | 10.29195 | 10.36 | -0.068213984 | -0.083790282 | 10.191828 | 10.176351 | 10.3835 | -0.207170228 | -0.191693494 |
| Gmds | 7.139504 | 7.142283 | 7.1284 | 0.013922367 | 0.011143364 | 7.225621 | 6.9636644 | 7.17094 | -0.207273225 | 0.054683334 |

| | | | | | | | | | | |
|---|---|---|---|---|---|---|---|---|---|---|
| Bfsp2 | 7.638735 | 7.539356 | 7.5578 | -0.018438095 | 0.080941373 | 8.2448482 | 7.5008679 | 7.70817 | -0.2073053 | 0.536675031 |
| Aldh2 | 11.04991 | 10.88927 | 11.047 | -0.15798129 | 0.002653528 | 11.229837 | 10.670813 | 10.8781 | -0.207331974 | 0.351691383 |
| Pcdhga1 | 2.539592 | 2.487145 | 2.3788 | 0.108335042 | 0.160781268 | 2.3694955 | 2.3238096 | 2.53122 | -0.207410427 | -0.161724602 |
| Nfkbid | 11.10805 | 11.28456 | 11.152 | 0.13206935 | -0.044435902 | 10.8054 | 10.727731 | 10.9352 | -0.20744359 | -0.129774512 |
| Slc12a8 | 4.57003 | 4.480576 | 4.4824 | -0.001783942 | 0.087670022 | 4.8250628 | 4.526241 | 4.73374 | -0.207494285 | 0.091327519 |
| Fcer1a | 4.338036 | 4.260857 | 3.9591 | 0.301714986 | 0.378893166 | 4.1195282 | 3.6646892 | 3.87225 | -0.207559572 | 0.247279417 |
| Nsun6 | 7.574598 | 7.66078 | 7.6368 | 0.023980916 | -0.062201177 | 7.3086599 | 7.3804023 | 7.58798 | -0.207573091 | -0.279315551 |
| 37135 | 12.54541 | 12.47548 | 12.566 | -0.090613352 | -0.020689968 | 12.698203 | 12.433041 | 12.6407 | -0.20761274 | 0.057549367 |
| Catsperd | 2.964729 | 2.886632 | 2.975 | -0.088402651 | -0.010306148 | 2.9273861 | 2.8660724 | 3.07373 | -0.207652778 | -0.146339045 |
| Gm10392 | 3.972722 | 3.835165 | 3.9911 | -0.1559123 | -0.01835579 | 3.8719669 | 3.6099563 | 3.81773 | -0.207770093 | 0.054240569 |
| Atg14 | 7.830803 | 7.818956 | 7.8891 | -0.07012592 | -0.058278893 | 7.9914174 | 7.6909412 | 7.89874 | -0.20780164 | 0.092674589 |
| Igflr1 | 6.729473 | 6.89973 | 7.0269 | -0.127195252 | -0.297452088 | 6.89018 | 6.727138 | 6.93523 | -0.208088209 | -0.045046283 |
| Fam168b | 12.38768 | 12.47039 | 12.374 | 0.096498784 | 0.013788364 | 12.394672 | 12.182287 | 12.3905 | -0.208217856 | 0.004166656 |
| Caskin2 | 7.768324 | 7.649644 | 7.6348 | 0.014887925 | 0.133567511 | 7.2413199 | 7.3167707 | 7.52508 | -0.208304488 | -0.283755206 |
| Entpd2 | 4.77286 | 4.520715 | 4.7824 | -0.261677061 | -0.009532166 | 4.4898309 | 4.2151613 | 4.42355 | -0.208386225 | 0.066283349 |
| Smc3 | 11.15338 | 11.14521 | 11.222 | -0.07634416 | -0.06817522 | 10.991146 | 10.992504 | 11.201 | -0.20848088 | -0.209838093 |
| Htatsf1 | 10.1506 | 10.0423 | 10.134 | -0.091869766 | 0.016430826 | 10.086344 | 9.9888398 | 10.1973 | -0.208482949 | -0.110978692 |
| Ptpn3 | 6.162923 | 6.042009 | 6.0339 | 0.008143195 | 0.129057874 | 5.9057285 | 5.6006363 | 5.80917 | -0.208531122 | 0.096561171 |
| Gamt | 4.083062 | 3.980368 | 4.0797 | -0.099315249 | 0.003378036 | 4.3480475 | 3.706696 | 3.91524 | -0.208539873 | 0.432811561 |
| Ighe | 2.786219 | 2.889547 | 3.1543 | -0.26471283 | -0.368040797 | 3.4227591 | 2.8641726 | 3.07274 | -0.208562887 | 0.350023656 |
| Zfyve28 | 4.843293 | 4.829758 | 5.1192 | -0.289460371 | -0.275925258 | 5.0022021 | 4.8641335 | 5.07271 | -0.208576493 | -0.070507901 |
| Adcy4 | 8.568907 | 8.258142 | 8.1904 | 0.067756668 | 0.378521826 | 7.8590801 | 7.431542 | 7.64027 | -0.208732668 | 0.218805415 |
| Hps5 | 9.87405 | 9.791391 | 9.8511 | -0.059737851 | 0.022920646 | 9.9571881 | 9.8694833 | 10.0783 | -0.208839562 | -0.121134757 |
| Zrsr1 | 8.696287 | 8.568085 | 8.6907 | -0.122590786 | 0.005611689 | 8.7400453 | 8.5380576 | 8.74691 | -0.20885267 | -0.006865007 |
| Cog6 | 8.738559 | 8.700832 | 8.7363 | -0.035468672 | 0.002258461 | 8.8015084 | 8.5823723 | 8.79127 | -0.208902204 | 0.010233953 |
| Ms4a2 | 4.153639 | 4.091439 | 4.0372 | 0.054249939 | 0.116449901 | 3.9047033 | 3.9507523 | 4.15967 | -0.208921803 | -0.254970808 |
| Vash1 | 4.850585 | 4.699434 | 4.7492 | -0.049805554 | 0.101345372 | 4.3770636 | 4.2897371 | 4.49879 | -0.209049757 | -0.121723242 |
| Ubn1 | 11.12832 | 11.15135 | 11.166 | -0.014814744 | -0.037840558 | 11.031166 | 10.989794 | 11.199 | -0.209156564 | -0.167784599 |
| Haghl | 7.110366 | 6.788748 | 7.1783 | -0.389538045 | -0.067919876 | 7.4575198 | 6.9632901 | 7.17263 | -0.209342228 | 0.284887487 |
| Fbln1 | 4.84383 | 4.919088 | 4.862 | 0.057122755 | -0.018135664 | 4.3924303 | 4.486318 | 4.69574 | -0.209417014 | -0.303304744 |
| Cerk | 10.9869 | 11.09145 | 11.061 | 0.030004936 | -0.074542184 | 11.095224 | 10.835748 | 11.0452 | -0.20949067 | 0.0499856 |
| Xrcc4 | 7.913052 | 7.900725 | 7.8798 | 0.020958839 | 0.033285769 | 7.9110724 | 7.8360434 | 8.04559 | -0.209549072 | -0.134520069 |

| | | | | | | | | | | |
|---|---|---|---|---|---|---|---|---|---|---|
| Trmu | 7.390052 | 7.420764 | 7.659 | -0.238257346 | -0.268969198 | 8.0999146 | 7.7789143 | 7.98859 | -0.209672492 | 0.111327819 |
| Zc4h2 | 6.332445 | 6.388967 | 6.3678 | 0.021152294 | -0.035370256 | 6.3631459 | 6.0655527 | 6.27531 | -0.209757715 | 0.087835484 |
| Unc79 | 2.602607 | 2.490257 | 2.473 | 0.01727296 | 0.129623454 | 2.4855307 | 2.4537649 | 2.66361 | -0.209840292 | -0.178074494 |
| Dbt | 8.585606 | 8.522697 | 8.589 | -0.06634065 | -0.003431474 | 8.55176 | 8.3756287 | 8.5855 | -0.209868136 | -0.033736841 |
| Golga3 | 10.37311 | 10.4473 | 10.37 | 0.07718431 | 0.002995136 | 10.315804 | 10.272443 | 10.4825 | -0.210024036 | -0.166663793 |
| Tuft1 | 6.457194 | 6.051889 | 6.0659 | -0.014043124 | 0.391262416 | 6.2203615 | 6.1340444 | 6.34412 | -0.210076848 | -0.12375976 |
| Olfr535 | 1.817858 | 1.823992 | 1.8141 | 0.009917781 | 0.003784742 | 1.866868 | 1.7912286 | 2.00136 | -0.210127478 | -0.134488077 |
| Flt3l | 9.517641 | 9.134694 | 9.4034 | -0.268692332 | 0.11425467 | 9.6394177 | 9.3060928 | 9.51633 | -0.21023822 | 0.123086695 |
| Gm6206 | 2.187508 | 2.561921 | 2.2917 | 0.270191912 | -0.104220213 | 2.2028291 | 2.0480985 | 2.25835 | -0.210251698 | -0.055521093 |
| Ppp3ca | 11.54119 | 11.34789 | 11.476 | -0.127998574 | 0.065298212 | 11.667722 | 11.255579 | 11.4659 | -0.210291782 | 0.201850726 |
| Olfr859 | 3.722228 | 3.769479 | 3.6823 | 0.08713809 | 0.039887659 | 3.7906293 | 3.6922745 | 3.90274 | -0.210467415 | -0.112112692 |
| Borcs5 | 7.346658 | 7.311555 | 7.3906 | -0.079075124 | -0.043972372 | 7.5890636 | 7.1663817 | 7.37694 | -0.210555857 | 0.2121261 |
| R3hcc1 | 7.925409 | 7.929427 | 7.9279 | 0.001528483 | -0.002488724 | 7.9802622 | 7.761789 | 7.97236 | -0.210566522 | 0.007906634 |
| Cstad | 2.446908 | 2.392502 | 2.5001 | -0.107557155 | -0.053151311 | 2.5179724 | 2.3200154 | 2.53062 | -0.210606576 | -0.01264952 |
| Ace2 | 3.041405 | 3.002613 | 3.0544 | -0.051805814 | -0.013014351 | 3.0822837 | 2.8147178 | 3.02534 | -0.210626769 | 0.056939105 |
| Ctsg | 2.847029 | 2.780586 | 2.8772 | -0.096659757 | -0.030216714 | 2.7854138 | 2.6576721 | 2.86845 | -0.210782149 | -0.083040396 |
| Rtkn2 | 2.810685 | 2.701813 | 2.8825 | -0.180664345 | -0.071791613 | 2.6878342 | 2.7144188 | 2.92521 | -0.210790216 | -0.237374759 |
| Olfr1173 | 4.822974 | 4.856841 | 4.8479 | 0.00896753 | -0.024899322 | 4.6929044 | 4.7783896 | 4.9892 | -0.210805828 | -0.296291002 |
| Calm2 | 11.5913 | 11.45622 | 11.592 | -0.135628804 | -0.000549148 | 11.587772 | 11.314149 | 11.525 | -0.210865668 | 0.062757124 |
| Grk6 | 12.65653 | 12.54276 | 12.622 | -0.079450978 | 0.034319716 | 12.876848 | 12.40731 | 12.6182 | -0.210872056 | 0.258666423 |
| Acat1 | 9.900959 | 9.765089 | 9.8511 | -0.085983183 | 0.04988689 | 9.8997623 | 9.5775171 | 9.78849 | -0.210977403 | 0.111267751 |
| Xpo7 | 10.48976 | 10.5113 | 10.566 | -0.054310122 | -0.075855848 | 10.441053 | 10.344622 | 10.5557 | -0.211030798 | -0.11459958 |
| Mical2 | 6.298322 | 6.181931 | 5.9805 | 0.201402493 | 0.317793921 | 5.9689968 | 5.8535956 | 6.06465 | -0.211057191 | -0.095656018 |
| 4930452B06Rik | 2.55182 | 2.434013 | 2.4891 | -0.055117848 | 0.062688386 | 2.6126525 | 2.3625835 | 2.57368 | -0.211092544 | 0.038976424 |
| Dsp | 3.554061 | 3.247055 | 3.4575 | -0.21045862 | 0.096547454 | 3.7963582 | 3.2923118 | 3.50345 | -0.211137937 | 0.292908443 |
| Acadsb | 8.846384 | 8.517564 | 8.5632 | -0.045634666 | 0.283186168 | 8.4840113 | 8.344774 | 8.55592 | -0.211145055 | -0.071907789 |
| Huwe1 | 13.24017 | 13.15263 | 13.201 | -0.048765422 | 0.038772844 | 13.13157 | 13.317854 | 13.529 | -0.211147278 | -0.397431502 |
| Tpgs2 | 7.866685 | 7.950195 | 7.9664 | -0.016178056 | -0.099687949 | 7.9513604 | 7.6677989 | 7.87898 | -0.211186075 | 0.072375415 |
| Alox5 | 4.916666 | 4.447313 | 4.6466 | -0.199284973 | 0.27006786 | 4.6909655 | 4.2482102 | 4.45944 | -0.211227134 | 0.231528174 |
| Ctps2 | 10.26343 | 10.30563 | 10.365 | -0.05911739 | -0.101317784 | 10.284201 | 10.16896 | 10.3802 | -0.211229912 | -0.095988281 |
| Klk8 | 9.05071 | 9.021682 | 9.0734 | -0.05174867 | -0.022720174 | 8.9178237 | 8.8001178 | 9.01136 | -0.21124439 | -0.093538526 |
| Casq2 | 1.310067 | 1.25416 | 1.4148 | -0.160659817 | -0.104752253 | 1.2618426 | 1.232289 | 1.44356 | -0.211275314 | -0.181721638 |

| | | | | | | | | | | |
|---|---|---|---|---|---|---|---|---|---|---|
| Gba2 | 9.696944 | 9.726912 | 9.7987 | -0.071830439 | -0.101797939 | 10.004749 | 9.6873342 | 9.89862 | -0.211283255 | 0.106131677 |
| Dusp8 | 4.443167 | 4.145203 | 4.2674 | -0.122174456 | 0.175789874 | 4.0038554 | 4.106142 | 4.31752 | -0.211375565 | -0.313662113 |
| Stard7 | 10.66571 | 10.69742 | 10.691 | 0.005943936 | -0.02576803 | 10.639241 | 10.487792 | 10.6992 | -0.211377536 | -0.059927898 |
| Kdm6a | 10.77259 | 10.7759 | 10.841 | -0.065019784 | -0.068328512 | 10.799597 | 10.729653 | 10.9411 | -0.211417962 | -0.141473967 |
| Crtam | 7.821271 | 7.834539 | 7.9831 | -0.148569801 | -0.161838284 | 7.816679 | 7.7643427 | 7.97584 | -0.211493782 | -0.159157506 |
| Smyd3 | 8.310361 | 8.24809 | 8.2212 | 0.026896309 | 0.089167063 | 8.1852967 | 8.0611076 | 8.27266 | -0.211555723 | -0.087366588 |
| Trav8d-1 | 3.72315 | 3.570888 | 3.7197 | -0.148854267 | 0.003407598 | 3.433169 | 3.4697471 | 3.68136 | -0.211608531 | -0.248186616 |
| Ahr | 8.579919 | 8.124572 | 8.0716 | 0.052931489 | 0.508279055 | 7.8561638 | 7.8948533 | 8.10658 | -0.211723155 | -0.250412623 |
| Sapcd1 | 3.837434 | 3.638409 | 3.7644 | -0.125975848 | 0.073048934 | 3.7916131 | 3.517619 | 3.72935 | -0.211726364 | 0.062267768 |
| Irf2bpl | 9.748619 | 9.686043 | 9.7717 | -0.085663936 | -0.023088148 | 9.628145 | 9.2769323 | 9.48867 | -0.211734016 | 0.139478765 |
| Trim46 | 4.974317 | 5.037088 | 4.8834 | 0.15369727 | 0.090925963 | 4.8797578 | 4.855809 | 5.06765 | -0.211843208 | -0.187894388 |
| Ighv1-19 | 7.648145 | 7.387865 | 7.3215 | 0.06638699 | 0.32666612 | 7.6124471 | 7.2305182 | 7.44243 | -0.211908369 | 0.170020556 |
| Usp11 | 9.609847 | 9.650633 | 9.6561 | -0.005490222 | -0.046275676 | 9.7848164 | 9.6873806 | 9.8993 | -0.211917844 | -0.114482045 |
| Bscl2 | 9.450774 | 9.335916 | 9.567 | -0.231120748 | -0.116262145 | 9.709451 | 9.3020622 | 9.51402 | -0.211957075 | 0.195431675 |
| Gm10260 | 3.323222 | 3.389893 | 3.4507 | -0.060785903 | -0.127457032 | 3.3144151 | 3.2788944 | 3.49086 | -0.211961369 | -0.176440654 |
| Tmeff1 | 3.188237 | 3.030644 | 3.2127 | -0.182079824 | -0.024487286 | 3.0882099 | 3.0516494 | 3.26378 | -0.212131564 | -0.175570984 |
| Gm10693 | 3.828668 | 3.687206 | 3.8494 | -0.162233077 | -0.020771904 | 3.9049093 | 3.6957101 | 3.90791 | -0.21220199 | -0.003002815 |
| Atf2 | 10.12953 | 10.06858 | 10.177 | -0.108548218 | -0.047598648 | 9.9790586 | 9.9240365 | 10.1364 | -0.212384981 | -0.157362935 |
| Atrn | 9.395986 | 9.460914 | 9.5013 | -0.040426849 | -0.105354812 | 9.3253814 | 9.3029609 | 9.5155 | -0.21253607 | -0.190115664 |
| Zfp12 | 8.675149 | 8.67477 | 8.7725 | -0.097732273 | -0.097352528 | 8.7240285 | 8.6148388 | 8.82739 | -0.212553405 | -0.103363721 |
| Clcn4 | 10.63329 | 10.64379 | 10.677 | -0.03338863 | -0.04388809 | 10.654302 | 10.505368 | 10.718 | -0.212642822 | -0.063709025 |
| Shisa8 | 3.181695 | 3.148056 | 3.3086 | -0.16058742 | -0.126949025 | 3.2484285 | 2.996574 | 3.20926 | -0.212683276 | 0.039171174 |
| Spag9 | 11.26753 | 11.40073 | 11.304 | 0.097060794 | -0.036140432 | 11.23371 | 11.207809 | 11.4205 | -0.21269776 | -0.186797523 |
| Glcci1 | 8.991277 | 8.981294 | 9.0895 | -0.108174372 | -0.098192018 | 8.9454762 | 8.8930874 | 9.10586 | -0.212768177 | -0.160379371 |
| Ldhb | 7.391744 | 7.317273 | 7.4839 | -0.166634616 | -0.092162996 | 7.1158959 | 7.0581906 | 7.27097 | -0.21278018 | -0.155074859 |
| Npat | 9.068311 | 9.067514 | 9.1359 | -0.068392548 | -0.067595373 | 8.8763607 | 9.0230897 | 9.23589 | -0.212798838 | -0.359527855 |
| Shb | 6.401705 | 6.205657 | 6.3854 | -0.179728922 | 0.016319374 | 6.0952843 | 5.9849534 | 6.19783 | -0.212876814 | -0.102545893 |
| Depdc7 | 4.331798 | 4.390804 | 4.3866 | 0.004233017 | -0.054772813 | 4.3607767 | 4.1960196 | 4.4089 | -0.212884744 | -0.048127652 |
| Commd8 | 10.29393 | 10.26795 | 10.293 | -0.02524 | 0.000740386 | 10.220628 | 10.091693 | 10.3047 | -0.212960917 | -0.08402623 |
| Trav10n | 3.332357 | 3.337785 | 3.3059 | 0.031931825 | 0.026504338 | 3.2530263 | 3.2423618 | 3.45534 | -0.21297824 | -0.202313683 |
| Pcm1 | 10.52776 | 10.53819 | 10.573 | -0.034583392 | -0.045011482 | 10.331635 | 10.327238 | 10.5402 | -0.212995072 | -0.208598303 |
| Mapk14 | 10.93782 | 10.9857 | 10.919 | 0.066458066 | 0.018572754 | 10.907796 | 10.726902 | 10.9402 | -0.21331447 | -0.0324208 |

| | | | | | | | | | | |
|---|---|---|---|---|---|---|---|---|---|---|
| Tbc1d15 | 10.44605 | 10.43957 | 10.504 | -0.06415711 | -0.057677186 | 10.352616 | 10.260987 | 10.4744 | -0.213365256 | -0.12173593 |
| Gpr162 | 5.134179 | 4.974323 | 5.1037 | -0.129382309 | 0.030474425 | 4.9735853 | 4.9373375 | 5.15078 | -0.213438289 | -0.177190575 |
| Vwa1 | 7.69226 | 7.158355 | 7.3381 | -0.179762312 | 0.354142158 | 7.0063536 | 6.5563005 | 6.76975 | -0.213450547 | 0.236602615 |
| Pard6b | 5.346879 | 5.364699 | 5.4163 | -0.051592698 | -0.069413395 | 5.4531797 | 5.0862643 | 5.29976 | -0.213492649 | 0.153422662 |
| Pitpnm3 | 3.117899 | 2.813445 | 2.9842 | -0.170705472 | 0.133748845 | 2.7175658 | 2.6521448 | 2.86566 | -0.213516649 | -0.14809563 |
| Gpr132 | 10.69398 | 10.81129 | 10.784 | 0.027424704 | -0.089879724 | 10.79565 | 10.760952 | 10.9746 | -0.21360965 | -0.178912197 |
| Stk38l | 7.764925 | 7.870373 | 7.8117 | 0.058711875 | -0.046736267 | 7.7648441 | 7.7038956 | 7.91753 | -0.213633357 | -0.152684886 |
| Limch1 | 4.364169 | 4.255003 | 4.4746 | -0.219571967 | -0.110406102 | 4.143997 | 4.1045227 | 4.31821 | -0.2136875 | -0.174213198 |
| Nenf | 6.292687 | 5.939787 | 6.1673 | -0.227516445 | 0.125383507 | 6.3303967 | 6.0150307 | 6.22873 | -0.213703206 | 0.101662786 |
| Smpd2 | 8.020668 | 7.913118 | 7.9927 | -0.079590512 | 0.02795928 | 8.3376641 | 7.9213352 | 8.13509 | -0.213758194 | 0.202570706 |
| Gm8935 | 3.893408 | 4.138344 | 4.0021 | 0.136274221 | -0.108662008 | 3.8514723 | 3.8441291 | 4.05791 | -0.213783802 | -0.206440621 |
| Adamdec1 | 3.21295 | 3.058369 | 3.1693 | -0.110897384 | 0.043683581 | 2.9165562 | 2.773418 | 2.98721 | -0.213794377 | -0.070656184 |
| Slc6a17 | 3.500248 | 3.551271 | 3.3707 | 0.180540083 | 0.129517048 | 3.3577068 | 3.3964279 | 3.61023 | -0.213800449 | -0.252521568 |
| Dnmt1 | 11.40538 | 11.42813 | 11.451 | -0.022851968 | -0.045606348 | 11.433627 | 11.34347 | 11.5573 | -0.21385787 | -0.123700443 |
| Neu3 | 7.061415 | 6.974002 | 7.1885 | -0.214521328 | -0.127108363 | 7.1313497 | 7.0175594 | 7.23142 | -0.213860962 | -0.100070729 |
| Wdr7 | 9.251023 | 9.174357 | 9.2039 | -0.029527424 | 0.047139145 | 9.1696331 | 8.9683543 | 9.18265 | -0.214294309 | -0.013015561 |
| Rdh1 | 4.202949 | 4.054678 | 4.1193 | -0.064665426 | 0.083606512 | 4.1010793 | 3.8859311 | 4.10025 | -0.214314988 | 0.000833206 |
| Vkorc1l1 | 8.797206 | 8.76233 | 8.9168 | -0.15442887 | -0.119552884 | 8.6700548 | 8.625059 | 8.8395 | -0.214443689 | -0.169447883 |
| Trav14d-2 | 4.752365 | 4.668031 | 4.7273 | -0.059249711 | 0.025083601 | 4.5131292 | 4.4758966 | 4.69042 | -0.214520316 | -0.177287685 |
| Trav14n-2 | 4.752365 | 4.668031 | 4.7273 | -0.059249711 | 0.025083601 | 4.5131292 | 4.4758966 | 4.69042 | -0.214520316 | -0.177287685 |
| Mfsd9 | 4.566717 | 4.434368 | 4.6309 | -0.196500913 | -0.064152549 | 4.4341303 | 4.2675713 | 4.48217 | -0.214602351 | -0.048043336 |
| Ube2q2 | 9.359358 | 9.380901 | 9.375 | 0.005901471 | -0.015641961 | 9.2410002 | 9.0936714 | 9.30828 | -0.214613544 | -0.067284742 |
| Ephb4 | 7.586684 | 7.260909 | 7.3863 | -0.125380451 | 0.200395107 | 6.6376397 | 6.5726136 | 6.78724 | -0.214627137 | -0.14960109 |
| Ncoa4 | 11.24866 | 11.28043 | 11.261 | 0.019247716 | -0.012520674 | 11.074958 | 10.910791 | 11.1255 | -0.214665746 | -0.050498896 |
| Ntpcr | 7.651886 | 7.265239 | 7.6104 | -0.34516247 | 0.041484537 | 7.7015361 | 7.3109063 | 7.52574 | -0.214829737 | 0.175800074 |
| Tor4a | 10.28758 | 10.26834 | 10.355 | -0.086560722 | -0.067321754 | 10.49541 | 10.075552 | 10.2906 | -0.21500836 | 0.204849919 |
| Rps27a-ps2 | 2.683035 | 2.426404 | 2.5941 | -0.167691785 | 0.088938488 | 2.524862 | 2.5141218 | 2.72913 | -0.215011834 | -0.204271681 |
| 9530077C05Rik | 3.084612 | 3.031281 | 3.0488 | -0.017532066 | 0.03579914 | 2.9834667 | 2.9353439 | 3.15049 | -0.215145453 | -0.167022687 |
| Pdik1l | 8.200038 | 8.259519 | 8.3036 | -0.044091472 | -0.103572388 | 8.1804082 | 7.9809614 | 8.19626 | -0.21529968 | -0.015852939 |
| Cyp20a1 | 7.758685 | 7.615973 | 7.7496 | -0.133631324 | 0.009080812 | 7.7811705 | 7.4737362 | 7.6892 | -0.215461864 | 0.09197244 |
| Zfp770 | 7.332267 | 7.422626 | 7.3851 | 0.037533508 | -0.052825461 | 7.6256776 | 7.3717129 | 7.58733 | -0.215621715 | 0.038342952 |
| Isca1 | 9.52719 | 9.478624 | 9.5039 | -0.025270073 | 0.023295937 | 9.5586837 | 9.2531824 | 9.46881 | -0.215628976 | 0.089872334 |

| | | | | | | | | | | |
|---|---|---|---|---|---|---|---|---|---|---|
| Pdzd8 | 10.16902 | 10.13885 | 10.173 | -0.033696485 | -0.003528225 | 9.9969938 | 9.9471668 | 10.163 | -0.215796828 | -0.165969876 |
| Spag4 | 3.38028 | 3.514363 | 3.4057 | 0.108626634 | -0.025456369 | 3.5669022 | 3.3739024 | 3.58971 | -0.215808716 | -0.022808912 |
| Zfp36l2 | 12.72832 | 12.72167 | 12.742 | -0.020427734 | -0.013784746 | 12.05953 | 12.220559 | 12.4364 | -0.215877588 | -0.376907033 |
| Dusp10 | 8.613488 | 8.59698 | 8.7145 | -0.117533312 | -0.101025533 | 8.412307 | 8.6156069 | 8.83157 | -0.215961465 | -0.419261368 |
| Lcp1 | 14.85252 | 14.84673 | 14.788 | 0.058738066 | 0.064536376 | 15.055414 | 14.617041 | 14.833 | -0.215976522 | 0.222396259 |
| Gm43110 | 4.082201 | 3.83566 | 4.0575 | -0.221796735 | 0.024744941 | 4.1349917 | 3.9297602 | 4.1458 | -0.216039846 | -0.010808283 |
| Rb1cc1 | 10.20012 | 10.22881 | 10.249 | -0.019754966 | -0.048442291 | 9.9964135 | 10.034443 | 10.2507 | -0.216239128 | -0.254269075 |
| Atxn10 | 10.17105 | 10.16113 | 10.213 | -0.051403551 | -0.041485772 | 10.194468 | 9.9806237 | 10.197 | -0.216411156 | -0.002566557 |
| Serhl | 7.701632 | 7.362571 | 7.6989 | -0.336371077 | 0.002689239 | 7.7969376 | 7.5511974 | 7.76768 | -0.216481358 | 0.029258881 |
| Itsn2 | 12.50388 | 12.53153 | 12.516 | 0.01557735 | -0.012071682 | 12.535191 | 12.474154 | 12.6908 | -0.216605816 | -0.1555685 |
| Csgalnact2 | 9.073725 | 9.056225 | 8.986 | 0.070191898 | 0.087692351 | 8.867277 | 8.6985342 | 8.91529 | -0.216760247 | -0.048017435 |
| Smc1a | 11.7064 | 11.57794 | 11.676 | -0.097722786 | 0.030736316 | 11.648862 | 11.565457 | 11.7825 | -0.217015878 | -0.133610316 |
| Rps6-ps3 | 3.019386 | 2.963747 | 3.0474 | -0.083651645 | -0.028012946 | 2.9103695 | 3.0228149 | 3.23985 | -0.217036967 | -0.329482368 |
| Rhbdd1 | 8.998676 | 8.91231 | 8.9958 | -0.083510389 | 0.002854928 | 9.1531166 | 8.9458723 | 9.16294 | -0.217068697 | -0.009824338 |
| Asxl2 | 10.82375 | 10.85197 | 10.88 | -0.027932422 | -0.056155808 | 10.639603 | 10.713541 | 10.9306 | -0.217076548 | -0.291014898 |
| Abcb7 | 10.30827 | 10.35972 | 10.29 | 0.069785868 | 0.018336034 | 10.30491 | 10.033331 | 10.2505 | -0.217199904 | 0.054378933 |
| Tdrd9 | 2.650581 | 2.608071 | 2.6406 | -0.032548953 | 0.009961334 | 2.7279824 | 2.4043326 | 2.62157 | -0.217235254 | 0.106414514 |
| Arfgef2 | 10.69263 | 10.66364 | 10.693 | -0.029552722 | -0.00055854 | 10.759774 | 10.479349 | 10.6966 | -0.21724756 | 0.063177722 |
| Wee1 | 6.930346 | 6.70674 | 7.0466 | -0.339843936 | -0.116238287 | 6.7051177 | 6.7639838 | 6.98128 | -0.217297669 | -0.276163761 |
| Pcyox1l | 5.66195 | 5.858531 | 5.7604 | 0.098091205 | -0.098490104 | 5.8890882 | 5.6251746 | 5.84252 | -0.217342469 | 0.046571114 |
| Diaph2 | 9.29461 | 9.162092 | 9.1427 | 0.019375462 | 0.15189338 | 9.0646375 | 9.0176545 | 9.235 | -0.217347224 | -0.170364275 |
| Cpq | 7.922722 | 7.430055 | 7.8703 | -0.440257492 | 0.052409065 | 6.9381735 | 7.1285307 | 7.34614 | -0.217604633 | -0.407961863 |
| Cdc27 | 10.06573 | 10.04571 | 10.043 | 0.003032092 | 0.023046297 | 10.007673 | 9.8788449 | 10.0966 | -0.217788157 | -0.088959615 |
| Zfp97 | 8.364514 | 8.357463 | 8.491 | -0.133533978 | -0.126482784 | 8.4845104 | 8.2587117 | 8.47652 | -0.217809528 | 0.00798914 |
| Hoga1 | 3.049208 | 2.839037 | 2.9705 | -0.131481883 | 0.078688757 | 2.8189887 | 2.717223 | 2.93508 | -0.217856285 | -0.116090607 |
| Fbh1 | 9.726023 | 9.625992 | 9.8007 | -0.174684789 | -0.074653494 | 9.7215027 | 9.7063416 | 9.92436 | -0.218014063 | -0.202852932 |
| Hivep2 | 11.24044 | 11.38778 | 11.254 | 0.133693714 | -0.013643312 | 11.066651 | 11.196954 | 11.415 | -0.218050266 | -0.34835293 |
| Sema6b | 6.915363 | 6.446212 | 6.4668 | -0.020561098 | 0.448589634 | 6.0453687 | 5.6708512 | 5.88902 | -0.218165083 | 0.15635251 |
| Oscp1 | 4.087229 | 4.125356 | 4.192 | -0.066614467 | -0.104741094 | 4.2064022 | 3.9702824 | 4.18851 | -0.21822813 | 0.017891699 |
| Adat2 | 6.038132 | 5.946055 | 6.1618 | -0.215720454 | -0.123643332 | 6.2073022 | 5.9145103 | 6.1328 | -0.218284877 | 0.074507083 |
| Myo5b | 4.652151 | 4.480185 | 4.2551 | 0.225127944 | 0.397094453 | 4.3874029 | 4.2985572 | 4.51686 | -0.218306173 | -0.129460495 |
| Enpp3 | 2.668655 | 2.506368 | 2.6807 | -0.174292471 | -0.012006062 | 2.6471446 | 2.4791913 | 2.69754 | -0.218344104 | -0.050390864 |

| | | | | | | | | | | |
|---|---|---|---|---|---|---|---|---|---|---|
| Neil1 | 8.404505 | 8.456472 | 8.4428 | 0.013674468 | -0.038292311 | 8.5853102 | 8.4617389 | 8.68009 | -0.218346386 | -0.09477507 |
| 39326 | 11.89514 | 11.86796 | 11.914 | -0.046050928 | -0.018876418 | 11.804398 | 11.699614 | 11.918 | -0.218400664 | -0.113616551 |
| Deup1 | 2.656691 | 2.556207 | 2.8125 | -0.2562898 | -0.155805597 | 2.5421025 | 2.4716484 | 2.69006 | -0.218409581 | -0.147955525 |
| Pithd1 | 8.649167 | 8.618176 | 8.6649 | -0.046753545 | -0.015762525 | 8.7695592 | 8.444645 | 8.6631 | -0.21845936 | 0.106454842 |
| Trub1 | 7.764902 | 7.808104 | 7.7676 | 0.040456869 | -0.002745026 | 8.0030319 | 7.7578948 | 7.97648 | -0.218589383 | 0.026547644 |
| Zfp82 | 7.289384 | 7.370171 | 7.3486 | 0.021564121 | -0.059222905 | 7.4407533 | 7.2378919 | 7.45649 | -0.21860032 | -0.015738907 |
| 39508 | 6.960478 | 6.533494 | 7.0986 | -0.565078322 | -0.138094511 | 6.6381366 | 6.2810416 | 6.49966 | -0.218617464 | 0.138477601 |
| Dennd1c | 10.75564 | 10.71453 | 10.836 | -0.121073806 | -0.07996448 | 11.242031 | 11.038277 | 11.2569 | -0.218627258 | -0.014873916 |
| Phlpp2 | 8.821646 | 8.878313 | 8.8986 | -0.020255847 | -0.076923425 | 8.7622533 | 8.6747631 | 8.89339 | -0.218629855 | -0.131139613 |
| Zfp458 | 5.743376 | 5.558936 | 5.844 | -0.285045625 | -0.10060538 | 5.7379396 | 5.6525224 | 5.87121 | -0.218687234 | -0.133270052 |
| Ipo8 | 10.23773 | 10.17067 | 10.216 | -0.045050945 | 0.021999751 | 10.275765 | 10.070027 | 10.2888 | -0.21880099 | -0.013063217 |
| Vps41 | 10.16598 | 10.10966 | 10.252 | -0.141835346 | -0.085524506 | 10.164822 | 10.022531 | 10.2413 | -0.218804513 | -0.076512721 |
| Alkbh8 | 9.252866 | 9.078542 | 9.1074 | -0.028853769 | 0.14546983 | 8.9551655 | 8.9056803 | 9.1246 | -0.218917596 | -0.169432411 |
| Pcdhgb4 | 4.040842 | 3.860538 | 4.1713 | -0.310761106 | -0.130457237 | 3.9609522 | 3.9388337 | 4.15778 | -0.218950293 | -0.196831749 |
| Pcdhga6 | 3.197691 | 3.196937 | 3.1651 | 0.031882389 | 0.032636916 | 3.1911248 | 2.9183674 | 3.13733 | -0.218961223 | 0.053796117 |
| Tnnt2 | 4.368168 | 4.493856 | 4.5914 | -0.097579305 | -0.223266798 | 4.2507096 | 4.4622625 | 4.68124 | -0.218976362 | -0.430529249 |
| Gtf2h3 | 8.749559 | 8.772818 | 8.7316 | 0.041170415 | 0.017912064 | 9.0509301 | 8.7916575 | 9.01064 | -0.218984585 | 0.040288014 |
| Kbtbd8 | 7.484114 | 7.681493 | 7.6948 | -0.013291682 | -0.210670339 | 7.4084649 | 7.3846788 | 7.60372 | -0.219039078 | -0.195253004 |
| Sec22c | 8.231494 | 8.323422 | 8.3291 | -0.005714388 | -0.097642447 | 8.3126835 | 8.044526 | 8.26358 | -0.219052837 | 0.049104668 |
| Mlx | 9.646032 | 9.507966 | 9.5436 | -0.035644345 | 0.102421808 | 9.8127297 | 9.4585739 | 9.67766 | -0.219087046 | 0.135068722 |
| D7Ertd443e | 2.262227 | 2.44856 | 2.3835 | 0.065085231 | -0.121247465 | 2.3596867 | 2.3201875 | 2.53937 | -0.21918593 | -0.179686658 |
| 2610044O15Rik8 | 8.655317 | 8.835742 | 8.7883 | 0.047466149 | -0.132958529 | 9.0934974 | 8.6696194 | 8.88884 | -0.219216683 | 0.204661284 |
| Tvp23b | 8.635713 | 8.502638 | 8.6252 | -0.122519756 | 0.010554717 | 8.5574294 | 8.4662857 | 8.68572 | -0.219436901 | -0.12829316 |
| Kdm3b | 11.45774 | 11.71486 | 11.767 | -0.052368618 | -0.309487718 | 11.604238 | 11.497275 | 11.7168 | -0.219506642 | -0.112544314 |
| Kcnj16 | 4.515674 | 4.341055 | 4.8937 | -0.552663394 | -0.378044097 | 4.2768876 | 4.0996076 | 4.31914 | -0.219528086 | -0.042248061 |
| P3h3 | 5.188861 | 5.089824 | 5.1686 | -0.078767275 | 0.020269451 | 4.9539946 | 4.9998246 | 5.21945 | -0.219625634 | -0.265455586 |
| Rec8 | 4.221265 | 4.263022 | 4.3449 | -0.081876838 | -0.123634402 | 4.0502571 | 3.9998769 | 4.21959 | -0.219709104 | -0.169328863 |
| Zfp623 | 6.747275 | 6.788039 | 6.8494 | -0.061373186 | -0.102137147 | 6.6516055 | 6.5335126 | 6.75342 | -0.219904707 | -0.101811841 |
| Il21 | 2.88705 | 2.739803 | 2.9705 | -0.23074154 | -0.083494 | 2.5887982 | 2.6686096 | 2.8886 | -0.219988702 | -0.299800117 |
| Trav16 | 5.291937 | 5.372282 | 5.3416 | 0.030696357 | -0.049648783 | 5.2144913 | 5.1424691 | 5.36252 | -0.220050961 | -0.148028728 |
| Il17rd | 4.678572 | 4.620426 | 4.8711 | -0.250720025 | -0.192574201 | 4.5201955 | 4.4473476 | 4.66743 | -0.220079284 | -0.147231426 |
| Ikzf5 | 9.739327 | 9.68236 | 9.7106 | -0.028204029 | 0.028762242 | 9.6897254 | 9.5405878 | 9.7607 | -0.2201094 | -0.070971829 |

| | | | | | | | | | | |
|---|---|---|---|---|---|---|---|---|---|---|
| Wasf1 | 4.025744 | 3.297909 | 3.8925 | -0.59455072 | 0.133285169 | 3.3871711 | 3.2444779 | 3.46459 | -0.220114863 | -0.077421706 |
| Tnrc6b | 11.4416 | 11.55325 | 11.45 | 0.102892994 | -0.008747862 | 11.277666 | 11.554801 | 11.7749 | -0.22012307 | -0.497257954 |
| Per3 | 6.505048 | 6.171248 | 6.5459 | -0.374653808 | -0.040854552 | 6.1343714 | 6.0514974 | 6.27172 | -0.220221595 | -0.137347648 |
| Acvr2b | 2.945567 | 3.010538 | 2.8917 | 0.118867493 | 0.053896523 | 2.9400573 | 2.8670956 | 3.0874 | -0.220309405 | -0.147347693 |
| Phf2 | 10.60058 | 10.5614 | 10.604 | -0.042568228 | -0.00338099 | 10.516056 | 10.33723 | 10.5576 | -0.220346882 | -0.041520285 |
| H2-Ke6 | 9.660616 | 9.659051 | 9.7725 | -0.11342666 | -0.111862427 | 10.080619 | 9.630509 | 9.85088 | -0.220367094 | 0.229742646 |
| Perp | 3.395377 | 3.206135 | 3.351 | -0.144830451 | 0.04441085 | 3.601175 | 3.2288872 | 3.44937 | -0.220481182 | 0.151806627 |
| Eda2r | 3.756508 | 3.833173 | 3.8455 | -0.012362164 | -0.089027364 | 3.6460615 | 3.4822562 | 3.70281 | -0.220550352 | -0.056745041 |
| Acoxl | 4.463567 | 4.477401 | 4.56 | -0.082625358 | -0.096458858 | 4.2513549 | 4.2838257 | 4.50442 | -0.22058989 | -0.253060713 |
| Mbtps2 | 8.757175 | 8.794363 | 8.9144 | -0.119994269 | -0.157182284 | 8.7922376 | 8.6474983 | 8.86813 | -0.220627186 | -0.075887954 |
| Noxo1 | 5.779949 | 5.37938 | 5.774 | -0.394608335 | 0.005961217 | 5.4387774 | 5.2034107 | 5.42416 | -0.220746173 | 0.01462049 |
| Rprd1b | 9.756356 | 9.698295 | 9.6994 | -0.001149787 | 0.05691127 | 9.6559853 | 9.4894631 | 9.71042 | -0.220955819 | -0.054433579 |
| Zfp939 | 4.411352 | 4.383365 | 4.4246 | -0.041234864 | -0.013247963 | 4.5343348 | 4.3404691 | 4.56144 | -0.220973029 | -0.027107386 |
| Meis3 | 7.415951 | 7.455224 | 7.5114 | -0.056149695 | -0.095422195 | 7.384401 | 7.2746978 | 7.4957 | -0.221000284 | -0.111297066 |
| Mex3d | 6.601549 | 6.627488 | 6.6551 | -0.027580244 | -0.053518903 | 6.6855429 | 6.3366184 | 6.55766 | -0.221042043 | 0.127882466 |
| Clasp2 | 9.970253 | 9.895584 | 9.981 | -0.085393912 | -0.010724973 | 9.808971 | 9.723034 | 9.94408 | -0.221050447 | -0.135113437 |
| Pdha1 | 11.19048 | 11.08009 | 11.174 | -0.09348394 | 0.016901618 | 11.074647 | 10.928799 | 11.1499 | -0.221073118 | -0.075224641 |
| Plagl1 | 4.810628 | 4.65097 | 4.9183 | -0.267367249 | -0.107709106 | 4.6823618 | 4.812595 | 5.03368 | -0.22108877 | -0.351321979 |
| Clec4a4 | 3.143371 | 3.185547 | 3.2754 | -0.089813011 | -0.131989148 | 3.4925298 | 3.0473086 | 3.26848 | -0.221174306 | 0.224046969 |
| Gsta4 | 3.765684 | 3.808645 | 3.6741 | 0.134587444 | 0.091626275 | 3.6978716 | 3.5463821 | 3.76761 | -0.221229812 | -0.069740317 |
| Mbp | 11.12633 | 11.19455 | 11.177 | 0.017628784 | -0.05058619 | 10.891755 | 10.88448 | 11.1058 | -0.221278014 | -0.214003045 |
| Triqk | 3.048475 | 2.783878 | 3.0506 | -0.26673503 | -0.00213778 | 2.8034603 | 2.696094 | 2.91743 | -0.221338569 | -0.113972257 |
| Stc1 | 7.128203 | 6.456066 | 6.4567 | -0.000654371 | 0.671482537 | 6.1665598 | 6.1983603 | 6.41972 | -0.221364606 | -0.253165085 |
| Rasa3 | 12.36713 | 12.41409 | 12.399 | 0.015411542 | -0.031551046 | 12.488179 | 12.200869 | 12.4223 | -0.221390146 | 0.065919171 |
| Anapc2 | 10.46137 | 10.42644 | 10.466 | -0.039651382 | -0.004719564 | 10.671922 | 10.488126 | 10.7096 | -0.221431278 | -0.037635001 |
| Ctnnd2 | 5.07477 | 4.681409 | 4.8941 | -0.212718303 | 0.180643401 | 5.1805429 | 4.8575948 | 5.07905 | -0.221451202 | 0.101496861 |
| Hist1h1a | 3.433361 | 3.388026 | 3.5716 | -0.183570002 | -0.138235145 | 3.5273482 | 3.3535321 | 3.57519 | -0.221653062 | -0.047836975 |
| Cbarp | 7.19302 | 6.920243 | 6.9424 | -0.022197205 | 0.250579574 | 6.7864965 | 6.7357554 | 6.95749 | -0.221730936 | -0.170989806 |
| Pfkfb1 | 5.448414 | 5.419842 | 5.5529 | -0.133062279 | -0.104490893 | 5.5582584 | 5.6084867 | 5.83036 | -0.221870406 | -0.272098763 |
| Hectd1 | 12.02958 | 11.98298 | 12.019 | -0.036201866 | 0.010393368 | 11.752913 | 11.825077 | 12.047 | -0.221957196 | -0.294121297 |
| Nat8l | 3.742759 | 3.820156 | 3.6843 | 0.135850119 | 0.058453021 | 3.9660813 | 3.5147384 | 3.7367 | -0.221965598 | 0.229377258 |
| Klc4 | 7.913939 | 7.985493 | 8.1126 | -0.127060732 | -0.198614494 | 8.1774002 | 7.8755766 | 8.09764 | -0.222067402 | 0.079756139 |

| | | | | | | | | | | |
|---|---|---|---|---|---|---|---|---|---|---|
| Elovl6 | 8.065769 | 7.610422 | 7.5583 | 0.052133135 | 0.50748079 | 7.5706739 | 7.5246743 | 7.74674 | -0.222068787 | -0.176069144 |
| Mmgt2 | 7.718259 | 7.572205 | 7.7158 | -0.143555167 | 0.002498193 | 7.9189651 | 7.5227609 | 7.74484 | -0.222082232 | 0.174121937 |
| Slc25a4 | 10.49979 | 10.40245 | 10.554 | -0.151574578 | -0.054232944 | 10.514068 | 10.276206 | 10.4983 | -0.222097854 | 0.015763618 |
| Ctcf | 11.82741 | 11.86635 | 11.876 | -0.009298822 | -0.048238484 | 11.850319 | 11.689583 | 11.9117 | -0.222124216 | -0.061388067 |
| P2rx2 | 2.057346 | 2.004433 | 2.1349 | -0.130468806 | -0.077554931 | 1.9936526 | 1.9056226 | 2.12784 | -0.222220905 | -0.134190877 |
| Traj12 | 3.221557 | 3.103007 | 3.1225 | -0.0195045 | 0.099045615 | 2.9982612 | 2.9797851 | 3.20217 | -0.222388779 | -0.203912714 |
| Insig2 | 8.301317 | 8.297347 | 8.3528 | -0.055453785 | -0.051484744 | 8.2920442 | 8.1801442 | 8.40255 | -0.222401701 | -0.110501692 |
| Pdcd4 | 11.93828 | 11.99883 | 12.083 | -0.08450362 | -0.14505194 | 11.338666 | 11.480771 | 11.7033 | -0.22252753 | -0.364631898 |
| Gm45062 | 10.56578 | 10.67162 | 10.697 | -0.02553965 | -0.131388094 | 10.53099 | 10.379579 | 10.6022 | -0.22266949 | -0.071259054 |
| Napg | 9.584376 | 9.66663 | 9.5761 | 0.090567182 | 0.008313231 | 9.6149852 | 9.6112275 | 9.83392 | -0.222689781 | -0.218932064 |
| Zfp507 | 9.441057 | 9.406362 | 9.4603 | -0.053975361 | -0.019280582 | 9.0333653 | 9.4607514 | 9.6835 | -0.22274589 | -0.650131982 |
| Bmp1 | 6.148007 | 6.136827 | 6.2022 | -0.065399635 | -0.054219404 | 5.8284898 | 5.7808862 | 6.00377 | -0.222879449 | -0.175275838 |
| Lcn4 | 6.305575 | 6.297118 | 6.3926 | -0.095518166 | -0.087061599 | 6.3911226 | 6.2159094 | 6.43891 | -0.223003677 | -0.047790421 |
| E330021D16Rik | 2.612679 | 2.523561 | 2.5838 | -0.060260003 | 0.02885789 | 2.5202092 | 2.4258369 | 2.64888 | -0.223040225 | -0.128667894 |
| Arrdc2 | 6.811757 | 6.75823 | 6.8528 | -0.09461457 | -0.041087509 | 7.0609609 | 6.759249 | 6.98231 | -0.223057298 | 0.07865467 |
| Acta2 | 7.603022 | 7.871017 | 7.7136 | 0.157376988 | -0.110618036 | 8.0552191 | 7.3186782 | 7.5418 | -0.223119354 | 0.513421619 |
| Fam131a | 5.335607 | 5.106297 | 5.4034 | -0.297118949 | -0.067808759 | 5.2532686 | 4.9457901 | 5.16892 | -0.223126753 | 0.084351738 |
| Ntn4 | 5.352243 | 5.0288 | 5.0851 | -0.05625594 | 0.267186656 | 4.6980689 | 4.6625186 | 4.88568 | -0.223160335 | -0.187609997 |
| Pcgf1 | 7.053287 | 6.989355 | 7.2147 | -0.22537328 | -0.161441506 | 7.3868017 | 7.1374337 | 7.36067 | -0.223236316 | 0.026131651 |
| Zc2hc1a | 5.50733 | 5.533321 | 5.6855 | -0.152200064 | -0.178190878 | 5.5082598 | 5.5234552 | 5.74674 | -0.223282174 | -0.238477514 |
| Ptbp2 | 9.061516 | 8.797268 | 8.9876 | -0.190335514 | 0.073912089 | 9.1273638 | 9.0173139 | 9.24074 | -0.223424493 | -0.113374602 |
| Igsf9b | 4.569112 | 4.376859 | 4.5466 | -0.169710815 | 0.022542632 | 4.5179121 | 4.437683 | 4.66125 | -0.223566542 | -0.143337484 |
| Ttll3 | 7.267644 | 7.278886 | 7.3729 | -0.094037427 | -0.105279375 | 7.8501957 | 7.5979553 | 7.82162 | -0.223667606 | 0.028572771 |
| Tmem132a | 5.260827 | 5.39236 | 5.2955 | 0.096850613 | -0.034682363 | 5.297229 | 4.8530835 | 5.07676 | -0.223678462 | 0.220467106 |
| Hist1h2ac | 4.747775 | 4.749797 | 4.9176 | -0.16781035 | -0.169832461 | 5.1110098 | 4.8584081 | 5.08211 | -0.223700522 | 0.028901123 |
| Ap1b1 | 11.46414 | 11.33925 | 11.414 | -0.07479202 | 0.050100596 | 11.461272 | 11.113902 | 11.3376 | -0.22372607 | 0.123644129 |
| Kif26b | 3.916104 | 3.982045 | 4.0746 | -0.092524042 | -0.158465335 | 4.0387282 | 3.9569363 | 4.18069 | -0.22374892 | -0.141957049 |
| Olfr374 | 5.141671 | 5.109406 | 5.5245 | -0.41508777 | -0.382823463 | 5.3629482 | 5.2346714 | 5.45848 | -0.223804275 | -0.095527527 |
| Rdh16f1 | 3.722547 | 3.515162 | 3.7099 | -0.19469448 | 0.012690826 | 3.5333952 | 3.2547419 | 3.47869 | -0.223943884 | 0.054709397 |
| Amt | 5.312704 | 5.200226 | 5.4233 | -0.223062719 | -0.110584858 | 5.3487741 | 5.1023837 | 5.32636 | -0.223977463 | 0.022412964 |
| Itpripl1 | 8.427781 | 8.333017 | 8.5904 | -0.257417836 | -0.162654318 | 8.4733482 | 8.1570202 | 8.381 | -0.223978433 | 0.092349543 |
| Parn | 9.586898 | 9.566491 | 9.5973 | -0.030849945 | -0.010442946 | 9.677847 | 9.4444613 | 9.6687 | -0.224235665 | 0.009150055 |

| | | | | | | | | | | |
|---|---|---|---|---|---|---|---|---|---|---|
| Tbc1d5 | 10.04576 | 10.14959 | 10.03 | 0.119971426 | 0.016144205 | 10.172291 | 9.9255647 | 10.15 | -0.224449659 | 0.022276739 |
| 1810010H24Rik | 2.90937 | 2.794908 | 2.9756 | -0.180741869 | -0.066279246 | 3.0780116 | 2.7215962 | 2.94605 | -0.224453015 | 0.131962457 |
| Ppp6r2 | 9.492812 | 9.5088 | 9.6106 | -0.101800919 | -0.11778959 | 9.8122113 | 9.4562151 | 9.68067 | -0.224453065 | 0.131543131 |
| Msantd4 | 8.506761 | 8.452589 | 8.5817 | -0.129101824 | -0.074930557 | 8.3574522 | 8.2721528 | 8.49663 | -0.224481514 | -0.139182114 |
| Cdk17 | 10.54943 | 10.70392 | 10.616 | 0.087965402 | -0.066523622 | 10.348308 | 10.404097 | 10.6287 | -0.224572286 | -0.280361539 |
| Neurl2 | 5.257803 | 4.981387 | 5.262 | -0.280658115 | -0.004242343 | 5.4160697 | 4.9190513 | 5.14366 | -0.224612674 | 0.272405676 |
| Dusp14 | 3.878606 | 3.893304 | 3.9692 | -0.075914713 | -0.090612738 | 3.9318489 | 3.9893319 | 4.21415 | -0.224814627 | -0.282297626 |
| Entpd4b | 8.326292 | 8.249291 | 8.4836 | -0.234335166 | -0.157333975 | 8.5440918 | 8.2328936 | 8.45794 | -0.225043686 | 0.086154527 |
| Stat5b | 11.14491 | 11.14354 | 11.184 | -0.040096776 | -0.038729848 | 11.160331 | 10.897254 | 11.1223 | -0.225064208 | 0.038012553 |
| Eya1 | 6.058359 | 6.278427 | 6.0881 | 0.190288347 | -0.029779605 | 5.9134787 | 6.0914311 | 6.31663 | -0.22520328 | -0.403155692 |
| Cyp4f18 | 10.01796 | 10.01759 | 9.9938 | 0.023746931 | 0.024114471 | 10.490356 | 9.7364339 | 9.96192 | -0.225482661 | 0.528439588 |
| Ncs1 | 3.987278 | 4.228555 | 3.8936 | 0.334972625 | 0.093695647 | 3.8288769 | 3.6731288 | 3.89873 | -0.225599236 | -0.069851224 |
| Gm19168 | 3.56167 | 3.517864 | 3.6176 | -0.099743764 | -0.055937328 | 3.8584527 | 3.5747784 | 3.80044 | -0.225662341 | 0.058011947 |
| Sh2b2 | 7.280766 | 7.104758 | 6.8639 | 0.24088064 | 0.41688833 | 6.728273 | 6.5504576 | 6.77622 | -0.225759356 | -0.047943892 |
| Kcnip2 | 3.702778 | 3.756442 | 3.9355 | -0.179062636 | -0.232726214 | 3.6666369 | 3.6634298 | 3.88924 | -0.225812507 | -0.222605392 |
| 2610301B20Rik | 6.775738 | 6.75242 | 6.8706 | -0.118204788 | -0.094886653 | 6.936018 | 6.6502949 | 6.87619 | -0.225898424 | 0.05982471 |
| Mtfr1 | 8.876078 | 8.896056 | 8.9506 | -0.0544979 | -0.074475493 | 8.7884313 | 8.6413539 | 8.86727 | -0.225919355 | -0.0788419 |
| Armh4 | 2.92694 | 2.912818 | 2.671 | 0.241854922 | 0.255976301 | 2.8944413 | 2.8123615 | 3.0384 | -0.226042055 | -0.143962254 |
| Fbxo3 | 9.55465 | 9.453685 | 9.6299 | -0.176202979 | -0.075237551 | 9.5825557 | 9.3741513 | 9.60024 | -0.226086734 | -0.017682262 |
| Acbd5 | 9.459066 | 9.489153 | 9.5967 | -0.107572745 | -0.137659744 | 9.3704992 | 9.3126239 | 9.53887 | -0.226244785 | -0.168369507 |
| Rbp1 | 5.499956 | 5.316341 | 5.5246 | -0.208245134 | -0.024629719 | 5.1772399 | 4.7906077 | 5.01693 | -0.22631743 | 0.160314763 |
| Sclt1 | 7.850372 | 7.779863 | 7.9166 | -0.136718213 | -0.066209248 | 7.9139676 | 7.7715165 | 7.998 | -0.226485705 | -0.084034593 |
| Cep290 | 6.249716 | 6.135241 | 6.3422 | -0.2069191 | -0.092443994 | 6.4954835 | 6.2277274 | 6.45434 | -0.226611928 | 0.041144186 |
| Bmp6 | 2.916732 | 2.989636 | 3.0639 | -0.074292186 | -0.147195991 | 2.7195407 | 2.754202 | 2.98085 | -0.226650536 | -0.26131177 |
| Pigp | 8.001439 | 7.877575 | 8.0197 | -0.142129254 | -0.018265996 | 7.9907066 | 7.6812719 | 7.90795 | -0.22667481 | 0.082759888 |
| Klhl12 | 8.359631 | 8.286637 | 8.2947 | -0.00807525 | 0.064919242 | 8.3984468 | 8.3019554 | 8.52866 | -0.226704946 | -0.130213542 |
| Trdv4 | 4.729246 | 4.559476 | 4.4204 | 0.139040218 | 0.308810891 | 4.0001546 | 4.1696484 | 4.39635 | -0.226705214 | -0.396199085 |
| Ptpra | 10.49661 | 10.42112 | 10.553 | -0.131808886 | -0.056324692 | 10.408954 | 10.317707 | 10.5444 | -0.22672442 | -0.135477196 |
| Colgalt1 | 10.94414 | 10.96716 | 11.059 | -0.091907352 | -0.1149295 | 11.349578 | 10.877172 | 11.1039 | -0.226732396 | 0.245673516 |
| Dtd1 | 8.163301 | 8.120239 | 8.2329 | -0.112683623 | -0.069620787 | 8.3466272 | 8.0392546 | 8.26607 | -0.226814703 | 0.080557962 |
| Rnf32 | 4.606898 | 4.781399 | 4.8664 | -0.085004158 | -0.259505562 | 4.6984244 | 4.6423748 | 4.86922 | -0.226841688 | -0.170792081 |
| Fkbp7 | 5.028563 | 4.782911 | 5.0032 | -0.220336061 | 0.025316447 | 4.797651 | 4.6080713 | 4.83495 | -0.226876947 | -0.037297315 |

| | | | | | | | | | | |
|---|---|---|---|---|---|---|---|---|---|---|
| Uap1 | 9.712493 | 9.575434 | 9.6784 | -0.102923386 | 0.034135702 | 9.5235888 | 9.4959848 | 9.72291 | -0.226925162 | -0.199321118 |
| Tada2b | 9.313769 | 9.19182 | 9.2814 | -0.089544235 | 0.032404589 | 9.0977759 | 8.8693755 | 9.09631 | -0.226930872 | 0.001469517 |
| Kdm4b | 9.70343 | 9.953387 | 9.7687 | 0.184666944 | -0.065290104 | 9.8796838 | 9.7365494 | 9.96351 | -0.226961086 | -0.083826653 |
| Bik | 3.439104 | 3.30627 | 3.4079 | -0.101581051 | 0.031253257 | 3.2782093 | 3.1847631 | 3.41174 | -0.226980573 | -0.133534341 |
| Pde7a | 10.79799 | 10.90315 | 10.925 | -0.022096834 | -0.127257844 | 11.036574 | 11.005397 | 11.2324 | -0.22704246 | -0.195865752 |
| 2700081O15Rik | 6.843495 | 6.630777 | 6.7598 | -0.129041215 | 0.083676624 | 6.5284163 | 6.3715724 | 6.59864 | -0.227065242 | -0.070221298 |
| Faap20 | 8.822679 | 8.760157 | 8.931 | -0.170842286 | -0.108320921 | 9.1723577 | 8.6451799 | 8.87239 | -0.227208324 | 0.299969487 |
| Fignl2 | 3.801059 | 3.758828 | 3.8183 | -0.059501681 | -0.017270479 | 3.5204858 | 3.4405818 | 3.66779 | -0.227208517 | -0.147304524 |
| 4930522L14Rik | 7.130875 | 6.908325 | 7.1818 | -0.273452638 | -0.050902191 | 7.3167197 | 6.9223302 | 7.14963 | -0.227296577 | 0.167092882 |
| Tgfbr2 | 12.48346 | 12.49603 | 12.506 | -0.01044867 | -0.023018978 | 12.350818 | 12.101066 | 12.3284 | -0.227323572 | 0.022427907 |
| Madcam1 | 4.987086 | 4.903401 | 4.7364 | 0.166954438 | 0.250639363 | 4.680282 | 4.6377245 | 4.86509 | -0.227367647 | -0.184810173 |
| BC053393 | 3.963545 | 3.871882 | 3.9782 | -0.106350588 | -0.014687749 | 3.9472213 | 3.6921262 | 3.91953 | -0.227400549 | 0.027694639 |
| Tmem135 | 8.99372 | 9.023197 | 9.0079 | 0.015300642 | -0.014176272 | 8.7444972 | 8.6661181 | 8.89375 | -0.22763133 | -0.149252217 |
| Foxp2 | 4.537667 | 4.492702 | 4.5915 | -0.098799507 | -0.053834567 | 4.6298794 | 4.5401137 | 4.76775 | -0.227638712 | -0.137873017 |
| Spef1 | 5.036804 | 5.135337 | 5.1687 | -0.03333729 | -0.131870144 | 5.2177967 | 4.9777364 | 5.20542 | -0.227681121 | 0.012379112 |
| Polr2a | 12.9469 | 12.97434 | 13.014 | -0.03949283 | -0.06692807 | 13.001345 | 12.816954 | 13.0447 | -0.227784524 | -0.043393203 |
| Lcp2 | 11.24179 | 11.31002 | 11.221 | 0.088867642 | 0.020635904 | 10.982272 | 11.08429 | 11.3122 | -0.227923664 | -0.329941736 |
| Zfp965 | 5.644642 | 5.677712 | 5.6811 | -0.003345411 | -0.036416274 | 5.6412816 | 5.3634855 | 5.59149 | -0.228002965 | 0.049793122 |
| Asrgl1 | 8.718506 | 8.584642 | 8.7198 | -0.135186803 | -0.001322602 | 9.0854945 | 8.6623166 | 8.89047 | -0.228156035 | 0.195021861 |
| Ncor1 | 12.68009 | 12.62823 | 12.647 | -0.018988746 | 0.032871532 | 12.539572 | 12.46412 | 12.6923 | -0.22818814 | -0.152736173 |
| Fbxw8 | 9.212616 | 9.074426 | 9.2034 | -0.128947116 | 0.009242786 | 9.1935095 | 8.9192185 | 9.14753 | -0.228315927 | 0.045975074 |
| Dennd5a | 10.67999 | 10.66003 | 10.737 | -0.077043468 | -0.057081366 | 10.714143 | 10.515604 | 10.7439 | -0.228339108 | -0.029800163 |
| Fam118a | 7.908081 | 7.849718 | 8.0113 | -0.161579933 | -0.103217217 | 8.1403675 | 7.7805494 | 8.0089 | -0.228355549 | 0.131462508 |
| Pcdhga11 | 2.996939 | 2.826597 | 2.8985 | -0.071923513 | 0.098419034 | 2.8066466 | 2.6579537 | 2.88634 | -0.228386737 | -0.079693885 |
| Dgkz | 12.02026 | 12.06846 | 12.187 | -0.118408724 | -0.166615088 | 12.261326 | 12.053156 | 12.2816 | -0.228405424 | -0.020235915 |
| Slc25a27 | 5.540732 | 5.37625 | 5.6164 | -0.240107084 | -0.075625128 | 5.4928511 | 5.5473792 | 5.77587 | -0.228488642 | -0.283016757 |
| B3galnt2 | 9.026931 | 9.132457 | 9.0246 | 0.107806809 | 0.002281514 | 9.1474194 | 8.8540534 | 9.08254 | -0.22849063 | 0.064875364 |
| Zfp386 | 8.552395 | 8.525227 | 8.717 | -0.191744515 | -0.164576885 | 8.6506527 | 8.4864993 | 8.71515 | -0.228646189 | -0.064492766 |
| Ighj3 | 6.37905 | 6.57708 | 6.2535 | 0.323560942 | 0.125530658 | 6.316832 | 6.1743463 | 6.40308 | -0.228732832 | -0.086247205 |
| Mdga2 | 4.888974 | 4.904917 | 4.9027 | 0.002250646 | -0.01369248 | 4.8956598 | 5.0734031 | 5.30215 | -0.228749314 | -0.406492669 |
| Kctd20 | 10.45898 | 10.43157 | 10.466 | -0.034176482 | -0.006766056 | 10.296159 | 10.254881 | 10.4836 | -0.228763396 | -0.187485097 |
| Zmynd11 | 10.90865 | 10.9492 | 11.041 | -0.091764178 | -0.132312096 | 11.019508 | 10.938164 | 11.1669 | -0.228774384 | -0.147429693 |

| | | | | | | | | | | |
|---|---|---|---|---|---|---|---|---|---|---|
| Cd163l1 | 7.825775 | 8.088639 | 7.9723 | 0.116305775 | -0.146558313 | 7.5197274 | 7.6387464 | 7.86753 | -0.228785602 | -0.347804631 |
| Tm6sf1 | 9.905251 | 9.957443 | 9.9557 | 0.001757306 | -0.05043375 | 10.351397 | 9.9357223 | 10.1646 | -0.228845817 | 0.186829053 |
| Calm3 | 11.4075 | 11.31192 | 11.399 | -0.086951548 | 0.008623746 | 11.368593 | 11.082999 | 11.3118 | -0.228848924 | 0.0567455 |
| Gm15443 | 4.67093 | 4.676252 | 4.8145 | -0.138253366 | -0.143575153 | 4.6836716 | 4.5858024 | 4.8148 | -0.229001993 | -0.131132816 |
| Fam205a4 | 2.140144 | 1.96207 | 2.2426 | -0.28048141 | -0.102407511 | 1.9024296 | 1.9275568 | 2.15657 | -0.229011538 | -0.254138802 |
| Sgsm1 | 4.4799 | 4.253417 | 4.3881 | -0.134689595 | 0.091793174 | 3.9909452 | 3.9727514 | 4.20181 | -0.229059022 | -0.210865233 |
| Zfp994 | 8.036446 | 7.984569 | 8.1676 | -0.183064111 | -0.131187407 | 8.0755401 | 7.9837086 | 8.21277 | -0.22906324 | -0.137231694 |
| Lmtk2 | 10.27946 | 10.33662 | 10.307 | 0.029200532 | -0.027966376 | 10.215412 | 10.095465 | 10.3245 | -0.229078914 | -0.109131758 |
| Vmn2r52 | 4.783269 | 4.848191 | 4.9851 | -0.136875377 | -0.201798128 | 4.9180376 | 4.8390626 | 5.06814 | -0.229081139 | -0.15010621 |
| Mccc1 | 8.600533 | 8.596255 | 8.7271 | -0.130805313 | -0.126527117 | 8.6006895 | 8.64689 | 8.87598 | -0.229085898 | -0.275286402 |
| Slc35a3 | 9.77301 | 9.604179 | 9.6963 | -0.092100553 | 0.076730551 | 9.6452895 | 9.516098 | 9.74521 | -0.229107876 | -0.099916314 |
| P3h4 | 4.597836 | 4.247953 | 4.4857 | -0.237710299 | 0.112172185 | 4.3225474 | 3.9600963 | 4.18921 | -0.229115659 | 0.133335475 |
| Gm15280 | 4.070559 | 3.973853 | 4.0204 | -0.046507944 | 0.050197407 | 4.0454365 | 3.934187 | 4.16331 | -0.229125798 | -0.117876327 |
| Usp54 | 7.307255 | 7.258354 | 7.3494 | -0.091084256 | -0.042183231 | 7.1873766 | 7.1094847 | 7.33865 | -0.229163527 | -0.151271683 |
| Acsf3 | 7.768174 | 7.77208 | 7.8259 | -0.053853868 | -0.057760299 | 8.0665914 | 7.665739 | 7.89494 | -0.229202194 | 0.171650209 |
| Atp7a | 8.411332 | 8.277627 | 8.0549 | 0.222699264 | 0.356404286 | 7.9982097 | 8.02113 | 8.25037 | -0.229237422 | -0.252157774 |
| Fbxo30 | 7.863385 | 7.700589 | 7.8044 | -0.103776236 | 0.059019816 | 7.252139 | 7.5326623 | 7.76192 | -0.229255354 | -0.509778585 |
| Scrn2 | 6.969895 | 6.783914 | 6.8169 | -0.033035012 | 0.152946837 | 6.9686501 | 6.6714468 | 6.90102 | -0.229577621 | 0.067625665 |
| Cdc73 | 10.25711 | 10.27591 | 10.23 | 0.04586929 | 0.027072332 | 10.109226 | 10.11822 | 10.3479 | -0.229646848 | -0.23864161 |
| Ifngr2 | 9.81379 | 9.941025 | 10.023 | -0.081871137 | -0.209106175 | 10.112643 | 9.7806166 | 10.0103 | -0.229663284 | 0.102363055 |
| Ggt5 | 8.58304 | 8.182464 | 8.3577 | -0.175260313 | 0.22531574 | 8.2554744 | 8.001416 | 8.23109 | -0.229673466 | 0.024384923 |
| Chaf1b | 7.346636 | 7.150284 | 7.4734 | -0.32308209 | -0.126730138 | 7.4330006 | 7.2436165 | 7.47341 | -0.22979826 | -0.040414108 |
| Zfp518a | 9.345326 | 9.368059 | 9.4071 | -0.038998789 | -0.061731838 | 9.3256771 | 9.0872468 | 9.31719 | -0.229942117 | 0.008488198 |
| Carnmt1 | 8.424565 | 8.398757 | 8.4695 | -0.07073963 | -0.044932267 | 8.3260354 | 8.2269118 | 8.45707 | -0.230158246 | -0.131034609 |
| Txlnb | 4.309378 | 4.473933 | 4.3822 | 0.09177582 | -0.072779353 | 4.2505678 | 4.2597281 | 4.48997 | -0.230245413 | -0.239405659 |
| Acsf2 | 10.41964 | 10.43905 | 10.548 | -0.109374816 | -0.128784194 | 10.935102 | 10.421116 | 10.6514 | -0.230255988 | 0.283730749 |
| Trim13 | 5.746239 | 5.512539 | 5.7143 | -0.201758122 | 0.031942299 | 5.3118062 | 5.4296105 | 5.6599 | -0.230287249 | -0.348091601 |
| Gpd2 | 9.948169 | 9.8111 | 9.9406 | -0.129533056 | 0.00753552 | 10.033403 | 9.7671 | 9.99739 | -0.23029092 | 0.036012056 |
| Camsap1 | 8.979877 | 9.024211 | 8.9105 | 0.113678304 | 0.069344015 | 8.9159446 | 8.7751795 | 9.00553 | -0.230352507 | -0.089587383 |
| Zfp605 | 6.843551 | 6.691251 | 6.8672 | -0.175928361 | -0.02362803 | 6.644339 | 6.6838528 | 6.91422 | -0.230370516 | -0.26988428 |
| Spast | 9.453872 | 9.413743 | 9.4803 | -0.066590245 | -0.02646119 | 9.2654725 | 9.1887594 | 9.41916 | -0.230395694 | -0.153682656 |
| Rtel1 | 9.615611 | 9.561416 | 9.6537 | -0.092302835 | -0.038108256 | 10.162984 | 9.8964949 | 10.1269 | -0.230409626 | 0.0360796 |

| | | | | | | | | | | |
|---|---|---|---|---|---|---|---|---|---|---|
| Dnajc1 | 9.634021 | 9.559135 | 9.4703 | 0.088853125 | 0.16373886 | 9.381393 | 9.4155418 | 9.64604 | -0.23049365 | -0.264642482 |
| Tnnt1 | 4.485241 | 4.493876 | 4.5942 | -0.10028218 | -0.108917816 | 4.5829074 | 4.3946661 | 4.62516 | -0.230495581 | -0.042254313 |
| Tmtc3 | 6.947997 | 6.79834 | 6.859 | -0.060697325 | 0.088959822 | 6.1978943 | 6.2036372 | 6.43417 | -0.230537461 | -0.236280361 |
| Egfl6 | 3.282772 | 3.326156 | 3.4442 | -0.118057556 | -0.161441963 | 3.3994012 | 3.1461398 | 3.37673 | -0.230590546 | 0.022670825 |
| Trabd2b | 3.568633 | 3.541475 | 3.5592 | -0.017735832 | 0.009421488 | 3.5314606 | 3.3173644 | 3.54798 | -0.230613012 | -0.016516862 |
| Tcf19 | 7.406667 | 7.142672 | 7.5022 | -0.359485469 | -0.095490315 | 7.4168172 | 7.4267745 | 7.65756 | -0.230782751 | -0.240740029 |
| Rad51d | 8.604098 | 8.735633 | 8.7715 | -0.035854788 | -0.167389482 | 8.7384985 | 8.4599366 | 8.69073 | -0.230793249 | 0.047768664 |
| Donson | 9.270058 | 9.262188 | 9.2843 | -0.022100482 | -0.014230949 | 9.4112383 | 9.2860288 | 9.51684 | -0.230814312 | -0.105604808 |
| Ppp3r1 | 11.25385 | 11.1969 | 11.255 | -0.057965354 | -0.001020026 | 11.156844 | 10.892023 | 11.1229 | -0.230862572 | 0.033958456 |
| BC017158 | 8.342461 | 8.168314 | 8.3376 | -0.169249543 | 0.004896952 | 8.5530561 | 8.2388128 | 8.46971 | -0.230897323 | 0.083346056 |
| Gm42688 | 7.593774 | 7.812711 | 7.8554 | -0.042731141 | -0.261667659 | 7.8166493 | 7.5482836 | 7.77927 | -0.230982101 | 0.037383678 |
| Dysf | 5.994794 | 6.240209 | 5.7858 | 0.454436554 | 0.209021868 | 6.0096968 | 5.6557742 | 5.88679 | -0.23101674 | 0.122905834 |
| Fbxo25 | 7.737844 | 7.852966 | 7.8649 | -0.011940309 | -0.127062787 | 7.7584933 | 7.7001331 | 7.93133 | -0.231201861 | -0.17284161 |
| Fnip1 | 9.209662 | 9.182175 | 9.3132 | -0.131065122 | -0.103577729 | 9.1285127 | 9.1222166 | 9.35347 | -0.231253059 | -0.224956955 |
| Trip13 | 6.313144 | 5.982293 | 6.2044 | -0.222089999 | 0.10876098 | 6.3795365 | 6.0005197 | 6.23177 | -0.23125505 | 0.147761764 |
| Wdr5b | 7.544753 | 7.284111 | 7.4616 | -0.177487947 | 0.083154488 | 7.8901588 | 7.4020851 | 7.63336 | -0.23127596 | 0.256797717 |
| Zfp760 | 6.091802 | 6.093362 | 6.2701 | -0.176781909 | -0.178342114 | 6.4587608 | 6.2705815 | 6.50186 | -0.231279665 | -0.043100355 |
| Slc6a6 | 10.99595 | 10.989 | 11.031 | -0.042478182 | -0.03552506 | 10.910922 | 10.867309 | 11.0987 | -0.231433896 | -0.187820801 |
| Map6 | 4.511645 | 4.497317 | 4.4889 | 0.008373096 | 0.022700925 | 4.159958 | 4.3541025 | 4.58556 | -0.231453141 | -0.425597602 |
| Prkcg | 2.477679 | 2.427422 | 2.4702 | -0.042742566 | 0.007514414 | 2.4641087 | 2.3887447 | 2.62031 | -0.231567984 | -0.156203921 |
| Mmaa | 6.95693 | 7.025652 | 7.0949 | -0.069220002 | -0.137941953 | 7.2276481 | 7.042438 | 7.27412 | -0.231680895 | -0.046470855 |
| Igkv8-24 | 7.53992 | 8.465157 | 7.5726 | 0.892596405 | -0.032640066 | 8.0373555 | 8.1340791 | 8.36579 | -0.231707568 | -0.328431188 |
| Akr1c12 | 3.645569 | 3.596233 | 3.7044 | -0.108156542 | -0.058820775 | 3.4622042 | 3.3333316 | 3.56505 | -0.231717226 | -0.102844676 |
| Wnt5a | 2.438675 | 2.557812 | 2.5228 | 0.035003801 | -0.084132922 | 2.5239856 | 2.2642031 | 2.49593 | -0.231725995 | 0.028056427 |
| Pms1 | 7.983033 | 8.0005 | 8.0613 | -0.060791211 | -0.078258276 | 8.017513 | 7.8195014 | 8.0513 | -0.231797767 | -0.033786157 |
| Micu1 | 9.377438 | 9.334429 | 9.4363 | -0.101894883 | -0.05888628 | 9.5067231 | 9.2866109 | 9.51848 | -0.231869239 | -0.011757059 |
| Syn3 | 2.774334 | 2.890293 | 2.8611 | 0.029242624 | -0.086716193 | 2.9374944 | 2.8326655 | 3.06456 | -0.231898568 | -0.127069663 |
| Ints10 | 8.838299 | 8.940901 | 8.9203 | 0.020643689 | -0.081958601 | 9.1035068 | 8.8059699 | 9.03799 | -0.232019653 | 0.0655173 |
| Trim32 | 7.537665 | 7.619875 | 7.6042 | 0.015723429 | -0.066486448 | 7.4402399 | 7.2262038 | 7.45835 | -0.232145552 | -0.018109471 |
| Vegfb | 6.734989 | 6.743874 | 6.8511 | -0.107206089 | -0.116090582 | 6.8736197 | 6.5202143 | 6.75238 | -0.232165708 | 0.12123972 |
| Spen | 10.51901 | 10.53929 | 10.47 | 0.069519572 | 0.049243604 | 10.290361 | 10.366981 | 10.5993 | -0.232269978 | -0.308890292 |
| Fzd6 | 4.709204 | 4.37972 | 4.6498 | -0.270110401 | 0.059373707 | 4.2676711 | 4.2050951 | 4.43739 | -0.232292012 | -0.16971597 |

| | | | | | | | | | | |
|---|---|---|---|---|---|---|---|---|---|---|
| mt-Nd1 | 15.04021 | 14.88114 | 15.094 | -0.212744038 | -0.053665898 | 14.934949 | 14.838478 | 15.0708 | -0.232314478 | -0.135843148 |
| Mphosph8 | 9.675968 | 9.623177 | 9.7033 | -0.080157916 | -0.027366945 | 9.5913695 | 9.4834718 | 9.71582 | -0.232348942 | -0.124451261 |
| St3gal5 | 6.723446 | 6.614837 | 6.6659 | -0.051068085 | 0.057540766 | 6.918491 | 6.4292159 | 6.6616 | -0.232383519 | 0.256891576 |
| Slc26a10 | 3.600571 | 3.520601 | 3.722 | -0.201386931 | -0.12141681 | 3.4669011 | 3.4513277 | 3.68378 | -0.232454872 | -0.216881418 |
| A930004D18Rik | 2.344238 | 2.310561 | 2.388 | -0.077448754 | -0.043772171 | 2.4386292 | 2.2693349 | 2.50185 | -0.232513135 | -0.063218829 |
| Lsm11 | 7.71402 | 7.738014 | 7.8149 | -0.076845419 | -0.100838655 | 7.8474964 | 7.4951918 | 7.72778 | -0.23259267 | 0.119711935 |
| Ankrd23 | 7.119602 | 7.433135 | 7.4977 | -0.064527191 | -0.378059421 | 7.7235674 | 7.4700037 | 7.70266 | -0.232657011 | 0.020906759 |
| Wls | 9.36495 | 9.191715 | 9.387 | -0.195250403 | -0.022015046 | 9.0881891 | 8.9778127 | 9.21051 | -0.232696183 | -0.12231975 |
| Ebf1 | 11.43417 | 11.64162 | 11.516 | 0.125911626 | -0.081538564 | 11.739113 | 11.331638 | 11.5644 | -0.23272412 | 0.174751094 |
| Gm26528 | 1.961068 | 1.96732 | 2.0003 | -0.033017932 | -0.039269871 | 1.9396476 | 1.9388742 | 2.17162 | -0.232743303 | -0.231969887 |
| Zfp870 | 6.931968 | 6.827751 | 7.062 | -0.234223715 | -0.130006931 | 6.947605 | 6.8196302 | 7.05252 | -0.232885133 | -0.104910348 |
| Arhgap44 | 3.628913 | 3.47183 | 3.6533 | -0.181439409 | -0.02435639 | 3.3865515 | 3.3204649 | 3.55336 | -0.232891914 | -0.166805295 |
| Gorasp1 | 7.258673 | 7.085548 | 7.362 | -0.276487951 | -0.103363087 | 7.5584438 | 7.0036491 | 7.23654 | -0.232892125 | 0.321902651 |
| Kank4 | 2.254542 | 2.325506 | 2.346 | -0.020490452 | -0.091453978 | 2.2439995 | 2.2039364 | 2.43683 | -0.23289367 | -0.192830573 |
| Creg1 | 8.489387 | 8.477921 | 8.5229 | -0.044943631 | -0.033477546 | 8.2389838 | 8.0159968 | 8.24892 | -0.232920774 | -0.009933779 |
| Fbxo40 | 3.015838 | 3.056628 | 3.0624 | -0.005766509 | -0.046556592 | 3.2518082 | 2.9565876 | 3.18973 | -0.233144481 | 0.062076209 |
| Nr2c1 | 7.402703 | 7.330117 | 7.4437 | -0.113601233 | -0.041015159 | 7.2300725 | 7.1387172 | 7.37187 | -0.233149633 | -0.141794268 |
| Wtip | 4.037915 | 3.87084 | 4.1216 | -0.250797606 | -0.08372255 | 3.81159 | 3.6955098 | 3.92875 | -0.233237754 | -0.117157547 |
| Fnbp1 | 12.28835 | 12.23574 | 12.246 | -0.010611276 | 0.041992614 | 12.130307 | 12.168692 | 12.4019 | -0.233257014 | -0.271641367 |
| Gm47860 | 3.004529 | 3.037142 | 3.0807 | -0.043601758 | -0.076214196 | 3.0405273 | 3.1121575 | 3.34552 | -0.233363897 | -0.304994082 |
| Rbm12 | 9.444874 | 9.370017 | 9.3571 | 0.012945142 | 0.087802275 | 9.3450949 | 9.2635688 | 9.49697 | -0.233398629 | -0.151872524 |
| E330009J07Rik | 8.98512 | 8.988991 | 9.0626 | -0.073593206 | -0.077463653 | 9.0148398 | 8.8220842 | 9.0555 | -0.233410813 | -0.040655229 |
| Tnfaip8 | 10.90679 | 11.00432 | 10.929 | 0.074876412 | -0.02264908 | 10.997978 | 10.686218 | 10.9197 | -0.233475694 | 0.078284336 |
| Nr2c2 | 10.9813 | 11.00547 | 11.085 | -0.079115828 | -0.10329437 | 10.943434 | 10.913564 | 11.1471 | -0.23353189 | -0.203662486 |
| C530008M17Rik | 3.913847 | 4.066012 | 4.0762 | -0.010194086 | -0.162358928 | 4.0221251 | 3.8807596 | 4.11433 | -0.233574114 | -0.092208637 |
| Orc1 | 4.899333 | 5.008443 | 4.9462 | 0.06227342 | -0.046836877 | 5.092941 | 5.1133355 | 5.34691 | -0.233575653 | -0.253970113 |
| Grb7 | 4.41175 | 4.459263 | 4.5401 | -0.080865138 | -0.128378584 | 4.760203 | 4.3003698 | 4.53397 | -0.233601625 | 0.226231661 |
| Bcl11b | 10.83379 | 10.82904 | 10.897 | -0.067579926 | -0.06283121 | 10.339861 | 10.507223 | 10.7408 | -0.23362582 | -0.400987057 |
| Wdr95 | 4.381224 | 4.210121 | 4.3257 | -0.115552787 | 0.055550072 | 4.4160652 | 4.1486864 | 4.38232 | -0.233638454 | 0.033740371 |
| Gatad2b | 11.26462 | 11.37725 | 11.315 | 0.06254505 | -0.050083638 | 11.212924 | 11.10384 | 11.3375 | -0.233673912 | -0.12458955 |
| Cep19 | 6.725559 | 6.699009 | 6.7369 | -0.037908396 | -0.011358576 | 6.5764922 | 6.5341086 | 6.76796 | -0.233854164 | -0.191470645 |
| B630019K06Rik | 4.138078 | 4.062839 | 4.1849 | -0.122091861 | -0.046853556 | 4.0192159 | 3.8678101 | 4.10167 | -0.233857462 | -0.082451658 |

| | | | | | | | | | | |
|---|---|---|---|---|---|---|---|---|---|---|
| Armh3 | 8.543127 | 8.534957 | 8.562 | -0.027040968 | -0.018870842 | 8.6404534 | 8.4130275 | 8.6469 | -0.233875277 | -0.006449432 |
| Zfp850 | 7.820511 | 7.835856 | 7.9279 | -0.092054006 | -0.107398453 | 7.6377049 | 7.6119558 | 7.84591 | -0.233951533 | -0.208202457 |
| Klhl15 | 7.814571 | 7.565708 | 7.753 | -0.187336939 | 0.061525604 | 7.6963856 | 7.5589449 | 7.79303 | -0.23408327 | -0.096642576 |
| Svip | 4.049056 | 3.945925 | 4.1353 | -0.189377516 | -0.08624644 | 3.9989544 | 3.8482965 | 4.08239 | -0.234090589 | -0.083432733 |
| Fdxacb1 | 6.964919 | 7.156853 | 7.2504 | -0.093562268 | -0.285495782 | 7.1272962 | 6.8364945 | 7.07068 | -0.23418689 | 0.056614749 |
| E130311K13Rik | 6.064795 | 5.960916 | 6.2043 | -0.243402 | -0.139523424 | 6.0405757 | 5.9433482 | 6.17757 | -0.234218306 | -0.136990843 |
| Olfr804 | 3.525291 | 3.424866 | 3.5842 | -0.159291611 | -0.058866996 | 3.6049807 | 3.5159585 | 3.75021 | -0.234254117 | -0.145231873 |
| Yipf2 | 7.628567 | 7.539259 | 7.6697 | -0.130392517 | -0.04108427 | 7.8404668 | 7.4633964 | 7.69778 | -0.234387774 | 0.142682571 |
| Dach1 | 3.108266 | 3.149861 | 3.3113 | -0.161455397 | -0.203050988 | 2.9581973 | 2.8399179 | 3.07435 | -0.234427851 | -0.116148415 |
| Ccdc157 | 6.867047 | 6.800036 | 6.8939 | -0.09389775 | -0.026887152 | 7.0723553 | 6.6492912 | 6.88379 | -0.234500101 | 0.188564042 |
| Trav13d-1 | 3.982067 | 3.756568 | 3.9548 | -0.19820908 | 0.027290275 | 3.7601961 | 3.7200707 | 3.95458 | -0.234506087 | -0.194380618 |
| Sec24a | 9.263997 | 9.219211 | 9.2698 | -0.050620171 | -0.005834112 | 9.1512911 | 9.1355833 | 9.37013 | -0.234545124 | -0.218837327 |
| Pnpla8 | 9.560499 | 9.605744 | 9.5525 | 0.053278089 | 0.008033109 | 9.3680322 | 9.3172626 | 9.55184 | -0.234579391 | -0.183809805 |
| Ttc16 | 2.40524 | 2.339404 | 2.409 | -0.069585584 | -0.003749019 | 2.3837231 | 2.2467998 | 2.48148 | -0.234679577 | -0.097756261 |
| Sema4b | 10.47303 | 10.57006 | 10.443 | 0.127064862 | 0.030040702 | 10.677694 | 10.159906 | 10.3947 | -0.234785252 | 0.283002593 |
| Spaca1 | 3.783318 | 3.895667 | 3.8915 | 0.00418023 | -0.108169454 | 3.9853439 | 3.7241721 | 3.95905 | -0.234881909 | 0.026289945 |
| Kansl1l | 8.251271 | 8.397132 | 8.4546 | -0.057456853 | -0.203317736 | 8.2707294 | 8.3042734 | 8.53916 | -0.234888682 | -0.268432687 |
| Arhgap22 | 7.292395 | 7.668614 | 7.4173 | 0.2512975 | -0.124922126 | 7.5449642 | 7.4648111 | 7.69979 | -0.234976201 | -0.154823131 |
| Tmem8b | 5.747959 | 5.79619 | 5.9689 | -0.172726497 | -0.220957654 | 5.8848832 | 5.6657039 | 5.90069 | -0.234991021 | -0.015811719 |
| Hilpda | 7.475621 | 7.296432 | 7.3612 | -0.064781388 | 0.114407494 | 7.1776323 | 7.1039148 | 7.33913 | -0.235213829 | -0.161496403 |
| Trim23 | 8.95475 | 8.784008 | 8.9375 | -0.153497783 | 0.017243932 | 8.8404621 | 8.6258027 | 8.86108 | -0.235281208 | -0.020621823 |
| Vpreb3 | 8.559799 | 8.616083 | 8.6163 | -0.000187346 | -0.056471819 | 9.1663809 | 8.1541671 | 8.38952 | -0.235355606 | 0.776858217 |
| Kdm5a | 11.87179 | 11.74693 | 11.715 | 0.031517436 | 0.156375138 | 11.67191 | 11.576743 | 11.8125 | -0.235714872 | -0.140547681 |
| Pbx1 | 7.771514 | 7.613214 | 7.8591 | -0.245862465 | -0.087562596 | 7.4242538 | 7.3625575 | 7.59834 | -0.235785825 | -0.174089513 |
| Ppcdc | 9.393964 | 9.39655 | 9.4954 | -0.098884718 | -0.101470672 | 9.999937 | 9.6149802 | 9.85084 | -0.235854896 | 0.149101947 |
| Lgals1 | 10.25751 | 10.34904 | 10.338 | 0.011151022 | -0.080371021 | 10.257459 | 10.224643 | 10.4605 | -0.235889503 | -0.203072917 |
| Ggt7 | 5.068521 | 5.132494 | 5.1437 | -0.011156812 | -0.075129923 | 5.1736231 | 4.9704615 | 5.20642 | -0.235959267 | -0.032797655 |
| Tfrc | 10.15212 | 10.10047 | 10.102 | -0.001243697 | 0.050405241 | 10.1735 | 10.14155 | 10.3776 | -0.236033088 | -0.204083542 |
| Lrrn4cl | 3.838925 | 3.847443 | 3.8204 | 0.027044544 | 0.0185268 | 3.7587745 | 3.6356056 | 3.87166 | -0.236058099 | -0.11288917 |
| Akr1c13 | 6.749148 | 6.675134 | 6.7637 | -0.088518412 | -0.014504206 | 6.4678755 | 6.3750534 | 6.61116 | -0.236110156 | -0.143288017 |
| Adm2 | 2.536753 | 2.493352 | 2.5194 | -0.026055628 | 0.017344877 | 2.4529734 | 2.3909162 | 2.62703 | -0.23611111 | -0.174053938 |
| Rad21 | 11.71714 | 11.72247 | 11.767 | -0.044704754 | -0.050032398 | 11.624496 | 11.509623 | 11.7459 | -0.23626849 | -0.121395601 |

| | | | | | | | | | | |
|---|---|---|---|---|---|---|---|---|---|---|
| Fermt2 | 7.099124 | 6.8394 | 6.7826 | 0.056812869 | 0.316536888 | 6.460053 | 6.2877215 | 6.52417 | -0.23644824 | -0.064116711 |
| Zfp715 | 9.217352 | 9.231318 | 9.2602 | -0.028863522 | -0.042828603 | 9.2337192 | 9.1005881 | 9.33705 | -0.236464016 | -0.103332963 |
| Stmn2 | 4.585046 | 4.371776 | 4.5387 | -0.166901573 | 0.046367512 | 4.4540786 | 4.085043 | 4.32151 | -0.236466018 | 0.132569617 |
| Zfp963 | 6.065791 | 6.017292 | 6.0773 | -0.060014981 | -0.011516058 | 6.0486223 | 6.0298918 | 6.26642 | -0.236529313 | -0.21779878 |
| Cd59a | 4.050964 | 3.928374 | 4.3855 | -0.457091073 | -0.334501087 | 3.8110021 | 3.9089594 | 4.1455 | -0.23653752 | -0.334494858 |
| Nadsyn1 | 7.865106 | 7.977441 | 8.0796 | -0.102202514 | -0.214537857 | 8.3844801 | 8.0259172 | 8.26248 | -0.236560775 | 0.122002187 |
| Gm20449 | 3.837512 | 3.817515 | 3.8561 | -0.038620746 | -0.01862369 | 4.0128933 | 3.8382455 | 4.07489 | -0.236649058 | -0.062001323 |
| Xlr3b | 6.540999 | 5.76591 | 6.585 | -0.819068995 | -0.043980428 | 6.2387349 | 6.0525272 | 6.28922 | -0.236695178 | -0.050487553 |
| Hcst | 9.052421 | 8.986239 | 9.0729 | -0.086685079 | -0.020502619 | 9.1252562 | 8.737715 | 8.97445 | -0.236738999 | 0.150802232 |
| Ercc4 | 8.941187 | 8.881736 | 8.9355 | -0.053802884 | 0.005648329 | 8.9428873 | 8.8391529 | 9.07595 | -0.236794731 | -0.133060307 |
| Kcnmb4 | 5.562973 | 5.439703 | 5.5381 | -0.098400106 | 0.024869816 | 5.2675441 | 5.2586899 | 5.49561 | -0.236920491 | -0.228066243 |
| Sike1 | 9.677457 | 9.637684 | 9.6027 | 0.03494458 | 0.074717633 | 9.7236712 | 9.5407291 | 9.77772 | -0.236987392 | -0.054045303 |
| Trav5-4 | 5.189315 | 5.215854 | 5.236 | -0.020160457 | -0.046699108 | 5.0900866 | 4.9941711 | 5.23126 | -0.237093185 | -0.141177684 |
| Plpbp | 9.603054 | 9.58543 | 9.3581 | 0.227328163 | 0.244951457 | 9.6767713 | 9.446762 | 9.68388 | -0.23711429 | -0.007105005 |
| Rad54b | 5.643432 | 5.352897 | 5.5506 | -0.197669907 | 0.092864728 | 5.3045559 | 5.5383068 | 5.77542 | -0.237114841 | -0.470865698 |
| Dgka | 13.06252 | 13.1459 | 13.152 | -0.005896432 | -0.089282048 | 13.102082 | 13.097785 | 13.3349 | -0.237160116 | -0.232863431 |
| Cldn15 | 3.444993 | 3.461858 | 3.4943 | -0.032484421 | -0.049349645 | 3.490788 | 3.4858601 | 3.72316 | -0.237302465 | -0.232374583 |
| Jade1 | 9.759678 | 9.744991 | 9.8128 | -0.067845355 | -0.053158528 | 9.4795833 | 9.4759441 | 9.71335 | -0.237405242 | -0.233766047 |
| Dtwd2 | 6.839901 | 6.740545 | 6.8076 | -0.067051517 | 0.032304562 | 6.7433368 | 6.5475176 | 6.78498 | -0.237463687 | -0.041644463 |
| Acap2 | 10.81817 | 10.83559 | 10.807 | 0.028551622 | 0.011138574 | 10.632155 | 10.571806 | 10.8093 | -0.237469696 | -0.177121306 |
| Pigc | 7.763539 | 8.13927 | 8.042 | 0.097241531 | -0.278489117 | 8.2232622 | 7.9921682 | 8.2298 | -0.237635779 | -0.006541721 |
| Rhobtb1 | 5.328265 | 5.129391 | 5.058 | 0.071344196 | 0.270218238 | 5.0511602 | 4.8832988 | 5.12096 | -0.237661352 | -0.069799954 |
| Mto1 | 8.273262 | 8.368438 | 8.3538 | 0.014681466 | -0.080494202 | 8.4484597 | 8.3026319 | 8.54035 | -0.237721002 | -0.091893247 |
| Rab40c | 8.469874 | 8.281123 | 8.367 | -0.085882198 | 0.102868906 | 8.3669843 | 8.1725578 | 8.41038 | -0.23782029 | -0.043393823 |
| Dut | 8.559954 | 8.455096 | 8.6228 | -0.167666317 | -0.06280826 | 8.6630453 | 8.5632561 | 8.80112 | -0.237864076 | -0.13807489 |
| Gm14276 | 4.35038 | 4.344195 | 4.4665 | -0.122321608 | -0.116136283 | 4.4742378 | 4.3282828 | 4.56618 | -0.237897984 | -0.091942977 |
| Itga7 | 5.313041 | 5.42965 | 5.5042 | -0.074584048 | -0.191193308 | 5.2862819 | 5.2528072 | 5.49084 | -0.238034934 | -0.204560293 |
| Irgq | 8.713259 | 8.766398 | 8.7642 | 0.00219398 | -0.05094539 | 8.7288495 | 8.5168447 | 8.75489 | -0.238047063 | -0.026042283 |
| Cdc42se2 | 11.80558 | 11.88951 | 11.913 | -0.023974954 | -0.107900934 | 11.806006 | 11.581986 | 11.8201 | -0.23809616 | -0.014075584 |
| Zik1 | 4.993625 | 4.940225 | 5.021 | -0.080734506 | -0.027333812 | 5.1734062 | 4.840414 | 5.07856 | -0.238147638 | 0.094844612 |
| Cp | 8.42538 | 8.247039 | 8.2866 | -0.039539577 | 0.138801793 | 7.7513446 | 7.5728549 | 7.81104 | -0.238184543 | -0.059694928 |
| Ldlrad3 | 6.790851 | 6.589218 | 6.6693 | -0.08007527 | 0.121557798 | 6.6696716 | 6.1981782 | 6.43655 | -0.23836744 | 0.233125981 |

| | | | | | | | | | | |
|---|---|---|---|---|---|---|---|---|---|---|
| Ankrd28 | 9.849532 | 9.498583 | 9.7579 | -0.259273855 | 0.091675084 | 9.7977121 | 9.6417477 | 9.88012 | -0.238376341 | -0.082411939 |
| Acadvl | 9.79905 | 9.71566 | 9.7961 | -0.080403348 | 0.002986182 | 9.9111195 | 9.5549828 | 9.79348 | -0.238498445 | 0.117638272 |
| Lrp2 | 3.65218 | 3.678318 | 3.7839 | -0.105550311 | -0.131688951 | 3.679199 | 3.8051279 | 4.04366 | -0.238527913 | -0.364456766 |
| SPATA24 | 2.415802 | 2.245279 | 2.038 | 0.207230924 | 0.377753934 | 2.1658159 | 2.2596608 | 2.49851 | -0.238849167 | -0.332694095 |
| Zfp646 | 10.92773 | 11.05413 | 11.005 | 0.049530558 | -0.07687469 | 11.020919 | 10.840241 | 11.0791 | -0.23888382 | -0.058206034 |
| Foxp1 | 12.23339 | 12.34711 | 12.339 | 0.008429696 | -0.105294832 | 12.403337 | 12.178016 | 12.4169 | -0.238905754 | -0.013584731 |
| Rptor | 10.00445 | 9.994413 | 10.02 | -0.025252012 | -0.015219885 | 9.9773489 | 9.9627441 | 10.2017 | -0.238924129 | -0.224319332 |
| Tsga10 | 6.409225 | 6.435032 | 6.3851 | 0.049900709 | 0.024093356 | 6.57242 | 6.2883011 | 6.52725 | -0.238953705 | 0.045165188 |
| Haus2 | 8.390304 | 8.059366 | 8.2394 | -0.180005812 | 0.150931637 | 8.1532437 | 7.990815 | 8.22992 | -0.239108554 | -0.076679834 |
| Slc39a4 | 6.563351 | 6.520976 | 6.6173 | -0.09629419 | -0.053918417 | 6.4281045 | 6.3750672 | 6.61422 | -0.239154051 | -0.186116693 |
| Tmem181a | 8.059522 | 8.011553 | 8.1611 | -0.149518504 | -0.101549345 | 8.0799158 | 8.019008 | 8.25817 | -0.239160134 | -0.178252306 |
| Bank1 | 12.45182 | 12.43102 | 12.393 | 0.038134468 | 0.058932848 | 13.041547 | 12.377954 | 12.6171 | -0.239164426 | 0.424428561 |
| Fra10ac1 | 7.832358 | 7.596742 | 7.7934 | -0.196656814 | 0.038959984 | 7.8385822 | 7.7402335 | 7.97945 | -0.239218662 | -0.140869915 |
| Rpgr | 6.300445 | 6.114281 | 6.1528 | -0.038553072 | 0.147610913 | 6.3247159 | 6.0038742 | 6.24317 | -0.239299513 | 0.081542163 |
| Ctbs | 7.400251 | 7.256129 | 7.3883 | -0.132180426 | 0.011941585 | 7.4055801 | 6.9557251 | 7.19504 | -0.239316633 | 0.210538297 |
| Alg14 | 6.686946 | 6.689798 | 6.6627 | 0.02712244 | 0.024270119 | 6.6830124 | 6.6264513 | 6.86577 | -0.239318768 | -0.182757643 |
| Slc39a8 | 5.597761 | 5.300483 | 5.5243 | -0.223791906 | 0.073486022 | 5.1782567 | 5.3351623 | 5.5745 | -0.239340889 | -0.396246513 |
| Itgb4 | 7.101556 | 7.077361 | 7.0961 | -0.01870641 | 0.005488243 | 6.4655759 | 6.4748267 | 6.71421 | -0.239386932 | -0.248637713 |
| Pten | 12.20584 | 12.2356 | 12.196 | 0.039783754 | 0.010024154 | 12.058816 | 11.886584 | 12.126 | -0.239420824 | -0.067189161 |
| Kcnj2 | 5.138805 | 5.162085 | 4.8295 | 0.332625613 | 0.309344791 | 5.3532402 | 4.5991673 | 4.83882 | -0.23965658 | 0.514416364 |
| Lox | 3.410341 | 3.258326 | 3.2938 | -0.035443379 | 0.11657098 | 3.1921651 | 3.0352121 | 3.27507 | -0.239857551 | -0.082904571 |
| Slc43a1 | 5.954888 | 5.874622 | 6.0054 | -0.130748834 | -0.050483328 | 6.2061778 | 5.8428791 | 6.08275 | -0.239873655 | 0.123425054 |
| Bmp8a | 2.339548 | 2.24862 | 2.3675 | -0.118846331 | -0.02791848 | 2.3440466 | 2.2064429 | 2.44639 | -0.239948562 | -0.102344908 |
| Ccdc61 | 7.27867 | 7.276399 | 7.3008 | -0.024385879 | -0.02211523 | 7.671992 | 7.1020596 | 7.34218 | -0.24011649 | 0.329815993 |
| Nek2 | 5.999431 | 5.96758 | 6.0687 | -0.101074182 | -0.069223145 | 5.9259867 | 6.0613231 | 6.30152 | -0.240197729 | -0.375534077 |
| Calu | 11.03692 | 10.90891 | 10.905 | 0.003779564 | 0.131784394 | 10.813008 | 10.579378 | 10.8197 | -0.240312808 | -0.006682133 |
| Tmem254b | 7.535877 | 7.531352 | 7.6018 | -0.070424099 | -0.06589854 | 7.8153402 | 7.2865353 | 7.52687 | -0.240331391 | 0.288473533 |
| Trac | 12.31267 | 12.19902 | 12.353 | -0.15390105 | -0.04025406 | 11.986771 | 12.081312 | 12.3217 | -0.240379546 | -0.334919911 |
| Gm49510 | 6.696463 | 6.494574 | 6.788 | -0.293412394 | -0.091523394 | 6.9839859 | 6.7911698 | 7.03168 | -0.24050777 | -0.047691611 |
| Pisd-ps2 | 6.358447 | 6.440739 | 6.5262 | -0.085427728 | -0.167720192 | 6.7657382 | 6.4276359 | 6.6682 | -0.240562078 | 0.097540226 |
| Rwdd2a | 4.283385 | 4.175663 | 4.3179 | -0.142222555 | -0.034500057 | 4.202304 | 4.1573957 | 4.39812 | -0.24071952 | -0.195811226 |
| Ntrk3 | 7.45024 | 7.285403 | 7.1367 | 0.148684597 | 0.313521557 | 6.9915513 | 7.0286034 | 7.26948 | -0.240874249 | -0.277926324 |

| | | | | | | | | | | |
|---|---|---|---|---|---|---|---|---|---|---|
| Fat1 | 5.6286 | 5.949828 | 5.6751 | 0.274706646 | -0.046520508 | 5.6198139 | 5.2859886 | 5.5269 | -0.240912715 | 0.092912612 |
| Haus1 | 6.571335 | 6.320879 | 6.5206 | -0.199709158 | 0.050747222 | 6.6832853 | 6.4159178 | 6.6569 | -0.240984883 | 0.026382602 |
| Itpkb | 11.62951 | 11.70102 | 11.733 | -0.031990078 | -0.103501224 | 11.505312 | 11.46315 | 11.7042 | -0.241091582 | -0.19892924 |
| Cobl | 3.828423 | 3.809059 | 4.0485 | -0.239438811 | -0.220075463 | 4.1142928 | 3.8445986 | 4.08579 | -0.241196185 | 0.028498029 |
| Rel | 11.68414 | 11.83778 | 11.725 | 0.112391774 | -0.041251822 | 11.599758 | 11.786091 | 12.0273 | -0.241241836 | -0.427574544 |
| Hid1 | 9.211105 | 9.241814 | 9.2572 | -0.015361131 | -0.046070121 | 8.7614105 | 8.8380459 | 9.07935 | -0.241307843 | -0.31794327 |
| Vmn2r96 | 5.041327 | 5.006575 | 5.1508 | -0.144208531 | -0.109457119 | 5.2911339 | 5.0583008 | 5.29963 | -0.241325337 | -0.008492165 |
| Fam204a | 9.126994 | 8.883759 | 9.142 | -0.258287737 | -0.015052867 | 9.0564188 | 8.8756726 | 9.11707 | -0.241392924 | -0.060646707 |
| Rab31 | 8.21538 | 8.086781 | 8.0548 | 0.03201736 | 0.160616798 | 8.244989 | 7.6149582 | 7.8564 | -0.241440111 | 0.388590725 |
| Taf1 | 10.89685 | 10.81384 | 10.889 | -0.075039962 | 0.007967318 | 10.808051 | 10.715757 | 10.9574 | -0.241600754 | -0.149306349 |
| Aamdc | 6.895297 | 6.781804 | 6.9252 | -0.14335953 | -0.02986666 | 7.0446384 | 6.69123 | 6.93292 | -0.241686421 | 0.111722015 |
| Cngb1 | 2.885173 | 2.986102 | 3.118 | -0.131849631 | -0.232778979 | 2.883643 | 2.6742418 | 2.91595 | -0.241711021 | -0.032309864 |
| Gm16523 | 2.963809 | 2.937887 | 3.0049 | -0.067011119 | -0.041089437 | 2.9183465 | 2.9449708 | 3.18682 | -0.241850501 | -0.268474775 |
| Ildr2 | 5.604947 | 5.205511 | 5.2002 | 0.005276171 | 0.404711662 | 4.8070777 | 4.7984493 | 5.0409 | -0.24245382 | -0.23382546 |
| Smim1 | 2.186292 | 2.133995 | 2.2945 | -0.160468007 | -0.108171566 | 2.3016007 | 2.0027558 | 2.2453 | -0.242549216 | 0.056295717 |
| Mfsd6 | 9.077412 | 8.853995 | 8.9885 | -0.13445898 | 0.088957957 | 8.7339403 | 8.7042233 | 8.947 | -0.242778446 | -0.213061456 |
| Dennd6a | 8.89967 | 8.943684 | 8.9981 | -0.054462475 | -0.098475714 | 8.7856317 | 8.581964 | 8.82484 | -0.242878247 | -0.039210536 |
| 2310057M21Rik | 7.587187 | 7.540105 | 7.5741 | -0.033952037 | 0.013130023 | 7.4449681 | 7.2687926 | 7.51173 | -0.242941179 | -0.066765698 |
| Sgip1 | 5.890637 | 5.902297 | 5.793 | 0.109345428 | 0.097685946 | 6.0270692 | 6.0082732 | 6.25127 | -0.243000898 | -0.224204925 |
| Slc18a2 | 3.172148 | 3.084364 | 3.1244 | -0.040000762 | 0.047782688 | 2.9263505 | 2.8827131 | 3.12572 | -0.243011707 | -0.199374234 |
| Zmat3 | 7.855224 | 7.901617 | 7.981 | -0.079352284 | -0.125744947 | 7.7166627 | 7.7098447 | 7.95287 | -0.243027993 | -0.236210021 |
| Fut10 | 4.131319 | 3.953938 | 3.9758 | -0.021833861 | 0.155547049 | 3.9882635 | 3.8364838 | 4.07951 | -0.243028231 | -0.091248561 |
| Zfp709 | 7.930963 | 7.823776 | 7.9498 | -0.126051672 | -0.018865059 | 7.6787172 | 7.6827868 | 7.92582 | -0.243036141 | -0.247105762 |
| Arhgef7 | 10.15929 | 10.08918 | 10.121 | -0.031585819 | 0.038523187 | 10.018772 | 9.9629982 | 10.2061 | -0.243060211 | -0.187286803 |
| Coq3 | 6.558058 | 6.478893 | 6.6204 | -0.141463704 | -0.062298336 | 6.5609113 | 6.4869483 | 6.73004 | -0.243089515 | -0.169126562 |
| Epor | 2.496289 | 2.307975 | 2.5665 | -0.258559796 | -0.0702458 | 2.3679314 | 2.2423401 | 2.48546 | -0.243119683 | -0.117528396 |
| Zfp113 | 8.199065 | 7.993714 | 8.0896 | -0.095907149 | 0.109443838 | 7.9068153 | 7.925971 | 8.16917 | -0.243196637 | -0.262352366 |
| Hs1bp3 | 7.492394 | 7.575787 | 7.7384 | -0.162591745 | -0.245984585 | 7.7823639 | 7.3460056 | 7.58923 | -0.243224219 | 0.193134082 |
| Tmem216 | 5.350818 | 5.166431 | 5.2604 | -0.093941958 | 0.090445344 | 5.4210833 | 5.2911171 | 5.53445 | -0.243332854 | -0.113366608 |
| Rdh5 | 5.47858 | 5.564822 | 5.6242 | -0.059344673 | -0.14558634 | 6.0150241 | 5.6698057 | 5.91317 | -0.243363228 | 0.101855183 |
| Il12a | 6.567941 | 6.636713 | 6.819 | -0.18227552 | -0.251047322 | 6.763465 | 6.538436 | 6.78181 | -0.243371707 | -0.018342687 |
| Zfp93 | 5.002415 | 4.828734 | 5.087 | -0.258245357 | -0.084564104 | 4.7119635 | 4.8407355 | 5.0842 | -0.243466894 | -0.372238831 |

| | | | | | | | | | | |
|---|---|---|---|---|---|---|---|---|---|---|
| Gm14403 | 3.239611 | 3.168752 | 3.2888 | -0.120001326 | -0.049142086 | 3.2043704 | 3.0394556 | 3.28295 | -0.243495069 | -0.078580353 |
| Fgd1 | 3.682374 | 3.634026 | 3.7259 | -0.091825593 | -0.043477393 | 3.6611839 | 3.6631798 | 3.90685 | -0.243668724 | -0.245664647 |
| Mpzl2 | 4.65433 | 4.842202 | 4.6429 | 0.199273988 | 0.011401938 | 4.8540043 | 4.5070538 | 4.75077 | -0.243718069 | 0.103232442 |
| Rad54l2 | 9.956165 | 9.866673 | 9.9504 | -0.083685575 | 0.005805627 | 9.8650156 | 9.7796272 | 10.0235 | -0.243833206 | -0.158444776 |
| Six5 | 3.700177 | 3.556591 | 3.8431 | -0.286540352 | -0.142954257 | 3.6297321 | 3.4664643 | 3.71037 | -0.243904712 | -0.080636875 |
| Arhgef3 | 11.90772 | 11.96728 | 11.967 | 2.818E-06 | -0.059563412 | 11.836363 | 11.695813 | 11.9398 | -0.243969102 | -0.103419442 |
| Susd3 | 8.476354 | 8.609266 | 8.5719 | 0.037352248 | -0.095559947 | 9.1090025 | 8.3099425 | 8.55394 | -0.244001456 | 0.555058549 |
| Vmn2r17 | 4.057341 | 4.01688 | 4.0223 | -0.005396063 | 0.035064771 | 4.0559145 | 3.9681577 | 4.21231 | -0.244155589 | -0.156398805 |
| Trav9-2 | 5.573179 | 5.433938 | 5.5285 | -0.094591331 | 0.04464988 | 5.2819111 | 5.2422544 | 5.48641 | -0.244159051 | -0.204502297 |
| Fam126a | 9.598092 | 9.632034 | 9.6645 | -0.032473487 | -0.066416276 | 9.5553122 | 9.3896582 | 9.63415 | -0.244494768 | -0.078840834 |
| Gm12184 | 2.834354 | 2.595071 | 2.6774 | -0.082331598 | 0.156952166 | 2.5768305 | 2.480241 | 2.72493 | -0.244687651 | -0.148098108 |
| Pspc1 | 9.251923 | 9.191523 | 9.2516 | -0.060081608 | 0.000318655 | 9.2237028 | 9.0058132 | 9.25065 | -0.244835533 | -0.026945994 |
| Kcnq3 | 3.733734 | 3.689818 | 3.7308 | -0.04095232 | 0.002964194 | 3.4097833 | 3.575185 | 3.82003 | -0.244844026 | -0.410245641 |
| Gm3373 | 2.30972 | 2.246652 | 2.1711 | 0.075545661 | 0.138613197 | 2.1237339 | 2.1073888 | 2.35228 | -0.244891384 | -0.22854627 |
| Coq9 | 8.414594 | 8.50608 | 8.4969 | 0.00922443 | -0.082261625 | 8.8183694 | 8.5370588 | 8.78218 | -0.245120608 | 0.036189966 |
| Otoa | 3.857926 | 3.834875 | 4.1889 | -0.354016044 | -0.33096462 | 4.1270378 | 3.9151606 | 4.16036 | -0.245203842 | -0.033326697 |
| Nipbl | 12.1767 | 12.22832 | 12.224 | 0.004672314 | -0.046952428 | 11.997102 | 11.997971 | 12.2432 | -0.245245268 | -0.24611489 |
| Gpatch2l | 9.839518 | 9.767735 | 9.7543 | 0.013439584 | 0.085222662 | 9.8967609 | 9.5816433 | 9.82694 | -0.245298199 | 0.069819349 |
| Ash2l | 9.554784 | 9.558529 | 9.6202 | -0.061651059 | -0.065395367 | 9.6014435 | 9.4125512 | 9.65792 | -0.245369693 | -0.056477421 |
| Hs3st3a1 | 3.065742 | 2.964479 | 3.0256 | -0.061109775 | 0.040152971 | 2.9427255 | 2.7372458 | 2.98273 | -0.245488559 | -0.040008913 |
| Chst7 | 4.485168 | 4.605282 | 4.3766 | 0.228702874 | 0.108588937 | 4.3888345 | 4.1439579 | 4.3895 | -0.245541899 | -0.000665324 |
| Pygl | 7.864371 | 7.582002 | 7.5629 | 0.019057223 | 0.301425506 | 7.9912183 | 7.5803318 | 7.8259 | -0.245569909 | 0.165316594 |
| Phf20l1 | 11.37589 | 11.26717 | 11.347 | -0.080040738 | 0.028681438 | 11.258202 | 11.326604 | 11.5722 | -0.24561523 | -0.314016562 |
| Gm14419 | 3.391234 | 3.133246 | 3.1245 | 0.008766525 | 0.266753882 | 3.027227 | 3.072894 | 3.31862 | -0.245730793 | -0.291397756 |
| Rag1 | 2.740673 | 2.776249 | 2.8007 | -0.024441668 | -0.060017921 | 2.8729525 | 2.672908 | 2.91865 | -0.245745179 | -0.045700608 |
| Ppm1d | 8.896875 | 8.994768 | 8.8973 | 0.097484763 | -0.000408622 | 8.7119177 | 8.6145368 | 8.86028 | -0.245747395 | -0.148366455 |
| Sarm1 | 3.97002 | 4.217339 | 4.0277 | 0.189688279 | -0.057630335 | 3.9138835 | 3.8991025 | 4.1449 | -0.245799562 | -0.231018569 |
| Pafah2 | 5.526657 | 5.71331 | 5.6671 | 0.046239357 | -0.140413907 | 5.5176425 | 5.3855689 | 5.6314 | -0.245828243 | -0.113754625 |
| Dtnb | 8.604953 | 8.573892 | 8.5539 | 0.0199721 | 0.051033045 | 8.7442522 | 8.4877446 | 8.73363 | -0.245883013 | 0.010624662 |
| Heca | 11.10074 | 11.16187 | 11.194 | -0.031813284 | -0.09294447 | 11.00511 | 10.914169 | 11.1601 | -0.24593492 | -0.154993319 |
| Cep164 | 9.075321 | 8.976571 | 9.0894 | -0.112793646 | -0.014044585 | 8.9299274 | 8.7869588 | 9.03295 | -0.245993944 | -0.103025326 |
| Bcat2 | 9.245726 | 9.315027 | 9.4271 | -0.112035001 | -0.181335704 | 9.6100226 | 9.2528052 | 9.49889 | -0.246080895 | 0.111136488 |

| | | | | | | | | | | |
|---|---|---|---|---|---|---|---|---|---|---|
| Atp2c1 | 10.45701 | 10.45264 | 10.369 | 0.083373054 | 0.087746226 | 10.488317 | 10.228545 | 10.4747 | -0.246170399 | 0.013602406 |
| Gm6741 | 4.429072 | 4.415768 | 4.1257 | 0.290026625 | 0.303330613 | 4.4176567 | 4.1992712 | 4.44545 | -0.246183334 | -0.027797851 |
| Prep | 9.695292 | 9.668395 | 9.7361 | -0.067700687 | -0.040803953 | 9.7427559 | 9.5428729 | 9.78911 | -0.246232534 | -0.046349557 |
| Yaf2 | 7.939708 | 7.885342 | 8.0551 | -0.169728846 | -0.115363521 | 7.9181086 | 7.59792 | 7.84421 | -0.246290693 | 0.073897957 |
| Kif1bp | 8.633363 | 8.697946 | 8.5869 | 0.111031349 | 0.046448976 | 8.6111827 | 8.4762383 | 8.72255 | -0.246311489 | -0.111367059 |
| St3gal3 | 7.764155 | 7.663737 | 7.8156 | -0.151856614 | -0.051437898 | 7.9095029 | 7.4971133 | 7.74359 | -0.246479303 | 0.165910368 |
| Mynn | 9.319946 | 9.380609 | 9.4622 | -0.081609696 | -0.14227295 | 9.4552663 | 9.1958724 | 9.44248 | -0.24661216 | 0.012781775 |
| Cap2 | 3.98955 | 3.920398 | 3.9104 | 0.009967355 | 0.079119355 | 3.7909143 | 3.6393983 | 3.88604 | -0.246636912 | -0.095120916 |
| Tbccd1 | 8.049444 | 7.95617 | 8.1002 | -0.143999766 | -0.050725972 | 8.0546076 | 7.8890616 | 8.13573 | -0.246666337 | -0.081120401 |
| Sptssa | 9.901327 | 9.845783 | 9.8959 | -0.050113428 | 0.005431015 | 9.9005575 | 9.578888 | 9.82566 | -0.246774192 | 0.074895309 |
| Kit | 8.923833 | 8.189703 | 8.5191 | -0.329365763 | 0.404764387 | 8.029126 | 7.9701334 | 8.21703 | -0.246892341 | -0.187899742 |
| Ms4a3 | 1.92295 | 1.763333 | 1.8437 | -0.080390795 | 0.079225849 | 1.890639 | 1.768477 | 2.01553 | -0.247056823 | -0.124894831 |
| Rprd1a | 7.884587 | 7.893703 | 7.9577 | -0.063984432 | -0.073101037 | 8.284077 | 8.0141169 | 8.26124 | -0.247127511 | 0.022832643 |
| Nhs | 3.168998 | 2.960357 | 3.0753 | -0.11498408 | 0.093657102 | 3.2675741 | 2.861717 | 3.10892 | -0.247201768 | 0.158655348 |
| Ighv1-31 | 3.56683 | 3.513416 | 3.4555 | 0.057939952 | 0.111354286 | 3.4921576 | 3.33556 | 3.58276 | -0.247204058 | -0.090606451 |
| Wasf2 | 12.12363 | 12.13555 | 12.138 | -0.00263514 | -0.014558712 | 12.024189 | 11.790907 | 12.0381 | -0.247221162 | -0.013939967 |
| Sh3d19 | 6.075306 | 5.893144 | 6.1197 | -0.226600745 | -0.044438883 | 5.8103012 | 5.8147719 | 6.06217 | -0.24739982 | -0.251870506 |
| Pogk | 8.757789 | 8.748656 | 8.6086 | 0.140015095 | 0.149147833 | 8.3652006 | 8.3690383 | 8.61644 | -0.247401801 | -0.251239517 |
| Echs1 | 9.167739 | 9.055229 | 9.2092 | -0.154000192 | -0.041489944 | 9.2945563 | 8.9007377 | 9.14822 | -0.247482361 | 0.146336324 |
| Nckap5l | 8.509206 | 8.494406 | 8.7002 | -0.205817836 | -0.191017306 | 8.1832878 | 8.4371166 | 8.6847 | -0.247581553 | -0.50141039 |
| Olfr586 | 4.124952 | 3.991891 | 4.0801 | -0.088168605 | 0.044892331 | 4.0541155 | 4.0922629 | 4.34006 | -0.247792508 | -0.285939947 |
| Ubqln2 | 9.634883 | 9.544963 | 9.673 | -0.12806227 | -0.038142113 | 9.5691922 | 9.2992139 | 9.54718 | -0.247962597 | 0.022015732 |
| Mtmr7 | 4.068715 | 3.801605 | 4.0352 | -0.233620293 | 0.033489372 | 4.0161301 | 3.8230987 | 4.07117 | -0.248070118 | -0.055038698 |
| Unc5c | 5.194287 | 4.9717 | 5.3665 | -0.394764582 | -0.172178217 | 4.9336853 | 4.7966596 | 5.04482 | -0.248157102 | -0.111131474 |
| Ankrd10 | 10.7199 | 10.84709 | 10.798 | 0.049495966 | -0.077694988 | 10.890312 | 10.73434 | 10.9825 | -0.248176486 | -0.092204832 |
| Neurl3 | 11.17443 | 11.18749 | 11.303 | -0.115383312 | -0.128442984 | 11.267918 | 11.05735 | 11.3057 | -0.248374638 | -0.037805705 |
| Cacnb3 | 10.11583 | 10.27225 | 10.357 | -0.085155053 | -0.241575242 | 10.215173 | 10.467136 | 10.7155 | -0.248412822 | -0.5003753 |
| Rev3l | 10.29737 | 10.31552 | 10.357 | -0.041727021 | -0.059877905 | 10.106271 | 10.155689 | 10.4042 | -0.248496536 | -0.297915096 |
| Wdr90 | 8.025625 | 8.265001 | 8.0454 | 0.219629689 | -0.019746568 | 8.5854675 | 8.2929542 | 8.54149 | -0.248539584 | 0.043973729 |
| Lrrc71 | 1.96907 | 1.928084 | 2.1059 | -0.177790228 | -0.13680404 | 2.1996607 | 1.9025216 | 2.1511 | -0.248582099 | 0.048556992 |
| Zfp982 | 4.371872 | 4.299943 | 4.6232 | -0.323232605 | -0.251304143 | 3.9267657 | 3.9628283 | 4.21148 | -0.248652437 | -0.284715116 |
| Pde4b | 11.1439 | 11.35635 | 11.157 | 0.199656866 | -0.01279661 | 11.148226 | 11.017495 | 11.2662 | -0.248715418 | -0.117984569 |

| | | | | | | | | | | |
|---|---|---|---|---|---|---|---|---|---|---|
| Gm21244 | 5.203672 | 5.210092 | 5.1887 | 0.02134286 | 0.01492233 | 5.2448047 | 5.4949953 | 5.74375 | -0.24875213 | -0.498942763 |
| Zfp568 | 8.408333 | 8.420197 | 8.3925 | 0.027669724 | 0.015805005 | 8.1575849 | 7.9787743 | 8.22756 | -0.248788927 | -0.06997828 |
| Recql4 | 5.986191 | 5.917579 | 5.9805 | -0.062969302 | 0.005642678 | 6.4848592 | 5.9854102 | 6.23425 | -0.248843215 | 0.250605789 |
| Acad10 | 7.56139 | 7.607602 | 7.7057 | -0.098086888 | -0.144299426 | 7.8093243 | 7.4738231 | 7.72272 | -0.248899289 | 0.086601964 |
| Oprd1 | 5.842849 | 5.900836 | 5.9297 | -0.028835951 | -0.086822169 | 5.6312713 | 5.9342451 | 6.18317 | -0.248928128 | -0.551901859 |
| Rapgef1 | 11.94109 | 11.90353 | 11.942 | -0.0387236 | -0.001158752 | 11.993516 | 11.772157 | 12.0211 | -0.248972434 | -0.02761374 |
| Hc | 4.100283 | 4.077752 | 3.9983 | 0.079449383 | 0.101980174 | 3.7153981 | 3.8439556 | 4.09298 | -0.249022936 | -0.377580377 |
| Rad50 | 9.855821 | 9.914965 | 9.9692 | -0.054241508 | -0.11338575 | 9.8246002 | 9.7713292 | 10.0205 | -0.249163167 | -0.195892225 |
| Fam160b1 | 9.545767 | 9.622356 | 9.6972 | -0.07479408 | -0.151383219 | 9.5335722 | 9.3857057 | 9.63494 | -0.249238239 | -0.101371739 |
| Zfp790 | 8.285534 | 8.36364 | 8.3411 | 0.022544785 | -0.055560836 | 8.2481587 | 8.1415909 | 8.39092 | -0.249325907 | -0.142758133 |
| Cars2 | 8.739668 | 8.665155 | 8.7617 | -0.096544651 | -0.022031124 | 9.0318165 | 8.764647 | 9.0141 | -0.249455395 | 0.017714185 |
| Snx4 | 10.28704 | 10.12909 | 10.299 | -0.170403023 | -0.012458416 | 10.144423 | 9.9408319 | 10.1903 | -0.249473541 | -0.04588205 |
| Zfp62 | 10.19564 | 10.3181 | 10.26 | 0.058206632 | -0.064254312 | 10.214831 | 10.11038 | 10.3599 | -0.249513562 | -0.145062292 |
| Itgb7 | 12.38256 | 12.39651 | 12.491 | -0.094136354 | -0.10808779 | 12.504863 | 12.305518 | 12.555 | -0.249521482 | -0.050176359 |
| Foxp3 | 9.147536 | 9.104221 | 9.151 | -0.046749109 | -0.003433716 | 8.8959548 | 9.0239702 | 9.27364 | -0.249666648 | -0.377681982 |
| Fam81a | 4.421828 | 4.209745 | 4.2437 | -0.033987042 | 0.178095914 | 4.3232927 | 4.213453 | 4.46313 | -0.249678479 | -0.139838809 |
| Ulk3 | 9.078919 | 9.06638 | 9.21 | -0.143570959 | -0.131031205 | 9.7535821 | 9.4282305 | 9.67802 | -0.249788295 | 0.075563316 |
| Rsu1 | 9.426199 | 9.264522 | 9.3685 | -0.103934142 | 0.05774245 | 9.3606467 | 8.9487761 | 9.19867 | -0.249897837 | 0.161972699 |
| Snrnp25 | 7.856835 | 7.850409 | 7.9578 | -0.107351251 | -0.100925081 | 8.2627657 | 7.7746002 | 8.0246 | -0.249995464 | 0.238169998 |
| Iqce | 8.614488 | 8.554095 | 8.6912 | -0.137152333 | -0.076759223 | 8.7460538 | 8.6186104 | 8.86861 | -0.250000636 | -0.122557159 |
| Rasa1 | 10.06055 | 10.11059 | 10.166 | -0.055824428 | -0.10586611 | 9.9505916 | 10.087318 | 10.3373 | -0.250023239 | -0.386749983 |
| Nek1 | 8.212481 | 8.329123 | 8.3133 | 0.01586449 | -0.100777219 | 8.131399 | 8.0336362 | 8.28373 | -0.250091112 | -0.152328377 |
| Ispd | 4.903954 | 4.910126 | 5.0261 | -0.115942799 | -0.12211433 | 4.8810414 | 4.682979 | 4.93308 | -0.250098507 | -0.052036066 |
| Wdr72 | 3.325365 | 3.197696 | 3.2655 | -0.067780178 | 0.05988847 | 3.1943335 | 3.2160523 | 3.46616 | -0.250106942 | -0.271825694 |
| Troap | 4.741246 | 4.875766 | 4.7613 | 0.114513983 | -0.020006102 | 4.9198442 | 4.8396846 | 5.08987 | -0.250188093 | -0.170028534 |
| Kcnh2 | 5.866628 | 5.850477 | 5.8389 | 0.011590917 | 0.027741956 | 5.8645008 | 5.4620771 | 5.71228 | -0.250202799 | 0.152220947 |
| Hmg20a | 10.16435 | 10.16694 | 10.186 | -0.018634405 | -0.021219128 | 10.122483 | 10.027755 | 10.278 | -0.250280827 | -0.155553378 |
| Cbr3 | 3.974976 | 3.755337 | 3.8926 | -0.137259536 | 0.082379816 | 4.1770709 | 3.6265763 | 3.87689 | -0.2503127 | 0.300181903 |
| Gm7336 | 5.813689 | 6.575547 | 6.2905 | 0.285028277 | -0.476829302 | 6.1739179 | 6.6858329 | 6.93636 | -0.250524505 | -0.762439456 |
| Spice1 | 8.487583 | 8.646353 | 8.6384 | 0.007941303 | -0.150828582 | 8.613997 | 8.5134081 | 8.76415 | -0.250741661 | -0.150152692 |
| Pou2f2 | 11.76732 | 11.50524 | 11.541 | -0.035342936 | 0.226746504 | 12.081 | 11.418216 | 11.669 | -0.250808568 | 0.411974822 |
| Sccpdh | 6.032329 | 5.822643 | 5.8824 | -0.059773134 | 0.149912215 | 5.7628592 | 5.6495464 | 5.90054 | -0.250990451 | -0.137677655 |

| | | | | | | | | | | |
|---|---|---|---|---|---|---|---|---|---|---|
| Dand5 | 5.269805 | 5.326917 | 5.4413 | -0.114347633 | -0.171459254 | 5.4851579 | 5.1720274 | 5.42312 | -0.251096119 | 0.06203442 |
| Ppa2 | 8.06286 | 7.983723 | 8.0275 | -0.043730401 | 0.035406608 | 8.0800099 | 7.8125979 | 8.06372 | -0.251120782 | 0.016291191 |
| Tmlhe | 6.300765 | 6.19332 | 6.3049 | -0.111602517 | -0.004157808 | 6.2113273 | 5.9635111 | 6.21483 | -0.251316074 | -0.003499869 |
| Gadd45a | 8.152432 | 7.885423 | 8.083 | -0.197609 | 0.069399669 | 7.6613145 | 7.3879501 | 7.63928 | -0.251333815 | 0.022030545 |
| Hist1h2ao | 2.930218 | 2.786029 | 2.9248 | -0.138794504 | 0.005394622 | 2.8340956 | 2.9291865 | 3.18073 | -0.251544991 | -0.346635927 |
| Speg | 4.509672 | 4.515924 | 4.3989 | 0.116986344 | 0.110734299 | 4.0991531 | 4.1520205 | 4.40359 | -0.251567983 | -0.304435363 |
| Brinp1 | 2.428454 | 2.313756 | 2.3582 | -0.044437402 | 0.07026064 | 2.4143734 | 2.2579584 | 2.50953 | -0.251576576 | -0.095161557 |
| Traj21 | 2.908801 | 2.865826 | 2.9491 | -0.083318539 | -0.040342985 | 2.830319 | 2.767854 | 3.01946 | -0.251601792 | -0.189136793 |
| Tmem106c | 7.476354 | 7.208789 | 7.4729 | -0.2641444 | 0.00342001 | 7.4181036 | 7.1553972 | 7.40711 | -0.251714846 | 0.010991534 |
| Lrrc57 | 7.900524 | 7.893487 | 7.9618 | -0.068354861 | -0.06131783 | 8.0034218 | 7.764337 | 8.01607 | -0.251728277 | -0.012643404 |
| Invs | 8.629028 | 8.64527 | 8.7371 | -0.091866829 | -0.108109027 | 8.7202095 | 8.5052482 | 8.75704 | -0.25179671 | -0.036835361 |
| Pigk | 9.990154 | 9.875028 | 10.023 | -0.147798404 | -0.03267261 | 9.8592291 | 9.6043351 | 9.85623 | -0.25189573 | 0.002998259 |
| Agbl3 | 4.902117 | 4.930263 | 5.0423 | -0.112013971 | -0.14015993 | 4.9907713 | 4.9318897 | 5.18381 | -0.251918387 | -0.193036727 |
| Wasl | 8.724123 | 8.642528 | 8.7207 | -0.078181088 | 0.003413432 | 8.5207244 | 8.5759171 | 8.82785 | -0.251931077 | -0.307123707 |
| Vwa8 | 9.036147 | 9.023179 | 9.1073 | -0.084157862 | -0.071189639 | 8.9974774 | 8.8326558 | 9.08459 | -0.251937262 | -0.087115656 |
| Dqx1 | 6.178936 | 6.380891 | 6.3499 | 0.031010073 | -0.170945086 | 6.7100611 | 6.5343897 | 6.7865 | -0.252107708 | -0.076436303 |
| Ccdc158 | 2.390977 | 2.489474 | 2.3765 | 0.112990688 | 0.014493483 | 2.4044871 | 2.3318755 | 2.58401 | -0.252139331 | -0.179527743 |
| Trim33 | 10.81686 | 10.70788 | 10.849 | -0.140954352 | -0.031974964 | 10.426647 | 10.467786 | 10.7199 | -0.252162854 | -0.293302246 |
| Polg2 | 7.553621 | 7.41141 | 7.6062 | -0.19476836 | -0.052557767 | 7.3900174 | 7.3683231 | 7.6205 | -0.25217647 | -0.230482205 |
| Fig4 | 9.102339 | 9.057798 | 9.1659 | -0.108082242 | -0.063541742 | 9.1628129 | 8.8069388 | 9.05916 | -0.252217763 | 0.103656389 |
| Acadm | 9.123263 | 9.209388 | 9.2725 | -0.063156014 | -0.14928058 | 9.1820436 | 9.0325775 | 9.28502 | -0.252442377 | -0.10297628 |
| Nek3 | 5.976174 | 5.975222 | 5.9893 | -0.01403874 | -0.013086749 | 6.0267895 | 5.8517477 | 6.10421 | -0.252466554 | -0.077424707 |
| Emc9 | 6.66829 | 6.63006 | 6.5638 | 0.066270467 | 0.104500979 | 6.7764382 | 6.4371585 | 6.6897 | -0.25254305 | 0.086736655 |
| Ppic | 7.313873 | 7.270525 | 7.419 | -0.148434152 | -0.105086 | 6.9511723 | 7.0162925 | 7.26886 | -0.252565074 | -0.317685259 |
| Zfp433 | 7.368953 | 6.911836 | 7.1647 | -0.252876689 | 0.204240669 | 7.2934892 | 7.1205386 | 7.37313 | -0.252596036 | -0.079645502 |
| Cyp2s1 | 6.192888 | 6.140879 | 6.2951 | -0.154260616 | -0.102251275 | 6.1187648 | 6.034861 | 6.28756 | -0.252701522 | -0.16879772 |
| Areg | 4.608801 | 4.785249 | 4.5743 | 0.210977559 | 0.034529306 | 4.7378463 | 4.7365777 | 4.98941 | -0.25282878 | -0.25156022 |
| Tubgcp2 | 9.382682 | 9.16816 | 9.3381 | -0.169958466 | 0.044562873 | 9.5168192 | 9.2393378 | 9.49225 | -0.252915908 | 0.024565473 |
| Eps8 | 8.102423 | 7.774877 | 7.4664 | 0.30844942 | 0.635995602 | 7.5203166 | 7.2286388 | 7.48175 | -0.253111382 | 0.03856641 |
| 1810014B01Rik | 6.091543 | 6.187403 | 6.006 | 0.181369616 | 0.085509982 | 6.4580743 | 6.1559155 | 6.40913 | -0.253215071 | 0.048943765 |
| Zfp946 | 6.915473 | 6.680299 | 6.8788 | -0.198527757 | 0.036646802 | 6.9111346 | 6.8117281 | 7.06512 | -0.253392586 | -0.15398603 |
| 37316 | 9.304152 | 9.158169 | 9.2561 | -0.097959288 | 0.048023219 | 9.2496031 | 8.9978021 | 9.2512 | -0.253393098 | -0.001592115 |

| | | | | | | | | | | |
|---|---|---|---|---|---|---|---|---|---|---|
| Xylt2 | 8.881212 | 8.856201 | 9.1107 | -0.254465674 | -0.229455271 | 8.9143498 | 8.758954 | 9.01238 | -0.253425821 | -0.098029992 |
| Gramd1b | 9.592084 | 9.520533 | 9.6656 | -0.145098376 | -0.073547398 | 9.1336518 | 9.6021905 | 9.85567 | -0.253474686 | -0.722013368 |
| Arhgef26 | 4.601528 | 4.621908 | 4.7754 | -0.153489425 | -0.17386959 | 4.0041769 | 4.0185011 | 4.27202 | -0.253519022 | -0.267843234 |
| 4930579G24Rik | 6.303875 | 6.215127 | 6.4064 | -0.191295557 | -0.102547708 | 6.2710567 | 6.1740537 | 6.42768 | -0.253621865 | -0.156618863 |
| Fads1 | 7.692555 | 7.420772 | 7.7461 | -0.325288917 | -0.053505666 | 7.2560544 | 7.1058667 | 7.3595 | -0.253628811 | -0.1034411 |
| Gm5134 | 4.636703 | 4.670578 | 4.6512 | 0.019356759 | -0.01451823 | 4.8645884 | 4.689444 | 4.9433 | -0.253860254 | -0.078715865 |
| Gm15448 | 5.764534 | 5.536315 | 5.8215 | -0.285151842 | -0.056932114 | 6.1812788 | 5.5951734 | 5.84912 | -0.253950648 | 0.332154697 |
| Ldlrap1 | 9.6736 | 9.859742 | 9.773 | 0.086730592 | -0.09941205 | 9.4743767 | 9.3982872 | 9.65268 | -0.254397166 | -0.178307641 |
| Dhcr7 | 8.816506 | 8.734463 | 8.9096 | -0.175180546 | -0.093137208 | 9.0409098 | 8.588879 | 8.84333 | -0.25445045 | 0.197580359 |
| Cd48 | 10.79439 | 10.8218 | 10.905 | -0.082950962 | -0.11036053 | 11.081799 | 10.709696 | 10.9642 | -0.254502988 | 0.117600849 |
| Dpyd | 4.289952 | 3.750201 | 4.1625 | -0.412258093 | 0.127493371 | 3.8895893 | 3.864743 | 4.11927 | -0.254523893 | -0.229677563 |
| Vtn | 6.042869 | 5.583536 | 5.6687 | -0.085164524 | 0.374168109 | 5.3478684 | 5.1411829 | 5.39607 | -0.254886674 | -0.048201116 |
| Cox7a2l | 12.50384 | 12.46144 | 12.54 | -0.078457288 | -0.036059844 | 12.514966 | 12.203327 | 12.4582 | -0.25489526 | 0.056743704 |
| Trav12-3 | 4.72699 | 4.815707 | 4.8895 | -0.073817807 | -0.162534694 | 4.6658431 | 4.5838403 | 4.83884 | -0.254995161 | -0.172992391 |
| Slx4ip | 7.933806 | 7.968377 | 8.0689 | -0.100524119 | -0.135095569 | 8.0154842 | 7.949852 | 8.20487 | -0.255013967 | -0.189381743 |
| Mxi1 | 7.655979 | 7.7213 | 7.8768 | -0.155532591 | -0.22085399 | 7.6065218 | 7.528985 | 7.78404 | -0.255057631 | -0.17752082 |
| Pcgf3 | 8.586957 | 8.511739 | 8.646 | -0.134270748 | -0.059052787 | 8.5008186 | 8.7206055 | 8.97571 | -0.255107796 | -0.474894697 |
| Hs3st3b1 | 8.403712 | 8.411396 | 8.3793 | 0.032093541 | 0.024409471 | 7.9097891 | 7.980132 | 8.23528 | -0.255147231 | -0.325490124 |
| Tfb2m | 8.130233 | 8.122119 | 8.1784 | -0.056314786 | -0.04820006 | 8.138443 | 8.2147381 | 8.46993 | -0.255187205 | -0.331482233 |
| Trav7-2 | 1.997721 | 2.271977 | 1.9894 | 0.282546806 | 0.008290595 | 3.1204246 | 2.0156532 | 2.271 | -0.255343735 | 0.849427724 |
| B230219D22Rik | 10.33598 | 10.27226 | 10.311 | -0.039164372 | 0.02455819 | 10.432896 | 10.182473 | 10.4379 | -0.255382586 | -0.004959652 |
| Cnbd2 | 6.616112 | 6.562921 | 6.6418 | -0.078907057 | -0.025716367 | 6.6835374 | 6.6632903 | 6.91876 | -0.255466379 | -0.235219312 |
| Slc6a13 | 3.201199 | 3.171407 | 3.3536 | -0.182151395 | -0.152359662 | 3.2059489 | 3.0120173 | 3.2675 | -0.255483433 | -0.061551781 |
| Csgalnact1 | 5.09768 | 4.778033 | 5.2801 | -0.50211614 | -0.182469117 | 4.6736299 | 4.4791367 | 4.73469 | -0.255550058 | -0.061056856 |
| Trbv20 | 6.489159 | 6.487103 | 6.5075 | -0.020361919 | -0.018305023 | 6.0415006 | 6.190624 | 6.44619 | -0.255569011 | -0.40469241 |
| Tmc3 | 3.538846 | 3.620423 | 3.6852 | -0.064825788 | -0.146402526 | 3.6524258 | 3.5082234 | 3.76392 | -0.255693901 | -0.111491536 |
| Hist1h3e | 3.253975 | 3.095901 | 3.1394 | -0.043456418 | 0.114617441 | 3.2555675 | 3.026091 | 3.28192 | -0.255824897 | -0.026348469 |
| Cep97 | 9.156091 | 9.11428 | 9.3635 | -0.249242214 | -0.207431102 | 9.0711075 | 8.9166785 | 9.17276 | -0.256078395 | -0.101649427 |
| Tmem255a | 2.620236 | 2.594557 | 2.7725 | -0.177943213 | -0.152264949 | 2.5686096 | 2.4772525 | 2.73336 | -0.25611135 | -0.164754218 |
| Bbof1 | 4.610299 | 4.604967 | 4.747 | -0.142064719 | -0.136733135 | 4.4505869 | 4.5856079 | 4.84182 | -0.256214303 | -0.391235299 |
| Zfp229 | 5.587819 | 5.741133 | 5.7894 | -0.048225167 | -0.201539664 | 5.8019996 | 5.7107473 | 5.9671 | -0.256356958 | -0.165104641 |
| Zfp128 | 5.891676 | 5.906334 | 6.0239 | -0.11754466 | -0.132202749 | 5.9184484 | 5.6485005 | 5.90493 | -0.256424708 | 0.013523172 |

| | | | | | | | | | | |
|---|---|---|---|---|---|---|---|---|---|---|
| Fhl3 | 7.511844 | 7.498794 | 7.6127 | -0.113925286 | -0.100875301 | 7.616099 | 7.2175666 | 7.47419 | -0.256620908 | 0.141911466 |
| Elmo1 | 11.36289 | 11.40133 | 11.438 | -0.036585056 | -0.075025354 | 11.438065 | 11.286425 | 11.5431 | -0.256686632 | -0.105046561 |
| Phtf1 | 9.017857 | 9.029216 | 9.053 | -0.023821999 | -0.035180596 | 9.0211892 | 8.9541596 | 9.21106 | -0.256897739 | -0.189868077 |
| Ppp1r3c | 3.681783 | 3.511327 | 3.6811 | -0.169731733 | 0.000723397 | 3.3670065 | 3.245867 | 3.50299 | -0.257125223 | -0.135985713 |
| Nf2 | 9.765209 | 9.863053 | 9.8748 | -0.011779176 | -0.10962258 | 9.8445233 | 9.6827505 | 9.93991 | -0.257157121 | -0.095384276 |
| Tcf7 | 13.49392 | 13.31066 | 13.352 | -0.040882466 | 0.142377966 | 13.141506 | 13.039384 | 13.2966 | -0.25717285 | -0.155051434 |
| Fyttd1 | 9.886715 | 9.909455 | 9.8954 | 0.014004998 | -0.00873486 | 9.6815877 | 9.733141 | 9.99051 | -0.257363992 | -0.308917346 |
| 2610005L07Rik | 9.255285 | 9.2085 | 9.2402 | -0.031741781 | 0.015042641 | 9.4718481 | 9.2019783 | 9.45952 | -0.25753879 | 0.01233102 |
| Hoxa3 | 3.296008 | 3.136991 | 3.3331 | -0.196089899 | -0.037072501 | 3.1075322 | 3.0636266 | 3.3212 | -0.257573462 | -0.213667868 |
| Aph1c | 5.080705 | 5.304501 | 5.1728 | 0.131728692 | -0.092066861 | 5.3625672 | 4.8104782 | 5.06808 | -0.257597346 | 0.294491674 |
| Fam83a | 2.671143 | 2.747208 | 2.7511 | -0.003912262 | -0.079977761 | 2.6805273 | 2.4976061 | 2.75543 | -0.257825763 | -0.074904553 |
| AW554918 | 8.174059 | 8.198955 | 8.287 | -0.088057732 | -0.112953767 | 8.2686767 | 8.1519279 | 8.40996 | -0.258030603 | -0.141281752 |
| mt-Nd6 | 8.999453 | 8.770913 | 8.94 | -0.169128575 | 0.059411881 | 8.353435 | 8.6201656 | 8.87824 | -0.258072439 | -0.524802953 |
| Fgf1 | 4.503073 | 4.408024 | 4.4572 | -0.049181469 | 0.045868164 | 4.3489364 | 4.0386848 | 4.29687 | -0.258184434 | 0.05206722 |
| Usp33 | 9.696993 | 9.691692 | 9.743 | -0.051269001 | -0.045968489 | 9.6082468 | 9.5150249 | 9.77324 | -0.258217642 | -0.164995731 |
| Iqcn | 2.202964 | 2.140996 | 2.2425 | -0.101491943 | -0.039523614 | 2.1223994 | 2.0044885 | 2.26271 | -0.258219743 | -0.140308883 |
| Fam210a | 9.022729 | 9.201777 | 8.9838 | 0.217976448 | 0.03892865 | 8.8664806 | 8.8381357 | 9.09636 | -0.258222695 | -0.229877793 |
| Gm43064 | 3.817132 | 3.503178 | 3.7695 | -0.266302124 | 0.047651701 | 3.4166746 | 3.4929985 | 3.75127 | -0.258272248 | -0.334596146 |
| Stox2 | 5.101611 | 4.894118 | 5.2415 | -0.347411367 | -0.139918328 | 4.6328221 | 4.7863627 | 5.04463 | -0.258272295 | -0.41181287 |
| Rock1 | 11.9416 | 11.93368 | 11.94 | -0.00616724 | 0.001761706 | 11.773578 | 11.717986 | 11.9763 | -0.25827539 | -0.202683128 |
| Apc2 | 3.174836 | 3.185997 | 3.2811 | -0.095072563 | -0.106233498 | 3.2103322 | 2.9890349 | 3.24742 | -0.25838418 | -0.037086906 |
| Pter | 4.590208 | 4.407311 | 4.6251 | -0.217780268 | -0.034883076 | 4.4339674 | 4.1380902 | 4.39651 | -0.258422951 | 0.037454284 |
| Lipc | 3.326511 | 3.402296 | 3.3506 | 0.051696848 | -0.024087739 | 3.3227179 | 3.2692067 | 3.52767 | -0.258467049 | -0.204955828 |
| Rabep1 | 10.24087 | 10.24003 | 10.317 | -0.076990873 | -0.07615698 | 10.00651 | 9.9973391 | 10.256 | -0.258632742 | -0.24946225 |
| Gpalpp1 | 9.989938 | 9.944921 | 9.9797 | -0.034754639 | 0.010262409 | 9.9794781 | 9.7670601 | 10.0259 | -0.258810237 | -0.046392173 |
| Dock8 | 12.62771 | 12.85079 | 12.696 | 0.154624612 | -0.068456458 | 12.629049 | 12.529936 | 12.7889 | -0.258976542 | -0.159863735 |
| Wdr60 | 4.996848 | 4.587766 | 4.7896 | -0.201789534 | 0.207292698 | 4.8468837 | 4.7382455 | 4.99729 | -0.259039623 | -0.15040147 |
| Gm28036 | 9.423976 | 9.608555 | 9.5506 | 0.057906022 | -0.126672828 | 9.4675659 | 9.3996354 | 9.65876 | -0.259129443 | -0.191198911 |
| Nedd9 | 11.71105 | 11.78151 | 11.733 | 0.048445116 | -0.022008418 | 11.654629 | 11.390042 | 11.6493 | -0.259256368 | 0.005330619 |
| Rps6kl1 | 3.181244 | 3.015692 | 3.0688 | -0.053121293 | 0.112430575 | 2.9707884 | 3.0078828 | 3.26717 | -0.259284474 | -0.296378885 |
| Ypel2 | 7.117897 | 6.982768 | 7.1207 | -0.13794988 | -0.002820482 | 6.6866461 | 6.6849006 | 6.94435 | -0.259448158 | -0.257702643 |
| Rpap2 | 8.255178 | 8.163377 | 8.3456 | -0.18223621 | -0.090435232 | 8.2098872 | 8.1192923 | 8.37874 | -0.259451764 | -0.168856828 |

| | | | | | | | | | | |
|---|---|---|---|---|---|---|---|---|---|---|
| Dnm1 | 4.431225 | 4.172323 | 4.1412 | 0.031077837 | 0.289979789 | 3.8141893 | 3.7148596 | 3.97452 | -0.259659619 | -0.160329871 |
| Ezh1 | 10.31703 | 10.34142 | 10.438 | -0.096700956 | -0.121089662 | 10.601481 | 10.377757 | 10.6375 | -0.259693264 | -0.035969642 |
| Ier3 | 9.786281 | 9.518393 | 9.5994 | -0.081002407 | 0.186886156 | 8.8042942 | 8.6407138 | 8.90043 | -0.259712842 | -0.096132447 |
| Nrg2 | 3.779114 | 3.684757 | 3.7306 | -0.045797013 | 0.048560542 | 3.6984444 | 3.3233992 | 3.58314 | -0.2597436 | 0.115301593 |
| Tle3 | 11.64576 | 11.43894 | 11.523 | -0.083608566 | 0.123209286 | 11.40345 | 11.319895 | 11.5797 | -0.259756134 | -0.176200981 |
| Fam213a | 7.792248 | 7.694899 | 7.6607 | 0.034176843 | 0.131526213 | 7.8555123 | 7.7037538 | 7.96366 | -0.25990996 | -0.108151381 |
| Zfp74 | 7.169569 | 7.175718 | 7.311 | -0.135238111 | -0.141387665 | 7.2834973 | 7.11421 | 7.37412 | -0.259912863 | -0.090625569 |
| Cep128 | 8.177609 | 8.182793 | 8.1596 | 0.023219686 | 0.018035769 | 8.2468301 | 7.9677983 | 8.22771 | -0.259915004 | 0.01911679 |
| Meis1 | 4.381912 | 4.272848 | 4.3697 | -0.096887168 | 0.01217714 | 4.4883995 | 4.1863787 | 4.4466 | -0.260217935 | 0.041802798 |
| Zfp101 | 7.435339 | 7.404301 | 7.2868 | 0.117517946 | 0.148555841 | 7.2443613 | 7.1204005 | 7.38068 | -0.26028316 | -0.136322331 |
| Dusp6 | 9.93225 | 9.540213 | 9.9471 | -0.406848565 | -0.014811264 | 9.4917474 | 8.9711682 | 9.23162 | -0.260452395 | 0.260126838 |
| Pcdhgc4 | 4.609398 | 4.239009 | 4.8647 | -0.625656554 | -0.255267566 | 4.4704876 | 4.3469101 | 4.60745 | -0.26053821 | -0.136960737 |
| Fam13a | 3.791117 | 3.589779 | 4.0581 | -0.468357475 | -0.267018962 | 3.5652897 | 3.5138407 | 3.77439 | -0.260544816 | -0.209095776 |
| Scn1b | 5.760374 | 5.394616 | 5.7406 | -0.345982278 | 0.019775882 | 5.2951421 | 5.1539482 | 5.41453 | -0.26057961 | -0.119385676 |
| Tmem94 | 9.870262 | 9.93252 | 9.9926 | -0.060100005 | -0.12235815 | 10.13416 | 9.7091005 | 9.96971 | -0.260612378 | 0.164447145 |
| Gm5281 | 4.653169 | 4.810801 | 4.6116 | 0.199195413 | 0.041563381 | 5.1062084 | 4.5967802 | 4.85755 | -0.260768231 | 0.24865994 |
| Pdzd4 | 6.162162 | 6.140369 | 6.3513 | -0.210928608 | -0.189135657 | 5.8310374 | 5.7790094 | 6.03982 | -0.260807817 | -0.208779766 |
| Trbv12-2 | 7.568488 | 7.623314 | 7.8283 | -0.204990348 | -0.259816662 | 7.4337703 | 7.5163664 | 7.77722 | -0.260857636 | -0.343453808 |
| Tbc1d31 | 7.076509 | 6.884092 | 7.1739 | -0.289854809 | -0.097437515 | 6.9222639 | 6.9772749 | 7.23819 | -0.26091725 | -0.315928196 |
| Armc9 | 6.963477 | 6.664014 | 6.6608 | 0.003223329 | 0.302685443 | 6.8569677 | 6.5951499 | 6.85609 | -0.260937049 | 0.000880735 |
| Snx33 | 8.161571 | 8.122019 | 8.2357 | -0.113671725 | -0.074120075 | 8.1164523 | 7.928123 | 8.18907 | -0.260951087 | -0.072621851 |
| Prkg2 | 3.796085 | 3.739507 | 4.001 | -0.26153532 | -0.204956586 | 3.5914281 | 3.573154 | 3.83416 | -0.261003269 | -0.242729196 |
| Dennd2d | 11.02079 | 10.94798 | 11.136 | -0.187705738 | -0.114888168 | 11.31733 | 11.078572 | 11.3396 | -0.261062288 | -0.022304419 |
| Atf7ip2 | 2.627151 | 2.537056 | 2.6054 | -0.068364851 | 0.021730271 | 2.6930446 | 2.6001656 | 2.86124 | -0.26107354 | -0.168194474 |
| Astl | 3.139355 | 2.976774 | 2.9598 | 0.016953444 | 0.179534797 | 3.0546789 | 2.8920914 | 3.1532 | -0.261112272 | -0.098524858 |
| 40422 | 6.422987 | 6.181952 | 6.4002 | -0.218278742 | 0.022755889 | 6.1208046 | 6.0257123 | 6.28686 | -0.261146697 | -0.166054324 |
| Mbd5 | 9.029544 | 8.769435 | 8.9925 | -0.223096748 | 0.037012344 | 8.7714335 | 8.6625185 | 8.92374 | -0.261221082 | -0.152306105 |
| Tenm3 | 4.889811 | 5.041033 | 4.9146 | 0.126445801 | -0.02477611 | 5.1690081 | 4.3908513 | 4.65216 | -0.261312324 | 0.516844465 |
| Ppp4r1 | 10.19608 | 10.20195 | 10.214 | -0.011795715 | -0.01766337 | 10.089996 | 9.905933 | 10.1672 | -0.261315807 | -0.077253061 |
| Ccdc34 | 6.382169 | 6.339181 | 6.5169 | -0.177673302 | -0.134685037 | 6.3152726 | 6.3207447 | 6.58207 | -0.261323924 | -0.266796002 |
| Sox12 | 4.989101 | 5.055672 | 5.3059 | -0.250206953 | -0.31677762 | 5.0256225 | 4.8496311 | 5.11096 | -0.261327535 | -0.0853362 |
| Ggcx | 6.332931 | 6.377453 | 6.4149 | -0.037490162 | -0.082012093 | 6.4093007 | 6.2398551 | 6.50141 | -0.261550082 | -0.092104536 |

| | | | | | | | | | | |
|---|---|---|---|---|---|---|---|---|---|---|
| Olfr365 | 4.839372 | 4.902311 | 4.8969 | 0.005427989 | -0.057510778 | 5.0374977 | 4.9941715 | 5.25581 | -0.261643244 | -0.218317016 |
| Notch2 | 12.29395 | 12.4515 | 12.369 | 0.082792324 | -0.074753196 | 12.412034 | 12.115349 | 12.3771 | -0.261757636 | 0.034927201 |
| Gprc5a | 4.116208 | 3.982904 | 4.0135 | -0.030563779 | 0.102739622 | 4.1402182 | 4.0171799 | 4.27898 | -0.261801927 | -0.138763671 |
| Cnot6 | 11.23843 | 11.2297 | 11.268 | -0.038508976 | -0.029780612 | 11.042309 | 10.977374 | 11.2392 | -0.26183108 | -0.196895393 |
| Msx1 | 3.885733 | 3.742758 | 3.937 | -0.194254753 | -0.051279631 | 3.4966188 | 3.4453274 | 3.70718 | -0.261848918 | -0.210557507 |
| B430305J03Rik | 3.192893 | 3.245585 | 3.3213 | -0.075686418 | -0.128378701 | 3.3031393 | 3.101881 | 3.36387 | -0.261990858 | -0.060732606 |
| Ccp110 | 7.868399 | 7.782729 | 7.867 | -0.084272972 | 0.001396449 | 7.8538075 | 7.816865 | 8.07892 | -0.26205922 | -0.225116684 |
| Mia3 | 10.93462 | 10.84802 | 10.888 | -0.039587422 | 0.04701692 | 10.851689 | 10.649784 | 10.9119 | -0.262099158 | -0.06019505 |
| Adm | 5.733044 | 5.673836 | 5.6869 | -0.013075274 | 0.046133354 | 5.2755083 | 5.4977395 | 5.75992 | -0.262180401 | -0.484411641 |
| Csnk1g1 | 9.980578 | 9.86591 | 10.088 | -0.221765287 | -0.107098126 | 9.9489195 | 9.8838474 | 10.1461 | -0.262224195 | -0.197152055 |
| Myh14 | 1.801409 | 1.806187 | 1.8208 | -0.014616951 | -0.019395125 | 1.728525 | 1.7862336 | 2.0486 | -0.262366542 | -0.320075132 |
| Gng7 | 2.766039 | 2.695476 | 2.7398 | -0.04433807 | 0.026224897 | 2.6664793 | 2.5899733 | 2.85237 | -0.262396627 | -0.18589054 |
| Josd1 | 8.857503 | 8.776805 | 8.867 | -0.090201842 | -0.009503268 | 8.7228914 | 8.5535687 | 8.81607 | -0.262498703 | -0.093176067 |
| Cfap74 | 6.987971 | 7.011773 | 6.9911 | 0.020718838 | -0.003082605 | 7.2272816 | 6.9688925 | 7.2314 | -0.262509812 | -0.004120644 |
| Cks1b | 7.194421 | 7.014626 | 7.2649 | -0.250283629 | -0.070488752 | 7.1408839 | 7.0829382 | 7.34549 | -0.262549639 | -0.204603948 |
| Adam10 | 11.47659 | 11.45506 | 11.476 | -0.021196396 | 0.00033508 | 11.173304 | 11.055181 | 11.3177 | -0.262556952 | -0.144433913 |
| Dnajc18 | 8.144777 | 8.222582 | 8.1538 | 0.068825322 | -0.00897886 | 8.1187392 | 8.143426 | 8.40602 | -0.26259156 | -0.287278395 |
| Rnft2 | 3.586898 | 3.482971 | 3.6127 | -0.129682782 | -0.025755227 | 3.8004416 | 3.527011 | 3.78979 | -0.262782551 | 0.010648133 |
| Cacna1g | 2.366293 | 2.22721 | 2.3277 | -0.100455094 | 0.038627737 | 2.4018783 | 2.1693771 | 2.43218 | -0.262806627 | -0.030305366 |
| Trav12-2 | 5.225327 | 5.244281 | 5.4204 | -0.176134404 | -0.195088685 | 5.234253 | 5.1140014 | 5.37683 | -0.262831141 | -0.142579536 |
| Msh6 | 9.73452 | 9.746033 | 9.7232 | 0.022796964 | 0.011283941 | 9.5178214 | 9.4902291 | 9.7534 | -0.263168209 | -0.235575894 |
| Gm47324 | 3.255454 | 3.273154 | 3.2689 | 0.004234627 | -0.01346591 | 3.3055409 | 3.2461657 | 3.50961 | -0.263442624 | -0.204067444 |
| Gm20518 | 5.203838 | 5.180461 | 5.0572 | 0.123254453 | 0.146631813 | 5.6173815 | 5.3162326 | 5.5797 | -0.263472094 | 0.037676856 |
| Mettl8 | 7.703879 | 7.91782 | 7.8693 | 0.048520103 | -0.165420253 | 7.7654328 | 7.6931549 | 7.95664 | -0.263483682 | -0.191205836 |
| Tmem14a | 4.172063 | 4.272972 | 4.2772 | -0.004238688 | -0.105147343 | 4.3706467 | 4.0606485 | 4.32419 | -0.263541978 | 0.046456142 |
| Cxadr | 4.530649 | 4.126716 | 4.908 | -0.781237827 | -0.37730453 | 4.4996552 | 4.2711338 | 4.53472 | -0.263588301 | -0.035066909 |
| Setd5 | 11.65812 | 11.54792 | 11.658 | -0.110386798 | -0.000191666 | 11.501963 | 11.462724 | 11.7263 | -0.263613926 | -0.224374461 |
| Cirbp | 10.59829 | 10.16733 | 10.568 | -0.40099899 | 0.029957154 | 10.786253 | 10.299312 | 10.5632 | -0.263855984 | 0.223084808 |
| Cast | 11.58881 | 11.55595 | 11.623 | -0.06743891 | -0.034585016 | 11.437452 | 11.313784 | 11.5776 | -0.263856306 | -0.140187352 |
| Cenpl | 7.179303 | 7.203268 | 7.2263 | -0.022989362 | -0.046955142 | 7.2501123 | 7.1798595 | 7.44374 | -0.263884314 | -0.193631499 |
| Zfp426 | 8.877176 | 8.837866 | 8.882 | -0.044093421 | -0.004783833 | 9.0111038 | 8.746388 | 9.01033 | -0.263944838 | 0.000771023 |
| Errfi1 | 8.740538 | 8.302782 | 8.3206 | -0.017796471 | 0.419959331 | 7.4149132 | 7.6652439 | 7.9292 | -0.263961023 | -0.5142917 |

| | | | | | | | | | | |
|---|---|---|---|---|---|---|---|---|---|---|
| Egfl7 | 9.576706 | 8.945795 | 9.5402 | -0.59436523 | 0.036545678 | 8.5202631 | 8.5171972 | 8.78121 | -0.264016454 | -0.260950569 |
| Snai2 | 3.732544 | 3.754263 | 3.8591 | -0.104798276 | -0.126517193 | 3.697737 | 3.6703144 | 3.93464 | -0.264320639 | -0.236898056 |
| Gsdme | 4.341337 | 4.043592 | 4.1708 | -0.127204176 | 0.170541169 | 4.2211164 | 3.8831152 | 4.14764 | -0.264525634 | 0.073475565 |
| Ncbp3 | 10.11457 | 10.07388 | 10.124 | -0.0499188 | -0.00922846 | 9.9606538 | 9.8558478 | 10.1204 | -0.264591602 | -0.159785592 |
| Fibin | 2.329906 | 2.261465 | 2.448 | -0.186553698 | -0.118112265 | 2.3292432 | 2.1662117 | 2.43089 | -0.264678196 | -0.101646618 |
| Dnah7a | 3.613114 | 3.288331 | 3.6122 | -0.323868064 | 0.000914201 | 3.3196554 | 3.2468076 | 3.51159 | -0.264786688 | -0.1919389 |
| Fam69a | 9.626094 | 9.630165 | 9.7766 | -0.146441045 | -0.150512359 | 9.7023415 | 9.4401399 | 9.70494 | -0.264795817 | -0.002594278 |
| Cdc23 | 9.507569 | 9.514416 | 9.6279 | -0.113494529 | -0.12034209 | 9.793912 | 9.4149041 | 9.67971 | -0.264809864 | 0.114198127 |
| Zfp597 | 9.230057 | 9.200072 | 9.1895 | 0.010611343 | 0.040596641 | 9.34424 | 8.9430168 | 9.20805 | -0.2650306 | 0.136192664 |
| Gm11223 | 2.602127 | 2.1516 | 2.3673 | -0.215694778 | 0.234831854 | 2.1640453 | 2.3785763 | 2.64365 | -0.265068752 | -0.479599758 |
| Tbc1d9b | 10.10977 | 10.09862 | 10.199 | -0.10038007 | -0.089236632 | 10.189171 | 9.9625157 | 10.2276 | -0.265125444 | -0.038469894 |
| Nnt | 8.566613 | 8.459086 | 8.6344 | -0.175357429 | -0.06783044 | 8.5141822 | 8.2708303 | 8.53603 | -0.265199731 | -0.021847835 |
| Soga1 | 6.849703 | 6.799472 | 6.6325 | 0.166971793 | 0.217203219 | 6.4429696 | 6.2607868 | 6.52607 | -0.265282801 | -0.083099991 |
| Lrrc75a | 6.534767 | 6.247619 | 6.531 | -0.283422013 | 0.003725833 | 6.5845654 | 5.945483 | 6.2108 | -0.265315786 | 0.373766709 |
| Pgm3 | 7.689128 | 7.805792 | 7.6848 | 0.120955254 | 0.004291501 | 7.7676522 | 7.5830341 | 7.84854 | -0.265509001 | -0.080890941 |
| Tmem254c | 8.054362 | 7.928067 | 8.192 | -0.263980509 | -0.137685961 | 7.7734447 | 7.7725419 | 8.03814 | -0.265602916 | -0.264700048 |
| Gm8741 | 1.997487 | 2.061592 | 2.2704 | -0.208769238 | -0.272874229 | 2.1257517 | 2.0498219 | 2.31556 | -0.265738319 | -0.189808475 |
| Nat8f1 | 4.809812 | 4.615154 | 4.9165 | -0.301354522 | -0.106696366 | 4.9082329 | 4.6730962 | 4.93889 | -0.265795334 | -0.03065867 |
| Jazf1 | 4.639246 | 4.547784 | 4.7056 | -0.157854708 | -0.06639287 | 4.7331109 | 4.492459 | 4.75839 | -0.265935384 | -0.025283435 |
| Cx3cl1 | 6.355793 | 6.553252 | 6.2164 | 0.336840673 | 0.139381634 | 6.0989643 | 6.0621435 | 6.3281 | -0.265960845 | -0.229140111 |
| Zfp235 | 6.85733 | 6.924512 | 6.9237 | 0.000801719 | -0.066379931 | 6.9056831 | 6.8723661 | 7.13834 | -0.265968971 | -0.232651979 |
| Gm15925 | 2.903101 | 2.899404 | 2.9848 | -0.085404031 | -0.081707145 | 3.0373292 | 2.8963547 | 3.16247 | -0.266112503 | -0.125137929 |
| Trbv5 | 7.029466 | 7.130213 | 7.2044 | -0.074174919 | -0.17492201 | 6.915956 | 6.8699734 | 7.13632 | -0.266344432 | -0.220361818 |
| Utp14b | 7.382048 | 7.322933 | 7.2208 | 0.102097929 | 0.161212931 | 7.3769619 | 7.0985714 | 7.36513 | -0.26656125 | 0.011829232 |
| Pea15a | 9.95737 | 9.969877 | 9.973 | -0.00314014 | -0.015647965 | 9.6939039 | 9.5381161 | 9.8048 | -0.266688659 | -0.110900782 |
| Odf2l | 7.017334 | 6.805179 | 6.9602 | -0.154974753 | 0.057180104 | 7.1550417 | 6.9645025 | 7.23124 | -0.266734461 | -0.076195292 |
| H3f3a | 13.72361 | 13.71445 | 13.749 | -0.03478888 | -0.025633066 | 13.963098 | 13.458669 | 13.7255 | -0.266838872 | 0.237589556 |
| Crtc1 | 8.865837 | 8.900213 | 8.9183 | -0.01809444 | -0.052470995 | 8.7672458 | 8.7306967 | 8.99757 | -0.26686902 | -0.230319847 |
| Smim13 | 7.706237 | 7.743785 | 7.69 | 0.0537851 | 0.016236996 | 7.5033641 | 7.2408406 | 7.50781 | -0.266966099 | -0.004442678 |
| Sesn3 | 12.11399 | 12.18728 | 12.108 | 0.079646456 | 0.006356526 | 11.964233 | 11.87406 | 12.1411 | -0.267026234 | -0.176853372 |
| Arid3a | 8.349112 | 8.248647 | 8.4345 | -0.185851309 | -0.085385466 | 8.2859582 | 8.0102403 | 8.27729 | -0.267046663 | 0.008671294 |
| Bpgm | 7.777479 | 7.580553 | 7.9361 | -0.355547398 | -0.158621657 | 7.8264823 | 7.4134096 | 7.68056 | -0.267145654 | 0.145927043 |

| | | | | | | | | | | |
|---|---|---|---|---|---|---|---|---|---|---|
| Abcb4 | 5.122183 | 5.056012 | 5.153 | -0.096961721 | -0.030789863 | 5.0283622 | 4.7925811 | 5.05978 | -0.267199747 | -0.031418686 |
| Mfng | 10.18298 | 10.33922 | 10.264 | 0.07526239 | -0.080973262 | 10.394374 | 9.9716795 | 10.2389 | -0.267224425 | 0.155470056 |
| Enpp1 | 9.064838 | 9.015257 | 9.0384 | -0.023151453 | 0.026429905 | 9.1655397 | 8.8026347 | 9.06986 | -0.26722556 | 0.095679446 |
| Rdh13 | 8.048046 | 7.884507 | 7.958 | -0.073515498 | 0.090023984 | 7.9505451 | 7.6933587 | 7.96074 | -0.267377075 | -0.010190666 |
| Prim2 | 7.819512 | 7.692149 | 7.9285 | -0.236367369 | -0.109004654 | 7.8971093 | 7.6957276 | 7.96311 | -0.267377664 | -0.065995938 |
| Epb41l3 | 5.770238 | 5.796283 | 5.7266 | 0.069636809 | 0.04359205 | 5.2824238 | 5.4024702 | 5.66985 | -0.267384236 | -0.387430635 |
| Kcnh3 | 4.527766 | 4.598215 | 4.4636 | 0.134594188 | 0.064145226 | 4.314268 | 4.1717508 | 4.4392 | -0.267448472 | -0.124931277 |
| Dexi | 7.550252 | 7.553009 | 7.6304 | -0.077343989 | -0.080100474 | 7.5923307 | 7.2467912 | 7.51429 | -0.267494661 | 0.078044881 |
| Pgrmc1 | 9.248313 | 9.081634 | 9.2553 | -0.173630762 | -0.00695169 | 9.0554402 | 8.8536169 | 9.12111 | -0.267496224 | -0.065672969 |
| Nt5dc3 | 8.799583 | 8.694889 | 8.5655 | 0.129418221 | 0.23411225 | 8.4897972 | 8.5169233 | 8.78446 | -0.267539462 | -0.294665518 |
| Brca1 | 7.523233 | 7.487259 | 7.7185 | -0.231210431 | -0.195236442 | 7.7196177 | 7.5362389 | 7.80398 | -0.267741615 | -0.084362806 |
| Arid1b | 11.1262 | 11.17855 | 11.175 | 0.003798562 | -0.048557658 | 11.13328 | 10.872035 | 11.1398 | -0.267756376 | -0.006511724 |
| Zer1 | 10.01161 | 10.00121 | 10.114 | -0.112468305 | -0.102077621 | 10.342242 | 9.9310352 | 10.1988 | -0.267781437 | 0.143424949 |
| 1810043G02Rik | 6.416627 | 6.378912 | 6.5435 | -0.164625227 | -0.126909608 | 6.924796 | 6.321604 | 6.58942 | -0.26782016 | 0.335371808 |
| Actr3b | 4.280757 | 4.283737 | 4.3672 | -0.083457039 | -0.086436526 | 4.3848906 | 4.1225189 | 4.3904 | -0.26787918 | -0.005507463 |
| Rnf225 | 5.296376 | 5.338454 | 5.6918 | -0.353318135 | -0.395395995 | 6.208178 | 5.5402714 | 5.80818 | -0.267907198 | 0.39999942 |
| Tubgcp5 | 8.897727 | 8.701524 | 8.8707 | -0.169148286 | 0.027055297 | 8.7733137 | 8.8379861 | 9.10622 | -0.268231069 | -0.332903463 |
| Tsnax | 10.50341 | 10.4254 | 10.498 | -0.072631562 | 0.005386138 | 10.578303 | 10.194356 | 10.4627 | -0.268384154 | 0.115562795 |
| Ctso | 10.48731 | 10.3502 | 10.492 | -0.14149443 | -0.00439125 | 10.871878 | 10.4605 | 10.729 | -0.268530464 | 0.142847975 |
| Hdac6 | 8.361775 | 8.177264 | 8.3724 | -0.195130338 | -0.010619649 | 8.5168481 | 8.2833552 | 8.5519 | -0.268540529 | -0.035047562 |
| Cd109 | 3.37669 | 3.065067 | 3.4223 | -0.357261001 | -0.045637972 | 3.1670285 | 2.9864215 | 3.25528 | -0.268857158 | -0.088250165 |
| Dhrs13 | 4.87244 | 4.857212 | 4.7999 | 0.057322618 | 0.072550492 | 4.7683015 | 4.687304 | 4.95618 | -0.268879127 | -0.187881705 |
| Foxj2 | 10.02511 | 10.07302 | 10.107 | -0.034284204 | -0.082193839 | 10.107251 | 9.7828058 | 10.0517 | -0.268885938 | 0.055558944 |
| Nde1 | 9.737291 | 9.755226 | 9.8282 | -0.072990439 | -0.090925034 | 9.8269667 | 9.6097656 | 9.8788 | -0.26903171 | -0.051830609 |
| Ap5m1 | 9.765657 | 9.657675 | 9.853 | -0.195328448 | -0.087346745 | 9.72999 | 9.4999902 | 9.7691 | -0.269109322 | -0.0391095 |
| Vps39 | 10.32192 | 10.35602 | 10.44 | -0.08414192 | -0.118240027 | 10.491407 | 10.301566 | 10.5707 | -0.269145794 | -0.079303949 |
| Rasa2 | 10.5538 | 10.52573 | 10.588 | -0.06246738 | -0.034399796 | 10.452417 | 10.285773 | 10.555 | -0.269270656 | -0.102626794 |
| Chd4 | 13.37745 | 13.32973 | 13.385 | -0.05491791 | -0.007197448 | 13.167294 | 13.131826 | 13.4011 | -0.269281 | -0.233813206 |
| Idh1 | 8.374867 | 8.339773 | 8.4744 | -0.134596437 | -0.099502169 | 8.0685668 | 8.0338034 | 8.30311 | -0.269304287 | -0.23454088 |
| Sdc2 | 5.385136 | 5.210199 | 5.4633 | -0.253097205 | -0.078159719 | 5.0140314 | 4.7881608 | 5.0575 | -0.269343842 | -0.043473215 |
| Gm21926 | 7.723859 | 7.547977 | 7.8054 | -0.257391014 | -0.08150925 | 7.8402061 | 7.5549277 | 7.82435 | -0.269422168 | 0.015856244 |
| Clip1 | 10.57805 | 10.68315 | 10.671 | 0.01222834 | -0.092865678 | 10.270736 | 10.412481 | 10.682 | -0.269561988 | -0.411306616 |

| | | | | | | | | | | |
|---|---|---|---|---|---|---|---|---|---|---|
| Slc29a1 | 9.198612 | 8.985202 | 9.1895 | -0.204286298 | 0.009124159 | 9.1136691 | 8.9003768 | 9.16995 | -0.269573975 | -0.056281728 |
| Cdc7 | 8.073179 | 8.179019 | 8.232 | -0.052948677 | -0.158788116 | 8.2412893 | 8.1498985 | 8.41947 | -0.269574725 | -0.178183914 |
| Lsp1 | 13.44317 | 13.45044 | 13.558 | -0.107420554 | -0.114690668 | 13.507273 | 13.314349 | 13.584 | -0.269652284 | -0.076728512 |
| Kntc1 | 7.104944 | 6.366469 | 6.772 | -0.405561112 | 0.332913638 | 6.5482903 | 6.908548 | 7.17821 | -0.269660587 | -0.629918318 |
| Cilp | 4.330703 | 4.29824 | 3.895 | 0.403218674 | 0.435682304 | 3.8802223 | 3.7519212 | 4.02164 | -0.269716006 | -0.141414897 |
| Tmem109 | 9.354076 | 9.41549 | 9.4327 | -0.017250888 | -0.078664614 | 9.5775963 | 9.0342328 | 9.30396 | -0.269730752 | 0.273632734 |
| Cyp4f37 | 3.566336 | 3.414148 | 3.2613 | 0.152827874 | 0.305015702 | 4.0248755 | 3.098061 | 3.36781 | -0.269752355 | 0.657062125 |
| 6330549D23Rik | 3.803978 | 3.767026 | 3.8239 | -0.056900593 | -0.019948545 | 3.7548114 | 3.5519126 | 3.82171 | -0.269794646 | -0.06689584 |
| Pard3 | 4.746659 | 4.461603 | 4.5331 | -0.071493601 | 0.213562384 | 4.3989086 | 4.3481675 | 4.61812 | -0.269949218 | -0.219208125 |
| Tmem206 | 8.432156 | 8.640001 | 8.5685 | 0.071524593 | -0.136321045 | 8.6182131 | 8.2091086 | 8.47908 | -0.269973843 | 0.13913066 |
| B4galnt2 | 3.03241 | 2.981657 | 2.9848 | -0.003168471 | 0.047584385 | 3.2167092 | 2.8978513 | 3.16789 | -0.270042324 | 0.048815632 |
| Nefh | 7.543301 | 7.335129 | 7.639 | -0.303834693 | -0.095662489 | 7.1957811 | 7.1569679 | 7.42706 | -0.270087559 | -0.23127435 |
| Zfp964 | 5.112191 | 5.078167 | 5.2714 | -0.193274922 | -0.159250748 | 5.1579024 | 5.0495803 | 5.31968 | -0.270103399 | -0.161781278 |
| Gm32687 | 7.105193 | 7.119214 | 7.1851 | -0.065898247 | -0.079919029 | 7.1685536 | 6.8392961 | 7.10943 | -0.270132921 | 0.059124517 |
| Map3k2 | 10.5705 | 10.64005 | 10.603 | 0.0366316 | -0.032920074 | 10.404939 | 10.3593 | 10.6295 | -0.270206408 | -0.224566582 |
| Gpr179 | 2.795796 | 2.785388 | 2.8344 | -0.049036117 | -0.038628743 | 2.707682 | 2.6630958 | 2.93338 | -0.270281873 | -0.225695636 |
| Eml1 | 6.755358 | 5.918363 | 6.2775 | -0.359095164 | 0.477899634 | 5.756214 | 5.7753589 | 6.0457 | -0.270341551 | -0.28948643 |
| Ufl1 | 9.539841 | 9.436652 | 9.5767 | -0.140020581 | -0.036831806 | 9.4284805 | 9.20578 | 9.47616 | -0.270375074 | -0.047674631 |
| Pltp | 9.063584 | 8.720833 | 9.1168 | -0.395943016 | -0.053191876 | 9.3141379 | 8.3465774 | 8.61702 | -0.27044127 | 0.697119185 |
| Zfp703 | 7.820268 | 7.664444 | 7.6748 | -0.010306447 | 0.145517471 | 7.6202028 | 7.1642306 | 7.43474 | -0.270504689 | 0.185467505 |
| Cux1 | 10.83131 | 10.7203 | 10.719 | 0.00160801 | 0.11261825 | 10.872154 | 10.493903 | 10.7644 | -0.270522602 | 0.107728611 |
| Cdk20 | 6.07623 | 6.01046 | 5.9799 | 0.030589067 | 0.096358187 | 6.2709898 | 5.7836801 | 6.05426 | -0.270578982 | 0.216730695 |
| Ggt1 | 7.562088 | 7.42988 | 7.2795 | 0.150385722 | 0.28259392 | 7.9894275 | 7.4868753 | 7.75747 | -0.270592327 | 0.231959911 |
| Exd2 | 8.712302 | 8.773487 | 8.8174 | -0.043894297 | -0.105079587 | 9.0583722 | 8.8616943 | 9.13232 | -0.27063005 | -0.073952238 |
| Zfp719 | 7.127876 | 7.126655 | 7.0829 | 0.043761717 | 0.044983042 | 7.0593682 | 6.868509 | 7.13922 | -0.270714985 | -0.079855701 |
| Snapc5 | 8.068701 | 7.860461 | 8.1573 | -0.296888884 | -0.08864929 | 8.095997 | 7.8729831 | 8.14374 | -0.270756252 | -0.047742375 |
| Prps2 | 10.94351 | 10.97735 | 10.964 | 0.012880398 | -0.020953146 | 10.972551 | 10.646512 | 10.9173 | -0.270800632 | 0.05523887 |
| Med23 | 9.662739 | 9.745049 | 9.723 | 0.022005238 | -0.060304406 | 9.7634969 | 9.6370004 | 9.90786 | -0.270859338 | -0.144362806 |
| Ulk1 | 8.91949 | 8.902871 | 9.0996 | -0.196711094 | -0.180091818 | 9.1538339 | 8.9252058 | 9.1961 | -0.270890514 | -0.042262468 |
| Gabbr1 | 9.231367 | 9.37082 | 9.4262 | -0.055357507 | -0.194810946 | 9.6470561 | 9.4622148 | 9.73312 | -0.270903522 | -0.086062226 |
| Tagln | 7.521151 | 7.817329 | 7.6024 | 0.214920894 | -0.081256987 | 7.5387765 | 6.8404366 | 7.11148 | -0.271048307 | 0.427291562 |
| Luc7l2 | 12.12166 | 12.06022 | 12.113 | -0.052446248 | 0.008993682 | 12.111441 | 12.144526 | 12.4157 | -0.271128962 | -0.304214469 |

| | | | | | | | | | | |
|---|---|---|---|---|---|---|---|---|---|---|
| Mst1 | 4.236656 | 4.138576 | 4.1423 | -0.0036999 | 0.094379493 | 4.2445564 | 4.1700646 | 4.44123 | -0.271164733 | -0.196672905 |
| Tyw1 | 9.203894 | 9.186027 | 9.2474 | -0.061385992 | -0.043518917 | 9.1966012 | 8.9229946 | 9.19417 | -0.271178863 | 0.002427745 |
| Hmgb2 | 10.61983 | 10.35679 | 10.659 | -0.30235853 | -0.03931947 | 10.440907 | 10.455765 | 10.7271 | -0.271375162 | -0.28623357 |
| Gmip | 11.54146 | 11.52809 | 11.665 | -0.136454894 | -0.123084488 | 11.878581 | 11.695656 | 11.9671 | -0.271434138 | -0.088509143 |
| Klhl29 | 5.196553 | 4.726911 | 5.0768 | -0.349893903 | 0.11974874 | 4.4645236 | 4.2902791 | 4.56176 | -0.27148072 | -0.09723621 |
| Gm49760 | 5.663137 | 5.464697 | 5.8627 | -0.397958465 | -0.199518183 | 5.6084164 | 5.4394562 | 5.71117 | -0.271714642 | -0.10275446 |
| C920009B18Rik | 8.004597 | 7.753024 | 7.9274 | -0.174328529 | 0.077244304 | 8.3615223 | 7.9841703 | 8.25595 | -0.271779772 | 0.105572231 |
| Gm5127 | 3.479859 | 3.492187 | 3.5113 | -0.019120936 | -0.031448818 | 3.4315696 | 3.256452 | 3.52824 | -0.271791263 | -0.096673626 |
| Nyx | 3.138025 | 3.118716 | 3.2066 | -0.087891556 | -0.068582464 | 3.1023546 | 3.1164155 | 3.38825 | -0.271831395 | -0.285892324 |
| Mamld1 | 5.612779 | 5.274976 | 5.5564 | -0.281448511 | 0.056354312 | 5.1227227 | 4.965365 | 5.23722 | -0.27185963 | -0.114501957 |
| Cryz | 5.546581 | 5.627931 | 5.6316 | -0.00364675 | -0.084996525 | 5.5410611 | 5.32287 | 5.59476 | -0.271890703 | -0.053699573 |
| Wrb | 6.732477 | 6.913875 | 6.7071 | 0.206804572 | 0.02540629 | 7.0143757 | 6.6585626 | 6.93061 | -0.272050276 | 0.083762852 |
| Hoxd9 | 4.212286 | 4.02206 | 4.3391 | -0.317089465 | -0.12686304 | 3.9431972 | 3.8287155 | 4.10085 | -0.272136799 | -0.157655088 |
| Hist2h4 | 5.049022 | 5.064504 | 5.1928 | -0.128252056 | -0.143733825 | 5.2921214 | 5.0130402 | 5.28537 | -0.272328808 | 0.006752482 |
| Sacm1l | 10.10095 | 10.10146 | 10.063 | 0.03859851 | 0.038089707 | 9.8845628 | 9.7882258 | 10.0607 | -0.272467974 | -0.176130945 |
| Tmem86a | 7.487266 | 7.018302 | 7.5989 | -0.580561671 | -0.111597185 | 7.0963514 | 6.6657683 | 6.93827 | -0.272499993 | 0.158083057 |
| Trak2 | 10.80249 | 10.84947 | 10.769 | 0.080240176 | 0.03325553 | 10.714074 | 10.511561 | 10.7841 | -0.27258728 | -0.070074736 |
| Gm14443 | 4.689131 | 4.76516 | 4.7482 | 0.016959681 | -0.059069572 | 4.742028 | 4.7344996 | 5.00712 | -0.272618337 | -0.265089962 |
| Intu | 4.453164 | 4.365052 | 4.3477 | 0.017386119 | 0.105498766 | 4.4341657 | 4.2593411 | 4.53205 | -0.272704345 | -0.097879787 |
| Klhdc8b | 5.240325 | 5.169456 | 5.2141 | -0.04460398 | 0.026264655 | 5.3606429 | 5.0830925 | 5.35581 | -0.272717244 | 0.004833142 |
| Thbd | 9.045131 | 8.650715 | 8.0227 | 0.627987575 | 1.022404101 | 7.5387867 | 7.3409576 | 7.61374 | -0.27278321 | -0.074954101 |
| Lpgat1 | 10.85179 | 10.97797 | 10.878 | 0.100116648 | -0.026062196 | 10.985923 | 10.754603 | 11.0276 | -0.272986188 | -0.041665807 |
| Eln | 7.94473 | 7.826131 | 7.7269 | 0.099195316 | 0.217794465 | 7.5173481 | 7.130635 | 7.40363 | -0.272993521 | 0.11371951 |
| Dag1 | 9.159117 | 9.168229 | 9.2366 | -0.068327281 | -0.077439332 | 8.9149282 | 8.7226479 | 8.99607 | -0.273421146 | -0.081140785 |
| Fam53b | 10.44852 | 10.57121 | 10.486 | 0.085125114 | -0.03756081 | 10.214348 | 10.314243 | 10.5877 | -0.273434756 | -0.373329644 |
| Cul3 | 11.3065 | 11.25967 | 11.306 | -0.045970812 | 0.000854846 | 11.037087 | 10.945194 | 11.2188 | -0.273579244 | -0.181686881 |
| Zmym5 | 10.16567 | 10.2199 | 10.376 | -0.15620101 | -0.210426898 | 10.243555 | 10.144874 | 10.4186 | -0.273700222 | -0.175019598 |
| Slc9a6 | 8.801897 | 8.79212 | 8.7399 | 0.052195427 | 0.061972324 | 8.6449064 | 8.2710751 | 8.54482 | -0.273744211 | 0.100087106 |
| Lta4h | 10.53693 | 10.3904 | 10.538 | -0.147104002 | -0.000566242 | 10.427619 | 10.249813 | 10.5238 | -0.273973724 | -0.096168039 |
| Stox1 | 3.276959 | 3.082897 | 3.3163 | -0.233385561 | -0.03932393 | 3.1888064 | 2.9798345 | 3.25384 | -0.274000774 | -0.065028931 |
| Slc35d1 | 7.285842 | 7.385178 | 7.2565 | 0.128676964 | 0.029340235 | 7.2074166 | 7.0952238 | 7.36925 | -0.274030596 | -0.161837815 |
| Gpc2 | 5.486046 | 5.484367 | 5.5873 | -0.102931717 | -0.101252058 | 5.6274024 | 5.4185236 | 5.69261 | -0.274087102 | -0.065208296 |

| | | | | | | | | | | |
|---|---|---|---|---|---|---|---|---|---|---|
| Osgin1 | 5.969855 | 5.674225 | 5.7679 | -0.093634407 | 0.201995135 | 6.0916061 | 5.5417318 | 5.81584 | -0.274112423 | 0.275761854 |
| Kcnk13 | 3.960558 | 3.394304 | 3.6644 | -0.270088131 | 0.29616589 | 3.4332322 | 3.2508199 | 3.52511 | -0.274293582 | -0.091881227 |
| Slc22a17 | 4.923818 | 4.847712 | 4.7613 | 0.086402472 | 0.162509254 | 4.7920245 | 4.6998959 | 4.97422 | -0.274328158 | -0.182199494 |
| Lactb | 8.744861 | 8.68121 | 8.8428 | -0.16159036 | -0.097938843 | 8.6849616 | 8.4596956 | 8.73409 | -0.274393165 | -0.049127173 |
| Gtf3c3 | 8.388974 | 8.421388 | 8.4665 | -0.045111579 | -0.077526494 | 8.4840878 | 8.4174849 | 8.69192 | -0.274431382 | -0.20782848 |
| Mxra7 | 4.911278 | 4.725451 | 4.999 | -0.273518993 | -0.087691692 | 4.8810867 | 4.4414629 | 4.71613 | -0.274668644 | 0.164955092 |
| Kifc1 | 7.305026 | 7.243686 | 7.4419 | -0.198200147 | -0.136859376 | 7.0819041 | 7.2177855 | 7.49247 | -0.274681726 | -0.410563199 |
| Nek7 | 10.95949 | 11.00181 | 11.071 | -0.069360532 | -0.111681094 | 10.891783 | 10.686737 | 10.9615 | -0.274751706 | -0.069705481 |
| Gm12161 | 3.854319 | 3.848476 | 3.9395 | -0.091039219 | -0.085196446 | 3.7872674 | 3.6704559 | 3.9456 | -0.275141223 | -0.158329728 |
| Aox1 | 3.318932 | 3.205191 | 3.4285 | -0.223306811 | -0.109565937 | 3.1054912 | 3.0584892 | 3.33389 | -0.275403304 | -0.228401289 |
| Ccdc38 | 2.261031 | 2.253958 | 2.3449 | -0.090931245 | -0.083858731 | 2.2332476 | 2.1466832 | 2.42218 | -0.275494515 | -0.1889302 |
| Ripor3 | 4.761861 | 4.577386 | 4.6497 | -0.072276919 | 0.112198816 | 4.3130698 | 4.1805745 | 4.45615 | -0.275571616 | -0.143076235 |
| E2f7 | 5.599403 | 5.364549 | 5.6783 | -0.313713322 | -0.078858657 | 5.5951871 | 5.5066603 | 5.78245 | -0.27579287 | -0.187266087 |
| Chsy3 | 3.798381 | 3.574914 | 3.7232 | -0.14832659 | 0.075141373 | 3.6882226 | 3.509031 | 3.78498 | -0.275947416 | -0.096755886 |
| Ikzf1 | 13.35733 | 13.29293 | 13.336 | -0.04334014 | 0.021062948 | 13.119469 | 12.938599 | 13.2148 | -0.276160444 | -0.095290152 |
| Trpc2 | 2.056069 | 2.116374 | 2.0664 | 0.050002077 | -0.01030261 | 2.0351354 | 1.8842058 | 2.16037 | -0.276165934 | -0.125236333 |
| Zbtb18 | 10.23756 | 10.47664 | 10.223 | 0.253463041 | 0.014379665 | 10.629243 | 10.652578 | 10.9288 | -0.276226641 | -0.299561954 |
| Mbd4 | 8.573451 | 8.781626 | 8.6735 | 0.108128587 | -0.100046389 | 8.852519 | 8.5479809 | 8.82427 | -0.276286112 | 0.028252066 |
| Adamts12 | 3.709301 | 3.704524 | 3.7347 | -0.030206798 | -0.02542931 | 3.6087867 | 3.4768614 | 3.75321 | -0.276347316 | -0.14442209 |
| Timeless | 9.09258 | 8.896507 | 9.1077 | -0.2112387 | -0.015165683 | 9.2289499 | 9.1872794 | 9.46375 | -0.276465918 | -0.234795465 |
| Tmem241 | 7.667571 | 7.53428 | 7.7491 | -0.214867194 | -0.081576048 | 7.6458403 | 7.3984941 | 7.67498 | -0.276485406 | -0.029139163 |
| Ffar4 | 4.421484 | 4.474244 | 4.5987 | -0.124491096 | -0.177251086 | 4.4189336 | 4.2793425 | 4.55583 | -0.276490102 | -0.136899019 |
| Phka1 | 7.862919 | 7.655617 | 7.8573 | -0.201634303 | 0.005668522 | 7.7791599 | 7.5417087 | 7.81826 | -0.276550665 | -0.039099443 |
| Ccdc91 | 7.798442 | 7.837657 | 7.9208 | -0.083102168 | -0.122317687 | 7.6350037 | 7.5780704 | 7.85462 | -0.276552787 | -0.21961943 |
| Arap3 | 8.324285 | 7.998571 | 8.1185 | -0.119949415 | 0.205763681 | 7.6754577 | 7.3986951 | 7.67538 | -0.276689593 | 7.29366E-05 |
| Hells | 8.637363 | 8.711605 | 8.7862 | -0.074603077 | -0.148845155 | 8.416933 | 8.7880937 | 9.06486 | -0.276761461 | -0.647922104 |
| Hcfc2 | 8.350697 | 8.401851 | 8.4426 | -0.040723033 | -0.091877115 | 8.1917838 | 8.1080741 | 8.3849 | -0.276823518 | -0.193113783 |
| Rhof | 10.61387 | 10.71177 | 10.709 | 0.002960732 | -0.094939438 | 10.595953 | 10.344249 | 10.6211 | -0.276850085 | -0.025146533 |
| Rab33b | 8.722462 | 8.707049 | 8.7572 | -0.050118622 | -0.034705682 | 8.5739547 | 8.2475138 | 8.52444 | -0.276929533 | 0.049511429 |
| Mtr | 10.15159 | 10.11584 | 10.154 | -0.038094513 | -0.002343777 | 10.09898 | 9.8902372 | 10.1672 | -0.27699587 | -0.068253271 |
| Plekha8 | 5.803847 | 5.99571 | 6.0372 | -0.041489247 | -0.233351623 | 5.5763882 | 5.6829436 | 5.95999 | -0.277042324 | -0.383597669 |
| Mcph1 | 7.617576 | 7.571677 | 7.6715 | -0.099794458 | -0.05389478 | 7.4698831 | 7.512543 | 7.78961 | -0.277069956 | -0.319729879 |

| | | | | | | | | | | |
|---|---|---|---|---|---|---|---|---|---|---|
| Ddb1 | 12.29881 | 12.24013 | 12.253 | -0.013022662 | 0.045663586 | 12.235213 | 12.074691 | 12.352 | -0.277302276 | -0.116780028 |
| Trbv14 | 6.973969 | 6.932465 | 6.9963 | -0.063862855 | -0.022358939 | 6.8837576 | 6.7588574 | 7.03617 | -0.277309583 | -0.152409405 |
| Trav7-4 | 5.711863 | 5.571686 | 5.7595 | -0.18784774 | -0.047670708 | 5.4404867 | 5.3224351 | 5.59977 | -0.277331364 | -0.159279755 |
| Tyro3 | 5.887362 | 6.222368 | 5.9396 | 0.282788539 | -0.052217588 | 5.8147661 | 5.7899419 | 6.0673 | -0.277360719 | -0.25253657 |
| Bckdk | 8.933901 | 8.783087 | 9.0079 | -0.224775293 | -0.073961369 | 9.1759628 | 8.7657852 | 9.04327 | -0.27748241 | 0.132695244 |
| Cln6 | 8.303604 | 8.114837 | 8.4506 | -0.335784905 | -0.147017658 | 8.336769 | 7.996196 | 8.27384 | -0.277643146 | 0.062929887 |
| Slc4a4 | 3.52179 | 3.471758 | 3.5748 | -0.103087951 | -0.053055964 | 3.5038062 | 3.2470452 | 3.5248 | -0.27775854 | -0.020997486 |
| Ptprr | 2.348166 | 2.430479 | 2.3897 | 0.04073278 | -0.041580162 | 2.3878196 | 2.2289289 | 2.50671 | -0.277781394 | -0.118890687 |
| Chil5 | 4.99101 | 4.955446 | 5.095 | -0.139602218 | -0.10403767 | 4.9253282 | 4.7674632 | 5.04526 | -0.277800638 | -0.119935613 |
| Zfpm2 | 4.797837 | 4.540916 | 4.854 | -0.313091819 | -0.056171227 | 4.2041103 | 4.206334 | 4.48435 | -0.278012707 | -0.280236372 |
| Rab3a | 5.552337 | 5.641535 | 5.7754 | -0.133833254 | -0.223030864 | 5.5772232 | 5.2768035 | 5.55484 | -0.278031945 | 0.022387689 |
| Tmem218 | 5.495093 | 5.329116 | 5.5834 | -0.254272614 | -0.088295287 | 5.4423344 | 5.1261711 | 5.40424 | -0.278068044 | 0.038095308 |
| Cytl1 | 2.267068 | 2.336094 | 2.5363 | -0.200235721 | -0.269261313 | 2.1419722 | 2.1433484 | 2.42166 | -0.278307063 | -0.279683222 |
| Ugt1a6a | 2.288211 | 2.408105 | 2.1617 | 0.246427728 | 0.126532882 | 2.2811658 | 2.2488362 | 2.52721 | -0.278378505 | -0.24604886 |
| Alg10b | 10.08025 | 10.08536 | 10.128 | -0.042232566 | -0.047343092 | 10.050218 | 9.860932 | 10.1394 | -0.278473097 | -0.089186821 |
| Gm7240 | 4.580644 | 4.53079 | 4.7159 | -0.185155711 | -0.13530191 | 4.7547976 | 4.409343 | 4.68783 | -0.278488577 | 0.066966052 |
| Prcp | 10.53881 | 10.53629 | 10.614 | -0.078195648 | -0.075675468 | 10.609912 | 10.199237 | 10.4777 | -0.278502557 | 0.132172428 |
| Trim16 | 5.191968 | 4.998684 | 4.9928 | 0.005863789 | 0.199147328 | 5.1092726 | 4.73175 | 5.01028 | -0.278526709 | 0.098995825 |
| Igkv14-126 | 8.830104 | 8.636183 | 8.4413 | 0.194844959 | 0.388765383 | 8.5218269 | 8.5983899 | 8.87693 | -0.27853549 | -0.355098474 |
| Hsf4 | 3.85052 | 3.784963 | 3.8909 | -0.105903235 | -0.040346858 | 3.9169374 | 3.7005093 | 3.97907 | -0.278556041 | -0.062127982 |
| Gm10767 | 4.123166 | 3.94512 | 4.0653 | -0.120203635 | 0.057841915 | 4.1471515 | 3.8297896 | 4.10837 | -0.278575843 | 0.038786042 |
| Dvl2 | 8.932243 | 8.893911 | 8.9772 | -0.083289554 | -0.044957617 | 9.0561307 | 8.7358013 | 9.01442 | -0.278620508 | 0.041708926 |
| Guf1 | 8.844076 | 8.664913 | 8.7494 | -0.084468811 | 0.094694752 | 8.8425472 | 8.7092885 | 8.98795 | -0.278657189 | -0.145398427 |
| Gprc5c | 5.938696 | 5.719847 | 5.6375 | 0.082311982 | 0.301161615 | 5.7300929 | 5.2681906 | 5.54693 | -0.278735239 | 0.18316704 |
| Dmtf1 | 9.848242 | 9.861941 | 10.01 | -0.148201642 | -0.161900385 | 9.9861021 | 9.9516251 | 10.2304 | -0.278788285 | -0.244311347 |
| Morc4 | 4.105364 | 4.287422 | 4.2974 | -0.009957645 | -0.192015197 | 4.0721889 | 3.8790975 | 4.15794 | -0.278841931 | -0.08575046 |
| Aktip | 7.571671 | 7.495417 | 7.6968 | -0.201398947 | -0.125144654 | 7.4806531 | 7.3066529 | 7.58551 | -0.278853108 | -0.104852933 |
| Klhl7 | 8.091355 | 8.112031 | 8.0929 | 0.019134672 | -0.001540902 | 8.0356021 | 7.7480971 | 8.02699 | -0.278891379 | 0.008613584 |
| Prkch | 10.98583 | 10.93708 | 11.023 | -0.086281206 | -0.03753536 | 10.528052 | 10.576918 | 10.8559 | -0.278972984 | -0.327839745 |
| Rfxank | 8.723714 | 8.663024 | 8.7383 | -0.07527408 | -0.014584032 | 9.0377432 | 8.6371893 | 8.91618 | -0.278987089 | 0.121566796 |
| Nek8 | 7.587212 | 7.35908 | 7.6607 | -0.301596168 | -0.073464618 | 8.1409641 | 7.4878289 | 7.7669 | -0.279066682 | 0.374068502 |
| 4833420G17Rik | 10.20012 | 10.19346 | 10.303 | -0.109819898 | -0.103159728 | 10.44999 | 10.281539 | 10.5606 | -0.279091619 | -0.110641307 |

| | | | | | | | | | | |
|---|---|---|---|---|---|---|---|---|---|---|
| Pdpk1 | 10.88935 | 10.91902 | 10.882 | 0.037060392 | 0.007393232 | 10.665703 | 10.611407 | 10.8906 | -0.279192096 | -0.224896861 |
| Ppp1ccb | 10.27081 | 10.20205 | 10.283 | -0.080830262 | -0.012069722 | 10.197913 | 9.9468077 | 10.2261 | -0.279323242 | -0.028217789 |
| Ylpm1 | 10.53216 | 10.64592 | 10.622 | 0.024278172 | -0.089487188 | 10.567326 | 10.548372 | 10.8278 | -0.279408556 | -0.260455092 |
| Cenpn | 5.663258 | 5.499968 | 5.851 | -0.351027501 | -0.187738369 | 5.738021 | 5.6939609 | 5.97349 | -0.27952688 | -0.235466844 |
| Pskh1 | 9.096872 | 8.964838 | 9.1649 | -0.20009797 | -0.06806478 | 9.0955767 | 8.7472902 | 9.02694 | -0.279650948 | 0.068635524 |
| Fus | 13.00257 | 12.63577 | 12.972 | -0.336691606 | 0.03010345 | 13.158341 | 12.903649 | 13.1833 | -0.279658814 | -0.024967528 |
| Trio | 9.653435 | 9.397955 | 9.7219 | -0.323934374 | -0.068454456 | 9.4730883 | 9.6369052 | 9.91667 | -0.27976603 | -0.443582938 |
| Cramp1l | 9.789247 | 9.892957 | 9.8682 | 0.024785796 | -0.078924608 | 9.7006526 | 9.6796881 | 9.95945 | -0.279766071 | -0.258801596 |
| BC030867 | 4.68245 | 4.544644 | 4.7792 | -0.234546063 | -0.096739217 | 4.7835023 | 4.7657509 | 5.04555 | -0.27979817 | -0.26204676 |
| Spry4 | 5.95133 | 5.470638 | 5.5532 | -0.082596329 | 0.398096022 | 5.1667348 | 5.0630394 | 5.34311 | -0.280071871 | -0.176376452 |
| Traip | 6.061784 | 6.149892 | 6.2802 | -0.130290489 | -0.218398462 | 6.1396822 | 6.0730353 | 6.35352 | -0.280485323 | -0.213838411 |
| Ago3 | 9.338454 | 9.231074 | 9.2961 | -0.065048182 | 0.042332003 | 9.0462054 | 9.2381557 | 9.51876 | -0.280599494 | -0.472549783 |
| Cyp2f2 | 4.693297 | 3.868018 | 4.4371 | -0.569088841 | 0.256190552 | 4.1880359 | 3.9627338 | 4.24362 | -0.280881224 | -0.055579185 |
| Atg16l2 | 9.521494 | 9.728794 | 9.7521 | -0.023312398 | -0.230612928 | 9.9918468 | 9.7195023 | 10.0004 | -0.280901159 | -0.008556691 |
| Cnpy4 | 8.275469 | 8.27374 | 8.3585 | -0.084777957 | -0.08304935 | 8.3201798 | 8.1241837 | 8.40516 | -0.280976939 | -0.084980806 |
| Ctdspl | 4.7446 | 4.711828 | 4.7853 | -0.073449058 | -0.040676611 | 4.606448 | 4.4346197 | 4.71562 | -0.280998919 | -0.109170575 |
| Pfkm | 7.353297 | 7.34553 | 7.5525 | -0.206969864 | -0.19920353 | 7.2597204 | 7.138516 | 7.41959 | -0.281069371 | -0.159864969 |
| Lonrf1 | 6.161294 | 5.908429 | 5.9597 | -0.051303131 | 0.201561346 | 5.9222091 | 5.7986711 | 6.07977 | -0.281099209 | -0.157561255 |
| Lrrc61 | 9.347055 | 9.258272 | 9.4703 | -0.212077909 | -0.1232954 | 9.6045385 | 9.1060123 | 9.38714 | -0.281130812 | 0.217395414 |
| Rcsd1 | 12.12427 | 12.20312 | 12.228 | -0.024651192 | -0.1034971 | 12.102168 | 11.881549 | 12.1628 | -0.281222266 | -0.060602873 |
| Etfbkmt | 6.518475 | 6.36639 | 6.5638 | -0.197435644 | -0.045351205 | 6.6867666 | 6.3211806 | 6.60251 | -0.281333588 | 0.084252399 |
| Ubr3 | 10.77754 | 10.64843 | 10.721 | -0.072772976 | 0.05632974 | 10.784689 | 10.5947 | 10.8761 | -0.281383792 | -0.091394876 |
| Pigh | 7.606964 | 7.624259 | 7.6551 | -0.030880406 | -0.048175389 | 7.8831872 | 7.4765656 | 7.75803 | -0.281463932 | 0.12515761 |
| Grhl1 | 4.922313 | 4.795347 | 4.9397 | -0.144375372 | -0.017409072 | 4.818343 | 4.7123318 | 4.99411 | -0.281773248 | -0.175762051 |
| Ptchd1 | 3.184465 | 3.04857 | 3.2202 | -0.171631378 | -0.035736134 | 2.9942744 | 2.8532738 | 3.13519 | -0.281912326 | -0.140911719 |
| Ptp4a2 | 12.97196 | 12.97851 | 13.03 | -0.051581642 | -0.05812624 | 12.97209 | 12.709872 | 12.9918 | -0.281944566 | -0.01972617 |
| Cyb5a | 9.716722 | 9.665338 | 9.731 | -0.065689185 | -0.014305022 | 9.5535506 | 9.3128162 | 9.59477 | -0.28194913 | -0.041214754 |
| Lamb1 | 7.264382 | 7.247982 | 6.943 | 0.305014466 | 0.321414377 | 6.8676708 | 6.4781586 | 6.76013 | -0.281975622 | 0.10753656 |
| Cox4i2 | 3.406437 | 3.021818 | 3.3528 | -0.331005899 | 0.053613195 | 2.9330122 | 2.8749044 | 3.15711 | -0.28220294 | -0.224095096 |
| Dusp9 | 3.800036 | 3.686594 | 3.7952 | -0.108588857 | 0.004853322 | 3.6107393 | 3.4984107 | 3.78081 | -0.282401294 | -0.170072696 |
| Ttc17 | 9.846649 | 9.815179 | 9.8743 | -0.059093998 | -0.027624186 | 9.8251885 | 9.6996867 | 9.98216 | -0.282474226 | -0.156972516 |
| Mnat1 | 7.857209 | 7.803943 | 7.8408 | -0.036881554 | 0.016385079 | 7.8216944 | 7.5951391 | 7.87767 | -0.282530487 | -0.055975148 |

| | | | | | | | | | | |
|---|---|---|---|---|---|---|---|---|---|---|
| Foxf1 | 5.907074 | 5.513423 | 5.7286 | -0.215211671 | 0.178439739 | 4.9192646 | 4.9756671 | 5.25827 | -0.282604503 | -0.33900698 |
| Ctnnal1 | 6.398163 | 6.359539 | 6.5195 | -0.159960122 | -0.121336208 | 6.4333364 | 6.130623 | 6.41341 | -0.282787446 | 0.019925886 |
| Mib1 | 9.591637 | 9.562663 | 9.6589 | -0.096218869 | -0.067245805 | 9.2263095 | 9.3896601 | 9.67257 | -0.282909219 | -0.446259915 |
| Gm13301 | 3.464648 | 3.301215 | 3.3237 | -0.022485388 | 0.140947499 | 3.2045962 | 2.9632098 | 3.2462 | -0.282989892 | -0.04160352 |
| Jakmip1 | 10.2176 | 10.25373 | 10.245 | 0.009118153 | -0.02700496 | 10.243019 | 9.9918382 | 10.275 | -0.283126086 | -0.031944938 |
| Ppip5k2 | 10.28332 | 10.31442 | 10.312 | 0.002886918 | -0.028208406 | 10.207336 | 10.105267 | 10.3884 | -0.283172216 | -0.18110325 |
| Pcbd2 | 6.882236 | 6.783843 | 6.8788 | -0.094960015 | 0.003433327 | 7.1711176 | 6.652884 | 6.93613 | -0.283248427 | 0.234985173 |
| Ptar1 | 9.616536 | 9.783705 | 9.6823 | 0.101364321 | -0.065804625 | 9.3551443 | 9.4983906 | 9.7818 | -0.283412597 | -0.426658978 |
| Sapcd2 | 4.477923 | 4.464029 | 4.6589 | -0.194847643 | -0.180954133 | 4.5566027 | 4.5363985 | 4.81987 | -0.283469613 | -0.263265407 |
| Trbv13-3 | 7.717258 | 7.603129 | 7.6586 | -0.055471133 | 0.058657348 | 7.4213326 | 7.4078543 | 7.69133 | -0.283477831 | -0.269999536 |
| Gm13071 | 3.172939 | 3.15062 | 3.2016 | -0.050987065 | -0.028668236 | 3.2452762 | 3.1461004 | 3.42961 | -0.283507635 | -0.184331791 |
| Adgrl1 | 9.341468 | 9.10011 | 9.556 | -0.455892386 | -0.214534639 | 8.7832395 | 9.0505227 | 9.33411 | -0.283591047 | -0.550874248 |
| Tspan6 | 5.826874 | 5.665089 | 5.8359 | -0.170779311 | -0.0089938 | 5.3676734 | 5.1877738 | 5.47152 | -0.283742837 | -0.103843256 |
| Pacsin3 | 2.809238 | 2.722793 | 2.9066 | -0.183759638 | -0.097314124 | 2.852282 | 2.7267017 | 3.01061 | -0.283904335 | -0.158324132 |
| Natd1 | 8.052004 | 8.065315 | 8.254 | -0.188693324 | -0.202003613 | 8.2318105 | 7.9615276 | 8.24551 | -0.28398443 | -0.013701585 |
| Gm12258 | 5.177697 | 5.186308 | 5.5374 | -0.351057751 | -0.359668616 | 6.0796022 | 5.5293562 | 5.81342 | -0.284064027 | 0.26618196 |
| Spindoc | 7.914006 | 7.82229 | 8.0162 | -0.193881937 | -0.102166294 | 8.2647573 | 7.8228913 | 8.10702 | -0.284124542 | 0.157741492 |
| Pf4 | 3.574629 | 3.199437 | 3.3554 | -0.155959042 | 0.219232747 | 2.9161284 | 2.9050178 | 3.18917 | -0.284149857 | -0.273039255 |
| Zfp451 | 9.766881 | 9.769974 | 9.7128 | 0.057221866 | 0.054128548 | 9.5316663 | 9.682636 | 9.96704 | -0.284405588 | -0.435375289 |
| Haus5 | 8.842978 | 8.770083 | 8.8742 | -0.104133643 | -0.031238434 | 9.4090002 | 8.8822375 | 9.16678 | -0.284543195 | 0.242219499 |
| Gm3752 | 2.166881 | 2.179684 | 2.2268 | -0.047141912 | -0.059944858 | 2.253639 | 2.1144596 | 2.39902 | -0.284556464 | -0.145377022 |
| Daglb | 8.198058 | 8.191944 | 8.1936 | -0.001703075 | 0.004410382 | 8.2987266 | 7.7289324 | 8.01352 | -0.284590797 | 0.285203344 |
| Akap13 | 13.31074 | 13.40729 | 13.317 | 0.090510212 | -0.006042472 | 13.113077 | 13.079584 | 13.3642 | -0.284655974 | -0.251162763 |
| Tspyl3 | 8.900054 | 8.841042 | 8.9949 | -0.153864919 | -0.094853117 | 9.0569151 | 8.6188324 | 8.90352 | -0.284688142 | 0.153394473 |
| Usp30 | 8.59875 | 8.390116 | 8.711 | -0.320914256 | -0.112280912 | 8.5183974 | 8.3220174 | 8.60673 | -0.28470794 | -0.088327887 |
| Tmem98 | 4.019158 | 3.83578 | 4.0981 | -0.262337017 | -0.078959257 | 3.7875214 | 3.5284554 | 3.81336 | -0.284908139 | -0.025842133 |
| Zfp282 | 8.335629 | 8.288864 | 8.4219 | -0.133004057 | -0.086238801 | 8.3044004 | 7.9891866 | 8.27412 | -0.284929954 | 0.03028389 |
| Hoxa7 | 4.399102 | 4.309306 | 4.4761 | -0.166815813 | -0.077019119 | 4.2656697 | 3.94323 | 4.22837 | -0.285138447 | 0.03730126 |
| Gramd2 | 4.20616 | 3.863266 | 3.9823 | -0.119041786 | 0.223851734 | 3.8890828 | 3.6732624 | 3.95841 | -0.285150758 | -0.069330307 |
| Iqgap1 | 13.87973 | 13.82501 | 13.844 | -0.019262784 | 0.035455308 | 13.84561 | 13.541458 | 13.8267 | -0.285228984 | 0.018922873 |
| Dnajb9 | 9.785912 | 9.562904 | 9.8663 | -0.30340938 | -0.08040104 | 8.829498 | 8.8861937 | 9.17145 | -0.285252321 | -0.341947965 |
| Zfp995 | 6.341306 | 6.290875 | 6.5743 | -0.283463081 | -0.233032025 | 6.3152633 | 6.1290363 | 6.41435 | -0.285316893 | -0.099089868 |

| | | | | | | | | | | |
|---|---|---|---|---|---|---|---|---|---|---|
| Mybl1 | 5.362069 | 5.465233 | 5.4878 | -0.022569501 | -0.125733575 | 5.6571924 | 5.3105481 | 5.59613 | -0.285582772 | 0.061061561 |
| Chmp4c | 3.933731 | 3.943488 | 4.0136 | -0.070122316 | -0.079879899 | 4.2271322 | 3.7931045 | 4.07878 | -0.285670911 | 0.148356737 |
| Xkr5 | 3.440346 | 3.383666 | 3.5201 | -0.136397623 | -0.079717629 | 3.462068 | 3.2528406 | 3.53859 | -0.285747394 | -0.076520025 |
| Hmgxb4 | 9.026421 | 8.932401 | 9.1626 | -0.230169743 | -0.136149668 | 9.1492401 | 8.849103 | 9.13502 | -0.285921736 | 0.014215413 |
| Polh | 8.616812 | 8.694434 | 8.7025 | -0.008072046 | -0.085694248 | 8.5390673 | 8.3736932 | 8.65962 | -0.285931204 | -0.120557102 |
| 1110008L16Rik | 7.08057 | 6.882074 | 7.1513 | -0.269262546 | -0.070766412 | 7.0161711 | 6.8357617 | 7.12176 | -0.285997777 | -0.105588446 |
| Cxcr4 | 12.68743 | 13.14636 | 12.785 | 0.361197586 | -0.097739946 | 12.428983 | 12.373998 | 12.6601 | -0.28606717 | -0.231081759 |
| Arrb2 | 10.8606 | 10.86606 | 10.974 | -0.107564222 | -0.113026774 | 11.009941 | 10.778415 | 11.0645 | -0.28607152 | -0.054545901 |
| Gm6136 | 3.874142 | 3.61076 | 3.4292 | 0.181574258 | 0.444956483 | 3.5266003 | 3.4120741 | 3.6983 | -0.286230501 | -0.171704333 |
| Uhmk1 | 11.7313 | 11.71095 | 11.748 | -0.037413302 | -0.017067462 | 11.673244 | 11.437962 | 11.7243 | -0.286385268 | -0.051102704 |
| Zfp950 | 8.726537 | 8.782311 | 8.7094 | 0.072868626 | 0.017094937 | 9.124158 | 8.8496292 | 9.13626 | -0.286633542 | -0.012104778 |
| Bicc1 | 5.963906 | 5.697301 | 5.6629 | 0.034352253 | 0.300957678 | 5.6050605 | 5.3322989 | 5.619 | -0.286699255 | -0.013937636 |
| Kdm4c | 9.67388 | 9.703696 | 9.7424 | -0.038662547 | -0.068478749 | 9.5768637 | 9.453798 | 9.74056 | -0.286757364 | -0.163691651 |
| Etfdh | 9.059152 | 9.047634 | 9.1361 | -0.088451442 | -0.076933367 | 9.0580666 | 8.8760308 | 9.163 | -0.286967812 | -0.104931964 |
| Fhl1 | 4.276572 | 4.382699 | 4.4886 | -0.105943437 | -0.212070274 | 4.1375904 | 4.0377632 | 4.32485 | -0.287085805 | -0.18725867 |
| Anapc1 | 11.49902 | 11.52998 | 11.482 | 0.048370182 | 0.01740267 | 11.4781 | 11.460197 | 11.7473 | -0.287137606 | -0.26923529 |
| Podnl1 | 6.861856 | 6.774856 | 7.019 | -0.244093936 | -0.157094505 | 6.6482457 | 6.654924 | 6.94212 | -0.287193728 | -0.293872011 |
| Celsr2 | 5.948305 | 5.910171 | 6.1178 | -0.207615459 | -0.169481822 | 6.0998137 | 5.6388203 | 5.92605 | -0.287234005 | 0.17375941 |
| Gm29609 | 5.4109 | 5.511198 | 5.2004 | 0.310829956 | 0.210531689 | 5.5928292 | 5.4340967 | 5.72133 | -0.287237519 | -0.128505011 |
| Suox | 8.18731 | 8.264903 | 8.3812 | -0.116261503 | -0.193853953 | 8.1284762 | 7.9457267 | 8.2331 | -0.287374819 | -0.104625268 |
| Dstn | 9.632898 | 9.495219 | 9.6128 | -0.117567919 | 0.020111094 | 9.5124336 | 9.1999601 | 9.48735 | -0.287391773 | 0.025081657 |
| Hsd17b4 | 9.330749 | 9.206178 | 9.3635 | -0.157320154 | -0.03274863 | 9.339274 | 8.9418965 | 9.22931 | -0.287414331 | 0.109963124 |
| Rab39b | 3.764273 | 3.588123 | 3.6869 | -0.098759228 | 0.077390962 | 3.6731886 | 3.4834551 | 3.77093 | -0.287470866 | -0.097737388 |
| Ighv14-4 | 7.120326 | 7.906134 | 7.1269 | 0.779277549 | -0.006530243 | 7.3209316 | 6.9437333 | 7.23146 | -0.287729124 | 0.089469226 |
| mt-Nd2 | 14.3711 | 14.25534 | 14.436 | -0.18032905 | -0.064576822 | 14.337799 | 14.175091 | 14.4629 | -0.287769088 | -0.125060304 |
| Lrch2 | 3.314341 | 3.244907 | 3.3477 | -0.102756224 | -0.033321748 | 3.0852316 | 2.9791869 | 3.26715 | -0.287960582 | -0.181915898 |
| Trbv19 | 7.501894 | 7.470954 | 7.5485 | -0.077544438 | -0.04660517 | 7.2632008 | 7.2014081 | 7.48937 | -0.287965528 | -0.22617276 |
| Fam205a2 | 3.088023 | 2.884345 | 3.1041 | -0.219764114 | -0.016085937 | 2.7325411 | 2.6787087 | 2.96675 | -0.288039289 | -0.234206918 |
| Zfp763 | 6.661195 | 6.638492 | 6.7405 | -0.102028388 | -0.079324855 | 6.5858597 | 6.4462972 | 6.73441 | -0.288117012 | -0.148554461 |
| Heatr9 | 5.766849 | 5.621525 | 5.6705 | -0.048970799 | 0.096353766 | 5.3564497 | 5.4862723 | 5.77461 | -0.288341134 | -0.418163741 |
| Gaa | 9.615368 | 9.702803 | 9.6416 | 0.061219876 | -0.026215546 | 9.768165 | 9.4226275 | 9.71124 | -0.28861129 | 0.05692617 |
| 6330403K07Rik | 3.211683 | 3.082903 | 3.287 | -0.204068109 | -0.075288678 | 2.997538 | 2.9287864 | 3.21758 | -0.2887905 | -0.220038848 |

| | | | | | | | | | | |
|---|---|---|---|---|---|---|---|---|---|---|
| Actn4 | 12.07318 | 12.03281 | 12.026 | 0.006375592 | 0.046744936 | 11.965477 | 11.700158 | 11.989 | -0.2888746 | -0.023555075 |
| Clec4b1 | 3.342211 | 3.238915 | 3.2183 | 0.020567678 | 0.123863781 | 3.3817268 | 2.9427537 | 3.2317 | -0.28894758 | 0.150025508 |
| Prss16 | 3.763926 | 3.579292 | 3.7708 | -0.191552719 | -0.006918802 | 4.0891464 | 3.669417 | 3.95837 | -0.288949011 | 0.130780362 |
| Xlr4a | 7.673632 | 7.682343 | 7.8685 | -0.186171738 | -0.194882877 | 7.5392229 | 7.5476925 | 7.83692 | -0.289230924 | -0.29770049 |
| Cacna1a | 5.536581 | 5.478034 | 5.5338 | -0.055738139 | 0.002809235 | 5.447853 | 5.4470979 | 5.73638 | -0.289283493 | -0.28852841 |
| Cenpp | 4.417895 | 4.300544 | 4.445 | -0.144481483 | -0.027130692 | 4.29248 | 4.2762807 | 4.56561 | -0.289333845 | -0.273134552 |
| Gm1821 | 7.494263 | 7.1594 | 7.1363 | 0.023064931 | 0.357927183 | 6.9327498 | 7.3473287 | 7.63673 | -0.289402196 | -0.703981111 |
| Ccdc142 | 5.013752 | 5.05992 | 5.0024 | 0.05751063 | 0.011342603 | 5.2867157 | 4.9034604 | 5.19288 | -0.28941764 | 0.093837642 |
| Kcna3 | 8.429066 | 8.388852 | 8.5185 | -0.129679032 | -0.089465406 | 8.4877081 | 8.4096856 | 8.69923 | -0.289539624 | -0.211517111 |
| Brwd3 | 9.241281 | 9.250733 | 9.2341 | 0.016662736 | 0.007210461 | 8.9695636 | 9.0564169 | 9.34612 | -0.289699758 | -0.376553121 |
| Cyp4f16 | 7.947836 | 7.802701 | 7.5912 | 0.211491716 | 0.356626032 | 8.426773 | 7.4451376 | 7.73491 | -0.28977428 | 0.691861142 |
| Dus2 | 7.548924 | 7.51029 | 7.7368 | -0.226477526 | -0.187843717 | 7.9856188 | 7.4607129 | 7.75051 | -0.289799919 | 0.235106026 |
| Pink1 | 10.17375 | 10.01351 | 10.27 | -0.256773737 | -0.096524431 | 10.017023 | 9.7302432 | 10.0201 | -0.289829821 | -0.00304957 |
| Dhx29 | 9.397256 | 9.316689 | 9.3676 | -0.050957608 | 0.029609477 | 9.3076446 | 9.162782 | 9.4527 | -0.289920248 | -0.145057596 |
| Gm10644 | 4.4707 | 4.453835 | 4.6273 | -0.17346844 | -0.156603792 | 4.7481631 | 4.3830821 | 4.67305 | -0.289970154 | 0.075110759 |
| Cadps2 | 5.382838 | 4.946419 | 5.3104 | -0.36397557 | 0.072443514 | 4.6409331 | 4.6845222 | 4.97479 | -0.290262818 | -0.333851941 |
| 5730507C01Rik | 3.453658 | 3.454016 | 3.549 | -0.095002835 | -0.095360149 | 3.6546563 | 3.3559961 | 3.64629 | -0.290290783 | 0.008369471 |
| Abl2 | 10.50392 | 10.41272 | 10.478 | -0.064921494 | 0.026280518 | 10.533432 | 10.291559 | 10.5819 | -0.290365064 | -0.048491897 |
| Pcdhgc3 | 7.532359 | 7.239703 | 7.1548 | 0.084912649 | 0.377568958 | 6.8693274 | 6.5239617 | 6.81453 | -0.290570843 | 0.054794889 |
| Smc4 | 12.34493 | 12.42143 | 12.525 | -0.103307056 | -0.179806512 | 12.069968 | 12.203599 | 12.4942 | -0.290586934 | -0.424217726 |
| Birc3 | 12.10662 | 11.91787 | 11.962 | -0.044538522 | 0.144206006 | 12.057788 | 11.841414 | 12.132 | -0.290603698 | -0.074229857 |
| Nanos1 | 6.658692 | 6.764681 | 6.7013 | 0.063399526 | -0.042589797 | 6.4785418 | 6.3258312 | 6.61649 | -0.290658081 | -0.137947447 |
| Exoc3l | 5.144138 | 4.849526 | 5.0277 | -0.178210017 | 0.116402134 | 4.7592981 | 4.7523153 | 5.04303 | -0.290715758 | -0.283732949 |
| Tiam2 | 4.88678 | 4.777472 | 4.7917 | -0.014182088 | 0.095126142 | 4.813979 | 4.5347362 | 4.82548 | -0.290740749 | -0.011498017 |
| Gm10521 | 6.752152 | 6.77825 | 6.8954 | -0.117181962 | -0.143280179 | 7.0307629 | 6.9537319 | 7.24463 | -0.290898523 | -0.213867458 |
| Itfg1 | 9.729219 | 9.62641 | 9.8031 | -0.176704899 | -0.07389516 | 9.7236426 | 9.5003113 | 9.79133 | -0.291015103 | -0.067683869 |
| Gid4 | 9.692957 | 9.686449 | 9.7433 | -0.056890085 | -0.050381678 | 9.5437408 | 9.3619563 | 9.65301 | -0.291048987 | -0.109264492 |
| Zfp664 | 9.685305 | 9.758162 | 9.6757 | 0.082449035 | 0.009592676 | 9.5298678 | 9.4400168 | 9.73107 | -0.291056117 | -0.201205137 |
| Nipsnap1 | 8.160333 | 7.981301 | 8.1961 | -0.214819387 | -0.035787854 | 8.2546151 | 7.8664762 | 8.15759 | -0.291112028 | 0.097026794 |
| Ccdc8 | 4.67068 | 4.639963 | 4.7484 | -0.108451512 | -0.077734228 | 4.7207553 | 4.6351748 | 4.92632 | -0.291141425 | -0.205560899 |
| Tbl1xr1 | 10.6285 | 10.72745 | 10.704 | 0.023362008 | -0.075585082 | 10.744661 | 10.564302 | 10.8555 | -0.29116026 | -0.110801507 |
| Grb10 | 6.378847 | 6.065576 | 6.2703 | -0.204708779 | 0.108562718 | 5.8835441 | 5.6807422 | 5.97201 | -0.291269132 | -0.088467235 |

| | | | | | | | | | | |
|---|---|---|---|---|---|---|---|---|---|---|
| Spidr | 8.577982 | 8.466579 | 8.5368 | -0.070230664 | 0.041171759 | 8.6643521 | 8.3174533 | 8.6089 | -0.291447227 | 0.055451605 |
| Sde2 | 9.711743 | 9.629087 | 9.7263 | -0.097218136 | -0.014561953 | 9.5250437 | 9.4151164 | 9.70666 | -0.291543731 | -0.181616407 |
| Cdnf | 4.840172 | 4.516623 | 4.5052 | 0.011470878 | 0.335019245 | 4.5924897 | 4.3721026 | 4.66369 | -0.291590222 | -0.071203133 |
| Ajuba | 4.298269 | 4.297786 | 4.3335 | -0.035679384 | -0.035196028 | 4.2364591 | 4.0384276 | 4.33003 | -0.291601546 | -0.093569996 |
| Pofut1 | 9.789761 | 9.847212 | 9.8508 | -0.003597303 | -0.061047956 | 9.9217893 | 9.6376469 | 9.92943 | -0.291783906 | -0.007641519 |
| Bptf | 12.19839 | 12.23412 | 12.237 | -0.002759098 | -0.03849076 | 12.117854 | 12.081544 | 12.3733 | -0.291799344 | -0.25548929 |
| Slc25a10 | 7.654397 | 7.5914 | 7.7917 | -0.200279098 | -0.1372822 | 7.7274577 | 7.3037743 | 7.59571 | -0.291933716 | 0.131749643 |
| Gm6361 | 5.894141 | 5.740908 | 5.7318 | 0.009082399 | 0.162314884 | 5.88677 | 5.7223746 | 6.01431 | -0.291939848 | -0.127544462 |
| Cd209f | 4.142868 | 3.276769 | 3.6951 | -0.418335553 | 0.447763448 | 3.087218 | 3.0021147 | 3.29409 | -0.291971933 | -0.206868644 |
| Slc4a7 | 10.58287 | 10.58023 | 10.562 | 0.018697458 | 0.021334984 | 10.590055 | 10.456105 | 10.7481 | -0.292036572 | -0.158086567 |
| P2ry10b | 8.840386 | 8.949679 | 8.9051 | 0.044590364 | -0.064702144 | 8.9264713 | 8.5363359 | 8.82843 | -0.292089368 | 0.098046005 |
| Zfp960 | 6.626449 | 6.473554 | 6.7079 | -0.234302252 | -0.08140729 | 6.6617978 | 6.3471038 | 6.63932 | -0.292219354 | 0.022474604 |
| Gulp1 | 3.816708 | 3.633108 | 3.8413 | -0.208162977 | -0.024562827 | 3.4706631 | 3.3716022 | 3.66394 | -0.292341846 | -0.193280923 |
| Slc25a25 | 9.1707 | 9.196208 | 9.0673 | 0.128891622 | 0.103384199 | 9.1125058 | 8.8913404 | 9.18374 | -0.292399047 | -0.071233582 |
| Nrbp2 | 5.056371 | 4.903656 | 4.9928 | -0.089192432 | 0.063522315 | 4.668083 | 4.6455241 | 4.93799 | -0.292469898 | -0.269910932 |
| Gm10273 | 3.103483 | 2.717528 | 4.358 | -1.640513588 | -1.254559236 | 3.2725522 | 2.9962638 | 3.28911 | -0.292841853 | -0.016553507 |
| Qpctl | 7.817105 | 7.693041 | 7.8644 | -0.171361653 | -0.047297155 | 7.9189532 | 7.4664582 | 7.75945 | -0.292994296 | 0.15950071 |
| Armcx5 | 7.863731 | 7.578494 | 7.8549 | -0.276380805 | 0.008855638 | 7.7216366 | 7.4639185 | 7.75694 | -0.293026165 | -0.035308076 |
| P2ry1 | 4.414119 | 4.273976 | 4.3976 | -0.123614646 | 0.016528332 | 4.1563583 | 4.0160748 | 4.30917 | -0.293090631 | -0.152807105 |
| Anp32e | 11.50456 | 11.40217 | 11.48 | -0.077714958 | 0.024673354 | 11.321348 | 11.253998 | 11.5471 | -0.293095112 | -0.225745547 |
| Fam91a1 | 9.117111 | 8.907156 | 9.0245 | -0.117367028 | 0.092588347 | 9.0006692 | 8.9031858 | 9.19632 | -0.293133387 | -0.195649944 |
| Rcor3 | 6.980398 | 7.024798 | 7.0614 | -0.036553312 | -0.080953194 | 6.8337386 | 6.6285799 | 6.92173 | -0.293145223 | -0.087986466 |
| Osbpl2 | 8.953384 | 8.915072 | 8.9826 | -0.067566083 | -0.029254033 | 9.0048981 | 8.7291933 | 9.02258 | -0.293381833 | -0.017677102 |
| Gm45844 | 3.358652 | 3.326447 | 3.4027 | -0.076210394 | -0.044005268 | 3.3810183 | 3.1916237 | 3.48515 | -0.293530127 | -0.104135523 |
| Rfx7 | 11.37984 | 11.40195 | 11.484 | -0.081852402 | -0.103967466 | 11.300115 | 11.195189 | 11.4887 | -0.293547938 | -0.188622212 |
| Usp22 | 10.23122 | 10.12013 | 10.334 | -0.213907909 | -0.10281931 | 10.195104 | 9.9352577 | 10.2289 | -0.293651685 | -0.033805288 |
| Nxn | 8.009568 | 7.788247 | 7.7768 | 0.011422791 | 0.232743509 | 7.6996185 | 7.4685663 | 7.76228 | -0.293717408 | -0.06266521 |
| Trav14-1 | 5.275595 | 5.032265 | 5.2957 | -0.263392511 | -0.02006279 | 5.2208246 | 5.0195666 | 5.31367 | -0.294108236 | -0.092850146 |
| Spata1 | 6.230165 | 5.923355 | 6.1618 | -0.238423182 | 0.06838698 | 6.1085209 | 5.9863564 | 6.28061 | -0.294249279 | -0.172084797 |
| Ehmt1 | 10.14314 | 10.09593 | 10.167 | -0.071301704 | -0.024095324 | 10.141986 | 9.9136653 | 10.2079 | -0.294282848 | -0.065962282 |
| Sncaip | 4.781598 | 4.43946 | 4.6919 | -0.252416584 | 0.089721438 | 4.4895893 | 4.3514873 | 4.64578 | -0.29428911 | -0.156187081 |
| Prdm8 | 2.794786 | 2.659491 | 2.8455 | -0.186038491 | -0.050743837 | 2.8360322 | 2.6769595 | 2.97129 | -0.294328143 | -0.135255371 |

| | | | | | | | | | | |
|---|---|---|---|---|---|---|---|---|---|---|
| Acad11 | 7.900723 | 7.961064 | 8.0599 | -0.098859802 | -0.159200787 | 8.0927765 | 7.6248636 | 7.9192 | -0.294331401 | 0.173581547 |
| Pam | 8.728358 | 8.360098 | 8.6807 | -0.320638293 | 0.047621707 | 7.9824966 | 7.9551309 | 8.24949 | -0.29436043 | -0.26699474 |
| Tnks2 | 11.95931 | 11.9415 | 11.971 | -0.029910014 | -0.012105362 | 11.804818 | 11.64514 | 11.9395 | -0.29436908 | -0.134691721 |
| Tsen2 | 6.574089 | 6.47735 | 6.6757 | -0.198349857 | -0.101610152 | 6.8603789 | 6.4807494 | 6.77516 | -0.294412757 | 0.0852168 |
| Ssh3 | 8.229192 | 7.902604 | 8.206 | -0.303386623 | 0.023201623 | 8.4242879 | 7.9973983 | 8.29192 | -0.294521768 | 0.132367835 |
| Klhl11 | 7.535156 | 7.473371 | 7.4739 | -0.000560967 | 0.061224725 | 7.4335629 | 7.3394461 | 7.63402 | -0.294571051 | -0.200454174 |
| Tbc1d9 | 8.497283 | 8.516061 | 8.3845 | 0.131512489 | 0.112734577 | 8.5178307 | 8.1032455 | 8.39784 | -0.294599506 | 0.119985707 |
| Rbm44 | 4.097719 | 3.790139 | 3.9304 | -0.140224866 | 0.167355342 | 3.8953096 | 3.7843361 | 4.07896 | -0.294620827 | -0.183647339 |
| Slc25a37 | 10.59835 | 10.72838 | 10.8 | -0.071126982 | -0.201159664 | 10.975157 | 10.563829 | 10.8585 | -0.294681638 | 0.116646202 |
| Zfp944 | 7.84989 | 7.875904 | 7.9337 | -0.057809095 | -0.083823004 | 8.09238 | 7.7193078 | 8.014 | -0.294688696 | 0.078383486 |
| Bsn | 5.126897 | 5.221306 | 5.2473 | -0.025999183 | -0.12040883 | 5.2565194 | 5.1234759 | 5.41823 | -0.294753444 | -0.161709937 |
| Mboat7 | 9.792216 | 9.727841 | 9.7916 | -0.063798696 | 0.000576024 | 10.057466 | 9.5713612 | 9.86617 | -0.294812799 | 0.191292028 |
| Rccd1 | 7.857047 | 7.670901 | 7.8875 | -0.216562757 | -0.030416275 | 8.254969 | 7.9568172 | 8.25171 | -0.294897036 | 0.003254749 |
| Ltn1 | 10.72805 | 10.802 | 10.746 | 0.056029586 | -0.01791931 | 10.652954 | 10.46796 | 10.7629 | -0.294905042 | -0.109910475 |
| Fndc3b | 8.302381 | 8.089491 | 7.9469 | 0.142552111 | 0.35544247 | 7.7034085 | 7.5409264 | 7.83586 | -0.294938459 | -0.132456333 |
| Prpf8 | 13.17685 | 13.14589 | 13.197 | -0.05069192 | -0.019725016 | 13.228906 | 12.977031 | 13.2721 | -0.295022336 | -0.043147024 |
| B3gnt7 | 6.49485 | 6.181787 | 6.2673 | -0.085534247 | 0.227528493 | 6.2311217 | 6.0148409 | 6.30992 | -0.295082368 | -0.078801549 |
| Rab3il1 | 7.818246 | 7.048388 | 7.3039 | -0.255548499 | 0.514309118 | 6.9751238 | 6.3706797 | 6.66577 | -0.295088632 | 0.30935547 |
| Plin3 | 8.889107 | 8.732423 | 9.0553 | -0.322923244 | -0.166238769 | 8.9600302 | 8.6584043 | 8.95355 | -0.295143136 | 0.006482774 |
| Radil | 2.97087 | 2.810628 | 2.8833 | -0.072689004 | 0.0875529 | 2.8420304 | 2.6870566 | 2.98234 | -0.295279883 | -0.140306095 |
| Naa30 | 9.754205 | 9.656173 | 9.6685 | -0.012327564 | 0.085704175 | 9.4484865 | 9.4492273 | 9.74455 | -0.29532585 | -0.296066699 |
| Stxbp5 | 9.47087 | 9.388659 | 9.4131 | -0.024456039 | 0.057754922 | 9.3476638 | 9.1008625 | 9.39622 | -0.295352484 | -0.048551277 |
| Fam228b | 3.033856 | 3.01979 | 2.9931 | 0.026692152 | 0.040758812 | 3.2147713 | 2.9431782 | 3.23853 | -0.295353046 | -0.023759947 |
| Mga | 11.35875 | 11.49042 | 11.478 | 0.012147558 | -0.11952455 | 11.396382 | 11.307536 | 11.603 | -0.295477446 | -0.206631597 |
| Adcy6 | 9.907293 | 9.909876 | 10.034 | -0.12459097 | -0.127174149 | 9.923521 | 9.8884032 | 10.184 | -0.29563506 | -0.260517331 |
| Lnpk | 8.44899 | 8.317268 | 8.3382 | -0.020974481 | 0.110746873 | 8.4172289 | 8.275131 | 8.5708 | -0.295665384 | -0.153567429 |
| Bcl7a | 9.187294 | 9.268898 | 9.2639 | 0.004972743 | -0.076631489 | 9.4904663 | 9.1221943 | 9.41803 | -0.295831382 | 0.072440598 |
| Tmcc1 | 9.017378 | 9.059322 | 9.1221 | -0.062811266 | -0.104754948 | 9.0599078 | 8.9226646 | 9.21855 | -0.295885746 | -0.158642552 |
| Gpr137b | 8.582843 | 8.850569 | 8.4643 | 0.386273567 | 0.118547055 | 8.4011274 | 8.2027509 | 8.49866 | -0.295913055 | -0.097536514 |
| Kcnq5 | 7.792512 | 7.737996 | 7.8536 | -0.1156058 | -0.061088988 | 7.7778632 | 7.5181659 | 7.81422 | -0.296058888 | -0.036361577 |
| Plekhh3 | 4.452893 | 4.275729 | 4.3959 | -0.120195802 | 0.056968887 | 4.2083318 | 4.0012068 | 4.29728 | -0.296077725 | -0.088952768 |
| St8sia1 | 9.632218 | 9.787318 | 9.7436 | 0.043747702 | -0.111351675 | 9.209325 | 9.3600739 | 9.6562 | -0.296126546 | -0.446875469 |

| | | | | | | | | | | |
|---|---|---|---|---|---|---|---|---|---|---|
| Rnf125 | 8.234144 | 7.988483 | 8.2084 | -0.219881523 | 0.025779389 | 7.6288672 | 7.7863564 | 8.08255 | -0.296191008 | -0.453680237 |
| Zfp125 | 6.973577 | 6.843745 | 7.0705 | -0.226717172 | -0.096885107 | 7.004266 | 6.8053168 | 7.1016 | -0.296281871 | -0.097332694 |
| Uggt2 | 5.985384 | 5.694741 | 5.8891 | -0.194366741 | 0.096275638 | 5.8314631 | 5.707767 | 6.00414 | -0.296374453 | -0.172678369 |
| 4930432K21Rik | 5.499988 | 5.453098 | 5.4086 | 0.044541309 | 0.091432143 | 5.521475 | 5.2345866 | 5.53111 | -0.296525618 | -0.00963717 |
| Idh2 | 9.259168 | 9.127314 | 9.2875 | -0.160185666 | -0.028331706 | 9.1379104 | 8.8836661 | 9.18019 | -0.296526118 | -0.042281828 |
| Zfp932 | 7.900298 | 7.931921 | 7.8343 | 0.097619294 | 0.065996933 | 8.0439366 | 7.899042 | 8.1956 | -0.296559183 | -0.151664573 |
| Cd84 | 11.11863 | 11.13507 | 11.227 | -0.091675372 | -0.108115418 | 11.159272 | 10.93157 | 11.2281 | -0.296572324 | -0.068870758 |
| Prkag1 | 9.872309 | 9.7475 | 10.036 | -0.288777622 | -0.163968684 | 10.107131 | 9.78253 | 10.0793 | -0.296770687 | 0.027830294 |
| Cped1 | 3.773308 | 3.729204 | 4.0855 | -0.35631653 | -0.312213232 | 3.5869708 | 3.5796801 | 3.87648 | -0.296804844 | -0.289514148 |
| Ezh2 | 9.385544 | 9.298258 | 9.5516 | -0.253381434 | -0.166095499 | 9.3610965 | 9.3549209 | 9.6518 | -0.296881404 | -0.290705796 |
| Zfp764 | 7.561842 | 7.474353 | 7.5033 | -0.028913344 | 0.058576224 | 7.5825332 | 7.304423 | 7.60149 | -0.297069532 | -0.018959262 |
| Phpt1 | 8.030248 | 8.079123 | 8.067 | 0.012092768 | -0.036782328 | 8.1536736 | 7.8554227 | 8.15257 | -0.297151102 | 0.001099775 |
| Atpaf1 | 7.922054 | 7.658348 | 7.9298 | -0.271459089 | -0.007752832 | 7.7980541 | 7.6374959 | 7.93489 | -0.297389928 | -0.136831732 |
| Ccdc107 | 7.407009 | 7.011019 | 7.2891 | -0.278055672 | 0.117934128 | 7.4868108 | 6.9951892 | 7.2927 | -0.29750638 | 0.194115219 |
| Taf4 | 9.620903 | 9.65757 | 9.6099 | 0.047679843 | 0.0110128 | 9.4780956 | 9.4402147 | 9.73779 | -0.297571492 | -0.259690584 |
| Zfyve16 | 8.782113 | 8.73685 | 8.7495 | -0.012638975 | 0.032623495 | 8.581702 | 8.6178238 | 8.9154 | -0.297573189 | -0.333694956 |
| Gm8750 | 3.129558 | 2.90834 | 3.2608 | -0.352438116 | -0.131219955 | 2.9932854 | 2.9492259 | 3.24697 | -0.297741302 | -0.253681793 |
| Kbtbd3 | 6.01554 | 6.118205 | 6.1202 | -0.001953967 | -0.104619769 | 6.2328547 | 6.009834 | 6.30769 | -0.29785787 | -0.074837106 |
| Galnt12 | 8.665883 | 8.699989 | 8.8139 | -0.113950402 | -0.14805613 | 8.6518086 | 8.4323321 | 8.73023 | -0.297897412 | -0.078420942 |
| Ehf | 6.68063 | 6.359141 | 6.5333 | -0.174156024 | 0.147332962 | 6.2038895 | 6.2254618 | 6.52352 | -0.298055107 | -0.319627446 |
| Zfp942 | 8.717426 | 8.724755 | 8.6899 | 0.034884879 | 0.027556136 | 8.7978377 | 8.602035 | 8.90031 | -0.298276201 | -0.102473443 |
| Snx32 | 5.82703 | 5.60856 | 5.9739 | -0.365304703 | -0.146835281 | 6.4097289 | 5.8910525 | 6.18937 | -0.298321357 | 0.220355042 |
| Jarid2 | 10.68763 | 10.68384 | 10.574 | 0.1096981 | 0.113492752 | 10.606321 | 10.430184 | 10.7286 | -0.298422282 | -0.122285237 |
| Fam135a | 5.054895 | 4.990708 | 5.2035 | -0.212833787 | -0.148646085 | 4.8643706 | 4.6872101 | 4.98576 | -0.29854544 | -0.121384943 |
| Csnk1g3 | 10.13881 | 10.15632 | 10.23 | -0.073961184 | -0.091477403 | 10.004558 | 9.7860836 | 10.0846 | -0.298552647 | -0.080078696 |
| Aldh4a1 | 8.637923 | 8.745765 | 8.7422 | 0.003589628 | -0.104252015 | 8.6856164 | 8.3169685 | 8.61555 | -0.298580946 | 0.070066956 |
| Camp | 3.95558 | 3.789834 | 3.9244 | -0.134524035 | 0.031222429 | 4.2871539 | 3.6882383 | 3.98683 | -0.298593752 | 0.300321784 |
| Gabarapl1 | 7.911891 | 7.713958 | 7.9974 | -0.283462898 | -0.085529907 | 7.4127465 | 7.4907889 | 7.78941 | -0.298618376 | -0.376660822 |
| 4921507P07Rik | 4.213051 | 4.051828 | 4.2725 | -0.220693558 | -0.059470211 | 4.0482976 | 3.9141876 | 4.21297 | -0.298786275 | -0.164676262 |
| Fah | 6.189353 | 6.129221 | 6.2768 | -0.147613714 | -0.087482001 | 6.1033678 | 5.8810101 | 6.17991 | -0.298897224 | -0.076539536 |
| Gm2a | 11.83803 | 11.78965 | 11.838 | -0.047882658 | 0.000501482 | 12.100041 | 11.44819 | 11.7472 | -0.298969438 | 0.352881074 |
| Tmco3 | 7.504311 | 7.736432 | 7.7439 | -0.007429223 | -0.23955043 | 7.6676149 | 7.5592714 | 7.85845 | -0.299177866 | -0.19083439 |

| | | | | | | | | | | |
|---|---|---|---|---|---|---|---|---|---|---|
| Slc12a6 | 11.74626 | 11.77828 | 11.929 | -0.150811294 | -0.182836952 | 12.115135 | 11.683681 | 11.9831 | -0.299462882 | 0.131991447 |
| Dbp | 7.964601 | 7.691792 | 8.2548 | -0.563057111 | -0.290247573 | 7.6706006 | 7.5990677 | 7.89862 | -0.299556155 | -0.228023218 |
| Xlr | 5.30917 | 5.302308 | 5.4471 | -0.144816464 | -0.137953617 | 5.2992782 | 5.1611335 | 5.46079 | -0.299658309 | -0.161513684 |
| Nrarp | 6.145626 | 5.74293 | 6.0027 | -0.259772443 | 0.142923342 | 5.915918 | 5.5594619 | 5.85921 | -0.299750817 | 0.056705264 |
| Thtpa | 7.735068 | 7.664828 | 7.8515 | -0.186697411 | -0.116456685 | 7.7481943 | 7.4723057 | 7.77211 | -0.299801568 | -0.023913029 |
| Eif2ak4 | 9.218185 | 9.02188 | 9.1931 | -0.171179931 | 0.025124933 | 9.31503 | 9.0397205 | 9.33967 | -0.299946914 | -0.02463746 |
| Igfbp5 | 7.942616 | 8.233764 | 7.7321 | 0.501640816 | 0.210492991 | 7.5866704 | 7.2371592 | 7.5374 | -0.300236433 | 0.049274797 |
| Zfp551 | 4.94473 | 4.887844 | 4.9561 | -0.068271847 | -0.011385459 | 4.8258375 | 4.6802117 | 4.98049 | -0.300281683 | -0.154655889 |
| Tmprss13 | 4.170995 | 4.115002 | 4.3039 | -0.188876702 | -0.132883865 | 4.3625687 | 4.0739182 | 4.37421 | -0.300288471 | -0.011638021 |
| BC025920 | 6.235931 | 6.18133 | 6.3247 | -0.143411624 | -0.088810416 | 6.7173286 | 6.4171486 | 6.71748 | -0.300334937 | -0.00015495 |
| Car7 | 4.878289 | 4.715874 | 4.9735 | -0.25765207 | -0.095236966 | 4.9800933 | 4.6479892 | 4.94834 | -0.300346747 | 0.031757345 |
| Abca6 | 3.405122 | 3.379213 | 3.4981 | -0.118913518 | -0.093004323 | 3.4175683 | 3.1013973 | 3.40185 | -0.300455542 | 0.015715496 |
| Fosl2 | 10.65436 | 10.11157 | 9.8307 | 0.280923621 | 0.823710914 | 9.7405119 | 9.5840067 | 9.88459 | -0.300579798 | -0.144074582 |
| Dgkd | 12.65506 | 12.6311 | 12.678 | -0.046843498 | -0.022889946 | 12.818011 | 12.493232 | 12.794 | -0.300810864 | 0.023967964 |
| Ciart | 7.652487 | 6.939619 | 7.6236 | -0.683950286 | 0.028918422 | 6.8521031 | 6.9708334 | 7.27167 | -0.300838901 | -0.419569174 |
| Tef | 9.664147 | 9.390823 | 9.757 | -0.366196424 | -0.092872691 | 9.5498327 | 9.4195443 | 9.72042 | -0.300878995 | -0.170590575 |
| Gpr55 | 7.620779 | 7.532892 | 7.6618 | -0.128896824 | -0.041010011 | 7.3369217 | 7.4098653 | 7.71076 | -0.300898436 | -0.373841994 |
| Drosha | 9.419009 | 9.330454 | 9.4979 | -0.16741212 | -0.078857701 | 9.4466975 | 9.2377381 | 9.53866 | -0.300925694 | -0.091966295 |
| Afap1l1 | 5.983711 | 5.912612 | 5.8427 | 0.069952172 | 0.141051389 | 5.4598255 | 5.2418316 | 5.54284 | -0.301004446 | -0.08301059 |
| Wipi1 | 5.672389 | 5.466361 | 5.6674 | -0.201068947 | 0.004958822 | 5.4085436 | 5.1774397 | 5.47852 | -0.301080156 | -0.069976317 |
| Btla | 12.41993 | 12.28681 | 12.507 | -0.220375432 | -0.087248932 | 12.622926 | 12.186743 | 12.4879 | -0.301128388 | 0.135054776 |
| Abcc4 | 8.892148 | 8.6639 | 8.8664 | -0.202500727 | 0.025746964 | 8.7152887 | 8.6125585 | 8.9137 | -0.301142386 | -0.198412284 |
| Gmpr | 4.648042 | 4.399182 | 4.7482 | -0.348980679 | -0.10012067 | 4.3881866 | 3.9815915 | 4.28284 | -0.30124915 | 0.105345927 |
| Enox2 | 8.380736 | 8.322282 | 8.4656 | -0.143346821 | -0.084892465 | 8.3430477 | 8.1402668 | 8.44159 | -0.301324769 | -0.098543835 |
| Bend3 | 7.71784 | 7.512997 | 7.6009 | -0.087877764 | 0.116965453 | 7.6140644 | 7.3381248 | 7.63954 | -0.30141996 | -0.025480382 |
| Zdhhc17 | 9.644707 | 9.616343 | 9.6996 | -0.083291484 | -0.054927652 | 9.7562954 | 9.559071 | 9.86051 | -0.30143629 | -0.104211897 |
| Zmynd8 | 10.34822 | 10.30642 | 10.456 | -0.150051398 | -0.108252469 | 10.351922 | 10.4041 | 10.7055 | -0.301447454 | -0.353625284 |
| Slc39a13 | 7.811954 | 7.906945 | 7.8687 | 0.038257784 | -0.056732571 | 8.0710105 | 7.7818592 | 8.08365 | -0.301795332 | -0.012644003 |
| Fmnl1 | 12.94617 | 13.01109 | 13.054 | -0.04294588 | -0.107856596 | 13.039804 | 12.795303 | 13.0971 | -0.301841536 | -0.057340303 |
| Ikzf2 | 9.420904 | 9.464767 | 9.6393 | -0.174525609 | -0.218389421 | 9.1711137 | 9.6205561 | 9.92244 | -0.301886058 | -0.751328459 |
| Tnni2 | 5.514097 | 5.141677 | 5.4797 | -0.338023935 | 0.034396485 | 5.3632002 | 5.1855012 | 5.48745 | -0.301945009 | -0.124245969 |
| Tbc1d14 | 11.18986 | 11.0964 | 11.183 | -0.086900452 | 0.00656557 | 11.151232 | 10.839796 | 11.142 | -0.302166002 | 0.009270166 |

| | | | | | | | | | | |
|---|---|---|---|---|---|---|---|---|---|---|
| Abcc5 | 9.107801 | 9.287043 | 9.3348 | -0.047798673 | -0.227040209 | 9.1734 | 9.1882438 | 9.49045 | -0.302207388 | -0.317051188 |
| Tnrc6c | 10.91355 | 10.89679 | 10.977 | -0.08007676 | -0.063315268 | 10.895084 | 10.789145 | 11.0914 | -0.302244648 | -0.196305215 |
| Fam193a | 10.38121 | 10.4708 | 10.436 | 0.034684118 | -0.054904328 | 10.222337 | 10.245819 | 10.5481 | -0.302262858 | -0.325744855 |
| Mtfr1l | 9.657533 | 9.578395 | 9.7542 | -0.175832684 | -0.096694451 | 9.7693489 | 9.3671937 | 9.66956 | -0.302365813 | 0.099789352 |
| Selenoi | 8.609365 | 8.684399 | 8.8235 | -0.139074062 | -0.21410741 | 8.7777393 | 8.5785232 | 8.88096 | -0.302431943 | -0.103215832 |
| Nmrk1 | 6.5222 | 6.572002 | 6.6152 | -0.043240206 | -0.0930424 | 6.6842199 | 6.5320836 | 6.83468 | -0.302599005 | -0.150462755 |
| Ctdspl2 | 9.894783 | 9.978442 | 9.9799 | -0.001426659 | -0.085085161 | 9.7600527 | 9.6911483 | 9.99392 | -0.302774367 | -0.233869999 |
| Neb | 6.922962 | 6.58068 | 6.5427 | 0.03793494 | 0.380217729 | 6.5429788 | 6.8256414 | 7.12857 | -0.302924987 | -0.585587593 |
| Zfp983 | 7.647998 | 7.59591 | 7.5694 | 0.026495949 | 0.078584307 | 7.840742 | 7.524003 | 7.82701 | -0.303005663 | 0.013733357 |
| Dnaaf5 | 8.584918 | 8.526966 | 8.5608 | -0.033836453 | 0.024115193 | 8.5173312 | 8.325842 | 8.62896 | -0.303119074 | -0.11162991 |
| Trav4d-4 | 4.43324 | 4.241071 | 4.3905 | -0.14947131 | 0.042697315 | 4.2294388 | 4.1278375 | 4.43099 | -0.303157444 | -0.201556158 |
| Trav4n-4 | 4.43324 | 4.241071 | 4.3905 | -0.14947131 | 0.042697315 | 4.2294388 | 4.1278375 | 4.43099 | -0.303157444 | -0.201556158 |
| Zfp382 | 6.514392 | 6.830934 | 6.6804 | 0.150508081 | -0.166033526 | 6.47006 | 6.5622205 | 6.86543 | -0.303208396 | -0.395368954 |
| Bin2 | 11.69122 | 11.62899 | 11.804 | -0.174947874 | -0.112717394 | 11.815893 | 11.630101 | 11.9336 | -0.303492008 | -0.117700263 |
| Coro7 | 11.28234 | 11.33963 | 11.405 | -0.065726544 | -0.12300996 | 11.660342 | 11.340956 | 11.6445 | -0.303560162 | 0.015825386 |
| Ocrl | 7.936073 | 7.589885 | 7.5487 | 0.041208388 | 0.387396275 | 7.5977184 | 7.5122205 | 7.81595 | -0.303727806 | -0.218229916 |
| Nfatc3 | 11.99783 | 12.09545 | 12.11 | -0.014644042 | -0.112268802 | 11.917503 | 11.766434 | 12.0702 | -0.303763886 | -0.152694339 |
| Ipo11 | 9.022782 | 8.933186 | 9.1595 | -0.226340743 | -0.136744431 | 8.8943548 | 8.8110899 | 9.11491 | -0.303815854 | -0.220551005 |
| Chst15 | 10.18142 | 10.30111 | 10.2 | 0.100693968 | -0.018995834 | 9.8835389 | 9.7270715 | 10.031 | -0.303896233 | -0.147428831 |
| Acvr1 | 6.476392 | 6.227971 | 6.2124 | 0.015525002 | 0.263945432 | 5.8140923 | 5.9974299 | 6.30156 | -0.304126276 | -0.487463893 |
| Zfp174 | 4.728103 | 4.748592 | 4.8196 | -0.070969528 | -0.091459372 | 4.7681658 | 4.4562976 | 4.76052 | -0.304218485 | 0.007649723 |
| Jrkl | 8.205626 | 8.083935 | 8.2719 | -0.187944301 | -0.066253634 | 8.4234925 | 8.1721718 | 8.4764 | -0.30422792 | -0.05290718 |
| Gpr137b-ps | 8.648893 | 8.59197 | 8.5463 | 0.04568921 | 0.102612224 | 8.5626861 | 8.1097505 | 8.41401 | -0.304260766 | 0.148674866 |
| Mgp | 5.168433 | 5.194274 | 5.0321 | 0.162158156 | 0.136317252 | 4.6003359 | 4.6309106 | 4.93541 | -0.304497563 | -0.33507232 |
| Ehhadh | 7.037584 | 7.12558 | 7.1503 | -0.024676089 | -0.112672075 | 7.0979665 | 6.8634251 | 7.16793 | -0.304506861 | -0.069965382 |
| Pcnx4 | 6.816517 | 6.763158 | 6.9394 | -0.1762121 | -0.122853156 | 6.8967437 | 6.6584016 | 6.96291 | -0.304511008 | -0.066168995 |
| Zfp317 | 9.432166 | 9.44683 | 9.4064 | 0.040458618 | 0.025794598 | 9.5814143 | 9.3740954 | 9.67871 | -0.304611915 | -0.09729302 |
| Mmp11 | 5.218488 | 5.038375 | 5.1199 | -0.081526126 | 0.098586624 | 5.0143204 | 4.71076 | 5.01544 | -0.304681418 | -0.001121018 |
| Hist3h2ba | 4.978137 | 4.921964 | 4.9669 | -0.044896061 | 0.01127617 | 5.1161962 | 4.7252031 | 5.0299 | -0.304696977 | 0.086296194 |
| Dkk3 | 4.594276 | 4.556521 | 4.6249 | -0.068384946 | -0.030629673 | 4.4097798 | 4.2821981 | 4.5869 | -0.304702901 | -0.177121194 |
| Nt5c3b | 9.126836 | 9.031966 | 9.2006 | -0.168638995 | -0.073769657 | 9.2634305 | 9.0003055 | 9.30503 | -0.304725514 | -0.041600532 |
| Lrp6 | 10.36983 | 10.27928 | 10.293 | -0.01364528 | 0.076905494 | 10.142704 | 10.032671 | 10.3374 | -0.304745497 | -0.194712666 |

| | | | | | | | | | | |
|---|---|---|---|---|---|---|---|---|---|---|
| Ice1 | 10.98038 | 11.01012 | 11.071 | -0.060997746 | -0.090738068 | 11.028953 | 10.903604 | 11.2084 | -0.304820216 | -0.179471354 |
| Ticam2 | 6.842569 | 6.584475 | 6.7103 | -0.125841129 | 0.132252476 | 6.9272508 | 6.4306917 | 6.73557 | -0.304874182 | 0.191684991 |
| Strbp | 10.74089 | 10.73905 | 10.79 | -0.050860274 | -0.049021484 | 10.84369 | 10.506465 | 10.8115 | -0.305054378 | 0.032170136 |
| Aaed1 | 9.340535 | 9.314813 | 9.5086 | -0.193758765 | -0.168036614 | 9.2069799 | 8.9397606 | 9.24509 | -0.305327015 | -0.038107697 |
| Apoe | 12.64835 | 12.28462 | 12.305 | -0.020476822 | 0.343251484 | 11.369655 | 11.291273 | 11.5966 | -0.30534954 | -0.226967395 |
| Tec | 10.31406 | 10.36167 | 10.466 | -0.104008862 | -0.151614326 | 10.671816 | 10.29681 | 10.6022 | -0.305418854 | 0.069587397 |
| Slc37a4 | 7.879391 | 7.851301 | 7.9581 | -0.106749076 | -0.078659481 | 8.4359255 | 7.8685341 | 8.17404 | -0.305508793 | 0.261882603 |
| Dbn1 | 5.983119 | 5.876107 | 5.96 | -0.083877664 | 0.023134171 | 6.2765208 | 5.7939381 | 6.09947 | -0.305536036 | 0.177046696 |
| Zzef1 | 11.25101 | 11.21964 | 11.309 | -0.089415258 | -0.058044436 | 11.337154 | 11.190491 | 11.4962 | -0.305693766 | -0.15902997 |
| Gdpgp1 | 7.302103 | 7.214977 | 7.2959 | -0.080896211 | 0.006230051 | 7.5624235 | 7.0961956 | 7.40201 | -0.305814778 | 0.160413073 |
| Gm3435 | 5.157095 | 5.001322 | 5.2946 | -0.293266263 | -0.13749356 | 5.5696915 | 5.2090454 | 5.51497 | -0.305924109 | 0.054721953 |
| Rcn3 | 7.676482 | 7.668892 | 7.7412 | -0.072283486 | -0.0646935 | 7.710094 | 7.3169127 | 7.62293 | -0.306015617 | 0.087165729 |
| Man2b2 | 9.87702 | 9.799636 | 9.8755 | -0.075830065 | 0.001553623 | 10.14056 | 9.5184907 | 9.82472 | -0.306233849 | 0.315835823 |
| Grik5 | 6.317615 | 6.298999 | 6.4249 | -0.125915324 | -0.107299276 | 6.336817 | 6.2887299 | 6.59497 | -0.306241813 | -0.258154678 |
| Nbdy | 6.828207 | 6.888643 | 6.9189 | -0.030231213 | -0.090666696 | 6.8445941 | 6.5959951 | 6.90225 | -0.30625577 | -0.057656691 |
| Noct | 12.98633 | 12.68913 | 13.032 | -0.343269004 | -0.046068948 | 12.483494 | 12.455815 | 12.7622 | -0.306408748 | -0.278729086 |
| Pogz | 10.07106 | 10.15999 | 10.174 | -0.013782842 | -0.102704882 | 10.16236 | 10.11279 | 10.4193 | -0.306508681 | -0.256938828 |
| Cables2 | 10.16456 | 10.17701 | 10.239 | -0.062141102 | -0.074593794 | 10.247172 | 9.9383945 | 10.245 | -0.306604188 | 0.002173609 |
| Osbpl6 | 3.519964 | 3.380256 | 3.6195 | -0.239220981 | -0.099513385 | 3.4596904 | 3.1791233 | 3.48573 | -0.306607582 | -0.026040457 |
| Ccdc148 | 4.990292 | 4.9649 | 4.7826 | 0.182318157 | 0.207710887 | 4.4000942 | 4.3464966 | 4.65313 | -0.306628759 | -0.25303122 |
| Popdc2 | 3.567209 | 3.570543 | 3.4736 | 0.096930457 | 0.093596503 | 3.3321686 | 3.3710147 | 3.67775 | -0.306733973 | -0.345580119 |
| Psrc1 | 4.895636 | 4.764483 | 5.0643 | -0.299779841 | -0.168627034 | 4.8556113 | 4.893534 | 5.20043 | -0.306895228 | -0.344817874 |
| Lrrc17 | 4.498664 | 4.552452 | 4.7123 | -0.159828467 | -0.213615977 | 4.6184745 | 4.3780938 | 4.68502 | -0.306923511 | -0.066542817 |
| Uggt1 | 11.53038 | 11.50226 | 11.556 | -0.053324978 | -0.025196888 | 11.551339 | 11.343151 | 11.6501 | -0.306985706 | -0.098797529 |
| Sh3tc1 | 8.747317 | 8.498558 | 8.9256 | -0.427041078 | -0.178281696 | 8.7287488 | 8.482791 | 8.78982 | -0.307033289 | -0.061075452 |
| Impact | 8.883738 | 8.453884 | 8.7807 | -0.326857466 | 0.102996421 | 8.780443 | 8.8998948 | 9.2073 | -0.307404709 | -0.426856509 |
| Plpp6 | 7.99117 | 7.939351 | 8.1267 | -0.187314083 | -0.135494937 | 8.0703681 | 7.6827634 | 7.99022 | -0.307455507 | 0.080149172 |
| Tcrg-C1 | 6.876511 | 6.646658 | 6.8433 | -0.196637057 | 0.033216081 | 6.4846382 | 6.4690804 | 6.77677 | -0.307684819 | -0.292127029 |
| Scg5 | 4.424724 | 4.40115 | 4.4297 | -0.028557252 | -0.004982847 | 4.3437276 | 4.1134848 | 4.42121 | -0.307720603 | -0.07747782 |
| Tcn2 | 9.42031 | 8.94294 | 9.3523 | -0.409400226 | 0.067969872 | 9.0015835 | 8.5969476 | 8.9048 | -0.307852684 | 0.096783157 |
| Pianp | 4.955334 | 4.645965 | 4.9553 | -0.309348465 | 2.10842E-05 | 4.5367629 | 4.3400345 | 4.64791 | -0.307879029 | -0.111150669 |
| B230217C12Rik | 3.360581 | 3.343613 | 3.4734 | -0.129776669 | -0.112807951 | 3.4155758 | 3.2408414 | 3.54873 | -0.307892789 | -0.133158333 |

| | | | | | | | | | | |
|---|---|---|---|---|---|---|---|---|---|---|
| Tbc1d12 | 6.284172 | 5.985837 | 6.2625 | -0.276689305 | 0.021646342 | 5.9023373 | 5.8451651 | 6.15306 | -0.307893778 | -0.250721572 |
| Clec3b | 4.654566 | 4.395231 | 4.4964 | -0.101184207 | 0.158150722 | 4.2377886 | 4.1388301 | 4.44675 | -0.307916594 | -0.208958011 |
| Zscan2 | 7.444921 | 7.498578 | 7.6333 | -0.134712375 | -0.188368787 | 7.3705634 | 7.174837 | 7.48281 | -0.307974142 | -0.112247801 |
| Tmem29 | 6.586395 | 6.658632 | 6.6782 | -0.019604638 | -0.091841382 | 6.7769166 | 6.4470996 | 6.7551 | -0.307998539 | 0.021818443 |
| Zfp78 | 5.332946 | 5.221993 | 5.3296 | -0.107574394 | 0.003378059 | 5.4874394 | 5.0502645 | 5.3585 | -0.308233732 | 0.128941199 |
| Sash1 | 7.657416 | 7.275437 | 7.4299 | -0.154414064 | 0.227565319 | 6.9438281 | 6.5754188 | 6.88366 | -0.308239138 | 0.060170129 |
| Ermard | 8.799482 | 8.715036 | 8.8358 | -0.120756741 | -0.036310493 | 9.1178541 | 8.906928 | 9.21518 | -0.308247283 | -0.097321157 |
| Eno2 | 7.1119 | 7.10515 | 7.0265 | 0.078695118 | 0.085444777 | 6.4893765 | 6.852213 | 7.1605 | -0.308285867 | -0.671122355 |
| Esrrg | 4.587684 | 4.254461 | 4.7451 | -0.490662741 | -0.157439389 | 4.6765057 | 4.3403942 | 4.64882 | -0.308421382 | 0.02769014 |
| Lancl3 | 3.12543 | 3.040423 | 3.1926 | -0.152218601 | -0.067212253 | 3.0852677 | 2.8357182 | 3.14424 | -0.308519882 | -0.058970307 |
| Hace1 | 8.589202 | 8.64662 | 8.6735 | -0.026859505 | -0.084277895 | 8.402154 | 8.238574 | 8.5472 | -0.308624757 | -0.145044821 |
| Maml2 | 10.02592 | 10.11922 | 10.193 | -0.073644868 | -0.166946412 | 9.9167437 | 9.8201771 | 10.1289 | -0.308676428 | -0.212109837 |
| Nckap1 | 8.500948 | 8.284105 | 8.4058 | -0.121714738 | 0.095128806 | 7.7688864 | 7.6583925 | 7.96708 | -0.308684411 | -0.198190549 |
| Cdyl | 8.461123 | 8.301658 | 8.3629 | -0.061203359 | 0.098262248 | 8.2069916 | 7.9903342 | 8.29925 | -0.308917157 | -0.092259763 |
| Mios | 8.913679 | 8.812693 | 8.9878 | -0.175095824 | -0.074109694 | 8.887958 | 8.6203272 | 8.92932 | -0.308995304 | -0.041364495 |
| AC149090.1 | 11.70857 | 11.56444 | 11.886 | -0.321640346 | -0.177505246 | 12.33561 | 11.883627 | 12.1928 | -0.309139646 | 0.142843114 |
| Shpk | 6.719386 | 6.622757 | 6.8772 | -0.254455889 | -0.157826945 | 6.9784172 | 6.5834776 | 6.89265 | -0.309174024 | 0.0857656 |
| Sgk1 | 10.30373 | 9.84833 | 10.416 | -0.568143871 | -0.112748087 | 9.4161455 | 9.408062 | 9.71725 | -0.30918753 | -0.301104049 |
| Alg6 | 5.871262 | 5.979187 | 5.8497 | 0.129493375 | 0.021568636 | 5.9867513 | 5.6172525 | 5.92648 | -0.309229107 | 0.060269671 |
| Slc25a32 | 8.193735 | 8.036497 | 8.1371 | -0.100590917 | 0.0566472 | 7.9959149 | 7.8196615 | 8.12893 | -0.309269532 | -0.133016092 |
| Atg9b | 3.65931 | 3.424975 | 3.6594 | -0.234446547 | -0.000111136 | 3.4153811 | 3.2489232 | 3.55846 | -0.309535229 | -0.143077381 |
| Magee1 | 5.978094 | 5.961951 | 5.9904 | -0.028432559 | -0.012289559 | 5.8555652 | 5.6967198 | 6.00649 | -0.309769032 | -0.150923669 |
| Lrch3 | 10.66966 | 10.81624 | 10.766 | 0.049913184 | -0.096666478 | 10.426948 | 10.404108 | 10.7139 | -0.309774328 | -0.286934618 |
| Rfk | 10.22701 | 10.26822 | 10.347 | -0.078789396 | -0.119995952 | 10.071219 | 9.8057648 | 10.1155 | -0.309776362 | -0.044321708 |
| Zfp788 | 7.939073 | 7.843329 | 7.9622 | -0.11887208 | -0.023127428 | 8.0065514 | 7.6525292 | 7.96252 | -0.309985862 | 0.044036309 |
| Gm13889 | 5.687634 | 5.1886 | 5.3923 | -0.203674914 | 0.29535966 | 4.7357239 | 4.6192103 | 4.92923 | -0.31001878 | -0.193505144 |
| Lig1 | 9.376436 | 8.986535 | 9.3992 | -0.412653344 | -0.022752168 | 9.2901161 | 9.3279462 | 9.63806 | -0.310114448 | -0.347944457 |
| Rab3gap2 | 10.06094 | 10.02577 | 10.05 | -0.02377129 | 0.011396344 | 10.238665 | 9.9040538 | 10.2142 | -0.310115113 | 0.024495595 |
| Vkorc1 | 7.271643 | 6.980461 | 7.2569 | -0.276443936 | 0.014738858 | 7.3965776 | 6.8545055 | 7.16469 | -0.310187837 | 0.231884278 |
| Tecpr1 | 11.16947 | 11.19699 | 11.279 | -0.082412064 | -0.109932362 | 11.260202 | 11.137541 | 11.4477 | -0.310204428 | -0.187543491 |
| Cep250 | 10.42478 | 10.36975 | 10.469 | -0.099637443 | -0.044603848 | 10.239198 | 10.213816 | 10.5241 | -0.310241938 | -0.284860334 |
| Phka2 | 8.742931 | 8.577185 | 8.8145 | -0.237285386 | -0.071539267 | 8.685141 | 8.7199348 | 9.03032 | -0.310384425 | -0.345178251 |

| | | | | | | | | | | |
|---|---|---|---|---|---|---|---|---|---|---|
| Hnrnpr | 11.34211 | 11.24221 | 11.3 | -0.057311098 | 0.042596006 | 11.32578 | 11.071191 | 11.3816 | -0.31043644 | -0.055848006 |
| Espl1 | 7.757728 | 7.397582 | 7.6549 | -0.257283259 | 0.102862972 | 7.4150056 | 7.525754 | 7.83621 | -0.310453246 | -0.421201592 |
| Rap2b | 9.484568 | 9.491371 | 9.5813 | -0.089930989 | -0.0967344 | 9.18131 | 9.103399 | 9.41389 | -0.310486689 | -0.232575744 |
| Lasp1 | 11.65242 | 11.693 | 11.654 | 0.038961108 | -0.00162564 | 11.468417 | 11.304865 | 11.6155 | -0.31058529 | -0.14703403 |
| Cygb | 6.743902 | 6.021252 | 6.4336 | -0.412334902 | 0.310314373 | 5.877365 | 5.4416248 | 5.75242 | -0.310791755 | 0.124948524 |
| Phf21b | 4.769769 | 4.881817 | 4.8231 | 0.058744136 | -0.053304192 | 4.4290257 | 4.7962591 | 5.10713 | -0.310874082 | -0.67810748 |
| Upf3b | 9.225161 | 9.175512 | 9.3803 | -0.20479136 | -0.155142681 | 9.2966043 | 9.1666807 | 9.47759 | -0.310910714 | -0.180987132 |
| Camk4 | 8.761204 | 8.771721 | 8.8093 | -0.037557419 | -0.048074281 | 8.3241449 | 8.3715465 | 8.68251 | -0.310961914 | -0.358363573 |
| Tln1 | 13.92319 | 13.84216 | 13.857 | -0.01519665 | 0.065833802 | 13.735161 | 13.569405 | 13.8804 | -0.311029324 | -0.145274064 |
| Trav6-3 | 4.795739 | 4.833526 | 4.8583 | -0.024784453 | -0.06257065 | 4.7458104 | 4.5034533 | 4.81469 | -0.3112405 | -0.068883416 |
| Mospd1 | 7.438482 | 7.212573 | 7.5771 | -0.364551913 | -0.138643567 | 7.2641344 | 7.0869049 | 7.39841 | -0.311506797 | -0.134277276 |
| Slc35f2 | 6.97444 | 6.812726 | 6.9576 | -0.144836195 | 0.016876976 | 7.0010695 | 6.7571616 | 7.06873 | -0.311570797 | -0.067662909 |
| Evc | 4.729668 | 4.59338 | 4.7632 | -0.169828065 | -0.033540465 | 4.3462943 | 4.3146993 | 4.62636 | -0.311665228 | -0.28007024 |
| Ramp2 | 7.581026 | 7.279327 | 7.7981 | -0.518733778 | -0.217034978 | 7.1205596 | 6.789441 | 7.10155 | -0.312111116 | 0.019007451 |
| Xrcc2 | 6.675796 | 6.732396 | 6.671 | 0.061372631 | 0.004772432 | 6.9893285 | 6.5429668 | 6.85513 | -0.312166099 | 0.134195526 |
| Senp7 | 10.58028 | 10.68349 | 10.64 | 0.04313353 | -0.060074196 | 10.503746 | 10.429589 | 10.7419 | -0.31228169 | -0.238125277 |
| Cdk8 | 8.869386 | 8.812024 | 8.9749 | -0.16292046 | -0.105558391 | 8.9396133 | 8.8290347 | 9.14133 | -0.312293737 | -0.201715088 |
| Zfp109 | 5.567033 | 5.505308 | 5.4905 | 0.014801552 | 0.076526027 | 5.4509772 | 5.141959 | 5.45436 | -0.312400965 | -0.003382697 |
| Mus81 | 8.441102 | 8.422511 | 8.5551 | -0.132564474 | -0.113973699 | 8.9131029 | 8.5023829 | 8.81484 | -0.312460357 | 0.098259682 |
| 2410004P03Rik | 3.548903 | 3.550676 | 3.6411 | -0.090424621 | -0.092197315 | 3.5153714 | 3.487908 | 3.80041 | -0.312506433 | -0.285043024 |
| D430019H16Rik | 4.183095 | 4.208684 | 4.233 | -0.024309808 | -0.049898875 | 4.0350245 | 3.8418473 | 4.1545 | -0.312655736 | -0.1194785 |
| Trbv31 | 7.172837 | 7.11464 | 7.1859 | -0.071242069 | -0.013044864 | 6.870075 | 6.7919915 | 7.10478 | -0.312793384 | -0.234709888 |
| Adam33 | 4.275913 | 4.239165 | 4.2433 | -0.00415122 | 0.032597571 | 3.8237073 | 4.0415542 | 4.35436 | -0.312804786 | -0.530651699 |
| Inpp5d | 12.32304 | 12.32071 | 12.282 | 0.039098116 | 0.041429964 | 12.408889 | 12.046746 | 12.3596 | -0.312839828 | 0.049302471 |
| Abcd4 | 8.019701 | 8.02058 | 8.2376 | -0.217045432 | -0.217923775 | 8.3214878 | 8.1120367 | 8.42495 | -0.312918025 | -0.103466926 |
| Zfp362 | 9.443057 | 9.462734 | 9.5888 | -0.126049289 | -0.14572606 | 9.5339835 | 9.1002021 | 9.41346 | -0.313259893 | 0.120521549 |
| Ctr9 | 10.10952 | 10.01556 | 10.059 | -0.042972712 | 0.05098477 | 10.060035 | 9.8500625 | 10.1634 | -0.313295528 | -0.103323129 |
| Rbak | 7.628506 | 7.54463 | 7.6699 | -0.125229338 | -0.04135328 | 7.7945863 | 7.5681709 | 7.88168 | -0.313507439 | -0.087092119 |
| Arl4d | 6.832586 | 6.635891 | 7.098 | -0.46215753 | -0.265463008 | 6.3462564 | 6.2101203 | 6.52363 | -0.313511647 | -0.177375555 |
| Oxct1 | 10.80448 | 10.48563 | 10.651 | -0.165769338 | 0.15308337 | 10.515859 | 10.378372 | 10.6919 | -0.3135751 | -0.176088473 |
| Zfp966 | 6.311738 | 6.579775 | 6.4737 | 0.106085645 | -0.161951395 | 6.6698196 | 6.4652767 | 6.77887 | -0.313591505 | -0.109048604 |
| Gsg1l | 5.489782 | 5.50636 | 5.6044 | -0.098016705 | -0.114594676 | 5.6755572 | 5.836753 | 6.15052 | -0.313770257 | -0.474966043 |

| | | | | | | | | | | |
|---|---|---|---|---|---|---|---|---|---|---|
| Nadk2 | 7.911153 | 7.799041 | 8.1407 | -0.341634106 | -0.229522803 | 7.9196326 | 7.7827178 | 8.09651 | -0.313794231 | -0.176879454 |
| Tmem110 | 8.044459 | 8.176507 | 8.1038 | 0.072698925 | -0.059349903 | 8.3382203 | 7.8412122 | 8.15516 | -0.31395127 | 0.183056872 |
| Nmb | 5.194572 | 5.200514 | 5.4333 | -0.232799193 | -0.238740883 | 5.2435758 | 4.9195556 | 5.23361 | -0.314050227 | 0.009970008 |
| Nbeal2 | 10.60396 | 10.60554 | 10.777 | -0.171817612 | -0.173394568 | 10.815833 | 10.614675 | 10.9288 | -0.314160252 | -0.113002278 |
| AA414768 | 4.860539 | 4.817441 | 4.9516 | -0.134195674 | -0.091098529 | 4.9100467 | 4.5447464 | 4.85894 | -0.314192683 | 0.051107654 |
| Zfp329 | 8.376158 | 8.251512 | 8.3982 | -0.146661429 | -0.022016179 | 8.8386838 | 8.4622189 | 8.77651 | -0.314295277 | 0.062169556 |
| Dnph1 | 4.074135 | 3.925929 | 4.0651 | -0.139178447 | 0.009027744 | 4.0262358 | 3.8839908 | 4.19848 | -0.314484608 | -0.172239672 |
| Slc22a21 | 4.365718 | 4.034891 | 4.0555 | -0.020563903 | 0.310263182 | 4.07215 | 3.780779 | 4.09527 | -0.314495683 | -0.023124687 |
| Cmtm4 | 6.936643 | 6.790481 | 7.0241 | -0.233657813 | -0.087496121 | 6.7863469 | 6.6372917 | 6.95191 | -0.314613464 | -0.165558316 |
| Mrnip | 5.419714 | 5.368285 | 5.615 | -0.246750978 | -0.195321625 | 5.5971141 | 5.3011857 | 5.61619 | -0.315003677 | -0.019075356 |
| Tbcel | 9.67826 | 9.682128 | 9.6817 | 0.00045552 | -0.003412933 | 9.5473715 | 9.4028699 | 9.71789 | -0.315020057 | -0.17051839 |
| Dnah11 | 3.272838 | 3.343078 | 3.1401 | 0.202977174 | 0.132737076 | 2.8764534 | 2.926843 | 3.24187 | -0.315031352 | -0.365420984 |
| Spsb1 | 8.069296 | 7.947107 | 7.9572 | -0.010126102 | 0.112063485 | 7.5192995 | 7.6691931 | 7.98448 | -0.315289336 | -0.465182926 |
| Ccng2 | 10.06052 | 10.15849 | 10.291 | -0.132608418 | -0.230577891 | 10.036336 | 9.8889628 | 10.2043 | -0.315320435 | -0.167947222 |
| Glud1 | 11.68412 | 11.67451 | 11.705 | -0.030910302 | -0.021308168 | 11.714561 | 11.336705 | 11.652 | -0.31533728 | 0.062518247 |
| Zmpste24 | 9.963581 | 9.963057 | 9.9129 | 0.050170136 | 0.050693968 | 9.9557098 | 9.6605906 | 9.97598 | -0.315389739 | -0.020270576 |
| Lrrn3 | 3.436043 | 3.449536 | 3.5418 | -0.092305491 | -0.105798636 | 3.5092912 | 3.2791449 | 3.59467 | -0.315521484 | -0.085375144 |
| Smim24 | 6.047593 | 5.943371 | 6.2434 | -0.300000643 | -0.195778802 | 6.2664368 | 5.9032222 | 6.21879 | -0.315569611 | 0.047645021 |
| Usp28 | 9.583435 | 9.417919 | 9.5933 | -0.175367343 | -0.009851597 | 9.3171582 | 9.3674505 | 9.68326 | -0.315811984 | -0.366104285 |
| Cdh13 | 5.546894 | 5.660104 | 5.9691 | -0.308996177 | -0.422205941 | 5.6544201 | 5.4397487 | 5.75556 | -0.315815447 | -0.101144054 |
| Exoc4 | 10.01126 | 9.98807 | 10.051 | -0.06339446 | -0.040204085 | 9.9467726 | 9.7538216 | 10.07 | -0.31618208 | -0.123231101 |
| Rarg | 8.816648 | 8.925568 | 8.9652 | -0.039679363 | -0.148599221 | 8.5979467 | 8.5925971 | 8.90878 | -0.316182787 | -0.310833201 |
| Il5ra | 8.147076 | 8.18616 | 8.1246 | 0.061550526 | 0.022465656 | 8.5523928 | 7.9834958 | 8.2997 | -0.316205321 | 0.252691624 |
| Iglc1 | 9.92684 | 10.09125 | 10.04 | 0.05149986 | -0.112915251 | 10.268992 | 10.030477 | 10.3468 | -0.316277553 | -0.077762274 |
| Ehd2 | 8.527663 | 8.566122 | 8.5637 | 0.002422101 | -0.036036573 | 8.3378582 | 7.837442 | 8.154 | -0.316554558 | 0.183861625 |
| Itga8 | 4.314704 | 3.868791 | 4.3015 | -0.432725231 | 0.013187535 | 4.0163591 | 3.6299719 | 3.94654 | -0.316567791 | 0.069819374 |
| Fpgt | 8.171561 | 8.155164 | 8.2056 | -0.050400428 | -0.034003421 | 8.1878076 | 7.9790931 | 8.2959 | -0.316809694 | -0.108095135 |
| mt-Atp8 | 13.78206 | 13.72682 | 13.874 | -0.147602048 | -0.092364454 | 13.562109 | 13.568599 | 13.8855 | -0.316859554 | -0.323349093 |
| Asb1 | 8.704512 | 8.527572 | 8.6191 | -0.091554153 | 0.08538632 | 8.6283469 | 8.2033549 | 8.5204 | -0.317042743 | 0.107949197 |
| Rchy1 | 9.849805 | 9.86644 | 9.9794 | -0.112943226 | -0.129578211 | 9.9643195 | 9.6397643 | 9.95707 | -0.317303956 | 0.007251285 |
| Nkd1 | 3.763709 | 3.563992 | 3.7273 | -0.163287852 | 0.036428432 | 3.6154715 | 3.3809022 | 3.69826 | -0.317352992 | -0.082783691 |
| Pcyt1b | 4.039571 | 3.92558 | 4.136 | -0.210417969 | -0.096426394 | 3.9645492 | 3.9800686 | 4.29745 | -0.317384888 | -0.332904247 |

| | | | | | | | | | | |
|---|---|---|---|---|---|---|---|---|---|---|
| Rpl14-ps1 | 11.11622 | 11.33291 | 11.306 | 0.026526162 | -0.190158286 | 11.033408 | 11.061364 | 11.3788 | -0.317387852 | -0.34534397 |
| Zfp975 | 5.417887 | 5.292149 | 5.4452 | -0.153024148 | -0.027285923 | 5.392547 | 5.2272551 | 5.5447 | -0.317440678 | -0.152148773 |
| Lama4 | 7.35484 | 7.022881 | 7.0825 | -0.059665035 | 0.272294074 | 6.8492276 | 6.6715663 | 6.98906 | -0.317498044 | -0.139836738 |
| Pde4a | 6.841441 | 6.922081 | 6.821 | 0.101091502 | 0.020451156 | 6.5779075 | 6.6488812 | 6.96641 | -0.317531696 | -0.388505351 |
| Ankrd6 | 5.540629 | 5.234652 | 5.5358 | -0.301184068 | 0.004793416 | 5.1992682 | 5.2150715 | 5.5327 | -0.317633393 | -0.333436737 |
| S100a10 | 10.9078 | 10.953 | 11.009 | -0.056359526 | -0.101562266 | 10.887383 | 10.752552 | 11.0705 | -0.317943002 | -0.183111178 |
| Robo4 | 8.118258 | 7.799463 | 7.9939 | -0.19446042 | 0.124333674 | 7.3852965 | 7.1241768 | 7.44213 | -0.317956076 | -0.056836351 |
| Slc18b1 | 3.145942 | 3.024809 | 3.1804 | -0.155632659 | -0.034499968 | 2.948271 | 2.8715843 | 3.18961 | -0.318026893 | -0.241340156 |
| Necab2 | 3.4936 | 3.218258 | 3.546 | -0.327746547 | -0.052405179 | 3.1046037 | 2.959047 | 3.27713 | -0.318081085 | -0.172524359 |
| Trip11 | 10.51888 | 10.48604 | 10.425 | 0.06109372 | 0.09392689 | 10.343947 | 10.278436 | 10.5966 | -0.318125956 | -0.252615171 |
| Rasip1 | 8.556659 | 8.051055 | 8.1471 | -0.096001887 | 0.409601879 | 7.3994622 | 7.2200439 | 7.5382 | -0.318155384 | -0.138737024 |
| Cers5 | 9.560385 | 9.560695 | 9.6371 | -0.076426506 | -0.076736033 | 9.6324793 | 9.4247378 | 9.74314 | -0.318398169 | -0.110656667 |
| Slc15a1 | 3.76341 | 3.626376 | 3.7536 | -0.127177875 | 0.009856948 | 3.8328802 | 3.5445713 | 3.86305 | -0.318480962 | -0.030172075 |
| Wdr6 | 10.35773 | 10.40018 | 10.485 | -0.084796934 | -0.127249716 | 10.391867 | 9.9879636 | 10.3065 | -0.318502244 | 0.085400737 |
| Igsf10 | 5.040822 | 4.939555 | 4.9882 | -0.048610424 | 0.052656452 | 4.8197356 | 4.8163935 | 5.13504 | -0.318648376 | -0.315306225 |
| Nprl3 | 8.74442 | 8.55783 | 8.832 | -0.274209937 | -0.087619991 | 9.2035219 | 8.7734889 | 9.09225 | -0.318756151 | 0.111276871 |
| Trav6-5 | 4.922374 | 4.859658 | 4.967 | -0.107349879 | -0.044634322 | 4.666281 | 4.6650051 | 4.9839 | -0.318892253 | -0.317616364 |
| Gtf3c1 | 10.64488 | 10.68415 | 10.754 | -0.070302902 | -0.109566998 | 10.645787 | 10.56292 | 10.8819 | -0.318954938 | -0.236087744 |
| Trbv2 | 7.320966 | 7.32912 | 7.325 | 0.004168156 | -0.003985836 | 7.2257938 | 7.0750259 | 7.39405 | -0.319026764 | -0.168258879 |
| Sh3glb2 | 9.412468 | 9.383569 | 9.4985 | -0.11498025 | -0.086080972 | 9.7504672 | 9.2719944 | 9.5912 | -0.319207675 | 0.15926518 |
| Zxdc | 8.873862 | 8.779939 | 8.8382 | -0.058287526 | 0.035635806 | 8.8984568 | 8.6682578 | 8.98759 | -0.319330998 | -0.089132046 |
| Git2 | 11.31927 | 11.38367 | 11.417 | -0.033820162 | -0.09821941 | 11.341672 | 11.245578 | 11.5651 | -0.319568708 | -0.223474716 |
| Efcab9 | 3.262616 | 3.333456 | 3.4584 | -0.12490254 | -0.195742762 | 3.4530302 | 3.2436399 | 3.56326 | -0.319616341 | -0.110226072 |
| Slc16a12 | 3.207613 | 3.185754 | 3.1902 | -0.004476912 | 0.017381797 | 3.0420255 | 2.895974 | 3.21569 | -0.3197158 | -0.173664274 |
| Fcna | 4.594809 | 4.161004 | 4.2742 | -0.113183151 | 0.320621992 | 3.6313673 | 3.7494567 | 4.06925 | -0.319794158 | -0.437883549 |
| Zfp974 | 6.998273 | 6.904995 | 7.0712 | -0.166160459 | -0.072881998 | 7.0977091 | 6.8682339 | 7.18805 | -0.319814732 | -0.090339556 |
| Gen1 | 6.947591 | 7.112547 | 7.1182 | -0.005642886 | -0.170599111 | 7.0777861 | 6.889098 | 7.20895 | -0.319851271 | -0.131163124 |
| Asxl3 | 2.743573 | 2.661999 | 2.712 | -0.05001429 | 0.031559585 | 2.4860158 | 2.5118128 | 2.83168 | -0.319866638 | -0.34566359 |
| Gpr180 | 7.792456 | 7.74107 | 7.827 | -0.085978989 | -0.034593082 | 7.7477716 | 7.479668 | 7.79958 | -0.319909387 | -0.051805729 |
| Kctd9 | 8.039684 | 7.980115 | 8.0905 | -0.11042381 | -0.050854379 | 7.7993934 | 7.6050195 | 7.92505 | -0.320025873 | -0.125651988 |
| Thada | 10.22062 | 10.22791 | 10.218 | 0.010034327 | 0.002746818 | 10.152553 | 10.153979 | 10.4741 | -0.320095032 | -0.321520439 |
| Myo7a | 8.61696 | 8.549518 | 8.4778 | 0.071686419 | 0.139128796 | 8.8322279 | 8.4133062 | 8.73351 | -0.320207639 | 0.098714042 |

| | | | | | | | | | | |
|---|---|---|---|---|---|---|---|---|---|---|
| Hivep3 | 8.616198 | 8.488995 | 8.6797 | -0.19074119 | -0.063538015 | 8.2838104 | 8.1974426 | 8.51772 | -0.320281843 | -0.233914038 |
| Reps1 | 9.819938 | 9.890347 | 9.9492 | -0.058807449 | -0.12921641 | 9.9400767 | 9.6899936 | 10.0104 | -0.320375825 | -0.070292799 |
| Zfp81 | 8.189653 | 8.229731 | 8.3435 | -0.113790195 | -0.153867623 | 8.1171671 | 8.0111989 | 8.33159 | -0.320390027 | -0.214421891 |
| Recql5 | 9.630662 | 9.444934 | 9.5786 | -0.133628208 | 0.052099594 | 9.9458719 | 9.6051785 | 9.92583 | -0.32064691 | 0.020046535 |
| Lrrc8a | 9.150455 | 9.148006 | 9.1935 | -0.045473837 | -0.043024334 | 9.0501033 | 8.8058276 | 9.12678 | -0.320948272 | -0.076672528 |
| Trpm7 | 11.07837 | 11.0134 | 11.074 | -0.060305506 | 0.004672358 | 10.906421 | 10.838923 | 11.1599 | -0.321006048 | -0.253508019 |
| Slc35f5 | 6.87943 | 6.264323 | 6.5207 | -0.256328342 | 0.358778623 | 6.667904 | 6.1402108 | 6.46131 | -0.321095217 | 0.206598021 |
| Fam149a | 5.538162 | 5.396453 | 5.6711 | -0.274632774 | -0.132923486 | 5.2354313 | 4.9176664 | 5.23876 | -0.321098415 | -0.003333504 |
| Numb | 9.171759 | 9.159834 | 9.1682 | -0.00832414 | 0.003600934 | 8.8083652 | 8.7723256 | 9.09346 | -0.321133309 | -0.285093704 |
| Tmem120b | 6.271197 | 6.5143 | 6.515 | -0.000652727 | -0.243756286 | 6.2518344 | 5.9938441 | 6.3155 | -0.321660891 | -0.06367055 |
| Slc16a2 | 4.936911 | 4.577713 | 4.9473 | -0.369628657 | -0.010430483 | 4.6693054 | 4.419383 | 4.74113 | -0.321751459 | -0.0718291 |
| Slc12a3 | 4.475679 | 4.630649 | 4.6618 | -0.03115054 | -0.186120417 | 4.6273167 | 4.6542303 | 4.9763 | -0.322070833 | -0.348984391 |
| Lmod1 | 3.80641 | 3.958489 | 4.0896 | -0.131143703 | -0.283222157 | 3.7408225 | 3.5326562 | 3.85493 | -0.322275008 | -0.114108682 |
| BC051142 | 5.894643 | 5.643701 | 5.6857 | -0.0420486 | 0.208894085 | 5.8409515 | 5.743194 | 6.06557 | -0.322377741 | -0.224620323 |
| Skint3 | 4.686175 | 4.619261 | 4.7988 | -0.179503125 | -0.112589216 | 5.1140622 | 4.4886931 | 4.81107 | -0.322378103 | 0.302990974 |
| Cmya5 | 4.738669 | 4.647182 | 4.8485 | -0.201282462 | -0.109796047 | 4.7568238 | 4.4764718 | 4.79896 | -0.322489036 | -0.042137079 |
| Lin54 | 8.680496 | 8.659957 | 8.7504 | -0.090451932 | -0.069913075 | 8.536962 | 8.4813082 | 8.8038 | -0.322494716 | -0.266840944 |
| Zbtb20 | 10.02956 | 9.787415 | 10.071 | -0.283917587 | -0.041773242 | 10.151296 | 10.226834 | 10.5493 | -0.322504254 | -0.398042587 |
| Ccser2 | 10.59514 | 10.5769 | 10.575 | 0.001569856 | 0.019813196 | 10.125802 | 10.220229 | 10.5429 | -0.322654512 | -0.417081242 |
| Smim10l1 | 8.825664 | 8.778033 | 8.891 | -0.112973688 | -0.065342644 | 8.6539043 | 8.4801538 | 8.80286 | -0.322705232 | -0.148954725 |
| Tox2 | 7.406699 | 6.888941 | 7.1145 | -0.225565059 | 0.292192998 | 6.4008271 | 6.251427 | 6.57428 | -0.322852274 | -0.173452256 |
| Asxl1 | 10.92539 | 10.83609 | 10.907 | -0.070410482 | 0.018888908 | 10.653783 | 10.639686 | 10.9629 | -0.323178322 | -0.309081266 |
| Muc3a | 5.078572 | 5.116477 | 5.2019 | -0.08542544 | -0.123329856 | 5.3076441 | 5.0098617 | 5.33304 | -0.323181893 | -0.025399576 |
| Gpr160 | 4.937196 | 5.05947 | 5.4342 | -0.374742752 | -0.497017098 | 5.1682259 | 5.062292 | 5.38558 | -0.323291877 | -0.21735791 |
| Kcnc3 | 5.790811 | 5.737059 | 5.7542 | -0.017165741 | 0.036585352 | 6.1838182 | 5.638672 | 5.962 | -0.323330889 | 0.221815397 |
| Trip4 | 9.524271 | 9.4571 | 9.4081 | 0.049022317 | 0.116193775 | 9.4694495 | 9.2002426 | 9.52374 | -0.323495934 | -0.054289068 |
| Lrp5 | 7.880352 | 7.781042 | 7.7673 | 0.013694422 | 0.113003919 | 7.7838591 | 7.2152213 | 7.53877 | -0.323544641 | 0.245093168 |
| Zfp955b | 7.342557 | 7.322874 | 7.2922 | 0.030660357 | 0.050342983 | 7.6308425 | 7.2304878 | 7.55405 | -0.323558164 | 0.076796524 |
| Sec22a | 7.174531 | 7.15075 | 7.1347 | 0.016087021 | 0.039867386 | 7.2798441 | 6.9677426 | 7.29147 | -0.323724812 | -0.011623329 |
| Carns1 | 8.706764 | 8.706982 | 8.7955 | -0.088558098 | -0.088775573 | 9.1225174 | 8.9543945 | 9.27848 | -0.324081194 | -0.155958322 |
| Sh3rf1 | 6.9642 | 6.86943 | 7.0406 | -0.171154588 | -0.076384441 | 6.7679892 | 6.6531138 | 6.97735 | -0.324240117 | -0.20936473 |
| Hmgcll1 | 3.835044 | 3.750079 | 3.9507 | -0.200587123 | -0.115621614 | 3.8605698 | 3.5362825 | 3.86054 | -0.324259833 | 2.75436E-05 |

| | | | | | | | | | | |
|---|---|---|---|---|---|---|---|---|---|---|
| Tpm1 | 9.47895 | 9.293036 | 9.4522 | -0.159141286 | 0.02677335 | 9.4259013 | 9.0452102 | 9.36968 | -0.324467771 | 0.056223324 |
| Rreb1 | 10.07475 | 9.960242 | 10.004 | -0.043409943 | 0.071095639 | 10.066242 | 9.7396561 | 10.0641 | -0.324474769 | 0.002110765 |
| AU040320 | 9.908105 | 10.01108 | 9.9609 | 0.050163624 | -0.052812645 | 10.093423 | 9.9313911 | 10.2559 | -0.324538421 | -0.162506027 |
| Glmn | 7.418849 | 7.385183 | 7.4776 | -0.092368161 | -0.058702529 | 7.6809241 | 7.2440866 | 7.56869 | -0.324608235 | 0.112229193 |
| Lyrm9 | 6.95994 | 6.951758 | 7.2947 | -0.342932463 | -0.3347509 | 7.6333857 | 7.2329205 | 7.55762 | -0.324696691 | 0.075768562 |
| Mgll | 6.025057 | 6.433254 | 6.2842 | 0.149068602 | -0.25912849 | 6.3453353 | 5.8988036 | 6.22369 | -0.324882792 | 0.121648876 |
| Adi1 | 8.898874 | 8.812444 | 9.0256 | -0.21313818 | -0.126708592 | 8.951116 | 8.5539485 | 8.87905 | -0.325103343 | 0.072064107 |
| Zfp777 | 9.355547 | 9.148267 | 9.2911 | -0.142800063 | 0.064480476 | 9.4204231 | 9.0624389 | 9.38768 | -0.325237396 | 0.032746812 |
| Lrrc23 | 3.876258 | 4.165178 | 3.948 | 0.217225605 | -0.071694228 | 3.7483561 | 3.6392215 | 3.96464 | -0.325418551 | -0.21628392 |
| Dyx1c1 | 5.035203 | 4.929247 | 5.2498 | -0.320590031 | -0.214633611 | 5.0386782 | 4.7983336 | 5.12392 | -0.325582641 | -0.085238094 |
| Med13l | 10.80355 | 10.84886 | 10.842 | 0.006795258 | -0.038521292 | 10.587926 | 10.531488 | 10.8571 | -0.325622526 | -0.269184322 |
| Pomk | 7.447163 | 7.541827 | 7.664 | -0.122206866 | -0.216870937 | 7.8007652 | 7.3416721 | 7.66746 | -0.325789211 | 0.133303923 |
| Rdx | 11.23433 | 11.21696 | 11.288 | -0.070738442 | -0.05337288 | 11.005715 | 10.831223 | 11.1572 | -0.325959794 | -0.151468135 |
| Itpkc | 7.778605 | 7.794928 | 7.7753 | 0.019592338 | 0.003269565 | 7.7355328 | 7.5904673 | 7.91644 | -0.325977095 | -0.180911586 |
| Slc35a1 | 8.137577 | 7.991913 | 7.9628 | 0.029067841 | 0.174731412 | 8.1076003 | 7.7592637 | 8.08534 | -0.326078585 | 0.022257994 |
| Hist1h2be | 4.865055 | 4.735481 | 5.0739 | -0.338430546 | -0.208856249 | 5.1721833 | 4.8412882 | 5.1675 | -0.326215242 | 0.004679766 |
| Cep170 | 10.02936 | 9.89076 | 9.8571 | 0.033627029 | 0.172226685 | 9.9816055 | 9.5855078 | 9.9118 | -0.326290841 | 0.069806823 |
| Siae | 9.37713 | 9.436349 | 9.5463 | -0.109979344 | -0.169198298 | 9.6063162 | 9.5442527 | 9.87056 | -0.326308611 | -0.264245112 |
| Miga1 | 8.7428 | 8.825863 | 8.7585 | 0.067348072 | -0.015714922 | 8.6949737 | 8.5631672 | 8.88957 | -0.326397857 | -0.194591359 |
| Phtf2 | 9.338294 | 9.333412 | 9.3857 | -0.05232444 | -0.047442439 | 9.377148 | 9.1758147 | 9.50236 | -0.326548335 | -0.125215035 |
| Kcnj1 | 3.904324 | 3.917852 | 4.0765 | -0.158637613 | -0.172166124 | 3.9563564 | 3.6565903 | 3.98339 | -0.326794873 | -0.02702879 |
| Sncg | 5.027024 | 4.805774 | 5.1598 | -0.353989166 | -0.132739372 | 4.5434357 | 4.4612286 | 4.78817 | -0.326939725 | -0.244732605 |
| Exoc5 | 10.43852 | 10.35745 | 10.305 | 0.05215609 | 0.13322829 | 10.173061 | 10.133067 | 10.4602 | -0.32714869 | -0.287154595 |
| Snai3 | 6.042331 | 5.956829 | 6.1335 | -0.17671984 | -0.091217353 | 6.011233 | 5.7190147 | 6.04627 | -0.327253491 | -0.035035138 |
| Prkra | 6.86588 | 6.706447 | 6.8083 | -0.101852576 | 0.057580805 | 6.88221 | 6.4083367 | 6.73584 | -0.327502935 | 0.146370385 |
| Crb3 | 5.105086 | 4.878977 | 5.1272 | -0.248208932 | -0.022100163 | 5.271588 | 4.7557053 | 5.08323 | -0.327527425 | 0.188355227 |
| As3mt | 8.026994 | 7.850646 | 8.1109 | -0.260205268 | -0.083856913 | 8.0488595 | 7.6962646 | 8.0238 | -0.327534193 | 0.025060699 |
| Engase | 7.19438 | 7.024363 | 7.1748 | -0.150407071 | 0.019610091 | 7.4539876 | 6.9450593 | 7.27265 | -0.327592477 | 0.181335747 |
| Gm4956 | 6.240818 | 6.0076 | 6.2444 | -0.236784729 | -0.003567119 | 5.9300639 | 5.7846095 | 6.11223 | -0.327624595 | -0.182170216 |
| Trp53inp2 | 8.075083 | 7.679181 | 8.0278 | -0.348635301 | 0.047266544 | 7.7097612 | 7.625159 | 7.9529 | -0.327742883 | -0.243140667 |
| Cr2 | 12.76343 | 12.73414 | 12.639 | 0.094921464 | 0.124209764 | 13.462039 | 12.616578 | 12.9446 | -0.32797992 | 0.517480421 |
| Esyt1 | 13.00575 | 12.99258 | 13.013 | -0.020216924 | -0.007050196 | 13.142631 | 12.714075 | 13.0423 | -0.328197186 | 0.100358977 |

| | | | | | | | | | | |
|---|---|---|---|---|---|---|---|---|---|---|
| Elfn2 | 2.434191 | 2.298859 | 2.4472 | -0.148383857 | -0.013051341 | 2.3651545 | 2.3235908 | 2.6518 | -0.328209519 | -0.286645816 |
| Adarb1 | 5.447106 | 5.467948 | 5.5776 | -0.109613952 | -0.130456075 | 5.4642839 | 5.0082121 | 5.33648 | -0.328270967 | 0.127800861 |
| Atp1b2 | 3.928218 | 3.765084 | 3.967 | -0.201897305 | -0.038763571 | 3.5225885 | 3.5506324 | 3.87903 | -0.328394295 | -0.356438159 |
| Ccdc122 | 5.2613 | 5.041981 | 5.2522 | -0.210185462 | 0.00913374 | 5.1908565 | 5.040701 | 5.36911 | -0.3284127 | -0.17825717 |
| Tmem63a | 9.900485 | 9.876748 | 9.8771 | -0.000311671 | 0.023425622 | 9.9732009 | 9.5984363 | 9.92687 | -0.328436142 | 0.046328501 |
| Slc35e3 | 7.888717 | 7.914401 | 7.8791 | 0.03531385 | 0.009630431 | 7.8711447 | 7.601701 | 7.9302 | -0.328502704 | -0.059059055 |
| Mtf2 | 10.07497 | 9.937869 | 10.093 | -0.155410302 | -0.01831211 | 9.8577017 | 9.8923585 | 10.2209 | -0.328546468 | -0.363203305 |
| Gmcl1 | 9.1611 | 9.127748 | 9.2571 | -0.129358851 | -0.096007043 | 8.9979004 | 8.8286841 | 9.15728 | -0.328594815 | -0.159378535 |
| Zfp182 | 8.020068 | 8.035635 | 8.1948 | -0.159200794 | -0.17476863 | 8.1362818 | 8.0693422 | 8.398 | -0.328656575 | -0.261716985 |
| Tusc3 | 8.619809 | 8.393765 | 8.5432 | -0.149450921 | 0.076593348 | 8.4395135 | 8.1750923 | 8.50382 | -0.328723902 | -0.064302637 |
| Tshr | 4.379307 | 4.047612 | 4.4245 | -0.376866095 | -0.045171018 | 4.1608494 | 3.7289376 | 4.05773 | -0.32879336 | 0.10311841 |
| Clcn1 | 2.429111 | 2.257954 | 2.281 | -0.023074984 | 0.148082441 | 2.5075346 | 2.3469012 | 2.67577 | -0.328871433 | -0.16823807 |
| 1500009L16Rik | 4.254871 | 4.071383 | 4.396 | -0.324648937 | -0.14116097 | 4.1440575 | 4.253136 | 4.58218 | -0.329046187 | -0.438124641 |
| Uaca | 7.296152 | 7.010763 | 7.3328 | -0.322026452 | -0.036637293 | 6.3505146 | 6.4206037 | 6.74979 | -0.329189782 | -0.399278911 |
| Arnt | 10.23881 | 10.16554 | 10.253 | -0.087627148 | -0.014355616 | 10.156185 | 9.9949877 | 10.3243 | -0.32928288 | -0.168085174 |
| Zfp248 | 4.949975 | 4.851004 | 5.02 | -0.168974765 | -0.070003626 | 4.9852434 | 4.8322222 | 5.1619 | -0.329677908 | -0.176656662 |
| Cntrob | 8.240114 | 8.439738 | 8.2752 | 0.164510233 | -0.035113835 | 8.7545957 | 8.3392461 | 8.66898 | -0.329733025 | 0.085616489 |
| Mxd3 | 5.094126 | 4.955032 | 5.2494 | -0.294365652 | -0.155271643 | 5.0618721 | 4.9992053 | 5.32914 | -0.329932878 | -0.267266145 |
| Mrgprb1 | 4.714785 | 4.441468 | 4.6113 | -0.169825912 | 0.10349104 | 4.1884296 | 4.10084 | 4.43091 | -0.330071975 | -0.242482366 |
| Gm20388 | 9.663062 | 9.875948 | 9.7515 | 0.124438578 | -0.088447693 | 9.6456731 | 9.4703282 | 9.80045 | -0.330121339 | -0.154776496 |
| Gdi1 | 11.12832 | 11.10067 | 11.201 | -0.10044039 | -0.07278613 | 11.674845 | 11.156083 | 11.4862 | -0.330149578 | 0.188612425 |
| Spin2c | 5.354113 | 5.23723 | 5.6101 | -0.372919397 | -0.256035838 | 5.2370294 | 4.9336578 | 5.26385 | -0.330189638 | -0.026817999 |
| BC024063 | 5.384605 | 5.375232 | 5.5707 | -0.195421524 | -0.186049 | 5.5101988 | 5.3191736 | 5.64937 | -0.330194573 | -0.139169404 |
| Akap1 | 6.989351 | 6.755668 | 6.9438 | -0.188129673 | 0.045553856 | 6.5531913 | 6.4860858 | 6.81634 | -0.330253647 | -0.263148088 |
| Dnmt3b | 7.3308 | 7.33482 | 7.4535 | -0.11872369 | -0.122743888 | 7.4542125 | 7.2467469 | 7.577 | -0.330257498 | -0.122791886 |
| Ltc4s | 4.516929 | 4.520525 | 4.6616 | -0.141033717 | -0.144629835 | 4.5895292 | 4.2372985 | 4.56778 | -0.330479046 | 0.021751669 |
| Mipol1 | 6.380413 | 6.073692 | 6.1829 | -0.109219383 | 0.197501286 | 6.1237183 | 6.0593914 | 6.38991 | -0.330520403 | -0.266193455 |
| Rtn4ip1 | 8.233827 | 8.130465 | 8.2418 | -0.111361313 | -0.007999345 | 8.4248068 | 8.0404615 | 8.37113 | -0.330668996 | 0.053676278 |
| Ckap5 | 10.45236 | 10.43638 | 10.497 | -0.060593308 | -0.044617422 | 10.433892 | 10.284187 | 10.6149 | -0.330729494 | -0.181025142 |
| Naglu | 8.903182 | 8.790022 | 8.9728 | -0.182758963 | -0.069598636 | 9.0993237 | 8.491899 | 8.82264 | -0.330738152 | 0.276686506 |
| Gm14399 | 3.932652 | 3.839997 | 3.8615 | -0.021504875 | 0.071149758 | 3.7144571 | 3.7284984 | 4.05938 | -0.330876736 | -0.344918098 |
| Arhgap4 | 12.26722 | 12.25442 | 12.349 | -0.094098696 | -0.081293842 | 12.74639 | 12.317402 | 12.6485 | -0.331133406 | 0.097855189 |

| | | | | | | | | | | |
|---|---|---|---|---|---|---|---|---|---|---|
| Cdr2 | 7.161462 | 7.305777 | 7.2901 | 0.015662216 | -0.128652613 | 6.924618 | 7.0340016 | 7.3652 | -0.331198368 | -0.440581947 |
| Gpr183 | 10.88382 | 10.93321 | 10.837 | 0.09644172 | 0.047055388 | 10.686045 | 10.356956 | 10.6882 | -0.331220568 | -0.002130954 |
| Aldh3a2 | 9.732397 | 9.758586 | 9.7845 | -0.025897406 | -0.05208708 | 9.917069 | 9.5322183 | 9.86351 | -0.331290043 | 0.053560662 |
| 6030468B19Rik | 4.028412 | 3.922842 | 4.0608 | -0.13793947 | -0.032369531 | 4.3723564 | 3.8992958 | 4.23073 | -0.331432837 | 0.141627787 |
| Mbtps1 | 11.1483 | 11.16006 | 11.273 | -0.112475938 | -0.124238194 | 11.247249 | 11.013857 | 11.3453 | -0.33148339 | -0.098090755 |
| Bmp2k | 11.45963 | 11.45603 | 11.485 | -0.029231962 | -0.0256297 | 11.41888 | 11.184953 | 11.5165 | -0.331532996 | -0.097606491 |
| Trbv13-1 | 7.821808 | 7.807232 | 7.9143 | -0.107036067 | -0.09246008 | 7.6327441 | 7.5102305 | 7.84177 | -0.331540335 | -0.209026737 |
| Zfp867 | 5.293231 | 5.165288 | 5.4292 | -0.263957988 | -0.136015074 | 5.4023901 | 5.1919052 | 5.52345 | -0.331543185 | -0.121058312 |
| Amot | 5.534012 | 5.347403 | 5.5606 | -0.213213044 | -0.026604654 | 5.3420452 | 5.1903036 | 5.52208 | -0.331777139 | -0.180035484 |
| Hdac4 | 9.215513 | 9.117845 | 9.1595 | -0.041614826 | 0.056053225 | 9.2344032 | 8.8644127 | 9.19622 | -0.331808746 | 0.038181681 |
| Arl10 | 5.717222 | 5.699623 | 5.8332 | -0.133607839 | -0.116009199 | 5.7001844 | 5.5454802 | 5.87767 | -0.332192525 | -0.177488307 |
| Clmn | 4.486195 | 4.678543 | 4.8811 | -0.202562253 | -0.394909598 | 4.4817285 | 4.4886772 | 4.82094 | -0.332264392 | -0.339213093 |
| Ap4m1 | 8.622903 | 8.722165 | 8.8133 | -0.091146444 | -0.190408466 | 9.0238473 | 8.7446375 | 9.07693 | -0.332295507 | -0.053085794 |
| Paqr8 | 6.66927 | 6.404065 | 6.479 | -0.074916111 | 0.190289406 | 6.2523866 | 5.9769777 | 6.30967 | -0.332695117 | -0.057286255 |
| Pask | 6.030427 | 5.518256 | 5.7165 | -0.198205838 | 0.313964523 | 5.8215651 | 5.7530059 | 6.08585 | -0.33284711 | -0.264287868 |
| Smc5 | 9.957422 | 9.953476 | 10.061 | -0.107125247 | -0.103179489 | 9.9707699 | 9.9130286 | 10.2459 | -0.332863309 | -0.275122042 |
| 4930402H24Rik | 8.542091 | 8.445756 | 8.557 | -0.111260588 | -0.014925548 | 8.3192715 | 8.1953895 | 8.52855 | -0.333165255 | -0.209283226 |
| Sirt1 | 9.815973 | 9.832501 | 9.9119 | -0.079408506 | -0.095936875 | 9.6806638 | 9.5799239 | 9.9134 | -0.33347939 | -0.232739422 |
| C1ql3 | 4.023802 | 3.782724 | 4.1267 | -0.343975519 | -0.102897727 | 3.4851092 | 3.4635628 | 3.79725 | -0.333689026 | -0.312142591 |
| Nyap1 | 4.34611 | 4.224832 | 4.2819 | -0.057047495 | 0.064229993 | 4.0573374 | 3.9076369 | 4.24136 | -0.333727586 | -0.184027163 |
| Trim37 | 8.54486 | 8.392924 | 8.6111 | -0.218201706 | -0.066265718 | 8.5574782 | 8.2793234 | 8.6131 | -0.333775082 | -0.055620338 |
| Sox7 | 6.125252 | 5.476337 | 5.8654 | -0.389040478 | 0.25987433 | 5.0553376 | 5.0557157 | 5.38962 | -0.333908329 | -0.334286442 |
| Mettl26 | 7.70982 | 7.551408 | 7.6435 | -0.09210435 | 0.066307129 | 8.0446156 | 7.3933554 | 7.72727 | -0.333914965 | 0.317345207 |
| Xlr5c | 3.151587 | 3.176196 | 3.0621 | 0.114137812 | 0.089528706 | 2.9497682 | 2.8236272 | 3.15761 | -0.333984429 | -0.207843424 |
| Nedd1 | 8.882035 | 8.991789 | 8.9309 | 0.06093315 | -0.048820689 | 8.9912587 | 8.6186747 | 8.95309 | -0.334413856 | 0.038170159 |
| Fgd6 | 8.563152 | 8.42904 | 8.5608 | -0.131722365 | 0.002389415 | 8.3713219 | 8.0990098 | 8.43349 | -0.334479075 | -0.062166949 |
| Prkca | 9.724896 | 9.793206 | 9.7506 | 0.042639889 | -0.025669488 | 9.3696995 | 9.4778097 | 9.81237 | -0.334559513 | -0.442669708 |
| Hoxd8 | 5.10506 | 5.161859 | 5.1 | 0.061866664 | 0.005067345 | 4.8491372 | 4.6181258 | 4.95284 | -0.334709915 | -0.103698501 |
| Mast2 | 9.368027 | 9.30085 | 9.4203 | -0.119448982 | -0.052272325 | 9.2199575 | 9.0349096 | 9.36975 | -0.334836865 | -0.149788971 |
| Tppp | 3.88785 | 3.746108 | 3.7107 | 0.035410279 | 0.177151767 | 3.4644948 | 3.4332064 | 3.76815 | -0.334940149 | -0.303651752 |
| Cuedc1 | 5.883378 | 5.807313 | 5.8733 | -0.065942609 | 0.010121943 | 5.7893212 | 5.5435463 | 5.87856 | -0.335011197 | -0.089236289 |
| Scd1 | 13.76597 | 13.58769 | 13.69 | -0.102112896 | 0.076173508 | 13.973782 | 13.317986 | 13.653 | -0.335046316 | 0.320749554 |

| | | | | | | | | | | |
|---|---|---|---|---|---|---|---|---|---|---|
| Hltf | 10.24732 | 10.25737 | 10.302 | -0.044809323 | -0.054858873 | 10.230429 | 9.9891774 | 10.3242 | -0.335051131 | -0.093799347 |
| Igsf8 | 8.076671 | 8.241171 | 7.9837 | 0.257514919 | 0.093014847 | 8.1004641 | 7.8169866 | 8.15211 | -0.335120729 | -0.051643308 |
| Zfp606 | 7.625446 | 7.612001 | 7.6944 | -0.082383426 | -0.068937867 | 7.552254 | 7.392249 | 7.72766 | -0.335415827 | -0.175410789 |
| Rida | 7.157056 | 7.024835 | 7.222 | -0.197166127 | -0.064944863 | 7.0718768 | 6.7762234 | 7.11176 | -0.335539287 | -0.039885805 |
| Tcaf2 | 6.198 | 6.03609 | 6.181 | -0.14493025 | 0.016979466 | 5.7540576 | 5.7214629 | 6.05713 | -0.335669106 | -0.30307442 |
| Pccb | 8.541406 | 8.483166 | 8.5157 | -0.032560141 | 0.025679561 | 8.6038011 | 8.2232664 | 8.55898 | -0.335714666 | 0.044819956 |
| Epg5 | 9.05226 | 9.054622 | 9.0236 | 0.031036503 | 0.02867418 | 8.9812778 | 8.8476428 | 9.18346 | -0.335819252 | -0.202184188 |
| Unc13b | 5.461058 | 5.308444 | 5.8209 | -0.512492441 | -0.359878654 | 5.8534327 | 5.2645415 | 5.60046 | -0.335913588 | 0.252977665 |
| Ccdc3 | 3.918657 | 3.932677 | 3.9532 | -0.020507185 | -0.034527711 | 3.6154864 | 3.4904151 | 3.82637 | -0.335957054 | -0.210885673 |
| Pqlc3 | 8.482177 | 8.356467 | 8.5893 | -0.232875498 | -0.107166383 | 8.5975634 | 8.2604854 | 8.59647 | -0.335988407 | 0.001089614 |
| Sh2d4a | 5.605201 | 5.458916 | 5.7062 | -0.247308044 | -0.101023036 | 5.3052686 | 5.1503253 | 5.48633 | -0.336002772 | -0.181059529 |
| Zfp612 | 4.075839 | 3.890954 | 4.1916 | -0.300613804 | -0.115729481 | 3.9194976 | 3.683777 | 4.0198 | -0.33602054 | -0.100299894 |
| Mettl9 | 9.198956 | 9.039879 | 9.1444 | -0.104532448 | 0.054544258 | 8.6981543 | 8.7092811 | 9.04556 | -0.33627542 | -0.347402254 |
| Trat1 | 9.280162 | 9.340878 | 9.5022 | -0.161359001 | -0.222075331 | 8.8304971 | 9.0166423 | 9.35296 | -0.336322165 | -0.522467279 |
| Arhgef9 | 5.661899 | 5.545383 | 5.7908 | -0.245431602 | -0.128915846 | 5.7236617 | 5.4466567 | 5.78366 | -0.337008126 | -0.060003157 |
| Zfp810 | 7.674418 | 7.488517 | 7.7235 | -0.234950183 | -0.049049705 | 7.7665754 | 7.4116057 | 7.74883 | -0.337227188 | 0.017742462 |
| Msc | 3.068338 | 3.056627 | 3.0898 | -0.033170702 | -0.021458971 | 3.1667314 | 2.9641875 | 3.30163 | -0.337444824 | -0.134900864 |
| Rufy1 | 9.294962 | 9.332184 | 9.2945 | 0.037680432 | 0.000458006 | 9.4600991 | 9.0850559 | 9.42256 | -0.337508104 | 0.037535075 |
| Prnd | 3.486816 | 3.492434 | 3.6619 | -0.169511435 | -0.175129397 | 3.713749 | 3.2540332 | 3.59173 | -0.33769918 | 0.12201657 |
| Rxylt1 | 7.655684 | 7.354343 | 7.6757 | -0.321365556 | -0.020024809 | 7.6394905 | 7.2534036 | 7.59111 | -0.337706464 | 0.048380444 |
| Pde4dip | 9.848022 | 9.790968 | 9.846 | -0.054987296 | 0.002066913 | 9.7821287 | 9.5296622 | 9.86753 | -0.337869276 | -0.085402794 |
| Arid3b | 9.623004 | 9.753015 | 9.6394 | 0.113622548 | -0.016388502 | 9.9389299 | 9.6414033 | 9.97984 | -0.338436672 | -0.040910052 |
| Pik3r1 | 11.09787 | 11.20783 | 11.133 | 0.074475592 | -0.035486704 | 10.669839 | 10.716129 | 11.0546 | -0.338471904 | -0.38476131 |
| Slc22a23 | 6.934254 | 6.860909 | 7.005 | -0.14413996 | -0.070795573 | 6.7602197 | 6.8680779 | 7.20659 | -0.338507956 | -0.446366161 |
| Pcdhgb6 | 5.679698 | 5.250945 | 5.4278 | -0.176872795 | 0.251880777 | 5.1680378 | 5.0012648 | 5.33989 | -0.338629043 | -0.171855957 |
| Grk4 | 5.901064 | 5.453801 | 5.7861 | -0.33225088 | 0.115011939 | 5.70181 | 5.7168918 | 6.05566 | -0.33876638 | -0.35384823 |
| Prkacb | 11.7544 | 11.71533 | 11.822 | -0.10666948 | -0.067591958 | 11.627147 | 11.390181 | 11.729 | -0.338800468 | -0.101834298 |
| Rwdd3 | 5.25745 | 5.069196 | 5.3146 | -0.245361813 | -0.057107787 | 5.3984886 | 5.1119418 | 5.45114 | -0.339193667 | -0.052646885 |
| Bex3 | 8.133452 | 8.077212 | 8.1864 | -0.109222673 | -0.052982111 | 8.1315748 | 7.8548319 | 8.19406 | -0.339225818 | -0.062482927 |
| Coq10a | 7.736801 | 7.596815 | 7.8823 | -0.285486448 | -0.145500569 | 8.0372027 | 7.5361781 | 7.87549 | -0.339308562 | 0.161716066 |
| Cep135 | 9.161711 | 9.072002 | 9.1893 | -0.11727936 | -0.027569954 | 9.1527351 | 8.9456917 | 9.28509 | -0.339401718 | -0.132358356 |
| Ell2 | 9.086049 | 9.222728 | 9.0077 | 0.21501994 | 0.078340033 | 8.9597519 | 8.763564 | 9.10299 | -0.339424896 | -0.143236933 |

| | | | | | | | | | | |
|---|---|---|---|---|---|---|---|---|---|---|
| Ash1l | 11.0961 | 10.99523 | 11.156 | -0.160802858 | -0.059929324 | 11.143514 | 11.104123 | 11.4436 | -0.339445656 | -0.300054981 |
| Ubxn8 | 8.257056 | 8.100423 | 8.1945 | -0.094077481 | 0.0625558 | 8.0922343 | 7.8742656 | 8.21372 | -0.339452477 | -0.121483755 |
| Creb1 | 11.30256 | 11.45804 | 11.321 | 0.137157636 | -0.018320016 | 11.138921 | 10.993825 | 11.3334 | -0.339619046 | -0.1945232 |
| Gm28900 | 3.913148 | 4.144522 | 4.1711 | -0.026554925 | -0.257929463 | 3.8103834 | 3.5654881 | 3.90511 | -0.339625196 | -0.094729972 |
| Ptgr2 | 8.953631 | 8.752227 | 8.9806 | -0.228348663 | -0.026944528 | 9.0005632 | 8.6401827 | 8.98007 | -0.339892249 | 0.020488185 |
| Rhoq | 9.457769 | 9.298527 | 9.4459 | -0.147373059 | 0.011869341 | 9.3692441 | 8.8795164 | 9.21948 | -0.339960484 | 0.1497673 |
| Ifnlr1 | 4.606107 | 4.644584 | 4.8602 | -0.215656709 | -0.254134535 | 4.8607695 | 4.5467106 | 4.88669 | -0.339978659 | -0.025919753 |
| Bhlhe40 | 10.98852 | 10.76822 | 10.733 | 0.034737892 | 0.255033698 | 9.9904367 | 10.077369 | 10.4174 | -0.340053815 | -0.426985816 |
| Usf3 | 10.66964 | 10.69069 | 10.725 | -0.034795972 | -0.055844182 | 10.720807 | 10.564545 | 10.9046 | -0.34008092 | -0.183818214 |
| Nrip2 | 2.933386 | 2.865411 | 3.0448 | -0.179348534 | -0.111373487 | 2.7953883 | 2.6252479 | 2.96548 | -0.340230321 | -0.170089956 |
| G2e3 | 8.157514 | 8.057597 | 8.1744 | -0.116838929 | -0.016922312 | 8.1123959 | 7.9437088 | 8.28403 | -0.340319531 | -0.171632436 |
| Brd9 | 9.995107 | 9.975806 | 10.09 | -0.114509778 | -0.095209167 | 10.23307 | 9.9507299 | 10.291 | -0.340319798 | -0.057979998 |
| Arhgap20 | 4.893834 | 4.689961 | 4.6262 | 0.063792994 | 0.267665638 | 4.5518209 | 4.35202 | 4.69246 | -0.340444043 | -0.140643106 |
| Arfgef3 | 4.428654 | 4.339415 | 4.4618 | -0.122435099 | -0.033195953 | 4.4422979 | 4.245571 | 4.58605 | -0.340482563 | -0.143755718 |
| Rasd1 | 5.924599 | 5.648403 | 5.8198 | -0.171347854 | 0.104847899 | 5.0691505 | 5.1649321 | 5.50552 | -0.340586038 | -0.436367612 |
| Zfp40 | 6.641905 | 6.394034 | 6.5727 | -0.178661956 | 0.069208754 | 6.5988305 | 6.3381261 | 6.67874 | -0.340609039 | -0.079904643 |
| Ednrb | 6.63751 | 6.261044 | 5.7503 | 0.510772362 | 0.887238254 | 6.0137471 | 5.4470484 | 5.78767 | -0.34062652 | 0.226072111 |
| Igkv5-45 | 5.395339 | 5.998206 | 5.3677 | 0.630479606 | 0.027613016 | 5.6820461 | 5.8268544 | 6.16769 | -0.340837584 | -0.485645928 |
| Kifc5b | 6.528615 | 6.64918 | 6.6131 | 0.03611623 | -0.084448596 | 6.6377383 | 6.4698344 | 6.81069 | -0.340854795 | -0.172950923 |
| Ubr1 | 10.72493 | 10.69593 | 10.765 | -0.068868762 | -0.039875822 | 10.712835 | 10.524394 | 10.8654 | -0.34095961 | -0.152518212 |
| Gstm4 | 6.067389 | 5.956807 | 6.2763 | -0.319458785 | -0.208876337 | 6.1706177 | 5.8625729 | 6.2036 | -0.341027522 | -0.03298272 |
| Rogdi | 9.55073 | 9.592032 | 9.8027 | -0.210664872 | -0.251966789 | 10.017047 | 9.4996175 | 9.84072 | -0.341106379 | 0.176322674 |
| Sdr39u1 | 8.36301 | 8.319998 | 8.4651 | -0.145097667 | -0.102085522 | 8.6750228 | 8.1791511 | 8.52035 | -0.341198428 | 0.154673211 |
| Rabgap1l | 11.18422 | 11.19565 | 11.241 | -0.045334502 | -0.056764998 | 11.069302 | 10.936401 | 11.2777 | -0.341266882 | -0.20836539 |
| Zfp931 | 6.262229 | 6.290789 | 6.2635 | 0.027291024 | -0.001269291 | 6.2656964 | 5.9466527 | 6.28805 | -0.341398904 | -0.022355238 |
| Bbs4 | 7.16899 | 7.050189 | 7.2863 | -0.236084835 | -0.117283836 | 7.2966171 | 7.1231015 | 7.46453 | -0.341424483 | -0.16790887 |
| Tyms | 7.62605 | 7.596204 | 7.7758 | -0.179611252 | -0.149764646 | 7.6262304 | 7.6538922 | 7.9954 | -0.341503585 | -0.369165355 |
| Zfp280b | 8.698719 | 8.859253 | 8.8328 | 0.026424554 | -0.134109387 | 8.6914814 | 8.5647563 | 8.9065 | -0.341746918 | -0.215021858 |
| Zfand4 | 4.717917 | 4.461503 | 4.9198 | -0.458335117 | -0.201921612 | 4.5755733 | 4.354827 | 4.69673 | -0.341907562 | -0.121161296 |
| Ssu2 | 2.969044 | 2.898006 | 3.0817 | -0.183739524 | -0.112702092 | 2.8632299 | 2.6979656 | 3.04003 | -0.342061392 | -0.176797085 |
| Cyp26b1 | 6.476711 | 5.86826 | 5.8508 | 0.017488012 | 0.625939465 | 5.3716877 | 4.7579952 | 5.10024 | -0.34224719 | 0.271445359 |
| Hist1h2bb | 3.843936 | 3.584035 | 3.7498 | -0.165728303 | 0.09417267 | 3.8318015 | 3.5099413 | 3.85221 | -0.34227022 | -0.020409966 |

| | | | | | | | | | | |
|---|---|---|---|---|---|---|---|---|---|---|
| Klf4 | 9.550499 | 8.83545 | 9.518 | -0.682574753 | 0.032474557 | 8.3120045 | 8.3792381 | 8.72152 | -0.342278837 | -0.409512438 |
| Dnm3 | 5.312707 | 4.942738 | 5.0292 | -0.086434569 | 0.283534423 | 4.8931957 | 4.73605 | 5.0784 | -0.342346871 | -0.185201149 |
| Osbpl1a | 5.549501 | 5.540768 | 5.5849 | -0.044082832 | -0.035349912 | 5.4344632 | 5.3977661 | 5.74019 | -0.342426721 | -0.305729599 |
| Gm14117 | 6.196058 | 6.039299 | 6.2503 | -0.210993771 | -0.054234679 | 6.1542297 | 5.8783063 | 6.22083 | -0.342525911 | -0.066602542 |
| Dlg4 | 8.336451 | 8.575596 | 8.4964 | 0.079239502 | -0.159905474 | 8.2004025 | 8.334839 | 8.67757 | -0.34273209 | -0.477168595 |
| Tmsb15b2 | 3.881455 | 3.96886 | 3.8913 | 0.077559326 | -0.009845339 | 3.9275596 | 3.7906573 | 4.13349 | -0.342832161 | -0.205929844 |
| Ccdc112 | 5.314296 | 5.534894 | 5.5062 | 0.028728972 | -0.191868252 | 5.5031688 | 5.219361 | 5.56223 | -0.342870773 | -0.059062958 |
| Zkscan7 | 7.163025 | 7.039758 | 7.1594 | -0.119654269 | 0.003612775 | 7.0157252 | 6.8801787 | 7.22367 | -0.343489256 | -0.207942732 |
| Atg10 | 6.876493 | 6.6308 | 6.7453 | -0.114491974 | 0.131201291 | 6.8021782 | 6.6623626 | 7.00586 | -0.343497761 | -0.203682134 |
| S100a16 | 6.523172 | 6.233871 | 6.5625 | -0.32862602 | -0.039325124 | 6.0604165 | 5.5980799 | 5.9416 | -0.343517994 | 0.118818601 |
| 38777 | 11.2906 | 11.23401 | 11.28 | -0.045953364 | 0.010642092 | 11.223871 | 10.930385 | 11.2742 | -0.343789822 | -0.050303794 |
| Pcbp3 | 6.497249 | 6.471346 | 6.7003 | -0.228966656 | -0.20306378 | 6.2315171 | 6.0388154 | 6.38267 | -0.34385638 | -0.151154703 |
| Nrgn | 5.252922 | 5.206205 | 5.3925 | -0.186264956 | -0.139548369 | 5.3596971 | 4.9114659 | 5.25552 | -0.344058538 | 0.104172667 |
| Pex13 | 8.35575 | 8.54637 | 8.322 | 0.224385664 | 0.033765049 | 8.3215789 | 8.2978226 | 8.64195 | -0.344128573 | -0.320372247 |
| Tube1 | 5.908009 | 5.772049 | 5.7814 | -0.009323447 | 0.126635846 | 5.9958179 | 5.8479846 | 6.19215 | -0.344166044 | -0.196332717 |
| Srgap2 | 9.783602 | 9.725419 | 9.7538 | -0.028411389 | 0.029771858 | 9.7061027 | 9.6704263 | 10.0148 | -0.344414587 | -0.308738233 |
| Ccdc18 | 5.388138 | 4.940217 | 5.2394 | -0.299170969 | 0.148749654 | 5.2307169 | 5.014524 | 5.35894 | -0.344415283 | -0.128222424 |
| Tlr1 | 10.66648 | 10.73491 | 10.724 | 0.011297468 | -0.057130014 | 11.224667 | 10.649742 | 10.9942 | -0.344449634 | 0.230476158 |
| Dnah6 | 5.038196 | 4.845925 | 4.7615 | 0.084450291 | 0.276720888 | 4.476742 | 4.3723698 | 4.717 | -0.344632368 | -0.240260117 |
| Rasgrp3 | 11.24258 | 11.4196 | 11.321 | 0.098768216 | -0.078250866 | 11.616467 | 11.130375 | 11.4751 | -0.344695108 | 0.141396921 |
| Tmem191c | 6.048786 | 6.070319 | 6.1768 | -0.10645255 | -0.127984694 | 6.1874748 | 5.9334959 | 6.27831 | -0.344816518 | -0.090837595 |
| Col4a3bp | 10.20025 | 10.18412 | 10.048 | 0.136081827 | 0.152206197 | 9.9291619 | 9.965769 | 10.3107 | -0.344945091 | -0.381552189 |
| Gm49405 | 6.303702 | 6.532422 | 6.6702 | -0.137806779 | -0.366526777 | 7.0425509 | 6.5843539 | 6.92946 | -0.345103184 | 0.11309383 |
| Abcc3 | 4.418909 | 4.181523 | 4.3956 | -0.214041961 | 0.023344231 | 4.1492727 | 4.0344075 | 4.3796 | -0.345191023 | -0.230325815 |
| Ephx2 | 4.682529 | 4.382526 | 4.691 | -0.30843558 | -0.008432429 | 4.1140984 | 4.0045041 | 4.34975 | -0.345243966 | -0.235649638 |
| Msrb3 | 6.854249 | 6.511917 | 6.6202 | -0.108275593 | 0.234055929 | 6.3313403 | 6.1425495 | 6.4878 | -0.345252414 | -0.156461677 |
| S1pr1 | 12.31423 | 12.20363 | 12.271 | -0.06781548 | 0.042783876 | 12.082815 | 11.816994 | 12.1623 | -0.345310496 | -0.079489393 |
| Bcar1 | 6.870319 | 6.453221 | 6.6596 | -0.206404771 | 0.210692286 | 6.1550211 | 6.0361468 | 6.3816 | -0.345453762 | -0.22657944 |
| Cotl1 | 12.85142 | 12.75428 | 12.796 | -0.041882636 | 0.05525864 | 12.923374 | 12.422184 | 12.7677 | -0.345539872 | 0.155649585 |
| Plekhm3 | 10.20503 | 10.41397 | 10.358 | 0.055744204 | -0.153200464 | 10.32597 | 10.116969 | 10.4626 | -0.345624472 | -0.136623855 |
| Tgfbrap1 | 9.542899 | 9.543335 | 9.6501 | -0.106811566 | -0.107246854 | 9.554975 | 9.3483073 | 9.69418 | -0.345873777 | -0.139206064 |
| Acer2 | 6.83945 | 6.630003 | 6.6879 | -0.057930172 | 0.151517594 | 6.0028568 | 6.2093288 | 6.55549 | -0.346164686 | -0.552636721 |

| | | | | | | | | | | |
|---|---|---|---|---|---|---|---|---|---|---|
| Mylip | 11.04728 | 10.85719 | 11.086 | -0.2286291 | -0.03854033 | 10.809809 | 10.542949 | 10.8893 | -0.346344008 | -0.079484083 |
| Cdk6 | 9.7157 | 9.713251 | 9.7585 | -0.045295196 | -0.04284582 | 9.7384902 | 9.5922352 | 9.93858 | -0.34634667 | -0.20009162 |
| Ccdc93 | 9.668811 | 9.66555 | 9.7621 | -0.096563753 | -0.093302862 | 9.6103858 | 9.3976275 | 9.74404 | -0.346414863 | -0.133656512 |
| Atrx | 12.40178 | 12.27535 | 12.279 | -0.00385333 | 0.122575688 | 12.133598 | 12.143458 | 12.49 | -0.346504954 | -0.356364986 |
| Kdelc1 | 6.632314 | 6.602617 | 6.6335 | -0.030891581 | -0.001195003 | 6.6214115 | 6.2882488 | 6.63483 | -0.346583197 | -0.01342044 |
| Ighv1-77 | 4.977077 | 5.470454 | 4.9493 | 0.521138006 | 0.027761568 | 5.2916215 | 5.1196791 | 5.46636 | -0.346682649 | -0.174740209 |
| Bdkrb2 | 3.955123 | 3.980951 | 3.9698 | 0.011178602 | -0.014649111 | 3.7959749 | 3.6580805 | 4.00484 | -0.346757979 | -0.208863525 |
| Trim45 | 5.336819 | 5.182582 | 5.3921 | -0.209472894 | -0.055236411 | 5.3777318 | 5.0882164 | 5.4351 | -0.34688458 | -0.057369184 |
| Tmem80 | 7.602331 | 7.5996 | 7.5614 | 0.03820055 | 0.040931936 | 8.0846875 | 7.6423444 | 7.98956 | -0.347219438 | 0.095123726 |
| Zbtb40 | 8.928438 | 9.115279 | 8.8949 | 0.22037316 | 0.0335316 | 9.2356281 | 8.8394209 | 9.18689 | -0.347473237 | 0.048733938 |
| Cobll1 | 9.858824 | 9.818157 | 9.8998 | -0.081673373 | -0.04100649 | 9.925287 | 9.5520033 | 9.89956 | -0.347555699 | 0.025727985 |
| Lrrc32 | 8.132303 | 8.031822 | 7.8952 | 0.136663257 | 0.237144909 | 7.9067171 | 7.7461629 | 8.09381 | -0.347651159 | -0.187096989 |
| Rffl | 8.654786 | 8.58686 | 8.6811 | -0.094250328 | -0.026324512 | 8.6717601 | 8.2922324 | 8.63996 | -0.347728479 | 0.031799159 |
| Arhgef15 | 7.10039 | 6.756433 | 6.9641 | -0.207639582 | 0.136317694 | 6.3409269 | 6.3273541 | 6.67513 | -0.347771393 | -0.334198506 |
| Zfp316 | 6.828358 | 6.624848 | 6.7776 | -0.152783377 | 0.05072625 | 6.7924577 | 6.3454936 | 6.69332 | -0.34782478 | 0.09913934 |
| Ddb2 | 9.439661 | 9.409726 | 9.5735 | -0.163738976 | -0.133803947 | 9.8089875 | 9.4702127 | 9.81834 | -0.348125285 | -0.00935048 |
| Aurka | 6.763301 | 6.667376 | 6.8934 | -0.226037163 | -0.130111735 | 6.7919848 | 6.726896 | 7.07521 | -0.348311245 | -0.28322246 |
| Rasgef1b | 11.04337 | 11.07859 | 10.846 | 0.232426784 | 0.197206768 | 10.920333 | 10.767142 | 11.1155 | -0.34833662 | -0.195146223 |
| Zfp783 | 7.357716 | 7.193594 | 7.3375 | -0.14386212 | 0.020259952 | 7.683322 | 7.4213325 | 7.76975 | -0.348422102 | -0.086432509 |
| Ptgds | 5.432561 | 5.036347 | 5.3581 | -0.32178704 | 0.074426509 | 4.6219708 | 4.3600222 | 4.70872 | -0.348701887 | -0.08675334 |
| Pds5a | 12.05661 | 12.02982 | 12.065 | -0.035550008 | -0.008756188 | 12.012912 | 11.800133 | 12.149 | -0.348843008 | -0.136063569 |
| Noxred1 | 6.717135 | 6.462788 | 6.6821 | -0.219312357 | 0.03503493 | 7.1350853 | 6.3585762 | 6.70754 | -0.348966352 | 0.427542732 |
| Clca3a1 | 7.400827 | 6.790482 | 7.14 | -0.349499883 | 0.260844606 | 6.0616183 | 6.6103549 | 6.95934 | -0.348981308 | -0.897717863 |
| Cnnm2 | 6.318923 | 6.334737 | 6.243 | 0.09173381 | 0.075920177 | 6.0403863 | 5.8861433 | 6.23528 | -0.349133958 | -0.19489087 |
| Ccne2 | 5.591293 | 5.395907 | 5.7299 | -0.334012272 | -0.138626044 | 5.4559796 | 5.4639609 | 5.8135 | -0.349543474 | -0.357524821 |
| Ttc38 | 8.854068 | 8.836185 | 8.8882 | -0.052002708 | -0.034119153 | 8.8156558 | 8.5306946 | 8.8804 | -0.349705214 | -0.064743972 |
| Ncoa1 | 10.17948 | 10.35032 | 10.287 | 0.063821278 | -0.10702331 | 10.283026 | 10.142265 | 10.492 | -0.349731824 | -0.208971184 |
| Rab11fip3 | 7.54406 | 7.402943 | 7.5228 | -0.119884375 | 0.021232401 | 7.3796605 | 7.21195 | 7.56184 | -0.349886934 | -0.182176435 |
| Cep70 | 7.073329 | 7.064688 | 7.226 | -0.161335781 | -0.152695371 | 7.1718965 | 6.9524138 | 7.30251 | -0.350099553 | -0.130616919 |
| Smc2 | 9.734802 | 9.741594 | 9.7653 | -0.02365804 | -0.0304503 | 9.6063806 | 9.5022086 | 9.85232 | -0.350113485 | -0.245941522 |
| Slc9a7 | 9.109091 | 9.332629 | 9.2052 | 0.127412321 | -0.096125294 | 9.4649133 | 9.0514473 | 9.40168 | -0.350235281 | 0.063230783 |
| Zfp740 | 11.31371 | 11.30553 | 11.344 | -0.038700618 | -0.030520104 | 11.257481 | 11.072174 | 11.4225 | -0.350310524 | -0.165003583 |

| | | | | | | | | | | |
|---|---|---|---|---|---|---|---|---|---|---|
| Acap3 | 8.366766 | 8.39575 | 8.4584 | -0.062641099 | -0.091624444 | 8.4226413 | 8.293468 | 8.64391 | -0.350437241 | -0.221263936 |
| Phf21a | 10.245 | 10.16603 | 10.217 | -0.050677908 | 0.028287614 | 10.209511 | 10.127052 | 10.4776 | -0.350525884 | -0.268067367 |
| AC134468.1 | 3.789844 | 3.867362 | 3.741 | 0.126406869 | 0.048888725 | 3.6182042 | 3.464537 | 3.81509 | -0.3505569 | -0.196889631 |
| Abtb1 | 8.81642 | 8.872588 | 8.9346 | -0.062004655 | -0.11817227 | 9.1243437 | 8.5932533 | 8.94389 | -0.350637843 | 0.180452527 |
| Myom1 | 4.972149 | 4.994489 | 5.0096 | -0.015147835 | -0.037487469 | 4.4710167 | 4.6982451 | 5.04888 | -0.35063872 | -0.577867127 |
| Sec23a | 8.882154 | 8.796339 | 8.8045 | -0.008157995 | 0.077656404 | 8.8116611 | 8.4763273 | 8.827 | -0.350670527 | -0.015336749 |
| Nfatc2 | 9.707061 | 9.735005 | 9.8103 | -0.075287781 | -0.103231822 | 9.4601967 | 9.3949194 | 9.74569 | -0.35077538 | -0.285498053 |
| Bicd1 | 6.272492 | 6.509068 | 6.2288 | 0.280289757 | 0.043714456 | 6.5755004 | 6.13181 | 6.48261 | -0.35080316 | 0.09288725 |
| Tmem106b | 10.25485 | 10.22058 | 10.362 | -0.141050824 | -0.106781152 | 10.214392 | 9.8638399 | 10.2147 | -0.35084449 | -0.000292553 |
| Hibch | 8.194212 | 8.089273 | 8.2449 | -0.155587611 | -0.05064902 | 8.1582746 | 7.9151477 | 8.26603 | -0.350885043 | -0.10775808 |
| Fads2 | 6.823575 | 6.601242 | 6.8657 | -0.264484588 | -0.0421519 | 6.4122201 | 5.8481785 | 6.19915 | -0.350966645 | 0.213074981 |
| Mgea5 | 11.98718 | 11.92222 | 12.046 | -0.124175754 | -0.059218892 | 11.984853 | 11.853734 | 12.2047 | -0.350999118 | -0.219880175 |
| Elac1 | 8.246862 | 8.189822 | 8.313 | -0.123214479 | -0.066174485 | 8.2613187 | 7.871109 | 8.22214 | -0.351029051 | 0.039180646 |
| Gna15 | 7.332368 | 7.323023 | 7.4074 | -0.084368644 | -0.075023734 | 7.2329902 | 7.1063939 | 7.45746 | -0.351068806 | -0.224472449 |
| Cyp7b1 | 5.050157 | 4.56828 | 4.5394 | 0.028878521 | 0.510755432 | 4.808351 | 4.1739347 | 4.52508 | -0.351145145 | 0.283271146 |
| Cst6 | 1.700541 | 1.761054 | 1.63 | 0.131065206 | 0.070552645 | 1.8339206 | 1.5227584 | 1.87394 | -0.351179806 | -0.040017585 |
| Tmem64 | 10.43386 | 10.33712 | 10.437 | -0.099603172 | -0.00286375 | 10.204206 | 9.9814245 | 10.3328 | -0.351335181 | -0.128553907 |
| Zfp11 | 5.548565 | 5.401326 | 5.5115 | -0.110133111 | 0.037105691 | 5.5912099 | 5.3871006 | 5.73863 | -0.351530956 | -0.147421659 |
| Gm19680 | 6.019085 | 6.059642 | 6.1016 | -0.041920385 | -0.082477026 | 5.9827702 | 5.5092803 | 5.86085 | -0.3515664 | 0.121923521 |
| 6720489N17Rik | 5.69919 | 5.681403 | 5.8005 | -0.119080886 | -0.101293585 | 5.7482694 | 5.4059797 | 5.75755 | -0.351567856 | -0.009278216 |
| Snrk | 10.57082 | 10.56126 | 10.619 | -0.058062006 | -0.048498322 | 10.142323 | 10.150011 | 10.5016 | -0.351612992 | -0.359300963 |
| Ibtk | 10.66109 | 10.73981 | 10.748 | -0.007818648 | -0.086543634 | 10.70137 | 10.410944 | 10.7626 | -0.351635264 | -0.061209364 |
| Sepsecs | 8.624155 | 8.593739 | 8.5968 | -0.003033546 | 0.027382766 | 8.6259131 | 8.4286049 | 8.78032 | -0.351717787 | -0.154409617 |
| Pawr | 5.160157 | 4.925728 | 4.9784 | -0.052663309 | 0.181765619 | 4.5259842 | 4.2565954 | 4.60835 | -0.351758706 | -0.082369982 |
| Ndc80 | 7.450011 | 7.500026 | 7.5717 | -0.071644601 | -0.121659211 | 7.4052085 | 7.3480141 | 7.7001 | -0.352089526 | -0.2948951 |
| C5ar2 | 5.093341 | 4.910266 | 5.0131 | -0.102795601 | 0.08027892 | 5.1039611 | 4.6674063 | 5.0195 | -0.352097495 | 0.084457397 |
| Ccdc180 | 4.7898 | 4.756765 | 4.8491 | -0.092322865 | -0.059287728 | 5.000774 | 4.6148606 | 4.96698 | -0.352122271 | 0.033791076 |
| Rhoj | 6.997857 | 6.512303 | 6.6595 | -0.147161978 | 0.33839263 | 6.0467136 | 5.9120471 | 6.2643 | -0.352249082 | -0.217582536 |
| Gm13378 | 2.755379 | 2.859838 | 2.7663 | 0.093541109 | -0.010918051 | 2.684641 | 2.6549017 | 3.00716 | -0.352262907 | -0.322523644 |
| Neo1 | 4.906074 | 4.87046 | 4.8749 | -0.004406606 | 0.031208011 | 4.8889514 | 4.6016497 | 4.95406 | -0.35240869 | -0.065107007 |
| Zfp445 | 10.71308 | 10.77502 | 10.792 | -0.017310858 | -0.079256782 | 10.98589 | 10.778644 | 11.1311 | -0.352448708 | -0.145202289 |
| Ptprf | 7.135882 | 6.985878 | 6.8638 | 0.122102689 | 0.272107242 | 6.4744466 | 6.2994609 | 6.65223 | -0.352773904 | -0.177788151 |

| | | | | | | | | | | |
|---|---|---|---|---|---|---|---|---|---|---|
| Myo10 | 7.871477 | 7.760755 | 7.8039 | -0.043163302 | 0.067559181 | 7.580104 | 7.1967112 | 7.54953 | -0.352823161 | 0.030569627 |
| Fbxo21 | 8.368947 | 8.475951 | 8.4951 | -0.019149747 | -0.126154354 | 8.5420796 | 8.1678325 | 8.52072 | -0.352889644 | 0.021357501 |
| G430095P16Rik | 5.157533 | 5.238858 | 5.2032 | 0.035660516 | -0.045664741 | 5.2512181 | 5.068088 | 5.42099 | -0.3529023 | -0.169772189 |
| Klhl5 | 9.187428 | 8.844516 | 9.2345 | -0.389996386 | -0.04708405 | 8.9316677 | 8.681444 | 9.03452 | -0.353077784 | -0.102854162 |
| Znrf3 | 8.341198 | 8.319717 | 8.3901 | -0.070392439 | -0.048911324 | 7.8841964 | 8.1034752 | 8.45672 | -0.353249086 | -0.572527837 |
| Mdm1 | 7.007631 | 6.837298 | 7.2442 | -0.40691506 | -0.236582103 | 7.1124326 | 6.7973672 | 7.15078 | -0.35341286 | -0.038347469 |
| Robo2 | 6.091357 | 5.243888 | 5.9041 | -0.66021344 | 0.187254767 | 5.2102379 | 4.7723037 | 5.12581 | -0.353501778 | 0.084432481 |
| Ints2 | 8.883865 | 9.016826 | 9.0061 | 0.010758729 | -0.122202273 | 9.1535859 | 8.8560556 | 9.20957 | -0.353517525 | -0.055987236 |
| Pls3 | 7.950402 | 7.629195 | 7.9196 | -0.290435124 | 0.030771841 | 7.2180054 | 7.0352553 | 7.38885 | -0.353593918 | -0.170843822 |
| Vat1 | 8.919844 | 9.18308 | 8.7848 | 0.398284043 | 0.135048804 | 8.2680679 | 7.8867753 | 8.24051 | -0.353730133 | 0.027562447 |
| Rad18 | 8.288856 | 8.215187 | 8.388 | -0.172783146 | -0.099113991 | 8.3507551 | 8.1433603 | 8.49712 | -0.353760847 | -0.146366072 |
| Zfp560 | 7.01046 | 6.726986 | 6.9728 | -0.245852167 | 0.037621967 | 6.974984 | 6.6627682 | 7.01673 | -0.353963687 | -0.041747838 |
| Lmntd2 | 6.085746 | 6.16042 | 6.2523 | -0.091865225 | -0.166539262 | 6.6332827 | 6.1910192 | 6.54504 | -0.354020704 | 0.088242822 |
| Slc25a16 | 8.206407 | 7.915669 | 8.1321 | -0.216416671 | 0.074321135 | 8.1387372 | 7.7423475 | 8.09648 | -0.354127966 | 0.04226173 |
| Ddc | 5.595335 | 5.585793 | 5.5632 | 0.022585284 | 0.032127303 | 5.5341146 | 5.4297921 | 5.78396 | -0.354167402 | -0.249844904 |
| Kif1c | 10.20559 | 10.16978 | 10.222 | -0.05204698 | -0.016234761 | 10.049713 | 9.7078814 | 10.0621 | -0.35418907 | -0.012357144 |
| Arhgap9 | 11.51482 | 11.46161 | 11.601 | -0.139325246 | -0.086116478 | 11.658451 | 11.234496 | 11.5887 | -0.3541964 | 0.069758103 |
| Pms2 | 8.731711 | 8.808623 | 8.9457 | -0.137073826 | -0.213986266 | 8.9019456 | 8.6713291 | 9.02567 | -0.35433693 | -0.123720401 |
| Bbs1 | 5.44404 | 5.332893 | 5.456 | -0.123122905 | -0.011975259 | 5.4821182 | 5.265309 | 5.6197 | -0.354391751 | -0.137582548 |
| Bmt2 | 9.636683 | 9.686233 | 9.7944 | -0.108136993 | -0.157687734 | 9.5988376 | 9.3942908 | 9.74883 | -0.354541773 | -0.149994925 |
| Pcbp4 | 6.644683 | 6.574834 | 6.6839 | -0.109110717 | -0.039261731 | 6.5758147 | 6.4666416 | 6.82131 | -0.354667769 | -0.245494709 |
| Dnajc28 | 5.346745 | 5.515162 | 5.4844 | 0.03079441 | -0.1376219 | 5.5026274 | 5.1539421 | 5.50879 | -0.354850572 | -0.006165299 |
| Ppp2r5a | 11.85455 | 11.88787 | 11.887 | 0.000633464 | -0.032685428 | 11.758227 | 11.584783 | 11.9397 | -0.35488723 | -0.181442999 |
| Glg1 | 11.76281 | 11.7597 | 11.782 | -0.021964722 | -0.01884885 | 11.656405 | 11.384264 | 11.7392 | -0.354914844 | -0.082773263 |
| Zfp59 | 7.448274 | 7.368537 | 7.367 | 0.001496329 | 0.081233628 | 7.3922604 | 7.1053334 | 7.46029 | -0.354953793 | -0.068026786 |
| Agbl5 | 7.785838 | 7.587297 | 7.8851 | -0.297776708 | -0.099236445 | 7.9916456 | 7.5339695 | 7.88903 | -0.355057765 | 0.102618302 |
| Arhgap28 | 6.523246 | 6.389407 | 6.4023 | -0.012926885 | 0.120911794 | 6.2261605 | 6.0408059 | 6.39619 | -0.35538889 | -0.170034337 |
| Zbtb46 | 7.961076 | 7.880318 | 7.9705 | -0.090145667 | -0.00938761 | 7.4612608 | 7.5489042 | 7.90453 | -0.355627947 | -0.443271344 |
| Bmx | 3.742853 | 3.641928 | 3.5957 | 0.0462039 | 0.147128724 | 3.6720328 | 3.2753976 | 3.63109 | -0.355693204 | 0.040942073 |
| Slx4 | 9.735449 | 9.83664 | 9.803 | 0.033661164 | -0.067530302 | 9.659304 | 9.4731839 | 9.82897 | -0.355788125 | -0.16966802 |
| Clock | 9.365929 | 9.267488 | 9.2865 | -0.019026378 | 0.079414865 | 9.1953751 | 9.0713448 | 9.42727 | -0.355925235 | -0.231894957 |
| Gm49378 | 4.734439 | 5.135663 | 5.1121 | 0.023568118 | -0.377655853 | 4.8411854 | 4.7711459 | 5.12708 | -0.355931876 | -0.285892391 |

| | | | | | | | | | | |
|---|---|---|---|---|---|---|---|---|---|---|
| 2510009E07Rik | 6.967263 | 7.106069 | 7.0633 | 0.042792288 | -0.096013059 | 6.863286 | 6.6821101 | 7.03808 | -0.355970079 | -0.174794167 |
| Sh3gl3 | 5.681033 | 5.257691 | 5.7352 | -0.477469771 | -0.054127731 | 4.8878755 | 4.8533255 | 5.20949 | -0.356165888 | -0.321615877 |
| Nudt17 | 7.78956 | 7.751358 | 7.9852 | -0.233839029 | -0.195636856 | 7.6086173 | 7.755367 | 8.11158 | -0.356213334 | -0.502962943 |
| 2810408A11Rik | 4.41893 | 4.300561 | 4.5635 | -0.262931652 | -0.144561989 | 4.3045881 | 4.0136586 | 4.36992 | -0.356256664 | -0.06532722 |
| Ttc3 | 10.66672 | 10.60082 | 10.688 | -0.087420162 | -0.021526865 | 10.225054 | 10.307191 | 10.6635 | -0.356279472 | -0.438416559 |
| Ift81 | 4.339756 | 4.397886 | 4.4824 | -0.084540137 | -0.142669756 | 4.3162337 | 4.2753191 | 4.63161 | -0.356291653 | -0.315376983 |
| Itgb1 | 11.77361 | 11.59091 | 11.739 | -0.148316078 | 0.03438072 | 11.312505 | 11.282043 | 11.6385 | -0.356488522 | -0.32602617 |
| Msh2 | 9.94695 | 9.951907 | 10.033 | -0.08144962 | -0.086405924 | 9.8033803 | 9.6341616 | 9.99077 | -0.356603979 | -0.187385258 |
| Sidt2 | 11.53399 | 11.49866 | 11.604 | -0.105095224 | -0.069758588 | 11.724547 | 11.424885 | 11.7815 | -0.356635432 | -0.056973609 |
| Map3k4 | 9.432836 | 9.530525 | 9.5723 | -0.041817578 | -0.139505837 | 9.3763584 | 9.2080059 | 9.56465 | -0.356644505 | -0.188291961 |
| Lrrc3 | 3.559286 | 3.536039 | 3.7482 | -0.212162913 | -0.188916365 | 3.5908858 | 3.2539374 | 3.61059 | -0.356652122 | -0.019703723 |
| Crebl2 | 8.881633 | 8.900569 | 8.9732 | -0.072656285 | -0.091592006 | 8.8220501 | 8.5835535 | 8.94026 | -0.35671148 | -0.118214891 |
| Gm17971 | 4.225373 | 4.165433 | 4.1866 | -0.021129062 | 0.038810495 | 4.5711336 | 4.2712669 | 4.62808 | -0.35681342 | -0.056946635 |
| Rnase4 | 6.496163 | 5.939746 | 6.4215 | -0.481764143 | 0.074653512 | 5.9422862 | 5.808833 | 6.16571 | -0.356878395 | -0.223425223 |
| Fam199x | 8.23401 | 8.370477 | 8.3458 | 0.024725699 | -0.111740787 | 8.113788 | 8.036798 | 8.39378 | -0.356978087 | -0.27998813 |
| Tacc1 | 11.40879 | 11.32871 | 11.469 | -0.140269868 | -0.060183148 | 11.371129 | 11.118801 | 11.4758 | -0.3570084 | -0.104681001 |
| Per1 | 10.83648 | 10.6168 | 10.729 | -0.112634894 | 0.107054294 | 10.639456 | 10.539312 | 10.8963 | -0.35703476 | -0.25689062 |
| Ripor2 | 12.61454 | 12.72996 | 12.697 | 0.032783452 | -0.082633452 | 12.656043 | 12.4052 | 12.7624 | -0.357155714 | -0.10631258 |
| Foxred2 | 8.2914 | 8.126902 | 8.369 | -0.242119943 | -0.077622032 | 8.4505536 | 8.0151602 | 8.37237 | -0.357206316 | 0.078187161 |
| Spg20 | 8.52271 | 8.466972 | 8.5894 | -0.122475627 | -0.066737389 | 8.3273674 | 8.1009765 | 8.45819 | -0.357217352 | -0.130826406 |
| Tlr5 | 5.013102 | 4.568753 | 4.9281 | -0.359389069 | 0.084959955 | 5.1512927 | 4.3794897 | 4.73677 | -0.357279581 | 0.414523449 |
| Ghdc | 7.640554 | 7.404714 | 7.5958 | -0.191080687 | 0.044759497 | 7.7503526 | 7.1639984 | 7.52145 | -0.357453244 | 0.228900957 |
| Cd207 | 4.322356 | 4.165927 | 4.3015 | -0.135530277 | 0.020899556 | 3.7216147 | 3.5696645 | 3.92715 | -0.357485735 | -0.205535538 |
| Retnla | 3.100675 | 3.179731 | 3.4017 | -0.221976038 | -0.301031488 | 2.9073844 | 2.9228648 | 3.28056 | -0.357695154 | -0.373175637 |
| Arhgef39 | 4.811545 | 4.921324 | 5.2379 | -0.316536735 | -0.426315371 | 4.9138051 | 4.8272376 | 5.18519 | -0.357956218 | -0.27138867 |
| Ppp1r15a | 12.56027 | 12.31255 | 12.586 | -0.273325502 | -0.025604648 | 11.708547 | 11.750272 | 12.1086 | -0.358287834 | -0.400012407 |
| Ccdc82 | 10.56274 | 10.51969 | 10.662 | -0.142113982 | -0.099067736 | 10.443175 | 10.362653 | 10.721 | -0.35831944 | -0.277797739 |
| Mcf2l | 5.333486 | 5.389275 | 5.5778 | -0.188489707 | -0.24427891 | 5.1750612 | 4.9604428 | 5.31888 | -0.358441039 | -0.143822647 |
| Elane | 2.490939 | 2.27069 | 2.445 | -0.174335002 | 0.045913916 | 2.586149 | 2.2221215 | 2.58067 | -0.358551759 | 0.005475741 |
| M1ap | 4.219958 | 4.226071 | 4.2061 | 0.020005091 | 0.013891879 | 4.3938339 | 4.1689834 | 4.52774 | -0.358760561 | -0.133910022 |
| Napepld | 5.62058 | 5.610686 | 5.4596 | 0.151116194 | 0.161010005 | 5.5374091 | 5.2875745 | 5.64634 | -0.358765925 | -0.108931274 |
| Foxn3 | 11.18296 | 11.23897 | 11.215 | 0.023990096 | -0.032012476 | 11.025395 | 10.938789 | 11.2976 | -0.358771792 | -0.27216592 |

| | | | | | | | | | | |
|---|---|---|---|---|---|---|---|---|---|---|
| Sirt5 | 7.994003 | 7.76365 | 8.0864 | -0.322708645 | -0.092356035 | 7.7156207 | 7.6158762 | 7.97467 | -0.358798147 | -0.259053646 |
| Tcf12 | 11.20029 | 11.12526 | 11.232 | -0.106327648 | -0.031294652 | 11.020189 | 10.806701 | 11.1656 | -0.358870978 | -0.145382456 |
| Pnmal2 | 4.293539 | 4.077944 | 4.4192 | -0.341294859 | -0.125699463 | 3.9335302 | 3.8367285 | 4.19563 | -0.358903605 | -0.262101959 |
| Atxn1 | 8.355551 | 8.077621 | 7.9535 | 0.12411671 | 0.402046499 | 8.0112195 | 8.0377385 | 8.39669 | -0.35894711 | -0.385466123 |
| Fbxo22 | 9.113731 | 9.06905 | 9.149 | -0.079979742 | -0.035298443 | 9.1520538 | 8.9922868 | 9.35134 | -0.359049016 | -0.199281988 |
| mt-Cytb | 15.5372 | 15.4101 | 15.566 | -0.155700382 | -0.02860058 | 15.387109 | 15.237318 | 15.5964 | -0.359077984 | -0.209286914 |
| Osgin2 | 7.79613 | 7.703809 | 7.8035 | -0.099643046 | -0.007321847 | 7.5742065 | 7.4319492 | 7.79129 | -0.35934382 | -0.217086478 |
| Slc35b2 | 8.990026 | 8.761649 | 8.9618 | -0.200120359 | 0.028256643 | 9.1730853 | 8.7023742 | 9.06174 | -0.359370449 | 0.11134056 |
| Asap2 | 7.864853 | 7.551288 | 7.6873 | -0.136020231 | 0.177545602 | 7.1617729 | 7.1797273 | 7.53915 | -0.359424166 | -0.377378593 |
| Cand1 | 11.07729 | 11.02494 | 11.144 | -0.119104176 | -0.066756274 | 10.959277 | 10.82468 | 11.1841 | -0.359436284 | -0.224838562 |
| Tedc1 | 7.40784 | 7.395263 | 7.517 | -0.121730517 | -0.109153596 | 7.8178553 | 7.1559465 | 7.51542 | -0.359469359 | 0.302439474 |
| B4gat1 | 6.953317 | 6.839092 | 7.1407 | -0.301608963 | -0.18738408 | 7.0620429 | 6.5768914 | 6.93649 | -0.359601411 | 0.125550067 |
| Lig4 | 7.306993 | 7.323531 | 7.5274 | -0.203837903 | -0.220376036 | 7.355029 | 7.140041 | 7.49975 | -0.359704971 | -0.144716959 |
| Nbea | 6.009096 | 5.624543 | 5.9465 | -0.321997987 | 0.062555502 | 5.5633908 | 5.4495539 | 5.80936 | -0.359808633 | -0.245971687 |
| Arhgap23 | 7.382096 | 7.050925 | 7.1303 | -0.079379918 | 0.25179067 | 6.6356123 | 6.5786453 | 6.93848 | -0.359831498 | -0.302864506 |
| Net1 | 10.6567 | 10.72377 | 10.878 | -0.154067696 | -0.221133054 | 10.380344 | 10.482578 | 10.8425 | -0.35990267 | -0.462136679 |
| Nsd1 | 12.22622 | 12.24113 | 12.293 | -0.051701802 | -0.066612618 | 12.055831 | 12.075657 | 12.4356 | -0.359924118 | -0.37975069 |
| Rbms3 | 5.978462 | 5.581816 | 5.5577 | 0.024120307 | 0.420766697 | 5.2533533 | 5.1849869 | 5.54493 | -0.359939764 | -0.291573379 |
| Ugdh | 9.110713 | 8.903782 | 9.1393 | -0.235484552 | -0.028553155 | 9.0639738 | 8.7341607 | 9.09412 | -0.359958025 | -0.030144931 |
| Micu3 | 6.711673 | 6.77846 | 6.8255 | -0.047002474 | -0.113789374 | 6.7909342 | 6.5460677 | 6.90604 | -0.359972459 | -0.115105953 |
| Trav14d-3-dv8 | 5.40857 | 5.227316 | 5.5829 | -0.355610889 | -0.174357305 | 5.196102 | 5.0954618 | 5.45551 | -0.360046095 | -0.259405918 |
| Egln3 | 5.415808 | 5.239227 | 5.3371 | -0.097917106 | 0.078663444 | 5.1984204 | 4.9171343 | 5.27726 | -0.360129102 | -0.078843006 |
| Plscr4 | 4.949253 | 4.541086 | 5.1067 | -0.565580543 | -0.157412932 | 4.3670759 | 4.3330915 | 4.69325 | -0.360160933 | -0.326176582 |
| Slc25a30 | 9.632577 | 9.415312 | 9.5961 | -0.180755925 | 0.036509809 | 9.5051939 | 9.2085749 | 9.5688 | -0.36022742 | -0.063608392 |
| Prrc2b | 12.13611 | 12.14511 | 12.208 | -0.063222892 | -0.072221134 | 12.009344 | 11.88122 | 12.2415 | -0.360241136 | -0.232116944 |
| Shroom2 | 4.659249 | 4.654384 | 4.795 | -0.14057528 | -0.135709886 | 4.2682942 | 4.2014792 | 4.56183 | -0.360355215 | -0.293540181 |
| Kcnk1 | 4.405019 | 4.144255 | 4.4089 | -0.264630589 | -0.003867375 | 4.3536465 | 4.0587771 | 4.41918 | -0.360405685 | -0.065536239 |
| Klhl8 | 6.48288 | 6.469931 | 6.4584 | 0.011549742 | 0.024498308 | 6.1471618 | 6.1055328 | 6.46598 | -0.360448286 | -0.318819285 |
| Bcas3 | 8.920551 | 8.861406 | 8.9059 | -0.044478875 | 0.014665904 | 8.8464374 | 8.5899482 | 8.95048 | -0.360527426 | -0.104038238 |
| Enpp5 | 6.63956 | 6.180861 | 6.7468 | -0.565907579 | -0.107208983 | 6.0737571 | 6.0021285 | 6.36274 | -0.360612584 | -0.288983919 |
| Klhl13 | 3.961512 | 3.808443 | 3.9776 | -0.169176113 | -0.016107415 | 3.5013224 | 3.3841867 | 3.74487 | -0.360684119 | -0.243548448 |
| Csrp2 | 7.180659 | 6.86825 | 6.9866 | -0.118367926 | 0.19404191 | 6.9873393 | 6.3174463 | 6.67821 | -0.360766786 | 0.309126268 |

| | | | | | | | | | | |
|---|---|---|---|---|---|---|---|---|---|---|
| Rgs7bp | 3.405338 | 3.395454 | 3.4673 | -0.07182211 | -0.061938133 | 3.1402479 | 3.0468163 | 3.40767 | -0.360849002 | -0.267417449 |
| 4930486L24Rik | 3.972979 | 3.580585 | 3.9196 | -0.338998338 | 0.053396571 | 3.9671506 | 3.6987576 | 4.05977 | -0.361011922 | -0.09261892 |
| Tlr11 | 6.604629 | 6.321527 | 6.7226 | -0.401033283 | -0.117930969 | 6.2973048 | 6.2378354 | 6.59897 | -0.361139412 | -0.301670042 |
| L2hgdh | 6.428963 | 6.400558 | 6.6601 | -0.259541507 | -0.231137012 | 6.5312343 | 6.3018418 | 6.66303 | -0.361183891 | -0.131791479 |
| Il11ra1 | 7.860716 | 7.782073 | 7.9663 | -0.184188075 | -0.105545527 | 7.6697764 | 7.2838517 | 7.64507 | -0.361221857 | 0.024702832 |
| Fam20a | 3.27432 | 3.21357 | 3.4975 | -0.283889177 | -0.223138745 | 3.2855944 | 3.1431061 | 3.50451 | -0.361401434 | -0.218913176 |
| Ppp1r9a | 3.897087 | 3.421981 | 3.7163 | -0.294309041 | 0.180797027 | 3.7395564 | 3.4041001 | 3.76572 | -0.36161852 | -0.026162244 |
| Galnt7 | 9.433175 | 9.365636 | 9.4828 | -0.117178517 | -0.049639224 | 8.9941277 | 9.0825622 | 9.44419 | -0.361623934 | -0.450058437 |
| Pcx | 5.952784 | 5.763407 | 6.0506 | -0.28723759 | -0.097860399 | 6.0402811 | 5.5876716 | 5.94937 | -0.361699238 | 0.090910269 |
| Rasd2 | 3.88268 | 3.473927 | 4.0466 | -0.572702375 | -0.16394918 | 3.4638336 | 3.2503704 | 3.61216 | -0.361788587 | -0.148325427 |
| Nfic | 9.276581 | 9.115472 | 9.3002 | -0.184757413 | -0.023648761 | 9.3091636 | 8.7818309 | 9.14372 | -0.361890721 | 0.165442051 |
| Adgre1 | 7.739947 | 7.615308 | 7.5964 | 0.018881669 | 0.14352019 | 7.5838539 | 7.1733678 | 7.53567 | -0.362306171 | 0.048179931 |
| Zfp69 | 4.877648 | 4.706009 | 4.8928 | -0.186838184 | -0.015199678 | 4.656518 | 4.59268 | 4.955 | -0.362323333 | -0.298485332 |
| Slc20a1 | 10.73184 | 10.64732 | 10.73 | -0.082829638 | 0.001688946 | 10.585043 | 10.590316 | 10.9527 | -0.36235216 | -0.367624658 |
| Ralgps2 | 11.96523 | 12.01526 | 12.053 | -0.03751859 | -0.08754122 | 12.437557 | 11.839895 | 12.2024 | -0.362554464 | 0.23510805 |
| Ifitm10 | 6.244856 | 5.912525 | 6.3082 | -0.395712827 | -0.063381448 | 6.0967865 | 5.6193317 | 5.98192 | -0.362587888 | 0.114866943 |
| 4930578C19Rik | 4.210469 | 4.062576 | 4.4311 | -0.368522679 | -0.220629022 | 3.9127516 | 3.7967437 | 4.15938 | -0.362638989 | -0.246631131 |
| Dhrs7 | 8.113372 | 7.848744 | 8.1204 | -0.271702147 | -0.007074576 | 8.0471135 | 7.6899755 | 8.05264 | -0.36266844 | -0.005530367 |
| Rasl11a | 4.153938 | 4.134249 | 4.2619 | -0.127661716 | -0.10797301 | 3.9168928 | 3.7591489 | 4.12189 | -0.362739796 | -0.20499592 |
| Jup | 8.245803 | 7.897409 | 8.1322 | -0.234785857 | 0.113608376 | 8.0469512 | 7.6492533 | 8.01201 | -0.362760744 | 0.034937084 |
| Ipmk | 8.74777 | 8.635936 | 8.767 | -0.131075617 | -0.019241337 | 8.4519557 | 8.3688008 | 8.73159 | -0.362794159 | -0.279639251 |
| Myo1g | 12.38809 | 12.3678 | 12.496 | -0.12831589 | -0.108028496 | 12.59802 | 12.281332 | 12.6442 | -0.362879608 | -0.046191562 |
| Arhgef6 | 11.48254 | 11.50189 | 11.687 | -0.185206204 | -0.204557382 | 11.600938 | 11.359276 | 11.7222 | -0.362929764 | -0.121268223 |
| Irx3 | 4.39292 | 4.076541 | 4.2257 | -0.14916924 | 0.167209476 | 3.9271877 | 3.7200265 | 4.08304 | -0.363009918 | -0.155848702 |
| Tmcc2 | 6.213042 | 6.169328 | 6.4791 | -0.309724523 | -0.266010194 | 5.9516102 | 5.8684291 | 6.23156 | -0.363128775 | -0.279947704 |
| A430033K04Rik | 6.746453 | 6.720847 | 6.8033 | -0.082446874 | -0.056840785 | 6.7312468 | 6.433015 | 6.79632 | -0.363302718 | -0.065070917 |
| Mettl7a1 | 6.861377 | 6.869028 | 7.0592 | -0.190127362 | -0.197778728 | 7.0361283 | 6.5424321 | 6.90585 | -0.36341311 | 0.130283082 |
| Sik1 | 11.22662 | 10.8594 | 11.048 | -0.188320142 | 0.178903214 | 10.545223 | 10.603369 | 10.9668 | -0.363434576 | -0.421581086 |
| Zfp148 | 10.90234 | 10.88851 | 10.971 | -0.082693232 | -0.068862734 | 10.672404 | 10.528062 | 10.8916 | -0.363570748 | -0.219228036 |
| Cetn4 | 4.615037 | 4.406228 | 4.4719 | -0.06564209 | 0.143167673 | 4.5601939 | 4.4442229 | 4.80822 | -0.363999138 | -0.248028167 |
| Fgf11 | 3.618136 | 3.39242 | 3.7336 | -0.341150965 | -0.115434451 | 3.4179628 | 3.2595597 | 3.62404 | -0.36448178 | -0.206078681 |
| Zhx3 | 7.808075 | 7.816547 | 7.7637 | 0.052827319 | 0.044355186 | 7.3232716 | 7.4383873 | 7.80327 | -0.364886875 | -0.480002503 |

| | | | | | | | | | | |
|---|---|---|---|---|---|---|---|---|---|---|
| Ighv1-9 | 8.207415 | 8.292192 | 7.9478 | 0.344390003 | 0.259612816 | 8.2079314 | 7.9485132 | 8.3135 | -0.364984284 | -0.105566087 |
| Actn2 | 6.235736 | 6.164762 | 6.3765 | -0.211696499 | -0.140722241 | 5.9670226 | 5.8350992 | 6.20021 | -0.365115085 | -0.233191714 |
| Ptpn9 | 8.841585 | 8.744884 | 8.8071 | -0.062250635 | 0.034450941 | 8.5223245 | 8.3266527 | 8.69185 | -0.365201778 | -0.169529947 |
| Vmac | 7.641503 | 7.494197 | 7.6806 | -0.186396836 | -0.039091252 | 7.7128842 | 7.0783389 | 7.4436 | -0.365258835 | 0.269286507 |
| Sfxn4 | 5.687754 | 5.622462 | 5.7423 | -0.119867818 | -0.054574978 | 6.0118371 | 5.4761493 | 5.84142 | -0.365275328 | 0.170412446 |
| Mpv17l | 5.055785 | 4.808241 | 4.9307 | -0.122468497 | 0.125076 | 5.0839233 | 4.798072 | 5.16367 | -0.365597665 | -0.079746396 |
| AI854703 | 4.881683 | 4.980958 | 4.7902 | 0.190776539 | 0.091501937 | 5.2379288 | 4.7622025 | 5.12789 | -0.365683615 | 0.110042743 |
| Mfsd8 | 6.835546 | 6.742616 | 6.9662 | -0.223548804 | -0.130618292 | 6.9042409 | 6.6631294 | 7.02882 | -0.365693483 | -0.12458195 |
| Gpihbp1 | 4.108698 | 4.122257 | 4.2576 | -0.135385104 | -0.148943956 | 4.2132402 | 3.8087654 | 4.17448 | -0.365710202 | 0.038764506 |
| Armcx2 | 8.813192 | 8.67526 | 8.8356 | -0.160326747 | -0.022395265 | 8.6538384 | 8.3784348 | 8.74423 | -0.365794485 | -0.090390882 |
| Paox | 6.725464 | 6.597709 | 6.6935 | -0.09577038 | 0.031984328 | 6.7659244 | 6.1797605 | 6.54565 | -0.365891008 | 0.220272925 |
| Adam12 | 6.018533 | 6.160931 | 5.9031 | 0.257834621 | 0.115435814 | 6.1573791 | 5.9416864 | 6.30758 | -0.365895639 | -0.150202953 |
| Fam210b | 7.99631 | 7.760626 | 8.0275 | -0.266890624 | -0.031206721 | 7.9690241 | 7.5159377 | 7.8819 | -0.365959492 | 0.087126976 |
| Adgrg3 | 6.4917 | 6.387607 | 6.1752 | 0.212416395 | 0.316509057 | 6.2684204 | 5.9410937 | 6.30713 | -0.366033679 | -0.038706993 |
| Chm | 9.883 | 9.659849 | 9.791 | -0.131175804 | 0.091974884 | 9.8161415 | 9.431906 | 9.79802 | -0.36611856 | 0.018116989 |
| Rbpj | 10.51439 | 10.44801 | 10.352 | 0.09594081 | 0.162320208 | 10.27028 | 10.195192 | 10.5613 | -0.366124714 | -0.291036693 |
| Homer3 | 6.200138 | 6.275579 | 6.2383 | 0.037301008 | -0.038140692 | 6.3831769 | 5.98729 | 6.35362 | -0.366334648 | 0.029552227 |
| Stam | 8.803267 | 8.836335 | 8.7452 | 0.091162794 | 0.05809473 | 8.5040806 | 8.3987848 | 8.76524 | -0.36645062 | -0.261154862 |
| Wdr26 | 12.34953 | 12.28973 | 12.332 | -0.0420154 | 0.017784624 | 12.151498 | 12.055778 | 12.4223 | -0.366524076 | -0.270803627 |
| Tsc22d2 | 9.603041 | 9.6961 | 9.797 | -0.100939085 | -0.193998953 | 9.453314 | 9.4102474 | 9.77695 | -0.366699382 | -0.323632805 |
| Abcg3 | 9.087185 | 9.204427 | 9.2369 | -0.032469143 | -0.149711319 | 9.2611691 | 8.7690944 | 9.1358 | -0.366701924 | 0.125372788 |
| Xxylt1 | 9.082921 | 8.837779 | 9.0415 | -0.203749439 | 0.041393019 | 8.7568021 | 8.4776311 | 8.84465 | -0.367022191 | -0.087851208 |
| Klhdc2 | 9.140739 | 8.941366 | 9.0775 | -0.136180251 | 0.063192836 | 8.9025708 | 8.6456065 | 9.01268 | -0.367076107 | -0.110111855 |
| Cma1 | 7.473655 | 7.485938 | 7.1986 | 0.287351729 | 0.275068901 | 6.8826224 | 6.6735283 | 7.04076 | -0.36723604 | -0.158141966 |
| Abcd3 | 8.5976 | 8.509748 | 8.6566 | -0.146820436 | -0.05896906 | 8.3618128 | 8.1421337 | 8.50968 | -0.367550835 | -0.147871769 |
| Tnfrsf11a | 7.68761 | 7.572445 | 7.6173 | -0.044831294 | 0.070333629 | 7.2956901 | 7.3571193 | 7.72478 | -0.367656017 | -0.429085184 |
| Gm15446 | 7.363583 | 7.435775 | 7.4171 | 0.01869307 | -0.053498686 | 7.6715422 | 7.216419 | 7.58411 | -0.367691207 | 0.087432019 |
| Sp3 | 11.65111 | 11.7022 | 11.69 | 0.012475306 | -0.038608118 | 11.486327 | 11.319115 | 11.6869 | -0.367828396 | -0.20061639 |
| mt-Nd4 | 15.22513 | 15.1246 | 15.283 | -0.158119912 | -0.057592786 | 15.084652 | 14.95459 | 15.3226 | -0.367975768 | -0.237913795 |
| Slc17a5 | 7.365064 | 7.23566 | 7.3517 | -0.11606359 | 0.013340718 | 7.3834383 | 6.9634831 | 7.33179 | -0.368311039 | 0.051644145 |
| Mindy1 | 8.927849 | 8.919785 | 8.9481 | -0.028318306 | -0.020254668 | 9.2649975 | 8.6847564 | 9.0531 | -0.368339752 | 0.211901268 |
| Mmp23 | 5.644624 | 5.66884 | 5.654 | 0.014839984 | -0.009375673 | 5.6616873 | 5.4843133 | 5.85268 | -0.368368575 | -0.190994623 |

| | | | | | | | | | | |
|---|---|---|---|---|---|---|---|---|---|---|
| Gm2446 | 5.657903 | 5.596071 | 5.4423 | 0.153733239 | 0.215565338 | 5.1388036 | 5.5536635 | 5.92215 | -0.368483238 | -0.783343107 |
| Zfp647 | 4.707919 | 4.543456 | 4.6127 | -0.069231667 | 0.095231379 | 4.7131455 | 4.3157577 | 4.68439 | -0.368633047 | 0.028754721 |
| Ighv8-12 | 7.551344 | 7.756674 | 7.3534 | 0.4032939 | 0.197964066 | 8.1110033 | 7.469821 | 7.83846 | -0.368635286 | 0.272546955 |
| Mettl15 | 6.32442 | 6.296099 | 6.3286 | -0.032483817 | -0.004163068 | 6.2361376 | 5.9926602 | 6.36134 | -0.368683658 | -0.125206226 |
| Frat2 | 7.105296 | 6.951353 | 7.1972 | -0.245808525 | -0.091864872 | 6.4881359 | 6.46542 | 6.83415 | -0.368734117 | -0.346018171 |
| Gm15429 | 5.760431 | 5.11555 | 5.6477 | -0.532174583 | 0.11270662 | 5.2302549 | 4.9702262 | 5.33911 | -0.368887164 | -0.108858471 |
| Xpo4 | 10.28017 | 10.13971 | 10.221 | -0.081527458 | 0.058937572 | 10.084546 | 9.9363165 | 10.3053 | -0.368970888 | -0.220741217 |
| Kmt2e | 12.29354 | 12.18732 | 12.274 | -0.087104086 | 0.019115236 | 12.100606 | 11.992759 | 12.3617 | -0.36898116 | -0.261134109 |
| Pla2g4b | 6.609207 | 6.816512 | 6.8755 | -0.058941072 | -0.266246205 | 7.134724 | 6.6406022 | 7.0097 | -0.369096821 | 0.125025053 |
| Palmd | 5.792062 | 5.426112 | 5.9028 | -0.476672143 | -0.110722187 | 5.2256002 | 5.0166216 | 5.38577 | -0.369144861 | -0.160166258 |
| Khdc1a | 5.115673 | 5.242384 | 5.3206 | -0.078183165 | -0.204893908 | 5.0233237 | 4.6812364 | 5.05087 | -0.369634123 | -0.027546728 |
| Zfp781 | 5.410502 | 5.361057 | 5.2768 | 0.084220024 | 0.13366497 | 5.349624 | 5.1664022 | 5.53606 | -0.369655005 | -0.186433193 |
| Pbx3 | 6.370028 | 6.000266 | 6.2288 | -0.228528592 | 0.141233097 | 5.9322642 | 5.74365 | 6.11331 | -0.369664724 | -0.181050478 |
| Pacs2 | 10.01386 | 9.921596 | 10.054 | -0.132691549 | -0.040426548 | 9.9420222 | 9.6903956 | 10.0602 | -0.3697964 | -0.118169814 |
| Ptch1 | 9.438706 | 9.425774 | 9.4913 | -0.065548965 | -0.052617064 | 9.0453043 | 8.9799984 | 9.3501 | -0.370105831 | -0.304799989 |
| Calm4 | 3.796914 | 3.825516 | 3.8516 | -0.026041986 | -0.054643954 | 3.6533774 | 3.5298909 | 3.9 | -0.370112459 | -0.246626047 |
| Npc1 | 10.01382 | 9.947112 | 10.069 | -0.121940332 | -0.055231669 | 9.9701204 | 9.6971016 | 10.0676 | -0.370527922 | -0.097509178 |
| Zbtb37 | 10.13741 | 10.10298 | 10.158 | -0.055406982 | -0.020975839 | 10.357883 | 10.159529 | 10.5301 | -0.37060228 | -0.172248001 |
| Arhgef18 | 11.87368 | 12.00089 | 12.01 | -0.008744486 | -0.135954062 | 11.879917 | 11.528934 | 11.8997 | -0.370792214 | -0.019809106 |
| Vwa3a | 2.653047 | 2.609665 | 2.6677 | -0.0580247 | -0.014642652 | 2.5201887 | 2.5386442 | 2.90952 | -0.370871176 | -0.389326624 |
| Chrnb4 | 4.898469 | 4.430736 | 4.8081 | -0.3773448 | 0.090387453 | 4.2095161 | 4.0228067 | 4.39374 | -0.370929962 | -0.184220612 |
| Aicda | 4.06065 | 4.554261 | 4.1191 | 0.435138354 | -0.058471899 | 4.1832052 | 4.0644738 | 4.43561 | -0.371134091 | -0.252402678 |
| Zfp27 | 6.851285 | 6.738864 | 7.0022 | -0.263299846 | -0.15087901 | 6.8202828 | 6.5879459 | 6.95917 | -0.371228146 | -0.138891236 |
| Tmem123 | 13.39154 | 13.34496 | 13.452 | -0.106668874 | -0.060083974 | 13.20855 | 13.047845 | 13.4191 | -0.37127506 | -0.210569745 |
| Lipa | 10.32197 | 10.57143 | 10.375 | 0.196309044 | -0.053153299 | 10.377525 | 9.9436085 | 10.3152 | -0.371572312 | 0.062344171 |
| Carmil3 | 5.095599 | 5.016251 | 5.0971 | -0.080822476 | -0.001474236 | 5.4123751 | 5.1080795 | 5.48012 | -0.372038686 | -0.067743076 |
| Rgs9 | 4.476894 | 4.296729 | 4.3825 | -0.085758002 | 0.094407379 | 4.218934 | 4.1640099 | 4.53612 | -0.372112376 | -0.317188228 |
| Adam6b | 4.364062 | 4.231065 | 4.2774 | -0.046314377 | 0.086683387 | 4.5709913 | 4.2006781 | 4.57284 | -0.372165407 | -0.001852209 |
| Hist1h1d | 5.58259 | 5.32918 | 5.6237 | -0.294567034 | -0.0411571 | 5.9219671 | 5.4049603 | 5.77715 | -0.372192365 | 0.144814407 |
| Efna1 | 5.72048 | 5.46949 | 5.9012 | -0.431685556 | -0.180695955 | 5.3481388 | 5.0274454 | 5.39985 | -0.372405438 | -0.051711947 |
| Alms1 | 7.328338 | 7.14712 | 7.5308 | -0.383715719 | -0.20249716 | 7.3659455 | 7.2281316 | 7.60065 | -0.372516333 | -0.234702486 |
| Tarbp1 | 8.485731 | 8.416676 | 8.4629 | -0.046261374 | 0.02279357 | 8.3879683 | 8.0529152 | 8.42547 | -0.372550581 | -0.037497456 |

| | | | | | | | | | | |
|---|---|---|---|---|---|---|---|---|---|---|
| Stx17 | 8.928534 | 9.019144 | 9.144 | -0.12488281 | -0.215492272 | 9.212075 | 8.8889001 | 9.26148 | -0.372583731 | -0.049408804 |
| Fign | 2.734085 | 2.648871 | 2.6397 | 0.009186563 | 0.094400525 | 2.7792957 | 2.5319529 | 2.90456 | -0.372609704 | -0.125266891 |
| Ndfip1 | 11.18197 | 11.07682 | 11.223 | -0.146136814 | -0.040982534 | 11.065809 | 10.784984 | 11.1576 | -0.372609986 | -0.091784734 |
| Hey1 | 4.376119 | 4.578158 | 4.6255 | -0.047376681 | -0.249415978 | 4.2916125 | 4.1042883 | 4.4771 | -0.372808785 | -0.185484559 |
| Slc19a2 | 6.22317 | 6.223616 | 6.2166 | 0.006973108 | 0.006527018 | 6.2422481 | 5.9633174 | 6.33636 | -0.373042987 | -0.094112306 |
| Dlg5 | 6.010765 | 5.742534 | 6.033 | -0.290465886 | -0.022235269 | 5.8525226 | 5.5936428 | 5.96681 | -0.373164241 | -0.11428443 |
| Zbtb14 | 8.928707 | 9.036054 | 9.1067 | -0.070625106 | -0.17797161 | 8.8503321 | 8.5725905 | 8.94578 | -0.37319181 | -0.095450176 |
| Kdm4a | 10.83033 | 10.81142 | 10.882 | -0.070946512 | -0.052030066 | 10.856447 | 10.512832 | 10.8861 | -0.373280162 | -0.029665441 |
| Wdhd1 | 8.586976 | 8.577166 | 8.8856 | -0.308482866 | -0.298672495 | 8.6645882 | 8.5896014 | 8.96294 | -0.373333884 | -0.298347075 |
| Dock6 | 7.718595 | 7.28784 | 7.7922 | -0.50433208 | -0.073577375 | 7.3158359 | 7.2711765 | 7.64458 | -0.373406661 | -0.328747264 |
| Gm10052 | 7.02216 | 6.2501 | 6.5419 | -0.291780812 | 0.480278565 | 4.1188997 | 5.17309 | 5.54651 | -0.373419633 | -1.427609975 |
| Iars2 | 10.26926 | 10.08633 | 10.252 | -0.165254813 | 0.01767573 | 10.185781 | 9.8466421 | 10.2201 | -0.37346428 | -0.034325736 |
| Eid1 | 9.984178 | 9.824344 | 9.974 | -0.149612973 | 0.010220769 | 9.9183708 | 9.5122527 | 9.88577 | -0.373514263 | 0.032603872 |
| Zfp3 | 6.115863 | 5.949685 | 6.3269 | -0.377244987 | -0.211067567 | 6.2591947 | 5.9494872 | 6.32306 | -0.373570323 | -0.063862873 |
| Adgrl2 | 7.295635 | 6.59462 | 7.1801 | -0.585441668 | 0.115573858 | 6.4297598 | 6.3282061 | 6.70178 | -0.37357371 | -0.272020012 |
| Dll4 | 7.79801 | 7.437 | 7.3883 | 0.048713254 | 0.409723516 | 7.1197095 | 6.8283529 | 7.20193 | -0.373581564 | -0.082224968 |
| Fbxo45 | 8.484864 | 8.404018 | 8.5339 | -0.129885399 | -0.049039215 | 8.2665623 | 8.0507334 | 8.42453 | -0.373794855 | -0.157965934 |
| BC052040 | 6.953308 | 6.898568 | 6.9974 | -0.098803687 | -0.044063611 | 6.8545688 | 6.6714974 | 7.04548 | -0.373981095 | -0.190909632 |
| Dtx3 | 8.688292 | 8.445495 | 8.7034 | -0.257866145 | -0.015069168 | 8.8805122 | 8.5439504 | 8.91793 | -0.373984559 | -0.037422699 |
| Ago1 | 10.98207 | 10.96866 | 11.086 | -0.117482444 | -0.104071828 | 10.888479 | 10.656064 | 11.0301 | -0.374068346 | -0.141653638 |
| Pisd-ps1 | 8.537458 | 8.443372 | 8.6055 | -0.162091675 | -0.068005669 | 9.2425765 | 8.5536331 | 8.92775 | -0.374115525 | 0.314827902 |
| Gm14584 | 5.196994 | 5.359188 | 5.4548 | -0.095623445 | -0.257817606 | 5.2036225 | 4.8543309 | 5.22856 | -0.374229073 | -0.024937439 |
| Ctse | 8.761712 | 8.50154 | 8.7232 | -0.221676741 | 0.03849462 | 8.7925982 | 8.4090323 | 8.78347 | -0.374436141 | 0.009129749 |
| Tpbgl | 5.447524 | 5.352744 | 5.4088 | -0.056054678 | 0.03872557 | 5.0407466 | 4.9050769 | 5.27994 | -0.374862406 | -0.239192713 |
| Ces2c | 3.88128 | 3.977599 | 4.0681 | -0.090491732 | -0.18681096 | 3.8905318 | 3.6892803 | 4.06423 | -0.374953723 | -0.173702186 |
| Irak3 | 8.780495 | 8.898184 | 8.8087 | 0.089481543 | -0.028207455 | 8.9202102 | 8.3673592 | 8.74261 | -0.375251845 | 0.177599077 |
| Nsf | 10.37801 | 10.27608 | 10.303 | -0.02712888 | 0.074798034 | 10.551995 | 10.160488 | 10.5358 | -0.375303216 | 0.016203364 |
| Baiap2 | 8.101402 | 7.99385 | 8.3551 | -0.361249217 | -0.253697612 | 8.1407459 | 7.8288947 | 8.20425 | -0.375355744 | -0.06350447 |
| Igkv18-36 | 3.176785 | 3.353535 | 3.3595 | -0.005996495 | -0.182746471 | 3.8554997 | 3.3982172 | 3.77381 | -0.375589464 | 0.081693014 |
| Lca5 | 6.396168 | 6.501755 | 6.4558 | 0.045952692 | -0.059634374 | 6.3006687 | 6.0367239 | 6.41245 | -0.375731036 | -0.111786309 |
| Wdr76 | 8.62679 | 8.52247 | 8.659 | -0.136573212 | -0.03225332 | 8.5740624 | 8.4245975 | 8.80052 | -0.375922314 | -0.226457434 |
| Nipal1 | 6.258216 | 6.353105 | 6.4501 | -0.096995842 | -0.19188391 | 6.1951205 | 6.0960472 | 6.47218 | -0.376135109 | -0.277061839 |

| | | | | | | | | | | |
|---|---|---|---|---|---|---|---|---|---|---|
| Fbxl3 | 10.12782 | 10.20624 | 10.208 | -0.001604426 | -0.080021461 | 10.17636 | 10.083641 | 10.4599 | -0.376236532 | -0.283516986 |
| Tspan5 | 7.866654 | 7.848952 | 7.8761 | -0.027149386 | -0.009447652 | 7.7632798 | 7.5390075 | 7.91528 | -0.376275696 | -0.152003333 |
| Mtmr1 | 10.18376 | 10.07914 | 10.186 | -0.106899457 | -0.002280168 | 9.8898778 | 9.6931785 | 10.0696 | -0.376429172 | -0.179729861 |
| Cisd3 | 7.752894 | 7.617464 | 7.8166 | -0.199093568 | -0.063664005 | 8.1716356 | 7.5610001 | 7.9375 | -0.376498699 | 0.234136775 |
| Mrgpre | 7.980245 | 7.940943 | 8.1335 | -0.192583658 | -0.153280953 | 8.3911638 | 7.6837777 | 8.06043 | -0.376647298 | 0.330738841 |
| Ttll7 | 4.971742 | 4.712561 | 4.8611 | -0.148502628 | 0.11067911 | 4.3546771 | 4.2110041 | 4.58771 | -0.376703481 | -0.233030456 |
| Pstk | 7.158784 | 7.142556 | 7.1567 | -0.014169466 | 0.002058622 | 7.1697206 | 6.9665959 | 7.3436 | -0.377004003 | -0.173879361 |
| Gtpbp10 | 7.715338 | 7.466094 | 7.7751 | -0.308990771 | -0.059747342 | 7.7659612 | 7.4786556 | 7.85582 | -0.377160429 | -0.089854881 |
| Ep300 | 11.82963 | 12.02596 | 12.025 | 0.000513028 | -0.195814932 | 11.633107 | 11.586785 | 11.9642 | -0.37738288 | -0.331060909 |
| Fv1 | 6.541323 | 6.367786 | 6.4692 | -0.101399952 | 0.072136684 | 6.521682 | 6.1498063 | 6.52735 | -0.37754225 | -0.005666626 |
| Ppp1r10 | 10.54317 | 10.50172 | 10.55 | -0.048622366 | -0.007173448 | 10.521086 | 10.297234 | 10.6749 | -0.377626568 | -0.153774395 |
| Ube2h | 11.38336 | 11.43334 | 11.417 | 0.016673828 | -0.033306668 | 11.173565 | 11.023061 | 11.4009 | -0.377821848 | -0.227317982 |
| Krba1 | 9.293066 | 9.228716 | 9.3969 | -0.168170128 | -0.103820736 | 9.3969002 | 9.1441065 | 9.52203 | -0.377928416 | -0.12513468 |
| Large1 | 6.368061 | 6.16262 | 6.375 | -0.21240646 | -0.006965296 | 5.6749883 | 5.6275682 | 6.00566 | -0.378091216 | -0.330671111 |
| D2hgdh | 8.980871 | 9.205317 | 9.2124 | -0.00711581 | -0.231562059 | 9.3158483 | 8.9419158 | 9.32003 | -0.37811682 | -0.004184393 |
| Vwf | 8.796305 | 7.972101 | 8.7587 | -0.786607776 | 0.037596494 | 7.1686228 | 7.536189 | 7.91451 | -0.378318431 | -0.745884615 |
| Pgm2 | 8.820984 | 8.607585 | 8.6285 | -0.020897259 | 0.192502028 | 8.6713672 | 8.219322 | 8.59767 | -0.37834428 | 0.073700944 |
| Ints4 | 10.42432 | 10.39852 | 10.58 | -0.181182014 | -0.15538663 | 10.595299 | 10.238791 | 10.6174 | -0.378611608 | -0.02210416 |
| Abhd18 | 6.650272 | 6.497407 | 6.7654 | -0.268025252 | -0.115160811 | 6.689332 | 6.4690101 | 6.84764 | -0.378629799 | -0.158307876 |
| Npr2 | 5.680714 | 5.490282 | 5.5802 | -0.089902213 | 0.100529404 | 5.2574873 | 5.333453 | 5.7121 | -0.378646846 | -0.454612498 |
| Ppp2r5d | 9.683479 | 9.786424 | 9.8337 | -0.047233878 | -0.150179219 | 9.8675884 | 9.5571221 | 9.93577 | -0.378651687 | -0.068185407 |
| C77080 | 5.979819 | 5.901613 | 6.0818 | -0.180138129 | -0.101932491 | 5.2902814 | 5.4091596 | 5.78785 | -0.378689912 | -0.497568074 |
| Tab3 | 8.905355 | 8.801546 | 8.9468 | -0.145282843 | -0.041473669 | 8.6286896 | 8.4497692 | 8.82865 | -0.378881193 | -0.199960763 |
| Prickle3 | 8.175503 | 8.076284 | 8.1767 | -0.100451917 | -0.001232636 | 8.2745133 | 7.9897745 | 8.36866 | -0.378882306 | -0.094143427 |
| Paqr4 | 5.758368 | 5.286861 | 5.6309 | -0.344080224 | 0.127426488 | 5.5739809 | 5.3893443 | 5.76825 | -0.378903347 | -0.194266815 |
| Rmnd1 | 8.857294 | 8.813423 | 8.9731 | -0.159661683 | -0.115790631 | 9.4116275 | 9.0581994 | 9.43728 | -0.379085075 | -0.025657041 |
| Zscan18 | 5.336551 | 5.224154 | 5.3355 | -0.111368244 | 0.001028715 | 5.2344348 | 5.1549589 | 5.53421 | -0.379256039 | -0.299780124 |
| Zfp799 | 6.794821 | 6.907302 | 7.1235 | -0.216226294 | -0.328707434 | 6.9756521 | 6.9001461 | 7.27943 | -0.379283004 | -0.303777023 |
| Trps1 | 8.497431 | 8.454215 | 8.2806 | 0.173643451 | 0.216860049 | 8.3687573 | 8.1374264 | 8.51673 | -0.379307698 | -0.147976805 |
| Wnt10a | 4.732213 | 4.697913 | 4.9535 | -0.255617614 | -0.221318194 | 4.3951561 | 4.3967296 | 4.77606 | -0.379327316 | -0.380900804 |
| Elovl7 | 5.823007 | 6.114592 | 6.0871 | 0.02753492 | -0.264049766 | 5.9079801 | 5.6198221 | 5.99933 | -0.379507743 | -0.091349825 |
| Ccdc80 | 4.990047 | 4.842397 | 5.2107 | -0.368327015 | -0.220677155 | 4.7391791 | 4.6388984 | 5.01844 | -0.379538895 | -0.279258223 |

| | | | | | | | | | | |
|---|---|---|---|---|---|---|---|---|---|---|
| Tspan33 | 6.942787 | 7.091542 | 7.2115 | -0.119975434 | -0.268731181 | 6.9975684 | 6.7253765 | 7.10499 | -0.379608541 | -0.107416621 |
| Gdpd1 | 4.349936 | 4.361881 | 4.572 | -0.210098226 | -0.222042649 | 4.0827295 | 3.8531989 | 4.23282 | -0.379623732 | -0.150093081 |
| Zfand1 | 8.267227 | 8.102155 | 8.3757 | -0.273568794 | -0.108496553 | 8.519832 | 8.0852428 | 8.46503 | -0.379784632 | 0.054804583 |
| Rps6ka3 | 11.79631 | 11.67367 | 11.758 | -0.084093426 | 0.03854453 | 11.367195 | 11.245029 | 11.6248 | -0.379805502 | -0.257639671 |
| Fam161a | 4.36631 | 4.105415 | 4.3537 | -0.248318869 | 0.01257623 | 4.2489418 | 4.0467807 | 4.42665 | -0.37986722 | -0.177706152 |
| Ngly1 | 9.805593 | 9.682909 | 9.8116 | -0.128675596 | -0.005990999 | 9.7145893 | 9.4477813 | 9.82769 | -0.379911688 | -0.113103753 |
| Ppp2r3a | 6.744168 | 6.622529 | 6.6957 | -0.073201709 | 0.048436528 | 6.4055504 | 6.2435309 | 6.6236 | -0.380067723 | -0.218048229 |
| Ppm1j | 6.679969 | 6.638056 | 6.8434 | -0.205349461 | -0.163436309 | 6.8354136 | 6.5480087 | 6.92813 | -0.380125621 | -0.092720658 |
| Zfp846 | 7.177871 | 7.062169 | 7.1897 | -0.127544948 | -0.011842422 | 7.1178917 | 6.7759209 | 7.15615 | -0.380228563 | -0.038257763 |
| Rfc1 | 10.66201 | 10.51946 | 10.738 | -0.218312796 | -0.07576851 | 10.627692 | 10.411136 | 10.7915 | -0.380322562 | -0.163766148 |
| Gpr83 | 7.077802 | 7.099241 | 7.1533 | -0.054091386 | -0.075530464 | 6.8687165 | 6.4953225 | 6.87572 | -0.380394674 | -0.007000736 |
| Poc1b | 8.748127 | 8.756502 | 8.719 | 0.037526525 | 0.029151536 | 8.9889258 | 8.3856453 | 8.7663 | -0.38065848 | 0.222622 |
| Zfp14 | 6.094526 | 6.164768 | 6.168 | -0.003221106 | -0.073463231 | 6.1027395 | 5.8529746 | 6.23385 | -0.380876257 | -0.13111133 |
| Cavin1 | 8.276484 | 8.153733 | 8.1644 | -0.010696525 | 0.112054387 | 7.5850036 | 7.4649858 | 7.8462 | -0.381209389 | -0.261191595 |
| Sema6c | 3.309817 | 3.156691 | 3.2516 | -0.094957369 | 0.058168163 | 3.1392315 | 2.9892209 | 3.37044 | -0.381219799 | -0.231209239 |
| Trav14d-1 | 5.570056 | 5.395373 | 5.6173 | -0.221926123 | -0.047242753 | 5.4544192 | 5.1683402 | 5.54959 | -0.381249187 | -0.095170214 |
| Trav14n-1 | 5.570056 | 5.395373 | 5.6173 | -0.221926123 | -0.047242753 | 5.4544192 | 5.1683402 | 5.54959 | -0.381249187 | -0.095170214 |
| Aldh7a1 | 7.250478 | 6.908317 | 7.0203 | -0.111956564 | 0.230204062 | 7.1893456 | 6.7714018 | 7.15271 | -0.381307376 | 0.036636436 |
| Ighv1-26 | 9.008301 | 9.285319 | 9.2085 | 0.07677037 | -0.200248281 | 9.2832192 | 8.8080265 | 9.18943 | -0.381406415 | 0.09378628 |
| Tstd3 | 7.376376 | 7.117049 | 7.4111 | -0.294036108 | -0.034708927 | 7.4171919 | 7.0013754 | 7.38297 | -0.381595523 | 0.034220966 |
| Nat8f4 | 6.476124 | 6.528639 | 6.6 | -0.071328596 | -0.123843758 | 6.7611494 | 6.2202496 | 6.60203 | -0.381782135 | 0.159117673 |
| Magi2 | 4.000607 | 3.882301 | 3.7703 | 0.11200227 | 0.230308069 | 3.9351391 | 3.6542855 | 4.0362 | -0.381917768 | -0.101064148 |
| Chrnb1 | 5.929455 | 5.789395 | 5.9852 | -0.195852591 | -0.055792838 | 6.1282534 | 5.7241538 | 6.10609 | -0.381933073 | 0.022166526 |
| Polr3gl | 8.51282 | 8.345847 | 8.5119 | -0.166021288 | 0.00095161 | 8.371513 | 8.1164108 | 8.49841 | -0.381995866 | -0.126893703 |
| Gnl3l | 10.04826 | 10.03435 | 10.045 | -0.010421544 | 0.00348335 | 10.047585 | 9.7896091 | 10.1716 | -0.381998942 | -0.124023131 |
| Gm9403 | 4.743241 | 4.636092 | 4.7275 | -0.091362562 | 0.015785866 | 4.7039164 | 4.493079 | 4.87509 | -0.382006325 | -0.171168894 |
| Stk38 | 11.34515 | 11.37863 | 11.432 | -0.05323748 | -0.086711686 | 11.477637 | 11.323564 | 11.7056 | -0.382082504 | -0.228010363 |
| Lrrc29 | 5.511827 | 5.108669 | 5.4479 | -0.339190377 | 0.063967012 | 5.5771476 | 5.0552891 | 5.43747 | -0.382179662 | 0.139678825 |
| Gm7694 | 5.627981 | 5.244041 | 5.334 | -0.089953412 | 0.293986481 | 5.2509411 | 4.9738363 | 5.35618 | -0.382345859 | -0.105241149 |
| Clcn2 | 6.055742 | 5.915796 | 6.127 | -0.211219788 | -0.071274108 | 6.4049751 | 6.1136178 | 6.49599 | -0.38237534 | -0.091017982 |
| Tcim | 6.075064 | 5.753703 | 5.8953 | -0.141628953 | 0.179731629 | 5.3703488 | 5.2253954 | 5.60783 | -0.38243541 | -0.237481974 |
| Pvr | 9.052836 | 8.951898 | 8.9464 | 0.005504024 | 0.106441659 | 8.6173591 | 8.6796439 | 9.06221 | -0.38256951 | -0.444854318 |

| | | | | | | | | | | |
|---|---|---|---|---|---|---|---|---|---|---|
| Slc48a1 | 8.880985 | 8.643111 | 8.9093 | -0.266175932 | -0.028301661 | 8.884907 | 8.3957284 | 8.77832 | -0.382596069 | 0.106582495 |
| Dapl1 | 8.930301 | 8.97729 | 9.1442 | -0.166955923 | -0.213944617 | 8.9572239 | 8.7847672 | 9.1674 | -0.382633517 | -0.210176841 |
| Actn1 | 12.02831 | 12.10176 | 12.037 | 0.065005116 | -0.008443906 | 11.646792 | 11.560255 | 11.9429 | -0.382665508 | -0.296127873 |
| Tpk1 | 7.495184 | 7.254249 | 7.4695 | -0.215272848 | 0.025661674 | 7.4241694 | 7.050798 | 7.4335 | -0.382704862 | -0.009333375 |
| Gm14325 | 6.510805 | 6.508695 | 6.7268 | -0.218154767 | -0.216044194 | 6.6169384 | 6.1959797 | 6.57892 | -0.38294462 | 0.038014136 |
| Pkd1 | 11.31253 | 11.25436 | 11.302 | -0.047471292 | 0.010704412 | 11.328196 | 10.951501 | 11.3346 | -0.383060544 | -0.006364925 |
| Trav6-7-dv9 | 5.571774 | 5.563299 | 5.6951 | -0.13183874 | -0.123363182 | 5.488796 | 5.2935123 | 5.67669 | -0.383181658 | -0.187897939 |
| Ttf2 | 8.858486 | 8.70982 | 8.9609 | -0.251045061 | -0.102379319 | 8.9799941 | 8.7523493 | 9.13558 | -0.383233196 | -0.155588387 |
| Ppm1b | 10.45244 | 10.44321 | 10.514 | -0.070734988 | -0.061511264 | 10.2847 | 10.062356 | 10.4457 | -0.383328584 | -0.160985404 |
| Zmat1 | 8.03016 | 8.035639 | 8.0207 | 0.014899576 | 0.009421123 | 8.1938194 | 7.7984602 | 8.18184 | -0.383378031 | 0.011981188 |
| Astn1 | 5.897159 | 5.563486 | 6.0033 | -0.439835145 | -0.106161531 | 4.8860265 | 5.464447 | 5.84785 | -0.383401273 | -0.961821815 |
| Znrf2 | 11.41327 | 11.3868 | 11.462 | -0.074702574 | -0.048226792 | 11.20258 | 10.973025 | 11.3565 | -0.383478948 | -0.153923774 |
| Styx | 9.235386 | 9.035856 | 9.21 | -0.174120994 | 0.025408676 | 8.9660183 | 8.9108735 | 9.29463 | -0.383758565 | -0.328613783 |
| Rps6ka1 | 11.08574 | 11.02578 | 11.205 | -0.179499678 | -0.119541228 | 11.128208 | 10.774919 | 11.1588 | -0.383838422 | -0.030549172 |
| Hcrtr2 | 5.544568 | 5.600465 | 5.5697 | 0.030778821 | -0.025118758 | 5.5685153 | 5.3230271 | 5.70706 | -0.384035738 | -0.13854752 |
| Aspn | 3.986937 | 3.857346 | 4.1679 | -0.31059046 | -0.180999316 | 3.7120917 | 3.4194645 | 3.80362 | -0.384150827 | -0.091523702 |
| Zdhhc23 | 7.618128 | 7.380008 | 7.7699 | -0.389893539 | -0.151773409 | 7.7406355 | 7.438544 | 7.82271 | -0.384169442 | -0.082077911 |
| Rprd2 | 10.87743 | 10.91841 | 10.795 | 0.123050408 | 0.08207021 | 10.712904 | 10.558185 | 10.9424 | -0.384195028 | -0.229475647 |
| Ryk | 6.12786 | 6.237746 | 6.2098 | 0.027971095 | -0.081915699 | 5.8715589 | 5.6985768 | 6.08284 | -0.384266528 | -0.211284421 |
| 38231 | 5.743621 | 5.460969 | 5.9842 | -0.523270412 | -0.240619013 | 5.248498 | 5.2450274 | 5.62931 | -0.384280747 | -0.380810202 |
| Tub | 3.995 | 4.094366 | 4.0258 | 0.068579542 | -0.030786133 | 3.8615886 | 3.7680913 | 4.15244 | -0.38435243 | -0.29085509 |
| Tmem198b | 7.278553 | 7.295357 | 7.3745 | -0.07917242 | -0.095976867 | 7.3082018 | 6.9247549 | 7.30911 | -0.384356302 | -0.000909364 |
| Hs2st1 | 8.113051 | 7.916807 | 8.0698 | -0.15301492 | 0.043229136 | 7.8169779 | 7.6032113 | 7.98761 | -0.38440113 | -0.170634523 |
| Ctsf | 5.03924 | 4.520038 | 4.9113 | -0.39127485 | 0.12792712 | 4.5759861 | 4.2310976 | 4.61562 | -0.384525583 | -0.039637087 |
| Fndc4 | 3.6952 | 3.83431 | 3.8239 | 0.010442685 | -0.128667027 | 3.600477 | 3.555096 | 3.93972 | -0.38462851 | -0.339247488 |
| Ppm1m | 9.772277 | 9.627138 | 9.8828 | -0.25565923 | -0.110519682 | 10.177417 | 9.5181875 | 9.90282 | -0.38463564 | 0.274593943 |
| Spc25 | 6.078556 | 6.03924 | 6.1648 | -0.125608921 | -0.086292673 | 6.081568 | 6.0371272 | 6.42185 | -0.384727169 | -0.340286291 |
| Usp24 | 12.14857 | 12.17779 | 12.141 | 0.036875384 | 0.007651218 | 12.063655 | 11.819149 | 12.2041 | -0.384985514 | -0.140480154 |
| Pbx4 | 5.092569 | 4.921962 | 5.1132 | -0.191260734 | -0.020653435 | 5.171804 | 4.7794698 | 5.16469 | -0.385216525 | 0.007117739 |
| Cysltr1 | 5.894023 | 5.735724 | 5.879 | -0.143304767 | 0.014994785 | 5.859115 | 5.4156865 | 5.80111 | -0.385423748 | 0.058004774 |
| Slc30a4 | 8.221006 | 7.993355 | 8.1038 | -0.110422013 | 0.117229091 | 7.9067301 | 7.5503846 | 7.93617 | -0.385780933 | -0.029435421 |
| Aldh9a1 | 9.916618 | 9.793812 | 9.9171 | -0.123319501 | -0.000513452 | 9.9521332 | 9.5264553 | 9.91243 | -0.385970753 | 0.039707094 |

| | | | | | | | | | | |
|---|---|---|---|---|---|---|---|---|---|---|
| Cep76 | 7.510776 | 7.483542 | 7.5119 | -0.028382395 | -0.0011484 | 7.5500934 | 7.1730703 | 7.55905 | -0.385976974 | -0.00895388 |
| Def8 | 8.200537 | 8.26283 | 8.1711 | 0.091699519 | 0.029406172 | 8.3350301 | 8.0243836 | 8.41039 | -0.386010257 | -0.075363747 |
| Them4 | 5.752313 | 5.74261 | 5.8664 | -0.123820416 | -0.11411791 | 5.9687971 | 5.6189427 | 6.00496 | -0.386018348 | -0.036163918 |
| Parp1 | 12.26775 | 12.27179 | 12.359 | -0.08724849 | -0.09129199 | 12.619382 | 12.003328 | 12.3894 | -0.386099804 | 0.229954203 |
| Ccdc167 | 7.805222 | 7.798984 | 7.9155 | -0.116533441 | -0.110295113 | 8.2621 | 7.6090521 | 7.99523 | -0.38617827 | 0.266869632 |
| Rap1gds1 | 11.1594 | 11.22369 | 11.218 | 0.005488712 | -0.058805532 | 11.145143 | 10.844557 | 11.2309 | -0.38631468 | -0.085728011 |
| Olfr270 | 6.216272 | 6.177734 | 6.2758 | -0.09810111 | -0.059563014 | 6.1117066 | 5.9619206 | 6.34831 | -0.386387015 | -0.236601051 |
| Trbv3 | 8.264652 | 8.288008 | 8.2906 | -0.002585443 | -0.025942386 | 7.963974 | 7.9605644 | 8.34715 | -0.386589876 | -0.38318025 |
| Fli1 | 11.22877 | 11.24006 | 11.319 | -0.078462324 | -0.08975363 | 11.138878 | 10.794576 | 11.1813 | -0.386730408 | -0.042428884 |
| Mospd2 | 8.391583 | 8.220696 | 8.3895 | -0.168768515 | 0.002118578 | 8.2537828 | 7.8536544 | 8.24076 | -0.387105563 | 0.013022772 |
| Gm44973 | 2.192719 | 2.350356 | 2.3603 | -0.009957709 | -0.167594773 | 2.2453291 | 2.1260443 | 2.5132 | -0.387153854 | -0.267869104 |
| Gm42372 | 6.452829 | 6.477235 | 6.5182 | -0.040957921 | -0.065363867 | 6.5487667 | 6.3601015 | 6.74733 | -0.387224854 | -0.198559653 |
| Ube2e3 | 8.299368 | 8.297172 | 8.425 | -0.127872815 | -0.125676331 | 7.9448583 | 7.9003205 | 8.2877 | -0.387383137 | -0.342845273 |
| Frrs1 | 7.722712 | 7.853724 | 7.845 | 0.008710658 | -0.122302251 | 8.1586823 | 7.7930354 | 8.18055 | -0.387515886 | -0.02186901 |
| Tmem72 | 8.733204 | 8.675646 | 8.8566 | -0.18098569 | -0.12342762 | 8.9103376 | 9.1606077 | 9.5482 | -0.387594652 | -0.637864829 |
| Col6a6 | 3.752622 | 3.660482 | 3.8482 | -0.187735076 | -0.095595217 | 3.4306183 | 3.3346326 | 3.7227 | -0.388064452 | -0.292078765 |
| Cd81 | 10.48768 | 10.42678 | 10.432 | -0.005674692 | 0.055223596 | 10.708353 | 10.017804 | 10.406 | -0.388221468 | 0.302327577 |
| Gstt2 | 7.976598 | 8.042829 | 8.1452 | -0.102389346 | -0.168619873 | 8.4282838 | 7.9267 | 8.31493 | -0.388234695 | 0.113349083 |
| Flnb | 10.71732 | 10.81042 | 10.725 | 0.085608172 | -0.007494234 | 10.381232 | 10.456686 | 10.8451 | -0.388445612 | -0.463899631 |
| Ccdc85a | 4.445392 | 4.422469 | 4.6008 | -0.178345076 | -0.155422153 | 3.9540659 | 3.8571808 | 4.24564 | -0.388457478 | -0.291572398 |
| Btnl1 | 4.044235 | 3.811716 | 3.9797 | -0.168029989 | 0.064488933 | 4.1584548 | 3.7100201 | 4.09874 | -0.388716013 | 0.059718651 |
| Gm15387 | 9.698862 | 9.539913 | 9.6539 | -0.113992383 | 0.044956012 | 9.3762767 | 9.2164511 | 9.60527 | -0.388820904 | -0.228995361 |
| Plcb2 | 10.87571 | 10.85854 | 10.958 | -0.099250302 | -0.082078342 | 11.20563 | 10.785007 | 11.1739 | -0.388862624 | 0.031760424 |
| Fam69b | 3.69542 | 3.593152 | 3.7261 | -0.132904769 | -0.030637151 | 3.6043297 | 3.4747027 | 3.8638 | -0.389099378 | -0.259472371 |
| Prox2 | 5.770253 | 5.649468 | 5.7954 | -0.145948165 | -0.025163658 | 5.8043898 | 5.4298898 | 5.81905 | -0.389159474 | -0.01465942 |
| Ift80 | 9.757939 | 9.989914 | 9.8841 | 0.105812209 | -0.126162473 | 9.5110349 | 9.4888803 | 9.87809 | -0.389205804 | -0.367051285 |
| Odf2 | 10.27872 | 10.25501 | 10.219 | 0.036149057 | 0.059864292 | 10.222133 | 10.043937 | 10.4334 | -0.389506251 | -0.21131024 |
| Atad5 | 8.073176 | 7.945602 | 8.1143 | -0.168731598 | -0.04115818 | 8.1163215 | 7.952536 | 8.34206 | -0.389525263 | -0.225739756 |
| Fgfr3 | 5.862951 | 5.668925 | 5.7019 | -0.032971061 | 0.161054315 | 5.1436392 | 5.1551247 | 5.54478 | -0.389656785 | -0.40114224 |
| Zfp473 | 5.075443 | 4.927297 | 4.9555 | -0.028190225 | 0.119956317 | 5.0099112 | 5.0315267 | 5.4212 | -0.389669487 | -0.411284958 |
| Itga9 | 7.857813 | 7.794661 | 7.4603 | 0.334366912 | 0.397518904 | 7.278188 | 7.0433989 | 7.43336 | -0.389965547 | -0.155176445 |
| Pde3a | 4.926539 | 4.878844 | 5.02 | -0.141107239 | -0.09341273 | 4.6856736 | 4.4226419 | 4.81262 | -0.389977745 | -0.12694609 |

| | | | | | | | | | | |
|---|---|---|---|---|---|---|---|---|---|---|
| Spata6 | 8.908346 | 8.941332 | 8.9934 | -0.052098729 | -0.085084971 | 8.8333057 | 8.6490363 | 9.03921 | -0.390173683 | -0.205904236 |
| Agmo | 3.870081 | 3.703029 | 3.9998 | -0.296765766 | -0.129713459 | 3.6558153 | 3.6506353 | 4.04088 | -0.390243125 | -0.385063114 |
| Cradd | 6.45822 | 6.520427 | 6.6207 | -0.100288005 | -0.162494493 | 6.6135353 | 6.2566837 | 6.64696 | -0.390273587 | -0.033421979 |
| Ptger4 | 9.380579 | 9.221574 | 9.3557 | -0.134128721 | 0.024876078 | 8.9395695 | 8.8078197 | 9.19814 | -0.3903214 | -0.258571602 |
| Kdm5b | 9.85251 | 9.881702 | 10.014 | -0.132784123 | -0.16197562 | 9.6336027 | 9.5817066 | 9.97239 | -0.3906834 | -0.338787256 |
| Pakap | 5.86746 | 5.979952 | 6.2469 | -0.266995511 | -0.379487872 | 5.6082151 | 5.5843967 | 5.9754 | -0.391007914 | -0.36718957 |
| Ighv10-3 | 6.969421 | 6.713893 | 6.6587 | 0.055207473 | 0.310735079 | 6.9644666 | 6.4589086 | 6.85006 | -0.391151522 | 0.114406479 |
| Gcat | 5.637441 | 5.479599 | 5.7163 | -0.236699872 | -0.078858386 | 5.578591 | 5.3166032 | 5.70781 | -0.391206102 | -0.12921833 |
| Arhgef11 | 10.05144 | 9.987648 | 10.078 | -0.090450261 | -0.026653942 | 9.9401224 | 9.6613578 | 10.0528 | -0.391438351 | -0.112673706 |
| Csad | 7.715078 | 7.367843 | 7.829 | -0.461162284 | -0.113926619 | 7.8084899 | 7.3777252 | 7.7693 | -0.391576568 | 0.039188154 |
| Plk4 | 6.848074 | 6.548377 | 6.8457 | -0.29737269 | 0.002324175 | 6.7760316 | 6.6613079 | 7.05326 | -0.391957098 | -0.277233424 |
| Myo1h | 4.971244 | 5.03614 | 5.2902 | -0.254077828 | -0.318973584 | 5.0315813 | 4.864644 | 5.25671 | -0.392065498 | -0.225128252 |
| Ext1 | 8.443234 | 8.445409 | 8.3355 | 0.109951597 | 0.107775928 | 8.2802451 | 7.9598695 | 8.35199 | -0.392122095 | -0.071746508 |
| Prex2 | 6.60965 | 6.047771 | 6.7393 | -0.69152834 | -0.129649646 | 6.014855 | 5.9711388 | 6.36365 | -0.392507847 | -0.348791643 |
| Clybl | 6.691473 | 6.571572 | 6.6917 | -0.120081946 | -0.000180898 | 6.6939558 | 6.2564501 | 6.64908 | -0.392627594 | 0.044878121 |
| Cdc42bpa | 6.481615 | 6.375986 | 6.4999 | -0.123869853 | -0.018241102 | 6.2061274 | 5.9016361 | 6.29433 | -0.392692656 | -0.0882014 |
| Grap | 11.37677 | 11.38635 | 11.471 | -0.084723064 | -0.09430549 | 11.515247 | 11.179887 | 11.5726 | -0.392704934 | -0.057344601 |
| Uprt | 7.856021 | 7.976 | 7.8769 | 0.099071632 | -0.020907591 | 7.7897923 | 7.4955764 | 7.88839 | -0.392809862 | -0.098593913 |
| Lin7c | 10.39868 | 10.34654 | 10.394 | -0.047080886 | 0.005061678 | 10.303693 | 9.9944682 | 10.3876 | -0.393098992 | -0.083873719 |
| Fgf2 | 4.618811 | 4.504359 | 4.2619 | 0.242498878 | 0.356951297 | 3.6738122 | 3.8105709 | 4.20374 | -0.393173113 | -0.529931827 |
| C1qtnf7 | 3.566768 | 3.25952 | 3.6535 | -0.394012081 | -0.086763966 | 3.1543384 | 3.0227612 | 3.41621 | -0.393452779 | -0.261875538 |
| Gm5537 | 5.737309 | 6.562836 | 6.245 | 0.317798643 | -0.507728678 | 6.1708299 | 6.2173497 | 6.61093 | -0.393575452 | -0.440095309 |
| Erbb3 | 6.877896 | 6.55656 | 6.9466 | -0.390066713 | -0.068730813 | 6.844165 | 6.4805324 | 6.87415 | -0.393615279 | -0.02998262 |
| Btrc | 8.578463 | 8.503848 | 8.6152 | -0.111374178 | -0.036759578 | 8.4938992 | 8.2376113 | 8.63131 | -0.393697711 | -0.137409881 |
| Lmbr1 | 5.761754 | 5.603635 | 5.7444 | -0.140752593 | 0.017366606 | 5.6438166 | 5.3380758 | 5.73196 | -0.393886352 | -0.088145588 |
| Akap8l | 10.49358 | 10.55295 | 10.621 | -0.067718746 | -0.127087074 | 10.946686 | 10.616459 | 11.0105 | -0.393996755 | -0.063769122 |
| Tcaim | 7.962217 | 7.879597 | 8.0641 | -0.184474621 | -0.101854637 | 7.99052 | 7.7589457 | 8.15304 | -0.394097972 | -0.162523683 |
| Pex3 | 8.317936 | 8.16904 | 8.2573 | -0.088214264 | 0.060681917 | 8.3527171 | 7.9759271 | 8.3702 | -0.394273759 | -0.017483765 |
| Myct1 | 5.702023 | 5.393953 | 5.9072 | -0.513277423 | -0.205207742 | 5.1523905 | 4.9007893 | 5.29518 | -0.394386448 | -0.142785312 |
| Retsat | 6.104318 | 5.744008 | 6.0641 | -0.320135436 | 0.040175143 | 5.9232424 | 5.5099502 | 5.90435 | -0.394395435 | 0.018896728 |
| Mthfsd | 7.951426 | 7.963856 | 8.0018 | -0.037970912 | -0.050401029 | 8.5426442 | 7.9380248 | 8.33248 | -0.394450799 | 0.210168623 |
| Klhl24 | 11.71211 | 11.76521 | 11.883 | -0.11783077 | -0.170929236 | 11.636569 | 11.314045 | 11.7085 | -0.394479448 | -0.071955548 |

| | | | | | | | | | | |
|---|---|---|---|---|---|---|---|---|---|---|
| Rcn2 | 9.029478 | 8.877116 | 9.0606 | -0.183443715 | -0.031081442 | 8.9926228 | 8.5541937 | 8.94875 | -0.394551712 | 0.043877384 |
| Gnpda2 | 8.138279 | 8.092491 | 8.2278 | -0.135298347 | -0.089510086 | 8.1075227 | 7.7915045 | 8.18618 | -0.394678287 | -0.078660041 |
| Cntn1 | 7.17017 | 7.240035 | 7.2058 | 0.034273504 | -0.035591048 | 6.9633726 | 6.7924861 | 7.18797 | -0.395485046 | -0.224598504 |
| Pard6g | 6.54226 | 6.525935 | 6.6522 | -0.126266501 | -0.109940827 | 6.239534 | 6.0989927 | 6.49462 | -0.395629931 | -0.255088696 |
| Pigg | 7.434278 | 7.099753 | 7.3199 | -0.220132031 | 0.114392403 | 7.5020985 | 7.0455087 | 7.44131 | -0.395800887 | 0.060788969 |
| Kidins220 | 10.88522 | 10.81342 | 10.84 | -0.026289292 | 0.045509308 | 10.979804 | 10.731678 | 11.1276 | -0.395918814 | -0.147792321 |
| Zbed6 | 8.795852 | 8.853294 | 8.6039 | 0.249387028 | 0.191945344 | 8.4967741 | 8.802379 | 9.19839 | -0.396012913 | -0.701617893 |
| Papss2 | 8.647372 | 8.321693 | 8.5707 | -0.249039658 | 0.076639443 | 7.7039933 | 7.8702242 | 8.26632 | -0.396097515 | -0.562328367 |
| Stap2 | 7.327615 | 7.00928 | 7.351 | -0.341705028 | -0.023369984 | 6.9778666 | 6.861047 | 7.25728 | -0.396235555 | -0.27941596 |
| Phkb | 9.054106 | 8.849886 | 9.0256 | -0.17571653 | 0.028503604 | 8.9404609 | 8.6776097 | 9.074 | -0.396392124 | -0.133540873 |
| Cbx2 | 6.567353 | 6.718904 | 6.6518 | 0.06710759 | -0.084444042 | 6.3329748 | 6.2470364 | 6.64349 | -0.396450038 | -0.310511608 |
| Prrg1 | 6.505728 | 6.637702 | 6.6451 | -0.007393296 | -0.139367577 | 6.314793 | 6.2301038 | 6.62739 | -0.397290215 | -0.312601046 |
| Scn4a | 8.046814 | 8.237638 | 8.3142 | -0.076572299 | -0.26739689 | 8.2043105 | 7.9626675 | 8.35996 | -0.397295575 | -0.155652619 |
| Ighv1-15 | 7.039382 | 7.038661 | 6.7502 | 0.288511012 | 0.289231767 | 6.9916466 | 6.4738989 | 6.87124 | -0.397338668 | 0.120409062 |
| Bmpr1a | 5.710645 | 5.394439 | 5.4748 | -0.080328675 | 0.23587747 | 5.4884074 | 4.972392 | 5.36974 | -0.397350488 | 0.118664896 |
| Adcy9 | 5.242775 | 4.983895 | 5.0903 | -0.106441012 | 0.152438987 | 4.7860085 | 4.6489629 | 5.04678 | -0.397817784 | -0.260772217 |
| Def6 | 10.22894 | 10.12003 | 10.242 | -0.122129643 | -0.013217322 | 10.352964 | 10.056818 | 10.4549 | -0.39808883 | -0.101942933 |
| Gabpb2 | 11.89149 | 11.88862 | 11.955 | -0.066310752 | -0.06343277 | 11.936292 | 11.726413 | 12.1247 | -0.398319852 | -0.188440877 |
| Chka | 8.761304 | 8.721378 | 8.8424 | -0.121004734 | -0.081078535 | 9.0341998 | 8.7062395 | 9.10461 | -0.398371316 | -0.070411075 |
| Abl1 | 10.51097 | 10.44331 | 10.61 | -0.166208534 | -0.09854278 | 10.603901 | 10.251902 | 10.6503 | -0.398402787 | -0.04640351 |
| Trbv4 | 7.371288 | 7.217215 | 7.3906 | -0.173352663 | -0.019278839 | 7.0896604 | 6.9531522 | 7.35159 | -0.398439364 | -0.261931169 |
| Col4a5 | 4.20094 | 3.955407 | 4.1875 | -0.232101105 | 0.013431862 | 3.9157336 | 3.722348 | 4.12081 | -0.398458709 | -0.205073122 |
| 5430403G16Rik | 6.963676 | 6.817934 | 6.9552 | -0.137279943 | 0.008462158 | 7.1384094 | 6.6754795 | 7.07395 | -0.398475235 | 0.064454672 |
| Smc6 | 11.43412 | 11.48504 | 11.451 | 0.033786328 | -0.017133518 | 11.210199 | 10.985001 | 11.3836 | -0.39859241 | -0.173393684 |
| Stxbp4 | 5.951064 | 6.009967 | 5.9003 | 0.109678313 | 0.050774507 | 6.1250284 | 5.9513233 | 6.35004 | -0.39871745 | -0.225012338 |
| Tsc1 | 9.977232 | 9.994412 | 10.083 | -0.088561807 | -0.105741988 | 10.015791 | 9.8395087 | 10.2383 | -0.398767441 | -0.222485411 |
| Car5b | 5.466755 | 5.438831 | 5.5665 | -0.127663362 | -0.099739514 | 5.2250746 | 5.5103549 | 5.90993 | -0.399570828 | -0.684851155 |
| Kif13b | 9.44069 | 9.501201 | 9.4731 | 0.028073388 | -0.032438122 | 9.300891 | 9.2618371 | 9.66151 | -0.399670321 | -0.36061648 |
| Dip2b | 10.8802 | 10.8885 | 10.885 | 0.003602978 | -0.004697296 | 10.747874 | 10.667881 | 11.0679 | -0.400009942 | -0.320016334 |
| Prune1 | 8.190049 | 8.081139 | 8.2082 | -0.127062293 | -0.018151932 | 8.2741835 | 7.786708 | 8.18675 | -0.400042055 | 0.087433426 |
| Crot | 9.213195 | 9.191819 | 9.2566 | -0.064787823 | -0.043412386 | 9.0876987 | 8.8260536 | 9.22611 | -0.400055446 | -0.13841039 |
| 5830411N06Rik | 8.359053 | 8.08373 | 8.3423 | -0.25859015 | 0.016733245 | 8.0753862 | 7.999185 | 8.39928 | -0.400094619 | -0.323893393 |

| | | | | | | | | | | |
|---|---|---|---|---|---|---|---|---|---|---|
| Zdhhc9 | 8.47391 | 8.403044 | 8.5344 | -0.131334153 | -0.060467857 | 8.5509867 | 8.0204243 | 8.42052 | -0.40009579 | 0.130466625 |
| Wnt11 | 5.023906 | 4.868361 | 4.9868 | -0.118485346 | 0.03706008 | 4.6991768 | 4.5230029 | 4.92338 | -0.400376544 | -0.22420264 |
| Me2 | 10.25714 | 10.19406 | 10.334 | -0.13973742 | -0.0766645 | 10.155659 | 9.8452562 | 10.2458 | -0.400497317 | -0.090094273 |
| Klhl36 | 7.782388 | 7.602535 | 7.8488 | -0.24624938 | -0.066396148 | 7.8949005 | 7.3202565 | 7.72104 | -0.400778661 | 0.173865338 |
| Sik2 | 8.882636 | 8.884654 | 9.0089 | -0.124251741 | -0.126269179 | 8.7334371 | 8.7563321 | 9.15712 | -0.400789952 | -0.423684973 |
| Helq | 8.114731 | 8.061015 | 8.1998 | -0.138776957 | -0.085061277 | 8.5040384 | 8.1409696 | 8.54215 | -0.401175883 | -0.038107036 |
| Btbd6 | 7.858385 | 7.931234 | 8.0851 | -0.153864759 | -0.226713889 | 7.7934527 | 7.4956482 | 7.89694 | -0.401292632 | -0.103488116 |
| Snai1 | 5.043242 | 4.763527 | 4.9694 | -0.205871226 | 0.073843467 | 4.5703641 | 4.3320659 | 4.73336 | -0.401298643 | -0.16300047 |
| Adgre5 | 12.35282 | 12.3778 | 12.398 | -0.02063295 | -0.045613106 | 12.427197 | 12.102222 | 12.5035 | -0.401310402 | -0.076335465 |
| B3gnt2 | 9.494231 | 9.708634 | 9.7921 | -0.083485761 | -0.297889473 | 9.3520038 | 9.256458 | 9.65781 | -0.401356005 | -0.305810116 |
| Spn | 11.96734 | 12.04651 | 12.135 | -0.088824914 | -0.167998246 | 11.828825 | 11.703818 | 12.1053 | -0.401482398 | -0.276474811 |
| Clcn5 | 8.361367 | 8.338984 | 8.4733 | -0.134293061 | -0.11191059 | 8.101622 | 7.8057696 | 8.2073 | -0.401529445 | -0.105676976 |
| Smo | 5.750787 | 5.550092 | 5.6907 | -0.140608759 | 0.060086297 | 5.6519002 | 5.3186019 | 5.72013 | -0.401531746 | -0.06823346 |
| Fam213b | 6.164138 | 5.95362 | 6.0586 | -0.10499831 | 0.105520012 | 6.1195323 | 5.3869328 | 5.78874 | -0.401803728 | 0.330795721 |
| Zfp366 | 9.10405 | 8.909229 | 8.9978 | -0.088544982 | 0.106276237 | 8.6731328 | 8.6224411 | 9.02431 | -0.401868335 | -0.351176613 |
| Evi5l | 7.966225 | 7.615712 | 7.7443 | -0.128627002 | 0.221886338 | 7.8799219 | 7.4643342 | 7.86624 | -0.40190986 | 0.01367785 |
| Hist1h2bg | 5.248758 | 5.183988 | 5.2896 | -0.105577262 | -0.04080656 | 5.3002419 | 4.9191961 | 5.32137 | -0.402170898 | -0.02112507 |
| Zfp367 | 8.322197 | 8.304244 | 8.3939 | -0.089687935 | -0.071734758 | 8.3768555 | 8.0442196 | 8.44654 | -0.402325269 | -0.069689414 |
| Smad1 | 8.643144 | 8.466084 | 8.7249 | -0.258772428 | -0.081712508 | 8.1205072 | 8.0007408 | 8.40311 | -0.402370675 | -0.28260428 |
| Scn7a | 4.024887 | 3.603734 | 3.9093 | -0.305567024 | 0.115586547 | 3.652287 | 3.4226722 | 3.82505 | -0.402375309 | -0.172760483 |
| Gm16867 | 7.804813 | 7.579215 | 7.8527 | -0.273467519 | -0.047869936 | 7.7521898 | 7.486012 | 7.8884 | -0.402383281 | -0.136205482 |
| Ero1l | 9.748806 | 9.656977 | 9.6805 | -0.023497413 | 0.068331709 | 9.3675065 | 9.2190704 | 9.62174 | -0.402667396 | -0.254231373 |
| Asb7 | 7.912516 | 7.731349 | 8.038 | -0.306668697 | -0.125500918 | 7.5607292 | 7.5753827 | 7.97806 | -0.402673906 | -0.417327445 |
| Kcna2 | 8.210688 | 8.317532 | 8.4148 | -0.097266693 | -0.204110651 | 7.9778417 | 8.0770269 | 8.47971 | -0.402684284 | -0.501869533 |
| Prss23 | 6.573794 | 6.845575 | 6.7206 | 0.125004569 | -0.146776197 | 6.4100423 | 5.9240912 | 6.32678 | -0.402686153 | 0.083264915 |
| Prickle1 | 5.867321 | 5.899021 | 5.9783 | -0.079311171 | -0.111010998 | 5.5345299 | 5.4893085 | 5.89207 | -0.402761213 | -0.357539845 |
| Myef2 | 9.110181 | 8.883556 | 9.0373 | -0.153735839 | 0.072889016 | 9.1297562 | 8.8522494 | 9.25504 | -0.402786822 | -0.125280063 |
| Pcdhgb7 | 4.395625 | 4.461017 | 4.4212 | 0.039825111 | -0.025567269 | 4.4296703 | 4.085343 | 4.48864 | -0.403301676 | -0.058974305 |
| Nav3 | 3.299815 | 3.004796 | 3.1418 | -0.136960759 | 0.158058537 | 3.1164922 | 2.9900115 | 3.39362 | -0.403606681 | -0.27712596 |
| Plp1 | 6.211459 | 6.245803 | 6.2544 | -0.008586377 | -0.042929436 | 6.4008968 | 6.1343062 | 6.53795 | -0.403642929 | -0.137052295 |
| Rnf19a | 9.982049 | 9.800234 | 9.9736 | -0.173374918 | 0.008440757 | 9.5493859 | 9.4989406 | 9.90263 | -0.403686049 | -0.35324072 |
| Ahdc1 | 9.672971 | 9.546151 | 9.5387 | 0.00747565 | 0.134295205 | 9.3511472 | 8.930212 | 9.33397 | -0.403754729 | 0.017180509 |

| | | | | | | | | | | |
|---|---|---|---|---|---|---|---|---|---|---|
| Stx6 | 9.541359 | 9.602056 | 9.6843 | -0.082234734 | -0.142932634 | 9.5770602 | 9.3302942 | 9.73408 | -0.403781113 | -0.157015099 |
| Mansc1 | 6.05636 | 5.697564 | 6.1431 | -0.445555492 | -0.08675949 | 5.6538572 | 5.4411872 | 5.84521 | -0.404022516 | -0.191352488 |
| Smim14 | 10.86179 | 10.85428 | 11.022 | -0.16796653 | -0.16045523 | 11.275971 | 10.543038 | 10.9472 | -0.404182642 | 0.328750153 |
| Glis2 | 5.586044 | 5.431095 | 5.3601 | 0.070980359 | 0.225928773 | 5.208068 | 5.0389361 | 5.44396 | -0.405026939 | -0.235895039 |
| Cdc42bpg | 6.760812 | 6.515538 | 6.7358 | -0.220265747 | 0.025008405 | 6.8288438 | 6.3114286 | 6.71662 | -0.405194469 | 0.112220712 |
| Syt7 | 4.211694 | 4.088893 | 4.2969 | -0.20796323 | -0.08516205 | 3.8354447 | 3.5843623 | 3.98959 | -0.405226857 | -0.154144503 |
| Bag4 | 8.883703 | 8.79126 | 8.8002 | -0.008942225 | 0.083500534 | 8.5869378 | 8.2880666 | 8.69354 | -0.405474833 | -0.10660363 |
| Smarcc2 | 11.2955 | 11.2169 | 11.349 | -0.132184302 | -0.053584808 | 11.422827 | 10.996596 | 11.4022 | -0.405593866 | 0.020637201 |
| Gm3604 | 5.567388 | 5.603166 | 5.5706 | 0.032572047 | -0.003206207 | 5.6630249 | 5.3412905 | 5.74704 | -0.405746933 | -0.084012514 |
| Purb | 12.18771 | 12.11808 | 12.181 | -0.062826218 | 0.006797524 | 12.140776 | 11.803291 | 12.2092 | -0.40588925 | -0.068404149 |
| Slc27a6 | 5.18195 | 4.88026 | 5.0987 | -0.218429424 | 0.083260578 | 4.7216495 | 4.5588985 | 4.96498 | -0.406086136 | -0.243335149 |
| Nsun3 | 8.083546 | 7.965512 | 8.1698 | -0.204323786 | -0.086289545 | 8.3436164 | 7.8546352 | 8.26082 | -0.406180088 | 0.082801091 |
| Zfp609 | 9.089103 | 9.159174 | 9.1426 | 0.016576917 | -0.053494005 | 8.9871089 | 8.8119502 | 9.21817 | -0.406215557 | -0.231056952 |
| Rassf2 | 12.41387 | 12.48907 | 12.442 | 0.046920518 | -0.028275266 | 12.36467 | 12.059297 | 12.4656 | -0.406342358 | -0.100969135 |
| Gm7334 | 4.907781 | 5.068697 | 5.1673 | -0.098626386 | -0.259541707 | 5.2346446 | 4.6573505 | 5.06374 | -0.406385946 | 0.170908128 |
| Zfp955a | 7.89659 | 7.955134 | 8.0951 | -0.139971107 | -0.198515644 | 7.8831515 | 7.6474233 | 8.05388 | -0.406454199 | -0.170726017 |
| Nipsnap2 | 9.084131 | 8.904081 | 9.1596 | -0.255524894 | -0.075474189 | 9.1901519 | 8.736665 | 9.14317 | -0.40650187 | 0.046984964 |
| Vps13c | 10.20289 | 10.15082 | 10.228 | -0.077048821 | -0.024972974 | 9.9023048 | 10.003125 | 10.4097 | -0.406564267 | -0.507384592 |
| Myo1b | 7.570642 | 7.292126 | 7.5366 | -0.244440127 | 0.034075957 | 6.7599068 | 6.5659391 | 6.97257 | -0.406632406 | -0.212664762 |
| Fchsd2 | 11.95715 | 12.00058 | 11.97 | 0.030431086 | -0.01299845 | 11.964097 | 11.757005 | 12.1641 | -0.407137326 | -0.200045022 |
| Wipf3 | 3.173358 | 3.197412 | 3.2942 | -0.096773471 | -0.120827524 | 3.0949038 | 2.9348632 | 3.34206 | -0.407200075 | -0.247159434 |
| Zfp526 | 8.319586 | 8.253171 | 8.278 | -0.024859614 | 0.041555887 | 8.2934181 | 7.9099721 | 8.31739 | -0.407415339 | -0.023969329 |
| Prkaca | 9.468256 | 9.324891 | 9.5118 | -0.186872023 | -0.043507684 | 9.3897809 | 9.006528 | 9.4142 | -0.407675612 | -0.024422712 |
| Mum1 | 10.14319 | 10.127 | 10.282 | -0.155456876 | -0.13926911 | 10.325348 | 9.9873512 | 10.3951 | -0.407699856 | -0.069703493 |
| Trmt2b | 9.383983 | 9.317944 | 9.4937 | -0.1757991 | -0.109759649 | 9.3294891 | 9.010674 | 9.41854 | -0.407868304 | -0.08905321 |
| 2210408I21Rik | 5.201012 | 5.084836 | 5.1755 | -0.090663324 | 0.025512792 | 5.0490404 | 4.8776262 | 5.2857 | -0.408070206 | -0.236656015 |
| Trav7d-4 | 4.204651 | 4.242826 | 4.2424 | 0.000445458 | -0.037729858 | 4.0415388 | 4.0067857 | 4.41511 | -0.408329085 | -0.373575943 |
| Ebpl | 7.011473 | 7.076722 | 7.1384 | -0.061693714 | -0.126942721 | 7.1199149 | 6.6876229 | 7.09597 | -0.408347473 | 0.023944538 |
| Dsn1 | 7.130574 | 7.109039 | 7.2363 | -0.127237179 | -0.10570268 | 7.3020799 | 7.0115643 | 7.4201 | -0.408537111 | -0.118021487 |
| Nlrc4 | 8.547003 | 8.626621 | 8.548 | 0.07864764 | -0.000970102 | 8.8216938 | 8.3284948 | 8.7372 | -0.408708329 | 0.084490636 |
| Mrs2 | 9.923078 | 9.827858 | 9.8244 | 0.003447391 | 0.098667981 | 10.06632 | 9.727898 | 10.1367 | -0.408843 | -0.070421056 |
| Dusp2 | 11.60743 | 11.64256 | 11.72 | -0.07763859 | -0.112767632 | 10.851457 | 10.83189 | 11.2408 | -0.408904928 | -0.389337539 |

| | | | | | | | | | | |
|---|---|---|---|---|---|---|---|---|---|---|
| Gal3st2c | 5.20993 | 5.072549 | 5.2587 | -0.186132472 | -0.048751357 | 5.1208197 | 4.9685274 | 5.37752 | -0.408993614 | -0.256701267 |
| Dlg3 | 7.437946 | 7.277435 | 7.4432 | -0.165765977 | -0.005254036 | 7.4128514 | 6.8286033 | 7.2376 | -0.408994487 | 0.175253648 |
| Jade3 | 7.379778 | 7.394414 | 7.3849 | 0.009563091 | -0.005073324 | 7.111964 | 7.0107802 | 7.41983 | -0.409049584 | -0.307865776 |
| Pbxip1 | 11.37558 | 11.35097 | 11.431 | -0.080490258 | -0.055879802 | 11.560101 | 11.119881 | 11.5292 | -0.409306496 | 0.030913277 |
| Primpol | 9.028914 | 8.954791 | 9.2412 | -0.286446389 | -0.212322713 | 9.3926585 | 8.8423589 | 9.25173 | -0.409368144 | 0.140931459 |
| Cd6 | 11.1387 | 11.1839 | 11.31 | -0.126092008 | -0.171296066 | 11.10466 | 10.942045 | 11.3517 | -0.409610326 | -0.246994997 |
| Ighv1-76 | 7.209557 | 8.132904 | 7.2712 | 0.86173647 | -0.061610437 | 7.2390051 | 6.7533504 | 7.16323 | -0.409875096 | 0.075779578 |
| Gdpd5 | 7.10394 | 6.926628 | 6.9783 | -0.051662033 | 0.125649881 | 6.9435737 | 6.5152179 | 6.92526 | -0.410044876 | 0.018310979 |
| Shprh | 10.11341 | 9.993575 | 10.053 | -0.05947258 | 0.060360169 | 9.9110999 | 9.6045791 | 10.0147 | -0.410130875 | -0.103610108 |
| Itih3 | 5.744234 | 5.280517 | 5.6805 | -0.399995469 | 0.063721887 | 4.9075197 | 4.7217031 | 5.13197 | -0.410267997 | -0.224451408 |
| Haus4 | 7.971437 | 8.006529 | 8.1448 | -0.138264035 | -0.173355632 | 8.1530766 | 7.8155271 | 8.22589 | -0.410365615 | -0.072816102 |
| Rab23 | 6.634713 | 6.689995 | 6.682 | 0.007986057 | -0.047295699 | 6.6335834 | 6.3176592 | 6.72832 | -0.410657325 | -0.094733214 |
| Ighv3-8 | 5.708834 | 6.314476 | 5.8467 | 0.467805013 | -0.137837163 | 6.2439943 | 5.7083102 | 6.11899 | -0.410680528 | 0.125003493 |
| Atp1a2 | 5.292624 | 5.18452 | 5.3864 | -0.201881047 | -0.093777365 | 4.8463445 | 4.6899806 | 5.10103 | -0.411051449 | -0.254687617 |
| Hrh1 | 3.578035 | 3.485176 | 3.6665 | -0.181338732 | -0.0884795 | 3.5495105 | 3.3201444 | 3.73129 | -0.411142723 | -0.181776679 |
| Card10 | 7.146313 | 6.679503 | 7.074 | -0.394504634 | 0.07230617 | 6.3519398 | 6.0908449 | 6.50212 | -0.411276151 | -0.150181181 |
| Nmnat3 | 7.094084 | 6.895594 | 7.1435 | -0.247901264 | -0.049412206 | 7.2910393 | 6.7944973 | 7.20585 | -0.41135689 | 0.085185133 |
| Nme4 | 5.24398 | 5.157378 | 5.3449 | -0.187548044 | -0.100946134 | 5.2076386 | 4.8965272 | 5.30789 | -0.411366268 | -0.100254865 |
| Map3k7 | 10.27165 | 10.23292 | 10.315 | -0.082025602 | -0.043291118 | 10.254312 | 10.187822 | 10.5995 | -0.411653866 | -0.345164184 |
| Zfp354c | 6.961666 | 6.954298 | 7.1443 | -0.189991225 | -0.182623042 | 6.8634529 | 6.6862578 | 7.09794 | -0.411681727 | -0.234486656 |
| Hemgn | 4.212486 | 3.992445 | 4.2297 | -0.23728964 | -0.017249151 | 4.1865736 | 3.8866027 | 4.29834 | -0.411736685 | -0.111765754 |
| Myh9 | 14.95625 | 14.93477 | 14.92 | 0.015047102 | 0.036527244 | 14.688112 | 14.44264 | 14.8544 | -0.411742036 | -0.166270127 |
| Rnf8 | 8.39565 | 8.214863 | 8.3561 | -0.141195864 | 0.039591102 | 8.481321 | 8.1575593 | 8.56931 | -0.41175241 | -0.087990678 |
| Aco1 | 9.172986 | 9.090513 | 9.2084 | -0.117880523 | -0.035407878 | 9.0172516 | 8.6915717 | 9.10341 | -0.411839112 | -0.086159155 |
| Fam83d | 4.452052 | 4.328624 | 4.5293 | -0.200690863 | -0.077262384 | 4.3154155 | 4.2301244 | 4.64201 | -0.411880771 | -0.326589612 |
| Mdc1 | 10.9073 | 11.03625 | 10.958 | 0.078318054 | -0.050629276 | 11.015089 | 10.720811 | 11.133 | -0.41214874 | -0.117870534 |
| Gpr34 | 6.631315 | 6.248492 | 6.7385 | -0.489958389 | -0.107135903 | 6.4458328 | 6.0647067 | 6.47721 | -0.412500643 | -0.031374531 |
| 2700049A03Rik | 8.271865 | 8.247134 | 8.4036 | -0.156422673 | -0.131692121 | 8.2484467 | 8.0067385 | 8.41931 | -0.412575162 | -0.170866886 |
| Pip4k2b | 10.86914 | 10.81408 | 10.895 | -0.080675692 | -0.025610226 | 10.662453 | 10.413703 | 10.8264 | -0.41269398 | -0.163944556 |
| Tll1 | 6.602423 | 6.205722 | 6.1973 | 0.008413397 | 0.405114238 | 5.5320551 | 5.251637 | 5.66434 | -0.412702018 | -0.132283854 |
| Lrrc20 | 7.667683 | 7.537851 | 7.7624 | -0.224517451 | -0.09468545 | 7.7270572 | 7.2375536 | 7.65034 | -0.412781706 | 0.076721941 |
| Plcg1 | 11.46971 | 11.41444 | 11.548 | -0.133421198 | -0.078147218 | 11.120085 | 11.116176 | 11.5292 | -0.413038004 | -0.40912919 |

| | | | | | | | | | | |
|---|---|---|---|---|---|---|---|---|---|---|
| Il1rl2 | 6.929831 | 6.954468 | 7.167 | -0.212491523 | -0.237128422 | 6.6859102 | 6.578333 | 6.99139 | -0.413061713 | -0.305484455 |
| Fto | 9.853222 | 9.886378 | 10.014 | -0.127130685 | -0.160286296 | 10.013492 | 9.5700962 | 9.98327 | -0.413173382 | 0.030222441 |
| Steap3 | 5.802055 | 5.485404 | 5.7601 | -0.274731288 | 0.041919837 | 5.7955504 | 5.1364934 | 5.54968 | -0.413187015 | 0.245870007 |
| Tgfb1 | 11.26971 | 11.25064 | 11.289 | -0.038025034 | -0.018960456 | 11.323758 | 10.876665 | 11.2899 | -0.413214978 | 0.033878067 |
| Ankmy2 | 8.573281 | 8.518463 | 8.7104 | -0.191961428 | -0.137143073 | 8.5935783 | 8.2113653 | 8.62477 | -0.413407073 | -0.03119412 |
| Tcerg1l | 3.894159 | 3.526894 | 3.9099 | -0.383024143 | -0.015759601 | 3.4023217 | 3.3546877 | 3.76865 | -0.413961231 | -0.366327199 |
| Ncmap | 4.619122 | 4.739066 | 4.9197 | -0.180630472 | -0.300574579 | 4.4752095 | 4.4048728 | 4.81892 | -0.414049216 | -0.343712578 |
| Cyb5d2 | 6.639079 | 6.611256 | 6.7349 | -0.12367052 | -0.095847135 | 6.6877727 | 6.3453606 | 6.75955 | -0.414185962 | -0.071773863 |
| 39692 | 7.344407 | 7.200591 | 7.4392 | -0.238601188 | -0.094785493 | 7.1601 | 6.9049659 | 7.31942 | -0.414455785 | -0.159321596 |
| Ttk | 5.587658 | 5.329605 | 5.6896 | -0.359971501 | -0.101919049 | 5.5126972 | 5.4745928 | 5.88906 | -0.414470454 | -0.376366045 |
| Cadm3 | 6.573682 | 6.464138 | 6.7795 | -0.315348591 | -0.205803894 | 6.3814546 | 6.3472955 | 6.76181 | -0.4145156 | -0.380356476 |
| Mfn1 | 8.862763 | 8.784009 | 8.8998 | -0.115814274 | -0.037060048 | 8.7548919 | 8.6362477 | 9.05092 | -0.414669542 | -0.296025411 |
| Dsel | 5.504181 | 5.451655 | 5.5735 | -0.12180977 | -0.069283335 | 5.3676275 | 5.0801118 | 5.49505 | -0.414939065 | -0.12742335 |
| Gm31166 | 4.537499 | 4.301255 | 4.6733 | -0.372078645 | -0.135835342 | 4.733134 | 4.2696777 | 4.6848 | -0.415120003 | 0.04833622 |
| Acpp | 7.463483 | 7.448135 | 7.6587 | -0.210555792 | -0.195207528 | 7.4334237 | 7.1539237 | 7.56906 | -0.415135497 | -0.135635537 |
| Hectd2 | 6.666478 | 6.404786 | 6.6844 | -0.279636582 | -0.017944289 | 6.1296537 | 6.2559535 | 6.67114 | -0.415186121 | -0.541485976 |
| Usp20 | 8.570727 | 8.442439 | 8.7853 | -0.342851222 | -0.214563207 | 8.6074097 | 8.2493073 | 8.6647 | -0.415391587 | -0.057289144 |
| Rhobtb3 | 4.043023 | 3.908187 | 4.1106 | -0.202443198 | -0.067607881 | 3.8472962 | 3.4361817 | 3.85187 | -0.415691997 | -0.00457749 |
| Sh3kbp1 | 12.19194 | 12.27214 | 12.277 | -0.004460384 | -0.084661264 | 12.21063 | 11.864285 | 12.28 | -0.415724008 | -0.069379236 |
| Bcl2l2 | 7.496165 | 7.243182 | 7.239 | 0.004175741 | 0.25715929 | 7.2938749 | 7.0397946 | 7.45571 | -0.41591706 | -0.161836692 |
| Phactr4 | 9.090469 | 9.184097 | 9.1393 | 0.044809583 | -0.048818408 | 8.79904 | 8.7351075 | 9.15105 | -0.415940663 | -0.352008188 |
| Ttc21b | 7.708443 | 7.643315 | 7.7885 | -0.145212024 | -0.080083822 | 7.631215 | 7.3193641 | 7.73532 | -0.415958048 | -0.104107178 |
| Farp1 | 6.578056 | 6.253256 | 6.6657 | -0.412462615 | -0.087662409 | 6.0984557 | 6.1562368 | 6.57243 | -0.41619242 | -0.473973442 |
| Syt11 | 9.024306 | 8.688421 | 8.8599 | -0.17152369 | 0.164360729 | 8.7710961 | 8.3406265 | 8.75689 | -0.416266391 | 0.014203202 |
| Gm14288 | 5.172248 | 5.75005 | 5.8883 | -0.138295943 | -0.716098111 | 5.7222153 | 5.2942175 | 5.71055 | -0.416329426 | 0.01166836 |
| Prag1 | 9.411961 | 9.257783 | 9.4329 | -0.175108938 | -0.020931197 | 9.4910312 | 8.8341804 | 9.25052 | -0.41633766 | 0.240513098 |
| Gucd1 | 9.942507 | 9.997701 | 10.04 | -0.042607684 | -0.097801097 | 10.089157 | 9.6384604 | 10.0548 | -0.416346811 | 0.034349421 |
| Ralgapb | 11.19625 | 11.16477 | 11.238 | -0.073419542 | -0.041944806 | 11.009889 | 10.851763 | 11.2681 | -0.416356744 | -0.258231511 |
| Rpa1 | 11.42799 | 11.32575 | 11.441 | -0.114855526 | -0.012611322 | 11.42217 | 11.035265 | 11.4519 | -0.41664877 | -0.029744694 |
| Slc25a36 | 10.25843 | 10.22847 | 10.359 | -0.130653685 | -0.100697728 | 10.409632 | 10.126475 | 10.5432 | -0.416769901 | -0.13361247 |
| AC154652.2 | 4.937272 | 4.854396 | 5.0063 | -0.151909225 | -0.069033186 | 4.8171506 | 4.7092686 | 5.12611 | -0.416842414 | -0.308960469 |
| Gm47654 | 4.903875 | 4.555762 | 4.7128 | -0.156991998 | 0.191121881 | 4.7863812 | 4.5785772 | 4.99575 | -0.417175082 | -0.209371118 |

| | | | | | | | | | | |
|---|---|---|---|---|---|---|---|---|---|---|
| Aurkb | 7.220249 | 7.022395 | 7.3505 | -0.328094445 | -0.130240666 | 7.3529736 | 7.2067818 | 7.62416 | -0.417380566 | -0.271188775 |
| Myo5a | 10.95348 | 10.86344 | 10.934 | -0.070409212 | 0.019639726 | 10.972967 | 10.608246 | 11.026 | -0.417708406 | -0.052986933 |
| 40787 | 11.08908 | 10.98474 | 11.081 | -0.096714726 | 0.007625896 | 11.086231 | 10.652895 | 11.0707 | -0.417759072 | 0.015576213 |
| Fam126b | 8.748284 | 8.469409 | 8.6943 | -0.224876428 | 0.053998243 | 8.5149917 | 8.2453321 | 8.66317 | -0.417834738 | -0.148175142 |
| Fam160a1 | 5.001301 | 4.923152 | 5.0182 | -0.09507097 | -0.016922442 | 4.7902237 | 4.7745167 | 5.19256 | -0.418044272 | -0.402337232 |
| Mfsd4b4 | 6.681566 | 6.530811 | 6.8169 | -0.286125635 | -0.1353713 | 6.5645868 | 6.3296918 | 6.74783 | -0.418142978 | -0.183247986 |
| Zmynd15 | 7.913355 | 7.914768 | 8.1259 | -0.211136124 | -0.212549005 | 7.7660072 | 7.8125642 | 8.23087 | -0.418303044 | -0.464860025 |
| Ghr | 5.407725 | 4.919797 | 5.4781 | -0.55830659 | -0.070377952 | 4.7929092 | 4.6495247 | 5.06799 | -0.418467813 | -0.275083281 |
| Pcyox1 | 8.622113 | 8.644056 | 8.7168 | -0.072702999 | -0.094646288 | 8.6781159 | 8.2644489 | 8.68336 | -0.418914417 | -0.005247488 |
| Ccdc17 | 6.30612 | 6.199205 | 6.452 | -0.252823455 | -0.145908786 | 6.8210198 | 6.2183724 | 6.63733 | -0.418960912 | 0.183686551 |
| Tram2 | 8.216911 | 8.29888 | 8.3367 | -0.037824175 | -0.119792974 | 8.3311261 | 7.8990463 | 8.31827 | -0.419222889 | 0.012856904 |
| Rab2b | 8.174395 | 7.999347 | 8.1798 | -0.180442515 | -0.005395381 | 8.324382 | 7.9059843 | 8.32524 | -0.419251803 | -0.000854125 |
| Hectd4 | 10.65309 | 10.83164 | 10.838 | -0.006683312 | -0.185232614 | 10.491598 | 10.573363 | 10.9929 | -0.419505894 | -0.501270626 |
| Tkfc | 7.189242 | 7.075366 | 7.2159 | -0.140537917 | -0.02666107 | 7.2302242 | 6.9266351 | 7.34627 | -0.419638876 | -0.1160497 |
| Ogfod1 | 8.832364 | 8.944264 | 8.8913 | 0.05295507 | -0.058945297 | 8.8764246 | 8.7353163 | 9.1551 | -0.419787518 | -0.278679233 |
| Tmed8 | 10.5795 | 10.5264 | 10.62 | -0.093395318 | -0.040297738 | 10.73648 | 10.28695 | 10.7069 | -0.419973074 | 0.029557314 |
| Pomt1 | 7.663632 | 7.631393 | 7.5961 | 0.035290221 | 0.067529235 | 7.9342185 | 7.5442412 | 7.96433 | -0.420089188 | -0.030111877 |
| Trp53bp2 | 9.030083 | 8.931254 | 9.0146 | -0.08331172 | 0.01551705 | 8.8761208 | 8.6718227 | 9.09232 | -0.420499709 | -0.216201638 |
| Tbc1d8b | 8.955133 | 8.963893 | 8.9148 | 0.049053812 | 0.040293562 | 8.7228228 | 8.4654758 | 8.88598 | -0.420506309 | -0.163159334 |
| Gm45716 | 5.167596 | 5.102824 | 5.2371 | -0.134261456 | -0.069490354 | 5.0382772 | 4.7954519 | 5.21615 | -0.4207014 | -0.177876108 |
| Cacna1b | 4.402134 | 4.069462 | 4.5287 | -0.459267443 | -0.126594776 | 4.051591 | 4.0430911 | 4.46393 | -0.420842967 | -0.412343137 |
| Pde8a | 7.459437 | 7.601153 | 7.3931 | 0.208083985 | 0.066367484 | 7.4022913 | 7.0204006 | 7.44139 | -0.420990763 | -0.03910012 |
| Ccni | 12.22219 | 12.13874 | 12.281 | -0.142238766 | -0.05878827 | 12.10588 | 11.780188 | 12.2012 | -0.421055728 | -0.095363444 |
| Dock9 | 9.755289 | 9.397219 | 9.5952 | -0.197988734 | 0.160082068 | 9.3502667 | 9.1857344 | 9.60695 | -0.421211178 | -0.256678943 |
| Wwc2 | 7.532989 | 7.221904 | 7.4436 | -0.221679091 | 0.089406579 | 7.2700594 | 6.8940145 | 7.31552 | -0.421507627 | -0.045462661 |
| Bhlhb9 | 8.085067 | 7.998838 | 8.1217 | -0.122899788 | -0.036670264 | 7.9264205 | 7.8058835 | 8.22742 | -0.421536173 | -0.3009991 |
| Abcc1 | 10.20405 | 10.0629 | 10.109 | -0.046549515 | 0.094600436 | 10.05934 | 9.7551749 | 10.1767 | -0.421567276 | -0.117402198 |
| Ptgdr2 | 3.981726 | 3.784523 | 4.0447 | -0.260165955 | -0.062962619 | 3.8181061 | 3.5587083 | 3.98062 | -0.421908027 | -0.162510263 |
| Mtmr12 | 10.14581 | 10.07205 | 10.153 | -0.081204548 | -0.007446648 | 9.7353338 | 9.6072321 | 10.0292 | -0.421938463 | -0.293836821 |
| Gm3739 | 4.993318 | 4.889773 | 5.105 | -0.215245265 | -0.111699624 | 4.7998019 | 4.6976368 | 5.11962 | -0.421985715 | -0.319820633 |
| Knstrn | 7.509079 | 7.271612 | 7.6261 | -0.354535797 | -0.117068447 | 7.2680913 | 7.2186578 | 7.6407 | -0.422037434 | -0.372603916 |
| Mapk3 | 9.920382 | 9.770948 | 10.089 | -0.317602568 | -0.16816837 | 10.082303 | 9.473338 | 9.8954 | -0.422060818 | 0.186903888 |

| | | | | | | | | | | |
|---|---|---|---|---|---|---|---|---|---|---|
| Sardh | 4.957989 | 4.964813 | 4.8863 | 0.07853093 | 0.071707128 | 4.7650963 | 4.5498945 | 4.97197 | -0.422075746 | -0.206873996 |
| Wscd2 | 5.643814 | 5.470567 | 5.6382 | -0.167652494 | 0.005594345 | 5.5476109 | 5.2946728 | 5.71694 | -0.422271051 | -0.169332931 |
| Disp1 | 7.676158 | 7.339395 | 7.6151 | -0.275703467 | 0.06106034 | 7.4552688 | 7.059306 | 7.48172 | -0.422416208 | -0.026453407 |
| Ppip5k1 | 8.472865 | 8.270749 | 8.3496 | -0.078877502 | 0.123239241 | 8.5480374 | 8.2081139 | 8.63081 | -0.422695262 | -0.082771706 |
| Pias3 | 8.63692 | 8.443274 | 8.6815 | -0.238235346 | -0.044589954 | 8.7211552 | 8.4598759 | 8.88292 | -0.423041523 | -0.161762204 |
| Impdh1 | 10.07671 | 9.924201 | 10.111 | -0.186871029 | -0.034358772 | 9.9781551 | 9.6704287 | 10.0937 | -0.423254825 | -0.115528414 |
| Afap1 | 7.882946 | 7.875232 | 7.8964 | -0.021180942 | -0.013467463 | 7.5376073 | 7.3434171 | 7.76671 | -0.423295185 | -0.229104994 |
| Qsox1 | 9.170392 | 9.148758 | 9.3761 | -0.227383987 | -0.205750581 | 9.0597961 | 8.7686007 | 9.19224 | -0.423635868 | -0.132440437 |
| C1qtnf1 | 7.300308 | 6.913587 | 7.2745 | -0.360939796 | 0.025780759 | 6.6153573 | 6.1676664 | 6.5914 | -0.423735083 | 0.023955827 |
| Kif15 | 7.466833 | 7.435852 | 7.7528 | -0.316942387 | -0.285962206 | 7.428912 | 7.6390017 | 8.06284 | -0.423835535 | -0.633925244 |
| Raph1 | 7.83918 | 7.491103 | 7.4839 | 0.007161661 | 0.3552385 | 7.3588969 | 7.2809598 | 7.70521 | -0.424253817 | -0.346316738 |
| Atp2b1 | 11.07033 | 10.80759 | 10.85 | -0.041940554 | 0.220796714 | 10.653352 | 10.586067 | 11.0103 | -0.424272142 | -0.356987332 |
| Egr3 | 11.09053 | 11.01221 | 11.135 | -0.123027962 | -0.044706404 | 10.680728 | 10.49667 | 10.921 | -0.424358646 | -0.240300631 |
| Bivm | 7.212527 | 7.127826 | 7.333 | -0.205190814 | -0.120490543 | 7.0147673 | 6.6756258 | 7.10013 | -0.424499483 | -0.08535797 |
| Slc43a2 | 10.0415 | 10.27911 | 10.224 | 0.055367616 | -0.18224311 | 10.280569 | 9.8303797 | 10.255 | -0.424611034 | 0.025578416 |
| Fsd2 | 5.91233 | 5.771297 | 6.0456 | -0.274305878 | -0.133272449 | 5.960106 | 5.6433018 | 6.06807 | -0.424768627 | -0.107964408 |
| Syndig1l | 3.37022 | 3.011835 | 3.7276 | -0.715793337 | -0.357408244 | 3.0189471 | 2.9384207 | 3.36326 | -0.424834834 | -0.344308501 |
| Fahd2a | 6.289513 | 6.091216 | 6.2422 | -0.150994586 | 0.047302445 | 6.4470251 | 5.8130419 | 6.23812 | -0.425077486 | 0.208905764 |
| Xlr3a | 3.437797 | 3.250246 | 3.6119 | -0.361674568 | -0.174123265 | 3.4481894 | 3.0684675 | 3.49355 | -0.425083453 | -0.04536159 |
| Frmd5 | 4.061413 | 3.895151 | 4.0808 | -0.185671987 | -0.019410048 | 4.0656537 | 3.7421186 | 4.16734 | -0.425219522 | -0.101684441 |
| Mindy2 | 10.64483 | 10.64689 | 10.752 | -0.104864222 | -0.106926702 | 10.591751 | 10.336569 | 10.7619 | -0.425299774 | -0.170117093 |
| Man1c1 | 7.401609 | 7.428524 | 7.4194 | 0.009142763 | -0.017772439 | 7.1028122 | 6.7627068 | 7.18804 | -0.425332329 | -0.08522695 |
| 9130023H24Rik | 6.641589 | 6.377663 | 6.7149 | -0.337196984 | -0.073271795 | 6.749578 | 6.2899154 | 6.71544 | -0.425524949 | 0.034137645 |
| Gm21092 | 6.451377 | 6.271472 | 6.3652 | -0.093723459 | 0.086181404 | 6.440369 | 6.2829131 | 6.70865 | -0.425735801 | -0.268279958 |
| Tnfaip3 | 12.00779 | 12.00224 | 11.983 | 0.019730202 | 0.025283534 | 11.382342 | 11.581678 | 12.0077 | -0.425975354 | -0.625311303 |
| Prdm10 | 8.831132 | 8.855951 | 8.8001 | 0.055853862 | 0.031034505 | 8.786515 | 8.5556632 | 8.98172 | -0.426052388 | -0.195200617 |
| Cyb5r1 | 8.165421 | 7.930286 | 8.0582 | -0.127903486 | 0.107231299 | 8.1633922 | 7.8046595 | 8.23076 | -0.426104094 | -0.067371406 |
| Ephx1 | 10.78224 | 10.7332 | 10.954 | -0.220383462 | -0.171340528 | 10.878962 | 10.444493 | 10.8706 | -0.426127002 | 0.008342364 |
| Abca3 | 10.57132 | 10.42688 | 10.642 | -0.214972846 | -0.070532412 | 10.57197 | 10.241149 | 10.6673 | -0.426161408 | -0.095340626 |
| Kantr | 7.226475 | 7.276429 | 7.1568 | 0.11966236 | 0.069708856 | 7.4856094 | 6.9292034 | 7.356 | -0.426799051 | 0.129606878 |
| Brca2 | 7.618469 | 7.072126 | 7.5609 | -0.488746935 | 0.057596032 | 7.6669687 | 7.6176893 | 8.04452 | -0.426826226 | -0.37754683 |
| Zfp157 | 8.016734 | 8.117956 | 8.1674 | -0.049478995 | -0.150700899 | 8.1657328 | 7.8026417 | 8.22976 | -0.427115394 | -0.064024285 |

| | | | | | | | | | | |
|---|---|---|---|---|---|---|---|---|---|---|
| Mastl | 5.491857 | 5.180007 | 5.4669 | -0.286932371 | 0.024917438 | 5.3646208 | 5.1840341 | 5.61116 | -0.427126489 | -0.246539781 |
| Aif1l | 4.568661 | 4.398533 | 4.4307 | -0.032192684 | 0.137935463 | 4.2556517 | 3.9977546 | 4.42496 | -0.427208518 | -0.16931143 |
| Arhgef40 | 7.941572 | 7.98431 | 7.8839 | 0.1004299 | 0.057691094 | 7.6328146 | 7.9010758 | 8.32851 | -0.427435012 | -0.695696282 |
| Trbv13-2 | 7.999954 | 7.991396 | 8.1252 | -0.133776437 | -0.12521846 | 7.8023221 | 7.7225066 | 8.15018 | -0.427670467 | -0.347854955 |
| Rnf13 | 9.95512 | 9.869016 | 9.9393 | -0.070289236 | 0.015814773 | 10.001563 | 9.5387348 | 9.96651 | -0.42777693 | 0.035051639 |
| Cdc42ep3 | 8.738183 | 8.718816 | 8.8419 | -0.123052505 | -0.103685194 | 8.5643312 | 8.377465 | 8.80525 | -0.427780528 | -0.240914385 |
| Zmym2 | 10.8618 | 10.76045 | 10.899 | -0.138621142 | -0.037272902 | 10.559136 | 10.470298 | 10.8982 | -0.427942419 | -0.339103932 |
| 9930012K11Rik | 6.195795 | 6.122935 | 6.2885 | -0.16560485 | -0.092743975 | 6.2041588 | 5.7010579 | 6.12921 | -0.428153644 | 0.074947247 |
| Acox1 | 9.417589 | 9.217347 | 9.4427 | -0.225358567 | -0.025116837 | 9.282818 | 8.971092 | 9.39925 | -0.428153798 | -0.116427778 |
| Mmd | 8.158284 | 8.069761 | 8.2745 | -0.204702857 | -0.116179634 | 7.9648765 | 7.7932052 | 8.22146 | -0.42825136 | -0.256580049 |
| Piezo2 | 4.613083 | 4.589865 | 4.5335 | 0.056361531 | 0.079579793 | 4.788936 | 4.4288687 | 4.85804 | -0.429166813 | -0.069099556 |
| Tnfrsf19 | 4.42399 | 4.582969 | 4.477 | 0.105963117 | -0.053016101 | 4.1815733 | 4.0034857 | 4.43275 | -0.429267954 | -0.251180373 |
| Polr3g | 6.877329 | 7.038264 | 7.1427 | -0.104391729 | -0.265327198 | 7.1572016 | 6.7346295 | 7.16394 | -0.429308781 | -0.006736645 |
| Spint2 | 9.742212 | 9.526349 | 9.7198 | -0.193451053 | 0.022412047 | 9.4835664 | 9.1673594 | 9.59681 | -0.429451657 | -0.113244723 |
| 40057 | 11.90459 | 11.68742 | 11.895 | -0.20766493 | 0.009508524 | 11.738572 | 11.374498 | 11.8041 | -0.429618072 | -0.065544179 |
| Jmy | 8.715478 | 8.769455 | 8.811 | -0.041498857 | -0.095475384 | 8.4896963 | 8.4714332 | 8.90116 | -0.429726419 | -0.411463296 |
| Scamp1 | 7.485994 | 6.998941 | 6.9752 | 0.023742586 | 0.510795298 | 6.6058262 | 6.293542 | 6.72366 | -0.430118272 | -0.117834101 |
| Atl1 | 5.261416 | 5.275572 | 5.3332 | -0.057604998 | -0.071760777 | 5.2222528 | 5.0794833 | 5.51049 | -0.431007767 | -0.288238331 |
| Tril | 3.489423 | 3.393956 | 3.7906 | -0.396623312 | -0.301157021 | 3.3241294 | 3.1555263 | 3.58714 | -0.431610651 | -0.263007573 |
| Gcdh | 8.302189 | 8.067592 | 8.2619 | -0.194307643 | 0.040288635 | 8.4864794 | 8.0730718 | 8.50502 | -0.431947372 | -0.01853979 |
| Zfp839 | 8.202108 | 8.115143 | 8.1805 | -0.065395241 | 0.021569377 | 8.2400033 | 7.7403605 | 8.17259 | -0.432231387 | 0.06741144 |
| Unc119 | 8.737856 | 8.535127 | 8.8499 | -0.314759527 | -0.11203084 | 8.9769733 | 8.4057441 | 8.838 | -0.432253042 | 0.138976113 |
| AC098880.2 | 6.969708 | 6.968615 | 7.109 | -0.140378127 | -0.139284808 | 6.881773 | 6.6741475 | 7.1065 | -0.432354767 | -0.224729286 |
| Il27ra | 11.32803 | 11.29665 | 11.344 | -0.047172372 | -0.015793952 | 11.439347 | 11.030106 | 11.4625 | -0.432403 | -0.023161958 |
| Mex3c | 9.598488 | 9.537222 | 9.6014 | -0.064215496 | -0.002949886 | 9.3120789 | 9.0817387 | 9.51443 | -0.432690492 | -0.202350303 |
| Usp2 | 5.353509 | 5.163213 | 4.7457 | 0.417558721 | 0.607854692 | 4.8237258 | 4.589709 | 5.02249 | -0.432782958 | -0.198766202 |
| Rgs13 | 4.195556 | 4.627601 | 4.355 | 0.272646242 | -0.159398848 | 4.135248 | 4.0320805 | 4.46496 | -0.432878909 | -0.329711368 |
| Afmid | 8.1006 | 8.126784 | 8.2064 | -0.079597917 | -0.105781832 | 8.4393065 | 7.8685472 | 8.30147 | -0.432920314 | 0.137839015 |
| Kank2 | 6.841862 | 6.432656 | 6.8237 | -0.391029808 | 0.018175789 | 6.1236785 | 5.9649635 | 6.39798 | -0.433019679 | -0.274304665 |
| Itga2 | 7.444797 | 7.140839 | 7.5915 | -0.450638827 | -0.146680214 | 7.0620073 | 6.921545 | 7.35465 | -0.433103136 | -0.29264081 |
| Slc25a29 | 6.430053 | 6.427236 | 6.4068 | 0.020434932 | 0.023252206 | 6.3614177 | 6.0946071 | 6.52776 | -0.433152576 | -0.166342039 |
| Ercc5 | 9.010579 | 8.874347 | 8.8726 | 0.001774067 | 0.138005994 | 8.921597 | 8.6279375 | 9.06113 | -0.433195771 | -0.139536208 |

| | | | | | | | | | | |
|---|---|---|---|---|---|---|---|---|---|---|
| Eme1 | 5.142342 | 5.189223 | 5.183 | 0.00626773 | -0.04061361 | 5.3576618 | 5.1598473 | 5.5935 | -0.433649036 | -0.235834617 |
| Mad2l1 | 8.415133 | 8.392182 | 8.6119 | -0.21973609 | -0.196784946 | 8.5273892 | 8.3275334 | 8.76133 | -0.433792942 | -0.233937149 |
| Pmm1 | 7.476245 | 7.453085 | 7.5271 | -0.074001689 | -0.050841297 | 7.6941172 | 7.0626765 | 7.49652 | -0.433847885 | 0.197592788 |
| Cep78 | 6.95441 | 6.921052 | 6.9807 | -0.059696405 | -0.026338805 | 7.115518 | 6.6227409 | 7.05669 | -0.433946998 | 0.058830092 |
| Wdr11 | 9.710435 | 9.605517 | 9.6922 | -0.086701212 | 0.018216919 | 9.8347367 | 9.4083494 | 9.84247 | -0.434117475 | -0.007730187 |
| Cep120 | 10.56987 | 10.51595 | 10.608 | -0.092130876 | -0.038215396 | 10.567989 | 10.169741 | 10.6044 | -0.434617318 | -0.036369003 |
| Smurf2 | 10.19324 | 10.07363 | 10.192 | -0.117973363 | 0.001641532 | 9.9230457 | 9.6778181 | 10.1126 | -0.434831643 | -0.189604043 |
| Tbc1d19 | 5.802318 | 5.566921 | 5.7725 | -0.205564298 | 0.029832384 | 5.5286412 | 5.5532459 | 5.98842 | -0.435173312 | -0.459777976 |
| Ankrd13b | 6.898292 | 6.657683 | 6.8761 | -0.218463925 | 0.022144534 | 6.4635505 | 6.3354156 | 6.77066 | -0.435248785 | -0.307113968 |
| Tnfrsf10b | 7.484873 | 7.193635 | 7.3195 | -0.125831717 | 0.165406739 | 7.1398424 | 6.9854546 | 7.42091 | -0.435456155 | -0.281068318 |
| Trpm4 | 6.716407 | 6.673052 | 6.7226 | -0.049527253 | -0.006171651 | 6.8472195 | 6.2772179 | 6.71269 | -0.435471087 | 0.134530543 |
| Uevld | 7.224941 | 7.276538 | 7.4317 | -0.155134268 | -0.206730662 | 7.4195323 | 7.0669286 | 7.50257 | -0.435640121 | -0.083036424 |
| Zfp318 | 11.55742 | 11.65741 | 11.705 | -0.047542232 | -0.147524938 | 11.771963 | 11.391998 | 11.8277 | -0.435747994 | -0.055782756 |
| Stradb | 8.212992 | 7.967285 | 8.1709 | -0.203606868 | 0.042099378 | 7.9627321 | 7.5346831 | 7.97055 | -0.435863652 | -0.007814645 |
| Peg3 | 3.150022 | 3.060322 | 3.2504 | -0.190096397 | -0.100396334 | 3.2480788 | 3.0478287 | 3.48387 | -0.436042366 | -0.235792223 |
| Chaf1a | 7.688032 | 7.597595 | 7.968 | -0.370388127 | -0.279950673 | 7.8213351 | 7.7318806 | 8.16807 | -0.436190656 | -0.346736074 |
| Gng11 | 6.705331 | 6.0974 | 6.691 | -0.593552662 | 0.01437859 | 5.8221278 | 5.4466725 | 5.88318 | -0.436505865 | -0.061050542 |
| Esco2 | 6.038446 | 5.96283 | 6.2709 | -0.308104465 | -0.232488241 | 5.8863841 | 6.0417382 | 6.47825 | -0.436510951 | -0.591865035 |
| Vcl | 8.968864 | 8.801152 | 8.6626 | 0.13856578 | 0.3062782 | 8.5231783 | 8.0855852 | 8.52213 | -0.436542198 | 0.00105091 |
| Mpzl1 | 5.450966 | 5.38533 | 5.4424 | -0.057049519 | 0.008587112 | 5.3408209 | 4.9340912 | 5.3708 | -0.43670899 | -0.0299793 |
| Map3k3 | 10.9027 | 10.93685 | 11.005 | -0.068476536 | -0.10262129 | 10.82394 | 10.484455 | 10.9212 | -0.436750496 | -0.097265131 |
| Hhex | 8.965008 | 8.89855 | 9.0617 | -0.163153747 | -0.096695705 | 9.0998721 | 8.3636813 | 8.80048 | -0.436799578 | 0.299391227 |
| Cenpu | 5.608914 | 5.31075 | 5.4854 | -0.174697111 | 0.123467059 | 5.5109014 | 5.3174304 | 5.75438 | -0.436945714 | -0.243474666 |
| Slc25a40 | 7.577648 | 7.553566 | 7.6197 | -0.066115109 | -0.042033104 | 7.5810042 | 7.4345782 | 7.87164 | -0.437061664 | -0.290635692 |
| Paqr7 | 8.179449 | 8.247778 | 8.3562 | -0.108437791 | -0.176766719 | 8.1000156 | 7.8250598 | 8.26229 | -0.437234169 | -0.162278378 |
| D930048N14Rik | 4.347171 | 4.261011 | 4.4536 | -0.192551049 | -0.106390663 | 4.6110213 | 3.9626149 | 4.39994 | -0.437329008 | 0.211077432 |
| Aga | 7.341637 | 7.230121 | 7.3347 | -0.104600416 | 0.006915959 | 7.4051928 | 6.9365978 | 7.37399 | -0.437387626 | 0.031207295 |
| Plbd1 | 10.62436 | 10.21645 | 10.441 | -0.22421704 | 0.18369642 | 10.647526 | 9.9034676 | 10.3409 | -0.437394088 | 0.306664058 |
| Dusp19 | 6.554654 | 6.634038 | 6.6801 | -0.046084134 | -0.125468062 | 6.8681137 | 6.3394009 | 6.7768 | -0.43739791 | 0.091314876 |
| Cand2 | 7.395958 | 7.12258 | 7.4028 | -0.280180172 | -0.00680243 | 7.2305484 | 6.8323482 | 7.2699 | -0.437554099 | -0.039353963 |
| Zfp7 | 8.471071 | 8.430439 | 8.6114 | -0.180985865 | -0.140354014 | 8.732303 | 8.3358771 | 8.77348 | -0.437606799 | -0.041180871 |
| Atxn3 | 9.515312 | 9.523285 | 9.5521 | -0.028821132 | -0.036794262 | 9.4156341 | 9.1362748 | 9.57391 | -0.437631841 | -0.158272477 |

| | | | | | | | | | | |
|---|---|---|---|---|---|---|---|---|---|---|
| Olfr147 | 6.09356 | 6.087042 | 6.18 | -0.092930534 | -0.086412833 | 6.4858303 | 6.1911951 | 6.62883 | -0.437639773 | -0.143004611 |
| Cdc42ep1 | 5.570924 | 5.275225 | 5.5298 | -0.254560817 | 0.041138423 | 5.2801236 | 4.8006644 | 5.23883 | -0.438165535 | 0.041293686 |
| Mks1 | 7.660293 | 7.626722 | 7.9143 | -0.287576895 | -0.254005494 | 8.2305332 | 7.5585402 | 7.99673 | -0.438185647 | 0.233807319 |
| 1700066M21Rik | 7.013504 | 7.110889 | 7.0105 | 0.100346796 | 0.002961186 | 7.2864632 | 6.8272496 | 7.26556 | -0.438308958 | 0.020904606 |
| Dis3l | 8.718704 | 8.740948 | 8.7147 | 0.026205271 | 0.003961564 | 8.7586615 | 8.2900803 | 8.72839 | -0.438312935 | 0.030268191 |
| Gm48111 | 6.634756 | 6.604538 | 6.8615 | -0.256937018 | -0.226718759 | 6.8396671 | 7.0508148 | 7.48928 | -0.438462211 | -0.649609909 |
| Sv2c | 4.193425 | 4.102542 | 4.2942 | -0.191631818 | -0.100749181 | 4.1755387 | 4.1551417 | 4.5938 | -0.438654198 | -0.418257215 |
| Clec10a | 7.894932 | 6.969947 | 7.5522 | -0.582253509 | 0.342732145 | 7.0324312 | 6.7163703 | 7.15524 | -0.438874193 | -0.122813327 |
| Xk | 7.247523 | 7.26517 | 7.347 | -0.081842757 | -0.099490059 | 7.3860882 | 6.8998286 | 7.33871 | -0.438881293 | 0.047378324 |
| Dhrs1 | 8.483538 | 8.256062 | 8.5161 | -0.26002572 | -0.032550564 | 8.5353942 | 7.95204 | 8.39112 | -0.43908387 | 0.144270367 |
| Dnase1l1 | 7.539708 | 7.325833 | 7.5196 | -0.193730465 | 0.020144485 | 7.6113721 | 7.1925015 | 7.63179 | -0.439283522 | -0.020413007 |
| Nusap1 | 8.036665 | 7.816026 | 8.1012 | -0.285196545 | -0.064557937 | 7.9251896 | 7.9312632 | 8.37067 | -0.439410689 | -0.44548431 |
| Mapre2 | 10.08794 | 10.05206 | 10.114 | -0.062040825 | -0.026156417 | 9.9906059 | 9.5716156 | 10.0112 | -0.439557654 | -0.02056738 |
| Crip2 | 7.488202 | 7.299466 | 7.5929 | -0.29338765 | -0.104652148 | 7.0218325 | 6.4153175 | 6.8549 | -0.439578806 | 0.166936205 |
| Olfr1388 | 4.917955 | 4.88102 | 4.9941 | -0.113115208 | -0.076180952 | 5.0201634 | 4.7977306 | 5.2374 | -0.43966776 | -0.21723495 |
| Svep1 | 5.734421 | 5.324851 | 5.1968 | 0.128072265 | 0.537641815 | 5.2464053 | 4.8441744 | 5.28399 | -0.439810662 | -0.037579799 |
| Thsd4 | 4.947998 | 4.899035 | 5.0019 | -0.102893393 | -0.053929642 | 4.7622529 | 4.5344186 | 4.97432 | -0.439899371 | -0.212064988 |
| Acss2 | 9.166173 | 9.295651 | 9.3086 | -0.012985782 | -0.142464525 | 8.9552929 | 8.8554627 | 9.29543 | -0.439962372 | -0.340132219 |
| D830030K20Rik | 5.26149 | 5.047583 | 5.3205 | -0.272924641 | -0.05901742 | 5.0175927 | 5.0743142 | 5.51438 | -0.440062624 | -0.49678408 |
| P4htm | 5.322851 | 5.480403 | 5.565 | -0.084630867 | -0.242182956 | 5.4530388 | 5.0869413 | 5.52702 | -0.440082919 | -0.073985418 |
| Specc1l | 10.90355 | 10.87818 | 10.925 | -0.04726243 | -0.021891202 | 10.759059 | 10.468333 | 10.9086 | -0.44028075 | -0.149554154 |
| Dap | 9.84783 | 9.761283 | 9.9942 | -0.232888646 | -0.146341491 | 9.9005157 | 9.5349723 | 9.97529 | -0.44031872 | -0.074775256 |
| Tmx4 | 9.123415 | 9.104713 | 9.1988 | -0.094076184 | -0.075374161 | 8.935262 | 8.6929939 | 9.13404 | -0.441045538 | -0.198777374 |
| Selp | 8.452411 | 7.979631 | 7.6873 | 0.292292424 | 0.765071763 | 7.4349492 | 6.8764843 | 7.31758 | -0.441098149 | 0.117366687 |
| Trove2 | 7.511561 | 7.219827 | 7.5065 | -0.286703602 | 0.005030135 | 6.8548869 | 6.8969658 | 7.33813 | -0.441169221 | -0.483248029 |
| Loxhd1 | 3.830024 | 3.739683 | 3.9138 | -0.174156893 | -0.083816072 | 3.9747704 | 3.5383323 | 3.97955 | -0.441222259 | -0.004784101 |
| Ogn | 3.996721 | 3.670894 | 4.0089 | -0.338023299 | -0.012196086 | 3.5102165 | 3.3937869 | 3.83504 | -0.441250876 | -0.324821289 |
| Gm7435 | 4.288532 | 4.193921 | 4.3316 | -0.137649221 | -0.043037921 | 4.13623 | 4.0733527 | 4.51465 | -0.441296565 | -0.378419331 |
| Bok | 5.31363 | 5.078132 | 5.1529 | -0.074802974 | 0.160695901 | 4.7655288 | 4.417737 | 4.85905 | -0.441308907 | -0.093517113 |
| Adamtsl2 | 3.904377 | 3.731387 | 3.9695 | -0.238131107 | -0.065140358 | 3.6337214 | 3.495345 | 3.93666 | -0.441311482 | -0.302935154 |
| Mospd3 | 8.446323 | 8.488433 | 8.6565 | -0.168030562 | -0.21014064 | 8.6143466 | 8.0270221 | 8.46843 | -0.441404945 | 0.145919544 |
| Sytl1 | 7.104715 | 6.965803 | 7.2654 | -0.299604339 | -0.160691555 | 6.8778205 | 6.6686818 | 7.11015 | -0.441463619 | -0.232324975 |

| | | | | | | | | | | |
|---|---|---|---|---|---|---|---|---|---|---|
| Klhl25 | 8.602859 | 8.699999 | 8.6945 | 0.005527109 | -0.09161332 | 8.6364276 | 8.2896749 | 8.73123 | -0.441550937 | -0.094798217 |
| Tcp11l2 | 10.87807 | 10.83466 | 10.92 | -0.084979466 | -0.041565954 | 10.873224 | 10.48052 | 10.9223 | -0.441763102 | -0.049059489 |
| Shld2 | 8.254631 | 8.077465 | 8.4139 | -0.336470351 | -0.159304549 | 8.3171767 | 7.8897264 | 8.33161 | -0.441887161 | -0.014436875 |
| Srpk2 | 10.34454 | 10.28495 | 10.396 | -0.111345962 | -0.051759026 | 10.175818 | 9.9741842 | 10.4161 | -0.441888915 | -0.240255207 |
| Dip2c | 6.637938 | 6.790211 | 6.694 | 0.09624525 | -0.056027809 | 6.3021774 | 6.1791525 | 6.62107 | -0.44191537 | -0.318890472 |
| Haus8 | 8.534962 | 8.54041 | 8.5838 | -0.043388382 | -0.048836192 | 8.7143833 | 8.3410839 | 8.78306 | -0.441974768 | -0.068675366 |
| Armcx1 | 6.007006 | 5.673563 | 5.9768 | -0.303235547 | 0.030207162 | 5.3552274 | 5.2369625 | 5.679 | -0.442037325 | -0.323772437 |
| Scn2b | 6.100302 | 6.085872 | 6.2783 | -0.192433248 | -0.178003025 | 6.0367981 | 5.8786614 | 6.32074 | -0.442078132 | -0.283941516 |
| Incenp | 9.150029 | 8.89239 | 9.266 | -0.373652986 | -0.116013986 | 9.1575639 | 9.0235397 | 9.46618 | -0.442641372 | -0.30861718 |
| Gk5 | 6.303461 | 6.177215 | 6.3841 | -0.20690879 | -0.080662882 | 6.1500501 | 6.0091208 | 6.45197 | -0.442848037 | -0.30191879 |
| Hmcn1 | 4.244514 | 4.121687 | 4.2087 | -0.087044208 | 0.035782765 | 3.358356 | 3.7256576 | 4.16858 | -0.442926997 | -0.810228589 |
| Fam13b | 10.14904 | 10.07487 | 10.213 | -0.138620818 | -0.06445429 | 9.9516354 | 9.8307813 | 10.2737 | -0.442956574 | -0.322102473 |
| Btd | 7.905295 | 7.861335 | 7.9754 | -0.114051096 | -0.070090167 | 7.9565162 | 7.557444 | 8.00043 | -0.442981447 | -0.043909208 |
| Pdlim3 | 3.806946 | 3.77226 | 3.7985 | -0.02622945 | 0.008456294 | 3.7723471 | 3.323416 | 3.76651 | -0.443095819 | 0.005835222 |
| Syp | 4.870556 | 4.689266 | 4.7695 | -0.080222551 | 0.101067434 | 4.6162083 | 4.5745407 | 5.01794 | -0.443403822 | -0.401736251 |
| Cln5 | 8.842757 | 8.748336 | 8.9866 | -0.238288401 | -0.143867387 | 9.0025213 | 8.504418 | 8.94792 | -0.443498381 | 0.054604988 |
| Cd28 | 10.48711 | 10.52832 | 10.668 | -0.139984126 | -0.18119116 | 10.170172 | 10.307031 | 10.7506 | -0.443573804 | -0.580433723 |
| Bnip3l | 11.11381 | 11.02484 | 11.257 | -0.23199054 | -0.14301991 | 11.097557 | 10.739288 | 11.1829 | -0.443609966 | -0.085341381 |
| Fam102b | 8.537376 | 8.563679 | 8.6595 | -0.095845852 | -0.122149187 | 8.3651419 | 8.0831615 | 8.52701 | -0.443845153 | -0.161864706 |
| Slc41a3 | 6.525141 | 6.523902 | 6.6419 | -0.117987104 | -0.11674813 | 6.6173926 | 6.1130389 | 6.55725 | -0.444213256 | 0.060140448 |
| Ctnnd1 | 9.236511 | 8.717137 | 8.9352 | -0.218090363 | 0.301283807 | 8.3748918 | 8.163657 | 8.60789 | -0.44423267 | -0.232997838 |
| Pitpnc1 | 10.63632 | 10.52553 | 10.481 | 0.0440614 | 0.154853024 | 10.279496 | 10.128533 | 10.5734 | -0.444828862 | -0.2938654 |
| Tdrd3 | 8.367483 | 8.180111 | 8.2569 | -0.076742366 | 0.110630091 | 8.2268947 | 7.9736138 | 8.41863 | -0.44501924 | -0.191738299 |
| Pla2g6 | 7.104247 | 7.140399 | 7.3153 | -0.174868976 | -0.211020516 | 7.4735682 | 7.0210244 | 7.46617 | -0.445145862 | 0.007398019 |
| Casp6 | 8.344711 | 8.166759 | 8.2721 | -0.105353333 | 0.072598513 | 8.4563132 | 7.8494804 | 8.29471 | -0.445233578 | 0.161599197 |
| Xkr6 | 4.367881 | 4.169853 | 4.3313 | -0.161473527 | 0.036554774 | 4.0764439 | 4.008962 | 4.45478 | -0.445816587 | -0.378334644 |
| Tmem35b | 6.641105 | 6.574571 | 6.597 | -0.022423243 | 0.044110375 | 6.9529571 | 6.3846847 | 6.83056 | -0.445874583 | 0.122397865 |
| Gpsm1 | 7.897677 | 7.883065 | 7.8564 | 0.026714422 | 0.04132621 | 7.9037801 | 7.6341606 | 8.08032 | -0.446157138 | -0.176537583 |
| Ivd | 9.850042 | 9.757731 | 9.8922 | -0.134437564 | -0.042127233 | 10.103512 | 9.6193011 | 10.0656 | -0.446318902 | 0.037891992 |
| Papss1 | 8.861324 | 8.9757 | 9.0241 | -0.048424788 | -0.162801757 | 8.8539831 | 8.5205239 | 8.967 | -0.446471795 | -0.113012547 |
| Fuz | 8.061459 | 7.973556 | 8.2445 | -0.270895457 | -0.182991887 | 8.5792396 | 7.9390309 | 8.38583 | -0.446799946 | 0.19340875 |
| Gm14410 | 5.854423 | 5.418203 | 5.6903 | -0.272066306 | 0.164152913 | 5.5736332 | 5.2930166 | 5.73982 | -0.446807576 | -0.166190978 |

| | | | | | | | | | | |
|---|---|---|---|---|---|---|---|---|---|---|
| Svil | 10.0505 | 9.93944 | 10.057 | -0.117149036 | -0.006090045 | 9.9126748 | 9.6251376 | 10.0722 | -0.447107202 | -0.159569987 |
| Dmrta1 | 4.943892 | 4.667722 | 4.9951 | -0.327366994 | -0.051197206 | 4.6131159 | 4.5340655 | 4.9817 | -0.447635911 | -0.36858559 |
| Ccdc14 | 6.695613 | 6.577889 | 6.7923 | -0.214377893 | -0.096653928 | 6.671623 | 6.5808856 | 7.02881 | -0.447928506 | -0.35719111 |
| Mut | 8.481758 | 8.407559 | 8.5806 | -0.17306837 | -0.098868798 | 8.5665833 | 8.1781255 | 8.62634 | -0.44821375 | -0.059756006 |
| Dhtkd1 | 6.785011 | 6.661703 | 6.5922 | 0.069513824 | 0.19282221 | 6.7413022 | 6.3461962 | 6.79474 | -0.448541732 | -0.053435717 |
| Ptprs | 8.60928 | 8.553816 | 8.7868 | -0.232962972 | -0.177498984 | 8.5895343 | 8.4904734 | 8.93916 | -0.448688393 | -0.34962748 |
| Map4k4 | 11.86253 | 11.85331 | 11.968 | -0.114251158 | -0.10502851 | 11.942364 | 11.632909 | 12.0816 | -0.448725878 | -0.139271563 |
| Sorcs2 | 7.127289 | 7.209999 | 7.1518 | 0.058208488 | -0.024501521 | 7.0546159 | 6.5328972 | 6.98176 | -0.448862021 | 0.072856689 |
| Fkbp9 | 7.530593 | 7.035356 | 7.4071 | -0.371695488 | 0.123541925 | 6.7874679 | 6.5023854 | 6.95147 | -0.449086586 | -0.164004137 |
| 1700073E17Rik | 3.561692 | 3.553145 | 3.5916 | -0.038490974 | -0.029943909 | 3.7319031 | 3.4270788 | 3.87629 | -0.449209671 | -0.144385422 |
| Retnlg | 5.905048 | 4.642364 | 4.4501 | 0.19224097 | 1.454925748 | 4.6166352 | 4.0934449 | 4.54282 | -0.449379159 | 0.07381111 |
| Hyi | 7.109039 | 6.979018 | 7.2383 | -0.259231809 | -0.129210977 | 7.1727219 | 6.8516099 | 7.30101 | -0.449401113 | -0.128289169 |
| Lmbrd2 | 8.968023 | 8.778911 | 8.8618 | -0.082878468 | 0.106233772 | 8.7853719 | 8.5242036 | 8.97366 | -0.44945251 | -0.188284289 |
| Plxnb1 | 5.974982 | 5.994689 | 5.9492 | 0.045489135 | 0.025781834 | 5.5657816 | 5.4538621 | 5.90364 | -0.44977675 | -0.337857186 |
| Zfp326 | 8.558971 | 8.492092 | 8.531 | -0.038934971 | 0.027944125 | 8.5092566 | 8.2118775 | 8.66177 | -0.449896487 | -0.1525174 |
| Nim1k | 4.459071 | 4.397321 | 4.5216 | -0.124315222 | -0.06256553 | 4.2567965 | 4.0377485 | 4.48799 | -0.450236933 | -0.231188903 |
| Socs5 | 8.915124 | 8.832917 | 8.8218 | 0.01108124 | 0.093288513 | 8.801304 | 8.4056443 | 8.85636 | -0.450720641 | -0.055060959 |
| mt-Nd5 | 14.87424 | 14.66879 | 14.889 | -0.220053506 | -0.01460147 | 14.632702 | 14.402234 | 14.8533 | -0.45108443 | -0.220615973 |
| Rnf38 | 10.84762 | 10.89508 | 10.958 | -0.062542716 | -0.110004868 | 10.827563 | 10.557713 | 11.0089 | -0.451232064 | -0.181382183 |
| Ganc | 9.735877 | 9.416338 | 9.6387 | -0.222357611 | 0.097181016 | 9.8608364 | 9.3065226 | 9.75783 | -0.451304268 | 0.103009485 |
| Angptl7 | 4.093471 | 3.724262 | 4.2116 | -0.487314043 | -0.11810498 | 3.7005227 | 3.5166319 | 3.96811 | -0.451481936 | -0.26759109 |
| Gapt | 7.253612 | 7.265564 | 7.2019 | 0.063659017 | 0.051706705 | 7.6901124 | 6.8996077 | 7.35173 | -0.452121062 | 0.338383646 |
| Il7 | 4.776763 | 4.392859 | 4.6165 | -0.223648327 | 0.160255905 | 4.5198747 | 4.145758 | 4.59821 | -0.452456012 | -0.078339344 |
| Akap7 | 6.162988 | 6.19967 | 6.3814 | -0.181773678 | -0.218455462 | 6.2325436 | 5.9786967 | 6.4313 | -0.45259863 | -0.198751814 |
| Fam120c | 7.63074 | 7.697485 | 7.688 | 0.009461422 | -0.057283641 | 7.5481433 | 7.2573193 | 7.70998 | -0.452664294 | -0.161840332 |
| Pola1 | 8.812909 | 8.659486 | 8.8291 | -0.169633274 | -0.016210899 | 8.8475205 | 8.5946114 | 9.04756 | -0.452949593 | -0.200040456 |
| Gys1 | 8.25105 | 8.190945 | 8.2856 | -0.094613074 | -0.034507757 | 8.4164301 | 7.9146845 | 8.36768 | -0.452993498 | 0.048752049 |
| Adgrg1 | 7.895556 | 7.479975 | 7.7922 | -0.312201862 | 0.103378548 | 6.9798548 | 6.7055434 | 7.15865 | -0.453110119 | -0.17879876 |
| Camsap2 | 9.512284 | 9.455086 | 9.5078 | -0.052741808 | 0.004456544 | 9.1485413 | 9.013632 | 9.46708 | -0.453447956 | -0.318538607 |
| Ttc23 | 4.993113 | 4.910265 | 5.2314 | -0.321158503 | -0.23831052 | 5.1560243 | 4.8900205 | 5.34357 | -0.453550156 | -0.187546352 |
| Ophn1 | 5.079843 | 5.036825 | 4.6763 | 0.360529751 | 0.403547385 | 4.5513412 | 4.3515118 | 4.80514 | -0.453623758 | -0.253794423 |
| Rtl6 | 8.172862 | 8.147679 | 8.2417 | -0.094029525 | -0.068847515 | 8.4277547 | 7.7751404 | 8.22913 | -0.453985567 | 0.198628754 |

| | | | | | | | | | | |
|---|---|---|---|---|---|---|---|---|---|---|
| Igha | 6.919865 | 7.187185 | 7.3743 | -0.18714246 | -0.45446315 | 7.9613339 | 6.7028507 | 7.15684 | -0.453989111 | 0.80449405 |
| Nbeal1 | 9.374185 | 9.411814 | 9.4386 | -0.026811187 | -0.064440137 | 9.1204332 | 9.0213328 | 9.47552 | -0.454190399 | -0.35508998 |
| Slc25a42 | 6.308277 | 6.324871 | 6.4101 | -0.085242902 | -0.101836694 | 6.4375117 | 6.0110654 | 6.46528 | -0.454212956 | -0.027766696 |
| Antxr1 | 6.37908 | 6.047593 | 6.2675 | -0.219946042 | 0.111540589 | 5.7861895 | 5.5466721 | 6.00097 | -0.454298152 | -0.214780734 |
| St8sia4 | 11.48011 | 11.57411 | 11.555 | 0.019377732 | -0.074614374 | 11.375286 | 11.045852 | 11.5004 | -0.454501096 | -0.125067242 |
| Lmnb1 | 10.45985 | 10.22394 | 10.389 | -0.164697614 | 0.071208292 | 10.410081 | 10.157304 | 10.6119 | -0.45457386 | -0.201797344 |
| Tgfbr1 | 9.644161 | 9.700894 | 9.7059 | -0.004979584 | -0.061712532 | 9.3263167 | 9.174283 | 9.629 | -0.454712124 | -0.302678437 |
| Trpv2 | 10.37537 | 10.38998 | 10.477 | -0.087458692 | -0.102069774 | 10.561826 | 10.062326 | 10.5173 | -0.454959609 | 0.044541117 |
| Tmem132c | 3.722904 | 3.556197 | 3.6943 | -0.138071343 | 0.028635674 | 3.842147 | 3.3450825 | 3.80019 | -0.455110001 | 0.041954486 |
| Kif13a | 7.108889 | 7.008974 | 7.3215 | -0.312491329 | -0.212576039 | 7.1864363 | 6.8554919 | 7.31105 | -0.455556737 | -0.124612316 |
| Decr1 | 7.944367 | 7.747496 | 8.0571 | -0.309650613 | -0.112780154 | 7.9797621 | 7.4894159 | 7.94504 | -0.455628272 | 0.034717898 |
| Scai | 9.320742 | 9.142211 | 9.296 | -0.153782852 | 0.02474885 | 9.3468368 | 9.0647118 | 9.52034 | -0.455629734 | -0.173504756 |
| Gal3st2b | 5.179606 | 5.173302 | 5.5922 | -0.418916911 | -0.412612608 | 4.8803473 | 4.7713032 | 5.22696 | -0.455656282 | -0.346612231 |
| Grhl2 | 3.137519 | 3.137292 | 3.1692 | -0.031890345 | -0.031663638 | 3.4337087 | 3.0949315 | 3.55098 | -0.456052093 | -0.117274948 |
| Mars2 | 8.953245 | 8.924708 | 8.984 | -0.059315357 | -0.030778222 | 9.2630152 | 8.790843 | 9.2471 | -0.456256574 | 0.015915637 |
| Zfp512 | 9.131377 | 8.873008 | 9.2036 | -0.330545777 | -0.072176859 | 9.1885768 | 8.8988386 | 9.35514 | -0.456304046 | -0.166565909 |
| Apold1 | 8.478303 | 8.018608 | 7.8323 | 0.186318899 | 0.646014 | 6.6325169 | 6.6612201 | 7.11755 | -0.456326174 | -0.485029337 |
| Smpd5 | 6.851366 | 6.637205 | 7.0867 | -0.449467004 | -0.235305284 | 7.0605475 | 6.5822918 | 7.03868 | -0.456383755 | 0.021871972 |
| Prickle2 | 5.217862 | 4.917268 | 5.1226 | -0.205291098 | 0.095302945 | 4.9472858 | 4.7606264 | 5.21722 | -0.45659086 | -0.26993139 |
| Il6ra | 11.27088 | 11.22551 | 11.224 | 0.001148662 | 0.046512428 | 10.977382 | 10.66677 | 11.1236 | -0.456796852 | -0.146184738 |
| Nos1 | 3.309272 | 3.023565 | 3.6323 | -0.608747343 | -0.323040053 | 3.7065126 | 3.1486524 | 3.60567 | -0.457017653 | 0.100842601 |
| Shcbp1l | 4.599283 | 4.692125 | 4.724 | -0.031900044 | -0.124742005 | 4.9588733 | 4.5330435 | 4.99007 | -0.457025825 | -0.031196042 |
| Plxna3 | 6.895383 | 6.776696 | 7.1062 | -0.329465138 | -0.210778152 | 7.3108347 | 6.643646 | 7.10099 | -0.457348026 | 0.209840696 |
| Ptgs1 | 5.677823 | 5.002786 | 5.5486 | -0.54582084 | 0.129215438 | 5.087118 | 4.5940448 | 5.05144 | -0.45739628 | 0.035676907 |
| Smpd1 | 8.730131 | 8.727933 | 8.887 | -0.159062158 | -0.156864323 | 8.8651932 | 8.2234717 | 8.6809 | -0.457426488 | 0.18429499 |
| Ate1 | 9.665519 | 9.671151 | 9.7622 | -0.091036557 | -0.096669147 | 9.6125994 | 9.2658147 | 9.72364 | -0.457826301 | -0.111041607 |
| Osgepl1 | 7.467076 | 7.363936 | 7.4306 | -0.066664626 | 0.036475351 | 7.6671588 | 7.145039 | 7.60303 | -0.457987006 | 0.064132733 |
| Ciita | 12.72753 | 12.62954 | 12.808 | -0.178834876 | -0.08085249 | 13.214538 | 12.441036 | 12.8992 | -0.45818358 | 0.315318255 |
| Fam114a1 | 7.181901 | 6.933641 | 7.2066 | -0.272932994 | -0.024673304 | 7.1113516 | 6.539182 | 6.99745 | -0.4582663 | 0.1139033 |
| Tcf7l1 | 6.342747 | 6.146408 | 6.3491 | -0.202665531 | -0.006326468 | 5.8942026 | 5.5963001 | 6.05479 | -0.458490301 | -0.160587809 |
| Mfhas1 | 9.24447 | 9.376954 | 9.3889 | -0.011962567 | -0.144446213 | 9.0857198 | 8.7929325 | 9.25142 | -0.458491054 | -0.165703692 |
| Rtl8c | 4.865587 | 4.602329 | 4.895 | -0.292701041 | -0.029442872 | 4.8078991 | 4.3706501 | 4.82922 | -0.458571199 | -0.021322264 |

| | | | | | | | | | | |
|---|---|---|---|---|---|---|---|---|---|---|
| Cyp4f13 | 8.582104 | 8.550178 | 8.6922 | -0.141980561 | -0.110054376 | 9.0132705 | 8.542475 | 9.00111 | -0.458640015 | 0.012155558 |
| Kif2c | 6.449791 | 6.301023 | 6.5559 | -0.254894715 | -0.106126869 | 6.4701195 | 6.3145398 | 6.77335 | -0.458807815 | -0.303228099 |
| Gpr182 | 8.160305 | 7.82405 | 7.8921 | -0.068055171 | 0.268200496 | 7.2246076 | 6.680547 | 7.13945 | -0.458898483 | 0.085162185 |
| Asf1b | 7.690415 | 7.526207 | 7.7886 | -0.262398408 | -0.098190361 | 7.6286128 | 7.4829948 | 7.9419 | -0.458903729 | -0.313285709 |
| Manea | 9.398944 | 9.468519 | 9.3782 | 0.090284664 | 0.020709794 | 9.5221642 | 8.9455793 | 9.40448 | -0.458905287 | 0.117679646 |
| Phf24 | 5.890703 | 5.592365 | 5.6861 | -0.093741714 | 0.204595712 | 5.5573011 | 5.4980735 | 5.95698 | -0.458908619 | -0.399680963 |
| Ctc1 | 8.774466 | 8.848522 | 8.8261 | 0.022384979 | -0.051670987 | 9.0844381 | 8.7358293 | 9.19487 | -0.459043642 | -0.110434845 |
| Col14a1 | 5.289521 | 5.142182 | 5.3675 | -0.225302051 | -0.077963595 | 4.8913252 | 4.6489614 | 5.10805 | -0.459085725 | -0.216721878 |
| Map4k2 | 11.33967 | 11.313 | 11.533 | -0.219507776 | -0.19283504 | 11.841368 | 11.425602 | 11.8848 | -0.45915428 | -0.043388417 |
| 4931406C07Rik | 7.125626 | 6.957301 | 7.1278 | -0.170521502 | -0.002196411 | 6.9822156 | 6.7098577 | 7.16915 | -0.459295945 | -0.186938021 |
| Gtf2ird1 | 6.946853 | 6.863331 | 7.0496 | -0.186224915 | -0.10270247 | 6.801442 | 6.5194312 | 6.97881 | -0.459378439 | -0.177367684 |
| Fech | 9.069601 | 8.958915 | 9.2347 | -0.275774704 | -0.165088031 | 8.9520224 | 8.530383 | 8.98996 | -0.45957616 | -0.037936842 |
| Tas1r3 | 4.684951 | 4.611985 | 4.6946 | -0.082624205 | -0.009657624 | 4.6437414 | 4.3271657 | 4.78678 | -0.459618175 | -0.143042503 |
| Map3k12 | 6.326127 | 6.421888 | 6.6076 | -0.185680439 | -0.281440986 | 6.2679449 | 6.1317658 | 6.59159 | -0.459822863 | -0.323643809 |
| Ptpn21 | 7.536173 | 7.412282 | 7.6448 | -0.232485961 | -0.108594974 | 7.2844215 | 7.1080327 | 7.56811 | -0.460082066 | -0.283693243 |
| Dscaml1 | 6.316232 | 6.415749 | 6.6083 | -0.192565672 | -0.292083076 | 5.9037012 | 6.0515825 | 6.51177 | -0.460185329 | -0.608066656 |
| Ckap4 | 9.016586 | 8.939766 | 8.8921 | 0.047692623 | 0.124513072 | 9.1470779 | 8.1679017 | 8.62812 | -0.460221174 | 0.518954968 |
| Prkar2a | 8.642484 | 8.391827 | 8.4128 | -0.020990265 | 0.229667201 | 8.1086083 | 8.0281122 | 8.48874 | -0.460625148 | -0.380129064 |
| Ighv2-5 | 6.074605 | 5.904958 | 6.1822 | -0.277205326 | -0.107558197 | 6.2035554 | 5.760273 | 6.22104 | -0.460770937 | -0.017488514 |
| Gpr155 | 8.603159 | 8.298458 | 8.6234 | -0.324953595 | -0.020252748 | 8.5482847 | 8.1378182 | 8.59875 | -0.460935089 | -0.050468617 |
| Gpat3 | 7.163287 | 6.870937 | 7.1051 | -0.234211796 | 0.058137702 | 7.3730402 | 6.6716345 | 7.13261 | -0.460971176 | 0.240434502 |
| Rcbtb2 | 8.855328 | 8.780178 | 9.0599 | -0.279693813 | -0.20454357 | 8.848861 | 8.5800463 | 9.04103 | -0.460983773 | -0.192169052 |
| Ptn | 5.170752 | 4.67702 | 5.0593 | -0.382235027 | 0.111496863 | 4.1782353 | 4.330349 | 4.79155 | -0.461200882 | -0.613314581 |
| Zbtb41 | 8.654273 | 8.609145 | 8.6667 | -0.057573419 | -0.012445679 | 8.5278092 | 8.2132473 | 8.67501 | -0.461766306 | -0.147204431 |
| Aak1 | 11.53975 | 11.45633 | 11.464 | -0.008030384 | 0.075387724 | 10.993378 | 11.065696 | 11.5276 | -0.461929034 | -0.534246908 |
| Tmem231 | 5.63331 | 5.259056 | 5.5577 | -0.298609651 | 0.07564478 | 5.7984425 | 5.2409132 | 5.70285 | -0.461932895 | 0.095596419 |
| Ska2 | 7.372176 | 6.961264 | 7.3248 | -0.363571329 | 0.047339986 | 6.9325843 | 7.0889783 | 7.55107 | -0.46209614 | -0.618490208 |
| Qser1 | 8.959971 | 8.911059 | 8.9535 | -0.042401089 | 0.006511142 | 8.8334049 | 8.5796819 | 9.0418 | -0.462120813 | -0.208397783 |
| Prrg3 | 3.781802 | 3.535751 | 3.7403 | -0.204558952 | 0.041492161 | 3.5338243 | 3.5086085 | 3.97091 | -0.462305233 | -0.437089434 |
| Fam13c | 4.82689 | 4.822539 | 5.0955 | -0.272988425 | -0.268637848 | 4.5107592 | 4.2779553 | 4.7404 | -0.46244813 | -0.229644324 |
| Plvap | 10.70876 | 10.23572 | 10.641 | -0.405560162 | 0.06747687 | 9.7927109 | 9.4368477 | 9.89935 | -0.462499036 | -0.10663586 |
| Acvr1b | 9.640384 | 9.538268 | 9.6573 | -0.119070465 | -0.016954463 | 9.7232583 | 9.1367201 | 9.59925 | -0.462525781 | 0.124012452 |

| | | | | | | | | | | |
|---|---|---|---|---|---|---|---|---|---|---|
| Snx13 | 9.48323 | 9.370077 | 9.4381 | -0.068002237 | 0.04515064 | 9.3091433 | 8.9787559 | 9.44148 | -0.462720751 | -0.132333342 |
| 2010315B03Rik | 7.065785 | 6.722204 | 6.998 | -0.275773848 | 0.067807337 | 7.0391701 | 6.6840352 | 7.14706 | -0.463023402 | -0.107888494 |
| Tulp4 | 10.48841 | 10.41083 | 10.442 | -0.031559498 | 0.046026074 | 10.376046 | 10.104231 | 10.5673 | -0.463117049 | -0.191301526 |
| Tns1 | 9.594432 | 9.625656 | 9.5653 | 0.060335026 | 0.029111168 | 8.6350764 | 8.7076746 | 9.17129 | -0.463618898 | -0.536217092 |
| Hsd11b1 | 9.248423 | 9.190178 | 9.3551 | -0.164883977 | -0.106639265 | 9.2856228 | 8.9008883 | 9.36483 | -0.463939703 | -0.079205226 |
| Etv1 | 4.124189 | 4.0517 | 4.1843 | -0.132607776 | -0.060119349 | 4.2018307 | 3.868792 | 4.33275 | -0.463953048 | -0.130914417 |
| Dclre1a | 6.456137 | 6.044244 | 6.4266 | -0.382342533 | 0.029550619 | 6.694751 | 6.2105854 | 6.67464 | -0.464052909 | 0.02011268 |
| Slco4a1 | 6.137759 | 5.833533 | 6.2662 | -0.432617377 | -0.128391891 | 5.9405176 | 5.686405 | 6.15059 | -0.464180802 | -0.21006817 |
| Map3k20 | 6.90713 | 6.715018 | 6.7965 | -0.081454446 | 0.110656983 | 6.7514828 | 6.2803604 | 6.7446 | -0.464236301 | 0.006886125 |
| Efr3b | 5.175416 | 5.139942 | 5.1507 | -0.010745755 | 0.024728081 | 5.2818557 | 5.0677177 | 5.53203 | -0.464307354 | -0.250169371 |
| Cdk1 | 7.364363 | 7.17192 | 7.4615 | -0.289624229 | -0.097180468 | 7.4021881 | 7.2987019 | 7.7634 | -0.464700568 | -0.361214342 |
| Cracr2a | 9.436426 | 9.238127 | 9.2696 | -0.031475059 | 0.166823064 | 9.1618592 | 9.1025251 | 9.56723 | -0.464701527 | -0.405367493 |
| Phf14 | 10.46758 | 10.2555 | 10.459 | -0.203697733 | 0.008384694 | 10.334962 | 10.005826 | 10.4706 | -0.464821838 | -0.135685838 |
| Plekha5 | 8.683499 | 8.665058 | 8.7901 | -0.125031231 | -0.106589945 | 8.5404998 | 8.3552938 | 8.82017 | -0.464874276 | -0.279668243 |
| Ski | 10.69718 | 10.61013 | 10.72 | -0.110006674 | -0.022955568 | 10.238351 | 10.077569 | 10.5426 | -0.465066166 | -0.304283607 |
| Zfp280c | 8.725238 | 8.609934 | 8.9095 | -0.299518777 | -0.184214227 | 8.8429798 | 8.5705301 | 9.03565 | -0.465118571 | -0.192668821 |
| Stk10 | 12.40534 | 12.50357 | 12.501 | 0.002559922 | -0.095676882 | 12.387614 | 12.000551 | 12.4657 | -0.465145394 | -0.078082356 |
| Extl3 | 10.0703 | 10.01964 | 10.148 | -0.128024202 | -0.077366373 | 9.7721008 | 9.5644191 | 10.0296 | -0.465173091 | -0.257491402 |
| Robo1 | 6.635483 | 6.26897 | 6.5297 | -0.260711503 | 0.105800715 | 6.0380245 | 5.726326 | 6.19196 | -0.465637072 | -0.15393853 |
| Gpr68 | 9.150041 | 9.341521 | 9.3322 | 0.009300934 | -0.18217865 | 9.046782 | 8.8520345 | 9.31779 | -0.465759388 | -0.2710119 |
| Cd44 | 11.4076 | 11.36782 | 11.128 | 0.240240448 | 0.28001512 | 10.835732 | 10.591272 | 11.0572 | -0.46590122 | -0.221440591 |
| Ptprk | 7.64897 | 7.259129 | 7.6126 | -0.353423663 | 0.036417593 | 6.6629878 | 6.3711007 | 6.83743 | -0.466330784 | -0.174443705 |
| Acaca | 10.01166 | 9.900161 | 9.9698 | -0.069598545 | 0.041898336 | 9.9756372 | 9.5474327 | 10.0138 | -0.46640166 | -0.038197127 |
| Chd3 | 12.70383 | 12.78733 | 12.846 | -0.059033672 | -0.142534464 | 12.615422 | 12.477598 | 12.944 | -0.46644412 | -0.328620431 |
| Cdca2 | 6.543929 | 6.463491 | 6.776 | -0.312472586 | -0.232035081 | 6.541864 | 6.5765164 | 7.04323 | -0.466710103 | -0.501362478 |
| Appl1 | 10.48853 | 10.31425 | 10.457 | -0.142949318 | 0.031333628 | 10.22462 | 9.9140079 | 10.3807 | -0.466732811 | -0.156120699 |
| Fam122b | 8.388015 | 8.258673 | 8.3902 | -0.131545591 | -0.002203455 | 8.4927264 | 7.9980796 | 8.4649 | -0.466821068 | 0.027825799 |
| Racgap1 | 9.428958 | 9.469763 | 9.5414 | -0.071658163 | -0.112463062 | 9.2967092 | 9.1417695 | 9.60865 | -0.466882958 | -0.311943271 |
| Acss1 | 10.29378 | 10.35812 | 10.435 | -0.076385198 | -0.140725934 | 10.369033 | 9.8621355 | 10.3293 | -0.467134559 | 0.039762779 |
| Coq10b | 8.935072 | 8.58155 | 8.7379 | -0.156361498 | 0.197161374 | 8.6453341 | 8.36307 | 8.83022 | -0.467150104 | -0.184886003 |
| Nexmif | 4.231953 | 3.973489 | 4.3516 | -0.378072612 | -0.119608162 | 3.7423 | 3.5984971 | 4.06612 | -0.467624776 | -0.323821789 |
| Lurap1 | 5.95716 | 5.800458 | 6.0195 | -0.219017254 | -0.062315647 | 6.2002075 | 5.5921119 | 6.05975 | -0.467638625 | 0.14045698 |

| | | | | | | | | | | |
|---|---|---|---|---|---|---|---|---|---|---|
| Vangl1 | 5.4326 | 5.322451 | 5.3992 | -0.076732644 | 0.033415364 | 5.3194787 | 4.7418246 | 5.20955 | -0.467725748 | 0.109928266 |
| 1110032A03Rik | 6.730694 | 6.406385 | 6.9232 | -0.516765964 | -0.192457508 | 6.7566432 | 6.3841742 | 6.85191 | -0.467734948 | -0.095265906 |
| Stum | 4.059294 | 4.026021 | 4.0912 | -0.065211799 | -0.031939154 | 4.0611003 | 3.6481464 | 4.11621 | -0.468068138 | -0.055114168 |
| Eml3 | 10.41148 | 10.33823 | 10.593 | -0.254429357 | -0.181179085 | 10.708443 | 10.332902 | 10.8014 | -0.468449282 | -0.092908298 |
| Rassf4 | 9.959936 | 9.847906 | 10.029 | -0.180747144 | -0.068717269 | 9.7396836 | 9.5736503 | 10.0425 | -0.46885519 | -0.302821885 |
| Npy1r | 4.781637 | 4.414314 | 4.8132 | -0.398872858 | -0.031549622 | 4.1229836 | 3.8798708 | 4.34874 | -0.468869187 | -0.225756388 |
| Sord | 7.004483 | 6.872674 | 7.1154 | -0.242766263 | -0.110957101 | 6.8390324 | 6.5048982 | 6.97381 | -0.468915047 | -0.134780898 |
| Lbh | 12.58476 | 12.58786 | 12.674 | -0.085889186 | -0.088988666 | 12.773135 | 12.08173 | 12.5507 | -0.468928694 | 0.222476259 |
| Nt5dc1 | 9.425831 | 9.459192 | 9.5155 | -0.05632786 | -0.089688959 | 9.543916 | 9.0707894 | 9.53977 | -0.468975962 | 0.00415064 |
| Ahrr | 4.586682 | 4.527251 | 4.5686 | -0.041374436 | 0.018056125 | 4.6223221 | 4.4444718 | 4.91374 | -0.469268295 | -0.291417948 |
| Serinc5 | 9.355563 | 9.432789 | 9.5558 | -0.12296232 | -0.200187985 | 9.3066009 | 9.0472303 | 9.51694 | -0.469713552 | -0.210342964 |
| Gga2 | 11.64183 | 11.75804 | 11.707 | 0.051047844 | -0.06516348 | 12.152645 | 11.364773 | 11.8346 | -0.469817968 | 0.318053226 |
| Cd200r1 | 6.927542 | 6.791038 | 6.7852 | 0.005806566 | 0.14231059 | 6.4354667 | 6.2847095 | 6.75464 | -0.469931669 | -0.319174468 |
| N4bp3 | 8.043387 | 7.998707 | 8.1035 | -0.104799877 | -0.060120026 | 8.3197026 | 7.4699456 | 7.94004 | -0.470097138 | 0.379659918 |
| Efnb1 | 7.012089 | 6.879554 | 6.9946 | -0.115056556 | 0.017478143 | 6.287414 | 6.1176189 | 6.58774 | -0.470121728 | -0.300326556 |
| Camk2g | 9.484324 | 9.287962 | 9.6771 | -0.389114575 | -0.192752518 | 9.3217635 | 9.1742174 | 9.64458 | -0.470360828 | -0.322814698 |
| Map3k9 | 7.301953 | 7.32212 | 7.3627 | -0.040616566 | -0.060784016 | 7.3509508 | 6.9494869 | 7.42053 | -0.471040839 | -0.069576939 |
| Pdcd5-ps | 3.70879 | 3.989671 | 4.3999 | -0.410275671 | -0.691156657 | 3.7114229 | 4.2147352 | 4.68599 | -0.471250985 | -0.974563335 |
| Gm28539 | 2.918708 | 3.677299 | 2.6944 | 0.982866225 | 0.224274475 | 3.2047774 | 2.78434 | 3.25576 | -0.471420506 | -0.050983086 |
| Ttll1 | 6.479305 | 6.460011 | 6.7507 | -0.290728436 | -0.271434832 | 6.5553548 | 6.2068544 | 6.6783 | -0.471450168 | -0.12294976 |
| Ankrd55 | 4.706248 | 4.753652 | 4.7664 | -0.012730621 | -0.060134415 | 4.7075588 | 4.384481 | 4.85593 | -0.471453547 | -0.148375769 |
| Stim1 | 10.60362 | 10.66455 | 10.675 | -0.01072939 | -0.071657976 | 10.499929 | 10.229198 | 10.7007 | -0.471550504 | -0.200818913 |
| Setx | 12.36147 | 12.36309 | 12.414 | -0.050517138 | -0.052137048 | 12.146946 | 12.030997 | 12.5027 | -0.471690496 | -0.355741411 |
| Dab2ip | 7.24635 | 6.96841 | 7.1911 | -0.222689938 | 0.055250422 | 7.2045391 | 6.6340923 | 7.1059 | -0.471810426 | 0.098636314 |
| 4930451G09Rik | 4.234024 | 4.160439 | 4.3392 | -0.178718286 | -0.10513307 | 4.2105686 | 3.9666149 | 4.43893 | -0.472313326 | -0.228359574 |
| Atg9a | 8.606069 | 8.430223 | 8.5564 | -0.126194924 | 0.049651134 | 8.6783971 | 8.2516971 | 8.72438 | -0.47268227 | -0.045982278 |
| Ssbp3 | 9.415615 | 9.423717 | 9.5589 | -0.135200229 | -0.143302456 | 9.1510329 | 9.0757002 | 9.54865 | -0.472951723 | -0.397619024 |
| Chst3 | 11.75698 | 11.76885 | 11.898 | -0.128919656 | -0.140783992 | 11.827297 | 11.331025 | 11.8041 | -0.473049826 | 0.023221785 |
| Homez | 6.982141 | 7.114226 | 7.3808 | -0.266622595 | -0.398707545 | 6.8459998 | 6.66003 | 7.13309 | -0.473057563 | -0.287087689 |
| Gm4631 | 8.951932 | 8.976823 | 9.1718 | -0.195003094 | -0.219894055 | 9.1320634 | 8.7608447 | 9.23397 | -0.473124793 | -0.101906085 |
| Depdc1b | 5.706516 | 5.690755 | 5.857 | -0.166246424 | -0.15048555 | 5.5661925 | 5.4446101 | 5.91801 | -0.473403837 | -0.351821401 |
| Gm15922 | 7.185902 | 7.090749 | 7.3751 | -0.284327353 | -0.189174644 | 7.5695436 | 6.9076915 | 7.38115 | -0.473459247 | 0.188392782 |

| | | | | | | | | | | |
|---|---|---|---|---|---|---|---|---|---|---|
| Pard3b | 5.988724 | 6.006468 | 6.0639 | -0.057388498 | -0.075132897 | 5.6704439 | 5.6527202 | 6.12629 | -0.473573769 | -0.455850073 |
| Ift140 | 10.41284 | 10.36655 | 10.455 | -0.088056882 | -0.041773946 | 10.837788 | 10.094632 | 10.5684 | -0.473784604 | 0.269371048 |
| Reps2 | 4.174982 | 3.682567 | 4.2268 | -0.544211809 | -0.051796833 | 3.6930196 | 3.3927086 | 3.8665 | -0.473793003 | -0.173482021 |
| Rgs3 | 10.22652 | 10.14888 | 10.386 | -0.237522682 | -0.1598858 | 10.066516 | 9.8950531 | 10.3692 | -0.474159997 | -0.302696967 |
| Prkd1 | 4.412559 | 4.113423 | 4.4283 | -0.314904727 | -0.015769434 | 3.9429844 | 3.7397662 | 4.21395 | -0.474181037 | -0.270962861 |
| Myo3b | 5.063978 | 5.04998 | 5.1391 | -0.089079904 | -0.075081432 | 5.1063987 | 4.8277177 | 5.30193 | -0.474216484 | -0.195535473 |
| Lactb2 | 8.00462 | 7.984455 | 7.8703 | 0.114200751 | 0.134366229 | 8.0061061 | 7.5154868 | 7.98974 | -0.474257987 | 0.016361358 |
| Galnt4 | 8.855599 | 8.732164 | 8.8904 | -0.158240417 | -0.034805126 | 9.0396118 | 8.4318961 | 8.90691 | -0.475010045 | 0.132705702 |
| Rasgrp2 | 10.7618 | 10.91247 | 10.956 | -0.043630988 | -0.194301898 | 11.171863 | 10.572498 | 11.0475 | -0.47503888 | 0.124325931 |
| Clspn | 7.231933 | 6.823318 | 7.4353 | -0.612021984 | -0.203407736 | 7.1755483 | 7.2442454 | 7.71955 | -0.47530113 | -0.543998234 |
| Sacs | 9.790106 | 9.88232 | 9.9047 | -0.022403838 | -0.114618529 | 9.7037165 | 9.5878083 | 10.0632 | -0.475375 | -0.359466754 |
| Lamb3 | 8.096637 | 8.163074 | 8.0843 | 0.078816977 | 0.012379666 | 8.4476816 | 7.8780687 | 8.35349 | -0.475423112 | 0.094189816 |
| Adgrf5 | 9.466243 | 9.123041 | 9.3819 | -0.258846783 | 0.084354845 | 8.2468042 | 8.3860489 | 8.86172 | -0.475669393 | -0.614914122 |
| Daam2 | 4.759656 | 4.5823 | 4.9839 | -0.401637931 | -0.224282314 | 4.5569743 | 4.4452579 | 4.92118 | -0.475918405 | -0.364202047 |
| Fam3a | 8.345947 | 8.09683 | 8.4875 | -0.390630214 | -0.141513521 | 8.5975371 | 7.9765328 | 8.45257 | -0.476038156 | 0.144966157 |
| Kmt2c | 11.99483 | 12.01036 | 12.058 | -0.047275572 | -0.062814066 | 11.781926 | 11.76173 | 12.2379 | -0.476143476 | -0.455947492 |
| Cd4 | 11.68948 | 11.70054 | 11.759 | -0.05867274 | -0.06974079 | 11.246296 | 11.269162 | 11.7453 | -0.476164908 | -0.499031105 |
| Itgb5 | 6.438538 | 6.160025 | 6.079 | 0.081061982 | 0.35957515 | 6.6707454 | 5.6326559 | 6.10884 | -0.476186828 | 0.561902627 |
| Tert | 5.735366 | 5.739106 | 5.9208 | -0.181730403 | -0.185470058 | 5.7656449 | 5.5013342 | 5.97782 | -0.476482024 | -0.21217126 |
| Farp2 | 5.448935 | 5.437584 | 5.5212 | -0.083620055 | -0.072268783 | 5.2267793 | 5.0578005 | 5.5348 | -0.477004404 | -0.308025555 |
| Gm48415 | 5.332886 | 5.395 | 5.7936 | -0.398564549 | -0.460678402 | 5.1508344 | 5.2135763 | 5.69065 | -0.477077882 | -0.539819829 |
| Hpgds | 6.777502 | 6.694471 | 6.6372 | 0.057277548 | 0.140307964 | 6.4091571 | 6.2006971 | 6.67787 | -0.477169352 | -0.268709316 |
| Pdia5 | 6.114232 | 5.949068 | 6.0491 | -0.100038809 | 0.065126089 | 5.6742268 | 5.3086045 | 5.78637 | -0.477765646 | -0.112143411 |
| Igfbp4 | 12.3049 | 11.57039 | 11.652 | -0.082049108 | 0.652458954 | 12.004624 | 11.143275 | 11.6211 | -0.477843474 | 0.383505265 |
| Bhlhe41 | 6.108513 | 6.062554 | 6.1704 | -0.107863427 | -0.061903979 | 5.6087992 | 5.572459 | 6.05045 | -0.477989151 | -0.44164902 |
| AU021092 | 5.662589 | 5.105404 | 5.7479 | -0.642464208 | -0.085278776 | 4.780477 | 4.5322028 | 5.01032 | -0.478113035 | -0.229838827 |
| Camkmt | 5.425695 | 5.564716 | 5.5933 | -0.028616409 | -0.167637667 | 5.6074274 | 5.1745617 | 5.65272 | -0.478162153 | -0.045296516 |
| Tmem19 | 8.797811 | 8.816843 | 8.975 | -0.158110923 | -0.177142807 | 8.8965256 | 8.4480123 | 8.92631 | -0.478294348 | -0.029780985 |
| Cnksr3 | 7.871246 | 7.667086 | 7.7144 | -0.047357244 | 0.15680291 | 7.1751612 | 6.9838802 | 7.46218 | -0.478297206 | -0.287016138 |
| Ighv3-6 | 9.0109 | 9.239581 | 9.0456 | 0.194024308 | -0.034657083 | 8.7726729 | 8.5820077 | 9.06031 | -0.478302103 | -0.287636958 |
| Plp2 | 10.07869 | 9.949321 | 10.154 | -0.204322761 | -0.074952265 | 10.24076 | 9.6273473 | 10.1058 | -0.47847214 | 0.134940211 |
| Cc2d2a | 5.373892 | 5.284179 | 5.5828 | -0.298596204 | -0.208883348 | 5.2048357 | 4.9128703 | 5.39135 | -0.478476698 | -0.186511255 |

| | | | | | | | | | | |
|---|---|---|---|---|---|---|---|---|---|---|
| Tubg2 | 4.721398 | 4.531518 | 4.7861 | -0.254569813 | -0.064689011 | 5.0946621 | 4.4647179 | 4.94322 | -0.478499079 | 0.15144515 |
| Tmem156 | 7.835587 | 7.846617 | 7.9109 | -0.064307616 | -0.075337835 | 8.0630373 | 7.5328622 | 8.0115 | -0.478637314 | 0.051537776 |
| Tmem245 | 10.87841 | 10.97386 | 10.903 | 0.071252078 | -0.024199078 | 10.973598 | 10.699624 | 11.1783 | -0.478716466 | -0.204742327 |
| Pkd2 | 6.671188 | 6.247606 | 6.4453 | -0.197656882 | 0.225925004 | 5.8961498 | 5.8604025 | 6.33934 | -0.47893803 | -0.443190708 |
| Gins2 | 6.220121 | 5.944471 | 6.0719 | -0.127464909 | 0.148184651 | 6.3771922 | 5.9867063 | 6.46567 | -0.478961236 | -0.088475289 |
| Ripor1 | 9.334565 | 9.358619 | 9.4055 | -0.046863266 | -0.07091753 | 9.1951468 | 8.7611685 | 9.24021 | -0.479039291 | -0.04506094 |
| Zfp934 | 6.783094 | 6.589383 | 6.8416 | -0.252209356 | -0.058498377 | 6.9835188 | 6.6075906 | 7.08684 | -0.479250058 | -0.103321863 |
| Enpep | 5.003286 | 4.673147 | 4.7726 | -0.099502365 | 0.230636342 | 4.3130076 | 4.3659983 | 4.8454 | -0.479404234 | -0.532394949 |
| 2810021J22Rik | 7.473918 | 7.267694 | 7.3816 | -0.113946373 | 0.092277093 | 7.590799 | 6.9570904 | 7.43674 | -0.479652568 | 0.154055994 |
| Gcfc2 | 7.649902 | 7.785156 | 7.8112 | -0.02607412 | -0.161328182 | 8.0233288 | 7.6613036 | 8.14117 | -0.479864667 | -0.117839492 |
| Zfp608 | 9.762858 | 9.836787 | 9.8086 | 0.028161127 | -0.045767482 | 9.7533178 | 9.194486 | 9.67441 | -0.479927646 | 0.078904209 |
| Wscd1 | 3.925251 | 3.874607 | 4.0909 | -0.21630159 | -0.165657224 | 3.8134646 | 3.6131928 | 4.09317 | -0.479979937 | -0.279708135 |
| Pald1 | 7.503599 | 7.155871 | 7.3991 | -0.243250583 | 0.104476803 | 6.8584976 | 6.2077269 | 6.68774 | -0.480015698 | 0.17075499 |
| Nedd4l | 9.019137 | 9.061412 | 9.0479 | 0.01346983 | -0.028804484 | 9.0215753 | 8.7538745 | 9.23399 | -0.480119109 | -0.212418325 |
| Tmem143 | 7.152877 | 7.149078 | 7.1501 | -0.001052481 | 0.002746342 | 7.390148 | 6.9552434 | 7.43553 | -0.480288504 | -0.045383937 |
| Rab3d | 8.262775 | 8.004494 | 8.1836 | -0.17908337 | 0.07919827 | 7.8040378 | 7.6669393 | 8.14725 | -0.480306122 | -0.343207601 |
| Tmem65 | 8.50276 | 8.573583 | 8.5099 | 0.063711548 | -0.007111512 | 7.9977784 | 7.7828959 | 8.26346 | -0.48056169 | -0.265679153 |
| C1qtnf9 | 4.262201 | 4.032152 | 4.5431 | -0.510948174 | -0.28089964 | 3.9597232 | 3.7060952 | 4.18667 | -0.480575064 | -0.226947058 |
| Gata3 | 8.34499 | 8.274061 | 8.4933 | -0.219282895 | -0.148354498 | 8.1690512 | 8.2295395 | 8.71021 | -0.480667096 | -0.541155385 |
| Cyb5d1 | 7.556164 | 7.391857 | 7.5501 | -0.158233579 | 0.006073396 | 7.5814516 | 7.167674 | 7.64853 | -0.48085952 | -0.067081954 |
| Usp46 | 7.217832 | 7.125562 | 7.0212 | 0.1043802 | 0.19665047 | 6.7721265 | 6.4042126 | 6.88523 | -0.481020685 | -0.113106737 |
| Zfhx2 | 8.693081 | 8.654762 | 8.8051 | -0.150362301 | -0.112043578 | 8.7272295 | 8.5027252 | 8.98419 | -0.481468773 | -0.256964497 |
| Ids | 10.11913 | 10.07846 | 10.197 | -0.118615722 | -0.077944943 | 10.010257 | 9.5553381 | 10.0369 | -0.481601021 | -0.026681668 |
| Alox8 | 4.237804 | 4.020507 | 4.285 | -0.264524618 | -0.047227927 | 4.0958457 | 3.8977693 | 4.37959 | -0.481816466 | -0.283740083 |
| Iffo2 | 8.179503 | 8.131022 | 8.2099 | -0.078881854 | -0.030401059 | 7.7702954 | 7.8526882 | 8.33456 | -0.481871409 | -0.564264265 |
| Lama5 | 8.581961 | 8.231922 | 8.2816 | -0.049706112 | 0.300333527 | 7.4567597 | 7.4374389 | 7.91933 | -0.481887579 | -0.462566746 |
| Gm48348 | 7.297046 | 7.165703 | 7.3093 | -0.143632388 | -0.012289412 | 7.5497759 | 7.2184247 | 7.70046 | -0.482030532 | -0.150679241 |
| Arhgap35 | 9.029767 | 8.731311 | 8.9396 | -0.208246345 | 0.09020984 | 8.5121623 | 8.3212185 | 8.80325 | -0.482031247 | -0.291087376 |
| Cntln | 6.798175 | 6.516365 | 6.7307 | -0.214321985 | 0.067488121 | 6.4955103 | 6.3335159 | 6.81577 | -0.482253293 | -0.320258953 |
| Slc27a1 | 7.358809 | 7.107622 | 7.4583 | -0.350672935 | -0.099486054 | 7.2600793 | 6.923301 | 7.40558 | -0.482274658 | -0.145496341 |
| Bbs2 | 7.597549 | 7.701695 | 7.7118 | -0.010083181 | -0.114229443 | 7.8248072 | 7.479581 | 7.96188 | -0.482295566 | -0.137069285 |
| Foxo4 | 8.76593 | 8.800022 | 8.8682 | -0.068161114 | -0.102252262 | 8.8444059 | 8.364538 | 8.84741 | -0.48287456 | -0.003006693 |

| | | | | | | | | | | |
|---|---|---|---|---|---|---|---|---|---|---|
| Maged1 | 8.357639 | 8.029746 | 8.2772 | -0.247483461 | 0.080409815 | 7.7266428 | 7.4365152 | 7.92002 | -0.483503102 | -0.193375505 |
| Cgnl1 | 6.226445 | 6.105104 | 6.2682 | -0.163090375 | -0.041749225 | 5.8818604 | 5.6221868 | 6.10574 | -0.483552936 | -0.223879303 |
| Cers4 | 9.484551 | 9.343725 | 9.4889 | -0.145224949 | -0.004398828 | 9.7716601 | 9.0778611 | 9.56159 | -0.48372399 | 0.210074976 |
| Arl5b | 8.966433 | 8.705196 | 8.8979 | -0.192726668 | 0.068510422 | 8.8948543 | 8.7623097 | 9.24651 | -0.484198783 | -0.351654142 |
| Adgrb2 | 4.638906 | 4.469336 | 4.4977 | -0.028388189 | 0.141181625 | 4.6555151 | 4.1040409 | 4.58863 | -0.484587652 | 0.066886564 |
| Dnhd1 | 6.499507 | 6.172013 | 6.7187 | -0.546719831 | -0.219226353 | 6.9416497 | 6.4998199 | 6.98441 | -0.484590731 | -0.042760905 |
| Heg1 | 10.73496 | 10.60783 | 10.683 | -0.074774592 | 0.052357132 | 10.477674 | 10.27286 | 10.7578 | -0.48489336 | -0.280078952 |
| Wdr47 | 8.009383 | 8.038387 | 7.9163 | 0.122134214 | 0.093130584 | 7.8724508 | 7.5793653 | 8.06428 | -0.484912447 | -0.191826957 |
| Ano10 | 8.770162 | 8.680786 | 8.837 | -0.156261862 | -0.066886552 | 8.8498615 | 8.3878644 | 8.87291 | -0.4850421 | -0.023044988 |
| Gdpd3 | 4.495367 | 4.45499 | 4.5509 | -0.095888959 | -0.055511674 | 4.5201884 | 4.2551622 | 4.74042 | -0.485256432 | -0.220230188 |
| Lilra6 | 6.834169 | 6.875897 | 6.6461 | 0.229787198 | 0.188058683 | 7.0407115 | 6.1753795 | 6.66068 | -0.485304431 | 0.380027634 |
| Fam234a | 8.52009 | 8.379744 | 8.6383 | -0.258563208 | -0.118216722 | 8.5237653 | 8.0453298 | 8.53075 | -0.48541904 | -0.006983528 |
| Fam168a | 9.858409 | 9.787447 | 9.8592 | -0.071765772 | -0.00080385 | 9.7783394 | 9.4060948 | 9.89152 | -0.48542509 | -0.113180461 |
| C2cd2 | 8.081196 | 8.070805 | 8.3269 | -0.256072485 | -0.245681365 | 8.2361388 | 7.5673588 | 8.05278 | -0.485425431 | 0.183354611 |
| Tgfa | 5.89741 | 5.657929 | 5.8505 | -0.192540448 | 0.046940961 | 5.1720445 | 5.299924 | 5.78569 | -0.485763838 | -0.613643317 |
| Chml | 7.799262 | 7.765891 | 7.8574 | -0.091554816 | -0.058184676 | 8.0090418 | 7.5482341 | 8.03401 | -0.485775255 | -0.024967521 |
| Prkcz | 6.666641 | 6.727279 | 6.7981 | -0.070864595 | -0.131502538 | 6.612674 | 6.3060782 | 6.79188 | -0.485796953 | -0.179201139 |
| Cdk2 | 9.572953 | 9.460609 | 9.6802 | -0.219616347 | -0.107271401 | 9.7809321 | 9.2847948 | 9.77063 | -0.485836292 | 0.010300974 |
| Mms22l | 7.874507 | 7.409228 | 7.623 | -0.213728812 | 0.251550327 | 7.9898101 | 7.4530379 | 7.93949 | -0.486448494 | 0.050323716 |
| Jkamp | 7.952329 | 8.103284 | 8.1437 | -0.040391286 | -0.191346088 | 8.1579249 | 7.7400211 | 8.22684 | -0.486814831 | -0.068910952 |
| Flna | 14.23864 | 14.27053 | 14.262 | 0.008234682 | -0.023650886 | 14.345408 | 13.899596 | 14.3866 | -0.486957604 | -0.041146126 |
| Atp5s | 5.627749 | 5.943167 | 5.8462 | 0.096967721 | -0.218450849 | 5.9348354 | 5.6034163 | 6.0904 | -0.48697974 | -0.155560641 |
| Herc1 | 12.11143 | 12.06076 | 12.118 | -0.056973306 | -0.006305596 | 11.856491 | 11.790123 | 12.2776 | -0.487455348 | -0.421086734 |
| Cd180 | 11.381 | 11.16418 | 11.378 | -0.214101568 | 0.002710484 | 11.732699 | 10.918073 | 11.4056 | -0.48757719 | 0.327049552 |
| 1190002N15Rik | 8.381277 | 8.282273 | 8.3343 | -0.051990578 | 0.047013843 | 7.9476338 | 7.752536 | 8.24015 | -0.487610783 | -0.292512972 |
| Sall2 | 6.321027 | 6.201745 | 6.4889 | -0.287125375 | -0.167844025 | 5.9573121 | 5.7693192 | 6.25721 | -0.48789352 | -0.299900589 |
| Cyp2d22 | 6.929835 | 6.993598 | 7.3035 | -0.309895445 | -0.373658134 | 7.0870289 | 6.6600644 | 7.14842 | -0.488356831 | -0.061392339 |
| Ankle1 | 4.688167 | 4.600331 | 4.7652 | -0.164849625 | -0.07701337 | 4.7518471 | 4.5726056 | 5.06179 | -0.489185002 | -0.309943519 |
| Cryzl2 | 6.417062 | 6.234298 | 6.4696 | -0.23531688 | -0.052552823 | 6.5768821 | 6.0253902 | 6.51474 | -0.489346668 | 0.062145239 |
| Rnf130 | 9.993586 | 9.866865 | 9.998 | -0.131121353 | -0.004400282 | 10.207343 | 9.3853395 | 9.87471 | -0.489373482 | 0.332629564 |
| Rnf103 | 8.470839 | 8.36499 | 8.4905 | -0.125529766 | -0.019680953 | 8.2529942 | 7.9105602 | 8.40033 | -0.489769671 | -0.147335682 |
| Gm48048 | 4.90686 | 5.197235 | 5.4464 | -0.249194733 | -0.539570661 | 5.2684467 | 4.4473383 | 4.93765 | -0.490308096 | 0.330800307 |

| | | | | | | | | | | |
|---|---|---|---|---|---|---|---|---|---|---|
| Agap1 | 5.911975 | 5.661318 | 6.0345 | -0.373211531 | -0.122555104 | 5.6187905 | 5.717274 | 6.20767 | -0.490392836 | -0.588876376 |
| Atp11c | 9.241574 | 9.041631 | 9.168 | -0.126371761 | 0.073571838 | 8.9754703 | 8.6964989 | 9.1872 | -0.490701891 | -0.211730433 |
| Dhx57 | 9.99011 | 9.961243 | 10.084 | -0.122334418 | -0.093467527 | 10.176364 | 9.9127682 | 10.4038 | -0.491014166 | -0.227418229 |
| Stk17b | 13.88798 | 13.92811 | 13.826 | 0.101921268 | 0.061796792 | 13.717116 | 13.348852 | 13.84 | -0.491137938 | -0.122874222 |
| Spred2 | 8.514423 | 8.330078 | 8.4385 | -0.108416924 | 0.07592774 | 8.0539149 | 7.8719227 | 8.36325 | -0.491329104 | -0.30933694 |
| D130040H23Rik | 7.11216 | 6.963061 | 7.2389 | -0.27585364 | -0.12675405 | 7.3570863 | 7.0103731 | 7.50204 | -0.491664964 | -0.144951792 |
| Havcr1 | 5.978197 | 6.066632 | 6.0931 | -0.02649099 | -0.114925394 | 5.3334974 | 5.2187876 | 5.71052 | -0.491729395 | -0.377019573 |
| Rasgef1a | 6.251327 | 6.002691 | 6.2325 | -0.229802161 | 0.018833865 | 5.2692442 | 5.3486746 | 5.84048 | -0.491801687 | -0.571232055 |
| Agtpbp1 | 9.523253 | 9.130659 | 9.4228 | -0.292189216 | 0.100405005 | 9.2592851 | 9.0683538 | 9.56025 | -0.491893825 | -0.300962621 |
| Nipal3 | 9.980694 | 10.0585 | 10.122 | -0.063871774 | -0.141674291 | 9.8884255 | 9.5666647 | 10.0589 | -0.492281656 | -0.170520845 |
| Wdfy3 | 7.6706 | 7.47827 | 7.475 | 0.003248005 | 0.195577919 | 7.0668876 | 6.7298292 | 7.2222 | -0.492367462 | -0.155308994 |
| Cyyr1 | 8.780007 | 8.408023 | 8.4406 | -0.032571661 | 0.339411856 | 7.3671693 | 7.3867859 | 7.87946 | -0.492670088 | -0.512286687 |
| Sspn | 4.923984 | 4.821394 | 5.1058 | -0.284383789 | -0.181793564 | 4.8111335 | 4.5010586 | 4.9941 | -0.493044733 | -0.18296983 |
| Phf20 | 9.948429 | 9.929893 | 10.052 | -0.122038064 | -0.103502284 | 10.060533 | 9.6742943 | 10.1679 | -0.493610673 | -0.107371836 |
| Adnp | 8.453674 | 8.496542 | 8.4646 | 0.031934333 | -0.010934286 | 7.9891547 | 8.1221926 | 8.61624 | -0.494044802 | -0.627082699 |
| Myb | 8.380225 | 8.244125 | 8.5744 | -0.33025311 | -0.194153455 | 8.0766761 | 8.2506496 | 8.7447 | -0.494054131 | -0.668027672 |
| Polg | 10.81688 | 10.86861 | 10.737 | 0.131972704 | 0.080245912 | 10.909162 | 10.489267 | 10.9834 | -0.494108406 | -0.074212572 |
| Srsf12 | 4.844932 | 4.80934 | 4.9768 | -0.167425596 | -0.131832954 | 4.6066884 | 4.563832 | 5.05805 | -0.494220921 | -0.451364533 |
| Sfxn5 | 6.399829 | 6.333947 | 6.2947 | 0.039285834 | 0.105167103 | 6.6318236 | 6.132418 | 6.62682 | -0.494397767 | 0.005007852 |
| Diras2 | 4.647973 | 4.714142 | 4.8856 | -0.17149813 | -0.237667137 | 4.6367341 | 4.2290602 | 4.72374 | -0.494677074 | -0.087003149 |
| Capn2 | 10.1321 | 10.01466 | 10.093 | -0.078659862 | 0.038778298 | 9.870468 | 9.6578268 | 10.1527 | -0.494830143 | -0.282188948 |
| Hist2h2be | 6.772727 | 6.812962 | 6.8916 | -0.078663606 | -0.118898333 | 7.0832801 | 6.4956998 | 6.99083 | -0.49512898 | 0.092451396 |
| Fcrls | 2.976572 | 2.874367 | 3.121 | -0.246597181 | -0.144391728 | 2.873277 | 2.7344027 | 3.22962 | -0.495213249 | -0.35633898 |
| Lims2 | 4.672459 | 4.255128 | 4.8796 | -0.624489164 | -0.207158798 | 4.2607874 | 4.3020681 | 4.7973 | -0.495236926 | -0.536517604 |
| Klhdc1 | 7.821454 | 7.828317 | 7.9566 | -0.128235634 | -0.13509845 | 7.7251583 | 7.57268 | 8.06815 | -0.495471083 | -0.342992731 |
| Gclm | 8.015768 | 7.668661 | 7.9456 | -0.276944435 | 0.07016214 | 7.7704315 | 7.4387735 | 7.93443 | -0.495653916 | -0.16399589 |
| Hspa12a | 5.92557 | 5.877829 | 6.0306 | -0.152772309 | -0.105031454 | 5.9790392 | 5.274589 | 5.77033 | -0.495745723 | 0.208704437 |
| Ccnf | 6.616313 | 6.621056 | 6.8568 | -0.235722922 | -0.24046621 | 6.688684 | 6.509151 | 7.00497 | -0.49581781 | -0.316284849 |
| Gucy1b1 | 5.168207 | 5.191544 | 5.2996 | -0.108102893 | -0.131440313 | 4.7912034 | 4.5528445 | 5.04866 | -0.495817987 | -0.257459083 |
| Stau2 | 6.595156 | 6.387487 | 6.4352 | -0.047698186 | 0.159970853 | 6.349301 | 6.03948 | 6.53545 | -0.495965078 | -0.186144116 |
| Fbxo10 | 6.201949 | 6.048766 | 6.2691 | -0.220356445 | -0.067173504 | 5.9930324 | 5.659253 | 6.15524 | -0.495986136 | -0.162206675 |
| Ptpn5 | 4.699409 | 4.701228 | 4.7786 | -0.077388852 | -0.079207977 | 4.6440238 | 4.4095986 | 4.90584 | -0.496239678 | -0.26181447 |

| | | | | | | | | | | |
|---|---|---|---|---|---|---|---|---|---|---|
| Enc1 | 8.621327 | 8.646596 | 8.7089 | -0.062288946 | -0.087558097 | 8.1185121 | 8.0740097 | 8.57028 | -0.49626849 | -0.451766078 |
| Dmpk | 5.816551 | 5.671887 | 5.8519 | -0.179994547 | -0.035330361 | 5.9948636 | 5.2866731 | 5.7831 | -0.496431477 | 0.211759034 |
| Lpcat1 | 10.32761 | 10.40066 | 10.372 | 0.028761552 | -0.044280352 | 10.696673 | 10.068446 | 10.5649 | -0.496465259 | 0.131762222 |
| Acbd4 | 7.71694 | 7.536032 | 7.6672 | -0.13117256 | 0.04973503 | 8.0416027 | 7.4210885 | 7.91762 | -0.496531461 | 0.123982724 |
| Gm21541 | 2.01839 | 1.740495 | 2.0428 | -0.302274042 | -0.024378867 | 1.8011617 | 1.7543221 | 2.25112 | -0.49680226 | -0.449962606 |
| Phc3 | 11.2076 | 11.17297 | 11.265 | -0.09219131 | -0.057562894 | 11.094342 | 10.9388 | 11.4358 | -0.497010936 | -0.341469211 |
| 36951 | 11.4037 | 11.35587 | 11.428 | -0.072285408 | -0.024458978 | 11.448593 | 10.733993 | 11.2311 | -0.49715521 | 0.217445164 |
| Ticrr | 6.912369 | 6.508208 | 6.7697 | -0.261470879 | 0.142690196 | 6.5828197 | 6.6392255 | 7.13663 | -0.497403128 | -0.553808853 |
| Smco4 | 7.151817 | 6.828069 | 7.1324 | -0.304370916 | 0.01937717 | 6.9403326 | 6.5370064 | 7.03447 | -0.497462863 | -0.094136606 |
| Hist1h1c | 8.252125 | 8.014035 | 8.2763 | -0.262289775 | -0.024199711 | 7.9016007 | 7.4582767 | 7.95643 | -0.498150569 | -0.054826571 |
| Tatdn3 | 6.383248 | 6.447022 | 6.464 | -0.016981364 | -0.080755748 | 6.6240457 | 6.2789425 | 6.77777 | -0.49882831 | -0.153725131 |
| Rdm1 | 7.938697 | 7.870195 | 8.0496 | -0.179363369 | -0.110860861 | 8.4941003 | 7.6449967 | 8.1441 | -0.499099673 | 0.350003968 |
| Herc2 | 11.87994 | 11.81336 | 11.911 | -0.097564392 | -0.030988058 | 11.639597 | 11.549435 | 12.0487 | -0.499273138 | -0.409110776 |
| Tspan12 | 5.242698 | 4.986855 | 5.3069 | -0.320047455 | -0.064204397 | 5.0472481 | 4.4915701 | 4.99089 | -0.499320712 | 0.056357311 |
| Ets1 | 14.10164 | 14.08745 | 14.193 | -0.105531274 | -0.091343688 | 13.90052 | 13.548772 | 14.0483 | -0.499512198 | -0.147764317 |
| Kifap3 | 6.735789 | 6.632213 | 6.6841 | -0.051841341 | 0.051734349 | 6.3735898 | 6.1306431 | 6.63035 | -0.499707375 | -0.256760643 |
| Cd83 | 12.1455 | 12.12516 | 12.025 | 0.100496536 | 0.120843518 | 11.750819 | 11.565227 | 12.065 | -0.499787996 | -0.31419581 |
| Sipa1l2 | 8.62643 | 8.702625 | 8.5487 | 0.153941561 | 0.077746233 | 8.5635589 | 8.1407128 | 8.64103 | -0.500312635 | -0.077466517 |
| Rnf152 | 4.828313 | 4.878434 | 4.9603 | -0.0818389 | -0.131959744 | 4.8041621 | 4.7150889 | 5.2158 | -0.500707391 | -0.411634214 |
| Fnbp1l | 9.136516 | 8.830668 | 9.1325 | -0.30179315 | 0.004054796 | 8.1735679 | 8.2096124 | 8.71078 | -0.501163045 | -0.537207529 |
| Scn3a | 4.349834 | 4.480378 | 4.5358 | -0.055379184 | -0.185922984 | 4.2955998 | 3.9923051 | 4.49385 | -0.501548866 | -0.198254242 |
| Gm4924 | 5.942177 | 5.790315 | 5.9497 | -0.15941626 | -0.007554087 | 5.9078899 | 5.5760192 | 6.07785 | -0.501830521 | -0.169959808 |
| Crocc | 7.346632 | 7.411514 | 7.2918 | 0.11970079 | 0.054818747 | 7.6367657 | 7.3001796 | 7.80209 | -0.501914893 | -0.165328836 |
| Rnd1 | 8.309367 | 7.797915 | 7.7629 | 0.035041417 | 0.546493539 | 7.0532896 | 7.0437626 | 7.546 | -0.502233322 | -0.492706337 |
| Acaa2 | 9.212874 | 9.224756 | 9.326 | -0.101240269 | -0.113121737 | 9.303241 | 8.7345376 | 9.23689 | -0.502354769 | 0.066348625 |
| Tspan18 | 4.819133 | 4.877 | 4.9805 | -0.10353318 | -0.161400491 | 4.7195266 | 4.465333 | 4.96771 | -0.502374232 | -0.248180658 |
| Plekhg5 | 7.001478 | 6.565197 | 7.0986 | -0.533378068 | -0.097096892 | 6.5006942 | 6.2574795 | 6.75986 | -0.502382749 | -0.259168046 |
| Smpd3 | 5.368176 | 5.248227 | 5.4363 | -0.188090726 | -0.068142322 | 5.6053211 | 4.9638382 | 5.46623 | -0.502391232 | 0.139091738 |
| Hlcs | 9.059058 | 9.000583 | 9.0967 | -0.0960745 | -0.037599112 | 9.1761973 | 8.6829268 | 9.18532 | -0.502395721 | -0.009125249 |
| Gstk1 | 6.38677 | 6.381728 | 6.4706 | -0.088847391 | -0.083806217 | 6.264175 | 5.9083559 | 6.41078 | -0.502427173 | -0.146608058 |
| Rab11fip5 | 7.689467 | 7.274822 | 7.6332 | -0.358375539 | 0.05627 | 7.5211148 | 6.6940142 | 7.19692 | -0.502905447 | 0.324195177 |
| Grap2 | 12.27981 | 12.35307 | 12.417 | -0.063925232 | -0.13719255 | 12.150837 | 11.932012 | 12.435 | -0.502960214 | -0.284134719 |

| | | | | | | | | | | |
|---|---|---|---|---|---|---|---|---|---|---|
| Ndst3 | 5.23927 | 4.945751 | 5.2097 | -0.263985106 | 0.029534549 | 4.559878 | 4.2396417 | 4.74296 | -0.503322372 | -0.183086041 |
| Zfp239 | 5.935848 | 5.78402 | 6.204 | -0.419942367 | -0.268114133 | 6.1760447 | 5.6432076 | 6.14691 | -0.503701558 | 0.029135455 |
| Zfp28 | 7.259632 | 6.836686 | 7.3536 | -0.516952429 | -0.094005826 | 7.0309783 | 6.9447179 | 7.44844 | -0.503717701 | -0.417457253 |
| Fbxl17 | 8.878742 | 8.802622 | 8.7436 | 0.058977264 | 0.135096587 | 8.7696635 | 8.4592645 | 8.963 | -0.503737742 | -0.193338678 |
| Slc9b2 | 4.848919 | 4.57371 | 4.9619 | -0.38817388 | -0.112964037 | 4.703536 | 4.5594006 | 5.0634 | -0.503998164 | -0.359862777 |
| Slc2a13 | 5.415038 | 4.856779 | 5.2518 | -0.394977877 | 0.163280905 | 4.562374 | 4.4395598 | 4.9443 | -0.504738134 | -0.381923885 |
| Rin3 | 11.35866 | 11.38505 | 11.34 | 0.044645214 | 0.018250766 | 11.370859 | 10.684708 | 11.1896 | -0.504931536 | 0.181220136 |
| Sox17 | 5.029405 | 4.604753 | 5.0161 | -0.411299916 | 0.013352135 | 4.4576018 | 4.3162061 | 4.82118 | -0.504971782 | -0.363576024 |
| Kremen1 | 9.44025 | 9.340265 | 9.4758 | -0.135541935 | -0.035556625 | 9.2706405 | 8.9826273 | 9.48793 | -0.505302631 | -0.217289444 |
| Bicral | 10.53905 | 10.6649 | 10.685 | -0.019961858 | -0.145804254 | 10.570592 | 10.180297 | 10.6856 | -0.50531127 | -0.11501635 |
| Dcaf6 | 8.432487 | 8.352287 | 8.5578 | -0.205493508 | -0.125293524 | 8.486778 | 8.1465587 | 8.65193 | -0.505371509 | -0.165152201 |
| Fndc10 | 5.832024 | 5.597441 | 5.8544 | -0.2569372 | -0.022354095 | 5.8182059 | 5.3333023 | 5.83879 | -0.505487644 | -0.020583973 |
| Plekhg1 | 7.921378 | 7.323758 | 7.7772 | -0.453451189 | 0.144168615 | 6.8082933 | 6.9615129 | 7.46734 | -0.505823029 | -0.659042707 |
| Ccdc186 | 9.074116 | 9.142186 | 9.1438 | -0.001659404 | -0.069729018 | 8.6013807 | 8.6922294 | 9.1985 | -0.506274608 | -0.597123311 |
| B3gntl1 | 7.536127 | 7.375417 | 7.6456 | -0.270179477 | -0.109469391 | 7.6581668 | 7.3444678 | 7.85134 | -0.50687298 | -0.193173975 |
| Wdr35 | 5.702696 | 5.681813 | 5.762 | -0.080196893 | -0.059313332 | 5.7436232 | 5.4578152 | 5.96471 | -0.506891888 | -0.221083892 |
| Ccdc88a | 9.394151 | 9.53079 | 9.4916 | 0.039181147 | -0.097457547 | 8.8326573 | 8.8027228 | 9.30969 | -0.506967867 | -0.477033394 |
| Plxna4 | 6.571873 | 6.042852 | 6.4939 | -0.451088569 | 0.077932301 | 5.5600894 | 5.5500769 | 6.05713 | -0.507048764 | -0.49703629 |
| Raver2 | 6.502218 | 6.397134 | 6.6195 | -0.222357863 | -0.117273594 | 6.2091227 | 5.9916276 | 6.49882 | -0.507191501 | -0.289696428 |
| Fgf10 | 3.701229 | 3.591434 | 3.744 | -0.152577761 | -0.042782914 | 3.2564063 | 3.1862099 | 3.69359 | -0.507383758 | -0.437187351 |
| Rtn1 | 6.721427 | 6.454213 | 6.676 | -0.221775748 | 0.045437889 | 6.4635587 | 6.0083711 | 6.51588 | -0.50751241 | -0.052324724 |
| Itga6 | 10.48875 | 10.46779 | 10.583 | -0.11491669 | -0.09395805 | 10.161734 | 9.9926091 | 10.5012 | -0.508593032 | -0.339468131 |
| Acyp1 | 7.579752 | 7.439207 | 7.6572 | -0.217947887 | -0.077402459 | 7.7370709 | 7.1419408 | 7.65107 | -0.50912797 | 0.086002163 |
| Mpp7 | 7.453176 | 7.638915 | 7.6503 | -0.011399121 | -0.197138052 | 7.4748385 | 7.3641165 | 7.8733 | -0.509180091 | -0.39845812 |
| Dab2 | 8.807118 | 7.898424 | 7.8201 | 0.078324934 | 0.987019024 | 7.2565933 | 7.0305545 | 7.53973 | -0.509180243 | -0.283141467 |
| Rab6b | 8.675543 | 8.659034 | 8.8019 | -0.142832782 | -0.126323704 | 8.3842904 | 8.1696556 | 8.67891 | -0.509254038 | -0.294619167 |
| Snap47 | 7.997946 | 7.968292 | 8.1202 | -0.15189865 | -0.122244627 | 8.0652471 | 7.5439119 | 8.05345 | -0.509534534 | 0.011800624 |
| Dock4 | 7.415659 | 7.166082 | 7.4117 | -0.245640048 | 0.003937236 | 6.6730919 | 6.756467 | 7.26602 | -0.509553682 | -0.592928761 |
| Isoc2b | 6.208098 | 6.004637 | 6.3313 | -0.326615576 | -0.123155113 | 6.2397254 | 5.8130539 | 6.32266 | -0.509606911 | -0.082935429 |
| Nudt4 | 9.961565 | 9.719275 | 9.8177 | -0.098459986 | 0.143830433 | 9.5067996 | 9.101103 | 9.61079 | -0.509684036 | -0.10398741 |
| Slc22a5 | 8.543085 | 8.209663 | 8.439 | -0.229367765 | 0.104053933 | 8.4032987 | 8.0074749 | 8.51738 | -0.509904003 | -0.114080218 |
| Dnajc10 | 10.45114 | 10.32768 | 10.472 | -0.14419577 | -0.020728572 | 10.390931 | 9.8927753 | 10.403 | -0.510206654 | -0.012051344 |

| | | | | | | | | | | |
|---|---|---|---|---|---|---|---|---|---|---|
| Zmym4 | 9.958713 | 9.899931 | 9.982 | -0.082111171 | -0.023328931 | 9.6237783 | 9.5458983 | 10.0565 | -0.510634002 | -0.432754002 |
| Tcaf1 | 5.629515 | 5.382593 | 5.6563 | -0.273711151 | -0.026789284 | 5.3168815 | 5.1529184 | 5.66372 | -0.510805768 | -0.346842678 |
| Fut8 | 8.909788 | 8.888645 | 8.9577 | -0.069032149 | -0.047889253 | 8.6531111 | 8.4453678 | 8.9562 | -0.510829445 | -0.303086086 |
| Crip3 | 4.866108 | 4.945162 | 5.1153 | -0.170157817 | -0.249212206 | 5.4093117 | 4.9323015 | 5.44351 | -0.511212042 | -0.034201834 |
| Chrm3 | 4.694452 | 4.405437 | 4.7507 | -0.345250637 | -0.056235774 | 4.1836555 | 3.9711689 | 4.48299 | -0.511820787 | -0.299334169 |
| Stard10 | 7.272225 | 7.110499 | 7.3537 | -0.243162687 | -0.081437358 | 7.2692381 | 6.728605 | 7.24116 | -0.512558565 | 0.028074565 |
| Blm | 7.873893 | 7.804634 | 8.1514 | -0.346792417 | -0.277533184 | 7.9384071 | 7.8014926 | 8.31438 | -0.512889622 | -0.375975121 |
| Nfat5 | 11.33023 | 11.28536 | 11.388 | -0.102298788 | -0.057426842 | 11.344082 | 11.213324 | 11.7265 | -0.513135552 | -0.382376899 |
| Arl13b | 8.150499 | 8.05222 | 8.0972 | -0.044960832 | 0.053318096 | 7.9837395 | 7.8216328 | 8.33478 | -0.513144882 | -0.351038163 |
| Efnb2 | 6.844477 | 6.843411 | 7.0989 | -0.255488591 | -0.25442192 | 6.4529358 | 6.4399971 | 6.95361 | -0.513613489 | -0.500674752 |
| Map6d1 | 3.911638 | 3.723586 | 4.0919 | -0.36831792 | -0.180266 | 3.835536 | 3.6022644 | 4.11641 | -0.514148616 | -0.280876926 |
| Tfpi | 7.88069 | 7.495754 | 7.923 | -0.427262269 | -0.042325701 | 7.2083151 | 6.8353015 | 7.34952 | -0.514218094 | -0.141204421 |
| Mns1 | 7.333979 | 7.176817 | 7.3515 | -0.174639925 | -0.017477589 | 7.2119429 | 7.015134 | 7.52947 | -0.514331863 | -0.317522895 |
| St3gal1 | 11.65374 | 11.67808 | 11.643 | 0.034999792 | 0.010653324 | 11.478092 | 11.167142 | 11.6815 | -0.514394452 | -0.203444849 |
| Pcmtd1 | 10.53311 | 10.41825 | 10.594 | -0.176048598 | -0.061193072 | 10.339885 | 10.06362 | 10.5781 | -0.514447151 | -0.238182725 |
| Col17a1 | 4.167688 | 4.01193 | 4.0427 | -0.030792466 | 0.124965081 | 4.0942173 | 3.6689763 | 4.18358 | -0.51460846 | -0.089367512 |
| Clu | 12.71264 | 12.11515 | 12.659 | -0.54379926 | 0.053694748 | 11.370583 | 11.206992 | 11.7216 | -0.514620462 | -0.351030036 |
| Ptpdc1 | 6.310781 | 6.144674 | 6.3414 | -0.196715832 | -0.030609024 | 6.5616681 | 6.0222061 | 6.53685 | -0.514639516 | 0.024822411 |
| Heph | 5.309144 | 4.816162 | 5.4159 | -0.599698539 | -0.106716409 | 4.6506952 | 4.4083641 | 4.92303 | -0.514670729 | -0.272339623 |
| Dynlt1a | 6.1029 | 6.088765 | 6.1351 | -0.046310358 | -0.032176233 | 6.2519132 | 6.0331525 | 6.54786 | -0.514707073 | -0.295946393 |
| Myo1d | 6.196498 | 6.019169 | 6.0578 | -0.038660601 | 0.138668165 | 5.7120076 | 5.4928664 | 6.00765 | -0.514779607 | -0.295638489 |
| Mecom | 6.177473 | 5.757046 | 6.4793 | -0.72229486 | -0.301866956 | 5.2937752 | 5.1590559 | 5.67422 | -0.515164999 | -0.380445677 |
| Kiz | 7.702719 | 7.549908 | 7.8268 | -0.276884811 | -0.124073829 | 7.8046582 | 7.4155469 | 7.93085 | -0.515306011 | -0.126194778 |
| Scrn3 | 7.348928 | 7.168201 | 7.4004 | -0.232237974 | -0.051510708 | 7.3394376 | 6.9035531 | 7.41902 | -0.515468232 | -0.07958366 |
| Plekhn1 | 9.488081 | 9.459333 | 9.6781 | -0.218760844 | -0.190012636 | 9.9013909 | 9.4144521 | 9.93008 | -0.515624444 | -0.028685619 |
| Thsd1 | 6.227319 | 5.821496 | 6.3016 | -0.480061108 | -0.074238292 | 5.7699067 | 5.2401722 | 5.7558 | -0.515627326 | 0.014107172 |
| Mef2c | 12.38023 | 12.43151 | 12.472 | -0.0407161 | -0.092004116 | 12.47996 | 12.011057 | 12.5267 | -0.515661846 | -0.04675914 |
| Gm20425 | 6.916872 | 6.796386 | 6.2874 | 0.508937693 | 0.629424289 | 5.3986002 | 5.696885 | 6.21261 | -0.515727387 | -0.81401216 |
| Abcg2 | 8.654778 | 8.138786 | 8.6494 | -0.510573859 | 0.005418204 | 7.3532927 | 7.2223275 | 7.73825 | -0.5159218 | -0.384956583 |
| Tpx2 | 8.093862 | 7.957096 | 8.2647 | -0.307595723 | -0.170829122 | 8.1183075 | 8.1735081 | 8.68979 | -0.516283326 | -0.571483929 |
| Tob2 | 11.13656 | 11.041 | 11.209 | -0.168474088 | -0.072917906 | 10.806075 | 10.455667 | 10.972 | -0.516315644 | -0.165908033 |
| Lgalsl | 6.643853 | 6.48458 | 6.7748 | -0.290217945 | -0.130945604 | 6.2290985 | 6.0598657 | 6.57619 | -0.516323261 | -0.347090504 |

| | | | | | | | | | | |
|---|---|---|---|---|---|---|---|---|---|---|
| Ap4e1 | 9.088304 | 8.996642 | 9.0711 | -0.074479097 | 0.01718249 | 8.9773627 | 8.771422 | 9.28795 | -0.516529292 | -0.310588524 |
| Kmt5b | 10.79276 | 10.76198 | 10.842 | -0.080125674 | -0.049347392 | 10.465038 | 10.330695 | 10.8476 | -0.516895823 | -0.382551928 |
| Adcy3 | 8.530966 | 8.407809 | 8.4683 | -0.060495529 | 0.06266108 | 8.4908866 | 8.0559595 | 8.57326 | -0.517298116 | -0.08237099 |
| Fam117b | 10.36636 | 10.3723 | 10.406 | -0.033386544 | -0.039327886 | 10.177845 | 9.8633278 | 10.3808 | -0.517434838 | -0.202917623 |
| Pla2g2d | 8.931841 | 9.341992 | 8.6709 | 0.671077017 | 0.260925868 | 7.2275344 | 7.3241292 | 7.84166 | -0.517535017 | -0.614129834 |
| Glul | 11.05241 | 10.82246 | 10.981 | -0.158639562 | 0.071308104 | 10.65061 | 10.235863 | 10.7535 | -0.517616698 | -0.10286911 |
| E2f8 | 6.641204 | 6.352725 | 6.8102 | -0.45745397 | -0.16897506 | 6.566121 | 6.5543371 | 7.07211 | -0.517772945 | -0.505988979 |
| Sorbs1 | 6.794011 | 6.755923 | 6.8007 | -0.044755663 | -0.006667638 | 6.6197262 | 6.1970379 | 6.71495 | -0.517915819 | -0.095227559 |
| Bmp7 | 5.95336 | 5.849112 | 6.1092 | -0.260115868 | -0.155868599 | 5.7504472 | 5.5048109 | 6.02325 | -0.518439029 | -0.272802807 |
| Lrsam1 | 7.135193 | 7.202694 | 7.602 | -0.399289902 | -0.466790368 | 7.4767132 | 7.0126712 | 7.53137 | -0.518694226 | -0.054652235 |
| Cd101 | 6.830586 | 6.765625 | 6.9013 | -0.135701544 | -0.070740712 | 6.4524174 | 6.3945143 | 6.91329 | -0.518777283 | -0.460874157 |
| Gpm6a | 5.638079 | 5.231608 | 5.5946 | -0.362997345 | 0.043473114 | 4.6908896 | 4.9234939 | 5.44237 | -0.518874855 | -0.751479114 |
| Smyd4 | 7.738128 | 7.740246 | 7.9593 | -0.219065533 | -0.221183855 | 7.7782104 | 7.3626415 | 7.88226 | -0.519615559 | -0.104046656 |
| Prune2 | 4.980031 | 4.548378 | 4.9546 | -0.406202987 | 0.025450283 | 4.5752041 | 4.333703 | 4.85373 | -0.520022086 | -0.278521002 |
| Cyp39a1 | 6.400357 | 6.513765 | 6.4705 | 0.043269862 | -0.070137673 | 6.8133973 | 5.9691312 | 6.48924 | -0.520112687 | 0.324153384 |
| Bub1 | 6.746576 | 6.449252 | 6.9924 | -0.543175423 | -0.245851689 | 6.7272688 | 6.6218006 | 7.14193 | -0.52012605 | -0.414657863 |
| Peak1 | 10.03327 | 9.96104 | 9.9576 | 0.003456699 | 0.075688659 | 9.6104425 | 9.6761113 | 10.1963 | -0.520162802 | -0.585831628 |
| Phf7 | 7.877574 | 7.612971 | 7.9438 | -0.330862911 | -0.066260037 | 8.0908925 | 7.4953893 | 8.01635 | -0.520958157 | 0.074545086 |
| Tspan32 | 10.56771 | 10.54624 | 10.712 | -0.165417908 | -0.143955802 | 10.950751 | 10.271392 | 10.7926 | -0.521195788 | 0.158163451 |
| Lrrc3b | 4.683485 | 4.278305 | 4.8567 | -0.578432269 | -0.173252966 | 4.1913675 | 3.8580558 | 4.37957 | -0.521516095 | -0.188204401 |
| Mapk11 | 9.867524 | 9.790824 | 9.8515 | -0.060642093 | 0.016057618 | 10.405436 | 9.5737131 | 10.0953 | -0.521608767 | 0.310113877 |
| Ankrd26 | 8.19595 | 8.241705 | 8.4087 | -0.16695227 | -0.21270677 | 8.0557764 | 8.0378592 | 8.55956 | -0.521698201 | -0.503780995 |
| mt-Nd4l | 12.44479 | 12.31669 | 12.518 | -0.201501436 | -0.073400464 | 12.238671 | 12.00714 | 12.5292 | -0.522010386 | -0.290479729 |
| Accs | 7.832909 | 7.806706 | 8.0389 | -0.232164524 | -0.205961356 | 8.0144739 | 7.6155729 | 8.13767 | -0.522097794 | -0.123196788 |
| Igkv8-30 | 8.354717 | 8.56338 | 8.3248 | 0.238612592 | 0.029949925 | 8.0108859 | 7.908525 | 8.43078 | -0.522258344 | -0.419897457 |
| Xpr1 | 10.02505 | 9.962275 | 10.108 | -0.145913842 | -0.08313885 | 9.909329 | 9.565435 | 10.0885 | -0.523094405 | -0.179200356 |
| Arhgap31 | 10.32942 | 10.22333 | 10.4 | -0.176482066 | -0.070398518 | 9.6777924 | 9.8133733 | 10.3365 | -0.523162264 | -0.658743174 |
| Ncoa2 | 11.17424 | 11.22249 | 11.282 | -0.059755218 | -0.108003844 | 11.048687 | 10.86172 | 11.3853 | -0.52355965 | -0.336592879 |
| Cd5l | 6.954463 | 7.821592 | 6.5961 | 1.225536734 | 0.358406983 | 7.2018416 | 5.8835845 | 6.40715 | -0.523563479 | 0.794693608 |
| Zfp951 | 7.562222 | 7.48206 | 7.6 | -0.117892908 | -0.037730903 | 7.6310323 | 7.3776942 | 7.9013 | -0.523603609 | -0.270265486 |
| Shroom3 | 5.356915 | 4.947355 | 5.4345 | -0.487117776 | -0.077558255 | 4.9724797 | 4.5667169 | 5.09042 | -0.52369915 | -0.11793635 |
| Capn11 | 3.619379 | 4.071901 | 3.8703 | 0.201638675 | -0.25088324 | 4.1489095 | 3.717792 | 4.24152 | -0.523732052 | -0.092614603 |

| | | | | | | | | | | |
|---|---|---|---|---|---|---|---|---|---|---|
| Ddx3x | 13.2262 | 12.95551 | 13.125 | -0.169021194 | 0.101667762 | 12.976314 | 12.558174 | 13.082 | -0.523843506 | -0.105703633 |
| Psd2 | 5.901231 | 5.893419 | 5.9651 | -0.071715041 | -0.063903271 | 5.8790014 | 5.6348989 | 6.15912 | -0.524221672 | -0.280119193 |
| Catsperg1 | 4.772376 | 4.505107 | 4.5481 | -0.043040352 | 0.22422862 | 4.7479717 | 4.5765183 | 5.10076 | -0.524239831 | -0.352786379 |
| Sorl1 | 13.22978 | 13.30308 | 13.254 | 0.048757888 | -0.024539318 | 13.263396 | 12.686722 | 13.211 | -0.524325364 | 0.052348768 |
| Tnk2 | 10.79515 | 10.71335 | 10.95 | -0.236455624 | -0.154655876 | 10.927925 | 10.393154 | 10.9175 | -0.524393548 | 0.010377292 |
| Matk | 7.630973 | 7.592651 | 7.6196 | -0.026899687 | 0.01142245 | 7.8168787 | 7.0994192 | 7.62385 | -0.524433143 | 0.193026363 |
| Depdc5 | 9.740263 | 9.839284 | 9.7424 | 0.096865777 | -0.00215531 | 9.8172202 | 9.3551126 | 9.87975 | -0.524640368 | -0.062532825 |
| Platr25 | 7.542216 | 7.599858 | 7.6556 | -0.055739278 | -0.113380429 | 7.573471 | 7.2232866 | 7.74798 | -0.524694159 | -0.174509799 |
| Fam149b | 8.478107 | 8.386575 | 8.6543 | -0.267706136 | -0.176174372 | 8.6613329 | 8.2646069 | 8.78942 | -0.524809097 | -0.128083114 |
| Dnajb14 | 8.455623 | 8.353328 | 8.3963 | -0.043019161 | 0.059276163 | 8.4205392 | 7.918124 | 8.44302 | -0.5248923 | -0.022477056 |
| Scn8a | 4.597888 | 4.565542 | 4.6339 | -0.068389637 | -0.036042896 | 4.7541073 | 4.4135994 | 4.93856 | -0.52496364 | -0.184455699 |
| Gja1 | 8.433846 | 7.953327 | 7.8694 | 0.083901812 | 0.564421579 | 7.3191299 | 7.1172176 | 7.64257 | -0.525352797 | -0.323440559 |
| Sntb1 | 8.633989 | 8.542199 | 8.7414 | -0.199228556 | -0.107438155 | 8.2392565 | 8.3727939 | 8.89828 | -0.525489932 | -0.659027348 |
| Gm29397 | 3.10959 | 3.190369 | 3.1595 | 0.030909363 | -0.049870094 | 2.9967329 | 2.9054412 | 3.4311 | -0.525653874 | -0.434362199 |
| Glipr1 | 8.804284 | 8.930567 | 8.9392 | -0.008676029 | -0.134959583 | 8.793092 | 8.3364217 | 8.86214 | -0.525721381 | -0.069051041 |
| Retreg1 | 10.96737 | 10.99364 | 11.046 | -0.05194844 | -0.07821246 | 10.974535 | 10.50205 | 11.0281 | -0.526005532 | -0.053520192 |
| Bpifb2 | 4.202028 | 3.528198 | 4.4582 | -0.930011642 | -0.256181735 | 3.2563806 | 3.2933538 | 3.81961 | -0.526256754 | -0.563229913 |
| Slc16a10 | 7.771091 | 7.978657 | 7.8768 | 0.10187728 | -0.105689269 | 7.8981966 | 7.375194 | 7.90161 | -0.526416915 | -0.003414242 |
| Cbx6 | 9.630313 | 9.670907 | 9.6565 | 0.014383284 | -0.026211144 | 9.3526149 | 9.1508958 | 9.67737 | -0.526478568 | -0.324759482 |
| Synrg | 10.97018 | 10.90455 | 11.001 | -0.095992642 | -0.03036432 | 10.797537 | 10.408783 | 10.9354 | -0.526609656 | -0.137856601 |
| Cd226 | 9.641831 | 9.772759 | 9.8745 | -0.101718215 | -0.232646068 | 9.3301661 | 9.3104187 | 9.83732 | -0.526896964 | -0.50714957 |
| Zscan20 | 8.609086 | 8.604584 | 8.7097 | -0.105101824 | -0.100600102 | 8.5279324 | 8.2174494 | 8.74436 | -0.526910468 | -0.216427444 |
| Eml2 | 8.539303 | 8.354346 | 8.6192 | -0.264836184 | -0.079879789 | 8.6244685 | 8.0836429 | 8.61061 | -0.526971222 | 0.013854414 |
| Fmod | 5.66917 | 5.303202 | 5.5883 | -0.28512426 | 0.080843525 | 5.1028998 | 4.6116486 | 5.13881 | -0.527163956 | -0.035912756 |
| Tmem8 | 9.096376 | 9.096315 | 9.272 | -0.175652504 | -0.175591095 | 9.3706554 | 8.8232859 | 9.35053 | -0.527241486 | 0.020128051 |
| Trim68 | 6.804193 | 6.909268 | 6.9964 | -0.087158326 | -0.192233568 | 7.2942586 | 6.763453 | 7.29097 | -0.527518535 | 0.00328703 |
| Pimreg | 4.968376 | 4.759497 | 5.0569 | -0.297353363 | -0.088474303 | 5.1230223 | 4.7721284 | 5.2997 | -0.527573453 | -0.176679506 |
| Melk | 6.544519 | 6.326675 | 6.6833 | -0.356619052 | -0.138775028 | 6.3863726 | 6.3551246 | 6.88276 | -0.527639122 | -0.496391149 |
| Sgo2a | 6.450624 | 6.117833 | 6.5578 | -0.439966005 | -0.107175034 | 6.28328 | 6.0617959 | 6.58966 | -0.527860431 | -0.306376352 |
| Gprc5b | 6.988243 | 7.137777 | 7.1283 | 0.009437254 | -0.14009682 | 6.6674889 | 6.443229 | 6.97143 | -0.528202474 | -0.30394253 |
| Drc1 | 4.647344 | 4.53108 | 4.4136 | 0.117457791 | 0.233721682 | 4.3666631 | 4.2927195 | 4.82108 | -0.528359383 | -0.454415773 |
| Smarca2 | 11.0054 | 10.92558 | 11.139 | -0.213260748 | -0.13343975 | 10.98318 | 10.642413 | 11.1709 | -0.528447112 | -0.187680733 |

| | | | | | | | | | | |
|---|---|---|---|---|---|---|---|---|---|---|
| Rasal1 | 8.567807 | 8.44223 | 8.2152 | 0.227061807 | 0.352639203 | 8.4017758 | 7.8064562 | 8.33494 | -0.528483243 | 0.066836369 |
| Sorbs2 | 6.669477 | 6.131179 | 6.48 | -0.34881182 | 0.189485599 | 5.8591972 | 5.5222156 | 6.05077 | -0.528557387 | -0.1915758 |
| Gpx3 | 7.643496 | 6.684231 | 7.255 | -0.570812254 | 0.388452847 | 6.5908812 | 6.247795 | 6.77638 | -0.528582385 | -0.185496124 |
| H2-Eb2 | 11.30462 | 11.16774 | 11.298 | -0.129802322 | 0.007086962 | 11.604047 | 10.864595 | 11.3936 | -0.529021872 | 0.210430426 |
| Ighv6-6 | 6.669511 | 6.97942 | 6.7493 | 0.230165422 | -0.079744009 | 7.3261797 | 6.437272 | 6.96632 | -0.529051566 | 0.359856163 |
| Pla2g4f | 5.540724 | 5.721479 | 5.8867 | -0.165175042 | -0.345930358 | 5.5670341 | 5.5990656 | 6.12841 | -0.529347021 | -0.561378512 |
| Erc1 | 8.472617 | 8.160222 | 8.4698 | -0.309551559 | 0.002843114 | 8.0694353 | 7.8264776 | 8.35599 | -0.529513174 | -0.286555504 |
| Slc25a35 | 6.353869 | 6.192192 | 6.4628 | -0.270592249 | -0.1089149 | 6.5300059 | 5.8901283 | 6.41976 | -0.529632278 | 0.110245359 |
| Lrba | 10.72278 | 10.70968 | 10.8 | -0.090630934 | -0.077534382 | 10.647466 | 10.33635 | 10.8662 | -0.52981 | -0.218694849 |
| F2rl3 | 5.417412 | 5.165586 | 5.325 | -0.159386193 | 0.092440202 | 5.3361232 | 4.7918546 | 5.32192 | -0.53006665 | 0.014201983 |
| Ap1s2 | 8.819782 | 8.792399 | 8.8673 | -0.074908459 | -0.047525855 | 8.805189 | 8.306598 | 8.83667 | -0.530069647 | -0.031478671 |
| Atrnl1 | 8.720642 | 8.533855 | 8.6069 | -0.073074946 | 0.11371277 | 8.5915939 | 8.0254341 | 8.55565 | -0.530219935 | 0.035939802 |
| Matn2 | 5.885999 | 5.729061 | 6.0351 | -0.306076592 | -0.14913823 | 5.6530189 | 5.3491715 | 5.87944 | -0.53026571 | -0.226418325 |
| Fbxo32 | 7.361587 | 7.265446 | 7.5052 | -0.239790567 | -0.143649248 | 7.2483573 | 6.9030451 | 7.43339 | -0.530344369 | -0.185032151 |
| Arl15 | 7.523327 | 7.405731 | 7.4663 | -0.060548793 | 0.057047182 | 7.289894 | 6.9016107 | 7.43215 | -0.530538741 | -0.142255418 |
| Rab11fip2 | 8.408504 | 8.190109 | 8.3327 | -0.142558272 | 0.075836705 | 8.2937896 | 7.9191324 | 8.44999 | -0.530856037 | -0.156198795 |
| Baz2b | 10.75879 | 10.73653 | 10.893 | -0.15600558 | -0.1337417 | 10.823889 | 10.41917 | 10.95 | -0.530867127 | -0.126148071 |
| Wdfy2 | 9.70335 | 9.653565 | 9.7843 | -0.130706287 | -0.080921353 | 9.6704733 | 9.2444143 | 9.77553 | -0.531117155 | -0.105058124 |
| Abhd4 | 7.939085 | 7.691689 | 8.1057 | -0.413983852 | -0.166587958 | 7.9578439 | 7.3971838 | 7.92838 | -0.531197865 | 0.029462205 |
| Tarsl2 | 7.19858 | 7.145767 | 6.9249 | 0.220857514 | 0.273670458 | 6.8795433 | 6.4389447 | 6.9704 | -0.531459037 | -0.090860455 |
| Lgals4 | 6.834137 | 6.734723 | 6.8303 | -0.095527701 | 0.003886505 | 7.2116583 | 6.7760186 | 7.30779 | -0.531769612 | -0.096129874 |
| Pbrm1 | 11.69529 | 11.57787 | 11.677 | -0.09910763 | 0.018315454 | 11.527609 | 11.191599 | 11.7234 | -0.53177252 | -0.195762511 |
| Mthfd2l | 5.401113 | 5.050746 | 5.2448 | -0.194067037 | 0.156300443 | 5.3885962 | 4.8731238 | 5.40496 | -0.531838104 | -0.016365735 |
| Fam184b | 4.238308 | 4.231975 | 4.3077 | -0.075675358 | -0.069342892 | 3.9317816 | 3.7361043 | 4.26802 | -0.531916278 | -0.336239058 |
| Il18r1 | 10.12699 | 9.897444 | 10.19 | -0.292671142 | -0.063128006 | 9.7908388 | 9.7997799 | 10.3318 | -0.531995293 | -0.540936388 |
| Gm48583 | 5.410779 | 5.596422 | 6.0848 | -0.488343299 | -0.673985959 | 5.9979213 | 5.4605466 | 5.9927 | -0.532158169 | 0.005216547 |
| Pip4p2 | 8.031858 | 7.958841 | 8.1687 | -0.209883165 | -0.136866387 | 8.1040974 | 7.531784 | 8.06408 | -0.53229204 | 0.040021368 |
| B3gnt8 | 8.408618 | 8.406345 | 8.6008 | -0.194492213 | -0.192218863 | 9.2857107 | 7.8824217 | 8.41489 | -0.532468527 | 0.870820456 |
| Pot1a | 8.480226 | 8.641857 | 8.6764 | -0.034512124 | -0.196143946 | 8.635355 | 8.2423698 | 8.77498 | -0.532612965 | -0.139627761 |
| Vamp1 | 9.097722 | 9.202275 | 9.2247 | -0.0224078 | -0.126961079 | 9.5188246 | 9.0540069 | 9.5867 | -0.532694477 | -0.067876746 |
| Mbnl3 | 8.485433 | 8.352658 | 8.447 | -0.094295574 | 0.038479941 | 8.2891904 | 8.0782348 | 8.61144 | -0.533207665 | -0.322252101 |
| Neil3 | 5.561772 | 5.461593 | 5.5754 | -0.113806425 | -0.013627624 | 5.3909005 | 5.4318087 | 5.96503 | -0.533226233 | -0.574134457 |

| | | | | | | | | | | |
|---|---|---|---|---|---|---|---|---|---|---|
| Kcnq1 | 4.116231 | 4.025288 | 4.1957 | -0.170401676 | -0.07945841 | 3.708397 | 3.4336795 | 3.96719 | -0.533507201 | -0.258789728 |
| Lix1l | 7.06321 | 6.732153 | 6.8474 | -0.115242424 | 0.215815328 | 6.52941 | 6.0997604 | 6.63362 | -0.533863669 | -0.104214061 |
| Slmap | 10.03536 | 9.914465 | 10.033 | -0.118101876 | 0.002793807 | 9.8126801 | 9.6274495 | 10.1613 | -0.53387428 | -0.348643661 |
| Rmnd5b | 9.329824 | 9.080259 | 9.3722 | -0.291909101 | -0.042344129 | 9.5030272 | 8.8503372 | 9.38426 | -0.533923014 | 0.118767017 |
| Habp4 | 7.313784 | 7.282977 | 7.5381 | -0.255121201 | -0.224314941 | 7.4978099 | 6.8067803 | 7.34079 | -0.534007315 | 0.157022323 |
| Wwtr1 | 7.46814 | 7.024455 | 7.2237 | -0.199260371 | 0.244424859 | 6.5389207 | 6.2305409 | 6.76483 | -0.534290551 | -0.225910691 |
| 5730409E04Rik | 5.366467 | 5.124306 | 5.3568 | -0.232476371 | 0.009684589 | 5.0803761 | 4.7036646 | 5.23796 | -0.534297892 | -0.157586412 |
| Sec16b | 4.731786 | 4.645749 | 4.8033 | -0.157557314 | -0.071520456 | 5.0028423 | 4.4613068 | 4.9958 | -0.534489774 | 0.007045764 |
| Ick | 6.895993 | 6.900531 | 7.0564 | -0.155825277 | -0.160363706 | 6.9290608 | 6.5852332 | 7.11987 | -0.534633604 | -0.190805997 |
| Slc35d2 | 7.569528 | 7.26502 | 7.5974 | -0.332352361 | -0.027844662 | 7.5311793 | 7.067533 | 7.60257 | -0.535038331 | -0.071392095 |
| Lyst | 11.63614 | 11.69239 | 11.677 | 0.015276072 | -0.040976864 | 11.608494 | 11.276684 | 11.8118 | -0.535150584 | -0.203341421 |
| Serac1 | 6.355046 | 6.009766 | 6.306 | -0.296239415 | 0.049040488 | 6.308511 | 5.786819 | 6.32213 | -0.535310954 | -0.013618889 |
| Asap1 | 10.66277 | 10.63289 | 10.718 | -0.084901842 | -0.055023942 | 10.588953 | 10.131143 | 10.6666 | -0.535422042 | -0.077612111 |
| Arpin | 7.046022 | 6.831976 | 7.107 | -0.275060315 | -0.061013472 | 6.8971304 | 6.6589813 | 7.19475 | -0.535768543 | -0.297619401 |
| Cyth3 | 8.974652 | 8.887215 | 8.8576 | 0.029573387 | 0.117010146 | 8.4057749 | 8.2910597 | 8.82683 | -0.5357694 | -0.421054204 |
| Sbf2 | 8.49706 | 8.280717 | 8.5825 | -0.301789636 | -0.085446698 | 7.888483 | 7.8685257 | 8.40482 | -0.536295277 | -0.516337973 |
| Dpysl3 | 8.206214 | 7.888823 | 7.8767 | 0.012110956 | 0.329501339 | 7.6444076 | 6.7848768 | 7.3215 | -0.53662669 | 0.322904155 |
| Mical1 | 11.25324 | 11.16105 | 11.392 | -0.231205362 | -0.139008976 | 11.753395 | 11.023051 | 11.5597 | -0.536640416 | 0.193703703 |
| Epm2a | 4.723115 | 4.500833 | 4.7681 | -0.267278832 | -0.044996638 | 4.5198963 | 4.2108994 | 4.74759 | -0.536688059 | -0.227691159 |
| Wbp1 | 8.32249 | 8.241436 | 8.5459 | -0.304459686 | -0.223406169 | 8.7069248 | 8.0522706 | 8.589 | -0.536729738 | 0.117924423 |
| Plcxd2 | 8.818082 | 8.792352 | 8.9742 | -0.181858536 | -0.156128963 | 8.6679456 | 8.5869035 | 9.12367 | -0.536769117 | -0.455726997 |
| Aff3 | 11.0064 | 10.93774 | 11.037 | -0.099509796 | -0.030854554 | 11.105403 | 10.643414 | 11.1804 | -0.536962506 | -0.074973604 |
| B4galt6 | 7.637483 | 7.246707 | 7.3803 | -0.133605332 | 0.257170659 | 7.194822 | 6.4312252 | 6.96858 | -0.537358415 | 0.226238376 |
| Dcaf4 | 8.097492 | 7.982472 | 8.2354 | -0.252973355 | -0.137952987 | 8.3454473 | 7.9004244 | 8.43826 | -0.537837254 | -0.092814359 |
| Tanc1 | 9.079202 | 9.030038 | 9.0489 | -0.01890807 | 0.030255713 | 8.5269534 | 8.61234 | 9.15029 | -0.53794923 | -0.623335854 |
| Zfp808 | 5.758031 | 5.625565 | 6.0186 | -0.393034847 | -0.260569066 | 5.7818328 | 5.4940822 | 6.03219 | -0.53810529 | -0.250354726 |
| AC124237.1 | 6.646393 | 6.503696 | 6.6821 | -0.178440699 | -0.035744459 | 6.6681101 | 6.4119166 | 6.95029 | -0.538369031 | -0.282175538 |
| Kdm6b | 10.86071 | 10.79866 | 10.732 | 0.066643812 | 0.12869001 | 10.580104 | 10.457213 | 10.9963 | -0.539114468 | -0.416223077 |
| Cdc14b | 7.744198 | 8.005351 | 8.1057 | -0.100349955 | -0.361502459 | 7.8875335 | 7.6199176 | 8.15919 | -0.539271582 | -0.271655634 |
| Akr1e1 | 6.57406 | 6.454005 | 6.8816 | -0.427595169 | -0.307539992 | 6.3705042 | 6.0097702 | 6.54923 | -0.539464754 | -0.17873079 |
| Ccdc30 | 5.581171 | 5.485859 | 5.6125 | -0.126652969 | -0.031341016 | 5.4877053 | 5.4289528 | 5.96857 | -0.539621269 | -0.480868768 |
| Slc4a8 | 9.133657 | 9.045195 | 9.2184 | -0.173207218 | -0.084745317 | 9.194591 | 9.1379354 | 9.67795 | -0.540013361 | -0.483357724 |

| | | | | | | | | | | |
|---|---|---|---|---|---|---|---|---|---|---|
| 5031439G07Rik | 10.90814 | 10.84771 | 10.817 | 0.030245604 | 0.09067674 | 10.648037 | 10.15284 | 10.693 | -0.540201343 | -0.045004779 |
| Zfp266 | 9.377271 | 9.214378 | 9.3901 | -0.175756089 | -0.012862341 | 9.141042 | 8.9412342 | 9.48183 | -0.540596153 | -0.340788312 |
| Hif3a | 5.472909 | 5.338363 | 5.5694 | -0.231030793 | -0.096485337 | 5.7929807 | 5.2698889 | 5.8107 | -0.540810069 | -0.017718304 |
| Ext2 | 9.516945 | 9.403659 | 9.5211 | -0.117442201 | -0.004156846 | 9.5547412 | 9.0214043 | 9.56223 | -0.540825583 | -0.007488605 |
| Grem1 | 5.345728 | 4.660153 | 4.9665 | -0.306395182 | 0.379179263 | 4.5430247 | 4.2296533 | 4.77049 | -0.540832797 | -0.227461406 |
| Dnajc6 | 6.011344 | 5.762974 | 5.8384 | -0.075377077 | 0.172992509 | 5.5250028 | 5.262622 | 5.80376 | -0.541137516 | -0.278756761 |
| Lrrc28 | 7.497814 | 7.375608 | 7.5718 | -0.196163179 | -0.073956492 | 7.5126205 | 7.0010899 | 7.54228 | -0.541193499 | -0.029662885 |
| Hddc3 | 6.691396 | 6.660609 | 6.7829 | -0.122324369 | -0.09153716 | 6.9157692 | 6.4304702 | 6.97177 | -0.54129739 | -0.055998459 |
| Slc22a15 | 7.037059 | 6.768536 | 6.9636 | -0.195103873 | 0.073419073 | 6.7879402 | 6.6270715 | 7.16867 | -0.541603177 | -0.380734454 |
| Dlgap5 | 7.462253 | 7.1289 | 7.5444 | -0.41552905 | -0.082175687 | 7.6831579 | 7.3667531 | 7.90845 | -0.541692311 | -0.225287492 |
| Kif22 | 7.215502 | 6.901058 | 7.3684 | -0.467326322 | -0.152881439 | 7.1372896 | 7.0859394 | 7.62786 | -0.541918646 | -0.490568523 |
| Kat6b | 9.469773 | 9.344324 | 9.4915 | -0.147140259 | -0.021691498 | 9.1957977 | 8.984356 | 9.52652 | -0.542159399 | -0.33071767 |
| Pde2a | 10.30527 | 10.24341 | 10.39 | -0.146599007 | -0.084739245 | 9.853895 | 9.4749189 | 10.0175 | -0.542561633 | -0.163585539 |
| Sgo1 | 5.566864 | 5.559942 | 5.9314 | -0.371492577 | -0.364571055 | 5.6429808 | 5.483436 | 6.02618 | -0.542744594 | -0.383199813 |
| Tle1 | 7.727084 | 7.778293 | 7.7376 | 0.04073533 | -0.010474401 | 7.6019958 | 7.0426124 | 7.58626 | -0.543651475 | 0.015731917 |
| Zfp512b | 9.811116 | 9.595215 | 9.9368 | -0.34158422 | -0.125682472 | 10.071259 | 9.5051853 | 10.0491 | -0.543899551 | 0.022174593 |
| Gstp3 | 6.97521 | 6.727014 | 7.0422 | -0.315221652 | -0.067026222 | 7.007464 | 6.4600908 | 7.00472 | -0.544633697 | 0.002739489 |
| Apbb2 | 6.865409 | 6.617849 | 6.9025 | -0.284685293 | -0.037125335 | 6.3708817 | 6.1270136 | 6.67183 | -0.544818536 | -0.300950451 |
| Slain2 | 10.59502 | 10.57647 | 10.597 | -0.020217226 | -0.00166281 | 10.46162 | 10.004414 | 10.5494 | -0.545019668 | -0.087813834 |
| Ints6l | 9.371518 | 9.345135 | 9.5137 | -0.168603146 | -0.142220529 | 9.6492048 | 9.2306535 | 9.77567 | -0.545019973 | -0.126468717 |
| Mmp9 | 7.200606 | 6.936383 | 6.538 | 0.398421867 | 0.662644456 | 7.2221799 | 5.9291242 | 6.47437 | -0.545242631 | 0.747813059 |
| Mpp5 | 8.710121 | 8.819061 | 8.7337 | 0.08535244 | -0.02358726 | 8.5504066 | 8.2726832 | 8.818 | -0.545318701 | -0.267595317 |
| C2cd2l | 8.55761 | 8.457177 | 8.5201 | -0.062918371 | 0.03751404 | 8.5721723 | 8.0158576 | 8.56127 | -0.54541028 | 0.010904435 |
| Aplp2 | 10.94094 | 10.55619 | 10.872 | -0.316304254 | 0.068442142 | 10.50229 | 10.279402 | 10.8253 | -0.545858736 | -0.322970008 |
| Tbx3 | 4.451767 | 4.39395 | 4.4604 | -0.066492418 | -0.008675325 | 4.1013278 | 3.8450527 | 4.39107 | -0.546022057 | -0.289746898 |
| Mterf2 | 6.74013 | 6.598 | 6.8621 | -0.264136548 | -0.122006266 | 6.9503881 | 6.5991157 | 7.14521 | -0.546094589 | -0.194822219 |
| Adamts10 | 10.09906 | 10.05195 | 10.242 | -0.190005545 | -0.142899813 | 10.452325 | 10.019097 | 10.5657 | -0.546625809 | -0.113397508 |
| Pdk4 | 5.360119 | 5.071405 | 5.6572 | -0.58581831 | -0.297104752 | 4.7558544 | 4.7901178 | 5.33689 | -0.546776815 | -0.581040167 |
| Cat | 10.93023 | 11.00633 | 11.134 | -0.127736446 | -0.203834138 | 10.979042 | 10.517939 | 11.0654 | -0.54747661 | -0.086374476 |
| Mettl27 | 6.46716 | 6.054331 | 6.7121 | -0.657805551 | -0.244976768 | 6.4031697 | 5.8596718 | 6.40717 | -0.547494793 | -0.003996912 |
| Dlg1 | 10.4139 | 10.37744 | 10.443 | -0.065238596 | -0.028783982 | 10.258555 | 10.049236 | 10.5968 | -0.54757083 | -0.338252539 |
| Ppfibp1 | 8.649402 | 8.400992 | 8.505 | -0.10401951 | 0.144390669 | 8.224726 | 7.8419235 | 8.38974 | -0.547817229 | -0.165014745 |

| | | | | | | | | | | |
|---|---|---|---|---|---|---|---|---|---|---|
| Hexb | 10.57252 | 10.52188 | 10.648 | -0.126010084 | -0.075367326 | 10.761524 | 10.018798 | 10.5666 | -0.547847494 | 0.194879416 |
| Prrt2 | 4.874736 | 4.975201 | 5.0758 | -0.100570127 | -0.201035246 | 4.5403517 | 4.7611946 | 5.30906 | -0.547869774 | -0.768712689 |
| Rcan3 | 8.375125 | 8.294977 | 8.552 | -0.256983116 | -0.176835625 | 8.1491313 | 7.8121248 | 8.36003 | -0.547903971 | -0.210897478 |
| Haao | 10.30635 | 10.13926 | 10.435 | -0.295760565 | -0.128679322 | 10.929271 | 10.158555 | 10.7069 | -0.548325175 | 0.222390407 |
| Ano6 | 10.6066 | 10.47006 | 10.643 | -0.172558662 | -0.036018204 | 10.458082 | 9.9831667 | 10.5315 | -0.548343223 | -0.073427731 |
| Pdgfd | 4.579614 | 4.433397 | 4.6262 | -0.192783105 | -0.046565762 | 4.2628777 | 3.9840536 | 4.53256 | -0.548508791 | -0.269684654 |
| Ttll5 | 8.113575 | 7.921871 | 8.0944 | -0.172551106 | 0.019153013 | 8.3536921 | 7.7946166 | 8.34334 | -0.548725057 | 0.010350467 |
| Crispld2 | 7.72723 | 7.524736 | 7.5909 | -0.066206694 | 0.136287339 | 7.8254882 | 7.1418578 | 7.69097 | -0.549109166 | 0.134521265 |
| Tspan7 | 9.123691 | 8.751155 | 8.8794 | -0.128286528 | 0.244249347 | 8.2243219 | 7.652337 | 8.20145 | -0.549117928 | 0.02286704 |
| Csrnp2 | 6.853833 | 7.018645 | 7.0619 | -0.043280933 | -0.208093371 | 6.8912365 | 6.538982 | 7.08818 | -0.549194628 | -0.19694011 |
| Prkar2b | 6.205137 | 6.069681 | 6.4262 | -0.356536042 | -0.221079861 | 6.2404455 | 5.6608596 | 6.21022 | -0.54935747 | 0.030228408 |
| Zfp683 | 4.557975 | 4.331007 | 4.6135 | -0.282508319 | -0.055540336 | 4.4139268 | 4.173966 | 4.72355 | -0.549584744 | -0.309623958 |
| Crtc3 | 10.13196 | 9.960589 | 10.047 | -0.086120724 | 0.085254831 | 9.8654069 | 9.5105312 | 10.0602 | -0.549696312 | -0.194820632 |
| Clasp1 | 10.34214 | 10.30186 | 10.437 | -0.134665108 | -0.094384342 | 10.413816 | 9.9882253 | 10.5381 | -0.549921278 | -0.124330517 |
| Manba | 8.881602 | 8.791645 | 9.1027 | -0.31103118 | -0.221074133 | 8.9959936 | 8.3795933 | 8.92978 | -0.550181688 | 0.066218557 |
| Bora | 7.125733 | 7.168976 | 7.4338 | -0.264831149 | -0.308073676 | 7.3237026 | 6.8934179 | 7.44368 | -0.550266973 | -0.119982228 |
| Ttbk1 | 5.990451 | 5.842917 | 5.9731 | -0.130232616 | 0.017301254 | 5.7967888 | 5.4681796 | 6.01884 | -0.550660041 | -0.222050819 |
| Zbtb4 | 10.69984 | 10.60179 | 10.75 | -0.147822748 | -0.049774772 | 10.691744 | 10.083921 | 10.6347 | -0.550738709 | 0.057084414 |
| Epn2 | 6.797342 | 6.702344 | 6.9072 | -0.204873202 | -0.109874485 | 6.1872681 | 6.2540711 | 6.805 | -0.550927951 | -0.617730949 |
| Fam107b | 12.57083 | 12.65462 | 12.579 | 0.075789008 | -0.007996976 | 12.287355 | 12.122705 | 12.6742 | -0.551497562 | -0.38684748 |
| Rapgef4 | 9.221486 | 9.225963 | 9.4933 | -0.267336718 | -0.271813039 | 9.4477456 | 9.1839072 | 9.73552 | -0.551607899 | -0.287769442 |
| Aqp3 | 6.765594 | 6.804 | 6.996 | -0.19204592 | -0.230451716 | 6.7017257 | 6.3806641 | 6.93244 | -0.551774132 | -0.230712496 |
| Plk1 | 7.047025 | 6.954306 | 7.1295 | -0.175147093 | -0.082427288 | 7.0296605 | 6.9500434 | 7.50183 | -0.551784916 | -0.472167821 |
| Zfp658 | 6.684188 | 6.558179 | 6.8911 | -0.332919396 | -0.206909893 | 6.2672983 | 6.1791889 | 6.73122 | -0.552034389 | -0.463924942 |
| Il4i1 | 10.8716 | 10.81539 | 10.84 | -0.025108072 | 0.031103744 | 10.936296 | 10.239215 | 10.7913 | -0.552076618 | 0.145003555 |
| Igsf9 | 6.944375 | 6.696496 | 6.8334 | -0.136914652 | 0.110964407 | 7.3393988 | 6.5722256 | 7.12464 | -0.552415967 | 0.21475716 |
| Lrrc8c | 10.65158 | 10.48926 | 10.569 | -0.079357892 | 0.082959078 | 10.34607 | 10.113138 | 10.6663 | -0.553119689 | -0.320187163 |
| Hyal2 | 8.064166 | 7.50786 | 7.7376 | -0.229756267 | 0.326549514 | 7.0323258 | 6.7041846 | 7.25776 | -0.553579948 | -0.225438685 |
| Fbxw10 | 5.499749 | 5.387755 | 5.3758 | 0.011955615 | 0.123950543 | 5.4546645 | 4.9250725 | 5.47886 | -0.553789853 | -0.024197857 |
| Ablim3 | 5.763722 | 5.374085 | 5.5546 | -0.180515565 | 0.209121777 | 4.9659472 | 4.8624813 | 5.41646 | -0.553975286 | -0.450509377 |
| D3Ertd751e | 7.301639 | 6.930517 | 7.2392 | -0.308639733 | 0.062481767 | 7.5361452 | 6.99546 | 7.54963 | -0.554167399 | -0.013482235 |
| Agfg2 | 8.410075 | 8.288007 | 8.4622 | -0.174213561 | -0.052144771 | 8.5269345 | 7.8546271 | 8.40882 | -0.554193849 | 0.118113488 |

| | | | | | | | | | | |
|---|---|---|---|---|---|---|---|---|---|---|
| Krt222 | 5.304867 | 5.192448 | 5.4567 | -0.264297773 | -0.151878622 | 5.3726479 | 4.8403925 | 5.39507 | -0.554682289 | -0.022426907 |
| Myzap | 7.320391 | 6.946121 | 7.1231 | -0.176968837 | 0.197301111 | 6.6122382 | 6.3571139 | 6.91244 | -0.555326409 | -0.300202083 |
| Slk | 10.88022 | 10.76392 | 10.82 | -0.056160926 | 0.06014734 | 10.573049 | 10.400497 | 10.9569 | -0.556358987 | -0.383807193 |
| Unc45b | 5.702172 | 5.306696 | 5.4743 | -0.167647293 | 0.227829417 | 4.7665839 | 4.9599546 | 5.51634 | -0.556388668 | -0.749759371 |
| Rflnb | 10.2295 | 10.45859 | 10.343 | 0.115778668 | -0.113311208 | 10.058261 | 9.5969474 | 10.1534 | -0.556411482 | -0.09509773 |
| Pcp4l1 | 4.091249 | 4.135209 | 4.3711 | -0.235855918 | -0.279815941 | 4.1820678 | 3.7648546 | 4.32211 | -0.557257595 | -0.140044449 |
| Igfbp7 | 10.55201 | 10.15051 | 10.51 | -0.358993057 | 0.042500126 | 9.6687536 | 9.272829 | 9.83029 | -0.557463747 | -0.161539082 |
| Ect2 | 7.341533 | 7.026581 | 7.3172 | -0.290642318 | 0.024309321 | 7.1420705 | 6.9426518 | 7.50018 | -0.557532755 | -0.358114137 |
| Ltf | 5.339868 | 5.140577 | 5.414 | -0.273450168 | -0.074158575 | 5.8173163 | 5.2316358 | 5.78928 | -0.557642581 | 0.028037935 |
| Thbs2 | 5.884054 | 6.016591 | 6.0683 | -0.051741615 | -0.184278832 | 5.8009265 | 5.3489985 | 5.90682 | -0.557819238 | -0.105891215 |
| Recql | 9.482141 | 9.40378 | 9.3224 | 0.081397241 | 0.159757868 | 9.5067973 | 9.1274713 | 9.68539 | -0.557918148 | -0.178592176 |
| Marveld1 | 8.741309 | 8.598986 | 8.9163 | -0.317298005 | -0.174975524 | 8.5749454 | 8.1449328 | 8.70319 | -0.558261279 | -0.128248692 |
| Fam124b | 4.875511 | 4.915937 | 4.9292 | -0.013294412 | -0.053720983 | 4.5196667 | 4.3918956 | 4.95041 | -0.558517021 | -0.430745969 |
| Scmh1 | 8.606534 | 8.635343 | 8.6299 | 0.005444581 | -0.023364361 | 8.4259501 | 8.2178979 | 8.77703 | -0.559136352 | -0.351084137 |
| Tjp2 | 7.778426 | 7.473956 | 7.757 | -0.282996576 | 0.021473438 | 7.2933727 | 7.0714291 | 7.63073 | -0.559303363 | -0.337359709 |
| Ttc7b | 8.48705 | 8.431301 | 8.4979 | -0.066633827 | -0.010885108 | 8.4672757 | 7.9456003 | 8.50492 | -0.559315747 | -0.037640362 |
| Aph1b | 5.483555 | 5.273612 | 5.8175 | -0.543936411 | -0.333993313 | 5.5189276 | 4.8245437 | 5.3839 | -0.55935417 | 0.135029696 |
| Ighv1-18 | 8.130869 | 8.582373 | 8.0155 | 0.56686479 | 0.115361433 | 8.2397867 | 7.7205311 | 8.28006 | -0.559526516 | -0.040271004 |
| Cplx2 | 5.532664 | 5.38736 | 5.4114 | -0.024021868 | 0.121282778 | 5.1783718 | 4.8243503 | 5.38391 | -0.559561721 | -0.205540201 |
| Ebf3 | 5.948812 | 5.442691 | 5.8178 | -0.37515426 | 0.130966111 | 4.9738644 | 4.8630805 | 5.4228 | -0.559717898 | -0.448933997 |
| Exoc3l2 | 5.638061 | 5.453448 | 5.4682 | -0.014730639 | 0.169882489 | 5.1906843 | 4.7468881 | 5.30673 | -0.559837252 | -0.116041089 |
| Ephb6 | 7.866877 | 7.580233 | 7.8672 | -0.286966715 | -0.000322973 | 7.6218848 | 7.3158261 | 7.87606 | -0.560230219 | -0.254171517 |
| Cip2a | 7.408223 | 7.177965 | 7.4739 | -0.295905576 | -0.065647397 | 7.4111307 | 7.2544438 | 7.81484 | -0.560394962 | -0.403708067 |
| Ska3 | 5.018873 | 5.034299 | 5.2782 | -0.243930008 | -0.259355452 | 5.0427705 | 5.0235913 | 5.58449 | -0.560901607 | -0.541722352 |
| Braf | 9.789707 | 9.675798 | 9.8057 | -0.129899322 | -0.015990294 | 9.7163832 | 9.4173992 | 9.9783 | -0.560903535 | -0.261919572 |
| Axin2 | 8.042757 | 8.004026 | 8.0186 | -0.014523722 | 0.02420716 | 7.5997233 | 7.6320785 | 8.19334 | -0.5612663 | -0.593621543 |
| Zfp956 | 8.365481 | 8.22833 | 8.5256 | -0.297262153 | -0.16011118 | 8.446134 | 7.9901653 | 8.55144 | -0.561273483 | -0.105304799 |
| Actg2 | 4.337214 | 4.351007 | 4.8396 | -0.488597113 | -0.502390292 | 4.3629069 | 4.0830046 | 4.64458 | -0.561578731 | -0.281676427 |
| Cd96 | 10.42015 | 10.41455 | 10.558 | -0.142980518 | -0.137383658 | 10.082039 | 9.9247502 | 10.4864 | -0.561611659 | -0.404323015 |
| Tdrkh | 6.232779 | 6.068442 | 6.2457 | -0.177224519 | -0.012887992 | 5.9852409 | 5.7307216 | 6.29235 | -0.561632517 | -0.307113215 |
| Srgap3 | 7.5531 | 7.388643 | 7.5639 | -0.175207817 | -0.010750847 | 7.0552155 | 6.7028833 | 7.26534 | -0.562454543 | -0.210122334 |
| Esm1 | 3.990421 | 3.919185 | 4.0189 | -0.099711511 | -0.028475844 | 4.011917 | 3.7162256 | 4.27902 | -0.562797372 | -0.267105989 |

| | | | | | | | | | | |
|---|---|---|---|---|---|---|---|---|---|---|
| Gfod1 | 7.742662 | 7.668986 | 7.7871 | -0.118115517 | -0.044439402 | 7.4765425 | 7.0932766 | 7.65646 | -0.563179007 | -0.179913124 |
| Pag1 | 11.78994 | 11.7745 | 11.767 | 0.007379688 | 0.022825762 | 11.671982 | 11.172547 | 11.7361 | -0.563527126 | -0.064091837 |
| Cacnb1 | 7.29709 | 7.46157 | 7.3912 | 0.070391066 | -0.094089228 | 7.2511485 | 6.9435983 | 7.50729 | -0.563691546 | -0.256141369 |
| Timp4 | 3.425084 | 3.092226 | 3.3216 | -0.229376217 | 0.10348194 | 3.073818 | 2.8791255 | 3.44317 | -0.564040758 | -0.369348244 |
| Arhgap24 | 7.742623 | 7.692068 | 7.7097 | -0.017603998 | 0.032951296 | 7.7457799 | 7.1477953 | 7.71195 | -0.564157629 | 0.033826956 |
| Nav1 | 8.671195 | 8.671079 | 8.7671 | -0.096056018 | -0.095939568 | 8.5213502 | 8.5509693 | 9.11523 | -0.564264097 | -0.593883172 |
| Gm2026 | 5.842943 | 5.577812 | 5.707 | -0.129174714 | 0.135956898 | 5.587573 | 5.1304773 | 5.69477 | -0.564293882 | -0.107198252 |
| Dmxl1 | 11.76304 | 11.83718 | 11.652 | 0.185498906 | 0.111356 | 11.785676 | 11.496457 | 12.0609 | -0.564393434 | -0.275174196 |
| Smtn | 6.965471 | 6.694808 | 6.991 | -0.296174742 | -0.025512541 | 6.8362633 | 6.2163615 | 6.78108 | -0.564722917 | 0.055178896 |
| Frmd6 | 8.511956 | 8.470857 | 8.6008 | -0.129903332 | -0.088804068 | 8.0278564 | 7.8255454 | 8.3909 | -0.565350348 | -0.363039298 |
| Ano1 | 6.219697 | 5.651372 | 6.1851 | -0.533735116 | 0.034589508 | 5.4609709 | 4.8015198 | 5.36741 | -0.565893061 | 0.093558064 |
| Ckap2 | 6.246406 | 6.003221 | 6.3472 | -0.343957543 | -0.10077274 | 6.1558549 | 6.0030788 | 6.56898 | -0.56589989 | -0.413123846 |
| Cd7 | 9.350745 | 9.362601 | 9.4864 | -0.123770243 | -0.135626201 | 9.4470508 | 8.8314411 | 9.39746 | -0.566023101 | 0.049586581 |
| Ly6g5b | 5.498181 | 5.329102 | 5.3646 | -0.035459292 | 0.133619606 | 5.0569112 | 4.8198093 | 5.38599 | -0.56618051 | -0.329078634 |
| Slc25a24 | 8.312363 | 8.102078 | 8.3495 | -0.24739034 | -0.037104988 | 8.0199511 | 7.6830796 | 8.24928 | -0.566199492 | -0.229328006 |
| Mpzl3 | 7.619557 | 7.609388 | 7.6199 | -0.010516803 | -0.000347914 | 7.6738661 | 7.2428265 | 7.80904 | -0.56620881 | -0.135169202 |
| Slamf1 | 9.629653 | 9.304908 | 9.5799 | -0.274980445 | 0.049764116 | 9.7314219 | 9.0218576 | 9.5881 | -0.566237596 | 0.143326662 |
| Sp6 | 5.539202 | 5.745139 | 5.6891 | 0.05601764 | -0.149919886 | 5.4339597 | 5.1943942 | 5.7611 | -0.566703787 | -0.327138287 |
| Zfp882 | 6.96542 | 6.97869 | 7.0779 | -0.099252996 | -0.112523678 | 7.1081422 | 6.4825379 | 7.04975 | -0.567216421 | 0.058387955 |
| Apbb1 | 5.354554 | 5.524836 | 5.6138 | -0.088998592 | -0.259280346 | 5.7180424 | 4.8601814 | 5.42771 | -0.567524152 | 0.290336794 |
| Tex9 | 7.441666 | 7.418636 | 7.6726 | -0.253999244 | -0.23096948 | 7.7707858 | 7.4007347 | 7.96834 | -0.567601583 | -0.197550545 |
| Mob3b | 6.983433 | 6.866265 | 7.1378 | -0.271542618 | -0.154373956 | 7.0221326 | 6.5797547 | 7.1474 | -0.567643998 | -0.125266062 |
| Tle2 | 4.763614 | 4.739021 | 4.8796 | -0.140580726 | -0.115987308 | 4.9075031 | 4.4302293 | 4.99791 | -0.567677973 | -0.090404152 |
| Gstm2 | 5.673841 | 5.359684 | 5.6226 | -0.262895678 | 0.05126117 | 5.4110865 | 4.879127 | 5.44693 | -0.567802121 | -0.035842624 |
| Hmga1b | 3.709042 | 3.442428 | 3.7458 | -0.303359721 | -0.036745039 | 2.90017 | 2.8419444 | 3.40975 | -0.567802772 | -0.509577223 |
| Tspoap1 | 6.903591 | 7.083332 | 6.9111 | 0.172266695 | -0.007474462 | 7.6280543 | 7.3681449 | 7.93606 | -0.56791824 | -0.308008825 |
| Ckb | 7.895159 | 7.615264 | 7.8946 | -0.27937159 | 0.00052351 | 8.2647407 | 7.2767736 | 7.84473 | -0.567952679 | 0.420014411 |
| Jam2 | 7.44115 | 7.050181 | 7.3908 | -0.340603072 | 0.050365504 | 6.673898 | 6.6059386 | 7.17411 | -0.568172765 | -0.50021337 |
| Rps6ka2 | 6.733109 | 6.836368 | 6.9636 | -0.127253139 | -0.230511984 | 6.7808259 | 6.4271196 | 6.99549 | -0.568368388 | -0.21466209 |
| Hipk3 | 10.92441 | 10.83344 | 10.862 | -0.028612592 | 0.062355056 | 10.57546 | 10.252663 | 10.8212 | -0.568539106 | -0.245742181 |
| Qpct | 5.948239 | 5.846737 | 5.8365 | 0.010273933 | 0.111776178 | 5.6684971 | 5.3230065 | 5.89179 | -0.568784357 | -0.223293722 |
| Mturn | 8.369788 | 8.27653 | 8.5134 | -0.236889125 | -0.143631346 | 8.2629334 | 7.8615455 | 8.43067 | -0.56912183 | -0.167733856 |

| | | | | | | | | | | |
|---|---|---|---|---|---|---|---|---|---|---|
| Zfp41 | 7.479487 | 7.517274 | 7.6246 | -0.10737344 | -0.145160558 | 7.4394359 | 7.0807099 | 7.65015 | -0.569438722 | -0.210712726 |
| Stap1 | 10.33628 | 10.25721 | 10.378 | -0.120772501 | -0.041704037 | 10.63068 | 10.04345 | 10.6131 | -0.569674925 | 0.017555111 |
| Rgs2 | 11.05873 | 10.75088 | 11.19 | -0.439617884 | -0.13176593 | 10.219195 | 10.044561 | 10.6144 | -0.569858871 | -0.395224633 |
| Hjurp | 10.53222 | 10.58897 | 10.721 | -0.131902546 | -0.188655524 | 10.651937 | 10.444053 | 11.014 | -0.569919606 | -0.362035556 |
| Endou | 5.558473 | 5.531749 | 5.6808 | -0.149017908 | -0.122294013 | 5.4317049 | 5.0498593 | 5.61981 | -0.569948896 | -0.18810331 |
| Mcpt4 | 8.414371 | 8.562954 | 8.3172 | 0.245796665 | 0.097213747 | 7.9523016 | 7.4086062 | 7.9786 | -0.569994959 | -0.026299607 |
| Ralgapa1 | 10.49318 | 10.42499 | 10.587 | -0.161580382 | -0.09339241 | 10.466603 | 10.161266 | 10.7313 | -0.570065858 | -0.264729237 |
| Fbxl20 | 10.27466 | 10.25007 | 10.465 | -0.215338212 | -0.190743801 | 10.21074 | 9.9331006 | 10.5037 | -0.570555942 | -0.292916206 |
| Ccsap | 6.86264 | 6.864936 | 6.9794 | -0.114487165 | -0.116783664 | 6.8343136 | 6.4717118 | 7.04235 | -0.570636859 | -0.208035071 |
| Birc5 | 7.324544 | 6.979799 | 7.4545 | -0.474731645 | -0.129986732 | 7.2454904 | 7.1862918 | 7.7577 | -0.571404287 | -0.512205693 |
| Otud7b | 10.2052 | 10.14589 | 10.124 | 0.021790804 | 0.081100888 | 9.9746739 | 9.6335499 | 10.205 | -0.571449222 | -0.230325234 |
| Zkscan8 | 8.92965 | 8.75432 | 8.9261 | -0.171741608 | 0.003588744 | 9.2492914 | 8.613284 | 9.18485 | -0.57156162 | 0.064445832 |
| Kdelc2 | 6.350095 | 6.017889 | 6.4152 | -0.397353442 | -0.065147601 | 5.9043891 | 5.576625 | 6.1482 | -0.571570478 | -0.243806363 |
| St5 | 6.524857 | 6.538124 | 6.5078 | 0.030351997 | 0.017085639 | 6.5614875 | 5.8762942 | 6.44815 | -0.571854319 | 0.113339021 |
| H2-Ob | 13.12133 | 13.10379 | 13.246 | -0.142610564 | -0.125074252 | 13.743883 | 12.964493 | 13.5364 | -0.571938538 | 0.20745122 |
| Uvssa | 9.368443 | 9.276792 | 9.4564 | -0.179614564 | -0.087963058 | 9.5787853 | 9.1881858 | 9.76018 | -0.571996577 | -0.181397076 |
| Adcy7 | 12.23169 | 12.2694 | 12.358 | -0.08901392 | -0.12672162 | 12.363905 | 11.949854 | 12.522 | -0.572146208 | -0.158094921 |
| Prkci | 8.623216 | 8.467174 | 8.7365 | -0.269340847 | -0.113299027 | 8.3574277 | 8.091081 | 8.66332 | -0.572242605 | -0.305895982 |
| Pcdh1 | 6.78787 | 6.530911 | 6.7131 | -0.182187534 | 0.074771516 | 6.2000085 | 5.9121818 | 6.4848 | -0.572620635 | -0.284793877 |
| Tpcn1 | 9.434976 | 9.501262 | 9.5705 | -0.069192397 | -0.135478897 | 9.3322604 | 8.9039948 | 9.47708 | -0.573081507 | -0.144815919 |
| Zfp2 | 5.345952 | 5.378515 | 5.3928 | -0.014238516 | -0.046801203 | 5.3781281 | 4.9627493 | 5.53599 | -0.573238348 | -0.15785956 |
| Dkk2 | 4.494727 | 4.718922 | 4.9006 | -0.181652789 | -0.405848304 | 4.8109378 | 4.3600889 | 4.93358 | -0.573490717 | -0.122641749 |
| Zfp667 | 6.429048 | 6.291606 | 6.485 | -0.193427932 | -0.055985908 | 6.3077315 | 5.8764094 | 6.44998 | -0.57356584 | -0.14224374 |
| Gabrr2 | 5.750646 | 5.61895 | 5.8272 | -0.208287538 | -0.076591444 | 5.6768624 | 5.2378591 | 5.81143 | -0.573569974 | -0.134566632 |
| Lancl1 | 8.751424 | 8.663504 | 8.8876 | -0.224099604 | -0.136179662 | 9.0986849 | 8.424096 | 8.998 | -0.57389925 | 0.100689604 |
| Aldh5a1 | 4.639756 | 4.55563 | 4.7137 | -0.158054608 | -0.073928624 | 4.7171776 | 4.3266307 | 4.90069 | -0.574061136 | -0.183514237 |
| Gng10 | 9.893964 | 9.603215 | 9.7439 | -0.140705432 | 0.150043834 | 10.025467 | 9.1166241 | 9.69089 | -0.574266993 | 0.334575853 |
| Arvcf | 5.421919 | 5.488786 | 5.7004 | -0.211573192 | -0.278440138 | 5.4162997 | 4.9228113 | 5.49739 | -0.574578833 | -0.08109045 |
| Rgs5 | 8.882373 | 8.704507 | 8.6421 | 0.062403813 | 0.24026995 | 8.3274373 | 7.6070573 | 8.18179 | -0.574730606 | 0.145649354 |
| Gm14391 | 7.411826 | 7.287164 | 7.4554 | -0.1682034 | -0.043541442 | 7.2561746 | 7.0232959 | 7.59834 | -0.575042315 | -0.342163591 |
| Zdhhc20 | 11.2883 | 11.10795 | 11.366 | -0.257589184 | -0.077239726 | 11.255959 | 10.783024 | 11.3585 | -0.575484818 | -0.102549681 |
| Steap4 | 8.2501 | 7.577508 | 7.1029 | 0.474632387 | 1.14722345 | 7.4756831 | 6.3818352 | 6.95764 | -0.575804283 | 0.518043548 |

| | | | | | | | | | | |
|---|---|---|---|---|---|---|---|---|---|---|
| Rrm2 | 8.644417 | 8.211471 | 8.7403 | -0.528828618 | -0.095882847 | 8.5361865 | 8.4856855 | 9.0616 | -0.575911689 | -0.525410715 |
| Akap11 | 10.42434 | 10.29448 | 10.33 | -0.035829612 | 0.094031532 | 10.006179 | 9.6337404 | 10.2098 | -0.576055427 | -0.2036163 |
| Igf2r | 10.6338 | 10.7177 | 10.721 | -0.003550848 | -0.087451626 | 10.35282 | 10.042968 | 10.6193 | -0.576377672 | -0.266525546 |
| Epcam | 6.181378 | 5.847017 | 6.1142 | -0.267140893 | 0.067220064 | 5.8971112 | 5.4954908 | 6.07189 | -0.576403636 | -0.174783238 |
| Endov | 8.77753 | 8.63219 | 8.8908 | -0.25857576 | -0.113235582 | 9.0709676 | 8.476854 | 9.05326 | -0.576410561 | 0.01770301 |
| Smad7 | 10.08765 | 10.17475 | 10.239 | -0.064511605 | -0.15160677 | 9.8076772 | 9.6426254 | 10.2191 | -0.576511657 | -0.411459857 |
| Wwp1 | 9.044922 | 8.909129 | 9.0664 | -0.157233417 | -0.021440779 | 8.6899219 | 8.3814272 | 8.9581 | -0.576671876 | -0.268177168 |
| Entpd4 | 9.009292 | 8.884387 | 9.0634 | -0.179044582 | -0.054139357 | 9.0018358 | 8.4644261 | 9.04124 | -0.576813732 | -0.039404049 |
| Gpam | 8.673229 | 8.267165 | 8.4958 | -0.2286211 | 0.177442258 | 8.3210927 | 7.7595847 | 8.3366 | -0.577015292 | -0.015507369 |
| Gm5512 | 2.895285 | 2.944997 | 2.8721 | 0.072866399 | 0.023154285 | 2.8907851 | 2.6480753 | 3.22512 | -0.577045481 | -0.334335671 |
| Kras | 10.4508 | 10.21582 | 10.419 | -0.202788666 | 0.032195452 | 10.498141 | 10.006757 | 10.584 | -0.577219841 | -0.085835949 |
| Ankrd44 | 12.24388 | 12.21307 | 12.313 | -0.099566248 | -0.06875416 | 12.145896 | 11.794541 | 12.3722 | -0.57766126 | -0.226306575 |
| Ide | 9.55073 | 9.411536 | 9.341 | 0.070577192 | 0.209771182 | 9.3627732 | 8.8233018 | 9.4011 | -0.577795361 | -0.038323942 |
| Abcd2 | 5.299569 | 4.918235 | 5.4525 | -0.534306881 | -0.152972891 | 5.8015587 | 4.8103014 | 5.38851 | -0.578212579 | 0.413044742 |
| Smpdl3a | 10.1665 | 9.950963 | 10.247 | -0.296354348 | -0.080812852 | 9.9052659 | 9.5410731 | 10.1194 | -0.578342836 | -0.214150021 |
| H1f0 | 8.470317 | 7.894933 | 8.5933 | -0.69839765 | -0.123013911 | 8.1237908 | 7.8765019 | 8.45548 | -0.578980491 | -0.331691687 |
| Arhgap32 | 5.901501 | 5.532536 | 5.6216 | -0.08910173 | 0.279862899 | 5.5229061 | 4.9361612 | 5.51542 | -0.57925826 | 0.007486603 |
| Bambi | 6.991313 | 6.719385 | 6.9324 | -0.212982091 | 0.058945336 | 6.3502709 | 6.0565021 | 6.63608 | -0.579574155 | -0.285805328 |
| Coq8a | 9.228532 | 9.233011 | 9.3814 | -0.148424857 | -0.152904121 | 9.7184581 | 8.8993764 | 9.47916 | -0.579784017 | 0.239297708 |
| Afdn | 8.195827 | 7.835786 | 8.1501 | -0.314284729 | 0.045756508 | 7.7667447 | 7.5222473 | 8.10208 | -0.579829059 | -0.335331637 |
| Prss12 | 6.602392 | 6.645685 | 6.694 | -0.048361769 | -0.091655448 | 6.0390053 | 5.8785104 | 6.4587 | -0.580190713 | -0.419695847 |
| Fat4 | 6.303459 | 5.935888 | 6.3368 | -0.400957904 | -0.033386721 | 5.2553071 | 5.3747429 | 5.95533 | -0.580589663 | -0.700025476 |
| Epb41l1 | 6.082938 | 5.950116 | 6.0431 | -0.092993373 | 0.039829127 | 5.5062235 | 5.3930629 | 5.97367 | -0.580606507 | -0.467445951 |
| Slc16a9 | 6.308143 | 5.897977 | 6.4005 | -0.502557904 | -0.092391895 | 5.6652478 | 5.1935365 | 5.77417 | -0.580629274 | -0.108917974 |
| Rnf144a | 10.40904 | 10.36171 | 10.6 | -0.237843288 | -0.190514678 | 10.534117 | 9.9122997 | 10.4933 | -0.581007727 | 0.040809534 |
| Zfp599 | 5.954554 | 5.872869 | 5.9786 | -0.105693656 | -0.024008457 | 5.7601603 | 5.3803287 | 5.96146 | -0.581135611 | -0.201304073 |
| Fktn | 7.283032 | 7.252004 | 7.2563 | -0.00430869 | 0.026719405 | 7.213492 | 6.81363 | 7.39549 | -0.581858598 | -0.181996663 |
| Swap70 | 12.66894 | 12.51571 | 12.748 | -0.231823734 | -0.07859551 | 12.892872 | 12.246514 | 12.8284 | -0.58186293 | 0.064494772 |
| Ankrd33b | 8.236587 | 8.231146 | 8.3681 | -0.136950028 | -0.131509755 | 7.7974749 | 7.8031798 | 8.38517 | -0.5819934 | -0.587698278 |
| Tonsl | 8.032703 | 7.933561 | 8.0513 | -0.117761505 | -0.018619623 | 8.1555842 | 7.8043324 | 8.38645 | -0.582120322 | -0.230868467 |
| Grm6 | 5.13741 | 5.146217 | 5.0919 | 0.054327119 | 0.045520355 | 5.4101736 | 4.7833412 | 5.36561 | -0.582268774 | 0.044563623 |
| Plxdc2 | 7.047083 | 6.451533 | 6.9775 | -0.525964587 | 0.069585926 | 6.038631 | 5.7220347 | 6.30623 | -0.584193235 | -0.267597001 |

| | | | | | | | | | | |
|---|---|---|---|---|---|---|---|---|---|---|
| Ccl22 | 11.29608 | 11.381 | 11.568 | -0.1869456 | -0.27186825 | 10.988661 | 10.694149 | 11.2784 | -0.584294156 | -0.289782367 |
| Snx14 | 8.908508 | 8.761965 | 9.0027 | -0.24075702 | -0.0942149 | 8.8520523 | 8.3567362 | 8.9411 | -0.584366616 | -0.089050439 |
| Mtus1 | 7.567822 | 7.124618 | 7.4516 | -0.32701627 | 0.116188038 | 6.8179918 | 6.1565562 | 6.74099 | -0.584435046 | 0.077000543 |
| Bphl | 7.137389 | 7.162224 | 7.3285 | -0.166258847 | -0.191094067 | 7.1422712 | 6.6211819 | 7.20573 | -0.584543781 | -0.063454474 |
| Exoc6b | 8.581887 | 8.468598 | 8.5963 | -0.127749152 | -0.014459848 | 8.4086201 | 8.0763572 | 8.66101 | -0.584649688 | -0.252386834 |
| Map3k1 | 11.93848 | 11.79773 | 11.905 | -0.10716917 | 0.033582296 | 11.907302 | 11.441381 | 12.0261 | -0.58468958 | -0.118769381 |
| Kynu | 8.2027 | 8.229197 | 8.3372 | -0.108032903 | -0.134529604 | 8.4757478 | 7.9380444 | 8.52296 | -0.584914651 | -0.047211224 |
| Pigb | 8.234471 | 8.061876 | 8.2841 | -0.222189505 | -0.049595332 | 8.436011 | 7.929858 | 8.51517 | -0.58531145 | -0.0791584 |
| Maml3 | 7.399249 | 7.414071 | 7.508 | -0.093922234 | -0.108744458 | 7.3545672 | 6.8701598 | 7.45569 | -0.585527964 | -0.101120526 |
| Maoa | 6.122435 | 5.848976 | 6.2623 | -0.413369692 | -0.139910868 | 5.4997867 | 5.2409635 | 5.82666 | -0.585700623 | -0.326877448 |
| Nrm | 9.552055 | 9.573278 | 9.7038 | -0.130509555 | -0.151732608 | 9.9150574 | 9.0273442 | 9.61357 | -0.586224331 | 0.301488909 |
| Rgs18 | 7.188366 | 7.141799 | 7.2573 | -0.115517121 | -0.068949832 | 7.2617268 | 6.4961046 | 7.08274 | -0.586638714 | 0.1789835 |
| Fmn1 | 5.811245 | 5.739456 | 5.7806 | -0.041113076 | 0.030675971 | 5.8905937 | 5.3029854 | 5.89117 | -0.588186931 | -0.000578591 |
| Spc24 | 6.004369 | 5.847141 | 6.0857 | -0.238556424 | -0.081328524 | 6.1549232 | 5.9453485 | 6.53397 | -0.588620293 | -0.379045625 |
| Rab30 | 8.903767 | 8.840049 | 9.0867 | -0.246614387 | -0.182896975 | 8.8726505 | 8.3745833 | 8.96338 | -0.588797047 | -0.090729869 |
| Osbpl3 | 8.060081 | 7.823011 | 8.2851 | -0.46206613 | -0.224996311 | 7.8518816 | 7.8433749 | 8.43228 | -0.588901909 | -0.580395203 |
| Rnf43 | 7.072447 | 6.946065 | 7.0268 | -0.080773132 | 0.045609316 | 7.0469763 | 6.6780748 | 7.26714 | -0.589065014 | -0.220163497 |
| Hist1h1b | 4.189662 | 3.962636 | 4.1614 | -0.198763146 | 0.0282626 | 4.2940559 | 3.8631586 | 4.45228 | -0.589118962 | -0.158221645 |
| Mylk | 8.948468 | 8.553685 | 9.0063 | -0.452597082 | -0.057813576 | 7.9392171 | 7.6911028 | 8.28034 | -0.589237471 | -0.341123161 |
| Pja2 | 10.04345 | 9.814267 | 10.029 | -0.214246936 | 0.014939307 | 9.8525236 | 9.3429164 | 9.93242 | -0.589507133 | -0.079899983 |
| Iqgap3 | 5.821148 | 5.741592 | 5.9632 | -0.221601594 | -0.142045483 | 5.837461 | 5.5153099 | 6.10596 | -0.590654417 | -0.268503308 |
| Rapgef3 | 6.63992 | 6.309534 | 6.4534 | -0.143866596 | 0.186519098 | 6.3617337 | 6.0440003 | 6.63543 | -0.591428912 | -0.273695454 |
| Inka2 | 5.778185 | 5.779179 | 5.9709 | -0.191765736 | -0.19275946 | 5.5094141 | 5.3840957 | 5.97568 | -0.591584914 | -0.466266539 |
| Izumo4 | 7.416408 | 7.2971 | 7.57 | -0.272866775 | -0.153559148 | 7.7417048 | 7.1774258 | 7.76965 | -0.59222467 | -0.02794567 |
| Slc38a1 | 11.56367 | 11.49424 | 11.617 | -0.122476558 | -0.053047748 | 11.61671 | 11.196254 | 11.7888 | -0.59250686 | -0.172051339 |
| Fam234b | 8.600608 | 8.58701 | 8.6637 | -0.07673082 | -0.063132707 | 8.5217194 | 8.0385022 | 8.63108 | -0.592582338 | -0.109365224 |
| Hvcn1 | 13.14095 | 12.99186 | 13.185 | -0.193129342 | -0.04403237 | 13.42338 | 12.453647 | 13.0478 | -0.594200242 | 0.375532834 |
| Gm2274 | 3.203353 | 3.447836 | 3.5408 | -0.092996659 | -0.337479542 | 3.0093436 | 3.4314001 | 4.02562 | -0.594215035 | -1.016271516 |
| Ppp1r12b | 8.644105 | 8.582653 | 8.5755 | 0.007109393 | 0.068561971 | 8.6647253 | 8.2139387 | 8.8084 | -0.594463751 | -0.143677147 |
| Ralgapa2 | 9.493284 | 9.503349 | 9.6029 | -0.099599117 | -0.10966379 | 9.5473927 | 9.3012525 | 9.89578 | -0.59453237 | -0.348392176 |
| Lmln | 4.887863 | 5.020169 | 5.1942 | -0.173991298 | -0.306297699 | 5.3195659 | 4.5935792 | 5.1882 | -0.594623084 | 0.131363643 |
| Rmnd5a | 10.90419 | 10.84299 | 10.933 | -0.090019708 | -0.028824496 | 10.632592 | 10.280376 | 10.8754 | -0.594979944 | -0.242763962 |

| | | | | | | | | | | |
|---|---|---|---|---|---|---|---|---|---|---|
| Traf3 | 10.7472 | 10.88403 | 10.883 | 0.001054954 | -0.135771344 | 10.737276 | 10.42891 | 11.024 | -0.595131938 | -0.286766374 |
| Papln | 4.717882 | 4.417228 | 4.9188 | -0.501539402 | -0.200885187 | 4.2555839 | 3.9857027 | 4.581 | -0.595301711 | -0.325420533 |
| Ampd1 | 6.248966 | 6.19456 | 6.152 | 0.042546946 | 0.096952612 | 6.3114809 | 5.792028 | 6.3876 | -0.595569101 | -0.076116208 |
| Gxylt2 | 5.330435 | 5.288904 | 5.3827 | -0.093761507 | -0.052230529 | 5.075541 | 4.8758858 | 5.47147 | -0.59558216 | -0.395926942 |
| Bace2 | 7.78575 | 7.154162 | 7.685 | -0.530887549 | 0.100701019 | 7.0576707 | 6.211079 | 6.80732 | -0.596238608 | 0.250353144 |
| Mrvi1 | 6.009723 | 6.006675 | 6.2161 | -0.209403174 | -0.206355878 | 5.8800781 | 5.4203733 | 6.01665 | -0.596272593 | -0.136567788 |
| Mex3b | 7.280059 | 7.266395 | 7.4022 | -0.135791689 | -0.122127114 | 7.0666769 | 6.8355144 | 7.43197 | -0.596455299 | -0.365292738 |
| Mapk12 | 7.455284 | 7.163532 | 7.4665 | -0.302941429 | -0.011189196 | 7.5339583 | 6.8658835 | 7.46251 | -0.596624357 | 0.071450526 |
| Nlrp10 | 5.592783 | 5.438086 | 5.5738 | -0.135728363 | 0.01896841 | 5.5996625 | 5.0430245 | 5.63976 | -0.596732355 | -0.040094353 |
| Tnfaip8l1 | 7.595434 | 7.416157 | 7.652 | -0.235891108 | -0.056614051 | 7.0477531 | 6.7619399 | 7.35877 | -0.596833827 | -0.311020664 |
| Foxm1 | 7.700164 | 7.617569 | 7.9243 | -0.30677816 | -0.224183259 | 7.9840546 | 7.6103987 | 8.20745 | -0.597053864 | -0.223397897 |
| Gm11688 | 4.655865 | 4.668871 | 4.791 | -0.122167032 | -0.13517325 | 4.5473091 | 4.4714307 | 5.06858 | -0.597147924 | -0.521269518 |
| Kdm1b | 9.418365 | 9.645893 | 9.506 | 0.139874906 | -0.087652909 | 9.4242056 | 9.0149685 | 9.61216 | -0.597187709 | -0.187950652 |
| Cd55b | 5.576944 | 5.473331 | 5.6561 | -0.182728914 | -0.079115926 | 5.7302374 | 5.3507787 | 5.94829 | -0.597514602 | -0.218055964 |
| Ptpn4 | 9.654551 | 9.605464 | 9.7698 | -0.164343336 | -0.11525596 | 9.5230714 | 9.3578587 | 9.95574 | -0.597878677 | -0.432665937 |
| Fbxl4 | 8.166586 | 7.966231 | 8.2753 | -0.309077023 | -0.108721863 | 8.4864718 | 7.7464792 | 8.34442 | -0.597943381 | 0.142049208 |
| Dab1 | 4.577231 | 4.410153 | 4.4352 | -0.025025588 | 0.142052292 | 4.3560859 | 3.7583514 | 4.35689 | -0.598539383 | -0.000804901 |
| Colq | 7.624656 | 7.416046 | 7.5967 | -0.180628715 | 0.027980849 | 7.6226709 | 6.936287 | 7.53519 | -0.598900015 | 0.087483855 |
| Plpp3 | 8.246804 | 7.872929 | 7.907 | -0.034047985 | 0.339826519 | 7.7190658 | 7.3737546 | 7.97268 | -0.598930312 | -0.253619046 |
| Fancb | 6.160471 | 6.303824 | 6.2438 | 0.060046362 | -0.083307021 | 6.2586693 | 5.7141191 | 6.31306 | -0.598940953 | -0.054390821 |
| Ak3 | 9.396765 | 9.29556 | 9.5553 | -0.259751387 | -0.158546059 | 9.2975308 | 8.7691458 | 9.36818 | -0.599038363 | -0.070653335 |
| Epha4 | 4.111211 | 4.072898 | 4.3348 | -0.261876842 | -0.223563688 | 3.8812355 | 3.604397 | 4.2035 | -0.599103188 | -0.322264683 |
| Gna12 | 9.413657 | 9.25495 | 9.531 | -0.276078614 | -0.117371604 | 9.112969 | 8.5005119 | 9.09965 | -0.599142631 | 0.013314499 |
| Eml5 | 8.521735 | 8.327675 | 8.4969 | -0.169235417 | 0.024824201 | 8.5123277 | 8.2825656 | 8.88172 | -0.599151675 | -0.369389555 |
| Il6st | 12.41741 | 12.06429 | 12.411 | -0.346704416 | 0.00641397 | 11.617712 | 11.540989 | 12.1402 | -0.599204636 | -0.522481813 |
| Clec4g | 3.539602 | 3.129827 | 3.4959 | -0.366042433 | 0.043732563 | 3.1444611 | 2.9875577 | 3.58708 | -0.599521113 | -0.442617648 |
| Plxdc1 | 6.197259 | 6.271714 | 6.4199 | -0.148184224 | -0.222638481 | 6.0185447 | 5.9106844 | 6.51076 | -0.600072808 | -0.492212558 |
| Cdkn2c | 7.215898 | 6.7561 | 7.1301 | -0.374045193 | 0.085752585 | 6.9294502 | 6.5036225 | 7.10374 | -0.600115722 | -0.174288062 |
| Fmo1 | 6.95097 | 6.502438 | 6.9781 | -0.475619236 | -0.027087457 | 6.7404635 | 6.3411999 | 6.94143 | -0.600228298 | -0.200964732 |
| Rftn2 | 7.61108 | 7.572447 | 7.7576 | -0.185199271 | -0.146566793 | 7.7623683 | 7.0561602 | 7.6564 | -0.600244026 | 0.105964081 |
| Slc35g1 | 7.599571 | 7.483572 | 7.6283 | -0.144772465 | -0.028773815 | 7.4314509 | 7.1397175 | 7.73996 | -0.600245413 | -0.308511991 |
| Tpp1 | 10.39504 | 10.37453 | 10.529 | -0.154885302 | -0.134374312 | 10.579989 | 9.9867753 | 10.5871 | -0.600368669 | -0.007155277 |

| | | | | | | | | | | |
|---|---|---|---|---|---|---|---|---|---|---|
| Pclaf | 8.808873 | 8.511887 | 8.8818 | -0.369879785 | -0.072893081 | 8.8012692 | 8.6505975 | 9.25112 | -0.600525595 | -0.449853944 |
| Gm4739 | 9.59102 | 9.428352 | 9.7037 | -0.275322257 | -0.112654235 | 9.4947316 | 9.3680036 | 9.96862 | -0.600613799 | -0.473885786 |
| Brwd1 | 11.77698 | 11.83329 | 11.851 | -0.017584224 | -0.073890382 | 11.854864 | 11.436177 | 12.0369 | -0.600753998 | -0.182066084 |
| Zfp827 | 7.598852 | 7.300961 | 7.6368 | -0.335838509 | -0.037947697 | 7.3912639 | 7.187325 | 7.78872 | -0.601393553 | -0.397454631 |
| Dpp10 | 5.448495 | 5.140781 | 5.2621 | -0.121334565 | 0.186378919 | 5.0761816 | 4.5928635 | 5.19442 | -0.601555623 | -0.118237581 |
| Gpr174 | 11.04213 | 11.06783 | 11.123 | -0.055647338 | -0.081351224 | 11.026367 | 10.539938 | 11.1416 | -0.60168331 | -0.115254453 |
| Rpgrip1 | 8.713254 | 8.967348 | 9.0658 | -0.098480195 | -0.352573989 | 9.2096173 | 8.6274597 | 9.22954 | -0.60208005 | -0.01992245 |
| Creb5 | 10.08469 | 9.888012 | 10.118 | -0.229883163 | -0.033204349 | 10.819951 | 10.064521 | 10.6668 | -0.602316387 | 0.153113829 |
| Zfp518b | 5.661905 | 5.642045 | 5.846 | -0.20390726 | -0.18404722 | 5.8793538 | 5.0702243 | 5.67254 | -0.602316969 | 0.20681251 |
| Man1a | 12.63559 | 12.56711 | 12.643 | -0.075839304 | -0.007352354 | 12.646789 | 11.97908 | 12.5817 | -0.60258102 | 0.065127414 |
| Ppfibp2 | 7.374137 | 7.218664 | 7.4805 | -0.26181592 | -0.106343558 | 7.2538739 | 6.8680273 | 7.47061 | -0.602584438 | -0.216737894 |
| Sfxn3 | 9.624059 | 9.713251 | 9.9394 | -0.226121663 | -0.315313488 | 10.026054 | 9.3189523 | 9.9219 | -0.602946851 | 0.104154479 |
| Tiam1 | 8.829782 | 8.614291 | 8.9815 | -0.367179917 | -0.151689332 | 8.6694838 | 8.3316179 | 8.93518 | -0.603564664 | -0.265698754 |
| Sit1 | 8.660556 | 8.550278 | 8.7731 | -0.22287128 | -0.112592945 | 8.6030417 | 7.9950636 | 8.59918 | -0.60411626 | 0.003861752 |
| Cdkn3 | 4.397796 | 4.339623 | 4.4428 | -0.103162133 | -0.044989154 | 4.2664486 | 4.086506 | 4.69085 | -0.604341119 | -0.424398515 |
| Myrip | 4.62884 | 4.292734 | 4.8911 | -0.59832449 | -0.262219146 | 4.4518864 | 4.0306036 | 4.63509 | -0.604482995 | -0.183200171 |
| Vps13b | 11.29751 | 11.23386 | 11.314 | -0.080342056 | -0.016691448 | 11.164427 | 10.819779 | 11.4243 | -0.604534956 | -0.259886869 |
| E130308A19Rik | 7.701686 | 7.732247 | 7.6408 | 0.091425931 | 0.060864566 | 7.6370176 | 7.156435 | 7.76122 | -0.604782565 | -0.124199919 |
| Pcdh7 | 7.470211 | 6.990696 | 7.4684 | -0.477708654 | 0.001806562 | 6.4077283 | 6.058042 | 6.6629 | -0.604860785 | -0.255174505 |
| Ralgps1 | 7.692774 | 7.570329 | 7.7652 | -0.194842717 | -0.072397274 | 7.6984111 | 7.2969247 | 7.9018 | -0.604872105 | -0.203385709 |
| Ift46 | 8.373455 | 8.006345 | 8.4861 | -0.479711935 | -0.112601176 | 8.5513233 | 7.9294028 | 8.53431 | -0.604909888 | 0.017010518 |
| Slc29a4 | 4.149235 | 4.26821 | 4.27 | -0.001745582 | -0.120720326 | 4.5721988 | 3.9425837 | 4.54754 | -0.60496092 | 0.024654129 |
| Golim4 | 8.609578 | 8.414221 | 8.4621 | -0.04791252 | 0.147444555 | 8.0755163 | 7.9798473 | 8.58532 | -0.605469947 | -0.509801036 |
| Nphp3 | 7.491634 | 7.345117 | 7.565 | -0.219877506 | -0.073360411 | 7.4631753 | 6.9892845 | 7.59505 | -0.605768153 | -0.131877442 |
| Slc12a7 | 11.01925 | 10.87536 | 11.006 | -0.130387272 | 0.01350296 | 10.798085 | 10.5999 | 11.206 | -0.606069668 | -0.407884221 |
| Cd99l2 | 7.964929 | 7.757714 | 7.9728 | -0.215133792 | -0.007919165 | 7.8909298 | 7.2435966 | 7.84971 | -0.606109214 | 0.041224018 |
| Aoc3 | 5.584787 | 5.505737 | 5.6705 | -0.164735171 | -0.085685034 | 5.4374915 | 4.8397808 | 5.44593 | -0.606150659 | -0.008439901 |
| Tmie | 8.128185 | 8.29981 | 8.3261 | -0.026302933 | -0.197927836 | 7.592864 | 7.4997601 | 8.10627 | -0.60650836 | -0.513404431 |
| Dchs1 | 8.20489 | 7.613861 | 7.6701 | -0.056281879 | 0.534746686 | 6.9020193 | 6.5876998 | 7.19431 | -0.606611725 | -0.292292268 |
| Il17re | 4.47035 | 4.349849 | 4.5928 | -0.242961263 | -0.122460425 | 4.2372976 | 4.0418891 | 4.64856 | -0.606668427 | -0.411259965 |
| Pdgfra | 6.579431 | 6.092887 | 6.2035 | -0.110591742 | 0.375952158 | 6.1228777 | 5.4748414 | 6.08163 | -0.606791087 | 0.041245203 |
| Sema3d | 7.185637 | 6.950642 | 6.6037 | 0.346931132 | 0.581926797 | 6.0482437 | 6.088471 | 6.69527 | -0.606796154 | -0.647023477 |

| | | | | | | | | | | |
|---|---|---|---|---|---|---|---|---|---|---|
| Gm48368 | 7.983523 | 7.94623 | 8.1514 | -0.205172546 | -0.167880419 | 7.9740643 | 7.7353465 | 8.34223 | -0.606883666 | -0.368165791 |
| Chd6 | 11.05474 | 11.06296 | 11.186 | -0.12270257 | -0.13092341 | 11.190426 | 10.763501 | 11.3717 | -0.608151068 | -0.181225859 |
| Abca5 | 5.605592 | 5.651881 | 5.7 | -0.048157736 | -0.094446995 | 5.6204114 | 5.0962084 | 5.70471 | -0.608496714 | -0.084293756 |
| Cpne5 | 5.917412 | 5.967353 | 6.0216 | -0.054247251 | -0.104187786 | 6.1890736 | 5.4725281 | 6.0811 | -0.608569217 | 0.107976344 |
| Rassf3 | 10.45046 | 10.41289 | 10.588 | -0.175534392 | -0.137955032 | 10.347253 | 9.8475551 | 10.4563 | -0.608713265 | -0.109015061 |
| Pik3r3 | 7.047991 | 6.89355 | 7.1574 | -0.263805705 | -0.109364491 | 6.7454249 | 6.4053861 | 7.01472 | -0.609334831 | -0.269295988 |
| Arl2bp | 10.17402 | 10.14628 | 10.2 | -0.053827817 | -0.026090232 | 10.193032 | 9.6282414 | 10.2377 | -0.609421079 | -0.044630399 |
| Kif18a | 6.660473 | 6.328763 | 6.7023 | -0.373532929 | -0.041823725 | 6.3975066 | 6.1359235 | 6.74566 | -0.609733602 | -0.348150529 |
| Mpp2 | 5.949605 | 5.452588 | 5.7717 | -0.319150497 | 0.17786613 | 5.3712619 | 5.1298558 | 5.73966 | -0.609806934 | -0.368400836 |
| Il1rl1 | 6.728069 | 6.425208 | 6.3423 | 0.082940595 | 0.385801621 | 6.4520314 | 6.1664606 | 6.77629 | -0.609832164 | -0.324261339 |
| Top2b | 12.60261 | 12.64361 | 12.718 | -0.074563236 | -0.115559202 | 12.535902 | 12.067659 | 12.6783 | -0.610615772 | -0.142372411 |
| Pdpr | 9.475586 | 9.31685 | 9.4913 | -0.174456227 | -0.015720852 | 9.3820262 | 8.9124991 | 9.5235 | -0.610997513 | -0.141470472 |
| Hdac7 | 11.42957 | 11.34397 | 11.555 | -0.211218972 | -0.125622716 | 11.580109 | 11.005305 | 11.6169 | -0.611639382 | -0.036834779 |
| Anln | 6.446865 | 6.226498 | 6.6257 | -0.39921848 | -0.178851839 | 6.4972444 | 6.3470489 | 6.95911 | -0.612062805 | -0.461867283 |
| Kyat1 | 6.958352 | 7.012907 | 7.0377 | -0.024800116 | -0.079355672 | 7.2739303 | 6.5584257 | 7.17058 | -0.612151526 | 0.103353098 |
| Sbk1 | 11.19086 | 10.99343 | 11.113 | -0.119859954 | 0.077567488 | 11.143102 | 10.496553 | 11.1087 | -0.612175952 | 0.034373487 |
| Slc16a7 | 7.517027 | 7.756822 | 7.3585 | 0.398274737 | 0.158480195 | 7.4226366 | 7.0771348 | 7.68975 | -0.612616991 | -0.26711515 |
| Kif4 | 6.873863 | 6.691634 | 7.2478 | -0.556147684 | -0.373918875 | 6.9490099 | 6.7123756 | 7.32516 | -0.612786875 | -0.376152533 |
| Gm28042 | 7.458005 | 7.480461 | 7.6089 | -0.1284858 | -0.150941629 | 7.9532363 | 7.3343812 | 7.94769 | -0.613310889 | 0.005544279 |
| Slc14a1 | 8.647368 | 8.342204 | 8.6074 | -0.265233743 | 0.039930229 | 8.8458396 | 8.1500602 | 8.76348 | -0.613415635 | 0.082363737 |
| Orai2 | 11.00228 | 11.04614 | 11.118 | -0.072157358 | -0.11601666 | 11.060743 | 10.553303 | 11.167 | -0.613659928 | -0.106219774 |
| Abhd14b | 6.497216 | 6.381991 | 6.5764 | -0.19437674 | -0.079152328 | 6.4603274 | 5.8605756 | 6.47476 | -0.614186193 | -0.014434408 |
| Etv5 | 7.345954 | 7.25625 | 7.3275 | -0.071290664 | 0.018413309 | 7.2517262 | 6.5354284 | 7.14962 | -0.61418768 | 0.102110175 |
| Prkdc | 10.37403 | 10.3823 | 10.456 | -0.07342293 | -0.081690914 | 10.343958 | 10.010656 | 10.6249 | -0.614238835 | -0.28093713 |
| Cep55 | 7.058755 | 6.790857 | 7.0164 | -0.225559988 | 0.042338222 | 7.0032607 | 6.6054829 | 7.21994 | -0.614457448 | -0.216679677 |
| Galm | 6.558962 | 6.612959 | 6.7619 | -0.148953247 | -0.202949517 | 6.4186468 | 6.0677303 | 6.68226 | -0.614525795 | -0.263609376 |
| Macf1 | 14.29519 | 14.22877 | 14.41 | -0.180780622 | -0.11436809 | 14.316715 | 13.994447 | 14.6094 | -0.614973662 | -0.292705541 |
| Gm10171 | 5.146518 | 5.226442 | 4.9558 | 0.270674764 | 0.19075042 | 4.6196569 | 4.480434 | 5.09643 | -0.615999193 | -0.476776263 |
| Gm10233 | 5.146518 | 5.226442 | 4.9558 | 0.270674764 | 0.19075042 | 4.6196569 | 4.480434 | 5.09643 | -0.615999193 | -0.476776263 |
| Car12 | 6.547521 | 6.125363 | 6.173 | -0.047619481 | 0.374538542 | 6.413327 | 5.8832989 | 6.4997 | -0.616398938 | -0.086370913 |
| Ldb2 | 6.989436 | 6.893997 | 6.8811 | 0.012876961 | 0.10831641 | 6.1566828 | 5.8044066 | 6.42117 | -0.616764625 | -0.264488455 |
| Vangl2 | 6.822105 | 6.734142 | 6.9485 | -0.214381467 | -0.126417734 | 6.6612644 | 6.1646576 | 6.78193 | -0.617270995 | -0.120664172 |

| | | | | | | | | | | |
|---|---|---|---|---|---|---|---|---|---|---|
| Dnmbp | 8.456229 | 8.565294 | 8.5403 | 0.024950981 | -0.084114655 | 8.315857 | 7.883674 | 8.50112 | -0.617443577 | -0.185260638 |
| Elk3 | 10.18779 | 10.16334 | 10.299 | -0.136001707 | -0.11155469 | 10.174705 | 9.5307756 | 10.1483 | -0.617476878 | 0.02645235 |
| Top2a | 10.31047 | 10.1096 | 10.491 | -0.381515908 | -0.180647664 | 10.28243 | 10.232004 | 10.8499 | -0.617893956 | -0.567467544 |
| Nxph1 | 6.405959 | 5.664962 | 6.2516 | -0.586647046 | 0.154350116 | 5.1644931 | 4.9172173 | 5.53577 | -0.61855446 | -0.371278648 |
| Cep72 | 7.270167 | 7.425757 | 7.5366 | -0.110828485 | -0.266418199 | 7.6526351 | 7.2746935 | 7.89341 | -0.61871536 | -0.240773776 |
| Sox18 | 6.268452 | 5.986451 | 6.2765 | -0.290007566 | -0.008005796 | 5.9129369 | 5.338372 | 5.95714 | -0.618763557 | -0.044198628 |
| Ube2j1 | 10.96308 | 10.97491 | 10.911 | 0.063725894 | 0.051897922 | 10.650876 | 10.349566 | 10.9684 | -0.61881421 | -0.317504295 |
| Kif20a | 7.736087 | 7.612432 | 7.7975 | -0.185025431 | -0.061370911 | 7.4402887 | 7.4477304 | 8.06771 | -0.619979814 | -0.627421439 |
| Cnst | 9.197197 | 9.008715 | 9.2131 | -0.20433903 | -0.015857232 | 8.8099271 | 8.6064167 | 9.22656 | -0.620143275 | -0.416632948 |
| Cdhr1 | 5.124916 | 5.097208 | 5.1725 | -0.075257663 | -0.047549559 | 4.8285271 | 4.8202889 | 5.44056 | -0.620269882 | -0.612031735 |
| Fcgrt | 9.052204 | 8.758086 | 9.0373 | -0.279198521 | 0.014919441 | 9.134733 | 8.1908826 | 8.81122 | -0.620341232 | 0.323509249 |
| Art4 | 6.482017 | 6.51205 | 6.6102 | -0.098105824 | -0.128138569 | 6.3150926 | 5.7446882 | 6.36521 | -0.620524673 | -0.050120245 |
| Phldb2 | 6.472043 | 5.932555 | 6.3763 | -0.443749425 | 0.095738958 | 5.4505076 | 5.5384044 | 6.15924 | -0.620832736 | -0.708729514 |
| Atcay | 5.189035 | 5.215169 | 5.4507 | -0.235560031 | -0.261694646 | 5.1042606 | 4.9417021 | 5.56307 | -0.621372866 | -0.458814357 |
| Plekhg2 | 11.07441 | 11.0161 | 11.101 | -0.085402992 | -0.027087706 | 11.216956 | 10.557465 | 11.1796 | -0.622178258 | 0.037313491 |
| Bach2 | 11.43493 | 11.45478 | 11.3 | 0.155251444 | 0.135401732 | 11.35742 | 11.021615 | 11.6439 | -0.622308054 | -0.286502348 |
| Hr | 5.054368 | 5.047435 | 5.1841 | -0.136696839 | -0.129763757 | 4.6468365 | 4.3723506 | 4.99572 | -0.623372439 | -0.348886533 |
| Ccdc88c | 11.78891 | 11.82327 | 11.982 | -0.15824545 | -0.192598788 | 11.820724 | 11.530372 | 12.1538 | -0.623465784 | -0.333112945 |
| Decr2 | 7.38726 | 7.421261 | 7.5422 | -0.120939366 | -0.154940337 | 7.5392173 | 7.1687461 | 7.79222 | -0.623474822 | -0.253003642 |
| Kcnab2 | 11.37878 | 11.17426 | 11.292 | -0.117365166 | 0.08714578 | 11.23382 | 10.672996 | 11.2965 | -0.623503962 | -0.062679293 |
| Chst2 | 7.93394 | 7.524951 | 7.8797 | -0.354741917 | 0.054247125 | 6.9030197 | 6.6103386 | 7.23426 | -0.623926265 | -0.331245171 |
| Dennd2c | 5.341757 | 5.21966 | 5.4285 | -0.208857894 | -0.08676115 | 5.2190357 | 4.7022445 | 5.3265 | -0.624259566 | -0.107468354 |
| Arhgap42 | 5.944242 | 5.909114 | 6.0063 | -0.097225006 | -0.062097072 | 5.6031005 | 5.0459416 | 5.67056 | -0.624622275 | -0.06746343 |
| Hadh | 9.117321 | 9.062733 | 9.2396 | -0.176874858 | -0.122286906 | 9.2086202 | 8.6132917 | 9.2387 | -0.625404608 | -0.030076073 |
| Mapkbp1 | 8.131143 | 7.965069 | 8.2328 | -0.267779811 | -0.101705937 | 8.3541127 | 7.9084898 | 8.53391 | -0.625418538 | -0.179795664 |
| Tanc2 | 6.880587 | 6.525916 | 6.8101 | -0.284221891 | 0.070449102 | 6.7920146 | 6.3901498 | 7.01656 | -0.626410575 | -0.224545842 |
| Col12a1 | 6.56361 | 5.90873 | 6.272 | -0.363296233 | 0.291584239 | 5.7046079 | 5.5710008 | 6.19803 | -0.627030376 | -0.493423364 |
| Mis18bp1 | 6.208177 | 5.984036 | 6.3286 | -0.344547141 | -0.120405966 | 5.9525619 | 6.0173241 | 6.64438 | -0.627053991 | -0.691816232 |
| Cryl1 | 7.477375 | 7.314077 | 7.5488 | -0.23474978 | -0.071451398 | 7.5778099 | 6.9020806 | 7.52919 | -0.62711238 | 0.048616955 |
| Slc2a8 | 6.246587 | 6.101832 | 6.3253 | -0.223453624 | -0.078698584 | 6.3142945 | 5.7136452 | 6.34103 | -0.627384061 | -0.026734792 |
| Zadh2 | 9.152239 | 8.988865 | 9.2662 | -0.277370312 | -0.11399609 | 9.1559782 | 8.5796483 | 9.20722 | -0.627573768 | -0.05124387 |
| Entpd5 | 10.5216 | 10.37863 | 10.541 | -0.162588846 | -0.019617154 | 10.484273 | 10.018321 | 10.6461 | -0.627826831 | -0.16187392 |

| | | | | | | | | | | |
|---|---|---|---|---|---|---|---|---|---|---|
| Mtss1l | 6.477121 | 6.351795 | 6.3766 | -0.024841027 | 0.10048497 | 6.1369227 | 5.9147124 | 6.54315 | -0.628439301 | -0.406228998 |
| Sntb2 | 9.626961 | 9.201832 | 9.3525 | -0.150623053 | 0.274505689 | 8.8734024 | 8.6362474 | 9.26481 | -0.628561461 | -0.391406463 |
| Iqgap2 | 10.04897 | 9.687497 | 10.026 | -0.338766624 | 0.022710601 | 9.5330271 | 9.3617715 | 9.99039 | -0.628619256 | -0.457363658 |
| Zfp607a | 8.069141 | 8.04737 | 8.2484 | -0.201063553 | -0.17929314 | 8.0281748 | 7.6769737 | 8.30591 | -0.628931484 | -0.27773043 |
| Eif2ak3 | 10.03436 | 10.02519 | 10.146 | -0.120991046 | -0.111821001 | 9.9384353 | 9.5380953 | 10.1678 | -0.629679423 | -0.22933946 |
| Appl2 | 8.958285 | 8.89328 | 9.1455 | -0.252187482 | -0.187182083 | 8.9651292 | 8.6043167 | 9.23517 | -0.630852481 | -0.270039961 |
| Zbtb38 | 9.481562 | 9.250641 | 9.5638 | -0.31312823 | -0.082207751 | 9.3722089 | 9.0081293 | 9.63903 | -0.630897127 | -0.266817556 |
| Myo18a | 10.89022 | 10.83081 | 10.857 | -0.026483154 | 0.032934338 | 10.886334 | 10.420762 | 11.0528 | -0.632029564 | -0.166457499 |
| Rassf8 | 5.253534 | 5.528293 | 5.2977 | 0.230596159 | -0.044162784 | 5.048329 | 4.6553039 | 5.28748 | -0.632180157 | -0.239155082 |
| Pglyrp1 | 8.834072 | 8.518885 | 8.874 | -0.35506707 | -0.039880844 | 8.6741014 | 7.823664 | 8.4559 | -0.632236343 | 0.218201017 |
| Sun1 | 9.650784 | 9.596103 | 9.8755 | -0.279375236 | -0.224694234 | 9.5777583 | 9.297243 | 9.92961 | -0.6323634 | -0.351848081 |
| Slc39a10 | 9.275287 | 9.209635 | 9.2713 | -0.061705161 | 0.003946918 | 8.9848803 | 8.4776351 | 9.11002 | -0.632385956 | -0.125140736 |
| Cxcr6 | 9.633497 | 9.461284 | 9.6133 | -0.152014682 | 0.020198515 | 8.9780039 | 8.9609876 | 9.59358 | -0.632595495 | -0.615579242 |
| AU041133 | 6.695647 | 6.479419 | 6.5645 | -0.085067052 | 0.131160158 | 6.6269378 | 6.1165812 | 6.74956 | -0.632976163 | -0.122619517 |
| Lama3 | 5.366057 | 5.191747 | 5.4096 | -0.217849987 | -0.043539214 | 4.9444302 | 4.7433698 | 5.37666 | -0.633293346 | -0.432232948 |
| Tmtc1 | 6.580053 | 6.088871 | 6.4229 | -0.334054787 | 0.157127468 | 5.6722214 | 5.4058385 | 6.03928 | -0.633441651 | -0.367058758 |
| Stk26 | 10.42126 | 10.40326 | 10.535 | -0.13155675 | -0.113558184 | 10.40376 | 9.8337794 | 10.4674 | -0.633592411 | -0.063611429 |
| Ear2 | 5.967915 | 5.914348 | 5.794 | 0.120391415 | 0.173958248 | 5.6611962 | 5.0837361 | 5.71835 | -0.634610225 | -0.057150171 |
| Usp53 | 8.410944 | 8.644581 | 8.588 | 0.056545302 | -0.177091138 | 8.1474902 | 7.9498162 | 8.58478 | -0.634962936 | -0.437288932 |
| Gpr137c | 6.660563 | 6.653703 | 6.6267 | 0.02695491 | 0.033814824 | 6.7459048 | 6.1704385 | 6.80565 | -0.635209425 | -0.05974316 |
| Kif16b | 9.461177 | 9.356185 | 9.5251 | -0.168920206 | -0.063928059 | 9.3454937 | 8.9508141 | 9.5867 | -0.63588741 | -0.241207876 |
| Fancm | 8.534104 | 8.814813 | 9.0287 | -0.213855777 | -0.494565 | 8.5603061 | 8.4205387 | 9.05659 | -0.636054893 | -0.496287497 |
| Ptgfrn | 7.634996 | 7.540802 | 7.5849 | -0.04410352 | 0.050090841 | 7.1482483 | 6.790274 | 7.42641 | -0.636134397 | -0.278160068 |
| Mctp1 | 6.426009 | 6.185896 | 6.3641 | -0.178206766 | 0.06190599 | 5.9800101 | 5.5677695 | 6.20394 | -0.636173995 | -0.223933419 |
| Apc | 10.98929 | 10.80345 | 10.964 | -0.160055874 | 0.025792032 | 10.547258 | 10.353525 | 10.9899 | -0.636367116 | -0.442634104 |
| Galnt6 | 11.13881 | 11.04951 | 11.113 | -0.063535558 | 0.025761038 | 10.951533 | 10.437778 | 11.0745 | -0.636688554 | -0.122933594 |
| Cacna2d4 | 7.343617 | 7.05079 | 7.4251 | -0.374338766 | -0.08151161 | 7.2735394 | 6.9658627 | 7.60299 | -0.637130511 | -0.329453783 |
| Ccdc69 | 8.664422 | 8.550882 | 8.6774 | -0.126529608 | -0.012989511 | 8.8990881 | 8.2049167 | 8.84208 | -0.637168126 | 0.057003331 |
| Nrep | 4.786611 | 4.796733 | 5.1103 | -0.313555056 | -0.323676678 | 4.9109033 | 4.2405486 | 4.87806 | -0.637507231 | 0.032847456 |
| Jmjd1c | 10.76722 | 10.73786 | 10.792 | -0.053649946 | -0.024285742 | 10.709218 | 10.306875 | 10.9444 | -0.637529712 | -0.235187197 |
| Rab11fip4 | 9.438324 | 9.461551 | 9.5289 | -0.067303068 | -0.090530458 | 9.1807275 | 8.8395392 | 9.47784 | -0.6383052 | -0.297116958 |
| Fmnl3 | 10.71267 | 10.53271 | 10.725 | -0.192765946 | -0.012804954 | 10.915219 | 10.147287 | 10.786 | -0.63867921 | 0.129252813 |

| | | | | | | | | | | |
|---|---|---|---|---|---|---|---|---|---|---|
| Tmem44 | 4.673905 | 4.395808 | 4.7533 | -0.357517552 | -0.07942124 | 4.5450549 | 4.1589365 | 4.79824 | -0.639299073 | -0.253180665 |
| Cacna1i | 10.51786 | 10.70353 | 10.641 | 0.062824034 | -0.122850038 | 10.729707 | 9.9672522 | 10.6067 | -0.639441405 | 0.123013164 |
| Pkhd1l1 | 3.862746 | 3.848936 | 4.0476 | -0.198667763 | -0.184857737 | 3.4122708 | 3.4101075 | 4.0498 | -0.639689774 | -0.637526456 |
| Yes1 | 7.008197 | 6.495211 | 6.8172 | -0.321996185 | 0.190990248 | 6.1991821 | 5.9854147 | 6.62527 | -0.639858319 | -0.426090865 |
| Ighv1-69 | 6.959177 | 7.148744 | 6.8368 | 0.311976231 | 0.122409062 | 7.3461056 | 6.8823896 | 7.52237 | -0.639979063 | -0.176263075 |
| Irf6 | 6.566507 | 6.488175 | 6.6255 | -0.137372965 | -0.059040734 | 6.3821255 | 5.9599842 | 6.60029 | -0.640301421 | -0.218160111 |
| 0610010F05Rik | 7.600446 | 7.045904 | 7.3304 | -0.284537022 | 0.270004843 | 7.2638553 | 6.6749248 | 7.31526 | -0.640339676 | -0.051409168 |
| Itpr3 | 12.16146 | 12.12172 | 12.268 | -0.146087866 | -0.106349364 | 12.430459 | 11.744222 | 12.3851 | -0.640918098 | 0.045318217 |
| Pou2f1 | 10.33585 | 10.37512 | 10.315 | 0.059726882 | 0.020461802 | 10.370713 | 10.04875 | 10.6899 | -0.641111813 | -0.319148811 |
| Thy1 | 12.58121 | 12.58267 | 12.735 | -0.152389896 | -0.153850226 | 12.516154 | 12.090854 | 12.7322 | -0.641337718 | -0.216038329 |
| Vsig10 | 7.557293 | 7.509631 | 7.8764 | -0.366816957 | -0.31915501 | 7.3705354 | 7.1711399 | 7.81263 | -0.641487105 | -0.442091611 |
| Scarb1 | 9.620958 | 9.234459 | 9.6791 | -0.444661666 | -0.058163197 | 9.2603126 | 8.5133332 | 9.15505 | -0.641721611 | 0.105257852 |
| Tmcc3 | 8.751226 | 8.792938 | 8.9973 | -0.204347935 | -0.246059559 | 8.6392185 | 8.5241091 | 9.16593 | -0.64182385 | -0.526714432 |
| Gab1 | 6.435139 | 6.020424 | 6.2613 | -0.240914919 | 0.173800026 | 5.8214222 | 5.0259314 | 5.66787 | -0.641943228 | 0.153547594 |
| Btbd8 | 6.825652 | 6.646014 | 6.9625 | -0.316495289 | -0.136857122 | 6.7460588 | 6.4702026 | 7.11245 | -0.642247076 | -0.366390856 |
| Il13ra2 | 3.80502 | 3.569583 | 3.922 | -0.352397422 | -0.116959993 | 3.4820336 | 3.3261344 | 3.96875 | -0.642620247 | -0.486721099 |
| Ggact | 7.346519 | 7.247519 | 7.3994 | -0.151851766 | -0.052852247 | 7.585173 | 7.0555936 | 7.69962 | -0.644029777 | -0.114450359 |
| Obscn | 5.743799 | 5.409428 | 5.9395 | -0.530074625 | -0.195703589 | 5.0046427 | 5.0759218 | 5.72024 | -0.644317111 | -0.715596249 |
| Rbl2 | 10.88326 | 10.86019 | 11.001 | -0.14096918 | -0.117893942 | 10.874077 | 10.418869 | 11.0633 | -0.644447298 | -0.189239672 |
| Nicn1 | 6.489351 | 6.417416 | 6.7031 | -0.285707559 | -0.213772519 | 6.5570459 | 5.8850799 | 6.53115 | -0.646068195 | 0.025897745 |
| Sspo | 6.249854 | 6.1796 | 6.4195 | -0.239876534 | -0.169622718 | 5.8626933 | 5.8408659 | 6.4871 | -0.646229829 | -0.624402403 |
| Dock5 | 8.115878 | 8.215593 | 8.2019 | 0.013726649 | -0.085988119 | 8.1990467 | 7.4575456 | 8.10444 | -0.646892216 | 0.094608861 |
| Zfp61 | 6.775995 | 6.753589 | 6.8945 | -0.140891055 | -0.118484858 | 6.9427462 | 6.514606 | 7.16188 | -0.647273169 | -0.219132951 |
| Podxl | 8.523081 | 8.235854 | 8.4077 | -0.171845337 | 0.115381071 | 7.926418 | 7.6838404 | 8.33124 | -0.64740342 | -0.404825755 |
| Sox13 | 6.273456 | 6.003392 | 6.1899 | -0.186528359 | 0.083535959 | 5.6280611 | 5.2596305 | 5.90783 | -0.648202183 | -0.279771665 |
| Shtn1 | 8.157553 | 7.989759 | 8.3257 | -0.335963113 | -0.168168813 | 7.9518423 | 7.4270449 | 8.07691 | -0.649865865 | -0.125068406 |
| Calcrl | 9.837002 | 9.738027 | 9.8163 | -0.078303192 | 0.02067145 | 9.3786458 | 8.9589507 | 9.60888 | -0.649925969 | -0.230230805 |
| Lad1 | 8.203781 | 8.459914 | 8.5699 | -0.1100158 | -0.366149389 | 8.1590419 | 8.0950961 | 8.74532 | -0.650222922 | -0.586277168 |
| Hacl1 | 7.182899 | 6.841032 | 7.1046 | -0.263594744 | 0.078272144 | 7.1569595 | 6.508439 | 7.15873 | -0.650288297 | -0.001767847 |
| Tnik | 9.990651 | 10.04979 | 10.158 | -0.107967394 | -0.167102511 | 9.6886368 | 9.5561868 | 10.2066 | -0.650368388 | -0.517918377 |
| Sh3bgrl2 | 6.574757 | 6.346211 | 6.5629 | -0.216705658 | 0.011840353 | 6.1284767 | 5.815026 | 6.46546 | -0.650430891 | -0.336980188 |
| Pacs1 | 11.04964 | 11.11547 | 11.174 | -0.05876806 | -0.12459976 | 11.081198 | 10.544068 | 11.1945 | -0.65046404 | -0.113333508 |

| | | | | | | | | | | |
|---|---|---|---|---|---|---|---|---|---|---|
| Kdm7a | 11.67255 | 11.641 | 11.65 | -0.0094469 | 0.022101984 | 11.47522 | 11.052772 | 11.7037 | -0.650920848 | -0.228473198 |
| Cav2 | 7.365825 | 7.113921 | 7.4779 | -0.363930597 | -0.112026586 | 6.5175948 | 6.1614526 | 6.81252 | -0.651063729 | -0.294921573 |
| Trim2 | 6.930451 | 6.791149 | 7.068 | -0.276885252 | -0.137582763 | 6.2155652 | 5.9934856 | 6.6447 | -0.651218509 | -0.429138909 |
| Filip1 | 5.230515 | 4.733133 | 5.1338 | -0.400634632 | 0.096747405 | 4.5433135 | 4.239204 | 4.89043 | -0.651225713 | -0.347116201 |
| Rapgef5 | 9.080129 | 8.761609 | 8.7376 | 0.024034861 | 0.342554791 | 8.5493768 | 7.9730872 | 8.62439 | -0.65130318 | -0.075013558 |
| Hecw2 | 5.817065 | 6.059684 | 6.1384 | -0.078716247 | -0.321335279 | 5.1658403 | 5.1762417 | 5.82778 | -0.651538542 | -0.661939969 |
| Stil | 6.146015 | 5.666331 | 6.1725 | -0.506183541 | -0.026499918 | 5.8836996 | 5.7262593 | 6.37787 | -0.651613225 | -0.494172896 |
| Emp3 | 10.49238 | 10.45158 | 10.57 | -0.118450648 | -0.07764665 | 10.603941 | 9.8598253 | 10.5115 | -0.651688345 | 0.092427464 |
| Ing1 | 9.844775 | 9.807518 | 9.8934 | -0.08586582 | -0.048608706 | 9.7127144 | 9.1450515 | 9.79716 | -0.652107519 | -0.084444657 |
| Scd2 | 11.13371 | 10.7809 | 11.231 | -0.450562916 | -0.09775345 | 10.876841 | 10.524944 | 11.1773 | -0.652369538 | -0.300472376 |
| Qrfp | 5.170097 | 5.05854 | 5.3515 | -0.292974883 | -0.181417681 | 5.3476604 | 4.6749616 | 5.32761 | -0.652650019 | 0.020048868 |
| Dkkl1 | 5.003716 | 4.841863 | 5.1126 | -0.270778392 | -0.108925295 | 4.9841719 | 4.5438476 | 5.19656 | -0.652711534 | -0.212387229 |
| Nuf2 | 6.843289 | 6.40691 | 6.8438 | -0.436920304 | -0.000541289 | 6.6695191 | 6.4753917 | 7.12851 | -0.653123226 | -0.45899578 |
| Icos | 9.654702 | 9.307036 | 9.8387 | -0.531701048 | -0.184034859 | 9.4109015 | 9.4300189 | 10.0837 | -0.653642877 | -0.672760207 |
| Add1 | 12.16377 | 12.0754 | 12.219 | -0.143745286 | -0.055380426 | 12.082057 | 11.496058 | 12.1502 | -0.65409701 | -0.06809757 |
| Ssh1 | 10.02347 | 10.0115 | 10.176 | -0.164794265 | -0.152826882 | 9.9457631 | 9.5238288 | 10.1792 | -0.655369292 | -0.233434966 |
| Ighv5-9-1 | 6.432252 | 6.216681 | 6.349 | -0.132364026 | 0.083207489 | 6.8058332 | 6.0927675 | 6.74823 | -0.655466924 | 0.057598842 |
| Klf12 | 6.583926 | 6.486291 | 6.6987 | -0.212421927 | -0.11478679 | 6.6553226 | 6.3531174 | 7.00863 | -0.655512324 | -0.353307117 |
| Rnf144b | 7.043573 | 6.769181 | 7.1851 | -0.415962128 | -0.14156992 | 6.6022837 | 6.1294052 | 6.7851 | -0.655694085 | -0.182815613 |
| Rdh12 | 8.250673 | 8.285739 | 8.317 | -0.031220726 | -0.066286648 | 8.5190898 | 7.7771058 | 8.43368 | -0.656576633 | 0.085407372 |
| Tctn1 | 8.30605 | 8.104742 | 8.4529 | -0.3481364 | -0.146828396 | 8.7775529 | 8.0652927 | 8.72194 | -0.656650799 | 0.055609434 |
| Polq | 7.153652 | 6.957764 | 7.2652 | -0.307471046 | -0.111583206 | 7.2692237 | 7.076296 | 7.7331 | -0.656806157 | -0.463878419 |
| Cdcp1 | 5.029172 | 4.906841 | 5.025 | -0.118191434 | 0.004139066 | 4.7332686 | 4.5574579 | 5.21483 | -0.657368293 | -0.481557641 |
| Pcp4 | 4.419158 | 4.155897 | 4.4009 | -0.245004047 | 0.018257004 | 4.3728606 | 3.7579445 | 4.41597 | -0.658021514 | -0.043105384 |
| Kctd12b | 5.97145 | 5.95989 | 6.168 | -0.208147177 | -0.196586953 | 5.6515827 | 5.3243173 | 5.98247 | -0.658148293 | -0.330882853 |
| Tesc | 7.257585 | 7.320743 | 7.4713 | -0.150600882 | -0.21375819 | 7.2226524 | 6.7153744 | 7.37369 | -0.658317616 | -0.151039625 |
| Shisa3 | 5.622729 | 5.034509 | 5.1271 | -0.092586869 | 0.495633635 | 5.4049696 | 4.5153091 | 5.17401 | -0.658698701 | 0.230961844 |
| Cbl | 12.27634 | 12.15534 | 12.239 | -0.083215672 | 0.03778133 | 12.120816 | 11.822314 | 12.4815 | -0.65917933 | -0.360677509 |
| Ikzf3 | 12.38115 | 12.33273 | 12.451 | -0.118491202 | -0.070071698 | 12.611997 | 12.049693 | 12.709 | -0.659355604 | -0.097051767 |
| Cd5 | 10.91536 | 10.948 | 11.15 | -0.201911458 | -0.234551542 | 10.903991 | 10.531586 | 11.1911 | -0.65955674 | -0.2871515 |
| Prkag2 | 8.194594 | 8.326592 | 8.4307 | -0.104076549 | -0.236074143 | 8.1944665 | 7.8698939 | 8.52961 | -0.65971718 | -0.335144572 |
| Lrg1 | 9.909638 | 9.562304 | 9.4946 | 0.067718558 | 0.415052315 | 9.0021958 | 7.9370524 | 8.59712 | -0.66006784 | 0.405075598 |

| | | | | | | | | | | |
|---|---|---|---|---|---|---|---|---|---|---|
| Tsc22d1 | 9.204576 | 8.912096 | 9.251 | -0.338868323 | -0.046388632 | 8.5710161 | 8.203254 | 8.86352 | -0.660270184 | -0.292508068 |
| Jcad | 7.412927 | 7.233332 | 7.4251 | -0.19180587 | -0.012210358 | 6.6600854 | 6.4070648 | 7.06798 | -0.660913434 | -0.407892841 |
| Vipr1 | 9.507721 | 9.66897 | 9.7386 | -0.069647348 | -0.230896165 | 9.3469545 | 9.0009384 | 9.66258 | -0.661646052 | -0.315629963 |
| Plcb4 | 7.7015 | 6.880154 | 7.7055 | -0.825303862 | -0.003958146 | 7.178627 | 6.8560465 | 7.51775 | -0.66170494 | -0.33912438 |
| Galnt16 | 4.869704 | 4.829619 | 5.0166 | -0.186958125 | -0.146873175 | 4.9372519 | 4.1883883 | 4.85042 | -0.662032893 | 0.086830733 |
| Gm9864 | 7.857421 | 7.824369 | 8.0619 | -0.237560696 | -0.204509106 | 7.7688919 | 7.4167758 | 8.07882 | -0.662047713 | -0.309931533 |
| Cd33 | 7.723615 | 7.139585 | 7.2687 | -0.129111266 | 0.45491883 | 6.8387652 | 6.1427705 | 6.80494 | -0.662172236 | 0.033822443 |
| Tmem150c | 6.770845 | 6.882266 | 7.3026 | -0.420336121 | -0.531756713 | 7.0060186 | 6.7371755 | 7.39936 | -0.662186174 | -0.393343012 |
| 1700047I17Rik2 | 9.098323 | 8.855493 | 9.2212 | -0.365693987 | -0.122864011 | 9.1830226 | 8.7201282 | 9.38266 | -0.662529287 | -0.199634902 |
| Eif5a2 | 7.724327 | 7.514662 | 7.7062 | -0.191514993 | 0.018149577 | 7.5249067 | 7.027469 | 7.69004 | -0.662573319 | -0.165135589 |
| Stk11ip | 10.13431 | 9.871922 | 10.205 | -0.332820916 | -0.070432746 | 10.694649 | 9.7956845 | 10.4583 | -0.6625734 | 0.236390947 |
| Gcsam | 6.875327 | 6.704633 | 6.8009 | -0.096235514 | 0.074458889 | 6.6181775 | 5.9807089 | 6.64355 | -0.662842834 | -0.025374209 |
| Hba-a1 | 8.868188 | 8.374826 | 9.0176 | -0.642814321 | -0.149452399 | 8.8251227 | 8.2157234 | 8.87942 | -0.663692391 | -0.054293107 |
| Dgke | 8.780914 | 8.642838 | 8.8663 | -0.223475366 | -0.085399085 | 8.7769971 | 8.3485053 | 9.0123 | -0.663793816 | -0.235302019 |
| Pqlc1 | 7.779301 | 7.725328 | 7.7347 | -0.009381152 | 0.044592216 | 7.773111 | 7.214283 | 7.87849 | -0.664203993 | -0.105375962 |
| Klhl4 | 6.17162 | 5.82713 | 6.2107 | -0.38353623 | -0.039045719 | 5.5610966 | 5.2874147 | 5.95164 | -0.664221185 | -0.390539273 |
| Rgs11 | 6.482701 | 6.531344 | 6.5275 | 0.003858711 | -0.044784631 | 6.7326876 | 6.2231116 | 6.88794 | -0.664827477 | -0.155251447 |
| Ccdc66 | 8.157706 | 8.074922 | 8.2656 | -0.190655796 | -0.107871321 | 7.9552961 | 7.8018202 | 8.46668 | -0.664857705 | -0.511381785 |
| Sh3bp5 | 11.70847 | 11.63527 | 11.673 | -0.037994254 | 0.035206644 | 11.663759 | 11.016615 | 11.6823 | -0.66566134 | -0.018517657 |
| Mr1 | 7.822502 | 7.693688 | 7.9935 | -0.299804531 | -0.17099053 | 7.8489012 | 7.2668801 | 7.93275 | -0.665874266 | -0.083853134 |
| Atp13a3 | 11.18591 | 11.15019 | 11.16 | -0.009666338 | 0.0260593 | 10.944349 | 10.61854 | 11.2853 | -0.666768846 | -0.340960163 |
| Acsbg1 | 7.246636 | 7.106795 | 7.4578 | -0.351033185 | -0.211192052 | 7.1547163 | 6.7375268 | 7.40431 | -0.666782366 | -0.249592858 |
| Ap1m2 | 6.207129 | 6.063705 | 6.1649 | -0.1012373 | 0.042187189 | 6.2766876 | 5.5490036 | 6.21639 | -0.667382954 | 0.060301055 |
| Eno3 | 8.98785 | 8.889478 | 9.1939 | -0.304407765 | -0.206036407 | 9.1977745 | 8.7158239 | 9.38351 | -0.66768134 | -0.185730757 |
| Gm3325 | 5.795325 | 5.697665 | 5.8101 | -0.112445107 | -0.014785739 | 5.5904404 | 5.3485307 | 6.01647 | -0.667939581 | -0.426029869 |
| Rasgrp1 | 11.62302 | 11.65587 | 11.784 | -0.12773118 | -0.160587616 | 11.499408 | 11.051704 | 11.7199 | -0.668153192 | -0.220448997 |
| Pik3c2b | 10.21138 | 10.23609 | 10.374 | -0.13764807 | -0.162356325 | 10.516488 | 9.8023817 | 10.4708 | -0.668370824 | 0.045735145 |
| Fbln2 | 7.337238 | 7.055078 | 7.4036 | -0.348484418 | -0.066323633 | 6.6478211 | 6.1652943 | 6.83382 | -0.668526223 | -0.185999407 |
| Sema3g | 6.931256 | 6.944566 | 7.206 | -0.26146199 | -0.274772835 | 6.3644149 | 6.3930078 | 7.06203 | -0.66901862 | -0.697611517 |
| Il1r1 | 8.495026 | 8.069643 | 8.1393 | -0.069629302 | 0.35575389 | 7.3119686 | 6.8644793 | 7.53367 | -0.669192014 | -0.221702713 |
| Ttbk2 | 7.936534 | 7.642881 | 7.9082 | -0.265272619 | 0.028379988 | 7.5073352 | 7.1839667 | 7.85362 | -0.669654846 | -0.346286432 |
| Galnt1 | 11.79729 | 11.62404 | 11.803 | -0.179076778 | -0.005825454 | 11.727214 | 11.094976 | 11.7648 | -0.669856594 | -0.037618686 |

| | | | | | | | | | | |
|---|---|---|---|---|---|---|---|---|---|---|
| Cacna1e | 9.120306 | 9.176758 | 9.2329 | -0.056169343 | -0.112621818 | 9.6489589 | 9.0179962 | 9.68809 | -0.670089548 | -0.039126917 |
| Anxa3 | 9.08461 | 9.155524 | 9.2448 | -0.089310356 | -0.160224629 | 8.781615 | 8.7201179 | 9.39133 | -0.67121123 | -0.609714193 |
| Mycl | 7.188853 | 7.019905 | 7.296 | -0.276049648 | -0.107101623 | 7.2332317 | 6.3770465 | 7.04839 | -0.671340282 | 0.184844908 |
| Chst1 | 6.897959 | 6.41262 | 6.7167 | -0.304093307 | 0.181246125 | 5.9889237 | 5.5383958 | 6.21026 | -0.671862742 | -0.221334865 |
| Mmrn1 | 7.740421 | 7.418208 | 7.4646 | -0.046421808 | 0.275790962 | 6.3455745 | 6.4909623 | 7.16359 | -0.672625594 | -0.81801334 |
| C2cd5 | 9.737927 | 9.605722 | 9.8998 | -0.294086181 | -0.161882057 | 9.8432769 | 9.3800302 | 10.0527 | -0.672654873 | -0.209408118 |
| Camk2a | 6.089022 | 5.899624 | 5.9087 | -0.009049559 | 0.180347817 | 5.7409592 | 5.2325509 | 5.90649 | -0.673937436 | -0.165529118 |
| Nfib | 8.345167 | 7.936458 | 8.3568 | -0.42034449 | -0.011635972 | 7.4120195 | 7.1869807 | 7.86129 | -0.674311231 | -0.449272419 |
| Rtn4rl1 | 7.512628 | 7.549043 | 7.654 | -0.104914536 | -0.141329416 | 7.3043462 | 6.8408842 | 7.51612 | -0.675239919 | -0.211777934 |
| Tmem204 | 5.634343 | 5.500503 | 5.6308 | -0.130305126 | 0.003534039 | 5.2240417 | 4.8253605 | 5.50076 | -0.675399978 | -0.276718779 |
| Nynrin | 6.289132 | 5.878024 | 6.4307 | -0.552637219 | -0.141529193 | 5.8505965 | 5.3191264 | 5.99482 | -0.67569552 | -0.144225419 |
| Slc26a2 | 9.762114 | 9.603869 | 9.789 | -0.185160532 | -0.026915549 | 9.654216 | 9.1869857 | 9.86305 | -0.676064387 | -0.208834053 |
| Insr | 9.579236 | 9.403925 | 9.444 | -0.040119307 | 0.135192222 | 9.3966414 | 8.959172 | 9.63576 | -0.676584252 | -0.239114828 |
| Mccc2 | 7.377597 | 7.169908 | 7.4677 | -0.297774807 | -0.090085954 | 7.5819114 | 6.8829247 | 7.55979 | -0.676861756 | 0.022124969 |
| Fmo2 | 5.218455 | 4.49026 | 5.4998 | -1.009588711 | -0.281393341 | 4.0685907 | 3.8859198 | 4.5629 | -0.676980883 | -0.494310013 |
| Lingo4 | 4.15731 | 3.877105 | 4.2671 | -0.390030619 | -0.109825525 | 3.9879824 | 3.6164208 | 4.29468 | -0.678260045 | -0.30669839 |
| Xkrx | 8.730919 | 8.766502 | 8.9449 | -0.178426906 | -0.21401037 | 8.8983316 | 8.1742781 | 8.85301 | -0.678729482 | 0.045324019 |
| Fan1 | 7.212729 | 6.769589 | 7.1583 | -0.388743746 | 0.054396113 | 7.2139168 | 6.607942 | 7.28687 | -0.678929165 | -0.072954418 |
| Mmgt1 | 8.943547 | 8.669384 | 8.8796 | -0.210219851 | 0.063943238 | 8.7611477 | 8.2384272 | 8.91742 | -0.678989445 | -0.15626897 |
| P3h2 | 5.496966 | 5.523094 | 5.4091 | 0.114036557 | 0.08790855 | 4.7124295 | 4.6992948 | 5.37914 | -0.679843947 | -0.666709205 |
| Cacna1d | 6.663348 | 6.433331 | 6.6696 | -0.236268805 | -0.006252139 | 6.31485 | 6.0367878 | 6.71675 | -0.679966745 | -0.401904569 |
| Tmem26 | 6.441496 | 6.489378 | 6.3981 | 0.091232288 | 0.043349985 | 6.4509364 | 5.6460684 | 6.3262 | -0.680132497 | 0.124735483 |
| Slc44a1 | 9.404097 | 9.387022 | 9.5053 | -0.118321633 | -0.101247294 | 9.1399323 | 8.7055377 | 9.38588 | -0.680344917 | -0.245950325 |
| Mblac2 | 7.351333 | 7.127849 | 7.5326 | -0.40473714 | -0.181252845 | 7.4342503 | 6.718164 | 7.39872 | -0.680557901 | 0.035528375 |
| Dstyk | 8.524886 | 8.298004 | 8.4977 | -0.199709522 | 0.027171773 | 8.4720058 | 8.0608165 | 8.74208 | -0.681259486 | -0.270070187 |
| Ipp | 7.868533 | 7.631504 | 7.8435 | -0.212035679 | 0.024992733 | 7.5802594 | 7.1771513 | 7.85917 | -0.682023008 | -0.278914947 |
| Amz2 | 8.539428 | 8.308183 | 8.5759 | -0.267682693 | -0.036437187 | 8.4495321 | 8.0384529 | 8.72058 | -0.682128064 | -0.271048889 |
| St6gal1 | 12.13218 | 12.01541 | 12.208 | -0.192719466 | -0.075946356 | 12.235793 | 11.475736 | 12.1584 | -0.68263971 | 0.077417184 |
| Lamb2 | 8.031001 | 7.41215 | 8.032 | -0.619808667 | -0.000958099 | 7.3088209 | 6.9313687 | 7.61422 | -0.682851511 | -0.305399318 |
| Msh5 | 5.953013 | 6.018287 | 5.9516 | 0.066717605 | 0.001443644 | 6.3341029 | 5.5581473 | 6.24114 | -0.682996505 | 0.092959097 |
| Sptb | 5.200826 | 5.221921 | 5.3295 | -0.107569078 | -0.128664909 | 4.9425831 | 4.7704551 | 5.45394 | -0.68348817 | -0.511360093 |
| Srl | 4.87879 | 5.099744 | 5.387 | -0.287261535 | -0.508215176 | 4.5514464 | 4.6729001 | 5.35642 | -0.683519129 | -0.804972859 |

| | | | | | | | | | | |
|---|---|---|---|---|---|---|---|---|---|---|
| Itgb8 | 6.275799 | 6.262256 | 6.2144 | 0.047861877 | 0.061404612 | 5.8348787 | 5.7011146 | 6.38466 | -0.683549952 | -0.549785844 |
| Glce | 8.065441 | 7.894491 | 8.0832 | -0.188729589 | -0.01777944 | 7.822613 | 7.3221735 | 8.00615 | -0.683980181 | -0.183540645 |
| Tacc2 | 6.847307 | 6.663565 | 6.9254 | -0.261789547 | -0.078047403 | 6.7396946 | 6.1561074 | 6.84009 | -0.683986749 | -0.100399494 |
| Slc25a23 | 8.38158 | 8.356168 | 8.5248 | -0.168657446 | -0.143246121 | 8.2032694 | 7.6495451 | 8.33358 | -0.684030959 | -0.130306625 |
| Maco1 | 9.645075 | 9.375912 | 9.6839 | -0.307960038 | -0.038797436 | 9.442022 | 8.9442922 | 9.62844 | -0.684151076 | -0.186421224 |
| Dmd | 4.975854 | 4.732964 | 5.0531 | -0.320118957 | -0.07722902 | 4.7080791 | 4.503295 | 5.18777 | -0.684471496 | -0.479687402 |
| Serpinf1 | 5.500211 | 5.305094 | 5.4937 | -0.188613917 | 0.006503273 | 5.3145689 | 4.7972135 | 5.48281 | -0.685595344 | -0.168239884 |
| Gm10359 | 7.249848 | 7.163151 | 6.6994 | 0.463758678 | 0.550456012 | 7.198809 | 5.6358503 | 6.322 | -0.68614528 | 0.876813421 |
| Gm12671 | 7.249848 | 7.163151 | 6.6994 | 0.463758678 | 0.550456012 | 7.198809 | 5.6358503 | 6.322 | -0.68614528 | 0.876813421 |
| Kif26a | 5.381708 | 5.374969 | 5.7756 | -0.400619006 | -0.393879741 | 4.8058504 | 4.4696127 | 5.15647 | -0.686860442 | -0.350622724 |
| Nebl | 6.326057 | 6.246969 | 6.4441 | -0.197161683 | -0.118074399 | 6.3321988 | 5.666574 | 6.35374 | -0.687165176 | -0.021540419 |
| Gpr19 | 6.000326 | 5.757365 | 5.9411 | -0.183770484 | 0.05919111 | 5.9654242 | 5.4494951 | 6.13669 | -0.687198509 | -0.171269388 |
| Mrtfb | 9.548247 | 9.336966 | 9.5802 | -0.243195439 | -0.031914186 | 9.2264565 | 8.6886938 | 9.37623 | -0.687540024 | -0.149777259 |
| 2900026A02Rik | 7.772754 | 7.500311 | 7.8683 | -0.367968191 | -0.095524636 | 7.286507 | 6.7987198 | 7.48692 | -0.688203323 | -0.200416104 |
| Btbd11 | 8.47159 | 8.286245 | 8.6621 | -0.375857479 | -0.190512568 | 7.9460424 | 7.8958917 | 8.58463 | -0.688741805 | -0.638591151 |
| Agl | 8.349462 | 8.217337 | 8.4766 | -0.259287538 | -0.127162354 | 8.3426558 | 7.7053109 | 8.39407 | -0.688759611 | -0.051414783 |
| Ulk2 | 9.69787 | 9.663434 | 9.7948 | -0.131325995 | -0.096889694 | 9.4293058 | 8.9791879 | 9.66796 | -0.688775355 | -0.238657458 |
| Ddx25 | 5.865728 | 5.912669 | 5.9129 | -0.000225982 | -0.047167845 | 6.3304217 | 5.2632998 | 5.95209 | -0.688788977 | 0.378332901 |
| Cdh4 | 4.776993 | 4.363314 | 4.8501 | -0.48677205 | -0.073093062 | 4.2189132 | 4.0577765 | 4.74672 | -0.688944289 | -0.527807589 |
| Shcbp1 | 6.528347 | 6.212308 | 6.6016 | -0.389260484 | -0.07322162 | 6.222469 | 6.1401827 | 6.8293 | -0.689115085 | -0.606828763 |
| Galnt10 | 10.54123 | 10.57544 | 10.681 | -0.105618178 | -0.139825604 | 10.515743 | 9.9193955 | 10.6085 | -0.689124751 | -0.092777531 |
| Slc41a2 | 7.403857 | 7.452313 | 7.5178 | -0.065460395 | -0.113916324 | 7.1608849 | 6.7696105 | 7.45942 | -0.689812884 | -0.298538478 |
| Prdm9 | 7.86179 | 7.860451 | 8.0642 | -0.203716102 | -0.202377085 | 8.2036246 | 7.6281099 | 8.31794 | -0.689825564 | -0.114310852 |
| Slc12a2 | 8.644559 | 8.34344 | 8.4526 | -0.109123904 | 0.191994457 | 8.071807 | 7.6399792 | 8.33034 | -0.690363507 | -0.258535698 |
| Gxylt1 | 9.671534 | 9.580398 | 9.6144 | -0.033955884 | 0.057180261 | 9.4712056 | 9.118022 | 9.80841 | -0.690392324 | -0.337208733 |
| Pxdn | 6.000788 | 5.920674 | 6.0862 | -0.165505609 | -0.085391387 | 5.7127824 | 5.206272 | 5.89724 | -0.690968281 | -0.184457868 |
| Acvrl1 | 8.903559 | 8.750052 | 8.9275 | -0.177403871 | -0.023896291 | 8.5233698 | 8.1026255 | 8.79361 | -0.690980626 | -0.270236362 |
| Ankrd50 | 9.273504 | 9.06691 | 9.2876 | -0.220712041 | -0.014117704 | 8.950773 | 8.4554567 | 9.14849 | -0.693032245 | -0.197715919 |
| Aldoc | 6.270925 | 6.17977 | 6.2824 | -0.102627213 | -0.011472086 | 6.4587395 | 5.5759433 | 6.26913 | -0.693183604 | 0.18961259 |
| Zfp398 | 9.705141 | 9.670793 | 9.7968 | -0.126049924 | -0.091702537 | 9.6378108 | 9.1797503 | 9.87332 | -0.693565973 | -0.235505551 |
| Ighv11-2 | 8.091147 | 7.73901 | 7.6468 | 0.09220166 | 0.444338423 | 7.551291 | 7.2930193 | 7.98679 | -0.693768136 | -0.435496459 |
| Ccr8 | 6.669736 | 6.520641 | 6.6105 | -0.089857517 | 0.059237681 | 6.3128812 | 6.050224 | 6.74431 | -0.694084418 | -0.431427219 |

| | | | | | | | | | | |
|---|---|---|---|---|---|---|---|---|---|---|
| Ckap2l | 6.872805 | 6.616967 | 6.9886 | -0.371595878 | -0.115757171 | 6.7533221 | 6.4969509 | 7.19135 | -0.694396676 | -0.438025482 |
| Lpin1 | 9.106086 | 9.10324 | 9.2884 | -0.185190274 | -0.182344652 | 9.0230362 | 8.5145852 | 9.21003 | -0.695446165 | -0.186995239 |
| Zcchc14 | 6.053966 | 5.71738 | 5.9893 | -0.271886831 | 0.06469906 | 5.2390662 | 5.2093885 | 5.90491 | -0.695526315 | -0.665848665 |
| Ankrd13d | 6.986186 | 7.123566 | 6.9063 | 0.217293042 | 0.079912884 | 7.2703819 | 6.5270554 | 7.2234 | -0.69634685 | 0.046979665 |
| Hook1 | 9.110437 | 8.797201 | 9.047 | -0.249829206 | 0.063406776 | 8.8911252 | 8.5179607 | 9.21485 | -0.696888617 | -0.323724143 |
| Cmah | 12.51203 | 12.34211 | 12.537 | -0.19483932 | -0.024918008 | 12.537482 | 11.967668 | 12.6647 | -0.697053822 | -0.127239639 |
| Gria3 | 6.474333 | 6.441876 | 6.6021 | -0.160218056 | -0.127761003 | 6.1193658 | 5.797266 | 6.49502 | -0.697757452 | -0.375657668 |
| Kif14 | 6.453474 | 6.040858 | 6.3725 | -0.331631232 | 0.080984773 | 6.2933941 | 6.0931144 | 6.79134 | -0.69822248 | -0.497942754 |
| Samd12 | 4.887186 | 4.289241 | 4.8822 | -0.592934643 | 0.005010081 | 4.0750406 | 4.0094479 | 4.70907 | -0.699626203 | -0.634033521 |
| Coro2b | 5.954132 | 5.935812 | 6.2051 | -0.269259883 | -0.250939672 | 5.7387208 | 5.4598787 | 6.16027 | -0.700389739 | -0.421547624 |
| Ttl | 6.501146 | 6.531063 | 6.6324 | -0.101325037 | -0.131242028 | 6.3844665 | 5.8106521 | 6.51353 | -0.702874061 | -0.129059586 |
| Cenpe | 8.118291 | 7.955289 | 8.2413 | -0.286043168 | -0.123040757 | 7.9460782 | 7.7741384 | 8.4784 | -0.704259114 | -0.532319292 |
| Cavin2 | 7.025241 | 7.072798 | 7.2001 | -0.127272263 | -0.17482919 | 6.5244909 | 6.088482 | 6.7928 | -0.704318238 | -0.268309349 |
| Zcchc24 | 8.91394 | 8.705245 | 8.8535 | -0.148218355 | 0.060477132 | 8.6439266 | 7.8930689 | 8.59745 | -0.704378679 | 0.046479062 |
| Mfge8 | 11.4054 | 10.9197 | 11.4 | -0.48025891 | 0.005440062 | 10.499129 | 10.030267 | 10.735 | -0.704734883 | -0.235872701 |
| Hepacam2 | 6.409078 | 6.398715 | 6.6718 | -0.273098538 | -0.262735137 | 6.6479916 | 5.9358643 | 6.64062 | -0.704759293 | 0.007367974 |
| Myo9a | 9.224741 | 9.128568 | 9.4368 | -0.308187974 | -0.212014874 | 8.8262608 | 8.625627 | 9.33041 | -0.704781551 | -0.504147806 |
| Spire1 | 7.069196 | 6.897169 | 7.0944 | -0.197261822 | -0.025235052 | 6.8300672 | 6.4657108 | 7.17065 | -0.704939597 | -0.34058319 |
| Gucy1a1 | 5.931928 | 5.758868 | 6.0638 | -0.304963613 | -0.131903882 | 5.5852718 | 5.1082738 | 5.81445 | -0.706180817 | -0.229182816 |
| Kifc2 | 6.510283 | 6.349763 | 6.5223 | -0.17256913 | -0.012048995 | 6.7479757 | 6.0886553 | 6.79537 | -0.706714242 | -0.047393861 |
| Trpm1 | 5.654757 | 5.725796 | 5.728 | -0.002161139 | -0.073200429 | 5.5378418 | 5.1229084 | 5.82976 | -0.706853556 | -0.291920122 |
| 2810025M15Rik | 6.174567 | 6.205366 | 6.4114 | -0.206081306 | -0.236880607 | 5.9823596 | 5.54257 | 6.24971 | -0.707143874 | -0.267354282 |
| Inpp5f | 9.757145 | 9.76123 | 9.7019 | 0.059376372 | 0.055291327 | 9.667104 | 9.124282 | 9.83174 | -0.707458592 | -0.164636571 |
| Atp6v0a1 | 9.62344 | 9.50371 | 9.6938 | -0.190135884 | -0.070406486 | 9.9194049 | 9.1289316 | 9.83659 | -0.707656928 | 0.082816354 |
| Crybg2 | 6.515413 | 6.202084 | 6.5376 | -0.335494068 | -0.02216549 | 6.7580798 | 6.0564545 | 6.7659 | -0.709440645 | -0.00781536 |
| Whrn | 8.118267 | 8.064219 | 8.1903 | -0.126130828 | -0.072083326 | 8.233048 | 7.6654093 | 8.37556 | -0.710146746 | -0.142508067 |
| Foxo1 | 12.2331 | 12.20509 | 12.253 | -0.048025326 | -0.020012408 | 12.056762 | 11.486318 | 12.1965 | -0.7101586 | -0.139714926 |
| Inpp4b | 11.96263 | 11.82741 | 11.877 | -0.04965181 | 0.085569122 | 11.503292 | 11.376079 | 12.0868 | -0.710734012 | -0.583520843 |
| Bmpr2 | 9.095723 | 8.73311 | 8.8281 | -0.094975824 | 0.267636571 | 8.1845458 | 7.9409461 | 8.65305 | -0.712101422 | -0.468501694 |
| Tob1 | 8.794389 | 8.58178 | 9.0103 | -0.428499664 | -0.215890937 | 8.0706955 | 7.8704532 | 8.58288 | -0.712430101 | -0.51218787 |
| Gnpda1 | 8.968829 | 8.854224 | 9.0261 | -0.171833603 | -0.057228522 | 9.2202572 | 8.4261945 | 9.13903 | -0.712834127 | 0.081228559 |
| Ccdc125 | 8.195595 | 8.067488 | 8.2356 | -0.16806856 | -0.03996167 | 8.1060761 | 7.4844041 | 8.19731 | -0.712904011 | -0.091231932 |

| | | | | | | | | | | |
|---|---|---|---|---|---|---|---|---|---|---|
| Fmo5 | 7.19949 | 7.139027 | 7.3556 | -0.216575729 | -0.156113012 | 7.3645604 | 6.7286895 | 7.44241 | -0.713724414 | -0.077853476 |
| A430078G23Rik | 9.919486 | 9.972968 | 10.008 | -0.034922308 | -0.088404677 | 9.8339948 | 9.3443432 | 10.0585 | -0.714159207 | -0.224507605 |
| Fgfr2 | 5.086129 | 4.73761 | 5.1712 | -0.433581011 | -0.085061795 | 4.5706989 | 4.1463356 | 4.86177 | -0.715438503 | -0.291075153 |
| Katnal1 | 6.973955 | 6.961586 | 7.0174 | -0.055812073 | -0.043443761 | 7.0709854 | 6.4389295 | 7.1563 | -0.717371638 | -0.085315766 |
| Rasgrf2 | 7.272845 | 7.331342 | 7.4353 | -0.103966212 | -0.162463137 | 7.3235034 | 6.5951558 | 7.31276 | -0.717604007 | 0.010743607 |
| Fam129a | 10.30241 | 10.27299 | 10.408 | -0.134910516 | -0.105489794 | 9.915313 | 9.7427662 | 10.4614 | -0.718677276 | -0.546130486 |
| Izumo1r | 9.016749 | 8.874393 | 9.1854 | -0.310966618 | -0.168610985 | 8.9215519 | 8.3357327 | 9.05451 | -0.718776906 | -0.1329577 |
| Klf9 | 7.724298 | 7.493821 | 7.6138 | -0.120023453 | 0.110453983 | 6.8873923 | 6.6233031 | 7.3422 | -0.718898924 | -0.454809752 |
| Alas2 | 5.218078 | 4.639761 | 5.2205 | -0.580747715 | -0.002430461 | 4.9030206 | 4.526086 | 5.24558 | -0.719493633 | -0.34255904 |
| Trim24 | 9.851397 | 9.735973 | 9.9451 | -0.209125788 | -0.09370126 | 9.8546532 | 9.359377 | 10.0802 | -0.720865467 | -0.225589252 |
| Rab37 | 9.148054 | 8.91487 | 9.2702 | -0.355315397 | -0.122131013 | 9.1065358 | 8.5484074 | 9.27002 | -0.721613064 | -0.163484706 |
| Zbtb16 | 5.965949 | 5.966889 | 6.1389 | -0.172059222 | -0.172998793 | 5.4028376 | 5.1518542 | 5.87378 | -0.721922975 | -0.47093963 |
| Il17rb | 5.808978 | 5.547294 | 6.2293 | -0.682020375 | -0.420336457 | 5.8139333 | 5.3855823 | 6.10998 | -0.724392944 | -0.29604186 |
| Pfkfb2 | 7.775592 | 7.663392 | 7.7867 | -0.1233264 | -0.011126212 | 7.9437264 | 7.2739485 | 7.99923 | -0.72527971 | -0.055501862 |
| Taf9b | 8.429413 | 8.229883 | 8.5118 | -0.281878418 | -0.082348779 | 8.4820234 | 7.7838902 | 8.51055 | -0.726656795 | -0.028523614 |
| Stmn1 | 9.491508 | 9.255202 | 9.6775 | -0.422271868 | -0.18596602 | 9.3485617 | 9.1833963 | 9.91049 | -0.727091729 | -0.561926352 |
| Mphosph9 | 9.419271 | 9.384416 | 9.3336 | 0.050802231 | 0.085656593 | 9.4205936 | 9.0387855 | 9.76592 | -0.727134608 | -0.345326416 |
| Lpar5 | 7.10766 | 7.213351 | 7.2539 | -0.040522846 | -0.146214602 | 7.2169534 | 6.4631904 | 7.19089 | -0.727699164 | 0.026063856 |
| Cep89 | 7.798187 | 7.679591 | 7.7714 | -0.091849248 | 0.026746295 | 8.0990374 | 7.380061 | 8.10787 | -0.727813425 | -0.00883703 |
| Kif21b | 12.71358 | 12.76023 | 12.868 | -0.108083674 | -0.15473241 | 12.642274 | 12.167526 | 12.8966 | -0.729094072 | -0.25434643 |
| Klf11 | 7.21437 | 7.094696 | 7.4423 | -0.347562923 | -0.227889451 | 7.1944173 | 6.7403498 | 7.46945 | -0.729095614 | -0.275028063 |
| Zfp979 | 6.279242 | 5.752846 | 6.428 | -0.675131179 | -0.14873589 | 5.7389654 | 5.0996979 | 5.82897 | -0.729275776 | -0.090008296 |
| Ldlrad4 | 7.302012 | 7.286582 | 7.4236 | -0.137037296 | -0.12160661 | 7.0082502 | 6.6329567 | 7.36391 | -0.730955606 | -0.355662083 |
| Tesk2 | 6.922289 | 6.865496 | 7.0693 | -0.203781183 | -0.146988401 | 7.1233906 | 6.465694 | 7.19765 | -0.731957655 | -0.07426105 |
| Samd10 | 7.196094 | 6.959541 | 7.3685 | -0.408961553 | -0.172408255 | 7.65784 | 6.6576256 | 7.38983 | -0.732206987 | 0.268007336 |
| Reck | 6.735986 | 6.700835 | 6.9232 | -0.222359642 | -0.187208833 | 6.294509 | 6.1760379 | 6.90842 | -0.732385569 | -0.613914459 |
| Zfp521 | 7.34858 | 7.136301 | 7.4293 | -0.293027161 | -0.08074801 | 6.1512011 | 6.1365237 | 6.86896 | -0.732437848 | -0.717760441 |
| Phactr2 | 8.190151 | 7.854329 | 8.27 | -0.415672391 | -0.079849961 | 7.6046097 | 7.3602242 | 8.09286 | -0.732636468 | -0.488250966 |
| Dip2a | 7.494933 | 7.526549 | 7.5636 | -0.037045559 | -0.068661995 | 7.5379514 | 7.0174754 | 7.75042 | -0.732944661 | -0.212468684 |
| Tshz2 | 7.301874 | 6.940942 | 7.1245 | -0.183560983 | 0.177370738 | 6.2277985 | 6.0256863 | 6.75904 | -0.733350778 | -0.531238613 |
| Ccpg1 | 10.18636 | 10.10789 | 10.25 | -0.142238619 | -0.063764198 | 10.035391 | 9.4712797 | 10.2046 | -0.733368595 | -0.169256877 |
| Pdk2 | 7.149792 | 6.934239 | 7.305 | -0.37077191 | -0.155219516 | 7.1169756 | 6.4199881 | 7.15467 | -0.734682001 | -0.037694539 |

| | | | | | | | | | | |
|---|---|---|---|---|---|---|---|---|---|---|
| Nr5a2 | 6.162022 | 5.693952 | 5.9473 | -0.25330975 | 0.2147602 | 5.216579 | 4.9260696 | 5.66101 | -0.734943425 | -0.444433982 |
| Slc9a3r2 | 6.929738 | 6.910674 | 7.008 | -0.097352835 | -0.078289278 | 6.4062177 | 5.9619023 | 6.69717 | -0.73527172 | -0.290956347 |
| Itga1 | 8.446388 | 7.723107 | 8.2693 | -0.546234523 | 0.177046049 | 7.3216554 | 7.2832154 | 8.02113 | -0.737911139 | -0.699471146 |
| Sema4f | 7.923253 | 8.045611 | 8.1034 | -0.057764137 | -0.18012192 | 7.7147803 | 7.2974789 | 8.03552 | -0.738042489 | -0.320741107 |
| Ncaph | 7.419501 | 7.293587 | 7.6481 | -0.354499855 | -0.228586084 | 7.3653914 | 7.1167552 | 7.85626 | -0.739505029 | -0.490868879 |
| Cbr2 | 6.03194 | 4.949498 | 5.7555 | -0.805987958 | 0.276453514 | 4.6122412 | 4.4533652 | 5.19353 | -0.740162999 | -0.581287 |
| Oit1 | 6.721849 | 5.995206 | 6.9176 | -0.922436154 | -0.19579341 | 5.557667 | 5.1389735 | 5.87922 | -0.740244129 | -0.321550615 |
| Extl1 | 7.768714 | 7.58717 | 7.8006 | -0.213434086 | -0.031889352 | 7.5034136 | 7.0462598 | 7.78694 | -0.740682029 | -0.283528218 |
| Depdc1a | 5.970546 | 5.796917 | 6.1563 | -0.359379639 | -0.185750839 | 5.8726067 | 5.561776 | 6.30265 | -0.740871548 | -0.430040869 |
| Pitpnm2 | 10.46164 | 10.43461 | 10.501 | -0.066689346 | -0.039655448 | 10.452491 | 9.7800534 | 10.5211 | -0.741093573 | -0.068655792 |
| Rgs4 | 5.661433 | 5.515239 | 5.2769 | 0.238318989 | 0.384513172 | 5.3660164 | 4.6628076 | 5.40472 | -0.741911947 | -0.038703069 |
| Sdc1 | 7.527095 | 7.746502 | 7.8484 | -0.101858861 | -0.321265819 | 7.3615371 | 6.7847422 | 7.52689 | -0.742149449 | -0.165354498 |
| Carf | 7.196955 | 7.076859 | 7.5067 | -0.429834888 | -0.30973875 | 7.3643611 | 6.8864807 | 7.62882 | -0.742344086 | -0.264463667 |
| Cldn1 | 5.950869 | 5.839574 | 6.1953 | -0.355718202 | -0.244423572 | 5.7674494 | 5.2084149 | 5.95202 | -0.743608803 | -0.18457433 |
| Trim36 | 7.019635 | 6.914298 | 7.1967 | -0.28240292 | -0.177065438 | 6.9427611 | 6.071996 | 6.81584 | -0.743839174 | 0.126925962 |
| Map4k5 | 8.833095 | 8.634736 | 8.726 | -0.091262299 | 0.107096472 | 8.8708116 | 8.2159211 | 8.96049 | -0.744572086 | -0.089681572 |
| St6galnac2 | 5.637993 | 5.429899 | 5.8341 | -0.404199492 | -0.19610545 | 5.196596 | 4.9561901 | 5.70098 | -0.74479112 | -0.504385198 |
| Rassf6 | 5.719626 | 5.473472 | 5.8808 | -0.407322467 | -0.161168506 | 5.4261305 | 4.9135697 | 5.65909 | -0.745520404 | -0.232959626 |
| Amotl1 | 7.361251 | 6.952231 | 7.2812 | -0.32893488 | 0.080084791 | 6.5990438 | 6.2326534 | 6.9784 | -0.745745933 | -0.379355535 |
| Pde1c | 5.979433 | 5.518971 | 5.9066 | -0.38760741 | 0.072855077 | 5.2425746 | 4.7912289 | 5.53791 | -0.746678396 | -0.295332642 |
| Gm21985 | 7.216095 | 7.454023 | 7.4472 | 0.006786405 | -0.231141831 | 7.2258903 | 7.1668068 | 7.91411 | -0.747303454 | -0.688219998 |
| Ttc28 | 9.495168 | 9.39505 | 9.6116 | -0.216589892 | -0.116472237 | 9.0120619 | 8.6342256 | 9.38299 | -0.74876931 | -0.370932997 |
| Mme | 5.677766 | 5.312282 | 5.7374 | -0.425150221 | -0.059665992 | 5.6727699 | 4.8244348 | 5.57326 | -0.748827339 | 0.099507771 |
| Notch4 | 6.632849 | 6.777213 | 6.8679 | -0.090722124 | -0.235086471 | 6.7705966 | 5.950515 | 6.69959 | -0.749073521 | 0.071008082 |
| Emcn | 7.619284 | 7.252923 | 7.706 | -0.453048979 | -0.08668725 | 6.8485493 | 6.3573274 | 7.10754 | -0.750208765 | -0.258986902 |
| Gal3st2 | 6.230296 | 5.714431 | 6.2538 | -0.539385141 | -0.023520183 | 5.9393733 | 5.5676245 | 6.31784 | -0.750219276 | -0.378470482 |
| Scn4b | 6.390975 | 6.055418 | 6.6429 | -0.587502162 | -0.251944636 | 5.8901591 | 5.7175336 | 6.46783 | -0.750296656 | -0.57767123 |
| Cdca3 | 7.126712 | 7.100001 | 7.3719 | -0.271918317 | -0.245207844 | 7.3713533 | 6.9682918 | 7.71951 | -0.751220649 | -0.348159113 |
| Acot11 | 7.378507 | 7.335959 | 7.4857 | -0.149699287 | -0.107151372 | 7.4497473 | 6.9757773 | 7.72859 | -0.752807766 | -0.278837835 |
| 2610528A11Rik | 5.466875 | 5.191385 | 5.5727 | -0.381341557 | -0.105851394 | 5.1385997 | 4.8427371 | 5.59556 | -0.752825247 | -0.456962673 |
| Ccr4 | 7.210771 | 7.007535 | 7.2089 | -0.2014002 | 0.001835334 | 6.841318 | 6.5432085 | 7.29617 | -0.752958096 | -0.45484865 |
| Sort1 | 7.375634 | 7.184142 | 7.4562 | -0.272042837 | -0.080551232 | 7.1740856 | 6.5301394 | 7.28389 | -0.753747354 | -0.10980122 |

| | | | | | | | | | | |
|---|---|---|---|---|---|---|---|---|---|---|
| Mtmr10 | 7.839093 | 7.637456 | 7.7668 | -0.129317807 | 0.07231938 | 7.4367767 | 7.010369 | 7.76415 | -0.753784241 | -0.327376568 |
| Gsn | 11.32192 | 11.1604 | 11.27 | -0.109664302 | 0.051855024 | 11.159169 | 10.295941 | 11.0499 | -0.753967241 | 0.109260556 |
| Bcor | 11.02605 | 11.04185 | 11.099 | -0.05686099 | -0.072657114 | 10.935539 | 10.339002 | 11.0935 | -0.754474058 | -0.157936884 |
| Dennd5b | 10.44272 | 10.30353 | 10.545 | -0.24164797 | -0.102453238 | 10.614907 | 9.9239163 | 10.6789 | -0.754951817 | -0.063961048 |
| Dlc1 | 8.103749 | 7.431744 | 7.6575 | -0.225709896 | 0.446295632 | 7.0994468 | 6.6346684 | 7.38968 | -0.755011779 | -0.290233422 |
| Mtss1 | 10.9376 | 10.84182 | 11.01 | -0.168477136 | -0.072697372 | 10.930725 | 10.171769 | 10.9272 | -0.75548081 | 0.003474789 |
| S100a8 | 9.611917 | 7.915791 | 7.266 | 0.649788519 | 2.345914625 | 8.3540844 | 6.9202775 | 7.67585 | -0.755575409 | 0.678231461 |
| Ptpn7 | 10.60944 | 10.39048 | 10.653 | -0.26227276 | -0.043318004 | 10.59174 | 9.9525312 | 10.7082 | -0.755668339 | -0.116459796 |
| Cdca8 | 7.457455 | 7.052879 | 7.462 | -0.409141919 | -0.004565307 | 7.3605272 | 6.9735373 | 7.72927 | -0.755733918 | -0.368744057 |
| Nuak1 | 6.884376 | 6.83896 | 7.0025 | -0.163542041 | -0.118126304 | 6.5419696 | 5.988102 | 6.74498 | -0.756876663 | -0.203009061 |
| Chst4 | 8.113374 | 7.507378 | 8.1388 | -0.631375889 | -0.025380423 | 6.679849 | 5.9729549 | 6.73011 | -0.757159381 | -0.050265305 |
| Syne3 | 9.188352 | 9.079068 | 9.2194 | -0.140341522 | -0.031057757 | 9.000241 | 8.3747935 | 9.13252 | -0.757731392 | -0.132283927 |
| Cadm1 | 7.267988 | 7.174382 | 7.0653 | 0.109074931 | 0.20268097 | 6.6875358 | 6.1067726 | 6.86588 | -0.759106253 | -0.178343068 |
| Mid2 | 6.272097 | 5.972379 | 6.3914 | -0.418989493 | -0.11927195 | 5.4139389 | 5.2005788 | 5.95977 | -0.759186816 | -0.545826672 |
| Lrig1 | 8.998171 | 8.837712 | 9.0741 | -0.236378603 | -0.075919104 | 8.2534493 | 8.2108035 | 8.97093 | -0.760127277 | -0.717481475 |
| Pif1 | 5.021644 | 4.954943 | 5.2106 | -0.255639947 | -0.188939179 | 4.9746622 | 4.7827687 | 5.54319 | -0.760420978 | -0.56852742 |
| Crybg3 | 8.713732 | 8.57314 | 8.6945 | -0.121338027 | 0.019253439 | 8.3572482 | 7.9756624 | 8.73621 | -0.76054718 | -0.378961414 |
| Lrrcc1 | 8.891054 | 8.606558 | 8.9866 | -0.380036123 | -0.095539932 | 8.802447 | 8.2506227 | 9.01135 | -0.760725541 | -0.208901264 |
| Chdh | 6.494417 | 6.58348 | 6.6824 | -0.098883789 | -0.187947108 | 6.0735294 | 5.8044461 | 6.56562 | -0.761169866 | -0.492086584 |
| Spry1 | 6.881696 | 6.659958 | 7.0465 | -0.386585814 | -0.164847948 | 6.0281396 | 5.6216335 | 6.38282 | -0.761185783 | -0.35467968 |
| Tnfrsf13b | 10.95642 | 10.72057 | 11.005 | -0.284081636 | -0.048234718 | 11.281035 | 10.16672 | 10.9281 | -0.761361676 | 0.352953933 |
| Capn3 | 6.202457 | 5.90318 | 6.4074 | -0.504206419 | -0.204929156 | 5.7387845 | 5.6377835 | 6.39931 | -0.761528248 | -0.66052731 |
| Kcnc1 | 6.435512 | 6.29912 | 6.4708 | -0.171659314 | -0.035266565 | 5.9946916 | 5.7435866 | 6.50646 | -0.762872805 | -0.511767761 |
| Abca8a | 5.56191 | 5.253593 | 5.73 | -0.476386962 | -0.168069764 | 5.0598156 | 4.3925341 | 5.1563 | -0.76376369 | -0.096482152 |
| Arhgef4 | 7.445501 | 7.164039 | 7.617 | -0.452950972 | -0.171489179 | 7.2072113 | 6.6384831 | 7.40349 | -0.765002533 | -0.19627434 |
| Gm6882 | 5.424898 | 5.365497 | 5.3983 | -0.032762677 | 0.02663903 | 5.5983948 | 4.803516 | 5.56862 | -0.765100216 | 0.02977863 |
| Igsf3 | 6.599002 | 6.748699 | 6.6058 | 0.142866716 | -0.006829798 | 6.4162437 | 6.1159117 | 6.88164 | -0.765726307 | -0.465394321 |
| Gstm1 | 7.714408 | 7.449504 | 7.9148 | -0.465261546 | -0.200358246 | 7.2319352 | 6.768373 | 7.53568 | -0.767309603 | -0.303747412 |
| Cdc20 | 7.574977 | 7.392725 | 7.6898 | -0.297089296 | -0.114837944 | 7.7491445 | 7.1108422 | 7.87818 | -0.767341514 | -0.12903921 |
| Plk2 | 10.03678 | 9.924435 | 9.9384 | -0.013999184 | 0.098343999 | 9.0091433 | 8.8030042 | 9.57079 | -0.76778131 | -0.561642174 |
| Itm2a | 9.098839 | 9.003369 | 9.196 | -0.192639393 | -0.097169858 | 8.7033539 | 8.1928047 | 8.9612 | -0.768396062 | -0.257846893 |
| Rorc | 6.767644 | 6.560189 | 6.9421 | -0.381878761 | -0.174424297 | 6.3572234 | 6.0400529 | 6.80975 | -0.769697513 | -0.452526948 |

| | | | | | | | | | | |
|---|---|---|---|---|---|---|---|---|---|---|
| Tspyl4 | 7.619222 | 7.628315 | 7.6936 | -0.065259325 | -0.07435241 | 7.5993669 | 7.2548892 | 8.02511 | -0.770216823 | -0.425739168 |
| Diaph3 | 7.088942 | 6.994816 | 7.0242 | -0.029340695 | 0.064786021 | 6.8826882 | 6.7052769 | 7.47664 | -0.771359242 | -0.593947908 |
| Nfia | 7.568593 | 7.275572 | 7.6139 | -0.338373695 | -0.045352695 | 6.6511966 | 6.5221451 | 7.2939 | -0.771759652 | -0.642708126 |
| Dpy19l4 | 8.715714 | 8.592361 | 8.7996 | -0.207198328 | -0.083845627 | 8.6468145 | 8.1804048 | 8.95266 | -0.7722521 | -0.305842373 |
| Etl4 | 6.20815 | 5.758891 | 6.2866 | -0.527662908 | -0.078403656 | 6.0381751 | 5.5797367 | 6.35204 | -0.77229904 | -0.313860604 |
| Cdkl1 | 6.159935 | 5.676862 | 6.0303 | -0.353398047 | 0.129674596 | 6.1828512 | 5.1332948 | 5.90617 | -0.77287157 | 0.276684787 |
| Themis | 10.66156 | 10.71724 | 10.821 | -0.10413313 | -0.159806618 | 10.347726 | 9.9921429 | 10.7668 | -0.774704112 | -0.419121274 |
| Fancd2 | 6.852038 | 6.500979 | 6.995 | -0.494016907 | -0.142957515 | 6.830638 | 6.4487461 | 7.22496 | -0.776211842 | -0.394319955 |
| Ngp | 5.918092 | 5.644569 | 6.0362 | -0.39162026 | -0.118097552 | 6.3221426 | 5.3690798 | 6.14544 | -0.776355805 | 0.176707016 |
| Ttn | 6.438636 | 5.629304 | 6.2786 | -0.64932082 | 0.160011398 | 5.347284 | 5.4224846 | 6.19994 | -0.777452782 | -0.852653392 |
| Ank | 8.581985 | 8.628807 | 8.823 | -0.194143627 | -0.240965471 | 8.6447683 | 7.917157 | 8.69564 | -0.778483297 | -0.050872043 |
| Fry | 9.938417 | 9.877281 | 9.9676 | -0.090334385 | -0.029197844 | 9.7604096 | 9.1901924 | 9.96905 | -0.778860401 | -0.208643185 |
| Il23r | 5.763511 | 5.508009 | 5.759 | -0.25096442 | 0.004537413 | 5.4589509 | 5.2202897 | 5.99931 | -0.779020627 | -0.540359392 |
| Bpifb3 | 4.023251 | 3.226846 | 4.2336 | -1.006795563 | -0.210390331 | 3.169165 | 3.1729399 | 3.95228 | -0.779343632 | -0.783118586 |
| Fzd4 | 7.627744 | 7.054912 | 7.6106 | -0.555677091 | 0.017155017 | 6.6033941 | 6.1668978 | 6.9468 | -0.779898252 | -0.343401925 |
| Colgalt2 | 4.74816 | 4.672377 | 4.8026 | -0.130271394 | -0.054488513 | 4.4132128 | 4.065266 | 4.84556 | -0.780296316 | -0.43234955 |
| Ctsl | 10.65949 | 10.00823 | 10.584 | -0.575988476 | 0.075266734 | 9.121099 | 8.8528511 | 9.63346 | -0.78061244 | -0.512364602 |
| Ppm1l | 8.181136 | 8.09646 | 8.1636 | -0.067147951 | 0.01752862 | 8.0925294 | 7.3930894 | 8.17552 | -0.782435422 | -0.082995386 |
| Ncapd2 | 10.20494 | 9.801819 | 10.253 | -0.451259463 | -0.048142156 | 10.21809 | 9.7746135 | 10.5571 | -0.782482435 | -0.339006085 |
| Ctla4 | 9.521034 | 9.104611 | 9.5297 | -0.425101211 | -0.008678672 | 9.3067757 | 9.2516197 | 10.0346 | -0.783029531 | -0.727873488 |
| Tmem108 | 7.744897 | 7.706558 | 7.8096 | -0.103082612 | -0.064743925 | 8.2048765 | 7.134432 | 7.91781 | -0.783376354 | 0.287068111 |
| Tk1 | 7.936546 | 7.643854 | 8.1089 | -0.465068725 | -0.172376364 | 8.4014974 | 7.6784226 | 8.46237 | -0.783951569 | -0.060876737 |
| N4bp2 | 10.52252 | 10.41601 | 10.52 | -0.103870046 | 0.002640996 | 10.422679 | 9.7523974 | 10.5365 | -0.784072322 | -0.113790963 |
| Zfp462 | 5.11897 | 4.517834 | 4.9843 | -0.466438133 | 0.134698259 | 4.721565 | 4.0896398 | 4.87419 | -0.784547144 | -0.152621991 |
| Myl9 | 6.061277 | 5.895758 | 6.2885 | -0.392759058 | -0.227239404 | 6.025315 | 5.2502347 | 6.03498 | -0.784740908 | -0.009660604 |
| Lats2 | 10.53532 | 10.45415 | 10.513 | -0.05929755 | 0.021879964 | 10.350643 | 9.8400058 | 10.6249 | -0.784915573 | -0.274278683 |
| Gdf11 | 9.09457 | 9.220229 | 9.2829 | -0.062683607 | -0.188342539 | 9.2189844 | 8.3742064 | 9.15956 | -0.78535705 | 0.059420994 |
| Ncapg | 7.478792 | 7.239777 | 7.5693 | -0.329546976 | -0.09053211 | 7.2836881 | 7.1092118 | 7.89518 | -0.785966379 | -0.611490031 |
| Insm1 | 5.883238 | 5.966605 | 6.0745 | -0.107860126 | -0.191226736 | 5.4296826 | 5.3060636 | 6.09338 | -0.787317175 | -0.6636982 |
| Tfdp2 | 8.433087 | 8.18464 | 8.5404 | -0.355764437 | -0.1073166 | 8.1183271 | 7.6852271 | 8.47276 | -0.787536878 | -0.35443682 |
| Arhgap5 | 8.807002 | 8.735157 | 8.9962 | -0.26104531 | -0.189199857 | 8.2591324 | 8.0708816 | 8.85849 | -0.787603776 | -0.599352999 |
| Tbc1d4 | 10.53539 | 10.46863 | 10.721 | -0.25216733 | -0.185409604 | 10.219741 | 10.084258 | 10.8724 | -0.788147754 | -0.652665292 |

| | | | | | | | | | | |
|---|---|---|---|---|---|---|---|---|---|---|
| Ttc39a | 6.535788 | 6.581989 | 6.7161 | -0.134143658 | -0.180344518 | 6.397342 | 6.0387134 | 6.82727 | -0.78856122 | -0.42993258 |
| Bace1 | 6.773159 | 6.678573 | 6.7652 | -0.086619315 | 0.007966924 | 6.6523631 | 5.8773421 | 6.66612 | -0.788781275 | -0.013760349 |
| Zfp46 | 9.391167 | 8.893149 | 8.9959 | -0.102714434 | 0.395303438 | 8.2262786 | 7.9520397 | 8.74142 | -0.789375756 | -0.515136828 |
| Chn2 | 6.987916 | 6.790012 | 7.1622 | -0.372223038 | -0.174319155 | 6.5467489 | 6.3084316 | 7.09943 | -0.79099393 | -0.552676636 |
| Plekha7 | 6.897177 | 6.737265 | 6.7475 | -0.01026097 | 0.149651173 | 6.5992149 | 6.3457852 | 7.1373 | -0.791518056 | -0.538088382 |
| Akap5 | 5.474288 | 5.547982 | 5.6467 | -0.098700602 | -0.172394008 | 5.688594 | 5.1765467 | 5.96827 | -0.791725353 | -0.279678088 |
| Gm28693 | 5.036076 | 4.983092 | 5.0278 | -0.044667963 | 0.008316197 | 5.3034795 | 4.8795308 | 5.67148 | -0.791949013 | -0.368000265 |
| Arhgap33 | 6.19937 | 6.205942 | 6.3992 | -0.193271495 | -0.199843497 | 6.6577937 | 5.8270017 | 6.61903 | -0.792026327 | 0.038765709 |
| Mlh3 | 8.180696 | 8.038468 | 8.2832 | -0.244736465 | -0.102508079 | 8.2589828 | 7.56202 | 8.35496 | -0.792937221 | -0.095974379 |
| Cldn11 | 5.443665 | 4.871193 | 5.5036 | -0.632372313 | -0.059900632 | 4.4093235 | 4.1830958 | 4.97625 | -0.793149661 | -0.566921972 |
| Armcx4 | 8.225108 | 7.89676 | 7.8112 | 0.085523236 | 0.413871534 | 7.3636646 | 6.8448485 | 7.63823 | -0.793380287 | -0.274564176 |
| Lepr | 8.502852 | 7.879615 | 7.9523 | -0.072643074 | 0.550593904 | 7.4054074 | 6.8380232 | 7.63167 | -0.793645009 | -0.226260831 |
| Siglech | 9.160825 | 8.65913 | 9.208 | -0.548897608 | -0.047201852 | 7.7960179 | 7.5256632 | 8.31955 | -0.793888225 | -0.523533554 |
| Gm47924 | 7.329437 | 7.256501 | 7.5047 | -0.248178927 | -0.175243786 | 7.3816011 | 7.3147071 | 8.10982 | -0.795114109 | -0.728220071 |
| Synpo2 | 9.226573 | 9.176593 | 9.3315 | -0.154926481 | -0.104946577 | 8.5662055 | 8.6178127 | 9.41412 | -0.796305955 | -0.847913183 |
| Rgcc | 7.767356 | 7.561622 | 7.8536 | -0.291977347 | -0.086243377 | 7.4819158 | 7.0702504 | 7.86729 | -0.797036886 | -0.385371527 |
| Lypd6b | 7.036147 | 7.247645 | 7.2676 | -0.019990563 | -0.231488849 | 6.8313137 | 6.5979819 | 7.39556 | -0.797573282 | -0.564241454 |
| Ptprg | 6.758167 | 6.762635 | 6.5503 | 0.2123291 | 0.207860674 | 6.0975795 | 5.8670928 | 6.66553 | -0.798438999 | -0.567952342 |
| Tox | 9.532799 | 9.288391 | 9.6406 | -0.352208357 | -0.107800151 | 8.9961172 | 8.7812563 | 9.57972 | -0.798459329 | -0.583598387 |
| Zfp395 | 8.910003 | 8.804702 | 9.0205 | -0.215769744 | -0.110468185 | 8.7530343 | 8.200889 | 8.99962 | -0.798731215 | -0.246585984 |
| Man2a2 | 11.04338 | 11.03382 | 11.233 | -0.199009702 | -0.189448352 | 11.039028 | 10.408312 | 11.208 | -0.7997244 | -0.169008493 |
| Ighv1-66 | 5.626157 | 5.949984 | 5.9607 | -0.010682342 | -0.334508954 | 6.1929207 | 5.898583 | 6.69832 | -0.7997327 | -0.505394968 |
| Nrn1 | 6.498898 | 6.306858 | 6.6602 | -0.353359802 | -0.161320518 | 6.2860068 | 5.6280958 | 6.43103 | -0.802939003 | -0.14502801 |
| B3galt5 | 4.114619 | 3.984811 | 4.3425 | -0.357736316 | -0.227928727 | 3.9577458 | 3.6052299 | 4.40832 | -0.803085442 | -0.450569559 |
| Prc1 | 7.6465 | 7.300178 | 7.7474 | -0.447225739 | -0.100903426 | 7.6115719 | 7.2594302 | 8.06253 | -0.803104344 | -0.450962557 |
| Rrm2b | 9.620938 | 9.589438 | 9.6847 | -0.095217726 | -0.063717896 | 9.7100055 | 8.9916745 | 9.79604 | -0.804361253 | -0.086030283 |
| Cenpa | 8.843543 | 8.873977 | 9.0279 | -0.153942151 | -0.18437587 | 8.9589969 | 8.4568978 | 9.26184 | -0.804942701 | -0.302843683 |
| Ccr10 | 4.569665 | 4.338973 | 4.7263 | -0.387314656 | -0.156622737 | 4.3595181 | 3.8897151 | 4.6947 | -0.804989165 | -0.335186121 |
| Creb3l2 | 8.78155 | 8.517768 | 8.8135 | -0.295731416 | -0.031949833 | 8.1678011 | 7.7764434 | 8.5819 | -0.805454004 | -0.41409631 |
| Pank1 | 6.733594 | 7.024282 | 6.8747 | 0.149626181 | -0.141062407 | 6.8852803 | 6.2223335 | 7.02837 | -0.806038923 | -0.143092135 |
| Dsg2 | 9.282127 | 8.458464 | 9.2716 | -0.813118766 | 0.01054458 | 7.8744002 | 7.4494039 | 8.25577 | -0.806367495 | -0.381371232 |
| Cpt1a | 10.15429 | 10.11063 | 10.341 | -0.230417263 | -0.186757402 | 10.642215 | 9.8696544 | 10.6771 | -0.807489239 | -0.034928566 |

| | | | | | | | | | | |
|---|---|---|---|---|---|---|---|---|---|---|
| Ncapg2 | 8.334427 | 7.927975 | 8.515 | -0.587064291 | -0.180613062 | 8.1836056 | 7.903344 | 8.71171 | -0.808361801 | -0.5281002 |
| Klhl3 | 7.28309 | 7.19343 | 7.2941 | -0.100623571 | -0.010963337 | 6.8113057 | 6.5324825 | 7.34137 | -0.808888304 | -0.530065083 |
| Nr1d1 | 7.387132 | 6.827809 | 7.6296 | -0.801834523 | -0.242510883 | 6.9540688 | 6.5149599 | 7.32565 | -0.810686437 | -0.371577574 |
| Phyhd1 | 6.607439 | 6.0657 | 6.5282 | -0.462488972 | 0.079250167 | 5.9507167 | 5.5300264 | 6.3408 | -0.810772058 | -0.390081778 |
| Nlrc3 | 10.2471 | 10.18071 | 10.343 | -0.161930705 | -0.095541497 | 10.099861 | 9.6800153 | 10.4908 | -0.810792452 | -0.390946612 |
| Ddhd2 | 8.854479 | 8.689653 | 8.8738 | -0.184116977 | -0.019290744 | 8.9130234 | 8.2227113 | 9.03422 | -0.81150419 | -0.121192082 |
| Ptpn13 | 6.825514 | 6.557105 | 6.8874 | -0.330251325 | -0.061841945 | 6.4906754 | 6.1161837 | 6.9282 | -0.812014862 | -0.437523166 |
| Tmem163 | 7.356182 | 7.269704 | 7.5003 | -0.230548835 | -0.144070681 | 7.6969269 | 6.8599112 | 7.67197 | -0.812058934 | 0.024956716 |
| Rhbdf1 | 6.879967 | 6.707827 | 6.7014 | 0.006426612 | 0.178566871 | 7.1144518 | 6.2742207 | 7.08628 | -0.812062555 | 0.02816848 |
| St6galnac3 | 6.702586 | 6.455689 | 6.6514 | -0.195741632 | 0.051155313 | 6.7668422 | 5.9855819 | 6.7983 | -0.812721283 | -0.031460995 |
| Susd1 | 9.61502 | 9.416504 | 9.7286 | -0.312060774 | -0.113544457 | 10.155792 | 9.0539898 | 9.86684 | -0.812854234 | 0.288948089 |
| Aff4 | 11.52163 | 11.52637 | 11.558 | -0.031691136 | -0.036436918 | 11.123466 | 10.883471 | 11.6967 | -0.813192732 | -0.573197845 |
| Slc36a4 | 7.764287 | 7.540067 | 7.761 | -0.220964384 | 0.003255723 | 7.5989502 | 6.9159073 | 7.72931 | -0.813404631 | -0.130361734 |
| Folr2 | 6.49376 | 5.978089 | 6.3227 | -0.344640298 | 0.171030207 | 5.0362805 | 5.1026775 | 5.91628 | -0.813601624 | -0.879998617 |
| Setbp1 | 8.703349 | 8.676099 | 8.7713 | -0.095239812 | -0.067989187 | 8.6320859 | 8.0262694 | 8.84101 | -0.814741158 | -0.208924702 |
| Rora | 8.189253 | 8.187986 | 8.2725 | -0.084527945 | -0.083261028 | 7.652909 | 7.6422148 | 8.45752 | -0.81530607 | -0.804611852 |
| Ngfr | 6.898321 | 6.687393 | 6.61 | 0.077425683 | 0.28835354 | 6.6467857 | 5.7171962 | 6.53265 | -0.815451287 | 0.114138235 |
| Tmem176b | 10.65909 | 10.47657 | 10.695 | -0.218327686 | -0.035800258 | 10.823705 | 9.7607922 | 10.5769 | -0.816152195 | 0.246760759 |
| Zc3h6 | 8.864332 | 8.428337 | 8.9603 | -0.531979842 | -0.095984685 | 8.8773922 | 8.2641119 | 9.08062 | -0.816505401 | -0.203225049 |
| Cfh | 8.373715 | 7.741194 | 7.8469 | -0.105713742 | 0.526808036 | 7.3496683 | 6.6577892 | 7.47454 | -0.816755007 | -0.124875957 |
| Tnfsf11 | 6.445343 | 5.979188 | 6.1586 | -0.17938105 | 0.286773682 | 5.6873983 | 5.3091766 | 6.12836 | -0.819178506 | -0.440956836 |
| Hebp1 | 6.372663 | 6.11129 | 6.4075 | -0.296212245 | -0.034839077 | 6.0374623 | 5.3408331 | 6.16102 | -0.820186593 | -0.123557316 |
| Mvb12b | 8.191567 | 8.07783 | 8.3492 | -0.271401808 | -0.157664854 | 8.0237505 | 7.3405921 | 8.16108 | -0.820490037 | -0.137331723 |
| Zfp831 | 10.53801 | 10.55123 | 10.639 | -0.088076494 | -0.101297874 | 10.417383 | 9.8164555 | 10.6375 | -0.821058168 | -0.22013034 |
| Shroom4 | 6.727488 | 6.10557 | 6.5 | -0.394417063 | 0.227501439 | 5.6991158 | 5.2882148 | 6.10931 | -0.821092551 | -0.41019152 |
| Mxd4 | 10.54948 | 10.66174 | 10.789 | -0.127126476 | -0.239381606 | 10.352997 | 9.7739742 | 10.5952 | -0.821239893 | -0.242216976 |
| Eef2k | 8.095851 | 8.171228 | 8.3907 | -0.219516533 | -0.29489309 | 8.2848481 | 7.6096433 | 8.43207 | -0.822426714 | -0.147221919 |
| Abca2 | 10.93225 | 10.92728 | 11.198 | -0.27077532 | -0.265807334 | 11.076677 | 10.477143 | 11.3004 | -0.823216938 | -0.223683004 |
| Bcam | 7.955686 | 7.687239 | 7.9539 | -0.266702536 | 0.001744862 | 7.158445 | 6.715902 | 7.53933 | -0.823425976 | -0.380882958 |
| Hpgd | 6.0162 | 5.997814 | 6.0548 | -0.056950814 | -0.03856468 | 5.4071948 | 4.6956937 | 5.51971 | -0.824013326 | -0.11251226 |
| Xcr1 | 7.837825 | 7.544985 | 7.8534 | -0.308429777 | -0.015590263 | 7.4121613 | 6.7061207 | 7.53019 | -0.824070813 | -0.118030227 |
| Adh1 | 6.473845 | 6.302154 | 6.5869 | -0.284708802 | -0.113017424 | 5.6712368 | 5.3292208 | 6.15361 | -0.82439174 | -0.482375728 |

| | | | | | | | | | | |
|---|---|---|---|---|---|---|---|---|---|---|
| Dpy19l3 | 9.120242 | 8.915534 | 8.893 | 0.022566267 | 0.227274026 | 9.2202788 | 8.3554315 | 9.18041 | -0.824982746 | 0.03986455 |
| Adam22 | 5.0318 | 4.952048 | 5.1977 | -0.245685951 | -0.165934447 | 4.6171889 | 4.4761121 | 5.30124 | -0.825128089 | -0.684051208 |
| Rasal2 | 7.302553 | 7.180775 | 7.3452 | -0.16444097 | -0.042663442 | 6.7055715 | 6.3438937 | 7.17088 | -0.826982586 | -0.465304789 |
| Thra | 7.662707 | 7.438187 | 7.8703 | -0.432113317 | -0.207592679 | 7.1646924 | 6.8290857 | 7.65808 | -0.82899021 | -0.493383415 |
| Plcl1 | 6.366322 | 6.463725 | 6.3688 | 0.094956874 | -0.002445777 | 6.4015381 | 5.682996 | 6.512 | -0.829004593 | -0.110462531 |
| Elmsan1 | 11.54933 | 11.47393 | 11.493 | -0.018921394 | 0.056480238 | 11.517376 | 11.061912 | 11.8917 | -0.829820184 | -0.374355778 |
| Caprin2 | 6.915169 | 6.79609 | 7.139 | -0.34290969 | -0.223830705 | 6.8584068 | 6.5984769 | 7.42831 | -0.829835331 | -0.569905424 |
| Nav2 | 6.970186 | 6.686117 | 7.0144 | -0.328241636 | -0.044172541 | 6.602204 | 6.5679028 | 7.39851 | -0.830602456 | -0.796301261 |
| Ecm2 | 6.337396 | 5.643953 | 6.4384 | -0.794487317 | -0.101044537 | 5.2281135 | 5.0888308 | 5.92314 | -0.834310197 | -0.695027526 |
| Ubash3b | 9.028912 | 8.628972 | 8.9593 | -0.33031076 | 0.069629349 | 8.4861179 | 8.1012539 | 8.93694 | -0.835682195 | -0.450818217 |
| Hmmr | 7.127174 | 7.042275 | 7.3886 | -0.346302297 | -0.261403978 | 6.9330952 | 6.8377774 | 7.67352 | -0.835745345 | -0.740427537 |
| Angptl2 | 7.262329 | 6.874924 | 7.2429 | -0.367942768 | 0.019461914 | 6.1995429 | 5.6886455 | 6.52487 | -0.836221291 | -0.325323868 |
| Esam | 8.340049 | 7.979038 | 8.3111 | -0.332105519 | 0.028904866 | 7.5653856 | 7.037052 | 7.87431 | -0.837258813 | -0.308925251 |
| Fbxl7 | 5.693272 | 5.332774 | 5.6877 | -0.354975095 | 0.005522947 | 5.0977447 | 4.4723532 | 5.31125 | -0.838892912 | -0.213501411 |
| Tmem62 | 8.095114 | 8.142265 | 8.2024 | -0.060147504 | -0.107297925 | 8.1822067 | 7.8392058 | 8.67852 | -0.839312342 | -0.496311438 |
| Gprin3 | 7.844811 | 7.839016 | 8.0584 | -0.219381114 | -0.213586178 | 7.6831393 | 7.3141125 | 8.15363 | -0.839516068 | -0.470489259 |
| Bmp2 | 6.108547 | 5.72955 | 5.9566 | -0.227051821 | 0.151945157 | 5.4246043 | 5.028942 | 5.86869 | -0.839743757 | -0.444081487 |
| Deptor | 7.524926 | 7.084908 | 7.4249 | -0.339991241 | 0.10002619 | 7.1001013 | 6.2655056 | 7.1055 | -0.839992469 | -0.005396836 |
| Adam19 | 10.41747 | 10.32362 | 10.525 | -0.201738936 | -0.107891318 | 10.333429 | 9.7778962 | 10.6188 | -0.840858812 | -0.285325745 |
| Afp | 7.019665 | 6.825813 | 7.026 | -0.200165214 | -0.006312638 | 6.9540018 | 6.1379887 | 6.98077 | -0.84277667 | -0.026763635 |
| Phf19 | 7.334838 | 7.149551 | 7.5618 | -0.412204648 | -0.226917516 | 7.172453 | 6.7768065 | 7.61961 | -0.842807263 | -0.447160764 |
| Cit | 7.03829 | 6.971496 | 7.3397 | -0.368218141 | -0.301424574 | 7.3099571 | 6.740641 | 7.58588 | -0.845242332 | -0.275926231 |
| Egr1 | 12.26987 | 11.66237 | 12.11 | -0.447851836 | 0.159654694 | 10.497793 | 10.236842 | 11.0874 | -0.85060387 | -0.589652192 |
| Zbtb10 | 8.969554 | 8.716766 | 9.0247 | -0.307975768 | -0.055187133 | 8.5238669 | 8.2898032 | 9.14377 | -0.853967608 | -0.619903995 |
| Ago4 | 7.081011 | 6.859756 | 7.2787 | -0.418986784 | -0.197731509 | 7.1054287 | 6.3976978 | 7.25262 | -0.854924333 | -0.147193407 |
| Camk1d | 10.3848 | 10.28036 | 10.397 | -0.116763112 | -0.01232895 | 10.199372 | 9.5519129 | 10.4069 | -0.854959768 | -0.207501147 |
| Dock1 | 7.611239 | 7.351782 | 7.5691 | -0.217356998 | 0.042099546 | 7.038585 | 6.3999815 | 7.25665 | -0.856665344 | -0.218061767 |
| Eri2 | 7.626968 | 7.337174 | 7.6361 | -0.298887124 | -0.009092581 | 7.4481603 | 6.8934334 | 7.75247 | -0.859034157 | -0.304307297 |
| Rab11b-ps2 | 6.035546 | 6.166034 | 6.1687 | -0.002640668 | -0.133128996 | 5.8211009 | 5.5604204 | 6.41967 | -0.859246004 | -0.598565478 |
| Thsd7a | 6.278862 | 5.900842 | 6.2063 | -0.305419845 | 0.072600523 | 5.4567278 | 4.9841962 | 5.84352 | -0.859319668 | -0.38678802 |
| Fabp4 | 7.397205 | 7.472508 | 7.4819 | -0.009396077 | -0.084699341 | 6.4699647 | 6.1450509 | 7.00515 | -0.860103502 | -0.535189688 |
| 2610008E11Rik | 8.523264 | 8.559136 | 8.8228 | -0.263649938 | -0.29952184 | 8.6537097 | 7.9892678 | 8.85022 | -0.86095094 | -0.19650906 |

| | | | | | | | | | | |
|---|---|---|---|---|---|---|---|---|---|---|
| S100a9 | 10.30945 | 8.605824 | 7.8436 | 0.762267955 | 2.46589086 | 9.0775172 | 7.5115767 | 8.37279 | -0.861217034 | 0.704723446 |
| Arhgef12 | 9.214392 | 8.7674 | 9.0448 | -0.277430963 | 0.16956024 | 8.4024678 | 8.1667008 | 9.02867 | -0.861971301 | -0.626204266 |
| Igkv4-61 | 5.678327 | 6.135855 | 5.7151 | 0.420780308 | -0.03674714 | 6.7864253 | 6.0162561 | 6.88 | -0.863740629 | -0.093571364 |
| Atp2a3 | 13.06176 | 12.95044 | 13.115 | -0.16406613 | -0.05274575 | 13.403751 | 12.377813 | 13.2419 | -0.86409628 | 0.161841441 |
| Clstn1 | 8.340874 | 8.241827 | 8.3781 | -0.136267103 | -0.037219813 | 8.0683853 | 7.4954677 | 8.36037 | -0.864899578 | -0.291981945 |
| Igf1r | 8.689734 | 8.440768 | 8.6727 | -0.231925516 | 0.017040412 | 8.3504107 | 7.6564123 | 8.5216 | -0.865189904 | -0.171191544 |
| Spock2 | 6.961703 | 6.775697 | 7.1519 | -0.376207023 | -0.190200734 | 6.326513 | 6.0249951 | 6.89042 | -0.865428158 | -0.563910199 |
| Pxylp1 | 7.42964 | 7.406747 | 7.5272 | -0.120478189 | -0.097585127 | 7.1611845 | 6.7548017 | 7.62114 | -0.866341342 | -0.459958553 |
| Knl1 | 7.636601 | 7.282562 | 7.5321 | -0.249547502 | 0.104491499 | 7.4084776 | 7.0103003 | 7.87857 | -0.868265327 | -0.47008805 |
| Gm10182 | 6.293377 | 6.24009 | 6.3321 | -0.091965155 | -0.038678428 | 6.0195707 | 6.1316692 | 7.00072 | -0.869052779 | -0.981151293 |
| Magi1 | 6.18752 | 5.740549 | 6.258 | -0.517421014 | -0.070449777 | 5.4253802 | 5.2161048 | 6.08545 | -0.869343545 | -0.660068118 |
| Ighv1-54 | 6.179787 | 6.46402 | 6.061 | 0.403062993 | 0.118829864 | 6.9544131 | 6.077875 | 6.94764 | -0.86976448 | 0.006773573 |
| Lrmp | 11.34602 | 11.20973 | 11.526 | -0.316706956 | -0.18041529 | 11.70712 | 10.833398 | 11.7043 | -0.870908986 | 0.002812313 |
| Hdac11 | 6.459817 | 6.376827 | 6.6913 | -0.314448039 | -0.23145854 | 6.607958 | 5.8409938 | 6.71264 | -0.871645799 | -0.104681656 |
| Ptprm | 7.223694 | 6.744829 | 7.0912 | -0.34637568 | 0.132489158 | 6.2358245 | 6.0316296 | 6.90568 | -0.874054441 | -0.669859527 |
| Clec1a | 7.129589 | 6.576807 | 7.0571 | -0.480251066 | 0.072531472 | 6.1046885 | 5.7530461 | 6.62782 | -0.874769938 | -0.523127497 |
| Ar | 8.23628 | 7.82564 | 8.3554 | -0.529784158 | -0.11914391 | 7.2030533 | 6.9597469 | 7.8347 | -0.874952611 | -0.631646164 |
| Prr11 | 6.521726 | 6.343876 | 6.7813 | -0.43745984 | -0.259609835 | 6.4917314 | 6.2119033 | 7.08724 | -0.875340564 | -0.595512484 |
| Megf9 | 7.519468 | 7.351219 | 7.5111 | -0.159910522 | 0.00833784 | 7.434251 | 6.5705885 | 7.44827 | -0.87768557 | -0.014023099 |
| Tshz1 | 9.304194 | 9.231488 | 9.4997 | -0.268207285 | -0.195501671 | 9.1881073 | 8.508932 | 9.39223 | -0.883300905 | -0.204125598 |
| Ramp1 | 8.195971 | 8.178731 | 8.3835 | -0.204781774 | -0.187540981 | 8.1778078 | 7.402454 | 8.28678 | -0.884322787 | -0.108968975 |
| Parpbp | 5.661911 | 5.465149 | 5.9081 | -0.442919861 | -0.246157369 | 5.5568446 | 5.1845517 | 6.06964 | -0.885090437 | -0.512797548 |
| Hlf | 5.440566 | 5.330628 | 5.9531 | -0.622486134 | -0.512548152 | 5.0292028 | 4.5800106 | 5.46552 | -0.885506009 | -0.436313788 |
| Mast4 | 9.438711 | 9.349736 | 9.5491 | -0.199391199 | -0.110415855 | 9.4538942 | 8.686962 | 9.57306 | -0.886096674 | -0.119164439 |
| Tek | 7.096058 | 6.708988 | 7.1145 | -0.405549253 | -0.018478426 | 6.3583649 | 5.7511523 | 6.63794 | -0.886783732 | -0.279571128 |
| Abhd15 | 6.631691 | 6.736599 | 6.7598 | -0.023190664 | -0.128098099 | 6.5263308 | 5.7269179 | 6.61383 | -0.886907997 | -0.087495115 |
| Abi2 | 7.8171 | 7.805891 | 7.8398 | -0.03387581 | -0.022667658 | 7.4483471 | 7.0420759 | 7.93031 | -0.888233564 | -0.48196243 |
| Prkcb | 11.903 | 11.87075 | 12.009 | -0.138143288 | -0.105892096 | 12.013313 | 11.093232 | 11.9815 | -0.88825912 | 0.031821754 |
| Rnf150 | 6.702602 | 6.287304 | 6.5733 | -0.28595556 | 0.129342597 | 6.3505208 | 5.6208661 | 6.51149 | -0.890622184 | -0.160967482 |
| Meox2 | 6.253803 | 5.925064 | 6.3736 | -0.448573653 | -0.119834931 | 5.772903 | 5.2107467 | 6.10182 | -0.891073001 | -0.328916653 |
| Cyp1b1 | 9.177288 | 8.696564 | 8.6324 | 0.064144092 | 0.544867808 | 8.2137269 | 7.5780431 | 8.47023 | -0.892182477 | -0.256498604 |
| Rgs16 | 7.659912 | 7.109483 | 7.303 | -0.193519308 | 0.35690944 | 6.4489608 | 6.281578 | 7.17478 | -0.893204679 | -0.725821915 |

| | | | | | | | | | | |
|---|---|---|---|---|---|---|---|---|---|---|
| Erg | 7.928627 | 7.384132 | 7.854 | -0.469897207 | 0.074597602 | 6.7664866 | 6.3213885 | 7.21491 | -0.893523236 | -0.448425188 |
| Arhgap21 | 8.111227 | 8.302618 | 8.2489 | 0.053739045 | -0.137651245 | 7.9642157 | 7.6403487 | 8.53438 | -0.894031442 | -0.570164426 |
| Fam214a | 9.970218 | 9.864657 | 10.156 | -0.291633344 | -0.1860725 | 9.9976806 | 9.2769093 | 10.1746 | -0.8977355 | -0.176964166 |
| Cd24a | 12.05673 | 11.79432 | 12.049 | -0.255034162 | 0.007373664 | 12.23185 | 11.050091 | 11.9488 | -0.898667876 | 0.283091731 |
| Ccna2 | 8.367257 | 8.123373 | 8.5868 | -0.463465247 | -0.219580906 | 8.2877972 | 8.0346346 | 8.93392 | -0.899284004 | -0.646121474 |
| Tm7sf3 | 8.417741 | 8.105146 | 8.3666 | -0.261456459 | 0.051138506 | 8.3578478 | 7.3554161 | 8.2553 | -0.899888796 | 0.102542904 |
| Lifr | 9.363142 | 8.988141 | 9.3191 | -0.330928092 | 0.044072604 | 8.3480895 | 7.6755035 | 8.57566 | -0.900157406 | -0.227571415 |
| Ube2c | 7.015681 | 6.91772 | 7.156 | -0.238285553 | -0.140324449 | 7.0324285 | 6.5300158 | 7.43087 | -0.900857456 | -0.39844475 |
| Mical3 | 8.937328 | 8.821523 | 9.0288 | -0.207242084 | -0.091436994 | 8.6661887 | 8.2209565 | 9.12234 | -0.901380456 | -0.456148272 |
| Cdon | 8.836042 | 8.967112 | 8.9052 | 0.061943243 | -0.069127325 | 8.5465351 | 8.1057014 | 9.0077 | -0.902001109 | -0.46116737 |
| Col15a1 | 9.914406 | 9.670218 | 9.7641 | -0.09386518 | 0.150322178 | 8.9473997 | 7.9997623 | 8.90298 | -0.903222054 | 0.04441534 |
| Golm1 | 9.666103 | 9.561929 | 9.8014 | -0.239460572 | -0.135287021 | 9.2657797 | 8.7967718 | 9.70026 | -0.903491759 | -0.434483824 |
| Meox1 | 8.484976 | 7.896379 | 8.248 | -0.351656773 | 0.236940101 | 7.4109653 | 6.6453179 | 7.54957 | -0.904249341 | -0.138601945 |
| Zmiz1 | 11.25738 | 11.11656 | 11.239 | -0.122741228 | 0.018074694 | 11.037738 | 10.351118 | 11.2557 | -0.904551458 | -0.217931318 |
| Slc9a9 | 8.972076 | 8.61165 | 8.959 | -0.34739083 | 0.013034744 | 8.6108679 | 8.0348715 | 8.93978 | -0.904907106 | -0.328910718 |
| Ceacam1 | 9.675674 | 9.730061 | 9.8028 | -0.072724754 | -0.127110967 | 9.8304049 | 8.8001744 | 9.70642 | -0.906245733 | 0.123984777 |
| Lhfpl2 | 7.197917 | 6.650198 | 7.2632 | -0.613038119 | -0.065319852 | 5.9352686 | 5.4376374 | 6.34474 | -0.907101557 | -0.409470344 |
| Scara5 | 5.993336 | 5.755633 | 6.231 | -0.475366141 | -0.23766272 | 5.3703997 | 5.0023566 | 5.90975 | -0.907394928 | -0.539351846 |
| Eya2 | 7.562382 | 7.350724 | 7.6806 | -0.329827377 | -0.118168753 | 7.2745791 | 6.7934135 | 7.70244 | -0.90902833 | -0.427862794 |
| Rps6ka5 | 9.130415 | 8.9607 | 9.2264 | -0.265726699 | -0.096012128 | 8.8805868 | 8.2903764 | 9.19953 | -0.909149206 | -0.31893887 |
| Itsn1 | 8.502503 | 7.932872 | 8.247 | -0.31415274 | 0.255478989 | 7.8033776 | 7.2280101 | 8.13831 | -0.910298617 | -0.334931138 |
| Pgk1-rs7 | 5.177778 | 4.512046 | 4.4505 | 0.06153902 | 0.727270858 | 4.0346638 | 3.6827023 | 4.59322 | -0.910516416 | -0.558554919 |
| Slco5a1 | 7.782522 | 7.837525 | 8.0395 | -0.201980835 | -0.256983727 | 7.463209 | 7.3888453 | 8.29937 | -0.910525057 | -0.836161381 |
| Arhgap29 | 9.204614 | 8.786263 | 8.8648 | -0.078530121 | 0.339820029 | 8.1305094 | 7.6533368 | 8.56501 | -0.911674769 | -0.434502173 |
| Jag1 | 7.572591 | 7.475087 | 7.4855 | -0.010379114 | 0.087125661 | 6.8771267 | 6.4353212 | 7.3474 | -0.912078148 | -0.470272581 |
| Arhgap18 | 8.615045 | 8.492048 | 8.7911 | -0.29900387 | -0.176007322 | 8.5460792 | 7.7656566 | 8.67885 | -0.913197561 | -0.132774922 |
| Sox4 | 8.40467 | 8.239944 | 8.5877 | -0.347765243 | -0.183038816 | 7.859198 | 7.3165793 | 8.23218 | -0.915602357 | -0.372983623 |
| Mpo | 4.955932 | 4.380179 | 4.5577 | -0.177485369 | 0.398267957 | 4.8734481 | 4.266573 | 5.18525 | -0.918681399 | -0.311806282 |
| Igfbp3 | 8.836448 | 8.640485 | 8.7049 | -0.064383731 | 0.131579522 | 8.6143502 | 7.713006 | 8.63172 | -0.918711004 | -0.017366721 |
| Abcc9 | 7.832 | 7.298145 | 7.501 | -0.202848449 | 0.331005967 | 6.8344417 | 6.0790441 | 6.99901 | -0.919969981 | -0.164572351 |
| Synj2 | 8.677104 | 8.47702 | 8.7953 | -0.318279369 | -0.118195176 | 8.3055778 | 7.8004165 | 8.72083 | -0.92041106 | -0.415249732 |
| Ehbp1 | 5.978902 | 6.104916 | 6.018 | 0.086946509 | -0.039067214 | 5.6896334 | 5.3445505 | 6.26868 | -0.924128277 | -0.579045368 |

| | | | | | | | | | | |
|---|---|---|---|---|---|---|---|---|---|---|
| Add3 | 12.68276 | 12.67948 | 12.815 | -0.135975584 | -0.132696052 | 12.734029 | 11.853887 | 12.7795 | -0.925600066 | -0.045458067 |
| Gm1043 | 7.374385 | 7.240805 | 7.4863 | -0.245462288 | -0.111882685 | 7.354416 | 6.7649884 | 7.69195 | -0.926964532 | -0.337536931 |
| Gad1 | 6.139534 | 5.907885 | 6.1847 | -0.276803567 | -0.045154337 | 6.176837 | 5.4249475 | 6.35238 | -0.927428006 | -0.175538457 |
| Atp1b1 | 9.885589 | 9.852 | 9.9667 | -0.114707079 | -0.081118288 | 9.6659255 | 8.9596442 | 9.88736 | -0.927717611 | -0.221436307 |
| She | 7.09056 | 6.634576 | 7.0765 | -0.441875622 | 0.014108441 | 6.2062944 | 5.9910961 | 6.91964 | -0.928547232 | -0.713348899 |
| Ankrd35 | 5.46765 | 5.30829 | 5.6295 | -0.321204073 | -0.16184354 | 5.2800405 | 5.1008131 | 6.03083 | -0.930019154 | -0.750791773 |
| Ltk | 6.226628 | 6.183499 | 6.3791 | -0.195605265 | -0.152475927 | 6.9791134 | 5.8281397 | 6.75898 | -0.930840765 | 0.22013289 |
| Ccnb2 | 7.260799 | 6.971227 | 7.3862 | -0.415016496 | -0.125443755 | 7.1271352 | 6.7649949 | 7.69724 | -0.932242925 | -0.57010259 |
| Cd40lg | 8.107745 | 8.069542 | 8.2762 | -0.206656859 | -0.168453735 | 7.5597402 | 7.3187723 | 8.25109 | -0.932317846 | -0.691349954 |
| Tspan9 | 8.163567 | 8.140193 | 8.3045 | -0.164307369 | -0.140933532 | 7.7605426 | 7.2924964 | 8.22656 | -0.934059166 | -0.46601291 |
| Mrip-ps | 5.335138 | 5.13704 | 5.4403 | -0.30322688 | -0.10512874 | 5.1505593 | 4.4170873 | 5.35122 | -0.934131132 | -0.20065911 |
| Pik3cg | 10.92529 | 10.89505 | 10.96 | -0.065258964 | -0.035020018 | 11.073048 | 10.051463 | 10.9858 | -0.934348629 | 0.087236508 |
| Fam198b | 5.417072 | 5.515297 | 5.7176 | -0.202315875 | -0.300541321 | 5.1820101 | 4.5673592 | 5.50403 | -0.936671802 | -0.32202093 |
| Synm | 6.224975 | 5.759625 | 6.0751 | -0.315456116 | 0.149894528 | 5.5300232 | 5.0440474 | 5.98108 | -0.937029842 | -0.451054033 |
| Syt15 | 7.499008 | 7.246929 | 7.6287 | -0.381749891 | -0.129671765 | 6.2009408 | 5.8020292 | 6.7392 | -0.937171848 | -0.538260195 |
| Ces2e | 5.867631 | 5.33901 | 5.803 | -0.464009857 | 0.064611124 | 4.9071573 | 4.5285197 | 5.46588 | -0.937360984 | -0.558723368 |
| Shank3 | 8.455783 | 7.700323 | 8.0828 | -0.382480996 | 0.372979315 | 7.1606548 | 6.6503617 | 7.58832 | -0.93795512 | -0.427662052 |
| Spag5 | 7.106374 | 6.768162 | 7.23 | -0.461875671 | -0.123664099 | 6.9348581 | 6.7394428 | 7.68084 | -0.941399099 | -0.745983811 |
| Cav1 | 9.032032 | 8.641614 | 9.154 | -0.512345741 | -0.121927797 | 8.1259251 | 7.5378483 | 8.47949 | -0.941638502 | -0.353561726 |
| Cdc25b | 9.414098 | 9.496761 | 9.6796 | -0.182792337 | -0.26545524 | 9.6155723 | 8.8755488 | 9.8218 | -0.946253972 | -0.206230395 |
| Kif23 | 8.214717 | 8.159727 | 8.3451 | -0.185368495 | -0.130378745 | 7.9514701 | 7.7042095 | 8.65064 | -0.946428544 | -0.699167993 |
| Alcam | 8.684631 | 8.545309 | 8.709 | -0.163672788 | -0.024351269 | 8.6875061 | 7.8449018 | 8.79251 | -0.947603813 | -0.104999525 |
| Slc16a5 | 7.993024 | 7.955335 | 8.0875 | -0.132212749 | -0.094523731 | 7.9337465 | 7.0899754 | 8.03789 | -0.947916169 | -0.10414503 |
| Lyz1 | 9.311097 | 8.880925 | 9.2565 | -0.375538767 | 0.054633304 | 8.1911524 | 7.5274857 | 8.47563 | -0.948147903 | -0.284481159 |
| Eogt | 8.209356 | 7.93778 | 8.2541 | -0.316270495 | -0.044694557 | 8.2248811 | 7.3911717 | 8.33978 | -0.948608137 | -0.11489878 |
| Crat | 8.356815 | 8.299744 | 8.4035 | -0.103727207 | -0.046655928 | 8.830216 | 7.4439659 | 8.39274 | -0.948773808 | 0.437476236 |
| Tmod2 | 6.510349 | 6.403881 | 6.6758 | -0.2718896 | -0.165421223 | 6.0617332 | 5.7800609 | 6.72912 | -0.949056415 | -0.667384083 |
| Gm14303 | 5.47242 | 5.564017 | 6.6462 | -1.082181003 | -1.17377854 | 6.2020767 | 5.553286 | 6.50292 | -0.949638715 | -0.300847969 |
| Prox1 | 6.601848 | 6.401399 | 6.4276 | -0.026238701 | 0.174210591 | 5.7469506 | 5.4257643 | 6.37652 | -0.950756005 | -0.629569657 |
| Dscam | 6.410843 | 6.555416 | 6.798 | -0.242573012 | -0.387146248 | 6.3974733 | 5.7997946 | 6.75411 | -0.954311792 | -0.356633133 |
| Snx22 | 6.418017 | 6.378234 | 6.4585 | -0.080259952 | -0.040477381 | 6.4215158 | 5.6200162 | 6.57543 | -0.955418745 | -0.153919163 |
| Evi5 | 7.971092 | 7.747803 | 7.8138 | -0.066006705 | 0.157282305 | 7.547832 | 6.9931597 | 7.95056 | -0.957401601 | -0.40272923 |

| | | | | | | | | | | |
|---|---|---|---|---|---|---|---|---|---|---|
| Psd3 | 7.864965 | 7.490018 | 7.6968 | -0.206751435 | 0.168196141 | 7.6189108 | 6.8509749 | 7.80896 | -0.957989626 | -0.190053721 |
| Ptpn14 | 7.931348 | 7.86036 | 7.8977 | -0.03734535 | 0.033642046 | 7.4626703 | 6.8699623 | 7.83212 | -0.962155107 | -0.369447148 |
| Ccnb1 | 7.548569 | 7.19748 | 7.7452 | -0.54775919 | -0.19667026 | 7.6575981 | 7.0963612 | 8.05859 | -0.962228164 | -0.400991259 |
| Irs2 | 10.19341 | 9.893671 | 10.136 | -0.242195043 | 0.057548403 | 9.4888951 | 9.0734707 | 10.0359 | -0.962414459 | -0.546989991 |
| Fgd5 | 7.858844 | 7.541608 | 7.8526 | -0.311005701 | 0.006230389 | 6.9875501 | 6.58781 | 7.55237 | -0.964558691 | -0.564818581 |
| Hba-a2 | 9.383591 | 8.861363 | 9.3876 | -0.526226763 | -0.003998419 | 9.1480607 | 8.4300201 | 9.39571 | -0.965687038 | -0.247646432 |
| Dglucy | 8.072224 | 7.980345 | 8.2259 | -0.245542739 | -0.153664129 | 8.2114787 | 7.2185105 | 8.18504 | -0.966529505 | 0.02643878 |
| Flt3 | 9.408226 | 9.240108 | 9.5851 | -0.344975136 | -0.176856822 | 9.2040818 | 8.582234 | 9.5516 | -0.969363047 | -0.347515236 |
| Nhsl1 | 6.040106 | 5.852482 | 6.0636 | -0.211100868 | -0.023476953 | 5.796507 | 5.2428283 | 6.21396 | -0.971135191 | -0.417456437 |
| Depp1 | 7.196961 | 7.253316 | 6.8572 | 0.396161223 | 0.339805745 | 6.8798478 | 6.0346388 | 7.0061 | -0.971456428 | -0.126247457 |
| Sema6d | 8.187456 | 8.041474 | 8.1674 | -0.125942836 | 0.02003865 | 7.3194789 | 7.188219 | 8.15968 | -0.971457796 | -0.840197908 |
| Slco2b1 | 9.371611 | 8.635521 | 9.5427 | -0.907200604 | -0.171110131 | 7.8931112 | 7.3404069 | 8.31281 | -0.972407247 | -0.419702938 |
| Xylt1 | 9.926266 | 10.03496 | 10.05 | -0.014952108 | -0.123644167 | 9.863158 | 9.1449531 | 10.1176 | -0.972658212 | -0.254453347 |
| Bub1b | 8.377466 | 8.069094 | 8.3371 | -0.267964908 | 0.040407216 | 8.37791 | 7.7812033 | 8.76448 | -0.983275948 | -0.386569236 |
| Ackr4 | 4.143106 | 3.966213 | 4.6821 | -0.715873458 | -0.538980355 | 3.8255345 | 3.5738103 | 4.56128 | -0.987470329 | -0.73574612 |
| Rb1 | 10.25554 | 10.08039 | 10.36 | -0.279622767 | -0.104470031 | 10.289072 | 9.4327187 | 10.4204 | -0.987705742 | -0.131352773 |
| Nrp1 | 8.915131 | 8.468337 | 8.9109 | -0.442540341 | 0.00425327 | 8.2401916 | 7.8255728 | 8.81337 | -0.987797875 | -0.573179088 |
| Irf4 | 11.95358 | 11.78032 | 11.975 | -0.19508206 | -0.021822792 | 11.297204 | 10.769832 | 11.7593 | -0.989490312 | -0.462117944 |
| Cd34 | 9.715123 | 9.031939 | 9.5972 | -0.565267983 | 0.117915579 | 8.6416131 | 7.8538543 | 8.84393 | -0.990079098 | -0.202320363 |
| Btnl9 | 5.509028 | 5.27323 | 5.6029 | -0.329639863 | -0.093842704 | 5.148204 | 4.7240853 | 5.71512 | -0.991031558 | -0.566912875 |
| Nhej1 | 6.113839 | 6.271587 | 5.8852 | 0.386368038 | 0.228619088 | 5.6655892 | 5.3684547 | 6.36002 | -0.991566953 | -0.694432448 |
| Tmem176a | 10.20924 | 10.06708 | 10.333 | -0.265849593 | -0.123683161 | 10.426067 | 9.2520323 | 10.2436 | -0.991572823 | 0.182461935 |
| Kif11 | 7.501341 | 7.226857 | 7.6895 | -0.462600274 | -0.188116333 | 7.2813438 | 7.036324 | 8.0291 | -0.992775705 | -0.747755981 |
| Aspm | 7.016581 | 7.017123 | 7.1933 | -0.176140651 | -0.176683113 | 7.037742 | 6.5213703 | 7.51601 | -0.994636518 | -0.47826489 |
| Dennd3 | 10.2239 | 10.1153 | 10.277 | -0.161890793 | -0.053286766 | 10.288224 | 9.3636346 | 10.3589 | -0.995241392 | -0.07065166 |
| Notch3 | 7.621643 | 7.396918 | 7.418 | -0.021086586 | 0.203638388 | 6.9721794 | 6.0492261 | 7.04601 | -0.996787913 | -0.073834655 |
| Kif18b | 6.376472 | 6.160392 | 6.5958 | -0.435405237 | -0.219325623 | 6.2129819 | 5.9471214 | 6.94519 | -0.998072622 | -0.732212059 |
| Aldh6a1 | 8.044412 | 7.713753 | 7.96 | -0.246284441 | 0.084374685 | 7.8643377 | 6.9911762 | 7.98935 | -0.998178311 | -0.125016879 |
| Spns3 | 7.176634 | 7.129387 | 7.2703 | -0.140896032 | -0.093648852 | 7.5510398 | 6.1211278 | 7.11988 | -0.998756812 | 0.431155118 |
| Sgms1 | 10.02752 | 10.11052 | 10.24 | -0.12962086 | -0.212612821 | 10.011368 | 9.1010436 | 10.1005 | -0.999411018 | -0.089086354 |
| Lclat1 | 8.936326 | 8.789263 | 8.9498 | -0.160555036 | -0.013491666 | 8.8000026 | 8.0919609 | 9.09144 | -0.999482378 | -0.291440639 |
| Bmf | 9.073552 | 8.962652 | 9.3698 | -0.407196321 | -0.296297097 | 8.920692 | 8.1451105 | 9.14488 | -0.999766017 | -0.224184479 |

| | | | | | | | | | | |
|---|---|---|---|---|---|---|---|---|---|---|
| Lrrc75b | 6.81763 | 6.562221 | 6.603 | -0.040801995 | 0.214606435 | 6.5286653 | 5.6797785 | 6.67966 | -0.999884016 | -0.150997189 |
| Edaradd | 8.043842 | 7.831706 | 8.036 | -0.204292368 | 0.007843538 | 8.2411002 | 7.2020238 | 8.20315 | -1.001124305 | 0.037952056 |
| Susd2 | 8.73828 | 7.967947 | 8.869 | -0.901082169 | -0.130749642 | 7.8589882 | 7.047615 | 8.05073 | -1.003110363 | -0.191737078 |
| Hmgb3 | 7.603615 | 7.084087 | 7.6369 | -0.552781192 | -0.033253617 | 7.2091884 | 6.6181144 | 7.62303 | -1.004917372 | -0.413843355 |
| Cntnap1 | 5.918797 | 5.594917 | 6.0906 | -0.495686708 | -0.171806568 | 5.5577036 | 5.1085258 | 6.1141 | -1.005572768 | -0.556394944 |
| Snx29 | 9.215009 | 9.077338 | 9.2439 | -0.16659537 | -0.02892421 | 9.3413394 | 8.2820561 | 9.29011 | -1.008050275 | 0.051233029 |
| Gfra2 | 7.47177 | 6.657747 | 7.0423 | -0.384551326 | 0.429471544 | 6.5005657 | 5.5876759 | 6.59595 | -1.008270988 | -0.095381147 |
| Cx3cr1 | 7.344354 | 7.189585 | 7.4311 | -0.241522547 | -0.086752936 | 7.4731107 | 6.2245591 | 7.23379 | -1.009233809 | 0.239317694 |
| H6pd | 9.510107 | 9.349963 | 9.4761 | -0.126184203 | 0.033960495 | 9.3435068 | 8.5069221 | 9.51642 | -1.009493896 | -0.172909197 |
| Creg2 | 6.141444 | 5.510501 | 5.7551 | -0.244593406 | 0.38634982 | 5.378387 | 4.7976227 | 5.80808 | -1.010456712 | -0.429692331 |
| Trim5 | 9.201508 | 8.779611 | 9.3205 | -0.540903682 | -0.119007138 | 9.3960865 | 8.3838137 | 9.3943 | -1.010483071 | 0.001789723 |
| Cpd | 8.98492 | 8.395923 | 8.96 | -0.564113939 | 0.024883183 | 8.395883 | 7.7347305 | 8.74657 | -1.011838914 | -0.350686419 |
| Tet1 | 7.714499 | 7.726115 | 8.0057 | -0.279619616 | -0.291235686 | 7.6103995 | 7.0577559 | 8.0714 | -1.013645878 | -0.461002304 |
| Crisp3 | 7.838277 | 7.939592 | 8.8534 | -0.913782627 | -1.015097916 | 9.4135423 | 7.4881175 | 8.50179 | -1.013677215 | 0.911747575 |
| Casc4 | 8.279417 | 7.901554 | 8.2124 | -0.310881938 | 0.066980492 | 8.0637043 | 7.3111346 | 8.32696 | -1.015825109 | -0.263255406 |
| Myh11 | 7.48829 | 7.492645 | 7.7624 | -0.269785241 | -0.274140499 | 7.1125806 | 6.4399909 | 7.45657 | -1.016575651 | -0.343985899 |
| Pip5k1b | 6.768743 | 6.603569 | 6.814 | -0.210463715 | -0.045289443 | 6.7553605 | 5.8446145 | 6.86164 | -1.017028089 | -0.10628203 |
| Lrrc8b | 8.522258 | 8.245807 | 8.5619 | -0.31612242 | -0.039671434 | 8.1879596 | 7.4941327 | 8.51363 | -1.019497644 | -0.325670758 |
| Pcdh12 | 6.192091 | 5.88563 | 6.1194 | -0.233817096 | 0.072643851 | 5.5060162 | 4.7763421 | 5.79673 | -1.020383881 | -0.290709714 |
| Sned1 | 7.876346 | 7.83497 | 7.9178 | -0.08282613 | -0.041449661 | 6.9019255 | 6.8625012 | 7.883 | -1.020502801 | -0.981078481 |
| Serpina1b | 8.115134 | 7.194588 | 8.0996 | -0.904963511 | 0.015583371 | 6.5564289 | 6.1901647 | 7.21082 | -1.020652649 | -0.654388543 |
| Plekhg3 | 8.556313 | 8.375557 | 8.5491 | -0.173497484 | 0.007259057 | 8.5912498 | 7.6594142 | 8.68229 | -1.022879037 | -0.091043462 |
| Map2k6 | 7.182754 | 7.310453 | 7.5071 | -0.196641591 | -0.324340889 | 7.2856977 | 6.3273511 | 7.3518 | -1.024449475 | -0.066102868 |
| Galnt15 | 7.440926 | 6.877082 | 7.5897 | -0.712643551 | -0.148799757 | 6.3080415 | 5.9319962 | 6.9566 | -1.024602526 | -0.648557308 |
| 37865 | 7.205088 | 6.291633 | 6.4759 | -0.184291686 | 0.72916363 | 6.6988857 | 5.7119018 | 6.744 | -1.032094865 | -0.045110972 |
| Tgfbr3 | 9.628713 | 9.340467 | 9.6152 | -0.274766627 | 0.01348004 | 8.736743 | 8.3472719 | 9.37955 | -1.032273144 | -0.642802044 |
| Acacb | 7.133457 | 6.114876 | 7.1847 | -1.069798756 | -0.051218392 | 6.2168511 | 5.540246 | 6.57323 | -1.032986415 | -0.356381288 |
| Tbc1d16 | 7.332907 | 7.180761 | 7.4491 | -0.268341527 | -0.11619608 | 7.1111196 | 6.2013279 | 7.23488 | -1.033552535 | -0.123760807 |
| Col6a5 | 5.919542 | 5.301896 | 5.7709 | -0.468979431 | 0.148666027 | 4.3686846 | 4.1127873 | 5.14704 | -1.034252304 | -0.778355065 |
| Cpa3 | 7.414323 | 7.405085 | 7.3325 | 0.072569604 | 0.081807004 | 6.5721851 | 6.0620467 | 7.09772 | -1.035672765 | -0.525534393 |
| Ptprb | 8.612876 | 8.186276 | 8.553 | -0.366688213 | 0.059911712 | 7.7509072 | 7.3253088 | 8.36153 | -1.036222572 | -0.610624201 |
| Parp16 | 7.09753 | 7.010375 | 7.2185 | -0.208097036 | -0.120942418 | 7.0964825 | 6.1733361 | 7.20958 | -1.036243222 | -0.113096907 |

| | | | | | | | | | | |
|---|---|---|---|---|---|---|---|---|---|---|
| Nedd4 | 11.04859 | 10.78155 | 11.054 | -0.272465356 | -0.005423692 | 10.43967 | 9.7850913 | 10.822 | -1.036877969 | -0.38229932 |
| Hbb-bt | 7.434882 | 6.871312 | 7.5289 | -0.657614907 | -0.094044734 | 7.1946129 | 6.5340627 | 7.57342 | -1.039356703 | -0.378806484 |
| Cecr2 | 8.257337 | 7.937808 | 8.2322 | -0.294409602 | 0.025118552 | 8.1835841 | 7.2820424 | 8.32262 | -1.040573452 | -0.139031773 |
| Gas6 | 8.100652 | 7.310408 | 7.6844 | -0.373990208 | 0.416253704 | 7.0402583 | 6.2574913 | 7.29891 | -1.041422379 | -0.258655372 |
| Cplane1 | 8.815854 | 8.972137 | 9.0406 | -0.068475611 | -0.224759073 | 8.9590822 | 8.4386252 | 9.4801 | -1.041475522 | -0.521018543 |
| Ushbp1 | 7.67087 | 6.993215 | 7.7858 | -0.792548144 | -0.114893322 | 6.3860618 | 5.955185 | 6.99807 | -1.04288289 | -0.612006119 |
| Ampd3 | 8.403142 | 8.143815 | 8.3349 | -0.191094927 | 0.068231258 | 8.3695876 | 7.2647843 | 8.31336 | -1.048570899 | 0.056232332 |
| Gm8221 | 7.751719 | 7.785469 | 7.8731 | -0.087630598 | -0.121380359 | 6.7566079 | 6.8192677 | 7.86925 | -1.049982613 | -1.112642398 |
| Itih5 | 9.037248 | 8.583584 | 8.979 | -0.395451065 | 0.058212559 | 8.6326184 | 7.6939668 | 8.74891 | -1.054942265 | -0.116290623 |
| Timp2 | 10.04275 | 9.75662 | 10.031 | -0.274828283 | 0.011302966 | 9.3003354 | 8.6303247 | 9.68841 | -1.058084995 | -0.388074344 |
| Mmrn2 | 9.338197 | 8.769685 | 9.199 | -0.429323274 | 0.139188784 | 8.3855825 | 7.5531973 | 8.61386 | -1.060663581 | -0.228278346 |
| Gm49369 | 6.015572 | 5.863753 | 6.1576 | -0.293833535 | -0.142014359 | 5.3846873 | 5.13516 | 6.19814 | -1.062982171 | -0.813454889 |
| Hbb-bs | 10.20151 | 9.706058 | 10.292 | -0.586312697 | -0.090857722 | 9.9530159 | 9.1989531 | 10.2649 | -1.065904319 | -0.311841512 |
| Apol7c | 10.9953 | 11.03977 | 11.27 | -0.229978772 | -0.27444374 | 10.193015 | 10.256446 | 11.3224 | -1.065966375 | -1.129397513 |
| Fxyd6 | 5.539748 | 5.36402 | 5.787 | -0.423027868 | -0.247300375 | 4.6226333 | 4.6361601 | 5.7078 | -1.071639276 | -1.085166077 |
| Adcy5 | 7.131993 | 6.731684 | 7.1198 | -0.388156088 | 0.012153515 | 6.6062659 | 5.2824838 | 6.35779 | -1.075304638 | 0.248477515 |
| Crmp1 | 6.680749 | 6.363474 | 6.7723 | -0.408821166 | -0.091545657 | 6.3030856 | 5.8918703 | 6.96744 | -1.075573872 | -0.664358573 |
| Cd209e | 6.761639 | 4.885922 | 5.5868 | -0.700832129 | 1.174884199 | 4.4568254 | 4.0600539 | 5.13766 | -1.077607853 | -0.680836336 |
| Fam171a1 | 6.817628 | 6.433371 | 7.0353 | -0.601969451 | -0.21771198 | 5.7990554 | 5.1239731 | 6.20202 | -1.07804678 | -0.402964462 |
| Rxra | 9.102891 | 8.698394 | 9.1238 | -0.425365986 | -0.020868928 | 8.4685412 | 7.7602057 | 8.84563 | -1.08542885 | -0.37709334 |
| Gm45234 | 5.082862 | 4.748758 | 5.6495 | -0.900787167 | -0.566682978 | 5.2196104 | 4.6252851 | 5.71097 | -1.085681689 | -0.491356408 |
| Sgsh | 9.894594 | 9.763047 | 9.9974 | -0.234331371 | -0.10278484 | 10.000886 | 8.8273623 | 9.91516 | -1.087799057 | 0.085724246 |
| Arnt2 | 6.084953 | 6.086586 | 6.2461 | -0.159492079 | -0.161125228 | 6.225708 | 5.5433171 | 6.63393 | -1.090616959 | -0.408226052 |
| Arrb1 | 10.5091 | 10.32003 | 10.71 | -0.389659636 | -0.200581888 | 10.18688 | 9.3959636 | 10.4873 | -1.091368583 | -0.300452524 |
| Cd36 | 8.512335 | 8.435919 | 8.6103 | -0.174411788 | -0.097995681 | 8.3327791 | 7.4978352 | 8.59081 | -1.092978834 | -0.258034944 |
| Ahnak | 12.5183 | 12.23175 | 12.498 | -0.266114474 | 0.020439592 | 11.491826 | 11.336805 | 12.431 | -1.094190226 | -0.939168524 |
| Fut7 | 8.131552 | 7.151401 | 8.1164 | -0.964992893 | 0.015157661 | 6.5607111 | 6.0848032 | 7.17966 | -1.094858444 | -0.618950538 |
| Hfe | 7.230261 | 6.612831 | 7.2624 | -0.649531827 | -0.032101683 | 6.9533596 | 5.9106639 | 7.00715 | -1.09648204 | -0.053786334 |
| Tie1 | 8.842431 | 8.307729 | 8.7979 | -0.490208357 | 0.044494039 | 7.8658006 | 7.0023018 | 8.09909 | -1.096783752 | -0.23328494 |
| Trf | 9.468953 | 8.788368 | 9.1317 | -0.343323927 | 0.337261078 | 7.8897314 | 7.5343812 | 8.63199 | -1.097606636 | -0.742256431 |
| Klf7 | 9.803862 | 9.593206 | 9.7058 | -0.112604047 | 0.098052519 | 9.6702978 | 8.8253311 | 9.92452 | -1.099188209 | -0.254221467 |
| L1cam | 8.503253 | 8.401059 | 8.7473 | -0.346253438 | -0.244059752 | 8.6714942 | 7.8913887 | 8.99453 | -1.103142033 | -0.32303657 |

| | | | | | | | | | | |
|---|---|---|---|---|---|---|---|---|---|---|
| Nr4a2 | 7.107215 | 6.67123 | 7.0248 | -0.35361842 | 0.082366688 | 6.8207702 | 6.4369564 | 7.54186 | -1.104903883 | -0.721090039 |
| Igf1 | 7.078246 | 6.552904 | 6.9707 | -0.417782304 | 0.107558949 | 5.9206074 | 5.5280559 | 6.63442 | -1.106368636 | -0.713817161 |
| C6 | 5.681951 | 4.775869 | 5.4072 | -0.631317261 | 0.274764761 | 4.5615925 | 3.9904628 | 5.09687 | -1.10640799 | -0.535278339 |
| Cpe | 8.55583 | 8.062511 | 8.2972 | -0.234640628 | 0.258678847 | 8.2551773 | 7.3084442 | 8.42044 | -1.111998389 | -0.165265319 |
| Ltbp4 | 8.736816 | 8.451718 | 8.7811 | -0.32933429 | -0.044235491 | 8.2764162 | 7.1946071 | 8.30852 | -1.11391308 | -0.032103934 |
| Zdhhc2 | 7.113981 | 6.835069 | 7.0623 | -0.227233279 | 0.051678596 | 6.8416597 | 6.2274913 | 7.34231 | -1.114821037 | -0.500652603 |
| Mmp15 | 7.57715 | 7.283816 | 7.7523 | -0.468450387 | -0.175116396 | 7.0877602 | 6.0917345 | 7.20922 | -1.117488161 | -0.121462475 |
| Chp2 | 5.578807 | 4.907692 | 5.9063 | -0.998560665 | -0.327445611 | 4.4526586 | 4.0803464 | 5.20084 | -1.120498431 | -0.748186258 |
| Cdh2 | 6.908936 | 6.709063 | 7.0856 | -0.376493452 | -0.176620033 | 6.3940983 | 5.7772093 | 6.89837 | -1.121158591 | -0.504269592 |
| Adam11 | 7.831285 | 7.880617 | 8.1334 | -0.252772308 | -0.302103367 | 8.0067359 | 7.4811018 | 8.60958 | -1.128475252 | -0.602841108 |
| St8sia6 | 9.632854 | 9.722432 | 9.8294 | -0.106947655 | -0.196525415 | 9.0420308 | 8.4786928 | 9.60774 | -1.129045308 | -0.565707343 |
| Bbs9 | 8.694339 | 8.572261 | 8.8341 | -0.261845024 | -0.139767179 | 8.7932909 | 7.7353312 | 8.8681 | -1.132770785 | -0.07481114 |
| Cpeb2 | 8.37583 | 8.135973 | 8.4291 | -0.293168274 | -0.053311515 | 7.7071105 | 7.1235153 | 8.26042 | -1.136901352 | -0.553306164 |
| Arhgap19 | 7.887306 | 7.559996 | 7.9961 | -0.436058655 | -0.108748157 | 7.5207903 | 6.9156251 | 8.05403 | -1.138403398 | -0.53323821 |
| Sap18 | 8.835421 | 8.377068 | 9.0109 | -0.633877137 | -0.175524358 | 8.6880919 | 7.8902249 | 9.0311 | -1.140878461 | -0.343011464 |
| Stac2 | 8.190961 | 8.123591 | 8.2109 | -0.087283105 | -0.019912777 | 8.3529527 | 7.040073 | 8.19344 | -1.153362063 | 0.159517643 |
| Slc2a3 | 10.02769 | 9.814412 | 10.11 | -0.296029877 | -0.082748962 | 9.9720191 | 8.9348233 | 10.0891 | -1.154315916 | -0.117120072 |
| Abca1 | 11.92772 | 11.5356 | 11.849 | -0.31371897 | 0.078398584 | 11.3348 | 10.788247 | 11.9486 | -1.160392938 | -0.613839167 |
| Mgl2 | 6.447231 | 5.499705 | 6.043 | -0.543286938 | 0.404239946 | 5.7366617 | 4.8915416 | 6.05456 | -1.16302011 | -0.317900092 |
| Sytl2 | 8.383597 | 8.188842 | 8.5558 | -0.366959595 | -0.172204414 | 7.9058937 | 7.4541085 | 8.62167 | -1.167556748 | -0.71577155 |
| Gm2223 | 6.828271 | 6.430408 | 6.4861 | -0.055736822 | 0.342126533 | 5.985819 | 5.5113879 | 6.68247 | -1.17108266 | -0.696651557 |
| Cd300ld3 | 6.931642 | 6.327248 | 7.0117 | -0.684415246 | -0.080021197 | 6.3565335 | 5.2150538 | 6.38706 | -1.172002978 | -0.030523373 |
| Ryr2 | 6.665039 | 6.397993 | 6.4958 | -0.097775362 | 0.169270732 | 6.4273576 | 5.8388476 | 7.01782 | -1.17896785 | -0.590457852 |
| Nr4a1 | 12.19678 | 12.0992 | 12.139 | -0.039309376 | 0.058267358 | 11.596113 | 10.924096 | 12.1052 | -1.181093574 | -0.509075938 |
| Slc26a11 | 8.389089 | 8.315701 | 8.5462 | -0.230472526 | -0.157084231 | 8.4604755 | 7.4584763 | 8.64017 | -1.181693446 | -0.179694192 |
| Shank1 | 7.539914 | 7.462568 | 7.3464 | 0.116206774 | 0.193552666 | 7.9294695 | 6.9329994 | 8.11613 | -1.18312752 | -0.186657334 |
| Cd93 | 9.35159 | 8.942133 | 9.1433 | -0.20113468 | 0.208322717 | 8.9571921 | 7.7289964 | 8.91985 | -1.190848851 | 0.037346889 |
| Fbln5 | 6.580008 | 6.168132 | 6.8362 | -0.668116105 | -0.256240646 | 5.8732615 | 5.419592 | 6.6131 | -1.193505117 | -0.739835674 |
| Dclk1 | 8.822568 | 8.532799 | 8.6613 | -0.128529941 | 0.161239346 | 7.6687916 | 6.70799 | 7.90549 | -1.197496596 | -0.236694996 |
| Scube3 | 5.817583 | 5.782798 | 5.8846 | -0.101806565 | -0.06702093 | 5.4156681 | 5.065741 | 6.26547 | -1.199727232 | -0.84980021 |
| Spns2 | 9.051466 | 8.751821 | 9.0695 | -0.317713631 | -0.018068893 | 8.677017 | 7.5411257 | 8.74139 | -1.200260332 | -0.064369074 |
| Cenpf | 7.746145 | 7.299979 | 7.8026 | -0.502600208 | -0.056434158 | 7.2807747 | 6.9617564 | 8.163 | -1.201246115 | -0.88222788 |

| | | | | | | | | | | |
|---|---|---|---|---|---|---|---|---|---|---|
| Prelp | 6.71324 | 6.105885 | 6.688 | -0.582098017 | 0.02525688 | 5.9815186 | 5.1034852 | 6.3073 | -1.203816796 | -0.325783436 |
| Syne1 | 10.43241 | 10.43962 | 10.506 | -0.06633645 | -0.073548572 | 10.080231 | 9.4852414 | 10.6952 | -1.209998905 | -0.615009726 |
| Rpl10a-ps1 | 11.25459 | 12.68849 | 12.668 | 0.020152234 | -1.413750841 | 11.450411 | 11.406272 | 12.6346 | -1.228325663 | -1.184186764 |
| Sult1a1 | 7.9291 | 6.802135 | 8.0796 | -1.277424528 | -0.150460151 | 6.0494733 | 5.6608352 | 6.89007 | -1.229239225 | -0.840601162 |
| E2f2 | 9.391881 | 9.230687 | 9.4655 | -0.234765448 | -0.073571377 | 9.5831741 | 8.3537615 | 9.58843 | -1.234671325 | -0.005258704 |
| Pgap1 | 10.45627 | 10.36364 | 10.57 | -0.20657366 | -0.113946082 | 10.418778 | 9.2620505 | 10.4976 | -1.235553796 | -0.078826556 |
| Selenop | 12.7694 | 12.03048 | 12.688 | -0.65779735 | 0.081122342 | 11.653362 | 10.950303 | 12.1872 | -1.23692454 | -0.533864991 |
| Serpina1e | 9.129853 | 8.320456 | 9.2332 | -0.912779395 | -0.103383341 | 7.4144434 | 6.9805378 | 8.21957 | -1.239032389 | -0.805126762 |
| Epas1 | 10.47764 | 9.768385 | 10.31 | -0.541879379 | 0.167371674 | 9.2002654 | 8.4491134 | 9.68973 | -1.240614981 | -0.489462953 |
| Cd300ld4 | 6.834554 | 5.994394 | 6.803 | -0.808588193 | 0.0315725 | 5.0874544 | 4.6802329 | 5.92186 | -1.241630013 | -0.834408497 |
| Cpm | 9.341149 | 9.182611 | 9.5001 | -0.317506315 | -0.158968307 | 9.3491797 | 8.1944848 | 9.43623 | -1.24174357 | -0.087048687 |
| Tjp1 | 8.216515 | 7.75362 | 8.2952 | -0.541591518 | -0.078696861 | 6.9786215 | 6.40356 | 7.65403 | -1.250468783 | -0.675407265 |
| Antxr2 | 9.5028 | 9.25074 | 9.52 | -0.269223613 | -0.017163572 | 9.3479906 | 8.3398756 | 9.59211 | -1.252237028 | -0.244122026 |
| Jam3 | 8.131768 | 7.394105 | 8.1497 | -0.75561795 | -0.017955684 | 6.7315159 | 6.3472572 | 7.60815 | -1.260890803 | -0.876632173 |
| Hmgcs2 | 6.248163 | 6.467772 | 6.5157 | -0.047903253 | -0.267512149 | 6.3404678 | 5.1108022 | 6.38081 | -1.270009479 | -0.040343883 |
| Kitl | 8.646929 | 8.151701 | 8.5546 | -0.4028672 | 0.092360462 | 7.2114312 | 6.7500187 | 8.02206 | -1.272046021 | -0.810633535 |
| Il9r | 9.387329 | 9.389302 | 9.6551 | -0.265772749 | -0.267745758 | 9.9091835 | 8.8411334 | 10.1162 | -1.275105582 | -0.20705547 |
| H2-M2 | 9.461627 | 9.341593 | 9.5326 | -0.190967521 | -0.070933453 | 8.4868648 | 8.3574302 | 9.6337 | -1.276273012 | -1.146838369 |
| Kansl2-ps | 4.452194 | 4.407852 | 3.8286 | 0.579228485 | 0.62357024 | 3.7105532 | 3.3965146 | 4.67375 | -1.277231294 | -0.96319268 |
| Kctd12 | 11.87312 | 11.48391 | 11.9 | -0.415839052 | -0.026621192 | 11.307443 | 10.203187 | 11.4867 | -1.283559502 | -0.179303105 |
| Hdac9 | 9.043129 | 8.795776 | 9.1838 | -0.388072811 | -0.140720676 | 9.1289586 | 8.0055296 | 9.28979 | -1.284256295 | -0.160827298 |
| Cyp4b1 | 6.807638 | 6.088876 | 6.6454 | -0.556493605 | 0.162267825 | 5.9258872 | 5.3027674 | 6.59876 | -1.295987726 | -0.672867896 |
| Abca9 | 6.377601 | 5.513055 | 6.4105 | -0.897470501 | -0.032924392 | 5.8085425 | 4.8641546 | 6.17015 | -1.305995674 | -0.361607794 |
| Pcdh17 | 8.562725 | 8.024523 | 8.6531 | -0.628544509 | -0.090342562 | 7.0508256 | 6.4446446 | 7.76084 | -1.316191277 | -0.710010294 |
| Mki67 | 10.60074 | 10.22666 | 10.748 | -0.521535461 | -0.147458514 | 10.226645 | 9.9129287 | 11.2427 | -1.329813402 | -1.016096921 |
| Adgrl4 | 8.259614 | 7.900627 | 8.2429 | -0.342277112 | 0.016709909 | 7.1787965 | 6.3803517 | 7.71213 | -1.331781941 | -0.533337161 |
| Ltbp2 | 9.67862 | 8.795843 | 9.5247 | -0.728837769 | 0.153939551 | 8.0713469 | 7.4193372 | 8.75557 | -1.336237716 | -0.684228003 |
| Marco | 10.14733 | 9.737026 | 9.5869 | 0.150086921 | 0.56039407 | 8.3367174 | 7.7828421 | 9.12289 | -1.340051934 | -0.786176692 |
| Itgae | 9.961941 | 9.933325 | 10.091 | -0.157749982 | -0.129133878 | 9.6615358 | 8.7711463 | 10.1116 | -1.340501882 | -0.450112349 |
| Pxdc1 | 10.30953 | 10.0398 | 10.217 | -0.176719759 | 0.093012986 | 10.705497 | 8.8897872 | 10.2459 | -1.356116909 | 0.45959317 |
| Cd300ld5 | 7.122109 | 6.048568 | 6.5837 | -0.535102061 | 0.538438883 | 4.6795464 | 4.4999659 | 5.87621 | -1.376246801 | -1.19666633 |
| Enpp2 | 11.76192 | 10.93372 | 11.785 | -0.851375126 | -0.023177884 | 10.114545 | 9.3753147 | 10.7675 | -1.392175865 | -0.652945929 |

| | | | | | | | | | | |
|---|---|---|---|---|---|---|---|---|---|---|
| Ifi27 | 9.819412 | 9.679 | 9.8155 | -0.136490216 | 0.003922214 | 10.172305 | 8.641434 | 10.0354 | -1.393954627 | 0.136916553 |
| Stab2 | 8.121038 | 7.352115 | 7.6091 | -0.257031439 | 0.511891024 | 6.7411877 | 5.931754 | 7.33881 | -1.407052133 | -0.597618417 |
| Sh3pxd2a | 10.7458 | 10.52766 | 10.773 | -0.245001998 | -0.026865992 | 10.622883 | 9.4141619 | 10.823 | -1.408881859 | -0.200161013 |
| Klhl14 | 7.163232 | 7.116631 | 7.3867 | -0.270030523 | -0.223429772 | 7.2777128 | 5.9540585 | 7.38447 | -1.430406651 | -0.106752356 |
| Cd55 | 12.21529 | 11.95182 | 12.313 | -0.361532126 | -0.098060622 | 12.357947 | 10.934305 | 12.3659 | -1.43159319 | -0.007951113 |
| Cxcl12 | 10.49491 | 9.768501 | 10.152 | -0.38377089 | 0.342638491 | 9.5861137 | 8.2310536 | 9.68067 | -1.449615807 | -0.094555768 |
| Fosb | 11.06765 | 10.45673 | 10.967 | -0.510714422 | 0.100202664 | 9.5232488 | 8.6530714 | 10.107 | -1.453930014 | -0.583752577 |
| Cd209a | 7.858611 | 7.099564 | 7.3312 | -0.231649047 | 0.527398533 | 6.8115212 | 5.675512 | 7.14749 | -1.471981947 | -0.335972815 |
| Trim7 | 10.0628 | 9.904409 | 10.099 | -0.194330377 | -0.035935335 | 10.412912 | 8.9644529 | 10.4535 | -1.489087879 | -0.040628595 |
| Adam23 | 9.334968 | 9.193938 | 9.4278 | -0.233860287 | -0.092830532 | 8.8822117 | 8.0680279 | 9.56081 | -1.492784543 | -0.678600757 |
| Sele | 8.130573 | 7.391393 | 7.3022 | 0.089205677 | 0.828385956 | 6.4718869 | 5.532272 | 7.04235 | -1.510078235 | -0.570463315 |
| Postn | 7.824335 | 7.561429 | 7.8341 | -0.272702629 | -0.009796996 | 6.8719906 | 6.0726286 | 7.59138 | -1.518756122 | -0.719394132 |
| F8 | 9.246659 | 8.388893 | 8.9539 | -0.564957041 | 0.292808882 | 7.2456963 | 6.9947182 | 8.51674 | -1.52202436 | -1.271046248 |
| Ces2g | 7.804442 | 7.147468 | 7.9083 | -0.760843182 | -0.103869446 | 6.9301075 | 5.6606572 | 7.20539 | -1.54473351 | -0.27528323 |
| Sema5a | 8.047589 | 7.608886 | 8.1029 | -0.494001935 | -0.055298913 | 7.8520931 | 6.4901508 | 8.05133 | -1.561181417 | -0.199239189 |
| Ccr6 | 9.828348 | 9.603289 | 9.8242 | -0.220905246 | 0.004154247 | 10.05829 | 8.2631622 | 9.84201 | -1.578847795 | 0.216280514 |
| Clec14a | 7.946348 | 7.289115 | 8.2243 | -0.935211443 | -0.27797778 | 6.8382458 | 5.8100647 | 7.39293 | -1.582864406 | -0.554683316 |
| Inmt | 5.893502 | 4.882475 | 6.4763 | -1.593858517 | -0.582831544 | 4.270191 | 4.0767149 | 5.66997 | -1.593258344 | -1.39978224 |
| Aqp1 | 10.7963 | 10.43513 | 10.813 | -0.378081851 | -0.01690327 | 9.9594648 | 8.4562405 | 10.055 | -1.598761675 | -0.095537354 |
| Olfml2a | 7.662784 | 7.218499 | 8.0344 | -0.815909244 | -0.371624559 | 6.7662652 | 5.5954387 | 7.19561 | -1.600173288 | -0.429346806 |
| Cd209d | 7.580597 | 6.146743 | 7.2519 | -1.105138218 | 0.32871598 | 5.691241 | 5.0052316 | 6.6073 | -1.602069546 | -0.916060134 |
| Mrc1 | 8.796611 | 7.739781 | 8.0694 | -0.329638095 | 0.727192046 | 7.0839224 | 5.7699838 | 7.41376 | -1.643772747 | -0.329834227 |
| Rps13-ps2 | 9.975725 | 10.43638 | 9.3446 | 1.091804226 | 0.631146718 | 10.439884 | 9.7044237 | 11.3508 | -1.646414949 | -0.910954579 |
| Gm10925 | 12.27748 | 12.36163 | 13.315 | -0.95298292 | -1.037124145 | 11.041849 | 11.673921 | 13.3548 | -1.680903555 | -2.312975656 |
| Cxcl1 | 9.329473 | 8.214999 | 8.9628 | -0.747812804 | 0.366661168 | 7.2905201 | 6.5438134 | 8.25941 | -1.715595565 | -0.968888859 |
| Timd4 | 9.739617 | 7.756181 | 9.4395 | -1.683308409 | 0.300127229 | 7.1943101 | 6.7506143 | 8.51004 | -1.759427994 | -1.315732188 |
| Dusp4 | 9.016038 | 8.383805 | 8.93 | -0.546156951 | 0.086075822 | 8.7640091 | 7.2947086 | 9.05668 | -1.761967667 | -0.292667184 |
| Glycam1 | 15.47465 | 14.52045 | 15.536 | -1.015783516 | -0.061580736 | 13.356268 | 12.363636 | 14.1337 | -1.770109544 | -0.777477953 |
| Gm28437 | 11.2831 | 11.09951 | 12.215 | -1.115923906 | -0.932334407 | 10.189628 | 10.507379 | 12.3194 | -1.812008244 | -2.129759609 |
| C130074G19Rik | 8.803886 | 8.113578 | 9.3051 | -1.191513456 | -0.501205189 | 7.4970284 | 6.5983412 | 8.47437 | -1.876031951 | -0.977344735 |
| Il22ra2 | 7.71593 | 6.620648 | 7.5921 | -0.971497082 | 0.123784731 | 5.8440823 | 4.7541235 | 6.692 | -1.937872789 | -0.847913995 |
| Sparcl1 | 8.541081 | 7.98311 | 8.6493 | -0.666209085 | -0.108237934 | 7.1026835 | 6.3391028 | 8.30323 | -1.964129984 | -1.200549245 |

| | | | | | | | | | | |
|---|---|---|---|---|---|---|---|---|---|---|
| Ccl21a | 12.35612 | 11.46362 | 12.429 | -0.965794588 | -0.073289936 | 10.735363 | 9.3328743 | 11.3267 | -1.993856264 | -0.59136808 |
| Fcer2a | 12.81319 | 12.60314 | 12.987 | -0.383561458 | -0.173518628 | 13.455853 | 11.119305 | 13.188 | -2.068654194 | 0.267893602 |
| Rps12l1 | 7.392482 | 3.913346 | 6.6705 | -2.757163935 | 0.721971744 | 5.4610536 | 4.9088085 | 7.04529 | -2.136484111 | -1.584239105 |
| Cd300lg | 8.983935 | 8.007884 | 9.2618 | -1.253874087 | -0.27782329 | 7.3529984 | 6.070053 | 8.39552 | -2.325467929 | -1.042522487 |
| Gm28661 | 9.683229 | 7.996176 | 9.6887 | -1.692513699 | -0.005460614 | 8.1739086 | 7.7260729 | 10.2509 | -2.524816585 | -2.076980867 |
| Cd209b | 10.16226 | 7.984128 | 9.8071 | -1.822970188 | 0.35516603 | 7.6627516 | 5.7015854 | 9.15168 | -3.4500962 | -1.488929971 |
